# HybridMine: pipeline for allele inheritance and gene copy number prediction in industrial yeast hybrids

**DOI:** 10.1101/2020.05.05.079186

**Authors:** Soukaina Timouma, Jean-Marc Schwartz, Daniela Delneri

**Author notes:** To whom correspondence should be addressed: Daniela Delneri,; correspondence may also be addressed to Soukaina Timouma:; and Jean-Marc Schwartz.

## Abstract

*Saccharomyces pastorianus* is an allopolyploid sterile yeast hybrid used in brewing to produce lager-style beers. The development of new yeast strains with valuable industrial traits such as improved maltose utilization or balanced flavour profiles are now a major ambition and challenge in craft brewing and distilling industries. Genome-scale computational approaches are opening opportunities to model and predict favourable combination of traits for strain development. However, mining the genome of these complex hybrids is not currently an easy task, due to the high level of redundancy and presence of homologous. Moreover, no genome annotation for these industrial strains have been published. Here, we developed HybridMine, a new user-friendly, open-source tool for functional annotation of hybrid aneuploid genomes of any species by predicting parental alleles including paralogs. As proof of principle, we carried out a comprehensive structural and functional annotation of complex yeast hybrids to enable system biology prediction studies. HybridMine is developed in Python, Perl and Bash programming languages and is available at https://github.com/Sookie-S/HybridMine.

## INTRODUCTION

Genome-scale computational approaches are opening opportunities in the beer processing market. *S. pastorianus* is an allopolyploid sterile hybrid of the mesophilic *Saccharomyces cerevisiae* and the cold tolerant *Saccharomyces eubayanus.* These aneuploid hybrid strains have the beneficial properties of both parents, such as a strong ability to ferment at low temperature (bottom-fermenting lager yeast) and under stressful conditions, such as anaerobiosis, high hydrostatic pressure and high gravity sugar solutions [1]. *S. pastorianus* strains carry multiple copies of *S. cerevisiae*-like, *S. eubayanus*-like and hybrid-gene alleles, which encode for different protein isoforms. This may lead to agonistic or antagonistic competition for substrates and varying biochemical activities resulting in novel phenotypes and unique cellular metabolism. Moreover, different chimeric protein complexes can be established in the hybrids producing a plethora of phenotypes [2]. In these environmental conditions, the fermenting process produces complex metabolites that lead to unique flavours and aromas, appreciated in beer beverage. *S. pastorianus,* originally named *S. carlsbergensis,* has been isolated from lager fermentation environments [3]. The analysis of transposon sequence distribution in the genome of different *S. pastorianus* strains suggested the presence of two genomically distinct groups [4], that may have arisen from different hybridization events (Figure 1) [5]. One theory supports the hypothesis that an initial hybridization event occurred between a diploid *S. cerevisiae* and a diploid *S. eubayanus* (Figure 1A), leading to a tetraploid hybrid progenitor, that evolved to give the Group II strains. This progenitor in parallel underwent chromosomal deletions of the *S. cerevisiae* sub-genome, leading to the Group I strains [6]. Another hypothesis states that an initial spontaneous hybridization event occurred between a haploid *S. cerevisiae* strain and a diploid *S. eubayanus* strain during the Middle Ages (Figure 1B) [7]. This event led to a progenitor of the Group I strain, which evolved through further reduction of the *S. cerevisiae* genome content, to produce the extant Group I strains, which are approximately triploid in nature. The *S. cerevisiae* parent of the Group I yeasts is related to yeast strains used for Ale beer production in Europe. In parallel, the Group I progenitor strain underwent a second hybridization event with a *S. cerevisiae* strain related to Stout fermentation, and evolved to give Group II strains with an approximate tetraploid genome. It appears that the Group II strains have 2-3 times more *S. cerevisiae* genomic content than the Group I strains. Therefore Group II yeasts have a rather complex genome containing Ale-like (*S. cerevisiae),* Stout-like (*S. cerevisiae,* British Isles) and lager-like (*S. eubayanus)* gene content. The geographical and brewery location are linked to the grouping of *S. pastorianus* strains. The Group I encompasses both Saaz-type strains from Czech Republic breweries and Carlsberg type strains from Denmark breweries (Figure 1C). The Group II, referred as Frohberg-type, includes strains found in two Canadian breweries, in Heineken and Oranjeboom breweries in the Netherlands, and in non-Carlsberg breweries in Denmark (Figure 1C) [8].

**Figure 1:**
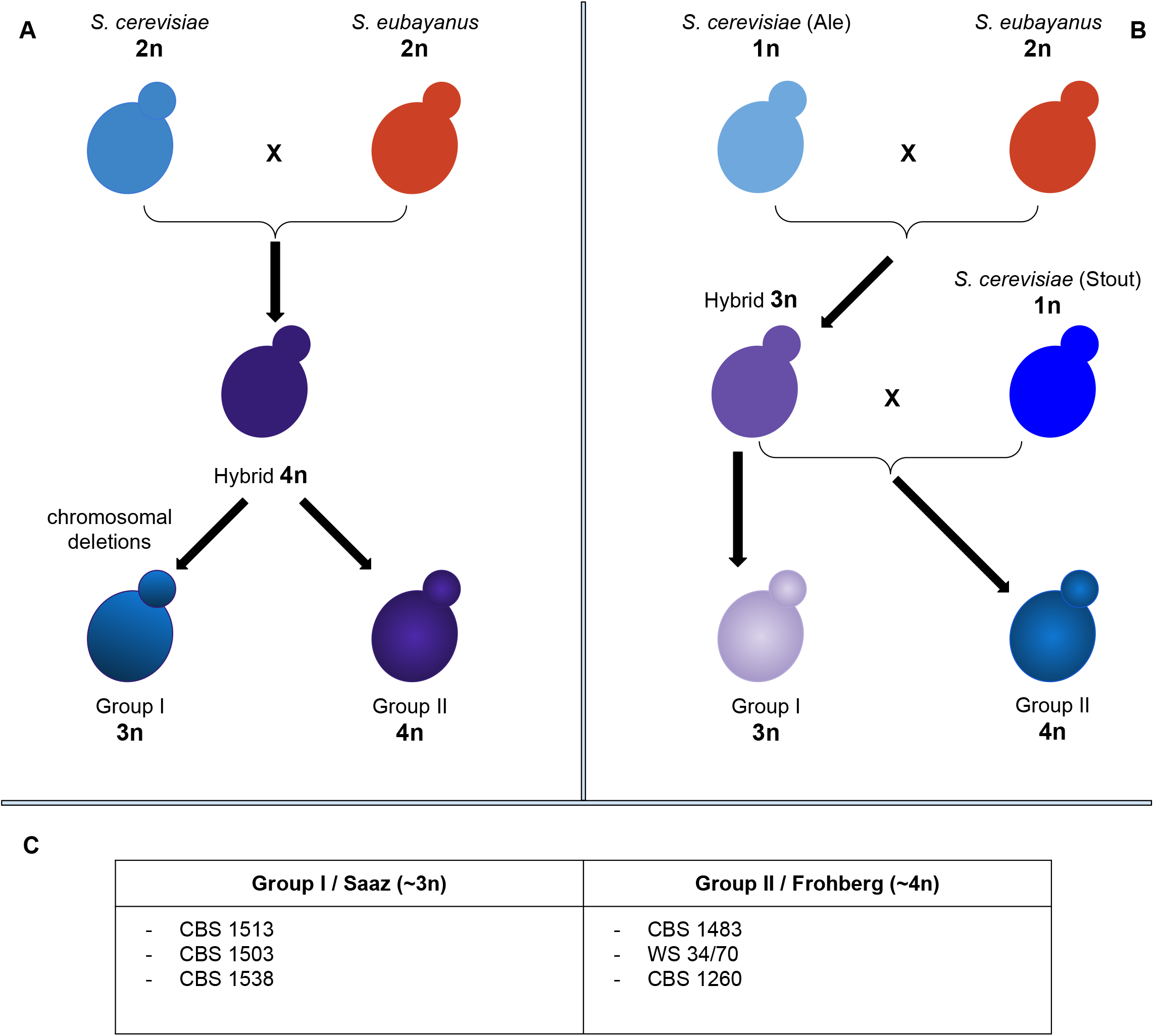
Proposed origins of Group I and II *Saccharomyces pastorianus.* **Panel A**: A hybridization event occurred between a diploid *S. cerevisiae* and a diploid *S. eubayanus*, followed by a different amount of genome reduction. **Panel B**: A hybridization event between a haploid *S. cerevisiae* and a diploid *S. eubayanus* lead to a triploid progenitor. Subsequently, a second hybridization event occurred between the triploid progenitor and a haploid *S. cerevisiae.* **Panel C**: Strains belonging to the Group I and II **(adapted from Alsammar *et al.,* 2020).**

Several lineages of *S. eubayanus* have been isolated from Nothofagus trees in Patagonia, and more recently in East Asia (Tibet) [9], while *S. cerevisiae* was isolated in Europe. The silk road, that connected Asia to Europe for trading purposes, can explain how the hybridization occurred between those two species. Before the discovery of *S. eubayanus,* the non-*S. cerevisiae* portion of the genome of *S. pastorianus* was considered as being *S. uvarum* and/or *S. bayanus* genome, that are closely related to *S. eubayanus* [10]. Moreover, *Saccharomyces* group yeast went through a whole genome duplication event (WGD), about 100 million years ago [11]. The WGD has important consequences as the organism doubles its genetic content leaving one copy of each gene free from constraints and able to evolve. Although the majority of paralogous will simply accumulate mutations and become pseudogenes (non-functionalization), some can acquire new functions (neo-functionalization) or can share the original function between them (sub-functionalization). Thereby, orthology relationships resulting from the WGD events are complicated as it leads to a 2:1 synteny relationship between genomic regions in post-WGD and non-WGD species [12]. We know now that almost all eukaryotic sequences show signs of ancient duplications, either WGDs or segmental duplications [12].

Mining the complex genome of these hybrids is therefore difficult. *Saccharomyces pastorianus* popularity is growing as one of the world’s most important industrial organism [13] and several R&D departments of the brewery industries worldwide are now focusing on strain improvement [14]. The strains *S. pastorianus* CBS 1503 (known as *S. monacensis),* CBS 1513 (known as *S. carlsbergensis),* CBS 1538 and WS 34/70 (known as *weihenstephan* strain) [15,16,17] used in the beer market, have been recently sequenced and assembled, but no annotation has been published, hampering biotechnological processes. It is in fact becoming essential to develop analytical and predictive tools to allow tailor-made improvements of specific yeast traits such as ethanol tolerance, maltose utilization and flavour profile. Functional annotation tools such as Blast2GO [18] are computationally intensive and come with a costly license. eggNOG-mapper [19] is not ideal for hybrid genomes as it transfers annotations by searching orthologs in a wide taxa group, hampering the discrimination of parental alleles. Finally, both Blast2GO and eggNOG-mapper are not designed to take into account aneuploidy and paralogous genes are discarded. Here we developed HybridMine, an open-source computational tool is built differently as it is specific for annotating any hybrid genomes by predicting parental alleles including paralogs. Using this tool, as proof of principle, the genome of four *S. pastorianus* strains have been functionally annotated showing a significant correlation between predictive and expected parental allele content.

## MATERIAL AND METHODS

### Genome sequences

The genome assembly for the strains *S. pastorianus* CBS 1503 [7,8], CBS 1513 [7,8], CBS 1538 [7,20,21,22] and WS 34/70 [7,20,21,22] have been downloaded from National Center for Biotechnology Information (NCBI). *S. cerevisiae* S288C genome has been used as a reference to annotate the *S. cerevisiae*-like genome content in the *S. pastorianus* strains. The last released *S. cerevisiae* genome has been downloaded from the Saccharomyces Genome Database (SGD). *S. eubayanus* FM1318 strain has been used as a reference to annotate the *S. eubayanus-like* genome content in *S. pastorianus*. Its genome assembly and annotation provided by the Tokyo Institute of Technology have been taken from NCBI database. Complementary information about the four *S. pastorianus strains* and the link to their repository are given in Supplementary Table 1.

### Genome structural annotation

The Yeast Genome Annotation Pipeline (YGAP) [23] has been used to predict the position of potential open reading frames (ORFs) and tRNAs in the yeast strains genome. YGAP is a structural annotation system that uses homology and synteny information from other yeast species present in the Yeast Gene Order Browser database, based on the hypothesis that the genes intron/exon structure is conserved through evolution (2 orthologous genes might have a similar intron/exon structure). The pipeline has been chosen as it is suitable for species that went through the Whole-Genome Duplication event.

### Pipeline architecture

The script for the HybridMine architecture has been developed in Bash language. The alignments have been done using BLAST 2.6.0+ program [24]. Information were extracted from the BLAST outputs using Perl. To identify orthologs, the parental alleles and paralogs in the hybrid genome, three scripts have been developed in Python 3.6 (see Results session).

### Statistical analysis

The difference between observed and expected number of parental alleles obtained in four *S. pastorianus* strains has been tested using the Chi square test. A p-value of less than 0.01 was considered statistically significant. Statistical analysis was performed using Python 3.6 packages Scipy and Stats models.

### Generation of annotation files

Bio::Tools::GFF module in the BioPerl bundle has been used to convert YGAP output GenBank files to GFF3 files for the four *S. pastorianus* strains. A Python 3.6 script has been developed to replace the fake gene IDs (generated by YGAP) by the parental gene name predicted by HybridMine.

## RESULTS

### Allele inheritance prediction pipeline

We developed a pipeline based on homology search using BLAST algorithms to identify the parental alleles in hybrid organisms. Divergence in orthologous genes is only considered to be due to speciation which allows direct functional inference. The main execution file (bash script) launching the pipeline runs on a Linux local machine and requires as inputs three FASTA files containing all the ORFs sequences of the hybrid strain to annotate, the first parent *(i.e.* Parent A) and the second parent (*i.e.* Parent B). Initially, BLAST databases are built to match query genomes. To determine the best hits, Nucleotide-Nucleotide blastn commands are run stringently (*i.e.* expectation value threshold for saving hits set at 0.05, default 10). Seven different blastn commands have been run to identify best bidirectional hits and paralogs (Figure 2A). For each run, the best alignments are written in a blast output file. Subsequently, a Perl script parses each output blast files (seven in total) and employs regular expressions in order to catch the query’s best hits in the database. The parser catches the e-value, the associated sequence identity and gap percentage for each best hit. As output, the script generates seven files containing the best hits for each gene in the queries. Those files are used as inputs of a Python 3.6 script that determines the 1:1 orthologs by finding the best bidirectional hits. Our script transforms each ORFs of the query genome into a Python object, defined by the following attributes: ID, best hit, best hit e-value, best paralog, best paralog e-value, best bidirectional hit, and best bidirectional hit e-value. A best hit is only considered if it shares more than 80% identity with the ORF of the query. A best bidirectional hit occurs when two ORFs are reciprocally found as best hits (Figure 2B). The two orthology output files generated (containing the 1:1 orthologs between the hybrid strain and the parent A, and those between the hybrid and the parent B) are then used as input in the next step of the pipeline. Another Python script determines which is the most likely parental allele the 1:1 ortholog evolved from. In the instance where a hybrid’s ORF has both a 1:1 ortholog in parent A and in parent B, the one that shares the highest percentage of identity is kept as parental allele. The only case in which a parental origin of an ortholog cannot be assigned is when the sequence of the orthologs in parent A and parent B are the same. For example, this can occur for tRNAs since they are extremely conserved and share 100% identity between the two parents. Once the origin of the alleles in the hybrid are identified, they are given the ID of the parental gene. The last Python script of our pipeline determines the groups of paralog genes (including the parental alleles in the hybrid genome that are paralogs). When two pairs of paralogs share one gene in common, they are grouped in as common paralogs.

**Figure 2:**
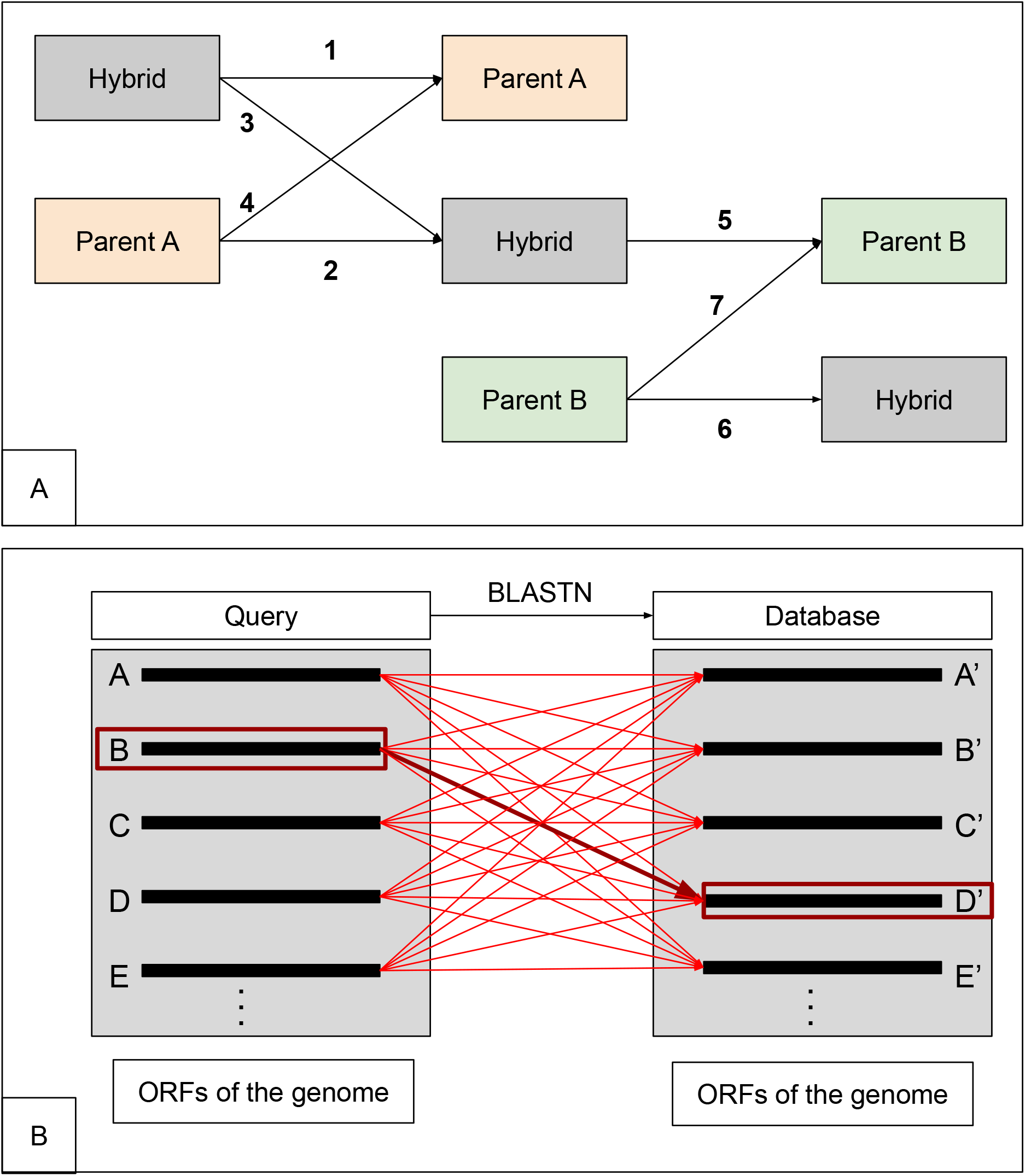
HybridMine usage. **Step 1:** Users to download HybridMine from its GitHub repository. **Step 2:** Users to add the three input fasta files required (ORFs of the hybrid organism, parent A and parent B) in the Data directory and rename them. **Step 3:** Users to run the main execution file “pipeline.sh” from the Script directory in a Unix terminal.

### HybridMine usage

HybridMine package is composed by two folders, “Script” and “Data”, which needs to stay colocated in the same directory when downloaded (Step 1 in Figure 3). The user then places in the Data directory the 3 fasta format files, containing the ORFs of the hybrid to annotate, the ORFs of the parent A and the ORFs of the parent B (Step 2 in Figure 3). To be recognized by HybridMine, the user has to rename the files, by adding “_orf.fasta” after the name of the organism *(i.e.* “[Hybrid]_orf.fasta”, “[ParentA]_orf.fasta” and “[ParentB]_orf.fasta”; see step 2, Figure 3). The user then launch the main execution file (pipeline.sh) from the Script directory, specifying as input the name of the hybrid to annotate and its two parental organism (*i.e.* command line: bash pipeline.sh [Hybrid] [ParentA] [ParentB]; see step 3, Figure 3). The documentation is presented in github (https://github.com/Sookie-S/HybridMine/blob/master/README.md), and the developed pipeline can be run locally on Unix OS.

**Figure 3:**
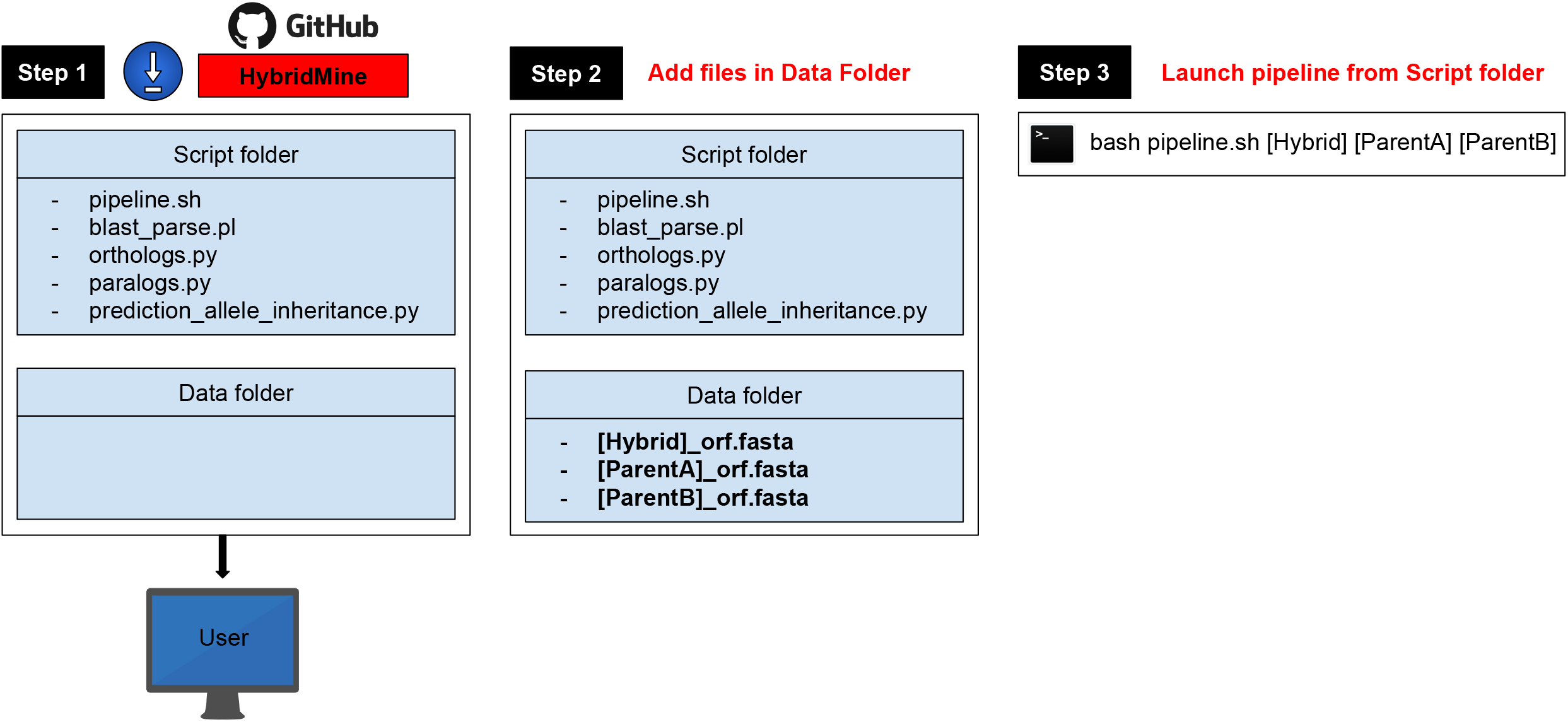
**Panel A**: The hybrid genome is blasted against the parental genomes (line 1 and 5) and reciprocally (line 2 and 6) to identify the orthologs. Each genome is searched against itself (line 3, 4 and 7) in order to get the best paralogs when they exist. **Panel B:** each ORF of the query genome is blasted against the database genome (red lines). When two ORFs are shown as reciprocal best hit (bold red arrows), the so called “best bidirectional hit” is identified. For example ORF B (boxed in red) of the hybrid has as best hit the ORF D’ (boxed in red) of the parent A and vice versa.

### Application and performances of HybridMine in *S. pastorianus* hybrids

As proof of principle, HybridMine was used to functionally annotate four *S. pastorianus* hybrid strains. The tool predicted approximately ⅔ *S. eubayanus-like* content and ⅓ *S. cerevisiae*-like content in the *S. pastorianus* hybrids CBS 1503, CBS 1513 and CBS 1538 belonging to the Group I; and approximately ½ *S. eubayanus-like* content and ½ *S. cerevisiae-like* content in the hybrid WS 34/70 that belongs to the group II (Table 1). Such predictions correlate with the expected genome content for these hybrids, approximately triploid and tetraploid for Group I and Group II strains, respectively (Figure 4). Lastly, we have generated GFF3 genome annotation files for the four *S. pastorianus* strains, that contain the parental gene IDs attributed by HybridMine.

**Figure 4:**
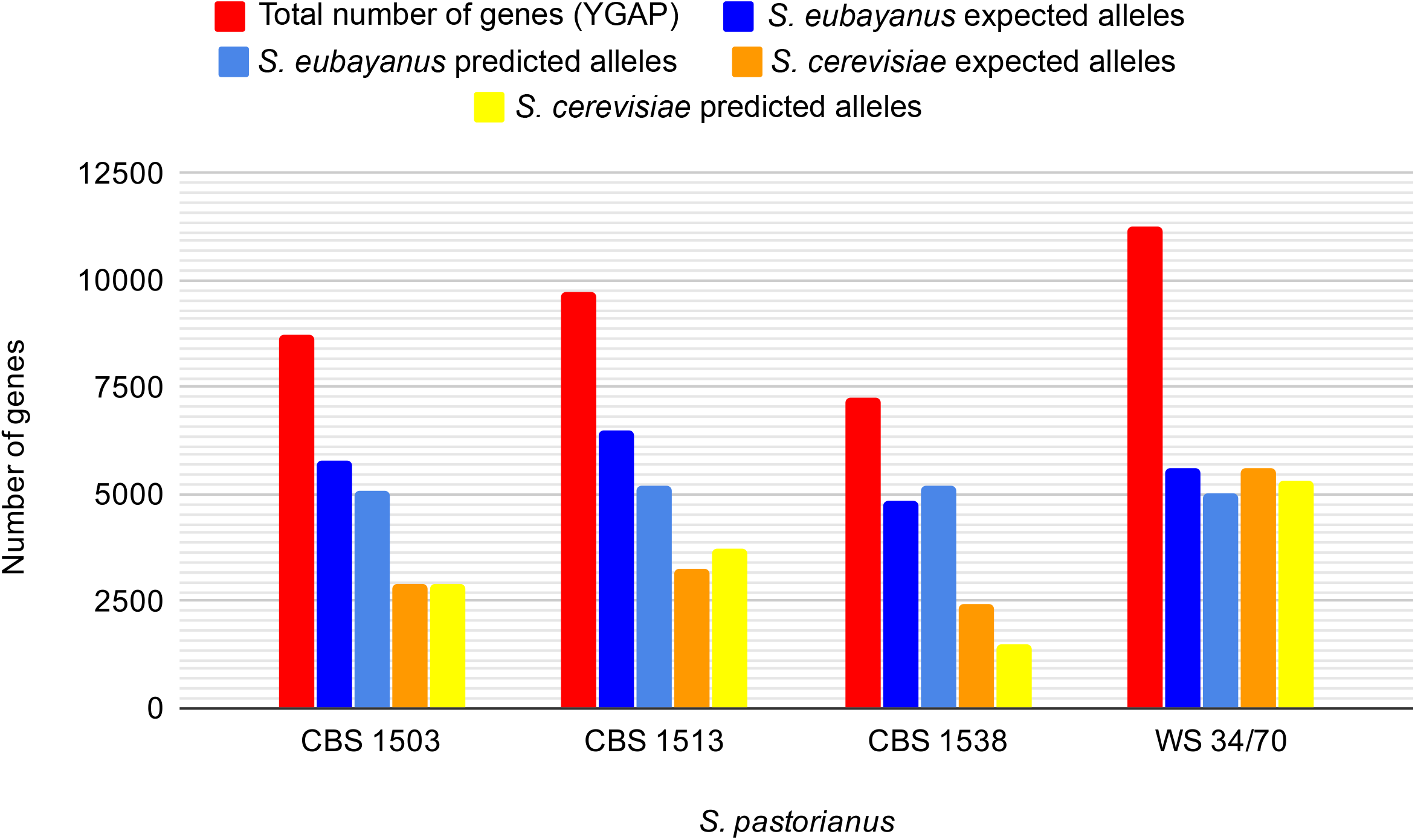
Histogram showing the amount of predicted versus expected *S. cerevisiae*-like and *S. eubayanus-like* genome content present in *S. pastorianus* strains CBS 1503, CBS 1513, CBS 1538 and WS 34/70, with 10% error bars.

**Table 1:** Prediction of the total number of *S. cerevisiae-like* and *S. eubayanus-like* alleles among the total number of genes predicted by YGAP.

| *S. pastorianus* strain annotated in this study | CBS 1538 | CBS 1503 | CBS 1513 | WS 34/70 |
| --- | --- | --- | --- | --- |
| Total number of genes predicted by YGAP | 7288 | 8714 | 9728 | 11265 |
| Number of predicted alleles in *S. eubayanus* | 5186 | 5075 | 5218 | 5024 |
| Number of predicted alleles in *S. cerevisiae* | 1517 | 2936 | 3739 | 5312 |
| 1:1 orthologs with same sequence identity in both parents | 9 | 10 | 9 | 7 |
| ORFs with less than 80% sequence identity with a predicted parental allele | 9 | 3 | 4 | 2 |
| *S. pastorianus* genes with no parental allele predicted | 567 | 690 | 758 | 920 |
| Number of different paralogs | 2814 | 3056 | 3316 | 3652 |

We determined whether the difference between the observed and the expected frequencies of the *S. eubayanus*-like and *S. cerevisiae*-like alleles was statistically significant in these four *S. pastorianus* strains. Here, the null hypothesis is that there is a difference between expected and observed data. We chose the value of 0.01 for the alpha risk to reject the null hypothesis when it is actually true. We obtained a chi square value of 865.44, which is superior to the critical value 18.48 of this test, and a p-value of 1.3 x 10^-182^, which is inferior to the risk alpha and approximable to zero. Hence, the values predicted are significantly concordant with the values expected.

Performance wise, HybridMine is a very fast tool. Our tests with three initial input containing up to 11,000 ORFs each, showed execution times lower than one minute on an Intel^®^ Core™ i9-7900X CPU @ 3.30GHz × 20 computer. The execution time was 34.358s for the largest hybrid predictions, *Saccharomyces pastorianus* WS34/76.

## DISCUSSION

To be able to mine the genome of yeast hybrids is becoming a major need in brewing and distilling industries. Until recently, due to sterile and complex nature of the yeast hybrids, the generation of improved yeast strains for beer making has been mainly carried out via industrial directional selection rather than breeding strategies. Computational predictive approaches are also rarely employed due to the lack of molecular data on these hybrid strains. The sequencing of large hybrids genome has now become more accurate with the development of the long-read third-generation sequencing technologies such as Nanopore [25,26]. Although annotation tools are in place, computational methods that specifically predict the parental alleles in hybrid genomes are lacking. The identification of parental alleles in hybrid genome is crucial to make accurate functional annotation and assigning the sequences to the right biological function. For example, correctly identifying the *S. cerevisiae*-like and *S. eubayanus*-like genes in *S. pastorianus* is important to study the transcriptome plasticity and the cis/trans regulation in this natural hybrid. Moreover, it can inform how the the genetic redundancy of *S. cerevisiae*-like and *S. eubayanus*-like ortholog alleles impact on the phenotype. The functional annotation of the genome also allows *in silico* design of new beneficial industrial strains using predictive approaches. Accurate annotation in these hybrids facilitate the study of their evolution as some of them are still undergoing genome reduction to acquire more stability. The generation of the functional annotation of four key industrial yeast strains using HybridMine can now be used for further comparative genomic analyses of industrial yeast strains, including studies on gene retention and gene loss, molecular adaptation processes and evolution of transcriptional plasticity.

Importantly, HybridMine has a general application since it can be used to map orthologs and paralogs of any hybrid organism of known parental species. Natural or artificial hybridization between strains or species is a common phenomenon that occurs in almost all sexually reproducing group of organisms, including bacteria, yeast, plants and animals [26]. It has been established that there is at least 25% of plant species and 10% of animal species involved in hybridization with other species [27]. Hence our tool have a broader application for any hybrid organisms.

## AVAILABILITY

### Code availability

The computational resources described in this paper and the genome annotations are available in GitHub (https://github.com/Sookie-S/HybridMine).

### Data availability

HybridMine has been used to predict the parental alleles and paralogs in four *S. pastorianus* strains: CBS 1503 (known as *S. monacensis),* CBS 1513 (known as *S. carlsbergensis),* CBS 1538 and WS 34/70 (known as *weihenstephan).* The predictions are downloadable from the Supplementary Files 1-4.

The genome annotations generated in GFF3 format are available in GitHub (https://github.com/Sookie-S/HybridMine/tree/master/Annotation_files) and can also be downloaded from Dryad (https://doi.org/10.5061/dryad.3n5tb2rdf).

## SUPPLEMENTARY DATA

Supplementary Table 1 - *S. pastorianus* genome sources

Supplementary File 1 - *Saccharomyces pastorianus* CBS 1503 parental alleles and paralogs prediction

Supplementary File 2 - *Saccharomyces pastorianus* CBS 1513 parental alleles and paralogs prediction

Supplementary File 3 - *Saccharomyces pastorianus* CBS 1538 parental alleles and paralogs prediction

Supplementary File 4 - *Saccharomyces pastorianus* CBS WS34/70 parental alleles and paralogs prediction

## ACKNOWLEDGEMENTS

Author contributions: D.D and J.M.S conceived the project; D.D. and J.M.S supervised the research; S.T. developed the experimental pipeline, wrote the code for HybridMine, and annotated the genomes. All the authors analysed the data and wrote the manuscript.

## REFERENCES

[1] Monerawela C., Bond U. Brewing up a storm: The genomes of lager yeasts and how they evolved. Biotechnology Advances. 2017; 35:512–519.

[2] Piatkowska E.M., Naseeb S., Knight D., Delneri D. Chimeric protein complexes in hybrid species generate novel phenotypes. PLoS Genetics. 2013; 9:e1003836.

[3] Dunn B. and Sherlock G. Reconstruction of the genome origins and evolution of the hybrid lager yeast *saccharomyces pastorianus*. Genome Research. 2008; 18:1610–1623.

[4] Liti G., Peruffo A., James S.A., Roberts I.N., Louis E.J. Inferences of evolutionary relationships from a population survey of ltr-retrotransposons and telomeric-associated sequences in the *saccharomyces* sensu stricto complex. Yeast. 2005; 22:177–192.

[5] Monerawela C., James T.C., Wolfe K.H., Bond U. Loss of lager specific genes and subtelomeric regions define two different *saccharomyces cerevisiae* lineages for *saccharomyces pastorianus* group i and ii strains. FEMS Yeast Research. 2015; 15:fou008.

[6] Alsammar H., Delneri D. An update on the diversity, ecology and biogeography of the *Saccharomyces* genus. FEMS Yeast Research. 2020; 20:foaa013.

[7] Bond U., Neal C., Donnelly D., James T.C. Aneuploidy and copy number breakpoints in the genome of lager yeasts mapped by microarray hybridisation. Current Genetics. 2004; 45:360–370.

[8] Noonan G.J. New brewing lager beer: The most comprehensive book for home- and microbrewers. Brewers Publications, Boulder, CO. 1996.

[9] Bing J., Han P.J., Liu W.Q., Wang Q.M., Bai F.Y. Evidence for a Far East Asian origin of lager beer yeast. Current Biology. 2014; 24:380–381.

[10] Vaughan-Martini A., Martini A. A Taxonomic Key for the Genus *Saccharomyces*. Systematic and Applied Microbiology. 1993; 16:113–119.

[11] Wolfe K. and Shields D. Molecular evidence for an ancient duplication of the entire yeast genome. Nature. 1997; 387:708–713.

[12] Wolfe K.H. Origin of the yeast whole-genome duplication. PLoS Biology. 2015; 13:e1002221.

[13] Gibson B., Liti G. *Saccharomyces pastorianus*: genomic insights inspiring innovation for industry. Yeast. 2014; 32:17–27.

[14] Bloomberg business, Beer Processing Market Worth $815.4 Billion by 2025. Exclusive Report by Markets and Markets™, July 2019.

[15] Okuno M., Kajitani R., Ryusui R., Morimoto H., Kodama Y., Itoh T. Next-generation sequencing analysis of lager brewing yeast strains reveals the evolutionary history of interspecies hybridization. DNA Research. 2016; 23:67–80.

[16] Walther A., Hesselbart A., Wendland J. Genome Sequence of *Saccharomyces carlsbergensis*, the World’s First Pure Culture Lager Yeast. G3: Genes, Genomes, Genetics. 2014; 4:783–793.

[17] Hewitt S.K., Donaldson I.J., Lovell S.C., Delneri D. Sequencing and characterisation of rearrangements in three *S. pastorianus* strains reveals the presence of chimeric genes and gives evidence of breakpoint reuse. PLoS one. 2014; 9:e92203.

[18] Conesa A., Götz S., Garcia-Gomez J.M., Terol J., Talon M., Robles M. Blast2GO: a universal tool for annotation, visualization and analysis in functional genomics research. Bioinformatics. 2005; 21:3674–3676.

[19] Huerta-Cepas J., Forslund K., Coelho L.P., Szklarczyk D., Jensen L.J., von Mering C., Bork P. Fast genome-wide functional annotation through orthology assignment by eggNOGmapper. Molecular Biology and Evolution. 2017; 34:2115–2122.

[20] Nakao Y., Kanamori T., Itoh T., Kodama Y., Rainieri S., Nakamura N., Shimonaga T., Hattori M., Ashikari T. Genome sequence of the lager brewing yeast, an interspecies hybrid. DNA Research. 2009; 16:115–129.

[21] van den Broek M., Bolat I., Nijkamp J.F., Ramos E., Luttik M.A., Koopman F., Geertman J.M., de Ridder D., Pronk J.T., Daran J.M. Chromosomal Copy Number Variation in Saccharomyces pastorianus Is Evidence for Extensive Genome Dynamics in Industrial Lager Brewing Strains. Applied and environmental microbiology. 2015: 81:6253–6267.

[22] De León-Medina P.M., Elizondo-González R., Damas-Buenrostro L.C., Geertman J.M., Van den Broek M., Galán-Wong L.J., Ortiz-López R., Pereyra-Alférez B. Genome annotation of a *Saccharomyces* sp. lager brewer’s yeast. Genomics data. 2015; 9:25–29.

[23] Proux-Wéra E., Armisén D., Byrne K.P. and Wolfe K.H. A pipeline for automated annotation of yeast genome sequences by a conserved-synteny approach. BMC Bioinformatics. 2012; 13:237.

[24] Altschul S.F., Gish W., Miller W., Myers E.W., Lipman D.J. Basic local alignment search tool. Journal of Molecular Biology. 1990; 215:403–410.

[25] McGinty R.J., Rubinstein R.G., Neil A.J., Dominska M., Kiktev D., Petes T.D. Nanopore sequencing of complex genomic rearrangements in yeast reveals mechanisms of repeat-mediated double-strand break repair. Genome Research. 2017; 27:2072–2082.

[26] Li W., Floyd Averette A., Desnos-Ollivier M., Ni M., Dromer F. and Heitman J. Genetic Diversity and Genomic Plasticity of *Cryptococcus neoformans* AD Hybrid Strains. G3: Genes Genomes Genetics. 2012; 2:83–97.

[27] Mallet J. Hybridization as an invasion of the genome. Trends in ecology and evolution. 2005; 20:229–237.

