## Supplementary Table 1 for "HybridMine: pipeline for allele inheritance and gene copy number prediction in industrial yeast hybrids"

Table 1 presents all the synonyms known for each *S. pastorianus* hybrid of the study, retrieved in the National Catalog of Yeast Collection, and a link to the repository where the genome assembly can be downloaded.

| **Saccharomyces pastorianus strain** | **CBS 1503** | **CBS 1538** | **CBS 1513** | **WS 34/70** |
|---|---|---|---|---|
| **Group strain** | Group I | Group I | Group I | Group II |
| **Equivalent names (from NCYC and litterature)** | NCYC 2801,<br>DBVPG 6261,<br>IFO 10610,<br>IGC 4581,<br>NRRL Y-1525<br>*S. monacensis* | NCYC 392,<br>ATCC 12752,<br>CCRC 21971,<br>DBVPG 6047,<br>IFO 0613,<br>IGC 4601,<br>NRRL Y-1551,<br>CLIB 281,<br>IFO 11024,<br>IFO 1132,<br>IFO 1943,<br>AS 2.2402 | NCYC 396,<br>CCRC 21423,<br>DBVPG 6033,<br>IFO 1167,<br>IGC 4457,<br>JCM 7256,<br>MUCL 38889,<br>NRRL Y-12693,<br>CLIB 176,<br>IFO 11023,<br>*S. carlsbergensis* | Weihenstephan<br>WS 34/70 |
| **Habitat** | from Perugia, Italy | Unknown | from Perugia, Italy | ? |
| **Sources<br>-<br>Repository** | Link to NCBI | Link to NCBI | Link to NCBI | Link to NCBI |

**Table 1: Equivalent names for the *S. pastorianus* strains collected from the National Catalog of Yeast Collection (NCYC) and litterature and link to their genome assembly.**
