## Supplementary file 1 for "HybridMine: pipeline for allele inheritance and gene copy number prediction in industrial yeast hybrids"

### Overview

genes alleles inherited from Saccharomyces eubayanus
genes alleles inherited from Saccharomyces cerevisiae
genes alleles below 80% identity
genes alleles copy with same identity in both parents
list of paralogs in Saccharomyces pastorianus CBS1503

#### Sheet 1: *genes alleles inherited from Saccharomyces eubayanus*

| *Saccharomyces pastorianus CBS1503 predicted ORF* | *Saccharomyces eubayanus parental allele* | Gene identities | Gene gaps |
|---|---|---|---|
| SPGP0NI00250 | XP_018223918.1 | 100 | 0 |
| SPGP0V00100 | XP_018222345.1 | 99 | 0 |
| SPGP0IC00260 | XP_018222781.1 | 99 | 0 |
| SPGP0OR00140 | XP_018218830.1 | 99 | 0 |
| SPGP0FC00110 | XP_018219052.1 | 99 | 1 |
| SPGP0WS00120 | XP_018222749.1 | 99 | 0 |
| SPGP0FW00190 | XP_018219577.1 | 99 | 0 |
| SPGP0LX00160 | XP_018219150.1 | 99 | 0 |
| SPGP0VH00130 | XP_018219684.1 | 99 | 0 |
| SPGP0DU00220 | XP_018222572.1 | 99 | 0 |
| SPGP0UH00140 | XP_018220410.1 | 98 | 0 |
| SPGP0K00100 | XP_018218750.1 | 99 | 0 |
| SPGP0KL00170 | XP_018221097.1 | 99 | 0 |
| SPGP0KZ00100 | XP_018223950.1 | 99 | 0 |
| SPGP0HA00150 | XP_018219873.1 | 99 | 0 |
| SPGP0OX00260 | XP_018219719.1 | 99 | 0 |
| SPGP0AA00220 | XP_018221073.1 | 99 | 0 |
| SPGP0TO00130 | XP_018222807.1 | 99 | 0 |
| SPGP0AC00110 | XP_018220415.1 | 99 | 0 |
| SPGP0K00770 | XP_018219509.1 | 100 | 0 |
| SPGP0HG00110 | XP_018220458.1 | 99 | 0 |
| SPGP0LX00140 | XP_018219148.1 | 99 | 0 |
| SPGP0NV00110 | XP_018223448.1 | 99 | 0 |
| SPGP0NY00140 | XP_018219623.1 | 99 | 0 |
| SPGP0NT00250 | XP_018222459.1 | 99 | 0 |
| SPGP0EY00120 | XP_018219090.1 | 99 | 0 |
| SPGP0TM00190 | XP_018219883.1 | 99 | 0 |
| SPGP0IR00140 | XP_018219763.1 | 99 | 0 |

Create PDF in your applications with the Pdfcrowd HTML to PDF API PDFCROWD

| | | | |
|---|---|---|---|
| SPGP0D01670 | XP_018221795.1 | 99 | 0 |
| SPGP0WQ00270 | XP_018222279.1 | 99 | 0 |
| SPGP0MO00260 | XP_018219110.1 | 99 | 0 |
| SPGP0KZ00150 | XP_018223945.1 | 99 | 0 |
| SPGP0DQ00150 | XP_018219674.1 | 99 | 0 |
| SPGP0LA00200 | XP_018220849.1 | 99 | 0 |
| SPGP0TQ00180 | XP_018220613.1 | 99 | 1 |
| SPGP0OD00110 | XP_018221044.1 | 99 | 0 |
| SPGP0WA00210 | XP_018219845.1 | 99 | 0 |
| SPGP0GF00150 | XP_018219646.1 | 99 | 0 |
| SPGP0OE00120 | XP_018219608.1 | 98 | 2 |
| SPGP0FM00120 | XP_018222148.1 | 99 | 0 |
| SPGP0OR00160 | XP_018218832.1 | 99 | 0 |
| SPGP0DP00140 | XP_018220648.1 | 99 | 0 |
| SPGP0OF00350 | XP_018222716.1 | 99 | 0 |
| SPGP0NY00120 | XP_018219621.1 | 99 | 0 |
| SPGP0TC00180 | XP_018219695.1 | 99 | 0 |
| SPGP0EZ00130 | XP_018219318.1 | 100 | 0 |
| SPGP0OI00730 | XP_018218877.1 | 99 | 0 |
| SPGP0GF00350 | XP_018219626.1 | 99 | 0 |
| SPGP0OE00290 | XP_018219592.1 | 99 | 0 |
| SPGP0MO00170 | XP_018219102.1 | 99 | 0 |
| SPGP0VM00230 | XP_018220671.1 | 100 | 0 |
| SPGP0RT00120 | XP_018223994.1 | 99 | 0 |
| SPGP0WW00100 | XP_018222072.1 | 99 | 0 |
| SPGP0MF00260 | XP_018222194.1 | 99 | 0 |
| SPGP0WI00160 | XP_018222124.1 | 99 | 0 |
| SPGP0EF00100 | XP_018220019.1 | 99 | 0 |
| SPGP0IH00110 | XP_018222239.1 | 99 | 0 |
| SPGP0HB00200 | XP_018219184.1 | 99 | 0 |
| SPGP0IE00170 | XP_018222759.1 | 99 | 0 |
| SPGP0QP00190 | XP_018219160.1 | 99 | 0 |
| SPGP0KZ00310 | XP_018223931.1 | 99 | 0 |
| SPGP0HU00130 | XP_018220429.1 | 99 | 0 |
| SPGP0GR00160 | XP_018222744.1 | 99 | 0 |
| SPGP0VU00140 | XP_018222738.1 | 99 | 0 |
| SPGP0HZ00350 | XP_018219792.1 | 99 | 0 |
| SPGP0OI00600 | XP_018218890.1 | 99 | 0 |
| SPGP0GI00190 | XP_018220691.1 | 99 | 0 |
| SPGP0CF00180 | XP_018222492.1 | 99 | 0 |

Create PDF in your applications with the Pdfcrowd HTML to PDF API PDFCROWD

| | | | |
|---|---|---|---|
| SPGP0GV00160 | XP_018218865.1 | 99 | 0 |
| SPGP0IK00120 | XP_018222109.1 | 99 | 0 |
| SPGP0GF00240 | XP_018219637.1 | 99 | 0 |
| SPGP0PA00200 | XP_018220464.1 | 99 | 0 |
| SPGP0RI00380 | XP_018220517.1 | 99 | 0 |
| SPGP0NI00270 | XP_018223920.1 | 99 | 0 |
| SPGP0OR00180 | XP_018218834.1 | 99 | 0 |
| SPGP0NG00220 | XP_018222766.1 | 99 | 0 |
| SPGP0KZ00200 | XP_018223941.1 | 99 | 0 |
| SPGP0JF00110 | XP_018222309.1 | 99 | 0 |
| SPGP0KZ00120 | XP_018223948.1 | 99 | 0 |
| SPGP0UV00160 | XP_018218803.1 | 99 | 0 |
| SPGP0IR00200 | XP_018219757.1 | 98 | 1 |
| SPGP0TQ00260 | XP_018220621.1 | 99 | 0 |
| SPGP0TQ00420 | XP_018220637.1 | 99 | 0 |
| SPGP0UV00170 | XP_018218804.1 | 99 | 0 |
| SPGP0CH00100 | XP_018219957.1 | 99 | 0 |
| SPGP0PE00240 | XP_018222369.1 | 99 | 0 |
| SPGP0GF00300 | XP_018219631.1 | 99 | 0 |
| SPGP0WI00310 | XP_018222139.1 | 98 | 0 |
| SPGP0HG00130 | XP_018220456.1 | 99 | 0 |
| SPGP0EH00110 | XP_018222262.1 | 99 | 0 |
| SPGP0RQ00170 | XP_018221128.1 | 99 | 0 |
| SPGP0HZ00170 | XP_018219774.1 | 99 | 0 |
| SPGP0NY00160 | XP_018219625.1 | 99 | 0 |
| SPGP0RC00290 | XP_018222104.1 | 99 | 0 |
| SPGP0MO00160 | XP_018219101.1 | 99 | 1 |
| SPGP0DQ00130 | XP_018219672.1 | 99 | 0 |
| SPGP0RI00320 | XP_018220510.1 | 87 | 0 |
| SPGP0JF00160 | XP_018222314.1 | 99 | 0 |
| SPGP0MO00220 | XP_018219107.1 | 99 | 0 |
| SPGP0WB00140 | XP_018222676.1 | 99 | 0 |
| SPGP0MO00280 | XP_018219112.1 | 99 | 0 |
| SPGP0FB00100 | XP_018220424.1 | 99 | 0 |
| SPGP0NZ00100 | XP_018222517.1 | 98 | 1 |
| SPGP0XC00100 | XP_018220586.1 | 99 | 0 |
| SPGP0VI00100 | XP_018222303.1 | 99 | 0 |
| SPGP0RB00130 | XP_018218997.1 | 99 | 0 |
| SPGP0QF00190 | XP_018224027.1 | 99 | 0 |
| SPGP0FC00130 | XP_018219054.1 | 99 | 0 |

Create PDF in your applications with the Pdfcrowd HTML to PDF API PDFCROWD

| | | | |
|---|---|---|---|
| SPGP0HY00130 | XP_018223990.1 | 99 | 0 |
| SPGP0IC00150 | XP_018222792.1 | 99 | 0 |
| SPGP0UR00110 | XP_018222343.1 | 99 | 0 |
| SPGP0VN00240 | XP_018219814.1 | 99 | 0 |
| SPGP0FW00250 | XP_018219571.1 | 99 | 0 |
| SPGP0DU00210 | XP_018222573.1 | 99 | 0 |
| SPGP0HR00110 | XP_018222179.1 | 98 | 1 |
| SPGP0TC00160 | XP_018219697.1 | 99 | 0 |
| SPGP0VO00100 | XP_018223435.1 | 99 | 0 |
| SPGP0SW00110 | XP_018220563.1 | 98 | 0 |
| SPGP0MG00120 | XP_018219616.1 | 99 | 0 |
| SPGP0HO00350 | XP_018219968.1 | 99 | 0 |
| SPGP0C03250 | XP_018220049.1 | 99 | 0 |
| SPGP0UZ00150 | XP_018223430.1 | 99 | 0 |
| SPGP0MO00150 | XP_018219100.1 | 99 | 0 |
| SPGP0WW00120 | XP_018222074.1 | 99 | 0 |
| SPGP0FE00140 | XP_018220536.1 | 100 | 0 |
| SPGP0EM00110 | XP_018220722.1 | 99 | 1 |
| SPGP0WI00300 | XP_018222138.1 | 99 | 0 |
| SPGP0JN00150 | XP_018219074.1 | 99 | 0 |
| SPGP0WY00210 | XP_018220435.1 | 99 | 0 |
| SPGP0QT00110 | XP_018220012.1 | 99 | 0 |
| SPGP0GD00100 | XP_018220834.1 | 99 | 0 |
| SPGP0LW00110 | XP_018222615.1 | 99 | 0 |
| SPGP0IC00220 | XP_018222785.1 | 99 | 0 |
| SPGP0NF00150 | XP_018222631.1 | 99 | 0 |
| SPGP0IH00120 | XP_018222238.1 | 99 | 0 |
| SPGP0VM00280 | XP_018220666.1 | 99 | 0 |
| SPGP0UQ00120 | XP_018220689.1 | 98 | 1 |
| SPGP0VZ00250 | XP_018223889.1 | 99 | 0 |
| SPGP0LH00100 | XP_018221036.1 | 99 | 0 |
| SPGP0GV00100 | XP_018218860.1 | 99 | 0 |
| SPGP0JF00260 | XP_018222324.1 | 99 | 0 |
| SPGP0HZ00210 | XP_018219778.1 | 99 | 0 |
| SPGP0MT00140 | XP_018219140.1 | 99 | 0 |
| SPGP0NF00140 | XP_018222632.1 | 99 | 0 |
| SPGP0LE00140 | XP_018220764.1 | 100 | 0 |
| SPGP0OI00480 | XP_018218902.1 | 99 | 0 |
| SPGP0XC00210 | XP_018220597.1 | 99 | 0 |
| SPGP0OL00130 | XP_018222285.1 | 99 | 0 |

Create PDF in your applications with the Pdfcrowd HTML to PDF API PDFCROWD

| | | | |
|---|---|---|---|
| SPGP0UN00230 | XP_018220775.1 | 99 | 0 |
| SPGP0MD00150 | XP_018222390.1 | 99 | 0 |
| SPGP0DN00140 | XP_018218990.1 | 100 | 0 |
| SPGP0LU00280 | XP_018223998.1 | 99 | 0 |
| SPGP0UH00100 | XP_018220406.1 | 99 | 0 |
| SPGP0CD00130 | XP_018222530.1 | 99 | 0 |
| SPGP0EM00260 | XP_018220709.1 | 99 | 0 |
| SPGP0TE00210 | XP_018219903.1 | 99 | 0 |
| SPGP0CY00110 | XP_018223952.1 | 99 | 0 |
| SPGP0LS00110 | XP_018222562.1 | 99 | 0 |
| SPGP0U00110 | XP_018224122.1 | 99 | 0 |
| SPGP0IR00180 | XP_018219759.1 | 100 | 0 |
| SPGP0TQ00240 | XP_018220619.1 | 99 | 0 |
| SPGP0FM00160 | XP_018222152.1 | 99 | 0 |
| SPGP0RP00100 | XP_018222609.1 | 99 | 0 |
| SPGP0ER00100 | XP_018218983.1 | 99 | 0 |
| SPGP0QV00220 | XP_018219998.1 | 99 | 0 |
| SPGP0VG00110 | XP_018223869.1 | 99 | 0 |
| SPGP0NQ00120 | XP_018220753.1 | 100 | 0 |
| SPGP0LK00140 | XP_018221067.1 | 98 | 1 |
| SPGP0OI00290 | XP_018218921.1 | 99 | 0 |
| SPGP0K00150 | XP_018218748.1 | 97 | 3 |
| SPGP0QT00100 | XP_018220013.1 | 99 | 0 |
| SPGP0TK00130 | XP_018222727.1 | 99 | 0 |
| SPGP0MO00390 | XP_018219123.1 | 99 | 0 |
| SPGP0OI00310 | XP_018218919.1 | 99 | 0 |
| SPGP0HT00110 | XP_018219003.1 | 99 | 0 |
| SPGP0DQ00110 | XP_018219670.1 | 99 | 0 |
| SPGP0NQ00170 | XP_018220748.1 | 99 | 0 |
| SPGP0SL00130 | XP_018220483.1 | 99 | 0 |
| SPGP0OF00280 | XP_018222709.1 | 99 | 0 |
| SPGP0NT00220 | XP_018222462.1 | 99 | 0 |
| SPGP0LE00110 | XP_018220768.1 | 99 | 0 |
| SPGP0MC00150 | XP_018222667.1 | 99 | 0 |
| SPGP0OZ00190 | XP_018222406.1 | 99 | 0 |
| SPGP0UN00220 | XP_018220776.1 | 99 | 0 |
| SPGP0KZ00320 | XP_018223930.1 | 99 | 0 |
| SPGP0FM00230 | XP_018222159.1 | 99 | 0 |
| SPGP0TQ00140 | XP_018220609.1 | 99 | 0 |
| SPGP0JF00180 | XP_018222316.1 | 100 | 0 |

Create PDF in your applications with the Pdfcrowd HTML to PDF API PDFCROWD

| | | | |
|---|---|---|---|
| SPGP0SP00100 | XP_018223960.1 | 99 | 0 |
| SPGP0OA00120 | XP_018218947.1 | 99 | 0 |
| SPGP0DW00130 | XP_018218953.1 | 99 | 0 |
| SPGP0UR00130 | XP_018222341.1 | 99 | 0 |
| SPGP0FW00230 | XP_018219573.1 | 100 | 0 |
| SPGP0HZ00340 | XP_018219791.1 | 99 | 0 |
| SPGP0WB00130 | XP_018222677.1 | 99 | 0 |
| SPGP0SP00130 | XP_018223963.1 | 100 | 0 |
| SPGP0GF00310 | XP_018219630.1 | 99 | 0 |
| SPGP0NK00180 | XP_018220447.1 | 99 | 0 |
| SPGP0LP00120 | XP_018222382.1 | 99 | 0 |
| SPGP0FM00260 | XP_018222161.1 | 98 | 1 |
| SPGP0IX00110 | XP_018221050.1 | 99 | 0 |
| SPGP0GR00150 | XP_018222743.1 | 100 | 0 |
| SPGP0LG00130 | XP_018222645.1 | 99 | 1 |
| SPGP0WW00140 | XP_018222076.1 | 99 | 0 |
| SPGP0XC00180 | XP_018220594.1 | 99 | 0 |
| SPGP0WG00210 | XP_018221017.1 | 100 | 0 |
| SPGP0VM00110 | XP_018220683.1 | 99 | 0 |
| SPGP0OE00160 | XP_018219604.1 | 99 | 0 |
| SPGP0MZ00100 | XP_018224017.1 | 99 | 0 |
| SPGP0NQ00240 | XP_018220741.1 | 99 | 0 |
| SPGP0UB00130 | XP_018219038.1 | 100 | 0 |
| SPGP0JW00210 | XP_018220983.1 | 99 | 0 |
| SPGP0WI00290 | XP_018222137.1 | 99 | 0 |
| SPGP0GL00120 | XP_018222613.1 | 99 | 0 |
| SPGP0ET00130 | XP_018222444.1 | 99 | 0 |
| SPGP0KO00110 | XP_018219703.1 | 99 | 0 |
| SPGP0TM00230 | XP_018219887.1 | 99 | 0 |
| SPGP0WQ00110 | XP_018222264.1 | 99 | 0 |
| SPGP0LH00160 | XP_018221031.1 | 99 | 0 |
| SPGP0HB00120 | XP_018219192.1 | 99 | 0 |
| SPGP0AH00100 | XP_018219888.1 | 99 | 0 |
| SPGP0WA00100 | XP_018219856.1 | 99 | 0 |
| SPGP0PE00330 | XP_018222378.1 | 99 | 0 |
| SPGP0MO00190 | XP_018219104.1 | 99 | 0 |
| SPGP0AB00100 | XP_018219707.1 | 99 | 0 |
| SPGP0RX00120 | XP_018222813.1 | 99 | 0 |
| SPGP0WG00110 | XP_018221027.1 | 99 | 0 |
| SPGP0RX00130 | XP_018222814.1 | 100 | 0 |

Create PDF in your applications with the Pdfcrowd HTML to PDF API PDFCROWD

| | | | |
|---|---|---|---|
| SPGP0NF00120 | XP_018222634.1 | 99 | 0 |
| SPGP0IX00100 | XP_018221051.1 | 99 | 0 |
| SPGP0DI00100 | XP_018219152.1 | 99 | 0 |
| SPGP0XC00230 | XP_018220599.1 | 99 | 0 |
| SPGP0BM00110 | XP_018222754.1 | 99 | 0 |
| SPGP0VM00270 | XP_018220667.1 | 99 | 0 |
| SPGP0TE00160 | XP_018219898.1 | 99 | 0 |
| SPGP0OI00750 | XP_018218875.1 | 99 | 0 |
| SPGP0RA00130 | XP_018218843.1 | 99 | 1 |
| SPGP0UN00210 | XP_018220777.1 | 100 | 0 |
| SPGP0UH00120 | XP_018220408.1 | 96 | 0 |
| SPGP0KU00110 | XP_018221004.1 | 98 | 2 |
| SPGP0DD00110 | XP_018221000.1 | 99 | 0 |
| SPGP0XC00160 | XP_018220592.1 | 99 | 0 |
| SPGP0OI00380 | XP_018218912.1 | 99 | 0 |
| SPGP0PZ00330 | XP_018222413.1 | 99 | 0 |
| SPGP0PE00180 | XP_018222362.1 | 99 | 0 |
| SPGP0NJ00100 | XP_018223418.1 | 99 | 0 |
| SPGP0MF00220 | XP_018222198.1 | 99 | 0 |
| SPGP0TK00150 | XP_018222729.1 | 99 | 0 |
| SPGP0BK00240 | XP_018221447.1 | 99 | 0 |
| SPGP0KN00200 | XP_018220576.1 | 98 | 2 |
| SPGP0KT00120 | XP_018222244.1 | 99 | 0 |
| SPGP0RC00240 | XP_018222099.1 | 99 | 0 |
| SPGP0CK00110 | XP_018219962.1 | 99 | 0 |
| SPGP0RQ00120 | XP_018221123.1 | 99 | 0 |
| SPGP0SM00160 | XP_018221111.1 | 99 | 0 |
| SPGP0NV00130 | XP_018223450.1 | 99 | 0 |
| SPGP0PZ00140 | XP_018222432.1 | 99 | 0 |
| SPGP0HT00170 | XP_018219009.1 | 99 | 0 |
| SPGP0OI00580 | XP_018218892.1 | 99 | 0 |
| SPGP0HL00110 | XP_018220997.1 | 99 | 0 |
| SPGP0NT00160 | XP_018222468.1 | 99 | 0 |
| SPGP0GX00150 | XP_018222539.1 | 99 | 0 |
| SPGP0IO00120 | XP_018222452.1 | 99 | 0 |
| SPGP0NU00130 | XP_018219058.1 | 99 | 0 |
| SPGP0CC00100 | XP_018220403.1 | 99 | 0 |
| SPGP0WM00100 | XP_018222724.1 | 99 | 0 |
| SPGP0EW00130 | XP_018222292.1 | 99 | 0 |
| SPGP0PL00220 | XP_018219744.1 | 99 | 0 |

Create PDF in your applications with the Pdfcrowd HTML to PDF API PDFCROWD

| | | | |
|---|---|---|---|
| SPGP0GH00160 | XP_018218962.1 | 99 | 0 |
| SPGP0LE00200 | XP_018220758.1 | 99 | 0 |
| SPGP0XF00110 | XP_018220854.1 | 99 | 0 |
| SPGP0LP00100 | XP_018222380.1 | 100 | 0 |
| SPGP0CX00200 | XP_018220789.1 | 99 | 0 |
| SPGP0KC00110 | XP_018222565.1 | 99 | 0 |
| SPGP0VO00140 | XP_018223439.1 | 99 | 0 |
| SPGP0LE00170 | XP_018220761.1 | 99 | 0 |
| SPGP0PO00100 | XP_018219001.1 | 99 | 0 |
| SPGP0VM00250 | XP_018220669.1 | 100 | 0 |
| SPGP0SY00110 | XP_018220801.1 | 99 | 0 |
| SPGP0IK00150 | XP_018222112.1 | 99 | 0 |
| SPGP0ET00140 | XP_018222445.1 | 99 | 0 |
| SPGP0WI00270 | XP_018222135.1 | 99 | 0 |
| SPGP0DZ00120 | XP_018220830.1 | 99 | 0 |
| SPGP0GL00100 | XP_018222611.1 | 99 | 0 |
| SPGP0PA00180 | XP_018220466.1 | 99 | 0 |
| SPGP0OW00100 | XP_018218807.1 | 99 | 0 |
| SPGP0HO00210 | XP_018219981.1 | 99 | 0 |
| SPGP0RI00100 | XP_018220488.1 | 99 | 0 |
| SPGP0LV00120 | XP_018223373.1 | 100 | 0 |
| SPGP0K01490 | XP_018218894.1 | 97 | 0 |
| SPGP0VZ00210 | XP_018223885.1 | 98 | 1 |
| SPGP0LH00140 | XP_018221033.1 | 99 | 0 |
| SPGP0KG00160 | XP_018222630.1 | 99 | 0 |
| SPGP0AH00120 | XP_018219890.1 | 98 | 1 |
| SPGP0IK00140 | XP_018222111.1 | 99 | 0 |
| SPGP0NI00120 | XP_018223908.1 | 99 | 0 |
| SPGP0FW00180 | XP_018219578.1 | 99 | 0 |
| SPGP0CH00110 | XP_018219956.1 | 99 | 0 |
| SPGP0OF00110 | XP_018222692.1 | 99 | 0 |
| SPGP0UN00270 | XP_018220771.1 | 99 | 0 |
| SPGP0VR00100 | XP_018220704.1 | 99 | 0 |
| SPGP0OT00150 | XP_018219134.1 | 99 | 0 |
| SPGP0HG00100 | XP_018220459.1 | 99 | 0 |
| SPGP0LS00150 | XP_018222559.1 | 99 | 0 |
| SPGP0QP00130 | XP_018219166.1 | 99 | 0 |
| SPGP0BR00120 | XP_018222221.1 | 99 | 0 |
| SPGP0WY00100 | XP_018220446.1 | 99 | 0 |
| SPGP0PK00100 | XP_018220423.1 | 99 | 0 |

Create PDF in your applications with the Pdfcrowd HTML to PDF API PDFCROWD

| | | | |
|---|---|---|---|
| SPGP0LK00180 | XP_018221063.1 | 99 | 1 |
| SPGP0KT00130 | XP_018222245.1 | 100 | 0 |
| SPGP0RQ00150 | XP_018221126.1 | 99 | 0 |
| SPGP0SM00140 | XP_018221109.1 | 99 | 0 |
| SPGP0VH00120 | XP_018219683.1 | 99 | 0 |
| SPGP0TX00100 | XP_018222347.1 | 99 | 0 |
| SPGP0HO00110 | XP_018219991.1 | 99 | 0 |
| SPGP0CR00130 | XP_018222211.1 | 98 | 0 |
| SPGP0OG00100 | XP_018220487.1 | 99 | 0 |
| SPGP0HL00130 | XP_018220995.1 | 99 | 0 |
| SPGP0LG00200 | XP_018222652.1 | 100 | 0 |
| SPGP0VU00130 | XP_018222737.1 | 99 | 0 |
| SPGP0IJ00100 | XP_018218823.1 | 99 | 0 |
| SPGP0AQ00100 | XP_018222241.1 | 100 | 0 |
| SPGP0DO00100 | XP_018219967.1 | 99 | 0 |
| SPGP0AP00100 | XP_018219059.1 | 99 | 0 |
| SPGP0EM00170 | XP_018220716.1 | 99 | 0 |
| SPGP0GX00130 | XP_018222537.1 | 99 | 0 |
| SPGP0TQ00100 | XP_018220605.1 | 99 | 0 |
| SPGP0CC00120 | XP_018220405.1 | 99 | 0 |
| SPGP0WG00120 | XP_018221026.1 | 99 | 0 |
| SPGP0IC00200 | XP_018222787.1 | 100 | 0 |
| SPGP0HZ00300 | XP_018219787.1 | 99 | 0 |
| SPGP0NG00130 | XP_018222775.1 | 99 | 0 |
| SPGP0EW00110 | XP_018222290.1 | 99 | 0 |
| SPGP0OI00590 | XP_018218891.1 | 99 | 0 |
| SPGP0CH00120 | XP_018219955.1 | 99 | 0 |
| SPGP0ER00130 | XP_018218980.1 | 99 | 0 |
| SPGP0C02620 | XP_018221216.1 | 87 | 0 |
| SPGP0QK00100 | XP_018220382.1 | 99 | 0 |
| SPGP0FM00220 | XP_018222158.1 | 99 | 0 |
| SPGP0ID00150 | XP_018222145.1 | 99 | 0 |
| SPGP0PL00200 | XP_018219746.1 | 99 | 0 |
| SPGP0IN00100 | XP_018219125.1 | 99 | 0 |
| SPGP0WA00130 | XP_018219853.1 | 99 | 0 |
| SPGP0LG00100 | XP_018222642.1 | 99 | 0 |
| SPGP0NT00210 | XP_018222463.1 | 99 | 0 |
| SPGP0KL00150 | XP_018221099.1 | 99 | 0 |
| SPGP0LU00210 | XP_018224005.1 | 99 | 0 |
| SPGP0LI00140 | XP_018222249.1 | 99 | 0 |

Create PDF in your applications with the Pdfcrowd HTML to PDF API PDFCROWD

| | | | |
|---|---|---|---|
| SPGP0HA00120 | XP_018219870.1 | 99 | 0 |
| SPGP0JW00150 | XP_018220977.1 | 99 | 1 |
| SPGP0PE00340 | XP_018222379.1 | 99 | 0 |
| SPGP0WI00250 | XP_018222133.1 | 99 | 0 |
| SPGP0WJ00120 | XP_018221058.1 | 99 | 0 |
| SPGP0WX00100 | XP_018219953.1 | 99 | 0 |
| SPGP0NQ00200 | XP_018220745.1 | 99 | 0 |
| SPGP0OC00100 | XP_018222531.1 | 100 | 0 |
| SPGP0DU00180 | XP_018222576.1 | 99 | 0 |
| SPGP0HO00230 | XP_018219979.1 | 99 | 0 |
| SPGP0UB00170 | XP_018219042.1 | 99 | 0 |
| SPGP0LT00100 | XP_018221091.1 | 100 | 0 |
| SPGP0MO00240 | XP_018219109.1 | 100 | 0 |
| SPGP0KC00100 | XP_018222566.1 | 100 | 0 |
| SPGP0CF00110 | XP_018222499.1 | 99 | 0 |
| SPGP0DJ00120 | XP_018219919.1 | 99 | 0 |
| SPGP0NT00280 | XP_018222456.1 | 99 | 0 |
| SPGP0UH00170 | XP_018220413.1 | 99 | 0 |
| SPGP0RC00260 | XP_018222101.1 | 99 | 0 |
| SPGP0RX00170 | XP_018222818.1 | 99 | 0 |
| SPGP0PZ00320 | XP_018222414.1 | 99 | 0 |
| SPGP0KN00230 | XP_018220579.1 | 99 | 0 |
| SPGP0DS00100 | XP_018220390.1 | 99 | 0 |
| SPGP0UN00250 | XP_018220773.1 | 99 | 0 |
| SPGP0GF00190 | XP_018219642.1 | 99 | 0 |
| SPGP0DB00110 | XP_018222188.1 | 99 | 0 |
| SPGP0EK00130 | XP_018220811.1 | 99 | 0 |
| SPGP0LU00110 | XP_018224015.1 | 99 | 0 |
| SPGP0LA00190 | XP_018220848.1 | 99 | 0 |
| SPGP0VR00120 | XP_018220702.1 | 99 | 0 |
| SPGP0JJ00110 | XP_018219064.1 | 99 | 1 |
| SPGP0CZ00100 | XP_018219905.1 | 99 | 0 |
| SPGP0MB00110 | XP_018222173.1 | 97 | 2 |
| SPGP0GW00100 | XP_018220829.1 | 99 | 0 |
| SPGP0MF00190 | XP_018222201.1 | 99 | 0 |
| SPGP0BI01870 | XP_018223693.1 | 99 | 0 |
| SPGP0JP00170 | XP_018218854.1 | 99 | 0 |
| SPGP0JC00120 | XP_018219862.1 | 99 | 0 |
| SPGP0EO00130 | XP_018222480.1 | 99 | 0 |
| SPGP0KY00120 | XP_018223903.1 | 99 | 0 |

Create PDF in your applications with the Pdfcrowd HTML to PDF API PDFCROWD

| | | | |
|---|---|---|---|
| SPGP0VJ00100 | XP_018218821.1 | 99 | 0 |
| SPGP0RC00300 | XP_018222105.1 | 99 | 0 |
| SPGP0JH00100 | XP_018218942.1 | 99 | 0 |
| SPGP0EK00210 | XP_018220804.1 | 99 | 0 |
| SPGP0HO00330 | XP_018219970.1 | 99 | 0 |
| SPGP0VZ00160 | XP_018223881.1 | 99 | 0 |
| SPGP0SL00110 | XP_018220485.1 | 99 | 0 |
| SPGP0DO00120 | XP_018219965.1 | 100 | 0 |
| SPGP0HT00220 | XP_018219013.1 | 99 | 0 |
| SPGP0UL00210 | XP_018219668.1 | 100 | 0 |
| SPGP0OX00220 | XP_018219723.1 | 99 | 1 |
| SPGP0US00110 | XP_018219155.1 | 99 | 0 |
| SPGP0EZ00190 | XP_018219324.1 | 99 | 0 |
| SPGP0SL00200 | XP_018220476.1 | 99 | 0 |
| SPGP0MO00290 | XP_018219113.1 | 99 | 0 |
| SPGP0WG00140 | XP_018221024.1 | 99 | 0 |
| SPGP0OF00260 | XP_018222707.1 | 99 | 0 |
| SPGP0LO00110 | XP_018223458.1 | 99 | 0 |
| SPGP0JW00190 | XP_018220981.1 | 99 | 0 |
| SPGP0OI00630 | XP_018218887.1 | 99 | 0 |
| SPGP0ER00110 | XP_018218982.1 | 99 | 0 |
| SPGP0HB00130 | XP_018219191.1 | 99 | 0 |
| SPGP0HO00150 | XP_018219987.1 | 99 | 0 |
| SPGP0TC00110 | XP_018219702.1 | 99 | 0 |
| SPGP0SH00250 | XP_018223983.1 | 99 | 0 |
| SPGP0LG00120 | XP_018222644.1 | 99 | 0 |
| SPGP0TO00140 | XP_018222808.1 | 100 | 0 |
| SPGP0TQ00370 | XP_018220632.1 | 99 | 0 |
| SPGP0NQ00180 | XP_018220747.1 | 99 | 0 |
| SPGP0OC00110 | XP_018222532.1 | 99 | 0 |
| SPGP0PE00320 | XP_018222377.1 | 99 | 0 |
| SPGP0WI00230 | XP_018222131.1 | 99 | 0 |
| SPGP0PZ00340 | XP_018222412.1 | 99 | 0 |
| SPGP0VO00180 | XP_018223442.1 | 99 | 0 |
| SPGP0OC00120 | XP_018222533.1 | 99 | 0 |
| SPGP0EW00150 | XP_018222294.1 | 99 | 0 |
| SPGP0SH00150 | XP_018223973.1 | 99 | 0 |
| SPGP0WQ00160 | XP_018222269.1 | 99 | 0 |
| SPGP0CX00120 | XP_018220797.1 | 99 | 0 |
| SPGP0WJ00110 | XP_018221059.1 | 99 | 0 |

Create PDF in your applications with the Pdfcrowd HTML to PDF API PDFCROWD

| | | | |
|---|---|---|---|
| SPGP0IA00100 | XP_018220585.1 | 99 | 0 |
| SPGP0NG00120 | XP_018222776.1 | 99 | 1 |
| SPGP0CZ00160 | XP_018219911.1 | 99 | 1 |
| SPGP0MD00180 | XP_018222393.1 | 99 | 0 |
| SPGP0NI00160 | XP_018223912.1 | 99 | 0 |
| SPGP0JF00240 | XP_018222322.1 | 99 | 0 |
| SPGP0RX00190 | XP_018222820.1 | 99 | 0 |
| SPGP0LG00140 | XP_018222646.1 | 99 | 0 |
| SPGP0AG00120 | XP_018223462.1 | 86 | 0 |
| SPGP0PE00150 | XP_018222359.1 | 99 | 0 |
| SPGP0IJ00120 | XP_018218824.1 | 99 | 0 |
| SPGP0OF00200 | XP_018222701.1 | 99 | 0 |
| SPGP0VR00130 | XP_018220701.1 | 99 | 0 |
| SPGP0XB00110 | XP_018219021.1 | 99 | 1 |
| SPGP0GF00170 | XP_018219644.1 | 99 | 0 |
| SPGP0NO00120 | XP_018222689.1 | 99 | 0 |
| SPGP0OI00400 | XP_018218910.1 | 99 | 0 |
| SPGP0VN00160 | XP_018219806.1 | 99 | 0 |
| SPGP0HZ00110 | XP_018219768.1 | 99 | 0 |
| SPGP0VZ00130 | XP_018223878.1 | 99 | 0 |
| SPGP0VM00120 | XP_018220682.1 | 99 | 0 |
| SPGP0GW00120 | XP_018220827.1 | 99 | 0 |
| SPGP0MY00110 | XP_018222623.1 | 99 | 0 |
| SPGP0EO00110 | XP_018222478.1 | 99 | 0 |
| SPGP0JH00120 | XP_018218940.1 | 99 | 0 |
| SPGP0NR00110 | XP_018222568.1 | 100 | 0 |
| SPGP0XC00260 | XP_018220602.1 | 99 | 0 |
| SPGP0NT00150 | XP_018222469.1 | 99 | 0 |
| SPGP0WI00150 | XP_018222123.1 | 99 | 0 |
| SPGP0HJ00170 | XP_018224043.1 | 97 | 0 |
| SPGP0OL00150 | XP_018222283.1 | 99 | 0 |
| SPGP0NQ00210 | XP_018220744.1 | 99 | 0 |
| SPGP0MP00100 | XP_018222751.1 | 99 | 0 |
| SPGP0VZ00180 | XP_018223883.1 | 99 | 0 |
| SPGP0EY00100 | XP_018219088.1 | 99 | 0 |
| SPGP0UL00180 | XP_018219665.1 | 99 | 0 |
| SPGP0ED00110 | XP_018222599.1 | 99 | 0 |
| SPGP0OB00100 | XP_018219793.1 | 99 | 0 |
| SPGP0LA00180 | XP_018220847.1 | 99 | 0 |
| SPGP0JW00170 | XP_018220979.1 | 99 | 0 |

Create PDF in your applications with the Pdfcrowd HTML to PDF API PDFCROWD

| | | | |
|---|---|---|---|
| SPGP0KS00110 | XP_018220461.1 | 99 | 0 |
| SPGP0OX00290 | XP_018219716.1 | 99 | 0 |
| SPGP0KY00110 | XP_018223902.1 | 99 | 0 |
| SPGP0RI00270 | XP_018220505.1 | 100 | 0 |
| SPGP0WG00160 | XP_018221022.1 | 99 | 0 |
| SPGP0UH00110 | XP_018220407.1 | 99 | 0 |
| SPGP0SB00130 | XP_018222301.1 | 99 | 0 |
| SPGP0HD00120 | XP_018222516.1 | 99 | 0 |
| SPGP0Y00120 | XP_018222683.1 | 99 | 0 |
| SPGP0FM00150 | XP_018222151.1 | 99 | 0 |
| SPGP0FX00110 | XP_018222823.1 | 99 | 0 |
| SPGP0UL00200 | XP_018219667.1 | 99 | 0 |
| SPGP0KU00120 | XP_018221003.1 | 99 | 0 |
| SPGP0QS00100 | XP_018223896.1 | 99 | 0 |
| SPGP0GI00130 | XP_018220697.1 | 99 | 0 |
| SPGP0HY00120 | XP_018223989.1 | 99 | 0 |
| SPGP0QP00110 | XP_018219168.1 | 99 | 0 |
| SPGP0IC00190 | XP_018222788.1 | 99 | 0 |
| SPGP0BK01260 | XP_018221348.1 | 99 | 0 |
| SPGP0DZ00110 | XP_018220831.1 | 99 | 0 |
| SPGP0HX00180 | XP_018221007.1 | 99 | 0 |
| SPGP0OF00390 | XP_018222720.1 | 99 | 0 |
| SPGP0TL00150 | XP_018219083.1 | 99 | 0 |
| SPGP0OH00110 | XP_018220985.1 | 98 | 1 |
| SPGP0OB00120 | XP_018219795.1 | 99 | 0 |
| SPGP0RE00150 | XP_018220399.1 | 99 | 0 |
| SPGP0OI00300 | XP_018218920.1 | 99 | 0 |
| SPGP0SS00100 | XP_018219049.1 | 99 | 0 |
| SPGP0DP00100 | XP_018220644.1 | 99 | 0 |
| SPGP0SW00150 | XP_018220559.1 | 99 | 0 |
| SPGP0BK00130 | XP_018221991.1 | 98 | 0 |
| SPGP0QJ00110 | XP_018219934.1 | 99 | 0 |
| SPGP0JN00200 | XP_018219079.1 | 99 | 0 |
| SPGP0OI00200 | XP_018218927.1 | 99 | 0 |
| SPGP0HO00270 | XP_018219975.1 | 99 | 0 |
| SPGP0OE00110 | XP_018219609.1 | 99 | 0 |
| SPGP0PU00100 | XP_018223959.1 | 99 | 0 |
| SPGP0ST00180 | XP_018220541.1 | 99 | 0 |
| SPGP0CF00140 | XP_018222496.1 | 99 | 0 |

Create PDF in your applications with the Pdfcrowd HTML to PDF API PDFCROWD

| | | | |
|---|---|---|---|
| SPGP0CZ00210 | XP_018219916.1 | 99 | 0 |
| SPGP0RI00180 | XP_018220496.1 | 99 | 0 |
| SPGP0UF00130 | XP_018221037.1 | 99 | 0 |
| SPGP0RX00200 | XP_018222821.1 | 99 | 0 |
| SPGP0OT00130 | XP_018219132.1 | 99 | 0 |
| SPGP0RC00220 | XP_018222097.1 | 99 | 0 |
| SPGP0LN00120 | XP_018223924.1 | 99 | 0 |
| SPGP0HB00100 | XP_018219194.1 | 98 | 0 |
| SPGP0TE00150 | XP_018219897.1 | 99 | 0 |
| SPGP0DK00110 | XP_018219068.1 | 99 | 0 |
| SPGP0DS00140 | XP_018220394.1 | 99 | 0 |
| SPGP0MF00100 | XP_018222210.1 | 98 | 1 |
| SPGP0XB00130 | XP_018219023.1 | 99 | 0 |
| SPGP0NF00100 | XP_018222636.1 | 100 | 0 |
| SPGP0HZ00280 | XP_018219785.1 | 99 | 0 |
| SPGP0FX00120 | XP_018222824.1 | 99 | 0 |
| SPGP0TC00120 | XP_018219701.1 | 97 | 2 |
| SPGP0LU00150 | XP_018224011.1 | 99 | 0 |
| SPGP0JP00110 | XP_018218848.1 | 99 | 1 |
| SPGP0VN00140 | XP_018219804.1 | 99 | 0 |
| SPGP0GB00120 | XP_018222657.1 | 98 | 0 |
| SPGP0VM00140 | XP_018220680.1 | 99 | 0 |
| SPGP0NG00170 | XP_018222771.1 | 99 | 0 |
| SPGP0PL00250 | XP_018219741.1 | 99 | 0 |
| SPGP0BT00110 | XP_018220735.1 | 99 | 0 |
| SPGP0NQ00230 | XP_018220742.1 | 99 | 0 |
| SPGP0GN00120 | XP_018222586.1 | 100 | 0 |
| SPGP0KI00110 | XP_018220840.1 | 99 | 0 |
| SPGP0WI00170 | XP_018222125.1 | 99 | 0 |
| SPGP0IV00110 | XP_018220639.1 | 99 | 0 |
| SPGP0OX00270 | XP_018219718.1 | 99 | 0 |
| SPGP0RI00210 | XP_018220499.1 | 99 | 0 |
| SPGP0WG00180 | XP_018221020.1 | 99 | 0 |
| SPGP0HD00100 | XP_018222514.1 | 97 | 2 |
| SPGP0DC00120 | XP_018222639.1 | 99 | 0 |
| SPGP0GN00110 | XP_018222587.1 | 99 | 0 |
| SPGP0FM00170 | XP_018222153.1 | 99 | 0 |
| SPGP0TM00120 | XP_018219876.1 | 100 | 0 |
| SPGP0OI00670 | XP_018218883.1 | 99 | 0 |
| SPGP0PZ00240 | XP_018222422.1 | 99 | 0 |

Create PDF in your applications with the Pdfcrowd HTML to PDF API PDFCROWD

| | | | |
|---|---|---|---|
| SPGP0FM00280 | XP_018222163.1 | 100 | 0 |
| SPGP0TC00150 | XP_018219698.1 | 99 | 0 |
| SPGP0HX00140 | XP_018221010.1 | 99 | 0 |
| SPGP0F01730 | XP_018223161.1 | 99 | 0 |
| SPGP0PV00120 | XP_018218977.1 | 99 | 0 |
| SPGP0TS00110 | XP_018219590.1 | 99 | 0 |
| SPGP0RC00120 | XP_018222087.1 | 99 | 0 |
| SPGP0NE00100 | XP_018221072.1 | 96 | 0 |
| SPGP0UL00150 | XP_018219662.1 | 99 | 0 |
| SPGP0RE00120 | XP_018220396.1 | 99 | 0 |
| SPGP0IW00110 | XP_018220015.1 | 99 | 0 |
| SPGP0TF00110 | XP_018222186.1 | 99 | 0 |
| SPGP0LP00110 | XP_018222381.1 | 99 | 0 |
| SPGP0UL00170 | XP_018219664.1 | 99 | 0 |
| SPGP0DP00160 | XP_018220650.1 | 99 | 0 |
| SPGP0KM00100 | XP_018222224.1 | 99 | 0 |
| SPGP0VO00130 | XP_018223438.1 | 99 | 0 |
| SPGP0KZ00330 | XP_018223929.1 | 99 | 0 |
| SPGP0LA00140 | XP_018220844.1 | 99 | 0 |
| SPGP0QV00240 | XP_018219996.1 | 99 | 0 |
| SPGP0WY00150 | XP_018220441.1 | 99 | 0 |
| SPGP0JF00230 | XP_018222321.1 | 99 | 0 |
| SPGP0JF00200 | XP_018222318.1 | 99 | 0 |
| SPGP0LN00100 | XP_018223922.1 | 99 | 0 |
| SPGP0Y00150 | XP_018222686.1 | 99 | 0 |
| SPGP0PA00110 | XP_018220473.1 | 99 | 0 |
| SPGP0XB00150 | XP_018219025.1 | 99 | 0 |
| SPGP0RI00400 | XP_018220519.1 | 99 | 0 |
| SPGP0RH00120 | XP_018220417.1 | 99 | 0 |
| SPGP0RQ00180 | XP_018221129.1 | 99 | 0 |
| SPGP0SH00100 | XP_018223968.1 | 99 | 0 |
| SPGP0AA00140 | XP_018221081.1 | 99 | 0 |
| SPGP0NL00100 | XP_018218984.1 | 99 | 0 |
| SPGP0AB00110 | XP_018219706.1 | 99 | 0 |
| SPGP0HZ00150 | XP_018219772.1 | 99 | 0 |
| SPGP0TQ00280 | XP_018220623.1 | 99 | 0 |
| SPGP0LS00160 | XP_018222558.1 | 99 | 0 |
| SPGP0GK00140 | XP_018219710.1 | 100 | 0 |
| SPGP0NG00150 | XP_018222773.1 | 99 | 0 |
| SPGP0HL00170 | XP_018220991.1 | 99 | 0 |

Create PDF in your applications with the Pdfcrowd HTML to PDF API PDFCROWD

| | | | |
|---|---|---|---|
| SPGP0BI02540 | XP_018223628.1 | 100 | 0 |
| SPGP0WI00190 | XP_018222127.1 | 99 | 0 |
| SPGP0HB00140 | XP_018219190.1 | 99 | 0 |
| SPGP0MF00120 | XP_018222208.1 | 99 | 0 |
| SPGP0QS00130 | XP_018223899.1 | 99 | 0 |
| SPGP0IE00110 | XP_018222764.1 | 99 | 0 |
| SPGP0SW00130 | XP_018220561.1 | 99 | 0 |
| SPGP0OZ00150 | XP_018222402.1 | 99 | 0 |
| SPGP0OU00100 | XP_018219195.1 | 99 | 0 |
| SPGP0NT00240 | XP_018222460.1 | 99 | 0 |
| SPGP0RI00230 | XP_018220501.1 | 99 | 0 |
| SPGP0JM00100 | XP_018222589.1 | 99 | 0 |
| SPGP0DC00100 | XP_018222637.1 | 99 | 0 |
| SPGP0NT00260 | XP_018222458.1 | 99 | 0 |
| SPGP0ID00110 | XP_018222141.1 | 99 | 0 |
| SPGP0NV00180 | XP_018223456.1 | 99 | 0 |
| SPGP0GI00170 | XP_018220693.1 | 99 | 0 |
| SPGP0TC00170 | XP_018219696.1 | 99 | 0 |
| SPGP0PK00110 | XP_018220422.1 | 99 | 0 |
| SPGP0OI00460 | XP_018218904.1 | 99 | 0 |
| SPGP0QP00150 | XP_018219164.1 | 99 | 0 |
| SPGP0WQ00180 | XP_018222271.1 | 99 | 0 |
| SPGP0VN00340 | XP_018219824.1 | 99 | 0 |
| SPGP0OE00210 | XP_018219599.1 | 99 | 0 |
| SPGP0QV00270 | XP_018219993.1 | 99 | 0 |
| SPGP0RC00100 | XP_018222085.1 | 99 | 0 |
| SPGP0TQ00230 | XP_018220618.1 | 99 | 0 |
| SPGP0TL00110 | XP_018219087.1 | 99 | 0 |
| SPGP0XC00220 | XP_018220598.1 | 99 | 0 |
| SPGP0CW00110 | XP_018222505.1 | 99 | 0 |
| SPGP0OF00240 | XP_018222705.1 | 99 | 0 |
| SPGP0WJ00160 | XP_018221054.1 | 99 | 0 |
| SPGP0SO00100 | XP_018219045.1 | 99 | 0 |
| SPGP0AA00170 | XP_018221078.1 | 99 | 0 |
| SPGP0CF00160 | XP_018222494.1 | 99 | 0 |
| SPGP0ST00140 | XP_018220545.1 | 99 | 0 |
| SPGP0WJ00170 | XP_018221053.1 | 99 | 0 |
| SPGP0PE00310 | XP_018222376.1 | 99 | 0 |
| SPGP0MG00150 | XP_018219613.1 | 99 | 0 |
| SPGP0DN00160 | XP_018218988.1 | 99 | 0 |

Create PDF in your applications with the Pdfcrowd HTML to PDF API PDFCROWD

| | | | |
|---|---|---|---|
| SPGP0GV00130 | XP_018218863.1 | 99 | 0 |
| SPGP0BE00130 | XP_018223321.1 | 99 | 0 |
| SPGP0HZ00230 | XP_018219780.1 | 99 | 0 |
| SPGP0FC00140 | XP_018219055.1 | 99 | 0 |
| SPGP0N00740 | XP_018219261.1 | 99 | 0 |
| SPGP0JJ00100 | XP_018219063.1 | 99 | 0 |
| SPGP0PA00170 | XP_018220467.1 | 99 | 0 |
| SPGP0PA00190 | XP_018220465.1 | 99 | 0 |
| SPGP0VN00100 | XP_018219800.1 | 99 | 0 |
| SPGP0FM00320 | XP_018222167.1 | 99 | 0 |
| SPGP0KO00100 | XP_018219704.1 | 99 | 0 |
| SPGP0AA00120 | XP_018221083.1 | 99 | 0 |
| SPGP0SH00210 | XP_018223979.1 | 99 | 0 |
| SPGP0HU00120 | XP_018220428.1 | 99 | 0 |
| SPGP0NQ00270 | XP_018220738.1 | 99 | 0 |
| SPGP0JN00180 | XP_018219077.1 | 99 | 0 |
| SPGP0IE00140 | XP_018222762.1 | 99 | 0 |
| SPGP0QV00130 | XP_018220007.1 | 99 | 0 |
| SPGP0OI00800 | XP_018218870.1 | 99 | 0 |
| SPGP0OZ00170 | XP_018222404.1 | 99 | 0 |
| SPGP0FT00130 | XP_018222733.1 | 99 | 0 |
| SPGP0RX00160 | XP_018222817.1 | 99 | 0 |
| SPGP0XC00130 | XP_018220589.1 | 99 | 0 |
| SPGP0PZ00280 | XP_018222418.1 | 99 | 0 |
| SPGP0VZ00150 | XP_018223880.1 | 99 | 0 |
| SPGP0FM00130 | XP_018222149.1 | 99 | 0 |
| SPGP0WA00120 | XP_018219854.1 | 99 | 0 |
| SPGP0VN00280 | XP_018219818.1 | 99 | 0 |
| SPGP0KQ00150 | XP_018219925.1 | 96 | 0 |
| SPGP0MC00180 | XP_018222669.1 | 100 | 0 |
| SPGP0UR00220 | XP_018222332.1 | 99 | 0 |
| SPGP0Y00140 | XP_018222685.1 | 99 | 0 |
| SPGP0RQ00100 | XP_018221122.1 | 99 | 0 |
| SPGP0CX00130 | XP_018220796.1 | 99 | 0 |
| SPGP0QP00170 | XP_018219162.1 | 99 | 0 |
| SPGP0JP00140 | XP_018218851.1 | 99 | 0 |
| SPGP0CW00130 | XP_018222506.1 | 99 | 0 |
| SPGP0EU00140 | XP_018222662.1 | 99 | 0 |
| SPGP0JQ00110 | XP_018222440.1 | 99 | 0 |
| SPGP0SH00220 | XP_018223980.1 | 99 | 0 |

Create PDF in your applications with the Pdfcrowd HTML to PDF API PDFCROWD

| | | | |
|---|---|---|---|
| SPGP0PE00170 | XP_018222361.1 | 99 | 0 |
| SPGP0KS00100 | XP_018220462.1 | 99 | 0 |
| SPGP0VP00130 | XP_018219948.1 | 99 | 0 |
| SPGP0WW00110 | XP_018222073.1 | 99 | 0 |
| SPGP0LJ00100 | XP_018219142.1 | 99 | 0 |
| SPGP0WS00110 | XP_018222748.1 | 99 | 0 |
| SPGP0EK00170 | XP_018220808.1 | 99 | 0 |
| SPGP0SK00110 | XP_018222280.1 | 99 | 0 |
| SPGP0GF00330 | XP_018219628.1 | 99 | 0 |
| SPGP0RI00130 | XP_018220491.1 | 99 | 0 |
| SPGP0NO00130 | XP_018222690.1 | 99 | 0 |
| SPGP0OB00160 | XP_018219799.1 | 99 | 0 |
| SPGP0HZ00250 | XP_018219782.1 | 99 | 0 |
| SPGP0FE00290 | XP_018220522.1 | 99 | 0 |
| SPGP0OX00160 | XP_018219729.1 | 99 | 0 |
| SPGP0KA00110 | XP_018222227.1 | 99 | 0 |
| SPGP0Y00110 | XP_018222682.1 | 99 | 0 |
| SPGP0OF00330 | XP_018222714.1 | 99 | 0 |
| SPGP0JJ00120 | XP_018219065.1 | 100 | 0 |
| SPGP0TT00130 | XP_018220835.1 | 99 | 0 |
| SPGP0GI00140 | XP_018220696.1 | 99 | 0 |
| SPGP0IR00150 | XP_018219762.1 | 99 | 0 |
| SPGP0NV00120 | XP_018223449.1 | 99 | 0 |
| SPGP0DT00130 | XP_018221085.1 | 99 | 0 |
| SPGP0NR00100 | XP_018222567.1 | 99 | 0 |
| SPGP0PL00120 | XP_018219754.1 | 99 | 0 |
| SPGP0PZ00130 | XP_018222433.1 | 99 | 0 |
| SPGP0QV00250 | XP_018219995.1 | 99 | 0 |
| SPGP0LS00120 | XP_018222561.1 | 99 | 0 |
| SPGP0GW00210 | XP_018220820.1 | 99 | 0 |
| SPGP0OR00200 | XP_018218836.1 | 99 | 0 |
| SPGP0LU00100 | XP_018224016.1 | 99 | 0 |
| SPGP0HU00140 | XP_018220430.1 | 99 | 0 |
| SPGP0VB00130 | XP_018221046.1 | 99 | 0 |
| SPGP0C01860 | XP_018220184.1 | 99 | 0 |
| SPGP0OX00190 | XP_018219726.1 | 100 | 0 |
| SPGP0NK00140 | XP_018220451.1 | 99 | 0 |
| SPGP0JF00210 | XP_018222319.1 | 99 | 0 |
| SPGP0FE00100 | XP_018220540.1 | 99 | 0 |
| SPGP0IE00160 | XP_018222760.1 | 99 | 0 |

Create PDF in your applications with the Pdfcrowd HTML to PDF API PDFCROWD

| | | | |
|---|---|---|---|
| SPGP0TM00180 | XP_018219882.1 | 99 | 0 |
| SPGP0EK00160 | XP_018220809.1 | 99 | 0 |
| SPGP0PJ00100 | XP_018222115.1 | 99 | 0 |
| SPGP0TE00170 | XP_018219899.1 | 99 | 0 |
| SPGP0SU00120 | XP_018218972.1 | 99 | 0 |
| SPGP0DC00140 | XP_018222641.1 | 99 | 0 |
| SPGP0NG00110 | XP_018222777.1 | 99 | 0 |
| SPGP0KQ00130 | XP_018219927.1 | 99 | 0 |
| SPGP0MD00200 | XP_018222395.1 | 99 | 0 |
| SPGP0SA00100 | XP_018222216.1 | 99 | 0 |
| SPGP0TW00110 | XP_018218844.1 | 99 | 0 |
| SPGP0KN00190 | XP_018220575.1 | 99 | 0 |
| SPGP0HX00100 | XP_018221014.1 | 99 | 0 |
| SPGP0WQ00140 | XP_018222267.1 | 99 | 0 |
| SPGP0II00100 | XP_018222555.1 | 99 | 0 |
| SPGP0OE00250 | XP_018219596.1 | 100 | 0 |
| SPGP0TQ00270 | XP_018220622.1 | 99 | 0 |
| SPGP0GB00100 | XP_018222655.1 | 99 | 0 |
| SPGP0GR00110 | XP_018222739.1 | 99 | 0 |
| SPGP0NX00110 | XP_018222475.1 | 99 | 0 |
| SPGP0FW00100 | XP_018219586.1 | 99 | 0 |
| SPGP0DX00110 | XP_018219327.1 | 99 | 0 |
| SPGP0EM00230 | XP_018220712.1 | 99 | 0 |
| SPGP0OR00110 | XP_018218827.1 | 99 | 0 |
| SPGP0UR00230 | XP_018222331.1 | 99 | 0 |
| SPGP0VP00150 | XP_018219946.1 | 99 | 0 |
| SPGP0HO00140 | XP_018219988.1 | 98 | 0 |
| SPGP0SM00150 | XP_018221110.1 | 99 | 0 |
| SPGP0EK00150 | XP_018220810.1 | 99 | 0 |
| SPGP0BU00110 | XP_018222250.1 | 99 | 0 |
| SPGP0QF00110 | XP_018224019.1 | 99 | 0 |
| SPGP0IR00190 | XP_018219758.1 | 99 | 0 |
| SPGP0KL00220 | XP_018221093.1 | 99 | 0 |
| SPGP0HZ00100 | XP_018219767.1 | 100 | 0 |
| SPGP0QP00180 | XP_018219161.1 | 99 | 0 |
| SPGP0HZ00270 | XP_018219784.1 | 99 | 0 |
| SPGP0MF00180 | XP_018222202.1 | 99 | 0 |
| SPGP0RE00140 | XP_018220398.1 | 99 | 0 |
| SPGP0GG00100 | XP_018220657.1 | 99 | 0 |
| SPGP0NV00160 | XP_018223454.1 | 99 | 0 |

Create PDF in your applications with the Pdfcrowd HTML to PDF API PDFCROWD

| | | | |
|---|---|---|---|
| SPGP0RI00390 | XP_018220518.1 | 99 | 0 |
| SPGP0RI00350 | XP_018220514.1 | 99 | 0 |
| SPGP0IR00130 | XP_018219764.1 | 99 | 0 |
| SPGP0XC00110 | XP_018220587.1 | 99 | 0 |
| SPGP0VN00410 | XP_018219831.1 | 99 | 0 |
| SPGP0LI00130 | XP_018222248.1 | 99 | 0 |
| SPGP0TV00100 | XP_018222351.1 | 99 | 0 |
| SPGP0PZ00150 | XP_018222431.1 | 99 | 0 |
| SPGP0HU00160 | XP_018220432.1 | 99 | 0 |
| SPGP0OI00330 | XP_018218917.1 | 99 | 0 |
| SPGP0MO00340 | XP_018219118.1 | 99 | 0 |
| SPGP0VS00120 | XP_018222522.1 | 99 | 0 |
| SPGP0OI00260 | XP_018218924.1 | 99 | 0 |
| SPGP0EX00140 | XP_018223423.1 | 99 | 0 |
| SPGP0NK00130 | XP_018220452.1 | 99 | 0 |
| SPGP0VZ00120 | XP_018223877.1 | 99 | 0 |
| SPGP0HA00110 | XP_018219869.1 | 99 | 0 |
| SPGP0PL00160 | XP_018219750.1 | 100 | 0 |
| SPGP0GG00110 | XP_018220656.1 | 99 | 0 |
| SPGP0PJ00120 | XP_018222117.1 | 99 | 0 |
| SPGP0TV00110 | XP_018222352.1 | 99 | 0 |
| SPGP0OI00470 | XP_018218903.1 | 99 | 0 |
| SPGP0RK00150 | XP_018224037.1 | 99 | 0 |
| SPGP0HT00100 | XP_018219002.1 | 99 | 0 |
| SPGP0WS00100 | XP_018222747.1 | 99 | 0 |
| SPGP0RC00170 | XP_018222092.1 | 99 | 0 |
| SPGP0RI00140 | XP_018220492.1 | 99 | 1 |
| SPGP0VN00250 | XP_018219815.1 | 99 | 0 |
| SPGP0GH00200 | XP_018218958.1 | 99 | 0 |
| SPGP0TG00100 | XP_018220660.1 | 99 | 0 |
| SPGP0HB00150 | XP_018219189.1 | 98 | 0 |
| SPGP0GI00180 | XP_018220692.1 | 99 | 0 |
| SPGP0MS00120 | XP_018222328.1 | 99 | 0 |
| SPGP0VE00130 | XP_018220965.1 | 99 | 0 |
| SPGP0QW00150 | XP_018222607.1 | 99 | 0 |
| SPGP0GK00120 | XP_018219712.1 | 99 | 0 |
| SPGP0GR00130 | XP_018222741.1 | 99 | 0 |
| SPGP0FW00160 | XP_018219580.1 | 99 | 0 |
| SPGP0UM00100 | XP_018222569.1 | 98 | 2 |
| SPGP0GF00220 | XP_018219639.1 | 99 | 1 |

Create PDF in your applications with the Pdfcrowd HTML to PDF API PDFCROWD

| | | | |
|---|---|---|---|
| SPGP0EU00100 | XP_018222658.1 | 99 | 0 |
| SPGP0CD00120 | XP_018222529.1 | 99 | 0 |
| SPGP0OR00130 | XP_018218829.1 | 99 | 0 |
| SPGP0NG00100 | XP_018222778.1 | 99 | 0 |
| SPGP0HO00160 | XP_018219986.1 | 99 | 0 |
| SPGP0EZ00160 | XP_018219321.1 | 99 | 0 |
| SPGP0MS00110 | XP_018222327.1 | 99 | 0 |
| SPGP0PG00110 | XP_018219178.1 | 99 | 0 |
| SPGP0EK00180 | XP_018220807.1 | 99 | 0 |
| SPGP0FE00250 | XP_018220526.1 | 99 | 0 |
| SPGP0RI00330 | XP_018220512.1 | 85 | 0 |
| SPGP0VZ00300 | XP_018223894.1 | 99 | 0 |
| SPGP0HZ00220 | XP_018219779.1 | 99 | 0 |
| SPGP0OI00550 | XP_018218895.1 | 99 | 0 |
| SPGP0ST00130 | XP_018220546.1 | 99 | 0 |
| SPGP0JF00130 | XP_018222311.1 | 99 | 0 |
| SPGP0LI00100 | XP_018222246.1 | 99 | 0 |
| SPGP0TB00110 | XP_018222798.1 | 99 | 0 |
| SPGP0JR00140 | XP_018219027.1 | 99 | 0 |
| SPGP0DT00100 | XP_018221088.1 | 99 | 0 |
| SPGP0NK00110 | XP_018220454.1 | 99 | 0 |
| SPGP0GF00100 | XP_018219651.1 | 99 | 0 |
| SPGP0MO00120 | XP_018219097.1 | 99 | 0 |
| SPGP0IE00120 | XP_018222763.1 | 99 | 0 |
| SPGP0RX00100 | XP_018222811.1 | 99 | 0 |
| SPGP0OI00410 | XP_018218909.1 | 99 | 0 |
| SPGP0WA00140 | XP_018219852.1 | 99 | 0 |
| SPGP0UO00120 | XP_018222184.1 | 98 | 1 |
| SPGP0FA00110 | XP_018219181.1 | 99 | 0 |
| SPGP0EQ00100 | XP_018220664.1 | 99 | 0 |
| SPGP0GT00100 | XP_018222508.1 | 99 | 0 |
| SPGP0WT00110 | XP_018220652.1 | 99 | 0 |
| SPGP0RC00190 | XP_018222094.1 | 99 | 0 |
| SPGP0PA00100 | XP_018220474.1 | 100 | 0 |
| SPGP0JW00200 | XP_018220982.1 | 99 | 0 |
| SPGP0KZ00240 | XP_018223938.1 | 100 | 0 |
| SPGP0UR00210 | XP_018222333.1 | 99 | 0 |
| SPGP0FW00140 | XP_018219582.1 | 99 | 0 |
| SPGP0UZ00160 | XP_018223431.1 | 99 | 0 |
| SPGP0VQ00130 | XP_018222546.1 | 99 | 0 |

Create PDF in your applications with the Pdfcrowd HTML to PDF API PDFCROWD

| | | | |
|---|---|---|---|
| SPGP0VN00350 | XP_018219825.1 | 99 | 0 |
| SPGP0JT00100 | XP_018219859.1 | 99 | 1 |
| SPGP0GW00150 | XP_018220825.1 | 99 | 0 |
| SPGP0KZ00280 | XP_018223934.1 | 99 | 0 |
| SPGP0EK00110 | XP_018220813.1 | 99 | 0 |
| SPGP0OW00160 | XP_018218813.1 | 99 | 0 |
| SPGP0U00320 | XP_018224104.1 | 100 | 0 |
| SPGP0HY00110 | XP_018223988.1 | 99 | 0 |
| SPGP0WY00200 | XP_018220436.1 | 99 | 0 |
| SPGP0VH00160 | XP_018219687.1 | 99 | 0 |
| SPGP0WQ00150 | XP_018222268.1 | 99 | 0 |
| SPGP0HZ00240 | XP_018219781.1 | 99 | 0 |
| SPGP0VO00210 | XP_018223444.1 | 99 | 0 |
| SPGP0OI00530 | XP_018218897.1 | 99 | 0 |
| SPGP0QF00160 | XP_018224024.1 | 99 | 0 |
| SPGP0OE00170 | XP_018219603.1 | 99 | 0 |
| SPGP0BE02360 | XP_018223102.1 | 99 | 0 |
| SPGP0MO00100 | XP_018219095.1 | 99 | 0 |
| SPGP0FM00210 | XP_018222157.1 | 98 | 0 |
| SPGP0LA00210 | XP_018220850.1 | 99 | 0 |
| SPGP0DP00110 | XP_018220645.1 | 99 | 0 |
| SPGP0EX00160 | XP_018223425.1 | 99 | 0 |
| SPGP0SW00160 | XP_018220558.1 | 100 | 0 |
| SPGP0GR00120 | XP_018222740.1 | 100 | 0 |
| SPGP0MB00140 | XP_018222176.1 | 99 | 0 |
| SPGP0WY00180 | XP_018220438.1 | 99 | 0 |
| SPGP0JH00110 | XP_018218941.1 | 99 | 0 |
| SPGP0OI00280 | XP_018218922.1 | 99 | 0 |
| SPGP0HA00130 | XP_018219871.1 | 99 | 0 |
| SPGP0HE00100 | XP_018220658.1 | 99 | 0 |
| SPGP0MT00110 | XP_018219137.1 | 99 | 0 |
| SPGP0QW00110 | XP_018222603.1 | 99 | 0 |
| SPGP0PZ00370 | XP_018222409.1 | 99 | 0 |
| SPGP0LX00110 | XP_018219145.1 | 99 | 0 |
| SPGP0KG00130 | XP_018222627.1 | 99 | 0 |
| SPGP0TQ00290 | XP_018220624.1 | 99 | 0 |
| SPGP0PZ00170 | XP_018222428.1 | 97 | 2 |
| SPGP0OF00150 | XP_018222696.1 | 99 | 0 |
| SPGP0OF00400 | XP_018222721.1 | 100 | 0 |
| SPGP0KZ00110 | XP_018223949.1 | 100 | 0 |

Create PDF in your applications with the Pdfcrowd HTML to PDF API PDFCROWD

| | | | |
|---|---|---|---|
| SPGP0WQ00200 | XP_018222273.1 | 99 | 0 |
| SPGP0WG00170 | XP_018221021.1 | 99 | 0 |
| SPGP0GF00340 | XP_018219627.1 | 99 | 0 |
| SPGP0WH00110 | XP_018219921.1 | 99 | 0 |
| SPGP0LK00110 | XP_018221070.1 | 99 | 0 |
| SPGP0JT00120 | XP_018219861.1 | 99 | 0 |
| SPGP0OI00270 | XP_018218923.1 | 99 | 0 |
| SPGP0MO00130 | XP_018219098.1 | 99 | 0 |
| SPGP0OZ00120 | XP_018222399.1 | 99 | 0 |
| SPGP0NY00150 | XP_018219624.1 | 99 | 0 |
| SPGP0NV00100 | XP_018223447.1 | 99 | 0 |
| SPGP0PG00130 | XP_018219176.1 | 99 | 0 |
| SPGP0HT00140 | XP_018219006.1 | 99 | 0 |
| SPGP0QV00210 | XP_018219999.1 | 99 | 0 |
| SPGP0EM00250 | XP_018220710.1 | 99 | 0 |
| SPGP0MO00270 | XP_018219111.1 | 99 | 0 |
| SPGP0FW00260 | XP_018219570.1 | 99 | 0 |
| SPGP0QF00180 | XP_018224026.1 | 99 | 0 |
| SPGP0CD00100 | XP_018222527.1 | 100 | 0 |
| SPGP0EM00160 | XP_018220717.1 | 99 | 0 |
| SPGP0GW00170 | XP_018220824.1 | 99 | 0 |
| SPGP0EF00110 | XP_018220018.1 | 99 | 0 |
| SPGP0IC00120 | XP_018222795.1 | 99 | 0 |
| SPGP0UZ00100 | XP_018223426.1 | 99 | 0 |
| SPGP0JN00160 | XP_018219075.1 | 99 | 0 |
| SPGP0DO00130 | XP_018219964.1 | 99 | 0 |
| SPGP0RJ00100 | XP_018222501.1 | 99 | 0 |
| SPGP0UN00170 | XP_018220781.1 | 99 | 0 |
| SPGP0GR00170 | XP_018222745.1 | 99 | 0 |
| SPGP0JF00100 | XP_018222308.1 | 99 | 0 |
| SPGP0BL00110 | XP_018222180.1 | 99 | 0 |
| SPGP0VL00110 | XP_018219128.1 | 99 | 0 |
| SPGP0LX00170 | XP_018219151.1 | 99 | 1 |
| SPGP0FW00200 | XP_018219576.1 | 99 | 0 |
| SPGP0OK00110 | XP_018219864.1 | 99 | 0 |
| SPGP0CX00190 | XP_018220790.1 | 99 | 0 |
| SPGP0TM00200 | XP_018219884.1 | 99 | 0 |
| SPGP0PE00230 | XP_018222368.1 | 99 | 0 |
| SPGP0PE00210 | XP_018222365.1 | 99 | 0 |
| SPGP0LK00130 | XP_018221068.1 | 100 | 0 |

Create PDF in your applications with the Pdfcrowd HTML to PDF API PDFCROWD

| | | | |
|---|---|---|---|
| SPGP0HL00160 | XP_018220992.1 | 99 | 0 |
| SPGP0TQ00410 | XP_018220636.1 | 99 | 0 |
| SPGP0VG00120 | XP_018223870.1 | 99 | 0 |
| SPGP0FE00230 | XP_018220528.1 | 99 | 0 |
| SPGP0OZ00140 | XP_018222401.1 | 99 | 0 |
| SPGP0PB00100 | XP_018222215.1 | 99 | 0 |
| SPGP0HX00110 | XP_018221013.1 | 99 | 0 |
| SPGP0MO00310 | XP_018219115.1 | 99 | 0 |
| SPGP0BK02300 | XP_018221247.1 | 99 | 0 |
| SPGP0DN00190 | XP_018218985.1 | 99 | 0 |
| SPGP0GR00180 | XP_018222746.1 | 99 | 0 |
| SPGP0RB00120 | XP_018218998.1 | 99 | 0 |
| SPGP0TQ00170 | XP_018220612.1 | 99 | 0 |
| SPGP0OE00130 | XP_018219607.1 | 100 | 0 |
| SPGP0KF00110 | XP_018219033.1 | 99 | 0 |
| SPGP0FE00170 | XP_018220533.1 | 99 | 0 |
| SPGP0BK00210 | XP_018221449.1 | 99 | 0 |
| SPGP0VN00230 | XP_018219813.1 | 99 | 0 |
| SPGP0GF00120 | XP_018219649.1 | 100 | 0 |
| SPGP0DP00130 | XP_018220647.1 | 99 | 0 |
| SPGP0IH00100 | XP_018222240.1 | 99 | 0 |
| SPGP0JN00140 | XP_018219073.1 | 99 | 0 |
| SPGP0KZ00300 | XP_018223932.1 | 100 | 0 |
| SPGP0MD00140 | XP_018222389.1 | 99 | 0 |
| SPGP0KB00130 | XP_018221115.1 | 99 | 0 |
| SPGP0MC00130 | XP_018222665.1 | 99 | 0 |
| SPGP0EH00100 | XP_018222261.1 | 99 | 0 |
| SPGP0HB00170 | XP_018219187.1 | 99 | 0 |
| SPGP0OI00490 | XP_018218901.1 | 99 | 0 |
| SPGP0PE00190 | XP_018222363.1 | 99 | 0 |
| SPGP0OI00190 | XP_018218928.1 | 99 | 0 |
| SPGP0OK00130 | XP_018219866.1 | 99 | 0 |
| SPGP0LK00150 | XP_018221066.1 | 99 | 0 |
| SPGP0DX00120 | XP_018219328.1 | 99 | 0 |
| SPGP0WT00120 | XP_018220653.1 | 99 | 0 |
| SPGP0HT00190 | XP_018219010.1 | 99 | 0 |
| SPGP0GH00170 | XP_018218961.1 | 99 | 1 |
| SPGP0UR00120 | XP_018222342.1 | 99 | 0 |
| SPGP0RB00100 | XP_018219000.1 | 99 | 0 |

Create PDF in your applications with the Pdfcrowd HTML to PDF API PDFCROWD

| | | | |
|---|---|---|---|
| SPGP0OE00260 | XP_018219595.1 | 99 | 0 |
| SPGP0TQ00330 | XP_018220628.1 | 99 | 0 |
| SPGP0FE00150 | XP_018220535.1 | 99 | 0 |
| SPGP0HU00100 | XP_018220426.1 | 99 | 0 |
| SPGP0IC00140 | XP_018222793.1 | 100 | 0 |
| SPGP0EM00290 | XP_018220706.1 | 99 | 0 |
| SPGP0EZ00100 | XP_018219315.1 | 99 | 0 |
| SPGP0VN00210 | XP_018219811.1 | 99 | 0 |
| SPGP0EM00120 | XP_018220721.1 | 100 | 0 |
| SPGP0IC00160 | XP_018222791.1 | 99 | 0 |
| SPGP0VB00100 | XP_018221049.1 | 99 | 0 |
| SPGP0VM00160 | XP_018220678.1 | 99 | 0 |
| SPGP0XA00100 | XP_018222801.1 | 99 | 0 |
| SPGP0VM00290 | XP_018220665.1 | 99 | 0 |
| SPGP0LH00110 | XP_018221035.1 | 99 | 0 |
| SPGP0VZ00290 | XP_018223893.1 | 99 | 0 |
| SPGP0OB00130 | XP_018219796.1 | 100 | 0 |
| SPGP0FW00170 | XP_018219579.1 | 99 | 0 |
| SPGP0HZ00180 | XP_018219775.1 | 99 | 0 |
| SPGP0UK00100 | XP_018219034.1 | 99 | 0 |
| SPGP0DD00100 | XP_018221001.1 | 99 | 0 |
| SPGP0MF00230 | XP_018222197.1 | 98 | 0 |
| SPGP0LK00170 | XP_018221064.1 | 99 | 0 |
| SPGP0NT00190 | XP_018222465.1 | 100 | 0 |
| SPGP0M00890 | XP_018220933.1 | 99 | 0 |
| SPGP0PL00230 | XP_018219743.1 | 99 | 0 |
| SPGP0FE00200 | XP_018220530.1 | 99 | 0 |
| SPGP0DW00150 | XP_018218951.1 | 99 | 0 |
| SPGP0XF00100 | XP_018220855.1 | 99 | 0 |
| SPGP0LP00130 | XP_018222383.1 | 99 | 0 |
| SPGP0FM00270 | XP_018222162.1 | 99 | 0 |
| SPGP0VM00100 | XP_018220684.1 | 99 | 0 |
| SPGP0LU00240 | XP_018224002.1 | 99 | 0 |
| SPGP0HO00200 | XP_018219982.1 | 99 | 0 |
| SPGP0EE00150 | XP_018220734.1 | 99 | 0 |
| SPGP0BM00100 | XP_018222753.1 | 99 | 0 |
| SPGP0AH00130 | XP_018219891.1 | 99 | 0 |
| SPGP0MA00100 | XP_018218969.1 | 99 | 0 |
| SPGP0UN00110 | XP_018220787.1 | 99 | 0 |
| SPGP0TL00120 | XP_018219086.1 | 99 | 0 |

Create PDF in your applications with the Pdfcrowd HTML to PDF API PDFCROWD

| | | | |
|---|---|---|---|
| SPGP0LE00190 | XP_018220759.1 | 99 | 0 |
| SPGP0MD00160 | XP_018222391.1 | 99 | 0 |
| SPGP0KW00110 | XP_018220727.1 | 99 | 0 |
| SPGP0UN00260 | XP_018220772.1 | 99 | 0 |
| SPGP0NF00130 | XP_018222633.1 | 99 | 0 |
| SPGP0MF00210 | XP_018222199.1 | 99 | 0 |
| SPGP0EZ00120 | XP_018219317.1 | 99 | 0 |
| SPGP0EW00100 | XP_018222289.1 | 99 | 0 |
| SPGP0JN00130 | XP_018219072.1 | 100 | 0 |
| SPGP0JP00220 | XP_018218859.1 | 99 | 0 |
| SPGP0SB00110 | XP_018222299.1 | 99 | 0 |
| SPGP0HZ00330 | XP_018219790.1 | 99 | 0 |
| SPGP0TX00130 | XP_018222350.1 | 99 | 0 |
| SPGP0PG00180 | XP_018219171.1 | 100 | 0 |
| SPGP0OI00450 | XP_018218905.1 | 99 | 0 |
| SPGP0PL00210 | XP_018219745.1 | 99 | 0 |
| SPGP0OW00180 | XP_018218815.1 | 99 | 0 |
| SPGP0OE00200 | XP_018219600.1 | 99 | 0 |
| SPGP0IN00110 | XP_018219126.1 | 99 | 0 |
| SPGP0VH00150 | XP_018219686.1 | 100 | 0 |
| SPGP0IH00160 | XP_018222234.1 | 99 | 0 |
| SPGP0SS00120 | XP_018219047.1 | 99 | 0 |
| SPGP0WX00110 | XP_018219952.1 | 99 | 0 |
| SPGP0OW00110 | XP_018218808.1 | 99 | 0 |
| SPGP0HO00220 | XP_018219980.1 | 99 | 0 |
| SPGP0GF00230 | XP_018219638.1 | 99 | 0 |
| SPGP0PZ00250 | XP_018222421.1 | 99 | 0 |
| SPGP0KN00220 | XP_018220578.1 | 99 | 0 |
| SPGP0VR00110 | XP_018220703.1 | 99 | 0 |
| SPGP0KG00150 | XP_018222629.1 | 99 | 1 |
| SPGP0MH00100 | XP_018222671.1 | 99 | 0 |
| SPGP0PE00250 | XP_018222370.1 | 99 | 0 |
| SPGP0LS00140 | XP_018222560.1 | 99 | 0 |
| SPGP0VG00140 | XP_018223872.1 | 99 | 0 |
| SPGP0OW00170 | XP_018218814.1 | 99 | 0 |
| SPGP0UR00100 | XP_018222344.1 | 98 | 2 |
| SPGP0FT00140 | XP_018222734.1 | 99 | 0 |
| SPGP0OF00270 | XP_018222708.1 | 100 | 0 |
| SPGP0LG00210 | XP_018222653.1 | 99 | 0 |
| SPGP0SL00100 | XP_018220486.1 | 99 | 1 |

Create PDF in your applications with the Pdfcrowd HTML to PDF API PDFCROWD

| | | | |
|---|---|---|---|
| SPGP0RI00240 | XP_018220502.1 | 99 | 0 |
| SPGP0WG00200 | XP_018221018.1 | 99 | 0 |
| SPGP0EW00160 | XP_018222295.1 | 99 | 0 |
| SPGP0LG00110 | XP_018222643.1 | 99 | 0 |
| SPGP0DQ00190 | XP_018219678.1 | 99 | 0 |
| SPGP0BK00290 | XP_018221981.1 | 100 | 0 |
| SPGP0TQ00390 | XP_018220634.1 | 99 | 0 |
| SPGP0TQ00220 | XP_018220617.1 | 99 | 0 |
| SPGP0BU00140 | XP_018222253.1 | 100 | 0 |
| SPGP0KL00140 | XP_018221100.1 | 99 | 0 |
| SPGP0IC00180 | XP_018222789.1 | 99 | 0 |
| SPGP0RC00310 | XP_018222106.1 | 100 | 0 |
| SPGP0SH00160 | XP_018223974.1 | 99 | 0 |
| SPGP0EE00110 | XP_018220730.1 | 99 | 0 |
| SPGP0VN00150 | XP_018219805.1 | 100 | 0 |
| SPGP0LU00200 | XP_018224006.1 | 99 | 0 |
| SPGP0OW00130 | XP_018218810.1 | 99 | 0 |
| SPGP0RC00250 | XP_018222100.1 | 99 | 0 |
| SPGP0HX00190 | XP_018221006.1 | 99 | 0 |
| SPGP0WF00100 | XP_018222552.1 | 99 | 0 |
| SPGP0UL00130 | XP_018219660.1 | 99 | 0 |
| SPGP0DU00100 | XP_018222584.1 | 99 | 0 |
| SPGP0GF00180 | XP_018219643.1 | 99 | 0 |
| SPGP0OI00660 | XP_018218884.1 | 99 | 0 |
| SPGP0EO00100 | XP_018222477.1 | 99 | 0 |
| SPGP0EK00100 | XP_018220815.1 | 100 | 0 |
| SPGP0UL00190 | XP_018219666.1 | 99 | 0 |
| SPGP0VS00140 | XP_018222524.1 | 99 | 0 |
| SPGP0KY00100 | XP_018223901.1 | 100 | 0 |
| SPGP0OE00190 | XP_018219601.1 | 99 | 0 |
| SPGP0SL00210 | XP_018220475.1 | 99 | 0 |
| SPGP0MO00360 | XP_018219120.1 | 99 | 0 |
| SPGP0VN00290 | XP_018219819.1 | 99 | 0 |
| SPGP0CF00130 | XP_018222497.1 | 99 | 0 |
| SPGP0KN00100 | XP_018220566.1 | 99 | 0 |
| SPGP0CW00140 | XP_018222507.1 | 99 | 0 |
| SPGP0EE00130 | XP_018220732.1 | 99 | 0 |
| SPGP0RI00110 | XP_018220489.1 | 99 | 0 |
| SPGP0JN00210 | XP_018219080.1 | 99 | 0 |
| SPGP0SH00140 | XP_018223972.1 | 99 | 0 |

Create PDF in your applications with the Pdfcrowd HTML to PDF API PDFCROWD

| | | | |
|---|---|---|---|
| SPGP0UB00140 | XP_018219039.1 | 99 | 0 |
| SPGP0LU00180 | XP_018224008.1 | 99 | 0 |
| SPGP0S00100 | XP_018219652.1 | 99 | 0 |
| SPGP0OI00770 | XP_018218873.1 | 99 | 0 |
| SPGP0KZ00370 | XP_018223925.1 | 99 | 0 |
| SPGP0LH00170 | XP_018221030.1 | 100 | 0 |
| SPGP0GF00160 | XP_018219645.1 | 99 | 0 |
| SPGP0TO00110 | XP_018222805.1 | 99 | 0 |
| SPGP0GW00130 | XP_018220826.1 | 99 | 0 |
| SPGP0JH00130 | XP_018218939.1 | 99 | 0 |
| SPGP0HL00100 | XP_018220998.1 | 99 | 0 |
| SPGP0UZ00140 | XP_018223429.1 | 99 | 0 |
| SPGP0ED00100 | XP_018222598.1 | 99 | 0 |
| SPGP0HT00250 | XP_018219016.1 | 99 | 0 |
| SPGP0VP00100 | XP_018219951.1 | 99 | 0 |
| SPGP0WA00220 | XP_018219844.1 | 99 | 0 |
| SPGP0JX00100 | XP_018218837.1 | 99 | 0 |
| SPGP0VZtrna23S | DI49_2374 | 100 | 0 |
| SPGP0FT00100 | XP_018222730.1 | 99 | 0 |
| SPGP0ID00140 | XP_018222144.1 | 99 | 0 |
| SPGP0OF00380 | XP_018222719.1 | 99 | 0 |
| SPGP0IV00100 | XP_018220638.1 | 99 | 0 |
| SPGP0RI00190 | XP_018220497.1 | 99 | 0 |
| SPGP0MD00110 | XP_018222385.1 | 99 | 0 |
| SPGP0HO00280 | XP_018219974.1 | 99 | 0 |
| SPGP0QS00120 | XP_018223898.1 | 100 | 0 |
| SPGP0UF00120 | XP_018221038.1 | 99 | 0 |
| SPGP0VI00120 | XP_018222305.1 | 99 | 0 |
| SPGP0OF00230 | XP_018222704.1 | 99 | 0 |
| SPGP0DK00100 | XP_018219067.1 | 99 | 0 |
| SPGP0PL00190 | XP_018219747.1 | 99 | 0 |
| SPGP0RH00110 | XP_018220418.1 | 99 | 0 |
| SPGP0HZ00140 | XP_018219771.1 | 99 | 0 |
| SPGP0JP00100 | XP_018218847.1 | 99 | 0 |
| SPGP0EO00140 | XP_018222481.1 | 97 | 0 |
| SPGP0SW00200 | XP_018220554.1 | 99 | 0 |
| SPGP0IO00110 | XP_018222453.1 | 99 | 0 |
| SPGP0PZ00380 | XP_018222408.1 | 99 | 0 |
| SPGP0RK00130 | XP_018224039.1 | 99 | 0 |
| SPGP0UZ00180 | XP_018223434.1 | 99 | 0 |

Create PDF in your applications with the Pdfcrowd HTML to PDF API PDFCROWD

| | | | |
|---|---|---|---|
| SPGP0CW00100 | XP_018222504.1 | 99 | 0 |
| SPGP0NT00170 | XP_018222467.1 | 99 | 0 |
| SPGP0DM00100 | XP_018218839.1 | 99 | 0 |
| SPGP0XC00250 | XP_018220601.1 | 100 | 0 |
| SPGP0BI03480 | XP_018223534.1 | 100 | 0 |
| SPGP0TE00120 | XP_018219895.1 | 99 | 0 |
| SPGP0GT00160 | XP_018222513.1 | 99 | 0 |
| SPGP0HX00150 | XP_018221009.1 | 99 | 0 |
| SPGP0QV00100 | XP_018220010.1 | 99 | 0 |
| SPGP0TQ00200 | XP_018220615.1 | 100 | 0 |
| SPGP0NT00110 | XP_018222473.1 | 99 | 0 |
| SPGP0UO00110 | XP_018222183.1 | 99 | 0 |
| SPGP0TX00120 | XP_018222349.1 | 99 | 0 |
| SPGP0UV00190 | XP_018218806.1 | 99 | 0 |
| SPGP0BU00120 | XP_018222251.1 | 100 | 0 |
| SPGP0KN00180 | XP_018220574.1 | 99 | 0 |
| SPGP0LK00200 | XP_018221061.1 | 100 | 0 |
| SPGP0OB00150 | XP_018219798.1 | 99 | 0 |
| SPGP0DN00150 | XP_018218989.1 | 99 | 0 |
| SPGP0EK00120 | XP_018220812.1 | 99 | 0 |
| SPGP0FM00310 | XP_018222166.1 | 98 | 1 |
| SPGP0PE00100 | XP_018222353.1 | 99 | 0 |
| SPGP0RC00270 | XP_018222102.1 | 99 | 0 |
| SPGP0GK00110 | XP_018219713.1 | 99 | 0 |
| SPGP0VP00160 | XP_018219944.1 | 99 | 0 |
| SPGP0IV00140 | XP_018220642.1 | 100 | 0 |
| SPGP0QV00120 | XP_018220008.1 | 100 | 0 |
| SPGP0MB00130 | XP_018222175.1 | 98 | 1 |
| SPGP0OJ00120 | XP_018219922.1 | 99 | 0 |
| SPGP0VE00110 | XP_018220963.1 | 99 | 0 |
| SPGP0DC00130 | XP_018222640.1 | 99 | 0 |
| SPGP0UB00190 | XP_018219044.1 | 99 | 0 |
| SPGP0CN00100 | XP_018219158.1 | 99 | 0 |
| SPGP0VP00140 | XP_018219947.1 | 99 | 0 |
| SPGP0Y00100 | XP_018222681.1 | 99 | 0 |
| SPGP0PL00130 | XP_018219753.1 | 99 | 0 |
| SPGP0OI00820 | XP_018218868.1 | 99 | 0 |
| SPGP0NT00130 | XP_018222471.1 | 99 | 0 |
| SPGP0KL00230 | XP_018221092.1 | 99 | 0 |
| SPGP0QW00140 | XP_018222606.1 | 99 | 0 |

Create PDF in your applications with the Pdfcrowd HTML to PDF API PDFCROWD

| | | | |
|---|---|---|---|
| SPGP0SP00140 | XP_018223964.1 | 99 | 0 |
| SPGP0C01340 | XP_018220230.1 | 99 | 0 |
| SPGP0PL00110 | XP_018219755.1 | 99 | 0 |
| SPGP0PL00170 | XP_018219749.1 | 99 | 0 |
| SPGP0WA00180 | XP_018219848.1 | 99 | 0 |
| SPGP0WV00110 | XP_018219958.1 | 99 | 0 |
| SPGP0QF00140 | XP_018224022.1 | 99 | 0 |
| SPGP0HU00170 | XP_018220433.1 | 99 | 0 |
| SPGP0WI00120 | XP_018222120.1 | 99 | 0 |
| SPGP0AA00200 | XP_018221075.1 | 99 | 0 |
| SPGP0WI00100 | XP_018222118.1 | 97 | 2 |
| SPGP0GR00140 | XP_018222742.1 | 99 | 0 |
| SPGP0WQ00210 | XP_018222274.1 | 99 | 0 |
| SPGP0BK00460 | XP_018221426.1 | 99 | 0 |
| SPGP0RN00170 | XP_018222826.1 | 99 | 0 |
| SPGP0HO00120 | XP_018219990.1 | 99 | 0 |
| SPGP0D03620 | XP_018221980.1 | 99 | 0 |
| SPGP0SL00150 | XP_018220481.1 | 99 | 0 |
| SPGP0BE03060 | XP_018223038.1 | 99 | 0 |
| SPGP0D00390 | XP_018221672.1 | 99 | 0 |
| SPGP0RX00140 | XP_018222815.1 | 99 | 0 |
| SPGP0K00230 | XP_018219562.1 | 99 | 0 |
| SPGP0BK00760 | XP_018221397.1 | 99 | 0 |
| SPGP0RK00110 | XP_018224041.1 | 100 | 0 |
| SPGP0LE00100 | XP_018220769.1 | 99 | 0 |
| SPGP0VN00320 | XP_018219822.1 | 99 | 0 |
| SPGP0C00850 | XP_018220279.1 | 99 | 0 |
| SPGP0BK00410 | XP_018221431.1 | 99 | 0 |
| SPGP0K02300 | XP_018219361.1 | 99 | 0 |
| SPGP0VO00150 | XP_018223440.1 | 99 | 0 |
| SPGP0N01140 | XP_018219301.1 | 99 | 0 |
| SPGP0BK01220 | XP_018221352.1 | 99 | 0 |
| SPGP0BK01810 | XP_018221296.1 | 99 | 0 |
| SPGP0BE04770 | XP_018222874.1 | 99 | 0 |
| SPGP0U00510 | XP_018224086.1 | 99 | 0 |
| SPGP0BE00480 | XP_018223286.1 | 99 | 0 |
| SPGP0D01440 | XP_018221772.1 | 99 | 0 |
| SPGP0D01910 | XP_018221817.1 | 99 | 0 |
| SPGP0BE01770 | XP_018223158.1 | 99 | 0 |
| SPGP0BE00850 | XP_018223249.1 | 99 | 0 |

Create PDF in your applications with the Pdfcrowd HTML to PDF API PDFCROWD

| | | | |
|---|---|---|---|
| SPGP0QF00150 | XP_018224023.1 | 99 | 0 |
| SPGP0BE00890 | XP_018223245.1 | 99 | 0 |
| SPGP0BI01540 | XP_018223726.1 | 99 | 0 |
| SPGP0D02930 | XP_018221915.1 | 98 | 0 |
| SPGP0N00760 | XP_018219263.1 | 100 | 0 |
| SPGP0P01510 | XP_018221508.1 | 99 | 0 |
| SPGP0BK01090 | XP_018221365.1 | 99 | 0 |
| SPGP0D01320 | XP_018221760.1 | 99 | 0 |
| SPGP0VQ00140 | XP_018222547.1 | 99 | 0 |
| SPGP0BE02620 | XP_018223076.1 | 99 | 0 |
| SPGP0P00320 | XP_018221621.1 | 99 | 0 |
| SPGP0WG00230 | XP_018221015.1 | 99 | 0 |
| SPGP0K02540 | XP_018219337.1 | 100 | 0 |
| SPGP0BK03030 | XP_018221180.1 | 99 | 0 |
| SPGP0ST00110 | XP_018220548.1 | 98 | 0 |
| SPGP0BE01850 | XP_018223150.1 | 99 | 0 |
| SPGP0N00830 | XP_018219270.1 | 99 | 0 |
| SPGP0RT00140 | XP_018223996.1 | 99 | 0 |
| SPGP0BK01320 | XP_018221342.1 | 99 | 0 |
| SPGP0NQ00140 | XP_018220751.1 | 99 | 0 |
| SPGP0DQ00210 | XP_018219680.1 | 99 | 0 |
| SPGP0M00960 | XP_018220940.1 | 99 | 0 |
| SPGP0P00180 | XP_018221635.1 | 99 | 0 |
| SPGP0WH00100 | XP_018219920.1 | 99 | 0 |
| SPGP0BK00230 | XP_018221448.1 | 99 | 0 |
| SPGP0BE01690 | XP_018223166.1 | 99 | 0 |
| SPGP0K02550 | XP_018219336.1 | 99 | 0 |
| SPGP0BK01580 | XP_018221318.1 | 99 | 0 |
| SPGP0U00280 | XP_018224108.1 | 99 | 0 |
| SPGP0BK00900 | XP_018221383.1 | 99 | 0 |
| SPGP0BI00410 | XP_018223837.1 | 99 | 0 |
| SPGP0Ltrna6G | DI49_4977 | 100 | 0 |
| SPGP0BI00140 | XP_018223864.1 | 99 | 0 |
| SPGP0BK03250 | XP_018221159.1 | 99 | 0 |
| SPGP0EV00100 | XP_018222488.1 | 88 | 0 |
| SPGP0KZtrna10G | DI49_0567 | 99 | 0 |
| SPGP0BI04170 | XP_018223469.1 | 99 | 0 |
| SPGP0LK00100 | XP_018221071.1 | 99 | 0 |
| SPGP0QW00100 | XP_018222602.1 | 99 | 0 |
| SPGP0C02730 | XP_018220100.1 | 99 | 0 |

Create PDF in your applications with the Pdfcrowd HTML to PDF API PDFCROWD

| | | | |
|---|---|---|---|
| SPGP0VZ00230 | XP_018223887.1 | 99 | 0 |
| SPGP0TQ00400 | XP_018220635.1 | 99 | 0 |
| SPGP0VI00110 | XP_018222304.1 | 99 | 0 |
| SPGP0HL00140 | XP_018220994.1 | 99 | 0 |
| SPGP0P00120 | XP_018221641.1 | 99 | 0 |
| SPGP0FA00120 | XP_018219180.1 | 99 | 0 |
| SPGP0BI02360 | XP_018223646.1 | 99 | 0 |
| SPGP0C01280 | XP_018220236.1 | 99 | 0 |
| SPGP0EZ00140 | XP_018219319.1 | 99 | 0 |
| SPGP0BI02810 | XP_018223602.1 | 99 | 0 |
| SPGP0C02880 | XP_018220085.1 | 99 | 0 |
| SPGP0C02550 | XP_018220116.1 | 99 | 1 |
| SPGP0QV00260 | XP_018219994.1 | 99 | 0 |
| SPGP0C00770 | XP_018220287.1 | 99 | 0 |
| SPGP0C00480 | XP_018220316.1 | 99 | 0 |
| SPGP0BE04710 | XP_018222880.1 | 99 | 0 |
| SPGP0BE02610 | XP_018223077.1 | 99 | 0 |
| SPGP0BK02070 | XP_018221270.1 | 99 | 0 |
| SPGP0BI03630 | XP_018223519.1 | 99 | 0 |
| SPGP0P01150 | XP_018221542.1 | 99 | 0 |
| SPGP0BI03090 | XP_018223574.1 | 99 | 0 |
| SPGP0UR00190 | XP_018222335.1 | 99 | 0 |
| SPGP0VH00110 | XP_018219682.1 | 99 | 0 |
| SPGP0BE00410 | XP_018223293.1 | 99 | 0 |
| SPGP0MO00300 | XP_018219114.1 | 99 | 0 |
| SPGP0P00670 | XP_018221587.1 | 99 | 0 |
| SPGP0BI03780 | XP_018223505.1 | 99 | 0 |
| SPGP0K01210 | XP_018219466.1 | 99 | 0 |
| SPGP0C00590 | XP_018220305.1 | 99 | 0 |
| SPGP0PG00120 | XP_018219177.1 | 99 | 0 |
| SPGP0JP00160 | XP_018218853.1 | 99 | 0 |
| SPGP0P00580 | XP_018221595.1 | 99 | 0 |
| SPGP0BK02310 | XP_018221246.1 | 99 | 0 |
| SPGP0BE04570 | XP_018222894.1 | 99 | 0 |
| SPGP0EM00270 | XP_018220708.1 | 99 | 0 |
| SPGP0JY00100 | XP_018222541.1 | 99 | 0 |
| SPGP0VP00280 | XP_018222852.1 | 97 | 0 |
| SPGP0BE04010 | XP_018222951.1 | 99 | 0 |
| SPGP0C00800 | XP_018220284.1 | 99 | 0 |
| SPGP0GH00140 | XP_018218964.1 | 99 | 0 |

Create PDF in your applications with the Pdfcrowd HTML to PDF API PDFCROWD

| | | | |
|---|---|---|---|
| SPGP0K01080 | XP_018219479.1 | 99 | 0 |
| SPGP0C00950 | XP_018220269.1 | 99 | 0 |
| SPGP0BE01410 | XP_018223193.1 | 99 | 0 |
| SPGP0D00890 | XP_018221720.1 | 99 | 0 |
| SPGP0P01980 | XP_018221461.1 | 96 | 1 |
| SPGP0BItrna20V | DI49_0484 | 100 | 0 |
| SPGP0N00220 | XP_018219210.1 | 96 | 2 |
| SPGP0K02110 | XP_018219380.1 | 99 | 0 |
| SPGP0K00420 | XP_018219543.1 | 99 | 0 |
| SPGP0K00930 | XP_018219494.1 | 99 | 0 |
| SPGP0K01480 | XP_018219440.1 | 99 | 0 |
| SPGP0RZ00110 | XP_018223380.1 | 99 | 0 |
| SPGP0P00850 | XP_018221569.1 | 99 | 0 |
| SPGP0BE02010 | XP_018223134.1 | 99 | 0 |
| SPGP0M01020 | XP_018220946.1 | 99 | 0 |
| SPGP0WI00200 | XP_018222128.1 | 99 | 0 |
| SPGP0K01330 | XP_018219454.1 | 99 | 0 |
| SPGP0BE03150 | XP_018223030.1 | 99 | 0 |
| SPGP0IE00100 | XP_018222765.1 | 99 | 0 |
| SPGP0C03040 | XP_018220070.1 | 99 | 0 |
| SPGP0P01180 | XP_018221539.1 | 99 | 0 |
| SPGP0K00680 | XP_018219518.1 | 99 | 0 |
| SPGP0N00590 | XP_018219247.1 | 99 | 0 |
| SPGP0BK01660 | XP_018221311.1 | 99 | 0 |
| SPGP0BK01530 | XP_018221323.1 | 99 | 0 |
| SPGP0FM00110 | XP_018222147.1 | 98 | 0 |
| SPGP0C02820 | XP_018220091.1 | 99 | 0 |
| SPGP0VG00130 | XP_018223871.1 | 99 | 0 |
| SPGP0RB00150 | XP_018218995.1 | 100 | 0 |
| SPGP0BE04260 | XP_018222927.1 | 99 | 0 |
| SPGP0JF00140 | XP_018222312.1 | 99 | 0 |
| SPGP0EA00120 | XP_018224029.1 | 99 | 0 |
| SPGP0D00240 | XP_018221657.1 | 99 | 0 |
| SPGP0EM00130 | XP_018220720.1 | 99 | 0 |
| SPGP0HL00200 | XP_018220989.1 | 99 | 0 |
| SPGP0N01170 | XP_018219304.1 | 99 | 0 |
| SPGP0LE00120 | XP_018220766.1 | 99 | 0 |
| SPGP0BI01730 | XP_018223707.1 | 99 | 0 |
| SPGP0RC00230 | XP_018222098.1 | 99 | 0 |
| SPGP0D00270 | XP_018221660.1 | 99 | 0 |

Create PDF in your applications with the Pdfcrowd HTML to PDF API PDFCROWD

| | | | |
|---|---|---|---|
| SPGP0OL00120 | XP_018222286.1 | 100 | 0 |
| SPGP0CF00170 | XP_018222493.1 | 99 | 0 |
| SPGP0K01520 | XP_018219436.1 | 99 | 0 |
| SPGP0P01720 | XP_018221487.1 | 99 | 0 |
| SPGP0LW00120 | XP_018222616.1 | 99 | 0 |
| SPGP0BE04100 | XP_018222943.1 | 99 | 0 |
| SPGP0BI03400 | XP_018223542.1 | 99 | 0 |
| SPGP0C01690 | XP_018220198.1 | 99 | 0 |
| SPGP0K02400 | XP_018219351.1 | 99 | 0 |
| SPGP0BK02750 | XP_018221206.1 | 100 | 0 |
| SPGP0SU00100 | XP_018218970.1 | 99 | 0 |
| SPGP0M00770 | XP_018220921.1 | 99 | 0 |
| SPGP0HQ00100 | XP_018223403.1 | 99 | 0 |
| SPGP0OW00140 | XP_018218811.1 | 98 | 1 |
| SPGP0U00720 | XP_018224066.1 | 99 | 0 |
| SPGP0VO00120 | XP_018223437.1 | 100 | 0 |
| SPGP0K02190 | XP_018219372.1 | 99 | 0 |
| SPGP0UV00180 | XP_018218805.1 | 99 | 0 |
| SPGP0FE00280 | XP_018220523.1 | 99 | 0 |
| SPGP0BE03720 | XP_018222979.1 | 99 | 0 |
| SPGP0P01440 | XP_018221514.1 | 99 | 0 |
| SPGP0WW00170 | XP_018222079.1 | 99 | 0 |
| SPGP0AK00100 | XP_018219739.1 | 99 | 0 |
| SPGP0HT00230 | XP_018219014.1 | 99 | 0 |
| SPGP0D00640 | XP_018221696.1 | 99 | 0 |
| SPGP0AC00120 | XP_018220416.1 | 100 | 0 |
| SPGP0M00510 | XP_018220895.1 | 100 | 0 |
| SPGP0BK01780 | XP_018221299.1 | 99 | 0 |
| SPGP0U00230 | XP_018224113.1 | 99 | 0 |
| SPGP0LW00130 | XP_018222617.1 | 99 | 0 |
| SPGP0N00140 | XP_018219203.1 | 98 | 0 |
| SPGP0BK00200 | XP_018221450.1 | 99 | 0 |
| SPGP0BI00730 | XP_018223805.1 | 99 | 0 |
| SPGP0VN00380 | XP_018219828.1 | 99 | 0 |
| SPGP0HX00130 | XP_018221011.1 | 93 | 7 |
| SPGP0BE01450 | XP_018223189.1 | 99 | 0 |
| SPGP0BE00530 | XP_018223281.1 | 100 | 0 |
| SPGP0FB00110 | XP_018220425.1 | 97 | 2 |
| SPGP0D02940 | XP_018221916.1 | 98 | 2 |
| SPGP0MJ00110 | XP_018222082.1 | 99 | 0 |

Create PDF in your applications with the Pdfcrowd HTML to PDF API PDFCROWD

| | | | |
|---|---|---|---|
| SPGP0BI01950 | XP_018223685.1 | 99 | 0 |
| SPGP0D03280 | XP_018221948.1 | 99 | 0 |
| SPGP0C02270 | XP_018220144.1 | 100 | 0 |
| SPGP0SH00280 | XP_018223985.1 | 99 | 0 |
| SPGP0QK00120 | XP_018220384.1 | 99 | 0 |
| SPGP0HZ00320 | XP_018219789.1 | 99 | 0 |
| SPGP0IC00130 | XP_018222794.1 | 100 | 0 |
| SPGP0LU00220 | XP_018224004.1 | 99 | 0 |
| SPGP0D02120 | XP_018221837.1 | 99 | 0 |
| SPGP0BI02680 | XP_018223614.1 | 100 | 0 |
| SPGP0BE02960 | XP_018223046.1 | 99 | 0 |
| SPGP0C02940 | XP_018220079.1 | 99 | 0 |
| SPGP0BI03550 | XP_018223527.1 | 99 | 0 |
| SPGP0KB00100 | XP_018221112.1 | 100 | 0 |
| SPGP0GF00210 | XP_018219640.1 | 99 | 0 |
| SPGP0BE04740 | XP_018222877.1 | 99 | 0 |
| SPGP0JN00170 | XP_018219076.1 | 99 | 0 |
| SPGP0D03540 | XP_018221972.1 | 99 | 0 |
| SPGP0BI00890 | XP_018223790.1 | 99 | 0 |
| SPGP0M00330 | XP_018220878.1 | 99 | 0 |
| SPGP0MF00170 | XP_018222203.1 | 99 | 0 |
| SPGP0C01360 | XP_018220228.1 | 99 | 0 |
| SPGP0D01110 | XP_018221741.1 | 98 | 1 |
| SPGP0BE04220 | XP_018222931.1 | 97 | 2 |
| SPGP0U00500 | XP_018224087.1 | 99 | 0 |
| SPGP0BI04210 | XP_018223465.1 | 99 | 0 |
| SPGP0UN00150 | XP_018220783.1 | 99 | 0 |
| SPGP0D00570 | XP_018221689.1 | 99 | 0 |
| SPGP0C00520 | XP_018220312.1 | 99 | 0 |
| SPGP0C03090 | XP_018220065.1 | 99 | 0 |
| SPGP0BI01800 | XP_018223700.1 | 99 | 0 |
| SPGP0C00860 | XP_018220278.1 | 100 | 0 |
| SPGP0BE04380 | XP_018222915.1 | 99 | 0 |
| SPGP0Ftrna13Q | DI49_1374 | 100 | 0 |
| SPGP0MO00330 | XP_018219117.1 | 99 | 0 |
| SPGP0BI01600 | XP_018223720.1 | 99 | 0 |
| SPGP0BI00600 | XP_018223818.1 | 99 | 0 |
| SPGP0VM00150 | XP_018220679.1 | 99 | 0 |
| SPGP0K00300 | XP_018219555.1 | 99 | 0 |

Create PDF in your applications with the Pdfcrowd HTML to PDF API PDFCROWD

| | | | |
|---|---|---|---|
| SPGP0BI02600 | XP_018223622.1 | 99 | 0 |
| SPGP0C01250 | XP_018220239.1 | 99 | 0 |
| SPGP0C01940 | XP_018220176.1 | 99 | 0 |
| SPGP0BK02980 | XP_018221185.1 | 99 | 0 |
| SPGP0SW00170 | XP_018220557.1 | 99 | 1 |
| SPGP0SN00110 | XP_018222593.1 | 98 | 1 |
| SPGP0K02510 | XP_018219340.1 | 99 | 0 |
| SPGP0C02250 | XP_018220146.1 | 99 | 0 |
| SPGP0BE00600 | XP_018223274.1 | 99 | 0 |
| SPGP0BE02420 | XP_018223096.1 | 99 | 0 |
| SPGP0OF00140 | XP_018222695.1 | 99 | 0 |
| SPGP0MC00120 | XP_018222664.1 | 99 | 0 |
| SPGP0C00540 | XP_018220310.1 | 99 | 0 |
| SPGP0BE02500 | XP_018223088.1 | 99 | 0 |
| SPGP0K01650 | XP_018219424.1 | 99 | 0 |
| SPGP0K02150 | XP_018219376.1 | 99 | 0 |
| SPGP0D01740 | XP_018221801.1 | 99 | 0 |
| SPGP0C02660 | XP_018220107.1 | 99 | 0 |
| SPGP0D03290 | XP_018221949.1 | 100 | 0 |
| SPGP0P01020 | XP_018221554.1 | 99 | 0 |
| SPGP0VZ00270 | XP_018223891.1 | 99 | 0 |
| SPGP0C02490 | XP_018220122.1 | 99 | 0 |
| SPGP0BI03490 | XP_018223533.1 | 98 | 0 |
| SPGP0BE03260 | XP_018223020.1 | 99 | 0 |
| SPGP0BE03740 | XP_018222977.1 | 99 | 0 |
| SPGP0D01020 | XP_018221733.1 | 99 | 0 |
| SPGP0BE01950 | XP_018223140.1 | 99 | 0 |
| SPGP0P01830 | XP_018221476.1 | 99 | 0 |
| SPGP0D02040 | XP_018221829.1 | 99 | 0 |
| SPGP0OT00100 | XP_018219129.1 | 99 | 0 |
| SPGP0BE03520 | XP_018222998.1 | 99 | 0 |
| SPGP0P00110 | XP_018221642.1 | 99 | 0 |
| SPGP0P00440 | XP_018221609.1 | 99 | 0 |
| SPGP0HI00100 | XP_018223349.1 | 99 | 0 |
| SPGP0U00190 | XP_018224117.1 | 99 | 0 |
| SPGP0JT00110 | XP_018219860.1 | 100 | 0 |
| SPGP0D02060 | XP_018221831.1 | 99 | 0 |
| SPGP0BI02280 | XP_018223654.1 | 99 | 0 |
| SPGP0BK00910 | XP_018221382.1 | 98 | 1 |
| SPGP0D03600 | XP_018221978.1 | 99 | 0 |

Create PDF in your applications with the Pdfcrowd HTML to PDF API PDFCROWD

| | | | |
|---|---|---|---|
| SPGP0BE03000 | XP_018223042.1 | 99 | 1 |
| SPGP0BK02930 | XP_018221190.1 | 98 | 1 |
| SPGP0BE03330 | XP_018223014.1 | 99 | 0 |
| SPGP0MT00120 | XP_018219138.1 | 100 | 0 |
| SPGP0GF00270 | XP_018219634.1 | 99 | 0 |
| SPGP0MD00130 | XP_018222388.1 | 100 | 0 |
| SPGP0P01530 | XP_018221506.1 | 99 | 0 |
| SPGP0LE00160 | XP_018220762.1 | 100 | 0 |
| SPGP0BE04800 | XP_018219681.1 | 99 | 0 |
| SPGP0BK00680 | XP_018221405.1 | 99 | 0 |
| SPGP0VP00180 | XP_018219943.1 | 99 | 0 |
| SPGP0BE03690 | XP_018222982.1 | 99 | 0 |
| SPGP0VO00220 | XP_018223445.1 | 99 | 0 |
| SPGP0BK00430 | XP_018221429.1 | 99 | 0 |
| SPGP0D01700 | XP_018221797.1 | 99 | 0 |
| SPGP0BI01380 | XP_018223743.1 | 99 | 0 |
| SPGP0GK00150 | XP_018219709.1 | 99 | 0 |
| SPGP0K00740 | XP_018219512.1 | 99 | 0 |
| SPGP0K00800 | XP_018219506.1 | 99 | 0 |
| SPGP0K00210 | XP_018219564.1 | 99 | 0 |
| SPGP0BE02120 | XP_018223125.1 | 99 | 0 |
| SPGP0BE00740 | XP_018223259.1 | 99 | 0 |
| SPGP0OI00710 | XP_018218879.1 | 99 | 0 |
| SPGP0U00530 | XP_018224084.1 | 99 | 0 |
| SPGP0BE00570 | XP_018223277.1 | 99 | 0 |
| SPGP0BK02690 | XP_018221211.1 | 99 | 0 |
| SPGP0HO00260 | XP_018219976.1 | 99 | 0 |
| SPGP0N00560 | XP_018219244.1 | 99 | 0 |
| SPGP0BE02350 | XP_018223103.1 | 99 | 0 |
| SPGP0C01000 | XP_018220264.1 | 99 | 0 |
| SPGP0BE02600 | XP_018223078.1 | 99 | 0 |
| SPGP0JO00130 | XP_018223353.1 | 99 | 0 |
| SPGP0BK01970 | XP_018221280.1 | 97 | 1 |
| SPGP0OF00360 | XP_018222717.1 | 99 | 0 |
| SPGP0BI01560 | XP_018223724.1 | 99 | 0 |
| SPGP0OK00120 | XP_018219865.1 | 99 | 0 |
| SPGP0NX00100 | XP_018222476.1 | 95 | 5 |
| SPGP0BE02400 | XP_018223098.1 | 99 | 0 |
| SPGP0BI01990 | XP_018223681.1 | 99 | 0 |
| SPGP0D02910 | XP_018221913.1 | 99 | 0 |

Create PDF in your applications with the Pdfcrowd HTML to PDF API PDFCROWD

| | | | |
|---|---|---|---|
| SPGP0N00780 | XP_018219265.1 | 99 | 1 |
| SPGP0UH00160 | XP_018220412.1 | 99 | 0 |
| SPGP0BI02210 | XP_018223661.1 | 99 | 0 |
| SPGP0D00110 | XP_018221644.1 | 95 | 0 |
| SPGP0CX00180 | XP_018220791.1 | 99 | 0 |
| SPGP0WQ00230 | XP_018222276.1 | 100 | 0 |
| SPGP0XD00100 | XP_018218816.1 | 100 | 0 |
| SPGP0BI00380 | XP_018223840.1 | 99 | 0 |
| SPGP0U00390 | XP_018224097.1 | 99 | 0 |
| SPGP0BK03010 | XP_018221182.1 | 99 | 0 |
| SPGP0BK01020 | XP_018221372.1 | 99 | 0 |
| SPGP0C01190 | XP_018220245.1 | 99 | 0 |
| SPGP0RT00110 | XP_018223993.1 | 100 | 0 |
| SPGP0DT00110 | XP_018221087.1 | 99 | 0 |
| SPGP0K02080 | XP_018219383.1 | 99 | 0 |
| SPGP0U00170 | XP_018224119.1 | 98 | 0 |
| SPGP0M00980 | XP_018220942.1 | 99 | 0 |
| SPGP0OX00170 | XP_018219728.1 | 99 | 0 |
| SPGP0OX00240 | XP_018219721.1 | 100 | 0 |
| SPGP0OF00290 | XP_018222710.1 | 99 | 0 |
| SPGP0BK02920 | XP_018221191.1 | 99 | 0 |
| SPGP0RX00180 | XP_018222819.1 | 99 | 0 |
| SPGP0BI00160 | XP_018223862.1 | 99 | 0 |
| SPGP0BK03270 | XP_018221157.1 | 99 | 0 |
| SPGP0K00580 | XP_018219526.1 | 99 | 0 |
| SPGP0LK00120 | XP_018221069.1 | 99 | 0 |
| SPGP0BE03320 | XP_018223015.1 | 99 | 0 |
| SPGP0NT00230 | XP_018222461.1 | 99 | 0 |
| SPGP0NI00260 | XP_018223919.1 | 99 | 0 |
| SPGP0K01200 | XP_018219467.1 | 99 | 0 |
| SPGP0K00720 | XP_018219514.1 | 99 | 0 |
| SPGP0C02710 | XP_018220102.1 | 99 | 0 |
| SPGP0D00180 | XP_018221651.1 | 99 | 0 |
| SPGP0M00180 | XP_018220863.1 | 99 | 0 |
| SPGP0BE02230 | XP_018223115.1 | 99 | 0 |
| SPGP0AP00130 | XP_018219062.1 | 99 | 0 |
| SPGP0BI04200 | XP_018223466.1 | 99 | 0 |
| SPGP0C00710 | XP_018220293.1 | 99 | 0 |
| SPGP0D01230 | XP_018221751.1 | 99 | 0 |
| SPGP0P01690 | XP_018221490.1 | 99 | 0 |

Create PDF in your applications with the Pdfcrowd HTML to PDF API PDFCROWD

| | | | |
|---|---|---|---|
| SPGP0BK01820 | XP_018221295.1 | 99 | 0 |
| SPGP0HB00180 | XP_018219186.1 | 98 | 0 |
| SPGP0VN00130 | XP_018219803.1 | 99 | 0 |
| SPGP0BK00440 | XP_018221428.1 | 99 | 0 |
| SPGP0BE01480 | XP_018223187.1 | 100 | 0 |
| SPGP0D03220 | XP_018221942.1 | 99 | 0 |
| SPGP0OI00140 | XP_018218933.1 | 99 | 0 |
| SPGP0C00660 | XP_018220298.1 | 99 | 0 |
| SPGP0BE03470 | XP_018223002.1 | 100 | 0 |
| SPGP0D00450 | XP_018221677.1 | 99 | 0 |
| SPGP0M00230 | XP_018220868.1 | 99 | 0 |
| SPGP0BK02050 | XP_018221272.1 | 99 | 0 |
| SPGP0D00250 | XP_018221658.1 | 99 | 0 |
| SPGP0MO00320 | XP_018219116.1 | 100 | 0 |
| SPGP0WI00140 | XP_018222122.1 | 98 | 0 |
| SPGP0C03490 | XP_018220025.1 | 99 | 0 |
| SPGP0HY00140 | XP_018223991.1 | 99 | 0 |
| SPGP0BE01070 | XP_018223227.1 | 99 | 0 |
| SPGP0D03190 | XP_018221939.1 | 99 | 1 |
| SPGP0BK02530 | XP_018221225.1 | 99 | 0 |
| SPGP0TE00220 | XP_018219904.1 | 99 | 0 |
| SPGP0BI02690 | XP_018223613.1 | 99 | 0 |
| SPGP0SM00120 | XP_018221107.1 | 99 | 0 |
| SPGP0BI02830 | XP_018223600.1 | 99 | 0 |
| SPGP0PL00140 | XP_018219752.1 | 99 | 0 |
| SPGP0OZ00160 | XP_018222403.1 | 99 | 0 |
| SPGP0GT00140 | XP_018222512.1 | 99 | 0 |
| SPGP0BE00470 | XP_018223287.1 | 99 | 0 |
| SPGP0BI03120 | XP_018223571.1 | 99 | 0 |
| SPGP0TK00110 | XP_018222725.1 | 100 | 0 |
| SPGP0BK02390 | XP_018221239.1 | 99 | 1 |
| SPGP0PG00140 | XP_018219175.1 | 99 | 0 |
| SPGP0D02180 | XP_018221843.1 | 99 | 0 |
| SPGP0BK02470 | XP_018221231.1 | 100 | 0 |
| SPGP0BI04150 | XP_018223471.1 | 99 | 0 |
| SPGP0JP00180 | XP_018218855.1 | 99 | 0 |
| SPGP0C03500 | XP_018220024.1 | 99 | 0 |
| SPGP0OD00100 | XP_018221045.1 | 99 | 0 |
| SPGP0NT00270 | XP_018222457.1 | 99 | 0 |
| SPGP0EY00140 | XP_018219092.1 | 99 | 0 |

Create PDF in your applications with the Pdfcrowd HTML to PDF API PDFCROWD

| | | | |
|---|---|---|---|
| SPGP0BE04590 | XP_018222892.1 | 99 | 0 |
| SPGP0KY00150 | XP_018223905.1 | 99 | 0 |
| SPGP0K01680 | XP_018219421.1 | 99 | 0 |
| SPGP0M00310 | XP_018220876.1 | 99 | 0 |
| SPGP0HT00130 | XP_018219005.1 | 99 | 0 |
| SPGP0D02350 | XP_018221860.1 | 99 | 0 |
| SPGP0SB00120 | XP_018222300.1 | 99 | 0 |
| SPGP0QP00160 | XP_018219163.1 | 99 | 0 |
| SPGP0BE04530 | XP_018222898.1 | 99 | 0 |
| SPGP0QS00110 | XP_018223897.1 | 99 | 0 |
| SPGP0C03380 | XP_018220036.1 | 99 | 0 |
| SPGP0OI00760 | XP_018218874.1 | 99 | 0 |
| SPGP0IH00140 | XP_018222236.1 | 99 | 0 |
| SPGP0BE01080 | XP_018223226.1 | 99 | 0 |
| SPGP0BE03920 | XP_018222959.1 | 98 | 0 |
| SPGP0BK01500 | XP_018221326.1 | 99 | 0 |
| SPGP0TM00210 | XP_018219885.1 | 99 | 0 |
| SPGP0BI00910 | XP_018223788.1 | 99 | 0 |
| SPGP0BE04370 | XP_018222916.1 | 99 | 0 |
| SPGP0K00910 | XP_018219496.1 | 99 | 0 |
| SPGP0N00200 | XP_018219208.1 | 99 | 0 |
| SPGP0BK01640 | XP_018221312.1 | 99 | 0 |
| SPGP0D02740 | XP_018221897.1 | 99 | 0 |
| SPGP0BK02240 | XP_018221253.1 | 99 | 0 |
| SPGP0PK00130 | XP_018220420.1 | 99 | 0 |
| SPGP0K01350 | XP_018219452.1 | 99 | 0 |
| SPGP0M01000 | XP_018220944.1 | 99 | 0 |
| SPGP0C02080 | XP_018220163.1 | 100 | 0 |
| SPGP0C03450 | XP_018220029.1 | 99 | 0 |
| SPGP0BK01540 | XP_018221322.1 | 99 | 0 |
| SPGP0N01010 | XP_018219288.1 | 99 | 0 |
| SPGP0C03540 | XP_018220020.1 | 99 | 0 |
| SPGP0M00970 | XP_018220941.1 | 99 | 0 |
| SPGP0QFtrna17L | DI49_5686 | 100 | 0 |
| SPGP0TQ00160 | XP_018220611.1 | 99 | 0 |
| SPGP0BI00820 | XP_018223796.1 | 99 | 0 |
| SPGP0EU00130 | XP_018222661.1 | 99 | 0 |
| SPGP0C03080 | XP_018220066.1 | 99 | 0 |
| SPGP0D03070 | XP_018221927.1 | 99 | 0 |
| SPGP0KZ00270 | XP_018223935.1 | 99 | 0 |

Create PDF in your applications with the Pdfcrowd HTML to PDF API PDFCROWD

| | | | |
|---|---|---|---|
| SPGP0D03170 | XP_018221937.1 | 99 | 0 |
| SPGP0BI01120 | XP_018223768.1 | 99 | 0 |
| SPGP0OT00110 | XP_018219130.1 | 99 | 1 |
| SPGP0C01910 | XP_018220179.1 | 99 | 1 |
| SPGP0BI01710 | XP_018223709.1 | 99 | 0 |
| SPGP0OE00100 | XP_018219610.1 | 99 | 0 |
| SPGP0RC00210 | XP_018222096.1 | 99 | 0 |
| SPGP0K01060 | XP_018219481.1 | 99 | 0 |
| SPGP0OL00100 | XP_018222288.1 | 99 | 0 |
| SPGP0D00210 | XP_018221654.1 | 99 | 0 |
| SPGP0BI03140 | XP_018223569.1 | 99 | 0 |
| SPGP0C02260 | XP_018220145.1 | 99 | 0 |
| SPGP0SK00100 | XP_018222281.1 | 99 | 0 |
| SPGP0TQ00300 | XP_018220625.1 | 99 | 0 |
| SPGP0M00750 | XP_018220919.1 | 99 | 0 |
| SPGP0K00620 | XP_018219524.1 | 99 | 0 |
| SPGP0D00540 | XP_018221686.1 | 99 | 0 |
| SPGP0PL00260 | XP_018219740.1 | 99 | 0 |
| SPGP0MY00100 | XP_018222622.1 | 99 | 0 |
| SPGP0BK02730 | XP_018221208.1 | 99 | 0 |
| SPGP0BE02290 | XP_018223109.1 | 99 | 0 |
| SPGP0P00870 | XP_018221567.1 | 99 | 0 |
| SPGP0BK00590 | XP_018221413.1 | 99 | 0 |
| SPGP0WB00110 | XP_018222679.1 | 99 | 0 |
| SPGP0SP00120 | XP_018223962.1 | 99 | 0 |
| SPGP0N01040 | XP_018219291.1 | 99 | 0 |
| SPGP0D00660 | XP_018221698.1 | 99 | 0 |
| SPGP0M00530 | XP_018220897.1 | 99 | 0 |
| SPGP0BE03300 | XP_018223016.1 | 99 | 0 |
| SPGP0LH00150 | XP_018221032.1 | 99 | 0 |
| SPGP0D02160 | XP_018221841.1 | 99 | 0 |
| SPGP0EZ00110 | XP_018219316.1 | 100 | 0 |
| SPGP0U00700 | XP_018224068.1 | 99 | 0 |
| SPGP0BE00200 | XP_018223314.1 | 99 | 0 |
| SPGP0M00720 | XP_018220916.1 | 99 | 0 |
| SPGP0MO00230 | XP_018219108.1 | 99 | 0 |
| SPGP0VN00220 | XP_018219812.1 | 99 | 0 |
| SPGP0Itrna27R | DI49_3064 | 100 | 0 |
| SPGP0BI02900 | XP_018223593.1 | 99 | 0 |
| SPGP0BI01970 | XP_018223683.1 | 100 | 0 |

Create PDF in your applications with the Pdfcrowd HTML to PDF API PDFCROWD

| | | | |
|---|---|---|---|
| SPGP0PE00280 | XP_018222373.1 | 100 | 0 |
| SPGP0HA00140 | XP_018219872.1 | 99 | 0 |
| SPGP0N00770 | XP_018219264.1 | 99 | 0 |
| SPGP0C02040 | XP_018220167.1 | 99 | 0 |
| SPGP0KA00100 | XP_018222228.1 | 99 | 0 |
| SPGP0BI03710 | XP_018223511.1 | 99 | 0 |
| SPGP0AG00100 | XP_018218752.1 | 99 | 0 |
| SPGP0BI03000 | XP_018223583.1 | 99 | 0 |
| SPGP0DP00170 | XP_018220651.1 | 99 | 0 |
| SPGP0BI04090 | XP_018223477.1 | 98 | 0 |
| SPGP0BI02730 | XP_018223609.1 | 99 | 0 |
| SPGP0D02100 | XP_018221835.1 | 99 | 0 |
| SPGP0BK03120 | XP_018221172.1 | 99 | 0 |
| SPGP0U00870 | XP_018224051.1 | 99 | 0 |
| SPGP0C02960 | XP_018220077.1 | 99 | 0 |
| SPGP0BI03820 | XP_018223502.1 | 99 | 0 |
| SPGP0IH00130 | XP_018222237.1 | 99 | 0 |
| SPGP0N00720 | XP_018219259.1 | 99 | 0 |
| SPGP0BE01240 | XP_018223210.1 | 99 | 0 |
| SPGP0BI00590 | XP_018223819.1 | 99 | 0 |
| SPGP0U00670 | XP_018224071.1 | 99 | 0 |
| SPGP0BE03870 | XP_018222964.1 | 99 | 0 |
| SPGP0BE03120 | XP_018223033.1 | 99 | 0 |
| SPGP0IE00190 | XP_018222757.1 | 99 | 0 |
| SPGP0BI02990 | XP_018223584.1 | 99 | 0 |
| SPGP0K02180 | XP_018219373.1 | 99 | 0 |
| SPGP0BI02440 | XP_018223638.1 | 99 | 0 |
| SPGP0BI02470 | XP_018223635.1 | 99 | 0 |
| SPGP0VP00120 | XP_018219949.1 | 99 | 0 |
| SPGP0BE00590 | XP_018223275.1 | 99 | 0 |
| SPGP0RI00260 | XP_018220504.1 | 99 | 0 |
| SPGP0KB00120 | XP_018221114.1 | 99 | 0 |
| SPGP0BE01310 | XP_018223203.1 | 99 | 0 |
| SPGP0JQ00100 | XP_018222439.1 | 99 | 0 |
| SPGP0U00730 | XP_018224065.1 | 100 | 0 |
| SPGP0HU00110 | XP_018220427.1 | 99 | 0 |
| SPGP0BI03670 | XP_018223515.1 | 99 | 0 |
| SPGP0BI00620 | XP_018223816.1 | 99 | 0 |
| SPGP0C01470 | XP_018220218.1 | 99 | 0 |
| SPGP0C03530 | XP_018220021.1 | 99 | 0 |

Create PDF in your applications with the Pdfcrowd HTML to PDF API PDFCROWD

| | | | |
|---|---|---|---|
| SPGP0WQ00220 | XP_018222275.1 | 99 | 0 |
| SPGP0C01180 | XP_018220246.1 | 99 | 0 |
| SPGP0JR00130 | XP_018219028.1 | 99 | 0 |
| SPGP0BI02660 | XP_018223616.1 | 99 | 0 |
| SPGP0BI02190 | XP_018223662.1 | 99 | 0 |
| SPGP0BE03600 | XP_018222991.1 | 99 | 0 |
| SPGP0WY00190 | XP_018220437.1 | 99 | 0 |
| SPGP0C01920 | XP_018220178.1 | 99 | 0 |
| SPGP0ET00110 | XP_018222442.1 | 99 | 0 |
| SPGP0D01920 | XP_018221818.1 | 98 | 1 |
| SPGP0BI04110 | XP_018223475.1 | 99 | 0 |
| SPGP0U00610 | XP_018224077.1 | 99 | 0 |
| SPGP0K01980 | XP_018219393.1 | 99 | 0 |
| SPGP0OX00110 | XP_018219734.1 | 99 | 0 |
| SPGP0C01600 | XP_018220207.1 | 99 | 0 |
| SPGP0BU00160 | XP_018222255.1 | 99 | 0 |
| SPGP0NQ00130 | XP_018220752.1 | 99 | 0 |
| SPGP0IK00100 | XP_018222107.1 | 99 | 0 |
| SPGP0BE00630 | XP_018223271.1 | 99 | 0 |
| SPGP0N00380 | XP_018219226.1 | 99 | 0 |
| SPGP0OT00160 | XP_018219135.1 | 100 | 0 |
| SPGP0BE00620 | XP_018223272.1 | 99 | 0 |
| SPGP0FW00120 | XP_018219584.1 | 99 | 0 |
| SPGP0P00420 | XP_018221611.1 | 99 | 0 |
| SPGP0SW00180 | XP_018220556.1 | 99 | 1 |
| SPGP0P00280 | XP_018221625.1 | 97 | 0 |
| SPGP0P00130 | XP_018221640.1 | 99 | 0 |
| SPGP0N01180 | XP_018219305.1 | 99 | 0 |
| SPGP0P00710 | XP_018221583.1 | 99 | 0 |
| SPGP0K01180 | XP_018219469.1 | 99 | 0 |
| SPGP0BE04900 | XP_018222861.1 | 98 | 0 |
| SPGP0BE00800 | XP_018223254.1 | 99 | 0 |
| SPGP0RZ00120 | XP_018223381.1 | 99 | 0 |
| SPGP0P01870 | XP_018221472.1 | 99 | 0 |
| SPGP0C02890 | XP_018220084.1 | 100 | 0 |
| SPGP0BK01210 | XP_018221353.1 | 99 | 0 |
| SPGP0RX00110 | XP_018222812.1 | 100 | 0 |
| SPGP0TQ00380 | XP_018220633.1 | 100 | 0 |
| SPGP0HL00210 | XP_018220988.1 | 99 | 0 |
| SPGP0PE00120 | XP_018222356.1 | 100 | 0 |

Create PDF in your applications with the Pdfcrowd HTML to PDF API PDFCROWD

| | | | |
|---|---|---|---|
| SPGP0BK01930 | XP_018221284.1 | 99 | 0 |
| SPGP0BE02990 | XP_018223043.1 | 99 | 0 |
| SPGP0BE01610 | XP_018223174.1 | 99 | 0 |
| SPGP0BE02020 | XP_018223133.1 | 99 | 0 |
| SPGP0BE02190 | XP_018223119.1 | 99 | 0 |
| SPGP0BE04750 | XP_018222876.1 | 99 | 0 |
| SPGP0XBtrna13F | DI49_5486 | 100 | 0 |
| SPGP0BI01360 | XP_018223745.1 | 100 | 0 |
| SPGP0N01250 | XP_018219310.1 | 99 | 0 |
| SPGP0CC00110 | XP_018220404.1 | 99 | 0 |
| SPGP0BI00200 | XP_018223858.1 | 100 | 0 |
| SPGP0P01550 | XP_018221504.1 | 99 | 0 |
| SPGP0D02680 | XP_018221891.1 | 99 | 0 |
| SPGP0N00410 | XP_018219229.1 | 97 | 2 |
| SPGP0GH00180 | XP_018218960.1 | 99 | 0 |
| SPGP0IE00180 | XP_018222758.1 | 99 | 0 |
| SPGP0BE02370 | XP_018223101.1 | 99 | 0 |
| SPGP0D01480 | XP_018221776.1 | 99 | 0 |
| SPGP0D01950 | XP_018221821.1 | 99 | 0 |
| SPGP0RA00110 | XP_018218841.1 | 99 | 0 |
| SPGP0BI01310 | XP_018223750.1 | 99 | 0 |
| SPGP0PF00100 | XP_018222571.1 | 99 | 0 |
| SPGP0OI00680 | XP_018218882.1 | 99 | 0 |
| SPGP0BE01150 | XP_018223219.1 | 99 | 0 |
| SPGP0D00870 | XP_018221718.1 | 99 | 0 |
| SPGP0D02970 | XP_018221918.1 | 99 | 0 |
| SPGP0OF00100 | XP_018222691.1 | 99 | 0 |
| SPGP0EY00130 | XP_018219091.1 | 98 | 2 |
| SPGP0BE00830 | XP_018223251.1 | 99 | 0 |
| SPGP0MO00140 | XP_018219099.1 | 99 | 0 |
| SPGP0BK02030 | XP_018221274.1 | 98 | 0 |
| SPGP0BI00360 | XP_018223842.1 | 99 | 0 |
| SPGP0BK03070 | XP_018221177.1 | 99 | 0 |
| SPGP0BK00660 | XP_018221407.1 | 99 | 0 |
| SPGP0BK03110 | XP_018221173.1 | 100 | 0 |
| SPGP0WQ00100 | XP_018222263.1 | 99 | 0 |
| SPGP0BK01410 | XP_018221333.1 | 99 | 0 |
| SPGP0D01940 | XP_018221820.1 | 99 | 0 |
| SPGP0D02020 | XP_018221827.1 | 99 | 0 |
| SPGP0P01950 | XP_018221464.1 | 96 | 0 |

Create PDF in your applications with the Pdfcrowd HTML to PDF API PDFCROWD

| | | | |
|---|---|---|---|
| SPGP0BK02230 | XP_018221254.1 | 99 | 0 |
| SPGP0BI00100 | XP_018223868.1 | 99 | 0 |
| SPGP0N00130 | XP_018219202.1 | 97 | 0 |
| SPGP0LU00160 | XP_018224010.1 | 99 | 0 |
| SPGP0D00620 | XP_018221694.1 | 99 | 0 |
| SPGP0BK03290 | XP_018221155.1 | 99 | 1 |
| SPGP0C00300 | XP_018220333.1 | 99 | 0 |
| SPGP0K01010 | XP_018219486.1 | 99 | 0 |
| SPGP0BI04030 | XP_018223483.1 | 99 | 0 |
| SPGP0K01500 | XP_018219438.1 | 99 | 0 |
| SPGP0N00950 | XP_018219282.1 | 99 | 0 |
| SPGP0C02770 | XP_018220096.1 | 99 | 0 |
| SPGP0BI03830 | XP_018223501.1 | 99 | 0 |
| SPGP0C01770 | XP_018220192.1 | 99 | 0 |
| SPGP0P00970 | XP_018221558.1 | 99 | 0 |
| SPGP0OI00210 | XP_018218926.1 | 99 | 0 |
| SPGP0C00730 | XP_018220291.1 | 99 | 0 |
| SPGP0BI03370 | XP_018223546.1 | 99 | 0 |
| SPGP0P01080 | XP_018221549.1 | 99 | 0 |
| SPGP0OH00120 | XP_018220986.1 | 99 | 0 |
| SPGP0P01060 | XP_018221551.1 | 99 | 0 |
| SPGP0OA00110 | XP_018218948.1 | 99 | 0 |
| SPGP0D00120 | XP_018221645.1 | 95 | 0 |
| SPGP0BI03210 | XP_018223562.1 | 99 | 0 |
| SPGP0C02590 | XP_018220112.1 | 99 | 0 |
| SPGP0M00250 | XP_018220870.1 | 99 | 0 |
| SPGP0VW00100 | XP_018218819.1 | 99 | 0 |
| SPGP0D00530 | XP_018221685.1 | 99 | 0 |
| SPGP0FE00190 | XP_018220531.1 | 99 | 0 |
| SPGP0BI03450 | XP_018223537.1 | 99 | 0 |
| SPGP0DU00150 | XP_018222579.1 | 99 | 0 |
| SPGP0BI03100 | XP_018223573.1 | 99 | 0 |
| SPGP0P00630 | XP_018221591.1 | 99 | 0 |
| SPGP0JP00210 | XP_018218858.1 | 99 | 0 |
| SPGP0K02250 | XP_018219366.1 | 99 | 0 |
| SPGP0BE01050 | XP_018223229.1 | 99 | 0 |
| SPGP0BE01500 | XP_018223185.1 | 99 | 0 |
| SPGP0SM00100 | XP_018221105.1 | 99 | 0 |
| SPGP0BI02850 | XP_018223598.1 | 99 | 0 |
| SPGP0D01610 | XP_018221789.1 | 99 | 0 |

Create PDF in your applications with the Pdfcrowd HTML to PDF API PDFCROWD

| | | | |
|---|---|---|---|
| SPGP0BE02890 | XP_018223053.1 | 99 | 0 |
| SPGP0TE00200 | XP_018219902.1 | 99 | 0 |
| SPGP0BI04120 | XP_018223474.1 | 98 | 0 |
| SPGP0MDtrna14L | DI49_2147 | 99 | 0 |
| SPGP0JZ00110 | XP_018223416.1 | 99 | 0 |
| SPGP0PG00160 | XP_018219173.1 | 99 | 0 |
| SPGP0VZ00100 | XP_018223875.1 | 99 | 0 |
| SPGP0M01160 | XP_018220960.1 | 99 | 0 |
| SPGP0C00190 | XP_018220344.1 | 99 | 0 |
| SPGP0BE00120 | XP_018223322.1 | 99 | 0 |
| SPGP0NT00290 | XP_018222455.1 | 99 | 0 |
| SPGP0K01130 | XP_018219474.1 | 99 | 0 |
| SPGP0D01300 | XP_018221758.1 | 99 | 1 |
| SPGP0K02530 | XP_018219338.1 | 99 | 0 |
| SPGP0M01120 | XP_018220956.1 | 99 | 0 |
| SPGP0LU00120 | XP_018224014.1 | 99 | 0 |
| SPGP0BK00780 | XP_018221395.1 | 100 | 0 |
| SPGP0D02370 | XP_018221862.1 | 99 | 0 |
| SPGP0K00970 | XP_018219490.1 | 99 | 0 |
| SPGP0BE00900 | XP_018223244.1 | 99 | 0 |
| SPGP0BE02480 | XP_018223090.1 | 99 | 0 |
| SPGP0KG00140 | XP_018222628.1 | 99 | 0 |
| SPGP0P01380 | XP_018221520.1 | 99 | 0 |
| SPGP0C00990 | XP_018220265.1 | 99 | 0 |
| SPGP0BK02740 | XP_018221207.1 | 99 | 0 |
| SPGP0BE03900 | XP_018222961.1 | 99 | 0 |
| SPGP0BI04160 | XP_018223470.1 | 99 | 0 |
| SPGP0N00510 | XP_018219239.1 | 99 | 0 |
| SPGP0BI02270 | XP_018223655.1 | 99 | 0 |
| SPGP0P01940 | XP_018221465.1 | 99 | 0 |
| SPGP0M00410 | XP_018220886.1 | 99 | 0 |
| SPGP0BK01620 | XP_018221314.1 | 99 | 0 |
| SPGP0BK03090 | XP_018221175.1 | 99 | 0 |
| SPGP0D02760 | XP_018221899.1 | 99 | 0 |
| SPGP0P01000 | XP_018221556.1 | 99 | 0 |
| SPGP0DP00120 | XP_018220646.1 | 99 | 0 |
| SPGP0BGtrna18P | DI49_4736 | 100 | 0 |
| SPGP0BI02700 | XP_018223612.1 | 99 | 0 |
| SPGP0P00890 | XP_018221565.1 | 99 | 0 |

Create PDF in your applications with the Pdfcrowd HTML to PDF API PDFCROWD

| | | | |
|---|---|---|---|
| SPGP0QF00100 | XP_018224018.1 | 99 | 0 |
| SPGP0FM00180 | XP_018222154.1 | 99 | 0 |
| SPGP0JR00110 | XP_018219030.1 | 99 | 0 |
| SPGP0MC00190 | XP_018222670.1 | 99 | 0 |
| SPGP0D02590 | XP_018221882.1 | 99 | 0 |
| SPGP0D00230 | XP_018221656.1 | 99 | 0 |
| SPGP0D00940 | XP_018221725.1 | 99 | 0 |
| SPGP0IC00170 | XP_018222790.1 | 99 | 0 |
| SPGP0BItrna16I | DI49_0395 | 100 | 0 |
| SPGP0BK01190 | XP_018221355.1 | 99 | 0 |
| SPGP0WW00180 | XP_018222080.1 | 100 | 0 |
| SPGP0RB00110 | XP_018218999.1 | 99 | 0 |
| SPGP0K00390 | XP_018219546.1 | 99 | 0 |
| SPGP0BI01940 | XP_018223686.1 | 99 | 0 |
| SPGP0SW00230 | XP_018220551.1 | 99 | 0 |
| SPGP0OF00370 | XP_018222718.1 | 99 | 0 |
| SPGP0K01370 | XP_018219450.1 | 99 | 0 |
| SPGP0U00470 | XP_018224090.1 | 99 | 0 |
| SPGP0N00310 | XP_018219219.1 | 99 | 0 |
| SPGP0VM00240 | XP_018220670.1 | 99 | 0 |
| SPGP0BKtrna7W | DI49_3006 | 100 | 0 |
| SPGP0BI01770 | XP_018223703.1 | 99 | 0 |
| SPGP0OL00160 | XP_018222282.1 | 99 | 0 |
| SPGP0C03470 | XP_018220027.1 | 99 | 0 |
| SPGP0BE03760 | XP_018222975.1 | 99 | 0 |
| SPGP0TQ00210 | XP_018220616.1 | 99 | 0 |
| SPGP0DB00100 | XP_018222189.1 | 99 | 0 |
| SPGP0DW00120 | XP_018218954.1 | 99 | 0 |
| SPGP0P01760 | XP_018221483.1 | 99 | 0 |
| SPGP0OF00160 | XP_018222697.1 | 99 | 0 |
| SPGP0N00620 | XP_018219249.1 | 99 | 0 |
| SPGP0TQ00320 | XP_018220627.1 | 99 | 0 |
| SPGP0IB00110 | XP_018222673.1 | 100 | 0 |
| SPGP0M00730 | XP_018220917.1 | 99 | 0 |
| SPGP0BI01760 | XP_018223704.1 | 99 | 0 |
| SPGP0PL00240 | XP_018219742.1 | 100 | 0 |
| SPGP0SU00140 | XP_018218974.1 | 99 | 0 |
| SPGP0BK02710 | XP_018221210.1 | 99 | 0 |
| SPGP0C01880 | XP_018220182.1 | 99 | 0 |
| SPGP0BE00490 | XP_018223285.1 | 99 | 0 |

Create PDF in your applications with the Pdfcrowd HTML to PDF API PDFCROWD

| | | | |
|---|---|---|---|
| SPGP0D03410 | XP_018221959.1 | 99 | 0 |
| SPGP0BI01470 | XP_018223734.1 | 99 | 0 |
| SPGP0BK00570 | XP_018221415.1 | 99 | 0 |
| SPGP0BI01660 | XP_018223714.1 | 99 | 0 |
| SPGP0BI01190 | XP_018223761.1 | 99 | 0 |
| SPGP0Itrna24A | DI49_5601 | 100 | 0 |
| SPGP0EZ00170 | XP_018219322.1 | 99 | 0 |
| SPGP0EM00280 | XP_018220707.1 | 99 | 0 |
| SPGP0U00270 | XP_018224109.1 | 99 | 0 |
| SPGP0D01540 | XP_018221782.1 | 99 | 0 |
| SPGP0NO00100 | XP_018222687.1 | 99 | 0 |
| SPGP0QF00120 | XP_018224020.1 | 100 | 0 |
| SPGP0VN00200 | XP_018219810.1 | 99 | 0 |
| SPGP0D02840 | XP_018221906.1 | 99 | 0 |
| SPGP0BE01780 | XP_018223157.1 | 99 | 0 |
| SPGP0BE03270 | XP_018223019.1 | 99 | 0 |
| SPGP0C01120 | XP_018220252.1 | 99 | 0 |
| SPGP0RT00130 | XP_018223995.1 | 99 | 0 |
| SPGP0K02470 | XP_018219344.1 | 99 | 0 |
| SPGP0TQ00150 | XP_018220610.1 | 99 | 0 |
| SPGP0BI03730 | XP_018223509.1 | 99 | 0 |
| SPGP0BK02960 | XP_018221187.1 | 99 | 0 |
| SPGP0OF00310 | XP_018222712.1 | 99 | 0 |
| SPGP0OR00190 | XP_018218835.1 | 99 | 0 |
| SPGP0HO00320 | XP_018219971.1 | 99 | 0 |
| SPGP0NQ00100 | XP_018220755.1 | 100 | 0 |
| SPGP0K02270 | XP_018219364.1 | 99 | 0 |
| SPGP0EZ00180 | XP_018219323.1 | 99 | 0 |
| SPGP0BI04080 | XP_018223478.1 | 99 | 0 |
| SPGP0KL00190 | XP_018221095.1 | 99 | 0 |
| SPGP0D01150 | XP_018221743.1 | 99 | 0 |
| SPGP0BI01450 | XP_018223736.1 | 99 | 0 |
| SPGP0BK02940 | XP_018221189.1 | 98 | 0 |
| SPGP0BI03840 | XP_018223500.1 | 99 | 0 |
| SPGP0C02900 | XP_018220083.1 | 99 | 0 |
| SPGP0WA00190 | XP_018219847.1 | 99 | 0 |
| SPGP0D02000 | XP_018221825.1 | 99 | 0 |
| SPGP0C02650 | XP_018220108.1 | 99 | 0 |
| SPGP0BE01260 | XP_018223208.1 | 99 | 0 |
| SPGP0U00850 | XP_018224053.1 | 99 | 0 |

Create PDF in your applications with the Pdfcrowd HTML to PDF API PDFCROWD

| | | | |
|---|---|---|---|
| SPGP0OF00180 | XP_018222699.1 | 99 | 0 |
| SPGP0BI02460 | XP_018223636.1 | 99 | 0 |
| SPGP0BE02560 | XP_018223082.1 | 99 | 0 |
| SPGP0QC00100 | XP_018221457.1 | 99 | 0 |
| SPGP0BE02840 | XP_018223057.1 | 99 | 0 |
| SPGP0SH00130 | XP_018223971.1 | 99 | 0 |
| SPGP0BK01950 | XP_018221282.1 | 99 | 0 |
| SPGP0OR00170 | XP_018218833.1 | 99 | 0 |
| SPGP0BE01190 | XP_018223215.1 | 99 | 0 |
| SPGP0BI00640 | XP_018223814.1 | 99 | 0 |
| SPGP0M01180 | XP_018220961.1 | 99 | 0 |
| SPGP0BI03440 | XP_018223538.1 | 99 | 0 |
| SPGP0BI01290 | XP_018223751.1 | 99 | 0 |
| SPGP0C01900 | XP_018220180.1 | 99 | 0 |
| SPGP0BI02640 | XP_018223618.1 | 99 | 0 |
| SPGP0K01660 | XP_018219423.1 | 99 | 0 |
| SPGP0D03300 | XP_018221950.1 | 99 | 0 |
| SPGP0C01660 | XP_018220201.1 | 99 | 0 |
| SPGP0BI03770 | XP_018223506.1 | 100 | 0 |
| SPGP0C02060 | XP_018220165.1 | 99 | 0 |
| SPGP0C01830 | XP_018220187.1 | 99 | 0 |
| SPGP0BK02150 | XP_018221262.1 | 99 | 0 |
| SPGP0BK00620 | XP_018221410.1 | 99 | 0 |
| SPGP0C00500 | XP_018220314.1 | 99 | 0 |
| SPGP0K01610 | XP_018219427.1 | 99 | 0 |
| SPGP0K01960 | XP_018219395.1 | 99 | 0 |
| SPGP0EE00140 | XP_018220733.1 | 99 | 0 |
| SPGP0RI00160 | XP_018220494.1 | 99 | 0 |
| SPGP0VN00420 | XP_018219832.1 | 99 | 0 |
| SPGP0BE04920 | XP_018222859.1 | 86 | 0 |
| SPGP0BE01220 | XP_018218943.1 | 99 | 0 |
| SPGP0LV00100 | XP_018223371.1 | 99 | 0 |
| SPGP0BE04250 | XP_018222928.1 | 99 | 0 |
| SPGP0PL00150 | XP_018219751.1 | 99 | 0 |
| SPGP0BK00800 | XP_018221393.1 | 99 | 0 |
| SPGP0U00440 | XP_018224093.1 | 99 | 0 |
| SPGP0GV00120 | XP_018218862.1 | 99 | 0 |
| SPGP0OI00350 | XP_018218915.1 | 99 | 0 |
| SPGP0U00210 | XP_018224115.1 | 99 | 1 |
| SPGP0BE01100 | XP_018223224.1 | 99 | 0 |

Create PDF in your applications with the Pdfcrowd HTML to PDF API PDFCROWD

| | | | |
|---|---|---|---|
| SPGP0C02850 | XP_018220088.1 | 99 | 0 |
| SPGP0C01300 | XP_018220234.1 | 99 | 0 |
| SPGP0OT00140 | XP_018219133.1 | 99 | 0 |
| SPGP0P00150 | XP_018221638.1 | 99 | 0 |
| SPGP0P00400 | XP_018221613.1 | 99 | 0 |
| SPGP0GK00130 | XP_018219711.1 | 99 | 1 |
| SPGP0KB00180 | XP_018221120.1 | 99 | 0 |
| SPGP0BK02520 | XP_018221226.1 | 99 | 0 |
| SPGP0HO00310 | XP_018219972.1 | 99 | 0 |
| SPGP0BK00970 | XP_018221376.1 | 99 | 0 |
| SPGP0BE03590 | XP_018222992.1 | 99 | 0 |
| SPGP0BK01470 | XP_018221328.1 | 99 | 0 |
| SPGP0IK00160 | XP_018222113.1 | 100 | 0 |
| SPGP0K00560 | XP_018219528.1 | 99 | 0 |
| SPGP0BK00790 | XP_018221394.1 | 100 | 0 |
| SPGP0NI00100 | XP_018223906.1 | 99 | 0 |
| SPGP0U00540 | XP_018224083.1 | 99 | 0 |
| SPGP0SL00180 | XP_018220478.1 | 99 | 0 |
| SPGP0NG00180 | XP_018222770.1 | 99 | 0 |
| SPGP0WB00100 | XP_018222680.1 | 99 | 0 |
| SPGP0BE01510 | XP_018223184.1 | 99 | 0 |
| SPGP0BI01810 | XP_018223699.1 | 99 | 0 |
| SPGP0BI01690 | XP_018223711.1 | 99 | 0 |
| SPGP0M00580 | XP_018220902.1 | 99 | 0 |
| SPGP0BI01340 | XP_018223747.1 | 99 | 0 |
| SPGP0EM00100 | XP_018220723.1 | 99 | 0 |
| SPGP0BE02780 | XP_018223062.1 | 99 | 0 |
| SPGP0P01570 | XP_018221502.1 | 99 | 0 |
| SPGP0K00700 | XP_018219516.1 | 99 | 0 |
| SPGP0N00430 | XP_018219231.1 | 99 | 0 |
| SPGP0MD00170 | XP_018222392.1 | 99 | 0 |
| SPGP0VZ00280 | XP_018223892.1 | 99 | 0 |
| SPGP0OX00210 | XP_018219724.1 | 99 | 0 |
| SPGP0KQ00120 | XP_018219928.1 | 99 | 0 |
| SPGP0JJ00130 | XP_018219066.1 | 99 | 0 |
| SPGP0BE02310 | XP_018223107.1 | 94 | 5 |
| SPGP0D01970 | XP_018221823.1 | 99 | 0 |
| SPGP0P02000 | XP_018221458.1 | 96 | 0 |
| SPGP0BI02880 | XP_018223595.1 | 99 | 0 |
| SPGP0D00330 | XP_018221666.1 | 99 | 0 |

Create PDF in your applications with the Pdfcrowd HTML to PDF API PDFCROWD

| | | | |
|---|---|---|---|
| SPGP0EX00110 | XP_018223420.1 | 99 | 0 |
| SPGP0OF00120 | XP_018222693.1 | 99 | 0 |
| SPGP0CD00110 | XP_018222528.1 | 99 | 0 |
| SPGP0BK02570 | XP_018221221.1 | 99 | 0 |
| SPGP0BI00340 | XP_018223844.1 | 99 | 0 |
| SPGP0P01520 | XP_018221507.1 | 99 | 0 |
| SPGP0BI01520 | XP_018223729.1 | 99 | 0 |
| SPGP0P01410 | XP_018221517.1 | 99 | 0 |
| SPGP0OI00520 | XP_018218898.1 | 99 | 0 |
| SPGP0C01980 | XP_018220172.1 | 99 | 0 |
| SPGP0C00700 | XP_018220294.1 | 99 | 0 |
| SPGP0BK01380 | XP_018221336.1 | 99 | 0 |
| SPGP0WG00220 | XP_018221016.1 | 99 | 0 |
| SPGP0BK02720 | XP_018221209.1 | 99 | 0 |
| SPGP0TL00140 | XP_018219084.1 | 99 | 0 |
| SPGP0EM00180 | XP_018220715.1 | 99 | 0 |
| SPGP0BI00120 | XP_018223866.1 | 99 | 0 |
| SPGP0U00130 | XP_018224121.1 | 94 | 1 |
| SPGP0BE02810 | XP_018223059.1 | 99 | 0 |
| SPGP0BI00550 | XP_018223823.1 | 99 | 0 |
| SPGP0UB00120 | XP_018219037.1 | 99 | 0 |
| SPGP0BK00380 | XP_018221434.1 | 99 | 0 |
| SPGP0BK02640 | XP_018221215.1 | 99 | 0 |
| SPGP0GY00130 | XP_018223399.1 | 99 | 0 |
| SPGP0JC00130 | XP_018219863.1 | 100 | 0 |
| SPGP0CZ00140 | XP_018219909.1 | 99 | 0 |
| SPGP0C02750 | XP_018220098.1 | 99 | 0 |
| SPGP0C02020 | XP_018220169.1 | 99 | 0 |
| SPGP0ID00120 | XP_018222142.1 | 97 | 0 |
| SPGP0QV00200 | XP_018220000.1 | 99 | 0 |
| SPGP0JA00100 | XP_018223406.1 | 99 | 0 |
| SPGP0M00270 | XP_018220872.1 | 99 | 0 |
| SPGP0BE04310 | XP_018222922.1 | 99 | 1 |
| SPGP0FM00300 | XP_018222165.1 | 99 | 0 |
| SPGP0LK00160 | XP_018221065.1 | 99 | 0 |
| SPGP0D02980 | XP_018221919.1 | 99 | 0 |
| SPGP0FP00100 | XP_018220839.1 | 99 | 0 |
| SPGP0BI01830 | XP_018223697.1 | 99 | 0 |
| SPGP0D00490 | XP_018221681.1 | 99 | 0 |
| SPGP0FM00360 | XP_018222171.1 | 99 | 0 |

Create PDF in your applications with the Pdfcrowd HTML to PDF API PDFCROWD

| | | | |
|---|---|---|---|
| SPGP0BI03470 | XP_018223535.1 | 99 | 0 |
| SPGP0TD00100 | XP_018218866.1 | 99 | 0 |
| SPGP0BK01980 | XP_018221279.1 | 99 | 0 |
| SPGP0BK02090 | XP_018221268.1 | 99 | 0 |
| SPGP0BE01030 | XP_018223231.1 | 100 | 0 |
| SPGP0BE01520 | XP_018223183.1 | 99 | 0 |
| SPGP0TM00100 | XP_018219874.1 | 99 | 0 |
| SPGP0ET00160 | XP_018222447.1 | 99 | 0 |
| SPGP0BI02870 | XP_018223596.1 | 99 | 0 |
| SPGP0C01590 | XP_018220208.1 | 99 | 0 |
| SPGP0UT00110 | XP_018222191.1 | 99 | 0 |
| SPGP0K00350 | XP_018219550.1 | 99 | 0 |
| SPGP0QK00110 | XP_018220383.1 | 99 | 0 |
| SPGP0N00980 | XP_018219285.1 | 99 | 0 |
| SPGP0D03470 | XP_018221965.1 | 99 | 0 |
| SPGP0C03120 | XP_018220062.1 | 99 | 0 |
| SPGP0K00820 | XP_018219504.1 | 99 | 0 |
| SPGP0EF00130 | XP_018220016.1 | 99 | 0 |
| SPGP0BI00860 | XP_018223792.1 | 99 | 0 |
| SPGP0BE04510 | XP_018222900.1 | 99 | 0 |
| SPGP0D01360 | XP_018221764.1 | 99 | 0 |
| SPGP0D02310 | XP_018221856.1 | 100 | 0 |
| SPGP0BI01030 | XP_018223776.1 | 99 | 0 |
| SPGP0K02440 | XP_018219347.1 | 99 | 0 |
| SPGP0C02460 | XP_018220125.1 | 99 | 0 |
| SPGP0C02220 | XP_018220149.1 | 99 | 0 |
| SPGP0BI00700 | XP_018223808.1 | 99 | 0 |
| SPGP0K00950 | XP_018219492.1 | 100 | 0 |
| SPGP0XD00120 | XP_018218818.1 | 99 | 0 |
| SPGP0CR00110 | XP_018222213.1 | 99 | 0 |
| SPGP0BE02520 | XP_018223086.1 | 99 | 0 |
| SPGP0BE00920 | XP_018223242.1 | 99 | 0 |
| SPGP0VU00110 | XP_018222735.1 | 99 | 0 |
| SPGP0P01500 | XP_018221509.1 | 99 | 0 |
| SPGP0BE00160 | XP_018223318.1 | 99 | 0 |
| SPGP0CX00150 | XP_018220794.1 | 99 | 0 |
| SPGP0ltrna16M | DI49_5351 | 100 | 0 |
| SPGP0C02420 | XP_018220129.1 | 99 | 0 |
| SPGP0BE03960 | XP_018222956.1 | 99 | 0 |
| SPGP0M00810 | XP_018220925.1 | 99 | 0 |

Create PDF in your applications with the Pdfcrowd HTML to PDF API PDFCROWD

| | | | |
|---|---|---|---|
| SPGP0BE02720 | XP_018223068.1 | 99 | 0 |
| SPGP0BI02250 | XP_018223657.1 | 99 | 0 |
| SPGP0BK01600 | XP_018221316.1 | 99 | 0 |
| SPGP0LU00250 | XP_018224001.1 | 99 | 0 |
| SPGP0BKtrna14M | DI49_4243 | 100 | 0 |
| SPGP0BK00390 | XP_018221433.1 | 99 | 0 |
| SPGP0N00610 | XP_018219248.1 | 99 | 0 |
| SPGP0C00310 | XP_018220332.1 | 99 | 0 |
| SPGP0P01710 | XP_018221488.1 | 99 | 0 |
| SPGP0D03040 | XP_018221924.1 | 99 | 0 |
| SPGP0BI01040 | XP_018223775.1 | 99 | 0 |
| SPGP0BI02340 | XP_018223648.1 | 100 | 0 |
| SPGP0OL00140 | XP_018222284.1 | 99 | 0 |
| SPGP0D00960 | XP_018221727.1 | 99 | 0 |
| SPGP0GF00280 | XP_018219633.1 | 99 | 0 |
| SPGP0P01290 | XP_018221529.1 | 99 | 0 |
| SPGP0GY00160 | XP_018223396.1 | 100 | 0 |
| SPGP0BI00740 | XP_018223804.1 | 99 | 0 |
| SPGP0K01850 | XP_018219404.1 | 99 | 0 |
| SPGP0TQ00120 | XP_018220607.1 | 99 | 0 |
| SPGP0U00220 | XP_018224114.1 | 99 | 1 |
| SPGP0P00210 | XP_018221632.1 | 99 | 0 |
| SPGP0K01390 | XP_018219448.1 | 99 | 0 |
| SPGP0RI00290 | XP_018220507.1 | 99 | 0 |
| SPGP0QV00230 | XP_018219997.1 | 99 | 0 |
| SPGP0WG00100 | XP_018221028.1 | 99 | 0 |
| SPGP0BK00630 | XP_018221409.1 | 99 | 0 |
| SPGP0N01110 | XP_018219298.1 | 99 | 0 |
| SPGP0P01620 | XP_018221497.1 | 99 | 0 |
| SPGP0WB00150 | XP_018222675.1 | 99 | 0 |
| SPGP0D03430 | XP_018221961.1 | 99 | 0 |
| SPGP0SP00160 | XP_018223966.1 | 99 | 0 |
| SPGP0C02200 | XP_018220151.1 | 99 | 0 |
| SPGP0N00660 | XP_018219253.1 | 99 | 0 |
| SPGP0P01030 | XP_018221553.1 | 99 | 0 |
| SPGP0M00600 | XP_018220904.1 | 100 | 0 |
| SPGP0P01740 | XP_018221485.1 | 99 | 0 |
| SPGP0M00710 | XP_018220915.1 | 99 | 0 |
| SPGP0C02500 | XP_018220121.1 | 99 | 0 |
| SPGP0C02360 | XP_018220135.1 | 99 | 0 |

Create PDF in your applications with the Pdfcrowd HTML to PDF API PDFCROWD

| | | | |
|---|---|---|---|
| SPGP0BK02770 | XP_018221204.1 | 98 | 0 |
| SPGP0KG00120 | XP_018222626.1 | 98 | 1 |
| SPGP0FW00210 | XP_018219575.1 | 99 | 0 |
| SPGP0C02680 | XP_018220105.1 | 98 | 2 |
| SPGP0SH00290 | XP_018223986.1 | 99 | 0 |
| SPGP0BK00550 | XP_018221417.1 | 99 | 0 |
| SPGP0BE03700 | XP_018222981.1 | 99 | 0 |
| SPGP0OE00150 | XP_018219605.1 | 99 | 0 |
| SPGP0U00630 | XP_018224075.1 | 99 | 0 |
| SPGP0D00260 | XP_018221659.1 | 99 | 0 |
| SPGP0NG00210 | XP_018222767.1 | 99 | 0 |
| SPGP0BI02510 | XP_018223631.1 | 99 | 0 |
| SPGP0U00200 | XP_018224116.1 | 99 | 0 |
| SPGP0WQ00120 | XP_018222265.1 | 99 | 0 |
| SPGP0BI00800 | XP_018223798.1 | 99 | 0 |
| SPGP0D00600 | XP_018221692.1 | 99 | 0 |
| SPGP0EZ00150 | XP_018219320.1 | 99 | 0 |
| SPGP0MG00160 | XP_018219612.1 | 99 | 0 |
| SPGP0BI03040 | XP_018223579.1 | 99 | 0 |
| SPGP0D01560 | XP_018221784.1 | 99 | 0 |
| SPGP0VN00260 | XP_018219816.1 | 99 | 0 |
| SPGP0BI02310 | XP_018223651.1 | 99 | 0 |
| SPGP0BK00990 | XP_018221375.1 | 99 | 0 |
| SPGP0K01310 | XP_018219456.1 | 99 | 0 |
| SPGP0BI02230 | XP_018223659.1 | 99 | 0 |
| SPGP0BI02090 | XP_018223672.1 | 99 | 0 |
| SPGP0OI00170 | XP_018218930.1 | 99 | 0 |
| SPGP0P00830 | XP_018221571.1 | 99 | 0 |
| SPGP0KL00160 | XP_018221098.1 | 99 | 0 |
| SPGP0C03400 | XP_018220034.1 | 99 | 0 |
| SPGP0WW00160 | XP_018222078.1 | 98 | 2 |
| SPGP0OX00120 | XP_018219733.1 | 100 | 0 |
| SPGP0N01020 | XP_018219289.1 | 99 | 0 |
| SPGP0D01170 | XP_018221745.1 | 99 | 0 |
| SPGP0K01970 | XP_018219394.1 | 99 | 0 |
| SPGP0UR00170 | XP_018222337.1 | 99 | 0 |
| SPGP0EW00180 | XP_018222297.1 | 99 | 0 |
| SPGP0P00930 | XP_018221561.1 | 99 | 0 |
| SPGP0VP00250 | XP_018218768.1 | 94 | 0 |
| SPGP0D02610 | XP_018221884.1 | 99 | 0 |

Create PDF in your applications with the Pdfcrowd HTML to PDF API PDFCROWD

| | | | |
|---|---|---|---|
| SPGP0BK02860 | XP_018221090.1 | 98 | 0 |
| SPGP0BE01200 | XP_018223214.1 | 99 | 0 |
| SPGP0IH00170 | XP_018222233.1 | 99 | 0 |
| SPGP0AP00110 | XP_018219060.1 | 99 | 0 |
| SPGP0U00830 | XP_018224055.1 | 99 | 0 |
| SPGP0C03030 | XP_018220071.1 | 99 | 0 |
| SPGP0BE00910 | XP_018223243.1 | 99 | 0 |
| SPGP0BI00660 | XP_018223812.1 | 99 | 0 |
| SPGP0BK01170 | XP_018221357.1 | 99 | 0 |
| SPGP0VM00130 | XP_018220681.1 | 100 | 0 |
| SPGP0BI02400 | XP_018223642.1 | 99 | 0 |
| SPGP0SH00110 | XP_018223969.1 | 99 | 0 |
| SPGP0OZ00130 | XP_018222400.1 | 99 | 0 |
| SPGP0UB00110 | XP_018219036.1 | 99 | 0 |
| SPGP0BE03350 | XP_018223012.1 | 99 | 0 |
| SPGP0BI03320 | XP_018223551.1 | 99 | 0 |
| SPGP0K01590 | XP_018219429.1 | 99 | 0 |
| SPGP0BE03420 | XP_018223005.1 | 99 | 0 |
| SPGP0BI03860 | XP_018223498.1 | 99 | 0 |
| SPGP0VS00150 | XP_018222525.1 | 99 | 0 |
| SPGP0C02920 | XP_018220081.1 | 99 | 0 |
| SPGP0K01800 | XP_018219409.1 | 99 | 0 |
| SPGP0BE03890 | XP_018222962.1 | 99 | 0 |
| SPGP0BE03750 | XP_018222976.1 | 99 | 0 |
| SPGP0OF00170 | XP_018222698.1 | 99 | 0 |
| SPGP0BE04490 | XP_018222902.1 | 99 | 0 |
| SPGP0OJ00110 | XP_018219923.1 | 99 | 0 |
| SPGP0D03230 | XP_018221943.1 | 99 | 0 |
| SPGP0UB00180 | XP_018219043.1 | 100 | 0 |
| SPGP0EU00110 | XP_018222659.1 | 99 | 0 |
| SPGP0ET00150 | XP_018222446.1 | 100 | 0 |
| SPGP0C01100 | XP_018220254.1 | 99 | 0 |
| SPGP0BE04160 | XP_018222937.1 | 99 | 0 |
| SPGP0BE00550 | XP_018223279.1 | 99 | 0 |
| SPGP0BE02280 | XP_018223110.1 | 99 | 0 |
| SPGP0BE02550 | XP_018223083.1 | 99 | 0 |
| SPGP0BK02170 | XP_018221260.1 | 99 | 0 |
| SPGP0K00530 | XP_018219531.1 | 99 | 0 |
| SPGP0Dtrna7F | DI49_2475 | 100 | 0 |
| SPGP0BE04440 | XP_018222908.1 | 99 | 0 |

Create PDF in your applications with the Pdfcrowd HTML to PDF API PDFCROWD

| | | | |
|---|---|---|---|
| SPGP0BK00820 | XP_018221391.1 | 99 | 1 |
| SPGP0BE01980 | XP_018223137.1 | 99 | 0 |
| SPGP0BE04270 | XP_018222926.1 | 99 | 0 |
| SPGP0VN00370 | XP_018219827.1 | 99 | 0 |
| SPGP0BE01130 | XP_018223221.1 | 99 | 0 |
| SPGP0D01030 | XP_018221734.1 | 99 | 0 |
| SPGP0BE02630 | XP_018223075.1 | 99 | 0 |
| SPGP0DJ00100 | XP_018219917.1 | 99 | 0 |
| SPGP0P00820 | XP_018221572.1 | 99 | 0 |
| SPGP0BI01550 | XP_018223725.1 | 99 | 0 |
| SPGP0M00130 | XP_018220859.1 | 99 | 0 |
| SPGP0BI03310 | XP_018223552.1 | 99 | 0 |
| SPGP0WI00110 | XP_018222119.1 | 98 | 0 |
| SPGP0P00170 | XP_018221636.1 | 99 | 0 |
| SPGP0DN00130 | XP_018218991.1 | 99 | 0 |
| SPGP0LE00180 | XP_018220760.1 | 99 | 0 |
| SPGP0BE00990 | XP_018223235.1 | 99 | 0 |
| SPGP0BI01090 | XP_018223770.1 | 99 | 0 |
| SPGP0KW00140 | XP_018220724.1 | 99 | 0 |
| SPGP0P01320 | XP_018221526.1 | 99 | 0 |
| SPGP0BE02850 | XP_018223056.1 | 99 | 0 |
| SPGP0RX00150 | XP_018222816.1 | 99 | 0 |
| SPGP0VD00100 | XP_018219833.1 | 99 | 0 |
| SPGP0K01540 | XP_018219434.1 | 98 | 1 |
| SPGP0K00280 | XP_018219557.1 | 99 | 0 |
| SPGP0TM00220 | XP_018219886.1 | 99 | 0 |
| SPGP0K00540 | XP_018219530.1 | 99 | 0 |
| SPGP0CZ00120 | XP_018219907.1 | 100 | 0 |
| SPGP0C00330 | XP_018220330.1 | 99 | 0 |
| SPGP0BI03430 | XP_018223539.1 | 99 | 0 |
| SPGP0BE03190 | XP_018223027.1 | 99 | 0 |
| SPGP0BE02150 | XP_018223123.1 | 99 | 0 |
| SPGP0BI04020 | XP_018223484.1 | 99 | 0 |
| SPGP0BI00310 | XP_018223847.1 | 99 | 0 |
| SPGP0K00810 | XP_018219505.1 | 99 | 0 |
| SPGP0K01230 | XP_018219464.1 | 99 | 0 |
| SPGP0FE00210 | XP_018220529.1 | 99 | 0 |
| SPGP0BK03020 | XP_018221181.1 | 98 | 0 |
| SPGP0C03330 | XP_018220041.1 | 99 | 0 |
| SPGP0P00450 | XP_018221608.1 | 99 | 0 |

Create PDF in your applications with the Pdfcrowd HTML to PDF API PDFCROWD

| | | | |
|---|---|---|---|
| SPGP0M00340 | XP_018220879.1 | 99 | 0 |
| SPGP0KN00210 | XP_018220577.1 | 99 | 0 |
| SPGP0K00610 | XP_018219525.1 | 99 | 0 |
| SPGP0P00940 | XP_018221560.1 | 99 | 0 |
| SPGP0N00450 | XP_018219233.1 | 99 | 0 |
| SPGP0BK02270 | XP_018221250.1 | 99 | 0 |
| SPGP0BI00870 | XP_018223791.1 | 99 | 0 |
| SPGP0K00830 | XP_018219503.1 | 99 | 0 |
| SPGP0GX00140 | XP_018222538.1 | 99 | 0 |
| SPGP0BE04690 | XP_018222882.1 | 99 | 0 |
| SPGP0Ctrna9P | DI49_4051 | 99 | 0 |
| SPGP0KW00130 | XP_018220725.1 | 97 | 0 |
| SPGP0BE04040 | XP_018222949.1 | 99 | 0 |
| SPGP0BE02330 | XP_018223105.1 | 99 | 0 |
| SPGP0D00300 | XP_018221663.1 | 99 | 0 |
| SPGP0D01990 | XP_018221824.1 | 99 | 0 |
| SPGP0BI03250 | XP_018223558.1 | 99 | 0 |
| SPGP0GE00100 | XP_018222601.1 | 97 | 0 |
| SPGP0BI01320 | XP_018223749.1 | 99 | 0 |
| SPGP0MF00160 | XP_018222204.1 | 99 | 0 |
| SPGP0C00120 | XP_018220351.1 | 99 | 0 |
| SPGP0BK01010 | XP_018221373.1 | 99 | 0 |
| SPGP0MO00370 | XP_018219121.1 | 99 | 0 |
| SPGP0D00400 | XP_018221673.1 | 99 | 0 |
| SPGP0RI00310 | XP_018220509.1 | 96 | 1 |
| SPGP0AA00180 | XP_018221077.1 | 98 | 2 |
| SPGP0BI00320 | XP_018223846.1 | 99 | 0 |
| SPGP0LU00130 | XP_018224013.1 | 100 | 0 |
| SPGP0U00480 | XP_018224089.1 | 99 | 0 |
| SPGP0MT00150 | XP_018219141.1 | 99 | 0 |
| SPGP0D03590 | XP_018221977.1 | 99 | 0 |
| SPGP0C00230 | XP_018220340.1 | 99 | 0 |
| SPGP0BE02950 | XP_018223047.1 | 99 | 0 |
| SPGP0DU00190 | XP_018222575.1 | 99 | 0 |
| SPGP0D01660 | XP_018221794.1 | 95 | 5 |
| SPGP0XD00110 | XP_018218817.1 | 99 | 0 |
| SPGP0D02860 | XP_018221908.1 | 99 | 0 |
| SPGP0BI00270 | XP_018223851.1 | 99 | 0 |
| SPGP0K00890 | XP_018219498.1 | 99 | 0 |

Create PDF in your applications with the Pdfcrowd HTML to PDF API PDFCROWD

| | | | |
|---|---|---|---|
| SPGP0BI00490 | XP_018223829.1 | 100 | 0 |
| SPGP0BI04130 | XP_018223473.1 | 99 | 0 |
| SPGP0D03080 | XP_018221928.1 | 99 | 0 |
| SPGP0C01700 | XP_018220197.1 | 99 | 0 |
| SPGP0RI00360 | XP_018220515.1 | 99 | 0 |
| SPGP0M00590 | XP_018220903.1 | 99 | 1 |
| SPGP0OX00150 | XP_018219730.1 | 99 | 0 |
| SPGP0M00290 | XP_018220874.1 | 100 | 0 |
| SPGP0VG00150 | XP_018223873.1 | 99 | 0 |
| SPGP0UJ00200 | XP_018223335.1 | 99 | 0 |
| SPGP0GD00110 | XP_018220833.1 | 99 | 0 |
| SPGP0TC00190 | XP_018219694.1 | 99 | 0 |
| SPGP0WB00120 | XP_018222678.1 | 99 | 0 |
| SPGP0BI03030 | XP_018223580.1 | 99 | 0 |
| SPGP0HB00190 | XP_018219185.1 | 98 | 0 |
| SPGP0M00370 | XP_018220882.1 | 99 | 0 |
| SPGP0GF00130 | XP_018219648.1 | 99 | 0 |
| SPGP0BI03410 | XP_018223541.1 | 99 | 0 |
| SPGP0MO00380 | XP_018219122.1 | 99 | 0 |
| SPGP0KT00100 | XP_018222242.1 | 99 | 0 |
| SPGP0CK00130 | XP_018219960.1 | 99 | 1 |
| SPGP0BK03410 | XP_018221143.1 | 99 | 0 |
| SPGP0BI01060 | XP_018223773.1 | 99 | 0 |
| SPGP0D01690 | XP_018221796.1 | 98 | 0 |
| SPGP0C01200 | XP_018220244.1 | 99 | 0 |
| SPGP0BE03840 | XP_018222967.1 | 100 | 0 |
| SPGP0BK03130 | XP_018221171.1 | 99 | 0 |
| SPGP0VD00140 | XP_018219837.1 | 87 | 2 |
| SPGP0BE02410 | XP_018223097.1 | 99 | 0 |
| SPGP0D02080 | XP_018221833.1 | 99 | 0 |
| SPGP0BI02500 | XP_018223632.1 | 99 | 0 |
| SPGP0D02330 | XP_018221858.1 | 99 | 0 |
| SPGP0N00120 | XP_018219201.1 | 97 | 0 |
| SPGP0JA00140 | XP_018223410.1 | 99 | 0 |
| SPGP0BE01540 | XP_018223181.1 | 99 | 0 |
| SPGP0BE01010 | XP_018223233.1 | 99 | 0 |
| SPGP0BU00130 | XP_018222252.1 | 100 | 0 |
| SPGP0ltrna28V | DI49_4248 | 100 | 0 |
| SPGP0VM00200 | XP_018220674.1 | 99 | 0 |
| SPGP0D01340 | XP_018221762.1 | 99 | 0 |

Create PDF in your applications with the Pdfcrowd HTML to PDF API PDFCROWD

| | | | |
|---|---|---|---|
| SPGP0K02570 | XP_018219334.1 | 100 | 0 |
| SPGP0C00810 | XP_018220283.1 | 99 | 0 |
| SPGP0P01900 | XP_018221469.1 | 99 | 1 |
| SPGP0BE00250 | XP_018223309.1 | 99 | 0 |
| SPGP0N00820 | XP_018219269.1 | 99 | 0 |
| SPGP0C02090 | XP_018220162.1 | 99 | 0 |
| SPGP0D02240 | XP_018221849.1 | 99 | 0 |
| SPGP0OM00110 | XP_018218950.1 | 99 | 0 |
| SPGP0BE00940 | XP_018223240.1 | 99 | 0 |
| SPGP0BE00760 | XP_018223257.1 | 97 | 3 |
| SPGP0C02130 | XP_018220158.1 | 99 | 0 |
| SPGP0TS00100 | XP_018219591.1 | 99 | 0 |
| SPGP0D02300 | XP_018221855.1 | 99 | 0 |
| SPGP0BE03940 | XP_018222958.1 | 98 | 0 |
| SPGP0OF00210 | XP_018222702.1 | 99 | 0 |
| SPGP0UJ00120 | XP_018223343.1 | 99 | 0 |
| SPGP0BI03340 | XP_018223549.1 | 99 | 1 |
| SPGP0EC00110 | XP_018222230.1 | 99 | 0 |
| SPGP0HO00170 | XP_018219985.1 | 99 | 0 |
| SPGP0BK00770 | XP_018221396.1 | 100 | 0 |
| SPGP0KL00130 | XP_018221101.1 | 99 | 0 |
| SPGP0BM00120 | XP_018222755.1 | 99 | 0 |
| SPGP0K01000 | XP_018219487.1 | 99 | 0 |
| SPGP0VB00120 | XP_018221047.1 | 99 | 0 |
| SPGP0UZ00110 | XP_018223427.1 | 99 | 0 |
| SPGP0BE04680 | XP_018222883.1 | 99 | 0 |
| SPGP0BE03410 | XP_018223006.1 | 99 | 0 |
| SPGP0C00460 | XP_018220318.1 | 99 | 0 |
| SPGP0EF00120 | XP_018220017.1 | 99 | 0 |
| SPGP0BK01800 | XP_018221297.1 | 99 | 0 |
| SPGP0D00480 | XP_018221680.1 | 99 | 0 |
| SPGP0BI04220 | XP_018223464.1 | 100 | 0 |
| SPGP0P00230 | XP_018221630.1 | 99 | 0 |
| SPGP0BK02280 | XP_018221249.1 | 99 | 0 |
| SPGP0BK01890 | XP_018221287.1 | 99 | 0 |
| SPGP0BK00530 | XP_018221419.1 | 99 | 0 |
| SPGP0D02790 | XP_018221902.1 | 99 | 0 |
| SPGP0PZ00160 | XP_018222430.1 | 99 | 0 |
| SPGP0D03450 | XP_018221963.1 | 99 | 0 |
| SPGP0VL00100 | XP_018219127.1 | 99 | 0 |

Create PDF in your applications with the Pdfcrowd HTML to PDF API PDFCROWD

| | | | |
|---|---|---|---|
| SPGP0P00370 | XP_018221616.1 | 99 | 0 |
| SPGP0K00240 | XP_018219561.1 | 99 | 0 |
| SPGP0P01010 | XP_018221555.1 | 99 | 0 |
| SPGP0M00840 | XP_018220928.1 | 99 | 0 |
| SPGP0D00610 | XP_018221693.1 | 100 | 0 |
| SPGP0IR00120 | XP_018219765.1 | 99 | 0 |
| SPGP0N01120 | XP_018219300.1 | 99 | 1 |
| SPGP0P00660 | XP_018221588.1 | 99 | 0 |
| SPGP0C03190 | XP_018220055.1 | 99 | 0 |
| SPGP0FM00190 | XP_018222155.1 | 99 | 0 |
| SPGP0BI01640 | XP_018223716.1 | 99 | 0 |
| SPGP0K02310 | XP_018219360.1 | 99 | 0 |
| SPGP0D01470 | XP_018221775.1 | 99 | 0 |
| SPGP0TT00120 | XP_018220836.1 | 99 | 0 |
| SPGP0K01730 | XP_018219416.1 | 99 | 0 |
| SPGP0BItrna19E | DI49_2698 | 100 | 0 |
| SPGP0BK00150 | XP_018221453.1 | 99 | 0 |
| SPGP0XF00130 | XP_018220853.1 | 99 | 0 |
| SPGP0RC00180 | XP_018222093.1 | 99 | 0 |
| SPGP0D02850 | XP_018221907.1 | 99 | 0 |
| SPGP0BE02090 | XP_018223128.1 | 99 | 0 |
| SPGP0P00330 | XP_018221620.1 | 99 | 0 |
| SPGP0PZ00360 | XP_018222410.1 | 99 | 0 |
| SPGP0EM00240 | XP_018220711.1 | 99 | 0 |
| SPGP0BK03390 | XP_018221145.1 | 99 | 0 |
| SPGP0LA00120 | XP_018220842.1 | 96 | 0 |
| SPGP0ST00170 | XP_018220542.1 | 99 | 0 |
| SPGP0K02350 | XP_018219356.1 | 99 | 0 |
| SPGP0BE01680 | XP_018223167.1 | 99 | 0 |
| SPGP0BE02450 | XP_018223093.1 | 99 | 0 |
| SPGP0LU00140 | XP_018224012.1 | 99 | 0 |
| SPGP0U00810 | XP_018224057.1 | 100 | 0 |
| SPGP0C00720 | XP_018220292.1 | 97 | 2 |
| SPGP0VP00270 | XP_018222851.1 | 92 | 0 |
| SPGP0VZ00220 | XP_018223886.1 | 99 | 0 |
| SPGP0C02110 | XP_018220160.1 | 99 | 0 |
| SPGP0JI00110 | XP_018222449.1 | 99 | 1 |
| SPGP0P00350 | XP_018221618.1 | 99 | 0 |
| SPGP0BI03720 | XP_018223510.1 | 100 | 0 |
| SPGP0BI00680 | XP_018223810.1 | 99 | 0 |

Create PDF in your applications with the Pdfcrowd HTML to PDF API PDFCROWD

| | | | |
|---|---|---|---|
| SPGP0HO00180 | XP_018219984.1 | 99 | 0 |
| SPGP0SH00170 | XP_018223975.1 | 99 | 0 |
| SPGP0BI02930 | XP_018223590.1 | 99 | 0 |
| SPGP0PL00180 | XP_018219748.1 | 99 | 0 |
| SPGP0BI02420 | XP_018223640.1 | 99 | 0 |
| SPGP0KN00110 | XP_018220567.1 | 99 | 0 |
| SPGP0D00750 | XP_018221707.1 | 99 | 0 |
| SPGP0KL00100 | XP_018221104.1 | 99 | 0 |
| SPGP0BI03880 | XP_018223496.1 | 99 | 0 |
| SPGP0UR00180 | XP_018222336.1 | 99 | 0 |
| SPGP0SW00190 | XP_018220555.1 | 99 | 0 |
| SPGP0EE00100 | XP_018220729.1 | 99 | 0 |
| SPGP0RI00120 | XP_018220490.1 | 99 | 0 |
| SPGP0C00400 | XP_018220323.1 | 99 | 0 |
| SPGP0K01920 | XP_018219398.1 | 99 | 0 |
| SPGP0D00460 | XP_018221678.1 | 97 | 0 |
| SPGP0OW00120 | XP_018218809.1 | 99 | 0 |
| SPGP0BK02060 | XP_018221271.1 | 99 | 0 |
| SPGP0BI03640 | XP_018223518.1 | 99 | 0 |
| SPGP0D03210 | XP_018221941.1 | 99 | 0 |
| SPGP0K02290 | XP_018219362.1 | 99 | 0 |
| SPGP0WY00130 | XP_018220443.1 | 99 | 0 |
| SPGP0P01140 | XP_018221543.1 | 99 | 0 |
| SPGP0BI03080 | XP_018223575.1 | 99 | 0 |
| SPGP0BK02110 | XP_018221266.1 | 99 | 0 |
| SPGP0FE00180 | XP_018220532.1 | 99 | 0 |
| SPGP0VE00150 | XP_018220967.1 | 99 | 0 |
| SPGP0BI03390 | XP_018223543.1 | 95 | 0 |
| SPGP0LV00140 | XP_018223375.1 | 99 | 0 |
| SPGP0BE04210 | XP_018222932.1 | 94 | 0 |
| SPGP0U00450 | XP_018224092.1 | 99 | 0 |
| SPGP0BK00840 | XP_018221389.1 | 99 | 0 |
| SPGP0OB00110 | XP_018219794.1 | 100 | 0 |
| SPGP0BE03880 | XP_018222963.1 | 99 | 0 |
| SPGP0BE00680 | XP_018223265.1 | 99 | 0 |
| SPGP0BK02780 | XP_018221203.1 | 99 | 0 |
| SPGP0GH00130 | XP_018218965.1 | 99 | 0 |
| SPGP0VZ00110 | XP_018223876.1 | 99 | 0 |
| SPGP0S00130 | XP_018219655.1 | 99 | 1 |
| SPGP0BI03330 | XP_018223550.1 | 99 | 0 |

Create PDF in your applications with the Pdfcrowd HTML to PDF API PDFCROWD

| | | | |
|---|---|---|---|
| SPGP0BK00490 | XP_018221423.1 | 99 | 0 |
| SPGP0BK02560 | XP_018221222.1 | 99 | 0 |
| SPGP0BK00730 | XP_018221400.1 | 99 | 0 |
| SPGP0IC00270 | XP_018222780.1 | 99 | 0 |
| SPGP0K02030 | XP_018219388.1 | 99 | 0 |
| SPGP0BE01420 | XP_018223192.1 | 99 | 0 |
| SPGP0BE02790 | XP_018223061.1 | 98 | 1 |
| SPGP0NI00140 | XP_018223910.1 | 99 | 0 |
| SPGP0K00520 | XP_018219532.1 | 99 | 0 |
| SPGP0K02430 | XP_018219348.1 | 99 | 0 |
| SPGP0D00350 | XP_018221668.1 | 100 | 0 |
| SPGP0BK02760 | XP_018221205.1 | 99 | 0 |
| SPGP0OI00790 | XP_018218871.1 | 99 | 0 |
| SPGP0BE02130 | XP_018223124.1 | 99 | 0 |
| SPGP0UJ00270 | XP_018223329.1 | 99 | 0 |
| SPGP0BE03080 | XP_018223037.1 | 99 | 0 |
| SPGP0HG00120 | XP_018220457.1 | 100 | 0 |
| SPGP0D02580 | XP_018221881.1 | 99 | 0 |
| SPGP0Ftrna5C | DI49_5567 | 100 | 0 |
| SPGP0D02620 | XP_018221885.1 | 99 | 0 |
| SPGP0N00470 | XP_018219235.1 | 98 | 2 |
| SPGP0K00290 | XP_018219556.1 | 99 | 0 |
| SPGP0GX00120 | XP_018222536.1 | 99 | 0 |
| SPGP0DQ00100 | XP_018219669.1 | 99 | 0 |
| SPGP0N00360 | XP_018219224.1 | 99 | 0 |
| SPGP0BK03490 | XP_018221138.1 | 99 | 0 |
| SPGP0BE03730 | XP_018222978.1 | 99 | 0 |
| SPGP0BE02680 | XP_018223072.1 | 99 | 0 |
| SPGP0DQ00160 | XP_018219675.1 | 99 | 0 |
| SPGP0VM00170 | XP_018220677.1 | 99 | 0 |
| SPGP0C03310 | XP_018220043.1 | 99 | 0 |
| SPGP0BI01650 | XP_018223715.1 | 99 | 0 |
| SPGP0MF00140 | XP_018222206.1 | 99 | 0 |
| SPGP0BK01030 | XP_018221371.1 | 99 | 0 |
| SPGP0FM00100 | XP_018222146.1 | 99 | 0 |
| SPGP0EE00120 | XP_018220731.1 | 99 | 0 |
| SPGP0GY00100 | XP_018223402.1 | 99 | 0 |
| SPGP0P01460 | XP_018221512.1 | 99 | 0 |
| SPGP0GH00210 | XP_018218957.1 | 100 | 0 |
| SPGP0BE04110 | XP_018222942.1 | 99 | 0 |

Create PDF in your applications with the Pdfcrowd HTML to PDF API PDFCROWD

| | | | |
|---|---|---|---|
| SPGP0P00650 | XP_018221589.1 | 100 | 0 |
| SPGP0HT00240 | XP_018219015.1 | 99 | 0 |
| SPGP0BI00300 | XP_018223848.1 | 99 | 0 |
| SPGP0U00310 | XP_018224105.1 | 99 | 0 |
| SPGP0BK00670 | XP_018221406.1 | 99 | 0 |
| SPGP0D02270 | XP_018221852.1 | 99 | 0 |
| SPGP0TS00120 | XP_018219589.1 | 99 | 0 |
| SPGP0BE03620 | XP_018222989.1 | 99 | 0 |
| SPGP0C01150 | XP_018220249.1 | 99 | 0 |
| SPGP0AA00190 | XP_018221076.1 | 99 | 0 |
| SPGP0M00100 | XP_018220856.1 | 98 | 0 |
| SPGP0BI01330 | XP_018223748.1 | 99 | 0 |
| SPGP0D01400 | XP_018221768.1 | 99 | 0 |
| SPGP0BI01400 | XP_018223741.1 | 99 | 0 |
| SPGP0JFtrna11L | DI49_2066 | 100 | 0 |
| SPGP0OX00130 | XP_018219732.1 | 100 | 0 |
| SPGP0BI03650 | XP_018223517.1 | 99 | 0 |
| SPGP0MFtrna6F | DI49_1939 | 100 | 0 |
| SPGP0K02240 | XP_018219367.1 | 99 | 0 |
| SPGP0BE00540 | XP_018223280.1 | 99 | 0 |
| SPGP0BI03150 | XP_018223568.1 | 99 | 0 |
| SPGP0C00130 | XP_018220350.1 | 98 | 0 |
| SPGP0CF00150 | XP_018222495.1 | 99 | 0 |
| SPGP0BI01070 | XP_018223772.1 | 99 | 0 |
| SPGP0K00250 | XP_018219560.1 | 99 | 0 |
| SPGP0VI00140 | XP_018222307.1 | 99 | 0 |
| SPGP0C01450 | XP_018220220.1 | 99 | 0 |
| SPGP0BK02970 | XP_018221186.1 | 99 | 0 |
| SPGP0BI01460 | XP_018223735.1 | 99 | 0 |
| SPGP0UJ00130 | XP_018223342.1 | 100 | 0 |
| SPGP0C01260 | XP_018220238.1 | 99 | 0 |
| SPGP0BK01590 | XP_018221317.1 | 99 | 0 |
| SPGP0MO00210 | XP_018219106.1 | 99 | 0 |
| SPGP0BE00440 | XP_018223290.1 | 99 | 0 |
| SPGP0PK00120 | XP_018220421.1 | 99 | 0 |
| SPGP0C02600 | XP_018220111.1 | 99 | 0 |
| SPGP0JA00160 | XP_018223411.1 | 99 | 0 |
| SPGP0HJ00150 | XP_018224045.1 | 96 | 0 |
| SPGP0BE01560 | XP_018223179.1 | 99 | 0 |
| SPGP0XC00270 | XP_018220603.1 | 99 | 0 |

Create PDF in your applications with the Pdfcrowd HTML to PDF API PDFCROWD

| | | | |
|---|---|---|---|
| SPGP0BI01590 | XP_018223721.1 | 95 | 4 |
| SPGP0C01080 | XP_018220256.1 | 99 | 0 |
| SPGP0TE00190 | XP_018219901.1 | 100 | 0 |
| SPGP0BI02740 | XP_018223608.1 | 99 | 0 |
| SPGP0BK01850 | XP_018221291.1 | 99 | 0 |
| SPGP0FX00130 | XP_018222825.1 | 99 | 0 |
| SPGP0P01920 | XP_018221467.1 | 97 | 0 |
| SPGP0C02520 | XP_018220119.1 | 99 | 0 |
| SPGP0BE00960 | XP_018223238.1 | 99 | 0 |
| SPGP0BE03510 | XP_018222999.1 | 99 | 0 |
| SPGP0BE00210 | XP_018223313.1 | 99 | 0 |
| SPGP0K02140 | XP_018219377.1 | 99 | 0 |
| SPGP0C00680 | XP_018220296.1 | 99 | 0 |
| SPGP0BE03660 | XP_018222985.1 | 99 | 0 |
| SPGP0D01730 | XP_018221800.1 | 99 | 0 |
| SPGP0BE03340 | XP_018223013.1 | 100 | 0 |
| SPGP0UJ00100 | XP_018223345.1 | 99 | 0 |
| SPGP0BI02330 | XP_018223649.1 | 100 | 0 |
| SPGP0D01330 | XP_018221761.1 | 99 | 0 |
| SPGP0BI03050 | XP_018223578.1 | 100 | 0 |
| SPGP0P00780 | XP_018221576.1 | 99 | 0 |
| SPGP0D01160 | XP_018221744.1 | 99 | 0 |
| SPGP0SN00120 | XP_018222592.1 | 95 | 1 |
| SPGP0C00350 | XP_018220328.1 | 99 | 0 |
| SPGP0K01020 | XP_018219485.1 | 99 | 0 |
| SPGP0NT00180 | XP_018222466.1 | 99 | 0 |
| SPGP0UZ00170 | XP_018223432.1 | 99 | 0 |
| SPGP0D03000 | XP_018221920.1 | 99 | 0 |
| SPGP0BK00940 | XP_018221379.1 | 99 | 0 |
| SPGP0TC00130 | XP_018219700.1 | 99 | 0 |
| SPGP0C00200 | XP_018220343.1 | 99 | 0 |
| SPGP0AP00120 | XP_018219061.1 | 100 | 0 |
| SPGP0OE00180 | XP_018219602.1 | 98 | 2 |
| SPGP0D00920 | XP_018221723.1 | 99 | 0 |
| SPGP0D00290 | XP_018221662.1 | 99 | 0 |
| SPGP0P01250 | XP_018221533.1 | 99 | 0 |
| SPGP0HO00250 | XP_018219977.1 | 99 | 0 |
| SPGP0BK01570 | XP_018221319.1 | 99 | 0 |
| SPGP0P00250 | XP_018221628.1 | 99 | 1 |
| SPGP0K01770 | XP_018219412.1 | 99 | 1 |

Create PDF in your applications with the Pdfcrowd HTML to PDF API PDFCROWD

| | | | |
|---|---|---|---|
| SPGP0KY00130 | XP_018223904.1 | 99 | 0 |
| SPGP0BK00690 | XP_018221404.1 | 99 | 0 |
| SPGP0EM00150 | XP_018220718.1 | 99 | 0 |
| SPGP0RI00250 | XP_018220503.1 | 100 | 0 |
| SPGP0OE00280 | XP_018219593.1 | 100 | 0 |
| SPGP0EW00170 | XP_018222296.1 | 99 | 0 |
| SPGP0P01070 | XP_018221550.1 | 99 | 0 |
| SPGP0K00880 | XP_018219499.1 | 99 | 0 |
| SPGP0BE04730 | XP_018222878.1 | 99 | 0 |
| SPGP0BI02130 | XP_018223668.1 | 99 | 0 |
| SPGP0GH00120 | XP_018218966.1 | 99 | 0 |
| SPGP0PZ00200 | XP_018222426.1 | 99 | 0 |
| SPGP0AK00110 | XP_018219738.1 | 99 | 0 |
| SPGP0D01520 | XP_018221780.1 | 99 | 0 |
| SPGP0JO00140 | XP_018223352.1 | 99 | 0 |
| SPGP0C03100 | XP_018220064.1 | 99 | 0 |
| SPGP0D00320 | XP_018221665.1 | 99 | 0 |
| SPGP0D02960 | XP_018221917.1 | 99 | 0 |
| SPGP0P01780 | XP_018221481.1 | 99 | 0 |
| SPGP0D02870 | XP_018221909.1 | 99 | 0 |
| SPGP0N00640 | XP_018219251.1 | 99 | 0 |
| SPGP0M00820 | XP_018220926.1 | 99 | 0 |
| SPGP0BI02220 | XP_018223660.1 | 100 | 0 |
| SPGP0U00780 | XP_018224060.1 | 99 | 0 |
| SPGP0NQ00160 | XP_018220749.1 | 99 | 0 |
| SPGP0K00550 | XP_018219529.1 | 100 | 0 |
| SPGP0BK03370 | XP_018221147.1 | 99 | 0 |
| SPGP0DT00120 | XP_018221086.1 | 99 | 0 |
| SPGP0PE00200 | XP_018222364.1 | 99 | 0 |
| SPGP0WA00110 | XP_018219855.1 | 99 | 0 |
| SPGP0OX00300 | XP_018219715.1 | 99 | 0 |
| SPGP0C00180 | XP_018220345.1 | 99 | 0 |
| SPGP0NT00100 | XP_018222474.1 | 99 | 0 |
| SPGP0OI00610 | XP_018218889.1 | 99 | 0 |
| SPGP0K01410 | XP_018219446.1 | 99 | 0 |
| SPGP0WI00130 | XP_018222121.1 | 99 | 0 |
| SPGP0BI00510 | XP_018223827.1 | 99 | 0 |
| SPGP0BK00510 | XP_018221421.1 | 99 | 0 |
| SPGP0BE00140 | XP_018223320.1 | 99 | 0 |
| SPGP0OI00150 | XP_018218932.1 | 99 | 0 |

Create PDF in your applications with the Pdfcrowd HTML to PDF API PDFCROWD

| | | | |
|---|---|---|---|
| SPGP0K00710 | XP_018219515.1 | 99 | 0 |
| SPGP0C02470 | XP_018220124.1 | 99 | 0 |
| SPGP0D02010 | XP_018221826.1 | 99 | 0 |
| SPGP0C03150 | XP_018220059.1 | 99 | 0 |
| SPGP0OI00420 | XP_018218908.1 | 99 | 0 |
| SPGP0BE04340 | XP_018222919.1 | 99 | 0 |
| SPGP0OH00130 | XP_018220987.1 | 99 | 0 |
| SPGP0SH00200 | XP_018223978.1 | 99 | 0 |
| SPGP0C01290 | XP_018220235.1 | 99 | 0 |
| SPGP0BE04280 | XP_018222925.1 | 99 | 0 |
| SPGP0P01680 | XP_018221491.1 | 99 | 0 |
| SPGP0QK00130 | XP_018220385.1 | 99 | 0 |
| SPGP0D01220 | XP_018221750.1 | 99 | 0 |
| SPGP0D00650 | XP_018221697.1 | 99 | 0 |
| SPGP0CF00100 | XP_018222500.1 | 99 | 0 |
| SPGP0BI02910 | XP_018223592.1 | 99 | 0 |
| SPGP0HT00120 | XP_018219004.1 | 99 | 0 |
| SPGP0C02210 | XP_018220150.1 | 99 | 0 |
| SPGP0BK01370 | XP_018221337.1 | 99 | 0 |
| SPGP0JA00130 | XP_018223409.1 | 99 | 0 |
| SPGP0K01840 | XP_018219405.1 | 99 | 1 |
| SPGP0BK02130 | XP_018221264.1 | 99 | 0 |
| SPGP0BE02470 | XP_018223091.1 | 99 | 0 |
| SPGP0K01900 | XP_018219400.1 | 99 | 0 |
| SPGP0BE04180 | XP_018222935.1 | 98 | 0 |
| SPGP0BK02540 | XP_018221224.1 | 99 | 0 |
| SPGP0VD00110 | XP_018219834.1 | 99 | 0 |
| SPGP0D03270 | XP_018221947.1 | 99 | 0 |
| SPGP0BE04450 | XP_018222907.1 | 99 | 0 |
| SPGP0C02070 | XP_018220164.1 | 99 | 0 |
| SPGP0UX00110 | XP_018222192.1 | 98 | 2 |
| SPGP0FC00120 | XP_018219053.1 | 99 | 0 |
| SPGP0UB00150 | XP_018219040.1 | 99 | 0 |
| SPGP0BE00510 | XP_018223283.1 | 99 | 0 |
| SPGP0C01560 | XP_018220210.1 | 99 | 0 |
| SPGP0SU00110 | XP_018218971.1 | 99 | 0 |
| SPGP0BE04230 | XP_018222930.1 | 99 | 0 |
| SPGP0LV00160 | XP_018223377.1 | 99 | 0 |
| SPGP0BE01600 | XP_018223175.1 | 99 | 0 |
| SPGP0SXtrna16S | DI49_0694 | 100 | 0 |

Create PDF in your applications with the Pdfcrowd HTML to PDF API PDFCROWD

| | | | |
|---|---|---|---|
| SPGP0BK00860 | XP_018221387.1 | 99 | 0 |
| SPGP0D01060 | XP_018221736.1 | 99 | 0 |
| SPGP0WY00110 | XP_018220445.1 | 100 | 0 |
| SPGP0TS00130 | XP_018219588.1 | 99 | 0 |
| SPGP0K00790 | XP_018219507.1 | 99 | 0 |
| SPGP0BI03350 | XP_018223548.1 | 99 | 0 |
| SPGP0D03480 | XP_018221966.1 | 99 | 0 |
| SPGP0S00110 | XP_018219653.1 | 99 | 1 |
| SPGP0D02730 | XP_018221896.1 | 99 | 0 |
| SPGP0OI00340 | XP_018218916.1 | 99 | 0 |
| SPGP0K02010 | XP_018219390.1 | 99 | 0 |
| SPGP0P01190 | XP_018221538.1 | 98 | 1 |
| SPGP0LB00100 | XP_018222553.1 | 97 | 3 |
| SPGP0RT00100 | XP_018223992.1 | 99 | 0 |
| SPGP0BE04360 | XP_018222917.1 | 99 | 0 |
| SPGP0K00500 | XP_018219534.1 | 100 | 0 |
| SPGP0D02600 | XP_018221883.1 | 99 | 0 |
| SPGP0FN00170 | XP_018223386.1 | 99 | 0 |
| SPGP0WY00140 | XP_018220442.1 | 99 | 0 |
| SPGP0GL00130 | XP_018222614.1 | 99 | 0 |
| SPGP0D00930 | XP_018221724.1 | 99 | 0 |
| SPGP0BE01720 | XP_018223163.1 | 100 | 0 |
| SPGP0BK01720 | XP_018221305.1 | 99 | 0 |
| SPGP0BI01390 | XP_018223742.1 | 99 | 0 |
| SPGP0BE02110 | XP_018223126.1 | 99 | 0 |
| SPGP0K01270 | XP_018219460.1 | 99 | 0 |
| SPGP0C03480 | XP_018220026.1 | 99 | 0 |
| SPGP0K02090 | XP_018219382.1 | 99 | 0 |
| SPGP0OI00640 | XP_018218886.1 | 99 | 0 |
| SPGP0BI02000 | XP_018223680.1 | 99 | 0 |
| SPGP0JO00100 | XP_018223356.1 | 99 | 0 |
| SPGP0BK01050 | XP_018221369.1 | 99 | 0 |
| SPGP0UH00180 | XP_018220414.1 | 98 | 0 |
| SPGP0GX00100 | XP_018222534.1 | 99 | 0 |
| SPGP0C03070 | XP_018220067.1 | 99 | 0 |
| SPGP0BK01200 | XP_018221354.1 | 99 | 0 |
| SPGP0BR00100 | XP_018222223.1 | 99 | 0 |
| SPGP0C02350 | XP_018220136.1 | 99 | 0 |
| SPGP0DS00120 | XP_018220392.1 | 100 | 0 |
| SPGP0BI01100 | XP_018223769.1 | 99 | 0 |

Create PDF in your applications with the Pdfcrowd HTML to PDF API PDFCROWD

| | | | |
|---|---|---|---|
| SPGP0BI01670 | XP_018223713.1 | 99 | 0 |
| SPGP0M00760 | XP_018220920.1 | 99 | 0 |
| SPGP0K02410 | XP_018219350.1 | 99 | 0 |
| SPGP0D03160 | XP_018221936.1 | 99 | 0 |
| SPGP0D02420 | XP_018221867.1 | 99 | 0 |
| SPGP0Dtrna11P | DI49_2389 | 99 | 0 |
| SPGP0OS00100 | XP_018219932.1 | 89 | 1 |
| SPGP0M00630 | XP_018220907.1 | 99 | 0 |
| SPGP0D00880 | XP_018221719.1 | 99 | 0 |
| SPGP0LU00170 | XP_018224009.1 | 99 | 0 |
| SPGP0BI00980 | XP_018223781.1 | 99 | 0 |
| SPGP0D01760 | XP_018221803.1 | 99 | 0 |
| SPGP0BI01580 | XP_018223722.1 | 99 | 0 |
| SPGP0EL00110 | XP_018223367.1 | 99 | 0 |
| SPGP0RE00110 | XP_018220395.1 | 99 | 0 |
| SPGP0RC00280 | XP_018222103.1 | 99 | 0 |
| SPGP0GY00110 | XP_018223401.1 | 99 | 0 |
| SPGP0RZ00130 | XP_018223382.1 | 99 | 0 |
| SPGP0FX00100 | XP_018222822.1 | 99 | 0 |
| SPGP0M00120 | XP_018220858.1 | 99 | 0 |
| SPGP0RI00220 | XP_018220500.1 | 99 | 0 |
| SPGP0U00370 | XP_018224099.1 | 99 | 0 |
| SPGP0BI02300 | XP_018223652.1 | 99 | 0 |
| SPGP0FN00140 | XP_018223389.1 | 99 | 0 |
| SPGP0C02030 | XP_018220168.1 | 99 | 0 |
| SPGP0C02830 | XP_018220090.1 | 99 | 0 |
| SPGP0BI01850 | XP_018223695.1 | 99 | 0 |
| SPGP0GF00250 | XP_018219636.1 | 99 | 0 |
| SPGP0D01090 | XP_018221739.1 | 99 | 0 |
| SPGP0C02050 | XP_018220166.1 | 99 | 0 |
| SPGP0EC00120 | XP_018222229.1 | 99 | 0 |
| SPGP0WW00150 | XP_018222077.1 | 99 | 0 |
| SPGP0BK02830 | XP_018221199.1 | 99 | 0 |
| SPGP0BI00180 | XP_018223860.1 | 99 | 0 |
| SPGP0BK00520 | XP_018221420.1 | 99 | 0 |
| SPGP0VNtrna17P | DI49_4699 | 100 | 0 |
| SPGP0K01760 | XP_018219413.1 | 99 | 0 |
| SPGP0OX00100 | XP_018219735.1 | 100 | 0 |
| SPGP0BE00290 | XP_018223305.1 | 99 | 0 |

Create PDF in your applications with the Pdfcrowd HTML to PDF API PDFCROWD

| | | | |
|---|---|---|---|
| SPGP0BE02200 | XP_018223118.1 | 99 | 0 |
| SPGP0GW00110 | XP_018220828.1 | 99 | 0 |
| SPGP0BE02650 | XP_018223074.1 | 99 | 0 |
| SPGP0BI01050 | XP_018223774.1 | 100 | 0 |
| SPGP0MJ00100 | XP_018222081.1 | 99 | 0 |
| SPGP0U00880 | XP_018224050.1 | 93 | 1 |
| SPGP0UR00150 | XP_018222339.1 | 99 | 0 |
| SPGP0LA00170 | XP_018220846.1 | 99 | 0 |
| SPGP0D03490 | XP_018221967.1 | 99 | 0 |
| SPGP0RC00200 | XP_018222095.1 | 99 | 0 |
| SPGP0N00710 | XP_018219258.1 | 99 | 0 |
| SPGP0BE00670 | XP_018223266.1 | 99 | 0 |
| SPGP0N00990 | XP_018219286.1 | 99 | 0 |
| SPGP0C01240 | XP_018220240.1 | 99 | 0 |
| SPGP0U00680 | XP_018224070.1 | 99 | 0 |
| SPGP0BI02580 | XP_018223624.1 | 99 | 0 |
| SPGP0D01380 | XP_018221766.1 | 99 | 1 |
| SPGP0C03140 | XP_018220060.1 | 99 | 0 |
| SPGP0GN00100 | XP_018222588.1 | 99 | 0 |
| SPGP0BE01320 | XP_018223202.1 | 99 | 0 |
| SPGP0BE04760 | XP_018222875.1 | 99 | 0 |
| SPGP0BI00720 | XP_018223806.1 | 98 | 2 |
| SPGP0BI03960 | XP_018223489.1 | 99 | 0 |
| SPGP0IV00130 | XP_018220641.1 | 99 | 0 |
| SPGP0C01170 | XP_018220247.1 | 99 | 0 |
| SPGP0WJ00180 | XP_018221052.1 | 99 | 0 |
| SPGP0BI02760 | XP_018223607.1 | 99 | 0 |
| SPGP0DP00150 | XP_018220649.1 | 99 | 0 |
| SPGP0D02770 | XP_018221900.1 | 99 | 0 |
| SPGP0BE00720 | XP_018223261.1 | 99 | 0 |
| SPGP0D01930 | XP_018221819.1 | 99 | 0 |
| SPGP0BI03890 | XP_018223495.1 | 99 | 0 |
| SPGP0QV00190 | XP_018220001.1 | 99 | 0 |
| SPGP0BItrna16Q | DI49_2421 | 100 | 0 |
| SPGP0TS00140 | XP_018219587.1 | 100 | 0 |
| SPGP0OX00250 | XP_018219720.1 | 99 | 0 |
| SPGP0BE01620 | XP_018223173.1 | 99 | 0 |
| SPGP0D03390 | XP_018221957.1 | 99 | 0 |
| SPGP0D03020 | XP_018221922.1 | 99 | 0 |
| SPGP0BK02290 | XP_018221248.1 | 97 | 1 |

Create PDF in your applications with the Pdfcrowd HTML to PDF API PDFCROWD

| | | | |
|---|---|---|---|
| SPGP0K01860 | XP_018219403.1 | 99 | 0 |
| SPGP0BI01270 | XP_018223753.1 | 99 | 0 |
| SPGP0UJ00160 | XP_018223339.1 | 99 | 0 |
| SPGP0K02460 | XP_018219345.1 | 99 | 0 |
| SPGP0HT00260 | XP_018219017.1 | 99 | 0 |
| SPGP0C03180 | XP_018220056.1 | 99 | 0 |
| SPGP0VH00140 | XP_018219685.1 | 99 | 0 |
| SPGP0HE00110 | XP_018220659.1 | 99 | 0 |
| SPGP0BE00230 | XP_018223311.1 | 99 | 0 |
| SPGP0TC00220 | XP_018219691.1 | 98 | 0 |
| SPGP0D01720 | XP_018221799.1 | 99 | 0 |
| SPGP0C00370 | XP_018220326.1 | 99 | 0 |
| SPGP0K01040 | XP_018219483.1 | 99 | 0 |
| SPGP0VS00130 | XP_018222523.1 | 99 | 0 |
| SPGP0OZ00110 | XP_018222398.1 | 99 | 0 |
| SPGP0GY00190 | XP_018223393.1 | 99 | 0 |
| SPGP0IK00110 | XP_018222108.1 | 99 | 0 |
| SPGP0XC00170 | XP_018220593.1 | 99 | 0 |
| SPGP0BK02340 | XP_018221243.1 | 99 | 0 |
| SPGP0BK01060 | XP_018221368.1 | 99 | 0 |
| SPGP0GT00130 | XP_018222511.1 | 99 | 0 |
| SPGP0MG00110 | XP_018219617.1 | 99 | 0 |
| SPGP0PE00130 | XP_018222357.1 | 99 | 0 |
| SPGP0P00270 | XP_018221626.1 | 98 | 0 |
| SPGP0TC00140 | XP_018219699.1 | 99 | 0 |
| SPGP0UQ00150 | XP_018220686.1 | 99 | 1 |
| SPGP0P01860 | XP_018221473.1 | 99 | 0 |
| SPGP0M00170 | XP_018220862.1 | 100 | 0 |
| SPGP0BE02030 | XP_018223132.1 | 99 | 0 |
| SPGP0D01890 | XP_018221815.1 | 99 | 0 |
| SPGP0BE03480 | XP_018223001.1 | 99 | 0 |
| SPGP0DQ00180 | XP_018219677.1 | 99 | 0 |
| SPGP0C03290 | XP_018220045.1 | 99 | 0 |
| SPGP0BKtrna10Y | DI49_2382 | 100 | 0 |
| SPGP0BK01690 | XP_018221308.1 | 99 | 0 |
| SPGP0P01270 | XP_018221531.1 | 99 | 0 |
| SPGP0K01820 | XP_018219407.1 | 99 | 0 |
| SPGP0K00310 | XP_018219554.1 | 99 | 0 |
| SPGP0WI00280 | XP_018222136.1 | 99 | 0 |
| SPGP0BI03980 | XP_018223487.1 | 99 | 0 |

Create PDF in your applications with the Pdfcrowd HTML to PDF API PDFCROWD

| | | | |
|---|---|---|---|
| SPGP0GH00100 | XP_018218968.1 | 99 | 0 |
| SPGP0PZ00220 | XP_018222424.1 | 99 | 0 |
| SPGP0P00900 | XP_018221564.1 | 99 | 0 |
| SPGP0BK03160 | XP_018221168.1 | 98 | 0 |
| SPGP0BE02100 | XP_018223127.1 | 99 | 0 |
| SPGP0UX00100 | XP_018222193.1 | 99 | 0 |
| SPGP0BI01440 | XP_018223737.1 | 100 | 0 |
| SPGP0QP00140 | XP_018219165.1 | 99 | 0 |
| SPGP0M00800 | XP_018220924.1 | 99 | 0 |
| SPGP0N00110 | XP_018219200.1 | 98 | 0 |
| SPGP0BE04790 | XP_018222872.1 | 99 | 0 |
| SPGP0D02810 | XP_018221904.1 | 99 | 0 |
| SPGP0TQ00310 | XP_018220626.1 | 99 | 0 |
| SPGP0BI01280 | XP_018223752.1 | 99 | 0 |
| SPGP0PV00100 | XP_018218976.1 | 99 | 0 |
| SPGP0NK00170 | XP_018220448.1 | 100 | 0 |
| SPGP0K01140 | XP_018219473.1 | 99 | 0 |
| SPGP0PA00130 | XP_018220471.1 | 99 | 0 |
| SPGP0BK03350 | XP_018221149.1 | 99 | 0 |
| SPGP0WA00170 | XP_018219849.1 | 99 | 0 |
| SPGP0VQ00120 | XP_018222544.1 | 96 | 0 |
| SPGP0VP00190 | XP_018219942.1 | 99 | 0 |
| SPGP0TO00160 | XP_018222810.1 | 99 | 0 |
| SPGP0K00900 | XP_018219497.1 | 99 | 0 |
| SPGP0BE03240 | XP_018223022.1 | 99 | 0 |
| SPGP0MO00400 | XP_018219124.1 | 100 | 0 |
| SPGP0D00730 | XP_018221705.1 | 99 | 0 |
| SPGP0BI00530 | XP_018223825.1 | 99 | 0 |
| SPGP0C02480 | XP_018220123.1 | 99 | 0 |
| SPGP0D01550 | XP_018221783.1 | 99 | 0 |
| SPGP0OI00100 | XP_018218937.1 | 99 | 0 |
| SPGP0BI03260 | XP_018223557.1 | 100 | 0 |
| SPGP0PE00260 | XP_018222371.1 | 99 | 0 |
| SPGP0M01030 | XP_018220947.1 | 100 | 0 |
| SPGP0C02180 | XP_018220153.1 | 99 | 0 |
| SPGP0BI00970 | XP_018223782.1 | 99 | 0 |
| SPGP0C02450 | XP_018220126.1 | 99 | 0 |
| SPGP0C01760 | XP_018220193.1 | 99 | 0 |
| SPGP0BI03590 | XP_018223523.1 | 99 | 0 |
| SPGP0KG00100 | XP_018222624.1 | 99 | 0 |

Create PDF in your applications with the Pdfcrowd HTML to PDF API PDFCROWD

| | | | |
|---|---|---|---|
| SPGP0BI03360 | XP_018223547.1 | 99 | 0 |
| SPGP0EU00150 | XP_018222663.1 | 99 | 0 |
| SPGP0BK03470 | XP_018221140.1 | 99 | 0 |
| SPGP0CF00120 | XP_018222498.1 | 99 | 0 |
| SPGP0OA00100 | XP_018218949.1 | 99 | 0 |
| SPGP0C01520 | XP_018220213.1 | 99 | 0 |
| SPGP0P00510 | XP_018221602.1 | 99 | 0 |
| SPGP0BI03200 | XP_018223563.1 | 99 | 0 |
| SPGP0K02600 | XP_018219331.1 | 99 | 0 |
| SPGP0C02980 | XP_018220075.1 | 99 | 0 |
| SPGP0D00710 | XP_018221703.1 | 99 | 0 |
| SPGP0MO00200 | XP_018219105.1 | 99 | 0 |
| SPGP0D00520 | XP_018221684.1 | 98 | 0 |
| SPGP0C00580 | XP_018220306.1 | 100 | 0 |
| SPGP0BE01910 | XP_018223145.1 | 98 | 1 |
| SPGP0HL00120 | XP_018220996.1 | 99 | 0 |
| SPGP0D03250 | XP_018221945.1 | 99 | 0 |
| SPGP0BI02970 | XP_018223586.1 | 99 | 0 |
| SPGP0PLtrna9F | DI49_4603 | 99 | 0 |
| SPGP0WI00210 | XP_018222129.1 | 99 | 0 |
| SPGP0FE00300 | XP_018220521.1 | 99 | 0 |
| SPGP0JW00160 | XP_018220978.1 | 99 | 0 |
| SPGP0WY00170 | XP_018220439.1 | 99 | 0 |
| SPGP0D01040 | XP_018221735.1 | 99 | 0 |
| SPGP0SN00100 | XP_018222594.1 | 99 | 0 |
| SPGP0WJ00130 | XP_018221057.1 | 99 | 0 |
| SPGP0D02250 | XP_018221850.1 | 99 | 0 |
| SPGP0EL00120 | XP_018223366.1 | 99 | 0 |
| SPGP0IA00120 | XP_018220583.1 | 99 | 0 |
| SPGP0OR00120 | XP_018218828.1 | 99 | 0 |
| SPGP0K02070 | XP_018219384.1 | 100 | 0 |
| SPGP0M00420 | XP_018220887.1 | 99 | 0 |
| SPGP0BI02390 | XP_018223643.1 | 99 | 0 |
| SPGP0BE00280 | XP_018223306.1 | 99 | 0 |
| SPGP0C02390 | XP_018220132.1 | 99 | 0 |
| SPGP0LH00180 | XP_018221029.1 | 99 | 0 |
| SPGP0BK01460 | XP_018221329.1 | 99 | 0 |
| SPGP0D02660 | XP_018221889.1 | 99 | 0 |
| SPGP0TT00100 | XP_018220838.1 | 99 | 0 |
| SPGP0FN00150 | XP_018223388.1 | 100 | 0 |

Create PDF in your applications with the Pdfcrowd HTML to PDF API PDFCROWD

| | | | |
|---|---|---|---|
| SPGP0JF00220 | XP_018222320.1 | 99 | 0 |
| SPGP0K02040 | XP_018219387.1 | 99 | 0 |
| SPGP0LK00190 | XP_018221062.1 | 99 | 0 |
| SPGP0BK02990 | XP_018221184.1 | 99 | 0 |
| SPGP0C03460 | XP_018220028.1 | 99 | 0 |
| SPGP0BI02800 | XP_018223603.1 | 99 | 0 |
| SPGP0BI02020 | XP_018223678.1 | 99 | 0 |
| SPGP0ST00150 | XP_018220544.1 | 99 | 0 |
| SPGP0BE04050 | XP_018222948.1 | 99 | 0 |
| SPGP0P00540 | XP_018221599.1 | 99 | 0 |
| SPGP0LG00160 | XP_018222648.1 | 100 | 0 |
| SPGP0BK01070 | XP_018221367.1 | 100 | 0 |
| SPGP0K01250 | XP_018219462.1 | 99 | 0 |
| SPGP0BI01610 | XP_018223719.1 | 99 | 0 |
| SPGP0FS00110 | XP_018222722.1 | 99 | 0 |
| SPGP0M00550 | XP_018220899.1 | 99 | 0 |
| SPGP0P01420 | XP_018221516.1 | 99 | 0 |
| SPGP0BI01750 | XP_018223705.1 | 99 | 0 |
| SPGP0BK00330 | XP_018221439.1 | 99 | 0 |
| SPGP0BE02390 | XP_018223099.1 | 99 | 0 |
| SPGP0U00240 | XP_018224112.1 | 100 | 0 |
| SPGP0TO00120 | XP_018222806.1 | 99 | 0 |
| SPGP0BK01340 | XP_018221340.1 | 99 | 0 |
| SPGP0BI03900 | XP_018223494.1 | 100 | 0 |
| SPGP0K02280 | XP_018219363.1 | 99 | 0 |
| SPGP0GF00110 | XP_018219650.1 | 99 | 0 |
| SPGP0PE00110 | XP_018222355.1 | 99 | 0 |
| SPGP0D03530 | XP_018221971.1 | 99 | 0 |
| SPGP0UN00160 | XP_018220782.1 | 99 | 0 |
| SPGP0OYtrna4W | DI49_5211 | 100 | 0 |
| SPGP0BK01730 | XP_018221304.1 | 99 | 0 |
| SPGP0N01000 | XP_018219287.1 | 99 | 0 |
| SPGP0RD00100 | XP_018220655.1 | 99 | 0 |
| SPGP0BE01710 | XP_018223164.1 | 99 | 0 |
| SPGP0P00260 | XP_018221627.1 | 99 | 0 |
| SPGP0P01850 | XP_018221474.1 | 99 | 0 |
| SPGP0U00350 | XP_018224101.1 | 99 | 0 |
| SPGP0U00420 | XP_018224095.1 | 99 | 0 |
| SPGP0HZ00130 | XP_018219770.1 | 99 | 0 |
| SPGP0BK00300 | XP_018221442.1 | 99 | 0 |

Create PDF in your applications with the Pdfcrowd HTML to PDF API PDFCROWD

| | | | |
|---|---|---|---|
| SPGP0BE01790 | XP_018223156.1 | 100 | 0 |
| SPGP0RQ00210 | XP_018221131.1 | 99 | 0 |
| SPGP0N01080 | XP_018219295.1 | 99 | 0 |
| SPGP0BE01990 | XP_018223136.1 | 99 | 0 |
| SPGP0OR00100 | XP_018218826.1 | 99 | 0 |
| SPGP0BI03610 | XP_018223521.1 | 99 | 0 |
| SPGP0BI02960 | XP_018223587.1 | 99 | 0 |
| SPGP0BE02530 | XP_018223085.1 | 99 | 0 |
| SPGP0JH00140 | XP_018218938.1 | 100 | 0 |
| SPGP0C01510 | XP_018220214.1 | 99 | 0 |
| SPGP0N00900 | XP_018219277.1 | 99 | 0 |
| SPGP0XF00140 | XP_018220852.1 | 99 | 0 |
| SPGP0GW00190 | XP_018220822.1 | 99 | 0 |
| SPGP0GW00180 | XP_018220823.1 | 99 | 0 |
| SPGP0TE00110 | XP_018219894.1 | 99 | 0 |
| SPGP0BI02560 | XP_018223626.1 | 99 | 0 |
| SPGP0BE02750 | XP_018223065.1 | 99 | 0 |
| SPGP0D02390 | XP_018221864.1 | 99 | 0 |
| SPGP0D00220 | XP_018221655.1 | 99 | 0 |
| SPGP0BK00250 | XP_018221446.1 | 96 | 3 |
| SPGP0U00490 | XP_018224088.1 | 99 | 0 |
| SPGP0D02130 | XP_018221838.1 | 99 | 0 |
| SPGP0BI02890 | XP_018223594.1 | 99 | 0 |
| SPGP0Utrna8P | DI49_4914 | 100 | 0 |
| SPGP0BI01960 | XP_018223684.1 | 99 | 0 |
| SPGP0BF01880 | XP_018222854.1 | 88 | 0 |
| SPGP0BI03950 | XP_018223490.1 | 99 | 0 |
| SPGP0JA00120 | XP_018223408.1 | 99 | 0 |
| SPGP0D00500 | XP_018221682.1 | 99 | 0 |
| SPGP0D01180 | XP_018221746.1 | 99 | 0 |
| SPGP0BI03750 | XP_018223507.1 | 99 | 0 |
| SPGP0C00170 | XP_018220346.1 | 99 | 0 |
| SPGP0DN00120 | XP_018218992.1 | 99 | 0 |
| SPGP0BE02590 | XP_018223079.1 | 99 | 0 |
| SPGP0VS00110 | XP_018222521.1 | 100 | 0 |
| SPGP0BK01550 | XP_018221321.1 | 99 | 0 |
| SPGP0UJ00140 | XP_018223341.1 | 99 | 0 |
| SPGP0OI00360 | XP_018218914.1 | 99 | 0 |
| SPGP0CK00100 | XP_018219963.1 | 99 | 0 |
| SPGP0BE00700 | XP_018223263.1 | 99 | 0 |

Create PDF in your applications with the Pdfcrowd HTML to PDF API PDFCROWD

| | | | |
|---|---|---|---|
| SPGP0KB00190 | XP_018221121.1 | 92 | 6 |
| SPGP0C00390 | XP_018220324.1 | 99 | 0 |
| SPGP0HO00300 | XP_018219973.1 | 99 | 0 |
| SPGP0IC00240 | XP_018222783.1 | 99 | 0 |
| SPGP0JX00110 | XP_018218838.1 | 99 | 0 |
| SPGP0SH00190 | XP_018223977.1 | 99 | 0 |
| SPGP0GT00110 | XP_018222509.1 | 99 | 0 |
| SPGP0BKtrna22R | DI49_4323 | 100 | 0 |
| SPGP0VN00310 | XP_018219821.1 | 99 | 0 |
| SPGP0HZ00160 | XP_018219773.1 | 99 | 0 |
| SPGP0P00290 | XP_018221624.1 | 99 | 1 |
| SPGP0UQ00140 | XP_018220687.1 | 99 | 0 |
| SPGP0C00820 | XP_018220282.1 | 99 | 0 |
| SPGP0D01870 | XP_018221813.1 | 99 | 0 |
| SPGP0BI00150 | XP_018223863.1 | 100 | 0 |
| SPGP0P01210 | XP_018221536.1 | 99 | 1 |
| SPGP0N00350 | XP_018219223.1 | 100 | 0 |
| SPGP0K00330 | XP_018219552.1 | 99 | 0 |
| SPGP0BE04350 | XP_018222918.1 | 99 | 0 |
| SPGP0BI00450 | XP_018223833.1 | 99 | 0 |
| SPGP0GI00110 | XP_018220699.1 | 97 | 0 |
| SPGP0BE04240 | XP_018222929.1 | 99 | 0 |
| SPGP0M01150 | XP_018220959.1 | 100 | 0 |
| SPGP0EM00220 | XP_018220713.1 | 99 | 0 |
| SPGP0D00160 | XP_018221649.1 | 100 | 0 |
| SPGP0PZ00300 | XP_018222416.1 | 99 | 0 |
| SPGP0BE02700 | XP_018223070.1 | 99 | 0 |
| SPGP0KU00100 | XP_018221005.1 | 99 | 0 |
| SPGP0MF00240 | XP_018222196.1 | 99 | 0 |
| SPGP0KN00130 | XP_018220569.1 | 99 | 0 |
| SPGP0BK01790 | XP_018221298.1 | 98 | 1 |
| SPGP0D02830 | XP_018221905.1 | 99 | 0 |
| SPGP0N00680 | XP_018219255.1 | 100 | 0 |
| SPGP0NU00110 | XP_018219056.1 | 99 | 0 |
| SPGP0KB00110 | XP_018221113.1 | 99 | 0 |
| SPGP0M00680 | XP_018220912.1 | 100 | 0 |
| SPGP0K00260 | XP_018219559.1 | 99 | 0 |
| SPGP0GW00220 | XP_018220819.1 | 99 | 1 |
| SPGP0BK03330 | XP_018221151.1 | 99 | 0 |
| SPGP0BK02580 | XP_018221220.1 | 99 | 0 |

Create PDF in your applications with the Pdfcrowd HTML to PDF API PDFCROWD

| | | | |
|---|---|---|---|
| SPGP0BI01130 | XP_018223767.1 | 99 | 0 |
| SPGP0UJ00170 | XP_018223338.1 | 99 | 0 |
| SPGP0TBtrna8H | DI49_4134 | 100 | 0 |
| SPGP0BK01240 | XP_018221350.1 | 99 | 0 |
| SPGP0WC00100 | XP_018222802.1 | 99 | 0 |
| SPGP0OI00510 | XP_018218899.1 | 99 | 0 |
| SPGP0BK02820 | XP_018221200.1 | 99 | 0 |
| SPGP0TL00130 | XP_018219085.1 | 99 | 0 |
| SPGP0EX00150 | XP_018223424.1 | 99 | 0 |
| SPGP0C02100 | XP_018220161.1 | 99 | 0 |
| SPGP0BI00460 | XP_018223832.1 | 99 | 0 |
| SPGP0WA00150 | XP_018219851.1 | 99 | 0 |
| SPGP0N01220 | XP_018219307.1 | 99 | 0 |
| SPGP0K01290 | XP_018219458.1 | 100 | 0 |
| SPGP0C02860 | XP_018220087.1 | 99 | 0 |
| SPGP0D03440 | XP_018221962.1 | 99 | 0 |
| SPGP0UL00110 | XP_018219658.1 | 99 | 0 |
| SPGP0C02430 | XP_018220128.1 | 99 | 0 |
| SPGP0HT00210 | XP_018219012.1 | 99 | 0 |
| SPGP0EK00200 | XP_018220805.1 | 99 | 0 |
| SPGP0M00350 | XP_018220880.1 | 99 | 0 |
| SPGP0DN00100 | XP_018218994.1 | 97 | 2 |
| SPGP0K01050 | XP_018219482.1 | 99 | 0 |
| SPGP0FM00290 | XP_018222164.1 | 99 | 0 |
| SPGP0BK00110 | XP_018221455.1 | 99 | 0 |
| SPGP0BE03970 | XP_018222955.1 | 99 | 0 |
| SPGP0BI01080 | XP_018223771.1 | 99 | 0 |
| SPGP0BI00610 | XP_018223817.1 | 99 | 0 |
| SPGP0DU00120 | XP_018222582.1 | 99 | 1 |
| SPGP0C01620 | XP_018220205.1 | 99 | 0 |
| SPGP0D01200 | XP_018221748.1 | 100 | 0 |
| SPGP0OE00140 | XP_018219606.1 | 99 | 0 |
| SPGP0AA00150 | XP_018221080.1 | 100 | 0 |
| SPGP0BE01700 | XP_018223165.1 | 99 | 0 |
| SPGP0BE01930 | XP_018223143.1 | 100 | 0 |
| SPGP0KT00110 | XP_018222243.1 | 99 | 0 |
| SPGP0BE01390 | XP_018223195.1 | 100 | 0 |
| SPGP0BE04140 | XP_018222939.1 | 98 | 1 |
| SPGP0BK00640 | XP_018221408.1 | 99 | 0 |
| SPGP0C01160 | XP_018220248.1 | 99 | 0 |

Create PDF in your applications with the Pdfcrowd HTML to PDF API PDFCROWD

| | | | |
|---|---|---|---|
| SPGP0BUtrna28S | DI49_4419 | 100 | 0 |
| SPGP0K00640 | XP_018219522.1 | 99 | 0 |
| SPGP0JW00140 | XP_018220976.1 | 98 | 2 |
| SPGP0C03110 | XP_018220063.1 | 99 | 0 |
| SPGP0LW00140 | XP_018222618.1 | 99 | 0 |
| SPGP0BI03300 | XP_018223553.1 | 99 | 0 |
| SPGP0BE01640 | XP_018223171.1 | 99 | 0 |
| SPGP0K02170 | XP_018219374.1 | 99 | 0 |
| SPGP0KZ00260 | XP_018223936.1 | 99 | 0 |
| SPGP0XC00240 | XP_018220600.1 | 99 | 0 |
| SPGP0C03260 | XP_018220048.1 | 99 | 0 |
| SPGP0WJ00150 | XP_018221055.1 | 99 | 0 |
| SPGP0ST00160 | XP_018220543.1 | 99 | 0 |
| SPGP0K02450 | XP_018219346.1 | 99 | 0 |
| SPGP0BE02440 | XP_018223094.1 | 100 | 0 |
| SPGP0C01330 | XP_018220231.1 | 100 | 0 |
| SPGP0BK01150 | XP_018221359.1 | 99 | 0 |
| SPGP0NG00160 | XP_018222772.1 | 100 | 0 |
| SPGP0IJ00130 | XP_018218825.1 | 99 | 0 |
| SPGP0LO00100 | XP_018223457.1 | 99 | 0 |
| SPGP0OB00140 | XP_018219797.1 | 99 | 1 |
| SPGP0CX00140 | XP_018220795.1 | 99 | 0 |
| SPGP0N00540 | XP_018219242.1 | 99 | 0 |
| SPGP0UF00110 | XP_018221039.1 | 99 | 0 |
| SPGP0BE03860 | XP_018222965.1 | 99 | 0 |
| SPGP0K02050 | XP_018219386.1 | 99 | 0 |
| SPGP0P00560 | XP_018221597.1 | 99 | 0 |
| SPGP0BK01400 | XP_018221334.1 | 99 | 0 |
| SPGP0P00550 | XP_018221598.1 | 99 | 0 |
| SPGP0BK03000 | XP_018221183.1 | 99 | 0 |
| SPGP0BI00760 | XP_018223802.1 | 99 | 0 |
| SPGP0FN00130 | XP_018223390.1 | 99 | 0 |
| SPGP0D02640 | XP_018221887.1 | 100 | 0 |
| SPGP0BE00110 | XP_018223323.1 | 98 | 0 |
| SPGP0M00150 | XP_018220860.1 | 99 | 0 |
| SPGP0C03440 | XP_018220030.1 | 99 | 0 |
| SPGP0BI02040 | XP_018223676.1 | 99 | 0 |
| SPGP0D00860 | XP_018221717.1 | 99 | 0 |
| SPGP0BE02580 | XP_018223080.1 | 99 | 0 |
| SPGP0GF00290 | XP_018219632.1 | 99 | 0 |

Create PDF in your applications with the Pdfcrowd HTML to PDF API PDFCROWD

| | | | |
|---|---|---|---|
| SPGP0C00140 | XP_018220349.1 | 99 | 0 |
| SPGP0LE00150 | XP_018220763.1 | 99 | 0 |
| SPGP0K01780 | XP_018219411.1 | 99 | 0 |
| SPGP0BI02260 | XP_018223656.1 | 100 | 0 |
| SPGP0BE03390 | XP_018223008.1 | 99 | 0 |
| SPGP0BI01630 | XP_018223717.1 | 99 | 0 |
| SPGP0BE03640 | XP_018222987.1 | 99 | 0 |
| SPGP0BI02380 | XP_018223644.1 | 99 | 0 |
| SPGP0P01400 | XP_018221518.1 | 99 | 0 |
| SPGP0BK02450 | XP_018221233.1 | 99 | 0 |
| SPGP0BE03710 | XP_018222980.1 | 99 | 0 |
| SPGP0D00900 | XP_018221721.1 | 99 | 0 |
| SPGP0PQ00100 | XP_018219157.1 | 98 | 0 |
| SPGP0BK00320 | XP_018221440.1 | 99 | 0 |
| SPGP0U00600 | XP_018224078.1 | 99 | 0 |
| SPGP0UN00140 | XP_018220784.1 | 100 | 0 |
| SPGP0C01110 | XP_018220253.1 | 99 | 0 |
| SPGP0BK01710 | XP_018221306.1 | 99 | 0 |
| SPGP0M00160 | XP_018220861.1 | 99 | 0 |
| SPGP0D01210 | XP_018221749.1 | 99 | 0 |
| SPGP0VO00200 | XP_018223443.1 | 99 | 0 |
| SPGP0BI00940 | XP_018223785.1 | 99 | 0 |
| SPGP0PZ00290 | XP_018222417.1 | 99 | 0 |
| SPGP0HO00240 | XP_018219978.1 | 99 | 0 |
| SPGP0VN00120 | XP_018219802.1 | 99 | 0 |
| SPGP0FM00340 | XP_018222169.1 | 99 | 0 |
| SPGP0BK00340 | XP_018221438.1 | 99 | 0 |
| SPGP0BK01000 | XP_018221374.1 | 99 | 0 |
| SPGP0BI01890 | XP_018223691.1 | 99 | 0 |
| SPGP0BI02080 | XP_018223673.1 | 99 | 0 |
| SPGP0BI03940 | XP_018223491.1 | 99 | 0 |
| SPGP0BE02510 | XP_018223087.1 | 99 | 0 |
| SPGP0OX00180 | XP_018219727.1 | 99 | 0 |
| SPGP0D01960 | XP_018221822.1 | 99 | 0 |
| SPGP0C01570 | XP_018220209.1 | 99 | 1 |
| SPGP0BE02800 | XP_018223060.1 | 99 | 0 |
| SPGP0N00960 | XP_018219283.1 | 99 | 0 |
| SPGP0P00530 | XP_018221600.1 | 99 | 0 |
| SPGP0C00980 | XP_018220266.1 | 99 | 0 |
| SPGP0BK02870 | XP_018221196.1 | 98 | 0 |

Create PDF in your applications with the Pdfcrowd HTML to PDF API PDFCROWD

| | | | |
|---|---|---|---|
| SPGP0KW00120 | XP_018220726.1 | 99 | 0 |
| SPGP0LA00130 | XP_018220843.1 | 99 | 0 |
| SPGP0NG00140 | XP_018222774.1 | 99 | 0 |
| SPGP0WV00100 | XP_018219959.1 | 99 | 0 |
| SPGP0NQ00250 | XP_018220740.1 | 100 | 0 |
| SPGP0BI04060 | XP_018223480.1 | 99 | 0 |
| SPGP0BE01120 | XP_018223222.1 | 99 | 0 |
| SPGP0DJ00110 | XP_018219918.1 | 99 | 0 |
| SPGP0SH00270 | XP_018223984.1 | 98 | 0 |
| SPGP0UH00130 | XP_018220409.1 | 98 | 0 |
| SPGP0OI00540 | XP_018218896.1 | 99 | 0 |
| SPGP0VN00400 | XP_018219830.1 | 99 | 0 |
| SPGP0P01670 | XP_018221492.1 | 99 | 0 |
| SPGP0BE01090 | XP_018223225.1 | 99 | 0 |
| SPGP0ltrna31L | DI49_3110 | 100 | 0 |
| SPGP0BK02550 | XP_018221223.1 | 99 | 0 |
| SPGP0BE01890 | XP_018223147.1 | 99 | 0 |
| SPGP0D03350 | XP_018221954.1 | 99 | 0 |
| SPGP0C01390 | XP_018220225.1 | 100 | 0 |
| SPGP0IQ00100 | XP_018221043.1 | 99 | 0 |
| SPGP0K01100 | XP_018219477.1 | 99 | 0 |
| SPGP0HZ00260 | XP_018219783.1 | 99 | 0 |
| SPGP0SY00100 | XP_018220802.1 | 99 | 0 |
| SPGP0BE02460 | XP_018223092.1 | 99 | 1 |
| SPGP0NI00220 | XP_018223917.1 | 99 | 0 |
| SPGP0C00150 | XP_018220348.1 | 99 | 0 |
| SPGP0C00620 | XP_018220302.1 | 100 | 0 |
| SPGP0BE04600 | XP_018222891.1 | 99 | 0 |
| SPGP0BK00920 | XP_018221381.1 | 99 | 0 |
| SPGP0NT00200 | XP_018222464.1 | 99 | 0 |
| SPGP0FT00110 | XP_018222731.1 | 99 | 0 |
| SPGP0VD00160 | XP_018219839.1 | 99 | 0 |
| SPGP0K02420 | XP_018219349.1 | 99 | 0 |
| SPGP0BE00270 | XP_018223307.1 | 99 | 0 |
| SPGP0BI03230 | XP_018223560.1 | 99 | 0 |
| SPGP0P00720 | XP_018221582.1 | 99 | 0 |
| SPGP0BK02200 | XP_018221257.1 | 99 | 0 |
| SPGP0AA00130 | XP_018221082.1 | 99 | 0 |
| SPGP0SP00150 | XP_018223965.1 | 99 | 0 |

Create PDF in your applications with the Pdfcrowd HTML to PDF API PDFCROWD

| | | | |
|---|---|---|---|
| SPGP0C00530 | XP_018220311.1 | 99 | 0 |
| SPGP0P01200 | XP_018221537.1 | 99 | 0 |
| SPGP0CZ00110 | XP_018219906.1 | 99 | 0 |
| SPGP0PZ00260 | XP_018222420.1 | 99 | 0 |
| SPGP0OI00120 | XP_018218935.1 | 99 | 0 |
| SPGP0OI00650 | XP_018218885.1 | 99 | 0 |
| SPGP0C03300 | XP_018220044.1 | 100 | 0 |
| SPGP0UQ00160 | XP_018220685.1 | 99 | 0 |
| SPGP0C00840 | XP_018220280.1 | 99 | 0 |
| SPGP0D01850 | XP_018221811.1 | 99 | 0 |
| SPGP0SB00100 | XP_018222298.1 | 99 | 0 |
| SPGP0D00980 | XP_018221729.1 | 99 | 0 |
| SPGP0P01890 | XP_018221470.1 | 99 | 0 |
| SPGP0P01230 | XP_018221535.1 | 99 | 0 |
| SPGP0N00370 | XP_018219225.1 | 99 | 0 |
| SPGP0GF00140 | XP_018219647.1 | 99 | 0 |
| SPGP0BE03200 | XP_018223026.1 | 100 | 0 |
| SPGP0JP00190 | XP_018218856.1 | 99 | 0 |
| SPGP0BK03400 | XP_018221144.1 | 99 | 0 |
| SPGP0D01580 | XP_018221786.1 | 100 | 0 |
| SPGP0D03500 | XP_018221968.1 | 99 | 0 |
| SPGP0FL00110 | XP_018219050.1 | 98 | 0 |
| SPGP0BE04150 | XP_018222938.1 | 98 | 0 |
| SPGP0BI00630 | XP_018223815.1 | 99 | 0 |
| SPGP0KN00150 | XP_018220571.1 | 99 | 0 |
| SPGP0BI03160 | XP_018223567.1 | 100 | 0 |
| SPGP0P01090 | XP_018221548.1 | 99 | 0 |
| SPGP0M00990 | XP_018220943.1 | 99 | 0 |
| SPGP0N00150 | XP_018219204.1 | 99 | 0 |
| SPGP0OJ00100 | XP_018219924.1 | 99 | 0 |
| SPGP0BE03250 | XP_018223021.1 | 99 | 0 |
| SPGP0RI00300 | XP_018220508.1 | 98 | 0 |
| SPGP0P01040 | XP_018221552.1 | 99 | 0 |
| SPGP0M00930 | XP_018220937.1 | 99 | 0 |
| SPGP0D02890 | XP_018221911.1 | 99 | 0 |
| SPGP0BK03310 | XP_018221153.1 | 100 | 0 |
| SPGP0JR00120 | XP_018219029.1 | 99 | 1 |
| SPGP0P01450 | XP_018221513.1 | 99 | 0 |
| SPGP0C03160 | XP_018220058.1 | 99 | 0 |
| SPGP0BI00430 | XP_018223835.1 | 99 | 0 |

Create PDF in your applications with the Pdfcrowd HTML to PDF API PDFCROWD

| | | | |
|---|---|---|---|
| SPGP0BE00780 | XP_018223256.1 | 99 | 0 |
| SPGP0VN00300 | XP_018219820.1 | 99 | 0 |
| SPGP0D00560 | XP_018221688.1 | 99 | 0 |
| SPGP0D01880 | XP_018221814.1 | 99 | 1 |
| SPGP0D03090 | XP_018221929.1 | 99 | 0 |
| SPGP0BI02980 | XP_018223585.1 | 99 | 0 |
| SPGP0C01650 | XP_018220202.1 | 99 | 0 |
| SPGP0BI00570 | XP_018223821.1 | 99 | 0 |
| SPGP0P01330 | XP_018221525.1 | 99 | 0 |
| SPGP0BI02920 | XP_018223591.1 | 99 | 0 |
| SPGP0C02140 | XP_018220157.1 | 99 | 0 |
| SPGP0K01910 | XP_018219399.1 | 99 | 0 |
| SPGP0C02410 | XP_018220130.1 | 99 | 0 |
| SPGP0GF00260 | XP_018219635.1 | 99 | 0 |
| SPGP0M00400 | XP_018220885.1 | 99 | 0 |
| SPGP0UJ00210 | XP_018223334.1 | 99 | 0 |
| SPGP0BK00170 | XP_018221451.1 | 99 | 0 |
| SPGP0GI00120 | XP_018220698.1 | 99 | 0 |
| SPGP0BE00320 | XP_018223302.1 | 99 | 0 |
| SPGP0C00220 | XP_018220341.1 | 98 | 0 |
| SPGP0D00740 | XP_018221706.1 | 99 | 0 |
| SPGP0N00790 | XP_018219266.1 | 99 | 0 |
| SPGP0M00380 | XP_018220883.1 | 99 | 0 |
| SPGP0RC00110 | XP_018222086.1 | 99 | 0 |
| SPGP0N00870 | XP_018219274.1 | 99 | 0 |
| SPGP0BK02190 | XP_018221258.1 | 99 | 0 |
| SPGP0KN00140 | XP_018220570.1 | 99 | 0 |
| SPGP0D00910 | XP_018221722.1 | 99 | 0 |
| SPGP0BE00650 | XP_018223269.1 | 99 | 0 |
| SPGP0U00650 | XP_018224073.1 | 99 | 0 |
| SPGP0C01140 | XP_018220250.1 | 99 | 0 |
| SPGP0HO00100 | XP_018219992.1 | 99 | 0 |
| SPGP0BK03140 | XP_018221170.1 | 98 | 0 |
| SPGP0BI02070 | XP_018223674.1 | 99 | 0 |
| SPGP0EK00190 | XP_018220806.1 | 99 | 0 |
| SPGP0JW00120 | XP_018220974.1 | 99 | 0 |
| SPGP0RI00280 | XP_018220506.1 | 99 | 0 |
| SPGP0BI03920 | XP_018223492.1 | 99 | 0 |
| SPGP0P00840 | XP_018221570.1 | 100 | 0 |
| SPGP0N00920 | XP_018219279.1 | 99 | 0 |

Create PDF in your applications with the Pdfcrowd HTML to PDF API PDFCROWD

| | | | |
|---|---|---|---|
| SPGP0WQ00240 | XP_018222277.1 | 99 | 0 |
| SPGP0D01000 | XP_018221731.1 | 99 | 0 |
| SPGP0K02210 | XP_018219370.1 | 99 | 0 |
| SPGP0BE04120 | XP_018222941.1 | 99 | 0 |
| SPGP0D02150 | XP_018221840.1 | 99 | 0 |
| SPGP0BK01420 | XP_018221332.1 | 99 | 0 |
| SPGP0N00550 | XP_018219243.1 | 100 | 0 |
| SPGP0EY00110 | XP_018219089.1 | 99 | 0 |
| SPGP0BE01660 | XP_018223169.1 | 97 | 3 |
| SPGP0BE04290 | XP_018222924.1 | 99 | 0 |
| SPGP0BE03800 | XP_018222971.1 | 99 | 0 |
| SPGP0KQ00140 | XP_018219926.1 | 98 | 0 |
| SPGP0C00650 | XP_018220299.1 | 99 | 0 |
| SPGP0BE01750 | XP_018223160.1 | 99 | 0 |
| SPGP0FN00110 | XP_018223392.1 | 99 | 0 |
| SPGP0OY00100 | XP_018220968.1 | 99 | 0 |
| SPGP0K01740 | XP_018219415.1 | 99 | 0 |
| SPGP0C03420 | XP_018220032.1 | 99 | 0 |
| SPGP0BE02380 | XP_018223100.1 | 99 | 1 |
| SPGP0BT00140 | XP_018220737.1 | 100 | 0 |
| SPGP0BK01990 | XP_018221278.1 | 99 | 0 |
| SPGP0BE03180 | XP_018223028.1 | 99 | 0 |
| SPGP0BE02300 | XP_018223108.1 | 99 | 0 |
| SPGP0BK01230 | XP_018221351.1 | 99 | 0 |
| SPGP0BE04170 | XP_018222936.1 | 99 | 0 |
| SPGP0TQ00350 | XP_018220631.1 | 99 | 0 |
| SPGP0D00840 | XP_018221715.1 | 100 | 0 |
| SPGP0P01390 | XP_018221519.1 | 100 | 0 |
| SPGP0K01810 | XP_018219408.1 | 99 | 0 |
| SPGP0BEtrna3C | DI49_5400 | 100 | 0 |
| SPGP0BK01900 | XP_018221286.1 | 99 | 0 |
| SPGP0JL00100 | XP_018220580.1 | 99 | 0 |
| SPGP0D03340 | XP_018221953.1 | 99 | 0 |
| SPGP0K01560 | XP_018219432.1 | 99 | 0 |
| SPGP0BE04830 | XP_018222868.1 | 100 | 0 |
| SPGP0XC00140 | XP_018220590.1 | 99 | 0 |
| SPGP0C03390 | XP_018220035.1 | 99 | 0 |
| SPGP0IR00170 | XP_018219760.1 | 99 | 0 |
| SPGP0D03570 | XP_018221975.1 | 99 | 0 |
| SPGP0P01930 | XP_018221466.1 | 98 | 0 |

Create PDF in your applications with the Pdfcrowd HTML to PDF API PDFCROWD

| | | | |
|---|---|---|---|
| SPGP0BE03110 | XP_018223034.1 | 99 | 0 |
| SPGP0P01170 | XP_018221540.1 | 99 | 0 |
| SPGP0BK01290 | XP_018221345.1 | 99 | 0 |
| SPGP0K00160 | XP_018218749.1 | 100 | 0 |
| SPGP0N00520 | XP_018219240.1 | 99 | 0 |
| SPGP0TO00100 | XP_018222804.1 | 99 | 0 |
| SPGP0BK02430 | XP_018221235.1 | 99 | 0 |
| SPGP0D01460 | XP_018221774.1 | 99 | 0 |
| SPGP0SJ00100 | XP_018219858.1 | 98 | 0 |
| SPGP0C00240 | XP_018220339.1 | 99 | 0 |
| SPGP0UN00120 | XP_018220786.1 | 99 | 0 |
| SPGP0SB00140 | XP_018222302.1 | 99 | 0 |
| SPGP0CZ00180 | XP_018219913.1 | 97 | 2 |
| SPGP0BE00310 | XP_018223303.1 | 99 | 0 |
| SPGP0LU00190 | XP_018224007.1 | 99 | 0 |
| SPGP0PZ00120 | XP_018222434.1 | 99 | 0 |
| SPGP0BI00960 | XP_018223783.1 | 99 | 1 |
| SPGP0U00460 | XP_018224091.1 | 99 | 0 |
| SPGP0K00960 | XP_018219491.1 | 100 | 0 |
| SPGP0P00620 | XP_018221592.1 | 99 | 0 |
| SPGP0IC00230 | XP_018222784.1 | 100 | 0 |
| SPGP0BI01210 | XP_018223759.1 | 99 | 0 |
| SPGP0BK01160 | XP_018221358.1 | 99 | 0 |
| SPGP0P00990 | XP_018221557.1 | 99 | 0 |
| SPGP0C01850 | XP_018220185.1 | 99 | 0 |
| SPGP0C02740 | XP_018220099.1 | 99 | 0 |
| SPGP0VZ00140 | XP_018223879.1 | 99 | 0 |
| SPGP0BE02570 | XP_018223081.1 | 99 | 0 |
| SPGP0PZ00110 | XP_018222435.1 | 99 | 0 |
| SPGP0BK03100 | XP_018221174.1 | 99 | 0 |
| SPGP0D01630 | XP_018221791.1 | 99 | 0 |
| SPGP0PS00110 | XP_018223359.1 | 99 | 0 |
| SPGP0N00940 | XP_018219281.1 | 99 | 0 |
| SPGP0P00480 | XP_018221605.1 | 99 | 0 |
| SPGP0JW00180 | XP_018220980.1 | 99 | 0 |
| SPGP0LE00130 | XP_018220765.1 | 99 | 0 |
| SPGP0BE04670 | XP_018222884.1 | 99 | 0 |
| SPGP0BK02850 | XP_018221198.1 | 99 | 0 |
| SPGP0D02460 | XP_018221871.1 | 99 | 0 |
| SPGP0K01190 | XP_018219468.1 | 95 | 4 |

Create PDF in your applications with the Pdfcrowd HTML to PDF API PDFCROWD

| | | | |
|---|---|---|---|
| SPGP0BK02020 | XP_018221275.1 | 99 | 0 |
| SPGP0BI00290 | XP_018223849.1 | 99 | 0 |
| SPGP0TC00200 | XP_018219693.1 | 99 | 0 |
| SPGP0HB00160 | XP_018219188.1 | 98 | 0 |
| SPGP0LX00130 | XP_018219147.1 | 99 | 0 |
| SPGP0BI04040 | XP_018223482.1 | 99 | 0 |
| SPGP0K00860 | XP_018219500.1 | 99 | 0 |
| SPGP0P00340 | XP_018221619.1 | 99 | 0 |
| SPGP0HL00150 | XP_018220993.1 | 99 | 0 |
| SPGP0BI02520 | XP_018223630.1 | 99 | 0 |
| SPGP0C00470 | XP_018220317.1 | 99 | 0 |
| SPGP0UB00160 | XP_018219041.1 | 99 | 0 |
| SPGP0BI00540 | XP_018223824.1 | 99 | 0 |
| SPGP0HZ00200 | XP_018219777.1 | 99 | 1 |
| SPGP0BI03420 | XP_018223540.1 | 99 | 0 |
| SPGP0C01270 | XP_018220237.1 | 99 | 0 |
| SPGP0FE00160 | XP_018220534.1 | 99 | 0 |
| SPGP0BI00110 | XP_018223867.1 | 99 | 0 |
| SPGP0D03370 | XP_018221955.1 | 99 | 0 |
| SPGP0IV00150 | XP_018220643.1 | 99 | 0 |
| SPGP0C01800 | XP_018220189.1 | 99 | 0 |
| SPGP0BI02290 | XP_018223653.1 | 99 | 0 |
| SPGP0M01100 | XP_018220954.1 | 99 | 0 |
| SPGP0DU00170 | XP_018222577.1 | 99 | 0 |
| SPGP0K00990 | XP_018219488.1 | 99 | 0 |
| SPGP0OI00810 | XP_018218869.1 | 99 | 0 |
| SPGP0BE01110 | XP_018223223.1 | 100 | 0 |
| SPGP0MO00180 | XP_018219103.1 | 99 | 0 |
| SPGP0C00600 | XP_018220304.1 | 99 | 0 |
| SPGP0VE00140 | XP_018220966.1 | 99 | 0 |
| SPGP0BE04330 | XP_018222920.1 | 99 | 0 |
| SPGP0BE04620 | XP_018222889.1 | 99 | 0 |
| SPGP0TT00110 | XP_018220837.1 | 99 | 0 |
| SPGP0JZ00130 | XP_018223414.1 | 99 | 0 |
| SPGP0P00700 | XP_018221584.1 | 99 | 0 |
| SPGP0P01360 | XP_018221522.1 | 99 | 0 |
| SPGP0GI00150 | XP_018220695.1 | 99 | 0 |
| SPGP0BK02220 | XP_018221255.1 | 99 | 0 |
| SPGP0N01150 | XP_018219302.1 | 99 | 0 |
| SPGP0C00900 | XP_018220274.1 | 99 | 0 |

Create PDF in your applications with the Pdfcrowd HTML to PDF API PDFCROWD

| | | | |
|---|---|---|---|
| SPGP0HT00200 | XP_018219011.1 | 99 | 0 |
| SPGP0UJ00180 | XP_018223337.1 | 99 | 0 |
| SPGP0OI00720 | XP_018218878.1 | 99 | 0 |
| SPGP0RZ00100 | XP_018223379.1 | 100 | 0 |
| SPGP0BK03230 | XP_018221161.1 | 99 | 0 |
| SPGP0NG00190 | XP_018222769.1 | 99 | 0 |
| SPGP0KB00150 | XP_018221117.1 | 99 | 0 |
| SPGP0K01170 | XP_018219470.1 | 99 | 0 |
| SPGP0D01830 | XP_018221809.1 | 99 | 0 |
| SPGP0N00390 | XP_018219227.1 | 99 | 0 |
| SPGP0BI03220 | XP_018223561.1 | 99 | 0 |
| SPGP0ST00120 | XP_018220547.1 | 99 | 0 |
| SPGP0D02570 | XP_018221880.1 | 99 | 0 |
| SPGP0K00370 | XP_018219548.1 | 99 | 0 |
| SPGP0C03430 | XP_018220031.1 | 99 | 0 |
| SPGP0C02790 | XP_018220094.1 | 99 | 0 |
| SPGP0C03350 | XP_018220039.1 | 99 | 0 |
| SPGP0BE02740 | XP_018223066.1 | 99 | 0 |
| SPGP0M01110 | XP_018220955.1 | 99 | 1 |
| SPGP0BE00810 | XP_018223253.1 | 99 | 0 |
| SPGP0BE03230 | XP_018223023.1 | 99 | 0 |
| SPGP0ID00130 | XP_018222143.1 | 99 | 0 |
| SPGP0VM00180 | XP_018220676.1 | 99 | 0 |
| SPGP0BE02060 | XP_018223131.1 | 100 | 0 |
| SPGP0BI03180 | XP_018223565.1 | 99 | 0 |
| SPGP0KN00170 | XP_018220573.1 | 100 | 0 |
| SPGP0BK01750 | XP_018221302.1 | 99 | 0 |
| SPGP0HX00120 | XP_018221012.1 | 99 | 1 |
| SPGP0BK01280 | XP_018221346.1 | 100 | 0 |
| SPGP0BItrna12F | DI49_0089 | 100 | 0 |
| SPGP0EA00100 | XP_018224032.1 | 99 | 0 |
| SPGP0LU00270 | XP_018223999.1 | 99 | 0 |
| SPGP0NQ00110 | XP_018220754.1 | 100 | 0 |
| SPGP0BK00400 | XP_018221432.1 | 99 | 0 |
| SPGP0D00670 | XP_018221699.1 | 99 | 0 |
| SPGP0OE00270 | XP_018219594.1 | 99 | 0 |
| SPGP0BI01170 | XP_018223763.1 | 99 | 0 |
| SPGP0HJ00160 | XP_018224044.1 | 97 | 0 |
| SPGP0RC00160 | XP_018222091.1 | 99 | 0 |
| SPGP0BE03560 | XP_018222994.1 | 99 | 0 |

Create PDF in your applications with the Pdfcrowd HTML to PDF API PDFCROWD

| | | | |
|---|---|---|---|
| SPGP0M00560 | XP_018220900.1 | 99 | 0 |
| SPGP0P01610 | XP_018221498.1 | 99 | 0 |
| SPGP0C02160 | XP_018220155.1 | 99 | 0 |
| SPGP0M00640 | XP_018220908.1 | 100 | 0 |
| SPGP0BI02950 | XP_018223588.1 | 99 | 0 |
| SPGP0CX00100 | XP_018220799.1 | 99 | 0 |
| SPGP0OA00130 | XP_018218946.1 | 99 | 0 |
| SPGP0PA00120 | XP_018220472.1 | 99 | 0 |
| SPGP0BK00480 | XP_018221424.1 | 99 | 0 |
| SPGP0N01260 | XP_018219311.1 | 100 | 0 |
| SPGP0UN00130 | XP_018220785.1 | 99 | 0 |
| SPGP0D02230 | XP_018221848.1 | 100 | 0 |
| SPGP0P01790 | XP_018221480.1 | 99 | 0 |
| SPGP0GK00160 | XP_018219708.1 | 99 | 0 |
| SPGP0BE00560 | XP_018223278.1 | 99 | 0 |
| SPGP0DU00160 | XP_018222578.1 | 99 | 0 |
| SPGP0C02290 | XP_018220142.1 | 99 | 0 |
| SPGP0M00390 | XP_018220884.1 | 99 | 0 |
| SPGP0M00460 | XP_018220891.1 | 99 | 0 |
| SPGP0D01650 | XP_018221793.1 | 91 | 1 |
| SPGP0D02500 | XP_018221875.1 | 99 | 0 |
| SPGP0BE01140 | XP_018223220.1 | 100 | 0 |
| SPGP0BE00750 | XP_018223258.1 | 99 | 0 |
| SPGP0C02700 | XP_018220103.1 | 99 | 0 |
| SPGP0P00640 | XP_018221590.1 | 99 | 0 |
| SPGP0OF00250 | XP_018222706.1 | 99 | 0 |
| SPGP0AA00110 | XP_018221084.1 | 99 | 0 |
| SPGP0BI03020 | XP_018223581.1 | 99 | 0 |
| SPGP0BK00260 | XP_018221445.1 | 99 | 0 |
| SPGP0D01190 | XP_018221747.1 | 99 | 0 |
| SPGP0WY00160 | XP_018220440.1 | 100 | 0 |
| SPGP0D01790 | XP_018221806.1 | 99 | 1 |
| SPGP0BE00970 | XP_018223237.1 | 99 | 0 |
| SPGP0N00880 | XP_018219275.1 | 99 | 0 |
| SPGP0JW00100 | XP_018220972.1 | 100 | 0 |
| SPGP0BE04420 | XP_018222910.1 | 98 | 1 |
| SPGP0QF00130 | XP_018224021.1 | 99 | 0 |
| SPGP0N00670 | XP_018219254.1 | 100 | 0 |
| SPGP0BE00950 | XP_018223239.1 | 99 | 0 |
| SPGP0KZ00220 | XP_018223940.1 | 99 | 0 |

Create PDF in your applications with the Pdfcrowd HTML to PDF API PDFCROWD

| | | | |
|---|---|---|---|
| SPGP0K02230 | XP_018219368.1 | 100 | 0 |
| SPGP0D03330 | XP_018221952.1 | 100 | 0 |
| SPGP0VH00180 | XP_018219689.1 | 99 | 0 |
| SPGP0K00140 | XP_018218747.1 | 99 | 0 |
| SPGP0BE00330 | XP_018223301.1 | 99 | 0 |
| SPGP0C00290 | XP_018220334.1 | 99 | 0 |
| SPGP0BK02380 | XP_018221240.1 | 99 | 0 |
| SPGP0D01310 | XP_018221759.1 | 99 | 0 |
| SPGP0BE03670 | XP_018222984.1 | 99 | 0 |
| SPGP0BE03820 | XP_018222969.1 | 99 | 0 |
| SPGP0DU00110 | XP_018222583.1 | 99 | 0 |
| SPGP0U00380 | XP_018224098.1 | 99 | 0 |
| SPGP0UJ00240 | XP_018223331.1 | 99 | 0 |
| SPGP0C02380 | XP_018220133.1 | 97 | 3 |
| SPGP0BE00150 | XP_018223319.1 | 99 | 0 |
| SPGP0BK00810 | XP_018221392.1 | 99 | 0 |
| SPGP0BE04870 | XP_018222864.1 | 99 | 0 |
| SPGP0P00390 | XP_018221614.1 | 99 | 0 |
| SPGP0D03140 | XP_018221934.1 | 100 | 0 |
| SPGP0BEtrna20K | DI49_5564 | 100 | 0 |
| SPGP0BK01700 | XP_018221307.1 | 99 | 0 |
| SPGP0D03640 | XP_018221982.1 | 99 | 0 |
| SPGP0D00820 | XP_018221713.1 | 99 | 0 |
| SPGP0PE00220 | XP_018222366.1 | 99 | 0 |
| SPGP0P01370 | XP_018221521.1 | 99 | 0 |
| SPGP0K00490 | XP_018219535.1 | 99 | 0 |
| SPGP0BK01940 | XP_018221283.1 | 99 | 0 |
| SPGP0FS00100 | XP_018222723.1 | 100 | 0 |
| SPGP0P00570 | XP_018221596.1 | 100 | 0 |
| SPGP0BI03740 | XP_018223508.1 | 99 | 0 |
| SPGP0XC00190 | XP_018220595.1 | 99 | 0 |
| SPGP0EM00140 | XP_018220719.1 | 99 | 0 |
| SPGP0PX00100 | XP_018222550.1 | 99 | 0 |
| SPGP0BK00710 | XP_018221402.1 | 99 | 0 |
| SPGP0BK00950 | XP_018221378.1 | 99 | 0 |
| SPGP0RI00370 | XP_018220516.1 | 99 | 0 |
| SPGP0BE04810 | XP_018222870.1 | 100 | 0 |
| SPGP0D03550 | XP_018221973.1 | 99 | 0 |
| SPGP0BE03170 | XP_018223029.1 | 99 | 0 |
| SPGP0LJ00110 | XP_018219143.1 | 99 | 0 |

Create PDF in your applications with the Pdfcrowd HTML to PDF API PDFCROWD

| | | | |
|---|---|---|---|
| SPGP0BE03680 | XP_018222983.1 | 99 | 0 |
| SPGP0PA00150 | XP_018220469.1 | 99 | 0 |
| SPGP0U00250 | XP_018224111.1 | 99 | 0 |
| SPGP0P01110 | XP_018221546.1 | 99 | 0 |
| SPGP0BE02880 | XP_018223054.1 | 100 | 0 |
| SPGP0BK02410 | XP_018221237.1 | 99 | 0 |
| SPGP0VD00120 | XP_018219835.1 | 99 | 0 |
| SPGP0BK03480 | XP_018221139.1 | 99 | 0 |
| SPGP0BE01490 | XP_018223186.1 | 99 | 0 |
| SPGP0JU00100 | XP_018219857.1 | 98 | 0 |
| SPGP0C01020 | XP_018220262.1 | 99 | 0 |
| SPGP0XC00200 | XP_018220596.1 | 99 | 0 |
| SPGP0BK00360 | XP_018221436.1 | 99 | 0 |
| SPGP0HZ00190 | XP_018219776.1 | 99 | 0 |
| SPGP0UN00100 | XP_018220788.1 | 99 | 0 |
| SPGP0P01470 | XP_018221511.1 | 99 | 0 |
| SPGP0QU00100 | XP_018219019.1 | 99 | 0 |
| SPGP0D03520 | XP_018221970.1 | 99 | 0 |
| SPGP0RQ00140 | XP_018221125.1 | 99 | 0 |
| SPGP0K00630 | XP_018219523.1 | 99 | 0 |
| SPGP0N00500 | XP_018219238.1 | 100 | 0 |
| SPGP0D01370 | XP_018221765.1 | 99 | 0 |
| SPGP0P01630 | XP_018221496.1 | 99 | 0 |
| SPGP0BI00900 | XP_018223789.1 | 99 | 0 |
| SPGP0BE02080 | XP_018223129.1 | 99 | 0 |
| SPGP0BE02220 | XP_018223116.1 | 99 | 0 |
| SPGP0N00290 | XP_018219217.1 | 99 | 0 |
| SPGP0BU00180 | XP_018222257.1 | 99 | 0 |
| SPGP0BE01970 | XP_018223138.1 | 99 | 0 |
| SPGP0K01300 | XP_018219457.1 | 99 | 0 |
| SPGP0C02930 | XP_018220080.1 | 99 | 0 |
| SPGP0IC00210 | XP_018222786.1 | 99 | 0 |
| SPGP0C00160 | XP_018220347.1 | 99 | 0 |
| SPGP0BK01140 | XP_018221360.1 | 99 | 0 |
| SPGP0HU00150 | XP_018220431.1 | 99 | 0 |
| SPGP0K00780 | XP_018219508.1 | 99 | 0 |
| SPGP0KZ00350 | XP_018223927.1 | 99 | 0 |
| SPGP0BI04140 | XP_018223472.1 | 99 | 0 |
| SPGP0OI00620 | XP_018218888.1 | 100 | 0 |
| SPGP0CV00120 | XP_018223383.1 | 99 | 0 |

Create PDF in your applications with the Pdfcrowd HTML to PDF API PDFCROWD

| | | | |
|---|---|---|---|
| SPGP0VZ00170 | XP_018223882.1 | 99 | 0 |
| SPGP0C01400 | XP_018220224.1 | 99 | 0 |
| SPGP0C01750 | XP_018220194.1 | 99 | 0 |
| SPGP0JO00150 | XP_018223351.1 | 99 | 0 |
| SPGP0IO00130 | XP_018222451.1 | 100 | 0 |
| SPGP0SH00230 | XP_018223981.1 | 99 | 0 |
| SPGP0D02720 | XP_018221895.1 | 99 | 0 |
| SPGP0BE00390 | XP_018223295.1 | 99 | 0 |
| SPGP0PS00130 | XP_018223357.1 | 99 | 1 |
| SPGP0BK03260 | XP_018221158.1 | 99 | 0 |
| SPGP0C02330 | XP_018220138.1 | 99 | 0 |
| SPGP0NK00150 | XP_018220450.1 | 99 | 0 |
| SPGP0C00110 | XP_018220352.1 | 99 | 1 |
| SPGP0D03310 | XP_018221951.1 | 99 | 0 |
| SPGP0RK00170 | XP_018224036.1 | 100 | 0 |
| SPGP0MB00150 | XP_018222177.1 | 100 | 0 |
| SPGP0QV00110 | XP_018220009.1 | 99 | 0 |
| SPGP0OI00830 | XP_018218867.1 | 100 | 0 |
| SPGP0UL00100 | XP_018219657.1 | 99 | 0 |
| SPGP0BE00360 | XP_018223298.1 | 99 | 0 |
| SPGP0P00760 | XP_018221578.1 | 99 | 0 |
| SPGP0NT00120 | XP_018222472.1 | 99 | 0 |
| SPGP0BE04640 | XP_018222887.1 | 99 | 1 |
| SPGP0BK00960 | XP_018221377.1 | 99 | 0 |
| SPGP0QP00200 | XP_018219159.1 | 99 | 0 |
| SPGP0K02390 | XP_018219352.1 | 99 | 0 |
| SPGP0MB00120 | XP_018222174.1 | 98 | 0 |
| SPGP0BI03270 | XP_018223556.1 | 99 | 0 |
| SPGP0BE04890 | XP_018222862.1 | 99 | 0 |
| SPGP0D03060 | XP_018221926.1 | 100 | 0 |
| SPGP0NK00120 | XP_018220453.1 | 99 | 0 |
| SPGP0D02400 | XP_018221865.1 | 99 | 0 |
| SPGP0D01070 | XP_018221737.1 | 99 | 0 |
| SPGP0VU00120 | XP_018222736.1 | 99 | 0 |
| SPGP0OI00160 | XP_018218931.1 | 99 | 0 |
| SPGP0OI00690 | XP_018218881.1 | 99 | 0 |
| SPGP0K01150 | XP_018219472.1 | 99 | 0 |
| SPGP0C00880 | XP_018220276.1 | 99 | 0 |
| SPGP0OY00110 | XP_018220969.1 | 99 | 0 |
| SPGP0D01810 | XP_018221807.1 | 99 | 0 |

Create PDF in your applications with the Pdfcrowd HTML to PDF API PDFCROWD

| | | | |
|---|---|---|---|
| SPGP0D02540 | XP_018221878.1 | 99 | 0 |
| SPGP0N00330 | XP_018219221.1 | 99 | 0 |
| SPGP0N01230 | XP_018219308.1 | 99 | 0 |
| SPGP0N00190 | XP_018219207.1 | 99 | 0 |
| SPGP0LV00150 | XP_018223376.1 | 99 | 0 |
| SPGP0BK01770 | XP_018221300.1 | 99 | 0 |
| SPGP0D03610 | XP_018221979.1 | 99 | 0 |
| SPGP0C02340 | XP_018220137.1 | 99 | 0 |
| SPGP0BE02760 | XP_018223064.1 | 99 | 0 |
| SPGP0BE02940 | XP_018223048.1 | 99 | 0 |
| SPGP0M01130 | XP_018220957.1 | 99 | 0 |
| SPGP0C03210 | XP_018220053.1 | 99 | 0 |
| SPGP0C02440 | XP_018220127.1 | 99 | 0 |
| SPGP0BK02880 | XP_018221195.1 | 99 | 0 |
| SPGP0D02410 | XP_018221866.1 | 99 | 0 |
| SPGP0P01580 | XP_018221501.1 | 99 | 0 |
| SPGP0M00620 | XP_018220906.1 | 99 | 0 |
| SPGP0BK00580 | XP_018221414.1 | 100 | 0 |
| SPGP0IC00110 | XP_018222796.1 | 99 | 0 |
| SPGP0D00850 | XP_018221716.1 | 99 | 0 |
| SPGP0U00580 | XP_018224080.1 | 99 | 0 |
| SPGP0WG00150 | XP_018221023.1 | 100 | 0 |
| SPGP0D01750 | XP_018221802.1 | 99 | 0 |
| SPGP0N01060 | XP_018219293.1 | 99 | 0 |
| SPGP0BE03540 | XP_018222996.1 | 99 | 0 |
| SPGP0RC00140 | XP_018222089.1 | 99 | 0 |
| SPGP0TM00110 | XP_018219875.1 | 99 | 0 |
| SPGP0D00360 | XP_018221669.1 | 99 | 0 |
| SPGP0GK00100 | XP_018219714.1 | 99 | 0 |
| SPGP0BK03180 | XP_018221166.1 | 98 | 0 |
| SPGP0M00660 | XP_018220910.1 | 99 | 0 |
| SPGP0CX00110 | XP_018220798.1 | 99 | 0 |
| SPGP0EX00130 | XP_018223422.1 | 98 | 1 |
| SPGP0BK02600 | XP_018221218.1 | 99 | 0 |
| SPGP0BU00170 | XP_018222256.1 | 99 | 0 |
| SPGP0N01240 | XP_018219309.1 | 99 | 0 |
| SPGP0IB00100 | XP_018222672.1 | 100 | 0 |
| SPGP0GF00200 | XP_018219641.1 | 99 | 0 |
| SPGP0D01080 | XP_018221738.1 | 99 | 0 |

Create PDF in your applications with the Pdfcrowd HTML to PDF API PDFCROWD

| | | | |
|---|---|---|---|
| SPGP0DU00140 | XP_018222580.1 | 99 | 0 |
| SPGP0BI02110 | XP_018223670.1 | 99 | 0 |
| SPGP0D00790 | XP_018221711.1 | 99 | 0 |
| SPGP0M00910 | XP_018220935.1 | 99 | 0 |
| SPGP0BK02950 | XP_018221188.1 | 97 | 1 |
| SPGP0DW00140 | XP_018218952.1 | 100 | 0 |
| SPGP0BI01900 | XP_018223690.1 | 99 | 0 |
| SPGP0LV00170 | XP_018223378.1 | 99 | 0 |
| SPGP0P00500 | XP_018221603.1 | 99 | 0 |
| SPGP0UJ00190 | XP_018223336.1 | 99 | 0 |
| SPGP0D01770 | XP_018221804.1 | 99 | 0 |
| SPGP0K01950 | XP_018219396.1 | 99 | 0 |
| SPGP0BE02340 | XP_018223104.1 | 99 | 0 |
| SPGP0ST00100 | XP_018220549.1 | 99 | 0 |
| SPGP0C02810 | XP_018220092.1 | 99 | 0 |
| SPGP0C01030 | XP_018220261.1 | 99 | 0 |
| SPGP0BE02970 | XP_018223045.1 | 99 | 0 |
| SPGP0BI02630 | XP_018223619.1 | 99 | 0 |
| SPGP0BE01440 | XP_018223190.1 | 99 | 0 |
| SPGP0BE01330 | XP_018223201.1 | 99 | 0 |
| SPGP0BI03970 | XP_018223488.1 | 99 | 0 |
| SPGP0QS00140 | XP_018223900.1 | 99 | 0 |
| SPGP0U00590 | XP_018224079.1 | 99 | 0 |
| SPGP0FY00100 | XP_018222557.1 | 99 | 0 |
| SPGP0D01100 | XP_018221740.1 | 99 | 0 |
| SPGP0OI00370 | XP_018218913.1 | 99 | 0 |
| SPGP0K00510 | XP_018219533.1 | 99 | 0 |
| SPGP0C00270 | XP_018220336.1 | 99 | 0 |
| SPGP0K01260 | XP_018219461.1 | 99 | 0 |
| SPGP0BE01960 | XP_018223139.1 | 99 | 0 |
| SPGP0P00730 | XP_018221581.1 | 100 | 0 |
| SPGP0UN00280 | XP_018220770.1 | 99 | 0 |
| SPGP0UJ00220 | XP_018223333.1 | 99 | 0 |
| SPGP0TM00160 | XP_018219880.1 | 99 | 0 |
| SPGP0BE01270 | XP_018223207.1 | 99 | 0 |
| SPGP0D01710 | XP_018221798.1 | 99 | 0 |
| SPGP0C00410 | XP_018220322.1 | 99 | 0 |
| SPGP0VH00190 | XP_018219690.1 | 99 | 0 |
| SPGP0K01700 | XP_018219419.1 | 99 | 0 |
| SPGP0D03120 | XP_018221932.1 | 99 | 0 |

Create PDF in your applications with the Pdfcrowd HTML to PDF API PDFCROWD

| | | | |
|---|---|---|---|
| SPGP0BE04780 | XP_018222873.1 | 99 | 0 |
| SPGP0BI03530 | XP_018223529.1 | 99 | 0 |
| SPGP0K02610 | XP_018219330.1 | 99 | 0 |
| SPGP0Y00130 | XP_018222684.1 | 99 | 0 |
| SPGP0BE03370 | XP_018223010.1 | 99 | 0 |
| SPGP0P01350 | XP_018221523.1 | 99 | 0 |
| SPGP0HR00100 | XP_018222178.1 | 99 | 0 |
| SPGP0BE00170 | XP_018223317.1 | 99 | 0 |
| SPGP0JN00190 | XP_018219078.1 | 99 | 0 |
| SPGP0Itrna30R | DI49_2918 | 100 | 0 |
| SPGP0BK02440 | XP_018221234.1 | 99 | 0 |
| SPGP0DQ00170 | XP_018219676.1 | 100 | 0 |
| SPGP0BI03190 | XP_018223564.1 | 99 | 0 |
| SPGP0BE03210 | XP_018223025.1 | 99 | 0 |
| SPGP0OI00570 | XP_018218893.1 | 100 | 0 |
| SPGP0BI04000 | XP_018223486.1 | 99 | 0 |
| SPGP0M01090 | XP_018220953.1 | 99 | 0 |
| SPGP0BE04500 | XP_018222901.1 | 99 | 0 |
| SPGP0D00280 | XP_018221661.1 | 99 | 0 |
| SPGP0WQ00260 | XP_018222278.1 | 99 | 0 |
| SPGP0C03020 | XP_018220072.1 | 99 | 0 |
| SPGP0BI01780 | XP_018223702.1 | 99 | 0 |
| SPGP0BI02530 | XP_018223629.1 | 99 | 0 |
| SPGP0BK03150 | XP_018221169.1 | 99 | 0 |
| SPGP0QJ00120 | XP_018219933.1 | 99 | 0 |
| SPGP0P01130 | XP_018221544.1 | 99 | 0 |
| SPGP0Ktrna3P | DI49_5158 | 99 | 0 |
| SPGP0PL00100 | XP_018219756.1 | 99 | 0 |
| SPGP0BI00920 | XP_018223787.1 | 99 | 0 |
| SPGP0WJ00140 | XP_018221056.1 | 99 | 0 |
| SPGP0BE04550 | XP_018222896.1 | 99 | 0 |
| SPGP0BE00190 | XP_018223315.1 | 98 | 1 |
| SPGP0M01060 | XP_018220950.1 | 99 | 0 |
| SPGP0PA00140 | XP_018220470.1 | 100 | 0 |
| SPGP0BI01250 | XP_018223755.1 | 99 | 0 |
| SPGP0RQ00160 | XP_018221127.1 | 99 | 0 |
| SPGP0BK01120 | XP_018221362.1 | 99 | 0 |
| SPGP0Ftrna19V | DI49_1074 | 100 | 0 |
| SPGP0FE00130 | XP_018220537.1 | 99 | 0 |
| SPGP0M00860 | XP_018220930.1 | 99 | 0 |

Create PDF in your applications with the Pdfcrowd HTML to PDF API PDFCROWD

| | | | |
|---|---|---|---|
| SPGP0VO00110 | XP_018223436.1 | 99 | 0 |
| SPGP0BI00250 | XP_018223853.1 | 99 | 0 |
| SPGP0BE03430 | XP_018223004.1 | 99 | 0 |
| SPGP0AU00100 | XP_018222232.1 | 99 | 0 |
| SPGP0VB00110 | XP_018221048.1 | 98 | 0 |
| SPGP0OI00110 | XP_018218936.1 | 100 | 0 |
| SPGP0K00750 | XP_018219511.1 | 99 | 0 |
| SPGP0RI00340 | XP_018220513.1 | 99 | 0 |
| SPGP0SS00110 | XP_018219048.1 | 100 | 0 |
| SPGP0LW00150 | XP_018222619.1 | 99 | 0 |
| SPGP0KG00110 | XP_018222625.1 | 99 | 0 |
| SPGP0BK03460 | XP_018221141.1 | 99 | 0 |
| SPGP0BI00780 | XP_018223800.1 | 99 | 0 |
| SPGP0U00690 | XP_018224069.1 | 99 | 0 |
| SPGP0C02640 | XP_018220109.1 | 99 | 0 |
| SPGP0QV00170 | XP_018220003.1 | 99 | 0 |
| SPGP0FE00120 | XP_018220538.1 | 99 | 0 |
| SPGP0SW00210 | XP_018220553.1 | 99 | 0 |
| SPGP0C00640 | XP_018220300.1 | 99 | 0 |
| SPGP0BE01830 | XP_018223152.1 | 99 | 0 |
| SPGP0BI02120 | XP_018223669.1 | 99 | 0 |
| SPGP0VP00260 | XP_018222850.1 | 91 | 0 |
| SPGP0IV00120 | XP_018220640.1 | 99 | 0 |
| SPGP0HZ00120 | XP_018219769.1 | 99 | 0 |
| SPGP0SD00100 | XP_018222219.1 | 99 | 0 |
| SPGP0N00860 | XP_018219273.1 | 99 | 0 |
| SPGP0D00580 | XP_018221690.1 | 99 | 0 |
| SPGP0M00520 | XP_018220896.1 | 99 | 0 |
| SPGP0BI03520 | XP_018223530.1 | 99 | 0 |
| SPGP0P00740 | XP_018221580.1 | 99 | 0 |
| SPGP0C01810 | XP_018220188.1 | 99 | 0 |
| SPGP0NT00140 | XP_018222470.1 | 99 | 0 |
| SPGP0GY00170 | XP_018223395.1 | 100 | 0 |
| SPGP0C02150 | XP_018220156.1 | 99 | 0 |
| SPGP0BE02540 | XP_018223084.1 | 99 | 0 |
| SPGP0BK02790 | XP_018221202.1 | 99 | 0 |
| SPGP0BE04660 | XP_018222885.1 | 99 | 1 |
| SPGP0BI02780 | XP_018223605.1 | 100 | 0 |
| SPGP0C01840 | XP_018220186.1 | 99 | 0 |
| SPGP0D02220 | XP_018221847.1 | 99 | 0 |

Create PDF in your applications with the Pdfcrowd HTML to PDF API PDFCROWD

| | | | |
|---|---|---|---|
| SPGP0K02370 | XP_018219354.1 | 99 | 0 |
| SPGP0MB00100 | XP_018222172.1 | 99 | 0 |
| SPGP0D02630 | XP_018221886.1 | 99 | 0 |
| SPGP0EM00200 | XP_018220714.1 | 99 | 0 |
| SPGP0P01240 | XP_018221534.1 | 99 | 0 |
| SPGP0BI03070 | XP_018223576.1 | 99 | 0 |
| SPGP0BE01180 | XP_018223216.1 | 99 | 0 |
| SPGP0BK01960 | XP_018221281.1 | 99 | 0 |
| SPGP0JP00150 | XP_018218852.1 | 99 | 0 |
| SPGP0N01290 | XP_018219314.1 | 99 | 0 |
| SPGP0BE02920 | XP_018223050.1 | 99 | 0 |
| SPGP0KX00120 | XP_018223362.1 | 99 | 0 |
| SPGP0C02310 | XP_018220140.1 | 99 | 0 |
| SPGP0C03230 | XP_018220051.1 | 99 | 0 |
| SPGP0BE01760 | XP_018223159.1 | 99 | 0 |
| SPGP0D00340 | XP_018221667.1 | 99 | 0 |
| SPGP0M00540 | XP_018220898.1 | 100 | 0 |
| SPGP0C01440 | XP_018220221.1 | 96 | 1 |
| SPGP0NI00200 | XP_018223915.1 | 99 | 0 |
| SPGP0LU00230 | XP_018224003.1 | 99 | 0 |
| SPGP0BK02590 | XP_018221219.1 | 99 | 0 |
| SPGP0GI00160 | XP_018220694.1 | 99 | 0 |
| SPGP0K02620 | XP_018219329.1 | 99 | 0 |
| SPGP0D01490 | XP_018221777.1 | 98 | 0 |
| SPGP0BI01700 | XP_018223710.1 | 99 | 0 |
| SPGP0DD00120 | XP_018220999.1 | 99 | 0 |
| SPGP0QN00110 | XP_018222490.1 | 99 | 1 |
| SPGP0TQ00250 | XP_018220620.1 | 99 | 0 |
| SPGP0P01650 | XP_018221494.1 | 99 | 0 |
| SPGP0OF00190 | XP_018222700.1 | 98 | 0 |
| SPGP0UL00120 | XP_018219659.1 | 99 | 0 |
| SPGP0P00140 | XP_018221639.1 | 99 | 0 |
| SPGP0BItrna11F | DI49_0495 | 100 | 0 |
| SPGP0P01840 | XP_018221475.1 | 99 | 0 |
| SPGP0BI01620 | XP_018223718.1 | 99 | 0 |
| SPGP0BE01360 | XP_018223198.1 | 100 | 0 |
| SPGP0C02870 | XP_018220086.1 | 99 | 0 |
| SPGP0DN00180 | XP_018218986.1 | 99 | 0 |
| SPGP0BI00400 | XP_018223838.1 | 99 | 0 |
| SPGP0VQ00110 | XP_018222543.1 | 99 | 0 |

Create PDF in your applications with the Pdfcrowd HTML to PDF API PDFCROWD

| | | | |
|---|---|---|---|
| SPGP0IK00130 | XP_018222110.1 | 99 | 0 |
| SPGP0D01430 | XP_018221771.1 | 97 | 0 |
| SPGP0GT00120 | XP_018222510.1 | 99 | 0 |
| SPGP0DQ00140 | XP_018219673.1 | 99 | 0 |
| SPGP0VN00330 | XP_018219823.1 | 99 | 0 |
| SPGP0BI00480 | XP_018223830.1 | 99 | 0 |
| SPGP0KL00120 | XP_018221102.1 | 99 | 0 |
| SPGP0BE02980 | XP_018223044.1 | 99 | 0 |
| SPGP0BI03060 | XP_018223577.1 | 99 | 0 |
| SPGP0C02780 | XP_018220095.1 | 99 | 0 |
| SPGP0OF00220 | XP_018222703.1 | 99 | 0 |
| SPGP0C01780 | XP_018220191.1 | 99 | 0 |
| SPGP0BK02140 | XP_018221263.1 | 99 | 0 |
| SPGP0QA00100 | XP_018223997.1 | 99 | 0 |
| SPGP0LB00110 | XP_018222554.1 | 99 | 0 |
| SPGP0QF00170 | XP_018224025.1 | 99 | 0 |
| SPGP0BI00440 | XP_018223834.1 | 99 | 0 |
| SPGP0BE03460 | XP_018223003.1 | 99 | 0 |
| SPGP0BI03660 | XP_018223516.1 | 99 | 0 |
| SPGP0BE04720 | XP_018222879.1 | 99 | 0 |
| SPGP0D01410 | XP_018221769.1 | 99 | 0 |
| SPGP0LG00190 | XP_018222651.1 | 99 | 0 |
| SPGP0BI03510 | XP_018223531.1 | 99 | 0 |
| SPGP0K00480 | XP_018219536.1 | 99 | 0 |
| SPGP0DU00200 | XP_018222574.1 | 99 | 0 |
| SPGP0GL00110 | XP_018222612.1 | 99 | 0 |
| SPGP0C00780 | XP_018220286.1 | 99 | 0 |
| SPGP0BE01530 | XP_018223182.1 | 99 | 0 |
| SPGP0C01040 | XP_018220260.1 | 99 | 0 |
| SPGP0D03100 | XP_018221930.1 | 99 | 0 |
| SPGP0QQtrna17A | DI49_3564 | 100 | 0 |
| SPGP0K02580 | XP_018219333.1 | 99 | 0 |
| SPGP0U00360 | XP_018224100.1 | 99 | 0 |
| SPGP0BM00130 | XP_018222756.1 | 98 | 1 |
| SPGP0BE00370 | XP_018223297.1 | 99 | 0 |
| SPGP0GH00110 | XP_018218967.1 | 99 | 0 |
| SPGP0HB00110 | XP_018219193.1 | 98 | 0 |
| SPGP0BE03020 | XP_018223041.1 | 99 | 0 |
| SPGP0K01720 | XP_018219417.1 | 99 | 0 |
| SPGP0BE04850 | XP_018222866.1 | 99 | 0 |

Create PDF in your applications with the Pdfcrowd HTML to PDF API PDFCROWD

| | | | |
|---|---|---|---|
| SPGP0K01340 | XP_018219453.1 | 100 | 0 |
| SPGP0IC00250 | XP_018222782.1 | 100 | 0 |
| SPGP0NK00100 | XP_018220455.1 | 99 | 0 |
| SPGP0P00100 | XP_018221643.1 | 99 | 0 |
| SPGP0NV00140 | XP_018223452.1 | 98 | 1 |
| SPGP0NY00110 | XP_018219620.1 | 99 | 0 |
| SPGP0K01580 | XP_018219430.1 | 100 | 0 |
| SPGP0BK02360 | XP_018221242.1 | 99 | 0 |
| SPGP0C00920 | XP_018220272.1 | 99 | 0 |
| SPGP0BE04000 | XP_018222952.1 | 99 | 0 |
| SPGP0IR00110 | XP_018219766.1 | 99 | 0 |
| SPGP0LG00180 | XP_018222650.1 | 99 | 0 |
| SPGP0K00450 | XP_018219540.1 | 99 | 0 |
| SPGP0K00980 | XP_018219489.1 | 99 | 0 |
| SPGP0N00270 | XP_018219215.1 | 99 | 0 |
| SPGP0BE01160 | XP_018223218.1 | 99 | 0 |
| SPGP0LN00110 | XP_018223923.1 | 99 | 0 |
| SPGP0D02650 | XP_018221888.1 | 99 | 0 |
| SPGP0MP00110 | XP_018222752.1 | 99 | 0 |
| SPGP0M01070 | XP_018220951.1 | 99 | 0 |
| SPGP0D00150 | XP_018221648.1 | 99 | 0 |
| SPGP0BK03200 | XP_018221164.1 | 99 | 0 |
| SPGP0BK03170 | XP_018221167.1 | 99 | 0 |
| SPGP0DB00120 | XP_018222187.1 | 98 | 0 |
| SPGP0D02320 | XP_018221857.1 | 99 | 0 |
| SPGP0K00180 | XP_018219567.1 | 99 | 0 |
| SPGP0ER00140 | XP_018218979.1 | 99 | 0 |
| SPGP0RK00190 | XP_018224035.1 | 98 | 1 |
| SPGP0TG00110 | XP_018220661.1 | 99 | 0 |
| SPGP0FW00110 | XP_018219585.1 | 99 | 0 |
| SPGP0BK01100 | XP_018221364.1 | 99 | 0 |
| SPGP0SE00100 | XP_018223404.1 | 99 | 0 |
| SPGP0BK02500 | XP_018221228.1 | 99 | 0 |
| SPGP0LG00150 | XP_018222647.1 | 99 | 0 |
| SPGP0BI01350 | XP_018223746.1 | 99 | 1 |
| SPGP0IW00130 | XP_018220014.1 | 99 | 0 |
| SPGP0U00770 | XP_018224061.1 | 99 | 0 |
| SPGP0Htrna30L | DI49_1927 | 100 | 0 |
| SPGP0SS00130 | XP_018219046.1 | 99 | 0 |
| SPGP0K00760 | XP_018219510.1 | 99 | 0 |

Create PDF in your applications with the Pdfcrowd HTML to PDF API PDFCROWD

| | | | |
|---|---|---|---|
| SPGP0UJ00150 | XP_018223340.1 | 99 | 0 |
| SPGP0RE00170 | XP_018220400.1 | 99 | 0 |
| SPGP0JO00110 | XP_018223355.1 | 99 | 0 |
| SPGP0C03370 | XP_018220037.1 | 99 | 0 |
| SPGP0BK00270 | XP_018221444.1 | 97 | 2 |
| SPGP0D00590 | XP_018221691.1 | 99 | 0 |
| SPGP0UL00140 | XP_018219661.1 | 99 | 0 |
| SPGP0QV00150 | XP_018220005.1 | 99 | 0 |
| SPGP0IH00150 | XP_018222235.1 | 99 | 0 |
| SPGP0JN00120 | XP_018219071.1 | 100 | 0 |
| SPGP0BE01810 | XP_018219094.1 | 99 | 0 |
| SPGP0BI01480 | XP_018223733.1 | 99 | 0 |
| SPGP0KN00120 | XP_018220568.1 | 99 | 0 |
| SPGP0K01830 | XP_018219406.1 | 99 | 0 |
| SPGP0BI03500 | XP_018223532.1 | 99 | 0 |
| SPGP0AA00210 | XP_018221074.1 | 99 | 0 |
| SPGP0K02160 | XP_018219375.1 | 99 | 0 |
| SPGP0GY00150 | XP_018223397.1 | 99 | 0 |
| SPGP0C01310 | XP_018220233.1 | 98 | 1 |
| SPGP0C01640 | XP_018220203.1 | 99 | 1 |
| SPGP0UN00180 | XP_018220780.1 | 99 | 0 |
| SPGP0D02190 | XP_018221844.1 | 99 | 0 |
| SPGP0BE03360 | XP_018223011.1 | 99 | 0 |
| SPGP0BE02430 | XP_018223095.1 | 99 | 0 |
| SPGP0VD00170 | XP_018219840.1 | 99 | 0 |
| SPGP0NO00110 | XP_018222688.1 | 99 | 0 |
| SPGP0ON00100 | XP_018222487.1 | 90 | 0 |
| SPGP0P00380 | XP_018221615.1 | 100 | 0 |
| SPGP0D01250 | XP_018221753.1 | 99 | 0 |
| SPGP0K02130 | XP_018219378.1 | 100 | 0 |
| SPGP0VI00130 | XP_018222306.1 | 99 | 0 |
| SPGP0D02440 | XP_018221869.1 | 99 | 0 |
| SPGP0P01810 | XP_018221478.1 | 99 | 0 |
| SPGP0BI02350 | XP_018223647.1 | 99 | 0 |
| SPGP0VM00210 | XP_018220673.1 | 99 | 0 |
| SPGP0C01490 | XP_018220216.1 | 99 | 0 |
| SPGP0U00260 | XP_018224110.1 | 99 | 0 |
| SPGP0BI02100 | XP_018223671.1 | 99 | 0 |
| SPGP0K01620 | XP_018219426.1 | 99 | 0 |
| SPGP0K00410 | XP_018219544.1 | 99 | 0 |

Create PDF in your applications with the Pdfcrowd HTML to PDF API PDFCROWD

| | | | |
|---|---|---|---|
| SPGP0BE02730 | XP_018223067.1 | 99 | 0 |
| SPGP0MG00100 | XP_018219618.1 | 99 | 0 |
| SPGP0TG00120 | XP_018220662.1 | 99 | 0 |
| SPGP0K01440 | XP_018219443.1 | 99 | 0 |
| SPGP0K01790 | XP_018219410.1 | 99 | 0 |
| SPGP0K01110 | XP_018219476.1 | 99 | 0 |
| SPGP0BI01860 | XP_018223694.1 | 99 | 0 |
| SPGP0P01910 | XP_018221468.1 | 99 | 0 |
| SPGP0BE03050 | XP_018223039.1 | 98 | 1 |
| SPGP0BE03500 | XP_018223000.1 | 99 | 0 |
| SPGP0FZ00100 | XP_018222220.1 | 98 | 0 |
| SPGP0N00480 | XP_018219236.1 | 100 | 0 |
| SPGP0PG00100 | XP_018219179.1 | 99 | 0 |
| SPGP0Dtrna8F | DI49_2529 | 100 | 0 |
| SPGP0GD00120 | XP_018220832.1 | 99 | 0 |
| SPGP0D03110 | XP_018221931.1 | 99 | 0 |
| SPGP0BK02180 | XP_018221259.1 | 99 | 0 |
| SPGP0NU00120 | XP_018219057.1 | 99 | 0 |
| SPGP0BE02900 | XP_018223052.1 | 99 | 0 |
| SPGP0KX00100 | XP_018223364.1 | 99 | 0 |
| SPGP0M00740 | XP_018220918.1 | 99 | 0 |
| SPGP0K01990 | XP_018219392.1 | 99 | 0 |
| SPGP0KZ00290 | XP_018223933.1 | 99 | 0 |
| SPGP0BI01500 | XP_018223731.1 | 99 | 0 |
| SPGP0N00730 | XP_018219260.1 | 99 | 0 |
| SPGP0BL00100 | XP_018222181.1 | 99 | 0 |
| SPGP0N00650 | XP_018219252.1 | 99 | 0 |
| SPGP0C00360 | XP_018220327.1 | 99 | 0 |
| SPGP0U00860 | XP_018224052.1 | 99 | 0 |
| SPGP0JI00120 | XP_018222448.1 | 99 | 0 |
| SPGP0N01280 | XP_018219313.1 | 99 | 0 |
| SPGP0BE02930 | XP_018223049.1 | 99 | 0 |
| SPGP0K00730 | XP_018219513.1 | 99 | 0 |
| SPGP0M00480 | XP_018220893.1 | 100 | 0 |
| SPGP0EU00120 | XP_018222660.1 | 99 | 0 |
| SPGP0M00950 | XP_018220939.1 | 99 | 0 |
| SPGP0OA00150 | XP_018218944.1 | 99 | 0 |
| SPGP0C01500 | XP_018220215.1 | 99 | 0 |
| SPGP0BE02670 | XP_018223073.1 | 99 | 0 |
| SPGP0OR00150 | XP_018218831.1 | 99 | 0 |

Create PDF in your applications with the Pdfcrowd HTML to PDF API PDFCROWD

| | | | |
|---|---|---|---|
| SPGP0BK03340 | XP_018221150.1 | 99 | 0 |
| SPGP0HA00100 | XP_018219868.1 | 99 | 0 |
| SPGP0BK02910 | XP_018221192.1 | 97 | 1 |
| SPGP0DW00100 | XP_018218956.1 | 99 | 0 |
| SPGP0OI00440 | XP_018218906.1 | 99 | 0 |
| SPGP0KZ00170 | XP_018223943.1 | 99 | 0 |
| SPGP0VO00170 | XP_018223441.1 | 99 | 0 |
| SPGP0D03400 | XP_018221958.1 | 99 | 0 |
| SPGP0LS00100 | XP_018222563.1 | 99 | 0 |
| SPGP0XB00120 | XP_018219022.1 | 99 | 0 |
| SPGP0BI00420 | XP_018223836.1 | 99 | 0 |
| SPGP0BE00430 | XP_018223291.1 | 99 | 0 |
| SPGP0BE03130 | XP_018223032.1 | 100 | 0 |
| SPGP0BI01180 | XP_018223762.1 | 99 | 0 |
| SPGP0C03010 | XP_018220073.1 | 99 | 0 |
| SPGP0D00420 | XP_018221675.1 | 99 | 0 |
| SPGP0NI00210 | XP_018223916.1 | 99 | 0 |
| SPGP0C00260 | XP_018220337.1 | 99 | 0 |
| SPGP0EK00220 | XP_018220803.1 | 99 | 0 |
| SPGP0FE00270 | XP_018220524.1 | 99 | 0 |
| SPGP0US00100 | XP_018219156.1 | 99 | 0 |
| SPGP0BE01590 | XP_018223176.1 | 99 | 0 |
| SPGP0OI00320 | XP_018218918.1 | 99 | 0 |
| SPGP0VD00150 | XP_018219838.1 | 99 | 0 |
| SPGP0D03420 | XP_018221960.1 | 99 | 0 |
| SPGP0BE00420 | XP_018223292.1 | 99 | 0 |
| SPGP0BE00350 | XP_018223299.1 | 99 | 0 |
| SPGP0BK02000 | XP_018221277.1 | 99 | 0 |
| SPGP0TE00140 | XP_018219896.1 | 99 | 0 |
| SPGP0MO00350 | XP_018219119.1 | 99 | 0 |
| SPGP0BE04460 | XP_018222905.1 | 99 | 0 |
| SPGP0D03010 | XP_018221921.1 | 99 | 0 |
| SPGP0BE04520 | XP_018222899.1 | 99 | 0 |
| SPGP0TM00140 | XP_018219878.1 | 99 | 0 |
| SPGP0BE03990 | XP_018222953.1 | 100 | 0 |
| SPGP0RE00180 | XP_018220401.1 | 99 | 0 |
| SPGP0UN00240 | XP_018220774.1 | 99 | 0 |
| SPGP0NY00130 | XP_018219622.1 | 99 | 0 |
| SPGP0JP00130 | XP_018218850.1 | 99 | 0 |
| SPGP0M00900 | XP_018220934.1 | 99 | 0 |

Create PDF in your applications with the Pdfcrowd HTML to PDF API PDFCROWD

| | | | |
|---|---|---|---|
| SPGP0XC00150 | XP_018220591.1 | 99 | 0 |
| SPGP0BE01800 | XP_018223155.1 | 99 | 0 |
| SPGP0C02760 | XP_018220097.1 | 99 | 0 |
| SPGP0P01310 | XP_018221527.1 | 99 | 0 |
| SPGP0QK00150 | XP_018220387.1 | 99 | 0 |
| SPGP0K00470 | XP_018219537.1 | 99 | 0 |
| SPGP0BI01240 | XP_018223756.1 | 99 | 0 |
| SPGP0P00800 | XP_018221574.1 | 99 | 1 |
| SPGP0M01050 | XP_018220949.1 | 99 | 0 |
| SPGP0RN00160 | XP_018222827.1 | 99 | 0 |
| SPGP0C03060 | XP_018220068.1 | 99 | 0 |
| SPGP0VN00180 | XP_018219808.1 | 99 | 0 |
| SPGP0M00650 | XP_018220909.1 | 99 | 0 |
| SPGP0K00650 | XP_018219521.1 | 99 | 0 |
| SPGP0BK01360 | XP_018221338.1 | 100 | 0 |
| SPGP0MG00130 | XP_018219615.1 | 99 | 0 |
| SPGP0BE02240 | XP_018223114.1 | 99 | 0 |
| SPGP0M00570 | XP_018220901.1 | 99 | 0 |
| SPGP0C01430 | XP_018220222.1 | 99 | 0 |
| SPGP0C02190 | XP_018220152.1 | 99 | 0 |
| SPGP0C00280 | XP_018220335.1 | 99 | 0 |
| SPGP0S00120 | XP_018219654.1 | 99 | 0 |
| SPGP0BI00210 | XP_018223857.1 | 99 | 0 |
| SPGP0U00750 | XP_018224063.1 | 99 | 0 |
| SPGP0BE00660 | XP_018223268.1 | 99 | 0 |
| SPGP0Gtrna14N | DI49_5581 | 100 | 0 |
| SPGP0P00750 | XP_018221579.1 | 99 | 0 |
| SPGP0N00750 | XP_018219262.1 | 99 | 0 |
| SPGP0D00630 | XP_018221695.1 | 99 | 0 |
| SPGP0UL00160 | XP_018219663.1 | 99 | 0 |
| SPGP0BK00610 | XP_018221411.1 | 99 | 0 |
| SPGP0BE01290 | XP_018223205.1 | 99 | 0 |
| SPGP0JN00100 | XP_018219069.1 | 99 | 0 |
| SPGP0BI00350 | XP_018223843.1 | 99 | 0 |
| SPGP0HO00190 | XP_018219983.1 | 99 | 0 |
| SPGP0C01050 | XP_018220259.1 | 99 | 0 |
| SPGP0C01130 | XP_018220251.1 | 99 | 0 |
| SPGP0BI03560 | XP_018223526.1 | 99 | 0 |
| SPGP0BI03290 | XP_018223554.1 | 99 | 0 |
| SPGP0KB00170 | XP_018221119.1 | 99 | 0 |

Create PDF in your applications with the Pdfcrowd HTML to PDF API PDFCROWD

| | | | |
|---|---|---|---|
| SPGP0BI03910 | XP_018223493.1 | 99 | 0 |
| SPGP0BR00110 | XP_018222222.1 | 99 | 0 |
| SPGP0XC00280 | XP_018220604.1 | 99 | 0 |
| SPGP0BK00310 | XP_018221441.1 | 99 | 0 |
| SPGP0PZ00310 | XP_018222415.1 | 99 | 0 |
| SPGP0JZ00100 | XP_018223417.1 | 99 | 0 |
| SPGP0KZ00340 | XP_018223928.1 | 99 | 0 |
| SPGP0K02330 | XP_018219358.1 | 99 | 1 |
| SPGP0D01390 | XP_018221767.1 | 96 | 2 |
| SPGP0UO00100 | XP_018222182.1 | 99 | 0 |
| SPGP0MC00170 | XP_018222668.1 | 99 | 0 |
| SPGP0D01270 | XP_018221755.1 | 99 | 0 |
| SPGP0K02480 | XP_018219343.1 | 99 | 0 |
| SPGP0BI01020 | XP_018223777.1 | 99 | 0 |
| SPGP0C00910 | XP_018220273.1 | 100 | 0 |
| SPGP0C00630 | XP_018220301.1 | 97 | 3 |
| SPGP0BK01080 | XP_018221366.1 | 99 | 0 |
| SPGP0OI00180 | XP_018218929.1 | 99 | 0 |
| SPGP0BE04650 | XP_018222886.1 | 100 | 0 |
| SPGP0C00570 | XP_018220307.1 | 99 | 0 |
| SPGP0VE00100 | XP_018220962.1 | 99 | 0 |
| SPGP0BI02160 | XP_018223665.1 | 99 | 0 |
| SPGP0KQ00110 | XP_018219929.1 | 99 | 0 |
| SPGP0BE03290 | XP_018223017.1 | 100 | 0 |
| SPGP0OY00130 | XP_018220971.1 | 99 | 0 |
| SPGP0C01680 | XP_018220199.1 | 100 | 0 |
| SPGP0BE00880 | XP_018223246.1 | 99 | 0 |
| SPGP0WI00260 | XP_018222134.1 | 99 | 0 |
| SPGP0BE02160 | XP_018223122.1 | 99 | 1 |
| SPGP0P00470 | XP_018221606.1 | 99 | 0 |
| SPGP0N00850 | XP_018219272.1 | 99 | 0 |
| SPGP0K01460 | XP_018219442.1 | 99 | 0 |
| SPGP0BK00890 | XP_018221384.1 | 99 | 0 |
| SPGP0C03270 | XP_018220047.1 | 99 | 0 |
| SPGP0MF00200 | XP_018222200.1 | 98 | 1 |
| SPGP0D00380 | XP_018221671.1 | 99 | 0 |
| SPGP0CZ00170 | XP_018219912.1 | 99 | 0 |
| SPGP0OK00140 | XP_018219867.1 | 99 | 0 |
| SPGP0UW00100 | XP_018223369.1 | 99 | 0 |

Create PDF in your applications with the Pdfcrowd HTML to PDF API PDFCROWD

| | | | |
|---|---|---|---|
| SPGP0D01500 | XP_018221778.1 | 99 | 0 |
| SPGP0K00220 | XP_018219563.1 | 99 | 0 |
| SPGP0C01630 | XP_018220204.1 | 99 | 0 |
| SPGP0BK02100 | XP_018221267.1 | 99 | 0 |
| SPGP0BI03540 | XP_018223528.1 | 99 | 0 |
| SPGP0BKtrna25T | DI49_2415 | 100 | 0 |
| SPGP0BItrna18R | DI49_2410 | 100 | 0 |
| SPGP0BI03580 | XP_018223524.1 | 99 | 0 |
| SPGP0U00560 | XP_018224081.1 | 99 | 0 |
| SPGP0BK02660 | XP_018221213.1 | 99 | 0 |
| SPGP0SP00170 | XP_018223967.1 | 99 | 0 |
| SPGP0BI00990 | XP_018223780.1 | 99 | 1 |
| SPGP0D02920 | XP_018221914.1 | 99 | 0 |
| SPGP0UH00150 | XP_018220411.1 | 96 | 2 |
| SPGP0K00920 | XP_018219495.1 | 99 | 0 |
| SPGP0D00700 | XP_018221702.1 | 99 | 0 |
| SPGP0VQ00150 | XP_018222548.1 | 99 | 0 |
| SPGP0K00430 | XP_018219542.1 | 99 | 0 |
| SPGP0BI03700 | XP_018223512.1 | 99 | 0 |
| SPGP0GI00100 | XP_018220700.1 | 99 | 0 |
| SPGP0BI03280 | XP_018223555.1 | 99 | 0 |
| SPGP0D01570 | XP_018221785.1 | 99 | 0 |
| SPGP0CX00170 | XP_018220792.1 | 99 | 0 |
| SPGP0N00800 | XP_018219267.1 | 99 | 0 |
| SPGP0QW00130 | XP_018222605.1 | 99 | 0 |
| SPGP0P00490 | XP_018221604.1 | 100 | 0 |
| SPGP0BK01310 | XP_018221343.1 | 99 | 0 |
| SPGP0LG00170 | XP_018222649.1 | 99 | 1 |
| SPGP0BK03380 | XP_018221146.1 | 99 | 0 |
| SPGP0BI04180 | XP_018223468.1 | 99 | 0 |
| SPGP0BK03220 | XP_018221162.1 | 100 | 0 |
| SPGP0PE00160 | XP_018222360.1 | 99 | 0 |
| SPGP0MS00130 | XP_018222329.1 | 99 | 0 |
| SPGP0K00400 | XP_018219545.1 | 99 | 0 |
| SPGP0BE01350 | XP_018223199.1 | 99 | 0 |
| SPGP0K01090 | XP_018219478.1 | 99 | 0 |
| SPGP0D00470 | XP_018221679.1 | 99 | 0 |
| SPGP0RI00150 | XP_018220493.1 | 99 | 0 |
| SPGP0M01080 | XP_018220952.1 | 100 | 0 |
| SPGP0BI02370 | XP_018223645.1 | 99 | 0 |

Create PDF in your applications with the Pdfcrowd HTML to PDF API PDFCROWD

| | | | |
|---|---|---|---|
| SPGP0M00200 | XP_018220865.1 | 99 | 0 |
| SPGP0C00740 | XP_018220290.1 | 99 | 0 |
| SPGP0C00210 | XP_018220342.1 | 99 | 0 |
| SPGP0BE03790 | XP_018222972.1 | 99 | 0 |
| SPGP0BK03240 | XP_018221160.1 | 99 | 0 |
| SPGP0BE04090 | XP_018222944.1 | 98 | 0 |
| SPGP0K02520 | XP_018219339.1 | 99 | 0 |
| SPGP0C02690 | XP_018220104.1 | 99 | 0 |
| SPGP0FT00120 | XP_018222732.1 | 99 | 0 |
| SPGP0BE00400 | XP_018223294.1 | 99 | 0 |
| SPGP0BI03170 | XP_018223566.1 | 99 | 0 |
| SPGP0P00310 | XP_018221622.1 | 98 | 1 |
| SPGP0N00240 | XP_018219212.1 | 99 | 0 |
| SPGP0BU00150 | XP_018222254.1 | 100 | 0 |
| SPGP0BE04540 | XP_018222897.1 | 99 | 0 |
| SPGP0BI02820 | XP_018223601.1 | 99 | 0 |
| SPGP0D00720 | XP_018221704.1 | 99 | 0 |
| SPGP0BE00300 | XP_018223304.1 | 98 | 1 |
| SPGP0M00360 | XP_018220881.1 | 99 | 0 |
| SPGP0U00790 | XP_018224059.1 | 99 | 0 |
| SPGP0HZ00290 | XP_018219786.1 | 100 | 0 |
| SPGP0C01320 | XP_018220232.1 | 99 | 0 |
| SPGP0KB00140 | XP_018221116.1 | 99 | 0 |
| SPGP0C01610 | XP_018220206.1 | 99 | 0 |
| SPGP0BI02670 | XP_018223615.1 | 99 | 0 |
| SPGP0BK02490 | XP_018221229.1 | 99 | 0 |
| SPGP0C00960 | XP_018220268.1 | 100 | 0 |
| SPGP0P01800 | XP_018221479.1 | 99 | 0 |
| SPGP0P01990 | XP_018221460.1 | 98 | 0 |
| SPGP0N00230 | XP_018219211.1 | 99 | 0 |
| SPGP0BK02250 | XP_018221252.1 | 99 | 0 |
| SPGP0N00570 | XP_018219245.1 | 99 | 0 |
| SPGP0C03360 | XP_018220038.1 | 99 | 0 |
| SPGP0P00860 | XP_018221568.1 | 99 | 0 |
| SPGP0BE02000 | XP_018223135.1 | 99 | 0 |
| SPGP0N00250 | XP_018219213.1 | 99 | 1 |
| SPGP0K01320 | XP_018219455.1 | 99 | 1 |
| SPGP0RA00100 | XP_018218840.1 | 99 | 0 |
| SPGP0D02170 | XP_018221842.1 | 99 | 0 |
| SPGP0C01370 | XP_018220227.1 | 99 | 0 |

Create PDF in your applications with the Pdfcrowd HTML to PDF API PDFCROWD

| | | | |
|---|---|---|---|
| SPGP0D02710 | XP_018221894.1 | 99 | 0 |
| SPGP0N00580 | XP_018219246.1 | 99 | 0 |
| SPGP0BK01670 | XP_018221310.1 | 100 | 0 |
| SPGP0RB00140 | XP_018218996.1 | 99 | 0 |
| SPGP0TQ00190 | XP_018220614.1 | 99 | 0 |
| SPGP0JF00170 | XP_018222315.1 | 99 | 0 |
| SPGP0BE01900 | XP_018223146.1 | 99 | 0 |
| SPGP0BE00450 | XP_018223289.1 | 99 | 0 |
| SPGP0EA00110 | XP_018224030.1 | 99 | 0 |
| SPGP0BI01720 | XP_018223708.1 | 99 | 0 |
| SPGP0OF00130 | XP_018222694.1 | 99 | 0 |
| SPGP0BK00350 | XP_018221437.1 | 99 | 0 |
| SPGP0BK01840 | XP_018221292.1 | 99 | 0 |
| SPGP0P01730 | XP_018221486.1 | 99 | 0 |
| SPGP0M00780 | XP_018220922.1 | 99 | 0 |
| SPGP0BI02940 | XP_018223589.1 | 99 | 0 |
| SPGP0SU00130 | XP_018218973.1 | 99 | 0 |
| SPGP0K02200 | XP_018219371.1 | 99 | 0 |
| SPGP0BI00230 | XP_018223855.1 | 99 | 0 |
| SPGP0BI00710 | XP_018223807.1 | 99 | 0 |
| SPGP0OS00120 | XP_018219931.1 | 90 | 0 |
| SPGP0P00220 | XP_018221631.1 | 99 | 0 |
| SPGP0BI00520 | XP_018223826.1 | 99 | 0 |
| SPGP0K02590 | XP_018219332.1 | 99 | 0 |
| SPGP0C02580 | XP_018220113.1 | 99 | 0 |
| SPGP0BK00700 | XP_018221403.1 | 99 | 0 |
| SPGP0HZtrna8L | DI49_4635 | 100 | 0 |
| SPGP0WG00130 | XP_018221025.1 | 99 | 0 |
| SPGP0MJ00120 | XP_018222083.1 | 99 | 0 |
| SPGP0C02950 | XP_018220078.1 | 99 | 0 |
| SPGP0K01430 | XP_018219444.1 | 99 | 0 |
| SPGP0C01350 | XP_018220229.1 | 100 | 0 |
| SPGP0BI04050 | XP_018223481.1 | 99 | 0 |
| SPGP0D02480 | XP_018221873.1 | 99 | 0 |
| SPGP0P01540 | XP_018221505.1 | 99 | 0 |
| SPGP0BE04390 | XP_018222913.1 | 99 | 0 |
| SPGP0D01290 | XP_018221757.1 | 99 | 0 |
| SPGP0C01970 | XP_018220173.1 | 99 | 0 |
| SPGP0BI02140 | XP_018223667.1 | 99 | 0 |
| SPGP0CK00120 | XP_018219961.1 | 99 | 0 |

Create PDF in your applications with the Pdfcrowd HTML to PDF API PDFCROWD

| | | | |
|---|---|---|---|
| SPGP0C00760 | XP_018220288.1 | 99 | 0 |
| SPGP0BK01740 | XP_018221303.1 | 99 | 0 |
| SPGP0BE03280 | XP_018223018.1 | 99 | 0 |
| SPGP0C00550 | XP_018220309.1 | 99 | 0 |
| SPGP0VE00120 | XP_018220964.1 | 99 | 0 |
| SPGP0VP00200 | XP_018219941.1 | 99 | 0 |
| SPGP0BK01490 | XP_018221327.1 | 99 | 0 |
| SPGP0C00490 | XP_018220315.1 | 99 | 0 |
| SPGP0BE01370 | XP_018223197.1 | 99 | 0 |
| SPGP0P00790 | XP_018221575.1 | 100 | 0 |
| SPGP0BE04930 | XP_018222858.1 | 99 | 0 |
| SPGP0K01400 | XP_018219447.1 | 99 | 0 |
| SPGP0BI00170 | XP_018223861.1 | 99 | 0 |
| SPGP0VZ00240 | XP_018223888.1 | 99 | 0 |
| SPGP0MD00100 | XP_018222384.1 | 99 | 0 |
| SPGP0BE01670 | XP_018223168.1 | 99 | 0 |
| SPGP0BE00870 | XP_018223247.1 | 99 | 0 |
| SPGP0BK00420 | XP_018221430.1 | 99 | 0 |
| SPGP0P01490 | XP_018221510.1 | 99 | 0 |
| SPGP0HT00270 | XP_018219018.1 | 99 | 0 |
| SPGP0UR00160 | XP_018222338.1 | 99 | 0 |
| SPGP0JE00100 | XP_018221134.1 | 93 | 0 |
| SPGP0OI00700 | XP_018218880.1 | 99 | 0 |
| SPGP0D02670 | XP_018221890.1 | 99 | 0 |
| SPGP0BK02680 | XP_018221212.1 | 99 | 0 |
| SPGP0D01510 | XP_018221779.1 | 99 | 0 |
| SPGP0D01900 | XP_018221816.1 | 99 | 0 |
| SPGP0D02750 | XP_018221898.1 | 99 | 1 |
| SPGP0D01450 | XP_018221773.1 | 99 | 0 |
| SPGP0UZ00120 | XP_018223428.1 | 99 | 1 |
| SPGP0N01160 | XP_018219303.1 | 99 | 0 |
| SPGP0D02900 | XP_018221912.1 | 99 | 0 |
| SPGP0BI00390 | XP_018223839.1 | 99 | 0 |
| SPGP0KZ00130 | XP_018223947.1 | 99 | 0 |
| SPGP0CY00120 | XP_018223951.1 | 99 | 0 |
| SPGP0QK00170 | XP_018220388.1 | 99 | 0 |
| SPGP0BK01330 | XP_018221341.1 | 99 | 0 |
| SPGP0BE04060 | XP_018222947.1 | 99 | 0 |
| SPGP0OX00140 | XP_018219731.1 | 99 | 0 |
| SPGP0BK01560 | XP_018221320.1 | 99 | 0 |

Create PDF in your applications with the Pdfcrowd HTML to PDF API PDFCROWD

| | | | |
|---|---|---|---|
| SPGP0C02530 | XP_018220118.1 | 99 | 0 |
| SPGP0JI00100 | XP_018222450.1 | 99 | 0 |
| SPGP0N00260 | XP_018219214.1 | 99 | 0 |
| SPGP0C02720 | XP_018220101.1 | 99 | 1 |
| SPGP0M00190 | XP_018220864.1 | 99 | 0 |
| SPGP0UZtrna20S | DI49_0808 | 99 | 0 |
| SPGP0D03580 | XP_018221976.1 | 100 | 0 |
| SPGP0TQ00110 | XP_018220606.1 | 99 | 0 |
| SPGP0BK00850 | XP_018221388.1 | 99 | 0 |
| SPGP0PZ00210 | XP_018222425.1 | 99 | 0 |
| SPGP0C02540 | XP_018220117.1 | 99 | 0 |
| SPGP0Itrna23E | DI49_5287 | 100 | 0 |
| SPGP0M00220 | XP_018220867.1 | 99 | 0 |
| SPGP0BK02040 | XP_018221273.1 | 99 | 0 |
| SPGP0P01590 | XP_018221500.1 | 99 | 0 |
| SPGP0BE04700 | XP_018222881.1 | 99 | 0 |
| SPGP0AK00120 | XP_018219737.1 | 100 | 0 |
| SPGP0C02570 | XP_018220114.1 | 99 | 0 |
| SPGP0BE00460 | XP_018223288.1 | 99 | 0 |
| SPGP0BE04880 | XP_018222863.1 | 99 | 0 |
| SPGP0D03200 | XP_018221940.1 | 99 | 0 |
| SPGP0PG00150 | XP_018219174.1 | 99 | 0 |
| SPGP0P00590 | XP_018221594.1 | 95 | 4 |
| SPGP0K01160 | XP_018219471.1 | 99 | 0 |
| SPGP0BE04560 | XP_018222895.1 | 99 | 0 |
| SPGP0U00330 | XP_018224103.1 | 98 | 0 |
| SPGP0EA00130 | XP_018224028.1 | 100 | 0 |
| SPGP0C01010 | XP_018220263.1 | 99 | 0 |
| SPGP0DQ00200 | XP_018219679.1 | 100 | 0 |
| SPGP0D02340 | XP_018221859.1 | 99 | 0 |
| SPGP0BI04190 | XP_018223467.1 | 99 | 0 |
| SPGP0Dtrna13V | DI49_2588 | 100 | 0 |
| SPGP0C00940 | XP_018220270.1 | 99 | 0 |
| SPGP0BI01790 | XP_018223701.1 | 100 | 0 |
| SPGP0D03260 | XP_018221946.1 | 99 | 0 |
| SPGP0BI00190 | XP_018223859.1 | 99 | 0 |
| SPGP0MW00100 | XP_018219153.1 | 99 | 0 |
| SPGP0XC00120 | XP_018220588.1 | 99 | 0 |
| SPGP0N00210 | XP_018219209.1 | 99 | 0 |
| SPGP0JF00150 | XP_018222313.1 | 99 | 0 |

Create PDF in your applications with the Pdfcrowd HTML to PDF API PDFCROWD

| | | | |
|---|---|---|---|
| SPGP0K00690 | XP_018219517.1 | 99 | 0 |
| SPGP0N01030 | XP_018219290.1 | 99 | 0 |
| SPGP0N00930 | XP_018219280.1 | 99 | 0 |
| SPGP0M01010 | XP_018220945.1 | 99 | 0 |
| SPGP0BE03530 | XP_018222997.1 | 99 | 0 |
| SPGP0D02140 | XP_018221839.1 | 99 | 0 |
| SPGP0BK02890 | XP_018221194.1 | 98 | 0 |
| SPGP0RK00140 | XP_018224038.1 | 99 | 0 |
| SPGP0WQ00190 | XP_018222272.1 | 99 | 0 |
| SPGP0DN00110 | XP_018218993.1 | 98 | 0 |
| SPGP0BE01020 | XP_018223232.1 | 99 | 0 |
| SPGP0BE03650 | XP_018222986.1 | 99 | 0 |
| SPGP0OL00110 | XP_018222287.1 | 99 | 0 |
| SPGP0MF00150 | XP_018222205.1 | 99 | 0 |
| SPGP0U00300 | XP_018224106.1 | 99 | 0 |
| SPGP0KW00100 | XP_018220728.1 | 99 | 0 |
| SPGP0FW00240 | XP_018219572.1 | 99 | 0 |
| SPGP0PZ00270 | XP_018222419.1 | 99 | 0 |
| SPGP0C00250 | XP_018220338.1 | 99 | 0 |
| SPGP0BI02010 | XP_018223679.1 | 99 | 0 |
| SPGP0U00290 | XP_018224107.1 | 99 | 0 |
| SPGP0OF00340 | XP_018222715.1 | 100 | 0 |
| SPGP0U00710 | XP_018224067.1 | 99 | 0 |
| SPGP0M00110 | XP_018220857.1 | 99 | 0 |
| SPGP0BK01450 | XP_018221330.1 | 99 | 0 |
| SPGP0D00410 | XP_018221674.1 | 98 | 1 |
| SPGP0K01930 | XP_018219397.1 | 99 | 0 |
| SPGP0IC00100 | XP_018222797.1 | 100 | 0 |
| SPGP0D01530 | XP_018221781.1 | 99 | 0 |
| SPGP0C01410 | XP_018220223.1 | 99 | 0 |
| SPGP0WI00220 | XP_018222130.1 | 98 | 0 |
| SPGP0C02970 | XP_018220076.1 | 99 | 0 |
| SPGP0K01570 | XP_018219431.1 | 99 | 0 |
| SPGP0TO00150 | XP_018222809.1 | 99 | 0 |
| SPGP0HY00100 | XP_018223987.1 | 99 | 1 |
| SPGP0BE01250 | XP_018223209.1 | 99 | 0 |
| SPGP0FM00140 | XP_018222150.1 | 99 | 0 |
| SPGP0NI00130 | XP_018223909.1 | 99 | 0 |
| SPGP0NV00150 | XP_018223453.1 | 99 | 0 |
| SPGP0VP00110 | XP_018219950.1 | 99 | 0 |

Create PDF in your applications with the Pdfcrowd HTML to PDF API PDFCROWD

| | | | |
|---|---|---|---|
| SPGP0BE03630 | XP_018222988.1 | 99 | 0 |
| SPGP0WI00180 | XP_018222126.1 | 98 | 0 |
| SPGP0VN00170 | XP_018219807.1 | 99 | 0 |
| SPGP0K02020 | XP_018219389.1 | 99 | 0 |
| SPGP0C01950 | XP_018220175.1 | 99 | 0 |
| SPGP0BI02610 | XP_018223621.1 | 99 | 0 |
| SPGP0OI00740 | XP_018218876.1 | 99 | 0 |
| SPGP0TM00170 | XP_018219881.1 | 99 | 0 |
| SPGP0BE00610 | XP_018223273.1 | 99 | 0 |
| SPGP0K01640 | XP_018219425.1 | 99 | 0 |
| SPGP0BE02070 | XP_018223130.1 | 99 | 0 |
| SPGP0VZ00260 | XP_018223890.1 | 99 | 0 |
| SPGP0P00680 | XP_018221586.1 | 99 | 0 |
| SPGP0LH00130 | XP_018221034.1 | 100 | 0 |
| SPGP0BI00950 | XP_018223784.1 | 99 | 0 |
| SPGP0TE00180 | XP_018219900.1 | 99 | 0 |
| SPGP0D01260 | XP_018221754.1 | 99 | 0 |
| SPGP0P00430 | XP_018221610.1 | 99 | 0 |
| SPGP0JP00120 | XP_018218849.1 | 100 | 0 |
| SPGP0K01420 | XP_018219445.1 | 99 | 0 |
| SPGP0BI01160 | XP_018223764.1 | 99 | 0 |
| SPGP0DS00130 | XP_018220393.1 | 99 | 0 |
| SPGP0DO00110 | XP_018219966.1 | 99 | 0 |
| SPGP0BE03030 | XP_018223040.1 | 99 | 0 |
| SPGP0BK02810 | XP_018221201.1 | 99 | 0 |
| SPGP0M00880 | XP_018220932.1 | 99 | 0 |
| SPGP0BI00500 | XP_018223828.1 | 99 | 0 |
| SPGP0OZ00100 | XP_018222397.1 | 99 | 0 |
| SPGP0C03280 | XP_018220046.1 | 99 | 0 |
| SPGP0BK02510 | XP_018221227.1 | 99 | 0 |
| SPGP0P00880 | XP_018221566.1 | 99 | 0 |
| SPGP0P00950 | XP_018221559.1 | 100 | 0 |
| SPGP0K00850 | XP_018219501.1 | 99 | 0 |
| SPGP0BI01510 | XP_018223730.1 | 99 | 0 |
| SPGP0D02690 | XP_018221892.1 | 99 | 0 |
| SPGP0NI00110 | XP_018223907.1 | 99 | 0 |
| SPGP0U00520 | XP_018224085.1 | 99 | 0 |
| SPGP0LX00150 | XP_018219149.1 | 99 | 0 |
| SPGP0BI02860 | XP_018223597.1 | 99 | 0 |
| SPGP0MF00110 | XP_018222209.1 | 98 | 0 |

Create PDF in your applications with the Pdfcrowd HTML to PDF API PDFCROWD

| | | | |
|---|---|---|---|
| SPGP0BI01570 | XP_018223723.1 | 99 | 0 |
| SPGP0DU00130 | XP_018222581.1 | 99 | 0 |
| SPGP0D03030 | XP_018221923.1 | 99 | 0 |
| SPGP0M00870 | XP_018220931.1 | 99 | 0 |
| SPGP0BE00240 | XP_018223310.1 | 99 | 0 |
| SPGP0BI00370 | XP_018223841.1 | 99 | 0 |
| SPGP0BE00340 | XP_018223300.1 | 99 | 0 |
| SPGP0CY00100 | XP_018223953.1 | 99 | 0 |
| SPGP0UN00200 | XP_018220778.1 | 99 | 0 |
| SPGP0BK01350 | XP_018221339.1 | 99 | 0 |
| SPGP0C02300 | XP_018220141.1 | 99 | 0 |
| SPGP0C00970 | XP_018220267.1 | 99 | 0 |
| SPGP0BK03300 | XP_018221154.1 | 99 | 0 |
| SPGP0D00760 | XP_018221708.1 | 99 | 0 |
| SPGP0D00190 | XP_018221652.1 | 99 | 0 |
| SPGP0BI02570 | XP_018223625.1 | 99 | 0 |
| SPGP0KN00160 | XP_018220572.1 | 100 | 0 |
| SPGP0BI02430 | XP_018223639.1 | 100 | 0 |
| SPGP0C02560 | XP_018220115.1 | 99 | 0 |
| SPGP0RI00170 | XP_018220495.1 | 100 | 0 |
| SPGP0M00240 | XP_018220869.1 | 99 | 0 |
| SPGP0D00810 | XP_018221712.1 | 99 | 0 |
| SPGP0P00600 | XP_018221593.1 | 99 | 0 |
| SPGP0OT00120 | XP_018219131.1 | 99 | 0 |
| SPGP0D03180 | XP_018221938.1 | 99 | 0 |
| SPGP0SM00130 | XP_018221108.1 | 99 | 0 |
| SPGP0D02210 | XP_018221846.1 | 99 | 0 |
| SPGP0IQ00110 | XP_018221042.1 | 99 | 0 |
| SPGP0BI03130 | XP_018223570.1 | 99 | 0 |
| SPGP0N00530 | XP_018219241.1 | 99 | 0 |
| SPGP0PG00170 | XP_018219172.1 | 99 | 0 |
| SPGP0OF00320 | XP_018222713.1 | 99 | 0 |
| SPGP0P00240 | XP_018221629.1 | 99 | 0 |
| SPGP0EY00150 | XP_018219093.1 | 99 | 0 |
| SPGP0BE04580 | XP_018222893.1 | 99 | 0 |
| SPGP0K02500 | XP_018219341.1 | 100 | 0 |
| SPGP0BE01170 | XP_018223217.1 | 100 | 0 |
| SPGP0D02360 | XP_018221861.1 | 99 | 0 |
| SPGP0JZ00120 | XP_018223415.1 | 99 | 0 |
| SPGP0DQ00120 | XP_018219671.1 | 99 | 0 |

Create PDF in your applications with the Pdfcrowd HTML to PDF API PDFCROWD

| | | | |
|---|---|---|---|
| SPGP0IA00130 | XP_018220582.1 | 99 | 0 |
| SPGP0OX00280 | XP_018219717.1 | 99 | 0 |
| SPGP0BK01630 | XP_018221313.1 | 99 | 0 |
| SPGP0OI00430 | XP_018218907.1 | 99 | 0 |
| SPGP0D01600 | XP_018221788.1 | 99 | 0 |
| SPGP0VH00170 | XP_018219688.1 | 99 | 0 |
| SPGP0BK01180 | XP_018221356.1 | 99 | 0 |
| SPGP0C01540 | XP_018220212.1 | 99 | 0 |
| SPGP0K01360 | XP_018219451.1 | 99 | 0 |
| SPGP0D00200 | XP_018221653.1 | 99 | 0 |
| SPGP0KZ00160 | XP_018223944.1 | 99 | 0 |
| SPGP0P01770 | XP_018221482.1 | 99 | 0 |
| SPGP0N00630 | XP_018219250.1 | 99 | 0 |
| SPGP0XF00150 | XP_018220851.1 | 99 | 0 |
| SPGP0BE01570 | XP_018223178.1 | 99 | 0 |
| SPGP0GH00190 | XP_018218959.1 | 99 | 0 |
| SPGP0FW00220 | XP_018219574.1 | 99 | 0 |
| SPGP0SP00110 | XP_018223961.1 | 99 | 0 |
| SPGP0P01750 | XP_018221484.1 | 99 | 0 |
| SPGP0GF00320 | XP_018219629.1 | 99 | 0 |
| SPGP0LA00160 | XP_018220845.1 | 99 | 0 |
| SPGP0C02230 | XP_018220148.1 | 99 | 0 |
| SPGP0C01710 | XP_018220196.1 | 99 | 0 |
| SPGP0KZ00250 | XP_018223937.1 | 99 | 0 |
| SPGP0FM00250 | XP_018222160.1 | 99 | 0 |
| SPGP0P00460 | XP_018221607.1 | 99 | 0 |
| SPGP0BI03570 | XP_018223525.1 | 99 | 0 |
| SPGP0BI01980 | XP_018223682.1 | 99 | 0 |
| SPGP0PE00290 | XP_018222374.1 | 99 | 0 |
| SPGP0KL00110 | XP_018221103.1 | 100 | 0 |
| SPGP0WW00130 | XP_018222075.1 | 99 | 0 |
| SPGP0U00430 | XP_018224094.1 | 99 | 0 |
| SPGP0D01120 | XP_018221742.1 | 99 | 0 |
| SPGP0BI04100 | XP_018223476.1 | 100 | 0 |
| SPGP0D02110 | XP_018221836.1 | 100 | 0 |
| SPGP0BI03850 | XP_018223499.1 | 99 | 0 |
| SPGP0BK01430 | XP_018221331.1 | 99 | 0 |
| SPGP0C02910 | XP_018220082.1 | 99 | 0 |
| SPGP0QV00180 | XP_018220002.1 | 99 | 0 |
| SPGP0KS00120 | XP_018220460.1 | 99 | 0 |

Create PDF in your applications with the Pdfcrowd HTML to PDF API PDFCROWD

| | | | |
|---|---|---|---|
| SPGP0WA00160 | XP_018219850.1 | 98 | 0 |
| SPGP0U00840 | XP_018224054.1 | 99 | 0 |
| SPGP0BI00580 | XP_018223820.1 | 100 | 0 |
| SPGP0BI02450 | XP_018223637.1 | 99 | 0 |
| SPGP0BE00580 | XP_018223276.1 | 99 | 0 |
| SPGP0MC00140 | XP_018222666.1 | 99 | 0 |
| SPGP0BI00130 | XP_018223865.1 | 99 | 0 |
| SPGP0WP00100 | XP_018222226.1 | 99 | 0 |
| SPGP0C01930 | XP_018220177.1 | 100 | 0 |
| SPGP0JR00100 | XP_018219031.1 | 99 | 0 |
| SPGP0BI02180 | XP_018223663.1 | 99 | 0 |
| SPGP0ET00100 | XP_018222441.1 | 99 | 0 |
| SPGP0C00510 | XP_018220313.1 | 99 | 0 |
| SPGP0M00260 | XP_018220871.1 | 99 | 0 |
| SPGP0K00670 | XP_018219519.1 | 99 | 0 |
| SPGP0AH00110 | XP_018219889.1 | 99 | 0 |
| SPGP0P00410 | XP_018221612.1 | 99 | 0 |
| SPGP0OE00230 | XP_018219597.1 | 100 | 0 |
| SPGP0BE03580 | XP_018222993.1 | 99 | 0 |
| SPGP0K00570 | XP_018219527.1 | 99 | 0 |
| SPGP0SL00190 | XP_018220477.1 | 99 | 0 |
| SPGP0BE02180 | XP_018223120.1 | 99 | 0 |
| SPGP0BK03190 | XP_018221165.1 | 98 | 1 |
| SPGP0JP00200 | XP_018218857.1 | 99 | 0 |
| SPGP0BI01370 | XP_018223744.1 | 99 | 0 |
| SPGP0BK03050 | XP_018221179.1 | 99 | 0 |
| SPGP0N00400 | XP_018219228.1 | 99 | 0 |
| SPGP0BE02260 | XP_018223112.1 | 99 | 0 |
| SPGP0PV00130 | XP_018218978.1 | 99 | 0 |
| SPGP0CS00100 | XP_018220818.1 | 99 | 0 |
| SPGP0NQ00150 | XP_018220750.1 | 99 | 0 |
| SPGP0BK03060 | XP_018221178.1 | 98 | 0 |
| SPGP0M00430 | XP_018220888.1 | 98 | 0 |
| SPGP0P01960 | XP_018221463.1 | 97 | 0 |
| SPGP0N01270 | XP_018219312.1 | 99 | 0 |
| SPGP0BE04840 | XP_018222867.1 | 98 | 1 |
| SPGP0U00100 | XP_018224123.1 | 96 | 0 |
| SPGP0BK03280 | XP_018221156.1 | 99 | 0 |
| SPGP0CZ00150 | XP_018219910.1 | 99 | 0 |
| SPGP0C02670 | XP_018220106.1 | 99 | 0 |

Create PDF in your applications with the Pdfcrowd HTML to PDF API PDFCROWD

| | | | |
|---|---|---|---|
| SPGP0C01550 | XP_018220211.1 | 99 | 0 |
| SPGP0BI02620 | XP_018223620.1 | 99 | 0 |
| SPGP0BE01820 | XP_018223153.1 | 99 | 0 |
| SPGP0BI01490 | XP_018223732.1 | 99 | 0 |
| SPGP0BK02080 | XP_018221269.1 | 99 | 0 |
| SPGP0BI03110 | XP_018223572.1 | 99 | 0 |
| SPGP0TK00120 | XP_018222726.1 | 100 | 0 |
| SPGP0MDtrna3W | DI49_2157 | 100 | 0 |
| SPGP0BE04030 | XP_018222950.1 | 99 | 0 |
| SPGP0C01230 | XP_018220241.1 | 99 | 0 |
| SPGP0SM00110 | XP_018221106.1 | 99 | 0 |
| SPGP0RQ00130 | XP_018221124.1 | 100 | 0 |
| SPGP0NQ00190 | XP_018220746.1 | 99 | 0 |
| SPGP0HT00160 | XP_018219008.1 | 99 | 0 |
| SPGP0BK00830 | XP_018221390.1 | 99 | 0 |
| SPGP0BK02260 | XP_018221251.1 | 99 | 0 |
| SPGP0HX00170 | XP_018221008.1 | 99 | 0 |
| SPGP0BK01520 | XP_018221324.1 | 99 | 0 |
| SPGP0BI02240 | XP_018223658.1 | 99 | 0 |
| SPGP0BE03910 | XP_018222960.1 | 99 | 0 |
| SPGP0P01970 | XP_018221462.1 | 97 | 0 |
| SPGP0BI01910 | XP_018223689.1 | 99 | 0 |
| SPGP0BK01610 | XP_018221315.1 | 99 | 0 |
| SPGP0BE01730 | XP_018223162.1 | 100 | 0 |
| SPGP0DN00170 | XP_018218987.1 | 99 | 0 |
| SPGP0OZ00180 | XP_018222405.1 | 99 | 0 |
| SPGP0OI00390 | XP_018218911.1 | 100 | 0 |
| SPGP0D00950 | XP_018221726.1 | 99 | 0 |
| SPGP0C00790 | XP_018220285.1 | 99 | 0 |
| SPGP0P01280 | XP_018221530.1 | 99 | 0 |
| SPGP0K00380 | XP_018219547.1 | 99 | 0 |
| SPGP0K00110 | XP_018218751.1 | 99 | 0 |
| SPGP0PE00300 | XP_018222375.1 | 99 | 0 |
| SPGP0MT00130 | XP_018219139.1 | 99 | 0 |
| SPGP0K01380 | XP_018219449.1 | 99 | 0 |
| SPGP0BI01740 | XP_018223706.1 | 99 | 0 |
| SPGP0BE03770 | XP_018222974.1 | 99 | 0 |
| SPGP0K01530 | XP_018219435.1 | 99 | 0 |
| SPGP0EW00120 | XP_018222291.1 | 99 | 0 |

Create PDF in your applications with the Pdfcrowd HTML to PDF API PDFCROWD

| | | | |
|---|---|---|---|
| SPGP0NI00150 | XP_018223911.1 | 99 | 0 |
| SPGP0SU00150 | XP_018218975.1 | 98 | 1 |
| SPGP0JF00190 | XP_018222317.1 | 99 | 0 |
| SPGP0BK00560 | XP_018221416.1 | 99 | 0 |
| SPGP0KZ00230 | XP_018223939.1 | 99 | 0 |
| SPGP0BI00830 | XP_018223795.1 | 99 | 0 |
| SPGP0VN00270 | XP_018219817.1 | 99 | 0 |
| SPGP0OY00120 | XP_018220970.1 | 99 | 0 |
| SPGP0GV00140 | XP_018218864.1 | 99 | 0 |
| SPGP0OF00300 | XP_018222711.1 | 99 | 1 |
| SPGP0P01560 | XP_018221503.1 | 99 | 0 |
| SPGP0KL00180 | XP_018221096.1 | 99 | 0 |
| SPGP0JD00100 | XP_018223361.1 | 99 | 0 |
| SPGP0BE03380 | XP_018223009.1 | 99 | 0 |
| SPGP0D00690 | XP_018221701.1 | 99 | 0 |
| SPGP0BE01210 | XP_018223213.1 | 99 | 0 |
| SPGP0BE02830 | XP_018223058.1 | 98 | 0 |
| SPGP0D01240 | XP_018221752.1 | 99 | 0 |
| SPGP0U00820 | XP_018224056.1 | 99 | 0 |
| SPGP0BI01200 | XP_018223760.1 | 99 | 0 |
| SPGP0SH00120 | XP_018223970.1 | 99 | 0 |
| SPGP0P00920 | XP_018221562.1 | 99 | 0 |
| SPGP0C02240 | XP_018220147.1 | 99 | 0 |
| SPGP0BI03870 | XP_018223497.1 | 99 | 0 |
| SPGP0BI00650 | XP_018223813.1 | 99 | 0 |
| SPGP0BE01880 | XP_018223148.1 | 99 | 0 |
| SPGP0RE00130 | XP_018220397.1 | 99 | 0 |
| SPGP0BE04480 | XP_018222903.1 | 99 | 0 |
| SPGP0BI02650 | XP_018223617.1 | 99 | 0 |
| SPGP0ET00120 | XP_018222443.1 | 99 | 0 |
| SPGP0GH00150 | XP_018218963.1 | 99 | 0 |
| SPGP0JF00270 | XP_018222325.1 | 99 | 0 |
| SPGP0BE04470 | XP_018222904.1 | 99 | 0 |
| SPGP0K01600 | XP_018219428.1 | 99 | 0 |
| SPGP0LV00110 | XP_018223372.1 | 99 | 0 |
| SPGP0C00430 | XP_018220320.1 | 99 | 0 |
| SPGP0M00850 | XP_018220929.1 | 99 | 0 |
| SPGP0BI01420 | XP_018223739.1 | 99 | 0 |
| SPGP0CR00120 | XP_018222212.1 | 99 | 0 |
| SPGP0BE00930 | XP_018223241.1 | 99 | 0 |

Create PDF in your applications with the Pdfcrowd HTML to PDF API PDFCROWD

| | | | |
|---|---|---|---|
| SPGP0K02380 | XP_018219353.1 | 99 | 0 |
| SPGP0P00810 | XP_018221573.1 | 99 | 0 |
| SPGP0RE00190 | XP_018220402.1 | 99 | 0 |
| SPGP0D02430 | XP_018221868.1 | 99 | 0 |
| SPGP0BK00720 | XP_018221401.1 | 99 | 0 |
| SPGP0IK00170 | XP_018222114.1 | 99 | 0 |
| SPGP0K00170 | XP_018219568.1 | 99 | 0 |
| SPGP0D02800 | XP_018221903.1 | 100 | 0 |
| SPGP0D01860 | XP_018221812.1 | 99 | 0 |
| SPGP0BE03220 | XP_018223024.1 | 99 | 0 |
| SPGP0BE00520 | XP_018223282.1 | 99 | 0 |
| SPGP0C02320 | XP_018220139.1 | 99 | 0 |
| SPGP0C02630 | XP_018220110.1 | 99 | 0 |
| SPGP0C02400 | XP_018220131.1 | 99 | 0 |
| SPGP0BK00370 | XP_018221435.1 | 99 | 0 |
| SPGP0BI01680 | XP_018223712.1 | 99 | 0 |
| SPGP0N00420 | XP_018219230.1 | 99 | 0 |
| SPGP0P00770 | XP_018221577.1 | 99 | 0 |
| SPGP0BI02410 | XP_018223641.1 | 99 | 0 |
| SPGP0RC00130 | XP_018222088.1 | 99 | 0 |
| SPGP0GY00180 | XP_018223394.1 | 99 | 0 |
| SPGP0N00490 | XP_018219237.1 | 99 | 0 |
| SPGP0BI00330 | XP_018223845.1 | 99 | 0 |
| SPGP0TK00140 | XP_018222728.1 | 99 | 0 |
| SPGP0BI01530 | XP_018223727.1 | 99 | 0 |
| SPGP0N00890 | XP_018219276.1 | 100 | 0 |
| SPGP0BK01390 | XP_018221335.1 | 99 | 0 |
| SPGP0UJ00110 | XP_018223344.1 | 99 | 0 |
| SPGP0EX00120 | XP_018223421.1 | 100 | 0 |
| SPGP0BI01880 | XP_018223692.1 | 99 | 0 |
| SPGP0BE00100 | XP_018223460.1 | 90 | 0 |
| SPGP0BI03680 | XP_018223514.1 | 99 | 0 |
| SPGP0VG00160 | XP_018223874.1 | 99 | 0 |
| SPGP0KL00210 | XP_018221094.1 | 99 | 0 |
| SPGP0IE00150 | XP_018222761.1 | 100 | 0 |
| SPGP0BI01820 | XP_018223698.1 | 99 | 0 |
| SPGP0C01060 | XP_018220258.1 | 99 | 0 |
| SPGP0C00670 | XP_018220297.1 | 99 | 0 |
| SPGP0BI03460 | XP_018223536.1 | 100 | 0 |
| SPGP0C01210 | XP_018220243.1 | 99 | 0 |

Create PDF in your applications with the Pdfcrowd HTML to PDF API PDFCROWD

| | | | |
|---|---|---|---|
| SPGP0BE02320 | XP_018223106.1 | 99 | 0 |
| SPGP0EW00140 | XP_018222293.1 | 99 | 0 |
| SPGP0D02090 | XP_018221834.1 | 100 | 0 |
| SPGP0D00990 | XP_018221730.1 | 99 | 0 |
| SPGP0OI00220 | XP_018218925.1 | 100 | 0 |
| SPGP0TC00210 | XP_018219692.1 | 99 | 0 |
| SPGP0D01350 | XP_018221763.1 | 99 | 0 |
| SPGP0K00940 | XP_018219493.1 | 99 | 1 |
| SPGP0BK02420 | XP_018221236.1 | 99 | 0 |
| SPGP0UR00140 | XP_018222340.1 | 99 | 0 |
| SPGP0VO00230 | XP_018223446.1 | 99 | 0 |
| SPGP0M00300 | XP_018220875.1 | 99 | 0 |
| SPGP0BK02330 | XP_018221244.1 | 99 | 0 |
| SPGP0PA00160 | XP_018220468.1 | 99 | 0 |
| SPGP0K01750 | XP_018219414.1 | 99 | 0 |
| SPGP0P00910 | XP_018221563.1 | 99 | 0 |
| SPGP0US00120 | XP_018219154.1 | 99 | 0 |
| SPGP0D03050 | XP_018221925.1 | 99 | 0 |
| SPGP0C03510 | XP_018220023.1 | 99 | 0 |
| SPGP0SW00140 | XP_018220560.1 | 99 | 0 |
| SPGP0D00970 | XP_018221728.1 | 99 | 0 |
| SPGP0C02510 | XP_018220120.1 | 99 | 0 |
| SPGP0P00200 | XP_018221633.1 | 99 | 0 |
| SPGP0K00340 | XP_018219551.1 | 99 | 0 |
| SPGP0MD00190 | XP_018222394.1 | 98 | 2 |
| SPGP0N01100 | XP_018219297.1 | 99 | 0 |
| SPGP0BK00540 | XP_018221418.1 | 99 | 0 |
| SPGP0BE03100 | XP_018223035.1 | 99 | 0 |
| SPGP0FE00110 | XP_018220539.1 | 100 | 0 |
| SPGP0M00700 | XP_018220914.1 | 99 | 0 |
| SPGP0CH00130 | XP_018219954.1 | 99 | 0 |
| SPGP0FE00260 | XP_018220525.1 | 99 | 0 |
| SPGP0BK00600 | XP_018221412.1 | 99 | 0 |
| SPGP0GY00140 | XP_018223398.1 | 99 | 0 |
| SPGP0FU00100 | XP_018222654.1 | 98 | 0 |
| SPGP0D01620 | XP_018221790.1 | 99 | 0 |
| SPGP0TQ00130 | XP_018220608.1 | 99 | 0 |
| SPGP0PZ00390 | XP_018222407.1 | 99 | 0 |
| SPGP0BI00810 | XP_018223797.1 | 99 | 0 |
| SPGP0CT00100 | XP_018219569.1 | 99 | 0 |

Create PDF in your applications with the Pdfcrowd HTML to PDF API PDFCROWD

| | | | |
|---|---|---|---|
| SPGP0OI00130 | XP_018218934.1 | 100 | 0 |
| SPGP0FM00330 | XP_018222168.1 | 99 | 0 |
| SPGP0U00800 | XP_018224058.1 | 99 | 0 |
| SPGP0BE01550 | XP_018223180.1 | 99 | 0 |
| SPGP0WJ00100 | XP_018221060.1 | 100 | 0 |
| SPGP0BE01230 | XP_018223211.1 | 99 | 0 |
| SPGP0BI00670 | XP_018223811.1 | 99 | 0 |
| SPGP0C01960 | XP_018220174.1 | 99 | 0 |
| SPGP0C00830 | XP_018220281.1 | 100 | 0 |
| SPGP0D02200 | XP_018221845.1 | 99 | 0 |
| SPGP0M01040 | XP_018220948.1 | 99 | 0 |
| SPGP0BU00190 | XP_018222258.1 | 99 | 0 |
| SPGP0N00840 | XP_018219271.1 | 99 | 0 |
| SPGP0N00910 | XP_018219278.1 | 99 | 0 |
| SPGP0K01880 | XP_018219401.1 | 99 | 0 |
| SPGP0C02170 | XP_018220154.1 | 99 | 0 |
| SPGP0BE01940 | XP_018223141.1 | 99 | 0 |
| SPGP0RK00100 | XP_018224042.1 | 98 | 1 |
| SPGP0WY00120 | XP_018220444.1 | 99 | 0 |
| SPGP0UB00100 | XP_018219035.1 | 100 | 0 |
| SPGP0M00490 | XP_018220894.1 | 99 | 0 |
| SPGP0BK02160 | XP_018221261.1 | 99 | 0 |
| SPGP0BE04200 | XP_018222933.1 | 99 | 0 |
| SPGP0D02550 | XP_018221879.1 | 99 | 0 |
| SPGP0BE01340 | XP_018223200.1 | 99 | 0 |
| SPGP0K00320 | XP_018219553.1 | 99 | 0 |
| SPGP0OI00780 | XP_018218872.1 | 99 | 0 |
| SPGP0BE03090 | XP_018223036.1 | 99 | 0 |
| SPGP0IT00100 | XP_018222437.1 | 99 | 0 |
| SPGP0N00440 | XP_018219232.1 | 99 | 0 |
| SPGP0DS00110 | XP_018220391.1 | 99 | 0 |
| SPGP0BI00840 | XP_018223794.1 | 99 | 0 |
| SPGP0TX00110 | XP_018222348.1 | 99 | 0 |
| SPGP0D00170 | XP_018221650.1 | 99 | 0 |
| SPGP0QK00140 | XP_018220386.1 | 99 | 0 |
| SPGP0BK01870 | XP_018221289.1 | 99 | 0 |
| SPGP0FN00180 | XP_018223385.1 | 99 | 0 |
| SPGP0BE04410 | XP_018222911.1 | 99 | 0 |
| SPGP0FY00110 | XP_018222556.1 | 99 | 0 |
| SPGP0PZ00180 | XP_018222427.1 | 99 | 0 |

Create PDF in your applications with the Pdfcrowd HTML to PDF API PDFCROWD

| | | | |
|---|---|---|---|
| SPGP0GN00130 | XP_018222585.1 | 100 | 0 |
| SPGP0BI03600 | XP_018223522.1 | 99 | 0 |
| SPGP0N00100 | XP_018219199.1 | 97 | 0 |
| SPGP0LV00130 | XP_018223374.1 | 99 | 0 |
| SPGP0EO00120 | XP_018222479.1 | 99 | 0 |
| SPGP0P01100 | XP_018221547.1 | 99 | 0 |
| SPGP0BE01000 | XP_018223234.1 | 99 | 0 |
| SPGP0BE04070 | XP_018222946.1 | 99 | 0 |
| SPGP0C01090 | XP_018220255.1 | 99 | 0 |
| SPGP0K02560 | XP_018219335.1 | 99 | 0 |
| SPGP0C01890 | XP_018220181.1 | 99 | 0 |
| SPGP0C00750 | XP_018220289.1 | 99 | 0 |
| SPGP0BE01380 | XP_018223196.1 | 99 | 0 |
| SPGP0UF00100 | XP_018221041.1 | 100 | 0 |
| SPGP0D03150 | XP_018221935.1 | 99 | 0 |
| SPGP0JW00110 | XP_018220973.1 | 99 | 0 |
| SPGP0BI00220 | XP_018223856.1 | 99 | 0 |
| SPGP0K01240 | XP_018219463.1 | 99 | 0 |
| SPGP0C00560 | XP_018220308.1 | 99 | 0 |
| SPGP0C00320 | XP_018220331.1 | 99 | 0 |
| SPGP0K01030 | XP_018219484.1 | 99 | 0 |
| SPGP0C01070 | XP_018220257.1 | 99 | 0 |
| SPGP0TF00100 | XP_018222185.1 | 99 | 0 |
| SPGP0K00270 | XP_018219558.1 | 99 | 0 |
| SPGP0HZ00310 | XP_018219788.1 | 99 | 0 |
| SPGP0D03460 | XP_018221964.1 | 99 | 0 |
| SPGP0K00460 | XP_018219539.1 | 99 | 0 |
| SPGP0M00210 | XP_018220866.1 | 99 | 0 |
| SPGP0SW00220 | XP_018220552.1 | 99 | 0 |
| SPGP0ER00120 | XP_018218981.1 | 99 | 0 |
| SPGP0TG00130 | XP_018220663.1 | 97 | 3 |
| SPGP0BK03080 | XP_018221176.1 | 99 | 0 |
| SPGP0BK03360 | XP_018221148.1 | 99 | 0 |
| SPGP0D02450 | XP_018221870.1 | 99 | 0 |
| SPGP0C01720 | XP_018220195.1 | 99 | 0 |
| SPGP0N00340 | XP_018219222.1 | 99 | 0 |
| SPGP0K01280 | XP_018219459.1 | 99 | 0 |
| SPGP0BE00840 | XP_018223250.1 | 99 | 0 |
| SPGP0BK03500 | XP_018221137.1 | 99 | 0 |
| SPGP0D02780 | XP_018221901.1 | 99 | 0 |

Create PDF in your applications with the Pdfcrowd HTML to PDF API PDFCROWD

| | | | |
|---|---|---|---|
| SPGP0TL00160 | XP_018219082.1 | 98 | 1 |
| SPGP0BE01920 | XP_018223144.1 | 99 | 0 |
| SPGP0BK00470 | XP_018221425.1 | 99 | 0 |
| SPGP0BI00690 | XP_018223809.1 | 99 | 0 |
| SPGP0BI00560 | XP_018223822.1 | 99 | 0 |
| SPGP0K02340 | XP_018219357.1 | 99 | 0 |
| SPGP0WI00240 | XP_018222132.1 | 99 | 0 |
| SPGP0M00320 | XP_018220877.1 | 99 | 0 |
| SPGP0BE04190 | XP_018222934.1 | 99 | 0 |
| SPGP0EL00130 | XP_018223365.1 | 99 | 0 |
| SPGP0FE00310 | XP_018220520.1 | 99 | 0 |
| SPGP0ED00120 | XP_018222600.1 | 99 | 0 |
| SPGP0D00140 | XP_018221647.1 | 99 | 0 |
| SPGP0D02070 | XP_018221832.1 | 99 | 0 |
| SPGP0NQ00220 | XP_018220743.1 | 99 | 0 |
| SPGP0HT00150 | XP_018219007.1 | 99 | 0 |
| SPGP0BK00500 | XP_018221422.1 | 99 | 0 |
| SPGP0BE02770 | XP_018223063.1 | 99 | 0 |
| SPGP0BI03690 | XP_018223513.1 | 99 | 0 |
| SPGP0PS00120 | XP_018223358.1 | 99 | 0 |
| SPGP0BK00140 | XP_018221454.1 | 99 | 0 |
| SPGP0BI01840 | XP_018223696.1 | 99 | 0 |
| SPGP0K02000 | XP_018219391.1 | 100 | 0 |
| SPGP0NI00170 | XP_018223913.1 | 99 | 0 |
| SPGP0FN00160 | XP_018223387.1 | 99 | 0 |
| SPGP0JF00250 | XP_018222323.1 | 99 | 0 |
| SPGP0UP00100 | XP_018222502.1 | 99 | 0 |
| SPGP0N00460 | XP_018219234.1 | 100 | 0 |
| SPGP0KX00110 | XP_018223363.1 | 99 | 0 |
| SPGP0BI01260 | XP_018223754.1 | 99 | 0 |
| SPGP0GX00110 | XP_018222535.1 | 99 | 0 |
| SPGP0BE02690 | XP_018223071.1 | 99 | 0 |
| SPGP0BE02870 | XP_018223055.1 | 99 | 0 |
| SPGP0MF00130 | XP_018222207.1 | 99 | 0 |
| SPGP0BK01680 | XP_018221309.1 | 100 | 0 |
| SPGP0BK01760 | XP_018221301.1 | 99 | 0 |
| SPGP0BI02170 | XP_018223664.1 | 99 | 0 |
| SPGP0BE03980 | XP_018222954.1 | 99 | 0 |
| SPGP0XB00100 | XP_018219020.1 | 99 | 0 |
| SPGP0BE03610 | XP_018222990.1 | 99 | 0 |

Create PDF in your applications with the Pdfcrowd HTML to PDF API PDFCROWD

| | | | |
|---|---|---|---|
| SPGP0OH00100 | XP_018220984.1 | 99 | 0 |
| SPGP0CZ00130 | XP_018219908.1 | 99 | 0 |
| SPGP0BK01110 | XP_018221363.1 | 99 | 0 |
| SPGP0BK02460 | XP_018221232.1 | 99 | 0 |
| SPGP0K01710 | XP_018219418.1 | 100 | 0 |
| SPGP0AA00160 | XP_018221079.1 | 98 | 2 |
| SPGP0QJ00100 | XP_018219935.1 | 91 | 1 |
| SPGP0D02050 | XP_018221830.1 | 99 | 0 |
| SPGP0NI00180 | XP_018223914.1 | 99 | 0 |
| SPGP0BT00130 | XP_018220736.1 | 99 | 0 |
| SPGP0P01640 | XP_018221495.1 | 99 | 0 |
| SPGP0D01640 | XP_018221792.1 | 100 | 0 |
| SPGP0BI02590 | XP_018223623.1 | 99 | 0 |
| SPGP0JA00110 | XP_018223407.1 | 99 | 0 |
| SPGP0BE04320 | XP_018222921.1 | 99 | 0 |
| SPGP0BI02770 | XP_018223606.1 | 99 | 0 |
| SPGP0BE00730 | XP_018223260.1 | 99 | 0 |
| SPGP0BE00220 | XP_018223312.1 | 99 | 0 |
| SPGP0KZ00140 | XP_018223946.1 | 99 | 0 |
| SPGP0C00690 | XP_018220295.1 | 99 | 0 |
| SPGP0BI00470 | XP_018223831.1 | 99 | 0 |
| SPGP0BI01410 | XP_018223740.1 | 99 | 0 |
| SPGP0D02880 | XP_018221910.1 | 98 | 0 |
| SPGP0C03170 | XP_018220057.1 | 99 | 0 |
| SPGP0BI01000 | XP_018223779.1 | 99 | 0 |
| SPGP0C00340 | XP_018220329.1 | 99 | 0 |
| SPGP0BI01150 | XP_018223765.1 | 99 | 0 |
| SPGP0P01260 | XP_018221532.1 | 99 | 0 |
| SPGP0M00920 | XP_018220936.1 | 99 | 0 |
| SPGP0GW00200 | XP_018220821.1 | 99 | 0 |
| SPGP0WY00230 | XP_018220434.1 | 99 | 0 |
| SPGP0VD00130 | XP_018219836.1 | 99 | 0 |
| SPGP0BI01220 | XP_018223758.1 | 99 | 0 |
| SPGP0PZ00230 | XP_018222423.1 | 99 | 0 |
| SPGP0UQ00110 | XP_018220690.1 | 99 | 0 |
| SPGP0PZ00350 | XP_018222411.1 | 99 | 0 |
| SPGP0P01880 | XP_018221471.1 | 98 | 0 |
| SPGP0M00830 | XP_018220927.1 | 99 | 0 |
| SPGP0BI00850 | XP_018223793.1 | 99 | 0 |
| SPGP0WQ00170 | XP_018222270.1 | 99 | 0 |

Create PDF in your applications with the Pdfcrowd HTML to PDF API PDFCROWD

| | | | |
|---|---|---|---|
| SPGP0M00690 | XP_018220913.1 | 100 | 0 |
| SPGP0D00770 | XP_018221709.1 | 99 | 0 |
| SPGP0BI02710 | XP_018223611.1 | 99 | 0 |
| SPGP0BK01270 | XP_018221347.1 | 99 | 0 |
| SPGP0M00450 | XP_018220890.1 | 99 | 0 |
| SPGP0BK00750 | XP_018221398.1 | 99 | 0 |
| SPGP0S00140 | XP_018219656.1 | 99 | 0 |
| SPGP0U00760 | XP_018224062.1 | 99 | 0 |
| SPGP0BI02790 | XP_018223604.1 | 100 | 0 |
| SPGP0WQ00130 | XP_018222266.1 | 98 | 0 |
| SPGP0K01870 | XP_018219402.1 | 99 | 0 |
| SPGP0C02990 | XP_018220074.1 | 99 | 0 |
| SPGP0BK02120 | XP_018221265.1 | 99 | 0 |
| SPGP0OW00150 | XP_018218812.1 | 99 | 0 |
| SPGP0D03240 | XP_018221944.1 | 99 | 0 |
| SPGP0NY00100 | XP_018219619.1 | 99 | 0 |
| SPGP0BE00500 | XP_018223284.1 | 99 | 0 |
| SPGP0K00200 | XP_018219565.1 | 99 | 0 |
| SPGP0K01470 | XP_018219441.1 | 100 | 0 |
| SPGP0BK01300 | XP_018221344.1 | 99 | 0 |
| SPGP0BE01630 | XP_018223172.1 | 99 | 0 |
| SPGP0BK00870 | XP_018221386.1 | 99 | 0 |
| SPGP0D02260 | XP_018221851.1 | 99 | 0 |
| SPGP0BE02250 | XP_018223113.1 | 99 | 0 |
| SPGP0IA00110 | XP_018220584.1 | 99 | 0 |
| SPGP0PJ00110 | XP_018222116.1 | 99 | 0 |
| SPGP0BE03850 | XP_018222966.1 | 99 | 0 |
| SPGP0K02060 | XP_018219385.1 | 99 | 0 |
| SPGP0BI01230 | XP_018223757.1 | 99 | 0 |
| SPGP0C00930 | XP_018220271.1 | 99 | 0 |
| SPGP0C03240 | XP_018220050.1 | 99 | 0 |
| SPGP0BE00980 | XP_018223236.1 | 99 | 0 |
| SPGP0TM00150 | XP_018219879.1 | 99 | 0 |
| SPGP0BK01040 | XP_018221370.1 | 99 | 0 |
| SPGP0D03510 | XP_018221969.1 | 99 | 0 |
| SPGP0K00130 | XP_018218753.1 | 99 | 0 |
| SPGP0D02290 | XP_018221854.1 | 100 | 0 |
| SPGP0C01460 | XP_018220219.1 | 99 | 0 |
| SPGP0BI00930 | XP_018223786.1 | 100 | 0 |
| SPGP0D00130 | XP_018221646.1 | 99 | 0 |

Create PDF in your applications with the Pdfcrowd HTML to PDF API PDFCROWD

| | | | |
|---|---|---|---|
| SPGP0P00160 | XP_018221637.1 | 99 | 0 |
| SPGP0D00370 | XP_018221670.1 | 100 | 0 |
| SPGP0QP00100 | XP_018219169.1 | 100 | 0 |
| SPGP0C03200 | XP_018220054.1 | 99 | 0 |
| SPGP0BI03620 | XP_018223520.1 | 99 | 0 |
| SPGP0P01660 | XP_018221493.1 | 99 | 0 |
| SPGP0U00180 | XP_018224118.1 | 99 | 0 |
| SPGP0D01590 | XP_018221787.1 | 99 | 0 |
| SPGP0MG00140 | XP_018219614.1 | 99 | 0 |
| SPGP0BI00260 | XP_018223852.1 | 99 | 0 |
| SPGP0D02380 | XP_018221863.1 | 100 | 0 |
| SPGP0CR00100 | XP_018222214.1 | 99 | 0 |
| SPGP0BK01860 | XP_018221290.1 | 99 | 0 |
| SPGP0WG00190 | XP_018221019.1 | 99 | 0 |
| SPGP0U00640 | XP_018224074.1 | 99 | 0 |
| SPGP0C00100 | XP_018220353.1 | 99 | 0 |
| SPGP0BE03810 | XP_018222970.1 | 99 | 0 |
| SPGP0D03380 | XP_018221956.1 | 99 | 0 |
| SPGP0DC00110 | XP_018222638.1 | 99 | 0 |
| SPGP0BE00710 | XP_018223262.1 | 99 | 0 |
| SPGP0K01070 | XP_018219480.1 | 99 | 0 |
| SPGP0RI00200 | XP_018220498.1 | 99 | 0 |
| SPGP0ltrna32L | DI49_5687 | 100 | 0 |
| SPGP0FW00150 | XP_018219581.1 | 99 | 0 |
| SPGP0D02030 | XP_018221828.1 | 99 | 0 |
| SPGP0UQ00130 | XP_018220688.1 | 99 | 0 |
| SPGP0BK00880 | XP_018221385.1 | 99 | 0 |
| SPGP0BK00740 | XP_018221399.1 | 99 | 0 |
| SPGP0BE00860 | XP_018223248.1 | 99 | 0 |
| SPGP0BK02900 | XP_018221193.1 | 99 | 0 |
| SPGP0M01140 | XP_018220958.1 | 99 | 0 |
| SPGP0BE02710 | XP_018223069.1 | 99 | 0 |
| SPGP0KU00130 | XP_018221002.1 | 100 | 0 |
| SPGP0BI01430 | XP_018223738.1 | 99 | 0 |
| SPGP0C03130 | XP_018220061.1 | 99 | 0 |
| SPGP0N00690 | XP_018219256.1 | 99 | 0 |
| SPGP0N00160 | XP_018219205.1 | 99 | 0 |
| SPGP0IB00120 | XP_018222674.1 | 97 | 1 |
| SPGP0P00300 | XP_018221623.1 | 98 | 2 |
| SPGP0BK03320 | XP_018221152.1 | 99 | 0 |

Create PDF in your applications with the Pdfcrowd HTML to PDF API PDFCROWD

| | | | |
|---|---|---|---|
| SPGP0JO00120 | XP_018223354.1 | 99 | 0 |
| SPGP0BK01250 | XP_018221349.1 | 99 | 0 |
| SPGP0PK00140 | XP_018220419.1 | 99 | 0 |
| SPGP0PE00270 | XP_018222372.1 | 99 | 0 |
| SPGP0BE02210 | XP_018223117.1 | 100 | 0 |
| SPGP0KZ00180 | XP_018223942.1 | 99 | 0 |
| SPGP0D01280 | XP_018221756.1 | 99 | 0 |
| SPGP0BE01430 | XP_018223191.1 | 99 | 0 |
| SPGP0BE04400 | XP_018222912.1 | 99 | 0 |
| SPGP0QV00160 | XP_018220004.1 | 100 | 0 |
| SPGP0BI03380 | XP_018223545.1 | 99 | 0 |
| SPGP0BE01650 | XP_018223170.1 | 99 | 0 |
| SPGP0K02260 | XP_018219365.1 | 99 | 0 |
| SPGP0BE01280 | XP_018223206.1 | 100 | 0 |
| SPGP0BE00380 | XP_018223296.1 | 99 | 0 |
| SPGP0MJ00130 | XP_018222084.1 | 99 | 0 |
| SPGP0U00410 | XP_018224096.1 | 99 | 0 |
| SPGP0BK01510 | XP_018221325.1 | 99 | 0 |
| SPGP0BI01920 | XP_018223688.1 | 99 | 0 |
| SPGP0FN00120 | XP_018223391.1 | 99 | 0 |
| SPGP0BI02030 | XP_018223677.1 | 99 | 0 |
| SPGP0XB00140 | XP_018219024.1 | 99 | 0 |
| SPGP0P01600 | XP_018221499.1 | 99 | 1 |
| SPGP0GA00100 | XP_018222800.1 | 100 | 0 |
| SPGP0P01430 | XP_018221515.1 | 99 | 0 |
| SPGP0C02800 | XP_018220093.1 | 99 | 0 |
| SPGP0NF00110 | XP_018222635.1 | 99 | 0 |
| SPGP0K00360 | XP_018219549.1 | 99 | 0 |
| SPGP0RQ00190 | XP_018221130.1 | 99 | 0 |
| SPGP0BE04080 | XP_018222945.1 | 99 | 0 |
| SPGP0U00340 | XP_018224102.1 | 99 | 0 |
| SPGP0FM00350 | XP_018222170.1 | 99 | 0 |
| SPGP0KB00160 | XP_018221118.1 | 99 | 0 |
| SPGP0Dtrna12P | DI49_2548 | 98 | 0 |
| SPGP0VA00100 | XP_018222570.1 | 99 | 1 |
| SPGP0C02370 | XP_018220134.1 | 99 | 0 |
| SPGP0IM00100 | XP_018220565.1 | 100 | 0 |
| SPGP0BI02550 | XP_018223627.1 | 99 | 0 |
| SPGP0BE03140 | XP_018223031.1 | 99 | 0 |
| SPGP0D03560 | XP_018221974.1 | 99 | 0 |

Create PDF in your applications with the Pdfcrowd HTML to PDF API PDFCROWD

| | | | |
|---|---|---|---|
| SPGP0BI00770 | XP_018223801.1 | 99 | 0 |
| SPGP0UR00200 | XP_018222334.1 | 99 | 0 |
| SPGP0C01380 | XP_018220226.1 | 99 | 0 |
| SPGP0BK01880 | XP_018221288.1 | 99 | 0 |
| SPGP0D00510 | XP_018221683.1 | 99 | 0 |
| SPGP0BE04860 | XP_018222865.1 | 99 | 0 |
| SPGP0BE03550 | XP_018222995.1 | 99 | 0 |
| SPGP0VS00100 | XP_018222520.1 | 100 | 0 |
| SPGP0BK00930 | XP_018221380.1 | 99 | 0 |
| SPGP0BE00260 | XP_018223308.1 | 99 | 0 |
| SPGP0C00380 | XP_018220325.1 | 99 | 0 |
| SPGP0FM00200 | XP_018222156.1 | 98 | 0 |
| SPGP0K02490 | XP_018219342.1 | 99 | 0 |
| SPGP0D01840 | XP_018221810.1 | 100 | 0 |
| SPGP0HD00110 | XP_018222515.1 | 98 | 0 |
| SPGP0N01050 | XP_018219292.1 | 99 | 0 |
| SPGP0FL00100 | XP_018219051.1 | 99 | 0 |
| SPGP0N01090 | XP_018219296.1 | 99 | 0 |
| SPGP0QP00120 | XP_018219167.1 | 99 | 0 |
| SPGP0BI01140 | XP_018223766.1 | 99 | 0 |
| SPGP0OE00220 | XP_018219598.1 | 99 | 0 |
| SPGP0D00430 | XP_018221676.1 | 99 | 0 |
| SPGP0U00660 | XP_018224072.1 | 99 | 0 |
| SPGP0SW00100 | XP_018220564.1 | 99 | 0 |
| SPGP0CZ00200 | XP_018219915.1 | 99 | 0 |
| SPGP0ID00100 | XP_018222140.1 | 99 | 0 |
| SPGP0K01550 | XP_018219433.1 | 99 | 0 |
| SPGP0BI03010 | XP_018223582.1 | 99 | 0 |
| SPGP0K00660 | XP_018219520.1 | 99 | 0 |
| SPGP0SH00180 | XP_018223976.1 | 99 | 0 |
| SPGP0JW00130 | XP_018220975.1 | 99 | 0 |
| SPGP0TE00100 | XP_018219893.1 | 99 | 0 |
| SPGP0C02840 | XP_018220089.1 | 99 | 0 |
| SPGP0VN00110 | XP_018219801.1 | 99 | 0 |
| SPGP0CZ00190 | XP_018219914.1 | 99 | 0 |
| SPGP0M00280 | XP_018220873.1 | 99 | 0 |
| SPGP0C00440 | XP_018220319.1 | 99 | 0 |
| SPGP0VN00190 | XP_018219809.1 | 99 | 0 |
| SPGP0VN00360 | XP_018219826.1 | 99 | 0 |

Create PDF in your applications with the Pdfcrowd HTML to PDF API PDFCROWD

| | | | |
|---|---|---|---|
| SPGP0K01510 | XP_018219437.1 | 99 | 0 |
| SPGP0BE04820 | XP_018222869.1 | 99 | 0 |
| SPGP0M00610 | XP_018220905.1 | 99 | 0 |
| SPGP0M00790 | XP_018220923.1 | 99 | 0 |
| SPGP0C03220 | XP_018220052.1 | 99 | 0 |
| SPGP0C02280 | XP_018220143.1 | 99 | 0 |
| SPGP0EZ00200 | XP_018219325.1 | 99 | 0 |
| SPGP0MD00120 | XP_018222387.1 | 99 | 0 |
| SPGP0PZ00100 | XP_018222436.1 | 99 | 0 |
| SPGP0N00970 | XP_018219284.1 | 99 | 0 |
| SPGP0BI00280 | XP_018223850.1 | 99 | 0 |
| SPGP0D01420 | XP_018221770.1 | 99 | 0 |
| SPGP0BI00750 | XP_018223803.1 | 99 | 0 |
| SPGP0BI04070 | XP_018223479.1 | 99 | 0 |
| SPGP0NQ00260 | XP_018220739.1 | 99 | 0 |
| SPGP0SH00240 | XP_018223982.1 | 99 | 0 |
| SPGP0D02470 | XP_018221872.1 | 99 | 0 |
| SPGP0HL00180 | XP_018220990.1 | 99 | 0 |
| SPGP0P00520 | XP_018221601.1 | 99 | 0 |
| SPGP0D00780 | XP_018221710.1 | 99 | 0 |
| SPGP0GY00120 | XP_018223400.1 | 99 | 0 |
| SPGP0BE04610 | XP_018222890.1 | 99 | 0 |
| SPGP0BK02210 | XP_018221256.1 | 99 | 0 |
| SPGP0D01820 | XP_018221808.1 | 99 | 0 |
| SPGP0N00300 | XP_018219218.1 | 99 | 0 |
| SPGP0BE00820 | XP_018223252.1 | 99 | 0 |
| SPGP0VUtrna22S | DI49_5523 | 100 | 0 |
| SPGP0PE00140 | XP_018222358.1 | 99 | 0 |
| SPGP0BI02320 | XP_018223650.1 | 99 | 0 |
| SPGP0P00360 | XP_018221617.1 | 99 | 1 |
| SPGP0LU00260 | XP_018224000.1 | 99 | 0 |
| SPGP0Ctrna7Q | DI49_4357 | 100 | 0 |
| SPGP0FW00130 | XP_018219583.1 | 99 | 0 |
| SPGP0C00610 | XP_018220303.1 | 99 | 0 |
| SPGP0BE02910 | XP_018223051.1 | 99 | 0 |
| SPGP0SW00120 | XP_018220562.1 | 99 | 0 |
| SPGP0BK00160 | XP_018221452.1 | 99 | 0 |
| SPGP0UR00240 | XP_018222330.1 | 99 | 0 |
| SPGP0BE04630 | XP_018222888.1 | 99 | 0 |
| SPGP0FE00240 | XP_018220527.1 | 99 | 0 |

Create PDF in your applications with the Pdfcrowd HTML to PDF API PDFCROWD

| | | | |
|---|---|---|---|
| SPGP0HO00130 | XP_018219989.1 | 100 | 0 |
| SPGP0C01480 | XP_018220217.1 | 97 | 0 |
| SPGP0GB00110 | XP_018222656.1 | 99 | 0 |
| SPGP0BE01470 | XP_018223188.1 | 99 | 0 |
| SPGP0K02220 | XP_018219369.1 | 99 | 0 |
| SPGP0D01010 | XP_018221732.1 | 99 | 0 |
| SPGP0BE03830 | XP_018222968.1 | 99 | 0 |
| SPGP0MN00100 | XP_018222260.1 | 99 | 0 |
| SPGP0VM00260 | XP_018220668.1 | 99 | 0 |
| SPGP0TM00130 | XP_018219877.1 | 99 | 0 |
| SPGP0TW00120 | XP_018218845.1 | 99 | 0 |
| SPGP0D00830 | XP_018221714.1 | 99 | 0 |
| SPGP0RK00120 | XP_018224040.1 | 99 | 0 |
| SPGP0VM00220 | XP_018220672.1 | 99 | 0 |
| SPGP0IR00160 | XP_018219761.1 | 99 | 0 |
| SPGP0SL00160 | XP_018220480.1 | 99 | 0 |
| SPGP0P01160 | XP_018221541.1 | 99 | 0 |
| SPGP0BK02400 | XP_018221238.1 | 99 | 0 |
| SPGP0N01190 | XP_018219306.1 | 99 | 0 |
| SPGP0NV00170 | XP_018223455.1 | 99 | 0 |
| SPGP0JF00120 | XP_018222310.1 | 99 | 0 |
| SPGP0PS00100 | XP_018223360.1 | 99 | 0 |
| SPGP0BE01840 | XP_018223151.1 | 99 | 0 |
| SPGP0P00190 | XP_018221634.1 | 99 | 0 |
| SPGP0JO00160 | XP_018223350.1 | 99 | 0 |
| SPGP0NK00160 | XP_018220449.1 | 99 | 0 |
| SPGP0SW00240 | XP_018220550.1 | 93 | 6 |
| SPGP0LX00100 | XP_018219144.1 | 99 | 0 |
| SPGP0BE01860 | XP_018223149.1 | 97 | 2 |
| SPGP0FA00100 | XP_018219182.1 | 98 | 0 |
| SPGP0D00550 | XP_018221687.1 | 99 | 0 |
| SPGP0BI03790 | XP_018223504.1 | 99 | 0 |
| SPGP0BI01930 | XP_018223687.1 | 99 | 0 |
| SPGP0C00870 | XP_018220277.1 | 99 | 0 |
| SPGP0K02100 | XP_018219381.1 | 99 | 0 |
| SPGP0N00320 | XP_018219220.1 | 99 | 0 |
| SPGP0OX00230 | XP_018219722.1 | 99 | 0 |
| SPGP0K00440 | XP_018219541.1 | 99 | 0 |
| SPGP0P01820 | XP_018221477.1 | 99 | 0 |
| SPGP0Itrna26D | DI49_4675 | 100 | 0 |

Create PDF in your applications with the Pdfcrowd HTML to PDF API PDFCROWD

| | | | |
|---|---|---|---|
| SPGP0D00310 | XP_018221664.1 | 100 | 0 |
| SPGP0RC00150 | XP_018222090.1 | 99 | 0 |
| SPGP0EX00100 | XP_018223419.1 | 99 | 0 |
| SPGP0DX00100 | XP_018219326.1 | 99 | 0 |
| SPGP0M00470 | XP_018220892.1 | 99 | 0 |
| SPGP0U00620 | XP_018224076.1 | 99 | 0 |
| SPGP0NG00200 | XP_018222768.1 | 99 | 0 |
| SPGP0QV00140 | XP_018220006.1 | 99 | 0 |
| SPGP0BI00790 | XP_018223799.1 | 99 | 0 |
| SPGP0D01780 | XP_018221805.1 | 99 | 0 |
| SPGP0BE03780 | XP_018222973.1 | 99 | 0 |
| SPGP0BE04130 | XP_018222940.1 | 99 | 0 |
| SPGP0K00840 | XP_018219502.1 | 100 | 0 |
| SPGP0BK02010 | XP_018221276.1 | 99 | 0 |
| SPGP0K00190 | XP_018219566.1 | 99 | 0 |
| SPGP0UJ00250 | XP_018223330.1 | 99 | 0 |
| SPGP0BI01010 | XP_018223778.1 | 99 | 0 |
| SPGP0D03130 | XP_018221933.1 | 99 | 0 |
| SPGP0K02360 | XP_018219355.1 | 99 | 0 |
| SPGP0BE00180 | XP_018223316.1 | 99 | 0 |
| SPGP0KP00100 | XP_018223348.1 | 99 | 0 |
| SPGP0LI00120 | XP_018222247.1 | 99 | 0 |
| SPGP0C01870 | XP_018220183.1 | 99 | 0 |
| SPGP0BTtrna25R | DI49_3845 | 100 | 0 |
| SPGP0D00680 | XP_018221700.1 | 99 | 0 |
| SPGP0BK03420 | XP_018221142.1 | 99 | 0 |
| SPGP0BI00240 | XP_018223854.1 | 99 | 0 |
| SPGP0BK02320 | XP_018221245.1 | 99 | 0 |
| SPGP0TQ00340 | XP_018220630.1 | 99 | 0 |
| SPGP0C02010 | XP_018220170.1 | 99 | 0 |
| SPGP0BE04300 | XP_018222923.1 | 99 | 0 |
| SPGP0C01670 | XP_018220200.1 | 99 | 0 |
| SPGP0BE00790 | XP_018223255.1 | 99 | 0 |
| SPGP0BK02480 | XP_018221230.1 | 99 | 0 |
| SPGP0N00700 | XP_018219257.1 | 98 | 2 |
| SPGP0K01690 | XP_018219420.1 | 99 | 0 |
| SPGP0C00890 | XP_018220275.1 | 99 | 0 |
| SPGP0K01120 | XP_018219475.1 | 99 | 0 |
| SPGP0SL00140 | XP_018220482.1 | 99 | 0 |
| SPGP0BE01400 | XP_018223194.1 | 99 | 0 |

Create PDF in your applications with the Pdfcrowd HTML to PDF API PDFCROWD

| | | | |
|---|---|---|---|
| SPGP0P01340 | XP_018221524.1 | 99 | 0 |
| SPGP0N01070 | XP_018219294.1 | 99 | 0 |
| SPGP0QW00160 | XP_018222608.1 | 100 | 0 |
| SPGP0OI00500 | XP_018218900.1 | 99 | 0 |
| SPGP0M00670 | XP_018220911.1 | 100 | 0 |
| SPGP0BE01300 | XP_018223204.1 | 99 | 0 |
| SPGP0SL00120 | XP_018220484.1 | 99 | 0 |
| SPGP0BI02480 | XP_018223634.1 | 99 | 0 |
| SPGP0C02120 | XP_018220159.1 | 99 | 0 |
| SPGP0U00740 | XP_018224064.1 | 99 | 0 |
| SPGP0BK03210 | XP_018221163.1 | 99 | 0 |
| SPGP0UJ00230 | XP_018223332.1 | 100 | 0 |
| SPGP0P00690 | XP_018221585.1 | 99 | 0 |
| SPGP0C00420 | XP_018220321.1 | 99 | 0 |
| SPGP0D02280 | XP_018221853.1 | 99 | 0 |
| SPGP0BE00640 | XP_018223270.1 | 99 | 0 |
| SPGP0D02530 | XP_018221877.1 | 99 | 0 |
| SPGP0BK02370 | XP_018221241.1 | 99 | 0 |
| SPGP0N00280 | XP_018219216.1 | 99 | 0 |
| SPGP0BK01920 | XP_018221285.1 | 99 | 0 |
| SPGP0P01120 | XP_018221545.1 | 99 | 0 |
| SPGP0BK01130 | XP_018221361.1 | 99 | 0 |
| SPGP0AU00110 | XP_018222231.1 | 99 | 0 |
| SPGP0BI04010 | XP_018223485.1 | 99 | 1 |
| SPGP0UI00100 | XP_018219183.1 | 99 | 0 |
| SPGP0BI03240 | XP_018223559.1 | 100 | 0 |
| SPGP0HO00340 | XP_018219969.1 | 99 | 0 |
| SPGP0VQ00100 | XP_018222542.1 | 99 | 0 |
| SPGP0LX00120 | XP_018219146.1 | 99 | 0 |
| SPGP0D02700 | XP_018221893.1 | 99 | 0 |
| SPGP0DW00110 | XP_018218955.1 | 99 | 0 |
| SPGP0U00550 | XP_018224082.1 | 99 | 0 |
| SPGP0BK00450 | XP_018221427.1 | 99 | 0 |
| SPGP0OA00140 | XP_018218945.1 | 99 | 0 |
| SPGP0M00940 | XP_018220938.1 | 99 | 0 |
| SPGP0C01790 | XP_018220190.1 | 99 | 0 |
| SPGP0K02120 | XP_018219379.1 | 99 | 0 |
| SPGP0K02320 | XP_018219359.1 | 99 | 0 |
| SPGP0BE01580 | XP_018223177.1 | 100 | 0 |
| SPGP0K01670 | XP_018219422.1 | 99 | 0 |

Create PDF in your applications with the Pdfcrowd HTML to PDF API PDFCROWD

| | | | |
|---|---|---|---|
| SPGP0SX00100 | XP_018223325.1 | 99 | 0 |
| SPGP0BK01830 | XP_018221294.1 | 99 | 0 |
| SPGP0OX00200 | XP_018219725.1 | 99 | 0 |
| SPGP0BI02150 | XP_018223666.1 | 99 | 0 |
| SPGP0BE02270 | XP_018223111.1 | 99 | 0 |
| SPGP0BK02650 | XP_018221214.1 | 99 | 0 |
| SPGP0GV00110 | XP_018218861.1 | 98 | 0 |
| SPGP0UN00190 | XP_018220779.1 | 99 | 0 |
| SPGP0JN00110 | XP_018219070.1 | 99 | 0 |
| SPGP0UT00120 | XP_018222190.1 | 98 | 1 |
| SPGP0VM00190 | XP_018220675.1 | 99 | 0 |
| SPGP0QW00120 | XP_018222604.1 | 99 | 0 |
| SPGP0K01220 | XP_018219465.1 | 100 | 0 |
| SPGP0C03050 | XP_018220069.1 | 99 | 0 |
| SPGP0N00810 | XP_018219268.1 | 99 | 0 |
| SPGP0P01300 | XP_018221528.1 | 99 | 1 |
| SPGP0MO00110 | XP_018219096.1 | 100 | 0 |
| SPGP0D02490 | XP_018221874.1 | 100 | 0 |

#### Sheet 2: *genes alleles inherited from Saccharomyces cerevisiae*

| *Saccharomyces pastorianus CBS1503 predicted ORF* | *Saccharomyces cerevisiae parental allele* | Gene identities | Gene gaps |
|---|---|---|---|
| SPGP0B02470 | YHR196W | 99 | 0 |
| SPGP0A04130 | YDR197W | 99 | 0 |
| SPGP0L01790 | YOR296W | 99 | 0 |
| SPGP0I00890 | YJR063W | 99 | 0 |
| SPGP0BF01750 | YIL156W-B | 99 | 0 |
| SPGP0I02390 | YJL111W | 99 | 0 |
| SPGP0L00840 | YOR190W | 99 | 0 |
| SPGP0BG00490 | YCL021W-A | 99 | 0 |
| SPGP0H03340 | YMR290C | 99 | 0 |
| SPGP0BS00100 | YGL216W | 100 | 0 |
| SPGP0VT00100 | YPL262W | 99 | 0 |
| SPGP0G00470 | YKL161C | 99 | 0 |
| SPGP0H00720 | YMR017W | 99 | 0 |
| SPGP0U00120 | YAR035W | 83 | 2 |
| SPGP0A01500 | YDL093W | 99 | 0 |
| SPGP0BF01580 | YIL139C | 99 | 0 |
| SPGP0L01500 | YOR264W | 99 | 0 |
| SPGP0L00150 | YOR114W | 99 | 0 |

Create PDF in your applications with the Pdfcrowd HTML to PDF API PDFCROWD

| | | | |
|---|---|---|---|
| SPGP0H01570 | YMR110C | 99 | 0 |
| SPGP0BF01740 | YIL156W | 99 | 0 |
| SPGP0H03020 | YMR258C | 99 | 0 |
| SPGP0G00120 | YKL197C | 99 | 0 |
| SPGP0A04950 | YDR291W | 99 | 0 |
| SPGP0A01020 | YDL145C | 99 | 0 |
| SPGP0H02000 | YMR155W | 99 | 0 |
| SPGP0F03670 | YBL085W | 99 | 0 |
| SPGP0BG00110 | YCL064C | 100 | 0 |
| SPGP0X00480 | YOL117W | 99 | 0 |
| SPGP0L02440 | YOR369C | 100 | 0 |
| SPGP0B02500 | YHR199C | 99 | 0 |
| SPGP0F00660 | YBR240C | 99 | 0 |
| SPGP0H02910 | YMR244C-A | 99 | 0 |
| SPGP0A02470 | YDR018C | 99 | 0 |
| SPGP0L00440 | YOR147W | 99 | 0 |
| SPGP0G00690 | YKL137W | 99 | 0 |
| SPGP0A03920 | YDR175C | 99 | 0 |
| SPGP0J01230 | YOR014W | 99 | 0 |
| SPGP0J00310 | YOL084W | 99 | 0 |
| SPGP0BO00100 | YHR205W | 99 | 0 |
| SPGP0J00640 | YOL051W | 98 | 2 |
| SPGP0BG00390 | YCL032W | 99 | 0 |
| SPGP0BF01270 | YIL108W | 99 | 0 |
| SPGP0BG00820 | YCR019W | 99 | 0 |
| SPGP0UV00120 | YPL243W | 99 | 0 |
| SPGP0A00930 | YDL156W | 99 | 0 |
| SPGP0F00480 | YBR257W | 99 | 0 |
| SPGP0I02990 | YJL179W | 99 | 0 |
| SPGP0I02030 | YJL070C | 99 | 0 |
| SPGP0B00380 | YHL014C | 99 | 0 |
| SPGP0L02550 | YKL221W | 99 | 0 |
| SPGP0A02160 | YDL018C | 99 | 0 |
| SPGP0H02950 | YMR251W | 99 | 0 |
| SPGP0F01280 | YBR172C | 99 | 0 |
| SPGP0QX00200 | YMR310C | 99 | 0 |
| SPGP0F01820 | YBR115C | 99 | 0 |
| SPGP0H00920 | YMR037C | 99 | 0 |
| SPGP0F03430 | YBL056W | 99 | 0 |

Create PDF in your applications with the Pdfcrowd HTML to PDF API PDFCROWD

| | | | |
|---|---|---|---|
| SPGP0A02660 | YDR040C | 98 | 0 |
| SPGP0B00620 | YHR010W | 99 | 0 |
| SPGP0A01920 | YDL048C | 99 | 0 |
| SPGP0A02040 | YDL033C | 100 | 0 |
| SPGP0I01280 | YJR015W | 100 | 0 |
| SPGP0BF00210 | YIR008C | 99 | 0 |
| SPGP0F01770 | YBR121C | 99 | 0 |
| SPGP0J01010 | YOL010W | 100 | 0 |
| SPGP0F01710 | YBR128C | 99 | 0 |
| SPGP0I02680 | YJL144W | 98 | 0 |
| SPGP0B00450 | YHL007C | 99 | 0 |
| SPGP0F00190 | YBR288C | 99 | 0 |
| SPGP0A03910 | YDR174W | 100 | 0 |
| SPGP0I02630 | YJL138C | 99 | 0 |
| SPGP0B01850 | YHR134W | 100 | 0 |
| SPGP0F02880 | YBR002C | 99 | 0 |
| SPGP0SR00110 | YGL192W | 99 | 0 |
| SPGP0A00830 | YDL168W | 99 | 0 |
| SPGP0B01040 | YHR052W | 99 | 0 |
| SPGP0BF00600 | YIL034C | 100 | 0 |
| SPGP0L01560 | YOR271C | 100 | 0 |
| SPGP0NH00130 | YAL005C | 99 | 0 |
| SPGP0G00260 | YKL183W | 99 | 0 |
| SPGP0A05220 | YDR318W | 99 | 0 |
| SPGP0FK00210 | YGL202W | 99 | 0 |
| SPGP0I00770 | YJR076C | 99 | 0 |
| SPGP0L01290 | YOR241W | 99 | 0 |
| SPGP0B01270 | YHR076W | 99 | 0 |
| SPGP0J00850 | YOL026C | 99 | 0 |
| SPGP0B01010 | YHR049W | 99 | 0 |
| SPGP0F01690 | YBR130C | 99 | 0 |
| SPGP0I02460 | YJL122W | 99 | 0 |
| SPGP0B00880 | YHR036W | 99 | 0 |
| SPGP0A02180 | YDL015C | 99 | 0 |
| SPGP0A04680 | YDR261C | 99 | 0 |
| SPGP0G00250 | YKL184W | 99 | 0 |
| SPGP0ME00130 | YGL231C | 100 | 0 |
| SPGP0A04700 | YDR263C | 99 | 0 |
| SPGP0WU00150 | YER182W | 99 | 0 |
| SPGP0DL00130 | YER187W | 92 | 0 |

Create PDF in your applications with the Pdfcrowd HTML to PDF API PDFCROWD

| | | | |
|---|---|---|---|
| SPGP0L00470 | YOR150W | 99 | 0 |
| SPGP0G00570 | YKL149C | 99 | 0 |
| SPGP0BG00630 | YCL001W | 100 | 0 |
| SPGP0A01320 | YDL113C | 99 | 0 |
| SPGP0BF01800 | YIL161W | 99 | 0 |
| SPGP0B02170 | YHR166C | 99 | 0 |
| SPGP0F03450 | YBL058W | 98 | 0 |
| SPGP0L00300 | YOR130C | 99 | 0 |
| SPGP0H01380 | YMR090W | 99 | 0 |
| SPGP0A00330 | YDL223C | 99 | 0 |
| SPGP0I01550 | YJL013C | 99 | 0 |
| SPGP0A04190 | YDR205W | 99 | 0 |
| SPGP0A04650 | YDR258C | 100 | 0 |
| SPGP0I02720 | YJL148W | 98 | 1 |
| SPGP0NA00110 | YCR098C | 99 | 0 |
| SPGP0B02360 | YHR185C | 100 | 0 |
| SPGP0H01960 | YMR150C | 99 | 0 |
| SPGP0A00390 | YDL216C | 99 | 0 |
| SPGP0A00740 | YDL178W | 99 | 0 |
| SPGP0L01310 | YOR243C | 99 | 0 |
| SPGP0G00780 | YKL126W | 99 | 0 |
| SPGP0BF01190 | YIL101C | 99 | 0 |
| SPGP0H03120 | YMR268C | 99 | 0 |
| SPGP0L01300 | YOR242C | 99 | 0 |
| SPGP0I03320 | YJL217W | 98 | 0 |
| SPGP0BG00120 | YCL063W | 99 | 0 |
| SPGP0L00420 | YOR144C | 99 | 0 |
| SPGP0B02190 | YHR168W | 99 | 0 |
| SPGP0A03440 | YDR123C | 99 | 0 |
| SPGP0F02140 | YBR081C | 99 | 0 |
| SPGP0VT00130 | YPL258C | 99 | 0 |
| SPGP0H00640 | YMR009W | 98 | 0 |
| SPGP0B01780 | YHR127W | 99 | 0 |
| SPGP0J00410 | YOL073C | 99 | 0 |
| SPGP0H00940 | YMR039C | 99 | 1 |
| SPGP0I00170 | YJR138W | 99 | 0 |
| SPGP0L01160 | YOR227W | 99 | 0 |
| SPGP0I00620 | YJR094C | 100 | 0 |
| SPGP0A03210 | YDR098C | 99 | 0 |
| SPGP0BG00940 | YCR030C | 99 | 0 |

Create PDF in your applications with the Pdfcrowd HTML to PDF API PDFCROWD

| | | | |
|---|---|---|---|
| SPGP0I02820 | YJL160C | 99 | 0 |
| SPGP0X00290 | YOL138C | 99 | 0 |
| SPGP0B01710 | YHR119W | 99 | 0 |
| SPGP0L02300 | YOR353C | 99 | 0 |
| SPGP0L02050 | YOR326W | 99 | 0 |
| SPGP0I02420 | YJL116C | 99 | 0 |
| SPGP0H02920 | YMR246W | 99 | 0 |
| SPGP0I00540 | YJR101W | 99 | 0 |
| SPGP0J01760 | YOR075W | 99 | 0 |
| SPGP0X00120 | YOL156W | 98 | 0 |
| SPGP0F00830 | YBR221C | 99 | 0 |
| SPGP0F00340 | YBR272C | 99 | 0 |
| SPGP0B00320 | YHL020C | 99 | 0 |
| SPGP0A02640 | YDR036C | 99 | 0 |
| SPGP0G00790 | YKL125W | 99 | 0 |
| SPGP0G00480 | YKL160W | 100 | 0 |
| SPGP0B00730 | YHR021C | 100 | 0 |
| SPGP0H00200 | YML037C | 99 | 0 |
| SPGP0BF01390 | YIL120W | 99 | 0 |
| SPGP0I01590 | YJL020C | 98 | 1 |
| SPGP0F02660 | YBR025C | 99 | 0 |
| SPGP0BF01020 | YIL079C | 99 | 0 |
| SPGP0J01530 | YOR047C | 99 | 0 |
| SPGP0F03860 | YBL106C | 99 | 0 |
| SPGP0F01950 | YBR102C | 99 | 0 |
| SPGP0H00710 | YMR016C | 99 | 0 |
| SPGP0B01460 | YHR094C | 99 | 0 |
| SPGP0B00840 | YHR032W | 99 | 0 |
| SPGP0B00590 | YHR007C | 99 | 0 |
| SPGP0A01510 | YDL092W | 99 | 0 |
| SPGP0BF00400 | YIL010W | 99 | 0 |
| SPGP0LL00110 | YAL026C | 99 | 0 |
| SPGP0F03770 | YBL095W | 99 | 0 |
| SPGP0A00100 | YNL336W | 99 | 0 |
| SPGP0LL00200 | YAL034W-A | 99 | 0 |
| SPGP0A01310 | YDL114W | 99 | 0 |
| SPGP0F02080 | YBR085C-A | 100 | 0 |
| SPGP0BF01000 | YIL077C | 99 | 0 |
| SPGP0B02280 | YHR177W | 99 | 0 |
| SPGP0BF00780 | YIL052C | 98 | 0 |

Create PDF in your applications with the Pdfcrowd HTML to PDF API PDFCROWD

| | | | |
|---|---|---|---|
| SPGP0F02000 | YBR095C | 99 | 0 |
| SPGP0I03010 | YJL181W | 99 | 0 |
| SPGP0A05080 | YDR304C | 99 | 0 |
| SPGP0F01250 | YBR176W | 99 | 0 |
| SPGP0I01700 | YJL035C | 99 | 0 |
| SPGP0IJ00110 | YPL218W | 90 | 0 |
| SPGP0B01640 | YHR112C | 99 | 0 |
| SPGP0BG00310 | YCL040W | 99 | 0 |
| SPGP0I00410 | YJR112W-A | 99 | 0 |
| SPGP0A01010 | YDL146W | 99 | 0 |
| SPGP0G01130 | YKL085W | 99 | 0 |
| SPGP0F02460 | YBR047W | 99 | 0 |
| SPGP0I01940 | YJL060W | 99 | 0 |
| SPGP0A04480 | YDR239C | 99 | 0 |
| SPGP0I01010 | YJR051W | 99 | 0 |
| SPGP0F01840 | YBR112C | 99 | 1 |
| SPGP0A03450 | YDR124W | 99 | 0 |
| SPGP0H02760 | YMR230W | 99 | 0 |
| SPGP0L01430 | YOR256C | 99 | 0 |
| SPGP0B02010 | YHR150W | 99 | 0 |
| SPGP0F03020 | YBL014C | 99 | 0 |
| SPGP0X00200 | YOL147C | 99 | 0 |
| SPGP0I02010 | YJL068C | 98 | 0 |
| SPGP0I00910 | YJR061W | 99 | 0 |
| SPGP0I02940 | YJL173C | 98 | 0 |
| SPGP0B02450 | YHR194W | 99 | 0 |
| SPGP0A04150 | YDR200C | 99 | 0 |
| SPGP0L00820 | YOR188W | 99 | 0 |
| SPGP0L00790 | YOR185C | 99 | 0 |
| SPGP0A00380 | YDL217C | 99 | 0 |
| SPGP0FK00140 | YGL210W | 99 | 0 |
| SPGP0F01190 | YBR183W | 99 | 0 |
| SPGP0G00400 | YKL167C | 99 | 0 |
| SPGP0TI00110 | YAL062W | 99 | 0 |
| SPGP0I01900 | YJL056C | 99 | 0 |
| SPGP0H01160 | YMR066W | 99 | 0 |
| SPGP0L01490 | YOR262W | 99 | 0 |
| SPGP0A04080 | YDR191W | 99 | 0 |
| SPGP0L00170 | YOR116C | 99 | 0 |
| SPGP0H01590 | YMR112C | 99 | 0 |

Create PDF in your applications with the Pdfcrowd HTML to PDF API PDFCROWD

| | | | |
|---|---|---|---|
| SPGP0B02340 | YHR183W | 99 | 0 |
| SPGP0BFtrna15E | YDR278C | 100 | 0 |
| SPGP0G00100 | YKL201C | 99 | 0 |
| SPGP0H02020 | YMR157C | 99 | 0 |
| SPGP0A00480 | YDL207W | 99 | 0 |
| SPGP0F00130 | YBR295W | 99 | 0 |
| SPGP0A02410 | YDR012W | 99 | 0 |
| SPGP0F00640 | YBR242W | 100 | 0 |
| SPGP0A01130 | YDL133W | 99 | 0 |
| SPGP0L02340 | YOR357C | 100 | 0 |
| SPGP0J00620 | YOL052C-A | 99 | 0 |
| SPGP0A02480 | YDR019C | 99 | 0 |
| SPGP0J00330 | YOL082W | 99 | 0 |
| SPGP0A01680 | YDL076C | 99 | 0 |
| SPGP0BF01250 | YIL106W | 99 | 0 |
| SPGP0BF01640 | YIL146C | 99 | 0 |
| SPGP0BG00920 | YCR028C | 99 | 0 |
| SPGP0F03560 | YBL071W-A | 98 | 0 |
| SPGP0I01880 | YJL054W | 99 | 0 |
| SPGP0B00360 | YHL016C | 99 | 0 |
| SPGP0L02530 | YOR380W | 99 | 0 |
| SPGP0B00690 | YHR017W | 99 | 0 |
| SPGP0SR00130 | YGL190C | 99 | 0 |
| SPGP0I02000 | YJL066C | 100 | 0 |
| SPGP0B00680 | YHR016C | 99 | 0 |
| SPGP0A02050 | YDL031W | 99 | 0 |
| SPGP0F03410 | YBL054W | 99 | 0 |
| SPGP0A02680 | YDR042C | 99 | 0 |
| SPGP0BF01540 | YIL135C | 99 | 0 |
| SPGP0QX00180 | YMR312W | 99 | 0 |
| SPGP0H02980 | YMR253C | 99 | 0 |
| SPGP0J01570 | YOR052C | 99 | 0 |
| SPGP0F01670 | YBR132C | 99 | 0 |
| SPGP0F01750 | YBR123C | 99 | 0 |
| SPGP0A04900 | YDR285W | 99 | 0 |
| SPGP0B02070 | YHR156C | 99 | 0 |
| SPGP0B00470 | YHL004W | 99 | 0 |
| SPGP0H01370 | YMR089C | 99 | 0 |
| SPGP0A03030 | YDR079W | 100 | 0 |
| SPGP0TH00100 | YPL273W | 100 | 0 |

Create PDF in your applications with the Pdfcrowd HTML to PDF API PDFCROWD

| | | | |
|---|---|---|---|
| SPGP0I00280 | YJR127C | 99 | 0 |
| SPGP0B01870 | YHR136C | 99 | 0 |
| SPGP0BF00660 | YIL040W | 99 | 0 |
| SPGP0I01300 | YJR013W | 99 | 0 |
| SPGP0I00940 | YJR058C | 99 | 0 |
| SPGP0I00550 | YJR100C | 99 | 0 |
| SPGP0I00750 | YJR078W | 99 | 0 |
| SPGP0I02950 | YJL174W | 99 | 0 |
| SPGP0A02060 | YDL030W | 99 | 0 |
| SPGP0H01000 | YMR047C | 99 | 0 |
| SPGP0J01910 | YOR092W | 99 | 0 |
| SPGP0B01890 | YHR138C | 99 | 0 |
| SPGP0I00360 | YJR118C | 99 | 0 |
| SPGP0F02690 | YBR022W | 100 | 0 |
| SPGP0A03960 | YDR179C | 99 | 0 |
| SPGP0J00360 | YOL078W | 99 | 0 |
| SPGP0F03570 | YBL072C | 100 | 0 |
| SPGP0A04490 | YDR240C | 100 | 0 |
| SPGP0F03610 | YBL078C | 100 | 0 |
| SPGP0A02170 | YDL017W | 99 | 0 |
| SPGP0B00900 | YHR038W | 99 | 0 |
| SPGP0A00310 | YDL225W | 99 | 0 |
| SPGP0A05140 | YDR310C | 99 | 0 |
| SPGP0DL00100 | YER184C | 99 | 0 |
| SPGP0I01390 | YJR005C-A | 100 | 0 |
| SPGP0L00360 | YOR137C | 99 | 0 |
| SPGP0L00490 | YOR152C | 99 | 0 |
| SPGP0F02170 | YBR078W | 99 | 0 |
| SPGP0F00740 | YBR233W | 99 | 0 |
| SPGP0I01480 | YJL004C | 99 | 0 |
| SPGP0F02520 | YBR041W | 99 | 0 |
| SPGP0DL00110 | YER185W | 99 | 0 |
| SPGP0G00230 | YKL186C | 99 | 0 |
| SPGP0IY00190 | YGL250W | 99 | 0 |
| SPGP0I03220 | YJL206C | 99 | 0 |
| SPGP0BO00150 | YHR210C | 99 | 0 |
| SPGP0B00180 | YHL034C | 99 | 0 |
| SPGP0A01300 | YDL115C | 99 | 0 |
| SPGP0BF01530 | YIL134W | 99 | 0 |
| SPGP0L01550 | YOR270C | 99 | 0 |

Create PDF in your applications with the Pdfcrowd HTML to PDF API PDFCROWD

| | | | |
|---|---|---|---|
| SPGP0I01570 | YJL016W | 99 | 0 |
| SPGP0QG00160 | YAL047C | 99 | 0 |
| SPGP0H02290 | YMR184W | 99 | 0 |
| SPGP0F02090 | YBR085W | 99 | 0 |
| SPGP0H01280 | YMR078C | 99 | 0 |
| SPGP0F01530 | YBR148W | 99 | 0 |
| SPGP0H01900 | YMR144W | 99 | 0 |
| SPGP0BG00520 | YCL016C | 99 | 0 |
| SPGP0G00800 | YKL124W | 99 | 0 |
| SPGP0L00580 | YOR161C | 99 | 0 |
| SPGP0X00300 | YOL137W | 99 | 0 |
| SPGP0J01130 | YOR003W | 99 | 0 |
| SPGP0F02310 | YBR062C | 99 | 0 |
| SPGP0BF01260 | YIL107C | 99 | 0 |
| SPGP0A02450 | YDR016C | 99 | 0 |
| SPGP0A00410 | YDL214C | 99 | 0 |
| SPGP0L01360 | YOR249C | 99 | 0 |
| SPGP0BG00130 | YCL061C | 99 | 0 |
| SPGP0A00350 | YDL220C | 99 | 0 |
| SPGP0A03420 | YDR121W | 98 | 1 |
| SPGP0BF01440 | YIL125W | 99 | 0 |
| SPGP0BG00370 | YCL034W | 99 | 0 |
| SPGP0NA00100 | YCR099C | 99 | 0 |
| SPGP0J01810 | YOR080W | 99 | 0 |
| SPGP0H01890 | YMR143W | 99 | 0 |
| SPGP0J00430 | YOL071W | 99 | 0 |
| SPGP0X00240 | YOL143C | 99 | 0 |
| SPGP0I00590 | YJR096W | 99 | 0 |
| SPGP0L01180 | YOR229W | 98 | 1 |
| SPGP0J00230 | YOL092W | 99 | 0 |
| SPGP0L01980 | YOR317W | 99 | 0 |
| SPGP0H02900 | YMR244W | 99 | 0 |
| SPGP0A03270 | YDR105C | 99 | 0 |
| SPGP0AC00100 | YLL021W | 81 | 1 |
| SPGP0I00180 | YJR137C | 99 | 0 |
| SPGP0F02330 | YBR060C | 99 | 0 |
| SPGP0BF00390 | YIL009C-A | 99 | 0 |
| SPGP0H00220 | YML035C | 99 | 0 |
| SPGP0I02840 | YJL162C | 98 | 1 |
| SPGP0J01740 | YOR073W | 99 | 0 |

Create PDF in your applications with the Pdfcrowd HTML to PDF API PDFCROWD

| | | | |
|---|---|---|---|
| SPGP0L01830 | YOR299W | 99 | 0 |
| SPGP0F00580 | YBR248C | 99 | 0 |
| SPGP0H03050 | YMR261C | 99 | 0 |
| SPGP0L02320 | YOR355W | 99 | 0 |
| SPGP0G00710 | YKL134C | 99 | 0 |
| SPGP0B01690 | YHR117W | 99 | 0 |
| SPGP0LA00100 | YER188W | 90 | 0 |
| SPGP0VR00140 | YLR300W | 86 | 0 |
| SPGP0F01270 | YBR173C | 99 | 0 |
| SPGP0I02400 | YJL112W | 99 | 0 |
| SPGP0F01980 | YBR097W | 99 | 0 |
| SPGP0H00550 | YMR001C | 99 | 0 |
| SPGP0H00880 | YMR033W | 99 | 0 |
| SPGP0X00140 | YOL154W | 99 | 0 |
| SPGP0F00360 | YBR270C | 99 | 0 |
| SPGP0L02180 | YOR341W | 99 | 0 |
| SPGP0A02580 | YDR030C | 99 | 0 |
| SPGP0J00290 | YOL086W-A | 99 | 0 |
| SPGP0A00770 | YDL175C | 99 | 0 |
| SPGP0L01040 | YOR213C | 99 | 0 |
| SPGP0I01150 | YJR035W | 99 | 0 |
| SPGP0B01450 | YHR093W | 99 | 0 |
| SPGP0A03890 | YDR172W | 99 | 0 |
| SPGP0I02750 | YJL153C | 99 | 0 |
| SPGP0H00770 | YMR022W | 100 | 0 |
| SPGP0B00860 | YHR034C | 99 | 1 |
| SPGP0ME00150 | YGL229C | 99 | 0 |
| SPGP0A01530 | YDL090C | 99 | 0 |
| SPGP0A01660 | YDL078C | 99 | 0 |
| SPGP0A00120 | YNL334C | 98 | 0 |
| SPGP0A01290 | YDL116W | 99 | 0 |
| SPGP0B01400 | YHR088W | 99 | 0 |
| SPGP0F02060 | YBR087W | 99 | 0 |
| SPGP0J00810 | YOL031C | 99 | 0 |
| SPGP0I03030 | YJL184W | 99 | 0 |
| SPGP0A05060 | YDR302W | 99 | 0 |
| SPGP0J00990 | YOL012C | 99 | 0 |
| SPGP0B01560 | YHR104W | 99 | 0 |
| SPGP0B00230 | YHL029C | 99 | 0 |
| SPGP0F02950 | YBL006C | 99 | 0 |

Create PDF in your applications with the Pdfcrowd HTML to PDF API PDFCROWD

| | | | |
|---|---|---|---|
| SPGP0F02280 | YBR066C | 99 | 0 |
| SPGP0A04310 | YDR221W | 99 | 0 |
| SPGP0QQ00170 | YAR007C | 99 | 0 |
| SPGP0J00120 | YOL104C | 99 | 0 |
| SPGP0I01760 | YJL042W | 99 | 0 |
| SPGP0I01530 | YJL011C | 100 | 0 |
| SPGP0B00960 | YHR044C | 99 | 0 |
| SPGP0CB00170 | YIR027C | 99 | 0 |
| SPGP0BF00830 | YIL061C | 99 | 0 |
| SPGP0B02430 | YHR192W | 99 | 0 |
| SPGP0A04810 | YDR277C | 99 | 0 |
| SPGP0A04170 | YDR202C | 99 | 0 |
| SPGP0A04990 | YDR295C | 99 | 0 |
| SPGP0F02120 | YBR083W | 99 | 0 |
| SPGP0I01850 | YJL051W | 99 | 0 |
| SPGP0F03000 | YBL011W | 99 | 0 |
| SPGP0F03550 | YBL069W | 99 | 0 |
| SPGP0G00420 | YKL165C | 99 | 0 |
| SPGP0H01750 | YMR128W | 99 | 0 |
| SPGP0L00800 | YOR186W | 98 | 0 |
| SPGP0A00290 | YDL227C | 99 | 0 |
| SPGP0I02070 | YJL074C | 99 | 0 |
| SPGP0L00970 | YOR206W | 99 | 0 |
| SPGP0F03240 | YBL035C | 99 | 0 |
| SPGP0BG00430 | YCL028W | 99 | 1 |
| SPGP0QO00120 | YGL238W | 99 | 0 |
| SPGP0H02220 | YMR177W | 99 | 0 |
| SPGP0H01140 | YMR064W | 99 | 0 |
| SPGP0BG00400 | YCL031C | 99 | 0 |
| SPGP0L00110 | YOR110W | 99 | 0 |
| SPGP0I01180 | YJR032W | 99 | 0 |
| SPGP0BF00750 | YIL049W | 98 | 0 |
| SPGP0B02320 | YHR181W | 99 | 0 |
| SPGP0A01480 | YDL097C | 99 | 0 |
| SPGP0A02780 | YDR054C | 99 | 0 |
| SPGP0H03090 | YMR265C | 99 | 0 |
| SPGP0G00380 | YKL170W | 100 | 0 |
| SPGP0A02430 | YDR014W | 99 | 0 |
| SPGP0BG00160 | YCL057C-A | 99 | 0 |
| SPGP0F01090 | YBR193C | 99 | 0 |

Create PDF in your applications with the Pdfcrowd HTML to PDF API PDFCROWD

| | | | |
|---|---|---|---|
| SPGP0H02040 | YMR159C | 99 | 0 |
| SPGP0I03090 | YJL191W | 99 | 0 |
| SPGP0X00440 | YOL122C | 99 | 0 |
| SPGP0A00460 | YDL209C | 99 | 0 |
| SPGP0F01360 | YBR164C | 99 | 0 |
| SPGP0J00350 | YOL080C | 99 | 0 |
| SPGP0J00600 | YOL054W | 100 | 0 |
| SPGP0L01640 | YOR280C | 99 | 0 |
| SPGP0F01380 | YBR162W-A | 99 | 0 |
| SPGP0BG00720 | YCR009C | 99 | 0 |
| SPGP0H00800 | YMR025W | 99 | 0 |
| SPGP0BF01620 | YIL144W | 99 | 0 |
| SPGP0H00280 | YML028W | 100 | 0 |
| SPGP0L01260 | YOR238W | 100 | 0 |
| SPGP0A01600 | YDL084W | 99 | 0 |
| SPGP0B00420 | YHL010C | 99 | 0 |
| SPGP0G00520 | YKL155C | 99 | 0 |
| SPGP0L02510 | YOR377W | 99 | 0 |
| SPGP0B00670 | YHR015W | 99 | 0 |
| SPGP0F03400 | YBL052C | 99 | 0 |
| SPGP0A02700 | YDR044W | 100 | 0 |
| SPGP0B01130 | YHR063C | 99 | 0 |
| SPGP0BF00230 | YIR006C | 99 | 0 |
| SPGP0H00960 | YMR041C | 100 | 0 |
| SPGP0I02980 | YJL178C | 99 | 0 |
| SPGP0J01550 | YOR049C | 99 | 0 |
| SPGP0J01000 | YOL011W | 99 | 0 |
| SPGP0B02050 | YHR154W | 99 | 0 |
| SPGP0A03460 | YDR125C | 99 | 0 |
| SPGP0J00970 | YOL014W | 93 | 0 |
| SPGP0B00410 | YHL011C | 99 | 0 |
| SPGP0I02550 | YJL130C | 99 | 0 |
| SPGP0F02840 | YBR006W | 99 | 0 |
| SPGP0F01740 | YBR125C | 99 | 0 |
| SPGP0L01970 | YOR316C | 99 | 0 |
| SPGP0H00340 | YML022W | 99 | 0 |
| SPGP0A03630 | YDR144C | 99 | 0 |
| SPGP0I01920 | YJL058C | 99 | 0 |
| SPGP0H00690 | YMR014W | 99 | 0 |

Create PDF in your applications with the Pdfcrowd HTML to PDF API PDFCROWD

| | | | |
|---|---|---|---|
| SPGP0J01540 | YOR048C | 99 | 0 |
| SPGP0I02960 | YJL176C | 99 | 0 |
| SPGP0VT00170 | YPL255W | 100 | 0 |
| SPGP0B01230 | YHR072W-A | 100 | 0 |
| SPGP0UV00130 | YPL242C | 99 | 0 |
| SPGP0X00430 | YOL123W | 99 | 0 |
| SPGP0ML00110 | YAL054C | 99 | 0 |
| SPGP0H01670 | YMR120C | 99 | 0 |
| SPGP0A02150 | YDL019C | 99 | 0 |
| SPGP0A01490 | YDL095W | 99 | 0 |
| SPGP0BG00810 | YHR145C | 98 | 0 |
| SPGP0F03170 | YBL029W | 99 | 0 |
| SPGP0RF00120 | YAL058W | 99 | 0 |
| SPGP0F03630 | YBL080C | 99 | 0 |
| SPGP0F03490 | YBL061C | 98 | 0 |
| SPGP0A05160 | YDR312W | 100 | 0 |
| SPGP0I01370 | YJR007W | 99 | 0 |
| SPGP0L00340 | YOR134W | 99 | 0 |
| SPGP0DL00120 | YER186C | 96 | 1 |
| SPGP0H02240 | YMR179W | 99 | 0 |
| SPGP0F00710 | YBR235W | 99 | 0 |
| SPGP0L00510 | YOR154W | 99 | 0 |
| SPGP0H03380 | YMR294W | 100 | 0 |
| SPGP0BO00130 | YHR208W | 99 | 0 |
| SPGP0A04780 | YDR273W | 99 | 0 |
| SPGP0BF01510 | YIL132C | 99 | 0 |
| SPGP0F01210 | YBR181C | 99 | 0 |
| SPGP0I01510 | YJL008C | 100 | 0 |
| SPGP0F00380 | YBR268W | 99 | 0 |
| SPGP0H01740 | YMR127C | 99 | 0 |
| SPGP0A04160 | YDR201W | 99 | 0 |
| SPGP0I00870 | YJR065C | 99 | 0 |
| SPGP0L00930 | YOR201C | 99 | 0 |
| SPGP0A03310 | YDR109C | 99 | 0 |
| SPGP0BF01240 | YIL105C | 99 | 0 |
| SPGP0G00810 | YKL122C | 99 | 0 |
| SPGP0H01250 | YMR075W | 99 | 0 |
| SPGP0H01920 | YMR146C | 100 | 0 |
| SPGP0A03240 | YDR101C | 99 | 0 |
| SPGP0A00430 | YDL212W | 99 | 0 |

Create PDF in your applications with the Pdfcrowd HTML to PDF API PDFCROWD

| | | | |
|---|---|---|---|
| SPGP0BF01090 | YIL090W | 99 | 0 |
| SPGP0H02690 | YMR223W | 99 | 0 |
| SPGP0A03640 | YDR145W | 99 | 0 |
| SPGP0L01340 | YOR246C | 99 | 0 |
| SPGP0I00130 | YJR142W | 99 | 0 |
| SPGP0A04060 | YDR189W | 99 | 0 |
| SPGP0F01010 | YBR201W | 99 | 0 |
| SPGP0B02210 | YHR170W | 99 | 0 |
| SPGP0F00790 | YBR227C | 99 | 0 |
| SPGP0I00660 | YJR090C | 99 | 0 |
| SPGP0J01830 | YOR083W | 98 | 1 |
| SPGP0A03940 | YDR177W | 99 | 0 |
| SPGP0F01410 | YBR160W | 99 | 0 |
| SPGP0L02260 | YOR349W | 99 | 0 |
| SPGP0F00810 | YBR223C | 99 | 0 |
| SPGP0F01140 | YBR188C | 98 | 0 |
| SPGP0H01170 | YMR067C | 99 | 0 |
| SPGP0A04970 | YDR293C | 99 | 0 |
| SPGP0A03100 | YDR085C | 98 | 0 |
| SPGP0A02600 | YDR032C | 99 | 0 |
| SPGP0BG00140 | YCL059C | 99 | 0 |
| SPGP0B01670 | YHR115C | 99 | 0 |
| SPGP0G00730 | YKL132C | 99 | 0 |
| SPGP0H03110 | YMR267W | 100 | 0 |
| SPGP0I00210 | YJR135C | 99 | 0 |
| SPGP0BF00870 | YIL065C | 100 | 0 |
| SPGP0A01670 | YDL077C | 99 | 0 |
| SPGP0A02690 | YDR043C | 100 | 0 |
| SPGP0BG00960 | YCR032W | 99 | 0 |
| SPGP0BF01350 | YIL116W | 99 | 0 |
| SPGP0DG00110 | YPL278C | 100 | 0 |
| SPGP0SC00120 | YER174C | 99 | 0 |
| SPGP0F01960 | YBR101C | 99 | 0 |
| SPGP0H03060 | YMR262W | 99 | 0 |
| SPGP0A00690 | YDL183C | 99 | 0 |
| SPGP0J00720 | YOL042W | 99 | 0 |
| SPGP0F02500 | YBR043C | 99 | 0 |
| SPGP0B01950 | YHR143W-A | 100 | 0 |
| SPGP0I02740 | YJL151C | 99 | 0 |
| SPGP0QG00140 | YAL044W-A | 99 | 0 |

Create PDF in your applications with the Pdfcrowd HTML to PDF API PDFCROWD

| | | | |
|---|---|---|---|
| SPGP0I01170 | YJR033C | 99 | 0 |
| SPGP0H02930 | YMR247C | 99 | 0 |
| SPGP0H00600 | YMR006C | 99 | 0 |
| SPGP0A01850 | YDL056W | 99 | 0 |
| SPGP0J01490 | YOR043W | 99 | 0 |
| SPGP0X00160 | YOL152W | 99 | 0 |
| SPGP0H00570 | YMR003W | 99 | 0 |
| SPGP0G01140 | YKL084W | 99 | 0 |
| SPGP0B00800 | YHR028C | 99 | 0 |
| SPGP0ME00170 | YGL227W | 99 | 0 |
| SPGP0A03750 | YDR159W | 99 | 0 |
| SPGP0J01890 | YOR090C | 99 | 0 |
| SPGP0I03050 | YJL186W | 99 | 0 |
| SPGP0J00590 | YOL055C | 100 | 0 |
| SPGP0B01530 | YHR101C | 99 | 0 |
| SPGP0H00750 | YMR020W | 99 | 0 |
| SPGP0B01420 | YHR090C | 99 | 0 |
| SPGP0B01930 | YHR142W | 100 | 0 |
| SPGP0F02040 | YBR089C-A | 100 | 0 |
| SPGP0F03820 | YBL102W | 99 | 0 |
| SPGP0A01350 | YDL110C | 100 | 0 |
| SPGP0A03660 | YDR147W | 99 | 0 |
| SPGP0A05040 | YDR300C | 99 | 0 |
| SPGP0B00330 | YHL019C | 99 | 0 |
| SPGP0F02970 | YBL008W | 99 | 0 |
| SPGP0B01680 | YHR116W | 100 | 0 |
| SPGP0F01470 | YBR154C | 99 | 0 |
| SPGP0L02060 | YOR327C | 99 | 0 |
| SPGP0QQ00110 | YFL020C | 97 | 0 |
| SPGP0A03410 | YDR120C | 99 | 0 |
| SPGP0I01040 | YJR048W | 100 | 0 |
| SPGP0F03790 | YBL098W | 99 | 0 |
| SPGP0H00170 | YML042W | 99 | 0 |
| SPGP0F03700 | YBL088C | 99 | 0 |
| SPGP0BD00110 | YAL037W | 99 | 0 |
| SPGP0A04790 | YDR275W | 99 | 0 |
| SPGP0H01820 | YMR135C | 99 | 1 |
| SPGP0A05010 | YDR297W | 99 | 0 |
| SPGP0B02410 | YHR190W | 99 | 0 |
| SPGP0FK00240 | YGL198W | 99 | 0 |

Create PDF in your applications with the Pdfcrowd HTML to PDF API PDFCROWD

| | | | |
|---|---|---|---|
| SPGP0I00830 | YJR069C | 99 | 0 |
| SPGP0H00250 | YML031W | 99 | 0 |
| SPGP0BG00790 | YCR017C | 99 | 0 |
| SPGP0L00280 | YOR128C | 99 | 0 |
| SPGP0L00750 | YOR180C | 99 | 0 |
| SPGP0I02090 | YJL078C | 99 | 0 |
| SPGP0VT00110 | YPL260W | 99 | 0 |
| SPGP0RN00140 | YER164W | 92 | 0 |
| SPGP0I01740 | YJL039C | 99 | 0 |
| SPGP0QO00100 | YGL236C | 99 | 0 |
| SPGP0A01110 | YDL135C | 99 | 0 |
| SPGP0BG00420 | YCL029C | 99 | 0 |
| SPGP0OP00140 | YAR028W | 98 | 0 |
| SPGP0L01780 | YOR295W | 99 | 0 |
| SPGP0A00500 | YDL205C | 99 | 0 |
| SPGP0G00640 | YKL141W | 99 | 0 |
| SPGP0L00130 | YOR112W | 99 | 0 |
| SPGP0F00170 | YBR290W | 100 | 0 |
| SPGP0I02470 | YJL123C | 98 | 2 |
| SPGP0L01080 | YOR217W | 99 | 0 |
| SPGP0H02200 | YMR175W | 100 | 0 |
| SPGP0X00420 | YOL124C | 99 | 0 |
| SPGP0BF01610 | YIL143C | 99 | 0 |
| SPGP0A00440 | YDL211C | 98 | 0 |
| SPGP0RN00130 | YER165W | 99 | 0 |
| SPGP0A00990 | YDL148C | 99 | 0 |
| SPGP0F01340 | YBR166C | 99 | 0 |
| SPGP0I02120 | YJL082W | 99 | 0 |
| SPGP0L02280 | YOR351C | 99 | 0 |
| SPGP0A05250 | YDR320C-A | 99 | 0 |
| SPGP0J00370 | YOL077W-A | 100 | 0 |
| SPGP0G01100 | YKL088W | 99 | 0 |
| SPGP0L01880 | YOR304C-A | 100 | 0 |
| SPGP0F03120 | YBL024W | 99 | 0 |
| SPGP0BF01600 | YIL142W | 99 | 0 |
| SPGP0X00390 | YOL127W | 100 | 0 |
| SPGP0L01650 | YOR281C | 99 | 0 |
| SPGP0G00500 | YKL157W | 99 | 0 |
| SPGP0I02040 | YJL071W | 99 | 0 |
| SPGP0B00650 | YHR013C | 99 | 0 |

Create PDF in your applications with the Pdfcrowd HTML to PDF API PDFCROWD

| | | | |
|---|---|---|---|
| SPGP0BG00880 | YCR024C | 99 | 0 |
| SPGP0B00400 | YHL012W | 99 | 0 |
| SPGP0A02720 | YDR046C | 99 | 0 |
| SPGP0A01990 | YDL042C | 99 | 0 |
| SPGP0A02030 | YDL035C | 99 | 0 |
| SPGP0I00200 | YJR135W-A | 99 | 0 |
| SPGP0CB00290 | YIR039C | 99 | 0 |
| SPGP0J01020 | YOL009C | 99 | 0 |
| SPGP0F01450 | YBR156C | 99 | 0 |
| SPGP0B01330 | YHR081W | 100 | 0 |
| SPGP0H00310 | YML025C | 99 | 0 |
| SPGP0J01770 | YOR076C | 99 | 0 |
| SPGP0F02860 | YBR004C | 99 | 0 |
| SPGP0F01720 | YBR127C | 95 | 0 |
| SPGP0I00240 | YJR132W | 99 | 0 |
| SPGP0SR00150 | YGL187C | 100 | 0 |
| SPGP0QD00140 | YGL179C | 99 | 0 |
| SPGP0B00430 | YHL009C | 99 | 0 |
| SPGP0H02260 | YMR181C | 99 | 0 |
| SPGP0I00190 | YJR136C | 99 | 0 |
| SPGP0I02910 | YJL170C | 99 | 0 |
| SPGP0A00890 | YDL160C-A | 100 | 0 |
| SPGP0BG00220 | YCL051W | 99 | 0 |
| SPGP0BF01160 | YIL097W | 99 | 0 |
| SPGP0I00790 | YJR074W | 99 | 0 |
| SPGP0H00980 | YMR043W | 99 | 1 |
| SPGP0X00450 | YOL121C | 99 | 0 |
| SPGP0H03370 | YMR293C | 99 | 0 |
| SPGP0RF00130 | YAL056W | 99 | 0 |
| SPGP0B01740 | YHR122W | 99 | 0 |
| SPGP0B01110 | YHR061C | 99 | 0 |
| SPGP0A03740 | YDR158W | 99 | 0 |
| SPGP0B01210 | YHR071W | 99 | 0 |
| SPGP0A00910 | YDL159W | 99 | 0 |
| SPGP0BF00160 | YIR013C | 99 | 0 |
| SPGP0G00540 | YKL152C | 99 | 0 |
| SPGP0B01430 | YHR091C | 99 | 0 |
| SPGP0RF00100 | YAL060W | 99 | 0 |
| SPGP0A01470 | YDL098C | 99 | 0 |
| SPGP0BF01570 | YIL138C | 100 | 0 |

Create PDF in your applications with the Pdfcrowd HTML to PDF API PDFCROWD

| | | | |
|---|---|---|---|
| SPGP0I01680 | YJL033W | 99 | 0 |
| SPGP0H02590 | YMR213W | 99 | 0 |
| SPGP0BF00290 | YIL001W | 99 | 0 |
| SPGP0A05100 | YDR306C | 99 | 0 |
| SPGP0I01350 | YJR009C | 100 | 0 |
| SPGP0F00690 | YBR237W | 99 | 0 |
| SPGP0L02450 | YOR370C | 99 | 0 |
| SPGP0I03260 | YJL210W | 99 | 0 |
| SPGP0QQ00130 | YAR018C | 99 | 0 |
| SPGP0A00790 | YDL173W | 99 | 0 |
| SPGP0L00530 | YOR156C | 99 | 0 |
| SPGP0BO00110 | YHR206W | 99 | 0 |
| SPGP0F03470 | YBL059C-A | 99 | 0 |
| SPGP0F00750 | YBR231C | 99 | 0 |
| SPGP0H01490 | YMR101C | 99 | 0 |
| SPGP0F02560 | YBR037C | 99 | 0 |
| SPGP0H01560 | YMR109W | 99 | 0 |
| SPGP0H03390 | YMR295C | 99 | 1 |
| SPGP0X00540 | YOL111C | 99 | 0 |
| SPGP0IY00150 | YGL246C | 99 | 0 |
| SPGP0A04630 | YDR256C | 99 | 0 |
| SPGP0L01410 | YOR254C | 99 | 0 |
| SPGP0H00410 | YML015C | 99 | 0 |
| SPGP0B02270 | YHR176W | 99 | 0 |
| SPGP0F03210 | YBL032W | 99 | 0 |
| SPGP0I01220 | YJR024C | 99 | 0 |
| SPGP0H01270 | YMR077C | 100 | 0 |
| SPGP0G00330 | YKL175W | 99 | 0 |
| SPGP0G00830 | YKL120W | 99 | 0 |
| SPGP0A00450 | YDL210W | 99 | 0 |
| SPGP0BF01110 | YIL092W | 99 | 0 |
| SPGP0A03620 | YDR143C | 99 | 0 |
| SPGP0BF01400 | YIL121W | 99 | 0 |
| SPGP0J01850 | YOR085W | 99 | 0 |
| SPGP0A00320 | YDL224C | 99 | 0 |
| SPGP0BF00580 | YIL031W | 100 | 0 |
| SPGP0I00640 | YJR092W | 99 | 0 |
| SPGP0BG00860 | YCR021C | 99 | 0 |
| SPGP0A00340 | YDL222C | 99 | 0 |
| SPGP0G00900 | YKL110C | 99 | 0 |

Create PDF in your applications with the Pdfcrowd HTML to PDF API PDFCROWD

| | | | |
|---|---|---|---|
| SPGP0H01470 | YMR099C | 100 | 0 |
| SPGP0G00390 | YKL168C | 99 | 0 |
| SPGP0A03120 | YDR087C | 99 | 0 |
| SPGP0A02930 | YDR069C | 99 | 0 |
| SPGP0B02460 | YHR195W | 99 | 0 |
| SPGP0I00230 | YJR133W | 99 | 0 |
| SPGP0A01170 | YDL129W | 99 | 0 |
| SPGP0F00120 | YBR296C | 99 | 0 |
| SPGP0A02570 | YDR028C | 99 | 0 |
| SPGP0G00750 | YKL129C | 99 | 0 |
| SPGP0A00780 | YDL174C | 99 | 0 |
| SPGP0A04870 | YDR282C | 99 | 0 |
| SPGP0NH00140 | YAL007C | 100 | 0 |
| SPGP0I03380 | YOL159C | 99 | 0 |
| SPGP0H00430 | YML013W | 99 | 0 |
| SPGP0BF01370 | YIL118W | 100 | 0 |
| SPGP0A01900 | YDL051W | 99 | 0 |
| SPGP0I00300 | YJR125C | 99 | 0 |
| SPGP0H00740 | YMR019W | 99 | 0 |
| SPGP0I02530 | YJL128C | 99 | 0 |
| SPGP0J01700 | YOR067C | 99 | 0 |
| SPGP0F00320 | YBR274W | 99 | 0 |
| SPGP0A02100 | YDL025C | 99 | 0 |
| SPGP0L02490 | YOR374W | 99 | 0 |
| SPGP0I02790 | YJL157C | 99 | 0 |
| SPGP0QG00120 | YAL043C | 99 | 0 |
| SPGP0I01190 | YJR031C | 99 | 0 |
| SPGP0BF01080 | YIL089W | 99 | 0 |
| SPGP0F01000 | YBR201C-A | 100 | 0 |
| SPGP0F01940 | YBR103W | 99 | 0 |
| SPGP0H00510 | YML005W | 99 | 0 |
| SPGP0X00180 | YOL149W | 99 | 0 |
| SPGP0B00780 | YHR026W | 99 | 0 |
| SPGP0F02680 | YBR023C | 99 | 0 |
| SPGP0ME00110 | YGL233W | 99 | 0 |
| SPGP0A04200 | YDR206W | 99 | 0 |
| SPGP0I03070 | YJL189W | 100 | 0 |
| SPGP0F01420 | YBR159W | 99 | 0 |
| SPGP0F02730 | YBR018C | 100 | 0 |
| SPGP0B01630 | YHR111W | 99 | 0 |

Create PDF in your applications with the Pdfcrowd HTML to PDF API PDFCROWD

| | | | |
|---|---|---|---|
| SPGP0BF01120 | YIL093C | 100 | 0 |
| SPGP0F03840 | YBL104C | 99 | 0 |
| SPGP0B01910 | YHR140W | 99 | 0 |
| SPGP0F02020 | YBR093C | 99 | 0 |
| SPGP0J01310 | YOR022C | 99 | 0 |
| SPGP0B00270 | YHL025W | 99 | 0 |
| SPGP0I02640 | YJL139C | 99 | 0 |
| SPGP0BF00560 | YIL027C | 100 | 0 |
| SPGP0F02790 | YBR011C | 99 | 0 |
| SPGP0A02810 | YDR057W | 99 | 0 |
| SPGP0F01880 | YBR109C | 99 | 0 |
| SPGP0A04350 | YDR225W | 100 | 0 |
| SPGP0A04420 | YDR233C | 99 | 0 |
| SPGP0BG00190 | YCL055W | 99 | 0 |
| SPGP0F03200 | YBL031W | 99 | 0 |
| SPGP0F00470 | YBR258C | 99 | 0 |
| SPGP0A01770 | YDL065C | 99 | 0 |
| SPGP0I02300 | YJL101C | 99 | 0 |
| SPGP0I02250 | YJL096W | 99 | 0 |
| SPGP0B01000 | YHR048W | 99 | 0 |
| SPGP0A04320 | YDR222W | 99 | 0 |
| SPGP0ME00200 | YGL225W | 99 | 0 |
| SPGP0L01700 | YOR287C | 99 | 0 |
| SPGP0H00130 | YML048W | 99 | 0 |
| SPGP0A01280 | YDL117W | 99 | 0 |
| SPGP0B01700 | YHR118C | 99 | 0 |
| SPGP0L00730 | YOR178C | 99 | 0 |
| SPGP0H01060 | YMR055C | 100 | 0 |
| SPGP0BF01380 | YIL119C | 98 | 1 |
| SPGP0F01780 | YBR120C | 100 | 0 |
| SPGP0A02740 | YDR049W | 99 | 0 |
| SPGP0BF00910 | YIL069C | 99 | 0 |
| SPGP0I02520 | YJL127C-B | 100 | 0 |
| SPGP0J00630 | YOL052C | 99 | 0 |
| SPGP0I00810 | YJR072C | 99 | 0 |
| SPGP0OP00120 | YAR029W | 98 | 0 |
| SPGP0B02020 | YHR151C | 99 | 0 |
| SPGP0BG00440 | YCL027W | 99 | 0 |
| SPGP0G00170 | YKL192C | 99 | 0 |
| SPGP0H01100 | YMR060C | 99 | 0 |

Create PDF in your applications with the Pdfcrowd HTML to PDF API PDFCROWD

| | | | |
|---|---|---|---|
| SPGP0G00620 | YKL143W | 99 | 0 |
| SPGP0A00520 | YDL203C | 99 | 0 |
| SPGP0H01680 | YMR121C | 99 | 0 |
| SPGP0A01570 | YDL086W | 99 | 0 |
| SPGP0BF01590 | YIL140W | 99 | 0 |
| SPGP0A00420 | YDL213C | 99 | 0 |
| SPGP0F00150 | YBR293W | 99 | 0 |
| SPGP0BF01340 | YIL115C | 99 | 0 |
| SPGP0BG00620 | YCL002C | 99 | 0 |
| SPGP0F01020 | YBR200W | 99 | 0 |
| SPGP0I01230 | YJR022W | 100 | 0 |
| SPGP0H03360 | YMR292W | 100 | 0 |
| SPGP0L01060 | YOR215C | 99 | 0 |
| SPGP0QL00150 | YAL019W | 99 | 0 |
| SPGP0X00400 | YOL126C | 99 | 0 |
| SPGP0B00770 | YHR025W | 99 | 0 |
| SPGP0GU00130 | YPL266W | 99 | 0 |
| SPGP0F02800 | YBR010W | 100 | 0 |
| SPGP0I02140 | YJL084C | 99 | 0 |
| SPGP0H03030 | YMR259C | 99 | 0 |
| SPGP0H03130 | YMR269W | 100 | 0 |
| SPGP0NS00130 | YGL194C-A | 99 | 0 |
| SPGP0H02080 | YMR163C | 99 | 0 |
| SPGP0G01160 | YKL081W | 99 | 0 |
| SPGP0BG00970 | YCR033W | 99 | 0 |
| SPGP0L01660 | YOR283W | 99 | 0 |
| SPGP0I03300 | YJL214W | 99 | 0 |
| SPGP0BF00880 | YIL066C | 99 | 0 |
| SPGP0B02540 | YHR202W | 99 | 0 |
| SPGP0G00560 | YKL150W | 99 | 0 |
| SPGP0A02000 | YDL040C | 99 | 0 |
| SPGP0H02880 | YMR242C | 100 | 0 |
| SPGP0H03310 | YMR287C | 99 | 0 |
| SPGP0JC00110 | YNL012W | 81 | 0 |
| SPGP0F02870 | YBR003W | 99 | 0 |
| SPGP0A03190 | YDR096W | 99 | 0 |
| SPGP0F02990 | YBL010C | 99 | 0 |
| SPGP0B01520 | YHR100C | 99 | 0 |
| SPGP0J01510 | YOR045W | 99 | 0 |
| SPGP0J01040 | YOL007C | 100 | 0 |

Create PDF in your applications with the Pdfcrowd HTML to PDF API PDFCROWD

| | | | |
|---|---|---|---|
| SPGP0A03830 | YDR167W | 100 | 0 |
| SPGP0B00980 | YHR046C | 99 | 0 |
| SPGP0B00630 | YHR011W | 99 | 0 |
| SPGP0B00110 | YHL044W | 98 | 0 |
| SPGP0H00290 | YML027W | 99 | 0 |
| SPGP0L00670 | YOR172W | 99 | 0 |
| SPGP0H01630 | YMR116C | 99 | 0 |
| SPGP0I00260 | YJR130C | 99 | 0 |
| SPGP0A03990 | YDR181C | 99 | 0 |
| SPGP0J01180 | YOR008C | 98 | 2 |
| SPGP0H00650 | YMR010W | 99 | 0 |
| SPGP0H02110 | YMR166C | 99 | 0 |
| SPGP0B01580 | YHR106W | 99 | 0 |
| SPGP0H02280 | YMR183C | 99 | 0 |
| SPGP0A01730 | YDL070W | 99 | 0 |
| SPGP0BF01140 | YIL095W | 99 | 0 |
| SPGP0B01170 | YHR067W | 100 | 0 |
| SPGP0BF00700 | YIL044C | 99 | 0 |
| SPGP0A05290 | YDR323C | 99 | 0 |
| SPGP0LL00160 | YAL031C | 99 | 0 |
| SPGP0B01760 | YHR124W | 99 | 0 |
| SPGP0G00980 | YKL101W | 99 | 0 |
| SPGP0B02180 | YHR167W | 99 | 0 |
| SPGP0BF00180 | YIR011C | 99 | 0 |
| SPGP0I02730 | YJL149W | 99 | 0 |
| SPGP0A01450 | YDL100C | 99 | 0 |
| SPGP0F02610 | YBR031W | 99 | 0 |
| SPGP0H02610 | YMR215W | 99 | 0 |
| SPGP0BF01550 | YIL136W | 99 | 0 |
| SPGP0A05120 | YDR308C | 99 | 0 |
| SPGP0I01330 | YJR010C-A | 100 | 0 |
| SPGP0F02220 | YBR072W | 99 | 0 |
| SPGP0F01170 | YBR185C | 99 | 0 |
| SPGP0G00600 | YKL145W | 99 | 0 |
| SPGP0L00550 | YOR158W | 99 | 0 |
| SPGP0H00760 | YMR021C | 99 | 0 |
| SPGP0H02210 | YMR176W | 99 | 0 |
| SPGP0I03240 | YJL208C | 99 | 0 |
| SPGP0I01790 | YJL045W | 99 | 0 |

Create PDF in your applications with the Pdfcrowd HTML to PDF API PDFCROWD

| | | | |
|---|---|---|---|
| SPGP0F00670 | YBR239C | 99 | 0 |
| SPGP0A01950 | YDL045W-A | 99 | 0 |
| SPGP0J00830 | YOL028C | 99 | 0 |
| SPGP0I02930 | YJL172W | 99 | 0 |
| SPGP0QX00210 | YMR309C | 99 | 0 |
| SPGP0BF00790 | YIL053W | 99 | 0 |
| SPGP0A00620 | YDL192W | 99 | 0 |
| SPGP0F01520 | YBR149W | 99 | 0 |
| SPGP0IY00170 | YGL248W | 99 | 0 |
| SPGP0H00680 | YMR013C | 100 | 0 |
| SPGP0I03060 | YJL187C | 99 | 0 |
| SPGP0L01470 | YOR260W | 99 | 0 |
| SPGP0A03600 | YDR141C | 99 | 0 |
| SPGP0F03230 | YBL034C | 99 | 0 |
| SPGP0H01210 | YMR071C | 99 | 0 |
| SPGP0SC00110 | YER175C | 99 | 0 |
| SPGP0FK00180 | YGL206C | 99 | 0 |
| SPGP0BF01130 | YIL094C | 99 | 0 |
| SPGP0BF01280 | YIL109C | 99 | 0 |
| SPGP0QX00100 | YMR319C | 99 | 0 |
| SPGP0A04030 | YDR185C | 99 | 0 |
| SPGP0BF00260 | YIR003W | 99 | 0 |
| SPGP0A00220 | YDL235C | 99 | 0 |
| SPGP0L01450 | YOR258W | 99 | 0 |
| SPGP0BF01330 | YIL114C | 99 | 0 |
| SPGP0BF01420 | YIL123W | 99 | 0 |
| SPGP0SR00140 | YGL189C | 100 | 0 |
| SPGP0QG00130 | YAL044C | 100 | 0 |
| SPGP0IG00120 | YPL249C | 99 | 0 |
| SPGP0G00490 | YKL159C | 100 | 0 |
| SPGP0H01950 | YMR149W | 100 | 0 |
| SPGP0J01870 | YOR087W | 99 | 0 |
| SPGP0L00620 | YOR165W | 99 | 0 |
| SPGP0F03690 | YBL087C | 98 | 1 |
| SPGP0H02740 | YMR228W | 99 | 0 |
| SPGP0BG00930 | YCR028C-A | 100 | 0 |
| SPGP0A03060 | YDR081C | 99 | 0 |
| SPGP0G01030 | YKL095W | 99 | 0 |
| SPGP0B02110 | YHR160C | 99 | 0 |
| SPGP0I00250 | YJR131W | 99 | 0 |

Create PDF in your applications with the Pdfcrowd HTML to PDF API PDFCROWD

| | | | |
|---|---|---|---|
| SPGP0B02440 | YHR193C | 100 | 0 |
| SPGP0B01960 | YHR144C | 99 | 0 |
| SPGP0A02550 | YDR026C | 99 | 0 |
| SPGP0F00520 | YBR254C | 99 | 0 |
| SPGP0G00770 | YKL127W | 96 | 0 |
| SPGP0NH00160 | YAL009W | 99 | 0 |
| SPGP0J00580 | YOL056W | 99 | 0 |
| SPGP0BF01310 | YIL112W | 99 | 0 |
| SPGP0X00210 | YOL146W | 99 | 0 |
| SPGP0J01720 | YOR070C | 100 | 0 |
| SPGP0I02770 | YJL155C | 99 | 0 |
| SPGP0B01730 | YHR121W | 99 | 0 |
| SPGP0QG00100 | YAL041W | 99 | 0 |
| SPGP0F01920 | YBR105C | 99 | 0 |
| SPGP0H00100 | YML051W | 100 | 0 |
| SPGP0H00820 | YMR027W | 99 | 0 |
| SPGP0A03520 | YDR131C | 99 | 0 |
| SPGP0X00280 | YOL139C | 99 | 0 |
| SPGP0H00530 | YML002W | 99 | 0 |
| SPGP0H02890 | YMR243C | 99 | 0 |
| SPGP0B01300 | YHR079C | 99 | 0 |
| SPGP0B00760 | YHR024C | 100 | 0 |
| SPGP0F00140 | YBR294W | 99 | 0 |
| SPGP0I01380 | YJR006W | 99 | 0 |
| SPGP0F03520 | YBL066C | 99 | 0 |
| SPGP0H03150 | YMR271C | 99 | 0 |
| SPGP0J01070 | YOL004W | 99 | 0 |
| SPGP0F00430 | YBR262C | 100 | 0 |
| SPGP0A04860 | YDR281C | 99 | 0 |
| SPGP0FK00190 | YGL205W | 99 | 0 |
| SPGP0H00790 | YMR024W | 100 | 0 |
| SPGP0H01050 | YMR054W | 99 | 0 |
| SPGP0B01970 | YHR146W | 99 | 0 |
| SPGP0OP00110 | YAR031W | 96 | 0 |
| SPGP0J00140 | YOL102C | 99 | 0 |
| SPGP0B00370 | YHL015W | 99 | 0 |
| SPGP0B00250 | YHL027W | 99 | 0 |
| SPGP0F03380 | YBL050W | 99 | 0 |
| SPGP0I00450 | YJR109C | 99 | 0 |
| SPGP0VX00130 | YGL219C | 100 | 0 |

Create PDF in your applications with the Pdfcrowd HTML to PDF API PDFCROWD

| | | | |
|---|---|---|---|
| SPGP0BF01210 | YIL102C-A | 99 | 0 |
| SPGP0BF00540 | YIL024C | 99 | 0 |
| SPGP0I03000 | YJL180C | 99 | 0 |
| SPGP0I02600 | YJL134W | 99 | 0 |
| SPGP0BF00890 | YIL067C | 99 | 0 |
| SPGP0I02320 | YJL103C | 99 | 0 |
| SPGP0B01390 | YHR087W | 100 | 0 |
| SPGP0F01990 | YBR096W | 99 | 0 |
| SPGP0A05030 | YDR299W | 99 | 0 |
| SPGP0A04400 | YDR231C | 99 | 0 |
| SPGP0F03750 | YBL092W | 100 | 0 |
| SPGP0H02840 | YMR238W | 99 | 0 |
| SPGP0L01930 | YOR310C | 99 | 1 |
| SPGP0I01250 | YJR019C | 99 | 0 |
| SPGP0A01750 | YDL067C | 99 | 0 |
| SPGP0F03220 | YBL033C | 99 | 0 |
| SPGP0L00240 | YOR124C | 99 | 0 |
| SPGP0BF00670 | YIL041W | 99 | 0 |
| SPGP0A04340 | YDR224C | 100 | 0 |
| SPGP0B01020 | YHR050W | 99 | 0 |
| SPGP0I02540 | YJL129C | 99 | 0 |
| SPGP0SR00120 | YGL191W | 99 | 0 |
| SPGP0H01840 | YMR137C | 99 | 0 |
| SPGP0F03530 | YBL067C | 99 | 0 |
| SPGP0H01040 | YMR053C | 99 | 0 |
| SPGP0L00710 | YOR176W | 100 | 0 |
| SPGP0I01470 | YJL003W | 100 | 0 |
| SPGP0BG00320 | YCL039W | 99 | 0 |
| SPGP0A02760 | YDR051C | 99 | 0 |
| SPGP0I00800 | YJR073C | 99 | 0 |
| SPGP0OP00100 | YAR033W | 98 | 0 |
| SPGP0A01150 | YDL131W | 99 | 0 |
| SPGP0BG00460 | YCL026C-A | 100 | 0 |
| SPGP0A00540 | YDL201W | 99 | 0 |
| SPGP0A00400 | YDL215C | 99 | 0 |
| SPGP0BG00610 | YCL004W | 99 | 0 |
| SPGP0IW00100 | YML056C | 90 | 1 |
| SPGP0A02490 | YDR020C | 99 | 0 |
| SPGP0BF01360 | YIL117C | 99 | 0 |
| SPGP0F00990 | YBR202W | 99 | 0 |

Create PDF in your applications with the Pdfcrowd HTML to PDF API PDFCROWD

| | | | |
|---|---|---|---|
| SPGP0I02170 | YJL088W | 99 | 0 |
| SPGP0BF00360 | YIL007C | 99 | 0 |
| SPGP0F03340 | YBL045C | 99 | 0 |
| SPGP0L00600 | YOR163W | 99 | 0 |
| SPGP0BG00470 | YCL025C | 99 | 0 |
| SPGP0H02530 | YMR207C | 99 | 0 |
| SPGP0L01590 | YOR274W | 99 | 0 |
| SPGP0A04910 | YDR286C | 99 | 0 |
| SPGP0QL00170 | YAL017W | 99 | 0 |
| SPGP0B01220 | YHR072W | 99 | 0 |
| SPGP0H00490 | YML007W | 99 | 0 |
| SPGP0A03580 | YDR139C | 100 | 0 |
| SPGP0RY00140 | YER169W | 99 | 0 |
| SPGP0H01980 | YMR153W | 99 | 0 |
| SPGP0GU00110 | YPL264C | 99 | 0 |
| SPGP0L01620 | YOR278W | 99 | 0 |
| SPGP0A03010 | YDR077W | 99 | 0 |
| SPGP0F01390 | YBR162C | 99 | 0 |
| SPGP0L01680 | YOR285W | 99 | 0 |
| SPGP0X00340 | YOL132W | 99 | 0 |
| SPGP0H01310 | YMR081C | 99 | 0 |
| SPGP0BG00950 | YCR031C | 100 | 0 |
| SPGP0F01230 | YBR179C | 99 | 0 |
| SPGP0MK00110 | YKR105C | 82 | 1 |
| SPGP0L02240 | YOR348C | 99 | 0 |
| SPGP0I02440 | YJL118W | 99 | 0 |
| SPGP0I00520 | YJR103W | 99 | 0 |
| SPGP0CB00250 | YIR035C | 100 | 0 |
| SPGP0A03170 | YDR092W | 100 | 0 |
| SPGP0F00270 | YBR280C | 99 | 0 |
| SPGP0H03420 | YMR298W | 99 | 0 |
| SPGP0F02100 | YBR084C-A | 99 | 0 |
| SPGP0I02690 | YJL145W | 99 | 0 |
| SPGP0J01060 | YOL005C | 99 | 0 |
| SPGP0F01040 | YBR198C | 99 | 0 |
| SPGP0RF00110 | YAL059W | 99 | 0 |
| SPGP0J00930 | YOL018C | 99 | 0 |
| SPGP0L02020 | YOR322C | 99 | 0 |
| SPGP0PZ00190 | YGR225W | 80 | 1 |
| SPGP0A02950 | YDR071C | 100 | 0 |

Create PDF in your applications with the Pdfcrowd HTML to PDF API PDFCROWD

| | | | |
|---|---|---|---|
| SPGP0B00610 | YHR009C | 99 | 0 |
| SPGP0H00350 | YML021C | 99 | 0 |
| SPGP0F03420 | YBL055C | 99 | 0 |
| SPGP0J00610 | YOL053W | 99 | 0 |
| SPGP0H01610 | YMR114C | 99 | 0 |
| SPGP0F01640 | YBR136W | 99 | 0 |
| SPGP0B01750 | YHR123W | 100 | 0 |
| SPGP0QD00100 | YGL184C | 99 | 0 |
| SPGP0H00670 | YMR012W | 99 | 0 |
| SPGP0BG00260 | YCL048W | 99 | 0 |
| SPGP0A01710 | YDL073W | 99 | 0 |
| SPGP0H02300 | YMR185W | 99 | 0 |
| SPGP0A01940 | YDL046W | 99 | 0 |
| SPGP0A01430 | YDL102W | 99 | 0 |
| SPGP0BF00680 | YIL042C | 99 | 0 |
| SPGP0LL00140 | YAL029C | 99 | 0 |
| SPGP0H00780 | YMR023C | 99 | 0 |
| SPGP0X00410 | YOL125W | 99 | 0 |
| SPGP0I00440 | YJR110W | 99 | 0 |
| SPGP0I01910 | YJL057C | 99 | 0 |
| SPGP0J00440 | YOL070C | 99 | 0 |
| SPGP0BF00120 | YIR017C | 100 | 0 |
| SPGP0J00860 | YOL025W | 99 | 0 |
| SPGP0B02160 | YHR165C | 99 | 0 |
| SPGP0QQ00150 | YAR014C | 99 | 0 |
| SPGP0A01380 | YDL107W | 99 | 0 |
| SPGP0F02930 | YBL004W | 99 | 0 |
| SPGP0I01640 | YJL027C | 100 | 0 |
| SPGP0GO00100 | YAL040C | 99 | 0 |
| SPGP0I01310 | YJR012C | 99 | 0 |
| SPGP0H02150 | YMR170C | 99 | 0 |
| SPGP0I01810 | YJL047C | 99 | 0 |
| SPGP0F02240 | YBR070C | 99 | 0 |
| SPGP0A02080 | YDL028C | 99 | 0 |
| SPGP0A04090 | YDR192C | 99 | 0 |
| SPGP0J01280 | YOR020C | 100 | 0 |
| SPGP0I01020 | YJR050W | 99 | 0 |
| SPGP0H01180 | YMR068W | 100 | 0 |
| SPGP0BF00440 | YGR161W-B | 99 | 0 |
| SPGP0A04590 | YDR252W | 99 | 0 |

Create PDF in your applications with the Pdfcrowd HTML to PDF API PDFCROWD

| | | | |
|---|---|---|---|
| SPGP0IY00110 | YGL242C | 99 | 0 |
| SPGP0F02260 | YBR068C | 99 | 0 |
| SPGP0H00840 | YMR029C | 99 | 0 |
| SPGP0A04670 | YDR260C | 99 | 0 |
| SPGP0I00990 | YJR053W | 99 | 0 |
| SPGP0B02390 | YHR188C | 99 | 0 |
| SPGP0F03250 | YBL036C | 99 | 0 |
| SPGP0VX00100 | YGL222C | 99 | 0 |
| SPGP0H01450 | YMR097C | 99 | 0 |
| SPGP0G00370 | YKL171W | 99 | 0 |
| SPGP0IY00200 | YGL251C | 99 | 0 |
| SPGP0WU00100 | YER177W | 99 | 0 |
| SPGP0H02180 | YMR173W | 87 | 5 |
| SPGP0H01230 | YMR073C | 99 | 0 |
| SPGP0A03080 | YDR083W | 99 | 0 |
| SPGP0A04470 | YDR238C | 99 | 0 |
| SPGP0L00500 | YOR153W | 99 | 0 |
| SPGP0IG00100 | YPL250C | 99 | 0 |
| SPGP0CB00160 | YIR026C | 99 | 0 |
| SPGP0BG00760 | YCR014C | 99 | 0 |
| SPGP0L00740 | YOR179C | 99 | 0 |
| SPGP0L01900 | YOR306C | 99 | 0 |
| SPGP0F01130 | YBR189W | 100 | 0 |
| SPGP0H03320 | YMR288W | 99 | 0 |
| SPGP0A01210 | YDL125C | 100 | 0 |
| SPGP0BF00850 | YIL063C | 99 | 0 |
| SPGP0B02420 | YHR191C | 100 | 0 |
| SPGP0B02130 | YHR162W | 99 | 0 |
| SPGP0A03810 | YDR165W | 99 | 0 |
| SPGP0F00500 | YBR255C-A | 99 | 0 |
| SPGP0A02530 | YDR024W | 99 | 0 |
| SPGP0QX00190 | YMR311C | 99 | 0 |
| SPGP0NH00100 | YAL001C | 99 | 0 |
| SPGP0J00210 | YOL094C | 99 | 0 |
| SPGP0BG00900 | YCR026C | 99 | 0 |
| SPGP0BF01150 | YIL096C | 99 | 0 |
| SPGP0SR00160 | YGL186C | 100 | 0 |
| SPGP0I03310 | YJL216C | 96 | 0 |
| SPGP0A03570 | YDR138W | 99 | 0 |
| SPGP0X00230 | YOL144W | 97 | 2 |

Create PDF in your applications with the Pdfcrowd HTML to PDF API PDFCROWD

| | | | |
|---|---|---|---|
| SPGP0I00270 | YJR129C | 99 | 0 |
| SPGP0L01020 | YOR211C | 99 | 0 |
| SPGP0QX00220 | YMR308C | 99 | 0 |
| SPGP0X00550 | YOL110W | 99 | 0 |
| SPGP0A02140 | YDL020C | 99 | 0 |
| SPGP0J00750 | YOL039W | 100 | 0 |
| SPGP0B00740 | YHR022C | 97 | 0 |
| SPGP0F01890 | YBR108W | 98 | 0 |
| SPGP0I01320 | YJR011C | 99 | 0 |
| SPGP0H02870 | YMR241W | 99 | 0 |
| SPGP0A02250 | YDL007W | 99 | 0 |
| SPGP0A02770 | YDR052C | 99 | 0 |
| SPGP0J01050 | YOL006C | 99 | 0 |
| SPGP0L02040 | YOR324C | 99 | 0 |
| SPGP0F03810 | YBL101C | 99 | 0 |
| SPGP0F01460 | YBR155W | 99 | 0 |
| SPGP0H00360 | YML020W | 99 | 0 |
| SPGP0A04840 | YDR279W | 99 | 0 |
| SPGP0BF00810 | YIL056W | 99 | 0 |
| SPGP0F02770 | YBR014C | 99 | 0 |
| SPGP0BF00420 | YIL013C | 99 | 0 |
| SPGP0I02080 | YJL076W | 99 | 0 |
| SPGP0J01300 | YOR021C | 100 | 0 |
| SPGP0B00950 | YHR042W | 99 | 0 |
| SPGP0F02700 | YBR021W | 99 | 0 |
| SPGP0B00390 | YHL013C | 100 | 0 |
| SPGP0I01210 | YJR025C | 99 | 0 |
| SPGP0I00430 | YJR111C | 99 | 0 |
| SPGP0B00310 | YHL021C | 99 | 0 |
| SPGP0H00110 | YML050W | 100 | 0 |
| SPGP0I02340 | YJL105W | 99 | 0 |
| SPGP0A02850 | YDR061W | 99 | 0 |
| SPGP0F03880 | YNR076W | 99 | 0 |
| SPGP0F03730 | YBL091C | 99 | 0 |
| SPGP0A01800 | YDL061C | 100 | 0 |
| SPGP0L00260 | YOR126C | 99 | 0 |
| SPGP0F00610 | YBR245C | 99 | 0 |
| SPGP0VX00110 | YGL221C | 100 | 0 |
| SPGP0F02420 | YBR052C | 99 | 0 |
| SPGP0A03430 | YDR122W | 99 | 0 |

Create PDF in your applications with the Pdfcrowd HTML to PDF API PDFCROWD

| | | | |
|---|---|---|---|
| SPGP0G00130 | YKL196C | 99 | 0 |
| SPGP0I01260 | YJR017C | 99 | 0 |
| SPGP0A04360 | YDR226W | 99 | 0 |
| SPGP0I03160 | YJL199C | 99 | 0 |
| SPGP0BF01690 | YIL151C | 99 | 0 |
| SPGP0A02970 | YDR073W | 99 | 0 |
| SPGP0L00190 | YOR118W | 100 | 0 |
| SPGP0I01450 | YJL001W | 99 | 0 |
| SPGP0A00560 | YDL199C | 99 | 0 |
| SPGP0I01100 | YJR042W | 99 | 0 |
| SPGP0H03100 | YMR266W | 99 | 0 |
| SPGP0BF00650 | YIL039W | 99 | 0 |
| SPGP0H01190 | YMR069W | 99 | 0 |
| SPGP0I00850 | YJR067C | 99 | 0 |
| SPGP0BF00980 | YIL075C | 99 | 0 |
| SPGP0BG00480 | YCL024W | 99 | 0 |
| SPGP0A01620 | YDL082W | 99 | 0 |
| SPGP0L00990 | YOR208W | 99 | 0 |
| SPGP0F01350 | YBR165W | 99 | 0 |
| SPGP0IY00250 | YGL256W | 99 | 0 |
| SPGP0H00700 | YMR015C | 99 | 0 |
| SPGP0H02510 | YMR205C | 99 | 0 |
| SPGP0A04280 | YDR217C | 99 | 0 |
| SPGP0F00970 | YBR204C | 99 | 0 |
| SPGP0A04890 | YDR284C | 99 | 0 |
| SPGP0QL00110 | YAL023C | 99 | 0 |
| SPGP0L00610 | YOR164C | 99 | 0 |
| SPGP0A00230 | YDL234C | 99 | 0 |
| SPGP0B02090 | YHR158C | 99 | 0 |
| SPGP0F02030 | YBR091C | 99 | 0 |
| SPGP0H01770 | YMR130W | 99 | 0 |
| SPGP0BF00330 | YIL004C | 99 | 0 |
| SPGP0I00580 | YJR097W | 99 | 0 |
| SPGP0A03150 | YDR090C | 99 | 0 |
| SPGP0H01320 | YMR083W | 99 | 0 |
| SPGP0L01190 | YOR230W | 99 | 0 |
| SPGP0BG01010 | YCR037C | 99 | 0 |
| SPGP0G00320 | YKL176C | 99 | 0 |
| SPGP0L02220 | YOR346W | 99 | 0 |
| SPGP0B00340 | YHL018W | 99 | 0 |

Create PDF in your applications with the Pdfcrowd HTML to PDF API PDFCROWD

| | | | |
|---|---|---|---|
| SPGP0I00150 | YJR140C | 99 | 0 |
| SPGP0F03360 | YBL047C | 99 | 1 |
| SPGP0I00220 | YJR134C | 99 | 0 |
| SPGP0H00470 | YML009C | 100 | 0 |
| SPGP0F00260 | YBR281C | 99 | 0 |
| SPGP0G00210 | YKL188C | 99 | 0 |
| SPGP0B00940 | YHR041C | 99 | 0 |
| SPGP0RH00100 | YLL015W | 80 | 0 |
| SPGP0A01880 | YDL053C | 99 | 0 |
| SPGP0BF01100 | YIL091C | 99 | 0 |
| SPGP0I03040 | YJL185C | 99 | 0 |
| SPGP0BG00240 | YCL049C | 99 | 0 |
| SPGP0H03490 | YMR305C | 99 | 0 |
| SPGP0H00330 | YML023C | 99 | 0 |
| SPGP0G00240 | YKL185W | 99 | 0 |
| SPGP0QD00120 | YGL181W | 97 | 3 |
| SPGP0I02380 | YJL110C | 99 | 0 |
| SPGP0H00610 | YMR007W | 99 | 0 |
| SPGP0A00820 | YDL169C | 99 | 0 |
| SPGP0A02280 | YDL004W | 99 | 0 |
| SPGP0B00490 | YHL002W | 99 | 1 |
| SPGP0CB00270 | YIR037W | 99 | 0 |
| SPGP0BG00100 | YJL225C | 99 | 0 |
| SPGP0I01830 | YJL049W | 99 | 0 |
| SPGP0J01680 | YOR065W | 99 | 0 |
| SPGP0LL00120 | YAL027W | 99 | 0 |
| SPGP0A01970 | YDL044C | 99 | 0 |
| SPGP0B01540 | YHR102W | 99 | 0 |
| SPGP0B02140 | YHR163W | 99 | 0 |
| SPGP0I01930 | YJL059W | 99 | 0 |
| SPGP0A01390 | YDL106C | 99 | 0 |
| SPGP0J00880 | YOL023W | 99 | 0 |
| SPGP0J00420 | YOL072W | 99 | 0 |
| SPGP0B01820 | YHR132C | 99 | 0 |
| SPGP0J00870 | YOL024W | 100 | 0 |
| SPGP0H02650 | YMR219W | 99 | 0 |
| SPGP0NH00170 | YAL010C | 99 | 0 |
| SPGP0F02190 | YBR076W | 99 | 0 |
| SPGP0B01720 | YHR120W | 99 | 0 |
| SPGP0A03400 | YDR119W-A | 100 | 0 |

Create PDF in your applications with the Pdfcrowd HTML to PDF API PDFCROWD

| | | | |
|---|---|---|---|
| SPGP0F02630 | YBR029C | 99 | 0 |
| SPGP0H02170 | YMR172W | 99 | 0 |
| SPGP0I01750 | YJL041W | 100 | 0 |
| SPGP0A04070 | YDR190C | 99 | 0 |
| SPGP0I01000 | YJR052W | 99 | 0 |
| SPGP0I01660 | YJL030W | 99 | 0 |
| SPGP0B01810 | YHR131C | 99 | 0 |
| SPGP0I02480 | YJL124C | 100 | 0 |
| SPGP0IY00130 | YGL244W | 99 | 0 |
| SPGP0F00450 | YBR260C | 100 | 0 |
| SPGP0BF00480 | YIL018W | 100 | 0 |
| SPGP0A04610 | YDR254W | 99 | 0 |
| SPGP0BG00250 | YCL048W-A | 100 | 0 |
| SPGP0NW00100 | YPL282C | 99 | 0 |
| SPGP0VX00120 | YGL220W | 99 | 0 |
| SPGP0G00850 | YKL117W | 99 | 0 |
| SPGP0A03670 | YDR148C | 99 | 1 |
| SPGP0RY00120 | YER171W | 99 | 0 |
| SPGP0I00820 | YJR070C | 99 | 0 |
| SPGP0B00130 | YHL040C | 99 | 0 |
| SPGP0B00930 | YHR040W | 99 | 0 |
| SPGP0G00110 | YKL198C | 98 | 1 |
| SPGP0H02160 | YMR171C | 99 | 0 |
| SPGP0SV00100 | YCR097W | 91 | 0 |
| SPGP0B02330 | YHR182W | 99 | 0 |
| SPGP0F01490 | YBR152W | 99 | 0 |
| SPGP0BF01480 | YIL129C | 99 | 0 |
| SPGP0F03270 | YBL038W | 99 | 0 |
| SPGP0L00520 | YOR155C | 99 | 0 |
| SPGP0PM00120 | YGR289C | 98 | 0 |
| SPGP0L01670 | YOR284W | 99 | 0 |
| SPGP0A02510 | YDR022C | 100 | 0 |
| SPGP0BF00320 | YIL003W | 100 | 0 |
| SPGP0F03180 | YBL029C-A | 100 | 0 |
| SPGP0F02490 | YBR044C | 99 | 0 |
| SPGP0F00850 | YBR218C | 99 | 0 |
| SPGP0SR00100 | YGL194C | 99 | 0 |
| SPGP0B02150 | YHR164C | 99 | 0 |
| SPGP0B02400 | YHR189W | 99 | 0 |

Create PDF in your applications with the Pdfcrowd HTML to PDF API PDFCROWD

| | | | |
|---|---|---|---|
| SPGP0A01190 | YDL127W | 99 | 0 |
| SPGP0IY00220 | YGL253W | 99 | 0 |
| SPGP0J00270 | YOL088C | 99 | 0 |
| SPGP0BF01170 | YIL098C | 99 | 0 |
| SPGP0A01960 | YDL045C | 99 | 0 |
| SPGP0L01920 | YOR308C | 99 | 0 |
| SPGP0X00250 | YOL142W | 99 | 0 |
| SPGP0A00610 | YDL193W | 99 | 0 |
| SPGP0QX00240 | YMR306W | 99 | 0 |
| SPGP0G00660 | YKL139W | 99 | 0 |
| SPGP0H02500 | YMR204C | 99 | 0 |
| SPGP0A02270 | YDL005C | 98 | 1 |
| SPGP0B01260 | YHR075C | 100 | 0 |
| SPGP0A01830 | YDL058W | 99 | 0 |
| SPGP0A04940 | YDR289C | 99 | 0 |
| SPGP0A04210 | YDR207C | 99 | 0 |
| SPGP0H00860 | YMR031C | 99 | 0 |
| SPGP0A02750 | YDR050C | 99 | 0 |
| SPGP0BF00190 | YIR010W | 99 | 1 |
| SPGP0B00720 | YHR020W | 99 | 0 |
| SPGP0F00100 | YBR298C | 99 | 0 |
| SPGP0H01730 | YMR126C | 99 | 0 |
| SPGP0B01320 | YHR080C | 99 | 0 |
| SPGP0F00920 | YBR211C | 99 | 0 |
| SPGP0L00780 | YOR184W | 99 | 0 |
| SPGP0J01800 | YOR079C | 99 | 0 |
| SPGP0FK00150 | YGL209W | 99 | 0 |
| SPGP0A02020 | YDL036C | 99 | 0 |
| SPGP0F01440 | YBR157C | 99 | 0 |
| SPGP0F02750 | YBR016W | 99 | 1 |
| SPGP0A01760 | YDL066W | 100 | 0 |
| SPGP0J01320 | YOR023C | 99 | 0 |
| SPGP0A01030 | YDL144C | 99 | 0 |
| SPGP0BF00510 | YIL021W | 99 | 0 |
| SPGP0GU00140 | YPL267W | 99 | 0 |
| SPGP0H02310 | YMR186W | 99 | 0 |
| SPGP0B02520 | YHR200W | 100 | 0 |
| SPGP0J00240 | YOL091W | 99 | 0 |
| SPGP0I02360 | YJL107C | 99 | 0 |
| SPGP0A02590 | YDR031W | 99 | 0 |

Create PDF in your applications with the Pdfcrowd HTML to PDF API PDFCROWD

| | | | |
|---|---|---|---|
| SPGP0QG00170 | YAL048C | 99 | 0 |
| SPGP0F03350 | YBL046W | 97 | 2 |
| SPGP0BG00450 | YCL026C-B | 100 | 0 |
| SPGP0RN00120 | YER166W | 99 | 0 |
| SPGP0BF01710 | YIL153W | 99 | 0 |
| SPGP0F03710 | YBL089W | 99 | 0 |
| SPGP0F03260 | YBL037W | 99 | 0 |
| SPGP0H03210 | YMR277W | 100 | 0 |
| SPGP0L00200 | YOR119C | 99 | 0 |
| SPGP0F00590 | YBR247C | 99 | 0 |
| SPGP0ME00220 | YGL223C | 99 | 0 |
| SPGP0A04920 | YDR287W | 99 | 0 |
| SPGP0B01060 | YHR056C | 99 | 0 |
| SPGP0L00210 | YOR120W | 99 | 0 |
| SPGP0F02400 | YBR054W | 98 | 1 |
| SPGP0I02220 | YJL093C | 99 | 0 |
| SPGP0G00150 | YKL194C | 99 | 0 |
| SPGP0H03190 | YMR275C | 99 | 0 |
| SPGP0I03140 | YJL197W | 99 | 0 |
| SPGP0A02870 | YDR063W | 99 | 0 |
| SPGP0BG00540 | YCL012C | 100 | 0 |
| SPGP0A04440 | YDR235W | 99 | 0 |
| SPGP0B02490 | YHR198C | 100 | 0 |
| SPGP0BG00710 | YCR008W | 99 | 0 |
| SPGP0BF01670 | YIL149C | 99 | 0 |
| SPGP0A00580 | YDL197C | 99 | 0 |
| SPGP0BF00630 | YIL037C | 99 | 0 |
| SPGP0I01430 | YJR002W | 99 | 0 |
| SPGP0A00670 | YDL186W | 99 | 0 |
| SPGP0BF00380 | YIL009W | 99 | 0 |
| SPGP0J00220 | YOL093W | 99 | 0 |
| SPGP0I00760 | YJR077C | 99 | 0 |
| SPGP0A01640 | YDL080C | 99 | 0 |
| SPGP0QL00130 | YAL021C | 99 | 0 |
| SPGP0BG00580 | YCL008C | 99 | 0 |
| SPGP0BF01320 | YIL113W | 99 | 0 |
| SPGP0H02570 | YMR211W | 99 | 0 |
| SPGP0L01000 | YOR209C | 99 | 0 |
| SPGP0H01780 | YMR131C | 99 | 0 |
| SPGP0NM00110 | YGL258W | 99 | 0 |

Create PDF in your applications with the Pdfcrowd HTML to PDF API PDFCROWD

| | | | |
|---|---|---|---|
| SPGP0A00210 | YDL236W | 99 | 0 |
| SPGP0H01300 | YMR080C | 99 | 0 |
| SPGP0I01630 | YJL026W | 99 | 0 |
| SPGP0A03050 | YDR080W | 99 | 0 |
| SPGP0H02390 | YMR194W | 99 | 0 |
| SPGP0A03130 | YDR088C | 99 | 0 |
| SPGP0BG00990 | YCR035C | 99 | 0 |
| SPGP0I03270 | YJL212C | 99 | 0 |
| SPGP0A03200 | YDR097C | 99 | 0 |
| SPGP0H00400 | YML016C | 99 | 0 |
| SPGP0L01210 | YOR232W | 99 | 0 |
| SPGP0BF01680 | YIL150C | 99 | 0 |
| SPGP0QQ00160 | YAR008W | 99 | 0 |
| SPGP0G00950 | YKL105C | 99 | 0 |
| SPGP0L01740 | YOR291W | 99 | 0 |
| SPGP0H00450 | YML011C | 99 | 0 |
| SPGP0VT00140 | YPL257W | 92 | 0 |
| SPGP0F00240 | YBR283C | 99 | 0 |
| SPGP0G00270 | YKL182W | 99 | 0 |
| SPGP0G00580 | YKL148C | 99 | 0 |
| SPGP0B00920 | YHR039C-A | 100 | 0 |
| SPGP0I01080 | YJR044C | 100 | 0 |
| SPGP0I02710 | YJL147C | 99 | 0 |
| SPGP0LA00150 | YKL215C | 81 | 1 |
| SPGP0SR00170 | YGL185C | 99 | 0 |
| SPGP0A00800 | YDL171C | 99 | 0 |
| SPGP0H03300 | YMR286W | 99 | 0 |
| SPGP0A02260 | YDL006W | 99 | 0 |
| SPGP0CB00210 | YIR031C | 100 | 0 |
| SPGP0A05270 | YDR322W | 99 | 0 |
| SPGP0I01950 | YJL061W | 99 | 0 |
| SPGP0F02600 | YBR033W | 99 | 0 |
| SPGP0A01410 | YDL104C | 99 | 0 |
| SPGP0OP00150 | YAR027W | 99 | 0 |
| SPGP0B00210 | YHL031C | 99 | 0 |
| SPGP0B00140 | YHL039W | 99 | 0 |
| SPGP0A01650 | YDL079C | 99 | 0 |
| SPGP0LL00100 | YAL025C | 99 | 0 |
| SPGP0F03500 | YBL063W | 99 | 0 |
| SPGP0F02200 | YBR074W | 99 | 0 |

Create PDF in your applications with the Pdfcrowd HTML to PDF API PDFCROWD

| | | | |
|---|---|---|---|
| SPGP0B01290 | YHR078W | 99 | 0 |
| SPGP0NH00190 | YAL012W | 99 | 0 |
| SPGP0ED00130 | YEL070W | 82 | 1 |
| SPGP0A04580 | YDR251W | 99 | 0 |
| SPGP0VZ00190 | YDR342C | 88 | 0 |
| SPGP0H02340 | YMR189W | 99 | 0 |
| SPGP0A04050 | YDR188W | 99 | 0 |
| SPGP0I00980 | YJR054W | 99 | 0 |
| SPGP0F03370 | YBL049W | 100 | 0 |
| SPGP0H02660 | YMR220W | 99 | 0 |
| SPGP0I00600 | YJR095W | 100 | 0 |
| SPGP0A05180 | YDR314C | 99 | 0 |
| SPGP0B02220 | YHR171W | 99 | 0 |
| SPGP0X00110 | YJL221C | 99 | 0 |
| SPGP0L01380 | YOR251C | 99 | 0 |
| SPGP0F01070 | YBR195C | 99 | 0 |
| SPGP0G00760 | YKL128C | 95 | 0 |
| SPGP0F01550 | YBR146W | 99 | 0 |
| SPGP0BF00250 | YIR004W | 99 | 0 |
| SPGP0H01410 | YMR093W | 99 | 0 |
| SPGP0G00870 | YKL114C | 99 | 0 |
| SPGP0NH00210 | YAL014C | 99 | 0 |
| SPGP0B00120 | YHL043W | 99 | 0 |
| SPGP0J00820 | YOL030W | 99 | 1 |
| SPGP0JG00100 | YPL281C | 99 | 0 |
| SPGP0H02140 | YMR169C | 99 | 0 |
| SPGP0H02490 | YMR203W | 99 | 0 |
| SPGP0L01570 | YOR272W | 99 | 0 |
| SPGP0L01390 | YOR252W | 99 | 0 |
| SPGP0A04010 | YDR183W | 99 | 0 |
| SPGP0H00500 | YML006C | 99 | 0 |
| SPGP0A00240 | YDL233W | 99 | 0 |
| SPGP0L00540 | YOR157C | 99 | 0 |
| SPGP0F02830 | YBR007C | 99 | 0 |
| SPGP0J00950 | YOL016C | 99 | 0 |
| SPGP0L00650 | YOR168W | 99 | 0 |
| SPGP0A03850 | YDR169C | 99 | 0 |
| SPGP0A00630 | YDL191W | 99 | 0 |
| SPGP0RY00100 | YER173W | 99 | 0 |
| SPGP0A04240 | YDR212W | 99 | 0 |

Create PDF in your applications with the Pdfcrowd HTML to PDF API PDFCROWD

| | | | |
|---|---|---|---|
| SPGP0FK00230 | YGL200C | 99 | 0 |
| SPGP0I02210 | YJL092W | 99 | 0 |
| SPGP0J00250 | YOL090W | 99 | 0 |
| SPGP0J00520 | YOL062C | 99 | 0 |
| SPGP0G01040 | YKL094W | 99 | 0 |
| SPGP0H02720 | YMR226C | 100 | 0 |
| SPGP0L01940 | YOR311C | 99 | 0 |
| SPGP0NS00110 | YGL196W | 99 | 0 |
| SPGP0A03540 | YDR134C | 99 | 0 |
| SPGP0H00660 | YMR011W | 99 | 0 |
| SPGP0BF00970 | YIL074C | 99 | 0 |
| SPGP0G00720 | YKL133C | 99 | 0 |
| SPGP0X00270 | YOL140W | 99 | 0 |
| SPGP0H03450 | YMR301C | 99 | 0 |
| SPGP0ME00190 | YGL226W | 99 | 0 |
| SPGP0L02410 | YOR365C | 99 | 0 |
| SPGP0B01280 | YHR077C | 99 | 0 |
| SPGP0A02290 | YDL003W | 99 | 0 |
| SPGP0F01630 | YBR137W | 99 | 0 |
| SPGP0J01120 | YOR002W | 99 | 0 |
| SPGP0J01690 | YOR066W | 99 | 0 |
| SPGP0I01160 | YJR034W | 100 | 0 |
| SPGP0A01250 | YDL121C | 100 | 0 |
| SPGP0B01340 | YHR082C | 99 | 0 |
| SPGP0F03080 | YBL020W | 99 | 0 |
| SPGP0F01500 | YBR151W | 99 | 0 |
| SPGP0FK00130 | YGL211W | 99 | 0 |
| SPGP0A04800 | YDR276C | 100 | 0 |
| SPGP0I00380 | YJR116W | 100 | 0 |
| SPGP0L02000 | YOR320C | 99 | 0 |
| SPGP0X00570 | YOL108C | 99 | 0 |
| SPGP0BF00530 | YIL023C | 99 | 0 |
| SPGP0J01340 | YOR026W | 99 | 0 |
| SPGP0G01000 | YKL099C | 100 | 0 |
| SPGP0I02560 | YJL131C | 99 | 0 |
| SPGP0H02380 | YMR193W | 99 | 0 |
| SPGP0BF00240 | YIR005W | 100 | 0 |
| SPGP0J00180 | YOL097C | 99 | 0 |
| SPGP0BF00930 | YIL071C | 99 | 0 |
| SPGP0F03050 | YBL017C | 99 | 0 |

Create PDF in your applications with the Pdfcrowd HTML to PDF API PDFCROWD

| | | | |
|---|---|---|---|
| SPGP0H00150 | YML046W | 99 | 0 |
| SPGP0B01880 | YHR137W | 99 | 0 |
| SPGP0QG00150 | YAL046C | 99 | 0 |
| SPGP0A02090 | YDL027C | 99 | 0 |
| SPGP0F03650 | YBL082C | 99 | 0 |
| SPGP0J00460 | YOL068C | 99 | 0 |
| SPGP0J01150 | YOR005C | 99 | 0 |
| SPGP0BF01650 | YIL147C | 99 | 0 |
| SPGP0I00780 | YJR075W | 99 | 0 |
| SPGP0F03590 | YBL075C | 99 | 0 |
| SPGP0L00220 | YOR122C | 99 | 0 |
| SPGP0F02450 | YBR048W | 99 | 0 |
| SPGP0B01080 | YHR058C | 99 | 0 |
| SPGP0B02510 | YHR199C-A | 98 | 2 |
| SPGP0A02890 | YDR065W | 99 | 0 |
| SPGP0I03120 | YJL194W | 99 | 0 |
| SPGP0A01200 | YDL126C | 99 | 0 |
| SPGP0BG00730 | YCR010C | 100 | 0 |
| SPGP0MC00110 | YER001W | 80 | 2 |
| SPGP0I01340 | YJR010W | 99 | 0 |
| SPGP0I01410 | YJR004C | 99 | 0 |
| SPGP0I01140 | YJR036C | 99 | 0 |
| SPGP0J01610 | YOR058C | 99 | 0 |
| SPGP0I01580 | YJL019W | 99 | 0 |
| SPGP0BF00610 | YIL035C | 99 | 0 |
| SPGP0L01870 | YOR304W | 99 | 0 |
| SPGP0I02160 | YJL087C | 99 | 0 |
| SPGP0I02970 | YJL177W | 100 | 0 |
| SPGP0F03330 | YBL043W | 99 | 0 |
| SPGP0H01350 | YMR087W | 99 | 0 |
| SPGP0L00950 | YOR204W | 99 | 0 |
| SPGP0L00480 | YOR151C | 99 | 0 |
| SPGP0H01600 | YMR113W | 99 | 0 |
| SPGP0BF01660 | YIL148W | 99 | 0 |
| SPGP0BF00920 | YIL070C | 99 | 0 |
| SPGP0IY00210 | YGL252C | 99 | 0 |
| SPGP0A04850 | YDR280W | 99 | 0 |
| SPGP0BF01520 | YIL133C | 99 | 0 |
| SPGP0A00270 | YDL230W | 99 | 0 |
| SPGP0F01240 | YBR177C | 99 | 0 |

Create PDF in your applications with the Pdfcrowd HTML to PDF API PDFCROWD

| | | | |
|---|---|---|---|
| SPGP0F01850 | YBR111W-A | 99 | 0 |
| SPGP0L02360 | YOR359W | 99 | 0 |
| SPGP0I01540 | YJL012C | 99 | 0 |
| SPGP0H02800 | YMR234W | 99 | 0 |
| SPGP0A03110 | YDR086C | 100 | 0 |
| SPGP0G00680 | YKL138C | 99 | 0 |
| SPGP0A04230 | YDR211W | 99 | 0 |
| SPGP0H00850 | YMR030W | 99 | 0 |
| SPGP0BG00600 | YCL005W | 99 | 0 |
| SPGP0BF01300 | YIL111W | 99 | 0 |
| SPGP0G00670 | YKL138C-A | 99 | 0 |
| SPGP0BG00740 | YCR011C | 99 | 0 |
| SPGP0L02540 | YOR381W | 99 | 0 |
| SPGP0H00540 | YML001W | 99 | 0 |
| SPGP0H00900 | YMR035W | 99 | 0 |
| SPGP0G00930 | YKL107W | 99 | 0 |
| SPGP0F00220 | YBR285W | 99 | 0 |
| SPGP0A02300 | YDL002C | 99 | 0 |
| SPGP0L02080 | YOR329C | 99 | 0 |
| SPGP0CB00100 | YIR019C | 96 | 0 |
| SPGP0BG00280 | YCL045C | 99 | 0 |
| SPGP0J00390 | YOL076W | 99 | 0 |
| SPGP0J01590 | YOR056C | 99 | 0 |
| SPGP0A00860 | YDL165W | 99 | 0 |
| SPGP0I02280 | YJL099W | 99 | 0 |
| SPGP0A02240 | YDL007C-A | 100 | 0 |
| SPGP0CB00230 | YIR033W | 99 | 0 |
| SPGP0F01050 | YBR197C | 99 | 0 |
| SPGP0J01640 | YOR061W | 99 | 0 |
| SPGP0A00960 | YDL153C | 99 | 0 |
| SPGP0H00380 | YML018C | 99 | 0 |
| SPGP0B01500 | YHR098C | 99 | 0 |
| SPGP0I01970 | YJL062W-A | 100 | 0 |
| SPGP0H03400 | YMR296C | 99 | 0 |
| SPGP0VZ00200 | YDR345C | 90 | 0 |
| SPGP0I01600 | YJL023C | 99 | 0 |
| SPGP0A05210 | YDR317W | 99 | 0 |
| SPGP0H02400 | YMR194C-B | 100 | 0 |
| SPGP0I00960 | YJR056C | 99 | 0 |
| SPGP0ME00180 | YGL226C-A | 100 | 0 |

Create PDF in your applications with the Pdfcrowd HTML to PDF API PDFCROWD

| | | | |
|---|---|---|---|
| SPGP0F00410 | YBR264C | 99 | 0 |
| SPGP0I01620 | YJL025W | 99 | 0 |
| SPGP0B00510 | YHR001W | 99 | 0 |
| SPGP0H02680 | YMR222C | 99 | 0 |
| SPGP0X00260 | YOL141W | 99 | 0 |
| SPGP0B00560 | YHR005C | 99 | 0 |
| SPGP0A04410 | YDR232W | 99 | 0 |
| SPGP0A04570 | YDR249C | 100 | 0 |
| SPGP0I00970 | YJR055W | 99 | 0 |
| SPGP0A03680 | YDR150W | 98 | 0 |
| SPGP0F03100 | YBL022C | 99 | 0 |
| SPGP0WY00220 | YLR002C | 83 | 1 |
| SPGP0WU00130 | YER180C | 99 | 0 |
| SPGP0I02190 | YJL090C | 99 | 0 |
| SPGP0H02270 | YMR182C | 99 | 0 |
| SPGP0GU00180 | YPL271W | 99 | 0 |
| SPGP0J00960 | YOL015W | 99 | 0 |
| SPGP0BG00500 | YCL018W | 99 | 0 |
| SPGP0H01290 | YMR079W | 99 | 0 |
| SPGP0A00260 | YDL231C | 99 | 0 |
| SPGP0L01370 | YOR250C | 99 | 0 |
| SPGP0F01120 | YBR190W | 99 | 0 |
| SPGP0BF01560 | YIL137C | 99 | 0 |
| SPGP0L00680 | YOR173W | 99 | 0 |
| SPGP0L02130 | YOR336W | 99 | 0 |
| SPGP0A03140 | YDR089W | 99 | 0 |
| SPGP0H02120 | YMR167W | 99 | 0 |
| SPGP0F01430 | YBR158W | 99 | 0 |
| SPGP0H00560 | YMR002W | 97 | 3 |
| SPGP0H03260 | YMR282C | 99 | 0 |
| SPGP0A00650 | YDL189W | 99 | 0 |
| SPGP0A04260 | YDR214W | 99 | 0 |
| SPGP0J01220 | YOR013W | 99 | 0 |
| SPGP0H03290 | YMR285C | 99 | 0 |
| SPGP0J00500 | YOL064C | 99 | 0 |
| SPGP0BG00300 | YCL043C | 99 | 0 |
| SPGP0L01960 | YOR315W | 99 | 0 |
| SPGP0BF00370 | YIL008W | 100 | 0 |
| SPGP0L01170 | YOR228C | 99 | 0 |
| SPGP0L02430 | YOR368W | 99 | 0 |

Create PDF in your applications with the Pdfcrowd HTML to PDF API PDFCROWD

| | | | |
|---|---|---|---|
| SPGP0I00610 | YJR094W-A | 100 | 0 |
| SPGP0H02230 | YMR178W | 99 | 0 |
| SPGP0A01870 | YDL054C | 99 | 0 |
| SPGP0CB00190 | YIR029W | 99 | 0 |
| SPGP0A02520 | YDR023W | 99 | 0 |
| SPGP0A02310 | YDL001W | 99 | 0 |
| SPGP0BF01840 | YIL167W | 99 | 0 |
| SPGP0I01710 | YJL036W | 99 | 0 |
| SPGP0A01230 | YDL123W | 99 | 0 |
| SPGP0L00640 | YOR167C | 100 | 0 |
| SPGP0J01100 | YOL001W | 99 | 0 |
| SPGP0F00960 | YBR205W | 99 | 0 |
| SPGP0H03230 | YMR279C | 99 | 0 |
| SPGP0A02390 | YDR009W | 99 | 0 |
| SPGP0F02780 | YBR012C | 99 | 0 |
| SPGP0QG00180 | YAL049C | 99 | 0 |
| SPGP0A03870 | YDR170W-A | 99 | 0 |
| SPGP0A03300 | YDR108W | 99 | 0 |
| SPGP0FK00110 | YGL213C | 99 | 0 |
| SPGP0G00410 | YKL166C | 99 | 0 |
| SPGP0F01560 | YBR145W | 99 | 0 |
| SPGP0F03660 | YBL084C | 98 | 1 |
| SPGP0J00200 | YOL095C | 99 | 0 |
| SPGP0LR00100 | YBR299W | 99 | 0 |
| SPGP0X00350 | YOL131W | 99 | 0 |
| SPGP0BF01010 | YIL078W | 99 | 0 |
| SPGP0B01310 | YHR079C-A | 100 | 0 |
| SPGP0I02900 | YJL168C | 99 | 0 |
| SPGP0BF00220 | YIR007W | 99 | 0 |
| SPGP0BF01820 | YIL164C | 99 | 0 |
| SPGP0QQ00200 | YAR002W | 99 | 0 |
| SPGP0BG00750 | YCR012W | 99 | 0 |
| SPGP0A01220 | YDL124W | 99 | 0 |
| SPGP0F02470 | YBR046C | 99 | 0 |
| SPGP0A05020 | YDR298C | 99 | 0 |
| SPGP0F03140 | YBL026W | 100 | 0 |
| SPGP0I03100 | YJL192C | 99 | 0 |
| SPGP0F00630 | YBR243C | 100 | 0 |
| SPGP0I02580 | YJL133W | 99 | 0 |
| SPGP0F00290 | YBR278W | 100 | 0 |

Create PDF in your applications with the Pdfcrowd HTML to PDF API PDFCROWD

| | | | |
|---|---|---|---|
| SPGP0A00700 | YDL182W | 99 | 0 |
| SPGP0F02440 | YBR049C | 99 | 0 |
| SPGP0BF00590 | YIL033C | 100 | 0 |
| SPGP0A04770 | YDR272W | 99 | 0 |
| SPGP0L01510 | YOR265W | 99 | 0 |
| SPGP0F03310 | YBL041W | 99 | 0 |
| SPGP0I02410 | YJL115W | 99 | 0 |
| SPGP0H01620 | YMR115W | 99 | 0 |
| SPGP0H01330 | YMR085W | 99 | 0 |
| SPGP0BF00900 | YIL068C | 99 | 0 |
| SPGP0G00200 | YKL189W | 99 | 0 |
| SPGP0A00600 | YDL194W | 99 | 0 |
| SPGP0IY00230 | YGL254W | 100 | 0 |
| SPGP0BG00270 | YCL047C | 99 | 0 |
| SPGP0A03090 | YDR084C | 99 | 0 |
| SPGP0BF00310 | YIL002W-A | 100 | 0 |
| SPGP0A00250 | YDL232W | 100 | 0 |
| SPGP0F02300 | YBR064W | 99 | 0 |
| SPGP0A03180 | YDR093W | 99 | 0 |
| SPGP0H00270 | YML029W | 99 | 0 |
| SPGP0H02820 | YMR236W | 99 | 0 |
| SPGP0H00870 | YMR032W | 99 | 0 |
| SPGP0B02030 | YHR152W | 100 | 0 |
| SPGP0L01240 | YOR236W | 99 | 0 |
| SPGP0F00650 | YBR241C | 99 | 0 |
| SPGP0A03720 | YDR155C | 100 | 0 |
| SPGP0J00190 | YOL096C | 99 | 0 |
| SPGP0A01820 | YDL059C | 99 | 0 |
| SPGP0TK00100 | YER056C-A | 95 | 0 |
| SPGP0B02250 | YHR174W | 100 | 0 |
| SPGP0F02150 | YBR080C | 99 | 0 |
| SPGP0F00200 | YBR287W | 99 | 0 |
| SPGP0A02320 | YDR001C | 99 | 0 |
| SPGP0VT00190 | YPL253C | 99 | 0 |
| SPGP0G01120 | YKL086W | 99 | 0 |
| SPGP0A02220 | YDL010W | 99 | 0 |
| SPGP0J00770 | YOL036W | 99 | 0 |
| SPGP0A01790 | YDL063C | 99 | 0 |
| SPGP0L01840 | YOR301W | 99 | 0 |

Create PDF in your applications with the Pdfcrowd HTML to PDF API PDFCROWD

| | | | |
|---|---|---|---|
| SPGP0BG00770 | YCR015C | 99 | 0 |
| SPGP0H00160 | YML043C | 99 | 0 |
| SPGP0A00840 | YDL167C | 99 | 0 |
| SPGP0L01770 | YOR294W | 99 | 0 |
| SPGP0NM00130 | YDR542W | 99 | 0 |
| SPGP0G00920 | YKL108W | 99 | 1 |
| SPGP0A00940 | YDL155W | 99 | 0 |
| SPGP0I01990 | YJL065C | 99 | 0 |
| SPGP0I02290 | YJL100W | 99 | 0 |
| SPGP0B00170 | YHL035C | 99 | 0 |
| SPGP0B00520 | YHR001W-A | 99 | 0 |
| SPGP0QQ00100 | YAR023C | 99 | 0 |
| SPGP0G01070 | YKL091C | 99 | 0 |
| SPGP0F03060 | YBL018C | 99 | 0 |
| SPGP0A05230 | YDR319C | 99 | 0 |
| SPGP0BF01810 | YIL162W | 99 | 0 |
| SPGP0B00480 | YHL003C | 99 | 0 |
| SPGP0BF00730 | YIL047C | 99 | 0 |
| SPGP0A02910 | YDR067C | 99 | 0 |
| SPGP0A04540 | YDR246W | 100 | 0 |
| SPGP0B00540 | YHR003C | 99 | 0 |
| SPGP0A01690 | YDL075W | 99 | 0 |
| SPGP0L02560 | YKL222C | 99 | 0 |
| SPGP0I02860 | YJL164C | 99 | 0 |
| SPGP0QX00230 | YMR307W | 99 | 0 |
| SPGP0B01120 | YHR062C | 99 | 0 |
| SPGP0L02390 | YOR362C | 100 | 0 |
| SPGP0BF00140 | YIR015W | 99 | 0 |
| SPGP0CB00180 | YIR028W | 99 | 0 |
| SPGP0A00640 | YDL190C | 99 | 0 |
| SPGP0B02310 | YHR180W | 99 | 0 |
| SPGP0F02580 | YBR035C | 100 | 0 |
| SPGP0WU00110 | YER178W | 99 | 0 |
| SPGP0L02110 | YOR334W | 99 | 0 |
| SPGP0QL00180 | YAL016W | 99 | 0 |
| SPGP0GU00160 | YPL269W | 99 | 0 |
| SPGP0J00940 | YOL017W | 99 | 0 |
| SPGP0J01620 | YOR059C | 99 | 0 |
| SPGP0BG00560 | YCL010C | 99 | 0 |
| SPGP0H01030 | YMR052W | 99 | 0 |

Create PDF in your applications with the Pdfcrowd HTML to PDF API PDFCROWD

| | | | |
|---|---|---|---|
| SPGP0A01070 | YDL140C | 99 | 0 |
| SPGP0A00200 | YDL237W | 99 | 0 |
| SPGP0I01200 | YJR030C | 99 | 0 |
| SPGP0L00660 | YOR171C | 100 | 0 |
| SPGP0H01810 | YMR134W | 99 | 0 |
| SPGP0F00910 | YBR212W | 99 | 0 |
| SPGP0H03240 | YMR280C | 99 | 0 |
| SPGP0NS00120 | YGL195W | 99 | 0 |
| SPGP0I01960 | YJL062W | 99 | 0 |
| SPGP0H02100 | YMR165C | 99 | 0 |
| SPGP0H02450 | YMR199W | 99 | 0 |
| SPGP0BF00960 | YIL073C | 99 | 0 |
| SPGP0L02150 | YOR338W | 100 | 0 |
| SPGP0J01840 | YOR084W | 99 | 0 |
| SPGP0I00720 | YJR083C | 99 | 0 |
| SPGP0A04270 | YDR216W | 99 | 0 |
| SPGP0F02250 | YBR069C | 99 | 0 |
| SPGP0G00300 | YKL179C | 99 | 0 |
| SPGP0H03270 | YMR283C | 99 | 0 |
| SPGP0QX00110 | YMR318C | 99 | 0 |
| SPGP0TI00100 | YAL061W | 99 | 0 |
| SPGP0BF00130 | YIR016W | 98 | 0 |
| SPGP0BG00910 | YCR027C | 99 | 0 |
| SPGP0A04140 | YDR198C | 99 | 0 |
| SPGP0H03410 | YMR297W | 99 | 0 |
| SPGP0NH00180 | YAL011W | 99 | 0 |
| SPGP0J00560 | YOL058W | 99 | 0 |
| SPGP0BG00980 | YCR034W | 100 | 0 |
| SPGP0A02500 | YDR021W | 99 | 0 |
| SPGP0G00880 | YKL113C | 99 | 0 |
| SPGP0L00400 | YOR142W | 99 | 0 |
| SPGP0I00340 | YJR121W | 99 | 0 |
| SPGP0X00360 | YOL130W | 99 | 0 |
| SPGP0J01160 | YOR006C | 99 | 0 |
| SPGP0B01380 | YHR086W | 99 | 0 |
| SPGP0I02760 | YJL154C | 99 | 0 |
| SPGP0F00800 | YBR225W | 99 | 0 |
| SPGP0G00350 | YKL173W | 99 | 0 |
| SPGP0L01520 | YOR266W | 99 | 0 |
| SPGP0A03320 | YDR110W | 99 | 0 |

Create PDF in your applications with the Pdfcrowd HTML to PDF API PDFCROWD

| | | | |
|---|---|---|---|
| SPGP0TR00100 | YAL039C | 99 | 0 |
| SPGP0H00990 | YMR044W | 99 | 0 |
| SPGP0A01740 | YDL069C | 100 | 0 |
| SPGP0J01380 | YOR031W | 99 | 0 |
| SPGP0F03680 | YBL086C | 99 | 0 |
| SPGP0F00940 | YBR208C | 99 | 0 |
| SPGP0A00880 | YDL161W | 100 | 0 |
| SPGP0J00380 | YOL077C | 99 | 0 |
| SPGP0BG00360 | YCL035C | 99 | 0 |
| SPGP0H00590 | YMR005W | 99 | 0 |
| SPGP0A00490 | YDL206W | 99 | 0 |
| SPGP0BF01770 | YIL158W | 99 | 0 |
| SPGP0F03800 | YBL099W | 100 | 0 |
| SPGP0B01240 | YHR073W | 99 | 0 |
| SPGP0QG00110 | YAL042W | 99 | 0 |
| SPGP0H00190 | YML038C | 99 | 0 |
| SPGP0X00330 | YOL133W | 99 | 0 |
| SPGP0I02350 | YJL106W | 99 | 0 |
| SPGP0BF00280 | YIR001C | 99 | 0 |
| SPGP0I00110 | YJR144W | 99 | 0 |
| SPGP0A03780 | YDR162C | 99 | 0 |
| SPGP0A01240 | YDL122W | 99 | 0 |
| SPGP0F02270 | YBR067C | 97 | 3 |
| SPGP0UV00100 | YPL245W | 99 | 0 |
| SPGP0I00140 | YJR141W | 99 | 0 |
| SPGP0I00920 | YJR060W | 99 | 0 |
| SPGP0A05000 | YDR296W | 99 | 0 |
| SPGP0J01190 | YOR009W | 96 | 0 |
| SPGP0I00420 | YJR112W | 99 | 0 |
| SPGP0I01890 | YJL055W | 100 | 0 |
| SPGP0I03200 | YJL204C | 99 | 0 |
| SPGP0F02430 | YBR050C | 98 | 1 |
| SPGP0B00710 | YHR019C | 99 | 0 |
| SPGP0J01580 | YOR054C | 99 | 0 |
| SPGP0F01860 | YBR111C | 99 | 0 |
| SPGP0A01550 | YDL088C | 99 | 0 |
| SPGP0BF00690 | YIL043C | 99 | 0 |
| SPGP0A00720 | YDL180W | 99 | 0 |
| SPGP0L00160 | YOR115C | 99 | 0 |
| SPGP0F01180 | YBR184W | 99 | 0 |

Create PDF in your applications with the Pdfcrowd HTML to PDF API PDFCROWD

| | | | |
|---|---|---|---|
| SPGP0A04750 | YDR268W | 99 | 0 |
| SPGP0L01530 | YOR267C | 97 | 2 |
| SPGP0I01290 | YJR014W | 100 | 0 |
| SPGP0L02290 | YOR352W | 99 | 0 |
| SPGP0G00740 | YKL130C | 99 | 0 |
| SPGP0BG00170 | YCL057W | 99 | 0 |
| SPGP0L00910 | YOR197W | 99 | 0 |
| SPGP0L02210 | YOR344C | 99 | 0 |
| SPGP0A03160 | YDR091C | 99 | 0 |
| SPGP0B00290 | YHL023C | 99 | 0 |
| SPGP0I02370 | YJL109C | 99 | 0 |
| SPGP0BF00410 | YIL011W | 94 | 3 |
| SPGP0A03710 | YDR153C | 99 | 0 |
| SPGP0G00290 | YKL180W | 99 | 0 |
| SPGP0J00170 | YOL098C | 99 | 0 |
| SPGP0SC00100 | YER176W | 99 | 0 |
| SPGP0A00130 | YNL333W | 99 | 0 |
| SPGP0H01660 | YMR119W | 99 | 0 |
| SPGP0A02340 | YDR003W | 99 | 0 |
| SPGP0F00770 | YBR229C | 99 | 0 |
| SPGP0A02200 | YDL013W | 99 | 0 |
| SPGP0J00710 | YOL043C | 99 | 0 |
| SPGP0L01090 | YOR219C | 99 | 0 |
| SPGP0F03040 | YBL016W | 99 | 0 |
| SPGP0L01860 | YOR303W | 99 | 0 |
| SPGP0L01750 | YOR292C | 99 | 0 |
| SPGP0A02900 | YDR066C | 99 | 0 |
| SPGP0H01440 | YMR096W | 99 | 0 |
| SPGP0H00180 | YML041C | 99 | 0 |
| SPGP0VT00180 | YPL254W | 99 | 0 |
| SPGP0I03250 | YJL209W | 99 | 0 |
| SPGP0I00560 | YJR099W | 99 | 0 |
| SPGP0F01540 | YBR147W | 99 | 0 |
| SPGP0A04980 | YDR294C | 99 | 0 |
| SPGP0I02890 | YJL167W | 99 | 0 |
| SPGP0G00530 | YKL154W | 99 | 0 |
| SPGP0BF00740 | YIL048W | 99 | 0 |
| SPGP0J01440 | YOR037W | 99 | 0 |
| SPGP0B00190 | YHL033C | 98 | 0 |
| SPGP0QD00110 | YGL183C | 99 | 0 |

Create PDF in your applications with the Pdfcrowd HTML to PDF API PDFCROWD

| | | | |
|---|---|---|---|
| SPGP0A02120 | YDL022W | 99 | 0 |
| SPGP0A04450 | YDR236C | 99 | 0 |
| SPGP0B01100 | YHR060W | 100 | 0 |
| SPGP0F02760 | YBR015C | 99 | 0 |
| SPGP0L00350 | YOR136W | 99 | 0 |
| SPGP0A04560 | YDR248C | 99 | 0 |
| SPGP0MU00100 | YGL175C | 99 | 0 |
| SPGP0H01120 | YMR062C | 99 | 0 |
| SPGP0A00190 | YDL238C | 99 | 0 |
| SPGP0QL00160 | YAL018C | 99 | 0 |
| SPGP0I01050 | YJR047C | 100 | 0 |
| SPGP0A04690 | YDR262W | 99 | 0 |
| SPGP0I00930 | YJR059W | 99 | 0 |
| SPGP0J00920 | YOL019W | 99 | 0 |
| SPGP0A01050 | YDL142C | 99 | 0 |
| SPGP0H02320 | YMR187C | 99 | 0 |
| SPGP0B02480 | YHR197W | 99 | 0 |
| SPGP0A03530 | YDR132C | 99 | 0 |
| SPGP0L02520 | YOR378W | 99 | 0 |
| SPGP0H01830 | YMR136W | 99 | 0 |
| SPGP0F00930 | YBR210W | 99 | 0 |
| SPGP0A01420 | YDL103C | 99 | 0 |
| SPGP0NS00100 | YGL197W | 99 | 0 |
| SPGP0H02470 | YMR201C | 99 | 0 |
| SPGP0L00720 | YOR177C | 99 | 0 |
| SPGP0A00550 | YDL200C | 99 | 0 |
| SPGP0BG00530 | YCL014W | 99 | 0 |
| SPGP0A04290 | YDR218C | 99 | 0 |
| SPGP0H01870 | YMR140W | 100 | 0 |
| SPGP0WU00120 | YER179W | 99 | 0 |
| SPGP0I00350 | YJR119C | 99 | 0 |
| SPGP0L01250 | YOR237W | 99 | 0 |
| SPGP0I02230 | YJL094C | 99 | 0 |
| SPGP0H02780 | YMR232W | 99 | 0 |
| SPGP0L02400 | YOR363C | 99 | 0 |
| SPGP0F01300 | YBR170C | 99 | 0 |
| SPGP0BF00110 | YIR018W | 99 | 0 |
| SPGP0X00130 | YOL155C | 91 | 3 |
| SPGP0G00190 | YKL190W | 99 | 0 |
| SPGP0H03250 | YMR281W | 99 | 0 |

Create PDF in your applications with the Pdfcrowd HTML to PDF API PDFCROWD

| | | | |
|---|---|---|---|
| SPGP0J00540 | YOL060C | 99 | 0 |
| SPGP0A02560 | YDR027C | 99 | 0 |
| SPGP0F01330 | YBR167C | 99 | 0 |
| SPGP0CB00150 | YIR025W | 99 | 0 |
| SPGP0I00320 | YJR123W | 99 | 0 |
| SPGP0A03340 | YDR113C | 99 | 0 |
| SPGP0A03380 | YDR118W | 99 | 0 |
| SPGP0J01630 | YOR060C | 99 | 0 |
| SPGP0BF00350 | YIL006W | 99 | 0 |
| SPGP0J01140 | YOR004W | 99 | 0 |
| SPGP0L02470 | YOR372C | 99 | 1 |
| SPGP0A02360 | YDR005C | 99 | 0 |
| SPGP0I00290 | YJR126C | 99 | 0 |
| SPGP0I00490 | YJR106W | 99 | 0 |
| SPGP0F00900 | YBR213W | 99 | 0 |
| SPGP0A01610 | YDL083C | 100 | 0 |
| SPGP0D00440 | YOL120C | 87 | 3 |
| SPGP0A05240 | YDR320C | 99 | 0 |
| SPGP0BF01790 | YIL160C | 99 | 0 |
| SPGP0A02820 | YDR058C | 99 | 0 |
| SPGP0H03220 | YMR278W | 99 | 0 |
| SPGP0B02100 | YHR159W | 99 | 0 |
| SPGP0I02880 | YJL166W | 100 | 0 |
| SPGP0H01500 | YMR102C | 99 | 0 |
| SPGP0I02660 | YJL141C | 99 | 0 |
| SPGP0IY00100 | YGL241W | 99 | 0 |
| SPGP0B01350 | YHR083W | 99 | 0 |
| SPGP0I01460 | YJL002C | 99 | 0 |
| SPGP0B00830 | YHR031C | 99 | 0 |
| SPGP0F02510 | YBR042C | 99 | 0 |
| SPGP0I01870 | YJL053W | 99 | 0 |
| SPGP0H02410 | YMR195W | 99 | 1 |
| SPGP0H01430 | YMR095C | 99 | 0 |
| SPGP0I02620 | YJL137C | 99 | 0 |
| SPGP0B02290 | YHR178W | 99 | 0 |
| SPGP0L00180 | YOR117W | 100 | 0 |
| SPGP0F02480 | YBR045C | 99 | 0 |
| SPGP0A04730 | YDR266C | 99 | 0 |
| SPGP0I02150 | YJL085W | 99 | 0 |
| SPGP0G00970 | YKL103C | 99 | 0 |

Create PDF in your applications with the Pdfcrowd HTML to PDF API PDFCROWD

| | | | |
|---|---|---|---|
| SPGP0L01580 | YOR273C | 99 | 0 |
| SPGP0BF00940 | YIL072W | 99 | 0 |
| SPGP0A05150 | YDR311W | 99 | 0 |
| SPGP0CB00140 | YIR024C | 99 | 0 |
| SPGP0L00390 | YOR141C | 99 | 0 |
| SPGP0J00730 | YOL041C | 100 | 0 |
| SPGP0H02640 | YMR218C | 99 | 0 |
| SPGP0H02860 | YMR240C | 99 | 0 |
| SPGP0A03260 | YDR104C | 99 | 0 |
| SPGP0A03690 | YDR151C | 100 | 0 |
| SPGP0BF01860 | YIL171W | 99 | 0 |
| SPGP0H00910 | YMR036C | 99 | 0 |
| SPGP0J00480 | YOL066C | 99 | 0 |
| SPGP0A04710 | YDR264C | 99 | 0 |
| SPGP0J00150 | YOL101C | 99 | 0 |
| SPGP0L01890 | YOR305W | 100 | 0 |
| SPGP0A01860 | YDL055C | 99 | 0 |
| SPGP0BF01040 | YIL084C | 99 | 0 |
| SPGP0H01080 | YMR058W | 99 | 0 |
| SPGP0A00660 | YDL188C | 99 | 0 |
| SPGP0A00730 | YDL179W | 100 | 0 |
| SPGP0VO00190 | YCR077C | 83 | 1 |
| SPGP0B02200 | YHR169W | 99 | 0 |
| SPGP0A02880 | YDR064W | 99 | 0 |
| SPGP0L01110 | YOR221C | 99 | 0 |
| SPGP0F00250 | YBR282W | 99 | 0 |
| SPGP0H01420 | YMR094W | 99 | 0 |
| SPGP0H00140 | YML047C | 100 | 0 |
| SPGP0L01200 | YOR231W | 99 | 0 |
| SPGP0H03070 | YMR263W | 99 | 0 |
| SPGP0A03280 | YDR106W | 99 | 0 |
| SPGP0F01110 | YBR191W | 99 | 0 |
| SPGP0J01460 | YOR039W | 99 | 0 |
| SPGP0GJ00100 | YHR204W | 99 | 0 |
| SPGP0J00280 | YOL087C | 99 | 0 |
| SPGP0LL00180 | YAL033W | 99 | 0 |
| SPGP0J00550 | YOL059W | 100 | 0 |
| SPGP0BG00850 | YCR020W-B | 99 | 0 |
| SPGP0I00680 | YJR088C | 99 | 0 |
| SPGP0F01590 | YBR141C | 99 | 0 |

Create PDF in your applications with the Pdfcrowd HTML to PDF API PDFCROWD

| | | | |
|---|---|---|---|
| SPGP0A04510 | YDR243C | 99 | 0 |
| SPGP0B01160 | YHR066W | 99 | 0 |
| SPGP0F00510 | YBR255W | 99 | 0 |
| SPGP0F02740 | YBR017C | 99 | 0 |
| SPGP0A04430 | YDR234W | 99 | 0 |
| SPGP0I03080 | YJL190C | 99 | 0 |
| SPGP0A04500 | YDR242W | 99 | 0 |
| SPGP0BF00270 | YIR002C | 99 | 0 |
| SPGP0MU00120 | YGL173C | 99 | 0 |
| SPGP0BG00780 | YCR016W | 100 | 0 |
| SPGP0F03160 | YBL028C | 99 | 0 |
| SPGP0I00880 | YJR064W | 99 | 0 |
| SPGP0I01070 | YJR045C | 99 | 0 |
| SPGP0I03330 | YJL218W | 99 | 0 |
| SPGP0J01400 | YOR033C | 99 | 0 |
| SPGP0Jtrna1P | YOL013W-A | 99 | 0 |
| SPGP0J00900 | YOL021C | 99 | 0 |
| SPGP0I01240 | YJR021C | 99 | 0 |
| SPGP0L00890 | YOR195W | 99 | 0 |
| SPGP0BF01700 | YIL152W | 99 | 1 |
| SPGP0GU00120 | YPL265W | 99 | 0 |
| SPGP0PM00100 | YGR288W | 99 | 0 |
| SPGP0I00670 | YJR089W | 99 | 0 |
| SPGP0F00870 | YBR216C | 99 | 0 |
| SPGP0H03280 | YMR284W | 100 | 0 |
| SPGP0L00700 | YOR175C | 99 | 0 |
| SPGP0H02700 | YMR224C | 99 | 0 |
| SPGP0A03470 | YDR126W | 99 | 0 |
| SPGP0H01850 | YMR138W | 99 | 0 |
| SPGP0ML00100 | YAL055W | 98 | 0 |
| SPGP0FK00160 | YGL208W | 99 | 0 |
| SPGP0L02380 | YOR361C | 99 | 0 |
| SPGP0F01100 | YBR192W | 99 | 0 |
| SPGP0A03770 | YDR161W | 99 | 0 |
| SPGP0L01270 | YOR239W | 99 | 0 |
| SPGP0A02380 | YDR007W | 100 | 0 |
| SPGP0BF00170 | YIR012W | 99 | 0 |
| SPGP0QX00150 | YMR315W | 99 | 0 |
| SPGP0A03360 | YDR116C | 99 | 0 |
| SPGP0L01850 | YOR302W | 100 | 0 |

Create PDF in your applications with the Pdfcrowd HTML to PDF API PDFCROWD

| | | | |
|---|---|---|---|
| SPGP0BG00680 | YCR005C | 99 | 0 |
| SPGP0X00320 | YOL135C | 99 | 0 |
| SPGP0A03970 | YDR179W-A | 99 | 0 |
| SPGP0F00180 | YBR289W | 98 | 1 |
| SPGP0J00130 | YOL103W | 99 | 0 |
| SPGP0G00310 | YKL178C | 99 | 0 |
| SPGP0A01630 | YDL081C | 100 | 0 |
| SPGP0F02810 | YBR009C | 99 | 0 |
| SPGP0BF01220 | YIL103W | 99 | 0 |
| SPGP0F00980 | YBR203W | 99 | 0 |
| SPGP0I03390 | YAL068C | 98 | 0 |
| SPGP0CB00130 | YIR023W | 99 | 0 |
| SPGP0A02540 | YDR025W | 97 | 3 |
| SPGP0A01890 | YDL052C | 99 | 0 |
| SPGP0A01040 | YDL143W | 99 | 0 |
| SPGP0A02010 | YDL039C | 99 | 0 |
| SPGP0I01800 | YJL046W | 99 | 0 |
| SPGP0H02550 | YMR209C | 99 | 0 |
| SPGP0B00850 | YHR033W | 99 | 0 |
| SPGP0F00570 | YBR249C | 99 | 0 |
| SPGP0B01370 | YHR085W | 99 | 0 |
| SPGP0B01800 | YHR129C | 99 | 0 |
| SPGP0F02530 | YBR040W | 99 | 0 |
| SPGP0NH00110 | YAL002W | 99 | 0 |
| SPGP0F00300 | YBR276C | 99 | 0 |
| SPGP0L00120 | YOR111W | 99 | 0 |
| SPGP0F02350 | YBR058C-A | 99 | 0 |
| SPGP0F03440 | YBL057C | 99 | 0 |
| SPGP0A04220 | YDR208W | 99 | 0 |
| SPGP0RE00160 | YLL039C | 84 | 0 |
| SPGP0A03560 | YDR137W | 99 | 0 |
| SPGP0I01490 | YJL005W | 99 | 0 |
| SPGP0A05170 | YDR313C | 100 | 0 |
| SPGP0L01320 | YOR244W | 99 | 0 |
| SPGP0NC00100 | YBR301W | 99 | 0 |
| SPGP0BG00660 | YCR003W | 99 | 0 |
| SPGP0H01020 | YMR049C | 99 | 0 |
| SPGP0L01720 | YOR289W | 99 | 0 |
| SPGP0H02620 | YMR216C | 99 | 0 |
| SPGP0F01290 | YBR171W | 99 | 0 |

Create PDF in your applications with the Pdfcrowd HTML to PDF API PDFCROWD

| | | | |
|---|---|---|---|
| SPGP0IY00160 | YGL247W | 99 | 0 |
| SPGP0A04930 | YDR288W | 99 | 0 |
| SPGP0A01840 | YDL057W | 99 | 0 |
| SPGP0L00560 | YOR159C | 99 | 0 |
| SPGP0H02330 | YMR188C | 99 | 0 |
| SPGP0RN00110 | YER167W | 99 | 0 |
| SPGP0A03490 | YDR128W | 99 | 0 |
| SPGP0X00510 | YOL114C | 99 | 0 |
| SPGP0A00710 | YDL181W | 99 | 0 |
| SPGP0G00700 | YKL135C | 99 | 0 |
| SPGP0F02160 | YBR079C | 99 | 0 |
| SPGP0H00240 | YML032C | 99 | 0 |
| SPGP0L01130 | YOR223W | 99 | 0 |
| SPGP0A02860 | YDR062W | 99 | 0 |
| SPGP0H01400 | YMR092C | 99 | 0 |
| SPGP0F01620 | YBR138C | 99 | 0 |
| SPGP0F02910 | YBL002W | 100 | 0 |
| SPGP0F02540 | YBR039W | 99 | 0 |
| SPGP0L00860 | YOR192C | 99 | 0 |
| SPGP0F00840 | YBR220C | 99 | 0 |
| SPGP0B01440 | YHR092C | 99 | 0 |
| SPGP0BF00520 | YIL022W | 99 | 0 |
| SPGP0J01110 | YOR001W | 99 | 0 |
| SPGP0H02420 | YMR196W | 99 | 0 |
| SPGP0G01110 | YKL087C | 99 | 0 |
| SPGP0J01710 | YOR069W | 99 | 0 |
| SPGP0H01240 | YMR074C | 100 | 0 |
| SPGP0F01570 | YBR143C | 99 | 0 |
| SPGP0I02050 | YJL072C | 99 | 0 |
| SPGP0I02700 | YJL146W | 99 | 0 |
| SPGP0H02960 | YMR251W-A | 100 | 0 |
| SPGP0I02800 | YJL158C | 100 | 0 |
| SPGP0F01650 | YBR135W | 99 | 0 |
| SPGP0MC00160 | YER006W | 86 | 1 |
| SPGP0I00510 | YJR104C | 100 | 0 |
| SPGP0B00810 | YHR029C | 99 | 0 |
| SPGP0BF01050 | YIL085C | 99 | 0 |
| SPGP0BG00210 | YCL052C | 99 | 0 |
| SPGP0B01140 | YHR064C | 99 | 0 |

Create PDF in your applications with the Pdfcrowd HTML to PDF API PDFCROWD

| | | | |
|---|---|---|---|
| SPGP0J01420 | YOR035C | 99 | 0 |
| SPGP0F00530 | YBR253W | 99 | 0 |
| SPGP0QM00100 | YNR053C | 85 | 0 |
| SPGP0F02720 | YBR019C | 99 | 0 |
| SPGP0BF00430 | YIL014W | 99 | 0 |
| SPGP0A04520 | YDR244W | 99 | 0 |
| SPGP0A01540 | YDL089W | 99 | 0 |
| SPGP0B02370 | YHR186C | 99 | 0 |
| SPGP0A01580 | YDL085C-A | 100 | 0 |
| SPGP0B02350 | YHR184W | 99 | 0 |
| SPGP0F01200 | YBR182C | 99 | 0 |
| SPGP0I01090 | YJR043C | 99 | 0 |
| SPGP0H00460 | YML010W | 99 | 0 |
| SPGP0BF01720 | YIL154C | 99 | 0 |
| SPGP0F00110 | YBR297W | 92 | 1 |
| SPGP0L00870 | YOR193W | 99 | 0 |
| SPGP0UV00110 | YPL244C | 99 | 0 |
| SPGP0F02340 | YBR059C | 99 | 0 |
| SPGP0F00890 | YBR214W | 99 | 0 |
| SPGP0I00700 | YJR085C | 100 | 0 |
| SPGP0GU00100 | YPL263C | 99 | 0 |
| SPGP0QQ00180 | YAR003W | 99 | 0 |
| SPGP0H02430 | YMR197C | 100 | 0 |
| SPGP0A04380 | YDR228C | 99 | 1 |
| SPGP0A01460 | YDL099W | 99 | 0 |
| SPGP0H01520 | YMR105C | 99 | 0 |
| SPGP0F01080 | YBR194W | 99 | 0 |
| SPGP0I02270 | YJL098W | 99 | 0 |
| SPGP0WU00160 | YER183C | 99 | 0 |
| SPGP0BF00150 | YIR014W | 99 | 0 |
| SPGP0L01100 | YOR220W | 99 | 0 |
| SPGP0I00120 | YJR143C | 99 | 0 |
| SPGP0QX00170 | YMR313C | 99 | 0 |
| SPGP0F01030 | YBR199W | 99 | 0 |
| SPGP0I01400 | YJR005W | 99 | 0 |
| SPGP0G00460 | YKL162C | 99 | 0 |
| SPGP0A02440 | YDR014W-A | 99 | 0 |
| SPGP0J00670 | YOL047C | 99 | 0 |
| SPGP0B00910 | YHR039C | 99 | 0 |
| SPGP0BF01200 | YIL102C | 100 | 0 |

Create PDF in your applications with the Pdfcrowd HTML to PDF API PDFCROWD

| | | | |
|---|---|---|---|
| SPGP0BG00870 | YCR023C | 99 | 0 |
| SPGP0J01670 | YOR064C | 100 | 0 |
| SPGP0B01600 | YHR108W | 99 | 0 |
| SPGP0BG00410 | YCL030C | 99 | 0 |
| SPGP0CB00110 | YIR021W | 99 | 0 |
| SPGP0BF00500 | YIL020C | 99 | 0 |
| SPGP0F01870 | YBR110W | 99 | 0 |
| SPGP0B01770 | YHR126C | 99 | 0 |
| SPGP0A01060 | YDL141W | 99 | 0 |
| SPGP0L00290 | YOR129C | 99 | 0 |
| SPGP0BJ00100 | YAR050W | 88 | 1 |
| SPGP0I02100 | YJL080C | 99 | 0 |
| SPGP0I03180 | YJL201W | 99 | 0 |
| SPGP0B00150 | YHL038C | 99 | 0 |
| SPGP0I01360 | YJR008W | 99 | 0 |
| SPGP0FK00120 | YGL212W | 99 | 0 |
| SPGP0B00870 | YHR035W | 99 | 0 |
| SPGP0J01090 | YOL002C | 99 | 0 |
| SPGP0H03000 | YMR256C | 100 | 0 |
| SPGP0J00660 | YOL048C | 99 | 0 |
| SPGP0F02380 | YBR056W | 99 | 0 |
| SPGP0B01180 | YHR068W | 99 | 0 |
| SPGP0G00430 | YKL164C | 99 | 0 |
| SPGP0FK00220 | YGL201C | 99 | 0 |
| SPGP0UY00100 | YOR391C | 99 | 0 |
| SPGP0F03460 | YBL059W | 99 | 0 |
| SPGP0X00150 | YOL153C | 99 | 0 |
| SPGP0H02370 | YMR192W | 99 | 0 |
| SPGP0L00140 | YOR113W | 98 | 1 |
| SPGP0F01480 | YBR153W | 99 | 0 |
| SPGP0L02100 | YOR332W | 99 | 0 |
| SPGP0F03640 | YBL081W | 99 | 0 |
| SPGP0G00280 | YKL181W | 99 | 0 |
| SPGP0I01500 | YJL006C | 99 | 0 |
| SPGP0H01010 | YMR048W | 98 | 1 |
| SPGP0A01370 | YDL108W | 99 | 0 |
| SPGP0BG00640 | YCL001W-A | 100 | 0 |
| SPGP0BF01850 | YIL169C | 97 | 0 |
| SPGP0A00360 | YDL219W | 99 | 0 |
| SPGP0L00460 | YOR149C | 99 | 0 |

Create PDF in your applications with the Pdfcrowd HTML to PDF API PDFCROWD

| | | | |
|---|---|---|---|
| SPGP0A02630 | YDR035W | 100 | 0 |
| SPGP0J00100 | YJR029W | 99 | 0 |
| SPGP0F02130 | YBR082C | 99 | 0 |
| SPGP0F00780 | YBR228W | 99 | 0 |
| SPGP0L02190 | YOR342C | 99 | 0 |
| SPGP0A03230 | YDR100W | 100 | 0 |
| SPGP0ME00120 | YGL232W | 99 | 0 |
| SPGP0F00280 | YBR279W | 100 | 0 |
| SPGP0J00110 | YOL105C | 98 | 1 |
| SPGP0H02350 | YMR190C | 99 | 0 |
| SPGP0L01480 | YOR261C | 99 | 0 |
| SPGP0H02600 | YMR214W | 99 | 0 |
| SPGP0L01330 | YOR245C | 99 | 0 |
| SPGP0BF01470 | YIL128W | 99 | 0 |
| SPGP0X00530 | YOL112W | 99 | 0 |
| SPGP0L02350 | YOR358W | 100 | 0 |
| SPGP0L01150 | YOR226C | 99 | 0 |
| SPGP0H00260 | YML030W | 100 | 0 |
| SPGP0L00430 | YOR145C | 99 | 0 |
| SPGP0J01240 | YOR016C | 99 | 0 |
| SPGP0J01790 | YOR078W | 99 | 0 |
| SPGP0F00860 | YBR217W | 100 | 0 |
| SPGP0L02170 | YOR340C | 99 | 0 |
| SPGP0H02970 | YMR252C | 99 | 0 |
| SPGP0L02090 | YOR330C | 99 | 0 |
| SPGP0F03720 | YBL090W | 99 | 0 |
| SPGP0B00700 | YHR018C | 99 | 0 |
| SPGP0F02620 | YBR030W | 99 | 0 |
| SPGP0J00780 | YOL034W | 99 | 0 |
| SPGP0J00320 | YOL083W | 99 | 0 |
| SPGP0I00530 | YJR102C | 100 | 0 |
| SPGP0I03190 | YJL203W | 99 | 0 |
| SPGP0BF01070 | YIL088C | 99 | 0 |
| SPGP0A00170 | YDL240W | 99 | 0 |
| SPGP0I01110 | YJR041C | 99 | 0 |
| SPGP0B01610 | YHR109W | 99 | 0 |
| SPGP0A04550 | YDR247W | 99 | 0 |
| SPGP0F00550 | YBR251W | 99 | 0 |
| SPGP0A02990 | YDR075W | 100 | 0 |
| SPGP0BF00460 | YIL016W | 99 | 0 |

Create PDF in your applications with the Pdfcrowd HTML to PDF API PDFCROWD

| | | | |
|---|---|---|---|
| SPGP0A01560 | YDL087C | 99 | 0 |
| SPGP0B02230 | YHR172W | 99 | 0 |
| SPGP0BF01460 | YIL127C | 99 | 0 |
| SPGP0I03370 | YOL158C | 99 | 0 |
| SPGP0F03070 | YBL019W | 99 | 0 |
| SPGP0Itrna18M | YJL007C | 100 | 0 |
| SPGP0H01720 | YMR125W | 99 | 0 |
| SPGP0H01090 | YMR059W | 99 | 0 |
| SPGP0L00850 | YOR191W | 99 | 0 |
| SPGP0BF00840 | YIL062C | 99 | 0 |
| SPGP0J01170 | YOR007C | 99 | 0 |
| SPGP0A04100 | YDR194C | 99 | 0 |
| SPGP0F02900 | YBL001C | 100 | 0 |
| SPGP0A05110 | YDR307W | 99 | 0 |
| SPGP0A03730 | YDR156W | 99 | 0 |
| SPGP0F02320 | YBR061C | 99 | 0 |
| SPGP0H01540 | YMR107W | 100 | 0 |
| SPGP0I02020 | YJL069C | 99 | 0 |
| SPGP0L01910 | YOR307C | 99 | 0 |
| SPGP0H02060 | YMR161W | 99 | 0 |
| SPGP0G01020 | YKL096W | 100 | 0 |
| SPGP0KZ00210 | YDR406W | 83 | 1 |
| SPGP0H02010 | YMR156C | 99 | 0 |
| SPGP0F01320 | YBR168W | 99 | 0 |
| SPGP0L01810 | YOR298W | 99 | 0 |
| SPGP0X00490 | YOL116W | 99 | 0 |
| SPGP0J01480 | YOR042W | 99 | 0 |
| SPGP0A02460 | YDR017C | 99 | 0 |
| SPGP0B01660 | YHR114W | 99 | 0 |
| SPGP0J00650 | YOL049W | 99 | 0 |
| SPGP0L01950 | YOR313C | 99 | 0 |
| SPGP0J00300 | YOL086C | 99 | 0 |
| SPGP0BF00200 | YIR009W | 100 | 0 |
| SPGP0A00900 | YDL160C | 99 | 0 |
| SPGP0L01120 | YOR222W | 99 | 0 |
| SPGP0H03430 | YMR299C | 100 | 0 |
| SPGP0I01120 | YJR040W | 99 | 0 |
| SPGP0B00890 | YHR037W | 99 | 0 |
| SPGP0F01810 | YBR117C | 99 | 0 |
| SPGP0I02670 | YJL143W | 99 | 0 |

Create PDF in your applications with the Pdfcrowd HTML to PDF API PDFCROWD

| | | | |
|---|---|---|---|
| SPGP0B02040 | YHR153C | 99 | 0 |
| SPGP0A01000 | YDL147W | 99 | 0 |
| SPGP0A05300 | YDR324C | 100 | 0 |
| SPGP0QQ00120 | YAR019C | 99 | 0 |
| SPGP0A03900 | YDR173C | 99 | 1 |
| SPGP0BF00640 | YIL038C | 99 | 0 |
| SPGP0A02210 | YDL012C | 99 | 0 |
| SPGP0F02390 | YBR055C | 99 | 0 |
| SPGP0A00160 | YDL241W | 98 | 0 |
| SPGP0F01790 | YBR119W | 99 | 0 |
| SPGP0H02940 | YMR250W | 99 | 0 |
| SPGP0FK00200 | YGL203C | 98 | 1 |
| SPGP0A01090 | YDL138W | 99 | 0 |
| SPGP0F02360 | YBR058C | 99 | 0 |
| SPGP0G00910 | YKL109W | 99 | 0 |
| SPGP0A04180 | YDR204W | 100 | 0 |
| SPGP0H03180 | YMR274C | 100 | 0 |
| SPGP0A04660 | YDR259C | 99 | 0 |
| SPGP0VT00160 | YPL256C | 99 | 0 |
| SPGP0A04880 | YDR283C | 99 | 0 |
| SPGP0L00330 | YDR385W | 99 | 0 |
| SPGP0H01390 | YMR091C | 99 | 0 |
| SPGP0A04250 | YDR213W | 99 | 0 |
| SPGP0A02610 | YDR033W | 100 | 0 |
| SPGP0VT00120 | YPL259C | 99 | 0 |
| SPGP0F01260 | YBR175W | 99 | 0 |
| SPGP0A03220 | YDR099W | 99 | 0 |
| SPGP0H01970 | YMR152W | 99 | 0 |
| SPGP0A00750 | YDL177C | 99 | 0 |
| SPGP0J01200 | YOR010C | 95 | 0 |
| SPGP0ME00140 | YGL230C | 99 | 0 |
| SPGP0I03150 | YJL198W | 99 | 0 |
| SPGP0H00120 | YML049C | 99 | 0 |
| SPGP0H00210 | YML036W | 100 | 0 |
| SPGP0I02810 | YJL159W | 96 | 1 |
| SPGP0IY00180 | YGL249W | 99 | 0 |
| SPGP0B01470 | YHR096C | 99 | 0 |
| SPGP0A05280 | YDR322C-A | 100 | 0 |
| SPGP0L02370 | YOR360C | 99 | 0 |
| SPGP0J00790 | YOL033W | 99 | 0 |

Create PDF in your applications with the Pdfcrowd HTML to PDF API PDFCROWD

| | | | |
|---|---|---|---|
| SPGP0J00840 | YOL027C | 99 | 0 |
| SPGP0BF01490 | YIL130W | 98 | 1 |
| SPGP0H00420 | YML014W | 99 | 0 |
| SPGP0BF00770 | YIL051C | 99 | 0 |
| SPGP0A03650 | YDR146C | 99 | 0 |
| SPGP0B01980 | YHR147C | 99 | 0 |
| SPGP0B00580 | YHR006W | 99 | 0 |
| SPGP0H02790 | YMR233W | 100 | 0 |
| SPGP0LL00210 | YAL035W | 99 | 0 |
| SPGP0A00110 | YNL335W | 99 | 0 |
| SPGP0I01270 | YJR016C | 99 | 0 |
| SPGP0F01580 | YBR142W | 99 | 0 |
| SPGP0BG00180 | YCL056C | 99 | 0 |
| SPGP0A04530 | YDR245W | 99 | 0 |
| SPGP0F02670 | YBR024W | 100 | 0 |
| SPGP0I01730 | YJL038C | 99 | 0 |
| SPGP0I01780 | YJL044C | 99 | 0 |
| SPGP0CB00120 | YIR022W | 99 | 0 |
| SPGP0QQ00140 | YAR015W | 99 | 0 |
| SPGP0BF00860 | YIL064W | 99 | 0 |
| SPGP0G00610 | YKL144C | 99 | 0 |
| SPGP0F02920 | YBL003C | 100 | 0 |
| SPGP0A04120 | YDR196C | 99 | 0 |
| SPGP0BG00890 | YCR024C-A | 100 | 0 |
| SPGP0A05090 | YDR305C | 99 | 0 |
| SPGP0H01700 | YMR123W | 99 | 0 |
| SPGP0L00830 | YOR189W | 99 | 0 |
| SPGP0G01150 | YKL082C | 99 | 0 |
| SPGP0L01140 | YOR224C | 100 | 0 |
| SPGP0RA00140 | YPL195W | 84 | 0 |
| SPGP0H02030 | YMR158W | 99 | 0 |
| SPGP0X00470 | YOL119C | 99 | 0 |
| SPGP0G00840 | YKL119C | 99 | 0 |
| SPGP0A02400 | YDR011W | 99 | 0 |
| SPGP0BG00830 | YCR020C | 99 | 0 |
| SPGP0BG00380 | YCL033C | 99 | 0 |
| SPGP0BF00820 | YIL057C | 100 | 0 |
| SPGP0A00920 | YDL157C | 99 | 0 |
| SPGP0A02070 | YDL029W | 99 | 0 |
| SPGP0ME00100 | YGL234W | 99 | 0 |

Create PDF in your applications with the Pdfcrowd HTML to PDF API PDFCROWD

| | | | |
|---|---|---|---|
| SPGP0F01830 | YBR114W | 99 | 0 |
| SPGP0H00930 | YMR038C | 99 | 0 |
| SPGP0A03930 | YDR176W | 99 | 0 |
| SPGP0A01140 | YDL132W | 99 | 0 |
| SPGP0H00890 | YMR034C | 99 | 0 |
| SPGP0B00440 | YHL008C | 87 | 2 |
| SPGP0A03860 | YDR170C | 99 | 0 |
| SPGP0A02190 | YDL014W | 99 | 0 |
| SPGP0B01860 | YHR135C | 99 | 0 |
| SPGP0B00350 | YHL017W | 99 | 0 |
| SPGP0G01050 | YKL093W | 99 | 0 |
| SPGP0BF01180 | YIL099W | 99 | 0 |
| SPGP0A02670 | YDR041W | 100 | 0 |
| SPGP0A01910 | YDL049C | 99 | 0 |
| SPGP0I00900 | YJR062C | 99 | 0 |
| SPGP0L00100 | YOR109W | 99 | 0 |
| SPGP0F03600 | YBL076C | 99 | 0 |
| SPGP0L00310 | YOR131C | 99 | 0 |
| SPGP0I02430 | YJL117W | 99 | 0 |
| SPGP0I03230 | YJL207C | 99 | 0 |
| SPGP0I02510 | YJL127C | 99 | 0 |
| SPGP0A01330 | YDL112W | 99 | 0 |
| SPGP0I01560 | YJL014W | 99 | 0 |
| SPGP0BF01430 | YIL124W | 99 | 0 |
| SPGP0F02070 | YBR086C | 99 | 0 |
| SPGP0H01880 | YMR142C | 100 | 0 |
| SPGP0F00760 | YBR230C | 99 | 0 |
| SPGP0H01910 | YMR145C | 99 | 0 |
| SPGP0H03440 | YMR300C | 99 | 0 |
| SPGP0L02500 | YOR375C | 99 | 0 |
| SPGP0F01800 | YPR080W | 99 | 0 |
| SPGP0J00400 | YOL075C | 99 | 0 |
| SPGP0A04740 | YDR267C | 100 | 0 |
| SPGP0I00160 | YJR139C | 99 | 0 |
| SPGP0L01990 | YOR319W | 100 | 0 |
| SPGP0I02830 | YJL161W | 99 | 0 |
| SPGP0J01750 | YOR074C | 99 | 0 |
| SPGP0L02310 | YOR354C | 99 | 0 |
| SPGP0H03040 | YMR260C | 99 | 0 |
| SPGP0L00920 | YOR198C | 100 | 0 |

Create PDF in your applications with the Pdfcrowd HTML to PDF API PDFCROWD

| | | | |
|---|---|---|---|
| SPGP0H03170 | YMR273C | 99 | 1 |
| SPGP0I00570 | YJR098C | 99 | 0 |
| SPGP0F00350 | YBR271W | 99 | 0 |
| SPGP0F00820 | YBR222C | 99 | 0 |
| SPGP0BF01030 | YIL083C | 99 | 0 |
| SPGP0H00230 | YML034W | 99 | 0 |
| SPGP0A01100 | YDL136W | 100 | 0 |
| SPGP0QX00140 | YMR316W | 99 | 0 |
| SPGP0A01520 | YDL091C | 99 | 0 |
| SPGP0H02770 | YMR231W | 99 | 0 |
| SPGP0BF00470 | YIL017C | 99 | 0 |
| SPGP0B00300 | YHL022C | 99 | 0 |
| SPGP0F02650 | YBR026C | 99 | 0 |
| SPGP0J00980 | YOL013C | 99 | 0 |
| SPGP0J01520 | YOR046C | 99 | 0 |
| SPGP0I02490 | YJL125C | 99 | 0 |
| SPGP0G01090 | YKL089W | 99 | 0 |
| SPGP0A02110 | YDL024C | 99 | 0 |
| SPGP0B01650 | YHR113W | 99 | 0 |
| SPGP0F02210 | YBR073W | 99 | 0 |
| SPGP0H01150 | YMR065W | 99 | 0 |
| SPGP0F03030 | YBL015W | 99 | 0 |
| SPGP0G00590 | YKL146W | 99 | 0 |
| SPGP0BF00800 | YIL055C | 99 | 0 |
| SPGP0B00240 | YHL028W | 97 | 2 |
| SPGP0A00300 | YDL226C | 99 | 0 |
| SPGP0F02940 | YBL005W | 99 | 0 |
| SPGP0BG00700 | YCR007C | 100 | 0 |
| SPGP0L00810 | YOR187W | 99 | 0 |
| SPGP0L00760 | YOR181W | 99 | 0 |
| SPGP0I02060 | YJL073W | 99 | 0 |
| SPGP0F02290 | YBR065C | 99 | 0 |
| SPGP0I01690 | YJL034W | 99 | 0 |
| SPGP0H02580 | YMR212C | 99 | 0 |
| SPGP0L02330 | YOR356W | 99 | 0 |
| SPGP0H01580 | YMR111C | 99 | 0 |
| SPGP0BG00350 | YCL036W | 99 | 0 |
| SPGP0H00810 | YMR026C | 99 | 0 |
| SPGP0A02790 | YDR055W | 99 | 0 |
| SPGP0A02420 | YDR013W | 99 | 0 |

Create PDF in your applications with the Pdfcrowd HTML to PDF API PDFCROWD

| | | | |
|---|---|---|---|
| SPGP0H02050 | YMR160W | 99 | 0 |
| SPGP0B02530 | YHR201C | 99 | 0 |
| SPGP0B02300 | YHR179W | 99 | 0 |
| SPGP0B02080 | YHR157W | 99 | 0 |
| SPGP0J00340 | YOL081W | 99 | 0 |
| SPGP0F02890 | YBR001C | 99 | 0 |
| SPGP0H01360 | YMR088C | 99 | 0 |
| SPGP0IG00110 | YPL249C-A | 100 | 0 |
| SPGP0F01760 | YBR122C | 99 | 0 |
| SPGP0A03020 | YDR078C | 99 | 0 |
| SPGP0F02850 | YBR005W | 99 | 0 |
| SPGP0J01900 | YOR091W | 99 | 0 |
| SPGP0H00950 | YMR040W | 100 | 0 |
| SPGP0A01160 | YDL130W | 99 | 0 |
| SPGP0A03950 | YDR178W | 99 | 0 |
| SPGP0J01560 | YOR051C | 98 | 0 |
| SPGP0B01900 | YHR139C | 99 | 0 |
| SPGP0X00100 | YJL222W | 99 | 0 |
| SPGP0A01270 | YDL119C | 99 | 0 |
| SPGP0B00460 | YHL006C | 98 | 0 |
| SPGP0A03250 | YDR103W | 99 | 0 |
| SPGP0H00370 | YML019W | 99 | 0 |
| SPGP0B02120 | YHR161C | 99 | 0 |
| SPGP0F02980 | YBL009W | 99 | 0 |
| SPGP0I00740 | YJR080C | 99 | 0 |
| SPGP0A04620 | YDR255C | 100 | 0 |
| SPGP0BF00300 | YIL002C | 99 | 0 |
| SPGP0F03620 | YBL079W | 99 | 0 |
| SPGP0A03510 | YDR130C | 99 | 1 |
| SPGP0L00370 | YOR138C | 99 | 0 |
| SPGP0A03790 | YDR163W | 99 | 0 |
| SPGP0I03210 | YJL205C | 99 | 0 |
| SPGP0BO00140 | YHR209W | 99 | 0 |
| SPGP0BF01830 | YIL166C | 99 | 0 |
| SPGP0F03480 | YBL060W | 99 | 0 |
| SPGP0B01990 | YHR148W | 99 | 0 |
| SPGP0A00760 | YDL176W | 99 | 0 |
| SPGP0BF01450 | YIL126W | 99 | 0 |
| SPGP0F02110 | YBR084W | 99 | 0 |
| SPGP0B00790 | YHR027C | 99 | 0 |

Create PDF in your applications with the Pdfcrowd HTML to PDF API PDFCROWD

| | | | |
|---|---|---|---|
| SPGP0I03110 | YJL193W | 99 | 0 |
| SPGP0I00650 | YJR091C | 99 | 0 |
| SPGP0L00410 | YOR143C | 99 | 0 |
| SPGP0H02560 | YMR210W | 99 | 0 |
| SPGP0L00590 | YOR162C | 99 | 0 |
| SPGP0H01930 | YMR147W | 99 | 0 |
| SPGP0G00220 | YKL187C | 99 | 0 |
| SPGP0BF00720 | YIL046W | 99 | 0 |
| SPGP0J01030 | YOL008W | 99 | 0 |
| SPGP0L01350 | YOR247W | 99 | 0 |
| SPGP0L01420 | YOR255W | 99 | 0 |
| SPGP0L01070 | YOR216C | 99 | 0 |
| SPGP0A03040 | YDR079C-A | 99 | 0 |
| SPGP0J01820 | YOR081C | 99 | 0 |
| SPGP0GU00170 | YPL270W | 99 | 0 |
| SPGP0BF01230 | YIL104C | 100 | 0 |
| SPGP0I02850 | YJL163C | 99 | 0 |
| SPGP0L02070 | YOR328W | 99 | 0 |
| SPGP0F01970 | YBR098W | 99 | 0 |
| SPGP0A00680 | YDL185W | 98 | 0 |
| SPGP0F00370 | YBR269C | 100 | 0 |
| SPGP0F03830 | YBL103C | 99 | 0 |
| SPGP0A03880 | YDR171W | 99 | 0 |
| SPGP0G00860 | YKL116C | 99 | 0 |
| SPGP0F00160 | YBR291C | 90 | 0 |
| SPGP0ME00160 | YGL228W | 99 | 0 |
| SPGP0UA00100 | YGR286C | 99 | 0 |
| SPGP0A02130 | YDL021W | 99 | 0 |
| SPGP0B02000 | YHR149C | 99 | 0 |
| SPGP0I03020 | YJL183W | 99 | 0 |
| SPGP0LL00220 | YAL036C | 99 | 0 |
| SPGP0B01410 | YHR089C | 99 | 0 |
| SPGP0B00970 | YHR045W | 99 | 0 |
| SPGP0F02050 | YBR088C | 100 | 0 |
| SPGP0J00800 | YOL032W | 99 | 0 |
| SPGP0J00260 | YOL089C | 99 | 0 |
| SPGP0F02710 | YBR020W | 100 | 0 |
| SPGP0F01900 | YBR107C | 99 | 0 |
| SPGP0J01500 | YOR044W | 99 | 0 |

Create PDF in your applications with the Pdfcrowd HTML to PDF API PDFCROWD

| | | | |
|---|---|---|---|
| SPGP0A04300 | YDR219C | 99 | 0 |
| SPGP0H01130 | YMR063W | 99 | 0 |
| SPGP0I01770 | YJL043W | 99 | 0 |
| SPGP0G00650 | YKL140W | 100 | 0 |
| SPGP0F01910 | YBR106W | 99 | 0 |
| SPGP0I01520 | YJL010C | 99 | 0 |
| SPGP0F03540 | YBL068W | 99 | 0 |
| SPGP0X00520 | YOL113W | 99 | 0 |
| SPGP0F03010 | YBL013W | 99 | 0 |
| SPGP0F02960 | YBL007C | 99 | 0 |
| SPGP0I02110 | YJL081C | 99 | 0 |
| SPGP0J01260 | YOR018W | 99 | 0 |
| SPGP0C03340 | YML026C | 91 | 1 |
| SPGP0L00900 | YOR196C | 99 | 0 |
| SPGP0BF01630 | YIL145C | 99 | 0 |
| SPGP0B01150 | YHR065C | 99 | 0 |
| SPGP0A00510 | YDL204W | 99 | 0 |
| SPGP0H03470 | YMR303C | 96 | 0 |
| SPGP0A00470 | YDL208W | 100 | 0 |
| SPGP0B02060 | YHR155W | 99 | 0 |
| SPGP0H02070 | YMR162C | 99 | 0 |
| SPGP0X00380 | YOL128C | 99 | 0 |
| SPGP0G00510 | YKL156W | 99 | 0 |
| SPGP0B00660 | YHR014W | 99 | 0 |
| SPGP0L00690 | YOR174W | 99 | 0 |
| SPGP0H00970 | YMR042W | 100 | 0 |
| SPGP0I00950 | YJR057W | 99 | 0 |
| SPGP0J01730 | YOR071C | 99 | 0 |
| SPGP0I02500 | YJL126W | 99 | 0 |
| SPGP0F03390 | YBL051C | 99 | 0 |
| SPGP0I00710 | YJR084W | 99 | 0 |
| SPGP0CB00280 | YIR038C | 99 | 0 |
| SPGP0A02710 | YDR045C | 100 | 0 |
| SPGP0A04640 | YDR257C | 100 | 0 |
| SPGP0H02750 | YMR229C | 99 | 0 |
| SPGP0B01200 | YHR070W | 99 | 0 |
| SPGP0OI00240 | YPL109C | 82 | 0 |
| SPGP0BF01500 | YIL131C | 99 | 0 |
| SPGP0RQ00110 | YKR094C | 95 | 0 |
| SPGP0A03370 | YDR117C | 99 | 0 |

Create PDF in your applications with the Pdfcrowd HTML to PDF API PDFCROWD

| | | | |
|---|---|---|---|
| SPGP0F00560 | YBR250W | 99 | 0 |
| SPGP0F02550 | YBR038W | 99 | 0 |
| SPGP0B02240 | YHR173C | 100 | 0 |
| SPGP0H02250 | YMR180C | 99 | 0 |
| SPGP0BO00120 | YHR207C | 99 | 0 |
| SPGP0BG00340 | YCL037C | 100 | 0 |
| SPGP0I00630 | YJR093C | 99 | 0 |
| SPGP0IY00140 | YGL245W | 99 | 0 |
| SPGP0F00720 | YBR234C | 99 | 0 |
| SPGP0H01260 | YMR076C | 99 | 0 |
| SPGP0L02460 | YOR371C | 99 | 0 |
| SPGP0F01220 | YBR180W | 99 | 0 |
| SPGP0L01400 | YOR253W | 99 | 0 |
| SPGP0H00440 | YML012W | 99 | 0 |
| SPGP0I00330 | YJR122W | 99 | 0 |
| SPGP0A04960 | YDR292C | 99 | 0 |
| SPGP0F00230 | YBR284W | 99 | 0 |
| SPGP0I02870 | YJL165C | 99 | 0 |
| SPGP0F03290 | YBL039W-B | 99 | 0 |
| SPGP0H03160 | YMR272C | 99 | 0 |
| SPGP0DG00100 | YPL277C | 99 | 0 |
| SPGP0F03850 | YBL105C | 99 | 0 |
| SPGP0F00310 | YBR275C | 99 | 0 |
| SPGP0A01340 | YDL111C | 99 | 0 |
| SPGP0J01880 | YOR089C | 99 | 0 |
| SPGP0H01460 | YMR098C | 99 | 0 |
| SPGP0A02920 | YDR068W | 99 | 0 |
| SPGP0X00170 | YOL151W | 98 | 0 |
| SPGP0I01030 | YJR049C | 99 | 0 |
| SPGP0A01780 | YDL064W | 100 | 0 |
| SPGP0B01920 | YHR141C | 100 | 0 |
| SPGP0B00990 | YHR047C | 99 | 0 |
| SPGP0X00580 | YOL107W | 99 | 0 |
| SPGP0F00460 | YBR259W | 99 | 0 |
| SPGP0I02780 | YJL156C | 99 | 0 |
| SPGP0OE00240 | YNL301C | 83 | 5 |
| SPGP0I00460 | YJR108W | 100 | 0 |
| SPGP0A02800 | YDR056C | 99 | 0 |
| SPGP0F03780 | YBL097W | 99 | 0 |
| SPGP0G00160 | YKL193C | 99 | 0 |

Create PDF in your applications with the Pdfcrowd HTML to PDF API PDFCROWD

| | | | |
|---|---|---|---|
| SPGP0G00630 | YKL142W | 99 | 0 |
| SPGP0B00280 | YHL024W | 99 | 0 |
| SPGP0NH00150 | YAL008W | 99 | 0 |
| SPGP0A04370 | YDR227W | 99 | 0 |
| SPGP0F01400 | YBR161W | 99 | 0 |
| SPGP0QO00110 | YGL237C | 99 | 1 |
| SPGP0I02130 | YJL083W | 99 | 0 |
| SPGP0A01120 | YDL134C | 98 | 0 |
| SPGP0A00530 | YDL202W | 99 | 0 |
| SPGP0A00980 | YDL149W | 99 | 0 |
| SPGP0IY00240 | YGL255W | 95 | 1 |
| SPGP0L02270 | YOR350C | 99 | 0 |
| SPGP0B01830 | YHR132W-A | 99 | 0 |
| SPGP0H02090 | YMR164C | 97 | 2 |
| SPGP0H01640 | YMR117C | 99 | 0 |
| SPGP0H00320 | YML024W | 99 | 0 |
| SPGP0A03000 | YDR076W | 99 | 0 |
| SPGP0L00570 | YOR160W | 99 | 0 |
| SPGP0J01450 | YOR038C | 99 | 0 |
| SPGP0L02010 | YOR321W | 99 | 0 |
| SPGP0BG00800 | YCR018C | 99 | 0 |
| SPGP0H00730 | YMR018W | 99 | 0 |
| SPGP0BD00100 | YAL038W | 99 | 0 |
| SPGP0QX00160 | YMR314W | 99 | 0 |
| SPGP0B00640 | YHR012W | 99 | 0 |
| SPGP0A03980 | YDR180W | 99 | 0 |
| SPGP0L01760 | YOR293W | 99 | 0 |
| SPGP0B01590 | YHR107C | 99 | 0 |
| SPGP0H01480 | YMR100W | 99 | 0 |
| SPGP0A01980 | YDL043C | 99 | 0 |
| SPGP0A02730 | YDR047W | 99 | 0 |
| SPGP0A01720 | YDL072C | 99 | 0 |
| SPGP0BG00230 | YCL050C | 99 | 0 |
| SPGP0B00600 | YHR008C | 99 | 0 |
| SPGP0G00990 | YKL100C | 99 | 0 |
| SPGP0BG00150 | YCL058C | 99 | 0 |
| SPGP0I02920 | YJL171C | 99 | 0 |
| SPGP0BF01410 | YIL122W | 99 | 0 |
| SPGP0F02570 | YBR036C | 100 | 0 |
| SPGP0H01940 | YMR148W | 100 | 0 |

Create PDF in your applications with the Pdfcrowd HTML to PDF API PDFCROWD

| | | | |
|---|---|---|---|
| SPGP0A03610 | YDR142C | 100 | 0 |
| SPGP0L01460 | YOR259C | 100 | 0 |
| SPGP0I00840 | YJR068W | 99 | 0 |
| SPGP0F03740 | YBL091C-A | 99 | 0 |
| SPGP0H01200 | YMR070W | 97 | 2 |
| SPGP0F01160 | YBR186W | 99 | 0 |
| SPGP0F00700 | YBR236C | 99 | 0 |
| SPGP0L01030 | YOR212W | 99 | 0 |
| SPGP0I00860 | YJR066W | 99 | 0 |
| SPGP0J01860 | YOR086C | 99 | 0 |
| SPGP0MU00110 | YGL174W | 99 | 0 |
| SPGP0BG00840 | YCR020C-A | 99 | 0 |
| SPGP0L00940 | YOR202W | 99 | 0 |
| SPGP0XF00120 | YKL205W | 84 | 0 |
| SPGP0FK00170 | YGL207W | 99 | 0 |
| SPGP0L00980 | YOR207C | 99 | 0 |
| SPGP0DG00120 | YOR390W | 99 | 0 |
| SPGP0F03870 | YBL107C | 99 | 0 |
| SPGP0H02830 | YMR237W | 99 | 0 |
| SPGP0F00880 | YBR215W | 99 | 0 |
| SPGP0F00330 | YBR273C | 99 | 0 |
| SPGP0L02480 | YOR373W | 99 | 1 |
| SPGP0F01930 | YBR104W | 100 | 0 |
| SPGP0X00190 | YOL148C | 99 | 0 |
| SPGP0H00520 | YML004C | 99 | 0 |
| SPGP0H00830 | YMR028W | 99 | 0 |
| SPGP0A03760 | YDR160W | 99 | 0 |
| SPGP0A04110 | YDR195W | 99 | 0 |
| SPGP0L00320 | YOR132W | 99 | 0 |
| SPGP0F02010 | YBR094W | 99 | 0 |
| SPGP0F03190 | YBL030C | 99 | 0 |
| SPGP0I02610 | YJL136C | 99 | 0 |
| SPGP0I00730 | YJR082C | 99 | 0 |
| SPGP0A02230 | YDL008W | 99 | 0 |
| SPGP0J01470 | YOR040W | 99 | 0 |
| SPGP0BF00570 | YIL030C | 99 | 0 |
| SPGP0H01710 | YMR124W | 99 | 0 |
| SPGP0F03760 | YBL093C | 99 | 0 |
| SPGP0G00180 | YKL191W | 99 | 0 |
| SPGP0I02330 | YJL104W | 100 | 0 |

Create PDF in your applications with the Pdfcrowd HTML to PDF API PDFCROWD

| | | | |
|---|---|---|---|
| SPGP0L01050 | YOR214C | 99 | 0 |
| SPGP0I01980 | YJL063C | 99 | 0 |
| SPGP0L01220 | YOR233W | 99 | 0 |
| SPGP0H01070 | YMR056C | 99 | 0 |
| SPGP0L00250 | YOR125C | 99 | 0 |
| SPGP0H01990 | YMR154C | 99 | 0 |
| SPGP0RN00100 | YER168C | 99 | 0 |
| SPGP0ME00210 | YGL224C | 99 | 0 |
| SPGP0H02520 | YMR206W | 100 | 0 |
| SPGP0A04330 | YDR223W | 99 | 0 |
| SPGP0QL00140 | YAL020C | 99 | 0 |
| SPGP0A02980 | YDR074W | 99 | 0 |
| SPGP0L01610 | YOR276W | 99 | 0 |
| SPGP0ND00100 | YPL246C | 99 | 0 |
| SPGP0H03140 | YMR270C | 99 | 0 |
| SPGP0H02540 | YMR208W | 99 | 0 |
| SPGP0H00480 | YML008C | 99 | 0 |
| SPGP0QG00190 | YAL051W | 99 | 0 |
| SPGP0L02030 | YOR323C | 99 | 0 |
| SPGP0F03090 | YBL021C | 100 | 0 |
| SPGP0BZ00100 | YAL063C | 87 | 0 |
| SPGP0B01570 | YHR105W | 99 | 0 |
| SPGP0CB00240 | YIR034C | 99 | 0 |
| SPGP0A01700 | YDL074C | 99 | 0 |
| SPGP0A01440 | YDL101C | 99 | 0 |
| SPGP0LL00170 | YAL032C | 100 | 0 |
| SPGP0TZ00100 | YOR388C | 99 | 0 |
| SPGP0NM00120 | YGL258W-A | 97 | 0 |
| SPGP0A04600 | YDR253C | 99 | 0 |
| SPGP0A05130 | YDR309C | 99 | 0 |
| SPGP0I01650 | YJL029C | 99 | 0 |
| SPGP0A04020 | YDR184C | 99 | 0 |
| SPGP0F02590 | YBR034C | 99 | 0 |
| SPGP0B01360 | YHR084W | 99 | 0 |
| SPGP0F00730 | YBR233W-A | 99 | 0 |
| SPGP0L01440 | YOR257W | 100 | 0 |
| SPGP0F02230 | YBR071W | 99 | 0 |
| SPGP0F02820 | YBR008C | 99 | 0 |
| SPGP0F00680 | YBR238C | 99 | 0 |
| SPGP0H01220 | YMR072W | 100 | 0 |

Create PDF in your applications with the Pdfcrowd HTML to PDF API PDFCROWD

| | | | |
|---|---|---|---|
| SPGP0L01010 | YOR210W | 100 | 0 |
| SPGP0B01790 | YHR128W | 100 | 0 |
| SPGP0IG00130 | YPL248C | 99 | 0 |
| SPGP0A01930 | YDL047W | 99 | 0 |
| SPGP0L01630 | YOR279C | 99 | 0 |
| SPGP0H03200 | YMR276W | 99 | 0 |
| SPGP0BF01730 | YIL155C | 99 | 0 |
| SPGP0F02640 | YBR028C | 99 | 0 |
| SPGP0H03330 | YMR289W | 100 | 0 |
| SPGP0A03070 | YDR082W | 99 | 0 |
| SPGP0F01660 | YBR133C | 99 | 0 |
| SPGP0A03820 | YDR166C | 99 | 0 |
| SPGP0A03390 | YDR119W | 99 | 0 |
| SPGP0A01810 | YDL060W | 99 | 0 |
| SPGP0H02850 | YMR239C | 99 | 0 |
| SPGP0BF00550 | YIL026C | 99 | 0 |
| SPGP0J00760 | YOL038W | 100 | 0 |
| SPGP0A02840 | YDR060W | 99 | 0 |
| SPGP0L01540 | YOR269W | 99 | 0 |
| SPGP0LL00150 | YAL030W | 99 | 0 |
| SPGP0B01030 | YHR051W | 99 | 0 |
| SPGP0TA00100 | YAL065C | 96 | 0 |
| SPGP0A04000 | YDR182W | 99 | 0 |
| SPGP0L00270 | YOR127W | 99 | 0 |
| SPGP0A00570 | YDL198C | 99 | 0 |
| SPGP0G01010 | YKL098W | 100 | 0 |
| SPGP0A04760 | YDR270W | 99 | 0 |
| SPGP0F00620 | YBR244W | 99 | 0 |
| SPGP0IY00120 | YGL243W | 99 | 0 |
| SPGP0A02940 | YDR070C | 100 | 0 |
| SPGP0BG00590 | YCL007C | 100 | 0 |
| SPGP0J01290 | YOR020W-A | 100 | 0 |
| SPGP0L02230 | YOR347C | 99 | 0 |
| SPGP0J00690 | YOL045W | 99 | 0 |
| SPGP0QG00200 | YAL053W | 99 | 0 |
| SPGP0BG01000 | YCR036W | 99 | 0 |
| SPGP0B01550 | YHR103W | 99 | 0 |
| SPGP0QD00130 | YGL180W | 99 | 0 |
| SPGP0J01650 | YOR062C | 99 | 0 |
| SPGP0CB00260 | YIR036C | 100 | 0 |

Create PDF in your applications with the Pdfcrowd HTML to PDF API PDFCROWD

| | | | |
|---|---|---|---|
| SPGP0F00420 | YBR263W | 99 | 0 |
| SPGP0A03800 | YDR164C | 99 | 0 |
| SPGP0I01670 | YJL031C | 99 | 0 |
| SPGP0LL00190 | YAL034C | 99 | 0 |
| SPGP0I03290 | YJL213W | 99 | 0 |
| SPGP0I02200 | YJL091C | 99 | 0 |
| SPGP0A04040 | YDR186C | 99 | 0 |
| SPGP0A00280 | YDL229W | 99 | 0 |
| SPGP0A03590 | YDR140W | 100 | 0 |
| SPGP0I00310 | YJR124C | 99 | 0 |
| SPGP0F00490 | YBR256C | 99 | 0 |
| SPGP0F02180 | YBR077C | 99 | 0 |
| SPGP0RY00110 | YER172C | 99 | 0 |
| SPGP0H02190 | YMR174C | 100 | 0 |
| SPGP0BF01780 | YIL159W | 99 | 0 |
| SPGP0H02990 | YMR255W | 99 | 0 |
| SPGP0X00220 | YOL145C | 99 | 0 |
| SPGP0BF00450 | YIL015W | 99 | 0 |
| SPGP0G00440 | YKL163W | 97 | 0 |
| SPGP0B01940 | YHR143W | 99 | 0 |
| SPGP0J01330 | YOR025W | 99 | 0 |
| SPGP0I00400 | YJR113C | 99 | 0 |
| SPGP0B00750 | YHR023W | 99 | 0 |
| SPGP0BF00620 | YIL036W | 99 | 0 |
| SPGP0J01780 | YOR077W | 99 | 0 |
| SPGP0A05070 | YDR303C | 99 | 0 |
| SPGP0B01050 | YHR055C | 100 | 0 |
| SPGP0J01660 | YOR063W | 100 | 0 |
| SPGP0F02410 | YBR053C | 99 | 0 |
| SPGP0HW00100 | YGL178W | 99 | 0 |
| SPGP0F02370 | YBR057C | 100 | 0 |
| SPGP0I03170 | YJL200C | 99 | 0 |
| SPGP0J01250 | YOR017W | 99 | 0 |
| SPGP0I01440 | YJR001W | 99 | 0 |
| SPGP0A00590 | YDL195W | 99 | 0 |
| SPGP0A04460 | YDR237W | 99 | 0 |
| SPGP0I02180 | YJL089W | 99 | 0 |
| SPGP0A01180 | YDL128W | 99 | 0 |
| SPGP0J01410 | YOR034C | 99 | 0 |
| SPGP0G00140 | YKL195W | 99 | 0 |

Create PDF in your applications with the Pdfcrowd HTML to PDF API PDFCROWD

| | | | |
|---|---|---|---|
| SPGP0F00600 | YBR246W | 99 | 0 |
| SPGP0I02590 | YJL133C-A | 99 | 0 |
| SPGP0H01110 | YMR061W | 99 | 0 |
| SPGP0QL00100 | YAL024C | 99 | 0 |
| SPGP0G00960 | YKL104C | 99 | 0 |
| SPGP0F01150 | YBR187W | 99 | 0 |
| SPGP0BG00570 | YCL009C | 99 | 0 |
| SPGP0BF01290 | YIL110W | 99 | 0 |
| SPGP0I02310 | YJL102W | 99 | 0 |
| SPGP0L00630 | YOR166C | 99 | 0 |
| SPGP0CB00200 | YIR030C | 99 | 0 |
| SPGP0B01840 | YHR133C | 99 | 0 |
| SPGP0A01400 | YDL105W | 99 | 0 |
| SPGP0A00970 | YDL150W | 99 | 0 |
| SPGP0F03280 | YBL039C | 99 | 0 |
| SPGP0A02960 | YDR072C | 99 | 0 |
| SPGP0H02730 | YMR227C | 99 | 0 |
| SPGP0H02670 | YMR221C | 99 | 0 |
| SPGP0A00140 | YFL058W | 99 | 0 |
| SPGP0LL00130 | YAL028W | 99 | 0 |
| SPGP0F00440 | YBR261C | 99 | 0 |
| SPGP0I01610 | YJL024C | 99 | 0 |
| SPGP0B01070 | YHR057C | 99 | 0 |
| SPGP0IR00100 | YNL112W | 87 | 1 |
| SPGP0H03460 | YMR302C | 99 | 0 |
| SPGP0B00220 | YHL030W | 99 | 0 |
| SPGP0H01550 | YMR108W | 99 | 0 |
| SPGP0A05260 | YDR321W | 99 | 0 |
| SPGP0WU00140 | YER180C-A | 99 | 0 |
| SPGP0H01800 | YMR133W | 99 | 0 |
| SPGP0I01720 | YJL037W | 99 | 0 |
| SPGP0BF01060 | YIL087C | 99 | 0 |
| SPGP0BF00490 | YIL019W | 99 | 0 |
| SPGP0H00580 | YMR004W | 99 | 0 |
| SPGP0H02710 | YMR225C | 100 | 0 |
| SPGP0L01690 | YOR286W | 99 | 0 |
| SPGP0G00360 | YKL172W | 99 | 0 |
| SPGP0RY00130 | YER170W | 99 | 0 |
| SPGP0F00390 | YBR267W | 99 | 0 |
| SPGP0QL00120 | YAL022C | 99 | 0 |

Create PDF in your applications with the Pdfcrowd HTML to PDF API PDFCROWD

| | | | |
|---|---|---|---|
| SPGP0J01080 | YOL003C | 99 | 0 |
| SPGP0F01510 | YBR150C | 99 | 0 |
| SPGP0L02120 | YOR335C | 99 | 0 |
| SPGP0F01600 | YBR140C | 99 | 0 |
| SPGP0QQ00190 | YAR002C-A | 99 | 0 |
| SPGP0A03330 | YDR111C | 99 | 0 |
| SPGP0J00450 | YOL069W | 99 | 0 |
| SPGP0I00390 | YJR115W | 100 | 0 |
| SPGP0J01350 | YOR027W | 99 | 0 |
| SPGP0QX00120 | YMR317W | 99 | 0 |
| SPGP0B01250 | YHR074W | 99 | 0 |
| SPGP0A05050 | YDR301W | 99 | 0 |
| SPGP0L00230 | YOR123C | 99 | 0 |
| SPGP0I01420 | YJR003C | 99 | 0 |
| SPGP0I01130 | YJR039W | 99 | 0 |
| SPGP0F01310 | YBR169C | 99 | 0 |
| SPGP0H02480 | YMR202W | 100 | 0 |
| SPGP0J00490 | YOL065C | 99 | 0 |
| SPGP0G00940 | YKL106W | 99 | 0 |
| SPGP0H02810 | YMR235C | 99 | 0 |
| SPGP0F00950 | YBR207W | 99 | 0 |
| SPGP0CB00220 | YIR032C | 99 | 0 |
| SPGP0J00740 | YOL040C | 100 | 0 |
| SPGP0B01620 | YHR110W | 100 | 0 |
| SPGP0A00810 | YDL170W | 99 | 0 |
| SPGP0HT00280 | YPL006W | 83 | 1 |
| SPGP0H00620 | YMR008C | 99 | 1 |
| SPGP0B01510 | YHR099W | 99 | 0 |
| SPGP0X00500 | YOL115W | 99 | 0 |
| SPGP0J00910 | YOL020W | 99 | 0 |
| SPGP0B00200 | YHL032C | 99 | 0 |
| SPGP0F00540 | YBR252W | 100 | 0 |
| SPGP0A05190 | YDR315C | 100 | 0 |
| SPGP0G01060 | YKL092C | 100 | 0 |
| SPGP0F03110 | YBL023C | 99 | 0 |
| SPGP0NH00200 | YAL013W | 99 | 0 |
| SPGP0F01700 | YBR129C | 99 | 0 |
| SPGP0BG00510 | YCL017C | 99 | 0 |
| SPGP0A03550 | YDR135C | 99 | 0 |
| SPGP0A03840 | YDR168W | 99 | 0 |

Create PDF in your applications with the Pdfcrowd HTML to PDF API PDFCROWD

| | | | |
|---|---|---|---|
| SPGP0J00530 | YOL061W | 99 | 0 |
| SPGP0H01690 | YMR122W-A | 100 | 0 |
| SPGP0L02140 | YOR337W | 99 | 0 |
| SPGP0X00310 | YOL136C | 99 | 0 |
| SPGP0H03350 | YMR291W | 99 | 0 |
| SPGP0I00370 | YJR117W | 99 | 0 |
| SPGP0J01370 | YOR030W | 99 | 0 |
| SPGP0F03580 | YBL074C | 99 | 0 |
| SPGP0X00560 | YOL109W | 99 | 0 |
| SPGP0B01090 | YHR059W | 99 | 0 |
| SPGP0I03130 | YJL196C | 99 | 0 |
| SPGP0BF00710 | YIL045W | 99 | 0 |
| SPGP0J00700 | YOL044W | 99 | 0 |
| SPGP0I02570 | YJL132W | 99 | 0 |
| SPGP0BF01760 | YIL157C | 99 | 0 |
| SPGP0H02460 | YMR200W | 99 | 0 |
| SPGP0F03320 | YBL042C | 99 | 0 |
| SPGP0H01340 | YMR086W | 99 | 0 |
| SPGP0L00960 | YOR205C | 99 | 0 |
| SPGP0XB00160 | YPR002W | 81 | 0 |
| SPGP0J00890 | YOL022C | 100 | 0 |
| SPGP0H03080 | YMR264W | 100 | 0 |
| SPGP0A00950 | YDL154W | 99 | 0 |
| SPGP0F00210 | YBR286W | 99 | 0 |
| SPGP0A02330 | YDR002W | 100 | 0 |
| SPGP0BG01020 | YCR038C | 99 | 0 |
| SPGP0BG00290 | YCL044C | 99 | 0 |
| SPGP0NM00100 | YGL257C | 98 | 0 |
| SPGP0A00870 | YDL164C | 99 | 0 |
| SPGP0B00530 | YHR002W | 99 | 0 |
| SPGP0B00260 | YHL026C | 99 | 0 |
| SPGP0F01680 | YBR131W | 99 | 0 |
| SPGP0I02650 | YJL140W | 99 | 0 |
| SPGP0L01800 | YOR297C | 99 | 0 |
| SPGP0A05200 | YDR316W | 99 | 0 |
| SPGP0G00820 | YKL121W | 99 | 0 |
| SPGP0G01080 | YKL090W | 99 | 0 |
| SPGP0B00160 | YHL036W | 99 | 0 |
| SPGP0F03130 | YBL025W | 99 | 0 |

Create PDF in your applications with the Pdfcrowd HTML to PDF API PDFCROWD

| | | | |
|---|---|---|---|
| SPGP0H02130 | YMR168C | 99 | 0 |
| SPGP0GU00190 | YPL272C | 99 | 0 |
| SPGP0I02260 | YJL097W | 99 | 0 |
| SPGP0G00550 | YKL151C | 99 | 0 |
| SPGP0A03350 | YDR115W | 99 | 0 |
| SPGP0L01730 | YOR290C | 99 | 0 |
| SPGP0BG00650 | YCR002C | 99 | 0 |
| SPGP0J00510 | YOL063C | 99 | 0 |
| SPGP0L02420 | YOR367W | 99 | 0 |
| SPGP0H01790 | YMR132C | 100 | 0 |
| SPGP0FK00100 | YGL215W | 99 | 0 |
| SPGP0J01390 | YOR032C | 99 | 0 |
| SPGP0F03300 | YBL040C | 99 | 0 |
| SPGP0I02240 | YJL095W | 99 | 0 |
| SPGP0BG00330 | YCL038C | 99 | 0 |
| SPGP0FH00100 | YGL262W | 82 | 0 |
| SPGP0VT00200 | YPL252C | 100 | 0 |
| SPGP0J00160 | YOL100W | 99 | 0 |
| SPGP0I02450 | YJL121C | 99 | 0 |
| SPGP0H03010 | YMR257C | 99 | 0 |
| SPGP0A02350 | YDR004W | 99 | 0 |
| SPGP0B01190 | YHR069C | 99 | 0 |
| SPGP0A03700 | YDR152W | 100 | 0 |
| SPGP0A00850 | YDL166C | 99 | 0 |
| SPGP0B00550 | YHR004C | 99 | 0 |
| SPGP0H03480 | YMR304W | 99 | 0 |
| SPGP0I01820 | YJL048C | 99 | 0 |
| SPGP0F03510 | YBL064C | 99 | 0 |
| SPGP0BG00550 | YCL011C | 99 | 0 |
| SPGP0J01430 | YOR036W | 99 | 0 |
| SPGP0H02630 | YMR217W | 99 | 0 |
| SPGP0EP00100 | YGL176C | 99 | 0 |
| SPGP0H02440 | YMR198W | 99 | 0 |
| SPGP0H01860 | YMR139W | 100 | 0 |
| SPGP0L01710 | YOR288C | 99 | 0 |
| SPGP0J00570 | YOL057W | 99 | 0 |
| SPGP0F01610 | YBR139W | 99 | 0 |
| SPGP0X00370 | YOL129W | 99 | 0 |
| SPGP0G00340 | YKL174C | 99 | 0 |
| SPGP0I00480 | YJR107W | 99 | 0 |

Create PDF in your applications with the Pdfcrowd HTML to PDF API PDFCROWD

| | | | |
|---|---|---|---|
| SPGP0F00400 | YBR265W | 99 | 0 |
| SPGP0H02360 | YMR191W | 99 | 0 |
| SPGP0BF00990 | YIL076W | 99 | 0 |
| SPGP0BG00200 | YCL054W | 99 | 0 |
| SPGP0H01650 | YMR118C | 99 | 0 |
| SPGP0L00380 | YOR140W | 99 | 0 |
| SPGP0A01260 | YDL120W | 99 | 0 |
| SPGP0A02370 | YDR006C | 99 | 0 |
| SPGP0L01600 | YOR275C | 99 | 0 |
| SPGP0EC00100 | YGR009C | 85 | 1 |
| SPGP0B02260 | YHR175W | 99 | 0 |
| SPGP0BG00690 | YCR006C | 99 | 0 |
| SPGP0A03290 | YDR107C | 99 | 0 |
| SPGP0BF00760 | YIL050W | 99 | 0 |
| SPGP0A01590 | YDL085W | 99 | 0 |
| SPGP0A00180 | YDL239C | 99 | 0 |
| SPGP0A04390 | YDR229W | 100 | 0 |
| SPGP0GU00150 | YPL268W | 99 | 0 |
| SPGP0A03480 | YDR127W | 99 | 0 |
| SPGP0H01510 | YMR104C | 99 | 0 |
| SPGP0I00690 | YJR086W | 99 | 0 |
| SPGP0J01600 | YOR057W | 99 | 0 |
| SPGP0L01820 | YOR298C-A | 99 | 0 |
| SPGP0F01060 | YBR196C | 99 | 0 |
| SPGP0G01170 | YKL080W | 98 | 0 |
| SPGP0A04720 | YDR265W | 99 | 0 |
| SPGP0A00370 | YDL218W | 99 | 0 |
| SPGP0B00820 | YHR030C | 99 | 0 |
| SPGP0I01860 | YJL052W | 99 | 0 |
| SPGP0B01480 | YHR097C | 99 | 0 |
| SPGP0B00570 | YHR005C-A | 99 | 0 |
| SPGP0L02160 | YOR339C | 100 | 0 |
| SPGP0L00880 | YOR194C | 99 | 0 |
| SPGP0B02380 | YHR187W | 100 | 0 |
| SPGP0BG00670 | YCR004C | 99 | 0 |
| SPGP0H01530 | YMR106C | 99 | 0 |
| SPGP0A01080 | YDL139C | 99 | 0 |
| SPGP0J00470 | YOL067C | 99 | 0 |
| SPGP0I01840 | YJL050W | 99 | 0 |
| SPGP0J01270 | YOR019W | 99 | 0 |

Create PDF in your applications with the Pdfcrowd HTML to PDF API PDFCROWD

| | | | |
|---|---|---|---|
| SPGP0A01360 | YDL109C | 99 | 0 |
| SPGP0A02620 | YDR034C | 99 | 0 |
| SPGP0L00450 | YOR148C | 98 | 0 |
| SPGP0I00500 | YJR105W | 99 | 0 |
| SPGP0BF00340 | YIL005W | 99 | 0 |
| SPGP0F01370 | YBR163W | 99 | 0 |

#### Sheet 3: *genes alleles below 80% identity*

| *Saccharomyces pastorianus CBS1503 predicted ORF* | Parentale allele ID | Parent | Gene identities | Gene gaps |
|---|---|---|---|---|
| SPGP0VO00160 | YCR075C | *Saccharomyces cerevisiae* | 78 | 1 |
| SPGP0KL00200 | YKR064W | *Saccharomyces cerevisiae* | 78 | 1 |
| SPGP0HJ00130 | XP_018224047.1 | *Saccharomyces eubayanus* | 78 | 2 |

#### Sheet 4: *genes alleles copy with same identity in both parents*

| *Saccharomyces pastorianus CBS1503 predicted ORF* | Parental allele ID | Parent | Gene identities | Gene gaps |
|---|---|---|---|---|
| SPGP0MF00250 | YGL030W | *Saccharomyces cerevisiae* | 99 | 0 |
| SPGP0MF00250 | XP_018222195.1 | *Saccharomyces eubayanus* | 99 | 0 |
| SPGP0Itrna26K | YDR445C | *Saccharomyces cerevisiae* | 100 | 0 |
| SPGP0Itrna26K | DI49_5468 | *Saccharomyces eubayanus* | 100 | 0 |
| SPGP0JF00280 | YPR132W | *Saccharomyces cerevisiae* | 99 | 0 |
| SPGP0JF00280 | XP_018222326.1 | *Saccharomyces eubayanus* | 99 | 0 |
| SPGP0TVtrna20G | YNL285W | *Saccharomyces cerevisiae* | 100 | 0 |
| SPGP0TVtrna20G | DI49_2297 | *Saccharomyces eubayanus* | 100 | 0 |
| SPGP0HJ00140 | YDR531W | *Saccharomyces cerevisiae* | 83 | 0 |
| SPGP0HJ00140 | XP_018224046.1 | *Saccharomyces eubayanus* | 83 | 0 |
| SPGP0SV00110 | YCR039C | *Saccharomyces cerevisiae* | 100 | 0 |
| SPGP0SV00110 | XP_018223412.1 | *Saccharomyces eubayanus* | 100 | 0 |
| SPGP0Dtrna17T | YHR180W-A | *Saccharomyces cerevisiae* | 100 | 0 |
| SPGP0Dtrna17T | DI49_5553 | *Saccharomyces eubayanus* | 100 | 0 |
| SPGP0Ftrna14I | YNL017C | *Saccharomyces cerevisiae* | 100 | 0 |
| SPGP0Ftrna14I | DI49_5614 | *Saccharomyces eubayanus* | 100 | 0 |
| SPGP0Itrna29R | YGR164W | *Saccharomyces cerevisiae* | 100 | 0 |
| SPGP0Itrna29R | DI49_4020 | *Saccharomyces eubayanus* | 100 | 0 |
| SPGP0Itrna32G | YPR159C-A | *Saccharomyces cerevisiae* | 100 | 0 |
| SPGP0Itrna32G | DI49_5603 | *Saccharomyces eubayanus* | 100 | 0 |

#### Sheet 5: *list of paralogs in Saccharomyces pastorianus CBS1503*

Create PDF in your applications with the Pdfcrowd HTML to PDF API PDFCROWD

| List of paralogs | *S. eubayanus-like content* | *S. pastorianus content* | *S. cerevisiae-like content* |
|---|---|---|---|
| SPGP0CVtrna3N; DI49_5581; SPGP0CV00100; SPGP0Jtrna1N; YOL054W; SPGP0Jtrna2N; SPGP0Mtrna4N; SPGP0EKtrna6N; DI49_3921; SPGP0Ntrna5N; SPGP0LJtrna7N; DI49_5582; DI49_1403; SPGP0MStrna9N; DI49_2081; DI49_3381; SPGP0Dtrna11N; DI49_2351; DI49_4738; YCR008W; SPGP0BGtrna13N | DI49_5581; DI49_3921; DI49_5582; DI49_1403; DI49_2081; DI49_3381; DI49_2351; DI49_4738 | SPGP0CVtrna3N; SPGP0CV00100; SPGP0Jtrna1N; SPGP0Jtrna2N; SPGP0Mtrna4N; SPGP0EKtrna6N; SPGP0Ntrna5N; SPGP0LJtrna7N; SPGP0MStrna9N; SPGP0Dtrna11N; SPGP0BGtrna13N | YOL054W; YCR008W |
| DI49_0756; SPGP0CV00110 | DI49_0756 | SPGP0CV00110 | |
| DI49_1621; DI49_1620 | DI49_1621; DI49_1620 | | |
| DI49_1622; SPGP0CW00120 | DI49_1622 | SPGP0CW00120 | |
| DI49_3910; DI49_2609; DI49_3909 | DI49_3910; DI49_2609; DI49_3909 | | |
| DI49_3907; DI49_3908 | DI49_3907; DI49_3908 | | |
| DI49_3905; DI49_3906 | DI49_3905; DI49_3906 | | |
| DI49_3903; SPGP0CX00160; YDL075W; SPGP0BE03400 | DI49_3903 | SPGP0CX00160; SPGP0BE03400 | YDL075W |
| DI49_3901; DI49_3902 | DI49_3901; DI49_3902 | | |
| DI49_0595; DI49_3900 | DI49_0595; DI49_3900 | | |
| DI49_0593; DI49_0594 | DI49_0593; DI49_0594 | | |
| DI49_4770; DI49_4769 | DI49_4770; DI49_4769 | | |
| DI49_4772; DI49_4771 | DI49_4772; DI49_4771 | | |
| DI49_4774; DI49_4773 | DI49_4774; DI49_4773 | | |
| DI49_4776; DI49_4775 | DI49_4776; DI49_4775 | | |
| DI49_4778; DI49_4777 | DI49_4778; DI49_4777 | | |
| DI49_4780; DI49_0833; DI49_0834 | DI49_4780; DI49_0833; DI49_0834 | | |
| SPGP0DA00100; YNR076W; DI49_1930; YBR301W; YPL246C | DI49_1930 | SPGP0DA00100 | YNR076W; YBR301W; YPL246C |
| DI49_1928; DI49_1820; DI49_1821 | DI49_1928; DI49_1820; DI49_1821 | | |
| SPGP0DBtrna2L; DI49_4635; DI49_1366 | DI49_4635; DI49_1366 | SPGP0DBtrna2L | |
| DI49_1368; DI49_1367 | DI49_1368; DI49_1367 | | |
| DI49_1370; DI49_1369 | DI49_1370; DI49_1369 | | |
| DI49_3335; DI49_3336 | DI49_3335; DI49_3336 | | |
| YJR029W; SPGP0GZ00100; DI49_4730 | DI49_4730 | SPGP0GZ00100 | YJR029W |
| SPGP0Jtrna2T; SPGP0Itrna26T; YOL105C; YOR013W; YDL007C-A; YHR006W; SPGP0NAtrna12T; SPGP0BFtrna21T; YIL102C | | SPGP0Jtrna2T; SPGP0Itrna26T; SPGP0NAtrna12T; SPGP0BFtrna21T | YOL105C; YOR013W; YDL007C-A; YHR006W; YIL102C |
| YOL103W; YOL104C | | | YOL103W; YOL104C |
| YOL101C; DI49_2305 | DI49_2305 | | YOL101C |
| YOL100W; DI49_2306 | DI49_2306 | | YOL100W |
| YOL098C; DI49_2307 | DI49_2307 | | YOL098C |
| YOL097C; DI49_2308 | DI49_2308 | | YOL097C |
| YOL096C; DI49_2309 | DI49_2309 | | YOL096C |
| YOL095C; DI49_2310 | DI49_2310 | | YOL095C |
| YOL094C; DI49_2311; YOL093W; DI49_2312 | DI49_2311; DI49_2312 | | YOL094C; YOL093W |
| YOL091W; DI49_2314 | DI49_2314 | | YOL091W |

Create PDF in your applications with the Pdfcrowd HTML to PDF API PDFCROWD

| | | | |
|---|---|---|---|
| YOL090W; DI49_2315 | DI49_2315 | | YOL090W |
| YOL089C; DI49_2316; YOL088C; DI49_2317 | DI49_2316; DI49_2317 | | YOL089C; YOL088C |
| YOL086W-A; YOL087C | | | YOL086W-A; YOL087C |
| YOL084W; YOL086C | | | YOL084W; YOL086C |
| YOL082W; YOL083W | | | YOL082W; YOL083W |
| YOL080C; YOL081W; DI49_2324; DI49_2325 | DI49_2324; DI49_2325 | | YOL080C; YOL081W |
| YOL077W-A; YOL078W | | | YOL077W-A; YOL078W |
| YOL076W; DI49_2329 | DI49_2329 | | YOL076W |
| YOL075C; DI49_2330; YOL073C; DI49_2331 | DI49_2330; DI49_2331 | | YOL075C; YOL073C |
| YOL071W; YOL072W | | | YOL071W; YOL072W |
| YOL069W; DI49_2335 | DI49_2335 | | YOL069W |
| YOL068C; DI49_2336; YOL067C; DI49_2337 | DI49_2336; DI49_2337 | | YOL068C; YOL067C |
| YOL065C; YOL066C | | | YOL065C; YOL066C |
| YOL063C; YOL064C | | | YOL063C; YOL064C |
| YOL061W; DI49_2343 | DI49_2343 | | YOL061W |
| YOL060C; DI49_2344 | DI49_2344 | | YOL060C |
| YOL059W; DI49_2345 | DI49_2345 | | YOL059W |
| YOL058W; DI49_2346; YOL057W; DI49_2347 | DI49_2346; DI49_2347 | | YOL058W; YOL057W |
| YOL055C; DI49_2348; YPL258C | DI49_2348 | | YOL055C; YPL258C |
| SPGP0Jtrna3G; SPGP0Itrna32G; DI49_0764; SPGP0Atrna2G; YDR262W; DI49_4046; SPGP0PZtrna18G; DI49_2184; SPGP0QPtrna19G; SPGP0Dtrna16G; SPGP0BEtrna21G; DI49_0870; DI49_1663; SPGP0BGtrna23G; YCR016W; SPGP0WFtrna25G; DI49_3364; DI49_0441; SPGP0BKtrna28G; DI49_3059; SPGP0BKtrna29G; SPGP0Htrna30G; YML043C; SPGP0Itrna31G; YJR051W | DI49_0764; DI49_4046; DI49_2184; DI49_0870; DI49_1663; DI49_3364; DI49_0441; DI49_3059 | SPGP0Jtrna3G; SPGP0Itrna32G; SPGP0Atrna2G; SPGP0PZtrna18G; SPGP0QPtrna19G; SPGP0Dtrna16G; SPGP0BEtrna21G; SPGP0BGtrna23G; SPGP0WFtrna25G; SPGP0BKtrna28G; SPGP0BKtrna29G; SPGP0Htrna30G; SPGP0Itrna31G | YDR262W; YCR016W; YML043C; YJR051W |
| YOL052C-A; YOL053W | | | YOL052C-A; YOL053W |
| YOL051W; DI49_2355; YOL049W; DI49_2356 | DI49_2355; DI49_2356 | | YOL051W; YOL049W |
| YOL047C; YOL048C | | | YOL047C; YOL048C |
| YOL045W; DI49_2357; YOL044W; DI49_2358 | DI49_2357; DI49_2358 | | YOL045W; YOL044W |
| YOL042W; YOL043C | | | YOL042W; YOL043C |
| YOL040C; DI49_2362 | DI49_2362 | | YOL040C |
| YOL039W; DI49_2363; YOL038W; DI49_2364 | DI49_2363; DI49_2364 | | YOL039W; YOL038W |
| YOL034W; DI49_2366; YOL033W; DI49_2367 | DI49_2366; DI49_2367 | | YOL034W; YOL033W |
| YOL031C; YOL032W | | | YOL031C; YOL032W |
| YOL028C; YOL030W | | | YOL028C; YOL030W |
| YOL026C; YOL027C | | | YOL026C; YOL027C |
| YOL025W; DI49_2374; SPGP0EMtrna5S; DI49_3829; DI49_0521 | DI49_2374; DI49_3829; DI49_0521 | SPGP0EMtrna5S | YOL025W |
| YOL023W; DI49_2377; YOL022C; DI49_2378 | DI49_2377; DI49_2378 | | YOL023W; YOL022C |
| YOL021C; DI49_2380; YOL020W; DI49_2381 | DI49_2380; DI49_2381 | | YOL021C; YOL020W |
| YOL019W; SPGP0Itrna12Y; SPGP0Atrna1Y; YJR056C | | SPGP0Itrna12Y; SPGP0Atrna1Y | YOL019W; YJR056C |

Create PDF in your applications with the Pdfcrowd HTML to PDF API PDFCROWD

| | | | |
|---|---|---|---|
| YOL017W; YOL018C | | | YOL017W; YOL018C |
| YOL015W; YOL016C | | | YOL015W; YOL016C |
| YOL013W-A; SPGP0Jtrna2P; SPGP0Htrna19P; SPGP0Ltrna4P; SPGP0NHtrna10P | | SPGP0Jtrna2P; SPGP0Htrna19P; SPGP0Ltrna4P; SPGP0NHtrna10P | YOL013W-A |
| YOL013C; DI49_2390 | DI49_2390 | | YOL013C |
| YOL012C; DI49_2391 | DI49_2391 | | YOL012C |
| YOL011W; DI49_2392; YOL010W; DI49_2393 | DI49_2392; DI49_2393 | | YOL011W; YOL010W |
| YOL008W; YOL009C | | | YOL008W; YOL009C |
| YOL006C; YOL007C; DI49_2397; DI49_2398 | DI49_2397; DI49_2398 | | YOL006C; YOL007C |
| YOL004W; DI49_2399 | DI49_2399 | | YOL004W |
| YOL003C; DI49_2400 | DI49_2400 | | YOL003C |
| YOL002C; DI49_2401; YOL001W; DI49_2402 | DI49_2401; DI49_2402 | | YOL002C; YOL001W |
| YOR002W; DI49_2404 | DI49_2404 | | YOR002W |
| YOR003W; DI49_2405; YOR004W; DI49_2406 | DI49_2405; DI49_2406 | | YOR003W; YOR004W |
| YOR006C; DI49_2408 | DI49_2408 | | YOR006C |
| YOR007C; DI49_2409 | DI49_2409 | | YOR007C |
| SPGP0Jtrna3R; DI49_2410; YOR008C; SPGP0Atrna2R; SPGP0LKtrna9R; DI49_3398; SPGP0RNtrna13R; DI49_1335; SPGP0Dtrna10R; DI49_2411; SPGP0STtrna14R; DI49_3638 | DI49_2410; DI49_3398; DI49_1335; DI49_2411; DI49_3638 | SPGP0Jtrna3R; SPGP0Atrna2R; SPGP0LKtrna9R; SPGP0RNtrna13R; SPGP0Dtrna10R; SPGP0STtrna14R | YOR008C |
| YOR010C; DI49_2413; SPGP0J01210; DI49_2414 | DI49_2413; DI49_2414 | SPGP0J01210 | YOR010C |
| YOR016C; DI49_2418 | DI49_2418 | | YOR016C |
| YOR017W; DI49_2419; YOR018W; DI49_2420 | DI49_2419; DI49_2420 | | YOR017W; YOR018W |
| YOR020C; DI49_4866; YOR020W-A; DI49_4867 | DI49_4866; DI49_4867 | | YOR020C; YOR020W-A |
| YOR022C; YOR021C | | | YOR022C; YOR021C |
| YOR025W; YOR023C | | | YOR025W; YOR023C |
| YOR027W; YOR026W; YOR030W | | | YOR027W; YOR026W; YOR030W |
| YOR032C; YOR031W | | | YOR032C; YOR031W |
| YOR034C; YOR033C | | | YOR034C; YOR033C |
| YOR036W; YOR035C | | | YOR036W; YOR035C |
| YOR038C; DI49_4883 | DI49_4883 | | YOR038C |
| YOR039W; DI49_4884; YOR040W; DI49_4885 | DI49_4884; DI49_4885 | | YOR039W; YOR040W |
| YOR043W; DI49_4887; YOR044W; DI49_4888 | DI49_4887; DI49_4888 | | YOR043W; YOR044W |
| YOR046C; DI49_4890 | DI49_4890 | | YOR046C |
| YOR047C; DI49_4891 | DI49_4891 | | YOR047C |
| YOR048C; DI49_4892; YOR049C; DI49_4893 | DI49_4892; DI49_4893 | | YOR048C; YOR049C |
| YOR052C; YOR051C | | | YOR052C; YOR051C |
| YOR056C; YOR054C | | | YOR056C; YOR054C |
| YOR058C; DI49_4899 | DI49_4899 | | YOR058C |
| SPGP0Jtrna2K; SPGP0Atrna1K; YOR059C; YGL202W; SPGP0FKtrna6K; YDL054C | | SPGP0Jtrna2K; SPGP0Atrna1K; SPGP0FKtrna6K | YOR059C; YGL202W; YDL054C |
| YOR061W; YOR060C | | | YOR061W; YOR060C |

Create PDF in your applications with the Pdfcrowd HTML to PDF API PDFCROWD

| | | | |
|---|---|---|---|
| YOR063W; DI49_4906; YOR064C; DI49_4907 | DI49_4906; DI49_4907 | | YOR063W; YOR064C |
| YOR066W; DI49_4909 | DI49_4909 | | YOR066W |
| YOR067C; DI49_4910 | DI49_4910 | | YOR067C |
| YOR069W; DI49_4911 | DI49_4911 | | YOR069W |
| YOR070C; DI49_4912 | DI49_4912 | | YOR070C |
| YOR071C; YOR192C; DI49_3747; DI49_3748 | DI49_3747; DI49_3748 | | YOR071C; YOR192C |
| YOR073W; DI49_4915 | DI49_4915 | | YOR073W |
| YOR074C; SPGP0N01130 | | SPGP0N01130 | YOR074C |
| YOR075W; DI49_4916; YOR076C; DI49_4917 | DI49_4916; DI49_4917 | | YOR075W; YOR076C |
| YOR078W; YOR077W | | | YOR078W; YOR077W |
| YOR080W; YOR079C | | | YOR080W; YOR079C |
| YOR083W; YOR081C | | | YOR083W; YOR081C |
| YOR085W; YOR084W | | | YOR085W; YOR084W |
| YOR087W; DI49_4926; YOR089C; DI49_4927 | DI49_4926; DI49_4927 | | YOR087W; YOR089C |
| YOR091W; YOR090C | | | YOR091W; YOR090C |
| YPL277C; YOR092W | | | YPL277C; YOR092W |
| YOR390W; YPL278C | | | YOR390W; YPL278C |
| DI49_4781; DI49_5593 | DI49_4781; DI49_5593 | | |
| DI49_4783; DI49_4782 | DI49_4783; DI49_4782 | | |
| DI49_5503; DI49_5502 | DI49_5503; DI49_5502 | | |
| YER185W; YER184C | | | YER185W; YER184C |
| YER187W; YER186C | | | YER187W; YER186C |
| DI49_5266; SPGP0DL00140 | DI49_5266 | SPGP0DL00140 | |
| DI49_5424; DI49_5425 | DI49_5424; DI49_5425 | | |
| DI49_5422; DI49_1959; DI49_5421; DI49_2045 | DI49_5422; DI49_1959; DI49_5421; DI49_2045 | | |
| DI49_5419; DI49_5420 | DI49_5419; DI49_5420 | | |
| DI49_5417; DI49_5418 | DI49_5417; DI49_5418 | | |
| DI49_3986; DI49_5416 | DI49_3986; DI49_5416 | | |
| DI49_3984; DI49_3985 | DI49_3984; DI49_3985 | | |
| DI49_5679; DI49_3983 | DI49_5679; DI49_3983 | | |
| SPGP0K00120; DI49_5681 | DI49_5681 | SPGP0K00120 | |
| DI49_5682; YFL058W; DI49_5683; SPGP0A00150 | DI49_5682; DI49_5683 | SPGP0A00150 | YFL058W |
| DI49_5685; DI49_5684 | DI49_5685; DI49_5684 | | |
| DI49_5191; YOR380W; DI49_5190; YOR381W | DI49_5191; DI49_5190 | | YOR380W; YOR381W |
| DI49_5188; YOR375C; YOR377W | DI49_5188 | | YOR375C; YOR377W |
| DI49_5187; YOR374W; DI49_5186 | DI49_5187; DI49_5186 | | YOR374W |
| DI49_5184; DI49_5185 | DI49_5184; DI49_5185 | | |
| DI49_5182; YOR369C; DI49_5181; YOR370C | DI49_5182; DI49_5181 | | YOR369C; YOR370C |
| DI49_5179; DI49_5180 | DI49_5179; DI49_5180 | | |
| DI49_5177; DI49_5178 | DI49_5177; DI49_5178 | | |

Create PDF in your applications with the Pdfcrowd HTML to PDF API PDFCROWD

| | | | |
|---|---|---|---|
| DI49_5175; YOR361C; YOR362C | DI49_5175 | | YOR361C; YOR362C |
| DI49_5174; YOR360C | DI49_5174 | | YOR360C |
| DI49_5173; YOR359W; DI49_5172 | DI49_5173; DI49_5172 | | YOR359W |
| DI49_5170; YOR356W | DI49_5170 | | YOR356W |
| DI49_5169; YOR355W; DI49_5168 | DI49_5169; DI49_5168 | | YOR355W |
| DI49_5166; DI49_5167 | DI49_5166; DI49_5167 | | |
| DI49_5164; DI49_5165 | DI49_5164; DI49_5165 | | |
| DI49_5161; YOR348C; SPGP0L02250 | DI49_5161 | SPGP0L02250 | YOR348C |
| DI49_5160; YOR347C | DI49_5160 | | YOR347C |
| DI49_5159; YOR346W | DI49_5159 | | YOR346W |
| DI49_5158; DI49_2389; DI49_5157; DI49_4051 | DI49_5158; DI49_2389; DI49_5157; DI49_4051 | | |
| SPGP0Ktrna1M; DI49_4243; SPGP0Ctrna6M; DI49_4242; SPGP0VZtrna11M; DI49_0528; SPGP0BKtrna13M; DI49_2990 | DI49_4243; DI49_4242; DI49_0528; DI49_2990 | SPGP0Ktrna1M; SPGP0Ctrna6M; SPGP0VZtrna11M; SPGP0BKtrna13M | |
| DI49_5154; YOR342C | DI49_5154 | | YOR342C |
| DI49_5153; YOR341W | DI49_5153 | | YOR341W |
| DI49_5152; YOR340C; DI49_5151 | DI49_5152; DI49_5151 | | YOR340C |
| DI49_5149; YOR337W; DI49_5148 | DI49_5149; DI49_5148 | | YOR337W |
| SPGP0K00600; SPGP0K00590; YOR335C; YOR336W | | SPGP0K00600; SPGP0K00590 | YOR335C; YOR336W |
| DI49_5146; YOR330C; DI49_5145; YOR332W | DI49_5146; DI49_5145 | | YOR330C; YOR332W |
| DI49_5143; YOR327C | DI49_5143 | | YOR327C |
| DI49_5142; YOR326W; DI49_5141 | DI49_5142; DI49_5141 | | YOR326W |
| DI49_5139; YOR322C; YOR323C | DI49_5139 | | YOR322C; YOR323C |
| DI49_5138; YOR321W; DI49_5137 | DI49_5138; DI49_5137 | | YOR321W |
| DI49_5135; DI49_5136 | DI49_5135; DI49_5136 | | |
| DI49_5133; DI49_5134 | DI49_5133; DI49_5134 | | |
| DI49_5131; YMR242C; DI49_4327 | DI49_5131; DI49_4327 | | YMR242C |
| DI49_5130; YOR311C; YOR313C | DI49_5130 | | YOR311C; YOR313C |
| DI49_5129; YOR310C; DI49_5128 | DI49_5129; DI49_5128 | | YOR310C |
| DI49_5126; YOR306C; DI49_5125 | DI49_5126; DI49_5125 | | YOR306C |
| DI49_5123; YOR304W; DI49_5122; YOR304C-A | DI49_5123; DI49_5122 | | YOR304W; YOR304C-A |
| DI49_5120; SPGP0K00870 | DI49_5120 | SPGP0K00870 | |
| DI49_5118; DI49_5119 | DI49_5118; DI49_5119 | | |
| DI49_5116; DI49_5117 | DI49_5116; DI49_5117 | | |
| DI49_5114; DI49_5115 | DI49_5114; DI49_5115 | | |
| DI49_5112; DI49_4314; DI49_5111; YOR293W; YMR230W | DI49_5112; DI49_4314; DI49_5111 | | YOR293W; YMR230W |
| DI49_5109; YOR290C; YOR291W | DI49_5109 | | YOR290C; YOR291W |
| SPGP0Ktrna2A; DI49_3564; DI49_5107; SPGP0Ltrna3A; SPGP0GRtrna7A; DI49_1482; SPGP0Utrna8A; DI49_0066; DI49_3563; SPGP0PEtrna15A; DI49_2116 | DI49_3564; DI49_5107; DI49_1482; DI49_0066; DI49_3563; DI49_2116 | SPGP0Ktrna2A; SPGP0Ltrna3A; SPGP0GRtrna7A; SPGP0Utrna8A; SPGP0PEtrna15A | |
| DI49_5105; DI49_5106 | DI49_5105; DI49_5106 | | |

Create PDF in your applications with the Pdfcrowd HTML to PDF API PDFCROWD

| | | | |
|---|---|---|---|
| DI49_5103; DI49_5104 | DI49_5103; DI49_5104 | | |
| DI49_5101; YOR283W; YOR284W | DI49_5101 | | YOR283W; YOR284W |
| DI49_5100; YOR281C | DI49_5100 | | YOR281C |
| DI49_5099; YOR280C; DI49_5098 | DI49_5099; DI49_5098 | | YOR280C |
| DI49_5096; YOR276W; DI49_5095; YOR278W | DI49_5096; DI49_5095 | | YOR276W; YOR278W |
| DI49_5093; YOR273C; YOR274W | DI49_5093 | | YOR273C; YOR274W |
| DI49_5092; YOR272W | DI49_5092 | | YOR272W |
| DI49_5091; YOR271C | DI49_5091 | | YOR271C |
| DI49_5090; YOR270C; DI49_5089 | DI49_5090; DI49_5089 | | YOR270C |
| DI49_5087; YOR267C; DI49_5086; YOR269W | DI49_5087; DI49_5086 | | YOR267C; YOR269W |
| DI49_5084; YOR264W; YOR265W | DI49_5084 | | YOR264W; YOR265W |
| DI49_5083; YOR262W | DI49_5083 | | YOR262W |
| DI49_5082; YOR261C | DI49_5082 | | YOR261C |
| DI49_5081; YOR260W | DI49_5081 | | YOR260W |
| DI49_5080; YOR259C; DI49_5079 | DI49_5080; DI49_5079 | | YOR259C |
| DI49_5077; DI49_5078 | DI49_5077; DI49_5078 | | |
| DI49_5075; YOR254C; DI49_5074 | DI49_5075; DI49_5074 | | YOR254C |
| DI49_5072; DI49_5073 | DI49_5072; DI49_5073 | | |
| DI49_5070; DI49_5071 | DI49_5070; DI49_5071 | | |
| DI49_5068; DI49_5069 | DI49_5068; DI49_5069 | | |
| DI49_5066; YOR244W; DI49_5065; YOR245C | DI49_5066; DI49_5065 | | YOR244W; YOR245C |
| DI49_5063; YOR241W; SPGP0K01450; YOR242C | DI49_5063 | SPGP0K01450 | YOR241W; YOR242C |
| DI49_5061; DI49_5062 | DI49_5061; DI49_5062 | | |
| DI49_5322; SPGP0OI00560; DI49_5058; SPGP0L01230; DI49_5321 | DI49_5322; DI49_5058; DI49_5321 | SPGP0OI00560; SPGP0L01230 | |
| DI49_5056; YOR231W; DI49_5055; YOR232W | DI49_5056; DI49_5055 | | YOR231W; YOR232W |
| DI49_5053; YOR228C; YOR229W | DI49_5053 | | YOR228C; YOR229W |
| DI49_5052; YOR227W; DI49_5051 | DI49_5052; DI49_5051 | | YOR227W |
| DI49_5049; DI49_5050 | DI49_5049; DI49_5050 | | |
| DI49_5047; YOR221C | DI49_5047 | | YOR221C |
| DI49_5046; YOR220W | DI49_5046 | | YOR220W |
| SPGP0K01630; YOR219C | | SPGP0K01630 | YOR219C |
| DI49_5045; YOR217W; DI49_5044 | DI49_5045; DI49_5044 | | YOR217W |
| DI49_5042; DI49_5043; YOR215C; YOR216C | DI49_5042; DI49_5043 | | YOR215C; YOR216C |
| DI49_5040; YOR212W; YOR213C | DI49_5040 | | YOR212W; YOR213C |
| DI49_5039; YOR211C; DI49_5038 | DI49_5039; DI49_5038 | | YOR211C |
| DI49_5036; DI49_5037 | DI49_5036; DI49_5037 | | |
| DI49_5034; DI49_5035; YOR207C; YOR208W | DI49_5034; DI49_5035 | | YOR207C; YOR208W |
| DI49_5032; YOR204W; DI49_5031; YOR205C; YNL112W; DI49_4621 | DI49_5032; DI49_5031; DI49_4621 | | YOR204W; YOR205C; YNL112W |
| DI49_5029; YOR198C; DI49_5028; YOR201C | DI49_5029; DI49_5028 | | YOR198C; YOR201C |

Create PDF in your applications with the Pdfcrowd HTML to PDF API PDFCROWD

| | | | |
|---|---|---|---|
| DI49_5026; DI49_5027 | DI49_5026; DI49_5027 | | |
| DI49_5024; DI49_5025 | DI49_5024; DI49_5025 | | |
| DI49_5022; YOR191W | DI49_5022 | | YOR191W |
| DI49_5021; YOR190W; DI49_5020 | DI49_5021; DI49_5020 | | YOR190W |
| DI49_5019; YOR187W; DI49_5018 | DI49_5019; DI49_5018 | | YOR187W |
| DI49_5016; DI49_5017; DI49_5015; DI49_3800; DI49_3799 | DI49_5016; DI49_5017; DI49_5015; DI49_3800; DI49_3799 | | |
| DI49_5013; DI49_5014 | DI49_5013; DI49_5014 | | |
| DI49_5011; DI49_5012 | DI49_5011; DI49_5012 | | |
| DI49_5009; DI49_4248; YOR177C; SPGP0KUtrna11V; DI49_3338; YGL225W; DI49_4247; SPGP0Ctrna10V; DI49_4138; DI49_3304; SPGP0PEtrna15V; DI49_2126; DI49_1560; DI49_3781; SPGP0XAtrna22V; DI49_1542; SPGP0BKtrna23V; SPGP0Htrna25V; SPGP0Htrna26V; YMR077C; YMR164C; SPGP0Htrna27V; YJL037W | DI49_5009; DI49_4248; DI49_3338; DI49_4247; DI49_4138; DI49_3304; DI49_2126; DI49_1560; DI49_3781; DI49_1542 | SPGP0KUtrna11V; SPGP0Ctrna10V; SPGP0PEtrna15V; SPGP0XAtrna22V; SPGP0BKtrna23V; SPGP0Htrna25V; SPGP0Htrna26V; SPGP0Htrna27V | YOR177C; YGL225W; YMR077C; YMR164C; YJL037W |
| DI49_5007; DI49_5008 | DI49_5007; DI49_5008 | | |
| DI49_5005; DI49_5006 | DI49_5005; DI49_5006 | | |
| DI49_5003; YOR168W | DI49_5003 | | YOR168W |
| DI49_5002; YOR167C; DI49_5001 | DI49_5002; DI49_5001 | | YOR167C |
| DI49_4999; YOR164C; DI49_4998 | DI49_4999; DI49_4998 | | YOR164C |
| DI49_4996; DI49_4997 | DI49_4996; DI49_4997 | | |
| DI49_4994; YOR160W; DI49_4993; YOR161C | DI49_4994; DI49_4993 | | YOR160W; YOR161C |
| DI49_4991; DI49_4992 | DI49_4991; DI49_4992 | | |
| DI49_4989; DI49_4990 | DI49_4989; DI49_4990 | | |
| DI49_4987; YOR153W; DI49_4986; YOR154W | DI49_4987; DI49_4986 | | YOR153W; YOR154W |
| DI49_4984; YOR150W | DI49_4984 | | YOR150W |
| DI49_4983; YOR149C; DI49_4982 | DI49_4983; DI49_4982 | | YOR149C |
| DI49_4980; YOR145C; DI49_4979 | DI49_4980; DI49_4979 | | YOR145C |
| SPGP0Ktrna5G; DI49_4977; YOR143C; DI49_0567; DI49_0565 | DI49_4977; DI49_0567; DI49_0565 | SPGP0Ktrna5G | YOR143C |
| DI49_4976; YOR142W | DI49_4976 | | YOR142W |
| DI49_4975; YOR141C; DI49_4974 | DI49_4975; DI49_4974 | | YOR141C |
| DI49_4972; DI49_4973 | DI49_4972; DI49_4973 | | |
| DI49_4970; DI49_4971 | DI49_4970; DI49_4971 | | |
| DI49_4968; YOR132W; DI49_4967 | DI49_4968; DI49_4967 | | YOR132W |
| DI49_4965; YOR130C; DI49_4964 | DI49_4965; DI49_4964 | | YOR130C |
| DI49_4962; YOR127W; DI49_4961 | DI49_4962; DI49_4961 | | YOR127W |
| DI49_4959; YOR124C; DI49_4958 | DI49_4959; DI49_4958 | | YOR124C |
| DI49_4956; DI49_4957 | DI49_4956; DI49_4957 | | |
| DI49_4954; DI49_4955 | DI49_4954; DI49_4955 | | |
| DI49_4952; YOR116C; DI49_4951 | DI49_4952; DI49_4951 | | YOR116C |
| DI49_4949; YOR113W | DI49_4949 | | YOR113W |
| DI49_4948; YOR112W; DI49_4947 | DI49_4948; DI49_4947 | | YOR112W |
| DI49_4945; YOR109W; DI49_3745; YOR110W | DI49_4945; DI49_3745 | | YOR109W; YOR110W |

Create PDF in your applications with the Pdfcrowd HTML to PDF API PDFCROWD

| | | | |
|---|---|---|---|
| DI49_3750; DI49_3749 | DI49_3750; DI49_3749 | | |
| DI49_3752; DI49_3751 | DI49_3752; DI49_3751 | | |
| DI49_4524; DI49_4523 | DI49_4524; DI49_4523 | | |
| DI49_4526; DI49_4525 | DI49_4526; DI49_4525 | | |
| DI49_4528; DI49_4527 | DI49_4528; DI49_4527 | | |
| DI49_4530; DI49_4529 | DI49_4530; DI49_4529 | | |
| DI49_4532; DI49_4531 | DI49_4532; DI49_4531 | | |
| DI49_4534; DI49_4533 | DI49_4534; DI49_4533 | | |
| DI49_3485; DI49_3484 | DI49_3485; DI49_3484 | | |
| DI49_3487; DI49_3486 | DI49_3487; DI49_3486 | | |
| DI49_3428; SPGP0BK02840; DI49_3427; DI49_2887 | DI49_3428; DI49_3427; DI49_2887 | SPGP0BK02840 | |
| DI49_3425; DI49_3426 | DI49_3425; DI49_3426 | | |
| DI49_1708; DI49_1709 | DI49_1708; DI49_1709 | | |
| DI49_1706; DI49_1707 | DI49_1706; DI49_1707 | | |
| DI49_1704; DI49_1705 | DI49_1704; DI49_1705 | | |
| DI49_1702; DI49_1703 | DI49_1702; DI49_1703 | | |
| DI49_1700; DI49_1701 | DI49_1700; DI49_1701 | | |
| DI49_1698; DI49_1699 | DI49_1698; DI49_1699 | | |
| DI49_5385; DI49_1697 | DI49_5385; DI49_1697 | | |
| DI49_5383; DI49_5384 | DI49_5383; DI49_5384 | | |
| DI49_5381; DI49_1215; DI49_5380; DI49_1214 | DI49_5381; DI49_1215; DI49_5380; DI49_1214 | | |
| DI49_4943; DI49_4942 | DI49_4943; DI49_4942 | | |
| SPGP0DX00130; DI49_4944 | DI49_4944 | SPGP0DX00130 | |
| YOR114W; DI49_4950; YOR115C | DI49_4950 | | YOR114W; YOR115C |
| YOR117W; DI49_4953; YOR118W | DI49_4953 | | YOR117W; YOR118W |
| YOR120W; YOR119C | | | YOR120W; YOR119C |
| YOR123C; YOR122C | | | YOR123C; YOR122C |
| YOR125C; DI49_4960; YOR126C | DI49_4960 | | YOR125C; YOR126C |
| YOR128C; DI49_4963; YOR129C | DI49_4963 | | YOR128C; YOR129C |
| SPGP0Ltrna3D; DI49_4675; YOR131C; SPGP0Ptrna4D; DI49_2768; SPGP0Ptrna5D; SPGP0Atrna1D; SPGP0A02830; DI49_3666; DI49_1987; SPGP0KWtrna9D; DI49_3834; SPGP0LItrna10D; DI49_1993; SPGP0LKtrna11D; DI49_3409; SPGP0Ctrna6D; SPGP0BEtrna12D; SPGP0VNtrna15D; DI49_4676; SPGP0BFtrna13D; YIL009C-A; SPGP0BFtrna14D; DI49_3066; SPGP0BKtrna19D; SPGP0BKtrna21D; SPGP0BK02620; SPGP0Ftrna16D; SPGP0Htrna22D; YJR055W; SPGP0Itrna24D; SPGP0Itrna25D | DI49_4675; DI49_2768; DI49_3666; DI49_1987; DI49_3834; DI49_1993; DI49_3409; DI49_4676; DI49_3066 | SPGP0Ltrna3D; SPGP0Ptrna4D; SPGP0Ptrna5D; SPGP0Atrna1D; SPGP0A02830; SPGP0KWtrna9D; SPGP0LItrna10D; SPGP0LKtrna11D; SPGP0Ctrna6D; SPGP0BEtrna12D; SPGP0VNtrna15D; SPGP0BFtrna13D; SPGP0BFtrna14D; SPGP0BKtrna19D; SPGP0BKtrna21D; SPGP0BK02620; SPGP0Ftrna16D; SPGP0Htrna22D; SPGP0Itrna24D; SPGP0Itrna25D | YOR131C; YIL009C-A; YJR055W |
| YDR385W; DI49_4969; YOR134W | DI49_4969 | | YDR385W; YOR134W |
| YOR137C; YOR136W | | | YOR137C; YOR136W |
| YOR140W; YOR138C | | | YOR140W; YOR138C |
| YOR147W; DI49_4981; YOR148C | DI49_4981 | | YOR147W; YOR148C |

Create PDF in your applications with the Pdfcrowd HTML to PDF API PDFCROWD

| | | | |
|---|---|---|---|
| YOR151C; DI49_4985; YOR152C | DI49_4985 | | YOR151C; YOR152C |
| YOR156C; YOR155C | | | YOR156C; YOR155C |
| YOR158W; YOR157C | | | YOR158W; YOR157C |
| YOR163W; YOR162C | | | YOR163W; YOR162C |
| YOR165W; DI49_5000; YOR166C | DI49_5000 | | YOR165W; YOR166C |
| YOR171C; DI49_5004; YOR172W | DI49_5004 | | YOR171C; YOR172W |
| YOR174W; YOR173W | | | YOR174W; YOR173W |
| YOR176W; YOR175C | | | YOR176W; YOR175C |
| YOR179C; YOR178C | | | YOR179C; YOR178C |
| YOR181W; YOR180C; YOR184W | | | YOR181W; YOR180C; YOR184W |
| YOR186W; YOR185C | | | YOR186W; YOR185C |
| YOR188W; SPGP0K01890; YOR189W | | SPGP0K01890 | YOR188W; YOR189W |
| SPGP0Ltrna3M; DI49_5351; YOR193W; SPGP0GYtrna5M; DI49_1363; SPGP0OItrna9M; DI49_5350; SPGP0BGtrna10M; DI49_3049; YJR043C | DI49_5351; DI49_1363; DI49_5350; DI49_3049 | SPGP0Ltrna3M; SPGP0GYtrna5M; SPGP0OItrna9M; SPGP0BGtrna10M | YOR193W; YJR043C |
| YOR195W; YOR194C | | | YOR195W; YOR194C |
| YOR197W; YOR196C | | | YOR197W; YOR196C |
| YOR210W; YOR209C | | | YOR210W; YOR209C |
| YOR222W; DI49_5048; YOR223W | DI49_5048 | | YOR222W; YOR223W |
| YOR226C; YOR224C | | | YOR226C; YOR224C |
| YOR236W; DI49_5060; YOR237W | DI49_5060 | | YOR236W; YOR237W |
| YOR239W; YOR238W | | | YOR239W; YOR238W |
| YOR247W; YOR246C | | | YOR247W; YOR246C |
| YOR250C; YOR249C | | | YOR250C; YOR249C |
| YOR252W; YOR251C | | | YOR252W; YOR251C |
| YOR255W; DI49_5076; YOR256C | DI49_5076 | | YOR255W; YOR256C |
| YOR258W; YOR257W | | | YOR258W; YOR257W |
| YOR286W; YOR285W | | | YOR286W; YOR285W |
| YOR288C; YOR287C | | | YOR288C; YOR287C |
| YOR294W; DI49_5113; YOR295W | DI49_5113 | | YOR294W; YOR295W |
| YOR297C; YOR296W | | | YOR297C; YOR296W |
| YOR298C-A; YOR298W | | | YOR298C-A; YOR298W |
| YOR301W; YOR299W | | | YOR301W; YOR299W |
| YOR303W; YOR302W | | | YOR303W; YOR302W |
| YOR307C; DI49_5127; YOR308C | DI49_5127 | | YOR307C; YOR308C |
| YOR316C; YOR315W | | | YOR316C; YOR315W |
| YOR319W; YOR317W | | | YOR319W; YOR317W |
| YOR328W; DI49_5144; YOR329C | DI49_5144 | | YOR328W; YOR329C |
| YOR338W; DI49_5150; YOR339C | DI49_5150 | | YOR338W; YOR339C |
| SPGP0L02200; YDR170W-A; YDR171W | | SPGP0L02200 | YDR170W-A; YDR171W |
| SPGP0Ltrna4M; YJL007C; SPGP0AItrna7M; YMR159C; | | SPGP0Ltrna4M; SPGP0AItrna7M | YJL007C; YMR159C; YJL029C |

Create PDF in your applications with the Pdfcrowd HTML to PDF API PDFCROWD

| | | | |
|---|---|---|---|
| YJL029C | | | |
| YOR344C; DI49_5156 | DI49_5156 | | YOR344C |
| YOR350C; YOR349W | | | YOR350C; YOR349W |
| YOR352W; YOR351C | | | YOR352W; YOR351C |
| YOR354C; YOR353C | | | YOR354C; YOR353C |
| YOR357C; DI49_5171; YOR358W | DI49_5171 | | YOR357C; YOR358W |
| YOR365C; YOR363C | | | YOR365C; YOR363C |
| YOR368W; YOR367W | | | YOR368W; YOR367W |
| YOR372C; YOR371C | | | YOR372C; YOR371C |
| YKL222C; YKL221W | | | YKL222C; YKL221W |
| SPGP0DZ00100; SPGP0L02570 | | SPGP0DZ00100; SPGP0L02570 | |
| DI49_3943; DI49_4014; SPGP0DZ00130; DI49_4013 | DI49_3943; DI49_4014; DI49_4013 | SPGP0DZ00130 | |
| DI49_0674; DI49_0676 | DI49_0674; DI49_0676 | | |
| DI49_0672; DI49_0673 | DI49_0672; DI49_0673 | | |
| YGR009C; DI49_5686; SPGP0GGtrna6L; DI49_5397; SPGP0GStrna7L; DI49_1624; SPGP0KXtrna13L; DI49_0734; DI49_2147; DI49_2148; DI49_0670 | DI49_5686; DI49_5397; DI49_1624; DI49_0734; DI49_2147; DI49_2148; DI49_0670 | SPGP0GGtrna6L; SPGP0GStrna7L; SPGP0KXtrna13L | YGR009C |
| DI49_1972; DI49_1973 | DI49_1972; DI49_1973 | | |
| DI49_1325; DI49_1324 | DI49_1325; DI49_1324 | | |
| YEL070W; SPGP0BK00120 | | SPGP0BK00120 | YEL070W |
| SPGP0ED00140; DI49_2618; DI49_3165 | DI49_2618; DI49_3165 | SPGP0ED00140 | |
| SPGP0EEtrna3E; DI49_5287; DI49_3838; SPGP0Mtrna2E; SPGP0Ptrna4E; SPGP0FXtrna6E; DI49_1880; DI49_2203; SPGP0LItrna10E; DI49_1991; SPGP0MBtrna11E; SPGP0MC00100; SPGP0NGtrna12E; YAL001C; SPGP0Ctrna9E; SPGP0BEtrna13E; SPGP0BFtrna14E; YCL021W-A; SPGP0WBtrna18E; DI49_1543; SPGP0Ftrna17E; SPGP0Gtrna21E; SPGP0Htrna22E | DI49_5287; DI49_3838; DI49_1880; DI49_2203; DI49_1991; DI49_1543 | SPGP0EEtrna3E; SPGP0Mtrna2E; SPGP0Ptrna4E; SPGP0FXtrna6E; SPGP0LItrna10E; SPGP0MBtrna11E; SPGP0MC00100; SPGP0NGtrna12E; SPGP0Ctrna9E; SPGP0BEtrna13E; SPGP0BFtrna14E; SPGP0WBtrna18E; SPGP0Ftrna17E; SPGP0Gtrna21E; SPGP0Htrna22E | YAL001C; YCL021W-A |
| DI49_3840; DI49_3839 | DI49_3840; DI49_3839 | | |
| DI49_3842; DI49_3841 | DI49_3842; DI49_3841 | | |
| DI49_4040; DI49_3843; DI49_1995 | DI49_4040; DI49_3843; DI49_1995 | | |
| SPGP0EFtrna3Y; DI49_2382; DI49_4037; DI49_1685; DI49_4369; DI49_1670; DI49_2383; SPGP0BItrna8Y; SPGP0BKtrna9Y; DI49_3068; DI49_2969 | DI49_2382; DI49_4037; DI49_1685; DI49_4369; DI49_1670; DI49_2383; DI49_3068; DI49_2969 | SPGP0EFtrna3Y; SPGP0BItrna8Y; SPGP0BKtrna9Y | |
| DI49_2008; DI49_4036 | DI49_2008; DI49_4036 | | |
| DI49_3182; DI49_2009 | DI49_3182; DI49_2009 | | |
| DI49_3184; YKL197C | DI49_3184 | | YKL197C |
| DI49_3186; YKL196C; SPGP0M00140; YKL195W | DI49_3186 | SPGP0M00140 | YKL196C; YKL195W |
| DI49_3188; DI49_3187 | DI49_3188; DI49_3187 | | |
| DI49_3190; DI49_3189 | DI49_3190; DI49_3189 | | |
| SPGP0Mtrna3L; DI49_5687; DI49_3193; DI49_3915; SPGP0OJtrna15L; DI49_4787; YOR391C; SPGP0BEtrna18L; DI49_0231; SPGP0BKtrna28L; DI49_3010; SPGP0Ftrna23L; SPGP0Gtrna29L; YKL189W; SPGP0Ftrna24L; YJL006C | DI49_5687; DI49_3193; DI49_3915; DI49_4787; DI49_0231; DI49_3010 | SPGP0Mtrna3L; SPGP0OJtrna15L; SPGP0BEtrna18L; SPGP0BKtrna28L; SPGP0Ftrna23L; SPGP0Gtrna29L; SPGP0Ftrna24L | YOR391C; YKL189W; YJL006C |
| DI49_3195; YKL187C; DI49_3196; YKL186C | DI49_3195; DI49_3196 | | YKL187C; YKL186C |

Create PDF in your applications with the Pdfcrowd HTML to PDF API PDFCROWD

| | | | |
|---|---|---|---|
| DI49_3198; DI49_3197 | DI49_3198; DI49_3197 | | |
| DI49_3200; YKL182W | DI49_3200 | | YKL182W |
| DI49_3201; YKL181W | DI49_3201 | | YKL181W |
| DI49_3202; YKL180W | DI49_3202 | | YKL180W |
| DI49_3203; YKL179C | DI49_3203 | | YKL179C |
| DI49_3204; YKL178C; DI49_3205; YKL176C | DI49_3204; DI49_3205 | | YKL178C; YKL176C |
| DI49_3207; DI49_3206 | DI49_3207; DI49_3206 | | |
| DI49_3209; DI49_3208 | DI49_3209; DI49_3208 | | |
| DI49_3211; DI49_3210 | DI49_3211; DI49_3210 | | |
| DI49_3213; DI49_3212 | DI49_3213; DI49_3212 | | |
| DI49_3215; YKL165C | DI49_3215 | | YKL165C |
| DI49_3217; DI49_2869; SPGP0M00440; YJL158C; YJL160C | DI49_3217; DI49_2869 | SPGP0M00440 | YJL158C; YJL160C |
| DI49_3220; DI49_3219 | DI49_3220; DI49_3219 | | |
| DI49_3222; DI49_3221 | DI49_3222; DI49_3221 | | |
| SPGP0M00500; YKL157W | | SPGP0M00500 | YKL157W |
| DI49_3224; YKL156W; DI49_3225; YKL155C | DI49_3224; DI49_3225 | | YKL156W; YKL155C |
| SPGP0Mtrna4R; DI49_4323; SPGP0Ptrna5R; SPGP0GBtrna6R; DI49_1387; SPGP0HRtrna7R; DI49_1917; SPGP0Ctrna8R; SPGP0PEtrna11R; DI49_2129; DI49_2082; SPGP0BEtrna16R; SPGP0BItrna19R; SPGP0BKtrna21R | DI49_4323; DI49_1387; DI49_1917; DI49_2129; DI49_2082 | SPGP0Mtrna4R; SPGP0Ptrna5R; SPGP0GBtrna6R; SPGP0HRtrna7R; SPGP0Ctrna8R; SPGP0PEtrna11R; SPGP0BEtrna16R; SPGP0BItrna19R; SPGP0BKtrna21R | |
| DI49_3228; YKL152C; DI49_3229; YKL151C | DI49_3228; DI49_3229 | | YKL152C; YKL151C |
| DI49_3231; DI49_3230 | DI49_3231; DI49_3230 | | |
| DI49_3233; YKL146W | DI49_3233 | | YKL146W |
| DI49_3234; YKL145W; DI49_3235; YKL144C | DI49_3234; DI49_3235 | | YKL145W; YKL144C |
| DI49_3237; YKL142W; DI49_3238; YKL141W | DI49_3237; DI49_3238 | | YKL142W; YKL141W |
| DI49_3240; DI49_3239 | DI49_3240; DI49_3239 | | |
| DI49_3242; YKL138C; DI49_3243; YKL137W | DI49_3242; DI49_3243 | | YKL138C; YKL137W |
| DI49_3245; DI49_3244 | DI49_3245; DI49_3244 | | |
| DI49_3247; DI49_3246 | DI49_3247; DI49_3246 | | |
| DI49_3249; YKL129C; DI49_3250; YKL128C | DI49_3249; DI49_3250 | | YKL129C; YKL128C |
| DI49_3252; YKL127W; DI49_3253; YKL126W | DI49_3252; DI49_3253 | | YKL127W; YKL126W |
| DI49_3255; YKL124W; DI49_3256; YKL122C | DI49_3255; DI49_3256 | | YKL124W; YKL122C |
| DI49_3258; YKL120W; DI49_3259; YKL119C | DI49_3258; DI49_3259 | | YKL120W; YKL119C |
| DI49_3261; DI49_5601; DI49_5413; SPGP0AAtrna9A; DI49_4721; SPGP0Btrna6A; DI49_1682; SPGP0Ctrna10A; DI49_4332; SPGP0Ctrna11A; DI49_4110; SPGP0QPtrna16A; DI49_5600; SPGP0Dtrna13A; SPGP0TXtrna18A; SPGP0BEtrna19A; DI49_1030; SPGP0BKtrna20A; DI49_2904; YKL117W; SPGP0Htrna22A; YMR250W; SPGP0Htrna23A; YJL115W | DI49_3261; DI49_5601; DI49_5413; DI49_4721; DI49_1682; DI49_4332; DI49_4110; DI49_5600; DI49_1030; DI49_2904 | SPGP0AAtrna9A; SPGP0Btrna6A; SPGP0Ctrna10A; SPGP0Ctrna11A; SPGP0QPtrna16A; SPGP0Dtrna13A; SPGP0TXtrna18A; SPGP0BEtrna19A; SPGP0BKtrna20A; SPGP0Htrna22A; SPGP0Htrna23A | YKL117W; YMR250W; YJL115W |
| DI49_3263; YKL114C | DI49_3263 | | YKL114C |
| DI49_3264; YKL113C | DI49_3264 | | YKL113C |

Create PDF in your applications with the Pdfcrowd HTML to PDF API PDFCROWD

| | | | |
|---|---|---|---|
| DI49_3265; SPGP0G00890; DI49_3266; YKL110C | DI49_3265; DI49_3266 | SPGP0G00890 | YKL110C |
| DI49_3268; DI49_3267 | DI49_3268; DI49_3267 | | |
| DI49_3270; YKL106W; DI49_3271; YKL105C | DI49_3270; DI49_3271 | | YKL106W; YKL105C |
| DI49_3273; YKL103C | DI49_3273 | | YKL103C |
| DI49_3274; YKL101W; DI49_3275; YKL100C | DI49_3274; DI49_3275 | | YKL101W; YKL100C |
| DI49_3277; DI49_3276 | DI49_3277; DI49_3276 | | |
| DI49_3279; DI49_3278 | DI49_3279; DI49_3278 | | |
| DI49_3281; YKL094W; DI49_3282; YKL093W | DI49_3281; DI49_3282 | | YKL094W; YKL093W |
| DI49_3284; DI49_3283 | DI49_3284; DI49_3283 | | |
| DI49_3286; YKL089W | DI49_3286 | | YKL089W |
| DI49_3287; YKL088W; DI49_3288; YKL087C | DI49_3287; DI49_3288 | | YKL088W; YKL087C |
| DI49_3290; DI49_3289 | DI49_3290; DI49_3289 | | |
| DI49_3292; YKL082C; SPGP0M01170; YKL081W | DI49_3292 | SPGP0M01170 | YKL082C; YKL081W |
| DI49_3928; DI49_3293 | DI49_3928; DI49_3293 | | |
| DI49_3925; DI49_3926 | DI49_3925; DI49_3926 | | |
| SPGP0EK00140; DI49_3924 | DI49_3924 | SPGP0EK00140 | |
| DI49_3919; DI49_3920 | DI49_3919; DI49_3920 | | |
| DI49_3917; DI49_3918 | DI49_3917; DI49_3918 | | |
| SPGP0EL00100; DI49_3914 | DI49_3914 | SPGP0EL00100 | |
| DI49_0738; YCL016C | DI49_0738 | | YCL016C |
| DI49_0737; YCL017C | DI49_0737 | | YCL017C |
| DI49_3827; DI49_3828 | DI49_3827; DI49_3828 | | |
| DI49_3825; DI49_3826 | DI49_3825; DI49_3826 | | |
| DI49_3823; DI49_3824 | DI49_3823; DI49_3824 | | |
| DI49_3821; DI49_3822 | DI49_3821; DI49_3822 | | |
| DI49_3820; SPGP0EM00190 | DI49_3820 | SPGP0EM00190 | |
| DI49_3819; SPGP0EM00210 | DI49_3819 | SPGP0EM00210 | |
| DI49_3817; DI49_3818 | DI49_3817; DI49_3818 | | |
| DI49_3815; DI49_3816 | DI49_3815; DI49_3816 | | |
| DI49_3813; DI49_3814 | DI49_3813; DI49_3814 | | |
| DI49_2242; DI49_3812 | DI49_2242; DI49_3812 | | |
| DI49_2244; DI49_2243 | DI49_2244; DI49_2243 | | |
| DI49_2246; DI49_2245 | DI49_2246; DI49_2245 | | |
| DI49_3766; YGL176C | DI49_3766 | | YGL176C |
| DI49_5411; DI49_5412 | DI49_5411; DI49_5412 | | |
| DI49_5409; DI49_5410 | DI49_5409; DI49_5410 | | |
| DI49_4809; DI49_4808 | DI49_4809; DI49_4808 | | |
| DI49_4811; DI49_4810 | DI49_4811; DI49_4810 | | |
| DI49_4813; DI49_4812 | DI49_4813; DI49_4812 | | |
| SPGP0N00170; DI49_3489; YHL033C; DI49_3490 | DI49_3489; DI49_3490 | SPGP0N00170 | YHL033C |

Create PDF in your applications with the Pdfcrowd HTML to PDF API PDFCROWD

| | | | |
|---|---|---|---|
| SPGP0N00180; YHL032C | | SPGP0N00180 | YHL032C |
| DI49_4816; YHL031C | DI49_4816 | | YHL031C |
| DI49_4817; YHL030W | DI49_4817 | | YHL030W |
| DI49_4818; YHL029C; DI49_4819; YHL028W | DI49_4818; DI49_4819 | | YHL029C; YHL028W |
| DI49_4821; DI49_4820 | DI49_4821; DI49_4820 | | |
| DI49_4823; YHL024W | DI49_4823 | | YHL024W |
| DI49_4824; YHL023C | DI49_4824 | | YHL023C |
| SPGP0Ntrna1H; DI49_4134; DI49_4826; SPGP0IDtrna4H; DI49_1876; SPGP0OFtrna6H; DI49_1425; SPGP0Ctrna5H; DI49_4133; SPGP0OYtrna7H; DI49_2160; SPGP0TB00100; YMR044W | DI49_4134; DI49_4826; DI49_1876; DI49_1425; DI49_4133; DI49_2160 | SPGP0Ntrna1H; SPGP0IDtrna4H; SPGP0OFtrna6H; SPGP0Ctrna5H; SPGP0OYtrna7H; SPGP0TB00100 | YMR044W |
| DI49_4828; YHL020C; DI49_4829; YHL019C | DI49_4828; DI49_4829 | | YHL020C; YHL019C |
| DI49_4831; YHL017W | DI49_4831 | | YHL017W |
| DI49_4832; YHL016C | DI49_4832 | | YHL016C |
| DI49_4833; YHL015W; DI49_4834; YHL014C | DI49_4833; DI49_4834 | | YHL015W; YHL014C |
| DI49_4836; DI49_4835 | DI49_4836; DI49_4835 | | |
| DI49_4838; YHL010C; DI49_4839; YHL009C | DI49_4838; DI49_4839 | | YHL010C; YHL009C |
| DI49_4841; YHL008C | DI49_4841 | | YHL008C |
| DI49_4842; YHL007C; DI49_4843; YHL006C | DI49_4842; DI49_4843 | | YHL007C; YHL006C |
| DI49_4845; YHL003C | DI49_4845 | | YHL003C |
| DI49_4846; YHL002W | DI49_4846 | | YHL002W |
| DI49_4847; DI49_3367; DI49_4848; SPGP0B00500; YHR001W; DI49_3366 | DI49_4847; DI49_3367; DI49_4848; DI49_3366 | SPGP0B00500 | YHR001W |
| DI49_4850; YHR002W | DI49_4850 | | YHR002W |
| DI49_4851; YHR003C; DI49_4852; YHR004C | DI49_4851; DI49_4852 | | YHR003C; YHR004C |
| DI49_4854; DI49_4853 | DI49_4854; DI49_4853 | | |
| DI49_4856; DI49_2415; SPGP0Ptrna6T; DI49_2683; SPGP0LFtrna9T; SPGP0MGtrna10T; DI49_4465; SPGP0Dtrna14T; DI49_2254; SPGP0Dtrna15T; DI49_2301; SPGP0D01680; SPGP0BEtrna19T; DI49_1043; SPGP0VNtrna22T; DI49_4691; SPGP0BItrna24T | DI49_4856; DI49_2415; DI49_2683; DI49_4465; DI49_2254; DI49_2301; DI49_1043; DI49_4691 | SPGP0Ptrna6T; SPGP0LFtrna9T; SPGP0MGtrna10T; SPGP0Dtrna14T; SPGP0Dtrna15T; SPGP0D01680; SPGP0BEtrna19T; SPGP0VNtrna22T; SPGP0BItrna24T | |
| DI49_4858; YHR008C | DI49_4858 | | YHR008C |
| DI49_4859; YHR009C | DI49_4859 | | YHR009C |
| SPGP0N00600; YHR010W; DI49_4860; YHR011W | DI49_4860 | SPGP0N00600 | YHR010W; YHR011W |
| DI49_4862; YHR013C; DI49_4863; YHR014W | DI49_4862; DI49_4863 | | YHR013C; YHR014W |
| DI49_4865; DI49_4419; SPGP0Ptrna7S; DI49_2817; SPGP0Utrna10S; SPGP0KVtrna12S; SPGP0NFtrna13S; SPGP0Ctrna11S; SPGP0TNtrna17S; SPGP0BEtrna19S; SPGP0BE02820; SPGP0BItrna25S; DI49_0176; SPGP0WYtrna26S; DI49_3537; SPGP0BKtrna27S; DI49_3053; DI49_2003 | DI49_4865; DI49_4419; DI49_2817; DI49_0176; DI49_3537; DI49_3053; DI49_2003 | SPGP0Ptrna7S; SPGP0Utrna10S; SPGP0KVtrna12S; SPGP0NFtrna13S; SPGP0Ctrna11S; SPGP0TNtrna17S; SPGP0BEtrna19S; SPGP0BE02820; SPGP0BItrna25S; SPGP0WYtrna26S; SPGP0BKtrna27S | |
| DI49_4869; DI49_4868 | DI49_4869; DI49_4868 | | |
| DI49_4871; DI49_4870 | DI49_4871; DI49_4870 | | |
| DI49_4873; DI49_4872 | DI49_4873; DI49_4872 | | |
| DI49_4875; DI49_4874 | DI49_4875; DI49_4874 | | |
| DI49_4877; DI49_4876 | DI49_4877; DI49_4876 | | |

Create PDF in your applications with the Pdfcrowd HTML to PDF API PDFCROWD

| | | | |
|---|---|---|---|
| DI49_4879; DI49_4878 | DI49_4879; DI49_4878 | | |
| DI49_4881; DI49_4880 | DI49_4881; DI49_4880 | | |
| DI49_4895; DI49_4894 | DI49_4895; DI49_4894 | | |
| DI49_4897; DI49_4896 | DI49_4897; DI49_4896 | | |
| SPGP0Ntrna4K; DI49_5564; DI49_4901; SPGP0INtrna9K; SPGP0JFtrna10K; DI49_2068; SPGP0KLtrna11K; DI49_3442; DI49_3876 | DI49_5564; DI49_4901; DI49_2068; DI49_3442; DI49_3876 | SPGP0Ntrna4K; SPGP0INtrna9K; SPGP0JFtrna10K; SPGP0KLtrna11K | |
| DI49_4903; DI49_4902 | DI49_4903; DI49_4902 | | |
| DI49_4905; DI49_4904 | DI49_4905; DI49_4904 | | |
| SPGP0Ntrna5P; DI49_4914; SPGP0OQtrna13P; DI49_5253; DI49_4699; DI49_4688 | DI49_4914; DI49_5253; DI49_4699; DI49_4688 | SPGP0Ntrna5P; SPGP0OQtrna13P | |
| DI49_4919; DI49_4918 | DI49_4919; DI49_4918 | | |
| DI49_4921; DI49_4920 | DI49_4921; DI49_4920 | | |
| SPGP0N01210; SPGP0N01200 | | SPGP0N01210; SPGP0N01200 | |
| DI49_4923; DI49_4922 | DI49_4923; DI49_4922 | | |
| DI49_4929; DI49_4928 | DI49_4929; DI49_4928 | | |
| DI49_2205; DI49_4930 | DI49_2205; DI49_4930 | | |
| DI49_2207; DI49_2206 | DI49_2207; DI49_2206 | | |
| DI49_2209; DI49_2208 | DI49_2209; DI49_2208 | | |
| DI49_2211; DI49_2210 | DI49_2211; DI49_2210 | | |
| DI49_1391; DI49_1390 | DI49_1391; DI49_1390 | | |
| DI49_1393; DI49_1392 | DI49_1393; DI49_1392 | | |
| DI49_1395; DI49_1394 | DI49_1395; DI49_1394 | | |
| DI49_2036; DI49_2253 | DI49_2036; DI49_2253 | | |
| DI49_2038; DI49_2037 | DI49_2038; DI49_2037 | | |
| DI49_2040; DI49_2039 | DI49_2040; DI49_2039 | | |
| DI49_2042; DI49_5546; DI49_2043; DI49_5547 | DI49_2042; DI49_5546; DI49_2043; DI49_5547 | | |
| DI49_0797; DI49_2044 | DI49_0797; DI49_2044 | | |
| DI49_0799; DI49_0798 | DI49_0799; DI49_0798 | | |
| DI49_0801; DI49_0800 | DI49_0801; DI49_0800 | | |
| DI49_0803; DI49_0802 | DI49_0803; DI49_0802 | | |
| DI49_5524; DI49_5523; SPGP0Ptrna8S; DI49_2723; DI49_0808; DI49_0809; SPGP0BEtrna18S; DI49_1141; YIL057C; SPGP0VU00100 | DI49_5524; DI49_5523; DI49_2723; DI49_0808; DI49_0809; DI49_1141 | SPGP0Ptrna8S; SPGP0BEtrna18S; SPGP0VU00100 | YIL057C |
| DI49_5526; YBR117C; DI49_5527 | DI49_5526; DI49_5527 | | YBR117C |
| DI49_5529; DI49_5528 | DI49_5529; DI49_5528 | | |
| DI49_4932; DI49_4931 | DI49_4932; DI49_4931 | | |
| DI49_4934; DI49_4636; DI49_4935; DI49_4637 | DI49_4934; DI49_4636; DI49_4935; DI49_4637 | | |
| DI49_4937; DI49_4936 | DI49_4937; DI49_4936 | | |
| DI49_4939; DI49_4938 | DI49_4939; DI49_4938 | | |
| DI49_4941; DI49_4940 | DI49_4941; DI49_4940 | | |
| DI49_5625; DI49_5626 | DI49_5625; DI49_5626 | | |

Create PDF in your applications with the Pdfcrowd HTML to PDF API PDFCROWD

| | | | |
|---|---|---|---|
| DI49_3519; DI49_5624 | DI49_3519; DI49_5624 | | |
| SPGP0FC00100; DI49_3520 | DI49_3520 | SPGP0FC00100 | |
| DI49_5488; DI49_5487 | DI49_5488; DI49_5487 | | |
| DI49_5490; DI49_5489 | DI49_5490; DI49_5489 | | |
| SPGP0O00110; SPGP0O00100 | | SPGP0O00110; SPGP0O00100 | |
| DI49_3636; DI49_3637 | DI49_3636; DI49_3637 | | |
| DI49_3634; DI49_3635 | DI49_3634; DI49_3635 | | |
| DI49_3632; DI49_3633 | DI49_3632; DI49_3633 | | |
| DI49_3630; DI49_3631 | DI49_3630; DI49_3631 | | |
| DI49_3628; DI49_3629 | DI49_3628; DI49_3629 | | |
| DI49_3626; DI49_3627 | DI49_3626; DI49_3627 | | |
| DI49_3625; SPGP0FE00220 | DI49_3625 | SPGP0FE00220 | |
| DI49_3623; DI49_3624 | DI49_3623; DI49_3624 | | |
| DI49_3621; DI49_3622 | DI49_3621; DI49_3622 | | |
| DI49_3619; DI49_3620 | DI49_3619; DI49_3620 | | |
| DI49_3617; DI49_3618 | DI49_3617; DI49_3618 | | |
| YGL215W; YGL262W | | | YGL215W; YGL262W |
| YGL212W; YGL213C | | | YGL212W; YGL213C |
| YGL210W; YGL211W | | | YGL210W; YGL211W |
| YGL208W; YGL209W | | | YGL208W; YGL209W |
| YGL206C; YGL207W | | | YGL206C; YGL207W |
| SPGP0FKtrna2H; SPGP0Htrna9H; YGL203C; SPGP0Btrna3H; YHL022C | | SPGP0FKtrna2H; SPGP0Htrna9H; SPGP0Btrna3H | YGL203C; YHL022C |
| SPGP0FKtrna7K; SPGP0Itrna26K; YGL200C; SPGP0GAtrna8K; SPGP0KYtrna13K; DI49_0541; SPGP0LGtrna15K; DI49_1372; DI49_2156; SPGP0Ctrna12K; DI49_4194; SPGP0PFtrna17K; SPGP0SHtrna18K; DI49_0613; SPGP0UBtrna19K; DI49_5469; SPGP0BFtrna21K; SPGP0BGtrna22K; DI49_1808; SPGP0BKtrna24K; DI49_3004; YMR107W; SPGP0Htrna25K; YJL011C | DI49_0541; DI49_1372; DI49_2156; DI49_4194; DI49_0613; DI49_5469; DI49_1808; DI49_3004 | SPGP0FKtrna7K; SPGP0Itrna26K; SPGP0GAtrna8K; SPGP0KYtrna13K; SPGP0LGtrna15K; SPGP0Ctrna12K; SPGP0PFtrna17K; SPGP0SHtrna18K; SPGP0UBtrna19K; SPGP0BFtrna21K; SPGP0BGtrna22K; SPGP0BKtrna24K; SPGP0Htrna25K | YGL200C; YMR107W; YJL011C |
| SPGP0FLtrna2F; DI49_5486; DI49_5485; DI49_2529; DI49_2530; DI49_0089; DI49_0088 | DI49_5486; DI49_5485; DI49_2529; DI49_2530; DI49_0089; DI49_0088 | SPGP0FLtrna2F | |
| DI49_1883; DI49_5484 | DI49_1883; DI49_5484 | | |
| DI49_1885; DI49_1884 | DI49_1885; DI49_1884 | | |
| DI49_1887; DI49_1886 | DI49_1887; DI49_1886 | | |
| DI49_1889; DI49_1888 | DI49_1889; DI49_1888 | | |
| DI49_1891; DI49_1890 | DI49_1891; DI49_1890 | | |
| DI49_1893; DI49_1892 | DI49_1893; DI49_1892 | | |
| DI49_1895; DI49_1894 | DI49_1895; DI49_1894 | | |
| DI49_1897; DI49_1896 | DI49_1897; DI49_1896 | | |
| DI49_1899; DI49_1898 | DI49_1899; DI49_1898 | | |
| DI49_1901; DI49_1900 | DI49_1901; DI49_1900 | | |
| DI49_1903; DI49_1902 | DI49_1903; DI49_1902 | | |

Create PDF in your applications with the Pdfcrowd HTML to PDF API PDFCROWD

| | | | |
|---|---|---|---|
| DI49_1905; DI49_1904 | DI49_1905; DI49_1904 | | |
| DI49_1907; DI49_1244; DI49_1908; DI49_1243 | DI49_1907; DI49_1244; DI49_1908; DI49_1243 | | |
| DI49_2815; DI49_2816 | DI49_2815; DI49_2816 | | |
| DI49_2813; YIR034C; YIR035C | DI49_2813 | | YIR034C; YIR035C |
| DI49_2812; YIR033W; DI49_2811 | DI49_2812; DI49_2811 | | YIR033W |
| DI49_2809; DI49_2810 | DI49_2809; DI49_2810 | | |
| DI49_2807; YIR028W; DI49_2806 | DI49_2807; DI49_2806 | | YIR028W |
| DI49_2804; DI49_2805 | DI49_2804; DI49_2805 | | |
| DI49_2802; DI49_2803 | DI49_2802; DI49_2803 | | |
| DI49_2800; DI49_2801 | DI49_2800; DI49_2801 | | |
| DI49_2798; YIR018W | DI49_2798 | | YIR018W |
| DI49_2797; YIR017C; DI49_2796; YIR016W | DI49_2797; DI49_2796 | | YIR017C; YIR016W |
| DI49_2794; DI49_2795 | DI49_2794; DI49_2795 | | |
| DI49_2792; YIR012W | DI49_2792 | | YIR012W |
| DI49_2791; YIR011C | DI49_2791 | | YIR011C |
| DI49_2790; YIR010W; DI49_2789; YIR009W | DI49_2790; DI49_2789 | | YIR010W; YIR009W |
| DI49_2787; YIR007W | DI49_2787 | | YIR007W |
| DI49_2785; YIR006C; DI49_2784; YIR005W | DI49_2785; DI49_2784 | | YIR006C; YIR005W |
| DI49_2782; DI49_2783 | DI49_2782; DI49_2783 | | |
| DI49_2780; YIR001C; DI49_2779; YIL001W | DI49_2780; DI49_2779 | | YIR001C; YIL001W |
| DI49_2777; YIL002W-A | DI49_2777 | | YIL002W-A |
| DI49_2776; YIL003W; DI49_2775; YIL004C | DI49_2776; DI49_2775 | | YIL003W; YIL004C |
| DI49_2773; DI49_2774 | DI49_2773; DI49_2774 | | |
| DI49_2771; DI49_2772 | DI49_2771; DI49_2772 | | |
| DI49_2766; DI49_2767 | DI49_2766; DI49_2767 | | |
| SPGP0P00610; DI49_2765 | DI49_2765 | SPGP0P00610 | |
| DI49_2762; YIL015W; DI49_2761; YIL016W | DI49_2762; DI49_2761 | | YIL015W; YIL016W |
| DI49_2759; YIL018W; DI49_1690; DI49_1689 | DI49_2759; DI49_1690; DI49_1689 | | YIL018W |
| DI49_2758; YIL019W; DI49_2757; YIL020C | DI49_2758; DI49_2757 | | YIL019W; YIL020C |
| DI49_2755; YIL022W; DI49_2754; YIL023C | DI49_2755; DI49_2754 | | YIL022W; YIL023C |
| DI49_2752; DI49_2753 | DI49_2752; DI49_2753 | | |
| DI49_2750; DI49_2751 | DI49_2750; DI49_2751 | | |
| DI49_2748; YIL030C; DI49_2747; YIL031W | DI49_2748; DI49_2747 | | YIL030C; YIL031W |
| DI49_2745; YIL034C | DI49_2745 | | YIL034C |
| DI49_2744; YIL035C; DI49_2743; YIL036W | DI49_2744; DI49_2743 | | YIL035C; YIL036W |
| DI49_2741; YIL038C | DI49_2741 | | YIL038C |
| DI49_2740; YIL039W; DI49_2739; YIL040W | DI49_2740; DI49_2739 | | YIL039W; YIL040W |
| DI49_2737; DI49_2738 | DI49_2737; DI49_2738 | | |
| DI49_2735; DI49_2736 | DI49_2735; DI49_2736 | | |
| DI49_2733; YIL046W | DI49_2733 | | YIL046W |

Create PDF in your applications with the Pdfcrowd HTML to PDF API PDFCROWD

| | | | |
|---|---|---|---|
| DI49_2732; YIL047C | DI49_2732 | | YIL047C |
| DI49_2731; YIL048W | DI49_2731 | | YIL048W |
| DI49_2730; YIL049W; DI49_2729; YIL050W | DI49_2730; DI49_2729 | | YIL049W; YIL050W |
| SPGP0P00960; YIL052C | | SPGP0P00960 | YIL052C |
| DI49_2727; YIL053W; SPGP0P00980 | DI49_2727 | SPGP0P00980 | YIL053W |
| DI49_2725; YIL056W | DI49_2725 | | YIL056W |
| DI49_2721; YIL062C | DI49_2721 | | YIL062C |
| DI49_2720; YIL063C; SPGP0P01050; YIL064W | DI49_2720 | SPGP0P01050 | YIL063C; YIL064W |
| DI49_2718; YIL066C | DI49_2718 | | YIL066C |
| DI49_2717; YIL067C | DI49_2717 | | YIL067C |
| DI49_2716; YIL068C | DI49_2716 | | YIL068C |
| DI49_2715; DI49_1478; DI49_1479; YIL069C | DI49_2715; DI49_1478; DI49_1479 | | YIL069C |
| DI49_2714; YIL070C; DI49_2713; YIL071C | DI49_2714; DI49_2713 | | YIL070C; YIL071C |
| DI49_2711; DI49_2712 | DI49_2711; DI49_2712 | | |
| DI49_2709; YIL075C; DI49_2708; YIL076W | DI49_2709; DI49_2708 | | YIL075C; YIL076W |
| DI49_2706; YIL077C | DI49_2706 | | YIL077C |
| DI49_2705; YIL078W; DI49_2704; YIL079C | DI49_2705; DI49_2704 | | YIL078W; YIL079C |
| SPGP0P01220; YIL083C; DI49_2702; YIL084C | DI49_2702 | SPGP0P01220 | YIL083C; YIL084C |
| DI49_2700; DI49_2701 | DI49_2700; DI49_2701 | | |
| SPGP0Ptrna5E; DI49_2698; DI49_2697 | DI49_2698; DI49_2697 | SPGP0Ptrna5E | |
| DI49_2695; YIL091C; DI49_2694; YIL092W | DI49_2695; DI49_2694 | | YIL091C; YIL092W |
| DI49_2692; YIL094C | DI49_2692 | | YIL094C |
| DI49_2691; YIL095W | DI49_2691 | | YIL095W |
| SPGP0Ptrna4I; SPGP0Ftrna14I; DI49_2689; DI49_3947; SPGP0Atrna1I; SPGP0LXtrna7I; DI49_5588; DI49_1849; SPGP0URtrna9I; DI49_2084; DI49_3294; SPGP0BFtrna12I; YIL096C; SPGP0VRtrna13I; DI49_3807; SPGP0WD00100; SPGP0WUtrna19I; YER182W; SPGP0BItrna17I; DI49_0301; SPGP0BItrna18I | DI49_2689; DI49_3947; DI49_5588; DI49_1849; DI49_2084; DI49_3294; DI49_3807; DI49_0301 | SPGP0Ptrna4I; SPGP0Ftrna14I; SPGP0Atrna1I; SPGP0LXtrna7I; SPGP0URtrna9I; SPGP0BFtrna12I; SPGP0VRtrna13I; SPGP0WD00100; SPGP0WUtrna19I; SPGP0BItrna17I; SPGP0BItrna18I | YIL096C; YER182W |
| DI49_2687; DI49_2688 | DI49_2687; DI49_2688 | | |
| DI49_2685; YIL101C | DI49_2685 | | YIL101C |
| DI49_2681; DI49_2682; YIL103W; YIL104C | DI49_2681; DI49_2682 | | YIL103W; YIL104C |
| DI49_2679; DI49_2680 | DI49_2679; DI49_2680 | | |
| DI49_2677; YIL108W | DI49_2677 | | YIL108W |
| DI49_2676; YIL109C | DI49_2676 | | YIL109C |
| DI49_2675; YIL110W; SPGP0P01480; YIL111W | DI49_2675 | SPGP0P01480 | YIL110W; YIL111W |
| DI49_2673; DI49_2674 | DI49_2673; DI49_2674 | | |
| DI49_2671; YIL115C; DI49_2670; YIL116W | DI49_2671; DI49_2670 | | YIL115C; YIL116W |
| DI49_2668; DI49_2669 | DI49_2668; DI49_2669 | | |
| DI49_2666; YIL120W; DI49_2665; YIL121W | DI49_2666; DI49_2665 | | YIL120W; YIL121W |
| DI49_2663; YIL123W | DI49_2663 | | YIL123W |
| DI49_2662; YIL124W | DI49_2662 | | YIL124W |

Create PDF in your applications with the Pdfcrowd HTML to PDF API PDFCROWD

| | | | |
|---|---|---|---|
| DI49_2661; YIL125W | DI49_2661 | | YIL125W |
| DI49_2660; YIL126W; DI49_2659; YIL127C | DI49_2660; DI49_2659 | | YIL126W; YIL127C |
| DI49_2657; YIL129C; DI49_2656; YIL130W | DI49_2657; DI49_2656 | | YIL129C; YIL130W |
| DI49_2654; DI49_2655 | DI49_2654; DI49_2655 | | |
| DI49_2652; SPGP0P01700 | DI49_2652 | SPGP0P01700 | |
| DI49_2650; DI49_2651 | DI49_2650; DI49_2651 | | |
| DI49_2648; DI49_2649 | DI49_2648; DI49_2649 | | |
| DI49_2646; DI49_2647 | DI49_2646; DI49_2647 | | |
| DI49_2644; YIL143C; DI49_2643; YIL144W | DI49_2644; DI49_2643 | | YIL143C; YIL144W |
| DI49_2641; YIL146C | DI49_2641 | | YIL146C |
| DI49_2640; YIL147C | DI49_2640 | | YIL147C |
| DI49_2639; YIL148W; YLL039C; DI49_3494; YKR094C; DI49_3464; DI49_3686; DI49_3687 | DI49_2639; DI49_3494; DI49_3464; DI49_3686; DI49_3687 | | YIL148W; YLL039C; YKR094C |
| DI49_2638; YIL149C | DI49_2638 | | YIL149C |
| DI49_2637; YIL150C | DI49_2637 | | YIL150C |
| DI49_2636; YIL151C | DI49_2636 | | YIL151C |
| DI49_2635; YIL152W; DI49_2634; YIL153W | DI49_2635; DI49_2634 | | YIL152W; YIL153W |
| DI49_2632; DI49_2633 | DI49_2632; DI49_2633 | | |
| DI49_2630; DI49_2631 | DI49_2630; DI49_2631 | | |
| DI49_2628; DI49_2629 | DI49_2628; DI49_2629 | | |
| DI49_2626; YIL160C | DI49_2626 | | YIL160C |
| DI49_2625; YIL161W | DI49_2625 | | YIL161W |
| DI49_2624; YIL162W; DI49_2622; YIL164C | DI49_2624; DI49_2622 | | YIL162W; YIL164C |
| DI49_0767; YCR017C; DI49_0766; YCR018C | DI49_0767; DI49_0766 | | YCR017C; YCR018C |
| DI49_0762; YCR012W; YCR014C | DI49_0762 | | YCR012W; YCR014C |
| DI49_0761; YCR011C; DI49_0760 | DI49_0761; DI49_0760 | | YCR011C |
| DI49_3953; SPGP0NZ00110; DI49_5378; DI49_4789 | DI49_3953; DI49_5378; DI49_4789 | SPGP0NZ00110 | |
| SPGP0FQ00100; DI49_0075; DI49_1458 | DI49_0075; DI49_1458 | SPGP0FQ00100 | |
| DI49_1465; DI49_1457 | DI49_1465; DI49_1457 | | |
| DI49_1467; DI49_1466 | DI49_1467; DI49_1466 | | |
| DI49_1469; DI49_1468 | DI49_1469; DI49_1468 | | |
| DI49_4437; DI49_1385 | DI49_4437; DI49_1385 | | |
| DI49_4435; DI49_4436 | DI49_4435; DI49_4436 | | |
| DI49_4433; DI49_4434 | DI49_4433; DI49_4434 | | |
| DI49_4431; DI49_4432 | DI49_4431; DI49_4432 | | |
| DI49_4429; DI49_4430 | DI49_4429; DI49_4430 | | |
| DI49_4427; DI49_4428 | DI49_4427; DI49_4428 | | |
| DI49_4425; DI49_4426 | DI49_4425; DI49_4426 | | |
| DI49_4423; DI49_4424 | DI49_4423; DI49_4424 | | |
| DI49_4421; DI49_4422 | DI49_4421; DI49_4422 | | |

Create PDF in your applications with the Pdfcrowd HTML to PDF API PDFCROWD

| | | | |
|---|---|---|---|
| DI49_1565; SPGP0FW00270 | DI49_1565 | SPGP0FW00270 | |
| SPGP0FXtrna1W; DI49_5211; DI49_1569; DI49_3302 | DI49_5211; DI49_1569; DI49_3302 | SPGP0FXtrna1W | |
| DI49_1679; DI49_1570 | DI49_1679; DI49_1570 | | |
| DI49_1963; DI49_1678 | DI49_1963; DI49_1678 | | |
| DI49_1389; DI49_1388 | DI49_1389; DI49_1388 | | |
| DI49_3945; DI49_3946 | DI49_3945; DI49_3946 | | |
| SPGP0GE00110; DI49_1327 | DI49_1327 | SPGP0GE00110 | |
| DI49_4504; DI49_4505 | DI49_4504; DI49_4505 | | |
| DI49_4502; DI49_4503 | DI49_4502; DI49_4503 | | |
| DI49_4500; DI49_4501 | DI49_4500; DI49_4501 | | |
| DI49_4498; DI49_4499 | DI49_4498; DI49_4499 | | |
| DI49_4496; DI49_4497 | DI49_4496; DI49_4497 | | |
| DI49_4494; DI49_4495 | DI49_4494; DI49_4495 | | |
| DI49_4492; DI49_4493 | DI49_4492; DI49_4493 | | |
| DI49_4490; DI49_4491 | DI49_4490; DI49_4491 | | |
| DI49_4488; DI49_4489 | DI49_4488; DI49_4489 | | |
| DI49_4486; DI49_4487 | DI49_4486; DI49_4487 | | |
| DI49_4484; DI49_4485 | DI49_4484; DI49_4485 | | |
| DI49_4482; DI49_4483 | DI49_4482; DI49_4483 | | |
| DI49_4480; DI49_4479; DI49_3759; DI49_1633 | DI49_4480; DI49_4479; DI49_3759; DI49_1633 | | |
| DI49_5395; DI49_5396 | DI49_5395; DI49_5396 | | |
| DI49_5393; DI49_5394 | DI49_5393; DI49_5394 | | |
| DI49_5391; DI49_5392 | DI49_5391; DI49_5392 | | |
| DI49_5389; DI49_5390 | DI49_5389; DI49_5390 | | |
| DI49_5387; DI49_5388 | DI49_5387; DI49_5388 | | |
| DI49_3804; DI49_5386 | DI49_3804; DI49_5386 | | |
| DI49_3802; DI49_3803 | DI49_3802; DI49_3803 | | |
| DI49_3797; DI49_3798 | DI49_3797; DI49_3798 | | |
| DI49_3795; DI49_3796 | DI49_3795; DI49_3796 | | |
| DI49_4568; YHR204W | DI49_4568 | | YHR204W |
| DI49_4566; DI49_4567 | DI49_4566; DI49_4567 | | |
| DI49_4564; DI49_4565 | DI49_4564; DI49_4565 | | |
| DI49_4562; DI49_4563 | DI49_4562; DI49_4563 | | |
| DI49_1338; DI49_1337 | DI49_1338; DI49_1337 | | |
| DI49_1340; DI49_1339 | DI49_1340; DI49_1339 | | |
| DI49_1712; DI49_1713 | DI49_1712; DI49_1713 | | |
| DI49_1710; DI49_1711 | DI49_1710; DI49_1711 | | |
| SPGP0GQ00100; YJL225C; DI49_4506; YCL064C | DI49_4506 | SPGP0GQ00100 | YJL225C; YCL064C |
| DI49_4508; DI49_4507 | DI49_4508; DI49_4507 | | |
| DI49_4510; DI49_4509 | DI49_4510; DI49_4509 | | |

Create PDF in your applications with the Pdfcrowd HTML to PDF API PDFCROWD

| | | | |
|---|---|---|---|
| YNL336W; SPGP0WO00100; YNL335W; DI49_1969 | DI49_1969 | SPGP0WO00100 | YNL336W; YNL335W |
| YNL333W; YNL334C | | | YNL333W; YNL334C |
| YDL240W; YDL241W | | | YDL240W; YDL241W |
| YDL238C; YDL239C | | | YDL238C; YDL239C |
| YDL236W; YDL237W | | | YDL236W; YDL237W |
| YDL234C; YDL235C | | | YDL234C; YDL235C |
| YDL232W; YDL233W | | | YDL232W; YDL233W |
| YDL230W; YDL231C | | | YDL230W; YDL231C |
| YDL227C; DI49_0855 | DI49_0855 | | YDL227C |
| YDL226C; DI49_0856 | DI49_0856 | | YDL226C |
| YDL225W; DI49_0857; YDL224C | DI49_0857 | | YDL225W; YDL224C |
| YDL222C; YDL223C | | | YDL222C; YDL223C |
| YDL219W; YDL220C | | | YDL219W; YDL220C |
| YDL217C; YDL218W | | | YDL217C; YDL218W |
| YDL215C; DI49_0866; YDL214C; DI49_0865 | DI49_0866; DI49_0865 | | YDL215C; YDL214C |
| YDL212W; DI49_0869; YDL211C; DI49_0868 | DI49_0869; DI49_0868 | | YDL212W; YDL211C |
| YDL210W; DI49_0872; YDL209C | DI49_0872 | | YDL210W; YDL209C |
| YDL207W; DI49_0875; YDL206W | DI49_0875 | | YDL207W; YDL206W |
| YDL204W; YDL205C | | | YDL204W; YDL205C |
| YDL202W; YDL203C | | | YDL202W; YDL203C |
| YDL200C; YDL201W | | | YDL200C; YDL201W |
| YDL198C; DI49_0884; YDL197C | DI49_0884 | | YDL198C; YDL197C |
| YDL194W; DI49_0887; YDL193W | DI49_0887 | | YDL194W; YDL193W |
| YDL191W; YDL136W | | | YDL191W; YDL136W |
| YDL190C; DI49_0891 | DI49_0891 | | YDL190C |
| YDL189W; DI49_0892 | DI49_0892 | | YDL189W |
| YDL188C; YDL134C; YDL186W | | | YDL188C; YDL134C; YDL186W |
| YDL183C; DI49_0896 | DI49_0896 | | YDL183C |
| YDL182W; YDL131W; YDL181W | | | YDL182W; YDL131W; YDL181W |
| YDL179W; DI49_0900; DI49_0899 | DI49_0900; DI49_0899 | | YDL179W |
| YDL178W; DI49_0901; YDL177C | DI49_0901 | | YDL178W; YDL177C |
| YDL175C; YDL176W | | | YDL175C; YDL176W |
| YDL173W; YDL174C | | | YDL173W; YDL174C |
| YDL170W; YDL171C; DI49_0907; DI49_0906 | DI49_0907; DI49_0906 | | YDL170W; YDL171C |
| YDL168W; DI49_0910; DI49_0909 | DI49_0910; DI49_0909 | | YDL168W |
| YDL167C; DI49_0911; YDL166C | DI49_0911 | | YDL167C; YDL166C |
| YDL164C; DI49_0914; YDL161W | DI49_0914 | | YDL164C; YDL161W |
| YDL160C; DI49_0917; DI49_0916 | DI49_0917; DI49_0916 | | YDL160C |
| YDL159W; DI49_0918 | DI49_0918 | | YDL159W |
| YDL157C; DI49_0919 | DI49_0919 | | YDL157C |

Create PDF in your applications with the Pdfcrowd HTML to PDF API PDFCROWD

| | | | |
|---|---|---|---|
| YDL156W; DI49_0920 | DI49_0920 | | YDL156W |
| YDL155W; DI49_0921; YDL154W | DI49_0921 | | YDL155W; YDL154W |
| YDL150W; DI49_0924; YDL149W; DI49_0923 | DI49_0924; DI49_0923 | | YDL150W; YDL149W |
| YDL147W; DI49_0927; DI49_0926 | DI49_0927; DI49_0926 | | YDL147W |
| YDL146W; DI49_0928 | DI49_0928 | | YDL146W |
| YDL145C; DI49_0929; YDL144C | DI49_0929 | | YDL145C; YDL144C |
| YDL142C; YDL143W | | | YDL142C; YDL143W |
| YDL140C; DI49_0934; YDL139C | DI49_0934 | | YDL140C; YDL139C |
| YDL135C; DI49_0937 | DI49_0937 | | YDL135C |
| YDL133W; DI49_0939 | DI49_0939 | | YDL133W |
| YDL132W; DI49_0940 | DI49_0940 | | YDL132W |
| YDL130W; SPGP0BE03930; YDL129W; DI49_0942 | DI49_0942 | SPGP0BE03930 | YDL130W; YDL129W |
| YDL127W; YDL128W | | | YDL127W; YDL128W |
| YDL125C; YDL126C; DI49_0946; DI49_0945 | DI49_0946; DI49_0945 | | YDL125C; YDL126C |
| YDL123W; YDL124W | | | YDL123W; YDL124W |
| YDL121C; YDL122W; DI49_0949; DI49_0948 | DI49_0949; DI49_0948 | | YDL121C; YDL122W |
| YDL119C; DI49_0952; YDL117W; DI49_0951 | DI49_0952; DI49_0951 | | YDL119C; YDL117W |
| YDL115C; DI49_0955; DI49_0954 | DI49_0955; DI49_0954 | | YDL115C |
| YDL114W; DI49_0956; YDL113C | DI49_0956 | | YDL114W; YDL113C |
| YDL111C; DI49_0959; DI49_0958 | DI49_0959; DI49_0958 | | YDL111C |
| YDL110C; DI49_0960; YDL109C | DI49_0960 | | YDL110C; YDL109C |
| YDL107W; YDL108W | | | YDL107W; YDL108W |
| YDL105W; DI49_0965; YDL104C | DI49_0965 | | YDL105W; YDL104C |
| YDL102W; YDL103C; DI49_0967; DI49_0966 | DI49_0967; DI49_0966 | | YDL102W; YDL103C |
| YDL100C; DI49_0970; YDL099W; DI49_0969 | DI49_0970; DI49_0969 | | YDL100C; YDL099W |
| YDL097C; DI49_0973; DI49_0972 | DI49_0973; DI49_0972 | | YDL097C |
| YDL095W; DI49_0974; YDL093W | DI49_0974 | | YDL095W; YDL093W |
| YDL091C; DI49_0977; YDL090C; DI49_0976 | DI49_0977; DI49_0976 | | YDL091C; YDL090C |
| YDL088C; YDL089W | | | YDL088C; YDL089W |
| YDL086W; YDL087C; DI49_0980; DI49_0979 | DI49_0980; DI49_0979 | | YDL086W; YDL087C |
| YDL085W; DI49_0983; DI49_0982 | DI49_0983; DI49_0982 | | YDL085W |
| YDL084W; DI49_0984 | DI49_0984 | | YDL084W |
| YDL083C; SPGP0BE03490 | | SPGP0BE03490 | YDL083C |
| YDL082W; DI49_0985; SPGP0C01740; DI49_4228 | DI49_0985; DI49_4228 | SPGP0C01740 | YDL082W |
| YDL081C; DI49_0986 | DI49_0986 | | YDL081C |
| YDL080C; DI49_0987; YDL079C | DI49_0987 | | YDL080C; YDL079C |
| YDL077C; DI49_0989 | DI49_0989 | | YDL077C |
| YDL076C; DI49_0990 | DI49_0990 | | YDL076C |
| YDL074C; DI49_0992 | DI49_0992 | | YDL074C |
| YDL073W; DI49_0993; YDL072C | DI49_0993 | | YDL073W; YDL072C |

Create PDF in your applications with the Pdfcrowd HTML to PDF API PDFCROWD

| | | | |
|---|---|---|---|
| YDL069C; YDL070W | | | YDL069C; YDL070W |
| YDL066W; DI49_0998; YDL065C | DI49_0998 | | YDL066W; YDL065C |
| YDL063C; DI49_1000 | DI49_1000 | | YDL063C |
| YDL061C; DI49_1001; YDL060W; DI49_3888; DI49_3887 | DI49_1001; DI49_3888; DI49_3887 | | YDL061C; YDL060W |
| YDL058W; DI49_1004; YDL057W; DI49_1003 | DI49_1004; DI49_1003 | | YDL058W; YDL057W |
| YDL055C; DI49_1007 | DI49_1007 | | YDL055C |
| YDL052C; YDL053C | | | YDL052C; YDL053C |
| YDL049C; DI49_1013 | DI49_1013 | | YDL049C |
| YDL048C; DI49_1014 | DI49_1014 | | YDL048C |
| YDL047W; DI49_1015; YDL046W; SPGP0BE03160 | DI49_1015 | SPGP0BE03160 | YDL047W; YDL046W |
| YDL045C; DI49_1018; DI49_1017 | DI49_1018; DI49_1017 | | YDL045C |
| YDL044C; DI49_1019; YDL043C | DI49_1019 | | YDL044C; YDL043C |
| YDL040C; DI49_1022; YDL039C | DI49_1022 | | YDL040C; YDL039C |
| YDL035C; YDL036C | | | YDL035C; YDL036C |
| YDL031W; DI49_1025; YDL030W; SPGP0BE03040 | DI49_1025 | SPGP0BE03040 | YDL031W; YDL030W |
| YDL028C; DI49_1027; YDL027C; SPGP0BE03010 | DI49_1027 | SPGP0BE03010 | YDL028C; YDL027C |
| YDL024C; YDL025C | | | YDL024C; YDL025C |
| YDL022W; DI49_1032; YDL021W | DI49_1032 | | YDL022W; YDL021W |
| YDL019C; DI49_1035; YDL018C; DI49_1034 | DI49_1035; DI49_1034 | | YDL019C; YDL018C |
| YDL015C; YDL017W | | | YDL015C; YDL017W |
| YDL013W; YDL014W; DI49_1040; DI49_1039 | DI49_1040; DI49_1039 | | YDL013W; YDL014W |
| YDL010W; DI49_1042; YDL008W; SPGP0BE02860 | DI49_1042 | SPGP0BE02860 | YDL010W; YDL008W |
| YDL007W; DI49_1047 | DI49_1047 | | YDL007W |
| YDL006W; DI49_1048 | DI49_1048 | | YDL006W |
| YDL005C; DI49_1049; YDL004W | DI49_1049 | | YDL005C; YDL004W |
| YDL002C; DI49_1052; YDL001W; DI49_1051 | DI49_1052; DI49_1051 | | YDL002C; YDL001W |
| YDR002W; DI49_1055; YDR003W | DI49_1055 | | YDR002W; YDR003W |
| YDR005C; YDR004W | | | YDR005C; YDR004W |
| YDR007W; DI49_1060; YDR009W | DI49_1060 | | YDR007W; YDR009W |
| YDR012W; YBR031W; YDR013W | | | YDR012W; YBR031W; YDR013W |
| YDR014W-A; YDR014W | | | YDR014W-A; YDR014W |
| YDR017C; DI49_0215; DI49_0214 | DI49_0215; DI49_0214 | | YDR017C |
| YDR018C; DI49_0216; YDR019C | DI49_0216 | | YDR018C; YDR019C |
| YDR021W; DI49_0219; DI49_0218 | DI49_0219; DI49_0218 | | YDR021W |
| YDR022C; DI49_0220 | DI49_0220 | | YDR022C |
| SPGP0Atrna1V; DI49_1074; DI49_1073; SPGP0BItrna21V; YBR044C | DI49_1074; DI49_1073 | SPGP0Atrna1V; SPGP0BItrna21V | YBR044C |
| YDR023W; DI49_0222; YDR024W | DI49_0222 | | YDR023W; YDR024W |
| YDR026C; DI49_0224 | DI49_0224 | | YDR026C |
| YDR027C; DI49_0225 | DI49_0225 | | YDR027C |
| YDR028C; DI49_0226; YDR030C | DI49_0226 | | YDR028C; YDR030C |

Create PDF in your applications with the Pdfcrowd HTML to PDF API PDFCROWD

| | | | |
|---|---|---|---|
| YDR032C; DI49_0229 | DI49_0229 | | YDR032C |
| YDR033W; DI49_0230; YDR034C | DI49_0230 | | YDR033W; YDR034C |
| SPGP0Atrna1Q; DI49_2421; SPGP0Atrna3Q; DI49_0779; DI49_3679; SPGP0Btrna4Q; SPGP0OFtrna9Q; DI49_1450; SPGP0BGtrna12Q; SPGP0BItrna15Q; DI49_0293; SPGP0BI02750 | DI49_2421; DI49_0779; DI49_3679; DI49_1450; DI49_0293 | SPGP0Atrna1Q; SPGP0Atrna3Q; SPGP0Btrna4Q; SPGP0OFtrna9Q; SPGP0BGtrna12Q; SPGP0BItrna15Q; SPGP0BI02750 | |
| YDR035W; DI49_0233 | DI49_0233 | | YDR035W |
| YDR036C; DI49_0234 | DI49_0234 | | YDR036C |
| SPGP0A02650; DI49_1093; YDR040C; SPGP0BI02720; DI49_1092 | DI49_1093; DI49_1092 | SPGP0A02650; SPGP0BI02720 | YDR040C |
| YDR042C; YDR041W; DI49_0237; DI49_0236 | DI49_0237; DI49_0236 | | YDR042C; YDR041W |
| YDR044W; DI49_0240; YDR045C; DI49_0239 | DI49_0240; DI49_0239 | | YDR044W; YDR045C |
| YDR047W; DI49_0243; YDR049W | DI49_0243 | | YDR047W; YDR049W |
| YDR051C; DI49_0246 | DI49_0246 | | YDR051C |
| YDR052C; DI49_0247 | DI49_0247 | | YDR052C |
| YDR054C; DI49_0248 | DI49_0248 | | YDR054C |
| YDR055W; DI49_0249; YDR056C | DI49_0249 | | YDR055W; YDR056C |
| YDR058C; DI49_0252 | DI49_0252 | | YDR058C |
| SPGP0Atrna1R; SPGP0Itrna29R; DI49_4019; SPGP0Ftrna17R; SPGP0Gtrna24R; SPGP0Htrna26R; YMR239C; SPGP0Itrna28R | DI49_4019 | SPGP0Atrna1R; SPGP0Itrna29R; SPGP0Ftrna17R; SPGP0Gtrna24R; SPGP0Htrna26R; SPGP0Itrna28R | YMR239C |
| YDR061W; DI49_0257 | DI49_0257 | | YDR061W |
| YDR062W; DI49_0258 | DI49_0258 | | YDR062W |
| YDR063W; DI49_0259 | DI49_0259 | | YDR063W |
| YDR064W; SPGP0BI02490; YDR065W | | SPGP0BI02490 | YDR064W; YDR065W |
| YDR067C; DI49_0263; YDR068W; DI49_0262 | DI49_0263; DI49_0262 | | YDR067C; YDR068W |
| YDR070C; YDR069C; DI49_0265; DI49_0264 | DI49_0265; DI49_0264 | | YDR070C; YDR069C |
| YDR072C; DI49_0268; YDR073W; DI49_0267 | DI49_0268; DI49_0267 | | YDR072C; YDR073W |
| YDR075W; DI49_0271 | DI49_0271 | | YDR075W |
| YDR076W; DI49_0272 | DI49_0272 | | YDR076W |
| YDR077W; DI49_0273; YDR078C | DI49_0273 | | YDR077W; YDR078C |
| YDR079C-A; YDR079W | | | YDR079C-A; YDR079W |
| YDR081C; YDR080W; DI49_0277; DI49_0276 | DI49_0277; DI49_0276 | | YDR081C; YDR080W |
| YDR083W; DI49_0280; YDR084C; DI49_0279 | DI49_0280; DI49_0279 | | YDR083W; YDR084C |
| YDR086C; YDR085C; DI49_0282; DI49_0281 | DI49_0282; DI49_0281 | | YDR086C; YDR085C |
| YDR088C; DI49_0285; DI49_0284 | DI49_0285; DI49_0284 | | YDR088C |
| YDR089W; DI49_0287; YDR090C | DI49_0287 | | YDR089W; YDR090C |
| YDR092W; YDR091C | | | YDR092W; YDR091C |
| YDR096W; DI49_0291 | DI49_0291 | | YDR096W |
| YDR097C; DI49_0292; YDR098C | DI49_0292 | | YDR097C; YDR098C |
| YDR099W; DI49_0295; YDR100W | DI49_0295 | | YDR099W; YDR100W |
| YDR103W; YDR101C; DI49_0297; DI49_0296 | DI49_0297; DI49_0296 | | YDR103W; YDR101C |

Create PDF in your applications with the Pdfcrowd HTML to PDF API PDFCROWD

| | | | |
|---|---|---|---|
| YDR105C; YDR104C | | | YDR105C; YDR104C |
| YDR107C; DI49_0303 | DI49_0303 | | YDR107C |
| YDR108W; DI49_0304; YDR109C | DI49_0304 | | YDR108W; YDR109C |
| YDR111C; YDR110W | | | YDR111C; YDR110W |
| YDR115W; YDR113C | | | YDR115W; YDR113C |
| YDR117C; YDR116C | | | YDR117C; YDR116C |
| YDR119W; DI49_0312; YDR119W-A; DI49_0311 | DI49_0312; DI49_0311 | | YDR119W; YDR119W-A |
| YDR121W; YDR120C | | | YDR121W; YDR120C |
| YDR123C; DI49_0317; YDR124W | DI49_0317 | | YDR123C; YDR124W |
| YDR126W; DI49_0320 | DI49_0320 | | YDR126W |
| YDR127W; DI49_0321; YDR128W | DI49_0321 | | YDR127W; YDR128W |
| YDR130C; SPGP0A03500 | | SPGP0A03500 | YDR130C |
| YDR132C; YDR131C | | | YDR132C; YDR131C |
| YDR135C; DI49_0328; YDR137W | DI49_0328 | | YDR135C; YDR137W |
| YDR139C; YDR138W | | | YDR139C; YDR138W |
| YDR141C; DI49_0333; DI49_0332 | DI49_0333; DI49_0332 | | YDR141C |
| YDR142C; DI49_0334 | DI49_0334 | | YDR142C |
| YDR143C; DI49_0335; YDR144C | DI49_0335 | | YDR143C; YDR144C |
| YDR146C; YDR145W; DI49_0337; DI49_0336 | DI49_0337; DI49_0336 | | YDR146C; YDR145W |
| YDR148C; DI49_0340; YDR150W; DI49_0339 | DI49_0340; DI49_0339 | | YDR148C; YDR150W |
| YDR152W; YDR151C | | | YDR152W; YDR151C |
| YDR155C; DI49_0344; YDR156W | DI49_0344 | | YDR155C; YDR156W |
| YDR159W; YDR158W | | | YDR159W; YDR158W |
| YDR161W; YDR160W | | | YDR161W; YDR160W |
| YDR163W; YDR162C; DI49_0350; DI49_0349 | DI49_0350; DI49_0349 | | YDR163W; YDR162C |
| YDR165W; DI49_0353; DI49_0352 | DI49_0353; DI49_0352 | | YDR165W |
| YDR166C; DI49_0354; YDR167W | DI49_0354 | | YDR166C; YDR167W |
| YDR169C; DI49_0357 | DI49_0357 | | YDR169C |
| YDR170C; DI49_0359 | DI49_0359 | | YDR170C |
| YDR173C; DI49_0363 | DI49_0363 | | YDR173C |
| YDR174W; DI49_0364; YDR175C | DI49_0364 | | YDR174W; YDR175C |
| YDR177W; YDR176W; DI49_0366; DI49_0365 | DI49_0366; DI49_0365 | | YDR177W; YDR176W |
| YDR179C; YDR178W | | | YDR179C; YDR178W |
| YDR180W; DI49_0371; YDR181C; DI49_0370 | DI49_0371; DI49_0370 | | YDR180W; YDR181C |
| YDR183W; YDR182W | | | YDR183W; YDR182W |
| YDR185C; DI49_0376; DI49_0375 | DI49_0376; DI49_0375 | | YDR185C |
| YDR186C; DI49_0377 | DI49_0377 | | YDR186C |
| YDR188W; DI49_0378 | DI49_0378 | | YDR188W |
| YDR189W; DI49_0379 | DI49_0379 | | YDR189W |
| YDR190C; DI49_0380; YDR191W | DI49_0380 | | YDR190C; YDR191W |

Create PDF in your applications with the Pdfcrowd HTML to PDF API PDFCROWD

| | | | |
|---|---|---|---|
| YDR194C; DI49_0382; SPGP0BI01300 | DI49_0382 | SPGP0BI01300 | YDR194C |
| YDR195W; DI49_0383; YDR196C | DI49_0383 | | YDR195W; YDR196C |
| YDR198C; DI49_0386; YDR200C; DI49_0385 | DI49_0386; DI49_0385 | | YDR198C; YDR200C |
| YDR202C; DI49_0389 | DI49_0389 | | YDR202C |
| YDR204W; DI49_0390; YDR205W | DI49_0390 | | YDR204W; YDR205W |
| YDR207C; YDR206W | | | YDR207C; YDR206W |
| SPGP0Atrna2I; DI49_0395; YDR211W | DI49_0395 | SPGP0Atrna2I | YDR211W |
| YDR213W; DI49_0398; YDR214W; DI49_3741; DI49_3742 | DI49_0398; DI49_3741; DI49_3742 | | YDR213W; YDR214W |
| YDR217C; YDR216W | | | YDR217C; YDR216W |
| YDR219C; YDR218C | | | YDR219C; YDR218C |
| YDR222W; YDR221W; DI49_0403; DI49_0402 | DI49_0403; DI49_0402 | | YDR222W; YDR221W |
| YDR224C; DI49_0406; DI49_0180; DI49_0405 | DI49_0406; DI49_0180; DI49_0405 | | YDR224C |
| YDR225W; YBL003C | | | YDR225W; YBL003C |
| YDR226W; DI49_0408; YDR227W | DI49_0408 | | YDR226W; YDR227W |
| YDR229W; DI49_0411 | DI49_0411 | | YDR229W |
| YDR231C; DI49_0412 | DI49_0412 | | YDR231C |
| YDR232W; DI49_0413 | DI49_0413 | | YDR232W |
| YDR233C; DI49_0414 | DI49_0414 | | YDR233C |
| YDR234W; DI49_0416; DI49_0415 | DI49_0416; DI49_0415 | | YDR234W |
| YDR235W; DI49_0417 | DI49_0417 | | YDR235W |
| YDR236C; DI49_0418; YDR237W | DI49_0418 | | YDR236C; YDR237W |
| YDR239C; YDR238C | | | YDR239C; YDR238C |
| YDR242W; SPGP0BI00880; YDR243C | | SPGP0BI00880 | YDR242W; YDR243C |
| YDR245W; DI49_0426; YDR246W | DI49_0426 | | YDR245W; YDR246W |
| YDR248C; DI49_0429; YDR249C; DI49_0428 | DI49_0429; DI49_0428 | | YDR248C; YDR249C |
| YDR252W; YDR251W | | | YDR252W; YDR251W |
| YDR254W; YDR253C | | | YDR254W; YDR253C |
| YDR256C; DI49_0436; YDR257C; DI49_0435 | DI49_0436; DI49_0435 | | YDR256C; YDR257C |
| YDR259C; YDR258C | | | YDR259C; YDR258C |
| YDR261C; YDR260C | | | YDR261C; YDR260C |
| YDR264C; YDR263C | | | YDR264C; YDR263C |
| YDR266C; YDR265W | | | YDR266C; YDR265W |
| YDR268W; DI49_0449 | DI49_0449 | | YDR268W |
| YDR270W; DI49_0450; YDR272W | DI49_0450 | | YDR270W; YDR272W |
| YDR275W; YDR273W | | | YDR275W; YDR273W |
| YDR277C; DI49_0455 | DI49_0455 | | YDR277C |
| SPGP0A04820; SPGP0A04830; YGR161W-B | | SPGP0A04820; SPGP0A04830 | YGR161W-B |
| SPGP0Atrna1E; YDR278C; YDR279W; YIL089W | | SPGP0Atrna1E | YDR278C; YDR279W; YIL089W |
| YDR281C; YDR280W; DI49_0458; DI49_0457 | DI49_0458; DI49_0457 | | YDR281C; YDR280W |
| YDR283C; DI49_0461; YDR284C; DI49_0460 | DI49_0461; DI49_0460 | | YDR283C; YDR284C |

Create PDF in your applications with the Pdfcrowd HTML to PDF API PDFCROWD

| | | | |
|---|---|---|---|
| YDR286C; YDR285W | | | YDR286C; YDR285W |
| YDR288W; YDR287W; DI49_0465; DI49_0464 | DI49_0465; DI49_0464 | | YDR288W; YDR287W |
| YDR291W; DI49_0468; DI49_0467 | DI49_0468; DI49_0467 | | YDR291W |
| YDR292C; DI49_0469 | DI49_0469 | | YDR292C |
| YDR293C; DI49_0470 | DI49_0470 | | YDR293C |
| YDR294C; DI49_0471 | DI49_0471 | | YDR294C |
| YDR295C; DI49_0472; YDR296W | DI49_0472 | | YDR295C; YDR296W |
| YDR298C; DI49_0475 | DI49_0475 | | YDR298C |
| YDR299W; DI49_0476 | DI49_0476 | | YDR299W |
| YDR300C; DI49_0477 | DI49_0477 | | YDR300C |
| YDR301W; DI49_0478; YDR302W | DI49_0478 | | YDR301W; YDR302W |
| YDR304C; YDR303C | | | YDR304C; YDR303C |
| YDR306C; YDR305C | | | YDR306C; YDR305C |
| YDR307W; SPGP0Btrna7V; SPGP0Atrna2V; DI49_2588 | DI49_2588 | SPGP0Btrna7V; SPGP0Atrna2V | YDR307W |
| YDR309C; YDR308C | | | YDR309C; YDR308C |
| YDR311W; YDR310C | | | YDR311W; YDR310C |
| YDR313C; YDR312W; YHR066W; YHR067W | | | YDR313C; YDR312W; YHR066W; YHR067W |
| YDR315C; YDR314C | | | YDR315C; YDR314C |
| SPGP0Atrna1F; DI49_0495; YDR317W | DI49_0495 | SPGP0Atrna1F | YDR317W |
| YDR319C; YDR318W | | | YDR319C; YDR318W |
| YDR320C-A; DI49_0500; DI49_0499 | DI49_0500; DI49_0499 | | YDR320C-A |
| YDR321W; DI49_0501 | DI49_0501 | | YDR321W |
| YDR322W; DI49_0502 | DI49_0502 | | YDR322W |
| YDR322C-A; DI49_0503; YDR323C | DI49_0503 | | YDR322C-A; YDR323C |
| SPGP0GR00100; YDR324C; DI49_0505; DI49_0504 | DI49_0505; DI49_0504 | SPGP0GR00100 | YDR324C |
| DI49_1476; DI49_1475 | DI49_1476; DI49_1475 | | |
| DI49_1626; DI49_1625 | DI49_1626; DI49_1625 | | |
| DI49_1628; DI49_1627 | DI49_1628; DI49_1627 | | |
| DI49_1629; SPGP0GT00150 | DI49_1629 | SPGP0GT00150 | |
| YPL264C; YPL263C | | | YPL264C; YPL263C |
| YPL266W; YPL265W | | | YPL266W; YPL265W |
| YPL268W; YPL267W | | | YPL268W; YPL267W |
| YPL270W; YPL269W | | | YPL270W; YPL269W |
| YPL272C; YPL271W | | | YPL272C; YPL271W |
| DI49_5288; SPGP0GU00200 | DI49_5288 | SPGP0GU00200 | |
| DI49_5290; DI49_5289 | DI49_5290; DI49_5289 | | |
| DI49_5292; DI49_5291 | DI49_5292; DI49_5291 | | |
| DI49_5293; SPGP0GV00150 | DI49_5293 | SPGP0GV00150 | |
| DI49_3941; DI49_4017; DI49_3940 | DI49_3941; DI49_4017; DI49_3940 | | |
| SPGP0GW00140; DI49_3939 | DI49_3939 | SPGP0GW00140 | |

Create PDF in your applications with the Pdfcrowd HTML to PDF API PDFCROWD

| | | | |
|---|---|---|---|
| SPGP0GW00160; DI49_3938 | DI49_3938 | SPGP0GW00160 | |
| DI49_3936; DI49_3937 | DI49_3936; DI49_3937 | | |
| DI49_3934; DI49_3935 | DI49_3934; DI49_3935 | | |
| DI49_3932; DI49_3933 | DI49_3932; DI49_3933 | | |
| DI49_1653; DI49_1652 | DI49_1653; DI49_1652 | | |
| DI49_1655; DI49_1654 | DI49_1655; DI49_1654 | | |
| DI49_1657; DI49_1656 | DI49_1657; DI49_1656 | | |
| DI49_0777; YCR024C-A; DI49_0776; YCR026C | DI49_0777; DI49_0776 | | YCR024C-A; YCR026C |
| DI49_0774; DI49_0775; YCR023C; YCR024C | DI49_0774; DI49_0775 | | YCR023C; YCR024C |
| DI49_0772; DI49_0773 | DI49_0772; DI49_0773 | | |
| DI49_0770; YCR019W; YCR020C | DI49_0770 | | YCR019W; YCR020C |
| DI49_4732; DI49_4731 | DI49_4732; DI49_4731 | | |
| DI49_4734; DI49_4733 | DI49_4734; DI49_4733 | | |
| SPGP0HAtrna7P; DI49_4736; DI49_5638; SPGP0RZtrna14P; YCR007C | DI49_4736; DI49_5638 | SPGP0HAtrna7P; SPGP0RZtrna14P | YCR007C |
| DI49_5636; DI49_5637 | DI49_5636; DI49_5637 | | |
| DI49_5634; DI49_5635 | DI49_5634; DI49_5635 | | |
| DI49_5632; DI49_5633 | DI49_5632; DI49_5633 | | |
| DI49_5630; DI49_5631 | DI49_5630; DI49_5631 | | |
| DI49_5628; DI49_5629 | DI49_5628; DI49_5629 | | |
| DI49_1631; DI49_1630 | DI49_1631; DI49_1630 | | |
| DI49_3760; DI49_4723; DI49_3761; DI49_4718 | DI49_3760; DI49_4723; DI49_3761; DI49_4718 | | |
| DI49_3554; DI49_3555 | DI49_3554; DI49_3555 | | |
| DI49_3552; DI49_3553 | DI49_3552; DI49_3553 | | |
| SPGP0HJ00100; DI49_0719 | DI49_0719 | SPGP0HJ00100 | |
| SPGP0HJ00120; SPGP0HJ00110 | | SPGP0HJ00120; SPGP0HJ00110 | |
| SPGP0HJ00140; SPGP0HJ00130 | | SPGP0HJ00140; SPGP0HJ00130 | |
| DI49_0688; DI49_0689 | DI49_0688; DI49_0689 | | |
| SPGP0HK00100; DI49_0687 | DI49_0687 | SPGP0HK00100 | |
| DI49_0076; DI49_0077; DI49_0001; DI49_3333 | DI49_0076; DI49_0077; DI49_0001; DI49_3333 | | |
| SPGP0U00140; SPGP0WQ00250; SPGP0U00150; DI49_2025 | DI49_2025 | SPGP0U00140; SPGP0WQ00250; SPGP0U00150 | |
| DI49_0072; SPGP0U00160 | DI49_0072 | SPGP0U00160 | |
| DI49_0070; DI49_0071 | DI49_0070; DI49_0071 | | |
| DI49_0068; YAR014C | DI49_0068 | | YAR014C |
| DI49_0064; DI49_0065 | DI49_0064; DI49_0065 | | |
| DI49_0062; YAR007C | DI49_0062 | | YAR007C |
| DI49_0061; YAR003W | DI49_0061 | | YAR003W |
| DI49_0060; YAR002C-A; DI49_0059; YAR002W | DI49_0060; DI49_0059 | | YAR002C-A; YAR002W |
| DI49_0057; DI49_0058 | DI49_0057; DI49_0058 | | |
| DI49_0055; DI49_3507; YAL005C; DI49_3508 | DI49_0055; DI49_3507; DI49_3508 | | YAL005C |

Create PDF in your applications with the Pdfcrowd HTML to PDF API PDFCROWD

| | | | |
|---|---|---|---|
| DI49_0053; YAL007C; DI49_0052; YAL008W | DI49_0053; DI49_0052 | | YAL007C; YAL008W |
| DI49_0050; DI49_0051 | DI49_0050; DI49_0051 | | |
| DI49_0048; YAL012W | DI49_0048 | | YAL012W |
| SPGP0U00400; YAL013W | | SPGP0U00400 | YAL013W |
| DI49_0047; YAL014C; DI49_0046; DI49_0546 | DI49_0047; DI49_0046; DI49_0546 | | YAL014C |
| DI49_0044; DI49_0045; YAL016W; YNR053C | DI49_0044; DI49_0045 | | YAL016W; YNR053C |
| DI49_0042; YAL019W; YAL018C | DI49_0042 | | YAL019W; YAL018C |
| DI49_0041; YAL020C | DI49_0041 | | YAL020C |
| DI49_0040; YAL021C | DI49_0040 | | YAL021C |
| DI49_0039; YAL022C | DI49_0039 | | YAL022C |
| DI49_0038; YAL023C | DI49_0038 | | YAL023C |
| DI49_0037; YAL024C; DI49_0036 | DI49_0037; DI49_0036 | | YAL024C |
| DI49_0034; DI49_0035; YAL026C; YAL027W | DI49_0034; DI49_0035 | | YAL026C; YAL027W |
| DI49_0032; YAL029C; SPGP0U00570; YAL030W | DI49_0032 | SPGP0U00570 | YAL029C; YAL030W |
| DI49_0030; DI49_0031 | DI49_0030; DI49_0031 | | |
| DI49_0028; YAL034C | DI49_0028 | | YAL034C |
| DI49_0027; YAL034W-A | DI49_0027 | | YAL034W-A |
| DI49_0026; YAL035W | DI49_0026 | | YAL035W |
| DI49_0025; YAL036C; DI49_0024; DI49_0562 | DI49_0025; DI49_0024; DI49_0562 | | YAL036C |
| DI49_0022; DI49_0023 | DI49_0022; DI49_0023 | | |
| DI49_0020; YAL041W | DI49_0020 | | YAL041W |
| DI49_0019; YAL042W | DI49_0019 | | YAL042W |
| DI49_0018; YAL043C; DI49_0017; YAL044C | DI49_0018; DI49_0017 | | YAL043C; YAL044C |
| DI49_0015; YAL046C; DI49_0014; YAL047C | DI49_0015; DI49_0014 | | YAL046C; YAL047C |
| DI49_0012; DI49_0013 | DI49_0012; DI49_0013 | | |
| DI49_0010; DI49_0011 | DI49_0010; DI49_0011 | | |
| DI49_0008; DI49_0009 | DI49_0008; DI49_0009 | | |
| DI49_0006; DI49_0007; YAL056W; YLL015W | DI49_0006; DI49_0007 | | YAL056W; YLL015W |
| DI49_0004; DI49_0005 | DI49_0004; DI49_0005 | | |
| DI49_0002; YAL062W; SPGP0TJ00100 | DI49_0002 | SPGP0TJ00100 | YAL062W |
| DI49_3331; DI49_3332 | DI49_3331; DI49_3332 | | |
| DI49_3329; DI49_3330 | DI49_3329; DI49_3330 | | |
| DI49_3327; DI49_3328 | DI49_3327; DI49_3328 | | |
| DI49_3325; DI49_3326 | DI49_3325; DI49_3326 | | |
| DI49_3324; SPGP0HL00190 | DI49_3324 | SPGP0HL00190 | |
| DI49_4011; DI49_3323 | DI49_4011; DI49_3323 | | |
| DI49_4009; DI49_4010 | DI49_4009; DI49_4010 | | |
| DI49_4007; DI49_4008 | DI49_4007; DI49_4008 | | |
| DI49_4005; DI49_3969; DI49_4004; DI49_3968 | DI49_4005; DI49_3969; DI49_4004; DI49_3968 | | |
| DI49_4002; DI49_4003 | DI49_4002; DI49_4003 | | |

Create PDF in your applications with the Pdfcrowd HTML to PDF API PDFCROWD

| | | | |
|---|---|---|---|
| DI49_4000; DI49_4001 | DI49_4000; DI49_4001 | | |
| DI49_3998; DI49_3999 | DI49_3998; DI49_3999 | | |
| DI49_3996; DI49_3997 | DI49_3996; DI49_3997 | | |
| DI49_3994; DI49_3995 | DI49_3994; DI49_3995 | | |
| SPGP0HO00290; DI49_3993 | DI49_3993 | SPGP0HO00290 | |
| DI49_3991; DI49_3992 | DI49_3991; DI49_3992 | | |
| DI49_3989; DI49_3990 | DI49_3989; DI49_3990 | | |
| DI49_3987; DI49_3988 | DI49_3987; DI49_3988 | | |
| DI49_5433; DI49_1918 | DI49_5433; DI49_1918 | | |
| DI49_5435; DI49_5434 | DI49_5435; DI49_5434 | | |
| DI49_5437; DI49_1649; DI49_5438; DI49_1650 | DI49_5437; DI49_1649; DI49_5438; DI49_1650 | | |
| DI49_5440; DI49_5439 | DI49_5440; DI49_5439 | | |
| DI49_5441; SPGP0HT00180 | DI49_5441 | SPGP0HT00180 | |
| DI49_5443; DI49_5442 | DI49_5443; DI49_5442 | | |
| DI49_5445; DI49_5444 | DI49_5445; DI49_5444 | | |
| DI49_5447; DI49_5446 | DI49_5447; DI49_5446 | | |
| DI49_5449; DI49_5448 | DI49_5449; DI49_5448 | | |
| DI49_3521; YPL006W | DI49_3521 | | YPL006W |
| DI49_3523; DI49_3522 | DI49_3523; DI49_3522 | | |
| DI49_3525; DI49_3524 | DI49_3525; DI49_3524 | | |
| DI49_3527; DI49_3526 | DI49_3527; DI49_3526 | | |
| DI49_2099; DI49_5598; YGL178W; DI49_0730 | DI49_2099; DI49_5598; DI49_0730 | | YGL178W |
| DI49_3349; DI49_3350 | DI49_3349; DI49_3350 | | |
| DI49_3347; DI49_3348 | DI49_3347; DI49_3348 | | |
| DI49_3345; DI49_3346 | DI49_3345; DI49_3346 | | |
| DI49_3344; SPGP0HX00160 | DI49_3344 | SPGP0HX00160 | |
| DI49_3342; DI49_3343 | DI49_3342; DI49_3343 | | |
| DI49_0631; DI49_0630 | DI49_0631; DI49_0630 | | |
| DI49_0633; DI49_0632 | DI49_0633; DI49_0632 | | |
| DI49_4622; DI49_0634 | DI49_4622; DI49_0634 | | |
| DI49_4624; DI49_4623 | DI49_4624; DI49_4623 | | |
| DI49_4626; DI49_4625 | DI49_4626; DI49_4625 | | |
| DI49_4628; DI49_4627 | DI49_4628; DI49_4627 | | |
| DI49_4630; DI49_4629 | DI49_4630; DI49_4629 | | |
| DI49_4632; DI49_4631 | DI49_4632; DI49_4631 | | |
| DI49_4634; DI49_4633 | DI49_4634; DI49_4633 | | |
| DI49_4639; DI49_4638 | DI49_4639; DI49_4638 | | |
| DI49_4641; DI49_4640 | DI49_4641; DI49_4640 | | |
| DI49_4643; DI49_4642 | DI49_4643; DI49_4642 | | |
| DI49_4645; DI49_4644 | DI49_4645; DI49_4644 | | |

Create PDF in your applications with the Pdfcrowd HTML to PDF API PDFCROWD

| | | | |
|---|---|---|---|
| DI49_4647; DI49_4646 | DI49_4647; DI49_4646 | | |
| DI49_3685; DI49_4648 | DI49_3685; DI49_4648 | | |
| DI49_3683; DI49_3684 | DI49_3683; DI49_3684 | | |
| SPGP0IA00140; DI49_3682 | DI49_3682 | SPGP0IA00140 | |
| DI49_1405; DI49_1404 | DI49_1405; DI49_1404 | | |
| DI49_1535; YGL189C; DI49_1534; YGL187C | DI49_1535; DI49_1534 | | YGL189C; YGL187C |
| DI49_1532; DI49_1533 | DI49_1532; DI49_1533 | | |
| DI49_1530; DI49_1531 | DI49_1530; DI49_1531 | | |
| DI49_1528; DI49_1529 | DI49_1528; DI49_1529 | | |
| DI49_1526; DI49_1527 | DI49_1526; DI49_1527 | | |
| DI49_1524; DI49_1525 | DI49_1524; DI49_1525 | | |
| DI49_1522; DI49_0101; DI49_1521 | DI49_1522; DI49_0101; DI49_1521 | | |
| DI49_1519; DI49_1520 | DI49_1519; DI49_1520 | | |
| DI49_1878; DI49_1877 | DI49_1878; DI49_1877 | | |
| DI49_1882; DI49_1881 | DI49_1882; DI49_1881 | | |
| DI49_1502; SPGP0ID00160 | DI49_1502 | SPGP0ID00160 | |
| DI49_1500; DI49_1501 | DI49_1500; DI49_1501 | | |
| DI49_1499; SPGP0IE00130 | DI49_1499 | SPGP0IE00130 | |
| DI49_1497; DI49_1498 | DI49_1497; DI49_1498 | | |
| DI49_1495; DI49_1496 | DI49_1495; DI49_1496 | | |
| YPL250C; DI49_1494 | DI49_1494 | | YPL250C |
| YPL249C; YPL249C-A | | | YPL249C; YPL249C-A |
| DI49_1983; YPL248C | DI49_1983 | | YPL248C |
| DI49_1981; DI49_1982 | DI49_1981; DI49_1982 | | |
| DI49_1979; DI49_1980 | DI49_1979; DI49_1980 | | |
| DI49_1977; DI49_1978 | DI49_1977; DI49_1978 | | |
| DI49_1676; DI49_1976 | DI49_1676; DI49_1976 | | |
| YPL218W; DI49_5250 | DI49_5250 | | YPL218W |
| DI49_5252; DI49_5251 | DI49_5252; DI49_5251 | | |
| DI49_1842; DI49_1841 | DI49_1842; DI49_1841 | | |
| DI49_1844; DI49_1843 | DI49_1844; DI49_1843 | | |
| DI49_1846; DI49_1845 | DI49_1846; DI49_1845 | | |
| DI49_1848; DI49_1847 | DI49_1848; DI49_1847 | | |
| DI49_5562; DI49_3663 | DI49_5562; DI49_3663 | | |
| SPGP0IO00100; DI49_2577; DI49_2217; DI49_2578 | DI49_2577; DI49_2217; DI49_2578 | SPGP0IO00100 | |
| DI49_2215; DI49_2216 | DI49_2215; DI49_2216 | | |
| YJL222W; YBL017C; YCR099C; YBL018C | | | YJL222W; YBL017C; YCR099C; YBL018C |
| YJL221C; SPGP0I03360; YBR299W; DI49_1687; SPGP0AG00130; DI49_1909; DI49_1598; DI49_1599; SPGP0BF01870; SPGP0CU00100 | DI49_1687; DI49_1909; DI49_1598; DI49_1599 | SPGP0I03360; SPGP0AG00130; SPGP0BF01870; SPGP0CU00100 | YJL221C; YBR299W |
| YOL156W; SPGP0I03350; YIL171W | | SPGP0I03350 | YOL156W; YIL171W |

Create PDF in your applications with the Pdfcrowd HTML to PDF API PDFCROWD

| | | | |
|---|---|---|---|
| YOL155C; YIL169C; YOL154W | | | YOL155C; YIL169C; YOL154W |
| YOL152W; DI49_2257 | DI49_2257 | | YOL152W |
| YOL151W; DI49_2258 | DI49_2258 | | YOL151W |
| YOL149W; DI49_2259 | DI49_2259 | | YOL149W |
| YOL148C; DI49_2260 | DI49_2260 | | YOL148C |
| YOL147C; DI49_2261; YOL146W; DI49_2262 | DI49_2261; DI49_2262 | | YOL147C; YOL146W |
| YOL144W; YOL145C | | | YOL144W; YOL145C |
| YOL142W; DI49_2266; YOL141W; DI49_2267 | DI49_2266; DI49_2267 | | YOL142W; YOL141W |
| YOL139C; DI49_2269; YOL138C; DI49_2270 | DI49_2269; DI49_2270 | | YOL139C; YOL138C |
| YOL136C; DI49_2272; YOL135C; DI49_2273 | DI49_2272; DI49_2273 | | YOL136C; YOL135C |
| YOL132W; DI49_2275; YOL131W; DI49_2276 | DI49_2275; DI49_2276 | | YOL132W; YOL131W |
| YOL129W; DI49_2278 | DI49_2278 | | YOL129W |
| YOL128C; DI49_2279 | DI49_2279 | | YOL128C |
| YOL127W; DI49_2280 | DI49_2280 | | YOL127W |
| YOL126C; DI49_2281; YOL125W; DI49_2282 | DI49_2281; DI49_2282 | | YOL126C; YOL125W |
| YOL123W; DI49_2284 | DI49_2284 | | YOL123W |
| YOL122C; DI49_2285 | DI49_2285 | | YOL122C |
| YOL121C; DI49_4447; DI49_4446; DI49_2286 | DI49_4447; DI49_4446; DI49_2286 | | YOL121C |
| SPGP0X00460; YOL120C; YNL301C | | SPGP0X00460 | YOL120C; YNL301C |
| YOL119C; DI49_2287; YOL117W; DI49_2288 | DI49_2287; DI49_2288 | | YOL119C; YOL117W |
| YOL115W; DI49_2290; YOL114C; DI49_2291 | DI49_2290; DI49_2291 | | YOL115W; YOL114C |
| YOL112W; YOL113W | | | YOL112W; YOL113W |
| YOL110W; DI49_2295; YOL109W; DI49_2296 | DI49_2295; DI49_2296 | | YOL110W; YOL109W |
| YOL108C; SPGP0TVtrna20G; SPGP0OEtrna14G; DI49_4461; DI49_2298; SPGP0Dtrna15G; DI49_2107 | DI49_4461; DI49_2298; DI49_2107 | SPGP0TVtrna20G; SPGP0OEtrna14G; SPGP0Dtrna15G | YOL108C |
| DI49_3379; YOL107W | DI49_3379 | | YOL107W |
| DI49_4619; DI49_4620 | DI49_4619; DI49_4620 | | |
| DI49_4617; DI49_4618 | DI49_4617; DI49_4618 | | |
| DI49_4615; DI49_4616 | DI49_4615; DI49_4616 | | |
| DI49_4613; DI49_4614 | DI49_4613; DI49_4614 | | |
| DI49_2201; DI49_4612 | DI49_2201; DI49_4612 | | |
| DI49_3739; DI49_3738 | DI49_3739; DI49_3738 | | |
| SPGP0IVtrna9L; SPGP0VRtrna22L; DI49_3805 | DI49_3805 | SPGP0IVtrna9L; SPGP0VRtrna22L | |
| SPGP0IVtrna10L; DI49_3110; DI49_3109; YJR100C | DI49_3110; DI49_3109 | SPGP0IVtrna10L | YJR100C |
| DI49_3744; SPGP0IVtrna6I | DI49_3744 | SPGP0IVtrna6I | |
| DI49_4035; YML056C | DI49_4035 | | YML056C |
| DI49_4034; SPGP0IW00120 | DI49_4034 | SPGP0IW00120 | |
| DI49_3387; DI49_3388 | DI49_3387; DI49_3388 | | |
| YGL242C; YGL241W | | | YGL242C; YGL241W |
| YGL244W; YGL243W | | | YGL244W; YGL243W |

Create PDF in your applications with the Pdfcrowd HTML to PDF API PDFCROWD

| | | | |
|---|---|---|---|
| YGL246C; YGL245W<br>YGL248W; YGL247W | | | YGL246C; YGL245W<br>YGL248W; YGL247W |
| YGL250W; YGL249W | | | YGL250W; YGL249W |
| YGL252C; YGL251C | | | YGL252C; YGL251C |
| YGL254W; YGL253W | | | YGL254W; YGL253W |
| YGL256W; YGL255W | | | YGL256W; YGL255W |
| DI49_1415; DI49_1414 | DI49_1415; DI49_1414 | | |
| DI49_1417; DI49_1416 | DI49_1417; DI49_1416 | | |
| DI49_1419; DI49_1418 | DI49_1419; DI49_1418 | | |
| DI49_0785; YCR033W | DI49_0785 | | YCR033W |
| DI49_0786; YCR034W | DI49_0786 | | YCR034W |
| DI49_0787; YCR035C; DI49_0788; YCR036W | DI49_0787; DI49_0788 | | YCR035C; YCR036W |
| DI49_0789; SPGP0JA00150 | DI49_0789 | SPGP0JA00150 | |
| YNL012W; SPGP0JC00100 | | SPGP0JC00100 | YNL012W |
| DI49_4725; DI49_4724 | DI49_4725; DI49_4724 | | |
| DI49_2821; DI49_0731 | DI49_2821; DI49_0731 | | |
| DI49_2057; DI49_2056 | DI49_2057; DI49_2056 | | |
| DI49_2059; DI49_2058 | DI49_2059; DI49_2058 | | |
| DI49_2061; DI49_2060 | DI49_2061; DI49_2060 | | |
| DI49_2063; DI49_2062 | DI49_2063; DI49_2062 | | |
| DI49_2065; DI49_2064 | DI49_2065; DI49_2064 | | |
| SPGP0JFtrna1C; DI49_5400; DI49_2071; SPGP0SUtrna2C; DI49_5401 | DI49_5400; DI49_2071; DI49_5401 | SPGP0JFtrna1C; SPGP0SUtrna2C | |
| DI49_2073; DI49_2072 | DI49_2073; DI49_2072 | | |
| DI49_2075; DI49_2074 | DI49_2075; DI49_2074 | | |
| SPGP0JF00280; SPGP0OT00170; YPL281C | | SPGP0JF00280; SPGP0OT00170 | YPL281C |
| DI49_5370; DI49_5371 | DI49_5370; DI49_5371 | | |
| DI49_5368; DI49_5369 | DI49_5368; DI49_5369 | | |
| DI49_2214; DI49_5367 | DI49_2214; DI49_5367 | | |
| DI49_2212; DI49_2213 | DI49_2212; DI49_2213 | | |
| DI49_5499; DI49_5498 | DI49_5499; DI49_5498 | | |
| DI49_5501; DI49_5500 | DI49_5501; DI49_5500 | | |
| DI49_5504; DI49_1714 | DI49_5504; DI49_1714 | | |
| DI49_5506; DI49_5505 | DI49_5506; DI49_5505 | | |
| DI49_5508; DI49_5507 | DI49_5508; DI49_5507 | | |
| DI49_5510; DI49_5509 | DI49_5510; DI49_5509 | | |
| DI49_5512; DI49_5511 | DI49_5512; DI49_5511 | | |
| DI49_5514; DI49_5513 | DI49_5514; DI49_5513 | | |
| DI49_0726; YCL029C; YCL028W | DI49_0726 | | YCL029C; YCL028W |
| DI49_0725; YCL030C | DI49_0725 | | YCL030C |
| DI49_0724; YCL031C | DI49_0724 | | YCL031C |

Create PDF in your applications with the Pdfcrowd HTML to PDF API PDFCROWD

| | | | |
|---|---|---|---|
| DI49_0723; YCL032W; DI49_0722 | DI49_0723; DI49_0722 | | YCL032W |
| DI49_0720; DI49_0721 | DI49_0720; DI49_0721 | | |
| DI49_5274; SPGP0JO00170; YCL036W; YCL035C | DI49_5274 | SPGP0JO00170 | YCL036W; YCL035C |
| DI49_5276; DI49_5275 | DI49_5276; DI49_5275 | | |
| DI49_5278; DI49_5277 | DI49_5278; DI49_5277 | | |
| DI49_5280; DI49_5279 | DI49_5280; DI49_5279 | | |
| DI49_5282; DI49_5281 | DI49_5282; DI49_5281 | | |
| DI49_5284; DI49_5283 | DI49_5284; DI49_5283 | | |
| DI49_5286; DI49_5285 | DI49_5286; DI49_5285 | | |
| DI49_5463; DI49_2204 | DI49_5463; DI49_2204 | | |
| DI49_5461; DI49_5462 | DI49_5461; DI49_5462 | | |
| DI49_5459; DI49_5460 | DI49_5459; DI49_5460 | | |
| DI49_3424; SPGP0AA00100 | DI49_3424 | SPGP0AA00100 | |
| DI49_3422; DI49_3423 | DI49_3422; DI49_3423 | | |
| DI49_3420; DI49_3421 | DI49_3420; DI49_3421 | | |
| DI49_3418; DI49_3419 | DI49_3418; DI49_3419 | | |
| DI49_3416; DI49_3417 | DI49_3416; DI49_3417 | | |
| DI49_3414; DI49_3415 | DI49_3414; DI49_3415 | | |
| DI49_3308; DI49_3307 | DI49_3308; DI49_3307 | | |
| DI49_3310; DI49_3309 | DI49_3310; DI49_3309 | | |
| DI49_3312; DI49_3311 | DI49_3312; DI49_3311 | | |
| DI49_3314; DI49_3313 | DI49_3314; DI49_3313 | | |
| DI49_3316; DI49_3315 | DI49_3316; DI49_3315 | | |
| DI49_3318; DI49_3317 | DI49_3318; DI49_3317 | | |
| DI49_5265; DI49_5264 | DI49_5265; DI49_5264 | | |
| DI49_0795; DI49_1659 | DI49_0795; DI49_1659 | | |
| DI49_0793; DI49_0794 | DI49_0793; DI49_0794 | | |
| DI49_1971; DI49_0792 | DI49_1971; DI49_0792 | | |
| DI49_3453; DI49_1970 | DI49_3453; DI49_1970 | | |
| DI49_3455; DI49_3454 | DI49_3455; DI49_3454 | | |
| DI49_3457; DI49_3456 | DI49_3457; DI49_3456 | | |
| DI49_3459; DI49_3458 | DI49_3459; DI49_3458 | | |
| DI49_3461; DI49_3460 | DI49_3461; DI49_3460 | | |
| DI49_4560; DI49_4561 | DI49_4560; DI49_4561 | | |
| DI49_5466; SPGP0KF00100 | DI49_5466 | SPGP0KF00100 | |
| DI49_1352; DI49_1351 | DI49_1352; DI49_1351 | | |
| DI49_1354; YJR048W; DI49_1355 | DI49_1354; DI49_1355 | | YJR048W |
| DI49_1357; DI49_1356 | DI49_1357; DI49_1356 | | |
| DI49_3954; SPGP0KI00100; YBR012C | DI49_3954 | SPGP0KI00100 | YBR012C |
| DI49_3445; SPGP0KJ00100 | DI49_3445 | SPGP0KJ00100 | |

Create PDF in your applications with the Pdfcrowd HTML to PDF API PDFCROWD

| | | | |
|---|---|---|---|
| DI49_3440; DI49_3441 | DI49_3440; DI49_3441 | | |
| DI49_3438; DI49_3439 | DI49_3438; DI49_3439 | | |
| DI49_3436; DI49_3437 | DI49_3436; DI49_3437 | | |
| SPGP0KL00200; DI49_3435 | DI49_3435 | SPGP0KL00200 | |
| DI49_3433; DI49_3434 | DI49_3433; DI49_3434 | | |
| DI49_1967; DI49_3432 | DI49_1967; DI49_3432 | | |
| YLL021W; SPGP0KM00110 | | SPGP0KM00110 | YLL021W |
| DI49_3511; DI49_3510 | DI49_3511; DI49_3510 | | |
| YHL044W; SPGP0B00100 | | SPGP0B00100 | YHL044W |
| YHL040C; YHL043W | | | YHL040C; YHL043W |
| YHL038C; YHL039W | | | YHL038C; YHL039W |
| YHL035C; YHL036W | | | YHL035C; YHL036W |
| YHL026C; YHL027W | | | YHL026C; YHL027W |
| YHL012W; YHL013C | | | YHL012W; YHL013C |
| YHR005C-A; YHR005C | | | YHR005C-A; YHR005C |
| YHR015W; DI49_2422 | DI49_2422 | | YHR015W |
| YHR016C; DI49_2423 | DI49_2423 | | YHR016C |
| YHR017W; DI49_2424 | DI49_2424 | | YHR017W |
| YHR018C; DI49_2425 | DI49_2425 | | YHR018C |
| YHR019C; DI49_2426 | DI49_2426 | | YHR019C |
| YHR020W; DI49_2427 | DI49_2427 | | YHR020W |
| YHR021C; SPGP0D01800; YHR022C | | SPGP0D01800 | YHR021C; YHR022C |
| YHR024C; YHR023W; DI49_2429; DI49_2430 | DI49_2429; DI49_2430 | | YHR024C; YHR023W |
| YHR026W; DI49_2432 | DI49_2432 | | YHR026W |
| YHR027C; DI49_2433 | DI49_2433 | | YHR027C |
| YHR028C; DI49_2434; YHR029C | DI49_2434 | | YHR028C; YHR029C |
| YHR031C; YHR030C; DI49_2436 | DI49_2436 | | YHR031C; YHR030C |
| YHR033W; DI49_2439; YHR034C; DI49_2440 | DI49_2439; DI49_2440 | | YHR033W; YHR034C |
| YHR036W; DI49_2442 | DI49_2442 | | YHR036W |
| YHR037W; DI49_2443; YHR038W; DI49_2444 | DI49_2443; DI49_2444 | | YHR037W; YHR038W |
| YHR039C-A; SPGP0D01980 | | SPGP0D01980 | YHR039C-A |
| YHR040W; DI49_2446; YHR041C; DI49_2447 | DI49_2446; DI49_2447 | | YHR040W; YHR041C |
| YHR044C; YHR042W | | | YHR044C; YHR042W |
| YHR046C; YHR045W | | | YHR046C; YHR045W |
| YHR048W; YHR047C | | | YHR048W; YHR047C |
| YHR050W; YHR049W | | | YHR050W; YHR049W |
| YHR052W; DI49_2457; YHR055C; DI49_2458 | DI49_2457; DI49_2458 | | YHR052W; YHR055C |
| YHR057C; YHR056C | | | YHR057C; YHR056C |
| YHR059W; DI49_2462; YHR060W; DI49_2463 | DI49_2462; DI49_2463 | | YHR059W; YHR060W |
| YHR062C; YHR061C | | | YHR062C; YHR061C |

Create PDF in your applications with the Pdfcrowd HTML to PDF API PDFCROWD

| | | | |
|---|---|---|---|
| YHR064C; DI49_2467 | DI49_2467 | | YHR064C |
| YHR065C; DI49_2468 | DI49_2468 | | YHR065C |
| YHR069C; DI49_2472; YHR070W | DI49_2472 | | YHR069C; YHR070W |
| SPGP0Btrna3F; SPGP0Btrna4F; YHR072W; YHR126C | | SPGP0Btrna3F; SPGP0Btrna4F | YHR072W; YHR126C |
| YHR073W; YHR072W-A | | | YHR073W; YHR072W-A |
| YHR075C; DI49_2480; YHR076W; DI49_2481 | DI49_2480; DI49_2481 | | YHR075C; YHR076W |
| YHR078W; YHR077C; DI49_2483; DI49_2484 | DI49_2483; DI49_2484 | | YHR078W; YHR077C |
| YHR079C-A; YHR079C | | | YHR079C-A; YHR079C |
| YHR081W; YHR080C | | | YHR081W; YHR080C |
| YHR083W; YHR082C | | | YHR083W; YHR082C |
| YHR085W; YHR084W; DI49_2492 | DI49_2492 | | YHR085W; YHR084W |
| YHR087W; DI49_2494; YHR088W; DI49_2495 | DI49_2494; DI49_2495 | | YHR087W; YHR088W |
| YHR090C; DI49_2497; YHR091C; DI49_2498 | DI49_2497; DI49_2498 | | YHR090C; YHR091C |
| YHR093W; YHR092C; SPGP0D02520; YJL214W | | SPGP0D02520 | YHR093W; YHR092C; YJL214W |
| YHR096C; YHR094C; YHR097C; DI49_2500; DI49_2501; DI49_2502; YDR345C; DI49_0523 | DI49_2500; DI49_2501; DI49_2502; DI49_0523 | | YHR096C; YHR094C; YHR097C; YDR345C |
| YHR098C; DI49_2503 | DI49_2503 | | YHR098C |
| YHR099W; DI49_2504; YHR100C; DI49_2505 | DI49_2504; DI49_2505 | | YHR099W; YHR100C |
| YHR102W; DI49_2507; YHR103W; DI49_2508 | DI49_2507; DI49_2508 | | YHR102W; YHR103W |
| YHR105W; YHR104W | | | YHR105W; YHR104W |
| YHR107C; DI49_2512; YHR108W; DI49_2513 | DI49_2512; DI49_2513 | | YHR107C; YHR108W |
| YHR110W; YHR109W | | | YHR110W; YHR109W |
| YHR112C; YHR111W | | | YHR112C; YHR111W |
| YHR114W; DI49_2519; YHR115C; DI49_2520 | DI49_2519; DI49_2520 | | YHR114W; YHR115C |
| YHR117W; YHR116W | | | YHR117W; YHR116W |
| YHR119W; DI49_2524; YHR120W; DI49_2525 | DI49_2524; DI49_2525 | | YHR119W; YHR120W |
| YHR122W; YHR121W | | | YHR122W; YHR121W |
| YHR124W; DI49_2528 | DI49_2528 | | YHR124W |
| YHR128W; DI49_2532 | DI49_2532 | | YHR128W |
| YHR129C; DI49_2533; YHR131C; DI49_2534 | DI49_2533; DI49_2534 | | YHR129C; YHR131C |
| YHR132W-A; DI49_2536; YHR133C; DI49_2537 | DI49_2536; DI49_2537 | | YHR132W-A; YHR133C |
| YHR135C; YHR134W | | | YHR135C; YHR134W |
| YHR137W; YHR136C | | | YHR137W; YHR136C |
| YHR139C; YHR138C | | | YHR139C; YHR138C |
| YHR141C; SPGP0D02990; DI49_4575; DI49_4574 | DI49_4575; DI49_4574 | SPGP0D02990 | YHR141C |
| YHR142W; DI49_2544; YHR143W; DI49_2545 | DI49_2544; DI49_2545 | | YHR142W; YHR143W |
| YHR144C; YHR143W-A | | | YHR144C; YHR143W-A |
| YHR146W; SPGP0Btrna6P | | SPGP0Btrna6P | YHR146W |
| YHR148W; DI49_2551 | DI49_2551 | | YHR148W |
| YHR149C; DI49_2552; YHR150W; DI49_2553 | DI49_2552; DI49_2553 | | YHR149C; YHR150W |

Create PDF in your applications with the Pdfcrowd HTML to PDF API PDFCROWD

| | | | |
|---|---|---|---|
| YHR152W; YHR151C | | | YHR152W; YHR151C |
| YHR154W; DI49_2557 | DI49_2557 | | YHR154W |
| YHR155W; DI49_2558; YHR156C; DI49_2559 | DI49_2558; DI49_2559 | | YHR155W; YHR156C |
| YHR158C; YHR157W | | | YHR158C; YHR157W |
| YHR160C; YHR159W | | | YHR160C; YHR159W |
| YHR162W; YHR161C | | | YHR162W; YHR161C |
| YHR164C; DI49_2567; DI49_2568 | DI49_2567; DI49_2568 | | YHR164C |
| YHR165C; DI49_2569 | DI49_2569 | | YHR165C |
| YHR166C; DI49_2570; YHR167W; DI49_2571 | DI49_2570; DI49_2571 | | YHR166C; YHR167W |
| YHR169W; DI49_2573 | DI49_2573 | | YHR169W |
| YHR170W; DI49_2574; YHR171W; DI49_2575 | DI49_2574; DI49_2575 | | YHR170W; YHR171W |
| YHR173C; YHR172W | | | YHR173C; YHR172W |
| YHR175W; YHR174W | | | YHR175W; YHR174W |
| YHR177W; YHR176W | | | YHR177W; YHR176W |
| YHR179W; DI49_2581; YHR180W | DI49_2581 | | YHR179W; YHR180W |
| YHR181W; DI49_2583; YHR182W; DI49_2584 | DI49_2583; DI49_2584 | | YHR181W; YHR182W |
| YHR184W; YHR183W | | | YHR184W; YHR183W |
| YHR186C; DI49_2589; YHR187W; DI49_2590 | DI49_2589; DI49_2590 | | YHR186C; YHR187W |
| YHR189W; YHR188C | | | YHR189W; YHR188C |
| YHR191C; YHR190W | | | YHR191C; YHR190W |
| YHR193C; DI49_2596 | DI49_2596 | | YHR193C |
| YHR194W; DI49_2597; YHR195W; DI49_2598 | DI49_2597; DI49_2598 | | YHR194W; YHR195W |
| YHR197W; DI49_2600; YHR198C; DI49_2601 | DI49_2600; DI49_2601 | | YHR197W; YHR198C |
| YHR199C-A; YHR199C | | | YHR199C-A; YHR199C |
| YHR201C; YHR200W | | | YHR201C; YHR200W |
| SPGP0B02550; SPGP0I00100; DI49_3664; YJR144W | DI49_3664 | SPGP0B02550; SPGP0I00100 | YJR144W |
| DI49_3668; DI49_3667 | DI49_3668; DI49_3667 | | |
| DI49_3670; DI49_3669 | DI49_3670; DI49_3669 | | |
| DI49_3672; DI49_3671 | DI49_3672; DI49_3671 | | |
| DI49_3674; DI49_3673 | DI49_3674; DI49_3673 | | |
| DI49_3676; DI49_3675 | DI49_3676; DI49_3675 | | |
| DI49_3678; DI49_3677 | DI49_3678; DI49_3677 | | |
| DI49_4557; DI49_4558 | DI49_4557; DI49_4558 | | |
| SPGP0KQ00100; DI49_3163; DI49_4794; SPGP0OS00130; DI49_5568; DI49_3162 | DI49_3163; DI49_4794; DI49_5568; DI49_3162 | SPGP0KQ00100; SPGP0OS00130 | |
| DI49_4792; DI49_4793 | DI49_4792; DI49_4793 | | |
| DI49_4790; DI49_4791 | DI49_4790; DI49_4791 | | |
| DI49_3558; DI49_4210; DI49_3557; YMR121C; YMR122W-A | DI49_3558; DI49_4210; DI49_3557 | | YMR121C; YMR122W-A |
| DI49_1985; DI49_3556 | DI49_1985; DI49_3556 | | |
| DI49_1989; DI49_3836; DI49_3341 | DI49_1989; DI49_3836; DI49_3341 | | |

Create PDF in your applications with the Pdfcrowd HTML to PDF API PDFCROWD

| | | | |
|---|---|---|---|
| DI49_3832; DI49_3833 | DI49_3832; DI49_3833 | | |
| DI49_0736; YCL018W | DI49_0736 | | YCL018W |
| DI49_0543; DI49_0542 | DI49_0543; DI49_0542 | | |
| SPGP0KY00140; DI49_0544 | DI49_0544 | SPGP0KY00140 | |
| DI49_0591; DI49_0545 | DI49_0591; DI49_0545 | | |
| DI49_0589; DI49_0590 | DI49_0589; DI49_0590 | | |
| DI49_0587; DI49_0588 | DI49_0587; DI49_0588 | | |
| DI49_0585; DI49_0586 | DI49_0585; DI49_0586 | | |
| DI49_0583; DI49_0584 | DI49_0583; DI49_0584 | | |
| DI49_0582; SPGP0KZ00190 | DI49_0582 | SPGP0KZ00190 | |
| DI49_0581; YDR406W | DI49_0581 | | YDR406W |
| DI49_0579; DI49_0580 | DI49_0579; DI49_0580 | | |
| DI49_0577; DI49_0578 | DI49_0577; DI49_0578 | | |
| DI49_0575; DI49_0576 | DI49_0575; DI49_0576 | | |
| DI49_0573; DI49_0574 | DI49_0573; DI49_0574 | | |
| DI49_0571; DI49_0572 | DI49_0571; DI49_0572 | | |
| DI49_0569; DI49_0570 | DI49_0569; DI49_0570 | | |
| SPGP0LA00110; YER188W | | SPGP0LA00110 | YER188W |
| DI49_3168; DI49_3167 | DI49_3168; DI49_3167 | | |
| YKL215C; DI49_3169 | DI49_3169 | | YKL215C |
| DI49_3171; DI49_3170 | DI49_3171; DI49_3170 | | |
| DI49_3173; DI49_3172 | DI49_3173; DI49_3172 | | |
| DI49_3175; DI49_3174 | DI49_3175; DI49_3174 | | |
| DI49_1675; DI49_1674 | DI49_1675; DI49_1674 | | |
| DI49_3879; DI49_3880 | DI49_3879; DI49_3880 | | |
| DI49_3874; DI49_3875 | DI49_3874; DI49_3875 | | |
| DI49_3872; DI49_3873 | DI49_3872; DI49_3873 | | |
| DI49_3870; DI49_3871 | DI49_3870; DI49_3871 | | |
| DI49_3868; DI49_3869 | DI49_3868; DI49_3869 | | |
| SPGP0LF00100; DI49_3161; DI49_3160 | DI49_3161; DI49_3160 | SPGP0LF00100 | |
| SPGP0LGtrna8Q; DI49_1374; DI49_1375 | DI49_1374; DI49_1375 | SPGP0LGtrna8Q | |
| DI49_1377; DI49_1376 | DI49_1377; DI49_1376 | | |
| DI49_1379; DI49_1378 | DI49_1379; DI49_1378 | | |
| DI49_1381; DI49_1380 | DI49_1381; DI49_1380 | | |
| DI49_1383; DI49_1382 | DI49_1383; DI49_1382 | | |
| DI49_3372; DI49_1384 | DI49_3372; DI49_1384 | | |
| SPGP0LH00120; DI49_3371 | DI49_3371 | SPGP0LH00120 | |
| DI49_3369; DI49_3370 | DI49_3369; DI49_3370 | | |
| DI49_3407; DI49_3408 | DI49_3407; DI49_3408 | | |
| DI49_3405; DI49_3406 | DI49_3405; DI49_3406 | | |

Create PDF in your applications with the Pdfcrowd HTML to PDF API PDFCROWD

| | | | |
|---|---|---|---|
| DI49_3403; DI49_3404 | DI49_3403; DI49_3404 | | |
| DI49_3401; DI49_3402 | DI49_3401; DI49_3402 | | |
| YAL032C; YAL031C | | | YAL032C; YAL031C |
| DI49_0564; DI49_0563 | DI49_0564; DI49_0563 | | |
| DI49_0837; DI49_0836 | DI49_0837; DI49_0836 | | |
| DI49_2140; SPGP0MVtrna13G; SPGP0LPtrna11G; DI49_5594 | DI49_2140; DI49_5594 | SPGP0MVtrna13G; SPGP0LPtrna11G | |
| DI49_2142; DI49_2141 | DI49_2142; DI49_2141 | | |
| SPGP0LS00130; DI49_1684 | DI49_1684 | SPGP0LS00130 | |
| DI49_1680; DI49_1681 | DI49_1680; DI49_1681 | | |
| DI49_0659; DI49_3431 | DI49_0659; DI49_3431 | | |
| DI49_0657; DI49_0658 | DI49_0657; DI49_0658 | | |
| DI49_0655; DI49_0656 | DI49_0655; DI49_0656 | | |
| DI49_0653; DI49_0654 | DI49_0653; DI49_0654 | | |
| DI49_0651; DI49_0652 | DI49_0651; DI49_0652 | | |
| DI49_0649; DI49_0650 | DI49_0649; DI49_0650 | | |
| DI49_0647; DI49_0648 | DI49_0647; DI49_0648 | | |
| DI49_0645; DI49_0646 | DI49_0645; DI49_0646 | | |
| DI49_0643; DI49_0644 | DI49_0643; DI49_0644 | | |
| DI49_0641; DI49_0642 | DI49_0641; DI49_0642 | | |
| DI49_0744; YCL008C; DI49_0745; YCL007C | DI49_0744; DI49_0745 | | YCL008C; YCL007C |
| DI49_0747; DI49_0746 | DI49_0747; DI49_0746 | | |
| DI49_0749; YCL001W; DI49_0750; YCL001W-A | DI49_0749; DI49_0750 | | YCL001W; YCL001W-A |
| DI49_1341; SPGP0LW00100 | DI49_1341 | SPGP0LW00100 | |
| DI49_1343; DI49_1342 | DI49_1343; DI49_1342 | | |
| DI49_1345; DI49_3566; DI49_1346; DI49_3565; DI49_3506; DI49_1411 | DI49_1345; DI49_3566; DI49_1346; DI49_3565; DI49_3506; DI49_1411 | | |
| DI49_5585; DI49_5584 | DI49_5585; DI49_5584 | | |
| DI49_5590; DI49_5589; DI49_2086 | DI49_5590; DI49_5589; DI49_2086 | | |
| DI49_5592; DI49_5591 | DI49_5592; DI49_5591 | | |
| DI49_5398; SPGP0LZ00100 | DI49_5398 | SPGP0LZ00100 | |
| SPGP0AG00110; DI49_5680 | DI49_5680 | SPGP0AG00110 | |
| DI49_0079; DI49_1597 | DI49_0079; DI49_1597 | | |
| DI49_1911; DI49_1910 | DI49_1911; DI49_1910 | | |
| DI49_1913; DI49_1912 | DI49_1913; DI49_1912 | | |
| DI49_1396; YER001W | DI49_1396 | | YER001W |
| DI49_1398; DI49_1397 | DI49_1398; DI49_1397 | | |
| YER006W; DI49_1399 | DI49_1399 | | YER006W |
| DI49_1401; DI49_1400 | DI49_1401; DI49_1400 | | |
| DI49_2144; DI49_1402 | DI49_2144; DI49_1402 | | |
| DI49_2150; DI49_2149 | DI49_2150; DI49_2149 | | |

Create PDF in your applications with the Pdfcrowd HTML to PDF API PDFCROWD

| | | | |
|---|---|---|---|
| DI49_2152; DI49_2151 | DI49_2152; DI49_2151 | | |
| DI49_2154; DI49_2153 | DI49_2154; DI49_2153 | | |
| DI49_2158; DI49_2157 | DI49_2158; DI49_2157 | | |
| YGL233W; YGL234W | | | YGL233W; YGL234W |
| YGL231C; YGL232W | | | YGL231C; YGL232W |
| YGL229C; YGL230C | | | YGL229C; YGL230C |
| YGL227W; YGL228W | | | YGL227W; YGL228W |
| YGL226W; YGL226C-A | | | YGL226W; YGL226C-A |
| YGL223C; YGL224C | | | YGL223C; YGL224C |
| DI49_1952; DI49_1953 | DI49_1952; DI49_1953 | | |
| DI49_1950; DI49_1951 | DI49_1950; DI49_1951 | | |
| DI49_1948; DI49_1949 | DI49_1948; DI49_1949 | | |
| DI49_1946; DI49_1947 | DI49_1946; DI49_1947 | | |
| DI49_1944; DI49_1945 | DI49_1944; DI49_1945 | | |
| DI49_1942; DI49_1943 | DI49_1942; DI49_1943 | | |
| DI49_1940; DI49_1941 | DI49_1940; DI49_1941 | | |
| DI49_1938; DI49_1939 | DI49_1938; DI49_1939 | | |
| DI49_1936; DI49_2112; DI49_4472; DI49_2113 | DI49_1936; DI49_2112; DI49_4472; DI49_2113 | | |
| DI49_4470; DI49_4471 | DI49_4470; DI49_4471 | | |
| DI49_4468; DI49_4469 | DI49_4468; DI49_4469 | | |
| DI49_1815; DI49_1814 | DI49_1815; DI49_1814 | | |
| DI49_1817; DI49_1816 | DI49_1817; DI49_1816 | | |
| YKR105C; SPGP0MK00100 | | SPGP0MK00100 | YKR105C |
| SPGP0MK00130; SPGP0MK00120 | | SPGP0MK00130; SPGP0MK00120 | |
| DI49_4753; DI49_4752 | DI49_4753; DI49_4752 | | |
| DI49_4755; DI49_4754; YAL055W | DI49_4755; DI49_4754 | | YAL055W |
| DI49_2007; YAL054C | DI49_2007 | | YAL054C |
| DI49_5532; DI49_5531 | DI49_5532; DI49_5531 | | |
| DI49_5534; DI49_5533 | DI49_5534; DI49_5533 | | |
| DI49_5536; DI49_5535 | DI49_5536; DI49_5535 | | |
| DI49_5538; DI49_5537 | DI49_5538; DI49_5537 | | |
| DI49_5540; DI49_5539 | DI49_5540; DI49_5539 | | |
| DI49_5542; DI49_5541 | DI49_5542; DI49_5541 | | |
| DI49_5544; DI49_5543 | DI49_5544; DI49_5543 | | |
| SPGP0MO00250; DI49_5545 | DI49_5545 | SPGP0MO00250 | |
| DI49_5549; DI49_5548 | DI49_5549; DI49_5548 | | |
| DI49_5551; DI49_5550 | DI49_5551; DI49_5550 | | |
| SPGP0MOtrna11T; SPGP0Dtrna17T; DI49_5554; SPGP0NTtrna13T; DI49_2219; DI49_2049; SPGP0SBtrna18T | DI49_5554; DI49_2219; DI49_2049 | SPGP0MOtrna11T; SPGP0Dtrna17T; SPGP0NTtrna13T; SPGP0SBtrna18T | |
| DI49_5556; DI49_5555 | DI49_5556; DI49_5555 | | |

Create PDF in your applications with the Pdfcrowd HTML to PDF API PDFCROWD

| | | | |
|---|---|---|---|
| DI49_5558; DI49_5557 | DI49_5558; DI49_5557 | | |
| DI49_5560; DI49_5559 | DI49_5560; DI49_5559 | | |
| DI49_1488; DI49_5561 | DI49_1488; DI49_5561 | | |
| SPGP0MS00100; SPGP0MT00100; DI49_2078; DI49_5576 | DI49_2078; DI49_5576 | SPGP0MS00100; SPGP0MT00100 | |
| DI49_5578; DI49_5577 | DI49_5578; DI49_5577 | | |
| DI49_5580; DI49_5579 | DI49_5580; DI49_5579 | | |
| YGL174W; YGL175C | | | YGL174W; YGL175C |
| DI49_1350; DI49_1349 | DI49_1350; DI49_1349 | | |
| YCR098C; DI49_4420 | DI49_4420 | | YCR098C |
| DI49_1365; DI49_3411 | DI49_1365; DI49_3411 | | |
| DI49_1361; DI49_1362 | DI49_1361; DI49_1362 | | |
| DI49_1358; DI49_3055; DI49_1516; YJR047C | DI49_1358; DI49_3055; DI49_1516 | | YJR047C |
| DI49_1514; DI49_1515 | DI49_1514; DI49_1515 | | |
| DI49_1512; DI49_1513 | DI49_1512; DI49_1513 | | |
| DI49_1510; DI49_1511 | DI49_1510; DI49_1511 | | |
| DI49_1508; DI49_0112; DI49_1507; DI49_0111; YBL072C; YBL074C | DI49_1508; DI49_0112; DI49_1507; DI49_0111 | | YBL072C; YBL074C |
| DI49_1505; DI49_1506 | DI49_1505; DI49_1506 | | |
| SPGP0NH00120; DI49_0056 | DI49_0056 | SPGP0NH00120 | |
| YAL010C; YAL009W | | | YAL010C; YAL009W |
| DI49_0548; DI49_0547 | DI49_0548; DI49_0547 | | |
| DI49_0550; DI49_0549 | DI49_0550; DI49_0549 | | |
| DI49_0552; DI49_0551 | DI49_0552; DI49_0551 | | |
| DI49_0554; DI49_0553 | DI49_0554; DI49_0553 | | |
| DI49_0555; SPGP0NI00190 | DI49_0555 | SPGP0NI00190 | |
| DI49_0557; DI49_0556 | DI49_0557; DI49_0556 | | |
| SPGP0NI00240; SPGP0NI00230 | | SPGP0NI00240; SPGP0NI00230 | |
| DI49_0559; DI49_0558 | DI49_0559; DI49_0558 | | |
| DI49_0796; DI49_0560 | DI49_0796; DI49_0560 | | |
| DI49_3550; DI49_3551 | DI49_3550; DI49_3551 | | |
| DI49_3548; DI49_3549 | DI49_3548; DI49_3549 | | |
| DI49_3546; DI49_3547 | DI49_3546; DI49_3547 | | |
| DI49_3544; DI49_3545 | DI49_3544; DI49_3545 | | |
| SPGP0NK00190; DI49_3543 | DI49_3543 | SPGP0NK00190 | |
| YGL257C; DI49_5414 | DI49_5414 | | YGL257C |
| YGL258W-A; YGL258W | | | YGL258W-A; YGL258W |
| DI49_1420; YDR542W | DI49_1420 | | YDR542W |
| DI49_1422; DI49_1421 | DI49_1422; DI49_1421 | | |
| DI49_4593; DI49_1423 | DI49_4593; DI49_1423 | | |
| DI49_4591; DI49_4592 | DI49_4591; DI49_4592 | | |
| DI49_3864; DI49_3865 | DI49_3864; DI49_3865 | | |

Create PDF in your applications with the Pdfcrowd HTML to PDF API PDFCROWD

| | | | |
|---|---|---|---|
| DI49_3862; DI49_3863 | DI49_3862; DI49_3863 | | |
| DI49_3860; DI49_3859; DI49_3858 | DI49_3860; DI49_3859; DI49_3858 | | |
| DI49_3856; DI49_3857 | DI49_3856; DI49_3857 | | |
| DI49_3854; DI49_3855 | DI49_3854; DI49_3855 | | |
| DI49_3852; DI49_3853 | DI49_3852; DI49_3853 | | |
| DI49_3850; DI49_3851 | DI49_3850; DI49_3851 | | |
| DI49_3848; DI49_3849 | DI49_3848; DI49_3849 | | |
| DI49_1692; DI49_1691 | DI49_1692; DI49_1691 | | |
| YGL196W; YGL197W | | | YGL196W; YGL197W |
| YGL194C-A; YGL195W | | | YGL194C-A; YGL195W |
| DI49_2238; DI49_2239 | DI49_2238; DI49_2239 | | |
| DI49_2236; DI49_2237 | DI49_2236; DI49_2237 | | |
| DI49_2234; DI49_2235 | DI49_2234; DI49_2235 | | |
| DI49_2232; DI49_2233 | DI49_2232; DI49_2233 | | |
| DI49_2230; DI49_2231 | DI49_2230; DI49_2231 | | |
| DI49_2228; DI49_2229 | DI49_2228; DI49_2229 | | |
| DI49_2226; DI49_2227 | DI49_2226; DI49_2227 | | |
| DI49_2224; DI49_2225 | DI49_2224; DI49_2225 | | |
| DI49_2222; DI49_2223 | DI49_2222; DI49_2223 | | |
| DI49_5491; SPGP0NU00100 | DI49_5491 | SPGP0NU00100 | |
| DI49_5493; DI49_5492 | DI49_5493; DI49_5492 | | |
| DI49_0827; DI49_0826 | DI49_0827; DI49_0826 | | |
| DI49_0829; DI49_0828 | DI49_0829; DI49_0828 | | |
| DI49_0832; DI49_0831 | DI49_0832; DI49_0831 | | |
| YPL282C; DI49_1412; DI49_2241; YFL020C; YAR019C | DI49_1412; DI49_2241 | | YPL282C; YFL020C; YAR019C |
| SPGP0NX00120; DI49_2240 | DI49_2240 | SPGP0NX00120 | |
| DI49_4474; DI49_4473 | DI49_4474; DI49_4473 | | |
| DI49_4476; DI49_4475 | DI49_4476; DI49_4475 | | |
| DI49_4478; DI49_4477 | DI49_4478; DI49_4477 | | |
| DI49_5376; DI49_5377 | DI49_5376; DI49_5377 | | |
| DI49_5374; DI49_5375 | DI49_5374; DI49_5375 | | |
| DI49_4649; DI49_5373 | DI49_4649; DI49_5373 | | |
| DI49_4651; DI49_4650 | DI49_4651; DI49_4650 | | |
| DI49_4653; DI49_4652 | DI49_4653; DI49_4652 | | |
| DI49_4655; DI49_4654 | DI49_4655; DI49_4654 | | |
| DI49_4459; DI49_4460 | DI49_4459; DI49_4460 | | |
| DI49_4457; DI49_4458 | DI49_4457; DI49_4458 | | |
| DI49_4455; DI49_4456 | DI49_4455; DI49_4456 | | |
| DI49_4453; DI49_4454 | DI49_4453; DI49_4454 | | |
| DI49_4451; DI49_4452 | DI49_4451; DI49_4452 | | |

Create PDF in your applications with the Pdfcrowd HTML to PDF API PDFCROWD

| | | | |
|---|---|---|---|
| DI49_4449; DI49_4450 | DI49_4449; DI49_4450 | | |
| DI49_4444; DI49_4445 | DI49_4444; DI49_4445 | | |
| DI49_1427; DI49_1426 | DI49_1427; DI49_1426 | | |
| DI49_1429; DI49_1428 | DI49_1429; DI49_1428 | | |
| DI49_1431; DI49_1430 | DI49_1431; DI49_1430 | | |
| DI49_1433; DI49_1432 | DI49_1433; DI49_1432 | | |
| DI49_1435; DI49_1434 | DI49_1435; DI49_1434 | | |
| DI49_1437; DI49_1436 | DI49_1437; DI49_1436 | | |
| DI49_1439; DI49_1438 | DI49_1439; DI49_1438 | | |
| DI49_1441; DI49_1440 | DI49_1441; DI49_1440 | | |
| DI49_1443; DI49_1442 | DI49_1443; DI49_1442 | | |
| DI49_1445; DI49_1444 | DI49_1445; DI49_1444 | | |
| DI49_1447; DI49_1446 | DI49_1447; DI49_1446 | | |
| DI49_1452; DI49_1451 | DI49_1452; DI49_1451 | | |
| DI49_1454; DI49_1453 | DI49_1454; DI49_1453 | | |
| DI49_1456; DI49_1455 | DI49_1456; DI49_1455 | | |
| DI49_3319; DI49_3584 | DI49_3319; DI49_3584 | | |
| DI49_3321; DI49_3320 | DI49_3321; DI49_3320 | | |
| DI49_5366; DI49_3322 | DI49_5366; DI49_3322 | | |
| DI49_5364; DI49_5365 | DI49_5364; DI49_5365 | | |
| DI49_5362; DI49_5363 | DI49_5362; DI49_5363 | | |
| DI49_5360; DI49_5361 | DI49_5360; DI49_5361 | | |
| DI49_5358; DI49_5359 | DI49_5358; DI49_5359 | | |
| DI49_5356; DI49_5357 | DI49_5356; DI49_5357 | | |
| DI49_5354; DI49_5355 | DI49_5354; DI49_5355 | | |
| YPL109C; SPGP0OI00230 | | SPGP0OI00230 | YPL109C |
| DI49_5353; SPGP0OI00250 | DI49_5353 | SPGP0OI00250 | |
| DI49_5348; DI49_5349 | DI49_5348; DI49_5349 | | |
| DI49_5346; DI49_5347 | DI49_5346; DI49_5347 | | |
| DI49_5344; DI49_5345 | DI49_5344; DI49_5345 | | |
| DI49_5342; DI49_5343 | DI49_5342; DI49_5343 | | |
| DI49_5340; DI49_5341 | DI49_5340; DI49_5341 | | |
| DI49_5338; DI49_5339 | DI49_5338; DI49_5339 | | |
| DI49_5336; DI49_5337 | DI49_5336; DI49_5337 | | |
| DI49_5334; DI49_5335 | DI49_5334; DI49_5335 | | |
| DI49_5332; DI49_5333 | DI49_5332; DI49_5333 | | |
| DI49_5330; DI49_5331 | DI49_5330; DI49_5331 | | |
| DI49_5328; DI49_5329 | DI49_5328; DI49_5329 | | |
| DI49_5326; DI49_5327 | DI49_5326; DI49_5327 | | |

Create PDF in your applications with the Pdfcrowd HTML to PDF API PDFCROWD

| | | | |
|---|---|---|---|
| DI49_5324; DI49_5325 | DI49_5324; DI49_5325 | | |
| DI49_5319; DI49_5320 | DI49_5319; DI49_5320 | | |
| DI49_5317; DI49_5318 | DI49_5317; DI49_5318 | | |
| DI49_5315; DI49_5316 | DI49_5315; DI49_5316 | | |
| DI49_5313; DI49_5314 | DI49_5313; DI49_5314 | | |
| DI49_5311; DI49_5312 | DI49_5311; DI49_5312 | | |
| DI49_5309; DI49_5310 | DI49_5309; DI49_5310 | | |
| DI49_5307; DI49_5308 | DI49_5307; DI49_5308 | | |
| DI49_5305; DI49_5306 | DI49_5305; DI49_5306 | | |
| DI49_5303; DI49_5304 | DI49_5303; DI49_5304 | | |
| DI49_5301; DI49_5302 | DI49_5301; DI49_5302 | | |
| DI49_5299; DI49_5300 | DI49_5299; DI49_5300 | | |
| DI49_5297; DI49_5298 | DI49_5297; DI49_5298 | | |
| DI49_5295; DI49_5296 | DI49_5295; DI49_5296 | | |
| DI49_4393; SPGP0AM00100; SPGP0WL00100 | DI49_4393 | SPGP0AM00100; SPGP0WL00100 | |
| DI49_4391; YMR304W | DI49_4391 | | YMR304W |
| DI49_4390; DI49_2320; DI49_2321; YMR303C | DI49_4390; DI49_2320; DI49_2321 | | YMR303C |
| DI49_4389; YMR302C | DI49_4389 | | YMR302C |
| DI49_4388; YMR301C | DI49_4388 | | YMR301C |
| DI49_4387; YMR300C; DI49_4386 | DI49_4387; DI49_4386 | | YMR300C |
| DI49_4384; YMR297W; DI49_4383 | DI49_4384; DI49_4383 | | YMR297W |
| DI49_4381; DI49_4382 | DI49_4381; DI49_4382 | | |
| DI49_4379; DI49_4380 | DI49_4379; DI49_4380 | | |
| DI49_4377; YMR290C; DI49_4376 | DI49_4377; DI49_4376 | | YMR290C |
| DI49_4374; YMR287C; DI49_4373 | DI49_4374; DI49_4373 | | YMR287C |
| DI49_4371; DI49_4372 | DI49_4371; DI49_4372 | | |
| DI49_4367; DI49_4368 | DI49_4367; DI49_4368 | | |
| DI49_4365; DI49_4366 | DI49_4365; DI49_4366 | | |
| DI49_4363; DI49_4364; YMR278W; YMR279C | DI49_4363; DI49_4364 | | YMR278W; YMR279C |
| DI49_4361; YMR275C; DI49_4360; YMR276W | DI49_4361; DI49_4360 | | YMR275C; YMR276W |
| DI49_4358; YMR272C; YMR273C | DI49_4358 | | YMR272C; YMR273C |
| DI49_4357; SPGP0Htrna17Q; SPGP0C00450 | DI49_4357 | SPGP0Htrna17Q; SPGP0C00450 | |
| DI49_4355; DI49_4356 | DI49_4355; DI49_4356 | | |
| DI49_4353; YMR269W; DI49_4352; YMR270C | DI49_4353; DI49_4352 | | YMR269W; YMR270C |
| DI49_4350; DI49_4351 | DI49_4350; DI49_4351 | | |
| DI49_4348; YMR264W; DI49_4347 | DI49_4348; DI49_4347 | | YMR264W |
| DI49_4345; YMR261C; YMR262W | DI49_4345 | | YMR261C; YMR262W |
| DI49_4344; YMR260C; DI49_4343 | DI49_4344; DI49_4343 | | YMR260C |
| DI49_4341; YMR257C; YMR258C | DI49_4341 | | YMR257C; YMR258C |
| DI49_4340; YMR256C; DI49_4339 | DI49_4340; DI49_4339 | | YMR256C |

Create PDF in your applications with the Pdfcrowd HTML to PDF API PDFCROWD

| | | | |
|---|---|---|---|
| DI49_4337; DI49_4338; YMR253C; YMR255W | DI49_4337; DI49_4338 | | YMR253C; YMR255W |
| DI49_4335; DI49_4336 | DI49_4335; DI49_4336 | | |
| DI49_4330; DI49_4331; YMR246W; YMR247C | DI49_4330; DI49_4331 | | YMR246W; YMR247C |
| DI49_4328; YMR243C; YMR244W | DI49_4328 | | YMR243C; YMR244W |
| DI49_4326; YMR241W; DI49_4325 | DI49_4326; DI49_4325 | | YMR241W |
| DI49_4322; YMR238W | DI49_4322 | | YMR238W |
| DI49_4321; YMR237W; DI49_4320 | DI49_4321; DI49_4320 | | YMR237W |
| DI49_4318; YMR234W; DI49_4317; YMR235C | DI49_4318; DI49_4317 | | YMR234W; YMR235C |
| DI49_4315; YMR231W | DI49_4315 | | YMR231W |
| DI49_4313; YMR229C | DI49_4313 | | YMR229C |
| DI49_4312; YMR228W | DI49_4312 | | YMR228W |
| DI49_4311; YMR227C | DI49_4311 | | YMR227C |
| DI49_4310; YMR226C; DI49_4309 | DI49_4310; DI49_4309 | | YMR226C |
| DI49_4307; DI49_4308; YMR224C; YMR225C | DI49_4307; DI49_4308 | | YMR224C; YMR225C |
| DI49_4305; YMR221C; YMR222C | DI49_4305 | | YMR221C; YMR222C |
| DI49_4304; YMR220W; DI49_4303 | DI49_4304; DI49_4303 | | YMR220W |
| DI49_4301; YMR217W; YMR218C | DI49_4301 | | YMR217W; YMR218C |
| DI49_4300; YMR216C; DI49_4299 | DI49_4300; DI49_4299 | | YMR216C |
| DI49_4297; DI49_4298 | DI49_4297; DI49_4298 | | |
| DI49_4295; DI49_4296 | DI49_4295; DI49_4296 | | |
| DI49_4293; DI49_4294 | DI49_4293; DI49_4294 | | |
| DI49_4291; YMR207C; DI49_4290 | DI49_4291; DI49_4290 | | YMR207C |
| DI49_4288; YMR204C | DI49_4288 | | YMR204C |
| DI49_4287; YMR203W | DI49_4287 | | YMR203W |
| DI49_4286; YMR202W; DI49_4285 | DI49_4286; DI49_4285 | | YMR202W |
| DI49_4283; YMR199W; DI49_4282; YMR200W | DI49_4283; DI49_4282 | | YMR199W; YMR200W |
| DI49_4280; YMR196W; DI49_4279 | DI49_4280; DI49_4279 | | YMR196W |
| DI49_4277; YMR194W; DI49_4276; YMR194C-B | DI49_4277; DI49_4276 | | YMR194W; YMR194C-B |
| DI49_4274; DI49_4275 | DI49_4274; DI49_4275 | | |
| DI49_4272; YMR189W; DI49_4271 | DI49_4272; DI49_4271 | | YMR189W |
| DI49_4269; SPGP0UV00150; DI49_5230; YMR186W | DI49_4269; DI49_5230 | SPGP0UV00150 | YMR186W |
| DI49_4268; YMR185W; DI49_4267 | DI49_4268; DI49_4267 | | YMR185W |
| DI49_4265; DI49_4266 | DI49_4265; DI49_4266 | | |
| DI49_4263; DI49_4264 | DI49_4263; DI49_4264 | | |
| DI49_4261; DI49_4262 | DI49_4261; DI49_4262 | | |
| DI49_4259; YMR176W; SPGP0C01420 | DI49_4259 | SPGP0C01420 | YMR176W |
| DI49_4257; DI49_4258 | DI49_4257; DI49_4258 | | |
| DI49_4255; YMR171C; YMR172W | DI49_4255 | | YMR171C; YMR172W |
| DI49_4254; DI49_4253; DI49_4252 | DI49_4254; DI49_4253; DI49_4252 | | |
| DI49_4250; YMR166C; YMR167W | DI49_4250 | | YMR166C; YMR167W |

Create PDF in your applications with the Pdfcrowd HTML to PDF API PDFCROWD

| | | | |
|---|---|---|---|
| DI49_4249; YMR165C; SPGP0C01530 | DI49_4249 | SPGP0C01530 | YMR165C |
| DI49_4245; YMR161W; DI49_4244 | DI49_4245; DI49_4244 | | YMR161W |
| DI49_4240; DI49_4241 | DI49_4240; DI49_4241 | | |
| DI49_4238; DI49_4239 | DI49_4238; DI49_4239 | | |
| DI49_4236; DI49_4237 | DI49_4236; DI49_4237 | | |
| DI49_4234; DI49_4235 | DI49_4234; DI49_4235 | | |
| DI49_4232; DI49_4233 | DI49_4232; DI49_4233 | | |
| DI49_4230; YMR145C | DI49_4230 | | YMR145C |
| DI49_4229; YMR144W | DI49_4229 | | YMR144W |
| SPGP0C01730; YMR143W | | SPGP0C01730 | YMR143W |
| DI49_4226; DI49_4227 | DI49_4226; DI49_4227 | | |
| DI49_4224; DI49_4225 | DI49_4224; DI49_4225 | | |
| DI49_4222; DI49_4223 | DI49_4222; DI49_4223 | | |
| DI49_4221; SPGP0C01820 | DI49_4221 | SPGP0C01820 | |
| DI49_4219; DI49_4220 | DI49_4219; DI49_4220 | | |
| DI49_4217; YMR128W | DI49_4217 | | YMR128W |
| DI49_4216; YMR127C; DI49_4215 | DI49_4216; DI49_4215 | | YMR127C |
| DI49_4213; YMR124W; DI49_4212 | DI49_4213; DI49_4212 | | YMR124W |
| DI49_4209; YMR120C; DI49_4208 | DI49_4209; DI49_4208 | | YMR120C |
| DI49_4206; SPGP0Ctrna12L | DI49_4206 | SPGP0Ctrna12L | |
| SPGP0C01990; YMR116C; YMR117C | | SPGP0C01990 | YMR116C; YMR117C |
| SPGP0C02000; YMR115W | | SPGP0C02000 | YMR115W |
| DI49_4203; YMR114C | DI49_4203 | | YMR114C |
| DI49_4202; YMR113W; DI49_4201 | DI49_4202; DI49_4201 | | YMR113W |
| DI49_4199; DI49_4200 | DI49_4199; DI49_4200 | | |
| DI49_4197; DI49_4198; YMR109W; YMR110C | DI49_4197; DI49_4198 | | YMR109W; YMR110C |
| DI49_4192; DI49_4193 | DI49_4192; DI49_4193 | | |
| DI49_4190; YMR101C; DI49_4189 | DI49_4190; DI49_4189 | | YMR101C |
| DI49_4186; YMR098C; DI49_4185 | DI49_4186; DI49_4185 | | YMR098C |
| DI49_4183; DI49_4184 | DI49_4183; DI49_4184 | | |
| DI49_4181; YMR093W; DI49_4180 | DI49_4181; DI49_4180 | | YMR093W |
| DI49_4178; DI49_2707 | DI49_4178; DI49_2707 | | |
| DI49_4177; YMR089C; YMR090W | DI49_4177 | | YMR089C; YMR090W |
| DI49_4176; YMR088C; DI49_4175 | DI49_4176; DI49_4175 | | YMR088C |
| DI49_4173; YMR085W | DI49_4173 | | YMR085W |
| DI49_4172; YMR083W; DI49_4171 | DI49_4172; DI49_4171 | | YMR083W |
| DI49_4169; DI49_4170 | DI49_4169; DI49_4170 | | |
| DI49_4167; DI49_4168 | DI49_4167; DI49_4168 | | |
| DI49_4165; YMR076C | DI49_4165 | | YMR076C |
| DI49_4164; YMR075W; DI49_4163 | DI49_4164; DI49_4163 | | YMR075W |

Create PDF in your applications with the Pdfcrowd HTML to PDF API PDFCROWD

| | | | |
|---|---|---|---|
| DI49_4161; DI49_4162 | DI49_4161; DI49_4162 | | |
| DI49_4159; DI49_4160; YMR071C; YMR072W | DI49_4159; DI49_4160 | | YMR071C; YMR072W |
| DI49_4157; YMR068W; DI49_4156; YMR069W | DI49_4157; DI49_4156 | | YMR068W; YMR069W |
| DI49_4154; YMR065W; DI49_4153; YMR066W | DI49_4154; DI49_4153 | | YMR065W; YMR066W |
| DI49_4151; DI49_4152 | DI49_4151; DI49_4152 | | |
| DI49_4149; DI49_4150; YMR061W; YMR062C | DI49_4149; DI49_4150 | | YMR061W; YMR062C |
| DI49_4147; YMR058W; YMR059W | DI49_4147 | | YMR058W; YMR059W |
| DI49_4146; YMR056C; DI49_4145 | DI49_4146; DI49_4145 | | YMR056C |
| DI49_4143; YMR053C; DI49_4142 | DI49_4143; DI49_4142 | | YMR053C |
| DI49_4140; DI49_3481; SPGP0C02610; SPGP0QK00160 | DI49_4140; DI49_3481 | SPGP0C02610; SPGP0QK00160 | |
| DI49_4136; DI49_4137 | DI49_4136; DI49_4137 | | |
| DI49_4131; DI49_4132 | DI49_4131; DI49_4132 | | |
| DI49_4129; SPGP0Ctrna5F | DI49_4129 | SPGP0Ctrna5F | |
| DI49_4127; DI49_4128 | DI49_4127; DI49_4128 | | |
| DI49_4125; DI49_4126 | DI49_4125; DI49_4126 | | |
| DI49_4123; DI49_4124 | DI49_4123; DI49_4124 | | |
| DI49_4121; DI49_4122 | DI49_4121; DI49_4122 | | |
| DI49_4119; DI49_4120 | DI49_4119; DI49_4120 | | |
| DI49_4117; DI49_4118; YMR030W; YMR031C | DI49_4117; DI49_4118 | | YMR030W; YMR031C |
| DI49_4115; YMR027W; DI49_4114; YMR028W | DI49_4115; DI49_4114 | | YMR027W; YMR028W |
| DI49_4112; YMR024W; YMR025W | DI49_4112 | | YMR024W; YMR025W |
| DI49_4108; YMR021C; YMR022W | DI49_4108 | | YMR021C; YMR022W |
| DI49_4107; YMR020W; DI49_4106 | DI49_4107; DI49_4106 | | YMR020W |
| DI49_4104; DI49_4105 | DI49_4104; DI49_4105 | | |
| DI49_4102; YMR015C; YMR016C | DI49_4102 | | YMR015C; YMR016C |
| DI49_4101; YMR014W | DI49_4101 | | YMR014W |
| DI49_4100; YMR013C | DI49_4100 | | YMR013C |
| DI49_4099; YMR012W | DI49_4099 | | YMR012W |
| SPGP0C03000; YMR011W | | SPGP0C03000 | YMR011W |
| DI49_4097; YMR010W; DI49_4096 | DI49_4097; DI49_4096 | | YMR010W |
| DI49_4094; YMR006C; DI49_4093; YMR007W | DI49_4094; DI49_4093 | | YMR006C; YMR007W |
| DI49_4091; DI49_4092 | DI49_4091; DI49_4092 | | |
| DI49_4089; YMR001C | DI49_4089 | | YMR001C |
| DI49_4088; YML001W | DI49_4088 | | YML001W |
| DI49_4087; YML002W | DI49_4087 | | YML002W |
| DI49_4086; YML004C | DI49_4086 | | YML004C |
| DI49_4085; YML005W | DI49_4085 | | YML005W |
| DI49_4083; YML006C | DI49_4083 | | YML006C |
| DI49_4082; YML007W | DI49_4082 | | YML007W |
| DI49_4081; YML008C | DI49_4081 | | YML008C |

Create PDF in your applications with the Pdfcrowd HTML to PDF API PDFCROWD

| | | | |
|---|---|---|---|
| DI49_4080; YML009C | DI49_4080 | | YML009C |
| DI49_4079; YML010W; DI49_4078 | DI49_4079; DI49_4078 | | YML010W |
| DI49_4076; DI49_4077 | DI49_4076; DI49_4077 | | |
| DI49_4074; DI49_4075 | DI49_4074; DI49_4075 | | |
| DI49_4072; DI49_4073 | DI49_4072; DI49_4073 | | |
| DI49_4070; DI49_4071 | DI49_4070; DI49_4071 | | |
| DI49_4068; YML021C; DI49_4067 | DI49_4068; DI49_4067 | | YML021C |
| SPGP0C03320; DI49_0614; DI49_4064; DI49_0615 | DI49_0614; DI49_4064; DI49_0615 | SPGP0C03320 | |
| DI49_4062; YML026C; DI49_0617; DI49_0618 | DI49_4062; DI49_0617; DI49_0618 | | YML026C |
| DI49_4060; DI49_4061; YML028W; YML027W | DI49_4060; DI49_4061 | | YML028W; YML027W |
| DI49_4058; YML031W; YML030W | DI49_4058 | | YML031W; YML030W |
| DI49_4057; YML032C | DI49_4057 | | YML032C |
| SPGP0C03410; YML034W | | SPGP0C03410 | YML034W |
| DI49_4055; YML035C; DI49_4054 | DI49_4055; DI49_4054 | | YML035C |
| DI49_4052; YML038C; YML037C | DI49_4052 | | YML038C; YML037C |
| DI49_4050; YML041C | DI49_4050 | | YML041C |
| DI49_4049; YML042W; DI49_4048 | DI49_4049; DI49_4048 | | YML042W |
| DI49_4044; YML048W; SPGP0C03520; YML047C | DI49_4044 | SPGP0C03520 | YML048W; YML047C |
| DI49_4041; YML051W; YML050W | DI49_4041 | | YML051W; YML050W |
| SPGP0OK00100; DI49_4786 | DI49_4786 | SPGP0OK00100 | |
| DI49_4727; DI49_4726 | DI49_4727; DI49_4726 | | |
| DI49_4729; DI49_4728 | DI49_4729; DI49_4728 | | |
| DI49_2034; DI49_2035 | DI49_2034; DI49_2035 | | |
| DI49_2032; DI49_2033 | DI49_2032; DI49_2033 | | |
| DI49_2030; DI49_2031 | DI49_2030; DI49_2031 | | |
| SPGP0OM00100; DI49_2029 | DI49_2029 | SPGP0OM00100 | |
| SPGP0OM00120; DI49_5379 | DI49_5379 | SPGP0OM00120 | |
| YAR033W; DI49_2252 | DI49_2252 | | YAR033W |
| YAR029W; YAR031W | | | YAR029W; YAR031W |
| YAR028W; YAR027W | | | YAR028W; YAR027W |
| SPGP0OPtrna15S; SPGP0Itrna30S; SPGP0Ftrna24S; YBL005W; SPGP0Htrna29S; SPGP0I01060 | | SPGP0OPtrna15S; SPGP0Itrna30S; SPGP0Ftrna24S; SPGP0Htrna29S; SPGP0I01060 | YBL005W |
| SPGP0OPtrna16L; DI49_1927; SPGP0BGtrna21L; YMR119W | DI49_1927 | SPGP0OPtrna16L; SPGP0BGtrna21L | YMR119W |
| DI49_5255; DI49_5254 | DI49_5255; DI49_5254 | | |
| DI49_5257; DI49_5256 | DI49_5257; DI49_5256 | | |
| DI49_5259; DI49_5258 | DI49_5259; DI49_5258 | | |
| DI49_5261; DI49_5260 | DI49_5261; DI49_5260 | | |
| DI49_5263; DI49_5262 | DI49_5263; DI49_5262 | | |
| SPGP0OS00110; DI49_4797 | DI49_4797 | SPGP0OS00110 | |
| DI49_5570; DI49_5569 | DI49_5570; DI49_5569 | | |

Create PDF in your applications with the Pdfcrowd HTML to PDF API PDFCROWD

| | | | |
|---|---|---|---|
| DI49_5572; DI49_5571 | DI49_5572; DI49_5571 | | |
| DI49_5574; DI49_5573 | DI49_5574; DI49_5573 | | |
| DI49_4804; YBR093C; SPGP0OV00100 | DI49_4804 | SPGP0OV00100 | YBR093C |
| DI49_5235; DI49_5234 | DI49_5235; DI49_5234 | | |
| DI49_5237; DI49_5236 | DI49_5237; DI49_5236 | | |
| DI49_5239; DI49_5238 | DI49_5239; DI49_5238 | | |
| DI49_5241; DI49_5240 | DI49_5241; DI49_5240 | | |
| DI49_4589; DI49_5242 | DI49_4589; DI49_5242 | | |
| DI49_4587; DI49_4588 | DI49_4587; DI49_4588 | | |
| DI49_4585; DI49_4586 | DI49_4585; DI49_4586 | | |
| DI49_4583; DI49_4584 | DI49_4583; DI49_4584 | | |
| DI49_4581; DI49_4582 | DI49_4581; DI49_4582 | | |
| DI49_4579; DI49_4580 | DI49_4579; DI49_4580 | | |
| DI49_4577; DI49_4578 | DI49_4577; DI49_4578 | | |
| DI49_4572; DI49_4573 | DI49_4572; DI49_4573 | | |
| DI49_4570; DI49_4571 | DI49_4570; DI49_4571 | | |
| DI49_3300; DI49_4569 | DI49_3300; DI49_4569 | | |
| DI49_2162; DI49_2161 | DI49_2162; DI49_2161 | | |
| DI49_2164; DI49_2163 | DI49_2164; DI49_2163 | | |
| DI49_2166; DI49_2165 | DI49_2166; DI49_2165 | | |
| DI49_2168; DI49_2167 | DI49_2168; DI49_2167 | | |
| DI49_3571; DI49_2169 | DI49_3571; DI49_2169 | | |
| DI49_3569; DI49_3570 | DI49_3569; DI49_3570 | | |
| DI49_3567; DI49_3568 | DI49_3567; DI49_3568 | | |
| DI49_3561; DI49_3562 | DI49_3561; DI49_3562 | | |
| DI49_1958; DI49_3560 | DI49_1958; DI49_3560 | | |
| DI49_2111; DI49_2109 | DI49_2111; DI49_2109 | | |
| DI49_2118; DI49_2117 | DI49_2118; DI49_2117 | | |
| DI49_2120; DI49_2119 | DI49_2120; DI49_2119 | | |
| DI49_2122; DI49_2121 | DI49_2122; DI49_2121 | | |
| DI49_2131; DI49_2130 | DI49_2131; DI49_2130 | | |
| DI49_2133; DI49_2132 | DI49_2133; DI49_2132 | | |
| DI49_2135; DI49_2134 | DI49_2135; DI49_2134 | | |
| DI49_2137; DI49_2136 | DI49_2137; DI49_2136 | | |
| DI49_1695; SPGP0PF00110; DI49_5623 | DI49_1695; DI49_5623 | SPGP0PF00110 | |
| DI49_5621; DI49_5622 | DI49_5621; DI49_5622 | | |
| DI49_5619; DI49_5620 | DI49_5619; DI49_5620 | | |
| DI49_5617; DI49_5618 | DI49_5617; DI49_5618 | | |
| DI49_5615; DI49_5616 | DI49_5615; DI49_5616 | | |
| DI49_1851; DI49_1850 | DI49_1851; DI49_1850 | | |

Create PDF in your applications with the Pdfcrowd HTML to PDF API PDFCROWD

| | | | |
|---|---|---|---|
| DI49_3517; DI49_3518 | DI49_3517; DI49_3518 | | |
| DI49_3515; DI49_3516 | DI49_3515; DI49_3516 | | |
| DI49_4611; DI49_3514 | DI49_4611; DI49_3514 | | |
| DI49_4609; DI49_4610 | DI49_4609; DI49_4610 | | |
| DI49_4607; DI49_4608 | DI49_4607; DI49_4608 | | |
| DI49_4605; DI49_4606 | DI49_4605; DI49_4606 | | |
| DI49_4603; DI49_2475; DI49_4602; DI49_2476 | DI49_4603; DI49_2475; DI49_4602; DI49_2476 | | |
| DI49_4600; DI49_4601 | DI49_4600; DI49_4601 | | |
| DI49_4598; DI49_4599 | DI49_4598; DI49_4599 | | |
| DI49_4596; DI49_4597 | DI49_4596; DI49_4597 | | |
| DI49_4594; DI49_4595 | DI49_4594; DI49_4595 | | |
| SPGP0PM00110; SPGP0D00100; YGR289C; DI49_5494 | DI49_5494 | SPGP0PM00110; SPGP0D00100 | YGR289C |
| DI49_5496; DI49_5495 | DI49_5496; DI49_5495 | | |
| DI49_5432; DI49_5497 | DI49_5432; DI49_5497 | | |
| DI49_0728; DI49_0729 | DI49_0728; DI49_0729 | | |
| DI49_0601; DI49_0727 | DI49_0601; DI49_0727 | | |
| SPGP0PV00110; DI49_5406 | DI49_5406 | SPGP0PV00110 | |
| DI49_5408; DI49_5407 | DI49_5408; DI49_5407 | | |
| DI49_1984; SPGP0PW00100 | DI49_1984 | SPGP0PW00100 | |
| DI49_2199; DI49_2200 | DI49_2199; DI49_2200 | | |
| DI49_2197; DI49_2198 | DI49_2197; DI49_2198 | | |
| DI49_2195; DI49_2196 | DI49_2195; DI49_2196 | | |
| DI49_2192; DI49_2194 | DI49_2192; DI49_2194 | | |
| YGR225W; DI49_2191 | DI49_2191 | | YGR225W |
| DI49_2189; DI49_2190 | DI49_2189; DI49_2190 | | |
| DI49_2187; DI49_2188 | DI49_2187; DI49_2188 | | |
| DI49_2182; DI49_2183 | DI49_2182; DI49_2183 | | |
| DI49_2180; DI49_2181 | DI49_2180; DI49_2181 | | |
| DI49_2178; DI49_2179 | DI49_2178; DI49_2179 | | |
| DI49_2176; DI49_2177 | DI49_2176; DI49_2177 | | |
| DI49_2174; DI49_2175 | DI49_2174; DI49_2175 | | |
| DI49_2172; DI49_2173 | DI49_2172; DI49_2173 | | |
| DI49_2170; DI49_2171 | DI49_2170; DI49_2171 | | |
| DI49_2621; DI49_0640 | DI49_2621; DI49_0640 | | |
| YGL183C; YGL184C | | | YGL183C; YGL184C |
| YGL180W; YGL181W | | | YGL180W; YGL181W |
| DI49_0661; YGL179C | DI49_0661 | | YGL179C |
| DI49_0663; DI49_0662 | DI49_0663; DI49_0662 | | |
| DI49_0665; DI49_3697; DI49_0666; DI49_3698 | DI49_0665; DI49_3697; DI49_0666; DI49_3698 | | |
| DI49_0668; DI49_0667 | DI49_0668; DI49_0667 | | |

Create PDF in your applications with the Pdfcrowd HTML to PDF API PDFCROWD

| | | | |
|---|---|---|---|
| YAL049C; YAL048C | | | YAL049C; YAL048C |
| YAL053W; YAL051W | | | YAL053W; YAL051W |
| DI49_4799; DI49_4800 | DI49_4799; DI49_4800 | | |
| DI49_3476; DI49_4798 | DI49_3476; DI49_4798 | | |
| DI49_3478; DI49_3477 | DI49_3478; DI49_3477 | | |
| DI49_3480; DI49_3479 | DI49_3480; DI49_3479 | | |
| DI49_1606; DI49_3482; YGL236C | DI49_1606; DI49_3482 | | YGL236C |
| YGL238W; YGL237C | | | YGL238W; YGL237C |
| DI49_5611; DI49_5612 | DI49_5611; DI49_5612 | | |
| DI49_5609; DI49_5610 | DI49_5609; DI49_5610 | | |
| DI49_5607; DI49_5608 | DI49_5607; DI49_5608 | | |
| DI49_5605; DI49_5606 | DI49_5605; DI49_5606 | | |
| DI49_5602; DI49_2105; YOR388C | DI49_5602; DI49_2105 | | YOR388C |
| YAR015W; DI49_0069 | DI49_0069 | | YAR015W |
| YAR008W; DI49_0063 | DI49_0063 | | YAR008W |
| SPGP0QR00110; SPGP0QR00100 | | SPGP0QR00110; SPGP0QR00100 | |
| DI49_0536; DI49_0535 | DI49_0536; DI49_0535 | | |
| DI49_0538; DI49_0537 | DI49_0538; DI49_0537 | | |
| DI49_4033; DI49_0539 | DI49_4033; DI49_0539 | | |
| DI49_5450; DI49_4032 | DI49_5450; DI49_4032 | | |
| DI49_4029; DI49_4030; DI49_2437 | DI49_4029; DI49_4030; DI49_2437 | | |
| DI49_4027; DI49_4028 | DI49_4027; DI49_4028 | | |
| DI49_4025; DI49_4026 | DI49_4025; DI49_4026 | | |
| DI49_4023; DI49_4024 | DI49_4023; DI49_4024 | | |
| DI49_4021; DI49_4022 | DI49_4021; DI49_4022 | | |
| DI49_4016; DI49_2435; DI49_4015 | DI49_4016; DI49_2435; DI49_4015 | | |
| DI49_1328; DI49_4012 | DI49_1328; DI49_4012 | | |
| DI49_1330; DI49_1329 | DI49_1330; DI49_1329 | | |
| DI49_1332; DI49_1331 | DI49_1332; DI49_1331 | | |
| DI49_1334; DI49_1333 | DI49_1334; DI49_1333 | | |
| YMR318C; YMR319C | | | YMR318C; YMR319C |
| SPGP0QX00130; YMR317W | | SPGP0QX00130 | YMR317W |
| YMR315W; YMR316W | | | YMR315W; YMR316W |
| YMR313C; YMR314W | | | YMR313C; YMR314W |
| YMR311C; YMR312W | | | YMR311C; YMR312W |
| YMR309C; YMR310C | | | YMR309C; YMR310C |
| YMR307W; YMR308C | | | YMR307W; YMR308C |
| DI49_5267; YMR306W | DI49_5267 | | YMR306W |
| DI49_5270; DI49_5268 | DI49_5270; DI49_5268 | | |
| DI49_5431; YPL195W | DI49_5431 | | YPL195W |

Create PDF in your applications with the Pdfcrowd HTML to PDF API PDFCROWD

| | | | |
|---|---|---|---|
| DI49_5429; DI49_5430 | DI49_5429; DI49_5430 | | |
| DI49_5427; DI49_5428 | DI49_5427; DI49_5428 | | |
| DI49_1819; DI49_5426 | DI49_1819; DI49_5426 | | |
| DI49_1823; DI49_1822 | DI49_1823; DI49_1822 | | |
| DI49_1825; DI49_1824 | DI49_1825; DI49_1824 | | |
| DI49_1827; DI49_1826 | DI49_1827; DI49_1826 | | |
| DI49_1829; DI49_1828 | DI49_1829; DI49_1828 | | |
| DI49_1831; DI49_1830 | DI49_1831; DI49_1830 | | |
| DI49_1833; DI49_1832 | DI49_1833; DI49_1832 | | |
| DI49_1835; DI49_1834 | DI49_1835; DI49_1834 | | |
| DI49_1837; DI49_1836 | DI49_1837; DI49_1836 | | |
| DI49_1839; DI49_1838 | DI49_1839; DI49_1838 | | |
| DI49_3756; DI49_1840 | DI49_3756; DI49_1840 | | |
| DI49_3492; DI49_3491 | DI49_3492; DI49_3491 | | |
| DI49_3496; DI49_3495 | DI49_3496; DI49_3495 | | |
| YAL059W; YAL060W | | | YAL059W; YAL060W |
| DI49_3512; DI49_3513 | DI49_3512; DI49_3513 | | |
| DI49_3586; DI49_3585 | DI49_3586; DI49_3585 | | |
| DI49_3588; DI49_3587 | DI49_3588; DI49_3587 | | |
| DI49_3590; DI49_3589 | DI49_3590; DI49_3589 | | |
| DI49_3592; DI49_3591 | DI49_3592; DI49_3591 | | |
| DI49_3594; DI49_3593 | DI49_3594; DI49_3593 | | |
| DI49_3596; DI49_3595 | DI49_3596; DI49_3595 | | |
| DI49_3598; DI49_3597 | DI49_3598; DI49_3597 | | |
| DI49_3600; DI49_3599 | DI49_3600; DI49_3599 | | |
| DI49_3602; DI49_3601 | DI49_3602; DI49_3601 | | |
| DI49_3604; DI49_3603 | DI49_3604; DI49_3603 | | |
| DI49_3606; DI49_3605 | DI49_3606; DI49_3605 | | |
| DI49_3609; DI49_3607 | DI49_3609; DI49_3607 | | |
| DI49_3611; DI49_3610 | DI49_3611; DI49_3610 | | |
| DI49_3613; DI49_3612 | DI49_3613; DI49_3612 | | |
| DI49_3615; DI49_3614 | DI49_3615; DI49_3614 | | |
| DI49_1617; DI49_3616 | DI49_1617; DI49_3616 | | |
| DI49_0685; DI49_0686 | DI49_0685; DI49_0686 | | |
| DI49_0683; DI49_0684 | DI49_0683; DI49_0684 | | |
| DI49_0681; DI49_0682 | DI49_0681; DI49_0682 | | |
| DI49_0680; SPGP0RK00160 | DI49_0680 | SPGP0RK00160 | |
| DI49_0679; SPGP0RK00180 | DI49_0679 | SPGP0RK00180 | |
| DI49_1974; DI49_1975 | DI49_1974; DI49_1975 | | |
| YER168C; SPGP0RM00100 | | SPGP0RM00100 | YER168C |

Create PDF in your applications with the Pdfcrowd HTML to PDF API PDFCROWD

| | | | |
|---|---|---|---|
| YER166W; YER167W | | | YER166W; YER167W |
| YER164W; YER165W | | | YER164W; YER165W |
| DI49_1573; SPGP0RN00150 | DI49_1573 | SPGP0RN00150 | |
| DI49_3466; DI49_3465 | DI49_3466; DI49_3465 | | |
| DI49_3468; DI49_3467 | DI49_3468; DI49_3467 | | |
| DI49_3470; DI49_3469 | DI49_3470; DI49_3469 | | |
| SPGP0RQ00200; DI49_3471 | DI49_3471 | SPGP0RQ00200 | |
| DI49_0635; DI49_3472 | DI49_0635; DI49_3472 | | |
| DI49_0637; DI49_0636 | DI49_0637; DI49_0636 | | |
| DI49_0639; DI49_0638 | DI49_0639; DI49_0638 | | |
| DI49_1554; DI49_1553 | DI49_1554; DI49_1553 | | |
| DI49_1556; DI49_1555 | DI49_1556; DI49_1555 | | |
| DI49_1558; DI49_1557 | DI49_1558; DI49_1557 | | |
| DI49_1562; DI49_1561 | DI49_1562; DI49_1561 | | |
| DI49_1564; DI49_1563 | DI49_1564; DI49_1563 | | |
| YER172C; YER173W | | | YER172C; YER173W |
| YER170W; YER171W | | | YER170W; YER171W |
| DI49_0751; YCR002C | DI49_0751 | | YCR002C |
| DI49_0752; YCR003W | DI49_0752 | | YCR003W |
| DI49_0753; YCR004C; DI49_0754; YCR005C | DI49_0753; DI49_0754 | | YCR004C; YCR005C |
| DI49_2047; DI49_2046 | DI49_2047; DI49_2046 | | |
| YER176W; DI49_2050 | DI49_2050 | | YER176W |
| YER174C; YER175C | | | YER174C; YER175C |
| DI49_0782; YCR030C | DI49_0782 | | YCR030C |
| DI49_2256; DI49_2255 | DI49_2256; DI49_2255 | | |
| DI49_2264; DI49_2263 | DI49_2264; DI49_2263 | | |
| DI49_2293; DI49_2292 | DI49_2293; DI49_2292 | | |
| DI49_2303; DI49_2302 | DI49_2303; DI49_2302 | | |
| DI49_2319; DI49_2318 | DI49_2319; DI49_2318 | | |
| DI49_2323; DI49_2322 | DI49_2323; DI49_2322 | | |
| DI49_2327; DI49_2326 | DI49_2327; DI49_2326 | | |
| DI49_2333; DI49_2332 | DI49_2333; DI49_2332 | | |
| DI49_2339; DI49_2338 | DI49_2339; DI49_2338 | | |
| DI49_2341; DI49_2340 | DI49_2341; DI49_2340 | | |
| DI49_2353; DI49_2352 | DI49_2353; DI49_2352 | | |
| SPGP0D01140; SPGP0D01130 | | SPGP0D01140; SPGP0D01130 | |
| DI49_2360; DI49_2359 | DI49_2360; DI49_2359 | | |
| DI49_2369; DI49_2368 | DI49_2369; DI49_2368 | | |
| DI49_2371; DI49_2370 | DI49_2371; DI49_2370 | | |
| DI49_2373; DI49_2372 | DI49_2373; DI49_2372 | | |

Create PDF in your applications with the Pdfcrowd HTML to PDF API PDFCROWD

| | | | |
|---|---|---|---|
| DI49_2375; SPGP0Dtrna14S | DI49_2375 | SPGP0Dtrna14S | |
| DI49_2385; DI49_2384 | DI49_2385; DI49_2384 | | |
| DI49_2387; DI49_2386 | DI49_2387; DI49_2386 | | |
| DI49_2395; DI49_2394 | DI49_2395; DI49_2394 | | |
| DI49_2417; YOR014W | DI49_2417 | | YOR014W |
| DI49_2438; YHR032W | DI49_2438 | | YHR032W |
| DI49_2449; DI49_2448 | DI49_2449; DI49_2448 | | |
| DI49_2451; DI49_2450 | DI49_2451; DI49_2450 | | |
| DI49_2453; DI49_2452 | DI49_2453; DI49_2452 | | |
| DI49_2455; DI49_2454 | DI49_2455; DI49_2454 | | |
| DI49_2460; DI49_2459 | DI49_2460; DI49_2459 | | |
| DI49_2465; DI49_2464 | DI49_2465; DI49_2464 | | |
| DI49_2469; DI49_0490; DI49_2470; DI49_0489 | DI49_2469; DI49_0490; DI49_2470; DI49_0489 | | |
| DI49_2473; DI49_2883; DI49_2474; YJL141C | DI49_2473; DI49_2883; DI49_2474 | | YJL141C |
| DI49_2478; DI49_2477 | DI49_2478; DI49_2477 | | |
| DI49_2486; DI49_2485 | DI49_2486; DI49_2485 | | |
| DI49_2488; DI49_2487 | DI49_2488; DI49_2487 | | |
| DI49_2490; DI49_2489 | DI49_2490; DI49_2489 | | |
| DI49_2493; YHR086W | DI49_2493 | | YHR086W |
| DI49_2510; DI49_2509 | DI49_2510; DI49_2509 | | |
| DI49_2515; DI49_2514 | DI49_2515; DI49_2514 | | |
| DI49_2517; DI49_2516 | DI49_2517; DI49_2516 | | |
| DI49_2522; DI49_2521 | DI49_2522; DI49_2521 | | |
| DI49_2527; DI49_2526 | DI49_2527; DI49_2526 | | |
| DI49_2539; DI49_2538 | DI49_2539; DI49_2538 | | |
| SPGP0D02950; DI49_2540 | DI49_2540 | SPGP0D02950 | |
| DI49_2542; DI49_2541 | DI49_2542; DI49_2541 | | |
| DI49_2547; DI49_2546 | DI49_2547; DI49_2546 | | |
| DI49_2549; DI49_2548 | DI49_2549; DI49_2548 | | |
| DI49_2555; DI49_2554 | DI49_2555; DI49_2554 | | |
| DI49_2561; DI49_2560 | DI49_2561; DI49_2560 | | |
| DI49_2563; DI49_2562 | DI49_2563; DI49_2562 | | |
| DI49_2565; DI49_2564 | DI49_2565; DI49_2564 | | |
| SPGP0D03320; DI49_2576 | DI49_2576 | SPGP0D03320 | |
| SPGP0D03360; DI49_2579 | DI49_2579 | SPGP0D03360 | |
| DI49_2586; DI49_2585 | DI49_2586; DI49_2585 | | |
| DI49_2592; DI49_2591 | DI49_2592; DI49_2591 | | |
| DI49_2594; DI49_2593 | DI49_2594; DI49_2593 | | |
| DI49_2603; DI49_2602 | DI49_2603; DI49_2602 | | |
| DI49_2605; DI49_2604 | DI49_2605; DI49_2604 | | |

Create PDF in your applications with the Pdfcrowd HTML to PDF API PDFCROWD

| | | | |
|---|---|---|---|
| DI49_2607; DI49_2606 | DI49_2607; DI49_2606 | | |
| SPGP0SF00100; SPGP0BN00100; DI49_0610; YHR205W | DI49_0610 | SPGP0SF00100; SPGP0BN00100 | YHR205W |
| DI49_0620; DI49_0619 | DI49_0620; DI49_0619 | | |
| DI49_0622; DI49_0621 | DI49_0622; DI49_0621 | | |
| DI49_0624; DI49_0623 | DI49_0624; DI49_0623 | | |
| DI49_0626; DI49_0625 | DI49_0626; DI49_0625 | | |
| DI49_0627; SPGP0SH00260 | DI49_0627 | SPGP0SH00260 | |
| DI49_0629; DI49_0628 | DI49_0629; DI49_0628 | | |
| DI49_2028; DI49_4719 | DI49_2028; DI49_4719 | | |
| DI49_3583; DI49_2027 | DI49_3583; DI49_2027 | | |
| DI49_3581; DI49_3582 | DI49_3581; DI49_3582 | | |
| DI49_3579; DI49_3580 | DI49_3579; DI49_3580 | | |
| DI49_3577; DI49_3578 | DI49_3577; DI49_3578 | | |
| DI49_3575; SPGP0SL00170 | DI49_3575 | SPGP0SL00170 | |
| DI49_3573; DI49_3574 | DI49_3573; DI49_3574 | | |
| DI49_3446; DI49_3572; DI49_3657; DI49_3656 | DI49_3446; DI49_3572; DI49_3657; DI49_3656 | | |
| DI49_3448; DI49_3447 | DI49_3448; DI49_3447 | | |
| DI49_3450; DI49_3449 | DI49_3450; DI49_3449 | | |
| DI49_3452; DI49_3451 | DI49_3452; DI49_3451 | | |
| DI49_1718; DI49_1719 | DI49_1718; DI49_1719 | | |
| DI49_5479; DI49_1717 | DI49_5479; DI49_1717 | | |
| SPGP0AX00110; SPGP0AX00100 | | SPGP0AX00110; SPGP0AX00100 | |
| DI49_0603; DI49_0602 | DI49_0603; DI49_0602 | | |
| DI49_0605; DI49_0604 | DI49_0605; DI49_0604 | | |
| DI49_0607; DI49_0606 | DI49_0607; DI49_0606 | | |
| DI49_0609; DI49_0608 | DI49_0609; DI49_0608 | | |
| YGL192W; YGL194C | | | YGL192W; YGL194C |
| YGL190C; YGL191W | | | YGL190C; YGL191W |
| YGL185C; YGL186C | | | YGL185C; YGL186C |
| DI49_5482; DI49_5483 | DI49_5482; DI49_5483 | | |
| DI49_5480; DI49_5481 | DI49_5480; DI49_5481 | | |
| DI49_3647; SPGP0SS00140 | DI49_3647 | SPGP0SS00140 | |
| DI49_3645; DI49_3646 | DI49_3645; DI49_3646 | | |
| DI49_3643; DI49_3644 | DI49_3643; DI49_3644 | | |
| DI49_3641; DI49_3642 | DI49_3641; DI49_3642 | | |
| DI49_5403; DI49_5402 | DI49_5403; DI49_5402 | | |
| DI49_5405; DI49_5404 | DI49_5405; DI49_5404 | | |
| SPGP0SV00110; YCR097W | | SPGP0SV00110 | YCR097W |
| DI49_3661; DI49_3662 | DI49_3661; DI49_3662 | | |
| DI49_3659; DI49_3660 | DI49_3659; DI49_3660 | | |

Create PDF in your applications with the Pdfcrowd HTML to PDF API PDFCROWD

| | | | |
|---|---|---|---|
| DI49_3654; DI49_3655 | DI49_3654; DI49_3655 | | |
| DI49_3652; DI49_3653 | DI49_3652; DI49_3653 | | |
| DI49_3650; DI49_3651 | DI49_3650; DI49_3651 | | |
| DI49_3648; DI49_3649 | DI49_3648; DI49_3649 | | |
| DI49_0695; DI49_1360; DI49_3913 | DI49_0695; DI49_1360; DI49_3913 | | |
| YAL065C; DI49_3912 | DI49_3912 | | YAL065C |
| SPGP0TC00100; DI49_1536 | DI49_1536 | SPGP0TC00100 | |
| DI49_4555; DI49_4556 | DI49_4555; DI49_4556 | | |
| DI49_4553; DI49_4554 | DI49_4553; DI49_4554 | | |
| DI49_4551; DI49_4552 | DI49_4551; DI49_4552 | | |
| DI49_4549; DI49_4550 | DI49_4549; DI49_4550 | | |
| DI49_4547; DI49_4548 | DI49_4547; DI49_4548 | | |
| DI49_4545; DI49_4546 | DI49_4545; DI49_4546 | | |
| DI49_5294; SPGP0TC00230 | DI49_5294 | SPGP0TC00230 | |
| DI49_4758; DI49_4757 | DI49_4758; DI49_4757 | | |
| SPGP0TE00130; DI49_4759 | DI49_4759 | SPGP0TE00130 | |
| DI49_4761; DI49_4760 | DI49_4761; DI49_4760 | | |
| DI49_4763; DI49_0818; DI49_4764; DI49_0819 | DI49_4763; DI49_0818; DI49_4764; DI49_0819 | | |
| DI49_4766; DI49_4765 | DI49_4766; DI49_4765 | | |
| DI49_4768; DI49_4767 | DI49_4768; DI49_4767 | | |
| DI49_1926; DI49_1925 | DI49_1926; DI49_1925 | | |
| DI49_3763; DI49_3762 | DI49_3763; DI49_3762 | | |
| DI49_3765; DI49_3764 | DI49_3765; DI49_3764 | | |
| YAL061W; DI49_0003 | DI49_0003 | | YAL061W |
| DI49_1460; YER056C-A | DI49_1460 | | YER056C-A |
| DI49_1462; DI49_1461 | DI49_1462; DI49_1461 | | |
| DI49_1464; DI49_1459; SPGP0TL00100 | DI49_1464; DI49_1459 | SPGP0TL00100 | |
| DI49_5521; DI49_5522 | DI49_5521; DI49_5522 | | |
| DI49_5519; DI49_5520 | DI49_5519; DI49_5520 | | |
| DI49_5517; DI49_5518 | DI49_5517; DI49_5518 | | |
| DI49_4740; DI49_4739 | DI49_4740; DI49_4739 | | |
| DI49_4742; DI49_4741 | DI49_4742; DI49_4741 | | |
| DI49_4744; DI49_4743 | DI49_4744; DI49_4743 | | |
| DI49_4746; DI49_4745 | DI49_4746; DI49_4745 | | |
| DI49_4748; DI49_4747 | DI49_4748; DI49_4747 | | |
| DI49_4750; DI49_4749 | DI49_4750; DI49_4749 | | |
| DI49_1546; DI49_2906; DI49_1547 | DI49_1546; DI49_2906; DI49_1547 | | |
| DI49_1549; DI49_1548 | DI49_1549; DI49_1548 | | |
| DI49_1551; DI49_1550 | DI49_1551; DI49_1550 | | |
| DI49_3705; DI49_1552 | DI49_3705; DI49_1552 | | |

Create PDF in your applications with the Pdfcrowd HTML to PDF API PDFCROWD

| | | | |
|---|---|---|---|
| DI49_3707; DI49_3706 | DI49_3707; DI49_3706 | | |
| DI49_3709; DI49_3708 | DI49_3709; DI49_3708 | | |
| DI49_3711; DI49_3710 | DI49_3711; DI49_3710 | | |
| DI49_3713; DI49_3712 | DI49_3713; DI49_3712 | | |
| DI49_3715; DI49_3714 | DI49_3715; DI49_3714 | | |
| DI49_3717; DI49_3716 | DI49_3717; DI49_3716 | | |
| DI49_3719; DI49_3718 | DI49_3719; DI49_3718 | | |
| DI49_3721; DI49_3720 | DI49_3721; DI49_3720 | | |
| DI49_3723; DI49_3722 | DI49_3723; DI49_3722 | | |
| DI49_3725; DI49_3724 | DI49_3725; DI49_3724 | | |
| DI49_3727; DI49_3726 | DI49_3727; DI49_3726 | | |
| DI49_3730; DI49_3728 | DI49_3730; DI49_3728 | | |
| SPGP0TQ00360; DI49_3731 | DI49_3731 | SPGP0TQ00360 | |
| DI49_3733; DI49_3732 | DI49_3733; DI49_3732 | | |
| DI49_3735; DI49_3734 | DI49_3735; DI49_3734 | | |
| DI49_3737; DI49_3736 | DI49_3737; DI49_3736 | | |
| DI49_4442; YAL039C | DI49_4442 | | YAL039C |
| DI49_4440; DI49_4441 | DI49_4440; DI49_4441 | | |
| DI49_4438; DI49_4439 | DI49_4438; DI49_4439 | | |
| DI49_3951; DI49_3952 | DI49_3951; DI49_3952 | | |
| DI49_3949; DI49_3950 | DI49_3949; DI49_3950 | | |
| SPGP0TW00100; DI49_2108 | DI49_2108 | SPGP0TW00100 | |
| DI49_5272; DI49_5271 | DI49_5272; DI49_5271 | | |
| DI49_2102; DI49_2101 | DI49_2102; DI49_2101 | | |
| DI49_5471; DI49_5470 | DI49_5471; DI49_5470 | | |
| DI49_5473; DI49_5472 | DI49_5473; DI49_5472 | | |
| DI49_5475; DI49_5474 | DI49_5475; DI49_5474 | | |
| DI49_5477; DI49_5476 | DI49_5477; DI49_5476 | | |
| DI49_3377; DI49_5478 | DI49_3377; DI49_5478 | | |
| DI49_3374; DI49_3375 | DI49_3374; DI49_3375 | | |
| DI49_3500; DI49_3373 | DI49_3500; DI49_3373 | | |
| DI49_3502; DI49_3501 | DI49_3502; DI49_3501 | | |
| DI49_3504; DI49_3503 | DI49_3504; DI49_3503 | | |
| DI49_5627; DI49_3509 | DI49_5627; DI49_3509 | | |
| DI49_0714; YCL044C | DI49_0714 | | YCL044C |
| DI49_0713; YCL045C; DI49_0712 | DI49_0713; DI49_0712 | | YCL045C |
| DI49_0710; DI49_0711; YCL048W; YCL047C | DI49_0710; DI49_0711 | | YCL048W; YCL047C |
| DI49_0708; DI49_0709; YCL050C; YCL049C | DI49_0708; DI49_0709 | | YCL050C; YCL049C |
| DI49_0706; YCL054W; YCL052C | DI49_0706 | | YCL054W; YCL052C |

Create PDF in your applications with the Pdfcrowd HTML to PDF API PDFCROWD

| | | | |
|---|---|---|---|
| DI49_0705; YCL055W; DI49_0704 | DI49_0705; DI49_0704 | | YCL055W |
| DI49_0702; DI49_0703; YCL057W; YCL056C | DI49_0702; DI49_0703 | | YCL057W; YCL056C |
| DI49_0700; YCL059C; SPGP0UJ00260; YCL058C | DI49_0700 | SPGP0UJ00260 | YCL059C; YCL058C |
| SPGP0UJ00280; DI49_0699 | DI49_0699 | SPGP0UJ00280 | |
| DI49_4511; DI49_5467 | DI49_4511; DI49_5467 | | |
| DI49_4513; DI49_4512 | DI49_4513; DI49_4512 | | |
| DI49_4515; DI49_4514 | DI49_4515; DI49_4514 | | |
| DI49_4517; DI49_4516 | DI49_4517; DI49_4516 | | |
| DI49_4519; DI49_4518 | DI49_4519; DI49_4518 | | |
| DI49_4521; DI49_4520 | DI49_4521; DI49_4520 | | |
| DI49_1693; DI49_4522 | DI49_1693; DI49_4522 | | |
| DI49_3898; DI49_3899 | DI49_3898; DI49_3899 | | |
| DI49_3896; DI49_3897 | DI49_3896; DI49_3897 | | |
| DI49_3894; DI49_3895 | DI49_3894; DI49_3895 | | |
| DI49_3892; DI49_3893 | DI49_3892; DI49_3893 | | |
| DI49_3890; DI49_3891 | DI49_3890; DI49_3891 | | |
| DI49_3885; DI49_3886 | DI49_3885; DI49_3886 | | |
| DI49_3883; DI49_3884 | DI49_3883; DI49_3884 | | |
| DI49_3881; DI49_3882 | DI49_3881; DI49_3882 | | |
| DI49_1923; DI49_1922 | DI49_1923; DI49_1922 | | |
| DI49_1618; DI49_1924; DI49_3794 | DI49_1618; DI49_1924; DI49_3794 | | |
| DI49_3792; DI49_3793 | DI49_3792; DI49_3793 | | |
| DI49_3790; DI49_3791 | DI49_3790; DI49_3791 | | |
| DI49_2098; DI49_3789 | DI49_2098; DI49_3789 | | |
| DI49_2096; DI49_2097 | DI49_2096; DI49_2097 | | |
| DI49_2094; DI49_2095 | DI49_2094; DI49_2095 | | |
| DI49_2092; DI49_2093 | DI49_2092; DI49_2093 | | |
| DI49_2090; DI49_2091 | DI49_2090; DI49_2091 | | |
| DI49_2088; DI49_2089 | DI49_2088; DI49_2089 | | |
| DI49_5596; DI49_5597 | DI49_5596; DI49_5597 | | |
| SPGP0US00130; DI49_5595 | DI49_5595 | SPGP0US00130 | |
| DI49_1932; SPGP0UT00100 | DI49_1932 | SPGP0UT00100 | |
| YPL245W; DI49_1931 | DI49_1931 | | YPL245W |
| YPL243W; YPL244C | | | YPL243W; YPL244C |
| DI49_5232; DI49_5231 | DI49_5232; DI49_5231 | | |
| DI49_0741; YCL011C; SPGP0UW00110; YCL010C | DI49_0741 | SPGP0UW00110 | YCL011C; YCL010C |
| YAL037W; YAL038W | | | YAL037W; YAL038W |
| DI49_1934; DI49_1935 | DI49_1934; DI49_1935 | | |
| DI49_0805; DI49_0804 | DI49_0805; DI49_0804 | | |
| SPGP0UZ00130; DI49_0806 | DI49_0806 | SPGP0UZ00130 | |

Create PDF in your applications with the Pdfcrowd HTML to PDF API PDFCROWD

| | | | |
|---|---|---|---|
| DI49_0811; DI49_0810 | DI49_0811; DI49_0810 | | |
| DI49_1694; DI49_0813 | DI49_1694; DI49_0813 | | |
| DI49_3385; DI49_3386 | DI49_3385; DI49_3386 | | |
| DI49_3383; DI49_3384 | DI49_3383; DI49_3384 | | |
| SPGP0VC00100; SPGP0VBtrna20L | | SPGP0VC00100; SPGP0VBtrna20L | |
| DI49_4693; DI49_4692 | DI49_4693; DI49_4692 | | |
| DI49_4695; DI49_4694 | DI49_4695; DI49_4694 | | |
| DI49_4697; YBL019W; DI49_4698 | DI49_4697; DI49_4698 | | YBL019W |
| DI49_4700; SPGP0VDtrna16P | DI49_4700 | SPGP0VDtrna16P | |
| DI49_3296; DI49_3295 | DI49_3296; DI49_3295 | | |
| DI49_3298; DI49_3297 | DI49_3298; DI49_3297 | | |
| SPGP0VG00100; DI49_3299 | DI49_3299 | SPGP0VG00100 | |
| DI49_0507; DI49_0506 | DI49_0507; DI49_0506 | | |
| DI49_0509; DI49_0508 | DI49_0509; DI49_0508 | | |
| DI49_0511; DI49_0510 | DI49_0511; DI49_0510 | | |
| DI49_1320; YJL217W; DI49_1319; YJL218W | DI49_1320; DI49_1319 | | YJL217W; YJL218W |
| DI49_1317; YBR296C; DI49_1316; YBR295W | DI49_1317; DI49_1316 | | YBR296C; YBR295W |
| DI49_1314; YBR291C; DI49_5565 | DI49_1314; DI49_5565 | | YBR291C |
| DI49_1313; YBR290W | DI49_1313 | | YBR290W |
| DI49_1312; YBR289W | DI49_1312 | | YBR289W |
| DI49_1311; YBR288C | DI49_1311 | | YBR288C |
| DI49_1310; YBR287W; DI49_1309; YBR286W | DI49_1310; DI49_1309 | | YBR287W; YBR286W |
| DI49_1307; DI49_1308 | DI49_1307; DI49_1308 | | |
| DI49_1305; YBR282W | DI49_1305 | | YBR282W |
| DI49_1304; YBR281C; DI49_1303; YBR280C | DI49_1304; DI49_1303 | | YBR281C; YBR280C |
| DI49_1301; DI49_1302 | DI49_1301; DI49_1302 | | |
| DI49_1299; YBR275C | DI49_1299 | | YBR275C |
| DI49_1298; YBR274W; DI49_1297; YBR273C | DI49_1298; DI49_1297 | | YBR274W; YBR273C |
| DI49_1295; DI49_1296 | DI49_1295; DI49_1296 | | |
| DI49_1293; DI49_1294 | DI49_1293; DI49_1294 | | |
| DI49_1291; YBR267W | DI49_1291 | | YBR267W |
| DI49_1290; YBR265W; DI49_1289; YBR264C | DI49_1290; DI49_1289 | | YBR265W; YBR264C |
| DI49_1287; DI49_1288 | DI49_1287; DI49_1288 | | |
| DI49_1285; YBR260C; DI49_1284; YBR259W | DI49_1285; DI49_1284 | | YBR260C; YBR259W |
| DI49_1282; DI49_1283 | DI49_1282; DI49_1283 | | |
| DI49_1280; YBR255C-A; DI49_1279; YBR255W | DI49_1280; DI49_1279 | | YBR255C-A; YBR255W |
| DI49_1277; DI49_1278 | DI49_1277; DI49_1278 | | |
| DI49_1275; DI49_1276 | DI49_1275; DI49_1276 | | |
| DI49_1273; YBR249C | DI49_1273 | | YBR249C |
| DI49_1272; YBR248C; DI49_1271; YBR247C | DI49_1272; DI49_1271 | | YBR248C; YBR247C |

Create PDF in your applications with the Pdfcrowd HTML to PDF API PDFCROWD

| | | | |
|---|---|---|---|
| DI49_1269; YBR245C; DI49_1268; YBR244W | DI49_1269; DI49_1268 | | YBR245C; YBR244W |
| DI49_1266; YBR242W | DI49_1266 | | YBR242W |
| DI49_1265; YBR241C | DI49_1265 | | YBR241C |
| DI49_1263; YBR240C | DI49_1263 | | YBR240C |
| DI49_1262; YBR239C | DI49_1262 | | YBR239C |
| SPGP0BE00690; YBR238C | | SPGP0BE00690 | YBR238C |
| DI49_1260; YBR237W; DI49_1259; YBR236C | DI49_1260; DI49_1259 | | YBR237W; YBR236C |
| DI49_1257; YBR234C; DI49_1256; YBR233W-A | DI49_1257; DI49_1256 | | YBR234C; YBR233W-A |
| DI49_1254; DI49_1255 | DI49_1254; DI49_1255 | | |
| DI49_1253; SPGP0BE00770 | DI49_1253 | SPGP0BE00770 | |
| DI49_1251; DI49_1252; YBR229C; YBR228W | DI49_1251; DI49_1252 | | YBR229C; YBR228W |
| DI49_1249; YBR225W; DI49_1248; YBR223C | DI49_1249; DI49_1248 | | YBR225W; YBR223C |
| DI49_1246; YBR221C; DI49_1245; YBR220C | DI49_1246; DI49_1245 | | YBR221C; YBR220C |
| DI49_1241; YBR215W | DI49_1241 | | YBR215W |
| DI49_1240; YBR214W; DI49_1239; YBR213W | DI49_1240; DI49_1239 | | YBR214W; YBR213W |
| DI49_1237; DI49_1238 | DI49_1237; DI49_1238 | | |
| DI49_1233; YBR208C | DI49_1233 | | YBR208C |
| DI49_1232; YBR207W | DI49_1232 | | YBR207W |
| DI49_1231; YBR205W; DI49_1230; YBR204C | DI49_1231; DI49_1230 | | YBR205W; YBR204C |
| DI49_1228; YBR202W; DI49_1227; YBR201C-A | DI49_1228; DI49_1227 | | YBR202W; YBR201C-A |
| SPGP0BE01040; DI49_1226 | DI49_1226 | SPGP0BE01040 | |
| SPGP0BE01060; DI49_1224 | DI49_1224 | SPGP0BE01060 | |
| DI49_1221; YBR196C; DI49_1220; YBR195C | DI49_1221; DI49_1220 | | YBR196C; YBR195C |
| DI49_1218; DI49_1219 | DI49_1218; DI49_1219 | | |
| DI49_1216; YBR191W; YBR190W | DI49_1216 | | YBR191W; YBR190W |
| DI49_1212; DI49_1213 | DI49_1212; DI49_1213 | | |
| DI49_1210; DI49_1211 | DI49_1210; DI49_1211 | | |
| DI49_1208; DI49_1209 | DI49_1208; DI49_1209 | | |
| DI49_1206; DI49_5372 | DI49_1206; DI49_5372 | | |
| DI49_1204; DI49_1205 | DI49_1204; DI49_1205 | | |
| DI49_1202; DI49_1203 | DI49_1202; DI49_1203 | | |
| DI49_1200; YBR172C | DI49_1200 | | YBR172C |
| DI49_1199; YBR171W | DI49_1199 | | YBR171W |
| DI49_1198; YBR170C | DI49_1198 | | YBR170C |
| DI49_1197; YBR169C; DI49_1196; YBR168W | DI49_1197; DI49_1196 | | YBR169C; YBR168W |
| DI49_1194; YBR166C; DI49_1193; YBR165W | DI49_1194; DI49_1193 | | YBR166C; YBR165W |
| DI49_1191; YBR163W; DI49_1190; YBR164C | DI49_1191; DI49_1190 | | YBR163W; YBR164C |
| DI49_1188; DI49_1189 | DI49_1188; DI49_1189 | | |
| DI49_1186; YBR159W | DI49_1186 | | YBR159W |
| DI49_1185; YBR158W; DI49_1184; YBR157C | DI49_1185; DI49_1184 | | YBR158W; YBR157C |

Create PDF in your applications with the Pdfcrowd HTML to PDF API PDFCROWD

| | | | |
|---|---|---|---|
| DI49_1183; YBR155W | DI49_1183 | | YBR155W |
| DI49_1182; YBR154C; DI49_1181; YBR153W | DI49_1182; DI49_1181 | | YBR154C; YBR153W |
| DI49_1179; YBR151W; DI49_1178; YBR150C | DI49_1179; DI49_1178 | | YBR151W; YBR150C |
| DI49_1176; DI49_1177 | DI49_1176; DI49_1177 | | |
| DI49_1174; YBR146W; DI49_1173; YBR145W | DI49_1174; DI49_1173 | | YBR146W; YBR145W |
| DI49_1171; DI49_1172; SPGP0BK01910; DI49_2982 | DI49_1171; DI49_1172; DI49_2982 | SPGP0BK01910 | |
| DI49_1169; YBR140C | DI49_1169 | | YBR140C |
| DI49_1168; YBR139W; DI49_1167; YBR138C | DI49_1168; DI49_1167 | | YBR139W; YBR138C |
| DI49_1165; YBR136W; DI49_1164; YBR135W | DI49_1165; DI49_1164 | | YBR136W; YBR135W |
| DI49_1162; YBR132C; DI49_1161; YBR131W | DI49_1162; DI49_1161 | | YBR132C; YBR131W |
| DI49_1159; DI49_1160 | DI49_1159; DI49_1160 | | |
| DI49_1157; YBR127C | DI49_1157 | | YBR127C |
| SPGP0BE01740; DI49_1156 | DI49_1156 | SPGP0BE01740 | |
| DI49_1155; YBR125C | DI49_1155 | | YBR125C |
| DI49_1154; YBR123C; DI49_1153; YBR122C | DI49_1154; DI49_1153 | | YBR123C; YBR122C |
| DI49_1151; DI49_1152 | DI49_1151; DI49_1152 | | |
| DI49_5530; YPR080W; DI49_1148 | DI49_5530; DI49_1148 | | YPR080W |
| DI49_1146; YBR114W; DI49_1145 | DI49_1146; DI49_1145 | | YBR114W |
| SPGP0BE01870; DI49_1144; YBR112C; YBR111W-A | DI49_1144 | SPGP0BE01870 | YBR112C; YBR111W-A |
| DI49_1139; DI49_1140 | DI49_1139; DI49_1140 | | |
| DI49_1137; YBR106W; DI49_1135; YBR105C | DI49_1137; DI49_1135 | | YBR106W; YBR105C |
| DI49_1133; DI49_1134 | DI49_1133; DI49_1134 | | |
| DI49_1131; DI49_1132 | DI49_1131; DI49_1132 | | |
| DI49_1129; YBR097W; DI49_1128; YBR096W | DI49_1129; DI49_1128 | | YBR097W; YBR096W |
| DI49_1126; YBR094W | DI49_1126 | | YBR094W |
| SPGP0BE02040; SPGP0BE02050; DI49_1125; YBR091C | DI49_1125 | SPGP0BE02040; SPGP0BE02050 | YBR091C |
| DI49_1123; YBR088C; DI49_1122; YBR087W | DI49_1123; DI49_1122 | | YBR088C; YBR087W |
| DI49_1120; DI49_1121 | DI49_1120; DI49_1121 | | |
| DI49_1118; YBR085W | DI49_1118 | | YBR085W |
| SPGP0BE02140; DI49_0153; DI49_0152; YBR084C-A | DI49_0153; DI49_0152 | SPGP0BE02140 | YBR084C-A |
| DI49_1117; YBR084W | DI49_1117 | | YBR084W |
| DI49_1116; YBR083W | DI49_1116 | | YBR083W |
| SPGP0BE02170; YBR082C | | SPGP0BE02170 | YBR082C |
| DI49_1112; YBR081C | DI49_1112 | | YBR081C |
| DI49_1111; YBR080C | DI49_1111 | | YBR080C |
| DI49_1110; YBR079C | DI49_1110 | | YBR079C |
| DI49_1109; YBR078W; DI49_1108; YBR077C | DI49_1109; DI49_1108 | | YBR078W; YBR077C |
| DI49_1106; YBR074W; DI49_1105; YBR073W | DI49_1106; DI49_1105 | | YBR074W; YBR073W |
| DI49_1103; DI49_1104 | DI49_1103; DI49_1104 | | |
| DI49_1101; YBR069C | DI49_1101 | | YBR069C |

Create PDF in your applications with the Pdfcrowd HTML to PDF API PDFCROWD

| | | | |
|---|---|---|---|
| DI49_1100; YBR068C; DI49_0242; DI49_0241 | DI49_1100; DI49_0242; DI49_0241 | | YBR068C |
| DI49_1099; YBR067C; DI49_1098; YBR066C | DI49_1099; DI49_1098 | | YBR067C; YBR066C |
| DI49_1096; DI49_1097 | DI49_1096; DI49_1097 | | |
| DI49_1090; DI49_1091 | DI49_1090; DI49_1091 | | |
| DI49_1088; DI49_1089 | DI49_1088; DI49_1089 | | |
| DI49_1086; YBR056W | DI49_1086 | | YBR056W |
| DI49_1084; YBR055C | DI49_1084 | | YBR055C |
| DI49_1083; YBR054W; DI49_1082; YBR053C | DI49_1083; DI49_1082 | | YBR054W; YBR053C |
| DI49_1080; YBR050C | DI49_1080 | | YBR050C |
| DI49_1079; YBR049C | DI49_1079 | | YBR049C |
| SPGP0BE02490; YBR048W | | SPGP0BE02490 | YBR048W |
| DI49_1077; YBR047W | DI49_1077 | | YBR047W |
| DI49_1076; YBR046C; DI49_1075; YBR045C | DI49_1076; DI49_1075 | | YBR046C; YBR045C |
| DI49_1071; YBR042C | DI49_1071 | | YBR042C |
| DI49_1070; YBR041W; DI49_1069; YBR040W | DI49_1070; DI49_1069 | | YBR041W; YBR040W |
| DI49_1067; YBR038W | DI49_1067 | | YBR038W |
| DI49_1066; YBR037C | DI49_1066 | | YBR037C |
| DI49_1065; YBR036C; DI49_1064; YBR035C | DI49_1065; DI49_1064 | | YBR036C; YBR035C |
| SPGP0BE02640; DI49_1063 | DI49_1063 | SPGP0BE02640 | |
| SPGP0BE02660; YDR011W; DI49_1061 | DI49_1061 | SPGP0BE02660 | YDR011W |
| DI49_1059; YDR006C; DI49_1058 | DI49_1059; DI49_1058 | | YDR006C |
| DI49_1056; DI49_1057 | DI49_1056; DI49_1057 | | |
| DI49_1054; YDR001C; DI49_1053 | DI49_1054; DI49_1053 | | YDR001C |
| DI49_1037; DI49_1038 | DI49_1037; DI49_1038 | | |
| DI49_1028; DI49_1029 | DI49_1028; DI49_1029 | | |
| DI49_1023; DI49_1024 | DI49_1023; DI49_1024 | | |
| DI49_1021; YDL042C; DI49_1020 | DI49_1021; DI49_1020 | | YDL042C |
| DI49_1012; YDL051W; DI49_1011 | DI49_1012; DI49_1011 | | YDL051W |
| DI49_1009; DI49_1010 | DI49_1009; DI49_1010 | | |
| DI49_1006; YDL056W; DI49_1005 | DI49_1006; DI49_1005 | | YDL056W |
| SPGP0BE03310; YDL064W; DI49_0999 | DI49_0999 | SPGP0BE03310 | YDL064W |
| DI49_0997; YDL067C; DI49_0996 | DI49_0997; DI49_0996 | | YDL067C |
| DI49_0994; DI49_0995 | DI49_0994; DI49_0995 | | |
| DI49_0988; YDL078C; SPGP0BE03440 | DI49_0988 | SPGP0BE03440 | YDL078C |
| SPGP0BE03570; DI49_0978 | DI49_0978 | SPGP0BE03570 | |
| DI49_0964; YDL106C; DI49_0963 | DI49_0964; DI49_0963 | | YDL106C |
| DI49_0961; DI49_0962 | DI49_0961; DI49_0962 | | |
| DI49_0943; DI49_0944 | DI49_0943; DI49_0944 | | |
| DI49_0938; SPGP0BE04430 | DI49_0938 | SPGP0BE04430 | |
| DI49_0936; DI49_0889; DI49_0935; YDL192W; DI49_0888 | DI49_0936; DI49_0889; DI49_0935; DI49_0888 | | YDL192W |

Create PDF in your applications with the Pdfcrowd HTML to PDF API PDFCROWD

| | | | |
|---|---|---|---|
| DI49_0933; YDL141W; DI49_0932 | DI49_0933; DI49_0932 | | YDL141W |
| DI49_0930; DI49_0931 | DI49_0930; DI49_0931 | | |
| DI49_0913; YDL165W; DI49_0912 | DI49_0913; DI49_0912 | | YDL165W |
| DI49_0904; DI49_0905 | DI49_0904; DI49_0905 | | |
| DI49_0902; DI49_0903 | DI49_0902; DI49_0903 | | |
| DI49_0895; YDL185W; DI49_0894 | DI49_0895; DI49_0894 | | YDL185W |
| DI49_0886; YDL195W; DI49_0885 | DI49_0886; DI49_0885 | | YDL195W |
| DI49_0883; YDL199C; DI49_0882 | DI49_0883; DI49_0882 | | YDL199C |
| DI49_0880; DI49_0881 | DI49_0880; DI49_0881 | | |
| DI49_0878; DI49_0879 | DI49_0878; DI49_0879 | | |
| DI49_0876; DI49_0877 | DI49_0876; DI49_0877 | | |
| DI49_0874; YDL208W; DI49_0873 | DI49_0874; DI49_0873 | | YDL208W |
| DI49_0863; DI49_0864 | DI49_0863; DI49_0864 | | |
| DI49_0861; DI49_0862 | DI49_0861; DI49_0862 | | |
| DI49_0859; DI49_0860 | DI49_0859; DI49_0860 | | |
| DI49_4535; YDL229W; DI49_0853 | DI49_4535; DI49_0853 | | YDL229W |
| DI49_0851; DI49_0852 | DI49_0851; DI49_0852 | | |
| DI49_0849; DI49_0850 | DI49_0849; DI49_0850 | | |
| DI49_0847; DI49_0848 | DI49_0847; DI49_0848 | | |
| DI49_0845; DI49_0846 | DI49_0845; DI49_0846 | | |
| SPGP0BE04910; DI49_0844 | DI49_0844 | SPGP0BE04910 | |
| DI49_0841; DI49_0842 | DI49_0841; DI49_0842 | | |
| SPGP0BE04950; SPGP0BE04940 | | SPGP0BE04950; SPGP0BE04940 | |
| DI49_4536; SPGP0VH00100 | DI49_4536 | SPGP0VH00100 | |
| DI49_4538; DI49_4537 | DI49_4538; DI49_4537 | | |
| DI49_4540; DI49_4539 | DI49_4540; DI49_4539 | | |
| DI49_4542; DI49_4541 | DI49_4542; DI49_4541 | | |
| DI49_4544; DI49_4543 | DI49_4544; DI49_4543 | | |
| DI49_2052; DI49_2051 | DI49_2052; DI49_2051 | | |
| DI49_2054; DI49_2053 | DI49_2054; DI49_2053 | | |
| DI49_5248; DI49_2055 | DI49_5248; DI49_2055 | | |
| SPGP0VLtrna4C; DI49_5567; DI49_3788 | DI49_5567; DI49_3788 | SPGP0VLtrna4C | |
| DI49_3786; DI49_3787 | DI49_3786; DI49_3787 | | |
| DI49_3784; YGL201C; DI49_3783 | DI49_3784; DI49_3783 | | YGL201C |
| DI49_3779; DI49_3780 | DI49_3779; DI49_3780 | | |
| DI49_3777; DI49_3778 | DI49_3777; DI49_3778 | | |
| DI49_3775; DI49_3776 | DI49_3775; DI49_3776 | | |
| DI49_3773; DI49_3774 | DI49_3773; DI49_3774 | | |
| DI49_3771; DI49_3772 | DI49_3771; DI49_3772 | | |
| DI49_3769; DI49_1665; DI49_3768; DI49_1666 | DI49_3769; DI49_1665; DI49_3768; DI49_1666 | | |

Create PDF in your applications with the Pdfcrowd HTML to PDF API PDFCROWD

| | | | |
|---|---|---|---|
| DI49_4657; DI49_4656 | DI49_4657; DI49_4656 | | |
| DI49_4659; DI49_4658 | DI49_4659; DI49_4658 | | |
| DI49_4661; DI49_4660 | DI49_4661; DI49_4660 | | |
| DI49_4663; DI49_4662 | DI49_4663; DI49_4662 | | |
| DI49_4665; DI49_4664 | DI49_4665; DI49_4664 | | |
| DI49_4667; DI49_4666 | DI49_4667; DI49_4666 | | |
| DI49_4669; DI49_4668 | DI49_4669; DI49_4668 | | |
| DI49_4671; DI49_4670 | DI49_4671; DI49_4670 | | |
| DI49_4673; DI49_4672 | DI49_4673; DI49_4672 | | |
| DI49_4678; DI49_4677 | DI49_4678; DI49_4677 | | |
| DI49_4680; DI49_4679 | DI49_4680; DI49_4679 | | |
| DI49_4682; DI49_4681 | DI49_4682; DI49_4681 | | |
| DI49_4684; DI49_4683 | DI49_4684; DI49_4683 | | |
| DI49_0815; DI49_0814 | DI49_0815; DI49_0814 | | |
| DI49_0817; DI49_0816 | DI49_0817; DI49_0816 | | |
| DI49_0820; SPGP0VO00160 | DI49_0820 | SPGP0VO00160 | |
| YCR077C; DI49_0821 | DI49_0821 | | YCR077C |
| DI49_0823; DI49_0822 | DI49_0823; DI49_0822 | | |
| DI49_0825; DI49_0824 | DI49_0825; DI49_0824 | | |
| DI49_3966; DI49_3967 | DI49_3966; DI49_3967 | | |
| DI49_3963; DI49_3965 | DI49_3963; DI49_3965 | | |
| DI49_3962; SPGP0VP00170 | DI49_3962 | SPGP0VP00170 | |
| DI49_3960; DI49_3961 | DI49_3960; DI49_3961 | | |
| SPGP0VP00220; YJL216C | | SPGP0VP00220 | YJL216C |
| SPGP0VP00230; DI49_5194; SPGP0VP00240; YBR297W | DI49_5194 | SPGP0VP00230; SPGP0VP00240 | YBR297W |
| DI49_1662; DI49_1661 | DI49_1662; DI49_1661 | | |
| SPGP0BF00100; DI49_1667 | DI49_1667 | SPGP0BF00100 | |
| YIR014W; YIR015W | | | YIR014W; YIR015W |
| YIR003W; YIR004W | | | YIR003W; YIR004W |
| YIL006W; YIL005W | | | YIL006W; YIL005W |
| YIL008W; YIL007C | | | YIL008W; YIL007C |
| YIL011W; YIL010W | | | YIL011W; YIL010W |
| YIL014W; YIL013C | | | YIL014W; YIL013C |
| YIL026C; YIL024C | | | YIL026C; YIL024C |
| YIL042C; YIL043C; YIL044C | | | YIL042C; YIL043C; YIL044C |
| YIL055C; DI49_2726 | DI49_2726 | | YIL055C |
| SPGP0BF00950; YIL072W | | SPGP0BF00950 | YIL072W |
| YIL074C; YIL073C | | | YIL074C; YIL073C |
| YIL087C; YIL085C | | | YIL087C; YIL085C |

Create PDF in your applications with the Pdfcrowd HTML to PDF API PDFCROWD

| | | | |
|---|---|---|---|
| YIL098C; YIL097W | | | YIL098C; YIL097W |
| YIL106W; YIL105C | | | YIL106W; YIL105C |
| YIL113W; YIL112W | | | YIL113W; YIL112W |
| YIL118W; YIL117C | | | YIL118W; YIL117C |
| YIL132C; YIL131C | | | YIL132C; YIL131C |
| YIL134W; YIL133C | | | YIL134W; YIL133C |
| YIL136W; YIL135C | | | YIL136W; YIL135C |
| YIL138C; YIL137C | | | YIL138C; YIL137C |
| YIL140W; YIL139C | | | YIL140W; YIL139C |
| YIL155C; YIL154C | | | YIL155C; YIL154C |
| YIL156W-B; YIL156W | | | YIL156W-B; YIL156W |
| YIL158W; YIL157C | | | YIL158W; YIL157C |
| YIL167W; YIL166C | | | YIL167W; YIL166C |
| DI49_1601; YOL158C | DI49_1601 | | YOL158C |
| SPGP0BF01890; YOL159C | | SPGP0BF01890 | YOL159C |
| SPGP0BF01900; YAL068C; DI49_3810; DI49_1966 | DI49_3810; DI49_1966 | SPGP0BF01900 | YAL068C |
| SPGP0VR00150; YLR300W | | SPGP0VR00150 | YLR300W |
| DI49_1638; DI49_1637 | DI49_1638; DI49_1637 | | |
| DI49_1640; DI49_1639 | DI49_1640; DI49_1639 | | |
| DI49_1642; DI49_1641 | DI49_1642; DI49_1641 | | |
| YPL260W; YPL262W | | | YPL260W; YPL262W |
| SPGP0VTtrna5W; SPGP0Itrna9W; YPL257W; SPGP0Htrna8W; YMR052W | | SPGP0VTtrna5W; SPGP0Itrna9W; SPGP0Htrna8W | YPL257W; YMR052W |
| YPL256C; SPGP0VT00150 | | SPGP0VT00150 | YPL256C |
| YPL254W; YPL255W | | | YPL254W; YPL255W |
| YPL252C; YPL253C | | | YPL252C; YPL253C |
| DI49_1472; DI49_1471 | DI49_1472; DI49_1471 | | |
| DI49_1474; DI49_1473 | DI49_1474; DI49_1473 | | |
| YGL222C; DI49_5246 | DI49_5246 | | YGL222C |
| YGL220W; YGL221C | | | YGL220W; YGL221C |
| DI49_0512; YGL219C | DI49_0512 | | YGL219C |
| DI49_0514; DI49_0513 | DI49_0514; DI49_0513 | | |
| DI49_0516; DI49_0515 | DI49_0516; DI49_0515 | | |
| DI49_0518; DI49_0517 | DI49_0518; DI49_0517 | | |
| DI49_0525; DI49_0524 | DI49_0525; DI49_0524 | | |
| DI49_0530; DI49_0529 | DI49_0530; DI49_0529 | | |
| DI49_0532; DI49_0531 | DI49_0532; DI49_0531 | | |
| DI49_4717; DI49_0533 | DI49_4717; DI49_0533 | | |
| DI49_4715; DI49_4716 | DI49_4715; DI49_4716 | | |
| DI49_4713; DI49_4714 | DI49_4713; DI49_4714 | | |
| DI49_4711; DI49_4712 | DI49_4711; DI49_4712 | | |

Create PDF in your applications with the Pdfcrowd HTML to PDF API PDFCROWD

| | | | |
|---|---|---|---|
| DI49_4709; DI49_4710 | DI49_4709; DI49_4710 | | |
| SPGP0WA00200; DI49_0189; DI49_0188; YBR009C; YBR008C | DI49_0189; DI49_0188 | SPGP0WA00200 | YBR009C; YBR008C |
| DI49_4706; DI49_0190; DI49_4705; YBR010W | DI49_4706; DI49_0190; DI49_4705 | | YBR010W |
| YCL061C; YCL063W | | | YCL061C; YCL063W |
| YCL043C; DI49_0715; YCL040W | DI49_0715 | | YCL043C; YCL040W |
| YCL038C; YCL039W | | | YCL038C; YCL039W |
| YCL033C; YCL034W | | | YCL033C; YCL034W |
| YCL026C-B; YCL027W | | | YCL026C-B; YCL027W |
| YCL025C; DI49_0671; YCL024W | DI49_0671 | | YCL025C; YCL024W |
| YCL014W; DI49_0739; YCL012C | DI49_0739 | | YCL014W; YCL012C |
| YCL004W; YCL005W | | | YCL004W; YCL005W |
| YCR010C; YCR009C | | | YCR010C; YCR009C |
| YCR020W-B; YCR020C-A | | | YCR020W-B; YCR020C-A |
| YCR028C; DI49_0781; YCR028C-A | DI49_0781 | | YCR028C; YCR028C-A |
| YCR031C; DI49_2840; YCR032W; YJL191W | DI49_2840 | | YCR031C; YCR032W; YJL191W |
| YCR038C; YCR037C | | | YCR038C; YCR037C |
| DI49_1409; DI49_1410 | DI49_1409; DI49_1410 | | |
| DI49_3362; DI49_3363 | DI49_3362; DI49_3363 | | |
| DI49_3360; DI49_3361 | DI49_3360; DI49_3361 | | |
| DI49_3358; DI49_3359 | DI49_3358; DI49_3359 | | |
| DI49_3356; DI49_3357 | DI49_3356; DI49_3357 | | |
| DI49_3354; DI49_3355 | DI49_3354; DI49_3355 | | |
| DI49_3352; DI49_3353 | DI49_3352; DI49_3353 | | |
| DI49_4784; DI49_3351 | DI49_4784; DI49_3351 | | |
| DI49_1852; DI49_4785 | DI49_1852; DI49_4785 | | |
| DI49_1854; DI49_1853 | DI49_1854; DI49_1853 | | |
| SPGP0WItrna6W; DI49_3006; DI49_1857 | DI49_3006; DI49_1857 | SPGP0WItrna6W | |
| DI49_1859; DI49_1858 | DI49_1859; DI49_1858 | | |
| DI49_1861; DI49_1860 | DI49_1861; DI49_1860 | | |
| DI49_1863; DI49_1862 | DI49_1863; DI49_1862 | | |
| DI49_1865; DI49_1864 | DI49_1865; DI49_1864 | | |
| DI49_1867; DI49_1866 | DI49_1867; DI49_1866 | | |
| DI49_1869; DI49_1868 | DI49_1869; DI49_1868 | | |
| DI49_1871; DI49_1870 | DI49_1871; DI49_1870 | | |
| DI49_1873; DI49_1872 | DI49_1873; DI49_1872 | | |
| DI49_3397; DI49_1874 | DI49_3397; DI49_1874 | | |
| DI49_3395; DI49_3396 | DI49_3395; DI49_3396 | | |
| DI49_3393; DI49_3394 | DI49_3393; DI49_3394 | | |
| DI49_3391; DI49_3392 | DI49_3391; DI49_3392 | | |

Create PDF in your applications with the Pdfcrowd HTML to PDF API PDFCROWD

| | | | |
|---|---|---|---|
| DI49_3389; DI49_3390 | DI49_3389; DI49_3390 | | |
| DI49_2011; DI49_2010 | DI49_2011; DI49_2010 | | |
| DI49_2013; DI49_2012 | DI49_2013; DI49_2012 | | |
| DI49_2015; DI49_2014 | DI49_2015; DI49_2014 | | |
| DI49_2017; DI49_2016 | DI49_2017; DI49_2016 | | |
| DI49_2019; DI49_2018 | DI49_2019; DI49_2018 | | |
| DI49_2021; DI49_2020 | DI49_2021; DI49_2020 | | |
| DI49_2023; DI49_2022 | DI49_2023; DI49_2022 | | |
| DI49_1484; DI49_2026 | DI49_1484; DI49_2026 | | |
| DI49_1486; DI49_2710; SPGP0WT00100 | DI49_1486; DI49_2710 | SPGP0WT00100 | |
| DI49_3754; DI49_3753 | DI49_3754; DI49_3753 | | |
| YER178W; YER177W | | | YER178W; YER177W |
| YER180C; YER179W | | | YER180C; YER179W |
| DI49_0497; DI49_0498 | DI49_0497; DI49_0498 | | |
| DI49_0494; YDR316W; DI49_0493 | DI49_0494; DI49_0493 | | YDR316W |
| DI49_0491; DI49_0492 | DI49_0491; DI49_0492 | | |
| DI49_0487; DI49_0488 | DI49_0487; DI49_0488 | | |
| DI49_0485; DI49_0486 | DI49_0485; DI49_0486 | | |
| DI49_0483; DI49_0484 | DI49_0483; DI49_0484 | | |
| DI49_0481; DI49_0482 | DI49_0481; DI49_0482 | | |
| DI49_0479; DI49_0480 | DI49_0479; DI49_0480 | | |
| DI49_0474; YDR297W; DI49_0473 | DI49_0474; DI49_0473 | | YDR297W |
| DI49_0462; DI49_0463 | DI49_0462; DI49_0463 | | |
| DI49_0454; YDR276C; DI49_0453 | DI49_0454; DI49_0453 | | YDR276C |
| DI49_0451; DI49_0452 | DI49_0451; DI49_0452 | | |
| DI49_0448; YDR267C; DI49_0447 | DI49_0448; DI49_0447 | | YDR267C |
| DI49_0445; DI49_0446 | DI49_0445; DI49_0446 | | |
| DI49_0443; DI49_0444 | DI49_0443; DI49_0444 | | |
| DI49_0439; DI49_0440 | DI49_0439; DI49_0440 | | |
| DI49_0437; DI49_0438 | DI49_0437; DI49_0438 | | |
| DI49_0433; DI49_0434 | DI49_0433; DI49_0434 | | |
| DI49_0431; DI49_0432 | DI49_0431; DI49_0432 | | |
| DI49_0425; YDR244W; DI49_0424 | DI49_0425; DI49_0424 | | YDR244W |
| DI49_0422; YDR240C; DI49_0421 | DI49_0422; DI49_0421 | | YDR240C |
| DI49_0419; DI49_0420 | DI49_0419; DI49_0420 | | |
| DI49_0410; YDR228C; DI49_0409 | DI49_0410; DI49_0409 | | YDR228C |
| DI49_0407; DI49_0179 | DI49_0407; DI49_0179 | | |
| SPGP0BI01110; DI49_0401 | DI49_0401 | SPGP0BI01110 | |
| DI49_0399; DI49_0400 | DI49_0399; DI49_0400 | | |
| DI49_0397; YDR212W; DI49_0396 | DI49_0397; DI49_0396 | | YDR212W |

Create PDF in your applications with the Pdfcrowd HTML to PDF API PDFCROWD

| | | | |
|---|---|---|---|
| DI49_0394; YDR208W; DI49_0393 | DI49_0394; DI49_0393 | | YDR208W |
| DI49_0391; DI49_0392 | DI49_0391; DI49_0392 | | |
| DI49_0388; YDR201W; DI49_0387 | DI49_0388; DI49_0387 | | YDR201W |
| DI49_0373; DI49_0374 | DI49_0373; DI49_0374 | | |
| DI49_0368; DI49_0369 | DI49_0368; DI49_0369 | | |
| DI49_0362; YDR172W; DI49_0361 | DI49_0362; DI49_0361 | | YDR172W |
| DI49_0356; YDR168W; DI49_0355 | DI49_0356; DI49_0355 | | YDR168W |
| DI49_0347; DI49_0348 | DI49_0347; DI49_0348 | | |
| DI49_0345; DI49_0346 | DI49_0345; DI49_0346 | | |
| DI49_0343; YDR153C; DI49_0342 | DI49_0343; DI49_0342 | | YDR153C |
| DI49_0330; DI49_0331 | DI49_0330; DI49_0331 | | |
| DI49_0327; YDR134C; DI49_0326 | DI49_0327; DI49_0326 | | YDR134C |
| DI49_0324; DI49_0325 | DI49_0324; DI49_0325 | | |
| DI49_0322; DI49_0323 | DI49_0322; DI49_0323 | | |
| DI49_0319; YDR125C; DI49_0318 | DI49_0319; DI49_0318 | | YDR125C |
| DI49_0316; YDR122W; DI49_0315 | DI49_0316; DI49_0315 | | YDR122W |
| DI49_0313; DI49_0314 | DI49_0313; DI49_0314 | | |
| DI49_0309; DI49_0310 | DI49_0309; DI49_0310 | | |
| DI49_0307; DI49_0308 | DI49_0307; DI49_0308 | | |
| SPGP0BI02050; DI49_0306 | DI49_0306 | SPGP0BI02050 | |
| DI49_0299; DI49_0300 | DI49_0299; DI49_0300 | | |
| DI49_0290; YDR093W; SPGP0BI02200 | DI49_0290 | SPGP0BI02200 | YDR093W |
| DI49_0288; DI49_0289 | DI49_0288; DI49_0289 | | |
| DI49_0274; DI49_0275 | DI49_0274; DI49_0275 | | |
| DI49_0270; YDR074W; DI49_0269 | DI49_0270; DI49_0269 | | YDR074W |
| DI49_0256; YDR060W; DI49_0255 | DI49_0256; DI49_0255 | | YDR060W |
| DI49_0251; YDR057W; DI49_0250 | DI49_0251; DI49_0250 | | YDR057W |
| DI49_0245; YDR050C; DI49_0244 | DI49_0245; DI49_0244 | | YDR050C |
| DI49_0228; YDR031W; DI49_0227 | DI49_0228; DI49_0227 | | YDR031W |
| SPGP0BI02840; YDR025W | | SPGP0BI02840 | YDR025W |
| DI49_0212; DI49_0213 | DI49_0212; DI49_0213 | | |
| DI49_0210; DI49_0211 | DI49_0210; DI49_0211 | | |
| DI49_0208; YBR030W | DI49_0208 | | YBR030W |
| DI49_0207; YBR029C; DI49_0206; YBR028C | DI49_0207; DI49_0206 | | YBR029C; YBR028C |
| DI49_0204; YBR025C; DI49_0203; YBR024W | DI49_0204; DI49_0203 | | YBR025C; YBR024W |
| DI49_0201; DI49_0202 | DI49_0201; DI49_0202 | | |
| DI49_0199; DI49_0200 | DI49_0199; DI49_0200 | | |
| DI49_0197; DI49_0198 | DI49_0197; DI49_0198 | | |
| DI49_0195; DI49_0196 | DI49_0195; DI49_0196 | | |
| DI49_0193; DI49_0194 | DI49_0193; DI49_0194 | | |

Create PDF in your applications with the Pdfcrowd HTML to PDF API PDFCROWD

| | | | |
|---|---|---|---|
| DI49_0191; YBR011C | DI49_0191 | | YBR011C |
| DI49_0186; DI49_0187; YBR006W; YBR005W | DI49_0186; DI49_0187 | | YBR006W; YBR005W |
| DI49_0184; YBR003W; DI49_0183; YBR002C | DI49_0184; DI49_0183 | | YBR003W; YBR002C |
| DI49_0181; DI49_0182 | DI49_0181; DI49_0182 | | |
| DI49_0178; YBL004W | DI49_0178 | | YBL004W |
| DI49_0174; YBL007C; DI49_0173; YBL008W | DI49_0174; DI49_0173 | | YBL007C; YBL008W |
| DI49_0171; YBL010C; DI49_0170; YBL011W | DI49_0171; DI49_0170 | | YBL010C; YBL011W |
| DI49_0168; DI49_0169 | DI49_0168; DI49_0169 | | |
| DI49_0165; YBL015W; DI49_0164; YBL016W | DI49_0165; DI49_0164 | | YBL015W; YBL016W |
| DI49_0161; YBL020W; DI49_0160; YBL021C | DI49_0161; DI49_0160 | | YBL020W; YBL021C |
| DI49_0158; YBL023C | DI49_0158 | | YBL023C |
| DI49_0157; YBL024W; DI49_0156; YBL025W | DI49_0157; DI49_0156 | | YBL024W; YBL025W |
| DI49_0154; DI49_0155 | DI49_0154; DI49_0155 | | |
| DI49_0150; DI49_0151; YBL029W; YBL029C-A | DI49_0150; DI49_0151 | | YBL029W; YBL029C-A |
| DI49_0148; YBL031W; DI49_0147; YBL032W | DI49_0148; DI49_0147 | | YBL031W; YBL032W |
| DI49_0145; DI49_0146 | DI49_0145; DI49_0146 | | |
| DI49_0143; DI49_0144 | DI49_0143; DI49_0144 | | |
| DI49_0141; YBL038W | DI49_0141 | | YBL038W |
| DI49_0140; YBL039C; DI49_0139; YBL039W-B | DI49_0140; DI49_0139 | | YBL039C; YBL039W-B |
| DI49_0137; YBL041W; DI49_0136; YBL042C | DI49_0137; DI49_0136 | | YBL041W; YBL042C |
| DI49_0134; DI49_0135 | DI49_0134; DI49_0135 | | |
| DI49_0132; DI49_0133; YBL045C; YBL046W | DI49_0132; DI49_0133 | | YBL045C; YBL046W |
| DI49_0130; DI49_0131 | DI49_0130; DI49_0131 | | |
| DI49_0128; YBL051C | DI49_0128 | | YBL051C |
| DI49_0127; YBL052C; DI49_0126; YBL054W | DI49_0127; DI49_0126 | | YBL052C; YBL054W |
| DI49_0125; YBL056W; DI49_0124; YBL057C | DI49_0125; DI49_0124 | | YBL056W; YBL057C |
| SPGP0BI03800; DI49_0123 | DI49_0123 | SPGP0BI03800 | |
| DI49_0121; SPGP0BI03810 | DI49_0121 | SPGP0BI03810 | |
| DI49_0119; YBL063W | DI49_0119 | | YBL063W |
| DI49_0118; YBL064C | DI49_0118 | | YBL064C |
| DI49_0117; YBL066C; DI49_0116; YBL067C | DI49_0117; DI49_0116 | | YBL066C; YBL067C |
| DI49_0114; DI49_0115 | DI49_0114; DI49_0115 | | |
| DI49_0110; YBL076C | DI49_0110 | | YBL076C |
| DI49_0109; YBL078C | DI49_0109 | | YBL078C |
| DI49_0108; YBL079W | DI49_0108 | | YBL079W |
| DI49_0107; YBL080C; DI49_0106; YBL081W | DI49_0107; DI49_0106 | | YBL080C; YBL081W |
| DI49_0105; SPGP0BI03990 | DI49_0105 | SPGP0BI03990 | |
| DI49_0103; YBL085W; DI49_0102; YBL086C | DI49_0103; DI49_0102 | | YBL085W; YBL086C |
| DI49_0100; YBL088C; DI49_0099; YBL089W | DI49_0100; DI49_0099 | | YBL088C; YBL089W |
| DI49_0097; YBL091C; DI49_0096; YBL091C-A | DI49_0097; DI49_0096 | | YBL091C; YBL091C-A |

Create PDF in your applications with the Pdfcrowd HTML to PDF API PDFCROWD

| | | | |
|---|---|---|---|
| DI49_0094; YBL093C; DI49_0093; YBL095W | DI49_0094; DI49_0093 | | YBL093C; YBL095W |
| DI49_0091; DI49_0092 | DI49_0091; DI49_0092 | | |
| DI49_0086; DI49_0087 | DI49_0086; DI49_0087 | | |
| DI49_0084; YBL105C | DI49_0084 | | YBL105C |
| DI49_0083; YBL106C; DI49_0082; YBL107C | DI49_0083; DI49_0082 | | YBL106C; YBL107C |
| DI49_3978; SPGP0BItrna26L | DI49_3978 | SPGP0BItrna26L | |
| DI49_1804; DI49_3977 | DI49_1804; DI49_3977 | | |
| DI49_1806; DI49_1805 | DI49_1806; DI49_1805 | | |
| DI49_1810; DI49_1809 | DI49_1810; DI49_1809 | | |
| DI49_1812; DI49_1811 | DI49_1812; DI49_1811 | | |
| DI49_3972; DI49_1813 | DI49_3972; DI49_1813 | | |
| DI49_3542; DI49_3971 | DI49_3542; DI49_3971 | | |
| DI49_3540; DI49_3541 | DI49_3540; DI49_3541 | | |
| DI49_3535; DI49_3536 | DI49_3535; DI49_3536 | | |
| DI49_3533; DI49_3534 | DI49_3533; DI49_3534 | | |
| DI49_3531; DI49_3532 | DI49_3531; DI49_3532 | | |
| YLR002C; DI49_3530 | DI49_3530 | | YLR002C |
| DI49_5452; DI49_5451 | DI49_5452; DI49_5451 | | |
| DI49_5454; DI49_5453 | DI49_5454; DI49_5453 | | |
| DI49_5456; DI49_5455 | DI49_5456; DI49_5455 | | |
| DI49_3689; DI49_3688 | DI49_3689; DI49_3688 | | |
| DI49_3691; DI49_3690 | DI49_3691; DI49_3690 | | |
| DI49_3693; DI49_3692 | DI49_3693; DI49_3692 | | |
| DI49_3695; DI49_3694 | DI49_3695; DI49_3694 | | |
| DI49_3700; DI49_3699 | DI49_3700; DI49_3699 | | |
| DI49_3702; DI49_3701 | DI49_3702; DI49_3701 | | |
| DI49_3704; DI49_3703 | DI49_3704; DI49_3703 | | |
| DI49_5244; DI49_5243 | DI49_5244; DI49_5243 | | |
| YAR050W; SPGP0KR00100 | | SPGP0KR00100 | YAR050W |
| DI49_3181; DI49_3079; DI49_3180 | DI49_3181; DI49_3079; DI49_3180 | | |
| DI49_3179; YKL205W | DI49_3179 | | YKL205W |
| DI49_3177; DI49_3178 | DI49_3177; DI49_3178 | | |
| DI49_3166; SPGP0BK00100 | DI49_3166 | SPGP0BK00100 | |
| SPGP0BK00190; SPGP0BK00180 | | SPGP0BK00190; SPGP0BK00180 | |
| DI49_3159; SPGP0BK00220; DI49_3157; DI49_3156 | DI49_3159; DI49_3157; DI49_3156 | SPGP0BK00220 | |
| SPGP0BK00280; DI49_3155 | DI49_3155 | SPGP0BK00280 | |
| DI49_3153; DI49_2608 | DI49_3153; DI49_2608 | | |
| DI49_3151; DI49_3152 | DI49_3151; DI49_3152 | | |
| DI49_3149; DI49_3150 | DI49_3149; DI49_3150 | | |
| DI49_3147; YJR138W | DI49_3147 | | YJR138W |

Create PDF in your applications with the Pdfcrowd HTML to PDF API PDFCROWD

| | | | |
|---|---|---|---|
| DI49_3146; YJR137C | DI49_3146 | | YJR137C |
| DI49_3145; YJR136C; DI49_3144; YJR135W-A | DI49_3145; DI49_3144 | | YJR136C; YJR135W-A |
| DI49_3142; YJR134C | DI49_3142 | | YJR134C |
| DI49_3141; YJR133W | DI49_3141 | | YJR133W |
| DI49_3140; YJR132W; DI49_3139; YJR131W | DI49_3140; DI49_3139 | | YJR132W; YJR131W |
| DI49_3137; DI49_3138 | DI49_3137; DI49_3138 | | |
| DI49_3135; DI49_3136 | DI49_3135; DI49_3136 | | |
| DI49_3133; YJR124C | DI49_3133 | | YJR124C |
| DI49_3132; YJR123W; DI49_3131; YJR122W | DI49_3132; DI49_3131 | | YJR123W; YJR122W |
| DI49_3129; YJR119C | DI49_3129 | | YJR119C |
| DI49_3128; YJR118C | DI49_3128 | | YJR118C |
| DI49_3127; YJR117W; DI49_3126; YJR116W | DI49_3127; DI49_3126 | | YJR117W; YJR116W |
| DI49_3124; DI49_3125 | DI49_3124; DI49_3125 | | |
| DI49_3122; DI49_3123 | DI49_3122; DI49_3123 | | |
| DI49_3120; YJR110W | DI49_3120 | | YJR110W |
| DI49_3119; YJR109C; SPGP0BK00650; YJR108W | DI49_3119 | SPGP0BK00650 | YJR109C; YJR108W |
| DI49_3117; YJR107W; DI49_3116; YJR106W | DI49_3117; DI49_3116 | | YJR107W; YJR106W |
| DI49_3114; YJR104C; DI49_3113; YJR103W | DI49_3114; DI49_3113 | | YJR104C; YJR103W |
| DI49_3111; DI49_3112 | DI49_3111; DI49_3112 | | |
| DI49_3107; YJR098C; DI49_3106; YJR097W | DI49_3107; DI49_3106 | | YJR098C; YJR097W |
| DI49_3104; YJR095W | DI49_3104 | | YJR095W |
| DI49_3103; YJR094W-A; DI49_3102; YJR094C | DI49_3103; DI49_3102 | | YJR094W-A; YJR094C |
| DI49_3100; DI49_3101 | DI49_3100; DI49_3101 | | |
| DI49_3098; YJR090C; DI49_3097; YJR089W | DI49_3098; DI49_3097 | | YJR090C; YJR089W |
| DI49_3095; DI49_3096 | DI49_3095; DI49_3096 | | |
| DI49_3093; DI49_3094 | DI49_3093; DI49_3094 | | |
| DI49_3091; DI49_3092 | DI49_3091; DI49_3092 | | |
| DI49_3089; DI49_3090 | DI49_3089; DI49_3090 | | |
| DI49_3087; YJR076C | DI49_3087 | | YJR076C |
| DI49_3086; YJR075W; SPGP0BK00980; YJR074W | DI49_3086 | SPGP0BK00980 | YJR075W; YJR074W |
| DI49_3084; DI49_3085 | DI49_3084; DI49_3085 | | |
| DI49_3082; YJR069C | DI49_3082 | | YJR069C |
| DI49_3081; YJR068W; DI49_3080; YJR067C | DI49_3081; DI49_3080 | | YJR068W; YJR067C |
| DI49_3078; YJR065C | DI49_3078 | | YJR065C |
| DI49_3077; YJR064W; DI49_3076; YJR063W | DI49_3077; DI49_3076 | | YJR064W; YJR063W |
| DI49_3074; DI49_3075 | DI49_3074; DI49_3075 | | |
| DI49_3072; YJR059W; DI49_3071; YJR058C | DI49_3072; DI49_3071 | | YJR059W; YJR058C |
| SPGP0BKtrna20R; DI49_3064 | DI49_3064 | SPGP0BKtrna20R | |
| DI49_3063; YJR054W; DI49_3062; YJR053W | DI49_3063; DI49_3062 | | YJR054W; YJR053W |
| DI49_3057; DI49_3058 | DI49_3057; DI49_3058 | | |

Create PDF in your applications with the Pdfcrowd HTML to PDF API PDFCROWD

| | | | |
|---|---|---|---|
| DI49_3051; DI49_3052 | DI49_3051; DI49_3052 | | |
| DI49_3047; DI49_3048 | DI49_3047; DI49_3048 | | |
| DI49_3045; YJR039W | DI49_3045 | | YJR039W |
| DI49_3044; YJR036C; DI49_3043; YJR035W | DI49_3044; DI49_3043 | | YJR036C; YJR035W |
| DI49_3041; YJR033C | DI49_3041 | | YJR033C |
| DI49_3040; YJR032W; DI49_3039; YJR031C | DI49_3040; DI49_3039 | | YJR032W; YJR031C |
| DI49_3037; DI49_3038 | DI49_3037; DI49_3038 | | |
| DI49_3035; DI49_3036 | DI49_3035; DI49_3036 | | |
| DI49_3034; SPGP0BK01440 | DI49_3034 | SPGP0BK01440 | |
| DI49_3032; YJR016C; SPGP0BK01480; YJR015W | DI49_3032 | SPGP0BK01480 | YJR016C; YJR015W |
| DI49_3030; DI49_3031 | DI49_3030; DI49_3031 | | |
| DI49_3028; YJR011C; DI49_3027; YJR010C-A | DI49_3028; DI49_3027 | | YJR011C; YJR010C-A |
| DI49_3025; YJR009C | DI49_3025 | | YJR009C |
| DI49_3024; YJR008W | DI49_3024 | | YJR008W |
| DI49_3023; YJR007W; DI49_3022; YJR006W | DI49_3023; DI49_3022 | | YJR007W; YJR006W |
| DI49_3020; YJR005W | DI49_3020 | | YJR005W |
| DI49_3019; YJR004C; DI49_3018; YJR003C | DI49_3019; DI49_3018 | | YJR004C; YJR003C |
| DI49_3016; DI49_3017 | DI49_3016; DI49_3017 | | |
| DI49_3015; SPGP0BK01650 | DI49_3015 | SPGP0BK01650 | |
| DI49_3013; DI49_3014 | DI49_3013; DI49_3014 | | |
| DI49_3008; YJL008C | DI49_3008 | | YJL008C |
| DI49_3007; YJL010C | DI49_3007 | | YJL010C |
| DI49_3002; DI49_3003 | DI49_3002; DI49_3003 | | |
| DI49_3000; DI49_3001 | DI49_3000; DI49_3001 | | |
| DI49_2998; DI49_2999 | DI49_2998; DI49_2999 | | |
| DI49_2995; YJL024C; DI49_2994; YJL025W | DI49_2995; DI49_2994 | | YJL024C; YJL025W |
| DI49_2988; DI49_2989 | DI49_2988; DI49_2989 | | |
| DI49_2986; YJL034W; DI49_2985; YJL035C | DI49_2986; DI49_2985 | | YJL034W; YJL035C |
| DI49_2978; YJL039C; DI49_2977; YJL041W | DI49_2978; DI49_2977 | | YJL039C; YJL041W |
| DI49_2975; DI49_2976 | DI49_2975; DI49_2976 | | |
| DI49_2973; DI49_2974 | DI49_2973; DI49_2974 | | |
| DI49_2967; DI49_2968 | DI49_2967; DI49_2968 | | |
| DI49_2965; YJL050W | DI49_2965 | | YJL050W |
| DI49_2964; YJL051W | DI49_2964 | | YJL051W |
| DI49_2963; YJL052W; DI49_2962; YJL053W | DI49_2963; DI49_2962 | | YJL052W; YJL053W |
| DI49_2960; DI49_2961 | DI49_2960; DI49_2961 | | |
| DI49_2958; DI49_2959 | DI49_2958; DI49_2959 | | |
| DI49_2956; YJL059W | DI49_2956 | | YJL059W |
| DI49_2955; YJL060W; DI49_2954; YJL061W | DI49_2955; DI49_2954 | | YJL060W; YJL061W |
| DI49_2952; YJL062W-A; DI49_2951; YJL063C | DI49_2952; DI49_2951 | | YJL062W-A; YJL063C |

Create PDF in your applications with the Pdfcrowd HTML to PDF API PDFCROWD

| | | | |
|---|---|---|---|
| DI49_2949; DI49_2950 | DI49_2949; DI49_2950 | | |
| DI49_2947; DI49_2948 | DI49_2947; DI49_2948 | | |
| DI49_2945; YJL071W | DI49_2945 | | YJL071W |
| DI49_2944; YJL072C; DI49_2943; YJL073W | DI49_2944; DI49_2943 | | YJL072C; YJL073W |
| DI49_2941; DI49_2942 | DI49_2941; DI49_2942 | | |
| DI49_2939; DI49_2940 | DI49_2939; DI49_2940 | | |
| DI49_2937; DI49_2938; YJL080C; YJL081C | DI49_2937; DI49_2938 | | YJL080C; YJL081C |
| DI49_2935; YJL083W | DI49_2935 | | YJL083W |
| SPGP0BK02350; YJL084C | | SPGP0BK02350 | YJL084C |
| DI49_2934; YJL085W | DI49_2934 | | YJL085W |
| DI49_2933; YJL087C; DI49_2932; YJL088W | DI49_2933; DI49_2932 | | YJL087C; YJL088W |
| DI49_2930; YJL090C | DI49_2930 | | YJL090C |
| DI49_2929; YJL091C | DI49_2929 | | YJL091C |
| DI49_2928; YJL092W; DI49_2927; YJL093C | DI49_2928; DI49_2927 | | YJL092W; YJL093C |
| DI49_2925; YJL095W; DI49_2924; YJL096W | DI49_2925; DI49_2924 | | YJL095W; YJL096W |
| DI49_2922; YJL098W; DI49_2921; YJL099W | DI49_2922; DI49_2921 | | YJL098W; YJL099W |
| DI49_2919; YJL101C | DI49_2919 | | YJL101C |
| SPGP0BKtrna23R; DI49_2918; DI49_2917; YJL102W | DI49_2918; DI49_2917 | SPGP0BKtrna23R | YJL102W |
| DI49_2915; YJL104W; DI49_2914; YJL105W | DI49_2915; DI49_2914 | | YJL104W; YJL105W |
| DI49_2912; YJL107C | DI49_2912 | | YJL107C |
| DI49_2911; YJL109C; DI49_2910; YJL110C | DI49_2911; DI49_2910 | | YJL109C; YJL110C |
| SPGP0BK02610; YJL112W | | SPGP0BK02610 | YJL112W |
| DI49_2902; DI49_2903 | DI49_2902; DI49_2903 | | |
| DI49_2901; SPGP0BK02670 | DI49_2901 | SPGP0BK02670 | |
| SPGP0BK02700; DI49_2900 | DI49_2900 | SPGP0BK02700 | |
| DI49_2898; YJL125C; DI49_2897; YJL126W | DI49_2898; DI49_2897 | | YJL125C; YJL126W |
| DI49_2895; YJL127C-B | DI49_2895 | | YJL127C-B |
| DI49_2894; YJL128C | DI49_2894 | | YJL128C |
| DI49_2893; YJL129C | DI49_2893 | | YJL129C |
| DI49_2892; YJL130C | DI49_2892 | | YJL130C |
| DI49_2891; YJL131C | DI49_2891 | | YJL131C |
| SPGP0BK02800; YJL132W | | SPGP0BK02800 | YJL132W |
| DI49_2890; YJL133W | DI49_2890 | | YJL133W |
| DI49_2889; YJL133C-A; DI49_2888; YJL134W | DI49_2889; DI49_2888 | | YJL133C-A; YJL134W |
| DI49_2885; DI49_3430 | DI49_2885; DI49_3430 | | |
| DI49_2882; YJL143W; DI49_2881; YJL144W | DI49_2882; DI49_2881 | | YJL143W; YJL144W |
| DI49_2879; YJL146W; DI49_2878; YJL147C | DI49_2879; DI49_2878 | | YJL146W; YJL147C |
| DI49_2876; YJL149W; DI49_2875; YJL151C | DI49_2876; DI49_2875 | | YJL149W; YJL151C |
| DI49_2873; YJL154C | DI49_2873 | | YJL154C |
| DI49_2872; YJL155C; DI49_2871; YJL156C | DI49_2872; DI49_2871 | | YJL155C; YJL156C |

Create PDF in your applications with the Pdfcrowd HTML to PDF API PDFCROWD

| | | | |
|---|---|---|---|
| SPGP0BK03040; YKL164C; YKL163W; SPGP0G00450 | | SPGP0BK03040; SPGP0G00450 | YKL164C; YKL163W |
| DI49_2868; YJL159W; DI49_2867 | DI49_2868; DI49_2867 | | YJL159W |
| DI49_2865; YJL162C; DI49_2864; YJL163C | DI49_2865; DI49_2864 | | YJL162C; YJL163C |
| DI49_2862; YJL165C | DI49_2862 | | YJL165C |
| DI49_2861; YJL166W | DI49_2861 | | YJL166W |
| DI49_2860; YJL167W | DI49_2860 | | YJL167W |
| DI49_2859; YJL168C; DI49_2858; YJL170C | DI49_2859; DI49_2858 | | YJL168C; YJL170C |
| DI49_2856; YJL172W; DI49_2855; YJL173C | DI49_2856; DI49_2855 | | YJL172W; YJL173C |
| DI49_2853; YJL176C | DI49_2853 | | YJL176C |
| DI49_2852; YJL177W; DI49_2851; YJL178C | DI49_2852; DI49_2851 | | YJL177W; YJL178C |
| DI49_2849; YJL180C; DI49_2848; YJL181W | DI49_2849; DI49_2848 | | YJL180C; YJL181W |
| DI49_2846; DI49_2847 | DI49_2846; DI49_2847 | | |
| DI49_2844; YJL186W; DI49_2843; YJL187C | DI49_2844; DI49_2843 | | YJL186W; YJL187C |
| DI49_2841; YJL190C | DI49_2841 | | YJL190C |
| DI49_2839; YJL192C | DI49_2839 | | YJL192C |
| DI49_2838; YJL193W; DI49_2837; YJL194W | DI49_2838; DI49_2837 | | YJL193W; YJL194W |
| DI49_2835; DI49_2836 | DI49_2835; DI49_2836 | | |
| DI49_2832; YJL200C | DI49_2832 | | YJL200C |
| DI49_2831; YJL201W; DI49_2830; YJL203W | DI49_2831; DI49_2830 | | YJL201W; YJL203W |
| SPGP0BK03430; DI49_2829 | DI49_2829 | SPGP0BK03430 | |
| SPGP0BK03450; YJL207C; DI49_2828; YJL208C | DI49_2828 | SPGP0BK03450 | YJL207C; YJL208C |
| DI49_2826; YJL209W; DI49_2825; YJL210W | DI49_2826; DI49_2825 | | YJL209W; YJL210W |
| SPGP0BK03510; DI49_2824 | DI49_2824 | SPGP0BK03510 | |
| DI49_1920; SPGP0BLtrna24V | DI49_1920 | SPGP0BLtrna24V | |
| DI49_1490; DI49_1919 | DI49_1490; DI49_1919 | | |
| DI49_1492; DI49_1491 | DI49_1492; DI49_1491 | | |
| YHR207C; YHR206W | | | YHR207C; YHR206W |
| YHR209W; YHR208W | | | YHR209W; YHR208W |
| SPGP0BO00160; YAL063C; SPGP0BQ00100; YKL201C | | SPGP0BO00160; SPGP0BQ00100 | YAL063C; YKL201C |
| YBR293W; DI49_1315 | DI49_1315 | | YBR293W |
| YBR284W; YBR285W | | | YBR284W; YBR285W |
| YBR278W; YBR279W | | | YBR278W; YBR279W |
| YBR271W; YBR272C | | | YBR271W; YBR272C |
| YBR269C; YBR270C | | | YBR269C; YBR270C |
| YBR262C; YBR263W | | | YBR262C; YBR263W |
| YBR257W; YBR258C | | | YBR257W; YBR258C |
| YBR253W; YBR254C | | | YBR253W; YBR254C |
| YBR251W; YBR252W | | | YBR251W; YBR252W |
| YBR231C; YBR233W | | | YBR231C; YBR233W |
| YBR217W; YBR218C | | | YBR217W; YBR218C |

Create PDF in your applications with the Pdfcrowd HTML to PDF API PDFCROWD

| | | | |
|---|---|---|---|
| YBR211C; YBR212W | | | YBR211C; YBR212W |
| YBR200W; YBR201W | | | YBR200W; YBR201W |
| YBR198C; YBR199W | | | YBR198C; YBR199W |
| YBR193C; YBR194W | | | YBR193C; YBR194W |
| YBR188C; YBR189W | | | YBR188C; YBR189W |
| YBR186W; YBR187W | | | YBR186W; YBR187W |
| YBR184W; YBR185C | | | YBR184W; YBR185C |
| YBR182C; YBR183W | | | YBR182C; YBR183W |
| YBR180W; YBR181C | | | YBR180W; YBR181C |
| YBR177C; YBR179C | | | YBR177C; YBR179C |
| YBR175W; YBR176W | | | YBR175W; YBR176W |
| YBR162C; YBR162W-A | | | YBR162C; YBR162W-A |
| YBR160W; DI49_1187 | DI49_1187 | | YBR160W |
| YBR148W; YBR149W | | | YBR148W; YBR149W |
| YBR142W; YBR143C | | | YBR142W; YBR143C |
| YBR129C; YBR130C | | | YBR129C; YBR130C |
| YBR120C; YBR121C | | | YBR120C; YBR121C |
| YBR115C; DI49_1147 | DI49_1147 | | YBR115C |
| YBR110W; YBR111C | | | YBR110W; YBR111C |
| YBR108W; YBR109C | | | YBR108W; YBR109C |
| YBR103W; YBR104W | | | YBR103W; YBR104W |
| YBR101C; YBR102C | | | YBR101C; YBR102C |
| YBR085C-A; YBR086C | | | YBR085C-A; YBR086C |
| YBR071W; YBR072W | | | YBR071W; YBR072W |
| YBR064W; YBR065C | | | YBR064W; YBR065C |
| YBR061C; DI49_1094; YBR060C | DI49_1094 | | YBR061C; YBR060C |
| YBR058C-A; YBR059C | | | YBR058C-A; YBR059C |
| YBR057C; YBR058C | | | YBR057C; YBR058C |
| YBR033W; YBR034C | | | YBR033W; YBR034C |
| YBR022W; YBR023C | | | YBR022W; YBR023C |
| YBR020W; YBR021W | | | YBR020W; YBR021W |
| YBR018C; YBR019C | | | YBR018C; YBR019C |
| YBR016W; YBR017C | | | YBR016W; YBR017C |
| YBR014C; YBR015C | | | YBR014C; YBR015C |
| YBL001C; YBR001C | | | YBL001C; YBR001C |
| YBL014C; YBL013W | | | YBL014C; YBL013W |
| SPGP0F03150; YBL026W | | SPGP0F03150 | YBL026W |
| YBL034C; YBL033C | | | YBL034C; YBL033C |
| YBL036C; YBL035C | | | YBL036C; YBL035C |
| YBL049W; YBL047C | | | YBL049W; YBL047C |

Create PDF in your applications with the Pdfcrowd HTML to PDF API PDFCROWD

| | | | |
|---|---|---|---|
| YBL059W; YBL058W | | | YBL059W; YBL058W |
| YBL060W; YBL059C-A | | | YBL060W; YBL059C-A |
| YBL069W; YBL068W | | | YBL069W; YBL068W |
| YBL084C; YBL082C | | | YBL084C; YBL082C |
| YBL098W; YBL097W | | | YBL098W; YBL097W |
| SPGP0Ftrna10F; SPGP0Htrna14F; YBL101C; YMR042W | | SPGP0Ftrna10F; SPGP0Htrna14F | YBL101C; YMR042W |
| YBL103C; YBL102W | | | YBL103C; YBL102W |
| DI49_1964; DI49_1965 | DI49_1964; DI49_1965 | | |
| SPGP0BT00100; SPGP0BU00100; DI49_3844; DI49_1996 | DI49_3844; DI49_1996 | SPGP0BT00100; SPGP0BU00100 | |
| SPGP0BT00120; DI49_1997; DI49_3846; DI49_1998 | DI49_1997; DI49_3846; DI49_1998 | SPGP0BT00120 | |
| DI49_2000; DI49_1999 | DI49_2000; DI49_1999 | | |
| DI49_2005; DI49_2004 | DI49_2005; DI49_2004 | | |
| YKL193C; YKL194C | | | YKL193C; YKL194C |
| YKL191W; YKL192C | | | YKL191W; YKL192C |
| YKL184W; YKL185W | | | YKL184W; YKL185W |
| YKL174C; YKL175W | | | YKL174C; YKL175W |
| YKL172W; YKL173W | | | YKL172W; YKL173W |
| YKL170W; YKL171W | | | YKL170W; YKL171W |
| YKL167C; YKL168C | | | YKL167C; YKL168C |
| YKL161C; YKL162C | | | YKL161C; YKL162C |
| YKL159C; YKL160W | | | YKL159C; YKL160W |
| YKL149C; YKL150W | | | YKL149C; YKL150W |
| YKL139W; YKL140W | | | YKL139W; YKL140W |
| YKL134C; YKL135C | | | YKL134C; YKL135C |
| YKL132C; YKL133C | | | YKL132C; YKL133C |
| YKL108W; YKL109W | | | YKL108W; YKL109W |
| YKL098W; YKL099C | | | YKL098W; YKL099C |
| YKL095W; DI49_3280 | DI49_3280 | | YKL095W |
| YKL091C; YKL092C | | | YKL091C; YKL092C |
| YKL085W; YKL086W | | | YKL085W; YKL086W |
| YIR019C; YKL080W | | | YIR019C; YKL080W |
| YIR022W; YIR021W | | | YIR022W; YIR021W |
| YIR024C; YIR023W | | | YIR024C; YIR023W |
| YIR026C; YIR025W | | | YIR026C; YIR025W |
| YIR029W; DI49_2808; YIR030C | DI49_2808 | | YIR029W; YIR030C |
| YIR032C; YIR031C | | | YIR032C; YIR031C |
| YIR037W; YIR036C | | | YIR037W; YIR036C |
| YIR039C; YIR038C | | | YIR039C; YIR038C |
| DI49_3497; SPGP0CB00300 | DI49_3497 | SPGP0CB00300 | |
| DI49_3499; DI49_3498 | DI49_3499; DI49_3498 | | |

Create PDF in your applications with the Pdfcrowd HTML to PDF API PDFCROWD

| | | | |
|---|---|---|---|
| DI49_1645; DI49_1644 | DI49_1645; DI49_1644 | | |
| DI49_1647; DI49_1646 | DI49_1647; DI49_1646 | | |
| DI49_1615; DI49_1616 | DI49_1615; DI49_1616 | | |
| DI49_1613; DI49_1614 | DI49_1613; DI49_1614 | | |
| DI49_1611; DI49_1612 | DI49_1611; DI49_1612 | | |
| DI49_1609; DI49_1610 | DI49_1609; DI49_1610 | | |
| SPGP0CF00190; DI49_2277; SPGP0CG00100 | DI49_2277 | SPGP0CF00190; SPGP0CG00100 | |
| DI49_3975; DI49_3976 | DI49_3975; DI49_3976 | | |
| DI49_3973; DI49_3974 | DI49_3973; DI49_3974 | | |
| DI49_3981; DI49_3982 | DI49_3981; DI49_3982 | | |
| DI49_3979; DI49_3980 | DI49_3979; DI49_3980 | | |
| YML025C; SPGP0H00300 | | SPGP0H00300 | YML025C |
| YML023C; YML024W | | | YML023C; YML024W |
| YML020W; DI49_4069; YML019W | DI49_4069 | | YML020W; YML019W |
| SPGP0H00390; YML018C | | SPGP0H00390 | YML018C |
| YML015C; YML016C | | | YML015C; YML016C |
| YML013W; YML014W | | | YML013W; YML014W |
| YML011C; YML012W | | | YML011C; YML012W |
| YMR002W; DI49_4090; YMR003W | DI49_4090 | | YMR002W; YMR003W |
| YMR005W; YMR004W | | | YMR005W; YMR004W |
| SPGP0H00630; YMR008C | | SPGP0H00630 | YMR008C |
| YMR018W; YMR017W | | | YMR018W; YMR017W |
| YMR033W; YMR032W | | | YMR033W; YMR032W |
| YMR035W; YMR034C | | | YMR035W; YMR034C |
| YMR037C; YMR036C | | | YMR037C; YMR036C |
| YMR039C; YMR038C | | | YMR039C; YMR038C |
| YMR041C; YMR040W | | | YMR041C; YMR040W |
| YMR047C; DI49_4135; YMR048W | DI49_4135 | | YMR047C; YMR048W |
| YMR054W; DI49_4144; YMR055C | DI49_4144 | | YMR054W; YMR055C |
| YMR064W; YMR063W | | | YMR064W; YMR063W |
| YMR074C; YMR073C | | | YMR074C; YMR073C |
| YMR079W; YMR078C | | | YMR079W; YMR078C |
| YMR081C; YMR080C | | | YMR081C; YMR080C |
| YMR086W; DI49_4174; YMR087W | DI49_4174 | | YMR086W; YMR087W |
| YMR092C; YMR091C | | | YMR092C; YMR091C |
| YMR094W; DI49_4182; YMR095C | DI49_4182 | | YMR094W; YMR095C |
| YMR097C; YMR096W | | | YMR097C; YMR096W |
| YMR099C; DI49_4188; YMR100W | DI49_4188 | | YMR099C; YMR100W |
| YMR102C; DI49_4191; YMR104C | DI49_4191 | | YMR102C; YMR104C |
| YMR106C; YMR105C | | | YMR106C; YMR105C |

Create PDF in your applications with the Pdfcrowd HTML to PDF API PDFCROWD

| | | | |
|---|---|---|---|
| YMR112C; YMR111C | | | YMR112C; YMR111C |
| YMR125W; DI49_4214; YMR126C | DI49_4214 | | YMR125W; YMR126C |
| SPGP0H01760; DI49_4218; YMR130W | DI49_4218 | SPGP0H01760 | YMR130W |
| YMR132C; YMR131C | | | YMR132C; YMR131C |
| YMR134W; YMR133W | | | YMR134W; YMR133W |
| YMR136W; YMR135C | | | YMR136W; YMR135C |
| YMR138W; YMR137C | | | YMR138W; YMR137C |
| YMR140W; YMR139W | | | YMR140W; YMR139W |
| YMR146C; DI49_4231; YMR147W | DI49_4231 | | YMR146C; YMR147W |
| YMR149W; YMR148W | | | YMR149W; YMR148W |
| YMR152W; YMR150C | | | YMR152W; YMR150C |
| YMR154C; YMR153W | | | YMR154C; YMR153W |
| YMR156C; YMR155W | | | YMR156C; YMR155W |
| YMR158W; YMR157C | | | YMR158W; YMR157C |
| YMR162C; DI49_4246; YMR163C | DI49_4246 | | YMR162C; YMR163C |
| YMR169C; YMR170C | | | YMR169C; YMR170C |
| YMR174C; YMR173W | | | YMR174C; YMR173W |
| YMR177W; DI49_4260; YMR178W | DI49_4260 | | YMR177W; YMR178W |
| YMR180C; YMR179W | | | YMR180C; YMR179W |
| YMR182C; YMR181C | | | YMR182C; YMR181C |
| YMR184W; YMR183C | | | YMR184W; YMR183C |
| YMR187C; DI49_4270; YMR188C | DI49_4270 | | YMR187C; YMR188C |
| YMR190C; DI49_4273; YMR191W | DI49_4273 | | YMR190C; YMR191W |
| YMR193W; YMR192W | | | YMR193W; YMR192W |
| YMR197C; DI49_4281; YMR198W | DI49_4281 | | YMR197C; YMR198W |
| YMR205C; DI49_4289; YMR206W | DI49_4289 | | YMR205C; YMR206W |
| YMR208W; DI49_4292; YMR209C | DI49_4292 | | YMR208W; YMR209C |
| YMR211W; YMR210W | | | YMR211W; YMR210W |
| YMR213W; YMR212C | | | YMR213W; YMR212C |
| YMR215W; YMR214W | | | YMR215W; YMR214W |
| YMR232W; DI49_4316; YMR233W | DI49_4316 | | YMR232W; YMR233W |
| YMR251W-A; YMR251W | | | YMR251W-A; YMR251W |
| YMR265C; DI49_4349; YMR266W | DI49_4349 | | YMR265C; YMR266W |
| YMR268C; YMR267W | | | YMR268C; YMR267W |
| YMR281W; YMR280C | | | YMR281W; YMR280C |
| YMR283C; YMR282C | | | YMR283C; YMR282C |
| YMR284W; SPGP0Itrna13Y; SPGP0Htrna11Y; YJL046W | | SPGP0Itrna13Y; SPGP0Htrna11Y | YMR284W; YJL046W |
| YMR286W; YMR285C | | | YMR286W; YMR285C |
| YMR288W; DI49_4375; YMR289W | DI49_4375 | | YMR288W; YMR289W |

Create PDF in your applications with the Pdfcrowd HTML to PDF API PDFCROWD

| | | | |
|---|---|---|---|
| YMR291W; DI49_4378; YMR292W | DI49_4378 | | YMR291W; YMR292W |
| YMR294W; YMR293C | | | YMR294W; YMR293C |
| YMR296C; YMR295C | | | YMR296C; YMR295C |
| YMR298W; DI49_4385; YMR299C | DI49_4385 | | YMR298W; YMR299C |
| YMR305C; DI49_4392; SPGP0H03500 | DI49_4392 | SPGP0H03500 | YMR305C |
| DI49_1957; DI49_5599 | DI49_1957; DI49_5599 | | |
| DI49_1955; DI49_1956 | DI49_1955; DI49_1956 | | |
| DI49_3931; DI49_1954 | DI49_3931; DI49_1954 | | |
| YJR142W; YJR143C | | | YJR142W; YJR143C |
| YJR140C; YJR141W | | | YJR140C; YJR141W |
| YJR129C; YJR130C | | | YJR129C; YJR130C |
| YJR126C; YJR127C | | | YJR126C; YJR127C |
| YJR113C; YJR115W | | | YJR113C; YJR115W |
| YJR112W; YJR112W-A | | | YJR112W; YJR112W-A |
| YJR101W; YJR102C | | | YJR101W; YJR102C |
| YJR092W; YJR093C | | | YJR092W; YJR093C |
| YJR086W; YJR088C | | | YJR086W; YJR088C |
| YJR084W; YJR085C | | | YJR084W; YJR085C |
| YJR082C; YJR083C | | | YJR082C; YJR083C |
| YJR078W; YJR080C | | | YJR078W; YJR080C |
| YJR072C; YJR073C | | | YJR072C; YJR073C |
| YJR061W; YJR062C | | | YJR061W; YJR062C |
| YJR049C; YJR050W | | | YJR049C; YJR050W |
| YJR044C; YJR045C | | | YJR044C; YJR045C |
| YJR041C; YJR042W | | | YJR041C; YJR042W |
| YJR025C; YJR030C | | | YJR025C; YJR030C |
| YJR022W; YJR024C | | | YJR022W; YJR024C |
| YJR019C; YJR021C | | | YJR019C; YJR021C |
| YJR013W; YJR012C | | | YJR013W; YJR012C |
| YJR001W; YJR002W | | | YJR001W; YJR002W |
| YJL002C; YJL001W | | | YJL002C; YJL001W |
| YJL004C; YJL003W | | | YJL004C; YJL003W |
| YJL013C; YJL012C | | | YJL013C; YJL012C |
| YJL016W; YJL014W | | | YJL016W; YJL014W |
| YJL020C; YJL019W | | | YJL020C; YJL019W |
| YJL027C; YJL026W | | | YJL027C; YJL026W |
| YJL031C; YJL030W | | | YJL031C; YJL030W |
| YJL043W; YJL042W | | | YJL043W; YJL042W |
| YJL045W; YJL044C | | | YJL045W; YJL044C |
| YJL048C; YJL047C | | | YJL048C; YJL047C |

Create PDF in your applications with the Pdfcrowd HTML to PDF API PDFCROWD

| | | | |
|---|---|---|---|
| YJL055W; YJL054W | | | YJL055W; YJL054W |
| YJL057C; YJL056C | | | YJL057C; YJL056C |
| YJL066C; YJL065C | | | YJL066C; YJL065C |
| YJL069C; YJL068C | | | YJL069C; YJL068C |
| YJL076W; YJL074C | | | YJL076W; YJL074C |
| YJL082W; DI49_2936 | DI49_2936 | | YJL082W |
| YJL117W; YJL116C | | | YJL117W; YJL116C |
| YJL121C; YJL118W | | | YJL121C; YJL118W |
| YJL123C; YJL122W | | | YJL123C; YJL122W |
| YJL137C; YJL136C | | | YJL137C; YJL136C |
| YJL139C; YJL138C | | | YJL139C; YJL138C |
| YJL184W; YJL183W | | | YJL184W; YJL183W |
| YJL197W; YJL196C | | | YJL197W; YJL196C |
| YJL199C; YJL198W | | | YJL199C; YJL198W |
| YJL205C; YJL204C | | | YJL205C; YJL204C |
| SPGP0I03280; YJL212C | | SPGP0I03280 | YJL212C |

Create PDF in your applications with the Pdfcrowd HTML to PDF API PDFCROWD
