## Supplementary file 2 for "HybridMine: pipeline for allele inheritance and gene copy number prediction in industrial yeast hybrids"

### Overview


genes alleles inherited from Saccharomyces eubayanus
genes alleles inherited from Saccharomyces cerevisiae
genes alleles below 80% identity
genes alleles copy with same identity in both parents
list of paralogs in Saccharomyces pastorianus CBS1513


#### Sheet 1: *genes alleles inherited from Saccharomyces eubayanus*

| *Saccharomyces pastorianus CBS1513 predicted ORF* | *Saccharomyces eubayanus parental allele* | Gene identities | Gene gaps |
|---|---|---|---|
| SPGP0U01640 | XP_018218830.1 | 99 | 0 |
| SPGP0W00160 | XP_018223318.1 | 99 | 0 |
| SPGP0EX00100 | XP_018223869.1 | 99 | 0 |
| SPGP0AH00300 | XP_018222572.1 | 99 | 0 |
| SPGP0CZ01880 | XP_018221009.1 | 99 | 0 |
| SPGP0CZ02780 | XP_018221097.1 | 99 | 0 |
| SPGP0N01210 | XP_018220691.1 | 99 | 0 |
| SPGP0Y00680 | XP_018223941.1 | 99 | 0 |
| SPGP0CZ02530 | XP_018221073.1 | 99 | 0 |
| SPGP0CZ01080 | XP_018220933.1 | 99 | 0 |
| SPGP0Y01020 | XP_018223972.1 | 99 | 0 |
| SPGP0U00470 | XP_018218942.1 | 99 | 0 |
| SPGP0Y01360 | XP_018224005.1 | 99 | 0 |
| SPGP0DU01760 | XP_018220415.1 | 99 | 0 |
| SPGP0DU01320 | XP_018220458.1 | 99 | 0 |
| SPGP0U00690 | XP_018218923.1 | 99 | 0 |
| SPGP0AC00640 | XP_018219148.1 | 99 | 0 |
| SPGP0BF00240 | XP_018220394.1 | 99 | 0 |
| SPGP0DZ01040 | XP_018219443.1 | 99 | 0 |
| SPGP0ET00140 | XP_018218983.1 | 99 | 0 |
| SPGP0BC00290 | XP_018220825.1 | 99 | 0 |
| SPGP0N01500 | XP_018220664.1 | 99 | 0 |
| SPGP0BC00120 | XP_018220810.1 | 99 | 0 |
| SPGP0U00490 | XP_018218940.1 | 99 | 0 |
| SPGP0DU01160 | XP_018220474.1 | 100 | 0 |
| SPGP0AC01070 | XP_018219190.1 | 99 | 0 |
| SPGP0DZ01250 | XP_018219464.1 | 99 | 0 |
| SPGP0EG00100 | XP_018221945.1 | 99 | 0 |

Create PDF in your applications with the Pdfcrowd HTML to PDF API PDFCROWD

| | | | |
|---|---|---|---|
| SPGP0Y00730 | XP_018223945.1 | 99 | 0 |
| SPGP0DZ00920 | XP_018219432.1 | 99 | 0 |
| SPGP0CZ00210 | XP_018220849.1 | 99 | 0 |
| SPGP0DU01230 | XP_018220467.1 | 99 | 0 |
| SPGP0AC00670 | XP_018219151.1 | 99 | 1 |
| SPGP0DZ02270 | XP_018219564.1 | 99 | 0 |
| SPGP0CZ00270 | XP_018220854.1 | 99 | 0 |
| SPGP0Q03560 | XP_018222148.1 | 99 | 0 |
| SPGP0CZ02520 | XP_018221072.1 | 96 | 0 |
| SPGP0N01660 | XP_018220648.1 | 99 | 0 |
| SPGP0AH01160 | XP_018222492.1 | 99 | 0 |
| SPGP0DZ00860 | XP_018219426.1 | 99 | 0 |
| SPGP0U01150 | XP_018218877.1 | 99 | 0 |
| SPGP0N01970 | XP_018220617.1 | 99 | 0 |
| SPGP0N00630 | XP_018220745.1 | 99 | 0 |
| SPGP0N01420 | XP_018220671.1 | 100 | 0 |
| SPGP0ET00160 | XP_018218985.1 | 99 | 0 |
| SPGP0U01010 | XP_018218891.1 | 99 | 0 |
| SPGP0DZ01860 | XP_018219524.1 | 99 | 0 |
| SPGP0AC01010 | XP_018219184.1 | 99 | 0 |
| SPGP0DU00790 | XP_018220512.1 | 85 | 0 |
| SPGP0U01800 | XP_018218814.1 | 99 | 0 |
| SPGP0DX00150 | XP_018219946.1 | 99 | 0 |
| SPGP0CZ01570 | XP_018220981.1 | 99 | 0 |
| SPGP0DZ02060 | XP_018219543.1 | 99 | 0 |
| SPGP0U01700 | XP_018218824.1 | 99 | 0 |
| SPGP0Y00200 | XP_018223898.1 | 100 | 0 |
| SPGP0CZ00790 | XP_018220904.1 | 100 | 0 |
| SPGP0N02010 | XP_018220613.1 | 99 | 1 |
| SPGP0CZ02800 | XP_018221099.1 | 99 | 0 |
| SPGP0N01930 | XP_018220621.1 | 99 | 0 |
| SPGP0P01570 | XP_018221802.1 | 99 | 0 |
| SPGP0U00320 | XP_018218957.1 | 99 | 0 |
| SPGP0U01900 | XP_018218804.1 | 99 | 0 |
| SPGP0U01490 | XP_018218844.1 | 99 | 0 |
| SPGP0N01090 | XP_018220702.1 | 99 | 0 |
| SPGP0DU01340 | XP_018220456.1 | 99 | 0 |
| SPGP0DZ01690 | XP_018219507.1 | 99 | 0 |
| SPGP0DX00670 | XP_018219996.1 | 99 | 0 |
| SPGP0CZ03100 | XP_018221128.1 | 99 | 0 |

Create PDF in your applications with the Pdfcrowd HTML to PDF API PDFCROWD

| | | | |
|---|---|---|---|
| SPGP0DZ00330 | XP_018219376.1 | 99 | 0 |
| SPGP0N00320 | XP_018220776.1 | 99 | 0 |
| SPGP0Z02890 | XP_018219596.1 | 100 | 0 |
| SPGP0DZ01920 | XP_018219528.1 | 99 | 0 |
| SPGP0DU00360 | XP_018220554.1 | 99 | 0 |
| SPGP0CZ02030 | XP_018221024.1 | 99 | 0 |
| SPGP0DU01360 | XP_018220454.1 | 99 | 0 |
| SPGP0AC00190 | XP_018219104.1 | 99 | 0 |
| SPGP0AC00220 | XP_018219107.1 | 99 | 0 |
| SPGP0DX00360 | XP_018219967.1 | 99 | 0 |
| SPGP0AH00920 | XP_018222514.1 | 97 | 2 |
| SPGP0CZ02170 | XP_018221037.1 | 99 | 0 |
| SPGP0CZ00780 | XP_018220903.1 | 99 | 1 |
| SPGP0CZ01780 | XP_018221001.1 | 99 | 0 |
| SPGP0DZ02210 | XP_018219558.1 | 99 | 0 |
| SPGP0Y01580 | XP_018224027.1 | 99 | 0 |
| SPGP0Y01210 | XP_018223990.1 | 99 | 0 |
| SPGP0AC00770 | XP_018219160.1 | 99 | 0 |
| SPGP0AH00540 | XP_018222551.1 | 99 | 0 |
| SPGP0AH01140 | XP_018222494.1 | 99 | 0 |
| SPGP0N02200 | XP_018220594.1 | 99 | 0 |
| SPGP0U01170 | XP_018218875.1 | 99 | 0 |
| SPGP0T00300 | XP_018220358.1 | 99 | 0 |
| SPGP0DU00270 | XP_018220563.1 | 98 | 0 |
| SPGP0N01950 | XP_018220619.1 | 99 | 0 |
| SPGP0DZ01710 | XP_018219509.1 | 100 | 0 |
| SPGP0DU01550 | XP_018220435.1 | 99 | 0 |
| SPGP0U01790 | XP_018218815.1 | 99 | 0 |
| SPGP0DU00770 | XP_018220514.1 | 99 | 0 |
| SPGP0EE00150 | XP_018220834.1 | 99 | 0 |
| SPGP0DU00110 | XP_018220579.1 | 99 | 0 |
| SPGP0DX00170 | XP_018219948.1 | 99 | 0 |
| SPGP0N01240 | XP_018220689.1 | 98 | 1 |
| SPGP0N00460 | XP_018220761.1 | 99 | 0 |
| SPGP0N01780 | XP_018220636.1 | 99 | 0 |
| SPGP0DZ01240 | XP_018219463.1 | 99 | 0 |
| SPGP0DZ01530 | XP_018219492.1 | 100 | 0 |
| SPGP0ET00210 | XP_018218990.1 | 100 | 0 |
| SPGP0N01020 | XP_018220709.1 | 99 | 0 |
| SPGP0AH00730 | XP_018222532.1 | 99 | 0 |

Create PDF in your applications with the Pdfcrowd HTML to PDF API PDFCROWD

| | | | |
|---|---|---|---|
| SPGP0DZ02080 | XP_018219545.1 | 99 | 0 |
| SPGP0BF00420 | XP_018220410.1 | 99 | 0 |
| SPGP0N00120 | XP_018220796.1 | 99 | 0 |
| SPGP0CZ02470 | XP_018221067.1 | 98 | 1 |
| SPGP0AA00130 | XP_018223413.1 | 99 | 0 |
| SPGP0AC00610 | XP_018219145.1 | 99 | 0 |
| SPGP0E01710 | XP_018222760.1 | 99 | 0 |
| SPGP0N00870 | XP_018220722.1 | 99 | 1 |
| SPGP0U01300 | XP_018218863.1 | 99 | 0 |
| SPGP0DZ02330 | XP_018218748.1 | 97 | 3 |
| SPGP0CZ02010 | XP_018221022.1 | 99 | 0 |
| SPGP0DU01430 | XP_018220447.1 | 99 | 0 |
| SPGP0N02290 | XP_018220585.1 | 99 | 0 |
| SPGP0N01610 | XP_018220653.1 | 99 | 0 |
| SPGP0FT00110 | XP_018220800.1 | 99 | 0 |
| SPGP0BC00160 | XP_018220813.1 | 99 | 0 |
| SPGP0DX00340 | XP_018219965.1 | 100 | 0 |
| SPGP0N00600 | XP_018220748.1 | 99 | 0 |
| SPGP0DX00610 | XP_018219991.1 | 99 | 0 |
| SPGP0AH00900 | XP_018222516.1 | 99 | 0 |
| SPGP0DU01280 | XP_018220462.1 | 99 | 0 |
| SPGP0DZ00960 | XP_018219436.1 | 99 | 0 |
| SPGP0CZ03050 | XP_018221123.1 | 99 | 0 |
| SPGP0Y00560 | XP_018223930.1 | 99 | 0 |
| SPGP0DU00910 | XP_018220499.1 | 99 | 0 |
| SPGP0DU00720 | XP_018220519.1 | 99 | 0 |
| SPGP0CZ02650 | XP_018221085.1 | 99 | 0 |
| SPGP0Y01720 | XP_018224039.1 | 99 | 0 |
| SPGP0CZ00500 | XP_018220876.1 | 99 | 0 |
| SPGP0Y00930 | XP_018223963.1 | 100 | 0 |
| SPGP0DU00740 | XP_018220517.1 | 99 | 0 |
| SPGP0N02220 | XP_018220592.1 | 99 | 1 |
| SPGP0U00950 | XP_018218897.1 | 99 | 0 |
| SPGP0CZ02300 | XP_018221050.1 | 99 | 0 |
| SPGP0E01550 | XP_018222743.1 | 100 | 0 |
| SPGP0W00180 | XP_018223316.1 | 99 | 0 |
| SPGP0U00820 | XP_018218910.1 | 99 | 0 |
| SPGP0Y00160 | XP_018223894.1 | 99 | 0 |
| SPGP0BC00150 | XP_018220812.1 | 99 | 0 |
| SPGP0U00170 | XP_018218972.1 | 99 | 0 |

Create PDF in your applications with the Pdfcrowd HTML to PDF API PDFCROWD

| | | | |
|---|---|---|---|
| SPGP0CZ01960 | XP_018221017.1 | 100 | 0 |
| SPGP0N01070 | XP_018220704.1 | 99 | 0 |
| SPGP0EE00130 | XP_018220832.1 | 99 | 0 |
| SPGP0N00670 | XP_018220741.1 | 99 | 0 |
| SPGP0N01190 | XP_018220693.1 | 99 | 0 |
| SPGP0DZ00530 | XP_018219396.1 | 99 | 0 |
| SPGP0N00210 | XP_018220787.1 | 99 | 0 |
| SPGP0DZ01970 | XP_018219533.1 | 99 | 0 |
| SPGP0DZ00160 | XP_018219359.1 | 99 | 0 |
| SPGP0CZ02060 | XP_018221027.1 | 99 | 0 |
| SPGP0T00170 | XP_018220345.1 | 99 | 0 |
| SPGP0N01460 | XP_018220667.1 | 99 | 0 |
| SPGP0R00310 | XP_018221154.1 | 99 | 0 |
| SPGP0AH00130 | XP_018222589.1 | 99 | 0 |
| SPGP0DZ02300 | XP_018219567.1 | 99 | 0 |
| SPGP0CZ01350 | XP_018220960.1 | 99 | 0 |
| SPGP0CZ01820 | XP_018221004.1 | 98 | 2 |
| SPGP0EG00130 | XP_018221948.1 | 99 | 0 |
| SPGP0DZ01410 | XP_018219480.1 | 99 | 0 |
| SPGP0DZ00390 | XP_018219382.1 | 99 | 0 |
| SPGP0CZ02460 | XP_018221066.1 | 99 | 0 |
| SPGP0R00290 | XP_018221152.1 | 99 | 0 |
| SPGP0DU00990 | XP_018220491.1 | 99 | 0 |
| SPGP0Y00870 | XP_018223957.1 | 100 | 0 |
| SPGP0CZ02240 | XP_018221044.1 | 99 | 0 |
| SPGP0N01670 | XP_018220647.1 | 99 | 0 |
| SPGP0BC00180 | XP_018220815.1 | 99 | 0 |
| SPGP0AC01090 | XP_018219192.1 | 99 | 0 |
| SPGP0CZ01740 | XP_018220997.1 | 99 | 0 |
| SPGP0CZ02920 | XP_018221111.1 | 99 | 0 |
| SPGP0T00140 | XP_018220342.1 | 99 | 0 |
| SPGP0DU00930 | XP_018220497.1 | 99 | 0 |
| SPGP0CZ01020 | XP_018220927.1 | 99 | 0 |
| SPGP0DZ01160 | XP_018219455.1 | 99 | 1 |
| SPGP0N01990 | XP_018220615.1 | 100 | 0 |
| SPGP0CZ00560 | XP_018220882.1 | 99 | 0 |
| SPGP0CZ00290 | XP_018220856.1 | 98 | 0 |
| SPGP0N00490 | XP_018220758.1 | 99 | 0 |
| SPGP0DZ01700 | XP_018219508.1 | 99 | 0 |
| SPGP0CZ02580 | XP_018221078.1 | 99 | 0 |

Create PDF in your applications with the Pdfcrowd HTML to PDF API PDFCROWD

| | | | |
|---|---|---|---|
| SPGP0CZ01160 | XP_018220941.1 | 99 | 0 |
| SPGP0W00140 | XP_018223320.1 | 99 | 0 |
| SPGP0T00340 | XP_018220362.1 | 99 | 0 |
| SPGP0U00800 | XP_018218912.1 | 99 | 0 |
| SPGP0N01440 | XP_018220669.1 | 100 | 0 |
| SPGP0U00110 | XP_018218977.1 | 99 | 0 |
| SPGP0DZ02020 | XP_018219539.1 | 99 | 0 |
| SPGP0DU00730 | XP_018220518.1 | 99 | 0 |
| SPGP0AH01090 | XP_018222499.1 | 99 | 0 |
| SPGP0Y01370 | XP_018224006.1 | 99 | 0 |
| SPGP0DZ01950 | XP_018219531.1 | 99 | 0 |
| SPGP0DU01740 | XP_018220417.1 | 99 | 0 |
| SPGP0Y01650 | XP_018224034.1 | 99 | 0 |
| SPGP0CZ00200 | XP_018220848.1 | 99 | 0 |
| SPGP0CZ01370 | XP_018220961.1 | 99 | 0 |
| SPGP0DZ00150 | XP_018219358.1 | 99 | 1 |
| SPGP0DU01310 | XP_018220459.1 | 99 | 0 |
| SPGP0EX00270 | XP_018223885.1 | 98 | 1 |
| SPGP0DZ01420 | XP_018219481.1 | 99 | 0 |
| SPGP0CZ01390 | XP_018220963.1 | 99 | 0 |
| SPGP0AC00830 | XP_018219166.1 | 99 | 0 |
| SPGP0CZ02440 | XP_018221064.1 | 99 | 0 |
| SPGP0FB00130 | XP_018220807.1 | 99 | 0 |
| SPGP0DU01580 | XP_018220433.1 | 99 | 0 |
| SPGP0DU01680 | XP_018220423.1 | 99 | 0 |
| SPGP0CZ02050 | XP_018221026.1 | 99 | 0 |
| SPGP0BF00380 | XP_018220406.1 | 99 | 0 |
| SPGP0BC00170 | XP_018220814.1 | 99 | 0 |
| SPGP0CZ03080 | XP_018221126.1 | 99 | 0 |
| SPGP0U01340 | XP_018218859.1 | 99 | 0 |
| SPGP0AC00170 | XP_018219102.1 | 99 | 0 |
| SPGP0DX00370 | XP_018219968.1 | 99 | 0 |
| SPGP0Y01700 | XP_018224037.1 | 99 | 0 |
| SPGP0AC00280 | XP_018219112.1 | 99 | 0 |
| SPGP0EC00110 | XP_018220413.1 | 99 | 0 |
| SPGP0CZ01720 | XP_018220995.1 | 99 | 0 |
| SPGP0N02090 | XP_018220605.1 | 99 | 0 |
| SPGP0DZ02140 | XP_018219551.1 | 99 | 0 |
| SPGP0N00360 | XP_018220772.1 | 99 | 0 |

Create PDF in your applications with the Pdfcrowd HTML to PDF API PDFCROWD

| | | | |
|---|---|---|---|
| SPGP0DU01290 | XP_018220461.1 | 99 | 0 |
| SPGP0DZ01490 | XP_018219488.1 | 99 | 0 |
| SPGP0CZ02610 | XP_018221081.1 | 99 | 0 |
| SPGP0DZ00580 | XP_018219400.1 | 99 | 0 |
| SPGP0AH00650 | XP_018222540.1 | 99 | 0 |
| SPGP0AC00410 | XP_018219125.1 | 99 | 0 |
| SPGP0U00280 | XP_018218961.1 | 99 | 1 |
| SPGP0BF00370 | XP_018220405.1 | 99 | 0 |
| SPGP0U00750 | XP_018218917.1 | 99 | 0 |
| SPGP0DU01490 | XP_018220441.1 | 99 | 0 |
| SPGP0W00150 | XP_018223319.1 | 99 | 0 |
| SPGP0AH00590 | XP_018222546.1 | 99 | 0 |
| SPGP0U01080 | XP_018218884.1 | 99 | 0 |
| SPGP0CZ02380 | XP_018221058.1 | 99 | 0 |
| SPGP0DZ00520 | XP_018219395.1 | 99 | 0 |
| SPGP0DZ01820 | XP_018219520.1 | 99 | 0 |
| SPGP0DU00240 | XP_018220566.1 | 99 | 0 |
| SPGP0Y01530 | XP_018224022.1 | 99 | 0 |
| SPGP0U01880 | XP_018218806.1 | 99 | 0 |
| SPGP0DU01590 | XP_018220432.1 | 99 | 0 |
| SPGP0DU01040 | XP_018220486.1 | 99 | 1 |
| SPGP0DU01620 | XP_018220429.1 | 99 | 0 |
| SPGP0AC00340 | XP_018219118.1 | 99 | 0 |
| SPGP0DX00510 | XP_018219981.1 | 99 | 0 |
| SPGP0AC00750 | XP_018219158.1 | 99 | 0 |
| SPGP0CZ00150 | XP_018220844.1 | 99 | 0 |
| SPGP0DZ01550 | XP_018219494.1 | 99 | 0 |
| SPGP0BC00140 | XP_018220811.1 | 99 | 0 |
| SPGP0AH00460 | XP_018222559.1 | 99 | 0 |
| SPGP0AA00970 | XP_018223405.1 | 100 | 0 |
| SPGP0CZ01170 | XP_018220942.1 | 99 | 0 |
| SPGP0CZ00310 | XP_018220858.1 | 99 | 0 |
| SPGP0BC00190 | XP_018220816.1 | 99 | 0 |
| SPGP0N01100 | XP_018220701.1 | 99 | 0 |
| SPGP0BC00230 | XP_018220820.1 | 99 | 0 |
| SPGP0DX00390 | XP_018219970.1 | 99 | 0 |
| SPGP0DU00500 | XP_018220540.1 | 99 | 0 |
| SPGP0EX00220 | XP_018223881.1 | 99 | 0 |
| SPGP0DU01050 | XP_018220485.1 | 99 | 0 |
| SPGP0CZ01530 | XP_018220977.1 | 98 | 1 |

Create PDF in your applications with the Pdfcrowd HTML to PDF API PDFCROWD

| | | | |
|---|---|---|---|
| SPGP0EE00110 | XP_018220830.1 | 99 | 0 |
| SPGP0Y00250 | XP_018223903.1 | 99 | 0 |
| SPGP0CZ02960 | XP_018221115.1 | 99 | 0 |
| SPGP0CZ00700 | XP_018220895.1 | 100 | 0 |
| SPGP0N01900 | XP_018220624.1 | 99 | 0 |
| SPGP0CZ02630 | XP_018221083.1 | 99 | 0 |
| SPGP0Y01090 | XP_018223979.1 | 99 | 0 |
| SPGP0T00380 | XP_018220367.1 | 99 | 0 |
| SPGP0EP00100 | XP_018220335.1 | 99 | 0 |
| SPGP0AC00390 | XP_018219123.1 | 99 | 0 |
| SPGP0CZ02430 | XP_018221063.1 | 99 | 0 |
| SPGP0CZ01610 | XP_018220985.1 | 98 | 1 |
| SPGP0DZ00780 | XP_018219419.1 | 99 | 0 |
| SPGP0N01820 | XP_018220632.1 | 99 | 0 |
| SPGP0DU01720 | XP_018220419.1 | 99 | 0 |
| SPGP0U00730 | XP_018218919.1 | 99 | 0 |
| SPGP0Y01550 | XP_018224024.1 | 99 | 0 |
| SPGP0N00610 | XP_018220747.1 | 99 | 0 |
| SPGP0T00100 | XP_018220338.1 | 99 | 0 |
| SPGP0N00750 | XP_018220734.1 | 97 | 0 |
| SPGP0U01600 | XP_018218834.1 | 99 | 0 |
| SPGP0CZ02390 | XP_018221059.1 | 99 | 0 |
| SPGP0F04410 | XP_018222050.1 | 99 | 0 |
| SPGP0DU01020 | XP_018220488.1 | 99 | 0 |
| SPGP0DZ01910 | XP_018219527.1 | 99 | 0 |
| SPGP0CZ00320 | XP_018220859.1 | 99 | 0 |
| SPGP0CZ03110 | XP_018221129.1 | 99 | 0 |
| SPGP0CZ00870 | XP_018220912.1 | 100 | 0 |
| SPGP0DZ00380 | XP_018219381.1 | 99 | 0 |
| SPGP0DU00850 | XP_018220505.1 | 100 | 0 |
| SPGP0AH00400 | XP_018222564.1 | 99 | 0 |
| SPGP0DZ01840 | XP_018219522.1 | 99 | 0 |
| SPGP0DX00490 | XP_018219979.1 | 99 | 0 |
| SPGP0CZ01260 | XP_018220951.1 | 99 | 0 |
| SPGP0DZ00190 | XP_018219362.1 | 99 | 0 |
| SPGP0N01310 | XP_018220682.1 | 99 | 0 |
| SPGP0DZ00470 | XP_018219390.1 | 99 | 0 |
| SPGP0N02120 | XP_018220602.1 | 99 | 0 |
| SPGP0DZ02010 | XP_018219537.1 | 99 | 0 |
| SPGP0CZ00430 | XP_018220869.1 | 99 | 0 |

Create PDF in your applications with the Pdfcrowd HTML to PDF API PDFCROWD

| | | | |
|---|---|---|---|
| SPGP0AC00680 | XP_018219152.1 | 99 | 0 |
| SPGP0Y00890 | XP_018223959.1 | 96 | 3 |
| SPGP0AH00360 | XP_018222568.1 | 100 | 0 |
| SPGP0N01690 | XP_018220645.1 | 99 | 0 |
| SPGP0N01160 | XP_018220696.1 | 99 | 0 |
| SPGP0Y00230 | XP_018223901.1 | 100 | 0 |
| SPGP0CZ02100 | XP_018221030.1 | 100 | 0 |
| SPGP0EX00240 | XP_018223883.1 | 99 | 0 |
| SPGP0CZ02180 | XP_018221038.1 | 99 | 0 |
| SPGP0CZ01550 | XP_018220979.1 | 99 | 0 |
| SPGP0AA00610 | XP_018223373.1 | 100 | 0 |
| SPGP0AC00290 | XP_018219113.1 | 99 | 0 |
| SPGP0CZ00450 | XP_018220871.1 | 99 | 0 |
| SPGP0Y00180 | XP_018223896.1 | 99 | 0 |
| SPGP0DU00440 | XP_018220546.1 | 99 | 0 |
| SPGP0DZ00410 | XP_018219384.1 | 100 | 0 |
| SPGP0U00710 | XP_018218921.1 | 99 | 0 |
| SPGP0Y01570 | XP_018224026.1 | 99 | 0 |
| SPGP0U01460 | XP_018218847.1 | 99 | 0 |
| SPGP0BF00200 | XP_018220390.1 | 99 | 0 |
| SPGP0AH00570 | XP_018222548.1 | 99 | 0 |
| SPGP0DZ00550 | XP_018219397.1 | 99 | 0 |
| SPGP0DU01260 | XP_018220464.1 | 99 | 0 |
| SPGP0N00770 | XP_018220732.1 | 99 | 0 |
| SPGP0DU00570 | XP_018220533.1 | 99 | 0 |
| SPGP0N01770 | XP_018220637.1 | 99 | 0 |
| SPGP0U00620 | XP_018218927.1 | 99 | 0 |
| SPGP0U01620 | XP_018218832.1 | 99 | 0 |
| SPGP0N01570 | XP_018220657.1 | 99 | 0 |
| SPGP0DU00940 | XP_018220496.1 | 99 | 0 |
| SPGP0AC00480 | XP_018219132.1 | 99 | 0 |
| SPGP0CZ02160 | XP_018221036.1 | 99 | 0 |
| SPGP0Y00500 | XP_018223924.1 | 99 | 0 |
| SPGP0DZ02310 | XP_018219568.1 | 99 | 0 |
| SPGP0AH00890 | XP_018222517.1 | 98 | 1 |
| SPGP0AH00420 | XP_018222562.1 | 99 | 0 |
| SPGP0DZ01660 | XP_018219504.1 | 99 | 0 |
| SPGP0CZ01240 | XP_018220949.1 | 99 | 0 |
| SPGP0EX00210 | XP_018223880.1 | 99 | 0 |
| SPGP0U01450 | XP_018218848.1 | 99 | 1 |

Create PDF in your applications with the Pdfcrowd HTML to PDF API PDFCROWD

| | | | |
|---|---|---|---|
| SPGP0DZ00640 | XP_018219405.1 | 99 | 1 |
| SPGP0N01330 | XP_018220680.1 | 99 | 0 |
| SPGP0N00740 | XP_018220735.1 | 99 | 0 |
| SPGP0Y01290 | XP_018223998.1 | 99 | 0 |
| SPGP0CZ01230 | XP_018220948.1 | 99 | 0 |
| SPGP0DU00540 | XP_018220536.1 | 100 | 0 |
| SPGP0BC00270 | XP_018220824.1 | 99 | 0 |
| SPGP0DZ02120 | XP_018219549.1 | 99 | 0 |
| SPGP0AH00150 | XP_018222587.1 | 99 | 0 |
| SPGP0DZ01330 | XP_018219472.1 | 99 | 0 |
| SPGP0DX00260 | XP_018219957.1 | 99 | 0 |
| SPGP0DZ01780 | XP_018219516.1 | 99 | 0 |
| SPGP0AH01010 | XP_018222506.1 | 99 | 0 |
| SPGP0N00930 | XP_018220716.1 | 99 | 0 |
| SPGP0CZ01190 | XP_018220944.1 | 99 | 0 |
| SPGP0N01750 | XP_018220639.1 | 99 | 0 |
| SPGP0AC00160 | XP_018219101.1 | 99 | 1 |
| SPGP0AC00660 | XP_018219150.1 | 99 | 0 |
| SPGP0FT00120 | XP_018220801.1 | 99 | 0 |
| SPGP0Y01330 | XP_018224002.1 | 99 | 0 |
| SPGP0Y00770 | XP_018223949.1 | 100 | 0 |
| SPGP0Y00480 | XP_018223922.1 | 99 | 0 |
| SPGP0B00270 | XP_018221475.1 | 99 | 0 |
| SPGP0N02280 | XP_018220586.1 | 99 | 0 |
| SPGP0CZ02290 | XP_018221049.1 | 99 | 0 |
| SPGP0CZ00460 | XP_018220872.1 | 99 | 0 |
| SPGP0DZ01310 | XP_018219470.1 | 99 | 0 |
| SPGP0Y00980 | XP_018223968.1 | 99 | 0 |
| SPGP0ET00150 | XP_018218984.1 | 99 | 0 |
| SPGP0DZ00990 | XP_018219439.1 | 100 | 0 |
| SPGP0N01910 | XP_018220623.1 | 99 | 0 |
| SPGP0AH00470 | XP_018222558.1 | 99 | 0 |
| SPGP0N01350 | XP_018220678.1 | 99 | 0 |
| SPGP0N01300 | XP_018220683.1 | 99 | 0 |
| SPGP0N02160 | XP_018220598.1 | 99 | 0 |
| SPGP0T00200 | XP_018220348.1 | 99 | 0 |
| SPGP0Y01130 | XP_018223983.1 | 99 | 0 |
| SPGP0AC01040 | XP_018219187.1 | 99 | 0 |
| SPGP0R01560 | XP_018221273.1 | 99 | 0 |
| SPGP0AC00530 | XP_018219137.1 | 99 | 0 |

Create PDF in your applications with the Pdfcrowd HTML to PDF API PDFCROWD

| | | | |
|---|---|---|---|
| SPGP0AH01070 | XP_018222501.1 | 100 | 0 |
| SPGP0CZ01410 | XP_018220965.1 | 99 | 0 |
| SPGP0AC00980 | XP_018219181.1 | 99 | 0 |
| SPGP0DU00800 | XP_018220510.1 | 87 | 0 |
| SPGP0DZ00930 | XP_018219433.1 | 99 | 0 |
| SPGP0U00200 | XP_018218969.1 | 99 | 0 |
| SPGP0U01330 | XP_018218860.1 | 99 | 0 |
| SPGP0AH01030 | XP_018222505.1 | 99 | 0 |
| SPGP0CZ02340 | XP_018221054.1 | 99 | 0 |
| SPGP0AC00790 | XP_018219162.1 | 99 | 0 |
| SPGP0CZ02330 | XP_018221053.1 | 99 | 0 |
| SPGP0DZ02260 | XP_018219563.1 | 99 | 0 |
| SPGP0U01000 | XP_018218892.1 | 99 | 0 |
| SPGP0U01550 | XP_018218839.1 | 99 | 0 |
| SPGP0DZ01230 | XP_018219462.1 | 99 | 0 |
| SPGP0BF00300 | XP_018220399.1 | 99 | 0 |
| SPGP0DU00150 | XP_018220575.1 | 99 | 0 |
| SPGP0AH00740 | XP_018222531.1 | 100 | 0 |
| SPGP0ET00190 | XP_018218988.1 | 99 | 0 |
| SPGP0Y00460 | XP_018223920.1 | 99 | 0 |
| SPGP0Y01460 | XP_018224015.1 | 99 | 0 |
| SPGP0DU00980 | XP_018220492.1 | 99 | 1 |
| SPGP0DU00600 | XP_018220530.1 | 99 | 0 |
| SPGP0Z01040 | XP_018219780.1 | 99 | 0 |
| SPGP0N00660 | XP_018220742.1 | 99 | 0 |
| SPGP0CZ00390 | XP_018220865.1 | 99 | 0 |
| SPGP0AH00930 | XP_018222513.1 | 99 | 0 |
| SPGP0N01200 | XP_018220692.1 | 99 | 0 |
| SPGP0CZ00400 | XP_018220866.1 | 99 | 0 |
| SPGP0DZ01000 | XP_018219440.1 | 99 | 0 |
| SPGP0AH00450 | XP_018222560.1 | 99 | 0 |
| SPGP0CZ01750 | XP_018220998.1 | 99 | 0 |
| SPGP0DZ00680 | XP_018219409.1 | 99 | 0 |
| SPGP0AHtrna26S | DI49_2374 | 100 | 0 |
| SPGP0CZ01320 | XP_018220957.1 | 99 | 0 |
| SPGP0Y01420 | XP_018224011.1 | 99 | 0 |
| SPGP0DU01630 | XP_018220428.1 | 99 | 0 |
| SPGP0N00700 | XP_018220738.1 | 99 | 0 |
| SPGP0Y01250 | XP_018223994.1 | 99 | 0 |
| SPGP0CZ02110 | XP_018221031.1 | 99 | 0 |

Create PDF in your applications with the Pdfcrowd HTML to PDF API PDFCROWD

| | | | |
|---|---|---|---|
| SPGP0DZ01520 | XP_018219491.1 | 100 | 0 |
| SPGP0N00270 | XP_018220781.1 | 99 | 0 |
| SPGP0T00270 | XP_018220355.1 | 100 | 0 |
| SPGP0Y00310 | XP_018223908.1 | 99 | 0 |
| SPGP0N02250 | XP_018220589.1 | 99 | 0 |
| SPGP0DZ00430 | XP_018219386.1 | 99 | 0 |
| SPGP0DZ00560 | XP_018219398.1 | 99 | 0 |
| SPGP0BF00270 | XP_018220396.1 | 99 | 0 |
| SPGP0CZ01920 | XP_018221013.1 | 99 | 0 |
| SPGP0CZ01690 | XP_018220992.1 | 99 | 0 |
| SPGP0CZ00490 | XP_018220875.1 | 99 | 0 |
| SPGP0CZ00340 | XP_018220860.1 | 99 | 0 |
| SPGP0DZ00740 | XP_018219415.1 | 99 | 0 |
| SPGP0CZ03030 | XP_018221122.1 | 99 | 0 |
| SPGP0U01420 | XP_018218851.1 | 99 | 0 |
| SPGP0CZ01070 | XP_018220932.1 | 99 | 0 |
| SPGP0AH01050 | XP_018222503.1 | 99 | 0 |
| SPGP0T00210 | XP_018220349.1 | 99 | 0 |
| SPGP0Y01100 | XP_018223980.1 | 99 | 0 |
| SPGP0CZ02310 | XP_018221051.1 | 99 | 0 |
| SPGP0U00480 | XP_018218941.1 | 99 | 0 |
| SPGP0U01570 | XP_018218837.1 | 99 | 0 |
| SPGP0AC00580 | XP_018219142.1 | 99 | 0 |
| SPGP0DZ01990 | XP_018219535.1 | 99 | 0 |
| SPGP0Y00440 | XP_018223918.1 | 100 | 0 |
| SPGP0AC00130 | XP_018219098.1 | 99 | 0 |
| SPGP0N00640 | XP_018220744.1 | 99 | 0 |
| SPGP0DZ01320 | XP_018219471.1 | 99 | 0 |
| SPGP0DU01390 | XP_018220451.1 | 99 | 0 |
| SPGP0Y01390 | XP_018224008.1 | 99 | 0 |
| SPGP0AC01110 | XP_018219194.1 | 98 | 0 |
| SPGP0CZ02480 | XP_018221068.1 | 100 | 0 |
| SPGP0N00920 | XP_018220717.1 | 99 | 0 |
| SPGP0CZ01590 | XP_018220983.1 | 99 | 0 |
| SPGP0Y00800 | XP_018223952.1 | 99 | 0 |
| SPGP0DX00660 | XP_018219995.1 | 99 | 0 |
| SPGP0Y01010 | XP_018223971.1 | 99 | 0 |
| SPGP0AH00430 | XP_018222561.1 | 99 | 0 |
| SPGP0CZ01770 | XP_018221000.1 | 99 | 0 |
| SPGP0U01580 | XP_018218836.1 | 99 | 0 |

Create PDF in your applications with the Pdfcrowd HTML to PDF API PDFCROWD

| | | | |
|---|---|---|---|
| SPGP0DX00330 | XP_018219964.1 | 99 | 0 |
| SPGP0Y01470 | XP_018224016.1 | 99 | 0 |
| SPGP0DU01610 | XP_018220430.1 | 99 | 0 |
| SPGP0AC00810 | XP_018219164.1 | 99 | 0 |
| SPGP0DZ00230 | XP_018219366.1 | 99 | 0 |
| SPGP0U01510 | XP_018218843.1 | 99 | 1 |
| SPGP0Y01760 | XP_018224043.1 | 97 | 0 |
| SPGP0CZ01060 | XP_018220931.1 | 99 | 0 |
| SPGP0U01740 | XP_018218821.1 | 99 | 0 |
| SPGP0BC00110 | XP_018220809.1 | 99 | 0 |
| SPGP0Y00900 | XP_018223960.1 | 99 | 0 |
| SPGP0T00240 | XP_018220352.1 | 99 | 1 |
| SPGP0DU01250 | XP_018220465.1 | 99 | 0 |
| SPGP0N01700 | XP_018220644.1 | 99 | 0 |
| SPGP0DX00250 | XP_018219956.1 | 99 | 0 |
| SPGP0DV00660 | XP_018219927.1 | 99 | 0 |
| SPGP0DZ02090 | XP_018219546.1 | 99 | 0 |
| SPGP0Y01480 | XP_018224017.1 | 99 | 0 |
| SPGP0EX00190 | XP_018223878.1 | 99 | 0 |
| SPGP0DZ00760 | XP_018219417.1 | 99 | 0 |
| SPGP0CZ01010 | XP_018220926.1 | 99 | 0 |
| SPGP0N00990 | XP_018220712.1 | 99 | 0 |
| SPGP0T00230 | XP_018220351.1 | 99 | 0 |
| SPGP0BC00320 | XP_018220827.1 | 99 | 0 |
| SPGP0CZ02910 | XP_018221110.1 | 99 | 0 |
| SPGP0CZ01130 | XP_018220938.1 | 99 | 0 |
| SPGP0DZ02100 | XP_018219547.1 | 99 | 0 |
| SPGP0AC00150 | XP_018219100.1 | 99 | 0 |
| SPGP0CZ00990 | XP_018220924.1 | 99 | 0 |
| SPGP0DZ01340 | XP_018219473.1 | 99 | 0 |
| SPGP0N01150 | XP_018220697.1 | 99 | 0 |
| SPGP0CZ00190 | XP_018220847.1 | 99 | 0 |
| SPGP0Y01030 | XP_018223973.1 | 99 | 0 |
| SPGP0DZ01370 | XP_018219476.1 | 100 | 0 |
| SPGP0DZ00610 | XP_018219402.1 | 99 | 0 |
| SPGP0DU00690 | XP_018220522.1 | 99 | 0 |
| SPGP0N01490 | XP_018221216.1 | 87 | 0 |
| SPGP0AH00830 | XP_018222522.1 | 99 | 0 |
| SPGP0U01720 | XP_018218823.1 | 99 | 0 |
| SPGP0EX00180 | XP_018223877.1 | 99 | 0 |

Create PDF in your applications with the Pdfcrowd HTML to PDF API PDFCROWD

| | | | |
|---|---|---|---|
| SPGP0U00890 | XP_018218903.1 | 99 | 0 |
| SPGP0N01720 | XP_018220642.1 | 100 | 0 |
| SPGP0U01840 | XP_018218810.1 | 99 | 0 |
| SPGP0DX00270 | XP_018219958.1 | 99 | 0 |
| SPGP0CZ02720 | XP_018221092.1 | 99 | 0 |
| SPGP0DZ00880 | XP_018219428.1 | 99 | 0 |
| SPGP0AH00350 | XP_018222569.1 | 98 | 2 |
| SPGP0CZ01040 | XP_018220929.1 | 99 | 0 |
| SPGP0N00400 | XP_018220768.1 | 99 | 0 |
| SPGP0U00910 | XP_018218901.1 | 99 | 0 |
| SPGP0DZ01220 | XP_018219461.1 | 99 | 0 |
| SPGP0Y01200 | XP_018223989.1 | 99 | 0 |
| SPGP0CZ02710 | XP_018221091.1 | 100 | 0 |
| SPGP0DX00310 | XP_018219962.1 | 99 | 0 |
| SPGP0Y00350 | XP_018223912.1 | 99 | 0 |
| SPGP0AH00860 | XP_018222519.1 | 99 | 0 |
| SPGP0N01620 | XP_018220652.1 | 99 | 0 |
| SPGP0N00350 | XP_018220773.1 | 99 | 0 |
| SPGP0CZ00810 | XP_018220906.1 | 99 | 0 |
| SPGP0CZ00600 | XP_018220886.1 | 99 | 0 |
| SPGP0CZ00610 | XP_018220887.1 | 99 | 0 |
| SPGP0DZ01940 | XP_018219530.1 | 99 | 0 |
| SPGP0DZ01070 | XP_018219446.1 | 99 | 0 |
| SPGP0N00480 | XP_018220759.1 | 99 | 0 |
| SPGP0AH00260 | XP_018222576.1 | 99 | 0 |
| SPGP0N01920 | XP_018220622.1 | 99 | 0 |
| SPGP0CZ02550 | XP_018221075.1 | 99 | 0 |
| SPGP0CZ01680 | XP_018220991.1 | 99 | 0 |
| SPGP0BF00290 | XP_018220398.1 | 99 | 0 |
| SPGP0BC00310 | XP_018220826.1 | 99 | 0 |
| SPGP0Y00940 | XP_018223964.1 | 99 | 0 |
| SPGP0DZ01830 | XP_018219521.1 | 99 | 0 |
| SPGP0AC00930 | XP_018219176.1 | 99 | 0 |
| SPGP0AC00780 | XP_018219161.1 | 99 | 0 |
| SPGP0U01910 | XP_018218803.1 | 99 | 0 |
| SPGP0DU01540 | XP_018220436.1 | 99 | 0 |
| SPGP0EP00120 | XP_018220337.1 | 99 | 0 |
| SPGP0DX00210 | XP_018219952.1 | 99 | 0 |
| SPGP0AC00310 | XP_018219115.1 | 99 | 0 |
| SPGP0AC00720 | XP_018219155.1 | 99 | 0 |

Create PDF in your applications with the Pdfcrowd HTML to PDF API PDFCROWD

| | | | |
|---|---|---|---|
| SPGP0U01870 | XP_018218807.1 | 99 | 0 |
| SPGP0DZ01280 | XP_018219467.1 | 99 | 0 |
| SPGP0N01640 | XP_018220650.1 | 99 | 0 |
| SPGP0CZ00770 | XP_018220902.1 | 99 | 0 |
| SPGP0AH00990 | XP_018222508.1 | 99 | 0 |
| SPGP0DZ00870 | XP_018219427.1 | 99 | 0 |
| SPGP0Y00640 | XP_018223938.1 | 100 | 0 |
| SPGP0DZ01790 | XP_018219517.1 | 99 | 0 |
| SPGP0U00870 | XP_018218905.1 | 99 | 0 |
| SPGP0AH00390 | XP_018222565.1 | 99 | 0 |
| SPGP0CZ01050 | XP_018220930.1 | 99 | 0 |
| SPGP0U00420 | XP_018218947.1 | 99 | 0 |
| SPGP0CZ02730 | XP_018221093.1 | 99 | 0 |
| SPGP0N00190 | XP_018220789.1 | 99 | 0 |
| SPGP0DU00550 | XP_018220535.1 | 99 | 0 |
| SPGP0AC00520 | XP_018219136.1 | 99 | 0 |
| SPGP0AC00120 | XP_018219097.1 | 99 | 0 |
| SPGP0N00370 | XP_018220771.1 | 99 | 0 |
| SPGP0U00310 | XP_018218958.1 | 99 | 0 |
| SPGP0EE00210 | XP_018220840.1 | 99 | 0 |
| SPGP0AC00950 | XP_018219178.1 | 99 | 0 |
| SPGP0CZ01890 | XP_018221010.1 | 99 | 0 |
| SPGP0CZ00480 | XP_018220874.1 | 100 | 0 |
| SPGP0DU01150 | XP_018220475.1 | 99 | 0 |
| SPGP0ET00130 | XP_018218982.1 | 99 | 0 |
| SPGP0DZ00220 | XP_018219365.1 | 99 | 0 |
| SPGP0DZ01960 | XP_018219532.1 | 99 | 0 |
| SPGP0CZ00220 | XP_018220850.1 | 99 | 0 |
| SPGP0AC00880 | XP_018219171.1 | 100 | 0 |
| SPGP0AH00530 | XP_018222552.1 | 99 | 0 |
| SPGP0Y00580 | XP_018223932.1 | 100 | 0 |
| SPGP0AC00440 | XP_018219128.1 | 99 | 0 |
| SPGP0AH01110 | XP_018222497.1 | 99 | 0 |
| SPGP0AH01120 | XP_018222496.1 | 99 | 0 |
| SPGP0U01270 | XP_018218865.1 | 99 | 0 |
| SPGP0N01760 | XP_018220638.1 | 99 | 0 |
| SPGP0T00290 | XP_018220357.1 | 99 | 0 |
| SPGP0DU00140 | XP_018220576.1 | 98 | 2 |
| SPGP0DU01520 | XP_018220438.1 | 99 | 0 |
| SPGP0DX00450 | XP_018219975.1 | 99 | 0 |

Create PDF in your applications with the Pdfcrowd HTML to PDF API PDFCROWD

| | | | |
|---|---|---|---|
| SPGP0U00270 | XP_018218962.1 | 99 | 0 |
| SPGP0N02130 | XP_018220601.1 | 100 | 0 |
| SPGP0Z00780 | XP_018219806.1 | 99 | 0 |
| SPGP0DU00890 | XP_018220501.1 | 99 | 0 |
| SPGP0DX00580 | XP_018219988.1 | 98 | 0 |
| SPGP0EX00130 | XP_018223872.1 | 99 | 0 |
| SPGP0N02170 | XP_018220597.1 | 99 | 0 |
| SPGP0AH00290 | XP_018222573.1 | 99 | 0 |
| SPGP0N02020 | XP_018220612.1 | 99 | 0 |
| SPGP0U01650 | XP_018218829.1 | 99 | 0 |
| SPGP0AH00760 | XP_018222529.1 | 99 | 0 |
| SPGP0AC00870 | XP_018219170.1 | 99 | 0 |
| SPGP0DZ00420 | XP_018219385.1 | 99 | 0 |
| SPGP0N00430 | XP_018220764.1 | 100 | 0 |
| SPGP0N00180 | XP_018220790.1 | 99 | 0 |
| SPGP0N00310 | XP_018220777.1 | 100 | 0 |
| SPGP0DN00100 | XP_018223462.1 | 86 | 0 |
| SPGP0DZ02000 | XP_018219536.1 | 99 | 0 |
| SPGP0DU01170 | XP_018220473.1 | 99 | 0 |
| SPGP0AC00270 | XP_018219111.1 | 99 | 0 |
| SPGP0DZ00200 | XP_018219363.1 | 99 | 0 |
| SPGP0N01010 | XP_018220710.1 | 99 | 0 |
| SPGP0DZ00240 | XP_018219367.1 | 99 | 0 |
| SPGP0U01020 | XP_018218890.1 | 99 | 0 |
| SPGP0DU01240 | XP_018220466.1 | 99 | 0 |
| SPGP0AH00780 | XP_018222527.1 | 100 | 0 |
| SPGP0N01960 | XP_018220618.1 | 99 | 0 |
| SPGP0AC01080 | XP_018219191.1 | 99 | 0 |
| SPGP0DU00160 | XP_018220574.1 | 99 | 0 |
| SPGP0AC01060 | XP_018219189.1 | 98 | 0 |
| SPGP0Y00110 | XP_018223889.1 | 99 | 0 |
| SPGP0Y00210 | XP_018223899.1 | 99 | 0 |
| SPGP0AH00160 | XP_018222586.1 | 100 | 0 |
| SPGP0EX00110 | XP_018223870.1 | 99 | 0 |
| SPGP0R01300 | XP_018221247.1 | 99 | 0 |
| SPGP0EG00140 | XP_018221949.1 | 100 | 0 |
| SPGP0AC00850 | XP_018219168.1 | 99 | 0 |
| SPGP0U00460 | XP_018218943.1 | 100 | 0 |
| SPGP0U00680 | XP_018218924.1 | 99 | 0 |
| SPGP0AH00750 | XP_018222530.1 | 99 | 0 |

Create PDF in your applications with the Pdfcrowd HTML to PDF API PDFCROWD

| | | | |
|---|---|---|---|
| SPGP0N00330 | XP_018220775.1 | 99 | 0 |
| SPGP0Y00150 | XP_018223893.1 | 99 | 0 |
| SPGP0DU00310 | XP_018220559.1 | 99 | 0 |
| SPGP0CZ02900 | XP_018221109.1 | 99 | 0 |
| SPGP0CZ02130 | XP_018221033.1 | 99 | 0 |
| SPGP0Y00600 | XP_018223934.1 | 99 | 0 |
| SPGP0U01860 | XP_018218808.1 | 99 | 0 |
| SPGP0CZ01840 | XP_018221006.1 | 99 | 0 |
| SPGP0DZ00260 | XP_018219369.1 | 99 | 0 |
| SPGP0CZ02680 | XP_018221088.1 | 99 | 0 |
| SPGP0CZ00960 | XP_018220921.1 | 99 | 0 |
| SPGP0EE00160 | XP_018220835.1 | 99 | 0 |
| SPGP0BC00330 | XP_018220828.1 | 99 | 0 |
| SPGP0U01810 | XP_018218813.1 | 99 | 0 |
| SPGP0CZ01290 | XP_018220954.1 | 99 | 0 |
| SPGP0U00500 | XP_018218939.1 | 99 | 0 |
| SPGP0CZ02200 | XP_018221040.1 | 99 | 0 |
| SPGP0CZ01300 | XP_018220955.1 | 99 | 1 |
| SPGP0AH00720 | XP_018222533.1 | 99 | 0 |
| SPGP0U01390 | XP_018218854.1 | 99 | 0 |
| SPGP0U00720 | XP_018218920.1 | 99 | 0 |
| SPGP0N01080 | XP_018220703.1 | 99 | 0 |
| SPGP0BF00350 | XP_018220403.1 | 99 | 0 |
| SPGP0CZ00890 | XP_018220914.1 | 99 | 0 |
| SPGP0U01050 | XP_018218887.1 | 99 | 0 |
| SPGP0DU01750 | XP_018220416.1 | 100 | 0 |
| SPGP0DU01380 | XP_018220452.1 | 99 | 0 |
| SPGP0DZ02190 | XP_018219556.1 | 99 | 0 |
| SPGP0CZ00410 | XP_018220867.1 | 99 | 0 |
| SPGP0U00980 | XP_018218894.1 | 100 | 0 |
| SPGP0DU01650 | XP_018220426.1 | 99 | 0 |
| SPGP0CZ00980 | XP_018220923.1 | 99 | 0 |
| SPGP0U00830 | XP_018218909.1 | 99 | 0 |
| SPGP0DZ00840 | XP_018219425.1 | 99 | 0 |
| SPGP0DU00290 | XP_018220561.1 | 99 | 0 |
| SPGP0DU00120 | XP_018220578.1 | 99 | 0 |
| SPGP0DX00160 | XP_018219947.1 | 99 | 0 |
| SPGP0N02270 | XP_018220587.1 | 99 | 0 |
| SPGP0Y00780 | XP_018223950.1 | 99 | 0 |

Create PDF in your applications with the Pdfcrowd HTML to PDF API PDFCROWD

| | | | |
|---|---|---|---|
| SPGP0CZ01990 | XP_018221020.1 | 99 | 0 |
| SPGP0N01050 | XP_018220706.1 | 99 | 0 |
| SPGP0Y01190 | XP_018223988.1 | 99 | 0 |
| SPGP0DZ00360 | XP_018219379.1 | 99 | 0 |
| SPGP0N00110 | XP_018220797.1 | 99 | 0 |
| SPGP0CZ02000 | XP_018221021.1 | 99 | 0 |
| SPGP0Y00570 | XP_018223931.1 | 99 | 0 |
| SPGP0DZ01190 | XP_018219458.1 | 100 | 0 |
| SPGP0U00970 | XP_018218895.1 | 99 | 0 |
| SPGP0W01730 | XP_018223162.1 | 100 | 0 |
| SPGP0DZ01670 | XP_018219505.1 | 99 | 0 |
| SPGP0EE00120 | XP_018220831.1 | 99 | 0 |
| SPGP0U01190 | XP_018218873.1 | 99 | 0 |
| SPGP0N01470 | XP_018220666.1 | 99 | 0 |
| SPGP0CZ00820 | XP_018220907.1 | 99 | 0 |
| SPGP0DU01670 | XP_018220424.1 | 99 | 0 |
| SPGP0EL00130 | XP_018219090.1 | 99 | 0 |
| SPGP0T00160 | XP_018220344.1 | 99 | 0 |
| SPGP0Z01030 | XP_018219781.1 | 99 | 0 |
| SPGP0FB00100 | XP_018220804.1 | 99 | 0 |
| SPGP0BF00390 | XP_018220407.1 | 99 | 0 |
| SPGP0T00280 | XP_018220356.1 | 99 | 0 |
| SPGP0N00820 | XP_018220727.1 | 99 | 0 |
| SPGP0DX00640 | XP_018219993.1 | 99 | 0 |
| SPGP0DZ01170 | XP_018219456.1 | 99 | 0 |
| SPGP0DZ00790 | XP_018219420.1 | 99 | 0 |
| SPGP0CZ00280 | XP_018220855.1 | 99 | 0 |
| SPGP0Y00240 | XP_018223902.1 | 99 | 0 |
| SPGP0AH00660 | XP_018222539.1 | 99 | 0 |
| SPGP0CZ01150 | XP_018220940.1 | 99 | 0 |
| SPGP0ET00110 | XP_018218980.1 | 99 | 0 |
| SPGP0CZ02260 | XP_018221046.1 | 99 | 0 |
| SPGP0Y01500 | XP_018224019.1 | 99 | 0 |
| SPGP0DU01070 | XP_018220483.1 | 99 | 0 |
| SPGP0S01150 | XP_018220230.1 | 99 | 0 |
| SPGP0DU00650 | XP_018220526.1 | 99 | 0 |
| SPGP0CZ02500 | XP_018221070.1 | 99 | 0 |
| SPGP0N01860 | XP_018220628.1 | 99 | 0 |
| SPGP0Y01040 | XP_018223974.1 | 99 | 0 |
| SPGP0N01290 | XP_018220684.1 | 99 | 0 |

Create PDF in your applications with the Pdfcrowd HTML to PDF API PDFCROWD

| | | | |
|---|---|---|---|
| SPGP0W00130 | XP_018223321.1 | 99 | 0 |
| SPGP0AC00100 | XP_018219095.1 | 99 | 0 |
| SPGP0DU01440 | XP_018220446.1 | 99 | 0 |
| SPGP0DZ01210 | XP_018219460.1 | 99 | 0 |
| SPGP0U00700 | XP_018218922.1 | 99 | 0 |
| SPGP0P01580 | XP_018221801.1 | 99 | 0 |
| SPGP0CZ01970 | XP_018221018.1 | 99 | 0 |
| SPGP0DZ00710 | XP_018219412.1 | 99 | 1 |
| SPGP0AH00640 | XP_018222541.1 | 99 | 0 |
| SPGP0EE00100 | XP_018220829.1 | 99 | 0 |
| SPGP0N01800 | XP_018220634.1 | 99 | 0 |
| SPGP0U01090 | XP_018218883.1 | 99 | 0 |
| SPGP0U01670 | XP_018218827.1 | 99 | 0 |
| SPGP0R03410 | XP_018221449.1 | 99 | 0 |
| SPGP0CZ00730 | XP_018220898.1 | 100 | 0 |
| SPGP0CZ01580 | XP_018220982.1 | 99 | 0 |
| SPGP0DV00270 | XP_018219891.1 | 99 | 0 |
| SPGP0DX00500 | XP_018219980.1 | 99 | 0 |
| SPGP0N00790 | XP_018220730.1 | 99 | 0 |
| SPGP0AH00180 | XP_018222584.1 | 99 | 0 |
| SPGP0DU01690 | XP_018220422.1 | 99 | 0 |
| SPGP0CZ02410 | XP_018221061.1 | 100 | 0 |
| SPGP0Y00550 | XP_018223929.1 | 99 | 0 |
| SPGP0AC00360 | XP_018219120.1 | 99 | 0 |
| SPGP0DX00570 | XP_018219987.1 | 99 | 0 |
| SPGP0DZ00500 | XP_018219393.1 | 99 | 0 |
| SPGP0FU00100 | XP_018219195.1 | 99 | 0 |
| SPGP0T00260 | XP_018220354.1 | 99 | 0 |
| SPGP0DU01030 | XP_018220487.1 | 99 | 0 |
| SPGP0N01560 | XP_018220658.1 | 99 | 0 |
| SPGP0Y00510 | XP_018223925.1 | 99 | 0 |
| SPGP0DX00520 | XP_018219982.1 | 99 | 0 |
| SPGP0CZ01850 | XP_018221007.1 | 99 | 0 |
| SPGP0AH00370 | XP_018222567.1 | 99 | 0 |
| SPGP0N02150 | XP_018220599.1 | 99 | 0 |
| SPGP0N00880 | XP_018220721.1 | 100 | 0 |
| SPGP0N00550 | XP_018220753.1 | 100 | 0 |
| SPGP0DZ02130 | XP_018219550.1 | 99 | 0 |
| SPGP0CZ02810 | XP_018221100.1 | 99 | 0 |
| SPGP0CZ01810 | XP_018221003.1 | 99 | 0 |

Create PDF in your applications with the Pdfcrowd HTML to PDF API PDFCROWD

| | | | |
|---|---|---|---|
| SPGP0AH01000 | XP_018222507.1 | 99 | 0 |
| SPGP0U00610 | XP_018218928.1 | 99 | 0 |
| SPGP0DU01010 | XP_018220489.1 | 99 | 0 |
| SPGP0N01540 | XP_018220660.1 | 99 | 0 |
| SPGP0DU00490 | XP_018220541.1 | 99 | 0 |
| SPGP0DU00630 | XP_018220528.1 | 99 | 0 |
| SPGP0Y00760 | XP_018223948.1 | 99 | 0 |
| SPGP0CZ00470 | XP_018220873.1 | 99 | 0 |
| SPGP0DU01730 | XP_018220418.1 | 99 | 0 |
| SPGP0AH00500 | XP_018222555.1 | 99 | 0 |
| SPGP0T00190 | XP_018220347.1 | 99 | 0 |
| SPGP0AC00500 | XP_018219134.1 | 99 | 0 |
| SPGP0CZ00940 | XP_018220919.1 | 99 | 0 |
| SPGP0DX00240 | XP_018219955.1 | 99 | 0 |
| SPGP0DX00200 | XP_018219951.1 | 99 | 0 |
| SPGP0AH00810 | XP_018222524.1 | 99 | 0 |
| SPGP0DX00560 | XP_018219986.1 | 99 | 0 |
| SPGP0CZ02150 | XP_018221035.1 | 99 | 0 |
| SPGP0DZ01630 | XP_018219501.1 | 99 | 0 |
| SPGP0U00380 | XP_018218951.1 | 99 | 0 |
| SPGP0AC00420 | XP_018219126.1 | 99 | 0 |
| SPGP0CZ01930 | XP_018221014.1 | 99 | 0 |
| SPGP0AH01040 | XP_018222504.1 | 99 | 0 |
| SPGP0DX00220 | XP_018219953.1 | 99 | 0 |
| SPGP0ET00200 | XP_018218989.1 | 99 | 0 |
| SPGP0N01580 | XP_018220656.1 | 99 | 0 |
| SPGP0DU00320 | XP_018220558.1 | 100 | 0 |
| SPGP0DZ00120 | XP_018219355.1 | 99 | 0 |
| SPGP0U01220 | XP_018218870.1 | 99 | 0 |
| SPGP0DZ00820 | XP_018219423.1 | 99 | 0 |
| SPGP0BF00110 | XP_018220382.1 | 99 | 0 |
| SPGP0Y00470 | XP_018223921.1 | 99 | 0 |
| SPGP0DZ00620 | XP_018219403.1 | 99 | 0 |
| SPGP0AC00560 | XP_018219140.1 | 99 | 0 |
| SPGP0AC00240 | XP_018219109.1 | 100 | 0 |
| SPGP0U01240 | XP_018218868.1 | 99 | 0 |
| SPGP0AH00680 | XP_018222537.1 | 99 | 0 |
| SPGP0U00900 | XP_018218902.1 | 99 | 0 |
| SPGP0DU00880 | XP_018220502.1 | 99 | 0 |
| SPGP0N01480 | XP_018220665.1 | 99 | 0 |

Create PDF in your applications with the Pdfcrowd HTML to PDF API PDFCROWD

| | | | |
|---|---|---|---|
| SPGP0BC00100 | XP_018220808.1 | 99 | 0 |
| SPGP0N02050 | XP_018220609.1 | 99 | 0 |
| SPGP0DU00450 | XP_018220545.1 | 99 | 0 |
| SPGP0DX00440 | XP_018219974.1 | 99 | 0 |
| SPGP0U00880 | XP_018218904.1 | 99 | 0 |
| SPGP0X00520 | XP_018219050.1 | 99 | 0 |
| SPGP0Q00470 | XP_018222454.1 | 99 | 0 |
| SPGP0L00690 | XP_018219294.1 | 99 | 0 |
| SPGP0Y01740 | XP_018224041.1 | 100 | 0 |
| SPGP0DU01090 | XP_018220481.1 | 99 | 0 |
| SPGP0Z02840 | XP_018219600.1 | 99 | 0 |
| SPGP0R01790 | XP_018221296.1 | 99 | 0 |
| SPGP0Z02510 | XP_018219634.1 | 99 | 0 |
| SPGP0S03390 | XP_018220016.1 | 99 | 0 |
| SPGP0Z01280 | XP_018219757.1 | 98 | 1 |
| SPGP0DX00600 | XP_018219990.1 | 99 | 0 |
| SPGP0N01870 | XP_018220627.1 | 99 | 0 |
| SPGP0DN01650 | XP_018223611.1 | 99 | 0 |
| SPGP0R02850 | XP_018221397.1 | 99 | 0 |
| SPGP0CZ00130 | XP_018220842.1 | 96 | 0 |
| SPGP0W00880 | XP_018223246.1 | 99 | 0 |
| SPGP0W04500 | XP_018222898.1 | 99 | 0 |
| SPGP0R01730 | XP_018221289.1 | 99 | 0 |
| SPGP0Z00620 | XP_018219822.1 | 99 | 0 |
| SPGP0Q04190 | XP_018222086.1 | 99 | 0 |
| SPGP0S00660 | XP_018220279.1 | 99 | 0 |
| SPGP0DN01520 | XP_018223599.1 | 99 | 0 |
| SPGP0DZ00180 | XP_018219361.1 | 99 | 0 |
| SPGP0R02640 | XP_018221376.1 | 99 | 0 |
| SPGP0B00470 | XP_018221495.1 | 99 | 0 |
| SPGP0CZ02770 | XP_018221096.1 | 99 | 0 |
| SPGP0DZ01200 | XP_018219459.1 | 99 | 0 |
| SPGP0AA00500 | XP_018223362.1 | 99 | 0 |
| SPGP0R00460 | XP_018221169.1 | 99 | 0 |
| SPGP0B01870 | XP_018221629.1 | 99 | 0 |
| SPGP0L01610 | XP_018219204.1 | 99 | 0 |
| SPGP0L00620 | XP_018219301.1 | 99 | 0 |
| SPGP0R01840 | XP_018221300.1 | 99 | 0 |
| SPGP0R01590 | XP_018221276.1 | 99 | 0 |
| SPGP0W04740 | XP_018222874.1 | 99 | 0 |

Create PDF in your applications with the Pdfcrowd HTML to PDF API PDFCROWD

| | | | |
|---|---|---|---|
| SPGP0T00110 | XP_018220339.1 | 99 | 0 |
| SPGP0Q01250 | XP_018222375.1 | 99 | 0 |
| SPGP0Q01450 | XP_018222353.1 | 99 | 0 |
| SPGP0S02510 | XP_018220102.1 | 99 | 0 |
| SPGP0DV00430 | XP_018219905.1 | 99 | 0 |
| SPGP0Q01920 | XP_018222306.1 | 99 | 0 |
| SPGP0DN02200 | XP_018223666.1 | 99 | 0 |
| SPGP0W00850 | XP_018223249.1 | 99 | 0 |
| SPGP0Y01540 | XP_018224023.1 | 99 | 0 |
| SPGP0R03450 | XP_018221452.1 | 99 | 0 |
| SPGP0S03100 | XP_018220044.1 | 100 | 0 |
| SPGP0B00650 | XP_018221512.1 | 99 | 0 |
| SPGP0Z01640 | XP_018219721.1 | 100 | 0 |
| SPGP0W01130 | XP_018223221.1 | 99 | 0 |
| SPGP0Q02690 | XP_018222231.1 | 99 | 0 |
| SPGP0P02000 | XP_018221760.1 | 99 | 0 |
| SPGP0Q01870 | XP_018222311.1 | 99 | 0 |
| SPGP0DN02620 | XP_018223707.1 | 99 | 0 |
| SPGP0S02460 | XP_018220107.1 | 99 | 0 |
| SPGP0Z01860 | XP_018219699.1 | 99 | 0 |
| SPGP0R01140 | XP_018221232.1 | 99 | 0 |
| SPGP0B00280 | XP_018221476.1 | 99 | 0 |
| SPGP0Q01670 | XP_018222331.1 | 99 | 0 |
| SPGP0W00620 | XP_018223272.1 | 99 | 0 |
| SPGP0L00240 | XP_018219337.1 | 100 | 0 |
| SPGP0Q01240 | XP_018222376.1 | 99 | 0 |
| SPGP0AH00770 | XP_018222528.1 | 99 | 0 |
| SPGP0DU00420 | XP_018220548.1 | 98 | 0 |
| SPGP0W01850 | XP_018223150.1 | 99 | 0 |
| SPGP0E01600 | XP_018222748.1 | 99 | 0 |
| SPGP0S00810 | XP_018220264.1 | 99 | 0 |
| SPGP0P00520 | XP_018221904.1 | 99 | 0 |
| SPGP0Y01270 | XP_018223996.1 | 99 | 0 |
| SPGP0Q03070 | XP_018222194.1 | 99 | 0 |
| SPGP0R02380 | XP_018221351.1 | 99 | 0 |
| SPGP0H00290 | XP_018224106.1 | 99 | 0 |
| SPGP0EG00150 | XP_018221950.1 | 99 | 0 |
| SPGP0W01860 | XP_018223149.1 | 97 | 2 |
| SPGP0B01930 | XP_018221635.1 | 99 | 0 |
| SPGP0W01680 | XP_018223167.1 | 99 | 0 |

Create PDF in your applications with the Pdfcrowd HTML to PDF API PDFCROWD

| | | | |
|---|---|---|---|
| SPGP0R03390 | XP_018221448.1 | 99 | 0 |
| SPGP0W01690 | XP_018223166.1 | 99 | 0 |
| SPGP0L00250 | XP_018219336.1 | 99 | 0 |
| SPGP0AC00840 | XP_018219167.1 | 99 | 0 |
| SPGP0P01150 | XP_018221842.1 | 99 | 0 |
| SPGP0DZtrna9G | DI49_4977 | 100 | 0 |
| SPGP0S00450 | XP_018220300.1 | 99 | 0 |
| SPGP0E00550 | XP_018222645.1 | 99 | 1 |
| SPGP0Y00190 | XP_018223897.1 | 99 | 0 |
| SPGP0Y00790 | XP_018223951.1 | 99 | 1 |
| SPGP0Z01010 | XP_018219783.1 | 99 | 0 |
| SPGP0EX00290 | XP_018223887.1 | 99 | 0 |
| SPGP0Z00910 | XP_018219793.1 | 99 | 0 |
| SPGP0Q01940 | XP_018222304.1 | 99 | 0 |
| SPGP0Z01790 | XP_018219706.1 | 99 | 0 |
| SPGP0DN01490 | XP_018223596.1 | 99 | 0 |
| SPGP0F04450 | XP_018222046.1 | 99 | 0 |
| SPGP0Q00550 | XP_018222446.1 | 100 | 0 |
| SPGP0N00160 | XP_018220792.1 | 99 | 0 |
| SPGP0DN00120 | XP_018218752.1 | 99 | 0 |
| SPGP0B01990 | XP_018221641.1 | 99 | 0 |
| SPGP0Z01360 | XP_018219749.1 | 99 | 0 |
| SPGP0DU00300 | XP_018220560.1 | 99 | 0 |
| SPGP0DN04040 | XP_018223846.1 | 99 | 0 |
| SPGP0H00500 | XP_018224086.1 | 99 | 0 |
| SPGP0P02290 | XP_018221734.1 | 99 | 0 |
| SPGP0Z01510 | XP_018219734.1 | 99 | 0 |
| SPGP0AA00600 | XP_018223372.1 | 99 | 0 |
| SPGP0DN01720 | XP_018223618.1 | 99 | 0 |
| SPGP0Q02840 | XP_018222217.1 | 99 | 0 |
| SPGP0E00470 | XP_018222637.1 | 99 | 0 |
| SPGP0N00300 | XP_018220778.1 | 99 | 0 |
| SPGP0F04900 | XP_018222004.1 | 99 | 0 |
| SPGP0Q00610 | XP_018222440.1 | 99 | 0 |
| SPGP0W02600 | XP_018223077.1 | 99 | 0 |
| SPGP0R01530 | XP_018221270.1 | 99 | 0 |
| SPGP0U01120 | XP_018218880.1 | 99 | 0 |
| SPGP0X00180 | XP_018219082.1 | 98 | 1 |
| SPGP0DN00730 | XP_018223519.1 | 99 | 0 |
| SPGP0B01470 | XP_018221590.1 | 99 | 0 |

Create PDF in your applications with the Pdfcrowd HTML to PDF API PDFCROWD

| | | | |
|---|---|---|---|
| SPGP0B00960 | XP_018221542.1 | 99 | 0 |
| SPGP0F04730 | XP_018222020.1 | 99 | 0 |
| SPGP0F05010 | XP_018221997.1 | 95 | 0 |
| SPGP0W04870 | XP_018222861.1 | 98 | 0 |
| SPGP0DZ00980 | XP_018219438.1 | 99 | 0 |
| SPGP0W03430 | XP_018223003.1 | 99 | 0 |
| SPGP0CZ00840 | XP_018220909.1 | 99 | 0 |
| SPGP0DN00580 | XP_018223505.1 | 99 | 0 |
| SPGP0DN00960 | XP_018223542.1 | 99 | 0 |
| SPGP0DZ01270 | XP_018219466.1 | 99 | 0 |
| SPGP0W02870 | XP_018223053.1 | 99 | 0 |
| SPGP0CZ00900 | XP_018220915.1 | 99 | 0 |
| SPGP0X00940 | XP_018219009.1 | 99 | 0 |
| SPGP0X01050 | XP_018218998.1 | 99 | 0 |
| SPGP0DU01300 | XP_018220460.1 | 99 | 0 |
| SPGP0L01270 | XP_018219237.1 | 99 | 0 |
| SPGP0E00860 | XP_018222674.1 | 95 | 4 |
| SPGP0E00520 | XP_018222642.1 | 99 | 0 |
| SPGP0W01900 | XP_018223146.1 | 99 | 0 |
| SPGP0DQ00100 | XP_018222852.1 | 97 | 0 |
| SPGP0X00540 | XP_018219048.1 | 100 | 0 |
| SPGP0DN02410 | XP_018223686.1 | 99 | 0 |
| SPGP0F04580 | XP_018222033.1 | 99 | 0 |
| SPGP0P03050 | XP_018221659.1 | 99 | 0 |
| SPGP0W02560 | XP_018223081.1 | 99 | 0 |
| SPGP0Q00330 | XP_018222468.1 | 99 | 0 |
| SPGP0U00250 | XP_018218964.1 | 99 | 0 |
| SPGP0W02030 | XP_018223132.1 | 99 | 0 |
| SPGP0P01080 | XP_018221849.1 | 99 | 0 |
| SPGP0S02680 | XP_018220085.1 | 99 | 0 |
| SPGP0AA00860 | XP_018223395.1 | 100 | 0 |
| SPGP0P01740 | XP_018221786.1 | 100 | 0 |
| SPGP0E01930 | XP_018222781.1 | 99 | 0 |
| SPGP0N01180 | XP_018220694.1 | 99 | 0 |
| SPGP0DN00920 | XP_018223538.1 | 99 | 0 |
| SPGP0Z02670 | XP_018219618.1 | 99 | 0 |
| SPGP0DZ00370 | XP_018219380.1 | 99 | 0 |
| SPGP0AC00200 | XP_018219105.1 | 99 | 0 |
| SPGP0L01570 | XP_018219207.1 | 99 | 0 |
| SPGP0E02560 | XP_018222842.1 | 99 | 0 |

Create PDF in your applications with the Pdfcrowd HTML to PDF API PDFCROWD

| | | | |
|---|---|---|---|
| SPGP0W01060 | XP_018223228.1 | 99 | 0 |
| SPGP0B01680 | XP_018221610.1 | 99 | 0 |
| SPGP0E02070 | XP_018222795.1 | 99 | 0 |
| SPGP0AA00680 | XP_018223380.1 | 99 | 0 |
| SPGP0W04190 | XP_018222931.1 | 97 | 2 |
| SPGP0E01200 | XP_018222710.1 | 99 | 0 |
| SPGP0Z01320 | XP_018219753.1 | 99 | 0 |
| SPGP0Z01630 | XP_018219722.1 | 99 | 0 |
| SPGP0P02890 | XP_018221675.1 | 99 | 0 |
| SPGP0E01770 | XP_018222765.1 | 99 | 0 |
| SPGP0AF00460 | XP_018221987.1 | 99 | 0 |
| SPGP0R02120 | XP_018221327.1 | 99 | 0 |
| SPGP0W03210 | XP_018223022.1 | 99 | 0 |
| SPGP0W03740 | XP_018222974.1 | 99 | 0 |
| SPGP0CZ03010 | XP_018221120.1 | 99 | 0 |
| SPGP0W03960 | XP_018222953.1 | 100 | 0 |
| SPGP0DN03730 | XP_018223815.1 | 100 | 0 |
| SPGP0DZ01680 | XP_018219506.1 | 99 | 0 |
| SPGP0CZ01500 | XP_018220974.1 | 99 | 0 |
| SPGP0AFtrna22V | DI49_2588 | 100 | 0 |
| SPGP0Q03570 | XP_018222147.1 | 98 | 0 |
| SPGP0S02620 | XP_018220091.1 | 99 | 0 |
| SPGP0B01460 | XP_018221589.1 | 99 | 0 |
| SPGP0EX00120 | XP_018223871.1 | 99 | 0 |
| SPGP0W04230 | XP_018222927.1 | 99 | 0 |
| SPGP0R01470 | XP_018221264.1 | 99 | 0 |
| SPGP0Z02170 | XP_018219668.1 | 100 | 0 |
| SPGP0W00430 | XP_018223291.1 | 99 | 0 |
| SPGP0P02090 | XP_018221751.1 | 99 | 0 |
| SPGP0L00220 | XP_018219339.1 | 99 | 0 |
| SPGP0L00730 | XP_018219290.1 | 99 | 0 |
| SPGP0AF00300 | XP_018221971.1 | 99 | 0 |
| SPGP0R00900 | XP_018221210.1 | 99 | 0 |
| SPGP0W02980 | XP_018223042.1 | 99 | 1 |
| SPGP0Q01770 | XP_018222321.1 | 99 | 0 |
| SPGP0R00870 | XP_018221207.1 | 99 | 0 |
| SPGP0DU00950 | XP_018220495.1 | 100 | 0 |
| SPGP0W04070 | XP_018222943.1 | 99 | 0 |
| SPGP0S01500 | XP_018220198.1 | 99 | 0 |
| SPGP0R00860 | XP_018221206.1 | 100 | 0 |

Create PDF in your applications with the Pdfcrowd HTML to PDF API PDFCROWD

| | | | |
|---|---|---|---|
| SPGP0AA00940 | XP_018223403.1 | 99 | 0 |
| SPGP0AC00260 | XP_018219110.1 | 99 | 0 |
| SPGP0Q02200 | XP_018222278.1 | 99 | 0 |
| SPGP0N00620 | XP_018220746.1 | 99 | 0 |
| SPGP0U01830 | XP_018218811.1 | 98 | 1 |
| SPGP0AH00560 | XP_018222549.1 | 99 | 0 |
| SPGP0W00480 | XP_018223286.1 | 99 | 0 |
| SPGP0P01510 | XP_018221807.1 | 99 | 0 |
| SPGP0DN02060 | XP_018223653.1 | 99 | 0 |
| SPGP0L01470 | XP_018219217.1 | 99 | 0 |
| SPGP0B00670 | XP_018221514.1 | 99 | 0 |
| SPGP0CZ02690 | XP_018221089.1 | 99 | 0 |
| SPGP0Z01460 | XP_018219739.1 | 99 | 0 |
| SPGP0X00880 | XP_018219014.1 | 99 | 0 |
| SPGP0AH00190 | XP_018222583.1 | 99 | 0 |
| SPGP0R00250 | XP_018221148.1 | 99 | 0 |
| SPGP0B01980 | XP_018221640.1 | 99 | 0 |
| SPGP0H00470 | XP_018224089.1 | 99 | 0 |
| SPGP0Q02060 | XP_018222292.1 | 99 | 0 |
| SPGP0R01830 | XP_018221299.1 | 99 | 0 |
| SPGP0DN01260 | XP_018223573.1 | 99 | 0 |
| SPGP0T00320 | XP_018220360.1 | 99 | 0 |
| SPGP0DV00590 | XP_018219921.1 | 99 | 0 |
| SPGP0E00690 | XP_018222659.1 | 99 | 0 |
| SPGP0DN02600 | XP_018223705.1 | 99 | 0 |
| SPGP0X00680 | XP_018219034.1 | 99 | 0 |
| SPGP0W01260 | XP_018223208.1 | 99 | 0 |
| SPGP0Z01370 | XP_018219748.1 | 99 | 0 |
| SPGP0Z00930 | XP_018219791.1 | 99 | 0 |
| SPGP0W01770 | XP_018223158.1 | 99 | 0 |
| SPGP0P00600 | XP_018221896.1 | 99 | 0 |
| SPGP0B00220 | XP_018221470.1 | 99 | 0 |
| SPGP0S03310 | XP_018220024.1 | 99 | 0 |
| SPGP0DN01420 | XP_018223589.1 | 99 | 0 |
| SPGP0DN03630 | XP_018223805.1 | 99 | 0 |
| SPGP0U00860 | XP_018218906.1 | 99 | 0 |
| SPGP0BF00150 | XP_018220386.1 | 99 | 0 |
| SPGP0S03490 | XP_018220007.1 | 99 | 0 |
| SPGP0W01470 | XP_018223188.1 | 99 | 0 |
| SPGP0DN01000 | XP_018223547.1 | 99 | 0 |

Create PDF in your applications with the Pdfcrowd HTML to PDF API PDFCROWD

| | | | |
|---|---|---|---|
| SPGP0E01570 | XP_018222745.1 | 99 | 0 |
| SPGP0W01450 | XP_018223189.1 | 99 | 0 |
| SPGP0N02060 | XP_018220608.1 | 99 | 0 |
| SPGP0AA00800 | XP_018223390.1 | 99 | 0 |
| SPGP0W00530 | XP_018223281.1 | 100 | 0 |
| SPGP0S02340 | XP_018220118.1 | 99 | 0 |
| SPGP0S03380 | XP_018220017.1 | 99 | 0 |
| SPGP0DZ01110 | XP_018219450.1 | 99 | 0 |
| SPGP0S02300 | XP_018220122.1 | 99 | 0 |
| SPGP0E02370 | XP_018222825.1 | 99 | 0 |
| SPGP0BF00130 | XP_018220384.1 | 99 | 0 |
| SPGP0Z00950 | XP_018219789.1 | 99 | 0 |
| SPGP0Q03330 | XP_018222170.1 | 99 | 0 |
| SPGP0Z03000 | XP_018219585.1 | 99 | 0 |
| SPGP0DN04180 | XP_018223860.1 | 99 | 0 |
| SPGP0DN00680 | XP_018223514.1 | 99 | 0 |
| SPGP0Q00520 | XP_018222449.1 | 99 | 0 |
| SPGP0Q00830 | XP_018222418.1 | 99 | 0 |
| SPGP0Q01470 | XP_018222351.1 | 99 | 0 |
| SPGP0E01560 | XP_018222744.1 | 99 | 0 |
| SPGP0U00190 | XP_018218970.1 | 99 | 0 |
| SPGP0CZ00170 | XP_018220845.1 | 99 | 0 |
| SPGP0Z02760 | XP_018219608.1 | 98 | 2 |
| SPGP0P02620 | XP_018221701.1 | 99 | 0 |
| SPGP0W03770 | XP_018222971.1 | 99 | 0 |
| SPGP0Y00990 | XP_018223969.1 | 99 | 0 |
| SPGP0E00610 | XP_018222651.1 | 99 | 0 |
| SPGP0N01790 | XP_018220635.1 | 99 | 0 |
| SPGP0L00130 | XP_018219348.1 | 99 | 0 |
| SPGP0N00250 | XP_018220783.1 | 99 | 0 |
| SPGP0EX00200 | XP_018223879.1 | 99 | 0 |
| SPGP0S00330 | XP_018220312.1 | 99 | 0 |
| SPGP0E01990 | XP_018222787.1 | 100 | 0 |
| SPGP0DN00350 | XP_018223485.1 | 99 | 1 |
| SPGP0Q00780 | XP_018222423.1 | 99 | 0 |
| SPGP0S01110 | XP_018220234.1 | 99 | 0 |
| SPGP0AA00290 | XP_018223342.1 | 99 | 0 |
| SPGP0H00700 | XP_018224067.1 | 99 | 0 |
| SPGP0F04440 | XP_018222047.1 | 99 | 0 |
| SPGP0E01690 | XP_018222758.1 | 99 | 0 |

Create PDF in your applications with the Pdfcrowd HTML to PDF API PDFCROWD

| | | | |
|---|---|---|---|
| SPGP0N00890 | XP_018220720.1 | 99 | 0 |
| SPGP0P00240 | XP_018221930.1 | 99 | 0 |
| SPGP0X00710 | XP_018219031.1 | 99 | 0 |
| SPGP0W00760 | XP_018223257.1 | 98 | 2 |
| SPGP0DN02760 | XP_018223720.1 | 99 | 0 |
| SPGP0Z00310 | XP_018219847.1 | 99 | 0 |
| SPGP0CZtrna1T | DI49_3176 | 100 | 0 |
| SPGP0R00630 | XP_018221185.1 | 99 | 0 |
| SPGP0S01200 | XP_018220225.1 | 100 | 0 |
| SPGP0DU00170 | XP_018220573.1 | 100 | 0 |
| SPGP0L00210 | XP_018219340.1 | 99 | 0 |
| SPGP0S02060 | XP_018220146.1 | 99 | 0 |
| SPGP0W02410 | XP_018223096.1 | 99 | 0 |
| SPGP0W00790 | XP_018223255.1 | 99 | 0 |
| SPGP0Gtrna30R | DI49_3064 | 100 | 0 |
| SPGP0P00790 | XP_018221877.1 | 99 | 0 |
| SPGP0DX00400 | XP_018219971.1 | 99 | 0 |
| SPGP0Y01640 | XP_018224033.1 | 99 | 0 |
| SPGP0L00820 | XP_018219281.1 | 99 | 0 |
| SPGP0R02230 | XP_018221336.1 | 99 | 0 |
| SPGP0N01940 | XP_018220620.1 | 99 | 0 |
| SPGP0R02800 | XP_018221392.1 | 99 | 0 |
| SPGP0N00240 | XP_018220784.1 | 100 | 0 |
| SPGP0Q01110 | XP_018222390.1 | 99 | 0 |
| SPGP0L00500 | XP_018219311.1 | 100 | 0 |
| SPGP0P02790 | XP_018221684.1 | 98 | 0 |
| SPGP0U01610 | XP_018218833.1 | 99 | 0 |
| SPGP0P02300 | XP_018221733.1 | 99 | 0 |
| SPGP0W02970 | XP_018223043.1 | 99 | 0 |
| SPGP0W03490 | XP_018222998.1 | 99 | 0 |
| SPGP0B01790 | XP_018221621.1 | 99 | 0 |
| SPGP0AA00360 | XP_018223349.1 | 99 | 0 |
| SPGP0Q04200 | XP_018222085.1 | 99 | 0 |
| SPGP0N00440 | XP_018220763.1 | 99 | 0 |
| SPGP0R02700 | XP_018221382.1 | 98 | 1 |
| SPGP0S01140 | XP_018220231.1 | 100 | 0 |
| SPGP0B01310 | XP_018221574.1 | 99 | 1 |
| SPGP0E01700 | XP_018222759.1 | 99 | 0 |
| SPGP0R00680 | XP_018221190.1 | 98 | 1 |

Create PDF in your applications with the Pdfcrowd HTML to PDF API PDFCROWD

| | | | |
|---|---|---|---|
| SPGP0W03300 | XP_018223014.1 | 99 | 0 |
| SPGP0Z02530 | XP_018219632.1 | 99 | 0 |
| SPGP0Q00380 | XP_018222463.1 | 99 | 0 |
| SPGP0DN00290 | XP_018223479.1 | 99 | 0 |
| SPGP0W00550 | XP_018223279.1 | 99 | 0 |
| SPGP0H00210 | XP_018224114.1 | 99 | 1 |
| SPGP0L01400 | XP_018219224.1 | 99 | 0 |
| SPGP0W04560 | XP_018222892.1 | 99 | 0 |
| SPGP0R02940 | XP_018221405.1 | 99 | 0 |
| SPGP0P01680 | XP_018221792.1 | 100 | 0 |
| SPGP0DZ00720 | XP_018219413.1 | 99 | 0 |
| SPGP0W02640 | XP_018223074.1 | 99 | 0 |
| SPGP0DN02470 | XP_018223692.1 | 99 | 0 |
| SPGP0P01530 | XP_018221806.1 | 99 | 1 |
| SPGP0F04280 | XP_018222063.1 | 99 | 0 |
| SPGP0Z01560 | XP_018219729.1 | 99 | 0 |
| SPGP0R02660 | XP_018221378.1 | 99 | 0 |
| SPGP0P02130 | XP_018221747.1 | 99 | 0 |
| SPGP0B00410 | XP_018221489.1 | 99 | 0 |
| SPGP0Z01760 | XP_018219709.1 | 99 | 0 |
| SPGP0S03240 | XP_018220031.1 | 99 | 0 |
| SPGP0Q03920 | XP_018222113.1 | 100 | 0 |
| SPGP0L01630 | XP_018219202.1 | 97 | 0 |
| SPGP0Z00200 | XP_018219863.1 | 99 | 0 |
| SPGP0DN02230 | XP_018223669.1 | 99 | 0 |
| SPGP0B01890 | XP_018221631.1 | 99 | 0 |
| SPGP0Q00680 | XP_018222433.1 | 99 | 0 |
| SPGP0R01860 | XP_018221302.1 | 99 | 0 |
| SPGP0T00130 | XP_018220341.1 | 98 | 0 |
| SPGP0Z01700 | XP_018219715.1 | 99 | 0 |
| SPGP0DX00460 | XP_018219976.1 | 99 | 0 |
| SPGP0AA00410 | XP_018223353.1 | 99 | 0 |
| SPGP0Q01270 | XP_018222373.1 | 100 | 0 |
| SPGP0S02530 | XP_018220100.1 | 99 | 0 |
| SPGP0AH00170 | XP_018222585.1 | 99 | 0 |
| SPGP0P02680 | XP_018221695.1 | 99 | 0 |
| SPGP0N01410 | XP_018220672.1 | 99 | 0 |
| SPGP0L00320 | XP_018219329.1 | 99 | 0 |
| SPGP0S03120 | XP_018220042.1 | 100 | 0 |
| SPGP0DZ01080 | XP_018219447.1 | 99 | 0 |

Create PDF in your applications with the Pdfcrowd HTML to PDF API PDFCROWD

| | | | |
|---|---|---|---|
| SPGP0EX00160 | XP_018223875.1 | 99 | 0 |
| SPGP0DN03070 | XP_018223751.1 | 99 | 0 |
| SPGP0R00400 | XP_018221163.1 | 99 | 0 |
| SPGP0W01110 | XP_018223223.1 | 100 | 0 |
| SPGP0DN02140 | XP_018223661.1 | 99 | 0 |
| SPGP0B01090 | XP_018221554.1 | 99 | 0 |
| SPGP0Q02230 | XP_018222276.1 | 100 | 0 |
| SPGP0R01310 | XP_018221248.1 | 97 | 1 |
| SPGP0F04350 | XP_018222056.1 | 99 | 0 |
| SPGP0R00640 | XP_018221186.1 | 99 | 0 |
| SPGP0W00600 | XP_018223274.1 | 99 | 0 |
| SPGP0AH00710 | XP_018222534.1 | 99 | 0 |
| SPGP0DZ00130 | XP_018219356.1 | 99 | 0 |
| SPGP0R02590 | XP_018221372.1 | 99 | 0 |
| SPGP0S01000 | XP_018220245.1 | 99 | 0 |
| SPGP0Y01240 | XP_018223993.1 | 99 | 0 |
| SPGP0CZ02670 | XP_018221087.1 | 99 | 0 |
| SPGP0N01890 | XP_018220625.1 | 99 | 0 |
| SPGP0Q01760 | XP_018222322.1 | 99 | 0 |
| SPGP0DZ00400 | XP_018219383.1 | 99 | 0 |
| SPGP0H00160 | XP_018224119.1 | 98 | 0 |
| SPGP0W03690 | XP_018222979.1 | 99 | 0 |
| SPGP0Z01570 | XP_018219728.1 | 99 | 0 |
| SPGP0X01090 | XP_018218994.1 | 97 | 2 |
| SPGP0R01880 | XP_018221304.1 | 99 | 0 |
| SPGP0W01800 | XP_018223155.1 | 99 | 0 |
| SPGP0B01320 | XP_018221575.1 | 100 | 0 |
| SPGP0DN01920 | XP_018223639.1 | 100 | 0 |
| SPGP0H00690 | XP_018224068.1 | 99 | 0 |
| SPGP0Y01340 | XP_018224003.1 | 99 | 0 |
| SPGP0Htrna22P | DI49_4914 | 100 | 0 |
| SPGP0DU01100 | XP_018220480.1 | 99 | 0 |
| SPGP0Z01410 | XP_018219744.1 | 99 | 0 |
| SPGP0DZ00350 | XP_018219378.1 | 100 | 0 |
| SPGP0E02530 | XP_018222839.1 | 99 | 0 |
| SPGP0E02310 | XP_018222819.1 | 99 | 0 |
| SPGP0S00470 | XP_018220298.1 | 99 | 0 |
| SPGP0Qtrna17L | DI49_2147 | 99 | 0 |
| SPGP0DZ01900 | XP_018219526.1 | 99 | 0 |
| SPGP0E00780 | XP_018222667.1 | 99 | 0 |

Create PDF in your applications with the Pdfcrowd HTML to PDF API PDFCROWD

| | | | |
|---|---|---|---|
| SPGP0E00570 | XP_018222647.1 | 99 | 0 |
| SPGP0W03290 | XP_018223015.1 | 99 | 0 |
| SPGP0Q00400 | XP_018222461.1 | 99 | 0 |
| SPGP0Y00450 | XP_018223919.1 | 99 | 0 |
| SPGP0U00450 | XP_018218944.1 | 99 | 0 |
| SPGP0DZ01760 | XP_018219514.1 | 99 | 0 |
| SPGP0Z00280 | XP_018219844.1 | 99 | 0 |
| SPGP0B01300 | XP_018221573.1 | 99 | 0 |
| SPGP0W02220 | XP_018223115.1 | 99 | 0 |
| SPGP0Q04020 | XP_018222103.1 | 99 | 0 |
| SPGP0X00390 | XP_018219062.1 | 99 | 0 |
| SPGP0U01380 | XP_018218855.1 | 99 | 0 |
| SPGP0E01120 | XP_018222702.1 | 99 | 0 |
| SPGP0W04320 | XP_018222918.1 | 99 | 0 |
| SPGP0Z00970 | XP_018219787.1 | 99 | 0 |
| SPGP0AA00660 | XP_018223378.1 | 99 | 0 |
| SPGP0Q01460 | XP_018222352.1 | 99 | 0 |
| SPGP0X00380 | XP_018219063.1 | 99 | 0 |
| SPGP0N00140 | XP_018220794.1 | 99 | 0 |
| SPGP0S00780 | XP_018220267.1 | 99 | 0 |
| SPGP0U01320 | XP_018218861.1 | 98 | 0 |
| SPGP0R02370 | XP_018221350.1 | 99 | 0 |
| SPGP0Z00810 | XP_018219803.1 | 99 | 0 |
| SPGP0P02210 | XP_018221741.1 | 98 | 1 |
| SPGP0U00560 | XP_018218933.1 | 99 | 0 |
| SPGP0Z00710 | XP_018219813.1 | 99 | 0 |
| SPGP0N01370 | XP_018220676.1 | 99 | 0 |
| SPGP0X00470 | XP_018219054.1 | 99 | 0 |
| SPGP0DN04060 | XP_018223848.1 | 99 | 0 |
| SPGP0W03440 | XP_018223002.1 | 100 | 0 |
| SPGP0R02810 | XP_018221393.1 | 99 | 0 |
| SPGP0CZ00860 | XP_018220911.1 | 100 | 0 |
| SPGP0P03060 | XP_018221658.1 | 99 | 0 |
| SPGP0DN01740 | XP_018223620.1 | 99 | 0 |
| SPGP0Q03830 | XP_018222122.1 | 98 | 0 |
| SPGP0X00280 | XP_018219073.1 | 99 | 0 |
| SPGP0Q02860 | XP_018222215.1 | 99 | 0 |
| SPGP0E01730 | XP_018222762.1 | 99 | 0 |
| SPGP0L01280 | XP_018219236.1 | 100 | 0 |
| SPGP0N00910 | XP_018220718.1 | 99 | 0 |

Create PDF in your applications with the Pdfcrowd HTML to PDF API PDFCROWD

| | | | |
|---|---|---|---|
| SPGP0B01520 | XP_018221594.1 | 96 | 0 |
| SPGP0U01140 | XP_018218878.1 | 99 | 0 |
| SPGP0Q01390 | XP_018222360.1 | 99 | 0 |
| SPGP0CZ02020 | XP_018221023.1 | 100 | 0 |
| SPGP0Q00980 | XP_018222403.1 | 99 | 0 |
| SPGP0AH00950 | XP_018222512.1 | 99 | 0 |
| SPGP0R03320 | XP_018221442.1 | 99 | 0 |
| SPGP0F04370 | XP_018222054.1 | 99 | 0 |
| SPGP0BF00360 | XP_018220404.1 | 99 | 0 |
| SPGP0R01210 | XP_018221239.1 | 99 | 1 |
| SPGP0Q00640 | XP_018222437.1 | 99 | 0 |
| SPGP0S01430 | XP_018220205.1 | 99 | 0 |
| SPGP0R01130 | XP_018221231.1 | 100 | 0 |
| SPGP0DN00210 | XP_018223471.1 | 99 | 0 |
| SPGP0P00470 | XP_018221908.1 | 99 | 0 |
| SPGP0DV00210 | XP_018219885.1 | 99 | 0 |
| SPGP0Q03220 | XP_018222180.1 | 99 | 0 |
| SPGP0X00960 | XP_018219007.1 | 99 | 0 |
| SPGP0Y01710 | XP_018224038.1 | 99 | 0 |
| SPGP0Y00280 | XP_018223905.1 | 99 | 0 |
| SPGP0DZ00800 | XP_018219421.1 | 99 | 0 |
| SPGP0R03350 | XP_018221444.1 | 97 | 2 |
| SPGP0N01630 | XP_018220651.1 | 99 | 0 |
| SPGP0P03070 | XP_018221657.1 | 99 | 0 |
| SPGP0S03460 | XP_018220010.1 | 99 | 0 |
| SPGP0FT00130 | XP_018220802.1 | 99 | 0 |
| SPGP0Z01210 | XP_018219764.1 | 99 | 0 |
| SPGP0Q03270 | XP_018222175.1 | 98 | 1 |
| SPGP0Q04220 | XP_018222083.1 | 99 | 0 |
| SPGP0W04380 | XP_018222911.1 | 99 | 0 |
| SPGP0U01180 | XP_018218874.1 | 99 | 0 |
| SPGP0B00780 | XP_018221525.1 | 99 | 0 |
| SPGP0AA00880 | XP_018223397.1 | 99 | 0 |
| SPGP0DU01180 | XP_018220472.1 | 99 | 0 |
| SPGP0P00400 | XP_018221915.1 | 98 | 0 |
| SPGP0R02320 | XP_018221345.1 | 99 | 0 |
| SPGP0CZ02350 | XP_018221055.1 | 99 | 0 |
| SPGP0B00930 | XP_018221539.1 | 99 | 0 |
| SPGP0DU00580 | XP_018220532.1 | 99 | 0 |
| SPGP0AC01050 | XP_018219188.1 | 98 | 0 |

Create PDF in your applications with the Pdfcrowd HTML to PDF API PDFCROWD

| | | | |
|---|---|---|---|
| SPGP0W04340 | XP_018222916.1 | 99 | 0 |
| SPGP0B01660 | XP_018221608.1 | 99 | 0 |
| SPGP0R01360 | XP_018221253.1 | 99 | 0 |
| SPGP0B00720 | XP_018221519.1 | 100 | 0 |
| SPGP0R01670 | XP_018221284.1 | 99 | 0 |
| SPGP0AC00540 | XP_018219138.1 | 100 | 0 |
| SPGP0CZ03070 | XP_018221125.1 | 99 | 0 |
| SPGP0W03720 | XP_018222976.1 | 99 | 0 |
| SPGP0DZ00900 | XP_018219430.1 | 100 | 0 |
| SPGP0R02490 | XP_018221362.1 | 99 | 0 |
| SPGP0W03230 | XP_018223020.1 | 99 | 0 |
| SPGP0DN03710 | XP_018223813.1 | 99 | 0 |
| SPGP0DN03240 | XP_018223768.1 | 99 | 0 |
| SPGP0DU00970 | XP_018220493.1 | 99 | 0 |
| SPGP0B01400 | XP_018221583.1 | 99 | 0 |
| SPGP0DN03540 | XP_018223796.1 | 99 | 0 |
| SPGP0E00710 | XP_018222661.1 | 99 | 0 |
| SPGP0S02880 | XP_018220066.1 | 99 | 0 |
| SPGP0DN02710 | XP_018223715.1 | 99 | 0 |
| SPGP0W04710 | XP_018222877.1 | 99 | 0 |
| SPGP0P00270 | XP_018221927.1 | 99 | 0 |
| SPGP0R01410 | XP_018221258.1 | 99 | 0 |
| SPGP0Q04010 | XP_018222104.1 | 99 | 0 |
| SPGP0AA00810 | XP_018223391.1 | 99 | 0 |
| SPGP0Q01060 | XP_018222395.1 | 99 | 0 |
| SPGP0L00710 | XP_018219292.1 | 99 | 0 |
| SPGP0AC00460 | XP_018219130.1 | 99 | 1 |
| SPGP0S01720 | XP_018220179.1 | 99 | 1 |
| SPGP0R01200 | XP_018221238.1 | 99 | 0 |
| SPGP0AF00360 | XP_018221977.1 | 99 | 0 |
| SPGP0DN01140 | XP_018223561.1 | 99 | 0 |
| SPGP0DN01220 | XP_018223569.1 | 99 | 0 |
| SPGP0S02070 | XP_018220145.1 | 99 | 0 |
| SPGP0E00190 | XP_018222609.1 | 99 | 0 |
| SPGP0Z00560 | XP_018219828.1 | 99 | 0 |
| SPGP0R02840 | XP_018221396.1 | 100 | 0 |
| SPGP0Q03800 | XP_018222125.1 | 99 | 0 |
| SPGP0Q02260 | XP_018222273.1 | 99 | 0 |
| SPGP0P02770 | XP_018221686.1 | 99 | 0 |
| SPGP0Z00160 | XP_018219867.1 | 99 | 0 |

Create PDF in your applications with the Pdfcrowd HTML to PDF API PDFCROWD

| | | | |
|---|---|---|---|
| SPGP0E00320 | XP_018222622.1 | 99 | 0 |
| SPGP0Q02950 | XP_018222206.1 | 99 | 0 |
| SPGP0CZ00520 | XP_018220878.1 | 99 | 0 |
| SPGP0CZ00250 | XP_018220853.1 | 99 | 0 |
| SPGP0N00510 | XP_018220757.1 | 99 | 0 |
| SPGP0DN02590 | XP_018223704.1 | 99 | 0 |
| SPGP0P03000 | XP_018221664.1 | 100 | 0 |
| SPGP0DN02040 | XP_018223651.1 | 99 | 0 |
| SPGP0B01240 | XP_018221567.1 | 99 | 0 |
| SPGP0R00350 | XP_018221158.1 | 99 | 0 |
| SPGP0B00200 | XP_018221468.1 | 99 | 0 |
| SPGP0L00860 | XP_018219277.1 | 99 | 0 |
| SPGP0DN04140 | XP_018223856.1 | 99 | 0 |
| SPGP0R00270 | XP_018221150.1 | 99 | 0 |
| SPGP0DZ01620 | XP_018219500.1 | 99 | 0 |
| SPGP0Z01450 | XP_018219740.1 | 99 | 0 |
| SPGP0R00230 | XP_018221146.1 | 99 | 0 |
| SPGP0S02660 | XP_018220087.1 | 99 | 0 |
| SPGP0FJ00100 | XP_018219929.1 | 99 | 0 |
| SPGP0CZ02120 | XP_018221032.1 | 99 | 0 |
| SPGP0CZ01120 | XP_018220937.1 | 99 | 0 |
| SPGP0Q02080 | XP_018222290.1 | 99 | 0 |
| SPGP0P03150 | XP_018221649.1 | 100 | 0 |
| SPGP0B00570 | XP_018221505.1 | 99 | 0 |
| SPGP0W00200 | XP_018223314.1 | 99 | 0 |
| SPGP0S02490 | XP_018220104.1 | 99 | 0 |
| SPGP0CZ00910 | XP_018220916.1 | 99 | 0 |
| SPGP0Q01680 | XP_018222330.1 | 99 | 0 |
| SPGP0Z00110 | XP_018219872.1 | 99 | 0 |
| SPGP0Q03680 | XP_018222137.1 | 99 | 0 |
| SPGP0L00990 | XP_018219264.1 | 99 | 0 |
| SPGP0W03890 | XP_018222959.1 | 98 | 0 |
| SPGP0S01850 | XP_018220167.1 | 99 | 0 |
| SPGP0Q03130 | XP_018222189.1 | 99 | 0 |
| SPGP0DN00990 | XP_018223546.1 | 99 | 0 |
| SPGP0Q02720 | XP_018222228.1 | 99 | 0 |
| SPGP0U00840 | XP_018218908.1 | 99 | 0 |
| SPGP0Z02480 | XP_018219637.1 | 99 | 0 |
| SPGP0W01490 | XP_018223186.1 | 99 | 0 |
| SPGP0W03510 | XP_018222996.1 | 99 | 0 |

Create PDF in your applications with the Pdfcrowd HTML to PDF API PDFCROWD

| | | | |
|---|---|---|---|
| SPGP0DN01360 | XP_018223583.1 | 99 | 0 |
| SPGP0DN01240 | XP_018223571.1 | 99 | 0 |
| SPGP0R00120 | XP_018221137.1 | 99 | 0 |
| SPGP0DN01630 | XP_018223609.1 | 99 | 0 |
| SPGP0P01020 | XP_018221855.1 | 99 | 0 |
| SPGP0S01670 | XP_018220184.1 | 99 | 0 |
| SPGP0W01320 | XP_018223202.1 | 99 | 0 |
| SPGP0BF00410 | XP_018220409.1 | 99 | 0 |
| SPGP0N01400 | XP_018220673.1 | 99 | 0 |
| SPGP0Z03060 | XP_018219579.1 | 99 | 0 |
| SPGP0R00490 | XP_018221172.1 | 99 | 0 |
| SPGP0U00150 | XP_018218974.1 | 99 | 0 |
| SPGP0E00950 | XP_018222685.1 | 99 | 0 |
| SPGP0DN00190 | XP_018223469.1 | 99 | 0 |
| SPGP0Q04260 | XP_018222080.1 | 100 | 0 |
| SPGP0Q00810 | XP_018222420.1 | 99 | 0 |
| SPGP0DN01020 | XP_018223549.1 | 99 | 1 |
| SPGP0E01500 | XP_018222738.1 | 99 | 0 |
| SPGP0L01040 | XP_018219259.1 | 99 | 0 |
| SPGP0E01300 | XP_018222720.1 | 99 | 0 |
| SPGP0H00710 | XP_018224066.1 | 99 | 0 |
| SPGP0H00660 | XP_018224071.1 | 99 | 0 |
| SPGP0W03840 | XP_018222964.1 | 99 | 0 |
| SPGP0P02390 | XP_018221724.1 | 99 | 0 |
| SPGP0H00580 | XP_018224079.1 | 99 | 0 |
| SPGP0E00630 | XP_018222653.1 | 99 | 0 |
| SPGP0E01170 | XP_018222707.1 | 99 | 0 |
| SPGP0DN01880 | XP_018223635.1 | 99 | 0 |
| SPGP0F04490 | XP_018222042.1 | 99 | 0 |
| SPGP0DU00860 | XP_018220504.1 | 99 | 0 |
| SPGP0W01310 | XP_018223203.1 | 99 | 0 |
| SPGP0P02470 | XP_018221716.1 | 99 | 0 |
| SPGP0H00720 | XP_018224065.1 | 100 | 0 |
| SPGP0DN01820 | XP_018223629.1 | 99 | 0 |
| SPGP0AA00270 | XP_018223340.1 | 99 | 0 |
| SPGP0L01120 | XP_018219251.1 | 99 | 0 |
| SPGP0S01280 | XP_018220218.1 | 99 | 0 |
| SPGP0Q02240 | XP_018222275.1 | 99 | 0 |
| SPGP0X00770 | XP_018219025.1 | 99 | 0 |
| SPGP0DX00650 | XP_018219994.1 | 99 | 0 |

Create PDF in your applications with the Pdfcrowd HTML to PDF API PDFCROWD

| | | | |
|---|---|---|---|
| SPGP0S00640 | XP_018220281.1 | 100 | 0 |
| SPGP0S00990 | XP_018220246.1 | 99 | 0 |
| SPGP0R02860 | XP_018221398.1 | 99 | 0 |
| SPGP0W03530 | XP_018222994.1 | 99 | 0 |
| SPGP0DN03800 | XP_018223822.1 | 99 | 0 |
| SPGP0S03040 | XP_018220050.1 | 99 | 0 |
| SPGP0P01400 | XP_018221818.1 | 98 | 1 |
| SPGP0B01230 | XP_018221566.1 | 99 | 0 |
| SPGP0FT00100 | XP_018220799.1 | 99 | 0 |
| SPGP0DN00460 | XP_018223494.1 | 100 | 0 |
| SPGP0DN02930 | XP_018223738.1 | 99 | 0 |
| SPGP0Y01660 | XP_018224035.1 | 98 | 1 |
| SPGP0E01270 | XP_018222717.1 | 99 | 0 |
| SPGP0X00590 | XP_018219043.1 | 100 | 0 |
| SPGP0Q03390 | XP_018222164.1 | 99 | 0 |
| SPGP0S00250 | XP_018220319.1 | 99 | 0 |
| SPGP0L00520 | XP_018219309.1 | 99 | 0 |
| SPGP0B01740 | XP_018221616.1 | 99 | 0 |
| SPGP0Z02080 | XP_018219677.1 | 99 | 0 |
| SPGP0Z02010 | XP_018219684.1 | 99 | 0 |
| SPGP0Z02730 | XP_018219612.1 | 99 | 0 |
| SPGP0W00630 | XP_018223271.1 | 99 | 0 |
| SPGP0F04630 | XP_018222029.1 | 100 | 0 |
| SPGP0L01380 | XP_018219226.1 | 99 | 0 |
| SPGP0W02910 | XP_018223049.1 | 99 | 0 |
| SPGP0AF00170 | XP_018221958.1 | 99 | 0 |
| SPGP0S01210 | XP_018220224.1 | 99 | 0 |
| SPGP0DX00280 | XP_018219959.1 | 99 | 0 |
| SPGP0Q01880 | XP_018222310.1 | 99 | 0 |
| SPGP0W00800 | XP_018223254.1 | 99 | 0 |
| SPGP0DU01640 | XP_018220427.1 | 99 | 0 |
| SPGP0B00240 | XP_018221472.1 | 99 | 0 |
| SPGP0W03390 | XP_018223005.1 | 99 | 0 |
| SPGP0R02400 | XP_018221353.1 | 99 | 0 |
| SPGP0N01810 | XP_018220633.1 | 100 | 0 |
| SPGP0E02380 | XP_018222826.1 | 99 | 0 |
| SPGP0Z02550 | XP_018219630.1 | 99 | 0 |
| SPGP0B01420 | XP_018221585.1 | 99 | 0 |
| SPGP0CZ01640 | XP_018220988.1 | 99 | 0 |
| SPGP0W02620 | XP_018223075.1 | 99 | 0 |

Create PDF in your applications with the Pdfcrowd HTML to PDF API PDFCROWD

| | | | |
|---|---|---|---|
| SPGP0L01430 | XP_018219221.1 | 99 | 0 |
| SPGP0DZ02040 | XP_018219541.1 | 99 | 0 |
| SPGP0Q00100 | XP_018219939.1 | 84 | 1 |
| SPGP0Q01410 | XP_018222358.1 | 99 | 0 |
| SPGP0W00840 | XP_018223250.1 | 99 | 0 |
| SPGP0W04540 | XP_018222894.1 | 99 | 0 |
| SPGP0W01610 | XP_018223174.1 | 99 | 0 |
| SPGP0W02020 | XP_018223133.1 | 99 | 0 |
| SPGP0T00150 | XP_018220343.1 | 99 | 0 |
| SPGP0DN02450 | XP_018223690.1 | 99 | 0 |
| SPGP0P02110 | XP_018221749.1 | 99 | 0 |
| SPGP0B00920 | XP_018221538.1 | 98 | 1 |
| SPGP0L00380 | XP_018219323.1 | 99 | 0 |
| SPGP0R02680 | XP_018221380.1 | 99 | 0 |
| SPGP0Q04040 | XP_018222101.1 | 99 | 0 |
| SPGP0B00430 | XP_018221491.1 | 99 | 0 |
| SPGP0DZ00650 | XP_018219406.1 | 99 | 0 |
| SPGP0Q00970 | XP_018222404.1 | 99 | 0 |
| SPGP0Q03900 | XP_018222115.1 | 99 | 0 |
| SPGP0Z00770 | XP_018219807.1 | 99 | 0 |
| SPGP0L01650 | XP_018219200.1 | 98 | 0 |
| SPGP0W04810 | XP_018222867.1 | 98 | 2 |
| SPGP0Q01210 | XP_018222379.1 | 99 | 0 |
| SPGP0U01100 | XP_018218882.1 | 99 | 0 |
| SPGP0W01150 | XP_018223219.1 | 99 | 0 |
| SPGP0Strna6W | DI49_3006 | 100 | 0 |
| SPGP0Itrna26A | DI49_3564 | 100 | 0 |
| SPGP0R00420 | XP_018221165.1 | 98 | 1 |
| SPGP0CZ01330 | XP_018220958.1 | 99 | 0 |
| SPGP0R00930 | XP_018221212.1 | 99 | 0 |
| SPGP0DN03500 | XP_018223792.1 | 99 | 0 |
| SPGP0EX00140 | XP_018223873.1 | 99 | 0 |
| SPGP0Z00480 | XP_018219835.1 | 99 | 0 |
| SPGP0EL00120 | XP_018219091.1 | 98 | 2 |
| SPGP0AC00730 | XP_018219156.1 | 99 | 0 |
| SPGP0S02890 | XP_018220065.1 | 99 | 0 |
| SPGP0AC00140 | XP_018219099.1 | 99 | 0 |
| SPGP0R01330 | XP_018221250.1 | 99 | 0 |
| SPGP0W00660 | XP_018223268.1 | 99 | 0 |
| SPGP0DN02900 | XP_018223735.1 | 99 | 0 |

Create PDF in your applications with the Pdfcrowd HTML to PDF API PDFCROWD

| | | | |
|---|---|---|---|
| SPGP0B00690 | XP_018221516.1 | 99 | 0 |
| SPGP0EG00110 | XP_018221946.1 | 99 | 0 |
| SPGP0W01340 | XP_018223200.1 | 99 | 0 |
| SPGP0W01820 | XP_018223153.1 | 99 | 0 |
| SPGP0B01970 | XP_018221639.1 | 99 | 0 |
| SPGP0W01390 | XP_018223195.1 | 100 | 0 |
| SPGP0DN01900 | XP_018223637.1 | 99 | 0 |
| SPGP0B01340 | XP_018221577.1 | 99 | 0 |
| SPGP0Q02740 | XP_018222226.1 | 99 | 0 |
| SPGP0Y01320 | XP_018224001.1 | 99 | 0 |
| SPGP0DZ02160 | XP_018219553.1 | 99 | 0 |
| SPGP0B02000 | XP_018221642.1 | 99 | 0 |
| SPGP0DN00300 | XP_018223480.1 | 99 | 0 |
| SPGP0E00560 | XP_018222646.1 | 99 | 0 |
| SPGP0P02690 | XP_018221694.1 | 99 | 0 |
| SPGP0DN04010 | XP_018223843.1 | 99 | 0 |
| SPGP0E02450 | XP_018222832.1 | 99 | 0 |
| SPGP0B01180 | XP_018221561.1 | 99 | 0 |
| SPGP0S00110 | XP_018220333.1 | 99 | 0 |
| SPGP0W01840 | XP_018223151.1 | 99 | 0 |
| SPGP0U00300 | XP_018218959.1 | 99 | 0 |
| SPGP0W00750 | XP_018223258.1 | 99 | 0 |
| SPGP0L01260 | XP_018219238.1 | 100 | 0 |
| SPGP0S00440 | XP_018220301.1 | 97 | 3 |
| SPGP0W04620 | XP_018222886.1 | 100 | 0 |
| SPGP0DN00530 | XP_018223501.1 | 99 | 0 |
| SPGP0S01580 | XP_018220192.1 | 99 | 0 |
| SPGP0E00720 | XP_018222662.1 | 99 | 0 |
| SPGP0W01530 | XP_018223182.1 | 99 | 0 |
| SPGP0U00630 | XP_018218926.1 | 99 | 0 |
| SPGP0Y01490 | XP_018224018.1 | 99 | 0 |
| SPGP0AC01030 | XP_018219186.1 | 98 | 0 |
| SPGP0S03480 | XP_018220008.1 | 100 | 0 |
| SPGP0DX00180 | XP_018219949.1 | 99 | 0 |
| SPGP0Q02030 | XP_018222295.1 | 99 | 0 |
| SPGP0CZ01620 | XP_018220986.1 | 99 | 0 |
| SPGP0U00410 | XP_018218948.1 | 99 | 0 |
| SPGP0P03190 | XP_018221645.1 | 95 | 0 |
| SPGP0DN01150 | XP_018223562.1 | 99 | 0 |
| SPGP0E01810 | XP_018222769.1 | 99 | 0 |

Create PDF in your applications with the Pdfcrowd HTML to PDF API PDFCROWD

| | | | |
|---|---|---|---|
| SPGP0Z01130 | XP_018219771.1 | 99 | 0 |
| SPGP0AH00200 | XP_018222582.1 | 99 | 1 |
| SPGP0U01440 | XP_018218849.1 | 100 | 0 |
| SPGP0W04240 | XP_018222926.1 | 99 | 0 |
| SPGP0BF00340 | XP_018220402.1 | 99 | 0 |
| SPGP0P02780 | XP_018221685.1 | 99 | 0 |
| SPGP0DU00590 | XP_018220531.1 | 99 | 0 |
| SPGP0AH00230 | XP_018222579.1 | 99 | 0 |
| SPGP0Q00480 | XP_018222453.1 | 99 | 0 |
| SPGP0CZ01440 | XP_018220968.1 | 99 | 0 |
| SPGP0W03560 | XP_018222992.1 | 99 | 0 |
| SPGP0CZ00880 | XP_018220913.1 | 100 | 0 |
| SPGP0DN01760 | XP_018223622.1 | 99 | 0 |
| SPGP0E01240 | XP_018222714.1 | 99 | 0 |
| SPGP0S00400 | XP_018220305.1 | 99 | 0 |
| SPGP0N00690 | XP_018220739.1 | 99 | 0 |
| SPGP0S02110 | XP_018220141.1 | 99 | 0 |
| SPGP0Y00810 | XP_018223953.1 | 99 | 0 |
| SPGP0E02590 | XP_018222845.1 | 99 | 0 |
| SPGP0W03700 | XP_018222978.1 | 99 | 0 |
| SPGP0AG00160 | XP_018219935.1 | 91 | 1 |
| SPGP0DU00340 | XP_018220556.1 | 99 | 1 |
| SPGP0S03500 | XP_018220006.1 | 99 | 0 |
| SPGP0E00810 | XP_018222669.1 | 100 | 0 |
| SPGP0R03300 | XP_018221440.1 | 99 | 0 |
| SPGP0F04820 | XP_018222012.1 | 99 | 0 |
| SPGP0Z01400 | XP_018219745.1 | 99 | 0 |
| SPGP0Q00660 | XP_018222435.1 | 99 | 0 |
| SPGP0T00310 | XP_018220359.1 | 99 | 0 |
| SPGP0P00450 | XP_018221910.1 | 98 | 0 |
| SPGP0DV00230 | XP_018219887.1 | 99 | 0 |
| SPGP0X00900 | XP_018219012.1 | 99 | 0 |
| SPGP0W01230 | XP_018223211.1 | 99 | 0 |
| SPGP0DZ01350 | XP_018219474.1 | 99 | 0 |
| SPGP0U01160 | XP_018218876.1 | 99 | 0 |
| SPGP0CZ01310 | XP_018220956.1 | 99 | 0 |
| SPGP0B00170 | XP_018221465.1 | 99 | 0 |
| SPGP0AC00890 | XP_018219172.1 | 99 | 0 |
| SPGP0S03440 | XP_018220012.1 | 99 | 0 |

Create PDF in your applications with the Pdfcrowd HTML to PDF API PDFCROWD

| | | | |
|---|---|---|---|
| SPGP0Q01380 | XP_018222361.1 | 99 | 0 |
| SPGP0Y01520 | XP_018224021.1 | 99 | 0 |
| SPGP0DN00410 | XP_018223490.1 | 99 | 0 |
| SPGP0DN02550 | XP_018223700.1 | 99 | 0 |
| SPGP0W02470 | XP_018223090.1 | 99 | 0 |
| SPGP0U01060 | XP_018218886.1 | 99 | 0 |
| SPGP0S01330 | XP_018220213.1 | 99 | 0 |
| SPGP0B00730 | XP_018221520.1 | 99 | 0 |
| SPGP0W03850 | XP_018222963.1 | 99 | 0 |
| SPGP0X00110 | XP_018219088.1 | 99 | 0 |
| SPGP0DN00200 | XP_018223470.1 | 99 | 0 |
| SPGP0R02300 | XP_018221343.1 | 99 | 0 |
| SPGP0S01090 | XP_018220236.1 | 99 | 0 |
| SPGP0L01250 | XP_018219239.1 | 99 | 0 |
| SPGP0R00520 | XP_018221175.1 | 99 | 0 |
| SPGP0DN03260 | XP_018223769.1 | 99 | 0 |
| SPGP0H00570 | XP_018224080.1 | 99 | 0 |
| SPGP0DZ02180 | XP_018219555.1 | 99 | 0 |
| SPGP0B01110 | XP_018221556.1 | 99 | 0 |
| SPGP0N01680 | XP_018220646.1 | 99 | 0 |
| SPGP0AEtrna18P | DI49_4736 | 100 | 0 |
| SPGP0B01640 | XP_018221606.1 | 99 | 0 |
| SPGP0W04480 | XP_018222900.1 | 99 | 0 |
| SPGP0Ctrna20V | DI49_1074 | 100 | 0 |
| SPGP0X00720 | XP_018219030.1 | 99 | 0 |
| SPGP0P02320 | XP_018221731.1 | 99 | 0 |
| SPGP0W03250 | XP_018223018.1 | 99 | 0 |
| SPGP0P00740 | XP_018221882.1 | 99 | 0 |
| SPGP0Z02940 | XP_018219591.1 | 99 | 0 |
| SPGP0DU01080 | XP_018220482.1 | 99 | 0 |
| SPGP0AF00340 | XP_018221975.1 | 99 | 0 |
| SPGP0DNtrna5I | DI49_0395 | 100 | 0 |
| SPGP0DZ02230 | XP_018219560.1 | 99 | 0 |
| SPGP0DN02690 | XP_018223713.1 | 99 | 0 |
| SPGP0P00870 | XP_018221870.1 | 99 | 0 |
| SPGP0S03530 | XP_018220003.1 | 99 | 0 |
| SPGP0R03190 | XP_018221429.1 | 99 | 0 |
| SPGP0DU00390 | XP_018220551.1 | 99 | 0 |
| SPGP0W04150 | XP_018222935.1 | 99 | 0 |
| SPGP0E01280 | XP_018222718.1 | 99 | 0 |

Create PDF in your applications with the Pdfcrowd HTML to PDF API PDFCROWD

| | | | |
|---|---|---|---|
| SPGP0B01270 | XP_018221570.1 | 100 | 0 |
| SPGP0E01780 | XP_018222766.1 | 99 | 0 |
| SPGP0L01450 | XP_018219219.1 | 99 | 0 |
| SPGP0N01430 | XP_018220670.1 | 99 | 0 |
| SPGP0X00640 | XP_018219038.1 | 100 | 0 |
| SPGP0L00260 | XP_018219335.1 | 99 | 0 |
| SPGP0S03280 | XP_018220027.1 | 99 | 0 |
| SPGP0W03870 | XP_018222961.1 | 99 | 0 |
| SPGP0B00590 | XP_018221507.1 | 99 | 0 |
| SPGP0Z02990 | XP_018219586.1 | 99 | 0 |
| SPGP0DZ01180 | XP_018219457.1 | 99 | 0 |
| SPGP0T00120 | XP_018220340.1 | 99 | 0 |
| SPGP0DN03990 | XP_018223841.1 | 99 | 0 |
| SPGP0E01070 | XP_018222697.1 | 99 | 0 |
| SPGP0Y01080 | XP_018223978.1 | 99 | 0 |
| SPGP0S03000 | XP_018220054.1 | 99 | 0 |
| SPGP0W03000 | XP_018223041.1 | 99 | 0 |
| SPGP0E00850 | XP_018222673.1 | 100 | 0 |
| SPGP0R01430 | XP_018221260.1 | 99 | 0 |
| SPGP0S01690 | XP_018220182.1 | 99 | 0 |
| SPGP0W04730 | XP_018222875.1 | 99 | 0 |
| SPGP0DZ01730 | XP_018219511.1 | 99 | 0 |
| SPGP0S01410 | XP_018220207.1 | 99 | 0 |
| SPGP0P01060 | XP_018221851.1 | 99 | 0 |
| SPGP0DN02020 | XP_018223649.1 | 100 | 0 |
| SPGP0P01470 | XP_018221811.1 | 99 | 0 |
| SPGP0W00490 | XP_018223285.1 | 99 | 0 |
| SPGP0AH00960 | XP_018222511.1 | 99 | 0 |
| SPGP0B01580 | XP_018221600.1 | 99 | 0 |
| SPGP0DN02700 | XP_018223714.1 | 99 | 0 |
| SPGP0L00840 | XP_018219279.1 | 99 | 0 |
| SPGP0R00740 | XP_018221196.1 | 98 | 0 |
| SPGP0R00210 | XP_018221144.1 | 99 | 0 |
| SPGP0W01700 | XP_018223165.1 | 99 | 0 |
| SPGP0Y01610 | XP_018224030.1 | 99 | 0 |
| SPGP0AF00420 | XP_018221983.1 | 98 | 0 |
| SPGP0T00360 | XP_018220364.1 | 99 | 0 |
| SPGP0E00970 | XP_018222687.1 | 99 | 0 |
| SPGP0Q01660 | XP_018222332.1 | 99 | 0 |
| SPGP0Z01940 | XP_018219691.1 | 98 | 0 |

Create PDF in your applications with the Pdfcrowd HTML to PDF API PDFCROWD

| | | | |
|---|---|---|---|
| SPGP0Y01510 | XP_018224020.1 | 100 | 0 |
| SPGP0H00100 | XP_018224123.1 | 96 | 0 |
| SPGP0S02470 | XP_018220106.1 | 99 | 0 |
| SPGP0P01000 | XP_018221857.1 | 99 | 0 |
| SPGP0E02610 | XP_018222847.1 | 99 | 0 |
| SPGP0Z00240 | XP_018219860.1 | 100 | 0 |
| SPGP0Q03110 | XP_018222191.1 | 99 | 0 |
| SPGP0L00170 | XP_018219344.1 | 99 | 0 |
| SPGP0U00790 | XP_018218913.1 | 99 | 0 |
| SPGP0N02040 | XP_018220610.1 | 99 | 0 |
| SPGP0U00160 | XP_018218973.1 | 99 | 0 |
| SPGP0DN01390 | XP_018223586.1 | 99 | 0 |
| SPGP0AC00940 | XP_018219177.1 | 99 | 0 |
| SPGP0AH01180 | XP_018222490.1 | 99 | 1 |
| SPGP0ET00120 | XP_018218981.1 | 99 | 0 |
| SPGP0S02590 | XP_018220094.1 | 99 | 0 |
| SPGP0DZ00210 | XP_018219364.1 | 99 | 0 |
| SPGP0E00380 | XP_018222628.1 | 99 | 0 |
| SPGP0E00430 | XP_018222633.1 | 99 | 0 |
| SPGP0Q03370 | XP_018222166.1 | 98 | 1 |
| SPGP0R00670 | XP_018221189.1 | 98 | 0 |
| SPGP0Z03040 | XP_018219581.1 | 99 | 0 |
| SPGP0L00650 | XP_018219298.1 | 99 | 0 |
| SPGP0DN00170 | XP_018223467.1 | 99 | 0 |
| SPGP0P01320 | XP_018221825.1 | 99 | 0 |
| SPGP0DN01590 | XP_018223606.1 | 99 | 0 |
| SPGP0S02450 | XP_018220108.1 | 99 | 0 |
| SPGP0E01520 | XP_018222740.1 | 100 | 0 |
| SPGP0DN00130 | XP_018218753.1 | 99 | 0 |
| SPGP0S00360 | XP_018220309.1 | 99 | 0 |
| SPGP0E01090 | XP_018222699.1 | 99 | 0 |
| SPGP0R02340 | XP_018221347.1 | 99 | 0 |
| SPGP0R03510 | XP_018221457.1 | 99 | 0 |
| SPGP0DN00630 | XP_018223509.1 | 99 | 0 |
| SPGP0P01200 | XP_018221837.1 | 99 | 0 |
| SPGP0DU00750 | XP_018220516.1 | 99 | 0 |
| SPGP0DN00310 | XP_018223481.1 | 99 | 0 |
| SPGP0R03250 | XP_018221435.1 | 99 | 0 |
| SPGP0CZ01860 | XP_018221008.1 | 99 | 0 |
| SPGP0S02050 | XP_018220147.1 | 99 | 0 |

Create PDF in your applications with the Pdfcrowd HTML to PDF API PDFCROWD

| | | | |
|---|---|---|---|
| SPGP0Q00870 | XP_018222414.1 | 99 | 0 |
| SPGP0AA00780 | XP_018223388.1 | 100 | 0 |
| SPGP0H00740 | XP_018224063.1 | 99 | 0 |
| SPGP0AA00250 | XP_018223338.1 | 99 | 0 |
| SPGP0P00200 | XP_018221934.1 | 100 | 0 |
| SPGP0DZ02370 | XP_018218751.1 | 99 | 0 |
| SPGP0X00750 | XP_018219027.1 | 99 | 0 |
| SPGP0E02100 | XP_018222798.1 | 99 | 0 |
| SPGP0W00570 | XP_018223277.1 | 99 | 0 |
| SPGP0F04910 | XP_018222003.1 | 99 | 0 |
| SPGP0W02590 | XP_018223078.1 | 99 | 0 |
| SPGP0P01910 | XP_018221769.1 | 99 | 0 |
| SPGP0DU00130 | XP_018220577.1 | 99 | 0 |
| SPGP0DN02400 | XP_018223685.1 | 99 | 0 |
| SPGP0S01470 | XP_018220201.1 | 99 | 0 |
| SPGP0DN00590 | XP_018223506.1 | 100 | 0 |
| SPGP0DX00110 | XP_018219942.1 | 99 | 0 |
| SPGP0AA00430 | XP_018223355.1 | 99 | 0 |
| SPGP0E01380 | XP_018222727.1 | 99 | 0 |
| SPGP0Y01680 | XP_018224036.1 | 100 | 0 |
| SPGP0Z01120 | XP_018219772.1 | 99 | 0 |
| SPGP0Z00510 | XP_018219832.1 | 99 | 0 |
| SPGP0Z00880 | XP_018219796.1 | 100 | 0 |
| SPGP0Q04210 | XP_018222084.1 | 99 | 0 |
| SPGP0H00430 | XP_018224093.1 | 99 | 0 |
| SPGP0W01360 | XP_018223198.1 | 100 | 0 |
| SPGP0U00770 | XP_018218915.1 | 99 | 0 |
| SPGP0H00200 | XP_018224115.1 | 99 | 1 |
| SPGP0W01100 | XP_018223224.1 | 99 | 0 |
| SPGP0Z02710 | XP_018219614.1 | 99 | 0 |
| SPGP0R03460 | XP_018221453.1 | 99 | 0 |
| SPGP0S02650 | XP_018220088.1 | 99 | 0 |
| SPGP0W02420 | XP_018223095.1 | 99 | 0 |
| SPGP0F04610 | XP_018222031.1 | 99 | 0 |
| SPGP0Z01540 | XP_018219731.1 | 99 | 0 |
| SPGP0W02930 | XP_018223047.1 | 99 | 0 |
| SPGP0Q00290 | XP_018222472.1 | 99 | 0 |
| SPGP0DV00390 | XP_018219901.1 | 100 | 0 |
| SPGP0Z01740 | XP_018219711.1 | 99 | 1 |
| SPGP0DU01660 | XP_018220425.1 | 97 | 2 |

Create PDF in your applications with the Pdfcrowd HTML to PDF API PDFCROWD

| | | | |
|---|---|---|---|
| SPGP0E01750 | XP_018222763.1 | 99 | 0 |
| SPGP0S02900 | XP_018220064.1 | 99 | 0 |
| SPGP0DN02940 | XP_018223739.1 | 99 | 0 |
| SPGP0Z02180 | XP_018219667.1 | 99 | 0 |
| SPGP0X00490 | XP_018219052.1 | 99 | 1 |
| SPGP0R02510 | XP_018221364.1 | 99 | 0 |
| SPGP0R02140 | XP_018221328.1 | 99 | 0 |
| SPGP0F04690 | XP_018222024.1 | 99 | 0 |
| SPGP0R02820 | XP_018221394.1 | 100 | 0 |
| SPGP0N00850 | XP_018220724.1 | 99 | 0 |
| SPGP0L01410 | XP_018219223.1 | 100 | 0 |
| SPGP0Q00300 | XP_018222471.1 | 99 | 0 |
| SPGP0H00530 | XP_018224083.1 | 99 | 0 |
| SPGP0Q03140 | XP_018222188.1 | 99 | 0 |
| SPGP0W00690 | XP_018223264.1 | 99 | 0 |
| SPGP0F04390 | XP_018222052.1 | 99 | 0 |
| SPGP0S01100 | XP_018220235.1 | 99 | 0 |
| SPGP0R01970 | XP_018221312.1 | 99 | 0 |
| SPGP0Q01180 | XP_018222382.1 | 99 | 0 |
| SPGP0W01510 | XP_018223184.1 | 99 | 0 |
| SPGP0H00230 | XP_018224112.1 | 100 | 0 |
| SPGP0DN02540 | XP_018223699.1 | 99 | 0 |
| SPGP0DN03610 | XP_018223803.1 | 99 | 0 |
| SPGP0N00860 | XP_018220723.1 | 99 | 0 |
| SPGP0DN02370 | XP_018223682.1 | 99 | 0 |
| SPGP0P02170 | XP_018221743.1 | 99 | 0 |
| SPGP0Z02570 | XP_018219628.1 | 99 | 0 |
| SPGP0Q00420 | XP_018222459.1 | 99 | 0 |
| SPGP0W02770 | XP_018223062.1 | 99 | 0 |
| SPGP0R01780 | XP_018221295.1 | 99 | 0 |
| SPGP0Q03960 | XP_018222109.1 | 99 | 0 |
| SPGP0E02120 | XP_018222800.1 | 99 | 0 |
| SPGP0DN02960 | XP_018223741.1 | 99 | 0 |
| SPGP0X01040 | XP_018218999.1 | 99 | 0 |
| SPGP0H00330 | XP_018224102.1 | 99 | 0 |
| SPGP0X00160 | XP_018219084.1 | 99 | 0 |
| SPGP0X00350 | XP_018219066.1 | 99 | 0 |
| SPGP0W00390 | XP_018223295.1 | 99 | 0 |
| SPGP0W04830 | XP_018222865.1 | 99 | 0 |
| SPGP0Q01980 | XP_018222300.1 | 99 | 0 |

Create PDF in your applications with the Pdfcrowd HTML to PDF API PDFCROWD

| | | | |
|---|---|---|---|
| SPGP0Q01230 | XP_018222377.1 | 99 | 0 |
| SPGP0P02980 | XP_018221666.1 | 99 | 0 |
| SPGP0B00900 | XP_018221536.1 | 99 | 1 |
| SPGP0W03110 | XP_018223032.1 | 100 | 0 |
| SPGP0DN03030 | XP_018223748.1 | 99 | 0 |
| SPGP0DN03520 | XP_018223794.1 | 99 | 0 |
| SPGP0Q01000 | XP_018222401.1 | 99 | 0 |
| SPGP0E02260 | XP_018222814.1 | 100 | 0 |
| SPGP0W00380 | XP_018223296.1 | 99 | 0 |
| SPGP0AC00710 | XP_018219154.1 | 99 | 0 |
| SPGP0Q04030 | XP_018222102.1 | 99 | 0 |
| SPGP0DN01960 | XP_018223643.1 | 99 | 0 |
| SPGP0Q02180 | XP_018222280.1 | 99 | 0 |
| SPGP0R01350 | XP_018221252.1 | 99 | 0 |
| SPGP0DZ00170 | XP_018219360.1 | 99 | 0 |
| SPGP0W00640 | XP_018223270.1 | 99 | 0 |
| SPGP0L00100 | XP_018219351.1 | 99 | 0 |
| SPGP0B00700 | XP_018221517.1 | 99 | 0 |
| SPGP0U00940 | XP_018218898.1 | 99 | 0 |
| SPGP0Y00590 | XP_018223933.1 | 99 | 0 |
| SPGP0B00710 | XP_018221518.1 | 99 | 0 |
| SPGP0DU01140 | XP_018220476.1 | 99 | 0 |
| SPGP0Q02560 | XP_018222244.1 | 99 | 0 |
| SPGP0R00890 | XP_018221209.1 | 99 | 0 |
| SPGP0Q02760 | XP_018222224.1 | 99 | 0 |
| SPGP0N00940 | XP_018220715.1 | 99 | 0 |
| SPGP0DZ02340 | XP_018218747.1 | 99 | 0 |
| SPGP0S00430 | XP_018220302.1 | 100 | 0 |
| SPGP0X00650 | XP_018219037.1 | 99 | 0 |
| SPGP0DN02920 | XP_018223737.1 | 100 | 0 |
| SPGP0B00380 | XP_018221486.1 | 99 | 0 |
| SPGP0N01850 | XP_018220630.1 | 99 | 0 |
| SPGP0R00970 | XP_018221215.1 | 99 | 0 |
| SPGP0Q03580 | XP_018222146.1 | 99 | 0 |
| SPGP0X00890 | XP_018219013.1 | 99 | 0 |
| SPGP0S03560 | XP_018220000.1 | 99 | 0 |
| SPGP0FB00110 | XP_018220805.1 | 99 | 0 |
| SPGP0W04280 | XP_018222922.1 | 99 | 1 |
| SPGP0R03500 | XP_018221455.1 | 99 | 0 |
| SPGP0N00100 | XP_018220798.1 | 99 | 0 |

Create PDF in your applications with the Pdfcrowd HTML to PDF API PDFCROWD

| | | | |
|---|---|---|---|
| SPGP0S02280 | XP_018220124.1 | 99 | 0 |
| SPGP0P00360 | XP_018221919.1 | 99 | 0 |
| SPGP0EE00200 | XP_018220839.1 | 99 | 0 |
| SPGP0E01820 | XP_018222770.1 | 99 | 0 |
| SPGP0W01590 | XP_018223176.1 | 99 | 0 |
| SPGP0X01100 | XP_018218993.1 | 98 | 0 |
| SPGP0CZ02070 | XP_018221028.1 | 99 | 0 |
| SPGP0Q03320 | XP_018222171.1 | 98 | 0 |
| SPGP0U00580 | XP_018218931.1 | 99 | 0 |
| SPGP0Q01480 | XP_018222350.1 | 99 | 0 |
| SPGP0Q00460 | XP_018222455.1 | 99 | 0 |
| SPGP0AC00300 | XP_018219114.1 | 99 | 0 |
| SPGP0E01260 | XP_018222716.1 | 99 | 0 |
| SPGP0DN01780 | XP_018223624.1 | 99 | 0 |
| SPGP0Q00540 | XP_018222447.1 | 99 | 0 |
| SPGP0S00750 | XP_018220270.1 | 99 | 0 |
| SPGP0S02130 | XP_018220139.1 | 99 | 0 |
| SPGP0P00830 | XP_018221874.1 | 100 | 0 |
| SPGP0BF00120 | XP_018220383.1 | 99 | 0 |
| SPGP0AA00820 | XP_018223392.1 | 99 | 0 |
| SPGP0F04200 | XP_018222071.1 | 99 | 0 |
| SPGP0S03520 | XP_018220004.1 | 100 | 0 |
| SPGP0Q01430 | XP_018222356.1 | 100 | 0 |
| SPGP0W03640 | XP_018222984.1 | 99 | 0 |
| SPGP0AF00240 | XP_018221965.1 | 99 | 0 |
| SPGP0DN00780 | XP_018223524.1 | 99 | 0 |
| SPGP0W02090 | XP_018223128.1 | 99 | 0 |
| SPGP0Q00600 | XP_018222441.1 | 99 | 0 |
| SPGP0W04880 | XP_018222860.1 | 99 | 0 |
| SPGP0E00290 | XP_018222619.1 | 99 | 0 |
| SPGP0ET00170 | XP_018218986.1 | 99 | 0 |
| SPGP0H00260 | XP_018224109.1 | 99 | 0 |
| SPGP0P00430 | XP_018221912.1 | 99 | 0 |
| SPGP0DN03050 | XP_018223750.1 | 99 | 0 |
| SPGP0L00340 | XP_018219327.1 | 99 | 0 |
| SPGP0Z02660 | XP_018219619.1 | 99 | 0 |
| SPGP0L00140 | XP_018219347.1 | 99 | 0 |
| SPGP0DN03660 | XP_018223808.1 | 99 | 0 |
| SPGP0U01760 | XP_018218818.1 | 99 | 0 |
| SPGP0B00620 | XP_018221510.1 | 99 | 0 |

Create PDF in your applications with the Pdfcrowd HTML to PDF API PDFCROWD

| | | | |
|---|---|---|---|
| SPGP0L01080 | XP_018219255.1 | 100 | 0 |
| SPGP0W00540 | XP_018223280.1 | 99 | 0 |
| SPGP0W02890 | XP_018223051.1 | 100 | 0 |
| SPGP0Q00720 | XP_018222428.1 | 97 | 2 |
| SPGP0W02000 | XP_018223135.1 | 99 | 0 |
| SPGP0B00610 | XP_018221509.1 | 99 | 0 |
| SPGP0DX00320 | XP_018219963.1 | 99 | 0 |
| SPGP0P02250 | XP_018221737.1 | 99 | 0 |
| SPGP0DU01410 | XP_018220449.1 | 99 | 0 |
| SPGP0X00920 | XP_018219010.1 | 99 | 0 |
| SPGP0S02230 | XP_018220129.1 | 99 | 0 |
| SPGP0R02980 | XP_018221408.1 | 99 | 0 |
| SPGP0F04460 | XP_018222045.1 | 97 | 2 |
| SPGP0E00150 | XP_018222605.1 | 99 | 0 |
| SPGP0DN03200 | XP_018223764.1 | 99 | 0 |
| SPGP0P03160 | XP_018221648.1 | 99 | 0 |
| SPGP0B01620 | XP_018221604.1 | 100 | 0 |
| SPGP0Ytrna9M | DI49_4243 | 100 | 0 |
| SPGP0R03230 | XP_018221433.1 | 99 | 0 |
| SPGP0Q02970 | XP_018222204.1 | 99 | 0 |
| SPGP0Y00490 | XP_018223923.1 | 99 | 0 |
| SPGP0AEtrna25S | DI49_0808 | 95 | 0 |
| SPGP0E00480 | XP_018222638.1 | 99 | 0 |
| SPGP0DZ00940 | XP_018219434.1 | 98 | 1 |
| SPGP0R02180 | XP_018221331.1 | 99 | 0 |
| SPGP0S00190 | XP_018220325.1 | 99 | 0 |
| SPGP0W03270 | XP_018223016.1 | 99 | 0 |
| SPGP0Z02960 | XP_018219589.1 | 99 | 0 |
| SPGP0R00710 | XP_018221193.1 | 99 | 0 |
| SPGP0Q02440 | XP_018222255.1 | 99 | 0 |
| SPGP0B01190 | XP_018221562.1 | 99 | 0 |
| SPGP0R01980 | XP_018221313.1 | 99 | 0 |
| SPGP0W04130 | XP_018222937.1 | 99 | 0 |
| SPGP0R01630 | XP_018221280.1 | 97 | 1 |
| SPGP0CZ00740 | XP_018220899.1 | 99 | 0 |
| SPGP0DN02520 | XP_018223697.1 | 99 | 0 |
| SPGP0E00400 | XP_018222630.1 | 99 | 0 |
| SPGP0R02720 | XP_018221384.1 | 99 | 0 |
| SPGP0R00580 | XP_018221180.1 | 99 | 0 |
| SPGP0Z00520 | XP_018219831.1 | 99 | 0 |

Create PDF in your applications with the Pdfcrowd HTML to PDF API PDFCROWD

| | | | |
|---|---|---|---|
| SPGP0P00660 | XP_018221890.1 | 99 | 0 |
| SPGP0S02310 | XP_018220121.1 | 99 | 0 |
| SPGP0S02170 | XP_018220135.1 | 99 | 0 |
| SPGP0AA00510 | XP_018223363.1 | 99 | 0 |
| SPGP0R01180 | XP_018221236.1 | 99 | 0 |
| SPGP0S02730 | XP_018220080.1 | 99 | 0 |
| SPGP0P01550 | XP_018221804.1 | 99 | 0 |
| SPGP0P01450 | XP_018221813.1 | 99 | 0 |
| SPGP0AF00150 | XP_018221956.1 | 99 | 0 |
| SPGP0S01700 | XP_018220181.1 | 99 | 0 |
| SPGP0E00760 | XP_018222665.1 | 99 | 0 |
| SPGP0Z02620 | XP_018219623.1 | 99 | 0 |
| SPGP0DN02000 | XP_018223647.1 | 99 | 0 |
| SPGP0DZ00140 | XP_018219357.1 | 99 | 0 |
| SPGP0DV00440 | XP_018219906.1 | 99 | 0 |
| SPGP0H00620 | XP_018224075.1 | 99 | 0 |
| SPGP0E01790 | XP_018222767.1 | 99 | 0 |
| SPGP0DN01840 | XP_018223631.1 | 99 | 0 |
| SPGP0H00190 | XP_018224116.1 | 99 | 0 |
| SPGP0R00760 | XP_018221198.1 | 99 | 0 |
| SPGP0Q02340 | XP_018222265.1 | 99 | 0 |
| SPGP0Q02510 | XP_018222248.1 | 99 | 0 |
| SPGP0W03520 | XP_018222995.1 | 99 | 0 |
| SPGP0Q02280 | XP_018222271.1 | 99 | 0 |
| SPGP0H00440 | XP_018224092.1 | 99 | 0 |
| SPGP0R02360 | XP_018221349.1 | 99 | 0 |
| SPGP0DV00530 | XP_018219915.1 | 99 | 0 |
| SPGP0Z01920 | XP_018219693.1 | 99 | 0 |
| SPGP0R02620 | XP_018221375.1 | 99 | 0 |
| SPGP0AH00670 | XP_018222538.1 | 99 | 0 |
| SPGP0DN02120 | XP_018223659.1 | 99 | 0 |
| SPGP0Q03170 | XP_018222185.1 | 99 | 0 |
| SPGP0W01720 | XP_018223163.1 | 100 | 0 |
| SPGP0N02240 | XP_018220590.1 | 99 | 0 |
| SPGP0DN01200 | XP_018223567.1 | 100 | 0 |
| SPGP0CZ02790 | XP_018221098.1 | 99 | 0 |
| SPGP0AC00960 | XP_018219179.1 | 99 | 0 |
| SPGP0Q02190 | XP_018222279.1 | 99 | 0 |
| SPGP0ET00100 | XP_018218979.1 | 99 | 0 |
| SPGP0BFtrna21P | DI49_3505 | 100 | 0 |

Create PDF in your applications with the Pdfcrowd HTML to PDF API PDFCROWD

| | | | |
|---|---|---|---|
| SPGP0L00740 | XP_018219289.1 | 99 | 0 |
| SPGP0E00410 | XP_018222631.1 | 99 | 0 |
| SPGP0DZ00510 | XP_018219394.1 | 99 | 0 |
| SPGP0Q01610 | XP_018222337.1 | 99 | 0 |
| SPGP0Q02010 | XP_018222297.1 | 99 | 0 |
| SPGP0CZ01600 | XP_018220984.1 | 99 | 0 |
| SPGP0DN04120 | XP_018223854.1 | 99 | 0 |
| SPGP0L01330 | XP_018219231.1 | 99 | 0 |
| SPGP0CZ02700 | XP_018221090.1 | 100 | 0 |
| SPGP0Q00910 | XP_018222410.1 | 99 | 0 |
| SPGP0S00600 | XP_018220285.1 | 99 | 0 |
| SPGP0R02440 | XP_018221357.1 | 99 | 0 |
| SPGP0E00670 | XP_018222657.1 | 98 | 0 |
| SPGP0F04780 | XP_018222016.1 | 99 | 0 |
| SPGP0E02000 | XP_018222788.1 | 99 | 0 |
| SPGP0Q01010 | XP_018222400.1 | 99 | 0 |
| SPGP0S02520 | XP_018220101.1 | 99 | 1 |
| SPGP0B01590 | XP_018221601.1 | 99 | 0 |
| SPGP0Q00850 | XP_018222416.1 | 99 | 0 |
| SPGP0AA00230 | XP_018223336.1 | 99 | 0 |
| SPGP0P00220 | XP_018221932.1 | 99 | 0 |
| SPGP0Z00300 | XP_018219846.1 | 99 | 0 |
| SPGP0DV00620 | XP_018219923.1 | 99 | 0 |
| SPGP0U01890 | XP_018218805.1 | 99 | 0 |
| SPGP0N00710 | XP_018220737.1 | 100 | 0 |
| SPGP0S02440 | XP_018220109.1 | 99 | 0 |
| SPGP0Q04000 | XP_018222105.1 | 99 | 0 |
| SPGP0P01930 | XP_018221767.1 | 96 | 2 |
| SPGP0Y00290 | XP_018223906.1 | 99 | 0 |
| SPGP0Z02160 | XP_018219669.1 | 99 | 0 |
| SPGP0R01510 | XP_018221268.1 | 99 | 0 |
| SPGP0S00910 | XP_018220254.1 | 99 | 0 |
| SPGP0DN00840 | XP_018223530.1 | 99 | 0 |
| SPGP0DN00330 | XP_018223483.1 | 99 | 0 |
| SPGP0F04980 | XP_018221994.1 | 98 | 0 |
| SPGP0W02270 | XP_018223110.1 | 99 | 0 |
| SPGP0W01650 | XP_018223170.1 | 99 | 0 |
| SPGP0E01100 | XP_018222700.1 | 98 | 0 |
| SPGP0E01330 | XP_018222723.1 | 100 | 0 |
| SPGP0R00840 | XP_018221204.1 | 98 | 0 |

Create PDF in your applications with the Pdfcrowd HTML to PDF API PDFCROWD

| | | | |
|---|---|---|---|
| SPGP0DU01060 | XP_018220484.1 | 99 | 0 |
| SPGP0AC00110 | XP_018219096.1 | 100 | 0 |
| SPGP0AC00330 | XP_018219117.1 | 99 | 0 |
| SPGP0R03440 | XP_018221451.1 | 99 | 0 |
| SPGP0B01220 | XP_018221565.1 | 99 | 0 |
| SPGP0F04670 | XP_018222025.1 | 99 | 0 |
| SPGP0B01290 | XP_018221572.1 | 99 | 0 |
| SPGP0DN02810 | XP_018223725.1 | 99 | 0 |
| SPGP0Z01890 | XP_018219696.1 | 99 | 0 |
| SPGP0Q03860 | XP_018222119.1 | 98 | 0 |
| SPGP0R03140 | XP_018221424.1 | 99 | 0 |
| SPGP0X00250 | XP_018219076.1 | 99 | 0 |
| SPGP0E01660 | XP_018222754.1 | 99 | 0 |
| SPGP0DU01600 | XP_018220431.1 | 99 | 0 |
| SPGP0N01320 | XP_018220681.1 | 100 | 0 |
| SPGP0S02920 | XP_018220062.1 | 99 | 0 |
| SPGP0P01660 | XP_018221794.1 | 95 | 5 |
| SPGP0B00790 | XP_018221526.1 | 99 | 0 |
| SPGP0W02840 | XP_018223056.1 | 99 | 0 |
| SPGP0R02530 | XP_018221366.1 | 99 | 0 |
| SPGP0P02480 | XP_018221715.1 | 100 | 0 |
| SPGP0Z00500 | XP_018219833.1 | 99 | 0 |
| SPGP0Z02770 | XP_018219607.1 | 100 | 0 |
| SPGP0DV00220 | XP_018219886.1 | 99 | 0 |
| SPGP0W01350 | XP_018223199.1 | 99 | 0 |
| SPGP0DV00450 | XP_018219907.1 | 100 | 0 |
| SPGP0Z00490 | XP_018219834.1 | 99 | 0 |
| SPGP0S00140 | XP_018220330.1 | 99 | 0 |
| SPGP0S01160 | XP_018220229.1 | 100 | 0 |
| SPGP0F04930 | XP_018222002.1 | 99 | 0 |
| SPGP0W03160 | XP_018223027.1 | 99 | 0 |
| SPGP0DN00340 | XP_018223484.1 | 99 | 0 |
| SPGP0DN04050 | XP_018223847.1 | 99 | 0 |
| SPGP0Y00260 | XP_018223904.1 | 99 | 0 |
| SPGP0L00230 | XP_018219338.1 | 99 | 0 |
| SPGP0R01220 | XP_018221240.1 | 99 | 0 |
| SPGP0E01850 | XP_018222773.1 | 99 | 0 |
| SPGP0Z02590 | XP_018219626.1 | 99 | 0 |
| SPGP0Q03940 | XP_018222111.1 | 99 | 0 |

Create PDF in your applications with the Pdfcrowd HTML to PDF API PDFCROWD

| | | | |
|---|---|---|---|
| SPGP0Z00730 | XP_018219811.1 | 99 | 0 |
| SPGP0DN03570 | XP_018223799.1 | 99 | 0 |
| SPGP0E02240 | XP_018222812.1 | 100 | 0 |
| SPGP0W04660 | XP_018222882.1 | 99 | 0 |
| SPGP0Strna12P | DI49_4051 | 99 | 0 |
| SPGP0S02370 | XP_018220115.1 | 99 | 0 |
| SPGP0P01390 | XP_018221819.1 | 99 | 0 |
| SPGP0W04670 | XP_018222881.1 | 99 | 0 |
| SPGP0AC00990 | XP_018219182.1 | 98 | 0 |
| SPGP0W04450 | XP_018222903.1 | 99 | 0 |
| SPGP0AH00110 | XP_018222593.1 | 98 | 1 |
| SPGP0DN01110 | XP_018223558.1 | 99 | 0 |
| SPGP0P02150 | XP_018221745.1 | 99 | 0 |
| SPGP0W03130 | XP_018223030.1 | 99 | 0 |
| SPGP0E00120 | XP_018222602.1 | 99 | 0 |
| SPGP0BF00180 | XP_018220388.1 | 99 | 0 |
| SPGP0DZ01430 | XP_018219482.1 | 99 | 0 |
| SPGP0Q03030 | XP_018222198.1 | 99 | 0 |
| SPGP0DN03010 | XP_018223746.1 | 99 | 1 |
| SPGP0CZ01800 | XP_018221002.1 | 99 | 0 |
| SPGP0AC00370 | XP_018219121.1 | 99 | 0 |
| SPGP0X00780 | XP_018219024.1 | 99 | 0 |
| SPGP0Z00190 | XP_018219864.1 | 99 | 0 |
| SPGP0P02740 | XP_018221689.1 | 99 | 0 |
| SPGP0DZ01510 | XP_018219490.1 | 99 | 0 |
| SPGP0B01380 | XP_018221581.1 | 100 | 0 |
| SPGP0DN01940 | XP_018223641.1 | 99 | 0 |
| SPGP0B00740 | XP_018221521.1 | 99 | 0 |
| SPGP0R01370 | XP_018221254.1 | 99 | 0 |
| SPGP0W00110 | XP_018223323.1 | 98 | 0 |
| SPGP0L00120 | XP_018219349.1 | 99 | 0 |
| SPGP0AG00110 | XP_018219931.1 | 90 | 0 |
| SPGP0S03060 | XP_018220048.1 | 99 | 0 |
| SPGP0Itrna19Q | DI49_2421 | 100 | 0 |
| SPGP0AH00270 | XP_018222575.1 | 99 | 0 |
| SPGP0R00390 | XP_018221162.1 | 100 | 0 |
| SPGP0AH00980 | XP_018222509.1 | 99 | 0 |
| SPGP0Z00990 | XP_018219785.1 | 99 | 0 |
| SPGP0Q02700 | XP_018222230.1 | 99 | 0 |
| SPGP0DZ01590 | XP_018219498.1 | 99 | 0 |

Create PDF in your applications with the Pdfcrowd HTML to PDF API PDFCROWD

| | | | |
|---|---|---|---|
| SPGP0DN01370 | XP_018223584.1 | 99 | 0 |
| SPGP0W04030 | XP_018222947.1 | 99 | 0 |
| SPGP0S01830 | XP_018220169.1 | 99 | 0 |
| SPGP0P00260 | XP_018221928.1 | 99 | 0 |
| SPGP0F04880 | XP_018222006.1 | 99 | 0 |
| SPGP0Z01710 | XP_018219714.1 | 99 | 0 |
| SPGP0DZ01390 | XP_018219478.1 | 99 | 0 |
| SPGP0U00340 | XP_018218955.1 | 99 | 0 |
| SPGP0Z00100 | XP_018219873.1 | 99 | 0 |
| SPGP0W01570 | XP_018223178.1 | 99 | 0 |
| SPGP0DN02260 | XP_018223672.1 | 99 | 0 |
| SPGP0AA00220 | XP_018223335.1 | 99 | 0 |
| SPGP0E01150 | XP_018222705.1 | 99 | 0 |
| SPGP0DN01410 | XP_018223588.1 | 99 | 0 |
| SPGP0S00290 | XP_018220316.1 | 99 | 0 |
| SPGP0E00900 | XP_018222678.1 | 99 | 0 |
| SPGP0U00590 | XP_018218930.1 | 99 | 0 |
| SPGP0Z02370 | XP_018219648.1 | 99 | 0 |
| SPGP0S03090 | XP_018220045.1 | 99 | 0 |
| SPGP0DN04090 | XP_018223851.1 | 99 | 0 |
| SPGP0F04860 | XP_018222008.1 | 99 | 0 |
| SPGP0DN01700 | XP_018223616.1 | 99 | 0 |
| SPGP0S01760 | XP_018220175.1 | 99 | 0 |
| SPGP0P00100 | XP_018221944.1 | 99 | 0 |
| SPGP0R00200 | XP_018221143.1 | 99 | 0 |
| SPGP0DN03300 | XP_018223773.1 | 99 | 0 |
| SPGP0U01430 | XP_018218850.1 | 99 | 0 |
| SPGP0DV00570 | XP_018219919.1 | 99 | 0 |
| SPGP0S02150 | XP_018220137.1 | 99 | 0 |
| SPGP0Y00850 | XP_018223955.1 | 99 | 0 |
| SPGP0W03810 | XP_018222967.1 | 100 | 0 |
| SPGP0R00480 | XP_018221171.1 | 99 | 0 |
| SPGP0Z00460 | XP_018219837.1 | 87 | 2 |
| SPGP0F04220 | XP_018222069.1 | 99 | 0 |
| SPGP0S03540 | XP_018220002.1 | 99 | 0 |
| SPGP0N01380 | XP_018220675.1 | 99 | 0 |
| SPGP0N01650 | XP_018220649.1 | 99 | 0 |
| SPGP0DN01850 | XP_018223632.1 | 99 | 0 |
| SPGP0N01120 | XP_018220700.1 | 99 | 0 |
| SPGP0H00220 | XP_018224113.1 | 99 | 0 |

Create PDF in your applications with the Pdfcrowd HTML to PDF API PDFCROWD

| | | | |
|---|---|---|---|
| SPGP0DU00830 | XP_018220507.1 | 99 | 0 |
| SPGP0DZ00320 | XP_018219375.1 | 99 | 0 |
| SPGP0S01490 | XP_018220199.1 | 100 | 0 |
| SPGP0L01640 | XP_018219201.1 | 97 | 0 |
| SPGP0Q00620 | XP_018222439.1 | 99 | 0 |
| SPGP0F04290 | XP_018222062.1 | 99 | 0 |
| SPGP0BC00210 | XP_018220818.1 | 99 | 0 |
| SPGP0N00380 | XP_018220770.1 | 99 | 0 |
| SPGP0N00830 | XP_018220726.1 | 99 | 0 |
| SPGP0N01390 | XP_018220674.1 | 99 | 0 |
| SPGP0P00700 | XP_018221886.1 | 99 | 0 |
| SPGP0L01020 | XP_018219261.1 | 99 | 0 |
| SPGP0W00250 | XP_018223309.1 | 99 | 0 |
| SPGP0DN00450 | XP_018223493.1 | 99 | 0 |
| SPGP0W01200 | XP_018223214.1 | 99 | 0 |
| SPGP0U00390 | XP_018218950.1 | 99 | 0 |
| SPGP0AA00150 | XP_018223329.1 | 99 | 0 |
| SPGP0EO00110 | XP_018222853.1 | 96 | 0 |
| SPGP0F04240 | XP_018222067.1 | 99 | 0 |
| SPGP0L01220 | XP_018219242.1 | 99 | 0 |
| SPGP0P00140 | XP_018221940.1 | 99 | 0 |
| SPGP0Qtrna21L | DI49_1818 | 100 | 0 |
| SPGP0DU01470 | XP_018220443.1 | 99 | 0 |
| SPGP0AA00570 | XP_018223369.1 | 99 | 0 |
| SPGP0P00410 | XP_018221914.1 | 99 | 0 |
| SPGP0X00270 | XP_018219074.1 | 99 | 0 |
| SPGP0S03320 | XP_018220023.1 | 99 | 0 |
| SPGP0Z03030 | XP_018219582.1 | 99 | 0 |
| SPGP0CZ03090 | XP_018221127.1 | 99 | 0 |
| SPGP0DN03220 | XP_018223766.1 | 99 | 0 |
| SPGP0H00830 | XP_018224054.1 | 99 | 0 |
| SPGP0P03140 | XP_018221650.1 | 99 | 0 |
| SPGP0DX00550 | XP_018219985.1 | 99 | 0 |
| SPGP0CZ02940 | XP_018221113.1 | 99 | 0 |
| SPGP0B01600 | XP_018221602.1 | 99 | 0 |
| SPGP0E01670 | XP_018222755.1 | 99 | 0 |
| SPGP0DX00300 | XP_018219961.1 | 99 | 0 |
| SPGP0CZ02270 | XP_018221047.1 | 99 | 0 |
| SPGP0N00520 | XP_018220756.1 | 99 | 0 |
| SPGP0W00710 | XP_018223262.1 | 99 | 0 |

Create PDF in your applications with the Pdfcrowd HTML to PDF API PDFCROWD

| | | | |
|---|---|---|---|
| SPGP0E00940 | XP_018222683.1 | 99 | 0 |
| SPGP0S00170 | XP_018220327.1 | 99 | 0 |
| SPGP0F04840 | XP_018222010.1 | 99 | 0 |
| SPGP0P02360 | XP_018221727.1 | 99 | 0 |
| SPGP0AF00380 | XP_018221979.1 | 99 | 0 |
| SPGP0Z02900 | XP_018219595.1 | 99 | 0 |
| SPGP0P02830 | XP_018221680.1 | 99 | 0 |
| SPGP0S00270 | XP_018220318.1 | 99 | 0 |
| SPGP0R01800 | XP_018221297.1 | 99 | 0 |
| SPGP0W04440 | XP_018222904.1 | 99 | 0 |
| SPGP0R01610 | XP_018221278.1 | 99 | 0 |
| SPGP0CZ00760 | XP_018220901.1 | 99 | 0 |
| SPGP0R01710 | XP_018221287.1 | 99 | 0 |
| SPGP0W04110 | XP_018222939.1 | 98 | 1 |
| SPGP0R03220 | XP_018221432.1 | 99 | 0 |
| SPGP0Q00710 | XP_018222430.1 | 99 | 0 |
| SPGP0W03830 | XP_018222965.1 | 99 | 0 |
| SPGP0B00550 | XP_018221503.1 | 99 | 0 |
| SPGP0R00560 | XP_018221179.1 | 99 | 0 |
| SPGP0CZ01490 | XP_018220973.1 | 99 | 0 |
| SPGP0DN03950 | XP_018223837.1 | 99 | 0 |
| SPGP0E01720 | XP_018222761.1 | 100 | 0 |
| SPGP0P02700 | XP_018221693.1 | 100 | 0 |
| SPGP0Z01200 | XP_018219765.1 | 99 | 0 |
| SPGP0DN03140 | XP_018223758.1 | 99 | 0 |
| SPGP0B01450 | XP_018221588.1 | 99 | 0 |
| SPGP0Y01350 | XP_018224004.1 | 99 | 0 |
| SPGP0CZ00540 | XP_018220880.1 | 99 | 0 |
| SPGP0DN02720 | XP_018223716.1 | 99 | 0 |
| SPGP0S02710 | XP_018220082.1 | 99 | 0 |
| SPGP0P01850 | XP_018221775.1 | 99 | 0 |
| SPGP0P01430 | XP_018221815.1 | 99 | 0 |
| SPGP0DN02680 | XP_018223712.1 | 99 | 0 |
| SPGP0Z01900 | XP_018219695.1 | 99 | 0 |
| SPGP0Q01370 | XP_018222362.1 | 99 | 0 |
| SPGP0S02430 | XP_018220110.1 | 99 | 0 |
| SPGP0DZ00750 | XP_018219416.1 | 99 | 0 |
| SPGP0Btrna13E | DI49_2698 | 100 | 0 |
| SPGP0DZ01720 | XP_018219510.1 | 99 | 0 |
| SPGP0Z01380 | XP_018219747.1 | 99 | 0 |

Create PDF in your applications with the Pdfcrowd HTML to PDF API PDFCROWD

| | | | |
|---|---|---|---|
| SPGP0R00700 | XP_018221192.1 | 97 | 1 |
| SPGP0P01990 | XP_018221761.1 | 99 | 0 |
| SPGP0Q02570 | XP_018222243.1 | 99 | 0 |
| SPGP0B01780 | XP_018221620.1 | 99 | 0 |
| SPGP0DV00510 | XP_018219913.1 | 97 | 2 |
| SPGP0B00290 | XP_018221477.1 | 99 | 0 |
| SPGP0W00560 | XP_018223278.1 | 99 | 0 |
| SPGP0S02380 | XP_018220114.1 | 99 | 0 |
| SPGP0P01040 | XP_018221853.1 | 99 | 0 |
| SPGP0R01820 | XP_018221298.1 | 99 | 1 |
| SPGP0Q03150 | XP_018222187.1 | 98 | 0 |
| SPGP0Q03620 | XP_018222143.1 | 99 | 0 |
| SPGP0W01960 | XP_018223139.1 | 99 | 0 |
| SPGP0CZ01140 | XP_018220939.1 | 99 | 0 |
| SPGP0DN03580 | XP_018223800.1 | 99 | 0 |
| SPGP0W01740 | XP_018223161.1 | 99 | 0 |
| SPGP0CZ01630 | XP_018220987.1 | 99 | 0 |
| SPGP0N02260 | XP_018220588.1 | 99 | 0 |
| SPGP0AC00900 | XP_018219173.1 | 99 | 0 |
| SPGP0F04510 | XP_018222040.1 | 97 | 2 |
| SPGP0Z00290 | XP_018219845.1 | 99 | 0 |
| SPGP0AHtrna12Y | DI49_2382 | 100 | 0 |
| SPGP0U01480 | XP_018218845.1 | 99 | 0 |
| SPGP0DZ00450 | XP_018219388.1 | 99 | 0 |
| SPGP0E02440 | XP_018222831.1 | 99 | 0 |
| SPGP0S01640 | XP_018220187.1 | 99 | 0 |
| SPGP0EX00280 | XP_018223886.1 | 99 | 0 |
| SPGP0S02690 | XP_018220084.1 | 100 | 0 |
| SPGP0U00740 | XP_018218918.1 | 99 | 0 |
| SPGP0Wtrna4C | DI49_5400 | 100 | 0 |
| SPGP0Q02390 | XP_018222260.1 | 99 | 0 |
| SPGP0DN01080 | XP_018223555.1 | 99 | 0 |
| SPGP0DN00540 | XP_018223502.1 | 99 | 0 |
| SPGP0Q02610 | XP_018222239.1 | 99 | 0 |
| SPGP0DV00610 | XP_018219922.1 | 99 | 0 |
| SPGP0B01760 | XP_018221618.1 | 99 | 0 |
| SPGP0DN00640 | XP_018223510.1 | 100 | 0 |
| SPGP0AF00390 | XP_018221980.1 | 99 | 0 |
| SPGP0B00340 | XP_018221482.1 | 99 | 0 |
| SPGP0DU00710 | XP_018220520.1 | 99 | 0 |

Create PDF in your applications with the Pdfcrowd HTML to PDF API PDFCROWD

| | | | |
|---|---|---|---|
| SPGP0Z01530 | XP_018219732.1 | 100 | 0 |
| SPGP0Z03080 | XP_018219577.1 | 99 | 0 |
| SPGP0DN04100 | XP_018223852.1 | 99 | 0 |
| SPGP0U00130 | XP_018218976.1 | 99 | 0 |
| SPGP0Z01080 | XP_018219776.1 | 99 | 0 |
| SPGP0AF00310 | XP_018221972.1 | 99 | 0 |
| SPGP0CZ02850 | XP_018221104.1 | 99 | 0 |
| SPGP0AF00400 | XP_018221981.1 | 100 | 0 |
| SPGP0Q01620 | XP_018222336.1 | 99 | 0 |
| SPGP0AA00210 | XP_018223334.1 | 99 | 0 |
| SPGP0AA00740 | XP_018223384.1 | 99 | 0 |
| SPGP0Ftrna35L | DI49_1927 | 100 | 0 |
| SPGP0DN01620 | XP_018223608.1 | 99 | 0 |
| SPGP0R01540 | XP_018221271.1 | 99 | 0 |
| SPGP0DN00720 | XP_018223518.1 | 99 | 0 |
| SPGP0U01660 | XP_018218828.1 | 99 | 0 |
| SPGP0P01950 | XP_018221765.1 | 99 | 0 |
| SPGP0DN00860 | XP_018223532.1 | 99 | 0 |
| SPGP0AG00170 | XP_018219936.1 | 99 | 0 |
| SPGP0B00970 | XP_018221543.1 | 99 | 0 |
| SPGP0F05000 | XP_018221996.1 | 98 | 0 |
| SPGP0F04470 | XP_018222044.1 | 99 | 0 |
| SPGP0Q03210 | XP_018222181.1 | 99 | 0 |
| SPGP0Z01850 | XP_018219700.1 | 99 | 0 |
| SPGP0CZ01430 | XP_018220967.1 | 99 | 0 |
| SPGP0DN00970 | XP_018223543.1 | 95 | 0 |
| SPGP0DV00350 | XP_018219897.1 | 99 | 0 |
| SPGP0DU00840 | XP_018220506.1 | 99 | 0 |
| SPGP0E00530 | XP_018222643.1 | 99 | 0 |
| SPGP0Z00900 | XP_018219794.1 | 100 | 0 |
| SPGP0B01170 | XP_018221560.1 | 99 | 0 |
| SPGP0B00420 | XP_018221490.1 | 99 | 0 |
| SPGP0Q01720 | XP_018222326.1 | 99 | 0 |
| SPGP0Z00600 | XP_018219824.1 | 99 | 0 |
| SPGP0DZ01930 | XP_018219529.1 | 100 | 0 |
| SPGP0R03420 | XP_018221450.1 | 99 | 0 |
| SPGP0DZ01500 | XP_018219489.1 | 99 | 0 |
| SPGP0R03130 | XP_018221423.1 | 99 | 0 |
| SPGP0U00240 | XP_018218965.1 | 99 | 0 |
| SPGP0W02460 | XP_018223091.1 | 99 | 1 |

Create PDF in your applications with the Pdfcrowd HTML to PDF API PDFCROWD

| | | | |
|---|---|---|---|
| SPGP0F04650 | XP_018222027.1 | 99 | 0 |
| SPGP0F04600 | XP_018222032.1 | 99 | 0 |
| SPGP0EX00170 | XP_018223876.1 | 99 | 0 |
| SPGP0S01530 | XP_018220195.1 | 99 | 0 |
| SPGP0Z02920 | XP_018219593.1 | 100 | 0 |
| SPGP0E02390 | XP_018222827.1 | 99 | 0 |
| SPGP0AA00960 | XP_018223404.1 | 99 | 0 |
| SPGP0W00820 | XP_018223252.1 | 99 | 0 |
| SPGP0P01600 | XP_018221799.1 | 99 | 0 |
| SPGP0S02940 | XP_018220060.1 | 99 | 0 |
| SPGP0CZ01950 | XP_018221016.1 | 99 | 0 |
| SPGP0E01920 | XP_018222780.1 | 99 | 0 |
| SPGP0Z02750 | XP_018219609.1 | 99 | 0 |
| SPGP0Z02020 | XP_018219683.1 | 99 | 0 |
| SPGP0W02680 | XP_018223071.1 | 99 | 0 |
| SPGP0L01320 | XP_018219232.1 | 99 | 0 |
| SPGP0R00850 | XP_018221205.1 | 99 | 0 |
| SPGP0H00520 | XP_018224084.1 | 99 | 0 |
| SPGP0Q04270 | XP_018222079.1 | 99 | 0 |
| SPGP0AA00690 | XP_018223381.1 | 99 | 0 |
| SPGP0Z01220 | XP_018219763.1 | 99 | 0 |
| SPGP0W03480 | XP_018222999.1 | 99 | 0 |
| SPGP0DU01330 | XP_018220457.1 | 100 | 0 |
| SPGP0R02550 | XP_018221368.1 | 99 | 0 |
| SPGP0P00750 | XP_018221881.1 | 99 | 0 |
| SPGP0W03330 | XP_018223011.1 | 99 | 0 |
| SPGP0CZ02230 | XP_018221043.1 | 99 | 0 |
| SPGP0E02160 | XP_018222804.1 | 99 | 0 |
| SPGP0DZ01440 | XP_018219483.1 | 99 | 0 |
| SPGP0P00950 | XP_018221862.1 | 99 | 0 |
| SPGP0W04650 | XP_018222883.1 | 99 | 0 |
| SPGP0W00350 | XP_018223299.1 | 99 | 0 |
| SPGP0W04300 | XP_018222920.1 | 99 | 0 |
| SPGP0DN00240 | XP_018223474.1 | 98 | 0 |
| SPGP0Z02100 | XP_018219675.1 | 99 | 0 |
| SPGP0N01360 | XP_018220677.1 | 99 | 0 |
| SPGP0P02600 | XP_018221703.1 | 99 | 0 |
| SPGP0L00490 | XP_018219312.1 | 99 | 0 |
| SPGP0B00940 | XP_018221540.1 | 99 | 0 |
| SPGP0W03150 | XP_018223028.1 | 99 | 0 |

Create PDF in your applications with the Pdfcrowd HTML to PDF API PDFCROWD

| | | | |
|---|---|---|---|
| SPGP0DN02800 | XP_018223724.1 | 99 | 0 |
| SPGP0R00950 | XP_018221213.1 | 99 | 0 |
| SPGP0DU00870 | XP_018220503.1 | 100 | 0 |
| SPGP0N00780 | XP_018220731.1 | 99 | 0 |
| SPGP0DN03560 | XP_018223798.1 | 99 | 0 |
| SPGP0Q02630 | XP_018222237.1 | 99 | 0 |
| SPGP0AA00930 | XP_018223402.1 | 99 | 0 |
| SPGP0W04080 | XP_018222942.1 | 99 | 0 |
| SPGP0Q02140 | XP_018222284.1 | 99 | 0 |
| SPGP0Q02490 | XP_018222250.1 | 99 | 0 |
| SPGP0Z01690 | XP_018219716.1 | 99 | 0 |
| SPGP0R01390 | XP_018221256.1 | 99 | 0 |
| SPGP0W00170 | XP_018223317.1 | 99 | 0 |
| SPGP0P02650 | XP_018221698.1 | 99 | 0 |
| SPGP0P01050 | XP_018221852.1 | 99 | 0 |
| SPGP0W01600 | XP_018223175.1 | 99 | 0 |
| SPGP0Q03060 | XP_018222195.1 | 99 | 0 |
| SPGP0Z01390 | XP_018219746.1 | 99 | 0 |
| SPGP0X01030 | XP_018219000.1 | 99 | 0 |
| SPGP0CZ02560 | XP_018221076.1 | 99 | 0 |
| SPGP0B01360 | XP_018221579.1 | 99 | 0 |
| SPGP0W01080 | XP_018223226.1 | 99 | 0 |
| SPGP0DV00100 | XP_018219874.1 | 99 | 0 |
| SPGP0AA00590 | XP_018223371.1 | 99 | 0 |
| SPGP0W00680 | XP_018223265.1 | 99 | 0 |
| SPGP0AH00790 | XP_018221597.1 | 99 | 0 |
| SPGP0F04810 | XP_018222013.1 | 99 | 0 |
| SPGP0DN02740 | XP_018223718.1 | 99 | 0 |
| SPGP0DN02210 | XP_018223667.1 | 99 | 0 |
| SPGP0B01000 | XP_018221546.1 | 99 | 0 |
| SPGP0CZ02420 | XP_018221062.1 | 99 | 0 |
| SPGP0Qtrna18L | DI49_2066 | 100 | 0 |
| SPGP0R02760 | XP_018221388.1 | 99 | 0 |
| SPGP0N02100 | XP_018220604.1 | 99 | 0 |
| SPGP0W01550 | XP_018223180.1 | 99 | 0 |
| SPGP0DN01430 | XP_018223590.1 | 99 | 0 |
| SPGP0E00260 | XP_018222616.1 | 99 | 0 |
| SPGP0S02930 | XP_018220061.1 | 99 | 0 |
| SPGP0P01380 | XP_018221820.1 | 99 | 0 |
| SPGP0W02100 | XP_018223127.1 | 99 | 0 |

Create PDF in your applications with the Pdfcrowd HTML to PDF API PDFCROWD

| | | | |
|---|---|---|---|
| SPGP0T00220 | XP_018220350.1 | 98 | 0 |
| SPGP0Z01310 | XP_018219754.1 | 99 | 0 |
| SPGP0AH01130 | XP_018222495.1 | 99 | 0 |
| SPGP0DN03290 | XP_018223772.1 | 99 | 0 |
| SPGP0E02580 | XP_018222844.1 | 99 | 0 |
| SPGP0Q03450 | XP_018222159.1 | 99 | 0 |
| SPGP0P02400 | XP_018221723.1 | 99 | 0 |
| SPGP0S01260 | XP_018220220.1 | 99 | 0 |
| SPGP0S01060 | XP_018220239.1 | 99 | 0 |
| SPGP0P00650 | XP_018221891.1 | 99 | 0 |
| SPGP0AC00800 | XP_018219163.1 | 99 | 0 |
| SPGP0R02000 | XP_018221315.1 | 99 | 0 |
| SPGP0Q03730 | XP_018222132.1 | 99 | 0 |
| SPGP0DN02840 | XP_018223729.1 | 99 | 0 |
| SPGP0U01410 | XP_018218852.1 | 99 | 0 |
| SPGP0F04400 | XP_018222051.1 | 99 | 0 |
| SPGP0AC00210 | XP_018219106.1 | 99 | 0 |
| SPGP0B00110 | XP_018221458.1 | 96 | 0 |
| SPGP0DV00200 | XP_018219884.1 | 99 | 0 |
| SPGP0DN03750 | XP_018223817.1 | 99 | 0 |
| SPGP0CZ02400 | XP_018221060.1 | 100 | 0 |
| SPGP0S02410 | XP_018220111.1 | 99 | 0 |
| SPGP0E00800 | XP_018222668.1 | 99 | 0 |
| SPGP0W02010 | XP_018223134.1 | 99 | 0 |
| SPGP0Q00390 | XP_018222462.1 | 99 | 0 |
| SPGP0W04720 | XP_018222876.1 | 99 | 0 |
| SPGP0P00380 | XP_018221917.1 | 99 | 0 |
| SPGP0Z02690 | XP_018219616.1 | 99 | 0 |
| SPGP0S03180 | XP_018220036.1 | 99 | 0 |
| SPGP0W01950 | XP_018223140.1 | 99 | 0 |
| SPGP0N02110 | XP_018220603.1 | 99 | 0 |
| SPGP0DN02770 | XP_018223721.1 | 95 | 4 |
| SPGP0DN04200 | XP_018223862.1 | 99 | 0 |
| SPGP0S01800 | XP_018220171.1 | 99 | 0 |
| SPGP0AF00110 | XP_018221953.1 | 99 | 0 |
| SPGP0S02330 | XP_018220119.1 | 99 | 0 |
| SPGP0R03090 | XP_018221419.1 | 99 | 0 |
| SPGP0W02400 | XP_018223097.1 | 99 | 0 |
| SPGP0AA00550 | XP_018223367.1 | 99 | 0 |
| SPGP0DZ00340 | XP_018219377.1 | 99 | 0 |

Create PDF in your applications with the Pdfcrowd HTML to PDF API PDFCROWD

| | | | |
|---|---|---|---|
| SPGP0DU01450 | XP_018220445.1 | 100 | 0 |
| SPGP0P00160 | XP_018221938.1 | 99 | 0 |
| SPGP0X00210 | XP_018219080.1 | 99 | 0 |
| SPGP0W03630 | XP_018222985.1 | 99 | 0 |
| SPGP0P01590 | XP_018221800.1 | 99 | 0 |
| SPGP0W04790 | XP_018222869.1 | 99 | 0 |
| SPGP0S00150 | XP_018220329.1 | 99 | 0 |
| SPGP0F04310 | XP_018222060.1 | 99 | 0 |
| SPGP0W01070 | XP_018223227.1 | 99 | 0 |
| SPGP0X00150 | XP_018219085.1 | 99 | 0 |
| SPGP0P03120 | XP_018221652.1 | 99 | 0 |
| SPGP0P02160 | XP_018221744.1 | 99 | 0 |
| SPGP0P01290 | XP_018221828.1 | 99 | 0 |
| SPGP0AH00120 | XP_018222592.1 | 95 | 1 |
| SPGP0Q02930 | XP_018222208.1 | 99 | 0 |
| SPGP0Z00630 | XP_018219821.1 | 99 | 0 |
| SPGP0Z01730 | XP_018219712.1 | 99 | 0 |
| SPGP0R02670 | XP_018221379.1 | 99 | 0 |
| SPGP0DZ01060 | XP_018219445.1 | 99 | 0 |
| SPGP0L00790 | XP_018219284.1 | 99 | 0 |
| SPGP0R02710 | XP_018221383.1 | 99 | 0 |
| SPGP0AA00470 | XP_018223359.1 | 99 | 0 |
| SPGP0Z02250 | XP_018219660.1 | 99 | 0 |
| SPGP0B00840 | XP_018221531.1 | 99 | 0 |
| SPGP0Q02940 | XP_018222207.1 | 99 | 0 |
| SPGP0DX00470 | XP_018219977.1 | 99 | 0 |
| SPGP0B01950 | XP_018221637.1 | 99 | 0 |
| SPGP0W02140 | XP_018223123.1 | 100 | 0 |
| SPGP0X00570 | XP_018219045.1 | 98 | 1 |
| SPGP0R02040 | XP_018221319.1 | 99 | 0 |
| SPGP0R01850 | XP_018221301.1 | 99 | 0 |
| SPGP0R01940 | XP_018221310.1 | 100 | 0 |
| SPGP0W04460 | XP_018222902.1 | 99 | 0 |
| SPGP0DV00160 | XP_018219880.1 | 99 | 0 |
| SPGP0Q04090 | XP_018222096.1 | 99 | 0 |
| SPGP0DN02080 | XP_018223655.1 | 99 | 0 |
| SPGP0R02930 | XP_018221404.1 | 99 | 0 |
| SPGP0DN02510 | XP_018223696.1 | 99 | 0 |
| SPGP0P01410 | XP_018221817.1 | 99 | 0 |
| SPGP0Y00540 | XP_018223928.1 | 99 | 0 |

Create PDF in your applications with the Pdfcrowd HTML to PDF API PDFCROWD

| | | | |
|---|---|---|---|
| SPGP0T00350 | XP_018220363.1 | 99 | 0 |
| SPGP0DZ00600 | XP_018219401.1 | 99 | 0 |
| SPGP0DZ01120 | XP_018219451.1 | 99 | 0 |
| SPGP0Z01770 | XP_018219708.1 | 99 | 0 |
| SPGP0B00530 | XP_018221501.1 | 99 | 0 |
| SPGP0L01150 | XP_018219248.1 | 99 | 0 |
| SPGP0DZ02290 | XP_018219566.1 | 99 | 0 |
| SPGP0B01480 | XP_018221591.1 | 99 | 0 |
| SPGP0W04700 | XP_018222878.1 | 99 | 0 |
| SPGP0DN02220 | XP_018223668.1 | 99 | 0 |
| SPGP0F04360 | XP_018222055.1 | 99 | 0 |
| SPGP0R01490 | XP_018221266.1 | 99 | 0 |
| SPGP0L00780 | XP_018219285.1 | 99 | 0 |
| SPGP0S02770 | XP_018220076.1 | 99 | 0 |
| SPGP0AA01000 | XP_018223408.1 | 99 | 0 |
| SPGP0Q01350 | XP_018222364.1 | 99 | 0 |
| SPGP0Q01600 | XP_018222338.1 | 99 | 0 |
| SPGP0AA00400 | XP_018223352.1 | 99 | 0 |
| SPGP0P02990 | XP_018221665.1 | 99 | 0 |
| SPGP0DZ02150 | XP_018219552.1 | 99 | 0 |
| SPGP0P01820 | XP_018221778.1 | 99 | 0 |
| SPGP0P01960 | XP_018221764.1 | 99 | 0 |
| SPGP0R00540 | XP_018221177.1 | 99 | 0 |
| SPGP0DN03970 | XP_018223839.1 | 99 | 0 |
| SPGP0Q02550 | XP_018222245.1 | 100 | 0 |
| SPGP0DN02130 | XP_018223660.1 | 100 | 0 |
| SPGP0DZ00290 | XP_018219372.1 | 99 | 0 |
| SPGP0R01230 | XP_018221241.1 | 99 | 0 |
| SPGP0Z01070 | XP_018219777.1 | 99 | 1 |
| SPGP0AH00630 | XP_018222542.1 | 99 | 0 |
| SPGP0AH00320 | XP_018222571.1 | 99 | 0 |
| SPGP0CZ02660 | XP_018221086.1 | 99 | 0 |
| SPGP0Q03600 | XP_018222145.1 | 99 | 0 |
| SPGP0Y01230 | XP_018223992.1 | 99 | 0 |
| SPGP0R01890 | XP_018221305.1 | 99 | 0 |
| SPGP0R00720 | XP_018221194.1 | 98 | 0 |
| SPGP0W01940 | XP_018223141.1 | 99 | 0 |
| SPGP0W01760 | XP_018223159.1 | 99 | 0 |
| SPGP0AC00920 | XP_018219175.1 | 99 | 0 |

Create PDF in your applications with the Pdfcrowd HTML to PDF API PDFCROWD

| | | | |
|---|---|---|---|
| SPGP0U01030 | XP_018218889.1 | 99 | 0 |
| SPGP0AA00700 | XP_018223382.1 | 99 | 0 |
| SPGP0Q03840 | XP_018222121.1 | 99 | 0 |
| SPGP0Ftrna11H | DI49_4134 | 100 | 0 |
| SPGP0E02300 | XP_018222818.1 | 99 | 0 |
| SPGP0E00450 | XP_018222635.1 | 99 | 0 |
| SPGP0U00570 | XP_018218932.1 | 99 | 0 |
| SPGP0Q03640 | XP_018222141.1 | 99 | 0 |
| SPGP0P01310 | XP_018221826.1 | 99 | 0 |
| SPGP0S02950 | XP_018220059.1 | 99 | 0 |
| SPGP0W04310 | XP_018222919.1 | 99 | 0 |
| SPGP0Y00170 | XP_018223895.1 | 99 | 0 |
| SPGP0B01370 | XP_018221580.1 | 99 | 0 |
| SPGP0W04250 | XP_018222925.1 | 99 | 0 |
| SPGP0AA00760 | XP_018223386.1 | 99 | 0 |
| SPGP0BF00140 | XP_018220385.1 | 99 | 0 |
| SPGP0DV00300 | XP_018219893.1 | 99 | 0 |
| SPGP0P02100 | XP_018221750.1 | 99 | 0 |
| SPGP0Q00560 | XP_018222445.1 | 99 | 0 |
| SPGP0Z01300 | XP_018219755.1 | 99 | 0 |
| SPGP0P01490 | XP_018221809.1 | 99 | 0 |
| SPGP0S01250 | XP_018220221.1 | 96 | 1 |
| SPGP0X00990 | XP_018219004.1 | 99 | 0 |
| SPGP0DN04160 | XP_018223858.1 | 100 | 0 |
| SPGP0Q01540 | XP_018222344.1 | 98 | 2 |
| SPGP0Z00720 | XP_018219812.1 | 99 | 0 |
| SPGP0DV00410 | XP_018219903.1 | 99 | 0 |
| SPGP0AA01010 | XP_018223409.1 | 99 | 0 |
| SPGP0X00290 | XP_018219072.1 | 100 | 0 |
| SPGP0AA00490 | XP_018223361.1 | 99 | 0 |
| SPGP0E01320 | XP_018222722.1 | 99 | 0 |
| SPGP0E01490 | XP_018222737.1 | 99 | 0 |
| SPGP0DN01600 | XP_018223607.1 | 99 | 0 |
| SPGP0R01060 | XP_018221224.1 | 99 | 0 |
| SPGP0CZ00350 | XP_018220861.1 | 99 | 0 |
| SPGP0Q03090 | XP_018222192.1 | 98 | 2 |
| SPGP0DU00260 | XP_018220564.1 | 99 | 0 |
| SPGP0DN00800 | XP_018223526.1 | 99 | 0 |
| SPGP0Q03260 | XP_018222176.1 | 99 | 0 |
| SPGP0S01510 | XP_018220197.1 | 99 | 0 |

Create PDF in your applications with the Pdfcrowd HTML to PDF API PDFCROWD

| | | | |
|---|---|---|---|
| SPGP0S00520 | XP_018220293.1 | 99 | 0 |
| SPGP0DV00370 | XP_018219899.1 | 99 | 0 |
| SPGP0E01140 | XP_018222704.1 | 99 | 0 |
| SPGP0P01360 | XP_018221822.1 | 99 | 0 |
| SPGP0S00120 | XP_018220332.1 | 99 | 0 |
| SPGP0Q04150 | XP_018222090.1 | 99 | 0 |
| SPGP0DU00190 | XP_018220571.1 | 99 | 0 |
| SPGP0P01970 | XP_018221763.1 | 99 | 0 |
| SPGP0DN03320 | XP_018223775.1 | 99 | 0 |
| SPGP0S02830 | XP_018220071.1 | 99 | 0 |
| SPGP0Z02970 | XP_018219588.1 | 99 | 0 |
| SPGP0E02470 | XP_018222834.1 | 99 | 0 |
| SPGP0R03150 | XP_018221425.1 | 99 | 0 |
| SPGP0W02330 | XP_018223104.1 | 99 | 0 |
| SPGP0AF00250 | XP_018221966.1 | 99 | 0 |
| SPGP0W02440 | XP_018223093.1 | 99 | 0 |
| SPGP0W01880 | XP_018223148.1 | 99 | 0 |
| SPGP0P00620 | XP_018221894.1 | 99 | 0 |
| SPGP0S02960 | XP_018220058.1 | 99 | 0 |
| SPGP0W00910 | XP_018223243.1 | 99 | 0 |
| SPGP0P01620 | XP_018221797.1 | 99 | 0 |
| SPGP0Z02000 | XP_018219685.1 | 99 | 0 |
| SPGP0B00310 | XP_018221479.1 | 99 | 0 |
| SPGP0L01300 | XP_018219234.1 | 100 | 0 |
| SPGP0E02140 | XP_018222802.1 | 99 | 0 |
| SPGP0Q03980 | XP_018222107.1 | 99 | 0 |
| SPGP0N02030 | XP_018220611.1 | 99 | 0 |
| SPGP0DU01480 | XP_018220442.1 | 99 | 0 |
| SPGP0E00240 | XP_018222614.1 | 99 | 0 |
| SPGP0F04530 | XP_018222038.1 | 99 | 0 |
| SPGP0AF00370 | XP_018221978.1 | 99 | 0 |
| SPGP0W04200 | XP_018222930.1 | 99 | 0 |
| SPGP0DX00530 | XP_018219983.1 | 99 | 0 |
| SPGP0S03110 | XP_018220043.1 | 99 | 0 |
| SPGP0W00440 | XP_018223290.1 | 99 | 0 |
| SPGP0W03060 | XP_018223037.1 | 99 | 0 |
| SPGP0R02570 | XP_018221370.1 | 99 | 0 |
| SPGP0P02440 | XP_018221719.1 | 99 | 0 |
| SPGP0W03310 | XP_018223013.1 | 100 | 0 |
| SPGP0AC00550 | XP_018219139.1 | 99 | 0 |

Create PDF in your applications with the Pdfcrowd HTML to PDF API PDFCROWD

| | | | |
|---|---|---|---|
| SPGP0S02870 | XP_018220067.1 | 99 | 0 |
| SPGP0P00970 | XP_018221860.1 | 99 | 1 |
| SPGP0E02200 | XP_018222808.1 | 100 | 0 |
| SPGP0R01000 | XP_018221218.1 | 99 | 0 |
| SPGP0S02160 | XP_018220136.1 | 99 | 0 |
| SPGP0Q01160 | XP_018222384.1 | 99 | 0 |
| SPGP0W04160 | XP_018222934.1 | 99 | 0 |
| SPGP0Q00270 | XP_018222474.1 | 99 | 0 |
| SPGP0S00830 | XP_018220262.1 | 99 | 0 |
| SPGP0L00110 | XP_018219350.1 | 99 | 0 |
| SPGP0P02660 | XP_018221697.1 | 99 | 0 |
| SPGP0W03170 | XP_018223026.1 | 100 | 0 |
| SPGP0L00300 | XP_018219331.1 | 99 | 0 |
| SPGP0F04550 | XP_018222036.1 | 99 | 0 |
| SPGP0DN03270 | XP_018223770.1 | 99 | 0 |
| SPGP0P01220 | XP_018221835.1 | 99 | 0 |
| SPGP0AF00260 | XP_018221967.1 | 99 | 0 |
| SPGP0Q00440 | XP_018222457.1 | 99 | 0 |
| SPGP0S02080 | XP_018220144.1 | 100 | 0 |
| SPGP0Z01520 | XP_018219733.1 | 100 | 0 |
| SPGP0DZ00700 | XP_018219411.1 | 99 | 0 |
| SPGP0CZ02620 | XP_018221082.1 | 99 | 0 |
| SPGP0E00330 | XP_018222623.1 | 99 | 0 |
| SPGP0Q02120 | XP_018222286.1 | 100 | 0 |
| SPGP0W00370 | XP_018223297.1 | 99 | 0 |
| SPGP0Q01290 | XP_018222371.1 | 99 | 0 |
| SPGP0BC00260 | XP_018220823.1 | 99 | 0 |
| SPGP0L00160 | XP_018219345.1 | 99 | 0 |
| SPGP0W00830 | XP_018223251.1 | 99 | 0 |
| SPGP0W02940 | XP_018223046.1 | 99 | 0 |
| SPGP0Z00860 | XP_018219798.1 | 99 | 0 |
| SPGP0B01010 | XP_018221547.1 | 99 | 0 |
| SPGP0W03220 | XP_018223021.1 | 99 | 0 |
| SPGP0Z02330 | XP_018219652.1 | 99 | 0 |
| SPGP0DZ00660 | XP_018219407.1 | 99 | 0 |
| SPGP0DU00900 | XP_018220500.1 | 99 | 0 |
| SPGP0P01180 | XP_018221839.1 | 99 | 0 |
| SPGP0DN02050 | XP_018223652.1 | 99 | 0 |
| SPGP0DN01450 | XP_018223592.1 | 99 | 0 |
| SPGP0Q01170 | XP_018222383.1 | 99 | 0 |

Create PDF in your applications with the Pdfcrowd HTML to PDF API PDFCROWD

| | | | |
|---|---|---|---|
| SPGP0CZ00920 | XP_018220917.1 | 99 | 0 |
| SPGP0S01890 | XP_018220163.1 | 100 | 0 |
| SPGP0S02630 | XP_018220090.1 | 99 | 0 |
| SPGP0Q03290 | XP_018222173.1 | 97 | 2 |
| SPGP0Q04160 | XP_018222089.1 | 99 | 0 |
| SPGP0P02230 | XP_018221739.1 | 99 | 0 |
| SPGP0S02840 | XP_018220070.1 | 99 | 0 |
| SPGP0X01110 | XP_018218992.1 | 99 | 0 |
| SPGP0S01870 | XP_018220165.1 | 99 | 0 |
| SPGP0R03100 | XP_018221420.1 | 99 | 0 |
| SPGP0H00640 | XP_018224073.1 | 99 | 0 |
| SPGP0Q01830 | XP_018222315.1 | 99 | 0 |
| SPGP0DN03700 | XP_018223812.1 | 99 | 0 |
| SPGP0W00290 | XP_018223305.1 | 99 | 0 |
| SPGP0L00980 | XP_018219265.1 | 99 | 1 |
| SPGP0W02190 | XP_018223118.1 | 99 | 0 |
| SPGP0S03580 | XP_018219998.1 | 99 | 0 |
| SPGP0Q03470 | XP_018222157.1 | 98 | 0 |
| SPGP0N01340 | XP_018220679.1 | 99 | 0 |
| SPGP0DN03600 | XP_018223802.1 | 99 | 0 |
| SPGP0Q04240 | XP_018222081.1 | 99 | 0 |
| SPGP0W03660 | XP_018222982.1 | 99 | 0 |
| SPGP0H00870 | XP_018224050.1 | 93 | 1 |
| SPGP0DU01000 | XP_018220490.1 | 99 | 0 |
| SPGP0Q01490 | XP_018222349.1 | 99 | 0 |
| SPGP0Q04100 | XP_018222095.1 | 99 | 0 |
| SPGP0L01050 | XP_018219258.1 | 99 | 0 |
| SPGP0DZ02050 | XP_018219542.1 | 99 | 0 |
| SPGP0P02380 | XP_018221725.1 | 99 | 0 |
| SPGP0U01470 | XP_018218846.1 | 99 | 0 |
| SPGP0W00670 | XP_018223266.1 | 99 | 0 |
| SPGP0L00770 | XP_018219286.1 | 99 | 0 |
| SPGP0S02190 | XP_018220133.1 | 97 | 3 |
| SPGP0W00960 | XP_018223238.1 | 99 | 0 |
| SPGP0H00670 | XP_018224070.1 | 99 | 0 |
| SPGP0E01160 | XP_018222706.1 | 99 | 0 |
| SPGP0S01860 | XP_018220166.1 | 99 | 0 |
| SPGP0Q01030 | XP_018222398.1 | 99 | 0 |
| SPGP0R03050 | XP_018221415.1 | 99 | 0 |
| SPGP0Q02250 | XP_018222274.1 | 99 | 0 |

Create PDF in your applications with the Pdfcrowd HTML to PDF API PDFCROWD

| | | | |
|---|---|---|---|
| SPGP0X00450 | XP_018219056.1 | 99 | 0 |
| SPGP0DZ01260 | XP_018219465.1 | 100 | 0 |
| SPGP0S00580 | XP_018220287.1 | 99 | 0 |
| SPGP0L01130 | XP_018219250.1 | 99 | 0 |
| SPGP0DN00400 | XP_018223489.1 | 99 | 0 |
| SPGP0R02960 | XP_018221407.1 | 99 | 0 |
| SPGP0S00980 | XP_018220247.1 | 99 | 0 |
| SPGP0B00520 | XP_018221500.1 | 99 | 0 |
| SPGP0DU00520 | XP_018220538.1 | 99 | 0 |
| SPGP0P02900 | XP_018221674.1 | 98 | 1 |
| SPGP0DN00490 | XP_018223497.1 | 99 | 0 |
| SPGP0BC00250 | XP_018220822.1 | 99 | 0 |
| SPGP0DN00470 | XP_018223495.1 | 99 | 0 |
| SPGP0AA00530 | XP_018223365.1 | 99 | 0 |
| SPGP0Z01650 | XP_018219720.1 | 99 | 0 |
| SPGP0X00980 | XP_018219005.1 | 99 | 0 |
| SPGP0X00230 | XP_018219078.1 | 99 | 0 |
| SPGP0X00580 | XP_018219044.1 | 99 | 0 |
| SPGP0S00130 | XP_018220331.1 | 99 | 0 |
| SPGP0W02300 | XP_018223107.1 | 94 | 5 |
| SPGP0Z01330 | XP_018219752.1 | 99 | 0 |
| SPGP0W03780 | XP_018222970.1 | 99 | 0 |
| SPGP0Z02490 | XP_018219636.1 | 99 | 0 |
| SPGP0E02210 | XP_018222809.1 | 99 | 0 |
| SPGP0F04750 | XP_018222019.1 | 99 | 0 |
| SPGP0W02830 | XP_018223057.1 | 99 | 0 |
| SPGP0DN02240 | XP_018223670.1 | 99 | 0 |
| SPGP0W00920 | XP_018223242.1 | 99 | 0 |
| SPGP0Q02910 | XP_018222210.1 | 98 | 1 |
| SPGP0N00560 | XP_018220752.1 | 99 | 0 |
| SPGP0Z00750 | XP_018219809.1 | 99 | 0 |
| SPGP0EE00170 | XP_018220836.1 | 99 | 0 |
| SPGP0Q03970 | XP_018222108.1 | 99 | 0 |
| SPGP0R02390 | XP_018221352.1 | 99 | 0 |
| SPGP0R02290 | XP_018221342.1 | 99 | 0 |
| SPGP0CZ00180 | XP_018220846.1 | 99 | 0 |
| SPGP0Z02270 | XP_018219658.1 | 99 | 0 |
| SPGP0R01260 | XP_018221243.1 | 99 | 0 |
| SPGP0P03100 | XP_018221654.1 | 99 | 0 |
| SPGP0R03210 | XP_018221431.1 | 99 | 0 |

Create PDF in your applications with the Pdfcrowd HTML to PDF API PDFCROWD

| | | | |
|---|---|---|---|
| SPGP0Z02680 | XP_018219617.1 | 99 | 0 |
| SPGP0CZ02980 | XP_018221117.1 | 99 | 0 |
| SPGP0Q01420 | XP_018222357.1 | 99 | 0 |
| SPGP0X00220 | XP_018219079.1 | 99 | 0 |
| SPGP0R01920 | XP_018221308.1 | 99 | 0 |
| SPGP0N01270 | XP_018220686.1 | 99 | 1 |
| SPGP0B00250 | XP_018221473.1 | 99 | 0 |
| SPGP0BF00430 | XP_018220411.1 | 100 | 0 |
| SPGP0DV00140 | XP_018219878.1 | 99 | 0 |
| SPGP0CZ00720 | XP_018220897.1 | 99 | 0 |
| SPGP0E01940 | XP_018222782.1 | 100 | 0 |
| SPGP0DN02530 | XP_018223698.1 | 99 | 0 |
| SPGP0Q04050 | XP_018222100.1 | 99 | 0 |
| SPGP0R02740 | XP_018221386.1 | 99 | 0 |
| SPGP0B00510 | XP_018221499.1 | 99 | 1 |
| SPGP0DZ01450 | XP_018219484.1 | 99 | 0 |
| SPGP0DZ01100 | XP_018219449.1 | 99 | 0 |
| SPGP0AH00970 | XP_018222510.1 | 99 | 0 |
| SPGP0L01170 | XP_018219247.1 | 99 | 0 |
| SPGP0Q03690 | XP_018222136.1 | 99 | 0 |
| SPGP0DN00380 | XP_018223487.1 | 99 | 0 |
| SPGP0CZ00580 | XP_018220884.1 | 99 | 0 |
| SPGP0DV00550 | XP_018219917.1 | 99 | 0 |
| SPGP0Q01330 | XP_018222366.1 | 99 | 0 |
| SPGP0R00450 | XP_018221168.1 | 98 | 0 |
| SPGP0W00610 | XP_018223273.1 | 99 | 0 |
| SPGP0Q00410 | XP_018222460.1 | 99 | 0 |
| SPGP0AC00820 | XP_018219165.1 | 99 | 0 |
| SPGP0H00180 | XP_018224117.1 | 99 | 0 |
| SPGP0CZ01980 | XP_018221019.1 | 99 | 0 |
| SPGP0CZ01450 | XP_018220969.1 | 99 | 0 |
| SPGP0AC00490 | XP_018219133.1 | 99 | 0 |
| SPGP0DN03080 | XP_018223752.1 | 99 | 0 |
| SPGP0DZ00270 | XP_018219370.1 | 99 | 0 |
| SPGP0AH00340 | XP_018222570.1 | 98 | 2 |
| SPGP0DN02390 | XP_018223684.1 | 99 | 0 |
| SPGP0X00820 | XP_018219020.1 | 99 | 0 |
| SPGP0AH00600 | XP_018222544.1 | 96 | 0 |
| SPGP0AGtrna32L | DI49_5686 | 100 | 0 |
| SPGP0Gtrna37L | DI49_5687 | 100 | 0 |

Create PDF in your applications with the Pdfcrowd HTML to PDF API PDFCROWD

| | | | |
|---|---|---|---|
| SPGP0CZ01670 | XP_018220990.1 | 99 | 0 |
| SPGP0W01920 | XP_018223144.1 | 99 | 0 |
| SPGP0DN03620 | XP_018223804.1 | 99 | 0 |
| SPGP0AHtrna16F | DI49_2529 | 100 | 0 |
| SPGP0CZ01180 | XP_018220943.1 | 99 | 0 |
| SPGP0E01510 | XP_018222739.1 | 99 | 0 |
| SPGP0B01200 | XP_018221563.1 | 99 | 0 |
| SPGP0E02410 | XP_018222828.1 | 99 | 0 |
| SPGP0DN04000 | XP_018223842.1 | 99 | 0 |
| SPGP0U00520 | XP_018218937.1 | 99 | 0 |
| SPGP0E00300 | XP_018222620.1 | 99 | 0 |
| SPGP0Q03660 | XP_018222139.1 | 98 | 0 |
| SPGP0AF00190 | XP_018221960.1 | 99 | 0 |
| SPGP0S01570 | XP_018220193.1 | 99 | 0 |
| SPGP0W03240 | XP_018223019.1 | 99 | 0 |
| SPGP0F04890 | XP_018222005.1 | 99 | 1 |
| SPGP0W04610 | XP_018222887.1 | 99 | 1 |
| SPGP0E00340 | XP_018222624.1 | 99 | 0 |
| SPGP0DV00670 | XP_018219928.1 | 99 | 0 |
| SPGP0E00730 | XP_018222663.1 | 99 | 0 |
| SPGP0U01540 | XP_018218840.1 | 99 | 0 |
| SPGP0DN02360 | XP_018223681.1 | 99 | 0 |
| SPGP0R00150 | XP_018221140.1 | 99 | 0 |
| SPGP0U01040 | XP_018218888.1 | 100 | 0 |
| SPGP0U01590 | XP_018218835.1 | 99 | 0 |
| SPGP0AF00290 | XP_018221970.1 | 99 | 0 |
| SPGP0P00230 | XP_018221931.1 | 99 | 0 |
| SPGP0Q02040 | XP_018222294.1 | 99 | 0 |
| SPGP0Q03050 | XP_018222196.1 | 99 | 0 |
| SPGP0DN01160 | XP_018223563.1 | 99 | 0 |
| SPGP0AF00440 | XP_018221985.1 | 98 | 0 |
| SPGP0CZ02210 | XP_018221041.1 | 98 | 2 |
| SPGP0Q03530 | XP_018222151.1 | 99 | 0 |
| SPGP0DN01660 | XP_018223612.1 | 99 | 0 |
| SPGP0EC00120 | XP_018220414.1 | 98 | 0 |
| SPGP0P01190 | XP_018221838.1 | 99 | 0 |
| SPGP0S02700 | XP_018220083.1 | 99 | 0 |
| SPGP0S02030 | XP_018220149.1 | 99 | 0 |
| SPGP0CZ00930 | XP_018220918.1 | 99 | 0 |
| SPGP0DN00390 | XP_018223488.1 | 99 | 0 |

Create PDF in your applications with the Pdfcrowd HTML to PDF API PDFCROWD

| | | | |
|---|---|---|---|
| SPGP0Q00570 | XP_018222444.1 | 99 | 0 |
| SPGP0B01720 | XP_018221614.1 | 99 | 0 |
| SPGP0DN00820 | XP_018223528.1 | 99 | 0 |
| SPGP0Z01490 | XP_018219736.1 | 99 | 0 |
| SPGP0F04430 | XP_018222048.1 | 99 | 0 |
| SPGP0CZ02370 | XP_018221057.1 | 99 | 0 |
| SPGP0S01610 | XP_018220189.1 | 99 | 0 |
| SPGP0Q00210 | XP_018222479.1 | 99 | 0 |
| SPGP0DV00310 | XP_018219894.1 | 99 | 0 |
| SPGP0W02070 | XP_018223130.1 | 99 | 0 |
| SPGP0Z02340 | XP_018219651.1 | 99 | 0 |
| SPGP0DN02340 | XP_018223679.1 | 99 | 0 |
| SPGP0P01010 | XP_018221856.1 | 100 | 0 |
| SPGP0Z02390 | XP_018219646.1 | 99 | 0 |
| SPGP0S01740 | XP_018220177.1 | 100 | 0 |
| SPGP0Z01020 | XP_018219782.1 | 99 | 0 |
| SPGP0P00710 | XP_018221885.1 | 99 | 0 |
| SPGP0E01410 | XP_018222730.1 | 99 | 0 |
| SPGP0R03170 | XP_018221427.1 | 99 | 0 |
| SPGP0L01360 | XP_018219228.1 | 99 | 0 |
| SPGP0CZ02950 | XP_018221114.1 | 99 | 0 |
| SPGP0P00640 | XP_018221892.1 | 99 | 0 |
| SPGP0W02520 | XP_018223085.1 | 99 | 0 |
| SPGP0E00890 | XP_018222677.1 | 99 | 0 |
| SPGP0W03860 | XP_018222962.1 | 99 | 0 |
| SPGP0Z01830 | XP_018219702.1 | 99 | 0 |
| SPGP0W00280 | XP_018223306.1 | 99 | 0 |
| SPGP0Z02060 | XP_018219679.1 | 100 | 0 |
| SPGP0U00360 | XP_018218953.1 | 100 | 0 |
| SPGP0P02610 | XP_018221702.1 | 99 | 0 |
| SPGP0H00840 | XP_018224053.1 | 99 | 0 |
| SPGP0Q02670 | XP_018222233.1 | 99 | 0 |
| SPGP0DN01670 | XP_018223613.1 | 99 | 0 |
| SPGP0R02830 | XP_018221395.1 | 100 | 0 |
| SPGP0R00220 | XP_018221145.1 | 99 | 0 |
| SPGP0Y00710 | XP_018223943.1 | 99 | 0 |
| SPGP0P02460 | XP_018221717.1 | 99 | 0 |
| SPGP0DN01560 | XP_018223603.1 | 99 | 0 |
| SPGP0DU00460 | XP_018220544.1 | 99 | 0 |
| SPGP0H00420 | XP_018224094.1 | 99 | 0 |

Create PDF in your applications with the Pdfcrowd HTML to PDF API PDFCROWD

| | | | |
|---|---|---|---|
| SPGP0B01570 | XP_018221599.1 | 99 | 0 |
| SPGP0DN01190 | XP_018223566.1 | 99 | 0 |
| SPGP0P00910 | XP_018221866.1 | 99 | 0 |
| SPGP0L01460 | XP_018219218.1 | 99 | 0 |
| SPGP0Z01260 | XP_018219759.1 | 100 | 0 |
| SPGP0Q02170 | XP_018222281.1 | 99 | 0 |
| SPGP0E02540 | XP_018222840.1 | 99 | 0 |
| SPGP0P02640 | XP_018221699.1 | 99 | 0 |
| SPGP0S00850 | XP_018220260.1 | 99 | 0 |
| SPGP0W03190 | XP_018223024.1 | 99 | 0 |
| SPGP0DN03790 | XP_018223821.1 | 99 | 0 |
| SPGP0E01060 | XP_018222696.1 | 99 | 0 |
| SPGP0AH01170 | XP_018222491.1 | 99 | 0 |
| SPGP0P01940 | XP_018221766.1 | 99 | 1 |
| SPGP0Z00680 | XP_018219816.1 | 99 | 0 |
| SPGP0Q02100 | XP_018222288.1 | 99 | 0 |
| SPGP0Z00130 | XP_018219870.1 | 99 | 0 |
| SPGP0R03290 | XP_018221439.1 | 99 | 0 |
| SPGP0R01020 | XP_018221220.1 | 99 | 0 |
| SPGP0CZ00440 | XP_018220870.1 | 99 | 0 |
| SPGP0DN02670 | XP_018223711.1 | 99 | 0 |
| SPGP0E02180 | XP_018222806.1 | 99 | 0 |
| SPGP0L00180 | XP_018219343.1 | 99 | 0 |
| SPGP0DN01750 | XP_018223621.1 | 99 | 0 |
| SPGP0DN02460 | XP_018223691.1 | 99 | 0 |
| SPGP0L00440 | XP_018219317.1 | 99 | 0 |
| SPGP0Etrna8W | DI49_5211 | 100 | 0 |
| SPGP0N02320 | XP_018220582.1 | 99 | 0 |
| SPGP0N01590 | XP_018220655.1 | 99 | 0 |
| SPGP0W01040 | XP_018223230.1 | 99 | 0 |
| SPGP0Z00350 | XP_018219851.1 | 99 | 0 |
| SPGP0E00100 | XP_018220372.1 | 99 | 0 |
| SPGP0B01850 | XP_018221627.1 | 99 | 0 |
| SPGP0B00260 | XP_018221474.1 | 99 | 0 |
| SPGP0Q01220 | XP_018222378.1 | 99 | 0 |
| SPGP0Z01880 | XP_018219697.1 | 99 | 0 |
| SPGP0DN02750 | XP_018223719.1 | 99 | 0 |
| SPGP0DN02250 | XP_018223671.1 | 99 | 0 |
| SPGP0Q00250 | XP_018222475.1 | 99 | 0 |
| SPGP0Y01000 | XP_018223970.1 | 99 | 0 |

Create PDF in your applications with the Pdfcrowd HTML to PDF API PDFCROWD

| | | | |
|---|---|---|---|
| SPGP0W01790 | XP_018223156.1 | 100 | 0 |
| SPGP0Z02810 | XP_018219603.1 | 99 | 0 |
| SPGP0Q01150 | XP_018222385.1 | 99 | 0 |
| SPGP0U01680 | XP_018218826.1 | 99 | 0 |
| SPGP0P01330 | XP_018221824.1 | 99 | 0 |
| SPGP0N02140 | XP_018220600.1 | 99 | 0 |
| SPGP0DN01400 | XP_018223587.1 | 99 | 0 |
| SPGP0Q01300 | XP_018222370.1 | 99 | 0 |
| SPGP0P00590 | XP_018221897.1 | 99 | 0 |
| SPGP0S02200 | XP_018220132.1 | 99 | 0 |
| SPGP0Q01810 | XP_018222317.1 | 99 | 0 |
| SPGP0L00390 | XP_018219322.1 | 99 | 0 |
| SPGP0DN02870 | XP_018223732.1 | 99 | 0 |
| SPGP0Q03410 | XP_018222162.1 | 99 | 0 |
| SPGP0AC00450 | XP_018219129.1 | 99 | 0 |
| SPGP0Z01060 | XP_018219778.1 | 99 | 0 |
| SPGP0E02620 | XP_018222848.1 | 98 | 0 |
| SPGP0DN00480 | XP_018223496.1 | 99 | 0 |
| SPGP0DN01470 | XP_018223594.1 | 99 | 0 |
| SPGP0E00170 | XP_018222607.1 | 99 | 0 |
| SPGP0E01080 | XP_018222698.1 | 99 | 0 |
| SPGP0W02740 | XP_018223065.1 | 99 | 0 |
| SPGP0N01140 | XP_018220698.1 | 99 | 0 |
| SPGP0R01170 | XP_018221235.1 | 99 | 0 |
| SPGP0F04520 | XP_018222039.1 | 99 | 0 |
| SPGP0DN00700 | XP_018223516.1 | 99 | 0 |
| SPGP0R03370 | XP_018221446.1 | 96 | 3 |
| SPGP0DQ00110 | XP_018222851.1 | 92 | 0 |
| SPGP0H00400 | XP_018224096.1 | 99 | 0 |
| SPGP0F04270 | XP_018222064.1 | 98 | 1 |
| SPGP0P00340 | XP_018221920.1 | 99 | 0 |
| SPGP0DN00710 | XP_018223517.1 | 99 | 0 |
| SPGP0DZ02380 | XP_018218750.1 | 99 | 0 |
| SPGP0R02900 | XP_018221401.1 | 99 | 0 |
| SPGP0P01830 | XP_018221777.1 | 98 | 0 |
| SPGP0Y01770 | XP_018224044.1 | 97 | 0 |
| SPGP0R00130 | XP_018221138.1 | 99 | 0 |
| SPGP0DN00610 | XP_018223507.1 | 99 | 0 |
| SPGP0U01400 | XP_018218853.1 | 99 | 0 |
| SPGP0P00120 | XP_018221942.1 | 99 | 0 |

Create PDF in your applications with the Pdfcrowd HTML to PDF API PDFCROWD

| | | | |
|---|---|---|---|
| SPGP0W02580 | XP_018223079.1 | 99 | 0 |
| SPGP0N00230 | XP_018220785.1 | 99 | 0 |
| SPGP0Q01730 | XP_018222325.1 | 99 | 0 |
| SPGP0N00540 | XP_018220754.1 | 100 | 0 |
| SPGP0P00770 | XP_018221879.1 | 100 | 0 |
| SPGP0W03650 | XP_018222983.1 | 99 | 0 |
| SPGP0Z00140 | XP_018219869.1 | 99 | 0 |
| SPGP0U00780 | XP_018218914.1 | 99 | 0 |
| SPGP0Z02470 | XP_018219638.1 | 99 | 0 |
| SPGP0F04770 | XP_018222017.1 | 99 | 0 |
| SPGP0S02640 | XP_018220089.1 | 99 | 0 |
| SPGP0W02760 | XP_018223063.1 | 99 | 0 |
| SPGP0CZ03020 | XP_018221121.1 | 92 | 6 |
| SPGP0E02190 | XP_018222807.1 | 99 | 0 |
| SPGP0S01650 | XP_018220186.1 | 99 | 0 |
| SPGP0E02230 | XP_018222811.1 | 99 | 0 |
| SPGP0E01950 | XP_018222783.1 | 99 | 0 |
| SPGP0Y00360 | XP_018223913.1 | 99 | 0 |
| SPGP0R02150 | XP_018221329.1 | 99 | 0 |
| SPGP0Y00320 | XP_018223909.1 | 99 | 0 |
| SPGP0Z02210 | XP_018219664.1 | 99 | 0 |
| SPGP0P02840 | XP_018221679.1 | 99 | 0 |
| SPGP0B01910 | XP_018221633.1 | 99 | 0 |
| SPGP0S01300 | XP_018220216.1 | 99 | 0 |
| SPGP0Q00310 | XP_018222470.1 | 99 | 0 |
| SPGP0W03790 | XP_018222969.1 | 99 | 0 |
| SPGP0R01900 | XP_018221306.1 | 99 | 0 |
| SPGP0DV00120 | XP_018219876.1 | 100 | 0 |
| SPGP0W04420 | XP_018222907.1 | 99 | 0 |
| SPGP0W01500 | XP_018223185.1 | 99 | 0 |
| SPGP0E01960 | XP_018222784.1 | 100 | 0 |
| SPGP0DN04210 | XP_018223863.1 | 100 | 0 |
| SPGP0DN03840 | XP_018223826.1 | 99 | 0 |
| SPGP0AC01100 | XP_018219193.1 | 98 | 0 |
| SPGP0AA01020 | XP_018223410.1 | 99 | 0 |
| SPGP0L01110 | XP_018219252.1 | 99 | 0 |
| SPGP0CZ01460 | XP_018220970.1 | 99 | 0 |
| SPGP0AC00510 | XP_018219135.1 | 100 | 0 |
| SPGP0H00170 | XP_018224118.1 | 99 | 0 |

Create PDF in your applications with the Pdfcrowd HTML to PDF API PDFCROWD

| | | | |
|---|---|---|---|
| SPGP0S01240 | XP_018220222.1 | 99 | 0 |
| SPGP0DN03910 | XP_018223833.1 | 99 | 0 |
| SPGP0W04210 | XP_018222929.1 | 99 | 0 |
| SPGP0Q01310 | XP_018222369.1 | 99 | 0 |
| SPGP0Q04140 | XP_018222091.1 | 99 | 0 |
| SPGP0Q02420 | XP_018222257.1 | 99 | 0 |
| SPGP0Q01640 | XP_018222334.1 | 99 | 0 |
| SPGP0Z02820 | XP_018219602.1 | 98 | 2 |
| SPGP0Z01340 | XP_018219751.1 | 99 | 0 |
| SPGP0W04010 | XP_018222949.1 | 99 | 0 |
| SPGP0DN03930 | XP_018223835.1 | 99 | 0 |
| SPGP0DN03390 | XP_018223782.1 | 99 | 0 |
| SPGP0R00500 | XP_018221173.1 | 100 | 0 |
| SPGP0DN03420 | XP_018223785.1 | 99 | 0 |
| SPGP0CZ01470 | XP_018220971.1 | 99 | 0 |
| SPGP0X00340 | XP_018219067.1 | 99 | 0 |
| SPGP0ltrna34S | DI49_5523 | 100 | 0 |
| SPGP0DN01860 | XP_018223633.1 | 99 | 0 |
| SPGP0X00740 | XP_018219028.1 | 99 | 0 |
| SPGP0R01270 | XP_018221244.1 | 99 | 0 |
| SPGP0BC00220 | XP_018220819.1 | 99 | 1 |
| SPGP0W00500 | XP_018223284.1 | 99 | 0 |
| SPGP0CZ00300 | XP_018220857.1 | 99 | 0 |
| SPGP0DZ00250 | XP_018219368.1 | 100 | 0 |
| SPGP0U00760 | XP_018218916.1 | 99 | 0 |
| SPGP0L00640 | XP_018219300.1 | 99 | 0 |
| SPGP0P02850 | XP_018221678.1 | 97 | 0 |
| SPGP0DU01710 | XP_018220420.1 | 99 | 0 |
| SPGP0DN03640 | XP_018223806.1 | 98 | 2 |
| SPGP0CZ01650 | XP_018220989.1 | 99 | 0 |
| SPGP0U00930 | XP_018218899.1 | 99 | 0 |
| SPGP0B01260 | XP_018221569.1 | 99 | 0 |
| SPGP0E02430 | XP_018222830.1 | 99 | 0 |
| SPGP0R00790 | XP_018221200.1 | 98 | 0 |
| SPGP0Q02660 | XP_018222234.1 | 99 | 0 |
| SPGP0DN02330 | XP_018223678.1 | 99 | 0 |
| SPGP0DU01220 | XP_018220468.1 | 99 | 0 |
| SPGP0R00880 | XP_018221208.1 | 99 | 0 |
| SPGP0S01930 | XP_018220159.1 | 99 | 0 |
| SPGP0R03240 | XP_018221434.1 | 99 | 0 |

Create PDF in your applications with the Pdfcrowd HTML to PDF API PDFCROWD

| | | | |
|---|---|---|---|
| SPGP0P00370 | XP_018221918.1 | 99 | 0 |
| SPGP0S03410 | XP_018220015.1 | 99 | 0 |
| SPGP0R01500 | XP_018221267.1 | 99 | 0 |
| SPGP0AA00890 | XP_018223398.1 | 99 | 0 |
| SPGP0L01310 | XP_018219233.1 | 99 | 0 |
| SPGP0U00260 | XP_018218963.1 | 99 | 0 |
| SPGP0E00160 | XP_018222606.1 | 99 | 0 |
| SPGP0Q00770 | XP_018222424.1 | 99 | 0 |
| SPGP0S03470 | XP_018220009.1 | 99 | 0 |
| SPGP0W01670 | XP_018223168.1 | 99 | 0 |
| SPGP0B00460 | XP_018221494.1 | 99 | 0 |
| SPGP0AC00320 | XP_018219116.1 | 100 | 0 |
| SPGP0S00680 | XP_018220277.1 | 99 | 0 |
| SPGP0W03940 | XP_018222955.1 | 99 | 0 |
| SPGP0DN02110 | XP_018223658.1 | 99 | 0 |
| SPGP0E00360 | XP_018222626.1 | 98 | 1 |
| SPGP0W04330 | XP_018222917.1 | 99 | 0 |
| SPGP0B00130 | XP_018221461.1 | 96 | 1 |
| SPGP0Y01180 | XP_018223987.1 | 99 | 0 |
| SPGP0Z01050 | XP_018219779.1 | 99 | 0 |
| SPGP0DN03210 | XP_018223765.1 | 99 | 0 |
| SPGP0DN02780 | XP_018223722.1 | 99 | 0 |
| SPGP0P02120 | XP_018221748.1 | 100 | 0 |
| SPGP0DN03000 | XP_018223745.1 | 100 | 0 |
| SPGP0Z02780 | XP_018219606.1 | 99 | 0 |
| SPGP0E01360 | XP_018222725.1 | 100 | 0 |
| SPGP0DN01130 | XP_018223560.1 | 99 | 0 |
| SPGP0DU00220 | XP_018220568.1 | 99 | 0 |
| SPGP0R02990 | XP_018221409.1 | 99 | 0 |
| SPGP0DU00210 | XP_018220569.1 | 99 | 0 |
| SPGP0Q02450 | XP_018222254.1 | 100 | 0 |
| SPGP0S03260 | XP_018220029.1 | 99 | 0 |
| SPGP0U00990 | XP_018218893.1 | 100 | 0 |
| SPGP0T00180 | XP_018220346.1 | 99 | 0 |
| SPGP0W02370 | XP_018223100.1 | 99 | 1 |
| SPGP0Q00230 | XP_018222477.1 | 99 | 0 |
| SPGP0S02910 | XP_018220063.1 | 99 | 0 |
| SPGP0E01180 | XP_018222708.1 | 100 | 0 |
| SPGP0DU01570 | XP_018220434.1 | 99 | 0 |
| SPGP0DZ00310 | XP_018219374.1 | 99 | 0 |

Create PDF in your applications with the Pdfcrowd HTML to PDF API PDFCROWD

| | | | |
|---|---|---|---|
| SPGP0AA00180 | XP_018223331.1 | 99 | 0 |
| SPGP0E00580 | XP_018222648.1 | 100 | 0 |
| SPGP0S02360 | XP_018220116.1 | 99 | 1 |
| SPGP0DN00510 | XP_018223499.1 | 99 | 0 |
| SPGP0DN02850 | XP_018223730.1 | 99 | 0 |
| SPGP0L00150 | XP_018219346.1 | 99 | 0 |
| SPGP0W02430 | XP_018223094.1 | 100 | 0 |
| SPGP0L01340 | XP_018219230.1 | 99 | 0 |
| SPGP0U01750 | XP_018218819.1 | 99 | 0 |
| SPGP0U01690 | XP_018218825.1 | 99 | 0 |
| SPGP0X00370 | XP_018219064.1 | 99 | 1 |
| SPGP0Z00870 | XP_018219797.1 | 99 | 1 |
| SPGP0N00130 | XP_018220795.1 | 99 | 0 |
| SPGP0S02240 | XP_018220128.1 | 99 | 0 |
| SPGP0Z02040 | XP_018219681.1 | 99 | 0 |
| SPGP0S02040 | XP_018220148.1 | 99 | 0 |
| SPGP0DU00480 | XP_018220542.1 | 99 | 0 |
| SPGP0R00610 | XP_018221183.1 | 99 | 0 |
| SPGP0W02150 | XP_018223122.1 | 99 | 1 |
| SPGP0S01840 | XP_018220168.1 | 99 | 0 |
| SPGP0E01220 | XP_018222712.1 | 99 | 0 |
| SPGP0DF00100 | XP_018219197.1 | 91 | 0 |
| SPGP0Z01810 | XP_018219704.1 | 99 | 0 |
| SPGP0R01750 | XP_018221291.1 | 99 | 0 |
| SPGP0CZ02820 | XP_018221101.1 | 99 | 0 |
| SPGP0Q02890 | XP_018222212.1 | 99 | 0 |
| SPGP0AA00980 | XP_018223406.1 | 99 | 0 |
| SPGP0CZ02040 | XP_018221025.1 | 99 | 0 |
| SPGP0DN03780 | XP_018223820.1 | 100 | 0 |
| SPGP0W03350 | XP_018223009.1 | 99 | 0 |
| SPGP0L00940 | XP_018219269.1 | 99 | 0 |
| SPGP0AF00220 | XP_018221963.1 | 99 | 0 |
| SPGP0DN03090 | XP_018223753.1 | 99 | 0 |
| SPGP0P02240 | XP_018221738.1 | 99 | 0 |
| SPGP0B00100 | XP_018222596.1 | 98 | 0 |
| SPGP0B01070 | XP_018221552.1 | 99 | 0 |
| SPGP0CZ00680 | XP_018220894.1 | 99 | 0 |
| SPGP0DU00350 | XP_018220555.1 | 99 | 0 |
| SPGP0Q04330 | XP_018222073.1 | 99 | 0 |
| SPGP0R01070 | XP_018221225.1 | 99 | 0 |

Create PDF in your applications with the Pdfcrowd HTML to PDF API PDFCROWD

| | | | |
|---|---|---|---|
| SPGP0DV00150 | XP_018219879.1 | 99 | 0 |
| SPGP0DN02090 | XP_018223656.1 | 100 | 0 |
| SPGP0L00430 | XP_018219318.1 | 100 | 0 |
| SPGP0S00870 | XP_018220258.1 | 99 | 0 |
| SPGP0DNtrna8V | DI49_0484 | 100 | 0 |
| SPGP0S00300 | XP_018220315.1 | 99 | 0 |
| SPGP0W03380 | XP_018223006.1 | 99 | 0 |
| SPGP0Gtrna24A | DI49_5601 | 100 | 0 |
| SPGP0R02460 | XP_018221359.1 | 99 | 0 |
| SPGP0DN03890 | XP_018223831.1 | 99 | 0 |
| SPGP0R03030 | XP_018221413.1 | 99 | 0 |
| SPGP0Q02470 | XP_018222252.1 | 100 | 0 |
| SPGP0E02130 | XP_018222801.1 | 99 | 0 |
| SPGP0E00820 | XP_018222670.1 | 99 | 0 |
| SPGP0R01040 | XP_018221222.1 | 99 | 0 |
| SPGP0B01140 | XP_018221558.1 | 99 | 0 |
| SPGP0W04360 | XP_018222913.1 | 99 | 0 |
| SPGP0W00990 | XP_018223235.1 | 99 | 0 |
| SPGP0W00330 | XP_018223301.1 | 99 | 0 |
| SPGP0S03070 | XP_018220047.1 | 99 | 0 |
| SPGP0Z01680 | XP_018219717.1 | 99 | 0 |
| SPGP0X01120 | XP_018218991.1 | 99 | 0 |
| SPGP0F04260 | XP_018222065.1 | 99 | 0 |
| SPGP0F04710 | XP_018222022.1 | 99 | 0 |
| SPGP0B00300 | XP_018221478.1 | 99 | 0 |
| SPGP0AA00260 | XP_018223339.1 | 99 | 0 |
| SPGP0DZ01640 | XP_018219502.1 | 100 | 0 |
| SPGP0W03820 | XP_018222966.1 | 99 | 0 |
| SPGP0CZ01220 | XP_018220947.1 | 100 | 0 |
| SPGP0R02600 | XP_018221373.1 | 99 | 0 |
| SPGP0Q02290 | XP_018222270.1 | 99 | 0 |
| SPGP0Q00840 | XP_018222417.1 | 99 | 0 |
| SPGP0W00190 | XP_018223315.1 | 98 | 1 |
| SPGP0DX00480 | XP_018219978.1 | 99 | 0 |
| SPGP0DN02270 | XP_018223673.1 | 99 | 0 |
| SPGP0Q01780 | XP_018222320.1 | 99 | 0 |
| SPGP0H00630 | XP_018224074.1 | 99 | 0 |
| SPGP0W03340 | XP_018223010.1 | 99 | 0 |
| SPGP0R03280 | XP_018221438.1 | 99 | 0 |
| SPGP0S02220 | XP_018220130.1 | 99 | 0 |

Create PDF in your applications with the Pdfcrowd HTML to PDF API PDFCROWD

| | | | |
|---|---|---|---|
| SPGP0Y01220 | XP_018223991.1 | 99 | 0 |
| SPGP0Q03780 | XP_018222127.1 | 99 | 0 |
| SPGP0B01030 | XP_018221549.1 | 99 | 0 |
| SPGP0Q03250 | XP_018222177.1 | 100 | 0 |
| SPGP0AA00650 | XP_018223377.1 | 99 | 0 |
| SPGP0P01370 | XP_018221821.1 | 99 | 0 |
| SPGP0Z00370 | XP_018219853.1 | 99 | 0 |
| SPGP0N02300 | XP_018220584.1 | 99 | 0 |
| SPGP0F04800 | XP_018222014.1 | 99 | 0 |
| SPGP0Q02780 | XP_018222223.1 | 99 | 0 |
| SPGP0S01780 | XP_018220173.1 | 99 | 0 |
| SPGP0F04330 | XP_018222058.1 | 100 | 0 |
| SPGP0P02260 | XP_018221736.1 | 99 | 0 |
| SPGP0R03470 | XP_018221454.1 | 99 | 0 |
| SPGP0X00620 | XP_018219040.1 | 99 | 0 |
| SPGP0E00770 | XP_018222666.1 | 99 | 0 |
| SPGP0Q03430 | XP_018222160.1 | 99 | 0 |
| SPGP0Q02320 | XP_018222267.1 | 99 | 0 |
| SPGP0DN04260 | XP_018223868.1 | 99 | 0 |
| SPGP0W01240 | XP_018223210.1 | 99 | 0 |
| SPGP0X00410 | XP_018219060.1 | 99 | 0 |
| SPGP0E02600 | XP_018222846.1 | 99 | 0 |
| SPGP0DZ00460 | XP_018219389.1 | 99 | 0 |
| SPGP0W04630 | XP_018222885.1 | 99 | 1 |
| SPGP0N01000 | XP_018220711.1 | 99 | 0 |
| SPGP0F04210 | XP_018222070.1 | 99 | 0 |
| SPGP0E00110 | XP_018222601.1 | 97 | 0 |
| SPGP0DU00810 | XP_018220509.1 | 96 | 1 |
| SPGP0R02430 | XP_018221356.1 | 99 | 0 |
| SPGP0Ptrna10P | DI49_2548 | 98 | 0 |
| SPGP0DV00560 | XP_018219918.1 | 99 | 0 |
| SPGP0W00420 | XP_018223292.1 | 99 | 0 |
| SPGP0U00960 | XP_018218896.1 | 99 | 0 |
| SPGP0Z00530 | XP_018219830.1 | 99 | 0 |
| SPGP0DQ00130 | XP_018218768.1 | 95 | 0 |
| SPGP0Z00800 | XP_018219804.1 | 99 | 0 |
| SPGP0P00320 | XP_018221922.1 | 99 | 0 |
| SPGP0X00140 | XP_018219086.1 | 99 | 0 |
| SPGP0B00880 | XP_018221535.1 | 99 | 0 |
| SPGP0B00440 | XP_018221492.1 | 99 | 0 |

Create PDF in your applications with the Pdfcrowd HTML to PDF API PDFCROWD

| | | | |
|---|---|---|---|
| SPGP0R01570 | XP_018221274.1 | 98 | 0 |
| SPGP0R01050 | XP_018221223.1 | 99 | 0 |
| SPGP0R02920 | XP_018221403.1 | 99 | 0 |
| SPGP0Y00300 | XP_018223907.1 | 99 | 0 |
| SPGP0P01810 | XP_018221779.1 | 99 | 0 |
| SPGP0P00180 | XP_018221936.1 | 99 | 0 |
| SPGP0DU00560 | XP_018220534.1 | 99 | 0 |
| SPGP0P00900 | XP_018221867.1 | 99 | 0 |
| SPGP0F04570 | XP_018222034.1 | 99 | 0 |
| SPGP0Y00410 | XP_018223917.1 | 99 | 0 |
| SPGP0X00260 | XP_018219075.1 | 99 | 0 |
| SPGP0P00490 | XP_018221906.1 | 99 | 0 |
| SPGP0Q03810 | XP_018222124.1 | 99 | 0 |
| SPGP0Q03400 | XP_018222163.1 | 100 | 0 |
| SPGP0W01990 | XP_018223136.1 | 99 | 0 |
| SPGP0E01640 | XP_018222752.1 | 99 | 0 |
| SPGP0DN01310 | XP_018223578.1 | 100 | 0 |
| SPGP0W01140 | XP_018223220.1 | 100 | 0 |
| SPGP0W02290 | XP_018223108.1 | 99 | 0 |
| SPGP0BF00210 | XP_018220391.1 | 99 | 0 |
| SPGP0DX00380 | XP_018219969.1 | 99 | 0 |
| SPGP0S02820 | XP_018220072.1 | 99 | 0 |
| SPGP0DV00320 | XP_018219895.1 | 99 | 0 |
| SPGP0DU01210 | XP_018220469.1 | 99 | 0 |
| SPGP0R02250 | XP_018221338.1 | 100 | 0 |
| SPGP0Y00950 | XP_018223965.1 | 99 | 0 |
| SPGP0Q00320 | XP_018222469.1 | 99 | 0 |
| SPGP0Z02450 | XP_018219640.1 | 99 | 0 |
| SPGP0W02120 | XP_018223124.1 | 100 | 0 |
| SPGP0S01320 | XP_018220214.1 | 99 | 0 |
| SPGP0Z02650 | XP_018219620.1 | 99 | 0 |
| SPGP0S01220 | XP_018220223.1 | 99 | 0 |
| SPGP0W00860 | XP_018223248.1 | 99 | 0 |
| SPGP0S02790 | XP_018220074.1 | 99 | 0 |
| SPGP0Ztrna28L | DI49_4635 | 100 | 0 |
| SPGP0Z03110 | XP_018219574.1 | 99 | 0 |
| SPGP0Q02000 | XP_018222298.1 | 99 | 0 |
| SPGP0Z01290 | XP_018219756.1 | 99 | 0 |
| SPGP0Z02230 | XP_018219662.1 | 99 | 0 |
| SPGP0E02040 | XP_018222792.1 | 99 | 0 |

Create PDF in your applications with the Pdfcrowd HTML to PDF API PDFCROWD

| | | | |
|---|---|---|---|
| SPGP0Z02380 | XP_018219647.1 | 99 | 0 |
| SPGP0E00230 | XP_018222613.1 | 99 | 0 |
| SPGP0R01340 | XP_018221251.1 | 99 | 0 |
| SPGP0DN03130 | XP_018223757.1 | 99 | 0 |
| SPGP0DV00420 | XP_018219904.1 | 99 | 0 |
| SPGP0P03020 | XP_018221662.1 | 99 | 0 |
| SPGP0R02780 | XP_018221390.1 | 99 | 0 |
| SPGP0E00130 | XP_018222603.1 | 99 | 0 |
| SPGP0DN03400 | XP_018223783.1 | 99 | 1 |
| SPGP0DV00630 | XP_018219924.1 | 99 | 0 |
| SPGP0Z00580 | XP_018219826.1 | 99 | 0 |
| SPGP0B01530 | XP_018221595.1 | 99 | 0 |
| SPGP0DZ01580 | XP_018219497.1 | 99 | 0 |
| SPGP0Y01120 | XP_018223982.1 | 99 | 0 |
| SPGP0P01560 | XP_018221803.1 | 99 | 0 |
| SPGP0W04000 | XP_018222950.1 | 99 | 0 |
| SPGP0R01520 | XP_018221269.1 | 99 | 0 |
| SPGP0Y00120 | XP_018223890.1 | 99 | 0 |
| SPGP0P00440 | XP_018221911.1 | 99 | 0 |
| SPGP0AH00690 | XP_018222536.1 | 99 | 0 |
| SPGP0B00660 | XP_018221513.1 | 99 | 0 |
| SPGP0DN03900 | XP_018223832.1 | 99 | 0 |
| SPGP0AA00480 | XP_018223360.1 | 99 | 0 |
| SPGP0L00660 | XP_018219297.1 | 99 | 0 |
| SPGP0Z00640 | XP_018219820.1 | 99 | 0 |
| SPGP0E02270 | XP_018222815.1 | 99 | 0 |
| SPGP0R00780 | XP_018221199.1 | 99 | 0 |
| SPGP0H00120 | XP_018224121.1 | 94 | 1 |
| SPGP0P02750 | XP_018221688.1 | 99 | 0 |
| SPGP0L00580 | XP_018219305.1 | 99 | 0 |
| SPGP0P00460 | XP_018221909.1 | 99 | 0 |
| SPGP0Q02310 | XP_018222268.1 | 99 | 0 |
| SPGP0N00340 | XP_018220774.1 | 99 | 0 |
| SPGP0Q02640 | XP_018222236.1 | 99 | 0 |
| SPGP0Q01080 | XP_018222393.1 | 99 | 0 |
| SPGP0P01440 | XP_018221814.1 | 99 | 1 |
| SPGP0P00250 | XP_018221929.1 | 99 | 0 |
| SPGP0S01910 | XP_018220161.1 | 99 | 0 |
| SPGP0DN01380 | XP_018223585.1 | 99 | 0 |
| SPGP0DN01440 | XP_018223591.1 | 99 | 0 |

Create PDF in your applications with the Pdfcrowd HTML to PDF API PDFCROWD

| | | | |
|---|---|---|---|
| SPGP0DZ00570 | XP_018219399.1 | 99 | 0 |
| SPGP0Z02500 | XP_018219635.1 | 99 | 0 |
| SPGP0AF00480 | XP_018221989.1 | 99 | 0 |
| SPGP0E01450 | XP_018222734.1 | 99 | 0 |
| SPGP0P01500 | XP_018221808.1 | 99 | 0 |
| SPGP0F04620 | XP_018222030.1 | 99 | 0 |
| SPGP0DV00480 | XP_018219910.1 | 99 | 0 |
| SPGP0W00320 | XP_018223302.1 | 99 | 0 |
| SPGP0W02900 | XP_018223050.1 | 99 | 0 |
| SPGP0P02570 | XP_018221706.1 | 99 | 0 |
| SPGP0L00970 | XP_018219266.1 | 99 | 0 |
| SPGP0Qtrna3W | DI49_2157 | 100 | 0 |
| SPGP0L00890 | XP_018219274.1 | 99 | 0 |
| SPGP0Q02130 | XP_018222285.1 | 99 | 0 |
| SPGP0DN01050 | XP_018223552.1 | 99 | 0 |
| SPGP0DN01710 | XP_018223617.1 | 99 | 0 |
| SPGP0E01430 | XP_018222732.1 | 99 | 0 |
| SPGP0E01340 | XP_018222724.1 | 99 | 0 |
| SPGP0W00650 | XP_018223269.1 | 99 | 0 |
| SPGP0E01190 | XP_018222709.1 | 99 | 0 |
| SPGP0S00920 | XP_018220253.1 | 99 | 0 |
| SPGP0EX00230 | XP_018223882.1 | 99 | 0 |
| SPGP0B01440 | XP_018221587.1 | 99 | 0 |
| SPGP0DN03450 | XP_018223788.1 | 99 | 0 |
| SPGP0R00470 | XP_018221170.1 | 98 | 0 |
| SPGP0S03200 | XP_018220034.1 | 99 | 0 |
| SPGP0DN02280 | XP_018223674.1 | 99 | 0 |
| SPGP0W02350 | XP_018223102.1 | 99 | 0 |
| SPGP0DN00440 | XP_018223492.1 | 99 | 0 |
| SPGP0H00610 | XP_018224076.1 | 99 | 0 |
| SPGP0DN03740 | XP_018223816.1 | 99 | 0 |
| SPGP0Q02220 | XP_018222277.1 | 99 | 0 |
| SPGP0E01110 | XP_018222701.1 | 99 | 0 |
| SPGP0P01170 | XP_018221840.1 | 99 | 0 |
| SPGP0N01530 | XP_018220661.1 | 99 | 0 |
| SPGP0Z00330 | XP_018219849.1 | 99 | 0 |
| SPGP0P01090 | XP_018221848.1 | 100 | 0 |
| SPGP0W01190 | XP_018223215.1 | 99 | 0 |
| SPGP0DN00570 | XP_018223504.1 | 99 | 0 |
| SPGP0F04380 | XP_018222053.1 | 99 | 0 |

Create PDF in your applications with the Pdfcrowd HTML to PDF API PDFCROWD

| | | | |
|---|---|---|---|
| SPGP0Z01350 | XP_018219750.1 | 100 | 0 |
| SPGP0DU01510 | XP_018220439.1 | 99 | 0 |
| SPGP0E02360 | XP_018222824.1 | 99 | 0 |
| SPGP0X00600 | XP_018219042.1 | 99 | 0 |
| SPGP0AH00800 | XP_018222525.1 | 99 | 0 |
| SPGP0DX00540 | XP_018219984.1 | 99 | 0 |
| SPGP0P03080 | XP_018221656.1 | 99 | 0 |
| SPGP0S00890 | XP_018220256.1 | 99 | 0 |
| SPGP0AA00300 | XP_018223343.1 | 99 | 0 |
| SPGP0P00680 | XP_018221888.1 | 99 | 0 |
| SPGP0Y00750 | XP_018223947.1 | 99 | 0 |
| SPGP0S01520 | XP_018220196.1 | 99 | 0 |
| SPGP0CW00100 | XP_018223054.1 | 100 | 0 |
| SPGP0P02420 | XP_018221721.1 | 99 | 0 |
| SPGP0DN01290 | XP_018223576.1 | 99 | 0 |
| SPGP0W03400 | XP_018223004.1 | 99 | 0 |
| SPGP0L00410 | XP_018219320.1 | 99 | 0 |
| SPGP0N01840 | XP_018220631.1 | 99 | 0 |
| SPGP0W04170 | XP_018222933.1 | 99 | 0 |
| SPGP0AH00840 | XP_018222521.1 | 100 | 0 |
| SPGP0N00680 | XP_018220740.1 | 100 | 0 |
| SPGP0DX00590 | XP_018219989.1 | 100 | 0 |
| SPGP0CZ00660 | XP_018220892.1 | 99 | 0 |
| SPGP0Q04310 | XP_018222075.1 | 99 | 0 |
| SPGP0S00320 | XP_018220313.1 | 99 | 0 |
| SPGP0Y01170 | XP_018223986.1 | 99 | 0 |
| SPGP0B01750 | XP_018221617.1 | 99 | 1 |
| SPGP0Z02400 | XP_018219645.1 | 99 | 0 |
| SPGP0B00180 | XP_018221466.1 | 98 | 0 |
| SPGP0DN03870 | XP_018223829.1 | 100 | 0 |
| SPGP0DN03330 | XP_018223776.1 | 99 | 0 |
| SPGP0Z02290 | XP_018219656.1 | 99 | 0 |
| SPGP0E00210 | XP_018222611.1 | 99 | 0 |
| SPGP0P01860 | XP_018221774.1 | 99 | 0 |
| SPGP0S02670 | XP_018220086.1 | 99 | 0 |
| SPGP0Z00420 | XP_018219858.1 | 98 | 0 |
| SPGP0P01790 | XP_018221781.1 | 99 | 0 |
| SPGP0Strna8Q | DI49_4357 | 100 | 0 |
| SPGP0L00350 | XP_018219326.1 | 99 | 0 |
| SPGP0B00320 | XP_018221480.1 | 99 | 0 |

Create PDF in your applications with the Pdfcrowd HTML to PDF API PDFCROWD

| | | | |
|---|---|---|---|
| SPGP0R00330 | XP_018221156.1 | 99 | 0 |
| SPGP0B01390 | XP_018221582.1 | 99 | 0 |
| SPGP0Q02410 | XP_018222258.1 | 99 | 0 |
| SPGP0Z01910 | XP_018219694.1 | 99 | 0 |
| SPGP0Q00670 | XP_018222434.1 | 99 | 0 |
| SPGP0P01240 | XP_018221833.1 | 99 | 0 |
| SPGP0H00780 | XP_018224059.1 | 99 | 0 |
| SPGP0Q01500 | XP_018222348.1 | 99 | 0 |
| SPGP0Q01850 | XP_018222313.1 | 99 | 0 |
| SPGP0P01300 | XP_018221827.1 | 99 | 0 |
| SPGP0Q03230 | XP_018222179.1 | 98 | 1 |
| SPGP0Y00960 | XP_018223966.1 | 99 | 0 |
| SPGP0H00650 | XP_018224072.1 | 99 | 0 |
| SPGP0S00690 | XP_018220276.1 | 99 | 0 |
| SPGP0H00300 | XP_018224105.1 | 99 | 0 |
| SPGP0CZ01200 | XP_018220945.1 | 99 | 0 |
| SPGP0W01000 | XP_018223234.1 | 99 | 0 |
| SPGP0Z00390 | XP_018219855.1 | 99 | 0 |
| SPGP0S02540 | XP_018220099.1 | 99 | 0 |
| SPGP0Etrna16Q | DI49_1374 | 100 | 0 |
| SPGP0DN04030 | XP_018223845.1 | 99 | 0 |
| SPGP0Ptrna11P | DI49_2389 | 99 | 0 |
| SPGP0P01760 | XP_018221784.1 | 99 | 0 |
| SPGP0DN02830 | XP_018223727.1 | 99 | 0 |
| SPGP0E00280 | XP_018222618.1 | 99 | 0 |
| SPGP0E00750 | XP_018222664.1 | 99 | 0 |
| SPGP0N01980 | XP_018220616.1 | 99 | 0 |
| SPGP0DN04240 | XP_018223866.1 | 99 | 0 |
| SPGP0N00420 | XP_018220765.1 | 99 | 0 |
| SPGP0N02180 | XP_018220596.1 | 99 | 0 |
| SPGP0DN01010 | XP_018223548.1 | 99 | 0 |
| SPGP0S01600 | XP_018220190.1 | 99 | 0 |
| SPGP0P00860 | XP_018221871.1 | 99 | 0 |
| SPGP0DZ01290 | XP_018219468.1 | 95 | 4 |
| SPGP0R01580 | XP_018221275.1 | 99 | 0 |
| SPGP0CZ03000 | XP_018221119.1 | 99 | 0 |
| SPGP0AC00970 | XP_018219180.1 | 99 | 0 |
| SPGP0S01710 | XP_018220180.1 | 99 | 0 |
| SPGP0Q03610 | XP_018222144.1 | 99 | 0 |
| SPGP0DU00680 | XP_018220523.1 | 99 | 0 |

Create PDF in your applications with the Pdfcrowd HTML to PDF API PDFCROWD

| | | | |
|---|---|---|---|
| SPGP0P03040 | XP_018221660.1 | 99 | 0 |
| SPGP0S00280 | XP_018220317.1 | 99 | 0 |
| SPGP0S00880 | XP_018220257.1 | 99 | 0 |
| SPGP0U00540 | XP_018218935.1 | 99 | 0 |
| SPGP0DN00940 | XP_018223540.1 | 99 | 0 |
| SPGP0W02710 | XP_018223068.1 | 99 | 0 |
| SPGP0AH00520 | XP_018222553.1 | 97 | 3 |
| SPGP0P00300 | XP_018221924.1 | 99 | 0 |
| SPGP0Y01780 | XP_018224045.1 | 96 | 0 |
| SPGP0DZ01740 | XP_018219512.1 | 99 | 0 |
| SPGP0DN04250 | XP_018223867.1 | 99 | 0 |
| SPGP0Q00750 | XP_018222426.1 | 99 | 0 |
| SPGP0P01870 | XP_018221773.1 | 99 | 0 |
| SPGP0DN00740 | XP_018223520.1 | 99 | 0 |
| SPGP0AH00250 | XP_018222577.1 | 99 | 0 |
| SPGP0CZ01420 | XP_018220966.1 | 99 | 0 |
| SPGP0N00580 | XP_018220750.1 | 99 | 0 |
| SPGP0Z00890 | XP_018219795.1 | 99 | 0 |
| SPGP0E00620 | XP_018222652.1 | 100 | 0 |
| SPGP0EE00180 | XP_018220837.1 | 99 | 0 |
| SPGP0R03010 | XP_018221411.1 | 99 | 0 |
| SPGP0W02570 | XP_018223080.1 | 99 | 0 |
| SPGP0B01880 | XP_018221630.1 | 99 | 0 |
| SPGP0W02720 | XP_018223067.1 | 99 | 0 |
| SPGP0BF00230 | XP_018220393.1 | 99 | 0 |
| SPGP0DN02170 | XP_018223663.1 | 99 | 0 |
| SPGP0H00800 | XP_018224057.1 | 100 | 0 |
| SPGP0B00750 | XP_018221522.1 | 99 | 0 |
| SPGP0N01170 | XP_018220695.1 | 99 | 0 |
| SPGP0X00830 | XP_018219019.1 | 99 | 0 |
| SPGP0Z00210 | XP_018219862.1 | 99 | 0 |
| SPGP0DZ01150 | XP_018219454.1 | 99 | 0 |
| SPGP0EE00190 | XP_018220838.1 | 99 | 0 |
| SPGP0X00700 | XP_018219032.1 | 99 | 0 |
| SPGP0DV00180 | XP_018219882.1 | 99 | 0 |
| SPGP0Z02430 | XP_018219642.1 | 99 | 0 |
| SPGP0DN01320 | XP_018223579.1 | 99 | 0 |
| SPGP0R03270 | XP_018221437.1 | 99 | 0 |
| SPGP0X00860 | XP_018219016.1 | 99 | 0 |

Create PDF in your applications with the Pdfcrowd HTML to PDF API PDFCROWD

| | | | |
|---|---|---|---|
| SPGP0DN01680 | XP_018223614.1 | 100 | 0 |
| SPGP0S03450 | XP_018220011.1 | 99 | 0 |
| SPGP0L01370 | XP_018219227.1 | 99 | 0 |
| SPGP0DU00330 | XP_018220557.1 | 99 | 1 |
| SPGP0Z01160 | XP_018219768.1 | 99 | 0 |
| SPGP0R02270 | XP_018221340.1 | 99 | 0 |
| SPGP0W03090 | XP_018223034.1 | 99 | 0 |
| SPGP0DU00430 | XP_018220547.1 | 99 | 0 |
| SPGP0E02060 | XP_018222794.1 | 100 | 0 |
| SPGP0L01200 | XP_018219244.1 | 99 | 0 |
| SPGP0S03150 | XP_018220039.1 | 99 | 0 |
| SPGP0AC00430 | XP_018219127.1 | 99 | 0 |
| SPGP0L01620 | XP_018219203.1 | 98 | 0 |
| SPGP0DN00250 | XP_018223475.1 | 99 | 0 |
| SPGP0W04690 | XP_018222879.1 | 99 | 0 |
| SPGP0W00270 | XP_018223307.1 | 99 | 0 |
| SPGP0W03200 | XP_018223023.1 | 99 | 0 |
| SPGP0Z00380 | XP_018219854.1 | 99 | 0 |
| SPGP0Z02110 | XP_018219674.1 | 99 | 0 |
| SPGP0R02050 | XP_018221320.1 | 99 | 0 |
| SPGP0DZ00810 | XP_018219422.1 | 99 | 0 |
| SPGP0W03670 | XP_018222981.1 | 99 | 0 |
| SPGP0W02060 | XP_018223131.1 | 100 | 0 |
| SPGP0DN03460 | XP_018223789.1 | 99 | 0 |
| SPGP0P01140 | XP_018221843.1 | 99 | 0 |
| SPGP0S03340 | XP_018220021.1 | 99 | 0 |
| SPGP0AH00140 | XP_018222588.1 | 99 | 0 |
| SPGP0DNtrna3F | DI49_0089 | 100 | 0 |
| SPGP0B01510 | XP_018221593.1 | 99 | 0 |
| SPGP0Q00930 | XP_018222408.1 | 99 | 0 |
| SPGP0Y01630 | XP_018224032.1 | 99 | 0 |
| SPGP0W02080 | XP_018223129.1 | 99 | 0 |
| SPGP0W04020 | XP_018222948.1 | 99 | 0 |
| SPGP0B01700 | XP_018221612.1 | 99 | 0 |
| SPGP0W02490 | XP_018223088.1 | 100 | 0 |
| SPGP0L00600 | XP_018219303.1 | 99 | 0 |
| SPGP0Y00100 | XP_018223888.1 | 99 | 0 |
| SPGP0W02250 | XP_018223112.1 | 100 | 0 |
| SPGP0E01050 | XP_018222695.1 | 99 | 0 |
| SPGP0DN03680 | XP_018223810.1 | 99 | 0 |

Create PDF in your applications with the Pdfcrowd HTML to PDF API PDFCROWD

| | | | |
|---|---|---|---|
| SPGP0Q02620 | XP_018222238.1 | 99 | 0 |
| SPGP0W04890 | XP_018222859.1 | 86 | 0 |
| SPGP0Q02330 | XP_018222266.1 | 98 | 0 |
| SPGP0DN03040 | XP_018223749.1 | 99 | 0 |
| SPGP0S01970 | XP_018220155.1 | 99 | 0 |
| SPGP0E01870 | XP_018222775.1 | 99 | 0 |
| SPGP0DN02480 | XP_018223693.1 | 99 | 0 |
| SPGP0E00920 | XP_018222681.1 | 99 | 0 |
| SPGP0Q03720 | XP_018222133.1 | 99 | 0 |
| SPGP0L00510 | XP_018219310.1 | 99 | 0 |
| SPGP0U00220 | XP_018218967.1 | 99 | 0 |
| SPGP0R00100 | XP_018221133.1 | 99 | 0 |
| SPGP0W01630 | XP_018223172.1 | 99 | 0 |
| SPGP0Y01410 | XP_018224010.1 | 99 | 0 |
| SPGP0DN01350 | XP_018223582.1 | 99 | 0 |
| SPGP0W00450 | XP_018223289.1 | 99 | 0 |
| SPGP0Q01090 | XP_018222392.1 | 99 | 0 |
| SPGP0S02580 | XP_018220095.1 | 99 | 0 |
| SPGP0S03190 | XP_018220035.1 | 99 | 0 |
| SPGP0DN00880 | XP_018223534.1 | 100 | 0 |
| SPGP0DN02890 | XP_018223734.1 | 99 | 0 |
| SPGP0F04940 | XP_018222001.1 | 98 | 0 |
| SPGP0DN01170 | XP_018223564.1 | 99 | 0 |
| SPGP0R03360 | XP_018221445.1 | 99 | 0 |
| SPGP0E02290 | XP_018222817.1 | 99 | 0 |
| SPGP0Z01480 | XP_018219737.1 | 100 | 0 |
| SPGP0H00590 | XP_018224078.1 | 99 | 0 |
| SPGP0DZ01770 | XP_018219515.1 | 99 | 0 |
| SPGP0R00650 | XP_018221187.1 | 99 | 0 |
| SPGP0N01040 | XP_018220707.1 | 99 | 0 |
| SPGP0N01510 | XP_018220663.1 | 97 | 3 |
| SPGP0Z03130 | XP_018219572.1 | 99 | 0 |
| SPGP0CZ01900 | XP_018221011.1 | 93 | 7 |
| SPGP0S01590 | XP_018220191.1 | 99 | 0 |
| SPGP0Q00900 | XP_018222411.1 | 99 | 0 |
| SPGP0H00320 | XP_018224103.1 | 98 | 0 |
| SPGP0U01560 | XP_018218838.1 | 99 | 0 |
| SPGP0AA00540 | XP_018223366.1 | 99 | 0 |
| SPGP0Y00140 | XP_018223892.1 | 99 | 0 |
| SPGP0AH00820 | XP_018222523.1 | 99 | 0 |

Create PDF in your applications with the Pdfcrowd HTML to PDF API PDFCROWD

| | | | |
|---|---|---|---|
| SPGP0Z01820 | XP_018219703.1 | 99 | 0 |
| SPGP0AF00100 | XP_018221952.1 | 100 | 0 |
| SPGP0Gtrna33R | DI49_2918 | 100 | 0 |
| SPGP0DN00220 | XP_018223472.1 | 99 | 0 |
| SPGP0Y01440 | XP_018224013.1 | 99 | 0 |
| SPGP0W02510 | XP_018223086.1 | 99 | 0 |
| SPGP0X01080 | XP_018218995.1 | 100 | 0 |
| SPGP0DU01530 | XP_018220437.1 | 99 | 0 |
| SPGP0X00330 | XP_018219068.1 | 99 | 0 |
| SPGP0P02010 | XP_018221759.1 | 99 | 0 |
| SPGP0E00880 | XP_018222676.1 | 99 | 0 |
| SPGP0Z01870 | XP_018219698.1 | 99 | 0 |
| SPGP0E00930 | XP_018222682.1 | 99 | 0 |
| SPGP0DN02010 | XP_018223648.1 | 100 | 0 |
| SPGP0CZ02860 | XP_018221105.1 | 99 | 0 |
| SPGP0E02500 | XP_018222837.1 | 99 | 0 |
| SPGP0Z01270 | XP_018219758.1 | 99 | 0 |
| SPGP0AA00320 | XP_018223345.1 | 99 | 0 |
| SPGP0R02110 | XP_018221326.1 | 99 | 0 |
| SPGP0F04300 | XP_018222061.1 | 99 | 0 |
| SPGP0H00450 | XP_018224091.1 | 99 | 0 |
| SPGP0P02910 | XP_018221673.1 | 99 | 0 |
| SPGP0B01100 | XP_018221555.1 | 99 | 0 |
| SPGP0AF00410 | XP_018221982.1 | 99 | 0 |
| SPGP0Ftrna16N | DI49_5581 | 100 | 0 |
| SPGP0L00470 | XP_018219314.1 | 99 | 0 |
| SPGP0S00340 | XP_018220311.1 | 99 | 0 |
| SPGP0F04660 | XP_018222026.1 | 96 | 2 |
| SPGP0W02790 | XP_018223060.1 | 100 | 0 |
| SPGP0S00310 | XP_018220314.1 | 99 | 0 |
| SPGP0R03180 | XP_018221428.1 | 99 | 0 |
| SPGP0R01660 | XP_018221283.1 | 99 | 0 |
| SPGP0Z01140 | XP_018219770.1 | 99 | 0 |
| SPGP0H00510 | XP_018224085.1 | 99 | 0 |
| SPGP0N01880 | XP_018220626.1 | 99 | 0 |
| SPGP0CZ00370 | XP_018220863.1 | 99 | 0 |
| SPGP0R02030 | XP_018221318.1 | 99 | 0 |
| SPGP0DN00620 | XP_018223508.1 | 99 | 0 |
| SPGP0N00900 | XP_018220719.1 | 99 | 0 |
| SPGP0CZ00640 | XP_018220890.1 | 99 | 0 |

Create PDF in your applications with the Pdfcrowd HTML to PDF API PDFCROWD

| | | | |
|---|---|---|---|
| SPGP0AH00550 | XP_018222550.1 | 99 | 0 |
| SPGP0N00840 | XP_018220725.1 | 97 | 0 |
| SPGP0R02420 | XP_018221355.1 | 99 | 0 |
| SPGP0DN03850 | XP_018223827.1 | 99 | 0 |
| SPGP0L01070 | XP_018219256.1 | 99 | 0 |
| SPGP0Q00880 | XP_018222413.1 | 99 | 0 |
| SPGP0CZ00420 | XP_018220868.1 | 99 | 0 |
| SPGP0S01790 | XP_018220172.1 | 99 | 0 |
| SPGP0S02610 | XP_018220092.1 | 99 | 0 |
| SPGP0U01290 | XP_018218864.1 | 99 | 0 |
| SPGP0P00480 | XP_018221907.1 | 99 | 0 |
| SPGP0DV00650 | XP_018219926.1 | 98 | 0 |
| SPGP0P01260 | XP_018221831.1 | 99 | 0 |
| SPGP0L00290 | XP_018219332.1 | 99 | 0 |
| SPGP0DZ01600 | XP_018219499.1 | 99 | 0 |
| SPGP0B00640 | XP_018221511.1 | 99 | 0 |
| SPGP0S00960 | XP_018220249.1 | 99 | 0 |
| SPGP0Q02430 | XP_018222256.1 | 99 | 0 |
| SPGP0B00480 | XP_018221496.1 | 99 | 0 |
| SPGP0L00910 | XP_018219272.1 | 99 | 0 |
| SPGP0W04900 | XP_018222858.1 | 99 | 0 |
| SPGP0Q01520 | XP_018222346.1 | 100 | 0 |
| SPGP0S02260 | XP_018220126.1 | 99 | 0 |
| SPGP0AH00220 | XP_018222580.1 | 99 | 0 |
| SPGP0T00250 | XP_018220353.1 | 99 | 0 |
| SPGP0Q03740 | XP_018222131.1 | 99 | 0 |
| SPGP0S03050 | XP_018220049.1 | 99 | 0 |
| SPGP0E01610 | XP_018222749.1 | 99 | 0 |
| SPGP0R00830 | XP_018221203.1 | 99 | 0 |
| SPGP0W01020 | XP_018223232.1 | 99 | 0 |
| SPGP0X00170 | XP_018219083.1 | 99 | 0 |
| SPGP0B01160 | XP_018221559.1 | 100 | 0 |
| SPGP0DN01460 | XP_018223593.1 | 99 | 0 |
| SPGP0Y00530 | XP_018223927.1 | 99 | 0 |
| SPGP0P02920 | XP_018221672.1 | 99 | 0 |
| SPGP0AA00710 | XP_018223383.1 | 99 | 0 |
| SPGP0Z02410 | XP_018219644.1 | 99 | 0 |
| SPGP0W00730 | XP_018223260.1 | 99 | 0 |
| SPGP0DU01460 | XP_018220444.1 | 99 | 0 |
| SPGP0DN01510 | XP_018223598.1 | 99 | 0 |

Create PDF in your applications with the Pdfcrowd HTML to PDF API PDFCROWD

| | | | |
|---|---|---|---|
| SPGP0BF00190 | XP_018220389.1 | 98 | 0 |
| SPGP0E01840 | XP_018222772.1 | 100 | 0 |
| SPGP0DV00490 | XP_018219911.1 | 99 | 1 |
| SPGP0Q03650 | XP_018222140.1 | 99 | 0 |
| SPGP0FB00120 | XP_018220806.1 | 99 | 0 |
| SPGP0Q03850 | XP_018222120.1 | 99 | 0 |
| SPGP0X00100 | XP_018219089.1 | 99 | 0 |
| SPGP0W04260 | XP_018222924.1 | 99 | 0 |
| SPGP0Q03360 | XP_018222167.1 | 99 | 0 |
| SPGP0S00500 | XP_018220295.1 | 99 | 0 |
| SPGP0Q00500 | XP_018222451.1 | 100 | 0 |
| SPGP0DN04220 | XP_018223864.1 | 99 | 0 |
| SPGP0DN00360 | XP_018223486.1 | 99 | 0 |
| SPGP0Q03190 | XP_018222183.1 | 97 | 1 |
| SPGP0Z00840 | XP_018219800.1 | 99 | 0 |
| SPGP0DN01540 | XP_018223601.1 | 99 | 0 |
| SPGP0Z03150 | XP_018219570.1 | 99 | 0 |
| SPGP0U01210 | XP_018218871.1 | 99 | 0 |
| SPGP0DN00760 | XP_018223522.1 | 99 | 0 |
| SPGP0W01620 | XP_018223173.1 | 99 | 0 |
| SPGP0DN02970 | XP_018223742.1 | 99 | 0 |
| SPGP0Itrna31V | DI49_4248 | 100 | 0 |
| SPGP0W00360 | XP_018223298.1 | 99 | 0 |
| SPGP0E00270 | XP_018222617.1 | 99 | 0 |
| SPGP0Q02990 | XP_018222202.1 | 99 | 0 |
| SPGP0BF00260 | XP_018220395.1 | 99 | 0 |
| SPGP0AC00760 | XP_018219159.1 | 99 | 0 |
| SPGP0P02550 | XP_018221708.1 | 99 | 0 |
| SPGP0B01540 | XP_018221596.1 | 99 | 0 |
| SPGP0DN03310 | XP_018223774.1 | 100 | 0 |
| SPGP0R03070 | XP_018221417.1 | 99 | 0 |
| SPGP0W02700 | XP_018223069.1 | 99 | 0 |
| SPGP0Q00590 | XP_018222442.1 | 99 | 0 |
| SPGP0P02020 | XP_018221758.1 | 99 | 1 |
| SPGP0Q00730 | XP_018222427.1 | 99 | 0 |
| SPGP0S01360 | XP_018220211.1 | 99 | 0 |
| SPGP0H00860 | XP_018224051.1 | 99 | 0 |
| SPGP0P00720 | XP_018221884.1 | 99 | 0 |
| SPGP0DU01700 | XP_018220421.1 | 99 | 0 |
| SPGP0S02860 | XP_018220068.1 | 99 | 0 |

Create PDF in your applications with the Pdfcrowd HTML to PDF API PDFCROWD

| | | | |
|---|---|---|---|
| SPGP0X00810 | XP_018219021.1 | 99 | 1 |
| SPGP0Y00650 | XP_018223939.1 | 99 | 0 |
| SPGP0CZ01730 | XP_018220996.1 | 99 | 0 |
| SPGP0Z02140 | XP_018219671.1 | 99 | 0 |
| SPGP0DN03550 | XP_018223797.1 | 99 | 0 |
| SPGP0L01240 | XP_018219240.1 | 99 | 0 |
| SPGP0P03180 | XP_018221646.1 | 99 | 0 |
| SPGP0AC00400 | XP_018219124.1 | 100 | 0 |
| SPGP0AC00570 | XP_018219141.1 | 99 | 0 |
| SPGP0S01170 | XP_018220228.1 | 99 | 0 |
| SPGP0Q04290 | XP_018222077.1 | 99 | 0 |
| SPGP0R02210 | XP_018221334.1 | 99 | 0 |
| SPGP0P02580 | XP_018221705.1 | 99 | 0 |
| SPGP0W03070 | XP_018223036.1 | 99 | 0 |
| SPGP0P00500 | XP_018221905.1 | 99 | 0 |
| SPGP0Q00920 | XP_018222409.1 | 99 | 0 |
| SPGP0L00720 | XP_018219291.1 | 99 | 0 |
| SPGP0W04220 | XP_018222928.1 | 99 | 0 |
| SPGP0W04590 | XP_018222889.1 | 99 | 0 |
| SPGP0R01740 | XP_018221290.1 | 99 | 0 |
| SPGP0BC00240 | XP_018220821.1 | 99 | 0 |
| SPGP0W03610 | XP_018222987.1 | 99 | 0 |
| SPGP0DZ00830 | XP_018219424.1 | 99 | 0 |
| SPGP0S02250 | XP_018220127.1 | 99 | 0 |
| SPGP0L00590 | XP_018219304.1 | 99 | 0 |
| SPGP0R00730 | XP_018221195.1 | 99 | 0 |
| SPGP0Z02130 | XP_018219672.1 | 99 | 0 |
| SPGP0Z02540 | XP_018219631.1 | 99 | 0 |
| SPGP0DN03440 | XP_018223787.1 | 99 | 0 |
| SPGP0L01660 | XP_018219199.1 | 97 | 0 |
| SPGP0W02780 | XP_018223061.1 | 99 | 1 |
| SPGP0R03040 | XP_018221414.1 | 100 | 0 |
| SPGP0B01280 | XP_018221571.1 | 99 | 0 |
| SPGP0Q03350 | XP_018222168.1 | 99 | 0 |
| SPGP0W04040 | XP_018222946.1 | 99 | 0 |
| SPGP0R01290 | XP_018221246.1 | 99 | 0 |
| SPGP0Q04120 | XP_018222093.1 | 99 | 0 |
| SPGP0W03080 | XP_018223035.1 | 99 | 0 |
| SPGP0E00990 | XP_018222689.1 | 99 | 0 |
| SPGP0F04250 | XP_018222066.1 | 99 | 0 |

Create PDF in your applications with the Pdfcrowd HTML to PDF API PDFCROWD

| | | | |
|---|---|---|---|
| SPGP0S01730 | XP_018220178.1 | 99 | 0 |
| SPGP0Z00450 | XP_018219838.1 | 99 | 0 |
| SPGP0N02080 | XP_018220606.1 | 99 | 0 |
| SPGP0Q02350 | XP_018222264.1 | 99 | 0 |
| SPGP0L00850 | XP_018219278.1 | 99 | 0 |
| SPGP0Q02600 | XP_018222240.1 | 99 | 0 |
| SPGP0CZ00360 | XP_018220862.1 | 100 | 0 |
| SPGP0DN02310 | XP_018223676.1 | 99 | 0 |
| SPGP0H00600 | XP_018224077.1 | 99 | 0 |
| SPGP0E00840 | XP_018222672.1 | 100 | 0 |
| SPGP0B00150 | XP_018221463.1 | 97 | 0 |
| SPGP0DZ01010 | XP_018219441.1 | 100 | 0 |
| SPGP0AC00590 | XP_018219143.1 | 99 | 0 |
| SPGP0R00160 | XP_018221141.1 | 99 | 0 |
| SPGP0X01020 | XP_018219001.1 | 99 | 0 |
| SPGP0AA00330 | XP_018223346.1 | 99 | 0 |
| SPGP0P01630 | XP_018221796.1 | 98 | 0 |
| SPGP0DZ00490 | XP_018219392.1 | 99 | 0 |
| SPGP0Z01670 | XP_018219718.1 | 99 | 0 |
| SPGP0Q01950 | XP_018222303.1 | 99 | 0 |
| SPGP0S00970 | XP_018220248.1 | 99 | 0 |
| SPGP0CZ01560 | XP_018220980.1 | 99 | 0 |
| SPGP0N01280 | XP_018220685.1 | 99 | 0 |
| SPGP0Q03000 | XP_018222201.1 | 99 | 0 |
| SPGP0Q00370 | XP_018222464.1 | 99 | 0 |
| SPGP0B01610 | XP_018221603.1 | 99 | 0 |
| SPGP0Q04110 | XP_018222094.1 | 99 | 0 |
| SPGP0L00760 | XP_018219287.1 | 99 | 0 |
| SPGP0W00950 | XP_018223239.1 | 99 | 0 |
| SPGP0P01110 | XP_018221846.1 | 99 | 0 |
| SPGP0CZ02600 | XP_018221080.1 | 100 | 0 |
| SPGP0N01060 | XP_018220705.1 | 99 | 0 |
| SPGP0U00600 | XP_018218929.1 | 99 | 0 |
| SPGP0DN00690 | XP_018223515.1 | 99 | 0 |
| SPGP0N02210 | XP_018220593.1 | 99 | 0 |
| SPGP0E01470 | XP_018222735.1 | 99 | 0 |
| SPGP0Q00890 | XP_018222412.1 | 99 | 0 |
| SPGP0DZ01850 | XP_018219523.1 | 99 | 0 |
| SPGP0Y00220 | XP_018223900.1 | 99 | 0 |
| SPGP0S00840 | XP_018220261.1 | 99 | 0 |

Create PDF in your applications with the Pdfcrowd HTML to PDF API PDFCROWD

| | | | |
|---|---|---|---|
| SPGP0E01680 | XP_018222756.1 | 98 | 1 |
| SPGP0AC00380 | XP_018219122.1 | 99 | 0 |
| SPGP0P02220 | XP_018221740.1 | 99 | 0 |
| SPGP0W03360 | XP_018223008.1 | 99 | 0 |
| SPGP0X00420 | XP_018219059.1 | 99 | 0 |
| SPGP0P00530 | XP_018221903.1 | 100 | 0 |
| SPGP0X00310 | XP_018219070.1 | 99 | 0 |
| SPGP0S00770 | XP_018220268.1 | 100 | 0 |
| SPGP0Qtrna8F | DI49_1939 | 100 | 0 |
| SPGP0Y01150 | XP_018223984.1 | 98 | 0 |
| SPGP0P00880 | XP_018221869.1 | 99 | 0 |
| SPGP0X01010 | XP_018219002.1 | 99 | 0 |
| SPGP0S01950 | XP_018220157.1 | 99 | 0 |
| SPGP0P02970 | XP_018221667.1 | 99 | 0 |
| SPGP0W03100 | XP_018223033.1 | 99 | 0 |
| SPGP0DU00640 | XP_018220527.1 | 99 | 0 |
| SPGP0S00710 | XP_018220274.1 | 99 | 0 |
| SPGP0P01610 | XP_018221798.1 | 99 | 0 |
| SPGP0AA00340 | XP_018223347.1 | 99 | 0 |
| SPGP0Z01950 | XP_018219690.1 | 98 | 2 |
| SPGP0E01290 | XP_018222719.1 | 99 | 0 |
| SPGP0Q00630 | XP_018222438.1 | 99 | 0 |
| SPGP0Y00920 | XP_018223962.1 | 99 | 0 |
| SPGP0L00450 | XP_018219316.1 | 100 | 0 |
| SPGP0R01240 | XP_018221242.1 | 99 | 0 |
| SPGP0R01080 | XP_018221226.1 | 99 | 0 |
| SPGP0AF00280 | XP_018221969.1 | 99 | 0 |
| SPGP0Z02350 | XP_018219650.1 | 99 | 0 |
| SPGP0Z02600 | XP_018219625.1 | 99 | 0 |
| SPGP0CZ02840 | XP_018221103.1 | 99 | 0 |
| SPGP0Q03240 | XP_018222178.1 | 99 | 0 |
| SPGP0R02560 | XP_018221369.1 | 99 | 0 |
| SPGP0R01160 | XP_018221234.1 | 99 | 0 |
| SPGP0CZ00620 | XP_018220888.1 | 98 | 0 |
| SPGP0N00410 | XP_018220766.1 | 99 | 0 |
| SPGP0N01260 | XP_018220687.1 | 99 | 0 |
| SPGP0BF00220 | XP_018220392.1 | 100 | 0 |
| SPGP0Q02730 | XP_018222227.1 | 99 | 0 |
| SPGP0DN04070 | XP_018223849.1 | 99 | 0 |
| SPGP0Z02090 | XP_018219676.1 | 100 | 0 |

Create PDF in your applications with the Pdfcrowd HTML to PDF API PDFCROWD

| | | | |
|---|---|---|---|
| SPGP0H00760 | XP_018224061.1 | 99 | 0 |
| SPGP0W03910 | XP_018222958.1 | 98 | 0 |
| SPGP0L01500 | XP_018219214.1 | 99 | 0 |
| SPGP0DN01090 | XP_018223556.1 | 99 | 0 |
| SPGP0W03180 | XP_018223025.1 | 99 | 0 |
| SPGP0W00930 | XP_018223241.1 | 99 | 0 |
| SPGP0Q01050 | XP_018222396.1 | 99 | 0 |
| SPGP0W04470 | XP_018222901.1 | 99 | 0 |
| SPGP0Z01960 | XP_018219689.1 | 99 | 0 |
| SPGP0P01280 | XP_018221829.1 | 99 | 0 |
| SPGP0S00940 | XP_018220251.1 | 99 | 0 |
| SPGP0L00900 | XP_018219273.1 | 99 | 0 |
| SPGP0Q02500 | XP_018222249.1 | 99 | 0 |
| SPGP0AH00480 | XP_018222557.1 | 99 | 0 |
| SPGP0AG00140 | XP_018219933.1 | 99 | 0 |
| SPGP0B00360 | XP_018221484.1 | 99 | 0 |
| SPGP0DN03830 | XP_018223825.1 | 99 | 0 |
| SPGP0S02750 | XP_018220078.1 | 99 | 1 |
| SPGP0W04680 | XP_018222880.1 | 99 | 0 |
| SPGP0S01010 | XP_018220244.1 | 99 | 0 |
| SPGP0W03120 | XP_018223031.1 | 99 | 0 |
| SPGP0CZ02360 | XP_018221056.1 | 99 | 0 |
| SPGP0B00600 | XP_018221508.1 | 99 | 0 |
| SPGP0W04520 | XP_018222896.1 | 99 | 0 |
| SPGP0CZ01250 | XP_018220950.1 | 99 | 0 |
| SPGP0Q01890 | XP_018222309.1 | 99 | 0 |
| SPGP0DU01200 | XP_018220470.1 | 100 | 0 |
| SPGP0DN03160 | XP_018223760.1 | 99 | 0 |
| SPGP0CZ01710 | XP_018220994.1 | 99 | 0 |
| SPGP0R00370 | XP_018221160.1 | 99 | 0 |
| SPGP0R00800 | XP_018221201.1 | 99 | 0 |
| SPGP0B01020 | XP_018221548.1 | 99 | 0 |
| SPGP0Z01780 | XP_018219707.1 | 99 | 0 |
| SPGP0B01080 | XP_018221553.1 | 99 | 0 |
| SPGP0AC00600 | XP_018219144.1 | 99 | 0 |
| SPGP0DZ01570 | XP_018219496.1 | 99 | 0 |
| SPGP0R00260 | XP_018221149.1 | 99 | 0 |
| SPGP0S03130 | XP_018220041.1 | 99 | 0 |
| SPGP0W00230 | XP_018223311.1 | 99 | 0 |
| SPGP0N00450 | XP_018220762.1 | 100 | 0 |

Create PDF in your applications with the Pdfcrowd HTML to PDF API PDFCROWD

| | | | |
|---|---|---|---|
| SPGP0DU00200 | XP_018220570.1 | 99 | 0 |
| SPGP0U00530 | XP_018218936.1 | 100 | 0 |
| SPGP0Q03760 | XP_018222129.1 | 99 | 0 |
| SPGP0DU00780 | XP_018220513.1 | 99 | 0 |
| SPGP0CZ01210 | XP_018220946.1 | 99 | 0 |
| SPGP0E00350 | XP_018222625.1 | 99 | 0 |
| SPGP0B00500 | XP_018221498.1 | 99 | 0 |
| SPGP0S00560 | XP_018220289.1 | 99 | 0 |
| SPGP0U01250 | XP_018218867.1 | 100 | 0 |
| SPGP0P00920 | XP_018221865.1 | 99 | 0 |
| SPGP0Q03490 | XP_018222155.1 | 99 | 0 |
| SPGP0R00320 | XP_018221155.1 | 99 | 1 |
| SPGP0Z02150 | XP_018219670.1 | 99 | 0 |
| SPGP0W01400 | XP_018223194.1 | 99 | 0 |
| SPGP0E01040 | XP_018222694.1 | 99 | 0 |
| SPGP0E01590 | XP_018222747.1 | 99 | 0 |
| SPGP0Z02070 | XP_018219678.1 | 99 | 0 |
| SPGP0DN01910 | XP_018223638.1 | 99 | 0 |
| SPGP0B01820 | XP_018221624.1 | 99 | 1 |
| SPGP0Z02700 | XP_018219615.1 | 99 | 0 |
| SPGP0CZ00710 | XP_018220896.1 | 99 | 0 |
| SPGP0P02500 | XP_018221713.1 | 99 | 0 |
| SPGP0E01620 | XP_018222750.1 | 99 | 0 |
| SPGP0E01630 | XP_018222751.1 | 99 | 0 |
| SPGP0S01960 | XP_018220156.1 | 99 | 0 |
| SPGP0R00440 | XP_018221167.1 | 99 | 0 |
| SPGP0W02530 | XP_018223084.1 | 99 | 0 |
| SPGP0U01230 | XP_018218869.1 | 99 | 0 |
| SPGP0P01100 | XP_018221847.1 | 99 | 0 |
| SPGP0L01000 | XP_018219263.1 | 100 | 0 |
| SPGP0DZ01810 | XP_018219519.1 | 99 | 0 |
| SPGP0E01420 | XP_018222731.1 | 99 | 0 |
| SPGP0S01380 | XP_018220209.1 | 99 | 1 |
| SPGP0Q04130 | XP_018222092.1 | 99 | 0 |
| SPGP0DU01270 | XP_018220463.1 | 99 | 0 |
| SPGP0X00870 | XP_018219015.1 | 99 | 0 |
| SPGP0W01170 | XP_018223217.1 | 100 | 0 |
| SPGP0B00870 | XP_018221534.1 | 99 | 0 |
| SPGP0E01010 | XP_018222691.1 | 99 | 0 |
| SPGP0W01180 | XP_018223216.1 | 99 | 0 |

Create PDF in your applications with the Pdfcrowd HTML to PDF API PDFCROWD

| | | | |
|---|---|---|---|
| SPGP0U00370 | XP_018218952.1 | 100 | 0 |
| SPGP0Q02680 | XP_018222232.1 | 99 | 0 |
| SPGP0L01190 | XP_018219245.1 | 99 | 0 |
| SPGP0W04570 | XP_018222891.1 | 99 | 0 |
| SPGP0E02020 | XP_018222790.1 | 99 | 0 |
| SPGP0H00850 | XP_018224052.1 | 99 | 0 |
| SPGP0Q02590 | XP_018222241.1 | 99 | 0 |
| SPGP0R01640 | XP_018221281.1 | 99 | 0 |
| SPGP0N02310 | XP_018220583.1 | 99 | 0 |
| SPGP0DN01550 | XP_018223602.1 | 99 | 0 |
| SPGP0P02560 | XP_018221707.1 | 99 | 0 |
| SPGP0W00120 | XP_018223322.1 | 99 | 0 |
| SPGP0R02890 | XP_018221400.1 | 99 | 0 |
| SPGP0W03580 | XP_018222990.1 | 99 | 0 |
| SPGP0Z03010 | XP_018219584.1 | 99 | 0 |
| SPGP0E00870 | XP_018222675.1 | 99 | 0 |
| SPGP0Q01800 | XP_018222318.1 | 99 | 0 |
| SPGP0AP00120 | XP_018219932.1 | 99 | 0 |
| SPGP0B01770 | XP_018221619.1 | 99 | 0 |
| SPGP0R01760 | XP_018221292.1 | 99 | 0 |
| SPGP0R02090 | XP_018221324.1 | 99 | 0 |
| SPGP0E02550 | XP_018222841.1 | 99 | 0 |
| SPGP0Gtrna36L | DI49_3110 | 100 | 0 |
| SPGP0AAtrna22S | DI49_0694 | 100 | 0 |
| SPGP0Z02830 | XP_018219601.1 | 99 | 0 |
| SPGP0Z01610 | XP_018219724.1 | 99 | 0 |
| SPGP0W01980 | XP_018223137.1 | 99 | 0 |
| SPGP0R03310 | XP_018221441.1 | 99 | 0 |
| SPGP0DN02650 | XP_018223710.1 | 99 | 0 |
| SPGP0CZ01760 | XP_018220999.1 | 99 | 0 |
| SPGP0S00210 | XP_018220323.1 | 99 | 0 |
| SPGP0Xtrna13F | DI49_5486 | 100 | 0 |
| SPGP0B00210 | XP_018221469.1 | 99 | 1 |
| SPGP0Z01090 | XP_018219775.1 | 99 | 0 |
| SPGP0Z02260 | XP_018219659.1 | 99 | 0 |
| SPGP0Q00360 | XP_018222465.1 | 100 | 0 |
| SPGP0CZ01380 | XP_018220962.1 | 99 | 0 |
| SPGP0Z00470 | XP_018219836.1 | 99 | 0 |
| SPGP0Q02370 | XP_018222262.1 | 99 | 0 |

Create PDF in your applications with the Pdfcrowd HTML to PDF API PDFCROWD

| | | | |
|---|---|---|---|
| SPGP0N01600 | XP_018220654.1 | 99 | 0 |
| SPGP0Q02880 | XP_018222213.1 | 99 | 0 |
| SPGP0W04060 | XP_018222944.1 | 98 | 0 |
| SPGP0W00580 | XP_018223276.1 | 99 | 0 |
| SPGP0CZ00380 | XP_018220864.1 | 99 | 0 |
| SPGP0W01560 | XP_018223179.1 | 99 | 0 |
| SPGP0L00400 | XP_018219321.1 | 99 | 0 |
| SPGP0P01540 | XP_018221805.1 | 99 | 0 |
| SPGP0Q01530 | XP_018222345.1 | 99 | 0 |
| SPGP0DZtrna9P | DI49_5158 | 99 | 0 |
| SPGP0S00930 | XP_018220252.1 | 99 | 0 |
| SPGP0R00140 | XP_018221139.1 | 99 | 0 |
| SPGP0CZ01030 | XP_018220928.1 | 99 | 0 |
| SPGP0Q03950 | XP_018222110.1 | 99 | 0 |
| SPGP0Q01970 | XP_018222301.1 | 99 | 0 |
| SPGP0DN02190 | XP_018223665.1 | 99 | 0 |
| SPGP0P01130 | XP_018221844.1 | 99 | 0 |
| SPGP0DN02990 | XP_018223744.1 | 99 | 0 |
| SPGP0Q03020 | XP_018222199.1 | 99 | 0 |
| SPGP0CZ02830 | XP_018221102.1 | 99 | 0 |
| SPGP0H00410 | XP_018224095.1 | 99 | 0 |
| SPGP0U01530 | XP_018218841.1 | 99 | 0 |
| SPGP0S02090 | XP_018220143.1 | 99 | 0 |
| SPGP0R02070 | XP_018221322.1 | 99 | 0 |
| SPGP0S00350 | XP_018220310.1 | 99 | 0 |
| SPGP0E00500 | XP_018222640.1 | 99 | 0 |
| SPGP0W02750 | XP_018223064.1 | 99 | 0 |
| SPGP0Z03170 | XP_018219569.1 | 99 | 0 |
| SPGP0W00520 | XP_018223282.1 | 99 | 0 |
| SPGP0AF00490 | XP_018221990.1 | 99 | 0 |
| SPGP0AH00510 | XP_018222554.1 | 99 | 0 |
| SPGP0DU01190 | XP_018220471.1 | 99 | 0 |
| SPGP0B00850 | XP_018221532.1 | 99 | 0 |
| SPGP0Z01420 | XP_018219743.1 | 99 | 0 |
| SPGP0DZ01400 | XP_018219479.1 | 99 | 0 |
| SPGP0DN01040 | XP_018223551.1 | 99 | 0 |
| SPGP0N00800 | XP_018220729.1 | 99 | 0 |
| SPGP0B01800 | XP_018221622.1 | 98 | 1 |
| SPGP0U01350 | XP_018218858.1 | 99 | 0 |
| SPGP0L00330 | XP_018219328.1 | 99 | 0 |

Create PDF in your applications with the Pdfcrowd HTML to PDF API PDFCROWD

| | | | |
|---|---|---|---|
| SPGP0DU00400 | XP_018220550.1 | 93 | 6 |
| SPGP0DN00260 | XP_018223476.1 | 100 | 0 |
| SPGP0Z00320 | XP_018219848.1 | 99 | 0 |
| SPGP0DN00850 | XP_018223531.1 | 99 | 0 |
| SPGP0N01740 | XP_018220640.1 | 99 | 0 |
| SPGP0AH00280 | XP_018222574.1 | 99 | 0 |
| SPGP0DN00420 | XP_018223491.1 | 99 | 0 |
| SPGP0Q00940 | XP_018222407.1 | 99 | 0 |
| SPGP0N00590 | XP_018220749.1 | 99 | 0 |
| SPGP0Y01400 | XP_018224009.1 | 99 | 0 |
| SPGP0Q02750 | XP_018222225.1 | 99 | 0 |
| SPGP0AA00100 | XP_018223325.1 | 99 | 0 |
| SPGP0P00550 | XP_018221901.1 | 99 | 0 |
| SPGP0X00660 | XP_018219036.1 | 99 | 0 |
| SPGP0P02950 | XP_018221669.1 | 99 | 0 |
| SPGP0DU00660 | XP_018220525.1 | 99 | 0 |
| SPGP0P02670 | XP_018221696.1 | 99 | 0 |
| SPGP0Z00170 | XP_018219866.1 | 99 | 0 |
| SPGP0B01670 | XP_018221609.1 | 99 | 0 |
| SPGP0Z02220 | XP_018219663.1 | 99 | 0 |
| SPGP0B01860 | XP_018221628.1 | 99 | 1 |
| SPGP0S02010 | XP_018220151.1 | 99 | 0 |
| SPGP0Q02810 | XP_018222220.1 | 99 | 0 |
| SPGP0X00550 | XP_018219047.1 | 99 | 0 |
| SPGP0DZ01980 | XP_018219534.1 | 100 | 0 |
| SPGP0EE00230 | XP_018220841.1 | 99 | 0 |
| SPGP0DU01350 | XP_018220455.1 | 99 | 0 |
| SPGP0Q01320 | XP_018222368.1 | 99 | 0 |
| SPGP0S01440 | XP_018220204.1 | 99 | 0 |
| SPGP0P02930 | XP_018221671.1 | 99 | 0 |
| SPGP0CZ00970 | XP_018220922.1 | 99 | 0 |
| SPGP0Ztrna16P | DI49_4699 | 100 | 0 |
| SPGP0DU01130 | XP_018220477.1 | 99 | 0 |
| SPGP0P01730 | XP_018221787.1 | 99 | 0 |
| SPGP0W03970 | XP_018222952.1 | 99 | 0 |
| SPGP0S03300 | XP_018220025.1 | 99 | 0 |
| SPGP0S01620 | XP_018220188.1 | 99 | 0 |
| SPGP0W03930 | XP_018222956.1 | 99 | 0 |
| SPGP0L01520 | XP_018219212.1 | 99 | 0 |
| SPGP0L01030 | XP_018219260.1 | 99 | 0 |

Create PDF in your applications with the Pdfcrowd HTML to PDF API PDFCROWD

| | | | |
|---|---|---|---|
| SPGP0U01370 | XP_018218856.1 | 99 | 0 |
| SPGP0Q01700 | XP_018222328.1 | 99 | 0 |
| SPGP0R00410 | XP_018221164.1 | 99 | 0 |
| SPGP0DU00820 | XP_018220508.1 | 98 | 0 |
| SPGP0Q03040 | XP_018222197.1 | 98 | 0 |
| SPGP0P02040 | XP_018221756.1 | 99 | 0 |
| SPGP0S00670 | XP_018220278.1 | 100 | 0 |
| SPGP0DN03810 | XP_018223823.1 | 99 | 0 |
| SPGP0Z02950 | XP_018219590.1 | 99 | 0 |
| SPGP0Z00660 | XP_018219818.1 | 99 | 0 |
| SPGP0X00730 | XP_018219029.1 | 99 | 1 |
| SPGP0R01150 | XP_018221233.1 | 99 | 0 |
| SPGP0Q01560 | XP_018222342.1 | 99 | 0 |
| SPGP0L00920 | XP_018219271.1 | 99 | 0 |
| SPGP0R00820 | XP_018221202.1 | 99 | 0 |
| SPGP0Q03990 | XP_018222106.1 | 100 | 0 |
| SPGP0B01410 | XP_018221584.1 | 99 | 0 |
| SPGP0E02350 | XP_018222823.1 | 99 | 0 |
| SPGP0AC00620 | XP_018219146.1 | 99 | 0 |
| SPGP0B01120 | XP_018221557.1 | 99 | 0 |
| SPGP0Q03700 | XP_018222135.1 | 99 | 0 |
| SPGP0W04820 | XP_018222866.1 | 99 | 0 |
| SPGP0DN01100 | XP_018223557.1 | 100 | 0 |
| SPGP0E01880 | XP_018222776.1 | 99 | 1 |
| SPGP0AG00100 | XP_018219930.1 | 97 | 0 |
| SPGP0S00100 | XP_018220334.1 | 99 | 0 |
| SPGP0S00540 | XP_018220291.1 | 99 | 0 |
| SPGP0U01310 | XP_018218862.1 | 99 | 0 |
| SPGP0P02720 | XP_018221691.1 | 99 | 0 |
| SPGP0L01600 | XP_018219205.1 | 99 | 0 |
| SPGP0CZ01830 | XP_018221005.1 | 99 | 0 |
| SPGP0W01420 | XP_018223192.1 | 99 | 0 |
| SPGP0DN01500 | XP_018223597.1 | 99 | 0 |
| SPGP0E01020 | XP_018222692.1 | 99 | 0 |
| SPGP0Z01250 | XP_018219760.1 | 99 | 0 |
| SPGP0B01330 | XP_018221576.1 | 99 | 0 |
| SPGP0W02180 | XP_018223119.1 | 99 | 0 |
| SPGP0DN01060 | XP_018223553.1 | 99 | 0 |
| SPGP0S03360 | XP_018220019.1 | 99 | 0 |
| SPGP0DN00900 | XP_018223536.1 | 99 | 0 |

Create PDF in your applications with the Pdfcrowd HTML to PDF API PDFCROWD

| | | | |
|---|---|---|---|
| SPGP0DN02880 | XP_018223733.1 | 99 | 0 |
| SPGP0Ptrna7F | DI49_2475 | 100 | 0 |
| SPGP0CZ02140 | XP_018221034.1 | 100 | 0 |
| SPGP0EL00100 | XP_018219093.1 | 99 | 0 |
| SPGP0F04790 | XP_018222015.1 | 99 | 0 |
| SPGP0Z02520 | XP_018219633.1 | 99 | 0 |
| SPGP0P01890 | XP_018221771.1 | 97 | 0 |
| SPGP0N00280 | XP_018220780.1 | 99 | 0 |
| SPGP0Z01620 | XP_018219723.1 | 99 | 1 |
| SPGP0AC00230 | XP_018219108.1 | 99 | 0 |
| SPGP0W04580 | XP_018222890.1 | 99 | 0 |
| SPGP0DN00670 | XP_018223513.1 | 99 | 0 |
| SPGP0Z00430 | XP_018219840.1 | 99 | 0 |
| SPGP0P02140 | XP_018221746.1 | 99 | 0 |
| SPGP0Q03540 | XP_018222150.1 | 99 | 0 |
| SPGP0E00980 | XP_018222688.1 | 99 | 0 |
| SPGP0P00760 | XP_018221880.1 | 99 | 0 |
| SPGP0N00290 | XP_018220779.1 | 99 | 0 |
| SPGP0B01730 | XP_018221615.1 | 100 | 0 |
| SPGP0R02790 | XP_018221391.1 | 99 | 0 |
| SPGP0S01120 | XP_018220233.1 | 98 | 1 |
| SPGP0H00350 | XP_018224100.1 | 99 | 0 |
| SPGP0X00850 | XP_018219017.1 | 99 | 0 |
| SPGP0Y00610 | XP_018223935.1 | 99 | 0 |
| SPGP0U01360 | XP_018218857.1 | 99 | 0 |
| SPGP0Z02420 | XP_018219643.1 | 99 | 0 |
| SPGP0P02540 | XP_018221709.1 | 99 | 0 |
| SPGP0W03030 | XP_018223039.1 | 98 | 1 |
| SPGP0H00250 | XP_018224110.1 | 99 | 0 |
| SPGP0DN03370 | XP_018223780.1 | 99 | 0 |
| SPGP0Y01750 | XP_018224042.1 | 98 | 1 |
| SPGP0DZ01470 | XP_018219486.1 | 99 | 0 |
| SPGP0CZ01940 | XP_018221015.1 | 99 | 0 |
| SPGP0B01710 | XP_018221613.1 | 99 | 0 |
| SPGP0DN02490 | XP_018223694.1 | 99 | 0 |
| SPGP0DV00470 | XP_018219909.1 | 99 | 0 |
| SPGP0U00400 | XP_018218949.1 | 99 | 0 |
| SPGP0Q04280 | XP_018222078.1 | 98 | 2 |
| SPGP0R01700 | XP_018221286.1 | 99 | 0 |
| SPGP0W00210 | XP_018223313.1 | 99 | 0 |

Create PDF in your applications with the Pdfcrowd HTML to PDF API PDFCROWD

| | | | |
|---|---|---|---|
| SPGP0L00800 | XP_018219283.1 | 99 | 0 |
| SPGP0L00370 | XP_018219324.1 | 99 | 0 |
| SPGP0AA00850 | XP_018223394.1 | 99 | 0 |
| SPGP0R02610 | XP_018221374.1 | 99 | 0 |
| SPGP0B00990 | XP_018221545.1 | 99 | 0 |
| SPGP0R01910 | XP_018221307.1 | 99 | 0 |
| SPGP0DN03590 | XP_018223801.1 | 99 | 0 |
| SPGP0W01160 | XP_018223218.1 | 99 | 0 |
| SPGP0Z00410 | XP_018219857.1 | 98 | 0 |
| SPGP0S01190 | XP_018220226.1 | 99 | 0 |
| SPGP0W01750 | XP_018223160.1 | 99 | 0 |
| SPGP0R02020 | XP_018221317.1 | 99 | 0 |
| SPGP0DN02420 | XP_018223687.1 | 99 | 0 |
| SPGP0DN02350 | XP_018223680.1 | 99 | 0 |
| SPGP0CZ02640 | XP_018221084.1 | 99 | 0 |
| SPGP0S03170 | XP_018220037.1 | 99 | 0 |
| SPGP0B00190 | XP_018221467.1 | 97 | 0 |
| SPGP0DZ01750 | XP_018219513.1 | 99 | 0 |
| SPGP0W01300 | XP_018223204.1 | 99 | 0 |
| SPGP0S01990 | XP_018220153.1 | 99 | 0 |
| SPGP0P02450 | XP_018221718.1 | 99 | 0 |
| SPGP0S02560 | XP_018220097.1 | 99 | 0 |
| SPGP0Q01910 | XP_018222307.1 | 99 | 0 |
| SPGP0B00350 | XP_018221483.1 | 99 | 0 |
| SPGP0R02520 | XP_018221365.1 | 99 | 0 |
| SPGP0Z00740 | XP_018219810.1 | 99 | 0 |
| SPGP0DN01330 | XP_018223580.1 | 99 | 0 |
| SPGP0DN01990 | XP_018223646.1 | 99 | 0 |
| SPGP0Q03820 | XP_018222123.1 | 99 | 0 |
| SPGP0Q02540 | XP_018222246.1 | 99 | 0 |
| SPGP0P03010 | XP_018221663.1 | 99 | 0 |
| SPGP0E02280 | XP_018222816.1 | 99 | 0 |
| SPGP0X00970 | XP_018219006.1 | 99 | 0 |
| SPGP0U00640 | XP_018218925.1 | 100 | 0 |
| SPGP0X00800 | XP_018219022.1 | 99 | 0 |
| SPGP0S00480 | XP_018220297.1 | 99 | 0 |
| SPGP0S00730 | XP_018220272.1 | 99 | 0 |
| SPGP0Etrna25R | DI49_4323 | 100 | 0 |
| SPGP0Y00400 | XP_018223916.1 | 99 | 0 |
| SPGP0CZ02510 | XP_018221071.1 | 99 | 0 |

Create PDF in your applications with the Pdfcrowd HTML to PDF API PDFCROWD

| | | | |
|---|---|---|---|
| SPGP0Q00960 | XP_018222405.1 | 99 | 0 |
| SPGP0FT00140 | XP_018220803.1 | 99 | 0 |
| SPGP0P00570 | XP_018221899.1 | 99 | 0 |
| SPGP0DZ00110 | XP_018219354.1 | 99 | 0 |
| SPGP0Q00790 | XP_018222422.1 | 99 | 0 |
| SPGP0U01110 | XP_018218881.1 | 99 | 0 |
| SPGP0CZ00830 | XP_018220908.1 | 100 | 0 |
| SPGP0DV00340 | XP_018219896.1 | 99 | 0 |
| SPGP0BF00330 | XP_018220401.1 | 99 | 0 |
| SPGP0W04430 | XP_018222905.1 | 99 | 0 |
| SPGP0Q02830 | XP_018222218.1 | 99 | 0 |
| SPGP0AA00380 | XP_018223350.1 | 99 | 0 |
| SPGP0CZ00950 | XP_018220920.1 | 99 | 0 |
| SPGP0DN01800 | XP_018223626.1 | 99 | 0 |
| SPGP0Q03080 | XP_018222193.1 | 99 | 0 |
| SPGP0Q00340 | XP_018222467.1 | 99 | 0 |
| SPGP0W00970 | XP_018223237.1 | 99 | 0 |
| SPGP0CZ01090 | XP_018220934.1 | 99 | 0 |
| SPGP0Z03100 | XP_018219575.1 | 99 | 0 |
| SPGP0DN03770 | XP_018223819.1 | 99 | 0 |
| SPGP0P01710 | XP_018221789.1 | 99 | 0 |
| SPGP0Ztrna14F | DI49_4603 | 99 | 0 |
| SPGP0Z02640 | XP_018219621.1 | 99 | 0 |
| SPGP0P00540 | XP_018221902.1 | 99 | 0 |
| SPGP0L01010 | XP_018219262.1 | 99 | 0 |
| SPGP0L01540 | XP_018219210.1 | 96 | 2 |
| SPGP0S02140 | XP_018220138.1 | 99 | 0 |
| SPGP0E00220 | XP_018222612.1 | 99 | 0 |
| SPGP0S01050 | XP_018220240.1 | 99 | 0 |
| SPGP0DN03120 | XP_018223756.1 | 99 | 0 |
| SPGP0Z00360 | XP_018219852.1 | 99 | 0 |
| SPGP0W02310 | XP_018223106.1 | 99 | 0 |
| SPGP0Q04060 | XP_018222099.1 | 99 | 0 |
| SPGP0Z01840 | XP_018219701.1 | 97 | 2 |
| SPGP0Q01690 | XP_018222329.1 | 99 | 0 |
| SPGP0W03950 | XP_018222954.1 | 99 | 0 |
| SPGP0E02090 | XP_018222797.1 | 100 | 0 |
| SPGP0Q03870 | XP_018222118.1 | 97 | 2 |
| SPGP0DNtrna4F | DI49_0495 | 100 | 0 |
| SPGP0E00310 | XP_018222621.1 | 99 | 0 |

Create PDF in your applications with the Pdfcrowd HTML to PDF API PDFCROWD

| | | | |
|---|---|---|---|
| SPGP0DZ02240 | XP_018219561.1 | 99 | 0 |
| SPGP0Q01580 | XP_018222340.1 | 99 | 0 |
| SPGP0AH00240 | XP_018222578.1 | 99 | 0 |
| SPGP0P02060 | XP_018221754.1 | 99 | 0 |
| SPGP0Q02980 | XP_018222203.1 | 99 | 0 |
| SPGP0Y01060 | XP_018223976.1 | 99 | 0 |
| SPGP0DV00360 | XP_018219898.1 | 99 | 0 |
| SPGP0CZ01280 | XP_018220953.1 | 99 | 0 |
| SPGP0Q02580 | XP_018222242.1 | 95 | 1 |
| SPGP0S02000 | XP_018220152.1 | 99 | 0 |
| SPGP0Z02310 | XP_018219654.1 | 99 | 0 |
| SPGP0E01400 | XP_018222729.1 | 99 | 0 |
| SPGP0W03470 | XP_018223000.1 | 99 | 0 |
| SPGP0B01650 | XP_018221607.1 | 99 | 0 |
| SPGP0DN00790 | XP_018223525.1 | 99 | 0 |
| SPGP0Z01600 | XP_018219725.1 | 99 | 0 |
| SPGP0Q02050 | XP_018222293.1 | 99 | 0 |
| SPGP0Q04080 | XP_018222097.1 | 99 | 0 |
| SPGP0E00200 | XP_018222610.1 | 99 | 0 |
| SPGP0W00900 | XP_018223244.1 | 99 | 0 |
| SPGP0R01680 | XP_018221285.1 | 99 | 0 |
| SPGP0CZ02570 | XP_018221077.1 | 98 | 2 |
| SPGP0Q04180 | XP_018222087.1 | 99 | 0 |
| SPGP0W01440 | XP_018223190.1 | 99 | 0 |
| SPGP0DN01070 | XP_018223554.1 | 99 | 0 |
| SPGP0E01000 | XP_018222690.1 | 99 | 0 |
| SPGP0S02350 | XP_018220117.1 | 99 | 0 |
| SPGP0S00860 | XP_018220259.1 | 99 | 0 |
| SPGP0DN02300 | XP_018223675.1 | 99 | 0 |
| SPGP0W02230 | XP_018223114.1 | 99 | 0 |
| SPGP0Q01710 | XP_018222327.1 | 99 | 0 |
| SPGP0P02730 | XP_018221690.1 | 99 | 0 |
| SPGP0P00290 | XP_018221925.1 | 99 | 0 |
| SPGP0Q02790 | XP_018222222.1 | 99 | 0 |
| SPGP0Q00860 | XP_018222415.1 | 99 | 0 |
| SPGP0P00690 | XP_018221887.1 | 100 | 0 |
| SPGP0Z01720 | XP_018219713.1 | 99 | 0 |
| SPGP0Z01240 | XP_018219761.1 | 99 | 0 |
| SPGP0DN00650 | XP_018223511.1 | 99 | 0 |
| SPGP0DN00750 | XP_018223521.1 | 99 | 0 |

Create PDF in your applications with the Pdfcrowd HTML to PDF API PDFCROWD

| | | | |
|---|---|---|---|
| SPGP0DZ02170 | XP_018219554.1 | 99 | 0 |
| SPGP0S00950 | XP_018220250.1 | 99 | 0 |
| SPGP0H00550 | XP_018224081.1 | 99 | 0 |
| SPGP0F04830 | XP_018222011.1 | 99 | 0 |
| SPGP0Q02160 | XP_018222282.1 | 99 | 0 |
| SPGP0B00820 | XP_018221529.1 | 99 | 0 |
| SPGP0R02240 | XP_018221337.1 | 99 | 0 |
| SPGP0Z02120 | XP_018219673.1 | 99 | 0 |
| SPGP0DZ01480 | XP_018219487.1 | 99 | 0 |
| SPGP0E00830 | XP_018222671.1 | 99 | 0 |
| SPGP0AA00440 | XP_018223356.1 | 99 | 0 |
| SPGP0Y00630 | XP_018223937.1 | 99 | 0 |
| SPGP0Q01570 | XP_018222341.1 | 99 | 0 |
| SPGP0E02420 | XP_018222829.1 | 99 | 1 |
| SPGP0R01620 | XP_018221279.1 | 99 | 0 |
| SPGP0P02520 | XP_018221711.1 | 99 | 0 |
| SPGP0U01850 | XP_018218809.1 | 99 | 0 |
| SPGP0W02360 | XP_018223101.1 | 99 | 0 |
| SPGP0R01280 | XP_018221245.1 | 99 | 0 |
| SPGP0W02280 | XP_018223109.1 | 99 | 0 |
| SPGP0DN03350 | XP_018223778.1 | 99 | 0 |
| SPGP0L01480 | XP_018219216.1 | 99 | 0 |
| SPGP0DZ02030 | XP_018219540.1 | 99 | 0 |
| SPGP0DX00230 | XP_018219954.1 | 99 | 0 |
| SPGP0Z01430 | XP_018219742.1 | 100 | 0 |
| SPGP0P03090 | XP_018221655.1 | 99 | 0 |
| SPGP0S00220 | XP_018220322.1 | 99 | 0 |
| SPGP0L00570 | XP_018219306.1 | 99 | 0 |
| SPGP0DZ00890 | XP_018219429.1 | 99 | 0 |
| SPGP0Q02090 | XP_018222289.1 | 99 | 0 |
| SPGP0L00680 | XP_018219295.1 | 99 | 0 |
| SPGP0Z02870 | XP_018219597.1 | 100 | 0 |
| SPGP0DV00500 | XP_018219912.1 | 99 | 0 |
| SPGP0AA00640 | XP_018223376.1 | 99 | 0 |
| SPGP0AGtrna27T | DI49_2415 | 100 | 0 |
| SPGP0CZ01520 | XP_018220976.1 | 98 | 1 |
| SPGP0Z03050 | XP_018219580.1 | 99 | 0 |
| SPGP0W04530 | XP_018222895.1 | 99 | 0 |
| SPGP0R01720 | XP_018221288.1 | 99 | 0 |
| SPGP0S02990 | XP_018220055.1 | 99 | 0 |

Create PDF in your applications with the Pdfcrowd HTML to PDF API PDFCROWD

| | | | |
|---|---|---|---|
| SPGP0E01580 | XP_018222746.1 | 100 | 0 |
| SPGP0CZ02760 | XP_018221095.1 | 99 | 0 |
| SPGP0ltrna35R | DI49_2410 | 100 | 0 |
| SPGP0R03260 | XP_018221436.1 | 99 | 0 |
| SPGP0B01050 | XP_018221551.1 | 99 | 0 |
| SPGP0E01540 | XP_018222742.1 | 99 | 0 |
| SPGP0S03350 | XP_018220020.1 | 99 | 0 |
| SPGP0Q01930 | XP_018222305.1 | 99 | 0 |
| SPGP0S02500 | XP_018220103.1 | 99 | 0 |
| SPGP0Y00970 | XP_018223967.1 | 99 | 0 |
| SPGP0E00650 | XP_018222655.1 | 99 | 0 |
| SPGP0CZ02090 | XP_018221029.1 | 99 | 0 |
| SPGP0CZ00240 | XP_018220852.1 | 99 | 0 |
| SPGP0Z00150 | XP_018219868.1 | 99 | 0 |
| SPGP0DZ01560 | XP_018219495.1 | 99 | 0 |
| SPGP0S02550 | XP_018220098.1 | 99 | 0 |
| SPGP0DN00660 | XP_018223512.1 | 99 | 0 |
| SPGP0R01600 | XP_018221277.1 | 99 | 0 |
| SPGP0S01660 | XP_018220185.1 | 99 | 0 |
| SPGP0DN04020 | XP_018223844.1 | 99 | 0 |
| SPGP0P01750 | XP_018221785.1 | 99 | 0 |
| SPGP0F04870 | XP_018222007.1 | 99 | 0 |
| SPGP0E02520 | XP_018222838.1 | 99 | 0 |
| SPGP0CZ02880 | XP_018221107.1 | 99 | 0 |
| SPGP0DN01280 | XP_018223575.1 | 99 | 0 |
| SPGP0DZ00300 | XP_018219373.1 | 99 | 0 |
| SPGP0E00540 | XP_018222644.1 | 99 | 0 |
| SPGP0B01840 | XP_018221626.1 | 98 | 0 |
| SPGP0Z00920 | XP_018219792.1 | 99 | 0 |
| SPGP0DN00950 | XP_018223541.1 | 99 | 0 |
| SPGP0E01830 | XP_018222771.1 | 99 | 0 |
| SPGP0N00170 | XP_018220791.1 | 99 | 0 |
| SPGP0T00390 | XP_018220368.1 | 99 | 0 |
| SPGP0AA00630 | XP_018223375.1 | 99 | 0 |
| SPGP0E01760 | XP_018222764.1 | 99 | 0 |
| SPGP0Z02320 | XP_018219653.1 | 99 | 0 |
| SPGP0Y01160 | XP_018223985.1 | 99 | 0 |
| SPGP0W03760 | XP_018222972.1 | 99 | 0 |
| SPGP0Q02850 | XP_018222216.1 | 99 | 0 |
| SPGP0DU01120 | XP_018220478.1 | 99 | 0 |

Create PDF in your applications with the Pdfcrowd HTML to PDF API PDFCROWD

| | | | |
|---|---|---|---|
| SPGP0U01130 | XP_018218879.1 | 99 | 0 |
| SPGP0Q01190 | XP_018222381.1 | 99 | 0 |
| SPGP0B00860 | XP_018221533.1 | 99 | 0 |
| SPGP0P00890 | XP_018221868.1 | 99 | 0 |
| SPGP0AF00140 | XP_018221955.1 | 99 | 0 |
| SPGP0CZ00850 | XP_018220910.1 | 99 | 0 |
| SPGP0N01550 | XP_018220659.1 | 99 | 0 |
| SPGP0S03230 | XP_018220032.1 | 99 | 0 |
| SPGP0X00950 | XP_018219008.1 | 99 | 0 |
| SPGP0R02350 | XP_018221348.1 | 99 | 0 |
| SPGP0P02590 | XP_018221704.1 | 99 | 0 |
| SPGP0W00300 | XP_018223304.1 | 98 | 1 |
| SPGP0DV00190 | XP_018219883.1 | 99 | 0 |
| SPGP0CZ00550 | XP_018220881.1 | 99 | 0 |
| SPGP0S01130 | XP_018220232.1 | 99 | 0 |
| SPGP0W02550 | XP_018223082.1 | 99 | 0 |
| SPGP0CZ02970 | XP_018221116.1 | 99 | 0 |
| SPGP0S01420 | XP_018220206.1 | 99 | 0 |
| SPGP0S01400 | XP_018220208.1 | 99 | 0 |
| SPGP0P00780 | XP_018221878.1 | 99 | 0 |
| SPGP0DN01690 | XP_018223615.1 | 99 | 0 |
| SPGP0P01770 | XP_018221783.1 | 99 | 0 |
| SPGP0Q01900 | XP_018222308.1 | 99 | 0 |
| SPGP0P00110 | XP_018221943.1 | 99 | 0 |
| SPGP0DN02070 | XP_018223654.1 | 99 | 0 |
| SPGP0AC01000 | XP_018219183.1 | 99 | 0 |
| SPGP0Q02650 | XP_018222235.1 | 99 | 0 |
| SPGP0H00750 | XP_018224062.1 | 99 | 0 |
| SPGP0L01560 | XP_018219208.1 | 99 | 0 |
| SPGP0S03250 | XP_018220030.1 | 99 | 0 |
| SPGP0B01690 | XP_018221611.1 | 99 | 0 |
| SPGP0E02490 | XP_018222836.1 | 99 | 0 |
| SPGP0S03160 | XP_018220038.1 | 99 | 0 |
| SPGP0S01070 | XP_018220238.1 | 99 | 0 |
| SPGP0W04180 | XP_018222932.1 | 94 | 0 |
| SPGP0X01000 | XP_018219003.1 | 99 | 0 |
| SPGP0Q01740 | XP_018222324.1 | 99 | 0 |
| SPGP0E01250 | XP_018222715.1 | 100 | 0 |
| SPGP0Z01580 | XP_018219727.1 | 99 | 0 |
| SPGP0P02880 | XP_018221676.1 | 99 | 0 |

Create PDF in your applications with the Pdfcrowd HTML to PDF API PDFCROWD

| | | | |
|---|---|---|---|
| SPGP0W03750 | XP_018222973.1 | 99 | 0 |
| SPGP0DN03720 | XP_018223814.1 | 99 | 0 |
| SPGP0E01970 | XP_018222785.1 | 99 | 0 |
| SPGP0Q03890 | XP_018222116.1 | 99 | 0 |
| SPGP0B02010 | XP_018221643.1 | 99 | 0 |
| SPGP0CZ01510 | XP_018220975.1 | 99 | 0 |
| SPGP0P00820 | XP_018221875.1 | 99 | 0 |
| SPGP0R01440 | XP_018221261.1 | 99 | 0 |
| SPGP0W04760 | XP_018222872.1 | 99 | 0 |
| SPGP0R01110 | XP_018221229.1 | 99 | 0 |
| SPGP0W00400 | XP_018223294.1 | 99 | 0 |
| SPGP0P02080 | XP_018221752.1 | 99 | 0 |
| SPGP0L00960 | XP_018219267.1 | 99 | 0 |
| SPGP0AH01150 | XP_018222493.1 | 99 | 0 |
| SPGP0AF00330 | XP_018221974.1 | 99 | 0 |
| SPGP0P01670 | XP_018221793.1 | 93 | 0 |
| SPGP0DZ00280 | XP_018219371.1 | 99 | 0 |
| SPGP0DN01480 | XP_018223595.1 | 99 | 0 |
| SPGP0DN03650 | XP_018223807.1 | 99 | 0 |
| SPGP0R00240 | XP_018221147.1 | 99 | 0 |
| SPGP0L00420 | XP_018219319.1 | 99 | 0 |
| SPGP0P01420 | XP_018221816.1 | 99 | 0 |
| SPGP0Q02070 | XP_018222291.1 | 99 | 0 |
| SPGP0W00740 | XP_018223259.1 | 99 | 0 |
| SPGP0DN01270 | XP_018223574.1 | 99 | 0 |
| SPGP0R00550 | XP_018221178.1 | 98 | 0 |
| SPGP0T00330 | XP_018220361.1 | 99 | 0 |
| SPGP0R03110 | XP_018221421.1 | 99 | 0 |
| SPGP0S02390 | XP_018220113.1 | 99 | 0 |
| SPGP0Z02800 | XP_018219604.1 | 99 | 0 |
| SPGP0Q01340 | XP_018222365.1 | 99 | 0 |
| SPGP0DN00520 | XP_018223500.1 | 99 | 0 |
| SPGP0L00360 | XP_018219325.1 | 99 | 0 |
| SPGP0Q03300 | XP_018222172.1 | 99 | 0 |
| SPGP0DN04190 | XP_018223861.1 | 99 | 0 |
| SPGP0DZ01870 | XP_018219525.1 | 99 | 0 |
| SPGP0Z01470 | XP_018219738.1 | 99 | 0 |
| SPGP0Q00510 | XP_018222450.1 | 99 | 0 |
| SPGP0Y00660 | XP_018223940.1 | 99 | 0 |

Create PDF in your applications with the Pdfcrowd HTML to PDF API PDFCROWD

| | | | |
|---|---|---|---|
| SPGP0E02320 | XP_018222820.1 | 99 | 0 |
| SPGP0DN03920 | XP_018223834.1 | 99 | 0 |
| SPGP0AF00200 | XP_018221961.1 | 99 | 0 |
| SPGP0X00320 | XP_018219069.1 | 99 | 0 |
| SPGP0U00290 | XP_018218960.1 | 99 | 0 |
| SPGP0Q01750 | XP_018222323.1 | 99 | 0 |
| SPGP0AA00280 | XP_018223341.1 | 99 | 0 |
| SPGP0DV00540 | XP_018219916.1 | 99 | 0 |
| SPGP0E01390 | XP_018222728.1 | 99 | 0 |
| SPGP0AA00420 | XP_018223354.1 | 99 | 0 |
| SPGP0S00570 | XP_018220288.1 | 99 | 0 |
| SPGP0E02080 | XP_018222796.1 | 99 | 0 |
| SPGP0R01870 | XP_018221303.1 | 99 | 0 |
| SPGP0Z01800 | XP_018219705.1 | 99 | 0 |
| SPGP0W02960 | XP_018223044.1 | 99 | 0 |
| SPGP0F04540 | XP_018222037.1 | 99 | 0 |
| SPGP0AA00460 | XP_018223358.1 | 99 | 0 |
| SPGP0H00460 | XP_018224090.1 | 99 | 0 |
| SPGP0DX00100 | XP_018219941.1 | 99 | 0 |
| SPGP0S00200 | XP_018220324.1 | 99 | 0 |
| SPGP0W01370 | XP_018223197.1 | 99 | 0 |
| SPGP0CZ01480 | XP_018220972.1 | 100 | 0 |
| SPGP0S00240 | XP_018220320.1 | 99 | 0 |
| SPGP0Z02850 | XP_018219599.1 | 99 | 0 |
| SPGP0W00310 | XP_018223303.1 | 99 | 0 |
| SPGP0DZ01130 | XP_018219452.1 | 99 | 0 |
| SPGP0L01290 | XP_018219235.1 | 98 | 2 |
| SPGP0W02610 | XP_018223076.1 | 99 | 0 |
| SPGP0Q02520 | XP_018222247.1 | 99 | 0 |
| SPGP0S01080 | XP_018220237.1 | 99 | 0 |
| SPGP0DU00610 | XP_018220529.1 | 99 | 0 |
| SPGP0W01480 | XP_018223187.1 | 100 | 0 |
| SPGP0W04510 | XP_018222897.1 | 99 | 0 |
| SPGP0W00870 | XP_018223247.1 | 99 | 0 |
| SPGP0P01690 | XP_018221791.1 | 99 | 0 |
| SPGP0Y00620 | XP_018223936.1 | 99 | 0 |
| SPGP0R02650 | XP_018221377.1 | 99 | 0 |
| SPGP0B00400 | XP_018221488.1 | 99 | 0 |
| SPGP0X00840 | XP_018219018.1 | 99 | 0 |
| SPGP0CZ02740 | XP_018221094.1 | 99 | 0 |

Create PDF in your applications with the Pdfcrowd HTML to PDF API PDFCROWD

| | | | |
|---|---|---|---|
| SPGP0Q03930 | XP_018222112.1 | 99 | 0 |
| SPGP0Z00230 | XP_018219861.1 | 99 | 0 |
| SPGP0E02030 | XP_018222791.1 | 99 | 0 |
| SPGP0CZ02540 | XP_018221074.1 | 99 | 0 |
| SPGP0B00490 | XP_018221497.1 | 99 | 0 |
| SPGP0DN01570 | XP_018223604.1 | 100 | 0 |
| SPGP0P00580 | XP_018221898.1 | 99 | 1 |
| SPGP0Q01260 | XP_018222374.1 | 99 | 0 |
| SPGP0BC00200 | XP_018220817.1 | 99 | 0 |
| SPGP0S02760 | XP_018220077.1 | 99 | 0 |
| SPGP0DZ01090 | XP_018219448.1 | 99 | 0 |
| SPGP0W02380 | XP_018223099.1 | 99 | 0 |
| SPGP0W01120 | XP_018223222.1 | 99 | 0 |
| SPGP0N02000 | XP_018220614.1 | 99 | 0 |
| SPGP0N01520 | XP_018220662.1 | 99 | 0 |
| SPGP0S03430 | XP_018220013.1 | 99 | 0 |
| SPGP0DN03760 | XP_018223818.1 | 99 | 0 |
| SPGP0DN00230 | XP_018223473.1 | 99 | 0 |
| SPGP0L01390 | XP_018219225.1 | 99 | 0 |
| SPGP0AH00700 | XP_018222535.1 | 99 | 0 |
| SPGP0S01560 | XP_018220194.1 | 99 | 0 |
| SPGP0W02390 | XP_018223098.1 | 99 | 0 |
| SPGP0B01810 | XP_018221623.1 | 98 | 2 |
| SPGP0R02910 | XP_018221402.1 | 99 | 0 |
| SPGP0AC00180 | XP_018219103.1 | 99 | 0 |
| SPGP0B00810 | XP_018221528.1 | 99 | 1 |
| SPGP0Q03200 | XP_018222182.1 | 99 | 0 |
| SPGP0AF00350 | XP_018221976.1 | 100 | 0 |
| SPGP0W04750 | XP_018222873.1 | 99 | 0 |
| SPGP0Z03140 | XP_018219571.1 | 99 | 0 |
| SPGP0W01710 | XP_018223164.1 | 99 | 0 |
| SPGP0N01250 | XP_018220688.1 | 99 | 0 |
| SPGP0H00380 | XP_018224097.1 | 99 | 0 |
| SPGP0DN02980 | XP_018223743.1 | 99 | 0 |
| SPGP0E01210 | XP_018222711.1 | 99 | 1 |
| SPGP0Q02870 | XP_018222214.1 | 99 | 0 |
| SPGP0Z00850 | XP_018219799.1 | 99 | 0 |
| SPGP0DN01730 | XP_018223619.1 | 99 | 0 |
| SPGP0CZ02280 | XP_018221048.1 | 98 | 0 |
| SPGP0R03000 | XP_018221410.1 | 99 | 0 |

Create PDF in your applications with the Pdfcrowd HTML to PDF API PDFCROWD

| | | | |
|---|---|---|---|
| SPGP0DN00280 | XP_018223478.1 | 99 | 0 |
| SPGP0Z02240 | XP_018219661.1 | 99 | 0 |
| SPGP0W04850 | XP_018222863.1 | 99 | 0 |
| SPGP0F04340 | XP_018222057.1 | 99 | 0 |
| SPGP0E01890 | XP_018222777.1 | 99 | 0 |
| SPGP0Q00650 | XP_018222436.1 | 99 | 0 |
| SPGP0Y01430 | XP_018224012.1 | 99 | 0 |
| SPGP0DU01370 | XP_018220453.1 | 99 | 0 |
| SPGP0Y01590 | XP_018224028.1 | 100 | 0 |
| SPGP0R00590 | XP_018221181.1 | 99 | 0 |
| SPGP0DZ01300 | XP_018219469.1 | 99 | 0 |
| SPGP0BF00280 | XP_018220397.1 | 99 | 0 |
| SPGP0AA00870 | XP_018223396.1 | 100 | 0 |
| SPGP0W00810 | XP_018223253.1 | 99 | 0 |
| SPGP0DU00470 | XP_018220543.1 | 99 | 0 |
| SPGP0R02330 | XP_018221346.1 | 100 | 0 |
| SPGP0S03210 | XP_018220033.1 | 99 | 0 |
| SPGP0F04850 | XP_018222009.1 | 99 | 0 |
| SPGP0L00830 | XP_018219280.1 | 99 | 0 |
| SPGP0DZ00910 | XP_018219431.1 | 99 | 0 |
| SPGP0CZ03060 | XP_018221124.1 | 100 | 0 |
| SPGP0R02480 | XP_018221361.1 | 99 | 0 |
| SPGP0P02310 | XP_018221732.1 | 99 | 0 |
| SPGP0W03730 | XP_018222975.1 | 99 | 0 |
| SPGP0S03030 | XP_018220051.1 | 99 | 0 |
| SPGP0DS00100 | XP_018219997.1 | 99 | 0 |
| SPGP0DZ02200 | XP_018219557.1 | 99 | 0 |
| SPGP0E02330 | XP_018222821.1 | 99 | 0 |
| SPGP0R01460 | XP_018221263.1 | 99 | 0 |
| SPGP0Q02270 | XP_018222272.1 | 99 | 0 |
| SPGP0W03620 | XP_018222986.1 | 99 | 0 |
| SPGP0DU00960 | XP_018220494.1 | 99 | 0 |
| SPGP0Q02960 | XP_018222205.1 | 99 | 0 |
| SPGP0DN01950 | XP_018223642.1 | 99 | 0 |
| SPGP0Z00570 | XP_018219827.1 | 99 | 0 |
| SPGP0Q00820 | XP_018222419.1 | 99 | 0 |
| SPGP0W02690 | XP_018223070.1 | 99 | 0 |
| SPGP0Z02790 | XP_018219605.1 | 99 | 0 |
| SPGP0P01350 | XP_018221823.1 | 99 | 0 |
| SPGP0DZ00440 | XP_018219387.1 | 99 | 0 |

Create PDF in your applications with the Pdfcrowd HTML to PDF API PDFCROWD

| | | | |
|---|---|---|---|
| SPGP0DN01250 | XP_018223572.1 | 99 | 0 |
| SPGP0Z01660 | XP_018219719.1 | 99 | 0 |
| SPGP0DN02610 | XP_018223706.1 | 99 | 0 |
| SPGP0DV00580 | XP_018219920.1 | 99 | 0 |
| SPGP0T00370 | XP_018220366.1 | 98 | 0 |
| SPGP0B00230 | XP_018221471.1 | 98 | 0 |
| SPGP0U00850 | XP_018218907.1 | 99 | 0 |
| SPGP0P01030 | XP_018221854.1 | 100 | 0 |
| SPGP0X00460 | XP_018219055.1 | 99 | 0 |
| SPGP0R01320 | XP_018221249.1 | 99 | 0 |
| SPGP0Z03070 | XP_018219578.1 | 99 | 0 |
| SPGP0Q03750 | XP_018222130.1 | 98 | 0 |
| SPGP0U00140 | XP_018218975.1 | 98 | 1 |
| SPGP0Y00860 | XP_018223956.1 | 99 | 0 |
| SPGP0Y00390 | XP_018223915.1 | 99 | 0 |
| SPGP0Q00530 | XP_018222448.1 | 99 | 0 |
| SPGP0DN01030 | XP_018223550.1 | 99 | 0 |
| SPGP0W02320 | XP_018223105.1 | 99 | 0 |
| SPGP0R02410 | XP_018221354.1 | 99 | 0 |
| SPGP0E00600 | XP_018222650.1 | 99 | 0 |
| SPGP0Q03460 | XP_018222158.1 | 99 | 0 |
| SPGP0DU00760 | XP_018220515.1 | 99 | 0 |
| SPGP0X00430 | XP_018219058.1 | 99 | 0 |
| SPGP0Q03160 | XP_018222186.1 | 99 | 0 |
| SPGP0W01410 | XP_018223193.1 | 99 | 0 |
| SPGP0H00770 | XP_018224060.1 | 99 | 0 |
| SPGP0W03600 | XP_018222988.1 | 99 | 0 |
| SPGP0Q03790 | XP_018222126.1 | 98 | 0 |
| SPGP0W01970 | XP_018223138.1 | 99 | 0 |
| SPGP0S00650 | XP_018220280.1 | 99 | 0 |
| SPGP0W03500 | XP_018222997.1 | 99 | 0 |
| SPGP0DU00100 | XP_018220580.1 | 99 | 0 |
| SPGP0DV00170 | XP_018219881.1 | 99 | 0 |
| SPGP0DX00410 | XP_018219972.1 | 99 | 0 |
| SPGP0DX00140 | XP_018219945.1 | 99 | 0 |
| SPGP0X00560 | XP_018219046.1 | 99 | 0 |
| SPGP0W04120 | XP_018222938.1 | 98 | 0 |
| SPGP0L00530 | XP_018219308.1 | 99 | 0 |
| SPGP0DN03410 | XP_018223784.1 | 99 | 0 |
| SPGP0DV00380 | XP_018219900.1 | 99 | 0 |

Create PDF in your applications with the Pdfcrowd HTML to PDF API PDFCROWD

| | | | |
|---|---|---|---|
| SPGP0S02720 | XP_018220081.1 | 99 | 0 |
| SPGP0Q04230 | XP_018222082.1 | 99 | 0 |
| SPGP0U00350 | XP_018218954.1 | 99 | 0 |
| SPGP0Z03090 | XP_018219576.1 | 99 | 0 |
| SPGP0DX00350 | XP_018219966.1 | 99 | 0 |
| SPGP0E02010 | XP_018222789.1 | 99 | 0 |
| SPGP0W04140 | XP_018222936.1 | 99 | 0 |
| SPGP0DN01180 | XP_018223565.1 | 99 | 0 |
| SPGP0DZ02070 | XP_018219544.1 | 99 | 0 |
| SPGP0R02160 | XP_018221330.1 | 99 | 0 |
| SPGP0DN03860 | XP_018223828.1 | 99 | 0 |
| SPGP0Q01040 | XP_018222397.1 | 99 | 0 |
| SPGP0AC00690 | XP_018219153.1 | 99 | 0 |
| SPGP0Q00430 | XP_018222458.1 | 99 | 0 |
| SPGP0DN02440 | XP_018223689.1 | 99 | 0 |
| SPGP0X00400 | XP_018219061.1 | 100 | 0 |
| SPGP0Q03910 | XP_018222114.1 | 99 | 0 |
| SPGP0DN03150 | XP_018223759.1 | 99 | 0 |
| SPGP0DZ01140 | XP_018219453.1 | 100 | 0 |
| SPGP0W04390 | XP_018222910.1 | 98 | 1 |
| SPGP0R01090 | XP_018221227.1 | 99 | 0 |
| SPGP0AH00490 | XP_018222556.1 | 99 | 0 |
| SPGP0E01030 | XP_018222693.1 | 99 | 0 |
| SPGP0DV00460 | XP_018219908.1 | 99 | 0 |
| SPGP0Q01200 | XP_018222380.1 | 100 | 0 |
| SPGP0W02800 | XP_018223059.1 | 99 | 0 |
| SPGP0B00680 | XP_018221515.1 | 99 | 0 |
| SPGP0W00470 | XP_018223287.1 | 99 | 0 |
| SPGP0P00310 | XP_018221923.1 | 99 | 0 |
| SPGP0W00240 | XP_018223310.1 | 99 | 0 |
| SPGP0DZ00690 | XP_018219410.1 | 99 | 0 |
| SPGP0Z00180 | XP_018219865.1 | 99 | 0 |
| SPGP0W00340 | XP_018223300.1 | 99 | 0 |
| SPGP0Y00700 | XP_018223942.1 | 99 | 0 |
| SPGP0DN02910 | XP_018223736.1 | 99 | 0 |
| SPGP0W01810 | XP_018219094.1 | 99 | 0 |
| SPGP0EG00160 | XP_018221951.1 | 99 | 0 |
| SPGP0CZ00650 | XP_018220891.1 | 99 | 0 |
| SPGP0Z02280 | XP_018219657.1 | 99 | 0 |
| SPGP0DN01930 | XP_018223640.1 | 99 | 0 |

Create PDF in your applications with the Pdfcrowd HTML to PDF API PDFCROWD

| | | | |
|---|---|---|---|
| SPGP0DN01580 | XP_018223605.1 | 100 | 0 |
| SPGP0S00460 | XP_018220299.1 | 99 | 0 |
| SPGP0E00510 | XP_018222641.1 | 99 | 0 |
| SPGP0Q02400 | XP_018222259.1 | 99 | 0 |
| SPGP0U01260 | XP_018218866.1 | 99 | 0 |
| SPGP0P00930 | XP_018221864.1 | 99 | 0 |
| SPGP0U00330 | XP_018218956.1 | 100 | 0 |
| SPGP0U00440 | XP_018218945.1 | 99 | 0 |
| SPGP0E01480 | XP_018222736.1 | 99 | 0 |
| SPGP0DN01790 | XP_018223625.1 | 99 | 0 |
| SPGP0Z00960 | XP_018219788.1 | 99 | 0 |
| SPGP0Q01020 | XP_018222399.1 | 99 | 0 |
| SPGP0AA00670 | XP_018223379.1 | 100 | 0 |
| SPGP0R03020 | XP_018221412.1 | 99 | 0 |
| SPGP0DU00180 | XP_018220572.1 | 100 | 0 |
| SPGP0E01370 | XP_018222726.1 | 100 | 0 |
| SPGP0S00790 | XP_018220266.1 | 99 | 0 |
| SPGP0N01730 | XP_018220641.1 | 99 | 0 |
| SPGP0AA00840 | XP_018223393.1 | 99 | 0 |
| SPGP0DN03820 | XP_018223824.1 | 99 | 0 |
| SPGP0Q00490 | XP_018222452.1 | 99 | 0 |
| SPGP0P02510 | XP_018221712.1 | 99 | 0 |
| SPGP0W01270 | XP_018223207.1 | 99 | 0 |
| SPGP0E01230 | XP_018222713.1 | 99 | 0 |
| SPGP0AA00560 | XP_018223368.1 | 99 | 0 |
| SPGP0Q00690 | XP_018222432.1 | 99 | 0 |
| SPGP0S02100 | XP_018220142.1 | 99 | 0 |
| SPGP0B01250 | XP_018221568.1 | 99 | 0 |
| SPGP0S03510 | XP_018220005.1 | 99 | 0 |
| SPGP0L01140 | XP_018219249.1 | 99 | 0 |
| SPGP0L01230 | XP_018219241.1 | 99 | 0 |
| SPGP0DV00260 | XP_018219890.1 | 98 | 1 |
| SPGP0H00680 | XP_018224069.1 | 99 | 0 |
| SPGP0DU01420 | XP_018220448.1 | 100 | 0 |
| SPGP0X00910 | XP_018219011.1 | 99 | 0 |
| SPGP0E02110 | XP_018222799.1 | 98 | 0 |
| SPGP0CZ01100 | XP_018220935.1 | 99 | 0 |
| SPGP0DN03940 | XP_018223836.1 | 99 | 0 |
| SPGP0W02950 | XP_018223045.1 | 99 | 0 |
| SPGP0R02310 | XP_018221344.1 | 99 | 0 |

Create PDF in your applications with the Pdfcrowd HTML to PDF API PDFCROWD

| | | | |
|---|---|---|---|
| SPGP0CZ02590 | XP_018221079.1 | 98 | 1 |
| SPGP0Q01790 | XP_018222319.1 | 99 | 0 |
| SPGP0R01550 | XP_018221272.1 | 99 | 0 |
| SPGP0W04490 | XP_018222899.1 | 99 | 0 |
| SPGP0DZ00970 | XP_018219437.1 | 99 | 0 |
| SPGP0P02330 | XP_018221730.1 | 99 | 0 |
| SPGP0W03710 | XP_018222977.1 | 99 | 0 |
| SPGP0W00590 | XP_018223275.1 | 99 | 0 |
| SPGP0L00460 | XP_018219315.1 | 99 | 0 |
| SPGP0P01720 | XP_018221788.1 | 99 | 0 |
| SPGP0DN03180 | XP_018223762.1 | 99 | 0 |
| SPGP0W01250 | XP_018223209.1 | 99 | 0 |
| SPGP0Z01970 | XP_018219688.1 | 99 | 0 |
| SPGP0BF00320 | XP_018220400.1 | 99 | 0 |
| SPGP0B01430 | XP_018221586.1 | 99 | 0 |
| SPGP0N00530 | XP_018220755.1 | 100 | 0 |
| SPGP0DZ02220 | XP_018219559.1 | 99 | 0 |
| SPGP0E02050 | XP_018222793.1 | 100 | 0 |
| SPGP0R01400 | XP_018221257.1 | 99 | 0 |
| SPGP0P01840 | XP_018221776.1 | 99 | 0 |
| SPGP0Q01070 | XP_018222394.1 | 98 | 2 |
| SPGP0B01350 | XP_018221578.1 | 99 | 0 |
| SPGP0L00700 | XP_018219293.1 | 99 | 0 |
| SPGP0B00120 | XP_018221460.1 | 98 | 0 |
| SPGP0DN03980 | XP_018223840.1 | 99 | 0 |
| SPGP0Y00720 | XP_018223944.1 | 99 | 0 |
| SPGP0W01780 | XP_018223157.1 | 99 | 0 |
| SPGP0AA01040 | XP_018223411.1 | 99 | 0 |
| SPGP0Z00550 | XP_018219829.1 | 99 | 0 |
| SPGP0CZ00230 | XP_018220851.1 | 99 | 0 |
| SPGP0CZ00530 | XP_018220879.1 | 99 | 0 |
| SPGP0DN02580 | XP_018223703.1 | 99 | 0 |
| SPGP0AH00410 | XP_018222563.1 | 99 | 0 |
| SPGP0L00870 | XP_018219276.1 | 100 | 0 |
| SPGP0R00530 | XP_018221176.1 | 99 | 0 |
| SPGP0CZ01110 | XP_018220936.1 | 99 | 0 |
| SPGP0DN02630 | XP_018223708.1 | 99 | 0 |
| SPGP0S02480 | XP_018220105.1 | 98 | 2 |
| SPGP0Y01260 | XP_018223995.1 | 99 | 0 |
| SPGP0AC00910 | XP_018219174.1 | 99 | 0 |

Create PDF in your applications with the Pdfcrowd HTML to PDF API PDFCROWD

| | | | |
|---|---|---|---|
| SPGP0Q01990 | XP_018222299.1 | 99 | 0 |
| SPGP0Q03120 | XP_018222190.1 | 98 | 1 |
| SPGP0DN00980 | XP_018223545.1 | 99 | 0 |
| SPGP0BF00160 | XP_018220387.1 | 99 | 0 |
| SPGP0DN01230 | XP_018223570.1 | 99 | 0 |
| SPGP0E00680 | XP_018222658.1 | 99 | 0 |
| SPGP0R01450 | XP_018221262.1 | 99 | 0 |
| SPGP0Q03340 | XP_018222169.1 | 99 | 0 |
| SPGP0DN00180 | XP_018223468.1 | 99 | 0 |
| SPGP0R01930 | XP_018221309.1 | 100 | 0 |
| SPGP0DV00640 | XP_018219925.1 | 96 | 0 |
| SPGP0E01530 | XP_018222741.1 | 99 | 0 |
| SPGP0Z00340 | XP_018219850.1 | 98 | 0 |
| SPGP0Z01150 | XP_018219769.1 | 99 | 0 |
| SPGP0AC00650 | XP_018219149.1 | 99 | 0 |
| SPGP0R02080 | XP_018221323.1 | 99 | 0 |
| SPGP0DU00700 | XP_018220521.1 | 99 | 0 |
| SPGP0U01780 | XP_018218816.1 | 100 | 0 |
| SPGP0CZ01270 | XP_018220952.1 | 100 | 0 |
| SPGP0Q00800 | XP_018222421.1 | 99 | 0 |
| SPGP0AA00790 | XP_018223389.1 | 99 | 0 |
| SPGP0AA00240 | XP_018223337.1 | 99 | 0 |
| SPGP0Z00590 | XP_018219825.1 | 99 | 0 |
| SPGP0W01520 | XP_018223183.1 | 99 | 0 |
| SPGP0R02060 | XP_018221321.1 | 99 | 0 |
| SPGP0DN04230 | XP_018223865.1 | 99 | 0 |
| SPGP0N00810 | XP_018220728.1 | 99 | 0 |
| SPGP0Q02380 | XP_018222261.1 | 99 | 0 |
| SPGP0S01350 | XP_018220212.1 | 99 | 0 |
| SPGP0S00630 | XP_018220282.1 | 99 | 0 |
| SPGP0R02870 | XP_018221399.1 | 99 | 0 |
| SPGP0P01900 | XP_018221770.1 | 99 | 0 |
| SPGP0B01900 | XP_018221632.1 | 99 | 0 |
| SPGP0P02820 | XP_018221681.1 | 99 | 0 |
| SPGP0P01800 | XP_018221780.1 | 99 | 0 |
| SPGP0N00470 | XP_018220760.1 | 99 | 0 |
| SPGP0Z02720 | XP_018219613.1 | 99 | 0 |
| SPGP0W02920 | XP_018223048.1 | 99 | 0 |
| SPGP0X00630 | XP_018219039.1 | 99 | 0 |
| SPGP0DX00290 | XP_018219960.1 | 99 | 1 |

Create PDF in your applications with the Pdfcrowd HTML to PDF API PDFCROWD

| | | | |
|---|---|---|---|
| SPGP0L01490 | XP_018219215.1 | 99 | 0 |
| SPGP0U00430 | XP_018218946.1 | 99 | 0 |
| SPGP0P01650 | XP_018221795.1 | 99 | 0 |
| SPGP0S01820 | XP_018220170.1 | 99 | 0 |
| SPGP0Z01170 | XP_018219767.1 | 100 | 0 |
| SPGP0DN04170 | XP_018223859.1 | 99 | 0 |
| SPGP0L01420 | XP_018219222.1 | 99 | 0 |
| SPGP0H00490 | XP_018224087.1 | 99 | 0 |
| SPGP0L01550 | XP_018219209.1 | 99 | 0 |
| SPGP0L01350 | XP_018219229.1 | 97 | 2 |
| SPGP0W04550 | XP_018222893.1 | 99 | 0 |
| SPGP0DN00270 | XP_018223477.1 | 98 | 0 |
| SPGP0E01650 | XP_018222753.1 | 99 | 0 |
| SPGP0U00230 | XP_018218966.1 | 99 | 0 |
| SPGP0Q01440 | XP_018222355.1 | 99 | 0 |
| SPGP0R02690 | XP_018221381.1 | 99 | 0 |
| SPGP0B00910 | XP_018221537.1 | 99 | 0 |
| SPGP0P01160 | XP_018221841.1 | 99 | 0 |
| SPGP0Z00760 | XP_018219808.1 | 99 | 0 |
| SPGP0P03110 | XP_018221653.1 | 99 | 0 |
| SPGP0U00100 | XP_018218978.1 | 99 | 0 |
| SPGP0W04800 | XP_018222868.1 | 100 | 0 |
| SPGP0R00920 | XP_018221211.1 | 99 | 0 |
| SPGP0EX00150 | XP_018223874.1 | 99 | 0 |
| SPGP0R00430 | XP_018221166.1 | 98 | 0 |
| SPGP0DN03510 | XP_018223793.1 | 99 | 0 |
| SPGP0DX00420 | XP_018219973.1 | 99 | 0 |
| SPGP0Q00200 | XP_018222480.1 | 99 | 0 |
| SPGP0DZ00100 | XP_018219353.1 | 99 | 0 |
| SPGP0R00690 | XP_018221191.1 | 99 | 0 |
| SPGP0W01380 | XP_018223196.1 | 99 | 0 |
| SPGP0W01830 | XP_018223152.1 | 99 | 0 |
| SPGP0Q01860 | XP_018222312.1 | 99 | 0 |
| SPGP0DN03020 | XP_018223747.1 | 99 | 0 |
| SPGP0Y01070 | XP_018223977.1 | 99 | 0 |
| SPGP0S01680 | XP_018220183.1 | 99 | 0 |
| SPGP0Z00940 | XP_018219790.1 | 99 | 0 |
| SPGP0Y01310 | XP_018224000.1 | 99 | 0 |
| SPGP0B00830 | XP_018221530.1 | 99 | 0 |
| SPGP0H00280 | XP_018224107.1 | 99 | 0 |

Create PDF in your applications with the Pdfcrowd HTML to PDF API PDFCROWD

| | | | |
|---|---|---|---|
| SPGP0Gtrna29D | DI49_4675 | 100 | 0 |
| SPGP0E02570 | XP_018222843.1 | 99 | 0 |
| SPGP0DN03100 | XP_018223754.1 | 99 | 0 |
| SPGP0Itrna29E | DI49_5287 | 100 | 0 |
| SPGP0AF00430 | XP_018221984.1 | 97 | 0 |
| SPGP0BF00400 | XP_018220408.1 | 99 | 0 |
| SPGP0W01930 | XP_018223143.1 | 100 | 0 |
| SPGP0S02290 | XP_018220123.1 | 99 | 0 |
| SPGP0Z00250 | XP_018219859.1 | 99 | 1 |
| SPGP0Y01300 | XP_018223999.1 | 99 | 0 |
| SPGP0DU00280 | XP_018220562.1 | 99 | 0 |
| SPGP0S01750 | XP_018220176.1 | 99 | 0 |
| SPGP0DN01770 | XP_018223623.1 | 99 | 0 |
| SPGP0X01060 | XP_018218997.1 | 99 | 0 |
| SPGP0S02120 | XP_018220140.1 | 99 | 0 |
| SPGP0Z01110 | XP_018219773.1 | 99 | 0 |
| SPGP0DU01400 | XP_018220450.1 | 99 | 0 |
| SPGP0P00420 | XP_018221913.1 | 99 | 0 |
| SPGP0AH00210 | XP_018222581.1 | 99 | 0 |
| SPGP0L01090 | XP_018219254.1 | 100 | 0 |
| SPGP0F04720 | XP_018222021.1 | 99 | 0 |
| SPGP0DX00620 | XP_018219992.1 | 99 | 0 |
| SPGP0X00240 | XP_018219077.1 | 99 | 0 |
| SPGP0F04640 | XP_018222028.1 | 99 | 0 |
| SPGP0AH00910 | XP_018222515.1 | 98 | 0 |
| SPGP0H00240 | XP_018224111.1 | 99 | 0 |
| SPGP0R00600 | XP_018221182.1 | 99 | 0 |
| SPGP0DN00910 | XP_018223537.1 | 99 | 0 |
| SPGP0W02730 | XP_018223066.1 | 99 | 0 |
| SPGP0ET00180 | XP_018218987.1 | 99 | 0 |
| SPGP0R03380 | XP_018221447.1 | 99 | 0 |
| SPGP0F04500 | XP_018222041.1 | 99 | 0 |
| SPGP0Ntrna10R | DI49_3845 | 100 | 0 |
| SPGP0DN02500 | XP_018223695.1 | 99 | 0 |
| SPGP0P03130 | XP_018221651.1 | 99 | 0 |
| SPGP0S00180 | XP_018220326.1 | 99 | 0 |
| SPGP0P02350 | XP_018221728.1 | 99 | 0 |
| SPGP0W03260 | XP_018223017.1 | 100 | 0 |
| SPGP0X01070 | XP_018218996.1 | 99 | 0 |
| SPGP0DU00920 | XP_018220498.1 | 99 | 0 |

Create PDF in your applications with the Pdfcrowd HTML to PDF API PDFCROWD

| | | | |
|---|---|---|---|
| SPGP0E02170 | XP_018222805.1 | 99 | 0 |
| SPGP0P00840 | XP_018221873.1 | 99 | 0 |
| SPGP0S00620 | XP_018220283.1 | 99 | 0 |
| SPGP0L00270 | XP_018219334.1 | 100 | 0 |
| SPGP0R02730 | XP_018221385.1 | 99 | 0 |
| SPGP0B00580 | XP_018221506.1 | 99 | 0 |
| SPGP0DZ00950 | XP_018219435.1 | 99 | 0 |
| SPGP0Y00340 | XP_018223911.1 | 99 | 0 |
| SPGP0R02010 | XP_018221316.1 | 99 | 0 |
| SPGP0R01420 | XP_018221259.1 | 99 | 0 |
| SPGP0CZ00510 | XP_018220877.1 | 99 | 0 |
| SPGP0W00460 | XP_018223288.1 | 99 | 0 |
| SPGP0P01460 | XP_018221812.1 | 99 | 0 |
| SPGP0DN02030 | XP_018223650.1 | 99 | 0 |
| SPGP0U01200 | XP_018218872.1 | 99 | 0 |
| SPGP0Q03500 | XP_018222154.1 | 99 | 0 |
| SPGP0S00550 | XP_018220290.1 | 99 | 0 |
| SPGP0P01070 | XP_018221850.1 | 99 | 0 |
| SPGP0U00810 | XP_018218911.1 | 100 | 0 |
| SPGP0E02480 | XP_018222835.1 | 99 | 0 |
| SPGP0N02230 | XP_018220591.1 | 99 | 0 |
| SPGP0S00610 | XP_018220284.1 | 99 | 0 |
| SPGP0DN01210 | XP_018223568.1 | 99 | 0 |
| SPGP0DV00240 | XP_018219888.1 | 99 | 0 |
| SPGP0CZ02450 | XP_018221065.1 | 99 | 0 |
| SPGP0E00420 | XP_018222632.1 | 99 | 0 |
| SPGP0E00390 | XP_018222629.1 | 99 | 1 |
| SPGP0DZ01020 | XP_018219442.1 | 99 | 0 |
| SPGP0N01450 | XP_018220668.1 | 99 | 0 |
| SPGP0DN04130 | XP_018223855.1 | 99 | 0 |
| SPGP0DN00160 | XP_018223466.1 | 99 | 0 |
| SPGP0P02030 | XP_018221757.1 | 99 | 0 |
| SPGP0X00790 | XP_018219023.1 | 99 | 0 |
| SPGP0W02820 | XP_018223058.1 | 98 | 0 |
| SPGP0Q03420 | XP_018222161.1 | 98 | 1 |
| SPGP0AH01080 | XP_018222500.1 | 99 | 0 |
| SPGP0E00640 | XP_018222654.1 | 98 | 0 |
| SPGP0Z02560 | XP_018219629.1 | 99 | 0 |
| SPGP0DN02950 | XP_018223740.1 | 99 | 0 |

Create PDF in your applications with the Pdfcrowd HTML to PDF API PDFCROWD

| | | | |
|---|---|---|---|
| SPGP0P00170 | XP_018221937.1 | 99 | 0 |
| SPGP0P00210 | XP_018221933.1 | 99 | 0 |
| SPGP0P01920 | XP_018221768.1 | 99 | 0 |
| SPGP0W04840 | XP_018222864.1 | 99 | 0 |
| SPGP0F04990 | XP_018221995.1 | 97 | 0 |
| SPGP0E00250 | XP_018222615.1 | 99 | 0 |
| SPGP0AC00740 | XP_018219157.1 | 99 | 0 |
| SPGP0R03080 | XP_018221418.1 | 99 | 0 |
| SPGP0S02600 | XP_018220093.1 | 99 | 0 |
| SPGP0DN03110 | XP_018223755.1 | 99 | 0 |
| SPGP0Z01500 | XP_018219735.1 | 100 | 0 |
| SPGP0AA00910 | XP_018223400.1 | 99 | 0 |
| SPGP0R02500 | XP_018221363.1 | 99 | 0 |
| SPGP0E00590 | XP_018222649.1 | 99 | 1 |
| SPGP0S02270 | XP_018220125.1 | 99 | 0 |
| SPGP0Ytrna21G | DI49_0567 | 99 | 0 |
| SPGP0Z01190 | XP_018219766.1 | 99 | 0 |
| SPGP0W02670 | XP_018223072.1 | 99 | 0 |
| SPGP0E00440 | XP_018222634.1 | 99 | 0 |
| SPGP0W01330 | XP_018223201.1 | 99 | 0 |
| SPGP0Z00700 | XP_018219814.1 | 99 | 0 |
| SPGP0Z02580 | XP_018219627.1 | 99 | 0 |
| SPGP0AH00100 | XP_018222594.1 | 99 | 0 |
| SPGP0AF00470 | XP_018221988.1 | 99 | 0 |
| SPGP0CZ01340 | XP_018220959.1 | 100 | 0 |
| SPGP0Q04170 | XP_018222088.1 | 99 | 0 |
| SPGP0CZ00750 | XP_018220900.1 | 99 | 0 |
| SPGP0R03060 | XP_018221416.1 | 99 | 0 |
| SPGP0AC00470 | XP_018219131.1 | 99 | 0 |
| SPGP0P00980 | XP_018221859.1 | 99 | 0 |
| SPGP0DN01970 | XP_018223644.1 | 99 | 0 |
| SPGP0EG00120 | XP_018221947.1 | 99 | 0 |
| SPGP0AA00310 | XP_018223344.1 | 99 | 0 |
| SPGP0DN02730 | XP_018223717.1 | 99 | 0 |
| SPGP0S00420 | XP_018220303.1 | 99 | 0 |
| SPGP0F04230 | XP_018222068.1 | 99 | 0 |
| SPGP0B00390 | XP_018221487.1 | 99 | 0 |
| SPGP0W01210 | XP_018223213.1 | 99 | 0 |
| SPGP0Strna9F | DI49_4130 | 99 | 0 |
| SPGP0F04480 | XP_018222043.1 | 99 | 0 |

Create PDF in your applications with the Pdfcrowd HTML to PDF API PDFCROWD

| | | | |
|---|---|---|---|
| SPGP0B00770 | XP_018221524.1 | 99 | 0 |
| SPGP0Q02300 | XP_018222269.1 | 99 | 0 |
| SPGP0W04050 | XP_018222945.1 | 99 | 0 |
| SPGP0AA00920 | XP_018223401.1 | 99 | 0 |
| SPGP0R02580 | XP_018221371.1 | 99 | 0 |
| SPGP0S03080 | XP_018220046.1 | 99 | 0 |
| SPGP0R00620 | XP_018221184.1 | 99 | 0 |
| SPGP0Q00350 | XP_018222466.1 | 99 | 0 |
| SPGP0S01770 | XP_018220174.1 | 99 | 0 |
| SPGP0P01980 | XP_018221762.1 | 99 | 0 |
| SPGP0Y00880 | XP_018223958.1 | 99 | 0 |
| SPGP0X00440 | XP_018219057.1 | 99 | 0 |
| SPGP0S03550 | XP_018220001.1 | 99 | 0 |
| SPGP0P02940 | XP_018221670.1 | 100 | 0 |
| SPGP0P02200 | XP_018221742.1 | 99 | 0 |
| SPGP0B01630 | XP_018221605.1 | 99 | 0 |
| SPGP0S01480 | XP_018220200.1 | 99 | 0 |
| SPGP0H00270 | XP_018224108.1 | 99 | 0 |
| SPGP0Z01930 | XP_018219692.1 | 99 | 0 |
| SPGP0N00390 | XP_018220769.1 | 99 | 0 |
| SPGP0W02210 | XP_018223116.1 | 99 | 0 |
| SPGP0W02880 | XP_018223052.1 | 99 | 0 |
| SPGP0P00150 | XP_018221939.1 | 99 | 1 |
| SPGP0DU00530 | XP_018220537.1 | 99 | 0 |
| SPGP0B00160 | XP_018221464.1 | 96 | 0 |
| SPGP0DN03230 | XP_018223767.1 | 99 | 0 |
| SPGP0P03170 | XP_018221647.1 | 99 | 0 |
| SPGP0B01960 | XP_018221638.1 | 99 | 0 |
| SPGP0CZ02930 | XP_018221112.1 | 100 | 0 |
| SPGP0W03980 | XP_018222951.1 | 99 | 0 |
| SPGP0P02370 | XP_018221726.1 | 99 | 0 |
| SPGP0S00160 | XP_018220328.1 | 99 | 0 |
| SPGP0R02190 | XP_018221332.1 | 99 | 0 |
| SPGP0AF00500 | XP_018221991.1 | 99 | 0 |
| SPGP0B01040 | XP_018221550.1 | 99 | 0 |
| SPGP0R01990 | XP_018221314.1 | 99 | 0 |
| SPGP0W04100 | XP_018222940.1 | 99 | 0 |
| SPGP0B00560 | XP_018221504.1 | 99 | 0 |
| SPGP0DN03490 | XP_018223791.1 | 99 | 0 |
| SPGP0CZ00570 | XP_018220883.1 | 99 | 0 |

Create PDF in your applications with the Pdfcrowd HTML to PDF API PDFCROWD

| | | | |
|---|---|---|---|
| SPGP0R01190 | XP_018221237.1 | 99 | 0 |
| SPGP0E01910 | XP_018222779.1 | 96 | 4 |
| SPGP0DN03380 | XP_018223781.1 | 99 | 0 |
| SPGP0P01700 | XP_018221790.1 | 99 | 0 |
| SPGP0DN03530 | XP_018223795.1 | 99 | 0 |
| SPGP0DV00520 | XP_018219914.1 | 99 | 0 |
| SPGP0U00550 | XP_018218934.1 | 100 | 0 |
| SPGP0R00340 | XP_018221157.1 | 99 | 0 |
| SPGP0AF00210 | XP_018221962.1 | 99 | 0 |
| SPGP0DN01890 | XP_018223636.1 | 99 | 0 |
| SPGP0P02860 | XP_018221677.1 | 99 | 0 |
| SPGP0Q03380 | XP_018222165.1 | 99 | 0 |
| SPGP0DN04110 | XP_018223853.1 | 99 | 0 |
| SPGP0Q02150 | XP_018222283.1 | 99 | 0 |
| SPGP0L01530 | XP_018219211.1 | 99 | 0 |
| SPGP0E00660 | XP_018222656.1 | 99 | 0 |
| SPGP0P01120 | XP_018221845.1 | 99 | 0 |
| SPGP0DN00140 | XP_018223464.1 | 100 | 0 |
| SPGP0AA00750 | XP_018223385.1 | 99 | 0 |
| SPGP0R00380 | XP_018221161.1 | 99 | 0 |
| SPGP0Q01590 | XP_018222339.1 | 99 | 0 |
| SPGP0Z01590 | XP_018219726.1 | 100 | 0 |
| SPGP0DN00320 | XP_018223482.1 | 99 | 0 |
| SPGP0DX00120 | XP_018219943.1 | 99 | 0 |
| SPGP0E01130 | XP_018222703.1 | 99 | 0 |
| SPGP0DN01530 | XP_018223600.1 | 99 | 0 |
| SPGP0DN03360 | XP_018223779.1 | 99 | 0 |
| SPGP0R03120 | XP_018221422.1 | 99 | 0 |
| SPGP0L01180 | XP_018219246.1 | 99 | 0 |
| SPGP0P02490 | XP_018221714.1 | 99 | 0 |
| SPGP0Z01000 | XP_018219784.1 | 99 | 0 |
| SPGP0AC00350 | XP_018219119.1 | 99 | 0 |
| SPGP0Q01820 | XP_018222316.1 | 100 | 0 |
| SPGP0Z01230 | XP_018219762.1 | 99 | 0 |
| SPGP0CZ03140 | XP_018221131.1 | 99 | 0 |
| SPGP0B01940 | XP_018221636.1 | 99 | 0 |
| SPGP0P03200 | XP_018221644.1 | 95 | 0 |
| SPGP0DN03280 | XP_018223771.1 | 99 | 0 |
| SPGP0DN02100 | XP_018223657.1 | 99 | 0 |
| SPGP0CZ00140 | XP_018220843.1 | 99 | 0 |

Create PDF in your applications with the Pdfcrowd HTML to PDF API PDFCROWD

| | | | |
|---|---|---|---|
| SPGP0W01890 | XP_018223147.1 | 99 | 0 |
| SPGP0S00390 | XP_018220306.1 | 100 | 0 |
| SPGP0L00480 | XP_018219313.1 | 99 | 0 |
| SPGP0Q00700 | XP_018222431.1 | 99 | 0 |
| SPGP0B00950 | XP_018221541.1 | 99 | 0 |
| SPGP0DZ01540 | XP_018219493.1 | 99 | 1 |
| SPGP0R00960 | XP_018221214.1 | 99 | 0 |
| SPGP0S00700 | XP_018220275.1 | 99 | 0 |
| SPGP0R01380 | XP_018221255.1 | 99 | 0 |
| SPGP0Q01510 | XP_018222347.1 | 99 | 0 |
| SPGP0Q01360 | XP_018222363.1 | 99 | 0 |
| SPGP0Q02710 | XP_018222229.1 | 99 | 0 |
| SPGP0Z00980 | XP_018219786.1 | 100 | 0 |
| SPGP0W02450 | XP_018223092.1 | 99 | 1 |
| SPGP0P00990 | XP_018221858.1 | 99 | 0 |
| SPGP0B00760 | XP_018221523.1 | 99 | 0 |
| SPGP0S03570 | XP_018219999.1 | 99 | 0 |
| SPGP0W03040 | XP_018223038.1 | 99 | 0 |
| SPGP0R03200 | XP_018221430.1 | 99 | 0 |
| SPGP0Y00840 | XP_018223954.1 | 99 | 0 |
| SPGP0Q00450 | XP_018222456.1 | 99 | 0 |
| SPGP0N01130 | XP_018220699.1 | 97 | 0 |
| SPGP0DN00870 | XP_018223533.1 | 98 | 0 |
| SPGP0W00410 | XP_018223293.1 | 99 | 0 |
| SPGP0AC01020 | XP_018219185.1 | 98 | 0 |
| SPGP0W02860 | XP_018223055.1 | 99 | 0 |
| SPGP0DN03170 | XP_018223761.1 | 99 | 0 |
| SPGP0R02950 | XP_018221406.1 | 99 | 0 |
| SPGP0S00370 | XP_018220308.1 | 99 | 0 |
| SPGP0Q02920 | XP_018222209.1 | 98 | 0 |
| SPGP0DN03880 | XP_018223830.1 | 99 | 0 |
| SPGP0E01900 | XP_018222778.1 | 99 | 0 |
| SPGP0S00510 | XP_018220294.1 | 99 | 0 |
| SPGP0Z02930 | XP_018219592.1 | 99 | 0 |
| SPGP0H00360 | XP_018224099.1 | 99 | 0 |
| SPGP0Z01100 | XP_018219774.1 | 99 | 0 |
| SPGP0CZ02490 | XP_018221069.1 | 99 | 0 |
| SPGP0DN02560 | XP_018223701.1 | 100 | 0 |
| SPGP0S00410 | XP_018220304.1 | 99 | 0 |
| SPGP0E01860 | XP_018222774.1 | 99 | 0 |

Create PDF in your applications with the Pdfcrowd HTML to PDF API PDFCROWD

| | | | |
|---|---|---|---|
| SPGP0R02770 | XP_018221389.1 | 99 | 0 |
| SPGP0W03800 | XP_018222968.1 | 99 | 0 |
| SPGP0B00540 | XP_018221502.1 | 99 | 0 |
| SPGP0DN03960 | XP_018223838.1 | 99 | 0 |
| SPGP0DZ02280 | XP_018219565.1 | 99 | 0 |
| SPGP0B01490 | XP_018221592.1 | 99 | 0 |
| SPGP0W04780 | XP_018222870.1 | 99 | 0 |
| SPGP0DU00380 | XP_018220552.1 | 99 | 0 |
| SPGP0L00810 | XP_018219282.1 | 99 | 0 |
| SPGP0S03270 | XP_018220028.1 | 99 | 0 |
| SPGP0R02200 | XP_018221333.1 | 99 | 0 |
| SPGP0Z01440 | XP_018219741.1 | 99 | 0 |
| SPGP0Q03630 | XP_018222142.1 | 97 | 0 |
| SPGP0AF00180 | XP_018221959.1 | 99 | 0 |
| SPGP0E00460 | XP_018222636.1 | 100 | 0 |
| SPGP0Q01650 | XP_018222333.1 | 99 | 0 |
| SPGP0AA00900 | XP_018223399.1 | 99 | 0 |
| SPGP0N02070 | XP_018220607.1 | 99 | 0 |
| SPGP0Z02740 | XP_018219610.1 | 99 | 0 |
| SPGP0AA00770 | XP_018223387.1 | 99 | 0 |
| SPGP0W02540 | XP_018223083.1 | 99 | 0 |
| SPGP0N00720 | XP_018220736.1 | 99 | 0 |
| SPGP0W03590 | XP_018222989.1 | 99 | 0 |
| SPGP0P01250 | XP_018221832.1 | 99 | 0 |
| SPGP0N00220 | XP_018220786.1 | 99 | 0 |
| SPGP0P01210 | XP_018221836.1 | 100 | 0 |
| SPGP0N00650 | XP_018220743.1 | 99 | 0 |
| SPGP0AA00580 | XP_018223370.1 | 97 | 0 |
| SPGP0X00690 | XP_018219033.1 | 99 | 0 |
| SPGP0Y01600 | XP_018224029.1 | 99 | 0 |
| SPGP0P00610 | XP_018221895.1 | 99 | 0 |
| SPGP0R01010 | XP_018221219.1 | 99 | 0 |
| SPGP0Z02030 | XP_018219682.1 | 99 | 0 |
| SPGP0DN01980 | XP_018223645.1 | 99 | 0 |
| SPGP0L01440 | XP_018219220.1 | 99 | 0 |
| SPGP0W01010 | XP_018223233.1 | 99 | 0 |
| SPGP0F04700 | XP_018222023.1 | 99 | 0 |
| SPGP0R02540 | XP_018221367.1 | 100 | 0 |
| SPGP0W03320 | XP_018223012.1 | 99 | 0 |
| SPGP0Q04300 | XP_018222076.1 | 99 | 0 |

Create PDF in your applications with the Pdfcrowd HTML to PDF API PDFCROWD

| | | | |
|---|---|---|---|
| SPGP0P00940 | XP_018221863.1 | 100 | 0 |
| SPGP0W04640 | XP_018222884.1 | 99 | 0 |
| SPGP0W04860 | XP_018222862.1 | 99 | 0 |
| SPGP0S01310 | XP_018220215.1 | 99 | 0 |
| SPGP0S00800 | XP_018220265.1 | 99 | 0 |
| SPGP0R02750 | XP_018221387.1 | 99 | 0 |
| SPGP0W03140 | XP_018223029.1 | 99 | 0 |
| SPGP0L00310 | XP_018219330.1 | 99 | 0 |
| SPGP0AF00270 | XP_018221968.1 | 99 | 0 |
| SPGP0Q02480 | XP_018222251.1 | 100 | 0 |
| SPGP0DN02180 | XP_018223664.1 | 99 | 0 |
| SPGP0Q01280 | XP_018222372.1 | 99 | 0 |
| SPGP0N00200 | XP_018220788.1 | 99 | 0 |
| SPGP0X00480 | XP_018219053.1 | 99 | 0 |
| SPGP0W01090 | XP_018223225.1 | 99 | 0 |
| SPGP0W02500 | XP_018223087.1 | 99 | 0 |
| SPGP0DN03690 | XP_018223811.1 | 99 | 0 |
| SPGP0Q03280 | XP_018222174.1 | 98 | 0 |
| SPGP0W01540 | XP_018223181.1 | 99 | 0 |
| SPGP0DZ00770 | XP_018219418.1 | 100 | 0 |
| SPGP0U01070 | XP_018218885.1 | 99 | 0 |
| SPGP0B00330 | XP_018221481.1 | 99 | 0 |
| SPGP0Y00370 | XP_018223914.1 | 99 | 0 |
| SPGP0R01100 | XP_018221228.1 | 99 | 0 |
| SPGP0S02180 | XP_018220134.1 | 99 | 0 |
| SPGP0S02780 | XP_018220075.1 | 99 | 0 |
| SPGP0E00140 | XP_018222604.1 | 99 | 0 |
| SPGP0H00340 | XP_018224101.1 | 99 | 0 |
| SPGP0DZ00730 | XP_018219414.1 | 99 | 0 |
| SPGP0W04290 | XP_018222921.1 | 99 | 0 |
| SPGP0DZ01050 | XP_018219444.1 | 99 | 0 |
| SPGP0DU00510 | XP_018220539.1 | 100 | 0 |
| SPGP0Y00740 | XP_018223946.1 | 99 | 0 |
| SPGP0DN02380 | XP_018223683.1 | 100 | 0 |
| SPGP0S02970 | XP_018220057.1 | 99 | 0 |
| SPGP0P00560 | XP_018221900.1 | 99 | 0 |
| SPGP0Q02900 | XP_018222211.1 | 98 | 0 |
| SPGP0N00570 | XP_018220751.1 | 99 | 0 |
| SPGP0R02280 | XP_018221341.1 | 99 | 0 |
| SPGP0R00300 | XP_018221153.1 | 100 | 0 |

Create PDF in your applications with the Pdfcrowd HTML to PDF API PDFCROWD

| | | | |
|---|---|---|---|
| SPGP0S00530 | XP_018220292.1 | 97 | 2 |
| SPGP0DZ02110 | XP_018219548.1 | 99 | 0 |
| SPGP0B01920 | XP_018221634.1 | 99 | 0 |
| SPGP0R01950 | XP_018221311.1 | 99 | 0 |
| SPGP0W04410 | XP_018222908.1 | 99 | 0 |
| SPGP0S02740 | XP_018220079.1 | 99 | 0 |
| SPGP0R01480 | XP_018221265.1 | 99 | 0 |
| SPGP0S01290 | XP_018220217.1 | 97 | 0 |
| SPGP0S02400 | XP_018220112.1 | 99 | 0 |
| SPGP0S01180 | XP_018220227.1 | 99 | 0 |
| SPGP0DV00400 | XP_018219902.1 | 99 | 0 |
| SPGP0H00790 | XP_018224058.1 | 99 | 0 |
| SPGP0E00370 | XP_018222627.1 | 99 | 0 |
| SPGP0DZ01380 | XP_018219477.1 | 99 | 0 |
| SPGP0AH00620 | XP_018222543.1 | 99 | 0 |
| SPGP0Q01120 | XP_018222389.1 | 99 | 0 |
| SPGP0DV00250 | XP_018219889.1 | 99 | 0 |
| SPGP0R00280 | XP_018221151.1 | 99 | 0 |
| SPGP0L00880 | XP_018219275.1 | 99 | 0 |
| SPGP0B01210 | XP_018221564.1 | 99 | 0 |
| SPGP0X00360 | XP_018219065.1 | 100 | 0 |
| SPGP0S02020 | XP_018220150.1 | 99 | 0 |
| SPGP0N00960 | XP_018220714.1 | 99 | 0 |
| SPGP0Q03510 | XP_018222153.1 | 99 | 0 |
| SPGP0DN04150 | XP_018223857.1 | 99 | 0 |
| SPGP0AF00450 | XP_018221986.1 | 98 | 0 |
| SPGP0X00610 | XP_018219041.1 | 99 | 0 |
| SPGP0W01280 | XP_018223206.1 | 100 | 0 |
| SPGP0E01310 | XP_018222721.1 | 99 | 0 |
| SPGP0CZ01700 | XP_018220993.1 | 99 | 0 |
| SPGP0DN00810 | XP_018223527.1 | 99 | 0 |
| SPGP0F04420 | XP_018222049.1 | 99 | 0 |
| SPGP0Q00990 | XP_018222402.1 | 99 | 0 |
| SPGP0DN02860 | XP_018223731.1 | 99 | 0 |
| SPGP0Q01550 | XP_018222343.1 | 99 | 0 |
| SPGP0R03160 | XP_018221426.1 | 99 | 0 |
| SPGP0W02240 | XP_018223113.1 | 99 | 0 |
| SPGP0Q00190 | XP_018222481.1 | 97 | 0 |
| SPGP0CZ01910 | XP_018221012.1 | 99 | 1 |
| SPGP0P00630 | XP_018221893.1 | 99 | 0 |

Create PDF in your applications with the Pdfcrowd HTML to PDF API PDFCROWD

| | | | |
|---|---|---|---|
| SPGP0DZ01460 | XP_018219485.1 | 99 | 0 |
| SPGP0S01020 | XP_018220243.1 | 99 | 0 |
| SPGP0W02340 | XP_018223103.1 | 99 | 0 |
| SPGP0S00740 | XP_018220271.1 | 99 | 0 |
| SPGP0Z00650 | XP_018219819.1 | 99 | 0 |
| SPGP0P03030 | XP_018221661.1 | 99 | 0 |
| SPGP0W02170 | XP_018223120.1 | 99 | 0 |
| SPGP0W00980 | XP_018223236.1 | 99 | 0 |
| SPGP0E02250 | XP_018222813.1 | 99 | 0 |
| SPGP0DZ02320 | XP_018218749.1 | 100 | 0 |
| SPGP0P00960 | XP_018221861.1 | 99 | 0 |
| SPGP0DV00110 | XP_018219875.1 | 99 | 0 |
| SPGP0R01030 | XP_018221221.1 | 99 | 0 |
| SPGP0S00820 | XP_018220263.1 | 99 | 0 |
| SPGP0Q01140 | XP_018222387.1 | 99 | 0 |
| SPGP0S02570 | XP_018220096.1 | 99 | 0 |
| SPGP0Q00280 | XP_018222473.1 | 99 | 0 |
| SPGP0CZ01000 | XP_018220925.1 | 99 | 0 |
| SPGP0L00190 | XP_018219342.1 | 99 | 0 |
| SPGP0DN03430 | XP_018223786.1 | 100 | 0 |
| SPGP0AC00860 | XP_018219169.1 | 100 | 0 |
| SPGP0DZ00670 | XP_018219408.1 | 99 | 0 |
| SPGP0S03020 | XP_018220052.1 | 99 | 0 |
| SPGP0DN01300 | XP_018223577.1 | 99 | 0 |
| SPGP0B00450 | XP_018221493.1 | 99 | 0 |
| SPGP0S01880 | XP_018220164.1 | 99 | 0 |
| SPGP0S02210 | XP_018220131.1 | 99 | 0 |
| SPGP0Q01100 | XP_018222391.1 | 99 | 0 |
| SPGP0AA00110 | XP_018223326.1 | 99 | 0 |
| SPGP0W02200 | XP_018223117.1 | 100 | 0 |
| SPGP0P00330 | XP_018221921.1 | 99 | 0 |
| SPGP0Y01280 | XP_018223997.1 | 96 | 3 |
| SPGP0E02220 | XP_018222810.1 | 99 | 0 |
| SPGP0S00590 | XP_018220286.1 | 99 | 0 |
| SPGP0P00670 | XP_018221889.1 | 99 | 0 |
| SPGP0CZ00670 | XP_018220893.1 | 100 | 0 |
| SPGP0AA00520 | XP_018223364.1 | 99 | 0 |
| SPGP0P00130 | XP_018221941.1 | 99 | 0 |
| SPGP0H00730 | XP_018224064.1 | 99 | 0 |
| SPGP0Z02460 | XP_018219639.1 | 99 | 1 |

Create PDF in your applications with the Pdfcrowd HTML to PDF API PDFCROWD

| | | | |
|---|---|---|---|
| SPGP0CZ02190 | XP_018221039.1 | 99 | 0 |
| SPGP0S02810 | XP_018220073.1 | 99 | 0 |
| SPGP0Y01380 | XP_018224007.1 | 99 | 0 |
| SPGP0P02340 | XP_018221729.1 | 99 | 0 |
| SPGP0L01100 | XP_018219253.1 | 99 | 0 |
| SPGP0W00720 | XP_018223261.1 | 99 | 0 |
| SPGP0Z02200 | XP_018219665.1 | 99 | 0 |
| SPGP0Z02440 | XP_018219641.1 | 99 | 0 |
| SPGP0N01710 | XP_018220643.1 | 99 | 0 |
| SPGP0S03010 | XP_018220053.1 | 99 | 0 |
| SPGP0CZ02990 | XP_018221118.1 | 99 | 0 |
| SPGP0DV00130 | XP_018219877.1 | 99 | 0 |
| SPGP0CZ00590 | XP_018220885.1 | 99 | 0 |
| SPGP0S01270 | XP_018220219.1 | 99 | 0 |
| SPGP0Q01630 | XP_018222335.1 | 99 | 0 |
| SPGP0Ftrna33K | DI49_5564 | 100 | 0 |
| SPGP0Ftrna6C | DI49_5567 | 100 | 0 |
| SPGP0R00510 | XP_018221174.1 | 99 | 0 |
| SPGP0DZ00630 | XP_018219404.1 | 99 | 0 |
| SPGP0Q00760 | XP_018222425.1 | 99 | 0 |
| SPGP0Y01560 | XP_018224025.1 | 99 | 0 |
| SPGP0W01910 | XP_018223145.1 | 98 | 1 |
| SPGP0P00850 | XP_018221872.1 | 99 | 0 |
| SPGP0DN01870 | XP_018223634.1 | 99 | 0 |
| SPGP0Y01110 | XP_018223981.1 | 99 | 0 |
| SPGP0DN02640 | XP_018223709.1 | 99 | 0 |
| SPGP0DN02320 | XP_018223677.1 | 99 | 0 |
| SPGP0Q00220 | XP_018222478.1 | 99 | 0 |
| SPGP0Q03670 | XP_018222138.1 | 99 | 0 |
| SPGP0U01730 | XP_018218822.1 | 100 | 0 |
| SPGP0Q01960 | XP_018222302.1 | 99 | 0 |
| SPGP0W01660 | XP_018223169.1 | 97 | 3 |
| SPGP0S01370 | XP_018220210.1 | 99 | 0 |
| SPGP0Z00610 | XP_018219823.1 | 99 | 0 |
| SPGP0Z00690 | XP_018219815.1 | 99 | 0 |
| SPGP0U00180 | XP_018218971.1 | 99 | 0 |
| SPGP0E02340 | XP_018222822.1 | 99 | 0 |
| SPGP0DN01120 | XP_018223559.1 | 100 | 0 |
| SPGP0N00760 | XP_018220733.1 | 99 | 0 |
| SPGP0Y00130 | XP_018223891.1 | 99 | 0 |

Create PDF in your applications with the Pdfcrowd HTML to PDF API PDFCROWD

| | | | |
|---|---|---|---|
| SPGP0DU00230 | XP_018220567.1 | 99 | 0 |
| SPGP0DN00830 | XP_018223529.1 | 99 | 0 |
| SPGP0DN00560 | XP_018223503.1 | 99 | 0 |
| SPGP0Q02020 | XP_018222296.1 | 99 | 0 |
| SPGP0AA00190 | XP_018223332.1 | 100 | 0 |
| SPGP0H00370 | XP_018224098.1 | 99 | 0 |
| SPGP0U01630 | XP_018218831.1 | 99 | 0 |
| SPGP0Q04340 | XP_018222072.1 | 99 | 0 |
| SPGP0B00980 | XP_018221544.1 | 99 | 0 |
| SPGP0R02260 | XP_018221339.1 | 99 | 0 |
| SPGP0Z00400 | XP_018219856.1 | 99 | 0 |
| SPGP0F04560 | XP_018222035.1 | 99 | 0 |
| SPGP0S00380 | XP_018220307.1 | 99 | 0 |
| SPGP0AA00990 | XP_018223407.1 | 99 | 0 |
| SPGP0DN03340 | XP_018223777.1 | 99 | 0 |
| SPGP0R02100 | XP_018221325.1 | 99 | 0 |
| SPGP0DN02430 | XP_018223688.1 | 99 | 0 |
| SPGP0DN03670 | XP_018223809.1 | 99 | 0 |
| SPGP0CZ03120 | XP_018221130.1 | 99 | 0 |
| SPGP0DN03190 | XP_018223763.1 | 99 | 0 |
| SPGP0E00960 | XP_018222686.1 | 99 | 0 |
| SPGP0AF00230 | XP_018221964.1 | 99 | 0 |
| SPGP0Q02800 | XP_018222221.1 | 99 | 0 |
| SPGP0P01880 | XP_018221772.1 | 99 | 0 |
| SPGP0Q01130 | XP_018222388.1 | 100 | 0 |
| SPGP0Q04320 | XP_018222074.1 | 99 | 0 |
| SPGP0B00370 | XP_018221485.1 | 99 | 0 |
| SPGP0Y00330 | XP_018223910.1 | 99 | 0 |
| SPGP0Z00120 | XP_018219871.1 | 99 | 0 |
| SPGP0Q02110 | XP_018222287.1 | 99 | 0 |
| SPGP0P02430 | XP_018221720.1 | 99 | 0 |
| SPGP0W04350 | XP_018222915.1 | 99 | 0 |
| SPGP0W04600 | XP_018222888.1 | 99 | 0 |
| SPGP0DZ01650 | XP_018219503.1 | 99 | 0 |
| SPGP0Itrna16M | DI49_5351 | 100 | 0 |
| SPGP0U00210 | XP_018218968.1 | 99 | 0 |
| SPGP0W01050 | XP_018223229.1 | 99 | 0 |
| SPGP0Z02190 | XP_018219666.1 | 99 | 0 |
| SPGP0P01230 | XP_018221834.1 | 100 | 0 |
| SPGP0W03570 | XP_018222991.1 | 99 | 0 |

Create PDF in your applications with the Pdfcrowd HTML to PDF API PDFCROWD

| | | | |
|---|---|---|---|
| SPGP0U01770 | XP_018218817.1 | 99 | 0 |
| SPGP0Y01050 | XP_018223975.1 | 99 | 0 |
| SPGP0S01460 | XP_018220202.1 | 99 | 0 |
| SPGP0Z01550 | XP_018219730.1 | 99 | 0 |
| SPGP0U00510 | XP_018218938.1 | 100 | 0 |
| SPGP0W01580 | XP_018223177.1 | 100 | 0 |
| SPGP0CZ02870 | XP_018221106.1 | 99 | 0 |
| SPGP0W00510 | XP_018223283.1 | 99 | 0 |
| SPGP0Z03120 | XP_018219573.1 | 100 | 0 |
| SPGP0DN02820 | XP_018223726.1 | 99 | 0 |
| SPGP0DN02570 | XP_018223702.1 | 99 | 0 |
| SPGP0F04760 | XP_018222018.1 | 99 | 0 |
| SPGP0P01480 | XP_018221810.1 | 100 | 0 |
| SPGP0EE00140 | XP_018220833.1 | 99 | 0 |
| SPGP0W02110 | XP_018223125.1 | 99 | 0 |
| SPGP0Z01750 | XP_018219710.1 | 100 | 0 |
| SPGP0R02450 | XP_018221358.1 | 99 | 0 |
| SPGP0CZ01400 | XP_018220964.1 | 99 | 0 |
| SPGP0DN01810 | XP_018223627.1 | 99 | 0 |
| SPGP0AA00390 | XP_018223351.1 | 99 | 0 |
| SPGP0S01920 | XP_018220160.1 | 99 | 0 |
| SPGP0W01640 | XP_018223171.1 | 99 | 0 |
| SPGP0DZ00480 | XP_018219391.1 | 100 | 0 |
| SPGP0R02470 | XP_018221360.1 | 99 | 0 |
| SPGP0CZ02220 | XP_018221042.1 | 99 | 0 |
| SPGP0CZ02320 | XP_018221052.1 | 99 | 0 |
| SPGP0S00490 | XP_018220296.1 | 99 | 0 |
| SPGP0P02070 | XP_018221753.1 | 99 | 0 |
| SPGP0Q03520 | XP_018222152.1 | 99 | 0 |
| SPGP0AA00170 | XP_018223330.1 | 99 | 0 |
| SPGP0DN04080 | XP_018223850.1 | 99 | 0 |
| SPGP0Z00440 | XP_018219839.1 | 99 | 0 |
| SPGP0X00300 | XP_018219071.1 | 100 | 0 |
| SPGP0Z00830 | XP_018219801.1 | 99 | 0 |
| SPGP0DN00500 | XP_018223498.1 | 99 | 0 |
| SPGP0Z02050 | XP_018219680.1 | 99 | 0 |
| SPGP0DN00150 | XP_018223465.1 | 99 | 0 |
| SPGP0AH01060 | XP_018222502.1 | 99 | 0 |
| SPGP0P02410 | XP_018221722.1 | 99 | 0 |

Create PDF in your applications with the Pdfcrowd HTML to PDF API PDFCROWD

| | | | |
|---|---|---|---|
| SPGP0Y00910 | XP_018223961.1 | 99 | 0 |
| SPGP0N01230 | XP_018220690.1 | 99 | 0 |
| SPGP0H00820 | XP_018224055.1 | 99 | 0 |
| SPGP0W04370 | XP_018222912.1 | 99 | 0 |
| SPGP0P01780 | XP_018221782.1 | 99 | 0 |
| SPGP0S00900 | XP_018220255.1 | 99 | 0 |
| SPGP0Q02460 | XP_018222253.1 | 100 | 0 |
| SPGP0Q01840 | XP_018222314.1 | 99 | 0 |
| SPGP0U01820 | XP_018218812.1 | 99 | 0 |
| SPGP0W00700 | XP_018223263.1 | 99 | 0 |
| SPGP0Z02860 | XP_018219598.1 | 99 | 0 |
| SPGP0Q03770 | XP_018222128.1 | 99 | 0 |
| SPGP0E00910 | XP_018222680.1 | 99 | 0 |
| SPGP0N02190 | XP_018220595.1 | 99 | 0 |
| SPGP0DN01340 | XP_018223581.1 | 99 | 0 |
| SPGP0L00930 | XP_018219270.1 | 99 | 0 |
| SPGP0L01060 | XP_018219257.1 | 98 | 2 |
| SPGP0DN01830 | XP_018223630.1 | 99 | 0 |
| SPGP0P02530 | XP_018221710.1 | 99 | 0 |
| SPGP0W01290 | XP_018223205.1 | 99 | 0 |
| SPGP0S00230 | XP_018220321.1 | 99 | 0 |
| SPGP0N00260 | XP_018220782.1 | 99 | 0 |
| SPGP0Q02360 | XP_018222263.1 | 99 | 0 |
| SPGP0L00540 | XP_018219307.1 | 99 | 0 |
| SPGP0P00190 | XP_018221935.1 | 99 | 0 |
| SPGP0W03880 | XP_018222960.1 | 99 | 0 |
| SPGP0B00140 | XP_018221462.1 | 97 | 0 |
| SPGP0DN03470 | XP_018223790.1 | 99 | 0 |
| SPGP0Q00950 | XP_018222406.1 | 99 | 0 |
| SPGP0L00670 | XP_018219296.1 | 99 | 0 |
| SPGP0S01900 | XP_018220162.1 | 99 | 0 |
| SPGP0AF00160 | XP_018221957.1 | 99 | 0 |
| SPGP0R02220 | XP_018221335.1 | 99 | 0 |
| SPGP0S00760 | XP_018220269.1 | 99 | 0 |
| SPGP0AH00580 | XP_018222547.1 | 99 | 0 |
| SPGP0DN00770 | XP_018223523.1 | 99 | 0 |
| SPGP0CZ01540 | XP_018220978.1 | 99 | 0 |
| SPGP0W01430 | XP_018223191.1 | 99 | 0 |
| SPGP0Q00240 | XP_018222476.1 | 95 | 5 |
| SPGP0Z01980 | XP_018219687.1 | 99 | 0 |

Create PDF in your applications with the Pdfcrowd HTML to PDF API PDFCROWD

| | | | |
|---|---|---|---|
| SPGP0P02800 | XP_018221683.1 | 99 | 0 |
| SPGP0DU01500 | XP_018220440.1 | 100 | 0 |
| SPGP0W04090 | XP_018222941.1 | 99 | 0 |
| SPGP0DU00670 | XP_018220524.1 | 99 | 0 |
| SPGP0Y01730 | XP_018224040.1 | 99 | 0 |
| SPGP0Q03180 | XP_018222184.1 | 99 | 0 |
| SPGP0EP00110 | XP_018220336.1 | 99 | 0 |
| SPGP0AA00200 | XP_018223333.1 | 99 | 0 |
| SPGP0DU00370 | XP_018220553.1 | 99 | 0 |
| SPGP0P01270 | XP_018221830.1 | 99 | 0 |
| SPGP0L00280 | XP_018219333.1 | 99 | 0 |
| SPGP0EC00100 | XP_018220412.1 | 99 | 0 |
| SPGP0AA00450 | XP_018223357.1 | 98 | 2 |
| SPGP0W03450 | XP_018223001.1 | 99 | 0 |
| SPGP0CZ02250 | XP_018221045.1 | 99 | 0 |
| SPGP0DU00410 | XP_018220549.1 | 99 | 0 |
| SPGP0S01940 | XP_018220158.1 | 99 | 0 |
| SPGP0S02320 | XP_018220120.1 | 99 | 0 |
| SPGP0P02050 | XP_018221755.1 | 99 | 0 |
| SPGP0AH01100 | XP_018222498.1 | 99 | 0 |
| SPGP0Z00790 | XP_018219805.1 | 100 | 0 |
| SPGP0N01030 | XP_018220708.1 | 99 | 0 |
| SPGP0Z02300 | XP_018219655.1 | 99 | 1 |
| SPGP0Q00580 | XP_018222443.1 | 99 | 0 |
| SPGP0W02660 | XP_018223073.1 | 99 | 0 |
| SPGP0H00810 | XP_018224056.1 | 99 | 0 |
| SPGP0P00730 | XP_018221883.1 | 99 | 0 |
| SPGP0DX00190 | XP_018219950.1 | 99 | 0 |
| SPGP0B01830 | XP_018221625.1 | 97 | 1 |
| SPGP0DN00930 | XP_018223539.1 | 99 | 0 |
| SPGP0Htrna32S | DI49_4419 | 100 | 0 |
| SPGP0W00260 | XP_018223308.1 | 99 | 0 |
| SPGP0Z02910 | XP_018219594.1 | 99 | 0 |
| SPGP0E02460 | XP_018222833.1 | 99 | 0 |
| SPGP0B01560 | XP_018221598.1 | 99 | 0 |
| SPGP0W00220 | XP_018223312.1 | 99 | 0 |
| SPGP0L00610 | XP_018219302.1 | 99 | 0 |
| SPGP0S01040 | XP_018220241.1 | 99 | 0 |
| SPGP0Q03480 | XP_018222156.1 | 98 | 0 |
| SPGP0P02960 | XP_018221668.1 | 99 | 0 |

Create PDF in your applications with the Pdfcrowd HTML to PDF API PDFCROWD

| | | | |
|---|---|---|---|
| SPGP0W01030 | XP_018223231.1 | 100 | 0 |
| SPGP0N00980 | XP_018220713.1 | 99 | 0 |
| SPGP0X00670 | XP_018219035.1 | 100 | 0 |
| SPGP0R00990 | XP_018221217.1 | 99 | 0 |
| SPGP0DN00890 | XP_018223535.1 | 99 | 0 |
| SPGP0AF00320 | XP_018221973.1 | 99 | 0 |
| SPGP0AH00850 | XP_018222520.1 | 100 | 0 |
| SPGP0H00310 | XP_018224104.1 | 100 | 0 |
| SPGP0Y01450 | XP_018224014.1 | 99 | 0 |
| SPGP0S02980 | XP_018220056.1 | 99 | 0 |
| SPGP0L00200 | XP_018219341.1 | 100 | 0 |
| SPGP0AA00350 | XP_018223348.1 | 99 | 0 |
| SPGP0S00720 | XP_018220273.1 | 100 | 0 |
| SPGP0DN02790 | XP_018223723.1 | 99 | 0 |
| SPGP0Z02360 | XP_018219649.1 | 100 | 0 |
| SPGP0B00800 | XP_018221527.1 | 99 | 0 |
| SPGP0DZ01360 | XP_018219475.1 | 99 | 0 |
| SPGP0W03550 | XP_018222993.1 | 99 | 0 |
| SPGP0Q04070 | XP_018222098.1 | 99 | 0 |
| SPGP0H00110 | XP_018224122.1 | 99 | 0 |
| SPGP0E00700 | XP_018222660.1 | 99 | 0 |
| SPGP0W03010 | XP_018223040.1 | 99 | 0 |
| SPGP0DN02160 | XP_018223662.1 | 99 | 0 |
| SPGP0X00130 | XP_018219087.1 | 99 | 0 |
| SPGP0Z00820 | XP_018219802.1 | 99 | 0 |
| SPGP0Q03710 | XP_018222134.1 | 99 | 0 |
| SPGP0R01120 | XP_018221230.1 | 99 | 0 |
| SPGP0Q03880 | XP_018222117.1 | 99 | 0 |
| SPGP0S02850 | XP_018220069.1 | 99 | 0 |
| SPGP0X00530 | XP_018219049.1 | 99 | 0 |
| SPGP0Z00670 | XP_018219817.1 | 99 | 0 |
| SPGP0P02710 | XP_018221692.1 | 99 | 0 |
| SPGP0P00280 | XP_018221926.1 | 100 | 0 |
| SPGP0E00180 | XP_018222608.1 | 100 | 0 |
| SPGP0U00920 | XP_018218900.1 | 99 | 0 |
| SPGP0AF00120 | XP_018221954.1 | 99 | 0 |
| SPGP0P02760 | XP_018221687.1 | 99 | 0 |
| SPGP0R00360 | XP_018221159.1 | 99 | 0 |
| SPGP0Q03010 | XP_018222200.1 | 98 | 1 |
| SPGP0L01210 | XP_018219243.1 | 100 | 0 |

Create PDF in your applications with the Pdfcrowd HTML to PDF API PDFCROWD

| | | | |
|---|---|---|---|
| SPGP0W00890 | XP_018223245.1 | 99 | 0 |
| SPGP0E01440 | XP_018222733.1 | 99 | 0 |
| SPGP0L01510 | XP_018219213.1 | 99 | 1 |
| SPGP0AA00620 | XP_018223374.1 | 99 | 0 |
| SPGP0Z02610 | XP_018219624.1 | 99 | 0 |
| SPGP0EL00110 | XP_018219092.1 | 99 | 0 |
| SPGP0DV00290 | XP_018219892.1 | 99 | 0 |
| SPGP0S03370 | XP_018220018.1 | 99 | 0 |
| SPGP0W02260 | XP_018223111.1 | 99 | 0 |
| SPGP0AG00150 | XP_018219934.1 | 99 | 0 |
| SPGP0DZ01800 | XP_018219518.1 | 99 | 0 |
| SPGP0S01980 | XP_018220154.1 | 99 | 0 |
| SPGP0X00510 | XP_018219051.1 | 99 | 0 |
| SPGP0E01980 | XP_018222786.1 | 99 | 0 |
| SPGP0CZ02890 | XP_018221108.1 | 99 | 0 |
| SPGP0Z01990 | XP_018219686.1 | 100 | 0 |
| SPGP0E01800 | XP_018222768.1 | 99 | 0 |
| SPGP0S01450 | XP_018220203.1 | 99 | 1 |
| SPGP0R01770 | XP_018221294.1 | 99 | 0 |
| SPGP0W04270 | XP_018222923.1 | 99 | 0 |
| SPGP0S03290 | XP_018220026.1 | 99 | 0 |
| SPGP0Y01620 | XP_018224031.1 | 99 | 0 |
| SPGP0DZ02250 | XP_018219562.1 | 99 | 0 |
| SPGP0R00660 | XP_018221188.1 | 98 | 1 |
| SPGP0CZ00800 | XP_018220905.1 | 99 | 0 |
| SPGP0Z02630 | XP_018219622.1 | 99 | 0 |
| SPGP0H00540 | XP_018224082.1 | 99 | 0 |
| SPGP0AC00630 | XP_018219147.1 | 99 | 0 |
| SPGP0L00950 | XP_018219268.1 | 99 | 0 |
| SPGP0W00780 | XP_018223256.1 | 99 | 0 |
| SPGP0AF00520 | XP_018221992.1 | 99 | 0 |
| SPGP0P02810 | XP_018221682.1 | 99 | 0 |
| SPGP0Q02820 | XP_018222219.1 | 99 | 0 |
| SPGP0E00490 | XP_018222639.1 | 99 | 0 |
| SPGP0P02280 | XP_018221735.1 | 99 | 0 |
| SPGP0Q01400 | XP_018222359.1 | 99 | 0 |
| SPGP0R00190 | XP_018221142.1 | 99 | 0 |
| SPGP0L00750 | XP_018219288.1 | 99 | 0 |
| SPGP0P02630 | XP_018221700.1 | 99 | 0 |
| SPGP0W03680 | XP_018222980.1 | 99 | 0 |

Create PDF in your applications with the Pdfcrowd HTML to PDF API PDFCROWD

| | | | |
|---|---|---|---|
| SPGP0W00940 | XP_018223240.1 | 99 | 0 |
| SPGP0Z03020 | XP_018219583.1 | 99 | 0 |

### Sheet 2: *genes alleles inherited from Saccharomyces cerevisiae*

| *Saccharomyces pastorianus CBS1513 predicted ORF* | *Saccharomyces cerevisiae parental allele* | Gene identities | Gene gaps |
|---|---|---|---|
| SPGP0DC00130 | YBL104C | 99 | 0 |
| SPGP0M03690 | YNR061C | 99 | 0 |
| SPGP0F01340 | YGR150C | 99 | 0 |
| SPGP0DD00220 | YIL156W-B | 99 | 0 |
| SPGP0C00910 | YDL153C | 99 | 0 |
| SPGP0AI00270 | YHR196W | 99 | 0 |
| SPGP0AI00160 | YHR206W | 99 | 0 |
| SPGP0I02270 | YER146W | 100 | 0 |
| SPGP0I00710 | YEL011W | 99 | 0 |
| SPGP0I01990 | YER114C | 99 | 0 |
| SPGP0AE00480 | YCL021W-A | 99 | 0 |
| SPGP0M03180 | YNR010W | 100 | 0 |
| SPGP0F00680 | YGR222W | 99 | 0 |
| SPGP0Q04720 | YGL216W | 100 | 0 |
| SPGP0F00930 | YGR196C | 99 | 0 |
| SPGP0C04580 | YDR256C | 99 | 0 |
| SPGP0DB02690 | YMR017W | 99 | 0 |
| SPGP0K00570 | YOL112W | 99 | 0 |
| SPGP0D00480 | YPR029C | 99 | 0 |
| SPGP0I01670 | YER081W | 100 | 0 |
| SPGP0M01040 | YNL227C | 99 | 0 |
| SPGP0F03010 | YGL038C | 99 | 0 |
| SPGP0DB02290 | YMR062C | 99 | 0 |
| SPGP0G01120 | YJR046W | 99 | 0 |
| SPGP0DB03210 | YML037C | 99 | 0 |
| SPGP0C05180 | YDR320C-A | 99 | 0 |
| SPGP0Q05020 | YGL249W | 99 | 0 |
| SPGP0DB01920 | YMR101C | 99 | 0 |
| SPGP0D00280 | YPR052C | 99 | 0 |
| SPGP0Q04990 | YGL246C | 99 | 0 |
| SPGP0I00690 | YEL013W | 99 | 0 |
| SPGP0K02500 | YOR097C | 99 | 0 |
| SPGP0DC03010 | YBR208C | 99 | 0 |

Create PDF in your applications with the Pdfcrowd HTML to PDF API PDFCROWD

| | | | |
|---|---|---|---|
| SPGP0DB02740 | YMR012W | 99 | 0 |
| SPGP0F01560 | YGR126W | 100 | 0 |
| SPGP0V00490 | YHL003C | 99 | 0 |
| SPGP0DB02030 | YMR090W | 99 | 0 |
| SPGP0M01240 | YNL206C | 99 | 0 |
| SPGP0G01870 | YJL047C | 99 | 0 |
| SPGP0G01690 | YJL026W | 99 | 0 |
| SPGP0D00560 | YPR021C | 99 | 0 |
| SPGP0K01620 | YOL003C | 99 | 0 |
| SPGP0V02000 | YHR150W | 99 | 0 |
| SPGP0DC01260 | YBR019C | 99 | 0 |
| SPGP0C03870 | YDR175C | 99 | 0 |
| SPGP0DC02700 | YBR175W | 99 | 0 |
| SPGP0AI00170 | YHR205W | 99 | 0 |
| SPGP0DB02650 | YMR021C | 99 | 0 |
| SPGP0K00420 | YOL128C | 99 | 0 |
| SPGP0I02430 | YER163C | 99 | 0 |
| SPGP0F01830 | YGR096W | 100 | 0 |
| SPGP0M00580 | YNL278W | 99 | 0 |
| SPGP0C02320 | YDR006C | 99 | 0 |
| SPGP0F00260 | YGR270W | 99 | 0 |
| SPGP0DB00520 | YMR243C | 99 | 0 |
| SPGP0U01950 | YPL243W | 99 | 0 |
| SPGP0C01450 | YDL093W | 99 | 0 |
| SPGP0G00610 | YJR100C | 99 | 0 |
| SPGP0K01560 | YOL009C | 99 | 0 |
| SPGP0D00640 | YPR011C | 99 | 0 |
| SPGP0DC01010 | YBL008W | 99 | 0 |
| SPGP0DB01210 | YMR175W | 100 | 0 |
| SPGP0V00390 | YHL014C | 99 | 0 |
| SPGP0DB00460 | YMR251W | 99 | 0 |
| SPGP0DD01500 | YIL016W | 99 | 0 |
| SPGP0DB00290 | YMR268C | 99 | 0 |
| SPGP0DB01170 | YMR179W | 99 | 0 |
| SPGP0DC00540 | YBL056W | 99 | 0 |
| SPGP0C02610 | YDR040C | 98 | 0 |
| SPGP0DB02340 | YMR056C | 99 | 0 |
| SPGP0C01870 | YDL048C | 99 | 0 |
| SPGP0M02160 | YNL100W | 100 | 0 |
| SPGP0G02670 | YJL136C | 99 | 0 |

Create PDF in your applications with the Pdfcrowd HTML to PDF API PDFCROWD

| | | | |
|---|---|---|---|
| SPGP0V01180 | YHR068W | 99 | 0 |
| SPGP0DD01750 | YIR008C | 99 | 0 |
| SPGP0DD01270 | YIL043C | 99 | 0 |
| SPGP0K01550 | YOL010W | 100 | 0 |
| SPGP0V01630 | YHR112C | 99 | 0 |
| SPGP0G02830 | YJL155C | 99 | 0 |
| SPGP0M02350 | YNL080C | 99 | 0 |
| SPGP0F01250 | YGR161C | 100 | 0 |
| SPGP0M02820 | YNL022C | 100 | 0 |
| SPGP0C01060 | YDL135C | 99 | 0 |
| SPGP0DC03760 | YBR288C | 99 | 0 |
| SPGP0M00730 | YNL260C | 100 | 0 |
| SPGP0D00400 | YPR036W-A | 100 | 0 |
| SPGP0DE00710 | YAR002C-A | 99 | 0 |
| SPGP0V00320 | YHL021C | 99 | 0 |
| SPGP0DE00240 | YAL046C | 99 | 0 |
| SPGP0DE00200 | YAL051W | 99 | 0 |
| SPGP0DC01100 | YBR002C | 99 | 0 |
| SPGP0Q04510 | YGL192W | 99 | 0 |
| SPGP0C00780 | YDL168W | 99 | 0 |
| SPGP0F00110 | YGR285C | 99 | 0 |
| SPGP0F01050 | YGR183C | 99 | 0 |
| SPGP0C05150 | YDR318W | 99 | 0 |
| SPGP0V01980 | YHR148W | 99 | 0 |
| SPGP0V01450 | YHR094C | 99 | 0 |
| SPGP0DE00650 | YAL005C | 99 | 0 |
| SPGP0EA00120 | YGR294W | 97 | 0 |
| SPGP0D00930 | YPL018W | 99 | 0 |
| SPGP0K00980 | YOL070C | 99 | 0 |
| SPGP0K00610 | YOL108C | 99 | 0 |
| SPGP0C03030 | YDR083W | 99 | 0 |
| SPGP0G00860 | YJR073C | 99 | 0 |
| SPGP0C03740 | YDR163W | 99 | 0 |
| SPGP0AE00830 | YCR020C-A | 99 | 0 |
| SPGP0F02580 | YGR009C | 99 | 0 |
| SPGP0DD00870 | YIL091C | 99 | 0 |
| SPGP0C02940 | YDR075W | 100 | 0 |
| SPGP0K01740 | YOR010C | 95 | 0 |
| SPGP0M02020 | YNL117W | 99 | 0 |
| SPGP0DC02030 | YBR103W | 99 | 0 |

Create PDF in your applications with the Pdfcrowd HTML to PDF API PDFCROWD

| | | | |
|---|---|---|---|
| SPGP0F03710 | YGL119W | 99 | 0 |
| SPGP0I01380 | YER054C | 99 | 0 |
| SPGP0DM00100 | YHR211W | 94 | 0 |
| SPGP0DC03240 | YBR235W | 99 | 0 |
| SPGP0V00580 | YHR005C-A | 99 | 0 |
| SPGP0Q04860 | YGL231C | 100 | 0 |
| SPGP0K01750 | YOR011W | 99 | 0 |
| SPGP0K00320 | YOL139C | 99 | 0 |
| SPGP0I02620 | YER182W | 99 | 0 |
| SPGP0U02160 | YPL264C | 99 | 0 |
| SPGP0I01460 | YER061C | 99 | 0 |
| SPGP0K02470 | YOR094W | 100 | 0 |
| SPGP0G03220 | YJL199C | 99 | 0 |
| SPGP0C00210 | YDL231C | 99 | 0 |
| SPGP0DD01260 | YIL044C | 99 | 0 |
| SPGP0I02040 | YER119C | 99 | 0 |
| SPGP0DB03170 | YML032C | 99 | 0 |
| SPGP0DC03630 | YBR274W | 99 | 0 |
| SPGP0K01810 | YOR019W | 99 | 0 |
| SPGP0V00910 | YHR039C | 99 | 0 |
| SPGP0DB00660 | YMR229C | 99 | 0 |
| SPGP0C04140 | YDR205W | 99 | 0 |
| SPGP0I00750 | YEL005C | 99 | 0 |
| SPGP0M01670 | YNL156C | 99 | 0 |
| SPGP0D00780 | YPL003W | 99 | 0 |
| SPGP0K01690 | YOR005C | 99 | 0 |
| SPGP0AE01260 | YCR072C | 99 | 0 |
| SPGP0F00440 | YGR249W | 99 | 0 |
| SPGP0AE01010 | YCR038C | 99 | 0 |
| SPGP0I00720 | YEL009C | 99 | 0 |
| SPGP0M00500 | YNL287W | 100 | 0 |
| SPGP0C00340 | YDL216C | 99 | 0 |
| SPGP0C03690 | YDR158W | 99 | 0 |
| SPGP0G00420 | YJR118C | 99 | 0 |
| SPGP0DC02990 | YBR205W | 99 | 0 |
| SPGP0DD00480 | YIL130W | 98 | 1 |
| SPGP0DD00780 | YIL101C | 99 | 0 |
| SPGP0Q04530 | YGL194C-A | 99 | 0 |
| SPGP0M01900 | YNL130C | 99 | 0 |
| SPGP0M00860 | YNL247W | 99 | 0 |

Create PDF in your applications with the Pdfcrowd HTML to PDF API PDFCROWD

| | | | |
|---|---|---|---|
| SPGP0DC03070 | YBR215W | 99 | 0 |
| SPGP0M01450 | YNL181W | 99 | 0 |
| SPGP0DB03000 | YML015C | 99 | 0 |
| SPGP0DD01080 | YIL066C | 99 | 0 |
| SPGP0G03380 | YJL217W | 98 | 0 |
| SPGP0C00290 | YDL222C | 99 | 0 |
| SPGP0AE00110 | YCL063W | 99 | 0 |
| SPGP0AE00380 | YCL032W | 99 | 0 |
| SPGP0DC03160 | YBR227C | 99 | 0 |
| SPGP0I01800 | YER093C-A | 97 | 0 |
| SPGP0D00190 | YPR063C | 100 | 0 |
| SPGP0C03390 | YDR123C | 99 | 0 |
| SPGP0DC02100 | YBR110W | 99 | 0 |
| SPGP0K01610 | YOL004W | 99 | 0 |
| SPGP0U02110 | YPL258C | 99 | 0 |
| SPGP0C04770 | YDR279W | 99 | 0 |
| SPGP0C04880 | YDR291W | 99 | 0 |
| SPGP0I00620 | YEL020C | 99 | 0 |
| SPGP0DC00850 | YBL025W | 99 | 0 |
| SPGP0M00400 | YNL298W | 99 | 0 |
| SPGP0C02510 | YDR027C | 99 | 0 |
| SPGP0I02300 | YER149C | 100 | 0 |
| SPGP0C03160 | YDR098C | 99 | 0 |
| SPGP0G03280 | YJL206C | 99 | 0 |
| SPGP0AE00930 | YCR030C | 99 | 0 |
| SPGP0DD01050 | YIL069C | 99 | 0 |
| SPGP0D00220 | YPR060C | 100 | 0 |
| SPGP0M00610 | YNL274C | 99 | 0 |
| SPGP0M01350 | YNL192W | 99 | 0 |
| SPGP0C04820 | YDR284C | 99 | 0 |
| SPGP0V01700 | YHR119W | 99 | 0 |
| SPGP0AE00350 | YCL035C | 99 | 0 |
| SPGP0AE00600 | YCL004W | 99 | 0 |
| SPGP0K01090 | YOL059W | 100 | 0 |
| SPGP0AE00170 | YCL056C | 99 | 0 |
| SPGP0K02300 | YOR075W | 99 | 0 |
| SPGP0G01480 | YJR003C | 99 | 0 |
| SPGP0AE01180 | YCR061W | 99 | 0 |
| SPGP0D00890 | YPL014W | 99 | 0 |
| SPGP0DB00880 | YMR207C | 99 | 0 |

Create PDF in your applications with the Pdfcrowd HTML to PDF API PDFCROWD

| | | | |
|---|---|---|---|
| SPGP0K02730 | YOR123C | 99 | 0 |
| SPGP0K02040 | YOR044W | 99 | 0 |
| SPGP0DC01090 | YBR001C | 99 | 0 |
| SPGP0C02590 | YDR036C | 99 | 0 |
| SPGP0M02100 | YNL107W | 100 | 0 |
| SPGP0K00390 | YOL131W | 99 | 0 |
| SPGP0DC01320 | YBR025C | 99 | 0 |
| SPGP0K02070 | YOR047C | 99 | 0 |
| SPGP0DC00110 | YBL106C | 99 | 0 |
| SPGP0DC02020 | YBR102C | 99 | 0 |
| SPGP0M03070 | YNL003C | 100 | 0 |
| SPGP0DD00670 | YIL111W | 99 | 0 |
| SPGP0DE00550 | YAL016W | 99 | 0 |
| SPGP0I00380 | YEL044W | 100 | 0 |
| SPGP0F03780 | YGL126W | 99 | 0 |
| SPGP0F04080 | YGL159W | 99 | 0 |
| SPGP0DC01890 | YBR085C-A | 100 | 0 |
| SPGP0G01680 | YJL025W | 99 | 0 |
| SPGP0DC02200 | YBR121C | 99 | 0 |
| SPGP0C04760 | YDR277C | 99 | 0 |
| SPGP0DE00450 | YAL026C | 99 | 0 |
| SPGP0DC00200 | YBL095W | 99 | 0 |
| SPGP0F02420 | YGR029W | 100 | 0 |
| SPGP0V00460 | YHL007C | 99 | 0 |
| SPGP0DE00500 | YAL021C | 99 | 0 |
| SPGP0F02530 | YGR015C | 99 | 0 |
| SPGP0DE00820 | YAR028W | 98 | 0 |
| SPGP0F02240 | YGR049W | 99 | 0 |
| SPGP0F02320 | YGR040W | 99 | 0 |
| SPGP0DD01180 | YIL052C | 99 | 0 |
| SPGP0I00120 | YEL071W | 99 | 0 |
| SPGP0F03580 | YGL104C | 99 | 0 |
| SPGP0F01450 | YGR138C | 99 | 0 |
| SPGP0I01550 | YER070W | 99 | 0 |
| SPGP0F01530 | YGR129W | 99 | 0 |
| SPGP0DD01860 | YIR019C | 98 | 1 |
| SPGP0DC00340 | YBL080C | 99 | 0 |
| SPGP0C00680 | YDL179W | 100 | 0 |
| SPGP0G02900 | YJL162C | 98 | 1 |
| SPGP0DC01050 | YBL004W | 99 | 0 |

Create PDF in your applications with the Pdfcrowd HTML to PDF API PDFCROWD

| | | | |
|---|---|---|---|
| SPGP0C05010 | YDR304C | 99 | 0 |
| SPGP0F04160 | YGL169W | 99 | 0 |
| SPGP0DC01680 | YBR064W | 99 | 0 |
| SPGP0C03480 | YDR132C | 99 | 0 |
| SPGP0Q04760 | YGL222C | 99 | 0 |
| SPGP0M03760 | YNR070W | 99 | 0 |
| SPGP0I01250 | YER044C | 100 | 0 |
| SPGP0I01910 | YER105C | 99 | 0 |
| SPGP0DB01320 | YMR164C | 97 | 2 |
| SPGP0V01840 | YHR134W | 100 | 0 |
| SPGP0DB00500 | YMR244C-A | 99 | 0 |
| SPGP0M02890 | YNL029C | 99 | 0 |
| SPGP0DB02900 | YML005W | 99 | 0 |
| SPGP0M02580 | YNL055C | 99 | 0 |
| SPGP0C00110 | YDL241W | 98 | 0 |
| SPGP0G02630 | YJL132W | 99 | 0 |
| SPGP0DB00650 | YMR230W | 100 | 0 |
| SPGP0M00260 | YNL313C | 99 | 0 |
| SPGP0G01380 | YJR011C | 99 | 0 |
| SPGP0C03260 | YDR109C | 99 | 0 |
| SPGP0G02070 | YJL068C | 98 | 0 |
| SPGP0G01610 | YJL013C | 99 | 0 |
| SPGP0I01070 | YER026C | 99 | 0 |
| SPGP0K00110 | YOL159C-A | 99 | 0 |
| SPGP0I01780 | YER092W | 99 | 0 |
| SPGP0C04210 | YDR214W | 99 | 0 |
| SPGP0K00800 | YOL089C | 99 | 0 |
| SPGP0AI00290 | YHR194W | 99 | 0 |
| SPGP0K00550 | YOL114C | 99 | 0 |
| SPGP0G00850 | YJR074W | 99 | 0 |
| SPGP0G00190 | YJR142W | 99 | 0 |
| SPGP0Q04670 | YGL210W | 99 | 0 |
| SPGP0C01230 | YDL117W | 99 | 0 |
| SPGP0F00660 | YGR224W | 99 | 0 |
| SPGP0I00870 | YER007C-A | 99 | 0 |
| SPGP0M01020 | YNL230C | 99 | 0 |
| SPGP0I01690 | YER083C | 99 | 0 |
| SPGP0DB02250 | YMR066W | 99 | 0 |
| SPGP0F02120 | YGR063C | 99 | 0 |
| SPGP0C01260 | YDL114W | 99 | 0 |

Create PDF in your applications with the Pdfcrowd HTML to PDF API PDFCROWD

| | | | |
|---|---|---|---|
| SPGP0G01100 | YJR048W | 100 | 0 |
| SPGP0F02310 | YGR041W | 99 | 0 |
| SPGP0C01440 | YDL095W | 99 | 0 |
| SPGP0D00210 | YPR061C | 99 | 0 |
| SPGP0DB03230 | YML041C | 99 | 0 |
| SPGP0I02520 | YER172C | 99 | 0 |
| SPGP0C05160 | YDR319C | 99 | 0 |
| SPGP0DD01470 | YIL019W | 99 | 0 |
| SPGP0C04900 | YDR293C | 99 | 0 |
| SPGP0DC01300 | YBR023C | 99 | 0 |
| SPGP0DB01900 | YMR104C | 99 | 0 |
| SPGP0DD00610 | YIL117C | 99 | 0 |
| SPGP0AE00290 | YCL043C | 99 | 0 |
| SPGP0I01840 | YER098W | 99 | 0 |
| SPGP0C04830 | YDR285W | 99 | 0 |
| SPGP0G01810 | YJL041W | 99 | 0 |
| SPGP0DB02010 | YMR092C | 99 | 0 |
| SPGP0C00980 | YDL144C | 99 | 0 |
| SPGP0C03890 | YDR177W | 99 | 0 |
| SPGP0F02400 | YGR031W | 99 | 0 |
| SPGP0K02430 | YOR090C | 99 | 0 |
| SPGP0DC01240 | YBR017C | 99 | 0 |
| SPGP0C01080 | YDL133W | 99 | 0 |
| SPGP0K01540 | YOL011W | 99 | 0 |
| SPGP0C01270 | YDL113C | 99 | 0 |
| SPGP0F01810 | YGR098C | 99 | 0 |
| SPGP0I00780 | YEL002C | 99 | 0 |
| SPGP0DD00330 | YIL146C | 99 | 0 |
| SPGP0G03060 | YJL180C | 99 | 0 |
| SPGP0DD01690 | YIR002C | 99 | 0 |
| SPGP0AE00910 | YCR028C | 99 | 0 |
| SPGP0I00150 | YEL069C | 98 | 0 |
| SPGP0G00670 | YJR094W-A | 100 | 0 |
| SPGP0M01960 | YNL124W | 99 | 0 |
| SPGP0Q04490 | YGL190C | 99 | 0 |
| SPGP0M02590 | YNL054W | 99 | 1 |
| SPGP0C04470 | YDR244W | 99 | 0 |
| SPGP0M00810 | YNL252C | 99 | 0 |
| SPGP0C02300 | YDR004W | 99 | 0 |
| SPGP0C02650 | YDR044W | 100 | 0 |

Create PDF in your applications with the Pdfcrowd HTML to PDF API PDFCROWD

| | | | |
|---|---|---|---|
| SPGP0F00760 | YGR213C | 99 | 0 |
| SPGP0F03110 | YGL050W | 99 | 0 |
| SPGP0F01580 | YGR124W | 99 | 0 |
| SPGP0C02000 | YDL031W | 99 | 0 |
| SPGP0K00140 | YJL221C | 98 | 0 |
| SPGP0F02710 | YGL005C | 99 | 0 |
| SPGP0DC00560 | YBL054W | 99 | 0 |
| SPGP0C02630 | YDR042C | 99 | 0 |
| SPGP0M02170 | YNL099C | 99 | 0 |
| SPGP0DC03040 | YBR212W | 99 | 0 |
| SPGP0K02380 | YOR084W | 99 | 0 |
| SPGP0K01490 | YOL016C | 99 | 0 |
| SPGP0M01600 | YNL162W-A | 99 | 0 |
| SPGP0D00870 | YPL012W | 99 | 0 |
| SPGP0V01610 | YHR110W | 100 | 0 |
| SPGP0D00570 | YPR020W | 99 | 0 |
| SPGP0F02760 | YGL011C | 99 | 0 |
| SPGP0DC01930 | YBR089C-A | 100 | 0 |
| SPGP0M00710 | YNL262W | 99 | 0 |
| SPGP0F03310 | YGL075C | 99 | 0 |
| SPGP0DC03090 | YBR217W | 99 | 0 |
| SPGP0DB02040 | YMR089C | 99 | 0 |
| SPGP0K00230 | YOL148C | 99 | 0 |
| SPGP0C02980 | YDR079W | 100 | 0 |
| SPGP0DE00260 | YAL044C | 100 | 0 |
| SPGP0C04570 | YDR255C | 100 | 0 |
| SPGP0C03010 | YDR081C | 99 | 0 |
| SPGP0F04150 | YGL167C | 99 | 0 |
| SPGP0K02560 | YOR104W | 99 | 0 |
| SPGP0M01150 | YNL215W | 99 | 0 |
| SPGP0V01470 | YHR097C | 99 | 0 |
| SPGP0DB02310 | YMR060C | 99 | 0 |
| SPGP0DC00680 | YBL039W-B | 99 | 0 |
| SPGP0F02480 | YGR021W | 100 | 0 |
| SPGP0C05170 | YDR320C | 99 | 0 |
| SPGP0M02750 | YNL037C | 99 | 0 |
| SPGP0DE00190 | YAL053W | 99 | 0 |
| SPGP0G03010 | YJL174W | 99 | 0 |
| SPGP0DB02410 | YMR047C | 99 | 0 |
| SPGP0K02450 | YOR092W | 99 | 0 |

Create PDF in your applications with the Pdfcrowd HTML to PDF API PDFCROWD

| | | | |
|---|---|---|---|
| SPGP0M02140 | YNL102W | 99 | 0 |
| SPGP0C03910 | YDR179C | 99 | 0 |
| SPGP0DD01980 | YIR032C | 99 | 0 |
| SPGP0M02930 | YNL019C | 100 | 0 |
| SPGP0K00900 | YOL078W | 99 | 0 |
| SPGP0DC00400 | YBL072C | 100 | 0 |
| SPGP0C03020 | YDR082W | 99 | 0 |
| SPGP0I00430 | YEL039C | 99 | 0 |
| SPGP0DC03260 | YBR237W | 99 | 0 |
| SPGP0C02120 | YDL017W | 99 | 0 |
| SPGP0V00900 | YHR038W | 99 | 0 |
| SPGP0V00700 | YHR018C | 99 | 0 |
| SPGP0DB02170 | YMR074C | 100 | 0 |
| SPGP0G02320 | YJL097W | 99 | 0 |
| SPGP0I02640 | YER184C | 99 | 0 |
| SPGP0Q04910 | YGL237C | 99 | 1 |
| SPGP0DC01810 | YBR078W | 99 | 0 |
| SPGP0U02140 | YPL262W | 99 | 0 |
| SPGP0DC03210 | YBR233W | 99 | 0 |
| SPGP0DD00580 | YIL120W | 99 | 0 |
| SPGP0I01440 | YER059W | 99 | 0 |
| SPGP0DD01210 | YIL049W | 98 | 0 |
| SPGP0I01080 | YER027C | 99 | 0 |
| SPGP0DC02770 | YBR183W | 99 | 0 |
| SPGP0V01080 | YHR058C | 99 | 0 |
| SPGP0C00270 | YDL224C | 99 | 0 |
| SPGP0I02060 | YER122C | 99 | 0 |
| SPGP0DC03650 | YBR276C | 99 | 0 |
| SPGP0V00950 | YHR042W | 99 | 0 |
| SPGP0K01830 | YOR020W-A | 100 | 0 |
| SPGP0F03130 | YGL054C | 99 | 0 |
| SPGP0C00550 | YDL194W | 99 | 0 |
| SPGP0M01610 | YNL162W | 99 | 0 |
| SPGP0DC01040 | YBL005W | 99 | 0 |
| SPGP0G03240 | YJL201W | 99 | 0 |
| SPGP0V02020 | YHR152W | 100 | 0 |
| SPGP0DC01730 | YBR069C | 99 | 0 |
| SPGP0DD01320 | YIL038C | 99 | 0 |
| SPGP0AE01240 | YCR069W | 99 | 0 |
| SPGP0G00950 | YJR063W | 99 | 0 |

Create PDF in your applications with the Pdfcrowd HTML to PDF API PDFCROWD

| | | | |
|---|---|---|---|
| SPGP0AE00510 | YCL016C | 99 | 0 |
| SPGP0G00440 | YJR116W | 100 | 0 |
| SPGP0DE00840 | YAR029W | 98 | 0 |
| SPGP0K01670 | YOR003W | 99 | 0 |
| SPGP0F00830 | YGR206W | 98 | 0 |
| SPGP0M01920 | YNL128W | 99 | 0 |
| SPGP0C02400 | YDR016C | 99 | 0 |
| SPGP0G01410 | YJR009C | 100 | 0 |
| SPGP0I02100 | YER126C | 100 | 0 |
| SPGP0DB01660 | YMR128W | 99 | 0 |
| SPGP0Q04820 | YGL227W | 99 | 0 |
| SPGP0DD01060 | YIL068C | 99 | 0 |
| SPGP0AE00120 | YCL061C | 99 | 0 |
| SPGP0C00300 | YDL220C | 99 | 0 |
| SPGP0DC01910 | YBR087W | 99 | 0 |
| SPGP0C03370 | YDR121W | 98 | 1 |
| SPGP0DD00530 | YIL125W | 99 | 0 |
| SPGP0C01990 | YDL033C | 100 | 0 |
| SPGP0I01860 | YER100W | 99 | 0 |
| SPGP0F01470 | YGR135W | 99 | 0 |
| SPGP0F00490 | YGR244C | 99 | 0 |
| SPGP0DB01520 | YMR143W | 100 | 0 |
| SPGP0DC03150 | YBR225W | 99 | 0 |
| SPGP0K00280 | YOL143C | 99 | 0 |
| SPGP0G00650 | YJR096W | 99 | 0 |
| SPGP0C02820 | YDR063W | 99 | 0 |
| SPGP0I00930 | YER013W | 99 | 0 |
| SPGP0DC00300 | YBL085W | 99 | 0 |
| SPGP0DC00870 | YBL023C | 99 | 0 |
| SPGP0C02790 | YDR060W | 99 | 0 |
| SPGP0DB02460 | YMR040W | 100 | 0 |
| SPGP0G03260 | YJL204C | 99 | 0 |
| SPGP0G01710 | YJL029C | 99 | 0 |
| SPGP0G03320 | YJL210W | 99 | 0 |
| SPGP0DB03190 | YML035C | 99 | 0 |
| SPGP0M01370 | YNL190W | 99 | 0 |
| SPGP0D00240 | YPR057W | 99 | 0 |
| SPGP0AE00370 | YCL033C | 99 | 0 |
| SPGP0DB01040 | YMR193W | 99 | 0 |

Create PDF in your applications with the Pdfcrowd HTML to PDF API PDFCROWD

| | | | |
|---|---|---|---|
| SPGP0C00690 | YDL178W | 99 | 0 |
| SPGP0V01920 | YHR142W | 100 | 0 |
| SPGP0V01680 | YHR117W | 99 | 0 |
| SPGP0C00180 | YDL234C | 99 | 0 |
| SPGP0DC02650 | YBR169C | 99 | 0 |
| SPGP0F03600 | YGL106W | 99 | 0 |
| SPGP0M00470 | YNL290W | 100 | 0 |
| SPGP0AE01190 | YCR063W | 99 | 0 |
| SPGP0G00400 | YJR121W | 99 | 0 |
| SPGP0M00340 | YNL305C | 99 | 0 |
| SPGP0DB00310 | YMR266W | 99 | 0 |
| SPGP0D00360 | YPR042C | 99 | 0 |
| SPGP0C02530 | YDR030C | 99 | 0 |
| SPGP0F02970 | YGL033W | 99 | 0 |
| SPGP0F03540 | YGL099W | 99 | 0 |
| SPGP0DC02220 | YBR123C | 99 | 0 |
| SPGP0U01500 | YPL195W | 81 | 1 |
| SPGP0K02050 | YOR045W | 99 | 0 |
| SPGP0G02810 | YJL153C | 99 | 0 |
| SPGP0K02710 | YOR120W | 99 | 0 |
| SPGP0I00260 | YEL056W | 99 | 0 |
| SPGP0M00530 | YNL283C | 99 | 0 |
| SPGP0V01280 | YHR078W | 99 | 0 |
| SPGP0C01900 | YDL045W-A | 99 | 0 |
| SPGP0G01800 | YJL039C | 99 | 0 |
| SPGP0DD01700 | YIR003W | 99 | 0 |
| SPGP0K02280 | YOR073W | 99 | 0 |
| SPGP0DE00520 | YAL019W | 99 | 0 |
| SPGP0M01000 | YNL232W | 99 | 0 |
| SPGP0C01610 | YDL078C | 99 | 0 |
| SPGP0C01240 | YDL116W | 99 | 0 |
| SPGP0M03560 | YNR048W | 99 | 0 |
| SPGP0K00760 | YOL093W | 99 | 0 |
| SPGP0K01060 | YOL062C | 99 | 0 |
| SPGP0F02510 | YGR017W | 99 | 0 |
| SPGP0M00130 | YNL328C | 99 | 0 |
| SPGP0DD01840 | YIR017C | 100 | 0 |
| SPGP0F01510 | YGR131W | 99 | 0 |
| SPGP0G02380 | YJL103C | 99 | 0 |
| SPGP0M00840 | YNL249C | 99 | 0 |

Create PDF in your applications with the Pdfcrowd HTML to PDF API PDFCROWD

| | | | |
|---|---|---|---|
| SPGP0G02920 | YJL164C | 99 | 0 |
| SPGP0G02430 | YJL109C | 99 | 0 |
| SPGP0V00240 | YHL029C | 99 | 0 |
| SPGP0C03460 | YDR130C | 99 | 1 |
| SPGP0I01270 | YER045C | 99 | 0 |
| SPGP0F02460 | YGR024C | 100 | 0 |
| SPGP0F02160 | YGR059W | 99 | 0 |
| SPGP0V01860 | YHR136C | 99 | 0 |
| SPGP0I00900 | YER010C | 100 | 0 |
| SPGP0G01360 | YJR013W | 99 | 0 |
| SPGP0K02330 | YOR078W | 99 | 0 |
| SPGP0M00200 | YNL320W | 99 | 0 |
| SPGP0G02650 | YJL133C-A | 99 | 0 |
| SPGP0K00660 | YOL104C | 99 | 0 |
| SPGP0M01490 | YNL176C | 99 | 0 |
| SPGP0F02130 | YGR062C | 99 | 0 |
| SPGP0F04140 | YGL166W | 99 | 0 |
| SPGP0DB02320 | YMR059W | 99 | 0 |
| SPGP0G01590 | YJL011C | 100 | 0 |
| SPGP0C03200 | YDR103W | 99 | 0 |
| SPGP0AE00970 | YCR034W | 100 | 0 |
| SPGP0I01050 | YER024W | 99 | 0 |
| SPGP0M01940 | YNL126W | 99 | 0 |
| SPGP0C04090 | YDR198C | 99 | 0 |
| SPGP0AI00310 | YHR192W | 99 | 0 |
| SPGP0K00820 | YOL087C | 99 | 0 |
| SPGP0Q04420 | YGL181W | 97 | 3 |
| SPGP0K01850 | YOR022C | 99 | 0 |
| SPGP0C04920 | YDR295C | 99 | 0 |
| SPGP0I00630 | YEL019C | 99 | 0 |
| SPGP0Z02880 | YNL301C | 83 | 5 |
| SPGP0M02680 | YNL045W | 99 | 0 |
| SPGP0DB03030 | YML018C | 99 | 0 |
| SPGP0G00870 | YJR072C | 99 | 0 |
| SPGP0M00150 | YNL326C | 99 | 0 |
| SPGP0C00240 | YDL227C | 99 | 0 |
| SPGP0D00800 | YPL005W | 99 | 0 |
| SPGP0I02470 | YER167W | 99 | 0 |
| SPGP0F00640 | YGR227W | 99 | 0 |
| SPGP0G02540 | YJL124C | 100 | 0 |

Create PDF in your applications with the Pdfcrowd HTML to PDF API PDFCROWD

| | | | |
|---|---|---|---|
| SPGP0G01780 | YJL037W | 99 | 0 |
| SPGP0Q04480 | YGL189C | 100 | 0 |
| SPGP0I00650 | YEL017C-A | 100 | 0 |
| SPGP0DC02820 | YBR188C | 98 | 0 |
| SPGP0M01550 | YNL167C | 99 | 0 |
| SPGP0V01740 | YHR123W | 100 | 0 |
| SPGP0I01700 | YER085C | 99 | 0 |
| SPGP0DB02270 | YMR064W | 99 | 0 |
| SPGP0C04080 | YDR197W | 99 | 0 |
| SPGP0I02500 | YER170W | 99 | 0 |
| SPGP0DB03250 | YML043C | 99 | 0 |
| SPGP0I02260 | YER145C | 97 | 3 |
| SPGP0Q04840 | YGL229C | 99 | 0 |
| SPGP0D00740 | YPR002W | 99 | 0 |
| SPGP0M02710 | YNL041C | 99 | 0 |
| SPGP0K01990 | YOR038C | 99 | 0 |
| SPGP0M03570 | YNR049C | 100 | 0 |
| SPGP0C01430 | YDL097C | 99 | 0 |
| SPGP0DB01880 | YMR106C | 99 | 0 |
| SPGP0DC01220 | YBR015C | 99 | 0 |
| SPGP0DD00630 | YIL115C | 99 | 0 |
| SPGP0M01260 | YNL202W | 99 | 0 |
| SPGP0C04630 | YDR261C | 99 | 0 |
| SPGP0DC02790 | YBR185C | 99 | 0 |
| SPGP0DB02070 | YMR086W | 99 | 0 |
| SPGP0DC02690 | YBR173C | 99 | 0 |
| SPGP0D00520 | YPR025C | 99 | 0 |
| SPGP0G02160 | YJL080C | 99 | 0 |
| SPGP0G00120 | YJR150C | 99 | 0 |
| SPGP0K01520 | YOL013C | 99 | 0 |
| SPGP0DB01780 | YMR116C | 99 | 0 |
| SPGP0DC02780 | YBR184W | 99 | 0 |
| SPGP0C02490 | YDR025W | 97 | 3 |
| SPGP0DC03270 | YBR238C | 99 | 0 |
| SPGP0DC02580 | YBR162W-A | 99 | 0 |
| SPGP0G01830 | YJL043W | 99 | 0 |
| SPGP0K00400 | YOL130W | 99 | 0 |
| SPGP0DB02610 | YMR025W | 99 | 0 |
| SPGP0V02070 | YHR157W | 99 | 0 |
| SPGP0DD00350 | YIL144W | 99 | 0 |

Create PDF in your applications with the Pdfcrowd HTML to PDF API PDFCROWD

| | | | |
|---|---|---|---|
| SPGP0M03510 | YNR043W | 99 | 0 |
| SPGP0F02230 | YGR052W | 99 | 0 |
| SPGP0C01550 | YDL084W | 99 | 0 |
| SPGP0K01290 | YOL039W | 100 | 0 |
| SPGP0V00430 | YHL010C | 99 | 0 |
| SPGP0DB01250 | YMR171C | 99 | 0 |
| SPGP0AE00410 | YCL029C | 99 | 0 |
| SPGP0M02570 | YNL056W | 99 | 0 |
| SPGP0F01280 | YGR157W | 99 | 0 |
| SPGP0C02670 | YDR046C | 99 | 0 |
| SPGP0DC00570 | YBL052C | 99 | 0 |
| SPGP0DD01540 | YIL013C | 99 | 0 |
| SPGP0V01130 | YHR063C | 99 | 0 |
| SPGP0DB00780 | YMR217W | 99 | 0 |
| SPGP0V00670 | YHR014W | 99 | 0 |
| SPGP0C01100 | YDL131W | 99 | 0 |
| SPGP0K02360 | YOR081C | 99 | 0 |
| SPGP0G00360 | YJR125C | 99 | 0 |
| SPGP0F00850 | YGR204W | 99 | 0 |
| SPGP0V01670 | YHR116W | 100 | 0 |
| SPGP0M02310 | YNL084C | 99 | 0 |
| SPGP0G01210 | YJR035W | 99 | 0 |
| SPGP0K01510 | YOL014W | 93 | 0 |
| SPGP0K00210 | YOL151W | 98 | 0 |
| SPGP0F02930 | YGL029W | 100 | 0 |
| SPGP0G01520 | YJL002C | 99 | 0 |
| SPGP0DC02230 | YBR125C | 99 | 0 |
| SPGP0G03390 | YJL218W | 99 | 0 |
| SPGP0DB03530 | YML072C | 99 | 0 |
| SPGP0C03070 | YDR087C | 99 | 0 |
| SPGP0DB00120 | YMR285C | 99 | 0 |
| SPGP0DB03430 | YML062C | 99 | 0 |
| SPGP0DE00620 | YAL009W | 99 | 0 |
| SPGP0C03580 | YDR144C | 99 | 0 |
| SPGP0V01490 | YHR098C | 99 | 0 |
| SPGP0DB02720 | YMR014W | 99 | 0 |
| SPGP0K02080 | YOR048C | 99 | 0 |
| SPGP0DB02450 | YMR041C | 100 | 0 |
| SPGP0G03020 | YJL176C | 99 | 0 |
| SPGP0M02910 | YNL031C | 100 | 0 |

Create PDF in your applications with the Pdfcrowd HTML to PDF API PDFCROWD

| | | | |
|---|---|---|---|
| SPGP0DB03390 | YML058W-A | 100 | 0 |
| SPGP0D00300 | YPR049C | 99 | 0 |
| SPGP0F00620 | YGR230W | 99 | 0 |
| SPGP0K00180 | YOL154W | 99 | 0 |
| SPGP0K00470 | YOL123W | 99 | 0 |
| SPGP0K01180 | YOL051W | 98 | 2 |
| SPGP0M03040 | YNL006W | 99 | 0 |
| SPGP0DE00180 | YAL054C | 99 | 0 |
| SPGP0M01640 | YNL159C | 99 | 0 |
| SPGP0I02670 | YER187W | 92 | 0 |
| SPGP0DC02070 | YBR107C | 99 | 0 |
| SPGP0F02660 | YGR001C | 99 | 0 |
| SPGP0G02500 | YJL118W | 99 | 0 |
| SPGP0DB02230 | YMR068W | 100 | 0 |
| SPGP0DC03200 | YBR231C | 99 | 0 |
| SPGP0DC00800 | YBL029W | 99 | 0 |
| SPGP0V02120 | YHR162W | 99 | 0 |
| SPGP0V00720 | YHR020W | 99 | 0 |
| SPGP0I00910 | YER011W | 94 | 0 |
| SPGP0DB00270 | YMR270C | 99 | 0 |
| SPGP0C00250 | YDL226C | 99 | 0 |
| SPGP0C00520 | YDL198C | 99 | 0 |
| SPGP0F01910 | YGR088W | 99 | 0 |
| SPGP0DC03670 | YBR279W | 100 | 0 |
| SPGP0C04110 | YDR201W | 100 | 0 |
| SPGP0C04560 | YDR254W | 99 | 0 |
| SPGP0M01630 | YNL160W | 99 | 0 |
| SPGP0G00930 | YJR065C | 99 | 0 |
| SPGP0AE01220 | YCR067C | 99 | 0 |
| SPGP0F01700 | YGR110W | 99 | 0 |
| SPGP0DD00730 | YIL105C | 99 | 0 |
| SPGP0I00410 | YEL041W | 99 | 0 |
| SPGP0D00440 | YPR033C | 99 | 0 |
| SPGP0M00540 | YNL282W | 100 | 0 |
| SPGP0G02570 | YJL127C | 99 | 0 |
| SPGP0AI00340 | YHR189W | 99 | 0 |
| SPGP0V01500 | YHR099W | 99 | 0 |
| SPGP0G00460 | YJR113C | 99 | 0 |
| SPGP0DD00740 | YIL104C | 100 | 0 |
| SPGP0C00380 | YDL212W | 99 | 0 |

Create PDF in your applications with the Pdfcrowd HTML to PDF API PDFCROWD

| | | | |
|---|---|---|---|
| SPGP0V00880 | YHR036W | 99 | 0 |
| SPGP0C04400 | YDR236C | 99 | 0 |
| SPGP0C01690 | YDL069C | 100 | 0 |
| SPGP0F03250 | YGL066W | 99 | 1 |
| SPGP0DD00880 | YIL090W | 99 | 0 |
| SPGP0K02170 | YOR060C | 99 | 0 |
| SPGP0I02080 | YER124C | 99 | 0 |
| SPGP0F01360 | YGR148C | 99 | 0 |
| SPGP0C02800 | YDR061W | 99 | 0 |
| SPGP0F02560 | YGR012W | 99 | 0 |
| SPGP0M02970 | YNL014W | 99 | 0 |
| SPGP0C04010 | YDR189W | 99 | 0 |
| SPGP0I00520 | YEL029C | 99 | 0 |
| SPGP0DC01880 | YBR085W | 99 | 0 |
| SPGP0DC02940 | YBR201W | 99 | 0 |
| SPGP0F01490 | YGR133W | 99 | 0 |
| SPGP0DB01240 | YMR172W | 99 | 0 |
| SPGP0C01340 | YDL106C | 99 | 0 |
| SPGP0DD00390 | YIL139C | 99 | 0 |
| SPGP0DC00320 | YBL082C | 99 | 0 |
| SPGP0AE00330 | YCL037C | 100 | 0 |
| SPGP0DC00620 | YBL046W | 97 | 2 |
| SPGP0DB02240 | YMR067C | 99 | 0 |
| SPGP0DB00990 | YMR197C | 100 | 0 |
| SPGP0G01730 | YJL031C | 99 | 0 |
| SPGP0V01530 | YHR102W | 99 | 0 |
| SPGP0DB01430 | YMR153W | 99 | 0 |
| SPGP0C03050 | YDR085C | 98 | 0 |
| SPGP0C02550 | YDR032C | 99 | 0 |
| SPGP0C04650 | YDR263C | 99 | 0 |
| SPGP0V01660 | YHR115C | 99 | 0 |
| SPGP0DC02480 | YBR153W | 99 | 0 |
| SPGP0AE00310 | YCL039W | 99 | 0 |
| SPGP0DB01060 | YMR191W | 99 | 0 |
| SPGP0I00460 | YEL036C | 99 | 0 |
| SPGP0G00270 | YJR135C | 99 | 0 |
| SPGP0DC02670 | YBR171W | 99 | 0 |
| SPGP0DD00790 | YIL099W | 99 | 0 |
| SPGP0DD01090 | YIL065C | 100 | 0 |
| SPGP0C02330 | YDR007W | 100 | 0 |

Create PDF in your applications with the Pdfcrowd HTML to PDF API PDFCROWD

| | | | |
|---|---|---|---|
| SPGP0F00270 | YGR268C | 99 | 0 |
| SPGP0F01970 | YGR082W | 99 | 0 |
| SPGP0AE01200 | YCR065W | 99 | 0 |
| SPGP0G00380 | YJR123W | 99 | 0 |
| SPGP0I00680 | YEL015W | 99 | 0 |
| SPGP0DB00330 | YMR264W | 100 | 0 |
| SPGP0DB00680 | YMR227C | 99 | 0 |
| SPGP0DB00350 | YMR262W | 99 | 0 |
| SPGP0D00260 | YPR055W | 99 | 0 |
| SPGP0V01940 | YHR143W-A | 100 | 0 |
| SPGP0K01940 | YOR033C | 99 | 0 |
| SPGP0G02800 | YJL151C | 99 | 0 |
| SPGP0DC02240 | YBR126C | 99 | 0 |
| SPGP0G01230 | YJR033C | 99 | 0 |
| SPGP0M00770 | YNL256W | 99 | 0 |
| SPGP0DB00480 | YMR247C | 99 | 0 |
| SPGP0C02110 | YDL018C | 99 | 0 |
| SPGP0DC01280 | YBR021W | 99 | 0 |
| SPGP0D00600 | YPR017C | 99 | 0 |
| SPGP0K02030 | YOR043W | 99 | 0 |
| SPGP0K02770 | YOR127W | 99 | 0 |
| SPGP0V00800 | YHR028C | 99 | 0 |
| SPGP0F02550 | YGR013W | 99 | 0 |
| SPGP0DC02570 | YBR162C | 99 | 0 |
| SPGP0M00450 | YNL292W | 99 | 0 |
| SPGP0K02270 | YOR071C | 99 | 0 |
| SPGP0D01040 | YPL030W | 99 | 0 |
| SPGP0K01130 | YOL055C | 100 | 0 |
| SPGP0M03500 | YNR041C | 99 | 0 |
| SPGP0M03230 | YNR015W | 99 | 0 |
| SPGP0F03550 | YGL100W | 99 | 0 |
| SPGP0I00280 | YEL054C | 100 | 0 |
| SPGP0DC00150 | YBL102W | 99 | 0 |
| SPGP0K01900 | YOR028C | 99 | 0 |
| SPGP0K01080 | YOL060C | 99 | 0 |
| SPGP0C01300 | YDL110C | 100 | 0 |
| SPGP0M02430 | YNL072W | 99 | 0 |
| SPGP0C04970 | YDR300C | 99 | 0 |
| SPGP0DE00870 | YAR035W | 99 | 0 |
| SPGP0DD01820 | YIR015W | 99 | 0 |

Create PDF in your applications with the Pdfcrowd HTML to PDF API PDFCROWD

| | | | |
|---|---|---|---|
| SPGP0DC00520 | YBL058W | 98 | 0 |
| SPGP0I00950 | YER015W | 99 | 0 |
| SPGP0K01010 | YOL067C | 99 | 0 |
| SPGP0G02410 | YJL106W | 99 | 0 |
| SPGP0DE00540 | YAL017W | 99 | 0 |
| SPGP0M02370 | YNL078W | 99 | 0 |
| SPGP0M03120 | YNR003C | 100 | 0 |
| SPGP0C03440 | YDR128W | 99 | 0 |
| SPGP0C03190 | YDR101C | 99 | 0 |
| SPGP0I01210 | YER040W | 99 | 0 |
| SPGP0C04220 | YDR216W | 99 | 0 |
| SPGP0V01800 | YHR131C | 99 | 0 |
| SPGP0DB02400 | YMR048W | 98 | 1 |
| SPGP0I00920 | YER012W | 99 | 0 |
| SPGP0F02040 | YGR074W | 99 | 0 |
| SPGP0D00320 | YPR047W | 99 | 0 |
| SPGP0DB03240 | YML042W | 99 | 0 |
| SPGP0DC00270 | YBL088C | 99 | 0 |
| SPGP0C03220 | YDR105C | 99 | 0 |
| SPGP0AE00950 | YCR032W | 99 | 0 |
| SPGP0AI00370 | YHR186C | 99 | 0 |
| SPGP0V00980 | YHR046C | 99 | 0 |
| SPGP0C03990 | YDR186C | 99 | 0 |
| SPGP0F00720 | YGR217W | 99 | 0 |
| SPGP0M01470 | YNL178W | 99 | 0 |
| SPGP0F03340 | YGL078C | 99 | 0 |
| SPGP0C01590 | YDL080C | 99 | 0 |
| SPGP0K00840 | YOL086C | 99 | 0 |
| SPGP0C04940 | YDR297W | 99 | 0 |
| SPGP0I02280 | YER147C | 99 | 0 |
| SPGP0K00590 | YOL110W | 99 | 0 |
| SPGP0AI00330 | YHR190W | 99 | 0 |
| SPGP0Q04570 | YGL198W | 99 | 0 |
| SPGP0M00950 | YNL238W | 99 | 0 |
| SPGP0M03240 | YNR016C | 99 | 0 |
| SPGP0M00220 | YNL317W | 99 | 0 |
| SPGP0DD00270 | YIL152W | 99 | 1 |
| SPGP0I00510 | YEL030W | 99 | 0 |
| SPGP0AI00450 | YHR178W | 99 | 0 |
| SPGP0G00810 | YJR078W | 99 | 0 |

Create PDF in your applications with the Pdfcrowd HTML to PDF API PDFCROWD

| | | | |
|---|---|---|---|
| SPGP0U02130 | YPL260W | 99 | 0 |
| SPGP0DC01690 | YBR065C | 99 | 0 |
| SPGP0I00670 | YEL016C | 99 | 0 |
| SPGP0G03100 | YJL185C | 99 | 0 |
| SPGP0Q04970 | YGL244W | 99 | 0 |
| SPGP0DC02800 | YBR186W | 99 | 0 |
| SPGP0I01720 | YER087W | 99 | 0 |
| SPGP0DB02380 | YMR052W | 99 | 0 |
| SPGP0C04060 | YDR195W | 99 | 0 |
| SPGP0F01950 | YGR084C | 99 | 0 |
| SPGP0I02290 | YER148W | 100 | 0 |
| SPGP0I02560 | YER176W | 99 | 0 |
| SPGP0C05120 | YDR315C | 100 | 0 |
| SPGP0K01640 | YOL001W | 99 | 0 |
| SPGP0F02910 | YGL027C | 99 | 0 |
| SPGP0C05020 | YDR305C | 99 | 0 |
| SPGP0M00780 | YNL255C | 99 | 0 |
| SPGP0M01280 | YNL200C | 99 | 0 |
| SPGP0DC01200 | YBR012C | 99 | 0 |
| SPGP0K01160 | YOL052C-A | 99 | 0 |
| SPGP0Q04600 | YGL202W | 99 | 0 |
| SPGP0C04610 | YDR259C | 99 | 0 |
| SPGP0I00580 | YEL023C | 99 | 0 |
| SPGP0G01880 | YJL048C | 99 | 0 |
| SPGP0F01930 | YGR086C | 99 | 0 |
| SPGP0C04600 | YDR258C | 100 | 0 |
| SPGP0G00540 | YJR107W | 99 | 0 |
| SPGP0DB01390 | YMR157C | 99 | 0 |
| SPGP0G01450 | YJR005C-A | 100 | 0 |
| SPGP0G00100 | YJL219W | 98 | 0 |
| SPGP0DC02620 | YBR166C | 99 | 0 |
| SPGP0K01500 | YOL015W | 99 | 0 |
| SPGP0DE00470 | YAL024C | 99 | 0 |
| SPGP0AE00780 | YCR017C | 99 | 0 |
| SPGP0D00980 | YPL023C | 99 | 0 |
| SPGP0DB02050 | YMR088C | 99 | 0 |
| SPGP0I01110 | YER030W | 99 | 0 |
| SPGP0C02470 | YDR023W | 99 | 0 |
| SPGP0DC00860 | YBL024W | 99 | 0 |
| SPGP0DD00370 | YIL142W | 99 | 0 |

Create PDF in your applications with the Pdfcrowd HTML to PDF API PDFCROWD

| | | | |
|---|---|---|---|
| SPGP0F04020 | YGL153W | 99 | 0 |
| SPGP0V02050 | YHR155W | 99 | 0 |
| SPGP0I01980 | YER113C | 99 | 0 |
| SPGP0C03540 | YDR140W | 100 | 0 |
| SPGP0K01320 | YOL034W | 99 | 0 |
| SPGP0DB01500 | YMR145C | 99 | 0 |
| SPGP0F01060 | YGR181W | 99 | 0 |
| SPGP0AE00430 | YCL027W | 99 | 0 |
| SPGP0AI00320 | YHR191C | 100 | 0 |
| SPGP0C01940 | YDL042C | 99 | 0 |
| SPGP0C01980 | YDL035C | 99 | 0 |
| SPGP0DD01560 | YIL010W | 99 | 0 |
| SPGP0DC00590 | YBL050W | 99 | 0 |
| SPGP0F02670 | YGL001C | 99 | 0 |
| SPGP0F04110 | YGL162W | 99 | 0 |
| SPGP0G00970 | YJR061W | 99 | 0 |
| SPGP0K02340 | YOR079C | 99 | 0 |
| SPGP0K02690 | YOR118W | 100 | 0 |
| SPGP0M02830 | YNL023C | 99 | 0 |
| SPGP0DC01970 | YBR095C | 99 | 0 |
| SPGP0V01650 | YHR114W | 99 | 0 |
| SPGP0V01120 | YHR062C | 99 | 0 |
| SPGP0V01320 | YHR081W | 100 | 0 |
| SPGP0M02550 | YNL059C | 99 | 0 |
| SPGP0DC01120 | YBR004C | 99 | 0 |
| SPGP0K00270 | YOL144W | 97 | 2 |
| SPGP0DB01840 | YMR110C | 99 | 0 |
| SPGP0C04590 | YDR257C | 100 | 0 |
| SPGP0DB03510 | YML070W | 99 | 0 |
| SPGP0DE00220 | YAL048C | 99 | 0 |
| SPGP0G01500 | YJR001W | 99 | 0 |
| SPGP0DB00100 | YMR287C | 99 | 0 |
| SPGP0F04120 | YGL163C | 99 | 0 |
| SPGP0K02520 | YOR099W | 99 | 0 |
| SPGP0DB01150 | YMR181C | 99 | 0 |
| SPGP0M02880 | YNL027W | 99 | 0 |
| SPGP0M02330 | YNL082W | 99 | 0 |
| SPGP0DD00210 | YIL157C | 99 | 0 |
| SPGP0G02970 | YJL170C | 99 | 0 |

Create PDF in your applications with the Pdfcrowd HTML to PDF API PDFCROWD

| | | | |
|---|---|---|---|
| SPGP0C00840 | YDL160C-A | 100 | 0 |
| SPGP0DC03190 | YBR230C | 99 | 0 |
| SPGP0AE01400 | YCR088W | 99 | 0 |
| SPGP0DB02430 | YMR043W | 99 | 1 |
| SPGP0F00600 | YGR232W | 99 | 0 |
| SPGP0DC03220 | YBR233W-A | 99 | 0 |
| SPGP0V00520 | YHR001W | 99 | 0 |
| SPGP0V00290 | YHL024W | 99 | 0 |
| SPGP0DE00480 | YAL023C | 99 | 0 |
| SPGP0V01110 | YHR061C | 99 | 0 |
| SPGP0AE00890 | YCR026C | 99 | 0 |
| SPGP0C00860 | YDL159W | 99 | 0 |
| SPGP0DD00810 | YIL097W | 99 | 0 |
| SPGP0F00970 | YGR192C | 99 | 0 |
| SPGP0DD01800 | YIR013C | 99 | 0 |
| SPGP0F02640 | YGR003W | 99 | 0 |
| SPGP0C02900 | YDR071C | 100 | 0 |
| SPGP0DC02050 | YBR105C | 99 | 0 |
| SPGP0M02960 | YNL015W | 100 | 0 |
| SPGP0Q04740 | YGL220W | 99 | 0 |
| SPGP0D01030 | YPL029W | 99 | 0 |
| SPGP0G02880 | YJL160C | 99 | 0 |
| SPGP0DB03480 | YML067C | 99 | 0 |
| SPGP0V00740 | YHR022C | 97 | 0 |
| SPGP0G00910 | YJR067C | 99 | 0 |
| SPGP0U02180 | YPL266W | 99 | 0 |
| SPGP0DC01590 | YBR055C | 99 | 0 |
| SPGP0DB01810 | YMR113W | 99 | 0 |
| SPGP0F03890 | YGL138C | 99 | 0 |
| SPGP0U01980 | YPL246C | 99 | 0 |
| SPGP0C00740 | YDL173W | 99 | 0 |
| SPGP0DB01220 | YMR174C | 100 | 0 |
| SPGP0DY00100 | YNL334C | 98 | 0 |
| SPGP0D00910 | YPL016W | 99 | 0 |
| SPGP0I00160 | YEL066W | 99 | 0 |
| SPGP0M02220 | YNL094W | 99 | 0 |
| SPGP0G01050 | YJR053W | 99 | 0 |
| SPGP0Q05030 | YGL250W | 99 | 0 |
| SPGP0K01870 | YOR025W | 99 | 0 |
| SPGP0C02420 | YDR018C | 99 | 0 |

Create PDF in your applications with the Pdfcrowd HTML to PDF API PDFCROWD

| | | | |
|---|---|---|---|
| SPGP0I01400 | YER056C | 97 | 0 |
| SPGP0I01150 | YER034W | 99 | 0 |
| SPGP0DX00630 | YML078W | 87 | 0 |
| SPGP0K00580 | YOL111C | 99 | 0 |
| SPGP0DC02110 | YBR111C | 99 | 0 |
| SPGP0DC00750 | YBL033C | 99 | 0 |
| SPGP0DD01360 | YIL034C | 100 | 0 |
| SPGP0F01720 | YGR108W | 99 | 0 |
| SPGP0DC01420 | YBR037C | 99 | 0 |
| SPGP0DD01020 | YIL072W | 99 | 0 |
| SPGP0I02170 | YER134C | 100 | 0 |
| SPGP0M00560 | YNL280C | 99 | 0 |
| SPGP0AI00240 | YHR199C | 99 | 0 |
| SPGP0F00410 | YGR252W | 99 | 0 |
| SPGP0DC01940 | YBR091C | 99 | 0 |
| SPGP0D00420 | YPR035W | 100 | 0 |
| SPGP0C00400 | YDL210W | 99 | 0 |
| SPGP0C02860 | YDR067C | 99 | 0 |
| SPGP0DD01790 | YIR012W | 99 | 0 |
| SPGP0DD00570 | YIL121W | 99 | 0 |
| SPGP0DB00560 | YMR239C | 99 | 0 |
| SPGP0C01360 | YDL104C | 99 | 0 |
| SPGP0DD01380 | YIL031W | 100 | 0 |
| SPGP0DC03120 | YBR221C | 99 | 0 |
| SPGP0M01210 | YNL209W | 99 | 0 |
| SPGP0M03010 | YNL009W | 99 | 0 |
| SPGP0F03640 | YGL111W | 99 | 0 |
| SPGP0DB01940 | YMR099C | 100 | 0 |
| SPGP0G01970 | YJL057C | 99 | 0 |
| SPGP0G01750 | YJL034W | 99 | 0 |
| SPGP0C02880 | YDR069C | 99 | 0 |
| SPGP0G00290 | YJR133W | 99 | 0 |
| SPGP0M02500 | YNL065W | 99 | 0 |
| SPGP0C01120 | YDL129W | 99 | 0 |
| SPGP0V01640 | YHR113W | 99 | 0 |
| SPGP0DC02610 | YBR165W | 99 | 0 |
| SPGP0I00440 | YEL038W | 99 | 0 |
| SPGP0DD00770 | YIL102C | 100 | 0 |
| SPGP0C00730 | YDL174C | 99 | 0 |
| SPGP0C01810 | YDL055C | 99 | 0 |

Create PDF in your applications with the Pdfcrowd HTML to PDF API PDFCROWD

| | | | |
|---|---|---|---|
| SPGP0C02360 | YDR012W | 99 | 0 |
| SPGP0K00120 | YOL159C | 97 | 0 |
| SPGP0DB02980 | YML013W | 99 | 0 |
| SPGP0I02630 | YER183C | 99 | 0 |
| SPGP0F01990 | YGR080W | 99 | 0 |
| SPGP0C02280 | YDR002W | 100 | 0 |
| SPGP0DB02670 | YMR019W | 99 | 0 |
| SPGP0DC00960 | YBL014C | 99 | 0 |
| SPGP0I00660 | YEL017W | 99 | 0 |
| SPGP0F00290 | YGR266W | 99 | 0 |
| SPGP0DB01340 | YMR162C | 99 | 0 |
| SPGP0C01760 | YDL060W | 99 | 0 |
| SPGP0U01990 | YPL247C | 99 | 0 |
| SPGP0F03090 | YGL048C | 99 | 0 |
| SPGP0G02850 | YJL157C | 99 | 0 |
| SPGP0F02170 | YGR058W | 98 | 2 |
| SPGP0DC02260 | YBR129C | 99 | 0 |
| SPGP0G01250 | YJR031C | 99 | 0 |
| SPGP0F01120 | YGR174C | 98 | 0 |
| SPGP0F00200 | YGR276C | 99 | 0 |
| SPGP0DC02040 | YBR104W | 100 | 0 |
| SPGP0DB02870 | YML001W | 99 | 0 |
| SPGP0DC01840 | YBR081C | 99 | 0 |
| SPGP0DC03050 | YBR213W | 99 | 0 |
| SPGP0C02130 | YDL015C | 99 | 0 |
| SPGP0V00780 | YHR026W | 99 | 0 |
| SPGP0K02750 | YOR125C | 99 | 0 |
| SPGP0F00470 | YGR246C | 99 | 0 |
| SPGP0F02110 | YGR065C | 99 | 0 |
| SPGP0M01790 | YNL141W | 99 | 0 |
| SPGP0K01880 | YOR026W | 99 | 0 |
| SPGP0I00390 | YEL043W | 99 | 0 |
| SPGP0DC02540 | YBR159W | 99 | 0 |
| SPGP0T00400 | YPL273W | 99 | 0 |
| SPGP0M00430 | YNL294C | 99 | 0 |
| SPGP0K02320 | YOR077W | 99 | 0 |
| SPGP0DD01730 | YIR006C | 99 | 0 |
| SPGP0G00200 | YJR141W | 99 | 0 |
| SPGP0DB02590 | YMR027W | 99 | 0 |
| SPGP0M03210 | YNR013C | 99 | 0 |

Create PDF in your applications with the Pdfcrowd HTML to PDF API PDFCROWD

| | | | |
|---|---|---|---|
| SPGP0C01280 | YDL112W | 99 | 0 |
| SPGP0C04990 | YDR302W | 99 | 0 |
| SPGP0V01350 | YHR084W | 99 | 0 |
| SPGP0M03390 | YNR032W | 99 | 0 |
| SPGP0V01820 | YHR132W-A | 99 | 0 |
| SPGP0DC00500 | YBL059C-A | 99 | 0 |
| SPGP0G02020 | YJL062W | 99 | 0 |
| SPGP0V00280 | YHL025W | 99 | 0 |
| SPGP0F02430 | YGR028W | 99 | 0 |
| SPGP0V00190 | YHL034C | 99 | 0 |
| SPGP0DE00560 | YAL015C | 99 | 0 |
| SPGP0V01850 | YHR135C | 99 | 0 |
| SPGP0G02700 | YJL139C | 99 | 0 |
| SPGP0Q04800 | YGL226W | 99 | 0 |
| SPGP0C03420 | YDR126W | 99 | 0 |
| SPGP0V01570 | YHR106W | 99 | 0 |
| SPGP0V01220 | YHR072W-A | 100 | 0 |
| SPGP0I01090 | YER028C | 98 | 1 |
| SPGP0D00960 | YPL021W | 99 | 0 |
| SPGP0F03300 | YGL073W | 99 | 0 |
| SPGP0I00940 | YER014W | 99 | 0 |
| SPGP0G00680 | YJR094C | 100 | 0 |
| SPGP0G03090 | YJL184W | 99 | 0 |
| SPGP0G02470 | YJL115W | 99 | 0 |
| SPGP0F04180 | YGL171W | 99 | 0 |
| SPGP0C01720 | YDL065C | 99 | 0 |
| SPGP0D00350 | YPR043W | 100 | 0 |
| SPGP0K00700 | YOL100W | 99 | 0 |
| SPGP0F01140 | YGR172C | 99 | 0 |
| SPGP0DE00350 | YAL035W | 99 | 0 |
| SPGP0C00370 | YDL213C | 99 | 0 |
| SPGP0U02040 | YPL252C | 100 | 0 |
| SPGP0C04270 | YDR222W | 99 | 0 |
| SPGP0AE00190 | YCL054W | 99 | 0 |
| SPGP0Q04790 | YGL225W | 99 | 0 |
| SPGP0K00860 | YOL083W | 99 | 0 |
| SPGP0M01770 | YNL145W | 99 | 0 |
| SPGP0I01260 | YER044C-A | 100 | 0 |
| SPGP0C00190 | YDL233W | 99 | 0 |
| SPGP0M00970 | YNL236W | 99 | 0 |

Create PDF in your applications with the Pdfcrowd HTML to PDF API PDFCROWD

| | | | |
|---|---|---|---|
| SPGP0G02690 | YJL138C | 99 | 0 |
| SPGP0Q04920 | YGL238W | 99 | 0 |
| SPGP0M00790 | YNL254C | 99 | 0 |
| SPGP0DB03070 | YML022W | 99 | 0 |
| SPGP0AI00470 | YHR176W | 99 | 0 |
| SPGP0DB02620 | YMR024W | 100 | 0 |
| SPGP0G00830 | YJR076C | 99 | 0 |
| SPGP0M00180 | YNL322C | 99 | 0 |
| SPGP0Q00140 | YGR289C | 98 | 0 |
| SPGP0DC01360 | YBR030W | 99 | 0 |
| SPGP0DC02190 | YBR120C | 100 | 0 |
| SPGP0I01750 | YER089C | 99 | 0 |
| SPGP0DC02860 | YBR193C | 99 | 0 |
| SPGP0C04320 | YDR227W | 99 | 0 |
| SPGP0G03120 | YJL187C | 99 | 0 |
| SPGP0M00390 | YNL299W | 99 | 0 |
| SPGP0M01510 | YNL173C | 99 | 0 |
| SPGP0M03450 | YNR036C | 100 | 0 |
| SPGP0C00990 | YDL143W | 99 | 0 |
| SPGP0I02540 | YER174C | 99 | 0 |
| SPGP0DB03290 | YML049C | 99 | 0 |
| SPGP0F02300 | YGR042W | 99 | 0 |
| SPGP0C05100 | YDR313C | 100 | 0 |
| SPGP0I00840 | YER005W | 99 | 0 |
| SPGP0DD00380 | YIL140W | 99 | 0 |
| SPGP0G00340 | YJR127C | 99 | 0 |
| SPGP0M00760 | YNL257C | 99 | 0 |
| SPGP0C03560 | YDR142C | 100 | 0 |
| SPGP0Q04620 | YGL205W | 99 | 0 |
| SPGP0DC02930 | YBR200W | 99 | 0 |
| SPGP0DB02210 | YMR070W | 97 | 2 |
| SPGP0AE01470 | YCR095C | 99 | 0 |
| SPGP0AE01410 | YCR089W | 99 | 0 |
| SPGP0AE00810 | YCR019W | 99 | 0 |
| SPGP0DB03100 | YML025C | 99 | 0 |
| SPGP0G02200 | YJL084C | 99 | 0 |
| SPGP0G00690 | YJR093C | 99 | 0 |
| SPGP0G00160 | YJR145C | 99 | 0 |
| SPGP0DB01410 | YMR155W | 99 | 0 |
| SPGP0M01030 | YNL229C | 99 | 0 |

Create PDF in your applications with the Pdfcrowd HTML to PDF API PDFCROWD

| | | | |
|---|---|---|---|
| SPGP0DC02550 | YBR160W | 99 | 0 |
| SPGP0DB01740 | YMR120C | 99 | 0 |
| SPGP0F01370 | YGR147C | 99 | 0 |
| SPGP0DB01020 | YMR194C-B | 100 | 0 |
| SPGP0I01130 | YER032W | 99 | 0 |
| SPGP0C02450 | YDR021W | 99 | 0 |
| SPGP0G03030 | YJL177W | 100 | 0 |
| SPGP0V02030 | YHR153C | 99 | 0 |
| SPGP0AE00960 | YCR033W | 99 | 0 |
| SPGP0M01390 | YNL188W | 99 | 0 |
| SPGP0DB02480 | YMR038C | 99 | 0 |
| SPGP0DB01560 | YMR138W | 99 | 0 |
| SPGP0DC01110 | YBR003W | 99 | 0 |
| SPGP0F02690 | YGL003C | 99 | 0 |
| SPGP0C01950 | YDL040C | 99 | 0 |
| SPGP0K00780 | YOL091W | 99 | 0 |
| SPGP0DD01580 | YIL009W | 99 | 0 |
| SPGP0Q04930 | YGL240W | 100 | 0 |
| SPGP0AE00160 | YCL057W | 99 | 0 |
| SPGP0V01100 | YHR060W | 100 | 0 |
| SPGP0Q00180 | YGR286C | 99 | 0 |
| SPGP0M00660 | YNL268W | 99 | 0 |
| SPGP0M02530 | YNL062C | 99 | 0 |
| SPGP0DD00180 | YIL160C | 99 | 0 |
| SPGP0V00120 | YHL044W | 98 | 0 |
| SPGP0F02750 | YGL010W | 99 | 0 |
| SPGP0K00250 | YOL146W | 99 | 0 |
| SPGP0DB03120 | YML027W | 99 | 0 |
| SPGP0DE00440 | YAL027W | 99 | 0 |
| SPGP0G01560 | YJL006C | 99 | 0 |
| SPGP0F02890 | YGL025C | 99 | 0 |
| SPGP0DB00160 | YMR281W | 99 | 0 |
| SPGP0DC00360 | YBL078C | 100 | 0 |
| SPGP0F00190 | YGR277C | 99 | 0 |
| SPGP0M03660 | YNR058W | 100 | 0 |
| SPGP0F02440 | YGR027C | 99 | 0 |
| SPGP0DC01950 | YBR093C | 99 | 0 |
| SPGP0F03850 | YGL134W | 99 | 0 |
| SPGP0DB01140 | YMR183C | 99 | 0 |
| SPGP0DD00230 | YIL156W | 99 | 0 |

Create PDF in your applications with the Pdfcrowd HTML to PDF API PDFCROWD

| | | | |
|---|---|---|---|
| SPGP0F04100 | YGL161C | 99 | 0 |
| SPGP0F00580 | YGR234W | 99 | 0 |
| SPGP0V00500 | YHL002W | 99 | 1 |
| SPGP0K00430 | YOL127W | 100 | 0 |
| SPGP0M01230 | YNL207W | 99 | 0 |
| SPGP0M00320 | YNL307C | 99 | 0 |
| SPGP0DD01780 | YIR011C | 99 | 0 |
| SPGP0F01350 | YGR149W | 99 | 0 |
| SPGP0G02790 | YJL149W | 99 | 0 |
| SPGP0Q04590 | YGL201C | 99 | 0 |
| SPGP0C00560 | YDL193W | 99 | 0 |
| SPGP0G02860 | YJL158C | 100 | 0 |
| SPGP0V00760 | YHR024C | 100 | 0 |
| SPGP0DE00600 | YAL011W | 99 | 0 |
| SPGP0C01580 | YDL081C | 100 | 0 |
| SPGP0DC01760 | YBR072W | 99 | 0 |
| SPGP0F01740 | YGR105W | 100 | 0 |
| SPGP0D00680 | YPR008W | 99 | 0 |
| SPGP0F00320 | YGR262C | 99 | 0 |
| SPGP0G01850 | YJL045W | 99 | 0 |
| SPGP0DC03280 | YBR239C | 99 | 0 |
| SPGP0M02290 | YNL086W | 99 | 0 |
| SPGP0D00850 | YPL010W | 100 | 0 |
| SPGP0G02990 | YJL172W | 99 | 0 |
| SPGP0M02470 | YNL068C | 99 | 0 |
| SPGP0I01170 | YER036C | 99 | 0 |
| SPGP0M02650 | YNL048W | 99 | 0 |
| SPGP0C00570 | YDL192W | 99 | 0 |
| SPGP0C00970 | YDL145C | 99 | 0 |
| SPGP0I01510 | YER066W | 100 | 0 |
| SPGP0DB02860 | YMR001C | 99 | 0 |
| SPGP0AE01080 | YCR046C | 99 | 0 |
| SPGP0F00430 | YGR250C | 99 | 0 |
| SPGP0DD01000 | YIL074C | 99 | 0 |
| SPGP0AI00220 | YHR200W | 100 | 0 |
| SPGP0I02150 | YER131W | 100 | 0 |
| SPGP0I02550 | YER175C | 99 | 0 |
| SPGP0Q04630 | YGL206C | 99 | 0 |
| SPGP0DE00410 | YAL030W | 99 | 0 |
| SPGP0DD00840 | YIL094C | 99 | 0 |

Create PDF in your applications with the Pdfcrowd HTML to PDF API PDFCROWD

| | | | |
|---|---|---|---|
| SPGP0M03720 | YNR064C | 100 | 0 |
| SPGP0DD00690 | YIL109C | 99 | 0 |
| SPGP0G02330 | YJL098W | 99 | 0 |
| SPGP0C01160 | YDL125C | 100 | 0 |
| SPGP0F03430 | YGL087C | 100 | 0 |
| SPGP0C02170 | YDL010W | 99 | 0 |
| SPGP0M01980 | YNL122C | 99 | 0 |
| SPGP0D00390 | YPR037C | 100 | 0 |
| SPGP0F00330 | YGR261C | 99 | 0 |
| SPGP0K02570 | YOR106W | 99 | 0 |
| SPGP0K00410 | YOL129W | 99 | 0 |
| SPGP0M01180 | YNL212W | 99 | 0 |
| SPGP0C01150 | YDL126C | 99 | 0 |
| SPGP0K01820 | YOR020C | 100 | 0 |
| SPGP0M02480 | YNL067W | 99 | 0 |
| SPGP0F03590 | YGL105W | 99 | 0 |
| SPGP0C01570 | YDL082W | 99 | 0 |
| SPGP0AE01040 | YCR042C | 99 | 0 |
| SPGP0G01980 | YJL058C | 99 | 0 |
| SPGP0DE00880 | YAR042W | 99 | 0 |
| SPGP0AE00130 | YCL059C | 99 | 0 |
| SPGP0D00650 | YPR010C-A | 99 | 0 |
| SPGP0DB01460 | YMR149W | 100 | 0 |
| SPGP0I00740 | YEL006W | 99 | 0 |
| SPGP0DD00430 | YIL135C | 99 | 0 |
| SPGP0C01380 | YDL102W | 99 | 0 |
| SPGP0DC02630 | YBR167C | 99 | 0 |
| SPGP0F03660 | YGL113W | 99 | 0 |
| SPGP0F03480 | YGL093W | 99 | 0 |
| SPGP0G02560 | YJL126W | 99 | 0 |
| SPGP0DB02150 | YMR076C | 99 | 0 |
| SPGP0G01950 | YJL055W | 100 | 0 |
| SPGP0AE00920 | YCR028C-A | 100 | 0 |
| SPGP0G01770 | YJL036W | 99 | 0 |
| SPGP0AI00490 | YHR174W | 100 | 0 |
| SPGP0C03780 | YDR167W | 100 | 0 |
| SPGP0G00310 | YJR131W | 99 | 0 |
| SPGP0V01950 | YHR144C | 99 | 0 |
| SPGP0V00420 | YHL011C | 99 | 0 |
| SPGP0F02590 | YGR008C | 99 | 0 |

Create PDF in your applications with the Pdfcrowd HTML to PDF API PDFCROWD

| | | | |
|---|---|---|---|
| SPGP0V01620 | YHR111W | 99 | 0 |
| SPGP0I00420 | YEL040W | 99 | 0 |
| SPGP0AE00680 | YCR006C | 99 | 0 |
| SPGP0C01830 | YDL053C | 99 | 0 |
| SPGP0DC00940 | YBL016W | 99 | 0 |
| SPGP0G00570 | YJR104C | 100 | 0 |
| SPGP0F00300 | YGR264C | 99 | 0 |
| SPGP0C04540 | YDR252W | 99 | 0 |
| SPGP0F02250 | YGR048W | 99 | 0 |
| SPGP0F01100 | YGR177C | 99 | 0 |
| SPGP0K02630 | YOR112W | 99 | 0 |
| SPGP0C02150 | YDL013W | 99 | 0 |
| SPGP0DC00650 | YBL042C | 99 | 0 |
| SPGP0DB03310 | YML051W | 100 | 0 |
| SPGP0DB00800 | YMR215W | 99 | 0 |
| SPGP0DB00370 | YMR260C | 99 | 0 |
| SPGP0V01300 | YHR079C-A | 100 | 0 |
| SPGP0C01460 | YDL092W | 99 | 0 |
| SPGP0V01290 | YHR079C | 99 | 0 |
| SPGP0K01330 | YOL033W | 99 | 0 |
| SPGP0AI00300 | YHR193C | 100 | 0 |
| SPGP0F01130 | YGR173W | 99 | 0 |
| SPGP0V01590 | YHR108W | 99 | 0 |
| SPGP0DB00260 | YMR271C | 99 | 0 |
| SPGP0M02270 | YNL088W | 99 | 0 |
| SPGP0C01970 | YDL036C | 99 | 0 |
| SPGP0K02550 | YOR103C | 98 | 0 |
| SPGP0M03270 | YNR019W | 99 | 0 |
| SPGP0AI00420 | YHR181W | 99 | 0 |
| SPGP0V01370 | YHR086W | 99 | 0 |
| SPGP0DE00850 | YAR031W | 96 | 0 |
| SPGP0K00680 | YOL102C | 99 | 0 |
| SPGP0DC02850 | YBR192W | 99 | 0 |
| SPGP0V00260 | YHL027W | 99 | 0 |
| SPGP0I01930 | YER107C | 99 | 0 |
| SPGP0C03400 | YDR124W | 99 | 0 |
| SPGP0C03150 | YDR097C | 99 | 0 |
| SPGP0F01180 | YGR169C | 99 | 0 |
| SPGP0M03490 | YNR040W | 99 | 0 |
| SPGP0DC01980 | YBR096W | 99 | 0 |

Create PDF in your applications with the Pdfcrowd HTML to PDF API PDFCROWD

| | | | |
|---|---|---|---|
| SPGP0M03050 | YNL005C | 99 | 0 |
| SPGP0DD01410 | YIL026C | 99 | 0 |
| SPGP0G02450 | YJL111W | 99 | 0 |
| SPGP0C01700 | YDL067C | 99 | 0 |
| SPGP0DE00380 | YAL033W | 99 | 0 |
| SPGP0K02740 | YOR124C | 99 | 0 |
| SPGP0Q00160 | YGR288W | 99 | 0 |
| SPGP0I01940 | YER109C | 99 | 0 |
| SPGP0C04290 | YDR224C | 100 | 0 |
| SPGP0U02060 | YPL254W | 99 | 0 |
| SPGP0K00880 | YOL081W | 99 | 0 |
| SPGP0M01750 | YNL147W | 100 | 0 |
| SPGP0DB01000 | YMR196W | 99 | 0 |
| SPGP0I00530 | YEL028W | 99 | 0 |
| SPGP0K00690 | YOL101C | 99 | 0 |
| SPGP0C00170 | YDL235C | 99 | 0 |
| SPGP0M00910 | YNL242W | 99 | 0 |
| SPGP0I00790 | YEL001C | 99 | 0 |
| SPGP0I00320 | YEL050C | 99 | 0 |
| SPGP0AI00410 | YHR182W | 99 | 0 |
| SPGP0I02700 | YPR204W | 99 | 0 |
| SPGP0G00140 | YJR148W | 99 | 0 |
| SPGP0DB03050 | YML020W | 99 | 0 |
| SPGP0F03910 | YGL140C | 99 | 0 |
| SPGP0K01930 | YOR032C | 99 | 0 |
| SPGP0DC01380 | YBR033W | 99 | 0 |
| SPGP0I01610 | YER075C | 99 | 0 |
| SPGP0M00880 | YNL245C | 100 | 0 |
| SPGP0DC02840 | YBR191W | 99 | 0 |
| SPGP0Q04390 | YGL178W | 99 | 0 |
| SPGP0G03140 | YJL190C | 99 | 0 |
| SPGP0C04380 | YDR234W | 99 | 0 |
| SPGP0D01050 | YPL031C | 99 | 0 |
| SPGP0F01680 | YGR112W | 99 | 0 |
| SPGP0C00710 | YDL176W | 99 | 0 |
| SPGP0V00360 | YHL017W | 99 | 0 |
| SPGP0M03420 | YNR034W | 99 | 0 |
| SPGP0DC03710 | YBR283C | 99 | 0 |
| SPGP0C01850 | YDL051W | 99 | 0 |

Create PDF in your applications with the Pdfcrowd HTML to PDF API PDFCROWD

| | | | |
|---|---|---|---|
| SPGP0DZ02350 | YFL058W | 88 | 1 |
| SPGP0D00580 | YPR019W | 99 | 0 |
| SPGP0K01680 | YOR004W | 99 | 0 |
| SPGP0F02950 | YGL031C | 100 | 0 |
| SPGP0DC02960 | YBR202W | 99 | 0 |
| SPGP0Q04640 | YGL207W | 99 | 0 |
| SPGP0F00880 | YGR201C | 99 | 0 |
| SPGP0M01530 | YNL169C | 100 | 0 |
| SPGP0DB02890 | YML004C | 99 | 0 |
| SPGP0C04840 | YDR286C | 99 | 0 |
| SPGP0M00820 | YNL251C | 99 | 0 |
| SPGP0M02260 | YNL090W | 99 | 0 |
| SPGP0DC01990 | YBR097W | 99 | 0 |
| SPGP0V02010 | YHR151C | 99 | 0 |
| SPGP0C01210 | YDL120W | 99 | 0 |
| SPGP0G02110 | YJL072C | 99 | 0 |
| SPGP0I02490 | YER169W | 99 | 0 |
| SPGP0DB01760 | YMR118C | 99 | 0 |
| SPGP0C04120 | YDR202C | 99 | 0 |
| SPGP0G01890 | YJL049W | 99 | 0 |
| SPGP0DB02090 | YMR083W | 99 | 0 |
| SPGP0C02430 | YDR019C | 99 | 0 |
| SPGP0DD01610 | YIL006W | 99 | 0 |
| SPGP0F03560 | YGL101W | 99 | 0 |
| SPGP0G03370 | YJL216C | 96 | 0 |
| SPGP0EX00250 | YDR342C | 88 | 0 |
| SPGP0Q04890 | YGL234W | 99 | 0 |
| SPGP0Q05110 | YGL258W | 99 | 0 |
| SPGP0DB02100 | YMR081C | 99 | 0 |
| SPGP0K01040 | YOL064C | 99 | 0 |
| SPGP0V00690 | YHR017W | 99 | 0 |
| SPGP0DD02010 | YIR035C | 100 | 0 |
| SPGP0DE00290 | YAL041W | 99 | 0 |
| SPGP0C02380 | YDR014W | 99 | 0 |
| SPGP0G01090 | YJR049C | 99 | 0 |
| SPGP0F03170 | YGL058W | 99 | 1 |
| SPGP0DC03680 | YBR280C | 99 | 0 |
| SPGP0G01540 | YJL004C | 99 | 0 |
| SPGP0F00370 | YGR256W | 99 | 0 |
| SPGP0DB00490 | YMR246W | 99 | 0 |

Create PDF in your applications with the Pdfcrowd HTML to PDF API PDFCROWD

| | | | |
|---|---|---|---|
| SPGP0K02650 | YOR114W | 99 | 0 |
| SPGP0AE00470 | YCL024W | 99 | 0 |
| SPGP0K01470 | YOL018C | 99 | 0 |
| SPGP0DB01540 | YMR140W | 100 | 0 |
| SPGP0C04710 | YDR270W | 99 | 0 |
| SPGP0M02040 | YNL115C | 99 | 0 |
| SPGP0DB03060 | YML021C | 99 | 0 |
| SPGP0F02490 | YGR020C | 99 | 0 |
| SPGP0M02510 | YNL064C | 99 | 0 |
| SPGP0F03800 | YGL128C | 99 | 0 |
| SPGP0K01150 | YOL053W | 99 | 0 |
| SPGP0M00630 | YNL272C | 99 | 0 |
| SPGP0DB01800 | YMR114C | 99 | 0 |
| SPGP0DC02320 | YBR136W | 99 | 0 |
| SPGP0DB02640 | YMR022W | 100 | 0 |
| SPGP0DW00120 | YCR102C | 99 | 0 |
| SPGP0DB00140 | YMR283C | 99 | 0 |
| SPGP0DB00630 | YMR232W | 99 | 0 |
| SPGP0M02990 | YNL011C | 100 | 0 |
| SPGP0M03640 | YNR056C | 99 | 0 |
| SPGP0DD00590 | YIL119C | 98 | 1 |
| SPGP0V01160 | YHR066W | 99 | 0 |
| SPGP0DE00670 | YAL002W | 99 | 0 |
| SPGP0DD00250 | YIL154C | 99 | 0 |
| SPGP0F03870 | YGL136C | 99 | 0 |
| SPGP0DB01120 | YMR185W | 99 | 0 |
| SPGP0C01890 | YDL046W | 99 | 0 |
| SPGP0F02260 | YGR047C | 99 | 0 |
| SPGP0AE01110 | YCR051W | 99 | 0 |
| SPGP0DD01280 | YIL042C | 99 | 0 |
| SPGP0V00560 | YHR004C | 99 | 0 |
| SPGP0DB02630 | YMR023C | 99 | 0 |
| SPGP0K00450 | YOL125W | 99 | 0 |
| SPGP0G00500 | YJR110W | 99 | 0 |
| SPGP0M03160 | YNR008W | 99 | 0 |
| SPGP0K01100 | YOL058W | 99 | 0 |
| SPGP0F02600 | YGR007W | 99 | 0 |
| SPGP0K01400 | YOL025W | 99 | 0 |
| SPGP0V02150 | YHR165C | 99 | 0 |
| SPGP0DC02090 | YBR109C | 99 | 0 |

Create PDF in your applications with the Pdfcrowd HTML to PDF API PDFCROWD

| | | | |
|---|---|---|---|
| SPGP0C00540 | YDL195W | 99 | 0 |
| SPGP0DC00790 | YBL029C-A | 100 | 0 |
| SPGP0G02840 | YJL156C | 99 | 0 |
| SPGP0M03290 | YNR021W | 99 | 0 |
| SPGP0D00130 | YPR070W | 100 | 0 |
| SPGP0M03020 | YNL008C | 99 | 0 |
| SPGP0DB01260 | YMR170C | 99 | 0 |
| SPGP0I01370 | YER053C-A | 100 | 0 |
| SPGP0DC01740 | YBR070C | 99 | 0 |
| SPGP0V01780 | YHR128W | 100 | 0 |
| SPGP0AE01420 | YCR090C | 99 | 0 |
| SPGP0C02030 | YDL028C | 99 | 0 |
| SPGP0D01080 | YPL034W | 99 | 0 |
| SPGP0DC01130 | YBR005W | 99 | 0 |
| SPGP0F01760 | YGR103W | 99 | 1 |
| SPGP0I01410 | YER056C-A | 100 | 0 |
| SPGP0I00890 | YER009W | 100 | 0 |
| SPGP0G02150 | YJL078C | 99 | 0 |
| SPGP0AE00300 | YCL040W | 99 | 0 |
| SPGP0G02210 | YJL085W | 99 | 0 |
| SPGP0C03360 | YDR120C | 99 | 0 |
| SPGP0DC03520 | YBR262C | 100 | 0 |
| SPGP0DC00710 | YBL037W | 99 | 0 |
| SPGP0F03940 | YGL143C | 99 | 0 |
| SPGP0DB02840 | YMR003W | 99 | 0 |
| SPGP0AI00350 | YHR188C | 99 | 0 |
| SPGP0C00810 | YDL165W | 99 | 0 |
| SPGP0I02130 | YER129W | 99 | 0 |
| SPGP0Q05040 | YGL251C | 99 | 0 |
| SPGP0C01630 | YDL076C | 99 | 0 |
| SPGP0C01140 | YDL127W | 99 | 0 |
| SPGP0DB02390 | YMR049C | 99 | 0 |
| SPGP0I02400 | YER159C | 99 | 0 |
| SPGP0AI00200 | YHR202W | 99 | 0 |
| SPGP0C04420 | YDR238C | 99 | 0 |
| SPGP0CZ00690 | YKL157W | 84 | 0 |
| SPGP0DC01440 | YBR039W | 99 | 0 |
| SPGP0DD00410 | YIL137C | 99 | 0 |
| SPGP0F03460 | YGL091C | 99 | 0 |
| SPGP0DD01920 | YIR026C | 99 | 0 |

Create PDF in your applications with the Pdfcrowd HTML to PDF API PDFCROWD

| | | | |
|---|---|---|---|
| SPGP0I00400 | YEL042W | 100 | 0 |
| SPGP0M01250 | YNL204C | 99 | 0 |
| SPGP0V00710 | YHR019C | 99 | 0 |
| SPGP0G01930 | YJL053W | 99 | 0 |
| SPGP0DD01110 | YIL063C | 99 | 0 |
| SPGP0C03760 | YDR165W | 99 | 0 |
| SPGP0DD00170 | YIL161W | 99 | 0 |
| SPGP0C00760 | YDL170W | 99 | 0 |
| SPGP0DC01030 | YBL006C | 99 | 0 |
| SPGP0DD00820 | YIL096C | 99 | 0 |
| SPGP0DC00920 | YBL018C | 99 | 0 |
| SPGP0I01470 | YER062C | 100 | 0 |
| SPGP0C03520 | YDR138W | 99 | 0 |
| SPGP0G00330 | YJR129C | 99 | 0 |
| SPGP0G00550 | YJR106W | 99 | 0 |
| SPGP0AI00440 | YHR179W | 99 | 0 |
| SPGP0F02150 | YGR060W | 99 | 0 |
| SPGP0F01160 | YGR170W | 99 | 0 |
| SPGP0I01030 | YER022W | 99 | 0 |
| SPGP0F02540 | YGR014W | 99 | 0 |
| SPGP0DC03450 | YBR255C-A | 99 | 0 |
| SPGP0DC02080 | YBR108W | 98 | 1 |
| SPGP0DD01420 | YIL024C | 99 | 0 |
| SPGP0DB00820 | YMR213W | 99 | 0 |
| SPGP0F00150 | YGR281W | 99 | 0 |
| SPGP0DB00540 | YMR241W | 99 | 0 |
| SPGP0C02200 | YDL007W | 99 | 0 |
| SPGP0M02230 | YNL093W | 99 | 0 |
| SPGP0C02720 | YDR052C | 99 | 0 |
| SPGP0DB03330 | YML053C | 98 | 1 |
| SPGP0K01590 | YOL006C | 99 | 0 |
| SPGP0V01960 | YHR146W | 99 | 0 |
| SPGP0G02960 | YJL168C | 99 | 0 |
| SPGP0DD00400 | YIL138C | 100 | 0 |
| SPGP0M02250 | YNL091W | 99 | 0 |
| SPGP0F01150 | YGR171C | 99 | 0 |
| SPGP0M02700 | YNL042W | 99 | 0 |
| SPGP0DC02500 | YBR155W | 99 | 0 |
| SPGP0AE01310 | YCR076C | 99 | 0 |
| SPGP0DB03490 | YML068W | 99 | 0 |

Create PDF in your applications with the Pdfcrowd HTML to PDF API PDFCROWD

| | | | |
|---|---|---|---|
| SPGP0DC01210 | YBR014C | 99 | 0 |
| SPGP0M03250 | YNR017W | 100 | 0 |
| SPGP0K02460 | YOR093C | 99 | 0 |
| SPGP0G02060 | YJL066C | 100 | 0 |
| SPGP0F00940 | YGR195W | 100 | 0 |
| SPGP0V01310 | YHR080C | 99 | 0 |
| SPGP0DE00860 | YAR033W | 99 | 0 |
| SPGP0DE00400 | YAL031C | 99 | 0 |
| SPGP0DB02420 | YMR044W | 99 | 0 |
| SPGP0M02420 | YNL073W | 99 | 0 |
| SPGP0M02540 | YNL061W | 99 | 0 |
| SPGP0C03280 | YDR111C | 99 | 0 |
| SPGP0DE00740 | YAR008W | 99 | 0 |
| SPGP0C01540 | YDL085W | 99 | 0 |
| SPGP0F03500 | YGL095C | 99 | 0 |
| SPGP0I00490 | YEL032W | 99 | 0 |
| SPGP0I02070 | YER123W | 100 | 0 |
| SPGP0DC02370 | YBR141C | 99 | 0 |
| SPGP0G02300 | YJL095W | 99 | 0 |
| SPGP0F03330 | YGL077C | 100 | 0 |
| SPGP0DC03360 | YBR247C | 99 | 0 |
| SPGP0F02050 | YGR072W | 99 | 0 |
| SPGP0C01750 | YDL061C | 100 | 0 |
| SPGP0DC03340 | YBR245C | 99 | 0 |
| SPGP0Q04750 | YGL221C | 100 | 0 |
| SPGP0K02760 | YOR126C | 99 | 0 |
| SPGP0U02000 | YPL248C | 99 | 0 |
| SPGP0DC01630 | YBR058C-A | 99 | 0 |
| SPGP0I01580 | YER073W | 99 | 0 |
| SPGP0C03380 | YDR122W | 99 | 0 |
| SPGP0C03800 | YDR169C | 99 | 0 |
| SPGP0F03350 | YGL079W | 100 | 0 |
| SPGP0I00980 | YER017C | 99 | 0 |
| SPGP0C00480 | YDL202W | 99 | 0 |
| SPGP0G01570 | YJL008C | 100 | 0 |
| SPGP0C00150 | YDL237W | 99 | 0 |
| SPGP0M00280 | YNL311C | 99 | 0 |
| SPGP0M00930 | YNL240C | 99 | 0 |
| SPGP0AI00430 | YHR180W | 99 | 0 |
| SPGP0D00950 | YPL020C | 99 | 0 |

Create PDF in your applications with the Pdfcrowd HTML to PDF API PDFCROWD

| | | | |
|---|---|---|---|
| SPGP0D00830 | YPL008W | 99 | 0 |
| SPGP0C00610 | YDL188C | 99 | 0 |
| SPGP0F03930 | YGL142C | 99 | 0 |
| SPGP0K01910 | YOR030W | 99 | 0 |
| SPGP0DC03640 | YBR275C | 99 | 0 |
| SPGP0C03820 | YDR170W-A | 99 | 0 |
| SPGP0K00130 | YOL158C | 98 | 0 |
| SPGP0Q04500 | YGL191W | 99 | 0 |
| SPGP0I01630 | YER077C | 99 | 0 |
| SPGP0DC03530 | YBR263W | 99 | 0 |
| SPGP0M02240 | YNL092W | 99 | 0 |
| SPGP0DC02970 | YBR203W | 99 | 0 |
| SPGP0M01880 | YNL132W | 99 | 0 |
| SPGP0G03300 | YJL208C | 99 | 0 |
| SPGP0M01730 | YNL149C | 100 | 0 |
| SPGP0DD00990 | YIL075C | 99 | 0 |
| SPGP0I02580 | YER178W | 99 | 0 |
| SPGP0M00420 | YNL295W | 99 | 0 |
| SPGP0DB01090 | YMR188C | 99 | 0 |
| SPGP0G00300 | YJR132W | 99 | 0 |
| SPGP0DD01900 | YIR024C | 99 | 0 |
| SPGP0M03590 | YNR051C | 99 | 0 |
| SPGP0G03160 | YJL192C | 99 | 0 |
| SPGP0DB02710 | YMR015C | 99 | 0 |
| SPGP0M01080 | YNL222W | 99 | 0 |
| SPGP0M00800 | YNL253W | 99 | 0 |
| SPGP0AE01450 | YCR093W | 99 | 0 |
| SPGP0DC02980 | YBR204C | 99 | 0 |
| SPGP0G03230 | YJL200C | 99 | 0 |
| SPGP0I01960 | YER111C | 99 | 0 |
| SPGP0DB01850 | YMR109W | 99 | 0 |
| SPGP0C04100 | YDR200C | 99 | 0 |
| SPGP0Q04660 | YGL209W | 99 | 0 |
| SPGP0C03120 | YDR092W | 100 | 0 |
| SPGP0G02230 | YJL088W | 99 | 0 |
| SPGP0V01560 | YHR105W | 99 | 0 |
| SPGP0I02410 | YER161C | 99 | 0 |
| SPGP0C02410 | YDR017C | 99 | 0 |
| SPGP0DD01630 | YIL004C | 100 | 0 |
| SPGP0G00640 | YJR097W | 99 | 0 |

Create PDF in your applications with the Pdfcrowd HTML to PDF API PDFCROWD

| | | | |
|---|---|---|---|
| SPGP0M02770 | YNL035C | 99 | 0 |
| SPGP0DB03080 | YML023C | 99 | 0 |
| SPGP0C01880 | YDL047W | 99 | 0 |
| SPGP0Q05130 | YDR542W | 99 | 0 |
| SPGP0F00550 | YGR237C | 99 | 0 |
| SPGP0K01350 | YOL031C | 99 | 0 |
| SPGP0DC00280 | YBL087C | 100 | 0 |
| SPGP0M00940 | YNL239W | 99 | 0 |
| SPGP0V00350 | YHL018W | 99 | 0 |
| SPGP0K00740 | YOL095C | 99 | 0 |
| SPGP0G00210 | YJR140C | 99 | 0 |
| SPGP0M03110 | YNR002C | 100 | 0 |
| SPGP0DD01650 | YIL002W-A | 100 | 0 |
| SPGP0F00910 | YGR198W | 99 | 0 |
| SPGP0F02770 | YGL012W | 99 | 0 |
| SPGP0AE00760 | YCR015C | 100 | 0 |
| SPGP0K02670 | YOR116C | 99 | 0 |
| SPGP0DB02910 | YML006C | 99 | 0 |
| SPGP0M02180 | YNL098C | 99 | 0 |
| SPGP0DC01190 | YBR011C | 99 | 0 |
| SPGP0AE00490 | YCL018W | 99 | 0 |
| SPGP0C01660 | YDL073W | 99 | 0 |
| SPGP0M02060 | YNL112W | 99 | 0 |
| SPGP0I00210 | YEL061C | 99 | 0 |
| SPGP0C02920 | YDR073W | 99 | 0 |
| SPGP0F00560 | YGR236C | 100 | 0 |
| SPGP0K00290 | YOL142W | 99 | 0 |
| SPGP0F02860 | YGL021W | 99 | 0 |
| SPGP0M00510 | YNL286W | 99 | 0 |
| SPGP0DC01400 | YBR035C | 100 | 0 |
| SPGP0G02750 | YJL145W | 99 | 0 |
| SPGP0G02440 | YJL110C | 99 | 0 |
| SPGP0DB00610 | YMR234W | 99 | 0 |
| SPGP0V01020 | YHR050W | 99 | 0 |
| SPGP0V01140 | YHR064C | 99 | 0 |
| SPGP0F03820 | YGL130W | 99 | 0 |
| SPGP0DB01370 | YMR159C | 99 | 0 |
| SPGP0DC02560 | YBR161W | 99 | 0 |
| SPGP0K00220 | YOL149W | 99 | 0 |
| SPGP0F00540 | YGR238C | 99 | 0 |

Create PDF in your applications with the Pdfcrowd HTML to PDF API PDFCROWD

| | | | |
|---|---|---|---|
| SPGP0V00230 | YHL030W | 99 | 0 |
| SPGP0C01920 | YDL044C | 99 | 0 |
| SPGP0DE00390 | YAL032C | 100 | 0 |
| SPGP0V00540 | YHR002W | 99 | 0 |
| SPGP0DE00420 | YAL029C | 99 | 0 |
| SPGP0DD01300 | YIL040W | 99 | 0 |
| SPGP0K02580 | YOR107W | 99 | 0 |
| SPGP0F02620 | YGR005C | 99 | 0 |
| SPGP0F03690 | YGL116W | 99 | 0 |
| SPGP0K01420 | YOL023W | 99 | 0 |
| SPGP0K00960 | YOL072W | 99 | 0 |
| SPGP0F01310 | YGR154C | 99 | 0 |
| SPGP0V01810 | YHR132C | 99 | 0 |
| SPGP0F03700 | YGL117W | 99 | 0 |
| SPGP0G02820 | YJL154C | 99 | 0 |
| SPGP0I01850 | YER099C | 99 | 0 |
| SPGP0DE00640 | YAL007C | 99 | 0 |
| SPGP0D00110 | YPR072W | 99 | 0 |
| SPGP0DC01790 | YBR076W | 99 | 0 |
| SPGP0C03350 | YDR119W-A | 100 | 0 |
| SPGP0K01980 | YOR037W | 99 | 0 |
| SPGP0I01350 | YER052C | 99 | 0 |
| SPGP0F02610 | YGR006W | 99 | 0 |
| SPGP0I02440 | YER164W | 99 | 0 |
| SPGP0DC00480 | YBL061C | 98 | 0 |
| SPGP0DC00730 | YBL035C | 99 | 0 |
| SPGP0AI00140 | YHR208W | 99 | 0 |
| SPGP0DC01750 | YBR071W | 99 | 0 |
| SPGP0M02730 | YNL039W | 99 | 0 |
| SPGP0G02170 | YJL081C | 99 | 0 |
| SPGP0C04030 | YDR191W | 99 | 0 |
| SPGP0C03340 | YDR119W | 99 | 0 |
| SPGP0G01700 | YJL027C | 100 | 0 |
| SPGP0D00760 | YPL001W | 99 | 0 |
| SPGP0K02540 | YOR101W | 97 | 0 |
| SPGP0Q05000 | YGL247W | 99 | 0 |
| SPGP0DC03470 | YBR257W | 99 | 0 |
| SPGP0DC03500 | YBR260C | 100 | 0 |
| SPGP0M01560 | YNL166C | 99 | 0 |
| SPGP0DD01480 | YIL018W | 99 | 0 |

Create PDF in your applications with the Pdfcrowd HTML to PDF API PDFCROWD

| | | | |
|---|---|---|---|
| SPGP0F03960 | YGL145W | 99 | 0 |
| SPGP0AE00630 | YCL001W-A | 100 | 0 |
| SPGP0AE01380 | YCR086W | 99 | 0 |
| SPGP0C00830 | YDL161W | 100 | 0 |
| SPGP0F00390 | YGR254W | 99 | 0 |
| SPGP0M03680 | YNR060W | 99 | 0 |
| SPGP0M01420 | YNL185C | 99 | 0 |
| SPGP0I02510 | YER171W | 99 | 0 |
| SPGP0I01000 | YER019W | 99 | 0 |
| SPGP0Q04870 | YGL232W | 99 | 0 |
| SPGP0C02680 | YDR047W | 99 | 0 |
| SPGP0DB02330 | YMR058W | 99 | 0 |
| SPGP0AE01030 | YCR097W | 100 | 0 |
| SPGP0I02110 | YER127W | 99 | 0 |
| SPGP0M03140 | YNR006W | 100 | 0 |
| SPGP0DB01230 | YMR173W | 87 | 5 |
| SPGP0DC01460 | YBR041W | 99 | 0 |
| SPGP0C03940 | YDR181C | 99 | 0 |
| SPGP0DD00470 | YIL131C | 99 | 0 |
| SPGP0M01070 | YNL223W | 99 | 0 |
| SPGP0AE00390 | YCL031C | 99 | 0 |
| SPGP0M01270 | YNL201C | 99 | 0 |
| SPGP0DB02110 | YMR080C | 99 | 0 |
| SPGP0DC03100 | YBR218C | 99 | 0 |
| SPGP0G00730 | YJR089W | 99 | 0 |
| SPGP0C03270 | YDR110W | 99 | 0 |
| SPGP0D00150 | YPR068C | 99 | 0 |
| SPGP0K01760 | YOR013W | 99 | 0 |
| SPGP0V00840 | YHR032W | 99 | 0 |
| SPGP0DC02910 | YBR198C | 99 | 0 |
| SPGP0DD00800 | YIL098C | 99 | 0 |
| SPGP0V00340 | YHL019C | 99 | 0 |
| SPGP0G01910 | YJL051W | 99 | 0 |
| SPGP0F01270 | YGR158C | 99 | 0 |
| SPGP0K01370 | YOL028C | 99 | 0 |
| SPGP0AE00530 | YCL012C | 100 | 0 |
| SPGP0DB00910 | YMR204C | 99 | 0 |
| SPGP0C02990 | YDR079C-A | 99 | 0 |
| SPGP0C02220 | YDL005C | 98 | 1 |
| SPGP0DC00900 | YBL020W | 99 | 0 |

Create PDF in your applications with the Pdfcrowd HTML to PDF API PDFCROWD

| | | | |
|---|---|---|---|
| SPGP0M00740 | YNL259C | 100 | 0 |
| SPGP0DD01400 | YIL027C | 100 | 0 |
| SPGP0DB00840 | YMR211W | 99 | 0 |
| SPGP0C02700 | YDR050C | 99 | 0 |
| SPGP0DI00100 | YBR298C | 99 | 0 |
| SPGP0D00510 | YPR026W | 99 | 0 |
| SPGP0M01050 | YNL225C | 99 | 0 |
| SPGP0C04930 | YDR296W | 99 | 0 |
| SPGP0V01550 | YHR104W | 99 | 0 |
| SPGP0I00360 | YEL046C | 99 | 0 |
| SPGP0C02190 | YDL007C-A | 100 | 0 |
| SPGP0G01660 | YJL023C | 99 | 0 |
| SPGP0DC01230 | YBR016W | 99 | 1 |
| SPGP0Q04400 | YGL179C | 99 | 0 |
| SPGP0C03110 | YDR091C | 99 | 0 |
| SPGP0K02110 | YOR052C | 99 | 0 |
| SPGP0K02440 | YOR091W | 99 | 0 |
| SPGP0F02290 | YGR043C | 99 | 0 |
| SPGP0V01880 | YHR138C | 99 | 0 |
| SPGP0V01330 | YHR082C | 99 | 0 |
| SPGP0C03730 | YDR162C | 99 | 0 |
| SPGP0T00420 | YOR391C | 99 | 0 |
| SPGP0F02200 | YGR055W | 99 | 0 |
| SPGP0D00690 | YPR007C | 99 | 0 |
| SPGP0G02940 | YJL166W | 100 | 0 |
| SPGP0C05050 | YDR308C | 99 | 0 |
| SPGP0V00680 | YLR410W-B | 99 | 0 |
| SPGP0DB03270 | YML047C | 100 | 0 |
| SPGP0U02240 | YPL272C | 99 | 0 |
| SPGP0AE00440 | YCL026C-B | 100 | 0 |
| SPGP0C02830 | YDR064W | 99 | 0 |
| SPGP0C01740 | YDL063C | 99 | 0 |
| SPGP0M00850 | YNL248C | 99 | 0 |
| SPGP0DC02350 | YBR139W | 99 | 0 |
| SPGP0G02360 | YJL101C | 99 | 0 |
| SPGP0D00970 | YPL022W | 99 | 0 |
| SPGP0DB00200 | YMR277W | 100 | 0 |
| SPGP0M03550 | YNR047W | 99 | 0 |
| SPGP0F01650 | YGR117C | 99 | 0 |

Create PDF in your applications with the Pdfcrowd HTML to PDF API PDFCROWD

| | | | |
|---|---|---|---|
| SPGP0Q04770 | YGL223C | 99 | 0 |
| SPGP0F02740 | YGL009C | 99 | 0 |
| SPGP0M03630 | YNR055C | 99 | 0 |
| SPGP0DC01610 | YBR057C | 100 | 0 |
| SPGP0U02020 | YPL249C-A | 99 | 0 |
| SPGP0DD00970 | YIL077C | 99 | 0 |
| SPGP0C00130 | YDL239C | 99 | 0 |
| SPGP0G02710 | YJL140W | 99 | 0 |
| SPGP0AI00260 | YHR197W | 99 | 0 |
| SPGP0G02580 | YJL127C-B | 100 | 0 |
| SPGP0DB01270 | YMR169C | 99 | 0 |
| SPGP0DD01330 | YIL037C | 99 | 0 |
| SPGP0F03950 | YGL144C | 99 | 0 |
| SPGP0DC03750 | YBR287W | 99 | 0 |
| SPGP0K01970 | YOR036W | 99 | 0 |
| SPGP0I01650 | YER079W | 99 | 0 |
| SPGP0C00620 | YDL186W | 99 | 0 |
| SPGP0G03360 | YJL214W | 99 | 0 |
| SPGP0M01710 | YNL152W | 99 | 0 |
| SPGP0C02930 | YDR074W | 99 | 0 |
| SPGP0M01860 | YNL134C | 99 | 0 |
| SPGP0G00820 | YJR077C | 99 | 0 |
| SPGP0G01940 | YJL054W | 99 | 0 |
| SPGP0DD01240 | YIL046W | 99 | 0 |
| SPGP0G01510 | YJL001W | 99 | 0 |
| SPGP0M03360 | YNR029C | 99 | 0 |
| SPGP0AE00570 | YCL008C | 99 | 0 |
| SPGP0K00830 | YOL086W-A | 99 | 0 |
| SPGP0DB03180 | YML034W | 99 | 0 |
| SPGP0G00980 | YJR060W | 99 | 0 |
| SPGP0G00320 | YJR130C | 99 | 0 |
| SPGP0D00230 | YPR058W | 99 | 0 |
| SPGP0AE01000 | YCR037C | 99 | 0 |
| SPGP0G03180 | YJL194W | 99 | 0 |
| SPGP0C04340 | YDR229W | 100 | 0 |
| SPGP0C04800 | YDR282C | 99 | 0 |
| SPGP0C00160 | YDL236W | 99 | 0 |
| SPGP0I01900 | YER104W | 99 | 0 |
| SPGP0G00180 | YJR143C | 99 | 0 |
| SPGP0DD01310 | YIL039W | 99 | 0 |

Create PDF in your applications with the Pdfcrowd HTML to PDF API PDFCROWD

| | | | |
|---|---|---|---|
| SPGP0F00590 | YGR233C | 99 | 0 |
| SPGP0I01190 | YER038C | 99 | 0 |
| SPGP0F00840 | YGR205W | 99 | 0 |
| SPGP0C04690 | YDR267C | 100 | 0 |
| SPGP0C03170 | YDR099W | 99 | 0 |
| SPGP0K02530 | YOR100C | 99 | 0 |
| SPGP0DC00220 | YBL092W | 100 | 0 |
| SPGP0AE00980 | YCR035C | 99 | 0 |
| SPGP0Q04370 | YGL175C | 99 | 0 |
| SPGP0G03330 | YJL212C | 99 | 0 |
| SPGP0DD00290 | YIL150C | 99 | 0 |
| SPGP0M01410 | YNL186W | 99 | 1 |
| SPGP0AE00700 | YCR008W | 99 | 0 |
| SPGP0G01270 | YJR025C | 99 | 0 |
| SPGP0G01580 | YJL010C | 99 | 0 |
| SPGP0DB00980 | YMR198W | 99 | 0 |
| SPGP0DB00450 | YMR251W-A | 100 | 0 |
| SPGP0G02780 | YJL148W | 99 | 0 |
| SPGP0DB01580 | YMR136W | 99 | 0 |
| SPGP0F03440 | YGL089C | 98 | 0 |
| SPGP0I01890 | YER103W | 99 | 0 |
| SPGP0M02000 | YNL119W | 99 | 0 |
| SPGP0I00230 | YEL059C-A | 99 | 0 |
| SPGP0I02050 | YER120W | 99 | 0 |
| SPGP0I00350 | YEL047C | 99 | 0 |
| SPGP0DC03110 | YBR220C | 99 | 0 |
| SPGP0K01440 | YOL021C | 99 | 0 |
| SPGP0DB00670 | YMR228W | 99 | 0 |
| SPGP0DC02290 | YBR132C | 99 | 0 |
| SPGP0G02770 | YJL147C | 99 | 0 |
| SPGP0Q04450 | YGL185C | 99 | 0 |
| SPGP0K02610 | YOR110W | 99 | 0 |
| SPGP0M00690 | YNL264C | 100 | 0 |
| SPGP0DB00110 | YMR286W | 99 | 0 |
| SPGP0V00370 | YHL016C | 99 | 0 |
| SPGP0F01080 | YGR179C | 99 | 0 |
| SPGP0F01330 | YGR152C | 99 | 0 |
| SPGP0I00180 | YEL064C | 99 | 0 |
| SPGP0DD01970 | YIR031C | 100 | 0 |
| SPGP0V02110 | YHR161C | 99 | 0 |

Create PDF in your applications with the Pdfcrowd HTML to PDF API PDFCROWD

| | | | |
|---|---|---|---|
| SPGP0DB00180 | YMR279C | 99 | 0 |
| SPGP0F03670 | YGL114W | 99 | 0 |
| SPGP0K01140 | YOL054W | 100 | 0 |
| SPGP0F02070 | YGR070W | 99 | 0 |
| SPGP0K01890 | YOR027W | 99 | 0 |
| SPGP0I01450 | YER060W | 99 | 0 |
| SPGP0V00150 | YHL039W | 99 | 0 |
| SPGP0C05130 | YDR316W | 99 | 0 |
| SPGP0V00210 | YHL032C | 99 | 0 |
| SPGP0DC00470 | YBL063W | 99 | 0 |
| SPGP0DC01780 | YBR074W | 99 | 0 |
| SPGP0DE00590 | YAL012W | 99 | 0 |
| SPGP0I01330 | YER050C | 99 | 0 |
| SPGP0F03470 | YGL092W | 99 | 0 |
| SPGP0G03040 | YJL178C | 99 | 0 |
| SPGP0M01360 | YNL191W | 99 | 0 |
| SPGP0P02870 | YOL120C | 87 | 3 |
| SPGP0DB01080 | YMR189W | 99 | 0 |
| SPGP0I00820 | YER003C | 99 | 0 |
| SPGP0C03830 | YDR171W | 99 | 0 |
| SPGP0G01040 | YJR054W | 99 | 0 |
| SPGP0F03220 | YGL063W | 99 | 0 |
| SPGP0DB00750 | YMR220W | 99 | 0 |
| SPGP0DE00660 | YAL003W | 99 | 1 |
| SPGP0G02000 | YJL060W | 99 | 0 |
| SPGP0DZ01890 | YOR335C | 87 | 0 |
| SPGP0C03320 | YDR117C | 99 | 0 |
| SPGP0C03630 | YDR150W | 98 | 0 |
| SPGP0V01060 | YHR056C | 99 | 0 |
| SPGP0M01190 | YNL211C | 100 | 0 |
| SPGP0M02050 | YNL113W | 99 | 0 |
| SPGP0I01480 | YER063W | 100 | 0 |
| SPGP0C01670 | YDL072C | 99 | 0 |
| SPGP0I01730 | YER087C-B | 100 | 0 |
| SPGP0F03900 | YGL139W | 99 | 0 |
| SPGP0M00870 | YNL246W | 99 | 0 |
| SPGP0M00650 | YNL270C | 99 | 0 |
| SPGP0DB03370 | YML057W | 99 | 0 |
| SPGP0F00130 | YGR283C | 99 | 0 |
| SPGP0T00410 | YPL274W | 99 | 0 |

Create PDF in your applications with the Pdfcrowd HTML to PDF API PDFCROWD

| | | | |
|---|---|---|---|
| SPGP0DD00450 | YIL133C | 99 | 0 |
| SPGP0F00450 | YGR248W | 99 | 0 |
| SPGP0DD00900 | YIL088C | 99 | 0 |
| SPGP0Q04850 | YGL230C | 99 | 0 |
| SPGP0K01360 | YOL030W | 99 | 1 |
| SPGP0C04240 | YDR218C | 99 | 0 |
| SPGP0DB02350 | YMR055C | 100 | 0 |
| SPGP0AE01390 | YCR087W | 94 | 6 |
| SPGP0M03190 | YNR011C | 99 | 0 |
| SPGP0M02620 | YNL051W | 99 | 0 |
| SPGP0DB00620 | YMR233W | 100 | 0 |
| SPGP0C00850 | YDL160C | 99 | 0 |
| SPGP0M03670 | YNR059W | 99 | 0 |
| SPGP0I02460 | YER166W | 99 | 0 |
| SPGP0C03960 | YDR183W | 99 | 0 |
| SPGP0DB00430 | YMR253C | 99 | 0 |
| SPGP0C01390 | YDL101C | 99 | 0 |
| SPGP0DB02490 | YMR037C | 99 | 0 |
| SPGP0D00660 | YPR010C | 99 | 0 |
| SPGP0M01290 | YNL199C | 99 | 0 |
| SPGP0F01540 | YGR128C | 99 | 0 |
| SPGP0F01850 | YGR094W | 99 | 0 |
| SPGP0AE01150 | YCR057C | 99 | 0 |
| SPGP0C03310 | YDR116C | 99 | 0 |
| SPGP0G00250 | YJR136C | 99 | 0 |
| SPGP0C04310 | YDR226W | 99 | 0 |
| SPGP0D00170 | YPR066W | 99 | 0 |
| SPGP0K01700 | YOR006C | 99 | 0 |
| SPGP0G00510 | YJR109C | 99 | 0 |
| SPGP0M03170 | YNR009W | 99 | 0 |
| SPGP0DC01070 | YBL002W | 100 | 0 |
| SPGP0C04190 | YDR212W | 99 | 0 |
| SPGP0I01310 | YER048W-A | 99 | 0 |
| SPGP0G02270 | YJL092W | 99 | 0 |
| SPGP0I02390 | YER158C | 98 | 1 |
| SPGP0DB02130 | YMR078C | 99 | 0 |
| SPGP0C03490 | YDR134C | 99 | 0 |
| SPGP0DB02750 | YMR011W | 99 | 0 |
| SPGP0M00300 | YNL309W | 99 | 0 |
| SPGP0V02060 | YHR156C | 99 | 0 |

Create PDF in your applications with the Pdfcrowd HTML to PDF API PDFCROWD

| | | | |
|---|---|---|---|
| SPGP0DD00100 | YJL222W | 99 | 0 |
| SPGP0I00480 | YEL034W | 99 | 0 |
| SPGP0G02280 | YJL093C | 99 | 0 |
| SPGP0DB01110 | YMR186W | 99 | 0 |
| SPGP0DB01440 | YMR152W | 99 | 0 |
| SPGP0DC00440 | YBL067C | 99 | 0 |
| SPGP0DD01460 | YIL020C | 99 | 0 |
| SPGP0V01270 | YHR077C | 99 | 0 |
| SPGP0C02240 | YDL003W | 99 | 0 |
| SPGP0F03030 | YGL040C | 99 | 0 |
| SPGP0DB00860 | YMR209C | 99 | 0 |
| SPGP0C02750 | YDR056C | 99 | 0 |
| SPGP0F01110 | YGR175C | 99 | 0 |
| SPGP0K01580 | YOL007C | 100 | 0 |
| SPGP0C04910 | YDR294C | 99 | 0 |
| SPGP0DB01700 | YMR124W | 99 | 0 |
| SPGP0C00410 | YDL209C | 99 | 0 |
| SPGP0DC02460 | YBR151W | 99 | 0 |
| SPGP0F03750 | YGL123W | 99 | 0 |
| SPGP0G01640 | YJL019W | 99 | 0 |
| SPGP0V01000 | YHR048W | 99 | 0 |
| SPGP0DD01430 | YIL023C | 99 | 0 |
| SPGP0C03130 | YDR093W | 99 | 0 |
| SPGP0DE00130 | YAL060W | 99 | 0 |
| SPGP0K02420 | YOR089C | 99 | 0 |
| SPGP0K02130 | YOR056C | 99 | 0 |
| SPGP0V00570 | YHR005C | 99 | 0 |
| SPGP0M02740 | YNL038W | 99 | 0 |
| SPGP0I02350 | YER154W | 99 | 0 |
| SPGP0G02620 | YJL131C | 99 | 0 |
| SPGP0M03430 | YNR034W-A | 99 | 0 |
| SPGP0F02180 | YGR057C | 99 | 0 |
| SPGP0DC02410 | YBR146W | 99 | 0 |
| SPGP0K00720 | YOL097C | 99 | 0 |
| SPGP0DD01030 | YIL071C | 99 | 0 |
| SPGP0K00340 | YOL137W | 99 | 0 |
| SPGP0V00660 | YHR013C | 99 | 0 |
| SPGP0U02220 | YPL270W | 99 | 0 |
| SPGP0DB01640 | YMR130W | 99 | 0 |
| SPGP0DB02700 | YMR016C | 99 | 0 |

Create PDF in your applications with the Pdfcrowd HTML to PDF API PDFCROWD

| | | | |
|---|---|---|---|
| SPGP0F02650 | YGR002C | 99 | 0 |
| SPGP0DD01880 | YIR022W | 99 | 0 |
| SPGP0DC02330 | YBR137W | 99 | 0 |
| SPGP0DD01720 | YIR005W | 100 | 0 |
| SPGP0G02340 | YJL099W | 99 | 0 |
| SPGP0DC03320 | YBR243C | 100 | 0 |
| SPGP0G01490 | YJR002W | 99 | 0 |
| SPGP0G00840 | YJR075W | 99 | 0 |
| SPGP0D00990 | YPL024W | 99 | 0 |
| SPGP0DC03300 | YBR241C | 99 | 0 |
| SPGP0DC01670 | YBR062C | 99 | 0 |
| SPGP0C03790 | YDR168W | 99 | 0 |
| SPGP0DC01530 | YBR048W | 99 | 0 |
| SPGP0F01690 | YGR111W | 99 | 0 |
| SPGP0AI00230 | YHR199C-A | 98 | 2 |
| SPGP0K02720 | YOR122C | 99 | 0 |
| SPGP0C00440 | YDL206W | 99 | 0 |
| SPGP0I00800 | YER001W | 99 | 0 |
| SPGP0F03080 | YGL047W | 99 | 0 |
| SPGP0D01070 | YPL033C | 99 | 0 |
| SPGP0K02700 | YOR119C | 99 | 0 |
| SPGP0F04010 | YGL151W | 99 | 0 |
| SPGP0K01950 | YOR034C | 99 | 0 |
| SPGP0K02150 | YOR058C | 99 | 0 |
| SPGP0C03860 | YDR174W | 100 | 0 |
| SPGP0M01840 | YNL136W | 99 | 0 |
| SPGP0F03970 | YGL146C | 100 | 0 |
| SPGP0DC00670 | YBL040C | 99 | 0 |
| SPGP0C02260 | YDL001W | 99 | 0 |
| SPGP0F01620 | YGR120C | 99 | 0 |
| SPGP0AE01280 | YCR073W-A | 99 | 0 |
| SPGP0M00460 | YNL291C | 99 | 0 |
| SPGP0AI00380 | YHR185C | 100 | 0 |
| SPGP0BK00100 | YMR182C | 99 | 0 |
| SPGP0F01290 | YGR156W | 99 | 0 |
| SPGP0Q04440 | YGL184C | 99 | 0 |
| SPGP0C01790 | YDL057W | 99 | 0 |
| SPGP0C04780 | YDR280W | 99 | 0 |
| SPGP0M01590 | YNL163C | 99 | 0 |
| SPGP0DB02830 | YMR004W | 99 | 0 |

Create PDF in your applications with the Pdfcrowd HTML to PDF API PDFCROWD

| | | | |
|---|---|---|---|
| SPGP0DD01670 | YIL001W | 99 | 0 |
| SPGP0G01000 | YJR058C | 99 | 0 |
| SPGP0I02320 | YER151C | 99 | 0 |
| SPGP0C00220 | YDL230W | 99 | 0 |
| SPGP0C05090 | YDR312W | 100 | 0 |
| SPGP0I01920 | YER106W | 99 | 0 |
| SPGP0DC02120 | YBR111W-A | 100 | 0 |
| SPGP0G02080 | YJL069C | 99 | 0 |
| SPGP0M01330 | YNL194C | 99 | 0 |
| SPGP0C02960 | YDR077W | 99 | 0 |
| SPGP0G01070 | YJR051W | 99 | 0 |
| SPGP0F00860 | YGR203W | 99 | 0 |
| SPGP0DB02060 | YMR087W | 99 | 0 |
| SPGP0F00990 | YGR189C | 96 | 4 |
| SPGP0AE00590 | YCL005W | 99 | 0 |
| SPGP0Q04880 | YGL233W | 99 | 0 |
| SPGP0U01710 | YPL218W | 90 | 0 |
| SPGP0K02590 | YOR108W | 99 | 0 |
| SPGP0C01930 | YDL043C | 99 | 0 |
| SPGP0DB02510 | YMR035W | 99 | 0 |
| SPGP0D00430 | YPR034W | 99 | 0 |
| SPGP0D00340 | YPR045C | 99 | 0 |
| SPGP0DB01870 | YMR107W | 100 | 0 |
| SPGP0DC02270 | YBR130C | 99 | 0 |
| SPGP0I00330 | YEL049W | 99 | 0 |
| SPGP0DC02520 | YBR157C | 99 | 0 |
| SPGP0F02020 | YGR076C | 100 | 0 |
| SPGP0C02010 | YDL030W | 99 | 0 |
| SPGP0DC03130 | YBR222C | 99 | 0 |
| SPGP0C03600 | YDR146C | 99 | 0 |
| SPGP0G00700 | YJR092W | 99 | 0 |
| SPGP0F01500 | YGR132C | 99 | 0 |
| SPGP0F02940 | YGL030W | 100 | 0 |
| SPGP0I01880 | YER102W | 99 | 0 |
| SPGP0DB01300 | YMR166C | 99 | 0 |
| SPGP0AI00210 | YHR201C | 99 | 0 |
| SPGP0DD01990 | YIR033W | 99 | 0 |
| SPGP0K02180 | YOR061W | 99 | 0 |
| SPGP0F03650 | YGL112C | 99 | 0 |
| SPGP0DC02950 | YBR201C-A | 100 | 0 |

Create PDF in your applications with the Pdfcrowd HTML to PDF API PDFCROWD

| | | | |
|---|---|---|---|
| SPGP0DD00850 | YIL093C | 100 | 0 |
| SPGP0EX00260 | YDR345C | 90 | 0 |
| SPGP0DE00460 | YAL025C | 99 | 0 |
| SPGP0V00270 | YHL026C | 99 | 0 |
| SPGP0DE00570 | YAL014C | 99 | 0 |
| SPGP0K02090 | YOR049C | 99 | 0 |
| SPGP0M03080 | YNL002C | 100 | 0 |
| SPGP0C05190 | YDR321W | 99 | 0 |
| SPGP0F03200 | YGL061C | 99 | 0 |
| SPGP0K00940 | YOL075C | 99 | 0 |
| SPGP0Q04810 | YGL226C-A | 100 | 0 |
| SPGP0M02070 | YNL111C | 100 | 0 |
| SPGP0DC03540 | YBR264C | 100 | 0 |
| SPGP0C01620 | YDL077C | 99 | 0 |
| SPGP0C03650 | YDR152W | 100 | 0 |
| SPGP0C03300 | YDR115W | 99 | 0 |
| SPGP0DE00360 | YAL034W-A | 99 | 0 |
| SPGP0K00300 | YOL141W | 99 | 0 |
| SPGP0M03790 | YNR074C | 99 | 0 |
| SPGP0C01650 | YDL074C | 99 | 0 |
| SPGP0F03920 | YGL141W | 99 | 0 |
| SPGP0I02340 | YER153C | 99 | 0 |
| SPGP0DB02580 | YMR028W | 99 | 0 |
| SPGP0C04520 | YDR249C | 100 | 0 |
| SPGP0M02630 | YNL050C | 99 | 0 |
| SPGP0G02130 | YJL074C | 99 | 0 |
| SPGP0I02660 | YER186C | 96 | 1 |
| SPGP0M00670 | YNL267W | 99 | 0 |
| SPGP0DU01560 | YLR002C | 83 | 1 |
| SPGP0I00550 | YEL026W | 100 | 0 |
| SPGP0F00350 | YGR258C | 99 | 0 |
| SPGP0F04070 | YGL158W | 99 | 0 |
| SPGP0DD00920 | YIL085C | 99 | 0 |
| SPGP0Q04830 | YGL228W | 99 | 0 |
| SPGP0M01100 | YNL220W | 99 | 0 |
| SPGP0ED00110 | YPL283C | 96 | 0 |
| SPGP0C00650 | YDL182W | 99 | 0 |
| SPGP0C00870 | YDL157C | 99 | 0 |
| SPGP0DC03490 | YBR259W | 99 | 0 |
| SPGP0D00840 | YPL009C | 99 | 0 |

Create PDF in your applications with the Pdfcrowd HTML to PDF API PDFCROWD

| | | | |
|---|---|---|---|
| SPGP0F00790 | YGR210C | 99 | 0 |
| SPGP0G00280 | YJR134C | 99 | 0 |
| SPGP0DB01620 | YMR132C | 100 | 0 |
| SPGP0K00640 | YJR029W | 99 | 0 |
| SPGP0U02010 | YPL249C | 99 | 0 |
| SPGP0C04550 | YDR253C | 99 | 0 |
| SPGP0V00530 | YHR001W-A | 99 | 0 |
| SPGP0C03090 | YDR089W | 99 | 0 |
| SPGP0F01870 | YGR092W | 99 | 0 |
| SPGP0M00230 | YNL316C | 99 | 0 |
| SPGP0K02400 | YOR086C | 99 | 0 |
| SPGP0F00140 | YGR282C | 99 | 0 |
| SPGP0DB02850 | YMR002W | 97 | 3 |
| SPGP0DB00150 | YMR282C | 99 | 0 |
| SPGP0EQ00110 | YPL276W | 100 | 0 |
| SPGP0G02250 | YJL090C | 99 | 0 |
| SPGP0I02120 | YER128W | 99 | 0 |
| SPGP0DB01680 | YMR126C | 99 | 0 |
| SPGP0Q05100 | YGL257C | 98 | 0 |
| SPGP0M02640 | YNL049C | 99 | 0 |
| SPGP0DD01190 | YIL051C | 99 | 0 |
| SPGP0DB01180 | YMR178W | 99 | 0 |
| SPGP0C02730 | YDR054C | 100 | 0 |
| SPGP0C01820 | YDL054C | 99 | 0 |
| SPGP0DD01370 | YIL033C | 100 | 0 |
| SPGP0DD01950 | YIR029W | 99 | 0 |
| SPGP0DD00130 | YIL167W | 99 | 0 |
| SPGP0C01180 | YDL123W | 99 | 0 |
| SPGP0V01040 | YHR052W | 99 | 0 |
| SPGP0D00550 | YPR022C | 99 | 0 |
| SPGP0V01510 | YHR100C | 99 | 0 |
| SPGP0M01140 | YNL216W | 99 | 0 |
| SPGP0C02340 | YDR009W | 99 | 0 |
| SPGP0F03730 | YGL121C | 99 | 0 |
| SPGP0C05230 | YDR324C | 100 | 0 |
| SPGP0C00360 | YDL214C | 99 | 0 |
| SPGP0G01620 | YJL014W | 99 | 0 |
| SPGP0DB00220 | YMR275C | 99 | 0 |
| SPGP0DE00110 | YAL062W | 99 | 0 |
| SPGP0K01280 | YOL040C | 100 | 0 |

Create PDF in your applications with the Pdfcrowd HTML to PDF API PDFCROWD

| | | | |
|---|---|---|---|
| SPGP0M02450 | YNL070W | 100 | 0 |
| SPGP0V00640 | YHR011W | 99 | 0 |
| SPGP0DD00960 | YIL078W | 99 | 0 |
| SPGP0F00700 | YGR220C | 99 | 0 |
| SPGP0U02200 | YPL268W | 99 | 0 |
| SPGP0K01260 | YOL042W | 99 | 0 |
| SPGP0DB01650 | YMR129W | 99 | 0 |
| SPGP0V00960 | YHR044C | 99 | 0 |
| SPGP0DC02310 | YBR135W | 99 | 0 |
| SPGP0DD01740 | YIR007W | 99 | 0 |
| SPGP0D01000 | YPL026C | 98 | 1 |
| SPGP0F02340 | YGR037C | 100 | 0 |
| SPGP0C00460 | YDL204W | 99 | 0 |
| SPGP0M00990 | YNL233W | 99 | 0 |
| SPGP0V02090 | YHR159W | 99 | 0 |
| SPGP0DE00140 | YAL059W | 99 | 0 |
| SPGP0DC01650 | YBR060C | 99 | 0 |
| SPGP0F01670 | YGR113W | 99 | 0 |
| SPGP0F01600 | YGR122W | 99 | 0 |
| SPGP0C00790 | YDL167C | 99 | 0 |
| SPGP0G02640 | YJL133W | 99 | 0 |
| SPGP0M00190 | YNL321W | 99 | 0 |
| SPGP0M02410 | YNL074C | 99 | 0 |
| SPGP0DC00990 | YBL010C | 99 | 0 |
| SPGP0F04030 | YGL154C | 99 | 0 |
| SPGP0DC03660 | YBR278W | 100 | 0 |
| SPGP0F03990 | YGL148W | 99 | 0 |
| SPGP0K01460 | YOL019W | 99 | 0 |
| SPGP0M01820 | YNL138W | 99 | 0 |
| SPGP0DD01200 | YIL050W | 99 | 0 |
| SPGP0DB02810 | YMR006C | 99 | 0 |
| SPGP0M00330 | YNL306W | 99 | 0 |
| SPGP0M00440 | YNL293W | 99 | 0 |
| SPGP0C03840 | YDR172W | 99 | 0 |
| SPGP0C03510 | YDR137W | 99 | 0 |
| SPGP0DD01640 | YIL003W | 100 | 0 |
| SPGP0C04300 | YDR225W | 100 | 0 |
| SPGP0DC00260 | YBL089W | 99 | 0 |
| SPGP0F03190 | YGL060W | 99 | 0 |

Create PDF in your applications with the Pdfcrowd HTML to PDF API PDFCROWD

| | | | |
|---|---|---|---|
| SPGP0C00200 | YDL232W | 100 | 0 |
| SPGP0F03140 | YGL055W | 99 | 0 |
| SPGP0AE00210 | YCL051W | 99 | 0 |
| SPGP0DD00550 | YIL123W | 99 | 0 |
| SPGP0F00800 | YGR209C | 98 | 0 |
| SPGP0M01310 | YNL196C | 99 | 0 |
| SPGP0DB02800 | YMR007W | 99 | 0 |
| SPGP0F03410 | YGL085W | 99 | 0 |
| SPGP0DB03140 | YML029W | 99 | 0 |
| SPGP0G01010 | YJR057W | 99 | 0 |
| SPGP0F02140 | YGR061C | 99 | 0 |
| SPGP0DB02540 | YMR032W | 99 | 0 |
| SPGP0AE00740 | YCR012W | 99 | 0 |
| SPGP0C03670 | YDR155C | 100 | 0 |
| SPGP0M01440 | YNL182C | 99 | 0 |
| SPGP0I00270 | YEL055C | 99 | 0 |
| SPGP0DD00120 | YIL169C | 97 | 0 |
| SPGP0F02030 | YGR075C | 100 | 0 |
| SPGP0DC00880 | YBL022C | 99 | 0 |
| SPGP0I00770 | YEL003W | 99 | 0 |
| SPGP0C05210 | YDR322C-A | 100 | 0 |
| SPGP0G00490 | YJR111C | 99 | 0 |
| SPGP0G00470 | YJR112W-A | 99 | 0 |
| SPGP0DB01890 | YMR105C | 99 | 0 |
| SPGP0F01040 | YGR184C | 99 | 0 |
| SPGP0I00310 | YEL051W | 100 | 0 |
| SPGP0C00890 | YDL155W | 99 | 0 |
| SPGP0G02730 | YJL143W | 99 | 0 |
| SPGP0M00520 | YNL284C | 100 | 0 |
| SPGP0DB00410 | YMR256C | 100 | 0 |
| SPGP0M01850 | YNL135C | 100 | 0 |
| SPGP0G00720 | YJR090C | 99 | 0 |
| SPGP0K01450 | YOL020W | 99 | 0 |
| SPGP0F02380 | YGR033C | 99 | 0 |
| SPGP0Q04940 | YGL241W | 99 | 0 |
| SPGP0DB00300 | YMR267W | 100 | 0 |
| SPGP0C03290 | YDR113C | 99 | 0 |
| SPGP0DB01720 | YMR122W-A | 100 | 0 |
| SPGP0C01400 | YDL100C | 99 | 0 |
| SPGP0K02200 | YOR063W | 100 | 0 |

Create PDF in your applications with the Pdfcrowd HTML to PDF API PDFCROWD

| | | | |
|---|---|---|---|
| SPGP0F03630 | YGL110C | 99 | 0 |
| SPGP0M02810 | YNL021W | 99 | 0 |
| SPGP0M03310 | YNR023W | 99 | 0 |
| SPGP0C04890 | YDR292C | 99 | 0 |
| SPGP0V01250 | YHR075C | 100 | 0 |
| SPGP0I01790 | YER093C | 99 | 0 |
| SPGP0V00180 | YHL035C | 99 | 0 |
| SPGP0DD01120 | YIL062C | 99 | 0 |
| SPGP0M01540 | YNL168C | 99 | 0 |
| SPGP0U01940 | YPL242C | 99 | 0 |
| SPGP0DC00460 | YBL064C | 99 | 0 |
| SPGP0M02950 | YNL016W | 99 | 0 |
| SPGP0I00240 | YEL058W | 99 | 0 |
| SPGP0DE00800 | YAR023C | 99 | 0 |
| SPGP0DB01010 | YMR195W | 99 | 1 |
| SPGP0V00250 | YHL028W | 97 | 2 |
| SPGP0DE00330 | YAL037W | 99 | 0 |
| SPGP0I01870 | YER101C | 99 | 0 |
| SPGP0G02600 | YJL129C | 99 | 0 |
| SPGP0DD00710 | YIL107C | 99 | 0 |
| SPGP0DB00830 | YMR212C | 99 | 0 |
| SPGP0K02210 | YOR064C | 100 | 0 |
| SPGP0K00920 | YOL077C | 99 | 0 |
| SPGP0F03260 | YGL067W | 99 | 0 |
| SPGP0I00860 | YER007W | 99 | 0 |
| SPGP0V00550 | YHR003C | 99 | 0 |
| SPGP0C01640 | YDL075W | 99 | 0 |
| SPGP0Q04540 | YGL195W | 99 | 0 |
| SPGP0F03490 | YGL094C | 99 | 0 |
| SPGP0C02090 | YDL020C | 99 | 0 |
| SPGP0DC00630 | YBL045C | 99 | 0 |
| SPGP0D00750 | YPR001W | 99 | 0 |
| SPGP0C00590 | YDL190C | 99 | 0 |
| SPGP0Q00170 | YGR287C | 99 | 0 |
| SPGP0G01080 | YJR050W | 99 | 0 |
| SPGP0M00830 | YNL250W | 99 | 0 |
| SPGP0DC03510 | YBR261C | 99 | 0 |
| SPGP0DC03170 | YBR228W | 99 | 0 |
| SPGP0DD00940 | YIL083C | 99 | 0 |
| SPGP0F03570 | YGL103W | 99 | 0 |

Create PDF in your applications with the Pdfcrowd HTML to PDF API PDFCROWD

| | | | |
|---|---|---|---|
| SPGP0DD00490 | YIL129C | 99 | 0 |
| SPGP0DB03300 | YML050W | 100 | 0 |
| SPGP0AC01140 | YKR105C | 82 | 1 |
| SPGP0K00850 | YOL084W | 99 | 0 |
| SPGP0C00320 | YDL218W | 99 | 0 |
| SPGP0C00630 | YDL185W | 98 | 0 |
| SPGP0I02420 | YER162C | 99 | 0 |
| SPGP0AI00280 | YHR195W | 99 | 0 |
| SPGP0K01790 | YOR017W | 99 | 0 |
| SPGP0G00260 | YJR135W-A | 99 | 0 |
| SPGP0V00820 | YHR030C | 99 | 0 |
| SPGP0DC02130 | YBR112C | 99 | 1 |
| SPGP0C01420 | YDL098C | 99 | 0 |
| SPGP0C04530 | YDR251W | 99 | 0 |
| SPGP0DB00170 | YMR280C | 99 | 0 |
| SPGP0M01120 | YNL218W | 99 | 0 |
| SPGP0I01820 | YER095W | 99 | 0 |
| SPGP0DD01010 | YIL073C | 99 | 0 |
| SPGP0AE01350 | YCR082W | 99 | 0 |
| SPGP0G00780 | YJR083C | 99 | 0 |
| SPGP0F03760 | YGL124C | 99 | 0 |
| SPGP0G02310 | YJL096W | 99 | 0 |
| SPGP0AE00360 | YCL034W | 99 | 0 |
| SPGP0Q04730 | YGL219C | 100 | 0 |
| SPGP0DD02000 | YIR034C | 99 | 0 |
| SPGP0AE01170 | YCR060W | 100 | 0 |
| SPGP0I01010 | YER020W | 99 | 0 |
| SPGP0M00250 | YNL314W | 99 | 0 |
| SPGP0DD01040 | YIL070C | 99 | 0 |
| SPGP0AE00900 | YCR027C | 99 | 0 |
| SPGP0DB01400 | YMR156C | 99 | 0 |
| SPGP0F03060 | YGL044C | 100 | 0 |
| SPGP0DD01930 | YIR027C | 99 | 0 |
| SPGP0DB02140 | YMR077C | 100 | 0 |
| SPGP0M02390 | YNL076W | 99 | 0 |
| SPGP0G01600 | YJL012C | 99 | 0 |
| SPGP0G01350 | YJR014W | 100 | 0 |
| SPGP0C00450 | YDL205C | 99 | 0 |
| SPGP0AE00230 | YCL049C | 99 | 0 |
| SPGP0I00370 | YEL045C | 99 | 0 |

Create PDF in your applications with the Pdfcrowd HTML to PDF API PDFCROWD

| | | | |
|---|---|---|---|
| SPGP0AE00710 | YCR009C | 99 | 0 |
| SPGP0C04950 | YDR298C | 99 | 0 |
| SPGP0EQ00100 | YPL275W | 99 | 0 |
| SPGP0AE00220 | YCL050C | 99 | 0 |
| SPGP0DB01830 | YMR111C | 99 | 0 |
| SPGP0AE00550 | YCL010C | 99 | 0 |
| SPGP0C04790 | YDR281C | 99 | 0 |
| SPGP0C01320 | YDL108W | 99 | 0 |
| SPGP0I01390 | YER055C | 99 | 0 |
| SPGP0DE00310 | YAL039C | 99 | 0 |
| SPGP0DC00290 | YBL086C | 99 | 0 |
| SPGP0F00250 | YGR271W | 99 | 0 |
| SPGP0DD02030 | YIR037W | 99 | 0 |
| SPGP0F02220 | YGR053C | 99 | 0 |
| SPGP0I02690 | YBL111C | 99 | 0 |
| SPGP0F00220 | YGR274C | 99 | 0 |
| SPGP0C01800 | YDL056W | 99 | 0 |
| SPGP0DE00280 | YAL042W | 99 | 0 |
| SPGP0DE00780 | YAR019C | 99 | 0 |
| SPGP0V00620 | YHR009C | 99 | 0 |
| SPGP0F02410 | YGR030C | 99 | 0 |
| SPGP0K00370 | YOL133W | 99 | 0 |
| SPGP0F02360 | YGR035C | 99 | 0 |
| SPGP0V01390 | YHR088W | 99 | 0 |
| SPGP0K01240 | YOL044W | 99 | 0 |
| SPGP0DD01680 | YIR001C | 99 | 0 |
| SPGP0I00340 | YEL048C | 99 | 0 |
| SPGP0M00550 | YNL281W | 100 | 0 |
| SPGP0F02680 | YGL002W | 99 | 0 |
| SPGP0K02220 | YOR065W | 99 | 0 |
| SPGP0I00100 | YEL073C | 100 | 0 |
| SPGP0F02080 | YGR068C | 99 | 0 |
| SPGP0DB00900 | YMR205C | 99 | 0 |
| SPGP0G01440 | YJR006W | 99 | 0 |
| SPGP0F00240 | YGR271C-A | 99 | 0 |
| SPGP0DC01490 | YBR044C | 99 | 0 |
| SPGP0DE00160 | YAL056W | 99 | 0 |
| SPGP0K01730 | YOR009W | 96 | 0 |
| SPGP0G00480 | YJR112W | 99 | 0 |
| SPGP0M00700 | YNL263C | 99 | 0 |

Create PDF in your applications with the Pdfcrowd HTML to PDF API PDFCROWD

| | | | |
|---|---|---|---|
| SPGP0DD01220 | YIL048W | 99 | 0 |
| SPGP0DC01550 | YBR050C | 99 | 1 |
| SPGP0F01660 | YGR116W | 99 | 0 |
| SPGP0I01060 | YER025W | 99 | 0 |
| SPGP0K02120 | YOR054C | 99 | 0 |
| SPGP0G01280 | YJR024C | 99 | 0 |
| SPGP0F04050 | YGL156W | 99 | 0 |
| SPGP0U02080 | YPL256C | 99 | 0 |
| SPGP0F00780 | YGR211W | 99 | 0 |
| SPGP0I01500 | YER065C | 99 | 0 |
| SPGP0C00670 | YDL180W | 99 | 0 |
| SPGP0M03620 | YNR054C | 100 | 0 |
| SPGP0V01930 | YHR143W | 99 | 0 |
| SPGP0V02130 | YHR163W | 99 | 0 |
| SPGP0M01690 | YNL154C | 99 | 0 |
| SPGP0M01800 | YNL139C | 99 | 0 |
| SPGP0DB02160 | YMR075W | 99 | 0 |
| SPGP0M00310 | YNL308C | 99 | 0 |
| SPGP0C03530 | YDR139C | 100 | 0 |
| SPGP0M03810 | YNR076W | 98 | 0 |
| SPGP0C01200 | YDL121C | 100 | 0 |
| SPGP0Q04960 | YGL243W | 99 | 0 |
| SPGP0G03290 | YJL207C | 99 | 0 |
| SPGP0DC00240 | YBL091C | 99 | 0 |
| SPGP0C05140 | YDR317W | 99 | 0 |
| SPGP0G03200 | YJL197W | 99 | 0 |
| SPGP0DB03470 | YML066C | 99 | 0 |
| SPGP0C01770 | YDL059C | 99 | 0 |
| SPGP0C00120 | YDL240W | 99 | 0 |
| SPGP0G00920 | YJR066W | 99 | 0 |
| SPGP0F00820 | YGR207C | 99 | 0 |
| SPGP0F00530 | YGR239C | 99 | 0 |
| SPGP0C04180 | YDR211W | 99 | 0 |
| SPGP0DC01700 | YBR066C | 99 | 0 |
| SPGP0G01030 | YJR055W | 99 | 0 |
| SPGP0I02360 | YER155C | 99 | 0 |
| SPGP0DU01770 | YLL021W | 81 | 1 |
| SPGP0M03730 | YNR065C | 99 | 0 |
| SPGP0C03660 | YDR153C | 99 | 0 |
| SPGP0DD00140 | YIL166C | 99 | 0 |

Create PDF in your applications with the Pdfcrowd HTML to PDF API PDFCROWD

| | | | |
|---|---|---|---|
| SPGP0D01090 | YPL036W | 99 | 0 |
| SPGP0I00290 | YEL053C | 99 | 0 |
| SPGP0DB03450 | YML064C | 99 | 0 |
| SPGP0K02510 | YOR098C | 99 | 0 |
| SPGP0C02580 | YDR035W | 100 | 0 |
| SPGP0DD00860 | YIL092W | 99 | 0 |
| SPGP0D00470 | YPR030W | 99 | 0 |
| SPGP0D00380 | YPR040W | 99 | 0 |
| SPGP0K01110 | YOL057W | 99 | 0 |
| SPGP0M01130 | YNL217W | 100 | 0 |
| SPGP0I00570 | YEL024W | 99 | 0 |
| SPGP0DB03020 | YML017W | 99 | 0 |
| SPGP0G01290 | YJR022W | 100 | 0 |
| SPGP0DD01760 | YIR009W | 100 | 0 |
| SPGP0DB00920 | YMR203W | 99 | 0 |
| SPGP0C02850 | YDR066C | 99 | 0 |
| SPGP0DB01970 | YMR096W | 99 | 0 |
| SPGP0DB00760 | YMR219W | 99 | 0 |
| SPGP0G03310 | YJL209W | 99 | 0 |
| SPGP0G00740 | YJR088C | 99 | 0 |
| SPGP0DE00610 | YAL010C | 99 | 0 |
| SPGP0F01710 | YGR109C | 99 | 0 |
| SPGP0DB00360 | YMR261C | 99 | 0 |
| SPGP0DD00510 | YIL127C | 99 | 0 |
| SPGP0F01390 | YGR145W | 99 | 0 |
| SPGP0DC02420 | YBR147W | 99 | 0 |
| SPGP0M01010 | YNL231C | 99 | 0 |
| SPGP0C02050 | YDL025C | 99 | 0 |
| SPGP0DB00690 | YMR226C | 100 | 0 |
| SPGP0G00880 | YJR070C | 99 | 0 |
| SPGP0M03330 | YNR026C | 99 | 0 |
| SPGP0V01720 | YHR121W | 99 | 0 |
| SPGP0V01230 | YHR073W | 99 | 0 |
| SPGP0DC01800 | YBR077C | 99 | 0 |
| SPGP0G02140 | YJL076W | 99 | 0 |
| SPGP0K00260 | YOL145C | 99 | 0 |
| SPGP0DC01450 | YBR040W | 99 | 0 |
| SPGP0U01960 | YPL244C | 99 | 0 |
| SPGP0DB02280 | YMR063W | 99 | 0 |
| SPGP0V00200 | YHL033C | 98 | 0 |

Create PDF in your applications with the Pdfcrowd HTML to PDF API PDFCROWD

| | | | |
|---|---|---|---|
| SPGP0K02230 | YOR066W | 99 | 0 |
| SPGP0I00500 | YEL031W | 99 | 0 |
| SPGP0M01830 | YNL137C | 99 | 0 |
| SPGP0Q04430 | YGL183C | 99 | 0 |
| SPGP0G02240 | YJL089W | 99 | 0 |
| SPGP0C02070 | YDL022W | 99 | 0 |
| SPGP0Ftrna30T | YPR108W-A | 100 | 0 |
| SPGP0F03240 | YGL065C | 99 | 0 |
| SPGP0F03390 | YGL083W | 99 | 0 |
| SPGP0AE01430 | YCR091W | 99 | 0 |
| SPGP0F02870 | YGL022W | 99 | 0 |
| SPGP0I01710 | YER086W | 99 | 0 |
| SPGP0DE00530 | YAL018C | 99 | 0 |
| SPGP0G01110 | YJR047C | 100 | 0 |
| SPGP0AE00450 | YCL026C-A | 100 | 0 |
| SPGP0K01720 | YOR008C | 98 | 2 |
| SPGP0I02190 | YER137C | 100 | 0 |
| SPGP0G01060 | YJR052W | 99 | 0 |
| SPGP0G00240 | YJR137C | 99 | 0 |
| SPGP0DC03020 | YBR210W | 99 | 0 |
| SPGP0DB02990 | YML014W | 99 | 0 |
| SPGP0AE01370 | YCR084C | 99 | 0 |
| SPGP0DB03340 | YML054C | 99 | 0 |
| SPGP0C00960 | YDL146W | 99 | 0 |
| SPGP0AE00520 | YCL014W | 99 | 0 |
| SPGP0I02590 | YER179W | 99 | 0 |
| SPGP0C01560 | YDL083C | 100 | 0 |
| SPGP0C04510 | YDR248C | 99 | 0 |
| SPGP0DB02190 | YMR072W | 100 | 0 |
| SPGP0G00410 | YJR119C | 99 | 0 |
| SPGP0G02290 | YJL094C | 99 | 0 |
| SPGP0DC00180 | YBL098W | 99 | 0 |
| SPGP0G01990 | YJL059W | 99 | 0 |
| SPGP0DD01150 | YIL056W | 99 | 0 |
| SPGP0G01190 | YJR039W | 99 | 0 |
| SPGP0AE00850 | YCR021C | 99 | 0 |
| SPGP0DC03060 | YBR214W | 99 | 0 |
| SPGP0F00170 | YGR279C | 99 | 0 |
| SPGP0K02820 | YOR132W | 99 | 0 |
| SPGP0C02910 | YDR072C | 99 | 0 |

Create PDF in your applications with the Pdfcrowd HTML to PDF API PDFCROWD

| | | | |
|---|---|---|---|
| SPGP0G02950 | YJL167W | 99 | 0 |
| SPGP0DD01910 | YIR025W | 99 | 0 |
| SPGP0DC00980 | YBL011W | 99 | 0 |
| SPGP0G01370 | YJR012C | 99 | 0 |
| SPGP0C03330 | YDR118W | 99 | 0 |
| SPGP0DB00590 | YMR236W | 99 | 0 |
| SPGP0AE00690 | YCR007C | 100 | 0 |
| SPGP0V00310 | YHL022C | 99 | 0 |
| SPGP0M02940 | YNL018C | 99 | 0 |
| SPGP0DB01420 | YMR154C | 99 | 0 |
| SPGP0AE00200 | YCL052C | 99 | 0 |
| SPGP0C00430 | YDL207W | 99 | 0 |
| SPGP0DD00680 | YIL110W | 99 | 0 |
| SPGP0M02440 | YNL071W | 99 | 0 |
| SPGP0C02310 | YDR005C | 99 | 0 |
| SPGP0DC02720 | YBR177C | 99 | 0 |
| SPGP0DD00930 | YIL084C | 99 | 0 |
| SPGP0DB00770 | YMR218C | 99 | 0 |
| SPGP0DC02250 | YBR128C | 99 | 0 |
| SPGP0G01530 | YJL003W | 100 | 0 |
| SPGP0Q04350 | YGL173C | 98 | 0 |
| SPGP0DB01630 | YMR131C | 99 | 0 |
| SPGP0G00590 | YJR102C | 100 | 0 |
| SPGP0DE00300 | YAL040C | 99 | 0 |
| SPGP0C02770 | YDR058C | 99 | 0 |
| SPGP0Gtrna15M | YJL007C | 100 | 0 |
| SPGP0V00600 | YHR007C | 99 | 0 |
| SPGP0DD00420 | YIL136W | 99 | 0 |
| SPGP0DE00760 | YAR015W | 99 | 0 |
| SPGP0DE00810 | YAR027W | 99 | 0 |
| SPGP0F01550 | YGR127W | 100 | 0 |
| SPGP0DC03690 | YBR281C | 99 | 0 |
| SPGP0DE00700 | YAR002W | 99 | 0 |
| SPGP0V01730 | YHR122W | 99 | 0 |
| SPGP0F01230 | YGR163W | 99 | 0 |
| SPGP0G01470 | YJR004C | 99 | 0 |
| SPGP0M02850 | YNL024C-A | 99 | 0 |
| SPGP0C00420 | YDL208W | 100 | 0 |
| SPGP0DB02820 | YMR005W | 99 | 0 |
| SPGP0DB00570 | YMR238W | 99 | 0 |

Create PDF in your applications with the Pdfcrowd HTML to PDF API PDFCROWD

| | | | |
|---|---|---|---|
| SPGP0F02840 | YGL019W | 99 | 0 |
| SPGP0M03100 | YNR001C | 99 | 0 |
| SPGP0M00100 | YNL331C | 99 | 0 |
| SPGP0V00830 | YHR031C | 99 | 0 |
| SPGP0DC01470 | YBR042C | 99 | 0 |
| SPGP0DD02050 | YIR039C | 99 | 0 |
| SPGP0DC03730 | YBR285W | 99 | 0 |
| SPGP0DC00610 | YBL047C | 99 | 1 |
| SPGP0F01640 | YGR118W | 99 | 0 |
| SPGP0G02050 | YJL065C | 99 | 0 |
| SPGP0G02680 | YJL137C | 99 | 0 |
| SPGP0F03070 | YGL045W | 99 | 0 |
| SPGP0C03080 | YDR088C | 99 | 0 |
| SPGP0K02680 | YOR117W | 100 | 0 |
| SPGP0I01520 | YER067W | 100 | 0 |
| SPGP0N01110 | YLR300W | 86 | 0 |
| SPGP0V01910 | YHR141C | 100 | 0 |
| SPGP0F03860 | YGL135W | 99 | 0 |
| SPGP0F00520 | YGR240C | 99 | 0 |
| SPGP0M02150 | YNL101W | 99 | 0 |
| SPGP0K01410 | YOL024W | 100 | 0 |
| SPGP0C00280 | YDL223C | 99 | 0 |
| SPGP0I00300 | YEL052W | 99 | 0 |
| SPGP0C02480 | YDR024W | 99 | 0 |
| SPGP0C03880 | YDR176W | 99 | 0 |
| SPGP0C03550 | YDR141C | 99 | 0 |
| SPGP0AE00840 | YCR020W-B | 99 | 0 |
| SPGP0M01570 | YNL165W | 99 | 0 |
| SPGP0C01500 | YDL088C | 99 | 0 |
| SPGP0C01480 | YDL090C | 99 | 0 |
| SPGP0C05080 | YDR311W | 99 | 0 |
| SPGP0Q04780 | YGL224C | 100 | 0 |
| SPGP0M01680 | YNL155W | 99 | 0 |
| SPGP0M00480 | YNL289W | 100 | 0 |
| SPGP0I02090 | YER125W | 98 | 0 |
| SPGP0DB00580 | YMR237W | 99 | 0 |
| SPGP0AE00180 | YCL055W | 99 | 0 |
| SPGP0G00900 | YJR068W | 99 | 0 |
| SPGP0DC01560 | YBR052C | 99 | 0 |
| SPGP0F00510 | YGR241C | 99 | 0 |

Create PDF in your applications with the Pdfcrowd HTML to PDF API PDFCROWD

| | | | |
|---|---|---|---|
| SPGP0V00140 | YHL040C | 99 | 0 |
| SPGP0F03370 | YGL081W | 99 | 0 |
| SPGP0G00600 | YJR101W | 99 | 0 |
| SPGP0DB02500 | YMR036C | 99 | 0 |
| SPGP0C03210 | YDR104C | 99 | 0 |
| SPGP0C03640 | YDR151C | 100 | 0 |
| SPGP0I02380 | YER157W | 99 | 0 |
| SPGP0D00670 | YPR009W | 99 | 0 |
| SPGP0F02700 | YGL004C | 99 | 0 |
| SPGP0G01310 | YJR019C | 99 | 0 |
| SPGP0DC01170 | YBR009C | 99 | 0 |
| SPGP0DD00160 | YIL162W | 99 | 0 |
| SPGP0C04660 | YDR264C | 99 | 0 |
| SPGP0C04700 | YDR268W | 99 | 0 |
| SPGP0Q05010 | YGL248W | 99 | 0 |
| SPGP0C04070 | YDR196C | 99 | 0 |
| SPGP0D00450 | YPR032W | 99 | 0 |
| SPGP0G00760 | YJR085C | 100 | 0 |
| SPGP0F01000 | YGR188C | 99 | 0 |
| SPGP0G01550 | YJL005W | 99 | 0 |
| SPGP0DB02370 | YMR053C | 99 | 0 |
| SPGP0M03370 | YNR030W | 99 | 0 |
| SPGP0DB01990 | YMR094W | 99 | 0 |
| SPGP0AE01130 | YCR053W | 99 | 0 |
| SPGP0C04850 | YDR287W | 99 | 0 |
| SPGP0DW00110 | YCR100C | 99 | 0 |
| SPGP0I01570 | YER072W | 100 | 0 |
| SPGP0DB01360 | YMR160W | 99 | 0 |
| SPGP0M02600 | YNL053W | 99 | 0 |
| SPGP0M02190 | YNL097C | 99 | 0 |
| SPGP0DB00340 | YMR263W | 99 | 0 |
| SPGP0AE01230 | YCR068W | 99 | 0 |
| SPGP0DB03460 | YML065W | 99 | 0 |
| SPGP0K02780 | YOR128C | 99 | 0 |
| SPGP0V01210 | YHR072W | 99 | 0 |
| SPGP0DC01820 | YBR079C | 99 | 0 |
| SPGP0F01010 | YGR187C | 98 | 1 |
| SPGP0DC00420 | YBL069W | 99 | 0 |
| SPGP0K00240 | YOL147C | 99 | 0 |

Create PDF in your applications with the Pdfcrowd HTML to PDF API PDFCROWD

| | | | |
|---|---|---|---|
| SPGP0DE00370 | YAL034C | 99 | 0 |
| SPGP0V00220 | YHL031C | 99 | 0 |
| SPGP0M03060 | YNL004W | 100 | 0 |
| SPGP0M01890 | YNL131W | 100 | 0 |
| SPGP0K00730 | YOL096C | 99 | 0 |
| SPGP0K02250 | YOR069W | 99 | 0 |
| SPGP0M00960 | YNL237W | 99 | 0 |
| SPGP0G02260 | YJL091C | 99 | 0 |
| SPGP0V00160 | YHL038C | 99 | 0 |
| SPGP0C04460 | YDR243C | 99 | 0 |
| SPGP0DC03440 | YBR255W | 99 | 0 |
| SPGP0M03710 | YNR063W | 99 | 0 |
| SPGP0M03090 | YNL001W | 99 | 0 |
| SPGP0Y00670 | YDR406W | 83 | 1 |
| SPGP0AI00390 | YHR184W | 99 | 0 |
| SPGP0DC02140 | YBR114W | 99 | 0 |
| SPGP0G02190 | YJL083W | 99 | 0 |
| SPGP0DC03550 | YBR265W | 99 | 0 |
| SPGP0AE00820 | YCR020C | 99 | 0 |
| SPGP0I02600 | YER180C | 99 | 0 |
| SPGP0DC03430 | YBR254C | 99 | 0 |
| SPGP0F04000 | YGL150C | 99 | 0 |
| SPGP0DC01480 | YBR043C | 99 | 0 |
| SPGP0G00940 | YJR064W | 99 | 0 |
| SPGP0F02850 | YGL020C | 99 | 0 |
| SPGP0DD00980 | YIL076W | 99 | 0 |
| SPGP0DD00950 | YIL079C | 99 | 0 |
| SPGP0F03980 | YGL147C | 99 | 0 |
| SPGP0G01130 | YJR045C | 99 | 0 |
| SPGP0F03520 | YGL097W | 99 | 0 |
| SPGP0M02870 | YNL026W | 99 | 0 |
| SPGP0K01020 | YOL066C | 99 | 0 |
| SPGP0G00220 | YJR139C | 99 | 0 |
| SPGP0M01160 | YNL214W | 99 | 0 |
| SPGP0DB02920 | YML007W | 99 | 0 |
| SPGP0DB00130 | YMR284W | 100 | 0 |
| SPGP0DD01170 | YIL053W | 99 | 0 |
| SPGP0Q05090 | YGL256W | 99 | 0 |
| SPGP0Q04650 | YGL208W | 99 | 0 |
| SPGP0M00570 | YNL279W | 99 | 0 |

Create PDF in your applications with the Pdfcrowd HTML to PDF API PDFCROWD

| | | | |
|---|---|---|---|
| SPGP0I01360 | YER053C | 99 | 0 |
| SPGP0F00980 | YGR191W | 99 | 0 |
| SPGP0C01010 | YDL141W | 99 | 0 |
| SPGP0K00520 | YOL117W | 99 | 0 |
| SPGP0AE00150 | YCL057C-A | 99 | 0 |
| SPGP0DC00160 | YBL101C | 99 | 0 |
| SPGP0AE00860 | YCR023C | 99 | 0 |
| SPGP0F01880 | YGR091W | 100 | 0 |
| SPGP0G01170 | YJR041C | 99 | 0 |
| SPGP0I02450 | YER165W | 99 | 0 |
| SPGP0M00290 | YNL310C | 99 | 0 |
| SPGP0K02800 | YOR130C | 99 | 0 |
| SPGP0G02090 | YJL070C | 99 | 0 |
| SPGP0G00150 | YJR147W | 99 | 0 |
| SPGP0AE00670 | YCR005C | 99 | 0 |
| SPGP0C00490 | YDL201W | 99 | 0 |
| SPGP0AE00260 | YCL047C | 99 | 0 |
| SPGP0DB01190 | YMR177W | 99 | 0 |
| SPGP0DC02440 | YBR149W | 99 | 0 |
| SPGP0DB00710 | YMR224C | 99 | 0 |
| SPGP0Q05060 | YGL253W | 99 | 0 |
| SPGP0I01530 | YER068W | 99 | 0 |
| SPGP0DD01890 | YIR023W | 99 | 0 |
| SPGP0F01210 | YGR166W | 99 | 0 |
| SPGP0C04680 | YDR266C | 99 | 0 |
| SPGP0C01040 | YDL138W | 99 | 0 |
| SPGP0C03810 | YDR170C | 99 | 0 |
| SPGP0F02000 | YGR079W | 99 | 0 |
| SPGP0F02190 | YGR056W | 99 | 0 |
| SPGP0V00850 | YHR033W | 99 | 0 |
| SPGP0F01630 | YGR119C | 99 | 0 |
| SPGP0DC02170 | YPR080W | 99 | 0 |
| SPGP0M03470 | YNR038W | 99 | 0 |
| SPGP0DC00380 | YBL075C | 99 | 0 |
| SPGP0DB01710 | YMR123W | 99 | 0 |
| SPGP0DC01580 | YBR054W | 98 | 1 |
| SPGP0G02520 | YJL122W | 99 | 0 |
| SPGP0G02030 | YJL062W-A | 100 | 0 |
| SPGP0F01940 | YGR085C | 99 | 0 |
| SPGP0DE00120 | YAL061W | 99 | 0 |

Create PDF in your applications with the Pdfcrowd HTML to PDF API PDFCROWD

| | | | |
|---|---|---|---|
| SPGP0F00900 | YGR199W | 99 | 0 |
| SPGP0C03710 | YDR160W | 99 | 0 |
| SPGP0F00340 | YGR260W | 99 | 0 |
| SPGP0V01970 | YHR147C | 99 | 0 |
| SPGP0DB00210 | YMR276W | 99 | 0 |
| SPGP0V02100 | YHR160C | 99 | 0 |
| SPGP0C04170 | YDR208W | 99 | 0 |
| SPGP0C01710 | YDL066W | 100 | 0 |
| SPGP0F03840 | YGL133W | 99 | 0 |
| SPGP0I02200 | YER139C | 99 | 0 |
| SPGP0M00170 | YNL323W | 99 | 0 |
| SPGP0AH01020 | YFL034C-A | 80 | 4 |
| SPGP0C03570 | YDR143C | 99 | 0 |
| SPGP0DD00200 | YIL158W | 99 | 0 |
| SPGP0K00750 | YOL094C | 99 | 0 |
| SPGP0AE01360 | YCR083W | 99 | 0 |
| SPGP0G00710 | YJR091C | 99 | 0 |
| SPGP0M00350 | YNL304W | 99 | 0 |
| SPGP0C00140 | YDL238C | 99 | 0 |
| SPGP0I02010 | YER116C | 99 | 0 |
| SPGP0DE00790 | YFL020C | 97 | 0 |
| SPGP0G00960 | YJR062C | 99 | 0 |
| SPGP0DC01540 | YBR049C | 99 | 0 |
| SPGP0Q04520 | YGL194C | 99 | 0 |
| SPGP0M03220 | YNR014W | 100 | 0 |
| SPGP0C04490 | YDR246W | 100 | 0 |
| SPGP0DB01510 | YMR144W | 99 | 0 |
| SPGP0M01400 | YNL187W | 99 | 0 |
| SPGP0G00620 | YJR099W | 99 | 0 |
| SPGP0K01650 | YOR001W | 99 | 0 |
| SPGP0FK00100 | R0020C | 99 | 0 |
| SPGP0D00610 | YPR016C | 99 | 0 |
| SPGP0C03230 | YDR106W | 99 | 0 |
| SPGP0DB01130 | YMR184W | 99 | 0 |
| SPGP0DC01150 | YBR007C | 99 | 0 |
| SPGP0I02160 | YER132C | 99 | 0 |
| SPGP0C04720 | YDR272W | 99 | 0 |
| SPGP0C02060 | YDL024C | 99 | 0 |
| SPGP0G01260 | YJR030C | 99 | 0 |
| SPGP0AE01050 | YCR043C | 100 | 0 |

Create PDF in your applications with the Pdfcrowd HTML to PDF API PDFCROWD

| | | | |
|---|---|---|---|
| SPGP0DB02520 | YMR034C | 99 | 0 |
| SPGP0AE00420 | YCL028W | 99 | 1 |
| SPGP0AE00730 | YCR011C | 99 | 0 |
| SPGP0K01430 | YOL022C | 100 | 0 |
| SPGP0D00730 | YPR003C | 99 | 0 |
| SPGP0F01020 | YGR186W | 99 | 0 |
| SPGP0C02890 | YDR070C | 100 | 0 |
| SPGP0DC00390 | YBL074C | 99 | 0 |
| SPGP0C02540 | YDR031W | 99 | 0 |
| SPGP0DC03080 | YBR216C | 99 | 0 |
| SPGP0Q04950 | YGL242C | 99 | 0 |
| SPGP0DB00960 | YMR200W | 99 | 0 |
| SPGP0DC02340 | YBR138C | 99 | 0 |
| SPGP0V01690 | YHR118C | 99 | 0 |
| SPGP0C04870 | YDR289C | 99 | 0 |
| SPGP0V01430 | YHR092C | 99 | 0 |
| SPGP0DB01380 | YMR158W | 99 | 0 |
| SPGP0AE00270 | YCL045C | 99 | 0 |
| SPGP0C02760 | YDR057W | 99 | 0 |
| SPGP0DC02390 | YBR143C | 99 | 0 |
| SPGP0C01780 | YDL058W | 99 | 0 |
| SPGP0DB00380 | YMR259C | 99 | 0 |
| SPGP0K02010 | YOR040W | 99 | 0 |
| SPGP0DC01180 | YBR010W | 100 | 0 |
| SPGP0V01760 | YHR126C | 99 | 0 |
| SPGP0F01030 | YGR185C | 99 | 0 |
| SPGP0G02530 | YJL123C | 98 | 2 |
| SPGP0F02570 | YGR010W | 99 | 0 |
| SPGP0G01430 | YJR007W | 99 | 0 |
| SPGP0DC01770 | YBR073W | 99 | 0 |
| SPGP0F00730 | YGR216C | 99 | 0 |
| SPGP0M03800 | YNR075W | 100 | 0 |
| SPGP0Q04550 | YGL196W | 99 | 0 |
| SPGP0M01870 | YNL133C | 99 | 0 |
| SPGP0F02810 | YGL016W | 99 | 0 |
| SPGP0DC00640 | YBL043W | 99 | 0 |
| SPGP0C04410 | YDR237W | 99 | 0 |
| SPGP0DC03570 | YBR268W | 99 | 0 |
| SPGP0V00410 | YHL012W | 99 | 0 |
| SPGP0DC01710 | YBR067C | 97 | 3 |

Create PDF in your applications with the Pdfcrowd HTML to PDF API PDFCROWD

| | | | |
|---|---|---|---|
| SPGP0M02840 | YNL024C | 99 | 0 |
| SPGP0V01900 | YHR140W | 99 | 0 |
| SPGP0K01960 | YOR035C | 99 | 0 |
| SPGP0DC03420 | YBR253W | 99 | 0 |
| SPGP0M03610 | YNR053C | 99 | 0 |
| SPGP0DD01530 | YIL014W | 99 | 0 |
| SPGP0DB03410 | YML060W | 99 | 0 |
| SPGP0I00880 | YER008C | 99 | 0 |
| SPGP0I00610 | YEL020W-A | 100 | 0 |
| SPGP0DC02160 | YBR117C | 99 | 0 |
| SPGP0DC03410 | YBR252W | 100 | 0 |
| SPGP0F04130 | YGL164C | 100 | 0 |
| SPGP0F02830 | YGL018C | 99 | 0 |
| SPGP0M02490 | YNL066W | 99 | 0 |
| SPGP0DC02760 | YBR182C | 99 | 0 |
| SPGP0G03000 | YJL173C | 98 | 0 |
| SPGP0G01150 | YJR043C | 99 | 0 |
| SPGP0G02400 | YJL105W | 99 | 0 |
| SPGP0DB02360 | YMR054W | 99 | 0 |
| SPGP0DB02950 | YML010W | 99 | 0 |
| SPGP0M03480 | YNR039C | 99 | 0 |
| SPGP0M00720 | YNL261W | 100 | 0 |
| SPGP0I01020 | YER021W | 99 | 0 |
| SPGP0C04260 | YDR221W | 99 | 0 |
| SPGP0Q04700 | YGL213C | 99 | 0 |
| SPGP0M02720 | YNL040W | 99 | 0 |
| SPGP0AE01330 | YCR079W | 99 | 0 |
| SPGP0F03770 | YGL125W | 99 | 0 |
| SPGP0C00920 | YDL150W | 99 | 0 |
| SPGP0DD01450 | YIL021W | 99 | 0 |
| SPGP0Q05070 | YGL254W | 100 | 0 |
| SPGP0F00960 | YGR193C | 99 | 0 |
| SPGP0DD01230 | YIL047C | 99 | 0 |
| SPGP0C04000 | YDR188W | 99 | 0 |
| SPGP0DC02870 | YBR194W | 99 | 0 |
| SPGP0C01030 | YDL139C | 99 | 0 |
| SPGP0DC00140 | YBL103C | 99 | 0 |
| SPGP0DB01330 | YMR163C | 99 | 0 |
| SPGP0I01340 | YER051W | 99 | 0 |
| SPGP0C02950 | YDR076W | 99 | 0 |

Create PDF in your applications with the Pdfcrowd HTML to PDF API PDFCROWD

| | | | |
|---|---|---|---|
| SPGP0Q05080 | YGL255W | 95 | 1 |
| SPGP0M00270 | YNL312W | 100 | 0 |
| SPGP0F00210 | YGR275W | 99 | 0 |
| SPGP0DB01600 | YMR134W | 99 | 0 |
| SPGP0DC02890 | YBR196C | 99 | 0 |
| SPGP0DB02770 | YMR009W | 98 | 0 |
| SPGP0AE00650 | YCR003W | 99 | 0 |
| SPGP0G01840 | YJL044C | 99 | 0 |
| SPGP0DB00550 | YMR240C | 99 | 0 |
| SPGP0I02020 | YER117W | 99 | 0 |
| SPGP0DD00640 | YIL114C | 99 | 0 |
| SPGP0DB01750 | YMR119W | 99 | 0 |
| SPGP0C02370 | YDR013W | 99 | 0 |
| SPGP0F02100 | YGR066C | 100 | 0 |
| SPGP0M00680 | YNL265C | 99 | 0 |
| SPGP0DT00100 | YNL333W | 99 | 0 |
| SPGP0M01910 | YNL129W | 100 | 0 |
| SPGP0DD01870 | YIR021W | 99 | 0 |
| SPGP0DC03380 | YBR249C | 99 | 0 |
| SPGP0I00110 | YEL072W | 99 | 0 |
| SPGP0DB00730 | YMR222C | 99 | 0 |
| SPGP0K02480 | YOR095C | 99 | 0 |
| SPGP0D00720 | YPR004C | 99 | 0 |
| SPGP0K01660 | YOR002W | 99 | 0 |
| SPGP0F02720 | YGL006W | 99 | 0 |
| SPGP0G01420 | YJR008W | 99 | 0 |
| SPGP0DE00720 | YAR003W | 99 | 0 |
| SPGP0Q04690 | YGL212W | 99 | 0 |
| SPGP0V00870 | YHR035W | 99 | 0 |
| SPGP0V01400 | YHR089C | 99 | 0 |
| SPGP0F03160 | YGL057C | 99 | 0 |
| SPGP0DB00970 | YMR199W | 99 | 0 |
| SPGP0K01200 | YOL048C | 99 | 0 |
| SPGP0DC01600 | YBR056W | 99 | 0 |
| SPGP0DC03590 | YBR270C | 99 | 0 |
| SPGP0F03230 | YGL064C | 99 | 0 |
| SPGP0C03770 | YDR166C | 99 | 0 |
| SPGP0I01560 | YER071C | 99 | 0 |
| SPGP0C01730 | YDL064W | 100 | 0 |
| SPGP0DD01590 | YIL008W | 100 | 0 |

Create PDF in your applications with the Pdfcrowd HTML to PDF API PDFCROWD

| | | | |
|---|---|---|---|
| SPGP0G01670 | YJL024C | 99 | 0 |
| SPGP0G02010 | YJL061W | 99 | 0 |
| SPGP0M00110 | YNL330C | 99 | 0 |
| SPGP0C03590 | YDR145W | 99 | 0 |
| SPGP0AE01340 | YCR081W | 99 | 0 |
| SPGP0I02030 | YER118C | 99 | 0 |
| SPGP0V00300 | YHL023C | 99 | 0 |
| SPGP0K01780 | YOR016C | 99 | 0 |
| SPGP0DC01520 | YBR047W | 99 | 0 |
| SPGP0V00940 | YHR041C | 99 | 0 |
| SPGP0DB01490 | YMR146C | 100 | 0 |
| SPGP0C01190 | YDL122W | 99 | 0 |
| SPGP0I00760 | YEL004W | 99 | 0 |
| SPGP0M01460 | YNL180C | 99 | 0 |
| SPGP0F01790 | YGR100W | 99 | 0 |
| SPGP0AE01290 | YCR075C | 99 | 0 |
| SPGP0DC01390 | YBR034C | 99 | 0 |
| SPGP0C03180 | YDR100W | 100 | 0 |
| SPGP0DC03600 | YBR271W | 99 | 0 |
| SPGP0C05030 | YDR306C | 99 | 0 |
| SPGP0D00490 | YPR028W | 100 | 0 |
| SPGP0Q04980 | YGL245W | 99 | 0 |
| SPGP0G02740 | YJL144W | 98 | 0 |
| SPGP0C04620 | YDR260C | 99 | 0 |
| SPGP0AE01070 | YCR045C | 100 | 0 |
| SPGP0C04280 | YDR223W | 99 | 0 |
| SPGP0DY00110 | YNL335W | 99 | 0 |
| SPGP0DD00500 | YIL128W | 99 | 0 |
| SPGP0K00360 | YOL135C | 99 | 0 |
| SPGP0F02990 | YGL036W | 99 | 0 |
| SPGP0C02560 | YDR033W | 100 | 0 |
| SPGP0DB03150 | YML030W | 100 | 0 |
| SPGP0DB02180 | YMR073C | 99 | 0 |
| SPGP0F01430 | YGR141W | 99 | 0 |
| SPGP0F02090 | YGR067C | 99 | 0 |
| SPGP0C04810 | YDR283C | 99 | 0 |
| SPGP0V01410 | YHR090C | 99 | 0 |
| SPGP0C02740 | YDR055W | 99 | 0 |
| SPGP0DD01140 | YIL057C | 100 | 0 |
| SPGP0M01110 | YNL219C | 99 | 0 |

Create PDF in your applications with the Pdfcrowd HTML to PDF API PDFCROWD

| | | | |
|---|---|---|---|
| SPGP0F00310 | YGR263C | 99 | 0 |
| SPGP0C00940 | YDL148C | 99 | 0 |
| SPGP0M02660 | YNL047C | 99 | 0 |
| SPGP0I02570 | YER177W | 99 | 0 |
| SPGP0F01770 | YGR102C | 99 | 0 |
| SPGP0F00750 | YGR214W | 99 | 0 |
| SPGP0M03530 | YNR045W | 99 | 0 |
| SPGP0DC01860 | YBR083W | 99 | 0 |
| SPGP0C03920 | YDR179W-A | 99 | 0 |
| SPGP0F02790 | YGL014W | 99 | 0 |
| SPGP0AI00130 | YHR209W | 99 | 0 |
| SPGP0G02220 | YJL087C | 99 | 0 |
| SPGP0K01920 | YOR031W | 99 | 0 |
| SPGP0D00820 | YPL007C | 99 | 0 |
| SPGP0F02210 | YGR054W | 99 | 0 |
| SPGP0G01820 | YJL042W | 99 | 0 |
| SPGP0DC03400 | YBR251W | 99 | 0 |
| SPGP0F01400 | YGR144W | 99 | 0 |
| SPGP0DC02180 | YBR119W | 99 | 0 |
| SPGP0G02480 | YJL116C | 99 | 0 |
| SPGP0F01890 | YGR090W | 99 | 0 |
| SPGP0C04040 | YDR192C | 99 | 0 |
| SPGP0K01710 | YOR007C | 99 | 0 |
| SPGP0F03320 | YGL076C | 99 | 0 |
| SPGP0DC03140 | YBR223C | 99 | 0 |
| SPGP0DB02970 | YML012W | 99 | 0 |
| SPGP0DC00120 | YBL105C | 99 | 0 |
| SPGP0DC01080 | YBL001C | 100 | 0 |
| SPGP0C05040 | YDR307W | 99 | 0 |
| SPGP0DD01130 | YIL061C | 99 | 0 |
| SPGP0C05000 | YDR303C | 99 | 0 |
| SPGP0G03270 | YJL205C | 99 | 0 |
| SPGP0C00900 | YDL154W | 99 | 0 |
| SPGP0V02040 | YHR154W | 99 | 0 |
| SPGP0C03040 | YDR084C | 99 | 0 |
| SPGP0M02610 | YNL052W | 100 | 0 |
| SPGP0C01050 | YDL136W | 100 | 0 |
| SPGP0DB01350 | YMR161W | 99 | 0 |
| SPGP0C04020 | YDR190C | 100 | 0 |
| SPGP0K00560 | YOL113W | 99 | 0 |

Create PDF in your applications with the Pdfcrowd HTML to PDF API PDFCROWD

| | | | |
|---|---|---|---|
| SPGP0I02370 | YER156C | 99 | 0 |
| SPGP0K00620 | YOL107W | 99 | 0 |
| SPGP0C02290 | YDR003W | 99 | 0 |
| SPGP0C02970 | YDR078C | 99 | 0 |
| SPGP0I00220 | YEL060C | 99 | 0 |
| SPGP0V01440 | YHR093W | 99 | 0 |
| SPGP0Q05050 | YGL252C | 99 | 0 |
| SPGP0DB01950 | YMR098C | 99 | 0 |
| SPGP0DC02740 | YBR180W | 99 | 0 |
| SPGP0K00190 | YOL153C | 99 | 0 |
| SPGP0DB02020 | YMR091C | 99 | 0 |
| SPGP0G01860 | YJL046W | 99 | 0 |
| SPGP0AI00250 | YHR198C | 100 | 0 |
| SPGP0DC01290 | YBR022W | 100 | 0 |
| SPGP0DB01480 | YMR147W | 99 | 0 |
| SPGP0Q04710 | YGL215W | 99 | 0 |
| SPGP0DC02400 | YBR145W | 99 | 0 |
| SPGP0C02350 | YDR011W | 99 | 0 |
| SPGP0DB00530 | YMR242C | 100 | 0 |
| SPGP0C00770 | YDL169C | 99 | 0 |
| SPGP0K01570 | YOL008W | 99 | 0 |
| SPGP0M01930 | YNL127W | 99 | 0 |
| SPGP0D00810 | YPL006W | 99 | 0 |
| SPGP0DB00280 | YMR269W | 100 | 0 |
| SPGP0V01750 | YHR124W | 99 | 0 |
| SPGP0D00290 | YPR051W | 99 | 0 |
| SPGP0M02340 | YNL081C | 99 | 0 |
| SPGP0U00660 | YPL109C | 82 | 0 |
| SPGP0V00330 | YHL020C | 99 | 0 |
| SPGP0DE00250 | YAL044W-A | 99 | 0 |
| SPGP0V00890 | YHR037W | 99 | 0 |
| SPGP0D00500 | YPR027C | 100 | 0 |
| SPGP0DB00470 | YMR250W | 99 | 0 |
| SPGP0DC00950 | YBL015W | 99 | 0 |
| SPGP0DC03610 | YBR272C | 99 | 0 |
| SPGP0BF00310 | YLL039C | 85 | 1 |
| SPGP0C03750 | YDR164C | 99 | 0 |
| SPGP0DC01620 | YBR058C | 99 | 0 |
| SPGP0F00120 | YGR284C | 99 | 0 |
| SPGP0F02880 | YGL023C | 99 | 0 |

Create PDF in your applications with the Pdfcrowd HTML to PDF API PDFCROWD

| | | | |
|---|---|---|---|
| SPGP0C00510 | YDL199C | 99 | 0 |
| SPGP0DC02660 | YBR170C | 99 | 0 |
| SPGP0V00590 | YHR006W | 99 | 0 |
| SPGP0M03340 | YNR027W | 99 | 0 |
| SPGP0DC01500 | YBR045C | 99 | 0 |
| SPGP0F02470 | YGR023W | 97 | 1 |
| SPGP0C00880 | YDL156W | 99 | 0 |
| SPGP0K00650 | YOL105C | 98 | 1 |
| SPGP0G03210 | YJL198W | 99 | 0 |
| SPGP0M01660 | YNL157W | 99 | 0 |
| SPGP0M02520 | YNL063W | 99 | 0 |
| SPGP0D00940 | YPL019C | 99 | 0 |
| SPGP0DB03160 | YML031W | 99 | 0 |
| SPGP0V00920 | YHR039C-A | 100 | 0 |
| SPGP0K01800 | YOR018W | 99 | 0 |
| SPGP0D00120 | YPR071W | 99 | 0 |
| SPGP0DC01830 | YBR080C | 99 | 0 |
| SPGP0AE01270 | YCR073C | 99 | 0 |
| SPGP0DC03620 | YBR273C | 99 | 0 |
| SPGP0K01070 | YOL061W | 99 | 0 |
| SPGP0I02180 | YER136W | 100 | 0 |
| SPGP0G00430 | YJR117W | 99 | 0 |
| SPGP0C05060 | YDR309C | 99 | 0 |
| SPGP0C04350 | YDR231C | 99 | 0 |
| SPGP0C04640 | YDR262W | 99 | 0 |
| SPGP0DB02560 | YMR030W | 99 | 0 |
| SPGP0DC01900 | YBR086C | 99 | 0 |
| SPGP0DB03220 | YML038C | 99 | 0 |
| SPGP0I00850 | YER006W | 99 | 0 |
| SPGP0DD00650 | YIL113W | 99 | 0 |
| SPGP0I01830 | YER096W | 99 | 0 |
| SPGP0DB01030 | YMR194W | 100 | 0 |
| SPGP0C01310 | YDL109C | 99 | 0 |
| SPGP0G00170 | YJR144W | 99 | 0 |
| SPGP0C02600 | YDR037W | 99 | 0 |
| SPGP0DC00820 | YBL027W | 99 | 0 |
| SPGP0C02500 | YDR026C | 99 | 0 |
| SPGP0DB03200 | YML036W | 99 | 0 |
| SPGP0C00330 | YDL217C | 99 | 0 |

Create PDF in your applications with the Pdfcrowd HTML to PDF API PDFCROWD

| | | | |
|---|---|---|---|
| SPGP0Q04460 | YGL186C | 100 | 0 |
| SPGP0V01460 | YHR096C | 99 | 0 |
| SPGP0K01120 | YOL056W | 99 | 0 |
| SPGP0G01720 | YJL030W | 99 | 0 |
| SPGP0DD00830 | YIL095W | 99 | 0 |
| SPGP0G01220 | YJR034W | 100 | 0 |
| SPGP0M03130 | YNR004W | 100 | 0 |
| SPGP0M02130 | YNL103W | 99 | 0 |
| SPGP0M03400 | YNR032C-A | 99 | 0 |
| SPGP0K01310 | YOL036W | 99 | 0 |
| SPGP0K00380 | YOL132W | 99 | 0 |
| SPGP0D00100 | YPR073C | 99 | 0 |
| SPGP0F00770 | YGR212W | 99 | 0 |
| SPGP0G00230 | YJR138W | 99 | 0 |
| SPGP0AE01160 | YCR059C | 99 | 0 |
| SPGP0K01380 | YOL027C | 99 | 0 |
| SPGP0V00440 | YHL009C | 99 | 0 |
| SPGP0C02100 | YDL019C | 99 | 0 |
| SPGP0DC02210 | YBR122C | 99 | 0 |
| SPGP0Ktrna3P | YOL013W-A | 99 | 0 |
| SPGP0M00890 | YNL244C | 99 | 0 |
| SPGP0V00450 | YHL008C | 87 | 2 |
| SPGP0V00100 | YNL336W | 99 | 0 |
| SPGP0G01330 | YJR016C | 99 | 0 |
| SPGP0C04480 | YDR245W | 99 | 0 |
| SPGP0K01030 | YOL065C | 99 | 0 |
| SPGP0F01420 | YGR142W | 99 | 0 |
| SPGP0F01520 | YGR130C | 99 | 0 |
| SPGP0F04170 | YGL170C | 99 | 0 |
| SPGP0M03770 | YNR071C | 100 | 0 |
| SPGP0I01240 | YER043C | 99 | 0 |
| SPGP0K02020 | YOR042W | 99 | 0 |
| SPGP0AI00480 | YHR175W | 99 | 0 |
| SPGP0C03250 | YDR108W | 99 | 0 |
| SPGP0C01680 | YDL070W | 99 | 0 |
| SPGP0D01020 | YPL028W | 99 | 0 |
| SPGP0K00540 | YOL115W | 99 | 0 |
| SPGP0K00810 | YOL088C | 99 | 0 |
| SPGP0C01910 | YDL045C | 99 | 0 |
| SPGP0DC01060 | YBL003C | 100 | 0 |

Create PDF in your applications with the Pdfcrowd HTML to PDF API PDFCROWD

| | | | |
|---|---|---|---|
| SPGP0DC00100 | YBL107C | 99 | 0 |
| SPGP0AE00880 | YCR024C-A | 100 | 0 |
| SPGP0D00160 | YPR067W | 99 | 0 |
| SPGP0DB01290 | YMR167W | 99 | 0 |
| SPGP0F00920 | YGR197C | 99 | 0 |
| SPGP0F00690 | YGR221C | 99 | 0 |
| SPGP0C01070 | YDL134C | 98 | 0 |
| SPGP0K00600 | YOL109W | 99 | 0 |
| SPGP0F01820 | YGR097W | 99 | 0 |
| SPGP0M01090 | YNL221C | 99 | 0 |
| SPGP0DB02660 | YMR020W | 99 | 0 |
| SPGP0D00770 | YPL002C | 99 | 0 |
| SPGP0F01780 | YGR101W | 100 | 0 |
| SPGP0DB01930 | YMR100W | 99 | 0 |
| SPGP0DC01310 | YBR024W | 100 | 0 |
| SPGP0I01420 | YER057C | 100 | 0 |
| SPGP0DD00600 | YIL118W | 100 | 0 |
| SPGP0M00160 | YNL325C | 99 | 0 |
| SPGP0I00560 | YEL025C | 99 | 0 |
| SPGP0DB02000 | YMR093W | 99 | 0 |
| SPGP0K01630 | YOL002C | 99 | 0 |
| SPGP0AE00280 | YCL044C | 99 | 0 |
| SPGP0DC01270 | YBR020W | 100 | 0 |
| SPGP0F01240 | YGR162W | 99 | 0 |
| SPGP0C02690 | YDR049W | 99 | 0 |
| SPGP0DB01670 | YMR127C | 99 | 0 |
| SPGP0DB01200 | YMR176W | 99 | 0 |
| SPGP0F01260 | YGR159C | 99 | 0 |
| SPGP0DB01770 | YMR117C | 99 | 0 |
| SPGP0DB02080 | YMR085W | 99 | 0 |
| SPGP0DD01510 | YIL015W | 99 | 0 |
| SPGP0C01090 | YDL132W | 99 | 0 |
| SPGP0K00330 | YOL138C | 99 | 0 |
| SPGP0V01190 | YHR069C | 99 | 0 |
| SPGP0M01720 | YNL151C | 100 | 0 |
| SPGP0M02360 | YNL079C | 100 | 0 |
| SPGP0DC00310 | YBL084C | 98 | 1 |
| SPGP0DE00270 | YAL043C | 99 | 0 |
| SPGP0C02140 | YDL014W | 99 | 0 |
| SPGP0DC02010 | YBR101C | 99 | 0 |

Create PDF in your applications with the Pdfcrowd HTML to PDF API PDFCROWD

| | | | |
|---|---|---|---|
| SPGP0V01710 | YHR120W | 99 | 0 |
| SPGP0V01990 | YHR149C | 99 | 0 |
| SPGP0DC00690 | YBL039C | 99 | 0 |
| SPGP0C02620 | YDR041W | 100 | 0 |
| SPGP0DE00340 | YAL036C | 99 | 0 |
| SPGP0DB00440 | YMR252C | 99 | 0 |
| SPGP0C03000 | YDR080W | 99 | 0 |
| SPGP0DC00550 | YBL055C | 99 | 0 |
| SPGP0K02600 | YOR109W | 99 | 0 |
| SPGP0DC03250 | YBR236C | 99 | 0 |
| SPGP0V00930 | YHR040W | 99 | 0 |
| SPGP0U02170 | YPL265W | 99 | 0 |
| SPGP0G00990 | YJR059W | 99 | 0 |
| SPGP0AI00360 | YHR187W | 100 | 0 |
| SPGP0DD00540 | YIL124W | 99 | 0 |
| SPGP0F01460 | YGR136W | 99 | 0 |
| SPGP0AE01250 | YCR071C | 99 | 0 |
| SPGP0DB01530 | YMR142C | 99 | 0 |
| SPGP0Q04900 | YGL236C | 99 | 0 |
| SPGP0G00450 | YJR115W | 100 | 0 |
| SPGP0CZ00260 | YKL205W | 84 | 0 |
| SPGP0I00730 | YEL007W | 99 | 0 |
| SPGP0DD01070 | YIL067C | 99 | 0 |
| SPGP0DB01100 | YMR187C | 99 | 0 |
| SPGP0D00180 | YPR065W | 99 | 0 |
| SPGP0I01810 | YER094C | 99 | 0 |
| SPGP0R02880 | YOR192C-B | 99 | 0 |
| SPGP0F00480 | YGR245C | 99 | 0 |
| SPGP0C01330 | YDL107W | 99 | 0 |
| SPGP0F02960 | YGL032C | 99 | 0 |
| SPGP0F03530 | YGL098W | 99 | 0 |
| SPGP0DC00840 | YBL026W | 99 | 0 |
| SPGP0DB02530 | YMR033W | 99 | 0 |
| SPGP0C02520 | YDR028C | 99 | 0 |
| SPGP0DD00720 | YIL106W | 99 | 0 |
| SPGP0C03700 | YDR159W | 99 | 0 |
| SPGP0C00350 | YDL215C | 99 | 0 |
| SPGP0C01860 | YDL049C | 99 | 0 |
| SPGP0U02100 | YPL257W | 92 | 0 |
| SPGP0D00590 | YPR018W | 99 | 0 |

Create PDF in your applications with the Pdfcrowd HTML to PDF API PDFCROWD

| | | | |
|---|---|---|---|
| SPGP0DB01050 | YMR192W | 99 | 0 |
| SPGP0AE00610 | YCL002C | 99 | 0 |
| SPGP0AE00340 | YCL036W | 99 | 0 |
| SPGP0DB03010 | YML016C | 99 | 0 |
| SPGP0F03610 | YGL107C | 99 | 0 |
| SPGP0C02780 | YDR059C | 99 | 0 |
| SPGP0DC00250 | YBL090W | 99 | 0 |
| SPGP0DB00890 | YMR206W | 100 | 0 |
| SPGP0F01840 | YGR095C | 99 | 0 |
| SPGP0C01600 | YDL079C | 99 | 0 |
| SPGP0M02110 | YNL106C | 99 | 0 |
| SPGP0C01250 | YDL115C | 99 | 0 |
| SPGP0F02520 | YGR016W | 99 | 0 |
| SPGP0I01490 | YER064C | 99 | 0 |
| SPGP0DC03000 | YBR207W | 99 | 0 |
| SPGP0M01340 | YNL193W | 99 | 0 |
| SPGP0G01400 | YJR010W | 99 | 0 |
| SPGP0DB00640 | YMR231W | 99 | 0 |
| SPGP0DB00190 | YMR278W | 99 | 0 |
| SPGP0V00470 | YHL006C | 98 | 0 |
| SPGP0DE00510 | YAL020C | 99 | 0 |
| SPGP0K01050 | YOL063C | 99 | 0 |
| SPGP0G02120 | YJL073W | 99 | 0 |
| SPGP0DC02150 | YBR115C | 99 | 0 |
| SPGP0F01440 | YGR140W | 99 | 0 |
| SPGP0M01500 | YNL175C | 99 | 0 |
| SPGP0DD01850 | YIR018W | 99 | 0 |
| SPGP0K02060 | YOR046C | 99 | 0 |
| SPGP0G02910 | YJL163C | 99 | 0 |
| SPGP0D01010 | YPL027W | 98 | 0 |
| SPGP0G02550 | YJL125C | 99 | 0 |
| SPGP0G01200 | YJR036C | 99 | 0 |
| SPGP0V01870 | YHR137W | 99 | 0 |
| SPGP0F03790 | YGL127C | 99 | 0 |
| SPGP0V01520 | YHR101C | 99 | 0 |
| SPGP0DB02930 | YML008C | 99 | 0 |
| SPGP0DB02260 | YMR065W | 99 | 0 |
| SPGP0I01680 | YER082C | 99 | 0 |
| SPGP0M00210 | YNL318C | 99 | 0 |
| SPGP0F00160 | YGR280C | 99 | 0 |

Create PDF in your applications with the Pdfcrowd HTML to PDF API PDFCROWD

| | | | |
|---|---|---|---|
| SPGP0I02250 | YER144C | 100 | 0 |
| SPGP0M03410 | YNR033W | 99 | 0 |
| SPGP0I01040 | YER023W | 99 | 0 |
| SPGP0F00710 | YGR218W | 99 | 0 |
| SPGP0C04200 | YDR213W | 99 | 0 |
| SPGP0U02210 | YPL269W | 99 | 0 |
| SPGP0K00990 | YOL069W | 99 | 0 |
| SPGP0F03740 | YGL122C | 98 | 1 |
| SPGP0DB01310 | YMR165C | 99 | 0 |
| SPGP0C04750 | YDR276C | 100 | 0 |
| SPGP0F00670 | YGR223C | 99 | 0 |
| SPGP0F03210 | YGL062W | 99 | 0 |
| SPGP0DC00720 | YBL036C | 99 | 0 |
| SPGP0F01800 | YGR099W | 99 | 0 |
| SPGP0I02530 | YER173W | 99 | 0 |
| SPGP0DD00320 | YIL147C | 99 | 0 |
| SPGP0I02240 | YER143W | 100 | 0 |
| SPGP0DC01330 | YBR026C | 99 | 0 |
| SPGP0DB02600 | YMR026C | 99 | 0 |
| SPGP0DB01910 | YMR102C | 99 | 0 |
| SPGP0F00230 | YGR273C | 99 | 0 |
| SPGP0DC01250 | YBR018C | 100 | 0 |
| SPGP0DB02790 | YMR008C | 99 | 1 |
| SPGP0DB02200 | YMR071C | 99 | 0 |
| SPGP0D00530 | YPR024W | 99 | 0 |
| SPGP0D00200 | YPR062W | 100 | 0 |
| SPGP0Q04680 | YGL211W | 99 | 0 |
| SPGP0K01530 | YOL012C | 99 | 0 |
| SPGP0F01590 | YGR123C | 99 | 0 |
| SPGP0D00630 | YPR013C | 99 | 0 |
| SPGP0K01480 | YOL017W | 99 | 0 |
| SPGP0G01140 | YJR044C | 100 | 0 |
| SPGP0K02240 | YOR067C | 99 | 0 |
| SPGP0G01390 | YJR010C-A | 100 | 0 |
| SPGP0G00660 | YJR095W | 100 | 0 |
| SPGP0AE00400 | YCL030C | 99 | 0 |
| SPGP0M02560 | YNL058C | 99 | 1 |
| SPGP0M00750 | YNL258C | 99 | 0 |
| SPGP0C02640 | YDR043C | 100 | 0 |
| SPGP0DB00790 | YMR216C | 99 | 0 |

Create PDF in your applications with the Pdfcrowd HTML to PDF API PDFCROWD

| | | | |
|---|---|---|---|
| SPGP0F04090 | YGL160W | 99 | 0 |
| SPGP0C01110 | YDL130W | 99 | 0 |
| SPGP0K01600 | YOL005C | 99 | 0 |
| SPGP0C03900 | YDR178W | 99 | 0 |
| SPGP0V01890 | YHR139C | 99 | 0 |
| SPGP0K02390 | YOR085W | 99 | 0 |
| SPGP0C01220 | YDL119C | 99 | 0 |
| SPGP0F03620 | YGL108C | 99 | 0 |
| SPGP0M01650 | YNL158W | 99 | 0 |
| SPGP0V01600 | YHR109W | 99 | 0 |
| SPGP0DC01920 | YBR088C | 100 | 0 |
| SPGP0K00480 | YOL122C | 99 | 0 |
| SPGP0M02300 | YNL085W | 99 | 0 |
| SPGP0DC00810 | YBL028C | 99 | 0 |
| SPGP0DE00210 | YAL049C | 99 | 0 |
| SPGP0DC00410 | YBL071W-A | 98 | 0 |
| SPGP0DB01070 | YMR190C | 99 | 0 |
| SPGP0AE00240 | YCL048W-A | 100 | 0 |
| SPGP0M01970 | YNL123W | 99 | 0 |
| SPGP0C03060 | YDR086C | 100 | 0 |
| SPGP0U01970 | YPL245W | 99 | 0 |
| SPGP0M03350 | YNR028W | 99 | 0 |
| SPGP0DD01660 | YIL002C | 100 | 0 |
| SPGP0F00570 | YGR235C | 100 | 0 |
| SPGP0DC02000 | YBR098W | 99 | 0 |
| SPGP0M00620 | YNL273W | 99 | 0 |
| SPGP0DB02120 | YMR079W | 99 | 0 |
| SPGP0V00730 | YHR021C | 100 | 0 |
| SPGP0C03680 | YDR156W | 99 | 0 |
| SPGP0C04440 | YDR240C | 100 | 0 |
| SPGP0U02150 | YPL263C | 99 | 0 |
| SPGP0M03600 | YNR052C | 99 | 0 |
| SPGP0D00900 | YPL015C | 99 | 0 |
| SPGP0C00260 | YDL225W | 99 | 0 |
| SPGP0V01240 | YHR074W | 99 | 0 |
| SPGP0M02760 | YNL036W | 99 | 0 |
| SPGP0DD00520 | YIL126W | 99 | 0 |
| SPGP0DC01870 | YBR084W | 99 | 0 |
| SPGP0K01840 | YOR021C | 100 | 0 |
| SPGP0V00790 | YHR027C | 99 | 0 |

Create PDF in your applications with the Pdfcrowd HTML to PDF API PDFCROWD

| | | | |
|---|---|---|---|
| SPGP0DC02730 | YBR179C | 99 | 0 |
| SPGP0G03170 | YJL193W | 99 | 0 |
| SPGP0G03250 | YJL203W | 99 | 0 |
| SPGP0F01480 | YGR134W | 99 | 0 |
| SPGP0M01620 | YNL161W | 99 | 0 |
| SPGP0CZ03040 | YKR094C | 95 | 0 |
| SPGP0F02920 | YGL028C | 98 | 1 |
| SPGP0DB00850 | YMR210W | 99 | 0 |
| SPGP0I02650 | YER185W | 99 | 0 |
| SPGP0C01130 | YDL128W | 99 | 0 |
| SPGP0M01480 | YNL177C | 99 | 0 |
| SPGP0M01950 | YNL125C | 99 | 0 |
| SPGP0F01960 | YGR083C | 99 | 0 |
| SPGP0C01350 | YDL105W | 99 | 0 |
| SPGP0C02810 | YDR062W | 99 | 0 |
| SPGP0DB02470 | YMR039C | 99 | 1 |
| SPGP0D00250 | YPR056W | 99 | 0 |
| SPGP0I00470 | YEL035C | 99 | 0 |
| SPGP0DC02640 | YBR168W | 99 | 0 |
| SPGP0K02260 | YOR070C | 100 | 0 |
| SPGP0DB02960 | YML011C | 99 | 0 |
| SPGP0DD01570 | YIL009C-A | 99 | 0 |
| SPGP0DB00390 | YMR258C | 99 | 0 |
| SPGP0I00700 | YEL012W | 99 | 0 |
| SPGP0M01430 | YNL183C | 99 | 0 |
| SPGP0DB02730 | YMR013C | 100 | 0 |
| SPGP0V00970 | YHR045W | 99 | 0 |
| SPGP0F02270 | YGR046W | 99 | 0 |
| SPGP0F03270 | YGL068W | 99 | 0 |
| SPGP0K01340 | YOL032W | 99 | 0 |
| SPGP0DD01830 | YIR016W | 98 | 0 |
| SPGP0M00140 | YNL327W | 99 | 0 |
| SPGP0G02420 | YJL107C | 99 | 0 |
| SPGP0G02390 | YJL104W | 100 | 0 |
| SPGP0G02930 | YJL165C | 99 | 0 |
| SPGP0C03470 | YDR131C | 99 | 0 |
| SPGP0I01200 | YER039C | 99 | 0 |
| SPGP0C01530 | YDL085C-A | 100 | 0 |
| SPGP0G00390 | YJR122W | 99 | 0 |
| SPGP0K00530 | YOL116W | 99 | 0 |

Create PDF in your applications with the Pdfcrowd HTML to PDF API PDFCROWD

| | | | |
|---|---|---|---|
| SPGP0C04050 | YDR194C | 99 | 0 |
| SPGP0M03380 | YNR031C | 99 | 0 |
| SPGP0F00650 | YGR225W | 99 | 0 |
| SPGP0DC02060 | YBR106W | 99 | 0 |
| SPGP0F03380 | YGL082W | 99 | 0 |
| SPGP0C04980 | YDR301W | 99 | 0 |
| SPGP0M01580 | YNL164C | 99 | 0 |
| SPGP0Q00110 | YBR297W | 81 | 2 |
| SPGP0DC01020 | YBL007C | 99 | 0 |
| SPGP0I00640 | YEL018W | 99 | 0 |
| SPGP0DC02810 | YBR187W | 99 | 0 |
| SPGP0G03110 | YJL186W | 99 | 0 |
| SPGP0DD00340 | YIL145C | 99 | 0 |
| SPGP0V01150 | YHR065C | 99 | 0 |
| SPGP0D00880 | YPL013C | 100 | 0 |
| SPGP0DC01350 | YBR029C | 99 | 0 |
| SPGP0Q04610 | YGL203C | 98 | 1 |
| SPGP0C04860 | YDR288W | 99 | 0 |
| SPGP0F01920 | YGR087C | 99 | 0 |
| SPGP0M03260 | YNR018W | 100 | 0 |
| SPGP0F03400 | YGL084C | 99 | 0 |
| SPGP0G00130 | YJR149W | 99 | 0 |
| SPGP0DB01790 | YMR115W | 99 | 0 |
| SPGP0AE00790 | YCR018C | 99 | 0 |
| SPGP0AE01090 | YCR047C | 98 | 2 |
| SPGP0AE00620 | YCL001W | 100 | 0 |
| SPGP0M02090 | YNL108C | 99 | 0 |
| SPGP0C02660 | YDR045C | 100 | 0 |
| SPGP0DD01550 | YIL011W | 94 | 3 |
| SPGP0K02370 | YOR083W | 97 | 2 |
| SPGP0DE00170 | YAL055W | 98 | 0 |
| SPGP0F00610 | YGR231C | 100 | 0 |
| SPGP0C05200 | YDR322W | 99 | 0 |
| SPGP0K00200 | YOL152W | 99 | 0 |
| SPGP0DE00230 | YAL047C | 99 | 0 |
| SPGP0M01990 | YNL121C | 99 | 0 |
| SPGP0M02320 | YNL083W | 99 | 0 |
| SPGP0F03050 | YGL043W | 99 | 0 |
| SPGP0V00400 | YHL013C | 100 | 0 |
| SPGP0DD02040 | YIR038C | 99 | 0 |

Create PDF in your applications with the Pdfcrowd HTML to PDF API PDFCROWD

| | | | |
|---|---|---|---|
| SPGP0DE00490 | YAL022C | 99 | 0 |
| SPGP0DC03580 | YBR269C | 100 | 0 |
| SPGP0D00310 | YPR048W | 99 | 0 |
| SPGP0C00530 | YDL197C | 99 | 0 |
| SPGP0I01540 | YER069W | 99 | 0 |
| SPGP0G02890 | YJL161W | 99 | 0 |
| SPGP0V00750 | YHR023W | 99 | 0 |
| SPGP0F02630 | YGR004W | 99 | 0 |
| SPGP0I01430 | YER058W | 99 | 0 |
| SPGP0DB01160 | YMR180C | 99 | 0 |
| SPGP0D00920 | YPL017C | 99 | 0 |
| SPGP0M02900 | YNL030W | 99 | 0 |
| SPGP0DC00580 | YBL051C | 99 | 0 |
| SPGP0DB00510 | YMR244W | 99 | 0 |
| SPGP0M03540 | YNR046W | 100 | 0 |
| SPGP0K01860 | YOR023C | 99 | 0 |
| SPGP0DC00740 | YBL034C | 99 | 0 |
| SPGP0AE01210 | YCR066W | 99 | 0 |
| SPGP0F01730 | YGR106C | 99 | 0 |
| SPGP0DC03230 | YBR234C | 99 | 0 |
| SPGP0DB03520 | YML071C | 99 | 0 |
| SPGP0F02280 | YGR044C | 99 | 0 |
| SPGP0I01290 | YER047C | 99 | 0 |
| SPGP0K00440 | YOL126C | 99 | 0 |
| SPGP0DC00330 | YBL081W | 99 | 0 |
| SPGP0C02870 | YDR068W | 99 | 0 |
| SPGP0F00280 | YGR267C | 100 | 0 |
| SPGP0F02730 | YGL008C | 99 | 0 |
| SPGP0C01370 | YDL103C | 99 | 0 |
| SPGP0M01200 | YNL210W | 99 | 0 |
| SPGP0U02120 | YPL259C | 99 | 0 |
| SPGP0DC03720 | YBR284W | 99 | 0 |
| SPGP0I00450 | YEL037C | 99 | 0 |
| SPGP0DB00250 | YMR272C | 99 | 0 |
| SPGP0M00590 | YNL277W | 99 | 0 |
| SPGP0DC02530 | YBR158W | 99 | 0 |
| SPGP0K01170 | YOL052C | 99 | 0 |
| SPGP0D00270 | YPR054W | 99 | 0 |
| SPGP0DE00150 | YAL058W | 99 | 0 |
| SPGP0C01290 | YDL111C | 99 | 0 |

Create PDF in your applications with the Pdfcrowd HTML to PDF API PDFCROWD

| | | | |
|---|---|---|---|
| SPGP0F01410 | YGR143W | 99 | 0 |
| SPGP0DB00320 | YMR265C | 99 | 0 |
| SPGP0M01300 | YNL197C | 99 | 0 |
| SPGP0C00700 | YDL177C | 99 | 0 |
| SPGP0DB02880 | YML002W | 99 | 0 |
| SPGP0K02190 | YOR062C | 99 | 0 |
| SPGP0DB00700 | YMR225C | 100 | 0 |
| SPGP0M03200 | YNR012W | 99 | 0 |
| SPGP0V00990 | YHR047C | 99 | 0 |
| SPGP0F03100 | YGL049C | 99 | 0 |
| SPGP0V01830 | YHR133C | 99 | 0 |
| SPGP0V01340 | YHR083W | 99 | 0 |
| SPGP0DC00530 | YBL057C | 99 | 0 |
| SPGP0G01240 | YJR032W | 99 | 0 |
| SPGP0DD01810 | YIR014W | 99 | 0 |
| SPGP0C03450 | YDR129C | 99 | 0 |
| SPGP0I01220 | YER041W | 99 | 0 |
| SPGP0AI00460 | YHR177W | 99 | 0 |
| SPGP0G01790 | YJL038C | 99 | 0 |
| SPGP0M03460 | YNR037C | 99 | 0 |
| SPGP0DB02300 | YMR061W | 99 | 0 |
| SPGP0C03980 | YDR185C | 99 | 0 |
| SPGP0U02050 | YPL253C | 99 | 0 |
| SPGP0F00630 | YGR229C | 99 | 0 |
| SPGP0K00870 | YOL082W | 99 | 0 |
| SPGP0M01780 | YNL144C | 100 | 0 |
| SPGP0D00140 | YPR069C | 99 | 0 |
| SPGP0G02660 | YJL134W | 99 | 0 |
| SPGP0DC01000 | YBL009W | 99 | 0 |
| SPGP0DE00630 | YAL008W | 99 | 0 |
| SPGP0DC01370 | YBR031W | 99 | 0 |
| SPGP0F03290 | YGL071W | 99 | 0 |
| SPGP0DD01160 | YIL055C | 99 | 0 |
| SPGP0DD00360 | YIL143C | 99 | 0 |
| SPGP0I01120 | YER031C | 100 | 0 |
| SPGP0I02230 | YER142C | 99 | 0 |
| SPGP0C05110 | YDR314C | 99 | 0 |
| SPGP0D00860 | YPL011C | 99 | 0 |
| SPGP0K02350 | YOR080W | 99 | 0 |

Create PDF in your applications with the Pdfcrowd HTML to PDF API PDFCROWD

| | | | |
|---|---|---|---|
| SPGP0F02900 | YGL026C | 99 | 0 |
| SPGP0CZ00160 | YKL215C | 81 | 1 |
| SPGP0F01900 | YGR089W | 99 | 0 |
| SPGP0F02500 | YGR019W | 99 | 0 |
| SPGP0V00860 | YHR034C | 99 | 1 |
| SPGP0F01190 | YGR168C | 99 | 0 |
| SPGP0DE00750 | YAR014C | 99 | 0 |
| SPGP0C02460 | YDR022C | 100 | 0 |
| SPGP0Q04580 | YGL200C | 99 | 0 |
| SPGP0DB03350 | YML055W | 99 | 0 |
| SPGP0DB02680 | YMR018W | 99 | 0 |
| SPGP0DC01960 | YBR094W | 99 | 0 |
| SPGP0DD00190 | YIL159W | 99 | 0 |
| SPGP0DD01770 | YIR010W | 99 | 1 |
| SPGP0F02010 | YGR077C | 99 | 0 |
| SPGP0F00810 | YGR208W | 99 | 0 |
| SPGP0V00610 | YHR008C | 99 | 0 |
| SPGP0DC02830 | YBR189W | 100 | 0 |
| SPGP0DC00350 | YBL079W | 99 | 0 |
| SPGP0G02870 | YJL159W | 97 | 1 |
| SPGP0V00770 | YHR025W | 99 | 0 |
| SPGP0F03880 | YGL137W | 99 | 0 |
| SPGP0G02980 | YJL171C | 99 | 0 |
| SPGP0DB02440 | YMR042W | 100 | 0 |
| SPGP0I02210 | YER140W | 99 | 0 |
| SPGP0DD00560 | YIL122W | 99 | 0 |
| SPGP0U02190 | YPL267W | 99 | 0 |
| SPGP0DB01470 | YMR148W | 100 | 0 |
| SPGP0DC00760 | YBL032W | 99 | 0 |
| SPGP0DB00810 | YMR214W | 99 | 0 |
| SPGP0DC00970 | YBL013W | 99 | 0 |
| SPGP0I02140 | YER130C | 99 | 0 |
| SPGP0DC00230 | YBL091C-A | 99 | 0 |
| SPGP0F01750 | YGR104C | 99 | 0 |
| SPGP0C04450 | YDR242W | 99 | 0 |
| SPGP0K00460 | YOL124C | 99 | 0 |
| SPGP0M00380 | YNL300W | 99 | 0 |
| SPGP0AI00120 | YHR210C | 99 | 0 |
| SPGP0C04150 | YDR206W | 99 | 0 |
| SPGP0F00400 | YGR253C | 99 | 0 |

Create PDF in your applications with the Pdfcrowd HTML to PDF API PDFCROWD

| | | | |
|---|---|---|---|
| SPGP0M01220 | YNL208W | 99 | 0 |
| SPGP0G01960 | YJL056C | 99 | 0 |
| SPGP0G01740 | YJL033W | 99 | 0 |
| SPGP0DD00700 | YIL108W | 99 | 0 |
| SPGP0AE00320 | YCL038C | 99 | 0 |
| SPGP0DC02600 | YBR164C | 99 | 0 |
| SPGP0M00600 | YNL275W | 99 | 0 |
| SPGP0M00410 | YNL297C | 99 | 0 |
| SPGP0K02290 | YOR074C | 99 | 0 |
| SPGP0G00560 | YJR105W | 99 | 0 |
| SPGP0I00250 | YEL057C | 99 | 0 |
| SPGP0C00750 | YDL171C | 99 | 0 |
| SPGP0C00720 | YDL175C | 99 | 0 |
| SPGP0K01300 | YOL038W | 100 | 0 |
| SPGP0M01320 | YNL195C | 99 | 0 |
| SPGP0V01580 | YHR107C | 99 | 0 |
| SPGP0C03140 | YDR096W | 99 | 0 |
| SPGP0I01280 | YER046W | 99 | 0 |
| SPGP0AI00180 | YHR204W | 99 | 0 |
| SPGP0F00890 | YGR200C | 99 | 0 |
| SPGP0V01360 | YHR085W | 99 | 0 |
| SPGP0DC00510 | YBL059W | 100 | 0 |
| SPGP0DC01340 | YBR028C | 99 | 0 |
| SPGP0C02180 | YDL008W | 99 | 0 |
| SPGP0C03430 | YDR127W | 99 | 0 |
| SPGP0C04430 | YDR239C | 99 | 0 |
| SPGP0I01740 | YER088C | 99 | 0 |
| SPGP0G02460 | YJL112W | 99 | 0 |
| SPGP0M03440 | YNR035C | 100 | 0 |
| SPGP0M02670 | YNL046W | 99 | 0 |
| SPGP0D01060 | YPL032C | 99 | 0 |
| SPGP0U02070 | YPL255W | 100 | 0 |
| SPGP0M01760 | YNL146W | 100 | 0 |
| SPGP0K00890 | YOL080C | 99 | 0 |
| SPGP0I00590 | YEL022W | 99 | 0 |
| SPGP0G03070 | YJL181W | 99 | 0 |
| SPGP0M00900 | YNL243W | 99 | 0 |
| SPGP0F01980 | YGR081C | 99 | 0 |
| SPGP0M01740 | YNL148C | 100 | 0 |
| SPGP0I00600 | YEL021W | 100 | 0 |

Create PDF in your applications with the Pdfcrowd HTML to PDF API PDFCROWD

| | | | |
|---|---|---|---|
| SPGP0Q04380 | YGL176C | 99 | 0 |
| SPGP0AE00140 | YCL058C | 99 | 0 |
| SPGP0D00460 | YPR031W | 99 | 0 |
| SPGP0DC02590 | YBR163W | 99 | 0 |
| SPGP0I01140 | YER033C | 99 | 0 |
| SPGP0DB03280 | YML048W | 99 | 0 |
| SPGP0M01810 | YNL138W-A | 99 | 0 |
| SPGP0G00350 | YJR126C | 99 | 0 |
| SPGP0D00620 | YPR015C | 99 | 0 |
| SPGP0I01950 | YER110C | 99 | 0 |
| SPGP0DB00240 | YMR273C | 99 | 1 |
| SPGP0F01220 | YGR165W | 99 | 0 |
| SPGP0DB03110 | YML026C | 99 | 0 |
| SPGP0C02440 | YDR020C | 99 | 0 |
| SPGP0D00330 | YPR046W | 99 | 0 |
| SPGP0DB01570 | YMR137C | 99 | 0 |
| SPGP0K00790 | YOL090W | 99 | 0 |
| SPGP0G03050 | YJL179W | 99 | 0 |
| SPGP0I01660 | YER080W | 99 | 0 |
| SPGP0DB00720 | YMR223W | 99 | 0 |
| SPGP0K01390 | YOL026C | 99 | 0 |
| SPGP0DB01280 | YMR168C | 99 | 0 |
| SPGP0AE00460 | YCL025C | 99 | 0 |
| SPGP0F00180 | YGR278W | 99 | 0 |
| SPGP0M02080 | YNL110C | 99 | 0 |
| SPGP0DD00240 | YIL155C | 99 | 0 |
| SPGP0M02280 | YNL087W | 99 | 0 |
| SPGP0M02980 | YNL012W | 99 | 0 |
| SPGP0V00810 | YHR029C | 99 | 0 |
| SPGP0DE00430 | YAL028W | 99 | 0 |
| SPGP0F02450 | YGR026W | 99 | 0 |
| SPGP0DD01350 | YIL035C | 99 | 0 |
| SPGP0C03970 | YDR184C | 99 | 0 |
| SPGP0AE01460 | YCR094W | 99 | 0 |
| SPGP0I01160 | YER035W | 100 | 0 |
| SPGP0DC00700 | YBL038W | 99 | 0 |
| SPGP0C03720 | YDR161W | 99 | 0 |
| SPGP0G00520 | YJR108W | 100 | 0 |
| SPGP0C00390 | YDL211C | 98 | 0 |
| SPGP0DC00370 | YBL076C | 99 | 0 |

Create PDF in your applications with the Pdfcrowd HTML to PDF API PDFCROWD

| | | | |
|---|---|---|---|
| SPGP0DB03540 | YML073C | 98 | 0 |
| SPGP0K02310 | YOR076C | 99 | 0 |
| SPGP0F00420 | YGR251W | 99 | 0 |
| SPGP0G01760 | YJL035C | 99 | 0 |
| SPGP0D00410 | YPR036W | 99 | 0 |
| SPGP0DC02490 | YBR154C | 99 | 0 |
| SPGP0C01840 | YDL052C | 99 | 0 |
| SPGP0C00580 | YDL191W | 99 | 0 |
| SPGP0F03450 | YGL090W | 99 | 0 |
| SPGP0K01000 | YOL068C | 99 | 0 |
| SPGP0C02160 | YDL012C | 99 | 0 |
| SPGP0DB03320 | YML052W | 99 | 0 |
| SPGP0C02210 | YDL006W | 99 | 0 |
| SPGP0C03410 | YDR125C | 99 | 0 |
| SPGP0DB00950 | YMR201C | 99 | 0 |
| SPGP0C01520 | YDL086W | 99 | 0 |
| SPGP0C00600 | YDL189W | 99 | 0 |
| SPGP0DC02360 | YBR140C | 99 | 0 |
| SPGP0DC03370 | YBR248C | 99 | 0 |
| SPGP0K00670 | YOL103W | 99 | 0 |
| SPGP0I01590 | YER074W | 99 | 0 |
| SPGP0K00770 | YOL092W | 99 | 0 |
| SPGP0I00970 | YER016W | 99 | 0 |
| SPGP0F01200 | YGR167W | 99 | 1 |
| SPGP0G03080 | YJL183W | 99 | 0 |
| SPGP0M00920 | YNL241C | 99 | 0 |
| SPGP0DB03040 | YML019W | 99 | 0 |
| SPGP0DC03700 | YBR282W | 99 | 0 |
| SPGP0F03040 | YGL041W-A | 99 | 0 |
| SPGP0I01620 | YER076C | 99 | 0 |
| SPGP0C04390 | YDR235W | 99 | 0 |
| SPGP0Q04410 | YGL180W | 99 | 0 |
| SPGP0F01860 | YGR093W | 99 | 0 |
| SPGP0ER00130 | YGL262W | 83 | 0 |
| SPGP0G00370 | YJR124C | 99 | 0 |
| SPGP0DC03330 | YBR244W | 99 | 0 |
| SPGP0M01520 | YNL172W | 99 | 0 |
| SPGP0V01420 | YHR091C | 99 | 0 |
| SPGP0C02270 | YDR001C | 99 | 0 |
| SPGP0I01970 | YER112W | 99 | 0 |

Create PDF in your applications with the Pdfcrowd HTML to PDF API PDFCROWD

| | | | |
|---|---|---|---|
| SPGP0AI00150 | YHR207C | 99 | 0 |
| SPGP0AE00750 | YCR014C | 99 | 0 |
| SPGP0M00120 | YNL329C | 99 | 0 |
| SPGP0C04130 | YDR204W | 100 | 0 |
| SPGP0I00200 | YEL062W | 99 | 0 |
| SPGP0DD01600 | YIL007C | 99 | 0 |
| SPGP0AE00580 | YCL007C | 100 | 0 |
| SPGP0C03610 | YDR147W | 99 | 0 |
| SPGP0DC02380 | YBR142W | 99 | 0 |
| SPGP0F01320 | YGR153W | 99 | 0 |
| SPGP0C02390 | YDR014W-A | 99 | 0 |
| SPGP0K02640 | YOR113W | 98 | 1 |
| SPGP0DB01550 | YMR139W | 100 | 0 |
| SPGP0DD01290 | YIL041W | 99 | 0 |
| SPGP0V02140 | YHR164C | 99 | 0 |
| SPGP0K00970 | YOL071W | 99 | 0 |
| SPGP0AE00990 | YCR036W | 99 | 0 |
| SPGP0DB00600 | YMR235C | 99 | 0 |
| SPGP0V01170 | YHR067W | 100 | 0 |
| SPGP0DD02020 | YIR036C | 100 | 0 |
| SPGP0DD00260 | YIL153W | 99 | 0 |
| SPGP0M02380 | YNL077W | 99 | 0 |
| SPGP0F03180 | YGL059W | 99 | 0 |
| SPGP0C02020 | YDL029W | 99 | 0 |
| SPGP0G02720 | YJL141C | 99 | 0 |
| SPGP0F00950 | YGR194C | 99 | 0 |
| SPGP0F01300 | YGR155W | 99 | 0 |
| SPGP0M02920 | YNL032W | 99 | 0 |
| SPGP0M03000 | YNL010W | 99 | 0 |
| SPGP0M03030 | YNL007C | 98 | 2 |
| SPGP0V01790 | YHR129C | 99 | 0 |
| SPGP0K02000 | YOR039W | 99 | 0 |
| SPGP0DC00490 | YBL060W | 99 | 0 |
| SPGP0F03830 | YGL131C | 99 | 0 |
| SPGP0V00130 | YHL043W | 99 | 0 |
| SPGP0I00990 | YER018C | 99 | 0 |
| SPGP0I01100 | YER029C | 99 | 0 |
| SPGP0C00230 | YDL229W | 99 | 0 |
| SPGP0DC00930 | YBL017C | 99 | 0 |
| SPGP0C00800 | YDL166C | 99 | 0 |

Create PDF in your applications with the Pdfcrowd HTML to PDF API PDFCROWD

| | | | |
|---|---|---|---|
| SPGP0DB03380 | YML058W | 99 | 0 |
| SPGP0DD00460 | YIL132C | 99 | 0 |
| SPGP0AE00100 | YCL064C | 100 | 0 |
| SPGP0F00360 | YGR257C | 99 | 0 |
| SPGP0G01920 | YJL052W | 99 | 0 |
| SPGP0C00470 | YDL203C | 99 | 0 |
| SPGP0K01770 | YOR014W | 99 | 0 |
| SPGP0M03740 | YNR067C | 100 | 0 |
| SPGP0M00640 | YNL271C | 99 | 0 |
| SPGP0M01060 | YNL224C | 99 | 0 |
| SPGP0DC02510 | YBR156C | 99 | 0 |
| SPGP0C04360 | YDR232W | 99 | 0 |
| SPGP0AE01320 | YCR077C | 99 | 0 |
| SPGP0DB00230 | YMR274C | 100 | 0 |
| SPGP0M00240 | YNL315C | 99 | 0 |
| SPGP0DC03480 | YBR258C | 99 | 0 |
| SPGP0F03510 | YGL096W | 99 | 0 |
| SPGP0F01170 | YGR169C-A | 99 | 0 |
| SPGP0C01170 | YDL124W | 99 | 0 |
| SPGP0V01540 | YHR103W | 99 | 0 |
| SPGP0V01090 | YHR059W | 99 | 0 |
| SPGP0DB02550 | YMR031C | 99 | 0 |
| SPGP0K02100 | YOR051C | 98 | 0 |
| SPGP0C02230 | YDL004W | 99 | 0 |
| SPGP0C02710 | YDR051C | 99 | 0 |
| SPGP0M01380 | YNL189W | 99 | 0 |
| SPGP0DD01340 | YIL036W | 99 | 0 |
| SPGP0I00190 | YEL063C | 99 | 0 |
| SPGP0C02080 | YDL021W | 99 | 0 |
| SPGP0DC03350 | YBR246W | 99 | 0 |
| SPGP0U02030 | YPL250C | 99 | 0 |
| SPGP0EA00100 | YAL067C | 99 | 0 |
| SPGP0DC01140 | YBR006W | 99 | 0 |
| SPGP0I00540 | YEL027W | 100 | 0 |
| SPGP0I01640 | YER078C | 99 | 0 |
| SPGP0C05220 | YDR323C | 99 | 0 |
| SPGP0I01180 | YER037W | 99 | 0 |
| SPGP0DD01250 | YIL045W | 99 | 0 |
| SPGP0DB01450 | YMR150C | 99 | 0 |
| SPGP0F03000 | YGL037C | 99 | 0 |

Create PDF in your applications with the Pdfcrowd HTML to PDF API PDFCROWD

| | | | |
|---|---|---|---|
| SPGP0G03190 | YJL196C | 99 | 0 |
| SPGP0M02690 | YNL044W | 100 | 0 |
| SPGP0AE01440 | YCR092C | 99 | 0 |
| SPGP0DD01620 | YIL005W | 99 | 0 |
| SPGP0AE00770 | YCR016W | 100 | 0 |
| SPGP0F04190 | YGL172W | 99 | 0 |
| SPGP0CZ00100 | YER188W | 90 | 0 |
| SPGP0C01470 | YDL091C | 99 | 0 |
| SPGP0V01770 | YHR127W | 99 | 0 |
| SPGP0C04160 | YDR207C | 99 | 0 |
| SPGP0AE00560 | YCL009C | 99 | 0 |
| SPGP0Q05120 | YGL258W-A | 97 | 0 |
| SPGP0D00370 | YPR041W | 100 | 0 |
| SPGP0C00500 | YDL200C | 99 | 0 |
| SPGP0F03150 | YGL056C | 99 | 0 |
| SPGP0DB01690 | YMR125W | 99 | 0 |
| SPGP0K00350 | YOL136C | 99 | 0 |
| SPGP0G01630 | YJL016W | 99 | 0 |
| SPGP0DD01960 | YIR030C | 99 | 0 |
| SPGP0F03680 | YGL115W | 99 | 0 |
| SPGP0DB01820 | YMR112C | 99 | 0 |
| SPGP0M02010 | YNL118C | 99 | 0 |
| SPGP0G03350 | YJL213W | 99 | 0 |
| SPGP0K02660 | YOR115C | 99 | 0 |
| SPGP0DD00890 | YIL089W | 99 | 0 |
| SPGP0DD00280 | YIL151C | 99 | 0 |
| SPGP0F01090 | YGR178C | 99 | 0 |
| SPGP0DB00740 | YMR221C | 99 | 0 |
| SPGP0DC03290 | YBR240C | 99 | 0 |
| SPGP0V01070 | YHR057C | 99 | 0 |
| SPGP0F02350 | YGR036C | 99 | 0 |
| SPGP0AE00720 | YCR010C | 100 | 0 |
| SPGP0DB01860 | YMR108W | 99 | 0 |
| SPGP0K00950 | YOL073C | 99 | 0 |
| SPGP0G02760 | YJL146W | 99 | 0 |
| SPGP0I00830 | YER004W | 99 | 0 |
| SPGP0DD01390 | YIL030C | 99 | 0 |
| SPGP0DD00660 | YIL112W | 99 | 0 |
| SPGP0G02100 | YJL071W | 99 | 0 |
| SPGP0C03930 | YDR180W | 99 | 0 |

Create PDF in your applications with the Pdfcrowd HTML to PDF API PDFCROWD

| | | | |
|---|---|---|---|
| SPGP0C03620 | YDR148C | 99 | 1 |
| SPGP0M03150 | YNR007C | 100 | 0 |
| SPGP0DB03500 | YML069W | 99 | 0 |
| SPGP0DD00910 | YIL087C | 99 | 0 |
| SPGP0DD00440 | YIL134W | 99 | 0 |
| SPGP0DC02920 | YBR199W | 99 | 0 |
| SPGP0C00820 | YDL164C | 99 | 0 |
| SPGP0DB02760 | YMR010W | 99 | 0 |
| SPGP0F00380 | YGR255C | 99 | 0 |
| SPGP0G00770 | YJR084W | 99 | 0 |
| SPGP0EW00100 | YPL282C | 99 | 0 |
| SPGP0K02410 | YOR087W | 99 | 0 |
| SPGP0AE01120 | YCR052W | 99 | 0 |
| SPGP0DC02450 | YBR150C | 99 | 0 |
| SPGP0DC00910 | YBL019W | 99 | 0 |
| SPGP0DD00150 | YIL164C | 99 | 0 |
| SPGP0F02330 | YGR038W | 100 | 0 |
| SPGP0DB03260 | YML046W | 100 | 0 |
| SPGP0C03100 | YDR090C | 99 | 0 |
| SPGP0K00930 | YOL076W | 99 | 0 |
| SPGP0D00710 | YPR005C | 99 | 0 |
| SPGP0C02250 | YDL002C | 99 | 0 |
| SPGP0M01170 | YNL213C | 100 | 0 |
| SPGP0U02230 | YPL271W | 99 | 0 |
| SPGP0G02370 | YJL102W | 99 | 0 |
| SPGP0DB02220 | YMR069W | 99 | 0 |
| SPGP0DC01660 | YBR061C | 99 | 0 |
| SPGP0G02610 | YJL130C | 99 | 0 |
| SPGP0Q04470 | YGL187C | 100 | 0 |
| SPGP0G02590 | YJL128C | 99 | 0 |
| SPGP0DC03740 | YBR286W | 99 | 0 |
| SPGP0M01700 | YNL153C | 99 | 0 |
| SPGP0DB00930 | YMR202W | 100 | 0 |
| SPGP0DD01710 | YIR004W | 99 | 0 |
| SPGP0M03520 | YNR044W | 95 | 1 |
| SPGP0F02980 | YGL035C | 99 | 0 |
| SPGP0I02310 | YER150W | 99 | 0 |
| SPGP0C01410 | YDL099W | 99 | 0 |
| SPGP0DC00210 | YBL093C | 99 | 0 |
| SPGP0DB01730 | YMR121C | 99 | 0 |

Create PDF in your applications with the Pdfcrowd HTML to PDF API PDFCROWD

| | | | |
|---|---|---|---|
| SPGP0K02140 | YOR057W | 99 | 0 |
| SPGP0DD00620 | YIL116W | 99 | 0 |
| SPGP0DB01590 | YMR135C | 99 | 1 |
| SPGP0M02030 | YNL116W | 99 | 0 |
| SPGP0F03810 | YGL129C | 99 | 0 |
| SPGP0DC03560 | YBR267W | 99 | 0 |
| SPGP0V01050 | YHR055C | 100 | 0 |
| SPGP0K00310 | YOL140W | 99 | 0 |
| SPGP0DC00450 | YBL066C | 99 | 0 |
| SPGP0F02060 | YGR071C | 99 | 0 |
| SPGP0DC02300 | YBR133C | 99 | 0 |
| SPGP0I01320 | YER049W | 99 | 0 |
| SPGP0M02860 | YNL025C | 99 | 0 |
| SPGP0I00810 | YER002W | 99 | 0 |
| SPGP0C03950 | YDR182W | 99 | 0 |
| SPGP0V00480 | YHL004W | 99 | 0 |
| SPGP0DE00580 | YAL013W | 99 | 0 |
| SPGP0F00460 | YGR247W | 99 | 0 |
| SPGP0C00660 | YDL181W | 99 | 0 |
| SPGP0C03500 | YDR135C | 99 | 0 |
| SPGP0DC01410 | YBR036C | 99 | 0 |
| SPGP0DC02680 | YBR172C | 99 | 0 |
| SPGP0C04370 | YDR233C | 99 | 0 |
| SPGP0FF00100 | YPL277C | 99 | 0 |
| SPGP0AE01100 | YCR048W | 99 | 1 |
| SPGP0DC02470 | YBR152W | 99 | 0 |
| SPGP0AE00500 | YCL017C | 99 | 0 |
| SPGP0V01260 | YHR076W | 99 | 0 |
| SPGP0DE00730 | YAR007C | 99 | 0 |
| SPGP0C04960 | YDR299W | 99 | 0 |
| SPGP0C04740 | YDR275W | 99 | 0 |
| SPGP0F02370 | YGR034W | 100 | 0 |
| SPGP0G01650 | YJL020C | 98 | 1 |
| SPGP0G01020 | YJR056C | 99 | 0 |
| SPGP0M02400 | YNL075W | 99 | 0 |
| SPGP0DC03310 | YBR242W | 100 | 0 |
| SPGP0K02160 | YOR059C | 99 | 0 |
| SPGP0DD01440 | YIL022W | 99 | 0 |
| SPGP0C00640 | YDL183C | 99 | 0 |
| SPGP0DC01850 | YBR082C | 99 | 0 |

Create PDF in your applications with the Pdfcrowd HTML to PDF API PDFCROWD

| | | | |
|---|---|---|---|
| SPGP0G02350 | YJL100W | 99 | 0 |
| SPGP0V01010 | YHR049W | 99 | 0 |
| SPGP0M00980 | YNL234W | 99 | 0 |
| SPGP0DC01640 | YBR059C | 99 | 0 |
| SPGP0K01270 | YOL041C | 100 | 0 |
| SPGP0DE00680 | YAL001C | 99 | 0 |
| SPGP0DC00660 | YBL041W | 99 | 0 |
| SPGP0DC03030 | YBR211C | 99 | 0 |
| SPGP0V02080 | YHR158C | 99 | 0 |
| SPGP0I02330 | YER152C | 99 | 0 |
| SPGP0F00870 | YGR202C | 99 | 0 |
| SPGP0K00710 | YOL098C | 99 | 0 |
| SPGP0G02490 | YJL117W | 99 | 0 |
| SPGP0DC00890 | YBL021C | 100 | 0 |
| SPGP0DB00400 | YMR257C | 99 | 0 |
| SPGP0K02620 | YOR111W | 99 | 0 |
| SPGP0K01190 | YOL049W | 99 | 0 |
| SPGP0DE00320 | YAL038W | 99 | 0 |
| SPGP0F01610 | YGR121C | 99 | 0 |
| SPGP0I01300 | YER048C | 99 | 0 |
| SPGP0G01320 | YJR017C | 99 | 0 |
| SPGP0DC00780 | YBL030C | 99 | 0 |
| SPGP0C01020 | YDL140C | 99 | 0 |
| SPGP0V00170 | YHL036W | 99 | 0 |
| SPGP0DC03180 | YBR229C | 99 | 0 |
| SPGP0F03280 | YGL070C | 99 | 0 |
| SPGP0C04230 | YDR217C | 99 | 0 |
| SPGP0DC01720 | YBR068C | 99 | 0 |
| SPGP0C00310 | YDL219W | 98 | 0 |
| SPGP0DC03460 | YBR256C | 99 | 0 |
| SPGP0F03720 | YGL120C | 99 | 0 |
| SPGP0F02780 | YGL013C | 99 | 0 |
| SPGP0G00790 | YJR082C | 99 | 0 |
| SPGP0V01200 | YHR070W | 99 | 0 |
| SPGP0AE00870 | YCR024C | 99 | 0 |
| SPGP0DD00300 | YIL149C | 99 | 0 |
| SPGP0DD01940 | YIR028W | 99 | 0 |
| SPGP0I01230 | YER042W | 99 | 0 |
| SPGP0V00380 | YHL015W | 99 | 0 |

Create PDF in your applications with the Pdfcrowd HTML to PDF API PDFCROWD

| | | | |
|---|---|---|---|
| SPGP0V00650 | YHR012W | 99 | 0 |
| SPGP0V01380 | YHR087W | 100 | 0 |
| SPGP0K01250 | YOL043C | 99 | 0 |
| SPGP0F04040 | YGL155W | 99 | 0 |
| SPGP0C03850 | YDR173C | 99 | 1 |
| SPGP0Q04360 | YGL174W | 99 | 0 |
| SPGP0Ctrna18E | YDR278C | 100 | 0 |
| SPGP0DC00770 | YBL031W | 99 | 0 |
| SPGP0D00790 | YPL004C | 99 | 0 |
| SPGP0DZ01610 | YOR302W | 96 | 0 |
| SPGP0F00740 | YGR215W | 100 | 0 |
| SPGP0F01380 | YGR146C | 99 | 0 |
| SPGP0DC02430 | YBR148W | 99 | 0 |
| SPGP0I02220 | YER141W | 100 | 0 |
| SPGP0G02510 | YJL121C | 99 | 0 |
| SPGP0C04730 | YDR273W | 99 | 0 |
| SPGP0F01070 | YGR180C | 99 | 0 |
| SPGP0DB03130 | YML028W | 100 | 0 |
| SPGP0DC03390 | YBR250W | 99 | 0 |
| SPGP0DB00420 | YMR255W | 99 | 0 |
| SPGP0M03300 | YNR022C | 100 | 0 |
| SPGP0V01030 | YHR051W | 99 | 0 |
| SPGP0I01760 | YER090W | 100 | 0 |
| SPGP0C04250 | YDR219C | 99 | 0 |
| SPGP0AI00400 | YHR183W | 99 | 0 |
| SPGP0DB03400 | YML059C | 99 | 0 |
| SPGP0F03420 | YGL086W | 99 | 0 |
| SPGP0G02180 | YJL082W | 99 | 0 |
| SPGP0C04500 | YDR247W | 99 | 0 |
| SPGP0AE01140 | YCR054C | 99 | 0 |
| SPGP0DB03440 | YML063W | 99 | 0 |
| SPGP0F01570 | YGR125W | 99 | 0 |
| SPGP0DD01490 | YIL017C | 99 | 0 |
| SPGP0G00800 | YJR080C | 99 | 0 |
| SPGP0G01340 | YJR015W | 100 | 0 |
| SPGP0G00580 | YJR103W | 99 | 0 |
| SPGP0AE00540 | YCL011C | 99 | 0 |
| SPGP0F02390 | YGR032W | 99 | 0 |
| SPGP0M03750 | YNR069C | 99 | 0 |
| SPGP0DB02570 | YMR029C | 99 | 0 |

Create PDF in your applications with the Pdfcrowd HTML to PDF API PDFCROWD

| | | | |
|---|---|---|---|
| SPGP0D00700 | YPR006C | 99 | 0 |
| SPGP0DC01570 | YBR053C | 99 | 0 |
| SPGP0V00630 | YHR010W | 99 | 0 |
| SPGP0C00930 | YDL149W | 99 | 0 |
| SPGP0DB00870 | YMR208W | 99 | 0 |
| SPGP0K01230 | YOL045W | 99 | 0 |
| SPGP0C05070 | YDR310C | 99 | 0 |
| SPGP0DB02940 | YML009C | 100 | 0 |
| SPGP0M00490 | YNL288W | 99 | 0 |
| SPGP0F04060 | YGL157W | 99 | 0 |
| SPGP0DC02900 | YBR197C | 99 | 0 |
| SPGP0K02490 | YOR096W | 99 | 1 |
| SPGP0C01960 | YDL039C | 99 | 0 |
| SPGP0DC00170 | YBL099W | 100 | 0 |
| SPGP0G00750 | YJR086W | 99 | 0 |
| SPGP0DW00100 | YCR098C | 99 | 0 |
| SPGP0DC01510 | YBR046C | 99 | 0 |
| SPGP0C02040 | YDL027C | 99 | 0 |
| SPGP0M03320 | YNR024W | 100 | 0 |
| SPGP0F03020 | YGL039W | 99 | 0 |
| SPGP0DC00430 | YBL068W | 99 | 0 |
| SPGP0DB01960 | YMR097C | 99 | 0 |
| SPGP0M03580 | YNR050C | 99 | 0 |
| SPGP0K00910 | YOL077W-A | 100 | 0 |
| SPGP0DC02280 | YBR131W | 99 | 0 |
| SPGP0I02610 | YER180C-A | 99 | 0 |
| SPGP0DD00760 | YIL102C-A | 99 | 0 |
| SPGP0AE00250 | YCL048W | 99 | 0 |
| SPGP0AE01300 | YCR075W-A | 100 | 0 |
| SPGP0C01000 | YDL142C | 99 | 0 |
| SPGP0DC00190 | YBL097W | 99 | 0 |
| SPGP0G01180 | YJR040W | 99 | 0 |
| SPGP0K02810 | YOR131C | 99 | 0 |
| SPGP0DB01980 | YMR095C | 99 | 0 |
| SPGP0C04330 | YDR228C | 99 | 1 |
| SPGP0I00130 | YEL070W | 99 | 0 |
| SPGP0K01210 | YOL047C | 99 | 0 |
| SPGP0DE00770 | YAR018C | 99 | 0 |
| SPGP0DC01160 | YBR008C | 99 | 0 |
| SPGP0F03360 | YGL080W | 99 | 0 |

Create PDF in your applications with the Pdfcrowd HTML to PDF API PDFCROWD

| | | | |
|---|---|---|---|
| SPGP0DC00600 | YBL049W | 100 | 0 |
| SPGP0G02040 | YJL063C | 99 | 0 |
| SPGP0G01460 | YJR005W | 99 | 0 |
| SPGP0F00500 | YGR243W | 100 | 0 |
| SPGP0C01490 | YDL089W | 99 | 0 |
| SPGP0C04670 | YDR265W | 99 | 0 |
| SPGP0DD00750 | YIL103W | 99 | 0 |
| SPGP0AE01060 | YCR044C | 99 | 0 |
| SPGP0AE00660 | YCR004C | 99 | 0 |
| SPGP0C03240 | YDR107C | 99 | 0 |
| SPGP0K02790 | YOR129C | 99 | 0 |
| SPGP0C02840 | YDR065W | 99 | 0 |
| SPGP0DC01430 | YBR038W | 99 | 0 |
| SPGP0G01160 | YJR042W | 99 | 0 |
| SPGP0I02000 | YER115C | 99 | 0 |
| SPGP0M02800 | YNL020C | 99 | 0 |
| SPGP0F02820 | YGL017W | 99 | 0 |
| SPGP0DD01100 | YIL064W | 99 | 0 |
| SPGP0C00950 | YDL147W | 99 | 0 |
| SPGP0DB01610 | YMR133W | 99 | 0 |
| SPGP0G01300 | YJR021C | 99 | 0 |
| SPGP0DC02710 | YBR176W | 99 | 0 |
| SPGP0M02120 | YNL104C | 99 | 0 |
| SPGP0G00890 | YJR069C | 99 | 0 |
| SPGP0DB03420 | YML061C | 99 | 0 |
| SPGP0D00540 | YPR023C | 99 | 0 |
| SPGP0G00630 | YJR098C | 99 | 0 |
| SPGP0AE00640 | YCR002C | 99 | 0 |
| SPGP0F02800 | YGL015C | 100 | 0 |
| SPGP0DC02880 | YBR195C | 99 | 0 |
| SPGP0C01510 | YDL087C | 99 | 0 |
| SPGP0I02480 | YER168C | 99 | 0 |
| SPGP0K00510 | YOL119C | 99 | 0 |
| SPGP0I00170 | YEL065W | 99 | 0 |
| SPGP0C02570 | YDR034C | 99 | 0 |

#### Sheet 3: *genes alleles below 80% identity*

| *Saccharomyces pastorianus CBS1513 predicted ORF* | Parental allele ID | Parent | Gene identities | Gene gaps |
|---|---|---|---|---|
| SPGP0CZ02750 | YKR064W | *Saccharomyces cerevisiae* | 78 | 1 |

Create PDF in your applications with the Pdfcrowd HTML to PDF API PDFCROWD

| | | | | |
|---|---|---|---|---|
| SPGP0DZ00590 | YOR188W | *Saccharomyces cerevisiae* | 79 | 2 |
| SPGP0DZ00850 | YOR219C | *Saccharomyces cerevisiae* | 78 | 1 |
| SPGP0Y01800 | XP_018224047.1 | *Saccharomyces eubayanus* | 78 | 2 |

### Sheet 4: *genes alleles copy with same identity in both parents*

| *Saccharomyces pastorianus CBS1513 predicted ORF* | Parental allele ID | Parent | Gene identities | Gene gaps |
|---|---|---|---|---|
| SPGP0Itrna24I | YNL017C | *Saccharomyces cerevisiae* | 100 | 0 |
| SPGP0Itrna24I | DI49_5614 | *Saccharomyces eubayanus* | 100 | 0 |
| SPGP0Itrna36K | YDR445C | *Saccharomyces cerevisiae* | 100 | 0 |
| SPGP0Itrna36K | DI49_5468 | *Saccharomyces eubayanus* | 100 | 0 |
| SPGP0Itrna34R | YGR164W | *Saccharomyces cerevisiae* | 100 | 0 |
| SPGP0Itrna34R | DI49_4020 | *Saccharomyces eubayanus* | 100 | 0 |
| SPGP0Ftrna34G | YNL285W | *Saccharomyces cerevisiae* | 100 | 0 |
| SPGP0Ftrna34G | DI49_2297 | *Saccharomyces eubayanus* | 100 | 0 |
| SPGP0Y01790 | YDR531W | *Saccharomyces cerevisiae* | 83 | 0 |
| SPGP0Y01790 | XP_018224046.1 | *Saccharomyces eubayanus* | 83 | 0 |
| SPGP0AA00120 | YCR039C | *Saccharomyces cerevisiae* | 99 | 0 |
| SPGP0AA00120 | XP_018223412.1 | *Saccharomyces eubayanus* | 99 | 0 |
| SPGP0AE01020 | YCR096C | *Saccharomyces cerevisiae* | 100 | 0 |
| SPGP0AE01020 | XP_018223459.1 | *Saccharomyces eubayanus* | 100 | 0 |
| SPGP0Itrna37G | YPR159C-A | *Saccharomyces cerevisiae* | 100 | 0 |
| SPGP0Itrna37G | DI49_5603 | *Saccharomyces eubayanus* | 100 | 0 |
| SPGP0AItrna28T | YHR180W-A | *Saccharomyces cerevisiae* | 100 | 0 |
| SPGP0AItrna28T | DI49_5553 | *Saccharomyces eubayanus* | 100 | 0 |

### Sheet 5: *list of paralogs in Saccharomyces pastorianus CBS1513*

| List of paralogs | *S. eubayanus-like content* | *S. pastorianus content* | *S. cerevisiae-like content* |
|---|---|---|---|
| DI49_1040; YDL014W; YDL013W | DI49_1040 | | YDL014W; YDL013W |
| SPGP0CY00100; SPGP0AJ00100; SPGP0CH00100; SPGP0BX00100; SPGP0BY00100 | | SPGP0CY00100; SPGP0AJ00100; SPGP0CH00100; SPGP0BX00100; SPGP0BY00100 | |
| YER188W; SPGP0I02680; SPGP0CZ00110 | | SPGP0I02680; SPGP0CZ00110 | YER188W |
| DI49_3167; SPGP0CZ00120 | DI49_3167 | SPGP0CZ00120 | |
| DI49_3169; DI49_3168 | DI49_3169; DI49_3168 | | |
| DI49_3170; YKL215C | DI49_3170 | | YKL215C |
| DI49_3172; DI49_3171 | DI49_3172; DI49_3171 | | |
| DI49_3174; DI49_3173 | DI49_3174; DI49_3173 | | |
| DI49_3176; DI49_3175 | DI49_3176; DI49_3175 | | |

Create PDF in your applications with the Pdfcrowd HTML to PDF API PDFCROWD

| | | | |
|---|---|---|---|
| DI49_3178; DI49_3177 | DI49_3178; DI49_3177 | | |
| YKL205W; DI49_3179 | DI49_3179 | | YKL205W |
| DI49_3181; DI49_3079; DI49_3182; YJR066W; DI49_3080 | DI49_3181; DI49_3079; DI49_3182; DI49_3080 | | YJR066W |
| DI49_3184; DI49_3183 | DI49_3184; DI49_3183 | | |
| DI49_3186; DI49_5581; SPGP0Ktrna2N; YOL054W; SPGP0Ktrna3N; SPGP0DVtrna5N; DI49_4738; SPGP0Ltrna4N; SPGP0Mtrna6N; YNL284C; SPGP0Mtrna7N; YNR002C; SPGP0Qtrna9N; SPGP0AHtrna14N; DI49_1650; SPGP0BCtrna15N; DI49_3923 | DI49_3186; DI49_5581; DI49_4738; DI49_1650; DI49_3923 | SPGP0Ktrna2N; SPGP0Ktrna3N; SPGP0DVtrna5N; SPGP0Ltrna4N; SPGP0Mtrna6N; SPGP0Mtrna7N; SPGP0Qtrna9N; SPGP0AHtrna14N; SPGP0BCtrna15N | YOL054W; YNL284C; YNR002C |
| DI49_3187; SPGP0CZ00330 | DI49_3187 | SPGP0CZ00330 | |
| DI49_3189; DI49_3188 | DI49_3189; DI49_3188 | | |
| DI49_3191; DI49_3190 | DI49_3191; DI49_3190 | | |
| DI49_3193; DI49_5687; SPGP0DCtrna4L; YBL107C; SPGP0DCtrna5L; SPGP0DNtrna7L; DI49_0082; SPGP0DN01610; SPGP0Mtrna11L; DI49_3917; SPGP0Qtrna19L; SPGP0Q03100; SPGP0Wtrna26L; SPGP0Ctrna30L; YJL006C | DI49_3193; DI49_5687; DI49_0082; DI49_3917 | SPGP0DCtrna4L; SPGP0DCtrna5L; SPGP0DNtrna7L; SPGP0DN01610; SPGP0Mtrna11L; SPGP0Qtrna19L; SPGP0Q03100; SPGP0Wtrna26L; SPGP0Ctrna30L | YBL107C; YJL006C |
| DI49_3195; DI49_3194 | DI49_3195; DI49_3194 | | |
| DI49_3197; DI49_3196 | DI49_3197; DI49_3196 | | |
| DI49_3199; DI49_3198 | DI49_3199; DI49_3198 | | |
| DI49_3201; DI49_3200 | DI49_3201; DI49_3200 | | |
| DI49_3203; DI49_3202 | DI49_3203; DI49_3202 | | |
| DI49_3205; DI49_3204 | DI49_3205; DI49_3204 | | |
| DI49_3207; DI49_3206 | DI49_3207; DI49_3206 | | |
| DI49_3209; DI49_3208 | DI49_3209; DI49_3208 | | |
| DI49_3211; DI49_3210 | DI49_3211; DI49_3210 | | |
| DI49_3213; DI49_3212 | DI49_3213; DI49_3212 | | |
| DI49_3215; DI49_3214 | DI49_3215; DI49_3214 | | |
| DI49_3217; DI49_2869; DI49_3219; YJL158C; DI49_2870 | DI49_3217; DI49_2869; DI49_3219; DI49_2870 | | YJL158C |
| DI49_3221; DI49_3220 | DI49_3221; DI49_3220 | | |
| DI49_3223; DI49_3222 | DI49_3223; DI49_3222 | | |
| DI49_3224; YKL157W | DI49_3224 | | YKL157W |
| DI49_3226; DI49_3225 | DI49_3226; DI49_3225 | | |
| DI49_3228; DI49_4323; SPGP0DNtrna7R; SPGP0Qtrna12R; SPGP0Qtrna13R; DI49_2082; SPGP0Rtrna16R; SPGP0Rtrna17R; DI49_2981; SPGP0Strna19R; SPGP0Wtrna22R; SPGP0Btrna21R | DI49_3228; DI49_4323; DI49_2082; DI49_2981 | SPGP0DNtrna7R; SPGP0Qtrna12R; SPGP0Qtrna13R; SPGP0Rtrna16R; SPGP0Rtrna17R; SPGP0Strna19R; SPGP0Wtrna22R; SPGP0Btrna21R | |
| DI49_3230; DI49_3229 | DI49_3230; DI49_3229 | | |
| DI49_3232; DI49_3231 | DI49_3232; DI49_3231 | | |
| DI49_3234; DI49_3233 | DI49_3234; DI49_3233 | | |
| DI49_3236; DI49_3235 | DI49_3236; DI49_3235 | | |
| DI49_3238; DI49_3237 | DI49_3238; DI49_3237 | | |
| DI49_3240; DI49_3239 | DI49_3240; DI49_3239 | | |
| DI49_3242; DI49_3241 | DI49_3242; DI49_3241 | | |

Create PDF in your applications with the Pdfcrowd HTML to PDF API PDFCROWD

| | | | |
|---|---|---|---|
| DI49_3244; DI49_3243 | DI49_3244; DI49_3243 | | |
| DI49_3246; DI49_3245 | DI49_3246; DI49_3245 | | |
| DI49_3248; DI49_3247 | DI49_3248; DI49_3247 | | |
| DI49_3250; DI49_3249 | DI49_3250; DI49_3249 | | |
| DI49_3252; DI49_4193; DI49_3253; DI49_4192 | DI49_3252; DI49_4193; DI49_3253; DI49_4192 | | |
| DI49_3255; DI49_3254 | DI49_3255; DI49_3254 | | |
| DI49_3257; DI49_3256 | DI49_3257; DI49_3256 | | |
| DI49_3259; DI49_3258 | DI49_3259; DI49_3258 | | |
| DI49_3261; DI49_5601; DI49_3413; SPGP0DBtrna3A; YMR247C; SPGP0DBtrna4A; YMR023C; SPGP0Ntrna8A; DI49_3766; DI49_5416; SPGP0Ptrna10A; SPGP0Qtrna12A; SPGP0Rtrna13A; SPGP0R00980; SPGP0Strna14A; DI49_4332; SPGP0Strna15A; DI49_4110; SPGP0Vtrna16A; SPGP0Wtrna17A; DI49_1030; SPGP0ACtrna18A; DI49_1682; SPGP0Ftrna23A; YJL115W | DI49_3261; DI49_5601; DI49_3413; DI49_3766; DI49_5416; DI49_4332; DI49_4110; DI49_1030; DI49_1682 | SPGP0DBtrna3A; SPGP0DBtrna4A; SPGP0Ntrna8A; SPGP0Ptrna10A; SPGP0Qtrna12A; SPGP0Rtrna13A; SPGP0R00980; SPGP0Strna14A; SPGP0Strna15A; SPGP0Vtrna16A; SPGP0Wtrna17A; SPGP0ACtrna18A; SPGP0Ftrna23A | YMR247C; YMR023C; YJL115W |
| DI49_3263; DI49_3262 | DI49_3263; DI49_3262 | | |
| DI49_3265; DI49_3264 | DI49_3265; DI49_3264 | | |
| DI49_3267; DI49_3266 | DI49_3267; DI49_3266 | | |
| DI49_3269; DI49_3268 | DI49_3269; DI49_3268 | | |
| DI49_3271; DI49_3270 | DI49_3271; DI49_3270 | | |
| DI49_3273; DI49_3272 | DI49_3273; DI49_3272 | | |
| DI49_3275; DI49_3274 | DI49_3275; DI49_3274 | | |
| DI49_3277; DI49_3276 | DI49_3277; DI49_3276 | | |
| DI49_3279; DI49_1406; DI49_3280 | DI49_3279; DI49_1406; DI49_3280 | | |
| DI49_3282; DI49_3281 | DI49_3282; DI49_3281 | | |
| DI49_3284; DI49_3283 | DI49_3284; DI49_3283 | | |
| DI49_3286; DI49_3285 | DI49_3286; DI49_3285 | | |
| DI49_3288; DI49_3287 | DI49_3288; DI49_3287 | | |
| DI49_3290; DI49_3289 | DI49_3290; DI49_3289 | | |
| DI49_3292; DI49_3291 | DI49_3292; DI49_3291 | | |
| DI49_3293; SPGP0CZ01360 | DI49_3293 | SPGP0CZ01360 | |
| DI49_3295; DI49_3294 | DI49_3295; DI49_3294 | | |
| DI49_3297; DI49_3296 | DI49_3297; DI49_3296 | | |
| DI49_3299; DI49_3298 | DI49_3299; DI49_3298 | | |
| SPGP0CZtrna1W; DI49_5211; DI49_3302; DI49_2157; DI49_2156 | DI49_5211; DI49_3302; DI49_2157; DI49_2156 | SPGP0CZtrna1W | |
| DI49_3304; DI49_4248; SPGP0CZtrna2V; DI49_3340; YMR163C; SPGP0DBtrna5V; YMR049C; SPGP0Ltrna9V; DI49_4839; DI49_5011; DI49_3781; SPGP0Qtrna12V; DI49_2123; SPGP0Qtrna13V; YGL226W; SPGP0Rtrna14V; DI49_4247; SPGP0Strna17V; DI49_4138; SPGP0Etrna25V; SPGP0Ftrna26V; YGR161C; DI49_1753; SPGP0Gtrna29V; YJL037W; SPGP0Itrna30V | DI49_3304; DI49_4248; DI49_3340; DI49_4839; DI49_5011; DI49_3781; DI49_2123; DI49_4247; DI49_4138; DI49_1753 | SPGP0CZtrna2V; SPGP0DBtrna5V; SPGP0Ltrna9V; SPGP0Qtrna12V; SPGP0Qtrna13V; SPGP0Rtrna14V; SPGP0Strna17V; SPGP0Etrna25V; SPGP0Ftrna26V; SPGP0Gtrna29V; SPGP0Itrna30V | YMR163C; YMR049C; YGL226W; YGR161C; YJL037W |
| SPGP0CZtrna1H; DI49_4134; DI49_3307; SPGP0Ltrna3H; SPGP0Strna6H; SPGP0Etrna8H; DI49_1425; SPGP0Etrna9H; DI49_1538; DI49_1773 | DI49_4134; DI49_3307; DI49_1425; DI49_1538; DI49_1773 | SPGP0CZtrna1H; SPGP0Ltrna3H; SPGP0Strna6H; SPGP0Etrna8H; SPGP0Etrna9H | |

Create PDF in your applications with the Pdfcrowd HTML to PDF API PDFCROWD

| | | | |
|---|---|---|---|
| DI49_3309; DI49_3308 | DI49_3309; DI49_3308 | | |
| DI49_3311; DI49_3310 | DI49_3311; DI49_3310 | | |
| DI49_3313; DI49_3312 | DI49_3313; DI49_3312 | | |
| DI49_3315; DI49_3314 | DI49_3315; DI49_3314 | | |
| DI49_3317; DI49_3316 | DI49_3317; DI49_3316 | | |
| DI49_3319; DI49_3318 | DI49_3319; DI49_3318 | | |
| DI49_3321; DI49_3320 | DI49_3321; DI49_3320 | | |
| DI49_3323; DI49_3322 | DI49_3323; DI49_3322 | | |
| SPGP0CZ01660; DI49_3324 | DI49_3324 | SPGP0CZ01660 | |
| DI49_3326; DI49_3325 | DI49_3326; DI49_3325 | | |
| DI49_3328; DI49_3327 | DI49_3328; DI49_3327 | | |
| DI49_3330; DI49_3329 | DI49_3330; DI49_3329 | | |
| DI49_3332; DI49_3331 | DI49_3332; DI49_3331 | | |
| DI49_3334; DI49_3333 | DI49_3334; DI49_3333 | | |
| DI49_3336; DI49_3335 | DI49_3336; DI49_3335 | | |
| DI49_3337; SPGP0CZ01790 | DI49_3337 | SPGP0CZ01790 | |
| DI49_3342; DI49_3341 | DI49_3342; DI49_3341 | | |
| DI49_3344; DI49_3343 | DI49_3344; DI49_3343 | | |
| DI49_3345; SPGP0CZ01870 | DI49_3345 | SPGP0CZ01870 | |
| DI49_3347; DI49_3346 | DI49_3347; DI49_3346 | | |
| DI49_3349; DI49_3348 | DI49_3349; DI49_3348 | | |
| DI49_3351; DI49_3350 | DI49_3351; DI49_3350 | | |
| DI49_3353; DI49_3352 | DI49_3353; DI49_3352 | | |
| DI49_3355; DI49_3354 | DI49_3355; DI49_3354 | | |
| DI49_3357; DI49_3356 | DI49_3357; DI49_3356 | | |
| DI49_3359; DI49_3358 | DI49_3359; DI49_3358 | | |
| DI49_3361; DI49_3360 | DI49_3361; DI49_3360 | | |
| DI49_3363; DI49_3362 | DI49_3363; DI49_3362 | | |
| SPGP0CZ02080; DI49_3364 | DI49_3364 | SPGP0CZ02080 | |
| DI49_3366; DI49_3365 | DI49_3366; DI49_3365 | | |
| DI49_3368; DI49_3367; DI49_4847; SPGP0V00510; YHR001W | DI49_3368; DI49_3367; DI49_4847 | SPGP0V00510 | YHR001W |
| DI49_3370; DI49_3369 | DI49_3370; DI49_3369 | | |
| DI49_3372; DI49_3371 | DI49_3372; DI49_3371 | | |
| DI49_3374; DI49_3373 | DI49_3374; DI49_3373 | | |
| DI49_3376; DI49_3375 | DI49_3376; DI49_3375 | | |
| DI49_3378; DI49_3377 | DI49_3378; DI49_3377 | | |
| DI49_3380; DI49_3379 | DI49_3380; DI49_3379 | | |
| SPGP0CZtrna2L; DI49_5686; DI49_3383; SPGP0Ntrna13L; DI49_3756; DI49_2147; SPGP0Rtrna22L; DI49_0670; SPGP0AAtrna29L; SPGP0Ytrna27L; DI49_1719 | DI49_5686; DI49_3383; DI49_3756; DI49_2147; DI49_0670; DI49_1719 | SPGP0CZtrna2L; SPGP0Ntrna13L; SPGP0Rtrna22L; SPGP0AAtrna29L; SPGP0Ytrna27L | |
| DI49_3385; DI49_3384 | DI49_3385; DI49_3384 | | |

Create PDF in your applications with the Pdfcrowd HTML to PDF API PDFCROWD

| | | | |
|---|---|---|---|
| DI49_3387; DI49_3386 | DI49_3387; DI49_3386 | | |
| DI49_3389; DI49_3388 | DI49_3389; DI49_3388 | | |
| DI49_3391; DI49_3390 | DI49_3391; DI49_3390 | | |
| DI49_3393; DI49_3392 | DI49_3393; DI49_3392 | | |
| DI49_3395; DI49_3394 | DI49_3395; DI49_3394 | | |
| DI49_3397; DI49_3396 | DI49_3397; DI49_3396 | | |
| SPGP0CZtrna2R; DI49_2410; DI49_3400; SPGP0DNtrna8R; SPGP0Ktrna6R; YOR008C; DI49_3638; SPGP0Ctrna24R; DI49_1572; YER161C | DI49_2410; DI49_3400; DI49_3638; DI49_1572 | SPGP0CZtrna2R; SPGP0DNtrna8R; SPGP0Ktrna6R; SPGP0Ctrna24R | YOR008C; YER161C |
| DI49_3402; DI49_3401 | DI49_3402; DI49_3401 | | |
| DI49_3404; DI49_3403 | DI49_3404; DI49_3403 | | |
| DI49_3406; DI49_3405 | DI49_3406; DI49_3405 | | |
| DI49_3408; DI49_3407 | DI49_3408; DI49_3407 | | |
| SPGP0CZtrna1D; DI49_4675; DI49_3411; SPGP0DBtrna2D; SPGP0DCtrna3D; SPGP0DDtrna4D; YIL009W; SPGP0DNtrna7D; SPGP0Ktrna6D; YOR131C; SPGP0DUtrna9D; DI49_3664; SPGP0Mtrna10D; SPGP0Ntrna11D; DI49_3834; SPGP0Q02530; DI49_1985; SPGP0Rtrna15D; DI49_2973; SPGP0Rtrna16D; SPGP0Rtrna17D; DI49_3068; SPGP0Strna18D; SPGP0Wtrna21D; SPGP0Ztrna22D; SPGP0Btrna19D; SPGP0Ctrna23D; YDR059C; YGR029W; YGR023W; YJR055W; SPGP0Gtrna27D; SPGP0Gtrna28D | DI49_4675; DI49_3411; DI49_3664; DI49_3834; DI49_1985; DI49_2973; DI49_3068 | SPGP0CZtrna1D; SPGP0DBtrna2D; SPGP0DCtrna3D; SPGP0DDtrna4D; SPGP0DNtrna7D; SPGP0Ktrna6D; SPGP0DUtrna9D; SPGP0Mtrna10D; SPGP0Ntrna11D; SPGP0Q02530; SPGP0Rtrna15D; SPGP0Rtrna16D; SPGP0Rtrna17D; SPGP0Strna18D; SPGP0Wtrna21D; SPGP0Ztrna22D; SPGP0Btrna19D; SPGP0Ctrna23D; SPGP0Gtrna27D; SPGP0Gtrna28D | YIL009W; YOR131C; YDR059C; YGR029W; YGR023W; YJR055W |
| DI49_3415; DI49_3414 | DI49_3415; DI49_3414 | | |
| DI49_3417; DI49_3416 | DI49_3417; DI49_3416 | | |
| DI49_3419; DI49_3418 | DI49_3419; DI49_3418 | | |
| DI49_3421; DI49_3420 | DI49_3421; DI49_3420 | | |
| DI49_3423; DI49_3422 | DI49_3423; DI49_3422 | | |
| DI49_3425; DI49_3424 | DI49_3425; DI49_3424 | | |
| DI49_3427; DI49_3426 | DI49_3427; DI49_3426 | | |
| DI49_3429; DI49_3428 | DI49_3429; DI49_3428 | | |
| DI49_3431; DI49_3430 | DI49_3431; DI49_3430 | | |
| DI49_3433; DI49_3432 | DI49_3433; DI49_3432 | | |
| SPGP0CZ02750; DI49_3434 | DI49_3434 | SPGP0CZ02750 | |
| DI49_3436; DI49_3435 | DI49_3436; DI49_3435 | | |
| DI49_3438; DI49_3437 | DI49_3438; DI49_3437 | | |
| DI49_3440; DI49_3439 | DI49_3440; DI49_3439 | | |
| DI49_3442; DI49_3441 | DI49_3442; DI49_3441 | | |
| DI49_3444; DI49_5564; DI49_3876; SPGP0ACtrna21K; DI49_5565; DI49_1775 | DI49_3444; DI49_5564; DI49_3876; DI49_5565; DI49_1775 | SPGP0ACtrna21K | |
| DI49_3446; DI49_3445 | DI49_3446; DI49_3445 | | |
| DI49_3448; DI49_3447 | DI49_3448; DI49_3447 | | |
| DI49_3450; DI49_3449 | DI49_3450; DI49_3449 | | |
| DI49_3452; DI49_3451 | DI49_3452; DI49_3451 | | |

Create PDF in your applications with the Pdfcrowd HTML to PDF API PDFCROWD

| | | | |
|---|---|---|---|
| DI49_3454; DI49_3453 | DI49_3454; DI49_3453 | | |
| DI49_3456; DI49_3455 | DI49_3456; DI49_3455 | | |
| DI49_3458; DI49_3457 | DI49_3458; DI49_3457 | | |
| DI49_3460; DI49_3459 | DI49_3460; DI49_3459 | | |
| DI49_3462; DI49_3461 | DI49_3462; DI49_3461 | | |
| YKR094C; DI49_2639; DI49_3464; DI49_3686; DI49_3685; YLL039C; DI49_3494 | DI49_2639; DI49_3464; DI49_3686; DI49_3685; DI49_3494 | | YKR094C; YLL039C |
| DI49_3466; DI49_3465 | DI49_3466; DI49_3465 | | |
| DI49_3468; DI49_3467 | DI49_3468; DI49_3467 | | |
| DI49_3470; DI49_3469 | DI49_3470; DI49_3469 | | |
| SPGP0CZ03130; DI49_3471 | DI49_3471 | SPGP0CZ03130 | |
| YMR287C; DI49_4374; YMR286W; DI49_4373 | DI49_4374; DI49_4373 | | YMR287C; YMR286W |
| YMR284W; YMR285C | | | YMR284W; YMR285C |
| YMR283C; SPGP0Gtrna14Y; SPGP0DBtrna1Y; YJL046W | | SPGP0Gtrna14Y; SPGP0DBtrna1Y | YMR283C; YJL046W |
| YMR281W; YMR282C | | | YMR281W; YMR282C |
| YMR279C; YMR280C | | | YMR279C; YMR280C |
| YMR277W; YMR278W | | | YMR277W; YMR278W |
| YMR275C; DI49_4361; YMR274C; DI49_4360 | DI49_4361; DI49_4360 | | YMR275C; YMR274C |
| YMR272C; DI49_4358 | DI49_4358 | | YMR272C |
| SPGP0DBtrna1Q; DI49_4357; YMR271C; SPGP0S00260 | DI49_4357 | SPGP0DBtrna1Q; SPGP0S00260 | YMR271C |
| YMR269W; DI49_4353; YMR268C; DI49_4352 | DI49_4353; DI49_4352 | | YMR269W; YMR268C |
| YMR266W; YMR267W | | | YMR266W; YMR267W |
| YMR264W; DI49_4348; YMR263W; DI49_4347 | DI49_4348; DI49_4347 | | YMR264W; YMR263W |
| YMR261C; DI49_4345 | DI49_4345 | | YMR261C |
| YMR260C; DI49_4344; YMR259C; DI49_4343 | DI49_4344; DI49_4343 | | YMR260C; YMR259C |
| YMR257C; DI49_4341 | DI49_4341 | | YMR257C |
| YMR256C; DI49_4340; YMR255W; DI49_4339 | DI49_4340; DI49_4339 | | YMR256C; YMR255W |
| YMR252C; YMR253C | | | YMR252C; YMR253C |
| YMR251W; YMR251W-A | | | YMR251W; YMR251W-A |
| YMR244C-A; YMR246W | | | YMR244C-A; YMR246W |
| YMR243C; DI49_4328 | DI49_4328 | | YMR243C |
| YMR242C; DI49_5131; DI49_5132; DI49_4327 | DI49_5131; DI49_5132; DI49_4327 | | YMR242C |
| YMR241W; DI49_4326; YMR240C; DI49_4325 | DI49_4326; DI49_4325 | | YMR241W; YMR240C |
| SPGP0DBtrna3R; SPGP0Itrna34R; SPGP0DBtrna4R; YML071C; SPGP0DCtrna5R; SPGP0Strna20R; DI49_4019; SPGP0Ctrna23R; SPGP0Ftrna27R; SPGP0Ftrna28R; YGR122W; SPGP0Gtrna31R; SPGP0Gtrna32R | DI49_4019 | SPGP0DBtrna3R; SPGP0Itrna34R; SPGP0DBtrna4R; SPGP0DCtrna5R; SPGP0Strna20R; SPGP0Ctrna23R; SPGP0Ftrna27R; SPGP0Ftrna28R; SPGP0Gtrna31R; SPGP0Gtrna32R | YML071C; YGR122W |
| YMR238W; DI49_4322 | DI49_4322 | | YMR238W |
| YMR237W; DI49_4321; YMR236W; DI49_4320 | DI49_4321; DI49_4320 | | YMR237W; YMR236W |
| YMR234W; DI49_4318; YMR233W; DI49_4317 | DI49_4318; DI49_4317 | | YMR234W; YMR233W |
| YMR231W; DI49_4315 | DI49_4315 | | YMR231W |
| YMR230W; DI49_4314; DI49_5112; DI49_5113 | DI49_4314; DI49_5112; DI49_5113 | | YMR230W |

Create PDF in your applications with the Pdfcrowd HTML to PDF API PDFCROWD

| | | | |
|---|---|---|---|
| YMR229C; DI49_4313 | DI49_4313 | | YMR229C |
| YMR228W; DI49_4312 | DI49_4312 | | YMR228W |
| YMR227C; DI49_4311 | DI49_4311 | | YMR227C |
| YMR226C; DI49_4310; YMR225C; DI49_4309 | DI49_4310; DI49_4309 | | YMR226C; YMR225C |
| YMR223W; YMR224C | | | YMR223W; YMR224C |
| YMR221C; DI49_4305 | DI49_4305 | | YMR221C |
| YMR220W; DI49_4304; YMR219W; DI49_4303 | DI49_4304; DI49_4303 | | YMR220W; YMR219W |
| YMR217W; DI49_4301 | DI49_4301 | | YMR217W |
| YMR216C; DI49_4300; YMR215W; DI49_4299 | DI49_4300; DI49_4299 | | YMR216C; YMR215W |
| YMR213W; YMR214W | | | YMR213W; YMR214W |
| YMR211W; YMR212C | | | YMR211W; YMR212C |
| YMR209C; YMR210W | | | YMR209C; YMR210W |
| YMR207C; DI49_4291; YMR206W; DI49_4290 | DI49_4291; DI49_4290 | | YMR207C; YMR206W |
| YMR204C; DI49_4288 | DI49_4288 | | YMR204C |
| YMR203W; DI49_4287 | DI49_4287 | | YMR203W |
| YMR202W; DI49_4286; SPGP0DB00940; DI49_4285 | DI49_4286; DI49_4285 | SPGP0DB00940 | YMR202W |
| YMR200W; YMR201C | | | YMR200W; YMR201C |
| YMR198W; YMR199W; DI49_4283; DI49_4282 | DI49_4283; DI49_4282 | | YMR198W; YMR199W |
| YMR196W; DI49_4280; YMR195W; DI49_4279 | DI49_4280; DI49_4279 | | YMR196W; YMR195W |
| YMR194W; DI49_4277; YMR193W; DI49_4276; YPL249C-A; YPL250C | DI49_4277; DI49_4276 | | YMR194W; YMR193W; YPL249C-A; YPL250C |
| YMR191W; YMR192W | | | YMR191W; YMR192W |
| YMR189W; DI49_4272; YMR188C; DI49_4271 | DI49_4272; DI49_4271 | | YMR189W; YMR188C |
| YMR186W; SPGP0U01920; DI49_4269; YPL242C | DI49_4269 | SPGP0U01920 | YMR186W; YPL242C |
| YMR185W; DI49_4268; YMR184W; DI49_4267 | DI49_4268; DI49_4267 | | YMR185W; YMR184W |
| YMR181C; DI49_4264; YMR180C; DI49_4263 | DI49_4264; DI49_4263 | | YMR181C; YMR180C |
| YMR178W; YMR179W | | | YMR178W; YMR179W |
| YMR176W; DI49_4259; YMR175W; SPGP0S01230 | DI49_4259 | SPGP0S01230 | YMR176W; YMR175W |
| YMR173W; YMR174C | | | YMR173W; YMR174C |
| YMR171C; DI49_4255 | DI49_4255 | | YMR171C |
| YMR170C; YMR169C; YMR168C | | | YMR170C; YMR169C; YMR168C |
| YMR166C; DI49_4250 | DI49_4250 | | YMR166C |
| YMR165C; DI49_4249; YMR164C; SPGP0S01340 | DI49_4249 | SPGP0S01340 | YMR165C; YMR164C |
| YMR161W; DI49_4245; YMR160W; DI49_4244 | DI49_4245; DI49_4244 | | YMR161W; YMR160W |
| SPGP0DBtrna1M; YJL007C; YMR158W; YJL029C | | SPGP0DBtrna1M | YJL007C; YMR158W; YJL029C |
| YMR156C; YMR157C | | | YMR156C; YMR157C |
| YMR154C; YMR155W | | | YMR154C; YMR155W |
| YMR152W; YMR153W | | | YMR152W; YMR153W |
| YMR149W; YMR150C | | | YMR149W; YMR150C |
| YMR147W; YMR148W | | | YMR147W; YMR148W |
| YMR145C; DI49_4230 | DI49_4230 | | YMR145C |

Create PDF in your applications with the Pdfcrowd HTML to PDF API PDFCROWD

| | | | |
|---|---|---|---|
| YMR144W; DI49_4229 | DI49_4229 | | YMR144W |
| YMR143W; SPGP0S01540 | | SPGP0S01540 | YMR143W |
| YMR142C; SPGP0S01550; YMR140W; DI49_0985; DI49_4228; YDL082W | DI49_0985; DI49_4228 | SPGP0S01550 | YMR142C; YMR140W; YDL082W |
| YMR138W; YMR139W | | | YMR138W; YMR139W |
| YMR136W; YMR137C | | | YMR136W; YMR137C |
| YMR134W; YMR135C | | | YMR134W; YMR135C |
| YMR132C; YMR133W | | | YMR132C; YMR133W |
| YMR130W; YMR131C | | | YMR130W; YMR131C |
| YMR128W; DI49_4217 | DI49_4217 | | YMR128W |
| YMR127C; DI49_4216; YMR126C; DI49_4215 | DI49_4216; DI49_4215 | | YMR127C; YMR126C |
| YMR124W; DI49_4213; YMR123W; DI49_4212 | DI49_4213; DI49_4212 | | YMR124W; YMR123W |
| YMR121C; DI49_3558; DI49_3557; DI49_4210 | DI49_3558; DI49_3557; DI49_4210 | | YMR121C |
| YMR120C; DI49_4209; YMR119W; DI49_4208 | DI49_4209; DI49_4208 | | YMR120C; YMR119W |
| YMR118C; DI49_1927; SPGP0Mtrna10L; SPGP0Ftrna33L | DI49_1927 | SPGP0Mtrna10L; SPGP0Ftrna33L | YMR118C |
| YMR116C; DI49_4204 | DI49_4204 | | YMR116C |
| YMR115W; SPGP0S01810 | | SPGP0S01810 | YMR115W |
| YMR114C; DI49_4203 | DI49_4203 | | YMR114C |
| YMR113W; DI49_4202; YMR112C; DI49_4201 | DI49_4202; DI49_4201 | | YMR113W; YMR112C |
| YMR110C; YMR111C | | | YMR110C; YMR111C |
| YMR108W; YMR109W | | | YMR108W; YMR109W |
| SPGP0DBtrna3K; SPGP0Itrna36K; YMR106C; SPGP0DDtrna4K; SPGP0Qtrna11K; SPGP0Strna14K; DI49_4194; SPGP0Ytrna18K; DI49_0541; DI49_0613; SPGP0Btrna15K; DI49_2750; SPGP0AEtrna23K; SPGP0AHtrna24K; SPGP0AWtrna26K; YPR073C; SPGP0Etrna27K; DI49_1540; SPGP0Ftrna32K; SPGP0Gtrna34K; YJL011C; SPGP0Itrna35K; YER134C | DI49_4194; DI49_0541; DI49_0613; DI49_2750; DI49_1540 | SPGP0DBtrna3K; SPGP0Itrna36K; SPGP0DDtrna4K; SPGP0Qtrna11K; SPGP0Strna14K; SPGP0Ytrna18K; SPGP0Btrna15K; SPGP0AEtrna23K; SPGP0AHtrna24K; SPGP0AWtrna26K; SPGP0Etrna27K; SPGP0Ftrna32K; SPGP0Gtrna34K; SPGP0Itrna35K | YMR106C; YPR073C; YJL011C; YER134C |
| YMR104C; YMR105C | | | YMR104C; YMR105C |
| YMR101C; DI49_4190; YMR100W; DI49_4189 | DI49_4190; DI49_4189 | | YMR101C; YMR100W |
| YMR098C; DI49_4186; YMR097C; DI49_4185 | DI49_4186; DI49_4185 | | YMR098C; YMR097C |
| YMR095C; YMR096W | | | YMR095C; YMR096W |
| YMR093W; DI49_4181; YMR092C; DI49_4180 | DI49_4181; DI49_4180 | | YMR093W; YMR092C |
| YMR090W; YMR091C | | | YMR090W; YMR091C |
| YMR088C; DI49_4176; YMR087W; DI49_4175 | DI49_4176; DI49_4175 | | YMR088C; YMR087W |
| YMR085W; DI49_4173 | DI49_4173 | | YMR085W |
| YMR083W; DI49_4172; YMR081C; DI49_4171 | DI49_4172; DI49_4171 | | YMR083W; YMR081C |
| YMR079W; YMR080C | | | YMR079W; YMR080C |
| YMR077C; YMR078C | | | YMR077C; YMR078C |
| YMR076C; DI49_4165 | DI49_4165 | | YMR076C |
| YMR075W; DI49_4164; YMR074C; DI49_4163 | DI49_4164; DI49_4163 | | YMR075W; YMR074C |
| YMR072W; YMR073C | | | YMR072W; YMR073C |

Create PDF in your applications with the Pdfcrowd HTML to PDF API PDFCROWD

| | | | |
|---|---|---|---|
| YMR070W; YMR071C | | | YMR070W; YMR071C |
| YMR068W; DI49_4157; YMR067C; DI49_4156 | DI49_4157; DI49_4156 | | YMR068W; YMR067C |
| YMR065W; DI49_4154; YMR064W; DI49_4153 | DI49_4154; DI49_4153 | | YMR065W; YMR064W |
| YMR062C; YMR063W | | | YMR062C; YMR063W |
| YMR060C; YMR061W | | | YMR060C; YMR061W |
| YMR058W; DI49_4147 | DI49_4147 | | YMR058W |
| YMR056C; DI49_4146; YMR055C; DI49_4145 | DI49_4146; DI49_4145 | | YMR056C; YMR055C |
| YMR053C; DI49_4143; YMR052W; DI49_4142 | DI49_4143; DI49_4142 | | YMR053C; YMR052W |
| YMR047C; DI49_4135 | DI49_4135 | | YMR047C |
| SPGP0DBtrna2H; SPGP0Itrna13H; YGL205W; SPGP0Vtrna7H; YHL022C; SPGP0Ftrna10H; SPGP0Itrna12H; YER026C | | SPGP0DBtrna2H; SPGP0Itrna13H; SPGP0Vtrna7H; SPGP0Ftrna10H; SPGP0Itrna12H | YGL205W; YHL022C; YER026C |
| YMR044W; DI49_4133; YMR043W | DI49_4133 | | YMR044W; YMR043W |
| SPGP0DBtrna1F; SPGP0Ftrna19F; YMR041C; YHR072W; SPGP0Vtrna11F; YHR126C; SPGP0Ctrna15F; YDR317W; YPR027C; SPGP0Dtrna18F; YGL029W | | SPGP0DBtrna1F; SPGP0Ftrna19F; SPGP0Vtrna11F; SPGP0Ctrna15F; SPGP0Dtrna18F | YMR041C; YHR072W; YHR126C; YDR317W; YPR027C; YGL029W |
| YMR039C; YMR040W | | | YMR039C; YMR040W |
| YMR037C; YMR038C | | | YMR037C; YMR038C |
| YMR035W; YMR036C | | | YMR035W; YMR036C |
| YMR033W; YMR034C | | | YMR033W; YMR034C |
| YMR031C; YMR032W | | | YMR031C; YMR032W |
| YMR029C; YMR030W | | | YMR029C; YMR030W |
| YMR027W; DI49_4115; YMR026C; DI49_4114 | DI49_4115; DI49_4114 | | YMR027W; YMR026C |
| YMR024W; DI49_4112 | DI49_4112 | | YMR024W |
| YMR021C; DI49_4108 | DI49_4108 | | YMR021C |
| YMR020W; DI49_4107; YMR019W; DI49_4106 | DI49_4107; DI49_4106 | | YMR020W; YMR019W |
| YMR017W; YMR018W | | | YMR017W; YMR018W |
| YMR015C; DI49_4102 | DI49_4102 | | YMR015C |
| YMR014W; DI49_4101 | DI49_4101 | | YMR014W |
| YMR013C; DI49_4100 | DI49_4100 | | YMR013C |
| YMR012W; DI49_4099 | DI49_4099 | | YMR012W |
| SPGP0DBtrna2E; DI49_5287; SPGP0DCtrna3E; YBR210W; SPGP0DDtrna5E; SPGP0Qtrna8E; SPGP0Qtrna9E; DI49_1914; DI49_1878; SPGP0Rtrna11E; DI49_2867; SPGP0Strna12E; DI49_5286; SPGP0Wtrna16E; DI49_0733; SPGP0Btrna14E; YCL021W-A; SPGP0Etrna20E; DI49_1408; DI49_1504; SPGP0Etrna22E; SPGP0Ftrna23E; SPGP0Ftrna24E; YGL054C; SPGP0Gtrna26E; SPGP0Itrna27E; YER012W; YER098W; SPGP0Itrna28E | DI49_5287; DI49_1914; DI49_1878; DI49_2867; DI49_5286; DI49_0733; DI49_1408; DI49_1504 | SPGP0DBtrna2E; SPGP0DCtrna3E; SPGP0DDtrna5E; SPGP0Qtrna8E; SPGP0Qtrna9E; SPGP0Rtrna11E; SPGP0Strna12E; SPGP0Wtrna16E; SPGP0Btrna14E; SPGP0Etrna20E; SPGP0Etrna22E; SPGP0Ftrna23E; SPGP0Ftrna24E; SPGP0Gtrna26E; SPGP0Itrna27E; SPGP0Itrna28E | YBR210W; YCL021W-A; YGL054C; YER012W; YER098W |
| YMR011W; SPGP0S02800 | | SPGP0S02800 | YMR011W |
| YMR010W; DI49_4097; YMR009W; DI49_4096 | DI49_4097; DI49_4096 | | YMR010W; YMR009W |
| YMR008C; DI49_4095; YMR007W | DI49_4095 | | YMR008C; YMR007W |
| YMR005W; YMR006C; DI49_4094; DI49_4093 | DI49_4094; DI49_4093 | | YMR005W; YMR006C |
| YMR003W; YMR004W | | | YMR003W; YMR004W |
| YMR001C; DI49_4089 | DI49_4089 | | YMR001C |

Create PDF in your applications with the Pdfcrowd HTML to PDF API PDFCROWD

| | | | |
|---|---|---|---|
| YML001W; DI49_4088 | DI49_4088 | | YML001W |
| YML002W; DI49_4087 | DI49_4087 | | YML002W |
| YML004C; DI49_4086 | DI49_4086 | | YML004C |
| YML005W; DI49_4085 | DI49_4085 | | YML005W |
| SPGP0DBtrna1S; SPGP0Itrna33S; SPGP0DCtrna2S; SPGP0DEtrna4S; YGR038W; SPGP0Gtrna31S | | SPGP0DBtrna1S; SPGP0Itrna33S; SPGP0DCtrna2S; SPGP0DEtrna4S; SPGP0Gtrna31S | YGR038W |
| YML006C; DI49_4083 | DI49_4083 | | YML006C |
| YML007W; DI49_4082 | DI49_4082 | | YML007W |
| YML008C; DI49_4081 | DI49_4081 | | YML008C |
| YML009C; DI49_4080 | DI49_4080 | | YML009C |
| YML010W; DI49_4079; YML011C; DI49_4078 | DI49_4079; DI49_4078 | | YML010W; YML011C |
| YML013W; YML012W | | | YML013W; YML012W |
| YML015C; YML014W | | | YML015C; YML014W |
| YML017W; DI49_4072; YML018C; DI49_4071 | DI49_4072; DI49_4071 | | YML017W; YML018C |
| YML020W; DI49_4069 | DI49_4069 | | YML020W |
| YML021C; DI49_4068; YML022W; DI49_4067 | DI49_4068; DI49_4067 | | YML021C; YML022W |
| SPGP0DB03090; DI49_0614; DI49_4065; DI49_0615 | DI49_0614; DI49_4065; DI49_0615 | SPGP0DB03090 | |
| YML025C; DI49_4064 | DI49_4064 | | YML025C |
| YML026C; SPGP0S03140; YML027W; DI49_4062; DI49_0617; DI49_0618 | DI49_4062; DI49_0617; DI49_0618 | SPGP0S03140 | YML026C; YML027W |
| YML029W; YML028W | | | YML029W; YML028W |
| YML031W; DI49_4058 | DI49_4058 | | YML031W |
| YML032C; DI49_4057; YML034W; DI49_4056 | DI49_4057; DI49_4056 | | YML032C; YML034W |
| YML036W; YML035C; DI49_4055; DI49_4054 | DI49_4055; DI49_4054 | | YML036W; YML035C |
| YML038C; DI49_4052 | DI49_4052 | | YML038C |
| SPGP0DBtrna1P; SPGP0Vtrna13P; YHR146W | | SPGP0DBtrna1P; SPGP0Vtrna13P | YHR146W |
| YML041C; DI49_4050 | DI49_4050 | | YML041C |
| YML042W; DI49_4049; YML043C; DI49_4048 | DI49_4049; DI49_4048 | | YML042W; YML043C |
| YML046W; SPGP0Itrna37G; SPGP0DCtrna2G; YBL014C; SPGP0DNtrna6G; DI49_0168; DI49_0443; SPGP0Ktrna4G; SPGP0Ptrna10G; DI49_2378; SPGP0Ptrna11G; SPGP0Qtrna13G; SPGP0Qtrna14G; SPGP0Rtrna16G; DI49_4046; SPGP0Wtrna19G; DI49_0870; SPGP0Xtrna20G; DI49_5463; SPGP0AAtrna23G; DI49_0766; SPGP0ACtrna24G; DI49_5604; SPGP0AEtrna27G; YCR016W; DI49_1672; SPGP0AH00610; SPGP0Ctrna26G; YDR262W; SPGP0Dtrna30G; YPR007C; SPGP0Etrna31G; SPGP0Ftrna32G; SPGP0Ftrna33G; SPGP0Gtrna35G; YJR051W; SPGP0Gtrna36G | DI49_0168; DI49_0443; DI49_2378; DI49_4046; DI49_0870; DI49_5463; DI49_0766; DI49_5604; DI49_1672 | SPGP0Itrna37G; SPGP0DCtrna2G; SPGP0DNtrna6G; SPGP0Ktrna4G; SPGP0Ptrna10G; SPGP0Ptrna11G; SPGP0Qtrna13G; SPGP0Qtrna14G; SPGP0Rtrna16G; SPGP0Wtrna19G; SPGP0Xtrna20G; SPGP0AAtrna23G; SPGP0ACtrna24G; SPGP0AEtrna27G; SPGP0AH00610; SPGP0Ctrna26G; SPGP0Dtrna30G; SPGP0Etrna31G; SPGP0Ftrna32G; SPGP0Ftrna33G; SPGP0Gtrna35G; SPGP0Gtrna36G | YML046W; YBL014C; YCR016W; YDR262W; YPR007C; YJR051W |
| YML048W; DI49_4044; YML049C; SPGP0S03330 | DI49_4044 | SPGP0S03330 | YML048W; YML049C |
| YML051W; DI49_4041 | DI49_4041 | | YML051W |
| YML052W; DI49_4040 | DI49_4040 | | YML052W |
| YML053C; DI49_4039 | DI49_4039 | | YML053C |
| SPGP0DBtrna2Y; SPGP0Gtrna13Y; YML054C; YOL019W; SPGP0Ctrna10Y; YJR056C | | SPGP0DBtrna2Y; SPGP0Gtrna13Y; SPGP0Ctrna10Y | YML054C; YOL019W; YJR056C |

Create PDF in your applications with the Pdfcrowd HTML to PDF API PDFCROWD

| | | | |
|---|---|---|---|
| SPGP0DB03360; SPGP0S03400 | | SPGP0DB03360; SPGP0S03400 | |
| YML057W; DI49_4035; YML058W; SPGP0S03420 | DI49_4035 | SPGP0S03420 | YML057W; YML058W |
| YML059C; DI49_4032 | DI49_4032 | | YML059C |
| YML060W; DI49_4031 | DI49_4031 | | YML060W |
| YML061C; DI49_4030; YML062C; DI49_2437; DI49_4029 | DI49_4030; DI49_2437; DI49_4029 | | YML061C; YML062C |
| YML064C; DI49_4027; YML065W; DI49_4026 | DI49_4027; DI49_4026 | | YML064C; YML065W |
| YML067C; YML066C | | | YML067C; YML066C |
| YML069W; YML068W | | | YML069W; YML068W |
| YML073C; DI49_4017; DI49_4378 | DI49_4017; DI49_4378 | | YML073C |
| YBL105C; DI49_0084 | DI49_0084 | | YBL105C |
| YBL104C; DI49_0085; YBL103C; DI49_0086 | DI49_0085; DI49_0086 | | YBL104C; YBL103C |
| YBL101C; YBL102W | | | YBL101C; YBL102W |
| YBL099W; DI49_0090; YBL098W; DI49_0091 | DI49_0090; DI49_0091 | | YBL099W; YBL098W |
| YBL095W; YBL097W | | | YBL095W; YBL097W |
| YBL092W; DI49_0095; YBL091C-A; DI49_0096 | DI49_0095; DI49_0096 | | YBL092W; YBL091C-A |
| YBL090W; YBL091C | | | YBL090W; YBL091C |
| YBL088C; DI49_0100 | DI49_0100 | | YBL088C |
| YBL087C; YER117W; YBL086C | | | YBL087C; YER117W; YBL086C |
| YBL084C; YBL085W | | | YBL084C; YBL085W |
| YBL081W; YBL082C | | | YBL081W; YBL082C |
| YBL079W; DI49_0108 | DI49_0108 | | YBL079W |
| YBL078C; DI49_0109 | DI49_0109 | | YBL078C |
| YBL076C; DI49_0110 | DI49_0110 | | YBL076C |
| YBL075C; SPGP0DN00430; YBL074C; DI49_0111 | DI49_0111 | SPGP0DN00430 | YBL075C; YBL074C |
| YBL071W-A; YBL072C; YER102W | | | YBL071W-A; YBL072C; YER102W |
| YBL068W; DI49_0115; YBL067C; DI49_0116 | DI49_0115; DI49_0116 | | YBL068W; YBL067C |
| YBL064C; DI49_0118 | DI49_0118 | | YBL064C |
| YBL063W; DI49_0119 | DI49_0119 | | YBL063W |
| YBL061C; DI49_0120; YBL060W; DI49_0121 | DI49_0120; DI49_0121 | | YBL061C; YBL060W |
| YBL059W; YBL059C-A | | | YBL059W; YBL059C-A |
| YBL057C; YBL058W | | | YBL057C; YBL058W |
| YBL055C; YBL056W | | | YBL055C; YBL056W |
| YBL052C; DI49_0127 | DI49_0127 | | YBL052C |
| YBL051C; DI49_0128; YBL050W; DI49_0129 | DI49_0128; DI49_0129 | | YBL051C; YBL050W |
| YBL047C; DI49_0131; YBL046W; DI49_0132 | DI49_0131; DI49_0132 | | YBL047C; YBL046W |
| YBL043W; YBL045C | | | YBL043W; YBL045C |
| YBL041W; DI49_0137; YBL040C; DI49_0138 | DI49_0137; DI49_0138 | | YBL041W; YBL040C |
| YBL039C; DI49_0140 | DI49_0140 | | YBL039C |
| YBL038W; DI49_0141 | DI49_0141 | | YBL038W |
| YBL037W; DI49_0142; YBL036C; DI49_0143 | DI49_0142; DI49_0143 | | YBL037W; YBL036C |

Create PDF in your applications with the Pdfcrowd HTML to PDF API PDFCROWD

| | | | |
|---|---|---|---|
| YBL034C; YBL035C | | | YBL034C; YBL035C |
| YBL032W; YBL033C | | | YBL032W; YBL033C |
| YBL030C; DI49_0149; YBL029C-A; DI49_0150 | DI49_0149; DI49_0150 | | YBL030C; YBL029C-A |
| YBL028C; YBL029W | | | YBL028C; YBL029W |
| SPGP0DC00830; YBL027W | | SPGP0DC00830 | YBL027W |
| YBL025W; YBL026W | | | YBL025W; YBL026W |
| YBL023C; DI49_0158 | DI49_0158 | | YBL023C |
| YBL022C; DI49_0159; YBL021C; DI49_0160 | DI49_0159; DI49_0160 | | YBL022C; YBL021C |
| YBL019W; DI49_4697; YBL018C | DI49_4697 | | YBL019W; YBL018C |
| YBL016W; YBL017C | | | YBL016W; YBL017C |
| SPGP0DCtrna1I; SPGP0Itrna24I; SPGP0DDtrna2I; YIL079C; SPGP0DNtrna3I; SPGP0DNtrna4I; SPGP0EEtrna8I; DI49_3949; SPGP0Mtrna7I; DI49_3807; SPGP0Qtrna11I; SPGP0Ztrna14I; SPGP0Ztrna15I; DI49_4700; DI49_5588; SPGP0ACtrna17I; DI49_5615; SPGP0Btrna12I; SPGP0Btrna13I; DI49_2704; SPGP0Ctrna18I; DI49_1592; SPGP0Ftrna21I; SPGP0BZtrna22I; YER182W | DI49_3949; DI49_3807; DI49_4700; DI49_5588; DI49_5615; DI49_2704; DI49_1592 | SPGP0DCtrna1I; SPGP0Itrna24I; SPGP0DDtrna2I; SPGP0DNtrna3I; SPGP0DNtrna4I; SPGP0EEtrna8I; SPGP0Mtrna7I; SPGP0Qtrna11I; SPGP0Ztrna14I; SPGP0Ztrna15I; SPGP0ACtrna17I; SPGP0Btrna12I; SPGP0Btrna13I; SPGP0Ctrna18I; SPGP0Ftrna21I; SPGP0BZtrna22I | YIL079C; YER182W |
| YBL011W; YBL013W | | | YBL011W; YBL013W |
| YBL009W; DI49_0172; YBL008W; DI49_0173 | DI49_0172; DI49_0173 | | YBL009W; YBL008W |
| YBL006C; YBL007C | | | YBL006C; YBL007C |
| YBL004W; DI49_0178 | DI49_0178 | | YBL004W |
| YBL003C; YDR225W | | | YBL003C; YDR225W |
| YBL002W; DI49_0406; YBL001C; DI49_0180; DI49_0181; YDR224C | DI49_0406; DI49_0180; DI49_0181 | | YBL002W; YBL001C; YDR224C |
| YBR002C; YBR001C | | | YBR002C; YBR001C |
| YBR004C; YBR003W | | | YBR004C; YBR003W |
| YBR006W; DI49_0187; YBR007C; DI49_0188 | DI49_0187; DI49_0188 | | YBR006W; YBR007C |
| YBR009C; YNL030W | | | YBR009C; YNL030W |
| YBR010W; DI49_0190; DI49_4706; YNL031C | DI49_0190; DI49_4706 | | YBR010W; YNL031C |
| YBR011C; DI49_0191 | DI49_0191 | | YBR011C |
| YBR012C; SPGP0EE00220 | | SPGP0EE00220 | YBR012C |
| SPGP0DCtrna2T; SPGP0Gtrna31T; YBR014C; SPGP0DDtrna3T; SPGP0DDtrna4T; YIL014W; SPGP0Ktrna5T; YOL105C; SPGP0Ktrna6T; YOR013W; YHR006W; YDL007C-A | | SPGP0DCtrna2T; SPGP0Gtrna31T; SPGP0DDtrna3T; SPGP0DDtrna4T; SPGP0Ktrna5T; SPGP0Ktrna6T | YBR014C; YIL014W; YOL105C; YOR013W; YHR006W; YDL007C-A |
| YBR016W; YBR015C | | | YBR016W; YBR015C |
| YBR018C; YBR017C | | | YBR018C; YBR017C |
| YBR020W; YBR019C | | | YBR020W; YBR019C |
| YBR022W; YBR021W | | | YBR022W; YBR021W |
| YBR024W; YBR023C | | | YBR024W; YBR023C |
| YBR026C; YBR025C | | | YBR026C; YBR025C |
| YBR029C; DI49_0207 | DI49_0207 | | YBR029C |
| YBR030W; DI49_0208 | DI49_0208 | | YBR030W |
| YBR031W; YDR012W; YBR033W; YDR013W | | | YBR031W; YDR012W; YBR033W; YDR013W |

Create PDF in your applications with the Pdfcrowd HTML to PDF API PDFCROWD

| | | | |
|---|---|---|---|
| YBR035C; YBR034C | | | YBR035C; YBR034C |
| YBR037C; DI49_1066 | DI49_1066 | | YBR037C |
| YBR038W; DI49_1067 | DI49_1067 | | YBR038W |
| YBR039W; DI49_1068; YBR040W | DI49_1068 | | YBR039W; YBR040W |
| YBR042C; DI49_1071 | DI49_1071 | | YBR042C |
| YBR043C; DI49_1072; YBR044C | DI49_1072 | | YBR043C; YBR044C |
| YBR045C; DI49_1074; SPGP0DNtrna7V; DI49_1073; SPGP0Wtrna19V | DI49_1074; DI49_1073 | SPGP0DNtrna7V; SPGP0Wtrna19V | YBR045C |
| YBR047W; DI49_1077 | DI49_1077 | | YBR047W |
| YBR048W; YDR025W; DI49_0223; SPGP0W02480 | DI49_0223 | SPGP0W02480 | YBR048W; YDR025W |
| YBR049C; DI49_1079 | DI49_1079 | | YBR049C |
| YBR050C; DI49_1080; YBR052C | DI49_1080 | | YBR050C; YBR052C |
| YBR054W; DI49_1083; DI49_1082 | DI49_1083; DI49_1082 | | YBR054W |
| YBR055C; DI49_1084 | DI49_1084 | | YBR055C |
| YBR056W; DI49_1086 | DI49_1086 | | YBR056W |
| SPGP0DCtrna2Q; DI49_1374; YBR057C; SPGP0Wtrna10Q; SPGP0Itrna18Q | DI49_1374 | SPGP0DCtrna2Q; SPGP0Wtrna10Q; SPGP0Itrna18Q | YBR057C |
| YBR058C-A; YBR058C | | | YBR058C-A; YBR058C |
| YBR060C; YBR059C | | | YBR060C; YBR059C |
| YBR062C; YBR061C | | | YBR062C; YBR061C |
| YBR065C; YBR064W | | | YBR065C; YBR064W |
| YBR067C; DI49_1099; DI49_1098 | DI49_1099; DI49_1098 | | YBR067C |
| YBR068C; DI49_1100 | DI49_1100 | | YBR068C |
| YBR069C; DI49_1101; YBR070C | DI49_1101 | | YBR069C; YBR070C |
| YBR072W; DI49_1104; YBR073W; DI49_1103 | DI49_1104; DI49_1103 | | YBR072W; YBR073W |
| YBR076W; YBR074W; DI49_1106; DI49_1105 | DI49_1106; DI49_1105 | | YBR076W; YBR074W |
| YBR078W; DI49_1109; DI49_1108 | DI49_1109; DI49_1108 | | YBR078W |
| YBR079C; DI49_1110 | DI49_1110 | | YBR079C |
| YBR080C; DI49_1111 | DI49_1111 | | YBR080C |
| YBR081C; DI49_1112 | DI49_1112 | | YBR081C |
| YBR082C; SPGP0W02160 | | SPGP0W02160 | YBR082C |
| YBR083W; DI49_1116 | DI49_1116 | | YBR083W |
| YBR084W; DI49_1117 | DI49_1117 | | YBR084W |
| YBR085W; DI49_1118; YBR085C-A | DI49_1118 | | YBR085W; YBR085C-A |
| YBR087W; YBR086C | | | YBR087W; YBR086C |
| YBR089C-A; YBR088C; DI49_1123; DI49_1122 | DI49_1123; DI49_1122 | | YBR089C-A; YBR088C |
| YBR093C; SPGP0W02050; DI49_4804; SPGP0W02040; DI49_1125 | DI49_4804; DI49_1125 | SPGP0W02050; SPGP0W02040 | YBR093C |
| YBR094W; DI49_1126 | DI49_1126 | | YBR094W |
| YBR095C; DI49_1127; YBR096W | DI49_1127 | | YBR095C; YBR096W |
| YBR098W; YBR097W; DI49_1129; DI49_1128 | DI49_1129; DI49_1128 | | YBR098W; YBR097W |
| YBR102C; DI49_1132; YBR103W; DI49_1131 | DI49_1132; DI49_1131 | | YBR102C; YBR103W |

Create PDF in your applications with the Pdfcrowd HTML to PDF API PDFCROWD

| | | | |
|---|---|---|---|
| YBR105C; YBR104W | | | YBR105C; YBR104W |
| YBR107C; YBR106W; DI49_1137; DI49_1135 | DI49_1137; DI49_1135 | | YBR107C; YBR106W |
| YBR109C; YBR108W | | | YBR109C; YBR108W |
| YBR111C; YBR110W | | | YBR111C; YBR110W |
| YBR112C; DI49_1144; SPGP0W01870 | DI49_1144 | SPGP0W01870 | YBR112C |
| YBR114W; DI49_1146; DI49_1145 | DI49_1146; DI49_1145 | | YBR114W |
| YBR115C; DI49_1147 | DI49_1147 | | YBR115C |
| YBR117C; DI49_5526; SPGP0EO00100 | DI49_5526 | SPGP0EO00100 | YBR117C |
| YPR080W; DI49_5530; YBR119W; DI49_1148 | DI49_5530; DI49_1148 | | YPR080W; YBR119W |
| YBR121C; DI49_1152; YBR122C; DI49_1151 | DI49_1152; DI49_1151 | | YBR121C; YBR122C |
| YBR125C; DI49_1155 | DI49_1155 | | YBR125C |
| YBR126C; DI49_1156 | DI49_1156 | | YBR126C |
| YBR128C; DI49_1158; YBR129C; DI49_1157 | DI49_1158; DI49_1157 | | YBR128C; YBR129C |
| YBR131W; YBR130C | | | YBR131W; YBR130C |
| YBR133C; DI49_1163; YBR135W | DI49_1163 | | YBR133C; YBR135W |
| YBR137W; YBR136W; DI49_1165; DI49_1164 | DI49_1165; DI49_1164 | | YBR137W; YBR136W |
| YBR139W; DI49_1168; DI49_1167 | DI49_1168; DI49_1167 | | YBR139W |
| YBR140C; DI49_1169; YBR141C | DI49_1169 | | YBR140C; YBR141C |
| YBR143C; DI49_1172; YBR145W; SPGP0R01690; DI49_1171 | DI49_1172; DI49_1171 | SPGP0R01690 | YBR143C; YBR145W |
| YBR147W; DI49_1175; YBR148W | DI49_1175 | | YBR147W; YBR148W |
| YBR150C; YBR149W | | | YBR150C; YBR149W |
| YBR152W; YBR151W; DI49_1179; DI49_1178 | DI49_1179; DI49_1178 | | YBR152W; YBR151W |
| YBR154C; DI49_1182; DI49_1181 | DI49_1182; DI49_1181 | | YBR154C |
| YBR155W; DI49_1183 | DI49_1183 | | YBR155W |
| YBR156C; SPGP0W01460; YBR157C | | SPGP0W01460 | YBR156C; YBR157C |
| YBR159W; DI49_1186 | DI49_1186 | | YBR159W |
| YBR160W; DI49_1187; YBR161W | DI49_1187 | | YBR160W; YBR161W |
| YBR162W-A; YBR162C | | | YBR162W-A; YBR162C |
| YBR164C; DI49_1192; YBR165W | DI49_1192 | | YBR164C; YBR165W |
| YBR167C; YBR166C; DI49_1194; DI49_1193 | DI49_1194; DI49_1193 | | YBR167C; YBR166C |
| YBR169C; DI49_1197; DI49_1196 | DI49_1197; DI49_1196 | | YBR169C |
| YBR170C; DI49_1198 | DI49_1198 | | YBR170C |
| YBR171W; DI49_1199 | DI49_1199 | | YBR171W |
| YBR172C; DI49_1200; YBR173C | DI49_1200 | | YBR172C; YBR173C |
| YBR176W; YBR175W | | | YBR176W; YBR175W |
| YBR179C; DI49_1205; YBR180W; DI49_1204 | DI49_1205; DI49_1204 | | YBR179C; YBR180W |
| YBR182C; SPGP0DC02750 | | SPGP0DC02750 | YBR182C |
| YBR184W; YBR183W | | | YBR184W; YBR183W |
| YBR186W; YBR185C | | | YBR186W; YBR185C |
| YBR188C; YBR187W | | | YBR188C; YBR187W |

Create PDF in your applications with the Pdfcrowd HTML to PDF API PDFCROWD

| | | | |
|---|---|---|---|
| YBR191W; DI49_1216 | DI49_1216 | | YBR191W |
| YBR192W; DI49_1217; YBR193C | DI49_1217 | | YBR192W; YBR193C |
| YBR195C; YBR194W | | | YBR195C; YBR194W |
| YBR197C; YBR196C | | | YBR197C; YBR196C |
| YBR199W; YBR198C | | | YBR199W; YBR198C |
| YBR201W; YBR200W; DI49_1225; DI49_1224 | DI49_1225; DI49_1224 | | YBR201W; YBR200W |
| YBR202W; DI49_1228; YBR203W; DI49_1227 | DI49_1228; DI49_1227 | | YBR202W; YBR203W |
| YBR205W; DI49_1231; DI49_1230 | DI49_1231; DI49_1230 | | YBR205W |
| YBR207W; DI49_1232 | DI49_1232 | | YBR207W |
| YBR208C; DI49_1233 | DI49_1233 | | YBR208C |
| SPGP0DCtrna1C; DI49_5567; SPGP0ACtrna5C; DI49_5568 | DI49_5567; DI49_5568 | SPGP0DCtrna1C; SPGP0ACtrna5C | |
| YBR212W; DI49_1238; YBR213W; DI49_1237 | DI49_1238; DI49_1237 | | YBR212W; YBR213W |
| YBR215W; DI49_1241; YBR216C | DI49_1241 | | YBR215W; YBR216C |
| YBR218C; DI49_1244; YBR220C; DI49_1243 | DI49_1244; DI49_1243 | | YBR218C; YBR220C |
| YBR222C; DI49_1247; YBR223C | DI49_1247 | | YBR222C; YBR223C |
| YBR227C; YBR225W; DI49_1249; DI49_1248 | DI49_1249; DI49_1248 | | YBR227C; YBR225W |
| YBR229C; DI49_1252; YBR230C; DI49_1251 | DI49_1252; DI49_1251 | | YBR229C; YBR230C |
| YBR233W; YBR231C | | | YBR233W; YBR231C |
| YBR234C; DI49_1257; YBR235W; DI49_1256 | DI49_1257; DI49_1256 | | YBR234C; YBR235W |
| YBR237W; DI49_1260; DI49_1259 | DI49_1260; DI49_1259 | | YBR237W |
| YBR238C; DI49_1261 | DI49_1261 | | YBR238C |
| YBR239C; DI49_1262 | DI49_1262 | | YBR239C |
| YBR240C; DI49_1263 | DI49_1263 | | YBR240C |
| YBR241C; DI49_1265 | DI49_1265 | | YBR241C |
| YBR242W; DI49_1266; YBR243C | DI49_1266 | | YBR242W; YBR243C |
| YBR245C; DI49_1269; YBR246W; DI49_1268 | DI49_1269; DI49_1268 | | YBR245C; YBR246W |
| YBR248C; DI49_1272; DI49_1271 | DI49_1272; DI49_1271 | | YBR248C |
| YBR249C; DI49_1273; YBR250W | DI49_1273 | | YBR249C; YBR250W |
| YBR252W; YBR251W | | | YBR252W; YBR251W |
| YBR254C; DI49_1278; YBR255W; DI49_1277 | DI49_1278; DI49_1277 | | YBR254C; YBR255W |
| YBR256C; DI49_1281; YBR257W | DI49_1281 | | YBR256C; YBR257W |
| YBR259W; YBR258C | | | YBR259W; YBR258C |
| YBR261C; YBR260C; DI49_1285; DI49_1284 | DI49_1285; DI49_1284 | | YBR261C; YBR260C |
| YBR263W; DI49_1288; YBR264C; DI49_1287 | DI49_1288; DI49_1287 | | YBR263W; YBR264C |
| YBR267W; DI49_1291; YBR268W | DI49_1291 | | YBR267W; YBR268W |
| YBR270C; YBR269C | | | YBR270C; YBR269C |
| YBR272C; YBR271W | | | YBR272C; YBR271W |
| YBR274W; DI49_1298; DI49_1297 | DI49_1298; DI49_1297 | | YBR274W |
| YBR275C; DI49_1299; YBR276C | DI49_1299 | | YBR275C; YBR276C |

Create PDF in your applications with the Pdfcrowd HTML to PDF API PDFCROWD

| | | | |
|---|---|---|---|
| YBR279W; YBR278W | | | YBR279W; YBR278W |
| YBR281C; DI49_1304; DI49_1303 | DI49_1304; DI49_1303 | | YBR281C |
| YBR282W; DI49_1305 | DI49_1305 | | YBR282W |
| YBR283C; DI49_1306; YBR284W | DI49_1306 | | YBR283C; YBR284W |
| YBR286W; YBR285W | | | YBR286W; YBR285W |
| YBR288C; DI49_1311 | DI49_1311 | | YBR288C |
| YJL222W; YNR065C; YNR067C | | | YJL222W; YNR065C; YNR067C |
| SPGP0DD00110; YJL219W; SPGP0K00150; SPGP0K00160; SPGP0BV00100 | | SPGP0DD00110; SPGP0K00150; SPGP0K00160; SPGP0BV00100 | YJL219W |
| YIL169C; SPGP0G00110; YIL167W; DI49_2820; SPGP0R00110; YJR150C | DI49_2820 | SPGP0G00110; SPGP0R00110 | YIL169C; YIL167W; YJR150C |
| YIL164C; YIL166C | | | YIL164C; YIL166C |
| YIL161W; DI49_2625 | DI49_2625 | | YIL161W |
| YIL160C; DI49_2626 | DI49_2626 | | YIL160C |
| YIL159W; DI49_2627; YIL158W; DI49_2628 | DI49_2627; DI49_2628 | | YIL159W; YIL158W |
| YIL156W-B; YIL157C | | | YIL156W-B; YIL157C |
| YIL155C; YIL156W | | | YIL155C; YIL156W |
| YIL153W; YIL154C | | | YIL153W; YIL154C |
| YIL151C; DI49_2636 | DI49_2636 | | YIL151C |
| YIL150C; DI49_2637 | DI49_2637 | | YIL150C |
| YIL149C; DI49_2638; SPGP0DD00310 | DI49_2638 | SPGP0DD00310 | YIL149C |
| YIL146C; DI49_2641; YIL145C; DI49_2642 | DI49_2641; DI49_2642 | | YIL146C; YIL145C |
| YIL143C; DI49_2644 | DI49_2644 | | YIL143C |
| YIL142W; DI49_2645; YIL140W; DI49_2646 | DI49_2645; DI49_2646 | | YIL142W; YIL140W |
| YIL138C; YIL139C | | | YIL138C; YIL139C |
| YIL136W; YIL137C | | | YIL136W; YIL137C |
| YIL134W; YIL135C | | | YIL134W; YIL135C |
| YIL132C; YIL133C | | | YIL132C; YIL133C |
| YIL130W; YIL131C | | | YIL130W; YIL131C |
| YIL128W; DI49_2658; YIL127C; DI49_2659 | DI49_2658; DI49_2659 | | YIL128W; YIL127C |
| YIL125W; DI49_2661 | DI49_2661 | | YIL125W |
| YIL124W; DI49_2662 | DI49_2662 | | YIL124W |
| YIL123W; DI49_2663 | DI49_2663 | | YIL123W |
| YIL122W; DI49_2664; YIL121W; DI49_2665 | DI49_2664; DI49_2665 | | YIL122W; YIL121W |
| YIL119C; YIL120W | | | YIL119C; YIL120W |
| YIL117C; YIL118W | | | YIL117C; YIL118W |
| YIL115C; DI49_2671; YIL114C; DI49_2672 | DI49_2671; DI49_2672 | | YIL115C; YIL114C |
| YIL112W; YIL113W | | | YIL112W; YIL113W |
| YIL110W; DI49_2675 | DI49_2675 | | YIL110W |
| YIL109C; DI49_2676 | DI49_2676 | | YIL109C |
| YIL108W; DI49_2677 | DI49_2677 | | YIL108W |

Create PDF in your applications with the Pdfcrowd HTML to PDF API PDFCROWD

| | | | |
|---|---|---|---|
| YIL107C; DI49_2678; YIL106W; DI49_2679 | DI49_2678; DI49_2679 | | YIL107C; YIL106W |
| YIL104C; YIL105C | | | YIL104C; YIL105C |
| YIL102C-A; YIL103W | | | YIL102C-A; YIL103W |
| YIL101C; DI49_2685; YIL099W; DI49_2686 | DI49_2685; DI49_2686 | | YIL101C; YIL099W |
| YIL097W; YIL098C | | | YIL097W; YIL098C |
| YIL095W; DI49_2691 | DI49_2691 | | YIL095W |
| YIL094C; DI49_2692 | DI49_2692 | | YIL094C |
| YIL093C; DI49_2693; YIL092W; DI49_2694 | DI49_2693; DI49_2694 | | YIL093C; YIL092W |
| YIL090W; YIL091C | | | YIL090W; YIL091C |
| SPGP0DDtrna4E; YDR278C; YIL088C; YDR279W | | SPGP0DDtrna4E | YDR278C; YIL088C; YDR279W |
| YIL085C; YIL087C | | | YIL085C; YIL087C |
| YIL083C; SPGP0B00890 | | SPGP0B00890 | YIL083C |
| YIL077C; DI49_2706; YIL076W | DI49_2706 | | YIL077C; YIL076W |
| YIL074C; YER081W; YIL073C | | | YIL074C; YER081W; YIL073C |
| YIL071C; YIL072W | | | YIL071C; YIL072W |
| YIL069C; YER074W; DI49_1479 | DI49_1479 | | YIL069C; YER074W |
| YIL068C; DI49_2716 | DI49_2716 | | YIL068C |
| YIL067C; DI49_2717 | DI49_2717 | | YIL067C |
| YIL066C; DI49_2718; YIL065C; DI49_2719 | DI49_2718; DI49_2719 | | YIL066C; YIL065C |
| YIL063C; DI49_2720 | DI49_2720 | | YIL063C |
| YIL062C; DI49_2721; YIL061C; DI49_2722 | DI49_2721; DI49_2722 | | YIL062C; YIL061C |
| SPGP0DDtrna3S; DI49_5523; YIL056W; SPGP0Wtrna18S; DI49_1141; SPGP0Xtrna20S; SPGP0X00120; SPGP0Btrna15S; DI49_2725; DI49_0808; YCR063W; SPGP0Dtrna27S; YPR071W; SPGP0Etrna29S; YER066W | DI49_5523; DI49_1141; DI49_2725; DI49_0808 | SPGP0DDtrna3S; SPGP0Wtrna18S; SPGP0Xtrna20S; SPGP0X00120; SPGP0Btrna15S; SPGP0Dtrna27S; SPGP0Etrna29S | YIL056W; YCR063W; YPR071W; YER066W |
| YIL053W; DI49_2727; DI49_1466; DI49_1467 | DI49_2727; DI49_1466; DI49_1467 | | YIL053W |
| YIL052C; SPGP0B01150 | | SPGP0B01150 | YIL052C |
| YIL051C; DI49_2728; YIL050W; DI49_2729 | DI49_2728; DI49_2729 | | YIL051C; YIL050W |
| YIL048W; DI49_2731 | DI49_2731 | | YIL048W |
| YIL047C; DI49_2732 | DI49_2732 | | YIL047C |
| YIL046W; DI49_2733 | DI49_2733 | | YIL046W |
| YIL045W; DI49_2734; YIL044C; DI49_2735 | DI49_2734; DI49_2735 | | YIL045W; YIL044C |
| YIL042C; YIL043C | | | YIL042C; YIL043C |
| YIL041W; DI49_2738; YIL040W; DI49_2739 | DI49_2738; DI49_2739 | | YIL041W; YIL040W |
| YIL038C; DI49_2741; YIL037C; DI49_2742 | DI49_2741; DI49_2742 | | YIL038C; YIL037C |
| YIL035C; DI49_2744 | DI49_2744 | | YIL035C |
| YIL034C; DI49_2745 | DI49_2745 | | YIL034C |
| YIL033C; DI49_2746; YIL031W; DI49_2747 | DI49_2746; DI49_2747 | | YIL033C; YIL031W |
| YIL027C; DI49_2751; YIL026C; DI49_2752 | DI49_2751; DI49_2752 | | YIL027C; YIL026C |
| YIL023C; YIL024C | | | YIL023C; YIL024C |
| YIL021W; DI49_2756; YIL020C; DI49_2757 | DI49_2756; DI49_2757 | | YIL021W; YIL020C |
| YIL018W; DI49_2759; YIL017C; SPGP0AH00380; | DI49_2759; DI49_2760; DI49_1689 | SPGP0AH00380 | YIL018W; YIL017C |

Create PDF in your applications with the Pdfcrowd HTML to PDF API PDFCROWD

| | | | |
|---|---|---|---|
| DI49_2760; DI49_1689 | | | |
| YIL015W; DI49_2762 | DI49_2762 | | YIL015W |
| SPGP0DD01520; YOR192C-B; YLR410W-B; SPGP0AE00800 | | SPGP0DD01520; SPGP0AE00800 | YOR192C-B; YLR410W-B |
| YIL011W; YIL013C | | | YIL011W; YIL013C |
| YIL009C-A; YIL010W | | | YIL009C-A; YIL010W |
| YIL007C; YIL008W | | | YIL007C; YIL008W |
| YIL005W; YIL006W | | | YIL005W; YIL006W |
| YIL003W; DI49_2776 | DI49_2776 | | YIL003W |
| YIL002W-A; DI49_2777 | DI49_2777 | | YIL002W-A |
| YIL002C; DI49_2778; YIL001W; DI49_2779 | DI49_2778; DI49_2779 | | YIL002C; YIL001W |
| YIR002C; DI49_2781; YIR003W; DI49_2782 | DI49_2781; DI49_2782 | | YIR002C; YIR003W |
| YIR005W; YIR004W | | | YIR005W; YIR004W |
| YIR007W; DI49_2787 | DI49_2787 | | YIR007W |
| YIR008C; DI49_2788; YIR009W; DI49_2789 | DI49_2788; DI49_2789 | | YIR008C; YIR009W |
| YIR011C; DI49_2791 | DI49_2791 | | YIR011C |
| YIR012W; DI49_2792; YIR013C; DI49_2793 | DI49_2792; DI49_2793 | | YIR012W; YIR013C |
| YIR015W; YIR014W | | | YIR015W; YIR014W |
| YIR017C; DI49_2797 | DI49_2797 | | YIR017C |
| YIR018W; DI49_2798; YIR019C; DI49_2799 | DI49_2798; DI49_2799 | | YIR018W; YIR019C |
| YIR022W; YIR021W | | | YIR022W; YIR021W |
| YIR024C; YIR023W | | | YIR024C; YIR023W |
| YIR026C; YIR025W | | | YIR026C; YIR025W |
| YIR028W; DI49_2807 | DI49_2807 | | YIR028W |
| YIR029W; DI49_2808; YIR030C; DI49_2809 | DI49_2808; DI49_2809 | | YIR029W; YIR030C |
| YIR032C; YIR031C | | | YIR032C; YIR031C |
| YIR034C; DI49_2813; YIR035C; DI49_2814 | DI49_2813; DI49_2814 | | YIR034C; YIR035C |
| YIR037W; YIR036C | | | YIR037W; YIR036C |
| YIR039C; YIR038C | | | YIR039C; YIR038C |
| SPGP0DE00100; SPGP0DE00900; SPGP0AI00110; YHR210C | | SPGP0DE00100; SPGP0DE00900; SPGP0AI00110 | YHR210C |
| YAL062W; DI49_0002; DI49_5188; DI49_5189 | DI49_0002; DI49_5188; DI49_5189 | | YAL062W |
| YAL061W; DI49_0003; YAL060W | DI49_0003 | | YAL061W; YAL060W |
| YAL058W; YAL059W | | | YAL058W; YAL059W |
| YAL055W; YAL056W | | | YAL055W; YAL056W |
| YAL053W; YAL054C | | | YAL053W; YAL054C |
| YAL049C; YAL051W | | | YAL049C; YAL051W |
| YAL047C; YAL048C | | | YAL047C; YAL048C |
| YAL044W-A; YAL046C; DI49_0015; DI49_0014 | DI49_0015; DI49_0014 | | YAL044W-A; YAL046C |
| YAL043C; DI49_0018; DI49_0017 | DI49_0018; DI49_0017 | | YAL043C |
| YAL042W; DI49_0019 | DI49_0019 | | YAL042W |

Create PDF in your applications with the Pdfcrowd HTML to PDF API PDFCROWD

| | | | |
|---|---|---|---|
| YAL041W; DI49_0020 | DI49_0020 | | YAL041W |
| YAL040C; DI49_0021; YAL039C | DI49_0021 | | YAL040C; YAL039C |
| YAL037W; YAL038W | | | YAL037W; YAL038W |
| YAL035W; DI49_0026 | DI49_0026 | | YAL035W |
| YAL034W-A; DI49_0027 | DI49_0027 | | YAL034W-A |
| YAL034C; DI49_0028 | DI49_0028 | | YAL034C |
| YAL033W; DI49_0029; YAL032C | DI49_0029 | | YAL033W; YAL032C |
| YAL030W; YAL031C | | | YAL030W; YAL031C |
| YAL028W; YAL029C; DI49_0032; SPGP0H00560 | DI49_0032 | SPGP0H00560 | YAL028W; YAL029C |
| YAL026C; DI49_0035; YAL025C; DI49_0034 | DI49_0035; DI49_0034 | | YAL026C; YAL025C |
| YAL023C; DI49_0038 | DI49_0038 | | YAL023C |
| YAL022C; SPGP0H00480 | | SPGP0H00480 | YAL022C |
| YAL021C; DI49_0040 | DI49_0040 | | YAL021C |
| YAL020C; DI49_0041 | DI49_0041 | | YAL020C |
| YAL019W; DI49_0042; YAL018C | DI49_0042 | | YAL019W; YAL018C |
| YAL016W; DI49_0045; YAL015C; DI49_0044 | DI49_0045; DI49_0044 | | YAL016W; YAL015C |
| YAL013W; SPGP0H00390 | | SPGP0H00390 | YAL013W |
| YAL012W; DI49_0048 | DI49_0048 | | YAL012W |
| YAL011W; DI49_0049; YAL010C | DI49_0049 | | YAL011W; YAL010C |
| YAL008W; YAL009W | | | YAL008W; YAL009W |
| SPGP0DEtrna2P; SPGP0Mtrna7P; SPGP0Mtrna6P; YNL042W; YNL035C | | SPGP0DEtrna2P; SPGP0Mtrna7P; SPGP0Mtrna6P | YNL042W; YNL035C |
| YAL005C; DI49_3507; DI49_0055; DI49_3508 | DI49_3507; DI49_0055; DI49_3508 | | YAL005C |
| YAL003W; DI49_0056; YAL002W | DI49_0056 | | YAL003W; YAL002W |
| SPGP0DE00690; YAL001C | | SPGP0DE00690 | YAL001C |
| YAR002C-A; DI49_0060; DI49_0059 | DI49_0060; DI49_0059 | | YAR002C-A |
| YAR003W; DI49_0061 | DI49_0061 | | YAR003W |
| YAR007C; DI49_0062 | DI49_0062 | | YAR007C |
| YAR008W; DI49_0063 | DI49_0063 | | YAR008W |
| SPGP0DEtrna5A; DI49_3564; DI49_3563; SPGP0Qtrna11A; DI49_2114; SPGP0Etrna21A; SPGP0Ftrna22A; YGR150C; SPGP0Htrna25A; DI49_0066; YER076C | DI49_3564; DI49_3563; DI49_2114; DI49_0066 | SPGP0DEtrna5A; SPGP0Qtrna11A; SPGP0Etrna21A; SPGP0Ftrna22A; SPGP0Htrna25A | YGR150C; YER076C |
| YAR014C; DI49_0068 | DI49_0068 | | YAR014C |
| YAR015W; DI49_0069; YAR018C | DI49_0069 | | YAR015W; YAR018C |
| YFL020C; DI49_4789; YAR023C; SPGP0H00150; DI49_3953; DI49_3954; SPGP0AH00880; DI49_1633; DI49_0072 | DI49_4789; DI49_3953; DI49_3954; DI49_1633; DI49_0072 | SPGP0H00150; SPGP0AH00880 | YFL020C; YAR023C |
| YAR027W; YAR028W; SPGP0DE00830; YAR031W | | SPGP0DE00830 | YAR027W; YAR028W; YAR031W |
| YAR033W; SPGP0F03120; YAR035W | | SPGP0F03120 | YAR033W; YAR035W |
| SPGP0DE00890; YAR042W; DI49_0076 | DI49_0076 | SPGP0DE00890 | YAR042W |
| SPGP0J00100; SPGP0BP00100; DI49_4806; SPGP0ED00120; DI49_3942; YPR204W | DI49_4806; DI49_3942 | SPGP0J00100; SPGP0BP00100; SPGP0ED00120 | YPR204W |
| SPGP0DH00100; YHR211W; SPGP0AI00100 | | SPGP0DH00100; SPGP0AI00100 | YHR211W |

Create PDF in your applications with the Pdfcrowd HTML to PDF API PDFCROWD

| | | | |
|---|---|---|---|
| YBR298C; DI49_3958; YBR297W | DI49_3958 | | YBR298C; YBR297W |
| SPGP0DJ00100; YGR294W | | SPGP0DJ00100 | YGR294W |
| SPGP0DK00100; YBL111C; SPGP0ED00100 | | SPGP0DK00100; SPGP0ED00100 | YBL111C |
| DI49_0079; SPGP0DQ00120; DI49_5194 | DI49_0079; DI49_5194 | SPGP0DQ00120 | |
| SPGP0DN00110; DI49_5680 | DI49_5680 | SPGP0DN00110 | |
| DI49_5681; SPGP0DZ02360 | DI49_5681 | SPGP0DZ02360 | |
| DI49_5682; YFL058W | DI49_5682 | | YFL058W |
| DI49_0088; DI49_0087 | DI49_0088; DI49_0087 | | |
| DI49_0093; DI49_0092 | DI49_0093; DI49_0092 | | |
| DI49_0098; DI49_0097 | DI49_0098; DI49_0097 | | |
| DI49_0101; DI49_1522; DI49_0102 | DI49_0101; DI49_1522; DI49_0102 | | |
| DI49_0104; DI49_0103 | DI49_0104; DI49_0103 | | |
| SPGP0DN00370; DI49_0105 | DI49_0105 | SPGP0DN00370 | |
| DI49_0107; YBL080C | DI49_0107 | | YBL080C |
| DI49_0113; DI49_0112; DI49_1508 | DI49_0113; DI49_0112; DI49_1508 | | |
| DI49_0122; SPGP0DN00550 | DI49_0122 | SPGP0DN00550 | |
| DI49_0124; DI49_0123 | DI49_0124; DI49_0123 | | |
| SPGP0DN00600; DI49_0125; YER089C | DI49_0125 | SPGP0DN00600 | YER089C |
| DI49_0134; DI49_0133 | DI49_0134; DI49_0133 | | |
| DI49_0136; DI49_0135 | DI49_0136; DI49_0135 | | |
| DI49_0145; DI49_0144 | DI49_0145; DI49_0144 | | |
| DI49_0147; DI49_0146 | DI49_0147; DI49_0146 | | |
| DI49_0152; DI49_0151 | DI49_0152; DI49_0151 | | |
| DI49_0154; DI49_0153; SPGP0W02130 | DI49_0154; DI49_0153 | SPGP0W02130 | |
| DI49_0156; DI49_0155 | DI49_0156; DI49_0155 | | |
| DI49_0162; DI49_4696; DI49_0164; DI49_4695 | DI49_0162; DI49_4696; DI49_0164; DI49_4695 | | |
| DI49_0170; DI49_0169 | DI49_0170; DI49_0169 | | |
| DI49_0175; DI49_0174 | DI49_0175; DI49_0174 | | |
| SPGP0DNtrna6S; DI49_4419; DI49_4863; DI49_2001; SPGP0Rtrna13S; SPGP0Strna14S; SPGP0Wtrna19S; SPGP0W02810; DI49_0593; DI49_0694; SPGP0Btrna16S; YCL064C; SPGP0Etrna28S | DI49_4419; DI49_4863; DI49_2001; DI49_0593; DI49_0694 | SPGP0DNtrna6S; SPGP0Rtrna13S; SPGP0Strna14S; SPGP0Wtrna19S; SPGP0W02810; SPGP0Btrna16S; SPGP0Etrna28S | YCL064C |
| DI49_0179; DI49_0407 | DI49_0179; DI49_0407 | | |
| DI49_0183; DI49_0182 | DI49_0183; DI49_0182 | | |
| DI49_0185; DI49_0184 | DI49_0185; DI49_0184 | | |
| SPGP0DNtrna7T; DI49_2415; DI49_0193; DI49_4854; SPGP0Ptrna12T; DI49_2299; SPGP0Rtrna17T; SPGP0Wtrna21T; DI49_1043; SPGP0Ztrna23T; DI49_4465; SPGP0Btrna19T; SPGP0B01500; SPGP0Btrna18T | DI49_2415; DI49_0193; DI49_4854; DI49_2299; DI49_1043; DI49_4465 | SPGP0DNtrna7T; SPGP0Ptrna12T; SPGP0Rtrna17T; SPGP0Wtrna21T; SPGP0Ztrna23T; SPGP0Btrna19T; SPGP0B01500; SPGP0Btrna18T | |
| DI49_0195; DI49_0194 | DI49_0195; DI49_0194 | | |
| DI49_0197; DI49_0196 | DI49_0197; DI49_0196 | | |
| DI49_0199; DI49_0198 | DI49_0199; DI49_0198 | | |
| DI49_0201; DI49_0200 | DI49_0201; DI49_0200 | | |

Create PDF in your applications with the Pdfcrowd HTML to PDF API PDFCROWD

| | | | |
|---|---|---|---|
| DI49_0203; DI49_0202 | DI49_0203; DI49_0202 | | |
| DI49_0205; DI49_0204 | DI49_0205; DI49_0204 | | |
| DI49_0209; DI49_1062; DI49_0210 | DI49_0209; DI49_1062; DI49_0210 | | |
| DI49_0212; DI49_0211 | DI49_0212; DI49_0211 | | |
| DI49_0214; DI49_0213 | DI49_0214; DI49_0213 | | |
| DI49_0216; YDR018C; DI49_0217; YDR019C | DI49_0216; DI49_0217 | | YDR018C; YDR019C |
| DI49_0219; YDR021W | DI49_0219 | | YDR021W |
| DI49_0220; YDR022C | DI49_0220 | | YDR022C |
| DI49_0222; YDR023W; YDR024W | DI49_0222 | | YDR023W; YDR024W |
| DI49_0224; YDR026C | DI49_0224 | | YDR026C |
| DI49_0225; YDR027C | DI49_0225 | | YDR027C |
| DI49_0226; YDR028C; DI49_0227; YDR030C | DI49_0226; DI49_0227 | | YDR028C; YDR030C |
| DI49_0229; YDR032C | DI49_0229 | | YDR032C |
| DI49_0230; YDR033W; DI49_0231; YDR034C | DI49_0230; DI49_0231 | | YDR033W; YDR034C |
| DI49_0233; YDR035W | DI49_0233 | | YDR035W |
| DI49_0234; YDR036C | DI49_0234 | | YDR036C |
| SPGP0DN01640; DI49_1093; DI49_0236; DI49_1092; YDR037W; YDR040C | DI49_1093; DI49_0236; DI49_1092 | SPGP0DN01640 | YDR037W; YDR040C |
| DI49_0238; DI49_0237 | DI49_0238; DI49_0237 | | |
| DI49_0240; YDR044W; DI49_0241; YDR045C | DI49_0240; DI49_0241 | | YDR044W; YDR045C |
| DI49_0243; YDR047W; DI49_0244; YDR049W | DI49_0243; DI49_0244 | | YDR047W; YDR049W |
| DI49_0246; YDR051C; DI49_0247; YDR052C | DI49_0246; DI49_0247 | | YDR051C; YDR052C |
| DI49_0249; YDR055W; DI49_0250; YDR056C | DI49_0249; DI49_0250 | | YDR055W; YDR056C |
| DI49_0252; YDR058C | DI49_0252 | | YDR058C |
| DI49_0256; YDR060W | DI49_0256 | | YDR060W |
| DI49_0257; YDR061W | DI49_0257 | | YDR061W |
| DI49_0258; YDR062W | DI49_0258 | | YDR062W |
| DI49_0259; YDR063W | DI49_0259 | | YDR063W |
| DI49_0260; YDR064W; DI49_0261; YDR065W | DI49_0260; DI49_0261 | | YDR064W; YDR065W |
| DI49_0263; YDR067C; DI49_0264; YDR068W | DI49_0263; DI49_0264 | | YDR067C; YDR068W |
| DI49_0266; DI49_0265 | DI49_0266; DI49_0265 | | |
| DI49_0268; YDR072C; DI49_0269; YDR073W | DI49_0268; DI49_0269 | | YDR072C; YDR073W |
| DI49_0271; YDR075W | DI49_0271 | | YDR075W |
| DI49_0272; YDR076W | DI49_0272 | | YDR076W |
| DI49_0273; YDR077W; DI49_0274; YDR078C | DI49_0273; DI49_0274 | | YDR077W; YDR078C |
| DI49_0276; DI49_0275 | DI49_0276; DI49_0275 | | |
| DI49_0278; DI49_0277 | DI49_0278; DI49_0277 | | |
| DI49_0280; YDR083W; DI49_0281; YDR084C | DI49_0280; DI49_0281 | | YDR083W; YDR084C |
| DI49_0283; DI49_0282 | DI49_0283; DI49_0282 | | |
| DI49_0285; YDR088C | DI49_0285 | | YDR088C |

Create PDF in your applications with the Pdfcrowd HTML to PDF API PDFCROWD

| | | | |
|---|---|---|---|
| DI49_0287; YDR089W; DI49_0288; YDR090C | DI49_0287; DI49_0288 | | YDR089W; YDR090C |
| SPGP0DN02150; DI49_0289 | DI49_0289 | SPGP0DN02150 | |
| DI49_0291; YDR096W | DI49_0291 | | YDR096W |
| DI49_0292; YDR097C; DI49_0293; YDR098C | DI49_0292; DI49_0293 | | YDR097C; YDR098C |
| DI49_0295; YDR099W; DI49_0296; YDR100W | DI49_0295; DI49_0296 | | YDR099W; YDR100W |
| DI49_0298; DI49_0297 | DI49_0298; DI49_0297 | | |
| DI49_0300; DI49_0299 | DI49_0300; DI49_0299 | | |
| DI49_0303; YDR107C | DI49_0303 | | YDR107C |
| DI49_0304; YDR108W; SPGP0DN02290; YDR109C | DI49_0304 | SPGP0DN02290 | YDR108W; YDR109C |
| DI49_0306; DI49_0305 | DI49_0306; DI49_0305 | | |
| DI49_0308; DI49_0307 | DI49_0308; DI49_0307 | | |
| DI49_0310; DI49_0309 | DI49_0310; DI49_0309 | | |
| DI49_0312; YDR119W; DI49_0313; YDR119W-A | DI49_0312; DI49_0313 | | YDR119W; YDR119W-A |
| DI49_0315; DI49_0314 | DI49_0315; DI49_0314 | | |
| DI49_0317; YDR123C; DI49_0318; YDR124W | DI49_0317; DI49_0318 | | YDR123C; YDR124W |
| DI49_0320; YDR126W | DI49_0320 | | YDR126W |
| DI49_0321; YDR127W; DI49_0322; YDR128W | DI49_0321; DI49_0322 | | YDR127W; YDR128W |
| DI49_0324; DI49_0323 | DI49_0324; DI49_0323 | | |
| DI49_0326; DI49_0325 | DI49_0326; DI49_0325 | | |
| DI49_0328; YDR135C; DI49_0329; YDR137W | DI49_0328; DI49_0329 | | YDR135C; YDR137W |
| DI49_0331; DI49_0330 | DI49_0331; DI49_0330 | | |
| DI49_0333; YDR141C | DI49_0333 | | YDR141C |
| DI49_0334; YDR142C | DI49_0334 | | YDR142C |
| DI49_0335; YDR143C; DI49_0336; YDR144C | DI49_0335; DI49_0336 | | YDR143C; YDR144C |
| DI49_0338; DI49_0337 | DI49_0338; DI49_0337 | | |
| DI49_0340; YDR148C; SPGP0DN02660; YDR150W | DI49_0340 | SPGP0DN02660 | YDR148C; YDR150W |
| DI49_0342; DI49_0341 | DI49_0342; DI49_0341 | | |
| DI49_0344; YDR155C; DI49_0345; YDR156W | DI49_0344; DI49_0345 | | YDR155C; YDR156W |
| DI49_0347; DI49_0346 | DI49_0347; DI49_0346 | | |
| DI49_0349; DI49_0348 | DI49_0349; DI49_0348 | | |
| DI49_0351; DI49_0350 | DI49_0351; DI49_0350 | | |
| DI49_0353; YDR165W | DI49_0353 | | YDR165W |
| DI49_0354; YDR166C; DI49_0355; YDR167W | DI49_0354; DI49_0355 | | YDR166C; YDR167W |
| DI49_0357; YDR169C | DI49_0357 | | YDR169C |
| DI49_0359; YDR170C | DI49_0359 | | YDR170C |
| SPGP0DNtrna5Q; DI49_2421; DI49_0361; DI49_3679; SPGP0Ptrna7Q; DI49_2420; SPGP0Vtrna9Q; SPGP0AAtrna11Q; SPGP0AEtrna15Q; SPGP0Ctrna12Q; SPGP0Ctrna14Q; YDR170W-A; SPGP0Etrna17Q; DI49_1450; YER048W-A | DI49_2421; DI49_0361; DI49_3679; DI49_2420; DI49_1450 | SPGP0DNtrna5Q; SPGP0Ptrna7Q; SPGP0Vtrna9Q; SPGP0AAtrna11Q; SPGP0AEtrna15Q; SPGP0Ctrna12Q; SPGP0Ctrna14Q; SPGP0Etrna17Q | YDR170W-A; YER048W-A |
| DI49_0363; YDR173C | DI49_0363 | | YDR173C |
| DI49_0364; YDR174W; DI49_0365; YDR175C | DI49_0364; DI49_0365 | | YDR174W; YDR175C |

Create PDF in your applications with the Pdfcrowd HTML to PDF API PDFCROWD

| | | | |
|---|---|---|---|
| DI49_0367; DI49_0366 | DI49_0367; DI49_0366 | | |
| DI49_0369; DI49_0368 | DI49_0369; DI49_0368 | | |
| DI49_0371; YDR180W; DI49_0372; YDR181C | DI49_0371; DI49_0372 | | YDR180W; YDR181C |
| DI49_0374; DI49_0373 | DI49_0374; DI49_0373 | | |
| DI49_0376; YDR185C | DI49_0376 | | YDR185C |
| DI49_0377; YDR186C | DI49_0377 | | YDR186C |
| DI49_0378; YDR188W | DI49_0378 | | YDR188W |
| DI49_0379; YDR189W | DI49_0379 | | YDR189W |
| DI49_0380; YDR190C; DI49_0381; YDR191W | DI49_0380; DI49_0381 | | YDR190C; YDR191W |
| DI49_0382; YDR194C | DI49_0382 | | YDR194C |
| DI49_0383; YDR195W; DI49_0384; YDR196C | DI49_0383; DI49_0384 | | YDR195W; YDR196C |
| DI49_0386; YDR198C; DI49_0387; YDR200C | DI49_0386; DI49_0387 | | YDR198C; YDR200C |
| DI49_0389; YDR202C | DI49_0389 | | YDR202C |
| DI49_0390; YDR204W; DI49_0391; YDR205W | DI49_0390; DI49_0391 | | YDR204W; YDR205W |
| DI49_0393; DI49_0392 | DI49_0393; DI49_0392 | | |
| DI49_0395; SPGP0Ntrna10I; DI49_0396; DI49_3742; SPGP0Ctrna19I; YDR211W | DI49_0395; DI49_0396; DI49_3742 | SPGP0Ntrna10I; SPGP0Ctrna19I | YDR211W |
| DI49_0398; YDR213W; DI49_0399; YDR214W | DI49_0398; DI49_0399 | | YDR213W; YDR214W |
| SPGP0DN03250; DI49_0400 | DI49_0400 | SPGP0DN03250 | |
| DI49_0402; DI49_0401 | DI49_0402; DI49_0401 | | |
| DI49_0404; DI49_0403 | DI49_0404; DI49_0403 | | |
| DI49_0408; YDR226W; DI49_0409; YDR227W | DI49_0408; DI49_0409 | | YDR226W; YDR227W |
| DI49_0411; YDR229W | DI49_0411 | | YDR229W |
| DI49_0412; YDR231C | DI49_0412 | | YDR231C |
| DI49_0413; YDR232W | DI49_0413 | | YDR232W |
| DI49_0414; YDR233C; DI49_0415 | DI49_0414; DI49_0415 | | YDR233C |
| DI49_0417; YDR235W | DI49_0417 | | YDR235W |
| DI49_0418; YDR236C; DI49_0419; YDR237W | DI49_0418; DI49_0419 | | YDR236C; YDR237W |
| DI49_0421; DI49_0420 | DI49_0421; DI49_0420 | | |
| SPGP0DNtrna4Y; DI49_2382; DI49_2381; DI49_3070; DI49_4369; SPGP0Strna9Y; DI49_4037; DI49_1685; DI49_1668 | DI49_2382; DI49_2381; DI49_3070; DI49_4369; DI49_4037; DI49_1685; DI49_1668 | SPGP0DNtrna4Y; SPGP0Strna9Y | |
| SPGP0DN03480; YDR242W; DI49_0424; YDR243C | DI49_0424 | SPGP0DN03480 | YDR242W; YDR243C |
| DI49_0426; YDR245W; DI49_0427; YDR246W | DI49_0426; DI49_0427 | | YDR245W; YDR246W |
| DI49_0429; YDR248C; DI49_0430; YDR249C | DI49_0429; DI49_0430 | | YDR248C; YDR249C |
| DI49_0432; DI49_0431 | DI49_0432; DI49_0431 | | |
| DI49_0434; DI49_0433 | DI49_0434; DI49_0433 | | |
| DI49_0436; YDR256C; DI49_0437; YDR257C | DI49_0436; DI49_0437 | | YDR256C; YDR257C |
| DI49_0439; DI49_0438 | DI49_0439; DI49_0438 | | |
| DI49_0441; DI49_0440 | DI49_0441; DI49_0440 | | |
| DI49_0445; DI49_0444 | DI49_0445; DI49_0444 | | |

Create PDF in your applications with the Pdfcrowd HTML to PDF API PDFCROWD

| | | | |
|---|---|---|---|
| DI49_0447; DI49_0446 | DI49_0447; DI49_0446 | | |
| DI49_0449; YDR268W | DI49_0449 | | YDR268W |
| DI49_0450; YDR270W; DI49_0451; YDR272W | DI49_0450; DI49_0451 | | YDR270W; YDR272W |
| DI49_0453; DI49_0452 | DI49_0453; DI49_0452 | | |
| DI49_0455; YDR277C | DI49_0455 | | YDR277C |
| SPGP0DNtrna6E; DI49_2698; DI49_0457; DI49_2699 | DI49_2698; DI49_0457; DI49_2699 | SPGP0DNtrna6E | |
| DI49_0459; DI49_0458 | DI49_0459; DI49_0458 | | |
| DI49_0461; YDR283C; DI49_0462; YDR284C | DI49_0461; DI49_0462 | | YDR283C; YDR284C |
| DI49_0464; DI49_0463 | DI49_0464; DI49_0463 | | |
| DI49_0466; DI49_0465 | DI49_0466; DI49_0465 | | |
| DI49_0468; YDR291W | DI49_0468 | | YDR291W |
| DI49_0469; YDR292C | DI49_0469 | | YDR292C |
| DI49_0470; YDR293C | DI49_0470 | | YDR293C |
| DI49_0471; YDR294C | DI49_0471 | | YDR294C |
| DI49_0472; YDR295C; DI49_0473; YDR296W | DI49_0472; DI49_0473 | | YDR295C; YDR296W |
| DI49_0475; YDR298C | DI49_0475 | | YDR298C |
| DI49_0476; YDR299W | DI49_0476 | | YDR299W |
| DI49_0477; YDR300C | DI49_0477 | | YDR300C |
| DI49_0478; YDR301W; DI49_0479; YDR302W | DI49_0478; DI49_0479 | | YDR301W; YDR302W |
| DI49_0481; DI49_0480 | DI49_0481; DI49_0480 | | |
| DI49_0483; DI49_0482 | DI49_0483; DI49_0482 | | |
| DI49_0485; DI49_0484 | DI49_0485; DI49_0484 | | |
| DI49_0487; DI49_0486 | DI49_0487; DI49_0486 | | |
| DI49_0489; DI49_0488 | DI49_0489; DI49_0488 | | |
| DI49_0491; DI49_0490 | DI49_0491; DI49_0490 | | |
| DI49_0493; DI49_0492 | DI49_0493; DI49_0492 | | |
| DI49_0495; SPGP0Mtrna5F; DI49_0496 | DI49_0495; DI49_0496 | SPGP0Mtrna5F | |
| DI49_0498; DI49_0497 | DI49_0498; DI49_0497 | | |
| DI49_0500; YDR320C-A | DI49_0500 | | YDR320C-A |
| DI49_0501; YDR321W | DI49_0501 | | YDR321W |
| DI49_0502; YDR322W | DI49_0502 | | YDR322W |
| DI49_0503; YDR322C-A; DI49_0504; YDR323C | DI49_0503; DI49_0504 | | YDR322C-A; YDR323C |
| SPGP0K00100; YDR542W; SPGP0V00110; YHL044W; YEL049W; YEL048C | | SPGP0K00100; SPGP0V00110 | YDR542W; YHL044W; YEL049W; YEL048C |
| YOL159C-A; DI49_3955; YOL159C; DI49_2570 | DI49_3955; DI49_2570 | | YOL159C-A; YOL159C |
| YJL221C; YGR287C; SPGP0Q00130 | | SPGP0Q00130 | YJL221C; YGR287C |
| YOL154W; SPGP0K00170 | | SPGP0K00170 | YOL154W |
| YOL152W; DI49_2257; DI49_2256 | DI49_2257; DI49_2256 | | YOL152W |
| YOL151W; DI49_2258 | DI49_2258 | | YOL151W |
| YOL149W; DI49_2259 | DI49_2259 | | YOL149W |
| YOL148C; DI49_2260 | DI49_2260 | | YOL148C |

Create PDF in your applications with the Pdfcrowd HTML to PDF API PDFCROWD

| | | | |
|---|---|---|---|
| YOL147C; DI49_2261; YOL146W | DI49_2261 | | YOL147C; YOL146W |
| YOL144W; YOL145C | | | YOL144W; YOL145C |
| YOL142W; DI49_2266; YOL141W | DI49_2266 | | YOL142W; YOL141W |
| YOL139C; DI49_2269; YOL138C | DI49_2269 | | YOL139C; YOL138C |
| YOL136C; DI49_2272; YOL135C; DI49_2271 | DI49_2272; DI49_2271 | | YOL136C; YOL135C |
| YOL132W; DI49_2275; YOL131W; DI49_2274 | DI49_2275; DI49_2274 | | YOL132W; YOL131W |
| YOL129W; DI49_2278 | DI49_2278 | | YOL129W |
| YOL128C; DI49_2279 | DI49_2279 | | YOL128C |
| YOL127W; DI49_2280 | DI49_2280 | | YOL127W |
| YOL126C; DI49_2281; YOL125W | DI49_2281 | | YOL126C; YOL125W |
| YOL123W; DI49_2284 | DI49_2284 | | YOL123W |
| YOL122C; DI49_2285 | DI49_2285 | | YOL122C |
| SPGP0K00490; SPGP0M00360; SPGP0K00500; DI49_4447; DI49_2286 | DI49_4447; DI49_2286 | SPGP0K00490; SPGP0M00360; SPGP0K00500 | |
| YOL117W; YOL119C; DI49_2287 | DI49_2287 | | YOL117W; YOL119C |
| YOL115W; DI49_2290; YOL114C; DI49_2289 | DI49_2290; DI49_2289 | | YOL115W; YOL114C |
| YOL112W; YOL113W | | | YOL112W; YOL113W |
| YOL110W; DI49_2295; YOL109W; DI49_2294 | DI49_2295; DI49_2294 | | YOL110W; YOL109W |
| YOL108C; SPGP0Ftrna34G; SPGP0Ptrna12G; DI49_2296; SPGP0Qtrna15G; SPGP0Ztrna22G; DI49_4461 | DI49_2296; DI49_4461 | SPGP0Ftrna34G; SPGP0Ptrna12G; SPGP0Qtrna15G; SPGP0Ztrna22G | YOL108C |
| SPGP0K00630; YOL107W | | SPGP0K00630 | YOL107W |
| YOL103W; YOL104C | | | YOL103W; YOL104C |
| YOL101C; DI49_2305 | DI49_2305 | | YOL101C |
| YOL100W; DI49_2306 | DI49_2306 | | YOL100W |
| YOL098C; DI49_2307 | DI49_2307 | | YOL098C |
| YOL097C; DI49_2308 | DI49_2308 | | YOL097C |
| YOL096C; DI49_2309 | DI49_2309 | | YOL096C |
| YOL095C; DI49_2310 | DI49_2310 | | YOL095C |
| YOL094C; DI49_2311; YOL093W | DI49_2311 | | YOL094C; YOL093W |
| YOL091W; DI49_2314; DI49_2313 | DI49_2314; DI49_2313 | | YOL091W |
| YOL090W; DI49_2315 | DI49_2315 | | YOL090W |
| YOL089C; DI49_2316; YOL088C | DI49_2316 | | YOL089C; YOL088C |
| YOL086W-A; YOL087C | | | YOL086W-A; YOL087C |
| YOL084W; YOL086C | | | YOL084W; YOL086C |
| YOL082W; YOL083W | | | YOL082W; YOL083W |
| YOL080C; YOL081W; DI49_2324; DI49_2323 | DI49_2324; DI49_2323 | | YOL080C; YOL081W |
| YOL077W-A; YOL078W | | | YOL077W-A; YOL078W |
| YOL076W; DI49_2329; DI49_2328 | DI49_2329; DI49_2328 | | YOL076W |
| YOL075C; DI49_2330; YOL073C | DI49_2330 | | YOL075C; YOL073C |
| YOL071W; YOL072W; DI49_2332; DI49_2331 | DI49_2332; DI49_2331 | | YOL071W; YOL072W |
| YOL069W; DI49_2335; DI49_2334 | DI49_2335; DI49_2334 | | YOL069W |

Create PDF in your applications with the Pdfcrowd HTML to PDF API PDFCROWD

| | | | |
|---|---|---|---|
| YOL068C; DI49_2336; YOL067C | DI49_2336 | | YOL068C; YOL067C |
| YOL065C; YOL066C | | | YOL065C; YOL066C |
| YOL063C; YOL064C | | | YOL063C; YOL064C |
| YOL061W; DI49_2343 | DI49_2343 | | YOL061W |
| YOL060C; DI49_2344 | DI49_2344 | | YOL060C |
| YOL059W; DI49_2345 | DI49_2345 | | YOL059W |
| YOL058W; DI49_2346; YOL057W | DI49_2346 | | YOL058W; YOL057W |
| YOL055C; DI49_2348; SPGP0P02270; YPL258C; YPL259C | DI49_2348 | SPGP0P02270 | YOL055C; YPL258C; YPL259C |
| YOL052C-A; YOL053W | | | YOL052C-A; YOL053W |
| YOL051W; DI49_2355; YOL049W | DI49_2355 | | YOL051W; YOL049W |
| YOL047C; YOL048C | | | YOL047C; YOL048C |
| YOL045W; DI49_2357; YOL044W; SPGP0P02180 | DI49_2357 | SPGP0P02180 | YOL045W; YOL044W |
| YOL042W; YOL043C | | | YOL042W; YOL043C |
| YOL040C; DI49_2362; DI49_2361 | DI49_2362; DI49_2361 | | YOL040C |
| YOL039W; DI49_2363; YOL038W | DI49_2363 | | YOL039W; YOL038W |
| YOL034W; DI49_2366; YOL033W; DI49_2365 | DI49_2366; DI49_2365 | | YOL034W; YOL033W |
| YOL031C; YOL032W | | | YOL031C; YOL032W |
| YOL028C; YOL030W | | | YOL028C; YOL030W |
| YOL026C; YOL027C | | | YOL026C; YOL027C |
| YOL025W; DI49_2374; SPGP0Ntrna9S; DI49_3829; SPGP0EXtrna10S; DI49_0521; SPGP0Ptrna11S; DI49_2373; DI49_1676 | DI49_2374; DI49_3829; DI49_0521; DI49_2373; DI49_1676 | SPGP0Ntrna9S; SPGP0EXtrna10S; SPGP0Ptrna11S | YOL025W |
| YOL023W; DI49_2377; YOL022C; DI49_2376 | DI49_2377; DI49_2376 | | YOL023W; YOL022C |
| YOL021C; DI49_2380; YOL020W | DI49_2380 | | YOL021C; YOL020W |
| YOL017W; YOL018C | | | YOL017W; YOL018C |
| YOL015W; YOL016C; DI49_2386; DI49_2385 | DI49_2386; DI49_2385 | | YOL015W; YOL016C |
| YOL013W-A; SPGP0Ktrna4P | | SPGP0Ktrna4P | YOL013W-A |
| YOL013C; DI49_2390 | DI49_2390 | | YOL013C |
| YOL012C; DI49_2391 | DI49_2391 | | YOL012C |
| YOL011W; DI49_2392; YOL010W | DI49_2392 | | YOL011W; YOL010W |
| YOL008W; YOL009C; DI49_2394; DI49_2393 | DI49_2394; DI49_2393 | | YOL008W; YOL009C |
| YOL006C; YOL007C | | | YOL006C; YOL007C |
| YOL004W; DI49_2399; DI49_2398 | DI49_2399; DI49_2398 | | YOL004W |
| YOL003C; DI49_2400 | DI49_2400 | | YOL003C |
| YOL002C; DI49_2401; YOL001W | DI49_2401 | | YOL002C; YOL001W |
| YOR002W; DI49_2404 | DI49_2404 | | YOR002W |
| YOR003W; DI49_2405; YOR004W | DI49_2405 | | YOR003W; YOR004W |
| YOR006C; DI49_2408 | DI49_2408 | | YOR006C |
| YOR007C; DI49_2409 | DI49_2409 | | YOR007C |
| YOR010C; DI49_2413; YER011W | DI49_2413 | | YOR010C; YER011W |
| YOR011W; DI49_2414 | DI49_2414 | | YOR011W |

Create PDF in your applications with the Pdfcrowd HTML to PDF API PDFCROWD

| | | | |
|---|---|---|---|
| YOR016C; DI49_2418 | DI49_2418 | | YOR016C |
| YOR017W; DI49_2419; YOR018W | DI49_2419 | | YOR017W; YOR018W |
| YOR020C; DI49_4866; YOR020W-A; DI49_4865 | DI49_4866; DI49_4865 | | YOR020C; YOR020W-A |
| YOR022C; YOR021C | | | YOR022C; YOR021C |
| YOR025W; YOR023C | | | YOR025W; YOR023C |
| YOR027W; YOR026W | | | YOR027W; YOR026W |
| YOR030W; YOR028C | | | YOR030W; YOR028C |
| YOR032C; YOR031W | | | YOR032C; YOR031W |
| YOR034C; YOR033C | | | YOR034C; YOR033C |
| YOR036W; YOR035C | | | YOR036W; YOR035C |
| YOR038C; DI49_4883 | DI49_4883 | | YOR038C |
| YOR039W; DI49_4884; YOR040W | DI49_4884 | | YOR039W; YOR040W |
| YOR043W; DI49_4887; YOR044W; DI49_4886 | DI49_4887; DI49_4886 | | YOR043W; YOR044W |
| YOR046C; DI49_4890; DI49_4889 | DI49_4890; DI49_4889 | | YOR046C |
| YOR047C; DI49_4891 | DI49_4891 | | YOR047C |
| YOR048C; DI49_4892; YOR049C | DI49_4892 | | YOR048C; YOR049C |
| YOR052C; YOR051C | | | YOR052C; YOR051C |
| YOR056C; YOR054C | | | YOR056C; YOR054C |
| YOR058C; DI49_4899 | DI49_4899 | | YOR058C |
| SPGP0Ktrna5K; SPGP0Ctrna22K; YOR059C; SPGP0Qtrna12K; YGL203C; YDL054C | | SPGP0Ktrna5K; SPGP0Ctrna22K; SPGP0Qtrna12K | YOR059C; YGL203C; YDL054C |
| YOR061W; YOR060C | | | YOR061W; YOR060C |
| YOR063W; DI49_4906; YOR064C; DI49_4905 | DI49_4906; DI49_4905 | | YOR063W; YOR064C |
| YOR066W; DI49_4909 | DI49_4909 | | YOR066W |
| YOR067C; DI49_4910 | DI49_4910 | | YOR067C |
| YOR069W; DI49_4911 | DI49_4911 | | YOR069W |
| YOR070C; DI49_4912 | DI49_4912 | | YOR070C |
| YOR071C; DI49_3747; DI49_3867; DI49_3866; DI49_3865; DI49_3746 | DI49_3747; DI49_3867; DI49_3866; DI49_3865; DI49_3746 | | YOR071C |
| YOR073W; DI49_4915 | DI49_4915 | | YOR073W |
| YOR074C; SPGP0L00630 | | SPGP0L00630 | YOR074C |
| YOR075W; DI49_4916; YOR076C | DI49_4916 | | YOR075W; YOR076C |
| YOR078W; YOR077W | | | YOR078W; YOR077W |
| YOR080W; YOR079C | | | YOR080W; YOR079C |
| YOR083W; YOR081C | | | YOR083W; YOR081C |
| YOR085W; YOR084W | | | YOR085W; YOR084W |
| YOR087W; DI49_4926; YOR089C | DI49_4926 | | YOR087W; YOR089C |
| YOR091W; YOR090C | | | YOR091W; YOR090C |
| YOR093C; DI49_4931; YOR094W | DI49_4931 | | YOR093C; YOR094W |
| YOR096W; DI49_4934; YOR097C; SPGP0M02200; SPGP0M02210 | DI49_4934 | SPGP0M02200; SPGP0M02210 | YOR096W; YOR097C |
| YOR099W; DI49_4937; YOR100C; DI49_4936 | DI49_4937; DI49_4936 | | YOR099W; YOR100C |

Create PDF in your applications with the Pdfcrowd HTML to PDF API PDFCROWD

| | | | |
|---|---|---|---|
| YOR103C; DI49_4940; YOR104W; DI49_4939 | DI49_4940; DI49_4939 | | YOR103C; YOR104W |
| YOR107W; DI49_4943; DI49_4942 | DI49_4943; DI49_4942 | | YOR107W |
| YOR108W; DI49_4944; YNL104C; YNL103W | DI49_4944 | | YOR108W; YNL104C; YNL103W |
| YOR109W; DI49_4945; YOR110W | DI49_4945 | | YOR109W; YOR110W |
| YOR112W; DI49_4948; DI49_4947 | DI49_4948; DI49_4947 | | YOR112W |
| YOR113W; DI49_4949 | DI49_4949 | | YOR113W |
| YOR114W; DI49_4950; YOR115C | DI49_4950 | | YOR114W; YOR115C |
| YOR117W; DI49_4953; YOR118W | DI49_4953 | | YOR117W; YOR118W |
| YOR120W; YOR119C | | | YOR120W; YOR119C |
| YOR123C; YOR122C | | | YOR123C; YOR122C |
| YOR125C; DI49_4960; YOR126C | DI49_4960 | | YOR125C; YOR126C |
| YOR128C; DI49_4963; YOR129C | DI49_4963 | | YOR128C; YOR129C |
| SPGP0K02830; DI49_0561; DI49_0562 | DI49_0561; DI49_0562 | SPGP0K02830 | |
| SPGP0DP00100; SPGP0FH00100; DI49_1599; SPGP0FI00100 | DI49_1599 | SPGP0DP00100; SPGP0FH00100; SPGP0FI00100 | |
| YNL333W; DI49_4016; DI49_2435 | DI49_4016; DI49_2435 | | YNL333W |
| DI49_3677; DI49_3678 | DI49_3677; DI49_3678 | | |
| DI49_3675; DI49_3676 | DI49_3675; DI49_3676 | | |
| DI49_3673; DI49_3674 | DI49_3673; DI49_3674 | | |
| DI49_3671; DI49_3672 | DI49_3671; DI49_3672 | | |
| DI49_3669; DI49_3670 | DI49_3669; DI49_3670 | | |
| DI49_3667; DI49_3668 | DI49_3667; DI49_3668 | | |
| DI49_3662; SPGP0DU00250 | DI49_3662 | SPGP0DU00250 | |
| DI49_3660; DI49_3661 | DI49_3660; DI49_3661 | | |
| DI49_3658; DI49_3659 | DI49_3658; DI49_3659 | | |
| DI49_3656; DI49_3657; YGR087C; YGR086C | DI49_3656; DI49_3657 | | YGR087C; YGR086C |
| DI49_3654; DI49_3655 | DI49_3654; DI49_3655 | | |
| DI49_3652; DI49_3653 | DI49_3652; DI49_3653 | | |
| DI49_3650; DI49_3651 | DI49_3650; DI49_3651 | | |
| DI49_3648; DI49_3649 | DI49_3648; DI49_3649 | | |
| DI49_3646; DI49_3647 | DI49_3646; DI49_3647 | | |
| DI49_3644; DI49_3645 | DI49_3644; DI49_3645 | | |
| DI49_3642; DI49_3643 | DI49_3642; DI49_3643 | | |
| DI49_3640; DI49_3641 | DI49_3640; DI49_3641 | | |
| DI49_3636; DI49_3637 | DI49_3636; DI49_3637 | | |
| DI49_3634; DI49_3635 | DI49_3634; DI49_3635 | | |
| DI49_3632; DI49_3633 | DI49_3632; DI49_3633 | | |
| DI49_3630; DI49_3631 | DI49_3630; DI49_3631 | | |
| DI49_3628; DI49_3629 | DI49_3628; DI49_3629 | | |
| DI49_3626; DI49_3627 | DI49_3626; DI49_3627 | | |

Create PDF in your applications with the Pdfcrowd HTML to PDF API PDFCROWD

| | | | |
|---|---|---|---|
| DI49_3625; SPGP0DU00620 | DI49_3625 | SPGP0DU00620 | |
| DI49_3623; DI49_3624 | DI49_3623; DI49_3624 | | |
| DI49_3621; DI49_3622 | DI49_3621; DI49_3622 | | |
| DI49_3619; DI49_3620 | DI49_3619; DI49_3620 | | |
| DI49_3617; DI49_3618 | DI49_3617; DI49_3618 | | |
| DI49_3615; DI49_3616 | DI49_3615; DI49_3616 | | |
| DI49_3613; DI49_3614 | DI49_3613; DI49_3614 | | |
| DI49_3611; DI49_3612 | DI49_3611; DI49_3612 | | |
| DI49_3609; DI49_3610 | DI49_3609; DI49_3610 | | |
| DI49_3606; DI49_3607 | DI49_3606; DI49_3607 | | |
| DI49_3604; DI49_3605 | DI49_3604; DI49_3605 | | |
| DI49_3602; DI49_3603 | DI49_3602; DI49_3603 | | |
| DI49_3600; DI49_3601 | DI49_3600; DI49_3601 | | |
| DI49_3598; DI49_3599 | DI49_3598; DI49_3599 | | |
| DI49_3596; DI49_3597 | DI49_3596; DI49_3597 | | |
| DI49_3594; DI49_3595 | DI49_3594; DI49_3595 | | |
| DI49_3592; DI49_3593 | DI49_3592; DI49_3593 | | |
| DI49_3590; DI49_3591 | DI49_3590; DI49_3591 | | |
| DI49_3588; DI49_3589 | DI49_3588; DI49_3589 | | |
| DI49_3586; DI49_3587 | DI49_3586; DI49_3587 | | |
| DI49_3584; DI49_3585 | DI49_3584; DI49_3585 | | |
| DI49_3582; DI49_3583 | DI49_3582; DI49_3583 | | |
| DI49_3580; DI49_3581 | DI49_3580; DI49_3581 | | |
| DI49_3578; DI49_3579 | DI49_3578; DI49_3579 | | |
| SPGP0DU01110; YGR214W; DI49_3575 | DI49_3575 | SPGP0DU01110 | YGR214W |
| DI49_3573; DI49_3574 | DI49_3573; DI49_3574 | | |
| DI49_3571; DI49_3572 | DI49_3571; DI49_3572 | | |
| DI49_3569; DI49_3570 | DI49_3569; DI49_3570 | | |
| DI49_3567; DI49_3568 | DI49_3567; DI49_3568 | | |
| DI49_3565; DI49_3566; YPL282C; DI49_0506; DI49_1345; DI49_1346; DI49_1756; DI49_1755 | DI49_3565; DI49_3566; DI49_0506; DI49_1345; DI49_1346; DI49_1756; DI49_1755 | | YPL282C |
| DI49_3561; DI49_3562 | DI49_3561; DI49_3562 | | |
| DI49_3559; DI49_3560 | DI49_3559; DI49_3560 | | |
| DI49_3555; DI49_3556 | DI49_3555; DI49_3556 | | |
| DI49_3553; DI49_3554 | DI49_3553; DI49_3554 | | |
| DI49_3551; DI49_3552 | DI49_3551; DI49_3552 | | |
| DI49_3549; DI49_3550 | DI49_3549; DI49_3550 | | |
| DI49_3547; DI49_3548 | DI49_3547; DI49_3548 | | |
| DI49_3545; DI49_3546 | DI49_3545; DI49_3546 | | |
| DI49_3543; DI49_3544 | DI49_3543; DI49_3544 | | |

Create PDF in your applications with the Pdfcrowd HTML to PDF API PDFCROWD

| | | | |
|---|---|---|---|
| DI49_3541; DI49_3542 | DI49_3541; DI49_3542 | | |
| DI49_3539; DI49_3540 | DI49_3539; DI49_3540 | | |
| DI49_3537; SPGP0DUtrna8S | DI49_3537 | SPGP0DUtrna8S | |
| DI49_3535; DI49_3536 | DI49_3535; DI49_3536 | | |
| DI49_3533; DI49_3534 | DI49_3533; DI49_3534 | | |
| DI49_3531; DI49_3532 | DI49_3531; DI49_3532 | | |
| YLR002C; DI49_3530 | DI49_3530 | | YLR002C |
| DI49_3528; DI49_3529 | DI49_3528; DI49_3529 | | |
| DI49_3526; DI49_3527 | DI49_3526; DI49_3527 | | |
| DI49_3524; DI49_3525 | DI49_3524; DI49_3525 | | |
| DI49_3522; DI49_3523 | DI49_3522; DI49_3523 | | |
| DI49_3520; DI49_3521 | DI49_3520; DI49_3521 | | |
| DI49_3518; DI49_3519 | DI49_3518; DI49_3519 | | |
| DI49_3516; DI49_3517 | DI49_3516; DI49_3517 | | |
| DI49_3514; DI49_3515 | DI49_3514; DI49_3515 | | |
| DI49_3512; DI49_3513 | DI49_3512; DI49_3513 | | |
| DI49_3510; DI49_3511 | DI49_3510; DI49_3511 | | |
| DI49_4740; YNR004W; DI49_4741; YNR006W | DI49_4740; DI49_4741 | | YNR004W; YNR006W |
| DI49_4743; DI49_4742 | DI49_4743; DI49_4742 | | |
| DI49_4745; DI49_4744 | DI49_4745; DI49_4744 | | |
| DI49_4747; YNR012W | DI49_4747 | | YNR012W |
| DI49_4748; YNR013C; DI49_4749; YNR014W | DI49_4748; DI49_4749 | | YNR013C; YNR014W |
| DI49_4751; YNR016C | DI49_4751 | | YNR016C |
| DI49_4752; YNR017W; DI49_4753; YNR018W | DI49_4752; DI49_4753 | | YNR017W; YNR018W |
| DI49_4755; DI49_4754; DI49_4756 | DI49_4755; DI49_4754; DI49_4756 | | |
| DI49_4758; DI49_4757 | DI49_4758; DI49_4757 | | |
| SPGP0DV00330; DI49_4759 | DI49_4759 | SPGP0DV00330 | |
| DI49_4761; DI49_4760 | DI49_4761; DI49_4760 | | |
| DI49_4763; YNR031C; DI49_4764; YNR032W | DI49_4763; DI49_4764 | | YNR031C; YNR032W |
| DI49_4766; DI49_4765 | DI49_4766; DI49_4765 | | |
| DI49_4768; DI49_4767 | DI49_4768; DI49_4767 | | |
| DI49_4770; DI49_4769 | DI49_4770; DI49_4769 | | |
| DI49_4772; DI49_4771 | DI49_4772; DI49_4771 | | |
| DI49_4774; DI49_4773 | DI49_4774; DI49_4773 | | |
| DI49_4776; DI49_4775 | DI49_4776; DI49_4775 | | |
| DI49_4778; DI49_4777 | DI49_4778; DI49_4777 | | |
| DI49_4780; YNR047W; DI49_4781; YNR048W; YCR091W; YCR092C | DI49_4780; DI49_4781 | | YNR047W; YNR048W; YCR091W; YCR092C |
| DI49_4783; YNR050C; DI49_4784; YNR051C | DI49_4783; DI49_4784 | | YNR050C; YNR051C |
| SPGP0DV00600; YNR053C | | SPGP0DV00600 | YNR053C |
| DI49_4786; YNR054C; DI49_4787 | DI49_4786; DI49_4787 | | YNR054C |

Create PDF in your applications with the Pdfcrowd HTML to PDF API PDFCROWD

| | | | |
|---|---|---|---|
| DI49_4790; YNR055C; DI49_4791; YNR056C | DI49_4790; DI49_4791 | | YNR055C; YNR056C |
| DI49_4793; DI49_4792 | DI49_4793; DI49_4792 | | |
| YCR098C; DI49_4420; YCR100C | DI49_4420 | | YCR098C; YCR100C |
| DI49_3960; YCR102C | DI49_3960 | | YCR102C |
| DI49_3962; DI49_3961 | DI49_3962; DI49_3961 | | |
| DI49_3964; SPGP0DX00130 | DI49_3964 | SPGP0DX00130 | |
| DI49_3966; DI49_3965 | DI49_3966; DI49_3965 | | |
| DI49_3968; DI49_3967 | DI49_3968; DI49_3967 | | |
| DI49_3970; DI49_3969; DI49_4005; DI49_4006 | DI49_3970; DI49_3969; DI49_4005; DI49_4006 | | |
| DI49_3972; DI49_3971 | DI49_3972; DI49_3971 | | |
| DI49_3974; DI49_3973 | DI49_3974; DI49_3973 | | |
| DI49_3976; DI49_3975 | DI49_3976; DI49_3975 | | |
| DI49_3978; DI49_3977 | DI49_3978; DI49_3977 | | |
| DI49_3980; DI49_3979 | DI49_3980; DI49_3979 | | |
| DI49_3982; DI49_3981 | DI49_3982; DI49_3981 | | |
| DI49_3984; DI49_3983 | DI49_3984; DI49_3983 | | |
| DI49_3986; DI49_3985 | DI49_3986; DI49_3985 | | |
| DI49_3988; DI49_3987 | DI49_3988; DI49_3987 | | |
| DI49_3990; DI49_3989 | DI49_3990; DI49_3989 | | |
| DI49_3992; DI49_3991 | DI49_3992; DI49_3991 | | |
| DI49_3993; SPGP0DX00430 | DI49_3993 | SPGP0DX00430 | |
| DI49_3995; DI49_3994 | DI49_3995; DI49_3994 | | |
| DI49_3997; DI49_3996 | DI49_3997; DI49_3996 | | |
| DI49_3999; DI49_3998 | DI49_3999; DI49_3998 | | |
| DI49_4001; DI49_4000 | DI49_4001; DI49_4000 | | |
| DI49_4003; DI49_4002 | DI49_4003; DI49_4002 | | |
| DI49_4008; DI49_4007 | DI49_4008; DI49_4007 | | |
| DI49_4010; DI49_4009 | DI49_4010; DI49_4009 | | |
| YML078W; DI49_4011 | DI49_4011 | | YML078W |
| DI49_4013; DI49_4012 | DI49_4013; DI49_4012 | | |
| DI49_4015; DI49_4014 | DI49_4015; DI49_4014 | | |
| YNL335W; YNL334C | | | YNL335W; YNL334C |
| DI49_4968; YOR132W; DI49_4967 | DI49_4968; DI49_4967 | | YOR132W |
| DI49_4965; YOR130C; DI49_4964 | DI49_4965; DI49_4964 | | YOR130C |
| DI49_4962; YOR127W; DI49_4961 | DI49_4962; DI49_4961 | | YOR127W |
| DI49_4959; YOR124C; DI49_4958 | DI49_4959; DI49_4958 | | YOR124C |
| DI49_4956; DI49_4957 | DI49_4956; DI49_4957 | | |
| DI49_4954; DI49_4955 | DI49_4954; DI49_4955 | | |
| DI49_4952; YOR116C; DI49_4951 | DI49_4952; DI49_4951 | | YOR116C |
| DI49_4933; YOR095C; DI49_4932 | DI49_4933; DI49_4932 | | YOR095C |

Create PDF in your applications with the Pdfcrowd HTML to PDF API PDFCROWD

| | | | |
|---|---|---|---|
| DI49_4930; YOR092W; DI49_4929 | DI49_4930; DI49_4929 | | YOR092W |
| DI49_4927; DI49_4928 | DI49_4927; DI49_4928 | | |
| DI49_4924; YOR086C; DI49_4923 | DI49_4924; DI49_4923 | | YOR086C |
| SPGP0L00550; DI49_4922 | DI49_4922 | SPGP0L00550 | |
| DI49_4921; SPGP0L00560 | DI49_4921 | SPGP0L00560 | |
| DI49_4919; DI49_4920 | DI49_4919; DI49_4920 | | |
| DI49_4917; DI49_4918 | DI49_4917; DI49_4918 | | |
| DI49_4908; YOR065W; DI49_4907 | DI49_4908; DI49_4907 | | YOR065W |
| DI49_4903; DI49_4904 | DI49_4903; DI49_4904 | | |
| DI49_4901; DI49_4902 | DI49_4901; DI49_4902 | | |
| DI49_4898; YOR057W; DI49_4897 | DI49_4898; DI49_4897 | | YOR057W |
| DI49_4895; DI49_4896 | DI49_4895; DI49_4896 | | |
| DI49_4893; DI49_4894 | DI49_4893; DI49_4894 | | |
| DI49_4882; YOR037W; DI49_4881 | DI49_4882; DI49_4881 | | YOR037W |
| DI49_4879; DI49_4880 | DI49_4879; DI49_4880 | | |
| DI49_4877; DI49_4878 | DI49_4877; DI49_4878 | | |
| DI49_4875; DI49_4876 | DI49_4875; DI49_4876 | | |
| DI49_4873; DI49_4874 | DI49_4873; DI49_4874 | | |
| DI49_4871; DI49_4872 | DI49_4871; DI49_4872 | | |
| DI49_4869; DI49_4870 | DI49_4869; DI49_4870 | | |
| DI49_4867; DI49_4868 | DI49_4867; DI49_4868 | | |
| DI49_4861; YHR012W; DI49_4860 | DI49_4861; DI49_4860 | | YHR012W |
| DI49_4859; YHR009C | DI49_4859 | | YHR009C |
| DI49_4858; YHR008C | DI49_4858 | | YHR008C |
| DI49_4857; YHR007C; DI49_4856 | DI49_4857; DI49_4856 | | YHR007C |
| DI49_4852; DI49_4853 | DI49_4852; DI49_4853 | | |
| DI49_4850; YHR002W; DI49_4849 | DI49_4850; DI49_4849 | | YHR002W |
| DI49_4846; YHL002W | DI49_4846 | | YHL002W |
| DI49_4845; YHL003C; DI49_4844 | DI49_4845; DI49_4844 | | YHL003C |
| DI49_4842; YHL007C; YHL006C | DI49_4842 | | YHL007C; YHL006C |
| DI49_4841; YHL008C | DI49_4841 | | YHL008C |
| DI49_4837; YHL011C; DI49_4836 | DI49_4837; DI49_4836 | | YHL011C |
| DI49_4834; DI49_4835 | DI49_4834; DI49_4835 | | |
| DI49_4832; YHL016C | DI49_4832 | | YHL016C |
| DI49_4831; YHL017W; DI49_4830 | DI49_4831; DI49_4830 | | YHL017W |
| DI49_4828; YHL020C; DI49_4827; YHL019C | DI49_4828; DI49_4827 | | YHL020C; YHL019C |
| DI49_4824; YHL023C | DI49_4824 | | YHL023C |
| DI49_4823; YHL024W; DI49_4822 | DI49_4823; DI49_4822 | | YHL024W |
| DI49_4820; DI49_4821 | DI49_4820; DI49_4821 | | |
| DI49_4818; YHL029C; YHL028W | DI49_4818 | | YHL029C; YHL028W |

Create PDF in your applications with the Pdfcrowd HTML to PDF API PDFCROWD

| | | | |
|---|---|---|---|
| DI49_4817; YHL030W | DI49_4817 | | YHL030W |
| DI49_4816; YHL031C | DI49_4816 | | YHL031C |
| SPGP0L01580; YHL032C | | SPGP0L01580 | YHL032C |
| SPGP0L01590; DI49_3489; YHL033C; DI49_3490 | DI49_3489; DI49_3490 | SPGP0L01590 | YHL033C |
| DI49_4814; YHL034C; DI49_4813 | DI49_4814; DI49_4813 | | YHL034C |
| DI49_4811; DI49_4812 | DI49_4811; DI49_4812 | | |
| DI49_4809; DI49_4810 | DI49_4809; DI49_4810 | | |
| DI49_4970; DI49_4808 | DI49_4970; DI49_4808 | | |
| DI49_4972; DI49_4971 | DI49_4972; DI49_4971 | | |
| DI49_4974; DI49_4973 | DI49_4974; DI49_4973 | | |
| DI49_4976; DI49_4975 | DI49_4976; DI49_4975 | | |
| DI49_4978; DI49_4977; DI49_0567; DI49_0568 | DI49_4978; DI49_4977; DI49_0567; DI49_0568 | | |
| DI49_4980; DI49_4979 | DI49_4980; DI49_4979 | | |
| DI49_4982; DI49_4981 | DI49_4982; DI49_4981 | | |
| DI49_4984; DI49_4983 | DI49_4984; DI49_4983 | | |
| DI49_4986; DI49_4985 | DI49_4986; DI49_4985 | | |
| DI49_4988; DI49_4987; YDR406W; DI49_0582 | DI49_4988; DI49_4987; DI49_0582 | | YDR406W |
| DI49_4990; DI49_4989 | DI49_4990; DI49_4989 | | |
| DI49_4992; DI49_4991 | DI49_4992; DI49_4991 | | |
| DI49_4994; DI49_4993 | DI49_4994; DI49_4993 | | |
| DI49_4996; DI49_4995 | DI49_4996; DI49_4995 | | |
| DI49_4998; DI49_4997 | DI49_4998; DI49_4997 | | |
| DI49_5000; DI49_4999 | DI49_5000; DI49_4999 | | |
| DI49_5002; DI49_3774; DI49_5003; DI49_3773 | DI49_5002; DI49_3774; DI49_5003; DI49_3773 | | |
| DI49_5005; DI49_5004 | DI49_5005; DI49_5004 | | |
| DI49_5007; DI49_5006 | DI49_5007; DI49_5006 | | |
| DI49_5009; DI49_5008 | DI49_5009; DI49_5008 | | |
| DI49_5013; DI49_5012 | DI49_5013; DI49_5012 | | |
| DI49_5015; DI49_5014; DI49_5016 | DI49_5015; DI49_5014; DI49_5016 | | |
| DI49_5018; DI49_5017; DI49_3800; DI49_3799 | DI49_5018; DI49_5017; DI49_3800; DI49_3799 | | |
| SPGP0DZ00590; DI49_5019 | DI49_5019 | SPGP0DZ00590 | |
| DI49_5021; DI49_5020 | DI49_5021; DI49_5020 | | |
| SPGP0DZtrna2M; DI49_5351; DI49_5024; DI49_3051; DI49_5350; SPGP0AAtrna10M; YJR043C; SPGP0Gtrna13M | DI49_5351; DI49_5024; DI49_3051; DI49_5350 | SPGP0DZtrna2M; SPGP0AAtrna10M; SPGP0Gtrna13M | YJR043C |
| DI49_5026; DI49_5025 | DI49_5026; DI49_5025 | | |
| DI49_5028; DI49_5027 | DI49_5028; DI49_5027 | | |
| DI49_5030; DI49_5029 | DI49_5030; DI49_5029 | | |
| DI49_5032; SPGP0Z01180; DI49_5033; YNL112W; DI49_4621 | DI49_5032; DI49_5033; DI49_4621 | SPGP0Z01180 | YNL112W |
| DI49_5035; DI49_5034 | DI49_5035; DI49_5034 | | |
| DI49_5037; DI49_5036 | DI49_5037; DI49_5036 | | |

Create PDF in your applications with the Pdfcrowd HTML to PDF API PDFCROWD

| | | |
|---|---|---|
| DI49_5039; DI49_5038 | DI49_5039; DI49_5038 | |
| DI49_5041; DI49_5040 | DI49_5041; DI49_5040 | |
| DI49_5043; DI49_5042 | DI49_5043; DI49_5042 | |
| DI49_5045; DI49_5044 | DI49_5045; DI49_5044 | |
| DI49_5046; SPGP0DZ00850 | DI49_5046 | SPGP0DZ00850 |
| DI49_5048; DI49_5047 | DI49_5048; DI49_5047 | |
| DI49_5050; DI49_5049 | DI49_5050; DI49_5049 | |
| DI49_5052; DI49_5051 | DI49_5052; DI49_5051 | |
| DI49_5054; DI49_5053 | DI49_5054; DI49_5053 | |
| DI49_5056; DI49_5055 | DI49_5056; DI49_5055 | |
| DI49_5058; DI49_5057 | DI49_5058; DI49_5057 | |
| DI49_5060; DI49_5059; DI49_5322; DI49_5321 | DI49_5060; DI49_5059; DI49_5322; DI49_5321 | |
| DI49_5062; DI49_5061 | DI49_5062; DI49_5061 | |
| DI49_5063; SPGP0DZ01030 | DI49_5063 | SPGP0DZ01030 |
| DI49_5065; DI49_5064 | DI49_5065; DI49_5064 | |
| DI49_5067; DI49_5066 | DI49_5067; DI49_5066 | |
| DI49_5069; DI49_5068 | DI49_5069; DI49_5068 | |
| DI49_5071; DI49_5070 | DI49_5071; DI49_5070 | |
| DI49_5073; DI49_5072 | DI49_5073; DI49_5072 | |
| DI49_5075; DI49_5074 | DI49_5075; DI49_5074 | |
| DI49_5077; DI49_5076 | DI49_5077; DI49_5076 | |
| DI49_5079; DI49_5078 | DI49_5079; DI49_5078 | |
| DI49_5081; DI49_5080 | DI49_5081; DI49_5080 | |
| DI49_5083; DI49_5082 | DI49_5083; DI49_5082 | |
| DI49_5085; DI49_5084 | DI49_5085; DI49_5084 | |
| DI49_5087; DI49_5086 | DI49_5087; DI49_5086 | |
| DI49_5089; DI49_5088 | DI49_5089; DI49_5088 | |
| DI49_5091; DI49_5090 | DI49_5091; DI49_5090 | |
| DI49_5093; DI49_5092 | DI49_5093; DI49_5092 | |
| DI49_5095; DI49_5094 | DI49_5095; DI49_5094 | |
| DI49_5097; DI49_5096 | DI49_5097; DI49_5096 | |
| DI49_5099; DI49_5098 | DI49_5099; DI49_5098 | |
| DI49_5101; DI49_5100 | DI49_5101; DI49_5100 | |
| DI49_5103; DI49_5102 | DI49_5103; DI49_5102 | |
| DI49_5105; DI49_5104 | DI49_5105; DI49_5104 | |
| DI49_5107; DI49_5106 | DI49_5107; DI49_5106 | |
| DI49_5109; DI49_2660; DI49_5110 | DI49_5109; DI49_2660; DI49_5110 | |
| DI49_5115; DI49_5114 | DI49_5115; DI49_5114 | |
| DI49_5117; DI49_5116 | DI49_5117; DI49_5116 | |
| DI49_5119; DI49_5118 | DI49_5119; DI49_5118 | |

Create PDF in your applications with the Pdfcrowd HTML to PDF API PDFCROWD

| | | | |
|---|---|---|---|
| YOR302W; DI49_5120 | DI49_5120 | | YOR302W |
| DI49_5123; YGL150C; DI49_5124 | DI49_5123; DI49_5124 | | YGL150C |
| DI49_5126; DI49_5125 | DI49_5126; DI49_5125 | | |
| DI49_5128; DI49_5127 | DI49_5128; DI49_5127 | | |
| DI49_5130; DI49_5129 | DI49_5130; DI49_5129 | | |
| DI49_5134; DI49_5133 | DI49_5134; DI49_5133 | | |
| DI49_5136; DI49_5135 | DI49_5136; DI49_5135 | | |
| DI49_5138; DI49_5137 | DI49_5138; DI49_5137 | | |
| DI49_5140; DI49_5139 | DI49_5140; DI49_5139 | | |
| DI49_5142; DI49_5141 | DI49_5142; DI49_5141 | | |
| DI49_5144; DI49_5143 | DI49_5144; DI49_5143 | | |
| DI49_5146; DI49_5145 | DI49_5146; DI49_5145 | | |
| SPGP0DZ01880; DI49_5147 | DI49_5147 | SPGP0DZ01880 | |
| DI49_5148; YOR335C | DI49_5148 | | YOR335C |
| DI49_5150; DI49_5149 | DI49_5150; DI49_5149 | | |
| DI49_5152; DI49_5151 | DI49_5152; DI49_5151 | | |
| DI49_5154; DI49_5153 | DI49_5154; DI49_5153 | | |
| DI49_5156; DI49_4243; SPGP0Rtrna5M; DI49_3010; SPGP0Strna7M; DI49_4242; DI49_0528 | DI49_5156; DI49_4243; DI49_3010; DI49_4242; DI49_0528 | SPGP0Rtrna5M; SPGP0Strna7M | |
| DI49_5158; DI49_2389; DI49_5159; DI49_4051; DI49_2388 | DI49_5158; DI49_2389; DI49_5159; DI49_4051; DI49_2388 | | |
| DI49_5161; DI49_5160 | DI49_5161; DI49_5160 | | |
| DI49_5164; DI49_5163 | DI49_5164; DI49_5163 | | |
| DI49_5166; DI49_5165 | DI49_5166; DI49_5165 | | |
| DI49_5168; DI49_5167 | DI49_5168; DI49_5167 | | |
| DI49_5170; DI49_5169 | DI49_5170; DI49_5169 | | |
| DI49_5172; DI49_5171 | DI49_5172; DI49_5171 | | |
| DI49_5174; DI49_5173 | DI49_5174; DI49_5173 | | |
| DI49_5176; DI49_5175 | DI49_5176; DI49_5175 | | |
| DI49_5178; DI49_5177 | DI49_5178; DI49_5177 | | |
| DI49_5180; DI49_5179 | DI49_5180; DI49_5179 | | |
| DI49_5182; DI49_5181 | DI49_5182; DI49_5181 | | |
| DI49_5184; DI49_5183 | DI49_5184; DI49_5183 | | |
| DI49_5186; DI49_5185 | DI49_5186; DI49_5185 | | |
| DI49_5191; DI49_5190 | DI49_5191; DI49_5190 | | |
| DI49_5685; DI49_5192 | DI49_5685; DI49_5192 | | |
| DI49_5683; DI49_5684 | DI49_5683; DI49_5684 | | |
| DI49_5679; YEL071W; YEL070W | DI49_5679 | | YEL071W; YEL070W |
| YAL067C; DI49_3158; SPGP0EA00110; DI49_3159 | DI49_3158; DI49_3159 | SPGP0EA00110 | YAL067C |
| YPL283C; SPGP0BI00100; YMR182C | | SPGP0BI00100 | YPL283C; YMR182C |
| DI49_3944; DI49_3943 | DI49_3944; DI49_3943 | | |

Create PDF in your applications with the Pdfcrowd HTML to PDF API PDFCROWD

| | | | |
|---|---|---|---|
| DI49_3946; DI49_3945 | DI49_3946; DI49_3945 | | |
| DI49_3951; DI49_3950 | DI49_3951; DI49_3950 | | |
| DI49_2572; DI49_2571 | DI49_2572; DI49_2571 | | |
| DI49_2574; DI49_2573 | DI49_2574; DI49_2573 | | |
| DI49_2576; DI49_2575 | DI49_2576; DI49_2575 | | |
| SPGP0EH00100; DI49_2501 | DI49_2501 | SPGP0EH00100 | |
| YNL331C; SPGP0C00100; YDL241W | | SPGP0C00100 | YNL331C; YDL241W |
| YNL330C; DI49_4422; YNL329C; DI49_4421 | DI49_4422; DI49_4421 | | YNL330C; YNL329C |
| YNL327W; YNL328C; DI49_4424; DI49_4423 | DI49_4424; DI49_4423 | | YNL327W; YNL328C |
| YNL325C; DI49_4427; DI49_4426 | DI49_4427; DI49_4426 | | YNL325C |
| YNL323W; DI49_4428 | DI49_4428 | | YNL323W |
| YNL322C; DI49_4429 | DI49_4429 | | YNL322C |
| YNL321W; DI49_4430 | DI49_4430 | | YNL321W |
| YNL320W; DI49_4431; YNL318C | DI49_4431 | | YNL320W; YNL318C |
| YNL316C; YNL317W | | | YNL316C; YNL317W |
| YNL314W; YNL315C | | | YNL314W; YNL315C |
| YNL312W; YNL313C | | | YNL312W; YNL313C |
| YNL310C; YNL311C | | | YNL310C; YNL311C |
| YNL308C; YNL309W | | | YNL308C; YNL309W |
| YNL306W; YNL307C; DI49_4443; DI49_4442 | DI49_4443; DI49_4442 | | YNL306W; YNL307C |
| YNL304W; DI49_4446; DI49_4445 | DI49_4446; DI49_4445 | | YNL304W |
| SPGP0M00370; YNL301C; YNL300W; YOL120C | | SPGP0M00370 | YNL301C; YNL300W; YOL120C |
| YNL298W; DI49_4450; YNL297C; DI49_4449 | DI49_4450; DI49_4449 | | YNL298W; YNL297C |
| YNL294C; YNL295W | | | YNL294C; YNL295W |
| YNL292W; DI49_4455; YNL291C | DI49_4455 | | YNL292W; YNL291C |
| YNL289W; DI49_4458; DI49_4457 | DI49_4458; DI49_4457 | | YNL289W |
| YNL288W; DI49_4459 | DI49_4459 | | YNL288W |
| YNL287W; DI49_4460; YNL286W | DI49_4460 | | YNL287W; YNL286W |
| YNL283C; DI49_4467; YNL282W | DI49_4467 | | YNL283C; YNL282W |
| YNL280C; DI49_4470; YNL279W; DI49_4469 | DI49_4470; DI49_4469 | | YNL280C; YNL279W |
| YNL277W; YNL278W | | | YNL277W; YNL278W |
| YNL274C; DI49_4475; YNL273W; DI49_4474 | DI49_4475; DI49_4474 | | YNL274C; YNL273W |
| YNL271C; DI49_4478 | DI49_4478 | | YNL271C |
| YNL270C; YNL268W; DI49_4480; DI49_4479 | DI49_4480; DI49_4479 | | YNL270C; YNL268W |
| YNL267W; DI49_4481 | DI49_4481 | | YNL267W |
| YNL265C; DI49_4482; YNL264C | DI49_4482 | | YNL265C; YNL264C |
| YNL262W; DI49_4485; YNL261W | DI49_4485 | | YNL262W; YNL261W |
| YNL259C; YNL260C | | | YNL259C; YNL260C |
| YNL257C; YNL258C | | | YNL257C; YNL258C |
| YNL255C; DI49_4492 | DI49_4492 | | YNL255C |

Create PDF in your applications with the Pdfcrowd HTML to PDF API PDFCROWD

| | | |
|---|---|---|
| YNL254C; DI49_4493; YNL253W | DI49_4493 | YNL254C; YNL253W |
| YNL251C; DI49_4496; DI49_4495 | DI49_4496; DI49_4495 | YNL251C |
| YNL250W; DI49_4497 | DI49_4497 | YNL250W |
| YNL249C; DI49_4498 | DI49_4498 | YNL249C |
| YNL248C; DI49_4499 | DI49_4499 | YNL248C |
| YNL247W; DI49_4500; YNL246W | DI49_4500 | YNL247W; YNL246W |
| YNL244C; DI49_4503; YNL243W; DI49_4502 | DI49_4503; DI49_4502 | YNL244C; YNL243W |
| YNL241C; DI49_4506; YNL240C | DI49_4506 | YNL241C; YNL240C |
| YNL238W; YNL239W | | YNL238W; YNL239W |
| YNL236W; YNL237W | | YNL236W; YNL237W |
| YNL233W; DI49_4513; DI49_4512 | DI49_4513; DI49_4512 | YNL233W |
| YNL232W; DI49_4514; YNL231C | DI49_4514 | YNL232W; YNL231C |
| YNL229C; YNL230C | | YNL229C; YNL230C |
| YNL225C; DI49_4519 | DI49_4519 | YNL225C |
| YNL224C; DI49_4520 | DI49_4520 | YNL224C |
| YNL223W; DI49_4521 | DI49_4521 | YNL223W |
| YNL222W; DI49_4522; YNL221C | DI49_4522 | YNL222W; YNL221C |
| YNL219C; YNL220W | | YNL219C; YNL220W |
| YNL217W; YNL218W | | YNL217W; YNL218W |
| YNL215W; YNL216W | | YNL215W; YNL216W |
| YNL213C; YNL214W | | YNL213C; YNL214W |
| YNL211C; YNL212W; DI49_4532; DI49_4531 | DI49_4532; DI49_4531 | YNL211C; YNL212W |
| YNL209W; YDL229W; YNL208W | | YNL209W; YDL229W; YNL208W |
| YNL206C; DI49_4538; YNL204C | DI49_4538 | YNL206C; YNL204C |
| YNL201C; DI49_4541; YNL200C | DI49_4541 | YNL201C; YNL200C |
| YNL197C; YNL199C | | YNL197C; YNL199C |
| YNL195C; YNL196C | | YNL195C; YNL196C |
| YNL193W; YNL194C | | YNL193W; YNL194C |
| YNL191W; DI49_4550 | DI49_4550 | YNL191W |
| YNL190W; DI49_4551 | DI49_4551 | YNL190W |
| YNL189W; DI49_4552 | DI49_4552 | YNL189W |
| YNL188W; DI49_4553; YNL187W | DI49_4553 | YNL188W; YNL187W |
| YNL185C; DI49_4556 | DI49_4556 | YNL185C |
| YNL183C; DI49_4557 | DI49_4557 | YNL183C |
| YNL182C; DI49_4558; YNL181W | DI49_4558 | YNL182C; YNL181W |
| YNL178W; DI49_4561; YNL177C | DI49_4561 | YNL178W; YNL177C |
| YNL175C; DI49_4564; YNL173C; DI49_4563 | DI49_4564; DI49_4563 | YNL175C; YNL173C |
| YNL169C; DI49_4567; YNL168C | DI49_4567 | YNL169C; YNL168C |
| YNL166C; YNL167C | | YNL166C; YNL167C |
| YNL164C; DI49_4572; DI49_4571 | DI49_4572; DI49_4571 | YNL164C |

Create PDF in your applications with the Pdfcrowd HTML to PDF API PDFCROWD

| | | | |
|---|---|---|---|
| YNL163C; DI49_4573; YNL162W-A | DI49_4573 | | YNL163C; YNL162W-A |
| YNL161W; YNL162W; SPGP0P00350; DI49_2543; YHR141C | DI49_2543 | SPGP0P00350 | YNL161W; YNL162W; YHR141C |
| YNL159C; YNL160W; DI49_4577; DI49_4576 | DI49_4577; DI49_4576 | | YNL159C; YNL160W |
| YNL157W; DI49_4580; YNL156C; DI49_4579 | DI49_4580; DI49_4579 | | YNL157W; YNL156C |
| YNL154C; DI49_4583; YNL153C; DI49_4582 | DI49_4583; DI49_4582 | | YNL154C; YNL153C |
| YNL151C; YNL152W | | | YNL151C; YNL152W |
| YNL148C; YNL149C | | | YNL148C; YNL149C |
| YNL146W; YNL147W | | | YNL146W; YNL147W |
| YNL144C; YNL145W | | | YNL144C; YNL145W |
| YNL139C; DI49_4595; YNL138W-A; DI49_4594 | DI49_4595; DI49_4594 | | YNL139C; YNL138W-A |
| YNL137C; DI49_4598 | DI49_4598 | | YNL137C |
| YNL136W; DI49_4599 | DI49_4599 | | YNL136W |
| YNL135C; DI49_4600; YNL134C | DI49_4600 | | YNL135C; YNL134C |
| YNL132W; DI49_4604 | DI49_4604 | | YNL132W |
| YNL131W; DI49_4605; YNL130C | DI49_4605 | | YNL131W; YNL130C |
| YNL128W; DI49_4608; DI49_4607 | DI49_4608; DI49_4607 | | YNL128W |
| YNL127W; DI49_4609 | DI49_4609 | | YNL127W |
| YNL126W; DI49_4610; YNL125C | DI49_4610 | | YNL126W; YNL125C |
| YNL123W; DI49_4613 | DI49_4613 | | YNL123W |
| YNL122C; DI49_4614; YNL121C | DI49_4614 | | YNL122C; YNL121C |
| YNL118C; DI49_4617; DI49_4616 | DI49_4617; DI49_4616 | | YNL118C |
| YNL117W; DI49_4618 | DI49_4618 | | YNL117W |
| YNL116W; DI49_4619; YNL115C | DI49_4619 | | YNL116W; YNL115C |
| YNL111C; DI49_4622 | DI49_4622 | | YNL111C |
| YNL110C; DI49_4623; YNL108C | DI49_4623 | | YNL110C; YNL108C |
| YNL106C; DI49_4626; DI49_4625 | DI49_4626; DI49_4625 | | YNL106C |
| YNL101W; DI49_4630; YNL100W | DI49_4630 | | YNL101W; YNL100W |
| YNL098C; DI49_4633; DI49_4632 | DI49_4633; DI49_4632 | | YNL098C |
| YNL097C; DI49_4634 | DI49_4634 | | YNL097C |
| YNL093W; YNL094W; DI49_4638; DI49_4637 | DI49_4638; DI49_4637 | | YNL093W; YNL094W |
| YNL091W; DI49_4641; YNL090W; DI49_4640 | DI49_4641; DI49_4640 | | YNL091W; YNL090W |
| YNL087W; YNL088W; DI49_4643; DI49_4642 | DI49_4643; DI49_4642 | | YNL087W; YNL088W |
| YNL085W; DI49_4646; DI49_4645 | DI49_4646; DI49_4645 | | YNL085W |
| YNL084C; DI49_4647 | DI49_4647 | | YNL084C |
| YNL083W; DI49_4648 | DI49_4648 | | YNL083W |
| YNL082W; DI49_4649 | DI49_4649 | | YNL082W |
| YNL081C; DI49_4650 | DI49_4650 | | YNL081C |
| YNL080C; DI49_4651 | DI49_4651 | | YNL080C |
| YNL079C; DI49_4652; YNL078W | DI49_4652 | | YNL079C; YNL078W |

Create PDF in your applications with the Pdfcrowd HTML to PDF API PDFCROWD

| | | | |
|---|---|---|---|
| YNL076W; YNL077W | | | YNL076W; YNL077W |
| YNL074C; YNL075W | | | YNL074C; YNL075W |
| YNL072W; YNL073W | | | YNL072W; YNL073W |
| YNL070W; DI49_4661 | DI49_4661 | | YNL070W |
| SPGP0M02460; DI49_4662; YNL068C | DI49_4662 | SPGP0M02460 | YNL068C |
| YNL066W; YNL067W; YGL148W | | | YNL066W; YNL067W; YGL148W |
| YNL064C; DI49_4667; YNL063W; DI49_4666 | DI49_4667; DI49_4666 | | YNL064C; YNL063W |
| YNL061W; DI49_4670 | DI49_4670 | | YNL061W |
| YNL059C; DI49_4671; YNL058C | DI49_4671 | | YNL059C; YNL058C |
| YNL055C; DI49_4674; DI49_4673 | DI49_4674; DI49_4673 | | YNL055C |
| YNL054W; DI49_4676; YNL053W | DI49_4676 | | YNL054W; YNL053W |
| YNL051W; YNL052W | | | YNL051W; YNL052W |
| YNL049C; DI49_4681; DI49_4680 | DI49_4681; DI49_4680 | | YNL049C |
| YNL048W; DI49_4682; YNL047C | DI49_4682 | | YNL048W; YNL047C |
| YNL045W; YNL046W | | | YNL045W; YNL046W |
| YNL040W; YNL041C; DI49_4689; DI49_4688 | DI49_4689; DI49_4688 | | YNL040W; YNL041C |
| YNL038W; YNL039W | | | YNL038W; YNL039W |
| YNL036W; YNL037C | | | YNL036W; YNL037C |
| SPGP0M02780; YNL018C | | SPGP0M02780 | YNL018C |
| SPGP0M02790; YNL019C | | SPGP0M02790 | YNL019C |
| YNL020C; DI49_4717; YNL021W | DI49_4717 | | YNL020C; YNL021W |
| YNL023C; DI49_4714 | DI49_4714 | | YNL023C |
| YNL024C; DI49_4713; YNL024C-A | DI49_4713 | | YNL024C; YNL024C-A |
| YNL026W; YNL025C | | | YNL026W; YNL025C |
| YNL029C; YNL027W | | | YNL029C; YNL027W |
| YNL032W; DI49_4705 | DI49_4705 | | YNL032W |
| YNL016W; DI49_4721; YNL015W | DI49_4721 | | YNL016W; YNL015W |
| YNL012W; SPGP0Z00220 | | SPGP0Z00220 | YNL012W |
| YNL011C; DI49_4724 | DI49_4724 | | YNL011C |
| YNL010W; DI49_4725 | DI49_4725 | | YNL010W |
| YNL009W; DI49_4726; YNL008C | DI49_4726 | | YNL009W; YNL008C |
| YNL006W; YNL007C; DI49_4728; DI49_4727 | DI49_4728; DI49_4727 | | YNL006W; YNL007C |
| YNL004W; DI49_4731; DI49_4730 | DI49_4731; DI49_4730 | | YNL004W |
| YNL003C; DI49_4732 | DI49_4732 | | YNL003C |
| YNL002C; DI49_4733; YNL001W | DI49_4733 | | YNL002C; YNL001W |
| SPGP0Mtrna8P; DI49_4736; DI49_0756; SPGP0AAtrna17P; YCR007C | DI49_4736; DI49_0756 | SPGP0Mtrna8P; SPGP0AAtrna17P | YCR007C |
| YNR008W; YNR007C | | | YNR008W; YNR007C |
| YNR010W; YNR009W | | | YNR010W; YNR009W |
| SPGP0M03280; YNR019W | | SPGP0M03280 | YNR019W |
| YNR022C; YNR021W | | | YNR022C; YNR021W |

Create PDF in your applications with the Pdfcrowd HTML to PDF API PDFCROWD

| | | | |
|---|---|---|---|
| YNR024W; YNR023W | | | YNR024W; YNR023W |
| YNR027W; YNR026C | | | YNR027W; YNR026C |
| YNR029C; YNR028W | | | YNR029C; YNR028W |
| YNR033W; YNR032C-A | | | YNR033W; YNR032C-A |
| YNR034W-A; YNR034W | | | YNR034W-A; YNR034W |
| YNR036C; YNR035C | | | YNR036C; YNR035C |
| YNR038W; YNR037C | | | YNR038W; YNR037C |
| YNR040W; YNR039C | | | YNR040W; YNR039C |
| YNR043W; YNR041C | | | YNR043W; YNR041C |
| YNR045W; YNR044W | | | YNR045W; YNR044W |
| YNR058W; SPGP0M03650 | | SPGP0M03650 | YNR058W |
| YNR060W; YNR059W | | | YNR060W; YNR059W |
| SPGP0M03700; YNR061C | | SPGP0M03700 | YNR061C |
| YNR064C; YNR063W | | | YNR064C; YNR063W |
| YNR070W; DI49_1724; YNR071C; DI49_1725 | DI49_1724; DI49_1725 | | YNR070W; YNR071C |
| YNR074C; SPGP0M03780; YNR075W; YEL069C; YEL066W | | SPGP0M03780 | YNR074C; YNR075W; YEL069C; YEL066W |
| DI49_5529; YNR076W | DI49_5529 | | YNR076W |
| DI49_5527; DI49_5528 | DI49_5527; DI49_5528 | | |
| DI49_4375; DI49_1600 | DI49_4375; DI49_1600 | | |
| DI49_4377; DI49_4376 | DI49_4377; DI49_4376 | | |
| YPL276W; YPL275W | | | YPL276W; YPL275W |
| SPGP0ER00110; SPGP0ER00100 | | SPGP0ER00110; SPGP0ER00100 | |
| YGL262W; SPGP0ER00120 | | SPGP0ER00120 | YGL262W |
| DI49_3908; DI49_3909 | DI49_3908; DI49_3909 | | |
| DI49_3906; DI49_3907 | DI49_3906; DI49_3907 | | |
| SPGP0N00150; YDL075W; DI49_3903; SPGP0W03370 | DI49_3903 | SPGP0N00150; SPGP0W03370 | YDL075W |
| DI49_3901; DI49_3902 | DI49_3901; DI49_3902 | | |
| DI49_3899; DI49_3900 | DI49_3899; DI49_3900 | | |
| DI49_3897; DI49_3898 | DI49_3897; DI49_3898 | | |
| DI49_3895; DI49_3896 | DI49_3895; DI49_3896 | | |
| DI49_3893; DI49_3894 | DI49_3893; DI49_3894 | | |
| DI49_3891; DI49_3892 | DI49_3891; DI49_3892 | | |
| DI49_3889; DI49_3890 | DI49_3889; DI49_3890 | | |
| DI49_3887; DI49_3888 | DI49_3887; DI49_3888 | | |
| DI49_3885; DI49_3886 | DI49_3885; DI49_3886 | | |
| DI49_3883; DI49_3884 | DI49_3883; DI49_3884 | | |
| DI49_3881; DI49_3882 | DI49_3881; DI49_3882 | | |
| DI49_3879; DI49_3880 | DI49_3879; DI49_3880 | | |
| DI49_3874; DI49_3875 | DI49_3874; DI49_3875 | | |
| DI49_3872; DI49_3873 | DI49_3872; DI49_3873 | | |

Create PDF in your applications with the Pdfcrowd HTML to PDF API PDFCROWD

| | | | |
|---|---|---|---|
| DI49_3870; DI49_3871 | DI49_3870; DI49_3871 | | |
| DI49_3868; DI49_3869 | DI49_3868; DI49_3869 | | |
| DI49_3863; DI49_3864 | DI49_3863; DI49_3864 | | |
| DI49_3861; DI49_3862 | DI49_3861; DI49_3862 | | |
| DI49_3859; DI49_3860; DI49_3858 | DI49_3859; DI49_3860; DI49_3858 | | |
| DI49_3856; DI49_3857 | DI49_3856; DI49_3857 | | |
| DI49_3854; DI49_3855 | DI49_3854; DI49_3855 | | |
| DI49_3852; DI49_3853 | DI49_3852; DI49_3853 | | |
| DI49_3850; DI49_3851 | DI49_3850; DI49_3851 | | |
| DI49_3848; DI49_3849 | DI49_3848; DI49_3849 | | |
| DI49_3846; DI49_3847 | DI49_3846; DI49_3847 | | |
| DI49_3845; SPGP0N00730 | DI49_3845 | SPGP0N00730 | |
| DI49_3843; DI49_1995; DI49_3842; YGR032W | DI49_3843; DI49_1995; DI49_3842 | | YGR032W |
| DI49_3840; DI49_3841 | DI49_3840; DI49_3841 | | |
| DI49_3838; DI49_3839 | DI49_3838; DI49_3839 | | |
| DI49_3836; DI49_1989; YGR027C | DI49_3836; DI49_1989 | | YGR027C |
| DI49_3832; DI49_3833 | DI49_3832; DI49_3833 | | |
| DI49_3827; DI49_3828 | DI49_3827; DI49_3828 | | |
| DI49_3825; DI49_3826 | DI49_3825; DI49_3826 | | |
| DI49_3823; DI49_3824 | DI49_3823; DI49_3824 | | |
| DI49_3821; DI49_3822 | DI49_3821; DI49_3822 | | |
| DI49_3820; SPGP0N00950 | DI49_3820 | SPGP0N00950 | |
| DI49_3819; SPGP0N00970 | DI49_3819 | SPGP0N00970 | |
| DI49_3817; DI49_3818 | DI49_3817; DI49_3818 | | |
| DI49_3815; DI49_3816 | DI49_3815; DI49_3816 | | |
| DI49_3813; DI49_3814 | DI49_3813; DI49_3814 | | |
| DI49_3811; DI49_3812 | DI49_3811; DI49_3812 | | |
| DI49_3809; DI49_3810 | DI49_3809; DI49_3810 | | |
| DI49_3805; SPGP0Ntrna12L; SPGP0Ntrna15L; DI49_3740 | DI49_3805; DI49_3740 | SPGP0Ntrna12L; SPGP0Ntrna15L | |
| DI49_3804; YLR300W | DI49_3804 | | YLR300W |
| DI49_3802; DI49_3803 | DI49_3802; DI49_3803 | | |
| DI49_3797; DI49_3798 | DI49_3797; DI49_3798 | | |
| DI49_3795; DI49_3796; DI49_3794 | DI49_3795; DI49_3796; DI49_3794 | | |
| DI49_3792; DI49_3793 | DI49_3792; DI49_3793 | | |
| DI49_3790; DI49_3791 | DI49_3790; DI49_3791 | | |
| DI49_3788; DI49_3789 | DI49_3788; DI49_3789 | | |
| DI49_3786; DI49_3787 | DI49_3786; DI49_3787 | | |
| DI49_3784; YGL201C; DI49_3783; DI49_1778 | DI49_3784; DI49_3783; DI49_1778 | | YGL201C |
| DI49_3779; DI49_3780 | DI49_3779; DI49_3780 | | |
| DI49_3777; DI49_3778 | DI49_3777; DI49_3778 | | |

Create PDF in your applications with the Pdfcrowd HTML to PDF API PDFCROWD

| | | | |
|---|---|---|---|
| DI49_3775; DI49_3776 | DI49_3775; DI49_3776 | | |
| DI49_3771; DI49_3772 | DI49_3771; DI49_3772 | | |
| DI49_3769; DI49_1665; DI49_3768; DI49_1663 | DI49_3769; DI49_1665; DI49_3768; DI49_1663 | | |
| DI49_3764; DI49_3765 | DI49_3764; DI49_3765 | | |
| DI49_3762; DI49_3763 | DI49_3762; DI49_3763 | | |
| DI49_3760; DI49_4723; DI49_3759; YNL014W; DI49_4722 | DI49_3760; DI49_4723; DI49_3759; DI49_4722 | | YNL014W |
| DI49_3754; DI49_3755 | DI49_3754; DI49_3755 | | |
| DI49_3752; DI49_3753 | DI49_3752; DI49_3753 | | |
| DI49_3750; DI49_3751 | DI49_3750; DI49_3751 | | |
| DI49_3748; DI49_3749 | DI49_3748; DI49_3749 | | |
| DI49_3744; DI49_3745 | DI49_3744; DI49_3745 | | |
| SPGP0Ntrna14L; DI49_3110; SPGP0Rtrna24L; DI49_3111; YJR100C | DI49_3110; DI49_3111 | SPGP0Ntrna14L; SPGP0Rtrna24L | YJR100C |
| DI49_3738; DI49_3739 | DI49_3738; DI49_3739 | | |
| DI49_3736; DI49_3737 | DI49_3736; DI49_3737 | | |
| DI49_3734; DI49_3735 | DI49_3734; DI49_3735 | | |
| DI49_3732; DI49_3733 | DI49_3732; DI49_3733 | | |
| DI49_3731; SPGP0N01830 | DI49_3731 | SPGP0N01830 | |
| DI49_3728; DI49_3730 | DI49_3728; DI49_3730 | | |
| DI49_3726; DI49_3727 | DI49_3726; DI49_3727 | | |
| DI49_3724; DI49_3725 | DI49_3724; DI49_3725 | | |
| DI49_3722; DI49_3723 | DI49_3722; DI49_3723 | | |
| DI49_3720; DI49_3721 | DI49_3720; DI49_3721 | | |
| DI49_3718; DI49_3719 | DI49_3718; DI49_3719 | | |
| DI49_3716; DI49_3717 | DI49_3716; DI49_3717 | | |
| DI49_3714; DI49_3715 | DI49_3714; DI49_3715 | | |
| DI49_3712; DI49_3713 | DI49_3712; DI49_3713 | | |
| DI49_3710; DI49_3711 | DI49_3710; DI49_3711 | | |
| DI49_3708; DI49_3709 | DI49_3708; DI49_3709 | | |
| DI49_3706; DI49_3707 | DI49_3706; DI49_3707 | | |
| DI49_3704; DI49_3705 | DI49_3704; DI49_3705 | | |
| DI49_3702; DI49_3703 | DI49_3702; DI49_3703 | | |
| DI49_3700; DI49_3701 | DI49_3700; DI49_3701 | | |
| DI49_3698; DI49_3699 | DI49_3698; DI49_3699 | | |
| DI49_3696; DI49_3697; DI49_0665; DI49_0666 | DI49_3696; DI49_3697; DI49_0665; DI49_0666 | | |
| DI49_3694; DI49_3695 | DI49_3694; DI49_3695 | | |
| DI49_3692; DI49_3693 | DI49_3692; DI49_3693 | | |
| DI49_3690; DI49_3691 | DI49_3690; DI49_3691 | | |
| DI49_3688; DI49_3689 | DI49_3688; DI49_3689 | | |
| DI49_3683; DI49_3684 | DI49_3683; DI49_3684 | | |

Create PDF in your applications with the Pdfcrowd HTML to PDF API PDFCROWD

| | | | |
|---|---|---|---|
| SPGP0N02330; DI49_3682 | DI49_3682 | SPGP0N02330 | |
| DI49_5410; DI49_5409 | DI49_5410; DI49_5409 | | |
| DI49_5412; DI49_5411 | DI49_5412; DI49_5411 | | |
| DI49_5414; DI49_5413 | DI49_5414; DI49_5413 | | |
| DI49_5418; DI49_5417 | DI49_5418; DI49_5417 | | |
| DI49_5420; DI49_5419 | DI49_5420; DI49_5419 | | |
| DI49_0508; DI49_0507 | DI49_0508; DI49_0507 | | |
| DI49_0510; DI49_0509 | DI49_0510; DI49_0509 | | |
| DI49_0512; DI49_0511 | DI49_0512; DI49_0511 | | |
| DI49_0514; DI49_0513 | DI49_0514; DI49_0513 | | |
| DI49_0516; DI49_0515 | DI49_0516; DI49_0515 | | |
| DI49_0518; DI49_0517 | DI49_0518; DI49_0517 | | |
| YDR345C; DI49_2500; DI49_0523; YHR094C; YHR096C; YHR097C | DI49_2500; DI49_0523 | | YDR345C; YHR094C; YHR096C; YHR097C |
| DI49_0525; DI49_0524 | DI49_0525; DI49_0524 | | |
| DI49_3915; SPGP0EZ00100; YJL216C | DI49_3915 | SPGP0EZ00100 | YJL216C |
| DI49_3919; DI49_3918 | DI49_3919; DI49_3918 | | |
| YPL277C; SPGP0FD00100 | | SPGP0FD00100 | YPL277C |
| R0020C; DI49_4794 | DI49_4794 | | R0020C |
| DI49_2568; DI49_2569; YHR165C; SPGP0W00100 | DI49_2568; DI49_2569 | SPGP0W00100 | YHR165C |
| DI49_2566; DI49_2567; YHR164C | DI49_2566; DI49_2567 | | YHR164C |
| DI49_2564; DI49_2565 | DI49_2564; DI49_2565 | | |
| DI49_2562; DI49_2563 | DI49_2562; DI49_2563 | | |
| DI49_2560; DI49_2561 | DI49_2560; DI49_2561 | | |
| DI49_2558; YHR155W; YHR156C | DI49_2558 | | YHR155W; YHR156C |
| DI49_2557; YHR154W; DI49_2556 | DI49_2557; DI49_2556 | | YHR154W |
| DI49_2554; DI49_2555 | DI49_2554; DI49_2555 | | |
| DI49_2552; YHR149C; YHR150W | DI49_2552 | | YHR149C; YHR150W |
| DI49_2551; YHR148W | DI49_2551 | | YHR148W |
| DI49_2550; YHR147C; DI49_2549 | DI49_2550; DI49_2549 | | YHR147C |
| DI49_2547; DI49_2548 | DI49_2547; DI49_2548 | | |
| DI49_2545; DI49_2546 | DI49_2545; DI49_2546 | | |
| DI49_2541; DI49_2542 | DI49_2541; DI49_2542 | | |
| DI49_2539; SPGP0P00390 | DI49_2539 | SPGP0P00390 | |
| DI49_2537; DI49_2538 | DI49_2537; DI49_2538 | | |
| DI49_2535; YHR132C; DI49_2534 | DI49_2535; DI49_2534 | | YHR132C |
| DI49_2532; YHR128W; DI49_2531 | DI49_2532; DI49_2531 | | YHR128W |
| SPGP0Ptrna6F; DI49_2529; DI49_1939; DI49_1938; DI49_1659 | DI49_2529; DI49_1939; DI49_1938; DI49_1659 | SPGP0Ptrna6F | |
| DI49_2528; YHR124W; SPGP0P00510 | DI49_2528 | SPGP0P00510 | YHR124W |
| DI49_2526; DI49_2527 | DI49_2526; DI49_2527 | | |

Create PDF in your applications with the Pdfcrowd HTML to PDF API PDFCROWD

| | | | |
|---|---|---|---|
| DI49_2524; YHR119W; DI49_2523; YHR120W | DI49_2524; DI49_2523 | | YHR119W; YHR120W |
| DI49_2521; DI49_2522 | DI49_2521; DI49_2522 | | |
| DI49_2519; YHR114W; DI49_2518; YHR115C | DI49_2519; DI49_2518 | | YHR114W; YHR115C |
| DI49_2516; DI49_2517 | DI49_2516; DI49_2517 | | |
| DI49_2514; DI49_2515 | DI49_2514; DI49_2515 | | |
| DI49_2512; YHR107C; DI49_2511; YHR108W | DI49_2512; DI49_2511 | | YHR107C; YHR108W |
| DI49_2509; DI49_2510 | DI49_2509; DI49_2510 | | |
| DI49_2507; YHR102W; DI49_2506; YHR103W | DI49_2507; DI49_2506 | | YHR102W; YHR103W |
| DI49_2504; YHR099W; YHR100C | DI49_2504 | | YHR099W; YHR100C |
| DI49_2503; YHR098C; DI49_2502 | DI49_2503; DI49_2502 | | YHR098C |
| SPGP0P00800; YHR092C; SPGP0P00810; DI49_2498; YDR342C; YHR093W; YJL214W | DI49_2498 | SPGP0P00800; SPGP0P00810 | YHR092C; YDR342C; YHR093W; YJL214W |
| DI49_2496; YHR089C; DI49_2495; YHR090C | DI49_2496; DI49_2495 | | YHR089C; YHR090C |
| DI49_2493; YHR086W | DI49_2493 | | YHR086W |
| DI49_2492; YHR085W | DI49_2492 | | YHR085W |
| DI49_2491; YHR084W; DI49_2490 | DI49_2491; DI49_2490 | | YHR084W |
| DI49_2488; DI49_2489 | DI49_2488; DI49_2489 | | |
| DI49_2486; DI49_2487 | DI49_2486; DI49_2487 | | |
| DI49_2484; DI49_2485 | DI49_2484; DI49_2485 | | |
| DI49_2482; DI49_2483 | DI49_2482; DI49_2483 | | |
| DI49_2480; YHR075C; YHR076W | DI49_2480 | | YHR075C; YHR076W |
| DI49_2479; YHR074W; DI49_2478 | DI49_2479; DI49_2478 | | YHR074W |
| DI49_2476; DI49_2477 | DI49_2476; DI49_2477 | | |
| DI49_2474; DI49_2475; DI49_4603; DI49_4602 | DI49_2474; DI49_2475; DI49_4603; DI49_4602 | | |
| DI49_2472; YHR069C; YHR070W | DI49_2472 | | YHR069C; YHR070W |
| DI49_2471; YHR068W; DI49_2470 | DI49_2471; DI49_2470 | | YHR068W |
| DI49_2468; YHR065C | DI49_2468 | | YHR065C |
| DI49_2467; YHR064C | DI49_2467 | | YHR064C |
| DI49_2466; YHR063C; DI49_2465 | DI49_2466; DI49_2465 | | YHR063C |
| DI49_2463; DI49_2464 | DI49_2463; DI49_2464 | | |
| DI49_2461; DI49_2462; YHR059W; YHR060W | DI49_2461; DI49_2462 | | YHR059W; YHR060W |
| DI49_2459; YHR056C; DI49_2458; YHR057C | DI49_2459; DI49_2458 | | YHR056C; YHR057C |
| DI49_2456; YHR051W; DI49_2455 | DI49_2456; DI49_2455 | | YHR051W |
| DI49_2453; DI49_2454 | DI49_2453; DI49_2454 | | |
| DI49_2451; DI49_2452 | DI49_2451; DI49_2452 | | |
| DI49_2449; DI49_2450 | DI49_2449; DI49_2450 | | |
| DI49_2447; DI49_2448 | DI49_2447; DI49_2448 | | |
| SPGP0P01340; YHR039C-A; DI49_2445 | DI49_2445 | SPGP0P01340 | YHR039C-A |
| DI49_2443; YHR037W; YHR038W | DI49_2443 | | YHR037W; YHR038W |
| DI49_2442; YHR036W | DI49_2442 | | YHR036W |
| DI49_2441; DI49_2440 | DI49_2441; DI49_2440 | | |

Create PDF in your applications with the Pdfcrowd HTML to PDF API PDFCROWD

| | | | |
|---|---|---|---|
| DI49_2439; YHR033W; YHR034C | DI49_2439 | | YHR033W; YHR034C |
| DI49_2438; YHR032W | DI49_2438 | | YHR032W |
| DI49_2436; YHR030C; YHR031C | DI49_2436 | | YHR030C; YHR031C |
| DI49_2434; YHR028C; YHR029C | DI49_2434 | | YHR028C; YHR029C |
| DI49_2433; YHR027C | DI49_2433 | | YHR027C |
| DI49_2432; YHR026W | DI49_2432 | | YHR026W |
| DI49_2431; YHR025W; DI49_2430 | DI49_2431; DI49_2430 | | YHR025W |
| SPGP0P01520; YHR021C; YHR022C | | SPGP0P01520 | YHR021C; YHR022C |
| DI49_2427; YHR020W | DI49_2427 | | YHR020W |
| DI49_2426; YHR019C | DI49_2426 | | YHR019C |
| DI49_2425; YHR018C | DI49_2425 | | YHR018C |
| DI49_2424; YHR017W; DI49_2423 | DI49_2424; DI49_2423 | | YHR017W |
| DI49_2417; YOR014W; DI49_2416 | DI49_2417; DI49_2416 | | YOR014W |
| DI49_2412; YOR009W; DI49_2411 | DI49_2412; DI49_2411 | | YOR009W |
| DI49_2407; YOR005C; DI49_2406 | DI49_2407; DI49_2406 | | YOR005C |
| DI49_2403; YOR001W; DI49_2402 | DI49_2403; DI49_2402 | | YOR001W |
| DI49_2396; DI49_2397 | DI49_2396; DI49_2397 | | |
| DI49_2383; DI49_2384 | DI49_2383; DI49_2384 | | |
| DI49_2371; DI49_2372 | DI49_2371; DI49_2372 | | |
| DI49_2369; DI49_2370 | DI49_2369; DI49_2370 | | |
| DI49_2367; DI49_2368 | DI49_2367; DI49_2368 | | |
| DI49_2359; DI49_2360 | DI49_2359; DI49_2360 | | |
| DI49_2356; SPGP0P02190 | DI49_2356 | SPGP0P02190 | |
| DI49_2354; YOL052C; DI49_2353 | DI49_2354; DI49_2353 | | YOL052C |
| DI49_2351; DI49_2352 | DI49_2351; DI49_2352 | | |
| DI49_2342; YOL062C; DI49_2341 | DI49_2342; DI49_2341 | | YOL062C |
| DI49_2339; DI49_2340 | DI49_2339; DI49_2340 | | |
| DI49_2337; DI49_2338 | DI49_2337; DI49_2338 | | |
| DI49_2326; DI49_2327 | DI49_2326; DI49_2327 | | |
| DI49_2321; DI49_2322 | DI49_2321; DI49_2322 | | |
| DI49_2319; DI49_2320; DI49_4390; DI49_4391 | DI49_2319; DI49_2320; DI49_4390; DI49_4391 | | |
| DI49_2317; DI49_2318 | DI49_2317; DI49_2318 | | |
| DI49_2304; YOL102C; DI49_2303 | DI49_2304; DI49_2303 | | YOL102C |
| DI49_2301; DI49_2302 | DI49_2301; DI49_2302 | | |
| DI49_2292; DI49_2293 | DI49_2292; DI49_2293 | | |
| DI49_2283; YOL124C; DI49_2282 | DI49_2283; DI49_2282 | | YOL124C |
| DI49_2277; DI49_1607; DI49_2276 | DI49_2277; DI49_1607; DI49_2276 | | |
| DI49_2268; YOL140W; DI49_2267 | DI49_2268; DI49_2267 | | YOL140W |
| DI49_2265; YOL143C; DI49_2264 | DI49_2265; DI49_2264 | | YOL143C |
| DI49_2262; DI49_2263 | DI49_2262; DI49_2263 | | |

Create PDF in your applications with the Pdfcrowd HTML to PDF API PDFCROWD

| | | | |
|---|---|---|---|
| DI49_2254; DI49_2255 | DI49_2254; DI49_2255 | | |
| SPGP0P03210; YGR289C; DI49_3910; SPGP0Q00150; YGR288W | DI49_3910 | SPGP0P03210; SPGP0Q00150 | YGR289C; YGR288W |
| DI49_3912; DI49_3911 | DI49_3912; DI49_3911 | | |
| DI49_3914; DI49_3913 | DI49_3914; DI49_3913 | | |
| YGR286C; SPGP0F00100 | | SPGP0F00100 | YGR286C |
| DI49_2246; YGR285C | DI49_2246 | | YGR285C |
| DI49_2245; YGR284C; DI49_2244; YGR283C | DI49_2245; DI49_2244 | | YGR284C; YGR283C |
| DI49_2242; YGR281W; DI49_2241; YGR280C | DI49_2242; DI49_2241 | | YGR281W; YGR280C |
| SPGP0Q00260; DI49_2240 | DI49_2240 | SPGP0Q00260 | |
| DI49_2238; YGR276C; DI49_2237; YGR275W | DI49_2238; DI49_2237 | | YGR276C; YGR275W |
| DI49_2235; DI49_2236 | DI49_2235; DI49_2236 | | |
| DI49_2233; YGR271W | DI49_2233 | | YGR271W |
| DI49_2232; YGR270W; DI49_2231; YGR268C | DI49_2232; DI49_2231 | | YGR270W; YGR268C |
| DI49_2229; DI49_2230 | DI49_2229; DI49_2230 | | |
| DI49_2227; YGR263C; DI49_2226; YGR262C | DI49_2227; DI49_2226 | | YGR263C; YGR262C |
| DI49_2224; YGR260W | DI49_2224 | | YGR260W |
| DI49_2223; YGR258C; DI49_2222; YGR257C | DI49_2223; DI49_2222 | | YGR258C; YGR257C |
| SPGP0Qtrna15T; SPGP0AItrna28T; DI49_2219; SPGP0Qtrna16T; DI49_2047; SPGP0ACtrna24T; DI49_5554; SPGP0AFtrna26T; YHR180W | DI49_2219; DI49_2047; DI49_5554 | SPGP0Qtrna15T; SPGP0AItrna28T; SPGP0Qtrna16T; SPGP0ACtrna24T; SPGP0AFtrna26T | YHR180W |
| DI49_2217; YGR253C | DI49_2217 | | YGR253C |
| DI49_2216; YGR252W; DI49_2215; YGR251W | DI49_2216; DI49_2215 | | YGR252W; YGR251W |
| DI49_2213; DI49_2214 | DI49_2213; DI49_2214 | | |
| DI49_2211; DI49_2212 | DI49_2211; DI49_2212 | | |
| DI49_2209; YGR246C | DI49_2209 | | YGR246C |
| DI49_2208; YGR245C; DI49_2207; YGR244C | DI49_2208; DI49_2207 | | YGR245C; YGR244C |
| DI49_2205; DI49_2206 | DI49_2205; DI49_2206 | | |
| DI49_2203; DI49_2204 | DI49_2203; DI49_2204 | | |
| DI49_2201; DI49_2202 | DI49_2201; DI49_2202 | | |
| DI49_2199; DI49_2200 | DI49_2199; DI49_2200 | | |
| DI49_2197; DI49_2198 | DI49_2197; DI49_2198 | | |
| DI49_2195; YGR231C; DI49_2194; YGR230W | DI49_2195; DI49_2194 | | YGR231C; YGR230W |
| DI49_2191; DI49_2192 | DI49_2191; DI49_2192 | | |
| DI49_2190; SPGP0Q00740 | DI49_2190 | SPGP0Q00740 | |
| DI49_2188; DI49_2189 | DI49_2188; DI49_2189 | | |
| DI49_2186; YGR218W | DI49_2186 | | YGR218W |
| DI49_2184; YGR217W | DI49_2184 | | YGR217W |
| DI49_2183; YGR216C; DI49_2182 | DI49_2183; DI49_2182 | | YGR216C |
| DI49_2180; DI49_2181; YGR213C | DI49_2180; DI49_2181 | | YGR213C |
| DI49_2178; DI49_2179 | DI49_2178; DI49_2179 | | |

Create PDF in your applications with the Pdfcrowd HTML to PDF API PDFCROWD

| | | |
|---|---|---|
| DI49_2176; YGR209C | DI49_2176 | YGR209C |
| DI49_2175; YGR208W; DI49_2174; YGR207C | DI49_2175; DI49_2174 | YGR208W; YGR207C |
| DI49_2172; YGR205W | DI49_2172 | YGR205W |
| DI49_2171; YGR204W; DI49_2170; YGR203W | DI49_2171; DI49_2170 | YGR204W; YGR203W |
| DI49_2168; DI49_2169 | DI49_2168; DI49_2169 | |
| DI49_2166; DI49_2167 | DI49_2166; DI49_2167 | |
| DI49_2164; YGR197C; DI49_2163; YGR196C | DI49_2164; DI49_2163 | YGR197C; YGR196C |
| DI49_2161; DI49_2162 | DI49_2161; DI49_2162 | |
| DI49_2159; DI49_3025 | DI49_2159; DI49_3025 | |
| DI49_2158; YGR191W | DI49_2158 | YGR191W |
| DI49_2154; YGR188C; DI49_2153; YGR187C | DI49_2154; DI49_2153 | YGR188C; YGR187C |
| DI49_2151; YGR185C | DI49_2151 | YGR185C |
| DI49_2150; YGR184C; DI49_2149; YGR183C | DI49_2150; DI49_2149 | YGR184C; YGR183C |
| DI49_2145; YGR180C; DI49_2144; YGR179C | DI49_2145; DI49_2144 | YGR180C; YGR179C |
| DI49_2142; YGR177C | DI49_2142 | YGR177C |
| DI49_2141; YGR175C | DI49_2141 | YGR175C |
| DI49_2140; YGR174C | DI49_2140 | YGR174C |
| DI49_2138; YGR173W | DI49_2138 | YGR173W |
| DI49_2137; YGR172C | DI49_2137 | YGR172C |
| DI49_2136; YGR171C | DI49_2136 | YGR171C |
| DI49_2135; YGR170W; DI49_2134; YGR169C-A | DI49_2135; DI49_2134 | YGR170W; YGR169C-A |
| DI49_2132; DI49_2133 | DI49_2132; DI49_2133 | |
| DI49_2130; DI49_2131 | DI49_2130; DI49_2131 | |
| DI49_2127; YGR163W | DI49_2127 | YGR163W |
| DI49_2126; YGR162W | DI49_2126 | YGR162W |
| DI49_2121; DI49_2122 | DI49_2121; DI49_2122 | |
| DI49_2119; YGR156W | DI49_2119 | YGR156W |
| DI49_2118; YGR155W; DI49_2117; YGR154C | DI49_2118; DI49_2117 | YGR155W; YGR154C |
| DI49_2112; DI49_1936; DI49_2111; DI49_1935; YGL032C | DI49_2112; DI49_1936; DI49_2111; DI49_1935 | YGL032C |
| DI49_2108; YGR145W | DI49_2108 | YGR145W |
| DI49_2107; YGR144W | DI49_2107 | YGR144W |
| DI49_2105; YGR143W; DI49_5602 | DI49_2105; DI49_5602 | YGR143W |
| DI49_2103; YGR142W; DI49_2102; YGR141W | DI49_2103; DI49_2102 | YGR142W; YGR141W |
| DI49_2100; DI49_2101 | DI49_2100; DI49_2101 | |
| DI49_2098; YGR136W | DI49_2098 | YGR136W |
| DI49_2097; YGR135W | DI49_2097 | YGR135W |
| DI49_2096; YGR134W | DI49_2096 | YGR134W |
| DI49_2095; YGR133W; DI49_2094; YGR132C | DI49_2095; DI49_2094 | YGR133W; YGR132C |
| DI49_2092; DI49_2093 | DI49_2092; DI49_2093 | |
| DI49_2090; DI49_2091 | DI49_2090; DI49_2091 | |

Create PDF in your applications with the Pdfcrowd HTML to PDF API PDFCROWD

| | | | |
|---|---|---|---|
| DI49_2088; DI49_2089 | DI49_2088; DI49_2089 | | |
| DI49_2086; YGR124W | DI49_2086 | | YGR124W |
| DI49_2084; YGR123C | DI49_2084 | | YGR123C |
| DI49_2079; YGR120C | DI49_2079 | | YGR120C |
| DI49_2078; YGR119C | DI49_2078 | | YGR119C |
| DI49_2077; DI49_5575; DI49_2076; DI49_5576; YGR118W; YGR117C | DI49_2077; DI49_5575; DI49_2076; DI49_5576 | | YGR118W; YGR117C |
| DI49_2074; DI49_2075 | DI49_2074; DI49_2075 | | |
| DI49_2072; YGR111W; DI49_2071; YGR110W | DI49_2072; DI49_2071 | | YGR111W; YGR110W |
| DI49_2069; YGR109C; DI49_2068; YGR108W | DI49_2069; DI49_2068 | | YGR109C; YGR108W |
| DI49_2066; SPGP0Ftrna34L; DI49_2065; YGR106C | DI49_2066; DI49_2065 | SPGP0Ftrna34L | YGR106C |
| DI49_2063; DI49_2064 | DI49_2063; DI49_2064 | | |
| DI49_2061; DI49_2062 | DI49_2061; DI49_2062 | | |
| DI49_2059; YGR100W; DI49_2058; YGR099W | DI49_2059; DI49_2058 | | YGR100W; YGR099W |
| DI49_2056; YGR097W; DI49_2055; YGR096W | DI49_2056; DI49_2055 | | YGR097W; YGR096W |
| DI49_2053; YGR094W; DI49_2052; YGR093W | DI49_2053; DI49_2052 | | YGR094W; YGR093W |
| DI49_2050; DI49_2051 | DI49_2050; DI49_2051 | | |
| DI49_2045; DI49_2046 | DI49_2045; DI49_2046 | | |
| DI49_2043; DI49_2044 | DI49_2043; DI49_2044 | | |
| DI49_2041; YGR084C; DI49_2040; YGR083C | DI49_2041; DI49_2040 | | YGR084C; YGR083C |
| DI49_2038; DI49_2039 | DI49_2038; DI49_2039 | | |
| DI49_2036; YGR079W; DI49_2035; YGR077C | DI49_2036; DI49_2035 | | YGR079W; YGR077C |
| DI49_2033; DI49_2034 | DI49_2033; DI49_2034 | | |
| DI49_2031; DI49_2032; YGR075C; YGR074W | DI49_2031; DI49_2032 | | YGR075C; YGR074W |
| DI49_2029; DI49_2030 | DI49_2029; DI49_2030 | | |
| DI49_2027; YGR068C | DI49_2027 | | YGR068C |
| DI49_2026; YGR067C; DI49_2025; YGR066C | DI49_2026; DI49_2025 | | YGR067C; YGR066C |
| DI49_2024; SPGP0Q02210; SPGP0H00130; SPGP0H00140 | DI49_2024 | SPGP0Q02210; SPGP0H00130; SPGP0H00140 | |
| DI49_2022; DI49_2023 | DI49_2022; DI49_2023 | | |
| DI49_2020; YGR060W; DI49_2019; YGR059W | DI49_2020; DI49_2019 | | YGR060W; YGR059W |
| DI49_2017; DI49_2018 | DI49_2017; DI49_2018 | | |
| DI49_2015; YGR055W | DI49_2015 | | YGR055W |
| DI49_2014; YGR054W; DI49_2013; YGR053C | DI49_2014; DI49_2013 | | YGR054W; YGR053C |
| DI49_2011; YGR049W; DI49_2010; YGR048W | DI49_2011; DI49_2010 | | YGR049W; YGR048W |
| DI49_2008; YGR046W; DI49_2007; YGR044C | DI49_2008; DI49_2007 | | YGR046W; YGR044C |
| DI49_2005; DI49_2006 | DI49_2005; DI49_2006 | | |
| DI49_2003; DI49_2004 | DI49_2003; DI49_2004 | | |
| DI49_1999; YGR036C; DI49_1998; YGR035C | DI49_1999; DI49_1998 | | YGR036C; YGR035C |
| DI49_1996; YGR033C | DI49_1996 | | YGR033C |
| DI49_1994; YGR031W; DI49_1993; YGR030C | DI49_1994; DI49_1993 | | YGR031W; YGR030C |
| DI49_1988; YGR026W; DI49_1987; YGR024C | DI49_1988; DI49_1987 | | YGR026W; YGR024C |

Create PDF in your applications with the Pdfcrowd HTML to PDF API PDFCROWD

| | | | |
|---|---|---|---|
| DI49_1983; YGR020C | DI49_1983 | | YGR020C |
| DI49_1982; YGR019W | DI49_1982 | | YGR019W |
| DI49_1981; YGR017W; DI49_1980; YGR016W | DI49_1981; DI49_1980 | | YGR017W; YGR016W |
| DI49_1978; DI49_1979 | DI49_1978; DI49_1979 | | |
| DI49_1976; DI49_1977 | DI49_1976; DI49_1977 | | |
| DI49_1974; YGR009C; DI49_1973; YGR008C | DI49_1974; DI49_1973 | | YGR009C; YGR008C |
| DI49_1971; DI49_1972 | DI49_1971; DI49_1972 | | |
| DI49_1969; YGR004W | DI49_1969 | | YGR004W |
| DI49_1968; YGR003W | DI49_1968 | | YGR003W |
| DI49_1967; YGR002C; SPGP0Q02770; YGR001C | DI49_1967 | SPGP0Q02770 | YGR002C; YGR001C |
| DI49_1965; DI49_1966 | DI49_1965; DI49_1966 | | |
| DI49_1963; DI49_1964 | DI49_1963; DI49_1964 | | |
| DI49_1961; YGL006W; DI49_1960 | DI49_1961; DI49_1960 | | YGL006W |
| DI49_1958; YGL009C; DI49_1957; YGL010W | DI49_1958; DI49_1957 | | YGL009C; YGL010W |
| DI49_1955; DI49_1956 | DI49_1955; DI49_1956 | | |
| DI49_1953; YGL014W; DI49_1952; YGL015C | DI49_1953; DI49_1952 | | YGL014W; YGL015C |
| DI49_1950; DI49_1951 | DI49_1950; DI49_1951 | | |
| DI49_1948; DI49_1949 | DI49_1948; DI49_1949 | | |
| DI49_1946; YGL021W | DI49_1946 | | YGL021W |
| DI49_1945; YGL022W | DI49_1945 | | YGL022W |
| DI49_1944; YGL023C; DI49_1943; YGL025C | DI49_1944; DI49_1943 | | YGL023C; YGL025C |
| DI49_1941; YGL027C | DI49_1941 | | YGL027C |
| DI49_1940; YGL028C | DI49_1940 | | YGL028C |
| DI49_1931; DI49_1932 | DI49_1931; DI49_1932 | | |
| DI49_1929; YGL038C; YGL039W | DI49_1929 | | YGL038C; YGL039W |
| DI49_1928; DI49_1820 | DI49_1928; DI49_1820 | | |
| SPGP0Qtrna20L; DI49_4635; DI49_1818; DI49_1817 | DI49_4635; DI49_1818; DI49_1817 | SPGP0Qtrna20L | |
| DI49_1926; YGL040C; DI49_1925; YGL041W-A | DI49_1926; DI49_1925 | | YGL040C; YGL041W-A |
| DI49_1923; YGL044C; DI49_1922; YGL045W | DI49_1923; DI49_1922 | | YGL044C; YGL045W |
| DI49_1919; YGL048C | DI49_1919 | | YGL048C |
| DI49_1918; YGL049C; DI49_1917; YGL050W | DI49_1918; DI49_1917 | | YGL049C; YGL050W |
| DI49_1912; YGL056C; DI49_1911; YGL057C | DI49_1912; DI49_1911 | | YGL056C; YGL057C |
| DI49_1909; DI49_1910 | DI49_1909; DI49_1910 | | |
| DI49_1908; SPGP0Q03310 | DI49_1908 | SPGP0Q03310 | |
| DI49_1906; DI49_1907 | DI49_1906; DI49_1907 | | |
| DI49_1904; YGL065C | DI49_1904 | | YGL065C |
| DI49_1903; YGL066W; DI49_1902; YGL067W | DI49_1903; DI49_1902 | | YGL066W; YGL067W |
| DI49_1900; YGL070C | DI49_1900 | | YGL070C |
| DI49_1899; YGL071W | DI49_1899 | | YGL071W |
| DI49_1898; YGL073W; DI49_1897; YGL075C | DI49_1898; DI49_1897 | | YGL073W; YGL075C |

Create PDF in your applications with the Pdfcrowd HTML to PDF API PDFCROWD

| | | | |
|---|---|---|---|
| DI49_1896; YGL077C | DI49_1896 | | YGL077C |
| DI49_1895; YGL078C | DI49_1895 | | YGL078C |
| DI49_1894; YGL079W; DI49_1893; YGL080W | DI49_1894; DI49_1893 | | YGL079W; YGL080W |
| DI49_1891; DI49_1892 | DI49_1891; DI49_1892 | | |
| DI49_1889; YGL084C | DI49_1889 | | YGL084C |
| DI49_1888; YGL085W; DI49_1887; YGL086W | DI49_1888; DI49_1887 | | YGL085W; YGL086W |
| DI49_1885; SPGP0Q03550 | DI49_1885 | SPGP0Q03550 | |
| DI49_1883; DI49_1884 | DI49_1883; DI49_1884 | | |
| DI49_1882; YGL093W | DI49_1882 | | YGL093W |
| DI49_1881; YGL094C; DI49_1880; YGL095C | DI49_1881; DI49_1880 | | YGL094C; YGL095C |
| DI49_1876; DI49_1877 | DI49_1876; DI49_1877 | | |
| DI49_1874; YGL098W | DI49_1874 | | YGL098W |
| DI49_1873; YGL099W | DI49_1873 | | YGL099W |
| DI49_1872; YGL100W; DI49_1871; YGL101W | DI49_1872; DI49_1871 | | YGL100W; YGL101W |
| DI49_1869; DI49_1870 | DI49_1869; DI49_1870 | | |
| DI49_1867; DI49_1868 | DI49_1867; DI49_1868 | | |
| DI49_1865; DI49_1866 | DI49_1865; DI49_1866 | | |
| DI49_1863; DI49_1864 | DI49_1863; DI49_1864 | | |
| DI49_1861; DI49_1862 | DI49_1861; DI49_1862 | | |
| DI49_1859; YGL115W | DI49_1859 | | YGL115W |
| DI49_1858; YGL116W; DI49_1857; YGL117W | DI49_1858; DI49_1857 | | YGL116W; YGL117W |
| DI49_1855; DI49_3006; SPGP0Rtrna5W | DI49_1855; DI49_3006 | SPGP0Rtrna5W | |
| DI49_1853; DI49_1854 | DI49_1853; DI49_1854 | | |
| DI49_1851; YGL123W; DI49_1850; YGL124C | DI49_1851; DI49_1850 | | YGL123W; YGL124C |
| DI49_1848; DI49_1849 | DI49_1848; DI49_1849 | | |
| DI49_1846; YGL128C | DI49_1846 | | YGL128C |
| DI49_1845; YGL129C; DI49_1844; YGL130W | DI49_1845; DI49_1844 | | YGL129C; YGL130W |
| DI49_1842; DI49_1843 | DI49_1842; DI49_1843 | | |
| DI49_1840; DI49_5249; DI49_5248; YGL135W | DI49_1840; DI49_5249; DI49_5248 | | YGL135W |
| DI49_1839; YGL136C | DI49_1839 | | YGL136C |
| DI49_1838; YGL137W; DI49_1837; YGL138C | DI49_1838; DI49_1837 | | YGL137W; YGL138C |
| DI49_1835; YGL140C | DI49_1835 | | YGL140C |
| DI49_1834; YGL141W | DI49_1834 | | YGL141W |
| DI49_1833; YGL142C; DI49_1832; YGL143C | DI49_1833; DI49_1832 | | YGL142C; YGL143C |
| DI49_1830; DI49_1831 | DI49_1830; DI49_1831 | | |
| DI49_1828; YGL147C; DI49_1827; DI49_4664; DI49_4663 | DI49_1828; DI49_1827; DI49_4664; DI49_4663 | | YGL147C |
| DI49_1825; YGL151W; DI49_1824; YGL153W | DI49_1825; DI49_1824 | | YGL151W; YGL153W |
| DI49_1822; DI49_1823 | DI49_1822; DI49_1823 | | |
| DI49_1819; YGL158W; YGL159W | DI49_1819 | | YGL158W; YGL159W |

Create PDF in your applications with the Pdfcrowd HTML to PDF API PDFCROWD

| | | | |
|---|---|---|---|
| DI49_1815; DI49_1816 | DI49_1815; DI49_1816 | | |
| SPGP0Q04250; DI49_1814 | DI49_1814 | SPGP0Q04250 | |
| DI49_1812; DI49_1813 | DI49_1812; DI49_1813 | | |
| DI49_1810; DI49_1811 | DI49_1810; DI49_1811 | | |
| DI49_1808; DI49_1809 | DI49_1808; DI49_1809 | | |
| DI49_1806; YGL170C | DI49_1806 | | YGL170C |
| DI49_1805; YGL171W | DI49_1805 | | YGL171W |
| DI49_1804; YGL172W | DI49_1804 | | YGL172W |
| YGL173C; DI49_1803; YGL174W; DI49_1802 | DI49_1803; DI49_1802 | | YGL173C; YGL174W |
| YGL176C; DI49_1800 | DI49_1800 | | YGL176C |
| YGL178W; DI49_1799; YGL179C; DI49_1798 | DI49_1799; DI49_1798 | | YGL178W; YGL179C |
| YGL181W; YGL180W | | | YGL181W; YGL180W |
| YGL184C; YGL183C | | | YGL184C; YGL183C |
| YGL186C; YGL185C | | | YGL186C; YGL185C |
| YGL189C; DI49_1790 | DI49_1790 | | YGL189C |
| YGL190C; DI49_1789; YGL191W; DI49_1788 | DI49_1789; DI49_1788 | | YGL190C; YGL191W |
| YGL194C; YGL192W | | | YGL194C; YGL192W |
| YGL195W; DI49_1784 | DI49_1784 | | YGL195W |
| YGL196W; DI49_1783; SPGP0Q04560; DI49_1782 | DI49_1783; DI49_1782 | SPGP0Q04560 | YGL196W |
| YGL200C; DI49_1780 | DI49_1780 | | YGL200C |
| YGL202W; DI49_1777 | DI49_1777 | | YGL202W |
| YGL207W; DI49_1771; YGL208W; DI49_1770 | DI49_1771; DI49_1770 | | YGL207W; YGL208W |
| YGL210W; YGL209W | | | YGL210W; YGL209W |
| YGL212W; DI49_1766 | DI49_1766 | | YGL212W |
| YGL213C; DI49_1765; YGL215W; DI49_1764 | DI49_1765; DI49_1764 | | YGL213C; YGL215W |
| YGL219C; YGL216W | | | YGL219C; YGL216W |
| YGL221C; DI49_1761 | DI49_1761 | | YGL221C |
| YGL222C; DI49_1760; YGL223C; DI49_1759 | DI49_1760; DI49_1759 | | YGL222C; YGL223C |
| YGL225W; DI49_1757 | DI49_1757 | | YGL225W |
| YGL227W; DI49_1752; YGL228W; DI49_1751 | DI49_1752; DI49_1751 | | YGL227W; YGL228W |
| YGL230C; YGL229C | | | YGL230C; YGL229C |
| YGL232W; YGL231C | | | YGL232W; YGL231C |
| YGL234W; DI49_1746 | DI49_1746 | | YGL234W |
| YGL236C; DI49_1745 | DI49_1745 | | YGL236C |
| YGL237C; DI49_1744 | DI49_1744 | | YGL237C |
| YGL238W; DI49_1743; YGL240W; DI49_1742 | DI49_1743; DI49_1742 | | YGL238W; YGL240W |
| YGL242C; YGL241W | | | YGL242C; YGL241W |
| YGL244W; YGL243W | | | YGL244W; YGL243W |
| YGL246C; YGL245W | | | YGL246C; YGL245W |
| YGL248W; YGL247W | | | YGL248W; YGL247W |

Create PDF in your applications with the Pdfcrowd HTML to PDF API PDFCROWD

| | | | |
|---|---|---|---|
| YGL250W; YGL249W | | | YGL250W; YGL249W |
| YGL252C; SPGP0F04920 | | SPGP0F04920 | YGL252C |
| YGL253W; DI49_1730; YGL254W; DI49_1729 | DI49_1730; DI49_1729 | | YGL253W; YGL254W |
| YGL256W; DI49_1594; YGL257C; DI49_1595 | DI49_1594; DI49_1595 | | YGL256W; YGL257C |
| YGL258W-A; YGL258W | | | YGL258W-A; YGL258W |
| DI49_2825; DI49_2824 | DI49_2825; DI49_2824 | | |
| DI49_2827; DI49_2826 | DI49_2827; DI49_2826 | | |
| SPGP0R00170; DI49_2828; YJL207C; YJL208C | DI49_2828 | SPGP0R00170 | YJL207C; YJL208C |
| DI49_2829; YJL204C; DI49_2830; YJL205C | DI49_2829; DI49_2830 | | YJL204C; YJL205C |
| DI49_2832; YJL200C | DI49_2832 | | YJL200C |
| DI49_2834; YJL198W; DI49_2835; YJL199C | DI49_2834; DI49_2835 | | YJL198W; YJL199C |
| DI49_2837; DI49_2836 | DI49_2837; DI49_2836 | | |
| DI49_2839; YJL192C | DI49_2839 | | YJL192C |
| DI49_2840; DI49_0783; DI49_0784; SPGP0AE00940; YCR032W; SPGP0G03150 | DI49_2840; DI49_0783; DI49_0784 | SPGP0AE00940; SPGP0G03150 | YCR032W |
| DI49_2841; YJL190C | DI49_2841 | | YJL190C |
| DI49_2842; SPGP0G03130; DI49_2843 | DI49_2842; DI49_2843 | SPGP0G03130 | |
| DI49_2845; DI49_2844; YJL186W; YJL187C | DI49_2845; DI49_2844 | | YJL186W; YJL187C |
| DI49_2847; DI49_2846 | DI49_2847; DI49_2846 | | |
| DI49_2849; YJL180C; YJL181W | DI49_2849 | | YJL180C; YJL181W |
| DI49_2850; YJL179W; DI49_2851 | DI49_2850; DI49_2851 | | YJL179W |
| DI49_2853; YJL176C; DI49_2854 | DI49_2853; DI49_2854 | | YJL176C |
| DI49_2856; YJL172W; YJL173C | DI49_2856 | | YJL172W; YJL173C |
| DI49_2857; YJL171C; DI49_2858 | DI49_2857; DI49_2858 | | YJL171C |
| DI49_2860; YJL167W | DI49_2860 | | YJL167W |
| DI49_2861; YJL166W | DI49_2861 | | YJL166W |
| DI49_2862; YJL165C; DI49_2863 | DI49_2862; DI49_2863 | | YJL165C |
| DI49_2865; YJL162C; YJL163C | DI49_2865 | | YJL162C; YJL163C |
| SPGP0R00570; YJL159W; YJL160C | | SPGP0R00570 | YJL159W; YJL160C |
| DI49_2872; YJL155C; YJL156C | DI49_2872 | | YJL155C; YJL156C |
| DI49_2873; YJL154C; DI49_2874 | DI49_2873; DI49_2874 | | YJL154C |
| DI49_2876; YJL149W; YJL151C | DI49_2876 | | YJL149W; YJL151C |
| DI49_2877; YJL148W; DI49_2878 | DI49_2877; DI49_2878 | | YJL148W |
| DI49_2880; YJL145W; DI49_2881 | DI49_2880; DI49_2881 | | YJL145W |
| DI49_2883; YJL141C | DI49_2883 | | YJL141C |
| DI49_2884; YJL140W; DI49_2885 | DI49_2884; DI49_2885 | | YJL140W |
| DI49_2887; SPGP0R00750 | DI49_2887 | SPGP0R00750 | |
| DI49_2888; SPGP0R00770 | DI49_2888 | SPGP0R00770 | |
| DI49_2890; YJL133W | DI49_2890 | | YJL133W |
| SPGP0R00810; YJL132W | | SPGP0R00810 | YJL132W |
| DI49_2891; YJL131C | DI49_2891 | | YJL131C |

Create PDF in your applications with the Pdfcrowd HTML to PDF API PDFCROWD

| | | | |
|---|---|---|---|
| DI49_2892; YJL130C | DI49_2892 | | YJL130C |
| DI49_2893; YJL129C | DI49_2893 | | YJL129C |
| DI49_2894; YJL128C | DI49_2894 | | YJL128C |
| DI49_2895; YJL127C-B; DI49_2896 | DI49_2895; DI49_2896 | | YJL127C-B |
| DI49_2898; YJL125C; DI49_2899; YJL126W | DI49_2898; DI49_2899 | | YJL125C; YJL126W |
| DI49_2900; SPGP0R00910 | DI49_2900 | SPGP0R00910 | |
| SPGP0R00940; DI49_2901 | DI49_2901 | SPGP0R00940 | |
| DI49_2903; DI49_2902 | DI49_2903; DI49_2902 | | |
| DI49_2908; YJL112W | DI49_2908 | | YJL112W |
| DI49_2909; YJL111W; DI49_2910 | DI49_2909; DI49_2910 | | YJL111W |
| DI49_2912; YJL107C | DI49_2912 | | YJL107C |
| DI49_2913; YJL106W | DI49_2913 | | YJL106W |
| DI49_2914; YJL105W | DI49_2914 | | YJL105W |
| DI49_2915; YJL104W; DI49_2916 | DI49_2915; DI49_2916 | | YJL104W |
| SPGP0Rtrna15R; DI49_2918; YJL102W | DI49_2918 | SPGP0Rtrna15R | YJL102W |
| DI49_2919; YJL101C; DI49_2920 | DI49_2919; DI49_2920 | | YJL101C |
| DI49_2922; YJL098W; DI49_2923; YJL099W | DI49_2922; DI49_2923 | | YJL098W; YJL099W |
| DI49_2925; YJL095W; DI49_2926; YJL096W | DI49_2925; DI49_2926 | | YJL095W; YJL096W |
| DI49_2928; YJL092W; YJL093C | DI49_2928 | | YJL092W; YJL093C |
| DI49_2929; YJL091C | DI49_2929 | | YJL091C |
| DI49_2930; YJL090C | DI49_2930 | | YJL090C |
| DI49_2931; YJL089W; DI49_2932 | DI49_2931; DI49_2932 | | YJL089W |
| DI49_2934; YJL085W | DI49_2934 | | YJL085W |
| SPGP0R01250; YJL084C | | SPGP0R01250 | YJL084C |
| DI49_2935; YJL083W | DI49_2935 | | YJL083W |
| DI49_2936; YJL082W; DI49_2937; YJL081C | DI49_2936; DI49_2937 | | YJL082W; YJL081C |
| DI49_2939; DI49_2938 | DI49_2939; DI49_2938 | | |
| DI49_2941; DI49_2940 | DI49_2941; DI49_2940 | | |
| DI49_2943; DI49_2942 | DI49_2943; DI49_2942 | | |
| DI49_2945; YJL071W; YJL072C | DI49_2945 | | YJL071W; YJL072C |
| DI49_2946; YJL070C; DI49_2947 | DI49_2946; DI49_2947 | | YJL070C |
| DI49_2949; DI49_2948 | DI49_2949; DI49_2948 | | |
| DI49_2951; DI49_2950 | DI49_2951; DI49_2950 | | |
| DI49_2953; YJL062W; DI49_2954 | DI49_2953; DI49_2954 | | YJL062W |
| DI49_2956; YJL059W | DI49_2956 | | YJL059W |
| DI49_2957; YJL058C; DI49_2958 | DI49_2957; DI49_2958 | | YJL058C |
| DI49_2960; DI49_2959 | DI49_2960; DI49_2959 | | |
| DI49_2962; DI49_2961 | DI49_2962; DI49_2961 | | |
| DI49_2964; YJL051W; DI49_2965 | DI49_2964; DI49_2965 | | YJL051W |
| DI49_2967; DI49_2966 | DI49_2967; DI49_2966 | | |

Create PDF in your applications with the Pdfcrowd HTML to PDF API PDFCROWD

| | | | |
|---|---|---|---|
| DI49_2969; DI49_2968 | DI49_2969; DI49_2968 | | |
| DI49_2975; DI49_2974 | DI49_2975; DI49_2974 | | |
| SPGP0R01650; DI49_2976 | DI49_2976 | SPGP0R01650 | |
| DI49_2984; YJL036W; DI49_2985 | DI49_2984; DI49_2985 | | YJL036W |
| DI49_2987; YJL033W; DI49_2988 | DI49_2987; DI49_2988 | | YJL033W |
| DI49_2990; DI49_2989 | DI49_2990; DI49_2989 | | |
| DI49_2993; YJL026W; DI49_2994; YJL027C | DI49_2993; DI49_2994 | | YJL026W; YJL027C |
| DI49_2997; YJL023C; SPGP0R01810 | DI49_2997 | SPGP0R01810 | YJL023C |
| DI49_2999; YJL019W; DI49_3000; YJL020C | DI49_2999; DI49_3000 | | YJL019W; YJL020C |
| DI49_3002; DI49_3001 | DI49_3002; DI49_3001 | | |
| DI49_3004; DI49_3003 | DI49_3004; DI49_3003 | | |
| DI49_3007; YJL010C | DI49_3007 | | YJL010C |
| DI49_3008; YJL008C | DI49_3008 | | YJL008C |
| DI49_3012; YJL005W; DI49_3013 | DI49_3012; DI49_3013 | | YJL005W |
| DI49_3015; DI49_3014 | DI49_3015; DI49_3014 | | |
| DI49_3016; SPGP0R01960 | DI49_3016 | SPGP0R01960 | |
| DI49_3018; DI49_3017 | DI49_3018; DI49_3017 | | |
| DI49_3020; YJR005W; DI49_3021 | DI49_3020; DI49_3021 | | YJR005W |
| DI49_3023; YJR007W; YJR006W | DI49_3023 | | YJR007W; YJR006W |
| DI49_3024; YJR008W | DI49_3024 | | YJR008W |
| DI49_3026; YJR010W; DI49_3027 | DI49_3026; DI49_3027 | | YJR010W |
| DI49_3029; DI49_3028; YJR011C; YJR010C-A | DI49_3029; DI49_3028 | | YJR011C; YJR010C-A |
| DI49_3031; YJR014W; SPGP0R02130 | DI49_3031 | SPGP0R02130 | YJR014W |
| DI49_3033; DI49_3032; YJR016C; YJR015W | DI49_3033; DI49_3032 | | YJR016C; YJR015W |
| SPGP0R02170; YJR021C; DI49_3035; YJR019C | DI49_3035 | SPGP0R02170 | YJR021C; YJR019C |
| DI49_3037; DI49_3036 | DI49_3037; DI49_3036 | | |
| DI49_3039; DI49_3038 | DI49_3039; DI49_3038 | | |
| DI49_3041; YJR033C; DI49_3042 | DI49_3041; DI49_3042 | | YJR033C |
| DI49_3044; YJR036C; YJR035W | DI49_3044 | | YJR036C; YJR035W |
| DI49_3045; YJR039W | DI49_3045 | | YJR039W |
| DI49_3046; YJR040W; DI49_3047 | DI49_3046; DI49_3047 | | YJR040W |
| DI49_3049; DI49_3048 | DI49_3049; DI49_3048 | | |
| DI49_3053; YJR046W | DI49_3053 | | YJR046W |
| DI49_3055; DI49_1358; DI49_3056; YEL034W; YJR047C | DI49_3055; DI49_1358; DI49_3056 | | YEL034W; YJR047C |
| DI49_3058; YJR050W; DI49_3059; YJR049C | DI49_3058; DI49_3059 | | YJR050W; YJR049C |
| DI49_3061; YJR052W; DI49_3062 | DI49_3061; DI49_3062 | | YJR052W |
| SPGP0Rtrna18R; DI49_3064; DI49_3065 | DI49_3064; DI49_3065 | SPGP0Rtrna18R | |
| DI49_3072; YJR059W; DI49_3073; YJR058C | DI49_3072; DI49_3073 | | YJR059W; YJR058C |
| DI49_3075; YJR062C; DI49_3076; YJR061W | DI49_3075; DI49_3076 | | YJR062C; YJR061W |

Create PDF in your applications with the Pdfcrowd HTML to PDF API PDFCROWD

| | | | |
|---|---|---|---|
| DI49_3078; YJR065C | DI49_3078 | | YJR065C |
| DI49_3082; YJR069C | DI49_3082 | | YJR069C |
| DI49_3083; YJR070C; DI49_3084 | DI49_3083; DI49_3084 | | YJR070C |
| SPGP0R02630; DI49_3085 | DI49_3085 | SPGP0R02630 | |
| DI49_3087; YJR076C | DI49_3087 | | YJR076C |
| DI49_3088; YJR077C; DI49_3089 | DI49_3088; DI49_3089 | | YJR077C |
| DI49_3091; DI49_3090 | DI49_3091; DI49_3090 | | |
| DI49_3093; DI49_3092 | DI49_3093; DI49_3092 | | |
| DI49_3095; DI49_3094 | DI49_3095; DI49_3094 | | |
| DI49_3097; DI49_3096 | DI49_3097; DI49_3096 | | |
| DI49_3099; YJR091C; DI49_3100 | DI49_3099; DI49_3100 | | YJR091C |
| DI49_3102; DI49_3101 | DI49_3102; DI49_3101 | | |
| DI49_3104; YJR095W; DI49_3105 | DI49_3104; DI49_3105 | | YJR095W |
| DI49_3107; YJR098C; DI49_3108; YJR097W | DI49_3107; DI49_3108 | | YJR098C; YJR097W |
| DI49_3113; DI49_3112 | DI49_3113; DI49_3112 | | |
| DI49_3115; YJR105W; DI49_3116 | DI49_3115; DI49_3116 | | YJR105W |
| DI49_3118; DI49_3117; YJR107W; YJR106W | DI49_3118; DI49_3117 | | YJR107W; YJR106W |
| DI49_3119; YJR109C; YJR108W | DI49_3119 | | YJR109C; YJR108W |
| DI49_3120; YJR110W | DI49_3120 | | YJR110W |
| DI49_3121; YJR111C; DI49_3122 | DI49_3121; DI49_3122 | | YJR111C |
| DI49_3124; DI49_3123 | DI49_3124; DI49_3123 | | |
| DI49_3126; DI49_3125 | DI49_3126; DI49_3125 | | |
| DI49_3128; YJR118C | DI49_3128 | | YJR118C |
| DI49_3129; YJR119C | DI49_3129 | | YJR119C |
| DI49_3130; YJR121W; DI49_3131 | DI49_3130; DI49_3131 | | YJR121W |
| DI49_3133; YJR124C | DI49_3133 | | YJR124C |
| DI49_3134; YJR125C; DI49_3135 | DI49_3134; DI49_3135 | | YJR125C |
| DI49_3137; DI49_3136 | DI49_3137; DI49_3136 | | |
| DI49_3139; DI49_3138 | DI49_3139; DI49_3138 | | |
| DI49_3141; YJR133W | DI49_3141 | | YJR133W |
| DI49_3142; YJR134C; DI49_3143 | DI49_3142; DI49_3143 | | YJR134C |
| DI49_3145; YJR136C; YJR135W-A | DI49_3145 | | YJR136C; YJR135W-A |
| DI49_3146; YJR137C | DI49_3146 | | YJR137C |
| DI49_3147; YJR138W | DI49_3147 | | YJR138W |
| DI49_3148; YJR139C; DI49_3149 | DI49_3148; DI49_3149 | | YJR139C |
| DI49_3151; DI49_3150 | DI49_3151; DI49_3150 | | |
| DI49_3153; DI49_3152 | DI49_3153; DI49_3152 | | |
| SPGP0R03340; DI49_2608; DI49_2609 | DI49_2608; DI49_2609 | SPGP0R03340 | |
| DI49_3156; DI49_3155 | DI49_3156; DI49_3155 | | |
| DI49_3160; SPGP0R03400 | DI49_3160 | SPGP0R03400 | |

Create PDF in your applications with the Pdfcrowd HTML to PDF API PDFCROWD

| | | | |
|---|---|---|---|
| SPGP0R03430; DI49_3161 | DI49_3161 | SPGP0R03430 | |
| DI49_3163; DI49_4795 | DI49_3163; DI49_4795 | | |
| DI49_3164; DI49_1722; DI49_3165; DI49_1723 | DI49_3164; DI49_1722; DI49_3165; DI49_1723 | | |
| SPGP0R03490; SPGP0AF00510; DI49_3166; DI49_2619 | DI49_3166; DI49_2619 | SPGP0R03490; SPGP0AF00510 | |
| SPGP0R03520; DI49_1322; DI49_2622 | DI49_1322; DI49_2622 | SPGP0R03520 | |
| DI49_4371; DI49_4372 | DI49_4371; DI49_4372 | | |
| DI49_4367; DI49_4368 | DI49_4367; DI49_4368 | | |
| DI49_4365; DI49_4366 | DI49_4365; DI49_4366 | | |
| DI49_4363; DI49_4364 | DI49_4363; DI49_4364 | | |
| DI49_4355; DI49_4356 | DI49_4355; DI49_4356 | | |
| DI49_4350; DI49_4351 | DI49_4350; DI49_4351 | | |
| DI49_4337; DI49_4338 | DI49_4337; DI49_4338 | | |
| DI49_4335; DI49_4336 | DI49_4335; DI49_4336 | | |
| DI49_4330; DI49_4331 | DI49_4330; DI49_4331 | | |
| DI49_4307; DI49_4308 | DI49_4307; DI49_4308 | | |
| DI49_4297; DI49_4298 | DI49_4297; DI49_4298 | | |
| DI49_4295; DI49_4296 | DI49_4295; DI49_4296 | | |
| DI49_4293; DI49_4294 | DI49_4293; DI49_4294 | | |
| DI49_4274; DI49_4275 | DI49_4274; DI49_4275 | | |
| DI49_4265; DI49_4266 | DI49_4265; DI49_4266 | | |
| DI49_4261; DI49_4262 | DI49_4261; DI49_4262 | | |
| DI49_4257; DI49_4258 | DI49_4257; DI49_4258 | | |
| DI49_4254; DI49_4253; DI49_4252 | DI49_4254; DI49_4253; DI49_4252 | | |
| DI49_4240; DI49_4241 | DI49_4240; DI49_4241 | | |
| DI49_4238; DI49_4239 | DI49_4238; DI49_4239 | | |
| DI49_4236; DI49_4237 | DI49_4236; DI49_4237 | | |
| DI49_4234; DI49_4235 | DI49_4234; DI49_4235 | | |
| DI49_4232; DI49_4233 | DI49_4232; DI49_4233 | | |
| DI49_4226; DI49_4227 | DI49_4226; DI49_4227 | | |
| DI49_4224; DI49_4225 | DI49_4224; DI49_4225 | | |
| DI49_4222; DI49_4223 | DI49_4222; DI49_4223 | | |
| DI49_4221; SPGP0S01630 | DI49_4221 | SPGP0S01630 | |
| DI49_4219; DI49_4220 | DI49_4219; DI49_4220 | | |
| DI49_4206; SPGP0Strna25L | DI49_4206 | SPGP0Strna25L | |
| DI49_4199; DI49_4200 | DI49_4199; DI49_4200 | | |
| DI49_4197; DI49_4198 | DI49_4197; DI49_4198 | | |
| DI49_4183; DI49_4184 | DI49_4183; DI49_4184 | | |
| DI49_4178; DI49_2707; DI49_2708 | DI49_4178; DI49_2707; DI49_2708 | | |
| DI49_4177; YMR089C | DI49_4177 | | YMR089C |
| DI49_4169; DI49_4170 | DI49_4169; DI49_4170 | | |

Create PDF in your applications with the Pdfcrowd HTML to PDF API PDFCROWD

| | | | |
|---|---|---|---|
| DI49_4167; DI49_4168 | DI49_4167; DI49_4168 | | |
| DI49_4161; DI49_4162 | DI49_4161; DI49_4162 | | |
| DI49_4159; DI49_4160 | DI49_4159; DI49_4160 | | |
| DI49_4151; DI49_4152 | DI49_4151; DI49_4152 | | |
| DI49_4149; DI49_4150 | DI49_4149; DI49_4150 | | |
| DI49_4140; DI49_3481; SPGP0BF00170 | DI49_4140; DI49_3481 | SPGP0BF00170 | |
| SPGP0S02420; DI49_2906; DI49_1546 | DI49_2906; DI49_1546 | SPGP0S02420 | |
| DI49_4136; DI49_4137 | DI49_4136; DI49_4137 | | |
| DI49_4131; DI49_4132 | DI49_4131; DI49_4132 | | |
| DI49_4129; DI49_4130 | DI49_4129; DI49_4130 | | |
| DI49_4127; DI49_4128 | DI49_4127; DI49_4128 | | |
| DI49_4125; DI49_4126 | DI49_4125; DI49_4126 | | |
| DI49_4123; DI49_4124 | DI49_4123; DI49_4124 | | |
| DI49_4121; DI49_4122 | DI49_4121; DI49_4122 | | |
| DI49_4119; DI49_4120 | DI49_4119; DI49_4120 | | |
| DI49_4117; DI49_4118 | DI49_4117; DI49_4118 | | |
| DI49_4104; DI49_4105 | DI49_4104; DI49_4105 | | |
| DI49_4091; DI49_4092 | DI49_4091; DI49_4092 | | |
| DI49_4076; DI49_4077 | DI49_4076; DI49_4077 | | |
| DI49_4074; DI49_4075 | DI49_4074; DI49_4075 | | |
| DI49_4060; DI49_4061 | DI49_4060; DI49_4061 | | |
| DI49_4024; DI49_4025 | DI49_4024; DI49_4025 | | |
| DI49_4022; DI49_4023 | DI49_4022; DI49_4023 | | |
| DI49_4380; DI49_4379 | DI49_4380; DI49_4379 | | |
| DI49_4382; DI49_4381 | DI49_4382; DI49_4381 | | |
| DI49_4384; DI49_4383 | DI49_4384; DI49_4383 | | |
| DI49_4386; DI49_4385 | DI49_4386; DI49_4385 | | |
| DI49_4388; DI49_4387 | DI49_4388; DI49_4387 | | |
| DI49_4393; DI49_4392 | DI49_4393; DI49_4392 | | |
| DI49_4395; DI49_4394 | DI49_4395; DI49_4394 | | |
| DI49_4397; DI49_4396 | DI49_4397; DI49_4396 | | |
| DI49_4399; DI49_4398 | DI49_4399; DI49_4398 | | |
| DI49_4401; DI49_4400 | DI49_4401; DI49_4400 | | |
| DI49_4403; DI49_4402 | DI49_4403; DI49_4402 | | |
| DI49_4406; DI49_4404 | DI49_4406; DI49_4404 | | |
| DI49_4408; DI49_4407 | DI49_4408; DI49_4407 | | |
| YPL274W; SPGP0U02260 | | SPGP0U02260 | YPL274W |
| YOR391C; SPGP0U02270; DI49_5408 | DI49_5408 | SPGP0U02270 | YOR391C |
| SPGP0U00120; DI49_5407 | DI49_5407 | SPGP0U00120 | |
| DI49_5405; DI49_5406 | DI49_5405; DI49_5406 | | |

Create PDF in your applications with the Pdfcrowd HTML to PDF API PDFCROWD

| | | | |
|---|---|---|---|
| DI49_5403; DI49_5404 | DI49_5403; DI49_5404 | | |
| DI49_5401; DI49_5402 | DI49_5401; DI49_5402 | | |
| DI49_5399; DI49_5400; SPGP0Utrna3C | DI49_5399; DI49_5400 | SPGP0Utrna3C | |
| DI49_5397; DI49_5398 | DI49_5397; DI49_5398 | | |
| DI49_5395; DI49_5396 | DI49_5395; DI49_5396 | | |
| DI49_5393; DI49_5394 | DI49_5393; DI49_5394 | | |
| DI49_5391; DI49_5392 | DI49_5391; DI49_5392 | | |
| DI49_5389; DI49_5390 | DI49_5389; DI49_5390 | | |
| DI49_5387; DI49_5388 | DI49_5387; DI49_5388 | | |
| DI49_5385; DI49_5386 | DI49_5385; DI49_5386 | | |
| DI49_5383; DI49_5384 | DI49_5383; DI49_5384 | | |
| DI49_5381; DI49_1215; DI49_5380; DI49_1214 | DI49_5381; DI49_1215; DI49_5380; DI49_1214 | | |
| DI49_5378; DI49_5379 | DI49_5378; DI49_5379 | | |
| DI49_5376; DI49_5377 | DI49_5376; DI49_5377 | | |
| DI49_5374; DI49_5375 | DI49_5374; DI49_5375 | | |
| DI49_5372; SPGP0W01220; DI49_5371; DI49_1206 | DI49_5372; DI49_5371; DI49_1206 | SPGP0W01220 | |
| DI49_5369; DI49_5370 | DI49_5369; DI49_5370 | | |
| DI49_5367; DI49_5368 | DI49_5367; DI49_5368 | | |
| DI49_5365; DI49_5366 | DI49_5365; DI49_5366 | | |
| DI49_5363; DI49_5364 | DI49_5363; DI49_5364 | | |
| DI49_5361; DI49_5362 | DI49_5361; DI49_5362 | | |
| DI49_5359; DI49_5360 | DI49_5359; DI49_5360 | | |
| DI49_5357; DI49_5358 | DI49_5357; DI49_5358 | | |
| DI49_5355; DI49_5356 | DI49_5355; DI49_5356 | | |
| SPGP0U00650; DI49_5354 | DI49_5354 | SPGP0U00650 | |
| SPGP0U00670; YPL109C | | SPGP0U00670 | YPL109C |
| DI49_5352; DI49_5353 | DI49_5352; DI49_5353 | | |
| DI49_5348; DI49_5349 | DI49_5348; DI49_5349 | | |
| DI49_5346; DI49_5347 | DI49_5346; DI49_5347 | | |
| DI49_5344; DI49_5345 | DI49_5344; DI49_5345 | | |
| DI49_5342; DI49_5343 | DI49_5342; DI49_5343 | | |
| DI49_5340; DI49_5341 | DI49_5340; DI49_5341 | | |
| DI49_5338; DI49_5339 | DI49_5338; DI49_5339 | | |
| DI49_5336; DI49_5337 | DI49_5336; DI49_5337 | | |
| DI49_5334; DI49_5335 | DI49_5334; DI49_5335 | | |
| DI49_5332; DI49_5333 | DI49_5332; DI49_5333 | | |
| DI49_5330; DI49_5331 | DI49_5330; DI49_5331 | | |
| DI49_5328; DI49_5329 | DI49_5328; DI49_5329 | | |
| DI49_5326; DI49_5327 | DI49_5326; DI49_5327 | | |
| DI49_5324; DI49_5325 | DI49_5324; DI49_5325 | | |

Create PDF in your applications with the Pdfcrowd HTML to PDF API PDFCROWD

| | | | |
|---|---|---|---|
| DI49_5319; DI49_5320 | DI49_5319; DI49_5320 | | |
| DI49_5317; DI49_5318 | DI49_5317; DI49_5318 | | |
| DI49_5315; DI49_5316 | DI49_5315; DI49_5316 | | |
| DI49_5313; DI49_5314 | DI49_5313; DI49_5314 | | |
| DI49_5311; DI49_5312 | DI49_5311; DI49_5312 | | |
| DI49_5309; DI49_5310 | DI49_5309; DI49_5310 | | |
| DI49_5307; DI49_5308 | DI49_5307; DI49_5308 | | |
| DI49_5305; DI49_5306 | DI49_5305; DI49_5306 | | |
| DI49_5303; DI49_5304 | DI49_5303; DI49_5304 | | |
| DI49_5301; DI49_5302 | DI49_5301; DI49_5302 | | |
| DI49_5299; DI49_5300 | DI49_5299; DI49_5300 | | |
| DI49_5297; DI49_5298 | DI49_5297; DI49_5298 | | |
| DI49_5295; DI49_5296 | DI49_5295; DI49_5296 | | |
| DI49_5293; DI49_5294 | DI49_5293; DI49_5294 | | |
| DI49_5292; SPGP0U01280 | DI49_5292 | SPGP0U01280 | |
| DI49_5290; DI49_5291 | DI49_5290; DI49_5291 | | |
| DI49_5288; DI49_5289 | DI49_5288; DI49_5289 | | |
| DI49_5284; DI49_5285 | DI49_5284; DI49_5285 | | |
| DI49_5282; DI49_5283 | DI49_5282; DI49_5283 | | |
| DI49_5280; DI49_5281 | DI49_5280; DI49_5281 | | |
| DI49_5278; DI49_5279 | DI49_5278; DI49_5279 | | |
| DI49_5276; DI49_5277 | DI49_5276; DI49_5277 | | |
| DI49_5274; DI49_5275 | DI49_5274; DI49_5275 | | |
| DI49_5272; DI49_5273 | DI49_5272; DI49_5273 | | |
| YPL195W; DI49_5271 | DI49_5271 | | YPL195W |
| DI49_5268; DI49_5270 | DI49_5268; DI49_5270 | | |
| DI49_5266; DI49_5267 | DI49_5266; DI49_5267 | | |
| DI49_5264; YJL164C; DI49_5263 | DI49_5264; DI49_5263 | | YJL164C |
| DI49_5261; DI49_5262 | DI49_5261; DI49_5262 | | |
| DI49_5259; DI49_5260 | DI49_5259; DI49_5260 | | |
| DI49_5257; DI49_5258 | DI49_5257; DI49_5258 | | |
| DI49_5255; DI49_5256 | DI49_5255; DI49_5256 | | |
| DI49_5253; DI49_5254 | DI49_5253; DI49_5254 | | |
| DI49_5251; DI49_5252 | DI49_5251; DI49_5252 | | |
| DI49_5250; YPL218W | DI49_5250 | | YPL218W |
| DI49_5245; DI49_5246 | DI49_5245; DI49_5246 | | |
| DI49_5243; DI49_5244 | DI49_5243; DI49_5244 | | |
| DI49_5241; DI49_5242 | DI49_5241; DI49_5242 | | |
| DI49_5239; DI49_5240 | DI49_5239; DI49_5240 | | |
| DI49_5237; DI49_5238 | DI49_5237; DI49_5238 | | |

Create PDF in your applications with the Pdfcrowd HTML to PDF API PDFCROWD

| | | | |
|---|---|---|---|
| DI49_5235; DI49_5236 | DI49_5235; DI49_5236 | | |
| DI49_5233; DI49_5234 | DI49_5233; DI49_5234 | | |
| DI49_5231; DI49_5232 | DI49_5231; DI49_5232 | | |
| YPL244C; YPL243W | | | YPL244C; YPL243W |
| YPL246C; YPL245W | | | YPL246C; YPL245W |
| YPL248C; YPL247C | | | YPL248C; YPL247C |
| YPL253C; YPL252C | | | YPL253C; YPL252C |
| YPL255W; YPL254W | | | YPL255W; YPL254W |
| SPGP0U02090; YPL256C | | SPGP0U02090 | YPL256C |
| SPGP0Utrna7W; SPGP0Ftrna10W; SPGP0Ftrna9W; YGR189C; YGL119W; SPGP0Gtrna11W | | SPGP0Utrna7W; SPGP0Ftrna10W; SPGP0Ftrna9W; SPGP0Gtrna11W | YGR189C; YGL119W |
| YPL262W; YPL260W | | | YPL262W; YPL260W |
| YPL264C; YPL263C | | | YPL264C; YPL263C |
| YPL266W; YPL265W | | | YPL266W; YPL265W |
| YPL268W; YPL267W | | | YPL268W; YPL267W |
| YPL270W; YPL269W | | | YPL270W; YPL269W |
| YPL272C; YPL271W | | | YPL272C; YPL271W |
| YNL336W; SPGP0DY00120; SPGP0CF00100 | | SPGP0DY00120; SPGP0CF00100 | YNL336W |
| YHL040C; YHL043W | | | YHL040C; YHL043W |
| YHL038C; YHL039W | | | YHL038C; YHL039W |
| YHL035C; YHL036W | | | YHL035C; YHL036W |
| YHL026C; YHL027W | | | YHL026C; YHL027W |
| YHL015W; DI49_4833; YHL014C | DI49_4833 | | YHL015W; YHL014C |
| YHL012W; YHL013C | | | YHL012W; YHL013C |
| YHL010C; DI49_4838; YHL009C | DI49_4838 | | YHL010C; YHL009C |
| YHR003C; DI49_4851; YHR004C | DI49_4851 | | YHR003C; YHR004C |
| YHR005C-A; YHR005C | | | YHR005C-A; YHR005C |
| YHR010W; SPGP0L01160; YHR011W | | SPGP0L01160 | YHR010W; YHR011W |
| YHR013C; DI49_4862; YHR014W | DI49_4862 | | YHR013C; YHR014W |
| YHR024C; YHR023W | | | YHR024C; YHR023W |
| YHR040W; DI49_2446; YHR041C | DI49_2446 | | YHR040W; YHR041C |
| YHR044C; YHR042W | | | YHR044C; YHR042W |
| YHR046C; YHR045W | | | YHR046C; YHR045W |
| YHR048W; YHR047C | | | YHR048W; YHR047C |
| YHR050W; YHR049W | | | YHR050W; YHR049W |
| YHR052W; DI49_2457; YHR055C | DI49_2457 | | YHR052W; YHR055C |
| YHR062C; YHR061C | | | YHR062C; YHR061C |
| YHR066W; YDR312W; YHR067W; YDR313C | | | YHR066W; YDR312W; YHR067W; YDR313C |
| YHR073W; YHR072W-A | | | YHR073W; YHR072W-A |
| YHR078W; YHR077C | | | YHR078W; YHR077C |
| YHR079C-A; YHR079C | | | YHR079C-A; YHR079C |

Create PDF in your applications with the Pdfcrowd HTML to PDF API PDFCROWD

| | | | |
|---|---|---|---|
| YHR081W; YHR080C | | | YHR081W; YHR080C |
| YHR083W; YHR082C | | | YHR083W; YHR082C |
| YHR087W; DI49_2494; YHR088W | DI49_2494 | | YHR087W; YHR088W |
| YHR105W; YHR104W | | | YHR105W; YHR104W |
| YHR110W; YHR109W | | | YHR110W; YHR109W |
| YHR112C; YHR111W | | | YHR112C; YHR111W |
| YHR117W; YHR116W | | | YHR117W; YHR116W |
| YHR122W; YHR121W | | | YHR122W; YHR121W |
| YHR129C; DI49_2533; YHR131C | DI49_2533 | | YHR129C; YHR131C |
| YHR132W-A; DI49_2536; YHR133C | DI49_2536 | | YHR132W-A; YHR133C |
| YHR135C; YHR134W | | | YHR135C; YHR134W |
| YHR137W; YHR136C | | | YHR137W; YHR136C |
| YHR139C; YHR138C | | | YHR139C; YHR138C |
| YHR142W; DI49_2544; YHR143W | DI49_2544 | | YHR142W; YHR143W |
| YHR144C; YHR143W-A | | | YHR144C; YHR143W-A |
| YHR152W; YHR151C | | | YHR152W; YHR151C |
| YHR158C; YHR157W | | | YHR158C; YHR157W |
| YHR160C; YHR159W | | | YHR160C; YHR159W |
| YHR162W; YHR161C | | | YHR162W; YHR161C |
| DI49_1319; DI49_1320; YJL217W; YJL218W | DI49_1319; DI49_1320 | | YJL217W; YJL218W |
| DI49_1317; DI49_1318 | DI49_1317; DI49_1318 | | |
| DI49_1315; DI49_1316 | DI49_1315; DI49_1316 | | |
| DI49_1313; DI49_1314 | DI49_1313; DI49_1314 | | |
| DI49_1310; YBR287W; DI49_1309 | DI49_1310; DI49_1309 | | YBR287W |
| DI49_1307; DI49_1308 | DI49_1307; DI49_1308 | | |
| DI49_1301; DI49_1302 | DI49_1301; DI49_1302 | | |
| DI49_1295; DI49_1296 | DI49_1295; DI49_1296 | | |
| DI49_1293; DI49_1294 | DI49_1293; DI49_1294 | | |
| DI49_1290; YBR265W; DI49_1289 | DI49_1290; DI49_1289 | | YBR265W |
| DI49_1282; DI49_1283 | DI49_1282; DI49_1283 | | |
| DI49_1280; YBR255C-A; DI49_1279 | DI49_1280; DI49_1279 | | YBR255C-A |
| DI49_1275; DI49_1276 | DI49_1275; DI49_1276 | | |
| DI49_1254; DI49_1255 | DI49_1254; DI49_1255 | | |
| DI49_1253; SPGP0W00770 | DI49_1253 | SPGP0W00770 | |
| DI49_1246; YBR221C; DI49_1245 | DI49_1246; DI49_1245 | | YBR221C |
| DI49_1240; YBR214W; DI49_1239 | DI49_1240; DI49_1239 | | YBR214W |
| DI49_1222; DI49_1223 | DI49_1222; DI49_1223 | | |
| DI49_1220; DI49_1221 | DI49_1220; DI49_1221 | | |
| DI49_1218; DI49_1219 | DI49_1218; DI49_1219 | | |

Create PDF in your applications with the Pdfcrowd HTML to PDF API PDFCROWD

| | | | |
|---|---|---|---|
| DI49_1212; DI49_1213 | DI49_1212; DI49_1213 | | |
| DI49_1210; DI49_1211 | DI49_1210; DI49_1211 | | |
| DI49_1208; DI49_1209 | DI49_1208; DI49_1209 | | |
| DI49_1202; DI49_1203 | DI49_1202; DI49_1203 | | |
| DI49_1191; YBR163W; DI49_1190 | DI49_1191; DI49_1190 | | YBR163W |
| DI49_1188; DI49_1189 | DI49_1188; DI49_1189 | | |
| DI49_1185; YBR158W; DI49_1184 | DI49_1185; DI49_1184 | | YBR158W |
| DI49_1176; DI49_1177 | DI49_1176; DI49_1177 | | |
| DI49_1174; YBR146W; DI49_1173 | DI49_1174; DI49_1173 | | YBR146W |
| DI49_1162; YBR132C; DI49_1161 | DI49_1162; DI49_1161 | | YBR132C |
| DI49_1159; DI49_1160 | DI49_1159; DI49_1160 | | |
| DI49_1154; YBR123C; DI49_1153 | DI49_1154; DI49_1153 | | YBR123C |
| DI49_1139; DI49_1140 | DI49_1139; DI49_1140 | | |
| DI49_1133; DI49_1134 | DI49_1133; DI49_1134 | | |
| DI49_1119; DI49_1121 | DI49_1119; DI49_1121 | | |
| DI49_1096; DI49_1097 | DI49_1096; DI49_1097 | | |
| DI49_1090; DI49_1091 | DI49_1090; DI49_1091 | | |
| DI49_1088; DI49_1089 | DI49_1088; DI49_1089 | | |
| DI49_1076; YBR046C; DI49_1075 | DI49_1076; DI49_1075 | | YBR046C |
| DI49_1070; YBR041W; DI49_1069 | DI49_1070; DI49_1069 | | YBR041W |
| DI49_1065; YBR036C; DI49_1064 | DI49_1065; DI49_1064 | | YBR036C |
| SPGP0W02630; DI49_1063 | DI49_1063 | SPGP0W02630 | |
| SPGP0W02650; YDR011W; DI49_1061 | DI49_1061 | SPGP0W02650 | YDR011W |
| DI49_1059; YDR006C; DI49_1058 | DI49_1059; DI49_1058 | | YDR006C |
| DI49_1056; DI49_1057 | DI49_1056; DI49_1057 | | |
| DI49_1054; YDR001C; DI49_1053 | DI49_1054; DI49_1053 | | YDR001C |
| DI49_1051; DI49_1052; YDL002C; YDL001W | DI49_1051; DI49_1052 | | YDL002C; YDL001W |
| DI49_1049; YDL005C; YDL004W | DI49_1049 | | YDL005C; YDL004W |
| DI49_1048; YDL006W | DI49_1048 | | YDL006W |
| DI49_1047; YDL007W | DI49_1047 | | YDL007W |
| SPGP0W02850; DI49_1042; YDL010W; YDL008W | DI49_1042 | SPGP0W02850 | YDL010W; YDL008W |
| DI49_1039; DI49_1041 | DI49_1039; DI49_1041 | | |
| DI49_1037; DI49_1038 | DI49_1037; DI49_1038 | | |
| DI49_1035; YDL019C; DI49_1034; YDL018C | DI49_1035; DI49_1034 | | YDL019C; YDL018C |
| DI49_1032; YDL022W; YDL021W | DI49_1032 | | YDL022W; YDL021W |
| DI49_1028; DI49_1029 | DI49_1028; DI49_1029 | | |
| SPGP0W02990; DI49_1027; YDL028C; YDL027C | DI49_1027 | SPGP0W02990 | YDL028C; YDL027C |
| DI49_1025; YDL031W; SPGP0W03020; YDL030W | DI49_1025 | SPGP0W03020 | YDL031W; YDL030W |
| DI49_1023; DI49_1024 | DI49_1023; DI49_1024 | | |
| DI49_1022; YDL040C; YDL039C | DI49_1022 | | YDL040C; YDL039C |

Create PDF in your applications with the Pdfcrowd HTML to PDF API PDFCROWD

| | | | |
|---|---|---|---|
| DI49_1021; YDL042C; DI49_1020 | DI49_1021; DI49_1020 | | YDL042C |
| DI49_1018; YDL045C; DI49_1017 | DI49_1018; DI49_1017 | | YDL045C |
| DI49_1015; YDL047W; YDL046W | DI49_1015 | | YDL047W; YDL046W |
| DI49_1014; YDL048C | DI49_1014 | | YDL048C |
| DI49_1013; YDL049C | DI49_1013 | | YDL049C |
| DI49_1012; YDL051W; DI49_1011 | DI49_1012; DI49_1011 | | YDL051W |
| DI49_1009; DI49_1010 | DI49_1009; DI49_1010 | | |
| DI49_1007; YDL055C | DI49_1007 | | YDL055C |
| DI49_1006; YDL056W; DI49_1005 | DI49_1006; DI49_1005 | | YDL056W |
| DI49_1003; DI49_1004; YDL058W; YDL057W | DI49_1003; DI49_1004 | | YDL058W; YDL057W |
| DI49_1001; YDL061C; YDL060W | DI49_1001 | | YDL061C; YDL060W |
| DI49_1000; YDL063C | DI49_1000 | | YDL063C |
| SPGP0W03280; YDL064W; DI49_0999 | DI49_0999 | SPGP0W03280 | YDL064W |
| DI49_0997; YDL067C; DI49_0996 | DI49_0997; DI49_0996 | | YDL067C |
| DI49_0994; DI49_0995 | DI49_0994; DI49_0995 | | |
| DI49_0992; YDL074C | DI49_0992 | | YDL074C |
| DI49_0990; YDL076C | DI49_0990 | | YDL076C |
| DI49_0989; YDL077C | DI49_0989 | | YDL077C |
| DI49_0988; YDL078C; SPGP0W03410 | DI49_0988 | SPGP0W03410 | YDL078C |
| DI49_0987; YDL080C; YDL079C | DI49_0987 | | YDL080C; YDL079C |
| DI49_0986; YDL081C | DI49_0986 | | YDL081C |
| SPGP0W03460; YDL083C | | SPGP0W03460 | YDL083C |
| DI49_0984; YDL084W | DI49_0984 | | YDL084W |
| DI49_0983; YDL085W; DI49_0982 | DI49_0983; DI49_0982 | | YDL085W |
| DI49_0980; YDL087C; DI49_0979; YDL086W | DI49_0980; DI49_0979 | | YDL087C; YDL086W |
| SPGP0W03540; DI49_0978 | DI49_0978 | SPGP0W03540 | |
| DI49_0976; DI49_0977; YDL091C; YDL090C | DI49_0976; DI49_0977 | | YDL091C; YDL090C |
| DI49_0974; YDL095W; YDL093W | DI49_0974 | | YDL095W; YDL093W |
| DI49_0973; YDL097C; DI49_0972 | DI49_0973; DI49_0972 | | YDL097C |
| DI49_0970; YDL100C; DI49_0969; YDL099W | DI49_0970; DI49_0969 | | YDL100C; YDL099W |
| DI49_0967; YDL103C; DI49_0966; YDL102W | DI49_0967; DI49_0966 | | YDL103C; YDL102W |
| DI49_0964; YDL106C; DI49_0963 | DI49_0964; DI49_0963 | | YDL106C |
| DI49_0961; DI49_0962 | DI49_0961; DI49_0962 | | |
| DI49_0959; YDL111C; DI49_0958 | DI49_0959; DI49_0958 | | YDL111C |
| DI49_0956; YDL114W; YDL113C | DI49_0956 | | YDL114W; YDL113C |
| DI49_0955; YDL115C; DI49_0954 | DI49_0955; DI49_0954 | | YDL115C |
| DI49_0952; YDL119C; DI49_0951; YDL117W | DI49_0952; DI49_0951 | | YDL119C; YDL117W |
| DI49_0949; YDL122W; DI49_0948; YDL121C | DI49_0949; DI49_0948 | | YDL122W; YDL121C |
| DI49_0946; YDL126C; DI49_0945; YDL125C | DI49_0946; DI49_0945 | | YDL126C; YDL125C |
| DI49_0943; DI49_0944 | DI49_0943; DI49_0944 | | |

Create PDF in your applications with the Pdfcrowd HTML to PDF API PDFCROWD

| | | | |
|---|---|---|---|
| DI49_0942; SPGP0W03900; YDL130W; YDL129W | DI49_0942 | SPGP0W03900 | YDL130W; YDL129W |
| DI49_0940; YDL132W | DI49_0940 | | YDL132W |
| DI49_0939; YDL133W | DI49_0939 | | YDL133W |
| DI49_0938; SPGP0W04400 | DI49_0938 | SPGP0W04400 | |
| DI49_0937; YDL135C | DI49_0937 | | YDL135C |
| DI49_0936; DI49_0889; DI49_0935; YDL192W; DI49_0888 | DI49_0936; DI49_0889; DI49_0935; DI49_0888 | | YDL192W |
| DI49_0934; YDL140C; YDL139C | DI49_0934 | | YDL140C; YDL139C |
| DI49_0933; YDL141W; DI49_0932 | DI49_0933; DI49_0932 | | YDL141W |
| DI49_0930; DI49_0931 | DI49_0930; DI49_0931 | | |
| DI49_0928; YDL146W | DI49_0928 | | YDL146W |
| DI49_0927; YDL147W; DI49_0926 | DI49_0927; DI49_0926 | | YDL147W |
| DI49_0924; YDL150W; DI49_0923; YDL149W | DI49_0924; DI49_0923 | | YDL150W; YDL149W |
| DI49_0921; YDL155W; YDL154W | DI49_0921 | | YDL155W; YDL154W |
| DI49_0920; YDL156W | DI49_0920 | | YDL156W |
| DI49_0919; YDL157C | DI49_0919 | | YDL157C |
| DI49_0918; YDL159W | DI49_0918 | | YDL159W |
| DI49_0917; YDL160C; DI49_0916 | DI49_0917; DI49_0916 | | YDL160C |
| DI49_0914; YDL164C; YDL161W | DI49_0914 | | YDL164C; YDL161W |
| DI49_0913; YDL165W; DI49_0912 | DI49_0913; DI49_0912 | | YDL165W |
| DI49_0910; YDL168W; DI49_0909 | DI49_0910; DI49_0909 | | YDL168W |
| DI49_0907; YDL171C; DI49_0906; YDL170W | DI49_0907; DI49_0906 | | YDL171C; YDL170W |
| DI49_0904; DI49_0905 | DI49_0904; DI49_0905 | | |
| DI49_0902; DI49_0903 | DI49_0902; DI49_0903 | | |
| DI49_0900; YDL179W; DI49_0899 | DI49_0900; DI49_0899 | | YDL179W |
| DI49_0896; YDL183C | DI49_0896 | | YDL183C |
| DI49_0895; YDL185W; DI49_0894 | DI49_0895; DI49_0894 | | YDL185W |
| DI49_0892; YDL189W | DI49_0892 | | YDL189W |
| DI49_0891; YDL190C | DI49_0891 | | YDL190C |
| DI49_0886; YDL195W; DI49_0885 | DI49_0886; DI49_0885 | | YDL195W |
| DI49_0883; YDL199C; DI49_0882 | DI49_0883; DI49_0882 | | YDL199C |
| DI49_0880; DI49_0881 | DI49_0880; DI49_0881 | | |
| DI49_0878; DI49_0879 | DI49_0878; DI49_0879 | | |
| DI49_0876; DI49_0877 | DI49_0876; DI49_0877 | | |
| DI49_0874; YDL208W; DI49_0873 | DI49_0874; DI49_0873 | | YDL208W |
| DI49_0868; DI49_0869; YDL212W; YDL211C | DI49_0868; DI49_0869 | | YDL212W; YDL211C |
| DI49_0866; YDL215C; DI49_0865; YDL214C | DI49_0866; DI49_0865 | | YDL215C; YDL214C |
| DI49_0863; DI49_0864 | DI49_0863; DI49_0864 | | |
| DI49_0861; DI49_0862 | DI49_0861; DI49_0862 | | |
| DI49_0859; DI49_0860 | DI49_0859; DI49_0860 | | |
| DI49_0857; YDL225W; YDL224C | DI49_0857 | | YDL225W; YDL224C |

Create PDF in your applications with the Pdfcrowd HTML to PDF API PDFCROWD

| | | | |
|---|---|---|---|
| DI49_0856; YDL226C | DI49_0856 | | YDL226C |
| DI49_0855; YDL227C | DI49_0855 | | YDL227C |
| SPGP0W04770; DI49_4535; DI49_0853; DI49_4534 | DI49_4535; DI49_0853; DI49_4534 | SPGP0W04770 | |
| DI49_0851; DI49_0852 | DI49_0851; DI49_0852 | | |
| DI49_0849; DI49_0850; YDL233W; YDL232W | DI49_0849; DI49_0850 | | YDL233W; YDL232W |
| DI49_0847; DI49_0848 | DI49_0847; DI49_0848 | | |
| DI49_0845; DI49_0846 | DI49_0845; DI49_0846 | | |
| DI49_0843; YDL240W; DI49_0842; YDL239C | DI49_0843; DI49_0842 | | YDL240W; YDL239C |
| SPGP0W04910; DI49_0841 | DI49_0841 | SPGP0W04910 | |
| DI49_5525; SPGP0W04920 | DI49_5525 | SPGP0W04920 | |
| DI49_5521; YPR069C; DI49_5520; YPR068C | DI49_5521; DI49_5520 | | YPR069C; YPR068C |
| DI49_5518; YPR066W; DI49_5517; YPR065W | DI49_5518; DI49_5517 | | YPR066W; YPR065W |
| SPGP0X00200; SPGP0X00190 | | SPGP0X00200; SPGP0X00190 | |
| DI49_5514; YPR061C; DI49_5513; YPR060C | DI49_5514; DI49_5513 | | YPR061C; YPR060C |
| DI49_5511; DI49_5512 | DI49_5511; DI49_5512 | | |
| DI49_5509; YPR055W | DI49_5509 | | YPR055W |
| DI49_5508; YPR054W | DI49_5508 | | YPR054W |
| DI49_5507; YPR052C; DI49_5506; YPR051W | DI49_5507; DI49_5506 | | YPR052C; YPR051W |
| DI49_5504; DI49_5505 | DI49_5504; DI49_5505 | | |
| DI49_5502; YPR046W; DI49_5501; YPR045C | DI49_5502; DI49_5501 | | YPR046W; YPR045C |
| DI49_5499; YPR042C | DI49_5499 | | YPR042C |
| DI49_5498; YPR041W; DI49_5497; YPR040W | DI49_5498; DI49_5497 | | YPR041W; YPR040W |
| DI49_5495; DI49_5496 | DI49_5495; DI49_5496 | | |
| DI49_5493; YPR035W; DI49_5492; YPR034W | DI49_5493; DI49_5492 | | YPR035W; YPR034W |
| DI49_5490; YPR032W | DI49_5490 | | YPR032W |
| DI49_5489; YPR031W; DI49_5488; YPR030W | DI49_5489; DI49_5488 | | YPR031W; YPR030W |
| SPGP0X00500; DI49_5487 | DI49_5487 | SPGP0X00500 | |
| DI49_5485; DI49_5486; SPGP0Xtrna12F | DI49_5485; DI49_5486 | SPGP0Xtrna12F | |
| DI49_5483; YPR026W; DI49_5482; YPR025C | DI49_5483; DI49_5482 | | YPR026W; YPR025C |
| DI49_5480; DI49_5481 | DI49_5480; DI49_5481 | | |
| DI49_5478; YPR021C; DI49_5477; YPR020W | DI49_5478; DI49_5477 | | YPR021C; YPR020W |
| DI49_5475; DI49_5476 | DI49_5475; DI49_5476 | | |
| DI49_5473; YPR016C | DI49_5473 | | YPR016C |
| DI49_5472; YPR015C; DI49_5471; YPR013C | DI49_5472; DI49_5471 | | YPR015C; YPR013C |
| DI49_5469; DI49_5470 | DI49_5469; DI49_5470 | | |
| DI49_5467; YPR010C | DI49_5467 | | YPR010C |
| DI49_5466; YPR009W | DI49_5466 | | YPR009W |
| DI49_5465; YPR008W | DI49_5465 | | YPR008W |
| DI49_5461; DI49_5462 | DI49_5461; DI49_5462 | | |
| DI49_5459; YPR003C | DI49_5459 | | YPR003C |

Create PDF in your applications with the Pdfcrowd HTML to PDF API PDFCROWD

| | | | |
|---|---|---|---|
| SPGP0X00760; YPR002W | | SPGP0X00760 | YPR002W |
| DI49_5456; YPR001W; DI49_5455; YPL001W | DI49_5456; DI49_5455 | | YPR001W; YPL001W |
| DI49_5453; DI49_5454 | DI49_5453; DI49_5454 | | |
| DI49_5451; DI49_5452 | DI49_5451; DI49_5452 | | |
| DI49_5449; DI49_5450 | DI49_5449; DI49_5450 | | |
| DI49_5447; YPL009C; DI49_5446; YPL010W | DI49_5447; DI49_5446 | | YPL009C; YPL010W |
| DI49_5444; YPL012W; DI49_5443; YPL013C | DI49_5444; DI49_5443 | | YPL012W; YPL013C |
| DI49_5441; YPL015C; SPGP0X00930 | DI49_5441 | SPGP0X00930 | YPL015C |
| DI49_5439; DI49_5440; YPL016W; YPL017C | DI49_5439; DI49_5440 | | YPL016W; YPL017C |
| DI49_5437; YPL019C; DI49_5436; YPL020C | DI49_5437; DI49_5436 | | YPL019C; YPL020C |
| DI49_5434; YPL022W | DI49_5434 | | YPL022W |
| DI49_5433; YPL023C | DI49_5433 | | YPL023C |
| DI49_5432; YPL024W; DI49_5431; YPL026C | DI49_5432; DI49_5431 | | YPL024W; YPL026C |
| DI49_5429; YPL028W | DI49_5429 | | YPL028W |
| DI49_5428; YPL029W; DI49_5427; YPL030W | DI49_5428; DI49_5427 | | YPL029W; YPL030W |
| DI49_5425; DI49_5426 | DI49_5425; DI49_5426 | | |
| DI49_5423; DI49_5424 | DI49_5423; DI49_5424 | | |
| DI49_0526; DI49_5422 | DI49_0526; DI49_5422 | | |
| DI49_0530; DI49_0529 | DI49_0530; DI49_0529 | | |
| DI49_0532; DI49_0531 | DI49_0532; DI49_0531 | | |
| DI49_0534; DI49_0533 | DI49_0534; DI49_0533 | | |
| DI49_0536; DI49_0535 | DI49_0536; DI49_0535 | | |
| DI49_0538; DI49_0537 | DI49_0538; DI49_0537 | | |
| DI49_0543; DI49_0542 | DI49_0543; DI49_0542 | | |
| SPGP0Y00270; DI49_0544 | DI49_0544 | SPGP0Y00270 | |
| DI49_0546; DI49_0545 | DI49_0546; DI49_0545 | | |
| DI49_0548; DI49_0547 | DI49_0548; DI49_0547 | | |
| DI49_0550; DI49_0549 | DI49_0550; DI49_0549 | | |
| DI49_0552; DI49_0551 | DI49_0552; DI49_0551 | | |
| DI49_0554; DI49_0553 | DI49_0554; DI49_0553 | | |
| DI49_0555; SPGP0Y00380 | DI49_0555 | SPGP0Y00380 | |
| DI49_0557; DI49_0556 | DI49_0557; DI49_0556 | | |
| SPGP0Y00430; SPGP0Y00420 | | SPGP0Y00430; SPGP0Y00420 | |
| DI49_0559; DI49_0558 | DI49_0559; DI49_0558 | | |
| DI49_0564; DI49_0563 | DI49_0564; DI49_0563 | | |
| DI49_0570; DI49_0569 | DI49_0570; DI49_0569 | | |
| DI49_0572; DI49_0571 | DI49_0572; DI49_0571 | | |
| DI49_0574; DI49_0573 | DI49_0574; DI49_0573 | | |
| DI49_0576; DI49_0575 | DI49_0576; DI49_0575 | | |
| DI49_0578; DI49_0577 | DI49_0578; DI49_0577 | | |

Create PDF in your applications with the Pdfcrowd HTML to PDF API PDFCROWD

| | | |
|---|---|---|
| DI49_0580; DI49_0579 | DI49_0580; DI49_0579 | |
| DI49_0583; SPGP0Y00690 | DI49_0583 | SPGP0Y00690 |
| DI49_0585; DI49_0584 | DI49_0585; DI49_0584 | |
| DI49_0587; DI49_0586 | DI49_0587; DI49_0586 | |
| DI49_0589; DI49_0588 | DI49_0589; DI49_0588 | |
| DI49_0591; DI49_0590 | DI49_0591; DI49_0590 | |
| DI49_0595; DI49_0594 | DI49_0595; DI49_0594 | |
| SPGP0Y00830; SPGP0Y00820 | | SPGP0Y00830; SPGP0Y00820 |
| DI49_0597; DI49_0596 | DI49_0597; DI49_0596 | |
| DI49_0599; DI49_0598 | DI49_0599; DI49_0598 | |
| DI49_0601; DI49_0600 | DI49_0601; DI49_0600 | |
| DI49_0603; DI49_0602 | DI49_0603; DI49_0602 | |
| DI49_0605; DI49_0604 | DI49_0605; DI49_0604 | |
| DI49_0607; DI49_0606 | DI49_0607; DI49_0606 | |
| DI49_0609; DI49_0608 | DI49_0609; DI49_0608 | |
| DI49_0611; DI49_0610 | DI49_0611; DI49_0610 | |
| DI49_0620; DI49_0619 | DI49_0620; DI49_0619 | |
| DI49_0622; DI49_0621 | DI49_0622; DI49_0621 | |
| DI49_0624; DI49_0623 | DI49_0624; DI49_0623 | |
| DI49_0626; DI49_0625 | DI49_0626; DI49_0625 | |
| DI49_0627; SPGP0Y01140 | DI49_0627 | SPGP0Y01140 |
| DI49_0629; DI49_0628 | DI49_0629; DI49_0628 | |
| DI49_0631; DI49_0630 | DI49_0631; DI49_0630 | |
| DI49_0633; DI49_0632 | DI49_0633; DI49_0632 | |
| DI49_0635; DI49_0634 | DI49_0635; DI49_0634 | |
| DI49_0637; DI49_0636 | DI49_0637; DI49_0636 | |
| DI49_0639; DI49_0638 | DI49_0639; DI49_0638 | |
| DI49_0641; DI49_0640 | DI49_0641; DI49_0640 | |
| DI49_0643; DI49_0642 | DI49_0643; DI49_0642 | |
| DI49_0645; DI49_0644 | DI49_0645; DI49_0644 | |
| DI49_0647; DI49_0646 | DI49_0647; DI49_0646 | |
| DI49_0649; DI49_0648 | DI49_0649; DI49_0648 | |
| DI49_0651; DI49_0650 | DI49_0651; DI49_0650 | |
| DI49_0653; DI49_0652 | DI49_0653; DI49_0652 | |
| DI49_0655; DI49_0654 | DI49_0655; DI49_0654 | |
| DI49_0657; DI49_0656 | DI49_0657; DI49_0656 | |
| DI49_0659; DI49_0658 | DI49_0659; DI49_0658 | |
| DI49_0661; DI49_0660 | DI49_0661; DI49_0660 | |
| DI49_0663; DI49_0662 | DI49_0663; DI49_0662 | |
| DI49_0668; DI49_0667 | DI49_0668; DI49_0667 | |

Create PDF in your applications with the Pdfcrowd HTML to PDF API PDFCROWD

| | | | |
|---|---|---|---|
| DI49_0672; DI49_0671; YCL025C; YCL024W | DI49_0672; DI49_0671 | | YCL025C; YCL024W |
| DI49_0674; DI49_0673 | DI49_0674; DI49_0673 | | |
| DI49_0676; DI49_0675 | DI49_0676; DI49_0675 | | |
| DI49_0678; DI49_0677 | DI49_0678; DI49_0677 | | |
| SPGP0Y01670; DI49_0679 | DI49_0679 | SPGP0Y01670 | |
| SPGP0Y01690; DI49_0680 | DI49_0680 | SPGP0Y01690 | |
| DI49_0682; DI49_0681 | DI49_0682; DI49_0681 | | |
| DI49_0684; DI49_0683 | DI49_0684; DI49_0683 | | |
| DI49_0686; DI49_0685 | DI49_0686; DI49_0685 | | |
| DI49_0688; DI49_0687 | DI49_0688; DI49_0687 | | |
| SPGP0Y01790; DI49_0689 | DI49_0689 | SPGP0Y01790 | |
| SPGP0Y01810; SPGP0Y01800 | | SPGP0Y01810; SPGP0Y01800 | |
| SPGP0Y01830; SPGP0Y01820 | | SPGP0Y01830; SPGP0Y01820 | |
| DI49_4735; YNR001C; DI49_4734 | DI49_4735; DI49_4734 | | YNR001C |
| SPGP0Z00260; DI49_4719 | DI49_4719 | SPGP0Z00260 | |
| SPGP0Z00270; DI49_4718 | DI49_4718 | SPGP0Z00270 | |
| DI49_4707; DI49_0189; DI49_4708 | DI49_4707; DI49_0189; DI49_4708 | | |
| DI49_4710; DI49_4709 | DI49_4710; DI49_4709 | | |
| DI49_4712; DI49_4711 | DI49_4712; DI49_4711 | | |
| DI49_4715; YNL022C; DI49_4716 | DI49_4715; DI49_4716 | | YNL022C |
| DI49_4698; SPGP0Ztrna15P | DI49_4698 | SPGP0Ztrna15P | |
| DI49_4693; DI49_4694 | DI49_4693; DI49_4694 | | |
| DI49_4691; DI49_4692 | DI49_4691; DI49_4692 | | |
| DI49_4686; YNL044W; DI49_4685 | DI49_4686; DI49_4685 | | YNL044W |
| DI49_4683; DI49_4684 | DI49_4683; DI49_4684 | | |
| DI49_4678; DI49_4679 | DI49_4678; DI49_4679 | | |
| DI49_4669; YNL062C; DI49_4668 | DI49_4669; DI49_4668 | | YNL062C |
| DI49_4660; YNL071W; DI49_4659 | DI49_4660; DI49_4659 | | YNL071W |
| DI49_4657; DI49_4658 | DI49_4657; DI49_4658 | | |
| DI49_4655; DI49_4656 | DI49_4655; DI49_4656 | | |
| DI49_4653; DI49_4654 | DI49_4653; DI49_4654 | | |
| DI49_4629; YNL102W; DI49_4628 | DI49_4629; DI49_4628 | | YNL102W |
| DI49_4612; YNL124W; DI49_4611 | DI49_4612; DI49_4611 | | YNL124W |
| DI49_4597; YNL138W; DI49_4596 | DI49_4597; DI49_4596 | | YNL138W |
| DI49_4592; DI49_4593 | DI49_4592; DI49_4593 | | |
| DI49_4590; DI49_4591 | DI49_4590; DI49_4591 | | |
| DI49_4588; DI49_4589 | DI49_4588; DI49_4589 | | |
| DI49_4586; DI49_4587 | DI49_4586; DI49_4587 | | |
| DI49_4584; DI49_4585 | DI49_4584; DI49_4585 | | |
| DI49_4574; DI49_4575 | DI49_4574; DI49_4575 | | |

Create PDF in your applications with the Pdfcrowd HTML to PDF API PDFCROWD

| | | | |
|---|---|---|---|
| DI49_4569; DI49_4570 | DI49_4569; DI49_4570 | | |
| DI49_4566; YNL172W; DI49_4565 | DI49_4566; DI49_4565 | | YNL172W |
| DI49_4560; YNL180C; DI49_4559 | DI49_4560; DI49_4559 | | YNL180C |
| DI49_4555; YNL186W; DI49_4554 | DI49_4555; DI49_4554 | | YNL186W |
| DI49_4549; YNL192W; DI49_4548 | DI49_4549; DI49_4548 | | YNL192W |
| DI49_4546; DI49_4547 | DI49_4546; DI49_4547 | | |
| DI49_4544; DI49_4545 | DI49_4544; DI49_4545 | | |
| DI49_4542; DI49_4543 | DI49_4542; DI49_4543 | | |
| DI49_4540; YNL202W; DI49_4539 | DI49_4540; DI49_4539 | | YNL202W |
| DI49_4537; YNL207W; DI49_4536 | DI49_4537; DI49_4536 | | YNL207W |
| DI49_4529; DI49_4530 | DI49_4529; DI49_4530 | | |
| DI49_4527; DI49_4528 | DI49_4527; DI49_4528 | | |
| DI49_4525; DI49_4526 | DI49_4525; DI49_4526 | | |
| DI49_4523; DI49_4524 | DI49_4523; DI49_4524 | | |
| DI49_4518; YNL227C; DI49_4517 | DI49_4518; DI49_4517 | | YNL227C |
| DI49_4515; DI49_4516 | DI49_4515; DI49_4516 | | |
| DI49_4510; DI49_4511 | DI49_4510; DI49_4511 | | |
| DI49_4508; DI49_4509 | DI49_4508; DI49_4509 | | |
| DI49_4505; YNL242W; DI49_4504 | DI49_4505; DI49_4504 | | YNL242W |
| DI49_4491; YNL256W; DI49_4490 | DI49_4491; DI49_4490 | | YNL256W |
| DI49_4488; DI49_4489 | DI49_4488; DI49_4489 | | |
| DI49_4486; DI49_4487 | DI49_4486; DI49_4487 | | |
| DI49_4484; YNL263C; DI49_4483 | DI49_4484; DI49_4483 | | YNL263C |
| DI49_4477; YNL272C; DI49_4476 | DI49_4477; DI49_4476 | | YNL272C |
| DI49_4472; DI49_4473 | DI49_4472; DI49_4473 | | |
| DI49_4454; YNL293W; DI49_4453 | DI49_4454; DI49_4453 | | YNL293W |
| DI49_4451; DI49_4452 | DI49_4451; DI49_4452 | | |
| DI49_4440; DI49_4441 | DI49_4440; DI49_4441 | | |
| SPGP0Z02980; DI49_4439 | DI49_4439 | SPGP0Z02980 | |
| DI49_4436; DI49_4437 | DI49_4436; DI49_4437 | | |
| DI49_4434; DI49_4435 | DI49_4434; DI49_4435 | | |
| DI49_4432; DI49_4433 | DI49_4432; DI49_4433 | | |
| DI49_0695; DI49_1360; YEL032W | DI49_0695; DI49_1360 | | YEL032W |
| DI49_0696; YCR038C; DI49_0789; DI49_5531 | DI49_0696; DI49_0789; DI49_5531 | | YCR038C |
| SPGP0AA00120; SPGP0AE01020; DI49_0791; YCR097W | DI49_0791 | SPGP0AA00120; SPGP0AE01020 | YCR097W |
| DI49_0699; SPGP0AA00140 | DI49_0699 | SPGP0AA00140 | |
| DI49_0700; YCL059C; DI49_0701; YCL058C | DI49_0700; DI49_0701 | | YCL059C; YCL058C |
| DI49_0703; YCL057W; DI49_0704; YCL056C | DI49_0703; DI49_0704 | | YCL057W; YCL056C |
| DI49_0706; YCL054W; DI49_0707; YCL052C | DI49_0706; DI49_0707 | | YCL054W; YCL052C |
| DI49_0709; YCL050C; DI49_0710; YCL049C | DI49_0709; DI49_0710 | | YCL050C; YCL049C |

Create PDF in your applications with the Pdfcrowd HTML to PDF API PDFCROWD

| | | | |
|---|---|---|---|
| DI49_0712; DI49_0711; YCL048W; YCL047C | DI49_0712; DI49_0711 | | YCL048W; YCL047C |
| DI49_0714; YCL044C | DI49_0714 | | YCL044C |
| DI49_0715; YCL043C | DI49_0715 | | YCL043C |
| DI49_0716; YCL040W | DI49_0716 | | YCL040W |
| DI49_0717; YCL039W; DI49_0718; YCL038C | DI49_0717; DI49_0718 | | YCL039W; YCL038C |
| SPGP0AA00370; YCL036W; DI49_0720; YCL035C | DI49_0720 | SPGP0AA00370 | YCL036W; YCL035C |
| DI49_0722; DI49_0721 | DI49_0722; DI49_0721 | | |
| DI49_0724; YCL031C | DI49_0724 | | YCL031C |
| DI49_0725; YCL030C | DI49_0725 | | YCL030C |
| DI49_0726; YCL029C; DI49_0727; YCL028W | DI49_0726; DI49_0727 | | YCL029C; YCL028W |
| DI49_0729; DI49_0728 | DI49_0729; DI49_0728 | | |
| DI49_0731; DI49_0730 | DI49_0731; DI49_0730 | | |
| DI49_0736; YCL018W | DI49_0736 | | YCL018W |
| DI49_0737; YCL017C | DI49_0737 | | YCL017C |
| DI49_0738; YCL016C | DI49_0738 | | YCL016C |
| DI49_0739; YCL014W; DI49_0740; YCL012C | DI49_0739; DI49_0740 | | YCL014W; YCL012C |
| DI49_0742; YCL010C | DI49_0742 | | YCL010C |
| DI49_0743; YCL009C | DI49_0743 | | YCL009C |
| DI49_0744; YCL008C; DI49_0745; YCL007C | DI49_0744; DI49_0745 | | YCL008C; YCL007C |
| DI49_0747; DI49_0746 | DI49_0747; DI49_0746 | | |
| DI49_0749; YCL001W; DI49_0750; YCL001W-A | DI49_0749; DI49_0750 | | YCL001W; YCL001W-A |
| DI49_0752; YCR003W | DI49_0752 | | YCR003W |
| DI49_0753; YCR004C; DI49_0754; YCR005C | DI49_0753; DI49_0754 | | YCR004C; YCR005C |
| SPGP0AA00730; SPGP0AA00720 | | SPGP0AA00730; SPGP0AA00720 | |
| DI49_0758; YCR008W; DI49_0759; YCR009C | DI49_0758; DI49_0759 | | YCR008W; YCR009C |
| DI49_0761; YCR011C | DI49_0761 | | YCR011C |
| DI49_0762; YCR012W; DI49_0763; YCR014C | DI49_0762; DI49_0763 | | YCR012W; YCR014C |
| SPGP0AA00830; DI49_0767 | DI49_0767 | SPGP0AA00830 | |
| DI49_0770; YCR019W; DI49_0771; YCR020C | DI49_0770; DI49_0771 | | YCR019W; YCR020C |
| DI49_0773; DI49_0772 | DI49_0773; DI49_0772 | | |
| DI49_0775; YCR023C; DI49_0776; YCR024C | DI49_0775; DI49_0776 | | YCR023C; YCR024C |
| DI49_0778; DI49_0777 | DI49_0778; DI49_0777 | | |
| DI49_0781; YCR028C; SPGP0AA00950; YCR028C-A | DI49_0781 | SPGP0AA00950 | YCR028C; YCR028C-A |
| DI49_0786; YCR034W | DI49_0786 | | YCR034W |
| DI49_0787; YCR035C; DI49_0788; YCR036W | DI49_0787; DI49_0788 | | YCR035C; YCR036W |
| DI49_5533; DI49_5532 | DI49_5533; DI49_5532 | | |
| DI49_5535; DI49_5534 | DI49_5535; DI49_5534 | | |
| DI49_5537; DI49_5536 | DI49_5537; DI49_5536 | | |
| DI49_5539; DI49_5538 | DI49_5539; DI49_5538 | | |
| DI49_5541; DI49_5540 | DI49_5541; DI49_5540 | | |

Create PDF in your applications with the Pdfcrowd HTML to PDF API PDFCROWD

| | | | |
|---|---|---|---|
| DI49_5543; DI49_5542 | DI49_5543; DI49_5542 | | |
| DI49_5545; DI49_5544 | DI49_5545; DI49_5544 | | |
| DI49_5546; DI49_2042; DI49_5547 | DI49_5546; DI49_2042; DI49_5547 | | |
| DI49_5549; DI49_5548 | DI49_5549; DI49_5548 | | |
| DI49_5551; DI49_5550 | DI49_5551; DI49_5550 | | |
| DI49_5556; DI49_5555 | DI49_5556; DI49_5555 | | |
| DI49_5558; DI49_5557 | DI49_5558; DI49_5557 | | |
| DI49_5560; DI49_5559 | DI49_5560; DI49_5559 | | |
| DI49_5562; DI49_5561 | DI49_5562; DI49_5561 | | |
| DI49_5570; DI49_5569 | DI49_5570; DI49_5569 | | |
| DI49_5572; DI49_5571 | DI49_5572; DI49_5571 | | |
| DI49_5574; DI49_5573 | DI49_5574; DI49_5573 | | |
| DI49_5578; DI49_5577 | DI49_5578; DI49_5577 | | |
| DI49_5580; DI49_5579 | DI49_5580; DI49_5579 | | |
| DI49_5582; SPGP0ACtrna12N | DI49_5582 | SPGP0ACtrna12N | |
| DI49_5584; DI49_5583 | DI49_5584; DI49_5583 | | |
| DI49_5586; DI49_5585 | DI49_5586; DI49_5585 | | |
| DI49_5590; DI49_5589 | DI49_5590; DI49_5589 | | |
| DI49_5592; DI49_5591 | DI49_5592; DI49_5591 | | |
| DI49_5594; DI49_5593 | DI49_5594; DI49_5593 | | |
| DI49_5595; SPGP0AC00700 | DI49_5595 | SPGP0AC00700 | |
| DI49_5597; DI49_5596 | DI49_5597; DI49_5596 | | |
| DI49_5599; DI49_5598 | DI49_5599; DI49_5598 | | |
| DI49_5606; DI49_5605 | DI49_5606; DI49_5605 | | |
| DI49_5608; DI49_5607 | DI49_5608; DI49_5607 | | |
| DI49_5610; DI49_5609 | DI49_5610; DI49_5609 | | |
| DI49_5612; DI49_5611 | DI49_5612; DI49_5611 | | |
| DI49_5617; DI49_5616 | DI49_5617; DI49_5616 | | |
| DI49_5619; DI49_5618 | DI49_5619; DI49_5618 | | |
| DI49_5621; DI49_5620 | DI49_5621; DI49_5620 | | |
| DI49_5623; DI49_5622 | DI49_5623; DI49_5622 | | |
| DI49_5625; DI49_5624 | DI49_5625; DI49_5624 | | |
| DI49_5627; DI49_5626 | DI49_5627; DI49_5626 | | |
| DI49_5629; DI49_5628 | DI49_5629; DI49_5628 | | |
| DI49_5631; DI49_5630 | DI49_5631; DI49_5630 | | |
| DI49_5633; DI49_5632 | DI49_5633; DI49_5632 | | |
| DI49_5635; DI49_5634 | DI49_5635; DI49_5634 | | |
| DI49_5637; DI49_5636 | DI49_5637; DI49_5636 | | |
| SPGP0AC01120; DI49_5638 | DI49_5638 | SPGP0AC01120 | |
| YKR105C; SPGP0AC01130 | | SPGP0AC01130 | YKR105C |

Create PDF in your applications with the Pdfcrowd HTML to PDF API PDFCROWD

| | | | |
|---|---|---|---|
| SPGP0AC01160; SPGP0AC01150 | | SPGP0AC01160; SPGP0AC01150 | |
| DI49_2630; DI49_2629 | DI49_2630; DI49_2629 | | |
| DI49_2632; DI49_2631 | DI49_2632; DI49_2631 | | |
| DI49_2634; DI49_2633 | DI49_2634; DI49_2633 | | |
| DI49_2640; YIL147C | DI49_2640 | | YIL147C |
| DI49_2648; DI49_2647 | DI49_2648; DI49_2647 | | |
| DI49_2650; DI49_2649 | DI49_2650; DI49_2649 | | |
| DI49_2652; DI49_2651 | DI49_2652; DI49_2651 | | |
| DI49_2654; DI49_2653 | DI49_2654; DI49_2653 | | |
| DI49_2656; DI49_2655 | DI49_2656; DI49_2655 | | |
| DI49_2667; DI49_2666 | DI49_2667; DI49_2666 | | |
| DI49_2669; DI49_2668 | DI49_2669; DI49_2668 | | |
| DI49_2674; DI49_2673 | DI49_2674; DI49_2673 | | |
| DI49_2681; DI49_2680 | DI49_2681; DI49_2680 | | |
| DI49_2683; DI49_2682 | DI49_2683; DI49_2682 | | |
| DI49_2688; DI49_2687 | DI49_2688; DI49_2687 | | |
| DI49_2696; DI49_2695 | DI49_2696; DI49_2695 | | |
| DI49_2701; DI49_2700 | DI49_2701; DI49_2700 | | |
| DI49_2710; DI49_1486; DI49_2711 | DI49_2710; DI49_1486; DI49_2711 | | |
| DI49_2713; DI49_2712 | DI49_2713; DI49_2712 | | |
| DI49_2715; DI49_1478 | DI49_2715; DI49_1478 | | |
| DI49_2726; YIL055C; SPGP0B01130 | DI49_2726 | SPGP0B01130 | YIL055C |
| DI49_2737; DI49_2736 | DI49_2737; DI49_2736 | | |
| DI49_2754; DI49_2753 | DI49_2754; DI49_2753 | | |
| DI49_2766; DI49_2765 | DI49_2766; DI49_2765 | | |
| DI49_2768; DI49_2767 | DI49_2768; DI49_2767 | | |
| SPGP0B01550; DI49_2770; DI49_2771; DI49_1642 | DI49_2770; DI49_2771; DI49_1642 | SPGP0B01550 | |
| DI49_2773; DI49_2772 | DI49_2773; DI49_2772 | | |
| DI49_2775; DI49_2774 | DI49_2775; DI49_2774 | | |
| DI49_2784; DI49_2783 | DI49_2784; DI49_2783 | | |
| DI49_2795; DI49_2794 | DI49_2795; DI49_2794 | | |
| DI49_2801; DI49_2800 | DI49_2801; DI49_2800 | | |
| DI49_2803; DI49_2802 | DI49_2803; DI49_2802 | | |
| DI49_2805; DI49_2804 | DI49_2805; DI49_2804 | | |
| DI49_2811; DI49_2810 | DI49_2811; DI49_2810 | | |
| DI49_2816; DI49_2815 | DI49_2816; DI49_2815 | | |
| YCL061C; YCL063W | | | YCL061C; YCL063W |
| YCL033C; YCL034W | | | YCL033C; YCL034W |
| YCL026C-B; YCL027W | | | YCL026C-B; YCL027W |

Create PDF in your applications with the Pdfcrowd HTML to PDF API PDFCROWD

| | | | |
|---|---|---|---|
| YCL004W; YCL005W | | | YCL004W; YCL005W |
| YCR018C; YCR017C | | | YCR018C; YCR017C |
| YCR020W-B; YCR020C-A | | | YCR020W-B; YCR020C-A |
| YCR026C; YCR024C-A | | | YCR026C; YCR024C-A |
| YCR043C; YCR042C | | | YCR043C; YCR042C |
| YCR045C; YCR044C | | | YCR045C; YCR044C |
| YCR047C; YCR046C | | | YCR047C; YCR046C |
| YCR051W; YCR048W | | | YCR051W; YCR048W |
| YCR053W; YCR052W | | | YCR053W; YCR052W |
| YCR057C; YCR054C | | | YCR057C; YCR054C |
| YCR060W; YCR059C | | | YCR060W; YCR059C |
| YCR066W; YCR065W | | | YCR066W; YCR065W |
| YCR068W; YCR067C | | | YCR068W; YCR067C |
| YCR071C; YCR069W | | | YCR071C; YCR069W |
| YCR073C; YCR072C | | | YCR073C; YCR072C |
| YCR075C; YCR073W-A | | | YCR075C; YCR073W-A |
| YCR076C; YCR075W-A | | | YCR076C; YCR075W-A |
| YCR079W; YCR077C | | | YCR079W; YCR077C |
| YCR082W; YCR081W | | | YCR082W; YCR081W |
| YCR084C; YCR083W | | | YCR084C; YCR083W |
| YCR087W; YCR086W | | | YCR087W; YCR086W |
| YCR089W; YCR088W | | | YCR089W; YCR088W |
| YCR094W; YCR093W | | | YCR094W; YCR093W |
| DI49_2577; DI49_2218; DI49_2578 | DI49_2577; DI49_2218; DI49_2578 | | |
| SPGP0AF00130; DI49_2579 | DI49_2579 | SPGP0AF00130 | |
| DI49_2581; YHR179W | DI49_2581 | | YHR179W |
| DI49_2583; YHR181W; DI49_2584 | DI49_2583; DI49_2584 | | YHR181W |
| DI49_2586; DI49_2585 | DI49_2586; DI49_2585 | | |
| DI49_2588; SPGP0Ctrna21V; SPGP0AItrna23V; YDR307W | DI49_2588 | SPGP0Ctrna21V; SPGP0AItrna23V | YDR307W |
| DI49_2589; YHR186C; DI49_2590 | DI49_2589; DI49_2590 | | YHR186C |
| DI49_2592; DI49_2591 | DI49_2592; DI49_2591 | | |
| DI49_2594; DI49_2593; YHR190W; YHR189W | DI49_2594; DI49_2593 | | YHR190W; YHR189W |
| DI49_2596; YHR193C; YHR192W | DI49_2596 | | YHR193C; YHR192W |
| DI49_2597; YHR194W; DI49_2598 | DI49_2597; DI49_2598 | | YHR194W |
| DI49_2600; YHR197W; DI49_2601; YHR196W | DI49_2600; DI49_2601 | | YHR197W; YHR196W |
| DI49_2603; DI49_2602 | DI49_2603; DI49_2602 | | |
| DI49_2605; DI49_2604; YHR200W; YHR199C-A | DI49_2605; DI49_2604 | | YHR200W; YHR199C-A |
| DI49_2607; DI49_2606 | DI49_2607; DI49_2606 | | |
| DI49_2611; YHR206W; DI49_2612 | DI49_2611; DI49_2612 | | YHR206W |
| DI49_2614; DI49_2613; YHR208W; YHR207C | DI49_2614; DI49_2613 | | YHR208W; YHR207C |

Create PDF in your applications with the Pdfcrowd HTML to PDF API PDFCROWD

| | | |
|---|---|---|
| DI49_2616; DI49_2615 | DI49_2616; DI49_2615 | |
| DI49_2618; SPGP0R03480; DI49_4412; DI49_1327 | DI49_2618; DI49_4412; DI49_1327 | SPGP0R03480 |
| DI49_4796; SPGP0AP00100 | DI49_4796 | SPGP0AP00100 |
| SPGP0AG00120; SPGP0AP00110 | | SPGP0AG00120; SPGP0AP00110 |
| SPGP0AG00130; DI49_4797; DI49_4798 | DI49_4797; DI49_4798 | SPGP0AG00130 |
| DI49_4800; DI49_4799 | DI49_4800; DI49_4799 | |
| DI49_1717; DI49_1718 | DI49_1717; DI49_1718 | |
| DI49_1713; DI49_1714 | DI49_1713; DI49_1714 | |
| DI49_1711; DI49_1712 | DI49_1711; DI49_1712 | |
| DI49_1709; DI49_1710 | DI49_1709; DI49_1710 | |
| DI49_1707; DI49_1708 | DI49_1707; DI49_1708 | |
| DI49_1705; DI49_1706 | DI49_1705; DI49_1706 | |
| DI49_1703; DI49_1704 | DI49_1703; DI49_1704 | |
| DI49_1701; DI49_1702 | DI49_1701; DI49_1702 | |
| DI49_1699; DI49_1700 | DI49_1699; DI49_1700 | |
| DI49_1697; DI49_1698 | DI49_1697; DI49_1698 | |
| DI49_1695; SPGP0AH00330; DI49_1694 | DI49_1695; DI49_1694 | SPGP0AH00330 |
| DI49_1692; DI49_1693 | DI49_1692; DI49_1693 | |
| DI49_1687; DI49_1688 | DI49_1687; DI49_1688 | |
| SPGP0AH00440; DI49_1684 | DI49_1684 | SPGP0AH00440 |
| DI49_1680; DI49_1681 | DI49_1680; DI49_1681 | |
| DI49_1678; DI49_1679 | DI49_1678; DI49_1679 | |
| DI49_1674; DI49_1675 | DI49_1674; DI49_1675 | |
| DI49_1670; DI49_1671 | DI49_1670; DI49_1671 | |
| DI49_1666; DI49_1667 | DI49_1666; DI49_1667 | |
| DI49_1661; DI49_1662 | DI49_1661; DI49_1662 | |
| DI49_1657; DI49_1658 | DI49_1657; DI49_1658 | |
| DI49_1655; DI49_1656 | DI49_1655; DI49_1656 | |
| DI49_1653; DI49_1654 | DI49_1653; DI49_1654 | |
| DI49_1648; DI49_1649 | DI49_1648; DI49_1649 | |
| DI49_1646; DI49_1647 | DI49_1646; DI49_1647 | |
| DI49_1644; DI49_1645; SPGP0E01460; DI49_1471 | DI49_1644; DI49_1645; DI49_1471 | SPGP0E01460 |
| DI49_1640; DI49_1641 | DI49_1640; DI49_1641 | |
| DI49_1638; DI49_1639 | DI49_1638; DI49_1639 | |
| DI49_1636; DI49_1637 | DI49_1636; DI49_1637 | |
| SPGP0AH00870; DI49_0075 | DI49_0075 | SPGP0AH00870 |
| DI49_1631; DI49_1632 | DI49_1631; DI49_1632 | |
| DI49_1629; DI49_1630 | DI49_1629; DI49_1630 | |
| DI49_1628; SPGP0AH00940 | DI49_1628 | SPGP0AH00940 |
| DI49_1626; DI49_1627 | DI49_1626; DI49_1627 | |

Create PDF in your applications with the Pdfcrowd HTML to PDF API PDFCROWD

| | | | |
|---|---|---|---|
| DI49_1624; DI49_1625 | DI49_1624; DI49_1625 | | |
| DI49_1622; DI49_1623 | DI49_1622; DI49_1623 | | |
| DI49_1621; YFL034C-A | DI49_1621 | | YFL034C-A |
| DI49_1619; DI49_1620 | DI49_1619; DI49_1620 | | |
| DI49_1617; DI49_1618 | DI49_1617; DI49_1618 | | |
| DI49_1615; DI49_1616 | DI49_1615; DI49_1616 | | |
| DI49_1613; DI49_1614 | DI49_1613; DI49_1614 | | |
| DI49_1611; DI49_1612 | DI49_1611; DI49_1612 | | |
| DI49_1609; DI49_1610 | DI49_1609; DI49_1610 | | |
| DI49_1606; DI49_3482; DI49_3483 | DI49_1606; DI49_3482; DI49_3483 | | |
| YHR205W; DI49_2610; YHR204W | DI49_2610 | | YHR205W; YHR204W |
| YHR202W; SPGP0AI00190; YJR145C; YJR144W | | SPGP0AI00190 | YHR202W; YJR145C; YJR144W |
| YHR198C; YHR199C | | | YHR198C; YHR199C |
| YHR187W; YHR188C | | | YHR187W; YHR188C |
| YHR185C; DI49_2587; YHR184W | DI49_2587 | | YHR185C; YHR184W |
| YHR182W; YHR183W | | | YHR182W; YHR183W |
| YHR178W; DI49_2580; YHR177W | DI49_2580 | | YHR178W; YHR177W |
| YHR175W; YHR176W | | | YHR175W; YHR176W |
| YDL237W; YDL238C | | | YDL237W; YDL238C |
| YDL235C; YDL236W | | | YDL235C; YDL236W |
| YDL230W; YDL231C | | | YDL230W; YDL231C |
| YDL222C; YDL223C | | | YDL222C; YDL223C |
| YDL219W; YDL220C | | | YDL219W; YDL220C |
| YDL217C; YDL218W | | | YDL217C; YDL218W |
| YDL210W; DI49_0872; YDL209C | DI49_0872 | | YDL210W; YDL209C |
| YDL207W; DI49_0875; YDL206W | DI49_0875 | | YDL207W; YDL206W |
| YDL204W; YDL205C | | | YDL204W; YDL205C |
| YDL202W; YDL203C | | | YDL202W; YDL203C |
| YDL200C; YDL201W | | | YDL200C; YDL201W |
| YDL198C; DI49_0884; YDL197C | DI49_0884 | | YDL198C; YDL197C |
| YDL194W; DI49_0887; YDL193W | DI49_0887 | | YDL194W; YDL193W |
| YDL191W; YDL136W | | | YDL191W; YDL136W |
| YDL188C; YDL134C; YDL186W | | | YDL188C; YDL134C; YDL186W |
| YDL182W; YDL131W; YDL181W | | | YDL182W; YDL131W; YDL181W |
| YDL178W; DI49_0901; YDL177C | DI49_0901 | | YDL178W; YDL177C |
| YDL175C; YDL176W | | | YDL175C; YDL176W |
| YDL173W; YDL174C | | | YDL173W; YDL174C |
| YDL167C; DI49_0911; YDL166C | DI49_0911 | | YDL167C; YDL166C |
| YDL145C; DI49_0929; YDL144C | DI49_0929 | | YDL145C; YDL144C |
| YDL142C; YDL143W | | | YDL142C; YDL143W |

Create PDF in your applications with the Pdfcrowd HTML to PDF API PDFCROWD

| | | |
|---|---|---|
| YDL127W; YDL128W | | YDL127W; YDL128W |
| YDL123W; YDL124W | | YDL123W; YDL124W |
| YDL110C; DI49_0960; YDL109C | DI49_0960 | YDL110C; YDL109C |
| YDL107W; YDL108W | | YDL107W; YDL108W |
| YDL105W; DI49_0965; YDL104C | DI49_0965 | YDL105W; YDL104C |
| YDL088C; YDL089W | | YDL088C; YDL089W |
| YDL073W; DI49_0993; YDL072C | DI49_0993 | YDL073W; YDL072C |
| YDL069C; YDL070W | | YDL069C; YDL070W |
| YDL066W; DI49_0998; YDL065C | DI49_0998 | YDL066W; YDL065C |
| YDL052C; YDL053C | | YDL052C; YDL053C |
| YDL044C; DI49_1019; YDL043C | DI49_1019 | YDL044C; YDL043C |
| YDL035C; YDL036C | | YDL035C; YDL036C |
| YDL024C; YDL025C | | YDL024C; YDL025C |
| YDL015C; YDL017W | | YDL015C; YDL017W |
| YDR002W; DI49_1055; YDR003W | DI49_1055 | YDR002W; YDR003W |
| YDR005C; YDR004W | | YDR005C; YDR004W |
| YDR007W; DI49_1060; YDR009W | DI49_1060 | YDR007W; YDR009W |
| YDR014W-A; YDR014W | | YDR014W-A; YDR014W |
| YDR017C; DI49_0215 | DI49_0215 | YDR017C |
| YDR042C; YDR041W | | YDR042C; YDR041W |
| YDR054C; DI49_0248 | DI49_0248 | YDR054C |
| YDR070C; YDR069C | | YDR070C; YDR069C |
| YDR079C-A; YDR079W | | YDR079C-A; YDR079W |
| YDR081C; YDR080W | | YDR081C; YDR080W |
| YDR086C; YDR085C | | YDR086C; YDR085C |
| YDR092W; YDR091C | | YDR092W; YDR091C |
| YDR103W; YDR101C | | YDR103W; YDR101C |
| YDR105C; YDR104C | | YDR105C; YDR104C |
| YDR111C; YDR110W | | YDR111C; YDR110W |
| YDR115W; YDR113C | | YDR115W; YDR113C |
| YDR117C; YDR116C | | YDR117C; YDR116C |
| YDR121W; YDR120C | | YDR121W; YDR120C |
| YDR130C; YDR129C | | YDR130C; YDR129C |
| YDR132C; YDR131C | | YDR132C; YDR131C |
| YDR139C; YDR138W | | YDR139C; YDR138W |
| YDR146C; YDR145W | | YDR146C; YDR145W |
| YDR152W; YDR151C | | YDR152W; YDR151C |
| YDR159W; YDR158W | | YDR159W; YDR158W |
| YDR161W; YDR160W | | YDR161W; YDR160W |
| YDR163W; YDR162C | | YDR163W; YDR162C |

Create PDF in your applications with the Pdfcrowd HTML to PDF API PDFCROWD

| | | | |
|---|---|---|---|
| YDR172W; DI49_0362 | DI49_0362 | | YDR172W |
| YDR177W; YDR176W | | | YDR177W; YDR176W |
| YDR179C; YDR178W | | | YDR179C; YDR178W |
| YDR183W; YDR182W | | | YDR183W; YDR182W |
| YDR207C; YDR206W | | | YDR207C; YDR206W |
| YDR217C; YDR216W | | | YDR217C; YDR216W |
| YDR219C; YDR218C | | | YDR219C; YDR218C |
| YDR222W; YDR221W | | | YDR222W; YDR221W |
| YDR234W; DI49_0416 | DI49_0416 | | YDR234W |
| YDR239C; YDR238C | | | YDR239C; YDR238C |
| YDR252W; YDR251W | | | YDR252W; YDR251W |
| YDR254W; YDR253C | | | YDR254W; YDR253C |
| YDR259C; YDR258C | | | YDR259C; YDR258C |
| YDR261C; YDR260C | | | YDR261C; YDR260C |
| YDR264C; YDR263C | | | YDR264C; YDR263C |
| YDR266C; YDR265W | | | YDR266C; YDR265W |
| YDR275W; YDR273W | | | YDR275W; YDR273W |
| YDR281C; YDR280W | | | YDR281C; YDR280W |
| YDR286C; YDR285W | | | YDR286C; YDR285W |
| YDR288W; YDR287W | | | YDR288W; YDR287W |
| YDR304C; YDR303C | | | YDR304C; YDR303C |
| YDR306C; YDR305C | | | YDR306C; YDR305C |
| YDR309C; YDR308C | | | YDR309C; YDR308C |
| YDR311W; YDR310C | | | YDR311W; YDR310C |
| YDR315C; YDR314C | | | YDR315C; YDR314C |
| YDR319C; YDR318W | | | YDR319C; YDR318W |
| SPGP0AS00100; SPGP0Q00120 | | SPGP0AS00100; SPGP0Q00120 | |
| SPGP0AT00100; DI49_3503 | DI49_3503 | SPGP0AT00100 | |
| SPGP0AT00110; DI49_3504 | DI49_3504 | SPGP0AT00110 | |
| SPGP0ATtrna20P; SPGP0AHtrna19P | | SPGP0ATtrna20P; SPGP0AHtrna19P | |
| SPGP0AT00120; DI49_3506 | DI49_3506 | SPGP0AT00120 | |
| YPR062W; DI49_5515 | DI49_5515 | | YPR062W |
| YPR057W; YPR058W | | | YPR057W; YPR058W |
| YPR048W; YPR049C | | | YPR048W; YPR049C |
| YPR036W-A; YPR037C | | | YPR036W-A; YPR037C |
| YPR028W; YPR029C | | | YPR028W; YPR029C |
| YPR023C; YPR024W | | | YPR023C; YPR024W |
| YPR018W; YPR019W | | | YPR018W; YPR019W |
| YPR010C-A; YPR011C | | | YPR010C-A; YPR011C |
| YPR005C; YPR006C | | | YPR005C; YPR006C |

Create PDF in your applications with the Pdfcrowd HTML to PDF API PDFCROWD

| | | | |
|---|---|---|---|
| YPL003W; YPL002C | | | YPL003W; YPL002C |
| YPL005W; YPL004C | | | YPL005W; YPL004C |
| YPL007C; YPL006W | | | YPL007C; YPL006W |
| YPL032C; YPL031C | | | YPL032C; YPL031C |
| YPL034W; YPL033C | | | YPL034W; YPL033C |
| SPGP0BA00100; SPGP0DG00100; DI49_3920 | DI49_3920 | SPGP0BA00100; SPGP0DG00100 | |
| DI49_3924; SPGP0BC00130 | DI49_3924 | SPGP0BC00130 | |
| DI49_3926; DI49_3925 | DI49_3926; DI49_3925 | | |
| DI49_3928; DI49_3927 | DI49_3928; DI49_3927 | | |
| DI49_3930; DI49_3929 | DI49_3930; DI49_3929 | | |
| DI49_3932; DI49_3931 | DI49_3932; DI49_3931 | | |
| DI49_3934; DI49_3933 | DI49_3934; DI49_3933 | | |
| DI49_3936; DI49_3935 | DI49_3936; DI49_3935 | | |
| SPGP0BC00280; DI49_3937 | DI49_3937 | SPGP0BC00280 | |
| SPGP0BC00300; DI49_3938 | DI49_3938 | SPGP0BC00300 | |
| DI49_3940; DI49_3939 | DI49_3940; DI49_3939 | | |
| SPGP0BF00100; DI49_3941 | DI49_3941 | SPGP0BF00100 | |
| DI49_3477; DI49_3476 | DI49_3477; DI49_3476 | | |
| DI49_3479; DI49_3478 | DI49_3479; DI49_3478 | | |
| DI49_3485; DI49_3484 | DI49_3485; DI49_3484 | | |
| DI49_3487; DI49_3486 | DI49_3487; DI49_3486 | | |
| SPGP0BF00250; DI49_3488 | DI49_3488 | SPGP0BF00250 | |
| DI49_3492; DI49_3491 | DI49_3492; DI49_3491 | | |
| DI49_3496; DI49_3495 | DI49_3496; DI49_3495 | | |
| DI49_3498; DI49_3497 | DI49_3498; DI49_3497 | | |
| DI49_3500; DI49_3499 | DI49_3500; DI49_3499 | | |
| DI49_3502; DI49_3501 | DI49_3502; DI49_3501 | | |
| DI49_3505; DI49_4914; SPGP0Ltrna5P | DI49_3505; DI49_4914 | SPGP0Ltrna5P | |
| DI49_1329; YEL064C; DI49_1330; YEL063C | DI49_1329; DI49_1330 | | YEL064C; YEL063C |
| DI49_1332; DI49_1331 | DI49_1332; DI49_1331 | | |
| DI49_1334; DI49_1333 | DI49_1334; DI49_1333 | | |
| DI49_1336; DI49_1335 | DI49_1336; DI49_1335 | | |
| DI49_1338; DI49_1337 | DI49_1338; DI49_1337 | | |
| DI49_1340; YEL053C | DI49_1340 | | YEL053C |
| DI49_1341; YEL052W; DI49_1342; YEL051W | DI49_1341; DI49_1342 | | YEL052W; YEL051W |
| DI49_1348; DI49_1347 | DI49_1348; DI49_1347 | | |
| DI49_1350; YEL043W; DI49_1351; YEL042W | DI49_1350; DI49_1351 | | YEL043W; YEL042W |
| DI49_1353; DI49_1352 | DI49_1353; DI49_1352 | | |
| DI49_1355; DI49_1354 | DI49_1355; DI49_1354 | | |
| DI49_1357; YEL036C; YEL035C | DI49_1357 | | YEL036C; YEL035C |

Create PDF in your applications with the Pdfcrowd HTML to PDF API PDFCROWD

| | | | |
|---|---|---|---|
| DI49_1361; YEL031W | DI49_1361 | | YEL031W |
| DI49_1362; YEL030W; DI49_1363; YEL029C | DI49_1362; DI49_1363 | | YEL030W; YEL029C |
| DI49_1365; YEL027W | DI49_1365 | | YEL027W |
| DI49_1366; YEL026W; DI49_1367; YEL025C | DI49_1366; DI49_1367 | | YEL026W; YEL025C |
| DI49_1369; YEL023C | DI49_1369 | | YEL023C |
| DI49_1370; YEL022W | DI49_1370 | | YEL022W |
| DI49_1372; YEL011W | DI49_1372 | | YEL011W |
| DI49_1373; YEL012W | DI49_1373 | | YEL012W |
| DI49_1375; YEL013W | DI49_1375 | | YEL013W |
| DI49_1376; YEL015W | DI49_1376 | | YEL015W |
| DI49_1377; YEL016C; DI49_1378 | DI49_1377; DI49_1378 | | YEL016C |
| DI49_1380; DI49_1379 | DI49_1380; DI49_1379 | | |
| DI49_1382; DI49_1381 | DI49_1382; DI49_1381 | | |
| DI49_1384; YEL021W; DI49_1385 | DI49_1384; DI49_1385 | | YEL021W |
| DI49_1387; YEL009C; DI49_1388 | DI49_1387; DI49_1388 | | YEL009C |
| DI49_1390; YEL006W; DI49_1391; YEL005C | DI49_1390; DI49_1391 | | YEL006W; YEL005C |
| DI49_1393; DI49_1392 | DI49_1393; DI49_1392 | | |
| DI49_1395; DI49_1394 | DI49_1395; DI49_1394 | | |
| DI49_1396; SPGP0E00740 | DI49_1396 | SPGP0E00740 | |
| DI49_1398; DI49_1397 | DI49_1398; DI49_1397 | | |
| SPGP0E00790; YER006W; DI49_1400; YER007W | DI49_1400 | SPGP0E00790 | YER006W; YER007W |
| DI49_1402; DI49_1401 | DI49_1402; DI49_1401 | | |
| DI49_1404; DI49_1403; YER008C; YER009W | DI49_1404; DI49_1403 | | YER008C; YER009W |
| DI49_1410; YER014W; DI49_1411; YER015W | DI49_1410; DI49_1411 | | YER014W; YER015W |
| DI49_1414; YER017C | DI49_1414 | | YER017C |
| DI49_1415; YER018C; DI49_1416; YER019W | DI49_1415; DI49_1416 | | YER018C; YER019W |
| DI49_1419; YER021W; DI49_1420; YER022W | DI49_1419; DI49_1420 | | YER021W; YER022W |
| DI49_1422; YER024W | DI49_1422 | | YER024W |
| DI49_1423; YER025W | DI49_1423 | | YER025W |
| DI49_1427; DI49_1426 | DI49_1427; DI49_1426 | | |
| DI49_1429; DI49_1428 | DI49_1429; DI49_1428 | | |
| DI49_1431; DI49_1430 | DI49_1431; DI49_1430 | | |
| DI49_1433; DI49_1432 | DI49_1433; DI49_1432 | | |
| DI49_1435; YER036C; DI49_1436; YER037W | DI49_1435; DI49_1436 | | YER036C; YER037W |
| DI49_1438; YER039C; DI49_1439; YER040W | DI49_1438; DI49_1439 | | YER039C; YER040W |
| DI49_1441; YER042W | DI49_1441 | | YER042W |
| DI49_1442; YER043C | DI49_1442 | | YER043C |
| DI49_1443; YER044C; DI49_1444; YER044C-A | DI49_1443; DI49_1444 | | YER044C; YER044C-A |
| DI49_1446; DI49_1445 | DI49_1446; DI49_1445 | | |
| DI49_1448; YER048C | DI49_1448 | | YER048C |

Create PDF in your applications with the Pdfcrowd HTML to PDF API PDFCROWD

| | | | |
|---|---|---|---|
| DI49_1452; DI49_1451 | DI49_1452; DI49_1451 | | |
| DI49_1454; YER052C | DI49_1454 | | YER052C |
| DI49_1455; YER053C | DI49_1455 | | YER053C |
| DI49_1456; YER053C-A; DI49_1457; YER054C | DI49_1456; DI49_1457 | | YER053C-A; YER054C |
| DI49_1459; YER056C; DI49_1463; DI49_1464; YER060W | DI49_1459; DI49_1463; DI49_1464 | | YER056C; YER060W |
| SPGP0E01350; YER056C-A; DI49_1460; YER057C | DI49_1460 | SPGP0E01350 | YER056C-A; YER057C |
| DI49_1462; YER059W | DI49_1462 | | YER059W |
| DI49_1465; YER061C; YER062C | DI49_1465 | | YER061C; YER062C |
| DI49_1469; YER065C | DI49_1469 | | YER065C |
| DI49_1473; YER069W | DI49_1473 | | YER069W |
| DI49_1474; YER070W | DI49_1474 | | YER070W |
| DI49_1475; YER071C | DI49_1475 | | YER071C |
| DI49_1476; YER072W | DI49_1476 | | YER072W |
| DI49_1477; YER073W | DI49_1477 | | YER073W |
| DI49_1482; YER077C | DI49_1482 | | YER077C |
| DI49_1483; YER078C | DI49_1483 | | YER078C |
| DI49_1484; YER079W | DI49_1484 | | YER079W |
| DI49_1485; YER080W | DI49_1485 | | YER080W |
| DI49_1487; YER082C | DI49_1487 | | YER082C |
| DI49_1488; YER083C; DI49_1489; YER085C | DI49_1488; DI49_1489 | | YER083C; YER085C |
| DI49_1491; YER087W; DI49_1492; YER087C-B | DI49_1491; DI49_1492 | | YER087W; YER087C-B |
| DI49_1495; DI49_1493 | DI49_1495; DI49_1493 | | |
| DI49_1497; SPGP0I01770; DI49_1498; YER092W | DI49_1497; DI49_1498 | SPGP0I01770 | YER092W |
| SPGP0E01740; DI49_1499 | DI49_1499 | SPGP0E01740 | |
| DI49_1501; YER095W; DI49_1502; YER096W | DI49_1501; DI49_1502 | | YER095W; YER096W |
| DI49_1506; DI49_1505 | DI49_1506; DI49_1505 | | |
| DI49_1509; YER103W; DI49_1510; YER104W | DI49_1509; DI49_1510 | | YER103W; YER104W |
| DI49_1512; DI49_1511 | DI49_1512; DI49_1511 | | |
| DI49_1514; YER109C | DI49_1514 | | YER109C |
| DI49_1515; YER110C | DI49_1515 | | YER110C |
| DI49_1516; YER111C; DI49_1517; YER112W | DI49_1516; DI49_1517 | | YER111C; YER112W |
| DI49_1519; YER114C; DI49_1520; YER115C | DI49_1519; DI49_1520 | | YER114C; YER115C |
| DI49_1523; YER118C; DI49_1524; YER119C | DI49_1523; DI49_1524 | | YER118C; YER119C |
| DI49_1526; YER122C | DI49_1526 | | YER122C |
| DI49_1527; YER123W; DI49_1528; YER124C | DI49_1527; DI49_1528 | | YER123W; YER124C |
| DI49_1530; YER126C | DI49_1530 | | YER126C |
| DI49_1531; YER127W; DI49_1532; YER128W | DI49_1531; DI49_1532 | | YER127W; YER128W |
| DI49_1534; DI49_1533 | DI49_1534; DI49_1533 | | |
| DI49_1536; YER132C | DI49_1536 | | YER132C |
| DI49_1542; YER136W; DI49_1543; YER137C | DI49_1542; DI49_1543 | | YER136W; YER137C |

Create PDF in your applications with the Pdfcrowd HTML to PDF API PDFCROWD

| | | | |
|---|---|---|---|
| DI49_1547; YER139C; DI49_1548; YER140W | DI49_1547; DI49_1548 | | YER139C; YER140W |
| DI49_1550; YER142C; DI49_1551; YER143W | DI49_1550; DI49_1551 | | YER142C; YER143W |
| DI49_1553; YER145C; DI49_1554; YER146W | DI49_1553; DI49_1554 | | YER145C; YER146W |
| DI49_1556; YER148W; DI49_1557; YER149C | DI49_1556; DI49_1557 | | YER148W; YER149C |
| DI49_1560; YER151C | DI49_1560 | | YER151C |
| DI49_1561; YER152C; DI49_1562; YER153C | DI49_1561; DI49_1562 | | YER152C; YER153C |
| DI49_1564; YER155C; DI49_1565; YER156C | DI49_1564; DI49_1565 | | YER155C; YER156C |
| DI49_1569; YER158C; DI49_1570; YER159C | DI49_1569; DI49_1570 | | YER158C; YER159C |
| SPGP0E02400; DI49_1573 | DI49_1573 | SPGP0E02400 | |
| DI49_1575; YER165W | DI49_1575 | | YER165W |
| DI49_1576; YER166W | DI49_1576 | | YER166W |
| DI49_1577; YER167W; DI49_1578; YER168C | DI49_1577; DI49_1578 | | YER167W; YER168C |
| DI49_1580; DI49_1579 | DI49_1580; DI49_1579 | | |
| DI49_1582; YER172C | DI49_1582 | | YER172C |
| DI49_1583; YER173W; SPGP0E02510; YER174C | DI49_1583 | SPGP0E02510 | YER173W; YER174C |
| DI49_1585; DI49_1584 | DI49_1585; DI49_1584 | | |
| DI49_1587; YER178W; DI49_1588; YER179W | DI49_1587; DI49_1588 | | YER178W; YER179W |
| DI49_1590; DI49_1589 | DI49_1590; DI49_1589 | | |
| YGR278W; YGR279C | | | YGR278W; YGR279C |
| YGR273C; YGR274C | | | YGR273C; YGR274C |
| YGR266W; YGR267C | | | YGR266W; YGR267C |
| SPGP0Ftrna29T; YPR108W-A; YGR255C; YGR089W | | SPGP0Ftrna29T | YPR108W-A; YGR255C; YGR089W |
| YGR249W; YGR250C | | | YGR249W; YGR250C |
| YGR247W; YGR248W | | | YGR247W; YGR248W |
| YGR241C; YGR243W | | | YGR241C; YGR243W |
| YGR239C; YGR240C | | | YGR239C; YGR240C |
| YGR237C; YGR238C | | | YGR237C; YGR238C |
| YGR235C; YGR236C | | | YGR235C; YGR236C |
| YGR233C; YGR234W | | | YGR233C; YGR234W |
| YGR227W; YGR229C | | | YGR227W; YGR229C |
| YGR224W; YGR225W | | | YGR224W; YGR225W |
| YGR222W; YGR223C | | | YGR222W; YGR223C |
| YGR220C; YGR221C | | | YGR220C; YGR221C |
| YGR211W; YGR212W | | | YGR211W; YGR212W |
| YGR201C; YGR202C | | | YGR201C; YGR202C |
| YGR199W; YGR200C | | | YGR199W; YGR200C |
| YGR194C; YGR195W | | | YGR194C; YGR195W |
| YGR192C; YJR009C | | | YGR192C; YJR009C |
| YGR168C; YGR169C | | | YGR168C; YGR169C |
| YGR166W; YGR167W | | | YGR166W; YGR167W |

Create PDF in your applications with the Pdfcrowd HTML to PDF API PDFCROWD

| | | | |
|---|---|---|---|
| YGR158C; YGR159C | | | YGR158C; YGR159C |
| YGR152C; DI49_2116 | DI49_2116 | | YGR152C |
| YGR148C; YGL031C; YGR147C | | | YGR148C; YGL031C; YGR147C |
| YGR138C; DI49_2099 | DI49_2099 | | YGR138C |
| YGR130C; YGR131W | | | YGR130C; YGR131W |
| YGR128C; YGR129W | | | YGR128C; YGR129W |
| YGR126W; YGR127W | | | YGR126W; YGR127W |
| YGR113W; YGR116W | | | YGR113W; YGR116W |
| YGR104C; YGR105W | | | YGR104C; YGR105W |
| YGR102C; YGR103W | | | YGR102C; YGR103W |
| YGR091W; YGR092W | | | YGR091W; YGR092W |
| YGR081C; YGR082W | | | YGR081C; YGR082W |
| YGR071C; YGR072W | | | YGR071C; YGR072W |
| YGR063C; YGR065C | | | YGR063C; YGR065C |
| YGR061C; DI49_2021 | DI49_2021 | | YGR061C |
| YGR057C; YGR058W | | | YGR057C; YGR058W |
| YGR042W; YGR043C | | | YGR042W; YGR043C |
| YGR040W; YGR041W | | | YGR040W; YGR041W |
| YGR014W; YGR015C | | | YGR014W; YGR015C |
| YGR012W; YGR013W | | | YGR012W; YGR013W |
| YGR006W; YGR007W | | | YGR006W; YGR007W |
| YGL002W; YGL001C | | | YGL002W; YGL001C |
| YGL004C; YGL003C | | | YGL004C; YGL003C |
| YGL008C; DI49_1959 | DI49_1959 | | YGL008C |
| YGL012W; YGL011C | | | YGL012W; YGL011C |
| YGL017W; YGL016W | | | YGL017W; YGL016W |
| YGL019W; YGL018C | | | YGL019W; YGL018C |
| YGL026C; DI49_1942 | DI49_1942 | | YGL026C |
| YGL035C; YGL033W | | | YGL035C; YGL033W |
| YGL037C; YGL036W | | | YGL037C; YGL036W |
| YGL047W; SPGP0Ftrna27V | | SPGP0Ftrna27V | YGL047W |
| YGL059W; YGL058W | | | YGL059W; YGL058W |
| YGL061C; YGL060W | | | YGL061C; YGL060W |
| YGL063W; YGL062W | | | YGL063W; YGL062W |
| YGL082W; YGL081W | | | YGL082W; YGL081W |
| YGL089C; YGL087C | | | YGL089C; YGL087C |
| YGL091C; YGL090W | | | YGL091C; YGL090W |
| YGL096W; SPGP0Ftrna25E | | SPGP0Ftrna25E | YGL096W |
| YGL104C; YGL103W | | | YGL104C; YGL103W |
| YGL106W; YGL105W | | | YGL106W; YGL105W |

Create PDF in your applications with the Pdfcrowd HTML to PDF API PDFCROWD

| | | | |
|---|---|---|---|
| YGL108C; YGL107C | | | YGL108C; YGL107C |
| YGL111W; YGL110C | | | YGL111W; YGL110C |
| YGL113W; YGL112C | | | YGL113W; YGL112C |
| YGL121C; YGL120C | | | YGL121C; YGL120C |
| YGL126W; YGL125W | | | YGL126W; YGL125W |
| YGL133W; YGL131C | | | YGL133W; YGL131C |
| YGL145W; YGL144C | | | YGL145W; YGL144C |
| YGL155W; YGL154C | | | YGL155W; YGL154C |
| YGL157W; YGL156W | | | YGL157W; YGL156W |
| YGL161C; YGL160W | | | YGL161C; YGL160W |
| YGL163C; YGL162W | | | YGL163C; YGL162W |
| YGL166W; YGL164C | | | YGL166W; YGL164C |
| YGL169W; YGL167C | | | YGL169W; YGL167C |
| DI49_1796; DI49_1797 | DI49_1796; DI49_1797 | | |
| DI49_1794; DI49_1795 | DI49_1794; DI49_1795 | | |
| DI49_1792; DI49_1793 | DI49_1792; DI49_1793 | | |
| DI49_1786; DI49_1787 | DI49_1786; DI49_1787 | | |
| DI49_1768; DI49_1769 | DI49_1768; DI49_1769 | | |
| DI49_1762; DI49_1763 | DI49_1762; DI49_1763 | | |
| DI49_1749; DI49_1750 | DI49_1749; DI49_1750 | | |
| SPGP0F04740; DI49_1748 | DI49_1748 | SPGP0F04740 | |
| DI49_1740; DI49_1741 | DI49_1740; DI49_1741 | | |
| DI49_1738; DI49_1739 | DI49_1738; DI49_1739 | | |
| DI49_1736; DI49_1737 | DI49_1736; DI49_1737 | | |
| DI49_1734; DI49_1735 | DI49_1734; DI49_1735 | | |
| DI49_1732; DI49_1733 | DI49_1732; DI49_1733 | | |
| SPGP0F04960; SPGP0F04950 | | SPGP0F04960; SPGP0F04950 | |
| YJR148W; YJR149W | | | YJR148W; YJR149W |
| YJR142W; YJR143C | | | YJR142W; YJR143C |
| YJR140C; YJR141W | | | YJR140C; YJR141W |
| YJR132W; DI49_3140; YJR131W | DI49_3140 | | YJR132W; YJR131W |
| YJR129C; YJR130C | | | YJR129C; YJR130C |
| YJR126C; YJR127C | | | YJR126C; YJR127C |
| YJR123W; DI49_3132; YJR122W | DI49_3132 | | YJR123W; YJR122W |
| YJR117W; DI49_3127; YJR116W | DI49_3127 | | YJR117W; YJR116W |
| YJR113C; YJR115W | | | YJR113C; YJR115W |
| YJR112W; YJR112W-A | | | YJR112W; YJR112W-A |
| YJR104C; DI49_3114; YJR103W | DI49_3114 | | YJR104C; YJR103W |
| YJR101W; YJR102C | | | YJR101W; YJR102C |

Create PDF in your applications with the Pdfcrowd HTML to PDF API PDFCROWD

| | | | |
|---|---|---|---|
| YJR094W-A; DI49_3103; YJR094C | DI49_3103 | | YJR094W-A; YJR094C |
| YJR092W; YJR093C | | | YJR092W; YJR093C |
| YJR090C; DI49_3098; YJR089W | DI49_3098 | | YJR090C; YJR089W |
| YJR086W; YJR088C | | | YJR086W; YJR088C |
| YJR084W; YJR085C | | | YJR084W; YJR085C |
| YJR082C; YJR083C | | | YJR082C; YJR083C |
| YJR078W; YJR080C | | | YJR078W; YJR080C |
| YJR075W; DI49_3086; YJR074W | DI49_3086 | | YJR075W; YJR074W |
| YJR072C; YJR073C | | | YJR072C; YJR073C |
| YJR068W; DI49_3081; YJR067C | DI49_3081 | | YJR068W; YJR067C |
| YJR064W; DI49_3077; YJR063W | DI49_3077 | | YJR064W; YJR063W |
| YJR054W; DI49_3063; YJR053W | DI49_3063 | | YJR054W; YJR053W |
| YJR045C; DI49_3052; YJR044C | DI49_3052 | | YJR045C; YJR044C |
| YJR041C; YJR042W | | | YJR041C; YJR042W |
| YJR032W; DI49_3040; YJR031C | DI49_3040 | | YJR032W; YJR031C |
| YJR025C; YJR030C | | | YJR025C; YJR030C |
| YJR022W; YJR024C | | | YJR022W; YJR024C |
| YJR013W; YJR012C | | | YJR013W; YJR012C |
| YJR004C; DI49_3019; YJR003C | DI49_3019 | | YJR004C; YJR003C |
| YJR001W; YJR002W | | | YJR001W; YJR002W |
| YJL002C; YJL001W | | | YJL002C; YJL001W |
| YJL004C; YJL003W | | | YJL004C; YJL003W |
| YJL013C; YJL012C | | | YJL013C; YJL012C |
| YJL016W; YJL014W | | | YJL016W; YJL014W |
| YJL024C; DI49_2995; YJL025W | DI49_2995 | | YJL024C; YJL025W |
| YJL031C; YJL030W | | | YJL031C; YJL030W |
| YJL034W; DI49_2986; YJL035C | DI49_2986 | | YJL034W; YJL035C |
| YJL039C; DI49_2978; YJL041W | DI49_2978 | | YJL039C; YJL041W |
| YJL043W; YJL042W | | | YJL043W; YJL042W |
| YJL045W; YJL044C | | | YJL045W; YJL044C |
| YJL048C; YJL047C | | | YJL048C; YJL047C |
| SPGP0G01900; YJL049W | | SPGP0G01900 | YJL049W |
| YJL052W; DI49_2963; YJL053W | DI49_2963 | | YJL052W; YJL053W |
| YJL055W; YJL054W | | | YJL055W; YJL054W |
| YJL057C; YJL056C | | | YJL057C; YJL056C |
| YJL060W; DI49_2955; YJL061W | DI49_2955 | | YJL060W; YJL061W |
| YJL062W-A; DI49_2952; YJL063C | DI49_2952 | | YJL062W-A; YJL063C |
| YJL066C; YJL065C | | | YJL066C; YJL065C |
| YJL069C; YJL068C | | | YJL069C; YJL068C |
| YJL074C; YJL073W | | | YJL074C; YJL073W |

Create PDF in your applications with the Pdfcrowd HTML to PDF API PDFCROWD

| | | | |
|---|---|---|---|
| YJL078C; YJL076W | | | YJL078C; YJL076W |
| YJL087C; DI49_2933; YJL088W | DI49_2933 | | YJL087C; YJL088W |
| YJL109C; DI49_2911; YJL110C | DI49_2911 | | YJL109C; YJL110C |
| YJL117W; YJL116C | | | YJL117W; YJL116C |
| YJL121C; YJL118W | | | YJL121C; YJL118W |
| YJL123C; YJL122W | | | YJL123C; YJL122W |
| YJL133C-A; DI49_2889; YJL134W | DI49_2889 | | YJL133C-A; YJL134W |
| YJL137C; YJL136C | | | YJL137C; YJL136C |
| YJL139C; YJL138C | | | YJL139C; YJL138C |
| YJL143W; DI49_2882; YJL144W | DI49_2882 | | YJL143W; YJL144W |
| YJL146W; DI49_2879; YJL147C | DI49_2879 | | YJL146W; YJL147C |
| YJL168C; DI49_2859; YJL170C | DI49_2859 | | YJL168C; YJL170C |
| YJL177W; DI49_2852; YJL178C | DI49_2852 | | YJL177W; YJL178C |
| YJL184W; YJL183W | | | YJL184W; YJL183W |
| YJL193W; DI49_2838; YJL194W | DI49_2838 | | YJL193W; YJL194W |
| YJL197W; YJL196C | | | YJL197W; YJL196C |
| YJL201W; DI49_2831; YJL203W | DI49_2831 | | YJL201W; YJL203W |
| YJL210W; YJL209W | | | YJL210W; YJL209W |
| SPGP0G03340; YJL212C | | SPGP0G03340 | YJL212C |
| DI49_0077; DI49_0001; SPGP0H00880 | DI49_0077; DI49_0001 | SPGP0H00880 | |
| DI49_0070; DI49_0071 | DI49_0070; DI49_0071 | | |
| DI49_0064; DI49_0065 | DI49_0064; DI49_0065 | | |
| DI49_0057; DI49_0058 | DI49_0057; DI49_0058 | | |
| DI49_0053; YAL007C; DI49_0052 | DI49_0053; DI49_0052 | | YAL007C |
| DI49_0050; DI49_0051 | DI49_0050; DI49_0051 | | |
| DI49_0047; YAL014C; DI49_0046 | DI49_0047; DI49_0046 | | YAL014C |
| DI49_0037; YAL024C; DI49_0036 | DI49_0037; DI49_0036 | | YAL024C |
| DI49_0030; DI49_0031 | DI49_0030; DI49_0031 | | |
| DI49_0025; YAL036C; DI49_0024 | DI49_0025; DI49_0024 | | YAL036C |
| DI49_0022; DI49_0023 | DI49_0022; DI49_0023 | | |
| DI49_0012; DI49_0013 | DI49_0012; DI49_0013 | | |
| DI49_0010; DI49_0011 | DI49_0010; DI49_0011 | | |
| DI49_0008; DI49_0009 | DI49_0008; DI49_0009 | | |
| DI49_0006; DI49_0007 | DI49_0006; DI49_0007 | | |
| DI49_0004; DI49_0005 | DI49_0004; DI49_0005 | | |
| YEL072W; YEL073C | | | YEL072W; YEL073C |
| YEL061C; YEL062W | | | YEL061C; YEL062W |
| YEL059C-A; YEL060C | | | YEL059C-A; YEL060C |
| YEL057C; YEL058W | | | YEL057C; YEL058W |
| YEL055C; YEL056W | | | YEL055C; YEL056W |

Create PDF in your applications with the Pdfcrowd HTML to PDF API PDFCROWD

| | | | |
|---|---|---|---|
| YEL046C; YEL047C | | | YEL046C; YEL047C |
| YEL044W; YEL045C | | | YEL044W; YEL045C |
| YEL040W; YEL041W | | | YEL040W; YEL041W |
| YEL038W; YEL039C | | | YEL038W; YEL039C |
| YEL020W-A; DI49_1383; YEL020C | DI49_1383 | | YEL020W-A; YEL020C |
| YEL018W; YEL019C | | | YEL018W; YEL019C |
| YEL017W; YEL017C-A | | | YEL017W; YEL017C-A |
| YEL007W; DI49_1389 | DI49_1389 | | YEL007W |
| YEL003W; YEL004W | | | YEL003W; YEL004W |
| YEL001C; YEL002C | | | YEL001C; YEL002C |
| YER002W; YER001W | | | YER002W; YER001W |
| YER004W; YER003C | | | YER004W; YER003C |
| YER016W; SPGP0I00960 | | SPGP0I00960 | YER016W |
| YER028C; YER027C | | | YER028C; YER027C |
| YER030W; YER029C | | | YER030W; YER029C |
| YER032W; YER031C | | | YER032W; YER031C |
| YER034W; YER033C | | | YER034W; YER033C |
| YER046W; YER045C | | | YER046W; YER045C |
| YER050C; YER049W | | | YER050C; YER049W |
| YER064C; DI49_1468 | DI49_1468 | | YER064C |
| YER068W; YER067W | | | YER068W; YER067W |
| YER075C; DI49_1480 | DI49_1480 | | YER075C |
| YER090W; DI49_1496 | DI49_1496 | | YER090W |
| YER093C-A; YER093C | | | YER093C-A; YER093C |
| YER100W; YER099C | | | YER100W; YER099C |
| YER106W; YER105C | | | YER106W; YER105C |
| YER130C; YER129W | | | YER130C; YER129W |
| YER163C; YER162C | | | YER163C; YER162C |
| YER170W; YER169W | | | YER170W; YER169W |
| YER176W; YER175C | | | YER176W; YER175C |
| YER180C-A; YER180C | | | YER180C-A; YER180C |
| YER184C; YER183C | | | YER184C; YER183C |
| YER186C; YER185W | | | YER186C; YER185W |

Create PDF in your applications with the Pdfcrowd HTML to PDF API PDFCROWD
