## Supplementary file 3 for "HybridMine: pipeline for allele inheritance and gene copy number prediction in industrial yeast hybrids"

### Overview


genes alleles inherited from Saccharomyces eubayanus
genes alleles inherited from Saccharomyces cerevisiae
genes alleles below 80% identity
genes alleles copy with same identity in both parents
list of paralogs in Saccharomyces pastorianus CBS1538


#### Sheet 1: *genes alleles inherited from Saccharomyces eubayanus*

| *Saccharomyces pastorianus CBS1538 predicted ORF* | *Saccharomyces eubayanus parental allele* | Gene identities | Gene gaps |
|---|---|---|---|
| SPGP0U01380 | XP_018222944.1 | 98 | 0 |
| SPGP0CU02160 | XP_018220618.1 | 99 | 0 |
| SPGP0CV00210 | XP_018222572.1 | 99 | 0 |
| SPGP0DF00140 | XP_018219194.1 | 98 | 0 |
| SPGP0V00490 | XP_018221736.1 | 99 | 0 |
| SPGP0L02060 | XP_018218830.1 | 99 | 0 |
| SPGP0V00940 | XP_018221693.1 | 100 | 0 |
| SPGP0AT00310 | XP_018219052.1 | 99 | 1 |
| SPGP0U00880 | XP_018222991.1 | 99 | 0 |
| SPGP0AT00510 | XP_018219072.1 | 100 | 0 |
| SPGP0IZ00100 | XP_018220338.1 | 99 | 0 |
| SPGP0DE00160 | XP_018223833.1 | 99 | 0 |
| SPGP0CX01390 | XP_018222157.1 | 98 | 0 |
| SPGP0CX01300 | XP_018222148.1 | 99 | 0 |
| SPGP0CP03190 | XP_018223521.1 | 99 | 0 |
| SPGP0CM00870 | XP_018223238.1 | 99 | 0 |
| SPGP0BE00240 | XP_018223354.1 | 99 | 0 |
| SPGP0GQ00110 | XP_018219558.1 | 99 | 0 |
| SPGP0CX01640 | XP_018222180.1 | 99 | 0 |
| SPGP0AJ00230 | XP_018221797.1 | 99 | 0 |
| SPGP0IO00100 | XP_018221983.1 | 98 | 0 |
| SPGP0CX01700 | XP_018222186.1 | 99 | 0 |
| SPGP0L00930 | XP_018218939.1 | 99 | 0 |
| SPGP0CW01760 | XP_018219384.1 | 100 | 0 |
| SPGP0AN00310 | XP_018223941.1 | 99 | 0 |
| SPGP0AF02090 | XP_018222467.1 | 99 | 0 |
| SPGP0H00720 | XP_018221073.1 | 99 | 0 |
| SPGP0CW02150 | XP_018219346.1 | 99 | 0 |

Create PDF in your applications with the Pdfcrowd HTML to PDF API PDFCROWD

| | | | |
|---|---|---|---|
| SPGP0L02320 | XP_018218804.1 | 99 | 0 |
| SPGP0CP03620 | XP_018223482.1 | 99 | 0 |
| SPGP0CW03400 | XP_018219224.1 | 99 | 0 |
| SPGP0CW01490 | XP_018219409.1 | 99 | 0 |
| SPGP0EP00150 | XP_018222748.1 | 99 | 0 |
| SPGP0EO00340 | XP_018221171.1 | 99 | 0 |
| SPGP0CP03350 | XP_018223506.1 | 100 | 0 |
| SPGP0AT00950 | XP_018219113.1 | 99 | 0 |
| SPGP0AF00620 | XP_018222316.1 | 100 | 0 |
| SPGP0AA01760 | XP_018220415.1 | 99 | 0 |
| SPGP0CM01690 | XP_018223157.1 | 99 | 0 |
| SPGP0CW00460 | XP_018219509.1 | 100 | 0 |
| SPGP0DA01590 | XP_018222598.1 | 99 | 0 |
| SPGP0CM00100 | XP_018223314.1 | 99 | 0 |
| SPGP0CM01390 | XP_018223187.1 | 100 | 0 |
| SPGP0U01560 | XP_018222926.1 | 99 | 0 |
| SPGP0CP01100 | XP_018223729.1 | 99 | 0 |
| SPGP0AJ00630 | XP_018221835.1 | 99 | 0 |
| SPGP0BE00130 | XP_018223365.1 | 99 | 0 |
| SPGP0AN00990 | XP_018224005.1 | 99 | 0 |
| SPGP0AT01300 | XP_018219148.1 | 99 | 0 |
| SPGP0CW01130 | XP_018219443.1 | 99 | 0 |
| SPGP0CW00170 | XP_018219536.1 | 99 | 0 |
| SPGP0AF01240 | XP_018222380.1 | 100 | 0 |
| SPGP0CX02170 | XP_018222231.1 | 99 | 0 |
| SPGP0CM02150 | XP_018223114.1 | 99 | 0 |
| SPGP0EO00280 | XP_018221165.1 | 98 | 1 |
| SPGP0IC00140 | XP_018222529.1 | 99 | 0 |
| SPGP0AJ00470 | XP_018221820.1 | 99 | 0 |
| SPGP0G00140 | XP_018222253.1 | 100 | 0 |
| SPGP0AF01630 | XP_018222421.1 | 99 | 0 |
| SPGP0V00550 | XP_018221731.1 | 99 | 0 |
| SPGP0CP00260 | XP_018223810.1 | 99 | 0 |
| SPGP0AJ00200 | XP_018221795.1 | 99 | 0 |
| SPGP0DA00840 | XP_018222669.1 | 100 | 0 |
| SPGP0CW01940 | XP_018219366.1 | 99 | 0 |
| SPGP0FE00210 | XP_018221973.1 | 99 | 0 |
| SPGP0CW00920 | XP_018219464.1 | 99 | 0 |
| SPGP0AJ01770 | XP_018221945.1 | 99 | 0 |
| SPGP0AN00360 | XP_018223945.1 | 99 | 0 |

Create PDF in your applications with the Pdfcrowd HTML to PDF API PDFCROWD

| | | | |
|---|---|---|---|
| SPGP0CW02710 | XP_018219292.1 | 99 | 0 |
| SPGP0CW02430 | XP_018219318.1 | 100 | 0 |
| SPGP0CU01370 | XP_018220696.1 | 99 | 0 |
| SPGP0CP00310 | XP_018223805.1 | 99 | 0 |
| SPGP0L00300 | XP_018219002.1 | 99 | 0 |
| SPGP0GL00200 | XP_018220849.1 | 99 | 0 |
| SPGP0Y02090 | XP_018220219.1 | 99 | 0 |
| SPGP0CM00420 | XP_018223282.1 | 99 | 0 |
| SPGP0IU00160 | XP_018220334.1 | 99 | 0 |
| SPGP0AT00420 | XP_018219063.1 | 99 | 0 |
| SPGP0Y00760 | XP_018220092.1 | 99 | 0 |
| SPGP0AJ00520 | XP_018221824.1 | 99 | 0 |
| SPGP0R00580 | XP_018219998.1 | 99 | 0 |
| SPGP0CP02480 | XP_018223593.1 | 99 | 0 |
| SPGP0U01710 | XP_018222911.1 | 99 | 0 |
| SPGP0DE00410 | XP_018223858.1 | 100 | 0 |
| SPGP0CX00810 | XP_018222100.1 | 99 | 0 |
| SPGP0Y00490 | XP_018220066.1 | 99 | 0 |
| SPGP0CM02090 | XP_018223120.1 | 99 | 0 |
| SPGP0AF00900 | XP_018222344.1 | 98 | 2 |
| SPGP0L00720 | XP_018218961.1 | 99 | 1 |
| SPGP0DF00170 | XP_018219191.1 | 99 | 0 |
| SPGP0DB00270 | XP_018223901.1 | 100 | 0 |
| SPGP0JA00120 | XP_018222244.1 | 99 | 0 |
| SPGP0CU01860 | XP_018220648.1 | 99 | 0 |
| SPGP0CP01290 | XP_018223710.1 | 99 | 0 |
| SPGP0CP01920 | XP_018223648.1 | 100 | 0 |
| SPGP0EO00150 | XP_018221152.1 | 99 | 0 |
| SPGP0AJ01300 | XP_018221901.1 | 99 | 0 |
| SPGP0CW00350 | XP_018219520.1 | 99 | 0 |
| SPGP0V00530 | XP_018221733.1 | 99 | 0 |
| SPGP0Y01010 | XP_018220116.1 | 99 | 1 |
| SPGP0JK00350 | XP_018223456.1 | 99 | 0 |
| SPGP0CW02620 | XP_018219301.1 | 99 | 0 |
| SPGP0IN00250 | XP_018221651.1 | 99 | 0 |
| SPGP0AJ00180 | XP_018221793.1 | 93 | 0 |
| SPGP0V00310 | XP_018221752.1 | 99 | 0 |
| SPGP0CU01630 | XP_018220671.1 | 100 | 0 |
| SPGP0CX01790 | XP_018222194.1 | 99 | 0 |
| SPGP0AA01040 | XP_018220486.1 | 99 | 1 |

Create PDF in your applications with the Pdfcrowd HTML to PDF API PDFCROWD

| | | | |
|---|---|---|---|
| SPGP0L01430 | XP_018218891.1 | 99 | 0 |
| SPGP0CW02850 | XP_018219278.1 | 99 | 0 |
| SPGP0IR00110 | XP_018223465.1 | 99 | 0 |
| SPGP0DF00240 | XP_018219184.1 | 99 | 0 |
| SPGP0L02640 | XP_018218773.1 | 99 | 0 |
| SPGP0AT01430 | XP_018219160.1 | 99 | 0 |
| SPGP0CW01530 | XP_018219405.1 | 99 | 1 |
| SPGP0CU00160 | XP_018220812.1 | 99 | 0 |
| SPGP0CW02940 | XP_018219269.1 | 99 | 0 |
| SPGP0U01070 | XP_018222972.1 | 99 | 0 |
| SPGP0AF00540 | XP_018222308.1 | 99 | 0 |
| SPGP0V01190 | XP_018221669.1 | 99 | 0 |
| SPGP0CM01610 | XP_018223165.1 | 99 | 0 |
| SPGP0CU01410 | XP_018220692.1 | 99 | 0 |
| SPGP0CM00630 | XP_018223261.1 | 99 | 0 |
| SPGP0CU01140 | XP_018220716.1 | 99 | 0 |
| SPGP0IG00120 | XP_018223346.1 | 99 | 0 |
| SPGP0CU01420 | XP_018220691.1 | 99 | 0 |
| SPGP0CM00410 | XP_018223283.1 | 99 | 0 |
| SPGP0Z00280 | XP_018222537.1 | 99 | 0 |
| SPGP0U01010 | XP_018222978.1 | 99 | 0 |
| SPGP0Y02690 | XP_018220278.1 | 100 | 0 |
| SPGP0FE00180 | XP_018221970.1 | 99 | 0 |
| SPGP0CU02490 | XP_018220585.1 | 99 | 0 |
| SPGP0CM01940 | XP_018223132.1 | 99 | 0 |
| SPGP0J01090 | XP_018219838.1 | 99 | 0 |
| SPGP0L02120 | XP_018218824.1 | 99 | 0 |
| SPGP0Y00670 | XP_018220083.1 | 99 | 0 |
| SPGP0EP00180 | XP_018222751.1 | 99 | 0 |
| SPGP0U00890 | XP_018222990.1 | 99 | 0 |
| SPGP0DE00180 | XP_018223835.1 | 99 | 0 |
| SPGP0R00260 | XP_018219968.1 | 99 | 0 |
| SPGP0AA01720 | XP_018220419.1 | 99 | 0 |
| SPGP0DE00350 | XP_018223852.1 | 99 | 0 |
| SPGP0K00240 | XP_018222766.1 | 99 | 0 |
| SPGP0U01900 | XP_018222891.1 | 99 | 0 |
| SPGP0DE00600 | XP_018223877.1 | 99 | 0 |
| SPGP0CW02490 | XP_018219312.1 | 99 | 0 |
| SPGP0U01260 | XP_018222955.1 | 99 | 0 |
| SPGP0CW03560 | XP_018219208.1 | 99 | 0 |

Create PDF in your applications with the Pdfcrowd HTML to PDF API PDFCROWD

| | | | |
|---|---|---|---|
| SPGP0CP03600 | XP_018223484.1 | 99 | 0 |
| SPGP0AF00550 | XP_018222309.1 | 99 | 0 |
| SPGP0CW01350 | XP_018219423.1 | 99 | 0 |
| SPGP0CW01700 | XP_018219390.1 | 99 | 0 |
| SPGP0L00910 | XP_018218941.1 | 99 | 0 |
| SPGP0CW02170 | XP_018219344.1 | 99 | 0 |
| SPGP0AQ00200 | XP_018223403.1 | 99 | 0 |
| SPGP0L02330 | XP_018218803.1 | 99 | 0 |
| SPGP0Y00530 | XP_018220070.1 | 99 | 0 |
| SPGP0AT00930 | XP_018219111.1 | 99 | 0 |
| SPGP0AN01020 | XP_018224008.1 | 99 | 0 |
| SPGP0CU02130 | XP_018220621.1 | 99 | 0 |
| SPGP0CW03420 | XP_018219222.1 | 99 | 0 |
| SPGP0JP00190 | XP_018218748.1 | 97 | 3 |
| SPGP0CU00390 | XP_018220790.1 | 99 | 0 |
| SPGP0AF01060 | XP_018222361.1 | 99 | 0 |
| SPGP0AJ00280 | XP_018221802.1 | 99 | 0 |
| SPGP0EO00320 | XP_018221169.1 | 99 | 0 |
| SPGP0U00310 | XP_018223040.1 | 99 | 0 |
| SPGP0Y02910 | XP_018220300.1 | 99 | 0 |
| SPGP0IC00160 | XP_018222527.1 | 100 | 0 |
| SPGP0V00140 | XP_018221769.1 | 99 | 0 |
| SPGP0AJ01580 | XP_018221926.1 | 100 | 0 |
| SPGP0R00150 | XP_018219957.1 | 99 | 0 |
| SPGP0AF01130 | XP_018222369.1 | 99 | 0 |
| SPGP0AE00400 | XP_018222519.1 | 99 | 0 |
| SPGP0AF01550 | XP_018222413.1 | 99 | 0 |
| SPGP0EP00130 | XP_018222746.1 | 99 | 0 |
| SPGP0AA01340 | XP_018220456.1 | 99 | 0 |
| SPGP0AN00200 | XP_018223932.1 | 100 | 0 |
| SPGP0CW00480 | XP_018219507.1 | 99 | 0 |
| SPGP0HC00100 | XP_018221781.1 | 99 | 0 |
| SPGP0R00560 | XP_018219996.1 | 99 | 0 |
| SPGP0CX00560 | XP_018222076.1 | 99 | 0 |
| SPGP0CW00150 | XP_018219539.1 | 99 | 0 |
| SPGP0AT01320 | XP_018219150.1 | 99 | 0 |
| SPGP0CP01120 | XP_018223726.1 | 99 | 0 |
| SPGP0CW00250 | XP_018219528.1 | 99 | 0 |
| SPGP0BE00110 | XP_018223367.1 | 99 | 0 |
| SPGP0CU02480 | XP_018220586.1 | 99 | 0 |

Create PDF in your applications with the Pdfcrowd HTML to PDF API PDFCROWD

| | | | |
|---|---|---|---|
| SPGP0CX02270 | XP_018222241.1 | 99 | 0 |
| SPGP0AA01980 | XP_018220395.1 | 99 | 0 |
| SPGP0CM01110 | XP_018223214.1 | 99 | 0 |
| SPGP0L00480 | XP_018218984.1 | 99 | 0 |
| SPGP0Y00910 | XP_018220107.1 | 99 | 0 |
| SPGP0V00100 | XP_018221773.1 | 99 | 0 |
| SPGP0DE00470 | XP_018223864.1 | 99 | 0 |
| SPGP0L02450 | XP_018218791.1 | 99 | 0 |
| SPGP0V00720 | XP_018221714.1 | 99 | 0 |
| SPGP0CM02120 | XP_018223117.1 | 100 | 0 |
| SPGP0AA00800 | XP_018220510.1 | 87 | 0 |
| SPGP0AF00600 | XP_018222314.1 | 99 | 0 |
| SPGP0CW02770 | XP_018219286.1 | 99 | 0 |
| SPGP0CM00400 | XP_018223284.1 | 99 | 0 |
| SPGP0U01780 | XP_018222903.1 | 99 | 0 |
| SPGP0AJ00720 | XP_018221844.1 | 99 | 0 |
| SPGP0CX02180 | XP_018222232.1 | 99 | 0 |
| SPGP0AF01200 | XP_018222376.1 | 99 | 0 |
| SPGP0Y02560 | XP_018220265.1 | 99 | 0 |
| SPGP0IU00140 | XP_018220332.1 | 99 | 0 |
| SPGP0AT00440 | XP_018219065.1 | 100 | 0 |
| SPGP0Y00700 | XP_018220086.1 | 99 | 0 |
| SPGP0AT00110 | XP_018219033.1 | 99 | 0 |
| SPGP0AJ00120 | XP_018221787.1 | 99 | 0 |
| SPGP0AJ00540 | XP_018221826.1 | 99 | 0 |
| SPGP0AT00330 | XP_018219054.1 | 99 | 0 |
| SPGP0J03820 | XP_018219569.1 | 99 | 0 |
| SPGP0AN00840 | XP_018223990.1 | 99 | 0 |
| SPGP0K00500 | XP_018222792.1 | 99 | 0 |
| SPGP0Y03170 | XP_018220325.1 | 99 | 0 |
| SPGP0CP02390 | XP_018223602.1 | 99 | 0 |
| SPGP0U00450 | XP_018223029.1 | 99 | 0 |
| SPGP0CP02020 | XP_018223638.1 | 99 | 0 |
| SPGP0AF00890 | XP_018222343.1 | 99 | 0 |
| SPGP0CW02010 | XP_018219359.1 | 99 | 0 |
| SPGP0L00800 | XP_018218953.1 | 100 | 0 |
| SPGP0L00210 | XP_018219010.1 | 99 | 0 |
| SPGP0DB00250 | XP_018223899.1 | 99 | 0 |
| SPGP0AN00560 | XP_018223963.1 | 100 | 0 |

Create PDF in your applications with the Pdfcrowd HTML to PDF API PDFCROWD

| | | | |
|---|---|---|---|
| SPGP0Y00150 | XP_018220032.1 | 99 | 0 |
| SPGP0CU01670 | XP_018220667.1 | 99 | 0 |
| SPGP0R00700 | XP_018220010.1 | 99 | 0 |
| SPGP0U01650 | XP_018222917.1 | 99 | 0 |
| SPGP0AA00270 | XP_018220563.1 | 98 | 0 |
| SPGP0HO00100 | XP_018224034.1 | 99 | 0 |
| SPGP0JK00160 | XP_018223438.1 | 99 | 0 |
| SPGP0AH00230 | XP_018223430.1 | 99 | 0 |
| SPGP0AA00910 | XP_018220499.1 | 99 | 0 |
| SPGP0AA00540 | XP_018220536.1 | 100 | 0 |
| SPGP0CU00980 | XP_018220732.1 | 99 | 0 |
| SPGP0AF00780 | XP_018222332.1 | 99 | 0 |
| SPGP0AH00180 | XP_018223426.1 | 99 | 0 |
| SPGP0AJ01910 | XP_018221957.1 | 99 | 0 |
| SPGP0AA01550 | XP_018220435.1 | 99 | 0 |
| SPGP0U00930 | XP_018222986.1 | 99 | 0 |
| SPGP0GG00170 | XP_018220834.1 | 99 | 0 |
| SPGP0AA01320 | XP_018220458.1 | 99 | 0 |
| SPGP0U00710 | XP_018223004.1 | 99 | 0 |
| SPGP0U01090 | XP_018222970.1 | 99 | 0 |
| SPGP0Y02820 | XP_018220291.1 | 99 | 0 |
| SPGP0Y00280 | XP_018220045.1 | 99 | 0 |
| SPGP0CX02240 | XP_018222238.1 | 99 | 0 |
| SPGP0GS00130 | XP_018222248.1 | 99 | 0 |
| SPGP0CM01630 | XP_018223163.1 | 100 | 0 |
| SPGP0CW03550 | XP_018219209.1 | 99 | 0 |
| SPGP0CX00930 | XP_018222112.1 | 99 | 0 |
| SPGP0FP00110 | XP_018223344.1 | 99 | 0 |
| SPGP0CU01450 | XP_018220689.1 | 98 | 1 |
| SPGP0U01030 | XP_018222976.1 | 99 | 0 |
| SPGP0CW00830 | XP_018219473.1 | 99 | 0 |
| SPGP0AT01180 | XP_018219136.1 | 99 | 0 |
| SPGP0CU02120 | XP_018220622.1 | 99 | 0 |
| SPGP0CP02250 | XP_018223615.1 | 99 | 0 |
| SPGP0CU01290 | XP_018220703.1 | 99 | 0 |
| SPGP0V00900 | XP_018221697.1 | 99 | 0 |
| SPGP0AJ00390 | XP_018221812.1 | 99 | 0 |
| SPGP0AT01450 | XP_018219162.1 | 99 | 0 |
| SPGP0K00660 | XP_018222808.1 | 100 | 0 |
| SPGP0CU02370 | XP_018220597.1 | 99 | 0 |

Create PDF in your applications with the Pdfcrowd HTML to PDF API PDFCROWD

| | | | |
|---|---|---|---|
| SPGP0FL00100 | XP_018222811.1 | 99 | 0 |
| SPGP0L00420 | XP_018218990.1 | 100 | 0 |
| SPGP0U00900 | XP_018222989.1 | 99 | 0 |
| SPGP0U00610 | XP_018223014.1 | 99 | 0 |
| SPGP0CP03180 | XP_018223522.1 | 99 | 0 |
| SPGP0EW00110 | XP_018223462.1 | 86 | 0 |
| SPGP0AA01380 | XP_018220452.1 | 99 | 0 |
| SPGP0AT00580 | XP_018219079.1 | 99 | 0 |
| SPGP0Y00510 | XP_018220068.1 | 99 | 0 |
| SPGP0CU01230 | XP_018220709.1 | 99 | 0 |
| SPGP0CP03660 | XP_018223478.1 | 99 | 0 |
| SPGP0K00430 | XP_018222785.1 | 99 | 0 |
| SPGP0IM00170 | XP_018219545.1 | 99 | 0 |
| SPGP0Y01380 | XP_018220153.1 | 99 | 0 |
| SPGP0DB00390 | XP_018223912.1 | 99 | 0 |
| SPGP0L01880 | XP_018218847.1 | 99 | 0 |
| SPGP0CM00730 | XP_018223252.1 | 99 | 0 |
| SPGP0JP00170 | XP_018219568.1 | 99 | 0 |
| SPGP0AA01820 | XP_018220410.1 | 98 | 0 |
| SPGP0CP03320 | XP_018223508.1 | 99 | 0 |
| SPGP0DE00480 | XP_018223865.1 | 99 | 0 |
| SPGP0L00490 | XP_018218983.1 | 99 | 0 |
| SPGP0AF01140 | XP_018222370.1 | 99 | 0 |
| SPGP0AJ01640 | XP_018221932.1 | 99 | 0 |
| SPGP0CU00760 | XP_018220753.1 | 100 | 0 |
| SPGP0AN00730 | XP_018223980.1 | 99 | 0 |
| SPGP0AN00220 | XP_018223934.1 | 99 | 0 |
| SPGP0EP00110 | XP_018222744.1 | 99 | 0 |
| SPGP0CP02760 | XP_018223565.1 | 99 | 0 |
| SPGP0CM01660 | XP_018223160.1 | 99 | 0 |
| SPGP0V00290 | XP_018221754.1 | 99 | 0 |
| SPGP0AT01270 | XP_018219145.1 | 99 | 0 |
| SPGP0AF00190 | XP_018222274.1 | 99 | 0 |
| SPGP0L01120 | XP_018218922.1 | 99 | 0 |
| SPGP0R00730 | XP_018220013.1 | 99 | 0 |
| SPGP0CM01130 | XP_018218943.1 | 99 | 0 |
| SPGP0Y00730 | XP_018220089.1 | 99 | 0 |
| SPGP0V01140 | XP_018221674.1 | 98 | 1 |
| SPGP0V00440 | XP_018221741.1 | 98 | 1 |
| SPGP0L02300 | XP_018218806.1 | 99 | 0 |

Create PDF in your applications with the Pdfcrowd HTML to PDF API PDFCROWD

| | | | |
|---|---|---|---|
| SPGP0L01150 | XP_018218919.1 | 99 | 0 |
| SPGP0CX00780 | XP_018222097.1 | 99 | 0 |
| SPGP0CP03080 | XP_018223532.1 | 99 | 0 |
| SPGP0GD00100 | XP_018223413.1 | 99 | 0 |
| SPGP0CW02750 | XP_018219288.1 | 99 | 0 |
| SPGP0CP03060 | XP_018223534.1 | 100 | 0 |
| SPGP0CU00810 | XP_018220748.1 | 99 | 0 |
| SPGP0CM02110 | XP_018223118.1 | 99 | 0 |
| SPGP0AF02040 | XP_018222462.1 | 99 | 0 |
| SPGP0FP00120 | XP_018223343.1 | 99 | 0 |
| SPGP0CW02320 | XP_018219329.1 | 99 | 0 |
| SPGP0AN00180 | XP_018223930.1 | 99 | 0 |
| SPGP0Z00240 | XP_018222541.1 | 99 | 0 |
| SPGP0AJ01100 | XP_018221881.1 | 99 | 0 |
| SPGP0CP01500 | XP_018223689.1 | 99 | 0 |
| SPGP0U01200 | XP_018222960.1 | 99 | 0 |
| SPGP0CM00310 | XP_018223293.1 | 99 | 0 |
| SPGP0Y03190 | XP_018220327.1 | 99 | 0 |
| SPGP0CP00910 | XP_018223748.1 | 99 | 0 |
| SPGP0U02080 | XP_018222873.1 | 99 | 0 |
| SPGP0HW00140 | XP_018224039.1 | 99 | 0 |
| SPGP0AF00870 | XP_018222341.1 | 99 | 0 |
| SPGP0L00760 | XP_018218957.1 | 99 | 0 |
| SPGP0AN00500 | XP_018223957.1 | 100 | 0 |
| SPGP0AA00740 | XP_018220517.1 | 99 | 0 |
| SPGP0Y02360 | XP_018220245.1 | 99 | 0 |
| SPGP0Y01920 | XP_018220203.1 | 99 | 1 |
| SPGP0CP02360 | XP_018223605.1 | 100 | 0 |
| SPGP0L02560 | XP_018218780.1 | 99 | 0 |
| SPGP0U00400 | XP_018223033.1 | 99 | 0 |
| SPGP0U01630 | XP_018222919.1 | 99 | 0 |
| SPGP0CX01140 | XP_018222133.1 | 99 | 0 |
| SPGP0EB00150 | XP_018223316.1 | 99 | 0 |
| SPGP0Y01050 | XP_018220120.1 | 99 | 0 |
| SPGP0AF00380 | XP_018222292.1 | 99 | 0 |
| SPGP0JK00100 | XP_018223431.1 | 99 | 0 |
| SPGP0U00670 | XP_018223008.1 | 99 | 0 |
| SPGP0V00350 | XP_018221748.1 | 100 | 0 |
| SPGP0CU00420 | XP_018220787.1 | 99 | 0 |
| SPGP0L01140 | XP_018218920.1 | 99 | 0 |

Create PDF in your applications with the Pdfcrowd HTML to PDF API PDFCROWD

| | | | |
|---|---|---|---|
| SPGP0CW03240 | XP_018219240.1 | 99 | 0 |
| SPGP0CX01530 | XP_018222170.1 | 99 | 0 |
| SPGP0CM01840 | XP_018223143.1 | 100 | 0 |
| SPGP0CU00210 | XP_018220808.1 | 99 | 0 |
| SPGP0CW02890 | XP_018219274.1 | 99 | 0 |
| SPGP0AJ01930 | XP_018221959.1 | 99 | 0 |
| SPGP0CP03410 | XP_018223501.1 | 99 | 0 |
| SPGP0GG00190 | XP_018220832.1 | 99 | 0 |
| SPGP0CU00880 | XP_018220741.1 | 99 | 0 |
| SPGP0Y02800 | XP_018220289.1 | 99 | 0 |
| SPGP0CX01180 | XP_018222137.1 | 99 | 0 |
| SPGP0CM01650 | XP_018223161.1 | 99 | 0 |
| SPGP0AT00920 | XP_018219110.1 | 99 | 0 |
| SPGP0CP01080 | XP_018223731.1 | 99 | 0 |
| SPGP0CX01450 | XP_018222162.1 | 99 | 0 |
| SPGP0AN00650 | XP_018223972.1 | 99 | 0 |
| SPGP0CW00850 | XP_018219471.1 | 99 | 0 |
| SPGP0CW02930 | XP_018219270.1 | 99 | 0 |
| SPGP0K00210 | XP_018222763.1 | 99 | 0 |
| SPGP0AT01610 | XP_018219178.1 | 99 | 0 |
| SPGP0DE00220 | XP_018223839.1 | 99 | 0 |
| SPGP0GH00110 | XP_018222813.1 | 99 | 0 |
| SPGP0FE00140 | XP_018221966.1 | 99 | 0 |
| SPGP0CU02450 | XP_018220589.1 | 99 | 0 |
| SPGP0CU02100 | XP_018220624.1 | 99 | 0 |
| SPGP0V00920 | XP_018221695.1 | 99 | 0 |
| SPGP0L02530 | XP_018218783.1 | 99 | 0 |
| SPGP0U00360 | XP_018223037.1 | 99 | 0 |
| SPGP0L02220 | XP_018218814.1 | 99 | 0 |
| SPGP0CX01330 | XP_018222151.1 | 99 | 0 |
| SPGP0CU00320 | XP_018220797.1 | 99 | 0 |
| SPGP0AT01470 | XP_018219164.1 | 99 | 0 |
| SPGP0AA01360 | XP_018220454.1 | 99 | 0 |
| SPGP0CP00860 | XP_018223752.1 | 99 | 0 |
| SPGP0CP03230 | XP_018223517.1 | 99 | 0 |
| SPGP0L01310 | XP_018218903.1 | 99 | 0 |
| SPGP0GJ00210 | XP_018223371.1 | 99 | 0 |
| SPGP0CM00230 | XP_018223301.1 | 99 | 0 |
| SPGP0Y02240 | XP_018220233.1 | 98 | 1 |
| SPGP0IU00190 | XP_018220337.1 | 99 | 0 |

Create PDF in your applications with the Pdfcrowd HTML to PDF API PDFCROWD

| | | | |
|---|---|---|---|
| SPGP0CU01810 | XP_018220653.1 | 99 | 0 |
| SPGP0DE00640 | XP_018223881.1 | 99 | 0 |
| SPGP0CP02310 | XP_018223609.1 | 99 | 0 |
| SPGP0L01220 | XP_018218912.1 | 99 | 0 |
| SPGP0CW00760 | XP_018219480.1 | 99 | 0 |
| SPGP0CU02170 | XP_018220617.1 | 99 | 0 |
| SPGP0U00370 | XP_018223036.1 | 99 | 0 |
| SPGP0AA01840 | XP_018220408.1 | 96 | 0 |
| SPGP0CX02080 | XP_018222223.1 | 99 | 0 |
| SPGP0AA00890 | XP_018220501.1 | 99 | 0 |
| SPGP0AJ01890 | XP_018221955.1 | 99 | 0 |
| SPGP0AT00120 | XP_018219034.1 | 99 | 0 |
| SPGP0AH00100 | XP_018223418.1 | 99 | 0 |
| SPGP0CX01830 | XP_018222198.1 | 99 | 0 |
| SPGP0Y02230 | XP_018220232.1 | 99 | 0 |
| SPGP0CW00110 | XP_018219543.1 | 99 | 0 |
| SPGP0CM00820 | XP_018223243.1 | 99 | 0 |
| SPGP0AJ01090 | XP_018221880.1 | 99 | 0 |
| SPGP0CX01490 | XP_018222166.1 | 98 | 1 |
| SPGP0Y02190 | XP_018220228.1 | 99 | 0 |
| SPGP0AT01400 | XP_018219157.1 | 99 | 0 |
| SPGP0Y01850 | XP_018220196.1 | 99 | 0 |
| SPGP0JK00300 | XP_018223450.1 | 99 | 0 |
| SPGP0AF01740 | XP_018222432.1 | 99 | 0 |
| SPGP0CP01220 | XP_018223716.1 | 99 | 0 |
| SPGP0U01480 | XP_018222934.1 | 99 | 0 |
| SPGP0V00620 | XP_018221724.1 | 99 | 0 |
| SPGP0R00500 | XP_018219991.1 | 99 | 0 |
| SPGP0CX02110 | XP_018222225.1 | 99 | 0 |
| SPGP0CW01300 | XP_018219427.1 | 99 | 0 |
| SPGP0L00350 | XP_018218997.1 | 99 | 0 |
| SPGP0AJ00250 | XP_018221799.1 | 99 | 0 |
| SPGP0AJ01570 | XP_018221925.1 | 99 | 0 |
| SPGP0CM01070 | XP_018223218.1 | 99 | 0 |
| SPGP0CW01240 | XP_018219433.1 | 99 | 0 |
| SPGP0L00230 | XP_018219009.1 | 99 | 0 |
| SPGP0Y02030 | XP_018220213.1 | 99 | 0 |
| SPGP0CU01310 | XP_018220701.1 | 99 | 0 |
| SPGP0CP00330 | XP_018223803.1 | 99 | 0 |
| SPGP0CP01140 | XP_018223724.1 | 99 | 0 |

Create PDF in your applications with the Pdfcrowd HTML to PDF API PDFCROWD

| | | | |
|---|---|---|---|
| SPGP0AU00230 | XP_018223078.1 | 99 | 0 |
| SPGP0CP02350 | XP_018223606.1 | 99 | 0 |
| SPGP0Z00260 | XP_018222539.1 | 99 | 0 |
| SPGP0AF01940 | XP_018222452.1 | 99 | 0 |
| SPGP0JK00210 | XP_018223442.1 | 99 | 0 |
| SPGP0AT00370 | XP_018219058.1 | 99 | 0 |
| SPGP0CP01520 | XP_018223687.1 | 99 | 0 |
| SPGP0U01930 | XP_018222888.1 | 99 | 0 |
| SPGP0H00600 | XP_018221085.1 | 99 | 0 |
| SPGP0CU01030 | XP_018220727.1 | 99 | 0 |
| SPGP0K00460 | XP_018222788.1 | 99 | 0 |
| SPGP0CP03430 | XP_018223499.1 | 99 | 0 |
| SPGP0DB00240 | XP_018223898.1 | 100 | 0 |
| SPGP0CW01930 | XP_018219367.1 | 99 | 0 |
| SPGP0V01200 | XP_018221668.1 | 100 | 0 |
| SPGP0AJ00580 | XP_018221830.1 | 99 | 0 |
| SPGP0U02060 | XP_018222875.1 | 99 | 0 |
| SPGP0CM00760 | XP_018223249.1 | 99 | 0 |
| SPGP0Y01070 | XP_018220122.1 | 99 | 0 |
| SPGP0CU00700 | XP_018220758.1 | 99 | 0 |
| SPGP0AN00520 | XP_018223959.1 | 96 | 3 |
| SPGP0Y01410 | XP_018220156.1 | 99 | 0 |
| SPGP0CW00680 | XP_018219488.1 | 99 | 0 |
| SPGP0AF02000 | XP_018222458.1 | 99 | 0 |
| SPGP0U00130 | XP_018223056.1 | 99 | 0 |
| SPGP0CX01160 | XP_018222135.1 | 99 | 0 |
| SPGP0U01610 | XP_018222921.1 | 99 | 0 |
| SPGP0IZ00210 | XP_018220349.1 | 99 | 0 |
| SPGP0JK00120 | XP_018223434.1 | 99 | 0 |
| SPGP0AF00360 | XP_018222290.1 | 99 | 0 |
| SPGP0CP02950 | XP_018223546.1 | 99 | 0 |
| SPGP0CM01860 | XP_018223140.1 | 99 | 0 |
| SPGP0CP00580 | XP_018223779.1 | 99 | 0 |
| SPGP0AJ00370 | XP_018221810.1 | 100 | 0 |
| SPGP0AJ01950 | XP_018221961.1 | 99 | 0 |
| SPGP0CP02620 | XP_018223579.1 | 99 | 0 |
| SPGP0AF01430 | XP_018222401.1 | 99 | 0 |
| SPGP0GG00210 | XP_018220830.1 | 99 | 0 |
| SPGP0CP01320 | XP_018223707.1 | 99 | 0 |
| SPGP0AA01240 | XP_018220466.1 | 99 | 0 |

Create PDF in your applications with the Pdfcrowd HTML to PDF API PDFCROWD

| | | | |
|---|---|---|---|
| SPGP0AE00120 | XP_018222494.1 | 99 | 0 |
| SPGP0R00750 | XP_018220015.1 | 99 | 0 |
| SPGP0CP01060 | XP_018223733.1 | 99 | 0 |
| SPGP0Z00190 | XP_018222546.1 | 99 | 0 |
| SPGP0CX01440 | XP_018222161.1 | 98 | 1 |
| SPGP0CM01670 | XP_018223159.1 | 99 | 0 |
| SPGP0FP00200 | XP_018223335.1 | 99 | 0 |
| SPGP0CP03110 | XP_018223529.1 | 99 | 0 |
| SPGP0V00930 | XP_018221694.1 | 99 | 0 |
| SPGP0K00650 | XP_018222807.1 | 99 | 0 |
| SPGP0Y00930 | XP_018220109.1 | 99 | 0 |
| SPGP0CU02470 | XP_018220587.1 | 99 | 0 |
| SPGP0DQ00120 | XP_018220023.1 | 99 | 0 |
| SPGP0AJ00740 | XP_018221846.1 | 99 | 0 |
| SPGP0CW02650 | XP_018219298.1 | 99 | 0 |
| SPGP0CP03210 | XP_018223519.1 | 99 | 0 |
| SPGP0AN00130 | XP_018223925.1 | 99 | 0 |
| SPGP0CX01310 | XP_018222149.1 | 99 | 0 |
| SPGP0GA00110 | XP_018223392.1 | 99 | 0 |
| SPGP0AT01490 | XP_018219166.1 | 99 | 0 |
| SPGP0Y01680 | XP_018220182.1 | 99 | 0 |
| SPGP0CM00780 | XP_018223247.1 | 99 | 0 |
| SPGP0CU01780 | XP_018220656.1 | 99 | 0 |
| SPGP0CW02060 | XP_018219355.1 | 99 | 0 |
| SPGP0JB00120 | XP_018222558.1 | 99 | 0 |
| SPGP0GL00190 | XP_018220848.1 | 99 | 0 |
| SPGP0L01330 | XP_018218901.1 | 99 | 0 |
| SPGP0CP01350 | XP_018223704.1 | 99 | 0 |
| SPGP0AJ00240 | XP_018221798.1 | 99 | 0 |
| SPGP0CM00950 | XP_018223230.1 | 99 | 0 |
| SPGP0Y03010 | XP_018220310.1 | 99 | 0 |
| SPGP0CU01870 | XP_018220647.1 | 99 | 0 |
| SPGP0L02580 | XP_018218778.1 | 99 | 0 |
| SPGP0CM00330 | XP_018223291.1 | 99 | 0 |
| SPGP0CU01280 | XP_018220704.1 | 99 | 0 |
| SPGP0CM01930 | XP_018223133.1 | 99 | 0 |
| SPGP0R00720 | XP_018220012.1 | 99 | 0 |
| SPGP0CM01330 | XP_018223192.1 | 99 | 0 |
| SPGP0AA01310 | XP_018220459.1 | 99 | 0 |
| SPGP0CP02160 | XP_018223624.1 | 99 | 0 |

Create PDF in your applications with the Pdfcrowd HTML to PDF API PDFCROWD

| | | | |
|---|---|---|---|
| SPGP0L01570 | XP_018218877.1 | 99 | 0 |
| SPGP0Y02650 | XP_018220274.1 | 99 | 0 |
| SPGP0AQ00150 | XP_018223407.1 | 99 | 0 |
| SPGP0CW00750 | XP_018219481.1 | 99 | 0 |
| SPGP0L01590 | XP_018218875.1 | 99 | 0 |
| SPGP0L01840 | XP_018218851.1 | 99 | 0 |
| SPGP0CX02060 | XP_018222221.1 | 99 | 0 |
| SPGP0CP02570 | XP_018223584.1 | 99 | 0 |
| SPGP0V00250 | XP_018221758.1 | 99 | 1 |
| SPGP0FP00220 | XP_018223333.1 | 99 | 0 |
| SPGP0U01280 | XP_018222953.1 | 100 | 0 |
| SPGP0AT00170 | XP_018219039.1 | 99 | 0 |
| SPGP0AA01680 | XP_018220423.1 | 99 | 0 |
| SPGP0AA01690 | XP_018220422.1 | 99 | 0 |
| SPGP0AN00260 | XP_018223938.1 | 100 | 0 |
| SPGP0AA01430 | XP_018220447.1 | 99 | 0 |
| SPGP0JA00130 | XP_018222245.1 | 100 | 0 |
| SPGP0IN00210 | XP_018221655.1 | 99 | 0 |
| SPGP0CP03530 | XP_018223490.1 | 99 | 0 |
| SPGP0CU00140 | XP_018220814.1 | 99 | 0 |
| SPGP0AT00670 | XP_018219087.1 | 99 | 0 |
| SPGP0CX01960 | XP_018222211.1 | 98 | 0 |
| SPGP0L02160 | XP_018218821.1 | 99 | 0 |
| SPGP0Y01300 | XP_018220145.1 | 99 | 0 |
| SPGP0AA00930 | XP_018220497.1 | 99 | 0 |
| SPGP0CW02820 | XP_018219281.1 | 99 | 0 |
| SPGP0V01180 | XP_018221670.1 | 100 | 0 |
| SPGP0AF01100 | XP_018222365.1 | 99 | 0 |
| SPGP0AJ01740 | XP_018221942.1 | 99 | 0 |
| SPGP0F00180 | XP_018221187.1 | 99 | 0 |
| SPGP0CP00630 | XP_018223774.1 | 100 | 0 |
| SPGP0CP01790 | XP_018223661.1 | 99 | 0 |
| SPGP0CP03020 | XP_018223538.1 | 99 | 0 |
| SPGP0CM02170 | XP_018223112.1 | 100 | 0 |
| SPGP0AA01030 | XP_018220487.1 | 99 | 0 |
| SPGP0Y02140 | XP_018220223.1 | 99 | 0 |
| SPGP0CM00590 | XP_018223265.1 | 99 | 0 |
| SPGP0EO00310 | XP_018221168.1 | 98 | 0 |
| SPGP0CU00570 | XP_018220772.1 | 99 | 0 |
| SPGP0U01760 | XP_018222905.1 | 99 | 0 |

Create PDF in your applications with the Pdfcrowd HTML to PDF API PDFCROWD

| | | | |
|---|---|---|---|
| SPGP0CX02120 | XP_018222226.1 | 99 | 0 |
| SPGP0Y00590 | XP_018220075.1 | 99 | 0 |
| SPGP0Y00480 | XP_018220065.1 | 99 | 0 |
| SPGP0U00280 | XP_018223042.1 | 99 | 1 |
| SPGP0CP02370 | XP_018223604.1 | 100 | 0 |
| SPGP0V00880 | XP_018221699.1 | 99 | 0 |
| SPGP0CM00520 | XP_018223272.1 | 99 | 0 |
| SPGP0K00450 | XP_018222787.1 | 100 | 0 |
| SPGP0CW03100 | XP_018219253.1 | 99 | 0 |
| SPGP0AF01230 | XP_018222379.1 | 99 | 0 |
| SPGP0CP00940 | XP_018223745.1 | 100 | 0 |
| SPGP0CP02080 | XP_018223632.1 | 99 | 0 |
| SPGP0CW01910 | XP_018219369.1 | 99 | 0 |
| SPGP0DE00530 | XP_018223870.1 | 99 | 0 |
| SPGP0IH00110 | XP_018220382.1 | 99 | 0 |
| SPGP0CV00120 | XP_018222565.1 | 99 | 0 |
| SPGP0Y01090 | XP_018220124.1 | 99 | 0 |
| SPGP0CX01260 | XP_018222145.1 | 99 | 0 |
| SPGP0CP02600 | XP_018223581.1 | 99 | 0 |
| SPGP0AF02020 | XP_018222460.1 | 99 | 0 |
| SPGP0AT01070 | XP_018219125.1 | 99 | 0 |
| SPGP0IZ00230 | XP_018220351.1 | 99 | 0 |
| SPGP0AA01590 | XP_018220432.1 | 99 | 0 |
| SPGP0AF02050 | XP_018222463.1 | 99 | 0 |
| SPGP0CM01880 | XP_018223138.1 | 99 | 0 |
| SPGP0AA01870 | XP_018220405.1 | 99 | 0 |
| SPGP0CX01570 | XP_018222173.1 | 97 | 2 |
| SPGP0F00140 | XP_018221183.1 | 99 | 0 |
| SPGP0CU00250 | XP_018220804.1 | 99 | 0 |
| SPGP0IN00170 | XP_018221659.1 | 99 | 0 |
| SPGP0AN00430 | XP_018223952.1 | 99 | 0 |
| SPGP0Y00320 | XP_018220049.1 | 99 | 0 |
| SPGP0AJ00900 | XP_018221862.1 | 99 | 0 |
| SPGP0EB00120 | XP_018223319.1 | 99 | 0 |
| SPGP0AJ00500 | XP_018221823.1 | 99 | 0 |
| SPGP0AF01410 | XP_018222399.1 | 99 | 0 |
| SPGP0CW01650 | XP_018219395.1 | 99 | 0 |
| SPGP0CU00840 | XP_018220745.1 | 99 | 0 |
| SPGP0EO00130 | XP_018221150.1 | 99 | 0 |
| SPGP0AE00100 | XP_018222492.1 | 99 | 0 |

Create PDF in your applications with the Pdfcrowd HTML to PDF API PDFCROWD

| | | | |
|---|---|---|---|
| SPGP0CP01040 | XP_018223735.1 | 99 | 0 |
| SPGP0AA01940 | XP_018220399.1 | 99 | 0 |
| SPGP0AJ01350 | XP_018221905.1 | 99 | 0 |
| SPGP0V00950 | XP_018221692.1 | 99 | 0 |
| SPGP0AN00100 | XP_018223922.1 | 99 | 0 |
| SPGP0EO00240 | XP_018221161.1 | 99 | 0 |
| SPGP0AE00170 | XP_018222499.1 | 99 | 0 |
| SPGP0Y02770 | XP_018220286.1 | 99 | 0 |
| SPGP0AT00880 | XP_018219107.1 | 99 | 0 |
| SPGP0CP01880 | XP_018223652.1 | 99 | 0 |
| SPGP0U00620 | XP_018223013.1 | 100 | 0 |
| SPGP0GZ00110 | XP_018222823.1 | 99 | 0 |
| SPGP0CW02180 | XP_018219343.1 | 99 | 0 |
| SPGP0K00340 | XP_018222776.1 | 99 | 1 |
| SPGP0FE00170 | XP_018221969.1 | 99 | 0 |
| SPGP0IH00240 | XP_018220394.1 | 99 | 0 |
| SPGP0CW02670 | XP_018219296.1 | 99 | 0 |
| SPGP0CP00790 | XP_018223759.1 | 99 | 0 |
| SPGP0CX00820 | XP_018222101.1 | 99 | 0 |
| SPGP0AJ00480 | XP_018221821.1 | 99 | 0 |
| SPGP0CV00330 | XP_018222584.1 | 99 | 0 |
| SPGP0DR00100 | XP_018219195.1 | 99 | 0 |
| SPGP0U00780 | XP_018223000.1 | 99 | 0 |
| SPGP0CW02240 | XP_018219337.1 | 100 | 0 |
| SPGP0J01670 | XP_018219781.1 | 99 | 0 |
| SPGP0CU01500 | XP_018220684.1 | 99 | 0 |
| SPGP0JB00140 | XP_018222560.1 | 99 | 0 |
| SPGP0AT00520 | XP_018219073.1 | 99 | 0 |
| SPGP0CM00920 | XP_018223233.1 | 99 | 0 |
| SPGP0U01180 | XP_018222962.1 | 99 | 0 |
| SPGP0CW01430 | XP_018219415.1 | 99 | 0 |
| SPGP0R00220 | XP_018219964.1 | 99 | 0 |
| SPGP0CU00610 | XP_018220768.1 | 99 | 0 |
| SPGP0Y00420 | XP_018220059.1 | 99 | 0 |
| SPGP0CM01910 | XP_018223135.1 | 99 | 0 |
| SPGP0FP00250 | XP_018223330.1 | 99 | 0 |
| SPGP0U00580 | XP_018223016.1 | 99 | 0 |
| SPGP0R00330 | XP_018219974.1 | 99 | 0 |
| SPGP0CU01300 | XP_018220702.1 | 99 | 0 |
| SPGP0U01980 | XP_018222883.1 | 99 | 0 |

Create PDF in your applications with the Pdfcrowd HTML to PDF API PDFCROWD

| | | | |
|---|---|---|---|
| SPGP0U01450 | XP_018222937.1 | 99 | 0 |
| SPGP0AA00500 | XP_018220540.1 | 99 | 0 |
| SPGP0CP02140 | XP_018223626.1 | 99 | 0 |
| SPGP0CP01820 | XP_018223658.1 | 99 | 0 |
| SPGP0Y02670 | XP_018220276.1 | 99 | 0 |
| SPGP0IN00110 | XP_018221665.1 | 99 | 0 |
| SPGP0AQ00170 | XP_018223405.1 | 100 | 0 |
| SPGP0L00440 | XP_018218988.1 | 99 | 0 |
| SPGP0J01680 | XP_018219780.1 | 99 | 0 |
| SPGP0V00270 | XP_018221756.1 | 99 | 0 |
| SPGP0CP02550 | XP_018223586.1 | 99 | 0 |
| SPGP0CX01860 | XP_018222201.1 | 99 | 0 |
| SPGP0Y01550 | XP_018220170.1 | 99 | 0 |
| SPGP0CM01560 | XP_018223170.1 | 99 | 0 |
| SPGP0CU00120 | XP_018220816.1 | 99 | 0 |
| SPGP0IM00130 | XP_018219549.1 | 99 | 0 |
| SPGP0CX00860 | XP_018222105.1 | 99 | 0 |
| SPGP0AN00820 | XP_018223988.1 | 99 | 0 |
| SPGP0AJ01760 | XP_018221944.1 | 99 | 0 |
| SPGP0R00280 | XP_018219970.1 | 99 | 0 |
| SPGP0CP02910 | XP_018223550.1 | 99 | 0 |
| SPGP0CP02850 | XP_018223556.1 | 99 | 0 |
| SPGP0AA01050 | XP_018220485.1 | 99 | 0 |
| SPGP0L00180 | XP_018219013.1 | 99 | 0 |
| SPGP0CM00480 | XP_018223276.1 | 99 | 0 |
| SPGP0CX02100 | XP_018222224.1 | 99 | 0 |
| SPGP0CW02370 | XP_018219324.1 | 99 | 0 |
| SPGP0BE00220 | XP_018223356.1 | 99 | 0 |
| SPGP0FE00120 | XP_018221964.1 | 99 | 0 |
| SPGP0AF01980 | XP_018222456.1 | 99 | 0 |
| SPGP0FO00120 | XP_018223458.1 | 99 | 0 |
| SPGP0L00290 | XP_018219003.1 | 99 | 0 |
| SPGP0Y01520 | XP_018220167.1 | 99 | 0 |
| SPGP0CU02330 | XP_018220601.1 | 100 | 0 |
| SPGP0CW01970 | XP_018219363.1 | 99 | 0 |
| SPGP0DB00290 | XP_018223903.1 | 99 | 0 |
| SPGP0CW03190 | XP_018219245.1 | 99 | 0 |
| SPGP0CP00490 | XP_018223788.1 | 99 | 0 |
| SPGP0CX01120 | XP_018222131.1 | 99 | 0 |

Create PDF in your applications with the Pdfcrowd HTML to PDF API PDFCROWD

| | | | |
|---|---|---|---|
| SPGP0U01840 | XP_018222897.1 | 99 | 0 |
| SPGP0CP01870 | XP_018223653.1 | 99 | 0 |
| SPGP0CP02120 | XP_018223628.1 | 100 | 0 |
| SPGP0IU00170 | XP_018220335.1 | 99 | 0 |
| SPGP0U00860 | XP_018222993.1 | 99 | 0 |
| SPGP0CP01150 | XP_018223723.1 | 99 | 0 |
| SPGP0U00460 | XP_018223028.1 | 99 | 0 |
| SPGP0AF00400 | XP_018222294.1 | 99 | 0 |
| SPGP0CW03220 | XP_018219242.1 | 99 | 0 |
| SPGP0AN00760 | XP_018223983.1 | 99 | 0 |
| SPGP0AT01050 | XP_018219123.1 | 99 | 0 |
| SPGP0CP00810 | XP_018223757.1 | 99 | 0 |
| SPGP0IN00140 | XP_018221662.1 | 99 | 0 |
| SPGP0CP00210 | XP_018223815.1 | 99 | 0 |
| SPGP0AF00910 | XP_018222345.1 | 99 | 0 |
| SPGP0CP03540 | XP_018223489.1 | 99 | 0 |
| SPGP0L01470 | XP_018218887.1 | 99 | 0 |
| SPGP0AJ01900 | XP_018221956.1 | 99 | 0 |
| SPGP0CU02020 | XP_018220632.1 | 99 | 0 |
| SPGP0V00340 | XP_018221749.1 | 99 | 0 |
| SPGP0CW01590 | XP_018219400.1 | 99 | 0 |
| SPGP0AA00990 | XP_018220491.1 | 99 | 0 |
| SPGP0AA00770 | XP_018220514.1 | 99 | 0 |
| SPGP0EM00100 | XP_018223059.1 | 99 | 0 |
| SPGP0AF00700 | XP_018222324.1 | 99 | 0 |
| SPGP0CW00470 | XP_018219508.1 | 99 | 0 |
| SPGP0AE00340 | XP_018222514.1 | 97 | 2 |
| SPGP0CU00820 | XP_018220747.1 | 99 | 0 |
| SPGP0IC00110 | XP_018222532.1 | 99 | 0 |
| SPGP0AF01210 | XP_018222377.1 | 99 | 0 |
| SPGP0CM01380 | XP_018223188.1 | 99 | 0 |
| SPGP0Y00290 | XP_018220046.1 | 99 | 0 |
| SPGP0CP01360 | XP_018223703.1 | 99 | 0 |
| SPGP0FP00190 | XP_018223336.1 | 99 | 0 |
| SPGP0AN00120 | XP_018223924.1 | 99 | 0 |
| SPGP0JP00160 | XP_018219567.1 | 99 | 0 |
| SPGP0CW00340 | XP_018219521.1 | 99 | 0 |
| SPGP0CP03150 | XP_018223525.1 | 99 | 0 |
| SPGP0CX01410 | XP_018222159.1 | 99 | 0 |
| SPGP0U00570 | XP_018223017.1 | 100 | 0 |

Create PDF in your applications with the Pdfcrowd HTML to PDF API PDFCROWD

| | | | |
|---|---|---|---|
| SPGP0AJ00100 | XP_018221785.1 | 99 | 0 |
| SPGP0V00450 | XP_018221740.1 | 99 | 0 |
| SPGP0JO00120 | XP_018222262.1 | 99 | 0 |
| SPGP0DF00180 | XP_018219190.1 | 99 | 0 |
| SPGP0AT01100 | XP_018219128.1 | 99 | 0 |
| SPGP0CU01360 | XP_018220697.1 | 99 | 0 |
| SPGP0DE00520 | XP_018223869.1 | 99 | 0 |
| SPGP0AF01500 | XP_018222408.1 | 99 | 0 |
| SPGP0CP00770 | XP_018223761.1 | 99 | 0 |
| SPGP0CX00800 | XP_018222099.1 | 99 | 0 |
| SPGP0CM00220 | XP_018223302.1 | 99 | 0 |
| SPGP0CW00960 | XP_018219460.1 | 99 | 0 |
| SPGP0CX01350 | XP_018222153.1 | 99 | 0 |
| SPGP0GA00150 | XP_018223388.1 | 100 | 0 |
| SPGP0CU00330 | XP_018220796.1 | 99 | 0 |
| SPGP0CU01520 | XP_018220682.1 | 99 | 0 |
| SPGP0JB00160 | XP_018222561.1 | 99 | 0 |
| SPGP0CV00350 | XP_018222586.1 | 100 | 0 |
| SPGP0CW01780 | XP_018219382.1 | 99 | 0 |
| SPGP0Y00610 | XP_018220077.1 | 99 | 0 |
| SPGP0AJ00730 | XP_018221845.1 | 99 | 0 |
| SPGP0CM00900 | XP_018223235.1 | 99 | 0 |
| SPGP0Y00440 | XP_018220061.1 | 99 | 0 |
| SPGP0CW01390 | XP_018219419.1 | 99 | 0 |
| SPGP0U01470 | XP_018222935.1 | 99 | 0 |
| SPGP0EO00380 | XP_018221175.1 | 99 | 0 |
| SPGP0U00430 | XP_018223030.1 | 99 | 0 |
| SPGP0U00700 | XP_018223005.1 | 99 | 0 |
| SPGP0GJ00180 | XP_018223374.1 | 99 | 0 |
| SPGP0U00660 | XP_018223009.1 | 99 | 0 |
| SPGP0AF01640 | XP_018222422.1 | 99 | 0 |
| SPGP0F00220 | XP_018221191.1 | 99 | 0 |
| SPGP0AJ01520 | XP_018221920.1 | 99 | 0 |
| SPGP0AT01340 | XP_018219152.1 | 99 | 0 |
| SPGP0IM00110 | XP_018219551.1 | 99 | 0 |
| SPGP0CV00150 | XP_018222568.1 | 100 | 0 |
| SPGP0CU02320 | XP_018220602.1 | 99 | 0 |
| SPGP0CW02860 | XP_018219277.1 | 99 | 0 |
| SPGP0CW02690 | XP_018219294.1 | 99 | 0 |
| SPGP0U00390 | XP_018223034.1 | 99 | 0 |

Create PDF in your applications with the Pdfcrowd HTML to PDF API PDFCROWD

| | | | |
|---|---|---|---|
| SPGP0CM00840 | XP_018223241.1 | 99 | 0 |
| SPGP0CX01040 | XP_018222123.1 | 99 | 0 |
| SPGP0U00990 | XP_018222980.1 | 99 | 0 |
| SPGP0V00210 | XP_018221762.1 | 99 | 0 |
| SPGP0V00660 | XP_018221720.1 | 99 | 0 |
| SPGP0AF00290 | XP_018222283.1 | 99 | 0 |
| SPGP0AJ01410 | XP_018221911.1 | 99 | 0 |
| SPGP0DE00660 | XP_018223883.1 | 99 | 0 |
| SPGP0R00790 | XP_018220018.1 | 99 | 0 |
| SPGP0GL00180 | XP_018220847.1 | 99 | 0 |
| SPGP0Y02050 | XP_018220215.1 | 99 | 0 |
| SPGP0HI00200 | XP_018219946.1 | 99 | 0 |
| SPGP0K00350 | XP_018222777.1 | 99 | 0 |
| SPGP0CM00360 | XP_018223288.1 | 99 | 0 |
| SPGP0AT01590 | XP_018219176.1 | 99 | 0 |
| SPGP0JK00280 | XP_018223448.1 | 99 | 0 |
| SPGP0EA00100 | XP_018222477.1 | 99 | 0 |
| SPGP0AF01960 | XP_018222454.1 | 99 | 0 |
| SPGP0V00840 | XP_018221703.1 | 99 | 0 |
| SPGP0Y03140 | XP_018220322.1 | 99 | 0 |
| SPGP0JB00130 | XP_018222559.1 | 99 | 0 |
| SPGP0JP00130 | XP_018219564.1 | 99 | 0 |
| SPGP0Y01020 | XP_018220117.1 | 99 | 0 |
| SPGP0CU02350 | XP_018220599.1 | 99 | 0 |
| SPGP0R00810 | XP_018220020.1 | 99 | 0 |
| SPGP0L00610 | XP_018218972.1 | 99 | 0 |
| SPGP0Y01690 | XP_018220183.1 | 99 | 0 |
| SPGP0CV00160 | XP_018222569.1 | 98 | 2 |
| SPGP0CX01220 | XP_018222141.1 | 99 | 0 |
| SPGP0U01860 | XP_018222895.1 | 99 | 0 |
| SPGP0CW01950 | XP_018219365.1 | 99 | 0 |
| SPGP0CU01950 | XP_018220639.1 | 99 | 0 |
| SPGP0CP03420 | XP_018223500.1 | 99 | 0 |
| SPGP0CP03520 | XP_018223491.1 | 99 | 0 |
| SPGP0EU00110 | XP_018223369.1 | 99 | 0 |
| SPGP0AN00830 | XP_018223989.1 | 99 | 0 |
| SPGP0L00810 | XP_018218952.1 | 100 | 0 |
| SPGP0AF00930 | XP_018222347.1 | 99 | 0 |
| SPGP0R00100 | XP_018219952.1 | 99 | 0 |
| SPGP0HS00100 | XP_018221774.1 | 99 | 0 |

Create PDF in your applications with the Pdfcrowd HTML to PDF API PDFCROWD

| | | | |
|---|---|---|---|
| SPGP0AJ00850 | XP_018221857.1 | 99 | 0 |
| SPGP0Y02730 | XP_018220282.1 | 99 | 0 |
| SPGP0JK00270 | XP_018223447.1 | 99 | 0 |
| SPGP0AJ00320 | XP_018221806.1 | 99 | 1 |
| SPGP0AT00270 | XP_018219049.1 | 99 | 0 |
| SPGP0CP01340 | XP_018223705.1 | 99 | 0 |
| SPGP0AE00360 | XP_018222516.1 | 99 | 0 |
| SPGP0IH00200 | XP_018220390.1 | 99 | 0 |
| SPGP0CW01620 | XP_018219397.1 | 99 | 0 |
| SPGP0AF01260 | XP_018222382.1 | 99 | 0 |
| SPGP0AU00350 | XP_018223090.1 | 99 | 0 |
| SPGP0EB00110 | XP_018223320.1 | 99 | 0 |
| SPGP0EO00170 | XP_018221154.1 | 99 | 0 |
| SPGP0AA01260 | XP_018220464.1 | 99 | 0 |
| SPGP0K00420 | XP_018222784.1 | 100 | 0 |
| SPGP0K00160 | XP_018222759.1 | 99 | 0 |
| SPGP0AE00140 | XP_018222496.1 | 99 | 0 |
| SPGP0AF01320 | XP_018222389.1 | 99 | 0 |
| SPGP0AF00130 | XP_018222268.1 | 99 | 0 |
| SPGP0Y01280 | XP_018220143.1 | 99 | 0 |
| SPGP0CX01200 | XP_018222139.1 | 98 | 0 |
| SPGP0R00110 | XP_018219953.1 | 99 | 0 |
| SPGP0CM01520 | XP_018223174.1 | 99 | 0 |
| SPGP0EO00360 | XP_018221173.1 | 100 | 0 |
| SPGP0CU01970 | XP_018220637.1 | 99 | 0 |
| SPGP0L01050 | XP_018218927.1 | 99 | 0 |
| SPGP0U02070 | XP_018222874.1 | 99 | 0 |
| SPGP0K00290 | XP_018222771.1 | 99 | 0 |
| SPGP0CP03550 | XP_018223488.1 | 99 | 0 |
| SPGP0L01420 | XP_018218892.1 | 99 | 0 |
| SPGP0CP00430 | XP_018223793.1 | 99 | 0 |
| SPGP0IM00150 | XP_018219547.1 | 99 | 0 |
| SPGP0Y02420 | XP_018220251.1 | 99 | 0 |
| SPGP0JO00140 | XP_018222264.1 | 99 | 0 |
| SPGP0AJ01060 | XP_018221877.1 | 99 | 0 |
| SPGP0AT01140 | XP_018219132.1 | 99 | 0 |
| SPGP0CP00750 | XP_018223763.1 | 99 | 0 |
| SPGP0Y00980 | XP_018220113.1 | 99 | 0 |
| SPGP0K00530 | XP_018222795.1 | 99 | 0 |
| SPGP0CU01090 | XP_018220721.1 | 100 | 0 |

Create PDF in your applications with the Pdfcrowd HTML to PDF API PDFCROWD

| | | | |
|---|---|---|---|
| SPGP0CW00940 | XP_018219462.1 | 99 | 0 |
| SPGP0AJ00440 | XP_018221817.1 | 99 | 0 |
| SPGP0AT01160 | XP_018219134.1 | 99 | 0 |
| SPGP0CX01190 | XP_018222138.1 | 99 | 0 |
| SPGP0CW01010 | XP_018219455.1 | 99 | 1 |
| SPGP0U00740 | XP_018223003.1 | 99 | 0 |
| SPGP0CU01540 | XP_018220680.1 | 99 | 0 |
| SPGP0AT00560 | XP_018219077.1 | 99 | 0 |
| SPGP0CU01890 | XP_018220645.1 | 99 | 0 |
| SPGP0F00150 | XP_018221184.1 | 99 | 0 |
| SPGP0AE00370 | XP_018222517.1 | 98 | 1 |
| SPGP0AN00920 | XP_018223998.1 | 99 | 0 |
| SPGP0CP02100 | XP_018223630.1 | 99 | 0 |
| SPGP0AT00780 | XP_018219097.1 | 99 | 0 |
| SPGP0AJ00990 | XP_018221871.1 | 99 | 0 |
| SPGP0J02020 | XP_018219747.1 | 99 | 0 |
| SPGP0U01490 | XP_018222933.1 | 99 | 0 |
| SPGP0CP01960 | XP_018223644.1 | 99 | 0 |
| SPGP0DB00190 | XP_018223893.1 | 99 | 0 |
| SPGP0L01870 | XP_018218848.1 | 99 | 1 |
| SPGP0AF01040 | XP_018222359.1 | 99 | 0 |
| SPGP0AA01140 | XP_018220476.1 | 99 | 0 |
| SPGP0AA01630 | XP_018220428.1 | 99 | 0 |
| SPGP0Y01500 | XP_018220165.1 | 99 | 0 |
| SPGP0V00190 | XP_018221764.1 | 99 | 0 |
| SPGP0CM01590 | XP_018223167.1 | 99 | 0 |
| SPGP0AJ01810 | XP_018221949.1 | 100 | 0 |
| SPGP0L01750 | XP_018218860.1 | 99 | 0 |
| SPGP0CU00950 | XP_018220735.1 | 99 | 0 |
| SPGP0CW01380 | XP_018219420.1 | 99 | 0 |
| SPGP0CX02000 | XP_018222215.1 | 99 | 0 |
| SPGP0CW01670 | XP_018219393.1 | 99 | 0 |
| SPGP0CW02840 | XP_018219279.1 | 99 | 0 |
| SPGP0Y02940 | XP_018220303.1 | 99 | 0 |
| SPGP0CX01060 | XP_018222125.1 | 99 | 0 |
| SPGP0IN00260 | XP_018221650.1 | 99 | 0 |
| SPGP0AA01850 | XP_018220407.1 | 99 | 0 |
| SPGP0CM01770 | XP_018223149.1 | 97 | 2 |
| SPGP0Y01200 | XP_018220135.1 | 99 | 0 |
| SPGP0J00590 | XP_018219891.1 | 99 | 0 |

Create PDF in your applications with the Pdfcrowd HTML to PDF API PDFCROWD

| | | | |
|---|---|---|---|
| SPGP0Y02270 | XP_018220236.1 | 99 | 0 |
| SPGP0Y02590 | XP_018220268.1 | 100 | 0 |
| SPGP0CU02060 | XP_018220628.1 | 99 | 0 |
| SPGP0AJ00650 | XP_018221837.1 | 99 | 0 |
| SPGP0K00330 | XP_018222775.1 | 99 | 0 |
| SPGP0CM01470 | XP_018223179.1 | 99 | 0 |
| SPGP0U01080 | XP_018222971.1 | 99 | 0 |
| SPGP0V00790 | XP_018221708.1 | 99 | 0 |
| SPGP0CV00360 | XP_018222587.1 | 99 | 0 |
| SPGP0CX01250 | XP_018222144.1 | 99 | 0 |
| SPGP0JK00330 | XP_018223454.1 | 99 | 0 |
| SPGP0X01820 | XP_018221625.1 | 94 | 2 |
| SPGP0CP03100 | XP_018223530.1 | 99 | 0 |
| SPGP0CV00140 | XP_018222567.1 | 99 | 0 |
| SPGP0IR00130 | XP_018223467.1 | 99 | 0 |
| SPGP0CP01110 | XP_018223727.1 | 99 | 0 |
| SPGP0CP03440 | XP_018223498.1 | 99 | 0 |
| SPGP0CP00730 | XP_018223765.1 | 99 | 0 |
| SPGP0GZ00120 | XP_018222824.1 | 99 | 0 |
| SPGP0AF00450 | XP_018222299.1 | 99 | 0 |
| SPGP0AA00150 | XP_018220575.1 | 99 | 0 |
| SPGP0CW02090 | XP_018219352.1 | 99 | 0 |
| SPGP0L00340 | XP_018218998.1 | 99 | 0 |
| SPGP0AA01970 | XP_018220396.1 | 99 | 0 |
| SPGP0IN00270 | XP_018221649.1 | 100 | 0 |
| SPGP0AA00730 | XP_018220518.1 | 99 | 0 |
| SPGP0AA01890 | XP_018220403.1 | 99 | 0 |
| SPGP0Y02710 | XP_018220280.1 | 99 | 0 |
| SPGP0Y01310 | XP_018220146.1 | 99 | 0 |
| SPGP0AF01250 | XP_018222381.1 | 99 | 0 |
| SPGP0AJ00940 | XP_018221866.1 | 99 | 0 |
| SPGP0EB00130 | XP_018223318.1 | 99 | 0 |
| SPGP0AA01280 | XP_018220462.1 | 99 | 0 |
| SPGP0CP02730 | XP_018223568.1 | 99 | 0 |
| SPGP0CP02900 | XP_018223551.1 | 99 | 0 |
| SPGP0EO00190 | XP_018221156.1 | 99 | 0 |
| SPGP0L02360 | XP_018218800.1 | 99 | 0 |
| SPGP0AN00170 | XP_018223929.1 | 99 | 0 |
| SPGP0IZ00140 | XP_018220342.1 | 99 | 0 |
| SPGP0BI00100 | XP_018223663.1 | 99 | 0 |

Create PDF in your applications with the Pdfcrowd HTML to PDF API PDFCROWD

| | | | |
|---|---|---|---|
| SPGP0V00910 | XP_018221696.1 | 99 | 0 |
| SPGP0GL00140 | XP_018220844.1 | 99 | 0 |
| SPGP0AU00150 | XP_018223072.1 | 99 | 0 |
| SPGP0GG00220 | XP_018220829.1 | 99 | 0 |
| SPGP0AJ00140 | XP_018221789.1 | 99 | 0 |
| SPGP0CM00560 | XP_018223268.1 | 99 | 0 |
| SPGP0AU00340 | XP_018223089.1 | 100 | 0 |
| SPGP0CM00700 | XP_018223255.1 | 99 | 0 |
| SPGP0Y02150 | XP_018220224.1 | 99 | 0 |
| SPGP0CW02220 | XP_018219339.1 | 99 | 0 |
| SPGP0CU01700 | XP_018220664.1 | 99 | 0 |
| SPGP0Y01120 | XP_018220127.1 | 99 | 0 |
| SPGP0AT00850 | XP_018219104.1 | 99 | 0 |
| SPGP0CP00530 | XP_018223784.1 | 99 | 0 |
| SPGP0CU01560 | XP_018220678.1 | 99 | 0 |
| SPGP0AT00540 | XP_018219075.1 | 99 | 0 |
| SPGP0X00260 | XP_018221475.1 | 99 | 0 |
| SPGP0AT00230 | XP_018219045.1 | 98 | 1 |
| SPGP0AT00470 | XP_018219068.1 | 99 | 0 |
| SPGP0AF01070 | XP_018222362.1 | 99 | 0 |
| SPGP0GA00190 | XP_018223384.1 | 99 | 0 |
| SPGP0CP00130 | XP_018223823.1 | 99 | 0 |
| SPGP0CP00920 | XP_018223747.1 | 99 | 0 |
| SPGP0Y02320 | XP_018220241.1 | 99 | 0 |
| SPGP0AT01220 | XP_018219140.1 | 99 | 0 |
| SPGP0V00580 | XP_018221728.1 | 99 | 0 |
| SPGP0R00460 | XP_018219987.1 | 99 | 0 |
| SPGP0L00640 | XP_018218969.1 | 99 | 0 |
| SPGP0AQ00130 | XP_018223409.1 | 99 | 0 |
| SPGP0CP01940 | XP_018223646.1 | 99 | 0 |
| SPGP0AN00610 | XP_018223968.1 | 99 | 0 |
| SPGP0CW00540 | XP_018219501.1 | 99 | 0 |
| SPGP0U00520 | XP_018223022.1 | 99 | 0 |
| SPGP0CW01640 | XP_018219396.1 | 99 | 0 |
| SPGP0CU02110 | XP_018220623.1 | 99 | 0 |
| SPGP0AA01610 | XP_018220430.1 | 99 | 0 |
| SPGP0Y00470 | XP_018220064.1 | 99 | 0 |
| SPGP0CX00730 | XP_018222092.1 | 99 | 0 |
| SPGP0L01510 | XP_018218883.1 | 99 | 0 |
| SPGP0CW03360 | XP_018219228.1 | 99 | 0 |

Create PDF in your applications with the Pdfcrowd HTML to PDF API PDFCROWD

| | | | |
|---|---|---|---|
| SPGP0K00310 | XP_018222773.1 | 99 | 0 |
| SPGP0F00250 | XP_018221194.1 | 98 | 0 |
| SPGP0V00690 | XP_018221717.1 | 99 | 0 |
| SPGP0CW03290 | XP_018219235.1 | 98 | 2 |
| SPGP0CW00620 | XP_018219494.1 | 99 | 0 |
| SPGP0Y03020 | XP_018220311.1 | 99 | 0 |
| SPGP0U01550 | XP_018222927.1 | 99 | 0 |
| SPGP0CM01280 | XP_018223197.1 | 99 | 0 |
| SPGP0CX01080 | XP_018222127.1 | 99 | 0 |
| SPGP0CM00880 | XP_018223237.1 | 99 | 0 |
| SPGP0K00220 | XP_018222764.1 | 99 | 0 |
| SPGP0Z00140 | XP_018222551.1 | 99 | 0 |
| SPGP0AE00270 | XP_018222508.1 | 99 | 0 |
| SPGP0FP00100 | XP_018223345.1 | 99 | 0 |
| SPGP0L02520 | XP_018218784.1 | 99 | 0 |
| SPGP0CP03490 | XP_018223493.1 | 99 | 0 |
| SPGP0IZ00200 | XP_018220348.1 | 99 | 0 |
| SPGP0CM01750 | XP_018223151.1 | 99 | 0 |
| SPGP0AT01080 | XP_018219126.1 | 99 | 0 |
| SPGP0GT00110 | XP_018219025.1 | 99 | 0 |
| SPGP0CP00560 | XP_018223781.1 | 99 | 0 |
| SPGP0CM02220 | XP_018223107.1 | 94 | 5 |
| SPGP0CV00380 | XP_018222589.1 | 99 | 0 |
| SPGP0EA00140 | XP_018222481.1 | 97 | 0 |
| SPGP0AT01330 | XP_018219151.1 | 99 | 1 |
| SPGP0Y01540 | XP_018220169.1 | 99 | 0 |
| SPGP0AT00760 | XP_018219095.1 | 99 | 0 |
| SPGP0AN00190 | XP_018223931.1 | 99 | 0 |
| SPGP0G00110 | XP_018222250.1 | 99 | 0 |
| SPGP0L00150 | XP_018219016.1 | 99 | 0 |
| SPGP0CP03460 | XP_018223496.1 | 99 | 0 |
| SPGP0CW03170 | XP_018219247.1 | 99 | 0 |
| SPGP0AE00190 | XP_018222501.1 | 99 | 0 |
| SPGP0CP01810 | XP_018223659.1 | 99 | 0 |
| SPGP0AF00970 | XP_018222351.1 | 99 | 0 |
| SPGP0GG00330 | XP_018220820.1 | 99 | 0 |
| SPGP0U01350 | XP_018222947.1 | 99 | 0 |
| SPGP0U01820 | XP_018222899.1 | 99 | 0 |
| SPGP0R00140 | XP_018219956.1 | 99 | 0 |
| SPGP0CP02430 | XP_018223598.1 | 99 | 0 |

Create PDF in your applications with the Pdfcrowd HTML to PDF API PDFCROWD

| | | | |
|---|---|---|---|
| SPGP0AJ00960 | XP_018221868.1 | 99 | 0 |
| SPGP0CU00960 | XP_018220734.1 | 97 | 0 |
| SPGP0X01540 | XP_018221597.1 | 99 | 0 |
| SPGP0R00380 | XP_018219979.1 | 99 | 0 |
| SPGP0K00120 | XP_018222754.1 | 99 | 0 |
| SPGP0Y01260 | XP_018220141.1 | 99 | 0 |
| SPGP0CW01460 | XP_018219412.1 | 99 | 1 |
| SPGP0CP00470 | XP_018223790.1 | 99 | 0 |
| SPGP0AU00320 | XP_018223087.1 | 99 | 0 |
| SPGP0DB00500 | XP_018223920.1 | 99 | 0 |
| SPGP0Y02460 | XP_018220255.1 | 99 | 0 |
| SPGP0AN01090 | XP_018224015.1 | 99 | 0 |
| SPGP0Z00130 | XP_018222552.1 | 99 | 0 |
| SPGP0AA00980 | XP_018220492.1 | 99 | 1 |
| SPGP0HC00120 | XP_018221783.1 | 99 | 0 |
| SPGP0CU01760 | XP_018220658.1 | 99 | 0 |
| SPGP0U01690 | XP_018222913.1 | 99 | 0 |
| SPGP0Y00990 | XP_018220114.1 | 99 | 0 |
| SPGP0CW03020 | XP_018219261.1 | 99 | 0 |
| SPGP0AA01160 | XP_018220474.1 | 100 | 0 |
| SPGP0CU02250 | XP_018220609.1 | 99 | 0 |
| SPGP0Y02280 | XP_018220237.1 | 99 | 0 |
| SPGP0L01240 | XP_018218910.1 | 99 | 0 |
| SPGP0F00110 | XP_018221180.1 | 99 | 0 |
| SPGP0DF00210 | XP_018219187.1 | 99 | 0 |
| SPGP0AA01250 | XP_018220465.1 | 99 | 0 |
| SPGP0R00400 | XP_018219981.1 | 99 | 0 |
| SPGP0GR00120 | XP_018221977.1 | 99 | 0 |
| SPGP0Y00780 | XP_018220094.1 | 99 | 0 |
| SPGP0CW01170 | XP_018219440.1 | 99 | 0 |
| SPGP0GD00110 | XP_018223412.1 | 99 | 0 |
| SPGP0CP01240 | XP_018223714.1 | 99 | 0 |
| SPGP0AA01670 | XP_018220424.1 | 99 | 0 |
| SPGP0DE00240 | XP_018223841.1 | 99 | 0 |
| SPGP0DB00480 | XP_018223918.1 | 100 | 0 |
| SPGP0CP02980 | XP_018223542.1 | 99 | 0 |
| SPGP0GG00290 | XP_018220824.1 | 99 | 0 |
| SPGP0CW03340 | XP_018219230.1 | 99 | 0 |
| SPGP0GJ00120 | XP_018223380.1 | 99 | 0 |
| SPGP0CW01610 | XP_018219398.1 | 99 | 0 |

Create PDF in your applications with the Pdfcrowd HTML to PDF API PDFCROWD

| | | | |
|---|---|---|---|
| SPGP0F00240 | XP_018221193.1 | 99 | 0 |
| SPGP0AA00850 | XP_018220505.1 | 100 | 0 |
| SPGP0AN01110 | XP_018224017.1 | 99 | 0 |
| SPGP0CW00640 | XP_018219492.1 | 100 | 0 |
| SPGP0Y00460 | XP_018220063.1 | 99 | 0 |
| SPGP0AJ01850 | XP_018221952.1 | 100 | 0 |
| SPGP0CM01440 | XP_018223182.1 | 99 | 0 |
| SPGP0CU00910 | XP_018220738.1 | 99 | 0 |
| SPGP0CM02070 | XP_018223122.1 | 99 | 1 |
| SPGP0Y00100 | XP_018220027.1 | 99 | 0 |
| SPGP0K00190 | XP_018222762.1 | 99 | 0 |
| SPGP0AN01210 | XP_018224027.1 | 99 | 0 |
| SPGP0CW03300 | XP_018219234.1 | 100 | 0 |
| SPGP0CM01000 | XP_018223225.1 | 99 | 0 |
| SPGP0HS00120 | XP_018221776.1 | 99 | 0 |
| SPGP0V00630 | XP_018221723.1 | 99 | 0 |
| SPGP0CP03260 | XP_018223514.1 | 99 | 0 |
| SPGP0CW00220 | XP_018219531.1 | 99 | 0 |
| SPGP0GT00130 | XP_018219023.1 | 99 | 0 |
| SPGP0AT01530 | XP_018219170.1 | 99 | 0 |
| SPGP0GH00150 | XP_018222817.1 | 99 | 0 |
| SPGP0U01750 | XP_018222907.1 | 99 | 0 |
| SPGP0U01110 | XP_018222968.1 | 99 | 0 |
| SPGP0AT00660 | XP_018219086.1 | 99 | 0 |
| SPGP0CW01740 | XP_018219386.1 | 99 | 0 |
| SPGP0AJ00360 | XP_018221809.1 | 99 | 0 |
| SPGP0U01590 | XP_018222923.1 | 99 | 0 |
| SPGP0AA00240 | XP_018220566.1 | 99 | 0 |
| SPGP0L02620 | XP_018218775.1 | 99 | 0 |
| SPGP0AF00990 | XP_018222353.1 | 99 | 0 |
| SPGP0CP00700 | XP_018223768.1 | 99 | 0 |
| SPGP0CM02200 | XP_018223109.1 | 99 | 0 |
| SPGP0BE00120 | XP_018223366.1 | 99 | 0 |
| SPGP0U01370 | XP_018222945.1 | 99 | 0 |
| SPGP0CP02280 | XP_018223612.1 | 99 | 0 |
| SPGP0AA00110 | XP_018220579.1 | 99 | 0 |
| SPGP0AA00440 | XP_018220546.1 | 99 | 0 |
| SPGP0CX01670 | XP_018222183.1 | 97 | 1 |
| SPGP0CP02450 | XP_018223596.1 | 99 | 0 |
| SPGP0CP02680 | XP_018223573.1 | 99 | 0 |

Create PDF in your applications with the Pdfcrowd HTML to PDF API PDFCROWD

| | | | |
|---|---|---|---|
| SPGP0U01880 | XP_018222893.1 | 99 | 0 |
| SPGP0AJ00980 | XP_018221870.1 | 99 | 0 |
| SPGP0R00160 | XP_018219958.1 | 99 | 0 |
| SPGP0AA01790 | XP_018220413.1 | 99 | 0 |
| SPGP0Y01610 | XP_018220175.1 | 99 | 0 |
| SPGP0HI00220 | XP_018219948.1 | 99 | 0 |
| SPGP0AU00330 | XP_018223088.1 | 100 | 0 |
| SPGP0CX00530 | XP_018222073.1 | 99 | 0 |
| SPGP0AJ01130 | XP_018221884.1 | 99 | 0 |
| SPGP0AT01240 | XP_018219142.1 | 99 | 0 |
| SPGP0L02040 | XP_018218832.1 | 99 | 0 |
| SPGP0CX01840 | XP_018222199.1 | 99 | 0 |
| SPGP0AF01340 | XP_018222391.1 | 99 | 0 |
| SPGP0CW00210 | XP_018219532.1 | 99 | 0 |
| SPGP0CP00900 | XP_018223749.1 | 99 | 0 |
| SPGP0K00170 | XP_018222760.1 | 99 | 0 |
| SPGP0L01610 | XP_018218873.1 | 99 | 0 |
| SPGP0AF00260 | XP_018222280.1 | 99 | 0 |
| SPGP0Y02440 | XP_018220253.1 | 99 | 0 |
| SPGP0AF02110 | XP_018222469.1 | 99 | 0 |
| SPGP0AU00360 | XP_018223091.1 | 99 | 1 |
| SPGP0AJ00310 | XP_018221805.1 | 99 | 0 |
| SPGP0CM01790 | XP_018223148.1 | 99 | 0 |
| SPGP0CU01740 | XP_018220660.1 | 99 | 0 |
| SPGP0AF01870 | XP_018222445.1 | 99 | 0 |
| SPGP0Y01190 | XP_018220134.1 | 99 | 0 |
| SPGP0L00900 | XP_018218942.1 | 99 | 0 |
| SPGP0CP01460 | XP_018223693.1 | 99 | 0 |
| SPGP0CU02210 | XP_018220613.1 | 99 | 1 |
| SPGP0JK00290 | XP_018223449.1 | 99 | 0 |
| SPGP0CW00200 | XP_018219533.1 | 99 | 0 |
| SPGP0FF00140 | XP_018223398.1 | 99 | 0 |
| SPGP0AF02100 | XP_018222468.1 | 99 | 0 |
| SPGP0CP00520 | XP_018223785.1 | 99 | 0 |
| SPGP0R00390 | XP_018219980.1 | 99 | 0 |
| SPGP0CP01250 | XP_018223713.1 | 99 | 0 |
| SPGP0AA01650 | XP_018220426.1 | 99 | 0 |
| SPGP0GH00180 | XP_018222820.1 | 99 | 0 |
| SPGP0DE00260 | XP_018223843.1 | 99 | 0 |

Create PDF in your applications with the Pdfcrowd HTML to PDF API PDFCROWD

| | | | |
|---|---|---|---|
| SPGP0R00550 | XP_018219995.1 | 99 | 0 |
| SPGP0GJ00150 | XP_018223377.1 | 99 | 0 |
| SPGP0CW02330 | XP_018219328.1 | 99 | 0 |
| SPGP0L02000 | XP_018218836.1 | 99 | 0 |
| SPGP0AT00250 | XP_018219047.1 | 99 | 0 |
| SPGP0GG00270 | XP_018220825.1 | 99 | 0 |
| SPGP0CX01720 | XP_018222188.1 | 99 | 0 |
| SPGP0Y01150 | XP_018220130.1 | 99 | 0 |
| SPGP0AN01100 | XP_018224016.1 | 99 | 0 |
| SPGP0AA00650 | XP_018220526.1 | 99 | 0 |
| SPGP0AN01130 | XP_018224019.1 | 99 | 0 |
| SPGP0Y00110 | XP_018220028.1 | 99 | 0 |
| SPGP0AA01390 | XP_018220451.1 | 99 | 0 |
| SPGP0Y00400 | XP_018220057.1 | 99 | 0 |
| SPGP0Y02980 | XP_018220307.1 | 99 | 0 |
| SPGP0L01930 | XP_018218843.1 | 99 | 1 |
| SPGP0AU00290 | XP_018223084.1 | 99 | 0 |
| SPGP0CM01460 | XP_018223180.1 | 99 | 0 |
| SPGP0AF01750 | XP_018222433.1 | 99 | 0 |
| SPGP0V01020 | XP_018221685.1 | 99 | 0 |
| SPGP0HB00160 | XP_018222867.1 | 98 | 2 |
| SPGP0IN00230 | XP_018221653.1 | 99 | 0 |
| SPGP0AN00660 | XP_018223973.1 | 99 | 0 |
| SPGP0AN00880 | XP_018223994.1 | 99 | 0 |
| SPGP0CU00200 | XP_018220809.1 | 99 | 0 |
| SPGP0CP01670 | XP_018223673.1 | 99 | 0 |
| SPGP0CP03280 | XP_018223512.1 | 99 | 0 |
| SPGP0AT00100 | XP_018219032.1 | 99 | 0 |
| SPGP0IZ00240 | XP_018220352.1 | 99 | 1 |
| SPGP0CX01500 | XP_018222167.1 | 99 | 0 |
| SPGP0CP00510 | XP_018223786.1 | 100 | 0 |
| SPGP0J00200 | XP_018219927.1 | 99 | 0 |
| SPGP0AT01510 | XP_018219168.1 | 99 | 0 |
| SPGP0CM02260 | XP_018223103.1 | 99 | 0 |
| SPGP0AA01490 | XP_018220441.1 | 99 | 0 |
| SPGP0DB00220 | XP_018223896.1 | 99 | 0 |
| SPGP0IG00110 | XP_018223347.1 | 99 | 0 |
| SPGP0CP01270 | XP_018223711.1 | 99 | 0 |
| SPGP0CP00100 | XP_018223826.1 | 99 | 0 |
| SPGP0AT01410 | XP_018219158.1 | 99 | 0 |

Create PDF in your applications with the Pdfcrowd HTML to PDF API PDFCROWD

| | | | |
|---|---|---|---|
| SPGP0CU01260 | XP_018220706.1 | 99 | 0 |
| SPGP0CW02520 | XP_018219309.1 | 99 | 0 |
| SPGP0CP00830 | XP_018223755.1 | 99 | 0 |
| SPGP0AJ00340 | XP_018221807.1 | 99 | 0 |
| SPGP0CW03130 | XP_018219250.1 | 99 | 0 |
| SPGP0EM00120 | XP_018223061.1 | 98 | 1 |
| SPGP0DE00230 | XP_018223840.1 | 99 | 0 |
| SPGP0AA00790 | XP_018220512.1 | 85 | 0 |
| SPGP0AF02180 | XP_018222475.1 | 99 | 0 |
| SPGP0CP02400 | XP_018223601.1 | 99 | 0 |
| SPGP0L00320 | XP_018219000.1 | 99 | 0 |
| SPGP0EO00350 | XP_018221172.1 | 99 | 0 |
| SPGP0AF00680 | XP_018222322.1 | 99 | 0 |
| SPGP0AN00410 | XP_018223950.1 | 99 | 0 |
| SPGP0CU01200 | XP_018220712.1 | 99 | 0 |
| SPGP0CX00680 | XP_018222087.1 | 99 | 0 |
| SPGP0Y02630 | XP_018220272.1 | 99 | 0 |
| SPGP0GG00240 | XP_018220827.1 | 99 | 0 |
| SPGP0AJ01420 | XP_018221912.1 | 99 | 0 |
| SPGP0CX01820 | XP_018222197.1 | 98 | 0 |
| SPGP0L02020 | XP_018218834.1 | 99 | 0 |
| SPGP0CW01790 | XP_018219381.1 | 99 | 0 |
| SPGP0CP02130 | XP_018223627.1 | 99 | 0 |
| SPGP0CM00170 | XP_018223307.1 | 99 | 0 |
| SPGP0AJ01390 | XP_018221909.1 | 99 | 0 |
| SPGP0V00560 | XP_018221730.1 | 99 | 0 |
| SPGP0CW03490 | XP_018219215.1 | 99 | 0 |
| SPGP0Y01440 | XP_018220159.1 | 99 | 0 |
| SPGP0CP03000 | XP_018223540.1 | 99 | 0 |
| SPGP0CW02140 | XP_018219347.1 | 99 | 0 |
| SPGP0AF00410 | XP_018222295.1 | 99 | 0 |
| SPGP0AU00280 | XP_018223083.1 | 99 | 0 |
| SPGP0Z00170 | XP_018222548.1 | 99 | 0 |
| SPGP0U01140 | XP_018222965.1 | 99 | 0 |
| SPGP0AF01820 | XP_018222440.1 | 99 | 0 |
| SPGP0L00310 | XP_018219001.1 | 99 | 0 |
| SPGP0L00920 | XP_018218940.1 | 99 | 0 |
| SPGP0CU01770 | XP_018220657.1 | 99 | 0 |
| SPGP0CW00310 | XP_018219524.1 | 99 | 0 |
| SPGP0AJ00110 | XP_018221786.1 | 100 | 0 |

Create PDF in your applications with the Pdfcrowd HTML to PDF API PDFCROWD

| | | | |
|---|---|---|---|
| SPGP0CW01410 | XP_018219417.1 | 99 | 0 |
| SPGP0CW01000 | XP_018219456.1 | 99 | 0 |
| SPGP0CV00220 | XP_018222573.1 | 99 | 0 |
| SPGP0CW00180 | XP_018219535.1 | 99 | 0 |
| SPGP0CU00530 | XP_018220776.1 | 99 | 0 |
| SPGP0U00540 | XP_018223020.1 | 99 | 0 |
| SPGP0DB00110 | XP_018223885.1 | 98 | 1 |
| SPGP0AA00690 | XP_018220522.1 | 99 | 0 |
| SPGP0AJ00270 | XP_018221801.1 | 99 | 0 |
| SPGP0GG00250 | XP_018220826.1 | 99 | 0 |
| SPGP0Y01430 | XP_018220158.1 | 99 | 0 |
| SPGP0AJ01800 | XP_018221948.1 | 99 | 0 |
| SPGP0CP02070 | XP_018223633.1 | 99 | 0 |
| SPGP0AA00630 | XP_018220528.1 | 99 | 0 |
| SPGP0AA00320 | XP_018220558.1 | 100 | 0 |
| SPGP0AH00150 | XP_018223423.1 | 99 | 0 |
| SPGP0AT01000 | XP_018219118.1 | 99 | 0 |
| SPGP0AE00430 | XP_018222522.1 | 99 | 0 |
| SPGP0CW01560 | XP_018219402.1 | 99 | 0 |
| SPGP0AF00980 | XP_018222352.1 | 99 | 0 |
| SPGP0AT00820 | XP_018219101.1 | 99 | 1 |
| SPGP0AE00260 | XP_018222507.1 | 99 | 0 |
| SPGP0AH00170 | XP_018223425.1 | 99 | 0 |
| SPGP0AF01730 | XP_018222431.1 | 99 | 0 |
| SPGP0AU00270 | XP_018223082.1 | 99 | 0 |
| SPGP0R00410 | XP_018219982.1 | 99 | 0 |
| SPGP0CW00650 | XP_018219491.1 | 100 | 0 |
| SPGP0JK00130 | XP_018223435.1 | 99 | 0 |
| SPGP0AA01950 | XP_018220398.1 | 99 | 0 |
| SPGP0U00210 | XP_018223049.1 | 99 | 0 |
| SPGP0CM01260 | XP_018223199.1 | 99 | 0 |
| SPGP0CU00220 | XP_018220807.1 | 99 | 0 |
| SPGP0CW03570 | XP_018219207.1 | 99 | 0 |
| SPGP0CW00260 | XP_018219527.1 | 99 | 0 |
| SPGP0CW03330 | XP_018219231.1 | 99 | 0 |
| SPGP0AJ01180 | XP_018221889.1 | 99 | 0 |
| SPGP0CP03220 | XP_018223518.1 | 99 | 0 |
| SPGP0JC00150 | XP_018220855.1 | 99 | 0 |
| SPGP0CM01040 | XP_018223221.1 | 99 | 0 |
| SPGP0CX00980 | XP_018222117.1 | 99 | 0 |

Create PDF in your applications with the Pdfcrowd HTML to PDF API PDFCROWD

| | | | |
|---|---|---|---|
| SPGP0CW02980 | XP_018219265.1 | 99 | 1 |
| SPGP0V01100 | XP_018221677.1 | 99 | 0 |
| SPGP0L00750 | XP_018218958.1 | 99 | 0 |
| SPGP0Y02300 | XP_018220239.1 | 99 | 0 |
| SPGP0X00220 | XP_018221471.1 | 98 | 0 |
| SPGP0AE00210 | XP_018222503.1 | 99 | 0 |
| SPGP0AF00740 | XP_018222328.1 | 99 | 0 |
| SPGP0CM01030 | XP_018223222.1 | 99 | 0 |
| SPGP0U01330 | XP_018222949.1 | 99 | 0 |
| SPGP0L00520 | XP_018218980.1 | 99 | 0 |
| SPGP0CP02790 | XP_018223562.1 | 99 | 0 |
| SPGP0R00670 | XP_018220007.1 | 99 | 0 |
| SPGP0CW01290 | XP_018219428.1 | 99 | 0 |
| SPGP0Y01650 | XP_018220179.1 | 99 | 1 |
| SPGP0AJ00160 | XP_018221791.1 | 99 | 0 |
| SPGP0L01690 | XP_018218865.1 | 99 | 0 |
| SPGP0DA01580 | XP_018222599.1 | 99 | 0 |
| SPGP0K00360 | XP_018222778.1 | 99 | 0 |
| SPGP0AA01750 | XP_018220416.1 | 100 | 0 |
| SPGP0CM00970 | XP_018223228.1 | 99 | 0 |
| SPGP0V00540 | XP_018221732.1 | 99 | 0 |
| SPGP0CW02100 | XP_018219351.1 | 99 | 0 |
| SPGP0U01530 | XP_018222929.1 | 99 | 0 |
| SPGP0CP00610 | XP_018223776.1 | 99 | 0 |
| SPGP0CU00670 | XP_018220761.1 | 99 | 0 |
| SPGP0CW02780 | XP_018219285.1 | 99 | 0 |
| SPGP0CM00150 | XP_018223309.1 | 99 | 0 |
| SPGP0FE00100 | XP_018221962.1 | 99 | 0 |
| SPGP0AF00650 | XP_018222319.1 | 99 | 0 |
| SPGP0R00200 | XP_018219962.1 | 99 | 0 |
| SPGP0AF00730 | XP_018222327.1 | 99 | 0 |
| SPGP0AF00470 | XP_018222301.1 | 99 | 0 |
| SPGP0BE00180 | XP_018223360.1 | 99 | 0 |
| SPGP0IS00120 | XP_018223094.1 | 100 | 0 |
| SPGP0IS00100 | XP_018223096.1 | 99 | 0 |
| SPGP0AF02010 | XP_018222459.1 | 99 | 0 |
| SPGP0AE00230 | XP_018222505.1 | 99 | 0 |
| SPGP0Y01460 | XP_018220161.1 | 99 | 0 |
| SPGP0R00680 | XP_018220008.1 | 100 | 0 |
| SPGP0CW00230 | XP_018219530.1 | 99 | 0 |

Create PDF in your applications with the Pdfcrowd HTML to PDF API PDFCROWD

| | | | |
|---|---|---|---|
| SPGP0DE00190 | XP_018223836.1 | 99 | 0 |
| SPGP0CM01210 | XP_018223204.1 | 99 | 0 |
| SPGP0CU01690 | XP_018220665.1 | 99 | 0 |
| SPGP0K00560 | XP_018222798.1 | 99 | 0 |
| SPGP0CP02200 | XP_018223620.1 | 99 | 0 |
| SPGP0Y02510 | XP_018220260.1 | 99 | 0 |
| SPGP0GA00160 | XP_018223387.1 | 99 | 0 |
| SPGP0ED00120 | XP_018223473.1 | 99 | 0 |
| SPGP0HZ00140 | XP_018219027.1 | 99 | 0 |
| SPGP0Y00170 | XP_018220034.1 | 99 | 0 |
| SPGP0CP02090 | XP_018223631.1 | 99 | 0 |
| SPGP0CP00550 | XP_018223782.1 | 99 | 0 |
| SPGP0AU00250 | XP_018223080.1 | 99 | 0 |
| SPGP0CP01950 | XP_018223645.1 | 99 | 0 |
| SPGP0AF00960 | XP_018222350.1 | 99 | 0 |
| SPGP0CW02660 | XP_018219297.1 | 99 | 0 |
| SPGP0CM01490 | XP_018223177.1 | 100 | 0 |
| SPGP0AF01330 | XP_018222390.1 | 99 | 0 |
| SPGP0AF01710 | XP_018222428.1 | 97 | 2 |
| SPGP0CM01200 | XP_018223205.1 | 99 | 0 |
| SPGP0CW00950 | XP_018219461.1 | 99 | 0 |
| SPGP0K00390 | XP_018222781.1 | 99 | 0 |
| SPGP0CX01930 | XP_018222208.1 | 99 | 0 |
| SPGP0CP00570 | XP_018223780.1 | 99 | 0 |
| SPGP0AA01150 | XP_018220475.1 | 99 | 0 |
| SPGP0CP03680 | XP_018223476.1 | 100 | 0 |
| SPGP0L01250 | XP_018218909.1 | 99 | 0 |
| SPGP0AT01440 | XP_018219161.1 | 99 | 0 |
| SPGP0AA01540 | XP_018220436.1 | 99 | 0 |
| SPGP0CP03170 | XP_018223523.1 | 99 | 0 |
| SPGP0JP00120 | XP_018219563.1 | 99 | 0 |
| SPGP0AT01380 | XP_018219155.1 | 99 | 0 |
| SPGP0Z00250 | XP_018222540.1 | 99 | 0 |
| SPGP0AT01640 | XP_018219181.1 | 99 | 0 |
| SPGP0CW00890 | XP_018219467.1 | 99 | 0 |
| SPGP0Y00390 | XP_018220056.1 | 99 | 0 |
| SPGP0EP00140 | XP_018222747.1 | 99 | 0 |
| SPGP0CU01220 | XP_018220710.1 | 99 | 0 |
| SPGP0AA01520 | XP_018220438.1 | 99 | 0 |
| SPGP0AT00530 | XP_018219074.1 | 99 | 0 |

Create PDF in your applications with the Pdfcrowd HTML to PDF API PDFCROWD

| | | | |
|---|---|---|---|
| SPGP0CP01760 | XP_018223664.1 | 99 | 0 |
| SPGP0BE00300 | XP_018223349.1 | 99 | 0 |
| SPGP0GJ00220 | XP_018223370.1 | 98 | 0 |
| SPGP0CP02220 | XP_018223618.1 | 99 | 0 |
| SPGP0CX02250 | XP_018222239.1 | 99 | 0 |
| SPGP0CM01990 | XP_018223129.1 | 99 | 0 |
| SPGP0L00500 | XP_018218982.1 | 99 | 0 |
| SPGP0EM00170 | XP_018223066.1 | 99 | 0 |
| SPGP0CW01310 | XP_018219426.1 | 99 | 0 |
| SPGP0R00230 | XP_018219965.1 | 100 | 0 |
| SPGP0IZ00170 | XP_018220345.1 | 99 | 0 |
| SPGP0CW00380 | XP_018219517.1 | 99 | 0 |
| SPGP0CX01630 | XP_018222179.1 | 98 | 1 |
| SPGP0CW01750 | XP_018219385.1 | 99 | 0 |
| SPGP0Y01240 | XP_018220139.1 | 99 | 0 |
| SPGP0AN01050 | XP_018224011.1 | 99 | 0 |
| SPGP0L02280 | XP_018218808.1 | 99 | 0 |
| SPGP0L01370 | XP_018218897.1 | 99 | 0 |
| SPGP0AF00670 | XP_018222321.1 | 99 | 0 |
| SPGP0AJ00410 | XP_018221814.1 | 99 | 1 |
| SPGP0F00260 | XP_018221195.1 | 99 | 0 |
| SPGP0AJ01020 | XP_018221874.1 | 100 | 0 |
| SPGP0Y02380 | XP_018220247.1 | 99 | 0 |
| SPGP0Y02930 | XP_018220302.1 | 100 | 0 |
| SPGP0L02420 | XP_018218794.1 | 99 | 0 |
| SPGP0AA01440 | XP_018220446.1 | 99 | 0 |
| SPGP0AJ00770 | XP_018221849.1 | 99 | 0 |
| SPGP0CW02810 | XP_018219282.1 | 99 | 0 |
| SPGP0F00190 | XP_018221188.1 | 97 | 1 |
| SPGP0V00850 | XP_018221702.1 | 99 | 0 |
| SPGP0L02370 | XP_018218799.1 | 99 | 0 |
| SPGP0CU00400 | XP_018220789.1 | 99 | 0 |
| SPGP0CW00980 | XP_018219458.1 | 100 | 0 |
| SPGP0Y00800 | XP_018220096.1 | 99 | 0 |
| SPGP0Y01400 | XP_018220155.1 | 99 | 0 |
| SPGP0CU01680 | XP_018220666.1 | 99 | 0 |
| SPGP0JK00240 | XP_018223444.1 | 99 | 0 |
| SPGP0K00510 | XP_018222793.1 | 100 | 0 |
| SPGP0V00520 | XP_018221734.1 | 99 | 0 |
| SPGP0CM02080 | XP_018223121.1 | 99 | 0 |

Create PDF in your applications with the Pdfcrowd HTML to PDF API PDFCROWD

| | | | |
|---|---|---|---|
| SPGP0AN01180 | XP_018224024.1 | 99 | 0 |
| SPGP0CW02460 | XP_018219315.1 | 99 | 0 |
| SPGP0AT00160 | XP_018219038.1 | 100 | 0 |
| SPGP0G00120 | XP_018222251.1 | 100 | 0 |
| SPGP0AJ00840 | XP_018221856.1 | 100 | 0 |
| SPGP0U01270 | XP_018222954.1 | 99 | 0 |
| SPGP0DB00150 | XP_018223889.1 | 99 | 0 |
| SPGP0Y02530 | XP_018220262.1 | 99 | 0 |
| SPGP0FF00190 | XP_018223393.1 | 99 | 0 |
| SPGP0ED00100 | XP_018223475.1 | 99 | 0 |
| SPGP0L00260 | XP_018219006.1 | 99 | 0 |
| SPGP0GH00160 | XP_018222818.1 | 99 | 0 |
| SPGP0CP02320 | XP_018223608.1 | 99 | 0 |
| SPGP0CP02110 | XP_018223629.1 | 99 | 0 |
| SPGP0AA00360 | XP_018220554.1 | 99 | 0 |
| SPGP0CP01970 | XP_018223643.1 | 99 | 0 |
| SPGP0AE00220 | XP_018222504.1 | 99 | 0 |
| SPGP0AF01080 | XP_018222363.1 | 99 | 0 |
| SPGP0AN00400 | XP_018223949.1 | 100 | 0 |
| SPGP0CP02440 | XP_018223597.1 | 99 | 0 |
| SPGP0AA00940 | XP_018220496.1 | 99 | 0 |
| SPGP0CU02150 | XP_018220619.1 | 99 | 0 |
| SPGP0AA01010 | XP_018220489.1 | 99 | 0 |
| SPGP0G00200 | XP_018222259.1 | 99 | 0 |
| SPGP0CM01220 | XP_018223203.1 | 99 | 0 |
| SPGP0AF01600 | XP_018222418.1 | 99 | 0 |
| SPGP0GL00210 | XP_018220850.1 | 99 | 0 |
| SPGP0JC00140 | XP_018220854.1 | 99 | 0 |
| SPGP0AA00450 | XP_018220545.1 | 99 | 0 |
| SPGP0CX01950 | XP_018222210.1 | 98 | 1 |
| SPGP0AF00770 | XP_018222331.1 | 99 | 0 |
| SPGP0CU00130 | XP_018220815.1 | 99 | 0 |
| SPGP0CP00160 | XP_018223820.1 | 100 | 0 |
| SPGP0AA01070 | XP_018220483.1 | 99 | 0 |
| SPGP0AT01020 | XP_018219120.1 | 99 | 0 |
| SPGP0CP01740 | XP_018223666.1 | 99 | 0 |
| SPGP0GH00120 | XP_018222814.1 | 100 | 0 |
| SPGP0CX02010 | XP_018222216.1 | 99 | 0 |
| SPGP0U01040 | XP_018222975.1 | 99 | 0 |
| SPGP0CU02190 | XP_018220615.1 | 100 | 0 |

Create PDF in your applications with the Pdfcrowd HTML to PDF API PDFCROWD

| | | | |
|---|---|---|---|
| SPGP0F02960 | XP_018221453.1 | 99 | 0 |
| SPGP0Y01990 | XP_018220210.1 | 99 | 0 |
| SPGP0AF01860 | XP_018222444.1 | 99 | 0 |
| SPGP0R00470 | XP_018219988.1 | 98 | 0 |
| SPGP0U00640 | XP_018223011.1 | 99 | 0 |
| SPGP0R00250 | XP_018219967.1 | 99 | 0 |
| SPGP0L02290 | XP_018218807.1 | 99 | 0 |
| SPGP0U01660 | XP_018222916.1 | 99 | 0 |
| SPGP0L02070 | XP_018218829.1 | 99 | 0 |
| SPGP0CM01350 | XP_018223190.1 | 99 | 0 |
| SPGP0CM00110 | XP_018223313.1 | 99 | 0 |
| SPGP0Y00560 | XP_018220073.1 | 99 | 0 |
| SPGP0CP03630 | XP_018223481.1 | 99 | 0 |
| SPGP0CW03430 | XP_018219221.1 | 99 | 0 |
| SPGP0CU00640 | XP_018220764.1 | 100 | 0 |
| SPGP0EP00160 | XP_018222749.1 | 99 | 0 |
| SPGP0EO00370 | XP_018221174.1 | 99 | 0 |
| SPGP0R00590 | XP_018219999.1 | 99 | 0 |
| SPGP0U01390 | XP_018222943.1 | 99 | 0 |
| SPGP0DE00510 | XP_018223868.1 | 99 | 0 |
| SPGP0L01110 | XP_018218923.1 | 99 | 0 |
| SPGP0IR00120 | XP_018223466.1 | 99 | 0 |
| SPGP0AT00790 | XP_018219098.1 | 99 | 0 |
| SPGP0U02090 | XP_018222872.1 | 99 | 0 |
| SPGP0CW03410 | XP_018219223.1 | 100 | 0 |
| SPGP0BE00140 | XP_018223364.1 | 99 | 0 |
| SPGP0CU00190 | XP_018220810.1 | 99 | 0 |
| SPGP0CX01600 | XP_018222176.1 | 99 | 0 |
| SPGP0Y02870 | XP_018220296.1 | 99 | 0 |
| SPGP0CM02310 | XP_018223098.1 | 99 | 0 |
| SPGP0CW00160 | XP_018219537.1 | 99 | 0 |
| SPGP0IC00130 | XP_018222530.1 | 99 | 0 |
| SPGP0CW02440 | XP_018219317.1 | 99 | 0 |
| SPGP0CP00720 | XP_018223766.1 | 99 | 0 |
| SPGP0CM02100 | XP_018223119.1 | 99 | 0 |
| SPGP0CM00990 | XP_018223226.1 | 99 | 0 |
| SPGP0CM00430 | XP_018223281.1 | 100 | 0 |
| SPGP0AN01200 | XP_018224026.1 | 99 | 0 |
| SPGP0CX01870 | XP_018222202.1 | 99 | 0 |
| SPGP0U01250 | XP_018222956.1 | 99 | 0 |

Create PDF in your applications with the Pdfcrowd HTML to PDF API PDFCROWD

| | | | |
|---|---|---|---|
| SPGP0U01700 | XP_018222912.1 | 99 | 0 |
| SPGP0Y00120 | XP_018220029.1 | 99 | 0 |
| SPGP0Y02430 | XP_018220252.1 | 99 | 0 |
| SPGP0L00710 | XP_018218962.1 | 99 | 0 |
| SPGP0L01440 | XP_018218890.1 | 99 | 0 |
| SPGP0CM02270 | XP_018223102.1 | 99 | 0 |
| SPGP0DE00490 | XP_018223866.1 | 99 | 0 |
| SPGP0Y03060 | XP_018220315.1 | 99 | 0 |
| SPGP0AF01120 | XP_018222368.1 | 99 | 0 |
| SPGP0AJ01250 | XP_018221896.1 | 99 | 0 |
| SPGP0L02350 | XP_018218801.1 | 99 | 0 |
| SPGP0CU00480 | XP_018220781.1 | 99 | 0 |
| SPGP0CU00150 | XP_018220813.1 | 99 | 0 |
| SPGP0CM00600 | XP_018223264.1 | 99 | 0 |
| SPGP0CU01400 | XP_018220693.1 | 99 | 0 |
| SPGP0CW00390 | XP_018219516.1 | 99 | 0 |
| SPGP0CP01720 | XP_018223668.1 | 99 | 0 |
| SPGP0CX02030 | XP_018222218.1 | 99 | 0 |
| SPGP0EP00120 | XP_018222745.1 | 99 | 0 |
| SPGP0AF01380 | XP_018222395.1 | 99 | 0 |
| SPGP0K00630 | XP_018222805.1 | 99 | 0 |
| SPGP0CW01840 | XP_018219376.1 | 99 | 0 |
| SPGP0Y00660 | XP_018220082.1 | 99 | 0 |
| SPGP0AA01730 | XP_018220418.1 | 99 | 0 |
| SPGP0J03480 | XP_018219600.1 | 99 | 0 |
| SPGP0AJ01610 | XP_018221929.1 | 99 | 0 |
| SPGP0Y00540 | XP_018220071.1 | 99 | 0 |
| SPGP0AT00940 | XP_018219112.1 | 99 | 0 |
| SPGP0Y02680 | XP_018220277.1 | 99 | 0 |
| SPGP0AF00490 | XP_018222303.1 | 99 | 0 |
| SPGP0AF01540 | XP_018222412.1 | 99 | 0 |
| SPGP0U00160 | XP_018223054.1 | 100 | 0 |
| SPGP0AT00810 | XP_018219100.1 | 99 | 0 |
| SPGP0AJ00290 | XP_018221803.1 | 99 | 0 |
| SPGP0CP00800 | XP_018223758.1 | 99 | 0 |
| SPGP0L00470 | XP_018218985.1 | 99 | 0 |
| SPGP0CW01980 | XP_018219362.1 | 99 | 0 |
| SPGP0IV00140 | XP_018222860.1 | 99 | 0 |
| SPGP0AT00430 | XP_018219064.1 | 99 | 1 |
| SPGP0AJ00880 | XP_018221860.1 | 99 | 0 |

Create PDF in your applications with the Pdfcrowd HTML to PDF API PDFCROWD

| | | | |
|---|---|---|---|
| SPGP0L02440 | XP_018218792.1 | 99 | 0 |
| SPGP0Y02410 | XP_018220250.1 | 99 | 0 |
| SPGP0ED00140 | XP_018223471.1 | 99 | 0 |
| SPGP0AA00570 | XP_018220533.1 | 99 | 0 |
| SPGP0DE00280 | XP_018223845.1 | 99 | 0 |
| SPGP0CM00490 | XP_018223275.1 | 99 | 0 |
| SPGP0CP02490 | XP_018223592.1 | 99 | 0 |
| SPGP0Y00330 | XP_018220050.1 | 99 | 0 |
| SPGP0AN00570 | XP_018223964.1 | 99 | 0 |
| SPGP0CP01530 | XP_018223686.1 | 99 | 0 |
| SPGP0GG00160 | XP_018220835.1 | 99 | 0 |
| SPGP0Y01720 | XP_018220186.1 | 99 | 0 |
| SPGP0GG00110 | XP_018220840.1 | 99 | 0 |
| SPGP0L01170 | XP_018218917.1 | 99 | 0 |
| SPGP0AF01220 | XP_018222378.1 | 99 | 0 |
| SPGP0AT00460 | XP_018219067.1 | 99 | 0 |
| SPGP0AF01560 | XP_018222414.1 | 99 | 0 |
| SPGP0U01950 | XP_018222886.1 | 100 | 0 |
| SPGP0AF00790 | XP_018222333.1 | 99 | 0 |
| SPGP0CU00170 | XP_018220811.1 | 99 | 0 |
| SPGP0Y02700 | XP_018220279.1 | 99 | 0 |
| SPGP0AF00570 | XP_018222311.1 | 99 | 0 |
| SPGP0AF00950 | XP_018222349.1 | 99 | 0 |
| SPGP0CW03600 | XP_018219205.1 | 99 | 0 |
| SPGP0CX00900 | XP_018222109.1 | 99 | 0 |
| SPGP0CX00660 | XP_018222085.1 | 99 | 0 |
| SPGP0AA01740 | XP_018220417.1 | 99 | 0 |
| SPGP0L01720 | XP_018218863.1 | 99 | 0 |
| SPGP0EP00170 | XP_018222750.1 | 99 | 0 |
| SPGP0CU02360 | XP_018220598.1 | 99 | 0 |
| SPGP0CW00800 | XP_018219476.1 | 99 | 0 |
| SPGP0CU01960 | XP_018220638.1 | 99 | 0 |
| SPGP0IM00160 | XP_018219546.1 | 99 | 0 |
| SPGP0DE00610 | XP_018223878.1 | 99 | 0 |
| SPGP0CV00250 | XP_018222576.1 | 99 | 0 |
| SPGP0HW00160 | XP_018224037.1 | 99 | 0 |
| SPGP0AT00590 | XP_018219080.1 | 99 | 0 |
| SPGP0CU01820 | XP_018220652.1 | 99 | 0 |
| SPGP0L02090 | XP_018218827.1 | 99 | 0 |
| SPGP0CP02810 | XP_018223560.1 | 99 | 0 |

Create PDF in your applications with the Pdfcrowd HTML to PDF API PDFCROWD

| | | | |
|---|---|---|---|
| SPGP0L01810 | XP_018218854.1 | 99 | 0 |
| SPGP0Y02600 | XP_018220269.1 | 99 | 0 |
| SPGP0AJ01590 | XP_018221927.1 | 99 | 0 |
| SPGP0V01130 | XP_018221675.1 | 99 | 0 |
| SPGP0AN00960 | XP_018224002.1 | 99 | 0 |
| SPGP0AF01440 | XP_018222402.1 | 99 | 0 |
| SPGP0CW03370 | XP_018219227.1 | 99 | 0 |
| SPGP0Y01800 | XP_018220193.1 | 99 | 0 |
| SPGP0V00110 | XP_018221772.1 | 99 | 0 |
| SPGP0U01790 | XP_018222902.1 | 99 | 0 |
| SPGP0Y01230 | XP_018220138.1 | 99 | 0 |
| SPGP0CX02260 | XP_018222240.1 | 99 | 0 |
| SPGP0CP00660 | XP_018223771.1 | 99 | 0 |
| SPGP0CW02400 | XP_018219321.1 | 99 | 0 |
| SPGP0CM02060 | XP_018223123.1 | 100 | 0 |
| SPGP0AJ00380 | XP_018221811.1 | 99 | 0 |
| SPGP0Y00860 | XP_018220102.1 | 99 | 0 |
| SPGP0AF00880 | XP_018222342.1 | 99 | 0 |
| SPGP0CM00750 | XP_018223250.1 | 99 | 0 |
| SPGP0EP00100 | XP_018222743.1 | 100 | 0 |
| SPGP0DE00110 | XP_018223828.1 | 99 | 0 |
| SPGP0U01740 | XP_018222908.1 | 99 | 0 |
| SPGP0AT00970 | XP_018219115.1 | 99 | 0 |
| SPGP0AA00550 | XP_018220535.1 | 99 | 0 |
| SPGP0CW01180 | XP_018218894.1 | 97 | 0 |
| SPGP0AN00530 | XP_018223960.1 | 99 | 0 |
| SPGP0CU01650 | XP_018220669.1 | 99 | 0 |
| SPGP0CP00870 | XP_018223751.1 | 99 | 0 |
| SPGP0CW01330 | XP_018219425.1 | 99 | 0 |
| SPGP0GJ00200 | XP_018223372.1 | 99 | 0 |
| SPGP0Y01840 | XP_018220195.1 | 99 | 0 |
| SPGP0CX01460 | XP_018222163.1 | 100 | 0 |
| SPGP0K00590 | XP_018222801.1 | 99 | 0 |
| SPGP0AF00510 | XP_018222305.1 | 99 | 0 |
| SPGP0CW03620 | XP_018219203.1 | 98 | 0 |
| SPGP0R00340 | XP_018219975.1 | 99 | 0 |
| SPGP0CM01640 | XP_018223162.1 | 100 | 0 |
| SPGP0GS00100 | XP_018222246.1 | 99 | 0 |
| SPGP0FP00160 | XP_018223339.1 | 99 | 0 |

Create PDF in your applications with the Pdfcrowd HTML to PDF API PDFCROWD

| | | | |
|---|---|---|---|
| SPGP0CX00960 | XP_018222115.1 | 99 | 0 |
| SPGP0CX00920 | XP_018222111.1 | 99 | 0 |
| SPGP0DF00190 | XP_018219189.1 | 98 | 0 |
| SPGP0AT01190 | XP_018219137.1 | 99 | 0 |
| SPGP0CP02880 | XP_018223553.1 | 99 | 0 |
| SPGP0U01680 | XP_018222914.1 | 100 | 0 |
| SPGP0CP02640 | XP_018223577.1 | 99 | 0 |
| SPGP0L02550 | XP_018218781.1 | 99 | 0 |
| SPGP0U00240 | XP_018223046.1 | 99 | 0 |
| SPGP0AU00140 | XP_018223071.1 | 99 | 0 |
| SPGP0AT00710 | XP_018219090.1 | 99 | 0 |
| SPGP0AJ01650 | XP_018221933.1 | 99 | 0 |
| SPGP0DE00630 | XP_018223880.1 | 99 | 0 |
| SPGP0CM01340 | XP_018223191.1 | 99 | 0 |
| SPGP0AT00900 | XP_018219109.1 | 100 | 0 |
| SPGP0AE00450 | XP_018222524.1 | 99 | 0 |
| SPGP0CU00690 | XP_018220759.1 | 99 | 0 |
| SPGP0AN00720 | XP_018223979.1 | 99 | 0 |
| SPGP0AF00180 | XP_018222273.1 | 99 | 0 |
| SPGP0CP03310 | XP_018223509.1 | 99 | 0 |
| SPGP0CU00540 | XP_018220775.1 | 99 | 0 |
| SPGP0CP03010 | XP_018223539.1 | 99 | 0 |
| SPGP0EO00270 | XP_018221164.1 | 99 | 0 |
| SPGP0CM00640 | XP_018223260.1 | 99 | 0 |
| SPGP0DE00130 | XP_018223830.1 | 99 | 0 |
| SPGP0AF02070 | XP_018222465.1 | 100 | 0 |
| SPGP0DE00250 | XP_018223842.1 | 99 | 0 |
| SPGP0L01390 | XP_018218895.1 | 99 | 0 |
| SPGP0AA00600 | XP_018220530.1 | 99 | 0 |
| SPGP0L00820 | XP_018218951.1 | 99 | 0 |
| SPGP0Y02470 | XP_018220256.1 | 99 | 0 |
| SPGP0CP01660 | XP_018223674.1 | 99 | 0 |
| SPGP0AT00340 | XP_018219055.1 | 99 | 0 |
| SPGP0L02230 | XP_018218813.1 | 99 | 0 |
| SPGP0CW00930 | XP_018219463.1 | 99 | 0 |
| SPGP0CU00870 | XP_018220742.1 | 99 | 0 |
| SPGP0GH00190 | XP_018222821.1 | 99 | 0 |
| SPGP0GG00200 | XP_018220831.1 | 99 | 0 |
| SPGP0CW02020 | XP_018219358.1 | 99 | 1 |
| SPGP0AA01230 | XP_018220467.1 | 99 | 0 |

Create PDF in your applications with the Pdfcrowd HTML to PDF API PDFCROWD

| | | | |
|---|---|---|---|
| SPGP0CP00590 | XP_018223778.1 | 99 | 0 |
| SPGP0AN00390 | XP_018223948.1 | 99 | 0 |
| SPGP0HB00180 | XP_018222869.1 | 99 | 0 |
| SPGP0CX01680 | XP_018222184.1 | 99 | 0 |
| SPGP0CW02920 | XP_018219271.1 | 99 | 0 |
| SPGP0Y02850 | XP_018220294.1 | 99 | 0 |
| SPGP0CP01070 | XP_018223732.1 | 99 | 0 |
| SPGP0K00110 | XP_018222753.1 | 99 | 0 |
| SPGP0CU01130 | XP_018220717.1 | 99 | 0 |
| SPGP0CW00840 | XP_018219472.1 | 99 | 0 |
| SPGP0CW03500 | XP_018219214.1 | 99 | 0 |
| SPGP0AF00160 | XP_018222271.1 | 99 | 0 |
| SPGP0AF01950 | XP_018222453.1 | 99 | 0 |
| SPGP0IZ00190 | XP_018220347.1 | 99 | 0 |
| SPGP0L01300 | XP_018218904.1 | 99 | 0 |
| SPGP0Y02250 | XP_018220234.1 | 99 | 0 |
| SPGP0CP02170 | XP_018223623.1 | 99 | 0 |
| SPGP0Y00250 | XP_018220042.1 | 100 | 0 |
| SPGP0L00430 | XP_018218989.1 | 99 | 0 |
| SPGP0CU02400 | XP_018220594.1 | 99 | 0 |
| SPGP0V00740 | XP_018221712.1 | 99 | 0 |
| SPGP0V01170 | XP_018221671.1 | 99 | 0 |
| SPGP0AA00880 | XP_018220502.1 | 99 | 0 |
| SPGP0AF00350 | XP_018222289.1 | 99 | 0 |
| SPGP0CU01840 | XP_018220650.1 | 99 | 0 |
| SPGP0CM00810 | XP_018223244.1 | 99 | 0 |
| SPGP0CW03180 | XP_018219246.1 | 99 | 0 |
| SPGP0AF00250 | XP_018222279.1 | 99 | 0 |
| SPGP0L00860 | XP_018218947.1 | 99 | 0 |
| SPGP0L02480 | XP_018218788.1 | 99 | 0 |
| SPGP0CU00560 | XP_018220773.1 | 99 | 0 |
| SPGP0CW01250 | XP_018219432.1 | 99 | 0 |
| SPGP0CP00620 | XP_018223775.1 | 99 | 0 |
| SPGP0CX00870 | XP_018222106.1 | 100 | 0 |
| SPGP0Y02000 | XP_018220211.1 | 99 | 0 |
| SPGP0DS00130 | XP_018220965.1 | 99 | 0 |
| SPGP0CX02130 | XP_018222227.1 | 99 | 0 |
| SPGP0CP02930 | XP_018223548.1 | 99 | 0 |
| SPGP0V00890 | XP_018221698.1 | 99 | 0 |
| SPGP0L01290 | XP_018218905.1 | 99 | 0 |

Create PDF in your applications with the Pdfcrowd HTML to PDF API PDFCROWD

| | | | |
|---|---|---|---|
| SPGP0AT00690 | XP_018219088.1 | 99 | 0 |
| SPGP0DB00200 | XP_018223894.1 | 99 | 0 |
| SPGP0U01520 | XP_018222930.1 | 99 | 0 |
| SPGP0CU00850 | XP_018220744.1 | 99 | 0 |
| SPGP0CP01990 | XP_018223641.1 | 99 | 0 |
| SPGP0Y02760 | XP_018220285.1 | 99 | 0 |
| SPGP0CP01900 | XP_018223650.1 | 99 | 0 |
| SPGP0AJ01380 | XP_018221908.1 | 99 | 0 |
| SPGP0EM00110 | XP_018223060.1 | 99 | 0 |
| SPGP0CW00860 | XP_018219470.1 | 99 | 0 |
| SPGP0V00400 | XP_018221743.1 | 99 | 0 |
| SPGP0JO00110 | XP_018222261.1 | 99 | 0 |
| SPGP0HN00100 | XP_018221986.1 | 98 | 0 |
| SPGP0CU00290 | XP_018220800.1 | 99 | 0 |
| SPGP0Y01570 | XP_018220171.1 | 99 | 0 |
| SPGP0CM01290 | XP_018223196.1 | 99 | 0 |
| SPGP0J03530 | XP_018219596.1 | 100 | 0 |
| SPGP0CW02250 | XP_018219336.1 | 99 | 0 |
| SPGP0CU01510 | XP_018220683.1 | 99 | 0 |
| SPGP0L01320 | XP_018218902.1 | 99 | 0 |
| SPGP0Y02350 | XP_018220244.1 | 99 | 0 |
| SPGP0HW00100 | XP_018224043.1 | 97 | 0 |
| SPGP0U01420 | XP_018222940.1 | 99 | 0 |
| SPGP0Y02660 | XP_018220275.1 | 99 | 0 |
| SPGP0CW01810 | XP_018219379.1 | 99 | 0 |
| SPGP0U00820 | XP_018222996.1 | 99 | 0 |
| SPGP0Y00410 | XP_018220058.1 | 99 | 0 |
| SPGP0AQ00180 | XP_018223404.1 | 99 | 0 |
| SPGP0CP02560 | XP_018223585.1 | 99 | 0 |
| SPGP0V01150 | XP_018221673.1 | 99 | 0 |
| SPGP0IM00120 | XP_018219550.1 | 99 | 0 |
| SPGP0Y01860 | XP_018220197.1 | 99 | 0 |
| SPGP0J01420 | XP_018219806.1 | 99 | 0 |
| SPGP0X00200 | XP_018221469.1 | 99 | 1 |
| SPGP0CM00300 | XP_018223294.1 | 99 | 0 |
| SPGP0CU00580 | XP_018220771.1 | 99 | 0 |
| SPGP0CX00850 | XP_018222104.1 | 99 | 0 |
| SPGP0K00490 | XP_018222791.1 | 99 | 0 |
| SPGP0DS00110 | XP_018220963.1 | 99 | 0 |
| SPGP0CX00540 | XP_018222074.1 | 99 | 0 |

Create PDF in your applications with the Pdfcrowd HTML to PDF API PDFCROWD

| | | | |
|---|---|---|---|
| SPGP0Y03110 | XP_018220319.1 | 99 | 0 |
| SPGP0Y00600 | XP_018220076.1 | 99 | 0 |
| SPGP0CV00110 | XP_018222564.1 | 99 | 0 |
| SPGP0CP00200 | XP_018223816.1 | 99 | 0 |
| SPGP0AT00380 | XP_018219059.1 | 99 | 0 |
| SPGP0L02260 | XP_018218810.1 | 99 | 0 |
| SPGP0AA01580 | XP_018220433.1 | 99 | 0 |
| SPGP0CU02000 | XP_018220634.1 | 99 | 0 |
| SPGP0L01130 | XP_018218921.1 | 99 | 0 |
| SPGP0EO00140 | XP_018221151.1 | 99 | 0 |
| SPGP0DF00160 | XP_018219192.1 | 99 | 0 |
| SPGP0AA01860 | XP_018220406.1 | 99 | 0 |
| SPGP0CW03250 | XP_018219239.1 | 99 | 0 |
| SPGP0AN01000 | XP_018224006.1 | 99 | 0 |
| SPGP0AU00100 | XP_018223067.1 | 99 | 0 |
| SPGP0R00800 | XP_018220019.1 | 99 | 0 |
| SPGP0CW00330 | XP_018219522.1 | 99 | 0 |
| SPGP0GQ00130 | XP_018219556.1 | 99 | 0 |
| SPGP0CU02290 | XP_018220605.1 | 99 | 0 |
| SPGP0CW03460 | XP_018219218.1 | 99 | 0 |
| SPGP0IO00110 | XP_018221984.1 | 97 | 0 |
| SPGP0L02570 | XP_018218779.1 | 99 | 0 |
| SPGP0CX01340 | XP_018222152.1 | 99 | 0 |
| SPGP0CU01000 | XP_018220730.1 | 99 | 0 |
| SPGP0Y02210 | XP_018220230.1 | 99 | 0 |
| SPGP0CP01180 | XP_018223720.1 | 99 | 0 |
| SPGP0U00790 | XP_018222999.1 | 99 | 0 |
| SPGP0Y01270 | XP_018220142.1 | 99 | 0 |
| SPGP0Y00430 | XP_018220060.1 | 99 | 0 |
| SPGP0CU00280 | XP_018220801.1 | 99 | 0 |
| SPGP0L01500 | XP_018218884.1 | 99 | 0 |
| SPGP0AA01170 | XP_018220473.1 | 99 | 0 |
| SPGP0AF01360 | XP_018222393.1 | 99 | 0 |
| SPGP0AU00220 | XP_018223077.1 | 99 | 0 |
| SPGP0CW01210 | XP_018219436.1 | 99 | 0 |
| SPGP0Y02040 | XP_018220214.1 | 99 | 0 |
| SPGP0DB00280 | XP_018223902.1 | 99 | 0 |
| SPGP0CP00930 | XP_018223746.1 | 99 | 1 |
| SPGP0BE00230 | XP_018223355.1 | 99 | 0 |
| SPGP0CW02290 | XP_018219332.1 | 99 | 0 |

Create PDF in your applications with the Pdfcrowd HTML to PDF API PDFCROWD

| | | | |
|---|---|---|---|
| SPGP0AF00640 | XP_018222318.1 | 99 | 0 |
| SPGP0CP01570 | XP_018223682.1 | 99 | 0 |
| SPGP0AN01160 | XP_018224022.1 | 99 | 0 |
| SPGP0Y00580 | XP_018220074.1 | 99 | 0 |
| SPGP0R00530 | XP_018219993.1 | 99 | 0 |
| SPGP0AJ00680 | XP_018221840.1 | 99 | 0 |
| SPGP0CP00220 | XP_018223814.1 | 99 | 0 |
| SPGP0L00550 | XP_018218977.1 | 99 | 0 |
| SPGP0U00470 | XP_018223027.1 | 99 | 0 |
| SPGP0Y01700 | XP_018220184.1 | 99 | 0 |
| SPGP0IC00120 | XP_018222531.1 | 100 | 0 |
| SPGP0DE00550 | XP_018223872.1 | 99 | 0 |
| SPGP0L01760 | XP_018218859.1 | 99 | 0 |
| SPGP0AA00160 | XP_018220574.1 | 99 | 0 |
| SPGP0Y00210 | XP_018220038.1 | 99 | 0 |
| SPGP0CU01980 | XP_018220636.1 | 99 | 0 |
| SPGP0AF01460 | XP_018222404.1 | 99 | 0 |
| SPGP0AE00150 | XP_018222497.1 | 99 | 0 |
| SPGP0EO00160 | XP_018221153.1 | 100 | 0 |
| SPGP0CX01400 | XP_018222158.1 | 99 | 0 |
| SPGP0FP00140 | XP_018223341.1 | 99 | 0 |
| SPGP0CW02120 | XP_018219349.1 | 100 | 0 |
| SPGP0CM01530 | XP_018223173.1 | 99 | 0 |
| SPGP0CP02190 | XP_018223621.1 | 99 | 0 |
| SPGP0AF00120 | XP_018222267.1 | 99 | 0 |
| SPGP0AA00120 | XP_018220578.1 | 99 | 0 |
| SPGP0CU01080 | XP_018220722.1 | 99 | 1 |
| SPGP0CU02420 | XP_018220592.1 | 99 | 0 |
| SPGP0CU01880 | XP_018220646.1 | 99 | 0 |
| SPGP0JB00170 | XP_018222562.1 | 99 | 0 |
| SPGP0Y00450 | XP_018220062.1 | 99 | 0 |
| SPGP0CW00510 | XP_018219504.1 | 99 | 0 |
| SPGP0CM00890 | XP_018223236.1 | 99 | 0 |
| SPGP0IZ00160 | XP_018220344.1 | 99 | 0 |
| SPGP0AQ00140 | XP_018223408.1 | 99 | 0 |
| SPGP0U00980 | XP_018222981.1 | 99 | 0 |
| SPGP0CX01050 | XP_018222124.1 | 99 | 0 |
| SPGP0AA01620 | XP_018220429.1 | 99 | 0 |
| SPGP0Y01350 | XP_018220150.1 | 99 | 0 |
| SPGP0CP00990 | XP_018223740.1 | 99 | 0 |

Create PDF in your applications with the Pdfcrowd HTML to PDF API PDFCROWD

| | | | |
|---|---|---|---|
| SPGP0L02210 | XP_018218815.1 | 99 | 0 |
| SPGP0U01810 | XP_018222900.1 | 99 | 0 |
| SPGP0CP00240 | XP_018223812.1 | 99 | 0 |
| SPGP0CP02720 | XP_018223569.1 | 99 | 0 |
| SPGP0AA00490 | XP_018220541.1 | 99 | 0 |
| SPGP0AA00140 | XP_018220576.1 | 98 | 2 |
| SPGP0AA00720 | XP_018220519.1 | 99 | 0 |
| SPGP0EB00100 | XP_018223321.1 | 99 | 0 |
| SPGP0L01040 | XP_018218928.1 | 99 | 0 |
| SPGP0EM00150 | XP_018223064.1 | 99 | 0 |
| SPGP0AF00310 | XP_018222285.1 | 99 | 0 |
| SPGP0CP00400 | XP_018223796.1 | 99 | 0 |
| SPGP0EA00130 | XP_018222480.1 | 99 | 0 |
| SPGP0AF00140 | XP_018222269.1 | 99 | 0 |
| SPGP0Z00100 | XP_018222555.1 | 99 | 0 |
| SPGP0CP01480 | XP_018223691.1 | 99 | 0 |
| SPGP0CM01230 | XP_018223202.1 | 99 | 0 |
| SPGP0CM00580 | XP_018223266.1 | 99 | 0 |
| SPGP0AT00220 | XP_018219044.1 | 99 | 0 |
| SPGP0EO00400 | XP_018221177.1 | 99 | 0 |
| SPGP0L02500 | XP_018218786.1 | 99 | 0 |
| SPGP0L00120 | XP_018219019.1 | 99 | 0 |
| SPGP0U00750 | XP_018223002.1 | 100 | 0 |
| SPGP0AF02150 | XP_018222473.1 | 99 | 0 |
| SPGP0CW02680 | XP_018219295.1 | 99 | 0 |
| SPGP0U00960 | XP_018222983.1 | 99 | 0 |
| SPGP0EO00250 | XP_018221162.1 | 100 | 0 |
| SPGP0Y02970 | XP_018220306.1 | 100 | 0 |
| SPGP0V01070 | XP_018221680.1 | 99 | 0 |
| SPGP0IN00180 | XP_018221658.1 | 99 | 0 |
| SPGP0F00210 | XP_018221190.1 | 98 | 1 |
| SPGP0V00680 | XP_018221718.1 | 99 | 0 |
| SPGP0AT01540 | XP_018219171.1 | 100 | 0 |
| SPGP0AF01290 | XP_018222385.1 | 99 | 0 |
| SPGP0CW01100 | XP_018219446.1 | 99 | 0 |
| SPGP0CP01410 | XP_018223698.1 | 99 | 0 |
| SPGP0CW03140 | XP_018219249.1 | 99 | 0 |
| SPGP0EA00110 | XP_018222478.1 | 99 | 0 |
| SPGP0Y01660 | XP_018220180.1 | 99 | 0 |
| SPGP0JK00170 | XP_018223439.1 | 99 | 0 |

Create PDF in your applications with the Pdfcrowd HTML to PDF API PDFCROWD

| | | | |
|---|---|---|---|
| SPGP0L01990 | XP_018218837.1 | 99 | 0 |
| SPGP0R00130 | XP_018219955.1 | 99 | 0 |
| SPGP0CW02510 | XP_018219310.1 | 99 | 0 |
| SPGP0AA01290 | XP_018220461.1 | 99 | 0 |
| SPGP0L02410 | XP_018218795.1 | 99 | 0 |
| SPGP0U00510 | XP_018223023.1 | 99 | 0 |
| SPGP0CX00830 | XP_018222102.1 | 99 | 0 |
| SPGP0HI00250 | XP_018219951.1 | 99 | 0 |
| SPGP0CM00550 | XP_018223269.1 | 99 | 0 |
| SPGP0AT00200 | XP_018219042.1 | 99 | 0 |
| SPGP0AJ00430 | XP_018221816.1 | 99 | 0 |
| SPGP0V00590 | XP_018221727.1 | 99 | 0 |
| SPGP0R00450 | XP_018219986.1 | 99 | 0 |
| SPGP0L01970 | XP_018218839.1 | 99 | 0 |
| SPGP0L00100 | XP_018219021.1 | 99 | 1 |
| SPGP0AF02130 | XP_018222471.1 | 99 | 0 |
| SPGP0CW02280 | XP_018219333.1 | 99 | 0 |
| SPGP0CW02340 | XP_018219327.1 | 99 | 0 |
| SPGP0GJ00130 | XP_018223379.1 | 100 | 0 |
| SPGP0CM00160 | XP_018223308.1 | 99 | 0 |
| SPGP0AF01480 | XP_018222406.1 | 99 | 0 |
| SPGP0CP01490 | XP_018223690.1 | 99 | 0 |
| SPGP0IN00200 | XP_018221656.1 | 99 | 0 |
| SPGP0CM01740 | XP_018223152.1 | 99 | 0 |
| SPGP0CM02210 | XP_018223108.1 | 99 | 0 |
| SPGP0AT00830 | XP_018219102.1 | 99 | 0 |
| SPGP0CU01900 | XP_018220644.1 | 99 | 0 |
| SPGP0AN00670 | XP_018223974.1 | 99 | 0 |
| SPGP0IC00100 | XP_018222533.1 | 99 | 0 |
| SPGP0AE00330 | XP_018222513.1 | 99 | 0 |
| SPGP0CP00440 | XP_018223792.1 | 99 | 0 |
| SPGP0Y00180 | XP_018220035.1 | 99 | 0 |
| SPGP0CU02220 | XP_018220612.1 | 99 | 0 |
| SPGP0F02910 | XP_018221449.1 | 99 | 0 |
| SPGP0K00470 | XP_018222789.1 | 99 | 0 |
| SPGP0GJ00110 | XP_018223381.1 | 99 | 0 |
| SPGP0AA00310 | XP_018220559.1 | 99 | 0 |
| SPGP0AJ00620 | XP_018221834.1 | 100 | 0 |
| SPGP0CU00520 | XP_018220777.1 | 100 | 0 |
| SPGP0CU01920 | XP_018220642.1 | 100 | 0 |

Create PDF in your applications with the Pdfcrowd HTML to PDF API PDFCROWD

| | | | |
|---|---|---|---|
| SPGP0L01640 | XP_018218870.1 | 99 | 0 |
| SPGP0L02140 | XP_018218823.1 | 99 | 0 |
| SPGP0CX02200 | XP_018222234.1 | 99 | 0 |
| SPGP0AF01270 | XP_018222383.1 | 99 | 0 |
| SPGP0AA01020 | XP_018220488.1 | 99 | 0 |
| SPGP0DB00350 | XP_018223908.1 | 99 | 0 |
| SPGP0CP02410 | XP_018223600.1 | 99 | 0 |
| SPGP0AJ00300 | XP_018221804.1 | 99 | 0 |
| SPGP0HI00210 | XP_018219947.1 | 99 | 0 |
| SPGP0CX01010 | XP_018222120.1 | 99 | 0 |
| SPGP0CW01550 | XP_018219403.1 | 99 | 0 |
| SPGP0AA00290 | XP_018220561.1 | 99 | 0 |
| SPGP0U00690 | XP_018223006.1 | 99 | 0 |
| SPGP0CP02040 | XP_018223636.1 | 99 | 0 |
| SPGP0CX00990 | XP_018222118.1 | 97 | 2 |
| SPGP0L01660 | XP_018218868.1 | 99 | 0 |
| SPGP0AJ01010 | XP_018221873.1 | 99 | 0 |
| SPGP0U01320 | XP_018222950.1 | 99 | 0 |
| SPGP0AE00250 | XP_018222506.1 | 99 | 0 |
| SPGP0CX00520 | XP_018222072.1 | 99 | 0 |
| SPGP0IS00110 | XP_018223095.1 | 99 | 0 |
| SPGP0CM01150 | XP_018223210.1 | 99 | 0 |
| SPGP0CP00150 | XP_018223821.1 | 99 | 0 |
| SPGP0CX01590 | XP_018222175.1 | 98 | 1 |
| SPGP0AT00630 | XP_018219083.1 | 99 | 0 |
| SPGP0CUtrna26S | DI49_2374 | 100 | 0 |
| SPGP0L01910 | XP_018218844.1 | 99 | 0 |
| SPGP0CP01770 | XP_018223662.1 | 99 | 0 |
| SPGP0AJ00780 | XP_018221850.1 | 99 | 0 |
| SPGP0CW02470 | XP_018219314.1 | 99 | 0 |
| SPGP0AF01510 | XP_018222409.1 | 99 | 0 |
| SPGP0AU00240 | XP_018223079.1 | 99 | 0 |
| SPGP0L02600 | XP_018218776.1 | 99 | 0 |
| SPGP0CM02290 | XP_018223100.1 | 99 | 1 |
| SPGP0F02660 | XP_018221426.1 | 99 | 0 |
| SPGP0F02970 | XP_018221454.1 | 99 | 0 |
| SPGP0J02740 | XP_018219675.1 | 99 | 0 |
| SPGP0Ftrna14M | DI49_5351 | 100 | 0 |
| SPGP0F01320 | XP_018221296.1 | 99 | 0 |
| SPGP0CP02210 | XP_018223619.1 | 99 | 0 |

Create PDF in your applications with the Pdfcrowd HTML to PDF API PDFCROWD

| | | | |
|---|---|---|---|
| SPGP0R00490 | XP_018219990.1 | 99 | 0 |
| SPGP0CU02070 | XP_018220627.1 | 99 | 0 |
| SPGP0CP02290 | XP_018223611.1 | 99 | 0 |
| SPGP0F02360 | XP_018221397.1 | 99 | 0 |
| SPGP0AF01850 | XP_018222443.1 | 99 | 0 |
| SPGP0J00920 | XP_018219859.1 | 99 | 1 |
| SPGP0AA01090 | XP_018220481.1 | 99 | 0 |
| SPGP0CV00390 | XP_018222591.1 | 99 | 0 |
| SPGP0Y01600 | XP_018220174.1 | 99 | 0 |
| SPGP0J02290 | XP_018219720.1 | 99 | 0 |
| SPGP0J02560 | XP_018219693.1 | 99 | 0 |
| SPGP0AA00200 | XP_018220570.1 | 99 | 0 |
| SPGP0CUtrna31L | DI49_3806 | 99 | 0 |
| SPGP0CW01830 | XP_018219377.1 | 99 | 0 |
| SPGP0F01770 | XP_018221339.1 | 99 | 0 |
| SPGP0F01950 | XP_018221357.1 | 99 | 0 |
| SPGP0U00380 | XP_018223035.1 | 99 | 0 |
| SPGP0CU00350 | XP_018220794.1 | 99 | 0 |
| SPGP0U00650 | XP_018223010.1 | 99 | 0 |
| SPGP0R00240 | XP_018219966.1 | 99 | 0 |
| SPGP0AA01700 | XP_018220421.1 | 99 | 0 |
| SPGP0CM01680 | XP_018223158.1 | 99 | 0 |
| SPGP0AN01170 | XP_018224023.1 | 99 | 0 |
| SPGP0F02950 | XP_018221452.1 | 99 | 0 |
| SPGP0CM00800 | XP_018223245.1 | 99 | 0 |
| SPGP0U01920 | XP_018222889.1 | 99 | 0 |
| SPGP0DE00370 | XP_018223854.1 | 99 | 0 |
| SPGP0V00230 | XP_018221760.1 | 99 | 0 |
| SPGP0AJ01140 | XP_018221885.1 | 99 | 0 |
| SPGP0L01620 | XP_018218872.1 | 99 | 0 |
| SPGP0X00270 | XP_018221476.1 | 99 | 0 |
| SPGP0X01780 | XP_018221621.1 | 99 | 0 |
| SPGP0CW03090 | XP_018219254.1 | 100 | 0 |
| SPGP0DA01370 | XP_018222618.1 | 99 | 0 |
| SPGP0L02670 | XP_018218770.1 | 97 | 0 |
| SPGP0AA00420 | XP_018220548.1 | 98 | 0 |
| SPGP0CM01760 | XP_018223150.1 | 99 | 0 |
| SPGP0Y02550 | XP_018220264.1 | 99 | 0 |
| SPGP0F02790 | XP_018221439.1 | 99 | 0 |
| SPGP0AN00900 | XP_018223996.1 | 99 | 0 |

Create PDF in your applications with the Pdfcrowd HTML to PDF API PDFCROWD

| | | | |
|---|---|---|---|
| SPGP0AT00360 | XP_018219057.1 | 99 | 0 |
| SPGP0J02690 | XP_018219680.1 | 99 | 0 |
| SPGP0X01920 | XP_018221635.1 | 99 | 0 |
| SPGP0X00750 | XP_018221523.1 | 99 | 0 |
| SPGP0F02890 | XP_018221448.1 | 99 | 0 |
| SPGP0CM01600 | XP_018223166.1 | 99 | 0 |
| SPGP0Y00550 | XP_018220072.1 | 99 | 0 |
| SPGP0CW03540 | XP_018219210.1 | 96 | 2 |
| SPGP0J03160 | XP_018219633.1 | 99 | 0 |
| SPGP0DN00100 | XP_018221992.1 | 99 | 0 |
| SPGP0CWtrna4G | DI49_4977 | 100 | 0 |
| SPGP0CU00620 | XP_018220766.1 | 99 | 0 |
| SPGP0AN01070 | XP_018224013.1 | 100 | 0 |
| SPGP0AF01680 | XP_018222426.1 | 99 | 0 |
| SPGP0DB00410 | XP_018223914.1 | 99 | 0 |
| SPGP0X01480 | XP_018221592.1 | 99 | 0 |
| SPGP0DB00130 | XP_018223887.1 | 99 | 0 |
| SPGP0X00280 | XP_018221477.1 | 99 | 0 |
| SPGP0AF00500 | XP_018222304.1 | 99 | 0 |
| SPGP0CU00300 | XP_018220799.1 | 99 | 0 |
| SPGP0Y00160 | XP_018220033.1 | 98 | 0 |
| SPGP0Y01900 | XP_018220201.1 | 99 | 0 |
| SPGP0H01550 | XP_018220994.1 | 99 | 0 |
| SPGP0AT00960 | XP_018219114.1 | 99 | 0 |
| SPGP0X01980 | XP_018221641.1 | 99 | 0 |
| SPGP0CU00260 | XP_018220803.1 | 99 | 0 |
| SPGP0J02000 | XP_018219749.1 | 99 | 0 |
| SPGP0AF01530 | XP_018222411.1 | 99 | 0 |
| SPGP0AN00950 | XP_018224001.1 | 99 | 0 |
| SPGP0J00400 | XP_018219908.1 | 99 | 0 |
| SPGP0CP02860 | XP_018223555.1 | 99 | 0 |
| SPGP0S00380 | XP_018224086.1 | 99 | 0 |
| SPGP0J02150 | XP_018219734.1 | 99 | 0 |
| SPGP0CN00860 | XP_018220886.1 | 99 | 0 |
| SPGP0AF00480 | XP_018222302.1 | 99 | 0 |
| SPGP0DA00420 | XP_018222712.1 | 99 | 0 |
| SPGP0DA00350 | XP_018222719.1 | 99 | 0 |
| SPGP0AJ01070 | XP_018221878.1 | 99 | 0 |
| SPGP0F00230 | XP_018221192.1 | 97 | 1 |
| SPGP0H01070 | XP_018221038.1 | 99 | 0 |

Create PDF in your applications with the Pdfcrowd HTML to PDF API PDFCROWD

| | | | |
|---|---|---|---|
| SPGP0L00390 | XP_018218993.1 | 98 | 0 |
| SPGP0L02470 | XP_018218789.1 | 99 | 0 |
| SPGP0X01150 | XP_018221559.1 | 99 | 0 |
| SPGP0CM01170 | XP_018223208.1 | 99 | 0 |
| SPGP0X01620 | XP_018221605.1 | 99 | 0 |
| SPGP0CW01990 | XP_018219361.1 | 99 | 0 |
| SPGP0CW01190 | XP_018219438.1 | 99 | 0 |
| SPGP0V00160 | XP_018221767.1 | 96 | 2 |
| SPGP0CW00900 | XP_018219466.1 | 99 | 0 |
| SPGP0Y02960 | XP_018220305.1 | 99 | 0 |
| SPGP0U00170 | XP_018223053.1 | 99 | 0 |
| SPGP0Y00770 | XP_018220093.1 | 99 | 0 |
| SPGP0DA01130 | XP_018222642.1 | 99 | 0 |
| SPGP0CU01240 | XP_018220708.1 | 99 | 0 |
| SPGP0CPtrna10Y | DI49_2382 | 100 | 0 |
| SPGP0HR00100 | XP_018222852.1 | 97 | 0 |
| SPGP0CN00730 | XP_018220898.1 | 100 | 0 |
| SPGP0Y02750 | XP_018220284.1 | 99 | 0 |
| SPGP0F00450 | XP_018221211.1 | 99 | 0 |
| SPGP0L00690 | XP_018218964.1 | 99 | 0 |
| SPGP0AF01990 | XP_018222457.1 | 99 | 0 |
| SPGP0F00490 | XP_018221214.1 | 99 | 0 |
| SPGP0Y00830 | XP_018220099.1 | 99 | 0 |
| SPGP0F00940 | XP_018221258.1 | 99 | 0 |
| SPGP0CM01320 | XP_018223193.1 | 99 | 0 |
| SPGP0DEtrna5V | DI49_0484 | 100 | 0 |
| SPGP0CW01800 | XP_018219380.1 | 99 | 0 |
| SPGP0F01540 | XP_018221317.1 | 99 | 0 |
| SPGP0X01410 | XP_018221585.1 | 99 | 0 |
| SPGP0S00430 | XP_018224091.1 | 99 | 0 |
| SPGP0F01010 | XP_018221265.1 | 99 | 0 |
| SPGP0J01400 | XP_018219808.1 | 99 | 0 |
| SPGP0J01150 | XP_018219832.1 | 99 | 0 |
| SPGP0H01130 | XP_018221033.1 | 99 | 0 |
| SPGP0CM01300 | XP_018223195.1 | 100 | 0 |
| SPGP0CX01090 | XP_018222128.1 | 99 | 0 |
| SPGP0JE00100 | XP_018222490.1 | 99 | 1 |
| SPGP0HN00110 | XP_018221987.1 | 99 | 0 |
| SPGP0CN00190 | XP_018220952.1 | 100 | 0 |

Create PDF in your applications with the Pdfcrowd HTML to PDF API PDFCROWD

| | | | |
|---|---|---|---|
| SPGP0CM00380 | XP_018223286.1 | 99 | 0 |
| SPGP0AA01270 | XP_018220463.1 | 99 | 0 |
| SPGP0CX01290 | XP_018222147.1 | 98 | 0 |
| SPGP0AT00130 | XP_018219035.1 | 100 | 0 |
| SPGP0DE00540 | XP_018223871.1 | 99 | 0 |
| SPGP0CW02380 | XP_018219323.1 | 99 | 0 |
| SPGP0F02200 | XP_018221381.1 | 99 | 0 |
| SPGP0J02810 | XP_018219668.1 | 100 | 0 |
| SPGP0J03370 | XP_018219612.1 | 99 | 0 |
| SPGP0CW03390 | XP_018219225.1 | 99 | 0 |
| SPGP0CU01100 | XP_018220720.1 | 99 | 0 |
| SPGP0Y03150 | XP_018220323.1 | 99 | 0 |
| SPGP0DE00140 | XP_018223831.1 | 99 | 0 |
| SPGP0F00430 | XP_018221210.1 | 99 | 0 |
| SPGP0Y02540 | XP_018220263.1 | 99 | 0 |
| SPGP0Y01870 | XP_018220198.1 | 99 | 0 |
| SPGP0Y02400 | XP_018220249.1 | 99 | 0 |
| SPGP0H01480 | XP_018221001.1 | 99 | 0 |
| SPGP0CU00830 | XP_018220746.1 | 99 | 0 |
| SPGP0H00330 | XP_018221111.1 | 99 | 0 |
| SPGP0L02250 | XP_018218811.1 | 98 | 1 |
| SPGP0L02310 | XP_018218805.1 | 99 | 0 |
| SPGP0GH00130 | XP_018222815.1 | 99 | 0 |
| SPGP0K00620 | XP_018222804.1 | 99 | 0 |
| SPGP0J02170 | XP_018219732.1 | 100 | 0 |
| SPGP0CW03470 | XP_018219217.1 | 99 | 0 |
| SPGP0X00660 | XP_018221514.1 | 99 | 0 |
| SPGP0H00930 | XP_018221052.1 | 99 | 0 |
| SPGP0F02090 | XP_018221371.1 | 99 | 0 |
| SPGP0CP02000 | XP_018223640.1 | 99 | 0 |
| SPGP0J02100 | XP_018219739.1 | 99 | 0 |
| SPGP0L00170 | XP_018219014.1 | 99 | 0 |
| SPGP0X01970 | XP_018221640.1 | 99 | 0 |
| SPGP0CW03060 | XP_018219257.1 | 98 | 2 |
| SPGP0H00870 | XP_018221058.1 | 99 | 0 |
| SPGP0AF01050 | XP_018222360.1 | 99 | 0 |
| SPGP0HO00120 | XP_018224032.1 | 99 | 0 |
| SPGP0AN00540 | XP_018223961.1 | 99 | 0 |
| SPGP0F02630 | XP_018221423.1 | 99 | 0 |
| SPGP0AF00330 | XP_018222287.1 | 99 | 0 |

Create PDF in your applications with the Pdfcrowd HTML to PDF API PDFCROWD

| | | | |
|---|---|---|---|
| SPGP0CP02470 | XP_018223594.1 | 99 | 0 |
| SPGP0X00140 | XP_018221463.1 | 97 | 0 |
| SPGP0H01360 | XP_018221011.1 | 93 | 7 |
| SPGP0CM01360 | XP_018223189.1 | 99 | 0 |
| SPGP0CPtrna25R | DI49_2410 | 100 | 0 |
| SPGP0GA00130 | XP_018223390.1 | 99 | 0 |
| SPGP0DA01290 | XP_018222626.1 | 98 | 1 |
| SPGP0H01510 | XP_018220998.1 | 99 | 0 |
| SPGP0R00780 | XP_018220017.1 | 99 | 0 |
| SPGP0AJ01460 | XP_018221916.1 | 98 | 2 |
| SPGP0AA01510 | XP_018220439.1 | 99 | 0 |
| SPGP0X01810 | XP_018221624.1 | 99 | 1 |
| SPGP0X01360 | XP_018221580.1 | 99 | 0 |
| SPGP0CW01890 | XP_018219371.1 | 99 | 0 |
| SPGP0JC00120 | XP_018220853.1 | 99 | 0 |
| SPGP0J01590 | XP_018219789.1 | 99 | 0 |
| SPGP0H00200 | XP_018221123.1 | 99 | 0 |
| SPGP0U01100 | XP_018222969.1 | 99 | 0 |
| SPGP0U02040 | XP_018222877.1 | 99 | 0 |
| SPGP0AJ01240 | XP_018221895.1 | 99 | 0 |
| SPGP0L00630 | XP_018218970.1 | 99 | 0 |
| SPGP0FE00200 | XP_018221972.1 | 99 | 0 |
| SPGP0DK00100 | XP_018222488.1 | 88 | 0 |
| SPGP0CW01680 | XP_018219392.1 | 99 | 0 |
| SPGP0AJ01630 | XP_018221931.1 | 99 | 0 |
| SPGP0DA01120 | XP_018222643.1 | 99 | 0 |
| SPGP0CX01880 | XP_018222203.1 | 99 | 0 |
| SPGP0F01550 | XP_018221318.1 | 99 | 0 |
| SPGP0U01510 | XP_018222931.1 | 97 | 2 |
| SPGP0X01230 | XP_018221567.1 | 99 | 0 |
| SPGP0S00220 | XP_018224071.1 | 99 | 0 |
| SPGP0CX01210 | XP_018222140.1 | 99 | 0 |
| SPGP0CP01390 | XP_018223700.1 | 99 | 0 |
| SPGP0CM00960 | XP_018223229.1 | 99 | 0 |
| SPGP0J00720 | XP_018219878.1 | 99 | 0 |
| SPGP0AT00990 | XP_018219117.1 | 99 | 0 |
| SPGP0AN00940 | XP_018224000.1 | 99 | 0 |
| SPGP0GQ00140 | XP_018219555.1 | 99 | 0 |
| SPGP0GLtrna6T | DI49_3176 | 100 | 0 |
| SPGP0CU00510 | XP_018220778.1 | 99 | 0 |

Create PDF in your applications with the Pdfcrowd HTML to PDF API PDFCROWD

| | | | |
|---|---|---|---|
| SPGP0CW02210 | XP_018219340.1 | 99 | 0 |
| SPGP0Y01030 | XP_018220118.1 | 99 | 0 |
| SPGP0DE00170 | XP_018223834.1 | 99 | 0 |
| SPGP0AC00100 | XP_018223460.1 | 90 | 0 |
| SPGP0DA00650 | XP_018222689.1 | 99 | 0 |
| SPGP0CX00910 | XP_018222110.1 | 99 | 0 |
| SPGP0F01740 | XP_018221336.1 | 99 | 0 |
| SPGP0CN00450 | XP_018220926.1 | 99 | 0 |
| SPGP0CN00500 | XP_018220921.1 | 99 | 0 |
| SPGP0Y02390 | XP_018220248.1 | 99 | 0 |
| SPGP0J01270 | XP_018219821.1 | 99 | 0 |
| SPGP0AJ00570 | XP_018221829.1 | 99 | 0 |
| SPGP0U00800 | XP_018222998.1 | 99 | 0 |
| SPGP0CP01730 | XP_018223667.1 | 99 | 0 |
| SPGP0CW00810 | XP_018219475.1 | 99 | 0 |
| SPGP0K00250 | XP_018222767.1 | 99 | 0 |
| SPGP0F00550 | XP_018221220.1 | 99 | 0 |
| SPGP0CP01860 | XP_018223654.1 | 99 | 0 |
| SPGP0Y02220 | XP_018220231.1 | 100 | 0 |
| SPGP0CU02140 | XP_018220620.1 | 99 | 0 |
| SPGP0J02760 | XP_018219673.1 | 99 | 0 |
| SPGP0CP02270 | XP_018223613.1 | 99 | 0 |
| SPGP0J03150 | XP_018219634.1 | 99 | 0 |
| SPGP0CW02420 | XP_018219319.1 | 99 | 0 |
| SPGP0V00960 | XP_018221691.1 | 99 | 0 |
| SPGP0H00770 | XP_018221068.1 | 100 | 0 |
| SPGP0AF01830 | XP_018222441.1 | 99 | 0 |
| SPGP0CW01450 | XP_018219413.1 | 99 | 0 |
| SPGP0J00940 | XP_018219857.1 | 98 | 0 |
| SPGP0IZ00120 | XP_018220340.1 | 99 | 0 |
| SPGP0U00970 | XP_018222982.1 | 99 | 0 |
| SPGP0JK00250 | XP_018223445.1 | 99 | 0 |
| SPGP0K00640 | XP_018222806.1 | 99 | 0 |
| SPGP0J02540 | XP_018219695.1 | 99 | 0 |
| SPGP0AT00140 | XP_018219036.1 | 99 | 0 |
| SPGP0CW01850 | XP_018219375.1 | 99 | 0 |
| SPGP0J02400 | XP_018219709.1 | 99 | 0 |
| SPGP0K00370 | XP_018222779.1 | 96 | 4 |
| SPGP0F01750 | XP_018221337.1 | 99 | 0 |
| SPGP0F01970 | XP_018221359.1 | 99 | 0 |

Create PDF in your applications with the Pdfcrowd HTML to PDF API PDFCROWD

| | | | |
|---|---|---|---|
| SPGP0CP01710 | XP_018223669.1 | 99 | 0 |
| SPGP0F01200 | XP_018221284.1 | 99 | 0 |
| SPGP0X00830 | XP_018221531.1 | 99 | 0 |
| SPGP0CP03270 | XP_018223513.1 | 99 | 0 |
| SPGP0J01280 | XP_018219820.1 | 99 | 0 |
| SPGP0CU00440 | XP_018220785.1 | 99 | 0 |
| SPGP0R00350 | XP_018219976.1 | 99 | 0 |
| SPGP0CX01660 | XP_018222182.1 | 99 | 0 |
| SPGP0AJ00950 | XP_018221867.1 | 99 | 0 |
| SPGP0CU02380 | XP_018220596.1 | 99 | 0 |
| SPGP0J01750 | XP_018219773.1 | 99 | 0 |
| SPGP0BE00250 | XP_018223353.1 | 99 | 0 |
| SPGP0H01090 | XP_018221036.1 | 99 | 0 |
| SPGP0CU01620 | XP_018220672.1 | 99 | 0 |
| SPGP0F02400 | XP_018221401.1 | 99 | 0 |
| SPGP0M00120 | XP_018221144.1 | 99 | 0 |
| SPGP0CP01580 | XP_018223681.1 | 99 | 0 |
| SPGP0IN00320 | XP_018221644.1 | 95 | 0 |
| SPGP0CM01250 | XP_018223200.1 | 99 | 0 |
| SPGP0L01350 | XP_018218899.1 | 99 | 0 |
| SPGP0J03490 | XP_018219599.1 | 99 | 0 |
| SPGP0J03630 | XP_018219586.1 | 99 | 0 |
| SPGP0J03140 | XP_018219635.1 | 99 | 0 |
| SPGP0FV00110 | XP_018223322.1 | 99 | 0 |
| SPGP0DA01350 | XP_018222620.1 | 99 | 0 |
| SPGP0CP03480 | XP_018223494.1 | 100 | 0 |
| SPGP0H00580 | XP_018221087.1 | 99 | 0 |
| SPGP0CW03110 | XP_018219252.1 | 99 | 0 |
| SPGP0AA00580 | XP_018220532.1 | 99 | 0 |
| SPGP0S00720 | XP_018224119.1 | 98 | 0 |
| SPGP0J02210 | XP_018219728.1 | 99 | 0 |
| SPGP0AA00230 | XP_018220567.1 | 99 | 0 |
| SPGP0F01400 | XP_018221304.1 | 99 | 0 |
| SPGP0X00770 | XP_018221525.1 | 99 | 0 |
| SPGP0H00470 | XP_018221097.1 | 99 | 0 |
| SPGP0CP00850 | XP_018223753.1 | 99 | 0 |
| SPGP0J02050 | XP_018219744.1 | 99 | 0 |
| SPGP0GH00170 | XP_018222819.1 | 99 | 0 |
| SPGP0CW00270 | XP_018219526.1 | 99 | 0 |
| SPGP0CN01020 | XP_018220870.1 | 99 | 0 |

Create PDF in your applications with the Pdfcrowd HTML to PDF API PDFCROWD

| | | | |
|---|---|---|---|
| SPGP0AA01180 | XP_018220472.1 | 99 | 0 |
| SPGP0U00600 | XP_018223015.1 | 99 | 0 |
| SPGP0AF02030 | XP_018222461.1 | 99 | 0 |
| SPGP0CW00410 | XP_018219514.1 | 99 | 0 |
| SPGP0J00950 | XP_018219844.1 | 99 | 0 |
| SPGP0CM02140 | XP_018223115.1 | 99 | 0 |
| SPGP0X00550 | XP_018221504.1 | 99 | 0 |
| SPGP0FE00190 | XP_018221971.1 | 99 | 0 |
| SPGP0U01640 | XP_018222918.1 | 99 | 0 |
| SPGP0ATtrna10F | DI49_5486 | 100 | 0 |
| SPGP0AF01620 | XP_018222420.1 | 99 | 0 |
| SPGP0Y00140 | XP_018220031.1 | 99 | 0 |
| SPGP0CX01980 | XP_018222213.1 | 99 | 0 |
| SPGP0F01880 | XP_018221350.1 | 99 | 0 |
| SPGP0J01450 | XP_018219803.1 | 99 | 0 |
| SPGP0L02180 | XP_018218818.1 | 99 | 0 |
| SPGP0AN00750 | XP_018223982.1 | 99 | 0 |
| SPGP0S00610 | XP_018224108.1 | 99 | 0 |
| SPGP0L00990 | XP_018218933.1 | 99 | 0 |
| SPGP0J01350 | XP_018219813.1 | 99 | 0 |
| SPGP0Y02890 | XP_018220298.1 | 99 | 0 |
| SPGP0AN00970 | XP_018224003.1 | 99 | 0 |
| SPGP0CP02800 | XP_018223561.1 | 99 | 0 |
| SPGP0DA00400 | XP_018222714.1 | 99 | 0 |
| SPGP0CN00880 | XP_018220884.1 | 99 | 0 |
| SPGP0DQ00100 | XP_018220025.1 | 99 | 0 |
| SPGP0CN01120 | XP_018220860.1 | 99 | 0 |
| SPGP0AJ01720 | XP_018221940.1 | 99 | 0 |
| SPGP0F00600 | XP_018221225.1 | 99 | 0 |
| SPGP0Y01810 | XP_018220194.1 | 99 | 0 |
| SPGP0J01970 | XP_018219752.1 | 99 | 0 |
| SPGP0AF01450 | XP_018222403.1 | 99 | 0 |
| SPGP0AE00310 | XP_018222512.1 | 99 | 0 |
| SPGP0AA00920 | XP_018220498.1 | 99 | 0 |
| SPGP0Y02110 | XP_018220221.1 | 96 | 1 |
| SPGP0AFtrna6W | DI49_2157 | 100 | 0 |
| SPGP0DA00750 | XP_018222678.1 | 99 | 0 |
| SPGP0AJ00710 | XP_018221843.1 | 99 | 0 |
| SPGP0F00660 | XP_018221231.1 | 100 | 0 |
| SPGP0J00550 | XP_018219894.1 | 99 | 0 |

Create PDF in your applications with the Pdfcrowd HTML to PDF API PDFCROWD

| | | | |
|---|---|---|---|
| SPGP0CM01920 | XP_018223134.1 | 99 | 0 |
| SPGP0AF00200 | XP_018222275.1 | 99 | 0 |
| SPGP0CM02130 | XP_018223116.1 | 99 | 0 |
| SPGP0X00820 | XP_018221530.1 | 99 | 0 |
| SPGP0CN00960 | XP_018220876.1 | 99 | 0 |
| SPGP0CM00740 | XP_018223251.1 | 99 | 0 |
| SPGP0CW02880 | XP_018219275.1 | 99 | 0 |
| SPGP0S00140 | XP_018224063.1 | 99 | 0 |
| SPGP0J01390 | XP_018219809.1 | 99 | 0 |
| SPGP0F01560 | XP_018221319.1 | 99 | 0 |
| SPGP0J01850 | XP_018219764.1 | 99 | 0 |
| SPGP0S00410 | XP_018224089.1 | 99 | 0 |
| SPGP0CN01040 | XP_018220868.1 | 99 | 0 |
| SPGP0V00570 | XP_018221729.1 | 99 | 0 |
| SPGP0CP03040 | XP_018223536.1 | 100 | 0 |
| SPGP0L01600 | XP_018218874.1 | 99 | 0 |
| SPGP0Y00850 | XP_018220101.1 | 99 | 1 |
| SPGP0F00960 | XP_018221260.1 | 99 | 0 |
| SPGP0CW01600 | XP_018219399.1 | 99 | 0 |
| SPGP0AJ01450 | XP_018221915.1 | 98 | 0 |
| SPGP0IH00160 | XP_018220387.1 | 99 | 0 |
| SPGP0X00920 | XP_018221539.1 | 99 | 0 |
| SPGP0J00650 | XP_018219885.1 | 99 | 0 |
| SPGP0CW01280 | XP_018219429.1 | 99 | 0 |
| SPGP0CP00650 | XP_018223772.1 | 99 | 0 |
| SPGP0CU01350 | XP_018220698.1 | 99 | 0 |
| SPGP0F01690 | XP_018221331.1 | 99 | 0 |
| SPGP0J01170 | XP_018219830.1 | 99 | 0 |
| SPGP0GG00180 | XP_018220833.1 | 99 | 0 |
| SPGP0CU01600 | XP_018220674.1 | 99 | 0 |
| SPGP0CW02410 | XP_018219320.1 | 99 | 0 |
| SPGP0F00890 | XP_018221253.1 | 99 | 0 |
| SPGP0AT01200 | XP_018219138.1 | 100 | 0 |
| SPGP0CP03610 | XP_018223483.1 | 99 | 0 |
| SPGP0CN00310 | XP_018220940.1 | 99 | 0 |
| SPGP0X00570 | XP_018221506.1 | 99 | 0 |
| SPGP0CW00550 | XP_018219500.1 | 99 | 0 |
| SPGP0CMtrna9Q | DI49_1374 | 100 | 0 |
| SPGP0DA00940 | XP_018222661.1 | 100 | 0 |
| SPGP0F00610 | XP_018221226.1 | 99 | 0 |

Create PDF in your applications with the Pdfcrowd HTML to PDF API PDFCROWD

| | | | |
|---|---|---|---|
| SPGP0GA00120 | XP_018223391.1 | 99 | 0 |
| SPGP0AJ01690 | XP_018221937.1 | 99 | 0 |
| SPGP0V00470 | XP_018221738.1 | 99 | 0 |
| SPGP0DA01540 | XP_018222601.1 | 97 | 0 |
| SPGP0DA01460 | XP_018222609.1 | 99 | 0 |
| SPGP0HZ00130 | XP_018219028.1 | 99 | 0 |
| SPGP0V01010 | XP_018221686.1 | 99 | 0 |
| SPGP0DA01330 | XP_018222622.1 | 99 | 0 |
| SPGP0Y01160 | XP_018220131.1 | 99 | 0 |
| SPGP0AJ00890 | XP_018221861.1 | 99 | 0 |
| SPGP0CU00720 | XP_018220757.1 | 99 | 0 |
| SPGP0IN00120 | XP_018221664.1 | 100 | 0 |
| SPGP0DE00290 | XP_018223846.1 | 99 | 0 |
| SPGP0L00740 | XP_018218959.1 | 99 | 0 |
| SPGP0H00910 | XP_018221054.1 | 99 | 0 |
| SPGP0EO00210 | XP_018221158.1 | 99 | 0 |
| SPGP0J03610 | XP_018219588.1 | 99 | 0 |
| SPGP0H01140 | XP_018221032.1 | 99 | 0 |
| SPGP0AJ00690 | XP_018221841.1 | 99 | 0 |
| SPGP0AF01030 | XP_018222358.1 | 99 | 0 |
| SPGP0CN00550 | XP_018220916.1 | 99 | 0 |
| SPGP0L02110 | XP_018218825.1 | 99 | 0 |
| SPGP0CP00890 | XP_018223750.1 | 99 | 0 |
| SPGP0X01830 | XP_018221626.1 | 98 | 0 |
| SPGP0AN00690 | XP_018223976.1 | 99 | 0 |
| SPGP0F02610 | XP_018221421.1 | 99 | 0 |
| SPGP0CW02990 | XP_018219264.1 | 99 | 0 |
| SPGP0CX02140 | XP_018222228.1 | 99 | 0 |
| SPGP0V00370 | XP_018221746.1 | 99 | 0 |
| SPGP0J03120 | XP_018219637.1 | 99 | 0 |
| SPGP0X00120 | XP_018221461.1 | 96 | 1 |
| SPGP0CP02580 | XP_018223583.1 | 99 | 0 |
| SPGP0CU01830 | XP_018220651.1 | 99 | 0 |
| SPGP0M00210 | XP_018221137.1 | 99 | 0 |
| SPGP0AF01780 | XP_018222436.1 | 99 | 0 |
| SPGP0IZ00250 | XP_018220353.1 | 99 | 0 |
| SPGP0CW01510 | XP_018219407.1 | 99 | 0 |
| SPGP0H00260 | XP_018221118.1 | 99 | 0 |
| SPGP0Itrna21D | DI49_4675 | 100 | 0 |
| SPGP0CX00600 | XP_018222080.1 | 100 | 0 |

Create PDF in your applications with the Pdfcrowd HTML to PDF API PDFCROWD

| | | | |
|---|---|---|---|
| SPGP0CM00180 | XP_018223306.1 | 99 | 0 |
| SPGP0AJ01260 | XP_018221897.1 | 99 | 0 |
| SPGP0CW03040 | XP_018219259.1 | 99 | 0 |
| SPGP0DA00340 | XP_018222720.1 | 99 | 0 |
| SPGP0CM00570 | XP_018223267.1 | 99 | 0 |
| SPGP0H01790 | XP_018220971.1 | 99 | 0 |
| SPGP0U01150 | XP_018222964.1 | 99 | 0 |
| SPGP0H01370 | XP_018221010.1 | 99 | 0 |
| SPGP0L02660 | XP_018218771.1 | 99 | 0 |
| SPGP0CX01510 | XP_018222168.1 | 99 | 0 |
| SPGP0Htrna9W | DI49_5211 | 100 | 0 |
| SPGP0S00300 | XP_018224079.1 | 99 | 0 |
| SPGP0J03510 | XP_018219597.1 | 100 | 0 |
| SPGP0CP02050 | XP_018223635.1 | 99 | 0 |
| SPGP0CM01580 | XP_018223168.1 | 99 | 0 |
| SPGP0CW02960 | XP_018219267.1 | 99 | 0 |
| SPGP0Htrna9H | DI49_4134 | 100 | 0 |
| SPGP0V00700 | XP_018221716.1 | 99 | 0 |
| SPGP0CM00980 | XP_018223227.1 | 99 | 0 |
| SPGP0J00700 | XP_018219880.1 | 99 | 0 |
| SPGP0GS00140 | XP_018222249.1 | 99 | 0 |
| SPGP0CW03120 | XP_018219251.1 | 99 | 0 |
| SPGP0Y02080 | XP_018220218.1 | 99 | 0 |
| SPGP0J01580 | XP_018219790.1 | 99 | 0 |
| SPGP0R00540 | XP_018219994.1 | 99 | 0 |
| SPGP0CW03610 | XP_018219204.1 | 99 | 0 |
| SPGP0Y02370 | XP_018220246.1 | 99 | 0 |
| SPGP0J02700 | XP_018219679.1 | 100 | 0 |
| SPGP0AJ00450 | XP_018221818.1 | 98 | 1 |
| SPGP0X01220 | XP_018221566.1 | 99 | 0 |
| SPGP0DA00890 | XP_018222665.1 | 99 | 0 |
| SPGP0L02650 | XP_018222485.1 | 99 | 0 |
| SPGP0J02650 | XP_018219684.1 | 99 | 0 |
| SPGP0X01960 | XP_018221639.1 | 99 | 0 |
| SPGP0CW03380 | XP_018219226.1 | 99 | 0 |
| SPGP0AJ01790 | XP_018221947.1 | 99 | 0 |
| SPGP0CX01370 | XP_018222155.1 | 99 | 0 |
| SPGP0K00270 | XP_018222769.1 | 99 | 0 |
| SPGP0CM00710 | XP_018223254.1 | 99 | 0 |
| SPGP0J00960 | XP_018219845.1 | 99 | 0 |

Create PDF in your applications with the Pdfcrowd HTML to PDF API PDFCROWD

| | | | |
|---|---|---|---|
| SPGP0H01120 | XP_018221034.1 | 100 | 0 |
| SPGP0AF01810 | XP_018222439.1 | 99 | 0 |
| SPGP0AT01630 | XP_018219180.1 | 99 | 0 |
| SPGP0F01910 | XP_018221353.1 | 99 | 0 |
| SPGP0CU02010 | XP_018220633.1 | 100 | 0 |
| SPGP0J02220 | XP_018219727.1 | 99 | 0 |
| SPGP0H01620 | XP_018220988.1 | 99 | 0 |
| SPGP0AT00320 | XP_018219053.1 | 99 | 0 |
| SPGP0AF01010 | XP_018222356.1 | 100 | 0 |
| SPGP0AT00480 | XP_018219069.1 | 99 | 0 |
| SPGP0J02830 | XP_018219666.1 | 99 | 0 |
| SPGP0CUtrna27R | DI49_3845 | 100 | 0 |
| SPGP0CW01520 | XP_018219406.1 | 99 | 0 |
| SPGP0AA01880 | XP_018220404.1 | 99 | 0 |
| SPGP0F01730 | XP_018221335.1 | 99 | 0 |
| SPGP0K00150 | XP_018222758.1 | 99 | 0 |
| SPGP0CP03250 | XP_018223515.1 | 99 | 0 |
| SPGP0S00600 | XP_018224107.1 | 99 | 0 |
| SPGP0U00340 | XP_018223038.1 | 99 | 0 |
| SPGP0Y02060 | XP_018220216.1 | 99 | 0 |
| SPGP0AJ00970 | XP_018221869.1 | 99 | 0 |
| SPGP0CP01550 | XP_018223684.1 | 99 | 0 |
| SPGP0F01060 | XP_018221270.1 | 99 | 0 |
| SPGP0DE00330 | XP_018223850.1 | 99 | 0 |
| SPGP0DE00620 | XP_018223879.1 | 99 | 0 |
| SPGP0U01960 | XP_018222885.1 | 99 | 1 |
| SPGP0CW01870 | XP_018219373.1 | 99 | 0 |
| SPGP0F02690 | XP_018221429.1 | 99 | 0 |
| SPGP0HW00150 | XP_018224038.1 | 99 | 0 |
| SPGP0J03750 | XP_018219574.1 | 99 | 0 |
| SPGP0J01610 | XP_018219787.1 | 99 | 0 |
| SPGP0CM00210 | XP_018223303.1 | 99 | 0 |
| SPGP0AT00720 | XP_018219091.1 | 98 | 2 |
| SPGP0X00950 | XP_018221542.1 | 99 | 0 |
| SPGP0CW01720 | XP_018219388.1 | 99 | 0 |
| SPGP0L00970 | XP_018218935.1 | 99 | 0 |
| SPGP0CU00500 | XP_018220779.1 | 99 | 0 |
| SPGP0F02420 | XP_018221403.1 | 99 | 0 |
| SPGP0AA00210 | XP_018220569.1 | 99 | 0 |
| SPGP0F01710 | XP_018221333.1 | 99 | 0 |

Create PDF in your applications with the Pdfcrowd HTML to PDF API PDFCROWD

| | | | |
|---|---|---|---|
| SPGP0X00710 | XP_018221519.1 | 100 | 0 |
| SPGP0CW02190 | XP_018219342.1 | 99 | 0 |
| SPGP0F00900 | XP_018221254.1 | 99 | 0 |
| SPGP0CP03640 | XP_018223480.1 | 99 | 0 |
| SPGP0AH00220 | XP_018223429.1 | 99 | 0 |
| SPGP0H00830 | XP_018221062.1 | 99 | 0 |
| SPGP0IU00150 | XP_018220333.1 | 99 | 0 |
| SPGP0CM00270 | XP_018223297.1 | 99 | 0 |
| SPGP0CW00700 | XP_018219486.1 | 99 | 0 |
| SPGP0CW00580 | XP_018219498.1 | 99 | 0 |
| SPGP0EO00300 | XP_018221167.1 | 99 | 0 |
| SPGP0Y02610 | XP_018220270.1 | 99 | 0 |
| SPGP0Y01790 | XP_018220192.1 | 99 | 0 |
| SPGP0DA00460 | XP_018222708.1 | 100 | 0 |
| SPGP0DA00930 | XP_018222662.1 | 99 | 0 |
| SPGP0X01130 | XP_018221558.1 | 99 | 0 |
| SPGP0L01060 | XP_018218926.1 | 99 | 0 |
| SPGP0X01020 | XP_018221549.1 | 99 | 0 |
| SPGP0DF00220 | XP_018219186.1 | 98 | 0 |
| SPGP0AE00420 | XP_018222521.1 | 100 | 0 |
| SPGP0H01300 | XP_018221017.1 | 100 | 0 |
| SPGP0HI00230 | XP_018219949.1 | 99 | 0 |
| SPGP0J02470 | XP_018219702.1 | 99 | 0 |
| SPGP0X01990 | XP_018221642.1 | 99 | 0 |
| SPGP0H01640 | XP_018220986.1 | 99 | 0 |
| SPGP0X01440 | XP_018221588.1 | 99 | 0 |
| SPGP0L00850 | XP_018218948.1 | 99 | 0 |
| SPGP0H00370 | XP_018221107.1 | 99 | 0 |
| SPGP0CP03360 | XP_018223505.1 | 99 | 0 |
| SPGP0CP02530 | XP_018223588.1 | 99 | 0 |
| SPGP0AF01160 | XP_018222372.1 | 99 | 0 |
| SPGP0CP02820 | XP_018223559.1 | 100 | 0 |
| SPGP0V00120 | XP_018221771.1 | 97 | 0 |
| SPGP0CW00130 | XP_018219541.1 | 99 | 0 |
| SPGP0CV00280 | XP_018222579.1 | 99 | 0 |
| SPGP0DS00160 | XP_018220968.1 | 99 | 0 |
| SPGP0AJ01700 | XP_018221938.1 | 99 | 0 |
| SPGP0J00390 | XP_018219909.1 | 99 | 0 |
| SPGP0AF01570 | XP_018222415.1 | 99 | 0 |
| SPGP0AA01300 | XP_018220460.1 | 99 | 0 |

Create PDF in your applications with the Pdfcrowd HTML to PDF API PDFCROWD

| | | | |
|---|---|---|---|
| SPGP0J00110 | XP_018219935.1 | 91 | 1 |
| SPGP0X01660 | XP_018221609.1 | 99 | 0 |
| SPGP0AFtrna15L | DI49_2147 | 99 | 0 |
| SPGP0AN01120 | XP_018224018.1 | 99 | 0 |
| SPGP0GP00110 | XP_018223416.1 | 99 | 0 |
| SPGP0R00190 | XP_018219961.1 | 99 | 0 |
| SPGP0IE00100 | XP_018221134.1 | 93 | 0 |
| SPGP0DE00580 | XP_018223875.1 | 99 | 0 |
| SPGP0AF00220 | XP_018222277.1 | 99 | 0 |
| SPGP0DA00280 | XP_018222725.1 | 100 | 0 |
| SPGP0J00570 | XP_018219892.1 | 99 | 0 |
| SPGP0CW00820 | XP_018219474.1 | 99 | 0 |
| SPGP0CN00150 | XP_018220956.1 | 99 | 0 |
| SPGP0J01650 | XP_018219783.1 | 99 | 0 |
| SPGP0F01390 | XP_018221303.1 | 99 | 0 |
| SPGP0J02890 | XP_018219660.1 | 99 | 0 |
| SPGP0F01500 | XP_018221313.1 | 99 | 0 |
| SPGP0CP00670 | XP_018223770.1 | 99 | 0 |
| SPGP0S00160 | XP_018224065.1 | 100 | 0 |
| SPGP0J01370 | XP_018219811.1 | 99 | 0 |
| SPGP0CN01060 | XP_018220866.1 | 99 | 0 |
| SPGP0X00720 | XP_018221520.1 | 99 | 0 |
| SPGP0U01170 | XP_018222963.1 | 99 | 0 |
| SPGP0IH00140 | XP_018220385.1 | 99 | 0 |
| SPGP0CW01260 | XP_018219431.1 | 99 | 0 |
| SPGP0CU01330 | XP_018220700.1 | 99 | 0 |
| SPGP0X01100 | XP_018221556.1 | 99 | 0 |
| SPGP0J01110 | XP_018219836.1 | 99 | 0 |
| SPGP0GJtrna8P | DI49_4736 | 100 | 0 |
| SPGP0FP00240 | XP_018223331.1 | 99 | 0 |
| SPGP0Y00870 | XP_018220103.1 | 99 | 0 |
| SPGP0CX02160 | XP_018222230.1 | 99 | 0 |
| SPGP0H00350 | XP_018221109.1 | 99 | 0 |
| SPGP0AJ01110 | XP_018221882.1 | 99 | 0 |
| SPGP0CN00330 | XP_018220938.1 | 99 | 0 |
| SPGP0AA01080 | XP_018220482.1 | 99 | 0 |
| SPGP0FF00100 | XP_018223402.1 | 99 | 0 |
| SPGP0Y02580 | XP_018220267.1 | 99 | 0 |
| SPGP0R00630 | XP_018220003.1 | 99 | 0 |

Create PDF in your applications with the Pdfcrowd HTML to PDF API PDFCROWD

| | | | |
|---|---|---|---|
| SPGP0Y00720 | XP_018220088.1 | 99 | 0 |
| SPGP0AJ00700 | XP_018221842.1 | 99 | 0 |
| SPGP0X01260 | XP_018221570.1 | 100 | 0 |
| SPGP0CV00290 | XP_018222580.1 | 99 | 0 |
| SPGP0CW03450 | XP_018219219.1 | 99 | 0 |
| SPGP0CU01640 | XP_018220670.1 | 99 | 0 |
| SPGP0DE00100 | XP_018223827.1 | 99 | 0 |
| SPGP0F02850 | XP_018221444.1 | 97 | 2 |
| SPGP0CMtrna23S | DI49_5523 | 100 | 0 |
| SPGP0H00340 | XP_018221110.1 | 99 | 0 |
| SPGP0AA00680 | XP_018220523.1 | 99 | 0 |
| SPGP0L00790 | XP_018218954.1 | 99 | 0 |
| SPGP0DA00570 | XP_018222697.1 | 99 | 0 |
| SPGP0HZ00110 | XP_018219030.1 | 99 | 0 |
| SPGP0AJ00560 | XP_018221828.1 | 99 | 0 |
| SPGP0AN00910 | XP_018223997.1 | 96 | 3 |
| SPGP0L01820 | XP_018218853.1 | 99 | 0 |
| SPGP0CM00390 | XP_018223285.1 | 99 | 0 |
| SPGP0Y01580 | XP_018220172.1 | 99 | 0 |
| SPGP0H01420 | XP_018221006.1 | 99 | 0 |
| SPGP0J02130 | XP_018219736.1 | 99 | 0 |
| SPGP0DB00230 | XP_018223897.1 | 99 | 0 |
| SPGP0CUtrna21A | DI49_5601 | 100 | 0 |
| SPGP0AF02060 | XP_018222464.1 | 99 | 0 |
| SPGP0HO00140 | XP_018224030.1 | 99 | 0 |
| SPGP0U00150 | XP_018223055.1 | 99 | 0 |
| SPGP0HO00110 | XP_018224033.1 | 99 | 0 |
| SPGP0DA00670 | XP_018222687.1 | 99 | 0 |
| SPGP0AN01140 | XP_018224020.1 | 100 | 0 |
| SPGP0Y01130 | XP_018220128.1 | 99 | 0 |
| SPGP0H00790 | XP_018221066.1 | 99 | 0 |
| SPGP0F02670 | XP_018221427.1 | 99 | 0 |
| SPGP0CU02240 | XP_018220610.1 | 99 | 0 |
| SPGP0IN00290 | XP_018221647.1 | 99 | 0 |
| SPGP0CW02030 | XP_018219357.1 | 98 | 0 |
| SPGP0L01410 | XP_018218893.1 | 100 | 0 |
| SPGP0J03260 | XP_018219623.1 | 99 | 0 |
| SPGP0DA01250 | XP_018222630.1 | 99 | 0 |
| SPGP0CW01960 | XP_018219364.1 | 99 | 0 |
| SPGP0H00530 | XP_018221092.1 | 99 | 0 |

Create PDF in your applications with the Pdfcrowd HTML to PDF API PDFCROWD

| | | | |
|---|---|---|---|
| SPGP0CP01030 | XP_018223736.1 | 99 | 0 |
| SPGP0F00200 | XP_018221189.1 | 98 | 0 |
| SPGP0H00240 | XP_018221120.1 | 99 | 0 |
| SPGP0J00980 | XP_018219847.1 | 99 | 0 |
| SPGP0X00100 | XP_018221458.1 | 96 | 0 |
| SPGP0X00450 | XP_018221494.1 | 99 | 0 |
| SPGP0AJ00530 | XP_018221825.1 | 99 | 0 |
| SPGP0CU01710 | XP_018220663.1 | 97 | 3 |
| SPGP0CU00730 | XP_018220756.1 | 99 | 0 |
| SPGP0F02940 | XP_018221451.1 | 99 | 0 |
| SPGP0AF00760 | XP_018222330.1 | 99 | 0 |
| SPGP0X00990 | XP_018221546.1 | 99 | 0 |
| SPGP0CW03630 | XP_018219202.1 | 97 | 0 |
| SPGP0DA00550 | XP_018222699.1 | 99 | 0 |
| SPGP0X01460 | XP_018221590.1 | 99 | 0 |
| SPGP0CX00950 | XP_018222114.1 | 99 | 0 |
| SPGP0DA01600 | XP_018221457.1 | 99 | 0 |
| SPGP0U00120 | XP_018223057.1 | 99 | 0 |
| SPGP0F02750 | XP_018221435.1 | 99 | 0 |
| SPGP0CN00920 | XP_018220880.1 | 99 | 0 |
| SPGP0F02880 | XP_018221447.1 | 99 | 0 |
| SPGP0J00760 | XP_018219874.1 | 99 | 0 |
| SPGP0EX00100 | XP_018218751.1 | 99 | 0 |
| SPGP0Y00260 | XP_018220043.1 | 99 | 0 |
| SPGP0CP01540 | XP_018223685.1 | 99 | 0 |
| SPGP0AJ01820 | XP_018221950.1 | 99 | 0 |
| SPGP0DA00610 | XP_018222693.1 | 99 | 0 |
| SPGP0J00180 | XP_018219929.1 | 99 | 0 |
| SPGP0HH00100 | XP_018222849.1 | 85 | 0 |
| SPGP0S00450 | XP_018224093.1 | 99 | 0 |
| SPGP0L01190 | XP_018218915.1 | 99 | 0 |
| SPGP0AE00160 | XP_018222498.1 | 99 | 0 |
| SPGP0S00680 | XP_018224115.1 | 99 | 1 |
| SPGP0CM01010 | XP_018223224.1 | 99 | 0 |
| SPGP0J02180 | XP_018219731.1 | 99 | 0 |
| SPGP0J02380 | XP_018219711.1 | 99 | 1 |
| SPGP0R00300 | XP_018219972.1 | 99 | 0 |
| SPGP0CM01900 | XP_018223136.1 | 99 | 0 |
| SPGP0CM02300 | XP_018223099.1 | 99 | 0 |
| SPGP0F02150 | XP_018221376.1 | 99 | 0 |

Create PDF in your applications with the Pdfcrowd HTML to PDF API PDFCROWD

| | | | |
|---|---|---|---|
| SPGP0J02720 | XP_018219677.1 | 99 | 0 |
| SPGP0L00540 | XP_018218978.1 | 99 | 0 |
| SPGP0F02330 | XP_018221394.1 | 100 | 0 |
| SPGP0V00990 | XP_018221688.1 | 99 | 0 |
| SPGP0S00350 | XP_018224083.1 | 99 | 0 |
| SPGP0S00590 | XP_018224106.1 | 99 | 0 |
| SPGP0J01760 | XP_018219772.1 | 99 | 0 |
| SPGP0V00430 | XP_018221742.1 | 99 | 0 |
| SPGP0F01490 | XP_018221312.1 | 99 | 0 |
| SPGP0CM01420 | XP_018223184.1 | 99 | 0 |
| SPGP0J03710 | XP_018219578.1 | 99 | 0 |
| SPGP0CP01400 | XP_018223699.1 | 99 | 0 |
| SPGP0X01250 | XP_018221569.1 | 99 | 0 |
| SPGP0J02580 | XP_018219691.1 | 98 | 0 |
| SPGP0J02850 | XP_018219664.1 | 99 | 0 |
| SPGP0EM00130 | XP_018223062.1 | 99 | 0 |
| SPGP0HB00170 | XP_018222868.1 | 100 | 0 |
| SPGP0CW01090 | XP_018219447.1 | 99 | 0 |
| SPGP0Y02720 | XP_018220281.1 | 100 | 0 |
| SPGP0S00620 | XP_018224109.1 | 99 | 0 |
| SPGP0F01240 | XP_018221287.1 | 99 | 0 |
| SPGP0AT00980 | XP_018219116.1 | 100 | 0 |
| SPGP0CP02670 | XP_018223574.1 | 99 | 0 |
| SPGP0JB00100 | XP_018222556.1 | 99 | 0 |
| SPGP0AT00450 | XP_018219066.1 | 99 | 0 |
| SPGP0AJ00910 | XP_018221863.1 | 100 | 0 |
| SPGP0CW03030 | XP_018219260.1 | 99 | 0 |
| SPGP0H01050 | XP_018221040.1 | 99 | 0 |
| SPGP0IN00100 | XP_018221666.1 | 99 | 0 |
| SPGP0BE00160 | XP_018223362.1 | 99 | 0 |
| SPGP0DA00900 | XP_018222664.1 | 99 | 0 |
| SPGP0Y03030 | XP_018220312.1 | 99 | 0 |
| SPGP0J02460 | XP_018219703.1 | 99 | 0 |
| SPGP0AH00120 | XP_018223420.1 | 99 | 0 |
| SPGP0HW00130 | XP_018224040.1 | 99 | 0 |
| SPGP0CP00950 | XP_018223744.1 | 99 | 0 |
| SPGP0J01630 | XP_018219785.1 | 99 | 0 |
| SPGP0F01860 | XP_018221348.1 | 99 | 0 |
| SPGP0CM00350 | XP_018223289.1 | 99 | 0 |
| SPGP0AT00700 | XP_018219089.1 | 99 | 0 |

Create PDF in your applications with the Pdfcrowd HTML to PDF API PDFCROWD

| | | | |
|---|---|---|---|
| SPGP0R00370 | XP_018219978.1 | 99 | 0 |
| SPGP0J03180 | XP_018219631.1 | 99 | 0 |
| SPGP0L01460 | XP_018218888.1 | 100 | 0 |
| SPGP0CP03590 | XP_018223485.1 | 99 | 1 |
| SPGP0Y01890 | XP_018220200.1 | 99 | 0 |
| SPGP0U01940 | XP_018222887.1 | 99 | 1 |
| SPGP0U01410 | XP_018222941.1 | 99 | 0 |
| SPGP0F02440 | XP_018221405.1 | 99 | 0 |
| SPGP0DE00310 | XP_018223848.1 | 99 | 0 |
| SPGP0J03230 | XP_018219626.1 | 99 | 0 |
| SPGP0X00690 | XP_018221517.1 | 99 | 0 |
| SPGP0L01360 | XP_018218898.1 | 99 | 0 |
| SPGP0F00330 | XP_018221201.1 | 99 | 0 |
| SPGP0X00730 | XP_018221521.1 | 99 | 0 |
| SPGP0X00590 | XP_018221508.1 | 99 | 0 |
| SPGP0CU01150 | XP_018220715.1 | 99 | 0 |
| SPGP0JP00200 | XP_018218747.1 | 99 | 0 |
| SPGP0AH00200 | XP_018223428.1 | 99 | 1 |
| SPGP0CW00970 | XP_018219459.1 | 99 | 0 |
| SPGP0AT00150 | XP_018219037.1 | 99 | 0 |
| SPGP0L00950 | XP_018218937.1 | 99 | 0 |
| SPGP0DB00400 | XP_018223913.1 | 99 | 0 |
| SPGP0F00500 | XP_018221215.1 | 99 | 0 |
| SPGP0FF00130 | XP_018223399.1 | 99 | 0 |
| SPGP0CX01230 | XP_018222142.1 | 97 | 0 |
| SPGP0R00600 | XP_018220000.1 | 99 | 0 |
| SPGP0AQ00160 | XP_018223406.1 | 99 | 0 |
| SPGP0DA00440 | XP_018222710.1 | 99 | 0 |
| SPGP0U01600 | XP_018222922.1 | 99 | 1 |
| SPGP0AE00440 | XP_018222523.1 | 99 | 0 |
| SPGP0AJ01540 | XP_018221922.1 | 99 | 0 |
| SPGP0CX01480 | XP_018222165.1 | 99 | 0 |
| SPGP0CU00680 | XP_018220760.1 | 99 | 0 |
| SPGP0JP00150 | XP_018219566.1 | 99 | 0 |
| SPGP0AJ01500 | XP_018221919.1 | 99 | 0 |
| SPGP0GG00120 | XP_018220839.1 | 99 | 0 |
| SPGP0K00280 | XP_018222770.1 | 99 | 0 |
| SPGP0AN00240 | XP_018223936.1 | 99 | 0 |
| SPGP0AN00710 | XP_018223978.1 | 99 | 0 |
| SPGP0J00160 | XP_018219931.1 | 90 | 0 |

Create PDF in your applications with the Pdfcrowd HTML to PDF API PDFCROWD

| | | | |
|---|---|---|---|
| SPGP0CP02510 | XP_018223590.1 | 99 | 0 |
| SPGP0CX01540 | XP_018222171.1 | 99 | 0 |
| SPGP0X01700 | XP_018221613.1 | 99 | 0 |
| SPGP0J00370 | XP_018219911.1 | 99 | 1 |
| SPGP0AF01890 | XP_018222447.1 | 99 | 0 |
| SPGP0V00240 | XP_018221759.1 | 99 | 0 |
| SPGP0DA01060 | XP_018222649.1 | 99 | 1 |
| SPGP0CX01940 | XP_018222209.1 | 98 | 0 |
| SPGP0AF01590 | XP_018222417.1 | 99 | 0 |
| SPGP0Y00750 | XP_018220091.1 | 99 | 0 |
| SPGP0IH00120 | XP_018220383.1 | 99 | 0 |
| SPGP0CM01480 | XP_018223178.1 | 99 | 0 |
| SPGP0FE00130 | XP_018221965.1 | 99 | 0 |
| SPGP0AN00860 | XP_018223992.1 | 99 | 0 |
| SPGP0S00360 | XP_018224084.1 | 99 | 0 |
| SPGP0DA01360 | XP_018222619.1 | 99 | 0 |
| SPGP0L00460 | XP_018218986.1 | 99 | 0 |
| SPGP0R00770 | XP_018220016.1 | 99 | 0 |
| SPGP0AF00240 | XP_018222278.1 | 99 | 0 |
| SPGP0J00510 | XP_018219897.1 | 99 | 0 |
| SPGP0L02490 | XP_018218787.1 | 99 | 0 |
| SPGP0J03300 | XP_018219619.1 | 99 | 0 |
| SPGP0Y01340 | XP_018220149.1 | 99 | 0 |
| SPGP0CP00280 | XP_018223808.1 | 99 | 0 |
| SPGP0H00990 | XP_018221046.1 | 99 | 0 |
| SPGP0F01520 | XP_018221315.1 | 99 | 0 |
| SPGP0X01290 | XP_018221573.1 | 99 | 0 |
| SPGP0G00170 | XP_018222256.1 | 99 | 0 |
| SPGP0X00600 | XP_018221509.1 | 99 | 0 |
| SPGP0CN01080 | XP_018220864.1 | 99 | 0 |
| SPGP0J01520 | XP_018219796.1 | 100 | 0 |
| SPGP0Y01140 | XP_018220129.1 | 99 | 0 |
| SPGP0CN00460 | XP_018220925.1 | 99 | 0 |
| SPGP0AU00110 | XP_018223068.1 | 99 | 0 |
| SPGP0CX00640 | XP_018222083.1 | 99 | 0 |
| SPGP0DA01500 | XP_018222605.1 | 99 | 0 |
| SPGP0F02390 | XP_018221400.1 | 99 | 0 |
| SPGP0Y00370 | XP_018220054.1 | 99 | 0 |
| SPGP0AA01100 | XP_018220480.1 | 99 | 0 |
| SPGP0CN00680 | XP_018220903.1 | 99 | 1 |

Create PDF in your applications with the Pdfcrowd HTML to PDF API PDFCROWD

| | | | |
|---|---|---|---|
| SPGP0CN00130 | XP_018220958.1 | 99 | 0 |
| SPGP0J01130 | XP_018219834.1 | 99 | 0 |
| SPGP0Ftrna13M | DI49_4243 | 100 | 0 |
| SPGP0F02730 | XP_018221433.1 | 99 | 0 |
| SPGP0HI00190 | XP_018219945.1 | 99 | 0 |
| SPGP0CP01690 | XP_018223671.1 | 99 | 0 |
| SPGP0DS00140 | XP_018220966.1 | 99 | 0 |
| SPGP0Y00890 | XP_018220105.1 | 98 | 2 |
| SPGP0AT01280 | XP_018219146.1 | 99 | 0 |
| SPGP0DA01170 | XP_018222638.1 | 99 | 0 |
| SPGP0CN00350 | XP_018220936.1 | 99 | 0 |
| SPGP0FF00160 | XP_018223396.1 | 100 | 0 |
| SPGP0CW01540 | XP_018219404.1 | 99 | 0 |
| SPGP0G00160 | XP_018222255.1 | 99 | 0 |
| SPGP0CU01990 | XP_018220635.1 | 99 | 0 |
| SPGP0F01380 | XP_018221302.1 | 99 | 0 |
| SPGP0AJ00760 | XP_018221848.1 | 100 | 0 |
| SPGP0F00650 | XP_018221230.1 | 99 | 0 |
| SPGP0AT01500 | XP_018219167.1 | 99 | 0 |
| SPGP0DE00120 | XP_018223829.1 | 100 | 0 |
| SPGP0F02870 | XP_018221446.1 | 96 | 3 |
| SPGP0U01770 | XP_018222904.1 | 99 | 0 |
| SPGP0J02440 | XP_018219705.1 | 99 | 0 |
| SPGP0J02110 | XP_018219738.1 | 99 | 0 |
| SPGP0CX00650 | XP_018222084.1 | 99 | 0 |
| SPGP0CN00670 | XP_018220904.1 | 100 | 0 |
| SPGP0Y01060 | XP_018220121.1 | 99 | 0 |
| SPGP0J02330 | XP_018219716.1 | 99 | 0 |
| SPGP0K00580 | XP_018222800.1 | 100 | 0 |
| SPGP0J03430 | XP_018219605.1 | 99 | 0 |
| SPGP0S00260 | XP_018224075.1 | 99 | 0 |
| SPGP0CV00410 | XP_018222593.1 | 98 | 1 |
| SPGP0DB00210 | XP_018223895.1 | 99 | 0 |
| SPGP0J01910 | XP_018219758.1 | 99 | 0 |
| SPGP0AF00100 | XP_018222265.1 | 99 | 0 |
| SPGP0CU00110 | XP_018220817.1 | 99 | 0 |
| SPGP0U00830 | XP_018222995.1 | 99 | 0 |
| SPGP0U00420 | XP_018223031.1 | 99 | 0 |
| SPGP0R00740 | XP_018220014.1 | 99 | 0 |
| SPGP0F01870 | XP_018221349.1 | 99 | 0 |

Create PDF in your applications with the Pdfcrowd HTML to PDF API PDFCROWD

| | | | |
|---|---|---|---|
| SPGP0F00390 | XP_018221206.1 | 100 | 0 |
| SPGP0CP01890 | XP_018223651.1 | 99 | 0 |
| SPGP0F02130 | XP_018221375.1 | 99 | 0 |
| SPGP0AA00300 | XP_018220560.1 | 99 | 0 |
| SPGP0X01870 | XP_018221630.1 | 99 | 0 |
| SPGP0F02650 | XP_018221425.1 | 99 | 0 |
| SPGP0F02120 | XP_018221374.1 | 99 | 0 |
| SPGP0CP01980 | XP_018223642.1 | 99 | 0 |
| SPGP0X01270 | XP_018221571.1 | 99 | 0 |
| SPGP0H00460 | XP_018221098.1 | 99 | 0 |
| SPGP0J03240 | XP_018219625.1 | 99 | 0 |
| SPGP0CP03580 | XP_018223486.1 | 99 | 0 |
| SPGP0DA01270 | XP_018222628.1 | 99 | 0 |
| SPGP0J02160 | XP_018219733.1 | 100 | 0 |
| SPGP0CW02740 | XP_018219289.1 | 99 | 0 |
| SPGP0IZ00130 | XP_018220341.1 | 98 | 0 |
| SPGP0CU00230 | XP_018220806.1 | 99 | 0 |
| SPGP0CW01660 | XP_018219394.1 | 99 | 0 |
| SPGP0AF00830 | XP_018222337.1 | 99 | 0 |
| SPGP0AF00430 | XP_018222297.1 | 99 | 0 |
| SPGP0S00330 | XP_018224081.1 | 99 | 0 |
| SPGP0H00510 | XP_018221094.1 | 99 | 0 |
| SPGP0H01660 | XP_018220984.1 | 99 | 0 |
| SPGP0AN00380 | XP_018223947.1 | 99 | 0 |
| SPGP0X00470 | XP_018221496.1 | 99 | 0 |
| SPGP0CU00460 | XP_018220783.1 | 99 | 0 |
| SPGP0Y00300 | XP_018220047.1 | 99 | 0 |
| SPGP0H00550 | XP_018221090.1 | 100 | 0 |
| SPGP0HB00190 | XP_018222870.1 | 99 | 0 |
| SPGP0CW03650 | XP_018219200.1 | 98 | 0 |
| SPGP0CN00950 | XP_018220877.1 | 99 | 0 |
| SPGP0L02100 | XP_018218826.1 | 99 | 0 |
| SPGP0AT00870 | XP_018219106.1 | 99 | 0 |
| SPGP0AF01420 | XP_018222400.1 | 99 | 0 |
| SPGP0IG00100 | XP_018223348.1 | 99 | 0 |
| SPGP0EO00110 | XP_018221148.1 | 99 | 0 |
| SPGP0DA00580 | XP_018222696.1 | 99 | 0 |
| SPGP0CN00900 | XP_018220882.1 | 99 | 0 |
| SPGP0Y02860 | XP_018220295.1 | 99 | 0 |
| SPGP0AF00520 | XP_018222306.1 | 99 | 0 |

Create PDF in your applications with the Pdfcrowd HTML to PDF API PDFCROWD

| | | | |
|---|---|---|---|
| SPGP0J00740 | XP_018219876.1 | 100 | 0 |
| SPGP0Y00240 | XP_018220041.1 | 99 | 0 |
| SPGP0AJ01370 | XP_018221907.1 | 99 | 0 |
| SPGP0AT00210 | XP_018219043.1 | 100 | 0 |
| SPGP0F02920 | XP_018221450.1 | 99 | 0 |
| SPGP0H01170 | XP_018221029.1 | 99 | 0 |
| SPGP0CU00470 | XP_018220782.1 | 99 | 0 |
| SPGP0DB00330 | XP_018223906.1 | 99 | 0 |
| SPGP0F01040 | XP_018221268.1 | 99 | 0 |
| SPGP0L02430 | XP_018218793.1 | 99 | 0 |
| SPGP0Y02450 | XP_018220254.1 | 99 | 0 |
| SPGP0CM02190 | XP_018223110.1 | 99 | 0 |
| SPGP0FP00150 | XP_018223340.1 | 99 | 0 |
| SPGP0J01950 | XP_018219754.1 | 99 | 0 |
| SPGP0K00180 | XP_018222761.1 | 100 | 0 |
| SPGP0F01850 | XP_018221347.1 | 99 | 0 |
| SPGP0CX00970 | XP_018222116.1 | 99 | 0 |
| SPGP0K00230 | XP_018222765.1 | 99 | 0 |
| SPGP0CU02390 | XP_018220595.1 | 99 | 0 |
| SPGP0CM01890 | XP_018223137.1 | 99 | 0 |
| SPGP0J01830 | XP_018219766.1 | 99 | 0 |
| SPGP0CU01120 | XP_018220718.1 | 99 | 0 |
| SPGP0X01210 | XP_018221565.1 | 99 | 0 |
| SPGP0X01280 | XP_018221572.1 | 99 | 0 |
| SPGP0CN01140 | XP_018220859.1 | 99 | 0 |
| SPGP0J02530 | XP_018219696.1 | 99 | 0 |
| SPGP0CX01000 | XP_018222119.1 | 98 | 0 |
| SPGP0CW00300 | XP_018219525.1 | 99 | 0 |
| SPGP0CW00870 | XP_018219469.1 | 99 | 0 |
| SPGP0AA01800 | XP_018220412.1 | 99 | 0 |
| SPGP0AT00550 | XP_018219076.1 | 99 | 0 |
| SPGP0CU01530 | XP_018220681.1 | 100 | 0 |
| SPGP0CM02320 | XP_018223097.1 | 99 | 0 |
| SPGP0X00780 | XP_018221526.1 | 99 | 0 |
| SPGP0J01140 | XP_018219833.1 | 99 | 0 |
| SPGP0GQ00120 | XP_018219557.1 | 99 | 0 |
| SPGP0J00640 | XP_018219886.1 | 99 | 0 |
| SPGP0J01990 | XP_018219750.1 | 100 | 0 |
| SPGP0CM02010 | XP_018223127.1 | 99 | 0 |
| SPGP0J00410 | XP_018219907.1 | 100 | 0 |

Create PDF in your applications with the Pdfcrowd HTML to PDF API PDFCROWD

| | | | |
|---|---|---|---|
| SPGP0IU00120 | XP_018220330.1 | 99 | 0 |
| SPGP0U00300 | XP_018223041.1 | 99 | 0 |
| SPGP0Z00150 | XP_018222550.1 | 99 | 0 |
| SPGP0DE00300 | XP_018223847.1 | 99 | 0 |
| SPGP0CN01070 | XP_018220865.1 | 99 | 0 |
| SPGP0AA00610 | XP_018220529.1 | 99 | 0 |
| SPGP0CW02230 | XP_018219338.1 | 99 | 0 |
| SPGP0CW01710 | XP_018219389.1 | 99 | 0 |
| SPGP0CW03200 | XP_018219244.1 | 99 | 0 |
| SPGP0F00750 | XP_018221240.1 | 99 | 0 |
| SPGP0CP00110 | XP_018223825.1 | 99 | 0 |
| SPGP0AF01520 | XP_018222410.1 | 99 | 0 |
| SPGP0CU00430 | XP_018220786.1 | 99 | 0 |
| SPGP0CN00930 | XP_018220879.1 | 99 | 0 |
| SPGP0F00860 | XP_018221250.1 | 99 | 0 |
| SPGP0F01260 | XP_018221289.1 | 99 | 0 |
| SPGP0CP00370 | XP_018223799.1 | 99 | 0 |
| SPGP0J03030 | XP_018219646.1 | 99 | 0 |
| SPGP0S00640 | XP_018224111.1 | 99 | 0 |
| SPGP0CW00520 | XP_018219503.1 | 99 | 0 |
| SPGP0U01990 | XP_018222882.1 | 99 | 0 |
| SPGP0J02270 | XP_018219722.1 | 99 | 0 |
| SPGP0CX01730 | XP_018222189.1 | 99 | 0 |
| SPGP0AJ00460 | XP_018221819.1 | 99 | 0 |
| SPGP0J03470 | XP_018219601.1 | 99 | 0 |
| SPGP0CW03010 | XP_018219262.1 | 99 | 0 |
| SPGP0AJ00930 | XP_018221865.1 | 99 | 0 |
| SPGP0IN00130 | XP_018221663.1 | 99 | 0 |
| SPGP0CP02830 | XP_018223558.1 | 99 | 0 |
| SPGP0J02870 | XP_018219662.1 | 99 | 0 |
| SPGP0M00180 | XP_018221140.1 | 99 | 0 |
| SPGP0DA01530 | XP_018222602.1 | 99 | 0 |
| SPGP0Gtrna22S | DI49_4419 | 100 | 0 |
| SPGP0CW00740 | XP_018219482.1 | 99 | 0 |
| SPGP0Y02260 | XP_018220235.1 | 99 | 0 |
| SPGP0CM00250 | XP_018223299.1 | 99 | 0 |
| SPGP0HW00110 | XP_018224042.1 | 98 | 1 |
| SPGP0AT01030 | XP_018219121.1 | 99 | 0 |
| SPGP0F01840 | XP_018221346.1 | 100 | 0 |
| SPGP0V00980 | XP_018221689.1 | 99 | 0 |

Create PDF in your applications with the Pdfcrowd HTML to PDF API PDFCROWD

| | | | |
|---|---|---|---|
| SPGP0AA00810 | XP_018220509.1 | 96 | 1 |
| SPGP0J03650 | XP_018219584.1 | 99 | 0 |
| SPGP0AT01230 | XP_018219141.1 | 99 | 0 |
| SPGP0U01430 | XP_018222939.1 | 98 | 1 |
| SPGP0F01510 | XP_018221314.1 | 99 | 0 |
| SPGP0AJ00350 | XP_018221808.1 | 99 | 0 |
| SPGP0F02460 | XP_018221407.1 | 99 | 0 |
| SPGP0CV00240 | XP_018222575.1 | 99 | 0 |
| SPGP0AA00520 | XP_018220538.1 | 99 | 0 |
| SPGP0AJ00190 | XP_018221794.1 | 95 | 5 |
| SPGP0AA00250 | XP_018220565.1 | 100 | 0 |
| SPGP0X01240 | XP_018221568.1 | 99 | 0 |
| SPGP0AE00280 | XP_018222509.1 | 99 | 0 |
| SPGP0CP01840 | XP_018223656.1 | 100 | 0 |
| SPGP0AJ01600 | XP_018221928.1 | 99 | 0 |
| SPGP0AA00760 | XP_018220515.1 | 99 | 0 |
| SPGP0J02300 | XP_018219719.1 | 99 | 0 |
| SPGP0CN00270 | XP_018220944.1 | 99 | 0 |
| SPGP0F00640 | XP_018221229.1 | 99 | 0 |
| SPGP0IN00310 | XP_018221645.1 | 95 | 0 |
| SPGP0AE00460 | XP_018222525.1 | 99 | 0 |
| SPGP0AJ01560 | XP_018221924.1 | 99 | 0 |
| SPGP0J00750 | XP_018219875.1 | 99 | 0 |
| SPGP0S00550 | XP_018224102.1 | 99 | 0 |
| SPGP0L01020 | XP_018218930.1 | 99 | 0 |
| SPGP0AA01120 | XP_018220478.1 | 99 | 0 |
| SPGP0X01400 | XP_018221584.1 | 99 | 0 |
| SPGP0J03010 | XP_018219648.1 | 99 | 0 |
| SPGP0X00340 | XP_018221483.1 | 99 | 0 |
| SPGP0DE00340 | XP_018223851.1 | 99 | 0 |
| SPGP0H00160 | XP_018221127.1 | 99 | 0 |
| SPGP0X00310 | XP_018221480.1 | 99 | 0 |
| SPGP0HR00110 | XP_018222851.1 | 94 | 0 |
| SPGP0J00680 | XP_018219882.1 | 99 | 0 |
| SPGP0M00130 | XP_018221143.1 | 99 | 0 |
| SPGP0H01340 | XP_018221013.1 | 99 | 0 |
| SPGP0J00290 | XP_018219919.1 | 99 | 0 |
| SPGP0DA01510 | XP_018222604.1 | 99 | 0 |
| SPGP0CX00760 | XP_018222095.1 | 99 | 0 |
| SPGP0U01120 | XP_018222967.1 | 100 | 0 |

Create PDF in your applications with the Pdfcrowd HTML to PDF API PDFCROWD

| | | | |
|---|---|---|---|
| SPGP0AA00830 | XP_018220507.1 | 99 | 0 |
| SPGP0DA00150 | XP_018222737.1 | 99 | 0 |
| SPGP0S00340 | XP_018224082.1 | 99 | 0 |
| SPGP0CW03640 | XP_018219201.1 | 97 | 0 |
| SPGP0J00350 | XP_018219913.1 | 97 | 2 |
| SPGP0L02340 | XP_018218802.1 | 98 | 0 |
| SPGP0Y02330 | XP_018220242.1 | 100 | 0 |
| SPGP0CW01220 | XP_018219435.1 | 99 | 0 |
| SPGP0V00860 | XP_018221701.1 | 99 | 0 |
| SPGP0IH00180 | XP_018220388.1 | 99 | 0 |
| SPGP0Y01470 | XP_018220162.1 | 99 | 0 |
| SPGP0L00830 | XP_018218950.1 | 99 | 0 |
| SPGP0FP00270 | XP_018223329.1 | 99 | 0 |
| SPGP0CW02790 | XP_018219284.1 | 99 | 0 |
| SPGP0S00120 | XP_018224061.1 | 99 | 0 |
| SPGP0J01330 | XP_018219815.1 | 99 | 0 |
| SPGP0CXtrna6L | DI49_1818 | 100 | 0 |
| SPGP0AJ00830 | XP_018221855.1 | 99 | 0 |
| SPGP0AJ00260 | XP_018221800.1 | 99 | 0 |
| SPGP0J03670 | XP_018219582.1 | 99 | 0 |
| SPGP0DA00520 | XP_018222702.1 | 99 | 0 |
| SPGP0JK00150 | XP_018223437.1 | 100 | 0 |
| SPGP0CN00110 | XP_018220960.1 | 99 | 0 |
| SPGP0Y02010 | XP_018220212.1 | 99 | 0 |
| SPGP0R00440 | XP_018219985.1 | 99 | 0 |
| SPGP0J01480 | XP_018219800.1 | 99 | 0 |
| SPGP0Ytrna10P | DI49_4051 | 99 | 0 |
| SPGP0CU01660 | XP_018220668.1 | 99 | 0 |
| SPGP0H00430 | XP_018221101.1 | 99 | 0 |
| SPGP0K00130 | XP_018222755.1 | 99 | 0 |
| SPGP0DS00120 | XP_018220964.1 | 99 | 0 |
| SPGP0H00980 | XP_018221047.1 | 99 | 0 |
| SPGP0CM00620 | XP_018223262.1 | 99 | 0 |
| SPGP0DA00700 | XP_018222683.1 | 99 | 0 |
| SPGP0F02280 | XP_018221389.1 | 99 | 0 |
| SPGP0J02680 | XP_018219681.1 | 99 | 0 |
| SPGP0F02810 | XP_018221441.1 | 99 | 0 |
| SPGP0CN00370 | XP_018220934.1 | 99 | 0 |
| SPGP0F01330 | XP_018221297.1 | 99 | 0 |

Create PDF in your applications with the Pdfcrowd HTML to PDF API PDFCROWD

| | | | |
|---|---|---|---|
| SPGP0X00540 | XP_018221503.1 | 99 | 0 |
| SPGP0J00820 | XP_018219868.1 | 99 | 0 |
| SPGP0F00670 | XP_018221232.1 | 99 | 0 |
| SPGP0J01470 | XP_018219801.1 | 99 | 0 |
| SPGP0F02720 | XP_018221432.1 | 99 | 0 |
| SPGP0X01170 | XP_018221561.1 | 99 | 0 |
| SPGP0X01730 | XP_018221616.1 | 99 | 0 |
| SPGP0CU02310 | XP_018220603.1 | 99 | 0 |
| SPGP0Y01170 | XP_018220132.1 | 99 | 0 |
| SPGP0V00180 | XP_018221765.1 | 99 | 0 |
| SPGP0CP01910 | XP_018223649.1 | 100 | 0 |
| SPGP0J01840 | XP_018219765.1 | 99 | 0 |
| SPGP0CW02640 | XP_018219300.1 | 99 | 0 |
| SPGP0AN00210 | XP_018223933.1 | 99 | 0 |
| SPGP0GA00140 | XP_018223389.1 | 99 | 0 |
| SPGP0HS00110 | XP_018221775.1 | 99 | 0 |
| SPGP0CM00770 | XP_018223248.1 | 99 | 0 |
| SPGP0F01700 | XP_018221332.1 | 99 | 0 |
| SPGP0F01100 | XP_018221274.1 | 98 | 0 |
| SPGP0V00220 | XP_018221761.1 | 99 | 0 |
| SPGP0CM02000 | XP_018223128.1 | 99 | 0 |
| SPGP0X01770 | XP_018221620.1 | 99 | 0 |
| SPGP0CX01100 | XP_018222129.1 | 99 | 0 |
| SPGP0U00560 | XP_018223018.1 | 99 | 0 |
| SPGP0AN00980 | XP_018224004.1 | 99 | 0 |
| SPGP0DA01210 | XP_018222634.1 | 99 | 0 |
| SPGP0F02100 | XP_018221372.1 | 99 | 0 |
| SPGP0DE00500 | XP_018223867.1 | 99 | 0 |
| SPGP0H00750 | XP_018221070.1 | 99 | 0 |
| SPGP0GL00120 | XP_018220842.1 | 96 | 0 |
| SPGP0J03440 | XP_018219604.1 | 99 | 0 |
| SPGP0CW02070 | XP_018219354.1 | 99 | 0 |
| SPGP0DEtrna3F | DI49_0495 | 100 | 0 |
| SPGP0X00670 | XP_018221515.1 | 99 | 0 |
| SPGP0CV00400 | XP_018222592.1 | 96 | 0 |
| SPGP0L01450 | XP_018218889.1 | 99 | 0 |
| SPGP0CP03560 | XP_018223487.1 | 99 | 0 |
| SPGP0F01620 | XP_018221325.1 | 99 | 0 |
| SPGP0J03220 | XP_018219627.1 | 99 | 0 |
| SPGP0AA00780 | XP_018220513.1 | 99 | 0 |

Create PDF in your applications with the Pdfcrowd HTML to PDF API PDFCROWD

| | | | |
|---|---|---|---|
| SPGP0Strna7P | DI49_4914 | 100 | 0 |
| SPGP0H01690 | XP_018220981.1 | 99 | 0 |
| SPGP0H00570 | XP_018221088.1 | 99 | 0 |
| SPGP0DB00120 | XP_018223886.1 | 99 | 0 |
| SPGP0Y00680 | XP_018220084.1 | 96 | 4 |
| SPGP0AA00750 | XP_018220516.1 | 99 | 0 |
| SPGP0X00410 | XP_018221490.1 | 99 | 0 |
| SPGP0AF00720 | XP_018222326.1 | 99 | 0 |
| SPGP0CP03400 | XP_018223502.1 | 99 | 0 |
| SPGP0Y00220 | XP_018220039.1 | 99 | 0 |
| SPGP0X01750 | XP_018221618.1 | 99 | 0 |
| SPGP0CP03300 | XP_018223510.1 | 100 | 0 |
| SPGP0H01540 | XP_018220995.1 | 99 | 0 |
| SPGP0GR00150 | XP_018221980.1 | 99 | 0 |
| SPGP0F01350 | XP_018221299.1 | 99 | 0 |
| SPGP0J02010 | XP_018219748.1 | 99 | 0 |
| SPGP0H00280 | XP_018221116.1 | 99 | 0 |
| SPGP0J01720 | XP_018219776.1 | 99 | 0 |
| SPGP0H00400 | XP_018221104.1 | 99 | 0 |
| SPGP0CP02960 | XP_018223545.1 | 99 | 0 |
| SPGP0Y02840 | XP_018220293.1 | 99 | 0 |
| SPGP0CP01300 | XP_018223709.1 | 99 | 0 |
| SPGP0S00390 | XP_018224087.1 | 99 | 0 |
| SPGP0F01070 | XP_018221271.1 | 99 | 0 |
| SPGP0H01150 | XP_018221031.1 | 99 | 0 |
| SPGP0DB00160 | XP_018223890.1 | 99 | 0 |
| SPGP0AU00130 | XP_018223070.1 | 99 | 0 |
| SPGP0CW00790 | XP_018219477.1 | 99 | 0 |
| SPGP0Y00640 | XP_018220080.1 | 99 | 0 |
| SPGP0J00100 | XP_018219936.1 | 99 | 0 |
| SPGP0AF01970 | XP_018222455.1 | 99 | 0 |
| SPGP0J02490 | XP_018219700.1 | 99 | 0 |
| SPGP0J02660 | XP_018219683.1 | 99 | 0 |
| SPGP0DS00150 | XP_018220967.1 | 99 | 0 |
| SPGP0CP02970 | XP_018223543.1 | 95 | 0 |
| SPGP0CW02270 | XP_018219334.1 | 99 | 0 |
| SPGP0CRtrna20A | DI49_3564 | 100 | 0 |
| SPGP0J01540 | XP_018219794.1 | 100 | 0 |
| SPGP0X01160 | XP_018221560.1 | 99 | 0 |
| SPGP0DA01020 | XP_018222653.1 | 99 | 0 |

Create PDF in your applications with the Pdfcrowd HTML to PDF API PDFCROWD

| | | | |
|---|---|---|---|
| SPGP0J01810 | XP_018219767.1 | 100 | 0 |
| SPGP0L00680 | XP_018218965.1 | 99 | 0 |
| SPGP0DE00590 | XP_018223876.1 | 99 | 0 |
| SPGP0CW00320 | XP_018219523.1 | 99 | 0 |
| SPGP0CP00960 | XP_018223743.1 | 99 | 0 |
| SPGP0L00400 | XP_018218992.1 | 99 | 0 |
| SPGP0JO00100 | XP_018222260.1 | 99 | 0 |
| SPGP0F01310 | XP_018221295.1 | 99 | 0 |
| SPGP0F00880 | XP_018221252.1 | 99 | 0 |
| SPGP0H01310 | XP_018221016.1 | 99 | 0 |
| SPGP0AT01120 | XP_018219130.1 | 99 | 1 |
| SPGP0AT01650 | XP_018219182.1 | 98 | 0 |
| SPGP0K00380 | XP_018222780.1 | 99 | 0 |
| SPGP0CN00580 | XP_018220913.1 | 100 | 0 |
| SPGP0CW02130 | XP_018219348.1 | 99 | 0 |
| SPGP0S00660 | XP_018224113.1 | 99 | 0 |
| SPGP0J01490 | XP_018219799.1 | 99 | 0 |
| SPGP0F00380 | XP_018221205.1 | 99 | 0 |
| SPGP0AT00490 | XP_018219070.1 | 99 | 0 |
| SPGP0X01200 | XP_018221564.1 | 99 | 0 |
| SPGP0CW01160 | XP_018219441.1 | 100 | 0 |
| SPGP0AA00340 | XP_018220556.1 | 99 | 1 |
| SPGP0L01550 | XP_018218879.1 | 99 | 0 |
| SPGP0CU02410 | XP_018220593.1 | 99 | 0 |
| SPGP0AA01330 | XP_018220457.1 | 100 | 0 |
| SPGP0H01020 | XP_018221043.1 | 99 | 0 |
| SPGP0CU01850 | XP_018220649.1 | 99 | 0 |
| SPGP0J02800 | XP_018219669.1 | 99 | 0 |
| SPGP0AT00290 | XP_018219051.1 | 99 | 0 |
| SPGP0CW03070 | XP_018219256.1 | 99 | 0 |
| SPGP0J03410 | XP_018219607.1 | 100 | 0 |
| SPGP0M00200 | XP_018221138.1 | 99 | 0 |
| SPGP0ED00110 | XP_018223474.1 | 98 | 0 |
| SPGP0CU01570 | XP_018220677.1 | 99 | 0 |
| SPGP0DQ00110 | XP_018220024.1 | 99 | 0 |
| SPGP0Y03070 | XP_018220316.1 | 99 | 0 |
| SPGP0CX01280 | XP_018222146.1 | 99 | 0 |
| SPGP0CU00990 | XP_018220731.1 | 99 | 0 |
| SPGP0Y02200 | XP_018220229.1 | 100 | 0 |
| SPGP0CX02230 | XP_018222237.1 | 99 | 0 |

Create PDF in your applications with the Pdfcrowd HTML to PDF API PDFCROWD

| | | | |
|---|---|---|---|
| SPGP0Y01390 | XP_018220154.1 | 99 | 0 |
| SPGP0U01400 | XP_018222942.1 | 99 | 0 |
| SPGP0AN01080 | XP_018224014.1 | 99 | 0 |
| SPGP0F01820 | XP_018221344.1 | 99 | 0 |
| SPGP0CP01850 | XP_018223655.1 | 99 | 0 |
| SPGP0AJ00800 | XP_018221852.1 | 99 | 0 |
| SPGP0L00620 | XP_018218971.1 | 99 | 0 |
| SPGP0L00190 | XP_018219012.1 | 99 | 0 |
| SPGP0AJ00660 | XP_018221838.1 | 99 | 0 |
| SPGP0J02030 | XP_018219746.1 | 99 | 0 |
| SPGP0J02090 | XP_018219740.1 | 99 | 0 |
| SPGP0J03600 | XP_018219589.1 | 99 | 0 |
| SPGP0X01350 | XP_018221579.1 | 99 | 0 |
| SPGP0L01860 | XP_018218849.1 | 100 | 0 |
| SPGP0FO00110 | XP_018223457.1 | 99 | 0 |
| SPGP0DA00730 | XP_018222680.1 | 99 | 0 |
| SPGP0AU00200 | XP_018223075.1 | 99 | 0 |
| SPGP0J02320 | XP_018219717.1 | 99 | 0 |
| SPGP0X00970 | XP_018221544.1 | 99 | 0 |
| SPGP0AFtrna14L | DI49_2066 | 100 | 0 |
| SPGP0V00170 | XP_018221766.1 | 99 | 1 |
| SPGP0DE00650 | XP_018223882.1 | 99 | 0 |
| SPGP0CU00310 | XP_018220798.1 | 99 | 0 |
| SPGP0JP00110 | XP_018219562.1 | 99 | 0 |
| SPGP0F03000 | XP_018221455.1 | 99 | 0 |
| SPGP0CM00440 | XP_018223280.1 | 99 | 0 |
| SPGP0X01420 | XP_018221586.1 | 99 | 0 |
| SPGP0AE00130 | XP_018222495.1 | 99 | 0 |
| SPGP0X01930 | XP_018221636.1 | 99 | 0 |
| SPGP0CM01190 | XP_018223206.1 | 100 | 0 |
| SPGP0Y02100 | XP_018220220.1 | 99 | 0 |
| SPGP0H00140 | XP_018221129.1 | 99 | 0 |
| SPGP0F00170 | XP_018221186.1 | 99 | 0 |
| SPGP0JA00110 | XP_018222243.1 | 99 | 0 |
| SPGP0H01410 | XP_018221007.1 | 99 | 0 |
| SPGP0X00330 | XP_018221482.1 | 99 | 0 |
| SPGP0CM00860 | XP_018223239.1 | 99 | 0 |
| SPGP0L01710 | XP_018218864.1 | 99 | 0 |
| SPGP0FP00130 | XP_018223342.1 | 100 | 0 |
| SPGP0U02050 | XP_018222876.1 | 99 | 0 |

Create PDF in your applications with the Pdfcrowd HTML to PDF API PDFCROWD

| | | | |
|---|---|---|---|
| SPGP0H01810 | XP_018220969.1 | 99 | 0 |
| SPGP0CX01900 | XP_018222205.1 | 99 | 0 |
| SPGP0CP01230 | XP_018223715.1 | 99 | 0 |
| SPGP0Y01320 | XP_018220147.1 | 99 | 0 |
| SPGP0AF01170 | XP_018222373.1 | 100 | 0 |
| SPGP0CP00190 | XP_018223817.1 | 99 | 0 |
| SPGP0X01680 | XP_018221611.1 | 99 | 0 |
| SPGP0H00850 | XP_018221060.1 | 100 | 0 |
| SPGP0Y00960 | XP_018220111.1 | 99 | 0 |
| SPGP0CW02800 | XP_018219283.1 | 99 | 0 |
| SPGP0J00330 | XP_018219915.1 | 99 | 0 |
| SPGP0J03330 | XP_018219616.1 | 99 | 0 |
| SPGP0DB00170 | XP_018223891.1 | 99 | 0 |
| SPGP0J02880 | XP_018219661.1 | 99 | 0 |
| SPGP0CP01170 | XP_018223721.1 | 95 | 4 |
| SPGP0CW01340 | XP_018219424.1 | 99 | 0 |
| SPGP0AA01810 | XP_018220411.1 | 96 | 2 |
| SPGP0AJ01860 | XP_018221953.1 | 99 | 0 |
| SPGP0AE00410 | XP_018222520.1 | 100 | 0 |
| SPGP0CW01200 | XP_018219437.1 | 99 | 0 |
| SPGP0Y01040 | XP_018220119.1 | 99 | 0 |
| SPGP0CM01730 | XP_018223153.1 | 99 | 0 |
| SPGP0J01310 | XP_018219817.1 | 99 | 0 |
| SPGP0DA00950 | XP_018222660.1 | 99 | 0 |
| SPGP0DA00770 | XP_018222676.1 | 99 | 0 |
| SPGP0U01580 | XP_018222924.1 | 99 | 0 |
| SPGP0AF00280 | XP_018222282.1 | 99 | 0 |
| SPGP0U00940 | XP_018222985.1 | 99 | 0 |
| SPGP0J01230 | XP_018219825.1 | 99 | 0 |
| SPGP0S00650 | XP_018224112.1 | 100 | 0 |
| SPGP0CN00170 | XP_018220954.1 | 99 | 0 |
| SPGP0Y02070 | XP_018220217.1 | 97 | 0 |
| SPGP0F02320 | XP_018221393.1 | 99 | 0 |
| SPGP0V00390 | XP_018221744.1 | 99 | 0 |
| SPGP0DS00100 | XP_018220962.1 | 99 | 0 |
| SPGP0J02370 | XP_018219712.1 | 99 | 0 |
| SPGP0F02180 | XP_018221379.1 | 99 | 0 |
| SPGP0F02830 | XP_018221981.1 | 100 | 0 |
| SPGP0IZ00150 | XP_018220343.1 | 99 | 0 |
| SPGP0F02220 | XP_018221383.1 | 99 | 0 |

Create PDF in your applications with the Pdfcrowd HTML to PDF API PDFCROWD

| | | | |
|---|---|---|---|
| SPGP0CN00390 | XP_018220932.1 | 99 | 0 |
| SPGP0CX01920 | XP_018222207.1 | 99 | 0 |
| SPGP0AA00510 | XP_018220539.1 | 100 | 0 |
| SPGP0F00160 | XP_018221185.1 | 99 | 0 |
| SPGP0CP01560 | XP_018223683.1 | 100 | 0 |
| SPGP0J00800 | XP_018219870.1 | 99 | 0 |
| SPGP0F00690 | XP_018221234.1 | 99 | 0 |
| SPGP0F02430 | XP_018221404.1 | 99 | 0 |
| SPGP0IH00130 | XP_018220384.1 | 99 | 0 |
| SPGP0AT01350 | XP_018219153.1 | 99 | 0 |
| SPGP0BE00260 | XP_018223352.1 | 99 | 0 |
| SPGP0J02410 | XP_018219708.1 | 99 | 0 |
| SPGP0H01250 | XP_018221022.1 | 99 | 0 |
| SPGP0U02030 | XP_018222878.1 | 99 | 0 |
| SPGP0AT00190 | XP_018219041.1 | 99 | 0 |
| SPGP0R00830 | XP_018220022.1 | 99 | 0 |
| SPGP0R00360 | XP_018219977.1 | 99 | 0 |
| SPGP0FP00230 | XP_018223332.1 | 100 | 0 |
| SPGP0CW03440 | XP_018219220.1 | 99 | 0 |
| SPGP0F02340 | XP_018221395.1 | 100 | 0 |
| SPGP0AJ01480 | XP_018221917.1 | 99 | 0 |
| SPGP0CW02580 | XP_018219305.1 | 99 | 0 |
| SPGP0F01120 | XP_018221276.1 | 99 | 0 |
| SPGP0J00460 | XP_018219902.1 | 99 | 0 |
| SPGP0J03200 | XP_018219629.1 | 99 | 0 |
| SPGP0L02030 | XP_018218833.1 | 99 | 0 |
| SPGP0DA01230 | XP_018222632.1 | 99 | 0 |
| SPGP0H00590 | XP_018221086.1 | 99 | 0 |
| SPGP0L00870 | XP_018218946.1 | 99 | 0 |
| SPGP0F02160 | XP_018221377.1 | 99 | 0 |
| SPGP0AA00190 | XP_018220571.1 | 99 | 0 |
| SPGP0X00650 | XP_018221513.1 | 99 | 0 |
| SPGP0CP01260 | XP_018223712.1 | 99 | 0 |
| SPGP0AF02160 | XP_018222474.1 | 99 | 0 |
| SPGP0CW02050 | XP_018219356.1 | 99 | 0 |
| SPGP0GJ00100 | XP_018223382.1 | 99 | 0 |
| SPGP0CX01020 | XP_018222121.1 | 99 | 0 |
| SPGP0L01000 | XP_018218932.1 | 99 | 0 |
| SPGP0CU00900 | XP_018220739.1 | 99 | 0 |
| SPGP0AJ01780 | XP_018221946.1 | 99 | 0 |

Create PDF in your applications with the Pdfcrowd HTML to PDF API PDFCROWD

| | | | |
|---|---|---|---|
| SPGP0CW00400 | XP_018219515.1 | 99 | 0 |
| SPGP0L01260 | XP_018218908.1 | 99 | 0 |
| SPGP0X00430 | XP_018221492.1 | 99 | 0 |
| SPGP0Y00840 | XP_018220100.1 | 99 | 0 |
| SPGP0Y00340 | XP_018220051.1 | 99 | 0 |
| SPGP0U01570 | XP_018222925.1 | 99 | 0 |
| SPGP0X00420 | XP_018221491.1 | 99 | 0 |
| SPGP0Y00200 | XP_018220037.1 | 99 | 0 |
| SPGP0V00330 | XP_018221750.1 | 99 | 0 |
| SPGP0J01940 | XP_018219755.1 | 99 | 0 |
| SPGP0DA01490 | XP_018222606.1 | 99 | 0 |
| SPGP0AF01470 | XP_018222405.1 | 99 | 0 |
| SPGP0L00280 | XP_018219004.1 | 99 | 0 |
| SPGP0H00800 | XP_018221065.1 | 99 | 0 |
| SPGP0J01360 | XP_018219812.1 | 99 | 0 |
| SPGP0J00450 | XP_018219903.1 | 99 | 0 |
| SPGP0ANtrna18G | DI49_0567 | 99 | 0 |
| SPGP0CP02940 | XP_018223547.1 | 99 | 0 |
| SPGP0DA00540 | XP_018222700.1 | 98 | 0 |
| SPGP0CP01020 | XP_018223737.1 | 100 | 0 |
| SPGP0AJ01330 | XP_018221904.1 | 99 | 0 |
| SPGP0F00590 | XP_018221224.1 | 99 | 0 |
| SPGP0X01740 | XP_018221617.1 | 99 | 1 |
| SPGP0CX01770 | XP_018222192.1 | 98 | 2 |
| SPGP0CM01500 | XP_018223176.1 | 99 | 0 |
| SPGP0CN01160 | XP_018220857.1 | 99 | 0 |
| SPGP0J00470 | XP_018219901.1 | 100 | 0 |
| SPGP0GG00310 | XP_018220822.1 | 99 | 0 |
| SPGP0J01870 | XP_018219762.1 | 99 | 0 |
| SPGP0GP00100 | XP_018223417.1 | 99 | 0 |
| SPGP0CW02730 | XP_018219290.1 | 99 | 0 |
| SPGP0Y00970 | XP_018220112.1 | 99 | 0 |
| SPGP0AJ01440 | XP_018221914.1 | 99 | 0 |
| SPGP0IH00220 | XP_018220392.1 | 100 | 0 |
| SPGP0F02210 | XP_018221382.1 | 98 | 1 |
| SPGP0X00910 | XP_018221538.1 | 98 | 1 |
| SPGP0CW02610 | XP_018219302.1 | 99 | 0 |
| SPGP0CP00760 | XP_018223762.1 | 99 | 0 |
| SPGP0AA01480 | XP_018220442.1 | 99 | 0 |
| SPGP0DA01410 | XP_018222614.1 | 99 | 0 |

Create PDF in your applications with the Pdfcrowd HTML to PDF API PDFCROWD

| | | | |
|---|---|---|---|
| SPGP0R00650 | XP_018220005.1 | 99 | 0 |
| SPGP0CM00340 | XP_018223290.1 | 99 | 0 |
| SPGP0CU02430 | XP_018220591.1 | 99 | 0 |
| SPGP0AT01210 | XP_018219139.1 | 99 | 0 |
| SPGP0AT00860 | XP_018219105.1 | 99 | 0 |
| SPGP0J01690 | XP_018219779.1 | 99 | 0 |
| SPGP0CW02260 | XP_018219335.1 | 99 | 0 |
| SPGP0Y00500 | XP_018220067.1 | 99 | 0 |
| SPGP0CM00290 | XP_018223295.1 | 99 | 0 |
| SPGP0AT00500 | XP_018219071.1 | 100 | 0 |
| SPGP0CW03050 | XP_018219258.1 | 99 | 0 |
| SPGP0Y01210 | XP_018220136.1 | 99 | 0 |
| SPGP0CP00680 | XP_018223769.1 | 99 | 0 |
| SPGP0CW02110 | XP_018219350.1 | 99 | 0 |
| SPGP0AJ01680 | XP_018221936.1 | 99 | 0 |
| SPGP0Y03050 | XP_018220314.1 | 99 | 0 |
| SPGP0M00140 | XP_018221142.1 | 99 | 0 |
| SPGP0CP00300 | XP_018223806.1 | 98 | 2 |
| SPGP0AA00560 | XP_018220534.1 | 99 | 0 |
| SPGP0Y01290 | XP_018220144.1 | 100 | 0 |
| SPGP0CN00640 | XP_018220907.1 | 99 | 0 |
| SPGP0F01800 | XP_018221342.1 | 99 | 0 |
| SPGP0DA01320 | XP_018222623.1 | 99 | 0 |
| SPGP0V00670 | XP_018221719.1 | 99 | 0 |
| SPGP0IZ00220 | XP_018220350.1 | 98 | 0 |
| SPGP0L00600 | XP_018218973.1 | 99 | 0 |
| SPGP0GG00300 | XP_018220823.1 | 99 | 0 |
| SPGP0J01500 | XP_018219798.1 | 99 | 0 |
| SPGP0CW00500 | XP_018219505.1 | 99 | 0 |
| SPGP0IV00130 | XP_018222859.1 | 86 | 0 |
| SPGP0CP01800 | XP_018223660.1 | 100 | 0 |
| SPGP0J02970 | XP_018219652.1 | 99 | 0 |
| SPGP0R00640 | XP_018220004.1 | 100 | 0 |
| SPGP0U00870 | XP_018222992.1 | 99 | 0 |
| SPGP0R00310 | XP_018219973.1 | 99 | 0 |
| SPGP0Y01750 | XP_018220188.1 | 99 | 0 |
| SPGP0CN00540 | XP_018220917.1 | 99 | 0 |
| SPGP0AU00310 | XP_018223086.1 | 99 | 0 |
| SPGP0F02570 | XP_018221417.1 | 99 | 0 |
| SPGP0Y00740 | XP_018220090.1 | 99 | 0 |

Create PDF in your applications with the Pdfcrowd HTML to PDF API PDFCROWD

| | | | |
|---|---|---|---|
| SPGP0CX00700 | XP_018222089.1 | 99 | 0 |
| SPGP0J03130 | XP_018219636.1 | 99 | 0 |
| SPGP0IV00120 | XP_018222858.1 | 99 | 0 |
| SPGP0Y01510 | XP_018220166.1 | 99 | 0 |
| SPGP0AA00380 | XP_018220552.1 | 99 | 0 |
| SPGP0CX02150 | XP_018222229.1 | 99 | 0 |
| SPGP0CX00570 | XP_018222077.1 | 99 | 0 |
| SPGP0L01800 | XP_018218855.1 | 99 | 0 |
| SPGP0J03190 | XP_018219630.1 | 99 | 0 |
| SPGP0CM00190 | XP_018223305.1 | 99 | 0 |
| SPGP0X01910 | XP_018221634.1 | 99 | 0 |
| SPGP0AU00180 | XP_018223074.1 | 99 | 0 |
| SPGP0CX00620 | XP_018222081.1 | 99 | 0 |
| SPGP0H01630 | XP_018220987.1 | 99 | 0 |
| SPGP0CW01400 | XP_018219418.1 | 100 | 0 |
| SPGP0AA01000 | XP_018220490.1 | 99 | 0 |
| SPGP0X00350 | XP_018221484.1 | 99 | 0 |
| SPGP0L01730 | XP_018218862.1 | 99 | 0 |
| SPGP0AJ01830 | XP_018221951.1 | 99 | 0 |
| SPGP0DA01000 | XP_018222655.1 | 99 | 0 |
| SPGP0CU00270 | XP_018220802.1 | 99 | 0 |
| SPGP0CU00590 | XP_018220770.1 | 99 | 0 |
| SPGP0CX01430 | XP_018222160.1 | 99 | 0 |
| SPGP0H00640 | XP_018221081.1 | 99 | 0 |
| SPGP0CW01360 | XP_018219422.1 | 99 | 0 |
| SPGP0AN00800 | XP_018223986.1 | 99 | 0 |
| SPGP0Z00160 | XP_018222549.1 | 99 | 0 |
| SPGP0L01490 | XP_018218885.1 | 99 | 0 |
| SPGP0Y02480 | XP_018220257.1 | 99 | 0 |
| SPGP0J03690 | XP_018219580.1 | 99 | 0 |
| SPGP0CW03260 | XP_018219238.1 | 100 | 0 |
| SPGP0AF00110 | XP_018222266.1 | 98 | 0 |
| SPGP0J00310 | XP_018219917.1 | 99 | 0 |
| SPGP0AH00190 | XP_018223427.1 | 99 | 0 |
| SPGP0Z00300 | XP_018222535.1 | 99 | 0 |
| SPGP0L02380 | XP_018218798.1 | 99 | 0 |
| SPGP0AA01830 | XP_018220409.1 | 98 | 0 |
| SPGP0AJ00550 | XP_018221827.1 | 99 | 0 |
| SPGP0CM01710 | XP_018223155.1 | 99 | 0 |
| SPGP0X01000 | XP_018221547.1 | 99 | 0 |

Create PDF in your applications with the Pdfcrowd HTML to PDF API PDFCROWD

| | | | |
|---|---|---|---|
| SPGP0AF01650 | XP_018222423.1 | 99 | 0 |
| SPGP0CPtrna12Q | DI49_2421 | 100 | 0 |
| SPGP0DA00200 | XP_018222733.1 | 99 | 0 |
| SPGP0V00480 | XP_018221737.1 | 99 | 0 |
| SPGP0F01570 | XP_018221320.1 | 99 | 0 |
| SPGP0CN00620 | XP_018220909.1 | 99 | 0 |
| SPGP0J02640 | XP_018219685.1 | 99 | 0 |
| SPGP0F01410 | XP_018221305.1 | 99 | 0 |
| SPGP0AF01110 | XP_018222366.1 | 99 | 0 |
| SPGP0F00320 | XP_018221200.1 | 99 | 0 |
| SPGP0BE00200 | XP_018223358.1 | 99 | 0 |
| SPGP0CX00890 | XP_018222108.1 | 99 | 0 |
| SPGP0F01900 | XP_018221352.1 | 99 | 0 |
| SPGP0X01690 | XP_018221612.1 | 99 | 0 |
| SPGP0GL00170 | XP_018220846.1 | 99 | 0 |
| SPGP0F00790 | XP_018221243.1 | 99 | 0 |
| SPGP0AA01370 | XP_018220453.1 | 99 | 0 |
| SPGP0F02060 | XP_018221368.1 | 99 | 0 |
| SPGP0CN01010 | XP_018220871.1 | 99 | 0 |
| SPGP0J01190 | XP_018219829.1 | 100 | 0 |
| SPGP0J03320 | XP_018219617.1 | 99 | 0 |
| SPGP0AF01020 | XP_018222357.1 | 99 | 0 |
| SPGP0U00630 | XP_018223012.1 | 99 | 0 |
| SPGP0CU01480 | XP_018220686.1 | 99 | 1 |
| SPGP0X00240 | XP_018221473.1 | 99 | 0 |
| SPGP0J00860 | XP_018219864.1 | 99 | 0 |
| SPGP0CN01100 | XP_018220862.1 | 100 | 0 |
| SPGP0L01030 | XP_018218929.1 | 99 | 0 |
| SPGP0AT01370 | XP_018219154.1 | 99 | 0 |
| SPGP0J02420 | XP_018219707.1 | 99 | 0 |
| SPGP0AE00290 | XP_018222510.1 | 99 | 0 |
| SPGP0CP00180 | XP_018223818.1 | 99 | 0 |
| SPGP0CX01170 | XP_018222136.1 | 99 | 0 |
| SPGP0X01790 | XP_018221622.1 | 98 | 1 |
| SPGP0J02340 | XP_018219715.1 | 99 | 0 |
| SPGP0L01830 | XP_018218852.1 | 99 | 0 |
| SPGP0J00140 | XP_018219932.1 | 89 | 1 |
| SPGP0AT01480 | XP_018219165.1 | 99 | 0 |
| SPGP0F01140 | XP_018221278.1 | 99 | 0 |
| SPGP0CP00230 | XP_018223813.1 | 99 | 0 |

Create PDF in your applications with the Pdfcrowd HTML to PDF API PDFCROWD

| | | | |
|---|---|---|---|
| SPGP0L00270 | XP_018219005.1 | 99 | 0 |
| SPGP0J03770 | XP_018219572.1 | 99 | 0 |
| SPGP0F01110 | XP_018221275.1 | 99 | 0 |
| SPGP0AA01190 | XP_018220471.1 | 99 | 0 |
| SPGP0Z00200 | XP_018222544.1 | 96 | 0 |
| SPGP0ANtrna17L | DI49_5686 | 100 | 0 |
| SPGP0CUtrna27L | DI49_5687 | 100 | 0 |
| SPGP0IN00160 | XP_018221660.1 | 99 | 0 |
| SPGP0AA00170 | XP_018220573.1 | 100 | 0 |
| SPGP0X00980 | XP_018221545.1 | 99 | 0 |
| SPGP0X00630 | XP_018221511.1 | 99 | 0 |
| SPGP0CW00590 | XP_018219497.1 | 99 | 0 |
| SPGP0AF00850 | XP_018222339.1 | 99 | 0 |
| SPGP0Y00360 | XP_018220053.1 | 99 | 0 |
| SPGP0F00630 | XP_018221228.1 | 99 | 0 |
| SPGP0Y00230 | XP_018220040.1 | 99 | 0 |
| SPGP0AF01180 | XP_018222374.1 | 99 | 0 |
| SPGP0CU00920 | XP_018220737.1 | 100 | 0 |
| SPGP0CW00450 | XP_018219510.1 | 99 | 0 |
| SPGP0H00710 | XP_018221074.1 | 99 | 0 |
| SPGP0CN00940 | XP_018220878.1 | 99 | 0 |
| SPGP0CN00240 | XP_018220947.1 | 100 | 0 |
| SPGP0AA00710 | XP_018220520.1 | 99 | 0 |
| SPGP0X00180 | XP_018221467.1 | 97 | 0 |
| SPGP0CM00120 | XP_018223312.1 | 99 | 0 |
| SPGP0DA01310 | XP_018222624.1 | 99 | 0 |
| SPGP0AF00580 | XP_018222312.1 | 99 | 0 |
| SPGP0DA00560 | XP_018222698.1 | 99 | 0 |
| SPGP0DA00920 | XP_018222663.1 | 99 | 0 |
| SPGP0H00810 | XP_018221064.1 | 99 | 0 |
| SPGP0AT01620 | XP_018219179.1 | 99 | 0 |
| SPGP0X01580 | XP_018221601.1 | 99 | 0 |
| SPGP0X01590 | XP_018221602.1 | 99 | 0 |
| SPGP0H00820 | XP_018221063.1 | 99 | 1 |
| SPGP0CP02780 | XP_018223563.1 | 99 | 0 |
| SPGP0AN00490 | XP_018223956.1 | 99 | 0 |
| SPGP0CP02920 | XP_018223549.1 | 99 | 1 |
| SPGP0Y02880 | XP_018220297.1 | 99 | 0 |
| SPGP0DA00690 | XP_018222685.1 | 99 | 0 |

Create PDF in your applications with the Pdfcrowd HTML to PDF API PDFCROWD

| | | | |
|---|---|---|---|
| SPGP0V01030 | XP_018221684.1 | 98 | 0 |
| SPGP0CP01000 | XP_018223739.1 | 99 | 0 |
| SPGP0AJ01310 | XP_018221902.1 | 99 | 0 |
| SPGP0F02680 | XP_018221428.1 | 99 | 0 |
| SPGP0DE00430 | XP_018223860.1 | 99 | 0 |
| SPGP0H01190 | XP_018221028.1 | 99 | 0 |
| SPGP0K00140 | XP_018222756.1 | 98 | 1 |
| SPGP0GP00120 | XP_018223415.1 | 99 | 0 |
| SPGP0H01430 | XP_018221005.1 | 99 | 0 |
| SPGP0CW00360 | XP_018219519.1 | 99 | 0 |
| SPGP0J02980 | XP_018219651.1 | 99 | 0 |
| SPGP0AF00320 | XP_018222286.1 | 100 | 0 |
| SPGP0L02240 | XP_018218812.1 | 99 | 0 |
| SPGP0DA01610 | XP_018222596.1 | 99 | 0 |
| SPGP0J01660 | XP_018219782.1 | 99 | 0 |
| SPGP0DA00230 | XP_018222730.1 | 99 | 0 |
| SPGP0CN01110 | XP_018220861.1 | 99 | 0 |
| SPGP0DA00140 | XP_018222738.1 | 99 | 0 |
| SPGP0CX00840 | XP_018222103.1 | 99 | 0 |
| SPGP0L02080 | XP_018218828.1 | 99 | 0 |
| SPGP0EA00120 | XP_018222479.1 | 99 | 0 |
| SPGP0AU00300 | XP_018223085.1 | 99 | 0 |
| SPGP0CW01570 | XP_018219401.1 | 99 | 0 |
| SPGP0DB00260 | XP_018223900.1 | 99 | 0 |
| SPGP0CN00850 | XP_018220887.1 | 99 | 0 |
| SPGP0F01790 | XP_018221341.1 | 99 | 0 |
| SPGP0CM00720 | XP_018223253.1 | 99 | 0 |
| SPGP0CU01720 | XP_018220662.1 | 99 | 0 |
| SPGP0J01120 | XP_018219835.1 | 99 | 0 |
| SPGP0AA00970 | XP_018220493.1 | 99 | 0 |
| SPGP0F00840 | XP_018221248.1 | 97 | 1 |
| SPGP0CP02380 | XP_018223603.1 | 99 | 0 |
| SPGP0J03640 | XP_018219585.1 | 99 | 0 |
| SPGP0K00100 | XP_018222752.1 | 99 | 0 |
| SPGP0JB00180 | XP_018222563.1 | 99 | 0 |
| SPGP0AN00870 | XP_018223993.1 | 100 | 0 |
| SPGP0GA00170 | XP_018223386.1 | 99 | 0 |
| SPGP0CV00370 | XP_018222588.1 | 99 | 0 |
| SPGP0GR00140 | XP_018221979.1 | 99 | 0 |
| SPGP0U01670 | XP_018222915.1 | 99 | 0 |

Create PDF in your applications with the Pdfcrowd HTML to PDF API PDFCROWD

| | | | |
|---|---|---|---|
| SPGP0CP03370 | XP_018223504.1 | 99 | 0 |
| SPGP0JE00110 | XP_018222491.1 | 99 | 0 |
| SPGP0CN00720 | XP_018220899.1 | 99 | 0 |
| SPGP0H01450 | XP_018221003.1 | 99 | 0 |
| SPGP0CW01770 | XP_018219383.1 | 99 | 0 |
| SPGP0EO00410 | XP_018221178.1 | 98 | 0 |
| SPGP0AJtrna13P | DI49_2548 | 98 | 0 |
| SPGP0AJtrna13V | DI49_2588 | 100 | 0 |
| SPGP0R00480 | XP_018219989.1 | 100 | 0 |
| SPGP0L00660 | XP_018218967.1 | 99 | 0 |
| SPGP0DE00390 | XP_018223856.1 | 99 | 0 |
| SPGP0J02360 | XP_018219713.1 | 99 | 0 |
| SPGP0CU01790 | XP_018220655.1 | 99 | 0 |
| SPGP0CM01620 | XP_018223164.1 | 99 | 0 |
| SPGP0X01840 | XP_018221627.1 | 99 | 0 |
| SPGP0U00810 | XP_018222997.1 | 99 | 0 |
| SPGP0X00250 | XP_018221474.1 | 99 | 0 |
| SPGP0R00660 | XP_018220006.1 | 99 | 0 |
| SPGP0CP01190 | XP_018223719.1 | 99 | 0 |
| SPGP0Y01770 | XP_018220190.1 | 99 | 0 |
| SPGP0CX01850 | XP_018222200.1 | 98 | 1 |
| SPGP0F02550 | XP_018221415.1 | 99 | 0 |
| SPGP0DB00440 | XP_018223916.1 | 99 | 0 |
| SPGP0AN00790 | XP_018223985.1 | 99 | 0 |
| SPGP0J03450 | XP_018219603.1 | 99 | 0 |
| SPGP0L01530 | XP_018218881.1 | 99 | 0 |
| SPGP0AN00680 | XP_018223975.1 | 99 | 0 |
| SPGP0CW02390 | XP_018219322.1 | 99 | 0 |
| SPGP0Y01360 | XP_018220151.1 | 99 | 0 |
| SPGP0F02410 | XP_018221402.1 | 99 | 0 |
| SPGP0H00100 | XP_018221131.1 | 99 | 0 |
| SPGP0H00450 | XP_018221099.1 | 99 | 0 |
| SPGP0AN01150 | XP_018224021.1 | 99 | 0 |
| SPGP0DA01570 | XP_018222600.1 | 99 | 0 |
| SPGP0X01390 | XP_018221583.1 | 99 | 0 |
| SPGP0X00370 | XP_018221486.1 | 99 | 0 |
| SPGP0X00840 | XP_018221532.1 | 99 | 0 |
| SPGP0J00660 | XP_018219884.1 | 99 | 0 |
| SPGP0EX00110 | XP_018218750.1 | 99 | 0 |
| SPGP0H01270 | XP_018221020.1 | 99 | 0 |

Create PDF in your applications with the Pdfcrowd HTML to PDF API PDFCROWD

| | | | |
|---|---|---|---|
| SPGP0F01080 | XP_018221272.1 | 99 | 0 |
| SPGP0IO00120 | XP_018221985.1 | 98 | 0 |
| SPGP0Y01940 | XP_018220205.1 | 99 | 0 |
| SPGP0C00120 | XP_018222854.1 | 88 | 0 |
| SPGP0DA00220 | XP_018222731.1 | 99 | 0 |
| SPGP0CP03330 | XP_018223507.1 | 99 | 0 |
| SPGP0DA01110 | XP_018222644.1 | 99 | 0 |
| SPGP0S00440 | XP_018224092.1 | 99 | 0 |
| SPGP0F01580 | XP_018221321.1 | 99 | 0 |
| SPGP0L01200 | XP_018218914.1 | 99 | 0 |
| SPGP0CN00600 | XP_018220911.1 | 100 | 0 |
| SPGP0R00210 | XP_018219963.1 | 99 | 0 |
| SPGP0DB00490 | XP_018223919.1 | 99 | 0 |
| SPGP0S00180 | XP_018224067.1 | 99 | 0 |
| SPGP0L00670 | XP_018218966.1 | 99 | 0 |
| SPGP0H00230 | XP_018221121.1 | 92 | 6 |
| SPGP0FI00150 | XP_018224045.1 | 96 | 0 |
| SPGP0CP01610 | XP_018223678.1 | 99 | 0 |
| SPGP0J00840 | XP_018219866.1 | 99 | 0 |
| SPGP0AT01570 | XP_018219174.1 | 99 | 0 |
| SPGP0F01670 | XP_018221329.1 | 99 | 0 |
| SPGP0AN00700 | XP_018223977.1 | 99 | 0 |
| SPGP0J02620 | XP_018219687.1 | 99 | 0 |
| SPGP0J02990 | XP_018219650.1 | 99 | 0 |
| SPGP0AA01500 | XP_018220440.1 | 100 | 0 |
| SPGP0CU01390 | XP_018220694.1 | 99 | 0 |
| SPGP0V00820 | XP_018221705.1 | 99 | 0 |
| SPGP0AF02120 | XP_018222470.1 | 99 | 0 |
| SPGP0CM01410 | XP_018223185.1 | 99 | 0 |
| SPGP0AT01310 | XP_018219149.1 | 99 | 0 |
| SPGP0DE00460 | XP_018223863.1 | 100 | 0 |
| SPGP0AT00410 | XP_018219062.1 | 99 | 0 |
| SPGP0J02190 | XP_018219730.1 | 99 | 0 |
| SPGP0AQ00120 | XP_018223410.1 | 99 | 0 |
| SPGP0H01800 | XP_018220970.1 | 99 | 0 |
| SPGP0F01300 | XP_018221294.1 | 99 | 0 |
| SPGP0CW01150 | XP_018219442.1 | 99 | 0 |
| SPGP0CU00740 | XP_018220755.1 | 100 | 0 |
| SPGP0AF02080 | XP_018222466.1 | 99 | 0 |
| SPGP0J03570 | XP_018219592.1 | 99 | 0 |

Create PDF in your applications with the Pdfcrowd HTML to PDF API PDFCROWD

| | | | |
|---|---|---|---|
| SPGP0CW03150 | XP_018219248.1 | 99 | 0 |
| SPGP0CX02210 | XP_018222235.1 | 99 | 0 |
| SPGP0F01890 | XP_018221351.1 | 99 | 0 |
| SPGP0CX01810 | XP_018222196.1 | 99 | 0 |
| SPGP0U01800 | XP_018222901.1 | 99 | 0 |
| SPGP0CP00250 | XP_018223811.1 | 99 | 0 |
| SPGP0V00320 | XP_018221751.1 | 99 | 0 |
| SPGP0J01710 | XP_018219777.1 | 99 | 1 |
| SPGP0F01160 | XP_018221280.1 | 97 | 1 |
| SPGP0CW02540 | XP_018219307.1 | 99 | 0 |
| SPGP0H00310 | XP_018221113.1 | 99 | 0 |
| SPGP0CN00590 | XP_018220912.1 | 100 | 0 |
| SPGP0GQ00100 | XP_018219559.1 | 99 | 0 |
| SPGP0CP03500 | XP_018223492.1 | 99 | 0 |
| SPGP0R00290 | XP_018219971.1 | 99 | 0 |
| SPGP0U00550 | XP_018223019.1 | 99 | 0 |
| SPGP0L01180 | XP_018218916.1 | 99 | 0 |
| SPGP0CX01110 | XP_018222130.1 | 98 | 0 |
| SPGP0F02300 | XP_018221391.1 | 99 | 1 |
| SPGP0AA00480 | XP_018220542.1 | 99 | 0 |
| SPGP0X00610 | XP_018221510.1 | 99 | 0 |
| SPGP0X00960 | XP_018221543.1 | 99 | 0 |
| SPGP0CM00140 | XP_018223310.1 | 99 | 0 |
| SPGP0F00410 | XP_018221208.1 | 99 | 0 |
| SPGP0Y00380 | XP_018220055.1 | 99 | 0 |
| SPGP0R00120 | XP_018219954.1 | 99 | 0 |
| SPGP0K00670 | XP_018222809.1 | 99 | 0 |
| SPGP0F02740 | XP_018221434.1 | 99 | 0 |
| SPGP0AJ01490 | XP_018221918.1 | 99 | 0 |
| SPGP0DE00560 | XP_018223873.1 | 99 | 0 |
| SPGP0Y00710 | XP_018220087.1 | 99 | 0 |
| SPGP0H00730 | XP_018221072.1 | 96 | 0 |
| SPGP0CP02710 | XP_018223570.1 | 99 | 0 |
| SPGP0L00960 | XP_018218936.1 | 100 | 0 |
| SPGP0DA00500 | XP_018222704.1 | 99 | 0 |
| SPGP0CN00980 | XP_018220874.1 | 100 | 0 |
| SPGP0R00610 | XP_018220001.1 | 99 | 0 |
| SPGP0EI00110 | XP_018223325.1 | 99 | 0 |
| SPGP0AE00300 | XP_018222511.1 | 99 | 0 |
| SPGP0Y02740 | XP_018220283.1 | 99 | 0 |

Create PDF in your applications with the Pdfcrowd HTML to PDF API PDFCROWD

| | | | |
|---|---|---|---|
| SPGP0H01460 | XP_018221002.1 | 99 | 0 |
| SPGP0X01560 | XP_018221599.1 | 99 | 0 |
| SPGP0H00840 | XP_018221061.1 | 100 | 0 |
| SPGP0GG00130 | XP_018220838.1 | 99 | 0 |
| SPGP0CW00710 | XP_018219485.1 | 99 | 0 |
| SPGP0J03420 | XP_018219606.1 | 99 | 0 |
| SPGP0J03720 | XP_018219577.1 | 99 | 0 |
| SPGP0AJ01660 | XP_018221934.1 | 100 | 0 |
| SPGP0AA01460 | XP_018220444.1 | 99 | 0 |
| SPGP0AU00370 | XP_018223092.1 | 99 | 1 |
| SPGP0AN00160 | XP_018223928.1 | 99 | 0 |
| SPGP0CM01540 | XP_018223172.1 | 99 | 0 |
| SPGP0HB00100 | XP_018222861.1 | 98 | 0 |
| SPGP0CW01860 | XP_018219374.1 | 99 | 0 |
| SPGP0J00120 | XP_018219934.1 | 99 | 0 |
| SPGP0J00430 | XP_018219905.1 | 99 | 0 |
| SPGP0ATtrna3C | DI49_5567 | 100 | 0 |
| SPGP0CU02340 | XP_018220600.1 | 99 | 0 |
| SPGP0CW01440 | XP_018219414.1 | 99 | 0 |
| SPGP0CP00410 | XP_018223795.1 | 99 | 0 |
| SPGP0J03310 | XP_018219618.1 | 99 | 0 |
| SPGP0CP01090 | XP_018223730.1 | 99 | 0 |
| SPGP0S00530 | XP_018224100.1 | 99 | 0 |
| SPGP0L02170 | XP_018218819.1 | 99 | 0 |
| SPGP0H00900 | XP_018221055.1 | 99 | 0 |
| SPGP0J01510 | XP_018219797.1 | 99 | 1 |
| SPGP0CU00340 | XP_018220795.1 | 99 | 0 |
| SPGP0Z00110 | XP_018222554.1 | 99 | 0 |
| SPGP0AF00840 | XP_018222338.1 | 99 | 0 |
| SPGP0BE00150 | XP_018223363.1 | 99 | 0 |
| SPGP0CX01030 | XP_018222122.1 | 98 | 0 |
| SPGP0Y01330 | XP_018220148.1 | 99 | 0 |
| SPGP0X01550 | XP_018221598.1 | 99 | 0 |
| SPGP0CN00290 | XP_018220942.1 | 99 | 0 |
| SPGP0CW01030 | XP_018219453.1 | 100 | 0 |
| SPGP0CP00340 | XP_018223802.1 | 99 | 0 |
| SPGP0CP03290 | XP_018223511.1 | 99 | 0 |
| SPGP0F00820 | XP_018221246.1 | 99 | 0 |
| SPGP0AA01710 | XP_018220420.1 | 99 | 0 |
| SPGP0CX00790 | XP_018222098.1 | 99 | 0 |

Create PDF in your applications with the Pdfcrowd HTML to PDF API PDFCROWD

| | | | |
|---|---|---|---|
| SPGP0Ftrna8W | DI49_3006 | 100 | 0 |
| SPGP0J01220 | XP_018219826.1 | 99 | 0 |
| SPGP0CN00470 | XP_018220924.1 | 99 | 0 |
| SPGP0DA01450 | XP_018222610.1 | 99 | 0 |
| SPGP0AJ00420 | XP_018221815.1 | 99 | 0 |
| SPGP0AA00350 | XP_018220555.1 | 99 | 0 |
| SPGP0CW01470 | XP_018219411.1 | 99 | 0 |
| SPGP0CP02610 | XP_018223580.1 | 99 | 0 |
| SPGP0Y03090 | XP_018220318.1 | 99 | 0 |
| SPGP0M00100 | XP_018221146.1 | 99 | 0 |
| SPGP0S00780 | XP_018224122.1 | 99 | 0 |
| SPGP0F02530 | XP_018221413.1 | 99 | 0 |
| SPGP0CX00590 | XP_018222079.1 | 99 | 0 |
| SPGP0X01600 | XP_018221603.1 | 99 | 0 |
| SPGP0DF00230 | XP_018219185.1 | 98 | 0 |
| SPGP0K00480 | XP_018222790.1 | 99 | 0 |
| SPGP0V00650 | XP_018221721.1 | 99 | 0 |
| SPGP0L00410 | XP_018218991.1 | 99 | 0 |
| SPGP0L00130 | XP_018219018.1 | 99 | 0 |
| SPGP0F01420 | XP_018221306.1 | 99 | 0 |
| SPGP0F02110 | XP_018221373.1 | 99 | 0 |
| SPGP0AF02140 | XP_018222472.1 | 99 | 0 |
| SPGP0AF00150 | XP_018222270.1 | 99 | 0 |
| SPGP0S00250 | XP_018224074.1 | 99 | 0 |
| SPGP0Y01710 | XP_018220185.1 | 99 | 0 |
| SPGP0F02780 | XP_018221438.1 | 99 | 0 |
| SPGP0AN00850 | XP_018223991.1 | 99 | 0 |
| SPGP0X01950 | XP_018221638.1 | 99 | 0 |
| SPGP0CW00430 | XP_018219512.1 | 99 | 0 |
| SPGP0J03380 | XP_018219610.1 | 99 | 0 |
| SPGP0F00740 | XP_018221239.1 | 99 | 1 |
| SPGP0CX01470 | XP_018222164.1 | 99 | 0 |
| SPGP0AT00180 | XP_018219040.1 | 99 | 0 |
| SPGP0CU00550 | XP_018220774.1 | 99 | 0 |
| SPGP0L01010 | XP_018218931.1 | 99 | 0 |
| SPGP0AF00420 | XP_018222296.1 | 99 | 0 |
| SPGP0AA01220 | XP_018220468.1 | 99 | 0 |
| SPGP0AT00390 | XP_018219060.1 | 99 | 0 |
| SPGP0AJ01870 | XP_018221954.1 | 99 | 0 |
| SPGP0L01770 | XP_018218858.1 | 99 | 0 |

Create PDF in your applications with the Pdfcrowd HTML to PDF API PDFCROWD

| | | | |
|---|---|---|---|
| SPGP0K00320 | XP_018222774.1 | 99 | 0 |
| SPGP0H00540 | XP_018221091.1 | 100 | 0 |
| SPGP0R00170 | XP_018219959.1 | 99 | 0 |
| SPGP0CU01210 | XP_018220711.1 | 99 | 0 |
| SPGP0CU00770 | XP_018220752.1 | 99 | 0 |
| SPGP0F00830 | XP_018221247.1 | 99 | 0 |
| SPGP0J00300 | XP_018219918.1 | 99 | 0 |
| SPGP0L01380 | XP_018218896.1 | 99 | 0 |
| SPGP0X00390 | XP_018221488.1 | 99 | 0 |
| SPGP0J00600 | XP_018219890.1 | 98 | 1 |
| SPGP0AT00840 | XP_018219103.1 | 99 | 0 |
| SPGP0F00580 | XP_018221223.1 | 99 | 0 |
| SPGP0AJ01290 | XP_018221900.1 | 99 | 0 |
| SPGP0CN00510 | XP_018220920.1 | 99 | 0 |
| SPGP0L01070 | XP_018218925.1 | 100 | 0 |
| SPGP0DB00450 | XP_018223917.1 | 99 | 0 |
| SPGP0DA00240 | XP_018222729.1 | 99 | 0 |
| SPGP0DA00710 | XP_018222682.1 | 99 | 0 |
| SPGP0AJ01120 | XP_018221883.1 | 99 | 0 |
| SPGP0F00360 | XP_018221203.1 | 99 | 0 |
| SPGP0AT01130 | XP_018219131.1 | 99 | 0 |
| SPGP0DB00320 | XP_018223905.1 | 99 | 0 |
| SPGP0J01080 | XP_018219839.1 | 99 | 0 |
| SPGP0CW01370 | XP_018219421.1 | 99 | 0 |
| SPGP0CM01020 | XP_018223223.1 | 100 | 0 |
| SPGP0H01590 | XP_018220990.1 | 99 | 0 |
| SPGP0V00360 | XP_018221747.1 | 99 | 0 |
| SPGP0V00610 | XP_018221725.1 | 99 | 0 |
| SPGP0CP01630 | XP_018223676.1 | 99 | 0 |
| SPGP0CX01130 | XP_018222132.1 | 99 | 0 |
| SPGP0AT01550 | XP_018219172.1 | 99 | 0 |
| SPGP0AN00580 | XP_018223965.1 | 99 | 0 |
| SPGP0AJ00640 | XP_018221836.1 | 100 | 0 |
| SPGP0J02600 | XP_018219689.1 | 99 | 0 |
| SPGP0CN00660 | XP_018220905.1 | 99 | 0 |
| SPGP0J00630 | XP_018219887.1 | 99 | 0 |
| SPGP0X00900 | XP_018221537.1 | 99 | 0 |
| SPGP0J00420 | XP_018219906.1 | 99 | 0 |
| SPGP0V00770 | XP_018221710.1 | 99 | 0 |
| SPGP0V00800 | XP_018221707.1 | 99 | 0 |

Create PDF in your applications with the Pdfcrowd HTML to PDF API PDFCROWD

| | | | |
|---|---|---|---|
| SPGP0H01230 | XP_018221024.1 | 99 | 0 |
| SPGP0CW00630 | XP_018219493.1 | 99 | 1 |
| SPGP0CX00750 | XP_018222094.1 | 99 | 0 |
| SPGP0AJ01670 | XP_018221935.1 | 99 | 0 |
| SPGP0J01930 | XP_018219756.1 | 99 | 0 |
| SPGP0J03020 | XP_018219647.1 | 99 | 0 |
| SPGP0F01360 | XP_018221300.1 | 99 | 0 |
| SPGP0S00700 | XP_018224117.1 | 99 | 0 |
| SPGP0U00480 | XP_018223026.1 | 100 | 0 |
| SPGP0F00870 | XP_018221251.1 | 99 | 0 |
| SPGP0CP00380 | XP_018223798.1 | 99 | 0 |
| SPGP0U00190 | XP_018223051.1 | 99 | 0 |
| SPGP0J03550 | XP_018219594.1 | 99 | 0 |
| SPGP0J00440 | XP_018219904.1 | 99 | 0 |
| SPGP0U01440 | XP_018222938.1 | 98 | 0 |
| SPGP0J01960 | XP_018219753.1 | 99 | 0 |
| SPGP0EM00160 | XP_018223065.1 | 99 | 0 |
| SPGP0DA01520 | XP_018222603.1 | 99 | 0 |
| SPGP0CP02740 | XP_018223567.1 | 100 | 0 |
| SPGP0CP00270 | XP_018223809.1 | 99 | 0 |
| SPGP0CN00280 | XP_018220943.1 | 99 | 0 |
| SPGP0F01180 | XP_018221282.1 | 99 | 0 |
| SPGP0J00230 | XP_018219924.1 | 99 | 0 |
| SPGP0J01770 | XP_018219771.1 | 99 | 0 |
| SPGP0F01980 | XP_018221360.1 | 99 | 0 |
| SPGP0AA00820 | XP_018220508.1 | 98 | 0 |
| SPGP0Y00630 | XP_018220079.1 | 99 | 0 |
| SPGP0H00860 | XP_018221059.1 | 99 | 0 |
| SPGP0CW03280 | XP_018219236.1 | 100 | 0 |
| SPGP0Z00290 | XP_018222536.1 | 99 | 0 |
| SPGP0L00580 | XP_018218975.1 | 98 | 1 |
| SPGP0F02980 | XP_018221991.1 | 98 | 0 |
| SPGP0AA00130 | XP_018220577.1 | 99 | 0 |
| SPGP0J00490 | XP_018219899.1 | 99 | 0 |
| SPGP0CX00580 | XP_018222078.1 | 98 | 2 |
| SPGP0X00940 | XP_018221541.1 | 99 | 0 |
| SPGP0CX00630 | XP_018222082.1 | 99 | 0 |
| SPGP0X00490 | XP_018221498.1 | 99 | 0 |
| SPGP0AF00810 | XP_018222335.1 | 99 | 0 |
| SPGP0J02940 | XP_018219655.1 | 99 | 1 |

Create PDF in your applications with the Pdfcrowd HTML to PDF API PDFCROWD

| | | | |
|---|---|---|---|
| SPGP0L01980 | XP_018218838.1 | 99 | 0 |
| SPGP0L00360 | XP_018218996.1 | 99 | 0 |
| SPGP0CP02500 | XP_018223591.1 | 99 | 0 |
| SPGP0F00310 | XP_018221199.1 | 99 | 0 |
| SPGP0EO00200 | XP_018221157.1 | 99 | 0 |
| SPGP0CW00730 | XP_018219483.1 | 99 | 0 |
| SPGP0CP02650 | XP_018223576.1 | 99 | 0 |
| SPGP0DA00190 | XP_018222734.1 | 99 | 0 |
| SPGP0CP01380 | XP_018223701.1 | 100 | 0 |
| SPGP0CN00340 | XP_018220937.1 | 99 | 0 |
| SPGP0FP00210 | XP_018223334.1 | 99 | 0 |
| SPGP0V00810 | XP_018221706.1 | 99 | 0 |
| SPGP0CW02970 | XP_018219266.1 | 99 | 0 |
| SPGP0CN00890 | XP_018220883.1 | 99 | 0 |
| SPGP0Vtrna9P | DI49_2389 | 99 | 0 |
| SPGP0H00920 | XP_018221053.1 | 99 | 0 |
| SPGP0J03000 | XP_018219649.1 | 100 | 0 |
| SPGP0CX00670 | XP_018222086.1 | 99 | 0 |
| SPGP0CP00740 | XP_018223764.1 | 99 | 0 |
| SPGP0X00230 | XP_018221472.1 | 99 | 0 |
| SPGP0CP02230 | XP_018223617.1 | 99 | 0 |
| SPGP0F02500 | XP_018221410.1 | 99 | 0 |
| SPGP0AE00180 | XP_018222500.1 | 99 | 0 |
| SPGP0DA01390 | XP_018222616.1 | 99 | 0 |
| SPGP0DA01380 | XP_018222617.1 | 99 | 0 |
| SPGP0J03210 | XP_018219628.1 | 99 | 0 |
| SPGP0AF01490 | XP_018222407.1 | 99 | 1 |
| SPGP0DA01430 | XP_018222612.1 | 99 | 0 |
| SPGP0X01720 | XP_018221615.1 | 100 | 0 |
| SPGP0EO00330 | XP_018221170.1 | 98 | 0 |
| SPGP0AA00840 | XP_018220506.1 | 99 | 0 |
| SPGP0DA00380 | XP_018222716.1 | 99 | 0 |
| SPGP0L02200 | XP_018218816.1 | 100 | 0 |
| SPGP0CM01100 | XP_018223215.1 | 99 | 0 |
| SPGP0CP00710 | XP_018223767.1 | 99 | 0 |
| SPGP0CW03210 | XP_018219243.1 | 100 | 0 |
| SPGP0Y00190 | XP_018220036.1 | 99 | 0 |
| SPGP0CM02030 | XP_018223125.1 | 99 | 0 |
| SPGP0R00710 | XP_018220011.1 | 99 | 0 |
| SPGP0CW01050 | XP_018219451.1 | 99 | 0 |

Create PDF in your applications with the Pdfcrowd HTML to PDF API PDFCROWD

| | | | |
|---|---|---|---|
| SPGP0J01210 | XP_018219827.1 | 99 | 0 |
| SPGP0R00430 | XP_018219984.1 | 99 | 0 |
| SPGP0Y02170 | XP_018220226.1 | 99 | 0 |
| SPGP0F00770 | XP_018221242.1 | 99 | 0 |
| SPGP0F00800 | XP_018221244.1 | 99 | 0 |
| SPGP0CU02040 | XP_018220631.1 | 99 | 0 |
| SPGP0CN00410 | XP_018220930.1 | 99 | 0 |
| SPGP0CU00890 | XP_018220740.1 | 100 | 0 |
| SPGP0CW01500 | XP_018219408.1 | 99 | 0 |
| SPGP0CU01580 | XP_018220676.1 | 99 | 0 |
| SPGP0F01230 | XP_018221286.1 | 99 | 0 |
| SPGP0AA00100 | XP_018220580.1 | 99 | 0 |
| SPGP0AJ00400 | XP_018221813.1 | 99 | 0 |
| SPGP0X00210 | XP_018221470.1 | 99 | 0 |
| SPGP0CU02440 | XP_018220590.1 | 99 | 0 |
| SPGP0L00110 | XP_018219020.1 | 99 | 0 |
| SPGP0X00170 | XP_018221466.1 | 98 | 0 |
| SPGP0CU02230 | XP_018220611.1 | 99 | 0 |
| SPGP0H00890 | XP_018221056.1 | 99 | 0 |
| SPGP0J00930 | XP_018219858.1 | 98 | 0 |
| SPGP0Y01480 | XP_018220163.1 | 100 | 0 |
| SPGP0Ytrna6Q | DI49_4357 | 100 | 0 |
| SPGP0CW02350 | XP_018219326.1 | 99 | 0 |
| SPGP0CV00260 | XP_018222577.1 | 99 | 0 |
| SPGP0Y01730 | XP_018220187.1 | 99 | 0 |
| SPGP0CU02050 | XP_018220630.1 | 99 | 0 |
| SPGP0F01640 | XP_018221327.1 | 99 | 0 |
| SPGP0X01380 | XP_018221582.1 | 99 | 0 |
| SPGP0AF01760 | XP_018222434.1 | 99 | 0 |
| SPGP0R00620 | XP_018220002.1 | 99 | 0 |
| SPGP0F02040 | XP_018221366.1 | 99 | 0 |
| SPGP0GL00130 | XP_018220843.1 | 99 | 0 |
| SPGP0V00460 | XP_018221739.1 | 99 | 0 |
| SPGP0F02510 | XP_018221411.1 | 99 | 0 |
| SPGP0H00670 | XP_018221078.1 | 99 | 0 |
| SPGP0U00530 | XP_018223021.1 | 99 | 0 |
| SPGP0CW00240 | XP_018219529.1 | 100 | 0 |
| SPGP0DA01070 | XP_018222648.1 | 100 | 0 |
| SPGP0Itrna11N | DI49_5581 | 100 | 0 |
| SPGP0CWtrna4P | DI49_5158 | 99 | 0 |

Create PDF in your applications with the Pdfcrowd HTML to PDF API PDFCROWD

| | | | |
|---|---|---|---|
| SPGP0H01700 | XP_018220980.1 | 99 | 0 |
| SPGP0AA00670 | XP_018220524.1 | 99 | 0 |
| SPGP0CN00300 | XP_018220941.1 | 99 | 0 |
| SPGP0U01970 | XP_018222884.1 | 99 | 0 |
| SPGP0CW00880 | XP_018219468.1 | 95 | 4 |
| SPGP0J03050 | XP_018219644.1 | 99 | 0 |
| SPGP0AT01290 | XP_018219147.1 | 99 | 0 |
| SPGP0AH00130 | XP_018223421.1 | 100 | 0 |
| SPGP0X01760 | XP_018221619.1 | 99 | 0 |
| SPGP0AT01420 | XP_018219159.1 | 99 | 0 |
| SPGP0CU00750 | XP_018220754.1 | 100 | 0 |
| SPGP0Y03080 | XP_018220317.1 | 99 | 0 |
| SPGP0Y02900 | XP_018220299.1 | 99 | 0 |
| SPGP0Z00120 | XP_018222553.1 | 97 | 3 |
| SPGP0AF00170 | XP_018222272.1 | 99 | 0 |
| SPGP0AT01560 | XP_018219173.1 | 100 | 0 |
| SPGP0AJ00870 | XP_018221859.1 | 99 | 0 |
| SPGP0AN00640 | XP_018223971.1 | 99 | 0 |
| SPGP0CMtrna4C | DI49_5400 | 100 | 0 |
| SPGP0DA00260 | XP_018222727.1 | 99 | 0 |
| SPGP0Ftrna21R | DI49_3064 | 100 | 0 |
| SPGP0J01530 | XP_018219795.1 | 99 | 0 |
| SPGP0IR00150 | XP_018223469.1 | 99 | 0 |
| SPGP0L02540 | XP_018218782.1 | 99 | 0 |
| SPGP0X01060 | XP_018221552.1 | 99 | 0 |
| SPGP0DA01030 | XP_018222652.1 | 100 | 0 |
| SPGP0GG00140 | XP_018220837.1 | 99 | 0 |
| SPGP0GT00140 | XP_018219022.1 | 99 | 0 |
| SPGP0GP00130 | XP_018223414.1 | 99 | 0 |
| SPGP0X00740 | XP_018221522.1 | 99 | 0 |
| SPGP0J00880 | XP_018219862.1 | 99 | 0 |
| SPGP0CW01020 | XP_018219454.1 | 99 | 0 |
| SPGP0X01190 | XP_018221563.1 | 99 | 0 |
| SPGP0H00270 | XP_018221117.1 | 99 | 0 |
| SPGP0CP02260 | XP_018223614.1 | 100 | 0 |
| SPGP0AA00330 | XP_018220557.1 | 99 | 1 |
| SPGP0GL00160 | XP_018220845.1 | 99 | 0 |
| SPGP0J02950 | XP_018219654.1 | 99 | 0 |
| SPGP0AA00430 | XP_018220547.1 | 99 | 0 |

Create PDF in your applications with the Pdfcrowd HTML to PDF API PDFCROWD

| | | | |
|---|---|---|---|
| SPGP0F01430 | XP_018221307.1 | 99 | 0 |
| SPGP0F00370 | XP_018221204.1 | 98 | 0 |
| SPGP0S00760 | XP_018224121.1 | 94 | 1 |
| SPGP0F01340 | XP_018221298.1 | 98 | 1 |
| SPGP0CW02500 | XP_018219311.1 | 99 | 0 |
| SPGP0J01020 | XP_018219851.1 | 99 | 0 |
| SPGP0CU01930 | XP_018220641.1 | 99 | 0 |
| SPGP0AA01780 | XP_018220414.1 | 98 | 0 |
| SPGP0U01000 | XP_018222979.1 | 99 | 0 |
| SPGP0U02020 | XP_018222879.1 | 99 | 0 |
| SPGP0J02590 | XP_018219690.1 | 99 | 0 |
| SPGP0J02480 | XP_018219701.1 | 97 | 2 |
| SPGP0J02750 | XP_018219674.1 | 99 | 0 |
| SPGP0EM00140 | XP_018223063.1 | 99 | 0 |
| SPGP0CM01970 | XP_018223131.1 | 100 | 0 |
| SPGP0V01080 | XP_018221679.1 | 99 | 0 |
| SPGP0HC00130 | XP_018221784.1 | 99 | 0 |
| SPGP0F00420 | XP_018221209.1 | 99 | 0 |
| SPGP0Y02340 | XP_018220243.1 | 99 | 0 |
| SPGP0CM00510 | XP_018223273.1 | 99 | 0 |
| SPGP0CP00290 | XP_018223807.1 | 99 | 0 |
| SPGP0CX02220 | XP_018222236.1 | 99 | 0 |
| SPGP0EDtrna4F | DI49_0089 | 100 | 0 |
| SPGP0AF01720 | XP_018222430.1 | 99 | 0 |
| SPGP0J03280 | XP_018219621.1 | 99 | 0 |
| SPGP0FI00160 | XP_018224044.1 | 97 | 0 |
| SPGP0DA00590 | XP_018222695.1 | 99 | 0 |
| SPGP0EO00220 | XP_018221159.1 | 99 | 0 |
| SPGP0CW01900 | XP_018219370.1 | 99 | 0 |
| SPGP0L01960 | XP_018218840.1 | 99 | 0 |
| SPGP0DA00720 | XP_018222681.1 | 99 | 0 |
| SPGP0CP02750 | XP_018223566.1 | 99 | 0 |
| SPGP0CM00460 | XP_018223278.1 | 99 | 0 |
| SPGP0CP02630 | XP_018223578.1 | 100 | 0 |
| SPGP0DB00340 | XP_018223907.1 | 99 | 0 |
| SPGP0AN01040 | XP_018224010.1 | 99 | 0 |
| SPGP0CM00660 | XP_018223258.1 | 99 | 0 |
| SPGP0X01520 | XP_018221595.1 | 99 | 0 |
| SPGP0H00880 | XP_018221057.1 | 99 | 0 |
| SPGP0AN00340 | XP_018223943.1 | 99 | 0 |

Create PDF in your applications with the Pdfcrowd HTML to PDF API PDFCROWD

| | | | |
|---|---|---|---|
| SPGP0S00210 | XP_018224070.1 | 99 | 0 |
| SPGP0CP01050 | XP_018223734.1 | 99 | 0 |
| SPGP0F02700 | XP_018221430.1 | 99 | 0 |
| SPGP0DA00480 | XP_018222706.1 | 99 | 0 |
| SPGP0L00700 | XP_018218963.1 | 99 | 0 |
| SPGP0AF00690 | XP_018222323.1 | 99 | 0 |
| SPGP0F02860 | XP_018221445.1 | 99 | 0 |
| SPGP0S00290 | XP_018224078.1 | 99 | 0 |
| SPGP0Y01220 | XP_018220137.1 | 99 | 0 |
| SPGP0L02010 | XP_018218835.1 | 99 | 0 |
| SPGP0CP00450 | XP_018223791.1 | 99 | 0 |
| SPGP0AJ01550 | XP_018221923.1 | 99 | 0 |
| SPGP0GH00100 | XP_018222812.1 | 100 | 0 |
| SPGP0AJ01620 | XP_018221930.1 | 99 | 0 |
| SPGP0CM01850 | XP_018223141.1 | 99 | 0 |
| SPGP0H01680 | XP_018220982.1 | 99 | 0 |
| SPGP0J02900 | XP_018219659.1 | 99 | 0 |
| SPGP0DB00180 | XP_018223892.1 | 99 | 0 |
| SPGP0AH00160 | XP_018223424.1 | 99 | 0 |
| SPGP0Ftrna18R | DI49_2918 | 100 | 0 |
| SPGP0AT00640 | XP_018219084.1 | 99 | 0 |
| SPGP0S00570 | XP_018224104.1 | 100 | 0 |
| SPGP0V00380 | XP_018221745.1 | 99 | 0 |
| SPGP0AF00340 | XP_018222288.1 | 99 | 0 |
| SPGP0U02010 | XP_018222880.1 | 99 | 0 |
| SPGP0L00370 | XP_018218995.1 | 99 | 0 |
| SPGP0U00950 | XP_018222984.1 | 99 | 0 |
| SPGP0CV00320 | XP_018222583.1 | 99 | 0 |
| SPGP0GG00150 | XP_018220836.1 | 99 | 0 |
| SPGP0CW01070 | XP_018219449.1 | 99 | 0 |
| SPGP0CN00780 | XP_018220894.1 | 99 | 0 |
| SPGP0CN00250 | XP_018220946.1 | 99 | 0 |
| SPGP0J01560 | XP_018219792.1 | 99 | 0 |
| SPGP0DA00100 | XP_018222742.1 | 99 | 0 |
| SPGP0F01630 | XP_018221326.1 | 99 | 0 |
| SPGP0EC00100 | XP_018221982.1 | 99 | 0 |
| SPGP0CN00430 | XP_018220928.1 | 99 | 0 |
| SPGP0F01190 | XP_018221283.1 | 99 | 0 |
| SPGP0J01780 | XP_018219770.1 | 99 | 0 |
| SPGP0DE00320 | XP_018223849.1 | 99 | 0 |

Create PDF in your applications with the Pdfcrowd HTML to PDF API PDFCROWD

| | | | |
|---|---|---|---|
| SPGP0S00370 | XP_018224085.1 | 99 | 0 |
| SPGP0AJ00170 | XP_018221792.1 | 99 | 0 |
| SPGP0CU01110 | XP_018220719.1 | 99 | 0 |
| SPGP0CU00410 | XP_018220788.1 | 99 | 0 |
| SPGP0F00130 | XP_018221182.1 | 99 | 0 |
| SPGP0F02170 | XP_018221378.1 | 99 | 0 |
| SPGP0CU01050 | XP_018220725.1 | 97 | 0 |
| SPGP0CM01830 | XP_018223144.1 | 99 | 0 |
| SPGP0X00560 | XP_018221505.1 | 99 | 0 |
| SPGP0R00420 | XP_018219983.1 | 99 | 0 |
| SPGP0AF01900 | XP_018222448.1 | 99 | 0 |
| SPGP0CM01400 | XP_018223186.1 | 99 | 0 |
| SPGP0K00440 | XP_018222786.1 | 99 | 0 |
| SPGP0J02730 | XP_018219676.1 | 100 | 0 |
| SPGP0BR00100 | XP_018218924.1 | 99 | 0 |
| SPGP0F01660 | XP_018221328.1 | 99 | 0 |
| SPGP0AJ00820 | XP_018221854.1 | 100 | 0 |
| SPGP0CU01250 | XP_018220707.1 | 99 | 0 |
| SPGP0J02920 | XP_018219657.1 | 99 | 0 |
| SPGP0CW00990 | XP_018219457.1 | 99 | 0 |
| SPGP0H00650 | XP_018221080.1 | 100 | 0 |
| SPGP0AA01600 | XP_018220431.1 | 99 | 0 |
| SPGP0Y00520 | XP_018220069.1 | 99 | 0 |
| SPGP0Y01960 | XP_018220207.1 | 99 | 0 |
| SPGP0CW03320 | XP_018219232.1 | 99 | 0 |
| SPGP0AN00150 | XP_018223927.1 | 99 | 0 |
| SPGP0ED00130 | XP_018223472.1 | 99 | 0 |
| SPGP0J03070 | XP_018219642.1 | 99 | 0 |
| SPGP0GA00230 | XP_018223383.1 | 99 | 0 |
| SPGP0DA01080 | XP_018222647.1 | 99 | 0 |
| SPGP0AF01770 | XP_018222435.1 | 99 | 0 |
| SPGP0IH00190 | XP_018220389.1 | 98 | 0 |
| SPGP0K00300 | XP_018222772.1 | 100 | 0 |
| SPGP0H01160 | XP_018221030.1 | 100 | 0 |
| SPGP0AA00870 | XP_018220503.1 | 100 | 0 |
| SPGP0AN01010 | XP_018224007.1 | 99 | 0 |
| SPGP0CX00770 | XP_018222096.1 | 99 | 0 |
| SPGP0AT00800 | XP_018219099.1 | 99 | 0 |
| SPGP0CP03470 | XP_018223495.1 | 99 | 0 |
| SPGP0AH00110 | XP_018223419.1 | 99 | 0 |

Create PDF in your applications with the Pdfcrowd HTML to PDF API PDFCROWD

| | | | |
|---|---|---|---|
| SPGP0AF01930 | XP_018222451.1 | 100 | 0 |
| SPGP0H00440 | XP_018221100.1 | 99 | 0 |
| SPGP0AN00470 | XP_018223954.1 | 99 | 0 |
| SPGP0CU00630 | XP_018220765.1 | 99 | 0 |
| SPGP0CW00670 | XP_018219489.1 | 99 | 0 |
| SPGP0Y02920 | XP_018220301.1 | 97 | 3 |
| SPGP0J03790 | XP_018219570.1 | 99 | 0 |
| SPGP0H00220 | XP_018221122.1 | 99 | 0 |
| SPGP0H01740 | XP_018220976.1 | 98 | 1 |
| SPGP0DA00970 | XP_018222658.1 | 99 | 0 |
| SPGP0H01210 | XP_018221026.1 | 99 | 0 |
| SPGP0X01880 | XP_018221631.1 | 99 | 0 |
| SPGP0CM01240 | XP_018223201.1 | 99 | 0 |
| SPGP0IN00150 | XP_018221661.1 | 99 | 0 |
| SPGP0X00870 | XP_018221535.1 | 99 | 0 |
| SPGP0CP00970 | XP_018223742.1 | 99 | 0 |
| SPGP0CUtrna21V | DI49_4248 | 100 | 0 |
| SPGP0L01160 | XP_018218918.1 | 99 | 0 |
| SPGP0CM00260 | XP_018223298.1 | 99 | 0 |
| SPGP0CPtrna15I | DI49_0395 | 100 | 0 |
| SPGP0GR00130 | XP_018221978.1 | 99 | 0 |
| SPGP0AJ01160 | XP_018221887.1 | 100 | 0 |
| SPGP0CM01060 | XP_018223219.1 | 99 | 0 |
| SPGP0X01080 | XP_018221554.1 | 99 | 0 |
| SPGP0FE00150 | XP_018221967.1 | 99 | 0 |
| SPGP0JK00320 | XP_018223453.1 | 99 | 0 |
| SPGP0HB00110 | XP_018222862.1 | 99 | 0 |
| SPGP0CW00490 | XP_018219506.1 | 99 | 0 |
| SPGP0V00730 | XP_018221713.1 | 99 | 0 |
| SPGP0DA00820 | XP_018222671.1 | 99 | 0 |
| SPGP0BE00280 | XP_018223350.1 | 99 | 0 |
| SPGP0AT01040 | XP_018219122.1 | 99 | 0 |
| SPGP0CU02090 | XP_018220625.1 | 99 | 0 |
| SPGP0AT01060 | XP_018219124.1 | 100 | 0 |
| SPGP0AA01470 | XP_018220443.1 | 99 | 0 |
| SPGP0CP03130 | XP_018223527.1 | 99 | 0 |
| SPGP0J02930 | XP_018219656.1 | 99 | 0 |
| SPGP0AA01060 | XP_018220484.1 | 99 | 0 |
| SPGP0CM00530 | XP_018223271.1 | 99 | 0 |
| SPGP0GJ00160 | XP_018223376.1 | 99 | 0 |

Create PDF in your applications with the Pdfcrowd HTML to PDF API PDFCROWD

| | | | |
|---|---|---|---|
| SPGP0J01000 | XP_018219849.1 | 99 | 0 |
| SPGP0F02520 | XP_018221412.1 | 99 | 0 |
| SPGP0AJ00610 | XP_018221833.1 | 99 | 0 |
| SPGP0J00810 | XP_018219869.1 | 99 | 0 |
| SPGP0V01060 | XP_018221681.1 | 99 | 0 |
| SPGP0CN00650 | XP_018220906.1 | 99 | 0 |
| SPGP0CW03660 | XP_018219199.1 | 97 | 0 |
| SPGP0AA00460 | XP_018220544.1 | 99 | 0 |
| SPGP0F02540 | XP_018221414.1 | 100 | 0 |
| SPGP0JK00140 | XP_018223436.1 | 99 | 0 |
| SPGP0H01240 | XP_018221023.1 | 100 | 0 |
| SPGP0U00920 | XP_018222987.1 | 99 | 0 |
| SPGP0F01000 | XP_018221264.1 | 99 | 0 |
| SPGP0S00670 | XP_018224114.1 | 98 | 1 |
| SPGP0CP02770 | XP_018223564.1 | 99 | 0 |
| SPGP0CW00440 | XP_018219511.1 | 99 | 0 |
| SPGP0Y01590 | XP_018220173.1 | 99 | 0 |
| SPGP0HI00100 | XP_018218768.1 | 94 | 0 |
| SPGP0J01740 | XP_018219774.1 | 99 | 0 |
| SPGP0Y01670 | XP_018220181.1 | 99 | 0 |
| SPGP0S00280 | XP_018224077.1 | 99 | 0 |
| SPGP0GZ00100 | XP_018222822.1 | 99 | 0 |
| SPGP0L00890 | XP_018218944.1 | 99 | 0 |
| SPGP0J02200 | XP_018219729.1 | 99 | 0 |
| SPGP0DB00360 | XP_018223909.1 | 99 | 0 |
| SPGP0F02490 | XP_018221409.1 | 99 | 0 |
| SPGP0CP01700 | XP_018223670.1 | 99 | 0 |
| SPGP0R00180 | XP_018219960.1 | 99 | 1 |
| SPGP0J02310 | XP_018219718.1 | 99 | 0 |
| SPGP0X01500 | XP_018221593.1 | 99 | 0 |
| SPGP0GJ00140 | XP_018223378.1 | 99 | 0 |
| SPGP0CP02010 | XP_018223639.1 | 100 | 0 |
| SPGP0CW02760 | XP_018219287.1 | 99 | 0 |
| SPGP0DA01470 | XP_018222608.1 | 100 | 0 |
| SPGP0CU00650 | XP_018220763.1 | 99 | 0 |
| SPGP0CW01420 | XP_018219416.1 | 99 | 0 |
| SPGP0X01470 | XP_018221591.1 | 99 | 0 |
| SPGP0DA00120 | XP_018222740.1 | 100 | 0 |
| SPGP0CPtrna26R | DI49_4323 | 100 | 0 |
| SPGP0Y01630 | XP_018220177.1 | 100 | 0 |

Create PDF in your applications with the Pdfcrowd HTML to PDF API PDFCROWD

| | | | |
|---|---|---|---|
| SPGP0X00150 | XP_018221464.1 | 96 | 0 |
| SPGP0HN00130 | XP_018221989.1 | 99 | 0 |
| SPGP0F01930 | XP_018221355.1 | 99 | 0 |
| SPGP0CW02910 | XP_018219272.1 | 99 | 0 |
| SPGP0AA01910 | XP_018220401.1 | 99 | 0 |
| SPGP0Y02520 | XP_018220261.1 | 99 | 0 |
| SPGP0IR00140 | XP_018223468.1 | 99 | 0 |
| SPGP0Y02160 | XP_018220225.1 | 100 | 0 |
| SPGP0L01210 | XP_018218913.1 | 99 | 0 |
| SPGP0CM01870 | XP_018223139.1 | 99 | 0 |
| SPGP0AF00270 | XP_018222281.1 | 99 | 0 |
| SPGP0H01000 | XP_018221045.1 | 99 | 0 |
| SPGP0DA00470 | XP_018222707.1 | 99 | 0 |
| SPGP0CN00230 | XP_018220948.1 | 99 | 0 |
| SPGP0CM01180 | XP_018223207.1 | 99 | 0 |
| SPGP0AT01110 | XP_018219129.1 | 99 | 0 |
| SPGP0AJ01040 | XP_018221876.1 | 99 | 0 |
| SPGP0F00730 | XP_018221238.1 | 99 | 0 |
| SPGP0AF00920 | XP_018222346.1 | 100 | 0 |
| SPGP0CW00600 | XP_018219496.1 | 99 | 0 |
| SPGP0F02480 | XP_018221408.1 | 99 | 0 |
| SPGP0AN00550 | XP_018223962.1 | 99 | 0 |
| SPGP0H00190 | XP_018221124.1 | 100 | 0 |
| SPGP0AF01670 | XP_018222425.1 | 99 | 0 |
| SPGP0CW02720 | XP_018219291.1 | 99 | 0 |
| SPGP0CX01620 | XP_018222178.1 | 99 | 0 |
| SPGP0CU01010 | XP_018220729.1 | 99 | 0 |
| SPGP0AT00570 | XP_018219078.1 | 99 | 0 |
| SPGP0J02500 | XP_018219699.1 | 99 | 0 |
| SPGP0CP01440 | XP_018223695.1 | 99 | 0 |
| SPGP0K00520 | XP_018222794.1 | 100 | 0 |
| SPGP0AA00860 | XP_018220504.1 | 99 | 0 |
| SPGP0CU01470 | XP_018220687.1 | 99 | 0 |
| SPGP0AT01250 | XP_018219143.1 | 99 | 0 |
| SPGP0AF01920 | XP_018222450.1 | 99 | 0 |
| SPGP0CU02270 | XP_018220607.1 | 99 | 0 |
| SPGP0CM01810 | XP_018223146.1 | 99 | 0 |
| SPGP0CP02180 | XP_018223622.1 | 99 | 0 |
| SPGP0U00490 | XP_018223025.1 | 99 | 0 |
| SPGP0J00850 | XP_018219865.1 | 99 | 0 |

Create PDF in your applications with the Pdfcrowd HTML to PDF API PDFCROWD

| | | | |
|---|---|---|---|
| SPGP0AF00560 | XP_018222310.1 | 99 | 0 |
| SPGP0F02260 | XP_018221387.1 | 99 | 0 |
| SPGP0J00130 | XP_018219933.1 | 99 | 0 |
| SPGP0J00240 | XP_018219923.1 | 99 | 0 |
| SPGP0F01600 | XP_018221323.1 | 99 | 0 |
| SPGP0U00230 | XP_018223047.1 | 99 | 0 |
| SPGP0U01850 | XP_018222896.1 | 99 | 0 |
| SPGP0EB00160 | XP_018223315.1 | 99 | 0 |
| SPGP0CN00210 | XP_018220950.1 | 99 | 0 |
| SPGP0CP00540 | XP_018223783.1 | 99 | 1 |
| SPGP0AA01200 | XP_018220470.1 | 100 | 0 |
| SPGP0DE00200 | XP_018223837.1 | 99 | 0 |
| SPGP0F02000 | XP_018221362.1 | 99 | 0 |
| SPGP0CW02310 | XP_018219330.1 | 99 | 0 |
| SPGP0AUtrna14V | DI49_1074 | 100 | 0 |
| SPGP0AA00530 | XP_018220537.1 | 99 | 0 |
| SPGP0Y00790 | XP_018220095.1 | 99 | 0 |
| SPGP0CW03230 | XP_018219241.1 | 99 | 0 |
| SPGP0J03090 | XP_018219640.1 | 99 | 0 |
| SPGP0EO00120 | XP_018221149.1 | 99 | 0 |
| SPGP0CM00130 | XP_018223311.1 | 99 | 0 |
| SPGP0CU00660 | XP_018220762.1 | 100 | 0 |
| SPGP0J01200 | XP_018219828.1 | 99 | 0 |
| SPGP0H00630 | XP_018221082.1 | 99 | 0 |
| SPGP0Y02490 | XP_018220258.1 | 99 | 0 |
| SPGP0CU00240 | XP_018220805.1 | 99 | 0 |
| SPGP0DA01300 | XP_018222625.1 | 99 | 0 |
| SPGP0H01580 | XP_018220991.1 | 99 | 0 |
| SPGP0L01670 | XP_018218867.1 | 100 | 0 |
| SPGP0X01450 | XP_018221589.1 | 99 | 0 |
| SPGP0EO00180 | XP_018221155.1 | 99 | 1 |
| SPGP0AF01400 | XP_018222398.1 | 99 | 0 |
| SPGP0J02790 | XP_018219670.1 | 99 | 0 |
| SPGP0CP01200 | XP_018223718.1 | 99 | 0 |
| SPGP0HB00140 | XP_018222865.1 | 99 | 0 |
| SPGP0J02710 | XP_018219678.1 | 99 | 0 |
| SPGP0H01760 | XP_018220974.1 | 99 | 0 |
| SPGP0F02640 | XP_018221424.1 | 99 | 0 |
| SPGP0CU00790 | XP_018220750.1 | 99 | 0 |
| SPGP0AN00600 | XP_018223967.1 | 99 | 0 |

Create PDF in your applications with the Pdfcrowd HTML to PDF API PDFCROWD

| | | | |
|---|---|---|---|
| SPGP0X01310 | XP_018221575.1 | 100 | 0 |
| SPGP0CU01940 | XP_018220640.1 | 99 | 0 |
| SPGP0CW02900 | XP_018219273.1 | 99 | 0 |
| SPGP0H01350 | XP_018221012.1 | 99 | 1 |
| SPGP0AN00480 | XP_018223955.1 | 99 | 0 |
| SPGP0GR00110 | XP_018221976.1 | 100 | 0 |
| SPGP0CX01580 | XP_018222174.1 | 98 | 0 |
| SPGP0H01570 | XP_018220992.1 | 99 | 0 |
| SPGP0M00170 | XP_018221141.1 | 99 | 0 |
| SPGP0AJ00750 | XP_018221847.1 | 99 | 0 |
| SPGP0CW03000 | XP_018219263.1 | 100 | 0 |
| SPGP0CUtrna25K | DI49_5564 | 100 | 0 |
| SPGP0J03590 | XP_018219590.1 | 99 | 0 |
| SPGP0J02510 | XP_018219698.1 | 99 | 0 |
| SPGP0L01630 | XP_018218871.1 | 99 | 0 |
| SPGP0V00710 | XP_018221715.1 | 100 | 0 |
| SPGP0AJ01150 | XP_018221886.1 | 99 | 0 |
| SPGP0EO00420 | XP_018221179.1 | 99 | 0 |
| SPGP0HW00180 | XP_018224036.1 | 100 | 0 |
| SPGP0DA00800 | XP_018222673.1 | 100 | 0 |
| SPGP0J00220 | XP_018219925.1 | 96 | 0 |
| SPGP0S00190 | XP_018224068.1 | 99 | 0 |
| SPGP0CM01080 | XP_018223217.1 | 100 | 0 |
| SPGP0X00860 | XP_018221534.1 | 99 | 0 |
| SPGP0CP00320 | XP_018223804.1 | 99 | 0 |
| SPGP0DA00630 | XP_018222691.1 | 99 | 0 |
| SPGP0ES00110 | XP_018222827.1 | 99 | 0 |
| SPGP0CM01090 | XP_018223216.1 | 99 | 0 |
| SPGP0F01470 | XP_018221311.1 | 99 | 0 |
| SPGP0CP00120 | XP_018223824.1 | 99 | 0 |
| SPGP0J01240 | XP_018219824.1 | 99 | 0 |
| SPGP0CU01460 | XP_018220688.1 | 99 | 0 |
| SPGP0F01170 | XP_018221281.1 | 99 | 0 |
| SPGP0CM02240 | XP_018223105.1 | 99 | 0 |
| SPGP0Z00230 | XP_018222542.1 | 99 | 0 |
| SPGP0J02910 | XP_018219658.1 | 99 | 0 |
| SPGP0H00180 | XP_018221125.1 | 99 | 0 |
| SPGP0CP01600 | XP_018223679.1 | 99 | 0 |
| SPGP0CN00570 | XP_018220914.1 | 99 | 0 |
| SPGP0J01790 | XP_018219769.1 | 99 | 0 |

Create PDF in your applications with the Pdfcrowd HTML to PDF API PDFCROWD

| | | | |
|---|---|---|---|
| SPGP0J01060 | XP_018219855.1 | 99 | 0 |
| SPGP0AT00620 | XP_018219082.1 | 98 | 1 |
| SPGP0J01050 | XP_018219854.1 | 99 | 0 |
| SPGP0FF00150 | XP_018223397.1 | 99 | 0 |
| SPGP0J01380 | XP_018219810.1 | 99 | 0 |
| SPGP0CX02050 | XP_018222220.1 | 98 | 0 |
| SPGP0V01040 | XP_018221683.1 | 99 | 0 |
| SPGP0L02150 | XP_018218822.1 | 100 | 0 |
| SPGP0AA00400 | XP_018220550.1 | 93 | 6 |
| SPGP0F01290 | XP_018221292.1 | 99 | 0 |
| SPGP0Y00650 | XP_018220081.1 | 99 | 0 |
| SPGP0HS00130 | XP_018221777.1 | 98 | 0 |
| SPGP0H01500 | XP_018220999.1 | 99 | 0 |
| SPGP0X01710 | XP_018221614.1 | 99 | 0 |
| SPGP0L02510 | XP_018218785.1 | 99 | 0 |
| SPGP0EO00260 | XP_018221163.1 | 99 | 0 |
| SPGP0CX00690 | XP_018222088.1 | 99 | 0 |
| SPGP0CU01800 | XP_018220654.1 | 99 | 0 |
| SPGP0J02520 | XP_018219697.1 | 99 | 0 |
| SPGP0J01730 | XP_018219775.1 | 99 | 0 |
| SPGP0J02260 | XP_018219723.1 | 99 | 1 |
| SPGP0DB00300 | XP_018223904.1 | 99 | 0 |
| SPGP0CX01610 | XP_018222177.1 | 100 | 0 |
| SPGP0AF01150 | XP_018222371.1 | 99 | 0 |
| SPGP0CU00490 | XP_018220780.1 | 99 | 0 |
| SPGP0H01670 | XP_018220983.1 | 99 | 0 |
| SPGP0H00420 | XP_018221102.1 | 99 | 0 |
| SPGP0U00260 | XP_018223044.1 | 99 | 0 |
| SPGP0J01320 | XP_018219816.1 | 99 | 0 |
| SPGP0AJ01400 | XP_018221910.1 | 98 | 0 |
| SPGP0JK00340 | XP_018223455.1 | 99 | 0 |
| SPGP0S00470 | XP_018224095.1 | 99 | 0 |
| SPGP0CW03510 | XP_018219213.1 | 99 | 1 |
| SPGP0F01590 | XP_018221322.1 | 99 | 0 |
| SPGP0CP02890 | XP_018223552.1 | 99 | 0 |
| SPGP0X00520 | XP_018221501.1 | 99 | 0 |
| SPGP0S00240 | XP_018224073.1 | 99 | 0 |
| SPGP0CW00770 | XP_018219479.1 | 99 | 0 |
| SPGP0Y01880 | XP_018220199.1 | 100 | 0 |
| SPGP0IR00160 | XP_018223470.1 | 99 | 0 |

Create PDF in your applications with the Pdfcrowd HTML to PDF API PDFCROWD

| | | | |
|---|---|---|---|
| SPGP0DA01040 | XP_018222651.1 | 99 | 0 |
| SPGP0CM01450 | XP_018223181.1 | 99 | 0 |
| SPGP0CP03090 | XP_018223531.1 | 99 | 0 |
| SPGP0CV00230 | XP_018222574.1 | 99 | 0 |
| SPGP0CN00810 | XP_018220891.1 | 99 | 0 |
| SPGP0CU00800 | XP_018220749.1 | 99 | 0 |
| SPGP0DA00490 | XP_018222705.1 | 99 | 0 |
| SPGP0L02460 | XP_018218790.1 | 98 | 0 |
| SPGP0Y00130 | XP_018220030.1 | 99 | 0 |
| SPGP0DA01100 | XP_018222645.1 | 99 | 1 |
| SPGP0J02860 | XP_018219663.1 | 99 | 0 |
| SPGP0CN00740 | XP_018220897.1 | 99 | 0 |
| SPGP0DF00150 | XP_018219193.1 | 98 | 0 |
| SPGP0X01850 | XP_018221628.1 | 99 | 1 |
| SPGP0F00710 | XP_018221236.1 | 99 | 0 |
| SPGP0K00400 | XP_018222782.1 | 100 | 0 |
| SPGP0CN00750 | XP_018220896.1 | 99 | 0 |
| SPGP0AA01350 | XP_018220455.1 | 99 | 0 |
| SPGP0CU01070 | XP_018220723.1 | 99 | 0 |
| SPGP0J03540 | XP_018219595.1 | 99 | 0 |
| SPGP0K00540 | XP_018222796.1 | 99 | 0 |
| SPGP0AA01420 | XP_018220448.1 | 100 | 0 |
| SPGP0DE00450 | XP_018223862.1 | 99 | 0 |
| SPGP0U01290 | XP_018222952.1 | 99 | 0 |
| SPGP0J02070 | XP_018219742.1 | 100 | 0 |
| SPGP0CU02520 | XP_018220582.1 | 99 | 0 |
| SPGP0S00460 | XP_018224094.1 | 99 | 0 |
| SPGP0F00400 | XP_018221207.1 | 99 | 0 |
| SPGP0AF01660 | XP_018222424.1 | 99 | 0 |
| SPGP0AJ00790 | XP_018221851.1 | 99 | 0 |
| SPGP0L01790 | XP_018218856.1 | 99 | 0 |
| SPGP0IN00280 | XP_018221648.1 | 99 | 0 |
| SPGP0DE00440 | XP_018223861.1 | 99 | 0 |
| SPGP0CP01620 | XP_018223677.1 | 99 | 0 |
| SPGP0CX01710 | XP_018222187.1 | 98 | 0 |
| SPGP0U01340 | XP_018222948.1 | 99 | 0 |
| SPGP0CW02480 | XP_018219313.1 | 99 | 0 |
| SPGP0AN01030 | XP_018224009.1 | 99 | 0 |
| SPGP0CM00470 | XP_018223277.1 | 99 | 0 |
| SPGP0JK00310 | XP_018223452.1 | 98 | 1 |

Create PDF in your applications with the Pdfcrowd HTML to PDF API PDFCROWD

| | | | |
|---|---|---|---|
| SPGP0U01540 | XP_018222928.1 | 99 | 0 |
| SPGP0GG00230 | XP_018220828.1 | 99 | 0 |
| SPGP0HZ00120 | XP_018219029.1 | 99 | 1 |
| SPGP0HW00200 | XP_018224035.1 | 98 | 1 |
| SPGP0CU01730 | XP_018220661.1 | 99 | 0 |
| SPGP0AJ00860 | XP_018221858.1 | 99 | 0 |
| SPGP0CU00100 | XP_018220818.1 | 99 | 0 |
| SPGP0GQ00160 | XP_018219553.1 | 99 | 0 |
| SPGP0F02240 | XP_018221385.1 | 99 | 0 |
| SPGP0S00150 | XP_018224064.1 | 99 | 0 |
| SPGP0H00610 | XP_018221084.1 | 99 | 0 |
| SPGP0H00960 | XP_018221049.1 | 99 | 0 |
| SPGP0CRtrna26L | DI49_1927 | 100 | 0 |
| SPGP0CP02840 | XP_018223557.1 | 100 | 0 |
| SPGP0L01480 | XP_018218886.1 | 99 | 0 |
| SPGP0AN00620 | XP_018223969.1 | 99 | 0 |
| SPGP0AN00590 | XP_018223966.1 | 99 | 0 |
| SPGP0AA00960 | XP_018220494.1 | 99 | 0 |
| SPGP0DA00790 | XP_018222674.1 | 93 | 5 |
| SPGP0CW00690 | XP_018219487.1 | 99 | 0 |
| SPGP0IQ00100 | XP_018220026.1 | 99 | 0 |
| SPGP0HB00120 | XP_018222863.1 | 99 | 0 |
| SPGP0J01890 | XP_018219760.1 | 99 | 0 |
| SPGP0AJ01210 | XP_018221892.1 | 99 | 0 |
| SPGP0CP01210 | XP_018223717.1 | 99 | 0 |
| SPGP0DA00130 | XP_018222739.1 | 99 | 0 |
| SPGP0X01330 | XP_018221577.1 | 99 | 0 |
| SPGP0AA00220 | XP_018220568.1 | 99 | 0 |
| SPGP0CPtrna19E | DI49_2698 | 100 | 0 |
| SPGP0Y01180 | XP_018220133.1 | 97 | 3 |
| SPGP0CM00910 | XP_018223234.1 | 99 | 0 |
| SPGP0J00790 | XP_018219871.1 | 99 | 0 |
| SPGP0AN00780 | XP_018223984.1 | 98 | 0 |
| SPGP0DA01150 | XP_018222640.1 | 99 | 0 |
| SPGP0HS00140 | XP_018221778.1 | 99 | 0 |
| SPGP0DA00660 | XP_018222688.1 | 99 | 0 |
| SPGP0J00190 | XP_018219928.1 | 99 | 0 |
| SPGP0DA01620 | XP_018222487.1 | 90 | 0 |
| SPGP0U01210 | XP_018222959.1 | 98 | 0 |

Create PDF in your applications with the Pdfcrowd HTML to PDF API PDFCROWD

| | | | |
|---|---|---|---|
| SPGP0CN00200 | XP_018220951.1 | 99 | 0 |
| SPGP0CP00140 | XP_018223822.1 | 99 | 0 |
| SPGP0CP01930 | XP_018223647.1 | 99 | 0 |
| SPGP0CU01190 | XP_018220713.1 | 99 | 0 |
| SPGP0CU01440 | XP_018220690.1 | 99 | 0 |
| SPGP0J01260 | XP_018219822.1 | 99 | 0 |
| SPGP0F01450 | XP_018221309.1 | 100 | 0 |
| SPGP0S00630 | XP_018224110.1 | 99 | 0 |
| SPGP0V01120 | XP_018221676.1 | 99 | 0 |
| SPGP0X00110 | XP_018221460.1 | 98 | 0 |
| SPGP0H01730 | XP_018220977.1 | 99 | 1 |
| SPGP0J01040 | XP_018219853.1 | 99 | 0 |
| SPGP0AT00600 | XP_018219081.1 | 99 | 0 |
| SPGP0CW01480 | XP_018219410.1 | 99 | 0 |
| SPGP0AJ01730 | XP_018221941.1 | 99 | 0 |
| SPGP0CP01450 | XP_018223694.1 | 99 | 0 |
| SPGP0CX02070 | XP_018222222.1 | 99 | 0 |
| SPGP0J02430 | XP_018219706.1 | 99 | 0 |
| SPGP0L00840 | XP_018218949.1 | 99 | 0 |
| SPGP0CW02300 | XP_018219331.1 | 99 | 0 |
| SPGP0F00700 | XP_018221235.1 | 99 | 0 |
| SPGP0J02280 | XP_018219721.1 | 100 | 0 |
| SPGP0FF00170 | XP_018223395.1 | 100 | 0 |
| SPGP0J01030 | XP_018219852.1 | 99 | 0 |
| SPGP0J02240 | XP_018219725.1 | 99 | 0 |
| SPGP0J00210 | XP_018219926.1 | 98 | 0 |
| SPGP0AJtrna8F | DI49_2475 | 100 | 0 |
| SPGP0CP02690 | XP_018223572.1 | 99 | 0 |
| SPGP0U00910 | XP_018222988.1 | 99 | 0 |
| SPGP0AF01580 | XP_018222416.1 | 99 | 0 |
| SPGP0CW00420 | XP_018219513.1 | 99 | 0 |
| SPGP0AA00280 | XP_018220562.1 | 99 | 0 |
| SPGP0L01900 | XP_018218845.1 | 99 | 0 |
| SPGP0AA01400 | XP_018220450.1 | 99 | 0 |
| SPGP0CX01750 | XP_018222191.1 | 99 | 0 |
| SPGP0F02760 | XP_018221436.1 | 99 | 0 |
| SPGP0F02030 | XP_018221365.1 | 99 | 0 |
| SPGP0L01280 | XP_018218906.1 | 99 | 0 |
| SPGP0CU02510 | XP_018220583.1 | 99 | 0 |
| SPGP0AT00400 | XP_018219061.1 | 100 | 0 |

Create PDF in your applications with the Pdfcrowd HTML to PDF API PDFCROWD

| | | | |
|---|---|---|---|
| SPGP0S00230 | XP_018224072.1 | 99 | 0 |
| SPGP0CW03530 | XP_018219211.1 | 99 | 0 |
| SPGP0J03110 | XP_018219638.1 | 99 | 0 |
| SPGP0L01340 | XP_018218900.1 | 99 | 0 |
| SPGP0H00390 | XP_018221105.1 | 99 | 0 |
| SPGP0F00530 | XP_018221218.1 | 99 | 0 |
| SPGP0X00500 | XP_018221499.1 | 99 | 1 |
| SPGP0FE00160 | XP_018221968.1 | 99 | 0 |
| SPGP0X00440 | XP_018221493.1 | 99 | 0 |
| SPGP0U00410 | XP_018223032.1 | 100 | 0 |
| SPGP0IR00100 | XP_018223464.1 | 100 | 0 |
| SPGP0CM01430 | XP_018223183.1 | 99 | 1 |
| SPGP0DA00110 | XP_018222741.1 | 99 | 0 |
| SPGP0CN00830 | XP_018220889.1 | 94 | 0 |
| SPGP0DA00300 | XP_018222724.1 | 99 | 0 |
| SPGP0AT01390 | XP_018219156.1 | 99 | 0 |
| SPGP0H01040 | XP_018221041.1 | 98 | 2 |
| SPGP0DA00870 | XP_018222667.1 | 99 | 0 |
| SPGP0X00890 | XP_018221536.1 | 99 | 1 |
| SPGP0AJ01940 | XP_018221960.1 | 99 | 0 |
| SPGP0DA00160 | XP_018222736.1 | 99 | 0 |
| SPGP0J00520 | XP_018219896.1 | 99 | 0 |
| SPGP0AJ01530 | XP_018221921.1 | 99 | 0 |
| SPGP0CM02160 | XP_018223113.1 | 99 | 0 |
| SPGP0HI00150 | XP_018219941.1 | 99 | 0 |
| SPGP0CN00760 | XP_018220895.1 | 100 | 0 |
| SPGP0J03740 | XP_018219575.1 | 99 | 0 |
| SPGP0F00990 | XP_018221263.1 | 99 | 0 |
| SPGP0HI00160 | XP_018219942.1 | 99 | 0 |
| SPGP0CP00170 | XP_018223819.1 | 99 | 0 |
| SPGP0CU02500 | XP_018220584.1 | 99 | 0 |
| SPGP0CN00490 | XP_018220922.1 | 99 | 0 |
| SPGP0J03580 | XP_018219591.1 | 99 | 0 |
| SPGP0F01020 | XP_018221266.1 | 99 | 0 |
| SPGP0U00270 | XP_018223043.1 | 99 | 0 |
| SPGP0CP00820 | XP_018223756.1 | 99 | 0 |
| SPGP0M00110 | XP_018221145.1 | 99 | 0 |
| SPGP0Jtrna1F | DI49_4603 | 99 | 0 |
| SPGP0DA01200 | XP_018222635.1 | 99 | 0 |
| SPGP0CM00450 | XP_018223279.1 | 99 | 0 |

Create PDF in your applications with the Pdfcrowd HTML to PDF API PDFCROWD

| | | | |
|---|---|---|---|
| SPGP0CP00350 | XP_018223801.1 | 99 | 0 |
| SPGP0GJ00170 | XP_018223375.1 | 99 | 0 |
| SPGP0Y01760 | XP_018220189.1 | 99 | 0 |
| SPGP0DA01340 | XP_018222621.1 | 99 | 0 |
| SPGP0DE00420 | XP_018223859.1 | 99 | 0 |
| SPGP0Y03120 | XP_018220320.1 | 99 | 0 |
| SPGP0BE00170 | XP_018223361.1 | 99 | 0 |
| SPGP0CN00700 | XP_018220901.1 | 99 | 0 |
| SPGP0CM00370 | XP_018223287.1 | 99 | 0 |
| SPGP0J00500 | XP_018219898.1 | 99 | 0 |
| SPGP0IU00130 | XP_018220331.1 | 99 | 0 |
| SPGP0AN01190 | XP_018224025.1 | 99 | 0 |
| SPGP0Y01370 | XP_018220152.1 | 99 | 0 |
| SPGP0H00940 | XP_018221051.1 | 99 | 0 |
| SPGP0F02590 | XP_018221419.1 | 99 | 0 |
| SPGP0IN00190 | XP_018221657.1 | 99 | 0 |
| SPGP0CW00610 | XP_018219495.1 | 99 | 0 |
| SPGP0DA00510 | XP_018222703.1 | 99 | 0 |
| SPGP0HO00150 | XP_018224029.1 | 99 | 0 |
| SPGP0CU00860 | XP_018220743.1 | 99 | 0 |
| SPGP0H00490 | XP_018221095.1 | 99 | 0 |
| SPGP0DA00170 | XP_018222735.1 | 99 | 0 |
| SPGP0Y01490 | XP_018220164.1 | 99 | 0 |
| SPGP0AJ01230 | XP_018221894.1 | 99 | 0 |
| SPGP0X00130 | XP_018221462.1 | 97 | 0 |
| SPGP0Y02500 | XP_018220259.1 | 99 | 0 |
| SPGP0CP01420 | XP_018223697.1 | 99 | 0 |
| SPGP0CP01640 | XP_018223675.1 | 100 | 0 |
| SPGP0V00970 | XP_018221690.1 | 99 | 0 |
| SPGP0CW00190 | XP_018219534.1 | 100 | 0 |
| SPGP0GZ00130 | XP_018222825.1 | 99 | 0 |
| SPGP0CM00930 | XP_018223232.1 | 99 | 0 |
| SPGP0CU02300 | XP_018220604.1 | 99 | 0 |
| SPGP0CXtrna2F | DI49_1939 | 100 | 0 |
| SPGP0L02630 | XP_018218774.1 | 100 | 0 |
| SPGP0JC00110 | XP_018220852.1 | 99 | 0 |
| SPGP0Y01980 | XP_018220209.1 | 99 | 1 |
| SPGP0AF00300 | XP_018222284.1 | 99 | 0 |
| SPGP0S00270 | XP_018224076.1 | 99 | 0 |
| SPGP0GQ00150 | XP_018219554.1 | 99 | 0 |

Create PDF in your applications with the Pdfcrowd HTML to PDF API PDFCROWD

| | | | |
|---|---|---|---|
| SPGP0AT00240 | XP_018219046.1 | 99 | 0 |
| SPGP0CU02080 | XP_018220626.1 | 99 | 0 |
| SPGP0X01300 | XP_018221574.1 | 99 | 1 |
| SPGP0S00510 | XP_018224098.1 | 99 | 0 |
| SPGP0X00160 | XP_018221465.1 | 98 | 0 |
| SPGP0F01280 | XP_018221291.1 | 99 | 0 |
| SPGP0Y02640 | XP_018220273.1 | 100 | 0 |
| SPGP0H00300 | XP_018221114.1 | 99 | 0 |
| SPGP0AF01700 | XP_018222427.1 | 99 | 0 |
| SPGP0V01090 | XP_018221678.1 | 97 | 0 |
| SPGP0Z00270 | XP_018222538.1 | 99 | 0 |
| SPGP0F01150 | XP_018221279.1 | 99 | 0 |
| SPGP0CM02280 | XP_018223101.1 | 99 | 0 |
| SPGP0F00810 | XP_018221245.1 | 99 | 0 |
| SPGP0CN00530 | XP_018220918.1 | 99 | 0 |
| SPGP0V01000 | XP_018221687.1 | 99 | 0 |
| SPGP0CM01570 | XP_018223169.1 | 97 | 3 |
| SPGP0CM00790 | XP_018223246.1 | 99 | 0 |
| SPGP0CX01150 | XP_018222134.1 | 99 | 0 |
| SPGP0F00520 | XP_018221217.1 | 99 | 0 |
| SPGP0J00360 | XP_018219912.1 | 99 | 0 |
| SPGP0J00830 | XP_018219867.1 | 99 | 0 |
| SPGP0AJ00670 | XP_018221839.1 | 99 | 0 |
| SPGP0F02070 | XP_018221369.1 | 99 | 0 |
| SPGP0J00910 | XP_018219860.1 | 100 | 0 |
| SPGP0L00380 | XP_018218994.1 | 98 | 0 |
| SPGP0CP03120 | XP_018223528.1 | 99 | 0 |
| SPGP0CPtrna23T | DI49_2415 | 100 | 0 |
| SPGP0CW01820 | XP_018219378.1 | 100 | 0 |
| SPGP0CV00170 | XP_018222570.1 | 99 | 1 |
| SPGP0F01920 | XP_018221354.1 | 99 | 0 |
| SPGP0X01040 | XP_018221551.1 | 99 | 0 |
| SPGP0CX01380 | XP_018222156.1 | 98 | 0 |
| SPGP0IZ00110 | XP_018220339.1 | 99 | 0 |
| SPGP0CP00640 | XP_018223773.1 | 99 | 0 |
| SPGP0L01650 | XP_018218869.1 | 99 | 0 |
| SPGP0Y00820 | XP_018220098.1 | 99 | 0 |
| SPGP0CW00120 | XP_018219542.1 | 99 | 0 |
| SPGP0DE00270 | XP_018223844.1 | 99 | 0 |
| SPGP0H01260 | XP_018221021.1 | 99 | 0 |

Create PDF in your applications with the Pdfcrowd HTML to PDF API PDFCROWD

| | | | |
|---|---|---|---|
| SPGP0X01610 | XP_018221604.1 | 100 | 0 |
| SPGP0Y01780 | XP_018220191.1 | 99 | 0 |
| SPGP0J03170 | XP_018219632.1 | 99 | 0 |
| SPGP0CX02190 | XP_018222233.1 | 99 | 0 |
| SPGP0J00770 | XP_018219873.1 | 99 | 0 |
| SPGP0S00100 | XP_018224059.1 | 99 | 0 |
| SPGP0X00290 | XP_018221478.1 | 99 | 0 |
| SPGP0AF00610 | XP_018222315.1 | 99 | 0 |
| SPGP0CP02990 | XP_018223541.1 | 99 | 0 |
| SPGP0CW00780 | XP_018219478.1 | 99 | 0 |
| SPGP0CN00160 | XP_018220955.1 | 99 | 1 |
| SPGP0AA00590 | XP_018220531.1 | 99 | 0 |
| SPGP0CN00220 | XP_018220949.1 | 99 | 0 |
| SPGP0DA00360 | XP_018222718.1 | 99 | 0 |
| SPGP0EO00230 | XP_018221160.1 | 99 | 0 |
| SPGP0AJ01000 | XP_018221872.1 | 99 | 0 |
| SPGP0H01060 | XP_018221039.1 | 99 | 0 |
| SPGP0L02400 | XP_018218796.1 | 99 | 0 |
| SPGP0CM01140 | XP_018223211.1 | 99 | 0 |
| SPGP0DA00210 | XP_018222732.1 | 99 | 0 |
| SPGP0V00150 | XP_018221768.1 | 99 | 0 |
| SPGP0DA00980 | XP_018222657.1 | 98 | 0 |
| SPGP0G00150 | XP_018222254.1 | 100 | 0 |
| SPGP0CU01750 | XP_018220659.1 | 99 | 0 |
| SPGP0V00830 | XP_018221704.1 | 99 | 0 |
| SPGP0CM00200 | XP_018223304.1 | 98 | 1 |
| SPGP0J00670 | XP_018219883.1 | 99 | 0 |
| SPGP0CN00910 | XP_018220881.1 | 99 | 0 |
| SPGP0J01620 | XP_018219786.1 | 100 | 0 |
| SPGP0Y01950 | XP_018220206.1 | 99 | 0 |
| SPGP0CN01030 | XP_018220869.1 | 99 | 0 |
| SPGP0AJ01750 | XP_018221943.1 | 99 | 0 |
| SPGP0Y01420 | XP_018220157.1 | 99 | 0 |
| SPGP0X00300 | XP_018221479.1 | 99 | 0 |
| SPGP0CW02700 | XP_018219293.1 | 99 | 0 |
| SPGP0AF01190 | XP_018222375.1 | 99 | 0 |
| SPGP0S00420 | XP_018224090.1 | 99 | 0 |
| SPGP0U00250 | XP_018223045.1 | 99 | 0 |
| SPGP0J01430 | XP_018219805.1 | 100 | 0 |
| SPGP0IU00110 | XP_018220329.1 | 99 | 0 |

Create PDF in your applications with the Pdfcrowd HTML to PDF API PDFCROWD

| | | | |
|---|---|---|---|
| SPGP0J03340 | XP_018219615.1 | 99 | 0 |
| SPGP0DE00150 | XP_018223832.1 | 99 | 0 |
| SPGP0DE00400 | XP_018223857.1 | 99 | 0 |
| SPGP0DA00600 | XP_018222694.1 | 99 | 0 |
| SPGP0AE00110 | XP_018222493.1 | 99 | 0 |
| SPGP0F02770 | XP_018221437.1 | 99 | 0 |
| SPGP0CP02520 | XP_018223589.1 | 99 | 0 |
| SPGP0CP02460 | XP_018223595.1 | 99 | 0 |
| SPGP0L00240 | XP_018219008.1 | 99 | 0 |
| SPGP0H00690 | XP_018221076.1 | 99 | 0 |
| SPGP0CU00450 | XP_018220784.1 | 100 | 0 |
| SPGP0Y01970 | XP_018220208.1 | 99 | 0 |
| SPGP0CM00650 | XP_018223259.1 | 99 | 0 |
| SPGP0H01080 | XP_018221037.1 | 99 | 0 |
| SPGP0AE00200 | XP_018222502.1 | 99 | 0 |
| SPGP0HO00130 | XP_018224031.1 | 99 | 0 |
| SPGP0IN00240 | XP_018221652.1 | 99 | 0 |
| SPGP0V00300 | XP_018221753.1 | 99 | 0 |
| SPGP0AF00460 | XP_018222300.1 | 99 | 0 |
| SPGP0H01220 | XP_018221025.1 | 99 | 0 |
| SPGP0H01520 | XP_018220997.1 | 99 | 0 |
| SPGP0AU00210 | XP_018223076.1 | 100 | 0 |
| SPGP0X01370 | XP_018221581.1 | 100 | 0 |
| SPGP0X01800 | XP_018221623.1 | 98 | 2 |
| SPGP0AN00280 | XP_018223940.1 | 99 | 0 |
| SPGP0H01290 | XP_018221018.1 | 99 | 0 |
| SPGP0AT00650 | XP_018219085.1 | 99 | 0 |
| SPGP0CV00310 | XP_018222582.1 | 99 | 1 |
| SPGP0HC00110 | XP_018221782.1 | 99 | 0 |
| SPGP0V01160 | XP_018221672.1 | 99 | 0 |
| SPGP0Y02790 | XP_018220288.1 | 99 | 0 |
| SPGP0J01880 | XP_018219761.1 | 99 | 0 |
| SPGP0DA00640 | XP_018222690.1 | 99 | 0 |
| SPGP0J00280 | XP_018219920.1 | 99 | 0 |
| SPGP0J01900 | XP_018219759.1 | 100 | 0 |
| SPGP0U01060 | XP_018222973.1 | 99 | 0 |
| SPGP0H01780 | XP_018220972.1 | 100 | 0 |
| SPGP0Y03000 | XP_018220309.1 | 99 | 0 |
| SPGP0CM00670 | XP_018223257.1 | 97 | 3 |
| SPGP0F00540 | XP_018221219.1 | 99 | 0 |

Create PDF in your applications with the Pdfcrowd HTML to PDF API PDFCROWD

| | | | |
|---|---|---|---|
| SPGP0H00290 | XP_018221115.1 | 99 | 0 |
| SPGP0CU01060 | XP_018220724.1 | 99 | 0 |
| SPGP0AJ00220 | XP_018221796.1 | 98 | 0 |
| SPGP0JK00110 | XP_018223432.1 | 99 | 0 |
| SPGP0AF01840 | XP_018222442.1 | 99 | 0 |
| SPGP0X01010 | XP_018221548.1 | 99 | 0 |
| SPGP0J02570 | XP_018219692.1 | 99 | 0 |
| SPGP0F01760 | XP_018221338.1 | 100 | 0 |
| SPGP0F01940 | XP_018221356.1 | 99 | 0 |
| SPGP0AN00630 | XP_018223970.1 | 99 | 0 |
| SPGP0X00480 | XP_018221497.1 | 99 | 0 |
| SPGP0R00270 | XP_018219969.1 | 99 | 0 |
| SPGP0AJ01190 | XP_018221890.1 | 99 | 0 |
| SPGP0AJ01270 | XP_018221898.1 | 99 | 1 |
| SPGP0J01550 | XP_018219793.1 | 99 | 0 |
| SPGP0DE00360 | XP_018223853.1 | 99 | 0 |
| SPGP0F00350 | XP_018221202.1 | 99 | 0 |
| SPGP0CN00610 | XP_018220910.1 | 99 | 0 |
| SPGP0FV00100 | XP_018223323.1 | 98 | 0 |
| SPGP0X00760 | XP_018221524.1 | 99 | 0 |
| SPGP0CW02160 | XP_018219345.1 | 99 | 0 |
| SPGP0JP00180 | XP_018218749.1 | 100 | 0 |
| SPGP0AA01570 | XP_018220434.1 | 99 | 0 |
| SPGP0CU00380 | XP_018220791.1 | 99 | 0 |
| SPGP0J02250 | XP_018219724.1 | 99 | 0 |
| SPGP0AF01090 | XP_018222364.1 | 99 | 0 |
| SPGP0AF00630 | XP_018222317.1 | 99 | 0 |
| SPGP0J03780 | XP_018219571.1 | 99 | 0 |
| SPGP0CU02280 | XP_018220606.1 | 99 | 0 |
| SPGP0F02270 | XP_018221388.1 | 99 | 0 |
| SPGP0CN00870 | XP_018220885.1 | 99 | 0 |
| SPGP0CP02870 | XP_018223554.1 | 99 | 0 |
| SPGP0DA00430 | XP_018222711.1 | 99 | 1 |
| SPGP0CW01880 | XP_018219372.1 | 99 | 0 |
| SPGP0CM02250 | XP_018223104.1 | 99 | 0 |
| SPGP0H00970 | XP_018221048.1 | 98 | 0 |
| SPGP0X01630 | XP_018221606.1 | 99 | 0 |
| SPGP0J02120 | XP_018219737.1 | 100 | 0 |
| SPGP0CM01160 | XP_018223209.1 | 99 | 0 |
| SPGP0CW00140 | XP_018219540.1 | 99 | 0 |

Create PDF in your applications with the Pdfcrowd HTML to PDF API PDFCROWD

| | | | |
|---|---|---|---|
| SPGP0CP03240 | XP_018223516.1 | 99 | 0 |
| SPGP0Ftrna21L | DI49_3110 | 100 | 0 |
| SPGP0AF00530 | XP_018222307.1 | 99 | 0 |
| SPGP0AN01060 | XP_018224012.1 | 99 | 0 |
| SPGP0J00540 | XP_018219895.1 | 99 | 0 |
| SPGP0S00560 | XP_018224103.1 | 98 | 0 |
| SPGP0HO00160 | XP_018224028.1 | 100 | 0 |
| SPGP0BE00190 | XP_018223359.1 | 99 | 0 |
| SPGP0F00120 | XP_018221181.1 | 98 | 0 |
| SPGP0IC00150 | XP_018222528.1 | 99 | 0 |
| SPGP0S00400 | XP_018224088.1 | 99 | 0 |
| SPGP0CN01050 | XP_018220867.1 | 99 | 0 |
| SPGP0CP03050 | XP_018223535.1 | 99 | 0 |
| SPGP0F00950 | XP_018221259.1 | 99 | 0 |
| SPGP0F00480 | XP_018221213.1 | 99 | 0 |
| SPGP0AA00470 | XP_018220543.1 | 99 | 0 |
| SPGP0F01460 | XP_018221310.1 | 100 | 0 |
| SPGP0J01410 | XP_018219807.1 | 99 | 0 |
| SPGP0F01680 | XP_018221330.1 | 99 | 0 |
| SPGP0CX00740 | XP_018222093.1 | 99 | 0 |
| SPGP0CW02830 | XP_018219280.1 | 99 | 0 |
| SPGP0CM01700 | XP_018223156.1 | 100 | 0 |
| SPGP0DF00200 | XP_018219188.1 | 98 | 0 |
| SPGP0CP00390 | XP_018223797.1 | 99 | 0 |
| SPGP0CN00180 | XP_018220953.1 | 99 | 0 |
| SPGP0Y02570 | XP_018220266.1 | 99 | 0 |
| SPGP0H00130 | XP_018221130.1 | 99 | 0 |
| SPGP0J03360 | XP_018219613.1 | 99 | 0 |
| SPGP0U01720 | XP_018222910.1 | 98 | 1 |
| SPGP0U01230 | XP_018222958.1 | 98 | 0 |
| SPGP0Y03160 | XP_018220324.1 | 99 | 0 |
| SPGP0V01210 | XP_018221667.1 | 99 | 0 |
| SPGP0F01440 | XP_018221308.1 | 99 | 0 |
| SPGP0JA00100 | XP_018222242.1 | 99 | 0 |
| SPGP0FF00110 | XP_018223401.1 | 99 | 0 |
| SPGP0DA00320 | XP_018222722.1 | 99 | 0 |
| SPGP0CP01750 | XP_018223665.1 | 99 | 0 |
| SPGP0AF01610 | XP_018222419.1 | 99 | 0 |
| SPGP0CM00320 | XP_018223292.1 | 99 | 0 |
| SPGP0L00730 | XP_018218960.1 | 99 | 0 |

Create PDF in your applications with the Pdfcrowd HTML to PDF API PDFCROWD

| | | | |
|---|---|---|---|
| SPGP0F02080 | XP_018221370.1 | 99 | 0 |
| SPGP0J02140 | XP_018219735.1 | 100 | 0 |
| SPGP0JP00100 | XP_018219561.1 | 99 | 0 |
| SPGP0CW01730 | XP_018219387.1 | 99 | 0 |
| SPGP0Y01000 | XP_018220115.1 | 99 | 0 |
| SPGP0H00620 | XP_018221083.1 | 99 | 0 |
| SPGP0J01920 | XP_018219757.1 | 98 | 1 |
| SPGP0F02620 | XP_018221422.1 | 99 | 0 |
| SPGP0AA00900 | XP_018220500.1 | 99 | 0 |
| SPGP0X01530 | XP_018221596.1 | 100 | 0 |
| SPGP0J02820 | XP_018219667.1 | 99 | 0 |
| SPGP0DA01280 | XP_018222627.1 | 99 | 0 |
| SPGP0J02670 | XP_018219682.1 | 99 | 0 |
| SPGP0F00850 | XP_018221249.1 | 99 | 0 |
| SPGP0AA00390 | XP_018220551.1 | 99 | 0 |
| SPGP0AT00280 | XP_018219050.1 | 99 | 0 |
| SPGP0DB00430 | XP_018223915.1 | 99 | 0 |
| SPGP0DA00810 | XP_018222672.1 | 100 | 0 |
| SPGP0AN00810 | XP_018223987.1 | 99 | 1 |
| SPGP0CX01320 | XP_018222150.1 | 99 | 0 |
| SPGP0Y00310 | XP_018220048.1 | 99 | 0 |
| SPGP0CW02950 | XP_018219268.1 | 99 | 0 |
| SPGP0J00730 | XP_018219877.1 | 99 | 0 |
| SPGP0CX01070 | XP_018222126.1 | 98 | 0 |
| SPGP0J00250 | XP_018219922.1 | 99 | 0 |
| SPGP0AA01640 | XP_018220427.1 | 99 | 0 |
| SPGP0CN01000 | XP_018220872.1 | 99 | 0 |
| SPGP0DA01480 | XP_018222607.1 | 99 | 0 |
| SPGP0GS00120 | XP_018222247.1 | 99 | 0 |
| SPGP0U00220 | XP_018223048.1 | 99 | 0 |
| SPGP0J00690 | XP_018219881.1 | 99 | 0 |
| SPGP0CU01170 | XP_018220714.1 | 99 | 0 |
| SPGP0H00170 | XP_018221126.1 | 99 | 0 |
| SPGP0DA00880 | XP_018222666.1 | 99 | 0 |
| SPGP0CM01980 | XP_018223130.1 | 99 | 0 |
| SPGP0J00480 | XP_018219900.1 | 99 | 0 |
| SPGP0K00260 | XP_018222768.1 | 99 | 0 |
| SPGP0F00560 | XP_018221221.1 | 99 | 0 |
| SPGP0J03730 | XP_018219576.1 | 99 | 0 |
| SPGP0IH00230 | XP_018220393.1 | 99 | 0 |

Create PDF in your applications with the Pdfcrowd HTML to PDF API PDFCROWD

| | | | |
|---|---|---|---|
| SPGP0J02770 | XP_018219672.1 | 99 | 0 |
| SPGP0U01460 | XP_018222936.1 | 99 | 0 |
| SPGP0CP02240 | XP_018223616.1 | 99 | 0 |
| SPGP0J03040 | XP_018219645.1 | 99 | 0 |
| SPGP0DA01440 | XP_018222611.1 | 99 | 0 |
| SPGP0CP02700 | XP_018223571.1 | 99 | 0 |
| SPGP0L01560 | XP_018218878.1 | 99 | 0 |
| SPGP0AF01390 | XP_018222397.1 | 99 | 0 |
| SPGP0CM00690 | XP_018223256.1 | 99 | 0 |
| SPGP0J02550 | XP_018219694.1 | 99 | 0 |
| SPGP0F01210 | XP_018221285.1 | 99 | 0 |
| SPGP0CU00780 | XP_018220751.1 | 99 | 0 |
| SPGP0F01960 | XP_018221358.1 | 99 | 0 |
| SPGP0Z00310 | XP_018222534.1 | 99 | 0 |
| SPGP0CX01690 | XP_018222185.1 | 99 | 0 |
| SPGP0U01910 | XP_018222890.1 | 99 | 0 |
| SPGP0CP02660 | XP_018223575.1 | 99 | 0 |
| SPGP0J01600 | XP_018219788.1 | 99 | 0 |
| SPGP0CN00400 | XP_018220931.1 | 99 | 0 |
| SPGP0CP03670 | XP_018223477.1 | 98 | 0 |
| SPGP0CN00320 | XP_018220939.1 | 99 | 0 |
| SPGP0CM00240 | XP_018223300.1 | 99 | 0 |
| SPGP0DB00380 | XP_018223911.1 | 99 | 0 |
| SPGP0X00700 | XP_018221518.1 | 99 | 0 |
| SPGP0L01680 | XP_018218866.1 | 99 | 0 |
| SPGP0AJ00920 | XP_018221864.1 | 99 | 0 |
| SPGP0CP02150 | XP_018223625.1 | 99 | 0 |
| SPGP0DA00410 | XP_018222713.1 | 99 | 0 |
| SPGP0AA00180 | XP_018220572.1 | 100 | 0 |
| SPGP0CX01990 | XP_018222214.1 | 99 | 0 |
| SPGP0L02190 | XP_018218817.1 | 99 | 0 |
| SPGP0AN00740 | XP_018223981.1 | 99 | 0 |
| SPGP0AA00950 | XP_018220495.1 | 100 | 0 |
| SPGP0CP01130 | XP_018223725.1 | 99 | 0 |
| SPGP0J00380 | XP_018219910.1 | 99 | 0 |
| SPGP0ES00100 | XP_018222826.1 | 99 | 0 |
| SPGP0BE00100 | XP_018223368.1 | 100 | 0 |
| SPGP0J03660 | XP_018219583.1 | 99 | 0 |
| SPGP0CW03080 | XP_018219255.1 | 100 | 0 |
| SPGP0AJ00130 | XP_018221788.1 | 99 | 0 |

Create PDF in your applications with the Pdfcrowd HTML to PDF API PDFCROWD

| | | | |
|---|---|---|---|
| SPGP0S00200 | XP_018224069.1 | 99 | 0 |
| SPGP0X01860 | XP_018221629.1 | 99 | 0 |
| SPGP0J00560 | XP_018219893.1 | 99 | 0 |
| SPGP0AF00210 | XP_018222276.1 | 100 | 0 |
| SPGP0K00570 | XP_018222799.1 | 98 | 0 |
| SPGP0S00170 | XP_018224066.1 | 99 | 0 |
| SPGP0CP03070 | XP_018223533.1 | 98 | 0 |
| SPGP0IH00150 | XP_018220386.1 | 99 | 0 |
| SPGP0J01160 | XP_018219831.1 | 99 | 0 |
| SPGP0CU01340 | XP_018220699.1 | 97 | 0 |
| SPGP0CU01610 | XP_018220673.1 | 99 | 0 |
| SPGP0F00970 | XP_018221261.1 | 99 | 0 |
| SPGP0AA01450 | XP_018220445.1 | 100 | 0 |
| SPGP0J02610 | XP_018219688.1 | 99 | 0 |
| SPGP0AA01920 | XP_018220400.1 | 99 | 0 |
| SPGP0F02800 | XP_018221440.1 | 99 | 0 |
| SPGP0J01440 | XP_018219804.1 | 99 | 0 |
| SPGP0Y03180 | XP_018220326.1 | 99 | 0 |
| SPGP0HZ00100 | XP_018219031.1 | 99 | 0 |
| SPGP0CP02030 | XP_018223637.1 | 99 | 0 |
| SPGP0X00190 | XP_018221468.1 | 99 | 0 |
| SPGP0AN00350 | XP_018223944.1 | 99 | 0 |
| SPGP0AQ00100 | XP_018223411.1 | 99 | 0 |
| SPGP0L00200 | XP_018219011.1 | 99 | 0 |
| SPGP0AF00660 | XP_018222320.1 | 99 | 0 |
| SPGP0AN00510 | XP_018223958.1 | 99 | 0 |
| SPGP0CP01310 | XP_018223708.1 | 99 | 0 |
| SPGP0EO00390 | XP_018221176.1 | 99 | 0 |
| SPGP0CX01890 | XP_018222204.1 | 99 | 0 |
| SPGP0AN00110 | XP_018223923.1 | 99 | 0 |
| SPGP0Y00810 | XP_018220097.1 | 99 | 0 |
| SPGP0AN00890 | XP_018223995.1 | 99 | 0 |
| SPGP0X01640 | XP_018221607.1 | 99 | 0 |
| SPGP0F02600 | XP_018221420.1 | 99 | 0 |
| SPGP0IZ00180 | XP_018220346.1 | 99 | 0 |
| SPGP0H00780 | XP_018221067.1 | 98 | 1 |
| SPGP0H00410 | XP_018221103.1 | 100 | 0 |
| SPGP0AF01790 | XP_018222437.1 | 99 | 0 |
| SPGP0F00980 | XP_018221262.1 | 99 | 0 |

Create PDF in your applications with the Pdfcrowd HTML to PDF API PDFCROWD

| | | | |
|---|---|---|---|
| SPGP0H00250 | XP_018221119.1 | 99 | 0 |
| SPGP0H00520 | XP_018221093.1 | 99 | 0 |
| SPGP0CU00970 | XP_018220733.1 | 99 | 0 |
| SPGP0AJ01920 | XP_018221958.1 | 99 | 0 |
| SPGP0CX01520 | XP_018222169.1 | 99 | 0 |
| SPGP0AA00700 | XP_018220521.1 | 99 | 0 |
| SPGP0DA01050 | XP_018222650.1 | 99 | 0 |
| SPGP0Y02830 | XP_018220292.1 | 97 | 2 |
| SPGP0J00710 | XP_018219879.1 | 99 | 0 |
| SPGP0L00570 | XP_018218976.1 | 99 | 0 |
| SPGP0CU01020 | XP_018220728.1 | 99 | 0 |
| SPGP0CUtrna20E | DI49_5287 | 100 | 0 |
| SPGP0Y02120 | XP_018220222.1 | 99 | 0 |
| SPGP0U01020 | XP_018222977.1 | 99 | 0 |
| SPGP0CW01040 | XP_018219452.1 | 99 | 0 |
| SPGP0H00680 | XP_018221077.1 | 98 | 2 |
| SPGP0AF01800 | XP_018222438.1 | 99 | 0 |
| SPGP0L00530 | XP_018218979.1 | 99 | 0 |
| SPGP0CX02040 | XP_018222219.1 | 99 | 0 |
| SPGP0X01320 | XP_018221576.1 | 99 | 0 |
| SPGP0CW03350 | XP_018219229.1 | 97 | 2 |
| SPGP0CU01490 | XP_018220685.1 | 99 | 0 |
| SPGP0L01780 | XP_018218857.1 | 99 | 0 |
| SPGP0CW01080 | XP_018219448.1 | 99 | 0 |
| SPGP0F01720 | XP_018221334.1 | 99 | 0 |
| SPGP0J02060 | XP_018219743.1 | 99 | 0 |
| SPGP0CP03140 | XP_018223526.1 | 99 | 0 |
| SPGP0F02230 | XP_018221384.1 | 99 | 0 |
| SPGP0CP03650 | XP_018223479.1 | 99 | 0 |
| SPGP0CN00840 | XP_018220888.1 | 98 | 0 |
| SPGP0F02450 | XP_018221406.1 | 99 | 0 |
| SPGP0CX02020 | XP_018222217.1 | 99 | 0 |
| SPGP0K00410 | XP_018222783.1 | 99 | 0 |
| SPGP0L01890 | XP_018218846.1 | 99 | 0 |
| SPGP0JP00140 | XP_018219565.1 | 99 | 0 |
| SPGP0L02590 | XP_018218777.1 | 99 | 0 |
| SPGP0L00940 | XP_018218938.1 | 99 | 0 |
| SPGP0Y00940 | XP_018220110.1 | 99 | 0 |
| SPGP0X00400 | XP_018221489.1 | 99 | 0 |
| SPGP0X01890 | XP_018221632.1 | 99 | 0 |

Create PDF in your applications with the Pdfcrowd HTML to PDF API PDFCROWD

| | | | |
|---|---|---|---|
| SPGP0Y02620 | XP_018220271.1 | 99 | 0 |
| SPGP0Y00900 | XP_018220106.1 | 99 | 0 |
| SPGP0H01330 | XP_018221014.1 | 99 | 0 |
| SPGP0AN00930 | XP_018223999.1 | 99 | 0 |
| SPGP0CU02200 | XP_018220614.1 | 99 | 0 |
| SPGP0AN00230 | XP_018223935.1 | 99 | 0 |
| SPGP0V00280 | XP_018221755.1 | 99 | 0 |
| SPGP0V00130 | XP_018221770.1 | 99 | 0 |
| SPGP0CW00100 | XP_018219544.1 | 99 | 0 |
| SPGP0CM00830 | XP_018223242.1 | 99 | 0 |
| SPGP0AJ01710 | XP_018221939.1 | 99 | 1 |
| SPGP0CW01920 | XP_018219368.1 | 100 | 0 |
| SPGP0X01670 | XP_018221610.1 | 99 | 0 |
| SPGP0CM01120 | XP_018223213.1 | 99 | 0 |
| SPGP0V00260 | XP_018221757.1 | 99 | 0 |
| SPGP0CV00300 | XP_018222581.1 | 99 | 0 |
| SPGP0CP00600 | XP_018223777.1 | 99 | 0 |
| SPGP0Y01110 | XP_018220126.1 | 99 | 0 |
| SPGP0F01530 | XP_018221316.1 | 99 | 0 |
| SPGP0F02290 | XP_018221390.1 | 99 | 0 |
| SPGP0S00110 | XP_018224060.1 | 99 | 0 |
| SPGP0H01400 | XP_018221008.1 | 99 | 0 |
| SPGP0S00500 | XP_018224097.1 | 99 | 0 |
| SPGP0F01610 | XP_018221324.1 | 99 | 0 |
| SPGP0F00270 | XP_018221196.1 | 98 | 0 |
| SPGP0CW01270 | XP_018219430.1 | 100 | 0 |
| SPGP0CP03030 | XP_018223537.1 | 99 | 0 |
| SPGP0J01100 | XP_018219837.1 | 87 | 2 |
| SPGP0L00450 | XP_018218987.1 | 99 | 0 |
| SPGP0F00910 | XP_018221255.1 | 99 | 0 |
| SPGP0Y00880 | XP_018220104.1 | 99 | 0 |
| SPGP0X01180 | XP_018221562.1 | 99 | 0 |
| SPGP0L01230 | XP_018218911.1 | 100 | 0 |
| SPGP0F02020 | XP_018221364.1 | 99 | 0 |
| SPGP0CP01510 | XP_018223688.1 | 99 | 0 |
| SPGP0F00620 | XP_018221227.1 | 99 | 0 |
| SPGP0J02450 | XP_018219704.1 | 99 | 0 |
| SPGP0CU01380 | XP_018220695.1 | 99 | 0 |
| SPGP0CN00380 | XP_018220933.1 | 99 | 0 |
| SPGP0J03700 | XP_018219579.1 | 99 | 0 |

Create PDF in your applications with the Pdfcrowd HTML to PDF API PDFCROWD

| | | | |
|---|---|---|---|
| SPGP0HB00150 | XP_018222866.1 | 99 | 0 |
| SPGP0AJ00590 | XP_018221831.1 | 99 | 0 |
| SPGP0AF01910 | XP_018222449.1 | 99 | 1 |
| SPGP0L00770 | XP_018218956.1 | 100 | 0 |
| SPGP0FE00110 | XP_018221963.1 | 99 | 0 |
| SPGP0AN00270 | XP_018223939.1 | 99 | 0 |
| SPGP0CX01360 | XP_018222154.1 | 99 | 0 |
| SPGP0Y02810 | XP_018220290.1 | 99 | 0 |
| SPGP0L01270 | XP_018218907.1 | 99 | 0 |
| SPGP0CU01270 | XP_018220705.1 | 99 | 0 |
| SPGP0AF01000 | XP_018222355.1 | 99 | 0 |
| SPGP0CP00840 | XP_018223754.1 | 99 | 0 |
| SPGP0L00140 | XP_018219017.1 | 99 | 0 |
| SPGP0X00360 | XP_018221485.1 | 99 | 0 |
| SPGP0J03270 | XP_018219622.1 | 99 | 0 |
| SPGP0DA01240 | XP_018222631.1 | 99 | 0 |
| SPGP0L00250 | XP_018219007.1 | 99 | 0 |
| SPGP0CX01650 | XP_018222181.1 | 99 | 0 |
| SPGP0FE00220 | XP_018221974.1 | 99 | 0 |
| SPGP0AF00750 | XP_018222329.1 | 99 | 0 |
| SPGP0U00110 | XP_018223058.1 | 98 | 0 |
| SPGP0FP00170 | XP_018223338.1 | 99 | 0 |
| SPGP0CP00780 | XP_018223760.1 | 99 | 0 |
| SPGP0U01500 | XP_018222932.1 | 94 | 0 |
| SPGP0EO00100 | XP_018221147.1 | 99 | 0 |
| SPGP0IN00220 | XP_018221654.1 | 99 | 0 |
| SPGP0Y00270 | XP_018220044.1 | 100 | 0 |
| SPGP0GR00100 | XP_018221975.1 | 99 | 0 |
| SPGP0AA01960 | XP_018220397.1 | 99 | 0 |
| SPGP0AJ01360 | XP_018221906.1 | 99 | 0 |
| SPGP0H01100 | XP_018221035.1 | 99 | 0 |
| SPGP0J02040 | XP_018219745.1 | 99 | 0 |
| SPGP0HB00130 | XP_018222864.1 | 99 | 0 |
| SPGP0CX00940 | XP_018222113.1 | 100 | 0 |
| SPGP0DA01400 | XP_018222615.1 | 99 | 0 |
| SPGP0DA00620 | XP_018222692.1 | 99 | 0 |
| SPGP0J03390 | XP_018219609.1 | 99 | 0 |
| SPGP0CN00420 | XP_018220929.1 | 99 | 0 |
| SPGP0Z00180 | XP_018222547.1 | 99 | 0 |
| SPGP0J00990 | XP_018219848.1 | 99 | 0 |

Create PDF in your applications with the Pdfcrowd HTML to PDF API PDFCROWD

| | | | |
|---|---|---|---|
| SPGP0CW02080 | XP_018219353.1 | 99 | 0 |
| SPGP0AT01600 | XP_018219177.1 | 99 | 0 |
| SPGP0Y01100 | XP_018220125.1 | 99 | 0 |
| SPGP0AJ01320 | XP_018221903.1 | 100 | 0 |
| SPGP0S00580 | XP_018224105.1 | 99 | 0 |
| SPGP0AT01460 | XP_018219163.1 | 99 | 0 |
| SPGP0J02840 | XP_018219665.1 | 99 | 0 |
| SPGP0CN01090 | XP_018220863.1 | 99 | 0 |
| SPGP0CN00630 | XP_018220908.1 | 100 | 0 |
| SPGP0F01250 | XP_018221288.1 | 99 | 0 |
| SPGP0JB00110 | XP_018222557.1 | 99 | 0 |
| SPGP0J03460 | XP_018219602.1 | 98 | 2 |
| SPGP0L00650 | XP_018218968.1 | 99 | 0 |
| SPGP0M00190 | XP_018221139.1 | 99 | 0 |
| SPGP0HN00120 | XP_018221988.1 | 99 | 0 |
| SPGP0HW00120 | XP_018224041.1 | 100 | 0 |
| SPGP0CN00710 | XP_018220900.1 | 99 | 0 |
| SPGP0FF00180 | XP_018223394.1 | 99 | 0 |
| SPGP0J01640 | XP_018219784.1 | 99 | 0 |
| SPGP0AT00730 | XP_018219092.1 | 99 | 0 |
| SPGP0H01320 | XP_018221015.1 | 99 | 0 |
| SPGP0AA00260 | XP_018220564.1 | 99 | 0 |
| SPGP0CW02870 | XP_018219276.1 | 100 | 0 |
| SPGP0GG00340 | XP_018220819.1 | 99 | 1 |
| SPGP0CP01470 | XP_018223692.1 | 99 | 0 |
| SPGP0F02710 | XP_018221431.1 | 99 | 0 |
| SPGP0DA00450 | XP_018222709.1 | 99 | 0 |
| SPGP0X00580 | XP_018221507.1 | 99 | 0 |
| SPGP0AA01130 | XP_018220477.1 | 99 | 0 |
| SPGP0AN00250 | XP_018223937.1 | 99 | 0 |
| SPGP0X00680 | XP_018221516.1 | 99 | 0 |
| SPGP0J00340 | XP_018219914.1 | 99 | 0 |
| SPGP0X01650 | XP_018221608.1 | 99 | 0 |
| SPGP0AF00390 | XP_018222293.1 | 99 | 0 |
| SPGP0F01810 | XP_018221343.1 | 99 | 0 |
| SPGP0AJ01080 | XP_018221879.1 | 99 | 0 |
| SPGP0F02050 | XP_018221367.1 | 100 | 0 |
| SPGP0S00130 | XP_018224062.1 | 99 | 0 |
| SPGP0J01340 | XP_018219814.1 | 99 | 0 |
| SPGP0AF00860 | XP_018222340.1 | 99 | 0 |

Create PDF in your applications with the Pdfcrowd HTML to PDF API PDFCROWD

| | | | |
|---|---|---|---|
| SPGP0JK00260 | XP_018223446.1 | 99 | 0 |
| SPGP0CN00120 | XP_018220959.1 | 100 | 0 |
| SPGP0AU00260 | XP_018223081.1 | 99 | 0 |
| SPGP0CP01680 | XP_018223672.1 | 99 | 0 |
| SPGP0U01300 | XP_018222951.1 | 99 | 0 |
| SPGP0F00930 | XP_018221257.1 | 99 | 0 |
| SPGP0AF00940 | XP_018222348.1 | 99 | 0 |
| SPGP0CP02340 | XP_018223607.1 | 99 | 0 |
| SPGP0L01950 | XP_018218841.1 | 99 | 0 |
| SPGP0J00270 | XP_018219921.1 | 99 | 0 |
| SPGP0X01030 | XP_018221550.1 | 99 | 0 |
| SPGP0CM00940 | XP_018223231.1 | 100 | 0 |
| SPGP0AF01370 | XP_018222394.1 | 98 | 2 |
| SPGP0CP00480 | XP_018223789.1 | 99 | 0 |
| SPGP0CV00190 | XP_018222571.1 | 99 | 0 |
| SPGP0AA00660 | XP_018220525.1 | 99 | 0 |
| SPGP0R00570 | XP_018219997.1 | 99 | 0 |
| SPGP0H01010 | XP_018221044.1 | 99 | 0 |
| SPGP0CV00130 | XP_018222566.1 | 100 | 0 |
| SPGP0AT00890 | XP_018219108.1 | 99 | 0 |
| SPGP0DA01010 | XP_018222654.1 | 98 | 0 |
| SPGP0AJ00150 | XP_018221790.1 | 99 | 0 |
| SPGP0DA01220 | XP_018222633.1 | 99 | 0 |
| SPGP0AF01280 | XP_018222384.1 | 99 | 0 |
| SPGP0DA01260 | XP_018222629.1 | 99 | 1 |
| SPGP0U01830 | XP_018222898.1 | 99 | 0 |
| SPGP0J00620 | XP_018219888.1 | 99 | 0 |
| SPGP0CW03310 | XP_018219233.1 | 99 | 0 |
| SPGP0H00740 | XP_018221071.1 | 99 | 0 |
| SPGP0CW02000 | XP_018219360.1 | 99 | 0 |
| SPGP0J03250 | XP_018219624.1 | 99 | 0 |
| SPGP0CW03270 | XP_018219237.1 | 99 | 0 |
| SPGP0DA01190 | XP_018222636.1 | 100 | 0 |
| SPGP0CX01560 | XP_018222172.1 | 99 | 0 |
| SPGP0DE00380 | XP_018223855.1 | 99 | 0 |
| SPGP0H00560 | XP_018221089.1 | 99 | 0 |
| SPGP0X00460 | XP_018221495.1 | 99 | 0 |
| SPGP0R00690 | XP_018220009.1 | 99 | 0 |
| SPGP0AU00380 | XP_018223093.1 | 99 | 0 |
| SPGP0CN00970 | XP_018220875.1 | 99 | 0 |

Create PDF in your applications with the Pdfcrowd HTML to PDF API PDFCROWD

| | | | |
|---|---|---|---|
| SPGP0CP01330 | XP_018223706.1 | 99 | 0 |
| SPGP0DA00310 | XP_018222723.1 | 100 | 0 |
| SPGP0AU00120 | XP_018223069.1 | 99 | 0 |
| SPGP0DA00370 | XP_018222717.1 | 99 | 0 |
| SPGP0V00760 | XP_018221711.1 | 99 | 0 |
| SPGP0G00190 | XP_018222258.1 | 99 | 0 |
| SPGP0J01800 | XP_018219768.1 | 99 | 0 |
| SPGP0Y00920 | XP_018220108.1 | 99 | 0 |
| SPGP0AT01660 | XP_018219183.1 | 99 | 0 |
| SPGP0CP03200 | XP_018223520.1 | 99 | 0 |
| SPGP0U00330 | XP_018223039.1 | 98 | 1 |
| SPGP0CU02460 | XP_018220588.1 | 99 | 0 |
| SPGP0CW02450 | XP_018219316.1 | 100 | 0 |
| SPGP0L02390 | XP_018218797.1 | 99 | 0 |
| SPGP0F01270 | XP_018221290.1 | 99 | 0 |
| SPGP0CW02590 | XP_018219304.1 | 99 | 0 |
| SPGP0CM01800 | XP_018223147.1 | 99 | 0 |
| SPGP0F01780 | XP_018221340.1 | 99 | 0 |
| SPGP0H01490 | XP_018221000.1 | 99 | 0 |
| SPGP0CM02040 | XP_018223124.1 | 99 | 0 |
| SPGP0AT00750 | XP_018219094.1 | 99 | 0 |
| SPGP0U00840 | XP_018222994.1 | 99 | 0 |
| SPGP0J00900 | XP_018219861.1 | 99 | 0 |
| SPGP0CP01830 | XP_018223657.1 | 99 | 0 |
| SPGP0L01850 | XP_018218850.1 | 99 | 0 |
| SPGP0L01580 | XP_018218876.1 | 99 | 0 |
| SPGP0L00980 | XP_018218934.1 | 100 | 0 |
| SPGP0CW02200 | XP_018219341.1 | 100 | 0 |
| SPGP0IN00300 | XP_018221646.1 | 99 | 0 |
| SPGP0U00200 | XP_018223050.1 | 99 | 0 |
| SPGP0CV00340 | XP_018222585.1 | 99 | 0 |
| SPGP0AF01300 | XP_018222387.1 | 99 | 0 |
| SPGP0X00320 | XP_018221481.1 | 99 | 0 |
| SPGP0CW01110 | XP_018219445.1 | 99 | 0 |
| SPGP0CX01970 | XP_018222212.1 | 99 | 0 |
| SPGP0Jtrna5L | DI49_4635 | 100 | 0 |
| SPGP0J00320 | XP_018219916.1 | 99 | 0 |
| SPGP0CW00570 | XP_018219499.1 | 99 | 0 |
| SPGP0CW01230 | XP_018219434.1 | 98 | 1 |
| SPGP0S00480 | XP_018224096.1 | 99 | 0 |

Create PDF in your applications with the Pdfcrowd HTML to PDF API PDFCROWD

| | | | |
|---|---|---|---|
| SPGP0DB00140 | XP_018223888.1 | 99 | 0 |
| SPGP0H01770 | XP_018220973.1 | 99 | 0 |
| SPGP0AT00350 | XP_018219056.1 | 99 | 0 |
| SPGP0CN00360 | XP_018220935.1 | 99 | 0 |
| SPGP0X01090 | XP_018221555.1 | 99 | 0 |
| SPGP0G00130 | XP_018222252.1 | 100 | 0 |
| SPGP0U01050 | XP_018222974.1 | 99 | 0 |
| SPGP0V00870 | XP_018221700.1 | 99 | 0 |
| SPGP0L01520 | XP_018218882.1 | 99 | 0 |
| SPGP0H00760 | XP_018221069.1 | 99 | 0 |
| SPGP0H00700 | XP_018221075.1 | 99 | 0 |
| SPGP0R00510 | XP_018219992.1 | 99 | 0 |
| SPGP0AJ01200 | XP_018221891.1 | 99 | 0 |
| SPGP0AJ01280 | XP_018221899.1 | 99 | 0 |
| SPGP0Y01080 | XP_018220123.1 | 99 | 0 |
| SPGP0F01130 | XP_018221277.1 | 99 | 0 |
| SPGP0GA00180 | XP_018223385.1 | 99 | 0 |
| SPGP0J02080 | XP_018219741.1 | 99 | 0 |
| SPGP0CM01270 | XP_018223198.1 | 100 | 0 |
| SPGP0U01360 | XP_018222946.1 | 99 | 0 |
| SPGP0AF00710 | XP_018222325.1 | 99 | 0 |
| SPGP0Y00350 | XP_018220052.1 | 99 | 0 |
| SPGP0AJ00600 | XP_018221832.1 | 100 | 0 |
| SPGP0CM01510 | XP_018223175.1 | 99 | 0 |
| SPGP0Y01530 | XP_018220168.1 | 99 | 0 |
| SPGP0FP00180 | XP_018223337.1 | 99 | 0 |
| SPGP0J01860 | XP_018219763.1 | 99 | 0 |
| SPGP0JO00130 | XP_018222263.1 | 99 | 0 |
| SPGP0CP01430 | XP_018223696.1 | 99 | 0 |
| SPGP0S00310 | XP_018224080.1 | 99 | 0 |
| SPGP0AJ00490 | XP_018221822.1 | 99 | 0 |
| SPGP0GH00140 | XP_018222816.1 | 99 | 0 |
| SPGP0DR00110 | XP_018219196.1 | 96 | 0 |
| SPGP0EO00290 | XP_018221166.1 | 98 | 0 |
| SPGP0U01620 | XP_018222920.1 | 99 | 0 |
| SPGP0U01190 | XP_018222961.1 | 99 | 0 |
| SPGP0F01370 | XP_018221301.1 | 99 | 0 |
| SPGP0F01830 | XP_018221345.1 | 99 | 0 |
| SPGP0AT00770 | XP_018219096.1 | 100 | 0 |
| SPGP0J03680 | XP_018219581.1 | 99 | 0 |

Create PDF in your applications with the Pdfcrowd HTML to PDF API PDFCROWD

| | | | |
|---|---|---|---|
| SPGP0CW00530 | XP_018219502.1 | 100 | 0 |
| SPGP0X01340 | XP_018221578.1 | 99 | 0 |
| SPGP0F02560 | XP_018221416.1 | 99 | 0 |
| SPGP0CP02540 | XP_018223587.1 | 99 | 0 |
| SPGP0F02010 | XP_018221363.1 | 99 | 0 |
| SPGP0X00510 | XP_018221500.1 | 99 | 0 |
| SPGP0F00510 | XP_018221216.1 | 99 | 0 |
| SPGP0X01430 | XP_018221587.1 | 99 | 0 |
| SPGP0H00150 | XP_018221128.1 | 99 | 0 |
| SPGP0J03350 | XP_018219614.1 | 99 | 0 |
| SPGP0CU00930 | XP_018220736.1 | 99 | 0 |
| SPGP0IM00100 | XP_018219552.1 | 99 | 0 |
| SPGP0CX01910 | XP_018222206.1 | 99 | 0 |
| SPGP0Jtrna3P | DI49_4699 | 100 | 0 |
| SPGP0S00540 | XP_018224101.1 | 99 | 0 |
| SPGP0CM00850 | XP_018223240.1 | 99 | 0 |
| SPGP0J01300 | XP_018219818.1 | 99 | 0 |
| SPGP0DA00760 | XP_018222677.1 | 99 | 0 |
| SPGP0CM02180 | XP_018223111.1 | 99 | 0 |
| SPGP0F00460 | XP_018221212.1 | 99 | 0 |
| SPGP0CP01160 | XP_018223722.1 | 99 | 0 |
| SPGP0AT01580 | XP_018219175.1 | 99 | 0 |
| SPGP0F02820 | XP_018221442.1 | 99 | 0 |
| SPGP0EB00140 | XP_018223317.1 | 99 | 0 |
| SPGP0F00680 | XP_018221233.1 | 99 | 0 |
| SPGP0U01870 | XP_018222894.1 | 99 | 0 |
| SPGP0J00870 | XP_018219863.1 | 99 | 0 |
| SPGP0GG00320 | XP_018220821.1 | 99 | 0 |
| SPGP0DA01180 | XP_018222637.1 | 99 | 0 |
| SPGP0AT01170 | XP_018219135.1 | 100 | 0 |
| SPGP0AHtrna17S | DI49_0808 | 99 | 0 |
| SPGP0R00820 | XP_018220021.1 | 99 | 0 |
| SPGP0J03290 | XP_018219620.1 | 99 | 0 |
| SPGP0F02350 | XP_018221396.1 | 100 | 0 |
| SPGP0Y02180 | XP_018220227.1 | 99 | 0 |
| SPGP0CX00720 | XP_018222091.1 | 99 | 0 |
| SPGP0V00780 | XP_018221709.1 | 99 | 0 |
| SPGP0Y01250 | XP_018220140.1 | 99 | 0 |
| SPGP0X00640 | XP_018221512.1 | 99 | 0 |
| SPGP0CN00820 | XP_018220890.1 | 99 | 0 |

Create PDF in your applications with the Pdfcrowd HTML to PDF API PDFCROWD

| | | | |
|---|---|---|---|
| SPGP0S00790 | XP_018224123.1 | 96 | 0 |
| SPGP0AF00590 | XP_018222313.1 | 99 | 0 |
| SPGP0AE00350 | XP_018222515.1 | 98 | 0 |
| SPGP0AN00330 | XP_018223942.1 | 99 | 0 |
| SPGP0AJ01030 | XP_018221875.1 | 99 | 0 |
| SPGP0CP01370 | XP_018223702.1 | 99 | 0 |
| SPGP0DA00680 | XP_018222686.1 | 99 | 0 |
| SPGP0CP01010 | XP_018223738.1 | 99 | 0 |
| SPGP0AU00160 | XP_018223073.1 | 99 | 0 |
| SPGP0H01610 | XP_018220989.1 | 99 | 0 |
| SPGP0H01560 | XP_018220993.1 | 99 | 0 |
| SPGP0F02580 | XP_018221418.1 | 99 | 0 |
| SPGP0CP00360 | XP_018223800.1 | 99 | 0 |
| SPGP0CN01170 | XP_018220856.1 | 98 | 0 |
| SPGP0CP00420 | XP_018223794.1 | 99 | 0 |
| SPGP0CW02530 | XP_018219308.1 | 99 | 0 |
| SPGP0AF01880 | XP_018222446.1 | 100 | 0 |
| SPGP0EItrna6S | DI49_0694 | 100 | 0 |
| SPGP0X00530 | XP_018221502.1 | 99 | 0 |
| SPGP0CW02600 | XP_018219303.1 | 100 | 0 |
| SPGP0J01290 | XP_018219819.1 | 99 | 0 |
| SPGP0S00690 | XP_018224116.1 | 99 | 0 |
| SPGP0JK00200 | XP_018223441.1 | 99 | 0 |
| SPGP0AT00260 | XP_018219048.1 | 100 | 0 |
| SPGP0CN00440 | XP_018220927.1 | 99 | 0 |
| SPGP0CU01550 | XP_018220679.1 | 99 | 0 |
| SPGP0CN01150 | XP_018220858.1 | 99 | 0 |
| SPGP0CM00280 | XP_018223296.1 | 99 | 0 |
| SPGP0X01070 | XP_018221553.1 | 99 | 0 |
| SPGP0Y03040 | XP_018220313.1 | 99 | 0 |
| SPGP0F01990 | XP_018221361.1 | 99 | 0 |
| SPGP0DE00210 | XP_018223838.1 | 99 | 0 |
| SPGP0AJtrna9F | DI49_2529 | 100 | 0 |
| SPGP0AT01520 | XP_018219169.1 | 100 | 0 |
| SPGP0J02350 | XP_018219714.1 | 99 | 0 |
| SPGP0X01900 | XP_018221633.1 | 99 | 0 |
| SPGP0F02310 | XP_018221392.1 | 99 | 0 |
| SPGP0V00200 | XP_018221763.1 | 99 | 0 |
| SPGP0Y02950 | XP_018220304.1 | 99 | 0 |
| SPGP0EI00100 | XP_018223326.1 | 100 | 0 |

Create PDF in your applications with the Pdfcrowd HTML to PDF API PDFCROWD

| | | | |
|---|---|---|---|
| SPGP0H01280 | XP_018221019.1 | 99 | 0 |
| SPGP0J01980 | XP_018219751.1 | 99 | 0 |
| SPGP0CN00790 | XP_018220893.1 | 100 | 0 |
| SPGP0X01510 | XP_018221594.1 | 95 | 4 |
| SPGP0DA00960 | XP_018222659.1 | 99 | 0 |
| SPGP0DA00740 | XP_018222679.1 | 99 | 0 |
| SPGP0F00920 | XP_018221256.1 | 99 | 0 |
| SPGP0U01130 | XP_018222966.1 | 99 | 0 |
| SPGP0CN00140 | XP_018220957.1 | 99 | 0 |
| SPGP0Y01930 | XP_018220204.1 | 99 | 0 |
| SPGP0H00660 | XP_018221079.1 | 99 | 1 |
| SPGP0BE00210 | XP_018223357.1 | 99 | 1 |
| SPGP0J02630 | XP_018219686.1 | 100 | 0 |
| SPGP0J03080 | XP_018219641.1 | 99 | 0 |
| SPGP0CU01910 | XP_018220643.1 | 99 | 0 |
| SPGP0CP01590 | XP_018223680.1 | 99 | 0 |
| SPGP0DA00390 | XP_018222715.1 | 100 | 0 |
| SPGP0F02380 | XP_018221399.1 | 99 | 0 |
| SPGP0CM00540 | XP_018223270.1 | 99 | 0 |
| SPGP0CP03450 | XP_018223497.1 | 99 | 0 |
| SPGP0J03760 | XP_018219573.1 | 100 | 0 |
| SPGP0G00180 | XP_018222257.1 | 99 | 0 |
| SPGP0F02370 | XP_018221398.1 | 99 | 0 |
| SPGP0AF00820 | XP_018222336.1 | 99 | 0 |
| SPGP0Y02290 | XP_018220238.1 | 99 | 0 |
| SPGP0DE00570 | XP_018223874.1 | 99 | 0 |
| SPGP0J01250 | XP_018219823.1 | 99 | 0 |
| SPGP0X01570 | XP_018221600.1 | 99 | 0 |
| SPGP0CN00990 | XP_018220873.1 | 99 | 0 |
| SPGP0H01530 | XP_018220996.1 | 99 | 0 |
| SPGP0CM01550 | XP_018223171.1 | 99 | 0 |
| SPGP0CW00370 | XP_018219518.1 | 99 | 0 |
| SPGP0L02270 | XP_018218809.1 | 99 | 0 |
| SPGP0JK00180 | XP_018223440.1 | 99 | 0 |
| SPGP0HI00170 | XP_018219943.1 | 99 | 0 |
| SPGP0IH00210 | XP_018220391.1 | 99 | 0 |
| SPGP0CX00880 | XP_018222107.1 | 99 | 0 |
| SPGP0Z00220 | XP_018222543.1 | 99 | 0 |
| SPGP0CV00420 | XP_018222594.1 | 99 | 0 |
| SPGP0GT00120 | XP_018219024.1 | 99 | 0 |

Create PDF in your applications with the Pdfcrowd HTML to PDF API PDFCROWD

| | | | |
|---|---|---|---|
| SPGP0K00550 | XP_018222797.1 | 100 | 0 |
| SPGP0DB00370 | XP_018223910.1 | 99 | 0 |
| SPGP0CW02360 | XP_018219325.1 | 99 | 0 |
| SPGP0IM00140 | XP_018219548.1 | 99 | 0 |
| SPGP0L01740 | XP_018218861.1 | 97 | 0 |
| SPGP0DA00250 | XP_018222728.1 | 99 | 0 |
| SPGP0AF01310 | XP_018222388.1 | 100 | 0 |
| SPGP0X00380 | XP_018221487.1 | 99 | 0 |
| SPGP0CP00980 | XP_018223741.1 | 99 | 0 |
| SPGP0J00610 | XP_018219889.1 | 99 | 0 |
| SPGP0AF00800 | XP_018222334.1 | 99 | 0 |
| SPGP0CW01690 | XP_018219391.1 | 100 | 0 |
| SPGP0AT01010 | XP_018219119.1 | 99 | 0 |
| SPGP0AT01090 | XP_018219127.1 | 99 | 0 |
| SPGP0BE00270 | XP_018223351.1 | 99 | 0 |
| SPGP0CM02230 | XP_018223106.1 | 99 | 0 |
| SPGP0J02960 | XP_018219653.1 | 99 | 1 |
| SPGP0CX01240 | XP_018222143.1 | 99 | 0 |
| SPGP0F02190 | XP_018221380.1 | 99 | 0 |
| SPGP0CX00550 | XP_018222075.1 | 99 | 0 |
| SPGP0S00710 | XP_018224118.1 | 99 | 0 |
| SPGP0J03560 | XP_018219593.1 | 100 | 0 |
| SPGP0X00810 | XP_018221529.1 | 99 | 0 |
| SPGP0J01700 | XP_018219778.1 | 99 | 0 |
| SPGP0Y00620 | XP_018220078.1 | 99 | 1 |
| SPGP0J02230 | XP_018219726.1 | 100 | 0 |
| SPGP0CU00370 | XP_018220792.1 | 99 | 0 |
| SPGP0CW00720 | XP_018219484.1 | 99 | 0 |
| SPGP0DA00530 | XP_018222701.1 | 99 | 0 |
| SPGP0AN00370 | XP_018223946.1 | 99 | 0 |
| SPGP0AF00370 | XP_018222291.1 | 99 | 0 |
| SPGP0H01650 | XP_018220985.1 | 98 | 1 |
| SPGP0CM01050 | XP_018223220.1 | 100 | 0 |
| SPGP0F00720 | XP_018221237.1 | 99 | 0 |
| SPGP0S00520 | XP_018224099.1 | 99 | 0 |
| SPGP0AA01660 | XP_018220425.1 | 97 | 2 |
| SPGP0CN00260 | XP_018220945.1 | 99 | 0 |
| SPGP0F00760 | XP_018221241.1 | 99 | 0 |
| SPGP0AJ01430 | XP_018221913.1 | 99 | 0 |

Create PDF in your applications with the Pdfcrowd HTML to PDF API PDFCROWD

| | | | |
|---|---|---|---|
| SPGP0J01070 | XP_018219856.1 | 99 | 0 |
| SPGP0CU01590 | XP_018220675.1 | 99 | 0 |
| SPGP0AA01530 | XP_018220437.1 | 99 | 0 |
| SPGP0CN00480 | XP_018220923.1 | 99 | 0 |
| SPGP0J03400 | XP_018219608.1 | 98 | 2 |
| SPGP0J02390 | XP_018219710.1 | 100 | 0 |
| SPGP0HI00240 | XP_018219950.1 | 99 | 0 |
| SPGP0AT00740 | XP_018219093.1 | 99 | 0 |
| SPGP0J03060 | XP_018219643.1 | 99 | 0 |
| SPGP0X01940 | XP_018221637.1 | 99 | 0 |
| SPGP0J03500 | XP_018219598.1 | 99 | 0 |
| SPGP0AT01150 | XP_018219133.1 | 99 | 0 |
| SPGP0V00600 | XP_018221726.1 | 99 | 0 |
| SPGP0J03100 | XP_018219639.1 | 99 | 1 |
| SPGP0CX00710 | XP_018222090.1 | 99 | 0 |
| SPGP0Y03130 | XP_018220321.1 | 99 | 0 |
| SPGP0Y02310 | XP_018220240.1 | 99 | 0 |
| SPGP0F00290 | XP_018221198.1 | 99 | 0 |
| SPGP0AF00440 | XP_018222298.1 | 99 | 0 |
| SPGP0CW02570 | XP_018219306.1 | 99 | 0 |
| SPGP0U00180 | XP_018223052.1 | 99 | 0 |
| SPGP0CM00500 | XP_018223274.1 | 99 | 0 |
| SPGP0L00590 | XP_018218974.1 | 99 | 0 |
| SPGP0DA01420 | XP_018222613.1 | 99 | 0 |
| SPGP0K00680 | XP_018222810.1 | 99 | 0 |
| SPGP0X00930 | XP_018221540.1 | 99 | 0 |
| SPGP0X00800 | XP_018221528.1 | 99 | 1 |
| SPGP0AN00420 | XP_018223951.1 | 99 | 1 |
| SPGP0H01720 | XP_018220978.1 | 99 | 0 |
| SPGP0AA00640 | XP_018220527.1 | 99 | 0 |
| SPGP0L02050 | XP_018218831.1 | 99 | 0 |
| SPGP0H00320 | XP_018221112.1 | 100 | 0 |
| SPGP0DA00990 | XP_018222656.1 | 99 | 0 |
| SPGP0U02000 | XP_018222881.1 | 99 | 0 |
| SPGP0CM02020 | XP_018223126.1 | 100 | 0 |
| SPGP0H00480 | XP_018221096.1 | 99 | 0 |
| SPGP0CW01060 | XP_018219450.1 | 99 | 0 |
| SPGP0J01570 | XP_018219791.1 | 99 | 0 |
| SPGP0CW00910 | XP_018219465.1 | 100 | 0 |
| SPGP0IU00180 | XP_018220336.1 | 99 | 0 |

Create PDF in your applications with the Pdfcrowd HTML to PDF API PDFCROWD

| | | | |
|---|---|---|---|
| SPGP0L00160 | XP_018219015.1 | 99 | 0 |
| SPGP0CM01820 | XP_018223145.1 | 98 | 1 |
| SPGP0F02250 | XP_018221386.1 | 99 | 0 |
| SPGP0AA00370 | XP_018220553.1 | 99 | 0 |
| SPGP0CV00270 | XP_018222578.1 | 99 | 0 |
| SPGP0J00780 | XP_018219872.1 | 99 | 0 |
| SPGP0L00780 | XP_018218955.1 | 99 | 0 |
| SPGP0U00760 | XP_018223001.1 | 99 | 0 |
| SPGP0AA00410 | XP_018220549.1 | 99 | 0 |
| SPGP0DA01090 | XP_018222646.1 | 99 | 0 |
| SPGP0CN00690 | XP_018220902.1 | 99 | 0 |
| SPGP0H01030 | XP_018221042.1 | 99 | 0 |
| SPGP0CU02180 | XP_018220616.1 | 99 | 0 |
| SPGP0AT01260 | XP_018219144.1 | 99 | 0 |
| SPGP0H01200 | XP_018221027.1 | 99 | 0 |
| SPGP0H01750 | XP_018220975.1 | 99 | 0 |
| SPGP0DA00270 | XP_018222726.1 | 100 | 0 |
| SPGP0JK00230 | XP_018223443.1 | 99 | 0 |
| SPGP0H00950 | XP_018221050.1 | 99 | 0 |
| SPGP0Y01910 | XP_018220202.1 | 99 | 0 |
| SPGP0DA00850 | XP_018222668.1 | 99 | 0 |
| SPGP0FF00120 | XP_018223400.1 | 99 | 0 |
| SPGP0X00850 | XP_018221533.1 | 99 | 0 |
| SPGP0JC00100 | XP_018220851.1 | 99 | 0 |
| SPGP0AA01410 | XP_018220449.1 | 99 | 0 |
| SPGP0AF01350 | XP_018222392.1 | 99 | 0 |
| SPGP0CX01780 | XP_018222193.1 | 99 | 0 |
| SPGP0CW03480 | XP_018219216.1 | 99 | 0 |
| SPGP0Y01450 | XP_018220160.1 | 99 | 0 |
| SPGP0F01090 | XP_018221273.1 | 99 | 0 |
| SPGP0K00600 | XP_018222802.1 | 99 | 0 |
| SPGP0CW00660 | XP_018219490.1 | 99 | 0 |
| SPGP0L01540 | XP_018218880.1 | 99 | 0 |
| SPGP0V00640 | XP_018221722.1 | 99 | 0 |
| SPGP0J01460 | XP_018219802.1 | 99 | 0 |
| SPGP0AJ01170 | XP_018221888.1 | 99 | 0 |
| SPGP0DA00830 | XP_018222670.1 | 99 | 0 |
| SPGP0J01010 | XP_018219850.1 | 98 | 0 |
| SPGP0U01890 | XP_018222892.1 | 99 | 0 |
| SPGP0Y01640 | XP_018220178.1 | 99 | 0 |

Create PDF in your applications with the Pdfcrowd HTML to PDF API PDFCROWD

| | | | |
|---|---|---|---|
| SPGP0CP00500 | XP_018223787.1 | 99 | 0 |
| SPGP0L00880 | XP_018218945.1 | 99 | 0 |
| SPGP0L00330 | XP_018218999.1 | 99 | 0 |
| SPGP0Y00690 | XP_018220085.1 | 99 | 0 |
| SPGP0CP02590 | XP_018223582.1 | 99 | 0 |
| SPGP0CN00560 | XP_018220915.1 | 99 | 0 |
| SPGP0H00360 | XP_018221108.1 | 99 | 0 |
| SPGP0DA00330 | XP_018222721.1 | 100 | 0 |
| SPGP0Y01620 | XP_018220176.1 | 99 | 0 |
| SPGP0AA01900 | XP_018220402.1 | 99 | 0 |
| SPGP0CP02060 | XP_018223634.1 | 99 | 0 |
| SPGP0CN00800 | XP_018220892.1 | 99 | 0 |
| SPGP0IU00100 | XP_018220328.1 | 99 | 0 |
| SPGP0AJ00810 | XP_018221853.1 | 99 | 0 |
| SPGP0F01030 | XP_018221267.1 | 99 | 0 |
| SPGP0Y02780 | XP_018220287.1 | 99 | 0 |
| SPGP0CU02260 | XP_018220608.1 | 99 | 0 |
| SPGP0CM01310 | XP_018223194.1 | 99 | 0 |
| SPGP0AN00440 | XP_018223953.1 | 99 | 0 |
| SPGP0HS00150 | XP_018221779.1 | 99 | 0 |
| SPGP0H01710 | XP_018220979.1 | 99 | 0 |
| SPGP0H00380 | XP_018221106.1 | 99 | 0 |
| SPGP0Y02990 | XP_018220308.1 | 99 | 0 |
| SPGP0DA01140 | XP_018222641.1 | 99 | 0 |
| SPGP0CP03160 | XP_018223524.1 | 99 | 0 |
| SPGP0CW01120 | XP_018219444.1 | 99 | 0 |
| SPGP0V01050 | XP_018221682.1 | 99 | 0 |
| SPGP0X01110 | XP_018221557.1 | 99 | 0 |
| SPGP0DA00780 | XP_018222675.1 | 99 | 0 |
| SPGP0AJ01220 | XP_018221893.1 | 99 | 0 |
| SPGP0J02780 | XP_018219671.1 | 99 | 0 |
| SPGP0X00790 | XP_018221527.1 | 99 | 0 |
| SPGP0AA01210 | XP_018220469.1 | 99 | 0 |
| SPGP0CX01740 | XP_018222190.1 | 98 | 1 |
| SPGP0H01380 | XP_018221009.1 | 99 | 0 |
| SPGP0HN00140 | XP_018221990.1 | 99 | 0 |
| SPGP0CU01040 | XP_018220726.1 | 99 | 0 |
| SPGP0F01050 | XP_018221269.1 | 99 | 0 |
| SPGP0V00510 | XP_018221735.1 | 99 | 0 |
| SPGP0AF02190 | XP_018222476.1 | 95 | 5 |

Create PDF in your applications with the Pdfcrowd HTML to PDF API PDFCROWD

| | | | |
|---|---|---|---|
| SPGP0AH00140 | XP_018223422.1 | 98 | 1 |
| SPGP0U00500 | XP_018223024.1 | 99 | 0 |
| SPGP0L00510 | XP_018218981.1 | 99 | 0 |
| SPGP0X02000 | XP_018221643.1 | 99 | 0 |
| SPGP0DA01160 | XP_018222639.1 | 99 | 0 |
| SPGP0CN00520 | XP_018220919.1 | 99 | 0 |
| SPGP0CW03520 | XP_018219212.1 | 99 | 0 |
| SPGP0CU00600 | XP_018220769.1 | 99 | 0 |
| SPGP0H01440 | XP_018221004.1 | 97 | 2 |
| SPGP0CM00610 | XP_018223263.1 | 99 | 0 |

### Sheet 2: *genes alleles inherited from Saccharomyces cerevisiae*

| *Saccharomyces pastorianus CBS1538 predicted ORF* | *Saccharomyces cerevisiae parental allele* | Gene identities | Gene gaps |
|---|---|---|---|
| SPGP0C00250 | YIL156W-B | 99 | 0 |
| SPGP0I02780 | YNL040W | 99 | 0 |
| SPGP0I00510 | YNL291C | 99 | 0 |
| SPGP0D00690 | YNR014W | 100 | 0 |
| SPGP0D00160 | YNR067C | 100 | 0 |
| SPGP0T00470 | YKL161C | 99 | 0 |
| SPGP0B00310 | YER054C | 99 | 0 |
| SPGP0T00860 | YKL116C | 99 | 0 |
| SPGP0I01300 | YNL206C | 99 | 0 |
| SPGP0C00960 | YIL077C | 99 | 0 |
| SPGP0B01580 | YEL071W | 99 | 0 |
| SPGP0E01110 | YKR031C | 99 | 0 |
| SPGP0E01430 | YKR067W | 99 | 0 |
| SPGP0B00850 | YER005W | 99 | 0 |
| SPGP0A01890 | YJL048C | 99 | 0 |
| SPGP0C00390 | YIL138C | 100 | 0 |
| SPGP0CR00500 | YAL012W | 99 | 0 |
| SPGP0I01190 | YNL217W | 100 | 0 |
| SPGP0IZ00300 | YMR310C | 99 | 0 |
| SPGP0CX00140 | YGL216W | 100 | 0 |
| SPGP0T00130 | YKL196C | 99 | 0 |
| SPGP0CX00110 | YGL221C | 100 | 0 |
| SPGP0CW03160 | YHR010W | 95 | 1 |
| SPGP0JF00120 | YER179W | 99 | 0 |
| SPGP0I02220 | YNL100W | 99 | 0 |

Create PDF in your applications with the Pdfcrowd HTML to PDF API PDFCROWD

| | | | |
|---|---|---|---|
| SPGP0B00140 | YER070W | 99 | 0 |
| SPGP0CR00760 | YAR031W | 96 | 0 |
| SPGP0I00790 | YNL260C | 100 | 0 |
| SPGP0E00790 | YKL003C | 100 | 0 |
| SPGP0A02260 | YJL089W | 99 | 0 |
| SPGP0I01990 | YNL128W | 99 | 0 |
| SPGP0CX00350 | YGL192W | 99 | 0 |
| SPGP0T00250 | YKL184W | 99 | 0 |
| SPGP0CR00250 | YAL036C | 99 | 0 |
| SPGP0CR00560 | YAL005C | 99 | 0 |
| SPGP0I02630 | YNL056W | 99 | 0 |
| SPGP0A01790 | YJL037W | 99 | 0 |
| SPGP0HU00100 | YGR294W | 97 | 0 |
| SPGP0A02040 | YJL062W-A | 100 | 0 |
| SPGP0IZ00270 | YMR307W | 99 | 0 |
| SPGP0C01420 | YIL023C | 99 | 0 |
| SPGP0C01130 | YIL057C | 100 | 0 |
| SPGP0I02090 | YNL117W | 99 | 0 |
| SPGP0IP00160 | YGL231C | 100 | 0 |
| SPGP0JF00150 | YER182W | 99 | 0 |
| SPGP0E00680 | YKL013C | 99 | 0 |
| SPGP0B01360 | YEL049W | 99 | 0 |
| SPGP0E00690 | YKL012W | 99 | 0 |
| SPGP0A01530 | YJL003W | 100 | 0 |
| SPGP0A01240 | YJR031C | 99 | 0 |
| SPGP0D00570 | YNR027W | 99 | 0 |
| SPGP0A00110 | YJR150C | 99 | 0 |
| SPGP0A00960 | YJR061W | 99 | 0 |
| SPGP0E00890 | YKR008W | 99 | 0 |
| SPGP0T00780 | YKL126W | 99 | 0 |
| SPGP0C00470 | YIL130W | 98 | 1 |
| SPGP0I00920 | YNL247W | 99 | 0 |
| SPGP0U01240 | YDL131W | 86 | 0 |
| SPGP0AI00270 | YJL217W | 98 | 0 |
| SPGP0E01560 | YKR081C | 99 | 0 |
| SPGP0A00670 | YJR094C | 100 | 0 |
| SPGP0DC00200 | YNL006W | 99 | 0 |
| SPGP0I02470 | YNL074C | 99 | 0 |
| SPGP0I01560 | YNL175C | 99 | 0 |
| SPGP0IF00100 | YGR288W | 99 | 0 |

Create PDF in your applications with the Pdfcrowd HTML to PDF API PDFCROWD

| | | | |
|---|---|---|---|
| SPGP0I01410 | YNL192W | 99 | 0 |
| SPGP0B00440 | YER044C | 100 | 0 |
| SPGP0E00960 | YKR016W | 99 | 0 |
| SPGP0I00370 | YNL307C | 99 | 0 |
| SPGP0B00620 | YER026C | 99 | 0 |
| SPGP0C01960 | YIR031C | 100 | 0 |
| SPGP0EL00150 | YGL246C | 99 | 0 |
| SPGP0I01380 | YNL195C | 99 | 0 |
| SPGP0DC00230 | YNL003C | 100 | 0 |
| SPGP0A01690 | YJL025W | 99 | 0 |
| SPGP0B01380 | YEL050C | 99 | 0 |
| SPGP0CR00360 | YAL026C | 99 | 0 |
| SPGP0AI00180 | YJL207C | 99 | 0 |
| SPGP0EL00220 | YGL253W | 99 | 0 |
| SPGP0A02450 | YJL109C | 99 | 0 |
| SPGP0IP00180 | YGL233W | 99 | 0 |
| SPGP0C00610 | YIL116W | 99 | 0 |
| SPGP0C01630 | YIL003W | 100 | 0 |
| SPGP0C01170 | YIL052C | 99 | 0 |
| SPGP0I00380 | YNL306W | 99 | 0 |
| SPGP0I02380 | YNL083W | 99 | 0 |
| SPGP0T00420 | YKL165C | 99 | 0 |
| SPGP0C00940 | YIL079C | 99 | 0 |
| SPGP0A00460 | YJR112W-A | 99 | 0 |
| SPGP0B01170 | YEL029C | 99 | 0 |
| SPGP0E00240 | YKL058W | 100 | 0 |
| SPGP0DD00190 | YNL029C | 99 | 0 |
| SPGP0I00830 | YNL256W | 99 | 0 |
| SPGP0I02420 | YNL079C | 100 | 0 |
| SPGP0I02640 | YNL055C | 99 | 0 |
| SPGP0I02340 | YNL087W | 99 | 0 |
| SPGP0A01370 | YJR011C | 99 | 0 |
| SPGP0A02080 | YJL068C | 98 | 0 |
| SPGP0A01610 | YJL013C | 99 | 0 |
| SPGP0D00360 | YNR047W | 99 | 0 |
| SPGP0A00320 | YJR129C | 99 | 0 |
| SPGP0C01090 | YIL064W | 99 | 0 |
| SPGP0B01280 | YEL041W | 99 | 0 |
| SPGP0A00710 | YJR090C | 99 | 0 |
| SPGP0D00140 | YNR069C | 99 | 0 |

Create PDF in your applications with the Pdfcrowd HTML to PDF API PDFCROWD

| | | | |
|---|---|---|---|
| SPGP0T00400 | YKL167C | 99 | 0 |
| SPGP0E01660 | YKR091W | 99 | 0 |
| SPGP0T00800 | YKL124W | 99 | 0 |
| SPGP0T00600 | YKL145W | 99 | 0 |
| SPGP0T00110 | YKL198C | 98 | 1 |
| SPGP0I00250 | YNL320W | 99 | 0 |
| SPGP0E01000 | YKR020W | 99 | 0 |
| SPGP0E00900 | YKR009C | 99 | 0 |
| SPGP0I01310 | YNL204C | 99 | 0 |
| SPGP0E01170 | YKR039W | 99 | 0 |
| SPGP0I02400 | YNL081C | 99 | 0 |
| SPGP0B00910 | YEL002C | 99 | 0 |
| SPGP0C00320 | YIL146C | 99 | 0 |
| SPGP0B01550 | YEL069C | 98 | 0 |
| SPGP0C00370 | YIL140W | 99 | 0 |
| SPGP0I02030 | YNL124W | 99 | 0 |
| SPGP0CX00370 | YGL190C | 99 | 0 |
| SPGP0DD00130 | YNL023C | 99 | 0 |
| SPGP0E00550 | YKL024C | 99 | 0 |
| SPGP0E00220 | YKL060C | 99 | 0 |
| SPGP0C00130 | YJL221C | 98 | 0 |
| SPGP0CX00130 | YGL219C | 100 | 0 |
| SPGP0I00110 | YNL335W | 99 | 0 |
| SPGP0IZ00320 | YMR312W | 99 | 0 |
| SPGP0T00230 | YKL186C | 99 | 0 |
| SPGP0B00160 | YER068W | 99 | 0 |
| SPGP0I00770 | YNL262W | 99 | 0 |
| SPGP0A02200 | YJL082W | 99 | 0 |
| SPGP0T00580 | YKL148C | 99 | 0 |
| SPGP0E01750 | YKR099W | 99 | 0 |
| SPGP0A00990 | YJR058C | 99 | 0 |
| SPGP0I02810 | YNL037C | 99 | 0 |
| SPGP0I02200 | YNL102W | 99 | 0 |
| SPGP0C00140 | YOL156W | 98 | 0 |
| SPGP0ES00150 | YER166W | 99 | 0 |
| SPGP0C01400 | YIL026C | 99 | 0 |
| SPGP0DW00100 | YBR299W | 99 | 0 |
| SPGP0FY00100 | YER184C | 99 | 0 |
| SPGP0HY00110 | YGL237C | 99 | 1 |
| SPGP0B01340 | YEL047C | 99 | 0 |

Create PDF in your applications with the Pdfcrowd HTML to PDF API PDFCROWD

| | | | |
|---|---|---|---|
| SPGP0CR00140 | YAL047C | 99 | 0 |
| SPGP0D00550 | YNR029C | 99 | 0 |
| SPGP0E00570 | YKL023W | 98 | 1 |
| SPGP0A00940 | YJR063W | 99 | 0 |
| SPGP0CR00750 | YAR029W | 98 | 0 |
| SPGP0C00700 | YIL107C | 99 | 0 |
| SPGP0A02330 | YJL096W | 99 | 0 |
| SPGP0U01220 | YDL130W | 86 | 2 |
| SPGP0U01730 | YDL188C | 83 | 1 |
| SPGP0I00940 | YNL245C | 100 | 0 |
| SPGP0C00520 | YIL125W | 99 | 0 |
| SPGP0HM00120 | YAL058W | 99 | 0 |
| SPGP0E01540 | YKR079C | 99 | 0 |
| SPGP0E01090 | YKR029C | 99 | 0 |
| SPGP0Y01830 | YMR143W | 97 | 0 |
| SPGP0I02600 | YNL061W | 99 | 0 |
| SPGP0T00960 | YKL104C | 99 | 0 |
| SPGP0I00890 | YNL250W | 99 | 0 |
| SPGP0A00640 | YJR096W | 99 | 0 |
| SPGP0DC00260 | YNR001C | 99 | 0 |
| SPGP0I01540 | YNL177C | 99 | 0 |
| SPGP0AI00150 | YJL204C | 99 | 0 |
| SPGP0A00230 | YJR137C | 99 | 0 |
| SPGP0I01430 | YNL190W | 99 | 0 |
| SPGP0B00420 | YER045C | 99 | 0 |
| SPGP0C01740 | YIR008C | 99 | 0 |
| SPGP0I00520 | YNL290W | 100 | 0 |
| SPGP0CU01320 | YLR300W | 86 | 0 |
| SPGP0E00940 | YKR014C | 100 | 0 |
| SPGP0I00390 | YNL305C | 99 | 0 |
| SPGP0I02360 | YNL085W | 99 | 0 |
| SPGP0E01320 | YKR056W | 99 | 0 |
| SPGP0C01980 | YIR033W | 99 | 0 |
| SPGP0A01200 | YJR035W | 99 | 0 |
| SPGP0I01810 | YNL147W | 100 | 0 |
| SPGP0C00580 | YIL119C | 98 | 1 |
| SPGP0I00630 | YNL278W | 99 | 0 |
| SPGP0HA00100 | YNL284C-B | 99 | 0 |
| SPGP0B01400 | YEL052W | 99 | 0 |
| SPGP0E01710 | YKR095W-A | 100 | 0 |

Create PDF in your applications with the Pdfcrowd HTML to PDF API PDFCROWD

| | | | |
|---|---|---|---|
| SPGP0A02430 | YJL106W | 99 | 0 |
| SPGP0C00630 | YIL114C | 99 | 0 |
| SPGP0I01770 | YNL152W | 99 | 0 |
| SPGP0ID00100 | YAL061W | 99 | 0 |
| SPGP0B00120 | YER072W | 100 | 0 |
| SPGP0B01110 | YEL023C | 99 | 0 |
| SPGP0I00270 | YNL317W | 99 | 0 |
| SPGP0I00850 | YNL254C | 99 | 0 |
| SPGP0A01160 | YJR041C | 99 | 0 |
| SPGP0A01350 | YJR013W | 99 | 0 |
| SPGP0E01020 | YKR022C | 99 | 0 |
| SPGP0A01590 | YJL011C | 100 | 0 |
| SPGP0I02240 | YNL098C | 99 | 0 |
| SPGP0A00340 | YJR126C | 99 | 0 |
| SPGP0T00770 | YKL127W | 96 | 0 |
| SPGP0D00300 | YNR053C | 99 | 0 |
| SPGP0I02010 | YNL126W | 99 | 0 |
| SPGP0C01210 | YIL048W | 99 | 0 |
| SPGP0CM01960 | YBR092C | 87 | 0 |
| SPGP0B01200 | YEL032W | 99 | 0 |
| SPGP0A01800 | YJL038C | 99 | 0 |
| SPGP0I00200 | YNL326C | 99 | 0 |
| SPGP0ES00160 | YER167W | 99 | 0 |
| SPGP0A00850 | YJR073C | 99 | 0 |
| SPGP0JF00160 | YER183C | 99 | 0 |
| SPGP0B00260 | YER058W | 99 | 0 |
| SPGP0T01110 | YKL087C | 99 | 0 |
| SPGP0I02720 | YNL047C | 99 | 0 |
| SPGP0IP00140 | YGL229C | 99 | 0 |
| SPGP0T00820 | YKL121W | 99 | 0 |
| SPGP0D00340 | YNR049C | 100 | 0 |
| SPGP0A02440 | YJL107C | 99 | 0 |
| SPGP0I01320 | YNL202W | 99 | 0 |
| SPGP0A01870 | YJL046W | 99 | 0 |
| SPGP0I02690 | YNL050C | 99 | 0 |
| SPGP0E01150 | YKR037C | 99 | 0 |
| SPGP0A02180 | YJL080C | 99 | 0 |
| SPGP0I02280 | YNL094W | 99 | 0 |
| SPGP0I00720 | YNL268W | 99 | 0 |
| SPGP0CX00480 | YGL176C | 99 | 0 |

Create PDF in your applications with the Pdfcrowd HTML to PDF API PDFCROWD

| | | | |
|---|---|---|---|
| SPGP0C00350 | YIL143C | 99 | 0 |
| SPGP0B00980 | YEL011W | 99 | 0 |
| SPGP0CR00540 | YAL008W | 99 | 0 |
| SPGP0C01710 | YIR005W | 100 | 0 |
| SPGP0I01480 | YNL185C | 99 | 0 |
| SPGP0E00200 | YKL062W | 99 | 0 |
| SPGP0T00170 | YKL192C | 99 | 0 |
| SPGP0E01290 | YKR053C | 99 | 0 |
| SPGP0T00210 | YKL188C | 99 | 0 |
| SPGP0I02820 | YNL036W | 99 | 0 |
| SPGP0B00180 | YER066W | 100 | 0 |
| SPGP0I01730 | YNL156C | 99 | 0 |
| SPGP0A02220 | YJL084C | 99 | 0 |
| SPGP0AI00280 | YJL218W | 99 | 0 |
| SPGP0CR00530 | YAL009W | 99 | 0 |
| SPGP0A01990 | YJL058C | 99 | 0 |
| SPGP0DD00210 | YNL031C | 100 | 0 |
| SPGP0JG00110 | YGL258W | 99 | 0 |
| SPGP0I02580 | YNL063W | 99 | 0 |
| SPGP0CY00100 | YAL054C | 99 | 0 |
| SPGP0I01700 | YNL159C | 99 | 0 |
| SPGP0A01750 | YJL033W | 99 | 0 |
| SPGP0A02520 | YJL118W | 99 | 0 |
| SPGP0T01160 | YKL081W | 99 | 0 |
| SPGP0B01320 | YEL045C | 99 | 0 |
| SPGP0A01570 | YJL008C | 100 | 0 |
| SPGP0A01280 | YJR022W | 100 | 0 |
| SPGP0CP03390 | YBL059C-A | 86 | 2 |
| SPGP0D00530 | YNR031C | 99 | 0 |
| SPGP0E00140 | YKL068W | 98 | 0 |
| SPGP0DZ00100 | YDR542W | 99 | 0 |
| SPGP0CRtrna17P | YOL013W-A | 96 | 1 |
| SPGP0T00810 | YKL122C | 99 | 0 |
| SPGP0A00150 | YJR145C | 99 | 0 |
| SPGP0A00620 | YJR098C | 99 | 0 |
| SPGP0T01140 | YKL084W | 99 | 0 |
| SPGP0C00870 | YIL090W | 99 | 0 |
| SPGP0CS00100 | YPR204W | 97 | 1 |
| SPGP0C00380 | YIL139C | 99 | 0 |
| SPGP0DC00240 | YNL002C | 100 | 0 |

Create PDF in your applications with the Pdfcrowd HTML to PDF API PDFCROWD

| | | | |
|---|---|---|---|
| SPGP0E01520 | YKR077W | 100 | 0 |
| SPGP0B00400 | YER047C | 99 | 0 |
| SPGP0A00260 | YJR135C | 99 | 0 |
| SPGP0E00950 | YKR015C | 99 | 0 |
| SPGP0C01080 | YIL065C | 100 | 0 |
| SPGP0CX00440 | YGL181W | 97 | 3 |
| SPGP0A00370 | YJR123W | 99 | 0 |
| SPGP0E00920 | YKR011C | 99 | 0 |
| SPGP0B00540 | YER034W | 99 | 0 |
| SPGP0I00610 | YNL280C | 99 | 0 |
| SPGP0A01220 | YJR033C | 99 | 0 |
| SPGP0CR00680 | YAR018C | 99 | 0 |
| SPGP0EL00190 | YGL250W | 99 | 0 |
| SPGP0I01830 | YNL145W | 99 | 0 |
| SPGP0CX00240 | YGL205W | 99 | 0 |
| SPGP0AY00120 | YPR202W | 99 | 0 |
| SPGP0I01220 | YNL214W | 99 | 0 |
| SPGP0HV00100 | YPL277C | 99 | 0 |
| SPGP0C00820 | YIL095W | 99 | 0 |
| SPGP0C00650 | YIL112W | 99 | 0 |
| SPGP0D00350 | YNR048W | 99 | 0 |
| SPGP0B01420 | YEL054C | 100 | 0 |
| SPGP0CR00780 | YAR035W | 99 | 0 |
| SPGP0B00740 | YER015W | 99 | 0 |
| SPGP0CW00560 | YOR302W | 96 | 0 |
| SPGP0A02140 | YJL074C | 99 | 0 |
| SPGP0A02410 | YJL104W | 100 | 0 |
| SPGP0B01130 | YEL025C | 99 | 0 |
| SPGP0CR00620 | YAR002C-A | 99 | 0 |
| SPGP0A01140 | YJR043C | 99 | 0 |
| SPGP0I00870 | YNL252C | 99 | 0 |
| SPGP0A01090 | YJR048W | 100 | 0 |
| SPGP0C00900 | YIL087C | 99 | 0 |
| SPGP0DD00220 | YNL032W | 99 | 0 |
| SPGP0CR00240 | YAL037W | 99 | 0 |
| SPGP0A00810 | YJR077C | 99 | 0 |
| SPGP0D00320 | YNR051C | 99 | 0 |
| SPGP0A00360 | YJR124C | 99 | 0 |
| SPGP0C01230 | YIL046W | 99 | 0 |
| SPGP0CX00290 | YGL198W | 99 | 0 |

Create PDF in your applications with the Pdfcrowd HTML to PDF API PDFCROWD

| | | | |
|---|---|---|---|
| SPGP0D00670 | YNR016C | 99 | 0 |
| SPGP0B01180 | YEL030W | 99 | 0 |
| SPGP0D00180 | YNR065C | 99 | 0 |
| SPGP0T01090 | YKL089W | 99 | 0 |
| SPGP0A01810 | YJL039C | 99 | 0 |
| SPGP0E01140 | YKR036C | 99 | 0 |
| SPGP0T00640 | YKL141W | 99 | 0 |
| SPGP0I01340 | YNL200C | 99 | 0 |
| SPGP0DD00110 | YNL021W | 99 | 0 |
| SPGP0ES00140 | YER165W | 99 | 0 |
| SPGP0E00490 | YKL032C | 98 | 2 |
| SPGP0C00360 | YIL142W | 99 | 0 |
| SPGP0E01100 | YKR030W | 99 | 0 |
| SPGP0T00150 | YKL194C | 99 | 0 |
| SPGP0C00480 | YIL129C | 99 | 0 |
| SPGP0I00600 | YNL281W | 100 | 0 |
| SPGP0B00210 | YER063W | 99 | 0 |
| SPGP0I01130 | YNL223W | 99 | 0 |
| SPGP0E00260 | YKL056C | 99 | 0 |
| SPGP0C02040 | YIR039C | 99 | 0 |
| SPGP0I02800 | YNL038W | 99 | 0 |
| SPGP0E01700 | YKR095W | 99 | 0 |
| SPGP0B01140 | YEL026W | 100 | 0 |
| SPGP0HM00130 | YAL056W | 99 | 0 |
| SPGP0B01350 | YEL048C | 99 | 0 |
| SPGP0C01790 | YIR013C | 99 | 0 |
| SPGP0A02500 | YJL116C | 99 | 0 |
| SPGP0I00400 | YNL304W | 99 | 0 |
| SPGP0A01770 | YJL035C | 99 | 0 |
| SPGP0E00740 | YKL007W | 99 | 0 |
| SPGP0DC00120 | YNL015W | 100 | 0 |
| SPGP0B00610 | YER027C | 99 | 0 |
| SPGP0A02600 | YJL127C-B | 100 | 0 |
| SPGP0A02360 | YJL099W | 99 | 0 |
| SPGP0A01550 | YJL005W | 99 | 0 |
| SPGP0E00410 | YKL040C | 99 | 0 |
| SPGP0I02850 | YNL033W | 99 | 0 |
| SPGP0T00630 | YKL142W | 99 | 0 |
| SPGP0B01300 | YEL043W | 99 | 0 |
| SPGP0C01520 | YIL014W | 99 | 0 |

Create PDF in your applications with the Pdfcrowd HTML to PDF API PDFCROWD

| | | | |
|---|---|---|---|
| SPGP0A01270 | YJR024C | 99 | 0 |
| SPGP0T00330 | YKL175W | 99 | 0 |
| SPGP0A00600 | YJR100C | 99 | 0 |
| SPGP0D00510 | YNR032C-A | 99 | 0 |
| SPGP0C00850 | YIL092W | 99 | 0 |
| SPGP0C00560 | YIL121W | 99 | 0 |
| SPGP0C01370 | YIL031W | 100 | 0 |
| SPGP0A00690 | YJR092W | 99 | 0 |
| SPGP0A00280 | YJR133W | 99 | 0 |
| SPGP0E01500 | YKR075C | 99 | 0 |
| SPGP0E01050 | YKR025W | 99 | 0 |
| SPGP0E00970 | YKR017C | 99 | 0 |
| SPGP0I01390 | YNL194C | 99 | 0 |
| SPGP0CZ00100 | YHR211W | 93 | 2 |
| SPGP0J03520 | YNL301C | 83 | 5 |
| SPGP0AI00310 | YOL159C | 99 | 0 |
| SPGP0CR00300 | YAL032C | 100 | 0 |
| SPGP0A00350 | YJR125C | 99 | 0 |
| SPGP0E00670 | YKL014C | 99 | 0 |
| SPGP0B01030 | YEL017W | 99 | 0 |
| SPGP0B00680 | YER020W | 99 | 0 |
| SPGP0I00670 | YNL273W | 99 | 0 |
| SPGP0AY00100 | YPL282C | 99 | 0 |
| SPGP0CR00660 | YAR014C | 99 | 0 |
| SPGP0I01860 | YNL141W | 99 | 0 |
| SPGP0AI00170 | YJL206C | 99 | 0 |
| SPGP0I00480 | YNL294C | 99 | 0 |
| SPGP0I01670 | YNL162W | 99 | 0 |
| SPGP0A00190 | YJR141W | 99 | 0 |
| SPGP0A02380 | YJL101C | 99 | 0 |
| SPGP0B01440 | YEL056W | 99 | 0 |
| SPGP0C00670 | YIL110W | 99 | 0 |
| SPGP0A02030 | YJL062W | 99 | 0 |
| SPGP0A02160 | YJL078C | 99 | 0 |
| SPGP0IP00120 | YGL227W | 99 | 0 |
| SPGP0I02430 | YNL078W | 99 | 0 |
| SPGP0A00830 | YJR075W | 99 | 0 |
| SPGP0C01780 | YIR012W | 99 | 0 |
| SPGP0C01270 | YIL042C | 99 | 0 |
| SPGP0A01120 | YJR045C | 99 | 0 |

Create PDF in your applications with the Pdfcrowd HTML to PDF API PDFCROWD

| | | | |
|---|---|---|---|
| SPGP0C00920 | YIL084C | 99 | 0 |
| SPGP0B00840 | YER006W | 99 | 0 |
| SPGP0CR00260 | YAL035W | 99 | 0 |
| SPGP0FK00100 | YGL225W | 99 | 0 |
| SPGP0C01250 | YIL044C | 99 | 0 |
| SPGP0AI00210 | YJL210W | 99 | 0 |
| SPGP0B00430 | YER044C-A | 100 | 0 |
| SPGP0B01230 | YEL036C | 99 | 0 |
| SPGP0D00650 | YNR018W | 100 | 0 |
| SPGP0C01040 | YIL069C | 99 | 0 |
| SPGP0B01510 | YEL063C | 99 | 0 |
| SPGP0I00440 | YNL299W | 99 | 0 |
| SPGP0D00460 | YNR036C | 100 | 0 |
| SPGP0I00820 | YNL257C | 99 | 0 |
| SPGP0C00620 | YIL115C | 99 | 0 |
| SPGP0D00700 | YNR013C | 99 | 0 |
| SPGP0C00310 | YIL147C | 99 | 0 |
| SPGP0E00450 | YKL034W | 99 | 0 |
| SPGP0CX00330 | YGL194C-A | 99 | 0 |
| SPGP0B00560 | YER032W | 99 | 0 |
| SPGP0I01450 | YNL188W | 99 | 0 |
| SPGP0CX00190 | YGL210W | 99 | 0 |
| SPGP0AI00250 | YJL214W | 99 | 0 |
| SPGP0E01260 | YKR050W | 99 | 0 |
| SPGP0E01340 | YKR058W | 99 | 0 |
| SPGP0DC00100 | YNL018C | 99 | 0 |
| SPGP0B00230 | YER061C | 99 | 0 |
| SPGP0I01440 | YNL189W | 99 | 0 |
| SPGP0I01110 | YNL225C | 99 | 0 |
| SPGP0CR00270 | YAL034W-A | 99 | 0 |
| SPGP0C01800 | YIR014W | 99 | 0 |
| SPGP0DT00100 | YER175C | 99 | 0 |
| SPGP0E01730 | YKR097W | 99 | 0 |
| SPGP0IZ00400 | YMR319C | 99 | 0 |
| SPGP0EL00250 | YGL256W | 99 | 0 |
| SPGP0T01120 | YKL086W | 99 | 0 |
| SPGP0I02150 | YNL108C | 99 | 0 |
| SPGP0C01770 | YIR011C | 99 | 0 |
| SPGP0A01710 | YJL027C | 100 | 0 |
| SPGP0CR00490 | YAL013W | 99 | 0 |

Create PDF in your applications with the Pdfcrowd HTML to PDF API PDFCROWD

| | | | |
|---|---|---|---|
| SPGP0I00100 | YNL336W | 97 | 0 |
| SPGP0A00880 | YJR069C | 99 | 0 |
| SPGP0E00390 | YKL042W | 99 | 0 |
| SPGP0A01860 | YJL045W | 99 | 0 |
| SPGP0B00480 | YER040W | 99 | 0 |
| SPGP0I02350 | YNL086W | 99 | 0 |
| SPGP0I00930 | YNL246W | 99 | 0 |
| SPGP0I02530 | YNL068C | 99 | 0 |
| SPGP0D00280 | YNR055C | 99 | 0 |
| SPGP0A00440 | YJR115W | 100 | 0 |
| SPGP0E00180 | YKL064W | 99 | 0 |
| SPGP0E00720 | YKL009W | 100 | 0 |
| SPGP0T00890 | YKL112W | 99 | 0 |
| SPGP0CX00230 | YGL206C | 99 | 0 |
| SPGP0C00830 | YIL094C | 99 | 0 |
| SPGP0A00660 | YJR094W-A | 100 | 0 |
| SPGP0C00680 | YIL109C | 99 | 0 |
| SPGP0A01070 | YJR050W | 99 | 0 |
| SPGP0T01170 | YKL080W | 98 | 0 |
| SPGP0T00490 | YKL159C | 100 | 0 |
| SPGP0I01820 | YNL146W | 100 | 0 |
| SPGP0A00300 | YJR131W | 99 | 0 |
| SPGP0E01070 | YKR027W | 99 | 0 |
| SPGP0E00380 | YKL043W | 100 | 0 |
| SPGP0A00330 | YJR127C | 99 | 0 |
| SPGP0I02300 | YNL092W | 99 | 0 |
| SPGP0E01380 | YKR062W | 99 | 0 |
| SPGP0CR00390 | YAL023C | 99 | 0 |
| SPGP0CR00640 | YAR007C | 99 | 0 |
| SPGP0I00650 | YNL275W | 99 | 0 |
| SPGP0E00650 | YKL016C | 99 | 0 |
| SPGP0I01150 | YNL221C | 99 | 0 |
| SPGP0I01650 | YNL163C | 99 | 0 |
| SPGP0I01870 | YNL139C | 99 | 0 |
| SPGP0C00690 | YIL108W | 99 | 0 |
| SPGP0A02010 | YJL060W | 99 | 0 |
| SPGP0A02100 | YJL070C | 99 | 0 |
| SPGP0IP00100 | YGL226W | 99 | 0 |
| SPGP0I00160 | YNL330C | 99 | 0 |

Create PDF in your applications with the Pdfcrowd HTML to PDF API PDFCROWD

| | | | |
|---|---|---|---|
| SPGP0CX00220 | YGL207W | 99 | 0 |
| SPGP0C01360 | YIL033C | 100 | 0 |
| SPGP0C01010 | YIL072W | 99 | 0 |
| SPGP0A01300 | YJR019C | 99 | 0 |
| SPGP0CR00290 | YAL033W | 99 | 0 |
| SPGP0T00190 | YKL190W | 99 | 0 |
| SPGP0A01100 | YJR047C | 100 | 0 |
| SPGP0A01450 | YJR005W | 99 | 0 |
| SPGP0A00130 | YJR148W | 99 | 0 |
| SPGP0B01210 | YEL034W | 99 | 0 |
| SPGP0A00270 | YJR134C | 99 | 0 |
| SPGP0C00660 | YIL111W | 99 | 0 |
| SPGP0D00490 | YNR034W | 99 | 0 |
| SPGP0B01530 | YEL065W | 99 | 0 |
| SPGP0I02290 | YNL093W | 99 | 0 |
| SPGP0T00370 | YKL171W | 99 | 0 |
| SPGP0D00400 | YNR043W | 99 | 0 |
| SPGP0A02240 | YJL087C | 99 | 0 |
| SPGP0E01120 | YKR034W | 100 | 0 |
| SPGP0D00720 | YNR011C | 99 | 0 |
| SPGP0C00600 | YIL117C | 99 | 0 |
| SPGP0I00880 | YNL251C | 99 | 0 |
| SPGP0D00170 | YNR066C | 98 | 0 |
| SPGP0T00990 | YKL100C | 99 | 0 |
| SPGP0CX00420 | YGL184C | 99 | 0 |
| SPGP0DT00160 | YER169W | 99 | 0 |
| SPGP0CR00730 | YAR028W | 98 | 0 |
| SPGP0AI00260 | YJL216C | 96 | 0 |
| SPGP0I01460 | YNL187W | 99 | 0 |
| SPGP0E01240 | YKR048C | 99 | 0 |
| SPGP0A00570 | YJR103W | 99 | 0 |
| SPGP0C02000 | YIR035C | 100 | 0 |
| SPGP0I01790 | YNL149C | 100 | 0 |
| SPGP0T00980 | YKL101W | 99 | 0 |
| SPGP0B00250 | YER059W | 99 | 0 |
| SPGP0C01860 | YIR021W | 99 | 0 |
| SPGP0IZ00390 | YMR318C | 99 | 0 |
| SPGP0DT00120 | YER173W | 99 | 0 |
| SPGP0EL00230 | YGL254W | 100 | 0 |
| SPGP0T01100 | YKL088W | 99 | 0 |

Create PDF in your applications with the Pdfcrowd HTML to PDF API PDFCROWD

| | | | |
|---|---|---|---|
| SPGP0A00490 | YJR110W | 99 | 0 |
| SPGP0C00730 | YIL104C | 100 | 0 |
| SPGP0C01830 | YIR017C | 100 | 0 |
| SPGP0A01730 | YJL030W | 99 | 0 |
| SPGP0B01220 | YEL035C | 99 | 0 |
| SPGP0A02640 | YJL131C | 99 | 0 |
| SPGP0A01880 | YJL047C | 99 | 0 |
| SPGP0C01480 | YIL017C | 99 | 0 |
| SPGP0E00610 | YKL019W | 99 | 0 |
| SPGP0I00910 | YNL248C | 99 | 0 |
| SPGP0CX00300 | YGL197W | 99 | 0 |
| SPGP0I02410 | YNL080C | 99 | 0 |
| SPGP0C01510 | YGR161W-B | 99 | 0 |
| SPGP0T00670 | YKL138C-A | 99 | 0 |
| SPGP0T00240 | YKL185W | 99 | 0 |
| SPGP0B01160 | YEL028W | 99 | 0 |
| SPGP0JF00100 | YER177W | 99 | 0 |
| SPGP0HV00110 | YPL278C | 100 | 0 |
| SPGP0B00460 | YER042W | 99 | 0 |
| SPGP0I02070 | YNL119W | 99 | 0 |
| SPGP0T01130 | YKL085W | 99 | 0 |
| SPGP0I01090 | YNL229C | 99 | 0 |
| SPGP0D00750 | YNR008W | 99 | 0 |
| SPGP0T00500 | YKL157W | 99 | 0 |
| SPGP0R00520 | YML078W | 87 | 0 |
| SPGP0IP00110 | YGL226C-A | 100 | 0 |
| SPGP0E00930 | YKR013W | 98 | 2 |
| SPGP0E01010 | YKR021W | 99 | 0 |
| SPGP0HY00100 | YGL236C | 99 | 0 |
| SPGP0C00810 | YIL096C | 99 | 0 |
| SPGP0C01900 | YIR025W | 99 | 0 |
| SPGP0C01100 | YIL063C | 99 | 0 |
| SPGP0C00770 | YIL101C | 99 | 0 |
| SPGP0E00630 | YKL018W | 99 | 0 |
| SPGP0C00500 | YIL127C | 99 | 0 |
| SPGP0I01630 | YNL165W | 99 | 0 |
| SPGP0A02340 | YJL097W | 99 | 0 |
| SPGP0CX00460 | YGL179C | 99 | 0 |
| SPGP0T00440 | YKL163W | 97 | 0 |
| SPGP0T00750 | YKL129C | 99 | 0 |

Create PDF in your applications with the Pdfcrowd HTML to PDF API PDFCROWD

| | | | |
|---|---|---|---|
| SPGP0CR00770 | YAR033W | 98 | 0 |
| SPGP0A01260 | YJR025C | 99 | 0 |
| SPGP0A02120 | YJL072C | 99 | 0 |
| SPGP0C01540 | YIL011W | 94 | 3 |
| SPGP0CR00650 | YAR008W | 99 | 0 |
| SPGP0A00870 | YJR070C | 99 | 0 |
| SPGP0C01070 | YIL066C | 99 | 0 |
| SPGP0D00410 | YNR041C | 99 | 0 |
| SPGP0B01080 | YEL020W-A | 100 | 0 |
| SPGP0C01290 | YIL040W | 99 | 0 |
| SPGP0A01470 | YJR003C | 99 | 0 |
| SPGP0B01000 | YEL013W | 99 | 0 |
| SPGP0A00250 | YJR135W-A | 99 | 0 |
| SPGP0C02010 | YIR036C | 100 | 0 |
| SPGP0T00680 | YKL138C | 99 | 0 |
| SPGP0D00740 | YNR009W | 99 | 0 |
| SPGP0I00860 | YNL253W | 99 | 0 |
| SPGP0DC00180 | YNL008C | 99 | 0 |
| SPGP0CX00400 | YGL186C | 100 | 0 |
| SPGP0C00400 | YIL137C | 99 | 0 |
| SPGP0B00520 | YER036C | 99 | 0 |
| SPGP0A02250 | YJL088W | 99 | 0 |
| SPGP0I02670 | YNL052W | 100 | 0 |
| SPGP0I02830 | YNL035C | 99 | 0 |
| SPGP0I01400 | YNL193W | 99 | 0 |
| SPGP0D00710 | YNR012W | 99 | 0 |
| SPGP0E01220 | YKR045C | 96 | 4 |
| SPGP0I01000 | YNL239W | 99 | 0 |
| SPGP0T00970 | YKL103C | 99 | 0 |
| SPGP0B00900 | YEL001C | 99 | 0 |
| SPGP0C01840 | YIR018W | 99 | 0 |
| SPGP0C01810 | YIR015W | 99 | 0 |
| SPGP0DT00140 | YER171W | 99 | 0 |
| SPGP0E00170 | YKL065C | 99 | 0 |
| SPGP0C01750 | YIR009W | 100 | 0 |
| SPGP0CR00100 | YAL053W | 99 | 0 |
| SPGP0A02460 | YJL110C | 99 | 0 |
| SPGP0E01680 | YKR093W | 99 | 0 |
| SPGP0GB00100 | YOR388C | 99 | 0 |
| SPGP0C02020 | YIR037W | 99 | 0 |

Create PDF in your applications with the Pdfcrowd HTML to PDF API PDFCROWD

| | | | |
|---|---|---|---|
| SPGP0B01290 | YEL042W | 100 | 0 |
| SPGP0C00710 | YIL106W | 99 | 0 |
| SPGP0E00160 | YKL065W-A | 99 | 0 |
| SPGP0B00220 | YER062C | 100 | 0 |
| SPGP0CR00520 | YAL010C | 99 | 0 |
| SPGP0IZ00380 | YMR317W | 99 | 0 |
| SPGP0CX00380 | YGL189C | 100 | 0 |
| SPGP0T00380 | YKL170W | 100 | 0 |
| SPGP0A01820 | YJL041W | 100 | 0 |
| SPGP0A00770 | YJR083C | 99 | 0 |
| SPGP0I00970 | YNL242W | 99 | 0 |
| SPGP0I02790 | YNL039W | 99 | 0 |
| SPGP0A01050 | YJR052W | 99 | 0 |
| SPGP0I02390 | YNL082W | 99 | 0 |
| SPGP0CW00280 | YOR335C | 87 | 0 |
| SPGP0T00650 | YKL140W | 100 | 0 |
| SPGP0D00240 | YNR059W | 99 | 0 |
| SPGP0A00400 | YJR119C | 99 | 0 |
| SPGP0I01620 | YNL166C | 99 | 0 |
| SPGP0C01470 | YIL018W | 99 | 0 |
| SPGP0B01490 | YEL061C | 99 | 0 |
| SPGP0A01310 | YJR017C | 99 | 0 |
| SPGP0GL00100 | YER188W | 90 | 0 |
| SPGP0D00230 | YNR060W | 99 | 0 |
| SPGP0T00350 | YKL173W | 99 | 0 |
| SPGP0CX00260 | YGL202W | 99 | 0 |
| SPGP0B00690 | YER019W | 99 | 0 |
| SPGP0GE00100 | YCR097W | 91 | 0 |
| SPGP0E00130 | YKL068W-A | 99 | 0 |
| SPGP0D00770 | YNR006W | 100 | 0 |
| SPGP0T00520 | YKL155C | 99 | 0 |
| SPGP0HY00120 | YGL238W | 99 | 0 |
| SPGP0T00260 | YKL183W | 99 | 0 |
| SPGP0GV00100 | YER176W | 99 | 0 |
| SPGP0CM01720 | YBR118W | 96 | 0 |
| SPGP0C00150 | YIL169C | 97 | 0 |
| SPGP0E01030 | YKR023W | 99 | 0 |
| SPGP0I00680 | YNL272C | 99 | 0 |
| SPGP0C00790 | YIL098C | 99 | 0 |
| SPGP0B00590 | YER029C | 99 | 0 |

Create PDF in your applications with the Pdfcrowd HTML to PDF API PDFCROWD

| | | | |
|---|---|---|---|
| SPGP0CR00350 | YAL027W | 99 | 0 |
| SPGP0I00800 | YNL259C | 100 | 0 |
| SPGP0FQ00110 | YBR298C | 99 | 0 |
| SPGP0A02320 | YJL095W | 99 | 0 |
| SPGP0E01640 | YKR089C | 99 | 0 |
| SPGP0T00460 | YKL162C | 99 | 0 |
| SPGP0C00980 | YIL075C | 99 | 0 |
| SPGP0C01440 | YIL021W | 99 | 0 |
| SPGP0A02070 | YJL066C | 100 | 0 |
| SPGP0AU00170 | YDR011W | 84 | 0 |
| SPGP0A02490 | YJL115W | 99 | 0 |
| SPGP0I02510 | YNL070W | 100 | 0 |
| SPGP0I01960 | YNL131W | 100 | 0 |
| SPGP0C01050 | YIL068C | 99 | 0 |
| SPGP0I00550 | YNL287W | 100 | 0 |
| SPGP0E01450 | YKR069W | 99 | 0 |
| SPGP0A01670 | YJL023C | 99 | 0 |
| SPGP0CR00310 | YAL031C | 99 | 0 |
| SPGP0CW02630 | YOR074C | 85 | 1 |
| SPGP0FK00120 | YGL223C | 99 | 0 |
| SPGP0A01410 | YJR008W | 99 | 0 |
| SPGP0E00810 | YKL001C | 100 | 0 |
| SPGP0C00890 | YIL088C | 99 | 0 |
| SPGP0C01320 | YIL037C | 99 | 0 |
| SPGP0A00910 | YJR066W | 99 | 0 |
| SPGP0C02030 | YIR038C | 99 | 0 |
| SPGP0C00640 | YIL113W | 99 | 0 |
| SPGP0EL00180 | YGL249W | 99 | 0 |
| SPGP0T01080 | YKL090W | 99 | 0 |
| SPGP0DC00160 | YNL010W | 99 | 0 |
| SPGP0C00420 | YIL135C | 99 | 0 |
| SPGP0D00760 | YNR007C | 100 | 0 |
| SPGP0B00500 | YER038C | 99 | 0 |
| SPGP0T01020 | YKL096W | 100 | 0 |
| SPGP0I01420 | YNL191W | 99 | 0 |
| SPGP0I01470 | YNL186W | 99 | 1 |
| SPGP0T00950 | YKL105C | 99 | 0 |
| SPGP0B00960 | YEL007W | 99 | 0 |
| SPGP0I00950 | YNL244C | 99 | 0 |
| SPGP0I02080 | YNL118C | 99 | 0 |

Create PDF in your applications with the Pdfcrowd HTML to PDF API PDFCROWD

| | | | |
|---|---|---|---|
| SPGP0E01200 | YKR043C | 99 | 0 |
| SPGP0B00920 | YEL003W | 100 | 0 |
| SPGP0CR00130 | YAL048C | 99 | 0 |
| SPGP0IZ00350 | YMR315W | 99 | 0 |
| SPGP0I01020 | YNL237W | 99 | 0 |
| SPGP0I01250 | YNL211C | 100 | 0 |
| SPGP0E00150 | YKL067W | 99 | 0 |
| SPGP0I01280 | YNL208W | 99 | 0 |
| SPGP0E00400 | YKL041W | 99 | 0 |
| SPGP0CX00410 | YGL185C | 99 | 0 |
| SPGP0I00750 | YNL264C | 100 | 0 |
| SPGP0CR00160 | YAL044W-A | 99 | 0 |
| SPGP0CU00710 | YLR367W | 80 | 4 |
| SPGP0B01270 | YEL040W | 99 | 0 |
| SPGP0CR00720 | YAR027W | 99 | 0 |
| SPGP0B00240 | YER060W | 99 | 0 |
| SPGP0A01080 | YJR049C | 99 | 0 |
| SPGP0I02370 | YNL084C | 99 | 0 |
| SPGP0CR00330 | YAL029C | 99 | 0 |
| SPGP0I01640 | YNL164C | 99 | 0 |
| SPGP0A02680 | YJL134W | 99 | 0 |
| SPGP0I01170 | YNL219C | 99 | 0 |
| SPGP0D00220 | YNR061C | 99 | 0 |
| SPGP0I02120 | YNL113W | 99 | 0 |
| SPGP0A00420 | YJR117W | 99 | 0 |
| SPGP0A00930 | YJR064W | 99 | 0 |
| SPGP0C01550 | YIL010W | 99 | 0 |
| SPGP0A01330 | YJR015W | 100 | 0 |
| SPGP0T00760 | YKL128C | 95 | 0 |
| SPGP0B01120 | YEL024W | 99 | 0 |
| SPGP0A00680 | YJR093C | 99 | 0 |
| SPGP0D00590 | YNR024W | 100 | 0 |
| SPGP0JM00100 | YPL274W | 99 | 0 |
| SPGP0A01840 | YJL043W | 99 | 0 |
| SPGP0C00590 | YIL118W | 100 | 0 |
| SPGP0Y01560 | YMR115W | 82 | 1 |
| SPGP0I02230 | YNL099C | 99 | 0 |
| SPGP0I02680 | YNL051W | 99 | 0 |
| SPGP0D00780 | YNR004W | 100 | 0 |
| SPGP0AY00130 | YER190W | 97 | 0 |

Create PDF in your applications with the Pdfcrowd HTML to PDF API PDFCROWD

| | | | |
|---|---|---|---|
| SPGP0A00240 | YJR136C | 99 | 0 |
| SPGP0Y00570 | YMR011W | 86 | 0 |
| SPGP0E01470 | YKR071C | 99 | 0 |
| SPGP0E00580 | YKL022C | 99 | 0 |
| SPGP0C00540 | YIL123W | 99 | 0 |
| SPGP0A02300 | YJL093C | 99 | 0 |
| SPGP0CR00370 | YAL025C | 99 | 0 |
| SPGP0E00360 | YKL046C | 99 | 0 |
| SPGP0B00710 | YER017C | 99 | 0 |
| SPGP0E01620 | YKR087C | 99 | 0 |
| SPGP0HM00100 | YAL060W | 98 | 0 |
| SPGP0T01000 | YKL099C | 100 | 0 |
| SPGP0D00480 | YNR034W-A | 99 | 0 |
| SPGP0C01020 | YIL071C | 99 | 0 |
| SPGP0A00510 | YJR108W | 100 | 0 |
| SPGP0I00150 | YNL331C | 99 | 0 |
| SPGP0CR00150 | YAL046C | 99 | 0 |
| SPGP0I00620 | YNL279W | 99 | 0 |
| SPGP0C01300 | YIL039W | 99 | 0 |
| SPGP0I00540 | YNL288W | 99 | 0 |
| SPGP0A01650 | YJL020C | 99 | 0 |
| SPGP0A01480 | YJR002W | 99 | 0 |
| SPGP0D00380 | YNR045W | 99 | 0 |
| SPGP0A01430 | YJR006W | 99 | 0 |
| SPGP0B00890 | YER001W | 99 | 0 |
| SPGP0C01990 | YIR034C | 99 | 0 |
| SPGP0C01340 | YIL035C | 99 | 0 |
| SPGP0I02740 | YNL045W | 99 | 0 |
| SPGP0A00210 | YJR139C | 99 | 0 |
| SPGP0B01590 | YEL072W | 99 | 0 |
| SPGP0C00440 | YIL133C | 99 | 0 |
| SPGP0DC00140 | YNL012W | 99 | 0 |
| SPGP0T00930 | YKL107W | 99 | 0 |
| SPGP0B00370 | YER049W | 99 | 0 |
| SPGP0L02130 | YPL218W | 90 | 0 |
| SPGP0E00860 | YKR005C | 99 | 0 |
| SPGP0DD00170 | YNL026W | 99 | 0 |
| SPGP0IZ00330 | YMR313C | 99 | 0 |
| SPGP0C01880 | YIR023W | 99 | 0 |
| SPGP0B00940 | YEL005C | 99 | 0 |

Create PDF in your applications with the Pdfcrowd HTML to PDF API PDFCROWD

| | | | |
|---|---|---|---|
| SPGP0B00510 | YER037W | 99 | 0 |
| SPGP0E01770 | YKR101W | 99 | 0 |
| SPGP0I00710 | YNL270C | 99 | 0 |
| SPGP0E00750 | YKL006C-A | 99 | 0 |
| SPGP0I01230 | YNL213C | 100 | 0 |
| SPGP0A01440 | YJR005C-A | 100 | 0 |
| SPGP0E00420 | YKL039W | 99 | 0 |
| SPGP0CX00390 | YGL187C | 100 | 0 |
| SPGP0B00360 | YER050C | 99 | 0 |
| SPGP0CR00410 | YAL021C | 99 | 0 |
| SPGP0E01480 | YKR072C | 99 | 0 |
| SPGP0A00430 | YJR116W | 100 | 0 |
| SPGP0C00750 | YIL102C-A | 99 | 0 |
| SPGP0B01330 | YEL046C | 99 | 0 |
| SPGP0C01150 | YIL055C | 99 | 0 |
| SPGP0T01050 | YKL093W | 99 | 0 |
| SPGP0A02570 | YJL125C | 99 | 0 |
| SPGP0A02510 | YJL117W | 99 | 0 |
| SPGP0CR00480 | YAL014C | 99 | 0 |
| SPGP0T00340 | YKL174C | 99 | 0 |
| SPGP0C01390 | YIL027C | 100 | 0 |
| SPGP0JG00120 | YGL258W-A | 97 | 0 |
| SPGP0I02130 | YNL111C | 100 | 0 |
| SPGP0T00690 | YKL137W | 99 | 0 |
| SPGP0D00200 | YNR063W | 99 | 0 |
| SPGP0C01570 | YIL009W | 99 | 0 |
| SPGP0AA01560 | YLR002C | 83 | 1 |
| SPGP0I02250 | YNL097C | 99 | 0 |
| SPGP0B00390 | YER048C | 99 | 0 |
| SPGP0D00800 | YNR002C | 100 | 0 |
| SPGP0T00550 | YKL151C | 99 | 0 |
| SPGP0E01060 | YKR026C | 99 | 0 |
| SPGP0JG00100 | YGL257C | 98 | 0 |
| SPGP0I02700 | YNL049C | 99 | 0 |
| SPGP0B01060 | YEL019C | 99 | 0 |
| SPGP0B00550 | YER033C | 99 | 0 |
| SPGP0B00880 | YER002W | 99 | 0 |
| SPGP0E00300 | YKL052C | 99 | 0 |
| SPGP0C01940 | YIR029W | 99 | 0 |
| SPGP0CR00210 | YAL040C | 99 | 0 |

Create PDF in your applications with the Pdfcrowd HTML to PDF API PDFCROWD

| | | | |
|---|---|---|---|
| SPGP0E00990 | YKR019C | 99 | 0 |
| SPGP0B00770 | YER012W | 99 | 0 |
| SPGP0GG00100 | YBR012C | 90 | 1 |
| SPGP0B00760 | YER013W | 99 | 0 |
| SPGP0E01600 | YKR085C | 99 | 0 |
| SPGP0ID00110 | YAL062W | 99 | 0 |
| SPGP0CR00170 | YAL044C | 100 | 0 |
| SPGP0DV00100 | YER168C | 99 | 0 |
| SPGP0I01920 | YNL135C | 100 | 0 |
| SPGP0C00950 | YIL078W | 99 | 0 |
| SPGP0A01980 | YJL057C | 99 | 0 |
| SPGP0A01630 | YJL016W | 99 | 0 |
| SPGP0I02170 | YNL106C | 99 | 0 |
| SPGP0I00580 | YNL283C | 99 | 0 |
| SPGP0E00770 | YKL005C | 99 | 0 |
| SPGP0FY00130 | YER187W | 92 | 0 |
| SPGP0A01180 | YJR039W | 99 | 0 |
| SPGP0A02660 | YJL133W | 99 | 0 |
| SPGP0B01020 | YEL016C | 99 | 0 |
| SPGP0I02660 | YNL053W | 99 | 0 |
| SPGP0A01460 | YJR004C | 99 | 0 |
| SPGP0T00200 | YKL189W | 99 | 0 |
| SPGP0A00760 | YJR084W | 99 | 0 |
| SPGP0C01180 | YIL051C | 99 | 0 |
| SPGP0A00170 | YJR143C | 99 | 0 |
| SPGP0C00460 | YIL131C | 99 | 0 |
| SPGP0T00910 | YKL109W | 99 | 0 |
| SPGP0E00840 | YKR003W | 99 | 0 |
| SPGP0E01280 | YKR052C | 99 | 0 |
| SPGP0IZ00310 | YMR311C | 99 | 0 |
| SPGP0E01400 | YKR064W | 99 | 0 |
| SPGP0B00280 | YER056C-A | 99 | 0 |
| SPGP0A01760 | YJL034W | 99 | 0 |
| SPGP0B01470 | YEL059C-A | 99 | 0 |
| SPGP0I02770 | YNL041C | 99 | 0 |
| SPGP0I01210 | YNL215W | 99 | 0 |
| SPGP0I00730 | YNL267W | 99 | 0 |
| SPGP0I02590 | YNL062C | 99 | 0 |
| SPGP0B00340 | YER052C | 99 | 0 |
| SPGP0CR00120 | YAL049C | 99 | 0 |

Create PDF in your applications with the Pdfcrowd HTML to PDF API PDFCROWD

| | | | |
|---|---|---|---|
| SPGP0CR00430 | YAL019W | 99 | 0 |
| SPGP0E01080 | YKR028W | 99 | 0 |
| SPGP0B00320 | YER053C-A | 100 | 0 |
| SPGP0E00440 | YKL035W | 99 | 0 |
| SPGP0B01310 | YEL044W | 100 | 0 |
| SPGP0C01110 | YIL062C | 99 | 0 |
| SPGP0I02110 | YNL115C | 99 | 0 |
| SPGP0I01600 | YNL168C | 99 | 0 |
| SPGP0I00190 | YNL327W | 99 | 0 |
| SPGP0CR00710 | YAR023C | 99 | 0 |
| SPGP0T01070 | YKL091C | 99 | 0 |
| SPGP0DD00200 | YNL030W | 99 | 0 |
| SPGP0A02530 | YJL121C | 99 | 0 |
| SPGP0A02620 | YJL129C | 99 | 0 |
| SPGP0T00360 | YKL172W | 99 | 0 |
| SPGP0CX00320 | YGL195W | 99 | 0 |
| SPGP0A00970 | YJR060W | 99 | 0 |
| SPGP0I00990 | YNL240C | 99 | 0 |
| SPGP0I02330 | YNL088W | 99 | 0 |
| SPGP0EL00210 | YGL252C | 99 | 0 |
| SPGP0B01070 | YEL020C | 99 | 0 |
| SPGP0CR00460 | YAL016W | 99 | 0 |
| SPGP0C01600 | YIL006W | 99 | 0 |
| SPGP0B01250 | YEL038W | 99 | 0 |
| SPGP0A00390 | YJR121W | 99 | 0 |
| SPGP0DD00120 | YNL022C | 100 | 0 |
| SPGP0D00620 | YNR021W | 99 | 0 |
| SPGP0C01820 | YIR016W | 98 | 0 |
| SPGP0C01000 | YIL073C | 99 | 0 |
| SPGP0E00530 | YKL026C | 100 | 0 |
| SPGP0T00300 | YKL179C | 99 | 0 |
| SPGP0I02570 | YNL064C | 99 | 0 |
| SPGP0I00300 | YNL314W | 99 | 0 |
| SPGP0E01630 | YKR088C | 99 | 0 |
| SPGP0C01920 | YIR027C | 99 | 0 |
| SPGP0I01330 | YNL201C | 99 | 0 |
| SPGP0E00320 | YKL050C | 99 | 0 |
| SPGP0T00880 | YKL113C | 99 | 0 |
| SPGP0T00430 | YKL164C | 99 | 0 |

Create PDF in your applications with the Pdfcrowd HTML to PDF API PDFCROWD

| | | | |
|---|---|---|---|
| SPGP0B00570 | YER031C | 100 | 0 |
| SPGP0E00850 | YKR004C | 99 | 0 |
| SPGP0CR00220 | YAL039C | 99 | 0 |
| SPGP0B00750 | YER014W | 99 | 0 |
| SPGP0E01580 | YKR083C | 99 | 0 |
| SPGP0E00620 | YKL018C-A | 99 | 0 |
| SPGP0I01900 | YNL137C | 99 | 0 |
| SPGP0CR00190 | YAL042W | 99 | 0 |
| SPGP0C01760 | YIR010W | 99 | 1 |
| SPGP0C01670 | YIR001C | 99 | 0 |
| SPGP0CR00590 | YAL001C | 99 | 0 |
| SPGP0I00500 | YNL292W | 99 | 0 |
| SPGP0A01960 | YJL055W | 100 | 0 |
| SPGP0AA01930 | YLL039C | 84 | 0 |
| SPGP0A02700 | YJL137C | 99 | 0 |
| SPGP0A00470 | YJR112W | 99 | 0 |
| SPGP0FY00110 | YER185W | 99 | 0 |
| SPGP0A00560 | YJR104C | 100 | 0 |
| SPGP0I01750 | YNL154C | 99 | 0 |
| SPGP0I01660 | YNL162W-A | 99 | 0 |
| SPGP0A00700 | YJR091C | 99 | 0 |
| SPGP0D00430 | YNR039C | 99 | 0 |
| SPGP0T00740 | YKL130C | 99 | 0 |
| SPGP0EL00120 | YGL243W | 99 | 0 |
| SPGP0B00830 | YER007W | 99 | 0 |
| SPGP0E00820 | YKR001C | 99 | 0 |
| SPGP0I00130 | YNL333W | 99 | 0 |
| SPGP0I02750 | YNL044W | 100 | 0 |
| SPGP0C00240 | YIL157C | 99 | 0 |
| SPGP0AI00200 | YJL209W | 99 | 0 |
| SPGP0E01360 | YKR060W | 99 | 0 |
| SPGP0A00180 | YJR142W | 99 | 0 |
| SPGP0I00570 | YNL284C | 100 | 0 |
| SPGP0E00780 | YKL004W | 99 | 0 |
| SPGP0I01070 | YNL231C | 99 | 0 |
| SPGP0E00460 | YKL033W-A | 99 | 0 |
| SPGP0I01590 | YNL169C | 100 | 0 |
| SPGP0CX00490 | YGL175C | 99 | 0 |
| SPGP0I02050 | YNL122C | 99 | 0 |
| SPGP0A02150 | YJL076W | 99 | 0 |

Create PDF in your applications with the Pdfcrowd HTML to PDF API PDFCROWD

| | | | |
|---|---|---|---|
| SPGP0A02550 | YJL123C | 98 | 2 |
| SPGP0A00950 | YJR062C | 99 | 0 |
| SPGP0C01680 | YIR002C | 99 | 0 |
| SPGP0C01350 | YIL034C | 100 | 0 |
| SPGP0CX00430 | YGL183C | 99 | 0 |
| SPGP0E01440 | YKR068C | 99 | 0 |
| SPGP0C01530 | YIL013C | 99 | 0 |
| SPGP0I00310 | YNL313C | 99 | 0 |
| SPGP0IF00110 | YGR289C | 99 | 0 |
| SPGP0AI00130 | YJL201W | 99 | 0 |
| SPGP0B01100 | YEL022W | 99 | 0 |
| SPGP0CR00440 | YAL018C | 99 | 0 |
| SPGP0DD00100 | YNL020C | 99 | 0 |
| SPGP0A01390 | YJR010W | 99 | 0 |
| SPGP0CX00470 | YGL178W | 99 | 0 |
| SPGP0D00600 | YNR023W | 99 | 0 |
| SPGP0C00280 | YIL150C | 99 | 0 |
| SPGP0A02310 | YJL094C | 99 | 0 |
| SPGP0I00280 | YNL316C | 99 | 0 |
| SPGP0E01610 | YKR086W | 99 | 0 |
| SPGP0I01690 | YNL160W | 99 | 0 |
| SPGP0C01690 | YIR003W | 99 | 0 |
| SPGP0I00430 | YNL300W | 99 | 0 |
| SPGP0E00870 | YKR006C | 99 | 0 |
| SPGP0I01610 | YNL167C | 99 | 0 |
| SPGP0CX00510 | YGL173C | 98 | 0 |
| SPGP0A00580 | YJR102C | 100 | 0 |
| SPGP0E00700 | YKL011C | 99 | 0 |
| SPGP0I00140 | YNL332W | 99 | 0 |
| SPGP0A01940 | YJL053W | 99 | 0 |
| SPGP0I02040 | YNL123W | 99 | 0 |
| SPGP0A01490 | YJR001W | 99 | 0 |
| SPGP0A00740 | YJR086W | 99 | 0 |
| SPGP0A01520 | YJL002C | 99 | 0 |
| SPGP0A00220 | YJR138W | 99 | 0 |
| SPGP0T00530 | YKL154W | 99 | 0 |
| SPGP0I02000 | YNL127W | 99 | 0 |
| SPGP0A00540 | YJR106W | 99 | 0 |
| SPGP0I02210 | YNL101W | 99 | 0 |
| SPGP0A01230 | YJR032W | 99 | 0 |

Create PDF in your applications with the Pdfcrowd HTML to PDF API PDFCROWD

| | | | |
|---|---|---|---|
| SPGP0T00730 | YKL132C | 99 | 0 |
| SPGP0A00720 | YJR089W | 99 | 0 |
| SPGP0D00450 | YNR037C | 99 | 0 |
| SPGP0D00110 | YNR074C | 99 | 0 |
| SPGP0A00590 | YJR101W | 99 | 0 |
| SPGP0I02480 | YNL073W | 99 | 0 |
| SPGP0HM00140 | YAL055W | 98 | 0 |
| SPGP0D00540 | YNR030W | 99 | 0 |
| SPGP0B00300 | YER055C | 99 | 0 |
| SPGP0B00650 | YER023W | 99 | 0 |
| SPGP0CR00470 | YAL015C | 99 | 0 |
| SPGP0C00260 | YIL156W | 99 | 0 |
| SPGP0CX00360 | YGL191W | 99 | 0 |
| SPGP0T00320 | YKL176C | 99 | 0 |
| SPGP0C01500 | YIL015W | 99 | 0 |
| SPGP0I01060 | YNL232W | 99 | 0 |
| SPGP0C01330 | YIL036W | 99 | 0 |
| SPGP0E00120 | YKL069W | 100 | 0 |
| SPGP0I00330 | YNL311C | 99 | 0 |
| SPGP0AI00110 | YJL199C | 99 | 0 |
| SPGP0A02400 | YJL103C | 99 | 0 |
| SPGP0D00660 | YNR017W | 100 | 0 |
| SPGP0I01200 | YNL216W | 99 | 0 |
| SPGP0CX00210 | YGL208W | 99 | 0 |
| SPGP0B00330 | YER053C | 99 | 0 |
| SPGP0I01720 | YNL157W | 99 | 0 |
| SPGP0B00640 | YER024W | 99 | 0 |
| SPGP0B01480 | YEL060C | 99 | 0 |
| SPGP0I00340 | YNL310C | 99 | 0 |
| SPGP0I01550 | YNL176C | 99 | 0 |
| SPGP0A00140 | YJR147W | 99 | 0 |
| SPGP0T00850 | YKL117W | 99 | 0 |
| SPGP0L01920 | YPL195W | 81 | 1 |
| SPGP0E01670 | YKR092C | 94 | 3 |
| SPGP0B00530 | YER035W | 100 | 0 |
| SPGP0I02310 | YNL091W | 99 | 0 |
| SPGP0E01490 | YKR074W | 99 | 0 |
| SPGP0B00790 | YER010C | 100 | 0 |
| SPGP0A01920 | YJL051W | 99 | 0 |
| SPGP0AN00290 | YDR406W | 81 | 1 |

Create PDF in your applications with the Pdfcrowd HTML to PDF API PDFCROWD

| | | | |
|---|---|---|---|
| SPGP0A02290 | YJL092W | 99 | 0 |
| SPGP0C00450 | YIL132C | 99 | 0 |
| SPGP0CP03510 | YBL075C | 85 | 0 |
| SPGP0CR00580 | YAL002W | 99 | 0 |
| SPGP0B01500 | YEL062W | 99 | 0 |
| SPGP0C00510 | YIL126W | 99 | 0 |
| SPGP0I02320 | YNL090W | 99 | 0 |
| SPGP0DC00220 | YNL004W | 100 | 0 |
| SPGP0T00510 | YKL156W | 99 | 0 |
| SPGP0B01040 | YEL017C-A | 100 | 0 |
| SPGP0E00430 | YKL038W | 99 | 0 |
| SPGP0A01210 | YJR034W | 100 | 0 |
| SPGP0T00710 | YKL134C | 99 | 0 |
| SPGP0D00470 | YNR035C | 100 | 0 |
| SPGP0E01720 | YKR096W | 99 | 0 |
| SPGP0A00290 | YJR132W | 99 | 0 |
| SPGP0C00230 | YIL158W | 99 | 0 |
| SPGP0AI00300 | YOL158C | 99 | 0 |
| SPGP0E01690 | YKR094C | 99 | 0 |
| SPGP0FQ00100 | YBR297W | 97 | 0 |
| SPGP0C01950 | YIR030C | 99 | 0 |
| SPGP0E01130 | YKR035W-A | 99 | 0 |
| SPGP0C00200 | YIL161W | 99 | 0 |
| SPGP0B00490 | YER039C | 99 | 0 |
| SPGP0B00670 | YER021W | 99 | 0 |
| SPGP0E00340 | YKL048C | 99 | 0 |
| SPGP0I02450 | YNL076W | 99 | 0 |
| SPGP0CX00340 | YGL194C | 99 | 0 |
| SPGP0C01910 | YIR026C | 99 | 0 |
| SPGP0I01080 | YNL230C | 99 | 0 |
| SPGP0I01880 | YNL138W-A | 99 | 0 |
| SPGP0E00350 | YKL047W | 99 | 0 |
| SPGP0C01640 | YIL002W-A | 100 | 0 |
| SPGP0I01940 | YNL133C | 99 | 0 |
| SPGP0C01310 | YIL038C | 99 | 0 |
| SPGP0DD00140 | YNL024C | 99 | 0 |
| SPGP0D00520 | YNR032W | 99 | 0 |
| SPGP0I00350 | YNL309W | 99 | 0 |
| SPGP0A01020 | YJR055W | 99 | 0 |
| SPGP0ES00130 | YER164W | 92 | 0 |

Create PDF in your applications with the Pdfcrowd HTML to PDF API PDFCROWD

| | | | |
|---|---|---|---|
| SPGP0C00990 | YIL074C | 99 | 0 |
| SPGP0I02550 | YNL066W | 99 | 0 |
| SPGP0D00390 | YNR044W | 95 | 1 |
| SPGP0B01450 | YEL057C | 99 | 0 |
| SPGP0A01000 | YJR057W | 99 | 0 |
| SPGP0A02420 | YJL105W | 99 | 0 |
| SPGP0C00190 | YIL162W | 99 | 0 |
| SPGP0T00700 | YKL135C | 99 | 0 |
| SPGP0A00750 | YJR085C | 100 | 0 |
| SPGP0I01360 | YNL197C | 99 | 0 |
| SPGP0CX00500 | YGL174W | 99 | 0 |
| SPGP0D00130 | YNR070W | 99 | 0 |
| SPGP0D00640 | YNR019W | 99 | 0 |
| SPGP0A02350 | YJL098W | 99 | 0 |
| SPGP0I02760 | YNL042W | 99 | 0 |
| SPGP0I00320 | YNL312W | 100 | 0 |
| SPGP0I00740 | YNL265C | 99 | 0 |
| SPGP0I00170 | YNL329C | 99 | 0 |
| SPGP0DD00180 | YNL027W | 99 | 0 |
| SPGP0E01650 | YKR090W | 99 | 0 |
| SPGP0I00210 | YNL325C | 99 | 0 |
| SPGP0AA01770 | YLL021W | 81 | 1 |
| SPGP0E00520 | YKL027W | 99 | 0 |
| SPGP0E00290 | YKL053C-A | 99 | 0 |
| SPGP0E00830 | YKR002W | 99 | 0 |
| SPGP0I01980 | YNL129W | 100 | 0 |
| SPGP0CR00280 | YAL034C | 99 | 0 |
| SPGP0A01900 | YJL049W | 99 | 0 |
| SPGP0C00430 | YIL134W | 99 | 0 |
| SPGP0C01030 | YIL070C | 99 | 0 |
| SPGP0B00110 | YER073W | 99 | 0 |
| SPGP0I02610 | YNL059C | 99 | 0 |
| SPGP0I01740 | YNL155W | 99 | 0 |
| SPGP0CX00170 | YGL212W | 99 | 0 |
| SPGP0T00280 | YKL181W | 99 | 0 |
| SPGP0CR00570 | YAL003W | 99 | 1 |
| SPGP0HV00130 | YOR391C | 99 | 0 |
| SPGP0I02160 | YNL107W | 100 | 0 |
| SPGP0A00500 | YJR109C | 99 | 0 |
| SPGP0C01450 | YIL020C | 99 | 0 |

Create PDF in your applications with the Pdfcrowd HTML to PDF API PDFCROWD

| | | | |
|---|---|---|---|
| SPGP0C01580 | YIL008W | 100 | 0 |
| SPGP0I02560 | YNL065W | 99 | 0 |
| SPGP0FK00110 | YGL224C | 99 | 0 |
| SPGP0C00340 | YIL144W | 99 | 0 |
| SPGP0A02470 | YJL111W | 99 | 0 |
| SPGP0IP00170 | YGL232W | 99 | 0 |
| SPGP0AJ00510 | YHR039C-A | 87 | 0 |
| SPGP0C00490 | YIL128W | 99 | 0 |
| SPGP0C00220 | YIL159W | 99 | 0 |
| SPGP0B00470 | YER041W | 99 | 0 |
| SPGP0C01720 | YIR006C | 99 | 0 |
| SPGP0I00560 | YNL286W | 99 | 0 |
| SPGP0C00760 | YIL102C | 100 | 0 |
| SPGP0I01530 | YNL178W | 99 | 0 |
| SPGP0E00190 | YKL063C | 99 | 0 |
| SPGP0I01240 | YNL212W | 99 | 0 |
| SPGP0C00880 | YIL089W | 99 | 0 |
| SPGP0IZ00280 | YMR308C | 99 | 0 |
| SPGP0C01930 | YIR028W | 99 | 0 |
| SPGP0CR00230 | YAL038W | 99 | 0 |
| SPGP0AI00140 | YJL203W | 99 | 0 |
| SPGP0EL00100 | YGL241W | 99 | 0 |
| SPGP0T01010 | YKL098W | 100 | 0 |
| SPGP0A01340 | YJR014W | 100 | 0 |
| SPGP0I00450 | YNL298W | 99 | 0 |
| SPGP0C01660 | YIL001W | 99 | 0 |
| SPGP0C01490 | YIL016W | 99 | 0 |
| SPGP0A01040 | YJR053W | 99 | 0 |
| SPGP0IZ00290 | YMR309C | 99 | 0 |
| SPGP0CX00150 | YGL215W | 99 | 0 |
| SPGP0C01590 | YIL007C | 99 | 0 |
| SPGP0E00310 | YKL051W | 99 | 0 |
| SPGP0B00600 | YER028C | 98 | 1 |
| SPGP0I01930 | YNL134C | 99 | 0 |
| SPGP0T00720 | YKL133C | 99 | 0 |
| SPGP0A00310 | YJR130C | 99 | 0 |
| SPGP0T00870 | YKL114C | 99 | 0 |
| SPGP0E00270 | YKL055C | 99 | 0 |
| SPGP0B01520 | YEL064C | 99 | 0 |
| SPGP0B01260 | YEL039C | 99 | 0 |

Create PDF in your applications with the Pdfcrowd HTML to PDF API PDFCROWD

| | | | |
|---|---|---|---|
| SPGP0B01390 | YEL051W | 100 | 0 |
| SPGP0T00570 | YKL149C | 99 | 0 |
| SPGP0B00860 | YER004W | 99 | 0 |
| SPGP0I00530 | YNL289W | 100 | 0 |
| SPGP0I01270 | YNL209W | 99 | 0 |
| SPGP0CR00510 | YAL011W | 99 | 0 |
| SPGP0AF01690 | YGR225W | 80 | 1 |
| SPGP0I01180 | YNL218W | 99 | 0 |
| SPGP0E00500 | YKL029C | 99 | 0 |
| SPGP0JF00110 | YER178W | 99 | 0 |
| SPGP0B01570 | YEL070W | 99 | 0 |
| SPGP0C00410 | YIL136W | 99 | 0 |
| SPGP0B00130 | YER071C | 99 | 0 |
| SPGP0E00760 | YKL006W | 99 | 0 |
| SPGP0A01780 | YJL036W | 99 | 0 |
| SPGP0A00780 | YJR082C | 99 | 0 |
| SPGP0C01120 | YIL061C | 99 | 0 |
| SPGP0C01430 | YIL022W | 99 | 0 |
| SPGP0E00600 | YKL020C | 99 | 0 |
| SPGP0I00760 | YNL263C | 99 | 0 |
| SPGP0A01250 | YJR030C | 99 | 0 |
| SPGP0A00120 | YJR149W | 99 | 0 |
| SPGP0D00560 | YNR028W | 99 | 0 |
| SPGP0CX00100 | YGL222C | 99 | 0 |
| SPGP0A00650 | YJR095W | 100 | 0 |
| SPGP0B00660 | YER022W | 99 | 0 |
| SPGP0IP00150 | YGL230C | 99 | 0 |
| SPGP0B00970 | YEL009C | 99 | 0 |
| SPGP0CX00270 | YGL201C | 99 | 0 |
| SPGP0AI00100 | YJL198W | 99 | 0 |
| SPGP0E01590 | YKR084C | 99 | 0 |
| SPGP0B00450 | YER043C | 99 | 0 |
| SPGP0A00160 | YJR144W | 99 | 0 |
| SPGP0I01910 | YNL136W | 99 | 0 |
| SPGP0I01510 | YNL181W | 99 | 0 |
| SPGP0E01370 | YKR061W | 99 | 0 |
| SPGP0B00630 | YER025W | 99 | 0 |
| SPGP0CR00180 | YAL043C | 99 | 0 |
| SPGP0I02490 | YNL072W | 99 | 0 |
| SPGP0JC00130 | YKL205W | 84 | 0 |

Create PDF in your applications with the Pdfcrowd HTML to PDF API PDFCROWD

| | | | |
|---|---|---|---|
| SPGP0I00840 | YNL255C | 99 | 0 |
| SPGP0EL00160 | YGL247W | 99 | 0 |
| SPGP0T01030 | YKL095W | 99 | 0 |
| SPGP0IP00190 | YGL234W | 99 | 0 |
| SPGP0A01320 | YJR016C | 99 | 0 |
| SPGP0E00100 | YKL071W | 99 | 0 |
| SPGP0I01570 | YNL173C | 99 | 0 |
| SPGP0A01060 | YJR051W | 99 | 0 |
| SPGP0B01410 | YEL053C | 99 | 0 |
| SPGP0I01800 | YNL148C | 100 | 0 |
| SPGP0T00610 | YKL144C | 99 | 0 |
| SPGP0FY00120 | YER186C | 96 | 1 |
| SPGP0T01150 | YKL082C | 99 | 0 |
| SPGP0D00680 | YNR015W | 99 | 0 |
| SPGP0B01540 | YEL066W | 99 | 0 |
| SPGP0A00480 | YJR111C | 99 | 0 |
| SPGP0E01160 | YKR038C | 99 | 0 |
| SPGP0T00410 | YKL166C | 99 | 0 |
| SPGP0T00840 | YKL119C | 99 | 0 |
| SPGP0I00780 | YNL261W | 100 | 0 |
| SPGP0B00780 | YER011W | 94 | 0 |
| SPGP0I02020 | YNL125C | 99 | 0 |
| SPGP0I00220 | YNL323W | 99 | 0 |
| SPGP0AI00220 | YJL212C | 99 | 0 |
| SPGP0A00800 | YJR078W | 99 | 0 |
| SPGP0E00560 | YKL023C-A | 100 | 0 |
| SPGP0E01420 | YKR066C | 99 | 1 |
| SPGP0I01160 | YNL220W | 99 | 0 |
| SPGP0JF00130 | YER180C | 99 | 0 |
| SPGP0T00100 | YKL201C | 99 | 0 |
| SPGP0I02190 | YNL103W | 99 | 0 |
| SPGP0B00150 | YER069W | 99 | 0 |
| SPGP0CR00790 | YAR042W | 99 | 0 |
| SPGP0A02270 | YJL090C | 99 | 0 |
| SPGP0I00660 | YNL274C | 99 | 0 |
| SPGP0A02050 | YJL063C | 99 | 0 |
| SPGP0HM00110 | YAL059W | 99 | 0 |
| SPGP0C01410 | YIL024C | 99 | 0 |
| SPGP0C01140 | YIL056W | 99 | 0 |
| SPGP0B01600 | YEL073C | 100 | 0 |

Create PDF in your applications with the Pdfcrowd HTML to PDF API PDFCROWD

| | | | |
|---|---|---|---|
| SPGP0E01740 | YKR098C | 99 | 0 |
| SPGP0FC00100 | YML045W-A | 100 | 0 |
| SPGP0A01500 | YJL001W | 99 | 0 |
| SPGP0A00980 | YJR059W | 99 | 0 |
| SPGP0C00530 | YIL124W | 99 | 0 |
| SPGP0A00920 | YJR065C | 99 | 0 |
| SPGP0A00630 | YJR097W | 99 | 0 |
| SPGP0Y01820 | YMR142C | 94 | 0 |
| SPGP0E00370 | YKL045W | 99 | 0 |
| SPGP0E00710 | YKL010C | 99 | 0 |
| SPGP0CR00450 | YAL017W | 99 | 0 |
| SPGP0E01570 | YKR082W | 99 | 0 |
| SPGP0C00720 | YIL105C | 99 | 0 |
| SPGP0DC00210 | YNL005C | 99 | 0 |
| SPGP0E01310 | YKR055W | 99 | 0 |
| SPGP0I02730 | YNL046W | 99 | 0 |
| SPGP0C01160 | YIL053W | 99 | 0 |
| SPGP0B00380 | YER048W-A | 99 | 0 |
| SPGP0C01970 | YIR032C | 99 | 0 |
| SPGP0I01710 | YNL158W | 99 | 0 |
| SPGP0EL00140 | YGL245W | 99 | 0 |
| SPGP0B00200 | YER064C | 99 | 0 |
| SPGP0I02460 | YNL075W | 99 | 0 |
| SPGP0AI00190 | YJL208C | 99 | 0 |
| SPGP0A02090 | YJL069C | 99 | 0 |
| SPGP0A01600 | YJL012C | 99 | 0 |
| SPGP0C01620 | YIL004C | 100 | 0 |
| SPGP0A01190 | YJR036C | 99 | 0 |
| SPGP0C00970 | YIL076W | 99 | 0 |
| SPGP0E00230 | YKL059C | 99 | 0 |
| SPGP0T00590 | YKL146W | 99 | 0 |
| SPGP0D00500 | YNR033W | 99 | 0 |
| SPGP0D00370 | YNR046W | 100 | 0 |
| SPGP0I00240 | YNL321W | 99 | 0 |
| SPGP0I01010 | YNL238W | 99 | 0 |
| SPGP0D00150 | YNR068C | 100 | 0 |
| SPGP0E01230 | YKR046C | 99 | 0 |
| SPGP0T00830 | YKL120W | 99 | 0 |
| SPGP0I00260 | YNL318C | 99 | 0 |
| SPGP0IP00130 | YGL228W | 99 | 0 |

Create PDF in your applications with the Pdfcrowd HTML to PDF API PDFCROWD

| | | | |
|---|---|---|---|
| SPGP0I02140 | YNL110C | 99 | 0 |
| SPGP0CX00180 | YGL211W | 99 | 0 |
| SPGP0I00120 | YNL334C | 99 | 0 |
| SPGP0GX00100 | R0020C | 99 | 0 |
| SPGP0CR00550 | YAL007C | 100 | 0 |
| SPGP0I00810 | YNL258C | 99 | 0 |
| SPGP0I01140 | YNL222W | 99 | 0 |
| SPGP0E00540 | YKL025C | 100 | 0 |
| SPGP0I01490 | YNL183C | 99 | 0 |
| SPGP0CX00120 | YGL220W | 99 | 0 |
| SPGP0T00160 | YKL193C | 99 | 0 |
| SPGP0T00220 | YKL187C | 99 | 0 |
| SPGP0B00170 | YER067W | 100 | 0 |
| SPGP0L01090 | YPL109C | 82 | 0 |
| SPGP0A02210 | YJL083W | 99 | 0 |
| SPGP0C01650 | YIL002C | 99 | 0 |
| SPGP0A01740 | YJL031C | 99 | 0 |
| SPGP0Ctrna14E | YDR278C | 100 | 0 |
| SPGP0A02000 | YJL059W | 99 | 0 |
| SPGP0A01290 | YJR021C | 99 | 0 |
| SPGP0A01560 | YJL006C | 99 | 0 |
| SPGP0A01540 | YJL004C | 99 | 0 |
| SPGP0I02540 | YNL067W | 99 | 0 |
| SPGP0C01730 | YIR007W | 99 | 0 |
| SPGP0T00790 | YKL125W | 99 | 0 |
| SPGP0E01550 | YKR080W | 100 | 0 |
| SPGP0B00410 | YER046W | 99 | 0 |
| SPGP0B01010 | YEL015W | 99 | 0 |
| SPGP0CR00200 | YAL041W | 99 | 0 |
| SPGP0E01330 | YKR057W | 99 | 0 |
| SPGP0C01560 | YIL009C-A | 99 | 0 |
| SPGP0IZ00260 | YMR306W | 95 | 0 |
| SPGP0AI00160 | YJL205C | 99 | 0 |
| SPGP0I00490 | YNL293W | 99 | 0 |
| SPGP0A01360 | YJR012C | 99 | 0 |
| SPGP0A00610 | YJR099W | 99 | 0 |
| SPGP0A01030 | YJR054W | 99 | 0 |
| SPGP0C01220 | YIL047C | 99 | 0 |
| SPGP0A01170 | YJR040W | 99 | 0 |

Create PDF in your applications with the Pdfcrowd HTML to PDF API PDFCROWD

| | | | |
|---|---|---|---|
| SPGP0C00910 | YIL085C | 99 | 0 |
| SPGP0D00310 | YNR052C | 99 | 0 |
| SPGP0C01200 | YIL049W | 98 | 0 |
| SPGP0A01580 | YJL010C | 99 | 0 |
| SPGP0DF00110 | YKR105C | 82 | 1 |
| SPGP0A02190 | YJL081C | 99 | 0 |
| SPGP0CX00160 | YGL213C | 99 | 0 |
| SPGP0C00330 | YIL145C | 99 | 0 |
| SPGP0I02060 | YNL121C | 99 | 0 |
| SPGP0E01210 | YKR044W | 99 | 0 |
| SPGP0T00140 | YKL195W | 99 | 0 |
| SPGP0B00990 | YEL012W | 99 | 0 |
| SPGP0E00210 | YKL061W | 100 | 0 |
| SPGP0I01120 | YNL224C | 99 | 0 |
| SPGP0B00190 | YER065C | 99 | 0 |
| SPGP0V01110 | YOL120C | 87 | 3 |
| SPGP0A02230 | YJL085W | 99 | 0 |
| SPGP0I00290 | YNL315C | 99 | 0 |
| SPGP0C00570 | YIL120W | 99 | 0 |
| SPGP0HV00140 | YPL281C | 99 | 0 |
| SPGP0A00380 | YJR122W | 99 | 0 |
| SPGP0C00860 | YIL091C | 99 | 0 |
| SPGP0E01530 | YKR078W | 99 | 0 |
| SPGP0I00640 | YNL277W | 99 | 0 |
| SPGP0E00910 | YKR010C | 99 | 0 |
| SPGP0E00660 | YKL015W | 99 | 0 |
| SPGP0I00470 | YNL295W | 99 | 0 |
| SPGP0A02390 | YJL102W | 99 | 0 |
| SPGP0A02170 | YJL079C | 99 | 0 |
| SPGP0A01150 | YJR042W | 99 | 0 |
| SPGP0A01720 | YJL029C | 99 | 0 |
| SPGP0C00930 | YIL083C | 99 | 0 |
| SPGP0J01820 | YNL112W | 86 | 1 |
| SPGP0I01260 | YNL210W | 99 | 0 |
| SPGP0A00820 | YJR076C | 99 | 0 |
| SPGP0D00330 | YNR050C | 99 | 0 |
| SPGP0D00190 | YNR064C | 100 | 0 |
| SPGP0I01050 | YNL233W | 99 | 0 |
| SPGP0T00940 | YKL106W | 99 | 0 |
| SPGP0I01030 | YNL236W | 99 | 0 |

Create PDF in your applications with the Pdfcrowd HTML to PDF API PDFCROWD

| | | | |
|---|---|---|---|
| SPGP0C00170 | YIL166C | 99 | 0 |
| SPGP0E00730 | YKL008C | 99 | 0 |
| SPGP0T00120 | YKL197C | 99 | 0 |
| SPGP0T00390 | YKL168C | 99 | 0 |
| SPGP0B01190 | YEL031W | 99 | 0 |
| SPGP0EL00240 | YGL255W | 95 | 1 |
| SPGP0E01270 | YKR051W | 99 | 0 |
| SPGP0I01780 | YNL151C | 100 | 0 |
| SPGP0I01100 | YNL227C | 99 | 0 |
| SPGP0CX00280 | YGL200C | 99 | 0 |
| SPGP0C01240 | YIL045W | 99 | 0 |
| SPGP0IZ00340 | YMR314W | 99 | 0 |
| SPGP0E01040 | YKR024C | 99 | 0 |
| SPGP0B01430 | YEL055C | 99 | 0 |
| SPGP0A01700 | YJL026W | 99 | 0 |
| SPGP0A02610 | YJL128C | 99 | 0 |
| SPGP0I01350 | YNL199C | 99 | 0 |
| SPGP0C00550 | YIL122W | 99 | 0 |
| SPGP0D00290 | YNR054C | 100 | 0 |
| SPGP0A00890 | YJR068W | 99 | 0 |
| SPGP0A01850 | YJL044C | 99 | 0 |
| SPGP0E01510 | YKR076W | 99 | 0 |
| SPGP0C00840 | YIL093C | 100 | 0 |
| SPGP0DD00150 | YNL024C-A | 99 | 0 |
| SPGP0E00640 | YKL017C | 99 | 0 |
| SPGP0CR00670 | YAR015W | 99 | 0 |
| SPGP0I01840 | YNL144C | 100 | 0 |
| SPGP0A02370 | YJL100W | 99 | 0 |
| SPGP0EL00200 | YGL251C | 99 | 0 |
| SPGP0A02110 | YJL071W | 99 | 0 |
| SPGP0E01410 | YKR065C | 99 | 0 |
| SPGP0A00840 | YJR074W | 99 | 0 |
| SPGP0A01130 | YJR044C | 100 | 0 |
| SPGP0T00180 | YKL191W | 99 | 0 |
| SPGP0B01240 | YEL037C | 99 | 0 |
| SPGP0B00810 | YER008C | 99 | 0 |
| SPGP0U00590 | YDL064W | 89 | 0 |
| SPGP0E01250 | YKR049C | 99 | 0 |
| SPGP0E00250 | YKL057C | 99 | 0 |
| SPGP0C01260 | YIL043C | 99 | 0 |

Create PDF in your applications with the Pdfcrowd HTML to PDF API PDFCROWD

| | | | |
|---|---|---|---|
| SPGP0CR00110 | YAL051W | 99 | 0 |
| SPGP0C01870 | YIR022W | 99 | 0 |
| SPGP0DT00110 | YER174C | 99 | 0 |
| SPGP0E01350 | YKR059W | 99 | 0 |
| SPGP0DC00250 | YNL001W | 99 | 0 |
| SPGP0A02580 | YJL126W | 99 | 0 |
| SPGP0I01680 | YNL161W | 99 | 0 |
| SPGP0CR00340 | YAL028W | 99 | 0 |
| SPGP0B00580 | YER030W | 99 | 0 |
| SPGP0A02630 | YJL130C | 99 | 0 |
| SPGP0A00450 | YJR113C | 99 | 0 |
| SPGP0D00270 | YNR056C | 99 | 0 |
| SPGP0C01700 | YIR004W | 99 | 0 |
| SPGP0E01390 | YKR063C | 99 | 0 |
| SPGP0CR00380 | YAL024C | 99 | 0 |
| SPGP0C00210 | YIL160C | 99 | 0 |
| SPGP0A00900 | YJR067C | 99 | 0 |
| SPGP0I01850 | YNL142W | 99 | 0 |
| SPGP0I02620 | YNL058C | 99 | 1 |
| SPGP0A02130 | YJL073W | 99 | 0 |
| SPGP0A00860 | YJR072C | 99 | 0 |
| SPGP0D00420 | YNR040W | 99 | 0 |
| SPGP0I02710 | YNL048W | 99 | 0 |
| SPGP0E00510 | YKL028W | 99 | 0 |
| SPGP0CR00320 | YAL030W | 99 | 0 |
| SPGP0W00100 | YAL065C | 96 | 0 |
| SPGP0A02020 | YJL061W | 99 | 0 |
| SPGP0A01110 | YJR046W | 99 | 0 |
| SPGP0T00290 | YKL180W | 99 | 0 |
| SPGP0B00350 | YER051W | 99 | 0 |
| SPGP0DD00160 | YNL025C | 99 | 0 |
| SPGP0D00730 | YNR010W | 100 | 0 |
| SPGP0DC00190 | YNL007C | 98 | 2 |
| SPGP0T00660 | YKL139W | 99 | 0 |
| SPGP0T01060 | YKL092C | 100 | 0 |
| SPGP0C01280 | YIL041W | 99 | 0 |
| SPGP0T00900 | YKL110C | 99 | 0 |
| SPGP0DT00130 | YER172C | 99 | 0 |
| SPGP0B00270 | YER057C | 100 | 0 |
| SPGP0I00230 | YNL322C | 99 | 0 |

Create PDF in your applications with the Pdfcrowd HTML to PDF API PDFCROWD

| | | | |
|---|---|---|---|
| SPGP0A02650 | YJL132W | 99 | 0 |
| SPGP0I00900 | YNL249C | 99 | 0 |
| SPGP0A01380 | YJR010C-A | 100 | 0 |
| SPGP0B00700 | YER018C | 99 | 0 |
| SPGP0E00590 | YKL021C | 99 | 0 |
| SPGP0D00250 | YNR058W | 100 | 0 |
| SPGP0C00800 | YIL097W | 99 | 0 |
| SPGP0CX00250 | YGL203C | 98 | 1 |
| SPGP0I00700 | YNL271C | 100 | 0 |
| SPGP0JK00220 | YCR077C | 83 | 1 |
| SPGP0A00730 | YJR088C | 99 | 0 |
| SPGP0CR00630 | YAR003W | 99 | 0 |
| SPGP0B00720 | YER016W | 99 | 0 |
| SPGP0A02480 | YJL112W | 99 | 0 |
| SPGP0C01060 | YIL067C | 99 | 0 |
| SPGP0A01680 | YJL024C | 99 | 0 |
| SPGP0EG00100 | YAL067C | 99 | 0 |
| SPGP0B01090 | YEL021W | 100 | 0 |
| SPGP0A00200 | YJR140C | 99 | 0 |
| SPGP0I01520 | YNL180C | 99 | 0 |
| SPGP0AI00240 | YJL213W | 99 | 0 |
| SPGP0I02500 | YNL071W | 99 | 0 |
| SPGP0A00410 | YJR118C | 99 | 0 |
| SPGP0A02670 | YJL133C-A | 99 | 0 |
| SPGP0C00160 | YIL167W | 99 | 0 |
| SPGP0DC00170 | YNL009W | 99 | 0 |
| SPGP0U00770 | YDL083C | 91 | 1 |
| SPGP0CR00700 | YFL020C | 97 | 0 |
| SPGP0IZ00360 | YMR316W | 99 | 0 |
| SPGP0DT00150 | YER170W | 99 | 0 |
| SPGP0I01290 | YNL207W | 99 | 0 |
| SPGP0D00790 | YNR003C | 100 | 0 |
| SPGP0E01760 | YKR100C | 99 | 0 |
| SPGP0T01040 | YKL094W | 99 | 0 |
| SPGP0I01890 | YNL138W | 99 | 0 |
| SPGP0A01830 | YJL042W | 99 | 0 |
| SPGP0I00960 | YNL243W | 99 | 0 |
| SPGP0E00330 | YKL049C | 100 | 0 |
| SPGP0A02710 | YJL138C | 99 | 0 |
| SPGP0D00580 | YNR026C | 99 | 0 |

Create PDF in your applications with the Pdfcrowd HTML to PDF API PDFCROWD

| | | | |
|---|---|---|---|
| SPGP0C00780 | YIL099W | 99 | 0 |
| SPGP0E01460 | YKR070W | 99 | 0 |
| SPGP0C01460 | YIL019W | 99 | 0 |
| SPGP0E00470 | YKL033W | 99 | 0 |
| SPGP0CR00610 | YAR002W | 99 | 0 |
| SPGP0C00180 | YIL164C | 99 | 0 |
| SPGP0GL00150 | YKL215C | 81 | 1 |
| SPGP0AJ00330 | YHR021C | 95 | 1 |
| SPGP0E00880 | YKR007W | 98 | 1 |
| SPGP0I01970 | YNL130C | 99 | 0 |
| SPGP0I02100 | YNL116W | 99 | 0 |
| SPGP0A02060 | YJL065C | 99 | 0 |
| SPGP0A01400 | YJR009C | 100 | 0 |
| SPGP0CR00690 | YAR019C | 99 | 0 |
| SPGP0HV00120 | YOR390W | 99 | 0 |
| SPGP0DC00150 | YNL011C | 99 | 0 |
| SPGP0B00950 | YEL006W | 99 | 0 |
| SPGP0T00920 | YKL108W | 99 | 1 |
| SPGP0EL00170 | YGL248W | 99 | 0 |
| SPGP0B00930 | YEL004W | 99 | 0 |
| SPGP0C01890 | YIR024C | 99 | 0 |
| SPGP0CR00400 | YAL022C | 99 | 0 |
| SPGP0C01380 | YIL030C | 99 | 0 |
| SPGP0T00540 | YKL152C | 99 | 0 |
| SPGP0A02690 | YJL136C | 99 | 0 |
| SPGP0DB00100 | YDR345C | 90 | 0 |
| SPGP0A02280 | YJL091C | 99 | 0 |
| SPGP0B01050 | YEL018W | 99 | 0 |
| SPGP0I00360 | YNL308C | 99 | 0 |
| SPGP0B01150 | YEL027W | 100 | 0 |
| SPGP0E00980 | YKR018C | 99 | 0 |
| SPGP0I00590 | YNL282W | 100 | 0 |
| SPGP0I01370 | YNL196C | 99 | 0 |
| SPGP0A01010 | YJR056C | 99 | 0 |
| SPGP0I01950 | YNL132W | 99 | 0 |
| SPGP0A01640 | YJL019W | 99 | 0 |
| SPGP0CP00460 | YDR242W | 81 | 0 |
| SPGP0A01420 | YJR007W | 99 | 0 |
| SPGP0T00560 | YKL150W | 99 | 0 |
| SPGP0F00100 | YJL159W | 92 | 0 |

Create PDF in your applications with the Pdfcrowd HTML to PDF API PDFCROWD

| | | | |
|---|---|---|---|
| SPGP0DC00130 | YNL014W | 99 | 0 |
| SPGP0CX00450 | YGL180W | 99 | 0 |
| SPGP0C00290 | YIL149C | 99 | 0 |
| SPGP0CR00420 | YAL020C | 99 | 0 |
| SPGP0C00740 | YIL103W | 99 | 0 |
| SPGP0Atrna3M | YJL007C | 100 | 0 |
| SPGP0GT00100 | YPR002W | 81 | 0 |
| SPGP0I02440 | YNL077W | 99 | 0 |
| SPGP0E00800 | YKL002W | 99 | 0 |
| SPGP0DC00110 | YNL016W | 99 | 0 |
| SPGP0CX00200 | YGL209W | 99 | 0 |
| SPGP0A01620 | YJL014W | 99 | 0 |
| SPGP0A01970 | YJL056C | 99 | 0 |
| SPGP0C01190 | YIL050W | 99 | 0 |
| SPGP0E01190 | YKR042W | 99 | 1 |
| SPGP0I01580 | YNL172W | 99 | 0 |
| SPGP0T00620 | YKL143W | 99 | 0 |
| SPGP0B00800 | YER009W | 100 | 0 |
| SPGP0I00180 | YNL328C | 99 | 0 |
| SPGP0CX00310 | YGL196W | 99 | 0 |
| SPGP0I02650 | YNL054W | 99 | 1 |
| SPGP0I00980 | YNL241C | 99 | 0 |
| SPGP0AI00120 | YJL200C | 99 | 0 |
| SPGP0JM00110 | YPL273W | 99 | 0 |
| SPGP0T00310 | YKL178C | 99 | 0 |
| SPGP0D00610 | YNR022C | 100 | 0 |
| SPGP0EA00150 | YGR286C | 99 | 0 |
| SPGP0A00790 | YJR080C | 99 | 0 |
| SPGP0A02560 | YJL124C | 100 | 0 |
| SPGP0A00550 | YJR105W | 99 | 0 |
| SPGP0A01950 | YJL054W | 99 | 0 |
| SPGP0D00100 | YNR075W | 100 | 0 |
| SPGP0EL00110 | YGL242C | 99 | 0 |
| SPGP0I01040 | YNL234W | 99 | 0 |
| SPGP0A02540 | YJL122W | 99 | 0 |
| SPGP0B01460 | YEL058W | 99 | 0 |
| SPGP0I02180 | YNL104C | 99 | 0 |
| SPGP0B00290 | YER056C | 97 | 0 |
| SPGP0JF00140 | YER180C-A | 99 | 0 |
| SPGP0I01760 | YNL153C | 99 | 0 |

Create PDF in your applications with the Pdfcrowd HTML to PDF API PDFCROWD

| | | | |
|---|---|---|---|
| SPGP0T00270 | YKL182W | 99 | 0 |
| SPGP0B00870 | YER003C | 99 | 0 |
| SPGP0A00530 | YJR107W | 99 | 0 |
| SPGP0D00440 | YNR038W | 99 | 0 |
| SPGP0A01930 | YJL052W | 99 | 0 |
| SPGP0C00270 | YIL151C | 99 | 0 |
| SPGP0EL00130 | YGL244W | 99 | 0 |
| SPGP0T00480 | YKL160W | 100 | 0 |
| SPGP0D00120 | YNR071C | 100 | 0 |
| SPGP0E00280 | YKL054C | 99 | 1 |
| SPGP0B00100 | YER074W | 99 | 0 |
| SPGP0A01910 | YJL050W | 99 | 0 |
| SPGP0A02590 | YJL127C | 99 | 0 |
| SPGP0I01500 | YNL182C | 99 | 0 |
| SPGP0C01610 | YIL005W | 99 | 0 |

#### Sheet 3: *genes alleles below 80% identity*

| *Saccharomyces pastorianus CBS1538 predicted ORF* | Parental allele ID | Parent | Gene identities | Gene gaps |
|---|---|---|---|---|
| SPGP0BE00290 | YCL036W | *Saccharomyces cerevisiae* | 78 | 1 |
| SPGP0E01780 | YPR196W | *Saccharomyces cerevisiae* | 79 | 0 |
| SPGP0CW01320 | YOR219C | *Saccharomyces cerevisiae* | 79 | 1 |
| SPGP0CM01370 | YBR156C | *Saccharomyces cerevisiae* | 77 | 2 |
| SPGP0CW02560 | YOR081C | *Saccharomyces cerevisiae* | 79 | 1 |
| SPGP0CW01580 | YOR188W | *Saccharomyces cerevisiae* | 79 | 2 |
| SPGP0L02610 | YPL269W | *Saccharomyces cerevisiae* | 78 | 1 |
| SPGP0JK00190 | YCR075C | *Saccharomyces cerevisiae* | 78 | 1 |
| SPGP0FI00130 | XP_018224047.1 | *Saccharomyces eubayanus* | 78 | 2 |

#### Sheet 4: *genes alleles copy with same identity in both parents*

| *Saccharomyces pastorianus CBS1538 predicted ORF* | Parental allele ID | Parent | Gene identities | Gene gaps |
|---|---|---|---|---|
| SPGP0Btrna9R | YGR164W | *Saccharomyces cerevisiae* | 100 | 0 |
| SPGP0Btrna9R | DI49_4020 | *Saccharomyces eubayanus* | 100 | 0 |
| SPGP0CU00360 | YDL075W | *Saccharomyces cerevisiae* | 93 | 0 |
| SPGP0CU00360 | XP_018223007.1 | *Saccharomyces eubayanus* | 93 | 0 |
| SPGP0Itrna14I | YNL017C | *Saccharomyces cerevisiae* | 100 | 0 |
| SPGP0Itrna14I | DI49_5614 | *Saccharomyces eubayanus* | 100 | 0 |
| SPGP0CNtrna24K | YDR445C | *Saccharomyces cerevisiae* | 100 | 0 |

Create PDF in your applications with the Pdfcrowd HTML to PDF API PDFCROWD

| | | | | |
|---|---|---|---|---|
| SPGP0CNtrna24K | DI49_5468 | *Saccharomyces eubayanus* | 100 | 0 |
| SPGP0FI00140 | YDR531W | *Saccharomyces cerevisiae* | 83 | 0 |
| SPGP0FI00140 | XP_018224046.1 | *Saccharomyces eubayanus* | 83 | 0 |
| SPGP0CX01800 | YGL030W | *Saccharomyces cerevisiae* | 99 | 0 |
| SPGP0CX01800 | XP_018222195.1 | *Saccharomyces eubayanus* | 99 | 0 |
| SPGP0CPtrna26G | YPR159C-A | *Saccharomyces cerevisiae* | 100 | 0 |
| SPGP0CPtrna26G | DI49_5603 | *Saccharomyces eubayanus* | 100 | 0 |
| SPGP0Itrna24G | YNL285W | *Saccharomyces cerevisiae* | 100 | 0 |
| SPGP0Itrna24G | DI49_2297 | *Saccharomyces eubayanus* | 100 | 0 |
| SPGP0CItrna20T | YHR180W-A | *Saccharomyces cerevisiae* | 100 | 0 |
| SPGP0CItrna20T | DI49_5553 | *Saccharomyces eubayanus* | 100 | 0 |

#### Sheet 5: *list of paralogs in Saccharomyces pastorianus CBS1538*

| List of paralogs | *S. eubayanus-like content* | *S. pastorianus content* | *S. cerevisiae-like content* |
|---|---|---|---|
| SPGP0CV00100; | | SPGP0CV00100; | |
| DI49_1689; DI49_1688 | DI49_1689; DI49_1688 | | |
| DI49_1691; DI49_1690 | DI49_1691; DI49_1690 | | |
| DI49_1693; DI49_1692 | DI49_1693; DI49_1692 | | |
| DI49_1695; SPGP0CV00180 | DI49_1695 | SPGP0CV00180 | |
| SPGP0CVtrna2K; SPGP0CNtrna24K; SPGP0CV00200; SPGP0CXtrna5K; YGL200C; SPGP0CXtrna6K; DI49_1808; SPGP0DAtrna7K; SPGP0DBtrna8K; DI49_0541; DI49_1540; SPGP0FFtrna10K; SPGP0Atrna1K; YJL011C; SPGP0Xtrna13K; DI49_2750; DI49_4196; SPGP0Btrna11K; DI49_2156; SPGP0Ctrna15K; DI49_0613; DI49_5469; SPGP0Ftrna22K; DI49_3250 | DI49_1808; DI49_0541; DI49_1540; DI49_2750; DI49_4196; DI49_2156; DI49_0613; DI49_5469; DI49_3250 | SPGP0CVtrna2K; SPGP0CNtrna24K; SPGP0CV00200; SPGP0CXtrna5K; SPGP0CXtrna6K; SPGP0DAtrna7K; SPGP0DBtrna8K; SPGP0FFtrna10K; SPGP0Atrna1K; SPGP0Xtrna13K; SPGP0Btrna11K; SPGP0Ctrna15K; SPGP0Ftrna22K | YGL200C; YJL011C |
| DI49_1698; DI49_1697 | DI49_1698; DI49_1697 | | |
| DI49_1700; DI49_1699 | DI49_1700; DI49_1699 | | |
| DI49_1702; DI49_1701 | DI49_1702; DI49_1701 | | |
| DI49_1704; DI49_1703 | DI49_1704; DI49_1703 | | |
| DI49_1706; DI49_1705 | DI49_1706; DI49_1705 | | |
| DI49_1708; DI49_1707 | DI49_1708; DI49_1707 | | |
| DI49_1710; DI49_1709 | DI49_1710; DI49_1709 | | |
| DI49_1712; DI49_1711 | DI49_1712; DI49_1711 | | |
| DI49_1714; YGL253W; DI49_1716; YGL254W | DI49_1714; DI49_1716 | | YGL253W; YGL254W |
| DI49_1718; DI49_1717 | DI49_1718; DI49_1717 | | |
| DI49_5168; DI49_1719 | DI49_5168; DI49_1719 | | |
| DI49_5166; DI49_5167 | DI49_5166; DI49_5167 | | |
| DI49_5164; DI49_5165 | DI49_5164; DI49_5165 | | |
| DI49_5161; DI49_5163 | DI49_5161; DI49_5163 | | |
| DI49_5159; DI49_5160 | DI49_5159; DI49_5160 | | |

Create PDF in your applications with the Pdfcrowd HTML to PDF API PDFCROWD

| | | | |
|---|---|---|---|
| DI49_5157; DI49_5158 | DI49_5157; DI49_5158 | | |
| SPGP0CWtrna4M; DI49_4243; DI49_5154; SPGP0DBtrna7M; DI49_0528; SPGP0Ytrna11M; DI49_4244; SPGP0Ftrna12M; DI49_3010 | DI49_4243; DI49_5154; DI49_0528; DI49_4244; DI49_3010 | SPGP0CWtrna4M; SPGP0DBtrna7M; SPGP0Ytrna11M; SPGP0Ftrna12M | |
| DI49_5152; DI49_5153 | DI49_5152; DI49_5153 | | |
| DI49_5150; DI49_5151 | DI49_5150; DI49_5151 | | |
| DI49_5148; DI49_5149 | DI49_5148; DI49_5149 | | |
| SPGP0CW00290; YOR335C | | SPGP0CW00290 | YOR335C |
| DI49_5146; DI49_5147 | DI49_5146; DI49_5147 | | |
| DI49_5144; DI49_5145 | DI49_5144; DI49_5145 | | |
| DI49_5142; DI49_5143 | DI49_5142; DI49_5143 | | |
| DI49_5140; DI49_5141 | DI49_5140; DI49_5141 | | |
| DI49_5138; DI49_5139 | DI49_5138; DI49_5139 | | |
| DI49_5136; DI49_5137 | DI49_5136; DI49_5137 | | |
| DI49_5134; DI49_5135 | DI49_5134; DI49_5135 | | |
| DI49_5132; DI49_5133 | DI49_5132; DI49_5133 | | |
| DI49_5130; DI49_5131 | DI49_5130; DI49_5131 | | |
| DI49_5128; DI49_5129 | DI49_5128; DI49_5129 | | |
| DI49_5126; DI49_5127 | DI49_5126; DI49_5127 | | |
| DI49_5124; DI49_5125 | DI49_5124; DI49_5125 | | |
| DI49_5122; DI49_5123 | DI49_5122; DI49_5123 | | |
| DI49_5120; YOR302W | DI49_5120 | | YOR302W |
| DI49_5118; DI49_5119 | DI49_5118; DI49_5119 | | |
| DI49_5116; DI49_5117 | DI49_5116; DI49_5117 | | |
| DI49_5114; DI49_5115 | DI49_5114; DI49_5115 | | |
| DI49_5112; DI49_4314; DI49_5111; DI49_4315 | DI49_5112; DI49_4314; DI49_5111; DI49_4315 | | |
| DI49_5109; DI49_2660 | DI49_5109; DI49_2660 | | |
| SPGP0CWtrna2A; DI49_3564; DI49_5107; SPGP0EPtrna4A; DI49_1482; DI49_3563; SPGP0AFtrna13A; DI49_2116 | DI49_3564; DI49_5107; DI49_1482; DI49_3563; DI49_2116 | SPGP0CWtrna2A; SPGP0EPtrna4A; SPGP0AFtrna13A | |
| DI49_5105; DI49_5106 | DI49_5105; DI49_5106 | | |
| DI49_5103; DI49_5104 | DI49_5103; DI49_5104 | | |
| DI49_5101; DI49_5102 | DI49_5101; DI49_5102 | | |
| DI49_5099; DI49_5100 | DI49_5099; DI49_5100 | | |
| DI49_5097; DI49_5098 | DI49_5097; DI49_5098 | | |
| DI49_5095; DI49_5096 | DI49_5095; DI49_5096 | | |
| DI49_5093; DI49_5094 | DI49_5093; DI49_5094 | | |
| DI49_5091; DI49_5092 | DI49_5091; DI49_5092 | | |
| DI49_5089; DI49_5090 | DI49_5089; DI49_5090 | | |
| DI49_5087; DI49_5088 | DI49_5087; DI49_5088 | | |
| DI49_5085; DI49_5086 | DI49_5085; DI49_5086 | | |
| DI49_5083; DI49_5084 | DI49_5083; DI49_5084 | | |
| DI49_5081; DI49_5082 | DI49_5081; DI49_5082 | | |

Create PDF in your applications with the Pdfcrowd HTML to PDF API PDFCROWD

| | | | |
|---|---|---|---|
| DI49_5079; DI49_5080 | DI49_5079; DI49_5080 | | |
| DI49_5077; DI49_5078 | DI49_5077; DI49_5078 | | |
| DI49_5075; DI49_5076 | DI49_5075; DI49_5076 | | |
| DI49_5073; DI49_5074 | DI49_5073; DI49_5074 | | |
| DI49_5071; DI49_5072 | DI49_5071; DI49_5072 | | |
| DI49_5069; DI49_5070 | DI49_5069; DI49_5070 | | |
| DI49_5067; DI49_5068 | DI49_5067; DI49_5068 | | |
| DI49_5065; DI49_5066 | DI49_5065; DI49_5066 | | |
| DI49_5063; DI49_5064 | DI49_5063; DI49_5064 | | |
| DI49_5062; SPGP0CW01140 | DI49_5062 | SPGP0CW01140 | |
| DI49_5060; DI49_5061 | DI49_5060; DI49_5061 | | |
| DI49_5058; DI49_5322; SPGP0L01400; DI49_5321 | DI49_5058; DI49_5322; DI49_5321 | SPGP0L01400 | |
| DI49_5056; DI49_5057 | DI49_5056; DI49_5057 | | |
| DI49_5054; DI49_5055 | DI49_5054; DI49_5055 | | |
| DI49_5052; DI49_5053 | DI49_5052; DI49_5053 | | |
| DI49_5050; DI49_5051 | DI49_5050; DI49_5051 | | |
| DI49_5048; DI49_5049 | DI49_5048; DI49_5049 | | |
| DI49_5046; DI49_5047 | DI49_5046; DI49_5047 | | |
| DI49_5045; SPGP0CW01320 | DI49_5045 | SPGP0CW01320 | |
| DI49_5043; DI49_5044 | DI49_5043; DI49_5044 | | |
| DI49_5041; DI49_5042 | DI49_5041; DI49_5042 | | |
| DI49_5039; DI49_5040 | DI49_5039; DI49_5040 | | |
| DI49_5037; DI49_5038 | DI49_5037; DI49_5038 | | |
| DI49_5035; DI49_5036 | DI49_5035; DI49_5036 | | |
| DI49_5033; DI49_5034 | DI49_5033; DI49_5034 | | |
| DI49_5031; DI49_5032; YNL112W; DI49_4621 | DI49_5031; DI49_5032; DI49_4621 | | YNL112W |
| DI49_5029; DI49_5030 | DI49_5029; DI49_5030 | | |
| DI49_5027; DI49_5028 | DI49_5027; DI49_5028 | | |
| DI49_5025; DI49_5026 | DI49_5025; DI49_5026 | | |
| SPGP0CWtrna5M; DI49_5351; DI49_5022; SPGP0DAtrna6M; DI49_1363; DI49_5350; SPGP0FFtrna9M; YJR043C; SPGP0Btrna10M; YEL028W; DI49_3051 | DI49_5351; DI49_5022; DI49_1363; DI49_5350; DI49_3051 | SPGP0CWtrna5M; SPGP0DAtrna6M; SPGP0FFtrna9M; SPGP0Btrna10M | YJR043C; YEL028W |
| DI49_5020; DI49_5021 | DI49_5020; DI49_5021 | | |
| DI49_5019; SPGP0CW01580 | DI49_5019 | SPGP0CW01580 | |
| DI49_5017; DI49_3800; DI49_5016; DI49_5015; DI49_3799 | DI49_5017; DI49_3800; DI49_5016; DI49_5015; DI49_3799 | | |
| DI49_5013; DI49_5014 | DI49_5013; DI49_5014 | | |
| DI49_5011; DI49_5012 | DI49_5011; DI49_5012 | | |
| DI49_5009; DI49_4248; SPGP0CWtrna3V; DI49_4839; SPGP0CXtrna4V; DI49_1922; DI49_1542; SPGP0GHtrna7V; DI49_1560; SPGP0Atrna1V; YJL037W; SPGP0IPtrna11V; YGL226W; SPGP0Ytrna8V; SPGP0Y00950; SPGP0Ytrna9V; DI49_4167; SPGP0Ytrna10V; SPGP0Y02020; | DI49_5009; DI49_4248; DI49_4839; DI49_1922; DI49_1542; DI49_1560; DI49_4167; DI49_2126; DI49_3302; DI49_3781 | SPGP0CWtrna3V; SPGP0CXtrna4V; SPGP0GHtrna7V; SPGP0Atrna1V; SPGP0IPtrna11V; SPGP0Ytrna8V; SPGP0Y00950; SPGP0Ytrna9V; SPGP0Ytrna10V; SPGP0Y02020; SPGP0AFtrna12V; SPGP0Etrna16V; SPGP0Ftrna17V; SPGP0Htrna18V; SPGP0Htrna19V | YJL037W; YGL226W; YKL068W-A |

Create PDF in your applications with the Pdfcrowd HTML to PDF API PDFCROWD

| | | | |
|---|---|---|---|
| SPGP0AFtrna12V; DI49_2126; YKL068W-A; SPGP0Etrna16V; SPGP0Ftrna17V; SPGP0Htrna18V; SPGP0Htrna19V; DI49_3302; DI49_3781 | | | |
| DI49_5007; DI49_5008 | DI49_5007; DI49_5008 | | |
| DI49_5005; DI49_5006 | DI49_5005; DI49_5006 | | |
| DI49_5003; DI49_5004 | DI49_5003; DI49_5004 | | |
| DI49_5001; DI49_5002 | DI49_5001; DI49_5002 | | |
| DI49_4999; DI49_5000 | DI49_4999; DI49_5000 | | |
| DI49_4997; DI49_4998 | DI49_4997; DI49_4998 | | |
| DI49_4995; DI49_4996 | DI49_4995; DI49_4996 | | |
| DI49_4993; DI49_4994 | DI49_4993; DI49_4994 | | |
| DI49_4991; DI49_4992 | DI49_4991; DI49_4992 | | |
| DI49_4989; DI49_4990 | DI49_4989; DI49_4990 | | |
| DI49_4987; SPGP0AN00300; DI49_4986; YDR406W; DI49_0582 | DI49_4987; DI49_4986; DI49_0582 | SPGP0AN00300 | YDR406W |
| DI49_4984; DI49_4985 | DI49_4984; DI49_4985 | | |
| DI49_4982; DI49_4983 | DI49_4982; DI49_4983 | | |
| DI49_4980; DI49_4981 | DI49_4980; DI49_4981 | | |
| DI49_4978; DI49_4979 | DI49_4978; DI49_4979 | | |
| DI49_4976; DI49_4977; DI49_0567; DI49_0568 | DI49_4976; DI49_4977; DI49_0567; DI49_0568 | | |
| DI49_4974; DI49_4975 | DI49_4974; DI49_4975 | | |
| DI49_4973; SPGP0CW02040 | DI49_4973 | SPGP0CW02040 | |
| DI49_4971; DI49_4972 | DI49_4971; DI49_4972 | | |
| DI49_4969; DI49_4970 | DI49_4969; DI49_4970 | | |
| DI49_4967; DI49_4968 | DI49_4967; DI49_4968 | | |
| DI49_4965; DI49_4675; SPGP0Jtrna5D; YJR055W; SPGP0Atrna2D; SPGP0Atrna4D; SPGP0GStrna7D; DI49_1993; SPGP0Xtrna8D; SPGP0Ytrna10D; DI49_4188; DI49_1987; SPGP0AAtrna12D; DI49_3664; SPGP0Ctrna13D; YIL009W; SPGP0Ftrna17D; DI49_2973; SPGP0Ftrna18D; SPGP0Ftrna19D; DI49_3068; SPGP0Htrna20D; DI49_3409; SPGP0CUtrna24D; DI49_3834 | DI49_4965; DI49_4675; DI49_1993; DI49_4188; DI49_1987; DI49_3664; DI49_2973; DI49_3068; DI49_3409; DI49_3834 | SPGP0Jtrna5D; SPGP0Atrna2D; SPGP0Atrna4D; SPGP0GStrna7D; SPGP0Xtrna8D; SPGP0Ytrna10D; SPGP0AAtrna12D; SPGP0Ctrna13D; SPGP0Ftrna17D; SPGP0Ftrna18D; SPGP0Ftrna19D; SPGP0Htrna20D; SPGP0CUtrna24D | YJR055W; YIL009W |
| DI49_4963; DI49_4964 | DI49_4963; DI49_4964 | | |
| DI49_4961; DI49_4962 | DI49_4961; DI49_4962 | | |
| DI49_4959; DI49_4960 | DI49_4959; DI49_4960 | | |
| DI49_4957; DI49_4958 | DI49_4957; DI49_4958 | | |
| DI49_4955; DI49_4956 | DI49_4955; DI49_4956 | | |
| DI49_4953; DI49_4954 | DI49_4953; DI49_4954 | | |
| DI49_4951; DI49_4952 | DI49_4951; DI49_4952 | | |
| DI49_4949; DI49_4950 | DI49_4949; DI49_4950 | | |
| DI49_4947; DI49_4948 | DI49_4947; DI49_4948 | | |
| DI49_4945; DI49_4946 | DI49_4945; DI49_4946 | | |
| DI49_4943; DI49_4944; YNL104C; YNL103W | DI49_4943; DI49_4944 | | YNL104C; YNL103W |
| DI49_4941; DI49_4942 | DI49_4941; DI49_4942 | | |

Create PDF in your applications with the Pdfcrowd HTML to PDF API PDFCROWD

| | | | |
|---|---|---|---|
| DI49_4939; DI49_4940 | DI49_4939; DI49_4940 | | |
| DI49_4937; DI49_4938 | DI49_4937; DI49_4938 | | |
| DI49_4935; DI49_4936 | DI49_4935; DI49_4936 | | |
| DI49_4933; DI49_4934; SPGP0I02260; SPGP0I02270 | DI49_4933; DI49_4934 | SPGP0I02260; SPGP0I02270 | |
| DI49_4931; DI49_4932 | DI49_4931; DI49_4932 | | |
| DI49_4929; DI49_4930 | DI49_4929; DI49_4930 | | |
| DI49_4927; DI49_4928 | DI49_4927; DI49_4928 | | |
| SPGP0CWtrna3N; DI49_5581; DI49_4924; SPGP0Jtrna1N; SPGP0GA00200; SPGP0GA00220; SPGP0Ttrna5N; SPGP0Vtrna6N; SPGP0ICtrna7N; DI49_1650; SPGP0AFtrna8N; DI49_2081; SPGP0Dtrna9N; SPGP0CNtrna12N; SPGP0CUtrna13N; DI49_3921; YNL284C | DI49_5581; DI49_4924; DI49_1650; DI49_2081; DI49_3921 | SPGP0CWtrna3N; SPGP0Jtrna1N; SPGP0GA00200; SPGP0GA00220; SPGP0Ttrna5N; SPGP0Vtrna6N; SPGP0ICtrna7N; SPGP0AFtrna8N; SPGP0Dtrna9N; SPGP0CNtrna12N; SPGP0CUtrna13N | YNL284C |
| DI49_4922; DI49_4923 | DI49_4922; DI49_4923 | | |
| SPGP0CW02560; SPGP0CW02550 | | SPGP0CW02560; SPGP0CW02550 | |
| DI49_4920; DI49_4921 | DI49_4920; DI49_4921 | | |
| DI49_4918; DI49_4919 | DI49_4918; DI49_4919 | | |
| DI49_4916; DI49_4917 | DI49_4916; DI49_4917 | | |
| DI49_4915; YOR074C | DI49_4915 | | YOR074C |
| DI49_4912; DI49_4914; SPGP0CWtrna5P; DI49_1636; DI49_2548; DI49_2549 | DI49_4912; DI49_4914; DI49_1636; DI49_2548; DI49_2549 | SPGP0CWtrna5P | |
| DI49_4910; DI49_4911 | DI49_4910; DI49_4911 | | |
| DI49_4908; DI49_4909 | DI49_4908; DI49_4909 | | |
| DI49_4906; DI49_4907 | DI49_4906; DI49_4907 | | |
| DI49_4904; DI49_4905 | DI49_4904; DI49_4905 | | |
| DI49_4902; DI49_4903 | DI49_4902; DI49_4903 | | |
| SPGP0CWtrna3K; DI49_5564; DI49_4899; SPGP0Utrna12K; DI49_1007; SPGP0AFtrna16K; DI49_2068; SPGP0ATtrna20K; DI49_5565; SPGP0Htrna23K; DI49_3442; DI49_3876 | DI49_5564; DI49_4899; DI49_1007; DI49_2068; DI49_5565; DI49_3442; DI49_3876 | SPGP0CWtrna3K; SPGP0Utrna12K; SPGP0AFtrna16K; SPGP0ATtrna20K; SPGP0Htrna23K | |
| DI49_4897; DI49_4898 | DI49_4897; DI49_4898 | | |
| DI49_4895; DI49_4896 | DI49_4895; DI49_4896 | | |
| DI49_4893; DI49_4894 | DI49_4893; DI49_4894 | | |
| DI49_4891; DI49_4892 | DI49_4891; DI49_4892 | | |
| DI49_4889; DI49_4890 | DI49_4889; DI49_4890 | | |
| DI49_4887; DI49_4888 | DI49_4887; DI49_4888 | | |
| DI49_4885; DI49_4886 | DI49_4885; DI49_4886 | | |
| DI49_4883; DI49_4884 | DI49_4883; DI49_4884 | | |
| DI49_4881; DI49_4882 | DI49_4881; DI49_4882 | | |
| DI49_4879; DI49_4880 | DI49_4879; DI49_4880 | | |
| DI49_4877; DI49_4878 | DI49_4877; DI49_4878 | | |
| DI49_4875; DI49_4876 | DI49_4875; DI49_4876 | | |
| DI49_4873; DI49_4874 | DI49_4873; DI49_4874 | | |
| DI49_4871; DI49_4872 | DI49_4871; DI49_4872 | | |

Create PDF in your applications with the Pdfcrowd HTML to PDF API PDFCROWD

| | | | |
|---|---|---|---|
| DI49_4869; DI49_4870 | DI49_4869; DI49_4870 | | |
| DI49_4867; DI49_4868 | DI49_4867; DI49_4868 | | |
| DI49_4865; DI49_4866 | DI49_4865; DI49_4866 | | |
| DI49_4863; DI49_4419; SPGP0DAtrna4S; DI49_0694; DI49_1047; SPGP0Strna8S; SPGP0Xtrna13S; DI49_0593; DI49_5465; SPGP0Ftrna21S; DI49_2003 | DI49_4863; DI49_4419; DI49_0694; DI49_1047; DI49_0593; DI49_5465; DI49_2003 | SPGP0DAtrna4S; SPGP0Strna8S; SPGP0Xtrna13S; SPGP0Ftrna21S | |
| DI49_4861; DI49_4862 | DI49_4861; DI49_4862 | | |
| YHR010W; DI49_0636; DI49_4859; DI49_0637 | DI49_0636; DI49_4859; DI49_0637 | | YHR010W |
| DI49_4857; DI49_4858 | DI49_4857; DI49_4858 | | |
| SPGP0CWtrna4T; DI49_2415; DI49_4854; SPGP0Jtrna1T; SPGP0Jtrna3T; DI49_4465; SPGP0Mtrna5T; SPGP0Utrna7T; DI49_1043; DI49_2299; SPGP0Xtrna10T; SPGP0X01490; SPGP0AJ00210; SPGP0AJtrna17T; DI49_0191 | DI49_2415; DI49_4854; DI49_4465; DI49_1043; DI49_2299; DI49_0191 | SPGP0CWtrna4T; SPGP0Jtrna1T; SPGP0Jtrna3T; SPGP0Mtrna5T; SPGP0Utrna7T; SPGP0Xtrna10T; SPGP0X01490; SPGP0AJ00210; SPGP0AJtrna17T | |
| DI49_4852; DI49_4853 | DI49_4852; DI49_4853 | | |
| DI49_4850; DI49_4851 | DI49_4850; DI49_4851 | | |
| DI49_4848; DI49_4849 | DI49_4848; DI49_4849 | | |
| DI49_4846; DI49_4847 | DI49_4846; DI49_4847 | | |
| DI49_4844; DI49_4845 | DI49_4844; DI49_4845 | | |
| DI49_4842; DI49_4843 | DI49_4842; DI49_4843 | | |
| DI49_4837; DI49_4838 | DI49_4837; DI49_4838 | | |
| DI49_4835; DI49_4836 | DI49_4835; DI49_4836 | | |
| DI49_4833; DI49_4834 | DI49_4833; DI49_4834 | | |
| DI49_4831; DI49_4832 | DI49_4831; DI49_4832 | | |
| DI49_4829; DI49_4830 | DI49_4829; DI49_4830 | | |
| DI49_4827; DI49_4828 | DI49_4827; DI49_4828 | | |
| SPGP0CWtrna1H; DI49_4134; DI49_4824; SPGP0CXtrna3H; DI49_1876; DI49_1538; DI49_3305 | DI49_4134; DI49_4824; DI49_1876; DI49_1538; DI49_3305 | SPGP0CWtrna1H; SPGP0CXtrna3H | |
| DI49_4822; DI49_4823 | DI49_4822; DI49_4823 | | |
| DI49_4820; DI49_4821 | DI49_4820; DI49_4821 | | |
| DI49_4818; DI49_4819 | DI49_4818; DI49_4819 | | |
| DI49_4816; DI49_4817 | DI49_4816; DI49_4817 | | |
| SPGP0CW03590; DI49_3489; DI49_4814; SPGP0AA01990 | DI49_3489; DI49_4814 | SPGP0CW03590; SPGP0AA01990 | |
| DI49_4812; DI49_4813 | DI49_4812; DI49_4813 | | |
| DI49_4810; DI49_4811 | DI49_4810; DI49_4811 | | |
| DI49_4808; DI49_4809 | DI49_4808; DI49_4809 | | |
| YGL221C; YGL222C | | | YGL221C; YGL222C |
| YGL219C; YGL220W | | | YGL219C; YGL220W |
| YGL215W; YGL216W | | | YGL215W; YGL216W |
| YGL212W; YGL213C | | | YGL212W; YGL213C |
| YGL210W; YGL211W | | | YGL210W; YGL211W |
| YGL208W; YGL209W | | | YGL208W; YGL209W |
| YGL206C; YGL207W | | | YGL206C; YGL207W |
| SPGP0CXtrna2H; SPGP0Etrna8H; YGL203C; SPGP0Btrna6H; | | SPGP0CXtrna2H; SPGP0Etrna8H; SPGP0Btrna6H | YGL203C; YKL067W |

Create PDF in your applications with the Pdfcrowd HTML to PDF API PDFCROWD

| | | | |
|---|---|---|---|
| YKL067W | | | |
| YGL202W; SPGP0Etrna21K; SPGP0CXtrna4K; YKR074W | | SPGP0Etrna21K; SPGP0CXtrna4K | YGL202W; YKR074W |
| YGL197W; YGL198W | | | YGL197W; YGL198W |
| YGL195W; YGL196W | | | YGL195W; YGL196W |
| YGL194C; YGL194C-A | | | YGL194C; YGL194C-A |
| YGL191W; YGL192W | | | YGL191W; YGL192W |
| YGL189C; DI49_1535; YGL187C; DI49_1536 | DI49_1535; DI49_1536 | | YGL189C; YGL187C |
| YGL185C; YGL186C | | | YGL185C; YGL186C |
| YGL183C; YGL184C | | | YGL183C; YGL184C |
| YGL180W; YGL181W | | | YGL180W; YGL181W |
| YGL178W; YGL179C | | | YGL178W; YGL179C |
| YGL175C; YGL176C | | | YGL175C; YGL176C |
| YGL173C; YGL174W | | | YGL173C; YGL174W |
| DI49_1805; DI49_1804 | DI49_1805; DI49_1804 | | |
| DI49_1810; DI49_1809 | DI49_1810; DI49_1809 | | |
| DI49_1812; DI49_1811 | DI49_1812; DI49_1811 | | |
| SPGP0CX00610; DI49_1813 | DI49_1813 | SPGP0CX00610 | |
| DI49_1815; DI49_1814 | DI49_1815; DI49_1814 | | |
| DI49_1817; DI49_1816 | DI49_1817; DI49_1816 | | |
| DI49_1819; DI49_1818 | DI49_1819; DI49_1818 | | |
| DI49_1821; DI49_1820; DI49_1928; DI49_1929 | DI49_1821; DI49_1820; DI49_1928; DI49_1929 | | |
| DI49_1823; DI49_1822 | DI49_1823; DI49_1822 | | |
| DI49_1825; DI49_1824 | DI49_1825; DI49_1824 | | |
| DI49_1827; DI49_1826 | DI49_1827; DI49_1826 | | |
| DI49_1829; DI49_1828; DI49_4664; DI49_4663 | DI49_1829; DI49_1828; DI49_4664; DI49_4663 | | |
| DI49_1831; DI49_1830 | DI49_1831; DI49_1830 | | |
| DI49_1833; DI49_1832 | DI49_1833; DI49_1832 | | |
| DI49_1835; DI49_1834 | DI49_1835; DI49_1834 | | |
| DI49_1837; DI49_1836 | DI49_1837; DI49_1836 | | |
| DI49_1839; DI49_1838 | DI49_1839; DI49_1838 | | |
| DI49_1841; DI49_1840; DI49_5249; DI49_5248 | DI49_1841; DI49_1840; DI49_5249; DI49_5248 | | |
| DI49_1843; DI49_1842 | DI49_1843; DI49_1842 | | |
| DI49_1845; DI49_1844 | DI49_1845; DI49_1844 | | |
| DI49_1847; DI49_1846 | DI49_1847; DI49_1846 | | |
| DI49_1849; DI49_1848 | DI49_1849; DI49_1848 | | |
| DI49_1851; DI49_1850 | DI49_1851; DI49_1850 | | |
| DI49_1853; DI49_1852 | DI49_1853; DI49_1852 | | |
| DI49_1855; DI49_1854 | DI49_1855; DI49_1854 | | |
| DI49_1857; DI49_3006; SPGP0Ytrna5W; DI49_4142 | DI49_1857; DI49_3006; DI49_4142 | SPGP0Ytrna5W | |
| DI49_1859; DI49_1858 | DI49_1859; DI49_1858 | | |

Create PDF in your applications with the Pdfcrowd HTML to PDF API PDFCROWD

| | | | |
|---|---|---|---|
| DI49_1861; DI49_1860 | DI49_1861; DI49_1860 | | |
| DI49_1863; DI49_1862 | DI49_1863; DI49_1862 | | |
| DI49_1865; DI49_1864 | DI49_1865; DI49_1864 | | |
| DI49_1867; DI49_1866 | DI49_1867; DI49_1866 | | |
| DI49_1869; DI49_1868 | DI49_1869; DI49_1868 | | |
| DI49_1871; DI49_1870 | DI49_1871; DI49_1870 | | |
| DI49_1873; DI49_1872 | DI49_1873; DI49_1872 | | |
| DI49_1878; DI49_1877 | DI49_1878; DI49_1877 | | |
| DI49_1880; DI49_5287; SPGP0CXtrna2E; SPGP0Ktrna4E; DI49_1504; DI49_5286; DI49_2867; SPGP0GStrna9E; DI49_1991; SPGP0GZtrna10E; SPGP0Ttrna8E; SPGP0Xtrna12E; SPGP0Ytrna13E; DI49_4099; SPGP0Ctrna15E; SPGP0BEtrna16E; DI49_0731; SPGP0CMtrna17E; SPGP0CNtrna18E | DI49_1880; DI49_5287; DI49_1504; DI49_5286; DI49_2867; DI49_1991; DI49_4099; DI49_0731 | SPGP0CXtrna2E; SPGP0Ktrna4E; SPGP0GStrna9E; SPGP0GZtrna10E; SPGP0Ttrna8E; SPGP0Xtrna12E; SPGP0Ytrna13E; SPGP0Ctrna15E; SPGP0BEtrna16E; SPGP0CMtrna17E; SPGP0CNtrna18E | |
| DI49_1882; DI49_1881 | DI49_1882; DI49_1881 | | |
| DI49_1883; SPGP0CX01270 | DI49_1883 | SPGP0CX01270 | |
| DI49_1885; DI49_1884 | DI49_1885; DI49_1884 | | |
| DI49_1887; DI49_1886 | DI49_1887; DI49_1886 | | |
| DI49_1889; DI49_1888 | DI49_1889; DI49_1888 | | |
| DI49_1891; DI49_1890 | DI49_1891; DI49_1890 | | |
| DI49_1893; DI49_1892 | DI49_1893; DI49_1892 | | |
| DI49_1895; DI49_1894 | DI49_1895; DI49_1894 | | |
| DI49_1897; DI49_1896 | DI49_1897; DI49_1896 | | |
| DI49_1899; DI49_1898 | DI49_1899; DI49_1898 | | |
| DI49_1901; DI49_1900 | DI49_1901; DI49_1900 | | |
| DI49_1903; DI49_1902 | DI49_1903; DI49_1902 | | |
| DI49_1905; DI49_1904 | DI49_1905; DI49_1904 | | |
| DI49_1907; DI49_1244; DI49_1908; DI49_1243 | DI49_1907; DI49_1244; DI49_1908; DI49_1243 | | |
| DI49_1909; SPGP0CX01550 | DI49_1909 | SPGP0CX01550 | |
| DI49_1911; DI49_1910 | DI49_1911; DI49_1910 | | |
| DI49_1913; DI49_1912 | DI49_1913; DI49_1912 | | |
| SPGP0CXtrna5R; DI49_4323; DI49_1917; SPGP0DAtrna6R; DI49_1385; SPGP0Xtrna11R; SPGP0Ytrna12R; DI49_4324; SPGP0AFtrna14R; DI49_2084; SPGP0AFtrna15R; DI49_2129; SPGP0Ftrna19R; SPGP0Ftrna20R; DI49_2981; SPGP0CMtrna23R; DI49_1112; SPGP0CNtrna24R; DI49_3226; DI49_0252 | DI49_4323; DI49_1917; DI49_1385; DI49_4324; DI49_2084; DI49_2129; DI49_2981; DI49_1112; DI49_3226; DI49_0252 | SPGP0CXtrna5R; SPGP0DAtrna6R; SPGP0Xtrna11R; SPGP0Ytrna12R; SPGP0AFtrna14R; SPGP0AFtrna15R; SPGP0Ftrna19R; SPGP0Ftrna20R; SPGP0CMtrna23R; SPGP0CNtrna24R | |
| DI49_1919; DI49_1918 | DI49_1919; DI49_1918 | | |
| DI49_1924; DI49_1923 | DI49_1924; DI49_1923 | | |
| DI49_1926; DI49_1925 | DI49_1926; DI49_1925 | | |
| DI49_1931; DI49_1930 | DI49_1931; DI49_1930 | | |
| SPGP0CX01760; DI49_1932 | DI49_1932 | SPGP0CX01760 | |
| DI49_1934; DI49_5687; SPGP0Jtrna4L; DI49_4787; SPGP0Atrna2L; YJL006C; SPGP0Ttrna10L; YKL189W; | DI49_1934; DI49_5687; DI49_4787; DI49_1084; DI49_3191; DI49_0231; DI49_3915 | SPGP0Jtrna4L; SPGP0Atrna2L; SPGP0Ttrna10L; SPGP0IStrna13L; SPGP0Dtrna16L; SPGP0CNtrna24L | YJL006C; YKL189W |

Create PDF in your applications with the Pdfcrowd HTML to PDF API PDFCROWD

| | | | |
|---|---|---|---|
| SPGP0IStrna13L; DI49_1084; SPGP0Dtrna16L; SPGP0CNtrna24L; DI49_3191; DI49_0231; DI49_3915 | | | |
| DI49_1936; DI49_2112; SPGP0CX01800; DI49_2113 | DI49_1936; DI49_2112; DI49_2113 | SPGP0CX01800 | |
| DI49_1939; DI49_5486; DI49_1940; SPGP0GTtrna5F; YPR002W; SPGP0AT00300 | DI49_1939; DI49_5486; DI49_1940 | SPGP0GTtrna5F; SPGP0AT00300 | YPR002W |
| DI49_1942; DI49_1941 | DI49_1942; DI49_1941 | | |
| DI49_1944; DI49_1943 | DI49_1944; DI49_1943 | | |
| DI49_1946; DI49_1945 | DI49_1946; DI49_1945 | | |
| DI49_1948; DI49_1947 | DI49_1948; DI49_1947 | | |
| DI49_1950; DI49_1949 | DI49_1950; DI49_1949 | | |
| DI49_1952; DI49_1951 | DI49_1952; DI49_1951 | | |
| DI49_1954; DI49_1953 | DI49_1954; DI49_1953 | | |
| DI49_1956; DI49_1955 | DI49_1956; DI49_1955 | | |
| DI49_1958; DI49_1957 | DI49_1958; DI49_1957 | | |
| DI49_1960; DI49_1959; DI49_5422; DI49_5421 | DI49_1960; DI49_1959; DI49_5422; DI49_5421 | | |
| DI49_1962; DI49_1961 | DI49_1962; DI49_1961 | | |
| DI49_1964; DI49_1963 | DI49_1964; DI49_1963 | | |
| DI49_1966; DI49_1965 | DI49_1966; DI49_1965 | | |
| DI49_1967; SPGP0CX02090 | DI49_1967 | SPGP0CX02090 | |
| DI49_1969; DI49_1968 | DI49_1969; DI49_1968 | | |
| DI49_1971; DI49_1970 | DI49_1971; DI49_1970 | | |
| DI49_1973; DI49_1972 | DI49_1973; DI49_1972 | | |
| DI49_1975; DI49_1974 | DI49_1975; DI49_1974 | | |
| DI49_1977; DI49_1976 | DI49_1977; DI49_1976 | | |
| DI49_1979; DI49_1978 | DI49_1979; DI49_1978 | | |
| DI49_1981; DI49_1980 | DI49_1981; DI49_1980 | | |
| DI49_1983; DI49_1982 | DI49_1983; DI49_1982 | | |
| SPGP0CX02280; DI49_1984 | DI49_1984 | SPGP0CX02280 | |
| YHR211W; YAL065C; YGR288W | | | YHR211W; YAL065C; YGR288W |
| DI49_1478; YER074W; DI49_1477; DI49_2715; YER073W; YIL069C | DI49_1478; DI49_1477; DI49_2715 | | YER074W; YER073W; YIL069C |
| DI49_1475; YER071C | DI49_1475 | | YER071C |
| DI49_1474; YER070W | DI49_1474 | | YER070W |
| DI49_1473; YER069W; DI49_1472; YER068W | DI49_1473; DI49_1472 | | YER069W; YER068W |
| SPGP0DA00180; DI49_1645; DI49_1644 | DI49_1645; DI49_1644 | SPGP0DA00180 | |
| SPGP0DAtrna3S; DI49_5523; SPGP0Xtrna12S; DI49_0808; DI49_0809; SPGP0Ctrna16S; SPGP0ATtrna20S; DI49_5524; DI49_1141 | DI49_5523; DI49_0808; DI49_0809; DI49_5524; DI49_1141 | SPGP0DAtrna3S; SPGP0Xtrna12S; SPGP0Ctrna16S; SPGP0ATtrna20S | |
| DI49_1469; YER065C | DI49_1469 | | YER065C |
| DI49_1468; YER064C; DI49_1467; YER063W | DI49_1468; DI49_1467 | | YER064C; YER063W |
| DI49_1465; YER061C | DI49_1465 | | YER061C |
| DI49_1464; DI49_1459; DI49_1463; YER056C; DI49_1458; YER060W; YER055C | DI49_1464; DI49_1459; DI49_1463; DI49_1458 | | YER056C; YER060W; YER055C |

Create PDF in your applications with the Pdfcrowd HTML to PDF API PDFCROWD

| | | | |
|---|---|---|---|
| DI49_1462; YER059W; DI49_1461; YER058W | DI49_1462; DI49_1461 | | YER059W; YER058W |
| SPGP0DA00290; YER056C-A | | SPGP0DA00290 | YER056C-A |
| DI49_1456; YER053C-A | DI49_1456 | | YER053C-A |
| DI49_1455; YER053C | DI49_1455 | | YER053C |
| DI49_1454; YER052C; DI49_1453; YER051W | DI49_1454; DI49_1453 | | YER052C; YER051W |
| DI49_1451; YER049W; DI49_1450; YER048W-A | DI49_1451; DI49_1450 | | YER049W; YER048W-A |
| DI49_1448; YER048C; DI49_1447; YER047C | DI49_1448; DI49_1447 | | YER048C; YER047C |
| DI49_1445; DI49_1446 | DI49_1445; DI49_1446 | | |
| DI49_1443; YER044C | DI49_1443 | | YER044C |
| DI49_1442; YER043C | DI49_1442 | | YER043C |
| DI49_1441; YER042W; DI49_1440; YER041W | DI49_1441; DI49_1440 | | YER042W; YER041W |
| DI49_1438; YER039C; DI49_1437; YER038C | DI49_1438; DI49_1437 | | YER039C; YER038C |
| DI49_1435; YER036C; DI49_1434; YER035W | DI49_1435; DI49_1434 | | YER036C; YER035W |
| DI49_1432; DI49_0132; DI49_1431; DI49_0133 | DI49_1432; DI49_0132; DI49_1431; DI49_0133 | | |
| DI49_1429; DI49_1430; YER031C; YER030W | DI49_1429; DI49_1430 | | YER031C; YER030W |
| DI49_1427; DI49_1428 | DI49_1427; DI49_1428 | | |
| DI49_1425; DI49_1426 | DI49_1425; DI49_1426 | | |
| DI49_1423; YER025W | DI49_1423 | | YER025W |
| DI49_1422; YER024W; DI49_1421; YER023W | DI49_1422; DI49_1421 | | YER024W; YER023W |
| DI49_1419; YER021W | DI49_1419 | | YER021W |
| DI49_1418; YER020W; DI49_1416; YER019W | DI49_1418; DI49_1416 | | YER020W; YER019W |
| DI49_1414; YER017C; DI49_1413; YER016W | DI49_1414; DI49_1413 | | YER017C; YER016W |
| DI49_1411; DI49_1412; YPL282C; SPGP0AY00110; YFL020C; YAR023C | DI49_1411; DI49_1412 | SPGP0AY00110 | YPL282C; YFL020C; YAR023C |
| DI49_1409; YER013W; DI49_1408; YER012W | DI49_1409; DI49_1408 | | YER013W; YER012W |
| DI49_1406; DI49_2413; DI49_1405; YER011W; YER010C; DI49_2414; DI49_3279; DI49_3278 | DI49_1406; DI49_2413; DI49_1405; DI49_2414; DI49_3279; DI49_3278 | | YER011W; YER010C |
| DI49_1403; YER008C; DI49_1402 | DI49_1403; DI49_1402 | | YER008C |
| DI49_1400; DI49_1401; SPGP0B00820; YER007W | DI49_1400; DI49_1401 | SPGP0B00820 | YER007W |
| DI49_1399; YER005W; DI49_1398; YER004W | DI49_1399; DI49_1398 | | YER005W; YER004W |
| DI49_1396; DI49_1397 | DI49_1396; DI49_1397 | | |
| DI49_1395; SPGP0DA00910 | DI49_1395 | SPGP0DA00910 | |
| DI49_1393; DI49_1394 | DI49_1393; DI49_1394 | | |
| DI49_1391; DI49_1392 | DI49_1391; DI49_1392 | | |
| DI49_1389; YEL007W; DI49_1388 | DI49_1389; DI49_1388 | | YEL007W |
| DI49_1383; YEL020W-A; DI49_1382 | DI49_1383; DI49_1382 | | YEL020W-A |
| DI49_1380; DI49_1381 | DI49_1380; DI49_1381 | | |
| DI49_1378; DI49_1379 | DI49_1378; DI49_1379 | | |
| DI49_1376; YEL015W | DI49_1376 | | YEL015W |
| DI49_1375; YEL013W | DI49_1375 | | YEL013W |
| SPGP0DAtrna2Q; DI49_1374; SPGP0Btrna5Q; SPGP0CN00100 | DI49_1374 | SPGP0DAtrna2Q; SPGP0Btrna5Q; SPGP0CN00100 | |

Create PDF in your applications with the Pdfcrowd HTML to PDF API PDFCROWD

| | | | |
|---|---|---|---|
| DI49_1373; YEL012W | DI49_1373 | | YEL012W |
| DI49_1372; YEL011W | DI49_1372 | | YEL011W |
| DI49_1370; YEL022W | DI49_1370 | | YEL022W |
| DI49_1369; YEL023C | DI49_1369 | | YEL023C |
| DI49_1368; YEL024W; DI49_1367; YEL025C | DI49_1368; DI49_1367 | | YEL024W; YEL025C |
| DI49_1365; YEL027W | DI49_1365 | | YEL027W |
| DI49_1361; YEL031W | DI49_1361 | | YEL031W |
| DI49_1360; YEL032W; DI49_0695 | DI49_1360; DI49_0695 | | YEL032W |
| DI49_1358; YEL034W; YEL035C; DI49_3056 | DI49_1358; DI49_3056 | | YEL034W; YEL035C |
| DI49_1357; YEL036C | DI49_1357 | | YEL036C |
| DI49_1356; YEL037C; DI49_1355; YEL038W | DI49_1356; DI49_1355 | | YEL037C; YEL038W |
| DI49_1353; DI49_1354 | DI49_1353; DI49_1354 | | |
| DI49_1351; DI49_1352 | DI49_1351; DI49_1352 | | |
| DI49_1349; DI49_1350 | DI49_1349; DI49_1350 | | |
| DI49_1347; DI49_1348 | DI49_1347; DI49_1348 | | |
| DI49_1345; DI49_3566; DI49_3565; DI49_3506 | DI49_1345; DI49_3566; DI49_3565; DI49_3506 | | |
| SPGP0DAtrna5G; SPGP0CPtrna26G; DI49_1343; DI49_0764; SPGP0Atrna1G; YJR051W; SPGP0Atrna2G; DI49_0870; SPGP0Vtrna9G; DI49_2378; SPGP0Vtrna10G; DI49_5463; DI49_1672; SPGP0Z00210; SPGP0Btrna7G; YEL050C; SPGP0AFtrna16G; DI49_2140; DI49_2186; SPGP0ATtrna19G; DI49_5604; SPGP0Ftrna20G; SPGP0CBtrna22G; SPGP0CCtrna23G; DI49_0441 | DI49_1343; DI49_0764; DI49_0870; DI49_2378; DI49_5463; DI49_1672; DI49_2140; DI49_2186; DI49_5604; DI49_0441 | SPGP0DAtrna5G; SPGP0CPtrna26G; SPGP0Atrna1G; SPGP0Atrna2G; SPGP0Vtrna9G; SPGP0Vtrna10G; SPGP0Z00210; SPGP0Btrna7G; SPGP0AFtrna16G; SPGP0ATtrna19G; SPGP0Ftrna20G; SPGP0CBtrna22G; SPGP0CCtrna23G | YJR051W; YEL050C |
| DI49_1341; YEL052W | DI49_1341 | | YEL052W |
| DI49_1340; YEL053C | DI49_1340 | | YEL053C |
| DI49_1339; YEL054C; DI49_1338; YEL055C | DI49_1339; DI49_1338 | | YEL054C; YEL055C |
| DI49_1336; DI49_1337 | DI49_1336; DI49_1337 | | |
| DI49_1334; DI49_1335 | DI49_1334; DI49_1335 | | |
| DI49_1332; DI49_1333 | DI49_1332; DI49_1333 | | |
| DI49_1330; DI49_1331 | DI49_1330; DI49_1331 | | |
| DI49_1328; DI49_1329 | DI49_1328; DI49_1329 | | |
| SPGP0DA01550; DI49_2618 | DI49_2618 | SPGP0DA01550 | |
| SPGP0DA01560; SPGP0F02990; DI49_1326; DI49_3166 | DI49_1326; DI49_3166 | SPGP0DA01560; SPGP0F02990 | |
| DI49_1324; DI49_1325 | DI49_1324; DI49_1325 | | |
| DI49_1322; DI49_2621 | DI49_1322; DI49_2621 | | |
| YDR345C; DI49_2500; DI49_0523; SPGP0HI00140; DI49_3960; DI49_2770; DI49_2771; YMR011W; DI49_2501; DI49_2502 | DI49_2500; DI49_0523; DI49_3960; DI49_2770; DI49_2771; DI49_2501; DI49_2502 | SPGP0HI00140 | YDR345C; YMR011W |
| DI49_0525; DI49_0524 | DI49_0525; DI49_0524 | | |
| DI49_0530; DI49_0529 | DI49_0530; DI49_0529 | | |
| DI49_0532; DI49_0531 | DI49_0532; DI49_0531 | | |
| DI49_0534; DI49_0533 | DI49_0534; DI49_0533 | | |
| DI49_0536; DI49_0535 | DI49_0536; DI49_0535 | | |
| DI49_0538; DI49_0537 | DI49_0538; DI49_0537 | | |

Create PDF in your applications with the Pdfcrowd HTML to PDF API PDFCROWD

| | | | |
|---|---|---|---|
| DI49_0543; DI49_0542 | DI49_0543; DI49_0542 | | |
| SPGP0DB00310; DI49_0544 | DI49_0544 | SPGP0DB00310 | |
| DI49_0546; DI49_0545 | DI49_0546; DI49_0545 | | |
| DI49_0548; DI49_0547 | DI49_0548; DI49_0547 | | |
| DI49_0550; DI49_0549 | DI49_0550; DI49_0549 | | |
| DI49_0552; DI49_0551 | DI49_0552; DI49_0551 | | |
| DI49_0554; DI49_0553 | DI49_0554; DI49_0553 | | |
| DI49_0555; SPGP0DB00420 | DI49_0555 | SPGP0DB00420 | |
| DI49_0557; DI49_0556 | DI49_0557; DI49_0556 | | |
| SPGP0DB00470; SPGP0DB00460 | | SPGP0DB00470; SPGP0DB00460 | |
| DI49_0559; DI49_0558; YDL130W; DI49_0942 | DI49_0559; DI49_0558; DI49_0942 | | YDL130W |
| YNL018C; SPGP0I02840; YNL033W | | SPGP0I02840 | YNL018C; YNL033W |
| SPGP0DCtrna2I; SPGP0Itrna14I; SPGP0Jtrna1I; DI49_4719; SPGP0Ktrna3I; SPGP0K00610; SPGP0GGtrna4I; DI49_3947; SPGP0Xtrna5I; SPGP0Xtrna6I; DI49_2704; YER182W; SPGP0AGtrna11I; DI49_0796; SPGP0Ctrna8I; SPGP0Ctrna9I; YIL079C; DI49_5588; SPGP0ATtrna13I; DI49_5615; DI49_0301; SPGP0CPtrna17I; DI49_0165; DI49_3807 | DI49_4719; DI49_3947; DI49_2704; DI49_0796; DI49_5588; DI49_5615; DI49_0301; DI49_0165; DI49_3807 | SPGP0DCtrna2I; SPGP0Itrna14I; SPGP0Jtrna1I; SPGP0Ktrna3I; SPGP0K00610; SPGP0GGtrna4I; SPGP0Xtrna5I; SPGP0Xtrna6I; SPGP0AGtrna11I; SPGP0Ctrna8I; SPGP0Ctrna9I; SPGP0ATtrna13I; SPGP0CPtrna17I | YER182W; YIL079C |
| YNL016W; DI49_4721; YNL015W | DI49_4721 | | YNL016W; YNL015W |
| YNL012W; SPGP0J00890 | | SPGP0J00890 | YNL012W |
| YNL011C; DI49_4724 | DI49_4724 | | YNL011C |
| YNL010W; DI49_4725 | DI49_4725 | | YNL010W |
| YNL009W; DI49_4726; YNL008C | DI49_4726 | | YNL009W; YNL008C |
| YNL006W; YNL007C; DI49_4728; DI49_4727 | DI49_4728; DI49_4727 | | YNL006W; YNL007C |
| YNL004W; DI49_4731; DI49_4730 | DI49_4731; DI49_4730 | | YNL004W |
| YNL003C; DI49_4732 | DI49_4732 | | YNL003C |
| YNL002C; DI49_4733; YNL001W | DI49_4733 | | YNL002C; YNL001W |
| SPGP0DCtrna6P; DI49_4736; SPGP0Jtrna1P; DI49_0754 | DI49_4736; DI49_0754 | SPGP0DCtrna6P; SPGP0Jtrna1P | |
| YNL020C; DI49_4717; YNL021W | DI49_4717 | | YNL020C; YNL021W |
| YNL023C; DI49_4714 | DI49_4714 | | YNL023C |
| YNL024C; DI49_4713; YNL024C-A | DI49_4713 | | YNL024C; YNL024C-A |
| YNL026W; YNL025C | | | YNL026W; YNL025C |
| YNL029C; YNL027W | | | YNL029C; YNL027W |
| YNL031C; DI49_4706; DI49_0190 | DI49_4706; DI49_0190 | | YNL031C |
| YNL032W; DI49_4705; DI49_0462 | DI49_4705; DI49_0462 | | YNL032W |
| DI49_0464; DI49_0463 | DI49_0464; DI49_0463 | | |
| DI49_0466; DI49_0465 | DI49_0466; DI49_0465 | | |
| DI49_0468; DI49_0467 | DI49_0468; DI49_0467 | | |
| DI49_0470; DI49_0469 | DI49_0470; DI49_0469 | | |
| DI49_0472; DI49_0471 | DI49_0472; DI49_0471 | | |
| DI49_0474; DI49_0473 | DI49_0474; DI49_0473 | | |
| DI49_0476; DI49_0475 | DI49_0476; DI49_0475 | | |

Create PDF in your applications with the Pdfcrowd HTML to PDF API PDFCROWD

| | | | |
|---|---|---|---|
| DI49_0478; DI49_0477 | DI49_0478; DI49_0477 | | |
| DI49_0480; DI49_0479 | DI49_0480; DI49_0479 | | |
| DI49_0482; DI49_0481 | DI49_0482; DI49_0481 | | |
| DI49_0484; DI49_2588; DI49_0485 | DI49_0484; DI49_2588; DI49_0485 | | |
| DI49_0487; DI49_0486 | DI49_0487; DI49_0486 | | |
| DI49_0489; DI49_0488 | DI49_0489; DI49_0488 | | |
| DI49_0491; DI49_0490; DI49_2469; DI49_2470 | DI49_0491; DI49_0490; DI49_2469; DI49_2470 | | |
| DI49_0493; DI49_0492 | DI49_0493; DI49_0492 | | |
| DI49_0495; SPGP0Itrna11F; DI49_0496 | DI49_0495; DI49_0496 | SPGP0Itrna11F | |
| DI49_0498; DI49_0497 | DI49_0498; DI49_0497 | | |
| DI49_0500; DI49_0499 | DI49_0500; DI49_0499 | | |
| DI49_0502; DI49_0501 | DI49_0502; DI49_0501 | | |
| DI49_0504; DI49_0503 | DI49_0504; DI49_0503 | | |
| DI49_0506; DI49_0505 | DI49_0506; DI49_0505 | | |
| DI49_0508; DI49_0507 | DI49_0508; DI49_0507 | | |
| DI49_0510; DI49_0509 | DI49_0510; DI49_0509 | | |
| DI49_0512; DI49_0511 | DI49_0512; DI49_0511 | | |
| DI49_0514; DI49_0513 | DI49_0514; DI49_0513 | | |
| DI49_0516; DI49_0515 | DI49_0516; DI49_0515 | | |
| DI49_0518; DI49_0517 | DI49_0518; DI49_0517 | | |
| SPGP0DEtrna5S; DI49_2374; DI49_0521; SPGP0Vtrna11S; DI49_2373; DI49_1676; SPGP0Ztrna14S; DI49_3829 | DI49_2374; DI49_0521; DI49_2373; DI49_1676; DI49_3829 | SPGP0DEtrna5S; SPGP0Vtrna11S; SPGP0Ztrna14S | |
| DI49_4801; DI49_3161; DI49_4800; SPGP0F02930 | DI49_4801; DI49_3161; DI49_4800 | SPGP0F02930 | |
| DI49_4798; DI49_4799 | DI49_4798; DI49_4799 | | |
| SPGP0J00150; DI49_4797 | DI49_4797 | SPGP0J00150 | |
| SPGP0J00170; DI49_4796 | DI49_4796 | SPGP0J00170 | |
| DI49_4793; DI49_4794 | DI49_4793; DI49_4794 | | |
| DI49_4791; DI49_4792; SPGP0D00260; YNR056C | DI49_4791; DI49_4792 | SPGP0D00260 | YNR056C |
| DI49_4789; SPGP0AE00380; DI49_3953; DI49_3952 | DI49_4789; DI49_3953; DI49_3952 | SPGP0AE00380 | |
| SPGP0J00260; YNR053C | | SPGP0J00260 | YNR053C |
| DI49_4785; YNR052C; DI49_4784; YNR051C | DI49_4785; DI49_4784 | | YNR052C; YNR051C |
| DI49_4782; DI49_4783 | DI49_4782; DI49_4783 | | |
| DI49_4780; YNR047W; DI49_4779; YNR046W | DI49_4780; DI49_4779 | | YNR047W; YNR046W |
| DI49_4777; DI49_4778 | DI49_4777; DI49_4778 | | |
| DI49_4775; YNR041C; DI49_4774; YNR040W | DI49_4775; DI49_4774 | | YNR041C; YNR040W |
| DI49_4772; DI49_4773 | DI49_4772; DI49_4773 | | |
| DI49_4770; DI49_4771 | DI49_4770; DI49_4771 | | |
| DI49_4768; DI49_4769 | DI49_4768; DI49_4769 | | |
| DI49_4766; DI49_4767 | DI49_4766; DI49_4767 | | |
| DI49_4764; DI49_4765 | DI49_4764; DI49_4765 | | |
| DI49_4762; YNR030W; DI49_4761; YNR029C | DI49_4762; DI49_4761 | | YNR030W; YNR029C |

Create PDF in your applications with the Pdfcrowd HTML to PDF API PDFCROWD

| | | | |
|---|---|---|---|
| SPGP0J00530; DI49_4760 | DI49_4760 | SPGP0J00530 | |
| DI49_4758; DI49_4759 | DI49_4758; DI49_4759 | | |
| DI49_4756; DI49_4757; DI49_4755 | DI49_4756; DI49_4757; DI49_4755 | | |
| DI49_4753; DI49_4754 | DI49_4753; DI49_4754 | | |
| DI49_4751; YNR016C; DI49_4750; YNR015W | DI49_4751; DI49_4750 | | YNR016C; YNR015W |
| DI49_4748; YNR013C | DI49_4748 | | YNR013C |
| DI49_4747; YNR012W | DI49_4747 | | YNR012W |
| DI49_4746; YNR011C; DI49_4745; YNR010W | DI49_4746; DI49_4745 | | YNR011C; YNR010W |
| DI49_4743; DI49_4744 | DI49_4743; DI49_4744 | | |
| DI49_4741; DI49_4742 | DI49_4741; DI49_4742 | | |
| DI49_4739; DI49_4740 | DI49_4739; DI49_4740 | | |
| DI49_4735; YNR001C; DI49_4734 | DI49_4735; DI49_4734 | | YNR001C |
| DI49_4723; YNL014W; DI49_4722; DI49_3760; DI49_3759 | DI49_4723; DI49_4722; DI49_3760; DI49_3759 | | YNL014W |
| SPGP0J00970; DI49_0189; DI49_4708; DI49_0188 | DI49_0189; DI49_4708; DI49_0188 | SPGP0J00970 | |
| DI49_4710; DI49_4709 | DI49_4710; DI49_4709 | | |
| DI49_4712; DI49_4711 | DI49_4712; DI49_4711 | | |
| DI49_4715; YNL022C; DI49_4716 | DI49_4715; DI49_4716 | | YNL022C |
| SPGP0Jtrna2P; DI49_4699; DI49_4698; SPGP0AMtrna14P | DI49_4699; DI49_4698 | SPGP0Jtrna2P; SPGP0AMtrna14P | |
| DI49_4696; DI49_0162; DI49_4695; DI49_0161 | DI49_4696; DI49_0162; DI49_4695; DI49_0161 | | |
| DI49_4693; DI49_4694 | DI49_4693; DI49_4694 | | |
| DI49_4691; DI49_4692 | DI49_4691; DI49_4692 | | |
| DI49_4689; YNL041C; DI49_4688; YNL040W | DI49_4689; DI49_4688 | | YNL041C; YNL040W |
| DI49_4686; YNL044W; DI49_4685 | DI49_4686; DI49_4685 | | YNL044W |
| DI49_4683; DI49_4684 | DI49_4683; DI49_4684 | | |
| DI49_4681; YNL049C; DI49_4680 | DI49_4681; DI49_4680 | | YNL049C |
| DI49_4678; DI49_4679 | DI49_4678; DI49_4679 | | |
| DI49_4676; YNL054W; YNL053W | DI49_4676 | | YNL054W; YNL053W |
| DI49_4674; YNL055C; DI49_4673 | DI49_4674; DI49_4673 | | YNL055C |
| DI49_4671; YNL059C; YNL058C | DI49_4671 | | YNL059C; YNL058C |
| DI49_4670; YNL061W | DI49_4670 | | YNL061W |
| DI49_4669; YNL062C; DI49_4668 | DI49_4669; DI49_4668 | | YNL062C |
| DI49_4666; DI49_4667; YNL064C; YNL063W | DI49_4666; DI49_4667 | | YNL064C; YNL063W |
| DI49_4661; YNL070W | DI49_4661 | | YNL070W |
| DI49_4660; YNL071W; DI49_4659 | DI49_4660; DI49_4659 | | YNL071W |
| DI49_4657; DI49_4658 | DI49_4657; DI49_4658 | | |
| DI49_4655; DI49_4656 | DI49_4655; DI49_4656 | | |
| DI49_4653; DI49_4654 | DI49_4653; DI49_4654 | | |
| DI49_4651; YNL080C | DI49_4651 | | YNL080C |
| DI49_4650; YNL081C | DI49_4650 | | YNL081C |

Create PDF in your applications with the Pdfcrowd HTML to PDF API PDFCROWD

| | | | |
|---|---|---|---|
| DI49_4649; YNL082W | DI49_4649 | | YNL082W |
| DI49_4648; YNL083W | DI49_4648 | | YNL083W |
| DI49_4647; YNL084C | DI49_4647 | | YNL084C |
| DI49_4646; YNL085W; DI49_4645 | DI49_4646; DI49_4645 | | YNL085W |
| DI49_4643; YNL088W; DI49_4642; YNL087W | DI49_4643; DI49_4642 | | YNL088W; YNL087W |
| DI49_4640; DI49_4641; YNL091W; YNL090W | DI49_4640; DI49_4641 | | YNL091W; YNL090W |
| DI49_4638; YNL094W; DI49_4637; YNL093W | DI49_4638; DI49_4637 | | YNL094W; YNL093W |
| DI49_4635; SPGP0CXtrna7L | DI49_4635 | SPGP0CXtrna7L | |
| DI49_4634; YNL097C | DI49_4634 | | YNL097C |
| DI49_4633; YNL098C; DI49_4632 | DI49_4633; DI49_4632 | | YNL098C |
| DI49_4630; YNL101W; YNL100W | DI49_4630 | | YNL101W; YNL100W |
| DI49_4629; YNL102W; DI49_4628 | DI49_4629; DI49_4628 | | YNL102W |
| DI49_4626; YNL106C; DI49_4625 | DI49_4626; DI49_4625 | | YNL106C |
| DI49_4623; YNL110C; YNL108C | DI49_4623 | | YNL110C; YNL108C |
| DI49_4622; YNL111C | DI49_4622 | | YNL111C |
| DI49_4619; YNL116W; YNL115C | DI49_4619 | | YNL116W; YNL115C |
| DI49_4618; YNL117W | DI49_4618 | | YNL117W |
| DI49_4617; YNL118C; DI49_4616 | DI49_4617; DI49_4616 | | YNL118C |
| DI49_4614; YNL122C; YNL121C | DI49_4614 | | YNL122C; YNL121C |
| DI49_4613; YNL123W | DI49_4613 | | YNL123W |
| DI49_4612; YNL124W; DI49_4611 | DI49_4612; DI49_4611 | | YNL124W |
| DI49_4609; YNL127W | DI49_4609 | | YNL127W |
| DI49_4608; YNL128W; DI49_4607 | DI49_4608; DI49_4607 | | YNL128W |
| DI49_4605; YNL131W; YNL130C | DI49_4605 | | YNL131W; YNL130C |
| DI49_4604; YNL132W | DI49_4604 | | YNL132W |
| DI49_4603; DI49_2475; DI49_4602; SPGP0Ytrna6F; DI49_4131; DI49_2476 | DI49_4603; DI49_2475; DI49_4602; DI49_4131; DI49_2476 | SPGP0Ytrna6F | |
| DI49_4600; YNL135C; YNL134C | DI49_4600 | | YNL135C; YNL134C |
| DI49_4599; YNL136W | DI49_4599 | | YNL136W |
| DI49_4598; YNL137C | DI49_4598 | | YNL137C |
| DI49_4597; YNL138W; DI49_4596 | DI49_4597; DI49_4596 | | YNL138W |
| DI49_4594; DI49_4595 | DI49_4594; DI49_4595 | | |
| DI49_4592; DI49_4593 | DI49_4592; DI49_4593 | | |
| DI49_4590; DI49_4591 | DI49_4590; DI49_4591 | | |
| DI49_4588; DI49_4589 | DI49_4588; DI49_4589 | | |
| DI49_4586; DI49_4587 | DI49_4586; DI49_4587 | | |
| DI49_4584; DI49_4585 | DI49_4584; DI49_4585 | | |
| DI49_4582; DI49_4583; YNL154C; YNL153C | DI49_4582; DI49_4583 | | YNL154C; YNL153C |
| DI49_4580; YNL157W; DI49_4579; YNL156C | DI49_4580; DI49_4579 | | YNL157W; YNL156C |
| DI49_4577; YNL160W; DI49_4576; DI49_0410; DI49_0409; YNL159C | DI49_4577; DI49_4576; DI49_0410; DI49_0409 | | YNL160W; YNL159C |

Create PDF in your applications with the Pdfcrowd HTML to PDF API PDFCROWD

| | | | |
|---|---|---|---|
| DI49_4574; DI49_4575 | DI49_4574; DI49_4575 | | |
| DI49_4572; YNL164C; DI49_4571 | DI49_4572; DI49_4571 | | YNL164C |
| DI49_4569; DI49_4570 | DI49_4569; DI49_4570 | | |
| DI49_4567; YNL169C; YNL168C | DI49_4567 | | YNL169C; YNL168C |
| DI49_4566; YNL172W; DI49_4565 | DI49_4566; DI49_4565 | | YNL172W |
| DI49_4563; DI49_4564; YNL175C; YNL173C | DI49_4563; DI49_4564 | | YNL175C; YNL173C |
| DI49_4561; YNL178W; YNL177C | DI49_4561 | | YNL178W; YNL177C |
| DI49_4560; YNL180C; DI49_4559 | DI49_4560; DI49_4559 | | YNL180C |
| DI49_4557; YNL183C | DI49_4557 | | YNL183C |
| DI49_4556; YNL185C | DI49_4556 | | YNL185C |
| DI49_4555; YNL186W; DI49_4554 | DI49_4555; DI49_4554 | | YNL186W |
| DI49_4552; YNL189W | DI49_4552 | | YNL189W |
| DI49_4551; YNL190W | DI49_4551 | | YNL190W |
| DI49_4550; YNL191W | DI49_4550 | | YNL191W |
| DI49_4549; YNL192W; DI49_4548 | DI49_4549; DI49_4548 | | YNL192W |
| DI49_4546; DI49_4547 | DI49_4546; DI49_4547 | | |
| DI49_4544; DI49_4545 | DI49_4544; DI49_4545 | | |
| DI49_4542; DI49_4543 | DI49_4542; DI49_4543 | | |
| DI49_4540; YNL202W; DI49_4539 | DI49_4540; DI49_4539 | | YNL202W |
| DI49_4537; YNL207W; DI49_4536 | DI49_4537; DI49_4536 | | YNL207W |
| DI49_4534; DI49_4535; SPGP0U02100; YAL060W; YNL209W; YNL208W | DI49_4534; DI49_4535 | SPGP0U02100 | YAL060W; YNL209W; YNL208W |
| DI49_4532; YNL212W; DI49_4531; YNL211C | DI49_4532; DI49_4531 | | YNL212W; YNL211C |
| DI49_4529; DI49_4530 | DI49_4529; DI49_4530 | | |
| DI49_4527; DI49_4528 | DI49_4527; DI49_4528 | | |
| DI49_4525; DI49_4526 | DI49_4525; DI49_4526 | | |
| DI49_4523; DI49_4524 | DI49_4523; DI49_4524 | | |
| DI49_4521; YNL223W | DI49_4521 | | YNL223W |
| DI49_4520; YNL224C | DI49_4520 | | YNL224C |
| DI49_4519; YNL225C | DI49_4519 | | YNL225C |
| DI49_4518; YNL227C; DI49_4517 | DI49_4518; DI49_4517 | | YNL227C |
| DI49_4515; DI49_4516 | DI49_4515; DI49_4516 | | |
| DI49_4513; YNL233W; DI49_4512 | DI49_4513; DI49_4512 | | YNL233W |
| DI49_4510; DI49_4511 | DI49_4510; DI49_4511 | | |
| DI49_4508; DI49_4509 | DI49_4508; DI49_4509 | | |
| DI49_4506; YNL241C; YNL240C | DI49_4506 | | YNL241C; YNL240C |
| DI49_4505; YNL242W; DI49_4504 | DI49_4505; DI49_4504 | | YNL242W |
| DI49_4502; DI49_4503; YNL244C; YNL243W | DI49_4502; DI49_4503 | | YNL244C; YNL243W |
| DI49_4500; YNL247W; YNL246W | DI49_4500 | | YNL247W; YNL246W |
| DI49_4499; YNL248C | DI49_4499 | | YNL248C |
| DI49_4498; YNL249C | DI49_4498 | | YNL249C |

Create PDF in your applications with the Pdfcrowd HTML to PDF API PDFCROWD

| | | | |
|---|---|---|---|
| DI49_4497; YNL250W | DI49_4497 | | YNL250W |
| DI49_4496; YNL251C; DI49_4495 | DI49_4496; DI49_4495 | | YNL251C |
| DI49_4493; YNL254C; YNL253W | DI49_4493 | | YNL254C; YNL253W |
| DI49_4492; YNL255C | DI49_4492 | | YNL255C |
| DI49_4491; YNL256W; DI49_4490 | DI49_4491; DI49_4490 | | YNL256W |
| DI49_4488; DI49_4489 | DI49_4488; DI49_4489 | | |
| DI49_4486; DI49_4487 | DI49_4486; DI49_4487 | | |
| DI49_4484; YNL263C; DI49_4483 | DI49_4484; DI49_4483 | | YNL263C |
| DI49_4481; YNL267W | DI49_4481 | | YNL267W |
| DI49_4480; YNL268W; DI49_4479; YNL270C | DI49_4480; DI49_4479 | | YNL268W; YNL270C |
| DI49_4478; YNL271C; SPGP0I00690 | DI49_4478 | SPGP0I00690 | YNL271C |
| DI49_4477; YNL272C; DI49_4476 | DI49_4477; DI49_4476 | | YNL272C |
| DI49_4474; DI49_4475; YNL274C; YNL273W | DI49_4474; DI49_4475 | | YNL274C; YNL273W |
| DI49_4472; DI49_4473 | DI49_4472; DI49_4473 | | |
| DI49_4470; YNL280C; DI49_4469; YNL279W | DI49_4470; DI49_4469 | | YNL280C; YNL279W |
| DI49_4467; YNL283C; YNL282W | DI49_4467 | | YNL283C; YNL282W |
| SPGP0Jtrna3G; SPGP0Itrna24G; DI49_4461; SPGP0Vtrna11G; DI49_2296; SPGP0AFtrna15G; DI49_2107 | DI49_4461; DI49_2296; DI49_2107 | SPGP0Jtrna3G; SPGP0Itrna24G; SPGP0Vtrna11G; SPGP0AFtrna15G | |
| DI49_4459; YNL288W | DI49_4459 | | YNL288W |
| DI49_4458; YNL289W; DI49_4457 | DI49_4458; DI49_4457 | | YNL289W |
| DI49_4455; YNL292W; YNL291C | DI49_4455 | | YNL292W; YNL291C |
| DI49_4454; YNL293W; DI49_4453 | DI49_4454; DI49_4453 | | YNL293W |
| DI49_4451; DI49_4452 | DI49_4451; DI49_4452 | | |
| DI49_4449; DI49_4450; YNL298W; SPGP0I00460 | DI49_4449; DI49_4450 | SPGP0I00460 | YNL298W |
| YNL301C; YOL120C; SPGP0I00420; YNL300W | | SPGP0I00420 | YNL301C; YOL120C; YNL300W |
| DI49_4447; DI49_2286; DI49_2285; SPGP0I00410 | DI49_4447; DI49_2286; DI49_2285 | SPGP0I00410 | |
| DI49_4446; YNL304W; DI49_4445 | DI49_4446; DI49_4445 | | YNL304W |
| DI49_4443; YNL307C; DI49_4442; YNL306W | DI49_4443; DI49_4442 | | YNL307C; YNL306W |
| DI49_4440; DI49_4441 | DI49_4440; DI49_4441 | | |
| SPGP0J03620; DI49_4439 | DI49_4439 | SPGP0J03620 | |
| DI49_4436; DI49_4437 | DI49_4436; DI49_4437 | | |
| DI49_4434; DI49_4435 | DI49_4434; DI49_4435 | | |
| DI49_4432; DI49_4433 | DI49_4432; DI49_4433 | | |
| DI49_4430; YNL321W | DI49_4430 | | YNL321W |
| DI49_4429; YNL322C | DI49_4429 | | YNL322C |
| DI49_4428; YNL323W | DI49_4428 | | YNL323W |
| DI49_4427; YNL325C; DI49_4426 | DI49_4427; DI49_4426 | | YNL325C |
| DI49_4424; YNL328C; DI49_4423; YNL327W | DI49_4424; DI49_4423 | | YNL328C; YNL327W |
| DI49_4421; DI49_4422 | DI49_4421; DI49_4422 | | |
| SPGP0J03810; SPGP0J03800 | | SPGP0J03810; SPGP0J03800 | |

Create PDF in your applications with the Pdfcrowd HTML to PDF API PDFCROWD

| | | | |
|---|---|---|---|
| SPGP0DF00100; DI49_4420 | DI49_4420 | SPGP0DF00100 | |
| SPGP0DF00120; YKR105C | | SPGP0DF00120 | YKR105C |
| DI49_5638; SPGP0DF00130 | DI49_5638 | SPGP0DF00130 | |
| DI49_5636; DI49_5637 | DI49_5636; DI49_5637 | | |
| DI49_5634; DI49_5635 | DI49_5634; DI49_5635 | | |
| DI49_5632; DI49_5633 | DI49_5632; DI49_5633 | | |
| DI49_5630; DI49_5631 | DI49_5630; DI49_5631 | | |
| DI49_5628; DI49_5629 | DI49_5628; DI49_5629 | | |
| SPGP0DJ00100; YER190W; DI49_2253 | DI49_2253 | SPGP0DJ00100 | YER190W |
| DI49_1489; DI49_2619 | DI49_1489; DI49_2619 | | |
| DI49_1491; DI49_1490 | DI49_1491; DI49_1490 | | |
| DI49_1493; DI49_1492 | DI49_1493; DI49_1492 | | |
| DI49_1496; DI49_1495 | DI49_1496; DI49_1495 | | |
| DI49_1498; DI49_1497 | DI49_1498; DI49_1497 | | |
| SPGP0K00200; DI49_1499 | DI49_1499 | SPGP0K00200 | |
| DI49_1501; DI49_1500 | DI49_1501; DI49_1500 | | |
| DI49_1506; DI49_1505 | DI49_1506; DI49_1505 | | |
| DI49_1508; DI49_0112; DI49_0111 | DI49_1508; DI49_0112; DI49_0111 | | |
| DI49_1509; YAL005C; DI49_1510 | DI49_1509; DI49_1510 | | YAL005C |
| DI49_1512; DI49_1511 | DI49_1512; DI49_1511 | | |
| DI49_1514; DI49_1513 | DI49_1514; DI49_1513 | | |
| DI49_1516; DI49_1515 | DI49_1516; DI49_1515 | | |
| DI49_1518; DI49_1517 | DI49_1518; DI49_1517 | | |
| DI49_1520; DI49_1519 | DI49_1520; DI49_1519 | | |
| DI49_1522; DI49_0101; DI49_1523; DI49_0100 | DI49_1522; DI49_0101; DI49_1523; DI49_0100 | | |
| DI49_1525; DI49_1524 | DI49_1525; DI49_1524 | | |
| DI49_1527; DI49_1526 | DI49_1527; DI49_1526 | | |
| DI49_1529; DI49_1528 | DI49_1529; DI49_1528 | | |
| DI49_1531; DI49_1530 | DI49_1531; DI49_1530 | | |
| DI49_1533; DI49_1532 | DI49_1533; DI49_1532 | | |
| DI49_1547; DI49_1546 | DI49_1547; DI49_1546 | | |
| DI49_1549; DI49_1548 | DI49_1549; DI49_1548 | | |
| DI49_1551; DI49_1550 | DI49_1551; DI49_1550 | | |
| DI49_4046; DI49_1552 | DI49_4046; DI49_1552 | | |
| DI49_4044; DI49_4045 | DI49_4044; DI49_4045 | | |
| DI49_4805; DI49_4804 | DI49_4805; DI49_4804 | | |
| DI49_3294; SPGP0DR00120 | DI49_3294 | SPGP0DR00120 | |
| DI49_3296; DI49_3295 | DI49_3296; DI49_3295 | | |
| DI49_3298; DI49_3297 | DI49_3298; DI49_3297 | | |
| DI49_3300; DI49_3299 | DI49_3300; DI49_3299 | | |

Create PDF in your applications with the Pdfcrowd HTML to PDF API PDFCROWD

| | | | |
|---|---|---|---|
| YER174C; YER175C | | | YER174C; YER175C |
| YER172C; YER173W | | | YER172C; YER173W |
| YER170W; YER171W | | | YER170W; YER171W |
| YER168C; YER169W | | | YER168C; YER169W |
| DI49_5452; YBR299W | DI49_5452 | | YBR299W |
| DI49_5450; DI49_5451 | DI49_5450; DI49_5451 | | |
| DI49_5448; DI49_5449 | DI49_5448; DI49_5449 | | |
| DI49_5446; DI49_5447 | DI49_5446; DI49_5447 | | |
| DI49_5444; DI49_5445 | DI49_5444; DI49_5445 | | |
| DI49_5442; DI49_5443 | DI49_5442; DI49_5443 | | |
| SPGP0L00220; DI49_5441 | DI49_5441 | SPGP0L00220 | |
| DI49_5439; DI49_5440 | DI49_5439; DI49_5440 | | |
| DI49_5437; DI49_1649; DI49_5436; DI49_1648 | DI49_5437; DI49_1649; DI49_5436; DI49_1648 | | |
| DI49_5434; DI49_5435 | DI49_5434; DI49_5435 | | |
| DI49_5432; DI49_5433 | DI49_5432; DI49_5433 | | |
| DI49_5430; DI49_5431 | DI49_5430; DI49_5431 | | |
| DI49_5428; DI49_5429 | DI49_5428; DI49_5429 | | |
| DI49_5426; DI49_5427 | DI49_5426; DI49_5427 | | |
| DI49_5424; DI49_5425 | DI49_5424; DI49_5425 | | |
| DI49_5419; DI49_5420 | DI49_5419; DI49_5420 | | |
| DI49_5417; DI49_5418 | DI49_5417; DI49_5418 | | |
| SPGP0Ltrna3A; DI49_5601; DI49_5414; SPGP0Atrna1A; YJL115W; YKL117W; DI49_1030; SPGP0Ytrna8A; DI49_4112; SPGP0JB00150; SPGP0AFtrna12A; SPGP0AJtrna14A; YHR021C; SPGP0ATtrna15A; SPGP0Etrna16A; YKR042W; SPGP0Ftrna17A; DI49_2906; SPGP0Htrna18A; DI49_3259; DI49_3766 | DI49_5601; DI49_5414; DI49_1030; DI49_4112; DI49_2906; DI49_3259; DI49_3766 | SPGP0Ltrna3A; SPGP0Atrna1A; SPGP0Ytrna8A; SPGP0JB00150; SPGP0AFtrna12A; SPGP0AJtrna14A; SPGP0ATtrna15A; SPGP0Etrna16A; SPGP0Ftrna17A; SPGP0Htrna18A | YJL115W; YKL117W; YHR021C; YKR042W |
| DI49_5412; DI49_5413 | DI49_5412; DI49_5413 | | |
| DI49_5410; DI49_5411 | DI49_5410; DI49_5411 | | |
| DI49_5408; DI49_5409 | DI49_5408; DI49_5409 | | |
| SPGP0L00560; DI49_5407 | DI49_5407 | SPGP0L00560 | |
| DI49_5405; DI49_5406 | DI49_5405; DI49_5406 | | |
| DI49_5403; DI49_5404 | DI49_5403; DI49_5404 | | |
| DI49_5401; DI49_5402 | DI49_5401; DI49_5402 | | |
| DI49_5399; DI49_5400; SPGP0AFtrna2C; DI49_2071; DI49_5567; DI49_5568; DI49_1233 | DI49_5399; DI49_5400; DI49_2071; DI49_5567; DI49_5568; DI49_1233 | SPGP0AFtrna2C | |
| DI49_5397; DI49_5398 | DI49_5397; DI49_5398 | | |
| DI49_5395; DI49_5396 | DI49_5395; DI49_5396 | | |
| DI49_5393; DI49_5394 | DI49_5393; DI49_5394 | | |
| DI49_5391; DI49_5392 | DI49_5391; DI49_5392 | | |
| DI49_5389; DI49_5390 | DI49_5389; DI49_5390 | | |
| DI49_5387; DI49_5388 | DI49_5387; DI49_5388 | | |
| DI49_5385; DI49_5386 | DI49_5385; DI49_5386 | | |

Create PDF in your applications with the Pdfcrowd HTML to PDF API PDFCROWD

| | | | |
|---|---|---|---|
| DI49_5383; DI49_5384 | DI49_5383; DI49_5384 | | |
| DI49_5381; DI49_1215; DI49_5380; DI49_1214 | DI49_5381; DI49_1215; DI49_5380; DI49_1214 | | |
| DI49_5378; DI49_5379 | DI49_5378; DI49_5379 | | |
| DI49_5376; DI49_5377 | DI49_5376; DI49_5377 | | |
| DI49_5374; DI49_5375 | DI49_5374; DI49_5375 | | |
| DI49_5371; DI49_5373 | DI49_5371; DI49_5373 | | |
| DI49_5369; DI49_5370 | DI49_5369; DI49_5370 | | |
| DI49_5367; DI49_5368 | DI49_5367; DI49_5368 | | |
| DI49_5365; DI49_5366 | DI49_5365; DI49_5366 | | |
| DI49_5363; DI49_5364 | DI49_5363; DI49_5364 | | |
| DI49_5361; DI49_5362 | DI49_5361; DI49_5362 | | |
| DI49_5359; DI49_5360 | DI49_5359; DI49_5360 | | |
| DI49_5357; DI49_5358 | DI49_5357; DI49_5358 | | |
| DI49_5355; DI49_5356 | DI49_5355; DI49_5356 | | |
| SPGP0L01080; DI49_5354 | DI49_5354 | SPGP0L01080 | |
| DI49_5352; YPL109C | DI49_5352 | | YPL109C |
| DI49_5348; DI49_5349 | DI49_5348; DI49_5349 | | |
| DI49_5346; DI49_5347 | DI49_5346; DI49_5347 | | |
| DI49_5344; DI49_5345 | DI49_5344; DI49_5345 | | |
| DI49_5342; DI49_5343 | DI49_5342; DI49_5343 | | |
| DI49_5340; DI49_5341 | DI49_5340; DI49_5341 | | |
| DI49_5338; DI49_5339 | DI49_5338; DI49_5339 | | |
| DI49_5336; DI49_5337 | DI49_5336; DI49_5337 | | |
| DI49_5334; DI49_5335 | DI49_5334; DI49_5335 | | |
| DI49_5332; DI49_5333 | DI49_5332; DI49_5333 | | |
| DI49_5330; DI49_5331 | DI49_5330; DI49_5331 | | |
| DI49_5328; DI49_5329 | DI49_5328; DI49_5329 | | |
| DI49_5326; DI49_5327 | DI49_5326; DI49_5327 | | |
| DI49_5324; DI49_5325 | DI49_5324; DI49_5325 | | |
| DI49_5319; DI49_5320 | DI49_5319; DI49_5320 | | |
| DI49_5317; DI49_5318 | DI49_5317; DI49_5318 | | |
| DI49_5315; DI49_5316 | DI49_5315; DI49_5316 | | |
| DI49_5313; DI49_5314 | DI49_5313; DI49_5314 | | |
| DI49_5311; DI49_5312 | DI49_5311; DI49_5312 | | |
| DI49_5309; DI49_5310 | DI49_5309; DI49_5310 | | |
| DI49_5307; DI49_5308 | DI49_5307; DI49_5308 | | |
| DI49_5305; DI49_5306 | DI49_5305; DI49_5306 | | |
| DI49_5303; DI49_5304 | DI49_5303; DI49_5304 | | |
| DI49_5301; DI49_5302 | DI49_5301; DI49_5302 | | |
| DI49_5299; DI49_5300 | DI49_5299; DI49_5300 | | |

Create PDF in your applications with the Pdfcrowd HTML to PDF API PDFCROWD

| | | | |
|---|---|---|---|
| DI49_5297; DI49_5298 | DI49_5297; DI49_5298 | | |
| DI49_5295; DI49_5296 | DI49_5295; DI49_5296 | | |
| DI49_5293; DI49_5294 | DI49_5293; DI49_5294 | | |
| DI49_5292; SPGP0L01700 | DI49_5292 | SPGP0L01700 | |
| DI49_5290; DI49_5291 | DI49_5290; DI49_5291 | | |
| DI49_5288; DI49_5289 | DI49_5288; DI49_5289 | | |
| DI49_5284; DI49_5285 | DI49_5284; DI49_5285 | | |
| DI49_5282; DI49_5283 | DI49_5282; DI49_5283 | | |
| DI49_5280; DI49_5281 | DI49_5280; DI49_5281 | | |
| DI49_5278; DI49_5279 | DI49_5278; DI49_5279 | | |
| DI49_5276; DI49_5277 | DI49_5276; DI49_5277 | | |
| DI49_5274; DI49_5275 | DI49_5274; DI49_5275 | | |
| DI49_5272; DI49_5273 | DI49_5272; DI49_5273 | | |
| YPL195W; DI49_5271 | DI49_5271 | | YPL195W |
| DI49_5268; DI49_5270 | DI49_5268; DI49_5270 | | |
| DI49_5266; DI49_5267 | DI49_5266; DI49_5267 | | |
| DI49_5264; DI49_5265 | DI49_5264; DI49_5265 | | |
| DI49_5262; DI49_5263 | DI49_5262; DI49_5263 | | |
| DI49_5260; DI49_5261 | DI49_5260; DI49_5261 | | |
| DI49_5258; DI49_5259 | DI49_5258; DI49_5259 | | |
| DI49_5256; DI49_5257 | DI49_5256; DI49_5257 | | |
| DI49_5254; DI49_5255 | DI49_5254; DI49_5255 | | |
| DI49_5252; DI49_5253 | DI49_5252; DI49_5253 | | |
| YPL218W; DI49_5251 | DI49_5251 | | YPL218W |
| DI49_5245; DI49_5246 | DI49_5245; DI49_5246 | | |
| DI49_5243; DI49_5244 | DI49_5243; DI49_5244 | | |
| DI49_5241; DI49_5242 | DI49_5241; DI49_5242 | | |
| DI49_5239; DI49_5240 | DI49_5239; DI49_5240 | | |
| DI49_5237; DI49_5238 | DI49_5237; DI49_5238 | | |
| DI49_5235; DI49_5236 | DI49_5235; DI49_5236 | | |
| DI49_5233; DI49_5234 | DI49_5233; DI49_5234 | | |
| DI49_5231; DI49_5232 | DI49_5231; DI49_5232 | | |
| DI49_5229; DI49_4269; DI49_5228; DI49_4270 | DI49_5229; DI49_4269; DI49_5228; DI49_4270 | | |
| DI49_5226; DI49_5227 | DI49_5226; DI49_5227 | | |
| DI49_5224; DI49_5225 | DI49_5224; DI49_5225 | | |
| DI49_5222; DI49_5223 | DI49_5222; DI49_5223 | | |
| DI49_5220; DI49_5221 | DI49_5220; DI49_5221 | | |
| DI49_5218; DI49_5219 | DI49_5218; DI49_5219 | | |
| DI49_5216; DI49_5217 | DI49_5216; DI49_5217 | | |
| DI49_5214; DI49_5215 | DI49_5214; DI49_5215 | | |

Create PDF in your applications with the Pdfcrowd HTML to PDF API PDFCROWD

| | | | |
|---|---|---|---|
| DI49_5212; DI49_2348; SPGP0V00500 | DI49_5212; DI49_2348 | SPGP0V00500 | |
| SPGP0Ltrna3W; DI49_5211; DI49_5210; SPGP0GZtrna4W; DI49_1569; DI49_1311 | DI49_5211; DI49_5210; DI49_1569; DI49_1311 | SPGP0Ltrna3W; SPGP0GZtrna4W | |
| DI49_5208; DI49_5209 | DI49_5208; DI49_5209 | | |
| DI49_5206; DI49_5207 | DI49_5206; DI49_5207 | | |
| DI49_5204; DI49_5205 | DI49_5204; DI49_5205 | | |
| DI49_5202; DI49_5203 | DI49_5202; DI49_5203 | | |
| DI49_5201; SPGP0L02610 | DI49_5201 | SPGP0L02610 | |
| DI49_5199; DI49_5200 | DI49_5199; DI49_5200 | | |
| DI49_5197; YPL274W; DI49_5196; YPL273W | DI49_5197; DI49_5196 | | YPL274W; YPL273W |
| DI49_2242; YDR542W | DI49_2242 | | YDR542W |
| DI49_2244; DI49_2243 | DI49_2244; DI49_2243 | | |
| DI49_2246; DI49_2245 | DI49_2246; DI49_2245 | | |
| SPGP0EA00160; YJL221C; DI49_1318; SPGP0IF00120; DI49_0718 | DI49_1318; DI49_0718 | SPGP0EA00160; SPGP0IF00120 | YJL221C |
| DI49_1316; DI49_1317 | DI49_1316; DI49_1317 | | |
| DI49_1314; DI49_1315 | DI49_1314; DI49_1315 | | |
| DI49_1312; DI49_1313 | DI49_1312; DI49_1313 | | |
| DI49_0094; DI49_2609 | DI49_0094; DI49_2609 | | |
| DI49_0092; DI49_0093 | DI49_0092; DI49_0093 | | |
| DI49_0090; DI49_0091 | DI49_0090; DI49_0091 | | |
| YAL067C; DI49_3158; SPGP0EH00100; DI49_3159 | DI49_3158; DI49_3159 | SPGP0EH00100 | YAL067C |
| DI49_0696; DI49_0789 | DI49_0696; DI49_0789 | | |
| DI49_2834; YJL198W; YJL199C; SPGP0AQ00110; DI49_0788 | DI49_2834; DI49_0788 | SPGP0AQ00110 | YJL198W; YJL199C |
| DI49_2832; YJL200C | DI49_2832 | | YJL200C |
| DI49_2831; YJL201W; DI49_2830; YJL203W | DI49_2831; DI49_2830 | | YJL201W; YJL203W |
| SPGP0M00150; DI49_2829 | DI49_2829 | SPGP0M00150 | |
| DI49_2828; YJL207C; DI49_2827; YJL208C | DI49_2828; DI49_2827 | | YJL207C; YJL208C |
| DI49_2825; DI49_2826 | DI49_2825; DI49_2826 | | |
| SPGP0EJ00100; SPGP0AB00100; YGL241W; SPGP0BD00100; DI49_0740 | DI49_0740 | SPGP0EJ00100; SPGP0AB00100; SPGP0BD00100 | YGL241W |
| YGL243W; YGL242C | | | YGL243W; YGL242C |
| YGL245W; YGL244W | | | YGL245W; YGL244W |
| YGL247W; YGL246C | | | YGL247W; YGL246C |
| YGL249W; YGL248W | | | YGL249W; YGL248W |
| YGL251C; YGL250W | | | YGL251C; YGL250W |
| YGL256W; YGL255W | | | YGL256W; YGL255W |
| DI49_1049; DI49_1048 | DI49_1049; DI49_1048 | | |
| DI49_1051; DI49_1050 | DI49_1051; DI49_1050 | | |
| DI49_1053; DI49_1052 | DI49_1053; DI49_1052 | | |
| DI49_2835; DI49_1054 | DI49_2835; DI49_1054 | | |

Create PDF in your applications with the Pdfcrowd HTML to PDF API PDFCROWD

| | | | |
|---|---|---|---|
| DI49_2837; DI49_2836 | DI49_2837; DI49_2836 | | |
| DI49_2839; DI49_2838 | DI49_2839; DI49_2838 | | |
| DI49_2841; DI49_2840 | DI49_2841; DI49_2840 | | |
| DI49_2843; DI49_2842 | DI49_2843; DI49_2842 | | |
| DI49_2845; DI49_2844 | DI49_2845; DI49_2844 | | |
| DI49_2847; DI49_2846 | DI49_2847; DI49_2846 | | |
| DI49_2849; DI49_2848 | DI49_2849; DI49_2848 | | |
| DI49_2851; DI49_2850 | DI49_2851; DI49_2850 | | |
| DI49_2853; DI49_2852 | DI49_2853; DI49_2852 | | |
| DI49_2855; DI49_2854 | DI49_2855; DI49_2854 | | |
| DI49_2857; DI49_2856 | DI49_2857; DI49_2856 | | |
| DI49_2859; DI49_2858 | DI49_2859; DI49_2858 | | |
| DI49_2861; DI49_2860 | DI49_2861; DI49_2860 | | |
| DI49_2863; DI49_2862 | DI49_2863; DI49_2862 | | |
| DI49_2865; DI49_2864 | DI49_2865; DI49_2864 | | |
| SPGP0EO00430; YKL164C; DI49_1479; YKL163W; SPGP0T00450 | DI49_1479 | SPGP0EO00430; SPGP0T00450 | YKL164C; YKL163W |
| DI49_1484; DI49_1483 | DI49_1484; DI49_1483 | | |
| DI49_1486; DI49_2710; DI49_1487; DI49_2711 | DI49_1486; DI49_2710; DI49_1487; DI49_2711 | | |
| SPGP0ER00100; YGR289C | | SPGP0ER00100 | YGR289C |
| SPGP0EStrna7R; DI49_2410; DI49_1572; DI49_3638; DI49_2411; SPGP0Etrna17R; YKR027W; DI49_0285 | DI49_2410; DI49_1572; DI49_3638; DI49_2411; DI49_0285 | SPGP0EStrna7R; SPGP0Etrna17R | YKR027W |
| SPGP0ES00120; DI49_1573 | DI49_1573 | SPGP0ES00120 | |
| YER165W; YER164W | | | YER165W; YER164W |
| YER167W; YER166W | | | YER167W; YER166W |
| DI49_0741; SPGP0EU00100 | DI49_0741 | SPGP0EU00100 | |
| DI49_0079; SPGP0EW00100 | DI49_0079 | SPGP0EW00100 | |
| DI49_5679; YEL071W; SPGP0FA00100; YEL072W | DI49_5679 | SPGP0FA00100 | YEL071W; YEL072W |
| YML045W-A; SPGP0FB00100 | | SPGP0FB00100 | YML045W-A |
| DI49_2589; DI49_5686; SPGP0Ytrna11L; DI49_4208; DI49_2147; DI49_2148; DI49_0670; SPGP0Htrna22L; DI49_3381; SPGP0CUtrna29L; DI49_3756 | DI49_2589; DI49_5686; DI49_4208; DI49_2147; DI49_2148; DI49_0670; DI49_3381; DI49_3756 | SPGP0Ytrna11L; SPGP0Htrna22L; SPGP0CUtrna29L | |
| DI49_2591; DI49_2590 | DI49_2591; DI49_2590 | | |
| DI49_2593; DI49_2592 | DI49_2593; DI49_2592 | | |
| DI49_2595; DI49_2594 | DI49_2595; DI49_2594 | | |
| DI49_2597; DI49_2596 | DI49_2597; DI49_2596 | | |
| DI49_2599; DI49_2598 | DI49_2599; DI49_2598 | | |
| DI49_2601; DI49_2600 | DI49_2601; DI49_2600 | | |
| DI49_0779; DI49_2421; DI49_4357; DI49_4358; DI49_3679; SPGP0AJtrna8Q; DI49_2422; DI49_0359; DI49_0293; SPGP0CPtrna11Q; SPGP0CP02330 | DI49_0779; DI49_2421; DI49_4357; DI49_4358; DI49_3679; DI49_2422; DI49_0359; DI49_0293 | SPGP0AJtrna8Q; SPGP0CPtrna11Q; SPGP0CP02330 | |
| DI49_0777; DI49_0778 | DI49_0777; DI49_0778 | | |
| DI49_0775; DI49_0776 | DI49_0775; DI49_0776 | | |

Create PDF in your applications with the Pdfcrowd HTML to PDF API PDFCROWD

| | | | |
|---|---|---|---|
| DI49_0773; DI49_0774 | DI49_0773; DI49_0774 | | |
| DI49_0771; DI49_0772 | DI49_0771; DI49_0772 | | |
| SPGP0FH00100; SPGP0C00100; SPGP0FI00100; YGR294W; DI49_2391; SPGP0AI00320; DI49_2403 | DI49_2391; DI49_2403 | SPGP0FH00100; SPGP0C00100; SPGP0FI00100; SPGP0AI00320 | YGR294W |
| SPGP0FI00120; SPGP0FI00110 | | SPGP0FI00120; SPGP0FI00110 | |
| SPGP0FI00140; SPGP0FI00130 | | SPGP0FI00140; SPGP0FI00130 | |
| DI49_0688; DI49_0689 | DI49_0688; DI49_0689 | | |
| YGL225W; SPGP0FJ00100 | | SPGP0FJ00100 | YGL225W |
| YGL223C; YGL224C | | | YGL223C; YGL224C |
| SPGP0FM00100; DI49_1553 | DI49_1553 | SPGP0FM00100 | |
| SPGP0FO00100; DI49_3025; YJR009C | DI49_3025 | SPGP0FO00100 | YJR009C |
| DI49_0837; DI49_0836 | DI49_0837; DI49_0836 | | |
| DI49_0714; DI49_0715 | DI49_0714; DI49_0715 | | |
| DI49_0712; DI49_0713 | DI49_0712; DI49_0713 | | |
| DI49_0710; DI49_0711 | DI49_0710; DI49_0711 | | |
| DI49_0708; DI49_0709 | DI49_0708; DI49_0709 | | |
| DI49_0706; DI49_0707 | DI49_0706; DI49_0707 | | |
| DI49_0704; DI49_0705 | DI49_0704; DI49_0705 | | |
| DI49_0702; DI49_0703 | DI49_0702; DI49_0703 | | |
| DI49_0700; DI49_0701 | DI49_0700; DI49_0701 | | |
| DI49_0699; SPGP0FP00260 | DI49_0699 | SPGP0FP00260 | |
| YBR297W; SPGP0FP00280 | | SPGP0FP00280 | YBR297W |
| SPGP0FR00100; YBR298C | | SPGP0FR00100 | YBR298C |
| DI49_1319; DI49_1320; YJL217W; YJL218W | DI49_1319; DI49_1320 | | YJL217W; YJL218W |
| YER184C; SPGP0FW00100 | | SPGP0FW00100 | YER184C |
| YER186C; YER185W | | | YER186C; YER185W |
| SPGP0FY00140; YER188W; SPGP0GA00100; SPGP0GL00110 | | SPGP0FY00140; SPGP0GA00100; SPGP0GL00110 | YER188W |
| DI49_0766; DI49_0767 | DI49_0766; DI49_0767 | | |
| DI49_0762; DI49_0763 | DI49_0762; DI49_0763 | | |
| DI49_0760; DI49_0761 | DI49_0760; DI49_0761 | | |
| DI49_0758; DI49_0759 | DI49_0758; DI49_0759 | | |
| YOR388C; DI49_0756 | DI49_0756 | | YOR388C |
| DI49_0790; DI49_0791 | DI49_0790; DI49_0791 | | |
| YBR012C; YCR097W | | | YBR012C; YCR097W |
| DI49_3950; DI49_3951 | DI49_3950; DI49_3951 | | |
| DI49_3945; DI49_3946 | DI49_3945; DI49_3946 | | |
| DI49_3943; DI49_4014; DI49_3942; DI49_4015 | DI49_3943; DI49_4014; DI49_3942; DI49_4015 | | |
| DI49_3940; DI49_3941; DI49_4017; DI49_4018 | DI49_3940; DI49_3941; DI49_4017; DI49_4018 | | |
| SPGP0GG00260; DI49_3939 | DI49_3939 | SPGP0GG00260 | |
| SPGP0GG00280; DI49_3938 | DI49_3938 | SPGP0GG00280 | |
| DI49_3936; DI49_3937 | DI49_3936; DI49_3937 | | |

Create PDF in your applications with the Pdfcrowd HTML to PDF API PDFCROWD

| | | | |
|---|---|---|---|
| DI49_3934; DI49_3935 | DI49_3934; DI49_3935 | | |
| DI49_3932; DI49_3933 | DI49_3932; DI49_3933 | | |
| DI49_3972; DI49_3971 | DI49_3972; DI49_3971 | | |
| DI49_3974; DI49_3973 | DI49_3974; DI49_3973 | | |
| DI49_3976; DI49_3975 | DI49_3976; DI49_3975 | | |
| DI49_3978; DI49_3977 | DI49_3978; DI49_3977 | | |
| DI49_3980; DI49_3979 | DI49_3980; DI49_3979 | | |
| DI49_3982; DI49_3981 | DI49_3982; DI49_3981 | | |
| DI49_3984; DI49_3983 | DI49_3984; DI49_3983 | | |
| DI49_3986; DI49_3985 | DI49_3986; DI49_3985 | | |
| DI49_3988; DI49_3987 | DI49_3988; DI49_3987 | | |
| DI49_3990; DI49_3989 | DI49_3990; DI49_3989 | | |
| DI49_3992; DI49_3991 | DI49_3992; DI49_3991 | | |
| DI49_3993; SPGP0R00320 | DI49_3993 | SPGP0R00320 | |
| DI49_3995; DI49_3994 | DI49_3995; DI49_3994 | | |
| DI49_3997; DI49_3996 | DI49_3997; DI49_3996 | | |
| DI49_3999; DI49_3998 | DI49_3999; DI49_3998 | | |
| DI49_4001; DI49_4000 | DI49_4001; DI49_4000 | | |
| DI49_4003; DI49_4002 | DI49_4003; DI49_4002 | | |
| DI49_4005; DI49_3969; DI49_4006; DI49_3970 | DI49_4005; DI49_3969; DI49_4006; DI49_3970 | | |
| DI49_4008; DI49_4007 | DI49_4008; DI49_4007 | | |
| DI49_4010; DI49_4009 | DI49_4010; DI49_4009 | | |
| YML078W; DI49_4011 | DI49_4011 | | YML078W |
| DI49_4013; DI49_4012 | DI49_4013; DI49_4012 | | |
| SPGP0Rtrna8R; SPGP0Btrna9R; DI49_4021; SPGP0Atrna2R; SPGP0Atrna3R; SPGP0Ttrna10R | DI49_4021 | SPGP0Rtrna8R; SPGP0Btrna9R; SPGP0Atrna2R; SPGP0Atrna3R; SPGP0Ttrna10R | |
| DI49_4023; DI49_4022 | DI49_4023; DI49_4022 | | |
| DI49_4025; DI49_4024 | DI49_4025; DI49_4024 | | |
| DI49_4027; DI49_4026 | DI49_4027; DI49_4026 | | |
| DI49_4029; DI49_4028 | DI49_4029; DI49_4028 | | |
| DI49_4031; DI49_4030 | DI49_4031; DI49_4030 | | |
| DI49_4033; DI49_4032 | DI49_4033; DI49_4032 | | |
| DI49_4035; DI49_4034 | DI49_4035; DI49_4034 | | |
| DI49_4036; SPGP0R00760 | DI49_4036 | SPGP0R00760 | |
| SPGP0Rtrna3Y; DI49_2382; DI49_4039; SPGP0Vtrna4Y; DI49_2381; DI49_4371; DI49_1687; DI49_1668; SPGP0Ftrna8Y; DI49_3070; DI49_0422 | DI49_2382; DI49_4039; DI49_2381; DI49_4371; DI49_1687; DI49_1668; DI49_3070; DI49_0422 | SPGP0Rtrna3Y; SPGP0Vtrna4Y; SPGP0Ftrna8Y | |
| DI49_4041; DI49_4040 | DI49_4041; DI49_4040 | | |
| DI49_4043; DI49_4042 | DI49_4043; DI49_4042 | | |
| DI49_1555; DI49_1554 | DI49_1555; DI49_1554 | | |
| DI49_1557; DI49_1556 | DI49_1557; DI49_1556 | | |

Create PDF in your applications with the Pdfcrowd HTML to PDF API PDFCROWD

| | | | |
|---|---|---|---|
| DI49_1562; DI49_1561 | DI49_1562; DI49_1561 | | |
| DI49_1564; DI49_1563 | DI49_1564; DI49_1563 | | |
| DI49_0752; DI49_0753 | DI49_0752; DI49_0753 | | |
| DI49_0750; DI49_0751 | DI49_0750; DI49_0751 | | |
| DI49_0748; DI49_0749 | DI49_0748; DI49_0749 | | |
| DI49_0746; DI49_0747 | DI49_0746; DI49_0747 | | |
| DI49_0744; SPGP0GJ00190 | DI49_0744 | SPGP0GJ00190 | |
| DI49_0742; DI49_0743 | DI49_0742; DI49_0743 | | |
| DI49_3168; DI49_3167 | DI49_3168; DI49_3167 | | |
| YKL215C; DI49_3169 | DI49_3169 | | YKL215C |
| DI49_3171; DI49_3170 | DI49_3171; DI49_3170 | | |
| DI49_3173; DI49_3172 | DI49_3173; DI49_3172 | | |
| DI49_3175; DI49_3174 | DI49_3175; DI49_3174 | | |
| DI49_0795; DI49_3176 | DI49_0795; DI49_3176 | | |
| DI49_0793; DI49_0794 | DI49_0793; DI49_0794 | | |
| DI49_5183; DI49_0792 | DI49_5183; DI49_0792 | | |
| DI49_5181; DI49_5182 | DI49_5181; DI49_5182 | | |
| DI49_5179; DI49_5180 | DI49_5179; DI49_5180 | | |
| DI49_5177; DI49_5178 | DI49_5177; DI49_5178 | | |
| DI49_0011; YAL051W; DI49_0012; YAL049C | DI49_0011; DI49_0012 | | YAL051W; YAL049C |
| DI49_0014; DI49_0013 | DI49_0014; DI49_0013 | | |
| DI49_0016; DI49_0015 | DI49_0016; DI49_0015 | | |
| DI49_0018; YAL043C | DI49_0018 | | YAL043C |
| DI49_0019; YAL042W | DI49_0019 | | YAL042W |
| DI49_0020; YAL041W | DI49_0020 | | YAL041W |
| DI49_0021; YAL040C; DI49_0022; YAL039C | DI49_0021; DI49_0022 | | YAL040C; YAL039C |
| DI49_0024; DI49_0023 | DI49_0024; DI49_0023 | | |
| DI49_0026; YAL035W | DI49_0026 | | YAL035W |
| DI49_0027; YAL034W-A | DI49_0027 | | YAL034W-A |
| DI49_0028; YAL034C | DI49_0028 | | YAL034C |
| DI49_0029; YAL033W; DI49_0030; YAL032C | DI49_0029; DI49_0030 | | YAL033W; YAL032C |
| SPGP0S00320; DI49_0031 | DI49_0031 | SPGP0S00320 | |
| DI49_0033; DI49_0032 | DI49_0033; DI49_0032 | | |
| DI49_0035; YAL026C; DI49_0036; YAL025C | DI49_0035; DI49_0036 | | YAL026C; YAL025C |
| DI49_0038; YAL023C | DI49_0038 | | YAL023C |
| DI49_0039; YAL022C | DI49_0039 | | YAL022C |
| DI49_0040; YAL021C | DI49_0040 | | YAL021C |
| DI49_0041; YAL020C | DI49_0041 | | YAL020C |
| DI49_0042; YAL019W; DI49_0043; YAL018C | DI49_0042; DI49_0043 | | YAL019W; YAL018C |
| DI49_0045; YAL016W; DI49_0046; YAL015C | DI49_0045; DI49_0046 | | YAL016W; YAL015C |

Create PDF in your applications with the Pdfcrowd HTML to PDF API PDFCROWD

| | | | |
|---|---|---|---|
| SPGP0S00490; YAL013W | | SPGP0S00490 | YAL013W |
| DI49_0048; YAL012W | DI49_0048 | | YAL012W |
| DI49_0049; YAL011W; DI49_0050; YAL010C | DI49_0049; DI49_0050 | | YAL011W; YAL010C |
| DI49_0052; DI49_0051 | DI49_0052; DI49_0051 | | |
| DI49_0055; DI49_3507 | DI49_0055; DI49_3507 | | |
| DI49_0056; YAL003W; DI49_0057; YAL002W | DI49_0056; DI49_0057 | | YAL003W; YAL002W |
| DI49_0059; DI49_0058 | DI49_0059; DI49_0058 | | |
| DI49_0061; YAR003W | DI49_0061 | | YAR003W |
| DI49_0062; YAR007C | DI49_0062 | | YAR007C |
| DI49_0063; YAR008W; DI49_0064 | DI49_0063; DI49_0064 | | YAR008W |
| DI49_0066; DI49_0065 | DI49_0066; DI49_0065 | | |
| DI49_0068; YAR014C | DI49_0068 | | YAR014C |
| DI49_0069; YAR015W; DI49_0070; YAR018C | DI49_0069; DI49_0070 | | YAR015W; YAR018C |
| DI49_0072; DI49_0071 | DI49_0072; DI49_0071 | | |
| SPGP0S00740; SPGP0S00730 | | SPGP0S00740; SPGP0S00730 | |
| DI49_0075; SPGP0AE00390 | DI49_0075 | SPGP0AE00390 | |
| SPGP0S00770; YAR035W | | SPGP0S00770 | YAR035W |
| DI49_0076; YAR042W; SPGP0CR00800 | DI49_0076 | SPGP0CR00800 | YAR042W |
| DI49_0077; SPGP0A00100; SPGP0S00800; YJR150C; DI49_3472; SPGP0H00110 | DI49_0077; DI49_3472 | SPGP0A00100; SPGP0S00800; SPGP0H00110 | YJR150C |
| YJR148W; YJR149W | | | YJR148W; YJR149W |
| YJR145C; DI49_2608; YJR144W; SPGP0F02840 | DI49_2608 | SPGP0F02840 | YJR145C; YJR144W |
| YJR142W; YJR143C | | | YJR142W; YJR143C |
| YJR140C; YJR141W | | | YJR140C; YJR141W |
| YJR138W; DI49_3147 | DI49_3147 | | YJR138W |
| YJR137C; DI49_3146 | DI49_3146 | | YJR137C |
| YJR136C; DI49_3145; YJR135W-A | DI49_3145 | | YJR136C; YJR135W-A |
| YJR134C; DI49_3142; DI49_3143 | DI49_3142; DI49_3143 | | YJR134C |
| YJR133W; DI49_3141 | DI49_3141 | | YJR133W |
| YJR132W; DI49_3140; YJR131W | DI49_3140 | | YJR132W; YJR131W |
| YJR129C; YJR130C | | | YJR129C; YJR130C |
| YJR126C; YJR127C | | | YJR126C; YJR127C |
| YJR124C; DI49_3133 | DI49_3133 | | YJR124C |
| YJR123W; DI49_3132; YJR122W | DI49_3132 | | YJR123W; YJR122W |
| YJR119C; DI49_3129 | DI49_3129 | | YJR119C |
| YJR118C; DI49_3128 | DI49_3128 | | YJR118C |
| YJR117W; DI49_3127; YJR116W | DI49_3127 | | YJR117W; YJR116W |
| YJR113C; YJR115W | | | YJR113C; YJR115W |
| YJR112W; YJR112W-A | | | YJR112W; YJR112W-A |
| YJR110W; DI49_3120 | DI49_3120 | | YJR110W |

Create PDF in your applications with the Pdfcrowd HTML to PDF API PDFCROWD

| | | | |
|---|---|---|---|
| YJR109C; DI49_3119; YJR108W | DI49_3119 | | YJR109C; YJR108W |
| YJR107W; DI49_3117; YJR106W; DI49_3118 | DI49_3117; DI49_3118 | | YJR107W; YJR106W |
| YJR104C; DI49_3114; YJR103W | DI49_3114 | | YJR104C; YJR103W |
| YJR101W; YJR102C | | | YJR101W; YJR102C |
| YJR100C; DI49_3110; SPGP0Atrna1L; DI49_3111; SPGP0CUtrna30L | DI49_3110; DI49_3111 | SPGP0Atrna1L; SPGP0CUtrna30L | YJR100C |
| YJR098C; DI49_3107; YJR097W; DI49_3108 | DI49_3107; DI49_3108 | | YJR098C; YJR097W |
| YJR095W; DI49_3104; DI49_3105 | DI49_3104; DI49_3105 | | YJR095W |
| YJR094W-A; DI49_3103; YJR094C; DI49_5500; DI49_5501 | DI49_3103; DI49_5500; DI49_5501 | | YJR094W-A; YJR094C |
| YJR092W; YJR093C | | | YJR092W; YJR093C |
| YJR090C; DI49_3098; YJR089W | DI49_3098 | | YJR090C; YJR089W |
| YJR086W; YJR088C | | | YJR086W; YJR088C |
| YJR084W; YJR085C | | | YJR084W; YJR085C |
| YJR082C; YJR083C | | | YJR082C; YJR083C |
| YJR078W; YJR080C | | | YJR078W; YJR080C |
| YJR076C; DI49_3087 | DI49_3087 | | YJR076C |
| YJR075W; DI49_3086; YJR074W | DI49_3086 | | YJR075W; YJR074W |
| YJR072C; YJR073C | | | YJR072C; YJR073C |
| YJR069C; DI49_3082 | DI49_3082 | | YJR069C |
| YJR068W; DI49_3081; YJR067C | DI49_3081 | | YJR068W; YJR067C |
| YJR065C; DI49_3078 | DI49_3078 | | YJR065C |
| YJR064W; DI49_3077; YJR063W | DI49_3077 | | YJR064W; YJR063W |
| YJR061W; YJR062C; DI49_3075; DI49_3076 | DI49_3075; DI49_3076 | | YJR061W; YJR062C |
| YJR059W; DI49_3072; YJR058C; DI49_3073 | DI49_3072; DI49_3073 | | YJR059W; YJR058C |
| SPGP0Atrna1Y; SPGP0Atrna2Y; YJR056C; YJL046W | | SPGP0Atrna1Y; SPGP0Atrna2Y | YJR056C; YJL046W |
| YJR054W; DI49_3063; YJR053W | DI49_3063 | | YJR054W; YJR053W |
| YJR049C; YJR050W; DI49_3058; DI49_3059 | DI49_3058; DI49_3059 | | YJR049C; YJR050W |
| YJR047C; DI49_3055 | DI49_3055 | | YJR047C |
| SPGP0Atrna1S; SPGP0CRtrna25S; SPGP0Btrna10S | | SPGP0Atrna1S; SPGP0CRtrna25S; SPGP0Btrna10S | |
| YJR046W; DI49_3053 | DI49_3053 | | YJR046W |
| YJR045C; DI49_3052; YJR044C | DI49_3052 | | YJR045C; YJR044C |
| YJR041C; YJR042W | | | YJR041C; YJR042W |
| YJR039W; DI49_3045 | DI49_3045 | | YJR039W |
| YJR036C; DI49_3044; YJR035W | DI49_3044 | | YJR036C; YJR035W |
| YJR033C; DI49_3041; DI49_3042 | DI49_3041; DI49_3042 | | YJR033C |
| YJR032W; DI49_3040; YJR031C | DI49_3040 | | YJR032W; YJR031C |
| YJR025C; YJR030C | | | YJR025C; YJR030C |
| YJR022W; YJR024C | | | YJR022W; YJR024C |
| YJR019C; YJR021C; SPGP0BX00100 | | SPGP0BX00100 | YJR019C; YJR021C |
| YJR016C; DI49_3032; YJR015W; DI49_3033 | DI49_3032; DI49_3033 | | YJR016C; YJR015W |
| YJR013W; YJR012C | | | YJR013W; YJR012C |

Create PDF in your applications with the Pdfcrowd HTML to PDF API PDFCROWD

| | | | |
|---|---|---|---|
| YJR011C; DI49_3028; YJR010C-A; DI49_3029 | DI49_3028; DI49_3029 | | YJR011C; YJR010C-A |
| YJR008W; DI49_3024 | DI49_3024 | | YJR008W |
| YJR007W; DI49_3023; YJR006W | DI49_3023 | | YJR007W; YJR006W |
| YJR005W; DI49_3020; DI49_3021 | DI49_3020; DI49_3021 | | YJR005W |
| YJR004C; DI49_3019; YJR003C | DI49_3019 | | YJR004C; YJR003C |
| YJR001W; YJR002W | | | YJR001W; YJR002W |
| SPGP0A01510; SPGP0F01480; YJL002C; DI49_3016 | DI49_3016 | SPGP0A01510; SPGP0F01480 | YJL002C |
| YJL004C; YJL003W | | | YJL004C; YJL003W |
| YJL008C; DI49_3008 | DI49_3008 | | YJL008C |
| YJL010C; DI49_3007 | DI49_3007 | | YJL010C |
| SPGP0Atrna1W; SPGP0Etrna7W; YKL071W | | SPGP0Atrna1W; SPGP0Etrna7W | YKL071W |
| YJL013C; YJL012C | | | YJL013C; YJL012C |
| YJL016W; YJL014W | | | YJL016W; YJL014W |
| YJL020C; YJL019W | | | YJL020C; YJL019W |
| YJL023C; DI49_2997; DI49_2998 | DI49_2997; DI49_2998 | | YJL023C |
| YJL024C; DI49_2995; YJL025W | DI49_2995 | | YJL024C; YJL025W |
| YJL027C; YJL026W; DI49_2993; DI49_2994 | DI49_2993; DI49_2994 | | YJL027C; YJL026W |
| YJL029C; YJL007C | | | YJL029C; YJL007C |
| YJL031C; YJL030W | | | YJL031C; YJL030W |
| YJL034W; DI49_2986; YJL035C | DI49_2986 | | YJL034W; YJL035C |
| YJL039C; DI49_2978; YJL041W | DI49_2978 | | YJL039C; YJL041W |
| YJL043W; YJL042W | | | YJL043W; YJL042W |
| YJL045W; YJL044C | | | YJL045W; YJL044C |
| YJL048C; YJL047C | | | YJL048C; YJL047C |
| YJL050W; DI49_2965; DI49_2966 | DI49_2965; DI49_2966 | | YJL050W |
| YJL051W; DI49_2964 | DI49_2964 | | YJL051W |
| YJL052W; DI49_2963; YJL053W | DI49_2963 | | YJL052W; YJL053W |
| YJL055W; YJL054W | | | YJL055W; YJL054W |
| YJL057C; YJL056C | | | YJL057C; YJL056C |
| YJL059W; DI49_2956 | DI49_2956 | | YJL059W |
| YJL060W; DI49_2955; YJL061W | DI49_2955 | | YJL060W; YJL061W |
| YJL062W-A; DI49_2952; YJL063C | DI49_2952 | | YJL062W-A; YJL063C |
| YJL066C; YJL065C | | | YJL066C; YJL065C |
| YJL069C; YJL068C | | | YJL069C; YJL068C |
| YJL071W; DI49_2945 | DI49_2945 | | YJL071W |
| YJL072C; DI49_2944; YJL073W | DI49_2944 | | YJL072C; YJL073W |
| YJL076W; YJL074C | | | YJL076W; YJL074C |
| YJL079C; DI49_2939; DI49_2940 | DI49_2939; DI49_2940 | | YJL079C |
| YJL080C; DI49_2938 | DI49_2938 | | YJL080C |
| YJL081C; DI49_2937 | DI49_2937 | | YJL081C |

Create PDF in your applications with the Pdfcrowd HTML to PDF API PDFCROWD

| | | | |
|---|---|---|---|
| YJL082W; DI49_2936 | DI49_2936 | | YJL082W |
| YJL083W; DI49_2935 | DI49_2935 | | YJL083W |
| YJL084C; SPGP0F00780 | | SPGP0F00780 | YJL084C |
| YJL085W; DI49_2934; YJL087C | DI49_2934 | | YJL085W; YJL087C |
| YJL089W; DI49_2931; DI49_2932 | DI49_2931; DI49_2932 | | YJL089W |
| YJL090C; DI49_2930 | DI49_2930 | | YJL090C |
| YJL091C; DI49_2929 | DI49_2929 | | YJL091C |
| YJL092W; DI49_2928; YJL093C | DI49_2928 | | YJL092W; YJL093C |
| YJL095W; DI49_2925; YJL096W; DI49_2926 | DI49_2925; DI49_2926 | | YJL095W; YJL096W |
| YJL098W; DI49_2922; YJL099W; DI49_2923 | DI49_2922; DI49_2923 | | YJL098W; YJL099W |
| YJL101C; DI49_2919; DI49_2920 | DI49_2919; DI49_2920 | | YJL101C |
| SPGP0Atrna4R; DI49_2918; YJL102W | DI49_2918 | SPGP0Atrna4R | YJL102W |
| YJL104W; DI49_2915; YJL105W; DI49_2916 | DI49_2915; DI49_2916 | | YJL104W; YJL105W |
| YJL107C; DI49_2912; SPGP0F00570 | DI49_2912 | SPGP0F00570 | YJL107C |
| YJL109C; DI49_2911; YJL110C | DI49_2911 | | YJL109C; YJL110C |
| YJL112W; DI49_2908 | DI49_2908 | | YJL112W |
| YJL117W; YJL116C | | | YJL117W; YJL116C |
| YJL121C; YJL118W | | | YJL121C; YJL118W |
| YJL123C; YJL122W | | | YJL123C; YJL122W |
| YJL125C; DI49_2898; YJL126W; DI49_2899 | DI49_2898; DI49_2899 | | YJL125C; YJL126W |
| YJL127C-B; DI49_2895; DI49_2896 | DI49_2895; DI49_2896 | | YJL127C-B |
| YJL128C; DI49_2894 | DI49_2894 | | YJL128C |
| YJL129C; DI49_2893 | DI49_2893 | | YJL129C |
| YJL130C; DI49_2892 | DI49_2892 | | YJL130C |
| YJL131C; DI49_2891 | DI49_2891 | | YJL131C |
| YJL132W; SPGP0F00340 | | SPGP0F00340 | YJL132W |
| YJL133W; DI49_2890 | DI49_2890 | | YJL133W |
| YJL133C-A; DI49_2889; YJL134W | DI49_2889 | | YJL133C-A; YJL134W |
| YJL137C; YJL136C | | | YJL137C; YJL136C |
| DI49_2602; YJL138C | DI49_2602 | | YJL138C |
| DI49_2604; DI49_2603 | DI49_2604; DI49_2603 | | |
| DI49_2606; DI49_2605 | DI49_2606; DI49_2605 | | |
| DI49_1995; DI49_3843; SPGP0G00100; DI49_1996; DI49_3842 | DI49_1995; DI49_3843; DI49_1996; DI49_3842 | SPGP0G00100 | |
| DI49_5455; DI49_5456 | DI49_5455; DI49_5456 | | |
| DI49_5453; DI49_5454 | DI49_5453; DI49_5454 | | |
| SPGP0GW00100; YER176W | | SPGP0GW00100 | YER176W |
| DI49_1565; R0020C | DI49_1565 | | R0020C |
| YNL284C-B; YGR161W-B; YKL201C; SPGP0B01370 | | SPGP0B01370 | YNL284C-B; YGR161W-B; YKL201C |
| YKL197C; DI49_3184; DI49_3183 | DI49_3184; DI49_3183 | | YKL197C |
| YKL196C; DI49_3186; YKL195W | DI49_3186 | | YKL196C; YKL195W |

Create PDF in your applications with the Pdfcrowd HTML to PDF API PDFCROWD

| | | | |
|---|---|---|---|
| YKL193C; YKL194C; DI49_3187; SPGP0CN01130 | DI49_3187 | SPGP0CN01130 | YKL193C; YKL194C |
| YKL191W; YKL192C | | | YKL191W; YKL192C |
| YKL187C; DI49_3195; YKL186C; DI49_3194 | DI49_3195; DI49_3194 | | YKL187C; YKL186C |
| YKL184W; YKL185W | | | YKL184W; YKL185W |
| YKL182W; DI49_3200 | DI49_3200 | | YKL182W |
| YKL181W; DI49_3201 | DI49_3201 | | YKL181W |
| YKL180W; DI49_3202 | DI49_3202 | | YKL180W |
| YKL179C; DI49_3203 | DI49_3203 | | YKL179C |
| YKL178C; DI49_3204; YKL176C | DI49_3204 | | YKL178C; YKL176C |
| YKL174C; YKL175W | | | YKL174C; YKL175W |
| YKL172W; YKL173W | | | YKL172W; YKL173W |
| YKL170W; YKL171W | | | YKL170W; YKL171W |
| YKL167C; YKL168C | | | YKL167C; YKL168C |
| YKL165C; DI49_3215 | DI49_3215 | | YKL165C |
| YKL161C; YKL162C | | | YKL161C; YKL162C |
| YKL159C; YKL160W | | | YKL159C; YKL160W |
| YKL156W; DI49_3224; YKL155C | DI49_3224 | | YKL156W; YKL155C |
| YKL152C; DI49_3228; YKL151C | DI49_3228 | | YKL152C; YKL151C |
| YKL149C; YKL150W | | | YKL149C; YKL150W |
| YKL146W; DI49_3233 | DI49_3233 | | YKL146W |
| YKL145W; DI49_3234; YKL144C | DI49_3234 | | YKL145W; YKL144C |
| YKL142W; DI49_3237; YKL141W | DI49_3237 | | YKL142W; YKL141W |
| YKL139W; YKL140W | | | YKL139W; YKL140W |
| YKL138C; DI49_3242; YKL137W; DI49_3241 | DI49_3242; DI49_3241 | | YKL138C; YKL137W |
| YKL134C; YKL135C; DI49_3244; DI49_3243 | DI49_3244; DI49_3243 | | YKL134C; YKL135C |
| YKL132C; YKL133C | | | YKL132C; YKL133C |
| YKL129C; DI49_3249; YKL128C; DI49_3248 | DI49_3249; DI49_3248 | | YKL129C; YKL128C |
| YKL126W; YKL127W | | | YKL126W; YKL127W |
| YKL124W; DI49_3255; YKL122C | DI49_3255 | | YKL124W; YKL122C |
| YKL120W; DI49_3258; YKL119C | DI49_3258 | | YKL120W; YKL119C |
| YKL114C; DI49_3263 | DI49_3263 | | YKL114C |
| YKL113C; DI49_3264 | DI49_3264 | | YKL113C |
| YKL112W; DI49_3265; YKL110C | DI49_3265 | | YKL112W; YKL110C |
| YKL108W; YKL109W | | | YKL108W; YKL109W |
| YKL106W; DI49_3270; YKL105C; DI49_3269 | DI49_3270; DI49_3269 | | YKL106W; YKL105C |
| YKL103C; DI49_3273 | DI49_3273 | | YKL103C |
| YKL101W; DI49_3274; YKL100C | DI49_3274 | | YKL101W; YKL100C |
| YKL098W; YKL099C; DI49_3276; DI49_3275 | DI49_3276; DI49_3275 | | YKL098W; YKL099C |
| YKL095W; DI49_3280 | DI49_3280 | | YKL095W |

Create PDF in your applications with the Pdfcrowd HTML to PDF API PDFCROWD

| | | | |
|---|---|---|---|
| YKL094W; DI49_3281; YKL093W | DI49_3281 | | YKL094W; YKL093W |
| YKL091C; YKL092C; DI49_3283; DI49_3282 | DI49_3283; DI49_3282 | | YKL091C; YKL092C |
| YKL089W; DI49_3286; DI49_3285 | DI49_3286; DI49_3285 | | YKL089W |
| YKL088W; DI49_3287; YKL087C | DI49_3287 | | YKL088W; YKL087C |
| YKL085W; YKL086W; DI49_3289; DI49_3288 | DI49_3289; DI49_3288 | | YKL085W; YKL086W |
| YKL082C; DI49_3292; YKL081W; DI49_3291 | DI49_3292; DI49_3291 | | YKL082C; YKL081W |
| DI49_0844; YKL080W | DI49_0844 | | YKL080W |
| DI49_0846; DI49_0845 | DI49_0846; DI49_0845 | | |
| DI49_0848; DI49_0847 | DI49_0848; DI49_0847 | | |
| DI49_0850; DI49_0849 | DI49_0850; DI49_0849 | | |
| DI49_0852; DI49_0851 | DI49_0852; DI49_0851 | | |
| DI49_2399; DI49_0853 | DI49_2399; DI49_0853 | | |
| DI49_2401; DI49_2400 | DI49_2401; DI49_2400 | | |
| SPGP0HE00100; DI49_2402 | DI49_2402 | SPGP0HE00100 | |
| DI49_5194; DI49_1596; SPGP0HI00110 | DI49_5194; DI49_1596 | SPGP0HI00110 | |
| SPGP0HI00130; YJL216C | | SPGP0HI00130 | YJL216C |
| DI49_3962; DI49_3961 | DI49_3962; DI49_3961 | | |
| DI49_3964; SPGP0HI00180 | DI49_3964 | SPGP0HI00180 | |
| DI49_3966; DI49_3965 | DI49_3966; DI49_3965 | | |
| DI49_3968; DI49_3967 | DI49_3968; DI49_3967 | | |
| SPGP0U00100; SPGP0HK00100 | | SPGP0U00100; SPGP0HK00100 | |
| SPGP0U00140; DI49_1042 | DI49_1042 | SPGP0U00140 | |
| DI49_1040; DI49_1041 | DI49_1040; DI49_1041 | | |
| DI49_1038; DI49_1039 | DI49_1038; DI49_1039 | | |
| DI49_1036; DI49_1037 | DI49_1036; DI49_1037 | | |
| DI49_1034; DI49_1035 | DI49_1034; DI49_1035 | | |
| DI49_1032; DI49_1033 | DI49_1032; DI49_1033 | | |
| DI49_1028; DI49_1029 | DI49_1028; DI49_1029 | | |
| SPGP0U00290; DI49_1027 | DI49_1027 | SPGP0U00290 | |
| DI49_1025; DI49_1026 | DI49_1025; DI49_1026 | | |
| DI49_1024; SPGP0U00320 | DI49_1024 | SPGP0U00320 | |
| SPGP0U00350; DI49_1023 | DI49_1023 | SPGP0U00350 | |
| DI49_1021; DI49_1022 | DI49_1021; DI49_1022 | | |
| DI49_1019; DI49_1020 | DI49_1019; DI49_1020 | | |
| DI49_1017; DI49_1018 | DI49_1017; DI49_1018 | | |
| DI49_1015; DI49_1016 | DI49_1015; DI49_1016 | | |
| DI49_1014; SPGP0U00440 | DI49_1014 | SPGP0U00440 | |
| DI49_1012; DI49_1013 | DI49_1012; DI49_1013 | | |
| DI49_1010; DI49_1011 | DI49_1010; DI49_1011 | | |
| DI49_1005; DI49_1006 | DI49_1005; DI49_1006 | | |

Create PDF in your applications with the Pdfcrowd HTML to PDF API PDFCROWD

| | | | |
|---|---|---|---|
| DI49_1003; DI49_1004 | DI49_1003; DI49_1004 | | |
| DI49_1001; DI49_3888; DI49_1000; DI49_3887 | DI49_1001; DI49_3888; DI49_1000; DI49_3887 | | |
| DI49_0999; DI49_0998; DI49_0997 | DI49_0999; DI49_0998; DI49_0997 | | |
| DI49_0995; DI49_0996 | DI49_0995; DI49_0996 | | |
| DI49_0993; DI49_0994 | DI49_0993; DI49_0994 | | |
| SPGP0U00680; SPGP0CU00360; DI49_0990; DI49_3903 | DI49_0990; DI49_3903 | SPGP0U00680; SPGP0CU00360 | |
| DI49_0988; DI49_0989 | DI49_0988; DI49_0989 | | |
| SPGP0U00730; SPGP0U00720 | | SPGP0U00730; SPGP0U00720 | |
| DI49_0986; DI49_2363; DI49_2362 | DI49_0986; DI49_2363; DI49_2362 | | |
| DI49_0985; YMR142C | DI49_0985 | | YMR142C |
| YDL083C; YMR143W; DI49_0984; DI49_4229 | DI49_0984; DI49_4229 | | YDL083C; YMR143W |
| DI49_0982; DI49_0983 | DI49_0982; DI49_0983 | | |
| DI49_0980; DI49_0981 | DI49_0980; DI49_0981 | | |
| DI49_0978; DI49_0979 | DI49_0978; DI49_0979 | | |
| DI49_0977; SPGP0U00850 | DI49_0977 | SPGP0U00850 | |
| DI49_0975; DI49_0976 | DI49_0975; DI49_0976 | | |
| DI49_0973; DI49_0974 | DI49_0973; DI49_0974 | | |
| DI49_0971; DI49_0972 | DI49_0971; DI49_0972 | | |
| DI49_0969; DI49_0970 | DI49_0969; DI49_0970 | | |
| DI49_0967; DI49_0968 | DI49_0967; DI49_0968 | | |
| DI49_0965; DI49_0966 | DI49_0965; DI49_0966 | | |
| DI49_0963; DI49_0964 | DI49_0963; DI49_0964 | | |
| DI49_0961; DI49_0962 | DI49_0961; DI49_0962 | | |
| DI49_0959; DI49_0960 | DI49_0959; DI49_0960 | | |
| DI49_0957; DI49_0958 | DI49_0957; DI49_0958 | | |
| DI49_0955; DI49_0956 | DI49_0955; DI49_0956 | | |
| DI49_0953; DI49_0954 | DI49_0953; DI49_0954 | | |
| DI49_0951; DI49_0952 | DI49_0951; DI49_0952 | | |
| DI49_0949; DI49_0950 | DI49_0949; DI49_0950 | | |
| SPGP0U01160; DI49_0948 | DI49_0948 | SPGP0U01160 | |
| DI49_0946; DI49_0947 | DI49_0946; DI49_0947 | | |
| DI49_0944; DI49_0945 | DI49_0944; DI49_0945 | | |
| DI49_0940; YDL131W | DI49_0940 | | YDL131W |
| DI49_0938; YDL188C; DI49_0937; DI49_0892 | DI49_0938; DI49_0937; DI49_0892 | | YDL188C |
| DI49_0935; DI49_0936; DI49_0889; DI49_0888 | DI49_0935; DI49_0936; DI49_0889; DI49_0888 | | |
| DI49_0934; SPGP0U01310 | DI49_0934 | SPGP0U01310 | |
| DI49_0932; DI49_0933 | DI49_0932; DI49_0933 | | |
| DI49_0930; DI49_0931 | DI49_0930; DI49_0931 | | |
| DI49_0928; DI49_0929 | DI49_0928; DI49_0929 | | |
| DI49_0926; DI49_0927 | DI49_0926; DI49_0927 | | |

Create PDF in your applications with the Pdfcrowd HTML to PDF API PDFCROWD

| | | |
|---|---|---|
| DI49_0924; DI49_0925 | DI49_0924; DI49_0925 | |
| DI49_0922; DI49_0923 | DI49_0922; DI49_0923 | |
| DI49_0920; DI49_0921 | DI49_0920; DI49_0921 | |
| DI49_0918; DI49_0919 | DI49_0918; DI49_0919 | |
| DI49_0916; DI49_0917 | DI49_0916; DI49_0917 | |
| DI49_0914; DI49_0915 | DI49_0914; DI49_0915 | |
| DI49_0912; DI49_0913 | DI49_0912; DI49_0913 | |
| DI49_0910; DI49_0911 | DI49_0910; DI49_0911 | |
| DI49_0908; DI49_0909 | DI49_0908; DI49_0909 | |
| DI49_0906; DI49_0907 | DI49_0906; DI49_0907 | |
| DI49_0904; DI49_0905 | DI49_0904; DI49_0905 | |
| DI49_0902; DI49_0903 | DI49_0902; DI49_0903 | |
| DI49_0900; DI49_0901 | DI49_0900; DI49_0901 | |
| DI49_0898; DI49_0899 | DI49_0898; DI49_0899 | |
| DI49_0896; DI49_0897 | DI49_0896; DI49_0897 | |
| DI49_0894; DI49_0895 | DI49_0894; DI49_0895 | |
| DI49_0886; DI49_0887 | DI49_0886; DI49_0887 | |
| DI49_0884; DI49_0885 | DI49_0884; DI49_0885 | |
| DI49_0882; DI49_0883 | DI49_0882; DI49_0883 | |
| DI49_0880; DI49_0881 | DI49_0880; DI49_0881 | |
| DI49_0878; DI49_0879 | DI49_0878; DI49_0879 | |
| DI49_0876; DI49_0877 | DI49_0876; DI49_0877 | |
| DI49_0874; DI49_0875 | DI49_0874; DI49_0875 | |
| DI49_0872; DI49_0873 | DI49_0872; DI49_0873 | |
| DI49_0868; DI49_0869 | DI49_0868; DI49_0869 | |
| DI49_0866; DI49_0867 | DI49_0866; DI49_0867 | |
| DI49_0864; DI49_0865 | DI49_0864; DI49_0865 | |
| DI49_0862; DI49_0863 | DI49_0862; DI49_0863 | |
| DI49_0860; DI49_0861 | DI49_0860; DI49_0861 | |
| DI49_0858; DI49_0859 | DI49_0858; DI49_0859 | |
| DI49_0856; DI49_0857 | DI49_0856; DI49_0857 | |
| YAL058W; YAL059W | | YAL058W; YAL059W |
| YAL055W; YAL056W | | YAL055W; YAL056W |
| DI49_2614; DI49_2613 | DI49_2614; DI49_2613 | |
| DI49_2616; DI49_2615 | DI49_2616; DI49_2615 | |
| DI49_0678; DI49_2617 | DI49_0678; DI49_2617 | |
| DI49_0676; DI49_0677 | DI49_0676; DI49_0677 | |
| DI49_0674; DI49_0675 | DI49_0674; DI49_0675 | |
| DI49_0672; DI49_0673 | DI49_0672; DI49_0673 | |
| DI49_1598; DI49_1599 | DI49_1598; DI49_1599 | |

Create PDF in your applications with the Pdfcrowd HTML to PDF API PDFCROWD

| | | |
|---|---|---|
| DI49_2393; DI49_2392 | DI49_2393; DI49_2392 | |
| DI49_2395; DI49_2394 | DI49_2395; DI49_2394 | |
| DI49_2397; DI49_2396 | DI49_2397; DI49_2396 | |
| DI49_2389; DI49_4051; DI49_2388; DI49_4052 | DI49_2389; DI49_4051; DI49_2388; DI49_4052 | |
| DI49_2386; DI49_2387 | DI49_2386; DI49_2387 | |
| DI49_2384; DI49_2385 | DI49_2384; DI49_2385 | |
| DI49_2376; DI49_2377 | DI49_2376; DI49_2377 | |
| DI49_2371; DI49_2372 | DI49_2371; DI49_2372 | |
| DI49_2369; DI49_2370 | DI49_2369; DI49_2370 | |
| DI49_2367; DI49_2368 | DI49_2367; DI49_2368 | |
| DI49_2365; DI49_2366 | DI49_2365; DI49_2366 | |
| DI49_2360; DI49_2361 | DI49_2360; DI49_2361 | |
| DI49_2358; DI49_2359 | DI49_2358; DI49_2359 | |
| SPGP0V00410; DI49_2357 | DI49_2357 | SPGP0V00410 |
| DI49_2356; SPGP0V00420 | DI49_2356 | SPGP0V00420 |
| DI49_2354; DI49_2355 | DI49_2354; DI49_2355 | |
| DI49_2352; DI49_2353 | DI49_2352; DI49_2353 | |
| DI49_2346; DI49_2347 | DI49_2346; DI49_2347 | |
| DI49_2344; DI49_2345 | DI49_2344; DI49_2345 | |
| DI49_2342; DI49_2343 | DI49_2342; DI49_2343 | |
| DI49_2340; DI49_2341 | DI49_2340; DI49_2341 | |
| DI49_2338; DI49_2339 | DI49_2338; DI49_2339 | |
| DI49_2336; DI49_2337 | DI49_2336; DI49_2337 | |
| DI49_2334; DI49_2335 | DI49_2334; DI49_2335 | |
| DI49_2332; DI49_2333 | DI49_2332; DI49_2333 | |
| DI49_2330; DI49_2331 | DI49_2330; DI49_2331 | |
| DI49_2328; DI49_2329 | DI49_2328; DI49_2329 | |
| DI49_2326; DI49_2327 | DI49_2326; DI49_2327 | |
| DI49_2324; DI49_2325 | DI49_2324; DI49_2325 | |
| DI49_2323; SPGP0V00750 | DI49_2323 | SPGP0V00750 |
| DI49_2321; DI49_2322 | DI49_2321; DI49_2322 | |
| DI49_2319; DI49_2320 | DI49_2319; DI49_2320 | |
| DI49_2317; DI49_2318 | DI49_2317; DI49_2318 | |
| DI49_2315; DI49_2316 | DI49_2315; DI49_2316 | |
| DI49_2313; DI49_2314 | DI49_2313; DI49_2314 | |
| DI49_2311; DI49_2312 | DI49_2311; DI49_2312 | |
| DI49_2309; DI49_2310 | DI49_2309; DI49_2310 | |
| DI49_2307; DI49_2308 | DI49_2307; DI49_2308 | |
| DI49_2305; DI49_2306 | DI49_2305; DI49_2306 | |
| DI49_2303; DI49_2304 | DI49_2303; DI49_2304 | |

Create PDF in your applications with the Pdfcrowd HTML to PDF API PDFCROWD

| | | | |
|---|---|---|---|
| DI49_2301; DI49_2302 | DI49_2301; DI49_2302 | | |
| DI49_2294; DI49_2295 | DI49_2294; DI49_2295 | | |
| DI49_2292; DI49_2293 | DI49_2292; DI49_2293 | | |
| DI49_2290; DI49_2291 | DI49_2290; DI49_2291 | | |
| DI49_2288; DI49_2289 | DI49_2288; DI49_2289 | | |
| DI49_2283; DI49_2284 | DI49_2283; DI49_2284 | | |
| DI49_2281; DI49_2282 | DI49_2281; DI49_2282 | | |
| DI49_2279; DI49_2280 | DI49_2279; DI49_2280 | | |
| DI49_2277; DI49_1607; YPL277C; YER177W | DI49_2277; DI49_1607 | | YPL277C; YER177W |
| YOR390W; YPL278C | | | YOR390W; YPL278C |
| YPL281C; YOR391C | | | YPL281C; YOR391C |
| DI49_0686; DI49_0687 | DI49_0686; DI49_0687 | | |
| DI49_0684; DI49_0685 | DI49_0684; DI49_0685 | | |
| DI49_0682; DI49_0683 | DI49_0682; DI49_0683 | | |
| SPGP0HW00170; DI49_0681 | DI49_0681 | SPGP0HW00170 | |
| SPGP0HW00190; DI49_0680 | DI49_0680 | SPGP0HW00190 | |
| YGL236C; DI49_0679 | DI49_0679 | | YGL236C |
| YGL238W; YGL237C | | | YGL238W; YGL237C |
| DI49_5461; DI49_5462 | DI49_5461; DI49_5462 | | |
| DI49_5459; DI49_5460 | DI49_5459; DI49_5460 | | |
| DI49_1646; DI49_1647 | DI49_1646; DI49_1647 | | |
| YAL062W; DI49_5188; DI49_2821; DI49_5189 | DI49_5188; DI49_2821; DI49_5189 | | YAL062W |
| DI49_0716; DI49_0717 | DI49_0716; DI49_0717 | | |
| DI49_3476; SPGP0IH00100 | DI49_3476 | SPGP0IH00100 | |
| DI49_3478; DI49_3477 | DI49_3478; DI49_3477 | | |
| DI49_3480; DI49_3479 | DI49_3480; DI49_3479 | | |
| SPGP0IH00170; DI49_3481 | DI49_3481 | SPGP0IH00170 | |
| DI49_3483; DI49_3482; DI49_1606 | DI49_3483; DI49_3482; DI49_1606 | | |
| DI49_3485; DI49_3484 | DI49_3485; DI49_3484 | | |
| DI49_3487; DI49_3486 | DI49_3487; DI49_3486 | | |
| SPGP0IK00100; DI49_3488 | DI49_3488 | SPGP0IK00100 | |
| DI49_5176; SPGP0IL00100 | DI49_5176 | SPGP0IL00100 | |
| DI49_5174; DI49_5175 | DI49_5174; DI49_5175 | | |
| DI49_5172; DI49_5173 | DI49_5172; DI49_5173 | | |
| DI49_5170; DI49_5171 | DI49_5170; DI49_5171 | | |
| DI49_2276; DI49_5169 | DI49_2276; DI49_5169 | | |
| DI49_2274; DI49_2275 | DI49_2274; DI49_2275 | | |
| DI49_2272; DI49_2273 | DI49_2272; DI49_2273 | | |
| DI49_2270; DI49_2271 | DI49_2270; DI49_2271 | | |
| DI49_2268; DI49_2269 | DI49_2268; DI49_2269 | | |

Create PDF in your applications with the Pdfcrowd HTML to PDF API PDFCROWD

| | | | |
|---|---|---|---|
| DI49_2266; DI49_2267 | DI49_2266; DI49_2267 | | |
| DI49_2264; DI49_2265 | DI49_2264; DI49_2265 | | |
| DI49_2262; DI49_2263 | DI49_2262; DI49_2263 | | |
| DI49_2260; DI49_2261 | DI49_2260; DI49_2261 | | |
| DI49_2258; DI49_2259 | DI49_2258; DI49_2259 | | |
| DI49_2256; DI49_2257 | DI49_2256; DI49_2257 | | |
| DI49_2254; DI49_2255 | DI49_2254; DI49_2255 | | |
| DI49_2610; SPGP0INtrna11T | DI49_2610 | SPGP0INtrna11T | |
| DI49_2612; DI49_2611 | DI49_2612; DI49_2611 | | |
| DI49_2624; YIL162W | DI49_2624 | | YIL162W |
| DI49_2625; YIL161W | DI49_2625 | | YIL161W |
| DI49_2626; YIL160C | DI49_2626 | | YIL160C |
| DI49_2627; YIL159W; DI49_2628; YIL158W | DI49_2627; DI49_2628 | | YIL159W; YIL158W |
| DI49_2630; DI49_2629 | DI49_2630; DI49_2629 | | |
| DI49_2632; DI49_2631 | DI49_2632; DI49_2631 | | |
| DI49_2634; DI49_2633 | DI49_2634; DI49_2633 | | |
| DI49_2636; YIL151C | DI49_2636 | | YIL151C |
| DI49_2637; YIL150C | DI49_2637 | | YIL150C |
| DI49_2638; YIL149C; SPGP0C00300 | DI49_2638 | SPGP0C00300 | YIL149C |
| DI49_2639; YKR094C; SPGP0H00210; DI49_3686; DI49_3685 | DI49_2639; DI49_3686; DI49_3685 | SPGP0H00210 | YKR094C |
| DI49_2640; YIL147C | DI49_2640 | | YIL147C |
| DI49_2641; YIL146C; DI49_2642; YIL145C | DI49_2641; DI49_2642 | | YIL146C; YIL145C |
| DI49_2644; YIL143C | DI49_2644 | | YIL143C |
| DI49_2645; YIL142W; DI49_2646; YIL140W | DI49_2645; DI49_2646 | | YIL142W; YIL140W |
| DI49_2648; DI49_2647 | DI49_2648; DI49_2647 | | |
| DI49_2650; DI49_2649 | DI49_2650; DI49_2649 | | |
| DI49_2652; DI49_2651 | DI49_2652; DI49_2651 | | |
| DI49_2654; DI49_2653 | DI49_2654; DI49_2653 | | |
| DI49_2656; DI49_2655 | DI49_2656; DI49_2655 | | |
| DI49_2658; YIL128W; DI49_2659; YIL127C | DI49_2658; DI49_2659 | | YIL128W; YIL127C |
| DI49_2661; YIL125W | DI49_2661 | | YIL125W |
| DI49_2662; YIL124W | DI49_2662 | | YIL124W |
| DI49_2663; YIL123W | DI49_2663 | | YIL123W |
| DI49_2664; YIL122W; DI49_2665; YIL121W | DI49_2664; DI49_2665 | | YIL122W; YIL121W |
| DI49_2667; DI49_2666 | DI49_2667; DI49_2666 | | |
| DI49_2669; DI49_2668 | DI49_2669; DI49_2668 | | |
| DI49_2671; YIL115C; DI49_2672; YIL114C | DI49_2671; DI49_2672 | | YIL115C; YIL114C |
| DI49_2674; DI49_2673 | DI49_2674; DI49_2673 | | |
| DI49_2675; YIL110W | DI49_2675 | | YIL110W |
| DI49_2676; YIL109C | DI49_2676 | | YIL109C |

Create PDF in your applications with the Pdfcrowd HTML to PDF API PDFCROWD

| | | | |
|---|---|---|---|
| DI49_2677; YIL108W | DI49_2677 | | YIL108W |
| DI49_2678; YIL107C; DI49_2679; YIL106W | DI49_2678; DI49_2679 | | YIL107C; YIL106W |
| DI49_2681; DI49_2680 | DI49_2681; DI49_2680 | | |
| DI49_2683; DI49_2682 | DI49_2683; DI49_2682 | | |
| DI49_2685; YIL101C; DI49_2686; YIL099W | DI49_2685; DI49_2686 | | YIL101C; YIL099W |
| DI49_2688; DI49_2687 | DI49_2688; DI49_2687 | | |
| DI49_2691; YIL095W | DI49_2691 | | YIL095W |
| DI49_2692; YIL094C | DI49_2692 | | YIL094C |
| DI49_2693; YIL093C; DI49_2694; YIL092W | DI49_2693; DI49_2694 | | YIL093C; YIL092W |
| DI49_2696; DI49_2695 | DI49_2696; DI49_2695 | | |
| SPGP0Xtrna11E; DI49_2698; DI49_2699; YDR278C; YIL088C; DI49_0455 | DI49_2698; DI49_2699; DI49_0455 | SPGP0Xtrna11E | YDR278C; YIL088C |
| DI49_2701; DI49_2700 | DI49_2701; DI49_2700 | | |
| SPGP0X00880; YIL083C | | SPGP0X00880 | YIL083C |
| DI49_2706; YIL077C; YIL076W | DI49_2706 | | YIL077C; YIL076W |
| DI49_2707; DI49_4178; DI49_2708; DI49_4179 | DI49_2707; DI49_4178; DI49_2708; DI49_4179 | | |
| DI49_2713; DI49_2712; YIL072W; YIL071C | DI49_2713; DI49_2712 | | YIL072W; YIL071C |
| DI49_2716; YIL068C | DI49_2716 | | YIL068C |
| DI49_2717; YIL067C | DI49_2717 | | YIL067C |
| DI49_2718; YIL066C; DI49_2719; YIL065C | DI49_2718; DI49_2719 | | YIL066C; YIL065C |
| DI49_2720; YIL063C | DI49_2720 | | YIL063C |
| DI49_2721; YIL062C; DI49_2722; YIL061C | DI49_2721; DI49_2722 | | YIL062C; YIL061C |
| DI49_2725; YIL056W; DI49_2726 | DI49_2725; DI49_2726 | | YIL056W |
| DI49_2727; YIL053W | DI49_2727 | | YIL053W |
| SPGP0X01140; YIL052C | | SPGP0X01140 | YIL052C |
| DI49_2728; YIL051C; DI49_2729; YIL050W | DI49_2728; DI49_2729 | | YIL051C; YIL050W |
| DI49_2731; YIL048W | DI49_2731 | | YIL048W |
| DI49_2732; YIL047C | DI49_2732 | | YIL047C |
| DI49_2733; YIL046W | DI49_2733 | | YIL046W |
| DI49_2734; YIL045W; DI49_2735; YIL044C | DI49_2734; DI49_2735 | | YIL045W; YIL044C |
| DI49_2737; DI49_2736 | DI49_2737; DI49_2736 | | |
| DI49_2739; DI49_2738; YIL041W; YIL040W | DI49_2739; DI49_2738 | | YIL041W; YIL040W |
| DI49_2741; YIL038C; DI49_2742; YIL037C | DI49_2741; DI49_2742 | | YIL038C; YIL037C |
| DI49_2744; YIL035C | DI49_2744 | | YIL035C |
| DI49_2745; YIL034C | DI49_2745 | | YIL034C |
| DI49_2746; YIL033C; DI49_2747; YIL031W | DI49_2746; DI49_2747 | | YIL033C; YIL031W |
| DI49_2752; DI49_2751; YIL027C; YIL026C | DI49_2752; DI49_2751 | | YIL027C; YIL026C |
| DI49_2754; DI49_2753 | DI49_2754; DI49_2753 | | |
| DI49_2756; YIL021W; DI49_2757; YIL020C | DI49_2756; DI49_2757 | | YIL021W; YIL020C |
| DI49_2759; YIL018W; DI49_2760; YIL017C | DI49_2759; DI49_2760 | | YIL018W; YIL017C |

Create PDF in your applications with the Pdfcrowd HTML to PDF API PDFCROWD

| | | | |
|---|---|---|---|
| DI49_2762; YIL015W | DI49_2762 | | YIL015W |
| DI49_2766; DI49_2765 | DI49_2766; DI49_2765 | | |
| DI49_2768; DI49_2767 | DI49_2768; DI49_2767 | | |
| DI49_2773; DI49_2772 | DI49_2773; DI49_2772 | | |
| DI49_2775; DI49_2774 | DI49_2775; DI49_2774 | | |
| DI49_2777; YIL002W-A | DI49_2777 | | YIL002W-A |
| DI49_2778; YIL002C; DI49_2779; YIL001W | DI49_2778; DI49_2779 | | YIL002C; YIL001W |
| DI49_2781; YIR002C; DI49_2782; YIR003W | DI49_2781; DI49_2782 | | YIR002C; YIR003W |
| DI49_2784; DI49_2783 | DI49_2784; DI49_2783 | | |
| DI49_2787; YIR007W | DI49_2787 | | YIR007W |
| DI49_2788; YIR008C; DI49_2789; YIR009W | DI49_2788; DI49_2789 | | YIR008C; YIR009W |
| DI49_2791; YIR011C | DI49_2791 | | YIR011C |
| DI49_2792; YIR012W; DI49_2793; YIR013C | DI49_2792; DI49_2793 | | YIR012W; YIR013C |
| DI49_2795; DI49_2794 | DI49_2795; DI49_2794 | | |
| DI49_2797; YIR017C | DI49_2797 | | YIR017C |
| DI49_2798; YIR018W; DI49_2799; SPGP0C01850 | DI49_2798; DI49_2799 | SPGP0C01850 | YIR018W |
| DI49_2801; DI49_2800 | DI49_2801; DI49_2800 | | |
| DI49_2803; DI49_2802 | DI49_2803; DI49_2802 | | |
| DI49_2805; DI49_2804 | DI49_2805; DI49_2804 | | |
| DI49_2807; YIR028W | DI49_2807 | | YIR028W |
| DI49_2808; YIR029W; DI49_2809; YIR030C | DI49_2808; DI49_2809 | | YIR029W; YIR030C |
| DI49_2811; DI49_2810 | DI49_2811; DI49_2810 | | |
| DI49_2813; YIR034C; DI49_2814; YIR035C | DI49_2813; DI49_2814 | | YIR034C; YIR035C |
| DI49_2816; DI49_2815 | DI49_2816; DI49_2815 | | |
| YGL227W; YGL226C-A | | | YGL227W; YGL226C-A |
| YGL229C; YGL228W | | | YGL229C; YGL228W |
| YGL231C; YGL230C | | | YGL231C; YGL230C |
| YGL233W; YGL232W | | | YGL233W; YGL232W |
| DI49_4048; YGL234W | DI49_4048 | | YGL234W |
| DI49_0082; SPGP0IRtrna12L | DI49_0082 | SPGP0IRtrna12L | |
| DI49_0084; DI49_0083 | DI49_0084; DI49_0083 | | |
| DI49_0086; DI49_0085 | DI49_0086; DI49_0085 | | |
| DI49_0088; DI49_0087 | DI49_0088; DI49_0087 | | |
| DI49_4367; DI49_1083 | DI49_4367; DI49_1083 | | |
| DI49_4369; DI49_4368 | DI49_4369; DI49_4368 | | |
| DI49_4373; DI49_4372 | DI49_4373; DI49_4372 | | |
| DI49_4375; DI49_4374 | DI49_4375; DI49_4374 | | |
| DI49_4377; DI49_4376 | DI49_4377; DI49_4376 | | |
| SPGP0IV00110; SPGP0IV00100 | | SPGP0IV00110; SPGP0IV00100 | |
| DI49_0842; DI49_0841 | DI49_0842; DI49_0841 | | |

Create PDF in your applications with the Pdfcrowd HTML to PDF API PDFCROWD

| | | |
|---|---|---|
| DI49_4049; DI49_0843 | DI49_4049; DI49_0843 | |
| DI49_4054; DI49_4053 | DI49_4054; DI49_4053 | |
| DI49_4056; DI49_4055 | DI49_4056; DI49_4055 | |
| DI49_4058; DI49_4057 | DI49_4058; DI49_4057 | |
| DI49_4060; DI49_4059 | DI49_4060; DI49_4059 | |
| DI49_4062; DI49_4061 | DI49_4062; DI49_4061 | |
| DI49_4064; DI49_4063; DI49_0617; DI49_0618 | DI49_4064; DI49_4063; DI49_0617; DI49_0618 | |
| DI49_4066; DI49_4065 | DI49_4066; DI49_4065 | |
| DI49_4068; DI49_4067 | DI49_4068; DI49_4067 | |
| DI49_4070; DI49_4069 | DI49_4070; DI49_4069 | |
| DI49_4072; DI49_4071 | DI49_4072; DI49_4071 | |
| DI49_4074; DI49_4073 | DI49_4074; DI49_4073 | |
| DI49_4076; DI49_4075 | DI49_4076; DI49_4075 | |
| DI49_4078; DI49_4077 | DI49_4078; DI49_4077 | |
| DI49_4080; DI49_4079 | DI49_4080; DI49_4079 | |
| DI49_4082; DI49_4081 | DI49_4082; DI49_4081 | |
| DI49_4085; DI49_4083 | DI49_4085; DI49_4083 | |
| DI49_4087; DI49_4086 | DI49_4087; DI49_4086 | |
| DI49_4089; DI49_4088 | DI49_4089; DI49_4088 | |
| DI49_4091; DI49_4090 | DI49_4091; DI49_4090 | |
| DI49_4093; DI49_4092 | DI49_4093; DI49_4092 | |
| DI49_4095; DI49_4094; DI49_4096 | DI49_4095; DI49_4094; DI49_4096 | |
| DI49_4101; DI49_4100 | DI49_4101; DI49_4100 | |
| DI49_4103; DI49_4102 | DI49_4103; DI49_4102 | |
| DI49_4105; DI49_4104 | DI49_4105; DI49_4104 | |
| DI49_4107; DI49_4106 | DI49_4107; DI49_4106 | |
| DI49_4109; DI49_4108 | DI49_4109; DI49_4108 | |
| DI49_4114; DI49_4113 | DI49_4114; DI49_4113 | |
| DI49_4116; DI49_4115 | DI49_4116; DI49_4115 | |
| DI49_4118; DI49_4117 | DI49_4118; DI49_4117 | |
| DI49_4120; DI49_4119 | DI49_4120; DI49_4119 | |
| DI49_4122; DI49_4121 | DI49_4122; DI49_4121 | |
| DI49_4124; DI49_4123 | DI49_4124; DI49_4123 | |
| DI49_4126; DI49_4125 | DI49_4126; DI49_4125 | |
| DI49_4128; DI49_4127 | DI49_4128; DI49_4127 | |
| DI49_4133; DI49_4132 | DI49_4133; DI49_4132 | |
| DI49_4135; SPGP0Ytrna7H | DI49_4135 | SPGP0Ytrna7H |
| DI49_4137; DI49_4136 | DI49_4137; DI49_4136 | |
| DI49_4144; DI49_4143 | DI49_4144; DI49_4143 | |
| DI49_4146; DI49_4145 | DI49_4146; DI49_4145 | |

Create PDF in your applications with the Pdfcrowd HTML to PDF API PDFCROWD

| | | | |
|---|---|---|---|
| DI49_4148; DI49_4147 | DI49_4148; DI49_4147 | | |
| DI49_4150; DI49_4149 | DI49_4150; DI49_4149 | | |
| DI49_4152; DI49_4151 | DI49_4152; DI49_4151 | | |
| DI49_4154; DI49_4153 | DI49_4154; DI49_4153 | | |
| DI49_4156; DI49_4155 | DI49_4156; DI49_4155 | | |
| DI49_4158; DI49_4157 | DI49_4158; DI49_4157 | | |
| DI49_4160; DI49_4159 | DI49_4160; DI49_4159 | | |
| DI49_4162; DI49_4161 | DI49_4162; DI49_4161 | | |
| DI49_4164; DI49_4163 | DI49_4164; DI49_4163 | | |
| DI49_4169; DI49_4168 | DI49_4169; DI49_4168 | | |
| DI49_4171; DI49_4170 | DI49_4171; DI49_4170 | | |
| DI49_4173; DI49_4172 | DI49_4173; DI49_4172 | | |
| DI49_4175; DI49_4174 | DI49_4175; DI49_4174 | | |
| DI49_4177; DI49_4176 | DI49_4177; DI49_4176 | | |
| DI49_4181; DI49_4180 | DI49_4181; DI49_4180 | | |
| DI49_4183; DI49_4182 | DI49_4183; DI49_4182 | | |
| DI49_4185; DI49_4184 | DI49_4185; DI49_4184 | | |
| DI49_4190; DI49_4189 | DI49_4190; DI49_4189 | | |
| DI49_4192; DI49_4191 | DI49_4192; DI49_4191 | | |
| DI49_4194; DI49_4193 | DI49_4194; DI49_4193 | | |
| DI49_4198; DI49_4197 | DI49_4198; DI49_4197 | | |
| DI49_4200; DI49_4199 | DI49_4200; DI49_4199 | | |
| DI49_4202; DI49_4201 | DI49_4202; DI49_4201 | | |
| YMR115W; DI49_4203 | DI49_4203 | | YMR115W |
| DI49_4205; DI49_4204 | DI49_4205; DI49_4204 | | |
| DI49_4210; DI49_3558; DI49_4211; DI49_3557 | DI49_4210; DI49_3558; DI49_4211; DI49_3557 | | |
| DI49_4213; DI49_4212 | DI49_4213; DI49_4212 | | |
| DI49_4215; DI49_4214 | DI49_4215; DI49_4214 | | |
| DI49_4217; DI49_4216 | DI49_4217; DI49_4216 | | |
| DI49_4219; DI49_4218 | DI49_4219; DI49_4218 | | |
| DI49_4221; DI49_4220 | DI49_4221; DI49_4220 | | |
| DI49_4222; SPGP0Y01740 | DI49_4222 | SPGP0Y01740 | |
| DI49_4224; DI49_4223 | DI49_4224; DI49_4223 | | |
| DI49_4226; DI49_4225 | DI49_4226; DI49_4225 | | |
| DI49_4228; DI49_4227 | DI49_4228; DI49_4227 | | |
| DI49_4231; DI49_4230 | DI49_4231; DI49_4230 | | |
| DI49_4233; DI49_4232 | DI49_4233; DI49_4232 | | |
| DI49_4235; DI49_4234 | DI49_4235; DI49_4234 | | |
| DI49_4237; DI49_4236 | DI49_4237; DI49_4236 | | |
| DI49_4239; DI49_4238 | DI49_4239; DI49_4238 | | |

Create PDF in your applications with the Pdfcrowd HTML to PDF API
PDFCROWD

| | | |
|---|---|---|
| DI49_4241; DI49_4240 | DI49_4241; DI49_4240 | |
| DI49_4246; DI49_4245 | DI49_4246; DI49_4245 | |
| DI49_4250; DI49_4249 | DI49_4250; DI49_4249 | |
| DI49_4252; DI49_4251 | DI49_4252; DI49_4251 | |
| DI49_4254; DI49_4253; DI49_4255 | DI49_4254; DI49_4253; DI49_4255 | |
| DI49_4257; DI49_4256 | DI49_4257; DI49_4256 | |
| SPGP0Y02130; DI49_4258 | DI49_4258 | SPGP0Y02130 |
| DI49_4260; DI49_4259 | DI49_4260; DI49_4259 | |
| DI49_4262; DI49_4261 | DI49_4262; DI49_4261 | |
| DI49_4264; DI49_4263 | DI49_4264; DI49_4263 | |
| DI49_4266; DI49_4265 | DI49_4266; DI49_4265 | |
| DI49_4268; DI49_4267 | DI49_4268; DI49_4267 | |
| DI49_4272; DI49_4271 | DI49_4272; DI49_4271 | |
| DI49_4274; DI49_4273 | DI49_4274; DI49_4273 | |
| DI49_4276; DI49_4275 | DI49_4276; DI49_4275 | |
| DI49_4278; DI49_4277 | DI49_4278; DI49_4277 | |
| DI49_4280; DI49_4279 | DI49_4280; DI49_4279 | |
| DI49_4282; DI49_4281 | DI49_4282; DI49_4281 | |
| DI49_4284; DI49_4283 | DI49_4284; DI49_4283 | |
| DI49_4286; DI49_4285 | DI49_4286; DI49_4285 | |
| DI49_4288; DI49_4287 | DI49_4288; DI49_4287 | |
| DI49_4290; DI49_4289 | DI49_4290; DI49_4289 | |
| DI49_4292; DI49_4291 | DI49_4292; DI49_4291 | |
| DI49_4294; DI49_4293 | DI49_4294; DI49_4293 | |
| DI49_4296; DI49_4295 | DI49_4296; DI49_4295 | |
| DI49_4298; DI49_4297 | DI49_4298; DI49_4297 | |
| DI49_4300; DI49_4299 | DI49_4300; DI49_4299 | |
| DI49_4302; DI49_4301 | DI49_4302; DI49_4301 | |
| DI49_4304; DI49_4303 | DI49_4304; DI49_4303 | |
| DI49_4306; DI49_4305 | DI49_4306; DI49_4305 | |
| DI49_4308; DI49_4307 | DI49_4308; DI49_4307 | |
| DI49_4310; DI49_4309 | DI49_4310; DI49_4309 | |
| DI49_4312; DI49_4311 | DI49_4312; DI49_4311 | |
| DI49_4317; DI49_4316 | DI49_4317; DI49_4316 | |
| DI49_4319; DI49_4318 | DI49_4319; DI49_4318 | |
| DI49_4321; DI49_4320 | DI49_4321; DI49_4320 | |
| DI49_4326; DI49_4325 | DI49_4326; DI49_4325 | |
| DI49_4328; DI49_4327 | DI49_4328; DI49_4327 | |
| DI49_4330; DI49_4329 | DI49_4330; DI49_4329 | |
| DI49_4332; DI49_4331 | DI49_4332; DI49_4331 | |

Create PDF in your applications with the Pdfcrowd HTML to PDF API PDFCROWD

| | | | |
|---|---|---|---|
| DI49_4334; SPGP0Ytrna9A | DI49_4334 | SPGP0Ytrna9A | |
| DI49_4336; DI49_4335 | DI49_4336; DI49_4335 | | |
| DI49_4338; DI49_4337 | DI49_4338; DI49_4337 | | |
| DI49_4340; DI49_4339 | DI49_4340; DI49_4339 | | |
| DI49_4342; DI49_4341 | DI49_4342; DI49_4341 | | |
| DI49_4344; DI49_4343 | DI49_4344; DI49_4343 | | |
| DI49_4346; DI49_4345 | DI49_4346; DI49_4345 | | |
| DI49_4348; DI49_4347 | DI49_4348; DI49_4347 | | |
| DI49_4350; DI49_4349 | DI49_4350; DI49_4349 | | |
| DI49_4352; DI49_4351 | DI49_4352; DI49_4351 | | |
| DI49_4354; DI49_4353 | DI49_4354; DI49_4353 | | |
| DI49_4356; DI49_4355 | DI49_4356; DI49_4355 | | |
| DI49_4360; DI49_4359 | DI49_4360; DI49_4359 | | |
| DI49_4362; DI49_4361 | DI49_4362; DI49_4361 | | |
| DI49_4364; DI49_4363 | DI49_4364; DI49_4363 | | |
| DI49_4366; DI49_4365 | DI49_4366; DI49_4365 | | |
| DI49_4379; DI49_4378 | DI49_4379; DI49_4378 | | |
| DI49_4381; DI49_4380 | DI49_4381; DI49_4380 | | |
| DI49_4383; DI49_4382 | DI49_4383; DI49_4382 | | |
| DI49_4385; DI49_4384 | DI49_4385; DI49_4384 | | |
| DI49_4387; DI49_4386 | DI49_4387; DI49_4386 | | |
| DI49_4389; DI49_4388 | DI49_4389; DI49_4388 | | |
| DI49_4391; DI49_4390 | DI49_4391; DI49_4390 | | |
| DI49_4393; DI49_4392 | DI49_4393; DI49_4392 | | |
| YMR307W; YMR306W | | | YMR307W; YMR306W |
| YMR309C; YMR308C | | | YMR309C; YMR308C |
| YMR311C; YMR310C | | | YMR311C; YMR310C |
| YMR313C; YMR312W | | | YMR313C; YMR312W |
| YMR315W; YMR314W | | | YMR315W; YMR314W |
| SPGP0IZ00370; YMR316W | | SPGP0IZ00370 | YMR316W |
| YMR318C; YMR317W | | | YMR318C; YMR317W |
| DI49_1985; YMR319C | DI49_1985 | | YMR319C |
| DI49_1989; DI49_3836; DI49_1678 | DI49_1989; DI49_3836; DI49_1678 | | |
| DI49_1680; DI49_1679 | DI49_1680; DI49_1679 | | |
| DI49_1682; DI49_1681 | DI49_1682; DI49_1681 | | |
| DI49_1685; DI49_1684 | DI49_1685; DI49_1684 | | |
| DI49_3178; DI49_3177 | DI49_3178; DI49_3177 | | |
| YKL205W; DI49_3179 | DI49_3179 | | YKL205W |
| DI49_3181; DI49_3079; YJR066W; DI49_3080 | DI49_3181; DI49_3079; DI49_3080 | | YJR066W |
| YER179W; YER178W | | | YER179W; YER178W |

Create PDF in your applications with the Pdfcrowd HTML to PDF API PDFCROWD

| | | | |
|---|---|---|---|
| YER180C-A; YER180C | | | YER180C-A; YER180C |
| YGL257C; YER183C | | | YGL257C; YER183C |
| YGL258W-A; YGL258W | | | YGL258W-A; YGL258W |
| DI49_1674; DI49_1675 | DI49_1674; DI49_1675 | | |
| DI49_1670; DI49_1671 | DI49_1670; DI49_1671 | | |
| DI49_1666; DI49_1667 | DI49_1666; DI49_1667 | | |
| DI49_1663; DI49_1665; DI49_3769; DI49_3768 | DI49_1663; DI49_1665; DI49_3769; DI49_3768 | | |
| DI49_1661; DI49_1662 | DI49_1661; DI49_1662 | | |
| DI49_1659; DI49_2529; SPGP0Ztrna7F; DI49_2530 | DI49_1659; DI49_2529; DI49_2530 | SPGP0Ztrna7F | |
| DI49_1657; DI49_1658 | DI49_1657; DI49_1658 | | |
| DI49_1655; DI49_1656 | DI49_1655; DI49_1656 | | |
| DI49_1653; DI49_1654 | DI49_1653; DI49_1654 | | |
| DI49_0810; DI49_1652 | DI49_0810; DI49_1652 | | |
| DI49_0813; DI49_0811 | DI49_0813; DI49_0811 | | |
| DI49_0815; DI49_0814 | DI49_0815; DI49_0814 | | |
| DI49_0817; DI49_0816 | DI49_0817; DI49_0816 | | |
| DI49_0819; DI49_0818 | DI49_0819; DI49_0818 | | |
| DI49_0820; SPGP0JK00190 | DI49_0820 | SPGP0JK00190 | |
| YCR077C; DI49_0821 | DI49_0821 | | YCR077C |
| DI49_0823; DI49_0822 | DI49_0823; DI49_0822 | | |
| DI49_0825; DI49_0824 | DI49_0825; DI49_0824 | | |
| DI49_0827; DI49_0826 | DI49_0827; DI49_0826 | | |
| DI49_0829; DI49_0828 | DI49_0829; DI49_0828 | | |
| DI49_0832; DI49_0831 | DI49_0832; DI49_0831 | | |
| DI49_0834; DI49_0833 | DI49_0834; DI49_0833 | | |
| DI49_2007; SPGP0JN00100; YPR204W; DI49_3931 | DI49_2007; DI49_3931 | SPGP0JN00100 | YPR204W |
| DI49_2009; DI49_2008 | DI49_2009; DI49_2008 | | |
| DI49_2011; DI49_2010 | DI49_2011; DI49_2010 | | |
| DI49_5186; DI49_5185 | DI49_5186; DI49_5185 | | |
| DI49_5191; DI49_5190 | DI49_5191; DI49_5190 | | |
| DI49_5685; DI49_5192 | DI49_5685; DI49_5192 | | |
| DI49_5683; DI49_5684 | DI49_5683; DI49_5684 | | |
| DI49_3677; DI49_3678 | DI49_3677; DI49_3678 | | |
| DI49_3675; DI49_3676 | DI49_3675; DI49_3676 | | |
| DI49_3673; DI49_3674 | DI49_3673; DI49_3674 | | |
| DI49_3671; DI49_3672 | DI49_3671; DI49_3672 | | |
| DI49_3669; DI49_3670 | DI49_3669; DI49_3670 | | |
| DI49_3667; DI49_3668 | DI49_3667; DI49_3668 | | |
| DI49_3662; DI49_3663 | DI49_3662; DI49_3663 | | |
| DI49_3660; DI49_3661 | DI49_3660; DI49_3661 | | |

Create PDF in your applications with the Pdfcrowd HTML to PDF API PDFCROWD

| | | |
|---|---|---|
| DI49_3658; DI49_3659 | DI49_3658; DI49_3659 | |
| DI49_3656; DI49_3657 | DI49_3656; DI49_3657 | |
| DI49_3654; DI49_3655 | DI49_3654; DI49_3655 | |
| DI49_3652; DI49_3653 | DI49_3652; DI49_3653 | |
| DI49_3650; DI49_3651 | DI49_3650; DI49_3651 | |
| DI49_3648; DI49_3649 | DI49_3648; DI49_3649 | |
| DI49_3646; DI49_3647 | DI49_3646; DI49_3647 | |
| DI49_3644; DI49_3645 | DI49_3644; DI49_3645 | |
| DI49_3642; DI49_3643 | DI49_3642; DI49_3643 | |
| DI49_3640; DI49_3641 | DI49_3640; DI49_3641 | |
| DI49_3636; DI49_3637 | DI49_3636; DI49_3637 | |
| DI49_3634; DI49_3635 | DI49_3634; DI49_3635 | |
| DI49_3632; DI49_3633 | DI49_3632; DI49_3633 | |
| DI49_3630; DI49_3631 | DI49_3630; DI49_3631 | |
| DI49_3628; DI49_3629 | DI49_3628; DI49_3629 | |
| DI49_3626; DI49_3627 | DI49_3626; DI49_3627 | |
| DI49_3625; SPGP0AA00620 | DI49_3625 | SPGP0AA00620 |
| DI49_3623; DI49_3624 | DI49_3623; DI49_3624 | |
| DI49_3621; DI49_3622 | DI49_3621; DI49_3622 | |
| DI49_3619; DI49_3620 | DI49_3619; DI49_3620 | |
| DI49_3617; DI49_3618 | DI49_3617; DI49_3618 | |
| DI49_3615; DI49_3616 | DI49_3615; DI49_3616 | |
| DI49_3613; DI49_3614 | DI49_3613; DI49_3614 | |
| DI49_3611; DI49_3612 | DI49_3611; DI49_3612 | |
| DI49_3609; DI49_3610 | DI49_3609; DI49_3610 | |
| DI49_3606; DI49_3607 | DI49_3606; DI49_3607 | |
| DI49_3604; DI49_3605 | DI49_3604; DI49_3605 | |
| DI49_3602; DI49_3603 | DI49_3602; DI49_3603 | |
| DI49_3600; DI49_3601 | DI49_3600; DI49_3601 | |
| DI49_3598; DI49_3599 | DI49_3598; DI49_3599 | |
| DI49_3596; DI49_3597 | DI49_3596; DI49_3597 | |
| DI49_3594; DI49_3595 | DI49_3594; DI49_3595 | |
| DI49_3592; DI49_3593 | DI49_3592; DI49_3593 | |
| DI49_3590; DI49_3591 | DI49_3590; DI49_3591 | |
| DI49_3588; DI49_3589; SPGP0AE00240; DI49_1622 | DI49_3588; DI49_3589; DI49_1622 | SPGP0AE00240 |
| DI49_3586; DI49_3587 | DI49_3586; DI49_3587 | |
| DI49_3584; DI49_3585 | DI49_3584; DI49_3585 | |
| DI49_3582; DI49_3583 | DI49_3582; DI49_3583 | |
| DI49_3580; DI49_3581 | DI49_3580; DI49_3581 | |
| DI49_3578; DI49_3579 | DI49_3578; DI49_3579 | |

Create PDF in your applications with the Pdfcrowd HTML to PDF API PDFCROWD

| | | | |
|---|---|---|---|
| SPGP0AA01110; DI49_2181; DI49_3575; DI49_2182 | DI49_2181; DI49_3575; DI49_2182 | SPGP0AA01110 | |
| DI49_3573; DI49_3574 | DI49_3573; DI49_3574 | | |
| DI49_3571; DI49_3572 | DI49_3571; DI49_3572 | | |
| DI49_3569; DI49_3570 | DI49_3569; DI49_3570 | | |
| DI49_3567; DI49_3568 | DI49_3567; DI49_3568 | | |
| DI49_3561; DI49_3562 | DI49_3561; DI49_3562 | | |
| DI49_3559; DI49_3560 | DI49_3559; DI49_3560 | | |
| DI49_3555; DI49_3556 | DI49_3555; DI49_3556 | | |
| DI49_3553; DI49_3554 | DI49_3553; DI49_3554 | | |
| DI49_3551; DI49_3552 | DI49_3551; DI49_3552 | | |
| DI49_3549; DI49_3550 | DI49_3549; DI49_3550 | | |
| DI49_3547; DI49_3548 | DI49_3547; DI49_3548 | | |
| DI49_3545; DI49_3546 | DI49_3545; DI49_3546 | | |
| DI49_3543; DI49_3544 | DI49_3543; DI49_3544 | | |
| DI49_3541; DI49_3542 | DI49_3541; DI49_3542 | | |
| DI49_3539; DI49_3540 | DI49_3539; DI49_3540 | | |
| DI49_3537; SPGP0AAtrna15S | DI49_3537 | SPGP0AAtrna15S | |
| DI49_3535; DI49_3536 | DI49_3535; DI49_3536 | | |
| DI49_3533; DI49_3534 | DI49_3533; DI49_3534 | | |
| DI49_3531; DI49_3532 | DI49_3531; DI49_3532 | | |
| YLR002C; DI49_3530 | DI49_3530 | | YLR002C |
| DI49_3528; DI49_3529 | DI49_3528; DI49_3529 | | |
| DI49_3526; DI49_3527 | DI49_3526; DI49_3527 | | |
| DI49_3524; DI49_3525 | DI49_3524; DI49_3525 | | |
| DI49_3522; DI49_3523 | DI49_3522; DI49_3523 | | |
| DI49_3520; DI49_3521 | DI49_3520; DI49_3521 | | |
| DI49_3518; DI49_3519 | DI49_3518; DI49_3519 | | |
| DI49_3516; DI49_3517 | DI49_3516; DI49_3517 | | |
| DI49_3514; DI49_3515 | DI49_3514; DI49_3515 | | |
| DI49_3512; DI49_3513 | DI49_3512; DI49_3513 | | |
| DI49_3510; DI49_3511 | DI49_3510; DI49_3511 | | |
| DI49_3509; YLL021W | DI49_3509 | | YLL021W |
| SPGP0AAtrna11P; SPGP0AEtrna12P; DI49_3504 | DI49_3504 | SPGP0AAtrna11P; SPGP0AEtrna12P | |
| DI49_3502; DI49_3503 | DI49_3502; DI49_3503 | | |
| DI49_3500; DI49_3501 | DI49_3500; DI49_3501 | | |
| DI49_3498; DI49_3499 | DI49_3498; DI49_3499 | | |
| DI49_3496; DI49_3497 | DI49_3496; DI49_3497 | | |
| DI49_3494; DI49_3495 | DI49_3494; DI49_3495 | | |
| DI49_3493; YLL039C | DI49_3493 | | YLL039C |
| DI49_3491; DI49_3492 | DI49_3491; DI49_3492 | | |

Create PDF in your applications with the Pdfcrowd HTML to PDF API PDFCROWD

| | | | |
|---|---|---|---|
| DI49_0839; YIL169C; YIL167W | DI49_0839 | | YIL169C; YIL167W |
| YER066W; YER067W | | | YER066W; YER067W |
| YER045C; YER046W | | | YER045C; YER046W |
| YER033C; YER034W | | | YER033C; YER034W |
| YER028C; YER029C | | | YER028C; YER029C |
| YER026C; YER027C | | | YER026C; YER027C |
| YER015W; SPGP0B00730 | | SPGP0B00730 | YER015W |
| YER002W; YER003C | | | YER002W; YER003C |
| YEL001C; YER001W | | | YEL001C; YER001W |
| YEL003W; YEL002C | | | YEL003W; YEL002C |
| YEL005C; YEL004W | | | YEL005C; YEL004W |
| YEL009C; DI49_1387 | DI49_1387 | | YEL009C |
| YEL016C; DI49_1377; YEL017W | DI49_1377 | | YEL016C; YEL017W |
| YEL018W; YEL017C-A | | | YEL018W; YEL017C-A |
| YEL020C; YEL019C | | | YEL020C; YEL019C |
| YEL021W; DI49_1384 | DI49_1384 | | YEL021W |
| YEL030W; DI49_1362 | DI49_1362 | | YEL030W |
| YEL040W; YEL039C | | | YEL040W; YEL039C |
| YEL042W; YEL041W | | | YEL042W; YEL041W |
| YEL044W; YEL043W | | | YEL044W; YEL043W |
| YEL046C; YEL045C | | | YEL046C; YEL045C |
| YEL048C; YEL047C | | | YEL048C; YEL047C |
| YEL057C; YEL056W | | | YEL057C; YEL056W |
| YEL059C-A; YEL058W | | | YEL059C-A; YEL058W |
| YEL061C; YEL060C | | | YEL061C; YEL060C |
| YEL063C; YEL062W | | | YEL063C; YEL062W |
| YEL065W; YEL064C | | | YEL065W; YEL064C |
| YEL069C; YNR074C; SPGP0B01560; YNR071C | | SPGP0B01560 | YEL069C; YNR074C; YNR071C |
| DI49_1608; YEL073C; DI49_0807 | DI49_1608; DI49_0807 | | YEL073C |
| DI49_1610; DI49_1609 | DI49_1610; DI49_1609 | | |
| DI49_1612; DI49_1611 | DI49_1612; DI49_1611 | | |
| DI49_1614; DI49_1613 | DI49_1614; DI49_1613 | | |
| DI49_1616; DI49_1615 | DI49_1616; DI49_1615 | | |
| DI49_1618; DI49_1617 | DI49_1618; DI49_1617 | | |
| DI49_1620; DI49_1619 | DI49_1620; DI49_1619 | | |
| DI49_1624; DI49_1623 | DI49_1624; DI49_1623 | | |
| DI49_1626; DI49_1625 | DI49_1626; DI49_1625 | | |
| DI49_1628; DI49_1627 | DI49_1628; DI49_1627 | | |
| DI49_1629; SPGP0AE00320 | DI49_1629 | SPGP0AE00320 | |
| DI49_1631; DI49_1630 | DI49_1631; DI49_1630 | | |

Create PDF in your applications with the Pdfcrowd HTML to PDF API PDFCROWD

| | | |
|---|---|---|
| DI49_1633; DI49_1632 | DI49_1633; DI49_1632 | |
| DI49_1638; DI49_1637 | DI49_1638; DI49_1637 | |
| DI49_1640; DI49_1639 | DI49_1640; DI49_1639 | |
| DI49_1642; DI49_1641 | DI49_1642; DI49_1641 | |
| DI49_2013; DI49_2012 | DI49_2013; DI49_2012 | |
| DI49_2015; DI49_2014 | DI49_2015; DI49_2014 | |
| DI49_2017; DI49_2016 | DI49_2017; DI49_2016 | |
| DI49_2019; DI49_2018 | DI49_2019; DI49_2018 | |
| DI49_2021; DI49_2020 | DI49_2021; DI49_2020 | |
| DI49_2023; DI49_2022 | DI49_2023; DI49_2022 | |
| SPGP0AF00230; SPGP0S00750; DI49_2025 | DI49_2025 | SPGP0AF00230; SPGP0S00750 |
| DI49_2027; DI49_2026 | DI49_2027; DI49_2026 | |
| DI49_2029; DI49_2028 | DI49_2029; DI49_2028 | |
| DI49_2031; DI49_2030 | DI49_2031; DI49_2030 | |
| DI49_2033; DI49_2032 | DI49_2033; DI49_2032 | |
| DI49_2035; DI49_2034 | DI49_2035; DI49_2034 | |
| DI49_2037; DI49_2036 | DI49_2037; DI49_2036 | |
| DI49_2039; DI49_2038 | DI49_2039; DI49_2038 | |
| DI49_2041; DI49_2040 | DI49_2041; DI49_2040 | |
| DI49_2043; DI49_2042; DI49_5546; DI49_5547 | DI49_2043; DI49_2042; DI49_5546; DI49_5547 | |
| DI49_2045; DI49_2044 | DI49_2045; DI49_2044 | |
| DI49_2047; DI49_2046 | DI49_2047; DI49_2046 | |
| DI49_2049; SPGP0CItrna20T; DI49_2583; SPGP0ATtrna19T; DI49_5554 | DI49_2049; DI49_2583; DI49_5554 | SPGP0CItrna20T; SPGP0ATtrna19T |
| DI49_2051; DI49_2050 | DI49_2051; DI49_2050 | |
| DI49_2053; DI49_2052 | DI49_2053; DI49_2052 | |
| DI49_2055; DI49_2054 | DI49_2055; DI49_2054 | |
| DI49_2057; DI49_2056 | DI49_2057; DI49_2056 | |
| DI49_2059; DI49_2058 | DI49_2059; DI49_2058 | |
| DI49_2061; DI49_2060 | DI49_2061; DI49_2060 | |
| DI49_2063; DI49_2062 | DI49_2063; DI49_2062 | |
| DI49_2065; DI49_2064 | DI49_2065; DI49_2064 | |
| DI49_2073; DI49_2072 | DI49_2073; DI49_2072 | |
| DI49_2075; DI49_2074 | DI49_2075; DI49_2074 | |
| DI49_2077; DI49_5575; DI49_2078; DI49_5576 | DI49_2077; DI49_5575; DI49_2078; DI49_5576 | |
| DI49_2086; DI49_5589; DI49_2087; DI49_5590 | DI49_2086; DI49_5589; DI49_2087; DI49_5590 | |
| DI49_2089; DI49_2088 | DI49_2089; DI49_2088 | |
| DI49_2091; DI49_2090 | DI49_2091; DI49_2090 | |
| DI49_2093; DI49_2092 | DI49_2093; DI49_2092 | |
| DI49_2095; DI49_2094 | DI49_2095; DI49_2094 | |

Create PDF in your applications with the Pdfcrowd HTML to PDF API PDFCROWD

| | | |
|---|---|---|
| DI49_2097; DI49_2096 | DI49_2097; DI49_2096 | |
| DI49_2099; DI49_5598; DI49_2100; DI49_5599 | DI49_2099; DI49_5598; DI49_2100; DI49_5599 | |
| DI49_2102; DI49_2101 | DI49_2102; DI49_2101 | |
| DI49_2105; DI49_5602 | DI49_2105; DI49_5602 | |
| DI49_2109; DI49_2108 | DI49_2109; DI49_2108 | |
| DI49_2118; DI49_2117 | DI49_2118; DI49_2117 | |
| DI49_2120; DI49_2119 | DI49_2120; DI49_2119 | |
| DI49_2122; DI49_2121 | DI49_2122; DI49_2121 | |
| DI49_2131; DI49_2130 | DI49_2131; DI49_2130 | |
| DI49_2133; DI49_2132 | DI49_2133; DI49_2132 | |
| DI49_2135; DI49_2134 | DI49_2135; DI49_2134 | |
| DI49_2137; DI49_2136 | DI49_2137; DI49_2136 | |
| DI49_2142; DI49_2141 | DI49_2142; DI49_2141 | |
| DI49_2144; DI49_2143 | DI49_2144; DI49_2143 | |
| DI49_2150; DI49_2149 | DI49_2150; DI49_2149 | |
| DI49_2152; DI49_2151 | DI49_2152; DI49_2151 | |
| DI49_2154; DI49_2153 | DI49_2154; DI49_2153 | |
| DI49_2158; DI49_2157 | DI49_2158; DI49_2157 | |
| DI49_2161; DI49_2160 | DI49_2161; DI49_2160 | |
| DI49_2163; DI49_2162 | DI49_2163; DI49_2162 | |
| DI49_2165; DI49_2164 | DI49_2165; DI49_2164 | |
| DI49_2167; DI49_2166 | DI49_2167; DI49_2166 | |
| DI49_2169; DI49_2168 | DI49_2169; DI49_2168 | |
| DI49_2171; DI49_2170 | DI49_2171; DI49_2170 | |
| DI49_2173; DI49_2172 | DI49_2173; DI49_2172 | |
| DI49_2175; DI49_2174 | DI49_2175; DI49_2174 | |
| DI49_2177; DI49_2176 | DI49_2177; DI49_2176 | |
| DI49_2179; DI49_2178 | DI49_2179; DI49_2178 | |
| DI49_2184; DI49_2183 | DI49_2184; DI49_2183 | |
| DI49_2188; DI49_2187 | DI49_2188; DI49_2187 | |
| DI49_2190; DI49_2189 | DI49_2190; DI49_2189 | |
| DI49_2191; YGR225W | DI49_2191 | YGR225W |
| DI49_2194; DI49_2192 | DI49_2194; DI49_2192 | |
| DI49_2196; DI49_2195 | DI49_2196; DI49_2195 | |
| DI49_2198; DI49_2197 | DI49_2198; DI49_2197 | |
| DI49_2200; DI49_2199 | DI49_2200; DI49_2199 | |
| DI49_2202; DI49_2201 | DI49_2202; DI49_2201 | |
| DI49_2204; DI49_2203 | DI49_2204; DI49_2203 | |
| DI49_2206; DI49_2205 | DI49_2206; DI49_2205 | |
| DI49_2208; DI49_2207 | DI49_2208; DI49_2207 | |

Create PDF in your applications with the Pdfcrowd HTML to PDF API PDFCROWD

| | | | |
|---|---|---|---|
| DI49_2210; DI49_2209 | DI49_2210; DI49_2209 | | |
| DI49_2212; DI49_2211 | DI49_2212; DI49_2211 | | |
| DI49_2214; DI49_2213 | DI49_2214; DI49_2213 | | |
| DI49_2216; DI49_2215 | DI49_2216; DI49_2215 | | |
| DI49_2218; DI49_2577; DI49_2219; DI49_2578 | DI49_2218; DI49_2577; DI49_2219; DI49_2578 | | |
| DI49_2221; DI49_2585; DI49_2222; DI49_2586 | DI49_2221; DI49_2585; DI49_2222; DI49_2586 | | |
| DI49_2224; DI49_2223 | DI49_2224; DI49_2223 | | |
| DI49_2226; DI49_2225 | DI49_2226; DI49_2225 | | |
| DI49_2228; DI49_2227 | DI49_2228; DI49_2227 | | |
| DI49_2230; DI49_2229 | DI49_2230; DI49_2229 | | |
| DI49_2232; DI49_2231 | DI49_2232; DI49_2231 | | |
| DI49_2234; DI49_2233 | DI49_2234; DI49_2233 | | |
| DI49_2236; DI49_2235 | DI49_2236; DI49_2235 | | |
| DI49_2238; DI49_2237 | DI49_2238; DI49_2237 | | |
| SPGP0AF02170; DI49_2239 | DI49_2239 | SPGP0AF02170 | |
| DI49_2241; DI49_2240 | DI49_2241; DI49_2240 | | |
| DI49_0798; DI49_0797 | DI49_0798; DI49_0797 | | |
| DI49_0800; DI49_0799 | DI49_0800; DI49_0799 | | |
| DI49_0802; DI49_0801 | DI49_0802; DI49_0801 | | |
| DI49_0804; DI49_0803 | DI49_0804; DI49_0803 | | |
| DI49_0806; DI49_0805 | DI49_0806; DI49_0805 | | |
| YJL205C; YJL204C | | | YJL205C; YJL204C |
| YJL210W; YJL209W | | | YJL210W; YJL209W |
| SPGP0AI00230; YJL212C | | SPGP0AI00230 | YJL212C |
| YJL214W; SPGP0AJ01050 | | SPGP0AJ01050 | YJL214W |
| YOL158C; DI49_1601 | DI49_1601 | | YOL158C |
| YOL159C; SPGP0C00110 | | SPGP0C00110 | YOL159C |
| DI49_2405; DI49_2404 | DI49_2405; DI49_2404 | | |
| DI49_2407; DI49_2406 | DI49_2407; DI49_2406 | | |
| DI49_2409; DI49_2408 | DI49_2409; DI49_2408 | | |
| DI49_2417; DI49_2416 | DI49_2417; DI49_2416 | | |
| DI49_2419; DI49_2418 | DI49_2419; DI49_2418 | | |
| DI49_2424; DI49_2423 | DI49_2424; DI49_2423 | | |
| DI49_2426; DI49_2425 | DI49_2426; DI49_2425 | | |
| DI49_2430; DI49_2429 | DI49_2430; DI49_2429 | | |
| DI49_2432; DI49_2431 | DI49_2432; DI49_2431 | | |
| DI49_2434; DI49_2433 | DI49_2434; DI49_2433 | | |
| DI49_2436; DI49_2435 | DI49_2436; DI49_2435 | | |
| DI49_2438; DI49_2437 | DI49_2438; DI49_2437 | | |
| DI49_2440; DI49_2439 | DI49_2440; DI49_2439 | | |

Create PDF in your applications with the Pdfcrowd HTML to PDF API PDFCROWD

| | | | |
|---|---|---|---|
| DI49_2442; DI49_2441 | DI49_2442; DI49_2441 | | |
| DI49_2444; DI49_2443 | DI49_2444; DI49_2443 | | |
| YHR039C-A; DI49_2445 | DI49_2445 | | YHR039C-A |
| DI49_2447; DI49_2446 | DI49_2447; DI49_2446 | | |
| DI49_2449; DI49_2448 | DI49_2449; DI49_2448 | | |
| DI49_2451; DI49_2450 | DI49_2451; DI49_2450 | | |
| DI49_2453; DI49_2452 | DI49_2453; DI49_2452 | | |
| DI49_2455; DI49_2454 | DI49_2455; DI49_2454 | | |
| DI49_2457; DI49_2456 | DI49_2457; DI49_2456 | | |
| DI49_2459; DI49_2458 | DI49_2459; DI49_2458 | | |
| DI49_2461; DI49_2460 | DI49_2461; DI49_2460 | | |
| DI49_2463; DI49_2462 | DI49_2463; DI49_2462 | | |
| DI49_2465; DI49_2464 | DI49_2465; DI49_2464 | | |
| DI49_2467; DI49_2466 | DI49_2467; DI49_2466 | | |
| DI49_2472; DI49_2471 | DI49_2472; DI49_2471 | | |
| DI49_2474; DI49_2473 | DI49_2474; DI49_2473 | | |
| DI49_2478; DI49_2477 | DI49_2478; DI49_2477 | | |
| DI49_2480; DI49_2479 | DI49_2480; DI49_2479 | | |
| DI49_2482; DI49_2481 | DI49_2482; DI49_2481 | | |
| DI49_2484; DI49_2483 | DI49_2484; DI49_2483 | | |
| DI49_2486; DI49_2485 | DI49_2486; DI49_2485 | | |
| DI49_2488; DI49_2487 | DI49_2488; DI49_2487 | | |
| DI49_2490; DI49_2489 | DI49_2490; DI49_2489 | | |
| DI49_2492; DI49_2491 | DI49_2492; DI49_2491 | | |
| DI49_2494; DI49_2493 | DI49_2494; DI49_2493 | | |
| DI49_2496; DI49_2495 | DI49_2496; DI49_2495 | | |
| DI49_2498; DI49_2497 | DI49_2498; DI49_2497 | | |
| DI49_2504; DI49_2503 | DI49_2504; DI49_2503 | | |
| DI49_2506; DI49_2505 | DI49_2506; DI49_2505 | | |
| DI49_2508; DI49_2507 | DI49_2508; DI49_2507 | | |
| DI49_2510; DI49_2509 | DI49_2510; DI49_2509 | | |
| DI49_2512; DI49_2511 | DI49_2512; DI49_2511 | | |
| DI49_2514; DI49_2513 | DI49_2514; DI49_2513 | | |
| DI49_2516; DI49_2515 | DI49_2516; DI49_2515 | | |
| DI49_2518; DI49_2517 | DI49_2518; DI49_2517 | | |
| DI49_2520; DI49_2519 | DI49_2520; DI49_2519 | | |
| DI49_2522; DI49_2521 | DI49_2522; DI49_2521 | | |
| DI49_2524; DI49_2523 | DI49_2524; DI49_2523 | | |
| DI49_2526; DI49_2525 | DI49_2526; DI49_2525 | | |
| SPGP0AJ01340; DI49_2527 | DI49_2527 | SPGP0AJ01340 | |

Create PDF in your applications with the Pdfcrowd HTML to PDF API PDFCROWD

| | | | |
|---|---|---|---|
| DI49_2532; DI49_2531 | DI49_2532; DI49_2531 | | |
| DI49_2534; DI49_2533 | DI49_2534; DI49_2533 | | |
| DI49_2536; DI49_2535 | DI49_2536; DI49_2535 | | |
| DI49_2538; DI49_2537 | DI49_2538; DI49_2537 | | |
| DI49_2540; DI49_2539 | DI49_2540; DI49_2539 | | |
| DI49_2541; SPGP0AJ01470 | DI49_2541 | SPGP0AJ01470 | |
| DI49_2543; DI49_2542 | DI49_2543; DI49_2542 | | |
| DI49_2544; SPGP0AJ01510 | DI49_2544 | SPGP0AJ01510 | |
| DI49_2546; DI49_2545 | DI49_2546; DI49_2545 | | |
| DI49_2551; DI49_2550 | DI49_2551; DI49_2550 | | |
| DI49_2553; DI49_2552 | DI49_2553; DI49_2552 | | |
| DI49_2555; DI49_2554 | DI49_2555; DI49_2554 | | |
| DI49_2557; DI49_2556 | DI49_2557; DI49_2556 | | |
| DI49_2559; DI49_2558 | DI49_2559; DI49_2558 | | |
| DI49_2561; DI49_2560 | DI49_2561; DI49_2560 | | |
| DI49_2563; DI49_2562 | DI49_2563; DI49_2562 | | |
| DI49_2565; DI49_2564 | DI49_2565; DI49_2564 | | |
| DI49_2567; DI49_2566 | DI49_2567; DI49_2566 | | |
| DI49_2569; DI49_2568 | DI49_2569; DI49_2568 | | |
| DI49_2571; DI49_2570 | DI49_2571; DI49_2570 | | |
| DI49_2573; DI49_2572 | DI49_2573; DI49_2572 | | |
| DI49_2575; DI49_2574 | DI49_2575; DI49_2574 | | |
| SPGP0AJ01840; DI49_2576 | DI49_2576 | SPGP0AJ01840 | |
| SPGP0AJ01880; DI49_2579 | DI49_2579 | SPGP0AJ01880 | |
| DI49_2581; DI49_2580 | DI49_2581; DI49_2580 | | |
| YOL156W; SPGP0BQ00100 | | SPGP0BQ00100 | YOL156W |
| YIL164C; YIL166C | | | YIL164C; YIL166C |
| YIL156W-B; YIL157C | | | YIL156W-B; YIL157C |
| YIL138C; YIL139C | | | YIL138C; YIL139C |
| YIL136W; YIL137C | | | YIL136W; YIL137C |
| YIL134W; YIL135C | | | YIL134W; YIL135C |
| YIL132C; YIL133C | | | YIL132C; YIL133C |
| YIL130W; YIL131C | | | YIL130W; YIL131C |
| YIL119C; YIL120W | | | YIL119C; YIL120W |
| YIL117C; YIL118W | | | YIL117C; YIL118W |
| YIL112W; YIL113W | | | YIL112W; YIL113W |
| YIL104C; YIL105C | | | YIL104C; YIL105C |
| YIL102C-A; YIL103W | | | YIL102C-A; YIL103W |
| SPGP0Ctrna12T; SPGP0Itrna22T; SPGP0Ctrna13T; YIL014W | | SPGP0Ctrna12T; SPGP0Itrna22T; SPGP0Ctrna13T | YIL014W |
| YIL097W; YIL098C | | | YIL097W; YIL098C |

Create PDF in your applications with the Pdfcrowd HTML to PDF API PDFCROWD

| | | | |
|---|---|---|---|
| YIL090W; YIL091C | | | YIL090W; YIL091C |
| YIL085C; YIL087C | | | YIL085C; YIL087C |
| YIL074C; YIL075C | | | YIL074C; YIL075C |
| YIL042C; YIL043C | | | YIL042C; YIL043C |
| YIL023C; YIL024C | | | YIL023C; YIL024C |
| YIL011W; YIL013C; YDR011W | | | YIL011W; YIL013C; YDR011W |
| YIL009C-A; YIL010W | | | YIL009C-A; YIL010W |
| YIL007C; YIL008W | | | YIL007C; YIL008W |
| YIL005W; YIL006W | | | YIL005W; YIL006W |
| YIL003W; DI49_2776 | DI49_2776 | | YIL003W |
| YIR005W; YIR004W | | | YIR005W; YIR004W |
| YIR015W; YIR014W | | | YIR015W; YIR014W |
| YIR022W; YIR021W | | | YIR022W; YIR021W |
| YIR024C; YIR023W | | | YIR024C; YIR023W |
| YIR026C; YIR025W | | | YIR026C; YIR025W |
| YIR032C; YIR031C | | | YIR032C; YIR031C |
| YIR037W; YIR036C | | | YIR037W; YIR036C |
| YIR039C; YIR038C | | | YIR039C; YIR038C |
| DI49_0562; SPGP0C02050 | DI49_0562 | SPGP0C02050 | |
| DI49_0564; DI49_0563 | DI49_0564; DI49_0563 | | |
| DI49_0570; DI49_0569 | DI49_0570; DI49_0569 | | |
| DI49_0572; DI49_0571 | DI49_0572; DI49_0571 | | |
| DI49_0574; DI49_0573 | DI49_0574; DI49_0573 | | |
| DI49_0576; DI49_0575 | DI49_0576; DI49_0575 | | |
| DI49_0578; DI49_0577 | DI49_0578; DI49_0577 | | |
| DI49_0580; DI49_0579 | DI49_0580; DI49_0579 | | |
| DI49_0583; SPGP0AN00320 | DI49_0583 | SPGP0AN00320 | |
| DI49_0585; DI49_0584 | DI49_0585; DI49_0584 | | |
| DI49_0587; DI49_0586 | DI49_0587; DI49_0586 | | |
| DI49_0589; DI49_0588 | DI49_0589; DI49_0588 | | |
| DI49_0591; DI49_0590 | DI49_0591; DI49_0590 | | |
| DI49_0595; DI49_0594 | DI49_0595; DI49_0594 | | |
| SPGP0AN00460; SPGP0AN00450 | | SPGP0AN00460; SPGP0AN00450 | |
| DI49_0597; DI49_0596 | DI49_0597; DI49_0596 | | |
| DI49_0599; DI49_0598 | DI49_0599; DI49_0598 | | |
| DI49_0601; DI49_0600 | DI49_0601; DI49_0600 | | |
| DI49_0603; DI49_0602 | DI49_0603; DI49_0602 | | |
| DI49_0605; DI49_0604 | DI49_0605; DI49_0604 | | |
| DI49_0607; DI49_0606 | DI49_0607; DI49_0606 | | |
| DI49_0609; DI49_0608 | DI49_0609; DI49_0608 | | |

Create PDF in your applications with the Pdfcrowd HTML to PDF API PDFCROWD

| | | |
|---|---|---|
| DI49_0611; DI49_0610 | DI49_0611; DI49_0610 | |
| DI49_0615; DI49_0614 | DI49_0615; DI49_0614 | |
| DI49_0620; DI49_0619 | DI49_0620; DI49_0619 | |
| DI49_0622; DI49_0621 | DI49_0622; DI49_0621 | |
| DI49_0624; DI49_0623 | DI49_0624; DI49_0623 | |
| DI49_0626; DI49_0625 | DI49_0626; DI49_0625 | |
| DI49_0627; SPGP0AN00770 | DI49_0627 | SPGP0AN00770 |
| DI49_0629; DI49_0628 | DI49_0629; DI49_0628 | |
| DI49_0631; DI49_0630 | DI49_0631; DI49_0630 | |
| DI49_0633; DI49_0632 | DI49_0633; DI49_0632 | |
| DI49_0635; DI49_0634 | DI49_0635; DI49_0634 | |
| DI49_0639; DI49_0638 | DI49_0639; DI49_0638 | |
| DI49_0641; DI49_0640 | DI49_0641; DI49_0640 | |
| DI49_0643; DI49_0642 | DI49_0643; DI49_0642 | |
| DI49_0645; DI49_0644 | DI49_0645; DI49_0644 | |
| DI49_0647; DI49_0646 | DI49_0647; DI49_0646 | |
| DI49_0649; DI49_0648 | DI49_0649; DI49_0648 | |
| DI49_0651; DI49_0650 | DI49_0651; DI49_0650 | |
| DI49_0653; DI49_0652 | DI49_0653; DI49_0652 | |
| DI49_0655; DI49_0654 | DI49_0655; DI49_0654 | |
| DI49_0657; DI49_0656 | DI49_0657; DI49_0656 | |
| DI49_0659; DI49_0658 | DI49_0659; DI49_0658 | |
| DI49_0661; DI49_0660 | DI49_0661; DI49_0660 | |
| DI49_0663; DI49_0662 | DI49_0663; DI49_0662 | |
| DI49_0665; DI49_3697; DI49_0666; DI49_3696 | DI49_0665; DI49_3697; DI49_0666; DI49_3696 | |
| DI49_0668; DI49_0667 | DI49_0668; DI49_0667 | |
| DI49_0786; DI49_0787 | DI49_0786; DI49_0787 | |
| DI49_0784; DI49_0785 | DI49_0784; DI49_0785 | |
| DI49_0782; DI49_0783 | DI49_0782; DI49_0783 | |
| DI49_0781; SPGP0AQ00190 | DI49_0781 | SPGP0AQ00190 |
| DI49_5467; DI49_5466 | DI49_5467; DI49_5466 | |
| DI49_5471; DI49_5470 | DI49_5471; DI49_5470 | |
| DI49_5473; DI49_5472 | DI49_5473; DI49_5472 | |
| DI49_5475; DI49_5474 | DI49_5475; DI49_5474 | |
| DI49_5477; DI49_5476 | DI49_5477; DI49_5476 | |
| DI49_5479; DI49_5478 | DI49_5479; DI49_5478 | |
| DI49_5481; DI49_5480 | DI49_5481; DI49_5480 | |
| DI49_5483; DI49_5482 | DI49_5483; DI49_5482 | |
| DI49_5485; DI49_5484 | DI49_5485; DI49_5484 | |
| DI49_5488; DI49_5487 | DI49_5488; DI49_5487 | |

Create PDF in your applications with the Pdfcrowd HTML to PDF API PDFCROWD

| | | | |
|---|---|---|---|
| DI49_5490; DI49_5489 | DI49_5490; DI49_5489 | | |
| DI49_5492; DI49_5491 | DI49_5492; DI49_5491 | | |
| DI49_5494; DI49_5493 | DI49_5494; DI49_5493 | | |
| DI49_5496; DI49_5495 | DI49_5496; DI49_5495 | | |
| DI49_5498; DI49_5497 | DI49_5498; DI49_5497 | | |
| DI49_5503; DI49_5502 | DI49_5503; DI49_5502 | | |
| DI49_5505; DI49_5504 | DI49_5505; DI49_5504 | | |
| DI49_5507; DI49_5506 | DI49_5507; DI49_5506 | | |
| DI49_5509; DI49_5508 | DI49_5509; DI49_5508 | | |
| DI49_5511; DI49_5510 | DI49_5511; DI49_5510 | | |
| DI49_5513; DI49_5512 | DI49_5513; DI49_5512 | | |
| DI49_5515; DI49_5514 | DI49_5515; DI49_5514 | | |
| SPGP0AT00610; DI49_5516 | DI49_5516 | SPGP0AT00610 | |
| DI49_5518; DI49_5517 | DI49_5518; DI49_5517 | | |
| DI49_5520; DI49_5519 | DI49_5520; DI49_5519 | | |
| DI49_5522; DI49_5521 | DI49_5522; DI49_5521 | | |
| DI49_5526; DI49_5525 | DI49_5526; DI49_5525 | | |
| DI49_5528; DI49_5527 | DI49_5528; DI49_5527 | | |
| DI49_5530; YBR118W; DI49_5531; DI49_1148 | DI49_5530; DI49_5531; DI49_1148 | | YBR118W |
| DI49_5533; DI49_5532 | DI49_5533; DI49_5532 | | |
| DI49_5535; DI49_5534 | DI49_5535; DI49_5534 | | |
| DI49_5537; DI49_5536 | DI49_5537; DI49_5536 | | |
| DI49_5539; DI49_5538 | DI49_5539; DI49_5538 | | |
| DI49_5541; DI49_5540 | DI49_5541; DI49_5540 | | |
| DI49_5543; DI49_5542 | DI49_5543; DI49_5542 | | |
| DI49_5545; DI49_5544 | DI49_5545; DI49_5544 | | |
| DI49_5549; DI49_5548 | DI49_5549; DI49_5548 | | |
| DI49_5551; DI49_5550 | DI49_5551; DI49_5550 | | |
| DI49_5556; DI49_5555 | DI49_5556; DI49_5555 | | |
| DI49_5558; DI49_5557 | DI49_5558; DI49_5557 | | |
| DI49_5560; DI49_5559 | DI49_5560; DI49_5559 | | |
| DI49_5562; DI49_5561 | DI49_5562; DI49_5561 | | |
| DI49_5570; DI49_5569 | DI49_5570; DI49_5569 | | |
| DI49_5572; DI49_5571 | DI49_5572; DI49_5571 | | |
| DI49_5574; DI49_5573 | DI49_5574; DI49_5573 | | |
| DI49_5578; DI49_5577 | DI49_5578; DI49_5577 | | |
| DI49_5580; DI49_5579 | DI49_5580; DI49_5579 | | |
| DI49_5582; SPGP0ATtrna10N | DI49_5582 | SPGP0ATtrna10N | |
| DI49_5584; DI49_5583 | DI49_5584; DI49_5583 | | |
| DI49_5586; DI49_5585 | DI49_5586; DI49_5585 | | |

Create PDF in your applications with the Pdfcrowd HTML to PDF API PDFCROWD

| | | | |
|---|---|---|---|
| DI49_5592; DI49_5591 | DI49_5592; DI49_5591 | | |
| DI49_5594; DI49_5593 | DI49_5594; DI49_5593 | | |
| DI49_5595; SPGP0AT01360 | DI49_5595 | SPGP0AT01360 | |
| DI49_5597; DI49_5596 | DI49_5597; DI49_5596 | | |
| DI49_5606; DI49_5605 | DI49_5606; DI49_5605 | | |
| DI49_5608; DI49_5607 | DI49_5608; DI49_5607 | | |
| DI49_5610; DI49_5609 | DI49_5610; DI49_5609 | | |
| DI49_5612; DI49_5611 | DI49_5612; DI49_5611 | | |
| DI49_5617; DI49_5616 | DI49_5617; DI49_5616 | | |
| DI49_5619; DI49_5618 | DI49_5619; DI49_5618 | | |
| DI49_5621; DI49_5620 | DI49_5621; DI49_5620 | | |
| DI49_5623; DI49_5622 | DI49_5623; DI49_5622 | | |
| DI49_5625; DI49_5624 | DI49_5625; DI49_5624 | | |
| DI49_5627; DI49_5626 | DI49_5627; DI49_5626 | | |
| DI49_1056; DI49_1055 | DI49_1056; DI49_1055 | | |
| DI49_1058; DI49_1057 | DI49_1058; DI49_1057 | | |
| DI49_1060; DI49_1059 | DI49_1060; DI49_1059 | | |
| DI49_1062; DI49_0209; SPGP0AU00190; DI49_0208 | DI49_1062; DI49_0209; DI49_0208 | SPGP0AU00190 | |
| DI49_1064; DI49_1063 | DI49_1064; DI49_1063 | | |
| DI49_1066; DI49_1065 | DI49_1066; DI49_1065 | | |
| DI49_1068; DI49_1067 | DI49_1068; DI49_1067 | | |
| DI49_1070; DI49_1069 | DI49_1070; DI49_1069 | | |
| DI49_1072; DI49_1071 | DI49_1072; DI49_1071 | | |
| DI49_1074; SPGP0CPtrna20V; DI49_1075; DI49_0220 | DI49_1074; DI49_1075; DI49_0220 | SPGP0CPtrna20V | |
| DI49_1077; DI49_1076 | DI49_1077; DI49_1076 | | |
| DI49_1079; DI49_1078; SPGP0CP02420; DI49_0222 | DI49_1079; DI49_1078; DI49_0222 | SPGP0CP02420 | |
| DI49_1081; DI49_1080 | DI49_1081; DI49_1080 | | |
| YNR075W; DI49_1082 | DI49_1082 | | YNR075W |
| YNR069C; YNR070W | | | YNR069C; YNR070W |
| YNR067C; YNR068C | | | YNR067C; YNR068C |
| YNR065C; YNR066C | | | YNR065C; YNR066C |
| YNR063W; YNR064C | | | YNR063W; YNR064C |
| YNR061C; SPGP0D00210 | | SPGP0D00210 | YNR061C |
| YNR059W; YNR060W | | | YNR059W; YNR060W |
| YNR054C; DI49_4786 | DI49_4786 | | YNR054C |
| YNR049C; YNR050C | | | YNR049C; YNR050C |
| YNR044W; YNR045W | | | YNR044W; YNR045W |
| YNR038W; YNR039C | | | YNR038W; YNR039C |
| YNR036C; YNR037C | | | YNR036C; YNR037C |
| YNR034W-A; YNR035C | | | YNR034W-A; YNR035C |

Create PDF in your applications with the Pdfcrowd HTML to PDF API PDFCROWD

| | | | |
|---|---|---|---|
| YNR033W; YNR034W | | | YNR033W; YNR034W |
| YNR032W; YNR032C-A | | | YNR032W; YNR032C-A |
| YNR027W; YNR028W | | | YNR027W; YNR028W |
| YNR024W; YNR026C | | | YNR024W; YNR026C |
| YNR022C; YNR023W | | | YNR022C; YNR023W |
| SPGP0D00630; YNR021W | | SPGP0D00630 | YNR021W |
| YNR018W; YNR019W | | | YNR018W; YNR019W |
| YNR008W; YNR009W | | | YNR008W; YNR009W |
| YNR006W; YNR007C | | | YNR006W; YNR007C |
| YNR003C; YNR004W | | | YNR003C; YNR004W |
| DI49_0738; DI49_0739 | DI49_0738; DI49_0739 | | |
| DI49_0736; DI49_0737 | DI49_0736; DI49_0737 | | |
| DI49_0734; SPGP0BEtrna19L | DI49_0734 | SPGP0BEtrna19L | |
| DI49_0729; DI49_0730 | DI49_0729; DI49_0730 | | |
| DI49_0727; DI49_0728 | DI49_0727; DI49_0728 | | |
| DI49_0725; DI49_0726 | DI49_0725; DI49_0726 | | |
| DI49_0723; DI49_0724 | DI49_0723; DI49_0724 | | |
| DI49_0721; DI49_0722 | DI49_0721; DI49_0722 | | |
| SPGP0BE00290; DI49_0720 | DI49_0720 | SPGP0BE00290 | |
| YKL069W; SPGP0E00110 | | SPGP0E00110 | YKL069W |
| YKL065C; YKL065W-A | | | YKL065C; YKL065W-A |
| YKL063C; YKL064W | | | YKL063C; YKL064W |
| YKL061W; YKL062W | | | YKL061W; YKL062W |
| YKL059C; YKL060C | | | YKL059C; YKL060C |
| YKL057C; DI49_3317 | DI49_3317 | | YKL057C |
| YKL056C; DI49_3318; YKL055C | DI49_3318 | | YKL056C; YKL055C |
| YKL053C-A; YKL054C; DI49_3320; DI49_3319 | DI49_3320; DI49_3319 | | YKL053C-A; YKL054C |
| YKL051W; YKL052C | | | YKL051W; YKL052C |
| YKL049C; YKL050C | | | YKL049C; YKL050C |
| YKL047W; DI49_3326; DI49_3325 | DI49_3326; DI49_3325 | | YKL047W |
| YKL046C; DI49_3327; YKL045W | DI49_3327 | | YKL046C; YKL045W |
| YKL042W; DI49_3330; YKL041W; DI49_3329 | DI49_3330; DI49_3329 | | YKL042W; YKL041W |
| YKL039W; DI49_3333; YKL038W | DI49_3333 | | YKL039W; YKL038W |
| YKL034W; DI49_3336; YKL033W-A | DI49_3336 | | YKL034W; YKL033W-A |
| SPGP0E00480; DI49_3340; YKL032C | DI49_3340 | SPGP0E00480 | YKL032C |
| YKL028W; DI49_3342; DI49_3341 | DI49_3342; DI49_3341 | | YKL028W |
| YKL027W; DI49_3343; YKL026C | DI49_3343 | | YKL027W; YKL026C |
| YKL024C; DI49_3346; YKL023C-A | DI49_3346 | | YKL024C; YKL023C-A |
| YKL022C; DI49_3349; DI49_3348 | DI49_3349; DI49_3348 | | YKL022C |
| YKL021C; DI49_3350 | DI49_3350 | | YKL021C |

Create PDF in your applications with the Pdfcrowd HTML to PDF API PDFCROWD

| | | | |
|---|---|---|---|
| YKL020C; DI49_3351; YKL019W | DI49_3351 | | YKL020C; YKL019W |
| YKL018W; DI49_3354; YKL017C | DI49_3354 | | YKL018W; YKL017C |
| YKL015W; DI49_3357 | DI49_3357 | | YKL015W |
| YKL014C; DI49_3358; YKL013C | DI49_3358 | | YKL014C; YKL013C |
| YKL011C; YKL012W | | | YKL011C; YKL012W |
| YKL009W; DI49_3363; YKL008C | DI49_3363 | | YKL009W; YKL008C |
| YKL006C-A; DI49_3366; DI49_3365 | DI49_3366; DI49_3365 | | YKL006C-A |
| YKL006W; DI49_3367; YKL005C | DI49_3367 | | YKL006W; YKL005C |
| YKL003C; YKL004W; DI49_3369; DI49_3368 | DI49_3369; DI49_3368 | | YKL003C; YKL004W |
| YKL001C; DI49_3372 | DI49_3372 | | YKL001C |
| YKR001C; DI49_3373 | DI49_3373 | | YKR001C |
| YKR002W; DI49_3374; YKR003W | DI49_3374 | | YKR002W; YKR003W |
| YKR005C; YKR004C | | | YKR005C; YKR004C |
| YKR007W; YKR006C | | | YKR007W; YKR006C |
| YKR009C; YKR008W | | | YKR009C; YKR008W |
| YKR010C; DI49_3383; YKR011C | DI49_3383 | | YKR010C; YKR011C |
| YKR014C; DI49_3386; DI49_3385 | DI49_3386; DI49_3385 | | YKR014C |
| YKR015C; DI49_3387 | DI49_3387 | | YKR015C |
| YKR016W; DI49_3388; YKR017C | DI49_3388 | | YKR016W; YKR017C |
| YKR019C; DI49_3391; YKR020W | DI49_3391 | | YKR019C; YKR020W |
| YKR022C; YKR021W | | | YKR022C; YKR021W |
| YKR024C; DI49_3396; YKR025W | DI49_3396 | | YKR024C; YKR025W |
| YKR029C; DI49_3402; YKR030W | DI49_3402 | | YKR029C; YKR030W |
| YKR034W; DI49_3405 | DI49_3405 | | YKR034W |
| YKR035W-A; DI49_3406; YKR036C | DI49_3406 | | YKR035W-A; YKR036C |
| YKR038C; YKR037C | | | YKR038C; YKR037C |
| YKR039W; DI49_3411 | DI49_3411 | | YKR039W |
| YKR044W; YKR043C; DI49_3415; DI49_3414 | DI49_3415; DI49_3414 | | YKR044W; YKR043C |
| YKR046C; DI49_3418; YKR048C; DI49_3417 | DI49_3418; DI49_3417 | | YKR046C; YKR048C |
| YKR050W; DI49_3421; YKR051W; DI49_3420 | DI49_3421; DI49_3420 | | YKR050W; YKR051W |
| YKR053C; YKR052C | | | YKR053C; YKR052C |
| YKR055W; SPGP0E01300; DI49_3425; DI49_3424 | DI49_3425; DI49_3424 | SPGP0E01300 | YKR055W |
| YKR057W; DI49_3428; YKR058W; DI49_3427 | DI49_3428; DI49_3427 | | YKR057W; YKR058W |
| YKR060W; DI49_3431 | DI49_3431 | | YKR060W |
| YKR061W; DI49_3432 | DI49_3432 | | YKR061W |
| YKR062W; DI49_3433 | DI49_3433 | | YKR062W |
| YKR063C; DI49_3434 | DI49_3434 | | YKR063C |
| YKR064W; SPGP0H00500; YKR065C | | SPGP0H00500 | YKR064W; YKR065C |
| YKR067W; DI49_3437; YKR068C; DI49_3436 | DI49_3437; DI49_3436 | | YKR067W; YKR068C |
| YKR070W; DI49_3440; YKR071C | DI49_3440 | | YKR070W; YKR071C |

Create PDF in your applications with the Pdfcrowd HTML to PDF API PDFCROWD

| | | | |
|---|---|---|---|
| YKR076W; YKR075C | | | YKR076W; YKR075C |
| YKR078W; YKR077W | | | YKR078W; YKR077W |
| YKR080W; DI49_3450; DI49_3449 | DI49_3450; DI49_3449 | | YKR080W |
| YKR081C; DI49_3451 | DI49_3451 | | YKR081C |
| YKR082W; DI49_3452; YKR083C | DI49_3452 | | YKR082W; YKR083C |
| YKR085C; YKR084C | | | YKR085C; YKR084C |
| YKR087C; YKR086W; DI49_3456; DI49_3455 | DI49_3456; DI49_3455 | | YKR087C; YKR086W |
| YKR089C; YKR088C | | | YKR089C; YKR088C |
| YKR091W; YKR090W | | | YKR091W; YKR090W |
| YKR093W; DI49_3463; DI49_3462 | DI49_3463; DI49_3462 | | YKR093W |
| YKR095W; DI49_3464; YKR095W-A | DI49_3464 | | YKR095W; YKR095W-A |
| YKR097W; DI49_3467; YKR098C; DI49_3466 | DI49_3467; DI49_3466 | | YKR097W; YKR098C |
| YKR100C; DI49_3470; YKR101W | DI49_3470 | | YKR100C; YKR101W |
| DI49_0291; SPGP0CD00100 | DI49_0291 | SPGP0CD00100 | |
| YJL159W; DI49_2869; DI49_2870 | DI49_2869; DI49_2870 | | YJL159W |
| DI49_2872; DI49_2871 | DI49_2872; DI49_2871 | | |
| DI49_2874; DI49_2873 | DI49_2874; DI49_2873 | | |
| DI49_2876; DI49_2875 | DI49_2876; DI49_2875 | | |
| DI49_2878; DI49_2877 | DI49_2878; DI49_2877 | | |
| DI49_2880; DI49_2879 | DI49_2880; DI49_2879 | | |
| DI49_2882; DI49_2881 | DI49_2882; DI49_2881 | | |
| DI49_2884; DI49_2883 | DI49_2884; DI49_2883 | | |
| SPGP0F00280; DI49_3430; DI49_2887; DI49_3429 | DI49_3430; DI49_2887; DI49_3429 | SPGP0F00280 | |
| DI49_2888; SPGP0F00300 | DI49_2888 | SPGP0F00300 | |
| DI49_2900; SPGP0F00440 | DI49_2900 | SPGP0F00440 | |
| SPGP0F00470; DI49_2901 | DI49_2901 | SPGP0F00470 | |
| DI49_2903; DI49_2902 | DI49_2903; DI49_2902 | | |
| DI49_2909; YJL111W; DI49_2910 | DI49_2909; DI49_2910 | | YJL111W |
| DI49_2942; DI49_2941 | DI49_2942; DI49_2941 | | |
| DI49_2946; YJL070C; DI49_2947 | DI49_2946; DI49_2947 | | YJL070C |
| DI49_2949; DI49_2948 | DI49_2949; DI49_2948 | | |
| DI49_2951; DI49_2950 | DI49_2951; DI49_2950 | | |
| DI49_2953; YJL062W; DI49_2954 | DI49_2953; DI49_2954 | | YJL062W |
| DI49_2957; YJL058C; DI49_2958 | DI49_2957; DI49_2958 | | YJL058C |
| DI49_2960; DI49_2959 | DI49_2960; DI49_2959 | | |
| DI49_2962; DI49_2961 | DI49_2962; DI49_2961 | | |
| DI49_2968; DI49_2967 | DI49_2968; DI49_2967 | | |
| DI49_2975; DI49_2974 | DI49_2975; DI49_2974 | | |
| DI49_2977; DI49_2976 | DI49_2977; DI49_2976 | | |
| SPGP0F01220; DI49_1172; DI49_1171 | DI49_1172; DI49_1171 | SPGP0F01220 | |

Create PDF in your applications with the Pdfcrowd HTML to PDF API PDFCROWD

| | | | |
|---|---|---|---|
| DI49_2984; YJL036W; DI49_2985 | DI49_2984; DI49_2985 | | YJL036W |
| DI49_2987; YJL033W; DI49_2988 | DI49_2987; DI49_2988 | | YJL033W |
| DI49_2990; DI49_2989 | DI49_2990; DI49_2989 | | |
| DI49_3000; DI49_2999 | DI49_3000; DI49_2999 | | |
| DI49_3002; DI49_3001 | DI49_3002; DI49_3001 | | |
| DI49_3004; DI49_3003 | DI49_3004; DI49_3003 | | |
| DI49_3012; YJL005W; DI49_3013 | DI49_3012; DI49_3013 | | YJL005W |
| DI49_3015; DI49_3014 | DI49_3015; DI49_3014 | | |
| DI49_3018; DI49_3017 | DI49_3018; DI49_3017 | | |
| DI49_3026; YJR010W; DI49_3027 | DI49_3026; DI49_3027 | | YJR010W |
| DI49_3031; DI49_3030 | DI49_3031; DI49_3030 | | |
| DI49_3035; DI49_3034 | DI49_3035; DI49_3034 | | |
| DI49_3037; DI49_3036 | DI49_3037; DI49_3036 | | |
| DI49_3039; DI49_3038 | DI49_3039; DI49_3038 | | |
| DI49_3046; YJR040W; DI49_3047 | DI49_3046; DI49_3047 | | YJR040W |
| DI49_3049; DI49_3048 | DI49_3049; DI49_3048 | | |
| DI49_3061; YJR052W; DI49_3062 | DI49_3061; DI49_3062 | | YJR052W |
| DI49_3064; SPGP0Atrna1R; DI49_3065 | DI49_3064; DI49_3065 | SPGP0Atrna1R | |
| DI49_3083; YJR070C; DI49_3084 | DI49_3083; DI49_3084 | | YJR070C |
| SPGP0F02140; DI49_3085 | DI49_3085 | SPGP0F02140 | |
| DI49_3088; YJR077C; DI49_3089 | DI49_3088; DI49_3089 | | YJR077C |
| DI49_3091; DI49_3090 | DI49_3091; DI49_3090 | | |
| DI49_3093; DI49_3092 | DI49_3093; DI49_3092 | | |
| DI49_3095; DI49_3094 | DI49_3095; DI49_3094 | | |
| DI49_3097; DI49_3096 | DI49_3097; DI49_3096 | | |
| DI49_3099; YJR091C; DI49_3100 | DI49_3099; DI49_3100 | | YJR091C |
| DI49_3102; DI49_3101 | DI49_3102; DI49_3101 | | |
| DI49_3113; DI49_3112 | DI49_3113; DI49_3112 | | |
| DI49_3115; YJR105W; DI49_3116 | DI49_3115; DI49_3116 | | YJR105W |
| DI49_3121; YJR111C; DI49_3122 | DI49_3121; DI49_3122 | | YJR111C |
| DI49_3124; DI49_3123 | DI49_3124; DI49_3123 | | |
| DI49_3126; DI49_3125 | DI49_3126; DI49_3125 | | |
| DI49_3130; YJR121W; DI49_3131 | DI49_3130; DI49_3131 | | YJR121W |
| DI49_3134; YJR125C; DI49_3135 | DI49_3134; DI49_3135 | | YJR125C |
| DI49_3137; DI49_3136 | DI49_3137; DI49_3136 | | |
| DI49_3139; DI49_3138 | DI49_3139; DI49_3138 | | |
| DI49_3148; YJR139C; DI49_3149 | DI49_3148; DI49_3149 | | YJR139C |
| DI49_3151; DI49_3150 | DI49_3151; DI49_3150 | | |
| DI49_3153; DI49_3152 | DI49_3153; DI49_3152 | | |

Create PDF in your applications with the Pdfcrowd HTML to PDF API PDFCROWD

| | | | |
|---|---|---|---|
| DI49_3156; DI49_3155 | DI49_3156; DI49_3155 | | |
| DI49_3160; SPGP0F02900 | DI49_3160 | SPGP0F02900 | |
| DI49_3163; DI49_3162 | DI49_3163; DI49_3162 | | |
| DI49_3165; DI49_3164 | DI49_3165; DI49_3164 | | |
| DI49_5353; SPGP0F03010 | DI49_5353 | SPGP0F03010 | |
| SPGP0BU00100; DI49_0337; DI49_0336 | DI49_0337; DI49_0336 | SPGP0BU00100 | |
| DI49_1998; DI49_1997 | DI49_1998; DI49_1997 | | |
| DI49_2000; DI49_1999 | DI49_2000; DI49_1999 | | |
| DI49_2005; DI49_2004 | DI49_2005; DI49_2004 | | |
| DI49_3471; SPGP0H00120 | DI49_3471 | SPGP0H00120 | |
| DI49_3469; YKR099W; DI49_3468 | DI49_3469; DI49_3468 | | YKR099W |
| DI49_3460; DI49_3461 | DI49_3460; DI49_3461 | | |
| DI49_3458; DI49_3459 | DI49_3458; DI49_3459 | | |
| DI49_3453; DI49_3454 | DI49_3453; DI49_3454 | | |
| DI49_3447; DI49_3448 | DI49_3447; DI49_3448 | | |
| DI49_3445; DI49_3446 | DI49_3445; DI49_3446 | | |
| DI49_3439; YKR069W; DI49_3438 | DI49_3439; DI49_3438 | | YKR069W |
| DI49_3422; DI49_3423 | DI49_3422; DI49_3423 | | |
| DI49_3407; DI49_3408 | DI49_3407; DI49_3408 | | |
| DI49_3404; YKR031C; DI49_3403 | DI49_3404; DI49_3403 | | YKR031C |
| DI49_3401; YKR028W; DI49_3400 | DI49_3401; DI49_3400 | | YKR028W |
| DI49_3398; YKR026C; DI49_3397 | DI49_3398; DI49_3397 | | YKR026C |
| DI49_3395; YKR023W; DI49_3394 | DI49_3395; DI49_3394 | | YKR023W |
| DI49_3392; DI49_3393 | DI49_3392; DI49_3393 | | |
| DI49_3390; YKR018C; DI49_3389 | DI49_3390; DI49_3389 | | YKR018C |
| DI49_3379; DI49_3380 | DI49_3379; DI49_3380 | | |
| DI49_3377; DI49_3378 | DI49_3377; DI49_3378 | | |
| DI49_3375; DI49_3376 | DI49_3375; DI49_3376 | | |
| DI49_3371; YKL002W; SPGP0H01110 | DI49_3371 | SPGP0H01110 | YKL002W |
| DI49_3364; SPGP0H01180 | DI49_3364 | SPGP0H01180 | |
| DI49_3362; YKL010C; DI49_3361 | DI49_3362; DI49_3361 | | YKL010C |
| DI49_3359; DI49_3360 | DI49_3359; DI49_3360 | | |
| DI49_3356; YKL016C; DI49_3355 | DI49_3356; DI49_3355 | | YKL016C |
| DI49_3353; YKL018C-A; DI49_3352 | DI49_3353; DI49_3352 | | YKL018C-A |
| DI49_3345; YKL025C; SPGP0H01390 | DI49_3345 | SPGP0H01390 | YKL025C |
| DI49_3338; YKL033W; DI49_3337 | DI49_3338; DI49_3337 | | YKL033W |
| DI49_3335; YKL035W; DI49_3334 | DI49_3335; DI49_3334 | | YKL035W |
| DI49_3332; YKL040C; DI49_3331 | DI49_3332; DI49_3331 | | YKL040C |
| DI49_3324; SPGP0H01600 | DI49_3324 | SPGP0H01600 | |
| DI49_3322; DI49_3323 | DI49_3322; DI49_3323 | | |

Create PDF in your applications with the Pdfcrowd HTML to PDF API PDFCROWD

| | | | |
|---|---|---|---|
| DI49_3316; YKL058W; DI49_3315 | DI49_3316; DI49_3315 | | YKL058W |
| DI49_3313; DI49_3314 | DI49_3313; DI49_3314 | | |
| DI49_3311; DI49_3312 | DI49_3311; DI49_3312 | | |
| DI49_3309; DI49_3310 | DI49_3309; DI49_3310 | | |
| DI49_3307; DI49_3308 | DI49_3307; DI49_3308 | | |
| DI49_1309; DI49_1310 | DI49_1309; DI49_1310 | | |
| DI49_1307; DI49_1308 | DI49_1307; DI49_1308 | | |
| DI49_1305; DI49_1306 | DI49_1305; DI49_1306 | | |
| DI49_1303; DI49_1304 | DI49_1303; DI49_1304 | | |
| DI49_1301; DI49_1302 | DI49_1301; DI49_1302 | | |
| DI49_1299; DI49_1300 | DI49_1299; DI49_1300 | | |
| DI49_1297; DI49_1298 | DI49_1297; DI49_1298 | | |
| DI49_1295; DI49_1296 | DI49_1295; DI49_1296 | | |
| DI49_1293; DI49_1294 | DI49_1293; DI49_1294 | | |
| DI49_1291; DI49_1292 | DI49_1291; DI49_1292 | | |
| DI49_1289; DI49_1290 | DI49_1289; DI49_1290 | | |
| DI49_1287; DI49_1288 | DI49_1287; DI49_1288 | | |
| DI49_1285; DI49_1286 | DI49_1285; DI49_1286 | | |
| DI49_1283; DI49_1284 | DI49_1283; DI49_1284 | | |
| DI49_1281; DI49_1282 | DI49_1281; DI49_1282 | | |
| DI49_1279; DI49_1280 | DI49_1279; DI49_1280 | | |
| DI49_1277; DI49_1278 | DI49_1277; DI49_1278 | | |
| DI49_1275; DI49_1276 | DI49_1275; DI49_1276 | | |
| DI49_1273; DI49_1274 | DI49_1273; DI49_1274 | | |
| DI49_1271; DI49_1272 | DI49_1271; DI49_1272 | | |
| DI49_1269; DI49_1270 | DI49_1269; DI49_1270 | | |
| DI49_1267; DI49_1268 | DI49_1267; DI49_1268 | | |
| DI49_1265; DI49_1266 | DI49_1265; DI49_1266 | | |
| DI49_1263; DI49_1264 | DI49_1263; DI49_1264 | | |
| DI49_1261; DI49_1262 | DI49_1261; DI49_1262 | | |
| DI49_1259; DI49_1260 | DI49_1259; DI49_1260 | | |
| DI49_1257; DI49_1258 | DI49_1257; DI49_1258 | | |
| DI49_1255; DI49_1256 | DI49_1255; DI49_1256 | | |
| SPGP0CM00680; DI49_1254 | DI49_1254 | SPGP0CM00680 | |
| DI49_1252; DI49_1253 | DI49_1252; DI49_1253 | | |
| DI49_1250; DI49_1251 | DI49_1250; DI49_1251 | | |
| DI49_1248; DI49_1249 | DI49_1248; DI49_1249 | | |
| DI49_1246; DI49_1247 | DI49_1246; DI49_1247 | | |
| DI49_1241; DI49_1242 | DI49_1241; DI49_1242 | | |
| DI49_1239; DI49_1240 | DI49_1239; DI49_1240 | | |

Create PDF in your applications with the Pdfcrowd HTML to PDF API PDFCROWD

| | | |
|---|---|---|
| DI49_1237; DI49_1238 | DI49_1237; DI49_1238 | |
| DI49_1231; DI49_1232 | DI49_1231; DI49_1232 | |
| DI49_1229; DI49_1230 | DI49_1229; DI49_1230 | |
| DI49_1227; DI49_1228 | DI49_1227; DI49_1228 | |
| DI49_1225; DI49_1226 | DI49_1225; DI49_1226 | |
| DI49_1223; DI49_1224 | DI49_1223; DI49_1224 | |
| DI49_1221; DI49_1222 | DI49_1221; DI49_1222 | |
| DI49_1219; DI49_1220 | DI49_1219; DI49_1220 | |
| DI49_1217; DI49_1218 | DI49_1217; DI49_1218 | |
| DI49_1212; DI49_1213 | DI49_1212; DI49_1213 | |
| DI49_1210; DI49_1211 | DI49_1210; DI49_1211 | |
| DI49_1208; DI49_1209 | DI49_1208; DI49_1209 | |
| DI49_1206; DI49_5372 | DI49_1206; DI49_5372 | |
| DI49_1204; DI49_1205 | DI49_1204; DI49_1205 | |
| DI49_1202; DI49_1203 | DI49_1202; DI49_1203 | |
| DI49_1200; DI49_1201 | DI49_1200; DI49_1201 | |
| DI49_1198; DI49_1199 | DI49_1198; DI49_1199 | |
| DI49_1196; DI49_1197 | DI49_1196; DI49_1197 | |
| DI49_1194; DI49_1195 | DI49_1194; DI49_1195 | |
| DI49_1192; DI49_1193 | DI49_1192; DI49_1193 | |
| DI49_1190; DI49_1191 | DI49_1190; DI49_1191 | |
| DI49_1188; DI49_1189 | DI49_1188; DI49_1189 | |
| DI49_1186; DI49_1187 | DI49_1186; DI49_1187 | |
| DI49_1184; DI49_1185 | DI49_1184; DI49_1185 | |
| DI49_1183; SPGP0CM01370 | DI49_1183 | SPGP0CM01370 |
| DI49_1181; DI49_1182 | DI49_1181; DI49_1182 | |
| DI49_1179; DI49_1180 | DI49_1179; DI49_1180 | |
| DI49_1177; DI49_1178 | DI49_1177; DI49_1178 | |
| DI49_1175; DI49_1176 | DI49_1175; DI49_1176 | |
| DI49_1173; DI49_1174 | DI49_1173; DI49_1174 | |
| DI49_1169; DI49_1170 | DI49_1169; DI49_1170 | |
| DI49_1167; DI49_1168 | DI49_1167; DI49_1168 | |
| DI49_1165; DI49_1166 | DI49_1165; DI49_1166 | |
| DI49_1163; DI49_1164 | DI49_1163; DI49_1164 | |
| DI49_1161; DI49_1162 | DI49_1161; DI49_1162 | |
| DI49_1159; DI49_1160 | DI49_1159; DI49_1160 | |
| DI49_1157; DI49_1158 | DI49_1157; DI49_1158 | |
| DI49_1155; DI49_1156 | DI49_1155; DI49_1156 | |
| DI49_1153; DI49_1154 | DI49_1153; DI49_1154 | |
| DI49_1151; DI49_1152 | DI49_1151; DI49_1152 | |

Create PDF in your applications with the Pdfcrowd HTML to PDF API PDFCROWD

| | | | |
|---|---|---|---|
| DI49_1146; DI49_1147 | DI49_1146; DI49_1147 | | |
| DI49_1144; DI49_1145 | DI49_1144; DI49_1145 | | |
| DI49_1143; SPGP0CM01780 | DI49_1143 | SPGP0CM01780 | |
| DI49_1139; DI49_1140 | DI49_1139; DI49_1140 | | |
| DI49_1137; DI49_1138 | DI49_1137; DI49_1138 | | |
| DI49_1134; DI49_1135 | DI49_1134; DI49_1135 | | |
| DI49_1132; DI49_1133 | DI49_1132; DI49_1133 | | |
| DI49_1130; DI49_1131 | DI49_1130; DI49_1131 | | |
| DI49_1128; DI49_1129 | DI49_1128; DI49_1129 | | |
| DI49_1126; DI49_1127 | DI49_1126; DI49_1127 | | |
| YBR092C; SPGP0CM01950; DI49_1125 | DI49_1125 | SPGP0CM01950 | YBR092C |
| DI49_1123; DI49_1124 | DI49_1123; DI49_1124 | | |
| DI49_1121; DI49_1122 | DI49_1121; DI49_1122 | | |
| DI49_1119; DI49_1120 | DI49_1119; DI49_1120 | | |
| SPGP0CM02050; DI49_0153; DI49_1117; DI49_0152 | DI49_0153; DI49_1117; DI49_0152 | SPGP0CM02050 | |
| DI49_1115; DI49_1116 | DI49_1115; DI49_1116 | | |
| DI49_1110; DI49_1111 | DI49_1110; DI49_1111 | | |
| DI49_1108; DI49_1109 | DI49_1108; DI49_1109 | | |
| DI49_1106; DI49_1107 | DI49_1106; DI49_1107 | | |
| DI49_1104; DI49_1105 | DI49_1104; DI49_1105 | | |
| DI49_1102; DI49_1103 | DI49_1102; DI49_1103 | | |
| DI49_1100; DI49_0242; DI49_1099; DI49_0241 | DI49_1100; DI49_0242; DI49_1099; DI49_0241 | | |
| DI49_1097; DI49_1098 | DI49_1097; DI49_1098 | | |
| DI49_1094; DI49_1096 | DI49_1094; DI49_1096 | | |
| DI49_1092; DI49_1093; SPGP0CP02300; DI49_0234 | DI49_1092; DI49_1093; DI49_0234 | SPGP0CP02300 | |
| DI49_1090; DI49_1091 | DI49_1090; DI49_1091 | | |
| DI49_1088; DI49_1089 | DI49_1088; DI49_1089 | | |
| DI49_3272; YKL104C; DI49_3271 | DI49_3272; DI49_3271 | | YKL104C |
| DI49_3267; DI49_3268 | DI49_3267; DI49_3268 | | |
| DI49_3262; YKL116C; DI49_3261 | DI49_3262; DI49_3261 | | YKL116C |
| DI49_3257; YKL121W; DI49_3256 | DI49_3257; DI49_3256 | | YKL121W |
| DI49_3254; YKL125W; DI49_3253 | DI49_3254; DI49_3253 | | YKL125W |
| DI49_3246; DI49_3247 | DI49_3246; DI49_3247 | | |
| DI49_3239; DI49_3240 | DI49_3239; DI49_3240 | | |
| DI49_3236; YKL143W; DI49_3235 | DI49_3236; DI49_3235 | | YKL143W |
| DI49_3232; YKL148C; DI49_3231 | DI49_3232; DI49_3231 | | YKL148C |
| DI49_3229; DI49_3230 | DI49_3229; DI49_3230 | | |
| SPGP0CN00770; YKL157W; DI49_3223 | DI49_3223 | SPGP0CN00770 | YKL157W |
| DI49_3221; DI49_3222 | DI49_3221; DI49_3222 | | |
| DI49_3219; DI49_3220 | DI49_3219; DI49_3220 | | |

Create PDF in your applications with the Pdfcrowd HTML to PDF API PDFCROWD

| | | | |
|---|---|---|---|
| DI49_3217; DI49_3218 | DI49_3217; DI49_3218 | | |
| DI49_3214; YKL166C; DI49_3213 | DI49_3214; DI49_3213 | | YKL166C |
| DI49_3211; DI49_3212 | DI49_3211; DI49_3212 | | |
| DI49_3209; DI49_3210 | DI49_3209; DI49_3210 | | |
| DI49_3207; DI49_3208 | DI49_3207; DI49_3208 | | |
| DI49_3205; DI49_3206 | DI49_3205; DI49_3206 | | |
| DI49_3199; YKL183W; DI49_3198 | DI49_3199; DI49_3198 | | YKL183W |
| DI49_3196; DI49_3197 | DI49_3196; DI49_3197 | | |
| DI49_3189; DI49_3190 | DI49_3189; DI49_3190 | | |
| SPGP0CN01180; DI49_3182 | DI49_3182 | SPGP0CN01180 | |
| DI49_0460; DI49_0461 | DI49_0460; DI49_0461 | | |
| DI49_0458; DI49_0459 | DI49_0458; DI49_0459 | | |
| DI49_0453; DI49_0454 | DI49_0453; DI49_0454 | | |
| DI49_0451; DI49_0452 | DI49_0451; DI49_0452 | | |
| DI49_0449; DI49_0450 | DI49_0449; DI49_0450 | | |
| DI49_0447; DI49_0448 | DI49_0447; DI49_0448 | | |
| DI49_0445; DI49_0446 | DI49_0445; DI49_0446 | | |
| DI49_0443; DI49_0444 | DI49_0443; DI49_0444 | | |
| DI49_0439; DI49_0440 | DI49_0439; DI49_0440 | | |
| DI49_0437; DI49_0438 | DI49_0437; DI49_0438 | | |
| DI49_0435; DI49_0436 | DI49_0435; DI49_0436 | | |
| DI49_0433; DI49_0434 | DI49_0433; DI49_0434 | | |
| DI49_0431; DI49_0432 | DI49_0431; DI49_0432 | | |
| DI49_0429; DI49_0430 | DI49_0429; DI49_0430 | | |
| DI49_0427; DI49_0428 | DI49_0427; DI49_0428 | | |
| DI49_0425; DI49_0426 | DI49_0425; DI49_0426 | | |
| YDR242W; DI49_0424 | DI49_0424 | | YDR242W |
| DI49_0420; DI49_0421 | DI49_0420; DI49_0421 | | |
| DI49_0418; DI49_0419 | DI49_0418; DI49_0419 | | |
| DI49_0416; DI49_0417 | DI49_0416; DI49_0417 | | |
| DI49_0414; DI49_0415 | DI49_0414; DI49_0415 | | |
| DI49_0412; DI49_0413 | DI49_0412; DI49_0413 | | |
| DI49_0407; DI49_0179; DI49_0178 | DI49_0407; DI49_0179; DI49_0178 | | |
| DI49_0406; DI49_0180; DI49_0405 | DI49_0406; DI49_0180; DI49_0405 | | |
| DI49_0403; DI49_0404 | DI49_0403; DI49_0404 | | |
| DI49_0401; DI49_0402 | DI49_0401; DI49_0402 | | |
| DI49_0400; SPGP0CP00690 | DI49_0400 | SPGP0CP00690 | |
| DI49_0398; DI49_0399 | DI49_0398; DI49_0399 | | |
| DI49_0396; DI49_0397 | DI49_0396; DI49_0397 | | |
| DI49_0394; DI49_0395 | DI49_0394; DI49_0395 | | |

Create PDF in your applications with the Pdfcrowd HTML to PDF API PDFCROWD

| | | |
|---|---|---|
| DI49_0392; DI49_0393 | DI49_0392; DI49_0393 | |
| DI49_0390; DI49_0391 | DI49_0390; DI49_0391 | |
| DI49_0388; DI49_0389 | DI49_0388; DI49_0389 | |
| DI49_0386; DI49_0387 | DI49_0386; DI49_0387 | |
| DI49_0384; DI49_0385 | DI49_0384; DI49_0385 | |
| DI49_0382; DI49_0383 | DI49_0382; DI49_0383 | |
| DI49_0381; SPGP0CP00880 | DI49_0381 | SPGP0CP00880 |
| DI49_0379; DI49_0380 | DI49_0379; DI49_0380 | |
| DI49_0377; DI49_0378 | DI49_0377; DI49_0378 | |
| DI49_0375; DI49_0376 | DI49_0375; DI49_0376 | |
| DI49_0373; DI49_0374 | DI49_0373; DI49_0374 | |
| DI49_0371; DI49_0372 | DI49_0371; DI49_0372 | |
| DI49_0369; DI49_0370 | DI49_0369; DI49_0370 | |
| DI49_0367; DI49_0368 | DI49_0367; DI49_0368 | |
| DI49_0365; DI49_0366 | DI49_0365; DI49_0366 | |
| DI49_0363; DI49_0364 | DI49_0363; DI49_0364 | |
| DI49_0361; DI49_0362 | DI49_0361; DI49_0362 | |
| DI49_0356; DI49_0357 | DI49_0356; DI49_0357 | |
| DI49_0354; DI49_0355 | DI49_0354; DI49_0355 | |
| DI49_0352; DI49_0353 | DI49_0352; DI49_0353 | |
| DI49_0350; DI49_0351 | DI49_0350; DI49_0351 | |
| DI49_0348; DI49_0349 | DI49_0348; DI49_0349 | |
| DI49_0346; DI49_0347 | DI49_0346; DI49_0347 | |
| DI49_0344; DI49_0345 | DI49_0344; DI49_0345 | |
| DI49_0342; DI49_0343 | DI49_0342; DI49_0343 | |
| SPGP0CP01280; DI49_0341 | DI49_0341 | SPGP0CP01280 |
| DI49_0339; DI49_0340 | DI49_0339; DI49_0340 | |
| DI49_0334; DI49_0335 | DI49_0334; DI49_0335 | |
| DI49_0332; DI49_0333 | DI49_0332; DI49_0333 | |
| DI49_0330; DI49_0331 | DI49_0330; DI49_0331 | |
| DI49_0328; DI49_0329 | DI49_0328; DI49_0329 | |
| DI49_0326; DI49_0327 | DI49_0326; DI49_0327 | |
| DI49_0324; DI49_0325 | DI49_0324; DI49_0325 | |
| DI49_0322; DI49_0323 | DI49_0322; DI49_0323 | |
| DI49_0320; DI49_0321 | DI49_0320; DI49_0321 | |
| DI49_0318; DI49_0319 | DI49_0318; DI49_0319 | |
| DI49_0316; DI49_0317 | DI49_0316; DI49_0317 | |
| DI49_0314; DI49_0315 | DI49_0314; DI49_0315 | |
| DI49_0312; DI49_0313 | DI49_0312; DI49_0313 | |
| DI49_0310; DI49_0311 | DI49_0310; DI49_0311 | |

Create PDF in your applications with the Pdfcrowd HTML to PDF API PDFCROWD

| | | |
|---|---|---|
| DI49_0308; DI49_0309 | DI49_0308; DI49_0309 | |
| DI49_0306; DI49_0307 | DI49_0306; DI49_0307 | |
| SPGP0CP01650; DI49_0305 | DI49_0305 | SPGP0CP01650 |
| DI49_0303; DI49_0304 | DI49_0303; DI49_0304 | |
| DI49_0299; DI49_0300 | DI49_0299; DI49_0300 | |
| DI49_0297; DI49_0298 | DI49_0297; DI49_0298 | |
| DI49_0295; DI49_0296 | DI49_0295; DI49_0296 | |
| DI49_0290; DI49_0292 | DI49_0290; DI49_0292 | |
| DI49_0289; SPGP0CP01780 | DI49_0289 | SPGP0CP01780 |
| DI49_0287; DI49_0288 | DI49_0287; DI49_0288 | |
| DI49_0283; DI49_0284 | DI49_0283; DI49_0284 | |
| DI49_0281; DI49_0282 | DI49_0281; DI49_0282 | |
| DI49_0279; DI49_0280 | DI49_0279; DI49_0280 | |
| DI49_0277; DI49_0278 | DI49_0277; DI49_0278 | |
| DI49_0275; DI49_0276 | DI49_0275; DI49_0276 | |
| DI49_0273; DI49_0274 | DI49_0273; DI49_0274 | |
| DI49_0271; DI49_0272 | DI49_0271; DI49_0272 | |
| DI49_0269; DI49_0270 | DI49_0269; DI49_0270 | |
| DI49_0267; DI49_0268 | DI49_0267; DI49_0268 | |
| DI49_0265; DI49_0266 | DI49_0265; DI49_0266 | |
| DI49_0263; DI49_0264 | DI49_0263; DI49_0264 | |
| DI49_0261; DI49_0262 | DI49_0261; DI49_0262 | |
| DI49_0259; DI49_0260 | DI49_0259; DI49_0260 | |
| DI49_0257; DI49_0258 | DI49_0257; DI49_0258 | |
| DI49_0255; DI49_0256 | DI49_0255; DI49_0256 | |
| DI49_0250; DI49_0251 | DI49_0250; DI49_0251 | |
| DI49_0248; DI49_0249 | DI49_0248; DI49_0249 | |
| DI49_0246; DI49_0247 | DI49_0246; DI49_0247 | |
| DI49_0244; DI49_0245 | DI49_0244; DI49_0245 | |
| DI49_0239; DI49_0240 | DI49_0239; DI49_0240 | |
| DI49_0237; DI49_0238 | DI49_0237; DI49_0238 | |
| DI49_0229; DI49_0230 | DI49_0229; DI49_0230 | |
| DI49_0227; DI49_0228 | DI49_0227; DI49_0228 | |
| DI49_0225; DI49_0226 | DI49_0225; DI49_0226 | |
| DI49_0218; DI49_0219 | DI49_0218; DI49_0219 | |
| DI49_0216; DI49_0217 | DI49_0216; DI49_0217 | |
| DI49_0214; DI49_0215 | DI49_0214; DI49_0215 | |
| DI49_0212; DI49_0213 | DI49_0212; DI49_0213 | |
| DI49_0210; DI49_0211 | DI49_0210; DI49_0211 | |
| DI49_0206; DI49_0207 | DI49_0206; DI49_0207 | |

Create PDF in your applications with the Pdfcrowd HTML to PDF API PDFCROWD

| | | | |
|---|---|---|---|
| DI49_0204; DI49_0205 | DI49_0204; DI49_0205 | | |
| DI49_0202; DI49_0203 | DI49_0202; DI49_0203 | | |
| DI49_0200; DI49_0201 | DI49_0200; DI49_0201 | | |
| DI49_0198; DI49_0199 | DI49_0198; DI49_0199 | | |
| DI49_0196; DI49_0197 | DI49_0196; DI49_0197 | | |
| DI49_0194; DI49_0195 | DI49_0194; DI49_0195 | | |
| DI49_0186; DI49_0187 | DI49_0186; DI49_0187 | | |
| DI49_0184; DI49_0185 | DI49_0184; DI49_0185 | | |
| DI49_0182; DI49_0183 | DI49_0182; DI49_0183 | | |
| DI49_0176; SPGP0CPtrna24S | DI49_0176 | SPGP0CPtrna24S | |
| DI49_0174; DI49_0175 | DI49_0174; DI49_0175 | | |
| DI49_0172; DI49_0173 | DI49_0172; DI49_0173 | | |
| DI49_0170; DI49_0171 | DI49_0170; DI49_0171 | | |
| DI49_0168; DI49_0169 | DI49_0168; DI49_0169 | | |
| DI49_0159; DI49_0160 | DI49_0159; DI49_0160 | | |
| DI49_0157; DI49_0158 | DI49_0157; DI49_0158 | | |
| DI49_0155; DI49_0156 | DI49_0155; DI49_0156 | | |
| DI49_0150; DI49_0151 | DI49_0150; DI49_0151 | | |
| DI49_0148; DI49_0149 | DI49_0148; DI49_0149 | | |
| DI49_0146; DI49_0147 | DI49_0146; DI49_0147 | | |
| DI49_0144; DI49_0145 | DI49_0144; DI49_0145 | | |
| DI49_0142; DI49_0143 | DI49_0142; DI49_0143 | | |
| DI49_0140; DI49_0141 | DI49_0140; DI49_0141 | | |
| DI49_0138; DI49_0139 | DI49_0138; DI49_0139 | | |
| DI49_0136; DI49_0137 | DI49_0136; DI49_0137 | | |
| DI49_0134; DI49_0135 | DI49_0134; DI49_0135 | | |
| DI49_0130; DI49_0131 | DI49_0130; DI49_0131 | | |
| DI49_0128; DI49_0129 | DI49_0128; DI49_0129 | | |
| DI49_0126; DI49_0127 | DI49_0126; DI49_0127 | | |
| DI49_0125; SPGP0CP03340 | DI49_0125 | SPGP0CP03340 | |
| DI49_0123; DI49_0124 | DI49_0123; DI49_0124 | | |
| YBL059C-A; SPGP0CP03380 | | SPGP0CP03380 | YBL059C-A |
| DI49_0120; DI49_0121 | DI49_0120; DI49_0121 | | |
| DI49_0118; DI49_0119 | DI49_0118; DI49_0119 | | |
| DI49_0116; DI49_0117 | DI49_0116; DI49_0117 | | |
| DI49_0114; DI49_0115 | DI49_0114; DI49_0115 | | |
| DI49_0110; YBL075C | DI49_0110 | | YBL075C |
| DI49_0108; DI49_0109 | DI49_0108; DI49_0109 | | |
| DI49_0106; DI49_0107 | DI49_0106; DI49_0107 | | |
| DI49_0105; SPGP0CP03570 | DI49_0105 | SPGP0CP03570 | |

Create PDF in your applications with the Pdfcrowd HTML to PDF API PDFCROWD

| | | | |
|---|---|---|---|
| DI49_0103; DI49_0104 | DI49_0103; DI49_0104 | | |
| DI49_0098; DI49_0099 | DI49_0098; DI49_0099 | | |
| DI49_0096; DI49_0097 | DI49_0096; DI49_0097 | | |
| SPGP0CQ00100; DI49_0095 | DI49_0095 | SPGP0CQ00100 | |
| YAL047C; YAL048C | | | YAL047C; YAL048C |
| YAL044W-A; YAL046C | | | YAL044W-A; YAL046C |
| YAL037W; YAL038W | | | YAL037W; YAL038W |
| YAL030W; YAL031C | | | YAL030W; YAL031C |
| YAL028W; YAL029C | | | YAL028W; YAL029C |
| YAL008W; YAL009W | | | YAL008W; YAL009W |
| YOL013W-A; SPGP0ltrna16P; SPGP0ltrna15P; YNL042W; YNL035C | | SPGP0ltrna16P; SPGP0ltrna15P | YOL013W-A; YNL042W; YNL035C |
| SPGP0CR00600; YAL001C | | SPGP0CR00600 | YAL001C |
| YAR002C-A; DI49_0060 | DI49_0060 | | YAR002C-A |
| YAR027W; YAR028W; SPGP0CR00740; YAR031W; YAR033W | | SPGP0CR00740 | YAR027W; YAR028W; YAR031W; YAR033W |
| DI49_3929; DI49_3930 | DI49_3929; DI49_3930 | | |
| DI49_3927; DI49_3928 | DI49_3927; DI49_3928 | | |
| DI49_3925; DI49_3926 | DI49_3925; DI49_3926 | | |
| SPGP0CU00180; DI49_3924 | DI49_3924 | SPGP0CU00180 | |
| DI49_3919; DI49_3920 | DI49_3919; DI49_3920 | | |
| DI49_3917; DI49_3918 | DI49_3917; DI49_3918 | | |
| DI49_3913; DI49_3914 | DI49_3913; DI49_3914 | | |
| DI49_3911; DI49_3912 | DI49_3911; DI49_3912 | | |
| DI49_3909; DI49_3910 | DI49_3909; DI49_3910 | | |
| DI49_3907; DI49_3908 | DI49_3907; DI49_3908 | | |
| DI49_3905; DI49_3906 | DI49_3905; DI49_3906 | | |
| DI49_3901; DI49_3902 | DI49_3901; DI49_3902 | | |
| DI49_3899; DI49_3900 | DI49_3899; DI49_3900 | | |
| DI49_3897; DI49_3898 | DI49_3897; DI49_3898 | | |
| DI49_3895; DI49_3896 | DI49_3895; DI49_3896 | | |
| DI49_3893; DI49_3894 | DI49_3893; DI49_3894 | | |
| DI49_3891; DI49_3892 | DI49_3891; DI49_3892 | | |
| DI49_3889; DI49_3890 | DI49_3889; DI49_3890 | | |
| DI49_3885; DI49_3886 | DI49_3885; DI49_3886 | | |
| DI49_3883; DI49_3884 | DI49_3883; DI49_3884 | | |
| DI49_3881; DI49_3882 | DI49_3881; DI49_3882 | | |
| DI49_3879; DI49_3880 | DI49_3879; DI49_3880 | | |
| DI49_3874; DI49_3875 | DI49_3874; DI49_3875 | | |
| DI49_3872; DI49_3873 | DI49_3872; DI49_3873 | | |
| DI49_3870; DI49_3871 | DI49_3870; DI49_3871 | | |

Create PDF in your applications with the Pdfcrowd HTML to PDF API PDFCROWD

| | | | |
|---|---|---|---|
| DI49_3868; DI49_3869 | DI49_3868; DI49_3869 | | |
| DI49_3867; DI49_3747; DI49_3866; DI49_3865; DI49_3746 | DI49_3867; DI49_3747; DI49_3866; DI49_3865; DI49_3746 | | |
| DI49_3863; DI49_3864 | DI49_3863; DI49_3864 | | |
| DI49_3861; DI49_3862 | DI49_3861; DI49_3862 | | |
| DI49_3859; DI49_3860; DI49_3858 | DI49_3859; DI49_3860; DI49_3858 | | |
| DI49_3856; DI49_3857 | DI49_3856; DI49_3857 | | |
| DI49_3854; DI49_3855 | DI49_3854; DI49_3855 | | |
| DI49_3852; DI49_3853 | DI49_3852; DI49_3853 | | |
| DI49_3850; DI49_3851 | DI49_3850; DI49_3851 | | |
| DI49_3848; DI49_3849 | DI49_3848; DI49_3849 | | |
| DI49_3846; DI49_3847 | DI49_3846; DI49_3847 | | |
| DI49_3845; SPGP0CU00940 | DI49_3845 | SPGP0CU00940 | |
| DI49_3840; DI49_3841 | DI49_3840; DI49_3841 | | |
| DI49_3838; DI49_3839 | DI49_3838; DI49_3839 | | |
| DI49_3832; DI49_3833 | DI49_3832; DI49_3833 | | |
| DI49_3827; DI49_3828 | DI49_3827; DI49_3828 | | |
| DI49_3825; DI49_3826 | DI49_3825; DI49_3826 | | |
| DI49_3823; DI49_3824 | DI49_3823; DI49_3824 | | |
| DI49_3821; DI49_3822 | DI49_3821; DI49_3822 | | |
| DI49_3820; SPGP0CU01160 | DI49_3820 | SPGP0CU01160 | |
| DI49_3819; SPGP0CU01180 | DI49_3819 | SPGP0CU01180 | |
| DI49_3817; DI49_3818 | DI49_3817; DI49_3818 | | |
| DI49_3815; DI49_3816 | DI49_3815; DI49_3816 | | |
| DI49_3813; DI49_3814 | DI49_3813; DI49_3814 | | |
| DI49_3811; DI49_3812 | DI49_3811; DI49_3812 | | |
| DI49_3809; DI49_3810 | DI49_3809; DI49_3810 | | |
| DI49_3805; SPGP0CUtrna28L; DI49_3806; DI49_3740 | DI49_3805; DI49_3806; DI49_3740 | SPGP0CUtrna28L | |
| DI49_3804; YLR300W | DI49_3804 | | YLR300W |
| DI49_3802; DI49_3803 | DI49_3802; DI49_3803 | | |
| DI49_3797; DI49_3798 | DI49_3797; DI49_3798 | | |
| DI49_3795; DI49_3796; DI49_3794 | DI49_3795; DI49_3796; DI49_3794 | | |
| DI49_3792; DI49_3793 | DI49_3792; DI49_3793 | | |
| DI49_3790; DI49_3791 | DI49_3790; DI49_3791 | | |
| DI49_3788; DI49_3789 | DI49_3788; DI49_3789 | | |
| DI49_3786; DI49_3787 | DI49_3786; DI49_3787 | | |
| DI49_3784; YGL201C; DI49_3783 | DI49_3784; DI49_3783 | | YGL201C |
| DI49_3779; DI49_3780 | DI49_3779; DI49_3780 | | |
| DI49_3777; DI49_3778 | DI49_3777; DI49_3778 | | |
| DI49_3775; DI49_3776 | DI49_3775; DI49_3776 | | |
| DI49_3773; DI49_3774 | DI49_3773; DI49_3774 | | |

Create PDF in your applications with the Pdfcrowd HTML to PDF API PDFCROWD

| | | | |
|---|---|---|---|
| DI49_3771; DI49_3772 | DI49_3771; DI49_3772 | | |
| DI49_3764; DI49_3765 | DI49_3764; DI49_3765 | | |
| DI49_3762; DI49_3763 | DI49_3762; DI49_3763 | | |
| DI49_3754; DI49_3755 | DI49_3754; DI49_3755 | | |
| DI49_3752; DI49_3753 | DI49_3752; DI49_3753 | | |
| DI49_3750; DI49_3751 | DI49_3750; DI49_3751 | | |
| DI49_3748; DI49_3749 | DI49_3748; DI49_3749 | | |
| DI49_3744; DI49_3745 | DI49_3744; DI49_3745 | | |
| DI49_3742; SPGP0CUtrna19I | DI49_3742 | SPGP0CUtrna19I | |
| DI49_3738; DI49_3739 | DI49_3738; DI49_3739 | | |
| DI49_3736; DI49_3737 | DI49_3736; DI49_3737 | | |
| DI49_3734; DI49_3735 | DI49_3734; DI49_3735 | | |
| DI49_3732; DI49_3733 | DI49_3732; DI49_3733 | | |
| DI49_3731; SPGP0CU02030 | DI49_3731 | SPGP0CU02030 | |
| DI49_3728; DI49_3730 | DI49_3728; DI49_3730 | | |
| DI49_3726; DI49_3727 | DI49_3726; DI49_3727 | | |
| DI49_3724; DI49_3725 | DI49_3724; DI49_3725 | | |
| DI49_3722; DI49_3723 | DI49_3722; DI49_3723 | | |
| DI49_3720; DI49_3721 | DI49_3720; DI49_3721 | | |
| DI49_3718; DI49_3719 | DI49_3718; DI49_3719 | | |
| DI49_3716; DI49_3717 | DI49_3716; DI49_3717 | | |
| DI49_3714; DI49_3715 | DI49_3714; DI49_3715 | | |
| DI49_3712; DI49_3713 | DI49_3712; DI49_3713 | | |
| DI49_3710; DI49_3711 | DI49_3710; DI49_3711 | | |
| DI49_3708; DI49_3709 | DI49_3708; DI49_3709 | | |
| DI49_3706; DI49_3707 | DI49_3706; DI49_3707 | | |
| DI49_3704; DI49_3705 | DI49_3704; DI49_3705 | | |
| DI49_3702; DI49_3703 | DI49_3702; DI49_3703 | | |
| DI49_3700; DI49_3701 | DI49_3700; DI49_3701 | | |
| DI49_3698; DI49_3699 | DI49_3698; DI49_3699 | | |
| DI49_3694; DI49_3695 | DI49_3694; DI49_3695 | | |
| DI49_3692; DI49_3693 | DI49_3692; DI49_3693 | | |
| DI49_3690; DI49_3691 | DI49_3690; DI49_3691 | | |
| DI49_3688; DI49_3689 | DI49_3688; DI49_3689 | | |
| DI49_3683; DI49_3684 | DI49_3683; DI49_3684 | | |
| SPGP0CU02530; DI49_3682 | DI49_3682 | SPGP0CU02530 | |
| YNL335W; YNL336W | | | YNL335W; YNL336W |
| YNL333W; YNL334C | | | YNL333W; YNL334C |
| YNL331C; YNL332W | | | YNL331C; YNL332W |
| YNL329C; YNL330C | | | YNL329C; YNL330C |

Create PDF in your applications with the Pdfcrowd HTML to PDF API PDFCROWD

| | | | |
|---|---|---|---|
| YNL320W; DI49_4431; YNL318C | DI49_4431 | | YNL320W; YNL318C |
| YNL316C; YNL317W | | | YNL316C; YNL317W |
| YNL314W; YNL315C | | | YNL314W; YNL315C |
| YNL312W; YNL313C | | | YNL312W; YNL313C |
| YNL310C; YNL311C | | | YNL310C; YNL311C |
| YNL308C; YNL309W | | | YNL308C; YNL309W |
| YNL294C; YNL295W | | | YNL294C; YNL295W |
| YNL287W; DI49_4460; YNL286W | DI49_4460 | | YNL287W; YNL286W |
| YNL277W; YNL278W | | | YNL277W; YNL278W |
| YNL265C; DI49_4482; YNL264C | DI49_4482 | | YNL265C; YNL264C |
| YNL262W; DI49_4485; YNL261W | DI49_4485 | | YNL262W; YNL261W |
| YNL259C; YNL260C | | | YNL259C; YNL260C |
| YNL257C; YNL258C | | | YNL257C; YNL258C |
| YNL238W; YNL239W | | | YNL238W; YNL239W |
| YNL236W; YNL237W | | | YNL236W; YNL237W |
| YNL232W; DI49_4514; YNL231C | DI49_4514 | | YNL232W; YNL231C |
| YNL229C; YNL230C | | | YNL229C; YNL230C |
| YNL222W; DI49_4522; YNL221C | DI49_4522 | | YNL222W; YNL221C |
| YNL219C; YNL220W | | | YNL219C; YNL220W |
| YNL217W; YNL218W | | | YNL217W; YNL218W |
| YNL215W; YNL216W | | | YNL215W; YNL216W |
| YNL213C; YNL214W | | | YNL213C; YNL214W |
| YNL206C; DI49_4538; YNL204C | DI49_4538 | | YNL206C; YNL204C |
| YNL201C; DI49_4541; YNL200C | DI49_4541 | | YNL201C; YNL200C |
| YNL197C; YNL199C | | | YNL197C; YNL199C |
| YNL195C; YNL196C | | | YNL195C; YNL196C |
| YNL193W; YNL194C | | | YNL193W; YNL194C |
| YNL188W; DI49_4553; YNL187W | DI49_4553 | | YNL188W; YNL187W |
| YNL182C; DI49_4558; YNL181W | DI49_4558 | | YNL182C; YNL181W |
| YNL166C; YNL167C | | | YNL166C; YNL167C |
| YNL163C; DI49_4573; YNL162W-A | DI49_4573 | | YNL163C; YNL162W-A |
| YNL161W; YNL162W | | | YNL161W; YNL162W |
| YNL151C; YNL152W | | | YNL151C; YNL152W |
| YNL148C; YNL149C | | | YNL148C; YNL149C |
| YNL146W; YNL147W | | | YNL146W; YNL147W |
| YNL144C; YNL145W | | | YNL144C; YNL145W |
| YNL141W; YNL142W | | | YNL141W; YNL142W |
| YNL138W-A; YNL139C | | | YNL138W-A; YNL139C |
| YNL126W; DI49_4610; YNL125C | DI49_4610 | | YNL126W; YNL125C |
| SPGP0ltrna23L; DI49_1927 | DI49_1927 | SPGP0ltrna23L | |

Create PDF in your applications with the Pdfcrowd HTML to PDF API PDFCROWD

| | | | |
|---|---|---|---|
| YNL079C; DI49_4652; YNL078W | DI49_4652 | | YNL079C; YNL078W |
| YNL076W; YNL077W | | | YNL076W; YNL077W |
| YNL074C; YNL075W | | | YNL074C; YNL075W |
| YNL072W; YNL073W | | | YNL072W; YNL073W |
| SPGP0I02520; DI49_4662; YNL068C | DI49_4662 | SPGP0I02520 | YNL068C |
| YNL066W; YNL067W | | | YNL066W; YNL067W |
| YNL051W; YNL052W | | | YNL051W; YNL052W |
| YNL048W; DI49_4682; YNL047C | DI49_4682 | | YNL048W; YNL047C |
| YNL045W; YNL046W | | | YNL045W; YNL046W |
| YNL038W; YNL039W | | | YNL038W; YNL039W |
| YNL036W; YNL037C | | | YNL036W; YNL037C |

Create PDF in your applications with the Pdfcrowd HTML to PDF API PDFCROWD
