## Supplementary file 4 for "HybridMine: pipeline for allele inheritance and gene copy number prediction in industrial yeast hybrids"

### Overview

genes alleles inherited from Saccharomyces eubayanus
genes alleles inherited from Saccharomyces cerevisiae
genes alleles below 80% identity
genes alleles copy with same identity in both parents
list of paralogs in Saccharomyces pastorianus WS3470

#### Sheet 1: *genes alleles inherited from Saccharomyces eubayanus*

| *Saccharomyces pastorianus WS34/70 predicted ORF* | *Saccharomyces eubayanus parental allele* | Gene identities | Gene gaps |
|---|---|---|---|
| SPGP0L02060 | XP_018218972.1 | 99 | 0 |
| SPGP0H03600 | XP_018219539.1 | 99 | 0 |
| SPGP0O01680 | XP_018221795.1 | 99 | 0 |
| SPGP0AI00650 | XP_018224104.1 | 100 | 0 |
| SPGP0I00130 | XP_018223321.1 | 99 | 0 |
| SPGP0AT00250 | XP_018219184.1 | 99 | 0 |
| SPGP0I03450 | XP_018223002.1 | 100 | 0 |
| SPGP0O01760 | XP_018221802.1 | 99 | 0 |
| SPGP0M03370 | XP_018221449.1 | 99 | 0 |
| SPGP0H03890 | XP_018219568.1 | 99 | 0 |
| SPGP0AT00390 | XP_018219170.1 | 99 | 0 |
| SPGP0S01050 | XP_018219747.1 | 99 | 0 |
| SPGP0AR00280 | XP_018219587.1 | 100 | 0 |
| SPGP0M00270 | XP_018221152.1 | 99 | 0 |
| SPGP0H03910 | XP_018218748.1 | 97 | 3 |
| SPGP0Q01190 | XP_018222743.1 | 100 | 0 |
| SPGP0BJ00120 | XP_018219164.1 | 98 | 1 |
| SPGP0E00840 | XP_018222413.1 | 99 | 0 |
| SPGP0AT00410 | XP_018219168.1 | 99 | 0 |
| SPGP0F02570 | XP_018221216.1 | 87 | 0 |
| SPGP0L01320 | XP_018219044.1 | 99 | 0 |
| SPGP0H03290 | XP_018219509.1 | 100 | 0 |
| SPGP0E00830 | XP_018222414.1 | 99 | 0 |
| SPGP0AG00950 | XP_018223405.1 | 100 | 0 |
| SPGP0L00870 | XP_018219087.1 | 99 | 0 |
| SPGP0W01630 | XP_018221607.1 | 99 | 0 |
| SPGP0W01440 | XP_018221589.1 | 99 | 0 |
| SPGP0AG00590 | XP_018223373.1 | 100 | 0 |

Create PDF in your applications with the Pdfcrowd HTML to PDF API PDFCROWD

| | | | |
|---|---|---|---|
| SPGP0Q01940 | XP_018222669.1 | 100 | 0 |
| SPGP0M03340 | XP_018221447.1 | 99 | 0 |
| SPGP0C04810 | XP_018220840.1 | 99 | 0 |
| SPGP0L02090 | XP_018218969.1 | 99 | 0 |
| SPGP0I01760 | XP_018223159.1 | 99 | 0 |
| SPGP0B01390 | XP_018223586.1 | 99 | 0 |
| SPGP0AT00190 | XP_018219190.1 | 99 | 0 |
| SPGP0BJ00100 | XP_018219166.1 | 99 | 0 |
| SPGP0AI00100 | XP_018224051.1 | 99 | 0 |
| SPGP0W01810 | XP_018221625.1 | 95 | 2 |
| SPGP0B00900 | XP_018223536.1 | 99 | 0 |
| SPGP0W00200 | XP_018221469.1 | 99 | 1 |
| SPGP0B05430 | XP_018223974.1 | 99 | 0 |
| SPGP0C01290 | XP_018220497.1 | 99 | 0 |
| SPGP0I03970 | XP_018222906.1 | 99 | 0 |
| SPGP0AT00380 | XP_018219171.1 | 100 | 0 |
| SPGP0Dtrna32S | DI49_2374 | 100 | 0 |
| SPGP0AL00660 | XP_018219927.1 | 98 | 0 |
| SPGP0AT00310 | XP_018219178.1 | 99 | 0 |
| SPGP0B00880 | XP_018223534.1 | 99 | 0 |
| SPGP0C04380 | XP_018220800.1 | 99 | 0 |
| SPGP0AT00280 | XP_018219181.1 | 99 | 0 |
| SPGP0Q01810 | XP_018222682.1 | 99 | 0 |
| SPGP0M00290 | XP_018221154.1 | 99 | 0 |
| SPGP0C02330 | XP_018220601.1 | 100 | 0 |
| SPGP0L00960 | XP_018219079.1 | 99 | 0 |
| SPGP0AT00220 | XP_018219187.1 | 99 | 0 |
| SPGP0AT00200 | XP_018219189.1 | 98 | 0 |
| SPGP0AF00450 | XP_018222560.1 | 99 | 0 |
| SPGP0F00400 | XP_018220319.1 | 99 | 0 |
| SPGP0S00400 | XP_018219811.1 | 99 | 0 |
| SPGP0AT00330 | XP_018219176.1 | 99 | 0 |
| SPGP0I02670 | XP_018223071.1 | 99 | 0 |
| SPGP0S00720 | XP_018219779.1 | 99 | 0 |
| SPGP0BJ00140 | XP_018219162.1 | 99 | 0 |
| SPGP0E02210 | XP_018222274.1 | 99 | 0 |
| SPGP0W01640 | XP_018221608.1 | 99 | 0 |
| SPGP0B03730 | XP_018223814.1 | 99 | 0 |
| SPGP0H01320 | XP_018219317.1 | 99 | 0 |
| SPGP0Q01530 | XP_018222711.1 | 99 | 1 |

Create PDF in your applications with the Pdfcrowd HTML to PDF API PDFCROWD

| | | | |
|---|---|---|---|
| SPGP0B06130 | XP_018224041.1 | 100 | 0 |
| SPGP0B02240 | XP_018223669.1 | 99 | 0 |
| SPGP0C01130 | XP_018220481.1 | 99 | 0 |
| SPGP0M00670 | XP_018221191.1 | 99 | 0 |
| SPGP0F00560 | XP_018220304.1 | 99 | 0 |
| SPGP0B02790 | XP_018223722.1 | 99 | 0 |
| SPGP0L00230 | XP_018219149.1 | 99 | 0 |
| SPGP0S00950 | XP_018219757.1 | 98 | 1 |
| SPGP0F03810 | XP_018219990.1 | 99 | 0 |
| SPGP0H00570 | XP_018219245.1 | 99 | 0 |
| SPGP0C02590 | XP_018220627.1 | 99 | 0 |
| SPGP0E01140 | XP_018222382.1 | 99 | 0 |
| SPGP0M02820 | XP_018221397.1 | 99 | 0 |
| SPGP0L00360 | XP_018219136.1 | 99 | 0 |
| SPGP0M01810 | XP_018221300.1 | 99 | 0 |
| SPGP0F02580 | XP_018220110.1 | 99 | 0 |
| SPGP0C01010 | XP_018220469.1 | 99 | 0 |
| SPGP0AF00970 | XP_018222510.1 | 99 | 0 |
| SPGP0B05640 | XP_018223994.1 | 99 | 0 |
| SPGP0B01520 | XP_018223599.1 | 99 | 0 |
| SPGP0H01760 | XP_018219361.1 | 99 | 0 |
| SPGP0M01970 | XP_018221315.1 | 99 | 0 |
| SPGP0I00460 | XP_018223288.1 | 99 | 0 |
| SPGP0F00690 | XP_018220291.1 | 99 | 0 |
| SPGP0Ctrna8L | DI49_3806 | 99 | 0 |
| SPGP0F02670 | XP_018220101.1 | 99 | 1 |
| SPGP0H01140 | XP_018219301.1 | 99 | 0 |
| SPGP0O00330 | XP_018221666.1 | 99 | 0 |
| SPGP0M01770 | XP_018221296.1 | 99 | 0 |
| SPGP0I00480 | XP_018223286.1 | 99 | 0 |
| SPGP0F01330 | XP_018220227.1 | 99 | 0 |
| SPGP0E00760 | XP_018222421.1 | 99 | 0 |
| SPGP0AR00240 | XP_018219583.1 | 99 | 0 |
| SPGP0Q02390 | XP_018222626.1 | 98 | 1 |
| SPGP0C01480 | XP_018220517.1 | 99 | 0 |
| SPGP0I01770 | XP_018223158.1 | 99 | 0 |
| SPGP0I00850 | XP_018223249.1 | 99 | 0 |
| SPGP0C01000 | XP_018220468.1 | 99 | 0 |
| SPGP0B00180 | XP_018223468.1 | 99 | 0 |
| SPGP0I00890 | XP_018223245.1 | 99 | 0 |

Create PDF in your applications with the Pdfcrowd HTML to PDF API PDFCROWD

| | | | |
|---|---|---|---|
| SPGP0B05970 | XP_018224027.1 | 99 | 0 |
| SPGP0L01600 | XP_018219016.1 | 99 | 0 |
| SPGP0I00790 | XP_018223255.1 | 99 | 0 |
| SPGP0O01310 | XP_018221758.1 | 99 | 1 |
| SPGP0Q00220 | XP_018222837.1 | 99 | 0 |
| SPGP0C01050 | XP_018220473.1 | 99 | 0 |
| SPGP0C00740 | XP_018220441.1 | 99 | 0 |
| SPGP0C03790 | XP_018220742.1 | 99 | 0 |
| SPGP0O01330 | XP_018221760.1 | 99 | 0 |
| SPGP0E01830 | XP_018222311.1 | 99 | 0 |
| SPGP0B02630 | XP_018223707.1 | 99 | 0 |
| SPGP0B04110 | XP_018223852.1 | 99 | 0 |
| SPGP0B00450 | XP_018223493.1 | 99 | 0 |
| SPGP0F02610 | XP_018220107.1 | 99 | 0 |
| SPGP0E01260 | XP_018222370.1 | 99 | 0 |
| SPGP0C03600 | XP_018220724.1 | 99 | 0 |
| SPGP0W00270 | XP_018221476.1 | 99 | 0 |
| SPGP0E01630 | XP_018222331.1 | 99 | 0 |
| SPGP0E04020 | XP_018222097.1 | 99 | 0 |
| SPGP0E01090 | XP_018222388.1 | 100 | 0 |
| SPGP0Strna26L | DI49_4635 | 100 | 0 |
| SPGP0H02640 | XP_018219445.1 | 99 | 0 |
| SPGP0P02950 | XP_018220856.1 | 98 | 0 |
| SPGP0O00340 | XP_018221667.1 | 99 | 0 |
| SPGP0Q02570 | XP_018222608.1 | 100 | 0 |
| SPGP0C01800 | XP_018220548.1 | 98 | 0 |
| SPGP0B01650 | XP_018223611.1 | 99 | 0 |
| SPGP0F00960 | XP_018220264.1 | 99 | 0 |
| SPGP0L01460 | XP_018219030.1 | 99 | 0 |
| SPGP0B05660 | XP_018223996.1 | 99 | 0 |
| SPGP0C00820 | XP_018220449.1 | 99 | 0 |
| SPGP0I02100 | XP_018223126.1 | 99 | 0 |
| SPGP0S01720 | XP_018219680.1 | 99 | 0 |
| SPGP0W01910 | XP_018221635.1 | 99 | 0 |
| SPGP0H03660 | XP_018219545.1 | 99 | 0 |
| SPGP0AF01080 | XP_018222500.1 | 99 | 0 |
| SPGP0B03700 | XP_018223811.1 | 99 | 0 |
| SPGP0I01690 | XP_018223166.1 | 99 | 0 |
| SPGP0H01510 | XP_018219336.1 | 99 | 0 |
| SPGP0H03330 | XP_018219513.1 | 99 | 0 |

Create PDF in your applications with the Pdfcrowd HTML to PDF API PDFCROWD

| | | | |
|---|---|---|---|
| SPGP0C02680 | XP_018220636.1 | 99 | 0 |
| SPGP0B04070 | XP_018223848.1 | 99 | 0 |
| SPGP0AI00690 | XP_018224108.1 | 99 | 0 |
| SPGP0H01110 | XP_018219298.1 | 99 | 0 |
| SPGP0F01800 | XP_018220185.1 | 99 | 0 |
| SPGP0O02180 | XP_018221842.1 | 99 | 0 |
| SPGP0Htrna41G | DI49_4977 | 100 | 0 |
| SPGP0W01720 | XP_018221616.1 | 99 | 0 |
| SPGP0AF00520 | XP_018222553.1 | 97 | 3 |
| SPGP0F04180 | XP_018219953.1 | 99 | 0 |
| SPGP0B01770 | XP_018223623.1 | 99 | 0 |
| SPGP0I00900 | XP_018223244.1 | 99 | 0 |
| SPGP0AG00770 | XP_018223389.1 | 99 | 0 |
| SPGP0B00330 | XP_018223483.1 | 99 | 0 |
| SPGP0AL00270 | XP_018219891.1 | 99 | 0 |
| SPGP0B00910 | XP_018223537.1 | 99 | 0 |
| SPGP0Ftrna16Q | DI49_4357 | 100 | 0 |
| SPGP0O01170 | XP_018221744.1 | 99 | 0 |
| SPGP0E02010 | XP_018222293.1 | 99 | 0 |
| SPGP0I01500 | XP_018223185.1 | 99 | 0 |
| SPGP0B04470 | XP_018223887.1 | 99 | 0 |
| SPGP0B06090 | XP_018224037.1 | 99 | 0 |
| SPGP0H00720 | XP_018219259.1 | 99 | 0 |
| SPGP0E01900 | XP_018222304.1 | 99 | 0 |
| SPGP0AG00670 | XP_018223381.1 | 99 | 0 |
| SPGP0L00730 | XP_018219100.1 | 99 | 0 |
| SPGP0P01540 | XP_018220994.1 | 99 | 0 |
| SPGP0B00120 | XP_018218752.1 | 99 | 0 |
| SPGP0L02850 | XP_018218896.1 | 99 | 0 |
| SPGP0O02150 | XP_018221839.1 | 99 | 0 |
| SPGP0L03120 | XP_018218869.1 | 99 | 0 |
| SPGP0E03760 | XP_018222123.1 | 99 | 0 |
| SPGP0S01450 | XP_018219707.1 | 99 | 0 |
| SPGP0P00810 | XP_018221063.1 | 99 | 1 |
| SPGP0P01430 | XP_018221004.1 | 98 | 2 |
| SPGP0E02900 | XP_018222205.1 | 99 | 0 |
| SPGP0AI00460 | XP_018224086.1 | 99 | 0 |
| SPGP0S01180 | XP_018219734.1 | 99 | 0 |
| SPGP0F02870 | XP_018220081.1 | 99 | 0 |
| SPGP0W00680 | XP_018221516.1 | 99 | 0 |

Create PDF in your applications with the Pdfcrowd HTML to PDF API PDFCROWD

| | | | |
|---|---|---|---|
| SPGP0F00150 | XP_018220344.1 | 99 | 0 |
| SPGP0C04050 | XP_018220768.1 | 99 | 0 |
| SPGP0B03930 | XP_018223834.1 | 99 | 0 |
| SPGP0E00570 | XP_018222440.1 | 99 | 0 |
| SPGP0C03170 | XP_018220685.1 | 99 | 0 |
| SPGP0Q01840 | XP_018222679.1 | 99 | 0 |
| SPGP0P00740 | XP_018221070.1 | 99 | 0 |
| SPGP0C03880 | XP_018220751.1 | 99 | 0 |
| SPGP0L03380 | XP_018218844.1 | 99 | 0 |
| SPGP0E03500 | XP_018222148.1 | 99 | 0 |
| SPGP0P00210 | XP_018221122.1 | 99 | 0 |
| SPGP0W00950 | XP_018221542.1 | 99 | 0 |
| SPGP0AL00180 | XP_018219882.1 | 99 | 0 |
| SPGP0W01620 | XP_018221606.1 | 99 | 0 |
| SPGP0P02000 | XP_018220950.1 | 99 | 0 |
| SPGP0C03220 | XP_018220690.1 | 99 | 0 |
| SPGP0F03940 | XP_018219977.1 | 99 | 0 |
| SPGP0B02510 | XP_018223695.1 | 99 | 0 |
| SPGP0H02560 | XP_018219438.1 | 99 | 0 |
| SPGP0M03240 | XP_018221438.1 | 99 | 0 |
| SPGP0AI00830 | XP_018224121.1 | 94 | 1 |
| SPGP0L03550 | XP_018218828.1 | 99 | 0 |
| SPGP0B00580 | XP_018223505.1 | 99 | 0 |
| SPGP0B00960 | XP_018223542.1 | 99 | 0 |
| SPGP0H02850 | XP_018219466.1 | 99 | 0 |
| SPGP0O03200 | XP_018221940.1 | 99 | 0 |
| SPGP0B04460 | XP_018223886.1 | 99 | 0 |
| SPGP0B01960 | XP_018223642.1 | 99 | 0 |
| SPGP0L02650 | XP_018218916.1 | 99 | 0 |
| SPGP0L01790 | XP_018218998.1 | 99 | 0 |
| SPGP0B05090 | XP_018223942.1 | 99 | 0 |
| SPGP0Q02230 | XP_018222642.1 | 99 | 0 |
| SPGP0H01400 | XP_018219325.1 | 99 | 0 |
| SPGP0C03420 | XP_018220708.1 | 99 | 0 |
| SPGP0W01950 | XP_018221639.1 | 99 | 0 |
| SPGP0B02420 | XP_018223686.1 | 99 | 0 |
| SPGP0I03990 | XP_018222951.1 | 99 | 0 |
| SPGP0P01290 | XP_018221017.1 | 100 | 0 |
| SPGP0L02140 | XP_018218964.1 | 99 | 0 |

Create PDF in your applications with the Pdfcrowd HTML to PDF API PDFCROWD

| | | | |
|---|---|---|---|
| SPGP0F04200 | XP_018219951.1 | 99 | 0 |
| SPGP0S00370 | XP_018219814.1 | 99 | 0 |
| SPGP0O03050 | XP_018221925.1 | 99 | 0 |
| SPGP0L02470 | XP_018218931.1 | 99 | 0 |
| SPGP0O02250 | XP_018221849.1 | 99 | 0 |
| SPGP0C01760 | XP_018220544.1 | 99 | 0 |
| SPGP0S01870 | XP_018219665.1 | 99 | 0 |
| SPGP0F03040 | XP_018220065.1 | 99 | 0 |
| SPGP0I01410 | XP_018223193.1 | 99 | 0 |
| SPGP0M00440 | XP_018221169.1 | 99 | 0 |
| SPGP0F03510 | XP_018220018.1 | 99 | 0 |
| SPGP0Itrna23R | DI49_4323 | 100 | 0 |
| SPGP0B00920 | XP_018223538.1 | 99 | 0 |
| SPGP0W00740 | XP_018221522.1 | 99 | 0 |
| SPGP0H01950 | XP_018219380.1 | 99 | 0 |
| SPGP0H01350 | XP_018219320.1 | 99 | 0 |
| SPGP0B04160 | XP_018223857.1 | 99 | 0 |
| SPGP0AF01140 | XP_018222494.1 | 99 | 0 |
| SPGP0B00230 | XP_018223473.1 | 99 | 0 |
| SPGP0Q00670 | XP_018222795.1 | 99 | 0 |
| SPGP0H00200 | XP_018219208.1 | 99 | 0 |
| SPGP0H03160 | XP_018219497.1 | 99 | 0 |
| SPGP0H00750 | XP_018219262.1 | 99 | 0 |
| SPGP0P01120 | XP_018221033.1 | 99 | 0 |
| SPGP0B01340 | XP_018223581.1 | 99 | 0 |
| SPGP0BI00130 | XP_018221948.1 | 99 | 0 |
| SPGP0M01710 | XP_018221289.1 | 99 | 0 |
| SPGP0E00530 | XP_018222444.1 | 99 | 0 |
| SPGP0C04340 | XP_018220796.1 | 99 | 0 |
| SPGP0E02640 | XP_018222232.1 | 99 | 0 |
| SPGP0M02300 | XP_018221346.1 | 100 | 0 |
| SPGP0P00900 | XP_018221054.1 | 99 | 0 |
| SPGP0S00470 | XP_018219804.1 | 99 | 0 |
| SPGP0I00940 | XP_018223240.1 | 99 | 0 |
| SPGP0F02770 | XP_018220091.1 | 99 | 0 |
| SPGP0S00280 | XP_018219823.1 | 99 | 0 |
| SPGP0P01990 | XP_018220951.1 | 99 | 0 |
| SPGP0O00780 | XP_018221709.1 | 99 | 0 |
| SPGP0L01060 | XP_018219069.1 | 99 | 0 |
| SPGP0O00830 | XP_018221713.1 | 99 | 0 |

Create PDF in your applications with the Pdfcrowd HTML to PDF API PDFCROWD

| | | | |
|---|---|---|---|
| SPGP0H02730 | XP_018219454.1 | 99 | 0 |
| SPGP0C00720 | XP_018220439.1 | 99 | 0 |
| SPGP0BB00180 | XP_018219852.1 | 99 | 0 |
| SPGP0Q01700 | XP_018222694.1 | 99 | 0 |
| SPGP0C04040 | XP_018220766.1 | 99 | 0 |
| SPGP0B04970 | XP_018223932.1 | 99 | 0 |
| SPGP0L01440 | XP_018219032.1 | 99 | 0 |
| SPGP0L01200 | XP_018219055.1 | 99 | 0 |
| SPGP0M00880 | XP_018221210.1 | 99 | 0 |
| SPGP0B00640 | XP_018223510.1 | 100 | 0 |
| SPGP0C03660 | XP_018220730.1 | 99 | 0 |
| SPGP0E01730 | XP_018222321.1 | 99 | 0 |
| SPGP0F03530 | XP_018220016.1 | 99 | 0 |
| SPGP0C03990 | XP_018220761.1 | 99 | 0 |
| SPGP0P00170 | XP_018221125.1 | 99 | 0 |
| SPGP0P03040 | XP_018220848.1 | 99 | 0 |
| SPGP0M00840 | XP_018221206.1 | 100 | 0 |
| SPGP0P01530 | XP_018220995.1 | 99 | 0 |
| SPGP0P01470 | XP_018221001.1 | 99 | 0 |
| SPGP0C03830 | XP_018220746.1 | 99 | 0 |
| SPGP0O00160 | XP_018221649.1 | 100 | 0 |
| SPGP0B05490 | XP_018223980.1 | 99 | 0 |
| SPGP0L03720 | XP_018218811.1 | 98 | 1 |
| SPGP0E01670 | XP_018222327.1 | 99 | 0 |
| SPGP0AQ00420 | XP_018221980.1 | 99 | 0 |
| SPGP0L03780 | XP_018218805.1 | 99 | 0 |
| SPGP0E04130 | XP_018222086.1 | 99 | 0 |
| SPGP0F03280 | XP_018220041.1 | 99 | 0 |
| SPGP0F00730 | XP_018220287.1 | 99 | 0 |
| SPGP0O01780 | XP_018221804.1 | 99 | 0 |
| SPGP0M00110 | XP_018221138.1 | 99 | 0 |
| SPGP0M01800 | XP_018221299.1 | 99 | 0 |
| SPGP0E01010 | XP_018222396.1 | 99 | 0 |
| SPGP0H02770 | XP_018219458.1 | 100 | 0 |
| SPGP0L01420 | XP_018219034.1 | 99 | 0 |
| SPGP0S01040 | XP_018219748.1 | 99 | 0 |
| SPGP0C01370 | XP_018220505.1 | 100 | 0 |
| SPGP0E02600 | XP_018222236.1 | 99 | 0 |
| SPGP0B00760 | XP_018223522.1 | 99 | 0 |
| SPGP0Q01070 | XP_018222755.1 | 99 | 0 |

Create PDF in your applications with the Pdfcrowd HTML to PDF API PDFCROWD

| | | | |
|---|---|---|---|
| SPGP0Q00400 | XP_018222821.1 | 99 | 0 |
| SPGP0I00630 | XP_018223271.1 | 99 | 0 |
| SPGP0B01420 | XP_018223589.1 | 99 | 0 |
| SPGP0C03410 | XP_018220707.1 | 99 | 0 |
| SPGP0AQ00170 | XP_018221955.1 | 99 | 0 |
| SPGP0B02720 | XP_018223715.1 | 99 | 0 |
| SPGP0F03630 | XP_018220007.1 | 99 | 0 |
| SPGP0B05930 | XP_018224023.1 | 99 | 0 |
| SPGP0I01450 | XP_018223189.1 | 99 | 0 |
| SPGP0M03410 | XP_018221452.1 | 99 | 0 |
| SPGP0AG00780 | XP_018223390.1 | 99 | 0 |
| SPGP0I00530 | XP_018223281.1 | 100 | 0 |
| SPGP0F03520 | XP_018220017.1 | 99 | 0 |
| SPGP0Q02180 | XP_018222647.1 | 99 | 0 |
| SPGP0O02940 | XP_018221916.1 | 98 | 2 |
| SPGP0S01200 | XP_018219732.1 | 100 | 0 |
| SPGP0C01670 | XP_018220535.1 | 99 | 0 |
| SPGP0H03820 | XP_018219561.1 | 99 | 0 |
| SPGP0L01810 | XP_018218996.1 | 99 | 0 |
| SPGP0H01860 | XP_018219371.1 | 99 | 0 |
| SPGP0C00130 | XP_018220384.1 | 99 | 0 |
| SPGP0S00620 | XP_018219789.1 | 99 | 0 |
| SPGP0W01930 | XP_018221637.1 | 99 | 0 |
| SPGP0P00390 | XP_018221104.1 | 99 | 0 |
| SPGP0B05740 | XP_018224004.1 | 99 | 0 |
| SPGP0B01680 | XP_018223614.1 | 100 | 0 |
| SPGP0S00910 | XP_018219761.1 | 99 | 0 |
| SPGP0AG00220 | XP_018223337.1 | 99 | 0 |
| SPGP0E01400 | XP_018222355.1 | 99 | 0 |
| SPGP0E02660 | XP_018222229.1 | 99 | 0 |
| SPGP0E01660 | XP_018222328.1 | 99 | 0 |
| SPGP0O00970 | XP_018221727.1 | 99 | 0 |
| SPGP0C02090 | XP_018220577.1 | 99 | 0 |
| SPGP0M02380 | XP_018221354.1 | 99 | 0 |
| SPGP0L02080 | XP_018218970.1 | 99 | 0 |
| SPGP0AQ00340 | XP_018221972.1 | 99 | 0 |
| SPGP0M02920 | XP_018221407.1 | 99 | 0 |
| SPGP0E04050 | XP_018222094.1 | 99 | 0 |
| SPGP0B03230 | XP_018223766.1 | 99 | 0 |
| SPGP0F00790 | XP_018220281.1 | 100 | 0 |

Create PDF in your applications with the Pdfcrowd HTML to PDF API PDFCROWD

| | | | |
|---|---|---|---|
| SPGP0F00300 | XP_018220329.1 | 99 | 0 |
| SPGP0E02920 | XP_018222203.1 | 99 | 0 |
| SPGP0B00470 | XP_018223495.1 | 99 | 0 |
| SPGP0AI00470 | XP_018224087.1 | 99 | 0 |
| SPGP0C00780 | XP_018220445.1 | 100 | 0 |
| SPGP0Btrna2V | DI49_0484 | 100 | 0 |
| SPGP0H01630 | XP_018219348.1 | 99 | 0 |
| SPGP0Q00850 | XP_018222777.1 | 99 | 0 |
| SPGP0P01600 | XP_018220989.1 | 99 | 0 |
| SPGP0Btrna2F | DI49_0495 | 100 | 0 |
| SPGP0E02590 | XP_018222237.1 | 99 | 0 |
| SPGP0M03380 | XP_018221450.1 | 99 | 0 |
| SPGP0W00230 | XP_018221472.1 | 99 | 0 |
| SPGP0F03590 | XP_018220011.1 | 99 | 0 |
| SPGP0W00940 | XP_018221541.1 | 99 | 0 |
| SPGP0M02650 | XP_018221380.1 | 99 | 0 |
| SPGP0B02770 | XP_018223720.1 | 99 | 0 |
| SPGP0H03310 | XP_018219511.1 | 99 | 0 |
| SPGP0H03810 | XP_018219560.1 | 99 | 0 |
| SPGP0M00870 | XP_018221209.1 | 99 | 0 |
| SPGP0C02340 | XP_018220602.1 | 99 | 0 |
| SPGP0H03760 | XP_018219555.1 | 99 | 0 |
| SPGP0L03240 | XP_018218858.1 | 99 | 0 |
| SPGP0AVtrna9T | DI49_3176 | 100 | 0 |
| SPGP0F01210 | XP_018220239.1 | 99 | 0 |
| SPGP0S00640 | XP_018219787.1 | 99 | 0 |
| SPGP0O01630 | XP_018221790.1 | 99 | 0 |
| SPGP0W01540 | XP_018221598.1 | 99 | 0 |
| SPGP0W01010 | XP_018221548.1 | 99 | 0 |
| SPGP0H02690 | XP_018219450.1 | 99 | 0 |
| SPGP0I02410 | XP_018223096.1 | 99 | 0 |
| SPGP0Q00160 | XP_018222842.1 | 99 | 0 |
| SPGP0AI00330 | XP_018224074.1 | 99 | 0 |
| SPGP0H01910 | XP_018219376.1 | 99 | 0 |
| SPGP0ACtrna7C | DI49_5567 | 100 | 0 |
| SPGP0C04310 | XP_018220793.1 | 100 | 0 |
| SPGP0Q01380 | XP_018222725.1 | 100 | 0 |
| SPGP0H00940 | XP_018219281.1 | 99 | 0 |
| SPGP0AQ00550 | XP_018221992.1 | 99 | 0 |
| SPGP0Q01980 | XP_018222666.1 | 99 | 0 |

Create PDF in your applications with the Pdfcrowd HTML to PDF API PDFCROWD

| | | | |
|---|---|---|---|
| SPGP0I01110 | XP_018223223.1 | 100 | 0 |
| SPGP0E00520 | XP_018222445.1 | 99 | 0 |
| SPGP0M01520 | XP_018221271.1 | 99 | 0 |
| SPGP0P00550 | XP_018221089.1 | 99 | 0 |
| SPGP0AG00610 | XP_018223375.1 | 99 | 0 |
| SPGP0F02440 | XP_018220122.1 | 99 | 0 |
| SPGP0L03500 | XP_018218833.1 | 99 | 0 |
| SPGP0M00380 | XP_018221163.1 | 99 | 0 |
| SPGP0F01540 | XP_018220208.1 | 99 | 0 |
| SPGP0O01030 | XP_018221733.1 | 99 | 0 |
| SPGP0C04650 | XP_018220825.1 | 99 | 0 |
| SPGP0S00300 | XP_018219821.1 | 99 | 0 |
| SPGP0O02050 | XP_018221829.1 | 99 | 0 |
| SPGP0B02980 | XP_018223742.1 | 99 | 0 |
| SPGP0I03500 | XP_018222998.1 | 99 | 0 |
| SPGP0E02330 | XP_018222262.1 | 99 | 0 |
| SPGP0O02240 | XP_018221848.1 | 100 | 0 |
| SPGP0AI00480 | XP_018224088.1 | 99 | 0 |
| SPGP0H00430 | XP_018219231.1 | 99 | 0 |
| SPGP0AG00340 | XP_018223349.1 | 99 | 0 |
| SPGP0B01570 | XP_018223604.1 | 100 | 0 |
| SPGP0Q01670 | XP_018222697.1 | 99 | 0 |
| SPGP0B03370 | XP_018223779.1 | 99 | 0 |
| SPGP0O02070 | XP_018221831.1 | 99 | 0 |
| SPGP0AF00950 | XP_018222512.1 | 99 | 0 |
| SPGP0Q01510 | XP_018222713.1 | 99 | 0 |
| SPGP0F02730 | XP_018220095.1 | 99 | 0 |
| SPGP0E02440 | XP_018222251.1 | 99 | 0 |
| SPGP0C02030 | XP_018220571.1 | 99 | 0 |
| SPGP0H01340 | XP_018219319.1 | 99 | 0 |
| SPGP0Q01040 | XP_018222759.1 | 99 | 0 |
| SPGP0M00650 | XP_018221189.1 | 98 | 0 |
| SPGP0M00120 | XP_018221139.1 | 99 | 0 |
| SPGP0M00660 | XP_018221190.1 | 98 | 1 |
| SPGP0I03310 | XP_018223014.1 | 99 | 0 |
| SPGP0F00500 | XP_018220310.1 | 99 | 0 |
| SPGP0S02180 | XP_018219634.1 | 99 | 0 |
| SPGP0B00290 | XP_018223479.1 | 99 | 0 |
| SPGP0E00290 | XP_018222468.1 | 99 | 0 |
| SPGP0L02040 | XP_018218974.1 | 99 | 0 |

Create PDF in your applications with the Pdfcrowd HTML to PDF API PDFCROWD

| | | | |
|---|---|---|---|
| SPGP0M01380 | XP_018221257.1 | 99 | 0 |
| SPGP0M01830 | XP_018221302.1 | 99 | 0 |
| SPGP0M02900 | XP_018221405.1 | 99 | 0 |
| SPGP0L03790 | XP_018218804.1 | 99 | 0 |
| SPGP0H02300 | XP_018219413.1 | 99 | 0 |
| SPGP0I03660 | XP_018222982.1 | 99 | 0 |
| SPGP0C01310 | XP_018220499.1 | 99 | 0 |
| SPGP0H03210 | XP_018219501.1 | 99 | 0 |
| SPGP0I01800 | XP_018223155.1 | 99 | 0 |
| SPGP0B04300 | XP_018223871.1 | 99 | 0 |
| SPGP0I00400 | XP_018223294.1 | 99 | 0 |
| SPGP0L01400 | XP_018219036.1 | 99 | 0 |
| SPGP0B03760 | XP_018223817.1 | 99 | 0 |
| SPGP0S01430 | XP_018219709.1 | 99 | 0 |
| SPGP0H02200 | XP_018219403.1 | 99 | 0 |
| SPGP0M00820 | XP_018221204.1 | 98 | 0 |
| SPGP0E00420 | XP_018222455.1 | 99 | 0 |
| SPGP0E00640 | XP_018222433.1 | 99 | 0 |
| SPGP0E03280 | XP_018222169.1 | 99 | 0 |
| SPGP0S01370 | XP_018219715.1 | 99 | 0 |
| SPGP0F03950 | XP_018219976.1 | 99 | 0 |
| SPGP0C02280 | XP_018220596.1 | 99 | 0 |
| SPGP0AG00390 | XP_018223353.1 | 99 | 0 |
| SPGP0Q02370 | XP_018222628.1 | 99 | 0 |
| SPGP0AR00510 | XP_018219609.1 | 99 | 0 |
| SPGP0P01080 | XP_018221036.1 | 99 | 0 |
| SPGP0M02160 | XP_018221332.1 | 99 | 0 |
| SPGP0O00630 | XP_018221695.1 | 99 | 0 |
| SPGP0C03040 | XP_018220672.1 | 99 | 0 |
| SPGP0L01660 | XP_018219010.1 | 99 | 0 |
| SPGP0I02290 | XP_018223108.1 | 99 | 0 |
| SPGP0B02370 | XP_018223681.1 | 99 | 0 |
| SPGP0B01080 | XP_018223555.1 | 99 | 0 |
| SPGP0Q00200 | XP_018222838.1 | 99 | 0 |
| SPGP0C01020 | XP_018220470.1 | 100 | 0 |
| SPGP0I00310 | XP_018223303.1 | 99 | 0 |
| SPGP0C00790 | XP_018220446.1 | 99 | 0 |
| SPGP0B02150 | XP_018223661.1 | 99 | 0 |
| SPGP0B02100 | XP_018223656.1 | 100 | 0 |
| SPGP0B04130 | XP_018223854.1 | 99 | 0 |

Create PDF in your applications with the Pdfcrowd HTML to PDF API PDFCROWD

| | | | |
|---|---|---|---|
| SPGP0I01340 | XP_018223200.1 | 99 | 0 |
| SPGP0F03480 | XP_018220021.1 | 99 | 0 |
| SPGP0B04050 | XP_018223846.1 | 99 | 0 |
| SPGP0AF00260 | XP_018222576.1 | 99 | 0 |
| SPGP0W01890 | XP_018221633.1 | 99 | 0 |
| SPGP0I01750 | XP_018223160.1 | 99 | 0 |
| SPGP0Q02510 | XP_018222614.1 | 99 | 0 |
| SPGP0F03210 | XP_018220048.1 | 99 | 0 |
| SPGP0F01150 | XP_018220245.1 | 99 | 0 |
| SPGP0Q01180 | XP_018222744.1 | 99 | 0 |
| SPGP0H01980 | XP_018219383.1 | 99 | 0 |
| SPGP0I02120 | XP_018223124.1 | 100 | 0 |
| SPGP0AI00790 | XP_018224119.1 | 98 | 0 |
| SPGP0O01370 | XP_018221764.1 | 99 | 0 |
| SPGP0C00800 | XP_018220447.1 | 99 | 0 |
| SPGP0H00710 | XP_018219258.1 | 99 | 0 |
| SPGP0M01850 | XP_018221304.1 | 99 | 0 |
| SPGP0I03530 | XP_018222995.1 | 99 | 0 |
| SPGP0B01010 | XP_018223548.1 | 99 | 0 |
| SPGP0H02340 | XP_018219417.1 | 99 | 0 |
| SPGP0E00210 | XP_018222475.1 | 99 | 0 |
| SPGP0P00460 | XP_018221097.1 | 99 | 0 |
| SPGP0L00280 | XP_018219144.1 | 99 | 0 |
| SPGP0C01120 | XP_018220480.1 | 99 | 0 |
| SPGP0B05480 | XP_018223979.1 | 99 | 0 |
| SPGP0S01080 | XP_018219744.1 | 99 | 0 |
| SPGP0H00690 | XP_018219256.1 | 99 | 0 |
| SPGP0L00530 | XP_018219119.1 | 99 | 0 |
| SPGP0B05520 | XP_018223983.1 | 99 | 0 |
| SPGP0O02310 | XP_018221855.1 | 99 | 0 |
| SPGP0C04750 | XP_018220834.1 | 99 | 0 |
| SPGP0I00190 | XP_018223315.1 | 98 | 2 |
| SPGP0B00970 | XP_018223543.1 | 95 | 0 |
| SPGP0C02440 | XP_018220612.1 | 99 | 0 |
| SPGP0E00360 | XP_018222461.1 | 99 | 0 |
| SPGP0B04830 | XP_018223919.1 | 99 | 0 |
| SPGP0O01190 | XP_018221746.1 | 99 | 0 |
| SPGP0H02860 | XP_018219467.1 | 99 | 0 |
| SPGP0H03340 | XP_018219514.1 | 99 | 0 |
| SPGP0BB00100 | XP_018219844.1 | 99 | 0 |

Create PDF in your applications with the Pdfcrowd HTML to PDF API PDFCROWD

| | | | |
|---|---|---|---|
| SPGP0I02220 | XP_018223115.1 | 99 | 0 |
| SPGP0E03960 | XP_018222103.1 | 99 | 0 |
| SPGP0F01250 | XP_018220235.1 | 99 | 0 |
| SPGP0L02770 | XP_018218904.1 | 99 | 0 |
| SPGP0I03980 | XP_018222952.1 | 99 | 0 |
| SPGP0P02720 | XP_018220878.1 | 99 | 0 |
| SPGP0P02090 | XP_018220941.1 | 99 | 0 |
| SPGP0L00750 | XP_018219098.1 | 99 | 0 |
| SPGP0B05300 | XP_018223962.1 | 99 | 0 |
| SPGP0P02290 | XP_018220921.1 | 99 | 0 |
| SPGP0M02340 | XP_018221350.1 | 99 | 0 |
| SPGP0Ftrna13Y | DI49_2382 | 100 | 0 |
| SPGP0B00930 | XP_018223539.1 | 99 | 0 |
| SPGP0E02040 | XP_018222290.1 | 99 | 0 |
| SPGP0B00240 | XP_018223474.1 | 98 | 0 |
| SPGP0M03140 | XP_018221428.1 | 99 | 0 |
| SPGP0L02450 | XP_018218933.1 | 99 | 0 |
| SPGP0AL00210 | XP_018219885.1 | 99 | 0 |
| SPGP0S00380 | XP_018219813.1 | 99 | 0 |
| SPGP0P00830 | XP_018221061.1 | 100 | 0 |
| SPGP0L01210 | XP_018219054.1 | 99 | 0 |
| SPGP0H02990 | XP_018219480.1 | 99 | 0 |
| SPGP0E02290 | XP_018222266.1 | 98 | 0 |
| SPGP0F02810 | XP_018220087.1 | 99 | 0 |
| SPGP0O00250 | XP_018221658.1 | 99 | 0 |
| SPGP0E04220 | XP_018222077.1 | 99 | 0 |
| SPGP0E03770 | XP_018222122.1 | 98 | 0 |
| SPGP0O03040 | XP_018221924.1 | 99 | 0 |
| SPGP0H02740 | XP_018219455.1 | 99 | 1 |
| SPGP0L01020 | XP_018219073.1 | 99 | 0 |
| SPGP0E01480 | XP_018222346.1 | 100 | 0 |
| SPGP0AF00930 | XP_018222513.1 | 99 | 0 |
| SPGP0C03540 | XP_018220718.1 | 99 | 0 |
| SPGP0C04030 | XP_018220765.1 | 99 | 0 |
| SPGP0B03910 | XP_018223832.1 | 99 | 0 |
| SPGP0Q01860 | XP_018222677.1 | 99 | 0 |
| SPGP0AF00360 | XP_018222568.1 | 100 | 0 |
| SPGP0M01050 | XP_018221225.1 | 99 | 0 |
| SPGP0AG00870 | XP_018223398.1 | 99 | 0 |
| SPGP0E01350 | XP_018222360.1 | 99 | 0 |

Create PDF in your applications with the Pdfcrowd HTML to PDF API PDFCROWD

| | | | |
|---|---|---|---|
| SPGP0E03740 | XP_018222125.1 | 99 | 0 |
| SPGP0P01230 | XP_018221023.1 | 100 | 0 |
| SPGP0C03890 | XP_018220752.1 | 99 | 0 |
| SPGP0F01400 | XP_018220221.1 | 96 | 1 |
| SPGP0C00390 | XP_018220407.1 | 99 | 0 |
| SPGP0M03260 | XP_018221440.1 | 99 | 0 |
| SPGP0M01190 | XP_018221239.1 | 99 | 1 |
| SPGP0E02430 | XP_018222252.1 | 100 | 0 |
| SPGP0M01110 | XP_018221231.1 | 100 | 0 |
| SPGP0C03450 | XP_018220711.1 | 99 | 0 |
| SPGP0W00550 | XP_018221504.1 | 99 | 0 |
| SPGP0S02160 | XP_018219636.1 | 99 | 0 |
| SPGP0L02670 | XP_018218914.1 | 99 | 0 |
| SPGP0H01640 | XP_018219349.1 | 99 | 0 |
| SPGP0B06180 | XP_018224046.1 | 83 | 0 |
| SPGP0P02740 | XP_018220876.1 | 99 | 0 |
| SPGP0F03840 | XP_018219987.1 | 99 | 0 |
| SPGP0W01970 | XP_018221641.1 | 99 | 0 |
| SPGP0Q00440 | XP_018222817.1 | 99 | 0 |
| SPGP0O02370 | XP_018221861.1 | 99 | 0 |
| SPGP0I00980 | XP_018223236.1 | 99 | 0 |
| SPGP0Q00150 | XP_018222843.1 | 99 | 0 |
| SPGP0P01010 | XP_018221043.1 | 99 | 0 |
| SPGP0E03210 | XP_018222175.1 | 98 | 1 |
| SPGP0BJ00130 | XP_018219163.1 | 99 | 0 |
| SPGP0E03080 | XP_018222188.1 | 99 | 0 |
| SPGP0B04570 | XP_018223897.1 | 99 | 0 |
| SPGP0C03050 | XP_018220673.1 | 99 | 0 |
| SPGP0H00590 | XP_018219247.1 | 99 | 0 |
| SPGP0L02900 | XP_018218891.1 | 99 | 0 |
| SPGP0L03070 | XP_018218874.1 | 99 | 0 |
| SPGP0W00770 | XP_018221525.1 | 99 | 0 |
| SPGP0S01850 | XP_018219667.1 | 99 | 0 |
| SPGP0E04240 | XP_018222075.1 | 99 | 0 |
| SPGP0O02930 | XP_018221915.1 | 98 | 0 |
| SPGP0H01420 | XP_018219327.1 | 99 | 0 |
| SPGP0I01080 | XP_018223226.1 | 99 | 0 |
| SPGP0F03060 | XP_018220063.1 | 99 | 0 |
| SPGP0M00460 | XP_018221171.1 | 99 | 0 |
| SPGP0I02720 | XP_018223066.1 | 99 | 0 |

Create PDF in your applications with the Pdfcrowd HTML to PDF API PDFCROWD

| | | | |
|---|---|---|---|
| SPGP0B03460 | XP_018223788.1 | 99 | 0 |
| SPGP0Itrna21T | DI49_2415 | 100 | 0 |
| SPGP0AF00120 | XP_018222592.1 | 95 | 1 |
| SPGP0W01200 | XP_018221565.1 | 99 | 0 |
| SPGP0AF01120 | XP_018222496.1 | 99 | 0 |
| SPGP0M01340 | XP_018221253.1 | 99 | 0 |
| SPGP0I03160 | XP_018223028.1 | 99 | 0 |
| SPGP0AI00670 | XP_018224106.1 | 99 | 0 |
| SPGP0H03500 | XP_018219528.1 | 99 | 0 |
| SPGP0B05440 | XP_018223975.1 | 99 | 0 |
| SPGP0M01730 | XP_018221291.1 | 99 | 0 |
| SPGP0B01320 | XP_018223579.1 | 99 | 0 |
| SPGP0AQ00530 | XP_018221991.1 | 99 | 0 |
| SPGP0H01010 | XP_018219288.1 | 99 | 0 |
| SPGP0S00450 | XP_018219806.1 | 99 | 0 |
| SPGP0B03120 | XP_018223755.1 | 99 | 0 |
| SPGP0I02460 | XP_018223091.1 | 99 | 1 |
| SPGP0B03550 | XP_018223796.1 | 99 | 0 |
| SPGP0Q02040 | XP_018222661.1 | 100 | 0 |
| SPGP0F03030 | XP_018220066.1 | 99 | 0 |
| SPGP0O00760 | XP_018221708.1 | 99 | 0 |
| SPGP0O03070 | XP_018221927.1 | 99 | 0 |
| SPGP0C02100 | XP_018220578.1 | 99 | 0 |
| SPGP0L00620 | XP_018219110.1 | 99 | 0 |
| SPGP0AG00790 | XP_018223391.1 | 99 | 0 |
| SPGP0Ctrna6R | DI49_3845 | 100 | 0 |
| SPGP0O03170 | XP_018221937.1 | 99 | 0 |
| SPGP0L03570 | XP_018218826.1 | 99 | 0 |
| SPGP0B03250 | XP_018223768.1 | 99 | 0 |
| SPGP0L00420 | XP_018219130.1 | 99 | 1 |
| SPGP0AG00400 | XP_018223354.1 | 99 | 0 |
| SPGP0B01140 | XP_018223561.1 | 99 | 0 |
| SPGP0H03000 | XP_018219481.1 | 99 | 0 |
| SPGP0F00710 | XP_018220289.1 | 99 | 0 |
| SPGP0B01220 | XP_018223569.1 | 99 | 0 |
| SPGP0F02210 | XP_018220145.1 | 99 | 0 |
| SPGP0Q02560 | XP_018222609.1 | 99 | 0 |
| SPGP0E00960 | XP_018222401.1 | 99 | 0 |
| SPGP0AQ00210 | XP_018221959.1 | 99 | 0 |
| SPGP0B05990 | XP_018224029.1 | 99 | 0 |

Create PDF in your applications with the Pdfcrowd HTML to PDF API PDFCROWD

| | | | |
|---|---|---|---|
| SPGP0C03940 | XP_018220757.1 | 99 | 0 |
| SPGP0O00310 | XP_018221664.1 | 100 | 0 |
| SPGP0W01220 | XP_018221567.1 | 99 | 0 |
| SPGP0AG00520 | XP_018223366.1 | 99 | 0 |
| SPGP0P01740 | XP_018220975.1 | 99 | 0 |
| SPGP0C01330 | XP_018220501.1 | 99 | 0 |
| SPGP0L01430 | XP_018219033.1 | 99 | 0 |
| SPGP0O01610 | XP_018221788.1 | 99 | 0 |
| SPGP0B00840 | XP_018223530.1 | 99 | 0 |
| SPGP0P01130 | XP_018221032.1 | 99 | 0 |
| SPGP0E00150 | XP_018222481.1 | 100 | 0 |
| SPGP0S01910 | XP_018219661.1 | 99 | 0 |
| SPGP0I00200 | XP_018223314.1 | 99 | 0 |
| SPGP0L01260 | XP_018219050.1 | 99 | 0 |
| SPGP0L02990 | XP_018218882.1 | 99 | 0 |
| SPGP0P02340 | XP_018220916.1 | 99 | 0 |
| SPGP0AR00160 | XP_018219575.1 | 99 | 0 |
| SPGP0Ntrna7R | DI49_3064 | 100 | 0 |
| SPGP0Q01090 | XP_018222753.1 | 99 | 0 |
| SPGP0B00740 | XP_018223520.1 | 99 | 0 |
| SPGP0L01830 | XP_018218994.1 | 98 | 0 |
| SPGP0L01340 | XP_018219042.1 | 99 | 0 |
| SPGP0Q01130 | XP_018222749.1 | 99 | 0 |
| SPGP0F01990 | XP_018220167.1 | 99 | 0 |
| SPGP0C03430 | XP_018220709.1 | 99 | 0 |
| SPGP0E02670 | XP_018222228.1 | 99 | 0 |
| SPGP0O02460 | XP_018221870.1 | 99 | 0 |
| SPGP0F00520 | XP_018220308.1 | 99 | 0 |
| SPGP0B01360 | XP_018223583.1 | 99 | 0 |
| SPGP0C02830 | XP_018220651.1 | 99 | 0 |
| SPGP0Q01690 | XP_018222695.1 | 99 | 0 |
| SPGP0BF00100 | XP_018222852.1 | 97 | 0 |
| SPGP0B01630 | XP_018223609.1 | 99 | 0 |
| SPGP0F02160 | XP_018220150.1 | 99 | 0 |
| SPGP0S01220 | XP_018219730.1 | 99 | 0 |
| SPGP0C01690 | XP_018220537.1 | 99 | 0 |
| SPGP0C03670 | XP_018220731.1 | 99 | 0 |
| SPGP0F00940 | XP_018220266.1 | 99 | 0 |
| SPGP0H02960 | XP_018219477.1 | 99 | 0 |

Create PDF in your applications with the Pdfcrowd HTML to PDF API PDFCROWD

| | | | |
|---|---|---|---|
| SPGP0C04630 | XP_018220824.1 | 99 | 0 |
| SPGP0M00470 | XP_018221172.1 | 99 | 0 |
| SPGP0C03840 | XP_018220747.1 | 99 | 0 |
| SPGP0S00930 | XP_018219759.1 | 100 | 0 |
| SPGP0H03450 | XP_018219525.1 | 99 | 0 |
| SPGP0P02390 | XP_018220911.1 | 100 | 0 |
| SPGP0M02410 | XP_018221357.1 | 99 | 0 |
| SPGP0Q01440 | XP_018222720.1 | 99 | 0 |
| SPGP0AG00210 | XP_018223336.1 | 99 | 0 |
| SPGP0I01560 | XP_018223179.1 | 99 | 0 |
| SPGP0C04550 | XP_018220816.1 | 99 | 0 |
| SPGP0AI00300 | XP_018224071.1 | 99 | 0 |
| SPGP0I03840 | XP_018222964.1 | 99 | 0 |
| SPGP0B05920 | XP_018224022.1 | 99 | 0 |
| SPGP0O02500 | XP_018221874.1 | 100 | 0 |
| SPGP0M00810 | XP_018221203.1 | 99 | 0 |
| SPGP0L03080 | XP_018218873.1 | 99 | 0 |
| SPGP0I02910 | XP_018223049.1 | 99 | 0 |
| SPGP0Q00870 | XP_018222775.1 | 99 | 0 |
| SPGP0B01890 | XP_018223635.1 | 99 | 0 |
| SPGP0AL00530 | XP_018219915.1 | 99 | 0 |
| SPGP0W01780 | XP_018221622.1 | 98 | 1 |
| SPGP0H02500 | XP_018219432.1 | 99 | 0 |
| SPGP0P01370 | XP_018221009.1 | 99 | 0 |
| SPGP0H01850 | XP_018219370.1 | 99 | 0 |
| SPGP0I01310 | XP_018223203.1 | 99 | 0 |
| SPGP0W00960 | XP_018221543.1 | 99 | 0 |
| SPGP0O00860 | XP_018221716.1 | 99 | 0 |
| SPGP0W00780 | XP_018221526.1 | 99 | 0 |
| SPGP0E00340 | XP_018222463.1 | 99 | 0 |
| SPGP0E02460 | XP_018222249.1 | 99 | 0 |
| SPGP0M02670 | XP_018221382.1 | 98 | 1 |
| SPGP0H00640 | XP_018219251.1 | 99 | 0 |
| SPGP0E02200 | XP_018222275.1 | 99 | 0 |
| SPGP0F03760 | XP_018219994.1 | 99 | 0 |
| SPGP0AG00980 | XP_018223408.1 | 99 | 0 |
| SPGP0F01140 | XP_018220246.1 | 99 | 0 |
| SPGP0AG00180 | XP_018223333.1 | 99 | 0 |
| SPGP0L03260 | XP_018218856.1 | 99 | 0 |
| SPGP0C00270 | XP_018220396.1 | 99 | 0 |

Create PDF in your applications with the Pdfcrowd HTML to PDF API PDFCROWD

| | | | |
|---|---|---|---|
| SPGP0S00660 | XP_018219785.1 | 99 | 0 |
| SPGP0H03830 | XP_018219562.1 | 99 | 0 |
| SPGP0O01930 | XP_018221818.1 | 97 | 2 |
| SPGP0W01560 | XP_018221600.1 | 99 | 0 |
| SPGP0W01210 | XP_018221566.1 | 99 | 0 |
| SPGP0C04370 | XP_018220799.1 | 99 | 0 |
| SPGP0Q00840 | XP_018222778.1 | 99 | 0 |
| SPGP0Q00750 | XP_018222787.1 | 100 | 0 |
| SPGP0B00460 | XP_018223494.1 | 100 | 0 |
| SPGP0P02970 | XP_018220854.1 | 99 | 0 |
| SPGP0L00610 | XP_018219111.1 | 99 | 0 |
| SPGP0C03810 | XP_018220744.1 | 99 | 0 |
| SPGP0F01320 | XP_018220228.1 | 99 | 0 |
| SPGP0B01710 | XP_018223617.1 | 99 | 0 |
| SPGP0C04540 | XP_018220815.1 | 99 | 0 |
| SPGP0M02000 | XP_018221318.1 | 99 | 0 |
| SPGP0Q01650 | XP_018222699.1 | 99 | 0 |
| SPGP0AG00360 | XP_018223350.1 | 99 | 0 |
| SPGP0P00530 | XP_018221091.1 | 100 | 0 |
| SPGP0E03260 | XP_018222171.1 | 99 | 0 |
| SPGP0F03490 | XP_018220020.1 | 99 | 0 |
| SPGP0B02540 | XP_018223698.1 | 99 | 0 |
| SPGP0S00590 | XP_018219792.1 | 99 | 0 |
| SPGP0S01680 | XP_018219684.1 | 99 | 0 |
| SPGP0H02590 | XP_018219441.1 | 100 | 0 |
| SPGP0B03750 | XP_018223816.1 | 99 | 0 |
| SPGP0F02090 | XP_018220157.1 | 99 | 0 |
| SPGP0F03010 | XP_018220068.1 | 99 | 0 |
| SPGP0M03100 | XP_018221424.1 | 99 | 0 |
| SPGP0B01280 | XP_018223575.1 | 99 | 0 |
| SPGP0F01360 | XP_018220224.1 | 99 | 0 |
| SPGP0H00900 | XP_018219277.1 | 99 | 0 |
| SPGP0H01180 | XP_018219305.1 | 99 | 0 |
| SPGP0B01510 | XP_018223598.1 | 99 | 0 |
| SPGP0P01110 | XP_018221034.1 | 100 | 0 |
| SPGP0B05680 | XP_018223998.1 | 99 | 0 |
| SPGP0AF00420 | XP_018222562.1 | 99 | 0 |
| SPGP0Q01230 | XP_018222739.1 | 99 | 0 |
| SPGP0H01360 | XP_018219321.1 | 99 | 0 |
| SPGP0M00100 | XP_018221137.1 | 99 | 0 |

Create PDF in your applications with the Pdfcrowd HTML to PDF API PDFCROWD

| | | | |
|---|---|---|---|
| SPGP0M02370 | XP_018221353.1 | 99 | 0 |
| SPGP0C04740 | XP_018220833.1 | 99 | 0 |
| SPGP0C02650 | XP_018220633.1 | 100 | 0 |
| SPGP0E01280 | XP_018222368.1 | 99 | 0 |
| SPGP0P01610 | XP_018220988.1 | 99 | 0 |
| SPGP0E01390 | XP_018222356.1 | 100 | 0 |
| SPGP0M01650 | XP_018221284.1 | 99 | 0 |
| SPGP0L02020 | XP_018218976.1 | 99 | 0 |
| SPGP0I01610 | XP_018223174.1 | 99 | 0 |
| SPGP0I02010 | XP_018223133.1 | 99 | 0 |
| SPGP0C02550 | XP_018220623.1 | 99 | 0 |
| SPGP0I01820 | XP_018223153.1 | 99 | 0 |
| SPGP0E03980 | XP_018222101.1 | 99 | 0 |
| SPGP0I00420 | XP_018223292.1 | 99 | 0 |
| SPGP0H02230 | XP_018219406.1 | 99 | 0 |
| SPGP0C00360 | XP_018220404.1 | 99 | 0 |
| SPGP0E00930 | XP_018222404.1 | 99 | 0 |
| SPGP0Q01050 | XP_018222758.1 | 99 | 0 |
| SPGP0B04480 | XP_018223888.1 | 99 | 0 |
| SPGP0O00370 | XP_018221670.1 | 100 | 0 |
| SPGP0B02190 | XP_018223664.1 | 99 | 0 |
| SPGP0S00630 | XP_018219788.1 | 99 | 0 |
| SPGP0I02600 | XP_018223077.1 | 99 | 0 |
| SPGP0I00560 | XP_018223278.1 | 99 | 0 |
| SPGP0M01510 | XP_018221270.1 | 99 | 0 |
| SPGP0E03840 | XP_018222115.1 | 99 | 0 |
| SPGP0M01500 | XP_018221269.1 | 99 | 0 |
| SPGP0F03870 | XP_018219984.1 | 99 | 0 |
| SPGP0Q02680 | XP_018222599.1 | 99 | 0 |
| SPGP0Q02350 | XP_018222630.1 | 99 | 0 |
| SPGP0H01660 | XP_018219351.1 | 99 | 0 |
| SPGP0L01130 | XP_018219062.1 | 99 | 0 |
| SPGP0L01640 | XP_018219012.1 | 99 | 0 |
| SPGP0M03150 | XP_018221429.1 | 99 | 0 |
| SPGP0F01050 | XP_018220255.1 | 99 | 0 |
| SPGP0H03590 | XP_018219537.1 | 99 | 0 |
| SPGP0C02500 | XP_018220618.1 | 99 | 0 |
| SPGP0L00820 | XP_018219091.1 | 98 | 2 |
| SPGP0P01810 | XP_018220968.1 | 99 | 0 |
| SPGP0B04150 | XP_018223856.1 | 99 | 0 |

Create PDF in your applications with the Pdfcrowd HTML to PDF API PDFCROWD

| | | | |
|---|---|---|---|
| SPGP0AF00100 | XP_018222594.1 | 99 | 0 |
| SPGP0O00590 | XP_018221691.1 | 99 | 0 |
| SPGP0C02640 | XP_018220632.1 | 99 | 0 |
| SPGP0C02310 | XP_018220599.1 | 99 | 0 |
| SPGP0M01550 | XP_018221274.1 | 98 | 0 |
| SPGP0P01580 | XP_018220990.1 | 99 | 0 |
| SPGP0CZ00100 | XP_018222854.1 | 98 | 0 |
| SPGP0O01350 | XP_018221762.1 | 99 | 0 |
| SPGP0O01420 | XP_018221769.1 | 99 | 0 |
| SPGP0C00860 | XP_018220454.1 | 99 | 0 |
| SPGP0I02140 | XP_018223123.1 | 100 | 0 |
| SPGP0C01930 | XP_018220561.1 | 99 | 0 |
| SPGP0I03210 | XP_018223023.1 | 99 | 0 |
| SPGP0B03020 | XP_018223746.1 | 99 | 1 |
| SPGP0M01350 | XP_018221254.1 | 99 | 0 |
| SPGP0H03740 | XP_018219553.1 | 99 | 0 |
| SPGP0B00300 | XP_018223480.1 | 99 | 0 |
| SPGP0L00510 | XP_018219121.1 | 99 | 0 |
| SPGP0B03680 | XP_018223809.1 | 99 | 0 |
| SPGP0I01480 | XP_018223187.1 | 100 | 0 |
| SPGP0P00820 | XP_018221062.1 | 99 | 0 |
| SPGP0B04020 | XP_018223843.1 | 99 | 0 |
| SPGP0O02330 | XP_018221857.1 | 99 | 0 |
| SPGP0F00260 | XP_018220333.1 | 99 | 0 |
| SPGP0H03050 | XP_018219486.1 | 99 | 0 |
| SPGP0P02170 | XP_018220933.1 | 99 | 0 |
| SPGP0B00750 | XP_018223521.1 | 99 | 0 |
| SPGP0E01740 | XP_018222320.1 | 99 | 0 |
| SPGP0F00590 | XP_018220301.1 | 97 | 3 |
| SPGP0B00530 | XP_018223501.1 | 99 | 0 |
| SPGP0F01720 | XP_018220192.1 | 99 | 0 |
| SPGP0AL00680 | XP_018219929.1 | 99 | 0 |
| SPGP0Q02030 | XP_018222662.1 | 99 | 0 |
| SPGP0B02040 | XP_018223650.1 | 99 | 0 |
| SPGP0C04440 | XP_018220806.1 | 99 | 0 |
| SPGP0F02830 | XP_018220085.1 | 99 | 0 |
| SPGP0L02520 | XP_018218926.1 | 99 | 0 |
| SPGP0E02050 | XP_018222289.1 | 99 | 0 |
| SPGP0B00990 | XP_018223546.1 | 99 | 0 |
| SPGP0W01020 | XP_018221549.1 | 99 | 0 |

Create PDF in your applications with the Pdfcrowd HTML to PDF API PDFCROWD

| | | | |
|---|---|---|---|
| SPGP0L00770 | XP_018219096.1 | 100 | 0 |
| SPGP0M02880 | XP_018221403.1 | 99 | 0 |
| SPGP0I00600 | XP_018223274.1 | 99 | 0 |
| SPGP0C00150 | XP_018220386.1 | 99 | 0 |
| SPGP0S01500 | XP_018219702.1 | 99 | 0 |
| SPGP0B00260 | XP_018223476.1 | 100 | 0 |
| SPGP0S00990 | XP_018219753.1 | 99 | 0 |
| SPGP0AL00230 | XP_018219887.1 | 99 | 0 |
| SPGP0L02300 | XP_018218948.1 | 99 | 0 |
| SPGP0P00850 | XP_018221059.1 | 99 | 0 |
| SPGP0C03590 | XP_018220723.1 | 99 | 0 |
| SPGP0B01150 | XP_018223562.1 | 99 | 0 |
| SPGP0C01300 | XP_018220498.1 | 99 | 0 |
| SPGP0S00800 | XP_018219771.1 | 99 | 0 |
| SPGP0B06030 | XP_018224033.1 | 99 | 0 |
| SPGP0L03330 | XP_018218849.1 | 100 | 0 |
| SPGP0O00530 | XP_018221685.1 | 99 | 0 |
| SPGP0AF00230 | XP_018222579.1 | 99 | 0 |
| SPGP0O03060 | XP_018221926.1 | 100 | 0 |
| SPGP0L03250 | XP_018218857.1 | 99 | 0 |
| SPGP0H01810 | XP_018219366.1 | 99 | 0 |
| SPGP0AF00460 | XP_018222559.1 | 99 | 0 |
| SPGP0AT00360 | XP_018219173.1 | 99 | 0 |
| SPGP0L03280 | XP_018218854.1 | 99 | 0 |
| SPGP0S01140 | XP_018219738.1 | 99 | 0 |
| SPGP0O02190 | XP_018221843.1 | 99 | 0 |
| SPGP0B03140 | XP_018223757.1 | 99 | 0 |
| SPGP0S01410 | XP_018219711.1 | 99 | 1 |
| SPGP0Q01080 | XP_018222754.1 | 99 | 0 |
| SPGP0I02870 | XP_018223053.1 | 99 | 0 |
| SPGP0C00480 | XP_018220416.1 | 100 | 0 |
| SPGP0Etrna40L | DI49_2147 | 99 | 0 |
| SPGP0I00160 | XP_018223318.1 | 99 | 0 |
| SPGP0W01770 | XP_018221621.1 | 99 | 0 |
| SPGP0W00530 | XP_018221502.1 | 99 | 0 |
| SPGP0E02410 | XP_018222254.1 | 100 | 0 |
| SPGP0F01230 | XP_018220237.1 | 99 | 0 |
| SPGP0S01070 | XP_018219745.1 | 99 | 0 |
| SPGP0B04340 | XP_018223875.1 | 99 | 0 |
| SPGP0S02140 | XP_018219638.1 | 99 | 0 |

Create PDF in your applications with the Pdfcrowd HTML to PDF API PDFCROWD

| | | | |
|---|---|---|---|
| SPGP0L02610 | XP_018218920.1 | 99 | 0 |
| SPGP0W00570 | XP_018221506.1 | 99 | 0 |
| SPGP0I00120 | XP_018223322.1 | 99 | 0 |
| SPGP0H03720 | XP_018219551.1 | 99 | 0 |
| SPGP0C00240 | XP_018220394.1 | 99 | 0 |
| SPGP0C04230 | XP_018220785.1 | 99 | 0 |
| SPGP0P00700 | XP_018221074.1 | 99 | 0 |
| SPGP0H01990 | XP_018219384.1 | 100 | 0 |
| SPGP0B04310 | XP_018223872.1 | 99 | 0 |
| SPGP0P01940 | XP_018220956.1 | 99 | 0 |
| SPGP0E03090 | XP_018222187.1 | 98 | 0 |
| SPGP0AL00490 | XP_018219911.1 | 99 | 1 |
| SPGP0W01990 | XP_018221643.1 | 99 | 0 |
| SPGP0E01680 | XP_018222326.1 | 99 | 0 |
| SPGP0S00680 | XP_018219783.1 | 99 | 0 |
| SPGP0W01190 | XP_018221564.1 | 99 | 0 |
| SPGP0P01030 | XP_018221041.1 | 98 | 2 |
| SPGP0E04030 | XP_018222096.1 | 99 | 0 |
| SPGP0L03540 | XP_018218829.1 | 99 | 0 |
| SPGP0I02470 | XP_018223090.1 | 99 | 0 |
| SPGP0C02990 | XP_018220667.1 | 99 | 0 |
| SPGP0AF00760 | XP_018222529.1 | 99 | 0 |
| SPGP0W00720 | XP_018221520.1 | 99 | 0 |
| SPGP0L02920 | XP_018218889.1 | 99 | 0 |
| SPGP0F04110 | XP_018219960.1 | 99 | 1 |
| SPGP0C04570 | XP_018220818.1 | 99 | 0 |
| SPGP0S00580 | XP_018219793.1 | 99 | 0 |
| SPGP0F01740 | XP_018220190.1 | 99 | 0 |
| SPGP0I03850 | XP_018222963.1 | 100 | 0 |
| SPGP0I01900 | XP_018223145.1 | 98 | 1 |
| SPGP0F03000 | XP_018220069.1 | 99 | 0 |
| SPGP0H01190 | XP_018219306.1 | 99 | 0 |
| SPGP0L03690 | XP_018218814.1 | 99 | 0 |
| SPGP0F01240 | XP_018220236.1 | 99 | 0 |
| SPGP0H00510 | XP_018219239.1 | 99 | 0 |
| SPGP0C02910 | XP_018220659.1 | 99 | 0 |
| SPGP0P02640 | XP_018220886.1 | 99 | 0 |
| SPGP0O00180 | XP_018221651.1 | 99 | 0 |
| SPGP0C02780 | XP_018220646.1 | 99 | 0 |
| SPGP0COtrna10P | DI49_4736 | 100 | 0 |

Create PDF in your applications with the Pdfcrowd HTML to PDF API PDFCROWD

| | | | |
|---|---|---|---|
| SPGP0AF01100 | XP_018222498.1 | 99 | 0 |
| SPGP0F00270 | XP_018220332.1 | 99 | 0 |
| SPGP0B01660 | XP_018223612.1 | 99 | 0 |
| SPGP0B05880 | XP_018224018.1 | 99 | 0 |
| SPGP0AG00160 | XP_018223331.1 | 99 | 0 |
| SPGP0B04560 | XP_018223896.1 | 99 | 0 |
| SPGP0L03520 | XP_018218831.1 | 99 | 0 |
| SPGP0C00900 | XP_018220458.1 | 99 | 0 |
| SPGP0S00430 | XP_018219808.1 | 99 | 0 |
| SPGP0O02590 | XP_018221882.1 | 99 | 0 |
| SPGP0Q00720 | XP_018222790.1 | 99 | 0 |
| SPGP0AG00910 | XP_018223402.1 | 99 | 0 |
| SPGP0F00220 | XP_018220337.1 | 99 | 0 |
| SPGP0O00740 | XP_018221706.1 | 99 | 0 |
| SPGP0O00290 | XP_018221662.1 | 99 | 0 |
| SPGP0B01300 | XP_018223577.1 | 99 | 0 |
| SPGP0C01830 | XP_018220551.1 | 99 | 0 |
| SPGP0Q01460 | XP_018222718.1 | 99 | 0 |
| SPGP0M01750 | XP_018221294.1 | 99 | 0 |
| SPGP0AI00500 | XP_018224090.1 | 99 | 0 |
| SPGP0M03320 | XP_018221445.1 | 99 | 0 |
| SPGP0Q00960 | XP_018222766.1 | 99 | 0 |
| SPGP0Ctrna4Q | DI49_2421 | 100 | 0 |
| SPGP0H00310 | XP_018219219.1 | 99 | 0 |
| SPGP0C03020 | XP_018220670.1 | 99 | 0 |
| SPGP0B02200 | XP_018223665.1 | 99 | 0 |
| SPGP0AG00420 | XP_018223356.1 | 99 | 0 |
| SPGP0L01380 | XP_018219038.1 | 100 | 0 |
| SPGP0F00760 | XP_018220284.1 | 99 | 0 |
| SPGP0F03420 | XP_018220027.1 | 99 | 0 |
| SPGP0O02770 | XP_018221900.1 | 99 | 0 |
| SPGP0AR00270 | XP_018219586.1 | 99 | 0 |
| SPGP0F00190 | XP_018220340.1 | 99 | 0 |
| SPGP0Q01900 | XP_018222673.1 | 100 | 0 |
| SPGP0F01830 | XP_018220182.1 | 99 | 0 |
| SPGP0H03670 | XP_018219546.1 | 99 | 0 |
| SPGP0L03290 | XP_018218853.1 | 99 | 0 |
| SPGP0O02270 | XP_018221851.1 | 99 | 0 |
| SPGP0L01550 | XP_018219021.1 | 99 | 1 |
| SPGP0I00490 | XP_018223285.1 | 99 | 0 |

Create PDF in your applications with the Pdfcrowd HTML to PDF API PDFCROWD

| | | | |
|---|---|---|---|
| SPGP0O02350 | XP_018221859.1 | 99 | 0 |
| SPGP0Q01300 | XP_018222733.1 | 99 | 0 |
| SPGP0L01580 | XP_018219018.1 | 99 | 0 |
| SPGP0AG00600 | XP_018223374.1 | 99 | 0 |
| SPGP0AL00560 | XP_018219918.1 | 99 | 0 |
| SPGP0O00120 | XP_018221645.1 | 95 | 0 |
| SPGP0B06000 | XP_018224030.1 | 99 | 0 |
| SPGP0F03410 | XP_018220028.1 | 99 | 0 |
| SPGP0E01890 | XP_018222305.1 | 99 | 0 |
| SPGP0Q01770 | XP_018222687.1 | 99 | 0 |
| SPGP0W01120 | XP_018221558.1 | 99 | 0 |
| SPGP0E01620 | XP_018222332.1 | 99 | 0 |
| SPGP0AR00140 | XP_018219573.1 | 100 | 0 |
| SPGP0B05900 | XP_018224020.1 | 100 | 0 |
| SPGP0C01500 | XP_018220519.1 | 99 | 0 |
| SPGP0H02260 | XP_018219409.1 | 99 | 0 |
| SPGP0I01780 | XP_018223157.1 | 99 | 0 |
| SPGP0Q00110 | XP_018222847.1 | 99 | 0 |
| SPGP0E00900 | XP_018222407.1 | 99 | 0 |
| SPGP0S02010 | XP_018219651.1 | 99 | 0 |
| SPGP0O01650 | XP_018221792.1 | 100 | 0 |
| SPGP0B02590 | XP_018223703.1 | 99 | 0 |
| SPGP0B05650 | XP_018223995.1 | 99 | 0 |
| SPGP0H01590 | XP_018219344.1 | 99 | 0 |
| SPGP0C02420 | XP_018220610.1 | 99 | 0 |
| SPGP0O02280 | XP_018221852.1 | 99 | 0 |
| SPGP0L02050 | XP_018218973.1 | 99 | 0 |
| SPGP0I03710 | XP_018222977.1 | 99 | 0 |
| SPGP0C04350 | XP_018220797.1 | 99 | 0 |
| SPGP0F02740 | XP_018220094.1 | 99 | 0 |
| SPGP0S00290 | XP_018219822.1 | 99 | 0 |
| SPGP0H01790 | XP_018219364.1 | 99 | 0 |
| SPGP0L03460 | XP_018218837.1 | 99 | 0 |
| SPGP0B02920 | XP_018223736.1 | 99 | 0 |
| SPGP0BB00130 | XP_018219847.1 | 99 | 0 |
| SPGP0O02010 | XP_018221825.1 | 99 | 0 |
| SPGP0C03930 | XP_018220756.1 | 99 | 0 |
| SPGP0C04610 | XP_018220822.1 | 99 | 0 |
| SPGP0B04950 | XP_018223930.1 | 99 | 0 |
| SPGP0I01540 | XP_018223181.1 | 99 | 0 |

Create PDF in your applications with the Pdfcrowd HTML to PDF API PDFCROWD

| | | | |
|---|---|---|---|
| SPGP0H03930 | XP_018218753.1 | 99 | 0 |
| SPGP0E01310 | XP_018222364.1 | 99 | 0 |
| SPGP0L02690 | XP_018218912.1 | 99 | 0 |
| SPGP0M00830 | XP_018221205.1 | 99 | 0 |
| SPGP0GZ00100 | XP_018221457.1 | 99 | 0 |
| SPGP0B00630 | XP_018223509.1 | 99 | 0 |
| SPGP0W01760 | XP_018221620.1 | 99 | 0 |
| SPGP0Q00890 | XP_018222773.1 | 99 | 0 |
| SPGP0I00370 | XP_018223297.1 | 99 | 0 |
| SPGP0P02010 | XP_018220949.1 | 99 | 0 |
| SPGP0P01390 | XP_018221008.1 | 99 | 0 |
| SPGP0BB00150 | XP_018219849.1 | 99 | 0 |
| SPGP0W00900 | XP_018221537.1 | 99 | 0 |
| SPGP0C02360 | XP_018220604.1 | 99 | 0 |
| SPGP0H03950 | XP_018218751.1 | 99 | 0 |
| SPGP0M02690 | XP_018221384.1 | 99 | 0 |
| SPGP0Q00640 | XP_018222798.1 | 99 | 0 |
| SPGP0H01170 | XP_018219304.1 | 99 | 0 |
| SPGP0B03890 | XP_018223830.1 | 99 | 0 |
| SPGP0I02590 | XP_018223078.1 | 99 | 0 |
| SPGP0F02060 | XP_018220160.1 | 99 | 0 |
| SPGP0L03200 | XP_018218862.1 | 99 | 0 |
| SPGP0S01600 | XP_018219692.1 | 99 | 0 |
| SPGP0S00600 | XP_018219791.1 | 99 | 0 |
| SPGP0H03390 | XP_018219519.1 | 99 | 0 |
| SPGP0H02400 | XP_018219423.1 | 99 | 0 |
| SPGP0W01500 | XP_018221594.1 | 93 | 6 |
| SPGP0AQ00100 | XP_018221950.1 | 99 | 0 |
| SPGP0F00290 | XP_018220330.1 | 99 | 0 |
| SPGP0H02520 | XP_018219434.1 | 98 | 1 |
| SPGP0I01120 | XP_018223222.1 | 99 | 0 |
| SPGP0F01610 | XP_018220201.1 | 99 | 0 |
| SPGP0B00590 | XP_018223506.1 | 100 | 0 |
| SPGP0F03580 | XP_018220012.1 | 99 | 0 |
| SPGP0S00180 | XP_018219832.1 | 99 | 0 |
| SPGP0L02350 | XP_018218943.1 | 100 | 0 |
| SPGP0AF00620 | XP_018222543.1 | 99 | 0 |
| SPGP0B05330 | XP_018223965.1 | 99 | 0 |
| SPGP0AF00190 | XP_018222583.1 | 99 | 0 |
| SPGP0S01010 | XP_018219751.1 | 99 | 0 |

Create PDF in your applications with the Pdfcrowd HTML to PDF API PDFCROWD

| | | | |
|---|---|---|---|
| SPGP0AG00300 | XP_018223345.1 | 99 | 0 |
| SPGP0AL00310 | XP_018219894.1 | 99 | 0 |
| SPGP0AI00530 | XP_018224093.1 | 99 | 0 |
| SPGP0M02910 | XP_018221406.1 | 99 | 0 |
| SPGP0P00510 | XP_018221093.1 | 99 | 0 |
| SPGP0L02660 | XP_018218915.1 | 99 | 0 |
| SPGP0I01930 | XP_018223141.1 | 99 | 0 |
| SPGP0B03650 | XP_018223806.1 | 98 | 2 |
| SPGP0M00340 | XP_018221159.1 | 99 | 0 |
| SPGP0I01100 | XP_018223224.1 | 99 | 0 |
| SPGP0F02800 | XP_018220088.1 | 99 | 0 |
| SPGP0C03610 | XP_018220725.1 | 99 | 0 |
| SPGP0P02280 | XP_018220922.1 | 99 | 0 |
| SPGP0M03120 | XP_018221426.1 | 99 | 0 |
| SPGP0E01770 | XP_018222317.1 | 99 | 0 |
| SPGP0H00960 | XP_018219283.1 | 99 | 0 |
| SPGP0AI00190 | XP_018224060.1 | 99 | 0 |
| SPGP0B01820 | XP_018223628.1 | 100 | 0 |
| SPGP0B03300 | XP_018223772.1 | 99 | 0 |
| SPGP0P00230 | XP_018221120.1 | 99 | 0 |
| SPGP0B01530 | XP_018223600.1 | 99 | 0 |
| SPGP0L02000 | XP_018218977.1 | 99 | 0 |
| SPGP0B01790 | XP_018223625.1 | 99 | 0 |
| SPGP0S01950 | XP_018219657.1 | 99 | 0 |
| SPGP0L01330 | XP_018219043.1 | 100 | 0 |
| SPGP0B02780 | XP_018223721.1 | 95 | 4 |
| SPGP0B02950 | XP_018223739.1 | 99 | 0 |
| SPGP0M02610 | XP_018221376.1 | 99 | 0 |
| SPGP0S00880 | XP_018219764.1 | 99 | 0 |
| SPGP0F00110 | XP_018220348.1 | 99 | 0 |
| SPGP0H01380 | XP_018219323.1 | 99 | 0 |
| SPGP0P01550 | XP_018220993.1 | 99 | 0 |
| SPGP0M00690 | XP_018221193.1 | 99 | 0 |
| SPGP0F03740 | XP_018219996.1 | 99 | 0 |
| SPGP0E00260 | XP_018222471.1 | 99 | 0 |
| SPGP0AI00430 | XP_018224083.1 | 99 | 0 |
| SPGP0B02250 | XP_018223670.1 | 99 | 0 |
| SPGP0E01130 | XP_018222383.1 | 99 | 0 |
| SPGP0S00790 | XP_018219772.1 | 99 | 0 |
| SPGP0L01940 | XP_018218983.1 | 99 | 0 |

Create PDF in your applications with the Pdfcrowd HTML to PDF API PDFCROWD

| | | | |
|---|---|---|---|
| SPGP0H02580 | XP_018219440.1 | 99 | 0 |
| SPGP0H00380 | XP_018219226.1 | 99 | 0 |
| SPGP0AI00730 | XP_018224113.1 | 99 | 0 |
| SPGP0I01510 | XP_018223184.1 | 99 | 0 |
| SPGP0AQ00330 | XP_018221971.1 | 99 | 0 |
| SPGP0B02550 | XP_018223699.1 | 99 | 0 |
| SPGP0I00180 | XP_018223316.1 | 99 | 0 |
| SPGP0W01240 | XP_018221569.1 | 99 | 0 |
| SPGP0I01840 | XP_018223151.1 | 99 | 0 |
| SPGP0AG00660 | XP_018223380.1 | 99 | 0 |
| SPGP0E00380 | XP_018222459.1 | 99 | 0 |
| SPGP0I02760 | XP_018223062.1 | 99 | 0 |
| SPGP0M01760 | XP_018221295.1 | 99 | 0 |
| SPGP0Q00790 | XP_018222783.1 | 99 | 0 |
| SPGP0C04450 | XP_018220807.1 | 99 | 0 |
| SPGP0F02640 | XP_018220104.1 | 99 | 0 |
| SPGP0O00350 | XP_018221668.1 | 99 | 0 |
| SPGP0B04170 | XP_018223858.1 | 100 | 0 |
| SPGP0L01090 | XP_018219066.1 | 99 | 0 |
| SPGP0M01560 | XP_018221275.1 | 99 | 0 |
| SPGP0Q02330 | XP_018222632.1 | 99 | 0 |
| SPGP0W01040 | XP_018221551.1 | 99 | 0 |
| SPGP0L02910 | XP_018218890.1 | 99 | 0 |
| SPGP0I03240 | XP_018223020.1 | 99 | 0 |
| SPGP0Q00240 | XP_018222835.1 | 99 | 0 |
| SPGP0I00380 | XP_018223296.1 | 99 | 0 |
| SPGP0AF01000 | XP_018222507.1 | 99 | 0 |
| SPGP0P01830 | XP_018220966.1 | 99 | 0 |
| SPGP0AL00170 | XP_018219881.1 | 99 | 0 |
| SPGP0C02560 | XP_018220624.1 | 99 | 0 |
| SPGP0I03570 | XP_018222992.1 | 99 | 0 |
| SPGP0B00420 | XP_018223491.1 | 99 | 0 |
| SPGP0AT00300 | XP_018219179.1 | 99 | 0 |
| SPGP0M01010 | XP_018221221.1 | 99 | 0 |
| SPGP0H01230 | XP_018219308.1 | 99 | 0 |
| SPGP0C04410 | XP_018220803.1 | 99 | 0 |
| SPGP0W00580 | XP_018221507.1 | 99 | 0 |
| SPGP0H02140 | XP_018219398.1 | 99 | 0 |
| SPGP0O01400 | XP_018221767.1 | 96 | 2 |

Create PDF in your applications with the Pdfcrowd HTML to PDF API PDFCROWD

| | | | |
|---|---|---|---|
| SPGP0L02830 | XP_018218898.1 | 99 | 0 |
| SPGP0C04170 | XP_018220779.1 | 99 | 0 |
| SPGP0C00840 | XP_018220452.1 | 99 | 0 |
| SPGP0M00780 | XP_018221201.1 | 99 | 0 |
| SPGP0B05200 | XP_018223953.1 | 99 | 0 |
| SPGP0E02520 | XP_018222244.1 | 99 | 0 |
| SPGP0C04420 | XP_018220804.1 | 99 | 0 |
| SPGP0I03410 | XP_018223004.1 | 99 | 0 |
| SPGP0B04940 | XP_018223929.1 | 99 | 0 |
| SPGP0L00570 | XP_018219115.1 | 99 | 0 |
| SPGP0H00340 | XP_018219222.1 | 99 | 0 |
| SPGP0B01740 | XP_018223620.1 | 99 | 0 |
| SPGP0O01180 | XP_018221745.1 | 99 | 0 |
| SPGP0Q01110 | XP_018222751.1 | 99 | 0 |
| SPGP0H02910 | XP_018219472.1 | 99 | 0 |
| SPGP0B05940 | XP_018224024.1 | 99 | 0 |
| SPGP0M00950 | XP_018221215.1 | 99 | 0 |
| SPGP0AG00880 | XP_018223399.1 | 99 | 0 |
| SPGP0L01880 | XP_018218989.1 | 99 | 0 |
| SPGP0F00100 | XP_018220349.1 | 99 | 0 |
| SPGP0L01630 | XP_018219013.1 | 99 | 0 |
| SPGP0I00500 | XP_018223284.1 | 99 | 0 |
| SPGP0B04840 | XP_018223920.1 | 99 | 0 |
| SPGP0F03620 | XP_018220008.1 | 100 | 0 |
| SPGP0E03570 | XP_018222142.1 | 97 | 0 |
| SPGP0F03700 | XP_018220000.1 | 99 | 0 |
| SPGP0M03460 | XP_018221455.1 | 99 | 0 |
| SPGP0M02860 | XP_018221401.1 | 99 | 0 |
| SPGP0L00790 | XP_018219094.1 | 99 | 0 |
| SPGP0E03320 | XP_018222165.1 | 99 | 0 |
| SPGP0O02980 | XP_018221919.1 | 99 | 0 |
| SPGP0S01380 | XP_018219714.1 | 99 | 0 |
| SPGP0F02280 | XP_018220138.1 | 99 | 0 |
| SPGP0C04800 | XP_018220839.1 | 99 | 0 |
| SPGP0B00200 | XP_018223470.1 | 99 | 0 |
| SPGP0C01060 | XP_018220474.1 | 100 | 0 |
| SPGP0P00870 | XP_018221057.1 | 99 | 0 |
| SPGP0E03170 | XP_018222179.1 | 98 | 1 |
| SPGP0C03570 | XP_018220721.1 | 100 | 0 |
| SPGP0L01820 | XP_018218995.1 | 99 | 0 |

Create PDF in your applications with the Pdfcrowd HTML to PDF API PDFCROWD

| | | | |
|---|---|---|---|
| SPGP0AI00850 | XP_018224122.1 | 99 | 0 |
| SPGP0W01000 | XP_018221547.1 | 99 | 0 |
| SPGP0O03000 | XP_018221920.1 | 99 | 0 |
| SPGP0H03850 | XP_018219564.1 | 99 | 0 |
| SPGP0L00580 | XP_018219114.1 | 99 | 0 |
| SPGP0P02580 | XP_018220893.1 | 100 | 0 |
| SPGP0E00500 | XP_018222447.1 | 99 | 0 |
| SPGP0E03510 | XP_018222147.1 | 98 | 0 |
| SPGP0E03870 | XP_018222112.1 | 99 | 0 |
| SPGP0S01160 | XP_018219736.1 | 99 | 0 |
| SPGP0E03700 | XP_018222129.1 | 99 | 0 |
| SPGP0AG00890 | XP_018223400.1 | 99 | 0 |
| SPGP0E02470 | XP_018222248.1 | 99 | 0 |
| SPGP0H03240 | XP_018219504.1 | 99 | 0 |
| SPGP0Q02460 | XP_018222619.1 | 99 | 0 |
| SPGP0B03510 | XP_018223792.1 | 99 | 0 |
| SPGP0W00590 | XP_018221508.1 | 99 | 0 |
| SPGP0B05710 | XP_018224001.1 | 99 | 0 |
| SPGP0B00570 | XP_018223504.1 | 99 | 0 |
| SPGP0S02120 | XP_018219640.1 | 99 | 0 |
| SPGP0L02630 | XP_018218918.1 | 99 | 0 |
| SPGP0C04360 | XP_018220798.1 | 99 | 0 |
| SPGP0C02060 | XP_018220574.1 | 99 | 0 |
| SPGP0H01970 | XP_018219382.1 | 99 | 0 |
| SPGP0L03190 | XP_018218863.1 | 99 | 0 |
| SPGP0B06140 | XP_018224042.1 | 98 | 1 |
| SPGP0I02890 | XP_018223051.1 | 99 | 0 |
| SPGP0H02570 | XP_018219439.1 | 100 | 0 |
| SPGP0B03340 | XP_018223776.1 | 99 | 0 |
| SPGP0B02870 | XP_018223731.1 | 99 | 0 |
| SPGP0H01620 | XP_018219347.1 | 99 | 0 |
| SPGP0AL00610 | XP_018219922.1 | 99 | 0 |
| SPGP0F02170 | XP_018220149.1 | 99 | 0 |
| SPGP0B03670 | XP_018223808.1 | 99 | 0 |
| SPGP0L03650 | XP_018218818.1 | 99 | 0 |
| SPGP0P00980 | XP_018221046.1 | 99 | 0 |
| SPGP0E00320 | XP_018222465.1 | 100 | 0 |
| SPGP0P01050 | XP_018221039.1 | 99 | 0 |
| SPGP0W00600 | XP_018221509.1 | 99 | 0 |
| SPGP0H00550 | XP_018219243.1 | 100 | 0 |

Create PDF in your applications with the Pdfcrowd HTML to PDF API PDFCROWD

| | | | |
|---|---|---|---|
| SPGP0L02940 | XP_018218887.1 | 99 | 0 |
| SPGP0B03080 | XP_018223751.1 | 99 | 0 |
| SPGP0S00550 | XP_018219796.1 | 100 | 0 |
| SPGP0Htrna18M | DI49_5351 | 100 | 0 |
| SPGP0S01660 | XP_018219686.1 | 100 | 0 |
| SPGP0F02370 | XP_018220129.1 | 99 | 0 |
| SPGP0AG00480 | XP_018223362.1 | 99 | 0 |
| SPGP0O02710 | XP_018221894.1 | 99 | 0 |
| SPGP0P02250 | XP_018220925.1 | 99 | 0 |
| SPGP0F03020 | XP_018220067.1 | 99 | 0 |
| SPGP0M00580 | XP_018221182.1 | 99 | 0 |
| SPGP0S00410 | XP_018219810.1 | 99 | 0 |
| SPGP0L03530 | XP_018218830.1 | 99 | 0 |
| SPGP0I02420 | XP_018223095.1 | 99 | 0 |
| SPGP0L02130 | XP_018218965.1 | 99 | 0 |
| SPGP0B05850 | XP_018224015.1 | 99 | 0 |
| SPGP0C03560 | XP_018220720.1 | 99 | 0 |
| SPGP0Htrna19M | DI49_4243 | 100 | 0 |
| SPGP0M03190 | XP_018221433.1 | 99 | 0 |
| SPGP0B04780 | XP_018223916.1 | 99 | 0 |
| SPGP0F02190 | XP_018220147.1 | 99 | 0 |
| SPGP0W00390 | XP_018221488.1 | 99 | 0 |
| SPGP0L00890 | XP_018219085.1 | 99 | 0 |
| SPGP0Q01750 | XP_018222689.1 | 99 | 0 |
| SPGP0H03170 | XP_018219498.1 | 99 | 0 |
| SPGP0M00570 | XP_018221181.1 | 98 | 0 |
| SPGP0P00840 | XP_018221060.1 | 100 | 0 |
| SPGP0B04320 | XP_018223873.1 | 99 | 0 |
| SPGP0S02190 | XP_018219633.1 | 99 | 0 |
| SPGP0AF00830 | XP_018222522.1 | 99 | 0 |
| SPGP0I02390 | XP_018223098.1 | 99 | 0 |
| SPGP0B03630 | XP_018223804.1 | 99 | 0 |
| SPGP0H02210 | XP_018219404.1 | 99 | 0 |
| SPGP0I03630 | XP_018222985.1 | 99 | 0 |
| SPGP0C02670 | XP_018220635.1 | 99 | 0 |
| SPGP0H00730 | XP_018219260.1 | 99 | 0 |
| SPGP0M01950 | XP_018221313.1 | 99 | 0 |
| SPGP0H03260 | XP_018219506.1 | 99 | 0 |
| SPGP0I00100 | XP_018223460.1 | 90 | 0 |
| SPGP0C01150 | XP_018220483.1 | 99 | 0 |

Create PDF in your applications with the Pdfcrowd HTML to PDF API PDFCROWD

| | | | |
|---|---|---|---|
| SPGP0B05160 | XP_018223949.1 | 100 | 0 |
| SPGP0F00200 | XP_018220339.1 | 99 | 0 |
| SPGP0AQ00190 | XP_018221957.1 | 99 | 0 |
| SPGP0AG00440 | XP_018223358.1 | 99 | 0 |
| SPGP0C03800 | XP_018220743.1 | 99 | 0 |
| SPGP0E04150 | XP_018222084.1 | 99 | 0 |
| SPGP0P02460 | XP_018220904.1 | 100 | 0 |
| SPGP0L02840 | XP_018218897.1 | 99 | 0 |
| SPGP0O02670 | XP_018221890.1 | 99 | 0 |
| SPGP0H03640 | XP_018219543.1 | 99 | 0 |
| SPGP0F02450 | XP_018220121.1 | 99 | 0 |
| SPGP0F02310 | XP_018220135.1 | 99 | 0 |
| SPGP0O00720 | XP_018221704.1 | 99 | 0 |
| SPGP0Q00610 | XP_018222800.1 | 100 | 0 |
| SPGP0H02540 | XP_018219436.1 | 99 | 0 |
| SPGP0O00900 | XP_018221720.1 | 99 | 0 |
| SPGP0I03670 | XP_018222981.1 | 99 | 0 |
| SPGP0AR00470 | XP_018219605.1 | 99 | 0 |
| SPGP0C01520 | XP_018220521.1 | 99 | 0 |
| SPGP0E00690 | XP_018222427.1 | 99 | 0 |
| SPGP0B05550 | XP_018223985.1 | 99 | 0 |
| SPGP0AI00340 | XP_018224075.1 | 99 | 0 |
| SPGP0O00170 | XP_018221650.1 | 99 | 0 |
| SPGP0C02390 | XP_018220607.1 | 99 | 0 |
| SPGP0Q00950 | XP_018222767.1 | 99 | 0 |
| SPGP0L03010 | XP_018218880.1 | 99 | 0 |
| SPGP0AI00760 | XP_018224116.1 | 99 | 0 |
| SPGP0F01640 | XP_018220198.1 | 99 | 0 |
| SPGP0E02300 | XP_018222265.1 | 99 | 0 |
| SPGP0O01210 | XP_018221748.1 | 100 | 0 |
| SPGP0C04560 | XP_018220817.1 | 99 | 0 |
| SPGP0E01340 | XP_018222361.1 | 99 | 0 |
| SPGP0E02240 | XP_018222271.1 | 99 | 0 |
| SPGP0E01610 | XP_018222333.1 | 99 | 0 |
| SPGP0M02330 | XP_018221349.1 | 99 | 0 |
| SPGP0L03480 | XP_018218835.1 | 99 | 0 |
| SPGP0B01980 | XP_018223644.1 | 99 | 0 |
| SPGP0B01070 | XP_018223554.1 | 99 | 0 |
| SPGP0AR00120 | XP_018219571.1 | 99 | 0 |
| SPGP0Q02470 | XP_018222618.1 | 99 | 0 |

Create PDF in your applications with the Pdfcrowd HTML to PDF API PDFCROWD

| | | | |
|---|---|---|---|
| SPGP0B02130 | XP_018223659.1 | 99 | 0 |
| SPGP0C01920 | XP_018220560.1 | 99 | 0 |
| SPGP0L01300 | XP_018219046.1 | 99 | 0 |
| SPGP0O01670 | XP_018221794.1 | 95 | 5 |
| SPGP0L00970 | XP_018219078.1 | 99 | 0 |
| SPGP0O01890 | XP_018221814.1 | 99 | 1 |
| SPGP0F01420 | XP_018220219.1 | 99 | 0 |
| SPGP0P00450 | XP_018221098.1 | 99 | 0 |
| SPGP0E00270 | XP_018222470.1 | 99 | 0 |
| SPGP0E02150 | XP_018222279.1 | 99 | 0 |
| SPGP0S01190 | XP_018219733.1 | 99 | 0 |
| SPGP0F04210 | XP_018219950.1 | 99 | 0 |
| SPGP0H01020 | XP_018219289.1 | 99 | 0 |
| SPGP0L01920 | XP_018218985.1 | 99 | 0 |
| SPGP0O02100 | XP_018221834.1 | 100 | 0 |
| SPGP0H02090 | XP_018219394.1 | 99 | 0 |
| SPGP0E01570 | XP_018222337.1 | 99 | 0 |
| SPGP0B00690 | XP_018223515.1 | 99 | 0 |
| SPGP0E01970 | XP_018222297.1 | 99 | 0 |
| SPGP0W01160 | XP_018221561.1 | 99 | 0 |
| SPGP0P01650 | XP_018220984.1 | 99 | 0 |
| SPGP0O02610 | XP_018221884.1 | 99 | 0 |
| SPGP0H03680 | XP_018219547.1 | 99 | 0 |
| SPGP0Strna18F | DI49_4603 | 99 | 0 |
| SPGP0H03410 | XP_018219521.1 | 99 | 0 |
| SPGP0P00540 | XP_018221090.1 | 100 | 0 |
| SPGP0H03140 | XP_018219495.1 | 99 | 0 |
| SPGP0M02320 | XP_018221348.1 | 99 | 0 |
| SPGP0E00870 | XP_018222410.1 | 99 | 0 |
| SPGP0I00910 | XP_018223243.1 | 99 | 0 |
| SPGP0Itrna5C | DI49_5400 | 100 | 0 |
| SPGP0S01590 | XP_018219693.1 | 99 | 0 |
| SPGP0I02950 | XP_018223045.1 | 99 | 0 |
| SPGP0B04200 | XP_018223861.1 | 99 | 0 |
| SPGP0AL00570 | XP_018219919.1 | 99 | 0 |
| SPGP0E02860 | XP_018222209.1 | 98 | 0 |
| SPGP0E01690 | XP_018222325.1 | 99 | 0 |
| SPGP0I02780 | XP_018223060.1 | 99 | 0 |
| SPGP0W01650 | XP_018221609.1 | 99 | 0 |
| SPGP0W00920 | XP_018221539.1 | 99 | 0 |

Create PDF in your applications with the Pdfcrowd HTML to PDF API PDFCROWD

| | | | |
|---|---|---|---|
| SPGP0F02970 | XP_018220072.1 | 99 | 0 |
| SPGP0W00290 | XP_018221478.1 | 99 | 0 |
| SPGP0C02380 | XP_018220606.1 | 99 | 0 |
| SPGP0P02930 | XP_018220858.1 | 99 | 0 |
| SPGP0BB00120 | XP_018219846.1 | 99 | 0 |
| SPGP0P02240 | XP_018220926.1 | 99 | 0 |
| SPGP0Q01920 | XP_018222671.1 | 99 | 0 |
| SPGP0AQ00380 | XP_018221976.1 | 100 | 0 |
| SPGP0E03940 | XP_018222105.1 | 99 | 0 |
| SPGP0C04200 | XP_018220782.1 | 99 | 0 |
| SPGP0S01620 | XP_018219690.1 | 98 | 2 |
| SPGP0E03110 | XP_018222185.1 | 99 | 0 |
| SPGP0C03380 | XP_018220704.1 | 99 | 0 |
| SPGP0W01740 | XP_018221618.1 | 99 | 0 |
| SPGP0C00400 | XP_018220408.1 | 96 | 0 |
| SPGP0B04100 | XP_018223851.1 | 99 | 0 |
| SPGP0B02690 | XP_018223712.1 | 99 | 0 |
| SPGP0I02270 | XP_018223110.1 | 99 | 0 |
| SPGP0S01840 | XP_018219668.1 | 100 | 0 |
| SPGP0H03530 | XP_018219531.1 | 99 | 0 |
| SPGP0S00980 | XP_018219754.1 | 99 | 0 |
| SPGP0F02550 | XP_018220111.1 | 99 | 0 |
| SPGP0I01650 | XP_018223170.1 | 99 | 0 |
| SPGP0M02310 | XP_018221347.1 | 99 | 0 |
| SPGP0E01270 | XP_018222369.1 | 99 | 0 |
| SPGP0W00430 | XP_018221492.1 | 99 | 0 |
| SPGP0Ttrna31L | DI49_3110 | 100 | 0 |
| SPGP0C03460 | XP_018220712.1 | 99 | 0 |
| SPGP0C01890 | XP_018220557.1 | 99 | 1 |
| SPGP0I01970 | XP_018223137.1 | 99 | 0 |
| SPGP0AF00600 | XP_018222544.1 | 96 | 0 |
| SPGP0AG00320 | XP_018223347.1 | 99 | 0 |
| SPGP0F03100 | XP_018220059.1 | 99 | 0 |
| SPGP0M00360 | XP_018221161.1 | 99 | 0 |
| SPGP0O01040 | XP_018221734.1 | 99 | 0 |
| SPGP0C03240 | XP_018220691.1 | 99 | 0 |
| SPGP0W01520 | XP_018221596.1 | 100 | 0 |
| SPGP0I00800 | XP_018223254.1 | 99 | 0 |
| SPGP0C03030 | XP_018220671.1 | 100 | 0 |
| SPGP0W01270 | XP_018221572.1 | 99 | 0 |

Create PDF in your applications with the Pdfcrowd HTML to PDF API PDFCROWD

| | | | |
|---|---|---|---|
| SPGP0P02920 | XP_018220859.1 | 99 | 0 |
| SPGP0I03790 | XP_018222969.1 | 99 | 0 |
| SPGP0H00810 | XP_018219268.1 | 99 | 0 |
| SPGP0S01560 | XP_018219696.1 | 99 | 0 |
| SPGP0E03800 | XP_018222119.1 | 98 | 0 |
| SPGP0C00430 | XP_018220412.1 | 99 | 0 |
| SPGP0M01890 | XP_018221308.1 | 99 | 0 |
| SPGP0L01620 | XP_018219014.1 | 99 | 0 |
| SPGP0C03130 | XP_018220681.1 | 100 | 0 |
| SPGP0L01910 | XP_018218986.1 | 99 | 0 |
| SPGP0S01970 | XP_018219655.1 | 99 | 1 |
| SPGP0B03280 | XP_018223770.1 | 99 | 0 |
| SPGP0I02830 | XP_018223056.1 | 99 | 0 |
| SPGP0H03840 | XP_018219563.1 | 99 | 0 |
| SPGP0M02230 | XP_018221339.1 | 99 | 0 |
| SPGP0H00370 | XP_018219225.1 | 99 | 0 |
| SPGP0AL00220 | XP_018219886.1 | 99 | 0 |
| SPGP0S01020 | XP_018219750.1 | 100 | 0 |
| SPGP0S01650 | XP_018219687.1 | 99 | 0 |
| SPGP0F02720 | XP_018220096.1 | 99 | 0 |
| SPGP0I02090 | XP_018223127.1 | 99 | 0 |
| SPGP0AL00450 | XP_018219907.1 | 100 | 0 |
| SPGP0AI00710 | XP_018224110.1 | 99 | 0 |
| SPGP0I03170 | XP_018223027.1 | 99 | 0 |
| SPGP0C02220 | XP_018220590.1 | 99 | 0 |
| SPGP0M01090 | XP_018221229.1 | 99 | 0 |
| SPGP0B00340 | XP_018223484.1 | 99 | 0 |
| SPGP0B04060 | XP_018223847.1 | 99 | 0 |
| SPGP0P02850 | XP_018220865.1 | 99 | 0 |
| SPGP0C01610 | XP_018220529.1 | 99 | 0 |
| SPGP0S00670 | XP_018219784.1 | 99 | 0 |
| SPGP0M00140 | XP_018221141.1 | 99 | 0 |
| SPGP0C00690 | XP_018220436.1 | 99 | 0 |
| SPGP0E00590 | XP_018222438.1 | 99 | 0 |
| SPGP0P02710 | XP_018220879.1 | 99 | 0 |
| SPGP0W01150 | XP_018221560.1 | 99 | 0 |
| SPGP0M01310 | XP_018221250.1 | 99 | 0 |
| SPGP0B03580 | XP_018223799.1 | 99 | 0 |
| SPGP0W01860 | XP_018221630.1 | 99 | 0 |
| SPGP0S02060 | XP_018219646.1 | 99 | 0 |

Create PDF in your applications with the Pdfcrowd HTML to PDF API PDFCROWD

| | | | |
|---|---|---|---|
| SPGP0C02540 | XP_018220622.1 | 99 | 0 |
| SPGP0BB00190 | XP_018219853.1 | 99 | 0 |
| SPGP0H03230 | XP_018219503.1 | 99 | 0 |
| SPGP0P01850 | XP_018220964.1 | 99 | 0 |
| SPGP0Ftrna20P | DI49_4051 | 99 | 0 |
| SPGP0B04190 | XP_018223860.1 | 99 | 0 |
| SPGP0O01940 | XP_018221819.1 | 99 | 0 |
| SPGP0AT00270 | XP_018219182.1 | 98 | 0 |
| SPGP0M01540 | XP_018221273.1 | 99 | 0 |
| SPGP0F03830 | XP_018219988.1 | 98 | 0 |
| SPGP0O00300 | XP_018221663.1 | 99 | 0 |
| SPGP0B01110 | XP_018223558.1 | 99 | 0 |
| SPGP0Q02310 | XP_018222634.1 | 99 | 0 |
| SPGP0S01820 | XP_018219670.1 | 99 | 0 |
| SPGP0AF00790 | XP_018221597.1 | 99 | 0 |
| SPGP0Q02630 | XP_018222602.1 | 99 | 0 |
| SPGP0Q01420 | XP_018222722.1 | 99 | 0 |
| SPGP0H03010 | XP_018219482.1 | 99 | 0 |
| SPGP0L02210 | XP_018218957.1 | 99 | 0 |
| SPGP0Q00910 | XP_018222771.1 | 99 | 0 |
| SPGP0O00570 | XP_018221689.1 | 99 | 0 |
| SPGP0H03090 | XP_018219490.1 | 99 | 0 |
| SPGP0I03590 | XP_018222989.1 | 99 | 0 |
| SPGP0O00400 | XP_018221673.1 | 99 | 0 |
| SPGP0C01410 | XP_018220509.1 | 96 | 1 |
| SPGP0C02600 | XP_018220628.1 | 99 | 0 |
| SPGP0P00860 | XP_018221058.1 | 99 | 0 |
| SPGP0I01730 | XP_018223162.1 | 100 | 0 |
| SPGP0L00310 | XP_018219141.1 | 99 | 0 |
| SPGP0E00470 | XP_018222450.1 | 99 | 0 |
| SPGP0DN00100 | XP_018220411.1 | 100 | 0 |
| SPGP0AL00700 | XP_018219931.1 | 90 | 0 |
| SPGP0B03180 | XP_018223761.1 | 99 | 0 |
| SPGP0O01460 | XP_018221773.1 | 99 | 0 |
| SPGP0H03520 | XP_018219530.1 | 99 | 0 |
| SPGP0S00360 | XP_018219815.1 | 99 | 0 |
| SPGP0F03200 | XP_018220049.1 | 99 | 0 |
| SPGP0C01170 | XP_018220485.1 | 99 | 0 |
| SPGP0O02860 | XP_018221908.1 | 99 | 0 |
| SPGP0AF00980 | XP_018222509.1 | 99 | 0 |

Create PDF in your applications with the Pdfcrowd HTML to PDF API PDFCROWD

| | | | |
|---|---|---|---|
| SPGP0B01370 | XP_018223584.1 | 99 | 0 |
| SPGP0Q00990 | XP_018222763.1 | 99 | 0 |
| SPGP0L00550 | XP_018219117.1 | 99 | 0 |
| SPGP0O03080 | XP_018221928.1 | 99 | 0 |
| SPGP0C01080 | XP_018220476.1 | 99 | 0 |
| SPGP0B05960 | XP_018224026.1 | 99 | 0 |
| SPGP0Q01170 | XP_018222745.1 | 99 | 0 |
| SPGP0C03630 | XP_018220727.1 | 99 | 0 |
| SPGP0L02760 | XP_018218905.1 | 99 | 0 |
| SPGP0F00120 | XP_018220347.1 | 99 | 0 |
| SPGP0H02940 | XP_018219475.1 | 99 | 0 |
| SPGP0P02470 | XP_018220903.1 | 99 | 1 |
| SPGP0O01280 | XP_018221755.1 | 99 | 0 |
| SPGP0E02980 | XP_018222197.1 | 98 | 0 |
| SPGP0H00270 | XP_018219215.1 | 99 | 0 |
| SPGP0L03300 | XP_018218852.1 | 99 | 0 |
| SPGP0P02200 | XP_018220930.1 | 99 | 0 |
| SPGP0M00630 | XP_018221187.1 | 99 | 0 |
| SPGP0AG00200 | XP_018223335.1 | 99 | 0 |
| SPGP0P02030 | XP_018220947.1 | 99 | 0 |
| SPGP0H03080 | XP_018219489.1 | 99 | 0 |
| SPGP0Q01850 | XP_018222678.1 | 99 | 0 |
| SPGP0S01360 | XP_018219716.1 | 99 | 0 |
| SPGP0O03120 | XP_018221932.1 | 99 | 0 |
| SPGP0L02290 | XP_018218949.1 | 99 | 0 |
| SPGP0B00220 | XP_018223472.1 | 99 | 0 |
| SPGP0F04090 | XP_018219962.1 | 99 | 0 |
| SPGP0F01780 | XP_018220187.1 | 99 | 0 |
| SPGP0S02040 | XP_018219648.1 | 99 | 0 |
| SPGP0P00890 | XP_018221055.1 | 99 | 0 |
| SPGP0W00340 | XP_018221483.1 | 99 | 0 |
| SPGP0C03550 | XP_018220719.1 | 99 | 0 |
| SPGP0I01590 | XP_018223176.1 | 99 | 0 |
| SPGP0E02180 | XP_018222277.1 | 99 | 0 |
| SPGP0B01700 | XP_018223616.1 | 99 | 0 |
| SPGP0C04520 | XP_018220813.1 | 99 | 0 |
| SPGP0P02270 | XP_018220923.1 | 99 | 0 |
| SPGP0AF00650 | XP_018222540.1 | 99 | 0 |
| SPGP0O03020 | XP_018221922.1 | 99 | 0 |
| SPGP0Htrna22P | DI49_4914 | 100 | 0 |

Create PDF in your applications with the Pdfcrowd HTML to PDF API PDFCROWD

| | | | |
|---|---|---|---|
| SPGP0H02870 | XP_018219468.1 | 95 | 4 |
| SPGP0B03640 | XP_018223805.1 | 99 | 0 |
| SPGP0S01100 | XP_018219742.1 | 100 | 0 |
| SPGP0AQ00510 | XP_018221989.1 | 99 | 0 |
| SPGP0I03810 | XP_018222967.1 | 100 | 0 |
| SPGP0I02510 | XP_018223086.1 | 99 | 0 |
| SPGP0B01860 | XP_018223632.1 | 99 | 0 |
| SPGP0C00640 | XP_018220432.1 | 99 | 0 |
| SPGP0Q00420 | XP_018222819.1 | 99 | 0 |
| SPGP0C01390 | XP_018220507.1 | 99 | 0 |
| SPGP0Q01250 | XP_018222737.1 | 99 | 0 |
| SPGP0P01210 | XP_018221025.1 | 99 | 0 |
| SPGP0H02280 | XP_018219411.1 | 100 | 0 |
| SPGP0E02450 | XP_018222250.1 | 99 | 0 |
| SPGP0E02320 | XP_018222263.1 | 99 | 0 |
| SPGP0H01920 | XP_018219377.1 | 99 | 0 |
| SPGP0H00120 | XP_018219201.1 | 97 | 0 |
| SPGP0F01590 | XP_018220203.1 | 99 | 1 |
| SPGP0F00340 | XP_018220325.1 | 99 | 0 |
| SPGP0M02150 | XP_018221331.1 | 99 | 0 |
| SPGP0H03880 | XP_018219567.1 | 99 | 0 |
| SPGP0O02480 | XP_018221872.1 | 99 | 0 |
| SPGP0P01060 | XP_018221038.1 | 99 | 0 |
| SPGP0B06120 | XP_018224040.1 | 99 | 0 |
| SPGP0C01180 | XP_018220486.1 | 99 | 1 |
| SPGP0F00770 | XP_018220283.1 | 99 | 0 |
| SPGP0S00750 | XP_018219776.1 | 99 | 0 |
| SPGP0H00740 | XP_018219261.1 | 99 | 0 |
| SPGP0I02740 | XP_018223064.1 | 99 | 0 |
| SPGP0C00550 | XP_018220423.1 | 99 | 0 |
| SPGP0O00690 | XP_018221701.1 | 99 | 0 |
| SPGP0AL00630 | XP_018219924.1 | 99 | 0 |
| SPGP0S01830 | XP_018219669.1 | 99 | 0 |
| SPGP0I01200 | XP_018223214.1 | 99 | 0 |
| SPGP0L02280 | XP_018218950.1 | 99 | 0 |
| SPGP0P01070 | XP_018221037.1 | 99 | 0 |
| SPGP0AG00130 | XP_018223329.1 | 99 | 0 |
| SPGP0H00540 | XP_018219242.1 | 99 | 0 |
| SPGP0S02100 | XP_018219642.1 | 99 | 0 |
| SPGP0I03830 | XP_018222965.1 | 99 | 0 |

Create PDF in your applications with the Pdfcrowd HTML to PDF API PDFCROWD

| | | | |
|---|---|---|---|
| SPGP0H01480 | XP_018219333.1 | 99 | 0 |
| SPGP0I01290 | XP_018223205.1 | 99 | 0 |
| SPGP0O02730 | XP_018221896.1 | 99 | 0 |
| SPGP0L03560 | XP_018218827.1 | 99 | 0 |
| SPGP0AG00970 | XP_018223407.1 | 99 | 0 |
| SPGP0Q01620 | XP_018222702.1 | 99 | 0 |
| SPGP0W01100 | XP_018221556.1 | 99 | 0 |
| SPGP0W01260 | XP_018221571.1 | 99 | 0 |
| SPGP0AI00130 | XP_018224054.1 | 99 | 0 |
| SPGP0M02810 | XP_018221396.1 | 100 | 0 |
| SPGP0AI00520 | XP_018224092.1 | 99 | 0 |
| SPGP0H00530 | XP_018219241.1 | 99 | 0 |
| SPGP0L02960 | XP_018218885.1 | 99 | 0 |
| SPGP0B01470 | XP_018223594.1 | 99 | 0 |
| SPGP0P00970 | XP_018221047.1 | 99 | 0 |
| SPGP0O02750 | XP_018221898.1 | 99 | 1 |
| SPGP0I00710 | XP_018223262.1 | 99 | 0 |
| SPGP0Q01800 | XP_018222683.1 | 99 | 0 |
| SPGP0I00570 | XP_018223277.1 | 99 | 0 |
| SPGP0P00300 | XP_018221113.1 | 99 | 0 |
| SPGP0O01860 | XP_018221811.1 | 99 | 0 |
| SPGP0AF00350 | XP_018222569.1 | 98 | 2 |
| SPGP0L01570 | XP_018219019.1 | 99 | 0 |
| SPGP0F00420 | XP_018220318.1 | 99 | 0 |
| SPGP0H02820 | XP_018219463.1 | 99 | 0 |
| SPGP0M01780 | XP_018221297.1 | 99 | 0 |
| SPGP0I02230 | XP_018223114.1 | 99 | 0 |
| SPGP0F01080 | XP_018220252.1 | 99 | 0 |
| SPGP0H00280 | XP_018219216.1 | 99 | 0 |
| SPGP0C02880 | XP_018220656.1 | 99 | 0 |
| SPGP0M01690 | XP_018221287.1 | 99 | 0 |
| SPGP0L00850 | XP_018219088.1 | 99 | 0 |
| SPGP0P01630 | XP_018220986.1 | 99 | 0 |
| SPGP0B05100 | XP_018223943.1 | 99 | 0 |
| SPGP0AF00810 | XP_018222524.1 | 99 | 0 |
| SPGP0E00670 | XP_018222430.1 | 99 | 0 |
| SPGP0Q01340 | XP_018222729.1 | 99 | 0 |
| SPGP0C03820 | XP_018220745.1 | 99 | 0 |
| SPGP0L02500 | XP_018218928.1 | 99 | 0 |

Create PDF in your applications with the Pdfcrowd HTML to PDF API PDFCROWD

| | | | |
|---|---|---|---|
| SPGP0Q01020 | XP_018222761.1 | 99 | 0 |
| SPGP0O00610 | XP_018221693.1 | 100 | 0 |
| SPGP0O00700 | XP_018221702.1 | 99 | 0 |
| SPGP0S00870 | XP_018219765.1 | 99 | 0 |
| SPGP0B03200 | XP_018223763.1 | 99 | 0 |
| SPGP0H01120 | XP_018219300.1 | 99 | 1 |
| SPGP0C01490 | XP_018220518.1 | 99 | 0 |
| SPGP0AX00100 | XP_018219859.1 | 99 | 1 |
| SPGP0AQ00130 | XP_018221952.1 | 100 | 0 |
| SPGP0B02730 | XP_018223716.1 | 99 | 0 |
| SPGP0H01750 | XP_018219360.1 | 99 | 0 |
| SPGP0O01480 | XP_018221775.1 | 99 | 0 |
| SPGP0AR00450 | XP_018219603.1 | 99 | 0 |
| SPGP0C01540 | XP_018220523.1 | 99 | 0 |
| SPGP0C04770 | XP_018220836.1 | 99 | 0 |
| SPGP0H02330 | XP_018219416.1 | 99 | 0 |
| SPGP0L01540 | XP_018219022.1 | 99 | 0 |
| SPGP0B05570 | XP_018223987.1 | 99 | 0 |
| SPGP0L00260 | XP_018219146.1 | 99 | 0 |
| SPGP0M03420 | XP_018221453.1 | 99 | 0 |
| SPGP0O01070 | XP_018221736.1 | 99 | 0 |
| SPGP0M02780 | XP_018221393.1 | 99 | 0 |
| SPGP0W00470 | XP_018221496.1 | 99 | 0 |
| SPGP0I02080 | XP_018223128.1 | 99 | 0 |
| SPGP0F01070 | XP_018220253.1 | 99 | 0 |
| SPGP0AR00360 | XP_018219595.1 | 99 | 0 |
| SPGP0E01360 | XP_018222359.1 | 99 | 0 |
| SPGP0S01570 | XP_018219695.1 | 99 | 0 |
| SPGP0P02730 | XP_018220877.1 | 99 | 0 |
| SPGP0W00280 | XP_018221477.1 | 99 | 0 |
| SPGP0E01190 | XP_018222377.1 | 99 | 0 |
| SPGP0Q02450 | XP_018222620.1 | 99 | 0 |
| SPGP0Q02100 | XP_018222655.1 | 99 | 0 |
| SPGP0C02290 | XP_018220597.1 | 99 | 0 |
| SPGP0L01890 | XP_018218988.1 | 99 | 0 |
| SPGP0I00240 | XP_018223310.1 | 99 | 0 |
| SPGP0P03110 | XP_018220842.1 | 96 | 0 |
| SPGP0C01740 | XP_018220542.1 | 99 | 0 |
| SPGP0F00810 | XP_018220279.1 | 99 | 0 |
| SPGP0AR00460 | XP_018219604.1 | 99 | 0 |

Create PDF in your applications with the Pdfcrowd HTML to PDF API PDFCROWD

| | | | |
|---|---|---|---|
| SPGP0C02840 | XP_018220652.1 | 99 | 0 |
| SPGP0E01410 | XP_018222353.1 | 99 | 0 |
| SPGP0BB00110 | XP_018219845.1 | 99 | 0 |
| SPGP0B04370 | XP_018223878.1 | 99 | 0 |
| SPGP0M01940 | XP_018221312.1 | 99 | 0 |
| SPGP0F00680 | XP_018220292.1 | 97 | 2 |
| SPGP0Q00290 | XP_018222831.1 | 99 | 0 |
| SPGP0H02410 | XP_018219424.1 | 99 | 0 |
| SPGP0AF00130 | XP_018222589.1 | 99 | 0 |
| SPGP0E04120 | XP_018222087.1 | 99 | 0 |
| SPGP0S00530 | XP_018219798.1 | 99 | 0 |
| SPGP0P01680 | XP_018220981.1 | 99 | 0 |
| SPGP0S00190 | XP_018219831.1 | 99 | 0 |
| SPGP0F02840 | XP_018220084.1 | 100 | 0 |
| SPGP0B00540 | XP_018223502.1 | 99 | 0 |
| SPGP0E01290 | XP_018222366.1 | 99 | 0 |
| SPGP0C02010 | XP_018220569.1 | 99 | 0 |
| SPGP0B04220 | XP_018223863.1 | 100 | 0 |
| SPGP0P02480 | XP_018220902.1 | 99 | 0 |
| SPGP0I00390 | XP_018223295.1 | 99 | 0 |
| SPGP0O02030 | XP_018221827.1 | 99 | 0 |
| SPGP0L03020 | XP_018218879.1 | 99 | 0 |
| SPGP0C00700 | XP_018220437.1 | 99 | 0 |
| SPGP0I02970 | XP_018223043.1 | 99 | 0 |
| SPGP0P00930 | XP_018221051.1 | 99 | 0 |
| SPGP0O00110 | XP_018221644.1 | 95 | 0 |
| SPGP0E02840 | XP_018222211.1 | 98 | 0 |
| SPGP0DM00100 | XP_018222488.1 | 88 | 0 |
| SPGP0Q01710 | XP_018222693.1 | 99 | 0 |
| SPGP0I03620 | XP_018222986.1 | 99 | 0 |
| SPGP0M02430 | XP_018221359.1 | 99 | 0 |
| SPGP0B04630 | XP_018223903.1 | 99 | 0 |
| SPGP0CStrna7L | DI49_1927 | 100 | 0 |
| SPGP0E00820 | XP_018222415.1 | 99 | 0 |
| SPGP0B00720 | XP_018223518.1 | 99 | 0 |
| SPGP0E03130 | XP_018222183.1 | 97 | 1 |
| SPGP0S01640 | XP_018219688.1 | 99 | 0 |
| SPGP0AL00760 | XP_018219936.1 | 99 | 0 |
| SPGP0E01510 | XP_018222343.1 | 99 | 0 |
| SPGP0F04190 | XP_018219952.1 | 99 | 0 |

Create PDF in your applications with the Pdfcrowd HTML to PDF API PDFCROWD

| | | | |
|---|---|---|---|
| SPGP0Q02220 | XP_018222643.1 | 99 | 0 |
| SPGP0AR00500 | XP_018219608.1 | 98 | 2 |
| SPGP0C01640 | XP_018220532.1 | 99 | 0 |
| SPGP0M03350 | XP_018221448.1 | 99 | 0 |
| SPGP0P01820 | XP_018220967.1 | 99 | 0 |
| SPGP0W00410 | XP_018221490.1 | 99 | 0 |
| SPGP0H01490 | XP_018219334.1 | 100 | 0 |
| SPGP0L01930 | XP_018218984.1 | 99 | 0 |
| SPGP0Htrna32A | DI49_3564 | 100 | 0 |
| SPGP0I02360 | XP_018223101.1 | 99 | 0 |
| SPGP0C01380 | XP_018220506.1 | 99 | 0 |
| SPGP0M02680 | XP_018221383.1 | 99 | 0 |
| SPGP0S00570 | XP_018219794.1 | 100 | 0 |
| SPGP0AF00660 | XP_018222539.1 | 99 | 0 |
| SPGP0P00960 | XP_018221048.1 | 98 | 0 |
| SPGP0H01030 | XP_018219290.1 | 99 | 0 |
| SPGP0AG00690 | XP_018223383.1 | 99 | 0 |
| SPGP0Q02120 | XP_018222653.1 | 99 | 0 |
| SPGP0E04200 | XP_018222079.1 | 99 | 0 |
| SPGP0I00820 | XP_018223252.1 | 99 | 0 |
| SPGP0B01760 | XP_018223622.1 | 99 | 0 |
| SPGP0M03160 | XP_018221430.1 | 99 | 0 |
| SPGP0B04350 | XP_018223876.1 | 99 | 0 |
| SPGP0I00430 | XP_018223291.1 | 99 | 0 |
| SPGP0L02510 | XP_018218927.1 | 99 | 0 |
| SPGP0M02850 | XP_018221400.1 | 99 | 0 |
| SPGP0AF00480 | XP_018222557.1 | 99 | 0 |
| SPGP0Q00820 | XP_018222780.1 | 99 | 0 |
| SPGP0Q01200 | XP_018222742.1 | 99 | 0 |
| SPGP0C01940 | XP_018220562.1 | 99 | 0 |
| SPGP0M00300 | XP_018221155.1 | 99 | 1 |
| SPGP0I01420 | XP_018223192.1 | 99 | 0 |
| SPGP0I03370 | XP_018223008.1 | 99 | 0 |
| SPGP0AL00510 | XP_018219913.1 | 97 | 2 |
| SPGP0C04500 | XP_018220811.1 | 99 | 0 |
| SPGP0H03280 | XP_018219508.1 | 99 | 0 |
| SPGP0E03420 | XP_018222156.1 | 98 | 0 |
| SPGP0B02810 | XP_018223724.1 | 99 | 0 |
| SPGP0Q00450 | XP_018222816.1 | 99 | 0 |
| SPGP0AL00640 | XP_018219925.1 | 96 | 0 |

Create PDF in your applications with the Pdfcrowd HTML to PDF API PDFCROWD

| | | | |
|---|---|---|---|
| SPGP0B04420 | XP_018223883.1 | 99 | 0 |
| SPGP0AI00770 | XP_018224117.1 | 99 | 0 |
| SPGP0C01880 | XP_018220556.1 | 99 | 1 |
| SPGP0L01230 | XP_018219052.1 | 99 | 1 |
| SPGP0C00890 | XP_018220457.1 | 100 | 0 |
| SPGP0O02580 | XP_018221881.1 | 99 | 0 |
| SPGP0H00130 | XP_018219202.1 | 97 | 0 |
| SPGP0Q00280 | XP_018222832.1 | 99 | 0 |
| SPGP0P01870 | XP_018220962.1 | 99 | 0 |
| SPGP0H03770 | XP_018219556.1 | 99 | 0 |
| SPGP0Q01730 | XP_018222691.1 | 99 | 0 |
| SPGP0F01220 | XP_018220238.1 | 99 | 0 |
| SPGP0M01210 | XP_018221241.1 | 99 | 0 |
| SPGP0I03700 | XP_018222978.1 | 99 | 0 |
| SPGP0S01770 | XP_018219675.1 | 99 | 0 |
| SPGP0M01910 | XP_018221310.1 | 100 | 0 |
| SPGP0C03090 | XP_018220677.1 | 99 | 0 |
| SPGP0B02180 | XP_018223663.1 | 99 | 0 |
| SPGP0E04080 | XP_018222091.1 | 99 | 0 |
| SPGP0F03450 | XP_018220024.1 | 99 | 0 |
| SPGP0H01650 | XP_018219350.1 | 99 | 0 |
| SPGP0I01880 | XP_018223147.1 | 99 | 0 |
| SPGP0S00200 | XP_018219830.1 | 99 | 0 |
| SPGP0H02290 | XP_018219412.1 | 99 | 0 |
| SPGP0I02210 | XP_018223116.1 | 99 | 0 |
| SPGP0E03520 | XP_018222146.1 | 99 | 0 |
| SPGP0C00880 | XP_018220456.1 | 99 | 0 |
| SPGP0P00330 | XP_018221110.1 | 99 | 0 |
| SPGP0E03070 | XP_018222189.1 | 99 | 0 |
| SPGP0C02320 | XP_018220600.1 | 99 | 0 |
| SPGP0O00660 | XP_018221698.1 | 99 | 0 |
| SPGP0F02360 | XP_018220130.1 | 99 | 0 |
| SPGP0I03300 | XP_018223015.1 | 99 | 0 |
| SPGP0I00140 | XP_018223320.1 | 99 | 0 |
| SPGP0Q01480 | XP_018222716.1 | 99 | 0 |
| SPGP0L01150 | XP_018219060.1 | 99 | 0 |
| SPGP0F02520 | XP_018220114.1 | 99 | 0 |
| SPGP0S01060 | XP_018219746.1 | 99 | 0 |
| SPGP0C01110 | XP_018220479.1 | 99 | 0 |
| SPGP0AR00300 | XP_018219589.1 | 99 | 0 |

Create PDF in your applications with the Pdfcrowd HTML to PDF API PDFCROWD

| | | | |
|---|---|---|---|
| SPGP0O01440 | XP_018221771.1 | 97 | 0 |
| SPGP0E04230 | XP_018222076.1 | 99 | 0 |
| SPGP0S00340 | XP_018219817.1 | 99 | 0 |
| SPGP0F01280 | XP_018220232.1 | 99 | 0 |
| SPGP0E03010 | XP_018222194.1 | 99 | 0 |
| SPGP0B03040 | XP_018223748.1 | 99 | 0 |
| SPGP0C00190 | XP_018220389.1 | 98 | 0 |
| SPGP0B03050 | XP_018223749.1 | 99 | 0 |
| SPGP0C03470 | XP_018220713.1 | 99 | 0 |
| SPGP0E01420 | XP_018222352.1 | 99 | 0 |
| SPGP0Q01830 | XP_018222680.1 | 99 | 0 |
| SPGP0Q02590 | XP_018222606.1 | 99 | 0 |
| SPGP0Q00900 | XP_018222772.1 | 100 | 0 |
| SPGP0I02620 | XP_018223075.1 | 99 | 0 |
| SPGP0C01280 | XP_018220496.1 | 99 | 0 |
| SPGP0Q01150 | XP_018222747.1 | 99 | 0 |
| SPGP0I00590 | XP_018223275.1 | 99 | 0 |
| SPGP0F00140 | XP_018220345.1 | 99 | 0 |
| SPGP0B02050 | XP_018223651.1 | 99 | 0 |
| SPGP0C02820 | XP_018220650.1 | 99 | 0 |
| SPGP0C03530 | XP_018220717.1 | 99 | 0 |
| SPGP0W00520 | XP_018221501.1 | 99 | 0 |
| SPGP0O01910 | XP_018221816.1 | 99 | 0 |
| SPGP0BI00100 | XP_018221945.1 | 99 | 0 |
| SPGP0F02890 | XP_018220079.1 | 99 | 0 |
| SPGP0B04410 | XP_018223882.1 | 99 | 0 |
| SPGP0H01820 | XP_018219367.1 | 99 | 0 |
| SPGP0I00540 | XP_018223280.1 | 99 | 0 |
| SPGP0F03080 | XP_018220061.1 | 99 | 0 |
| SPGP0C01750 | XP_018220543.1 | 99 | 0 |
| SPGP0BI00140 | XP_018221949.1 | 100 | 0 |
| SPGP0S01340 | XP_018219718.1 | 99 | 0 |
| SPGP0S01990 | XP_018219653.1 | 99 | 1 |
| SPGP0AF01130 | XP_018222495.1 | 99 | 0 |
| SPGP0Q00140 | XP_018222844.1 | 99 | 0 |
| SPGP0F01410 | XP_018220220.1 | 99 | 0 |
| SPGP0AG00920 | XP_018223403.1 | 99 | 0 |
| SPGP0M00620 | XP_018221186.1 | 99 | 0 |
| SPGP0B02910 | XP_018223735.1 | 99 | 0 |
| SPGP0B05750 | XP_018224005.1 | 99 | 0 |

Create PDF in your applications with the Pdfcrowd HTML to PDF API PDFCROWD

| | | | |
|---|---|---|---|
| SPGP0S00830 | XP_018219768.1 | 99 | 0 |
| SPGP0P01400 | XP_018221007.1 | 99 | 0 |
| SPGP0L01680 | XP_018219009.1 | 99 | 0 |
| SPGP0AG00270 | XP_018223342.1 | 100 | 0 |
| SPGP0M02790 | XP_018221394.1 | 100 | 0 |
| SPGP0E01110 | XP_018222385.1 | 99 | 0 |
| SPGP0L03710 | XP_018218812.1 | 99 | 0 |
| SPGP0E03670 | XP_018222132.1 | 99 | 0 |
| SPGP0B02850 | XP_018223729.1 | 99 | 0 |
| SPGP0S01120 | XP_018219740.1 | 99 | 0 |
| SPGP0L03490 | XP_018218834.1 | 99 | 0 |
| SPGP0L00670 | XP_018219106.1 | 99 | 0 |
| SPGP0W00100 | XP_018221458.1 | 96 | 0 |
| SPGP0I00580 | XP_018223276.1 | 99 | 0 |
| SPGP0L02720 | XP_018218909.1 | 99 | 0 |
| SPGP0B05820 | XP_018224012.1 | 99 | 0 |
| SPGP0AL00380 | XP_018219900.1 | 99 | 0 |
| SPGP0AL00430 | XP_018219905.1 | 99 | 0 |
| SPGP0P01270 | XP_018221019.1 | 99 | 0 |
| SPGP0H03730 | XP_018219552.1 | 99 | 0 |
| SPGP0I02000 | XP_018223134.1 | 99 | 0 |
| SPGP0E03790 | XP_018222120.1 | 99 | 0 |
| SPGP0H01530 | XP_018219338.1 | 99 | 0 |
| SPGP0M02170 | XP_018221333.1 | 99 | 0 |
| SPGP0I01940 | XP_018223140.1 | 99 | 0 |
| SPGP0B03990 | XP_018223840.1 | 99 | 0 |
| SPGP0C02350 | XP_018220603.1 | 99 | 0 |
| SPGP0H01930 | XP_018219378.1 | 100 | 0 |
| SPGP0H03220 | XP_018219502.1 | 100 | 0 |
| SPGP0M03180 | XP_018221432.1 | 99 | 0 |
| SPGP0H03040 | XP_018219485.1 | 99 | 0 |
| SPGP0B06100 | XP_018224038.1 | 99 | 0 |
| SPGP0AQ00140 | XP_018221953.1 | 99 | 0 |
| SPGP0C00570 | XP_018220425.1 | 97 | 2 |
| SPGP0AL00650 | XP_018219926.1 | 98 | 0 |
| SPGP0F02470 | XP_018220119.1 | 99 | 0 |
| SPGP0P01090 | XP_018221035.1 | 99 | 0 |
| SPGP0I03490 | XP_018222999.1 | 99 | 0 |
| SPGP0M02190 | XP_018221335.1 | 99 | 0 |
| SPGP0P01240 | XP_018221022.1 | 99 | 0 |

Create PDF in your applications with the Pdfcrowd HTML to PDF API PDFCROWD

| | | | |
|---|---|---|---|
| SPGP0AI00640 | XP_018224103.1 | 98 | 0 |
| SPGP0C03370 | XP_018220703.1 | 99 | 0 |
| SPGP0O01740 | XP_018221800.1 | 99 | 0 |
| SPGP0I01270 | XP_018223207.1 | 99 | 0 |
| SPGP0F00670 | XP_018220293.1 | 99 | 0 |
| SPGP0E03300 | XP_018222167.1 | 99 | 0 |
| SPGP0B02030 | XP_018223649.1 | 100 | 0 |
| SPGP0I01070 | XP_018223227.1 | 99 | 0 |
| SPGP0C02800 | XP_018220648.1 | 99 | 0 |
| SPGP0W01280 | XP_018221573.1 | 99 | 0 |
| SPGP0B02560 | XP_018223700.1 | 99 | 0 |
| SPGP0AI00540 | XP_018224094.1 | 99 | 0 |
| SPGP0O02040 | XP_018221828.1 | 99 | 0 |
| SPGP0L02980 | XP_018218883.1 | 99 | 0 |
| SPGP0B01450 | XP_018223592.1 | 99 | 0 |
| SPGP0E00300 | XP_018222467.1 | 99 | 0 |
| SPGP0W01820 | XP_018221626.1 | 98 | 0 |
| SPGP0S01400 | XP_018219712.1 | 99 | 0 |
| SPGP0M01900 | XP_018221309.1 | 100 | 0 |
| SPGP0B05120 | XP_018223945.1 | 99 | 0 |
| SPGP0AF00340 | XP_018222570.1 | 99 | 1 |
| SPGP0M02640 | XP_018221379.1 | 99 | 0 |
| SPGP0S01520 | XP_018219700.1 | 99 | 0 |
| SPGP0C02510 | XP_018220619.1 | 99 | 0 |
| SPGP0M00530 | XP_018221178.1 | 98 | 0 |
| SPGP0W01940 | XP_018221638.1 | 99 | 1 |
| SPGP0I02770 | XP_018223061.1 | 98 | 1 |
| SPGP0E00460 | XP_018222451.1 | 100 | 0 |
| SPGP0AI00440 | XP_018224084.1 | 99 | 0 |
| SPGP0AT00320 | XP_018219177.1 | 99 | 0 |
| SPGP0M02010 | XP_018221319.1 | 99 | 0 |
| SPGP0F00890 | XP_018220271.1 | 99 | 0 |
| SPGP0L02180 | XP_018218960.1 | 99 | 0 |
| SPGP0B02090 | XP_018223655.1 | 99 | 0 |
| SPGP0M02890 | XP_018221404.1 | 99 | 0 |
| SPGP0F04120 | XP_018219959.1 | 99 | 0 |
| SPGP0E04140 | XP_018222085.1 | 99 | 0 |
| SPGP0Itrna29S | DI49_4419 | 100 | 0 |
| SPGP0Q01360 | XP_018222727.1 | 99 | 0 |
| SPGP0S01440 | XP_018219708.1 | 99 | 0 |

Create PDF in your applications with the Pdfcrowd HTML to PDF API PDFCROWD

| | | | |
|---|---|---|---|
| SPGP0M02630 | XP_018221378.1 | 99 | 0 |
| SPGP0I03030 | XP_018223039.1 | 98 | 1 |
| SPGP0O00270 | XP_018221660.1 | 99 | 0 |
| SPGP0B02230 | XP_018223668.1 | 99 | 0 |
| SPGP0P02760 | XP_018220874.1 | 100 | 0 |
| SPGP0AX00120 | XP_018219861.1 | 99 | 0 |
| SPGP0H00980 | XP_018219285.1 | 99 | 0 |
| SPGP0E01030 | XP_018222394.1 | 98 | 2 |
| SPGP0L02230 | XP_018218955.1 | 99 | 0 |
| SPGP0M01660 | XP_018221285.1 | 99 | 0 |
| SPGP0L01520 | XP_018219024.1 | 99 | 0 |
| SPGP0Htrna30V | DI49_4248 | 100 | 0 |
| SPGP0AG00380 | XP_018223352.1 | 99 | 0 |
| SPGP0C01560 | XP_018220525.1 | 99 | 0 |
| SPGP0O00320 | XP_018221665.1 | 99 | 0 |
| SPGP0I00690 | XP_018223264.1 | 99 | 0 |
| SPGP0O01510 | XP_018221778.1 | 99 | 0 |
| SPGP0O01580 | XP_018221785.1 | 99 | 0 |
| SPGP0O02960 | XP_018221917.1 | 99 | 0 |
| SPGP0F03380 | XP_018220031.1 | 99 | 0 |
| SPGP0B02140 | XP_018223660.1 | 100 | 0 |
| SPGP0AL00400 | XP_018219902.1 | 99 | 0 |
| SPGP0L01760 | XP_018219001.1 | 99 | 0 |
| SPGP0Q02430 | XP_018222622.1 | 99 | 0 |
| SPGP0B04620 | XP_018223902.1 | 99 | 0 |
| SPGP0P00580 | XP_018221086.1 | 99 | 0 |
| SPGP0E03240 | XP_018222172.1 | 99 | 0 |
| SPGP0O01050 | XP_018221735.1 | 99 | 0 |
| SPGP0E04060 | XP_018222093.1 | 99 | 0 |
| SPGP0AL00440 | XP_018219906.1 | 99 | 0 |
| SPGP0B04910 | XP_018223926.1 | 99 | 0 |
| SPGP0F01470 | XP_018220213.1 | 99 | 0 |
| SPGP0O00980 | XP_018221728.1 | 99 | 0 |
| SPGP0C03980 | XP_018220760.1 | 99 | 0 |
| SPGP0AG00680 | XP_018223382.1 | 99 | 0 |
| SPGP0H02650 | XP_018219446.1 | 99 | 0 |
| SPGP0M02360 | XP_018221352.1 | 99 | 0 |
| SPGP0L00690 | XP_018219104.1 | 99 | 0 |
| SPGP0L00780 | XP_018219095.1 | 99 | 0 |
| SPGP0M03070 | XP_018221421.1 | 99 | 0 |

Create PDF in your applications with the Pdfcrowd HTML to PDF API PDFCROWD

| | | | |
|---|---|---|---|
| SPGP0Q00430 | XP_018222818.1 | 99 | 0 |
| SPGP0C04470 | XP_018220809.1 | 99 | 0 |
| SPGP0O02230 | XP_018221847.1 | 99 | 0 |
| SPGP0B04700 | XP_018223909.1 | 99 | 0 |
| SPGP0L02460 | XP_018218932.1 | 99 | 0 |
| SPGP0H03350 | XP_018219515.1 | 99 | 0 |
| SPGP0B03720 | XP_018223813.1 | 99 | 0 |
| SPGP0E03580 | XP_018222141.1 | 99 | 0 |
| SPGP0O02020 | XP_018221826.1 | 99 | 0 |
| SPGP0B04820 | XP_018223918.1 | 100 | 0 |
| SPGP0L02730 | XP_018218908.1 | 99 | 0 |
| SPGP0P02610 | XP_018220890.1 | 99 | 0 |
| SPGP0C03520 | XP_018220716.1 | 99 | 0 |
| SPGP0B05470 | XP_018223978.1 | 99 | 0 |
| SPGP0BF00130 | XP_018218768.1 | 94 | 0 |
| SPGP0E01300 | XP_018222365.1 | 99 | 0 |
| SPGP0I03640 | XP_018222984.1 | 99 | 0 |
| SPGP0C04780 | XP_018220837.1 | 99 | 0 |
| SPGP0O01230 | XP_018221750.1 | 99 | 0 |
| SPGP0M02990 | XP_018221413.1 | 99 | 0 |
| SPGP0S00970 | XP_018219755.1 | 99 | 0 |
| SPGP0O01840 | XP_018221809.1 | 99 | 0 |
| SPGP0L01730 | XP_018219004.1 | 99 | 0 |
| SPGP0L03000 | XP_018218881.1 | 99 | 0 |
| SPGP0P00910 | XP_018221053.1 | 99 | 0 |
| SPGP0L02860 | XP_018218895.1 | 99 | 0 |
| SPGP0E02820 | XP_018222213.1 | 99 | 0 |
| SPGP0S00390 | XP_018219812.1 | 99 | 0 |
| SPGP0AL00410 | XP_018219903.1 | 99 | 0 |
| SPGP0E03230 | XP_018222173.1 | 97 | 2 |
| SPGP0AG00990 | XP_018223409.1 | 99 | 0 |
| SPGP0C01960 | XP_018220564.1 | 99 | 0 |
| SPGP0F02930 | XP_018220075.1 | 99 | 0 |
| SPGP0L01030 | XP_018219072.1 | 100 | 0 |
| SPGP0F02300 | XP_018220136.1 | 99 | 0 |
| SPGP0M01040 | XP_018221224.1 | 99 | 0 |
| SPGP0M02580 | XP_018221374.1 | 99 | 0 |
| SPGP0H02010 | XP_018219386.1 | 99 | 0 |
| SPGP0AG00100 | XP_018223325.1 | 99 | 0 |
| SPGP0E03030 | XP_018222192.1 | 98 | 2 |

Create PDF in your applications with the Pdfcrowd HTML to PDF API PDFCROWD

| | | | |
|---|---|---|---|
| SPGP0L01220 | XP_018219053.1 | 99 | 0 |
| SPGP0C00760 | XP_018220443.1 | 99 | 0 |
| SPGP0W01870 | XP_018221631.1 | 99 | 0 |
| SPGP0W01700 | XP_018221614.1 | 99 | 0 |
| SPGP0E03200 | XP_018222176.1 | 99 | 0 |
| SPGP0AL00130 | XP_018219877.1 | 99 | 0 |
| SPGP0C04390 | XP_018220801.1 | 99 | 0 |
| SPGP0L02070 | XP_018218971.1 | 99 | 0 |
| SPGP0F01200 | XP_018220240.1 | 99 | 0 |
| SPGP0H03550 | XP_018219533.1 | 99 | 0 |
| SPGP0B01840 | XP_018223630.1 | 99 | 0 |
| SPGP0L00130 | XP_018219158.1 | 99 | 0 |
| SPGP0W00170 | XP_018221466.1 | 98 | 0 |
| SPGP0AF00640 | XP_018222541.1 | 99 | 0 |
| SPGP0B05350 | XP_018223967.1 | 99 | 0 |
| SPGP0E03150 | XP_018222181.1 | 99 | 0 |
| SPGP0F02980 | XP_018220071.1 | 99 | 0 |
| SPGP0S01300 | XP_018219722.1 | 99 | 0 |
| SPGP0C04530 | XP_018220814.1 | 99 | 0 |
| SPGP0I00840 | XP_018223250.1 | 99 | 0 |
| SPGP0AQ00280 | XP_018221966.1 | 99 | 0 |
| SPGP0AF00400 | XP_018222564.1 | 99 | 0 |
| SPGP0AI00280 | XP_018224069.1 | 99 | 0 |
| SPGP0S02310 | XP_018219622.1 | 99 | 0 |
| SPGP0S01930 | XP_018219659.1 | 99 | 0 |
| SPGP0C00300 | XP_018220399.1 | 99 | 0 |
| SPGP0M00980 | XP_018221218.1 | 99 | 0 |
| SPGP0B02080 | XP_018223654.1 | 99 | 0 |
| SPGP0H01050 | XP_018219292.1 | 99 | 0 |
| SPGP0O02920 | XP_018221914.1 | 99 | 0 |
| SPGP0M00320 | XP_018221157.1 | 99 | 0 |
| SPGP0E01120 | XP_018222384.1 | 99 | 0 |
| SPGP0C00180 | XP_018220388.1 | 99 | 0 |
| SPGP0W00910 | XP_018221538.1 | 98 | 1 |
| SPGP0B03690 | XP_018223810.1 | 99 | 0 |
| SPGP0B05620 | XP_018223992.1 | 99 | 0 |
| SPGP0Q00770 | XP_018222785.1 | 99 | 0 |
| SPGP0B05720 | XP_018224002.1 | 99 | 0 |
| SPGP0B02220 | XP_018223667.1 | 99 | 0 |
| SPGP0I02020 | XP_018223132.1 | 99 | 0 |

Create PDF in your applications with the Pdfcrowd HTML to PDF API PDFCROWD

| | | | |
|---|---|---|---|
| SPGP0C00750 | XP_018220442.1 | 99 | 0 |
| SPGP0B02830 | XP_018223726.1 | 99 | 0 |
| SPGP0F01300 | XP_018220230.1 | 99 | 0 |
| SPGP0O00940 | XP_018221724.1 | 99 | 0 |
| SPGP0AI00750 | XP_018224115.1 | 99 | 1 |
| SPGP0M01860 | XP_018221305.1 | 99 | 0 |
| SPGP0M00180 | XP_018221143.1 | 99 | 0 |
| SPGP0L00680 | XP_018219105.1 | 99 | 0 |
| SPGP0H03360 | XP_018219516.1 | 99 | 0 |
| SPGP0O01540 | XP_018221781.1 | 99 | 0 |
| SPGP0AL00520 | XP_018219914.1 | 99 | 0 |
| SPGP0W00750 | XP_018221523.1 | 99 | 0 |
| SPGP0Q00540 | XP_018222807.1 | 99 | 0 |
| SPGP0C02850 | XP_018220653.1 | 99 | 0 |
| SPGP0C01870 | XP_018220555.1 | 99 | 0 |
| SPGP0M01270 | XP_018221246.1 | 99 | 0 |
| SPGP0F02650 | XP_018220103.1 | 99 | 0 |
| SPGP0I03250 | XP_018223019.1 | 99 | 0 |
| SPGP0P02300 | XP_018220920.1 | 99 | 0 |
| SPGP0O03160 | XP_018221936.1 | 99 | 0 |
| SPGP0O02430 | XP_018221867.1 | 99 | 0 |
| SPGP0C00110 | XP_018220382.1 | 99 | 0 |
| SPGP0S00220 | XP_018219829.1 | 99 | 0 |
| SPGP0AQ00500 | XP_018221988.1 | 99 | 0 |
| SPGP0E03000 | XP_018222195.1 | 99 | 0 |
| SPGP0S00320 | XP_018219819.1 | 99 | 0 |
| SPGP0F02220 | XP_018220144.1 | 100 | 0 |
| SPGP0P02430 | XP_018220907.1 | 99 | 0 |
| SPGP0O00890 | XP_018221719.1 | 99 | 0 |
| SPGP0O00510 | XP_018221683.1 | 99 | 0 |
| SPGP0O01770 | XP_018221803.1 | 99 | 0 |
| SPGP0M01580 | XP_018221277.1 | 99 | 0 |
| SPGP0Q02600 | XP_018222605.1 | 99 | 0 |
| SPGP0H01070 | XP_018219294.1 | 99 | 0 |
| SPGP0AG00530 | XP_018223367.1 | 99 | 0 |
| SPGP0C00260 | XP_018220395.1 | 99 | 0 |
| SPGP0I00830 | XP_018223251.1 | 99 | 0 |
| SPGP0F00970 | XP_018220263.1 | 99 | 0 |
| SPGP0F00160 | XP_018220343.1 | 99 | 0 |

Create PDF in your applications with the Pdfcrowd HTML to PDF API PDFCROWD

| | | | |
|---|---|---|---|
| SPGP0H00920 | XP_018219279.1 | 99 | 0 |
| SPGP0S02000 | XP_018219652.1 | 99 | 0 |
| SPGP0H02900 | XP_018219471.1 | 99 | 0 |
| SPGP0B04240 | XP_018223865.1 | 99 | 0 |
| SPGP0B02060 | XP_018223652.1 | 99 | 0 |
| SPGP0P02330 | XP_018220917.1 | 99 | 0 |
| SPGP0B00400 | XP_018223489.1 | 99 | 0 |
| SPGP0B00150 | XP_018223465.1 | 99 | 0 |
| SPGP0F02780 | XP_018220090.1 | 99 | 0 |
| SPGP0E02760 | XP_018222220.1 | 99 | 0 |
| SPGP0C03510 | XP_018220715.1 | 99 | 0 |
| SPGP0E04100 | XP_018222089.1 | 99 | 0 |
| SPGP0B01750 | XP_018223621.1 | 99 | 0 |
| SPGP0O01100 | XP_018221739.1 | 99 | 0 |
| SPGP0F02990 | XP_018220070.1 | 99 | 0 |
| SPGP0M02470 | XP_018221363.1 | 99 | 0 |
| SPGP0E00600 | XP_018222437.1 | 99 | 0 |
| SPGP0M03060 | XP_018221420.1 | 99 | 0 |
| SPGP0E03660 | XP_018222133.1 | 99 | 0 |
| SPGP0F01910 | XP_018220174.1 | 99 | 0 |
| SPGP0C01070 | XP_018220475.1 | 99 | 0 |
| SPGP0AR00310 | XP_018219590.1 | 99 | 0 |
| SPGP0M02420 | XP_018221358.1 | 99 | 0 |
| SPGP0O02910 | XP_018221913.1 | 99 | 0 |
| SPGP0S01320 | XP_018219720.1 | 99 | 0 |
| SPGP0E00560 | XP_018222441.1 | 99 | 0 |
| SPGP0I00290 | XP_018223305.1 | 99 | 0 |
| SPGP0H00780 | XP_018219265.1 | 99 | 1 |
| SPGP0L00560 | XP_018219116.1 | 100 | 0 |
| SPGP0AG00650 | XP_018223379.1 | 100 | 0 |
| SPGP0Q00970 | XP_018222765.1 | 99 | 0 |
| SPGP0F03900 | XP_018219981.1 | 99 | 0 |
| SPGP0B03610 | XP_018223802.1 | 99 | 0 |
| SPGP0E04180 | XP_018222081.1 | 99 | 0 |
| SPGP0S00810 | XP_018219770.1 | 99 | 0 |
| SPGP0C03850 | XP_018220748.1 | 99 | 0 |
| SPGP0C01220 | XP_018220490.1 | 99 | 0 |
| SPGP0E02140 | XP_018222280.1 | 99 | 0 |
| SPGP0C03150 | XP_018220683.1 | 99 | 0 |
| SPGP0M02770 | XP_018221392.1 | 99 | 0 |

Create PDF in your applications with the Pdfcrowd HTML to PDF API PDFCROWD

| | | | |
|---|---|---|---|
| SPGP0E04040 | XP_018222095.1 | 99 | 0 |
| SPGP0M02800 | XP_018221395.1 | 100 | 0 |
| SPGP0H03630 | XP_018219542.1 | 99 | 0 |
| SPGP0O00950 | XP_018221725.1 | 99 | 0 |
| SPGP0F03710 | XP_018219999.1 | 99 | 0 |
| SPGP0I00670 | XP_018223266.1 | 99 | 0 |
| SPGP0H00990 | XP_018219286.1 | 99 | 0 |
| SPGP0I00960 | XP_018223238.1 | 99 | 0 |
| SPGP0S01490 | XP_018219703.1 | 99 | 0 |
| SPGP0P00630 | XP_018221081.1 | 99 | 0 |
| SPGP0S02240 | XP_018219628.1 | 99 | 0 |
| SPGP0AI00370 | XP_018224078.1 | 99 | 0 |
| SPGP0C00370 | XP_018220405.1 | 99 | 0 |
| SPGP0C04680 | XP_018220827.1 | 99 | 0 |
| SPGP0P01250 | XP_018221021.1 | 99 | 0 |
| SPGP0P01520 | XP_018220996.1 | 99 | 0 |
| SPGP0M03010 | XP_018221415.1 | 99 | 0 |
| SPGP0E02360 | XP_018222259.1 | 99 | 0 |
| SPGP0F04140 | XP_018219957.1 | 99 | 0 |
| SPGP0W00860 | XP_018221534.1 | 99 | 0 |
| SPGP0W00640 | XP_018221512.1 | 99 | 0 |
| SPGP0I01320 | XP_018223202.1 | 99 | 0 |
| SPGP0F04040 | XP_018219967.1 | 99 | 0 |
| SPGP0M02110 | XP_018221328.1 | 99 | 0 |
| SPGP0L00480 | XP_018219124.1 | 100 | 0 |
| SPGP0I01330 | XP_018223201.1 | 99 | 0 |
| SPGP0B04380 | XP_018223879.1 | 99 | 0 |
| SPGP0H00630 | XP_018219250.1 | 99 | 0 |
| SPGP0F03750 | XP_018219995.1 | 99 | 0 |
| SPGP0C03780 | XP_018220741.1 | 99 | 0 |
| SPGP0I01250 | XP_018223209.1 | 99 | 0 |
| SPGP0F01130 | XP_018220247.1 | 99 | 0 |
| SPGP0W00510 | XP_018221500.1 | 99 | 0 |
| SPGP0C00510 | XP_018220419.1 | 99 | 0 |
| SPGP0I01030 | XP_018223231.1 | 100 | 0 |
| SPGP0W01110 | XP_018221557.1 | 99 | 0 |
| SPGP0H02790 | XP_018219460.1 | 99 | 0 |
| SPGP0AL00580 | XP_018219920.1 | 99 | 0 |
| SPGP0H03540 | XP_018219532.1 | 99 | 0 |
| SPGP0I03870 | XP_018222961.1 | 99 | 0 |

Create PDF in your applications with the Pdfcrowd HTML to PDF API PDFCROWD

| | | | |
|---|---|---|---|
| SPGP0S02410 | XP_018219612.1 | 99 | 0 |
| SPGP0I01620 | XP_018223173.1 | 99 | 0 |
| SPGP0AF00580 | XP_018222547.1 | 99 | 0 |
| SPGP0H03870 | XP_018219566.1 | 99 | 0 |
| SPGP0E04170 | XP_018222082.1 | 99 | 0 |
| SPGP0O01080 | XP_018221737.1 | 99 | 0 |
| SPGP0I02300 | XP_018223107.1 | 94 | 5 |
| SPGP0E03360 | XP_018222161.1 | 98 | 1 |
| SPGP0L02270 | XP_018218951.1 | 99 | 0 |
| SPGP0B02440 | XP_018223688.1 | 99 | 0 |
| SPGP0W00320 | XP_018221481.1 | 99 | 0 |
| SPGP0S01610 | XP_018219691.1 | 98 | 0 |
| SPGP0O01730 | XP_018221799.1 | 99 | 0 |
| SPGP0AI00560 | XP_018224096.1 | 99 | 0 |
| SPGP0L03220 | XP_018218860.1 | 99 | 0 |
| SPGP0L02120 | XP_018218966.1 | 99 | 0 |
| SPGP0I00920 | XP_018223242.1 | 99 | 0 |
| SPGP0B01430 | XP_018223590.1 | 99 | 0 |
| SPGP0L01120 | XP_018219063.1 | 99 | 0 |
| SPGP0E00580 | XP_018222439.1 | 99 | 0 |
| SPGP0P00340 | XP_018221109.1 | 99 | 0 |
| SPGP0P03060 | XP_018220846.1 | 99 | 0 |
| SPGP0I03130 | XP_018223030.1 | 99 | 0 |
| SPGP0W00420 | XP_018221491.1 | 99 | 0 |
| SPGP0M00510 | XP_018221176.1 | 99 | 0 |
| SPGP0E00510 | XP_018222446.1 | 100 | 0 |
| SPGP0M01240 | XP_018221243.1 | 99 | 0 |
| SPGP0M02520 | XP_018221368.1 | 99 | 0 |
| SPGP0S02360 | XP_018219617.1 | 99 | 0 |
| SPGP0AF01180 | XP_018222490.1 | 99 | 1 |
| SPGP0E01380 | XP_018222357.1 | 99 | 0 |
| SPGP0S01530 | XP_018219699.1 | 99 | 0 |
| SPGP0S01730 | XP_018219679.1 | 100 | 0 |
| SPGP0B01380 | XP_018223585.1 | 99 | 0 |
| SPGP0C03180 | XP_018220686.1 | 99 | 1 |
| SPGP0W00240 | XP_018221473.1 | 99 | 0 |
| SPGP0P02880 | XP_018220862.1 | 100 | 0 |
| SPGP0H02480 | XP_018219430.1 | 100 | 0 |
| SPGP0E00770 | XP_018222420.1 | 99 | 0 |
| SPGP0C03070 | XP_018220675.1 | 99 | 0 |

Create PDF in your applications with the Pdfcrowd HTML to PDF API PDFCROWD

| | | | |
|---|---|---|---|
| SPGP0I01920 | XP_018223143.1 | 100 | 0 |
| SPGP0S01750 | XP_018219677.1 | 99 | 0 |
| SPGP0H01770 | XP_018219362.1 | 99 | 0 |
| SPGP0C03860 | XP_018220749.1 | 99 | 0 |
| SPGP0E04160 | XP_018222083.1 | 99 | 0 |
| SPGP0H02240 | XP_018219407.1 | 99 | 0 |
| SPGP0O00210 | XP_018221654.1 | 99 | 0 |
| SPGP0B05410 | XP_018223972.1 | 99 | 0 |
| SPGP0B00380 | XP_018223487.1 | 99 | 0 |
| SPGP0AX00140 | XP_018219862.1 | 99 | 0 |
| SPGP0S01000 | XP_018219752.1 | 99 | 0 |
| SPGP0M01640 | XP_018221283.1 | 99 | 0 |
| SPGP0C03730 | XP_018220736.1 | 99 | 0 |
| SPGP0AR00410 | XP_018219599.1 | 99 | 0 |
| SPGP0M00430 | XP_018221168.1 | 98 | 0 |
| SPGP0BJ00110 | XP_018219165.1 | 99 | 0 |
| SPGP0E00720 | XP_018222425.1 | 99 | 0 |
| SPGP0O00800 | XP_018221711.1 | 99 | 0 |
| SPGP0Q00120 | XP_018222846.1 | 99 | 0 |
| SPGP0B04670 | XP_018223906.1 | 99 | 0 |
| SPGP0F03850 | XP_018219986.1 | 99 | 0 |
| SPGP0E01320 | XP_018222363.1 | 99 | 0 |
| SPGP0Q02410 | XP_018222624.1 | 99 | 0 |
| SPGP0C01030 | XP_018220471.1 | 99 | 0 |
| SPGP0H03430 | XP_018219523.1 | 99 | 0 |
| SPGP0Q02140 | XP_018222651.1 | 99 | 0 |
| SPGP0H03150 | XP_018219496.1 | 99 | 0 |
| SPGP0F01490 | XP_018220212.1 | 99 | 0 |
| SPGP0E00980 | XP_018222399.1 | 99 | 0 |
| SPGP0E00450 | XP_018222452.1 | 99 | 0 |
| SPGP0ALtrna38L | DI49_5686 | 100 | 0 |
| SPGP0Itrna24L | DI49_5687 | 100 | 0 |
| SPGP0AF00300 | XP_018222572.1 | 99 | 0 |
| SPGP0C01680 | XP_018220536.1 | 100 | 0 |
| SPGP0F03330 | XP_018220036.1 | 99 | 0 |
| SPGP0I02700 | XP_018223068.1 | 99 | 0 |
| SPGP0M01060 | XP_018221226.1 | 99 | 0 |
| SPGP0I03270 | XP_018223017.1 | 100 | 0 |
| SPGP0B03160 | XP_018223759.1 | 99 | 0 |
| SPGP0B01600 | XP_018223607.1 | 99 | 0 |

Create PDF in your applications with the Pdfcrowd HTML to PDF API PDFCROWD

| | | | |
|---|---|---|---|
| SPGP0C02050 | XP_018220573.1 | 100 | 0 |
| SPGP0B04010 | XP_018223842.1 | 99 | 0 |
| SPGP0O02210 | XP_018221845.1 | 99 | 0 |
| SPGP0O01560 | XP_018221783.1 | 99 | 0 |
| SPGP0B01100 | XP_018223557.1 | 100 | 0 |
| SPGP0H03300 | XP_018219510.1 | 99 | 0 |
| SPGP0C01430 | XP_018220512.1 | 85 | 0 |
| SPGP0B05380 | XP_018223969.1 | 99 | 0 |
| SPGP0E01640 | XP_018222330.1 | 99 | 0 |
| SPGP0AR00490 | XP_018219607.1 | 100 | 0 |
| SPGP0L02490 | XP_018218929.1 | 99 | 0 |
| SPGP0B03400 | XP_018223782.1 | 99 | 0 |
| SPGP0F01710 | XP_018220193.1 | 99 | 0 |
| SPGP0F00600 | XP_018220300.1 | 99 | 0 |
| SPGP0F02910 | XP_018220077.1 | 99 | 0 |
| SPGP0B01000 | XP_018223547.1 | 99 | 0 |
| SPGP0H01150 | XP_018219302.1 | 99 | 0 |
| SPGP0AL00670 | XP_018219928.1 | 99 | 0 |
| SPGP0W00980 | XP_018221545.1 | 99 | 0 |
| SPGP0Q02020 | XP_018222663.1 | 99 | 0 |
| SPGP0P02440 | XP_018220906.1 | 99 | 0 |
| SPGP0S01940 | XP_018219658.1 | 99 | 0 |
| SPGP0M00130 | XP_018221140.1 | 99 | 0 |
| SPGP0B00360 | XP_018223486.1 | 99 | 0 |
| SPGP0O03110 | XP_018221931.1 | 99 | 0 |
| SPGP0E02000 | XP_018222294.1 | 99 | 0 |
| SPGP0W01580 | XP_018221602.1 | 99 | 0 |
| SPGP0B01800 | XP_018223626.1 | 99 | 0 |
| SPGP0E02800 | XP_018222215.1 | 99 | 0 |
| SPGP0AL00120 | XP_018219876.1 | 100 | 0 |
| SPGP0O00520 | XP_018221684.1 | 98 | 0 |
| SPGP0B05690 | XP_018223999.1 | 99 | 0 |
| SPGP0W00180 | XP_018221467.1 | 97 | 0 |
| SPGP0P02220 | XP_018220928.1 | 99 | 0 |
| SPGP0P02990 | XP_018220853.1 | 99 | 0 |
| SPGP0B03810 | XP_018223822.1 | 99 | 0 |
| SPGP0F04280 | XP_018219943.1 | 99 | 0 |
| SPGP0F02850 | XP_018220083.1 | 99 | 0 |
| SPGP0P02320 | XP_018220918.1 | 99 | 0 |
| SPGP0Q01950 | XP_018222668.1 | 99 | 0 |

Create PDF in your applications with the Pdfcrowd HTML to PDF API PDFCROWD

| | | | |
|---|---|---|---|
| SPGP0I03950 | XP_018222954.1 | 99 | 0 |
| SPGP0O03180 | XP_018221938.1 | 99 | 0 |
| SPGP0C02520 | XP_018220620.1 | 99 | 1 |
| SPGP0W01850 | XP_018221629.1 | 99 | 0 |
| SPGP0F01750 | XP_018220189.1 | 99 | 0 |
| SPGP0W00450 | XP_018221494.1 | 99 | 0 |
| SPGP0S02080 | XP_018219644.1 | 99 | 0 |
| SPGP0O02320 | XP_018221856.1 | 99 | 0 |
| SPGP0P01930 | XP_018220957.1 | 99 | 0 |
| SPGP0B03380 | XP_018223780.1 | 99 | 0 |
| SPGP0O02620 | XP_018221885.1 | 99 | 0 |
| SPGP0Q01540 | XP_018222710.1 | 99 | 0 |
| SPGP0Q01330 | XP_018222730.1 | 99 | 0 |
| SPGP0P01750 | XP_018220974.1 | 99 | 0 |
| SPGP0Q01240 | XP_018222738.1 | 99 | 0 |
| SPGP0F03110 | XP_018220058.1 | 99 | 0 |
| SPGP0F02050 | XP_018220161.1 | 99 | 0 |
| SPGP0B02260 | XP_018223671.1 | 99 | 0 |
| SPGP0L03110 | XP_018218870.1 | 99 | 0 |
| SPGP0F02200 | XP_018220146.1 | 99 | 0 |
| SPGP0P00290 | XP_018221114.1 | 99 | 0 |
| SPGP0F00990 | XP_018220261.1 | 99 | 0 |
| SPGP0S02330 | XP_018219620.1 | 99 | 0 |
| SPGP0AF00110 | XP_018222593.1 | 98 | 1 |
| SPGP0AG00830 | XP_018223394.1 | 99 | 0 |
| SPGP0P00590 | XP_018221085.1 | 99 | 0 |
| SPGP0E00180 | XP_018222478.1 | 99 | 0 |
| SPGP0F03430 | XP_018220026.1 | 99 | 0 |
| SPGP0I03860 | XP_018222962.1 | 99 | 0 |
| SPGP0I00280 | XP_018223306.1 | 99 | 0 |
| SPGP0L02250 | XP_018218953.1 | 99 | 0 |
| SPGP0M01320 | XP_018221251.1 | 99 | 0 |
| SPGP0C04010 | XP_018220763.1 | 99 | 0 |
| SPGP0W01320 | XP_018221577.1 | 99 | 0 |
| SPGP0AI00120 | XP_018224053.1 | 99 | 0 |
| SPGP0Q00710 | XP_018222791.1 | 99 | 0 |
| SPGP0E02630 | XP_018222233.1 | 99 | 0 |
| SPGP0H00300 | XP_018219218.1 | 99 | 0 |
| SPGP0I00680 | XP_018223265.1 | 99 | 0 |
| SPGP0AQ00270 | XP_018221965.1 | 99 | 0 |

Create PDF in your applications with the Pdfcrowd HTML to PDF API PDFCROWD

| | | | |
|---|---|---|---|
| SPGP0B01560 | XP_018223603.1 | 99 | 0 |
| SPGP0C04080 | XP_018220770.1 | 99 | 0 |
| SPGP0C04720 | XP_018220831.1 | 99 | 0 |
| SPGP0W01550 | XP_018221599.1 | 99 | 0 |
| SPGP0I02250 | XP_018223112.1 | 100 | 0 |
| SPGP0B03620 | XP_018223803.1 | 99 | 0 |
| SPGP0C02870 | XP_018220655.1 | 99 | 0 |
| SPGP0M01070 | XP_018221227.1 | 99 | 0 |
| SPGP0F03260 | XP_018220043.1 | 99 | 0 |
| SPGP0F04100 | XP_018219961.1 | 99 | 0 |
| SPGP0B05980 | XP_018224028.1 | 100 | 0 |
| SPGP0S00240 | XP_018219827.1 | 99 | 0 |
| SPGP0M00700 | XP_018221194.1 | 98 | 0 |
| SPGP0Q01680 | XP_018222696.1 | 99 | 0 |
| SPGP0AF01170 | XP_018222491.1 | 99 | 0 |
| SPGP0AF01010 | XP_018222506.1 | 99 | 0 |
| SPGP0P01440 | XP_018221003.1 | 99 | 0 |
| SPGP0M02540 | XP_018221370.1 | 99 | 0 |
| SPGP0H01900 | XP_018219375.1 | 99 | 0 |
| SPGP0B02610 | XP_018223705.1 | 99 | 0 |
| SPGP0M03250 | XP_018221439.1 | 99 | 0 |
| SPGP0Q00320 | XP_018222828.1 | 99 | 0 |
| SPGP0M01250 | XP_018221244.1 | 99 | 0 |
| SPGP0Q02620 | XP_018222603.1 | 99 | 0 |
| SPGP0P01500 | XP_018220998.1 | 99 | 0 |
| SPGP0S01240 | XP_018219728.1 | 99 | 0 |
| SPGP0C03650 | XP_018220729.1 | 99 | 0 |
| SPGP0F00180 | XP_018220341.1 | 98 | 0 |
| SPGP0CL00110 | XP_018219857.1 | 98 | 0 |
| SPGP0I01710 | XP_018223164.1 | 99 | 0 |
| SPGP0F00950 | XP_018220265.1 | 99 | 0 |
| SPGP0W01830 | XP_018221627.1 | 99 | 0 |
| SPGP0F03090 | XP_018220060.1 | 99 | 0 |
| SPGP0W00250 | XP_018221474.1 | 99 | 0 |
| SPGP0E01180 | XP_018222378.1 | 99 | 0 |
| SPGP0H03420 | XP_018219522.1 | 99 | 0 |
| SPGP0B02760 | XP_018223719.1 | 99 | 0 |
| SPGP0C01240 | XP_018220492.1 | 99 | 1 |
| SPGP0AF00850 | XP_018222520.1 | 100 | 0 |
| SPGP0P01330 | XP_018221013.1 | 99 | 0 |

Create PDF in your applications with the Pdfcrowd HTML to PDF API PDFCROWD

| | | | |
|---|---|---|---|
| SPGP0C02260 | XP_018220594.1 | 99 | 0 |
| SPGP0B00280 | XP_018223478.1 | 99 | 0 |
| SPGP0I01790 | XP_018223156.1 | 100 | 0 |
| SPGP0O01950 | XP_018221820.1 | 99 | 0 |
| SPGP0C02660 | XP_018220634.1 | 99 | 0 |
| SPGP0B05500 | XP_018223981.1 | 99 | 0 |
| SPGP0P01960 | XP_018220954.1 | 99 | 0 |
| SPGP0E01440 | XP_018222350.1 | 99 | 0 |
| SPGP0B01400 | XP_018223587.1 | 99 | 0 |
| SPGP0L00240 | XP_018219148.1 | 99 | 0 |
| SPGP0B03540 | XP_018223795.1 | 99 | 0 |
| SPGP0O02970 | XP_018221918.1 | 99 | 0 |
| SPGP0AR00330 | XP_018219592.1 | 99 | 0 |
| SPGP0Q02080 | XP_018222657.1 | 98 | 0 |
| SPGP0C01790 | XP_018220547.1 | 99 | 0 |
| SPGP0H01370 | XP_018219322.1 | 99 | 0 |
| SPGP0F01970 | XP_018220169.1 | 99 | 0 |
| SPGP0M02870 | XP_018221402.1 | 99 | 0 |
| SPGP0AL00460 | XP_018219908.1 | 99 | 0 |
| SPGP0E02160 | XP_018222278.1 | 99 | 0 |
| SPGP0S00730 | XP_018219778.1 | 99 | 0 |
| SPGP0H03790 | XP_018219558.1 | 99 | 0 |
| SPGP0B03480 | XP_018223790.1 | 99 | 0 |
| SPGP0M02750 | XP_018221390.1 | 99 | 0 |
| SPGP0I01680 | XP_018223167.1 | 99 | 0 |
| SPGP0C04620 | XP_018220823.1 | 99 | 0 |
| SPGP0I00610 | XP_018223273.1 | 99 | 0 |
| SPGP0I02810 | XP_018223058.1 | 98 | 0 |
| SPGP0C00680 | XP_018220435.1 | 99 | 0 |
| SPGP0B00820 | XP_018223528.1 | 99 | 0 |
| SPGP0C00350 | XP_018220403.1 | 99 | 0 |
| SPGP0Q00480 | XP_018222813.1 | 99 | 0 |
| SPGP0P00150 | XP_018221127.1 | 99 | 0 |
| SPGP0W00840 | XP_018221532.1 | 99 | 0 |
| SPGP0M03090 | XP_018221423.1 | 99 | 0 |
| SPGP0W00660 | XP_018221514.1 | 99 | 0 |
| SPGP0B03330 | XP_018223775.1 | 99 | 0 |
| SPGP0B00710 | XP_018223517.1 | 99 | 0 |
| SPGP0I01350 | XP_018223199.1 | 99 | 0 |
| SPGP0M01530 | XP_018221272.1 | 99 | 0 |

Create PDF in your applications with the Pdfcrowd HTML to PDF API PDFCROWD

| | | | |
|---|---|---|---|
| SPGP0I01230 | XP_018223211.1 | 99 | 0 |
| SPGP0O02790 | XP_018221902.1 | 99 | 0 |
| SPGP0F04060 | XP_018219965.1 | 100 | 0 |
| SPGP0H03480 | XP_018219526.1 | 99 | 0 |
| SPGP0L03630 | XP_018218821.1 | 99 | 0 |
| SPGP0E03690 | XP_018222130.1 | 98 | 0 |
| SPGP0S01210 | XP_018219731.1 | 99 | 0 |
| SPGP0B02400 | XP_018223684.1 | 99 | 0 |
| SPGP0M03080 | XP_018221422.1 | 99 | 0 |
| SPGP0E01020 | XP_018222395.1 | 99 | 0 |
| SPGP0Q02210 | XP_018222644.1 | 99 | 0 |
| SPGP0L00810 | XP_018219092.1 | 99 | 0 |
| SPGP0B05250 | XP_018223957.1 | 99 | 0 |
| SPGP0AF00560 | XP_018222549.1 | 99 | 0 |
| SPGP0AG00730 | XP_018223385.1 | 99 | 0 |
| SPGP0P00100 | XP_018221131.1 | 99 | 0 |
| SPGP0B02490 | XP_018223693.1 | 99 | 0 |
| SPGP0M00280 | XP_018221153.1 | 100 | 0 |
| SPGP0S00490 | XP_018219802.1 | 99 | 0 |
| SPGP0C01230 | XP_018220491.1 | 99 | 0 |
| SPGP0F04080 | XP_018219963.1 | 99 | 0 |
| SPGP0B02460 | XP_018223690.1 | 99 | 0 |
| SPGP0F02790 | XP_018220089.1 | 99 | 0 |
| SPGP0L00590 | XP_018219113.1 | 99 | 0 |
| SPGP0M01960 | XP_018221314.1 | 99 | 0 |
| SPGP0P00220 | XP_018221121.1 | 92 | 6 |
| SPGP0B01410 | XP_018223588.1 | 99 | 0 |
| SPGP0E00740 | XP_018222423.1 | 99 | 0 |
| SPGP0Q02690 | XP_018222598.1 | 99 | 0 |
| SPGP0B05460 | XP_018223977.1 | 99 | 0 |
| SPGP0W00400 | XP_018221489.1 | 99 | 0 |
| SPGP0P00360 | XP_018221107.1 | 99 | 0 |
| SPGP0F03600 | XP_018220010.1 | 99 | 0 |
| SPGP0H02130 | XP_018219397.1 | 99 | 0 |
| SPGP0AG00800 | XP_018223392.1 | 99 | 0 |
| SPGP0L03590 | XP_018218824.1 | 99 | 0 |
| SPGP0F03120 | XP_018220057.1 | 99 | 0 |
| SPGP0AX00160 | XP_018219864.1 | 99 | 0 |
| SPGP0B04860 | XP_018223922.1 | 99 | 0 |
| SPGP0C03580 | XP_018220722.1 | 99 | 1 |

Create PDF in your applications with the Pdfcrowd HTML to PDF API PDFCROWD

| | | | |
|---|---|---|---|
| SPGP0L02530 | XP_018218925.1 | 100 | 0 |
| SPGP0C04430 | XP_018220805.1 | 99 | 0 |
| SPGP0C02690 | XP_018220637.1 | 99 | 0 |
| SPGP0P01790 | XP_018220970.1 | 99 | 0 |
| SPGP0O00230 | XP_018221656.1 | 99 | 0 |
| SPGP0C03920 | XP_018220755.1 | 100 | 0 |
| SPGP0B03920 | XP_018223833.1 | 99 | 0 |
| SPGP0H00110 | XP_018219200.1 | 98 | 0 |
| SPGP0M01620 | XP_018221281.1 | 99 | 0 |
| SPGP0AR00430 | XP_018219601.1 | 99 | 0 |
| SPGP0L03800 | XP_018218803.1 | 99 | 0 |
| SPGP0C03970 | XP_018220759.1 | 99 | 0 |
| SPGP0I02860 | XP_018223054.1 | 100 | 0 |
| SPGP0Q00100 | XP_018222848.1 | 98 | 0 |
| SPGP0E03540 | XP_018222145.1 | 99 | 0 |
| SPGP0B04610 | XP_018223901.1 | 100 | 0 |
| SPGP0B05780 | XP_018224008.1 | 99 | 0 |
| SPGP0O00490 | XP_018221681.1 | 99 | 0 |
| SPGP0H03800 | XP_018219559.1 | 99 | 0 |
| SPGP0C02920 | XP_018220660.1 | 99 | 0 |
| SPGP0C04580 | XP_018220819.1 | 99 | 1 |
| SPGP0F02340 | XP_018220132.1 | 99 | 0 |
| SPGP0C03330 | XP_018220700.1 | 99 | 0 |
| SPGP0AF00720 | XP_018222533.1 | 99 | 0 |
| SPGP0O01010 | XP_018221731.1 | 99 | 0 |
| SPGP0H03130 | XP_018219494.1 | 99 | 0 |
| SPGP0S00920 | XP_018219760.1 | 99 | 0 |
| SPGP0M00770 | XP_018221200.1 | 99 | 0 |
| SPGP0M00860 | XP_018221208.1 | 99 | 0 |
| SPGP0C02070 | XP_018220575.1 | 99 | 0 |
| SPGP0I03040 | XP_018223038.1 | 99 | 0 |
| SPGP0L00380 | XP_018219134.1 | 99 | 0 |
| SPGP0L00650 | XP_018219108.1 | 99 | 0 |
| SPGP0M03200 | XP_018221434.1 | 99 | 0 |
| SPGP0C02470 | XP_018220615.1 | 100 | 0 |
| SPGP0W00970 | XP_018221544.1 | 99 | 0 |
| SPGP0H00450 | XP_018219233.1 | 99 | 0 |
| SPGP0S00480 | XP_018219803.1 | 99 | 0 |
| SPGP0AQ00240 | XP_018221962.1 | 99 | 0 |
| SPGP0F00640 | XP_018220296.1 | 99 | 0 |

Create PDF in your applications with the Pdfcrowd HTML to PDF API PDFCROWD

| | | | |
|---|---|---|---|
| SPGP0F02140 | XP_018220152.1 | 99 | 0 |
| SPGP0F03930 | XP_018219978.1 | 99 | 0 |
| SPGP0B04900 | XP_018223925.1 | 99 | 0 |
| SPGP0L02420 | XP_018218936.1 | 100 | 0 |
| SPGP0F03690 | XP_018220001.1 | 99 | 0 |
| SPGP0C01470 | XP_018220516.1 | 99 | 0 |
| SPGP0I03940 | XP_018222955.1 | 99 | 0 |
| SPGP0B00610 | XP_018223507.1 | 99 | 0 |
| SPGP0B05280 | XP_018223960.1 | 99 | 0 |
| SPGP0P00950 | XP_018221049.1 | 99 | 0 |
| SPGP0I00740 | XP_018223259.1 | 99 | 0 |
| SPGP0B03290 | XP_018223771.1 | 99 | 0 |
| SPGP0H00910 | XP_018219278.1 | 99 | 0 |
| SPGP0E01560 | XP_018222338.1 | 99 | 0 |
| SPGP0B03010 | XP_018223745.1 | 100 | 0 |
| SPGP0AR00480 | XP_018219606.1 | 99 | 0 |
| SPGP0O03140 | XP_018221934.1 | 100 | 0 |
| SPGP0E01990 | XP_018222295.1 | 99 | 0 |
| SPGP0B03830 | XP_018223824.1 | 99 | 0 |
| SPGP0C03160 | XP_018220684.1 | 99 | 0 |
| SPGP0O02090 | XP_018221833.1 | 99 | 0 |
| SPGP0S01510 | XP_018219701.1 | 97 | 2 |
| SPGP0M02950 | XP_018221409.1 | 99 | 0 |
| SPGP0B05760 | XP_018224006.1 | 99 | 0 |
| SPGP0P01190 | XP_018221027.1 | 99 | 0 |
| SPGP0B04400 | XP_018223881.1 | 99 | 0 |
| SPGP0W00300 | XP_018221479.1 | 99 | 0 |
| SPGP0H01270 | XP_018219312.1 | 99 | 0 |
| SPGP0H01890 | XP_018219374.1 | 99 | 0 |
| SPGP0B02110 | XP_018223657.1 | 99 | 0 |
| SPGP0I02320 | XP_018223105.1 | 99 | 0 |
| SPGP0H00890 | XP_018219276.1 | 100 | 0 |
| SPGP0L00640 | XP_018219109.1 | 100 | 0 |
| SPGP0C02160 | XP_018220584.1 | 99 | 0 |
| SPGP0F02500 | XP_018220116.1 | 99 | 1 |
| SPGP0C04190 | XP_018220781.1 | 99 | 0 |
| SPGP0S02350 | XP_018219618.1 | 99 | 0 |
| SPGP0B02860 | XP_018223730.1 | 99 | 0 |
| SPGP0H01610 | XP_018219346.1 | 99 | 0 |

Create PDF in your applications with the Pdfcrowd HTML to PDF API PDFCROWD

| | | | |
|---|---|---|---|
| SPGP0I02430 | XP_018223094.1 | 100 | 0 |
| SPGP0F01290 | XP_018220231.1 | 100 | 0 |
| SPGP0F01500 | XP_018220211.1 | 99 | 0 |
| SPGP0P01770 | XP_018220972.1 | 100 | 0 |
| SPGP0L03640 | XP_018218819.1 | 99 | 0 |
| SPGP0E00940 | XP_018222403.1 | 99 | 0 |
| SPGP0B03060 | XP_018223750.1 | 99 | 0 |
| SPGP0L03580 | XP_018218825.1 | 99 | 0 |
| SPGP0AI00240 | XP_018224065.1 | 100 | 0 |
| SPGP0C01810 | XP_018220549.1 | 99 | 0 |
| SPGP0AF00170 | XP_018222585.1 | 99 | 0 |
| SPGP0S00540 | XP_018219797.1 | 99 | 1 |
| SPGP0I03560 | XP_018222993.1 | 99 | 0 |
| SPGP0F02380 | XP_018220128.1 | 99 | 0 |
| SPGP0O02630 | XP_018221886.1 | 99 | 0 |
| SPGP0M01200 | XP_018221240.1 | 99 | 0 |
| SPGP0B05010 | XP_018223936.1 | 99 | 0 |
| SPGP0M00590 | XP_018221183.1 | 99 | 0 |
| SPGP0AQ00250 | XP_018221963.1 | 99 | 0 |
| SPGP0AR00230 | XP_018219582.1 | 99 | 0 |
| SPGP0C02400 | XP_018220608.1 | 99 | 0 |
| SPGP0W01300 | XP_018221575.1 | 100 | 0 |
| SPGP0Q00730 | XP_018222789.1 | 99 | 0 |
| SPGP0AI00620 | XP_018224101.1 | 99 | 0 |
| SPGP0P02890 | XP_018220861.1 | 99 | 0 |
| SPGP0C00520 | XP_018220420.1 | 99 | 0 |
| SPGP0AF00280 | XP_018222574.1 | 99 | 0 |
| SPGP0L03400 | XP_018218843.1 | 99 | 1 |
| SPGP0Ftrna8W | DI49_3006 | 100 | 0 |
| SPGP0M00680 | XP_018221192.1 | 97 | 1 |
| SPGP0M00490 | XP_018221174.1 | 99 | 0 |
| SPGP0S00260 | XP_018219825.1 | 99 | 0 |
| SPGP0AX00110 | XP_018219860.1 | 100 | 0 |
| SPGP0I02570 | XP_018223080.1 | 99 | 0 |
| SPGP0S01800 | XP_018219672.1 | 99 | 0 |
| SPGP0E00610 | XP_018222436.1 | 99 | 0 |
| SPGP0C02890 | XP_018220657.1 | 99 | 0 |
| SPGP0B01990 | XP_018223645.1 | 99 | 0 |
| SPGP0H00320 | XP_018219220.1 | 99 | 0 |
| SPGP0E04090 | XP_018222090.1 | 99 | 0 |

Create PDF in your applications with the Pdfcrowd HTML to PDF API PDFCROWD

| | | | |
|---|---|---|---|
| SPGP0I01630 | XP_018223172.1 | 99 | 0 |
| SPGP0AG00500 | XP_018223364.1 | 99 | 0 |
| SPGP0O01120 | XP_018221741.1 | 98 | 1 |
| SPGP0I03390 | XP_018223006.1 | 99 | 0 |
| SPGP0Q00460 | XP_018222815.1 | 99 | 0 |
| SPGP0M00720 | XP_018221196.1 | 98 | 0 |
| SPGP0L00160 | XP_018219155.1 | 99 | 0 |
| SPGP0E00970 | XP_018222400.1 | 99 | 0 |
| SPGP0E00250 | XP_018222472.1 | 99 | 0 |
| SPGP0B01190 | XP_018223566.1 | 99 | 0 |
| SPGP0H00950 | XP_018219282.1 | 99 | 0 |
| SPGP0S01350 | XP_018219717.1 | 99 | 0 |
| SPGP0I03680 | XP_018222980.1 | 99 | 0 |
| SPGP0H01600 | XP_018219345.1 | 99 | 0 |
| SPGP0O00910 | XP_018221721.1 | 99 | 0 |
| SPGP0L01860 | XP_018218991.1 | 99 | 0 |
| SPGP0E00280 | XP_018222469.1 | 99 | 0 |
| SPGP0L02710 | XP_018218910.1 | 99 | 0 |
| SPGP0I01960 | XP_018223138.1 | 99 | 0 |
| SPGP0M01870 | XP_018221306.1 | 99 | 0 |
| SPGP0P02370 | XP_018220913.1 | 100 | 0 |
| SPGP0C03200 | XP_018220688.1 | 99 | 0 |
| SPGP0O00550 | XP_018221687.1 | 99 | 0 |
| SPGP0O01220 | XP_018221749.1 | 99 | 0 |
| SPGP0E02250 | XP_018222270.1 | 99 | 0 |
| SPGP0E00800 | XP_018222417.1 | 99 | 0 |
| SPGP0C03120 | XP_018220680.1 | 99 | 0 |
| SPGP0C01260 | XP_018220494.1 | 99 | 0 |
| SPGP0AI00780 | XP_018224118.1 | 99 | 0 |
| SPGP0AQ00370 | XP_018221975.1 | 99 | 0 |
| SPGP0H02190 | XP_018219402.1 | 99 | 0 |
| SPGP0E00660 | XP_018222431.1 | 99 | 0 |
| SPGP0L01710 | XP_018219006.1 | 99 | 0 |
| SPGP0F00880 | XP_018220272.1 | 99 | 0 |
| SPGP0H03320 | XP_018219512.1 | 99 | 0 |
| SPGP0E01500 | XP_018222344.1 | 98 | 2 |
| SPGP0O01750 | XP_018221801.1 | 99 | 0 |
| SPGP0S00440 | XP_018219807.1 | 99 | 1 |
| SPGP0O01970 | XP_018221822.1 | 99 | 0 |
| SPGP0O01960 | XP_018221821.1 | 99 | 0 |

Create PDF in your applications with the Pdfcrowd HTML to PDF API PDFCROWD

| | | | |
|---|---|---|---|
| SPGP0E00240 | XP_018222473.1 | 99 | 0 |
| SPGP0B04930 | XP_018223928.1 | 99 | 0 |
| SPGP0L00370 | XP_018219135.1 | 100 | 0 |
| SPGP0L03430 | XP_018218840.1 | 99 | 0 |
| SPGP0AR00350 | XP_018219594.1 | 99 | 0 |
| SPGP0Q02200 | XP_018222645.1 | 99 | 1 |
| SPGP0B04580 | XP_018223898.1 | 100 | 0 |
| SPGP0L01360 | XP_018219040.1 | 99 | 0 |
| SPGP0B03560 | XP_018223797.1 | 99 | 0 |
| SPGP0C04460 | XP_018220808.1 | 99 | 0 |
| SPGP0C00590 | XP_018220427.1 | 99 | 0 |
| SPGP0Q00930 | XP_018222769.1 | 99 | 0 |
| SPGP0E01980 | XP_018222296.1 | 99 | 0 |
| SPGP0L03270 | XP_018218855.1 | 99 | 0 |
| SPGP0AG00260 | XP_018223341.1 | 99 | 0 |
| SPGP0C02790 | XP_018220647.1 | 99 | 0 |
| SPGP0H02040 | XP_018219389.1 | 99 | 0 |
| SPGP0M02730 | XP_018221388.1 | 99 | 0 |
| SPGP0C00660 | XP_018220434.1 | 99 | 0 |
| SPGP0E02480 | XP_018222247.1 | 99 | 0 |
| SPGP0I01370 | XP_018223197.1 | 99 | 0 |
| SPGP0O02400 | XP_018221864.1 | 99 | 0 |
| SPGP0M01280 | XP_018221247.1 | 99 | 0 |
| SPGP0B03270 | XP_018223769.1 | 99 | 0 |
| SPGP0B00800 | XP_018223526.1 | 99 | 0 |
| SPGP0C00330 | XP_018220401.1 | 99 | 0 |
| SPGP0AL00320 | XP_018219895.1 | 99 | 0 |
| SPGP0W01380 | XP_018221583.1 | 99 | 0 |
| SPGP0E02100 | XP_018222284.1 | 99 | 0 |
| SPGP0F03230 | XP_018220046.1 | 99 | 0 |
| SPGP0C02110 | XP_018220579.1 | 99 | 0 |
| SPGP0W00820 | XP_018221530.1 | 99 | 0 |
| SPGP0C00470 | XP_018220415.1 | 99 | 0 |
| SPGP0W00870 | XP_018221535.1 | 99 | 0 |
| SPGP0M01030 | XP_018221223.1 | 99 | 0 |
| SPGP0C03140 | XP_018220682.1 | 99 | 0 |
| SPGP0I01210 | XP_018223213.1 | 99 | 0 |
| SPGP0E03950 | XP_018222104.1 | 99 | 0 |
| SPGP0B02170 | XP_018223662.1 | 99 | 0 |
| SPGP0F00390 | XP_018220320.1 | 99 | 0 |

Create PDF in your applications with the Pdfcrowd HTML to PDF API PDFCROWD

| | | | |
|---|---|---|---|
| SPGP0P01560 | XP_018220992.1 | 99 | 0 |
| SPGP0C04400 | XP_018220802.1 | 99 | 0 |
| SPGP0Otrna8P | DI49_2548 | 98 | 0 |
| SPGP0B04790 | XP_018223917.1 | 99 | 0 |
| SPGP0L01000 | XP_018219075.1 | 99 | 0 |
| SPGP0E03750 | XP_018222124.1 | 99 | 0 |
| SPGP0E03340 | XP_018222163.1 | 100 | 0 |
| SPGP0P00470 | XP_018221096.1 | 99 | 0 |
| SPGP0L00410 | XP_018219131.1 | 99 | 0 |
| SPGP0B05270 | XP_018223959.1 | 98 | 0 |
| SPGP0AF00540 | XP_018222551.1 | 99 | 0 |
| SPGP0L00830 | XP_018219090.1 | 99 | 0 |
| SPGP0B04660 | XP_018223905.1 | 99 | 0 |
| SPGP0P00160 | XP_018221126.1 | 99 | 0 |
| SPGP0L02620 | XP_018218919.1 | 99 | 0 |
| SPGP0AG00750 | XP_018223387.1 | 99 | 0 |
| SPGP0M00260 | XP_018221151.1 | 99 | 0 |
| SPGP0S00110 | XP_018219839.1 | 99 | 0 |
| SPGP0E03410 | XP_018222157.1 | 98 | 0 |
| SPGP0H02380 | XP_018219421.1 | 99 | 0 |
| SPGP0P02100 | XP_018220940.1 | 99 | 0 |
| SPGP0O01200 | XP_018221747.1 | 99 | 0 |
| SPGP0L02240 | XP_018218954.1 | 99 | 0 |
| SPGP0E02080 | XP_018222286.1 | 100 | 0 |
| SPGP0AL00350 | XP_018219897.1 | 99 | 0 |
| SPGP0B01690 | XP_018223615.1 | 99 | 0 |
| SPGP0C03250 | XP_018220692.1 | 99 | 0 |
| SPGP0B05180 | XP_018223951.1 | 99 | 1 |
| SPGP0AF00890 | XP_018222517.1 | 98 | 1 |
| SPGP0P00380 | XP_018221105.1 | 99 | 0 |
| SPGP0F03660 | XP_018220004.1 | 100 | 0 |
| SPGP0H03110 | XP_018219492.1 | 100 | 0 |
| SPGP0AT00340 | XP_018219175.1 | 99 | 0 |
| SPGP0L00100 | XP_018219161.1 | 99 | 0 |
| SPGP0H03380 | XP_018219518.1 | 99 | 0 |
| SPGP0F03370 | XP_018220032.1 | 99 | 0 |
| SPGP0H03120 | XP_018219493.1 | 99 | 1 |
| SPGP0F03250 | XP_018220044.1 | 99 | 0 |
| SPGP0AX00180 | XP_018219866.1 | 99 | 0 |
| SPGP0I00860 | XP_018223248.1 | 99 | 0 |

Create PDF in your applications with the Pdfcrowd HTML to PDF API PDFCROWD

| | | | |
|---|---|---|---|
| SPGP0O03150 | XP_018221935.1 | 99 | 0 |
| SPGP0E03780 | XP_018222121.1 | 99 | 0 |
| SPGP0S00960 | XP_018219756.1 | 99 | 0 |
| SPGP0B03770 | XP_018223818.1 | 99 | 0 |
| SPGP0E01760 | XP_018222318.1 | 99 | 0 |
| SPGP0Q00690 | XP_018222793.1 | 100 | 0 |
| SPGP0Q02520 | XP_018222613.1 | 99 | 0 |
| SPGP0I03180 | XP_018223026.1 | 100 | 0 |
| SPGP0L01530 | XP_018219023.1 | 99 | 0 |
| SPGP0AF01050 | XP_018222503.1 | 99 | 0 |
| SPGP0F03050 | XP_018220064.1 | 99 | 0 |
| SPGP0M01600 | XP_018221279.1 | 99 | 0 |
| SPGP0B01030 | XP_018223550.1 | 99 | 0 |
| SPGP0AL00420 | XP_018219904.1 | 99 | 0 |
| SPGP0AR00180 | XP_018219577.1 | 99 | 0 |
| SPGP0AI00510 | XP_018224091.1 | 99 | 0 |
| SPGP0AG00370 | XP_018223351.1 | 99 | 0 |
| SPGP0B03740 | XP_018223815.1 | 99 | 0 |
| SPGP0H02670 | XP_018219448.1 | 99 | 0 |
| SPGP0B01200 | XP_018223567.1 | 100 | 0 |
| SPGP0P02070 | XP_018220943.1 | 99 | 0 |
| SPGP0F03220 | XP_018220047.1 | 99 | 0 |
| SPGP0E00920 | XP_018222405.1 | 99 | 0 |
| SPGP0L00520 | XP_018219120.1 | 99 | 0 |
| SPGP0P02130 | XP_018220937.1 | 99 | 0 |
| SPGP0O00960 | XP_018221726.1 | 99 | 0 |
| SPGP0L00320 | XP_018219140.1 | 99 | 0 |
| SPGP0O02890 | XP_018221911.1 | 99 | 0 |
| SPGP0W00650 | XP_018221513.1 | 99 | 0 |
| SPGP0AG00460 | XP_018223360.1 | 99 | 0 |
| SPGP0L01010 | XP_018219074.1 | 99 | 0 |
| SPGP0BB00210 | XP_018219855.1 | 99 | 0 |
| SPGP0I02400 | XP_018223097.1 | 99 | 0 |
| SPGP0CY00110 | XP_018222596.1 | 99 | 0 |
| SPGP0F02230 | XP_018220143.1 | 99 | 0 |
| SPGP0AL00370 | XP_018219899.1 | 99 | 0 |
| SPGP0O00560 | XP_018221688.1 | 99 | 0 |
| SPGP0O02870 | XP_018221909.1 | 99 | 0 |
| SPGP0H01390 | XP_018219324.1 | 99 | 0 |
| SPGP0AF00990 | XP_018222508.1 | 99 | 0 |

Create PDF in your applications with the Pdfcrowd HTML to PDF API PDFCROWD

| | | | |
|---|---|---|---|
| SPGP0O03090 | XP_018221929.1 | 99 | 0 |
| SPGP0B05860 | XP_018224016.1 | 99 | 0 |
| SPGP0B01440 | XP_018223591.1 | 99 | 0 |
| SPGP0C02450 | XP_018220613.1 | 99 | 1 |
| SPGP0M02560 | XP_018221372.1 | 99 | 0 |
| SPGP0C03750 | XP_018220738.1 | 99 | 0 |
| SPGP0L01390 | XP_018219037.1 | 99 | 0 |
| SPGP0F00660 | XP_018220294.1 | 99 | 0 |
| SPGP0I02240 | XP_018223113.1 | 99 | 0 |
| SPGP0E00620 | XP_018222435.1 | 99 | 0 |
| SPGP0L02640 | XP_018218917.1 | 99 | 0 |
| SPGP0S02170 | XP_018219635.1 | 99 | 0 |
| SPGP0Q01290 | XP_018222734.1 | 99 | 0 |
| SPGP0P02620 | XP_018220888.1 | 98 | 0 |
| SPGP0M03110 | XP_018221425.1 | 99 | 0 |
| SPGP0M01150 | XP_018221235.1 | 99 | 0 |
| SPGP0P02400 | XP_018220910.1 | 99 | 0 |
| SPGP0P02150 | XP_018220935.1 | 99 | 0 |
| SPGP0AG00190 | XP_018223334.1 | 99 | 0 |
| SPGP0L01610 | XP_018219015.1 | 99 | 0 |
| SPGP0I00320 | XP_018223302.1 | 99 | 0 |
| SPGP0C01450 | XP_018220514.1 | 99 | 0 |
| SPGP0B00320 | XP_018223482.1 | 99 | 0 |
| SPGP0I00760 | XP_018223257.1 | 97 | 3 |
| SPGP0P02670 | XP_018220883.1 | 99 | 0 |
| SPGP0Otrna7P | DI49_2389 | 99 | 0 |
| SPGP0H00870 | XP_018219274.1 | 99 | 0 |
| SPGP0E02090 | XP_018222285.1 | 99 | 0 |
| SPGP0M02960 | XP_018221410.1 | 99 | 0 |
| SPGP0W00140 | XP_018221463.1 | 97 | 0 |
| SPGP0O00920 | XP_018221722.1 | 99 | 0 |
| SPGP0E02260 | XP_018222269.1 | 99 | 0 |
| SPGP0W00190 | XP_018221468.1 | 99 | 0 |
| SPGP0I00650 | XP_018223269.1 | 99 | 0 |
| SPGP0AI00320 | XP_018224073.1 | 99 | 0 |
| SPGP0Q01550 | XP_018222709.1 | 99 | 0 |
| SPGP0W01420 | XP_018221587.1 | 99 | 0 |
| SPGP0L01900 | XP_018218987.1 | 99 | 0 |
| SPGP0B02290 | XP_018223674.1 | 99 | 0 |
| SPGP0F02000 | XP_018220166.1 | 99 | 0 |

Create PDF in your applications with the Pdfcrowd HTML to PDF API PDFCROWD

| | | | |
|---|---|---|---|
| SPGP0B01870 | XP_018223633.1 | 99 | 0 |
| SPGP0Q02240 | XP_018222641.1 | 99 | 0 |
| SPGP0B00440 | XP_018223492.1 | 99 | 0 |
| SPGP0W01250 | XP_018221570.1 | 100 | 0 |
| SPGP0S00820 | XP_018219769.1 | 99 | 0 |
| SPGP0H03560 | XP_018219534.1 | 100 | 0 |
| SPGP0B03210 | XP_018223764.1 | 99 | 0 |
| SPGP0W00360 | XP_018221485.1 | 99 | 0 |
| SPGP0M02070 | XP_018221325.1 | 99 | 0 |
| SPGP0B03850 | XP_018223826.1 | 99 | 0 |
| SPGP0C03290 | XP_018220696.1 | 99 | 0 |
| SPGP0Q02130 | XP_018222652.1 | 100 | 0 |
| SPGP0F00720 | XP_018220288.1 | 99 | 0 |
| SPGP0E01240 | XP_018222372.1 | 99 | 0 |
| SPGP0AL00150 | XP_018219879.1 | 99 | 0 |
| SPGP0C01040 | XP_018220472.1 | 99 | 0 |
| SPGP0I01190 | XP_018223215.1 | 99 | 0 |
| SPGP0B06170 | XP_018224045.1 | 96 | 0 |
| SPGP0P01710 | XP_018220978.1 | 99 | 0 |
| SPGP0P01360 | XP_018221010.1 | 99 | 0 |
| SPGP0AI00260 | XP_018224067.1 | 99 | 0 |
| SPGP0C00230 | XP_018220393.1 | 99 | 0 |
| SPGP0Q00370 | XP_018222824.1 | 99 | 0 |
| SPGP0AF00150 | XP_018222587.1 | 99 | 0 |
| SPGP0AQ00450 | XP_018221983.1 | 98 | 0 |
| SPGP0B00390 | XP_018223488.1 | 99 | 0 |
| SPGP0M03030 | XP_018221417.1 | 99 | 0 |
| SPGP0E03060 | XP_018222190.1 | 98 | 1 |
| SPGP0W01360 | XP_018221581.1 | 100 | 0 |
| SPGP0E00850 | XP_018222412.1 | 99 | 0 |
| SPGP0B02890 | XP_018223733.1 | 99 | 0 |
| SPGP0L02590 | XP_018218922.1 | 99 | 0 |
| SPGP0S02370 | XP_018219616.1 | 99 | 0 |
| SPGP0F00910 | XP_018220269.1 | 99 | 0 |
| SPGP0L00990 | XP_018219076.1 | 99 | 0 |
| SPGP0Q01890 | XP_018222674.1 | 97 | 1 |
| SPGP0AF00740 | XP_018222531.1 | 100 | 0 |
| SPGP0H00770 | XP_018219264.1 | 99 | 0 |
| SPGP0I01000 | XP_018223234.1 | 99 | 0 |
| SPGP0M01590 | XP_018221278.1 | 99 | 0 |

Create PDF in your applications with the Pdfcrowd HTML to PDF API PDFCROWD

| | | | |
|---|---|---|---|
| SPGP0P00440 | XP_018221099.1 | 99 | 0 |
| SPGP0C02620 | XP_018220631.1 | 99 | 0 |
| SPGP0B04600 | XP_018223900.1 | 99 | 0 |
| SPGP0H02270 | XP_018219410.1 | 99 | 0 |
| SPGP0H02250 | XP_018219408.1 | 99 | 0 |
| SPGP0M01680 | XP_018221286.1 | 99 | 0 |
| SPGP0C02120 | XP_018220580.1 | 99 | 0 |
| SPGP0W00210 | XP_018221470.1 | 99 | 0 |
| SPGP0F02620 | XP_018220106.1 | 99 | 0 |
| SPGP0I03260 | XP_018223018.1 | 99 | 0 |
| SPGP0B05040 | XP_018223939.1 | 99 | 0 |
| SPGP0AG00560 | XP_018223370.1 | 97 | 0 |
| SPGP0W01730 | XP_018221617.1 | 99 | 1 |
| SPGP0Q01400 | XP_018222724.1 | 99 | 0 |
| SPGP0S02070 | XP_018219645.1 | 99 | 0 |
| SPGP0M00740 | XP_018221198.1 | 99 | 0 |
| SPGP0Q00570 | XP_018222804.1 | 99 | 0 |
| SPGP0L02330 | XP_018218945.1 | 99 | 0 |
| SPGP0M01290 | XP_018221248.1 | 97 | 1 |
| SPGP0S01880 | XP_018219664.1 | 99 | 0 |
| SPGP0CL00100 | XP_018219858.1 | 97 | 1 |
| SPGP0L00340 | XP_018219138.1 | 100 | 0 |
| SPGP0C01100 | XP_018220478.1 | 99 | 0 |
| SPGP0C04110 | XP_018220773.1 | 99 | 0 |
| SPGP0H01410 | XP_018219326.1 | 99 | 0 |
| SPGP0C01200 | XP_018220488.1 | 99 | 0 |
| SPGP0AF00430 | XP_018222561.1 | 99 | 0 |
| SPGP0I00510 | XP_018223283.1 | 99 | 0 |
| SPGP0O01110 | XP_018221740.1 | 99 | 0 |
| SPGP0B03570 | XP_018223798.1 | 99 | 0 |
| SPGP0F02070 | XP_018220159.1 | 99 | 0 |
| SPGP0W01370 | XP_018221582.1 | 99 | 0 |
| SPGP0S01580 | XP_018219694.1 | 99 | 0 |
| SPGP0H02930 | XP_018219474.1 | 99 | 0 |
| SPGP0AR00370 | XP_018219596.1 | 100 | 0 |
| SPGP0C03010 | XP_018220669.1 | 100 | 0 |
| SPGP0P03100 | XP_018220843.1 | 99 | 0 |
| SPGP0C04000 | XP_018220762.1 | 100 | 0 |
| SPGP0B05340 | XP_018223966.1 | 99 | 0 |
| SPGP0Q00310 | XP_018222829.1 | 99 | 0 |

Create PDF in your applications with the Pdfcrowd HTML to PDF API PDFCROWD

| | | | |
|---|---|---|---|
| SPGP0F00820 | XP_018220278.1 | 100 | 0 |
| SPGP0I03230 | XP_018223021.1 | 99 | 0 |
| SPGP0F02690 | XP_018220099.1 | 99 | 0 |
| SPGP0E03720 | XP_018222127.1 | 99 | 0 |
| SPGP0Itrna15Q | DI49_1374 | 100 | 0 |
| SPGP0M01570 | XP_018221276.1 | 99 | 0 |
| SPGP0E01230 | XP_018222373.1 | 100 | 0 |
| SPGP0B04040 | XP_018223845.1 | 99 | 0 |
| SPGP0H03510 | XP_018219529.1 | 100 | 0 |
| SPGP0Q02170 | XP_018222648.1 | 100 | 0 |
| SPGP0O01570 | XP_018221784.1 | 99 | 0 |
| SPGP0Q00300 | XP_018222830.1 | 99 | 0 |
| SPGP0B00790 | XP_018223525.1 | 99 | 0 |
| SPGP0AF00140 | XP_018222588.1 | 99 | 0 |
| SPGP0W01610 | XP_018221605.1 | 99 | 0 |
| SPGP0C00910 | XP_018220459.1 | 99 | 0 |
| SPGP0O02470 | XP_018221871.1 | 99 | 0 |
| SPGP0M02240 | XP_018221340.1 | 99 | 0 |
| SPGP0P00240 | XP_018221119.1 | 99 | 0 |
| SPGP0M02710 | XP_018221386.1 | 99 | 0 |
| SPGP0L00250 | XP_018219147.1 | 99 | 0 |
| SPGP0L00170 | XP_018219154.1 | 99 | 0 |
| SPGP0AT00290 | XP_018219180.1 | 99 | 0 |
| SPGP0I01390 | XP_018223195.1 | 100 | 0 |
| SPGP0F01850 | XP_018220180.1 | 99 | 0 |
| SPGP0O02420 | XP_018221866.1 | 99 | 0 |
| SPGP0AQ00430 | XP_018221981.1 | 100 | 0 |
| SPGP0I02850 | XP_018223055.1 | 99 | 0 |
| SPGP0F00430 | XP_018220317.1 | 99 | 0 |
| SPGP0E00430 | XP_018222454.1 | 99 | 0 |
| SPGP0AL00300 | XP_018219893.1 | 99 | 0 |
| SPGP0B00860 | XP_018223532.1 | 99 | 0 |
| SPGP0B01130 | XP_018223560.1 | 99 | 0 |
| SPGP0E02120 | XP_018222282.1 | 99 | 0 |
| SPGP0S00740 | XP_018219777.1 | 99 | 1 |
| SPGP0B00940 | XP_018223540.1 | 99 | 0 |
| SPGP0F03350 | XP_018220034.1 | 99 | 0 |
| SPGP0W00800 | XP_018221528.1 | 99 | 1 |
| SPGP0C00940 | XP_018220462.1 | 99 | 0 |
| SPGP0P02360 | XP_018220914.1 | 99 | 0 |

Create PDF in your applications with the Pdfcrowd HTML to PDF API PDFCROWD

| | | | |
|---|---|---|---|
| SPGP0I03890 | XP_018222959.1 | 98 | 0 |
| SPGP0E00750 | XP_018222422.1 | 99 | 0 |
| SPGP0E02550 | XP_018222241.1 | 99 | 0 |
| SPGP0B04260 | XP_018223867.1 | 99 | 0 |
| SPGP0M01100 | XP_018221230.1 | 99 | 0 |
| SPGP0H02950 | XP_018219476.1 | 99 | 0 |
| SPGP0C03620 | XP_018220726.1 | 99 | 0 |
| SPGP0L03670 | XP_018218816.1 | 100 | 0 |
| SPGP0E03650 | XP_018222134.1 | 99 | 0 |
| SPGP0S01700 | XP_018219682.1 | 99 | 0 |
| SPGP0E00710 | XP_018222426.1 | 99 | 0 |
| SPGP0O02410 | XP_018221865.1 | 99 | 0 |
| SPGP0H02880 | XP_018219469.1 | 99 | 0 |
| SPGP0AF00250 | XP_018222577.1 | 99 | 0 |
| SPGP0M03040 | XP_018221418.1 | 99 | 0 |
| SPGP0H03070 | XP_018219488.1 | 99 | 0 |
| SPGP0C01650 | XP_018220533.1 | 99 | 0 |
| SPGP0C02230 | XP_018220591.1 | 99 | 0 |
| SPGP0L03310 | XP_018218851.1 | 99 | 0 |
| SPGP0L01290 | XP_018219047.1 | 99 | 0 |
| SPGP0C04160 | XP_018220778.1 | 99 | 0 |
| SPGP0C01990 | XP_018220567.1 | 99 | 0 |
| SPGP0Q01790 | XP_018222685.1 | 99 | 0 |
| SPGP0S00560 | XP_018219795.1 | 99 | 0 |
| SPGP0B00190 | XP_018223469.1 | 99 | 0 |
| SPGP0E03470 | XP_018222151.1 | 99 | 0 |
| SPGP0P00140 | XP_018221128.1 | 99 | 0 |
| SPGP0W01170 | XP_018221562.1 | 99 | 0 |
| SPGP0H02080 | XP_018219393.1 | 99 | 0 |
| SPGP0AG00850 | XP_018223396.1 | 100 | 0 |
| SPGP0B04360 | XP_018223877.1 | 99 | 0 |
| SPGP0E03820 | XP_018222117.1 | 99 | 0 |
| SPGP0M01990 | XP_018221317.1 | 99 | 0 |
| SPGP0E04000 | XP_018222099.1 | 99 | 0 |
| SPGP0AX00210 | XP_018219869.1 | 99 | 0 |
| SPGP0L02810 | XP_018218900.1 | 99 | 0 |
| SPGP0C01190 | XP_018220487.1 | 99 | 0 |
| SPGP0H01570 | XP_018219342.1 | 99 | 0 |
| SPGP0P02700 | XP_018220880.1 | 99 | 0 |
| SPGP0M00240 | XP_018221149.1 | 99 | 0 |

Create PDF in your applications with the Pdfcrowd HTML to PDF API PDFCROWD

| | | | |
|---|---|---|---|
| SPGP0I00930 | XP_018223241.1 | 99 | 0 |
| SPGP0F01860 | XP_018220179.1 | 99 | 1 |
| SPGP0P00260 | XP_018221117.1 | 99 | 0 |
| SPGP0B02930 | XP_018223737.1 | 100 | 0 |
| SPGP0O01720 | XP_018221798.1 | 99 | 0 |
| SPGP0I03770 | XP_018222971.1 | 99 | 0 |
| SPGP0AI00450 | XP_018224085.1 | 99 | 0 |
| SPGP0AI00290 | XP_018224070.1 | 99 | 0 |
| SPGP0H00480 | XP_018219236.1 | 100 | 0 |
| SPGP0H00390 | XP_018219227.1 | 99 | 0 |
| SPGP0O03030 | XP_018221923.1 | 99 | 0 |
| SPGP0F03130 | XP_018220056.1 | 99 | 0 |
| SPGP0H01280 | XP_018219313.1 | 99 | 0 |
| SPGP0H02680 | XP_018219449.1 | 99 | 0 |
| SPGP0H01700 | XP_018219355.1 | 99 | 0 |
| SPGP0F03640 | XP_018220006.1 | 99 | 0 |
| SPGP0AR00190 | XP_018219578.1 | 99 | 0 |
| SPGP0Q00880 | XP_018222774.1 | 99 | 0 |
| SPGP0O02570 | XP_018221880.1 | 99 | 0 |
| SPGP0P01620 | XP_018220987.1 | 99 | 0 |
| SPGP0C02760 | XP_018220644.1 | 99 | 0 |
| SPGP0H00560 | XP_018219244.1 | 99 | 0 |
| SPGP0F03300 | XP_018220039.1 | 99 | 0 |
| SPGP0AF01070 | XP_018222501.1 | 100 | 0 |
| SPGP0H00150 | XP_018219204.1 | 99 | 0 |
| SPGP0I03690 | XP_018222979.1 | 99 | 0 |
| SPGP0H00620 | XP_018219249.1 | 99 | 0 |
| SPGP0B00250 | XP_018223475.1 | 99 | 0 |
| SPGP0M01370 | XP_018221256.1 | 99 | 0 |
| SPGP0E00230 | XP_018222474.1 | 99 | 0 |
| SPGP0E03290 | XP_018222168.1 | 99 | 0 |
| SPGP0S01780 | XP_018219674.1 | 99 | 0 |
| SPGP0O03220 | XP_018221942.1 | 99 | 0 |
| SPGP0F03440 | XP_018220025.1 | 99 | 0 |
| SPGP0E03160 | XP_018222180.1 | 99 | 0 |
| SPGP0C04130 | XP_018220775.1 | 99 | 0 |
| SPGP0L00470 | XP_018219125.1 | 99 | 0 |
| SPGP0C00610 | XP_018220429.1 | 99 | 0 |
| SPGP0B06020 | XP_018224032.1 | 99 | 0 |

Create PDF in your applications with the Pdfcrowd HTML to PDF API PDFCROWD

| | | | |
|---|---|---|---|
| SPGP0I02070 | XP_018223129.1 | 99 | 0 |
| SPGP0O00670 | XP_018221699.1 | 99 | 0 |
| SPGP0I02490 | XP_018223088.1 | 100 | 0 |
| SPGP0AR00290 | XP_018219588.1 | 99 | 0 |
| SPGP0Itrna16V | DI49_1074 | 100 | 0 |
| SPGP0B06160 | XP_018224044.1 | 97 | 0 |
| SPGP0F00900 | XP_018220270.1 | 99 | 0 |
| SPGP0F01020 | XP_018220258.1 | 99 | 0 |
| SPGP0C03900 | XP_018220753.1 | 100 | 0 |
| SPGP0O01500 | XP_018221777.1 | 98 | 0 |
| SPGP0B03170 | XP_018223760.1 | 99 | 0 |
| SPGP0E00630 | XP_018222434.1 | 99 | 0 |
| SPGP0E00860 | XP_018222411.1 | 99 | 0 |
| SPGP0O00540 | XP_018221686.1 | 99 | 0 |
| SPGP0S00130 | XP_018219837.1 | 87 | 2 |
| SPGP0E01520 | XP_018222342.1 | 99 | 0 |
| SPGP0B02310 | XP_018223675.1 | 99 | 0 |
| SPGP0Q01820 | XP_018222681.1 | 99 | 0 |
| SPGP0F01890 | XP_018220176.1 | 99 | 0 |
| SPGP0B05840 | XP_018224014.1 | 99 | 0 |
| SPGP0C03770 | XP_018220740.1 | 100 | 0 |
| SPGP0W00310 | XP_018221480.1 | 99 | 0 |
| SPGP0H01250 | XP_018219310.1 | 99 | 0 |
| SPGP0C02960 | XP_018220664.1 | 99 | 0 |
| SPGP0H01330 | XP_018219318.1 | 100 | 0 |
| SPGP0F02540 | XP_018220112.1 | 99 | 0 |
| SPGP0M00850 | XP_018221207.1 | 99 | 0 |
| SPGP0P00400 | XP_018221103.1 | 99 | 0 |
| SPGP0B05800 | XP_018224010.1 | 99 | 0 |
| SPGP0AG00280 | XP_018223343.1 | 99 | 0 |
| SPGP0I00750 | XP_018223258.1 | 99 | 0 |
| SPGP0O02850 | XP_018221907.1 | 99 | 0 |
| SPGP0Q01560 | XP_018222708.1 | 100 | 0 |
| SPGP0L01690 | XP_018219008.1 | 99 | 0 |
| SPGP0B00650 | XP_018223511.1 | 99 | 0 |
| SPGP0P00990 | XP_018221045.1 | 99 | 0 |
| SPGP0I00700 | XP_018223263.1 | 99 | 0 |
| SPGP0B02900 | XP_018223734.1 | 99 | 0 |
| SPGP0Q01580 | XP_018222706.1 | 99 | 0 |
| SPGP0P02230 | XP_018220927.1 | 99 | 0 |

Create PDF in your applications with the Pdfcrowd HTML to PDF API PDFCROWD

| | | | |
|---|---|---|---|
| SPGP0P01410 | XP_018221006.1 | 99 | 0 |
| SPGP0C01910 | XP_018220559.1 | 99 | 0 |
| SPGP0W00160 | XP_018221465.1 | 99 | 0 |
| SPGP0E01930 | XP_018222301.1 | 99 | 0 |
| SPGP0O03100 | XP_018221930.1 | 99 | 0 |
| SPGP0B02710 | XP_018223714.1 | 99 | 0 |
| SPGP0H03780 | XP_018219557.1 | 99 | 0 |
| SPGP0S01550 | XP_018219697.1 | 99 | 0 |
| SPGP0L01960 | XP_018218981.1 | 99 | 0 |
| SPGP0M03400 | XP_018221451.1 | 99 | 0 |
| SPGP0P01350 | XP_018221011.1 | 93 | 7 |
| SPGP0P01480 | XP_018221000.1 | 99 | 0 |
| SPGP0P02210 | XP_018220929.1 | 99 | 0 |
| SPGP0P01150 | XP_018221030.1 | 100 | 0 |
| SPGP0E02710 | XP_018222224.1 | 99 | 0 |
| SPGP0F01620 | XP_018220200.1 | 99 | 0 |
| SPGP0M02050 | XP_018221323.1 | 99 | 0 |
| SPGP0M02590 | XP_018221375.1 | 99 | 0 |
| SPGP0B03870 | XP_018223828.1 | 99 | 0 |
| SPGP0C03100 | XP_018220678.1 | 99 | 0 |
| SPGP0M02500 | XP_018221366.1 | 99 | 0 |
| SPGP0H00670 | XP_018219254.1 | 100 | 0 |
| SPGP0O01020 | XP_018221732.1 | 99 | 0 |
| SPGP0E01250 | XP_018222371.1 | 99 | 0 |
| SPGP0B04520 | XP_018223892.1 | 99 | 0 |
| SPGP0C00450 | XP_018220414.1 | 98 | 0 |
| SPGP0W01690 | XP_018221613.1 | 99 | 0 |
| SPGP0L00900 | XP_018219084.1 | 99 | 0 |
| SPGP0P01730 | XP_018220976.1 | 98 | 1 |
| SPGP0M01700 | XP_018221288.1 | 99 | 0 |
| SPGP0H03920 | XP_018218747.1 | 99 | 0 |
| SPGP0F00850 | XP_018220275.1 | 99 | 0 |
| SPGP0W00500 | XP_018221499.1 | 99 | 1 |
| SPGP0L02820 | XP_018218899.1 | 99 | 0 |
| SPGP0C01850 | XP_018220553.1 | 99 | 0 |
| SPGP0O01320 | XP_018221759.1 | 99 | 0 |
| SPGP0I01740 | XP_018223161.1 | 99 | 0 |
| SPGP0M00900 | XP_018221211.1 | 99 | 0 |
| SPGP0P00280 | XP_018221115.1 | 99 | 0 |
| SPGP0L03440 | XP_018218839.1 | 99 | 0 |

Create PDF in your applications with the Pdfcrowd HTML to PDF API PDFCROWD

| | | | |
|---|---|---|---|
| SPGP0S01540 | XP_018219698.1 | 99 | 0 |
| SPGP0W01430 | XP_018221588.1 | 99 | 0 |
| SPGP0S00100 | XP_018219840.1 | 99 | 0 |
| SPGP0B02020 | XP_018223648.1 | 100 | 0 |
| SPGP0E00400 | XP_018222457.1 | 99 | 0 |
| SPGP0W01340 | XP_018221579.1 | 99 | 0 |
| SPGP0I00950 | XP_018223239.1 | 99 | 0 |
| SPGP0AI00200 | XP_018224061.1 | 99 | 0 |
| SPGP0S02390 | XP_018219614.1 | 99 | 0 |
| SPGP0S00940 | XP_018219758.1 | 99 | 0 |
| SPGP0C04730 | XP_018220832.1 | 99 | 0 |
| SPGP0P02820 | XP_018220868.1 | 99 | 0 |
| SPGP0M02080 | XP_018221326.1 | 99 | 0 |
| SPGP0I01020 | XP_018223232.1 | 99 | 0 |
| SPGP0AG00230 | XP_018223338.1 | 99 | 0 |
| SPGP0AQ00440 | XP_018221982.1 | 99 | 0 |
| SPGP0C02200 | XP_018220588.1 | 99 | 0 |
| SPGP0AG00540 | XP_018223368.1 | 99 | 0 |
| SPGP0W01840 | XP_018221628.1 | 99 | 1 |
| SPGP0B03710 | XP_018223812.1 | 99 | 0 |
| SPGP0F00460 | XP_018220314.1 | 99 | 0 |
| SPGP0B00620 | XP_018223508.1 | 99 | 0 |
| SPGP0F03970 | XP_018219974.1 | 99 | 0 |
| SPGP0AF00550 | XP_018222550.1 | 99 | 0 |
| SPGP0B05020 | XP_018223937.1 | 99 | 0 |
| SPGP0I03090 | XP_018223034.1 | 99 | 0 |
| SPGP0M00760 | XP_018221199.1 | 99 | 0 |
| SPGP0F00470 | XP_018220313.1 | 99 | 0 |
| SPGP0L02890 | XP_018218892.1 | 99 | 0 |
| SPGP0B02620 | XP_018223706.1 | 99 | 0 |
| SPGP0W00560 | XP_018221505.1 | 99 | 0 |
| SPGP0F02900 | XP_018220078.1 | 99 | 0 |
| SPGP0F02430 | XP_018220123.1 | 99 | 0 |
| SPGP0L02430 | XP_018218935.1 | 99 | 0 |
| SPGP0L02310 | XP_018218947.1 | 99 | 0 |
| SPGP0I01490 | XP_018223186.1 | 99 | 0 |
| SPGP0F00980 | XP_018220262.1 | 99 | 0 |
| SPGP0S01760 | XP_018219676.1 | 100 | 0 |
| SPGP0W00630 | XP_018221511.1 | 99 | 0 |
| SPGP0F02250 | XP_018220141.1 | 99 | 0 |

Create PDF in your applications with the Pdfcrowd HTML to PDF API PDFCROWD

| | | | |
|---|---|---|---|
| SPGP0O01380 | XP_018221765.1 | 99 | 0 |
| SPGP0B01060 | XP_018223553.1 | 99 | 0 |
| SPGP0B04770 | XP_018223915.1 | 99 | 0 |
| SPGP0L01110 | XP_018219064.1 | 99 | 1 |
| SPGP0I03350 | XP_018223010.1 | 99 | 0 |
| SPGP0L03610 | XP_018218823.1 | 99 | 0 |
| SPGP0M02350 | XP_018221351.1 | 99 | 0 |
| SPGP0M01410 | XP_018221260.1 | 99 | 0 |
| SPGP0AR00390 | XP_018219597.1 | 100 | 0 |
| SPGP0F03920 | XP_018219979.1 | 99 | 0 |
| SPGP0E00350 | XP_018222462.1 | 99 | 0 |
| SPGP0E03620 | XP_018222137.1 | 99 | 0 |
| SPGP0AF00750 | XP_018222530.1 | 99 | 0 |
| SPGP0L01750 | XP_018219002.1 | 99 | 0 |
| SPGP0H02760 | XP_018219457.1 | 99 | 0 |
| SPGP0F02880 | XP_018220080.1 | 99 | 0 |
| SPGP0W00760 | XP_018221524.1 | 99 | 0 |
| SPGP0C00630 | XP_018220431.1 | 99 | 0 |
| SPGP0P01900 | XP_018220960.1 | 99 | 0 |
| SPGP0L02740 | XP_018218907.1 | 99 | 0 |
| SPGP0B01460 | XP_018223593.1 | 99 | 0 |
| SPGP0B05360 | XP_018223968.1 | 99 | 0 |
| SPGP0B04920 | XP_018223927.1 | 99 | 0 |
| SPGP0M02260 | XP_018221342.1 | 99 | 0 |
| SPGP0L02930 | XP_018218888.1 | 100 | 0 |
| SPGP0O02360 | XP_018221860.1 | 99 | 0 |
| SPGP0O01710 | XP_018221797.1 | 99 | 0 |
| SPGP0O01240 | XP_018221751.1 | 99 | 0 |
| SPGP0C00930 | XP_018220461.1 | 99 | 0 |
| SPGP0E03590 | XP_018222140.1 | 99 | 0 |
| SPGP0Etrna11W | DI49_2157 | 100 | 0 |
| SPGP0C02750 | XP_018220643.1 | 99 | 0 |
| SPGP0B05140 | XP_018223947.1 | 99 | 0 |
| SPGP0L00150 | XP_018219156.1 | 99 | 0 |
| SPGP0B03790 | XP_018223820.1 | 100 | 0 |
| SPGP0E02060 | XP_018222288.1 | 99 | 0 |
| SPGP0L00400 | XP_018219132.1 | 99 | 0 |
| SPGP0L01720 | XP_018219005.1 | 99 | 0 |
| SPGP0O02440 | XP_018221868.1 | 99 | 0 |
| SPGP0B03880 | XP_018223829.1 | 100 | 0 |

Create PDF in your applications with the Pdfcrowd HTML to PDF API PDFCROWD

| | | | |
|---|---|---|---|
| SPGP0F02180 | XP_018220148.1 | 99 | 0 |
| SPGP0P00430 | XP_018221100.1 | 99 | 0 |
| SPGP0I02560 | XP_018223081.1 | 99 | 0 |
| SPGP0S00890 | XP_018219763.1 | 99 | 0 |
| SPGP0B05220 | XP_018223954.1 | 99 | 0 |
| SPGP0E04270 | XP_018222072.1 | 99 | 0 |
| SPGP0S02150 | XP_018219637.1 | 99 | 0 |
| SPGP0AR00110 | XP_018219570.1 | 99 | 0 |
| SPGP0AQ00390 | XP_018221977.1 | 99 | 0 |
| SPGP0C02080 | XP_018220576.1 | 98 | 2 |
| SPGP0B00510 | XP_018223499.1 | 99 | 0 |
| SPGP0AL00360 | XP_018219898.1 | 99 | 0 |
| SPGP0M03020 | XP_018221416.1 | 99 | 0 |
| SPGP0E03190 | XP_018222177.1 | 100 | 0 |
| SPGP0C00200 | XP_018220390.1 | 99 | 0 |
| SPGP0I00360 | XP_018223298.1 | 99 | 0 |
| SPGP0L02800 | XP_018218901.1 | 99 | 0 |
| SPGP0E02190 | XP_018222276.1 | 100 | 0 |
| SPGP0L00120 | XP_018219159.1 | 99 | 0 |
| SPGP0C04100 | XP_018220772.1 | 99 | 0 |
| SPGP0B05230 | XP_018223955.1 | 99 | 0 |
| SPGP0AF00500 | XP_018222555.1 | 99 | 0 |
| SPGP0E01840 | XP_018222310.1 | 99 | 0 |
| SPGP0I00300 | XP_018223304.1 | 98 | 1 |
| SPGP0Q02050 | XP_018222660.1 | 99 | 0 |
| SPGP0Strna18P | DI49_4699 | 100 | 0 |
| SPGP0B03320 | XP_018223774.1 | 100 | 0 |
| SPGP0C01550 | XP_018220524.1 | 99 | 0 |
| SPGP0E02500 | XP_018222246.1 | 99 | 0 |
| SPGP0B03100 | XP_018223753.1 | 99 | 0 |
| SPGP0AX00230 | XP_018219871.1 | 99 | 0 |
| SPGP0S02230 | XP_018219629.1 | 99 | 0 |
| SPGP0B05830 | XP_018224013.1 | 100 | 0 |
| SPGP0C00850 | XP_018220453.1 | 99 | 0 |
| SPGP0E03900 | XP_018222109.1 | 99 | 0 |
| SPGP0P00610 | XP_018221083.1 | 99 | 0 |
| SPGP0M00220 | XP_018221147.1 | 99 | 0 |
| SPGP0E03450 | XP_018222153.1 | 99 | 0 |
| SPGP0M00560 | XP_018221180.1 | 99 | 0 |
| SPGP0B03840 | XP_018223825.1 | 99 | 0 |

Create PDF in your applications with the Pdfcrowd HTML to PDF API PDFCROWD

| | | | |
|---|---|---|---|
| SPGP0Q00530 | XP_018222808.1 | 100 | 0 |
| SPGP0AF00240 | XP_018222578.1 | 99 | 0 |
| SPGP0P00120 | XP_018221130.1 | 99 | 0 |
| SPGP0I00550 | XP_018223279.1 | 99 | 0 |
| SPGP0H02600 | XP_018219442.1 | 99 | 0 |
| SPGP0P00690 | XP_018221075.1 | 99 | 0 |
| SPGP0B04540 | XP_018223894.1 | 99 | 0 |
| SPGP0F03670 | XP_018220003.1 | 99 | 0 |
| SPGP0M01170 | XP_018221237.1 | 99 | 0 |
| SPGP0I03220 | XP_018223022.1 | 99 | 0 |
| SPGP0B02880 | XP_018223732.1 | 99 | 0 |
| SPGP0H01800 | XP_018219365.1 | 99 | 0 |
| SPGP0F01880 | XP_018220177.1 | 100 | 0 |
| SPGP0H01550 | XP_018219340.1 | 99 | 0 |
| SPGP0O02830 | XP_018221905.1 | 99 | 0 |
| SPGP0Q00650 | XP_018222797.1 | 100 | 0 |
| SPGP0Q01990 | XP_018222665.1 | 99 | 0 |
| SPGP0AG00620 | XP_018223376.1 | 99 | 0 |
| SPGP0AQ00410 | XP_018221979.1 | 99 | 0 |
| SPGP0Q00860 | XP_018222776.1 | 99 | 1 |
| SPGP0H01040 | XP_018219291.1 | 99 | 0 |
| SPGP0F02290 | XP_018220137.1 | 99 | 0 |
| SPGP0Q00550 | XP_018222806.1 | 99 | 0 |
| SPGP0B05580 | XP_018223988.1 | 99 | 0 |
| SPGP0I02190 | XP_018223118.1 | 99 | 0 |
| SPGP0B05420 | XP_018223973.1 | 99 | 0 |
| SPGP0F02390 | XP_018220127.1 | 99 | 0 |
| SPGP0S01670 | XP_018219685.1 | 99 | 0 |
| SPGP0M00710 | XP_018221195.1 | 99 | 0 |
| SPGP0H02030 | XP_018219388.1 | 99 | 0 |
| SPGP0S02210 | XP_018219631.1 | 99 | 0 |
| SPGP0I02550 | XP_018223082.1 | 99 | 0 |
| SPGP0O02140 | XP_018221838.1 | 99 | 0 |
| SPGP0I01530 | XP_018223182.1 | 99 | 0 |
| SPGP0H00100 | XP_018219199.1 | 97 | 0 |
| SPGP0AQ00480 | XP_018221986.1 | 98 | 0 |
| SPGP0M03000 | XP_018221414.1 | 100 | 0 |
| SPGP0O00380 | XP_018221671.1 | 88 | 0 |
| SPGP0E02220 | XP_018222273.1 | 99 | 0 |
| SPGP0O00430 | XP_018221676.1 | 99 | 0 |

Create PDF in your applications with the Pdfcrowd HTML to PDF API PDFCROWD

| | | | |
|---|---|---|---|
| SPGP0H02460 | XP_018219428.1 | 99 | 0 |
| SPGP0I03610 | XP_018222987.1 | 99 | 0 |
| SPGP0I03510 | XP_018222997.1 | 99 | 0 |
| SPGP0F00580 | XP_018220302.1 | 100 | 0 |
| SPGP0B01970 | XP_018223643.1 | 99 | 0 |
| SPGP0B00680 | XP_018223514.1 | 99 | 0 |
| SPGP0AL00110 | XP_018219875.1 | 99 | 0 |
| SPGP0O01520 | XP_018221779.1 | 99 | 0 |
| SPGP0F00620 | XP_018220298.1 | 99 | 0 |
| SPGP0AI00740 | XP_018224114.1 | 99 | 1 |
| SPGP0S00150 | XP_018219835.1 | 99 | 0 |
| SPGP0I02340 | XP_018223103.1 | 99 | 0 |
| SPGP0F02110 | XP_018220155.1 | 99 | 0 |
| SPGP0O00650 | XP_018221697.1 | 99 | 0 |
| SPGP0E02390 | XP_018222256.1 | 99 | 0 |
| SPGP0Q02490 | XP_018222616.1 | 99 | 0 |
| SPGP0B03450 | XP_018223787.1 | 99 | 0 |
| SPGP0B02070 | XP_018223653.1 | 99 | 0 |
| SPGP0AI00360 | XP_018224077.1 | 99 | 0 |
| SPGP0Q01210 | XP_018222741.1 | 99 | 0 |
| SPGP0F02510 | XP_018220115.1 | 99 | 0 |
| SPGP0B03500 | XP_018223791.1 | 99 | 0 |
| SPGP0C03710 | XP_018220735.1 | 99 | 0 |
| SPGP0W01800 | XP_018221624.1 | 100 | 0 |
| SPGP0I00720 | XP_018223261.1 | 99 | 0 |
| SPGP0O01090 | XP_018221738.1 | 99 | 0 |
| SPGP0F02130 | XP_018220153.1 | 99 | 0 |
| SPGP0E02880 | XP_018222207.1 | 99 | 0 |
| SPGP0AG00310 | XP_018223346.1 | 99 | 0 |
| SPGP0F01120 | XP_018220248.1 | 99 | 0 |
| SPGP0P01690 | XP_018220980.1 | 99 | 0 |
| SPGP0H03100 | XP_018219491.1 | 100 | 0 |
| SPGP0B00670 | XP_018223513.1 | 99 | 0 |
| SPGP0L01410 | XP_018219035.1 | 100 | 0 |
| SPGP0P01140 | XP_018221031.1 | 99 | 0 |
| SPGP0L02200 | XP_018218958.1 | 99 | 0 |
| SPGP0E00330 | XP_018222464.1 | 99 | 0 |
| SPGP0C03360 | XP_018220702.1 | 99 | 0 |
| SPGP0W01590 | XP_018221603.1 | 99 | 0 |
| SPGP0S01810 | XP_018219671.1 | 99 | 0 |

Create PDF in your applications with the Pdfcrowd HTML to PDF API PDFCROWD

| | | | |
|---|---|---|---|
| SPGP0E00370 | XP_018222460.1 | 99 | 0 |
| SPGP0P02110 | XP_018220939.1 | 99 | 0 |
| SPGP0F01870 | XP_018220178.1 | 99 | 0 |
| SPGP0H01000 | XP_018219287.1 | 99 | 0 |
| SPGP0E03910 | XP_018222108.1 | 99 | 0 |
| SPGP0H02110 | XP_018219396.1 | 99 | 0 |
| SPGP0C02730 | XP_018220641.1 | 99 | 0 |
| SPGP0I02330 | XP_018223104.1 | 99 | 0 |
| SPGP0H01260 | XP_018219311.1 | 100 | 0 |
| SPGP0F01040 | XP_018220256.1 | 99 | 0 |
| SPGP0F04020 | XP_018219969.1 | 99 | 0 |
| SPGP0F02600 | XP_018220108.1 | 99 | 0 |
| SPGP0F00250 | XP_018220334.1 | 99 | 0 |
| SPGP0Q01220 | XP_018222740.1 | 100 | 0 |
| SPGP0C02140 | XP_018220582.1 | 99 | 0 |
| SPGP0M02030 | XP_018221321.1 | 99 | 0 |
| SPGP0W00150 | XP_018221464.1 | 96 | 0 |
| SPGP0F01310 | XP_018220229.1 | 100 | 0 |
| SPGP0M02390 | XP_018221355.1 | 99 | 0 |
| SPGP0B04690 | XP_018223908.1 | 99 | 0 |
| SPGP0H00850 | XP_018219272.1 | 99 | 0 |
| SPGP0O02380 | XP_018221862.1 | 99 | 0 |
| SPGP0O02650 | XP_018221888.1 | 99 | 0 |
| SPGP0O02120 | XP_018221836.1 | 100 | 0 |
| SPGP0L00290 | XP_018219143.1 | 99 | 0 |
| SPGP0C00120 | XP_018220383.1 | 99 | 0 |
| SPGP0AI00380 | XP_018224079.1 | 99 | 0 |
| SPGP0Q01870 | XP_018222676.1 | 99 | 0 |
| SPGP0F01350 | XP_018220225.1 | 100 | 0 |
| SPGP0F02410 | XP_018220125.1 | 99 | 0 |
| SPGP0F03790 | XP_018219992.1 | 99 | 0 |
| SPGP0H02160 | XP_018219400.1 | 99 | 0 |
| SPGP0L01160 | XP_018219059.1 | 99 | 0 |
| SPGP0F00830 | XP_018220277.1 | 99 | 0 |
| SPGP0AR00400 | XP_018219598.1 | 99 | 0 |
| SPGP0I01950 | XP_018223139.1 | 99 | 0 |
| SPGP0I03910 | XP_018222958.1 | 98 | 0 |
| SPGP0E02130 | XP_018222281.1 | 99 | 0 |
| SPGP0Etrna22F | DI49_1939 | 100 | 0 |
| SPGP0L01310 | XP_018219045.1 | 98 | 1 |

Create PDF in your applications with the Pdfcrowd HTML to PDF API PDFCROWD

| | | | |
|---|---|---|---|
| SPGP0I01040 | XP_018223230.1 | 99 | 0 |
| SPGP0Q01570 | XP_018222707.1 | 99 | 0 |
| SPGP0F01100 | XP_018220250.1 | 99 | 0 |
| SPGP0E03020 | XP_018222193.1 | 99 | 0 |
| SPGP0AL00160 | XP_018219880.1 | 99 | 0 |
| SPGP0S00710 | XP_018219780.1 | 99 | 0 |
| SPGP0F00860 | XP_018220274.1 | 99 | 0 |
| SPGP0W01410 | XP_018221586.1 | 99 | 0 |
| SPGP0AI00220 | XP_018224063.1 | 99 | 0 |
| SPGP0I02050 | XP_018223131.1 | 100 | 0 |
| SPGP0P02800 | XP_018220870.1 | 99 | 0 |
| SPGP0L00950 | XP_018219080.1 | 99 | 0 |
| SPGP0B05000 | XP_018223935.1 | 99 | 0 |
| SPGP0B02380 | XP_018223682.1 | 99 | 0 |
| SPGP0M01220 | XP_018221242.1 | 99 | 0 |
| SPGP0H03710 | XP_018219550.1 | 99 | 0 |
| SPGP0AT00240 | XP_018219185.1 | 98 | 0 |
| SPGP0F03360 | XP_018220033.1 | 100 | 0 |
| SPGP0M02530 | XP_018221369.1 | 99 | 0 |
| SPGP0Ntrna10R | DI49_2918 | 100 | 0 |
| SPGP0M01140 | XP_018221234.1 | 99 | 0 |
| SPGP0C00420 | XP_018220410.1 | 98 | 0 |
| SPGP0H00830 | XP_018219270.1 | 99 | 0 |
| SPGP0B01240 | XP_018223571.1 | 99 | 0 |
| SPGP0P02080 | XP_018220942.1 | 99 | 0 |
| SPGP0F00210 | XP_018220338.1 | 99 | 0 |
| SPGP0C01360 | XP_018220504.1 | 99 | 0 |
| SPGP0AF01030 | XP_018222505.1 | 99 | 0 |
| SPGP0C03190 | XP_018220687.1 | 99 | 0 |
| SPGP0C00220 | XP_018220392.1 | 100 | 0 |
| SPGP0E03880 | XP_018222111.1 | 99 | 0 |
| SPGP0H00790 | XP_018219266.1 | 99 | 0 |
| SPGP0B04080 | XP_018223849.1 | 99 | 0 |
| SPGP0P02870 | XP_018220863.1 | 99 | 0 |
| SPGP0I01600 | XP_018223175.1 | 99 | 0 |
| SPGP0P00760 | XP_018221068.1 | 100 | 0 |
| SPGP0L02380 | XP_018218940.1 | 99 | 0 |
| SPGP0F02680 | XP_018220100.1 | 99 | 0 |
| SPGP0C03960 | XP_018220758.1 | 99 | 0 |
| SPGP0B01170 | XP_018223564.1 | 99 | 0 |

Create PDF in your applications with the Pdfcrowd HTML to PDF API PDFCROWD

| | | | |
|---|---|---|---|
| SPGP0F00450 | XP_018220315.1 | 99 | 0 |
| SPGP0AF00960 | XP_018222511.1 | 99 | 0 |
| SPGP0B01090 | XP_018223556.1 | 99 | 0 |
| SPGP0I03190 | XP_018223025.1 | 99 | 0 |
| SPGP0L00200 | XP_018219152.1 | 99 | 0 |
| SPGP0L02370 | XP_018218941.1 | 99 | 0 |
| SPGP0AX00170 | XP_018219865.1 | 99 | 0 |
| SPGP0O00280 | XP_018221661.1 | 99 | 0 |
| SPGP0O02220 | XP_018221846.1 | 99 | 0 |
| SPGP0AL00730 | XP_018219933.1 | 99 | 0 |
| SPGP0F04220 | XP_018219949.1 | 99 | 0 |
| SPGP0AL00620 | XP_018219923.1 | 99 | 0 |
| SPGP0DV00100 | XP_018220218.1 | 100 | 0 |
| SPGP0B04750 | XP_018223914.1 | 99 | 0 |
| SPGP0I03120 | XP_018223031.1 | 99 | 0 |
| SPGP0H02180 | XP_018219401.1 | 99 | 0 |
| SPGP0P00880 | XP_018221056.1 | 99 | 0 |
| SPGP0C00600 | XP_018220428.1 | 99 | 0 |
| SPGP0M01430 | XP_018221262.1 | 99 | 0 |
| SPGP0Q02000 | XP_018222664.1 | 99 | 0 |
| SPGP0L01770 | XP_018219000.1 | 99 | 0 |
| SPGP0E03990 | XP_018222100.1 | 99 | 0 |
| SPGP0Q00350 | XP_018222826.1 | 99 | 0 |
| SPGP0AF00290 | XP_018222573.1 | 99 | 0 |
| SPGP0P02120 | XP_018220938.1 | 99 | 0 |
| SPGP0C02770 | XP_018220645.1 | 99 | 0 |
| SPGP0E02420 | XP_018222253.1 | 100 | 0 |
| SPGP0I00230 | XP_018223311.1 | 99 | 0 |
| SPGP0P02040 | XP_018220946.1 | 99 | 0 |
| SPGP0Q02260 | XP_018222639.1 | 99 | 0 |
| SPGP0B03410 | XP_018223783.1 | 99 | 0 |
| SPGP0C01440 | XP_018220513.1 | 99 | 0 |
| SPGP0O01260 | XP_018221753.1 | 99 | 0 |
| SPGP0C00950 | XP_018220463.1 | 99 | 0 |
| SPGP0P03030 | XP_018220849.1 | 99 | 0 |
| SPGP0C00710 | XP_018220438.1 | 99 | 0 |
| SPGP0O02840 | XP_018221906.1 | 99 | 0 |
| SPGP0O00730 | XP_018221705.1 | 99 | 0 |
| SPGP0F04130 | XP_018219958.1 | 99 | 0 |
| SPGP0B01050 | XP_018223552.1 | 99 | 0 |

Create PDF in your applications with the Pdfcrowd HTML to PDF API PDFCROWD

| | | | |
|---|---|---|---|
| SPGP0S01290 | XP_018219723.1 | 99 | 1 |
| SPGP0L03140 | XP_018218867.1 | 100 | 0 |
| SPGP0BB00160 | XP_018219850.1 | 98 | 0 |
| SPGP0E00780 | XP_018222419.1 | 99 | 0 |
| SPGP0E02510 | XP_018222245.1 | 100 | 0 |
| SPGP0E00990 | XP_018222398.1 | 99 | 0 |
| SPGP0S00310 | XP_018219820.1 | 99 | 0 |
| SPGP0B01720 | XP_018223618.1 | 99 | 0 |
| SPGP0B04980 | XP_018223933.1 | 99 | 0 |
| SPGP0P02630 | XP_018220887.1 | 99 | 0 |
| SPGP0E03630 | XP_018222136.1 | 99 | 0 |
| SPGP0S01740 | XP_018219678.1 | 99 | 0 |
| SPGP0B01920 | XP_018223638.1 | 99 | 0 |
| SPGP0AL00340 | XP_018219896.1 | 99 | 0 |
| SPGP0C02720 | XP_018220640.1 | 99 | 0 |
| SPGP0I03760 | XP_018222972.1 | 99 | 0 |
| SPGP0B02750 | XP_018223718.1 | 99 | 0 |
| SPGP0C04790 | XP_018220838.1 | 99 | 0 |
| SPGP0F03340 | XP_018220035.1 | 99 | 0 |
| SPGP0P01340 | XP_018221012.1 | 99 | 1 |
| SPGP0S02020 | XP_018219650.1 | 99 | 0 |
| SPGP0W00730 | XP_018221521.1 | 99 | 0 |
| SPGP0F01760 | XP_018220188.1 | 99 | 0 |
| SPGP0F02100 | XP_018220156.1 | 99 | 0 |
| SPGP0B04120 | XP_018223853.1 | 99 | 0 |
| SPGP0S01030 | XP_018219749.1 | 99 | 0 |
| SPGP0E03610 | XP_018222138.1 | 99 | 0 |
| SPGP0AL00480 | XP_018219910.1 | 99 | 0 |
| SPGP0F01160 | XP_018220244.1 | 99 | 0 |
| SPGP0I01550 | XP_018223180.1 | 99 | 0 |
| SPGP0F03770 | XP_018219993.1 | 99 | 0 |
| SPGP0AI00310 | XP_018224072.1 | 99 | 0 |
| SPGP0Q01320 | XP_018222731.1 | 99 | 0 |
| SPGP0P01670 | XP_018220982.1 | 99 | 0 |
| SPGP0BB00170 | XP_018219851.1 | 99 | 0 |
| SPGP0H00520 | XP_018219240.1 | 99 | 0 |
| SPGP0AX00250 | XP_018219873.1 | 99 | 0 |
| SPGP0P00670 | XP_018221077.1 | 98 | 2 |
| SPGP0B06050 | XP_018224035.1 | 98 | 1 |

Create PDF in your applications with the Pdfcrowd HTML to PDF API PDFCROWD

| | | | |
|---|---|---|---|
| SPGP0I00270 | XP_018223307.1 | 99 | 0 |
| SPGP0M00200 | XP_018221145.1 | 99 | 0 |
| SPGP0O00710 | XP_018221703.1 | 99 | 0 |
| SPGP0I02980 | XP_018223042.1 | 99 | 1 |
| SPGP0Q00510 | XP_018222810.1 | 99 | 0 |
| SPGP0I01180 | XP_018223216.1 | 99 | 0 |
| SPGP0L02260 | XP_018218952.1 | 100 | 0 |
| SPGP0P01180 | XP_018221028.1 | 99 | 0 |
| SPGP0W01060 | XP_018221552.1 | 99 | 0 |
| SPGP0AI00110 | XP_018224052.1 | 99 | 0 |
| SPGP0C02970 | XP_018220665.1 | 99 | 0 |
| SPGP0B02970 | XP_018223741.1 | 99 | 0 |
| SPGP0E02680 | XP_018222227.1 | 99 | 0 |
| SPGP0AI00140 | XP_018224055.1 | 99 | 0 |
| SPGP0AI00410 | XP_018224081.1 | 99 | 0 |
| SPGP0Q01660 | XP_018222698.1 | 99 | 0 |
| SPGP0H00440 | XP_018219232.1 | 99 | 0 |
| SPGP0C02150 | XP_018220583.1 | 99 | 0 |
| SPGP0B01580 | XP_018223605.1 | 100 | 0 |
| SPGP0B01550 | XP_018223602.1 | 99 | 0 |
| SPGP0AQ00350 | XP_018221973.1 | 99 | 0 |
| SPGP0F03170 | XP_018220052.1 | 99 | 0 |
| SPGP0L03760 | XP_018218807.1 | 99 | 0 |
| SPGP0F03680 | XP_018220002.1 | 99 | 0 |
| SPGP0AR00250 | XP_018219584.1 | 99 | 0 |
| SPGP0E01790 | XP_018222315.1 | 99 | 0 |
| SPGP0H01290 | XP_018219314.1 | 99 | 0 |
| SPGP0H02720 | XP_018219453.1 | 100 | 0 |
| SPGP0Q00180 | XP_018222840.1 | 99 | 0 |
| SPGP0H00190 | XP_018219207.1 | 99 | 0 |
| SPGP0H00660 | XP_018219253.1 | 99 | 0 |
| SPGP0C03260 | XP_018220693.1 | 99 | 0 |
| SPGP0L03160 | XP_018218865.1 | 99 | 0 |
| SPGP0Q00170 | XP_018222841.1 | 99 | 0 |
| SPGP0S00500 | XP_018219801.1 | 99 | 0 |
| SPGP0B01160 | XP_018223563.1 | 99 | 0 |
| SPGP0P02520 | XP_018220898.1 | 100 | 0 |
| SPGP0O00410 | XP_018221674.1 | 98 | 1 |
| SPGP0M01330 | XP_018221252.1 | 99 | 0 |
| SPGP0M01840 | XP_018221303.1 | 99 | 0 |

Create PDF in your applications with the Pdfcrowd HTML to PDF API PDFCROWD

| | | | |
|---|---|---|---|
| SPGP0C03270 | XP_018220694.1 | 99 | 0 |
| SPGP0H01440 | XP_018219329.1 | 99 | 0 |
| SPGP0B02660 | XP_018223710.1 | 99 | 0 |
| SPGP0P01490 | XP_018220999.1 | 99 | 0 |
| SPGP0B05610 | XP_018223991.1 | 99 | 0 |
| SPGP0AF00920 | XP_018222514.1 | 97 | 2 |
| SPGP0F00360 | XP_018220323.1 | 99 | 0 |
| SPGP0W00120 | XP_018221461.1 | 96 | 1 |
| SPGP0S00700 | XP_018219781.1 | 99 | 0 |
| SPGP0M01610 | XP_018221280.1 | 97 | 1 |
| SPGP0S00170 | XP_018219833.1 | 99 | 0 |
| SPGP0I01700 | XP_018223165.1 | 99 | 0 |
| SPGP0S02260 | XP_018219626.1 | 99 | 0 |
| SPGP0C02860 | XP_018220654.1 | 99 | 0 |
| SPGP0Q00700 | XP_018222792.1 | 99 | 0 |
| SPGP0W00260 | XP_018221475.1 | 99 | 0 |
| SPGP0E01170 | XP_018222379.1 | 99 | 0 |
| SPGP0S01980 | XP_018219654.1 | 99 | 0 |
| SPGP0B05770 | XP_018224007.1 | 99 | 0 |
| SPGP0Htrna23P | DI49_5158 | 99 | 0 |
| SPGP0F02820 | XP_018220086.1 | 99 | 0 |
| SPGP0B04390 | XP_018223880.1 | 99 | 0 |
| SPGP0F02530 | XP_018220113.1 | 99 | 0 |
| SPGP0B03970 | XP_018223838.1 | 99 | 0 |
| SPGP0F01170 | XP_018220243.1 | 99 | 0 |
| SPGP0I02350 | XP_018223102.1 | 99 | 0 |
| SPGP0AR00210 | XP_018219580.1 | 99 | 0 |
| SPGP0O02810 | XP_018221904.1 | 99 | 0 |
| SPGP0P01660 | XP_018220983.1 | 99 | 0 |
| SPGP0P00410 | XP_018221102.1 | 99 | 0 |
| SPGP0L00720 | XP_018219101.1 | 99 | 1 |
| SPGP0I02960 | XP_018223044.1 | 99 | 0 |
| SPGP0H02370 | XP_018219420.1 | 99 | 0 |
| SPGP0AI00550 | XP_018224095.1 | 99 | 0 |
| SPGP0F02010 | XP_018220165.1 | 99 | 0 |
| SPGP0AT00230 | XP_018219186.1 | 98 | 0 |
| SPGP0M02040 | XP_018221322.1 | 99 | 0 |
| SPGP0AL00140 | XP_018219878.1 | 99 | 0 |
| SPGP0AL00750 | XP_018219935.1 | 91 | 1 |
| SPGP0E02340 | XP_018222261.1 | 99 | 0 |

Create PDF in your applications with the Pdfcrowd HTML to PDF API PDFCROWD

| | | | |
|---|---|---|---|
| SPGP0M01440 | XP_018221263.1 | 99 | 0 |
| SPGP0H02920 | XP_018219473.1 | 99 | 0 |
| SPGP0AQ00520 | XP_018221990.1 | 99 | 0 |
| SPGP0AF00510 | XP_018222554.1 | 99 | 0 |
| SPGP0H02980 | XP_018219479.1 | 99 | 0 |
| SPGP0I01470 | XP_018223188.1 | 99 | 0 |
| SPGP0I03440 | XP_018223003.1 | 99 | 0 |
| SPGP0B00700 | XP_018223516.1 | 99 | 0 |
| SPGP0B01040 | XP_018223551.1 | 99 | 0 |
| SPGP0H03570 | XP_018219535.1 | 99 | 0 |
| SPGP0B05170 | XP_018223950.1 | 99 | 0 |
| SPGP0M00940 | XP_018221214.1 | 99 | 0 |
| SPGP0L03150 | XP_018218866.1 | 99 | 0 |
| SPGP0C00410 | XP_018220409.1 | 98 | 0 |
| SPGP0H02100 | XP_018219395.1 | 99 | 0 |
| SPGP0H03580 | XP_018219536.1 | 99 | 0 |
| SPGP0C01660 | XP_018220534.1 | 99 | 0 |
| SPGP0Q00340 | XP_018222827.1 | 99 | 0 |
| SPGP0W00380 | XP_018221487.1 | 99 | 0 |
| SPGP0I03930 | XP_018222956.1 | 99 | 0 |
| SPGP0H00250 | XP_018219213.1 | 99 | 1 |
| SPGP0C02240 | XP_018220592.1 | 99 | 0 |
| SPGP0B03960 | XP_018223837.1 | 99 | 0 |
| SPGP0AQ00290 | XP_018221967.1 | 99 | 0 |
| SPGP0B05240 | XP_018223956.1 | 99 | 0 |
| SPGP0I01060 | XP_018223228.1 | 99 | 0 |
| SPGP0I00440 | XP_018223290.1 | 99 | 0 |
| SPGP0L03370 | XP_018218845.1 | 99 | 0 |
| SPGP0P00790 | XP_018221065.1 | 99 | 0 |
| SPGP0E03550 | XP_018222144.1 | 99 | 0 |
| SPGP0W01470 | XP_018221592.1 | 99 | 0 |
| SPGP0O02390 | XP_018221863.1 | 100 | 0 |
| SPGP0C01900 | XP_018220558.1 | 100 | 0 |
| SPGP0F03720 | XP_018219998.1 | 99 | 0 |
| SPGP0P02860 | XP_018220864.1 | 99 | 0 |
| SPGP0C04710 | XP_018220830.1 | 99 | 0 |
| SPGP0AF00710 | XP_018222534.1 | 99 | 0 |
| SPGP0E02690 | XP_018222226.1 | 99 | 0 |
| SPGP0H01100 | XP_018219297.1 | 99 | 0 |
| SPGP0AG00580 | XP_018223372.1 | 99 | 0 |

Create PDF in your applications with the Pdfcrowd HTML to PDF API PDFCROWD

| | | | |
|---|---|---|---|
| SPGP0P02540 | XP_018220896.1 | 99 | 0 |
| SPGP0C00440 | XP_018220413.1 | 99 | 0 |
| SPGP0M00410 | XP_018221166.1 | 98 | 0 |
| SPGP0AT00260 | XP_018219183.1 | 99 | 0 |
| SPGP0B02360 | XP_018223680.1 | 99 | 0 |
| SPGP0F02630 | XP_018220105.1 | 98 | 2 |
| SPGP0H00800 | XP_018219267.1 | 99 | 0 |
| SPGP0B01910 | XP_018223637.1 | 99 | 0 |
| SPGP0L02680 | XP_018218913.1 | 99 | 0 |
| SPGP0B01260 | XP_018223573.1 | 99 | 0 |
| SPGP0H00400 | XP_018219228.1 | 99 | 0 |
| SPGP0B04210 | XP_018223862.1 | 99 | 0 |
| SPGP0P01780 | XP_018220971.1 | 99 | 0 |
| SPGP0AG01020 | XP_018223411.1 | 99 | 0 |
| SPGP0H01450 | XP_018219330.1 | 99 | 0 |
| SPGP0I03720 | XP_018222976.1 | 99 | 0 |
| SPGP0E00730 | XP_018222424.1 | 99 | 0 |
| SPGP0Q01100 | XP_018222752.1 | 99 | 0 |
| SPGP0AI00600 | XP_018224099.1 | 99 | 0 |
| SPGP0O02740 | XP_018221897.1 | 99 | 0 |
| SPGP0O00150 | XP_018221648.1 | 99 | 0 |
| SPGP0C04120 | XP_018220774.1 | 99 | 0 |
| SPGP0M00390 | XP_018221164.1 | 99 | 0 |
| SPGP0M00420 | XP_018221167.1 | 99 | 0 |
| SPGP0C01950 | XP_018220563.1 | 98 | 0 |
| SPGP0B01880 | XP_018223634.1 | 99 | 0 |
| SPGP0L03450 | XP_018218838.1 | 99 | 0 |
| SPGP0B05790 | XP_018224009.1 | 99 | 0 |
| SPGP0L01980 | XP_018218979.1 | 99 | 0 |
| SPGP0B04730 | XP_018223912.1 | 99 | 0 |
| SPGP0L01470 | XP_018219029.1 | 99 | 1 |
| SPGP0E01580 | XP_018222336.1 | 99 | 0 |
| SPGP0O00880 | XP_018221718.1 | 99 | 0 |
| SPGP0M01450 | XP_018221264.1 | 99 | 0 |
| SPGP0F01570 | XP_018220205.1 | 99 | 0 |
| SPGP0Q02060 | XP_018222659.1 | 99 | 0 |
| SPGP0F03960 | XP_018219975.1 | 99 | 0 |
| SPGP0AL00260 | XP_018219890.1 | 99 | 1 |
| SPGP0H01300 | XP_018219315.1 | 99 | 0 |
| SPGP0I02450 | XP_018223092.1 | 99 | 1 |

Create PDF in your applications with the Pdfcrowd HTML to PDF API PDFCROWD

| | | | |
|---|---|---|---|
| SPGP0E01210 | XP_018222375.1 | 99 | 0 |
| SPGP0W01390 | XP_018221584.1 | 99 | 0 |
| SPGP0E02570 | XP_018222239.1 | 99 | 0 |
| SPGP0BB00220 | XP_018219856.1 | 99 | 0 |
| SPGP0C02710 | XP_018220639.1 | 99 | 0 |
| SPGP0E01950 | XP_018222299.1 | 99 | 0 |
| SPGP0B03420 | XP_018223784.1 | 99 | 0 |
| SPGP0Q02280 | XP_018222637.1 | 99 | 0 |
| SPGP0L02410 | XP_018218937.1 | 99 | 0 |
| SPGP0P03010 | XP_018220851.1 | 99 | 0 |
| SPGP0I03400 | XP_018223005.1 | 99 | 0 |
| SPGP0C00970 | XP_018220465.1 | 99 | 0 |
| SPGP0B05870 | XP_018224017.1 | 99 | 0 |
| SPGP0L00110 | XP_018219160.1 | 99 | 0 |
| SPGP0AG00250 | XP_018223340.1 | 99 | 0 |
| SPGP0L00440 | XP_018219128.1 | 99 | 0 |
| SPGP0E01450 | XP_018222349.1 | 99 | 0 |
| SPGP0C01590 | XP_018220528.1 | 99 | 0 |
| SPGP0AF01110 | XP_018222497.1 | 99 | 0 |
| SPGP0B00730 | XP_018223519.1 | 99 | 0 |
| SPGP0E01100 | XP_018222387.1 | 99 | 0 |
| SPGP0E02730 | XP_018222223.1 | 99 | 0 |
| SPGP0P01420 | XP_018221005.1 | 99 | 0 |
| SPGP0H03060 | XP_018219487.1 | 99 | 0 |
| SPGP0M02480 | XP_018221364.1 | 99 | 0 |
| SPGP0I01050 | XP_018223229.1 | 99 | 0 |
| SPGP0M02940 | XP_018221408.1 | 99 | 0 |
| SPGP0C01620 | XP_018220530.1 | 99 | 0 |
| SPGP0W01310 | XP_018221576.1 | 99 | 0 |
| SPGP0O01530 | XP_018221780.1 | 99 | 0 |
| SPGP0B01590 | XP_018223606.1 | 99 | 0 |
| SPGP0I03460 | XP_018223001.1 | 99 | 0 |
| SPGP0C00620 | XP_018220430.1 | 99 | 0 |
| SPGP0M01390 | XP_018221258.1 | 99 | 0 |
| SPGP0M00600 | XP_018221184.1 | 99 | 0 |
| SPGP0C02000 | XP_018220568.1 | 99 | 0 |
| SPGP0C03440 | XP_018220710.1 | 99 | 0 |
| SPGP0Etrna41L | DI49_2066 | 100 | 0 |
| SPGP0O01820 | XP_018221807.1 | 99 | 0 |
| SPGP0B02480 | XP_018223692.1 | 99 | 0 |

Create PDF in your applications with the Pdfcrowd HTML to PDF API PDFCROWD

| | | | |
|---|---|---|---|
| SPGP0L03100 | XP_018218871.1 | 99 | 0 |
| SPGP0W00710 | XP_018221519.1 | 100 | 0 |
| SPGP0B05540 | XP_018223984.1 | 98 | 0 |
| SPGP0L02100 | XP_018218968.1 | 99 | 0 |
| SPGP0H01840 | XP_018219369.1 | 99 | 0 |
| SPGP0C04180 | XP_018220780.1 | 99 | 0 |
| SPGP0I00210 | XP_018223313.1 | 99 | 0 |
| SPGP0I03340 | XP_018223011.1 | 99 | 0 |
| SPGP0P01300 | XP_018221016.1 | 99 | 0 |
| SPGP0E02540 | XP_018222242.1 | 95 | 2 |
| SPGP0I01570 | XP_018223178.1 | 99 | 0 |
| SPGP0Q01760 | XP_018222688.1 | 99 | 0 |
| SPGP0S02270 | XP_018219625.1 | 99 | 0 |
| SPGP0L02160 | XP_018218962.1 | 99 | 0 |
| SPGP0CY00100 | XP_018222487.1 | 90 | 0 |
| SPGP0W01710 | XP_018221615.1 | 100 | 0 |
| SPGP0AG00170 | XP_018223332.1 | 100 | 0 |
| SPGP0P00730 | XP_018221071.1 | 99 | 0 |
| SPGP0H01730 | XP_018219358.1 | 99 | 1 |
| SPGP0B06070 | XP_018224036.1 | 100 | 0 |
| SPGP0Q01520 | XP_018222712.1 | 99 | 0 |
| SPGP0B02010 | XP_018223647.1 | 99 | 0 |
| SPGP0F01060 | XP_018220254.1 | 99 | 0 |
| SPGP0C04020 | XP_018220764.1 | 100 | 0 |
| SPGP0P02190 | XP_018220931.1 | 99 | 0 |
| SPGP0E02350 | XP_018222260.1 | 99 | 0 |
| SPGP0AI00160 | XP_018224057.1 | 100 | 0 |
| SPGP0E02790 | XP_018222216.1 | 99 | 0 |
| SPGP0H00460 | XP_018219234.1 | 100 | 0 |
| SPGP0H01710 | XP_018219356.1 | 99 | 0 |
| SPGP0AI00610 | XP_018224100.1 | 99 | 0 |
| SPGP0F03150 | XP_018220054.1 | 99 | 0 |
| SPGP0W00110 | XP_018221460.1 | 98 | 0 |
| SPGP0S00690 | XP_018219782.1 | 99 | 0 |
| SPGP0P01720 | XP_018220977.1 | 99 | 1 |
| SPGP0S00520 | XP_018219799.1 | 99 | 0 |
| SPGP0I02710 | XP_018223067.1 | 99 | 0 |
| SPGP0E03920 | XP_018222107.1 | 99 | 0 |
| SPGP0H02220 | XP_018219405.1 | 98 | 1 |
| SPGP0E02850 | XP_018222210.1 | 98 | 1 |

Create PDF in your applications with the Pdfcrowd HTML to PDF API PDFCROWD

| | | | |
|---|---|---|---|
| SPGP0Itrna18D | DI49_4675 | 100 | 0 |
| SPGP0I03100 | XP_018223033.1 | 99 | 0 |
| SPGP0M01130 | XP_018221233.1 | 99 | 0 |
| SPGP0O03210 | XP_018221941.1 | 99 | 0 |
| SPGP0B02500 | XP_018223694.1 | 99 | 0 |
| SPGP0F00700 | XP_018220290.1 | 99 | 0 |
| SPGP0Q00470 | XP_018222814.1 | 99 | 0 |
| SPGP0AG00640 | XP_018223378.1 | 99 | 0 |
| SPGP0S01460 | XP_018219706.1 | 99 | 0 |
| SPGP0F00840 | XP_018220276.1 | 99 | 0 |
| SPGP0F00550 | XP_018220305.1 | 99 | 0 |
| SPGP0O02290 | XP_018221853.1 | 99 | 0 |
| SPGP0S01310 | XP_018219721.1 | 100 | 0 |
| SPGP0L01870 | XP_018218990.1 | 100 | 0 |
| SPGP0H00860 | XP_018219273.1 | 99 | 0 |
| SPGP0H02070 | XP_018219392.1 | 99 | 0 |
| SPGP0L00330 | XP_018219139.1 | 99 | 0 |
| SPGP0B05630 | XP_018223993.1 | 100 | 0 |
| SPGP0AF00900 | XP_018222516.1 | 99 | 0 |
| SPGP0H03030 | XP_018219484.1 | 99 | 0 |
| SPGP0C04700 | XP_018220829.1 | 99 | 0 |
| SPGP0M02740 | XP_018221389.1 | 99 | 0 |
| SPGP0I00450 | XP_018223289.1 | 99 | 0 |
| SPGP0Otrna9F | DI49_2475 | 100 | 0 |
| SPGP0H02620 | XP_018219443.1 | 99 | 0 |
| SPGP0H00650 | XP_018219252.1 | 99 | 0 |
| SPGP0O00470 | XP_018221679.1 | 99 | 0 |
| SPGP0L03340 | XP_018218848.1 | 99 | 1 |
| SPGP0F00320 | XP_018220327.1 | 99 | 0 |
| SPGP0E00810 | XP_018222416.1 | 99 | 0 |
| SPGP0F04230 | XP_018219948.1 | 99 | 0 |
| SPGP0I01150 | XP_018223219.1 | 99 | 0 |
| SPGP0E00490 | XP_018222448.1 | 99 | 0 |
| SPGP0P00600 | XP_018221084.1 | 99 | 0 |
| SPGP0L01180 | XP_018219057.1 | 99 | 0 |
| SPGP0B05600 | XP_018223990.1 | 99 | 0 |
| SPGP0F04160 | XP_018219955.1 | 99 | 0 |
| SPGP0P01880 | XP_018220961.1 | 99 | 0 |
| SPGP0B04890 | XP_018223924.1 | 99 | 0 |
| SPGP0F01260 | XP_018220234.1 | 99 | 0 |

Create PDF in your applications with the Pdfcrowd HTML to PDF API PDFCROWD

| | | | |
|---|---|---|---|
| SPGP0B01850 | XP_018223631.1 | 99 | 0 |
| SPGP0Q02540 | XP_018222611.1 | 99 | 0 |
| SPGP0E04070 | XP_018222092.1 | 99 | 0 |
| SPGP0L01450 | XP_018219031.1 | 99 | 0 |
| SPGP0M00250 | XP_018221150.1 | 99 | 0 |
| SPGP0L01140 | XP_018219061.1 | 100 | 0 |
| SPGP0H03690 | XP_018219548.1 | 99 | 0 |
| SPGP0F02080 | XP_018220158.1 | 99 | 0 |
| SPGP0I03080 | XP_018223035.1 | 99 | 0 |
| SPGP0L01350 | XP_018219041.1 | 99 | 0 |
| SPGP0E00390 | XP_018222458.1 | 99 | 0 |
| SPGP0O02340 | XP_018221858.1 | 99 | 0 |
| SPGP0I03480 | XP_018223000.1 | 99 | 0 |
| SPGP0W00440 | XP_018221493.1 | 99 | 0 |
| SPGP0B03360 | XP_018223778.1 | 99 | 0 |
| SPGP0M01020 | XP_018221222.1 | 99 | 0 |
| SPGP0B00410 | XP_018223490.1 | 99 | 0 |
| SPGP0O02690 | XP_018221892.1 | 99 | 0 |
| SPGP0I03110 | XP_018223032.1 | 100 | 0 |
| SPGP0O02160 | XP_018221840.1 | 99 | 0 |
| SPGP0I02680 | XP_018223070.1 | 99 | 0 |
| SPGP0P02690 | XP_018220881.1 | 99 | 0 |
| SPGP0B04030 | XP_018223844.1 | 99 | 0 |
| SPGP0E02950 | XP_018222200.1 | 99 | 0 |
| SPGP0P03070 | XP_018220845.1 | 99 | 0 |
| SPGP0P00620 | XP_018221082.1 | 99 | 0 |
| SPGP0P02840 | XP_018220866.1 | 99 | 0 |
| SPGP0B03940 | XP_018223835.1 | 99 | 0 |
| SPGP0L00910 | XP_018219083.1 | 99 | 0 |
| SPGP0H01690 | XP_018219354.1 | 99 | 0 |
| SPGP0W00890 | XP_018221536.1 | 99 | 1 |
| SPGP0S00120 | XP_018219838.1 | 99 | 0 |
| SPGP0AQ00220 | XP_018221960.1 | 99 | 0 |
| SPGP0W01670 | XP_018221611.1 | 99 | 0 |
| SPGP0E02990 | XP_018222196.1 | 99 | 0 |
| SPGP0M03210 | XP_018221435.1 | 99 | 0 |
| SPGP0F04260 | XP_018219944.1 | 99 | 0 |
| SPGP0L02580 | XP_018218923.1 | 99 | 0 |
| SPGP0F01550 | XP_018220207.1 | 99 | 0 |
| SPGP0B01930 | XP_018223639.1 | 100 | 0 |

Create PDF in your applications with the Pdfcrowd HTML to PDF API PDFCROWD

| | | | |
|---|---|---|---|
| SPGP0AF00220 | XP_018222580.1 | 99 | 0 |
| SPGP0AF00730 | XP_018222532.1 | 99 | 0 |
| SPGP0C04210 | XP_018220783.1 | 99 | 0 |
| SPGP0C02270 | XP_018220595.1 | 99 | 0 |
| SPGP0W01900 | XP_018221634.1 | 99 | 0 |
| SPGP0W01450 | XP_018221590.1 | 99 | 0 |
| SPGP0C01160 | XP_018220484.1 | 99 | 0 |
| SPGP0AR00150 | XP_018219574.1 | 99 | 0 |
| SPGP0AI00590 | XP_018224098.1 | 99 | 0 |
| SPGP0L02400 | XP_018218938.1 | 100 | 0 |
| SPGP0P02160 | XP_018220934.1 | 99 | 0 |
| SPGP0F02710 | XP_018220097.1 | 99 | 0 |
| SPGP0B03780 | XP_018223819.1 | 99 | 0 |
| SPGP0M03430 | XP_018221454.1 | 99 | 0 |
| SPGP0H01670 | XP_018219352.1 | 99 | 0 |
| SPGP0L00660 | XP_018219107.1 | 99 | 0 |
| SPGP0AR00320 | XP_018219591.1 | 99 | 0 |
| SPGP0L00490 | XP_018219123.1 | 99 | 0 |
| SPGP0B01290 | XP_018223576.1 | 99 | 0 |
| SPGP0F01190 | XP_018220241.1 | 99 | 0 |
| SPGP0B04710 | XP_018223910.1 | 99 | 0 |
| SPGP0E02770 | XP_018222219.1 | 99 | 0 |
| SPGP0Q00600 | XP_018222801.1 | 99 | 0 |
| SPGP0M01360 | XP_018221255.1 | 99 | 0 |
| SPGP0C03870 | XP_018220750.1 | 99 | 0 |
| SPGP0Q02300 | XP_018222635.1 | 99 | 0 |
| SPGP0O02450 | XP_018221869.1 | 99 | 0 |
| SPGP0P02410 | XP_018220909.1 | 99 | 0 |
| SPGP0Q02440 | XP_018222621.1 | 99 | 0 |
| SPGP0P01220 | XP_018221024.1 | 99 | 0 |
| SPGP0P02530 | XP_018220897.1 | 99 | 0 |
| SPGP0M01470 | XP_018221266.1 | 99 | 0 |
| SPGP0AT00350 | XP_018219174.1 | 99 | 0 |
| SPGP0L02880 | XP_018218893.1 | 100 | 0 |
| SPGP0AG00470 | XP_018223361.1 | 99 | 0 |
| SPGP0Q00390 | XP_018222822.1 | 99 | 0 |
| SPGP0O01900 | XP_018221815.1 | 99 | 0 |
| SPGP0BB00200 | XP_018219854.1 | 99 | 0 |
| SPGP0F00240 | XP_018220335.1 | 99 | 0 |
| SPGP0H03490 | XP_018219527.1 | 99 | 0 |

Create PDF in your applications with the Pdfcrowd HTML to PDF API PDFCROWD

| | | | |
|---|---|---|---|
| SPGP0E00880 | XP_018222409.1 | 99 | 0 |
| SPGP0I01980 | XP_018223136.1 | 99 | 0 |
| SPGP0I03060 | XP_018223037.1 | 99 | 0 |
| SPGP0C00990 | XP_018220467.1 | 99 | 0 |
| SPGP0B05190 | XP_018223952.1 | 99 | 0 |
| SPGP0O00480 | XP_018221680.1 | 99 | 0 |
| SPGP0L00460 | XP_018219126.1 | 99 | 0 |
| SPGP0I00660 | XP_018223268.1 | 99 | 0 |
| SPGP0Q02190 | XP_018222646.1 | 99 | 0 |
| SPGP0M00450 | XP_018221170.1 | 98 | 0 |
| SPGP0L02360 | XP_018218942.1 | 99 | 0 |
| SPGP0Q01270 | XP_018222735.1 | 99 | 0 |
| SPGP0O01800 | XP_018221806.1 | 99 | 1 |
| SPGP0M02970 | XP_018221411.1 | 99 | 0 |
| SPGP0E02270 | XP_018222268.1 | 99 | 0 |
| SPGP0Btrna1F | DI49_0089 | 100 | 0 |
| SPGP0Btrna1E | DI49_2698 | 100 | 0 |
| SPGP0H00970 | XP_018219284.1 | 99 | 0 |
| SPGP0S01250 | XP_018219727.1 | 99 | 0 |
| SPGP0O02300 | XP_018221854.1 | 100 | 0 |
| SPGP0L01840 | XP_018218993.1 | 98 | 0 |
| SPGP0F03880 | XP_018219983.1 | 99 | 0 |
| SPGP0E02930 | XP_018222202.1 | 99 | 0 |
| SPGP0B00480 | XP_018223496.1 | 99 | 0 |
| SPGP0F01090 | XP_018220251.1 | 99 | 0 |
| SPGP0Q00500 | XP_018222811.1 | 99 | 0 |
| SPGP0O00580 | XP_018221690.1 | 99 | 0 |
| SPGP0C04830 | XP_018220841.1 | 99 | 0 |
| SPGP0M03270 | XP_018221441.1 | 99 | 0 |
| SPGP0Q00190 | XP_018222839.1 | 99 | 0 |
| SPGP0Q01640 | XP_018222700.1 | 98 | 0 |
| SPGP0F00350 | XP_018220324.1 | 99 | 0 |
| SPGP0Q00800 | XP_018222782.1 | 100 | 0 |
| SPGP0P01320 | XP_018221014.1 | 99 | 0 |
| SPGP0E02560 | XP_018222240.1 | 99 | 0 |
| SPGP0H03750 | XP_018219554.1 | 99 | 0 |
| SPGP0AF00590 | XP_018222546.1 | 99 | 0 |
| SPGP0E02400 | XP_018222255.1 | 99 | 0 |
| SPGP0B06010 | XP_018224031.1 | 99 | 0 |
| SPGP0M02210 | XP_018221337.1 | 99 | 0 |

Create PDF in your applications with the Pdfcrowd HTML to PDF API PDFCROWD

| | | | |
|---|---|---|---|
| SPGP0W01290 | XP_018221574.1 | 99 | 1 |
| SPGP0F00870 | XP_018220273.1 | 100 | 0 |
| SPGP0H02660 | XP_018219447.1 | 99 | 0 |
| SPGP0S02250 | XP_018219627.1 | 99 | 0 |
| SPGP0O00460 | XP_018221678.1 | 97 | 0 |
| SPGP0F01730 | XP_018220191.1 | 99 | 0 |
| SPGP0AQ00310 | XP_018221969.1 | 99 | 0 |
| SPGP0P02510 | XP_018220899.1 | 99 | 0 |
| SPGP0M01260 | XP_018221245.1 | 99 | 0 |
| SPGP0I02280 | XP_018223109.1 | 99 | 0 |
| SPGP0P01920 | XP_018220958.1 | 99 | 0 |
| SPGP0C02610 | XP_018220630.1 | 99 | 0 |
| SPGP0S01090 | XP_018219743.1 | 99 | 0 |
| SPGP0I00880 | XP_018223246.1 | 99 | 0 |
| SPGP0I02150 | XP_018223122.1 | 99 | 1 |
| SPGP0H02320 | XP_018219415.1 | 99 | 0 |
| SPGP0C00490 | XP_018220417.1 | 99 | 0 |
| SPGP0B01540 | XP_018223601.1 | 99 | 0 |
| SPGP0B05400 | XP_018223971.1 | 99 | 0 |
| SPGP0O00220 | XP_018221655.1 | 99 | 0 |
| SPGP0Q01600 | XP_018222704.1 | 99 | 0 |
| SPGP0AF00690 | XP_018222536.1 | 99 | 0 |
| SPGP0H03860 | XP_018219565.1 | 99 | 0 |
| SPGP0H01310 | XP_018219316.1 | 100 | 0 |
| SPGP0E00160 | XP_018222480.1 | 99 | 0 |
| SPGP0E03930 | XP_018222106.1 | 100 | 0 |
| SPGP0F00570 | XP_018220303.1 | 99 | 0 |
| SPGP0AX00190 | XP_018219867.1 | 99 | 0 |
| SPGP0B04740 | XP_018223913.1 | 99 | 0 |
| SPGP0AQ00490 | XP_018221987.1 | 99 | 0 |
| SPGP0P01260 | XP_018221020.1 | 99 | 0 |
| SPGP0M03170 | XP_018221431.1 | 99 | 0 |
| SPGP0C01340 | XP_018220502.1 | 99 | 0 |
| SPGP0M01480 | XP_018221267.1 | 99 | 0 |
| SPGP0I00470 | XP_018223287.1 | 99 | 0 |
| SPGP0Q01160 | XP_018222746.1 | 99 | 0 |
| SPGP0Dtrna24R | DI49_2410 | 100 | 0 |
| SPGP0O00450 | XP_018221677.1 | 99 | 0 |
| SPGP0B02270 | XP_018223672.1 | 99 | 0 |

Create PDF in your applications with the Pdfcrowd HTML to PDF API PDFCROWD

| | | | |
|---|---|---|---|
| SPGP0F02120 | XP_018220154.1 | 99 | 0 |
| SPGP0Q02380 | XP_018222627.1 | 99 | 0 |
| SPGP0AR00520 | XP_018219610.1 | 99 | 0 |
| SPGP0C01730 | XP_018220541.1 | 99 | 0 |
| SPGP0E01650 | XP_018222329.1 | 99 | 0 |
| SPGP0I00780 | XP_018223256.1 | 99 | 0 |
| SPGP0O01490 | XP_018221776.1 | 99 | 0 |
| SPGP0P01160 | XP_018221029.1 | 99 | 0 |
| SPGP0Q00230 | XP_018222836.1 | 99 | 0 |
| SPGP0F02700 | XP_018220098.1 | 99 | 0 |
| SPGP0B00660 | XP_018223512.1 | 99 | 0 |
| SPGP0B03860 | XP_018223827.1 | 99 | 0 |
| SPGP0L01270 | XP_018219049.1 | 99 | 0 |
| SPGP0Htrna13H | DI49_4134 | 100 | 0 |
| SPGP0W01600 | XP_018221604.1 | 100 | 0 |
| SPGP0C00830 | XP_018220450.1 | 99 | 0 |
| SPGP0M02400 | XP_018221356.1 | 99 | 0 |
| SPGP0P01800 | XP_018220969.1 | 99 | 0 |
| SPGP0F01810 | XP_018220184.1 | 99 | 0 |
| SPGP0F04290 | XP_018219942.1 | 99 | 0 |
| SPGP0Q02160 | XP_018222649.1 | 98 | 1 |
| SPGP0C01530 | XP_018220522.1 | 99 | 0 |
| SPGP0I01430 | XP_018223191.1 | 99 | 0 |
| SPGP0C04090 | XP_018220771.1 | 99 | 0 |
| SPGP0AL00240 | XP_018219888.1 | 99 | 0 |
| SPGP0H02150 | XP_018219399.1 | 99 | 0 |
| SPGP0E02020 | XP_018222292.1 | 99 | 0 |
| SPGP0B01670 | XP_018223613.1 | 99 | 0 |
| SPGP0L00210 | XP_018219151.1 | 99 | 1 |
| SPGP0B00950 | XP_018223541.1 | 99 | 0 |
| SPGP0H02970 | XP_018219478.1 | 99 | 0 |
| SPGP0B04250 | XP_018223866.1 | 99 | 0 |
| SPGP0L00700 | XP_018219103.1 | 99 | 0 |
| SPGP0C01250 | XP_018220493.1 | 99 | 0 |
| SPGP0P01950 | XP_018220955.1 | 99 | 1 |
| SPGP0C01630 | XP_018220531.1 | 99 | 0 |
| SPGP0C03110 | XP_018220679.1 | 99 | 0 |
| SPGP0P00800 | XP_018221064.1 | 99 | 1 |
| SPGP0E02970 | XP_018222198.1 | 99 | 0 |
| SPGP0S01270 | XP_018219725.1 | 99 | 0 |

Create PDF in your applications with the Pdfcrowd HTML to PDF API PDFCROWD

| | | | |
|---|---|---|---|
| SPGP0F02860 | XP_018220082.1 | 99 | 0 |
| SPGP0I01670 | XP_018223168.1 | 99 | 0 |
| SPGP0W00690 | XP_018221517.1 | 99 | 0 |
| SPGP0E01490 | XP_018222345.1 | 99 | 0 |
| SPGP0M00350 | XP_018221160.1 | 99 | 0 |
| SPGP0I02500 | XP_018223087.1 | 99 | 0 |
| SPGP0E01150 | XP_018222381.1 | 99 | 0 |
| SPGP0H01540 | XP_018219339.1 | 99 | 0 |
| SPGP0P00750 | XP_018221069.1 | 99 | 0 |
| SPGP0Q01310 | XP_018222732.1 | 99 | 0 |
| SPGP0B02520 | XP_018223696.1 | 99 | 0 |
| SPGP0H02490 | XP_018219431.1 | 99 | 0 |
| SPGP0M03230 | XP_018221437.1 | 99 | 0 |
| SPGP0P02550 | XP_018220895.1 | 100 | 0 |
| SPGP0O03230 | XP_018221944.1 | 99 | 0 |
| SPGP0B01950 | XP_018223641.1 | 99 | 0 |
| SPGP0B06110 | XP_018224039.1 | 99 | 0 |
| SPGP0H00240 | XP_018219212.1 | 99 | 0 |
| SPGP0E02380 | XP_018222257.1 | 99 | 0 |
| SPGP0F04240 | XP_018219947.1 | 99 | 0 |
| SPGP0AF00200 | XP_018222582.1 | 99 | 1 |
| SPGP0AL00190 | XP_018219883.1 | 99 | 0 |
| SPGP0L03420 | XP_018218841.1 | 99 | 0 |
| SPGP0AI00180 | XP_018224059.1 | 99 | 0 |
| SPGP0W01920 | XP_018221636.1 | 99 | 0 |
| SPGP0AR00260 | XP_018219585.1 | 99 | 0 |
| SPGP0Ltrna8F | DI49_5486 | 100 | 0 |
| SPGP0F01450 | XP_018220215.1 | 99 | 0 |
| SPGP0P00270 | XP_018221116.1 | 99 | 0 |
| SPGP0F01560 | XP_018220206.1 | 99 | 0 |
| SPGP0O02550 | XP_018221878.1 | 99 | 0 |
| SPGP0AL00550 | XP_018219917.1 | 99 | 0 |
| SPGP0B00100 | XP_018223462.1 | 86 | 0 |
| SPGP0S01860 | XP_018219666.1 | 99 | 0 |
| SPGP0F03500 | XP_018220019.1 | 99 | 0 |
| SPGP0C01780 | XP_018220546.1 | 99 | 0 |
| SPGP0B01900 | XP_018223636.1 | 99 | 0 |
| SPGP0E02610 | XP_018222235.1 | 99 | 0 |
| SPGP0F03390 | XP_018220030.1 | 99 | 0 |
| SPGP0L02110 | XP_018218967.1 | 99 | 0 |

Create PDF in your applications with the Pdfcrowd HTML to PDF API PDFCROWD

| | | | |
|---|---|---|---|
| SPGP0AF01150 | XP_018222493.1 | 99 | 0 |
| SPGP0F03310 | XP_018220038.1 | 99 | 0 |
| SPGP0H00230 | XP_018219211.1 | 99 | 0 |
| SPGP0B02700 | XP_018223713.1 | 99 | 0 |
| SPGP0Q02290 | XP_018222636.1 | 100 | 0 |
| SPGP0Q01490 | XP_018222715.1 | 100 | 0 |
| SPGP0Q01140 | XP_018222748.1 | 99 | 0 |
| SPGP0I02930 | XP_018223047.1 | 99 | 0 |
| SPGP0P01980 | XP_018220952.1 | 100 | 0 |
| SPGP0I01890 | XP_018223146.1 | 99 | 0 |
| SPGP0O00840 | XP_018221714.1 | 99 | 0 |
| SPGP0M01490 | XP_018221268.1 | 99 | 0 |
| SPGP0L00300 | XP_018219142.1 | 99 | 0 |
| SPGP0M01740 | XP_018221292.1 | 99 | 0 |
| SPGP0P03050 | XP_018220847.1 | 99 | 0 |
| SPGP0CV00100 | XP_018222858.1 | 99 | 0 |
| SPGP0E03890 | XP_018222110.1 | 99 | 0 |
| SPGP0E01910 | XP_018222303.1 | 99 | 0 |
| SPGP0B01480 | XP_018223595.1 | 99 | 0 |
| SPGP0C00960 | XP_018220464.1 | 99 | 0 |
| SPGP0M02700 | XP_018221385.1 | 99 | 0 |
| SPGP0F04000 | XP_018219971.1 | 99 | 0 |
| SPGP0L01280 | XP_018219048.1 | 100 | 0 |
| SPGP0C04220 | XP_018220784.1 | 100 | 0 |
| SPGP0O01790 | XP_018221805.1 | 99 | 0 |
| SPGP0B03110 | XP_018223754.1 | 99 | 0 |
| SPGP0F00740 | XP_018220286.1 | 99 | 0 |
| SPGP0E01960 | XP_018222298.1 | 99 | 0 |
| SPGP0L03210 | XP_018218861.1 | 98 | 0 |
| SPGP0H01470 | XP_018219332.1 | 99 | 0 |
| SPGP0Q01060 | XP_018222756.1 | 98 | 1 |
| SPGP0B00770 | XP_018223523.1 | 99 | 0 |
| SPGP0I01260 | XP_018223208.1 | 99 | 0 |
| SPGP0C03400 | XP_018220706.1 | 99 | 0 |
| SPGP0Btrna3I | DI49_0395 | 100 | 0 |
| SPGP0M03310 | XP_018221444.1 | 97 | 2 |
| SPGP0Btrna3G | DI49_0567 | 99 | 0 |
| SPGP0B00520 | XP_018223500.1 | 99 | 0 |
| SPGP0O02880 | XP_018221910.1 | 98 | 0 |
| SPGP0I00250 | XP_018223309.1 | 99 | 0 |

Create PDF in your applications with the Pdfcrowd HTML to PDF API PDFCROWD

| | | | |
|---|---|---|---|
| SPGP0S00900 | XP_018219762.1 | 99 | 0 |
| SPGP0B05050 | XP_018223940.1 | 99 | 0 |
| SPGP0Q00410 | XP_018222820.1 | 99 | 0 |
| SPGP0B04290 | XP_018223870.1 | 99 | 0 |
| SPGP0B04530 | XP_018223893.1 | 99 | 0 |
| SPGP0AG01000 | XP_018223410.1 | 99 | 0 |
| SPGP0W00990 | XP_018221546.1 | 99 | 0 |
| SPGP0E02580 | XP_018222238.1 | 99 | 0 |
| SPGP0F00370 | XP_018220322.1 | 99 | 0 |
| SPGP0AI00490 | XP_018224089.1 | 99 | 0 |
| SPGP0E01710 | XP_018222323.1 | 99 | 0 |
| SPGP0M02620 | XP_018221377.1 | 99 | 0 |
| SPGP0FW00100 | XP_018221133.1 | 80 | 1 |
| SPGP0L03700 | XP_018218813.1 | 99 | 0 |
| SPGP0E03180 | XP_018222178.1 | 99 | 0 |
| SPGP0O01550 | XP_018221782.1 | 99 | 0 |
| SPGP0B05590 | XP_018223989.1 | 99 | 0 |
| SPGP0S00650 | XP_018219786.1 | 100 | 0 |
| SPGP0B02990 | XP_018223743.1 | 99 | 0 |
| SPGP0P00190 | XP_018221123.1 | 99 | 0 |
| SPGP0O01390 | XP_018221766.1 | 99 | 1 |
| SPGP0F04300 | XP_018219941.1 | 99 | 0 |
| SPGP0E01780 | XP_018222316.1 | 100 | 0 |
| SPGP0H01880 | XP_018219373.1 | 99 | 0 |
| SPGP0M02090 | XP_018221327.1 | 98 | 0 |
| SPGP0P00560 | XP_018221088.1 | 99 | 0 |
| SPGP0F03910 | XP_018219980.1 | 99 | 0 |
| SPGP0B04490 | XP_018223889.1 | 99 | 0 |
| SPGP0H02710 | XP_018219452.1 | 99 | 0 |
| SPGP0F00510 | XP_018220309.1 | 99 | 0 |
| SPGP0H00420 | XP_018219230.1 | 99 | 0 |
| SPGP0H00930 | XP_018219280.1 | 99 | 0 |
| SPGP0H02810 | XP_018219462.1 | 99 | 0 |
| SPGP0B05670 | XP_018223997.1 | 96 | 3 |
| SPGP0AF00410 | XP_018222563.1 | 99 | 0 |
| SPGP0AF01090 | XP_018222499.1 | 99 | 0 |
| SPGP0H00680 | XP_018219255.1 | 100 | 0 |
| SPGP0M01820 | XP_018221301.1 | 99 | 0 |
| SPGP0F00230 | XP_018220336.1 | 99 | 0 |
| SPGP0B02680 | XP_018223711.1 | 99 | 0 |

Create PDF in your applications with the Pdfcrowd HTML to PDF API PDFCROWD

| | | | |
|---|---|---|---|
| SPGP0C01320 | XP_018220500.1 | 99 | 0 |
| SPGP0B04590 | XP_018223899.1 | 99 | 0 |
| SPGP0I00410 | XP_018223293.1 | 99 | 0 |
| SPGP0W00610 | XP_018221510.1 | 99 | 0 |
| SPGP0H02450 | XP_018219427.1 | 99 | 0 |
| SPGP0W00480 | XP_018221497.1 | 99 | 0 |
| SPGP0Q02360 | XP_018222629.1 | 99 | 1 |
| SPGP0H02750 | XP_018219456.1 | 99 | 0 |
| SPGP0C03000 | XP_018220668.1 | 99 | 0 |
| SPGP0B02330 | XP_018223677.1 | 99 | 0 |
| SPGP0AF00800 | XP_018222525.1 | 99 | 0 |
| SPGP0M00800 | XP_018221202.1 | 99 | 0 |
| SPGP0F03190 | XP_018220050.1 | 99 | 0 |
| SPGP0M02490 | XP_018221365.1 | 99 | 0 |
| SPGP0C02940 | XP_018220662.1 | 99 | 0 |
| SPGP0F03570 | XP_018220013.1 | 99 | 0 |
| SPGP0Q02500 | XP_018222615.1 | 99 | 0 |
| SPGP0H02360 | XP_018219419.1 | 99 | 0 |
| SPGP0L01490 | XP_018219027.1 | 99 | 0 |
| SPGP0F01700 | XP_018220194.1 | 99 | 0 |
| SPGP0I02110 | XP_018223125.1 | 99 | 0 |
| SPGP0O01300 | XP_018221757.1 | 99 | 0 |
| SPGP0C00810 | XP_018220448.1 | 100 | 0 |
| SPGP0L03680 | XP_018218815.1 | 99 | 0 |
| SPGP0C04670 | XP_018220826.1 | 99 | 0 |
| SPGP0Qtrna2W | DI49_5211 | 100 | 0 |
| SPGP0C02410 | XP_018220609.1 | 99 | 0 |
| SPGP0F02020 | XP_018220164.1 | 99 | 0 |
| SPGP0C04250 | XP_018220787.1 | 99 | 0 |
| SPGP0P00420 | XP_018221101.1 | 99 | 0 |
| SPGP0O01160 | XP_018221743.1 | 99 | 0 |
| SPGP0E03140 | XP_018222182.1 | 99 | 0 |
| SPGP0I02440 | XP_018223093.1 | 99 | 0 |
| SPGP0S01420 | XP_018219710.1 | 100 | 0 |
| SPGP0L03130 | XP_018218868.1 | 99 | 0 |
| SPGP0E02910 | XP_018222204.1 | 99 | 0 |
| SPGP0C02250 | XP_018220593.1 | 99 | 0 |
| SPGP0H01870 | XP_018219372.1 | 99 | 0 |
| SPGP0I00350 | XP_018223299.1 | 99 | 0 |
| SPGP0B03900 | XP_018223831.1 | 99 | 0 |

Create PDF in your applications with the Pdfcrowd HTML to PDF API PDFCROWD

| | | | |
|---|---|---|---|
| SPGP0P00770 | XP_018221067.1 | 98 | 1 |
| SPGP0H02470 | XP_018219429.1 | 99 | 0 |
| SPGP0AG00570 | XP_018223371.1 | 99 | 0 |
| SPGP0B02210 | XP_018223666.1 | 99 | 0 |
| SPGP0E01870 | XP_018222307.1 | 99 | 0 |
| SPGP0B03090 | XP_018223752.1 | 99 | 0 |
| SPGP0W00540 | XP_018221503.1 | 99 | 0 |
| SPGP0AF00770 | XP_018222528.1 | 99 | 0 |
| SPGP0AG00450 | XP_018223359.1 | 99 | 0 |
| SPGP0AG00860 | XP_018223397.1 | 99 | 0 |
| SPGP0W01490 | XP_018221593.1 | 99 | 0 |
| SPGP0O01620 | XP_018221789.1 | 99 | 0 |
| SPGP0L02440 | XP_018218934.1 | 100 | 0 |
| SPGP0I00810 | XP_018223253.1 | 99 | 0 |
| SPGP0BI00110 | XP_018221946.1 | 99 | 0 |
| SPGP0H01430 | XP_018219328.1 | 99 | 0 |
| SPGP0AG00290 | XP_018223344.1 | 99 | 0 |
| SPGP0F03070 | XP_018220062.1 | 99 | 0 |
| SPGP0P00480 | XP_018221095.1 | 99 | 0 |
| SPGP0B05910 | XP_018224021.1 | 99 | 0 |
| SPGP0I02940 | XP_018223046.1 | 99 | 0 |
| SPGP0AQ00400 | XP_018221978.1 | 99 | 0 |
| SPGP0AI00660 | XP_018224105.1 | 99 | 0 |
| SPGP0H00210 | XP_018219209.1 | 99 | 0 |
| SPGP0B01350 | XP_018223582.1 | 99 | 0 |
| SPGP0S01470 | XP_018219705.1 | 99 | 0 |
| SPGP0L00740 | XP_018219099.1 | 99 | 0 |
| SPGP0S00460 | XP_018219805.1 | 100 | 0 |
| SPGP0P00660 | XP_018221078.1 | 99 | 0 |
| SPGP0B00850 | XP_018223531.1 | 99 | 0 |
| SPGP0AL00720 | XP_018219932.1 | 99 | 0 |
| SPGP0E01860 | XP_018222308.1 | 99 | 0 |
| SPGP0O00820 | XP_018221712.1 | 99 | 0 |
| SPGP0O00790 | XP_018221710.1 | 98 | 0 |
| SPGP0E02230 | XP_018222272.1 | 99 | 0 |
| SPGP0S02380 | XP_018219615.1 | 99 | 0 |
| SPGP0C01770 | XP_018220545.1 | 99 | 0 |
| SPGP0C01140 | XP_018220482.1 | 99 | 0 |
| SPGP0O02000 | XP_018221824.1 | 99 | 0 |
| SPGP0O00620 | XP_018221694.1 | 99 | 0 |

Create PDF in your applications with the Pdfcrowd HTML to PDF API PDFCROWD

| | | | |
|---|---|---|---|
| SPGP0P02790 | XP_018220871.1 | 99 | 0 |
| SPGP0H02700 | XP_018219451.1 | 99 | 0 |
| SPGP0AL00740 | XP_018219934.1 | 99 | 0 |
| SPGP0O01980 | XP_018221823.1 | 99 | 0 |
| SPGP0H02020 | XP_018219387.1 | 99 | 0 |
| SPGP0S01330 | XP_018219719.1 | 99 | 0 |
| SPGP0B02470 | XP_018223691.1 | 99 | 0 |
| SPGP0P01450 | XP_018221002.1 | 99 | 0 |
| SPGP0I00620 | XP_018223272.1 | 99 | 0 |
| SPGP0H02840 | XP_018219465.1 | 100 | 0 |
| SPGP0M02460 | XP_018221362.1 | 99 | 0 |
| SPGP0I02180 | XP_018223119.1 | 99 | 0 |
| SPGP0AF00630 | XP_018222542.1 | 99 | 0 |
| SPGP0L01050 | XP_018219070.1 | 99 | 0 |
| SPGP0I03920 | XP_018222913.1 | 94 | 0 |
| SPGP0L01800 | XP_018218997.1 | 99 | 0 |
| SPGP0F00540 | XP_018220306.1 | 100 | 0 |
| SPGP0M01300 | XP_018221249.1 | 99 | 0 |
| SPGP0F01370 | XP_018220223.1 | 99 | 0 |
| SPGP0Q00520 | XP_018222809.1 | 99 | 0 |
| SPGP0H02800 | XP_018219461.1 | 99 | 0 |
| SPGP0B03220 | XP_018223765.1 | 99 | 0 |
| SPGP0L03620 | XP_018218822.1 | 100 | 0 |
| SPGP0E03480 | XP_018222150.1 | 99 | 0 |
| SPGP0S00770 | XP_018219774.1 | 99 | 0 |
| SPGP0L03090 | XP_018218872.1 | 99 | 0 |
| SPGP0E03400 | XP_018222158.1 | 99 | 0 |
| SPGP0C03350 | XP_018220701.1 | 99 | 0 |
| SPGP0I02900 | XP_018223050.1 | 99 | 0 |
| SPGP0P00780 | XP_018221066.1 | 99 | 0 |
| SPGP0W01790 | XP_018221623.1 | 98 | 2 |
| SPGP0Q00590 | XP_018222802.1 | 99 | 0 |
| SPGP0H02510 | XP_018219433.1 | 99 | 0 |
| SPGP0E03100 | XP_018222186.1 | 99 | 0 |
| SPGP0W00220 | XP_018221471.1 | 98 | 0 |
| SPGP0I03600 | XP_018222988.1 | 99 | 0 |
| SPGP0L01740 | XP_018219003.1 | 99 | 0 |
| SPGP0P02780 | XP_018220872.1 | 99 | 0 |
| SPGP0P02180 | XP_018220932.1 | 99 | 0 |
| SPGP0W01570 | XP_018221601.1 | 99 | 0 |

Create PDF in your applications with the Pdfcrowd HTML to PDF API PDFCROWD

| | | | |
|---|---|---|---|
| SPGP0H00160 | XP_018219205.1 | 99 | 0 |
| SPGP0H02550 | XP_018219437.1 | 99 | 0 |
| SPGP0F01390 | XP_018220222.1 | 99 | 0 |
| SPGP0S02290 | XP_018219624.1 | 99 | 0 |
| SPGP0P02940 | XP_018220857.1 | 99 | 0 |
| SPGP0P02450 | XP_018220905.1 | 99 | 0 |
| SPGP0I02060 | XP_018223130.1 | 99 | 0 |
| SPGP0F02240 | XP_018220142.1 | 99 | 0 |
| SPGP0B05310 | XP_018223963.1 | 100 | 0 |
| SPGP0O01270 | XP_018221754.1 | 99 | 0 |
| SPGP0B06040 | XP_018224034.1 | 99 | 0 |
| SPGP0AF01040 | XP_018222504.1 | 99 | 0 |
| SPGP0B01500 | XP_018223597.1 | 99 | 0 |
| SPGP0W01460 | XP_018221591.1 | 99 | 0 |
| SPGP0AR00170 | XP_018219576.1 | 99 | 0 |
| SPGP0Q01500 | XP_018222714.1 | 99 | 0 |
| SPGP0M00640 | XP_018221188.1 | 98 | 1 |
| SPGP0F00530 | XP_018220307.1 | 99 | 0 |
| SPGP0B02000 | XP_018223646.1 | 99 | 0 |
| SPGP0B01180 | XP_018223565.1 | 99 | 0 |
| SPGP0L03030 | XP_018218878.1 | 99 | 0 |
| SPGP0E01000 | XP_018222397.1 | 99 | 0 |
| SPGP0H03020 | XP_018219483.1 | 99 | 0 |
| SPGP0E04010 | XP_018222098.1 | 99 | 0 |
| SPGP0L00270 | XP_018219145.1 | 99 | 0 |
| SPGP0L00190 | XP_018219153.1 | 99 | 0 |
| SPGP0C04240 | XP_018220786.1 | 99 | 0 |
| SPGP0M02280 | XP_018221344.1 | 99 | 0 |
| SPGP0AQ00470 | XP_018221985.1 | 98 | 0 |
| SPGP0P02050 | XP_018220945.1 | 99 | 0 |
| SPGP0L01100 | XP_018219065.1 | 100 | 0 |
| SPGP0B03150 | XP_018223758.1 | 99 | 0 |
| SPGP0B04680 | XP_018223907.1 | 99 | 0 |
| SPGP0O00360 | XP_018221669.1 | 99 | 0 |
| SPGP0Q02340 | XP_018222631.1 | 99 | 0 |
| SPGP0AQ00300 | XP_018221968.1 | 99 | 0 |
| SPGP0L01650 | XP_018219011.1 | 99 | 0 |
| SPGP0I02790 | XP_018223059.1 | 99 | 0 |
| SPGP0H03900 | XP_018218749.1 | 100 | 0 |
| SPGP0Q00270 | XP_018222833.1 | 99 | 0 |

Create PDF in your applications with the Pdfcrowd HTML to PDF API PDFCROWD

| | | | |
|---|---|---|---|
| SPGP0C02530 | XP_018220621.1 | 99 | 0 |
| SPGP0C00870 | XP_018220455.1 | 99 | 0 |
| SPGP0B04510 | XP_018223891.1 | 99 | 0 |
| SPGP0I00340 | XP_018223300.1 | 99 | 0 |
| SPGP0P02380 | XP_018220912.1 | 100 | 0 |
| SPGP0O01360 | XP_018221763.1 | 99 | 0 |
| SPGP0P02600 | XP_018220891.1 | 99 | 0 |
| SPGP0I03520 | XP_018222996.1 | 99 | 0 |
| SPGP0E04110 | XP_018222088.1 | 99 | 0 |
| SPGP0I02730 | XP_018223065.1 | 99 | 0 |
| SPGP0Q01260 | XP_018222736.1 | 99 | 0 |
| SPGP0F02750 | XP_018220093.1 | 99 | 0 |
| SPGP0M02980 | XP_018221412.1 | 99 | 0 |
| SPGP0C02040 | XP_018220572.1 | 100 | 0 |
| SPGP0AL00470 | XP_018219909.1 | 99 | 0 |
| SPGP0F01920 | XP_018220173.1 | 99 | 0 |
| SPGP0O01290 | XP_018221756.1 | 99 | 0 |
| SPGP0AL00200 | XP_018219884.1 | 99 | 0 |
| SPGP0H03610 | XP_018219540.1 | 99 | 0 |
| SPGP0E02280 | XP_018222267.1 | 99 | 0 |
| SPGP0C01270 | XP_018220495.1 | 100 | 0 |
| SPGP0AG00820 | XP_018223393.1 | 99 | 0 |
| SPGP0B01730 | XP_018223619.1 | 99 | 0 |
| SPGP0AI00860 | XP_018224123.1 | 99 | 0 |
| SPGP0Itrna19A | DI49_5601 | 100 | 0 |
| SPGP0P02570 | XP_018220894.1 | 99 | 0 |
| SPGP0S01150 | XP_018219737.1 | 100 | 0 |
| SPGP0W01230 | XP_018221568.1 | 99 | 0 |
| SPGP0E03730 | XP_018222126.1 | 98 | 0 |
| SPGP0H02390 | XP_018219422.1 | 99 | 0 |
| SPGP0C03210 | XP_018220689.1 | 98 | 1 |
| SPGP0F02760 | XP_018220092.1 | 99 | 0 |
| SPGP0E04250 | XP_018222074.1 | 99 | 0 |
| SPGP0Q01910 | XP_018222672.1 | 100 | 0 |
| SPGP0L02600 | XP_018218921.1 | 99 | 0 |
| SPGP0I00110 | XP_018223323.1 | 98 | 0 |
| SPGP0Q00620 | XP_018222799.1 | 98 | 0 |
| SPGP0F03270 | XP_018220042.1 | 100 | 0 |
| SPGP0I01170 | XP_018223217.1 | 100 | 0 |
| SPGP0Q01740 | XP_018222690.1 | 99 | 0 |

Create PDF in your applications with the Pdfcrowd HTML to PDF API PDFCROWD

| | | | |
|---|---|---|---|
| SPGP0W01960 | XP_018221640.1 | 99 | 0 |
| SPGP0I00990 | XP_018223235.1 | 99 | 0 |
| SPGP0P01000 | XP_018221044.1 | 99 | 0 |
| SPGP0H01240 | XP_018219309.1 | 99 | 0 |
| SPGP0H00580 | XP_018219246.1 | 99 | 0 |
| SPGP0B02600 | XP_018223704.1 | 99 | 0 |
| SPGP0H03960 | XP_018218750.1 | 99 | 0 |
| SPGP0M02450 | XP_018221361.1 | 99 | 0 |
| SPGP0E01750 | XP_018222319.1 | 99 | 0 |
| SPGP0E02740 | XP_018222222.1 | 99 | 0 |
| SPGP0AI00680 | XP_018224107.1 | 99 | 0 |
| SPGP0B01330 | XP_018223580.1 | 99 | 0 |
| SPGP0L03180 | XP_018218864.1 | 99 | 0 |
| SPGP0M00970 | XP_018221217.1 | 99 | 0 |
| SPGP0L01510 | XP_018219025.1 | 99 | 0 |
| SPGP0L03510 | XP_018218832.1 | 99 | 0 |
| SPGP0E00890 | XP_018222408.1 | 99 | 0 |
| SPGP0C00320 | XP_018220400.1 | 99 | 0 |
| SPGP0E00680 | XP_018222428.1 | 97 | 2 |
| SPGP0M01720 | XP_018221290.1 | 99 | 0 |
| SPGP0AG00410 | XP_018223355.1 | 99 | 0 |
| SPGP0O01340 | XP_018221761.1 | 99 | 0 |
| SPGP0B05110 | XP_018223944.1 | 99 | 0 |
| SPGP0P03090 | XP_018220844.1 | 99 | 0 |
| SPGP0E00650 | XP_018222432.1 | 99 | 0 |
| SPGP0C03280 | XP_018220695.1 | 99 | 0 |
| SPGP0AF01160 | XP_018222492.1 | 99 | 0 |
| SPGP0W01090 | XP_018221555.1 | 99 | 0 |
| SPGP0B02280 | XP_018223673.1 | 99 | 0 |
| SPGP0B05320 | XP_018223964.1 | 99 | 0 |
| SPGP0F01660 | XP_018220196.1 | 99 | 0 |
| SPGP0L02970 | XP_018218884.1 | 99 | 0 |
| SPGP0M03130 | XP_018221427.1 | 99 | 0 |
| SPGP0B04880 | XP_018223923.1 | 99 | 0 |
| SPGP0S01900 | XP_018219662.1 | 99 | 0 |
| SPGP0C01510 | XP_018220520.1 | 99 | 0 |
| SPGP0F00130 | XP_018220346.1 | 99 | 0 |
| SPGP0O01640 | XP_018221791.1 | 99 | 0 |
| SPGP0C04600 | XP_018220821.1 | 99 | 0 |
| SPGP0O01590 | XP_018221786.1 | 100 | 0 |

Create PDF in your applications with the Pdfcrowd HTML to PDF API PDFCROWD

| | | | |
|---|---|---|---|
| SPGP0AF00160 | XP_018222586.1 | 100 | 0 |
| SPGP0C04270 | XP_018220789.1 | 99 | 0 |
| SPGP0S01230 | XP_018219729.1 | 99 | 0 |
| SPGP0Q02070 | XP_018222658.1 | 99 | 0 |
| SPGP0B03310 | XP_018223773.1 | 99 | 0 |
| SPGP0AG00960 | XP_018223406.1 | 99 | 0 |
| SPGP0F00490 | XP_018220311.1 | 99 | 0 |
| SPGP0L01850 | XP_018218992.1 | 99 | 0 |
| SPGP0C03690 | XP_018220733.1 | 99 | 0 |
| SPGP0I02580 | XP_018223079.1 | 99 | 0 |
| SPGP0H03440 | XP_018219524.1 | 99 | 0 |
| SPGP0C00920 | XP_018220460.1 | 99 | 0 |
| SPGP0C04140 | XP_018220776.1 | 99 | 0 |
| SPGP0L03230 | XP_018218859.1 | 99 | 0 |
| SPGP0O02510 | XP_018221875.1 | 99 | 0 |
| SPGP0I03000 | XP_018223041.1 | 99 | 0 |
| SPGP0I02920 | XP_018223048.1 | 99 | 0 |
| SPGP0E01700 | XP_018222324.1 | 99 | 0 |
| SPGP0Q02150 | XP_018222650.1 | 99 | 0 |
| SPGP0AL00540 | XP_018219916.1 | 99 | 0 |
| SPGP0H02440 | XP_018219426.1 | 99 | 0 |
| SPGP0H02530 | XP_018219435.1 | 99 | 0 |
| SPGP0W00790 | XP_018221527.1 | 99 | 0 |
| SPGP0I01520 | XP_018223183.1 | 99 | 1 |
| SPGP0M02660 | XP_018221381.1 | 99 | 0 |
| SPGP0C03640 | XP_018220728.1 | 99 | 0 |
| SPGP0E02620 | XP_018222234.1 | 99 | 0 |
| SPGP0I01160 | XP_018223218.1 | 99 | 0 |
| SPGP0Itrna19E | DI49_5287 | 100 | 0 |
| SPGP0P02060 | XP_018220944.1 | 99 | 0 |
| SPGP0F01510 | XP_018220210.1 | 99 | 0 |
| SPGP0P01280 | XP_018221018.1 | 99 | 0 |
| SPGP0S00610 | XP_018219790.1 | 99 | 0 |
| SPGP0L02220 | XP_018218956.1 | 99 | 0 |
| SPGP0W01510 | XP_018221595.1 | 99 | 0 |
| SPGP0F00280 | XP_018220331.1 | 99 | 0 |
| SPGP0F04150 | XP_018219956.1 | 99 | 0 |
| SPGP0Q02420 | XP_018222623.1 | 99 | 0 |
| SPGP0P02960 | XP_018220855.1 | 99 | 0 |

Create PDF in your applications with the Pdfcrowd HTML to PDF API PDFCROWD

| | | | |
|---|---|---|---|
| SPGP0AF00180 | XP_018222584.1 | 99 | 0 |
| SPGP0P00520 | XP_018221092.1 | 99 | 0 |
| SPGP0H00470 | XP_018219235.1 | 98 | 2 |
| SPGP0E03270 | XP_018222170.1 | 99 | 0 |
| SPGP0L01370 | XP_018219039.1 | 99 | 0 |
| SPGP0C02980 | XP_018220666.1 | 99 | 0 |
| SPGP0B01830 | XP_018223629.1 | 99 | 1 |
| SPGP0E00170 | XP_018222479.1 | 99 | 0 |
| SPGP0I03960 | XP_018222953.1 | 100 | 0 |
| SPGP0F01960 | XP_018220170.1 | 99 | 0 |
| SPGP0S00840 | XP_018219767.1 | 100 | 0 |
| SPGP0L03350 | XP_018218847.1 | 99 | 0 |
| SPGP0F03730 | XP_018219997.1 | 99 | 0 |
| SPGP0B00170 | XP_018223467.1 | 99 | 0 |
| SPGP0C02740 | XP_018220642.1 | 100 | 0 |
| SPGP0B04180 | XP_018223859.1 | 99 | 0 |
| SPGP0F01030 | XP_018220257.1 | 99 | 0 |
| SPGP0H00410 | XP_018219229.1 | 97 | 2 |
| SPGP0B01780 | XP_018223624.1 | 99 | 0 |
| SPGP0L02030 | XP_018218975.1 | 98 | 1 |
| SPGP0B00270 | XP_018223477.1 | 98 | 0 |
| SPGP0M00400 | XP_018221165.1 | 98 | 1 |
| SPGP0C02930 | XP_018220661.1 | 99 | 0 |
| SPGP0P00320 | XP_018221111.1 | 99 | 0 |
| SPGP0I01830 | XP_018223152.1 | 99 | 0 |
| SPGP0M00540 | XP_018221179.1 | 99 | 0 |
| SPGP0O02170 | XP_018221841.1 | 99 | 0 |
| SPGP0P00640 | XP_018221080.1 | 100 | 0 |
| SPGP0Q00780 | XP_018222784.1 | 100 | 0 |
| SPGP0O00200 | XP_018221653.1 | 99 | 0 |
| SPGP0L01990 | XP_018218978.1 | 99 | 0 |
| SPGP0E03310 | XP_018222166.1 | 98 | 1 |
| SPGP0H01220 | XP_018219307.1 | 99 | 0 |
| SPGP0AT00210 | XP_018219188.1 | 98 | 0 |
| SPGP0F03860 | XP_018219985.1 | 99 | 0 |
| SPGP0Q02320 | XP_018222633.1 | 99 | 0 |
| SPGP0Q01030 | XP_018222760.1 | 99 | 0 |
| SPGP0S01280 | XP_018219724.1 | 99 | 0 |
| SPGP0Q00250 | XP_018222834.1 | 99 | 0 |
| SPGP0B05510 | XP_018223982.1 | 99 | 0 |

Create PDF in your applications with the Pdfcrowd HTML to PDF API PDFCROWD

| | | | |
|---|---|---|---|
| SPGP0B04140 | XP_018223855.1 | 99 | 0 |
| SPGP0B04000 | XP_018223841.1 | 99 | 0 |
| SPGP0F02660 | XP_018220102.1 | 99 | 0 |
| SPGP0O01430 | XP_018221770.1 | 99 | 0 |
| SPGP0E00310 | XP_018222466.1 | 99 | 0 |
| SPGP0F00170 | XP_018220342.1 | 99 | 0 |
| SPGP0E01820 | XP_018222312.1 | 99 | 0 |
| SPGP0B03030 | XP_018223747.1 | 99 | 0 |
| SPGP0I03540 | XP_018222994.1 | 99 | 0 |
| SPGP0C02180 | XP_018220586.1 | 99 | 0 |
| SPGP0B02640 | XP_018223708.1 | 99 | 0 |
| SPGP0L00500 | XP_018219122.1 | 99 | 0 |
| SPGP0C02570 | XP_018220625.1 | 99 | 0 |
| SPGP0Q01120 | XP_018222750.1 | 99 | 0 |
| SPGP0B00490 | XP_018223497.1 | 99 | 0 |
| SPGP0W01880 | XP_018221632.1 | 99 | 0 |
| SPGP0P02770 | XP_018220873.1 | 99 | 0 |
| SPGP0L00760 | XP_018219097.1 | 99 | 0 |
| SPGP0S01390 | XP_018219713.1 | 99 | 0 |
| SPGP0C02460 | XP_018220614.1 | 99 | 0 |
| SPGP0L03060 | XP_018218875.1 | 99 | 0 |
| SPGP0AG00940 | XP_018223404.1 | 99 | 0 |
| SPGP0P02590 | XP_018220892.1 | 99 | 0 |
| SPGP0H01830 | XP_018219368.1 | 100 | 0 |
| SPGP0S01170 | XP_018219735.1 | 100 | 0 |
| SPGP0E03710 | XP_018222128.1 | 99 | 0 |
| SPGP0B04500 | XP_018223890.1 | 99 | 0 |
| SPGP0I03330 | XP_018223012.1 | 99 | 0 |
| SPGP0C03490 | XP_018220714.1 | 99 | 0 |
| SPGP0S00780 | XP_018219773.1 | 99 | 0 |
| SPGP0E04260 | XP_018222073.1 | 99 | 0 |
| SPGP0H01960 | XP_018219381.1 | 99 | 0 |
| SPGP0B06150 | XP_018224043.1 | 97 | 0 |
| SPGP0W01980 | XP_018221642.1 | 99 | 0 |
| SPGP0W01180 | XP_018221563.1 | 99 | 0 |
| SPGP0F02400 | XP_018220126.1 | 99 | 0 |
| SPGP0P01020 | XP_018221042.1 | 99 | 0 |
| SPGP0I02530 | XP_018223084.1 | 99 | 0 |
| SPGP0M02220 | XP_018221338.1 | 100 | 0 |
| SPGP0L02480 | XP_018218930.1 | 99 | 0 |

Create PDF in your applications with the Pdfcrowd HTML to PDF API PDFCROWD

| | | | |
|---|---|---|---|
| SPGP0L02950 | XP_018218886.1 | 99 | 0 |
| SPGP0AI00580 | XP_018224097.1 | 99 | 0 |
| SPGP0E00440 | XP_018222453.1 | 99 | 0 |
| SPGP0O02700 | XP_018221893.1 | 99 | 0 |
| SPGP0F00440 | XP_018220316.1 | 99 | 0 |
| SPGP0I01660 | XP_018223169.1 | 97 | 3 |
| SPGP0E02810 | XP_018222214.1 | 99 | 0 |
| SPGP0H00700 | XP_018219257.1 | 98 | 2 |
| SPGP0L00880 | XP_018219086.1 | 99 | 0 |
| SPGP0S00420 | XP_018219809.1 | 99 | 0 |
| SPGP0L02700 | XP_018218911.1 | 100 | 0 |
| SPGP0F04070 | XP_018219964.1 | 99 | 0 |
| SPGP0H02310 | XP_018219414.1 | 99 | 0 |
| SPGP0I02380 | XP_018223099.1 | 99 | 0 |
| SPGP0F00750 | XP_018220285.1 | 99 | 0 |
| SPGP0H03650 | XP_018219544.1 | 99 | 0 |
| SPGP0I00170 | XP_018223317.1 | 99 | 0 |
| SPGP0B01310 | XP_018223578.1 | 100 | 0 |
| SPGP0AF00840 | XP_018222521.1 | 100 | 0 |
| SPGP0B05150 | XP_018223948.1 | 99 | 0 |
| SPGP0AG00430 | XP_018223357.1 | 98 | 1 |
| SPGP0L01080 | XP_018219067.1 | 99 | 0 |
| SPGP0E01220 | XP_018222374.1 | 99 | 0 |
| SPGP0I03070 | XP_018223036.1 | 99 | 0 |
| SPGP0AR00200 | XP_018219579.1 | 99 | 0 |
| SPGP0B04720 | XP_018223911.1 | 99 | 0 |
| SPGP0O00750 | XP_018221707.1 | 99 | 0 |
| SPGP0W00490 | XP_018221498.1 | 99 | 0 |
| SPGP0B04960 | XP_018223931.1 | 99 | 0 |
| SPGP0L01590 | XP_018219017.1 | 99 | 0 |
| SPGP0O00130 | XP_018221646.1 | 99 | 0 |
| SPGP0C03390 | XP_018220705.1 | 99 | 0 |
| SPGP0I04010 | XP_018222950.1 | 98 | 0 |
| SPGP0W00330 | XP_018221482.1 | 99 | 0 |
| SPGP0O01660 | XP_018221793.1 | 95 | 0 |
| SPGP0C03910 | XP_018220754.1 | 100 | 0 |
| SPGP0O01880 | XP_018221813.1 | 99 | 0 |
| SPGP0F04050 | XP_018219966.1 | 99 | 0 |
| SPGP0L01700 | XP_018219007.1 | 99 | 0 |
| SPGP0I03740 | XP_018222974.1 | 99 | 0 |

Create PDF in your applications with the Pdfcrowd HTML to PDF API PDFCROWD

| | | | |
|---|---|---|---|
| SPGP0S02400 | XP_018219613.1 | 99 | 0 |
| SPGP0E01040 | XP_018222393.1 | 99 | 0 |
| SPGP0F01630 | XP_018220199.1 | 100 | 0 |
| SPGP0C04510 | XP_018220812.1 | 99 | 0 |
| SPGP0O02060 | XP_018221830.1 | 99 | 0 |
| SPGP0L03050 | XP_018218876.1 | 99 | 0 |
| SPGP0W01750 | XP_018221619.1 | 99 | 0 |
| SPGP0C00160 | XP_018220387.1 | 99 | 0 |
| SPGP0E02870 | XP_018222208.1 | 99 | 0 |
| SPGP0C03760 | XP_018220739.1 | 99 | 0 |
| SPGP0E02650 | XP_018222231.1 | 99 | 0 |
| SPGP0B02960 | XP_018223740.1 | 99 | 0 |
| SPGP0C03680 | XP_018220732.1 | 99 | 0 |
| SPGP0Q01930 | XP_018222670.1 | 99 | 0 |
| SPGP0AG00150 | XP_018223330.1 | 99 | 0 |
| SPGP0I01140 | XP_018223220.1 | 100 | 0 |
| SPGP0L02150 | XP_018218963.1 | 99 | 0 |
| SPGP0AQ00230 | XP_018221961.1 | 99 | 0 |
| SPGP0M03330 | XP_018221446.1 | 96 | 3 |
| SPGP0H00760 | XP_018219263.1 | 100 | 0 |
| SPGP0F02480 | XP_018220118.1 | 99 | 0 |
| SPGP0AF00780 | XP_018222527.1 | 100 | 0 |
| SPGP0L00140 | XP_018219157.1 | 99 | 0 |
| SPGP0P02900 | XP_018220860.1 | 99 | 0 |
| SPGP0AG00840 | XP_018223395.1 | 100 | 0 |
| SPGP0P00500 | XP_018221094.1 | 99 | 0 |
| SPGP0L01070 | XP_018219068.1 | 99 | 0 |
| SPGP0B01810 | XP_018223627.1 | 99 | 0 |
| SPGP0E00410 | XP_018222456.1 | 99 | 0 |
| SPGP0M01880 | XP_018221307.1 | 99 | 0 |
| SPGP0M00170 | XP_018221142.1 | 99 | 0 |
| SPGP0H01580 | XP_018219343.1 | 99 | 0 |
| SPGP0O02800 | XP_018221903.1 | 100 | 0 |
| SPGP0O01870 | XP_018221812.1 | 99 | 0 |
| SPGP0F02270 | XP_018220139.1 | 99 | 0 |
| SPGP0AI00720 | XP_018224111.1 | 99 | 0 |
| SPGP0M01080 | XP_018221228.1 | 99 | 0 |
| SPGP0I01850 | XP_018223149.1 | 99 | 0 |
| SPGP0Q00810 | XP_018222781.1 | 99 | 0 |
| SPGP0P02420 | XP_018220908.1 | 100 | 0 |

Create PDF in your applications with the Pdfcrowd HTML to PDF API PDFCROWD

| | | | |
|---|---|---|---|
| SPGP0H01060 | XP_018219293.1 | 99 | 0 |
| SPGP0F03800 | XP_018219991.1 | 99 | 0 |
| SPGP0F01110 | XP_018220249.1 | 99 | 0 |
| SPGP0AGtrna41S | DI49_0694 | 100 | 0 |
| SPGP0P00680 | XP_018221076.1 | 99 | 0 |
| SPGP0I03780 | XP_018222970.1 | 99 | 0 |
| SPGP0P02650 | XP_018220885.1 | 99 | 0 |
| SPGP0B02320 | XP_018223676.1 | 99 | 0 |
| SPGP0H03620 | XP_018219541.1 | 99 | 0 |
| SPGP0P01310 | XP_018221015.1 | 99 | 0 |
| SPGP0B00160 | XP_018223466.1 | 99 | 0 |
| SPGP0H02350 | XP_018219418.1 | 100 | 0 |
| SPGP0O01410 | XP_018221768.1 | 99 | 0 |
| SPGP0L01480 | XP_018219028.1 | 99 | 0 |
| SPGP0B02740 | XP_018223717.1 | 99 | 0 |
| SPGP0B05950 | XP_018224025.1 | 99 | 0 |
| SPGP0F02030 | XP_018220163.1 | 100 | 0 |
| SPGP0AQtrna33V | DI49_2588 | 100 | 0 |
| SPGP0M01000 | XP_018221220.1 | 99 | 0 |
| SPGP0H01940 | XP_018219379.1 | 99 | 0 |
| SPGP0P02020 | XP_018220948.1 | 99 | 0 |
| SPGP0H02050 | XP_018219390.1 | 99 | 0 |
| SPGP0M02550 | XP_018221371.1 | 99 | 0 |
| SPGP0L01780 | XP_018218999.1 | 99 | 0 |
| SPGP0F03990 | XP_018219973.1 | 99 | 0 |
| SPGP0C04590 | XP_018220820.1 | 99 | 0 |
| SPGP0F03890 | XP_018219982.1 | 99 | 0 |
| SPGP0B03350 | XP_018223777.1 | 99 | 0 |
| SPGP0P01040 | XP_018221040.1 | 99 | 0 |
| SPGP0O03010 | XP_018221921.1 | 99 | 0 |
| SPGP0B03190 | XP_018223762.1 | 99 | 0 |
| SPGP0C02370 | XP_018220605.1 | 99 | 0 |
| SPGP0F00310 | XP_018220328.1 | 99 | 0 |
| SPGP0E01810 | XP_018222313.1 | 99 | 0 |
| SPGP0S01110 | XP_018219741.1 | 99 | 0 |
| SPGP0I02310 | XP_018223106.1 | 99 | 0 |
| SPGP0I01400 | XP_018223194.1 | 99 | 0 |
| SPGP0F02150 | XP_018220151.1 | 99 | 0 |
| SPGP0M02270 | XP_018221343.1 | 99 | 0 |
| SPGP0S02130 | XP_018219639.1 | 99 | 1 |

Create PDF in your applications with the Pdfcrowd HTML to PDF API PDFCROWD

| | | | |
|---|---|---|---|
| SPGP0E02110 | XP_018222283.1 | 99 | 0 |
| SPGP0S00510 | XP_018219800.1 | 99 | 0 |
| SPGP0O02490 | XP_018221873.1 | 99 | 0 |
| SPGP0I02880 | XP_018223052.1 | 99 | 0 |
| SPGP0C00540 | XP_018220422.1 | 99 | 0 |
| SPGP0H03270 | XP_018219507.1 | 99 | 0 |
| SPGP0M02510 | XP_018221367.1 | 100 | 0 |
| SPGP0AF00530 | XP_018222552.1 | 99 | 0 |
| SPGP0F01790 | XP_018220186.1 | 99 | 0 |
| SPGP0M01160 | XP_018221236.1 | 99 | 0 |
| SPGP0I03820 | XP_018222966.1 | 99 | 0 |
| SPGP0E03430 | XP_018222155.1 | 99 | 0 |
| SPGP0I01580 | XP_018223177.1 | 100 | 0 |
| SPGP0B04230 | XP_018223864.1 | 99 | 0 |
| SPGP0AF00370 | XP_018222567.1 | 99 | 0 |
| SPGP0O02720 | XP_018221895.1 | 99 | 0 |
| SPGP0P02750 | XP_018220875.1 | 99 | 0 |
| SPGP0C00560 | XP_018220424.1 | 99 | 0 |
| SPGP0F02330 | XP_018220133.1 | 97 | 3 |
| SPGP0Q00660 | XP_018222796.1 | 99 | 0 |
| SPGP0C03320 | XP_018220699.1 | 97 | 0 |
| SPGP0E03640 | XP_018222135.1 | 99 | 0 |
| SPGP0AT00400 | XP_018219169.1 | 100 | 0 |
| SPGP0C01970 | XP_018220565.1 | 100 | 0 |
| SPGP0P02810 | XP_018220869.1 | 99 | 0 |
| SPGP0W01660 | XP_018221610.1 | 99 | 0 |
| SPGP0P00310 | XP_018221112.1 | 100 | 0 |
| SPGP0AL00590 | XP_018219921.1 | 99 | 0 |
| SPGP0W01030 | XP_018221550.1 | 99 | 0 |
| SPGP0L00220 | XP_018219150.1 | 99 | 0 |
| SPGP0E03600 | XP_018222139.1 | 98 | 0 |
| SPGP0I02520 | XP_018223085.1 | 99 | 0 |
| SPGP0AF00860 | XP_018222519.1 | 99 | 0 |
| SPGP0Q01350 | XP_018222728.1 | 99 | 0 |
| SPGP0W00850 | XP_018221533.1 | 99 | 0 |
| SPGP0COtrna9N | DI49_5581 | 100 | 0 |
| SPGP0B04330 | XP_018223874.1 | 99 | 0 |
| SPGP0L02190 | XP_018218959.1 | 99 | 0 |
| SPGP0B00830 | XP_018223529.1 | 99 | 0 |
| SPGP0C01710 | XP_018220539.1 | 100 | 0 |

Create PDF in your applications with the Pdfcrowd HTML to PDF API PDFCROWD

| | | | |
|---|---|---|---|
| SPGP0AF00320 | XP_018222571.1 | 99 | 0 |
| SPGP0F03550 | XP_018220015.1 | 99 | 0 |
| SPGP0B00140 | XP_018223464.1 | 100 | 0 |
| SPGP0L03660 | XP_018218817.1 | 99 | 0 |
| SPGP0B05560 | XP_018223986.1 | 99 | 0 |
| SPGP0B03390 | XP_018223781.1 | 99 | 0 |
| SPGP0L02790 | XP_018218902.1 | 99 | 0 |
| SPGP0W01330 | XP_018221578.1 | 99 | 0 |
| SPGP0F02490 | XP_018220117.1 | 99 | 0 |
| SPGP0AR00130 | XP_018219572.1 | 99 | 0 |
| SPGP0E02890 | XP_018222206.1 | 99 | 0 |
| SPGP0O01600 | XP_018221787.1 | 99 | 0 |
| SPGP0E01080 | XP_018222389.1 | 99 | 0 |
| SPGP0O02260 | XP_018221850.1 | 99 | 0 |
| SPGP0C00500 | XP_018220418.1 | 99 | 0 |
| SPGP0H03400 | XP_018219520.1 | 99 | 0 |
| SPGP0F01000 | XP_018220260.1 | 99 | 0 |
| SPGP0F03610 | XP_018220009.1 | 99 | 0 |
| SPGP0C00730 | XP_018220440.1 | 100 | 0 |
| SPGP0F02960 | XP_018220073.1 | 99 | 0 |
| SPGP0W00930 | XP_018221540.1 | 99 | 0 |
| SPGP0Q01410 | XP_018222723.1 | 100 | 0 |
| SPGP0P01910 | XP_018220959.1 | 100 | 0 |
| SPGP0B01620 | XP_018223608.1 | 99 | 0 |
| SPGP0AG00240 | XP_018223339.1 | 99 | 0 |
| SPGP0E01550 | XP_018222339.1 | 99 | 0 |
| SPGP0B03820 | XP_018223823.1 | 99 | 0 |
| SPGP0S01630 | XP_018219689.1 | 99 | 0 |
| SPGP0E03330 | XP_018222164.1 | 99 | 0 |
| SPGP0Q01470 | XP_018222717.1 | 99 | 0 |
| SPGP0AI00250 | XP_018224066.1 | 99 | 0 |
| SPGP0I03750 | XP_018222973.1 | 99 | 0 |
| SPGP0AF00670 | XP_018222538.1 | 99 | 0 |
| SPGP0P00650 | XP_018221079.1 | 99 | 1 |
| SPGP0I01240 | XP_018223210.1 | 99 | 0 |
| SPGP0W01080 | XP_018221554.1 | 99 | 0 |
| SPGP0AI00170 | XP_018224058.1 | 99 | 0 |
| SPGP0M02200 | XP_018221336.1 | 99 | 0 |
| SPGP0C02170 | XP_018220585.1 | 99 | 0 |
| SPGP0O02560 | XP_018221879.1 | 99 | 0 |

Create PDF in your applications with the Pdfcrowd HTML to PDF API PDFCROWD

| | | | |
|---|---|---|---|
| SPGP0I01720 | XP_018223163.1 | 100 | 0 |
| SPGP0S01960 | XP_018219656.1 | 99 | 0 |
| SPGP0AL00500 | XP_018219912.1 | 99 | 0 |
| SPGP0M00370 | XP_018221162.1 | 100 | 0 |
| SPGP0C00650 | XP_018220433.1 | 99 | 0 |
| SPGP0B02800 | XP_018223723.1 | 99 | 0 |
| SPGP0AI00700 | XP_018224109.1 | 99 | 0 |
| SPGP0M00230 | XP_018221148.1 | 99 | 0 |
| SPGP0M02760 | XP_018221391.1 | 99 | 1 |
| SPGP0B03660 | XP_018223807.1 | 99 | 0 |
| SPGP0Q00740 | XP_018222788.1 | 99 | 0 |
| SPGP0C00530 | XP_018220421.1 | 99 | 0 |
| SPGP0P01840 | XP_018220965.1 | 99 | 0 |
| SPGP0M01980 | XP_018221316.1 | 99 | 0 |
| SPGP0Q02670 | XP_018222600.1 | 99 | 0 |
| SPGP0F03820 | XP_018219989.1 | 100 | 0 |
| SPGP0W01680 | XP_018221612.1 | 99 | 0 |
| SPGP0C02950 | XP_018220663.1 | 97 | 3 |
| SPGP0O01000 | XP_018221730.1 | 99 | 0 |
| SPGP0E01470 | XP_018222347.1 | 99 | 0 |
| SPGP0BW00110 | XP_018219196.1 | 96 | 0 |
| SPGP0E00480 | XP_018222449.1 | 99 | 0 |
| SPGP0P02260 | XP_018220924.1 | 99 | 0 |
| SPGP0O01470 | XP_018221774.1 | 99 | 0 |
| SPGP0F01520 | XP_018220209.1 | 99 | 1 |
| SPGP0L00540 | XP_018219118.1 | 99 | 0 |
| SPGP0Q02580 | XP_018222607.1 | 99 | 0 |
| SPGP0L01560 | XP_018219020.1 | 99 | 0 |
| SPGP0C01860 | XP_018220554.1 | 99 | 0 |
| SPGP0AF00490 | XP_018222556.1 | 99 | 0 |
| SPGP0AF00570 | XP_018222548.1 | 99 | 0 |
| SPGP0B00350 | XP_018223485.1 | 99 | 1 |
| SPGP0E03440 | XP_018222154.1 | 99 | 0 |
| SPGP0B00980 | XP_018223545.1 | 99 | 0 |
| SPGP0L03740 | XP_018218809.1 | 99 | 0 |
| SPGP0I03580 | XP_018222991.1 | 99 | 0 |
| SPGP0Q00980 | XP_018222764.1 | 99 | 0 |
| SPGP0M01420 | XP_018221261.1 | 99 | 0 |
| SPGP0B05070 | XP_018223941.1 | 99 | 0 |
| SPGP0B02580 | XP_018223702.1 | 99 | 0 |

Create PDF in your applications with the Pdfcrowd HTML to PDF API PDFCROWD

| | | | |
|---|---|---|---|
| SPGP0F03650 | XP_018220005.1 | 99 | 0 |
| SPGP0S01130 | XP_018219739.1 | 99 | 0 |
| SPGP0M02830 | XP_018221398.1 | 99 | 0 |
| SPGP0M02020 | XP_018221320.1 | 99 | 0 |
| SPGP0F00480 | XP_018220312.1 | 99 | 0 |
| SPGP0C00380 | XP_018220406.1 | 99 | 0 |
| SPGP0B00870 | XP_018223533.1 | 98 | 0 |
| SPGP0P01200 | XP_018221026.1 | 99 | 0 |
| SPGP0L02170 | XP_018218961.1 | 99 | 1 |
| SPGP0M01790 | XP_018221298.1 | 99 | 1 |
| SPGP0B03980 | XP_018223839.1 | 99 | 0 |
| SPGP0P01570 | XP_018220991.1 | 99 | 0 |
| SPGP0L02340 | XP_018218944.1 | 99 | 0 |
| SPGP0F04030 | XP_018219968.1 | 99 | 0 |
| SPGP0H03180 | XP_018219499.1 | 99 | 0 |
| SPGP0I02750 | XP_018223063.1 | 99 | 0 |
| SPGP0M00500 | XP_018221175.1 | 99 | 0 |
| SPGP0H03250 | XP_018219505.1 | 99 | 0 |
| SPGP0I03800 | XP_018222968.1 | 99 | 0 |
| SPGP0S02110 | XP_018219641.1 | 99 | 0 |
| SPGP0E01600 | XP_018222334.1 | 99 | 0 |
| SPGP0S00230 | XP_018219828.1 | 99 | 0 |
| SPGP0C02480 | XP_018220616.1 | 99 | 0 |
| SPGP0I01380 | XP_018223196.1 | 99 | 0 |
| SPGP0ltrna28S | DI49_5523 | 100 | 0 |
| SPGP0P01760 | XP_018220973.1 | 99 | 0 |
| SPGP0I00330 | XP_018223301.1 | 99 | 0 |
| SPGP0P01860 | XP_018220963.1 | 99 | 0 |
| SPGP0L00840 | XP_018219089.1 | 99 | 0 |
| SPGP0E02750 | XP_018222221.1 | 99 | 0 |
| SPGP0M00520 | XP_018221177.1 | 98 | 0 |
| SPGP0M01920 | XP_018221311.1 | 99 | 0 |
| SPGP0AG00550 | XP_018223369.1 | 99 | 0 |
| SPGP0Q01880 | XP_018222675.1 | 99 | 0 |
| SPGP0Q01370 | XP_018222726.1 | 100 | 0 |
| SPGP0H00500 | XP_018219238.1 | 100 | 0 |
| SPGP0O00240 | XP_018221657.1 | 99 | 0 |
| SPGP0C01840 | XP_018220552.1 | 99 | 0 |
| SPGP0O02680 | XP_018221891.1 | 99 | 0 |
| SPGP0AR00440 | XP_018219602.1 | 98 | 2 |

Create PDF in your applications with the Pdfcrowd HTML to PDF API PDFCROWD

| | | | |
|---|---|---|---|
| SPGP0E00910 | XP_018222406.1 | 99 | 0 |
| SPGP0O02760 | XP_018221899.1 | 99 | 0 |
| SPGP0C01570 | XP_018220526.1 | 99 | 0 |
| SPGP0F02920 | XP_018220076.1 | 99 | 0 |
| SPGP0C02190 | XP_018220587.1 | 99 | 0 |
| SPGP0B04640 | XP_018223904.1 | 99 | 0 |
| SPGP0Q02110 | XP_018222654.1 | 98 | 0 |
| SPGP0I01360 | XP_018223198.1 | 100 | 0 |
| SPGP0E02960 | XP_018222199.1 | 99 | 0 |
| SPGP0E00950 | XP_018222402.1 | 99 | 0 |
| SPGP0F01670 | XP_018220195.1 | 99 | 0 |
| SPGP0B03000 | XP_018223744.1 | 99 | 0 |
| SPGP0H02780 | XP_018219459.1 | 99 | 0 |
| SPGP0C01980 | XP_018220566.1 | 99 | 0 |
| SPGP0E04210 | XP_018222078.1 | 98 | 2 |
| SPGP0W00370 | XP_018221486.1 | 99 | 0 |
| SPGP0I01910 | XP_018223144.1 | 99 | 0 |
| SPGP0P02490 | XP_018220901.1 | 99 | 0 |
| SPGP0H01720 | XP_018219357.1 | 99 | 0 |
| SPGP0P00920 | XP_018221052.1 | 99 | 0 |
| SPGP0E00540 | XP_018222443.1 | 99 | 0 |
| SPGP0C02490 | XP_018220617.1 | 99 | 0 |
| SPGP0E02830 | XP_018222212.1 | 99 | 0 |
| SPGP0F02940 | XP_018220074.1 | 99 | 0 |
| SPGP0O03190 | XP_018221939.1 | 99 | 1 |
| SPGP0F02590 | XP_018220109.1 | 99 | 0 |
| SPGP0I02540 | XP_018223083.1 | 99 | 0 |
| SPGP0B02820 | XP_018223725.1 | 99 | 0 |
| SPGP0Q01970 | XP_018222667.1 | 99 | 0 |
| SPGP0O02080 | XP_018221832.1 | 100 | 0 |
| SPGP0M02120 | XP_018221329.1 | 99 | 0 |
| SPGP0E01880 | XP_018222306.1 | 99 | 0 |
| SPGP0C01420 | XP_018220510.1 | 87 | 0 |
| SPGP0S00270 | XP_018219824.1 | 99 | 0 |
| SPGP0E03120 | XP_018222184.1 | 99 | 0 |
| SPGP0C03060 | XP_018220674.1 | 99 | 0 |
| SPGP0B02390 | XP_018223683.1 | 100 | 0 |
| SPGP0S02300 | XP_018219623.1 | 99 | 0 |
| SPGP0BI00120 | XP_018221947.1 | 99 | 0 |
| SPGP0M00310 | XP_018221156.1 | 99 | 0 |

Create PDF in your applications with the Pdfcrowd HTML to PDF API PDFCROWD

| | | | |
|---|---|---|---|
| SPGP0M00990 | XP_018221219.1 | 99 | 0 |
| SPGP0AI00390 | XP_018224080.1 | 99 | 0 |
| SPGP0H00880 | XP_018219275.1 | 99 | 0 |
| SPGP0I01010 | XP_018223233.1 | 99 | 0 |
| SPGP0M02720 | XP_018221387.1 | 99 | 0 |
| SPGP0I01090 | XP_018223225.1 | 99 | 0 |
| SPGP0S01480 | XP_018219704.1 | 99 | 0 |
| SPGP0P00570 | XP_018221087.1 | 99 | 0 |
| SPGP0Q00760 | XP_018222786.1 | 99 | 0 |
| SPGP0E00790 | XP_018222418.1 | 99 | 0 |
| SPGP0I02660 | XP_018223072.1 | 99 | 0 |
| SPGP0I02200 | XP_018223117.1 | 100 | 0 |
| SPGP0F00780 | XP_018220282.1 | 99 | 0 |
| SPGP0S00350 | XP_018219816.1 | 99 | 0 |
| SPGP0Q00920 | XP_018222770.1 | 99 | 0 |
| SPGP0O00500 | XP_018221682.1 | 99 | 0 |
| SPGP0Q02610 | XP_018222604.1 | 99 | 0 |
| SPGP0C04260 | XP_018220788.1 | 99 | 0 |
| SPGP0O01450 | XP_018221772.1 | 99 | 0 |
| SPGP0O00420 | XP_018221675.1 | 99 | 0 |
| SPGP0M01120 | XP_018221232.1 | 99 | 0 |
| SPGP0F01340 | XP_018220226.1 | 99 | 0 |
| SPGP0E01430 | XP_018222351.1 | 99 | 0 |
| SPGP0O02900 | XP_018221912.1 | 99 | 0 |
| SPGP0W00460 | XP_018221495.1 | 99 | 0 |
| SPGP0AG00740 | XP_018223386.1 | 99 | 0 |
| SPGP0E03390 | XP_018222159.1 | 99 | 0 |
| SPGP0AL00390 | XP_018219901.1 | 100 | 0 |
| SPGP0E00200 | XP_018222476.1 | 95 | 5 |
| SPGP0AI00270 | XP_018224068.1 | 99 | 0 |
| SPGP0E02310 | XP_018222264.1 | 99 | 0 |
| SPGP0C02210 | XP_018220589.1 | 99 | 0 |
| SPGP0C00290 | XP_018220398.1 | 99 | 0 |
| SPGP0I01300 | XP_018223204.1 | 99 | 0 |
| SPGP0C04760 | XP_018220835.1 | 99 | 0 |
| SPGP0F01600 | XP_018220202.1 | 99 | 0 |
| SPGP0Q02530 | XP_018222612.1 | 99 | 0 |
| SPGP0L00430 | XP_018219129.1 | 99 | 0 |
| SPGP0P02680 | XP_018220882.1 | 99 | 0 |

Create PDF in your applications with the Pdfcrowd HTML to PDF API PDFCROWD

| | | | |
|---|---|---|---|
| SPGP0I03280 | XP_018223016.1 | 99 | 0 |
| SPGP0E03050 | XP_018222191.1 | 99 | 0 |
| SPGP0P00130 | XP_018221129.1 | 99 | 0 |
| SPGP0C02430 | XP_018220611.1 | 99 | 0 |
| SPGP0E03370 | XP_018222160.1 | 99 | 0 |
| SPGP0B03430 | XP_018223785.1 | 99 | 0 |
| SPGP0AF00820 | XP_018222523.1 | 99 | 0 |
| SPGP0B05130 | XP_018223946.1 | 99 | 0 |
| SPGP0AG00490 | XP_018223363.1 | 99 | 0 |
| SPGP0P00350 | XP_018221108.1 | 99 | 0 |
| SPGP0H00140 | XP_018219203.1 | 98 | 0 |
| SPGP0S02200 | XP_018219632.1 | 99 | 0 |
| SPGP0F01980 | XP_018220168.1 | 99 | 0 |
| SPGP0Q02550 | XP_018222610.1 | 99 | 0 |
| SPGP0AG00760 | XP_018223388.1 | 100 | 0 |
| SPGP0P02140 | XP_018220936.1 | 99 | 0 |
| SPGP0AG00630 | XP_018223377.1 | 99 | 0 |
| SPGP0C00280 | XP_018220397.1 | 99 | 0 |
| SPGP0E01540 | XP_018222340.1 | 99 | 0 |
| SPGP0S00140 | XP_018219836.1 | 99 | 0 |
| SPGP0O00260 | XP_018221659.1 | 99 | 0 |
| SPGP0AF00700 | XP_018222535.1 | 99 | 0 |
| SPGP0C02580 | XP_018220626.1 | 99 | 0 |
| SPGP0B01490 | XP_018223596.1 | 99 | 0 |
| SPGP0H03200 | XP_018219500.1 | 99 | 0 |
| SPGP0O00190 | XP_018221652.1 | 99 | 0 |
| SPGP0E01330 | XP_018222362.1 | 99 | 0 |
| SPGP0AQ00200 | XP_018221958.1 | 99 | 0 |
| SPGP0Q02400 | XP_018222625.1 | 99 | 0 |
| SPGP0F02260 | XP_018220140.1 | 99 | 0 |
| SPGP0O00850 | XP_018221715.1 | 100 | 0 |
| SPGP0O01920 | XP_018221817.1 | 99 | 0 |
| SPGP0O00990 | XP_018221729.1 | 99 | 0 |
| SPGP0C02020 | XP_018220570.1 | 99 | 0 |
| SPGP0AI00210 | XP_018224062.1 | 99 | 0 |
| SPGP0E02370 | XP_018222258.1 | 99 | 0 |
| SPGP0E01050 | XP_018222392.1 | 99 | 0 |
| SPGP0I03650 | XP_018222983.1 | 99 | 0 |
| SPGP0P00720 | XP_018221072.1 | 96 | 0 |
| SPGP0I01280 | XP_018223206.1 | 100 | 0 |

Create PDF in your applications with the Pdfcrowd HTML to PDF API PDFCROWD

| | | | |
|---|---|---|---|
| SPGP0S02340 | XP_018219619.1 | 99 | 0 |
| SPGP0C00770 | XP_018220444.1 | 99 | 0 |
| SPGP0F00330 | XP_018220326.1 | 99 | 0 |
| SPGP0B03590 | XP_018223800.1 | 99 | 0 |
| SPGP0S02090 | XP_018219643.1 | 99 | 0 |
| SPGP0B05810 | XP_018224011.1 | 99 | 0 |
| SPGP0C01820 | XP_018220550.1 | 93 | 6 |
| SPGP0M00610 | XP_018221185.1 | 99 | 0 |
| SPGP0I00870 | XP_018223247.1 | 99 | 0 |
| SPGP0C03080 | XP_018220676.1 | 99 | 0 |
| SPGP0E03830 | XP_018222116.1 | 99 | 0 |
| SPGP0S02320 | XP_018219621.1 | 99 | 0 |
| SPGP0S01920 | XP_018219660.1 | 99 | 0 |
| SPGP0AG00330 | XP_018223348.1 | 99 | 0 |
| SPGP0F03400 | XP_018220029.1 | 99 | 0 |
| SPGP0M00330 | XP_018221158.1 | 99 | 0 |
| SPGP0I02650 | XP_018223073.1 | 99 | 0 |
| SPGP0P00940 | XP_018221050.1 | 99 | 0 |
| SPGP0I02170 | XP_018223120.1 | 99 | 0 |
| SPGP0B02840 | XP_018223727.1 | 99 | 0 |
| SPGP0H01520 | XP_018219337.1 | 100 | 0 |
| SPGP0F01820 | XP_018220183.1 | 99 | 0 |
| SPGP0Q02640 | XP_018222601.1 | 97 | 0 |
| SPGP0AQ00260 | XP_018221964.1 | 99 | 0 |
| SPGP0O00640 | XP_018221696.1 | 99 | 0 |
| SPGP0M00190 | XP_018221144.1 | 99 | 0 |
| SPGP0S01710 | XP_018219681.1 | 99 | 0 |
| SPGP0L02390 | XP_018218939.1 | 99 | 0 |
| SPGP0BB00140 | XP_018219848.1 | 99 | 0 |
| SPGP0I03010 | XP_018223040.1 | 99 | 0 |
| SPGP0W01070 | XP_018221553.1 | 99 | 0 |
| SPGP0B01230 | XP_018223570.1 | 99 | 0 |
| SPGP0C04290 | XP_018220791.1 | 99 | 0 |
| SPGP0S00250 | XP_018219826.1 | 99 | 0 |
| SPGP0S00330 | XP_018219818.1 | 99 | 0 |
| SPGP0P02660 | XP_018220884.1 | 99 | 0 |
| SPGP0B02350 | XP_018223679.1 | 99 | 0 |
| SPGP0M01460 | XP_018221265.1 | 99 | 0 |
| SPGP0S01790 | XP_018219673.1 | 99 | 0 |
| SPGP0AFtrna17F | DI49_2529 | 100 | 0 |

Create PDF in your applications with the Pdfcrowd HTML to PDF API PDFCROWD

| | | | |
|---|---|---|---|
| SPGP0B03440 | XP_018223786.1 | 100 | 0 |
| SPGP0H01780 | XP_018219363.1 | 99 | 0 |
| SPGP0F01650 | XP_018220197.1 | 99 | 0 |
| SPGP0S01890 | XP_018219663.1 | 99 | 0 |
| SPGP0B01270 | XP_018223574.1 | 99 | 0 |
| SPGP0I03360 | XP_018223009.1 | 99 | 0 |
| SPGP0Q02090 | XP_018222656.1 | 99 | 0 |
| SPGP0Q00940 | XP_018222768.1 | 99 | 0 |
| SPGP0H02830 | XP_018219464.1 | 99 | 0 |
| SPGP0P01640 | XP_018220985.1 | 98 | 1 |
| SPGP0M01400 | XP_018221259.1 | 99 | 0 |
| SPGP0AF00470 | XP_018222558.1 | 99 | 0 |
| SPGP0M02290 | XP_018221345.1 | 99 | 0 |
| SPGP0F01900 | XP_018220175.1 | 99 | 0 |
| SPGP0C04150 | XP_018220777.1 | 100 | 0 |
| SPGP0P01970 | XP_018220953.1 | 99 | 0 |
| SPGP0I02820 | XP_018223057.1 | 99 | 0 |
| SPGP0C00340 | XP_018220402.1 | 99 | 0 |
| SPGP0Q00490 | XP_018222812.1 | 100 | 0 |
| SPGP0W00670 | XP_018221515.1 | 99 | 0 |
| SPGP0W00830 | XP_018221531.1 | 99 | 0 |
| SPGP0O02780 | XP_018221901.1 | 99 | 0 |
| SPGP0E03680 | XP_018222131.1 | 99 | 0 |
| SPGP0C04330 | XP_018220795.1 | 98 | 0 |
| SPGP0L00800 | XP_018219093.1 | 99 | 0 |
| SPGP0Q01720 | XP_018222692.1 | 99 | 0 |
| SPGP0E03350 | XP_018222162.1 | 99 | 0 |
| SPGP0F01270 | XP_018220233.1 | 98 | 1 |
| SPGP0Q02270 | XP_018222638.1 | 99 | 0 |
| SPGP0O02540 | XP_018221877.1 | 99 | 0 |
| SPGP0F01580 | XP_018220204.1 | 99 | 0 |
| SPGP0H00260 | XP_018219214.1 | 99 | 0 |
| SPGP0P00370 | XP_018221106.1 | 99 | 0 |
| SPGP0E02780 | XP_018222218.1 | 99 | 0 |
| SPGP0AR00220 | XP_018219581.1 | 99 | 0 |
| SPGP0L03360 | XP_018218846.1 | 99 | 0 |
| SPGP0P00180 | XP_018221124.1 | 100 | 0 |
| SPGP0F03320 | XP_018220037.1 | 99 | 0 |
| SPGP0M02840 | XP_018221399.1 | 99 | 0 |
| SPGP0AX00150 | XP_018219863.1 | 99 | 0 |

Create PDF in your applications with the Pdfcrowd HTML to PDF API PDFCROWD

| | | | |
|---|---|---|---|
| SPGP0I01870 | XP_018223148.1 | 99 | 0 |
| SPGP0O01700 | XP_018221796.1 | 98 | 0 |
| SPGP0L01040 | XP_018219071.1 | 99 | 0 |
| SPGP0Itrna24K | DI49_5564 | 100 | 0 |
| SPGP0Q00130 | XP_018222845.1 | 99 | 0 |
| SPGP0H01160 | XP_018219303.1 | 99 | 0 |
| SPGP0B02940 | XP_018223738.1 | 99 | 0 |
| SPGP0F00650 | XP_018220295.1 | 99 | 0 |
| SPGP0B02570 | XP_018223701.1 | 100 | 0 |
| SPGP0I03200 | XP_018223024.1 | 99 | 0 |
| SPGP0B02650 | XP_018223709.1 | 99 | 0 |
| SPGP0L00390 | XP_018219133.1 | 99 | 0 |
| SPGP0B04270 | XP_018223868.1 | 99 | 0 |
| SPGP0O02200 | XP_018221844.1 | 99 | 0 |
| SPGP0B05890 | XP_018224019.1 | 99 | 0 |
| SPGP0B03800 | XP_018223821.1 | 99 | 0 |
| SPGP0E01920 | XP_018222302.1 | 99 | 0 |
| SPGP0M03280 | XP_018221442.1 | 99 | 0 |
| SPGP0L02320 | XP_018218946.1 | 99 | 0 |
| SPGP0C02300 | XP_018220598.1 | 99 | 0 |
| SPGP0C01460 | XP_018220515.1 | 99 | 0 |
| SPGP0E01200 | XP_018222376.1 | 99 | 0 |
| SPGP0O00140 | XP_018221647.1 | 99 | 0 |
| SPGP0B04450 | XP_018223885.1 | 98 | 1 |
| SPGP0O02110 | XP_018221835.1 | 99 | 0 |
| SPGP0AT00420 | XP_018219167.1 | 99 | 0 |
| SPGP0B04850 | XP_018223921.1 | 99 | 0 |
| SPGP0I02690 | XP_018223069.1 | 99 | 0 |
| SPGP0C03310 | XP_018220698.1 | 99 | 0 |
| SPGP0B00560 | XP_018223503.1 | 99 | 0 |
| SPGP0C01700 | XP_018220538.1 | 99 | 0 |
| SPGP0S02050 | XP_018219647.1 | 99 | 0 |
| SPGP0S01690 | XP_018219683.1 | 99 | 0 |
| SPGP0E00550 | XP_018222442.1 | 99 | 0 |
| SPGP0AQ00460 | XP_018221984.1 | 97 | 0 |
| SPGP0H00330 | XP_018219221.1 | 99 | 0 |
| SPGP0Q02250 | XP_018222640.1 | 99 | 0 |
| SPGP0B03130 | XP_018223756.1 | 99 | 0 |
| SPGP0AG00510 | XP_018223365.1 | 99 | 0 |
| SPGP0AQ00360 | XP_018221974.1 | 99 | 0 |

Create PDF in your applications with the Pdfcrowd HTML to PDF API PDFCROWD

| | | | |
|---|---|---|---|
| SPGP0M00910 | XP_018221212.1 | 99 | 0 |
| SPGP0B01250 | XP_018223572.1 | 99 | 0 |
| SPGP0F00920 | XP_018220268.1 | 100 | 0 |
| SPGP0Q01450 | XP_018222719.1 | 99 | 0 |
| SPGP0H03700 | XP_018219549.1 | 99 | 0 |
| SPGP0H00490 | XP_018219237.1 | 99 | 0 |
| SPGP0B05030 | XP_018223938.1 | 100 | 0 |
| SPGP0I00520 | XP_018223282.1 | 99 | 0 |
| SPGP0P02500 | XP_018220900.1 | 99 | 0 |
| SPGP0AG00900 | XP_018223401.1 | 99 | 0 |
| SPGP0C03300 | XP_018220697.1 | 99 | 0 |
| SPGP0P00250 | XP_018221118.1 | 99 | 0 |
| SPGP0L01970 | XP_018218980.1 | 99 | 0 |
| SPGP0L01250 | XP_018219051.1 | 99 | 0 |
| SPGP0AQ00320 | XP_018221970.1 | 99 | 0 |
| SPGP0AR00340 | XP_018219593.1 | 100 | 0 |
| SPGP0E03860 | XP_018222113.1 | 100 | 0 |
| SPGP0S00860 | XP_018219766.1 | 99 | 0 |
| SPGP0L03750 | XP_018218808.1 | 99 | 0 |
| SPGP0E03460 | XP_018222152.1 | 99 | 0 |
| SPGP0M03220 | XP_018221436.1 | 99 | 0 |
| SPGP0H01080 | XP_018219295.1 | 99 | 0 |
| SPGP0F04010 | XP_018219970.1 | 99 | 0 |
| SPGP0W00700 | XP_018221518.1 | 99 | 0 |
| SPGP0F03240 | XP_018220045.1 | 99 | 0 |
| SPGP0P01510 | XP_018220997.1 | 99 | 0 |
| SPGP0M03050 | XP_018221419.1 | 99 | 0 |
| SPGP0L01190 | XP_018219056.1 | 99 | 0 |
| SPGP0AG00720 | XP_018223384.1 | 99 | 0 |
| SPGP0AR00420 | XP_018219600.1 | 99 | 0 |
| SPGP0E01850 | XP_018222309.1 | 99 | 0 |
| SPGP0C00140 | XP_018220385.1 | 99 | 0 |
| SPGP0F02420 | XP_018220124.1 | 99 | 0 |
| SPGP0H01500 | XP_018219335.1 | 99 | 0 |
| SPGP0I00150 | XP_018223319.1 | 99 | 0 |
| SPGP0AT00370 | XP_018219172.1 | 99 | 0 |
| SPGP0H02890 | XP_018219470.1 | 99 | 0 |
| SPGP0E01160 | XP_018222380.1 | 100 | 0 |
| SPGP0B01940 | XP_018223640.1 | 99 | 0 |
| SPGP0C02700 | XP_018220638.1 | 99 | 0 |

Create PDF in your applications with the Pdfcrowd HTML to PDF API PDFCROWD

| | | | |
|---|---|---|---|
| SPGP0F01010 | XP_018220259.1 | 99 | 0 |
| SPGP0CBtrna33K | DI49_5468 | 100 | 0 |
| SPGP0S01260 | XP_018219726.1 | 100 | 0 |
| SPGP0O02130 | XP_018221837.1 | 99 | 0 |
| SPGP0F02350 | XP_018220131.1 | 99 | 0 |
| SPGP0H02000 | XP_018219385.1 | 99 | 0 |
| SPGP0B05450 | XP_018223976.1 | 99 | 0 |
| SPGP0H02060 | XP_018219391.1 | 100 | 0 |
| SPGP0M02440 | XP_018221360.1 | 99 | 0 |
| SPGP0H02420 | XP_018219425.1 | 99 | 0 |
| SPGP0O03130 | XP_018221933.1 | 99 | 0 |
| SPGP0P01700 | XP_018220979.1 | 99 | 0 |
| SPGP0I02610 | XP_018223076.1 | 99 | 0 |
| SPGP0AF01060 | XP_018222502.1 | 99 | 0 |
| SPGP0B05290 | XP_018223961.1 | 99 | 0 |
| SPGP0M00480 | XP_018221173.1 | 100 | 0 |
| SPGP0E03810 | XP_018222118.1 | 97 | 2 |
| SPGP0E01530 | XP_018222341.1 | 99 | 0 |
| SPGP0H00350 | XP_018219223.1 | 100 | 0 |
| SPGP0B01210 | XP_018223568.1 | 99 | 0 |
| SPGP0B00210 | XP_018223471.1 | 99 | 0 |
| SPGP0M02180 | XP_018221334.1 | 99 | 0 |
| SPGP0L00600 | XP_018219112.1 | 99 | 0 |
| SPGP0AF00680 | XP_018222537.1 | 99 | 0 |
| SPGP0B03520 | XP_018223793.1 | 99 | 0 |
| SPGP0B03530 | XP_018223794.1 | 99 | 0 |
| SPGP0AL00250 | XP_018219889.1 | 99 | 0 |
| SPGP0I00220 | XP_018223312.1 | 99 | 0 |
| SPGP0AI00630 | XP_018224102.1 | 99 | 0 |
| SPGP0L03040 | XP_018218877.1 | 99 | 0 |
| SPGP0M02250 | XP_018221341.1 | 99 | 0 |
| SPGP0B00810 | XP_018223527.1 | 99 | 0 |
| SPGP0W00810 | XP_018221529.1 | 99 | 0 |
| SPGP0L01170 | XP_018219058.1 | 99 | 0 |
| SPGP0I03730 | XP_018222975.1 | 99 | 0 |
| SPGP0H00220 | XP_018219210.1 | 96 | 2 |
| SPGP0B05260 | XP_018223958.1 | 99 | 0 |
| SPGP0Q01590 | XP_018222705.1 | 99 | 0 |
| SPGP0B02340 | XP_018223678.1 | 99 | 0 |
| SPGP0F00380 | XP_018220321.1 | 99 | 0 |

Create PDF in your applications with the Pdfcrowd HTML to PDF API PDFCROWD

| | | | |
|---|---|---|---|
| SPGP0L03470 | XP_018218836.1 | 99 | 0 |
| SPGP0I03880 | XP_018222960.1 | 99 | 0 |
| SPGP0I01640 | XP_018223171.1 | 99 | 0 |
| SPGP0W01140 | XP_018221559.1 | 100 | 0 |
| SPGP0B02410 | XP_018223685.1 | 99 | 0 |
| SPGP0I02260 | XP_018223111.1 | 99 | 0 |
| SPGP0H01560 | XP_018219341.1 | 100 | 0 |
| SPGP0P00710 | XP_018221073.1 | 99 | 0 |
| SPGP0O01130 | XP_018221742.1 | 99 | 0 |
| SPGP0Q00680 | XP_018222794.1 | 100 | 0 |
| SPGP0E01370 | XP_018222358.1 | 99 | 0 |
| SPGP0F01460 | XP_018220214.1 | 96 | 3 |
| SPGP0W00130 | XP_018221462.1 | 97 | 0 |
| SPGP0C04320 | XP_018220794.1 | 99 | 0 |
| SPGP0B04280 | XP_018223869.1 | 99 | 0 |
| SPGP0B00500 | XP_018223498.1 | 99 | 0 |
| SPGP0O00600 | XP_018221692.1 | 99 | 0 |
| SPGP0L00350 | XP_018219137.1 | 99 | 0 |
| SPGP0C03740 | XP_018220737.1 | 100 | 0 |
| SPGP0B02120 | XP_018223658.1 | 99 | 0 |
| SPGP0E01060 | XP_018222391.1 | 99 | 0 |
| SPGP0C00980 | XP_018220466.1 | 99 | 0 |
| SPGP0B00310 | XP_018223481.1 | 99 | 0 |
| SPGP0L02570 | XP_018218924.1 | 99 | 0 |
| SPGP0C01580 | XP_018220527.1 | 99 | 0 |
| SPGP0E02700 | XP_018222225.1 | 99 | 0 |
| SPGP0M02060 | XP_018221324.1 | 99 | 0 |
| SPGP0E02070 | XP_018222287.1 | 99 | 0 |
| SPGP0W01350 | XP_018221580.1 | 99 | 0 |
| SPGP0B04090 | XP_018223850.1 | 99 | 0 |
| SPGP0M00930 | XP_018221213.1 | 99 | 0 |
| SPGP0O02600 | XP_018221883.1 | 99 | 0 |
| SPGP0F03460 | XP_018220023.1 | 99 | 0 |
| SPGP0L03320 | XP_018218850.1 | 99 | 0 |
| SPGP0I01130 | XP_018223221.1 | 99 | 0 |
| SPGP0H00290 | XP_018219217.1 | 99 | 0 |
| SPGP0L00980 | XP_018219077.1 | 99 | 0 |
| SPGP0B04990 | XP_018223934.1 | 99 | 0 |
| SPGP0F01930 | XP_018220172.1 | 99 | 0 |
| SPGP0E01070 | XP_018222390.1 | 99 | 0 |

Create PDF in your applications with the Pdfcrowd HTML to PDF API PDFCROWD

| | | | |
|---|---|---|---|
| SPGP0Q00830 | XP_018222779.1 | 96 | 4 |
| SPGP0C01400 | XP_018220508.1 | 98 | 0 |
| SPGP0H00610 | XP_018219248.1 | 99 | 0 |
| SPGP0C01210 | XP_018220489.1 | 99 | 0 |
| SPGP0AX00240 | XP_018219872.1 | 99 | 0 |
| SPGP0O00930 | XP_018221723.1 | 99 | 0 |
| SPGP0L02780 | XP_018218903.1 | 99 | 0 |
| SPGP0B00780 | XP_018223524.1 | 99 | 0 |
| SPGP0C01090 | XP_018220477.1 | 99 | 0 |
| SPGP0F02460 | XP_018220120.1 | 99 | 0 |
| SPGP0P02350 | XP_018220915.1 | 99 | 0 |
| SPGP0B05390 | XP_018223970.1 | 99 | 0 |
| SPGP0Q01780 | XP_018222686.1 | 99 | 0 |
| SPGP0Q00360 | XP_018222825.1 | 99 | 0 |
| SPGP0C00580 | XP_018220426.1 | 99 | 0 |
| SPGP0B02430 | XP_018223687.1 | 99 | 0 |
| SPGP0AX00200 | XP_018219868.1 | 99 | 0 |
| SPGP0E01460 | XP_018222348.1 | 99 | 0 |
| SPGP0B02530 | XP_018223697.1 | 99 | 0 |
| SPGP0B04550 | XP_018223895.1 | 99 | 0 |
| SPGP0AI00420 | XP_018224082.1 | 99 | 0 |
| SPGP0M01180 | XP_018221238.1 | 99 | 0 |
| SPGP0I03150 | XP_018223029.1 | 99 | 0 |
| SPGP0O01850 | XP_018221810.1 | 100 | 0 |
| SPGP0F04170 | XP_018219954.1 | 99 | 0 |
| SPGP0Ftrna20F | DI49_4130 | 99 | 0 |
| SPGP0C04070 | XP_018220769.1 | 99 | 0 |
| SPGP0L03730 | XP_018218810.1 | 99 | 0 |
| SPGP0F00610 | XP_018220299.1 | 99 | 0 |
| SPGP0C04480 | XP_018220810.1 | 99 | 0 |
| SPGP0M02130 | XP_018221330.1 | 99 | 0 |
| SPGP0E01720 | XP_018222322.1 | 99 | 0 |
| SPGP0I01440 | XP_018223190.1 | 99 | 0 |
| SPGP0B03470 | XP_018223789.1 | 99 | 0 |
| SPGP0M00210 | XP_018221146.1 | 99 | 0 |
| SPGP0AL00100 | XP_018219874.1 | 99 | 0 |
| SPGP0W00350 | XP_018221484.1 | 99 | 0 |
| SPGP0C00210 | XP_018220391.1 | 99 | 0 |
| SPGP0AQ00150 | XP_018221954.1 | 99 | 0 |
| SPGP0O02660 | XP_018221889.1 | 99 | 0 |

Create PDF in your applications with the Pdfcrowd HTML to PDF API PDFCROWD

| | | | |
|---|---|---|---|
| SPGP0W01400 | XP_018221585.1 | 99 | 0 |
| SPGP0P02830 | XP_018220867.1 | 99 | 0 |
| SPGP0Q02480 | XP_018222617.1 | 99 | 0 |
| SPGP0C04300 | XP_018220792.1 | 99 | 0 |
| SPGP0AF00270 | XP_018222575.1 | 99 | 0 |
| SPGP0F00630 | XP_018220297.1 | 99 | 0 |
| SPGP0H00820 | XP_018219269.1 | 99 | 0 |
| SPGP0S02030 | XP_018219649.1 | 100 | 0 |
| SPGP0Q01430 | XP_018222721.1 | 100 | 0 |
| SPGP0H01460 | XP_018219331.1 | 99 | 0 |
| SPGP0M02570 | XP_018221373.1 | 99 | 0 |
| SPGP0O01250 | XP_018221752.1 | 99 | 0 |
| SPGP0P03020 | XP_018220850.1 | 99 | 0 |
| SPGP0F01440 | XP_018220216.1 | 99 | 0 |
| SPGP0AF00210 | XP_018222581.1 | 99 | 0 |
| SPGP0O01830 | XP_018221808.1 | 99 | 0 |
| SPGP0E02030 | XP_018222291.1 | 99 | 0 |
| SPGP0F00930 | XP_018220267.1 | 99 | 0 |
| SPGP0S02220 | XP_018219630.1 | 99 | 0 |
| SPGP0AX00220 | XP_018219870.1 | 99 | 0 |
| SPGP0I01990 | XP_018223135.1 | 99 | 0 |
| SPGP0F02040 | XP_018220162.1 | 99 | 0 |
| SPGP0E03220 | XP_018222174.1 | 98 | 0 |
| SPGP0AI00150 | XP_018224056.1 | 99 | 0 |
| SPGP0I00970 | XP_018223237.1 | 99 | 0 |
| SPGP0C04690 | XP_018220828.1 | 99 | 0 |
| SPGP0F03160 | XP_018220053.1 | 99 | 0 |
| SPGP0AF00910 | XP_018222515.1 | 98 | 0 |
| SPGP0O00390 | XP_018221672.1 | 99 | 0 |
| SPGP0Q01630 | XP_018222701.1 | 99 | 0 |
| SPGP0F03180 | XP_018220051.1 | 99 | 0 |
| SPGP0S00160 | XP_018219834.1 | 99 | 0 |
| SPGP0E01590 | XP_018222335.1 | 99 | 0 |
| SPGP0C03700 | XP_018220734.1 | 97 | 0 |
| SPGP0I00730 | XP_018223260.1 | 99 | 0 |
| SPGP0L00710 | XP_018219102.1 | 99 | 0 |
| SPGP0L01950 | XP_018218982.1 | 99 | 0 |
| SPGP0Q00560 | XP_018222805.1 | 99 | 0 |
| SPGP0AL00290 | XP_018219892.1 | 99 | 0 |
| SPGP0H01740 | XP_018219359.1 | 99 | 0 |

Create PDF in your applications with the Pdfcrowd HTML to PDF API PDFCROWD

| | | | |
|---|---|---|---|
| SPGP0O02640 | XP_018221887.1 | 100 | 0 |
| SPGP0O00870 | XP_018221717.1 | 99 | 0 |
| SPGP0I00640 | XP_018223270.1 | 99 | 0 |
| SPGP0AI00230 | XP_018224064.1 | 99 | 0 |
| SPGP0AQ00180 | XP_018221956.1 | 99 | 0 |
| SPGP0S00760 | XP_018219775.1 | 99 | 0 |
| SPGP0B05730 | XP_018224003.1 | 99 | 0 |
| SPGP0F00800 | XP_018220280.1 | 99 | 0 |
| SPGP0BF00110 | XP_018222851.1 | 92 | 0 |
| SPGP0H00840 | XP_018219271.1 | 99 | 0 |
| SPGP0B01120 | XP_018223559.1 | 100 | 0 |
| SPGP0E01800 | XP_018222314.1 | 99 | 0 |
| SPGP0E03850 | XP_018222114.1 | 99 | 1 |
| SPGP0Q00380 | XP_018222823.1 | 99 | 0 |
| SPGP0P02310 | XP_018220919.1 | 99 | 0 |
| SPGP0I02370 | XP_018223100.1 | 99 | 1 |
| SPGP0B03950 | XP_018223836.1 | 99 | 0 |
| SPGP0F04250 | XP_018219946.1 | 99 | 0 |
| SPGP0B00890 | XP_018223535.1 | 99 | 0 |
| SPGP0E02530 | XP_018222243.1 | 99 | 0 |
| SPGP0P03000 | XP_018220852.1 | 99 | 0 |
| SPGP0H02630 | XP_018219444.1 | 99 | 0 |
| SPGP0AF00390 | XP_018222565.1 | 99 | 0 |
| SPGP0F03140 | XP_018220055.1 | 99 | 0 |
| SPGP0L03770 | XP_018218806.1 | 99 | 0 |
| SPGP0E02940 | XP_018222201.1 | 99 | 0 |
| SPGP0B01020 | XP_018223549.1 | 99 | 1 |
| SPGP0H03370 | XP_018219517.1 | 99 | 0 |
| SPGP0F01840 | XP_018220181.1 | 99 | 0 |
| SPGP0C02900 | XP_018220658.1 | 99 | 0 |
| SPGP0B03600 | XP_018223801.1 | 99 | 0 |
| SPGP0AQ00110 | XP_018221951.1 | 99 | 0 |
| SPGP0L00920 | XP_018219082.1 | 98 | 1 |
| SPGP0Etrna44L | DI49_1818 | 100 | 0 |
| SPGP0E01940 | XP_018222300.1 | 99 | 0 |
| SPGP0I03320 | XP_018223013.1 | 100 | 0 |
| SPGP0L00450 | XP_018219127.1 | 99 | 0 |
| SPGP0E03560 | XP_018222143.1 | 99 | 0 |
| SPGP0C01350 | XP_018220503.1 | 100 | 0 |
| SPGP0E03970 | XP_018222102.1 | 99 | 0 |

Create PDF in your applications with the Pdfcrowd HTML to PDF API PDFCROWD

| | | | |
|---|---|---|---|
| SPGP0B03240 | XP_018223767.1 | 99 | 0 |
| SPGP0B05700 | XP_018224000.1 | 99 | 0 |
| SPGP0O00680 | XP_018221700.1 | 99 | 0 |
| SPGP0AI00350 | XP_018224076.1 | 99 | 0 |
| SPGP0H01680 | XP_018219353.1 | 99 | 0 |
| SPGP0I00260 | XP_018223308.1 | 99 | 0 |
| SPGP0Q01610 | XP_018222703.1 | 99 | 0 |

#### Sheet 2: *genes alleles inherited from Saccharomyces cerevisiae*

| *Saccharomyces pastorianus WS34/70 predicted ORF* | *Saccharomyces cerevisiae parental allele* | Gene identities | Gene gaps |
|---|---|---|---|
| SPGP0R01360 | YGL039W | 99 | 0 |
| SPGP0Y00480 | YGR150C | 99 | 0 |
| SPGP0AC01150 | YPR137W | 99 | 0 |
| SPGP0N03230 | YJL206C | 99 | 0 |
| SPGP0AP00400 | YHR196W | 99 | 0 |
| SPGP0AB01480 | YIL156W | 99 | 0 |
| SPGP0A04720 | YDR034C | 99 | 0 |
| SPGP0T01880 | YLR208W | 99 | 0 |
| SPGP0A04230 | YDR085C | 98 | 0 |
| SPGP0A03880 | YDR124W | 99 | 0 |
| SPGP0Z01610 | YKL192C | 100 | 0 |
| SPGP0AH00480 | YCL021W-A | 99 | 0 |
| SPGP0AA01770 | YPL067C | 99 | 0 |
| SPGP0AA00400 | YPL211W | 100 | 0 |
| SPGP0G02240 | YMR081C | 99 | 0 |
| SPGP0AT00130 | YPR198W | 99 | 0 |
| SPGP0Z01310 | YKL161C | 99 | 0 |
| SPGP0D03550 | YOL047C | 99 | 0 |
| SPGP0AO00500 | YLR417W | 99 | 0 |
| SPGP0V01670 | YER081W | 100 | 0 |
| SPGP0AP00450 | YHR200W | 100 | 0 |
| SPGP0N02790 | YJL156C | 99 | 0 |
| SPGP0Y00600 | YGR165W | 99 | 0 |
| SPGP0V01380 | YER054C | 99 | 0 |
| SPGP0AE00880 | YFL025C | 99 | 0 |
| SPGP0A05790 | YDL093W | 99 | 0 |
| SPGP0D00790 | YOR264W | 99 | 0 |
| SPGP0D04420 | YOL141W | 99 | 0 |

Create PDF in your applications with the Pdfcrowd HTML to PDF API PDFCROWD

| | | | |
|---|---|---|---|
| SPGP0AD00340 | YKR080W | 100 | 0 |
| SPGP0D00740 | YOR270C | 99 | 0 |
| SPGP0T01490 | YLR253W | 99 | 0 |
| SPGP0AE01040 | YFL041W | 99 | 0 |
| SPGP0BA00120 | YCR068W | 99 | 0 |
| SPGP0A02410 | YDR291W | 99 | 0 |
| SPGP0AM00200 | YER149C | 99 | 0 |
| SPGP0T00440 | YLR372W | 99 | 0 |
| SPGP0BL00140 | YCR054C | 99 | 0 |
| SPGP0Y00260 | YGR126W | 100 | 0 |
| SPGP0T00130 | YLR407W | 100 | 0 |
| SPGP0AH00100 | YCL064C | 100 | 0 |
| SPGP0K02620 | YNL206C | 99 | 0 |
| SPGP0AE00930 | YFL030W | 99 | 0 |
| SPGP0AE00300 | YFR036W | 100 | 0 |
| SPGP0J01230 | YBR177C | 99 | 0 |
| SPGP0T01080 | YLR301W | 99 | 0 |
| SPGP0AB00740 | YIL077C | 99 | 0 |
| SPGP0V00120 | YEL071W | 99 | 0 |
| SPGP0AY00190 | YOR369C | 100 | 0 |
| SPGP0G00650 | YMR244C-A | 99 | 0 |
| SPGP0AD00460 | YKR067W | 99 | 0 |
| SPGP0AW00100 | YHR150W | 99 | 0 |
| SPGP0K00490 | YNR034W-A | 99 | 0 |
| SPGP0AC01020 | YPR121W | 99 | 0 |
| SPGP0AC00460 | YPR056W | 100 | 0 |
| SPGP0T01700 | YLR227C | 99 | 0 |
| SPGP0AP00500 | YHR205W | 99 | 0 |
| SPGP0BA00190 | YCR075W-A | 100 | 0 |
| SPGP0G02790 | YMR021C | 99 | 0 |
| SPGP0AM00330 | YER163C | 99 | 0 |
| SPGP0T00860 | YLR325C | 100 | 0 |
| SPGP0V01980 | YER113C | 99 | 0 |
| SPGP0R00200 | YGR096W | 100 | 0 |
| SPGP0T00680 | YLR346C | 99 | 0 |
| SPGP0Y01040 | YGR213C | 99 | 0 |
| SPGP0AB00290 | YIL024C | 99 | 0 |
| SPGP0A01290 | YDR407C | 99 | 0 |
| SPGP0V01990 | YER114C | 99 | 0 |
| SPGP0Y01540 | YGR270W | 99 | 0 |

Create PDF in your applications with the Pdfcrowd HTML to PDF API PDFCROWD

| | | | |
|---|---|---|---|
| SPGP0AA00860 | YPL163C | 99 | 0 |
| SPGP0AD00710 | YKR041W | 99 | 0 |
| SPGP0AH00330 | YCL037C | 100 | 0 |
| SPGP0AH00800 | YCR019W | 99 | 0 |
| SPGP0AO00430 | YLR424W | 99 | 0 |
| SPGP0D04300 | YOL128C | 99 | 0 |
| SPGP0FJ00100 | YPR011C | 99 | 0 |
| SPGP0AK00740 | YGL187C | 100 | 0 |
| SPGP0A02180 | YDR314C | 99 | 0 |
| SPGP0X00370 | YHL014C | 99 | 0 |
| SPGP0K00670 | YNR015W | 99 | 0 |
| SPGP0AA01010 | YPL148C | 99 | 0 |
| SPGP0U00450 | YLL022C | 99 | 0 |
| SPGP0AB01420 | YIL150C | 99 | 0 |
| SPGP0N00940 | YJR063W | 99 | 0 |
| SPGP0N00490 | YJR110W | 99 | 0 |
| SPGP0J00440 | YBR260C | 99 | 0 |
| SPGP0U00490 | YLL018C | 99 | 0 |
| SPGP0G00610 | YMR251W | 99 | 0 |
| SPGP0AW00120 | YHR152W | 100 | 0 |
| SPGP0BD00140 | YMR310C | 99 | 0 |
| SPGP0A01260 | YDR410C | 99 | 0 |
| SPGP0A04680 | YDR040C | 98 | 0 |
| SPGP0T01620 | YLR239C | 99 | 0 |
| SPGP0X00610 | YHR010W | 99 | 0 |
| SPGP0A05380 | YDL048C | 99 | 0 |
| SPGP0A05270 | YDL033C | 100 | 0 |
| SPGP0K01690 | YNL100W | 100 | 0 |
| SPGP0J03060 | YBL020W | 99 | 0 |
| SPGP0U00470 | YLL019C | 99 | 0 |
| SPGP0J01760 | YBR121C | 99 | 0 |
| SPGP0A01150 | YDR423C | 99 | 0 |
| SPGP0A02980 | YDR225W | 100 | 0 |
| SPGP0A06400 | YDL160C-A | 100 | 0 |
| SPGP0U00900 | YLR026C | 99 | 0 |
| SPGP0Y00800 | YGR187C | 99 | 0 |
| SPGP0G04050 | YML114C | 99 | 0 |
| SPGP0V00710 | YEL011W | 99 | 0 |
| SPGP0K03130 | YNL260C | 100 | 0 |
| SPGP0AC00310 | YPR036W-A | 100 | 0 |

Create PDF in your applications with the Pdfcrowd HTML to PDF API PDFCROWD

| | | | |
|---|---|---|---|
| SPGP0X01330 | YHR083W | 99 | 0 |
| SPGP0G03700 | YML076C | 99 | 0 |
| SPGP0J02860 | YBR002C | 99 | 0 |
| SPGP0A06460 | YDL168W | 99 | 0 |
| SPGP0X01020 | YHR052W | 99 | 0 |
| SPGP0AO00340 | YLR435W | 99 | 0 |
| SPGP0U01720 | YLR114C | 98 | 0 |
| SPGP0D01820 | YOR149C | 99 | 0 |
| SPGP0D00730 | YOR271C | 99 | 0 |
| SPGP0AC01220 | YPR145W | 99 | 0 |
| SPGP0A00290 | YDR519W | 99 | 0 |
| SPGP0A00850 | YDR459C | 100 | 0 |
| SPGP0A02140 | YDR318W | 99 | 0 |
| SPGP0R02050 | YGL119W | 99 | 0 |
| SPGP0AB00330 | YIL031W | 100 | 0 |
| SPGP0AN00310 | YPL018W | 99 | 0 |
| SPGP0J02330 | YBR059C | 99 | 0 |
| SPGP0A03580 | YDR159W | 99 | 0 |
| SPGP0D03770 | YOL070C | 99 | 0 |
| SPGP0AN00550 | YPL045W | 99 | 0 |
| SPGP0D03680 | YOL061W | 99 | 0 |
| SPGP0Y01230 | YGR235C | 100 | 0 |
| SPGP0Y00880 | YGR196C | 99 | 0 |
| SPGP0AP00100 | YHR166C | 99 | 0 |
| SPGP0G02010 | YMR107W | 100 | 0 |
| SPGP0A01980 | YDR333C | 99 | 0 |
| SPGP0A06140 | YDL131W | 99 | 0 |
| SPGP0X01430 | YHR093W | 99 | 0 |
| SPGP0BV00110 | YCR082W | 99 | 0 |
| SPGP0J00860 | YBR216C | 99 | 0 |
| SPGP0R01550 | YGL062W | 99 | 0 |
| SPGP0R01880 | YGL099W | 99 | 0 |
| SPGP0K01830 | YNL117W | 99 | 0 |
| SPGP0Y01680 | YGR284C | 99 | 0 |
| SPGP0K01070 | YNL021W | 100 | 0 |
| SPGP0AC00710 | YPR085C | 99 | 0 |
| SPGP0Y01130 | YGR223C | 99 | 0 |
| SPGP0X00860 | YHR036W | 100 | 0 |
| SPGP0V00250 | YEL057C | 99 | 0 |
| SPGP0V00700 | YEL012W | 100 | 0 |

Create PDF in your applications with the Pdfcrowd HTML to PDF API PDFCROWD

| | | | |
|---|---|---|---|
| SPGP0A03630 | YDR152W | 99 | 0 |
| SPGP0A05150 | YDL015C | 99 | 0 |
| SPGP0Z01530 | YKL184W | 99 | 0 |
| SPGP0A00250 | YDR524C | 99 | 0 |
| SPGP0A02630 | YDR263C | 99 | 0 |
| SPGP0AM00520 | YER182W | 99 | 0 |
| SPGP0AA01530 | YPL093W | 99 | 0 |
| SPGP0AD01220 | YKL013C | 99 | 0 |
| SPGP0AA00500 | YPL200W | 99 | 0 |
| SPGP0AD01210 | YKL012W | 99 | 0 |
| SPGP0A01180 | YDR420W | 98 | 0 |
| SPGP0AA00850 | YPL164C | 99 | 0 |
| SPGP0D01810 | YOR150W | 99 | 0 |
| SPGP0K02250 | YNL162W | 99 | 0 |
| SPGP0D01350 | YOR201C | 99 | 0 |
| SPGP0K02520 | YNL193W | 99 | 0 |
| SPGP0R01770 | YGL087C | 100 | 0 |
| SPGP0A00120 | YDR540C | 99 | 0 |
| SPGP0AO00230 | YLR445W | 99 | 0 |
| SPGP0V00950 | YER015W | 99 | 0 |
| SPGP0Y01350 | YGR248W | 99 | 0 |
| SPGP0AC00600 | YPR072W | 99 | 0 |
| SPGP0AC00830 | YPR102C | 99 | 0 |
| SPGP0A04000 | YDR111C | 99 | 0 |
| SPGP0J01280 | YBR171W | 99 | 0 |
| SPGP0AT00120 | YPR199C | 99 | 0 |
| SPGP0X00740 | YHR024C | 99 | 0 |
| SPGP0U01910 | YLR136C | 99 | 0 |
| SPGP0D00880 | YOR254C | 99 | 0 |
| SPGP0X00210 | YHL030W | 99 | 0 |
| SPGP0A03140 | YDR205W | 99 | 0 |
| SPGP0T02210 | YLR172C | 99 | 0 |
| SPGP0D01170 | YOR221C | 99 | 0 |
| SPGP0R01080 | YGL009C | 99 | 0 |
| SPGP0AN00360 | YPL023C | 99 | 0 |
| SPGP0AA01470 | YPL098C | 99 | 0 |
| SPGP0A01370 | YDR398W | 99 | 0 |
| SPGP0AE00480 | YFR017C | 99 | 0 |
| SPGP0N00370 | YJR123W | 99 | 0 |
| SPGP0Y01360 | YGR249W | 99 | 0 |

Create PDF in your applications with the Pdfcrowd HTML to PDF API PDFCROWD

| | | | |
|---|---|---|---|
| SPGP0G03630 | YML069W | 99 | 0 |
| SPGP0BD00110 | YMR313C | 99 | 0 |
| SPGP0AJ00370 | YAL011W | 99 | 0 |
| SPGP0U01680 | YLR108C | 99 | 0 |
| SPGP0K03120 | YNL259C | 100 | 0 |
| SPGP0AD00750 | YKR036C | 99 | 0 |
| SPGP0K03690 | YNL323W | 99 | 0 |
| SPGP0A03590 | YDR158W | 99 | 0 |
| SPGP0AA00810 | YPL168W | 99 | 0 |
| SPGP0G02900 | YMR010W | 99 | 0 |
| SPGP0A00410 | YDR506C | 99 | 0 |
| SPGP0AB01230 | YIL130W | 98 | 1 |
| SPGP0Z00440 | YKL063C | 99 | 0 |
| SPGP0G01870 | YMR121C | 99 | 0 |
| SPGP0D01680 | YOR163W | 99 | 0 |
| SPGP0A04870 | YDR018C | 99 | 0 |
| SPGP0G00440 | YMR268C | 99 | 0 |
| SPGP0A05600 | YDL074C | 99 | 0 |
| SPGP0A05350 | YDL045W-A | 99 | 0 |
| SPGP0G03130 | YML015C | 99 | 0 |
| SPGP0D02600 | YOR061W | 99 | 0 |
| SPGP0G00410 | YMR271C | 100 | 0 |
| SPGP0AJ00310 | YAL002W | 99 | 0 |
| SPGP0J01010 | YBR200W | 99 | 0 |
| SPGP0A06930 | YDL222C | 99 | 0 |
| SPGP0AH00110 | YCL063W | 99 | 0 |
| SPGP0AH00380 | YCL032W | 99 | 0 |
| SPGP0A04910 | YDR014W | 99 | 0 |
| SPGP0T01430 | YLR260W | 98 | 1 |
| SPGP0K03340 | YNL284C | 99 | 0 |
| SPGP0D01860 | YOR144C | 99 | 0 |
| SPGP0CS00100 | YNL098C | 100 | 0 |
| SPGP0D00550 | YOR291W | 99 | 0 |
| SPGP0A00280 | YDR520C | 99 | 0 |
| SPGP0D00220 | YOR328W | 99 | 0 |
| SPGP0V00790 | YEL001C | 99 | 0 |
| SPGP0AS00260 | YPL260W | 99 | 0 |
| SPGP0AS00280 | YPL258C | 99 | 0 |
| SPGP0A01780 | YDR358W | 99 | 0 |

Create PDF in your applications with the Pdfcrowd HTML to PDF API PDFCROWD

| | | | |
|---|---|---|---|
| SPGP0AC00610 | YPR073C | 99 | 0 |
| SPGP0Z01420 | YKL172W | 99 | 0 |
| SPGP0AH00720 | YCR011C | 99 | 0 |
| SPGP0AB00530 | YIL052C | 98 | 0 |
| SPGP0G01900 | YMR118C | 99 | 0 |
| SPGP0AH00270 | YCL045C | 99 | 0 |
| SPGP0Z01130 | YKL140W | 99 | 0 |
| SPGP0U01180 | YLR056W | 99 | 0 |
| SPGP0N00670 | YJR094C | 99 | 1 |
| SPGP0D02200 | YOR107W | 99 | 0 |
| SPGP0N01950 | YJL060W | 99 | 0 |
| SPGP0G03410 | YML048W | 99 | 0 |
| SPGP0N01220 | YJR033C | 99 | 0 |
| SPGP0AB00660 | YIL069C | 99 | 0 |
| SPGP0A01350 | YDR400W | 99 | 0 |
| SPGP0K03250 | YNL274C | 99 | 0 |
| SPGP0K02510 | YNL192W | 99 | 0 |
| SPGP0AS00400 | YPL247C | 99 | 0 |
| SPGP0X01690 | YHR119W | 99 | 0 |
| SPGP0AA01220 | YPL125W | 99 | 0 |
| SPGP0K00800 | YNL001W | 99 | 0 |
| SPGP0R00750 | YGR031W | 99 | 0 |
| SPGP0AA00890 | YPL160W | 99 | 0 |
| SPGP0J00820 | YBR221C | 99 | 0 |
| SPGP0D03660 | YOL059W | 99 | 0 |
| SPGP0A02340 | YDR298C | 99 | 0 |
| SPGP0J00270 | YBR279W | 99 | 0 |
| SPGP0T00600 | YLR355C | 99 | 0 |
| SPGP0G01840 | YMR124W | 99 | 0 |
| SPGP0AB00650 | YIL068C | 99 | 0 |
| SPGP0A06340 | YDL154W | 99 | 0 |
| SPGP0G01590 | YMR150C | 100 | 0 |
| SPGP0D02480 | YOR075W | 99 | 0 |
| SPGP0AK00610 | YGL205W | 99 | 0 |
| SPGP0G01080 | YMR202W | 100 | 0 |
| SPGP0A03350 | YDR180W | 99 | 0 |
| SPGP0K03540 | YNL307C | 99 | 0 |
| SPGP0J01900 | YBR106W | 99 | 0 |
| SPGP0CN00110 | YCR094W | 99 | 0 |
| SPGP0J02290 | YBR063C | 99 | 0 |

Create PDF in your applications with the Pdfcrowd HTML to PDF API PDFCROWD

| | | | |
|---|---|---|---|
| SPGP0T01400 | YLR264W | 99 | 0 |
| SPGP0G00460 | YMR266W | 99 | 0 |
| SPGP0N01610 | YJL016W | 99 | 0 |
| SPGP0J00930 | YBR208C | 99 | 0 |
| SPGP0A06870 | YDL215C | 99 | 0 |
| SPGP0A01620 | YDR374W-A | 100 | 0 |
| SPGP0A06650 | YDL192W | 99 | 0 |
| SPGP0R01290 | YGL031C | 100 | 0 |
| SPGP0A04700 | YDR036C | 99 | 0 |
| SPGP0AB01440 | YIL152W | 99 | 0 |
| SPGP0G03340 | YML037C | 99 | 0 |
| SPGP0D03470 | YOL038W | 99 | 0 |
| SPGP0AN00530 | YPL042C | 99 | 0 |
| SPGP0U01650 | YLR105C | 99 | 0 |
| SPGP0AA01030 | YPL146C | 99 | 0 |
| SPGP0N02420 | YJL115W | 99 | 0 |
| SPGP0U00210 | YLL050C | 99 | 0 |
| SPGP0R01310 | YGL033W | 99 | 0 |
| SPGP0J02650 | YBR025C | 99 | 0 |
| SPGP0T00470 | YLR369W | 99 | 0 |
| SPGP0D02710 | YOR047C | 99 | 0 |
| SPGP0A06640 | YDL190C | 99 | 0 |
| SPGP0D02120 | YOR115C | 99 | 0 |
| SPGP0A03130 | YDR206W | 99 | 0 |
| SPGP0D03850 | YOL078W | 99 | 0 |
| SPGP0AV00100 | YKL219W | 99 | 0 |
| SPGP0AV00110 | YKL218C | 99 | 0 |
| SPGP0N01660 | YJL025W | 99 | 0 |
| SPGP0N00570 | YJR103W | 99 | 0 |
| SPGP0AC01350 | YPR159W | 99 | 0 |
| SPGP0Y00230 | YGR123C | 99 | 0 |
| SPGP0AN00490 | YPL038W | 99 | 0 |
| SPGP0U00800 | YLR015W | 99 | 0 |
| SPGP0AV00140 | YKL215C | 99 | 0 |
| SPGP0D03720 | YOL065C | 99 | 0 |
| SPGP0G01510 | YMR159C | 99 | 0 |
| SPGP0D02580 | YOR063W | 99 | 0 |
| SPGP0J01650 | YBR133C | 99 | 0 |
| SPGP0AJ00520 | YAL026C | 99 | 0 |
| SPGP0N01480 | YJR001W | 99 | 0 |

Create PDF in your applications with the Pdfcrowd HTML to PDF API PDFCROWD

| | | | |
|---|---|---|---|
| SPGP0G01330 | YMR179W | 99 | 0 |
| SPGP0G01140 | YMR197C | 100 | 0 |
| SPGP0AA01670 | YPL078C | 99 | 0 |
| SPGP0AE00950 | YFL033C | 99 | 0 |
| SPGP0N01230 | YJR032W | 99 | 0 |
| SPGP0J03500 | YBL066C | 99 | 0 |
| SPGP0R00880 | YGR015C | 99 | 0 |
| SPGP0K00210 | YNR063W | 99 | 0 |
| SPGP0V01420 | YER057C | 100 | 0 |
| SPGP0AS00200 | YPL267W | 99 | 0 |
| SPGP0V01390 | YER055C | 99 | 0 |
| SPGP0AA01710 | YPL074W | 99 | 0 |
| SPGP0AP00210 | YHR177W | 99 | 0 |
| SPGP0V01930 | YER107C | 99 | 0 |
| SPGP0R00670 | YGR040W | 99 | 0 |
| SPGP0AB01210 | YIL128W | 99 | 0 |
| SPGP0G02540 | YMR048W | 98 | 1 |
| SPGP0U02090 | YLR154C | 100 | 0 |
| SPGP0IV00100 | YIR004W | 100 | 0 |
| SPGP0A00110 | YDR541C | 99 | 0 |
| SPGP0D02680 | YOR051C | 98 | 0 |
| SPGP0K01440 | YNL072W | 99 | 0 |
| SPGP0K01190 | YNL045W | 99 | 0 |
| SPGP0A01840 | YDR351W | 99 | 0 |
| SPGP0D02410 | YOR083W | 99 | 0 |
| SPGP0N03170 | YJL199C | 99 | 0 |
| SPGP0J02910 | YBL004W | 99 | 0 |
| SPGP0J02100 | YBR084W | 99 | 0 |
| SPGP0X00910 | YHR040W | 99 | 0 |
| SPGP0G00880 | YMR222C | 99 | 0 |
| SPGP0D03140 | YOL001W | 99 | 0 |
| SPGP0AH00300 | YCL040W | 99 | 0 |
| SPGP0A01860 | YDR349C | 99 | 0 |
| SPGP0G01460 | YMR164C | 97 | 2 |
| SPGP0N01730 | YJL035C | 99 | 0 |
| SPGP0V01640 | YER078C | 99 | 0 |
| SPGP0T01190 | YLR288C | 99 | 0 |
| SPGP0A06280 | YDL146W | 99 | 0 |
| SPGP0J02260 | YBR067C | 94 | 6 |
| SPGP0R01960 | YGL108C | 99 | 0 |

Create PDF in your applications with the Pdfcrowd HTML to PDF API PDFCROWD

| | | | |
|---|---|---|---|
| SPGP0Z00390 | YKL058W | 100 | 0 |
| SPGP0J01840 | YBR111W-A | 99 | 0 |
| SPGP0T00270 | YLR390W-A | 99 | 0 |
| SPGP0K01060 | YNL022C | 100 | 0 |
| SPGP0AJ00140 | YAR031W | 96 | 0 |
| SPGP0D03970 | YOL091W | 99 | 0 |
| SPGP0AE01110 | YFL049W | 99 | 0 |
| SPGP0V02010 | YER116C | 99 | 0 |
| SPGP0K01590 | YNL087W | 99 | 0 |
| SPGP0K01220 | YNL048W | 99 | 0 |
| SPGP0G03330 | YML036W | 99 | 0 |
| SPGP0U01240 | YLR063W | 99 | 0 |
| SPGP0G00800 | YMR230W | 100 | 0 |
| SPGP0D02810 | YOR036W | 99 | 0 |
| SPGP0N01360 | YJR011C | 99 | 0 |
| SPGP0N01760 | YJL038C | 99 | 0 |
| SPGP0N02020 | YJL068C | 99 | 0 |
| SPGP0N01590 | YJL013C | 99 | 0 |
| SPGP0N03390 | YOL159C-A | 100 | 0 |
| SPGP0X01560 | YHR106W | 100 | 0 |
| SPGP0AO00150 | YLR453C | 99 | 0 |
| SPGP0A03410 | YDR175C | 99 | 0 |
| SPGP0Z00540 | YKL072W | 99 | 0 |
| SPGP0DZ00100 | YGR087C | 99 | 0 |
| SPGP0AP00380 | YHR194W | 99 | 0 |
| SPGP0R01340 | YGL037C | 99 | 0 |
| SPGP0N02220 | YJL092W | 99 | 0 |
| SPGP0AC01130 | YPR134W | 99 | 0 |
| SPGP0V00110 | YEL072W | 99 | 0 |
| SPGP0AA00340 | YPL217C | 99 | 0 |
| SPGP0D04220 | YOL120C | 99 | 0 |
| SPGP0AM00310 | YER161C | 99 | 0 |
| SPGP0G02220 | YMR085W | 99 | 0 |
| SPGP0AA01600 | YPL086C | 99 | 0 |
| SPGP0G02730 | YMR027W | 99 | 0 |
| SPGP0AK00560 | YGL210W | 100 | 0 |
| SPGP0X01350 | YHR085W | 99 | 0 |
| SPGP0Z01380 | YKL167C | 99 | 0 |
| SPGP0K02750 | YNL219C | 99 | 0 |
| SPGP0D01540 | YOR179C | 99 | 0 |

Create PDF in your applications with the Pdfcrowd HTML to PDF API PDFCROWD

| | | | |
|---|---|---|---|
| SPGP0AC00880 | YPR107C | 99 | 0 |
| SPGP0AP00430 | YHR199C | 99 | 0 |
| SPGP0V01690 | YER083C | 99 | 0 |
| SPGP0G02390 | YMR066W | 99 | 0 |
| SPGP0N01630 | YJL020C | 99 | 0 |
| SPGP0D00800 | YOR262W | 99 | 0 |
| SPGP0AH00660 | YCR004C | 99 | 0 |
| SPGP0V01820 | YER095W | 99 | 0 |
| SPGP0A04210 | YDR087C | 99 | 0 |
| SPGP0AD00230 | YKR091W | 99 | 0 |
| SPGP0J03700 | YBL090W | 99 | 0 |
| SPGP0A03250 | YDR191W | 99 | 0 |
| SPGP0A04740 | YDR032C | 100 | 0 |
| SPGP0D00940 | YOR247W | 99 | 0 |
| SPGP0AE01020 | YFL039C | 100 | 0 |
| SPGP0A02390 | YDR293C | 99 | 0 |
| SPGP0D00760 | YOR267C | 97 | 2 |
| SPGP0J02450 | YBR047W | 99 | 0 |
| SPGP0Z01670 | YKL198C | 98 | 1 |
| SPGP0BE00100 | YPR005C | 99 | 0 |
| SPGP0T00460 | YLR370C | 99 | 0 |
| SPGP0G03750 | YML081W | 99 | 0 |
| SPGP0AC00260 | YPR032W | 99 | 0 |
| SPGP0A05160 | YDL019C | 99 | 0 |
| SPGP0Y00550 | YGR158C | 99 | 0 |
| SPGP0V01840 | YER098W | 99 | 0 |
| SPGP0T00110 | YLR409C | 99 | 0 |
| SPGP0K02610 | YNL204C | 99 | 0 |
| SPGP0AP00390 | YHR195W | 99 | 0 |
| SPGP0A03390 | YDR177W | 99 | 0 |
| SPGP0A02740 | YDR252W | 99 | 0 |
| SPGP0Z00940 | YKL119C | 99 | 0 |
| SPGP0AD00440 | YKR069W | 99 | 0 |
| SPGP0A06160 | YDL133W | 99 | 0 |
| SPGP0AU00220 | YOR357C | 100 | 0 |
| SPGP0R00180 | YGR098C | 99 | 0 |
| SPGP0AC00400 | YPR048W | 99 | 0 |
| SPGP0U00650 | YLL001W | 99 | 0 |
| SPGP0U00120 | YLL061W | 99 | 0 |
| SPGP0AH00980 | YCR036W | 99 | 0 |

Create PDF in your applications with the Pdfcrowd HTML to PDF API PDFCROWD

| | | | |
|---|---|---|---|
| SPGP0T00800 | YLR330W | 98 | 0 |
| SPGP0AB01380 | YIL146C | 99 | 0 |
| SPGP0AE00770 | YFL014W | 100 | 0 |
| SPGP0IE00100 | YIL004C | 100 | 0 |
| SPGP0AH00310 | YCL039W | 99 | 0 |
| SPGP0AH00900 | YCR028C | 99 | 0 |
| SPGP0G00670 | YMR243C | 99 | 0 |
| SPGP0AA00130 | YPL239W | 99 | 0 |
| SPGP0AM00480 | YER178W | 99 | 0 |
| SPGP0N02580 | YJL132W | 99 | 0 |
| SPGP0T01890 | YLR207W | 99 | 0 |
| SPGP0X00350 | YHL016C | 99 | 0 |
| SPGP0U00430 | YLL024C | 98 | 0 |
| SPGP0AA01070 | YPL141C | 99 | 0 |
| SPGP0R00250 | YGR091W | 100 | 0 |
| SPGP0N00960 | YJR061W | 99 | 0 |
| SPGP0T01770 | YLR220W | 100 | 0 |
| SPGP0J00420 | YBR262C | 99 | 0 |
| SPGP0K03050 | YNL252C | 99 | 0 |
| SPGP0G00430 | YMR269W | 100 | 0 |
| SPGP0G01740 | YMR134W | 99 | 0 |
| SPGP0R01450 | YGL050W | 99 | 0 |
| SPGP0A03830 | YDR129C | 99 | 0 |
| SPGP0N03360 | YJL221C | 98 | 0 |
| SPGP0R01050 | YGL005C | 99 | 0 |
| SPGP0AT00160 | YPR193C | 100 | 0 |
| SPGP0AC00950 | YPR114W | 99 | 0 |
| SPGP0J03390 | YBL054W | 99 | 0 |
| SPGP0A04660 | YDR042C | 99 | 0 |
| SPGP0BD00120 | YMR312W | 99 | 0 |
| SPGP0T01600 | YLR241W | 99 | 0 |
| SPGP0AN00340 | YPL021W | 99 | 0 |
| SPGP0D03290 | YOL016C | 99 | 0 |
| SPGP0G00320 | YMR280C | 99 | 0 |
| SPGP0V01090 | YER028C | 98 | 1 |
| SPGP0D04100 | YOL107W | 99 | 0 |
| SPGP0AW00170 | YHR156C | 99 | 0 |
| SPGP0K03150 | YNL262W | 99 | 0 |
| SPGP0BR00100 | YCR042C | 99 | 0 |
| SPGP0A04300 | YDR079W | 100 | 0 |

Create PDF in your applications with the Pdfcrowd HTML to PDF API PDFCROWD

| | | | |
|---|---|---|---|
| SPGP0A05180 | YDL021W | 100 | 0 |
| SPGP0G01100 | YMR201C | 99 | 0 |
| SPGP0U00880 | YLR024C | 99 | 0 |
| SPGP0D03380 | YOL025W | 99 | 0 |
| SPGP0D00360 | YOR310C | 99 | 0 |
| SPGP0Y00530 | YGR156W | 100 | 0 |
| SPGP0G01700 | YMR138W | 99 | 0 |
| SPGP0T00300 | YLR388W | 100 | 0 |
| SPGP0Y00240 | YGR124W | 99 | 0 |
| SPGP0AH00510 | YCL016C | 99 | 0 |
| SPGP0AV00200 | YKL209C | 99 | 0 |
| SPGP0D01950 | YDR385W | 99 | 0 |
| SPGP0AC01240 | YPR147C | 99 | 0 |
| SPGP0V01890 | YER103W | 99 | 0 |
| SPGP0X00510 | YHR001W-A | 99 | 0 |
| SPGP0AO00360 | YLR432W | 99 | 0 |
| SPGP0D00140 | YOR338W | 99 | 0 |
| SPGP0T02240 | YLR167W | 100 | 0 |
| SPGP0K02960 | YNL243W | 99 | 0 |
| SPGP0N00990 | YJR058C | 99 | 0 |
| SPGP0R00830 | YGR021W | 99 | 0 |
| SPGP0K01120 | YNL037C | 99 | 0 |
| SPGP0A01120 | YDR427W | 99 | 0 |
| SPGP0D00980 | YOR243C | 99 | 0 |
| SPGP0AB00350 | YIL034C | 100 | 0 |
| SPGP0R02070 | YGL121C | 99 | 0 |
| SPGP0AN00540 | YPL043W | 99 | 0 |
| SPGP0N02960 | YJL174W | 99 | 0 |
| SPGP0J02350 | YBR058C | 99 | 0 |
| SPGP0N01150 | YJR042W | 99 | 0 |
| SPGP0G02550 | YMR047C | 99 | 0 |
| SPGP0D02330 | YOR092W | 99 | 0 |
| SPGP0X01870 | YHR138C | 99 | 0 |
| SPGP0K01710 | YNL102W | 99 | 0 |
| SPGP0G02030 | YMR105C | 99 | 0 |
| SPGP0R00900 | YGR013W | 99 | 0 |
| SPGP0N00410 | YJR118C | 99 | 0 |
| SPGP0X01180 | YHR070W | 99 | 0 |
| SPGP0A00870 | YDR457W | 99 | 0 |
| SPGP0J03550 | YBL072C | 99 | 0 |

Create PDF in your applications with the Pdfcrowd HTML to PDF API PDFCROWD

| | | | |
|---|---|---|---|
| SPGP0D02090 | YOR118W | 100 | 0 |
| SPGP0Y01110 | YGR221C | 99 | 0 |
| SPGP0K01050 | YNL023C | 99 | 0 |
| SPGP0BV00130 | YCR084C | 99 | 0 |
| SPGP0R01430 | YGL048C | 99 | 0 |
| SPGP0AA01180 | YPL129W | 99 | 0 |
| SPGP0A03650 | YDR150W | 97 | 1 |
| SPGP0V00270 | YEL055C | 99 | 0 |
| SPGP0AM00540 | YER184C | 99 | 0 |
| SPGP0AK00320 | YGL237C | 99 | 0 |
| SPGP0D01790 | YOR152C | 99 | 0 |
| SPGP0J02160 | YBR078W | 99 | 0 |
| SPGP0J00730 | YBR233W | 99 | 0 |
| SPGP0U01410 | YLR081W | 99 | 0 |
| SPGP0D04140 | YOL111C | 99 | 0 |
| SPGP0Z01550 | YKL186C | 99 | 0 |
| SPGP0AB00500 | YIL049W | 98 | 0 |
| SPGP0K02500 | YNL191W | 99 | 0 |
| SPGP0AB01000 | YIL107C | 99 | 0 |
| SPGP0AH00790 | YHR145C | 98 | 0 |
| SPGP0J01180 | YBR183W | 99 | 0 |
| SPGP0N00680 | YJR093C | 99 | 0 |
| SPGP0Y01370 | YGR250C | 99 | 0 |
| SPGP0J01460 | YBR154C | 99 | 0 |
| SPGP0J02180 | YBR076W | 99 | 0 |
| SPGP0AE01140 | YHR213W-B | 99 | 0 |
| SPGP0AD00400 | YKR074W | 99 | 0 |
| SPGP0N03090 | YJL190C | 99 | 0 |
| SPGP0A04020 | YDR109C | 99 | 0 |
| SPGP0N03320 | YJL216C | 96 | 0 |
| SPGP0BT00100 | YIR021W | 99 | 0 |
| SPGP0V00370 | YEL044W | 99 | 0 |
| SPGP0V00850 | YER006W | 99 | 0 |
| SPGP0G01280 | YMR184W | 99 | 0 |
| SPGP0Z00520 | YKL070W | 99 | 0 |
| SPGP0G02690 | YMR031C | 99 | 0 |
| SPGP0D01370 | YOR197W | 99 | 0 |
| SPGP0R01750 | YGL085W | 99 | 0 |
| SPGP0D02660 | YOR054C | 99 | 0 |
| SPGP0N00350 | YJR125C | 99 | 0 |

Create PDF in your applications with the Pdfcrowd HTML to PDF API PDFCROWD

| | | | |
|---|---|---|---|
| SPGP0G03610 | YML067C | 99 | 0 |
| SPGP0Y00860 | YGR194C | 99 | 0 |
| SPGP0J01070 | YBR194W | 99 | 0 |
| SPGP0AA00630 | YPL187W | 99 | 0 |
| SPGP0AD00730 | YKR038C | 99 | 0 |
| SPGP0CX00120 | YCR047C | 99 | 0 |
| SPGP0AA01650 | YPL081W | 99 | 0 |
| SPGP0Z00980 | YKL124W | 99 | 0 |
| SPGP0K03100 | YNL257C | 99 | 0 |
| SPGP0D03110 | YOR003W | 99 | 0 |
| SPGP0J03410 | YBL056W | 99 | 0 |
| SPGP0AC00250 | YPR031W | 99 | 0 |
| SPGP0G02960 | YMR004W | 99 | 0 |
| SPGP0R00890 | YGR014W | 99 | 0 |
| SPGP0T02230 | YLR168C | 99 | 0 |
| SPGP0A00760 | YDR468C | 99 | 0 |
| SPGP0D01150 | YOR223W | 99 | 0 |
| SPGP0A05370 | YDL047W | 99 | 0 |
| SPGP0D03460 | YOL036W | 99 | 0 |
| SPGP0D00710 | YOR273C | 99 | 0 |
| SPGP0N01390 | YJR009C | 100 | 0 |
| SPGP0A06860 | YDL214C | 99 | 0 |
| SPGP0AE00540 | YFR012W | 94 | 4 |
| SPGP0K02980 | YNL245C | 99 | 0 |
| SPGP0G01200 | YMR192W | 99 | 0 |
| SPGP0AH00120 | YCL061C | 99 | 0 |
| SPGP0A03310 | YDR184C | 99 | 0 |
| SPGP0A06920 | YDL220C | 99 | 0 |
| SPGP0A06100 | YDL128W | 99 | 0 |
| SPGP0A04970 | YDR005C | 99 | 0 |
| SPGP0A03910 | YDR121W | 99 | 1 |
| SPGP0AH00360 | YCL034W | 99 | 0 |
| SPGP0AJ00810 | YAL058W | 99 | 0 |
| SPGP0G03080 | YML010W | 99 | 0 |
| SPGP0D00200 | YOR330C | 99 | 0 |
| SPGP0A01710 | YDR365C | 99 | 0 |
| SPGP0A01080 | YDR432W | 99 | 0 |
| SPGP0K03360 | YNL287W | 99 | 0 |
| SPGP0Y00350 | YGR135W | 99 | 0 |
| SPGP0A02680 | YDR258C | 99 | 0 |

Create PDF in your applications with the Pdfcrowd HTML to PDF API PDFCROWD

| | | | |
|---|---|---|---|
| SPGP0D01880 | YOR142W | 99 | 0 |
| SPGP0G01660 | YMR143W | 99 | 0 |
| SPGP0AC00670 | YPR081C | 99 | 0 |
| SPGP0J00790 | YBR225W | 99 | 0 |
| SPGP0K01330 | YNL061W | 99 | 0 |
| SPGP0D04360 | YOL135C | 99 | 0 |
| SPGP0AA01660 | YPL079W | 99 | 0 |
| SPGP0D04440 | YOL143C | 99 | 0 |
| SPGP0N02050 | YJL071W | 99 | 0 |
| SPGP0T02140 | YLR179C | 99 | 0 |
| SPGP0Z01150 | YKL142W | 99 | 0 |
| SPGP0A05080 | YDL007W | 99 | 0 |
| SPGP0Z01400 | YKL170W | 99 | 0 |
| SPGP0G03380 | YML043C | 99 | 0 |
| SPGP0AB00510 | YIL050W | 99 | 0 |
| SPGP0G01920 | YMR116C | 100 | 0 |
| SPGP0T00960 | YLR313C | 99 | 0 |
| SPGP0AD01060 | YKR003W | 99 | 0 |
| SPGP0G01820 | YMR126C | 100 | 0 |
| SPGP0AB00630 | YIL066C | 99 | 0 |
| SPGP0AK00450 | YGL224C | 100 | 0 |
| SPGP0N03210 | YJL204C | 99 | 0 |
| SPGP0N01970 | YJL062W | 99 | 0 |
| SPGP0V01010 | YER020W | 99 | 0 |
| SPGP0Y01020 | YGR211W | 99 | 0 |
| SPGP0G00470 | YMR265C | 99 | 0 |
| SPGP0G03320 | YML035C | 99 | 0 |
| SPGP0D00460 | YOR299W | 99 | 0 |
| SPGP0K02490 | YNL190W | 99 | 0 |
| SPGP0G00190 | YMR293C | 99 | 0 |
| SPGP0K02130 | YNL149C | 100 | 0 |
| SPGP0A02350 | YDR297W | 99 | 0 |
| SPGP0AA01240 | YPL123C | 99 | 0 |
| SPGP0K00860 | YNL007C | 98 | 2 |
| SPGP0A06550 | YDL178W | 99 | 0 |
| SPGP0AZ00180 | YIR008C | 99 | 0 |
| SPGP0X01390 | YHR089C | 99 | 0 |
| SPGP0J01310 | YBR168W | 99 | 0 |
| SPGP0X01670 | YHR117W | 99 | 0 |

Create PDF in your applications with the Pdfcrowd HTML to PDF API PDFCROWD

| | | | |
|---|---|---|---|
| SPGP0U01930 | YLR138W | 99 | 0 |
| SPGP0Y01030 | YGR212W | 99 | 0 |
| SPGP0J00210 | YBR285W | 100 | 0 |
| SPGP0U01290 | YLR068W | 100 | 0 |
| SPGP0D03230 | YOL010W | 99 | 0 |
| SPGP0A05880 | YDL104C | 99 | 0 |
| SPGP0K03390 | YNL290W | 99 | 0 |
| SPGP0Z00700 | YKL090W | 99 | 0 |
| SPGP0G03150 | YML017W | 99 | 0 |
| SPGP0N02440 | YJL117W | 99 | 0 |
| SPGP0AB00180 | YIL014W | 99 | 0 |
| SPGP0JE00100 | YCR063W | 99 | 0 |
| SPGP0AK00670 | YGL196W | 99 | 0 |
| SPGP0T01380 | YLR266C | 99 | 0 |
| SPGP0AK00180 | YGL252C | 99 | 0 |
| SPGP0J01920 | YBR104W | 100 | 0 |
| SPGP0A03330 | YDR182W | 99 | 0 |
| SPGP0G03780 | YML083C | 99 | 0 |
| SPGP0AJ00550 | YAL030W | 99 | 0 |
| SPGP0A06850 | YDL213C | 99 | 0 |
| SPGP0A02530 | YDR277C | 99 | 0 |
| SPGP0A06670 | YDL194W | 99 | 0 |
| SPGP0U01670 | YLR107W | 99 | 0 |
| SPGP0A01060 | YDR435C | 99 | 0 |
| SPGP0A04760 | YDR030C | 99 | 0 |
| SPGP0AA01050 | YPL144W | 99 | 0 |
| SPGP0A06520 | YDL175C | 99 | 0 |
| SPGP0U00350 | YLL033W | 99 | 0 |
| SPGP0D02670 | YOR052C | 99 | 0 |
| SPGP0U00820 | YLR017W | 99 | 0 |
| SPGP0A00720 | YDR472W | 99 | 0 |
| SPGP0A03480 | YDR169C | 99 | 0 |
| SPGP0A07100 | YDL240W | 98 | 0 |
| SPGP0D02730 | YOR045W | 100 | 0 |
| SPGP0R00470 | YGR063C | 99 | 0 |
| SPGP0G00330 | YMR279C | 99 | 0 |
| SPGP0R02100 | YGL124C | 99 | 0 |
| SPGP0G04060 | YML115C | 99 | 0 |
| SPGP0J03740 | YBL093C | 99 | 0 |
| SPGP0K03330 | YNL283C | 99 | 0 |

Create PDF in your applications with the Pdfcrowd HTML to PDF API PDFCROWD

| | | | |
|---|---|---|---|
| SPGP0G00940 | YMR216C | 99 | 0 |
| SPGP0V00800 | YER001W | 99 | 0 |
| SPGP0AN00630 | YPL053C | 99 | 0 |
| SPGP0U00680 | YLR003C | 99 | 1 |
| SPGP0AB01580 | YIL167W | 99 | 0 |
| SPGP0AV00160 | YKL213C | 99 | 0 |
| SPGP0R01180 | YGL019W | 99 | 0 |
| SPGP0J01670 | YBR131W | 99 | 0 |
| SPGP0Y01580 | YGR274C | 99 | 0 |
| SPGP0G01310 | YMR181C | 99 | 0 |
| SPGP0AO00140 | YLR454W | 99 | 0 |
| SPGP0V00300 | YEL052W | 99 | 0 |
| SPGP0K02860 | YNL232W | 99 | 0 |
| SPGP0A05640 | YDL078C | 99 | 0 |
| SPGP0Y00180 | YGR118W | 99 | 0 |
| SPGP0Y00440 | YGR146C | 99 | 0 |
| SPGP0A06000 | YDL116W | 99 | 0 |
| SPGP0D03830 | YOL077C | 99 | 0 |
| SPGP0J01520 | YBR148W | 99 | 0 |
| SPGP0R01940 | YGL106W | 99 | 0 |
| SPGP0AP00150 | YHR171W | 99 | 0 |
| SPGP0J03290 | YBL041W | 99 | 0 |
| SPGP0J03560 | YBL074C | 99 | 0 |
| SPGP0D03420 | YOL031C | 99 | 0 |
| SPGP0A02820 | YDR243C | 99 | 0 |
| SPGP0V01400 | YER056C | 99 | 0 |
| SPGP0K00130 | YNR071C | 100 | 0 |
| SPGP0AA01620 | YPL084W | 99 | 0 |
| SPGP0U02070 | YLR152C | 99 | 0 |
| SPGP0V01570 | YER072W | 100 | 0 |
| SPGP0N00160 | YJR144W | 99 | 0 |
| SPGP0D02350 | YOR090C | 99 | 0 |
| SPGP0R02330 | YGL148W | 99 | 0 |
| SPGP0A01820 | YDR353W | 99 | 0 |
| SPGP0K01420 | YNL070W | 100 | 0 |
| SPGP0A02020 | YDR329C | 99 | 0 |
| SPGP0N03150 | YJL197W | 99 | 0 |
| SPGP0A02790 | YDR246W | 99 | 0 |
| SPGP0Y00760 | YGR183C | 100 | 0 |
| SPGP0Y00980 | YGR206W | 99 | 0 |

Create PDF in your applications with the Pdfcrowd HTML to PDF API PDFCROWD

| | | | |
|---|---|---|---|
| SPGP0Z00350 | YKL054C | 98 | 0 |
| SPGP0AN00180 | YPL005W | 99 | 0 |
| SPGP0J02930 | YBL006C | 99 | 0 |
| SPGP0V01660 | YER080W | 99 | 0 |
| SPGP0T01220 | YLR286C | 98 | 2 |
| SPGP0J02270 | YBR066C | 99 | 0 |
| SPGP0N01340 | YJR013W | 99 | 0 |
| SPGP0G00910 | YMR219W | 99 | 0 |
| SPGP0D04080 | YOL104C | 99 | 0 |
| SPGP0K02370 | YNL176C | 98 | 1 |
| SPGP0KQ00100 | YGL192W | 99 | 0 |
| SPGP0N01740 | YJL036W | 99 | 0 |
| SPGP0X00970 | YHR047C | 99 | 0 |
| SPGP0AA00880 | YPL161C | 99 | 0 |
| SPGP0N01570 | YJL011C | 100 | 0 |
| SPGP0AM00290 | YER158C | 99 | 0 |
| SPGP0X00940 | YHR044C | 98 | 0 |
| SPGP0J01260 | YBR173C | 99 | 0 |
| SPGP0N01100 | YJR047C | 100 | 0 |
| SPGP0R00540 | YGR056W | 99 | 0 |
| SPGP0K01910 | YNL126W | 99 | 0 |
| SPGP0J00980 | YBR202W | 99 | 0 |
| SPGP0AO00130 | YLR455W | 99 | 0 |
| SPGP0AA00950 | YPL154C | 100 | 0 |
| SPGP0X01370 | YHR087W | 100 | 0 |
| SPGP0AP00360 | YHR192W | 99 | 0 |
| SPGP0D03760 | YOL069W | 99 | 0 |
| SPGP0N02200 | YJL090C | 99 | 0 |
| SPGP0R01320 | YGL035C | 99 | 0 |
| SPGP0AC01110 | YPR133C | 99 | 0 |
| SPGP0V00130 | YEL070W | 99 | 0 |
| SPGP0Z01360 | YKL165C | 99 | 0 |
| SPGP0D04200 | YOL117W | 99 | 0 |
| SPGP0AD00120 | YKR101W | 99 | 0 |
| SPGP0Y01390 | YGR252W | 99 | 0 |
| SPGP0V00490 | YEL032W | 99 | 0 |
| SPGP0V01100 | YER029C | 99 | 0 |
| SPGP0K03710 | YNL326C | 99 | 0 |
| SPGP0G02750 | YMR025W | 99 | 0 |
| SPGP0G02200 | YMR087W | 99 | 0 |

Create PDF in your applications with the Pdfcrowd HTML to PDF API PDFCROWD

| | | | |
|---|---|---|---|
| SPGP0AE00250 | YFR041C | 99 | 0 |
| SPGP0D01310 | YOR206W | 99 | 0 |
| SPGP0AM00370 | YER167W | 99 | 0 |
| SPGP0AS00250 | YPL262W | 99 | 0 |
| SPGP0AH00420 | YCL028W | 99 | 0 |
| SPGP0D03270 | YOL014W | 93 | 0 |
| SPGP0R02180 | YGL133W | 99 | 0 |
| SPGP0N01750 | YJL037W | 100 | 0 |
| SPGP0X01650 | YHR115C | 99 | 0 |
| SPGP0V01430 | YER058W | 99 | 0 |
| SPGP0K02770 | YNL221C | 99 | 0 |
| SPGP0AN00430 | YPL031C | 99 | 0 |
| SPGP0D01520 | YOR181W | 99 | 0 |
| SPGP0AP00410 | YHR197W | 99 | 0 |
| SPGP0Z00670 | YKL087C | 99 | 0 |
| SPGP0AP00140 | YHR170W | 99 | 0 |
| SPGP0V01700 | YER085C | 99 | 0 |
| SPGP0G02410 | YMR064W | 99 | 0 |
| SPGP0J02200 | YBR073W | 99 | 0 |
| SPGP0AE01100 | YFL048C | 99 | 0 |
| SPGP0A03200 | YDR197W | 99 | 0 |
| SPGP0AJ00580 | YAL033W | 99 | 0 |
| SPGP0T01840 | YLR212C | 99 | 0 |
| SPGP0AJ00230 | YAR015W | 99 | 0 |
| SPGP0AK00390 | YGL229C | 99 | 0 |
| SPGP0AD00380 | YKR076W | 99 | 0 |
| SPGP0A04840 | YDR021W | 99 | 0 |
| SPGP0AN00480 | YPL037C | 99 | 0 |
| SPGP0X00390 | YHL012W | 99 | 0 |
| SPGP0K01160 | YNL041C | 99 | 0 |
| SPGP0AE01000 | YFL037W | 99 | 0 |
| SPGP0A05810 | YDL097C | 99 | 0 |
| SPGP0BE00120 | YPR007C | 99 | 0 |
| SPGP0Z00900 | YKL113C | 99 | 0 |
| SPGP0Z00960 | YKL121W | 99 | 0 |
| SPGP0V00260 | YEL056W | 99 | 0 |
| SPGP0AC00790 | YPR097W | 99 | 0 |
| SPGP0K02600 | YNL202W | 99 | 0 |
| SPGP0J00700 | YBR235W | 99 | 0 |
| SPGP0A05140 | YDL014W | 99 | 0 |

Create PDF in your applications with the Pdfcrowd HTML to PDF API PDFCROWD

| | | | |
|---|---|---|---|
| SPGP0BL00100 | YCR048W | 99 | 1 |
| SPGP0AO00100 | YLR459W | 99 | 0 |
| SPGP0A00580 | YDR487C | 100 | 0 |
| SPGP0A03370 | YDR179C | 99 | 0 |
| SPGP0N02470 | YJL122W | 99 | 0 |
| SPGP0N02110 | YJL080C | 99 | 0 |
| SPGP0T01000 | YLR309C | 99 | 0 |
| SPGP0A06990 | YDL229W | 99 | 0 |
| SPGP0AN00400 | YPL028W | 99 | 0 |
| SPGP0K00450 | YNR038W | 99 | 0 |
| SPGP0AA01580 | YPL088W | 99 | 0 |
| SPGP0AC00420 | YPR051W | 99 | 0 |
| SPGP0AU00200 | YOR355W | 99 | 0 |
| SPGP0AC01050 | YPR125W | 100 | 0 |
| SPGP0BA00150 | YCR072C | 99 | 0 |
| SPGP0U00100 | YLL063C | 99 | 0 |
| SPGP0J01370 | YBR162W-A | 99 | 0 |
| SPGP0AE00970 | YFL034C-A | 99 | 0 |
| SPGP0R02500 | YGL169W | 99 | 0 |
| SPGP0AB01360 | YIL144W | 99 | 0 |
| SPGP0CR00100 | YMR303C | 97 | 0 |
| SPGP0G01290 | YMR183C | 99 | 0 |
| SPGP0AH00760 | YCR016W | 100 | 0 |
| SPGP0AK00260 | YGL244W | 99 | 0 |
| SPGP0D01020 | YOR238W | 99 | 0 |
| SPGP0AE00530 | YFR012W-A | 99 | 0 |
| SPGP0A06190 | YDL137W | 100 | 0 |
| SPGP0K00630 | YNR019W | 99 | 0 |
| SPGP0AA01720 | YPL072W | 99 | 0 |
| SPGP0K01870 | YNL122C | 100 | 0 |
| SPGP0N00980 | YJR059W | 99 | 0 |
| SPGP0R00270 | YGR089W | 99 | 0 |
| SPGP0R01740 | YGL084C | 99 | 0 |
| SPGP0T00390 | YLR378C | 99 | 0 |
| SPGP0J03380 | YBL052C | 99 | 0 |
| SPGP0A04640 | YDR044W | 100 | 0 |
| SPGP0X01110 | YHR063C | 99 | 0 |
| SPGP0Z01490 | YKL180W | 100 | 0 |
| SPGP0A03810 | YDR131C | 99 | 0 |
| SPGP0BC00100 | YIR023W | 99 | 0 |

Create PDF in your applications with the Pdfcrowd HTML to PDF API PDFCROWD

| | | | |
|---|---|---|---|
| SPGP0X00650 | YHR014W | 99 | 0 |
| SPGP0T01660 | YLR233C | 99 | 0 |
| SPGP0D03310 | YOL018C | 99 | 0 |
| SPGP0D01900 | YOR140W | 99 | 0 |
| SPGP0D04510 | YOL151W | 98 | 0 |
| SPGP0AW00140 | YHR154W | 99 | 0 |
| SPGP0U01500 | YLR090W | 99 | 0 |
| SPGP0X00950 | YHR045W | 99 | 0 |
| SPGP0J03770 | YBL098W | 99 | 0 |
| SPGP0A00800 | YDR463W | 99 | 0 |
| SPGP0AS00150 | YPL272C | 99 | 0 |
| SPGP0G02670 | YMR033W | 100 | 0 |
| SPGP0J00400 | YBR264C | 99 | 0 |
| SPGP0J00150 | YBR291C | 99 | 0 |
| SPGP0R01270 | YGL029W | 100 | 0 |
| SPGP0G01120 | YMR199W | 99 | 0 |
| SPGP0J01730 | YBR125C | 99 | 0 |
| SPGP0AK00240 | YGL246C | 99 | 0 |
| SPGP0A04920 | YDR013W | 99 | 0 |
| SPGP0D01030 | YOR237W | 99 | 0 |
| SPGP0BH00110 | YCR097W | 100 | 0 |
| SPGP0N03340 | YJL218W | 99 | 0 |
| SPGP0G03660 | YML072C | 99 | 0 |
| SPGP0A00100 | YAL068C | 98 | 0 |
| SPGP0N01170 | YJR040W | 99 | 0 |
| SPGP0AJ00350 | YAL009W | 99 | 0 |
| SPGP0A03700 | YDR144C | 99 | 0 |
| SPGP0J03170 | YBL030C | 99 | 0 |
| SPGP0AO00300 | YLR438C-A | 99 | 0 |
| SPGP0U01760 | YLR118C | 99 | 0 |
| SPGP0X01610 | YHR111W | 100 | 0 |
| SPGP0V01870 | YER101C | 99 | 0 |
| SPGP0D00190 | YOR332W | 99 | 0 |
| SPGP0D02700 | YOR048C | 99 | 0 |
| SPGP0K02940 | YNL241C | 99 | 0 |
| SPGP0N02970 | YJL176C | 99 | 0 |
| SPGP0K00970 | YNL031C | 100 | 0 |
| SPGP0AN00520 | YPL041C | 99 | 0 |
| SPGP0T00530 | YLR362W | 99 | 0 |
| SPGP0G03520 | YML058W-A | 99 | 0 |

Create PDF in your applications with the Pdfcrowd HTML to PDF API PDFCROWD

| | | | |
|---|---|---|---|
| SPGP0AB00370 | YIL036W | 99 | 0 |
| SPGP0A01450 | YDR389W | 99 | 0 |
| SPGP0J02370 | YBR056W | 99 | 0 |
| SPGP0R01070 | YGL008C | 99 | 0 |
| SPGP0AN00270 | YPL014W | 99 | 0 |
| SPGP0A03670 | YDR147W | 99 | 0 |
| SPGP0D04540 | YOL154W | 99 | 0 |
| SPGP0D04250 | YOL123W | 99 | 0 |
| SPGP0AA00720 | YPL177C | 99 | 0 |
| SPGP0D01280 | YOR209C | 99 | 0 |
| SPGP0G00750 | YMR235C | 99 | 0 |
| SPGP0D01000 | YOR241W | 99 | 0 |
| SPGP0AB01390 | YIL147C | 99 | 0 |
| SPGP0AA01310 | YPL116W | 99 | 0 |
| SPGP0A00890 | YDR454C | 99 | 0 |
| SPGP0T02250 | YLR166C | 99 | 0 |
| SPGP0Y01170 | YGR229C | 99 | 0 |
| SPGP0AH00680 | YCR007C | 99 | 0 |
| SPGP0A02070 | YDR323C | 99 | 0 |
| SPGP0Y01070 | YGR216C | 99 | 0 |
| SPGP0AP00270 | YHR183W | 99 | 0 |
| SPGP0Y00960 | YGR204W | 99 | 0 |
| SPGP0AW00230 | YHR162W | 100 | 0 |
| SPGP0G00990 | YMR211W | 99 | 0 |
| SPGP0X00560 | YHR005C-A | 100 | 0 |
| SPGP0AM00560 | YER186C | 96 | 1 |
| SPGP0U01430 | YLR083C | 99 | 0 |
| SPGP0AA01110 | YPL137C | 99 | 0 |
| SPGP0G01470 | YMR163C | 99 | 0 |
| SPGP0AA00540 | YPL195W | 99 | 0 |
| SPGP0J02960 | YBL009W | 99 | 0 |
| SPGP0G00180 | YMR294W | 99 | 0 |
| SPGP0V01250 | YER044C | 100 | 0 |
| SPGP0R01510 | YGL058W | 99 | 1 |
| SPGP0AC00870 | YPR106W | 99 | 0 |
| SPGP0T00920 | YLR318W | 99 | 0 |
| SPGP0V00910 | YER011W | 93 | 3 |
| SPGP0D02190 | YOR108W | 99 | 0 |
| SPGP0J01200 | YBR181C | 99 | 0 |
| SPGP0D02990 | YOR017W | 99 | 0 |

Create PDF in your applications with the Pdfcrowd HTML to PDF API PDFCROWD

| | | | |
|---|---|---|---|
| SPGP0G03360 | YML041C | 99 | 0 |
| SPGP0X00700 | YHR020W | 99 | 0 |
| SPGP0J00370 | YBR268W | 99 | 0 |
| SPGP0G01810 | YMR127C | 99 | 0 |
| SPGP0A02720 | YDR254W | 99 | 0 |
| SPGP0Z00490 | YKL068W | 98 | 0 |
| SPGP0A06080 | YDL126C | 99 | 0 |
| SPGP0K02560 | YNL197C | 99 | 0 |
| SPGP0T00310 | YLR387C | 99 | 0 |
| SPGP0A01400 | YDR394W | 99 | 0 |
| SPGP0AE00560 | YFR010W | 99 | 0 |
| SPGP0G03090 | YML011C | 99 | 0 |
| SPGP0J01050 | YBR196C | 99 | 0 |
| SPGP0Z00970 | YKL122C | 99 | 0 |
| SPGP0K01350 | YNL063W | 99 | 0 |
| SPGP0AD00240 | YKR090W | 99 | 0 |
| SPGP0D04560 | YOL155C | 92 | 2 |
| SPGP0AE00600 | YFR006W | 99 | 0 |
| SPGP0K03160 | YNL263C | 99 | 0 |
| SPGP0X01490 | YHR099W | 99 | 0 |
| SPGP0U01870 | YLR132C | 99 | 0 |
| SPGP0A06840 | YDL212W | 99 | 0 |
| SPGP0Z00640 | YKL084W | 99 | 0 |
| SPGP0AB00830 | YIL090W | 99 | 0 |
| SPGP0A05310 | YDL042C | 99 | 0 |
| SPGP0AO00290 | YLR439W | 99 | 0 |
| SPGP0Y00460 | YGR148C | 99 | 0 |
| SPGP0AY00110 | YOR378W | 99 | 0 |
| SPGP0A06230 | YDL141W | 99 | 0 |
| SPGP0BO00100 | YCR091W | 99 | 0 |
| SPGP0AE00690 | YFL004W | 99 | 0 |
| SPGP0A03270 | YDR189W | 99 | 0 |
| SPGP0AC01300 | YPR154W | 98 | 1 |
| SPGP0A06970 | YDL226C | 99 | 0 |
| SPGP0A04400 | YDR069C | 99 | 0 |
| SPGP0J01000 | YBR201W | 99 | 0 |
| SPGP0K03400 | YNL291C | 99 | 0 |
| SPGP0Z00790 | YKL100C | 99 | 0 |
| SPGP0A04950 | YDR007W | 99 | 0 |
| SPGP0T01470 | YLR256W | 99 | 0 |

Create PDF in your applications with the Pdfcrowd HTML to PDF API PDFCROWD

| | | | |
|---|---|---|---|
| SPGP0K03380 | YNL289W | 100 | 0 |
| SPGP0AE00630 | YFR003C | 99 | 0 |
| SPGP0D00590 | YOR287C | 99 | 0 |
| SPGP0AH00340 | YCL036W | 99 | 0 |
| SPGP0DD00110 | YCR060W | 100 | 0 |
| SPGP0G02940 | YMR007W | 99 | 0 |
| SPGP0G02120 | YMR095C | 99 | 0 |
| SPGP0U00140 | YLL058W | 99 | 0 |
| SPGP0J00780 | YBR227C | 99 | 0 |
| SPGP0N00710 | YJR090C | 99 | 0 |
| SPGP0K02470 | YNL188W | 99 | 0 |
| SPGP0AC00650 | YPR079W | 99 | 0 |
| SPGP0K02380 | YNL177C | 99 | 0 |
| SPGP0J00800 | YBR223C | 99 | 0 |
| SPGP0N02070 | YJL073W | 99 | 0 |
| SPGP0AB00570 | YIL057C | 100 | 0 |
| SPGP0G01940 | YMR114C | 99 | 0 |
| SPGP0Z01170 | YKL144C | 99 | 0 |
| SPGP0J00740 | YBR231C | 99 | 0 |
| SPGP0AD01040 | YKR005C | 99 | 0 |
| SPGP0A01040 | YDR437W | 99 | 0 |
| SPGP0G01800 | YMR128W | 99 | 0 |
| SPGP0AB00610 | YIL064W | 99 | 0 |
| SPGP0J03360 | YBL050W | 99 | 0 |
| SPGP0N01910 | YJL055W | 100 | 0 |
| SPGP0U00770 | YLR012C | 99 | 0 |
| SPGP0G01110 | YMR200W | 99 | 0 |
| SPGP0AK00470 | YGL222C | 99 | 0 |
| SPGP0GB00100 | YMR006C | 99 | 0 |
| SPGP0G01880 | YMR120C | 99 | 0 |
| SPGP0D01330 | YOR204W | 99 | 0 |
| SPGP0A03540 | YDR163W | 99 | 0 |
| SPGP0A02330 | YDR299W | 98 | 2 |
| SPGP0AS00220 | YPL265W | 99 | 0 |
| SPGP0X01970 | YHR148W | 99 | 0 |
| SPGP0A06530 | YDL176W | 99 | 0 |
| SPGP0AA01260 | YPL121C | 100 | 0 |
| SPGP0K00840 | YNL005C | 99 | 0 |
| SPGP0N00260 | YJR135C | 99 | 0 |
| SPGP0AD00950 | YKR015C | 98 | 0 |

Create PDF in your applications with the Pdfcrowd HTML to PDF API PDFCROWD

| | | | |
|---|---|---|---|
| SPGP0AB00620 | YIL065C | 100 | 0 |
| SPGP0J00160 | YBR290W | 100 | 0 |
| SPGP0G04170 | YML128C | 99 | 0 |
| SPGP0Y01530 | YGR268C | 99 | 0 |
| SPGP0U00590 | YLL007C | 99 | 0 |
| SPGP0N02460 | YJL121C | 99 | 0 |
| SPGP0T00700 | YLR344W | 99 | 0 |
| SPGP0A01670 | YDR370C | 99 | 0 |
| SPGP0EI00100 | YCR065W | 99 | 0 |
| SPGP0AM00440 | YER174C | 99 | 0 |
| SPGP0AK00650 | YGL198W | 99 | 0 |
| SPGP0V00680 | YEL015W | 99 | 0 |
| SPGP0J01940 | YBR102C | 99 | 0 |
| SPGP0D01200 | YOR217W | 99 | 0 |
| SPGP0AH00290 | YCL043C | 99 | 0 |
| SPGP0U00570 | YLL009C | 99 | 0 |
| SPGP0A06830 | YDL211C | 98 | 0 |
| SPGP0X01930 | YHR143W-A | 99 | 0 |
| SPGP0N00530 | YJR107W | 99 | 0 |
| SPGP0N02750 | YJL151C | 99 | 0 |
| SPGP0R00680 | YGR038W | 100 | 0 |
| SPGP0K03090 | YNL256W | 99 | 0 |
| SPGP0G00630 | YMR247C | 99 | 0 |
| SPGP0U00370 | YLL031C | 99 | 0 |
| SPGP0A00650 | YDR480W | 99 | 0 |
| SPGP0D03940 | YOL088C | 99 | 0 |
| SPGP0U01150 | YLR053C | 100 | 0 |
| SPGP0A05470 | YDL058W | 99 | 0 |
| SPGP0A05450 | YDL056W | 99 | 0 |
| SPGP0D02750 | YOR043W | 99 | 0 |
| SPGP0AB00140 | YIL009C-A | 99 | 0 |
| SPGP0A00460 | YDR500C | 99 | 0 |
| SPGP0G00310 | YMR281W | 100 | 0 |
| SPGP0J03760 | YBL097W | 99 | 0 |
| SPGP0X00780 | YHR028C | 99 | 0 |
| SPGP0AN00610 | YPL051W | 100 | 0 |
| SPGP0AB01560 | YIL164C | 99 | 0 |
| SPGP0U01400 | YLR080W | 99 | 0 |
| SPGP0J01380 | YBR162C | 99 | 0 |

Create PDF in your applications with the Pdfcrowd HTML to PDF API PDFCROWD

| | | | |
|---|---|---|---|
| SPGP0AO00160 | YLR452C | 99 | 0 |
| SPGP0A00660 | YDR479C | 99 | 0 |
| SPGP0CM00100 | YMR304W | 99 | 0 |
| SPGP0D03620 | YOL055C | 99 | 0 |
| SPGP0T00640 | YLR351C | 99 | 0 |
| SPGP0A06540 | YDL177C | 99 | 0 |
| SPGP0J01610 | YBR138C | 99 | 0 |
| SPGP0Z01460 | YKL176C | 99 | 0 |
| SPGP0T01740 | YLR223C | 100 | 0 |
| SPGP0K00420 | YNR041C | 99 | 0 |
| SPGP0J02490 | YBR043C | 99 | 0 |
| SPGP0G01370 | YMR175W | 100 | 0 |
| SPGP0A07120 | YDL243C | 95 | 0 |
| SPGP0AP00170 | YHR173C | 100 | 0 |
| SPGP0D03810 | YOL075C | 99 | 0 |
| SPGP0Y00160 | YGR116W | 99 | 0 |
| SPGP0R01890 | YGL100W | 99 | 0 |
| SPGP0AJ00680 | YAL042W | 99 | 0 |
| SPGP0V00280 | YEL054C | 99 | 0 |
| SPGP0V01460 | YER061C | 99 | 0 |
| SPGP0D02880 | YOR028C | 99 | 0 |
| SPGP0A02800 | YDR245W | 99 | 0 |
| SPGP0A05940 | YDL110C | 99 | 0 |
| SPGP0K00150 | YNR069C | 99 | 0 |
| SPGP0A02320 | YDR300C | 99 | 0 |
| SPGP0AJ00130 | YAR035W | 99 | 0 |
| SPGP0A07010 | YDL231C | 99 | 0 |
| SPGP0U02050 | YLR150W | 99 | 0 |
| SPGP0K01400 | YNL068C | 99 | 0 |
| SPGP0K01150 | YNL040W | 99 | 0 |
| SPGP0R02350 | YGL151W | 99 | 0 |
| SPGP0G00150 | YMR297W | 99 | 0 |
| SPGP0J02950 | YBL008W | 99 | 0 |
| SPGP0Y00700 | YGR174C | 99 | 0 |
| SPGP0D02370 | YOR087W | 99 | 0 |
| SPGP0G03760 | YML081C-A | 99 | 0 |
| SPGP0N00870 | YJR070C | 99 | 0 |
| SPGP0AA00390 | YPL212C | 99 | 0 |
| SPGP0V00550 | YEL026W | 100 | 0 |
| SPGP0AO00110 | YLR457C | 99 | 0 |

Create PDF in your applications with the Pdfcrowd HTML to PDF API PDFCROWD

| | | | |
|---|---|---|---|
| SPGP0D00230 | YOR327C | 99 | 0 |
| SPGP0G01250 | YMR187C | 99 | 0 |
| SPGP0R01630 | YGL071W | 99 | 0 |
| SPGP0N01090 | YJR048W | 100 | 0 |
| SPGP0J02630 | YBR028C | 99 | 0 |
| SPGP0J01170 | YBR184W | 99 | 0 |
| SPGP0J02140 | YBR080C | 99 | 0 |
| SPGP0V00330 | YEL049W | 99 | 0 |
| SPGP0N01720 | YJL034W | 99 | 0 |
| SPGP0J03680 | YBL088C | 99 | 0 |
| SPGP0X00960 | YHR046C | 99 | 0 |
| SPGP0AC00760 | YPR093C | 99 | 0 |
| SPGP0D02790 | YOR038C | 99 | 0 |
| SPGP0K02390 | YNL178W | 99 | 0 |
| SPGP0AA00970 | YPL152W | 99 | 0 |
| SPGP0K02170 | YNL154C | 99 | 0 |
| SPGP0R01300 | YGL032C | 99 | 0 |
| SPGP0AM00180 | YER147C | 99 | 0 |
| SPGP0R01650 | YGL075C | 99 | 0 |
| SPGP0AD01180 | YKL009W | 100 | 0 |
| SPGP0N02260 | YJL096W | 100 | 0 |
| SPGP0Z01600 | YKL191W | 99 | 0 |
| SPGP0K03660 | YNL320W | 99 | 0 |
| SPGP0K00660 | YNR016C | 99 | 0 |
| SPGP0K03310 | YNL281W | 100 | 0 |
| SPGP0A02120 | YDR320C | 99 | 0 |
| SPGP0V00510 | YEL030W | 99 | 0 |
| SPGP0A00300 | YDR518W | 99 | 0 |
| SPGP0J01020 | YBR199W | 99 | 0 |
| SPGP0G02770 | YMR023C | 99 | 0 |
| SPGP0BE00140 | YPR009W | 99 | 0 |
| SPGP0Z00690 | YKL089W | 99 | 0 |
| SPGP0K02710 | YNL215W | 99 | 0 |
| SPGP0D01500 | YOR184W | 99 | 0 |
| SPGP0D03560 | YOL048C | 99 | 0 |
| SPGP0D01050 | YOR234C | 99 | 0 |
| SPGP0V01720 | YER087W | 99 | 0 |
| SPGP0N01770 | YJL039C | 99 | 0 |
| SPGP0AP00160 | YHR172W | 99 | 0 |
| SPGP0R00430 | YGR068C | 99 | 0 |

Create PDF in your applications with the Pdfcrowd HTML to PDF API PDFCROWD

| | | | |
|---|---|---|---|
| SPGP0T01440 | YLR259C | 99 | 0 |
| SPGP0A03220 | YDR195W | 99 | 0 |
| SPGP0DC00110 | YPL060W | 99 | 0 |
| SPGP0G00950 | YMR215W | 99 | 0 |
| SPGP0T01860 | YLR210W | 99 | 0 |
| SPGP0V00840 | YER005W | 99 | 0 |
| SPGP0K00530 | YNR030W | 99 | 0 |
| SPGP0D04480 | YOL147C | 99 | 0 |
| SPGP0D00900 | YOR252W | 99 | 0 |
| SPGP0D03410 | YOL030W | 99 | 1 |
| SPGP0AJ00430 | YAL017W | 99 | 0 |
| SPGP0K03080 | YNL255C | 99 | 0 |
| SPGP0AE00920 | YFL029C | 99 | 0 |
| SPGP0K02580 | YNL200C | 99 | 0 |
| SPGP0V02080 | YER124C | 99 | 0 |
| SPGP0D03600 | YOL052C-A | 100 | 0 |
| SPGP0A01850 | YDR350C | 99 | 0 |
| SPGP0T02110 | YLR182W | 99 | 0 |
| SPGP0A01520 | YDR382W | 100 | 0 |
| SPGP0AC00220 | YPR028W | 99 | 0 |
| SPGP0N01840 | YJL048C | 99 | 0 |
| SPGP0HR00100 | R0020C | 99 | 0 |
| SPGP0A05120 | YDL012C | 99 | 0 |
| SPGP0Y01490 | YGR263C | 99 | 0 |
| SPGP0Z01270 | YKL156W | 100 | 0 |
| SPGP0X01060 | YHR058C | 99 | 0 |
| SPGP0G02650 | YMR035W | 99 | 0 |
| SPGP0N01510 | YJL003W | 100 | 0 |
| SPGP0G03420 | YML049C | 99 | 0 |
| SPGP0D02310 | YOR094W | 100 | 0 |
| SPGP0R02340 | YGL150C | 99 | 0 |
| SPGP0AH00460 | YCL025C | 99 | 0 |
| SPGP0AM00350 | YER165W | 99 | 0 |
| SPGP0K03440 | YNL295W | 99 | 0 |
| SPGP0AD00500 | YKR063C | 99 | 0 |
| SPGP0Z00150 | YKL032C | 98 | 1 |
| SPGP0J03750 | YBL095W | 99 | 0 |
| SPGP0AU00180 | YOR353C | 99 | 0 |
| SPGP0A00860 | YDR458C | 99 | 0 |
| SPGP0ACtrna26T | YPR108W-A | 100 | 0 |

Create PDF in your applications with the Pdfcrowd HTML to PDF API PDFCROWD

| | | | |
|---|---|---|---|
| SPGP0X00690 | YHR019C | 99 | 0 |
| SPGP0K00430 | YNR040W | 99 | 0 |
| SPGP0A02110 | YDR320C-A | 99 | 0 |
| SPGP0T01030 | YLR307W | 99 | 0 |
| SPGP0A05430 | YDL054C | 99 | 0 |
| SPGP0V01110 | YER030W | 99 | 0 |
| SPGP0BL00120 | YCR052W | 99 | 0 |
| SPGP0AN00390 | YPL027W | 99 | 0 |
| SPGP0AE00220 | YFR044C | 99 | 0 |
| SPGP0AE00730 | YFL010C | 99 | 0 |
| SPGP0AB01340 | YIL142W | 99 | 0 |
| SPGP0R02360 | YGL153W | 99 | 0 |
| SPGP0AD00790 | YKR030W | 99 | 0 |
| SPGP0T01130 | YLR295C | 99 | 0 |
| SPGP0T01640 | YLR237W | 97 | 0 |
| SPGP0G02720 | YMR028W | 99 | 0 |
| SPGP0AH00870 | YCR024C-A | 100 | 0 |
| SPGP0A02470 | YDR284C | 99 | 0 |
| SPGP0AM00190 | YER148W | 99 | 0 |
| SPGP0AA01700 | YPL075W | 99 | 0 |
| SPGP0K00380 | YNR046W | 100 | 0 |
| SPGP0AP00550 | YHR210C | 99 | 0 |
| SPGP0A07190 | YDL248W | 89 | 1 |
| SPGP0AK00770 | YGL184C | 99 | 0 |
| SPGP0J00390 | YBR265W | 99 | 0 |
| SPGP0BA00170 | YCR073W-A | 99 | 0 |
| SPGP0U00160 | YLL056C | 99 | 0 |
| SPGP0A01270 | YDR409W | 99 | 0 |
| SPGP0U00390 | YLL028W | 99 | 0 |
| SPGP0A03870 | YDR125C | 99 | 0 |
| SPGP0R01010 | YGL001C | 99 | 0 |
| SPGP0R01790 | YGL090W | 99 | 0 |
| SPGP0BM00100 | YIR039C | 99 | 0 |
| SPGP0Y00570 | YGR161C | 99 | 0 |
| SPGP0BU00110 | YIR022W | 99 | 0 |
| SPGP0G00760 | YMR234W | 99 | 0 |
| SPGP0U01560 | YLR096W | 99 | 0 |
| SPGP0BC00120 | YIR025W | 99 | 0 |
| SPGP0A06310 | YDL149W | 99 | 0 |
| SPGP0G03230 | YML025C | 99 | 0 |

Create PDF in your applications with the Pdfcrowd HTML to PDF API PDFCROWD

| | | | |
|---|---|---|---|
| SPGP0D02470 | YOR076C | 99 | 0 |
| SPGP0J00170 | YBR289W | 98 | 2 |
| SPGP0J02840 | YBR004C | 99 | 0 |
| SPGP0G03510 | YML058W | 99 | 0 |
| SPGP0G00230 | YMR289W | 99 | 0 |
| SPGP0CX00110 | YCR046C | 99 | 0 |
| SPGP0G01980 | YMR110C | 99 | 0 |
| SPGP0N02870 | YJL164C | 99 | 0 |
| SPGP0G03640 | YML070W | 99 | 0 |
| SPGP0R01250 | YGL027C | 99 | 0 |
| SPGP0A00960 | YDR447C | 100 | 0 |
| SPGP0G01210 | YMR191W | 99 | 0 |
| SPGP0AK00220 | YGL248W | 99 | 0 |
| SPGP0V01040 | YER023W | 99 | 0 |
| SPGP0N01110 | YJR046W | 99 | 0 |
| SPGP0J02700 | YBR020W | 99 | 0 |
| SPGP0J03600 | YBL079W | 99 | 0 |
| SPGP0U01220 | YLR060W | 99 | 0 |
| SPGP0J03150 | YBL029W | 99 | 0 |
| SPGP0X01940 | YHR144C | 99 | 0 |
| SPGP0AO00320 | YLR437C | 98 | 1 |
| SPGP0AD00190 | YKR095W | 99 | 0 |
| SPGP0AC01200 | YPR143W | 99 | 0 |
| SPGP0N02920 | YJL170C | 99 | 0 |
| SPGP0K02920 | YNL239W | 100 | 0 |
| SPGP0A01360 | YDR399W | 99 | 0 |
| SPGP0U00780 | YLR013W | 100 | 0 |
| SPGP0T00510 | YLR363W-A | 100 | 0 |
| SPGP0D04130 | YOL110W | 99 | 0 |
| SPGP0AH00520 | YCL014W | 99 | 0 |
| SPGP0AB00390 | YIL038C | 99 | 0 |
| SPGP0J02820 | YBR006W | 99 | 0 |
| SPGP0G02570 | YMR043W | 99 | 1 |
| SPGP0AB01530 | YIL160C | 99 | 0 |
| SPGP0Y01200 | YGR232W | 99 | 0 |
| SPGP0A06880 | YDL216C | 99 | 0 |
| SPGP0A03690 | YDR145W | 98 | 2 |
| SPGP0Y00390 | YGR141W | 99 | 0 |
| SPGP0Z00270 | YKL046C | 99 | 0 |
| SPGP0A06250 | YDL143W | 99 | 0 |

Create PDF in your applications with the Pdfcrowd HTML to PDF API PDFCROWD

| | | | |
|---|---|---|---|
| SPGP0N03070 | YJL187C | 99 | 0 |
| SPGP0Y00280 | YGR128C | 99 | 0 |
| SPGP0AZ00130 | YIR013C | 100 | 0 |
| SPGP0G03740 | YML080W | 99 | 0 |
| SPGP0N02430 | YJL116C | 99 | 0 |
| SPGP0R00980 | YGR003W | 99 | 0 |
| SPGP0AY00170 | YOR371C | 99 | 0 |
| SPGP0AD01160 | YKL007W | 99 | 0 |
| SPGP0R00350 | YGR078C | 99 | 0 |
| SPGP0Y01150 | YGR225W | 99 | 0 |
| SPGP0Z01000 | YKL126W | 99 | 0 |
| SPGP0D02420 | YOR081C | 99 | 0 |
| SPGP0AO00490 | YLR418C | 99 | 0 |
| SPGP0AH00910 | YCR028C-A | 100 | 0 |
| SPGP0N03050 | YJL185C | 99 | 0 |
| SPGP0G00970 | YMR213W | 99 | 0 |
| SPGP0T01560 | YLR245C | 100 | 0 |
| SPGP0X00540 | YHR004C | 99 | 0 |
| SPGP0X01960 | YHR147C | 99 | 0 |
| SPGP0AB00930 | YIL101C | 99 | 0 |
| SPGP0J00130 | YBR294W | 99 | 0 |
| SPGP0V01140 | YER033C | 99 | 0 |
| SPGP0AY00180 | YOR370C | 99 | 0 |
| SPGP0R01820 | YGL093W | 99 | 0 |
| SPGP0AB00220 | YIL017C | 99 | 0 |
| SPGP0R01530 | YGL060W | 99 | 0 |
| SPGP0J01140 | YBR187W | 99 | 0 |
| SPGP0D01750 | YOR156C | 99 | 0 |
| SPGP0N00640 | YJR096W | 99 | 0 |
| SPGP0J01420 | YBR158W | 99 | 0 |
| SPGP0J03450 | YBL059C-A | 99 | 0 |
| SPGP0V00160 | YEL066W | 99 | 0 |
| SPGP0K01560 | YNL084C | 99 | 0 |
| SPGP0Y01550 | YGR271W | 99 | 0 |
| SPGP0X00720 | YHR022C | 97 | 0 |
| SPGP0T00120 | YLR408C | 100 | 0 |
| SPGP0A01690 | YDR368W | 99 | 0 |
| SPGP0AA00560 | YPL193W | 99 | 0 |
| SPGP0G00170 | YMR295C | 99 | 0 |
| SPGP0K02540 | YNL195C | 99 | 0 |

Create PDF in your applications with the Pdfcrowd HTML to PDF API PDFCROWD

| | | | |
|---|---|---|---|
| SPGP0D01990 | YOR129C | 99 | 0 |
| SPGP0FA00100 | YIL007C | 99 | 0 |
| SPGP0T00330 | YLR385C | 100 | 0 |
| SPGP0N00390 | YJR121W | 99 | 0 |
| SPGP0AA01630 | YPL083C | 99 | 0 |
| SPGP0AE00500 | YFR015C | 99 | 0 |
| SPGP0R00690 | YGR037C | 100 | 0 |
| SPGP0X01550 | YHR105W | 99 | 0 |
| SPGP0AE00620 | YFR004W | 99 | 0 |
| SPGP0D04010 | YOL096C | 99 | 0 |
| SPGP0AD00260 | YKR088C | 99 | 0 |
| SPGP0AA00220 | YPL229W | 96 | 4 |
| SPGP0A06820 | YDL210W | 99 | 0 |
| SPGP0U01690 | YLR109W | 99 | 0 |
| SPGP0T02270 | YLR164W | 99 | 0 |
| SPGP0D01190 | YOR219C | 99 | 0 |
| SPGP0AE00130 | YFR053C | 99 | 0 |
| SPGP0A05330 | YDL044C | 99 | 0 |
| SPGP0V02060 | YER122C | 99 | 0 |
| SPGP0AB00800 | YIL087C | 99 | 0 |
| SPGP0Z00450 | YKL064W | 99 | 0 |
| SPGP0E00130 | YGR288W | 99 | 0 |
| SPGP0AB00550 | YIL055C | 99 | 0 |
| SPGP0G01960 | YMR112C | 99 | 0 |
| SPGP0AC01320 | YPR156C | 99 | 0 |
| SPGP0A01760 | YDR361C | 99 | 0 |
| SPGP0AB01140 | YIL121W | 99 | 0 |
| SPGP0U00510 | YLL015W | 99 | 0 |
| SPGP0A04420 | YDR067C | 99 | 0 |
| SPGP0Y01260 | YGR238C | 99 | 0 |
| SPGP0V01480 | YER063W | 100 | 0 |
| SPGP0G02430 | YMR062C | 100 | 0 |
| SPGP0G04150 | YML126C | 100 | 0 |
| SPGP0A06180 | YDL135C | 99 | 0 |
| SPGP0Y00310 | YGR131W | 99 | 0 |
| SPGP0K01390 | YNL067W | 99 | 0 |
| SPGP0A06950 | YDL224C | 99 | 0 |
| SPGP0D02210 | YOR106W | 99 | 0 |
| SPGP0N00690 | YJR092W | 99 | 0 |
| SPGP0N02010 | YJL066C | 99 | 0 |

Create PDF in your applications with the Pdfcrowd HTML to PDF API PDFCROWD

| | | | |
|---|---|---|---|
| SPGP0AP00200 | YHR176W | 99 | 0 |
| SPGP0K00880 | YNL009W | 99 | 0 |
| SPGP0T00480 | YLR368W | 99 | 0 |
| SPGP0A05840 | YDL100C | 99 | 0 |
| SPGP0A05240 | YDL029W | 99 | 0 |
| SPGP0G02080 | YMR099C | 99 | 0 |
| SPGP0AD01020 | YKR007W | 99 | 1 |
| SPGP0AK00410 | YGL227W | 99 | 0 |
| SPGP0N01280 | YJR021C | 99 | 0 |
| SPGP0N01930 | YJL058C | 99 | 0 |
| SPGP0D02960 | YOR020C | 100 | 0 |
| SPGP0D00240 | YOR326W | 99 | 0 |
| SPGP0A03560 | YDR161W | 99 | 0 |
| SPGP0N00280 | YJR133W | 99 | 0 |
| SPGP0X00170 | YHL034C | 99 | 0 |
| SPGP0J00410 | YBR263W | 99 | 0 |
| SPGP0A02370 | YDR295C | 99 | 0 |
| SPGP0X01630 | YHR113W | 99 | 0 |
| SPGP0AD00920 | YKR017C | 99 | 0 |
| SPGP0K02530 | YNL194C | 100 | 0 |
| SPGP0A06510 | YDL174C | 99 | 0 |
| SPGP0N00880 | YJR069C | 99 | 0 |
| SPGP0N03380 | YOL159C | 99 | 0 |
| SPGP0BA00110 | YCR067C | 99 | 0 |
| SPGP0R00330 | YGR080W | 99 | 0 |
| SPGP0R00530 | YGR057C | 99 | 0 |
| SPGP0T00720 | YLR342W | 99 | 0 |
| SPGP0V00660 | YEL017W | 99 | 0 |
| SPGP0A03990 | YDR113C | 99 | 0 |
| SPGP0X01820 | YHR133C | 99 | 0 |
| SPGP0U01630 | YLR103C | 99 | 0 |
| SPGP0A06350 | YDL155W | 99 | 0 |
| SPGP0AM00500 | YER180C | 99 | 0 |
| SPGP0A06810 | YDL209C | 99 | 0 |
| SPGP0D01220 | YOR215C | 99 | 0 |
| SPGP0AD00350 | YKR079C | 99 | 0 |
| SPGP0A03890 | YDR123C | 99 | 0 |
| SPGP0T01420 | YLR262C | 99 | 0 |
| SPGP0AB01540 | YIL161W | 99 | 0 |
| SPGP0A00630 | YDR482C | 100 | 0 |

Create PDF in your applications with the Pdfcrowd HTML to PDF API PDFCROWD

| | | | |
|---|---|---|---|
| SPGP0N02800 | YJL157C | 99 | 0 |
| SPGP0Y00630 | YGR168C | 99 | 0 |
| SPGP0U00310 | YLL038C | 99 | 0 |
| SPGP0G01680 | YMR140W | 99 | 0 |
| SPGP0J01960 | YBR098W | 99 | 0 |
| SPGP0X00760 | YHR026W | 100 | 0 |
| SPGP0R02140 | YGL128C | 99 | 0 |
| SPGP0AH00140 | YCL058C | 99 | 0 |
| SPGP0N02450 | YJL118W | 99 | 0 |
| SPGP0G00480 | YMR264W | 99 | 0 |
| SPGP0J02670 | YBR023C | 99 | 0 |
| SPGP0Y01330 | YGR246C | 99 | 0 |
| SPGP0K02060 | YNL141W | 99 | 0 |
| SPGP0Y00130 | YGR111W | 99 | 0 |
| SPGP0A06760 | YDL204W | 99 | 0 |
| SPGP0G00390 | YMR273C | 99 | 1 |
| SPGP0K00170 | YNR067C | 100 | 0 |
| SPGP0J01410 | YBR159W | 99 | 0 |
| SPGP0T00620 | YLR353W | 99 | 0 |
| SPGP0D01510 | YOR182C | 99 | 0 |
| SPGP0J00610 | YBR244W | 100 | 0 |
| SPGP0A00640 | YDR481C | 99 | 0 |
| SPGP0R02210 | YGL136C | 99 | 0 |
| SPGP0J01630 | YBR136W | 99 | 0 |
| SPGP0AE00240 | YFR042W | 99 | 0 |
| SPGP0R02110 | YGL125W | 99 | 0 |
| SPGP0A00340 | YDR514C | 99 | 0 |
| SPGP0A07140 | YDL245C | 99 | 0 |
| SPGP0A03190 | YDR198C | 99 | 0 |
| SPGP0J03820 | YBL104C | 99 | 0 |
| SPGP0Y00140 | YGR112W | 99 | 0 |
| SPGP0Y00490 | YGR152C | 100 | 0 |
| SPGP0V01330 | YER050C | 99 | 0 |
| SPGP0U00970 | YLR034C | 99 | 0 |
| SPGP0A02860 | YDR238C | 99 | 0 |
| SPGP0V01440 | YER059W | 99 | 0 |
| SPGP0AP00220 | YHR178W | 99 | 0 |
| SPGP0J02130 | YBR081C | 99 | 0 |
| SPGP0Z01470 | YKL178C | 99 | 0 |
| SPGP0A01790 | YDR357C | 99 | 0 |

Create PDF in your applications with the Pdfcrowd HTML to PDF API PDFCROWD

| | | | |
|---|---|---|---|
| SPGP0N03280 | YJL212C | 99 | 0 |
| SPGP0AS00360 | YPL250C | 99 | 0 |
| SPGP0U02030 | YLR148W | 99 | 0 |
| SPGP0A00420 | YDR505C | 99 | 0 |
| SPGP0K01170 | YNL042W | 99 | 0 |
| SPGP0G01350 | YMR177W | 99 | 0 |
| SPGP0X00770 | YHR027C | 99 | 0 |
| SPGP0D03120 | YOR002W | 99 | 0 |
| SPGP0N02650 | YJL139C | 99 | 0 |
| SPGP0X01210 | YHR072W-A | 100 | 0 |
| SPGP0A01590 | YDR377W | 100 | 0 |
| SPGP0D01910 | YOR138C | 99 | 0 |
| SPGP0J00940 | YBR207W | 99 | 0 |
| SPGP0R01490 | YGL056C | 99 | 0 |
| SPGP0R00340 | YGR079W | 99 | 0 |
| SPGP0AB00430 | YIL042C | 99 | 0 |
| SPGP0R01640 | YGL073W | 99 | 0 |
| SPGP0R02370 | YGL154C | 99 | 0 |
| SPGP0N00480 | YJR111C | 99 | 0 |
| SPGP0V00310 | YEL051W | 100 | 0 |
| SPGP0A04120 | YDR098C | 99 | 0 |
| SPGP0N01700 | YJL031C | 99 | 0 |
| SPGP0A03730 | YDR141C | 99 | 0 |
| SPGP0A05530 | YDL065C | 99 | 0 |
| SPGP0D04040 | YOL100W | 99 | 0 |
| SPGP0AJ00610 | YAL035W | 99 | 0 |
| SPGP0AC00750 | YPR091C | 99 | 0 |
| SPGP0AM00120 | YER141W | 99 | 0 |
| SPGP0A01770 | YDR359C | 99 | 0 |
| SPGP0A02560 | YDR273W | 99 | 0 |
| SPGP0A03010 | YDR222W | 99 | 0 |
| SPGP0K02480 | YNL189W | 99 | 0 |
| SPGP0AA00910 | YPL158C | 98 | 1 |
| SPGP0D01290 | YOR208W | 99 | 0 |
| SPGP0G01520 | YMR158W | 99 | 0 |
| SPGP0K02150 | YNL152W | 99 | 0 |
| SPGP0R01670 | YGL077C | 100 | 0 |
| SPGP0U01570 | YLR097C | 100 | 0 |
| SPGP0D02650 | YOR056C | 99 | 0 |

Create PDF in your applications with the Pdfcrowd HTML to PDF API PDFCROWD

| | | | |
|---|---|---|---|
| SPGP0V01260 | YER044C-A | 99 | 0 |
| SPGP0V00450 | YEL036C | 99 | 0 |
| SPGP0K01280 | YNL054W | 99 | 1 |
| SPGP0T02260 | YLR165C | 99 | 0 |
| SPGP0A00320 | YDR516C | 99 | 0 |
| SPGP0Z00100 | YKL025C | 100 | 0 |
| SPGP0X01130 | YHR065C | 99 | 0 |
| SPGP0V00170 | YEL065W | 99 | 0 |
| SPGP0AE00210 | YFR045W | 99 | 0 |
| SPGP0BS00110 | YEL069C | 96 | 0 |
| SPGP0AE00800 | YFL017W-A | 99 | 0 |
| SPGP0K03680 | YNL322C | 99 | 0 |
| SPGP0K00640 | YNR018W | 100 | 0 |
| SPGP0J01770 | YBR120C | 100 | 0 |
| SPGP0V01750 | YER089C | 100 | 0 |
| SPGP0R00610 | YGR047C | 99 | 0 |
| SPGP0AU00270 | YOR362C | 100 | 0 |
| SPGP0D01070 | YOR232W | 99 | 0 |
| SPGP0J03230 | YBL036C | 99 | 0 |
| SPGP0K02730 | YNL217W | 100 | 0 |
| SPGP0X01150 | YHR067W | 99 | 0 |
| SPGP0N03260 | YJL209W | 99 | 0 |
| SPGP0T01800 | YLR216C | 99 | 0 |
| SPGP0AJ00270 | YAR003W | 99 | 0 |
| SPGP0Z01160 | YKL143W | 99 | 0 |
| SPGP0Z01620 | YKL193C | 99 | 0 |
| SPGP0D03240 | YOL011W | 99 | 0 |
| SPGP0AH00440 | YCL026C-B | 100 | 0 |
| SPGP0AD00690 | YKR043C | 99 | 0 |
| SPGP0A05720 | YDL086W | 99 | 0 |
| SPGP0A05070 | YDL006W | 99 | 0 |
| SPGP0D01700 | YOR161C | 99 | 0 |
| SPGP0Z00920 | YKL116C | 99 | 0 |
| SPGP0AB01080 | YIL115C | 99 | 0 |
| SPGP0T02130 | YLR180W | 99 | 0 |
| SPGP0AC00200 | YPR026W | 99 | 0 |
| SPGP0R00940 | YGR007W | 99 | 0 |
| SPGP0D01610 | YOR172W | 99 | 0 |
| SPGP0LN00100 | YCR095C | 99 | 0 |
| SPGP0AK00550 | YGL211W | 99 | 0 |

Create PDF in your applications with the Pdfcrowd HTML to PDF API PDFCROWD

| | | | |
|---|---|---|---|
| SPGP0A02510 | YDR280W | 99 | 0 |
| SPGP0G03110 | YML013W | 99 | 0 |
| SPGP0K01820 | YNL116W | 99 | 0 |
| SPGP0G03440 | YML051W | 99 | 0 |
| SPGP0J01390 | YBR161W | 99 | 0 |
| SPGP0D02290 | YOR096W | 99 | 1 |
| SPGP0AS00210 | YPL266W | 99 | 0 |
| SPGP0A04850 | YDR020C | 99 | 0 |
| SPGP0K03460 | YNL298W | 99 | 0 |
| SPGP0AD00520 | YKR061W | 99 | 0 |
| SPGP0D00650 | YOR280C | 99 | 0 |
| SPGP0T01050 | YLR305C | 99 | 0 |
| SPGP0J01990 | YBR095C | 99 | 0 |
| SPGP0G01550 | YMR155W | 99 | 0 |
| SPGP0K02830 | YNL229C | 99 | 0 |
| SPGP0AN00440 | YPL032C | 99 | 0 |
| SPGP0AC01080 | YPR129W | 99 | 0 |
| SPGP0AA01540 | YPL092W | 99 | 0 |
| SPGP0AK00690 | YGL194C-A | 99 | 0 |
| SPGP0V01130 | YER032W | 99 | 0 |
| SPGP0Z01030 | YKL129C | 99 | 0 |
| SPGP0G03710 | YML077W | 99 | 0 |
| SPGP0AE00710 | YFL008W | 99 | 0 |
| SPGP0AU00160 | YOR351C | 99 | 0 |
| SPGP0T01940 | YLR201C | 99 | 0 |
| SPGP0AH00950 | YCR033W | 99 | 0 |
| SPGP0D00630 | YOR283W | 99 | 0 |
| SPGP0T01110 | YLR298C | 99 | 0 |
| SPGP0N03310 | YJL214W | 99 | 0 |
| SPGP0G02620 | YMR038C | 99 | 0 |
| SPGP0A02450 | YDR286C | 100 | 0 |
| SPGP0AP00470 | YHR202W | 99 | 0 |
| SPGP0A06150 | YDL132W | 99 | 0 |
| SPGP0A02100 | YDR321W | 99 | 0 |
| SPGP0X00880 | YHR038W | 99 | 0 |
| SPGP0AA01090 | YPL139C | 99 | 0 |
| SPGP0AA01760 | YPL068C | 100 | 0 |
| SPGP0R01030 | YGL003C | 99 | 0 |
| SPGP0R00450 | YGR066C | 100 | 0 |
| SPGP0A03230 | YDR194C | 99 | 0 |

Create PDF in your applications with the Pdfcrowd HTML to PDF API PDFCROWD

| | | | |
|---|---|---|---|
| SPGP0T00850 | YLR326W | 99 | 0 |
| SPGP0U01780 | YLR120C | 99 | 0 |
| SPGP0J02970 | YBL010C | 99 | 0 |
| SPGP0AD00250 | YKR089C | 99 | 0 |
| SPGP0A03850 | YDR127W | 99 | 0 |
| SPGP0AK00200 | YGL250W | 99 | 0 |
| SPGP0Y00640 | YGR169C | 99 | 0 |
| SPGP0AH00160 | YCL057W | 99 | 0 |
| SPGP0G00740 | YMR236W | 99 | 0 |
| SPGP0A06370 | YDL157C | 99 | 0 |
| SPGP0U01540 | YLR094C | 99 | 0 |
| SPGP0K03200 | YNL268W | 99 | 0 |
| SPGP0D04170 | YOL114C | 99 | 0 |
| SPGP0R00230 | YGR093W | 99 | 0 |
| SPGP0X00100 | YHL044W | 98 | 0 |
| SPGP0AD00650 | YKR048C | 99 | 0 |
| SPGP0U00550 | YLL011W | 99 | 0 |
| SPGP0J00110 | YBR296C | 99 | 0 |
| SPGP0G03250 | YML027W | 99 | 0 |
| SPGP0N02810 | YJL158C | 100 | 0 |
| SPGP0R01230 | YGL025C | 99 | 0 |
| SPGP0G01160 | YMR195W | 99 | 1 |
| SPGP0X01050 | YHR057C | 100 | 0 |
| SPGP0A02190 | YDR313C | 100 | 0 |
| SPGP0A03500 | YDR167W | 99 | 0 |
| SPGP0N01130 | YJR044C | 100 | 0 |
| SPGP0G03990 | YML108W | 99 | 0 |
| SPGP0AB01060 | YIL113W | 99 | 0 |
| SPGP0R00790 | YGR027C | 99 | 0 |
| SPGP0A06480 | YDL170W | 99 | 0 |
| SPGP0AD00160 | YKR097W | 99 | 0 |
| SPGP0R02190 | YGL134W | 99 | 0 |
| SPGP0K02900 | YNL237W | 99 | 0 |
| SPGP0R02440 | YGL161C | 99 | 0 |
| SPGP0A01110 | YDR428C | 99 | 0 |
| SPGP0X00500 | YHR001W | 99 | 0 |
| SPGP0T00570 | YLR359W | 99 | 0 |
| SPGP0Z00410 | YKL060C | 99 | 0 |
| SPGP0Y01220 | YGR234W | 99 | 0 |
| SPGP0AB00450 | YIL044C | 99 | 0 |

Create PDF in your applications with the Pdfcrowd HTML to PDF API PDFCROWD

| | | | |
|---|---|---|---|
| SPGP0V00780 | YEL002C | 99 | 0 |
| SPGP0D04290 | YOL127W | 100 | 0 |
| SPGP0AE00570 | YFR009W | 99 | 0 |
| SPGP0A01960 | YDR335W | 99 | 0 |
| SPGP0V01780 | YER092W | 99 | 0 |
| SPGP0AP00110 | YHR167W | 99 | 0 |
| SPGP0X01920 | YHR143W | 99 | 0 |
| SPGP0AZ00150 | YIR011C | 99 | 0 |
| SPGP0N02740 | YJL149W | 99 | 0 |
| SPGP0G03960 | YML105C | 99 | 0 |
| SPGP0Y01600 | YGR276C | 99 | 0 |
| SPGP0D02060 | YOR122C | 99 | 0 |
| SPGP0R01000 | YGR001C | 99 | 0 |
| SPGP0J01400 | YBR160W | 99 | 0 |
| SPGP0A04170 | YDR091C | 100 | 0 |
| SPGP0A01440 | YDR390C | 99 | 0 |
| SPGP0X00520 | YHR002W | 99 | 0 |
| SPGP0J02210 | YBR072W | 99 | 0 |
| SPGP0V01160 | YER035W | 99 | 0 |
| SPGP0Z00750 | YKL095W | 99 | 0 |
| SPGP0J02230 | YBR070C | 99 | 0 |
| SPGP0J01160 | YBR185C | 99 | 0 |
| SPGP0AY00160 | YOR372C | 99 | 0 |
| SPGP0D01730 | YOR158W | 99 | 0 |
| SPGP0R01800 | YGL091C | 100 | 0 |
| SPGP0J00660 | YBR239C | 99 | 0 |
| SPGP0N00660 | YJR094W-A | 99 | 0 |
| SPGP0K01580 | YNL086W | 99 | 0 |
| SPGP0AN00230 | YPL010W | 99 | 0 |
| SPGP0R01810 | YGL092W | 99 | 0 |
| SPGP0N02940 | YJL172W | 99 | 0 |
| SPGP0K01610 | YNL091W | 99 | 0 |
| SPGP0K01340 | YNL062C | 99 | 0 |
| SPGP0AD01290 | YKL019W | 99 | 0 |
| SPGP0N02400 | YJL111W | 99 | 0 |
| SPGP0X00290 | YHL022C | 99 | 0 |
| SPGP0Z00550 | YKL073W | 99 | 0 |
| SPGP0AA00580 | YPL191C | 99 | 0 |
| SPGP0AE00740 | YFL011W | 99 | 0 |
| SPGP0AA00320 | YPL219W | 99 | 0 |

Create PDF in your applications with the Pdfcrowd HTML to PDF API PDFCROWD

| | | | |
|---|---|---|---|
| SPGP0N01040 | YJR053W | 99 | 0 |
| SPGP0A06270 | YDL145C | 99 | 0 |
| SPGP0V01510 | YER066W | 99 | 0 |
| SPGP0G02990 | YMR001C | 99 | 0 |
| SPGP0Y01240 | YGR236C | 99 | 0 |
| SPGP0D01320 | YOR205C | 99 | 0 |
| SPGP0AE00520 | YFR013W | 99 | 0 |
| SPGP0AE00640 | YFR002W | 99 | 0 |
| SPGP0A04620 | YDR046C | 99 | 0 |
| SPGP0A04190 | YDR089W | 99 | 0 |
| SPGP0AD00200 | YKR094C | 99 | 0 |
| SPGP0D04520 | YOL152W | 99 | 0 |
| SPGP0AM00450 | YER175C | 99 | 0 |
| SPGP0AK00600 | YGL206C | 99 | 0 |
| SPGP0G02180 | YMR089C | 99 | 0 |
| SPGP0AB00870 | YIL094C | 99 | 0 |
| SPGP0K00200 | YNR064C | 100 | 0 |
| SPGP0AB01020 | YIL109C | 99 | 0 |
| SPGP0V02050 | YER120W | 99 | 0 |
| SPGP0Y00650 | YGR169C-A | 99 | 0 |
| SPGP0U01840 | YLR129W | 99 | 0 |
| SPGP0AA00240 | YPL227C | 99 | 0 |
| SPGP0N02380 | YJL109C | 99 | 0 |
| SPGP0R01110 | YGL012W | 99 | 0 |
| SPGP0A01740 | YDR363W | 99 | 0 |
| SPGP0N02030 | YJL069C | 99 | 0 |
| SPGP0A03300 | YDR185C | 99 | 0 |
| SPGP0AB00820 | YIL089W | 99 | 0 |
| SPGP0T02150 | YLR178C | 99 | 0 |
| SPGP0D00840 | YOR258W | 99 | 0 |
| SPGP0AE00190 | YFR047C | 99 | 0 |
| SPGP0AC01340 | YPR158W | 100 | 0 |
| SPGP0G03850 | YML093W | 99 | 0 |
| SPGP0G03580 | YML064C | 99 | 0 |
| SPGP0J00430 | YBR261C | 99 | 0 |
| SPGP0K03610 | YNL314W | 99 | 0 |
| SPGP0Z00610 | YKL080W | 99 | 0 |
| SPGP0X00890 | YHR039C | 100 | 0 |
| SPGP0D01460 | YOR188W | 99 | 0 |
| SPGP0G02450 | YMR060C | 99 | 0 |

Create PDF in your applications with the Pdfcrowd HTML to PDF API PDFCROWD

| | | | |
|---|---|---|---|
| SPGP0A01010 | YDR440W | 99 | 0 |
| SPGP0Z01290 | YKL159C | 100 | 0 |
| SPGP0A01530 | YDR381C-A | 100 | 0 |
| SPGP0A05230 | YDL028C | 99 | 0 |
| SPGP0N01960 | YJL061W | 99 | 0 |
| SPGP0D01710 | YOR160W | 99 | 0 |
| SPGP0Z00650 | YKL085W | 99 | 0 |
| SPGP0AD01000 | YKR009C | 99 | 0 |
| SPGP0N02510 | YJL126W | 99 | 0 |
| SPGP0K03290 | YNL279W | 99 | 0 |
| SPGP0AE00380 | YFR030W | 99 | 0 |
| SPGP0D00690 | YOR275C | 99 | 0 |
| SPGP0AK00430 | YGL226W | 99 | 0 |
| SPGP0A04270 | YDR081C | 99 | 0 |
| SPGP0A04990 | YDR003W | 99 | 0 |
| SPGP0N00300 | YJR131W | 99 | 0 |
| SPGP0AH00450 | YCL026C-A | 99 | 0 |
| SPGP0D01110 | YOR228C | 99 | 0 |
| SPGP0X00630 | YHR012W | 99 | 0 |
| SPGP0U01160 | YLR054C | 99 | 0 |
| SPGP0D03370 | YOL024W | 100 | 0 |
| SPGP0AD00980 | YKR011C | 99 | 0 |
| SPGP0AC00560 | YPR068C | 99 | 0 |
| SPGP0AC00980 | YPR117W | 99 | 0 |
| SPGP0A00810 | YDR462W | 100 | 0 |
| SPGP0A05710 | YDL085C-A | 100 | 0 |
| SPGP0T00740 | YLR340W | 100 | 0 |
| SPGP0A01220 | YDR415C | 99 | 0 |
| SPGP0JU00100 | YGR086C | 99 | 0 |
| SPGP0AE00420 | YFR023W | 99 | 0 |
| SPGP0A00130 | YDR539W | 99 | 0 |
| SPGP0A00390 | YDR508C | 99 | 0 |
| SPGP0A04980 | YDR004W | 99 | 0 |
| SPGP0AD00370 | YKR077W | 100 | 0 |
| SPGP0Y01500 | YGR264C | 99 | 0 |
| SPGP0N00330 | YJR127C | 99 | 0 |
| SPGP0A02590 | YDR268W | 100 | 0 |
| SPGP0D03820 | YOL076W | 99 | 0 |
| SPGP0AN00380 | YPL026C | 98 | 1 |
| SPGP0AA01370 | YPL109C | 99 | 0 |

Create PDF in your applications with the Pdfcrowd HTML to PDF API PDFCROWD

| | | | |
|---|---|---|---|
| SPGP0K00990 | YNL029C | 99 | 0 |
| SPGP0AN00120 | YPR002W | 99 | 0 |
| SPGP0AB01520 | YIL159W | 99 | 0 |
| SPGP0R00600 | YGR048W | 99 | 0 |
| SPGP0N02780 | YJL155C | 99 | 0 |
| SPGP0T01260 | YLR281C | 99 | 0 |
| SPGP0J00580 | YBR247C | 99 | 0 |
| SPGP0U00330 | YLL035W | 99 | 0 |
| SPGP0A00610 | YDR484W | 99 | 0 |
| SPGP0D02150 | YOR112W | 99 | 0 |
| SPGP0AC01270 | YPR151C | 100 | 0 |
| SPGP0J01980 | YBR096W | 99 | 0 |
| SPGP0D04400 | YOL139C | 100 | 0 |
| SPGP0K00820 | YNL003C | 100 | 0 |
| SPGP0Z00430 | YKL062W | 99 | 0 |
| SPGP0G00350 | YMR277W | 99 | 0 |
| SPGP0J03720 | YBL091C-A | 99 | 0 |
| SPGP0X01290 | YHR079C-A | 100 | 0 |
| SPGP0A00730 | YDR471W | 99 | 0 |
| SPGP0V01310 | YER048W-A | 99 | 0 |
| SPGP0A06740 | YDL202W | 99 | 0 |
| SPGP0A02840 | YDR240C | 99 | 0 |
| SPGP0K00190 | YNR065C | 96 | 0 |
| SPGP0AZ00110 | YIR017C | 100 | 0 |
| SPGP0AB01290 | YIL136W | 99 | 0 |
| SPGP0D03170 | YOL004W | 99 | 0 |
| SPGP0A01160 | YDR422C | 99 | 0 |
| SPGP0D01530 | YOR180C | 99 | 0 |
| SPGP0A00990 | YDR443C | 99 | 0 |
| SPGP0J00630 | YBR242W | 100 | 0 |
| SPGP0U00990 | YLR036C | 100 | 0 |
| SPGP0K01630 | YNL093W | 99 | 0 |
| SPGP0A05050 | YDL004W | 99 | 0 |
| SPGP0K00410 | YNR043W | 99 | 0 |
| SPGP0D03790 | YOL072W | 99 | 0 |
| SPGP0A07160 | YJR159W | 99 | 0 |
| SPGP0R02120 | YGL126W | 99 | 0 |
| SPGP0Y00120 | YGR110W | 99 | 0 |
| SPGP0G00920 | YMR218C | 99 | 0 |
| SPGP0U01060 | YLR044C | 99 | 0 |

Create PDF in your applications with the Pdfcrowd HTML to PDF API PDFCROWD

| | | | |
|---|---|---|---|
| SPGP0R01280 | YGL030W | 100 | 0 |
| SPGP0U00290 | YLL040C | 99 | 0 |
| SPGP0AB01430 | YIL151C | 99 | 0 |
| SPGP0D04060 | YOL102C | 99 | 0 |
| SPGP0U02010 | YLR146W-A | 100 | 0 |
| SPGP0A00190 | YDR531W | 99 | 0 |
| SPGP0G01400 | YMR172W | 99 | 0 |
| SPGP0AB00210 | YIL016W | 99 | 0 |
| SPGP0R02170 | YGL131C | 99 | 0 |
| SPGP0J02980 | YBL011W | 99 | 0 |
| SPGP0AE01090 | YFL047W | 99 | 0 |
| SPGP0N03010 | YJL180C | 99 | 0 |
| SPGP0J03540 | YBL071W-A | 98 | 0 |
| SPGP0X00750 | YHR025W | 99 | 0 |
| SPGP0A01570 | YDR379W | 99 | 0 |
| SPGP0Y00250 | YGR125W | 99 | 0 |
| SPGP0J00960 | YBR204C | 99 | 0 |
| SPGP0G00250 | YMR287C | 99 | 0 |
| SPGP0R01470 | YGL054C | 99 | 0 |
| SPGP0Y00520 | YGR155W | 99 | 0 |
| SPGP0K01110 | YNL036W | 99 | 0 |
| SPGP0Y01290 | YGR241C | 99 | 0 |
| SPGP0A05130 | YDL013W | 99 | 0 |
| SPGP0R02390 | YGL156W | 99 | 0 |
| SPGP0G00720 | YMR238W | 99 | 0 |
| SPGP0A03710 | YDR143C | 99 | 0 |
| SPGP0AC00720 | YPR086W | 99 | 0 |
| SPGP0X00930 | YHR042W | 99 | 0 |
| SPGP0A05550 | YDL067C | 99 | 0 |
| SPGP0J03200 | YBL033C | 99 | 0 |
| SPGP0Z01660 | YKL197C | 99 | 0 |
| SPGP0D02040 | YOR124C | 99 | 0 |
| SPGP0V01940 | YER109C | 99 | 0 |
| SPGP0A02990 | YDR224C | 100 | 0 |
| SPGP0Z01590 | YKL190W | 100 | 0 |
| SPGP0AP00370 | YHR193C | 100 | 0 |
| SPGP0AM00140 | YER143W | 99 | 0 |
| SPGP0U00920 | YLR028C | 99 | 0 |
| SPGP0AA00930 | YPL156C | 99 | 0 |
| SPGP0D01270 | YOR210W | 100 | 0 |

Create PDF in your applications with the Pdfcrowd HTML to PDF API PDFCROWD

| | | | |
|---|---|---|---|
| SPGP0U01550 | YLR095C | 99 | 0 |
| SPGP0AA01590 | YPL087W | 100 | 0 |
| SPGP0Y01470 | YGR261C | 99 | 0 |
| SPGP0D02890 | YOR027W | 99 | 0 |
| SPGP0AH00890 | YCR027C | 99 | 0 |
| SPGP0N00130 | YJR148W | 99 | 0 |
| SPGP0V00190 | YEL063C | 99 | 0 |
| SPGP0V00470 | YEL034W | 99 | 0 |
| SPGP0R02250 | YGL140C | 99 | 0 |
| SPGP0U01750 | YLR117C | 99 | 0 |
| SPGP0AM00170 | YER146W | 99 | 0 |
| SPGP0AM00160 | YER145C | 99 | 0 |
| SPGP0D01010 | YOR239W | 99 | 0 |
| SPGP0N00850 | YJR073C | 99 | 0 |
| SPGP0AC00570 | YPR069C | 99 | 0 |
| SPGP0AA01800 | YPL064C | 99 | 0 |
| SPGP0AC00820 | YPR101W | 99 | 0 |
| SPGP0Y00100 | YGR108W | 99 | 0 |
| SPGP0V02140 | YER130C | 99 | 0 |
| SPGP0J01210 | YBR180W | 99 | 0 |
| SPGP0A04290 | YDR079C-A | 99 | 0 |
| SPGP0U01200 | YLR058C | 99 | 0 |
| SPGP0N03240 | YJL207C | 99 | 0 |
| SPGP0T01820 | YLR214W | 99 | 0 |
| SPGP0A05690 | YDL084W | 99 | 0 |
| SPGP0AK00190 | YGL251C | 99 | 0 |
| SPGP0K03260 | YNL275W | 99 | 0 |
| SPGP0AD00770 | YKR034W | 100 | 0 |
| SPGP0D03480 | YOL039W | 100 | 0 |
| SPGP0G02840 | YMR016C | 99 | 0 |
| SPGP0AB01100 | YIL117C | 99 | 0 |
| SPGP0V02040 | YER119C | 99 | 0 |
| SPGP0AC00770 | YPR094W | 99 | 0 |
| SPGP0AC00410 | YPR049C | 99 | 0 |
| SPGP0N03270 | YJL210W | 99 | 0 |
| SPGP0BK00100 | YGL178W | 99 | 0 |
| SPGP0D01300 | YOR207C | 99 | 0 |
| SPGP0K01800 | YNL113W | 99 | 0 |
| SPGP0Z01560 | YKL187C | 99 | 0 |
| SPGP0G03460 | YML053C | 96 | 4 |

Create PDF in your applications with the Pdfcrowd HTML to PDF API PDFCROWD

| | | | |
|---|---|---|---|
| SPGP0G03050 | YML007W | 99 | 1 |
| SPGP0K00540 | YNR029C | 99 | 0 |
| SPGP0AA00260 | YPL225W | 99 | 0 |
| SPGP0G02510 | YMR053C | 99 | 0 |
| SPGP0J01970 | YBR097W | 99 | 0 |
| SPGP0BC00180 | YIR031C | 100 | 0 |
| SPGP0T01350 | YLR270W | 99 | 0 |
| SPGP0N02060 | YJL072C | 99 | 0 |
| SPGP0AH00390 | YCL031C | 99 | 0 |
| SPGP0AD00540 | YKR059W | 99 | 0 |
| SPGP0AB01320 | YIL139C | 99 | 0 |
| SPGP0AV00180 | YKL211C | 99 | 0 |
| SPGP0AM00390 | YER169W | 99 | 0 |
| SPGP0G01570 | YMR153W | 99 | 0 |
| SPGP0AS00230 | YPL264C | 99 | 0 |
| SPGP0AN00460 | YPL034W | 99 | 0 |
| SPGP0T01070 | YLR303W | 99 | 0 |
| SPGP0V01150 | YER034W | 99 | 0 |
| SPGP0AH00770 | YCR017C | 99 | 0 |
| SPGP0Z01010 | YKL127W | 99 | 0 |
| SPGP0AJ00170 | YAR028W | 98 | 0 |
| SPGP0AE00260 | YFR040W | 99 | 0 |
| SPGP0AC01280 | YPR152C | 99 | 1 |
| SPGP0AK00590 | YGL207W | 99 | 0 |
| SPGP0N01870 | YJL051W | 99 | 0 |
| SPGP0G03870 | YML095C | 99 | 0 |
| SPGP0V01230 | YER042W | 99 | 0 |
| SPGP0T01170 | YLR290C | 99 | 0 |
| SPGP0D00610 | YOR285W | 99 | 0 |
| SPGP0T01680 | YLR229C | 99 | 0 |
| SPGP0T00590 | YLR356W | 99 | 0 |
| SPGP0AC00480 | YPR058W | 100 | 0 |
| SPGP0BY00110 | YIR035C | 100 | 0 |
| SPGP0J00350 | YBR270C | 99 | 0 |
| SPGP0T00880 | YLR323C | 99 | 0 |
| SPGP0AH00410 | YCL029C | 99 | 0 |
| SPGP0V00930 | YER013W | 99 | 0 |
| SPGP0AE00840 | YFL022C | 99 | 0 |
| SPGP0AA00750 | YPL174C | 99 | 0 |

Create PDF in your applications with the Pdfcrowd HTML to PDF API PDFCROWD

| | | | |
|---|---|---|---|
| SPGP0J02090 | YBR084C-A | 99 | 0 |
| SPGP0R01920 | YGL104C | 99 | 0 |
| SPGP0Z00340 | YKL053C-A | 99 | 0 |
| SPGP0Z00800 | YKL101W | 99 | 0 |
| SPGP0Z01480 | YKL179C | 99 | 0 |
| SPGP0G00590 | YMR252C | 99 | 0 |
| SPGP0D00430 | YOR303W | 99 | 0 |
| SPGP0AA01740 | YPL070W | 99 | 0 |
| SPGP0A06170 | YDL134C | 99 | 0 |
| SPGP0R01220 | YGL023C | 99 | 0 |
| SPGP0K00690 | YNR013C | 99 | 0 |
| SPGP0A04380 | YDR071C | 100 | 0 |
| SPGP0G03190 | YML021C | 99 | 0 |
| SPGP0G03140 | YML016C | 99 | 0 |
| SPGP0D03610 | YOL053W | 99 | 0 |
| SPGP0X00680 | YHR018C | 99 | 0 |
| SPGP0K03230 | YNL272C | 99 | 0 |
| SPGP0U00530 | YLL013C | 99 | 0 |
| SPGP0N02830 | YJL160C | 99 | 0 |
| SPGP0T00550 | YLR361C | 99 | 0 |
| SPGP0V01550 | YER070W | 99 | 0 |
| SPGP0T00220 | YLR396C | 99 | 0 |
| SPGP0X01730 | YHR123W | 99 | 0 |
| SPGP0A03520 | YDR165W | 99 | 0 |
| SPGP0K00900 | YNL011C | 100 | 0 |
| SPGP0R00740 | YGR032W | 99 | 0 |
| SPGP0AO00350 | YLR433C | 99 | 0 |
| SPGP0N00140 | YJR147W | 99 | 0 |
| SPGP0AM00260 | YER155C | 99 | 0 |
| SPGP0U01210 | YLR059C | 99 | 0 |
| SPGP0K03480 | YNL300W | 99 | 0 |
| SPGP0J03040 | YBL018C | 99 | 0 |
| SPGP0G01270 | YMR185W | 99 | 0 |
| SPGP0R00770 | YGR029W | 99 | 0 |
| SPGP0A05360 | YDL046W | 98 | 0 |
| SPGP0AV00150 | YKL214C | 99 | 0 |
| SPGP0Y01660 | YGR282C | 99 | 0 |
| SPGP0AK00810 | YGL179C | 99 | 0 |
| SPGP0A01460 | YDR388W | 99 | 0 |
| SPGP0BN00110 | YCR087W | 94 | 6 |

Create PDF in your applications with the Pdfcrowd HTML to PDF API PDFCROWD

| | | | |
|---|---|---|---|
| SPGP0X01230 | YHR074W | 99 | 0 |
| SPGP0D04270 | YOL125W | 99 | 0 |
| SPGP0K03670 | YNL321W | 99 | 0 |
| SPGP0J03440 | YBL059W | 99 | 0 |
| SPGP0K00740 | YNR008W | 99 | 0 |
| SPGP0J03110 | YBL025W | 99 | 0 |
| SPGP0X01900 | YHR141C | 100 | 0 |
| SPGP0V01760 | YER090W | 100 | 0 |
| SPGP0AT00140 | YPR196W | 99 | 0 |
| SPGP0R02310 | YGL146C | 99 | 0 |
| SPGP0AW00260 | YHR165C | 99 | 0 |
| SPGP0AJ00240 | YAR014C | 99 | 0 |
| SPGP0N00600 | YJR100C | 99 | 0 |
| SPGP0AZ00210 | YIR005W | 100 | 0 |
| SPGP0A01420 | YDR392W | 100 | 0 |
| SPGP0J03160 | YBL029C-A | 100 | 0 |
| SPGP0J02240 | YBR069C | 99 | 0 |
| SPGP0A02280 | YDR304C | 100 | 0 |
| SPGP0Y00820 | YGR189C | 96 | 4 |
| SPGP0D01470 | YOR187W | 99 | 0 |
| SPGP0N01830 | YJL047C | 99 | 0 |
| SPGP0N01250 | YJR030C | 99 | 0 |
| SPGP0AA00740 | YPL175W | 99 | 0 |
| SPGP0D04390 | YOL138C | 99 | 0 |
| SPGP0R01860 | YGL097W | 99 | 0 |
| SPGP0AY00200 | YOR368W | 99 | 0 |
| SPGP0X01000 | YHR050W | 99 | 0 |
| SPGP0A05800 | YDL095W | 99 | 0 |
| SPGP0AN00250 | YPL012W | 99 | 0 |
| SPGP0A00380 | YDR510W | 99 | 0 |
| SPGP0K01320 | YNL059C | 99 | 0 |
| SPGP0V01530 | YER068W | 99 | 0 |
| SPGP0Y01080 | YGR217W | 99 | 0 |
| SPGP0AA00530 | YPL196W | 99 | 0 |
| SPGP0D01040 | YOR236W | 100 | 0 |
| SPGP0K01520 | YNL080C | 99 | 0 |
| SPGP0A01030 | YDR438W | 99 | 0 |
| SPGP0K02230 | YNL160W | 99 | 0 |
| SPGP0G02980 | YMR002W | 97 | 2 |
| SPGP0T00370 | YLR381W | 99 | 0 |

Create PDF in your applications with the Pdfcrowd HTML to PDF API PDFCROWD

| | | | |
|---|---|---|---|
| SPGP0D02320 | YOR093C | 99 | 0 |
| SPGP0D02230 | YOR103C | 98 | 0 |
| SPGP0AE00490 | YFR016C | 99 | 0 |
| SPGP0G02970 | YMR003W | 99 | 0 |
| SPGP0AP00320 | YHR188C | 99 | 0 |
| SPGP0Y01430 | YGR256W | 99 | 0 |
| SPGP0Y01620 | YGR278W | 99 | 0 |
| SPGP0V00530 | YEL028W | 99 | 0 |
| SPGP0AE00110 | YFR055W | 99 | 0 |
| SPGP0Z01410 | YKL171W | 99 | 0 |
| SPGP0G02610 | YMR039C | 99 | 1 |
| SPGP0AM00470 | YER177W | 99 | 0 |
| SPGP0K02850 | YNL231C | 99 | 0 |
| SPGP0K00180 | YNR066C | 98 | 0 |
| SPGP0U01860 | YLR131C | 99 | 0 |
| SPGP0X01270 | YHR078W | 100 | 0 |
| SPGP0D01480 | YOR186W | 98 | 0 |
| SPGP0K02670 | YNL211C | 100 | 0 |
| SPGP0X01760 | YHR127W | 99 | 0 |
| SPGP0A02430 | YDR288W | 99 | 0 |
| SPGP0AD00550 | YKR058W | 99 | 0 |
| SPGP0R01130 | YGL014W | 99 | 0 |
| SPGP0AB00840 | YIL091C | 99 | 0 |
| SPGP0AC00100 | YPR016C | 100 | 0 |
| SPGP0T01990 | YLR195C | 99 | 0 |
| SPGP0AB00590 | YIL062C | 99 | 0 |
| SPGP0U01360 | YLR075W | 100 | 0 |
| SPGP0A03450 | YDR171W | 99 | 0 |
| SPGP0AC01360 | YPR160W | 99 | 0 |
| SPGP0AM00300 | YER159C | 99 | 0 |
| SPGP0A04600 | YDR049W | 99 | 0 |
| SPGP0X00580 | YHR007C | 99 | 0 |
| SPGP0D04050 | YOL101C | 99 | 0 |
| SPGP0AD00220 | YKR092C | 94 | 3 |
| SPGP0D04500 | YOL149W | 99 | 0 |
| SPGP0A06220 | YDL140C | 99 | 0 |
| SPGP0G02470 | YMR058W | 99 | 0 |
| SPGP0AM00400 | YER170W | 99 | 0 |
| SPGP0BC00130 | YIR026C | 99 | 0 |
| SPGP0A06430 | YDL165W | 100 | 0 |

Create PDF in your applications with the Pdfcrowd HTML to PDF API PDFCROWD

| | | | |
|---|---|---|---|
| SPGP0AP00240 | YHR180W | 99 | 0 |
| SPGP0A04250 | YDR083W | 99 | 0 |
| SPGP0AJ00750 | YAL049C | 99 | 0 |
| SPGP0A04460 | YDR063W | 99 | 0 |
| SPGP0A06070 | YDL125C | 100 | 0 |
| SPGP0AB00600 | YIL063C | 99 | 0 |
| SPGP0D00280 | YOR321W | 99 | 0 |
| SPGP0G00240 | YMR288W | 99 | 0 |
| SPGP0A02600 | YDR266C | 99 | 0 |
| SPGP0A05090 | YDL007C-A | 100 | 0 |
| SPGP0D02170 | YOR110W | 99 | 0 |
| SPGP0R00290 | YGR084C | 99 | 0 |
| SPGP0AC00540 | YPR066W | 99 | 0 |
| SPGP0AC00960 | YPR115W | 99 | 0 |
| SPGP0AB00890 | YIL096C | 99 | 0 |
| SPGP0A05730 | YDL087C | 99 | 0 |
| SPGP0T00940 | YLR315W | 100 | 0 |
| SPGP0Z00290 | YKL048C | 99 | 0 |
| SPGP0T00760 | YLR336C | 99 | 0 |
| SPGP0A01640 | YDR373W | 99 | 0 |
| SPGP0A03760 | YDR138W | 99 | 0 |
| SPGP0AD00810 | YKR028W | 99 | 0 |
| SPGP0Y01480 | YGR262C | 100 | 0 |
| SPGP0N00320 | YJR129C | 99 | 0 |
| SPGP0Y01690 | YGR285C | 99 | 0 |
| SPGP0AP00230 | YHR179W | 99 | 0 |
| SPGP0T01780 | YLR219W | 99 | 0 |
| SPGP0AN00100 | YPR004C | 99 | 0 |
| SPGP0AA01350 | YPL111W | 99 | 0 |
| SPGP0AB01500 | YIL157C | 99 | 0 |
| SPGP0R00370 | YGR076C | 99 | 0 |
| SPGP0V01030 | YER022W | 99 | 0 |
| SPGP0AE00100 | YFR057W | 100 | 0 |
| SPGP0J00490 | YBR255C-A | 99 | 0 |
| SPGP0J01880 | YBR108W | 98 | 1 |
| SPGP0AD00150 | YKR098C | 99 | 0 |
| SPGP0A03950 | YDR118W | 99 | 0 |
| SPGP0T02190 | YLR174W | 99 | 0 |
| SPGP0G00690 | YMR241W | 100 | 0 |
| SPGP0A00590 | YDR486C | 100 | 0 |

Create PDF in your applications with the Pdfcrowd HTML to PDF API PDFCROWD

| | | | |
|---|---|---|---|
| SPGP0A04570 | YDR052C | 99 | 0 |
| SPGP0T01540 | YLR247C | 99 | 0 |
| SPGP0AH00180 | YCL055W | 99 | 0 |
| SPGP0D00350 | YOR311C | 99 | 0 |
| SPGP0D03190 | YOL006C | 99 | 0 |
| SPGP0N01410 | YJR007W | 99 | 0 |
| SPGP0V01370 | YER053C-A | 100 | 0 |
| SPGP0D02490 | YOR074C | 99 | 0 |
| SPGP0A06720 | YDL200C | 99 | 0 |
| SPGP0T00580 | YLR357W | 100 | 0 |
| SPGP0K02790 | YNL223W | 99 | 0 |
| SPGP0Y00670 | YGR171C | 99 | 0 |
| SPGP0A00970 | YDR446W | 99 | 0 |
| SPGP0J00650 | YBR240C | 99 | 0 |
| SPGP0N02990 | YJL178C | 99 | 0 |
| SPGP0AJ00460 | YAL020C | 99 | 0 |
| SPGP0G03180 | YML020W | 99 | 0 |
| SPGP0G03620 | YML068W | 99 | 0 |
| SPGP0G01640 | YMR145C | 99 | 0 |
| SPGP0U00940 | YLR030W | 99 | 0 |
| SPGP0AC00190 | YPR025C | 99 | 0 |
| SPGP0D03250 | YOL012C | 100 | 0 |
| SPGP0J02760 | YBR014C | 99 | 0 |
| SPGP0D00210 | YOR329C | 99 | 0 |
| SPGP0K01810 | YNL115C | 99 | 0 |
| SPGP0J01830 | YBR112C | 99 | 1 |
| SPGP0N01860 | YJL050W | 99 | 0 |
| SPGP0U01000 | YLR037C | 100 | 0 |
| SPGP0D02140 | YOR113W | 99 | 1 |
| SPGP0A03290 | YDR186C | 99 | 0 |
| SPGP0T00250 | YLR393W | 100 | 0 |
| SPGP0R02150 | YGL129C | 99 | 0 |
| SPGP0G01390 | YMR173W | 87 | 5 |
| SPGP0X00300 | YHL021C | 99 | 0 |
| SPGP0BX00100 | YFR026C | 100 | 0 |
| SPGP0AY00100 | YOR380W | 99 | 0 |
| SPGP0N01070 | YJR050W | 99 | 0 |
| SPGP0A00840 | YDR460W | 99 | 0 |
| SPGP0K02090 | YNL145W | 99 | 0 |
| SPGP0A01950 | YDR336W | 99 | 0 |

Create PDF in your applications with the Pdfcrowd HTML to PDF API PDFCROWD

| | | | |
|---|---|---|---|
| SPGP0N00840 | YJR074W | 100 | 0 |
| SPGP0AJ00250 | YAR008W | 100 | 0 |
| SPGP0A01550 | YDR380W | 99 | 0 |
| SPGP0V01590 | YER074W | 99 | 0 |
| SPGP0J03300 | YBL042C | 99 | 0 |
| SPGP0X01040 | YHR056C | 99 | 0 |
| SPGP0T01750 | YLR222C | 99 | 0 |
| SPGP0AA01450 | YPL100W | 99 | 0 |
| SPGP0J03220 | YBL035C | 99 | 0 |
| SPGP0U01370 | YLR077W | 99 | 0 |
| SPGP0K01130 | YNL038W | 99 | 0 |
| SPGP0Y01270 | YGR239C | 99 | 0 |
| SPGP0AC00780 | YPR095C | 99 | 0 |
| SPGP0X00450 | YHL006C | 99 | 0 |
| SPGP0V00350 | YEL047C | 99 | 0 |
| SPGP0V00420 | YEL039C | 99 | 0 |
| SPGP0A01890 | YDR346C | 99 | 1 |
| SPGP0A03770 | YDR137W | 99 | 0 |
| SPGP0A05500 | YDL061C | 100 | 0 |
| SPGP0D03130 | YOR001W | 99 | 0 |
| SPGP0AK00480 | YGL221C | 100 | 0 |
| SPGP0D02020 | YOR126C | 99 | 0 |
| SPGP0A01730 | YDR363W-A | 99 | 0 |
| SPGP0A03900 | YDR122W | 99 | 0 |
| SPGP0Z01650 | YKL196C | 99 | 0 |
| SPGP0D02130 | YOR114W | 99 | 0 |
| SPGP0D01250 | YOR212W | 99 | 0 |
| SPGP0A02970 | YDR226W | 99 | 0 |
| SPGP0K02110 | YNL147W | 100 | 0 |
| SPGP0Y01410 | YGR254W | 99 | 0 |
| SPGP0N01550 | YJL008C | 100 | 0 |
| SPGP0Z01630 | YKL194C | 99 | 0 |
| SPGP0AB00420 | YIL041W | 99 | 0 |
| SPGP0D04190 | YOL116W | 99 | 0 |
| SPGP0AH00740 | YCR014C | 99 | 0 |
| SPGP0R02270 | YGL142C | 99 | 0 |
| SPGP0J01590 | YBR140C | 99 | 0 |
| SPGP0X00310 | YHL020C | 99 | 1 |
| SPGP0AE01080 | YFL046W | 99 | 0 |
| SPGP0N00860 | YJR072C | 99 | 0 |

Create PDF in your applications with the Pdfcrowd HTML to PDF API PDFCROWD

| | | | |
|---|---|---|---|
| SPGP0K01620 | YNL092W | 99 | 0 |
| SPGP0AA00940 | YPL155C | 99 | 0 |
| SPGP0T02060 | YLR188W | 99 | 0 |
| SPGP0AC00840 | YPR103W | 100 | 0 |
| SPGP0K01790 | YNL112W | 99 | 0 |
| SPGP0AB00720 | YIL075C | 99 | 0 |
| SPGP0D03570 | YOL049W | 99 | 0 |
| SPGP0N03220 | YJL205C | 99 | 0 |
| SPGP0T01590 | YLR242C | 99 | 0 |
| SPGP0J01350 | YBR164C | 99 | 0 |
| SPGP0A05670 | YDL082W | 99 | 0 |
| SPGP0K03240 | YNL273W | 99 | 0 |
| SPGP0A01500 | YDR384C | 99 | 0 |
| SPGP0AE00510 | YFR014C | 99 | 0 |
| SPGP0G02860 | YMR014W | 99 | 0 |
| SPGP0G03160 | YML018C | 99 | 0 |
| SPGP0AP00490 | YHR204W | 99 | 0 |
| SPGP0G00160 | YMR296C | 99 | 0 |
| SPGP0K03060 | YNL253W | 99 | 0 |
| SPGP0AE00860 | YFL023W | 100 | 0 |
| SPGP0AZ00190 | YIR007W | 99 | 0 |
| SPGP0G03030 | YML005W | 99 | 0 |
| SPGP0K01860 | YNL121C | 99 | 0 |
| SPGP0A07040 | YDL234C | 99 | 0 |
| SPGP0AE00170 | YFR049W | 100 | 0 |
| SPGP0Z01260 | YKL155C | 99 | 0 |
| SPGP0N02080 | YJL074C | 99 | 0 |
| SPGP0A04810 | YDR024W | 99 | 0 |
| SPGP0K03420 | YNL293W | 99 | 0 |
| SPGP0AD00560 | YKR057W | 99 | 0 |
| SPGP0D04310 | YOL129W | 99 | 0 |
| SPGP0T01370 | YLR267W | 99 | 0 |
| SPGP0A00360 | YDR512C | 100 | 0 |
| SPGP0AB01300 | YIL137C | 99 | 0 |
| SPGP0J02020 | YBR091C | 99 | 0 |
| SPGP0T02170 | YLR176C | 99 | 0 |
| SPGP0V01170 | YER036C | 99 | 0 |
| SPGP0G00860 | YMR224C | 99 | 0 |
| SPGP0N02180 | YJL088W | 99 | 0 |
| SPGP0AO00210 | YLR447C | 99 | 0 |

Create PDF in your applications with the Pdfcrowd HTML to PDF API PDFCROWD

| | | | |
|---|---|---|---|
| SPGP0D01260 | YOR211C | 99 | 0 |
| SPGP0N02170 | YJL087C | 99 | 0 |
| SPGP0N02520 | YJL127C | 99 | 0 |
| SPGP0AB00470 | YIL046W | 99 | 0 |
| SPGP0Z01070 | YKL134C | 99 | 0 |
| SPGP0A01480 | YDR386W | 100 | 0 |
| SPGP0AC01000 | YPR119W | 99 | 0 |
| SPGP0G03860 | YML094W | 100 | 0 |
| SPGP0X01700 | YHR120W | 99 | 0 |
| SPGP0G02230 | YMR083W | 99 | 0 |
| SPGP0AK00570 | YGL209W | 99 | 0 |
| SPGP0K00700 | YNR012W | 99 | 0 |
| SPGP0G03210 | YML023C | 99 | 0 |
| SPGP0T01150 | YLR292C | 99 | 0 |
| SPGP0D01090 | YOR230W | 99 | 0 |
| SPGP0X00460 | YHL004W | 99 | 0 |
| SPGP0AA01500 | YPL096W | 99 | 0 |
| SPGP0R00140 | YGR102C | 99 | 0 |
| SPGP0R00410 | YGR071C | 99 | 0 |
| SPGP0G04130 | YML123C | 99 | 0 |
| SPGP0J00330 | YBR272C | 99 | 0 |
| SPGP0A01240 | YDR412W | 100 | 0 |
| SPGP0AK00760 | YGL185C | 100 | 0 |
| SPGP0AE00320 | YFR034C | 99 | 0 |
| SPGP0N02500 | YJL125C | 99 | 0 |
| SPGP0Z00810 | YKL103C | 100 | 0 |
| SPGP0AE00820 | YFL020C | 99 | 0 |
| SPGP0J01820 | YBR114W | 99 | 0 |
| SPGP0AD00320 | YKR082W | 99 | 0 |
| SPGP0AH00750 | YCR015C | 99 | 0 |
| SPGP0G03350 | YML038C | 99 | 0 |
| SPGP0A06110 | YDL129W | 99 | 0 |
| SPGP0A05510 | YDL063C | 99 | 0 |
| SPGP0A05590 | YDL073W | 99 | 0 |
| SPGP0J00480 | YBR256C | 99 | 0 |
| SPGP0A04360 | YDR073W | 99 | 0 |
| SPGP0A05190 | YDL022W | 100 | 0 |
| SPGP0R01200 | YGL021W | 99 | 0 |
| SPGP0U00270 | YLL042C | 98 | 0 |
| SPGP0T00200 | YLR398C | 99 | 0 |

Create PDF in your applications with the Pdfcrowd HTML to PDF API PDFCROWD

| | | | |
|---|---|---|---|
| SPGP0N00590 | YJR101W | 99 | 0 |
| SPGP0J02570 | YBR035C | 99 | 0 |
| SPGP0N02700 | YJL145W | 99 | 0 |
| SPGP0Z01540 | YKL185W | 99 | 0 |
| SPGP0AA00410 | YPL210C | 100 | 0 |
| SPGP0A04690 | YDR037W | 99 | 0 |
| SPGP0G00700 | YMR240C | 99 | 0 |
| SPGP0U01270 | YLR066W | 100 | 0 |
| SPGP0U00760 | YLR011W | 99 | 0 |
| SPGP0AM00280 | YER157W | 99 | 0 |
| SPGP0AC01260 | YPR149W | 100 | 0 |
| SPGP0K01210 | YNL047C | 99 | 0 |
| SPGP0X01360 | YHR086W | 99 | 0 |
| SPGP0K02260 | YNL162W-A | 99 | 0 |
| SPGP0AE00340 | YFR032C-A | 99 | 0 |
| SPGP0BY00130 | YIR037W | 99 | 0 |
| SPGP0Y01640 | YGR280C | 99 | 0 |
| SPGP0G01890 | YMR119W | 99 | 0 |
| SPGP0T02080 | YLR186W | 99 | 0 |
| SPGP0G01070 | YMR203W | 99 | 0 |
| SPGP0A02130 | YDR319C | 99 | 0 |
| SPGP0D02560 | YOR065W | 99 | 0 |
| SPGP0KM00100 | YKL222C | 99 | 0 |
| SPGP0A05910 | YDL107W | 99 | 0 |
| SPGP0A02930 | YDR231C | 99 | 0 |
| SPGP0R02510 | YGL170C | 99 | 0 |
| SPGP0AW00240 | YHR163W | 100 | 0 |
| SPGP0J03460 | YBL060W | 99 | 0 |
| SPGP0V01740 | YER088C | 99 | 0 |
| SPGP0R02030 | YGL116W | 99 | 0 |
| SPGP0A05900 | YDL106C | 99 | 0 |
| SPGP0D03360 | YOL023W | 99 | 0 |
| SPGP0AB01310 | YIL138C | 100 | 0 |
| SPGP0R00400 | YGR072W | 99 | 0 |
| SPGP0J03190 | YBL032W | 99 | 0 |
| SPGP0X01800 | YHR132C | 99 | 0 |
| SPGP0J01750 | YBR122C | 99 | 0 |
| SPGP0Z00470 | YKL065W-A | 99 | 0 |
| SPGP0N00620 | YJR098C | 99 | 0 |
| SPGP0R02040 | YGL117W | 99 | 0 |

Create PDF in your applications with the Pdfcrowd HTML to PDF API PDFCROWD

| | | | |
|---|---|---|---|
| SPGP0R02450 | YGL162W | 99 | 0 |
| SPGP0T02200 | YLR173W | 99 | 0 |
| SPGP0V00830 | YER004W | 99 | 0 |
| SPGP0V01850 | YER099C | 99 | 0 |
| SPGP0V01470 | YER062C | 100 | 0 |
| SPGP0N03030 | YJL183W | 99 | 0 |
| SPGP0J03020 | YBL016W | 99 | 0 |
| SPGP0J02220 | YBR071W | 99 | 0 |
| SPGP0AJ00360 | YAL010C | 99 | 0 |
| SPGP0Y00660 | YGR170W | 99 | 0 |
| SPGP0G02910 | YMR009W | 99 | 0 |
| SPGP0CC00100 | YMR319C | 99 | 0 |
| SPGP0Y00840 | YGR192C | 99 | 0 |
| SPGP0A03930 | YDR119W-A | 100 | 0 |
| SPGP0D02800 | YOR037W | 99 | 0 |
| SPGP0A01940 | YDR337W | 100 | 0 |
| SPGP0V01120 | YER031C | 99 | 0 |
| SPGP0U01490 | YLR089C | 99 | 0 |
| SPGP0R00950 | YGR006W | 99 | 0 |
| SPGP0D02780 | YOR039W | 99 | 0 |
| SPGP0N01780 | YJL041W | 97 | 1 |
| SPGP0R01840 | YGL095C | 99 | 0 |
| SPGP0K03780 | YFL059W | 99 | 0 |
| SPGP0D00930 | YOR249C | 99 | 0 |
| SPGP0J00120 | YBR295W | 99 | 0 |
| SPGP0K01140 | YNL039W | 99 | 0 |
| SPGP0Z01500 | YKL181W | 99 | 0 |
| SPGP0K01650 | YNL095C | 99 | 0 |
| SPGP0K01540 | YNL082W | 99 | 0 |
| SPGP0K01300 | YNL056W | 99 | 0 |
| SPGP0D02910 | YOR025W | 99 | 0 |
| SPGP0A05340 | YDL045C | 99 | 0 |
| SPGP0N01680 | YJL029C | 99 | 0 |
| SPGP0N03130 | YJL194W | 99 | 0 |
| SPGP0D02930 | YOR022C | 99 | 0 |
| SPGP0D02240 | YOR101W | 97 | 0 |
| SPGP0D01930 | YOR136W | 99 | 0 |
| SPGP0X01780 | YHR129C | 99 | 0 |
| SPGP0AH00490 | YCL018W | 99 | 0 |

Create PDF in your applications with the Pdfcrowd HTML to PDF API PDFCROWD

| | | | |
|---|---|---|---|
| SPGP0X01250 | YHR076W | 99 | 0 |
| SPGP0X01520 | YHR102W | 99 | 0 |
| SPGP0U01800 | YLR125W | 99 | 0 |
| SPGP0N02340 | YJL104W | 100 | 0 |
| SPGP0BN00100 | YCR086W | 99 | 0 |
| SPGP0R00420 | YGR070W | 99 | 0 |
| SPGP0V00320 | YEL050C | 99 | 0 |
| SPGP0Z01330 | YKL162C-A | 100 | 0 |
| SPGP0X01950 | YHR146W | 99 | 0 |
| SPGP0V00650 | YEL017C-A | 100 | 0 |
| SPGP0Y01650 | YGR281W | 99 | 0 |
| SPGP0AA00280 | YPL223C | 99 | 0 |
| SPGP0N01530 | YJL005W | 99 | 0 |
| SPGP0K02440 | YNL185C | 99 | 0 |
| SPGP0AN00600 | YPL050C | 100 | 0 |
| SPGP0D00660 | YOR279C | 99 | 0 |
| SPGP0U01740 | YLR116W | 99 | 0 |
| SPGP0U01660 | YLR106C | 99 | 0 |
| SPGP0X00120 | YHL040C | 99 | 0 |
| SPGP0K02650 | YNL209W | 99 | 0 |
| SPGP0V01000 | YER019W | 99 | 0 |
| SPGP0K02300 | YNL166C | 99 | 0 |
| SPGP0R01150 | YGL016W | 99 | 0 |
| SPGP0Z00420 | YKL061W | 100 | 0 |
| SPGP0AB00860 | YIL093C | 100 | 0 |
| SPGP0Z01240 | YKL152C | 99 | 0 |
| SPGP0J02870 | YBR001C | 99 | 0 |
| SPGP0AP00260 | YHR182W | 99 | 0 |
| SPGP0Z00500 | YKL068W-A | 99 | 0 |
| SPGP0AJ00190 | YAR023C | 99 | 0 |
| SPGP0AD00970 | YKR013W | 98 | 2 |
| SPGP0K00760 | YNR006W | 99 | 0 |
| SPGP0K03650 | YNL318C | 99 | 0 |
| SPGP0A03340 | YDR181C | 99 | 0 |
| SPGP0J00680 | YBR237W | 99 | 0 |
| SPGP0A00500 | YDR496C | 99 | 0 |
| SPGP0G02490 | YMR055C | 99 | 0 |
| SPGP0A01050 | YDR436W | 99 | 0 |
| SPGP0D02220 | YOR104W | 99 | 0 |
| SPGP0Z00370 | YKL056C | 100 | 0 |

Create PDF in your applications with the Pdfcrowd HTML to PDF API PDFCROWD

| | | | |
|---|---|---|---|
| SPGP0A05280 | YDL035C | 99 | 0 |
| SPGP0A05820 | YDL098C | 99 | 0 |
| SPGP0J00840 | YBR218C | 99 | 0 |
| SPGP0K01380 | YNL066W | 99 | 0 |
| SPGP0N00720 | YJR089W | 99 | 0 |
| SPGP0Z01520 | YKL183W | 99 | 0 |
| SPGP0AU00280 | YOR363C | 99 | 0 |
| SPGP0G03810 | YML087C | 99 | 0 |
| SPGP0A04480 | YDR061W | 99 | 0 |
| SPGP0AJ00770 | YAL053W | 99 | 0 |
| SPGP0A04010 | YDR110W | 99 | 0 |
| SPGP0J01030 | YBR198C | 99 | 0 |
| SPGP0BA00200 | YCR076C | 99 | 0 |
| SPGP0T01390 | YLR265C | 99 | 0 |
| SPGP0AK00170 | YGL253W | 99 | 0 |
| SPGP0AC00940 | YPR113W | 99 | 0 |
| SPGP0AC00520 | YPR063C | 99 | 0 |
| SPGP0G04090 | YML118W | 99 | 0 |
| SPGP0AB00910 | YIL098C | 99 | 0 |
| SPGP0Y01460 | YGR260W | 99 | 0 |
| SPGP0G03720 | YML078W | 100 | 0 |
| SPGP0AB01510 | YIL158W | 99 | 0 |
| SPGP0G00130 | YMR299C | 99 | 0 |
| SPGP0D00400 | YOR305W | 99 | 0 |
| SPGP0AS00190 | YPL268W | 99 | 0 |
| SPGP0A02080 | YDR322C-A | 100 | 0 |
| SPGP0AA01330 | YPL113C | 99 | 0 |
| SPGP0AN00160 | YPL003W | 99 | 0 |
| SPGP0Z00680 | YKL088W | 99 | 0 |
| SPGP0BA00210 | YCR077C | 99 | 0 |
| SPGP0T01020 | YLR307C-A | 100 | 0 |
| SPGP0J02620 | YBR029C | 99 | 0 |
| SPGP0X01240 | YHR075C | 100 | 0 |
| SPGP0AE00120 | YFR054C | 99 | 0 |
| SPGP0A03970 | YDR116C | 99 | 0 |
| SPGP0D03580 | YOL051W | 98 | 2 |
| SPGP0A04590 | YDR050C | 100 | 0 |
| SPGP0T01520 | YLR249W | 99 | 0 |
| SPGP0FC00100 | YBR298C | 88 | 1 |
| SPGP0K03210 | YNL270C | 99 | 0 |

Create PDF in your applications with the Pdfcrowd HTML to PDF API PDFCROWD

| | | | |
|---|---|---|---|
| SPGP0K02810 | YNL225C | 99 | 0 |
| SPGP0Z00360 | YKL055C | 99 | 0 |
| SPGP0D02740 | YOR044W | 99 | 0 |
| SPGP0U01390 | YLR079W | 99 | 0 |
| SPGP0V01350 | YER052C | 99 | 0 |
| SPGP0A06700 | YDL198C | 99 | 0 |
| SPGP0D03700 | YOL063C | 99 | 0 |
| SPGP0A02880 | YDR236C | 99 | 0 |
| SPGP0Y00710 | YGR177C | 99 | 0 |
| SPGP0Y00450 | YGR147C | 99 | 0 |
| SPGP0T01970 | YLR197W | 99 | 0 |
| SPGP0AY00140 | YOR374W | 99 | 0 |
| SPGP0AB00780 | YIL084C | 99 | 0 |
| SPGP0AK00380 | YGL230C | 99 | 0 |
| SPGP0N01330 | YJR014W | 100 | 0 |
| SPGP0D00270 | YOR322C | 99 | 0 |
| SPGP0J02740 | YBR016W | 99 | 0 |
| SPGP0U00950 | YLR032W | 99 | 0 |
| SPGP0Y00470 | YGR149W | 99 | 0 |
| SPGP0N00800 | YJR078W | 99 | 0 |
| SPGP0BX00120 | YFR028C | 99 | 0 |
| SPGP0Y00780 | YGR185C | 99 | 0 |
| SPGP0A06260 | YDL144C | 99 | 0 |
| SPGP0J03090 | YBL023C | 99 | 0 |
| SPGP0K02880 | YNL234W | 100 | 0 |
| SPGP0A00950 | YDR448W | 99 | 0 |
| SPGP0J00670 | YBR238C | 99 | 0 |
| SPGP0R01870 | YGL098W | 99 | 0 |
| SPGP0G01440 | YMR166C | 99 | 0 |
| SPGP0AB00250 | YIL020C | 99 | 0 |
| SPGP0N02890 | YJL166W | 100 | 0 |
| SPGP0N01050 | YJR052W | 99 | 0 |
| SPGP0A03280 | YDR188W | 99 | 0 |
| SPGP0A03750 | YDR139C | 100 | 0 |
| SPGP0A07070 | YDL237W | 99 | 0 |
| SPGP0A03020 | YDR221W | 99 | 0 |
| SPGP0T01730 | YLR224W | 99 | 0 |
| SPGP0D03980 | YOL092W | 99 | 0 |
| SPGP0AA01400 | YPL106C | 99 | 0 |
| SPGP0V00920 | YER012W | 99 | 0 |

Create PDF in your applications with the Pdfcrowd HTML to PDF API PDFCROWD

| | | | |
|---|---|---|---|
| SPGP0J03320 | YBL045C | 99 | 0 |
| SPGP0AM00620 | YPR204W | 91 | 0 |
| SPGP0U01990 | YLR145W | 100 | 0 |
| SPGP0AA00460 | YPL204W | 99 | 0 |
| SPGP0AJ00450 | YAL019W | 99 | 0 |
| SPGP0A00480 | YDR498C | 99 | 0 |
| SPGP0Y01250 | YGR237C | 99 | 0 |
| SPGP0AE01070 | YFL045C | 99 | 0 |
| SPGP0V00400 | YEL041W | 99 | 0 |
| SPGP0J03240 | YBL037W | 99 | 0 |
| SPGP0AW00200 | YHR159W | 99 | 0 |
| SPGP0Y00170 | YGR117C | 99 | 0 |
| SPGP0AK00460 | YGL223C | 99 | 0 |
| SPGP0CO00100 | YNR003C | 100 | 0 |
| SPGP0T00400 | YLR377C | 99 | 0 |
| SPGP0G00490 | YMR263W | 100 | 0 |
| SPGP0U01510 | YLR091W | 99 | 0 |
| SPGP0AA00200 | YPL231W | 99 | 0 |
| SPGP0AB01120 | YIL119C | 98 | 1 |
| SPGP0T00770 | YLR335W | 99 | 0 |
| SPGP0D03060 | YOR008C | 97 | 3 |
| SPGP0AD00580 | YKR055W | 99 | 0 |
| SPGP0AU00110 | YOR346W | 99 | 0 |
| SPGP0J01780 | YBR119W | 99 | 0 |
| SPGP0AD01090 | YKL001C | 100 | 0 |
| SPGP0K00470 | YNR036C | 99 | 0 |
| SPGP0N03200 | YJL203W | 99 | 0 |
| SPGP0AB00380 | YIL037C | 99 | 0 |
| SPGP0R02290 | YGL144C | 99 | 0 |
| SPGP0G01320 | YMR180C | 99 | 0 |
| SPGP0AO00330 | YLR436C | 99 | 0 |
| SPGP0A06610 | YDL186W | 99 | 0 |
| SPGP0K02460 | YNL187W | 99 | 0 |
| SPGP0D01230 | YOR214C | 99 | 0 |
| SPGP0G04190 | YML130C | 99 | 0 |
| SPGP0R01380 | YGL041W-A | 99 | 0 |
| SPGP0AH00600 | YCL004W | 99 | 0 |
| SPGP0A00220 | YDR528W | 99 | 0 |
| SPGP0V01770 | YER091C | 99 | 0 |
| SPGP0N01490 | YJL001W | 99 | 0 |

Create PDF in your applications with the Pdfcrowd HTML to PDF API PDFCROWD

| | | | |
|---|---|---|---|
| SPGP0J01250 | YBR175W | 99 | 0 |
| SPGP0K01840 | YNL118C | 99 | 0 |
| SPGP0A05650 | YDL080C | 99 | 0 |
| SPGP0J01330 | YBR166C | 99 | 0 |
| SPGP0AH00570 | YCL008C | 99 | 0 |
| SPGP0D03920 | YOL086W-A | 99 | 0 |
| SPGP0A00740 | YDR470C | 99 | 0 |
| SPGP0K03220 | YNL271C | 99 | 0 |
| SPGP0D00180 | YOR334W | 99 | 0 |
| SPGP0X01910 | YHR142W | 99 | 0 |
| SPGP0G02800 | YMR020W | 99 | 0 |
| SPGP0AK00210 | YGL249W | 99 | 0 |
| SPGP0T00380 | YLR380W | 99 | 0 |
| SPGP0D01580 | YOR175C | 99 | 0 |
| SPGP0K03720 | YNL327W | 99 | 0 |
| SPGP0J02470 | YBR045C | 99 | 0 |
| SPGP0Y00510 | YGR154C | 99 | 0 |
| SPGP0AZ00170 | YIR009W | 99 | 0 |
| SPGP0A05760 | YDL090C | 99 | 0 |
| SPGP0A04830 | YDR022C | 99 | 0 |
| SPGP0A04340 | YDR075W | 100 | 0 |
| SPGP0D04330 | YOL131W | 99 | 0 |
| SPGP0T01310 | YLR274W | 99 | 0 |
| SPGP0A07060 | YDL236W | 99 | 0 |
| SPGP0AB01280 | YIL135C | 99 | 0 |
| SPGP0G02250 | YMR080C | 99 | 0 |
| SPGP0J00540 | YBR251W | 99 | 0 |
| SPGP0V02000 | YER115C | 99 | 0 |
| SPGP0V01190 | YER038C | 99 | 0 |
| SPGP0AC00730 | YPR088C | 99 | 0 |
| SPGP0AA01250 | YPL122C | 99 | 0 |
| SPGP0Z01050 | YKL132C | 99 | 0 |
| SPGP0AB00410 | YIL040W | 99 | 0 |
| SPGP0G03880 | YML096W | 99 | 0 |
| SPGP0AH00970 | YCR035C | 99 | 0 |
| SPGP0Y00930 | YGR201C | 99 | 0 |
| SPGP0A01380 | YDR397C | 100 | 0 |
| SPGP0A04130 | YDR097C | 99 | 0 |
| SPGP0AC00530 | YPR065W | 99 | 0 |
| SPGP0G04110 | YML120C | 99 | 0 |

Create PDF in your applications with the Pdfcrowd HTML to PDF API PDFCROWD

| | | | |
|---|---|---|---|
| SPGP0T00750 | YLR337C | 99 | 0 |
| SPGP0AA00590 | YPL190C | 98 | 1 |
| SPGP0AA01520 | YPL094C | 99 | 0 |
| SPGP0AA01200 | YPL127C | 99 | 0 |
| SPGP0R00160 | YGR100W | 99 | 0 |
| SPGP0J00310 | YBR274W | 99 | 0 |
| SPGP0U01090 | YLR047C | 99 | 0 |
| SPGP0N02560 | YJL130C | 99 | 0 |
| SPGP0AD01230 | YKL014C | 99 | 0 |
| SPGP0Z00830 | YKL105C | 99 | 0 |
| SPGP0AK00730 | YGL189C | 100 | 0 |
| SPGP0V00730 | YEL007W | 99 | 0 |
| SPGP0G02710 | YMR029C | 99 | 0 |
| SPGP0J01800 | YBR117C | 99 | 0 |
| SPGP0D04160 | YOL113W | 99 | 0 |
| SPGP0AS00290 | YPL257W | 92 | 0 |
| SPGP0AA00480 | YPL202C | 99 | 0 |
| SPGP0Z00890 | YKL112W | 99 | 0 |
| SPGP0R01780 | YGL089C | 99 | 0 |
| SPGP0AA01780 | YPL066W | 99 | 0 |
| SPGP0A06130 | YDL130W-A | 99 | 0 |
| SPGP0A05540 | YDL066W | 99 | 0 |
| SPGP0J00900 | YBR212W | 99 | 0 |
| SPGP0U00250 | YLL045C | 99 | 0 |
| SPGP0U00700 | YLR005W | 99 | 0 |
| SPGP0T00260 | YLR392C | 99 | 0 |
| SPGP0AH00960 | YCR034W | 100 | 0 |
| SPGP0J02590 | YBR033W | 99 | 0 |
| SPGP0N02720 | YJL147C | 99 | 0 |
| SPGP0G00270 | YMR285C | 99 | 0 |
| SPGP0J03660 | YBL086C | 100 | 0 |
| SPGP0U01250 | YLR064W | 99 | 0 |
| SPGP0AN00570 | YPL047W | 99 | 0 |
| SPGP0K02050 | YNL139C | 99 | 0 |
| SPGP0T00710 | YLR343W | 99 | 0 |
| SPGP0D02540 | YOR067C | 99 | 0 |
| SPGP0R02130 | YGL127C | 99 | 0 |
| SPGP0AA00690 | YPL180W | 99 | 0 |
| SPGP0J02940 | YBL007C | 99 | 0 |
| SPGP0G01050 | YMR205C | 99 | 0 |

Create PDF in your applications with the Pdfcrowd HTML to PDF API PDFCROWD

| | | | |
|---|---|---|---|
| SPGP0AW00220 | YHR161C | 99 | 1 |
| SPGP0A01100 | YDR429C | 100 | 0 |
| SPGP0V00410 | YEL040W | 99 | 0 |
| SPGP0AJ00710 | YAL044W-A | 99 | 0 |
| SPGP0X01830 | YHR134W | 100 | 0 |
| SPGP0R02010 | YGL114W | 99 | 0 |
| SPGP0D02070 | YOR120W | 99 | 0 |
| SPGP0AB01330 | YIL140W | 99 | 0 |
| SPGP0AK00720 | YGL190C | 99 | 0 |
| SPGP0R00930 | YGR009C | 99 | 0 |
| SPGP0AJ00180 | YAR027W | 99 | 0 |
| SPGP0R00920 | YGR010W | 99 | 0 |
| SPGP0A01880 | YDR347W | 99 | 0 |
| SPGP0X00130 | YHL039W | 99 | 0 |
| SPGP0A02160 | YDR316W | 99 | 0 |
| SPGP0R02470 | YGL164C | 100 | 0 |
| SPGP0J03480 | YBL063W | 99 | 0 |
| SPGP0D03040 | YOR010C | 95 | 0 |
| SPGP0U01480 | YLR088W | 99 | 0 |
| SPGP0AJ00380 | YAL012W | 99 | 0 |
| SPGP0J03000 | YBL014C | 99 | 0 |
| SPGP0AB00540 | YIL053W | 99 | 0 |
| SPGP0G01230 | YMR189W | 99 | 0 |
| SPGP0AB01090 | YIL116W | 99 | 0 |
| SPGP0R01560 | YGL063W | 99 | 0 |
| SPGP0J03350 | YBL049W | 100 | 0 |
| SPGP0J02750 | YBR015C | 99 | 0 |
| SPGP0K02280 | YNL164C | 99 | 0 |
| SPGP0LW00100 | YJL078C | 99 | 0 |
| SPGP0K00160 | YNR068C | 99 | 0 |
| SPGP0A05260 | YDL031W | 99 | 0 |
| SPGP0A02040 | YDR326C | 99 | 0 |
| SPGP0X01500 | YHR100C | 99 | 0 |
| SPGP0U01820 | YLR127C | 99 | 0 |
| SPGP0V01730 | YER087C-B | 100 | 0 |
| SPGP0N02360 | YJL106W | 99 | 0 |
| SPGP0AS00140 | YPL273W | 99 | 0 |
| SPGP0AU00240 | YOR359W | 99 | 0 |
| SPGP0U01530 | YLR093C | 99 | 0 |
| SPGP0GY00100 | YBR299W | 99 | 0 |

Create PDF in your applications with the Pdfcrowd HTML to PDF API PDFCROWD

| | | | |
|---|---|---|---|
| SPGP0AH00230 | YCL049C | 99 | 0 |
| SPGP0Z01020 | YKL128C | 99 | 0 |
| SPGP0V00670 | YEL016C | 99 | 0 |
| SPGP0V00890 | YER009W | 100 | 0 |
| SPGP0Y01670 | YGR283C | 99 | 0 |
| SPGP0AA00300 | YPL221W | 99 | 0 |
| SPGP0AE00150 | YFR051C | 99 | 0 |
| SPGP0A01070 | YDR434W | 99 | 0 |
| SPGP0A01680 | YDR369C | 99 | 0 |
| SPGP0D00640 | YOR281C | 99 | 0 |
| SPGP0K02320 | YNL168C | 99 | 0 |
| SPGP0K02630 | YNL207W | 99 | 0 |
| SPGP0AB01110 | YIL118W | 100 | 0 |
| SPGP0N02090 | YJL076W | 99 | 0 |
| SPGP0R01170 | YGL018C | 99 | 0 |
| SPGP0AB00880 | YIL095W | 99 | 0 |
| SPGP0G02280 | YMR077C | 100 | 0 |
| SPGP0K01680 | YNL099C | 99 | 0 |
| SPGP0K01250 | YNL051W | 99 | 0 |
| SPGP0D00720 | YOR272W | 99 | 0 |
| SPGP0K00770 | YNR004W | 100 | 0 |
| SPGP0AD00910 | YKR018C | 99 | 0 |
| SPGP0A03320 | YDR183W | 99 | 0 |
| SPGP0G00870 | YMR223W | 99 | 0 |
| SPGP0AP00280 | YHR184W | 99 | 0 |
| SPGP0T00560 | YLR360W | 100 | 0 |
| SPGP0FA00110 | YIL008W | 100 | 0 |
| SPGP0N00200 | YJR140C | 99 | 0 |
| SPGP0AE01120 | YFL050C | 99 | 0 |
| SPGP0G03290 | YML031W | 99 | 0 |
| SPGP0D02430 | YOR080W | 99 | 0 |
| SPGP0U01520 | YLR092W | 99 | 0 |
| SPGP0X00420 | YHL009C | 98 | 0 |
| SPGP0CH00100 | YKL024C | 99 | 0 |
| SPGP0AK00150 | YGL255W | 95 | 1 |
| SPGP0AM00430 | YER173W | 99 | 0 |
| SPGP0A06890 | YDL217C | 99 | 0 |
| SPGP0Z00760 | YKL096W | 100 | 0 |
| SPGP0AC00920 | YPR111W | 99 | 0 |
| SPGP0A00790 | YDR464W | 99 | 0 |

Create PDF in your applications with the Pdfcrowd HTML to PDF API PDFCROWD

| | | | |
|---|---|---|---|
| SPGP0R00280 | YGR088W | 99 | 0 |
| SPGP0A03790 | YDR134C | 97 | 3 |
| SPGP0AD00850 | YKR024C | 99 | 0 |
| SPGP0K03560 | YNL309W | 99 | 0 |
| SPGP0V01070 | YER026C | 99 | 0 |
| SPGP0T01500 | YLR251W | 99 | 0 |
| SPGP0D00420 | YOR304W | 99 | 0 |
| SPGP0N02230 | YJL093C | 100 | 0 |
| SPGP0K02840 | YNL230C | 99 | 0 |
| SPGP0D02610 | YOR060C | 99 | 0 |
| SPGP0AA01640 | YPL082C | 99 | 0 |
| SPGP0AN00140 | YPL001W | 99 | 0 |
| SPGP0D01400 | YOR194C | 99 | 0 |
| SPGP0K00240 | YNR060W | 99 | 0 |
| SPGP0X00730 | YHR023W | 99 | 0 |
| SPGP0R00620 | YGR046W | 99 | 0 |
| SPGP0A07110 | YDL241W | 99 | 0 |
| SPGP0AE00140 | YFR052W | 99 | 0 |
| SPGP0X01260 | YHR077C | 99 | 0 |
| SPGP0G03500 | YML057W | 99 | 0 |
| SPGP0R01370 | YGL040C | 100 | 0 |
| SPGP0Y00410 | YGR143W | 99 | 0 |
| SPGP0K02820 | YNL227C | 99 | 0 |
| SPGP0AM00320 | YER162C | 99 | 0 |
| SPGP0A02200 | YDR312W | 99 | 0 |
| SPGP0N01210 | YJR034W | 100 | 0 |
| SPGP0AS00310 | YPL256C | 99 | 0 |
| SPGP0U01620 | YLR102C | 99 | 0 |
| SPGP0J01490 | YBR151W | 99 | 0 |
| SPGP0N02950 | YJL173C | 99 | 0 |
| SPGP0J02790 | YBR010W | 100 | 0 |
| SPGP0G01600 | YMR149W | 99 | 0 |
| SPGP0D00290 | YOR320C | 99 | 0 |
| SPGP0N01310 | YJR016C | 99 | 0 |
| SPGP0X00980 | YHR048W | 99 | 0 |
| SPGP0N01030 | YJR054W | 99 | 0 |
| SPGP0AD00270 | YKR087C | 99 | 0 |
| SPGP0AJ00830 | YAL060W | 99 | 0 |
| SPGP0G00510 | YMR261C | 99 | 0 |
| SPGP0A06620 | YDL188C | 99 | 0 |

Create PDF in your applications with the Pdfcrowd HTML to PDF API PDFCROWD

| | | | |
|---|---|---|---|
| SPGP0Z00780 | YKL099C | 100 | 0 |
| SPGP0J00690 | YBR236C | 99 | 0 |
| SPGP0G00930 | YMR217W | 99 | 0 |
| SPGP0T00670 | YLR347C | 99 | 0 |
| SPGP0G01190 | YMR193W | 99 | 0 |
| SPGP0J02920 | YBL005W | 99 | 0 |
| SPGP0J01540 | YBR146W | 99 | 0 |
| SPGP0U02060 | YLR151C | 99 | 0 |
| SPGP0J02430 | YBR049C | 99 | 0 |
| SPGP0J02690 | YBR021W | 99 | 0 |
| SPGP0A03000 | YDR223W | 99 | 0 |
| SPGP0G03680 | YML074C | 100 | 0 |
| SPGP0V00460 | YEL035C | 99 | 0 |
| SPGP0D04380 | YOL137W | 99 | 0 |
| SPGP0A07050 | YDL235C | 99 | 0 |
| SPGP0AV00130 | YKL216W | 99 | 0 |
| SPGP0T01710 | YLR226W | 99 | 0 |
| SPGP0D03330 | YOL020W | 99 | 0 |
| SPGP0A01510 | YDR383C | 99 | 0 |
| SPGP0AJ00720 | YAL046C | 99 | 0 |
| SPGP0J03340 | YBL047C | 99 | 1 |
| SPGP0AA00190 | YPL232W | 99 | 0 |
| SPGP0G04100 | YML119W | 99 | 0 |
| SPGP0AB00400 | YIL039W | 99 | 0 |
| SPGP0J03630 | YBL082C | 99 | 0 |
| SPGP0U00930 | YLR029C | 99 | 0 |
| SPGP0U02020 | YLR147C | 100 | 0 |
| SPGP0N01820 | YJL046W | 99 | 0 |
| SPGP0D03750 | YOL068C | 100 | 0 |
| SPGP0D03440 | YOL033W | 99 | 0 |
| SPGP0BK00130 | YGL174W | 99 | 0 |
| SPGP0A00690 | YDR476C | 99 | 0 |
| SPGP0D03090 | YOR005C | 99 | 0 |
| SPGP0N01470 | YJR002W | 99 | 0 |
| SPGP0N00830 | YJR075W | 99 | 0 |
| SPGP0K00390 | YNR045W | 99 | 0 |
| SPGP0J00640 | YBR241C | 99 | 0 |
| SPGP0R02490 | YGL167C | 99 | 0 |
| SPGP0J02440 | YBR048W | 99 | 0 |

Create PDF in your applications with the Pdfcrowd HTML to PDF API PDFCROWD

| | | | |
|---|---|---|---|
| SPGP0AP00440 | YHR199C-A | 100 | 0 |
| SPGP0AE01060 | YFL044C | 99 | 0 |
| SPGP0R01410 | YGL045W | 100 | 0 |
| SPGP0AU00130 | YOR348C | 99 | 0 |
| SPGP0N00780 | YJR082C | 100 | 0 |
| SPGP0AD01110 | YKL003C | 100 | 0 |
| SPGP0AA00920 | YPL157W | 99 | 0 |
| SPGP0J01270 | YBR172C | 99 | 0 |
| SPGP0AD00140 | YKR099W | 99 | 0 |
| SPGP0AJ00540 | YAL029C | 99 | 0 |
| SPGP0D02080 | YOR119C | 99 | 0 |
| SPGP0AO00510 | YLR414C | 99 | 0 |
| SPGP0B04440 | YDR345C | 90 | 0 |
| SPGP0U00670 | YLR002C | 99 | 0 |
| SPGP0D02630 | YOR058C | 99 | 0 |
| SPGP0J00180 | YBR288C | 99 | 0 |
| SPGP0X01980 | YHR149C | 99 | 0 |
| SPGP0K03800 | YNL335W | 99 | 0 |
| SPGP0K02400 | YNL180C | 99 | 0 |
| SPGP0AA00990 | YPL150W | 99 | 0 |
| SPGP0D01210 | YOR216C | 99 | 0 |
| SPGP0AB00360 | YIL035C | 99 | 0 |
| SPGP0G02370 | YMR068W | 100 | 0 |
| SPGP0AO00310 | YLR438W | 99 | 0 |
| SPGP0AA01510 | YPL095C | 99 | 0 |
| SPGP0AE00470 | YFR018C | 99 | 0 |
| SPGP0K01530 | YNL081C | 99 | 0 |
| SPGP0U01470 | YLR087C | 99 | 0 |
| SPGP0A04580 | YDR051C | 99 | 0 |
| SPGP0AD00610 | YKR052C | 99 | 0 |
| SPGP0AB01400 | YIL148W | 99 | 0 |
| SPGP0AD00300 | YKR084C | 99 | 0 |
| SPGP0AE00900 | YFL027C | 99 | 0 |
| SPGP0K03000 | YNL247W | 99 | 0 |
| SPGP0A00670 | YDR478W | 99 | 0 |
| SPGP0Z01450 | YKL175W | 99 | 0 |
| SPGP0AC01140 | YPR135W | 99 | 0 |
| SPGP0J02550 | YBR037C | 99 | 0 |
| SPGP0AE00790 | YFL017C | 99 | 0 |
| SPGP0J03840 | YBL106C | 99 | 0 |

Create PDF in your applications with the Pdfcrowd HTML to PDF API PDFCROWD

| | | | |
|---|---|---|---|
| SPGP0D03840 | YOL077W-A | 99 | 0 |
| SPGP0AH00470 | YCL024W | 99 | 0 |
| SPGP0D00300 | YOR319W | 100 | 0 |
| SPGP0D04350 | YOL133W | 99 | 0 |
| SPGP0T01330 | YLR272C | 99 | 0 |
| SPGP0G02820 | YMR018W | 99 | 0 |
| SPGP0V02020 | YER117W | 99 | 0 |
| SPGP0AJ00500 | YAL024C | 99 | 0 |
| SPGP0D01630 | YOR168W | 99 | 0 |
| SPGP0R02160 | YGL130W | 99 | 0 |
| SPGP0AC00990 | YPR118W | 99 | 0 |
| SPGP0Z01700 | YKL204W | 99 | 0 |
| SPGP0N02130 | YJL082W | 99 | 0 |
| SPGP0J03510 | YBL067C | 99 | 0 |
| SPGP0A05980 | YDL114W | 99 | 0 |
| SPGP0AC01040 | YPR124W | 99 | 0 |
| SPGP0A04320 | YDR077W | 99 | 0 |
| SPGP0Z01100 | YKL138C | 100 | 0 |
| SPGP0AK00530 | YGL213C | 99 | 0 |
| SPGP0Y00950 | YGR203W | 99 | 0 |
| SPGP0G02700 | YMR030W | 99 | 0 |
| SPGP0AH00590 | YCL005W | 99 | 0 |
| SPGP0AD00630 | YKR050W | 99 | 0 |
| SPGP0AA01810 | YPL063W | 99 | 0 |
| SPGP0AA00570 | YPL192C | 99 | 0 |
| SPGP0L03600 | YPL218W | 90 | 0 |
| SPGP0AB01040 | YIL111W | 99 | 0 |
| SPGP0R00100 | YGR106C | 99 | 0 |
| SPGP0AO00410 | YLR426W | 99 | 0 |
| SPGP0U01830 | YLR128W | 99 | 0 |
| SPGP0T00910 | YLR319C | 99 | 0 |
| SPGP0T01870 | YLR209C | 99 | 0 |
| SPGP0A04060 | YDR105C | 99 | 0 |
| SPGP0Z01110 | YKL138C-A | 99 | 0 |
| SPGP0J00190 | YBR287W | 99 | 0 |
| SPGP0AD01250 | YKL016C | 99 | 0 |
| SPGP0G03000 | YML001W | 100 | 0 |
| SPGP0N02540 | YJL128C | 99 | 0 |
| SPGP0K03790 | YNL334C | 99 | 0 |
| SPGP0AK00280 | YGL242C | 99 | 0 |

Create PDF in your applications with the Pdfcrowd HTML to PDF API PDFCROWD

| | | | |
|---|---|---|---|
| SPGP0J01860 | YBR110W | 99 | 0 |
| SPGP0A06790 | YDL207W | 99 | 0 |
| SPGP0AS00320 | YPL255W | 100 | 0 |
| SPGP0Z00330 | YKL052C | 99 | 0 |
| SPGP0AM00130 | YER142C | 99 | 0 |
| SPGP0X01680 | YHR118C | 99 | 0 |
| SPGP0J02810 | YBR007C | 99 | 0 |
| SPGP0J01680 | YBR130C | 99 | 0 |
| SPGP0N02670 | YJL141C | 99 | 0 |
| SPGP0U00720 | YLR007W | 100 | 0 |
| SPGP0U00230 | YLL048C | 99 | 0 |
| SPGP0T00240 | YLR394W | 99 | 0 |
| SPGP0AD01150 | YKL006C-A | 100 | 0 |
| SPGP0N03350 | YOL156W | 97 | 0 |
| SPGP0K02690 | YNL213C | 99 | 0 |
| SPGP0A00540 | YDR492W | 100 | 0 |
| SPGP0A04440 | YDR065W | 99 | 0 |
| SPGP0N01430 | YJR005C-A | 100 | 0 |
| SPGP0D00260 | YOR323C | 99 | 0 |
| SPGP0G00210 | YMR291W | 100 | 0 |
| SPGP0AC01120 | YPR133W-A | 100 | 0 |
| SPGP0J03640 | YBL084C | 99 | 0 |
| SPGP0D00500 | YOR296W | 99 | 0 |
| SPGP0K00580 | YNR024W | 100 | 0 |
| SPGP0G02880 | YMR012W | 99 | 0 |
| SPGP0K00720 | YNR010W | 99 | 0 |
| SPGP0AA01100 | YPL138C | 99 | 0 |
| SPGP0AB01350 | YIL143C | 99 | 0 |
| SPGP0R00810 | YGR024C | 99 | 0 |
| SPGP0D01660 | YOR165W | 99 | 0 |
| SPGP0G01030 | YMR207C | 99 | 0 |
| SPGP0A06330 | YDL153C | 99 | 0 |
| SPGP0AC01180 | YPR140W | 100 | 0 |
| SPGP0G00680 | YMR242C | 100 | 0 |
| SPGP0A05860 | YDL102W | 99 | 0 |
| SPGP0A01470 | YDR387C | 99 | 0 |
| SPGP0J03420 | YBL057C | 99 | 0 |
| SPGP0J00990 | YBR201C-A | 99 | 0 |
| SPGP0Z00730 | YKL093W | 99 | 1 |
| SPGP0T00340 | YLR384C | 99 | 0 |

Create PDF in your applications with the Pdfcrowd HTML to PDF API PDFCROWD

| | | | |
|---|---|---|---|
| SPGP0N01640 | YJL023C | 99 | 0 |
| SPGP0G00620 | YMR250W | 99 | 0 |
| SPGP0A01990 | YDR332W | 99 | 0 |
| SPGP0R02410 | YGL158W | 99 | 0 |
| SPGP0J01480 | YBR152W | 99 | 0 |
| SPGP0J03490 | YBL064C | 99 | 0 |
| SPGP0AJ00400 | YAL014C | 99 | 0 |
| SPGP0J03470 | YBL061C | 99 | 0 |
| SPGP0D01970 | YOR131C | 99 | 0 |
| SPGP0R01540 | YGL061C | 99 | 0 |
| SPGP0Y00750 | YGR181W | 99 | 0 |
| SPGP0X01740 | YHR124W | 99 | 0 |
| SPGP0A01930 | YDR338C | 99 | 0 |
| SPGP0G00110 | YMR301C | 99 | 0 |
| SPGP0J02770 | YBR012C | 99 | 0 |
| SPGP0AK00420 | YGL226C-A | 100 | 0 |
| SPGP0K01780 | YNL111C | 99 | 0 |
| SPGP0A05630 | YDL077C | 99 | 0 |
| SPGP0AJ00600 | YAL034W-A | 99 | 0 |
| SPGP0AA00660 | YPL183W-A | 99 | 0 |
| SPGP0A02920 | YDR232W | 99 | 0 |
| SPGP0R01990 | YGL112C | 99 | 0 |
| SPGP0N02300 | YJL100W | 99 | 0 |
| SPGP0AA01570 | YPL089C | 99 | 1 |
| SPGP0Y01570 | YGR273C | 99 | 0 |
| SPGP0D02950 | YOR020W-A | 100 | 0 |
| SPGP0AN00640 | YPL054W | 99 | 0 |
| SPGP0V00610 | YEL020W-A | 100 | 0 |
| SPGP0AA01120 | YPL135W | 99 | 0 |
| SPGP0G02100 | YMR097C | 99 | 0 |
| SPGP0Y01450 | YGR258C | 99 | 0 |
| SPGP0V01300 | YER048C | 99 | 0 |
| SPGP0T02220 | YLR170C | 99 | 0 |
| SPGP0A01090 | YDR430C | 99 | 0 |
| SPGP0AM00270 | YER156C | 99 | 0 |
| SPGP0AP00330 | YHR189W | 99 | 0 |
| SPGP0K02340 | YNL172W | 99 | 0 |
| SPGP0AM00610 | YPL283C | 97 | 0 |
| SPGP0Y01440 | YGR257C | 99 | 0 |
| SPGP0BY00100 | YIR034C | 99 | 0 |

Create PDF in your applications with the Pdfcrowd HTML to PDF API PDFCROWD

| | | | |
|---|---|---|---|
| SPGP0CN00100 | YCR093W | 99 | 0 |
| SPGP0N00190 | YJR141W | 99 | 0 |
| SPGP0G00540 | YMR258C | 99 | 0 |
| SPGP0AM00340 | YER164W | 99 | 0 |
| SPGP0Z01230 | YKL151C | 100 | 0 |
| SPGP0AJ00480 | YAL022C | 99 | 0 |
| SPGP0Z00480 | YKL067W | 98 | 0 |
| SPGP0AJ00150 | YAR029W | 98 | 0 |
| SPGP0A04110 | YDR099W | 99 | 0 |
| SPGP0AD00280 | YKR086W | 99 | 0 |
| SPGP0AD00930 | YKR016W | 99 | 0 |
| SPGP0U01610 | YLR100W | 99 | 0 |
| SPGP0AN00220 | YPL009C | 99 | 0 |
| SPGP0D00920 | YOR250C | 99 | 0 |
| SPGP0G01760 | YMR132C | 100 | 0 |
| SPGP0AD00830 | YKR026C | 99 | 0 |
| SPGP0AD01200 | YKL011C | 99 | 0 |
| SPGP0D03860 | YOL080C | 99 | 0 |
| SPGP0D00160 | YOR336W | 99 | 0 |
| SPGP0N00220 | YJR138W | 99 | 0 |
| SPGP0D02380 | YOR086C | 99 | 0 |
| SPGP0BR00120 | YCR044C | 99 | 0 |
| SPGP0J02340 | YBR058C-A | 99 | 0 |
| SPGP0K01730 | YNL104C | 99 | 0 |
| SPGP0AJ00730 | YAL047C | 99 | 0 |
| SPGP0J00600 | YBR245C | 99 | 0 |
| SPGP0A02260 | YDR306C | 99 | 0 |
| SPGP0X00160 | YHL035C | 99 | 0 |
| SPGP0T01290 | YLR276C | 99 | 0 |
| SPGP0AC00860 | YPR105C | 99 | 0 |
| SPGP0AS00120 | YPL276W | 99 | 0 |
| SPGP0AH00250 | YCL048W | 99 | 0 |
| SPGP0A03070 | YDR214W | 99 | 0 |
| SPGP0V02120 | YER128W | 99 | 0 |
| SPGP0AA01480 | YPL097W | 99 | 0 |
| SPGP0AN00300 | YPL017C | 99 | 0 |
| SPGP0AK00130 | YGL257C | 99 | 0 |
| SPGP0K01230 | YNL049C | 99 | 0 |
| SPGP0Z01390 | YKL168C | 99 | 0 |
| SPGP0AD01080 | YKR001C | 99 | 0 |

Create PDF in your applications with the Pdfcrowd HTML to PDF API PDFCROWD

| | | | |
|---|---|---|---|
| SPGP0X00570 | YHR006W | 99 | 0 |
| SPGP0J01100 | YBR191W | 99 | 0 |
| SPGP0Z00200 | YKL038W | 99 | 0 |
| SPGP0A01650 | YDR372C | 99 | 0 |
| SPGP0D00440 | YOR302W | 100 | 0 |
| SPGP0A00270 | YDR522C | 99 | 0 |
| SPGP0K00910 | YNL012W | 99 | 0 |
| SPGP0K00260 | YNR058W | 100 | 0 |
| SPGP0G01340 | YMR178W | 99 | 0 |
| SPGP0A04560 | YDR054C | 100 | 0 |
| SPGP0AE00160 | YFR050C | 99 | 0 |
| SPGP0A04820 | YDR023W | 99 | 0 |
| SPGP0X00530 | YHR003C | 99 | 0 |
| SPGP0N00580 | YJR102C | 99 | 0 |
| SPGP0Y00430 | YGR145W | 100 | 0 |
| SPGP0A06060 | YDL123W | 99 | 0 |
| SPGP0R01570 | YGL064C | 99 | 0 |
| SPGP0AD00900 | YKR019C | 99 | 0 |
| SPGP0A02220 | YDR310C | 99 | 0 |
| SPGP0AC00160 | YPR022C | 99 | 0 |
| SPGP0Z00110 | YKL026C | 99 | 0 |
| SPGP0R00310 | YGR082W | 99 | 0 |
| SPGP0Z00660 | YKL086W | 99 | 0 |
| SPGP0R00640 | YGR043C | 99 | 0 |
| SPGP0K02720 | YNL216W | 99 | 0 |
| SPGP0J00500 | YBR255W | 99 | 0 |
| SPGP0G04070 | YML116W | 99 | 0 |
| SPGP0N02680 | YJL143W | 99 | 0 |
| SPGP0AE00670 | YFL002C | 99 | 0 |
| SPGP0N02930 | YJL171C | 99 | 0 |
| SPGP0G01620 | YMR147W | 99 | 0 |
| SPGP0N01370 | YJR010C-A | 100 | 0 |
| SPGP0J02900 | YBL003C | 100 | 0 |
| SPGP0N01010 | YJR056C | 99 | 0 |
| SPGP0A03240 | YDR192C | 99 | 0 |
| SPGP0AD00290 | YKR085C | 99 | 0 |
| SPGP0T00840 | YLR327C | 100 | 0 |
| SPGP0N01850 | YJL049W | 99 | 0 |
| SPGP0G00530 | YMR259C | 99 | 0 |
| SPGP0D03490 | YOL040C | 100 | 0 |

Create PDF in your applications with the Pdfcrowd HTML to PDF API PDFCROWD

| | | | |
|---|---|---|---|
| SPGP0D04000 | YOL095C | 99 | 0 |
| SPGP0A00910 | YDR452W | 99 | 0 |
| SPGP0T00650 | YLR350W | 99 | 0 |
| SPGP0X01190 | YHR071W | 99 | 0 |
| SPGP0AJ00700 | YAL044C | 100 | 0 |
| SPGP0V00440 | YEL037C | 99 | 0 |
| SPGP0AC00850 | YPR104C | 99 | 0 |
| SPGP0A07030 | YDL233W | 99 | 0 |
| SPGP0A03060 | YDR216W | 99 | 0 |
| SPGP0AB00750 | YIL078W | 99 | 0 |
| SPGP0Y00290 | YGR129W | 99 | 0 |
| SPGP0R00730 | YGR033C | 99 | 0 |
| SPGP0N00420 | YJR117W | 99 | 0 |
| SPGP0AJ00290 | YAR002W | 99 | 0 |
| SPGP0U02040 | YLR149C | 99 | 0 |
| SPGP0U00910 | YLR027C | 99 | 0 |
| SPGP0Y01210 | YGR233C | 99 | 0 |
| SPGP0J02460 | YBR046C | 99 | 0 |
| SPGP0BK00110 | YGL176C | 99 | 0 |
| SPGP0AN00500 | YPL039W | 99 | 0 |
| SPGP0A00240 | YDR525W-A | 100 | 0 |
| SPGP0AM00110 | YER140W | 99 | 0 |
| SPGP0X01720 | YHR122W | 99 | 0 |
| SPGP0V01060 | YER025W | 99 | 0 |
| SPGP0AD01130 | YKL005C | 99 | 0 |
| SPGP0Y00150 | YGR113W | 99 | 0 |
| SPGP0AB01160 | YIL123W | 99 | 0 |
| SPGP0A00510 | YDR495C | 99 | 0 |
| SPGP0A02520 | YDR279W | 99 | 0 |
| SPGP0J00620 | YBR243C | 100 | 0 |
| SPGP0N01790 | YJL042W | 99 | 0 |
| SPGP0N00760 | YJR084W | 99 | 0 |
| SPGP0AH00990 | YCR037C | 99 | 0 |
| SPGP0AK00370 | YGL231C | 100 | 0 |
| SPGP0N02590 | YJL133W | 99 | 0 |
| SPGP0J01290 | YBR170C | 99 | 0 |
| SPGP0J02010 | YBR093C | 99 | 0 |
| SPGP0K01660 | YNL096C | 99 | 0 |
| SPGP0K01460 | YNL074C | 99 | 0 |
| SPGP0X00800 | YHR030C | 99 | 0 |

Create PDF in your applications with the Pdfcrowd HTML to PDF API PDFCROWD

| | | | |
|---|---|---|---|
| SPGP0J00280 | YBR278W | 100 | 0 |
| SPGP0U01330 | YLR072W | 99 | 0 |
| SPGP0A06580 | YDL182W | 99 | 0 |
| SPGP0N01980 | YJL062W-A | 100 | 0 |
| SPGP0D01890 | YOR141C | 99 | 0 |
| SPGP0K02420 | YNL182C | 99 | 0 |
| SPGP0N00460 | YJR112W-A | 99 | 0 |
| SPGP0R01690 | YGL079W | 99 | 0 |
| SPGP0AB00340 | YIL033C | 100 | 0 |
| SPGP0A00260 | YDR523C | 99 | 0 |
| SPGP0AB00980 | YIL105C | 99 | 0 |
| SPGP0Y01310 | YGR244C | 99 | 0 |
| SPGP0V01560 | YER071C | 99 | 0 |
| SPGP0K01880 | YNL123W | 99 | 0 |
| SPGP0N01450 | YJR004C | 99 | 0 |
| SPGP0K01550 | YNL083W | 99 | 0 |
| SPGP0T02010 | YLR193C | 99 | 0 |
| SPGP0K03770 | YNL332W | 99 | 0 |
| SPGP0AC00340 | YPR041W | 100 | 0 |
| SPGP0G02590 | YMR041C | 100 | 0 |
| SPGP0D03340 | YOL021C | 99 | 0 |
| SPGP0BN00120 | YCR088W | 99 | 0 |
| SPGP0T01160 | YLR291C | 99 | 0 |
| SPGP0X01170 | YHR069C | 99 | 0 |
| SPGP0T01570 | YLR244C | 100 | 0 |
| SPGP0X00330 | YHL018W | 99 | 0 |
| SPGP0U01980 | YLR144C | 99 | 0 |
| SPGP0T01180 | YLR289W | 99 | 0 |
| SPGP0J03080 | YBL022C | 99 | 0 |
| SPGP0AB01240 | YIL131C | 99 | 0 |
| SPGP0G02270 | YMR078C | 99 | 0 |
| SPGP0G03970 | YML106W | 99 | 0 |
| SPGP0AB00110 | YIL002W-A | 100 | 0 |
| SPGP0A07020 | YDL232W | 100 | 0 |
| SPGP0AH00210 | YCL051W | 99 | 0 |
| SPGP0U00960 | YLR033W | 99 | 0 |
| SPGP0A05960 | YDL112W | 99 | 0 |
| SPGP0AO00260 | YLR442C | 99 | 0 |
| SPGP0G03270 | YML029W | 99 | 0 |
| SPGP0AC01060 | YPR127W | 99 | 0 |

Create PDF in your applications with the Pdfcrowd HTML to PDF API PDFCROWD

| | | | |
|---|---|---|---|
| SPGP0AK00510 | YGL216W | 100 | 0 |
| SPGP0G02680 | YMR032W | 99 | 0 |
| SPGP0D04370 | YOL136C | 99 | 0 |
| SPGP0D00850 | YOR257W | 100 | 0 |
| SPGP0D00320 | YOR316C | 99 | 0 |
| SPGP0AE01170 | YHR216W | 99 | 0 |
| SPGP0A03610 | YDR155C | 100 | 0 |
| SPGP0AN00110 | YPR003C | 99 | 0 |
| SPGP0R00120 | YGR104C | 99 | 0 |
| SPGP0AC00900 | YPR109W | 99 | 0 |
| SPGP0J02280 | YBR065C | 99 | 0 |
| SPGP0T02180 | YLR175W | 99 | 1 |
| SPGP0AD01270 | YKL018W | 99 | 0 |
| SPGP0BD00130 | YMR311C | 100 | 0 |
| SPGP0V00770 | YEL003W | 99 | 0 |
| SPGP0V01410 | YER056C-A | 100 | 0 |
| SPGP0AK00710 | YGL191W | 99 | 0 |
| SPGP0A01250 | YDR411C | 99 | 0 |
| SPGP0D02400 | YOR084W | 99 | 0 |
| SPGP0K00500 | YNR034W | 99 | 0 |
| SPGP0Z00870 | YKL109W | 99 | 0 |
| SPGP0AM00150 | YER144C | 99 | 0 |
| SPGP0A00530 | YDR493W | 100 | 0 |
| SPGP0K00110 | YNR074C | 99 | 0 |
| SPGP0AA01290 | YPL118W | 99 | 0 |
| SPGP0AC00330 | YPR040W | 99 | 0 |
| SPGP0A01300 | YDR406W | 99 | 0 |
| SPGP0A00550 | YDR490C | 99 | 0 |
| SPGP0R00390 | YGR074W | 99 | 0 |
| SPGP0AA00170 | YPL234C | 99 | 0 |
| SPGP0N01500 | YJL002C | 100 | 0 |
| SPGP0U00740 | YLR009W | 99 | 0 |
| SPGP0D04340 | YOL132W | 99 | 0 |
| SPGP0K02000 | YNL135C | 100 | 0 |
| SPGP0D00520 | YOR294W | 99 | 0 |
| SPGP0J03620 | YBL081W | 99 | 0 |
| SPGP0AA00760 | YPL173W | 99 | 0 |
| SPGP0AK00790 | YGL181W | 99 | 1 |
| SPGP0A06680 | YDL195W | 99 | 0 |
| SPGP0J02720 | YBR018C | 100 | 0 |

Create PDF in your applications with the Pdfcrowd HTML to PDF API PDFCROWD

| | | | |
|---|---|---|---|
| SPGP0K02870 | YNL233W | 99 | 0 |
| SPGP0R00870 | YGR016W | 99 | 0 |
| SPGP0G00450 | YMR267W | 99 | 0 |
| SPGP0J00710 | YBR234C | 99 | 0 |
| SPGP0J01510 | YBR149W | 99 | 0 |
| SPGP0D00870 | YOR255W | 99 | 0 |
| SPGP0Z00860 | YKL108W | 99 | 1 |
| SPGP0K02970 | YNL244C | 99 | 0 |
| SPGP0R01970 | YGL110C | 99 | 0 |
| SPGP0V01270 | YER045C | 99 | 0 |
| SPGP0V01180 | YER037W | 99 | 0 |
| SPGP0V01790 | YER093C | 99 | 0 |
| SPGP0R01390 | YGL043W | 99 | 0 |
| SPGP0J02660 | YBR024W | 100 | 0 |
| SPGP0A02010 | YDR330W | 99 | 0 |
| SPGP0Z00710 | YKL091C | 99 | 0 |
| SPGP0AB00170 | YIL013C | 99 | 0 |
| SPGP0G01540 | YMR156C | 99 | 0 |
| SPGP0G01010 | YMR209C | 99 | 0 |
| SPGP0AB01550 | YIL162W | 99 | 0 |
| SPGP0N02550 | YJL129C | 99 | 0 |
| SPGP0D03000 | YOR016C | 99 | 0 |
| SPGP0K00870 | YNL008C | 99 | 0 |
| SPGP0A00980 | YDR444W | 99 | 0 |
| SPGP0R01600 | YGL067W | 99 | 0 |
| SPGP0G02150 | YMR092C | 99 | 0 |
| SPGP0K01600 | YNL088W | 99 | 0 |
| SPGP0J02190 | YBR074W | 99 | 0 |
| SPGP0A05610 | YDL075W | 99 | 0 |
| SPGP0R02430 | YGL160W | 99 | 0 |
| SPGP0V00580 | YEL023C | 99 | 0 |
| SPGP0J02710 | YBR019C | 99 | 0 |
| SPGP0A03470 | YDR170C | 99 | 0 |
| SPGP0BD00170 | YMR307W | 99 | 0 |
| SPGP0AT00100 | YPR201W | 99 | 0 |
| SPGP0A05170 | YDL020C | 99 | 0 |
| SPGP0Y00970 | YGR205W | 99 | 0 |
| SPGP0G02160 | YMR091C | 99 | 0 |
| SPGP0X01540 | YHR104W | 100 | 0 |

Create PDF in your applications with the Pdfcrowd HTML to PDF API PDFCROWD

| | | | |
|---|---|---|---|
| SPGP0X01810 | YHR132W-A | 99 | 0 |
| SPGP0AM00220 | YER151C | 99 | 0 |
| SPGP0N02320 | YJL102W | 99 | 0 |
| SPGP0E00140 | YGR287C | 99 | 0 |
| SPGP0V00620 | YEL020C | 99 | 0 |
| SPGP0D02970 | YOR019W | 99 | 0 |
| SPGP0Y01000 | YGR208W | 99 | 0 |
| SPGP0AJ00590 | YAL034C | 99 | 0 |
| SPGP0V00630 | YEL019C | 99 | 0 |
| SPGP0AE00580 | YFR008W | 99 | 0 |
| SPGP0AJ00420 | YAL016W | 99 | 0 |
| SPGP0D03210 | YOL008W | 99 | 0 |
| SPGP0K02360 | YNL175C | 99 | 0 |
| SPGP0G03430 | YML050W | 99 | 0 |
| SPGP0K01290 | YNL055C | 99 | 0 |
| SPGP0D00860 | YOR256C | 99 | 0 |
| SPGP0K00290 | YNR055C | 99 | 0 |
| SPGP0N03180 | YJL200C | 99 | 0 |
| SPGP0G00830 | YMR227C | 99 | 0 |
| SPGP0AD00890 | YKR020W | 99 | 0 |
| SPGP0X00140 | YHL038C | 99 | 0 |
| SPGP0K02080 | YNL144C | 100 | 0 |
| SPGP0D01620 | YOR171C | 100 | 0 |
| SPGP0D02720 | YOR046C | 99 | 0 |
| SPGP0A00170 | YDR533C | 99 | 0 |
| SPGP0G02780 | YMR022W | 100 | 0 |
| SPGP0V02180 | YER136W | 99 | 0 |
| SPGP0N02390 | YJL110C | 99 | 0 |
| SPGP0BY00120 | YIR036C | 100 | 0 |
| SPGP0K01410 | YNL069C | 99 | 0 |
| SPGP0AB00700 | YIL073C | 99 | 0 |
| SPGP0N00240 | YJR136C | 99 | 0 |
| SPGP0U01960 | YLR142W | 99 | 0 |
| SPGP0K01940 | YNL129W | 100 | 0 |
| SPGP0G03560 | YML062C | 99 | 0 |
| SPGP0N00770 | YJR083C | 99 | 0 |
| SPGP0U00260 | YLL043W | 99 | 0 |
| SPGP0T00690 | YLR345W | 99 | 0 |
| SPGP0A04790 | YDR026C | 99 | 0 |
| SPGP0J02050 | YBR087W | 99 | 0 |

Create PDF in your applications with the Pdfcrowd HTML to PDF API PDFCROWD

| | | | |
|---|---|---|---|
| SPGP0K03010 | YNL248C | 100 | 0 |
| SPGP0T01270 | YLR278C | 99 | 0 |
| SPGP0AA01460 | YPL099C | 99 | 0 |
| SPGP0AN00320 | YPL019C | 99 | 0 |
| SPGP0AB01180 | YIL125W | 99 | 0 |
| SPGP0A00710 | YDR473C | 99 | 0 |
| SPGP0Z01370 | YKL166C | 99 | 0 |
| SPGP0J03210 | YBL034C | 99 | 0 |
| SPGP0AO00420 | YLR425W | 99 | 0 |
| SPGP0AE00360 | YFR031C-A | 99 | 0 |
| SPGP0AA00670 | YPL183C | 99 | 0 |
| SPGP0T01450 | YLR258W | 99 | 0 |
| SPGP0T01850 | YLR211C | 99 | 0 |
| SPGP0D01240 | YOR213C | 99 | 0 |
| SPGP0T01010 | YLR308W | 100 | 0 |
| SPGP0A06050 | YDL122W | 99 | 0 |
| SPGP0R01400 | YGL044C | 99 | 0 |
| SPGP0BC00140 | YIR027C | 99 | 0 |
| SPGP0G04030 | YML112W | 99 | 0 |
| SPGP0Z01510 | YKL182W | 99 | 0 |
| SPGP0AS00240 | YPL263C | 99 | 0 |
| SPGP0AE00180 | YFR048W | 99 | 0 |
| SPGP0K00950 | YNL034W | 97 | 0 |
| SPGP0BL00110 | YCR051W | 99 | 0 |
| SPGP0AE00980 | YFL034C-B | 99 | 0 |
| SPGP0K01080 | YNL020C | 99 | 0 |
| SPGP0N01350 | YJR012C | 99 | 0 |
| SPGP0A06770 | YDL205C | 99 | 0 |
| SPGP0V00900 | YER010C | 99 | 0 |
| SPGP0Y00730 | YGR179C | 99 | 0 |
| SPGP0AH00700 | YCR009C | 99 | 0 |
| SPGP0Y01280 | YGR240C | 99 | 0 |
| SPGP0AM00360 | YER166W | 99 | 0 |
| SPGP0A02240 | YDR308C | 99 | 0 |
| SPGP0AS00350 | YPL252C | 100 | 0 |
| SPGP0K00930 | YNL015W | 100 | 0 |
| SPGP0R00660 | YGR041W | 99 | 0 |
| SPGP0R01240 | YGL026C | 99 | 0 |
| SPGP0AD00130 | YKR100C | 99 | 0 |
| SPGP0J00520 | YBR253W | 99 | 0 |

Create PDF in your applications with the Pdfcrowd HTML to PDF API PDFCROWD

| | | | |
|---|---|---|---|
| SPGP0N02910 | YJL168C | 99 | 0 |
| SPGP0N02660 | YJL140W | 99 | 0 |
| SPGP0AB00760 | YIL079C | 99 | 0 |
| SPGP0G03540 | YML060W | 99 | 0 |
| SPGP0A03260 | YDR190C | 100 | 0 |
| SPGP0AJ00650 | YAL039C | 99 | 0 |
| SPGP0A05560 | YDL069C | 99 | 0 |
| SPGP0G00550 | YMR257C | 99 | 0 |
| SPGP0Z00130 | YKL028W | 99 | 0 |
| SPGP0A06940 | YDL223C | 99 | 0 |
| SPGP0A06410 | YDL161W | 100 | 0 |
| SPGP0R00570 | YGR053C | 99 | 0 |
| SPGP0R02520 | YGL171W | 99 | 0 |
| SPGP0AC01160 | YPR138C | 99 | 0 |
| SPGP0AE00780 | YFL016C | 99 | 0 |
| SPGP0D02440 | YOR079C | 99 | 0 |
| SPGP0AB00310 | YIL027C | 100 | 0 |
| SPGP0AD00310 | YKR083C | 99 | 0 |
| SPGP0T00630 | YLR352W | 99 | 0 |
| SPGP0AD01280 | YKL018C-A | 99 | 0 |
| SPGP0X01220 | YHR073W | 99 | 0 |
| SPGP0A03040 | YDR218C | 99 | 0 |
| SPGP0J00470 | YBR257W | 99 | 0 |
| SPGP0D03960 | YOL090W | 99 | 0 |
| SPGP0R00710 | YGR035C | 99 | 0 |
| SPGP0U01080 | YLR046C | 99 | 0 |
| SPGP0AB01490 | YIL156W-B | 98 | 0 |
| SPGP0V00340 | YEL048C | 99 | 0 |
| SPGP0T01790 | YLR218C | 100 | 0 |
| SPGP0N00400 | YJR119C | 99 | 0 |
| SPGP0AA01380 | YPL108W | 99 | 0 |
| SPGP0N00740 | YJR086W | 100 | 0 |
| SPGP0D02690 | YOR049C | 99 | 0 |
| SPGP0N01420 | YJR006W | 99 | 0 |
| SPGP0J01720 | YBR126C | 99 | 0 |
| SPGP0Y01560 | YGR271C-A | 99 | 0 |
| SPGP0J02480 | YBR044C | 99 | 0 |
| SPGP0J02380 | YBR055C | 99 | 0 |
| SPGP0D03050 | YOR009W | 96 | 0 |
| SPGP0A04860 | YDR019C | 99 | 0 |

Create PDF in your applications with the Pdfcrowd HTML to PDF API PDFCROWD

| | | | |
|---|---|---|---|
| SPGP0AW00210 | YHR160C | 99 | 0 |
| SPGP0AA00150 | YPL236C | 99 | 0 |
| SPGP0AH00130 | YCL059C | 99 | 0 |
| SPGP0V00390 | YEL042W | 99 | 0 |
| SPGP0G02260 | YMR079W | 99 | 0 |
| SPGP0J02420 | YBR050C | 99 | 1 |
| SPGP0K01640 | YNL094W | 99 | 0 |
| SPGP0U01350 | YLR074C | 99 | 0 |
| SPGP0J01850 | YBR111C | 99 | 0 |
| SPGP0N01270 | YJR024C | 99 | 0 |
| SPGP0J02030 | YBR089C-A | 100 | 0 |
| SPGP0X00820 | YHR032W | 99 | 0 |
| SPGP0AK00290 | YGL241W | 99 | 0 |
| SPGP0DY00100 | YPR013C | 99 | 0 |
| SPGP0AA01140 | YPL133C | 99 | 0 |
| SPGP0A06570 | YDL180W | 99 | 0 |
| SPGP0D03300 | YOL017W | 99 | 0 |
| SPGP0G02130 | YMR094W | 100 | 0 |
| SPGP0R00960 | YGR005C | 99 | 0 |
| SPGP0J03100 | YBL024W | 99 | 0 |
| SPGP0AB00960 | YIL103W | 99 | 0 |
| SPGP0T02030 | YLR191W | 100 | 0 |
| SPGP0K01570 | YNL085W | 99 | 0 |
| SPGP0V01290 | YER047C | 99 | 0 |
| SPGP0G02300 | YMR075W | 99 | 0 |
| SPGP0AA01550 | YPL091W | 99 | 0 |
| SPGP0A04540 | YDR056C | 99 | 0 |
| SPGP0A04090 | YDR101C | 99 | 0 |
| SPGP0AK00310 | YGL238W | 99 | 0 |
| SPGP0T01140 | YLR293C | 99 | 0 |
| SPGP0J00890 | YBR213W | 99 | 0 |
| SPGP0AK00270 | YGL243W | 99 | 0 |
| SPGP0D00890 | YOR253W | 99 | 0 |
| SPGP0D00310 | YOR317W | 99 | 0 |
| SPGP0Z01740 | YKL208W | 100 | 0 |
| SPGP0A02150 | YDR317W | 99 | 0 |
| SPGP0FZ00100 | YGR081C | 100 | 0 |
| SPGP0AD00180 | YKR095W-A | 100 | 0 |
| SPGP0G03950 | YML104C | 100 | 0 |
| SPGP0AA00870 | YPL162C | 100 | 0 |

Create PDF in your applications with the Pdfcrowd HTML to PDF API PDFCROWD

| | | | |
|---|---|---|---|
| SPGP0AD00870 | YKR022C | 99 | 0 |
| SPGP0D00120 | YOR340C | 99 | 0 |
| SPGP0K00280 | YNR056C | 99 | 0 |
| SPGP0G03470 | YML054C | 99 | 0 |
| SPGP0G02310 | YMR074C | 100 | 0 |
| SPGP0AA00820 | YPL167C | 99 | 0 |
| SPGP0AU00100 | YOR344C | 99 | 0 |
| SPGP0T02280 | YLR163C | 99 | 0 |
| SPGP0AO00450 | YLR422W | 99 | 0 |
| SPGP0D02830 | YOR034C | 99 | 0 |
| SPGP0Z01220 | YKL150W | 99 | 0 |
| SPGP0AJ00670 | YAL041W | 99 | 0 |
| SPGP0AD00590 | YKR054C | 99 | 0 |
| SPGP0X00280 | YHL023C | 99 | 0 |
| SPGP0A04890 | YDR016C | 99 | 0 |
| SPGP0BC00160 | YIR029W | 99 | 0 |
| SPGP0A02830 | YDR242W | 100 | 0 |
| SPGP0BP00100 | YGL262W | 83 | 0 |
| SPGP0R02280 | YGL143C | 99 | 0 |
| SPGP0Y00990 | YGR207C | 99 | 0 |
| SPGP0AB00160 | YIL011W | 94 | 3 |
| SPGP0K01990 | YNL134C | 99 | 0 |
| SPGP0A03620 | YDR153C | 99 | 0 |
| SPGP0AC00680 | YPR082C | 99 | 0 |
| SPGP0AC00130 | YPR019W | 99 | 0 |
| SPGP0D00960 | YOR245C | 99 | 0 |
| SPGP0D00970 | YOR244W | 99 | 0 |
| SPGP0BX00110 | YFR027W | 99 | 0 |
| SPGP0G03040 | YML006C | 99 | 0 |
| SPGP0AB00850 | YIL092W | 99 | 0 |
| SPGP0D04150 | YOL112W | 99 | 0 |
| SPGP0J03810 | YBL103C | 99 | 0 |
| SPGP0A00210 | YDR529C | 99 | 0 |
| SPGP0A02490 | YDR282C | 99 | 0 |
| SPGP0D03640 | YOL057W | 99 | 0 |
| SPGP0K00890 | YNL010W | 99 | 0 |
| SPGP0AN00510 | YPL040C | 99 | 0 |
| SPGP0G04000 | YML109W | 99 | 0 |
| SPGP0R01350 | YGL038C | 99 | 0 |
| SPGP0AC00350 | YPR042C | 99 | 0 |

Create PDF in your applications with the Pdfcrowd HTML to PDF API PDFCROWD

| | | | |
|---|---|---|---|
| SPGP0U00620 | YLL004W | 99 | 0 |
| SPGP0V00570 | YEL024W | 99 | 0 |
| SPGP0T00280 | YLR390W | 99 | 0 |
| SPGP0D00540 | YOR292C | 99 | 0 |
| SPGP0A04430 | YDR066C | 99 | 0 |
| SPGP0T01480 | YLR254C | 99 | 0 |
| SPGP0G02110 | YMR096W | 100 | 0 |
| SPGP0G00780 | YMR232W | 99 | 0 |
| SPGP0AD00420 | YKR071C | 99 | 0 |
| SPGP0V01210 | YER040W | 99 | 0 |
| SPGP0R00500 | YGR060W | 99 | 0 |
| SPGP0R00850 | YGR019W | 99 | 0 |
| SPGP0D01380 | YOR196C | 99 | 0 |
| SPGP0J01530 | YBR147W | 99 | 0 |
| SPGP0J00770 | YBR228W | 99 | 0 |
| SPGP0N00170 | YJR143C | 99 | 0 |
| SPGP0G04080 | YML117W | 99 | 0 |
| SPGP0A00490 | YDR497C | 99 | 0 |
| SPGP0G01780 | YMR130W | 99 | 0 |
| SPGP0AC00630 | YPR075C | 99 | 0 |
| SPGP0AB00490 | YIL048W | 99 | 0 |
| SPGP0U01140 | YLR052W | 99 | 0 |
| SPGP0D04460 | YOL145C | 99 | 0 |
| SPGP0GK00100 | YPR015C | 99 | 0 |
| SPGP0N02490 | YJL124C | 100 | 0 |
| SPGP0R02230 | YGL138C | 99 | 0 |
| SPGP0AB00150 | YIL010W | 99 | 0 |
| SPGP0G01560 | YMR154C | 99 | 0 |
| SPGP0X00180 | YHL033C | 98 | 0 |
| SPGP0K03110 | YNL258C | 99 | 0 |
| SPGP0Y00370 | YGR138C | 99 | 0 |
| SPGP0AA01130 | YPL134C | 100 | 0 |
| SPGP0AE01130 | YFL051C | 98 | 0 |
| SPGP0K02020 | YNL137C | 99 | 0 |
| SPGP0K01430 | YNL071W | 99 | 0 |
| SPGP0AK00780 | YGL183C | 99 | 0 |
| SPGP0X00550 | YHR005C | 99 | 0 |
| SPGP0BQ00100 | YMR305C | 99 | 0 |
| SPGP0J02730 | YBR017C | 99 | 0 |
| SPGP0Z00570 | YKL075C | 99 | 0 |

Create PDF in your applications with the Pdfcrowd HTML to PDF API PDFCROWD

| | | | |
|---|---|---|---|
| SPGP0R01580 | YGL065C | 100 | 0 |
| SPGP0N01600 | YJL014W | 99 | 0 |
| SPGP0G00840 | YMR226C | 99 | 0 |
| SPGP0AD00450 | YKR068C | 99 | 0 |
| SPGP0K03040 | YNL251C | 99 | 0 |
| SPGP0G02140 | YMR093W | 99 | 0 |
| SPGP0T00890 | YLR321C | 99 | 0 |
| SPGP0AA00360 | YPL215W | 99 | 0 |
| SPGP0A01320 | YDR404C | 100 | 0 |
| SPGP0Z00600 | YKL079W | 99 | 0 |
| SPGP0E00110 | YGR289C | 98 | 0 |
| SPGP0V00590 | YEL022W | 99 | 0 |
| SPGP0G00810 | YMR229C | 99 | 0 |
| SPGP0AJ00440 | YAL018C | 99 | 0 |
| SPGP0T02070 | YLR187W | 99 | 0 |
| SPGP0A03640 | YDR151C | 99 | 0 |
| SPGP0N02310 | YJL101C | 99 | 0 |
| SPGP0J01550 | YBR145W | 99 | 0 |
| SPGP0AA01690 | YPL076W | 99 | 0 |
| SPGP0AJ00110 | YHR211W | 94 | 1 |
| SPGP0A04150 | YDR093W | 99 | 0 |
| SPGP0Y00830 | YGR191W | 99 | 0 |
| SPGP0X01880 | YHR139C | 99 | 0 |
| SPGP0V02160 | YER132C | 99 | 0 |
| SPGP0A03800 | YDR132C | 99 | 0 |
| SPGP0G01720 | YMR136W | 99 | 0 |
| SPGP0D01420 | YOR192C | 99 | 0 |
| SPGP0J00920 | YBR210W | 99 | 0 |
| SPGP0AD01070 | YKR002W | 99 | 0 |
| SPGP0X00260 | YHL025W | 99 | 0 |
| SPGP0K01960 | YNL131W | 100 | 0 |
| SPGP0N01200 | YJR035W | 99 | 0 |
| SPGP0T01250 | YLR283W | 99 | 0 |
| SPGP0K03520 | YNL305C | 99 | 0 |
| SPGP0AM00490 | YER179W | 99 | 0 |
| SPGP0AA01280 | YPL119C | 99 | 0 |
| SPGP0K03030 | YNL250W | 99 | 0 |
| SPGP0AA01320 | YPL115C | 99 | 0 |
| SPGP0D03710 | YOL064C | 99 | 0 |
| SPGP0A06450 | YDL167C | 99 | 0 |

Create PDF in your applications with the Pdfcrowd HTML to PDF API PDFCROWD

| | | | |
|---|---|---|---|
| SPGP0A05200 | YDL024C | 99 | 0 |
| SPGP0Z01680 | YKL201C | 99 | 1 |
| SPGP0Z01350 | YKL164C | 92 | 4 |
| SPGP0N02240 | YJL094C | 99 | 0 |
| SPGP0AE00430 | YFR022W | 99 | 0 |
| SPGP0T01830 | YLR213C | 99 | 0 |
| SPGP0AC00810 | YPR100W | 100 | 0 |
| SPGP0K03630 | YNL316C | 99 | 0 |
| SPGP0D00480 | YOR298W | 99 | 0 |
| SPGP0AH00840 | YCR021C | 99 | 0 |
| SPGP0AS00110 | YPL282C | 99 | 0 |
| SPGP0D02620 | YOR059C | 99 | 0 |
| SPGP0AA01440 | YPL101W | 99 | 0 |
| SPGP0N02760 | YJL153C | 99 | 0 |
| SPGP0G04010 | YML110C | 99 | 0 |
| SPGP0A04780 | YDR027C | 99 | 0 |
| SPGP0AP00190 | YHR175W | 99 | 0 |
| SPGP0D01960 | YOR132W | 99 | 0 |
| SPGP0AK00300 | YGL240W | 100 | 0 |
| SPGP0T00970 | YLR312W-A | 99 | 0 |
| SPGP0X01380 | YHR088W | 99 | 0 |
| SPGP0A02000 | YDR331W | 99 | 0 |
| SPGP0A06030 | YDL120W | 99 | 0 |
| SPGP0U01190 | YLR057W | 99 | 0 |
| SPGP0A06040 | YDL121C | 99 | 0 |
| SPGP0N02640 | YJL138C | 99 | 0 |
| SPGP0AZ00220 | YIR003W | 99 | 0 |
| SPGP0T00610 | YLR354C | 100 | 0 |
| SPGP0AJ00510 | YAL025C | 100 | 0 |
| SPGP0AW00190 | YHR158C | 99 | 0 |
| SPGP0J01700 | YBR128C | 99 | 0 |
| SPGP0AS00370 | YPL249C-A | 100 | 0 |
| SPGP0A06960 | YDL225W | 99 | 0 |
| SPGP0A06780 | YDL206W | 99 | 0 |
| SPGP0D00990 | YOR242C | 99 | 0 |
| SPGP0K02800 | YNL224C | 99 | 0 |
| SPGP0AD00330 | YKR081C | 99 | 0 |
| SPGP0D03520 | YOL044W | 99 | 0 |
| SPGP0BK00140 | YGL173C | 99 | 0 |
| SPGP0U00400 | YLL027W | 99 | 0 |

Create PDF in your applications with the Pdfcrowd HTML to PDF API PDFCROWD

| | | | |
|---|---|---|---|
| SPGP0D02500 | YOR073W | 99 | 0 |
| SPGP0AJ00660 | YAL040C | 99 | 0 |
| SPGP0A04520 | YDR058C | 99 | 0 |
| SPGP0G03060 | YML008C | 99 | 0 |
| SPGP0Y00270 | YGR127W | 99 | 0 |
| SPGP0J00250 | YBR281C | 99 | 0 |
| SPGP0G00570 | YMR255W | 99 | 0 |
| SPGP0J03010 | YBL015W | 99 | 0 |
| SPGP0D03690 | YOL062C | 99 | 0 |
| SPGP0X00200 | YHL031C | 99 | 0 |
| SPGP0K00370 | YNR047W | 99 | 0 |
| SPGP0D03320 | YOL019W | 99 | 0 |
| SPGP0AU00190 | YOR354C | 99 | 0 |
| SPGP0G02950 | YMR005W | 100 | 0 |
| SPGP0G01220 | YMR190C | 99 | 0 |
| SPGP0AH00850 | YCR023C | 99 | 0 |
| SPGP0D02000 | YOR128C | 99 | 0 |
| SPGP0D03900 | YOL084W | 99 | 0 |
| SPGP0A00680 | YDR477W | 99 | 0 |
| SPGP0V01020 | YER021W | 99 | 0 |
| SPGP0X00810 | YHR031C | 99 | 0 |
| SPGP0A02910 | YDR233C | 99 | 0 |
| SPGP0J02500 | YBR042C | 99 | 0 |
| SPGP0J03830 | YBL105C | 99 | 0 |
| SPGP0J03250 | YBL038W | 99 | 0 |
| SPGP0AA01610 | YPL085W | 99 | 0 |
| SPGP0K01890 | YNL124W | 99 | 0 |
| SPGP0J02410 | YBR052C | 99 | 0 |
| SPGP0T00990 | YLR310C | 99 | 0 |
| SPGP0K01770 | YNL110C | 99 | 0 |
| SPGP0U01770 | YLR119W | 99 | 0 |
| SPGP0D03630 | YOL056W | 99 | 0 |
| SPGP0D02100 | YOR117W | 99 | 0 |
| SPGP0G04120 | YML121W | 99 | 0 |
| SPGP0D01850 | YOR145C | 99 | 0 |
| SPGP0X01620 | YHR112C | 99 | 0 |
| SPGP0D02270 | YOR098C | 99 | 0 |
| SPGP0U01940 | YLR139C | 99 | 0 |
| SPGP0AB00940 | YIL102C | 100 | 0 |
| SPGP0R02420 | YGL159W | 99 | 0 |

Create PDF in your applications with the Pdfcrowd HTML to PDF API PDFCROWD

| | | | |
|---|---|---|---|
| SPGP0K01700 | YNL101W | 99 | 0 |
| SPGP0AA00430 | YPL208W | 99 | 0 |
| SPGP0AC00140 | YPR020W | 99 | 0 |
| SPGP0K03810 | YNL336W | 97 | 0 |
| SPGP0Z00210 | YKL039W | 99 | 0 |
| SPGP0X00840 | YHR034C | 99 | 0 |
| SPGP0T01530 | YLR248W | 100 | 0 |
| SPGP0AN00130 | YPR001W | 99 | 0 |
| SPGP0K00610 | YNR021W | 99 | 0 |
| SPGP0A05740 | YDL088C | 99 | 0 |
| SPGP0N02690 | YJL144W | 97 | 0 |
| SPGP0A02210 | YDR311W | 99 | 0 |
| SPGP0AP00340 | YHR190W | 99 | 0 |
| SPGP0V00860 | YER007W | 99 | 0 |
| SPGP0BA00220 | YCR079W | 99 | 0 |
| SPGP0A02960 | YDR227W | 99 | 0 |
| SPGP0J01220 | YBR179C | 99 | 0 |
| SPGP0K03730 | YNL328C | 99 | 0 |
| SPGP0AD00670 | YKR045C | 96 | 4 |
| SPGP0D00100 | YOR342C | 99 | 0 |
| SPGP0X01100 | YHR062C | 100 | 0 |
| SPGP0G02330 | YMR072W | 100 | 0 |
| SPGP0A00940 | YDR449C | 99 | 0 |
| SPGP0D01670 | YOR164C | 99 | 0 |
| SPGP0R01710 | YGL081W | 98 | 1 |
| SPGP0AP00520 | YHR207C | 99 | 0 |
| SPGP0A05920 | YDL108W | 99 | 0 |
| SPGP0AD01100 | YKL002W | 99 | 0 |
| SPGP0AE00410 | YFR024C-A | 99 | 0 |
| SPGP0A04070 | YDR104C | 99 | 0 |
| SPGP0G03930 | YML102W | 99 | 0 |
| SPGP0T01930 | YLR203C | 99 | 0 |
| SPGP0AJ00300 | YAL001C | 99 | 0 |
| SPGP0D00810 | YOR261C | 98 | 1 |
| SPGP0R01040 | YGL004C | 99 | 0 |
| SPGP0G02640 | YMR036C | 99 | 0 |
| SPGP0N01290 | YJR019C | 99 | 0 |
| SPGP0AD00570 | YKR056W | 99 | 0 |
| SPGP0AN00280 | YPL015C | 99 | 0 |

Create PDF in your applications with the Pdfcrowd HTML to PDF API PDFCROWD

| | | | |
|---|---|---|---|
| SPGP0AC00110 | YPR017C | 99 | 0 |
| SPGP0A02620 | YDR264C | 99 | 0 |
| SPGP0AB00280 | YIL023C | 99 | 0 |
| SPGP0R00860 | YGR017W | 99 | 0 |
| SPGP0AA00450 | YPL206C | 99 | 0 |
| SPGP0J01080 | YBR193C | 99 | 0 |
| SPGP0AB01130 | YIL120W | 99 | 0 |
| SPGP0AE00390 | YFR029W | 99 | 0 |
| SPGP0K03760 | YNL331C | 100 | 0 |
| SPGP0K03190 | YNL267W | 99 | 0 |
| SPGP0X00660 | YHR015W | 99 | 0 |
| SPGP0X01710 | YHR121W | 99 | 0 |
| SPGP0G02060 | YMR101C | 99 | 0 |
| SPGP0K00560 | YNR027W | 99 | 0 |
| SPGP0AP00130 | YHR169W | 100 | 0 |
| SPGP0AC00370 | YPR045C | 99 | 0 |
| SPGP0U00640 | YLL002W | 99 | 0 |
| SPGP0Y01520 | YGR267C | 100 | 0 |
| SPGP0A00520 | YDR494W | 99 | 0 |
| SPGP0A04450 | YDR064W | 99 | 0 |
| SPGP0AE00660 | YFL001W | 99 | 0 |
| SPGP0G02810 | YMR019W | 99 | 0 |
| SPGP0G03400 | YML047C | 99 | 0 |
| SPGP0BL00130 | YCR053W | 99 | 0 |
| SPGP0R01610 | YGL068W | 99 | 0 |
| SPGP0AC00910 | YPR110C | 99 | 0 |
| SPGP0A02300 | YDR302W | 99 | 0 |
| SPGP0D01440 | YOR190W | 99 | 0 |
| SPGP0A03430 | YDR173C | 99 | 1 |
| SPGP0AA01160 | YPL131W | 99 | 0 |
| SPGP0Y00110 | YGR109C | 99 | 0 |
| SPGP0K01270 | YNL053W | 99 | 0 |
| SPGP0R00520 | YGR058W | 99 | 0 |
| SPGP0Y00300 | YGR130C | 99 | 0 |
| SPGP0AD00960 | YKR014C | 99 | 0 |
| SPGP0G03590 | YML065W | 99 | 0 |
| SPGP0AJ00470 | YAL021C | 99 | 0 |
| SPGP0R00260 | YGR090W | 99 | 0 |
| SPGP0AV00190 | YKL210W | 99 | 0 |
| SPGP0D04070 | YOL103W | 99 | 0 |

Create PDF in your applications with the Pdfcrowd HTML to PDF API PDFCROWD

| | | | |
|---|---|---|---|
| SPGP0J00750 | YBR230C | 100 | 0 |
| SPGP0D02250 | YOR100C | 99 | 0 |
| SPGP0AA00730 | YPL176C | 99 | 0 |
| SPGP0AB00670 | YIL070C | 100 | 0 |
| SPGP0AB00130 | YIL006W | 99 | 0 |
| SPGP0G01500 | YMR160W | 99 | 0 |
| SPGP0A01970 | YDR334W | 99 | 0 |
| SPGP0AK00110 | YGL258W-A | 97 | 0 |
| SPGP0K00830 | YNL004W | 100 | 0 |
| SPGP0N01620 | YJL019W | 99 | 0 |
| SPGP0AS00130 | YPL274W | 99 | 0 |
| SPGP0X00440 | YHL007C | 99 | 0 |
| SPGP0D00560 | YOR290C | 99 | 0 |
| SPGP0V01490 | YER064C | 99 | 0 |
| SPGP0G00370 | YMR275C | 99 | 0 |
| SPGP0D03160 | YOL003C | 99 | 0 |
| SPGP0V00540 | YEL027W | 100 | 0 |
| SPGP0D00530 | YOR293W | 99 | 0 |
| SPGP0G02630 | YMR037C | 99 | 0 |
| SPGP0AJ00220 | YAR018C | 99 | 0 |
| SPGP0K00330 | YNR051C | 99 | 0 |
| SPGP0A01340 | YDR402C | 99 | 0 |
| SPGP0AA00610 | YPL189W | 99 | 0 |
| SPGP0A02900 | YDR234W | 99 | 0 |
| SPGP0X01580 | YHR108W | 99 | 0 |
| SPGP0D01940 | YOR134W | 99 | 0 |
| SPGP0X01850 | YHR136C | 100 | 0 |
| SPGP0AB01260 | YIL133C | 99 | 0 |
| SPGP0V01650 | YER079W | 99 | 0 |
| SPGP0Y01510 | YGR266W | 99 | 0 |
| SPGP0D02300 | YOR095C | 99 | 0 |
| SPGP0AD00990 | YKR010C | 99 | 0 |
| SPGP0U01300 | YLR069C | 99 | 0 |
| SPGP0N02330 | YJL103C | 99 | 0 |
| SPGP0K01010 | YNL026W | 99 | 0 |
| SPGP0J01570 | YBR142W | 99 | 0 |
| SPGP0D02860 | YOR031W | 99 | 0 |
| SPGP0AJ00790 | YAL055W | 98 | 0 |
| SPGP0X00670 | YHR017W | 99 | 0 |
| SPGP0T02050 | YLR189C | 99 | 0 |

Create PDF in your applications with the Pdfcrowd HTML to PDF API PDFCROWD

| | | | |
|---|---|---|---|
| SPGP0Y00200 | YGR120C | 99 | 0 |
| SPGP0G00280 | YMR284W | 99 | 0 |
| SPGP0K01900 | YNL125C | 99 | 0 |
| SPGP0D03020 | YOR013W | 99 | 0 |
| SPGP0AH00530 | YCL012C | 100 | 0 |
| SPGP0A03170 | YDR201W | 100 | 0 |
| SPGP0D04450 | YOL144W | 99 | 0 |
| SPGP0K03500 | YNL302C | 99 | 0 |
| SPGP0AK00580 | YGL208W | 99 | 0 |
| SPGP0AU00260 | YOR361C | 99 | 0 |
| SPGP0J01090 | YBR192W | 99 | 0 |
| SPGP0Z00320 | YKL051W | 99 | 0 |
| SPGP0V01050 | YER024W | 99 | 0 |
| SPGP0V00220 | YEL060C | 99 | 0 |
| SPGP0X00240 | YHL027W | 99 | 0 |
| SPGP0V00230 | YEL059C-A | 99 | 0 |
| SPGP0T01580 | YLR243W | 99 | 0 |
| SPGP0K03570 | YNL310C | 99 | 0 |
| SPGP0D01160 | YOR222W | 99 | 0 |
| SPGP0G02170 | YMR090W | 99 | 0 |
| SPGP0AA01420 | YPL104W | 99 | 0 |
| SPGP0N02040 | YJL070C | 99 | 0 |
| SPGP0AD00390 | YKR075C | 99 | 0 |
| SPGP0T01810 | YLR215C | 99 | 0 |
| SPGP0T00980 | YLR312C | 99 | 0 |
| SPGP0AE00330 | YFR033C | 99 | 0 |
| SPGP0A05480 | YDL059C | 100 | 0 |
| SPGP0AD01310 | YKL021C | 99 | 0 |
| SPGP0N02620 | YJL136C | 99 | 0 |
| SPGP0T00180 | YLR401C | 99 | 0 |
| SPGP0D03070 | YOR007C | 99 | 0 |
| SPGP0AE00450 | YFR020W | 99 | 0 |
| SPGP0U01120 | YLR050C | 100 | 0 |
| SPGP0Z00930 | YKL117W | 99 | 0 |
| SPGP0A06090 | YDL127W | 99 | 0 |
| SPGP0AH00670 | YCR005C | 99 | 0 |
| SPGP0A02060 | YDR324C | 99 | 0 |
| SPGP0AS00170 | YPL270W | 99 | 0 |
| SPGP0Z00310 | YKL050C | 99 | 0 |
| SPGP0A00900 | YDR453C | 100 | 0 |

Create PDF in your applications with the Pdfcrowd HTML to PDF API PDFCROWD

| | | | |
|---|---|---|---|
| SPGP0AA01390 | YPL107W | 99 | 0 |
| SPGP0U01810 | YLR126C | 100 | 0 |
| SPGP0J00560 | YBR249C | 99 | 0 |
| SPGP0AO00270 | YLR441C | 100 | 0 |
| SPGP0T00360 | YLR382C | 99 | 0 |
| SPGP0BK00120 | YGL175C | 99 | 0 |
| SPGP0AM00410 | YER171W | 99 | 0 |
| SPGP0AS00390 | YPL248C | 99 | 0 |
| SPGP0A06900 | YDL218W | 99 | 0 |
| SPGP0D00820 | YOR260W | 99 | 0 |
| SPGP0AH00560 | YCL009C | 99 | 0 |
| SPGP0AK00660 | YGL197W | 99 | 0 |
| SPGP0N00440 | YJR115W | 96 | 2 |
| SPGP0N00910 | YJR066W | 99 | 0 |
| SPGP0AB01030 | YIL110W | 99 | 0 |
| SPGP0U00420 | YLL025W | 100 | 0 |
| SPGP0D02520 | YOR070C | 99 | 0 |
| SPGP0A01600 | YDR376W | 99 | 0 |
| SPGP0D02260 | YOR099W | 99 | 0 |
| SPGP0A03080 | YDR213W | 99 | 0 |
| SPGP0D02640 | YOR057W | 99 | 0 |
| SPGP0A00770 | YDR466W | 99 | 0 |
| SPGP0A06200 | YDL138W | 99 | 0 |
| SPGP0T00150 | YLR405W | 99 | 0 |
| SPGP0AB01370 | YIL145C | 99 | 0 |
| SPGP0AD00530 | YKR060W | 99 | 0 |
| SPGP0J03030 | YBL017C | 99 | 0 |
| SPGP0AB01450 | YIL153W | 99 | 0 |
| SPGP0G04020 | YML111W | 99 | 0 |
| SPGP0A01610 | YDR375C | 99 | 0 |
| SPGP0N00820 | YJR076C | 99 | 0 |
| SPGP0A01170 | YDR421W | 99 | 0 |
| SPGP0A00750 | YDR469W | 99 | 0 |
| SPGP0AM00550 | YER185W | 99 | 0 |
| SPGP0N00700 | YJR091C | 99 | 0 |
| SPGP0R02000 | YGL113W | 99 | 0 |
| SPGP0G01240 | YMR188C | 99 | 0 |
| SPGP0A02030 | YDR328C | 99 | 0 |
| SPGP0R01090 | YGL010W | 99 | 0 |
| SPGP0X00830 | YHR033W | 99 | 0 |

Create PDF in your applications with the Pdfcrowd HTML to PDF API PDFCROWD

| | | | |
|---|---|---|---|
| SPGP0Y00580 | YGR162W | 99 | 0 |
| SPGP0J02520 | YBR040W | 99 | 0 |
| SPGP0U01590 | YLR099C | 99 | 0 |
| SPGP0D03930 | YOL087C | 99 | 0 |
| SPGP0A00570 | YDR488C | 99 | 0 |
| SPGP0V00820 | YER003C | 100 | 0 |
| SPGP0K01750 | YNL107W | 99 | 0 |
| SPGP0K01020 | YNL025C | 100 | 0 |
| SPGP0Y01180 | YGR230W | 99 | 0 |
| SPGP0N01990 | YJL063C | 99 | 0 |
| SPGP0A01910 | YDR341C | 99 | 0 |
| SPGP0Y00910 | YGR199W | 99 | 0 |
| SPGP0D03650 | YOL058W | 99 | 0 |
| SPGP0D02160 | YOR111W | 99 | 0 |
| SPGP0AU00300 | YOR367W | 99 | 0 |
| SPGP0U01380 | YLR078C | 100 | 0 |
| SPGP0D01870 | YOR143C | 99 | 0 |
| SPGP0T00500 | YLR364W | 100 | 0 |
| SPGP0AO00390 | YLR429W | 99 | 0 |
| SPGP0X01440 | YHR094C | 99 | 0 |
| SPGP0G03920 | YML100W | 99 | 0 |
| SPGP0U01920 | YLR137W | 99 | 0 |
| SPGP0A03110 | YDR208W | 99 | 0 |
| SPGP0G02340 | YMR071C | 99 | 0 |
| SPGP0AB00920 | YIL099W | 99 | 0 |
| SPGP0AM00100 | YER139C | 99 | 0 |
| SPGP0V00750 | YEL005C | 99 | 0 |
| SPGP0AB01270 | YIL134W | 99 | 0 |
| SPGP0G02530 | YMR049C | 99 | 0 |
| SPGP0T01120 | YLR297W | 100 | 0 |
| SPGP0AA00800 | YPL169C | 99 | 0 |
| SPGP0K00850 | YNL006W | 99 | 0 |
| SPGP0K02220 | YNL159C | 99 | 0 |
| SPGP0G02870 | YMR013C | 99 | 0 |
| SPGP0R01720 | YGL082W | 99 | 0 |
| SPGP0Z01280 | YKL157W | 99 | 0 |
| SPGP0N03120 | YJL193W | 99 | 0 |
| SPGP0AN00580 | YPL048W | 99 | 0 |
| SPGP0A07080 | YDL238C | 99 | 0 |
| SPGP0A04050 | YDR106W | 99 | 0 |

Create PDF in your applications with the Pdfcrowd HTML to PDF API PDFCROWD

| | | | |
|---|---|---|---|
| SPGP0AA01150 | YPL132W | 99 | 0 |
| SPGP0T00950 | YLR314C | 99 | 0 |
| SPGP0K02040 | YNL138W-A | 100 | 0 |
| SPGP0D01650 | YOR166C | 99 | 0 |
| SPGP0D01120 | YOR227W | 99 | 0 |
| SPGP0D02570 | YOR064C | 100 | 0 |
| SPGP0G01650 | YMR144W | 99 | 0 |
| SPGP0N00610 | YJR099W | 99 | 0 |
| SPGP0R01730 | YGL083W | 99 | 0 |
| SPGP0A05850 | YDL101C | 99 | 0 |
| SPGP0G03910 | YML099C | 99 | 0 |
| SPGP0AJ00630 | YAL037W | 99 | 0 |
| SPGP0T01910 | YLR205C | 99 | 0 |
| SPGP0D00380 | YOR307C | 99 | 0 |
| SPGP0D00830 | YOR259C | 99 | 0 |
| SPGP0AK00230 | YGL247W | 99 | 0 |
| SPGP0AN00260 | YPL013C | 99 | 0 |
| SPGP0AK00700 | YGL194C | 100 | 0 |
| SPGP0AC00170 | YPR023C | 99 | 0 |
| SPGP0K03470 | YNL299W | 99 | 0 |
| SPGP0A05010 | YDR001C | 99 | 0 |
| SPGP0AD01120 | YKL004W | 99 | 0 |
| SPGP0N01890 | YJL053W | 99 | 0 |
| SPGP0J02640 | YBR026C | 100 | 0 |
| SPGP0A03840 | YDR128W | 99 | 0 |
| SPGP0AW00150 | YHR155W | 99 | 0 |
| SPGP0K00250 | YNR059W | 99 | 0 |
| SPGP0AH00550 | YCL010C | 99 | 0 |
| SPGP0BC00170 | YIR030C | 99 | 0 |
| SPGP0AD00760 | YKR035W-A | 99 | 0 |
| SPGP0J03850 | YBL107C | 99 | 0 |
| SPGP0G00290 | YMR283C | 99 | 0 |
| SPGP0AC00390 | YPR047W | 100 | 0 |
| SPGP0G03300 | YML032C | 99 | 0 |
| SPGP0U00170 | YLL055W | 99 | 0 |
| SPGP0A04470 | YDR062W | 99 | 0 |
| SPGP0J02530 | YBR039W | 99 | 0 |
| SPGP0A01310 | YDR405W | 99 | 0 |
| SPGP0CC00110 | YMR318C | 99 | 0 |
| SPGP0AH00830 | YCR020W-B | 99 | 0 |

Create PDF in your applications with the Pdfcrowd HTML to PDF API PDFCROWD

| | | | |
|---|---|---|---|
| SPGP0A02540 | YDR276C | 100 | 0 |
| SPGP0X01420 | YHR092C | 99 | 0 |
| SPGP0J03400 | YBL055C | 99 | 0 |
| SPGP0Z00880 | YKL110C | 99 | 0 |
| SPGP0G01150 | YMR196W | 99 | 0 |
| SPGP0D02530 | YOR069W | 99 | 0 |
| SPGP0Z00220 | YKL040C | 99 | 0 |
| SPGP0J01560 | YBR143C | 99 | 0 |
| SPGP0K01480 | YNL076W | 99 | 0 |
| SPGP0IU00100 | YPL245W | 99 | 0 |
| SPGP0U01450 | YLR085C | 99 | 0 |
| SPGP0A06590 | YDL183C | 99 | 0 |
| SPGP0AJ00490 | YAL023C | 99 | 0 |
| SPGP0N02480 | YJL123C | 99 | 1 |
| SPGP0D00600 | YOR286W | 99 | 0 |
| SPGP0T00790 | YLR332W | 99 | 0 |
| SPGP0AK00340 | YGL234W | 99 | 0 |
| SPGP0J02510 | YBR041W | 99 | 0 |
| SPGP0V00560 | YEL025C | 99 | 0 |
| SPGP0Z00280 | YKL047W | 99 | 0 |
| SPGP0N01140 | YJR043C | 99 | 0 |
| SPGP0A00560 | YDR489W | 99 | 0 |
| SPGP0K02570 | YNL199C | 99 | 0 |
| SPGP0V01960 | YER111C | 99 | 0 |
| SPGP0K01980 | YNL133C | 99 | 0 |
| SPGP0AO00250 | YLR443W | 99 | 0 |
| SPGP0AA00840 | YPL165C | 99 | 0 |
| SPGP0U00850 | YLR021W | 99 | 0 |
| SPGP0T00420 | YLR375W | 99 | 0 |
| SPGP0Y01190 | YGR231C | 100 | 0 |
| SPGP0AY00120 | YOR377W | 99 | 0 |
| SPGP0G03200 | YML022W | 99 | 0 |
| SPGP0K01040 | YNL024C | 99 | 0 |
| SPGP0X01890 | YHR140W | 99 | 0 |
| SPGP0D02820 | YOR035C | 99 | 0 |
| SPGP0A01810 | YDR354W | 99 | 0 |
| SPGP0K00510 | YNR032W | 99 | 0 |
| SPGP0V00760 | YEL004W | 99 | 0 |
| SPGP0Y01060 | YGR215W | 100 | 0 |
| SPGP0U00690 | YLR004C | 99 | 0 |

Create PDF in your applications with the Pdfcrowd HTML to PDF API PDFCROWD

| | | | |
|---|---|---|---|
| SPGP0J03530 | YBL069W | 99 | 0 |
| SPGP0AO00470 | YLR420W | 99 | 0 |
| SPGP0U01640 | YLR104W | 99 | 0 |
| SPGP0X00470 | YHL003C | 99 | 0 |
| SPGP0K00350 | YNR049C | 100 | 0 |
| SPGP0A07090 | YDL239C | 99 | 0 |
| SPGP0Z00400 | YKL059C | 99 | 0 |
| SPGP0V01610 | YER075C | 99 | 0 |
| SPGP0A04370 | YDR072C | 99 | 0 |
| SPGP0R01910 | YGL103W | 99 | 0 |
| SPGP0R01260 | YGL028C | 98 | 1 |
| SPGP0A00140 | YDR538W | 99 | 0 |
| SPGP0N02350 | YJL105W | 99 | 0 |
| SPGP0N00110 | YJR150C | 98 | 1 |
| SPGP0D02840 | YOR033C | 99 | 0 |
| SPGP0K01920 | YNL127W | 99 | 0 |
| SPGP0Z01200 | YKL148C | 99 | 0 |
| SPGP0N02210 | YJL091C | 99 | 0 |
| SPGP0J02070 | YBR085C-A | 100 | 0 |
| SPGP0AC00450 | YPR055W | 99 | 0 |
| SPGP0J00720 | YBR233W-A | 99 | 0 |
| SPGP0J00880 | YBR214W | 99 | 0 |
| SPGP0Y00220 | YGR122W | 99 | 0 |
| SPGP0AE00940 | YFL031W | 99 | 0 |
| SPGP0A02480 | YDR283C | 99 | 0 |
| SPGP0AC00280 | YPR034W | 99 | 0 |
| SPGP0AC00240 | YPR030W | 99 | 0 |
| SPGP0A04260 | YDR082W | 99 | 0 |
| SPGP0AB00260 | YIL021W | 99 | 0 |
| SPGP0T01210 | YLR287C | 99 | 0 |
| SPGP0D04470 | YOL146W | 99 | 0 |
| SPGP0K03070 | YNL254C | 99 | 0 |
| SPGP0R00300 | YGR083C | 99 | 0 |
| SPGP0CH00110 | YKL023C-A | 100 | 0 |
| SPGP0AB00480 | YIL047C | 99 | 0 |
| SPGP0G00710 | YMR239C | 100 | 0 |
| SPGP0X01300 | YHR080C | 99 | 0 |
| SPGP0N02290 | YJL099W | 99 | 0 |
| SPGP0AP00510 | YHR206W | 99 | 0 |
| SPGP0A05410 | YDL052C | 99 | 0 |

Create PDF in your applications with the Pdfcrowd HTML to PDF API PDFCROWD

| | | | |
|---|---|---|---|
| SPGP0V00200 | YEL062W | 99 | 0 |
| SPGP0Y01590 | YGR275W | 99 | 0 |
| SPGP0A01280 | YDR408C | 99 | 0 |
| SPGP0AH00620 | YCL001W | 100 | 0 |
| SPGP0J01430 | YBR157C | 99 | 0 |
| SPGP0AA00960 | YPL153C | 99 | 0 |
| SPGP0AA01230 | YPL124W | 99 | 0 |
| SPGP0A06800 | YDL208W | 100 | 0 |
| SPGP0AE00370 | YFR031C | 99 | 0 |
| SPGP0N02600 | YJL133C-A | 99 | 0 |
| SPGP0T00160 | YLR404W | 100 | 0 |
| SPGP0AE00290 | YFR037C | 99 | 0 |
| SPGP0K03740 | YNL329C | 99 | 0 |
| SPGP0J01580 | YBR141C | 99 | 0 |
| SPGP0G00770 | YMR233W | 99 | 0 |
| SPGP0A02730 | YDR253C | 100 | 0 |
| SPGP0Z00990 | YKL125W | 99 | 0 |
| SPGP0Z01320 | YKL162C | 99 | 0 |
| SPGP0J03180 | YBL031W | 99 | 0 |
| SPGP0A06750 | YDL203C | 99 | 0 |
| SPGP0AN00210 | YPL008W | 99 | 0 |
| SPGP0U00190 | YLL053C | 99 | 0 |
| SPGP0N01880 | YJL052W | 99 | 0 |
| SPGP0D03730 | YOL066C | 99 | 0 |
| SPGP0Y00560 | YGR159C | 99 | 0 |
| SPGP0G00600 | YMR251W-A | 100 | 0 |
| SPGP0AM00380 | YER168C | 99 | 0 |
| SPGP0Z00190 | YKL035W | 99 | 0 |
| SPGP0U00870 | YLR023C | 99 | 0 |
| SPGP0AB01470 | YIL155C | 99 | 0 |
| SPGP0N00930 | YJR064W | 99 | 0 |
| SPGP0U00220 | YLL049W | 99 | 0 |
| SPGP0U00440 | YLL023C | 99 | 0 |
| SPGP0AA01790 | YPL065W | 99 | 0 |
| SPGP0G02480 | YMR056C | 99 | 0 |
| SPGP0U01020 | YLR039C | 99 | 0 |
| SPGP0A03440 | YDR172W | 99 | 0 |
| SPGP0X01750 | YHR126C | 99 | 0 |
| SPGP0AD00510 | YKR062W | 99 | 0 |
| SPGP0U00580 | YLL008W | 99 | 0 |

Create PDF in your applications with the Pdfcrowd HTML to PDF API PDFCROWD

| | | | |
|---|---|---|---|
| SPGP0J03050 | YBL019W | 99 | 0 |
| SPGP0N01120 | YJR045C | 99 | 0 |
| SPGP0G04040 | YML113W | 98 | 1 |
| SPGP0N01400 | YJR008W | 99 | 0 |
| SPGP0R02020 | YGL115W | 99 | 0 |
| SPGP0U02080 | YLR153C | 99 | 1 |
| SPGP0R02530 | YGL172W | 99 | 0 |
| SPGP0AK00540 | YGL212W | 99 | 0 |
| SPGP0X00850 | YHR035W | 99 | 0 |
| SPGP0Y00210 | YGR121C | 99 | 0 |
| SPGP0A00160 | YDR534C | 91 | 3 |
| SPGP0A02950 | YDR228C | 99 | 0 |
| SPGP0U01710 | YLR113W | 99 | 0 |
| SPGP0R01500 | YGL057C | 99 | 0 |
| SPGP0G02520 | YMR052W | 99 | 0 |
| SPGP0G00560 | YMR256C | 100 | 0 |
| SPGP0BA00180 | YCR075C | 99 | 0 |
| SPGP0K01000 | YNL027W | 99 | 0 |
| SPGP0G01260 | YMR186W | 99 | 0 |
| SPGP0X01160 | YHR068W | 99 | 0 |
| SPGP0D04530 | YOL153C | 99 | 0 |
| SPGP0CC00120 | YMR317W | 99 | 0 |
| SPGP0N01180 | YJR039W | 99 | 0 |
| SPGP0D03670 | YOL060C | 99 | 0 |
| SPGP0X00640 | YHR013C | 99 | 0 |
| SPGP0AA00790 | YPL170W | 99 | 0 |
| SPGP0J00830 | YBR220C | 99 | 0 |
| SPGP0X01460 | YHR097C | 99 | 0 |
| SPGP0U01900 | YLR135W | 99 | 0 |
| SPGP0AA00600 | YPL189C-A | 100 | 0 |
| SPGP0EA00100 | YPL243W | 99 | 0 |
| SPGP0AB00900 | YIL097W | 99 | 0 |
| SPGP0A05020 | YDL001W | 99 | 0 |
| SPGP0V01450 | YER060W | 99 | 0 |
| SPGP0K00140 | YNR070W | 99 | 0 |
| SPGP0A03600 | YDR156W | 99 | 0 |
| SPGP0N03040 | YJL184W | 99 | 0 |
| SPGP0R02300 | YGL145W | 99 | 0 |
| SPGP0G00100 | YMR302C | 99 | 0 |

Create PDF in your applications with the Pdfcrowd HTML to PDF API PDFCROWD

| | | | |
|---|---|---|---|
| SPGP0K02200 | YNL157W | 99 | 0 |
| SPGP0R01700 | YGL080W | 99 | 0 |
| SPGP0N02190 | YJL089W | 99 | 0 |
| SPGP0AP00300 | YHR186C | 99 | 0 |
| SPGP0N01540 | YJL006C | 99 | 0 |
| SPGP0AE00750 | YFL012W | 100 | 0 |
| SPGP0Y01300 | YGR243W | 99 | 0 |
| SPGP0BV00100 | YCR081W | 99 | 0 |
| SPGP0K01370 | YNL065W | 99 | 0 |
| SPGP0AN00560 | YPL046C | 99 | 0 |
| SPGP0AC00120 | YPR018W | 99 | 0 |
| SPGP0AE00200 | YFR046C | 99 | 0 |
| SPGP0R00480 | YGR062C | 99 | 0 |
| SPGP0G01630 | YMR146C | 99 | 0 |
| SPGP0T00930 | YLR316C | 99 | 0 |
| SPGP0J02120 | YBR082C | 99 | 0 |
| SPGP0D01100 | YOR229W | 99 | 0 |
| SPGP0CH00130 | YKL022C | 99 | 0 |
| SPGP0G03390 | YML046W | 99 | 0 |
| SPGP0AJ00340 | YAL008W | 99 | 0 |
| SPGP0N00250 | YJR135W-A | 99 | 0 |
| SPGP0A04100 | YDR100W | 100 | 0 |
| SPGP0X00220 | YHL029C | 99 | 0 |
| SPGP0AB00810 | YIL088C | 99 | 0 |
| SPGP0J00230 | YBR283C | 99 | 0 |
| SPGP0AK00360 | YGL232W | 99 | 0 |
| SPGP0AK00250 | YGL245W | 99 | 0 |
| SPGP0Z00850 | YKL107W | 99 | 0 |
| SPGP0AN00240 | YPL011C | 99 | 0 |
| SPGP0AC00620 | YPR074C | 99 | 0 |
| SPGP0A02660 | YDR260C | 100 | 0 |
| SPGP0V00970 | YER016W | 99 | 0 |
| SPGP0A05890 | YDL105W | 99 | 0 |
| SPGP0AJ00740 | YAL048C | 98 | 0 |
| SPGP0A02380 | YDR294C | 99 | 0 |
| SPGP0AO00190 | YLR449W | 99 | 0 |
| SPGP0K00520 | YNR031C | 99 | 0 |
| SPGP0AA01210 | YPL126W | 99 | 0 |
| SPGP0AA01000 | YPL149W | 99 | 0 |
| SPGP0A05030 | YDL002C | 99 | 0 |

Create PDF in your applications with the Pdfcrowd HTML to PDF API PDFCROWD

| | | | |
|---|---|---|---|
| SPGP0G03280 | YML030W | 100 | 0 |
| SPGP0A04710 | YDR035W | 99 | 0 |
| SPGP0A04490 | YDR060W | 99 | 0 |
| SPGP0G02320 | YMR073C | 99 | 0 |
| SPGP0AN00450 | YPL033C | 99 | 0 |
| SPGP0R02460 | YGL163C | 99 | 0 |
| SPGP0A00440 | YDR502C | 100 | 0 |
| SPGP0K02930 | YNL240C | 99 | 0 |
| SPGP0A06910 | YDL219W | 99 | 0 |
| SPGP0A06600 | YDL185W | 98 | 0 |
| SPGP0X00410 | YHL010C | 99 | 0 |
| SPGP0K00780 | YNR002C | 100 | 0 |
| SPGP0R00700 | YGR036C | 99 | 0 |
| SPGP0J00240 | YBR282W | 99 | 0 |
| SPGP0A01200 | YDR418W | 100 | 0 |
| SPGP0AB00680 | YIL071C | 99 | 0 |
| SPGP0D03500 | YOL041C | 99 | 0 |
| SPGP0X00600 | YHR009C | 99 | 0 |
| SPGP0A03380 | YDR178W | 99 | 0 |
| SPGP0T02000 | YLR194C | 99 | 0 |
| SPGP0D03080 | YOR006C | 99 | 0 |
| SPGP0G01060 | YMR204C | 99 | 0 |
| SPGP0AP00180 | YHR174W | 99 | 0 |
| SPGP0R00560 | YGR054W | 99 | 0 |
| SPGP0D01450 | YOR189W | 99 | 0 |
| SPGP0R00910 | YGR012W | 99 | 0 |
| SPGP0D02050 | YOR123C | 99 | 0 |
| SPGP0D03450 | YOL034W | 99 | 0 |
| SPGP0D03890 | YOL083W | 99 | 0 |
| SPGP0R00150 | YGR101W | 100 | 0 |
| SPGP0Y01050 | YGR214W | 99 | 0 |
| SPGP0V00500 | YEL031W | 99 | 0 |
| SPGP0A03160 | YDR202C | 99 | 0 |
| SPGP0Z01210 | YKL149C | 99 | 0 |
| SPGP0T01650 | YLR234W | 99 | 0 |
| SPGP0BL00150 | YCR057C | 100 | 0 |
| SPGP0D03880 | YOL082W | 99 | 0 |
| SPGP0A01330 | YDR403W | 99 | 0 |
| SPGP0J03260 | YBL039C | 99 | 0 |
| SPGP0AA00140 | YPL237W | 99 | 0 |

Create PDF in your applications with the Pdfcrowd HTML to PDF API PDFCROWD

| | | | |
|---|---|---|---|
| SPGP0U00410 | YLL026W | 99 | 0 |
| SPGP0A03360 | YDR179W-A | 99 | 0 |
| SPGP0T01230 | YLR285W | 99 | 0 |
| SPGP0G02350 | YMR070W | 98 | 2 |
| SPGP0N01160 | YJR041C | 99 | 0 |
| SPGP0AN00200 | YPL007C | 99 | 0 |
| SPGP0Z00560 | YKL074C | 99 | 0 |
| SPGP0A02780 | YDR247W | 99 | 0 |
| SPGP0V01900 | YER104W | 99 | 0 |
| SPGP0AB01220 | YIL129C | 99 | 0 |
| SPGP0G00900 | YMR220W | 99 | 0 |
| SPGP0U00710 | YLR006C | 99 | 0 |
| SPGP0K01490 | YNL077W | 99 | 0 |
| SPGP0AA01340 | YPL112C | 99 | 0 |
| SPGP0EW00110 | YCR090C | 99 | 0 |
| SPGP0V01360 | YER053C | 99 | 0 |
| SPGP0A01830 | YDR352W | 99 | 0 |
| SPGP0V01630 | YER077C | 99 | 0 |
| SPGP0R02090 | YGL123W | 99 | 0 |
| SPGP0G03690 | YML075C | 99 | 0 |
| SPGP0R01930 | YGL105W | 99 | 0 |
| SPGP0N02630 | YJL137C | 99 | 0 |
| SPGP0G03240 | YML026C | 99 | 0 |
| SPGP0AA01750 | YPL069C | 100 | 0 |
| SPGP0D00780 | YOR265W | 100 | 0 |
| SPGP0T01510 | YLR250W | 99 | 0 |
| SPGP0K00810 | YNL002C | 99 | 0 |
| SPGP0X01090 | YHR061C | 99 | 0 |
| SPGP0A02940 | YDR229W | 99 | 0 |
| SPGP0A02250 | YDR307W | 99 | 0 |
| SPGP0U01110 | YLR049C | 99 | 0 |
| SPGP0AJ00280 | YAR002C-A | 99 | 0 |
| SPGP0A04240 | YDR084C | 99 | 0 |
| SPGP0D04410 | YOL140W | 99 | 0 |
| SPGP0K01260 | YNL052W | 100 | 0 |
| SPGP0AU00290 | YOR365C | 99 | 0 |
| SPGP0V02100 | YER126C | 100 | 0 |
| SPGP0X01320 | YHR082C | 99 | 0 |
| SPGP0G01490 | YMR161W | 99 | 0 |
| SPGP0U01070 | YLR045C | 99 | 0 |

Create PDF in your applications with the Pdfcrowd HTML to PDF API PDFCROWD

| | | | |
|---|---|---|---|
| SPGP0AC00430 | YPR052C | 99 | 0 |
| SPGP0AP00460 | YHR201C | 99 | 0 |
| SPGP0A06440 | YDL166C | 99 | 0 |
| SPGP0V00180 | YEL064C | 99 | 0 |
| SPGP0R00210 | YGR095C | 99 | 0 |
| SPGP0AE01050 | YFL042C | 99 | 0 |
| SPGP0D00750 | YOR269W | 99 | 0 |
| SPGP0AD00720 | YKR039W | 99 | 0 |
| SPGP0AD00700 | YKR042W | 99 | 1 |
| SPGP0Y01610 | YGR277C | 99 | 0 |
| SPGP0D01130 | YOR226C | 99 | 0 |
| SPGP0T00140 | YLR406C | 99 | 0 |
| SPGP0AP00420 | YHR198C | 100 | 0 |
| SPGP0T01090 | YLR300W | 99 | 0 |
| SPGP0AS00180 | YPL269W | 99 | 0 |
| SPGP0K00710 | YNR011C | 99 | 0 |
| SPGP0A02710 | YDR255C | 99 | 0 |
| SPGP0AA01680 | YPL077C | 99 | 0 |
| SPGP0N02850 | YJL162C | 99 | 1 |
| SPGP0D01410 | YOR193W | 99 | 0 |
| SPGP0T00320 | YLR386W | 99 | 0 |
| SPGP0D00580 | YOR288C | 99 | 0 |
| SPGP0T00870 | YLR324W | 99 | 0 |
| SPGP0A06470 | YDL169C | 99 | 0 |
| SPGP0AJ00560 | YAL031C | 99 | 0 |
| SPGP0AH00320 | YCL038C | 99 | 0 |
| SPGP0A04750 | YDR031W | 100 | 0 |
| SPGP0AA01020 | YPL147W | 99 | 0 |
| SPGP0AB01410 | YIL149C | 99 | 0 |
| SPGP0AC01030 | YPR122W | 99 | 0 |
| SPGP0V01830 | YER096W | 99 | 0 |
| SPGP0U00460 | YLL021W | 99 | 0 |
| SPGP0N02880 | YJL165C | 99 | 0 |
| SPGP0U00660 | YLR001C | 99 | 0 |
| SPGP0AW00130 | YHR153C | 99 | 0 |
| SPGP0A03420 | YDR174W | 99 | 0 |
| SPGP0A06290 | YDL147W | 99 | 0 |
| SPGP0T01630 | YLR238W | 99 | 0 |
| SPGP0J03070 | YBL021C | 100 | 0 |
| SPGP0N00560 | YJR104C | 100 | 0 |

Create PDF in your applications with the Pdfcrowd HTML to PDF API PDFCROWD

| | | | |
|---|---|---|---|
| SPGP0G03070 | YML009C | 100 | 0 |
| SPGP0AJ00210 | YAR019C | 99 | 0 |
| SPGP0K02190 | YNL156C | 99 | 0 |
| SPGP0BA00130 | YCR069W | 99 | 0 |
| SPGP0U01730 | YLR115W | 99 | 0 |
| SPGP0D01830 | YOR148C | 98 | 0 |
| SPGP0X01660 | YHR116W | 100 | 0 |
| SPGP0X00360 | YHL015W | 99 | 0 |
| SPGP0A00820 | YDR461C-A | 100 | 0 |
| SPGP0R01480 | YGL055W | 99 | 0 |
| SPGP0J02300 | YBR062C | 99 | 0 |
| SPGP0Y00870 | YGR195W | 100 | 0 |
| SPGP0Y00890 | YGR197C | 99 | 0 |
| SPGP0N00810 | YJR077C | 99 | 0 |
| SPGP0X01400 | YHR090C | 99 | 0 |
| SPGP0AJ00530 | YAL027W | 99 | 0 |
| SPGP0J00810 | YBR222C | 99 | 0 |
| SPGP0A02670 | YDR259C | 99 | 0 |
| SPGP0A06710 | YDL199C | 99 | 0 |
| SPGP0V00240 | YEL058W | 99 | 0 |
| SPGP0AU00120 | YOR347C | 99 | 0 |
| SPGP0A01190 | YDR419W | 100 | 0 |
| SPGP0R00820 | YGR023W | 97 | 1 |
| SPGP0A01660 | YDR371W | 99 | 0 |
| SPGP0AA00510 | YPL199C | 99 | 0 |
| SPGP0D01340 | YOR202W | 99 | 0 |
| SPGP0R01760 | YGL086W | 99 | 0 |
| SPGP0D00950 | YOR246C | 99 | 0 |
| SPGP0Y00190 | YGR119C | 99 | 0 |
| SPGP0AK00160 | YGL254W | 100 | 0 |
| SPGP0AS00270 | YPL259C | 99 | 0 |
| SPGP0D01690 | YOR162C | 99 | 0 |
| SPGP0D01840 | YOR147W | 99 | 0 |
| SPGP0N00650 | YJR095W | 100 | 0 |
| SPGP0N03370 | YOL158C | 98 | 0 |
| SPGP0T01950 | YLR200W | 99 | 0 |
| SPGP0D01800 | YOR151C | 99 | 0 |
| SPGP0AD00740 | YKR037C | 99 | 0 |
| SPGP0G02930 | YMR008C | 99 | 1 |
| SPGP0A02640 | YDR262W | 99 | 0 |

Create PDF in your applications with the Pdfcrowd HTML to PDF API PDFCROWD

| | | | |
|---|---|---|---|
| SPGP0R01210 | YGL022W | 99 | 0 |
| SPGP0AP00120 | YHR168W | 99 | 0 |
| SPGP0J02800 | YBR008C | 99 | 0 |
| SPGP0V00940 | YER014W | 99 | 0 |
| SPGP0V00720 | YEL009C | 99 | 0 |
| SPGP0T01610 | YLR240W | 99 | 0 |
| SPGP0J02060 | YBR086C | 99 | 0 |
| SPGP0N03160 | YJL198W | 99 | 0 |
| SPGP0A04900 | YDR014W-A | 99 | 0 |
| SPGP0K03350 | YNL286W | 99 | 0 |
| SPGP0A02360 | YDR296W | 99 | 0 |
| SPGP0AN00590 | YPL049C | 99 | 0 |
| SPGP0Z01120 | YKL139W | 99 | 0 |
| SPGP0Z01430 | YKL173W | 99 | 0 |
| SPGP0G01910 | YMR117C | 99 | 0 |
| SPGP0AB00520 | YIL051C | 99 | 0 |
| SPGP0U00200 | YLL051C | 99 | 0 |
| SPGP0N01940 | YJL059W | 99 | 0 |
| SPGP0J00550 | YBR250W | 99 | 0 |
| SPGP0A00400 | YDR507C | 99 | 0 |
| SPGP0A02500 | YDR281C | 99 | 0 |
| SPGP0X01450 | YHR096C | 99 | 0 |
| SPGP0K02990 | YNL246W | 99 | 0 |
| SPGP0R00760 | YGR030C | 99 | 0 |
| SPGP0BE00130 | YPR008W | 100 | 0 |
| SPGP0J00260 | YBR280C | 99 | 0 |
| SPGP0G03120 | YML014W | 99 | 0 |
| SPGP0A01410 | YDR393W | 99 | 0 |
| SPGP0Z01730 | YKL207W | 99 | 0 |
| SPGP0AJ00780 | YAL054C | 99 | 0 |
| SPGP0J03790 | YBL101C | 99 | 0 |
| SPGP0U00480 | YLL018C-A | 99 | 0 |
| SPGP0R00580 | YGR052W | 99 | 0 |
| SPGP0GO00100 | YNR032C-A | 100 | 0 |
| SPGP0Y01320 | YGR245C | 99 | 0 |
| SPGP0AC00890 | YPR108W | 99 | 0 |
| SPGP0N02410 | YJL112W | 99 | 0 |
| SPGP0AE00830 | YFL021W | 99 | 1 |
| SPGP0A03680 | YDR146C | 99 | 0 |
| SPGP0A03100 | YDR211W | 99 | 0 |

Create PDF in your applications with the Pdfcrowd HTML to PDF API PDFCROWD

| | | | |
|---|---|---|---|
| SPGP0V00520 | YEL029C | 99 | 0 |
| SPGP0N03190 | YJL201W | 99 | 0 |
| SPGP0DD00100 | YCR059C | 99 | 0 |
| SPGP0U01010 | YLR038C | 100 | 0 |
| SPGP0D01640 | YOR167C | 100 | 0 |
| SPGP0N00540 | YJR106W | 99 | 0 |
| SPGP0J03520 | YBL068W | 100 | 0 |
| SPGP0U00810 | YLR016C | 100 | 0 |
| SPGP0Dtrna16P | YOL013W-A | 99 | 0 |
| SPGP0AT00180 | YPR191W | 92 | 0 |
| SPGP0AH00150 | YCL057C-A | 99 | 0 |
| SPGP0V01920 | YER106W | 99 | 0 |
| SPGP0T00810 | YLR329W | 99 | 0 |
| SPGP0AB01200 | YIL127C | 99 | 0 |
| SPGP0K00100 | YNR075W | 100 | 0 |
| SPGP0Z00530 | YKL071W | 99 | 0 |
| SPGP0J01930 | YBR103W | 99 | 0 |
| SPGP0G01420 | YMR168C | 99 | 0 |
| SPGP0N00150 | YJR145C | 99 | 0 |
| SPGP0K01470 | YNL075W | 99 | 0 |
| SPGP0J02110 | YBR083W | 99 | 0 |
| SPGP0G00890 | YMR221C | 99 | 0 |
| SPGP0X00920 | YHR041C | 99 | 0 |
| SPGP0V00380 | YEL043W | 99 | 0 |
| SPGP0X00430 | YHL008C | 88 | 2 |
| SPGP0R01950 | YGL107C | 99 | 0 |
| SPGP0CE00100 | YMR316W | 99 | 0 |
| SPGP0G02380 | YMR067C | 99 | 0 |
| SPGP0A05520 | YDL064W | 99 | 0 |
| SPGP0A05780 | YDL092W | 99 | 0 |
| SPGP0A05570 | YDL070W | 99 | 0 |
| SPGP0A05830 | YDL099W | 99 | 0 |
| SPGP0AC00550 | YPR067W | 100 | 0 |
| SPGP0A04220 | YDR086C | 100 | 0 |
| SPGP0AD00170 | YKR096W | 99 | 0 |
| SPGP0D04030 | YOL098C | 99 | 0 |
| SPGP0J01110 | YBR190W | 99 | 0 |
| SPGP0Z00630 | YKL082C | 99 | 0 |
| SPGP0G01430 | YMR167W | 99 | 0 |
| SPGP0X01340 | YHR084W | 99 | 0 |

Create PDF in your applications with the Pdfcrowd HTML to PDF API PDFCROWD

| | | | |
|---|---|---|---|
| SPGP0D01570 | YOR176W | 99 | 0 |
| SPGP0K02740 | YNL218W | 99 | 0 |
| SPGP0R00190 | YGR097W | 99 | 0 |
| SPGP0AC01010 | YPR120C | 99 | 0 |
| SPGP0A04730 | YDR033W | 99 | 0 |
| SPGP0D04430 | YOL142W | 99 | 0 |
| SPGP0D00770 | YOR266W | 99 | 0 |
| SPGP0AE01030 | YFL040W | 99 | 0 |
| SPGP0U01460 | YLR086W | 99 | 0 |
| SPGP0D01140 | YOR224C | 100 | 0 |
| SPGP0AC00290 | YPR035W | 99 | 0 |
| SPGP0K03700 | YNL325C | 99 | 0 |
| SPGP0Z01720 | YKL206C | 99 | 0 |
| SPGP0AE00310 | YFR035C | 100 | 0 |
| SPGP0A02770 | YDR248C | 99 | 0 |
| SPGP0Z00950 | YKL120W | 99 | 0 |
| SPGP0K00480 | YNR035C | 99 | 0 |
| SPGP0DJ00100 | YCR061W | 99 | 0 |
| SPGP0AC00470 | YPR057W | 100 | 0 |
| SPGP0K03140 | YNL261W | 99 | 0 |
| SPGP0AW00110 | YHR151C | 99 | 0 |
| SPGP0G04180 | YML129C | 99 | 0 |
| SPGP0D01490 | YOR185C | 99 | 0 |
| SPGP0BR00110 | YCR043C | 100 | 0 |
| SPGP0J02850 | YBR003W | 99 | 0 |
| SPGP0G00640 | YMR246W | 99 | 0 |
| SPGP0BD00160 | YMR308C | 99 | 0 |
| SPGP0N00970 | YJR060W | 99 | 0 |
| SPGP0R00240 | YGR092W | 99 | 0 |
| SPGP0J00450 | YBR259W | 99 | 0 |
| SPGP0T00430 | YLR373C | 99 | 0 |
| SPGP0AK00400 | YGL228W | 99 | 0 |
| SPGP0AD00470 | YKR066C | 99 | 1 |
| SPGP0A03820 | YDR130C | 100 | 0 |
| SPGP0AC00210 | YPR027C | 99 | 0 |
| SPGP0A03400 | YDR176W | 99 | 0 |
| SPGP0AE00590 | YFR007W | 99 | 0 |
| SPGP0AH00400 | YCL030C | 99 | 0 |
| SPGP0G02660 | YMR034C | 99 | 0 |
| SPGP0BV00120 | YCR083W | 99 | 0 |

Create PDF in your applications with the Pdfcrowd HTML to PDF API PDFCROWD

| | | | |
|---|---|---|---|
| SPGP0A03920 | YDR120C | 99 | 0 |
| SPGP0K02160 | YNL153C | 99 | 0 |
| SPGP0AA01190 | YPL128C | 99 | 0 |
| SPGP0T01900 | YLR206W | 98 | 1 |
| SPGP0AO00370 | YLR431C | 99 | 0 |
| SPGP0X00340 | YHL017W | 99 | 0 |
| SPGP0X01640 | YHR114W | 99 | 0 |
| SPGP0J02780 | YBR011C | 99 | 0 |
| SPGP0A06010 | YDL117W | 99 | 0 |
| SPGP0A04670 | YDR041W | 100 | 0 |
| SPGP0R02060 | YGL120C | 99 | 0 |
| SPGP0J02320 | YBR060C | 99 | 0 |
| SPGP0A05390 | YDL049C | 99 | 0 |
| SPGP0G02000 | YMR108W | 99 | 0 |
| SPGP0J00850 | YBR217W | 100 | 0 |
| SPGP0J00870 | YBR215W | 99 | 0 |
| SPGP0HL00100 | YIL001W | 99 | 0 |
| SPGP0N00950 | YJR062C | 99 | 0 |
| SPGP0Y01120 | YGR222W | 99 | 0 |
| SPGP0D02180 | YOR109W | 99 | 0 |
| SPGP0X00590 | YHR008C | 100 | 0 |
| SPGP0AC01210 | YPR144C | 99 | 0 |
| SPGP0T00100 | YLR410W | 99 | 0 |
| SPGP0R02480 | YGL166W | 99 | 0 |
| SPGP0D01360 | YOR198C | 100 | 0 |
| SPGP0K02240 | YNL161W | 99 | 0 |
| SPGP0AD00660 | YKR046C | 100 | 0 |
| SPGP0Y01340 | YGR247W | 99 | 0 |
| SPGP0G02560 | YMR044W | 99 | 1 |
| SPGP0J01470 | YBR153W | 99 | 0 |
| SPGP0A04030 | YDR108W | 99 | 0 |
| SPGP0N02900 | YJL167W | 99 | 0 |
| SPGP0T00170 | YLR403W | 99 | 0 |
| SPGP0A00230 | YDR527W | 99 | 0 |
| SPGP0AH00370 | YCL033C | 99 | 0 |
| SPGP0Z01090 | YKL137W | 100 | 0 |
| SPGP0AB01170 | YIL124W | 99 | 0 |
| SPGP0Y00360 | YGR136W | 99 | 0 |
| SPGP0N00630 | YJR097W | 99 | 0 |
| SPGP0N00360 | YJR124C | 99 | 0 |

Create PDF in your applications with the Pdfcrowd HTML to PDF API PDFCROWD

| | | | |
|---|---|---|---|
| SPGP0G01670 | YMR142C | 99 | 0 |
| SPGP0BD00100 | YMR314W | 99 | 0 |
| SPGP0G03600 | YML066C | 99 | 0 |
| SPGP0AM00530 | YER183C | 99 | 0 |
| SPGP0AK00330 | YGL236C | 99 | 0 |
| SPGP0AH00240 | YCL048W-A | 100 | 0 |
| SPGP0N03060 | YJL186W | 99 | 0 |
| SPGP0D01780 | YOR153W | 99 | 0 |
| SPGP0J02150 | YBR079C | 99 | 0 |
| SPGP0Z00260 | YKL045W | 99 | 0 |
| SPGP0AH00630 | YCL001W-A | 99 | 0 |
| SPGP0AC00640 | YPR078C | 99 | 0 |
| SPGP0U00280 | YLL041C | 99 | 0 |
| SPGP0AD01190 | YKL010C | 99 | 0 |
| SPGP0AY00130 | YOR375C | 100 | 0 |
| SPGP0AE00550 | YFR011C | 99 | 0 |
| SPGP0D03430 | YOL032W | 99 | 0 |
| SPGP0K03530 | YNL306W | 99 | 0 |
| SPGP0AC00660 | YPR080W | 100 | 0 |
| SPGP0AE00440 | YFR021W | 100 | 0 |
| SPGP0K03370 | YNL288W | 99 | 0 |
| SPGP0A02690 | YDR257C | 99 | 0 |
| SPGP0T00540 | YLR361C-A | 98 | 0 |
| SPGP0J03570 | YBL075C | 99 | 0 |
| SPGP0AN00350 | YPL022W | 99 | 0 |
| SPGP0G01930 | YMR115W | 99 | 0 |
| SPGP0Z01140 | YKL141W | 99 | 0 |
| SPGP0AA01040 | YPL145C | 99 | 0 |
| SPGP0AB00640 | YIL067C | 99 | 0 |
| SPGP0G01850 | YMR123W | 99 | 0 |
| SPGP0AB00990 | YIL106W | 99 | 0 |
| SPGP0AK00440 | YGL225W | 99 | 0 |
| SPGP0AC00270 | YPR033C | 99 | 0 |
| SPGP0AS00410 | YPL246C | 99 | 0 |
| SPGP0AK00120 | YGL258W | 99 | 0 |
| SPGP0CX00100 | YCR045C | 100 | 0 |
| SPGP0A04880 | YDR017C | 99 | 0 |
| SPGP0J00200 | YBR286W | 99 | 0 |
| SPGP0G01580 | YMR152W | 99 | 0 |

Create PDF in your applications with the Pdfcrowd HTML to PDF API PDFCROWD

| | | | |
|---|---|---|---|
| SPGP0J01910 | YBR105C | 99 | 0 |
| SPGP0Z00230 | YKL041W | 99 | 0 |
| SPGP0AK00490 | YGL220W | 99 | 0 |
| SPGP0A05990 | YDL115C | 99 | 0 |
| SPGP0U00830 | YLR018C | 100 | 0 |
| SPGP0U00340 | YLL034C | 99 | 0 |
| SPGP0N00210 | YJR139C | 99 | 0 |
| SPGP0G03310 | YML034W | 99 | 0 |
| SPGP0A03120 | YDR207C | 99 | 0 |
| SPGP0J02580 | YBR034C | 99 | 0 |
| SPGP0KB00100 | YNL090W | 100 | 0 |
| SPGP0J00140 | YBR293W | 99 | 0 |
| SPGP0G02760 | YMR024W | 100 | 0 |
| SPGP0J01640 | YBR135W | 99 | 0 |
| SPGP0N01380 | YJR010W | 99 | 0 |
| SPGP0J02400 | YBR053C | 99 | 0 |
| SPGP0Y00770 | YGR184C | 99 | 0 |
| SPGP0AA00620 | YPL188W | 99 | 0 |
| SPGP0J01810 | YBR115C | 99 | 0 |
| SPGP0A00470 | YDR499W | 99 | 0 |
| SPGP0K01180 | YNL044W | 100 | 0 |
| SPGP0K01450 | YNL073W | 99 | 0 |
| SPGP0J01950 | YBR101C | 99 | 0 |
| SPGP0N03140 | YJL196C | 99 | 0 |
| SPGP0N01190 | YJR036C | 99 | 0 |
| SPGP0J02170 | YBR077C | 100 | 0 |
| SPGP0D03150 | YOL002C | 99 | 0 |
| SPGP0Z00140 | YKL029C | 99 | 0 |
| SPGP0A01870 | YDR348C | 99 | 0 |
| SPGP0V01340 | YER051W | 99 | 0 |
| SPGP0N00470 | YJR112W | 99 | 0 |
| SPGP0G02400 | YMR065W | 99 | 0 |
| SPGP0V01680 | YER082C | 100 | 0 |
| SPGP0T02120 | YLR181C | 99 | 0 |
| SPGP0X00900 | YHR039C-A | 100 | 0 |
| SPGP0J01660 | YBR132C | 99 | 0 |
| SPGP0Z01190 | YKL146W | 99 | 0 |
| SPGP0A03210 | YDR196C | 99 | 0 |
| SPGP0R01830 | YGL094C | 99 | 0 |
| SPGP0T01720 | YLR225C | 99 | 0 |

Create PDF in your applications with the Pdfcrowd HTML to PDF API PDFCROWD

| | | | |
|---|---|---|---|
| SPGP0K02180 | YNL155W | 99 | 0 |
| SPGP0G01990 | YMR109W | 99 | 0 |
| SPGP0X00230 | YHL028W | 97 | 2 |
| SPGP0AC01100 | YPR132W | 99 | 0 |
| SPGP0A02290 | YDR303C | 99 | 0 |
| SPGP0V00100 | YEL073C | 100 | 0 |
| SPGP0J02080 | YBR085W | 99 | 0 |
| SPGP0G01450 | YMR165C | 99 | 0 |
| SPGP0G02210 | YMR086W | 99 | 0 |
| SPGP0G02740 | YMR026C | 99 | 0 |
| SPGP0K02760 | YNL220W | 99 | 0 |
| SPGP0BC00200 | YIR033W | 99 | 0 |
| SPGP0D01550 | YOR178C | 99 | 0 |
| SPGP0J01340 | YBR165W | 99 | 0 |
| SPGP0AU00210 | YOR356W | 99 | 0 |
| SPGP0R00170 | YGR099W | 99 | 0 |
| SPGP0AA00770 | YPL172C | 99 | 0 |
| SPGP0D03260 | YOL013C | 99 | 0 |
| SPGP0A04200 | YDR088C | 99 | 0 |
| SPGP0AJ00410 | YAL015C | 100 | 0 |
| SPGP0AA01410 | YPL105C | 99 | 0 |
| SPGP0K03580 | YNL311C | 99 | 0 |
| SPGP0AE01010 | YFL038C | 99 | 0 |
| SPGP0DA00100 | YAL061W | 99 | 0 |
| SPGP0BE00110 | YPR006C | 99 | 0 |
| SPGP0Y00330 | YGR133W | 100 | 0 |
| SPGP0A02400 | YDR292C | 99 | 0 |
| SPGP0U01040 | YLR042C | 100 | 0 |
| SPGP0A05400 | YDL051W | 99 | 0 |
| SPGP0Z01080 | YKL135C | 99 | 0 |
| SPGP0AA00350 | YPL216W | 99 | 0 |
| SPGP0AC00510 | YPR062W | 100 | 0 |
| SPGP0A02750 | YDR251W | 99 | 0 |
| SPGP0AW00180 | YHR157W | 99 | 0 |
| SPGP0K00460 | YNR037C | 99 | 0 |
| SPGP0AN00410 | YPL029W | 99 | 0 |
| SPGP0U00130 | YLL060C | 99 | 0 |
| SPGP0BA00140 | YCR071C | 99 | 0 |
| SPGP0G01950 | YMR113W | 99 | 0 |
| SPGP0Y00790 | YGR186W | 99 | 0 |

Create PDF in your applications with the Pdfcrowd HTML to PDF API PDFCROWD

| | | | |
|---|---|---|---|
| SPGP0AE00760 | YFL013C | 99 | 0 |
| SPGP0AV00170 | YKL212W | 99 | 0 |
| SPGP0A04310 | YDR078C | 99 | 0 |
| SPGP0U00890 | YLR025W | 100 | 0 |
| SPGP0G00660 | YMR244W | 99 | 0 |
| SPGP0AA01060 | YPL143W | 99 | 0 |
| SPGP0K00600 | YNR022C | 100 | 0 |
| SPGP0T00450 | YLR371W | 99 | 0 |
| SPGP0G04200 | YML131W | 99 | 0 |
| SPGP0AA00650 | YPL184C | 99 | 0 |
| SPGP0G01530 | YMR157C | 99 | 0 |
| SPGP0A06120 | YDL130W | 99 | 0 |
| SPGP0T01670 | YLR231C | 99 | 0 |
| SPGP0B04430 | YDR342C | 88 | 0 |
| SPGP0K02210 | YNL158W | 99 | 0 |
| SPGP0V01080 | YER027C | 99 | 0 |
| SPGP0A06320 | YDL150W | 99 | 0 |
| SPGP0A05750 | YDL089W | 99 | 0 |
| SPGP0D04240 | YOL122C | 99 | 0 |
| SPGP0J03140 | YBL028C | 99 | 0 |
| SPGP0G01130 | YMR198W | 99 | 0 |
| SPGP0Y00500 | YGR153W | 99 | 0 |
| SPGP0T01920 | YLR204W | 99 | 0 |
| SPGP0X00400 | YHL011C | 99 | 0 |
| SPGP0V01880 | YER102W | 99 | 0 |
| SPGP0K00440 | YNR039C | 99 | 0 |
| SPGP0A04650 | YDR043C | 100 | 0 |
| SPGP0A03660 | YDR148C | 99 | 0 |
| SPGP0Y00610 | YGR166W | 99 | 0 |
| SPGP0G00200 | YMR292W | 100 | 0 |
| SPGP0A03180 | YDR200C | 99 | 0 |
| SPGP0DS00100 | YIL002C | 100 | 0 |
| SPGP0D03280 | YOL015W | 99 | 0 |
| SPGP0AT00170 | YPR192W | 99 | 0 |
| SPGP0G03260 | YML028W | 100 | 0 |
| SPGP0N02570 | YJL131C | 99 | 0 |
| SPGP0Y01100 | YGR220C | 99 | 0 |
| SPGP0D03740 | YOL067C | 99 | 0 |
| SPGP0K02590 | YNL201C | 99 | 0 |
| SPGP0AA00550 | YPL194W | 99 | 0 |

Create PDF in your applications with the Pdfcrowd HTML to PDF API PDFCROWD

| | | | |
|---|---|---|---|
| SPGP0AB01010 | YIL108W | 99 | 0 |
| SPGP0D04280 | YOL126C | 99 | 0 |
| SPGP0R00840 | YGR020C | 99 | 0 |
| SPGP0J01450 | YBR155W | 99 | 0 |
| SPGP0D00450 | YOR301W | 99 | 0 |
| SPGP0N01520 | YJL004C | 100 | 0 |
| SPGP0N03330 | YJL217W | 98 | 0 |
| SPGP0AB01190 | YIL126W | 99 | 0 |
| SPGP0X00710 | YHR021C | 100 | 0 |
| SPGP0D00330 | YOR315W | 99 | 0 |
| SPGP0Y00340 | YGR134W | 99 | 0 |
| SPGP0N00340 | YJR126C | 99 | 0 |
| SPGP0K02550 | YNL196C | 100 | 0 |
| SPGP0G03370 | YML042W | 99 | 0 |
| SPGP0G01000 | YMR210W | 99 | 0 |
| SPGP0J01060 | YBR195C | 99 | 0 |
| SPGP0U01420 | YLR082C | 100 | 0 |
| SPGP0D01760 | YOR155C | 99 | 0 |
| SPGP0K03170 | YNL264C | 99 | 0 |
| SPGP0AH00220 | YCL050C | 99 | 0 |
| SPGP0D02280 | YOR097C | 99 | 0 |
| SPGP0J01190 | YBR182C | 99 | 0 |
| SPGP0AE00680 | YFL003C | 99 | 0 |
| SPGP0K03510 | YNL304W | 99 | 0 |
| SPGP0A04940 | YDR009W | 99 | 0 |
| SPGP0A04410 | YDR068W | 99 | 0 |
| SPGP0K02270 | YNL163C | 99 | 0 |
| SPGP0AC00700 | YPR084W | 99 | 0 |
| SPGP0AB00560 | YIL056W | 99 | 0 |
| SPGP0AB00970 | YIL104C | 100 | 0 |
| SPGP0G01830 | YMR125W | 99 | 0 |
| SPGP0N01900 | YJL054W | 99 | 0 |
| SPGP0D00250 | YOR324C | 99 | 0 |
| SPGP0A06730 | YDL201W | 99 | 0 |
| SPGP0V02110 | YER127W | 99 | 0 |
| SPGP0AA01270 | YPL120W | 99 | 0 |
| SPGP0D00340 | YOR313C | 100 | 0 |
| SPGP0D03510 | YOL042W | 99 | 0 |
| SPGP0J00220 | YBR284W | 99 | 0 |
| SPGP0AB00190 | YIL014C-A | 99 | 0 |

Create PDF in your applications with the Pdfcrowd HTML to PDF API PDFCROWD

| | | | |
|---|---|---|---|
| SPGP0AK00640 | YGL200C | 99 | 0 |
| SPGP0AM00570 | YER187W | 92 | 0 |
| SPGP0A06660 | YDL193W | 99 | 0 |
| SPGP0V00290 | YEL053C | 99 | 0 |
| SPGP0J02390 | YBR054W | 98 | 1 |
| SPGP0U00360 | YLL032C | 99 | 0 |
| SPGP0N00270 | YJR134C | 99 | 0 |
| SPGP0X00870 | YHR037W | 99 | 0 |
| SPGP0A07130 | YJR156C | 99 | 0 |
| SPGP0A03490 | YDR168W | 99 | 0 |
| SPGP0A06560 | YDL179W | 99 | 0 |
| SPGP0G00300 | YMR282C | 99 | 0 |
| SPGP0D03220 | YOL009C | 99 | 0 |
| SPGP0AN00620 | YPL052W | 100 | 0 |
| SPGP0AB01570 | YIL166C | 99 | 0 |
| SPGP0N00500 | YJR109C | 99 | 0 |
| SPGP0R01620 | YGL070C | 99 | 0 |
| SPGP0K02430 | YNL183C | 99 | 0 |
| SPGP0G01300 | YMR182C | 99 | 0 |
| SPGP0AJ00620 | YAL036C | 99 | 0 |
| SPGP0D04090 | YOL105C | 99 | 0 |
| SPGP0J03280 | YBL040C | 99 | 0 |
| SPGP0A02810 | YDR244W | 99 | 0 |
| SPGP0D03990 | YOL094C | 99 | 0 |
| SPGP0G03900 | YML098W | 100 | 0 |
| SPGP0D03950 | YOL089C | 99 | 0 |
| SPGP0R02320 | YGL147C | 99 | 0 |
| SPGP0J01890 | YBR107C | 99 | 0 |
| SPGP0A00430 | YDR503C | 100 | 0 |
| SPGP0T02100 | YLR183C | 99 | 0 |
| SPGP0G02420 | YMR063W | 99 | 0 |
| SPGP0AN00470 | YPL036W | 99 | 0 |
| SPGP0AO00120 | YLR456W | 99 | 0 |
| SPGP0V01580 | YER073W | 99 | 0 |
| SPGP0R01330 | YGL036W | 99 | 0 |
| SPGP0N01560 | YJL010C | 99 | 0 |
| SPGP0A02310 | YDR301W | 99 | 0 |
| SPGP0D04210 | YOL119C | 99 | 0 |
| SPGP0K01510 | YNL079C | 100 | 0 |
| SPGP0N02120 | YJL081C | 99 | 0 |

Create PDF in your applications with the Pdfcrowd HTML to PDF API PDFCROWD

| | | | |
|---|---|---|---|
| SPGP0J01320 | YBR167C | 99 | 0 |
| SPGP0K02700 | YNL214W | 99 | 0 |
| SPGP0Y00850 | YGR193C | 99 | 0 |
| SPGP0N01000 | YJR057W | 99 | 0 |
| SPGP0A04770 | YDR028C | 99 | 0 |
| SPGP0D00910 | YOR251C | 99 | 0 |
| SPGP0X00380 | YHL013C | 99 | 0 |
| SPGP0AD00680 | YKR044W | 99 | 0 |
| SPGP0N03250 | YJL208C | 99 | 0 |
| SPGP0Z00910 | YKL114C | 99 | 0 |
| SPGP0A05420 | YDL053C | 99 | 0 |
| SPGP0AT00110 | YPR200C | 99 | 0 |
| SPGP0K02950 | YNL242W | 99 | 0 |
| SPGP0AA00370 | YPL214C | 99 | 0 |
| SPGP0K00650 | YNR017W | 100 | 0 |
| SPGP0A02090 | YDR322W | 99 | 0 |
| SPGP0AE00270 | YFR039C | 99 | 1 |
| SPGP0A05250 | YDL030W | 99 | 0 |
| SPGP0G01970 | YMR111C | 99 | 0 |
| SPGP0AJ00570 | YAL032C | 99 | 0 |
| SPGP0R00460 | YGR065C | 99 | 0 |
| SPGP0BA00160 | YCR073C | 99 | 0 |
| SPGP0AE00960 | YFL034W | 99 | 1 |
| SPGP0AH00200 | YCL052C | 99 | 0 |
| SPGP0D02770 | YOR040W | 99 | 0 |
| SPGP0AZ00140 | YIR012W | 99 | 0 |
| SPGP0G03670 | YML073C | 99 | 0 |
| SPGP0AE00910 | YFL028C | 99 | 0 |
| SPGP0AA01730 | YPL071C | 99 | 0 |
| SPGP0K00620 | YNR020C | 99 | 0 |
| SPGP0U00110 | YLL062C | 99 | 0 |
| SPGP0A03860 | YDR126W | 99 | 0 |
| SPGP0D01590 | YOR174W | 100 | 0 |
| SPGP0AM00210 | YER150W | 99 | 0 |
| SPGP0A06300 | YDL148C | 99 | 0 |
| SPGP0K03410 | YNL292W | 99 | 0 |
| SPGP0Z01250 | YKL154W | 99 | 0 |
| SPGP0D04120 | YOL109W | 99 | 0 |
| SPGP0N02860 | YJL163C | 99 | 0 |
| SPGP0AA01300 | YPL117C | 99 | 0 |

Create PDF in your applications with the Pdfcrowd HTML to PDF API PDFCROWD

| | | | |
|---|---|---|---|
| SPGP0K00940 | YNL016W | 99 | 0 |
| SPGP0EO00100 | YGR286C | 100 | 0 |
| SPGP0V01860 | YER100W | 99 | 0 |
| SPGP0X01600 | YHR110W | 100 | 0 |
| SPGP0A05290 | YDL036C | 99 | 0 |
| SPGP0J03370 | YBL051C | 99 | 0 |
| SPGP0R01060 | YGL006W | 99 | 0 |
| SPGP0T00520 | YLR363C | 99 | 0 |
| SPGP0X01080 | YHR060W | 99 | 0 |
| SPGP0BY00140 | YIR038C | 99 | 0 |
| SPGP0J02360 | YBR057C | 100 | 0 |
| SPGP0A05490 | YDL060W | 99 | 0 |
| SPGP0J00360 | YBR269C | 100 | 0 |
| SPGP0AA00710 | YPL178W | 99 | 0 |
| SPGP0A00880 | YDR456W | 99 | 0 |
| SPGP0A06690 | YDL197C | 99 | 0 |
| SPGP0Y01160 | YGR227W | 99 | 0 |
| SPGP0V01540 | YER069W | 99 | 0 |
| SPGP0AD01300 | YKL020C | 99 | 0 |
| SPGP0A01390 | YDR395W | 99 | 0 |
| SPGP0Z00770 | YKL098W | 99 | 0 |
| SPGP0AB00270 | YIL022W | 99 | 0 |
| SPGP0R01520 | YGL059W | 99 | 0 |
| SPGP0G02600 | YMR040W | 100 | 0 |
| SPGP0Y01380 | YGR251W | 99 | 0 |
| SPGP0D02980 | YOR018W | 99 | 0 |
| SPGP0N01800 | YJL043W | 99 | 0 |
| SPGP0AJ00690 | YAL043C | 99 | 0 |
| SPGP0A06980 | YDL227C | 99 | 0 |
| SPGP0R00720 | YGR034W | 100 | 0 |
| SPGP0D01980 | YOR130C | 99 | 0 |
| SPGP0Y00320 | YGR132C | 100 | 0 |
| SPGP0A00330 | YDR515W | 99 | 0 |
| SPGP0J03800 | YBL102W | 99 | 0 |
| SPGP0U01130 | YLR051C | 99 | 0 |
| SPGP0J01040 | YBR197C | 99 | 0 |
| SPGP0K03620 | YNL315C | 99 | 0 |
| SPGP0D01180 | YOR220W | 99 | 0 |
| SPGP0U01890 | YLR134W | 99 | 0 |
| SPGP0D01740 | YOR157C | 99 | 0 |

Create PDF in your applications with the Pdfcrowd HTML to PDF API PDFCROWD

| | | | |
|---|---|---|---|
| SPGP0A05300 | YDL040C | 99 | 0 |
| SPGP0J03580 | YBL076C | 99 | 0 |
| SPGP0A04630 | YDR045C | 100 | 0 |
| SPGP0AC01310 | YPR155C | 100 | 0 |
| SPGP0G03100 | YML012W | 99 | 0 |
| SPGP0Z00300 | YKL049C | 99 | 0 |
| SPGP0T01460 | YLR257W | 99 | 0 |
| SPGP0AD00360 | YKR078W | 99 | 0 |
| SPGP0N02000 | YJL065C | 99 | 0 |
| SPGP0AD01050 | YKR004C | 99 | 0 |
| SPGP0K03020 | YNL249C | 99 | 0 |
| SPGP0A05060 | YDL005C | 98 | 1 |
| SPGP0AA00230 | YPL228W | 99 | 0 |
| SPGP0A02650 | YDR261C | 99 | 0 |
| SPGP0Y00380 | YGR140W | 99 | 0 |
| SPGP0X01480 | YHR098C | 99 | 0 |
| SPGP0AP00540 | YHR209W | 99 | 0 |
| SPGP0Y00620 | YGR167W | 99 | 1 |
| SPGP0U00560 | YLL010C | 98 | 1 |
| SPGP0T00730 | YLR341W | 99 | 0 |
| SPGP0K02450 | YNL186W | 99 | 0 |
| SPGP0AO00170 | YLR451W | 99 | 0 |
| SPGP0AH00280 | YCL044C | 99 | 0 |
| SPGP0G00140 | YMR298W | 99 | 0 |
| SPGP0AM00510 | YER180C-A | 99 | 0 |
| SPGP0R01900 | YGL101W | 99 | 0 |
| SPGP0U00980 | YLR035C | 99 | 0 |
| SPGP0A05950 | YDL111C | 99 | 0 |
| SPGP0D02360 | YOR089C | 99 | 0 |
| SPGP0G02090 | YMR098C | 99 | 0 |
| SPGP0U00300 | YLL039C | 92 | 0 |
| SPGP0AC00180 | YPR024W | 99 | 0 |
| SPGP0N01020 | YJR055W | 99 | 0 |
| SPGP0N00290 | YJR132W | 99 | 0 |
| SPGP0K01500 | YNL078W | 99 | 0 |
| SPGP0G00360 | YMR276W | 99 | 0 |
| SPGP0J00100 | YBR297W | 97 | 0 |
| SPGP0N01080 | YJR049C | 99 | 0 |
| SPGP0J01600 | YBR139W | 99 | 0 |
| SPGP0G01360 | YMR176W | 99 | 0 |

Create PDF in your applications with the Pdfcrowd HTML to PDF API PDFCROWD

| | | | |
|---|---|---|---|
| SPGP0J03690 | YBL089W | 99 | 0 |
| SPGP0D03800 | YOL073C | 99 | 0 |
| SPGP0V01320 | YER049W | 99 | 0 |
| SPGP0A02870 | YDR237W | 99 | 0 |
| SPGP0R01440 | YGL049C | 99 | 0 |
| SPGP0D02760 | YOR042W | 99 | 0 |
| SPGP0X00790 | YHR029C | 100 | 0 |
| SPGP0AA00380 | YPL213W | 99 | 0 |
| SPGP0J00950 | YBR205W | 99 | 0 |
| SPGP0V00740 | YEL006W | 99 | 0 |
| SPGP0T01320 | YLR273C | 99 | 0 |
| SPGP0AM00460 | YER176W | 99 | 0 |
| SPGP0G02440 | YMR061W | 99 | 0 |
| SPGP0K02140 | YNL151C | 100 | 0 |
| SPGP0N02270 | YJL097W | 100 | 0 |
| SPGP0X00270 | YHL024W | 99 | 0 |
| SPGP0A01800 | YDR356W | 99 | 0 |
| SPGP0BC00110 | YIR024C | 99 | 0 |
| SPGP0K02890 | YNL236W | 99 | 0 |
| SPGP0G02190 | YMR088C | 99 | 0 |
| SPGP0BE00150 | YPR010C | 99 | 0 |
| SPGP0N00900 | YJR067C | 99 | 0 |
| SPGP0AA00310 | YPL220W | 99 | 0 |
| SPGP0AU00170 | YOR352W | 99 | 0 |
| SPGP0T01040 | YLR306W | 98 | 0 |
| SPGP0Y01630 | YGR279C | 99 | 0 |
| SPGP0AH00730 | YCR012W | 100 | 0 |
| SPGP0G00960 | YMR214W | 99 | 0 |
| SPGP0D04490 | YOL148C | 99 | 0 |
| SPGP0AP00350 | YHR191C | 100 | 0 |
| SPGP0V02090 | YER125W | 99 | 0 |
| SPGP0AC00230 | YPR029C | 99 | 0 |
| SPGP0K03450 | YNL297C | 99 | 0 |
| SPGP0U01100 | YLR048W | 99 | 0 |
| SPGP0AB00120 | YIL003W | 100 | 0 |
| SPGP0R00440 | YGR067C | 99 | 0 |
| SPGP0A05110 | YDL010W | 99 | 0 |
| SPGP0A05440 | YDL055C | 99 | 0 |
| SPGP0AE00720 | YFL009W | 99 | 0 |
| SPGP0AE00230 | YFR043C | 99 | 0 |

Create PDF in your applications with the Pdfcrowd HTML to PDF API PDFCROWD

| | | | |
|---|---|---|---|
| SPGP0J02830 | YBR005W | 99 | 0 |
| SPGP0T01100 | YLR299W | 99 | 0 |
| SPGP0AH01000 | YCR038C | 99 | 0 |
| SPGP0A02460 | YDR285W | 99 | 0 |
| SPGP0AO00480 | YLR419W | 99 | 0 |
| SPGP0J00380 | YBR267W | 99 | 0 |
| SPGP0AH00780 | YCR018C | 99 | 0 |
| SPGP0N02980 | YJL177W | 99 | 0 |
| SPGP0D03870 | YOL081W | 99 | 0 |
| SPGP0BN00130 | YCR089W | 99 | 0 |
| SPGP0D00700 | YOR274W | 99 | 0 |
| SPGP0Y00900 | YGR198W | 99 | 0 |
| SPGP0G03480 | YML055W | 99 | 0 |
| SPGP0Y00540 | YGR157W | 99 | 0 |
| SPGP0A01750 | YDR362C | 99 | 0 |
| SPGP0AB00460 | YIL045W | 99 | 0 |
| SPGP0V02170 | YER134C | 100 | 0 |
| SPGP0A06490 | YDL171C | 99 | 0 |
| SPGP0A06360 | YDL156W | 99 | 0 |
| SPGP0R00220 | YGR094W | 99 | 0 |
| SPGP0R00990 | YGR002C | 99 | 0 |
| SPGP0AZ00160 | YIR010W | 99 | 1 |
| SPGP0G01170 | YMR194C-B | 100 | 0 |
| SPGP0R00550 | YGR055W | 100 | 0 |
| SPGP0AH00940 | YCR032W | 99 | 0 |
| SPGP0A03510 | YDR166C | 99 | 0 |
| SPGP0A02760 | YDR249C | 100 | 0 |
| SPGP0A05320 | YDL043C | 99 | 0 |
| SPGP0A04610 | YDR047W | 99 | 0 |
| SPGP0U01790 | YLR121C | 99 | 0 |
| SPGP0G02890 | YMR011W | 99 | 0 |
| SPGP0U01970 | YLR143W | 100 | 0 |
| SPGP0K00960 | YNL032W | 99 | 0 |
| SPGP0Y00810 | YGR188C | 99 | 0 |
| SPGP0AH00920 | YCR030C | 99 | 0 |
| SPGP0V01520 | YER067W | 100 | 0 |
| SPGP0Z00460 | YKL065C | 99 | 0 |
| SPGP0Y01140 | YGR224W | 99 | 0 |
| SPGP0K00920 | YNL014W | 99 | 0 |

Create PDF in your applications with the Pdfcrowd HTML to PDF API PDFCROWD

| | | | |
|---|---|---|---|
| SPGP0G03020 | YML004C | 99 | 0 |
| SPGP0G04160 | YML127W | 99 | 0 |
| SPGP0V00360 | YEL046C | 99 | 0 |
| SPGP0N03300 | YJL213W | 99 | 0 |
| SPGP0N00380 | YJR122W | 99 | 0 |
| SPGP0J02040 | YBR088C | 100 | 0 |
| SPGP0AB01150 | YIL122W | 99 | 0 |
| SPGP0G01610 | YMR148W | 100 | 0 |
| SPGP0A00350 | YDR513W | 100 | 0 |
| SPGP0AE00280 | YFR038W | 99 | 0 |
| SPGP0N00890 | YJR068W | 99 | 0 |
| SPGP0A04180 | YDR090C | 99 | 0 |
| SPGP0AA00900 | YPL159C | 100 | 0 |
| SPGP0D04020 | YOL097C | 99 | 0 |
| SPGP0N01810 | YJL044C | 99 | 0 |
| SPGP0Z00240 | YKL042W | 99 | 0 |
| SPGP0U02000 | YLR146C | 99 | 0 |
| SPGP0Z00250 | YKL043W | 99 | 0 |
| SPGP0V01220 | YER041W | 99 | 0 |
| SPGP0R01100 | YGL011C | 100 | 0 |
| SPGP0D03390 | YOL027C | 99 | 0 |
| SPGP0AC00970 | YPR116W | 99 | 0 |
| SPGP0Z00740 | YKL094W | 99 | 0 |
| SPGP0AZ00200 | YIR006C | 99 | 1 |
| SPGP0A04930 | YDR011W | 99 | 0 |
| SPGP0AC01330 | YPR157W | 99 | 0 |
| SPGP0BA00100 | YCR066W | 99 | 0 |
| SPGP0G03840 | YML092C | 100 | 0 |
| SPGP0T00830 | YLR328W | 99 | 0 |
| SPGP0T01410 | YLR263W | 99 | 0 |
| SPGP0D02550 | YOR066W | 99 | 0 |
| SPGP0BH00140 | YCR100C | 99 | 0 |
| SPGP0A05770 | YDL091C | 99 | 0 |
| SPGP0Y01400 | YGR253C | 99 | 0 |
| SPGP0A03460 | YDR170W-A | 100 | 0 |
| SPGP0A05580 | YDL072C | 99 | 0 |
| SPGP0AD01030 | YKR006C | 100 | 0 |
| SPGP0AH00820 | YCR020C-A | 99 | 0 |
| SPGP0U00860 | YLR022C | 99 | 0 |
| SPGP0V02030 | YER118C | 99 | 0 |

Create PDF in your applications with the Pdfcrowd HTML to PDF API PDFCROWD

| | | | |
|---|---|---|---|
| SPGP0J03310 | YBL043W | 99 | 0 |
| SPGP0U00790 | YLR014C | 99 | 0 |
| SPGP0U01580 | YLR098C | 99 | 0 |
| SPGP0T00900 | YLR320W | 99 | 0 |
| SPGP0AE00810 | YFL018C | 99 | 0 |
| SPGP0Z01440 | YKL174C | 99 | 0 |
| SPGP0J00530 | YBR252W | 100 | 0 |
| SPGP0J03730 | YBL092W | 100 | 0 |
| SPGP0G00980 | YMR212C | 99 | 0 |
| SPGP0U00540 | YLL012W | 99 | 0 |
| SPGP0A02170 | YDR315C | 99 | 0 |
| SPGP0J01240 | YBR176W | 99 | 0 |
| SPGP0A05970 | YDL113C | 99 | 0 |
| SPGP0G00730 | YMR237W | 99 | 0 |
| SPGP0A05040 | YDL003W | 99 | 0 |
| SPGP0K00980 | YNL030W | 100 | 0 |
| SPGP0AA01360 | YPL110C | 99 | 0 |
| SPGP0A04280 | YDR080W | 99 | 0 |
| SPGP0U00320 | YLL036C | 99 | 0 |
| SPGP0J00590 | YBR246W | 99 | 0 |
| SPGP0K00680 | YNR014W | 100 | 0 |
| SPGP0X01770 | YHR128W | 100 | 0 |
| SPGP0G01690 | YMR139W | 99 | 0 |
| SPGP0P03130 | YER188W | 90 | 0 |
| SPGP0A04080 | YDR103W | 99 | 0 |
| SPGP0G00340 | YMR278W | 99 | 0 |
| SPGP0J03710 | YBL091C | 99 | 0 |
| SPGP0N00310 | YJR130C | 99 | 0 |
| SPGP0A02850 | YDR239C | 99 | 0 |
| SPGP0K00310 | YNR053C | 99 | 0 |
| SPGP0J01620 | YBR137W | 99 | 0 |
| SPGP0A07170 | YJR160C | 99 | 0 |
| SPGP0AB00950 | YIL102C-A | 99 | 0 |
| SPGP0U01050 | YLR043C | 99 | 0 |
| SPGP0Y00920 | YGR200C | 99 | 0 |
| SPGP0N02770 | YJL154C | 99 | 0 |
| SPGP0J02880 | YBL001C | 100 | 0 |
| SPGP0R00590 | YGR049W | 99 | 0 |
| SPGP0A00620 | YDR483W | 100 | 0 |
| SPGP0J02990 | YBL013W | 99 | 0 |

Create PDF in your applications with the Pdfcrowd HTML to PDF API PDFCROWD

| | | | |
|---|---|---|---|
| SPGP0D03350 | YOL022C | 99 | 0 |
| SPGP0Z00170 | YKL033W-A | 99 | 0 |
| SPGP0AK00680 | YGL195W | 99 | 0 |
| SPGP0A01580 | YDR378C | 100 | 0 |
| SPGP0J00970 | YBR203W | 99 | 0 |
| SPGP0A03720 | YDR142C | 99 | 0 |
| SPGP0K01200 | YNL046W | 99 | 0 |
| SPGP0N01710 | YJL033W | 99 | 0 |
| SPGP0AS00330 | YPL254W | 99 | 0 |
| SPGP0AP00530 | YHR208W | 99 | 0 |
| SPGP0X00250 | YHL026C | 99 | 0 |
| SPGP0K02120 | YNL148C | 100 | 0 |
| SPGP0N02250 | YJL095W | 99 | 0 |
| SPGP0N03020 | YJL181W | 99 | 0 |
| SPGP0J02000 | YBR094W | 99 | 0 |
| SPGP0D03780 | YOL071W | 99 | 0 |
| SPGP0D01060 | YOR233W | 99 | 0 |
| SPGP0J01360 | YBR163W | 99 | 0 |
| SPGP0G03820 | YML088W | 99 | 0 |
| SPGP0T01060 | YLR304C | 99 | 0 |
| SPGP0AU00150 | YOR350C | 99 | 0 |
| SPGP0X01280 | YHR079C | 99 | 0 |
| SPGP0A02570 | YDR272W | 99 | 0 |
| SPGP0G02850 | YMR015C | 99 | 0 |
| SPGP0AC00740 | YPR089W | 99 | 0 |
| SPGP0G03450 | YML052W | 99 | 0 |
| SPGP0N00120 | YJR149W | 99 | 0 |
| SPGP0AA00420 | YPL209C | 100 | 0 |
| SPGP0D00620 | YOR284W | 99 | 0 |
| SPGP0D04110 | YOL108C | 99 | 0 |
| SPGP0K00400 | YNR044W | 95 | 1 |
| SPGP0A05460 | YDL057W | 99 | 0 |
| SPGP0AE00700 | YFL007W | 99 | 0 |
| SPGP0J01440 | YBR156C | 99 | 0 |
| SPGP0T01690 | YLR228C | 99 | 0 |
| SPGP0G00220 | YMR290C | 99 | 0 |
| SPGP0A02440 | YDR287W | 99 | 0 |
| SPGP0Z01040 | YKL130C | 99 | 0 |
| SPGP0U00150 | YLL057C | 99 | 0 |
| SPGP0AE00870 | YFL024C | 99 | 0 |

Create PDF in your applications with the Pdfcrowd HTML to PDF API PDFCROWD

| | | | |
|---|---|---|---|
| SPGP0D04260 | YOL124C | 99 | 0 |
| SPGP0AB00440 | YIL043C | 99 | 0 |
| SPGP0AJ00760 | YAL051W | 99 | 0 |
| SPGP0X01120 | YHR064C | 99 | 0 |
| SPGP0AM00250 | YER154W | 99 | 0 |
| SPGP0AA01080 | YPL140C | 99 | 0 |
| SPGP0N02820 | YJL159W | 96 | 1 |
| SPGP0N02530 | YJL127C-B | 100 | 0 |
| SPGP0N02730 | YJL148W | 99 | 0 |
| SPGP0A03530 | YDR164C | 99 | 0 |
| SPGP0G02050 | YMR102C | 99 | 0 |
| SPGP0N01920 | YJL056C | 100 | 0 |
| SPGP0R01020 | YGL002W | 99 | 0 |
| SPGP0G02500 | YMR054W | 99 | 0 |
| SPGP0K02910 | YNL238W | 99 | 0 |
| SPGP0A01630 | YDR374C | 99 | 0 |
| SPGP0D02940 | YOR021C | 100 | 0 |
| SPGP0N01060 | YJR051W | 99 | 0 |
| SPGP0AZ00120 | YIR014W | 99 | 0 |
| SPGP0V01500 | YER065C | 99 | 0 |
| SPGP0AO00380 | YLR430W | 99 | 0 |
| SPGP0D00370 | YOR308C | 99 | 0 |
| SPGP0AA00640 | YPL186C | 99 | 0 |
| SPGP0A00830 | YDR461W | 100 | 0 |
| SPGP0N03000 | YJL179W | 99 | 0 |
| SPGP0R02220 | YGL137W | 99 | 0 |
| SPGP0AU00230 | YOR358W | 100 | 0 |
| SPGP0A00370 | YDR511W | 99 | 0 |
| SPGP0U01950 | YLR141W | 99 | 1 |
| SPGP0V01240 | YER043C | 99 | 0 |
| SPGP0D02110 | YOR116C | 99 | 0 |
| SPGP0A05220 | YDL027C | 99 | 0 |
| SPGP0A01430 | YDR391C | 99 | 0 |
| SPGP0AC00590 | YPR071W | 95 | 0 |
| SPGP0G01410 | YMR171C | 99 | 0 |
| SPGP0U01850 | YLR130C | 99 | 0 |
| SPGP0U01170 | YLR055C | 99 | 0 |
| SPGP0X01200 | YHR072W | 100 | 0 |
| SPGP0R01120 | YGL013C | 99 | 0 |
| SPGP0A01230 | YDR414C | 100 | 0 |

Create PDF in your applications with the Pdfcrowd HTML to PDF API PDFCROWD

| | | | |
|---|---|---|---|
| SPGP0AB00580 | YIL061C | 99 | 0 |
| SPGP0K03270 | YNL277W | 99 | 0 |
| SPGP0AA00100 | YPL242C | 99 | 0 |
| SPGP0J03610 | YBL080C | 99 | 0 |
| SPGP0A01020 | YDR439W | 100 | 0 |
| SPGP0AD00800 | YKR029C | 99 | 0 |
| SPGP0D01720 | YOR159C | 99 | 0 |
| SPGP0AP00250 | YHR181W | 100 | 0 |
| SPGP0A05870 | YDL103C | 99 | 0 |
| SPGP0AD01010 | YKR008W | 99 | 0 |
| SPGP0AC00300 | YPR036W | 99 | 0 |
| SPGP0A02610 | YDR265W | 99 | 0 |
| SPGP0D03590 | YOL052C | 99 | 0 |
| SPGP0AA00270 | YPL224C | 99 | 0 |
| SPGP0K02780 | YNL222W | 100 | 0 |
| SPGP0A05700 | YDL085W | 99 | 0 |
| SPGP0Y00680 | YGR172C | 99 | 0 |
| SPGP0U00520 | YLL014W | 99 | 0 |
| SPGP0A05100 | YDL008W | 99 | 0 |
| SPGP0D03200 | YOL007C | 99 | 0 |
| SPGP0D02030 | YOR125C | 99 | 0 |
| SPGP0A06500 | YDL173W | 99 | 1 |
| SPGP0AH00170 | YCL056C | 99 | 0 |
| SPGP0J03590 | YBL078C | 100 | 0 |
| SPGP0K02070 | YNL142W | 99 | 0 |
| SPGP0T01760 | YLR221C | 100 | 0 |
| SPGP0Y00740 | YGR180C | 99 | 0 |
| SPGP0R01660 | YGL076C | 99 | 0 |
| SPGP0D02340 | YOR091W | 99 | 0 |
| SPGP0J01870 | YBR109C | 99 | 0 |
| SPGP0A00600 | YDR485C | 99 | 0 |
| SPGP0AB00200 | YIL015W | 99 | 0 |
| SPGP0A01560 | YDR379C-A | 100 | 0 |
| SPGP0J00910 | YBR211C | 99 | 0 |
| SPGP0R02260 | YGL141W | 99 | 0 |
| SPGP0AD00860 | YKR023W | 99 | 0 |
| SPGP0K01100 | YNL035C | 99 | 0 |
| SPGP0R02380 | YGL155W | 99 | 0 |
| SPGP0AJ00330 | YAL007C | 100 | 0 |
| SPGP0AN00420 | YPL030W | 99 | 0 |

Create PDF in your applications with the Pdfcrowd HTML to PDF API PDFCROWD

| | | | |
|---|---|---|---|
| SPGP0N02280 | YJL098W | 99 | 0 |
| SPGP0G02290 | YMR076C | 99 | 0 |
| SPGP0A01700 | YDR367W | 99 | 0 |
| SPGP0AA00250 | YPL226W | 99 | 0 |
| SPGP0Z01640 | YKL195W | 99 | 0 |
| SPGP0K02100 | YNL146W | 100 | 0 |
| SPGP0AH00880 | YCR026C | 99 | 0 |
| SPGP0Z00380 | YKL057C | 99 | 0 |
| SPGP0V00980 | YER017C | 99 | 0 |
| SPGP0AK00500 | YGL219C | 100 | 0 |
| SPGP0N03110 | YJL192C | 99 | 0 |
| SPGP0D04180 | YOL115W | 99 | 0 |
| SPGP0A02580 | YDR270W | 99 | 0 |
| SPGP0A00920 | YDR451C | 99 | 0 |
| SPGP0AA01490 | YPL096C-A | 99 | 0 |
| SPGP0X01410 | YHR091C | 99 | 0 |
| SPGP0T01280 | YLR277C | 99 | 0 |
| SPGP0BC00190 | YIR032C | 99 | 0 |
| SPGP0T01360 | YLR268W | 99 | 0 |
| SPGP0U01310 | YLR070C | 99 | 0 |
| SPGP0A04390 | YDR070C | 100 | 0 |
| SPGP0AH00580 | YCL007C | 100 | 0 |
| SPGP0G00850 | YMR225C | 99 | 0 |
| SPGP0D00410 | YOR304C-A | 100 | 0 |
| SPGP0A02420 | YDR289C | 99 | 0 |
| SPGP0AC00490 | YPR060C | 100 | 0 |
| SPGP0J00340 | YBR271W | 99 | 0 |
| SPGP0Z00720 | YKL092C | 99 | 0 |
| SPGP0A00310 | YDR517W | 99 | 0 |
| SPGP0J02890 | YBL002W | 100 | 0 |
| SPGP0V01810 | YER094C | 99 | 0 |
| SPGP0AW00250 | YHR164C | 99 | 0 |
| SPGP0Y00590 | YGR163W | 99 | 0 |
| SPGP0X01790 | YHR131C | 99 | 0 |
| SPGP0T00230 | YLR395C | 100 | 0 |
| SPGP0A03550 | YDR162C | 99 | 0 |
| SPGP0J00460 | YBR258C | 99 | 0 |
| SPGP0U01260 | YLR065C | 99 | 0 |
| SPGP0G00260 | YMR286W | 99 | 0 |
| SPGP0D02920 | YOR023C | 99 | 0 |

Create PDF in your applications with the Pdfcrowd HTML to PDF API PDFCROWD

| | | | |
|---|---|---|---|
| SPGP0X00320 | YHL019C | 99 | 0 |
| SPGP0V01800 | YER093C-A | 97 | 0 |
| SPGP0K00790 | YNR001C | 99 | 0 |
| SPGP0N01670 | YJL026W | 99 | 0 |
| SPGP0J02560 | YBR036C | 100 | 0 |
| SPGP0A07000 | YDL230W | 99 | 0 |
| SPGP0J02250 | YBR068C | 99 | 0 |
| SPGP0AH00650 | YCR003W | 99 | 0 |
| SPGP0A00200 | YDR530C | 99 | 0 |
| SPGP0AD00480 | YKR065C | 99 | 0 |
| SPGP0R01850 | YGL096W | 99 | 0 |
| SPGP0R00780 | YGR028W | 99 | 0 |
| SPGP0X00110 | YHL043W | 99 | 0 |
| SPGP0K01310 | YNL058C | 99 | 1 |
| SPGP0Y01090 | YGR218W | 99 | 0 |
| SPGP0A01720 | YDR364C | 99 | 0 |
| SPGP0K00730 | YNR009W | 99 | 0 |
| SPGP0N01690 | YJL030W | 99 | 0 |
| SPGP0D00110 | YOR341W | 99 | 0 |
| SPGP0AO00280 | YLR440C | 99 | 0 |
| SPGP0D01920 | YOR137C | 99 | 0 |
| SPGP0A01920 | YDR339C | 100 | 0 |
| SPGP0A03740 | YDR140W | 100 | 0 |
| SPGP0V00640 | YEL018W | 99 | 0 |
| SPGP0K02660 | YNL210W | 99 | 0 |
| SPGP0R01140 | YGL015C | 99 | 0 |
| SPGP0AC00930 | YPR112C | 99 | 0 |
| SPGP0K03750 | YNL330C | 99 | 0 |
| SPGP0AD00410 | YKR072C | 99 | 0 |
| SPGP0K03640 | YNL317W | 99 | 0 |
| SPGP0AD00780 | YKR031C | 99 | 0 |
| SPGP0G02460 | YMR059W | 99 | 0 |
| SPGP0AJ00800 | YAL056W | 99 | 0 |
| SPGP0G03800 | YML086C | 99 | 0 |
| SPGP0AK00620 | YGL203C | 99 | 1 |
| SPGP0AA00290 | YPL222W | 99 | 0 |
| SPGP0A06380 | YDL159W | 99 | 0 |
| SPGP0G01790 | YMR129W | 99 | 0 |
| SPGP0U00500 | YLL016W | 99 | 0 |
| SPGP0AA00700 | YPL179W | 99 | 0 |

Create PDF in your applications with the Pdfcrowd HTML to PDF API PDFCROWD

| | | | |
|---|---|---|---|
| SPGP0AC00800 | YPR098C | 99 | 0 |
| SPGP0G03730 | YML079W | 99 | 0 |
| SPGP0D02450 | YOR078W | 99 | 0 |
| SPGP0G00380 | YMR274C | 100 | 0 |
| SPGP0G00120 | YMR300C | 99 | 0 |
| SPGP0AA00980 | YPL151C | 99 | 0 |
| SPGP0N00730 | YJR088C | 99 | 0 |
| SPGP0K00230 | YNR061C | 99 | 0 |
| SPGP0R00360 | YGR077C | 99 | 0 |
| SPGP0A03940 | YDR119W | 99 | 0 |
| SPGP0K03550 | YNL308C | 99 | 0 |
| SPGP0K03180 | YNL265C | 99 | 0 |
| SPGP0AH00190 | YCL054W | 99 | 0 |
| SPGP0T01550 | YLR246W | 99 | 0 |
| SPGP0A02890 | YDR235W | 99 | 0 |
| SPGP0Z00510 | YKL069W | 99 | 0 |
| SPGP0X01010 | YHR051W | 99 | 0 |
| SPGP0N01320 | YJR015W | 100 | 0 |
| SPGP0R01680 | YGL078C | 99 | 0 |
| SPGP0T00350 | YLR383W | 99 | 0 |
| SPGP0G01380 | YMR174C | 100 | 0 |
| SPGP0R01160 | YGL017W | 99 | 0 |
| SPGP0A01540 | YDR381W | 100 | 0 |
| SPGP0AA00490 | YPL201C | 99 | 0 |
| SPGP0X01070 | YHR059W | 99 | 0 |
| SPGP0AH00350 | YCL035C | 99 | 0 |
| SPGP0D02590 | YOR062C | 99 | 0 |
| SPGP0K00360 | YNR048W | 99 | 0 |
| SPGP0V00430 | YEL038W | 99 | 0 |
| SPGP0AD00880 | YKR021W | 99 | 0 |
| SPGP0Y01420 | YGR255C | 99 | 0 |
| SPGP0A02230 | YDR309C | 99 | 0 |
| SPGP0AH00430 | YCL027W | 99 | 0 |
| SPGP0R00970 | YGR004W | 99 | 0 |
| SPGP0AY00150 | YOR373W | 99 | 1 |
| SPGP0D02010 | YOR127W | 99 | 0 |
| SPGP0G02830 | YMR017W | 99 | 0 |
| SPGP0K01760 | YNL108C | 100 | 0 |
| SPGP0Z00820 | YKL104C | 99 | 0 |
| SPGP0N02840 | YJL161W | 99 | 0 |

Create PDF in your applications with the Pdfcrowd HTML to PDF API PDFCROWD

| | | | |
|---|---|---|---|
| SPGP0T01300 | YLR275W | 100 | 0 |
| SPGP0K03430 | YNL294C | 99 | 0 |
| SPGP0V02070 | YER123W | 99 | 0 |
| SPGP0T02160 | YLR177W | 99 | 0 |
| SPGP0G03170 | YML019W | 99 | 0 |
| SPGP0R01190 | YGL020C | 99 | 0 |
| SPGP0N02160 | YJL085W | 99 | 0 |
| SPGP0Z01060 | YKL133C | 99 | 0 |
| SPGP0K00550 | YNR028W | 99 | 0 |
| SPGP0A05680 | YDL083C | 99 | 0 |
| SPGP0V01280 | YER046W | 99 | 0 |
| SPGP0AN00330 | YPL020C | 99 | 0 |
| SPGP0K03280 | YNL278W | 99 | 0 |
| SPGP0J00320 | YBR273C | 99 | 0 |
| SPGP0G04140 | YML124C | 99 | 0 |
| SPGP0A01210 | YDR416W | 99 | 0 |
| SPGP0N00450 | YJR113C | 99 | 0 |
| SPGP0U00180 | YLL054C | 99 | 0 |
| SPGP0T00210 | YLR397C | 99 | 0 |
| SPGP0A03570 | YDR160W | 99 | 0 |
| SPGP0D01080 | YOR231W | 99 | 0 |
| SPGP0J03670 | YBL087C | 99 | 0 |
| SPGP0K02030 | YNL138W | 99 | 0 |
| SPGP0V00810 | YER002W | 99 | 0 |
| SPGP0G01040 | YMR206W | 99 | 0 |
| SPGP0K02290 | YNL165W | 99 | 0 |
| SPGP0AC01250 | YPR148C | 98 | 0 |
| SPGP0U00750 | YLR010C | 99 | 0 |
| SPGP0AO00460 | YLR421C | 99 | 0 |
| SPGP0D02900 | YOR026W | 99 | 0 |
| SPGP0A01130 | YDR425W | 99 | 0 |
| SPGP0A02700 | YDR256C | 99 | 0 |
| SPGP0K00750 | YNR007C | 99 | 0 |
| SPGP0X01530 | YHR103W | 99 | 0 |
| SPGP0AA00680 | YPL181W | 99 | 0 |
| SPGP0N02370 | YJL107C | 99 | 0 |
| SPGP0V00880 | YER008C | 99 | 0 |
| SPGP0D02510 | YOR071C | 99 | 0 |
| SPGP0AN00150 | YPL002C | 99 | 0 |
| SPGP0K02310 | YNL167C | 99 | 0 |

Create PDF in your applications with the Pdfcrowd HTML to PDF API PDFCROWD

| | | | |
|---|---|---|---|
| SPGP0K02640 | YNL208W | 99 | 0 |
| SPGP0AK00630 | YGL201C | 99 | 0 |
| SPGP0CH00120 | YKL023W | 98 | 1 |
| SPGP0AD00430 | YKR070W | 99 | 0 |
| SPGP0AB00240 | YIL019W | 99 | 0 |
| SPGP0G03530 | YML059C | 99 | 0 |
| SPGP0D03180 | YOL005C | 99 | 0 |
| SPGP0X01310 | YHR081W | 100 | 0 |
| SPGP0R00510 | YGR059W | 99 | 0 |
| SPGP0N02140 | YJL083W | 99 | 0 |
| SPGP0J01500 | YBR150C | 99 | 0 |
| SPGP0KH00100 | R0010W | 99 | 0 |
| SPGP0G01770 | YMR131C | 99 | 0 |
| SPGP0G00790 | YMR231W | 99 | 0 |
| SPGP0A03960 | YDR117C | 99 | 0 |
| SPGP0V01910 | YER105C | 99 | 0 |
| SPGP0A06240 | YDL142C | 100 | 0 |
| SPGP0Y00400 | YGR142W | 99 | 0 |
| SPGP0U00380 | YLL029W | 99 | 0 |
| SPGP0AB00790 | YIL085C | 99 | 0 |
| SPGP0N01300 | YJR017C | 99 | 0 |
| SPGP0AO00200 | YLR448W | 99 | 0 |
| SPGP0G00500 | YMR262W | 99 | 0 |
| SPGP0AB00300 | YIL026C | 99 | 0 |
| SPGP0U01030 | YLR040C | 99 | 0 |
| SPGP0A04530 | YDR057W | 99 | 0 |
| SPGP0T00660 | YLR348C | 99 | 0 |
| SPGP0A03030 | YDR219C | 99 | 0 |
| SPGP0AJ00820 | YAL059W | 99 | 0 |
| SPGP0AD01170 | YKL008C | 99 | 0 |
| SPGP0AA00470 | YPL203W | 100 | 0 |
| SPGP0J03330 | YBL046W | 97 | 2 |
| SPGP0AO00180 | YLR450W | 99 | 0 |
| SPGP0Z00160 | YKL033W | 99 | 0 |
| SPGP0Y00690 | YGR173W | 99 | 0 |
| SPGP0T01240 | YLR284C | 99 | 0 |
| SPGP0J02680 | YBR022W | 99 | 0 |
| SPGP0G02360 | YMR069W | 99 | 0 |
| SPGP0R00630 | YGR044C | 100 | 0 |

Create PDF in your applications with the Pdfcrowd HTML to PDF API PDFCROWD

| | | | |
|---|---|---|---|
| SPGP0AA00210 | YPL230W | 99 | 0 |
| SPGP0N03400 | YHL046C | 99 | 0 |
| SPGP0R01420 | YGL047W | 99 | 0 |
| SPGP0N00790 | YJR080C | 99 | 0 |
| SPGP0J01300 | YBR169C | 99 | 0 |
| SPGP0K02410 | YNL181W | 99 | 0 |
| SPGP0V01200 | YER039C | 99 | 0 |
| SPGP0K01850 | YNL119W | 99 | 0 |
| SPGP0AH00540 | YCL011C | 99 | 0 |
| SPGP0D00170 | YOR335C | 99 | 0 |
| SPGP0A06020 | YDL119C | 99 | 0 |
| SPGP0G03010 | YML002W | 99 | 0 |
| SPGP0AH00710 | YCR010C | 99 | 0 |
| SPGP0AC01230 | YPR145C-A | 99 | 0 |
| SPGP0AJ00390 | YAL013W | 99 | 0 |
| SPGP0A04350 | YDR074W | 99 | 0 |
| SPGP0A03090 | YDR212W | 99 | 0 |
| SPGP0T01980 | YLR196W | 99 | 0 |
| SPGP0D04320 | YOL130W | 99 | 0 |
| SPGP0Z00840 | YKL106W | 99 | 0 |
| SPGP0AB01050 | YIL112W | 99 | 0 |
| SPGP0AE00460 | YFR019W | 99 | 0 |
| SPGP0AC00380 | YPR046W | 99 | 0 |
| SPGP0AB00770 | YIL083C | 99 | 0 |
| SPGP0AE00890 | YFL026W | 99 | 0 |
| SPGP0K00590 | YNR023W | 99 | 0 |
| SPGP0R00110 | YGR105W | 100 | 0 |
| SPGP0AO00400 | YLR427W | 99 | 0 |
| SPGP0J00300 | YBR275C | 99 | 0 |
| SPGP0AD01240 | YKL015W | 99 | 0 |
| SPGP0X01840 | YHR135C | 99 | 0 |
| SPGP0A04140 | YDR096W | 99 | 0 |
| SPGP0U00240 | YLL046C | 99 | 0 |
| SPGP0G02040 | YMR104C | 99 | 0 |
| SPGP0G03890 | YML097C | 100 | 0 |
| SPGP0J03650 | YBL085W | 99 | 0 |
| SPGP0R00800 | YGR026W | 99 | 0 |
| SPGP0AU00250 | YOR360C | 99 | 0 |
| SPGP0J03430 | YBL058W | 98 | 0 |
| SPGP0FM00100 | YKL221W | 99 | 0 |

Create PDF in your applications with the Pdfcrowd HTML to PDF API PDFCROWD

| | | | |
|---|---|---|---|
| SPGP0X01030 | YHR055C | 99 | 0 |
| SPGP0A04550 | YDR055W | 99 | 0 |
| SPGP0AZ00100 | YIR018W | 99 | 0 |
| SPGP0AJ00120 | YAR042W | 99 | 0 |
| SPGP0N02710 | YJL146W | 99 | 0 |
| SPGP0R02400 | YGL157W | 99 | 0 |
| SPGP0AK00750 | YGL186C | 99 | 0 |
| SPGP0U01340 | YLR073C | 99 | 0 |
| SPGP0AT00150 | YPR194C | 99 | 0 |
| SPGP0K00320 | YNR052C | 99 | 0 |
| SPGP0BO00110 | YCR092C | 99 | 0 |
| SPGP0K01240 | YNL050C | 99 | 0 |
| SPGP0R01980 | YGL111W | 99 | 0 |
| SPGP0G00580 | YMR253C | 99 | 0 |
| SPGP0V00600 | YEL021W | 100 | 0 |
| SPGP0A00780 | YDR465C | 99 | 0 |
| SPGP0AK00140 | YGL256W | 99 | 0 |
| SPGP0K02330 | YNL169C | 99 | 0 |
| SPGP0AM00420 | YER172C | 99 | 0 |
| SPGP0A03780 | YDR135C | 99 | 0 |
| SPGP0AD00820 | YKR027W | 100 | 0 |
| SPGP0K01720 | YNL103W | 99 | 0 |
| SPGP0CI00100 | YPL055C | 99 | 0 |
| SPGP0AN00370 | YPL024W | 99 | 0 |
| SPGP0J00290 | YBR276C | 99 | 0 |
| SPGP0G03770 | YML082W | 99 | 0 |
| SPGP0D03530 | YOL045W | 99 | 0 |
| SPGP0G01860 | YMR122W-A | 100 | 0 |
| SPGP0G01750 | YMR133W | 99 | 0 |
| SPGP0D00150 | YOR337W | 99 | 0 |
| SPGP0AV00120 | YKL217W | 99 | 0 |
| SPGP0AJ00260 | YAR007C | 99 | 0 |
| SPGP0X00990 | YHR049W | 99 | 0 |
| SPGP0K03320 | YNL282W | 100 | 0 |
| SPGP0J00510 | YBR254C | 99 | 0 |
| SPGP0AP00290 | YHR185C | 100 | 0 |
| SPGP0AM00240 | YER153C | 99 | 0 |
| SPGP0J01710 | YBR127C | 99 | 0 |
| SPGP0G00520 | YMR260C | 99 | 0 |
| SPGP0Z01340 | YKL163W | 96 | 1 |

Create PDF in your applications with the Pdfcrowd HTML to PDF API PDFCROWD

| | | | |
|---|---|---|---|
| SPGP0X01140 | YHR066W | 99 | 0 |
| SPGP0AA00180 | YPL233W | 99 | 0 |
| SPGP0Z00580 | YKL077W | 99 | 0 |
| SPGP0N01240 | YJR031C | 99 | 0 |
| SPGP0X01510 | YHR101C | 99 | 0 |
| SPGP0AB01250 | YIL132C | 99 | 0 |
| SPGP0AM00230 | YER152C | 99 | 0 |
| SPGP0Z01580 | YKL189W | 99 | 0 |
| SPGP0J02610 | YBR030W | 99 | 0 |
| SPGP0A00450 | YDR501W | 100 | 0 |
| SPGP0J01130 | YBR188C | 98 | 0 |
| SPGP0D01770 | YOR154W | 99 | 0 |
| SPGP0N01460 | YJR003C | 99 | 0 |
| SPGP0AD00600 | YKR053C | 100 | 0 |
| SPGP0Z01710 | YKL205W | 99 | 0 |
| SPGP0U01440 | YLR084C | 99 | 0 |
| SPGP0G03980 | YML107C | 99 | 0 |
| SPGP0T01340 | YLR271W | 99 | 0 |
| SPGP0AC00360 | YPR043W | 99 | 0 |
| SPGP0T02020 | YLR192C | 100 | 0 |
| SPGP0AC01070 | YPR128C | 100 | 0 |
| SPGP0D00680 | YOR276W | 99 | 0 |
| SPGP0AK00520 | YGL215W | 99 | 0 |
| SPGP0AD01140 | YKL006W | 99 | 0 |
| SPGP0AC00440 | YPR054W | 99 | 0 |
| SPGP0N00920 | YJR065C | 99 | 0 |
| SPGP0AE01160 | YAR071W | 99 | 0 |
| SPGP0Z01570 | YKL188C | 99 | 0 |
| SPGP0R00130 | YGR103W | 99 | 1 |
| SPGP0D01390 | YOR195W | 99 | 0 |
| SPGP0AD01260 | YKL017C | 99 | 0 |
| SPGP0G03940 | YML103C | 99 | 0 |
| SPGP0X01860 | YHR137W | 99 | 0 |
| SPGP0J03270 | YBL039W-B | 99 | 0 |
| SPGP0A04330 | YDR076W | 99 | 0 |
| SPGP0AC01090 | YPR131C | 99 | 0 |
| SPGP0G03830 | YML091C | 99 | 0 |
| SPGP0AD00620 | YKR051W | 99 | 0 |
| SPGP0Y00940 | YGR202C | 99 | 0 |
| SPGP0X00480 | YHL002W | 100 | 0 |

Create PDF in your applications with the Pdfcrowd HTML to PDF API PDFCROWD

| | | | |
|---|---|---|---|
| SPGP0R02240 | YGL139W | 99 | 0 |
| SPGP0J03780 | YBL099W | 99 | 0 |
| SPGP0D02460 | YOR077W | 99 | 0 |
| SPGP0A05620 | YDL076C | 99 | 0 |
| SPGP0V01710 | YER086W | 99 | 0 |
| SPGP0K02680 | YNL212W | 99 | 0 |
| SPGP0A06630 | YDL189W | 99 | 0 |
| SPGP0D03010 | YOR014W | 99 | 0 |
| SPGP0AK00800 | YGL180W | 99 | 0 |
| SPGP0AO00220 | YLR446W | 99 | 0 |
| SPGP0X01570 | YHR107C | 99 | 0 |
| SPGP0X00150 | YHL036W | 99 | 0 |
| SPGP0Atrna16E | YDR278C | 100 | 0 |
| SPGP0D03100 | YOR004W | 100 | 0 |
| SPGP0N01580 | YJL012C | 99 | 0 |
| SPGP0AS00160 | YPL271W | 99 | 0 |
| SPGP0A03050 | YDR217C | 99 | 0 |
| SPGP0K02350 | YNL173C | 99 | 0 |
| SPGP0N00180 | YJR142W | 99 | 0 |
| SPGP0A00180 | YDR532C | 99 | 0 |
| SPGP0A04160 | YDR092W | 100 | 0 |
| SPGP0G02580 | YMR042W | 100 | 0 |
| SPGP0X00620 | YHR011W | 99 | 0 |
| SPGP0V02190 | YER137C | 99 | 0 |
| SPGP0N00230 | YJR137C | 99 | 0 |
| SPGP0K01950 | YNL130C | 99 | 0 |
| SPGP0G03570 | YML063W | 99 | 0 |
| SPGP0A03980 | YDR115W | 99 | 0 |
| SPGP0G03550 | YML061C | 99 | 0 |
| SPGP0G03790 | YML085C | 99 | 0 |
| SPGP0G01730 | YMR135C | 99 | 1 |
| SPGP0CF00100 | YCR002C | 99 | 0 |
| SPGP0D02390 | YOR085W | 99 | 0 |
| SPGP0AE00990 | YFL036W | 99 | 0 |
| SPGP0Y00420 | YGR144W | 100 | 0 |
| SPGP0Y00720 | YGR178C | 99 | 0 |
| SPGP0K03300 | YNL280C | 99 | 0 |
| SPGP0R00650 | YGR042W | 99 | 0 |
| SPGP0AH00860 | YCR024C | 99 | 0 |
| SPGP0AC00320 | YPR037C | 99 | 0 |

Create PDF in your applications with the Pdfcrowd HTML to PDF API PDFCROWD

| | | | |
|---|---|---|---|
| SPGP0N02610 | YJL134W | 99 | 0 |
| SPGP0AS00380 | YPL249C | 99 | 0 |
| SPGP0D01600 | YOR173W | 99 | 0 |
| SPGP0N00430 | YJR116W | 99 | 0 |
| SPGP0AB00320 | YIL030C | 99 | 0 |
| SPGP0Z00180 | YKL034W | 99 | 0 |
| SPGP0AJ00640 | YAL038W | 99 | 0 |
| SPGP0N00750 | YJR085C | 100 | 0 |
| SPGP0K00300 | YNR054C | 100 | 0 |
| SPGP0N02100 | YJL079C | 99 | 0 |
| SPGP0AB01070 | YIL114C | 99 | 0 |
| SPGP0A00700 | YDR475C | 99 | 0 |
| SPGP0D00130 | YOR339C | 99 | 0 |
| SPGP0K02010 | YNL136W | 99 | 0 |
| SPGP0AO00440 | YLR423C | 99 | 0 |
| SPGP0T00780 | YLR333C | 100 | 0 |
| SPGP0AC01190 | YPR141C | 99 | 0 |
| SPGP0G01480 | YMR162C | 99 | 0 |
| SPGP0AB00710 | YIL074C | 99 | 0 |
| SPGP0R01590 | YGL066W | 99 | 1 |
| SPGP0AA00780 | YPL171C | 99 | 0 |
| SPGP0R00380 | YGR075C | 99 | 0 |
| SPGP0Z01690 | YKL203C | 99 | 0 |
| SPGP0D00510 | YOR295W | 99 | 0 |
| SPGP0A06390 | YDL160C | 99 | 0 |
| SPGP0A01000 | YDR441C | 99 | 0 |
| SPGP0AC01290 | YPR153W | 99 | 0 |
| SPGP0U01880 | YLR133W | 99 | 0 |
| SPGP0Z01300 | YKL160W | 100 | 0 |
| SPGP0AD00210 | YKR093W | 99 | 0 |
| SPGP0AH00500 | YCL017C | 99 | 0 |
| SPGP0D01430 | YOR191W | 99 | 0 |
| SPGP0K01930 | YNL128W | 99 | 0 |
| SPGP0G00820 | YMR228W | 99 | 0 |
| SPGP0AH00610 | YCL002C | 100 | 0 |
| SPGP0Z00590 | YKL078W | 99 | 0 |
| SPGP0K01970 | YNL132W | 99 | 0 |
| SPGP0U00840 | YLR020C | 99 | 0 |
| SPGP0AH00690 | YCR008W | 99 | 0 |
| SPGP0AH00260 | YCL047C | 99 | 0 |

Create PDF in your applications with the Pdfcrowd HTML to PDF API PDFCROWD

| | | | |
|---|---|---|---|
| SPGP0K03600 | YNL313C | 99 | 0 |
| SPGP0AE00350 | YFR032C | 99 | 0 |
| SPGP0D00470 | YOR298C-A | 99 | 0 |
| SPGP0G01710 | YMR137C | 99 | 0 |
| SPGP0AS00340 | YPL253C | 99 | 0 |
| SPGP0AN00190 | YPL006W | 99 | 0 |
| SPGP0J01150 | YBR186W | 99 | 0 |
| SPGP0AD00640 | YKR049C | 99 | 0 |
| SPGP0A02270 | YDR305C | 100 | 0 |
| SPGP0Z00120 | YKL027W | 99 | 0 |
| SPGP0Gtrna17M | YJL007C | 100 | 0 |
| SPGP0A04510 | YDR059C | 100 | 0 |
| SPGP0U00730 | YLR008C | 99 | 0 |
| SPGP0AN00170 | YPL004C | 99 | 0 |
| SPGP0U01600 | YLR099W-A | 99 | 0 |
| SPGP0N01260 | YJR025C | 99 | 0 |
| SPGP0A06210 | YDL139C | 99 | 0 |
| SPGP0J01690 | YBR129C | 99 | 0 |
| SPGP0AK00100 | YDR542W | 99 | 0 |
| SPGP0U01230 | YLR061W | 99 | 0 |
| SPGP0A03150 | YDR204W | 99 | 0 |
| SPGP0A05930 | YDL109C | 99 | 0 |
| SPGP0V02150 | YER131W | 99 | 0 |
| SPGP0AN00290 | YPL016W | 99 | 1 |
| SPGP0AC00690 | YPR083W | 100 | 0 |
| SPGP0AE00650 | YFR001W | 100 | 0 |
| SPGP0AA00440 | YPL207W | 99 | 0 |
| SPGP0J00760 | YBR229C | 99 | 0 |
| SPGP0T00290 | YLR389C | 99 | 0 |
| SPGP0U00610 | YLL005C | 99 | 0 |
| SPGP0N00550 | YJR105W | 99 | 0 |
| SPGP0G02070 | YMR100W | 99 | 0 |
| SPGP0J02540 | YBR038W | 99 | 0 |
| SPGP0BH00130 | YCR098C | 99 | 0 |
| SPGP0X00190 | YHL032C | 99 | 0 |
| SPGP0T02040 | YLR190W | 99 | 0 |
| SPGP0G00420 | YMR270C | 99 | 0 |
| SPGP0D00490 | YOR297C | 99 | 0 |
| SPGP0V00690 | YEL013W | 99 | 0 |
| SPGP0V01970 | YER112W | 99 | 0 |

Create PDF in your applications with the Pdfcrowd HTML to PDF API PDFCROWD

| | | | |
|---|---|---|---|
| SPGP0A05210 | YDL025C | 99 | 0 |
| SPGP0J02310 | YBR061C | 99 | 0 |
| SPGP0A00150 | YDR536W | 99 | 0 |
| SPGP0AA01430 | YPL103C | 99 | 0 |
| SPGP0T00190 | YLR399C | 99 | 0 |
| SPGP0J00570 | YBR248C | 99 | 0 |
| SPGP0V00210 | YEL061C | 99 | 0 |
| SPGP0AH00810 | YCR020C | 99 | 0 |
| SPGP0G01180 | YMR194W | 100 | 0 |
| SPGP0K00340 | YNR050C | 99 | 0 |
| SPGP0D03910 | YOL086C | 99 | 0 |
| SPGP0AA00160 | YPL235W | 99 | 0 |
| SPGP0G03650 | YML071C | 99 | 0 |
| SPGP0N01440 | YJR005W | 99 | 0 |
| SPGP0AA00830 | YPL166W | 99 | 0 |
| SPGP0AD00840 | YKR025W | 99 | 0 |
| SPGP0G01020 | YMR208W | 99 | 0 |
| SPGP0U01320 | YLR071C | 99 | 0 |
| SPGP0D00390 | YOR306C | 99 | 0 |
| SPGP0BC00150 | YIR028W | 99 | 0 |
| SPGP0A05000 | YDR002W | 100 | 0 |
| SPGP0K00570 | YNR026C | 99 | 0 |
| SPGP0R02080 | YGL122C | 98 | 1 |
| SPGP0A04040 | YDR107C | 99 | 0 |
| SPGP0AA01170 | YPL130W | 99 | 0 |
| SPGP0D03400 | YOL028C | 99 | 0 |
| SPGP0U00630 | YLL003W | 99 | 0 |
| SPGP0N01650 | YJL024C | 99 | 0 |
| SPGP0V01950 | YER110C | 99 | 0 |
| SPGP0T00410 | YLR376C | 99 | 0 |
| SPGP0V02130 | YER129W | 99 | 0 |
| SPGP0X01590 | YHR109W | 99 | 0 |
| SPGP0D02870 | YOR030W | 99 | 0 |
| SPGP0A01490 | YOR133W | 99 | 0 |
| SPGP0D02850 | YOR032C | 99 | 0 |
| SPGP0AP00310 | YHR187W | 99 | 0 |
| SPGP0G02020 | YMR106C | 100 | 0 |
| SPGP0U00600 | YLL006W | 100 | 0 |
| SPGP0AB00730 | YIL076W | 99 | 0 |
| SPGP0AK00350 | YGL233W | 99 | 0 |

Create PDF in your applications with the Pdfcrowd HTML to PDF API

PDFCROWD

| | | | |
|---|---|---|---|
| SPGP0A02050 | YDR325W | 99 | 0 |
| SPGP0U01700 | YLR110C | 100 | 0 |
| SPGP0D00670 | YOR278W | 100 | 0 |
| SPGP0A06420 | YDL164C | 99 | 0 |
| SPGP0K01030 | YNL024C-A | 99 | 0 |
| SPGP0K01740 | YNL106C | 99 | 0 |
| SPGP0A01140 | YDR424C | 100 | 0 |
| SPGP0A02550 | YDR275W | 99 | 0 |
| SPGP0BD00150 | YMR309C | 99 | 0 |
| SPGP0T01960 | YLR199C | 99 | 0 |
| SPGP0AD00490 | YKR064W | 99 | 0 |
| SPGP0AC01170 | YPR139C | 99 | 0 |
| SPGP0AE00610 | YFR005C | 99 | 0 |
| SPGP0AB00690 | YIL072W | 99 | 0 |
| SPGP0Z01180 | YKL145W | 99 | 0 |
| SPGP0D00570 | YOR289W | 99 | 0 |
| SPGP0V01620 | YER076C | 99 | 0 |

#### Sheet 3: *genes alleles below 80% identity*

| *Saccharomyces pastorianus WS34/70 predicted ORF* | Parentale allele ID | Parent | Gene identities | Gene gaps |
|---|---|---|---|---|
| SPGP0M01230 | YJL084C | *Saccharomyces cerevisiae* | 79 | 2 |
| SPGP0B06190 | XP_018224047.1 | *Saccharomyces eubayanus* | 78 | 2 |

#### Sheet 4: *genes alleles copy with same identity in both parents*

| *Saccharomyces pastorianus WS34/70 predicted ORF* | Parental allele ID | Parent | Gene identities | Gene gaps |
|---|---|---|---|---|
| SPGP0Etrna28I | YNL017C | *Saccharomyces cerevisiae* | 100 | 0 |
| SPGP0Etrna28I | DI49_5614 | *Saccharomyces eubayanus* | 100 | 0 |
| SPGP0AG00110 | YCR040W | *Saccharomyces cerevisiae* | 100 | 0 |
| SPGP0AG00110 | XP_018223413.1 | *Saccharomyces eubayanus* | 100 | 0 |
| SPGP0Etrna44G | YNL285W | *Saccharomyces cerevisiae* | 100 | 0 |
| SPGP0Etrna44G | DI49_2297 | *Saccharomyces eubayanus* | 100 | 0 |
| SPGP0BH00100 | YCR096C | *Saccharomyces cerevisiae* | 100 | 0 |
| SPGP0BH00100 | XP_018223459.1 | *Saccharomyces eubayanus* | 100 | 0 |
| SPGP0Gtrna34R | YGR164W | *Saccharomyces cerevisiae* | 100 | 0 |
| SPGP0Gtrna34R | DI49_4020 | *Saccharomyces eubayanus* | 100 | 0 |
| SPGP0Gtrna40G | YPR159C-A | *Saccharomyces cerevisiae* | 100 | 0 |
| SPGP0Gtrna40G | DI49_5603 | *Saccharomyces eubayanus* | 100 | 0 |

Create PDF in your applications with the Pdfcrowd HTML to PDF API PDFCROWD

| | | | | |
|---|---|---|---|---|
| SPGP0Etrna34T | YHR180W-A | *Saccharomyces cerevisiae* | 100 | 0 |
| SPGP0Etrna34T | DI49_5553 | *Saccharomyces eubayanus* | 100 | 0 |

### Sheet 5: *list of paralogs in Saccharomyces pastorianus WS3470*

| List of paralogs | *S. eubayanus-like content* | *S. pastorianus content* | *S. cerevisiae-like content* |
|---|---|---|---|
| YCR045C; SPGP0CV00110 | | SPGP0CV00110 | YCR045C |
| YCR047C; YCR046C | | | YCR047C; YCR046C |
| DI49_1322; DI49_2252 | DI49_1322; DI49_2252 | | |
| YAL061W; DI49_0003; DI49_0004 | DI49_0003; DI49_0004 | | YAL061W |
| SPGP0DB00100; YPL273W; YPL272C | | SPGP0DB00100 | YPL273W; YPL272C |
| SPGP0DC00100; DI49_5398; YPL060W; DI49_5397 | DI49_5398; DI49_5397 | SPGP0DC00100 | YPL060W |
| YCR059C; DI49_5567; SPGP0Jtrna3C; SPGP0Ytrna4C; YGR110W; YPR121W | DI49_5567 | SPGP0Jtrna3C; SPGP0Ytrna4C | YCR059C; YGR110W; YPR121W |
| YBR297W; DI49_5194; SPGP0BF00150; SPGP0BF00140; SPGP0BF00160 | DI49_5194 | SPGP0BF00150; SPGP0BF00140; SPGP0BF00160 | YBR297W |
| YBR296C; DI49_1317; YBR295W; DI49_1316 | DI49_1317; DI49_1316 | | YBR296C; YBR295W |
| YBR293W; DI49_1315; YBR291C; DI49_1314 | DI49_1315; DI49_1314 | | YBR293W; YBR291C |
| YBR289W; YBR290W; DI49_1312 | DI49_1312 | | YBR289W; YBR290W |
| YBR287W; DI49_1310; YBR286W; DI49_1309 | DI49_1310; DI49_1309 | | YBR287W; YBR286W |
| YBR284W; YBR285W | | | YBR284W; YBR285W |
| YBR282W; DI49_1305 | DI49_1305 | | YBR282W |
| YBR281C; DI49_1304; YBR280C; DI49_1303 | DI49_1304; DI49_1303 | | YBR281C; YBR280C |
| YBR278W; YBR279W | | | YBR278W; YBR279W |
| YBR275C; DI49_1299 | DI49_1299 | | YBR275C |
| YBR274W; DI49_1298; YBR273C; DI49_1297 | DI49_1298; DI49_1297 | | YBR274W; YBR273C |
| YBR271W; YBR272C | | | YBR271W; YBR272C |
| YBR269C; YBR270C | | | YBR269C; YBR270C |
| YBR267W; DI49_1291 | DI49_1291 | | YBR267W |
| YBR265W; DI49_1290; YBR264C; DI49_1289 | DI49_1290; DI49_1289 | | YBR265W; YBR264C |
| YBR262C; YBR263W | | | YBR262C; YBR263W |
| YBR260C; DI49_1285; YBR259W; DI49_1284 | DI49_1285; DI49_1284 | | YBR260C; YBR259W |
| YBR257W; YBR258C | | | YBR257W; YBR258C |
| YBR255C-A; DI49_1280; YBR255W; DI49_1279 | DI49_1280; DI49_1279 | | YBR255C-A; YBR255W |
| YBR253W; YBR254C | | | YBR253W; YBR254C |
| YBR251W; YBR252W | | | YBR251W; YBR252W |
| YBR249C; DI49_1273 | DI49_1273 | | YBR249C |
| YBR248C; DI49_1272; YBR247C; DI49_1271 | DI49_1272; DI49_1271 | | YBR248C; YBR247C |
| YBR245C; DI49_1269; YBR244W; DI49_1268 | DI49_1269; DI49_1268 | | YBR245C; YBR244W |

Create PDF in your applications with the Pdfcrowd HTML to PDF API PDFCROWD

| | | | |
|---|---|---|---|
| YBR242W; DI49_1266 | DI49_1266 | | YBR242W |
| YBR241C; DI49_1265 | DI49_1265 | | YBR241C |
| YBR240C; DI49_1263 | DI49_1263 | | YBR240C |
| YBR239C; DI49_1262 | DI49_1262 | | YBR239C |
| YBR238C; DI49_1261 | DI49_1261 | | YBR238C |
| YBR237W; DI49_1260; YBR236C; DI49_1259 | DI49_1260; DI49_1259 | | YBR237W; YBR236C |
| YBR234C; DI49_1257; YBR233W-A; DI49_1256 | DI49_1257; DI49_1256 | | YBR234C; YBR233W-A |
| YBR231C; YBR233W | | | YBR231C; YBR233W |
| YBR229C; DI49_1252; YBR228W; DI49_1251 | DI49_1252; DI49_1251 | | YBR229C; YBR228W |
| YBR225W; DI49_1249; YBR223C; DI49_1248 | DI49_1249; DI49_1248 | | YBR225W; YBR223C |
| YBR221C; DI49_1246; YBR220C; DI49_1245 | DI49_1246; DI49_1245 | | YBR221C; YBR220C |
| YBR217W; YBR218C | | | YBR217W; YBR218C |
| YBR215W; DI49_1241 | DI49_1241 | | YBR215W |
| YBR214W; DI49_1240; YBR213W; DI49_1239 | DI49_1240; DI49_1239 | | YBR214W; YBR213W |
| YBR211C; YBR212W | | | YBR211C; YBR212W |
| SPGP0Jtrna17E; DI49_5287; SPGP0Ltrna5E; SPGP0Mtrna25E; DI49_2867; SPGP0Ntrna6E; SPGP0Ptrna12E; SPGP0Qtrna7E; SPGP0Qtrna8E; DI49_1502; SPGP0Qtrna9E; SPGP0Rtrna13E; YGL054C; SPGP0Vtrna10E; YER012W; YER098W; SPGP0Wtrna3E; SPGP0Ztrna21E; SPGP0AAtrna20E; DI49_0733; YCL021W-A; SPGP0AMtrna31E; SPGP0Ctrna4E; DI49_3838; SPGP0AZtrna22E; SPGP0Etrna27E; SPGP0Etrna28E; DI49_1914; SPGP0Ftrna23E; SPGP0Gtrna26E | DI49_5287; DI49_2867; DI49_1502; DI49_0733; DI49_3838; DI49_1914 | SPGP0Jtrna17E; SPGP0Ltrna5E; SPGP0Mtrna25E; SPGP0Ntrna6E; SPGP0Ptrna12E; SPGP0Qtrna7E; SPGP0Qtrna8E; SPGP0Qtrna9E; SPGP0Rtrna13E; SPGP0Vtrna10E; SPGP0Wtrna3E; SPGP0Ztrna21E; SPGP0AAtrna20E; SPGP0AMtrna31E; SPGP0Ctrna4E; SPGP0AZtrna22E; SPGP0Etrna27E; SPGP0Etrna28E; SPGP0Ftrna23E; SPGP0Gtrna26E | YGL054C; YER012W; YER098W; YCL021W-A |
| YBR208C; DI49_1233 | DI49_1233 | | YBR208C |
| YBR207W; DI49_1232 | DI49_1232 | | YBR207W |
| YBR205W; DI49_1231 | DI49_1231 | | YBR205W |
| YBR204C; DI49_1230; YBR203W; DI49_1229 | DI49_1230; DI49_1229 | | YBR204C; YBR203W |
| YBR201C-A; YBR202W | | | YBR201C-A; YBR202W |
| YBR200W; DI49_1225; YBR199W; DI49_1224 | DI49_1225; DI49_1224 | | YBR200W; YBR199W |
| YBR197C; YBR198C | | | YBR197C; YBR198C |
| YBR195C; YBR196C | | | YBR195C; YBR196C |
| YBR193C; YBR194W | | | YBR193C; YBR194W |
| YBR191W; DI49_1216; YBR190W | DI49_1216 | | YBR191W; YBR190W |
| YBR188C; SPGP0J01120 | | SPGP0J01120 | YBR188C |
| YBR186W; YBR187W | | | YBR186W; YBR187W |
| YBR184W; YBR185C | | | YBR184W; YBR185C |
| YBR182C; YBR183W | | | YBR182C; YBR183W |
| YBR180W; YBR181C; SPGP0AA01560; YPL089C | | SPGP0AA01560 | YBR180W; YBR181C; YPL089C |

Create PDF in your applications with the Pdfcrowd HTML to PDF API PDFCROWD

| | | | |
|---|---|---|---|
| YBR177C; YBR179C | | | YBR177C; YBR179C |
| YBR175W; YBR176W | | | YBR175W; YBR176W |
| YBR172C; DI49_1200 | DI49_1200 | | YBR172C |
| YBR171W; DI49_1199 | DI49_1199 | | YBR171W |
| YBR170C; DI49_1198 | DI49_1198 | | YBR170C |
| YBR169C; DI49_1197; YBR168W; DI49_1196 | DI49_1197; DI49_1196 | | YBR169C; YBR168W |
| YBR166C; DI49_1194; YBR165W; DI49_1193 | DI49_1194; DI49_1193 | | YBR166C; YBR165W |
| YBR163W; YBR164C; DI49_1191; DI49_1190 | DI49_1191; DI49_1190 | | YBR163W; YBR164C |
| YBR162C; YBR162W-A | | | YBR162C; YBR162W-A |
| YBR160W; DI49_1187 | DI49_1187 | | YBR160W |
| YBR159W; DI49_1186 | DI49_1186 | | YBR159W |
| YBR158W; DI49_1185; YBR157C; DI49_1184 | DI49_1185; DI49_1184 | | YBR158W; YBR157C |
| YBR155W; DI49_1183 | DI49_1183 | | YBR155W |
| YBR154C; DI49_1182; YBR153W; DI49_1181 | DI49_1182; DI49_1181 | | YBR154C; YBR153W |
| YBR151W; DI49_1179; YBR150C; DI49_1178 | DI49_1179; DI49_1178 | | YBR151W; YBR150C |
| YBR148W; YBR149W | | | YBR148W; YBR149W |
| YBR146W; DI49_1174; YBR145W; DI49_1173 | DI49_1174; DI49_1173 | | YBR146W; YBR145W |
| YBR142W; YBR143C | | | YBR142W; YBR143C |
| YBR140C; DI49_1169 | DI49_1169 | | YBR140C |
| YBR139W; DI49_1168; YBR138C; DI49_1167 | DI49_1168; DI49_1167 | | YBR139W; YBR138C |
| YBR136W; DI49_1165; YBR135W; DI49_1164 | DI49_1165; DI49_1164 | | YBR136W; YBR135W |
| YBR132C; DI49_1162; YBR131W; DI49_1161 | DI49_1162; DI49_1161 | | YBR132C; YBR131W |
| YBR129C; YBR130C | | | YBR129C; YBR130C |
| YBR127C; DI49_1157 | DI49_1157 | | YBR127C |
| YBR126C; DI49_1156 | DI49_1156 | | YBR126C |
| YBR125C; DI49_1155 | DI49_1155 | | YBR125C |
| SPGP0J01740; DI49_1154; YBR122C; DI49_1153 | DI49_1154; DI49_1153 | SPGP0J01740 | YBR122C |
| YBR120C; YBR121C | | | YBR120C; YBR121C |
| SPGP0J01790; YPR080W | | SPGP0J01790 | YPR080W |
| YBR117C; DI49_5526; DI49_5525 | DI49_5526; DI49_5525 | | YBR117C |
| YBR115C; DI49_1147 | DI49_1147 | | YBR115C |
| YBR114W; DI49_1146 | DI49_1146 | | YBR114W |
| YBR112C; DI49_1144; YBR111W-A; SPGP0I01860 | DI49_1144 | SPGP0I01860 | YBR112C; YBR111W-A |
| YBR110W; YBR111C | | | YBR110W; YBR111C |
| YBR108W; YBR109C | | | YBR108W; YBR109C |
| YBR106W; DI49_1137; YBR105C; DI49_1135 | DI49_1137; DI49_1135 | | YBR106W; YBR105C |
| YBR103W; YBR104W | | | YBR103W; YBR104W |
| YBR101C; YBR102C | | | YBR101C; YBR102C |
| YBR097W; DI49_1129; YBR096W; DI49_1128 | DI49_1129; DI49_1128 | | YBR097W; YBR096W |
| YBR094W; DI49_1126 | DI49_1126 | | YBR094W |

Create PDF in your applications with the Pdfcrowd HTML to PDF API PDFCROWD

| | | | |
|---|---|---|---|
| YBR093C; SPGP0I02040; YBR091C; SPGP0I02030; DI49_1125 | DI49_1125 | SPGP0I02040; SPGP0I02030 | YBR093C; YBR091C |
| YBR088C; DI49_1123; YBR087W; DI49_1122 | DI49_1123; DI49_1122 | | YBR088C; YBR087W |
| YBR085C-A; YBR086C | | | YBR085C-A; YBR086C |
| YBR084C-A; DI49_0153; DI49_0154; SPGP0I02130 | DI49_0153; DI49_0154 | SPGP0I02130 | YBR084C-A |
| YBR084W; DI49_1117 | DI49_1117 | | YBR084W |
| YBR083W; DI49_1116 | DI49_1116 | | YBR083W |
| YBR082C; SPGP0I02160 | | SPGP0I02160 | YBR082C |
| SPGP0Jtrna16D; DI49_4675; SPGP0DFtrna2D; SPGP0Ktrna3D; SPGP0Mtrna26D; DI49_2973; SPGP0Mtrna27D; SPGP0Mtrna28D; DI49_3068; YJR055W; SPGP0Ntrna9D; SPGP0Ntrna10D; SPGP0Ntrna11D; SPGP0Ptrna12D; DI49_3409; SPGP0Rtrna13D; YGR029W; YGR023W; SPGP0Strna20D; SPGP0Atrna15D; SPGP0Ttrna24D; SPGP0Utrna17D; YLR143W; SPGP0Wtrna4D; YIL009C-A; SPGP0Btrna1D; SPGP0ADtrna30D; YKR038C; SPGP0Ctrna6D; DI49_3666; SPGP0Ctrna7D; SPGP0Etrna32D; SPGP0E02490; DI49_1985; SPGP0Ftrna21D; SPGP0Gtrna29D; SPGP0Htrna31D; DI49_4967 | DI49_4675; DI49_2973; DI49_3068; DI49_3409; DI49_3666; DI49_1985; DI49_4967 | SPGP0Jtrna16D; SPGP0DFtrna2D; SPGP0Ktrna3D; SPGP0Mtrna26D; SPGP0Mtrna27D; SPGP0Mtrna28D; SPGP0Ntrna9D; SPGP0Ntrna10D; SPGP0Ntrna11D; SPGP0Ptrna12D; SPGP0Rtrna13D; SPGP0Strna20D; SPGP0Atrna15D; SPGP0Ttrna24D; SPGP0Utrna17D; SPGP0Wtrna4D; SPGP0Btrna1D; SPGP0ADtrna30D; SPGP0Ctrna6D; SPGP0Ctrna7D; SPGP0Etrna32D; SPGP0E02490; SPGP0Ftrna21D; SPGP0Gtrna29D; SPGP0Htrna31D | YJR055W; YGR029W; YGR023W; YLR143W; YIL009C-A; YKR038C |
| SPGP0Jtrna20R; SPGP0Gtrna34R; SPGP0DFtrna3R; SPGP0Ntrna8R; SPGP0Ntrna9R; SPGP0R01460; SPGP0Atrna19R; YDR058C; SPGP0Vtrna13R; SPGP0Ytrna21R; YGR165W; SPGP0Ztrna25R; YKL154W; SPGP0Ftrna27R; DI49_4019; SPGP0Gtrna33R; YML071C | DI49_4019 | SPGP0Jtrna20R; SPGP0Gtrna34R; SPGP0DFtrna3R; SPGP0Ntrna8R; SPGP0Ntrna9R; SPGP0R01460; SPGP0Atrna19R; SPGP0Vtrna13R; SPGP0Ytrna21R; SPGP0Ztrna25R; SPGP0Ftrna27R; SPGP0Gtrna33R | YDR058C; YGR165W; YKL154W; YML071C |
| YBR081C; DI49_1112 | DI49_1112 | | YBR081C |
| YBR080C; DI49_1111 | DI49_1111 | | YBR080C |
| YBR079C; DI49_1110 | DI49_1110 | | YBR079C |
| YBR078W; DI49_1109; YBR077C; DI49_1108 | DI49_1109; DI49_1108 | | YBR078W; YBR077C |
| YBR074W; DI49_1106; YBR073W; DI49_1105 | DI49_1106; DI49_1105 | | YBR074W; YBR073W |
| YBR071W; YBR072W | | | YBR071W; YBR072W |
| YBR069C; DI49_1101 | DI49_1101 | | YBR069C |
| YBR068C; DI49_1100 | DI49_1100 | | YBR068C |
| YBR067C; DI49_1099; YBR066C; DI49_1098 | DI49_1099; DI49_1098 | | YBR067C; YBR066C |
| YBR063C; YBR065C | | | YBR063C; YBR065C |
| YBR061C; DI49_1094; YBR060C | DI49_1094 | | YBR061C; YBR060C |
| YBR058C-A; YBR059C | | | YBR058C-A; YBR059C |
| YBR057C; YBR058C | | | YBR057C; YBR058C |
| YBR056W; DI49_1086 | DI49_1086 | | YBR056W |
| SPGP0Jtrna19L; DI49_5687; SPGP0Ktrna5L; SPGP0Ptrna15L; DI49_3191; SPGP0Ztrna27L; YKL190W; SPGP0Btrna1L; | DI49_5687; DI49_3191; DI49_0082; DI49_3917 | SPGP0Jtrna19L; SPGP0Ktrna5L; SPGP0Ptrna15L; SPGP0Ztrna27L; SPGP0Btrna1L; SPGP0B01610; | YKL190W; YGL192W |

Create PDF in your applications with the Pdfcrowd HTML to PDF API PDFCROWD

| | | | |
|---|---|---|---|
| DI49_0082; SPGP0B01610; SPGP0KOtrna22L; YGL192W; SPGP0Ctrna12L; DI49_3917; SPGP0AOtrna39L; SPGP0Etrna42L; SPGP0E03040 | | SPGP0KOtrna22L; SPGP0Ctrna12L; SPGP0AOtrna39L; SPGP0Etrna42L; SPGP0E03040 | |
| YBR055C; DI49_1084 | DI49_1084 | | YBR055C |
| YBR054W; DI49_1083; YBR053C; DI49_1082 | DI49_1083; DI49_1082 | | YBR054W; YBR053C |
| YBR050C; DI49_1080 | DI49_1080 | | YBR050C |
| YBR049C; DI49_1079 | DI49_1079 | | YBR049C |
| YBR048W; DI49_0223; SPGP0A04800; YDR024W; SPGP0I02480 | DI49_0223 | SPGP0A04800; SPGP0I02480 | YBR048W; YDR024W |
| YBR047W; DI49_1077 | DI49_1077 | | YBR047W |
| YBR046C; DI49_1076; YBR045C; DI49_1075 | DI49_1076; DI49_1075 | | YBR046C; YBR045C |
| YBR044C; DI49_1074; SPGP0Atrna13V; SPGP0Btrna1V; DI49_1073 | DI49_1074; DI49_1073 | SPGP0Atrna13V; SPGP0Btrna1V | YBR044C |
| YBR042C; DI49_1071 | DI49_1071 | | YBR042C |
| YBR041W; DI49_1070; YBR040W; DI49_1069 | DI49_1070; DI49_1069 | | YBR041W; YBR040W |
| YBR038W; DI49_1067 | DI49_1067 | | YBR038W |
| YBR037C; DI49_1066 | DI49_1066 | | YBR037C |
| YBR036C; DI49_1065; YBR035C; DI49_1064 | DI49_1065; DI49_1064 | | YBR036C; YBR035C |
| YBR033W; YBR034C | | | YBR033W; YBR034C |
| YBR030W; DI49_0208 | DI49_0208 | | YBR030W |
| YBR029C; DI49_0207; YBR028C | DI49_0207 | | YBR029C; YBR028C |
| YBR025C; DI49_0204; YBR024W; DI49_0205 | DI49_0204; DI49_0205 | | YBR025C; YBR024W |
| YBR022W; YBR023C | | | YBR022W; YBR023C |
| YBR020W; YBR021W | | | YBR020W; YBR021W |
| YBR018C; YBR019C | | | YBR018C; YBR019C |
| YBR016W; YBR017C | | | YBR016W; YBR017C |
| YBR014C; YBR015C | | | YBR014C; YBR015C |
| YBR012C; SPGP0C04820 | | SPGP0C04820 | YBR012C |
| YBR011C; DI49_0191 | DI49_0191 | | YBR011C |
| YBR010W; DI49_0190; YBR008C | DI49_0190 | | YBR010W; YBR008C |
| YBR006W; DI49_0187; YBR005W; DI49_0188 | DI49_0187; DI49_0188 | | YBR006W; YBR005W |
| YBR003W; DI49_0184; YBR002C; DI49_0185 | DI49_0184; DI49_0185 | | YBR003W; YBR002C |
| YBL001C; YBR001C | | | YBL001C; YBR001C |
| YBL003C; YDR225W; DI49_0407; DI49_0179 | DI49_0407; DI49_0179 | | YBL003C; YDR225W |
| YBL004W; DI49_0178 | DI49_0178 | | YBL004W |
| SPGP0Jtrna26S; SPGP0AJtrna31S; YBL005W; SPGP0DKtrna38S; SPGP0DL00100; SPGP0Ntrna12S; YGR038W; SPGP0Atrna22S; YDR261C; SPGP0Atrna23S; YDL007C-A; SPGP0Vtrna15S; SPGP0KCtrna7S; SPGP0AB00100; SPGP0KWtrna25S | | SPGP0Jtrna26S; SPGP0AJtrna31S; SPGP0DKtrna38S; SPGP0DL00100; SPGP0Ntrna12S; SPGP0Atrna22S; SPGP0Atrna23S; SPGP0Vtrna15S; SPGP0KCtrna7S; SPGP0AB00100; SPGP0KWtrna25S | YBL005W; YGR038W; YDR261C; YDL007C-A |

Create PDF in your applications with the Pdfcrowd HTML to PDF API PDFCROWD

| | | | |
|---|---|---|---|
| YBL007C; DI49_0174; YBL008W; DI49_0175 | DI49_0174; DI49_0175 | | YBL007C; YBL008W |
| YBL010C; DI49_0171; YBL011W | DI49_0171 | | YBL010C; YBL011W |
| YBL014C; YBL013W | | | YBL014C; YBL013W |
| SPGP0Jtrna17I; SPGP0Etrna28I; SPGP0Ktrna5I; SPGP0Ktrna6I; YNL035C; DI49_5586; SPGP0EUtrna26I; DI49_1590; SPGP0Qtrna14I; DI49_1543; DI49_4700; SPGP0Atrna16I; YDR106W; SPGP0Ttrna24I; SPGP0Wtrna7I; SPGP0Wtrna8I; DI49_2704; SPGP0Ytrna18I; SPGP0ABtrna23I; YIL096C; YPR143W; SPGP0ACtrna21I; YPR144C; SPGP0Btrna1I; SPGP0Btrna2I; YER182W; SPGP0Ctrna10I; SPGP0Ctrna11I; SPGP0AOtrna27I; SPGP0ATtrna4I; DI49_5613; SPGP0AXtrna20I | DI49_5586; DI49_1590; DI49_1543; DI49_4700; DI49_2704; DI49_5613 | SPGP0Jtrna17I; SPGP0Etrna28I; SPGP0Ktrna5I; SPGP0Ktrna6I; SPGP0EUtrna26I; SPGP0Qtrna14I; SPGP0Atrna16I; SPGP0Ttrna24I; SPGP0Wtrna7I; SPGP0Wtrna8I; SPGP0Ytrna18I; SPGP0ABtrna23I; SPGP0ACtrna21I; SPGP0Btrna1I; SPGP0Btrna2I; SPGP0Ctrna10I; SPGP0Ctrna11I; SPGP0AOtrna27I; SPGP0ATtrna4I; SPGP0AXtrna20I | YNL035C; YDR106W; YIL096C; YPR143W; YPR144C; YER182W |
| YBL015W; DI49_0165; YBL016W | DI49_0165 | | YBL015W; YBL016W |
| YBL018C; YBL017C; YNR065C; YNR064C | | | YBL018C; YBL017C; YNR065C; YNR064C |
| YBL020W; DI49_0161; YBL021C | DI49_0161 | | YBL020W; YBL021C |
| YBL023C; DI49_0158 | DI49_0158 | | YBL023C |
| YBL024W; DI49_0157; YBL025W | DI49_0157 | | YBL024W; YBL025W |
| SPGP0J03130; SPGP0J03120 | | SPGP0J03130; SPGP0J03120 | |
| YBL029W; DI49_0151; YBL029C-A; DI49_0152 | DI49_0151; DI49_0152 | | YBL029W; YBL029C-A |
| YBL031W; DI49_0148; YBL032W | DI49_0148 | | YBL031W; YBL032W |
| YBL034C; YBL033C | | | YBL034C; YBL033C |
| YBL036C; YBL035C | | | YBL036C; YBL035C |
| YBL038W; DI49_0141 | DI49_0141 | | YBL038W |
| YBL039C; DI49_0140; YBL039W-B | DI49_0140 | | YBL039C; YBL039W-B |
| YBL041W; DI49_0137; YBL042C; DI49_0138 | DI49_0137; DI49_0138 | | YBL041W; YBL042C |
| YBL045C; DI49_0133; YBL046W; DI49_0134 | DI49_0133; DI49_0134 | | YBL045C; YBL046W |
| YBL049W; YBL047C; DI49_0131 | DI49_0131 | | YBL049W; YBL047C |
| YBL051C; DI49_0128; DI49_0129 | DI49_0128; DI49_0129 | | YBL051C |
| YBL052C; DI49_0127; YBL054W | DI49_0127 | | YBL052C; YBL054W |
| YBL056W; DI49_0125; YBL057C; YER089C; SPGP0B00600 | DI49_0125 | SPGP0B00600 | YBL056W; YBL057C; YER089C |
| YBL059W; YBL058W | | | YBL059W; YBL058W |
| YBL060W; YBL059C-A | | | YBL060W; YBL059C-A |
| YBL063W; DI49_0119 | DI49_0119 | | YBL063W |
| YBL064C; DI49_0118 | DI49_0118 | | YBL064C |
| YBL066C; DI49_0117; YBL067C | DI49_0117 | | YBL066C; YBL067C |
| YBL069W; YBL068W; DI49_0115; DI49_0116 | DI49_0115; DI49_0116 | | YBL069W; YBL068W |
| YBL072C; YER102W; YBL074C | | | YBL072C; YER102W; YBL074C |

Create PDF in your applications with the Pdfcrowd HTML to PDF API PDFCROWD

| | | | |
|---|---|---|---|
| YBL076C; DI49_0110 | DI49_0110 | | YBL076C |
| YBL078C; DI49_0109 | DI49_0109 | | YBL078C |
| YBL079W; DI49_0108 | DI49_0108 | | YBL079W |
| YBL080C; DI49_0107; YBL081W | DI49_0107 | | YBL080C; YBL081W |
| YBL084C; YBL082C | | | YBL084C; YBL082C |
| YBL086C; YBL085W | | | YBL086C; YBL085W |
| YBL088C; DI49_0100; YBL089W | DI49_0100 | | YBL088C; YBL089W |
| YBL091C; DI49_0097; YBL091C-A; DI49_0098 | DI49_0097; DI49_0098 | | YBL091C; YBL091C-A |
| YBL093C; DI49_0094; YBL095W | DI49_0094 | | YBL093C; YBL095W |
| YBL098W; YBL097W | | | YBL098W; YBL097W |
| SPGP0Jtrna15F; SPGP0Gtrna21F; YBL101C | | SPGP0Jtrna15F; SPGP0Gtrna21F | YBL101C |
| YBL103C; YBL102W | | | YBL103C; YBL102W |
| YBL105C; DI49_0084 | DI49_0084 | | YBL105C |
| YBL106C; DI49_0083; YBL107C | DI49_0083 | | YBL106C; YBL107C |
| SPGP0DGtrna23A; DI49_5601; YCR061W; SPGP0Ltrna5A; DI49_5600; SPGP0Ltrna6A; SPGP0Ntrna7A; YJL115W; SPGP0Otrna12A; SPGP0Ptrna10A; DI49_3259; SPGP0Atrna14A; SPGP0Ttrna28A; SPGP0Xtrna13A; SPGP0Ytrna16A; YKL119C; SPGP0ADtrna31A; DI49_1682; SPGP0Ctrna4A; SPGP0Etrna35A; SPGP0Ftrna26A; DI49_4332; SPGP0Ftrna27A; DI49_4110; SPGP0BXtrna21A; YFR026C; SPGP0Gtrna29A; YMR247C; SPGP0Gtrna30A; YMR023C; DI49_1030 | DI49_5601; DI49_5600; DI49_3259; DI49_1682; DI49_4332; DI49_4110; DI49_1030 | SPGP0DGtrna23A; SPGP0Ltrna5A; SPGP0Ltrna6A; SPGP0Ntrna7A; SPGP0Otrna12A; SPGP0Ptrna10A; SPGP0Atrna14A; SPGP0Ttrna28A; SPGP0Xtrna13A; SPGP0Ytrna16A; SPGP0ADtrna31A; SPGP0Ctrna4A; SPGP0Etrna35A; SPGP0Ftrna26A; SPGP0Ftrna27A; SPGP0BXtrna21A; SPGP0Gtrna29A; SPGP0Gtrna30A | YCR061W; YJL115W; YKL119C; YFR026C; YMR247C; YMR023C |
| DI49_3506; SPGP0DP00100; YNR075W | DI49_3506 | SPGP0DP00100 | YNR075W |
| SPGP0K00120; SPGP0V00150; YNR071C | | SPGP0K00120; SPGP0V00150 | YNR071C |
| YNR069C; YNR070W | | | YNR069C; YNR070W |
| YNR067C; YNR068C | | | YNR067C; YNR068C |
| SPGP0K00220; YNR063W | | SPGP0K00220 | YNR063W |
| YNR060W; YNR061C | | | YNR060W; YNR061C |
| YNR058W; YNR059W | | | YNR058W; YNR059W |
| YNR056C; SPGP0K00270 | | SPGP0K00270 | YNR056C |
| YNR054C; DI49_4786; DI49_4787 | DI49_4786; DI49_4787 | | YNR054C |
| YNR053C; SPGP0AL00600 | | SPGP0AL00600 | YNR053C |
| YNR052C; DI49_4785; YNR051C | DI49_4785 | | YNR052C; YNR051C |
| YNR049C; YNR050C; DI49_4783; DI49_4784 | DI49_4783; DI49_4784 | | YNR049C; YNR050C |
| YNR047W; DI49_4780; YNR046W; DI49_4781; YCR091W; YCR092C | DI49_4780; DI49_4781 | | YNR047W; YNR046W; YCR091W; YCR092C |
| YNR044W; YNR045W | | | YNR044W; YNR045W |
| YNR041C; DI49_4775; YNR040W; DI49_4776 | DI49_4775; DI49_4776 | | YNR041C; YNR040W |
| YNR038W; YNR039C | | | YNR038W; YNR039C |

Create PDF in your applications with the Pdfcrowd HTML to PDF API PDFCROWD

| | | | |
|---|---|---|---|
| YNR036C; YNR037C | | | YNR036C; YNR037C |
| YNR034W-A; YNR035C | | | YNR034W-A; YNR035C |
| YNR032W; YNR034W | | | YNR032W; YNR034W |
| YNR030W; DI49_4762; YNR029C | DI49_4762 | | YNR030W; YNR029C |
| YNR027W; YNR028W | | | YNR027W; YNR028W |
| YNR024W; YNR026C | | | YNR024W; YNR026C |
| YNR022C; YNR023W | | | YNR022C; YNR023W |
| YNR020C; DI49_4755; DI49_4756 | DI49_4755; DI49_4756 | | YNR020C |
| YNR019W; DI49_4754; YNR018W | DI49_4754 | | YNR019W; YNR018W |
| YNR016C; DI49_4751; YNR015W | DI49_4751 | | YNR016C; YNR015W |
| YNR013C; DI49_4748; DI49_4749 | DI49_4748; DI49_4749 | | YNR013C |
| YNR012W; DI49_4747 | DI49_4747 | | YNR012W |
| YNR011C; DI49_4746; YNR010W | DI49_4746 | | YNR011C; YNR010W |
| YNR008W; YNR009W | | | YNR008W; YNR009W |
| YNR006W; YNR007C | | | YNR006W; YNR007C |
| YNR002C; YNR004W; DI49_4740; DI49_4741 | DI49_4740; DI49_4741 | | YNR002C; YNR004W |
| SPGP0Ktrna1P; DI49_4736; DI49_0756; YCR007C; SPGP0AXtrna19P | DI49_4736; DI49_0756 | SPGP0Ktrna1P; SPGP0AXtrna19P | YCR007C |
| YNR001C; DI49_4735; YNL001W | DI49_4735 | | YNR001C; YNL001W |
| YNL003C; DI49_4732 | DI49_4732 | | YNL003C |
| YNL004W; DI49_4731; YNL005C | DI49_4731 | | YNL004W; YNL005C |
| YNL007C; DI49_4728; YNL008C; DI49_4729 | DI49_4728; DI49_4729 | | YNL007C; YNL008C |
| YNL010W; DI49_4725 | DI49_4725 | | YNL010W |
| YNL011C; DI49_4724 | DI49_4724 | | YNL011C |
| YNL012W; SPGP0AX00130 | | SPGP0AX00130 | YNL012W |
| YNL014W; DI49_4723; YNL015W | DI49_4723 | | YNL014W; YNL015W |
| YNL034W; SPGP0K01090 | | SPGP0K01090 | YNL034W |
| YNL032W; DI49_4705 | DI49_4705 | | YNL032W |
| YNL031C; DI49_4706 | DI49_4706 | | YNL031C |
| YNL030W; DI49_0189; YNL029C; DI49_4707; DI49_4708 | DI49_0189; DI49_4707; DI49_4708 | | YNL030W; YNL029C |
| YNL026W; YNL027W | | | YNL026W; YNL027W |
| YNL024C-A; YNL025C | | | YNL024C-A; YNL025C |
| YNL023C; DI49_4714 | DI49_4714 | | YNL023C |
| YNL022C; DI49_4715; YNL021W; DI49_4716 | DI49_4715; DI49_4716 | | YNL022C; YNL021W |
| SPGP0Ktrna2T; SPGP0BHtrna30T; YNL036W; SPGP0Ktrna3T; YNL284C; SPGP0FPtrna4T; SPGP0FS00100; SPGP0GJtrna5T; YHR006W; YIL014C-A; SPGP0ABtrna28T; YIL102C; SPGP0RJtrna8T; SPGP0Dtrna22T; YCR098C | | SPGP0Ktrna2T; SPGP0BHtrna30T; SPGP0Ktrna3T; SPGP0FPtrna4T; SPGP0FS00100; SPGP0GJtrna5T; SPGP0ABtrna28T; SPGP0RJtrna8T; SPGP0Dtrna22T | YNL036W; YNL284C; YHR006W; YIL014C-A; YIL102C; YCR098C |

Create PDF in your applications with the Pdfcrowd HTML to PDF API PDFCROWD

| | | | |
|---|---|---|---|
| YNL038W; YNL037C | | | YNL038W; YNL037C |
| YNL040W; YNL039W | | | YNL040W; YNL039W |
| YNL042W; YNL041C; DI49_4689; DI49_4688 | DI49_4689; DI49_4688 | | YNL042W; YNL041C |
| YNL044W; DI49_4686; YNL045W; DI49_4685 | DI49_4686; DI49_4685 | | YNL044W; YNL045W |
| YNL047C; YNL046W | | | YNL047C; YNL046W |
| YNL049C; DI49_4681 | DI49_4681 | | YNL049C |
| YNL050C; DI49_4680; YNL051W | DI49_4680 | | YNL050C; YNL051W |
| YNL053W; YNL052W | | | YNL053W; YNL052W |
| YNL055C; DI49_4674; YNL056W; DI49_4673 | DI49_4674; DI49_4673 | | YNL055C; YNL056W |
| YNL059C; DI49_4671 | DI49_4671 | | YNL059C |
| YNL061W; DI49_4670 | DI49_4670 | | YNL061W |
| YNL062C; DI49_4669; YNL063W; DI49_4668 | DI49_4669; DI49_4668 | | YNL062C; YNL063W |
| YNL065W; SPGP0K01360 | | SPGP0K01360 | YNL065W |
| YNL067W; YGL147C; YNL068C; YGL148W; DI49_1827 | DI49_1827 | | YNL067W; YGL147C; YNL068C; YGL148W |
| YNL070W; DI49_4661 | DI49_4661 | | YNL070W |
| YNL071W; DI49_4660; YNL072W; DI49_4659 | DI49_4660; DI49_4659 | | YNL071W; YNL072W |
| YNL074C; YNL073W | | | YNL074C; YNL073W |
| YNL076W; YNL075W | | | YNL076W; YNL075W |
| YNL078W; YNL077W | | | YNL078W; YNL077W |
| YNL080C; DI49_4651 | DI49_4651 | | YNL080C |
| YNL081C; DI49_4650 | DI49_4650 | | YNL081C |
| YNL082W; DI49_4649 | DI49_4649 | | YNL082W |
| YNL083W; DI49_4648 | DI49_4648 | | YNL083W |
| YNL084C; DI49_4647 | DI49_4647 | | YNL084C |
| YNL085W; DI49_4646; YNL086W; DI49_4645 | DI49_4646; DI49_4645 | | YNL085W; YNL086W |
| YNL088W; DI49_4643; DI49_4642 | DI49_4643; DI49_4642 | | YNL088W |
| YNL091W; DI49_4641; YNL092W; DI49_4640 | DI49_4641; DI49_4640 | | YNL091W; YNL092W |
| YNL094W; DI49_4638 | DI49_4638 | | YNL094W |
| YNL095C; DI49_4637 | DI49_4637 | | YNL095C |
| YNL096C; DI49_4636 | DI49_4636 | | YNL096C |
| SPGP0Ktrna6L; DI49_1927; SPGP0DUtrna4L; SPGP0Rtrna18L; YGL159W; DI49_4635; DI49_4634; SPGP0Ttrna30L; YLR246W; SPGP0Ytrna21L; SPGP0ADtrna36L; YKR009C; SPGP0AJtrna25L; YAR023C; SPGP0Etrna43L; YMR118C; SPGP0Gtrna35L | DI49_1927; DI49_4635; DI49_4634 | SPGP0Ktrna6L; SPGP0DUtrna4L; SPGP0Rtrna18L; SPGP0Ttrna30L; SPGP0Ytrna21L; SPGP0ADtrna36L; SPGP0AJtrna25L; SPGP0Etrna43L; SPGP0Gtrna35L | YGL159W; YLR246W; YKR009C; YAR023C; YMR118C |
| SPGP0K01670; YNL098C; YNL099C | | SPGP0K01670 | YNL098C; YNL099C |
| YNL101W; DI49_4630 | DI49_4630 | | YNL101W |
| YNL102W; DI49_4629; YNL103W; DI49_4628 | DI49_4629; DI49_4628 | | YNL102W; YNL103W |
| YNL106C; DI49_4626; YNL107W; DI49_4625 | DI49_4626; DI49_4625 | | YNL106C; YNL107W |

Create PDF in your applications with the Pdfcrowd HTML to PDF API PDFCROWD

| | | | |
|---|---|---|---|
| YNL110C; DI49_4623 | DI49_4623 | | YNL110C |
| YNL111C; DI49_4622 | DI49_4622 | | YNL111C |
| YNL112W; SPGP0S00850; YNL113W; DI49_4621 | DI49_4621 | SPGP0S00850 | YNL112W; YNL113W |
| YNL116W; DI49_4619 | DI49_4619 | | YNL116W |
| YNL117W; DI49_4618 | DI49_4618 | | YNL117W |
| YNL118C; DI49_4617; YNL119W; DI49_4616 | DI49_4617; DI49_4616 | | YNL118C; YNL119W |
| YNL122C; DI49_4614 | DI49_4614 | | YNL122C |
| YNL123W; DI49_4613 | DI49_4613 | | YNL123W |
| YNL124W; DI49_4612; YNL125C; DI49_4611 | DI49_4612; DI49_4611 | | YNL124W; YNL125C |
| YNL127W; DI49_4609 | DI49_4609 | | YNL127W |
| YNL128W; DI49_4608; YNL129W; DI49_4607 | DI49_4608; DI49_4607 | | YNL128W; YNL129W |
| YNL131W; DI49_4605 | DI49_4605 | | YNL131W |
| YNL132W; DI49_4604 | DI49_4604 | | YNL132W |
| SPGP0Ktrna3F; SPGP0OVtrna4F; YNL133C; YLR459W | | SPGP0Ktrna3F; SPGP0OVtrna4F | YNL133C; YLR459W |
| YNL135C; DI49_4600 | DI49_4600 | | YNL135C |
| YNL136W; DI49_4599 | DI49_4599 | | YNL136W |
| YNL137C; DI49_4598 | DI49_4598 | | YNL137C |
| YNL138W; DI49_4597; YNL138W-A; DI49_4596 | DI49_4597; DI49_4596 | | YNL138W; YNL138W-A |
| YNL141W; YNL139C | | | YNL141W; YNL139C |
| YNL144C; YNL142W | | | YNL144C; YNL142W |
| YNL146W; YNL145W | | | YNL146W; YNL145W |
| YNL148C; YNL147W | | | YNL148C; YNL147W |
| YNL151C; YNL149C | | | YNL151C; YNL149C |
| YNL153C; YNL152W | | | YNL153C; YNL152W |
| YNL155W; YNL154C | | | YNL155W; YNL154C |
| YNL157W; DI49_4580; YNL158W; DI49_4579 | DI49_4580; DI49_4579 | | YNL157W; YNL158W |
| YNL160W; DI49_4577; YNL161W; DI49_4576 | DI49_4577; DI49_4576 | | YNL160W; YNL161W |
| YNL162W-A; YNL162W; SPGP0O02990; YHR141C | | SPGP0O02990 | YNL162W-A; YNL162W; YHR141C |
| YNL164C; DI49_4572; YNL165W; DI49_4571 | DI49_4572; DI49_4571 | | YNL164C; YNL165W |
| YNL167C; YNL166C | | | YNL167C; YNL166C |
| YNL169C; DI49_4567 | DI49_4567 | | YNL169C |
| YNL172W; DI49_4566; YNL173C; DI49_4565 | DI49_4566; DI49_4565 | | YNL172W; YNL173C |
| YNL176C; YNL175C | | | YNL176C; YNL175C |
| YNL178W; DI49_4561 | DI49_4561 | | YNL178W |
| YNL180C; DI49_4560; YNL181W; DI49_4559 | DI49_4560; DI49_4559 | | YNL180C; YNL181W |
| YNL183C; DI49_4557 | DI49_4557 | | YNL183C |
| YNL185C; DI49_4556 | DI49_4556 | | YNL185C |
| YNL186W; DI49_4555; YNL187W; DI49_4554 | DI49_4555; DI49_4554 | | YNL186W; YNL187W |

Create PDF in your applications with the Pdfcrowd HTML to PDF API PDFCROWD

| | | | |
|---|---|---|---|
| YNL189W; DI49_4552 | DI49_4552 | | YNL189W |
| YNL190W; DI49_4551 | DI49_4551 | | YNL190W |
| YNL191W; DI49_4550 | DI49_4550 | | YNL191W |
| YNL192W; DI49_4549; YNL193W; DI49_4548 | DI49_4549; DI49_4548 | | YNL192W; YNL193W |
| YNL195C; YNL194C | | | YNL195C; YNL194C |
| YNL197C; YNL196C | | | YNL197C; YNL196C |
| YNL200C; YNL199C | | | YNL200C; YNL199C |
| YNL202W; DI49_4540; YNL204C; DI49_4539 | DI49_4540; DI49_4539 | | YNL202W; YNL204C |
| YNL207W; DI49_4537; YNL208W; DI49_4536 | DI49_4537; DI49_4536 | | YNL207W; YNL208W |
| YNL210W; YNL209W; YDL229W; YDL230W | | | YNL210W; YNL209W; YDL229W; YDL230W |
| YNL212W; DI49_4532; YNL213C; DI49_4531 | DI49_4532; DI49_4531 | | YNL212W; YNL213C |
| YNL215W; YNL214W | | | YNL215W; YNL214W |
| YNL217W; YNL216W | | | YNL217W; YNL216W |
| YNL219C; YNL218W | | | YNL219C; YNL218W |
| YNL221C; YNL220W | | | YNL221C; YNL220W |
| YNL223W; DI49_4521 | DI49_4521 | | YNL223W |
| YNL224C; DI49_4520 | DI49_4520 | | YNL224C |
| YNL225C; DI49_4519 | DI49_4519 | | YNL225C |
| YNL227C; DI49_4518; YNL229C; DI49_4517 | DI49_4518; DI49_4517 | | YNL227C; YNL229C |
| YNL231C; YNL230C | | | YNL231C; YNL230C |
| YNL233W; DI49_4513; YNL234W; DI49_4512 | DI49_4513; DI49_4512 | | YNL233W; YNL234W |
| YNL237W; YNL236W | | | YNL237W; YNL236W |
| YNL239W; YNL238W | | | YNL239W; YNL238W |
| YNL241C; DI49_4506 | DI49_4506 | | YNL241C |
| YNL242W; DI49_4505; YNL243W; DI49_4504 | DI49_4505; DI49_4504 | | YNL242W; YNL243W |
| YNL245C; YNL244C | | | YNL245C; YNL244C |
| YNL247W; DI49_4500 | DI49_4500 | | YNL247W |
| YNL248C; DI49_4499 | DI49_4499 | | YNL248C |
| YNL249C; DI49_4498 | DI49_4498 | | YNL249C |
| YNL250W; DI49_4497 | DI49_4497 | | YNL250W |
| YNL251C; DI49_4496; YNL252C; DI49_4495 | DI49_4496; DI49_4495 | | YNL251C; YNL252C |
| YNL254C; DI49_4493 | DI49_4493 | | YNL254C |
| YNL255C; DI49_4492 | DI49_4492 | | YNL255C |
| YNL256W; DI49_4491; YNL257C; DI49_4490 | DI49_4491; DI49_4490 | | YNL256W; YNL257C |
| YNL259C; YNL258C | | | YNL259C; YNL258C |
| YNL261W; YNL260C | | | YNL261W; YNL260C |
| YNL263C; DI49_4484; YNL264C; DI49_4483 | DI49_4484; DI49_4483 | | YNL263C; YNL264C |
| YNL267W; DI49_4481 | DI49_4481 | | YNL267W |
| YNL268W; DI49_4480; YNL270C; DI49_4479 | DI49_4480; DI49_4479 | | YNL268W; YNL270C |
| YNL271C; SPGP0S02280; DI49_4478 | DI49_4478 | SPGP0S02280 | YNL271C |

Create PDF in your applications with the Pdfcrowd HTML to PDF API PDFCROWD

| | | | |
|---|---|---|---|
| YNL272C; DI49_4477; YNL273W; DI49_4476 | DI49_4477; DI49_4476 | | YNL272C; YNL273W |
| YNL275W; YNL274C | | | YNL275W; YNL274C |
| YNL278W; YNL277W | | | YNL278W; YNL277W |
| YNL280C; DI49_4470; YNL281W; DI49_4469 | DI49_4470; DI49_4469 | | YNL280C; YNL281W |
| YNL283C; DI49_4467 | DI49_4467 | | YNL283C |
| SPGP0Ktrna5G; SPGP0Etrna44G; YNL286W; DI49_2298; SPGP0Ytrna24G; SPGP0Dtrna33G; YOL109W | DI49_2298 | SPGP0Ktrna5G; SPGP0Etrna44G; SPGP0Ytrna24G; SPGP0Dtrna33G | YNL286W; YOL109W |
| YNL288W; DI49_4459 | DI49_4459 | | YNL288W |
| YNL289W; DI49_4458; YNL290W | DI49_4458 | | YNL289W; YNL290W |
| YNL292W; DI49_4455; DI49_4456 | DI49_4455; DI49_4456 | | YNL292W |
| YNL293W; DI49_4454; YNL294C | DI49_4454 | | YNL293W; YNL294C |
| YNL297C; YNL295W | | | YNL297C; YNL295W |
| YNL299W; YNL298W; DI49_4450; DI49_4451 | DI49_4450; DI49_4451 | | YNL299W; YNL298W |
| SPGP0K03490; YOL120C; SPGP0O00440; SPGP0AR00380; DI49_4448 | DI49_4448 | SPGP0K03490; SPGP0O00440; SPGP0AR00380 | YOL120C |
| YNL302C; DI49_4447; DI49_2286; SPGP0D04230 | DI49_4447; DI49_2286 | SPGP0D04230 | YNL302C |
| YNL304W; DI49_4446; YNL305C | DI49_4446 | | YNL304W; YNL305C |
| YNL307C; DI49_4443; YNL308C; DI49_4444 | DI49_4443; DI49_4444 | | YNL307C; YNL308C |
| YNL310C; YNL309W | | | YNL310C; YNL309W |
| SPGP0K03590; YNL311C | | SPGP0K03590 | YNL311C |
| YNL314W; YNL313C | | | YNL314W; YNL313C |
| YNL316C; YNL315C | | | YNL316C; YNL315C |
| YNL318C; YNL317W | | | YNL318C; YNL317W |
| YNL321W; DI49_4430 | DI49_4430 | | YNL321W |
| YNL322C; DI49_4429 | DI49_4429 | | YNL322C |
| YNL323W; DI49_4428 | DI49_4428 | | YNL323W |
| YNL325C; DI49_4427; YNL326C | DI49_4427 | | YNL325C; YNL326C |
| YNL328C; DI49_4424; YNL329C; DI49_4425 | DI49_4424; DI49_4425 | | YNL328C; YNL329C |
| YNL331C; YDL243C; DI49_4421 | DI49_4421 | | YNL331C; YDL243C |
| YNL332W; YJR156C; YFL059W | | | YNL332W; YJR156C; YFL059W |
| YNL335W; YNL334C | | | YNL335W; YNL334C |
| YIL002C; DI49_2778; DI49_2779 | DI49_2778; DI49_2779 | | YIL002C |
| SPGP0DT00100; YDL024C; YDL025C | | SPGP0DT00100 | YDL024C; YDL025C |
| SPGP0DU00100; YGR181W | | SPGP0DU00100 | YGR181W |
| SPGP0DU00110; YGR183C; DI49_4254 | DI49_4254 | SPGP0DU00110 | YGR183C |
| YPR013C; SPGP0OR00100; YPR004C | | SPGP0OR00100 | YPR013C; YPR004C |
| DI49_5604; YPR160W | DI49_5604 | | YPR160W |
| SPGP0Ltrna9G; SPGP0Gtrna40G; SPGP0Ltrna10G; DI49_5463; SPGP0Mtrna38G; DI49_2997; SPGP0Ntrna11G; YJR051W; SPGP0Ntrna12G; SPGP0Otrna16G; SPGP0Qtrna13G; DI49_1343; SPGP0GEtrna6G; YDL211C; SPGP0Vtrna14G; | DI49_5463; DI49_2997; DI49_1343; DI49_0168; DI49_0443; DI49_0766; DI49_4046 | SPGP0Ltrna9G; SPGP0Gtrna40G; SPGP0Ltrna10G; SPGP0Mtrna38G; SPGP0Ntrna11G; SPGP0Ntrna12G; SPGP0Otrna16G; SPGP0Qtrna13G; SPGP0GEtrna6G; SPGP0Vtrna14G; SPGP0Ytrna25G; SPGP0Ytrna26G; | YJR051W; YDL211C; YFR057W; YFR016C; YFR008W; YCR016W; YOL022C; YML046W |

Create PDF in your applications with the Pdfcrowd HTML to PDF API PDFCROWD

| | | | |
|---|---|---|---|
| SPGP0Ytrna25G; SPGP0Ytrna26G; SPGP0ACtrna35G; SPGP0Btrna1G; DI49_0168; DI49_0443; SPGP0KXtrna23G; YFR057W; SPGP0AEtrna7G; YFR016C; SPGP0AEtrna8G; YFR008W; SPGP0AFtrna28G; SPGP0AF00610; SPGP0AGtrna45G; DI49_0766; SPGP0AHtrna27G; YCR016W; SPGP0Dtrna31G; YOL022C; SPGP0Dtrna32G; SPGP0Etrna42G; SPGP0Etrna43G; DI49_4046; SPGP0CAtrna37G; YML046W | | SPGP0ACtrna35G; SPGP0Btrna1G; SPGP0KXtrna23G; SPGP0AEtrna7G; SPGP0AEtrna8G; SPGP0AFtrna28G; SPGP0AF00610; SPGP0AGtrna45G; SPGP0AHtrna27G; SPGP0Dtrna31G; SPGP0Dtrna32G; SPGP0Etrna42G; SPGP0Etrna43G; SPGP0CAtrna37G | |
| DI49_5602; YPR159W | DI49_5602 | | YPR159W |
| DI49_5598; DI49_2099; DI49_5597 | DI49_5598; DI49_2099; DI49_5597 | | |
| DI49_5595; DI49_5596 | DI49_5595; DI49_5596 | | |
| DI49_5594; SPGP0L00180 | DI49_5594 | SPGP0L00180 | |
| DI49_5592; DI49_5593 | DI49_5592; DI49_5593 | | |
| DI49_5590; DI49_5591 | DI49_5590; DI49_5591 | | |
| DI49_5588; DI49_5589 | DI49_5588; DI49_5589 | | |
| DI49_5584; DI49_5585; YPR141C | DI49_5584; DI49_5585 | | YPR141C |
| DI49_5582; YPR138C; YPR139C | DI49_5582 | | YPR138C; YPR139C |
| SPGP0Ltrna5N; DI49_5581; DI49_5580; SPGP0Otrna8N; DI49_2351; SPGP0Ptrna7N; SPGP0Ytrna10N; YGR121C; SPGP0Ztrna16N; YFL003C; DI49_1650; DI49_0758; YCR008W; SPGP0ALtrna18N; DI49_4738; SPGP0Ctrna4N; SPGP0AOtrna20N; SPGP0Dtrna13N; SPGP0Dtrna14N; SPGP0Etrna21N; SPGP0Htrna19N | DI49_5581; DI49_5580; DI49_2351; DI49_1650; DI49_0758; DI49_4738 | SPGP0Ltrna5N; SPGP0Otrna8N; SPGP0Ptrna7N; SPGP0Ytrna10N; SPGP0Ztrna16N; SPGP0ALtrna18N; SPGP0Ctrna4N; SPGP0AOtrna20N; SPGP0Dtrna13N; SPGP0Dtrna14N; SPGP0Etrna21N; SPGP0Htrna19N | YGR121C; YFL003C; YCR008W |
| DI49_5578; DI49_5579; YPR135W; YPR137W | DI49_5578; DI49_5579 | | YPR135W; YPR137W |
| DI49_5576; YPR133C; YPR133W-A | DI49_5576 | | YPR133C; YPR133W-A |
| DI49_5575; YPR132W; DI49_5574; DI49_2077; DI49_2076 | DI49_5575; DI49_5574; DI49_2077; DI49_2076 | | YPR132W |
| DI49_5572; DI49_5573 | DI49_5572; DI49_5573 | | |
| DI49_5570; DI49_5571; YPR127W; YPR128C | DI49_5570; DI49_5571 | | YPR127W; YPR128C |
| DI49_5568; DI49_5569 | DI49_5568; DI49_5569 | | |
| DI49_5566; YPR120C; DI49_5565 | DI49_5566; DI49_5565 | | YPR120C |
| DI49_5563; YPR118W | DI49_5563 | | YPR118W |
| DI49_5562; YPR117W; DI49_5561 | DI49_5562; DI49_5561 | | YPR117W |
| DI49_5559; DI49_5560; YPR115W; YPR116W | DI49_5559; DI49_5560 | | YPR115W; YPR116W |
| DI49_5557; YPR112C; DI49_5556; YPR113W | DI49_5557; DI49_5556 | | YPR112C; YPR113W |
| DI49_5554; DI49_5555 | DI49_5554; DI49_5555 | | |

Create PDF in your applications with the Pdfcrowd HTML to PDF API PDFCROWD

| | | | |
|---|---|---|---|
| DI49_5552; SPGP0Etrna34T; SPGP0AQtrna35T; SPGP0Etrna33T; DI49_2219; DI49_2047 | DI49_5552; DI49_2219; DI49_2047 | SPGP0Etrna34T; SPGP0AQtrna35T; SPGP0Etrna33T | |
| DI49_5550; DI49_5551 | DI49_5550; DI49_5551 | | |
| DI49_5548; YPR104C | DI49_5548 | | YPR104C |
| DI49_5547; YPR103W | DI49_5547 | | YPR103W |
| DI49_5546; DI49_2042; SPGP0L00630 | DI49_5546; DI49_2042 | SPGP0L00630 | |
| DI49_5544; DI49_5545 | DI49_5544; DI49_5545 | | |
| DI49_5542; DI49_5543; YPR097W; YPR098C | DI49_5542; DI49_5543 | | YPR097W; YPR098C |
| DI49_5540; YPR093C; DI49_5539; YPR094W | DI49_5540; DI49_5539 | | YPR093C; YPR094W |
| DI49_5537; YPR088C | DI49_5537 | | YPR088C |
| DI49_5536; YPR086W; DI49_5535 | DI49_5536; DI49_5535 | | YPR086W |
| DI49_5533; YPR083W; DI49_5532 | DI49_5533; DI49_5532 | | YPR083W |
| DI49_5530; SPGP0I01810; DI49_5529; DI49_1148 | DI49_5530; DI49_5529; DI49_1148 | SPGP0I01810 | |
| DI49_5527; DI49_5528 | DI49_5527; DI49_5528 | | |
| SPGP0Ltrna11S; DI49_5523; SPGP0L00860; SPGP0Qtrna13S; SPGP0Vtrna16S; SPGP0Wtrna5S; YIL057C; SPGP0ACtrna33S; DI49_1141 | DI49_5523; DI49_1141 | SPGP0Ltrna11S; SPGP0L00860; SPGP0Qtrna13S; SPGP0Vtrna16S; SPGP0Wtrna5S; SPGP0ACtrna33S | YIL057C |
| DI49_5521; YPR069C; DI49_5520; SPGP0AC00580 | DI49_5521; DI49_5520 | SPGP0AC00580 | YPR069C |
| DI49_5518; YPR066W; DI49_5517; YPR067W | DI49_5518; DI49_5517 | | YPR066W; YPR067W |
| SPGP0L00940; SPGP0L00930 | | SPGP0L00940; SPGP0L00930 | |
| DI49_5514; DI49_5515; YPR062W; YPR063C | DI49_5514; DI49_5515 | | YPR062W; YPR063C |
| DI49_5512; YPR058W; DI49_5511; YPR060C | DI49_5512; DI49_5511 | | YPR058W; YPR060C |
| DI49_5509; YPR055W | DI49_5509 | | YPR055W |
| DI49_5508; YPR054W | DI49_5508 | | YPR054W |
| DI49_5507; YPR052C; DI49_5506 | DI49_5507; DI49_5506 | | YPR052C |
| DI49_5504; DI49_5505 | DI49_5504; DI49_5505 | | |
| DI49_5502; YPR046W; DI49_5501 | DI49_5502; DI49_5501 | | YPR046W |
| DI49_5499; YPR042C | DI49_5499 | | YPR042C |
| DI49_5498; YPR041W; DI49_5497 | DI49_5498; DI49_5497 | | YPR041W |
| DI49_5495; DI49_5496 | DI49_5495; DI49_5496 | | |
| DI49_5493; YPR035W; DI49_5492 | DI49_5493; DI49_5492 | | YPR035W |
| DI49_5490; YPR032W; YPR033C | DI49_5490 | | YPR032W; YPR033C |
| DI49_5489; YPR031W; DI49_5488 | DI49_5489; DI49_5488 | | YPR031W |
| SPGP0L01240; DI49_5487 | DI49_5487 | SPGP0L01240 | |
| DI49_5485; DI49_5486; SPGP0Ltrna7F | DI49_5485; DI49_5486 | SPGP0Ltrna7F | |
| DI49_5483; YPR026W; DI49_5482; YPR027C | DI49_5483; DI49_5482 | | YPR026W; YPR027C |
| DI49_5480; DI49_5481 | DI49_5480; DI49_5481 | | |

Create PDF in your applications with the Pdfcrowd HTML to PDF API PDFCROWD

| | | | |
|---|---|---|---|
| DI49_5478; DI49_5479; YPR022C; YPR023C | DI49_5478; DI49_5479 | | YPR022C; YPR023C |
| DI49_5476; YPR019W; DI49_5475; YPR020W | DI49_5476; DI49_5475 | | YPR019W; YPR020W |
| DI49_5473; YPR016C; YPR017C | DI49_5473 | | YPR016C; YPR017C |
| DI49_5472; YPR015C; DI49_5471 | DI49_5472; DI49_5471 | | YPR015C |
| DI49_5469; DI49_5470 | DI49_5469; DI49_5470 | | |
| DI49_5467; YPR010C | DI49_5467 | | YPR010C |
| DI49_5466; YPR009W | DI49_5466 | | YPR009W |
| DI49_5465; YPR008W | DI49_5465 | | YPR008W |
| DI49_5461; DI49_5462 | DI49_5461; DI49_5462 | | |
| DI49_5459; YPR003C | DI49_5459 | | YPR003C |
| SPGP0L01500; YPR002W | | SPGP0L01500 | YPR002W |
| DI49_5456; YPR001W; DI49_5455; YPL001W | DI49_5456; DI49_5455 | | YPR001W; YPL001W |
| DI49_5453; DI49_5454 | DI49_5453; DI49_5454 | | |
| DI49_5451; DI49_5452 | DI49_5451; DI49_5452 | | |
| DI49_5449; DI49_5450 | DI49_5449; DI49_5450 | | |
| DI49_5447; YPL009C; DI49_5446; YPL010W | DI49_5447; DI49_5446 | | YPL009C; YPL010W |
| DI49_5444; YPL012W; DI49_5443; YPL013C | DI49_5444; DI49_5443 | | YPL012W; YPL013C |
| DI49_5441; YPL015C; SPGP0L01670 | DI49_5441 | SPGP0L01670 | YPL015C |
| DI49_5439; DI49_5440; YPL016W; YPL017C | DI49_5439; DI49_5440 | | YPL016W; YPL017C |
| DI49_5437; YPL019C; DI49_5436; YPL020C | DI49_5437; DI49_5436 | | YPL019C; YPL020C |
| DI49_5434; YPL022W | DI49_5434 | | YPL022W |
| DI49_5433; YPL023C | DI49_5433 | | YPL023C |
| DI49_5432; YPL024W; DI49_5431; YPL026C | DI49_5432; DI49_5431 | | YPL024W; YPL026C |
| DI49_5429; YPL028W | DI49_5429 | | YPL028W |
| DI49_5428; YPL029W; DI49_5427; YPL030W | DI49_5428; DI49_5427 | | YPL029W; YPL030W |
| DI49_5425; DI49_5426 | DI49_5425; DI49_5426 | | |
| DI49_5423; DI49_5424 | DI49_5423; DI49_5424 | | |
| DI49_5421; YPL037C | DI49_5421 | | YPL037C |
| DI49_5420; YPL038W | DI49_5420 | | YPL038W |
| DI49_5419; YPL039W; DI49_5418; YPL040C | DI49_5419; DI49_5418 | | YPL039W; YPL040C |
| DI49_5416; YPL042C | DI49_5416 | | YPL042C |
| DI49_5414; YPL043W | DI49_5414 | | YPL043W |
| DI49_5413; YPL045W | DI49_5413 | | YPL045W |
| DI49_5412; YPL046C; DI49_5411; YPL047W | DI49_5412; DI49_5411 | | YPL046C; YPL047W |
| DI49_5409; DI49_5410 | DI49_5409; DI49_5410 | | |
| DI49_5407; YPL051W; SPGP0L02010; YPL052W | DI49_5407 | SPGP0L02010 | YPL051W; YPL052W |
| DI49_5405; DI49_5406 | DI49_5405; DI49_5406 | | |
| DI49_5403; DI49_5404 | DI49_5403; DI49_5404 | | |
| DI49_5401; DI49_5402 | DI49_5401; DI49_5402 | | |
| DI49_5399; DI49_5400; SPGP0Etrna8C | DI49_5399; DI49_5400 | SPGP0Etrna8C | |

Create PDF in your applications with the Pdfcrowd HTML to PDF API PDFCROWD

| | | | |
|---|---|---|---|
| DI49_5395; DI49_5396 | DI49_5395; DI49_5396 | | |
| DI49_5393; DI49_5394 | DI49_5393; DI49_5394 | | |
| DI49_5391; YPL069C; YPL068C | DI49_5391 | | YPL069C; YPL068C |
| DI49_5390; YPL070W; DI49_5389 | DI49_5390; DI49_5389 | | YPL070W |
| DI49_5387; YPL074W; YPL072W | DI49_5387 | | YPL074W; YPL072W |
| DI49_5386; YPL075W; DI49_5385 | DI49_5386; DI49_5385 | | YPL075W |
| DI49_5383; DI49_5384 | DI49_5383; DI49_5384 | | |
| DI49_5381; DI49_1215; DI49_1214 | DI49_5381; DI49_1215; DI49_1214 | | |
| DI49_5380; YPL082C; DI49_5379; YPL081W | DI49_5380; DI49_5379 | | YPL082C; YPL081W |
| DI49_5377; DI49_5378 | DI49_5377; DI49_5378 | | |
| DI49_5375; DI49_5376 | DI49_5375; DI49_5376 | | |
| DI49_5373; DI49_5374 | DI49_5373; DI49_5374 | | |
| DI49_5371; YPL091W; DI49_5370 | DI49_5371; DI49_5370 | | YPL091W |
| DI49_5368; YPL094C; DI49_5367 | DI49_5368; DI49_5367 | | YPL094C |
| DI49_5365; DI49_5366 | DI49_5365; DI49_5366 | | |
| DI49_5363; DI49_5364 | DI49_5363; DI49_5364 | | |
| DI49_5361; DI49_5362 | DI49_5361; DI49_5362 | | |
| DI49_5359; DI49_5360 | DI49_5359; DI49_5360 | | |
| DI49_5357; YPL105C; YPL104W | DI49_5357 | | YPL105C; YPL104W |
| DI49_5356; YPL106C; DI49_5355 | DI49_5356; DI49_5355 | | YPL106C |
| SPGP0L02540; DI49_5354 | DI49_5354 | SPGP0L02540 | |
| SPGP0L02560; SPGP0L02550 | | SPGP0L02560; SPGP0L02550 | |
| DI49_5352; YPL111W | DI49_5352 | | YPL111W |
| SPGP0Ltrna3M; DI49_5351; DI49_3051; YJR043C; SPGP0Qtrna7M; DI49_1363; SPGP0Vtrna8M; SPGP0AAtrna11M; SPGP0AGtrna20M; SPGP0AHtrna10M; SPGP0Dtrna12M; DI49_5024 | DI49_5351; DI49_3051; DI49_1363; DI49_5024 | SPGP0Ltrna3M; SPGP0Qtrna7M; SPGP0Vtrna8M; SPGP0AAtrna11M; SPGP0AGtrna20M; SPGP0AHtrna10M; SPGP0Dtrna12M | YJR043C |
| DI49_5350; YPL112C | DI49_5350 | | YPL112C |
| DI49_5349; YPL113C | DI49_5349 | | YPL113C |
| DI49_5348; YPL115C | DI49_5348 | | YPL115C |
| DI49_5347; YPL116W; DI49_5346 | DI49_5347; DI49_5346 | | YPL116W |
| DI49_5344; DI49_5345 | DI49_5344; DI49_5345 | | |
| DI49_5342; DI49_5343 | DI49_5342; DI49_5343 | | |
| DI49_5340; YPL123C; DI49_5339 | DI49_5340; DI49_5339 | | YPL123C |
| DI49_5337; YPL126W; DI49_5336 | DI49_5337; DI49_5336 | | YPL126W |
| SPGP0L02750; DI49_5335 | DI49_5335 | SPGP0L02750 | |
| DI49_5332; YPL131W; YPL130W | DI49_5332 | | YPL131W; YPL130W |
| DI49_5331; YPL132W | DI49_5331 | | YPL132W |
| DI49_5330; YPL133C | DI49_5330 | | YPL133C |
| DI49_5329; YPL134C; DI49_5328 | DI49_5329; DI49_5328 | | YPL134C |

Create PDF in your applications with the Pdfcrowd HTML to PDF API PDFCROWD

| | | | |
|---|---|---|---|
| DI49_5326; YPL138C; YPL137C | DI49_5326 | | YPL138C; YPL137C |
| DI49_5325; YPL139C; DI49_5324 | DI49_5325; DI49_5324 | | YPL139C |
| SPGP0L02870; DI49_5059; DI49_5321 | DI49_5059; DI49_5321 | SPGP0L02870 | |
| DI49_5319; YPL146C; DI49_5318 | DI49_5319; DI49_5318 | | YPL146C |
| DI49_5316; DI49_5317 | DI49_5316; DI49_5317 | | |
| DI49_5314; DI49_5315; YPL150W; YPL149W | DI49_5314; DI49_5315 | | YPL150W; YPL149W |
| DI49_5312; YPL153C; YPL152W | DI49_5312 | | YPL153C; YPL152W |
| DI49_5311; YPL154C; DI49_5310 | DI49_5311; DI49_5310 | | YPL154C |
| DI49_5308; DI49_5309 | DI49_5308; DI49_5309 | | |
| DI49_5306; DI49_5307 | DI49_5306; DI49_5307 | | |
| DI49_5304; DI49_5305 | DI49_5304; DI49_5305 | | |
| DI49_5302; DI49_5303 | DI49_5302; DI49_5303 | | |
| DI49_5300; DI49_5301; YPL164C; YPL163C | DI49_5300; DI49_5301 | | YPL164C; YPL163C |
| DI49_5298; YPL167C; DI49_5297; YPL166W | DI49_5298; DI49_5297 | | YPL167C; YPL166W |
| DI49_5295; DI49_5296; YPL169C; YPL168W | DI49_5295; DI49_5296 | | YPL169C; YPL168W |
| DI49_5293; YPL172C; SPGP0L03170; YPL171C | DI49_5293 | SPGP0L03170 | YPL172C; YPL171C |
| DI49_5291; YPL175W; DI49_5290; YPL174C | DI49_5291; DI49_5290 | | YPL175W; YPL174C |
| DI49_5288; YPL178W; YPL177C | DI49_5288 | | YPL178W; YPL177C |
| DI49_5286; YPL179W; DI49_5285 | DI49_5286; DI49_5285 | | YPL179W |
| DI49_5283; DI49_5284 | DI49_5283; DI49_5284 | | |
| DI49_5281; YPL184C; DI49_5280; YPL183W-A | DI49_5281; DI49_5280 | | YPL184C; YPL183W-A |
| DI49_5278; YPL188W; DI49_5277 | DI49_5278; DI49_5277 | | YPL188W |
| DI49_5275; YPL190C; YPL189C-A | DI49_5275 | | YPL190C; YPL189C-A |
| DI49_5274; YPL191C; DI49_5273 | DI49_5274; DI49_5273 | | YPL191C |
| DI49_5271; YPL194W; YPL193W | DI49_5271 | | YPL194W; YPL193W |
| SPGP0L03390; YPL195W | | SPGP0L03390 | YPL195W |
| DI49_5270; YPL196W; DI49_5268 | DI49_5270; DI49_5268 | | YPL196W |
| DI49_5266; YPL201C; DI49_5265; YPL200W | DI49_5266; DI49_5265 | | YPL201C; YPL200W |
| DI49_5263; YPL204W; DI49_5262 | DI49_5263; DI49_5262 | | YPL204W |
| DI49_5260; DI49_5261 | DI49_5260; DI49_5261 | | |
| DI49_5258; DI49_5259 | DI49_5258; DI49_5259 | | |
| DI49_5256; YPL212C; DI49_5255 | DI49_5256; DI49_5255 | | YPL212C |
| DI49_5253; DI49_5254; YPL214C; YPL213W | DI49_5253; DI49_5254 | | YPL214C; YPL213W |
| DI49_5251; YPL217C; YPL218W; YPL216W | DI49_5251 | | YPL217C; YPL218W; YPL216W |
| DI49_5249; DI49_1840 | DI49_5249; DI49_1840 | | |
| DI49_5248; YPL221W; DI49_5246 | DI49_5248; DI49_5246 | | YPL221W |
| DI49_5244; DI49_5245 | DI49_5244; DI49_5245 | | |
| DI49_5242; YPL226W | DI49_5242 | | YPL226W |
| DI49_5241; YPL227C | DI49_5241 | | YPL227C |

Create PDF in your applications with the Pdfcrowd HTML to PDF API PDFCROWD

| | | | |
|---|---|---|---|
| DI49_5240; YPL228W; DI49_5239 | DI49_5240; DI49_5239 | | YPL228W |
| DI49_5237; YPL231W; DI49_5236; YPL230W | DI49_5237; DI49_5236 | | YPL231W; YPL230W |
| DI49_5234; YPL234C | DI49_5234 | | YPL234C |
| DI49_5233; YPL235W; DI49_5232 | DI49_5233; DI49_5232 | | YPL235W |
| DI49_5230; YPL239W | DI49_5230 | | YPL239W |
| SPGP0L03810; DI49_4269 | DI49_4269 | SPGP0L03810 | |
| SPGP0L03830; YPL242C | | SPGP0L03830 | YPL242C |
| YGR087C; DI49_3657; YPL243W; YLR134W; YLR135W; DI49_3658 | DI49_3657; DI49_3658 | | YGR087C; YPL243W; YLR134W; YLR135W |
| YCR065W; SPGP0EB00100 | | SPGP0EB00100 | YCR065W |
| DI49_2825; DI49_2824 | DI49_2825; DI49_2824 | | |
| DI49_2827; DI49_2826 | DI49_2827; DI49_2826 | | |
| SPGP0M00150; DI49_2828; YJL207C; YJL208C | DI49_2828 | SPGP0M00150 | YJL207C; YJL208C |
| DI49_2829; YJL204C; DI49_2830; YJL205C | DI49_2829; DI49_2830 | | YJL204C; YJL205C |
| DI49_2832; YJL200C | DI49_2832 | | YJL200C |
| SPGP0Mtrna29T; DI49_2415; SPGP0Otrna14T; DI49_2254; SPGP0Otrna15T; DI49_2301; SPGP0Otrna16T; SPGP0O01690; SPGP0Strna25T; DI49_4465; SPGP0Wtrna7T; SPGP0W01480; SPGP0Btrna1T; DI49_0193; SPGP0ALtrna31T; DI49_4856; SPGP0Htrna32T; DI49_1043 | DI49_2415; DI49_2254; DI49_2301; DI49_4465; DI49_0193; DI49_4856; DI49_1043 | SPGP0Mtrna29T; SPGP0Otrna14T; SPGP0Otrna15T; SPGP0Otrna16T; SPGP0O01690; SPGP0Strna25T; SPGP0Wtrna7T; SPGP0W01480; SPGP0Btrna1T; SPGP0ALtrna31T; SPGP0Htrna32T | |
| DI49_2834; YJL198W; DI49_2835; YJL199C | DI49_2834; DI49_2835 | | YJL198W; YJL199C |
| DI49_2837; DI49_2836 | DI49_2837; DI49_2836 | | |
| DI49_2839; YJL192C | DI49_2839 | | YJL192C |
| DI49_2840; DI49_0783; DI49_0784; YCR032W | DI49_2840; DI49_0783; DI49_0784 | | YCR032W |
| DI49_2841; YJL190C | DI49_2841 | | YJL190C |
| DI49_2842; SPGP0N03080; DI49_2843 | DI49_2842; DI49_2843 | SPGP0N03080 | |
| DI49_2845; DI49_2844; YJL186W; YJL187C | DI49_2845; DI49_2844 | | YJL186W; YJL187C |
| DI49_2847; DI49_2846 | DI49_2847; DI49_2846 | | |
| DI49_2849; YJL180C; YJL181W | DI49_2849 | | YJL180C; YJL181W |
| DI49_2850; YJL179W; DI49_2851 | DI49_2850; DI49_2851 | | YJL179W |
| DI49_2853; YJL176C; DI49_2854 | DI49_2853; DI49_2854 | | YJL176C |
| DI49_2856; YJL172W; YJL173C | DI49_2856 | | YJL172W; YJL173C |
| DI49_2857; YJL171C; DI49_2858 | DI49_2857; DI49_2858 | | YJL171C |
| DI49_2860; YJL167W | DI49_2860 | | YJL167W |
| DI49_2861; YJL166W | DI49_2861 | | YJL166W |
| DI49_2862; YJL165C | DI49_2862 | | YJL165C |
| DI49_2863; YJL164C; DI49_2864 | DI49_2863; DI49_2864 | | YJL164C |
| SPGP0M00550; YJL159W; YJL160C | | SPGP0M00550 | YJL159W; YJL160C |
| DI49_2869; YJL158C; DI49_2870 | DI49_2869; DI49_2870 | | YJL158C |

Create PDF in your applications with the Pdfcrowd HTML to PDF API PDFCROWD

| | | | |
|---|---|---|---|
| DI49_2872; YJL155C; DI49_2873; YJL156C | DI49_2872; DI49_2873 | | YJL155C; YJL156C |
| DI49_2875; DI49_2874 | DI49_2875; DI49_2874 | | |
| DI49_2877; YJL148W; DI49_2878 | DI49_2877; DI49_2878 | | YJL148W |
| DI49_2880; YJL145W; DI49_2881 | DI49_2880; DI49_2881 | | YJL145W |
| DI49_2883; YJL141C; DI49_2473; DI49_2474 | DI49_2883; DI49_2473; DI49_2474 | | YJL141C |
| DI49_2884; YJL140W; DI49_2885 | DI49_2884; DI49_2885 | | YJL140W |
| DI49_2887; SPGP0M00730; DI49_3430; DI49_3429 | DI49_2887; DI49_3430; DI49_3429 | SPGP0M00730 | |
| DI49_2888; SPGP0M00750 | DI49_2888 | SPGP0M00750 | |
| DI49_2890; YJL133W | DI49_2890 | | YJL133W |
| SPGP0M00790; YJL132W | | SPGP0M00790 | YJL132W |
| DI49_2891; YJL131C | DI49_2891 | | YJL131C |
| DI49_2892; YJL130C | DI49_2892 | | YJL130C |
| DI49_2893; YJL129C | DI49_2893 | | YJL129C |
| DI49_2894; YJL128C | DI49_2894 | | YJL128C |
| DI49_2895; YJL127C-B; DI49_2896 | DI49_2895; DI49_2896 | | YJL127C-B |
| DI49_2898; YJL125C; DI49_2899; YJL126W | DI49_2898; DI49_2899 | | YJL125C; YJL126W |
| DI49_2900; SPGP0M00890 | DI49_2900 | SPGP0M00890 | |
| SPGP0M00920; DI49_2901 | DI49_2901 | SPGP0M00920 | |
| DI49_2903; DI49_2902 | DI49_2903; DI49_2902 | | |
| SPGP0M00960; DI49_2904 | DI49_2904 | SPGP0M00960 | |
| DI49_2908; YJL112W | DI49_2908 | | YJL112W |
| DI49_2909; YJL111W; DI49_2910 | DI49_2909; DI49_2910 | | YJL111W |
| DI49_2912; YJL107C | DI49_2912 | | YJL107C |
| DI49_2913; YJL106W; DI49_2914 | DI49_2913; DI49_2914 | | YJL106W |
| DI49_2916; DI49_2915; YJL104W; YJL105W | DI49_2916; DI49_2915 | | YJL104W; YJL105W |
| SPGP0Mtrna29R; DI49_2918; YJL102W | DI49_2918 | SPGP0Mtrna29R | YJL102W |
| DI49_2919; YJL101C; DI49_2920 | DI49_2919; DI49_2920 | | YJL101C |
| DI49_2922; YJL098W; DI49_2923; YJL099W | DI49_2922; DI49_2923 | | YJL098W; YJL099W |
| DI49_2925; YJL095W; DI49_2926; YJL096W | DI49_2925; DI49_2926 | | YJL095W; YJL096W |
| DI49_2928; YJL092W; YJL093C | DI49_2928 | | YJL092W; YJL093C |
| DI49_2929; YJL091C | DI49_2929 | | YJL091C |
| DI49_2930; YJL090C | DI49_2930 | | YJL090C |
| DI49_2931; YJL089W; DI49_2932 | DI49_2931; DI49_2932 | | YJL089W |
| DI49_2934; YJL085W | DI49_2934 | | YJL085W |
| SPGP0M01230; SPGP0N02150 | | SPGP0M01230; SPGP0N02150 | |
| DI49_2935; YJL083W | DI49_2935 | | YJL083W |
| DI49_2936; YJL082W; DI49_2937; YJL081C | DI49_2936; DI49_2937 | | YJL082W; YJL081C |
| DI49_2939; YJL079C; DI49_2940 | DI49_2939; DI49_2940 | | YJL079C |

Create PDF in your applications with the Pdfcrowd HTML to PDF API PDFCROWD

| | | | |
|---|---|---|---|
| DI49_2942; DI49_2941 | DI49_2942; DI49_2941 | | |
| DI49_2944; YJL072C; YJL073W | DI49_2944 | | YJL072C; YJL073W |
| DI49_2945; YJL071W | DI49_2945 | | YJL071W |
| DI49_2946; YJL070C; DI49_2947 | DI49_2946; DI49_2947 | | YJL070C |
| DI49_2949; DI49_2948 | DI49_2949; DI49_2948 | | |
| DI49_2951; DI49_2950 | DI49_2951; DI49_2950 | | |
| DI49_2953; YJL062W; DI49_2954 | DI49_2953; DI49_2954 | | YJL062W |
| DI49_2956; YJL059W | DI49_2956 | | YJL059W |
| DI49_2957; YJL058C; DI49_2958 | DI49_2957; DI49_2958 | | YJL058C |
| DI49_2960; DI49_2959 | DI49_2960; DI49_2959 | | |
| DI49_2962; DI49_2961 | DI49_2962; DI49_2961 | | |
| DI49_2964; YJL051W | DI49_2964 | | YJL051W |
| DI49_2965; YJL050W; DI49_2966 | DI49_2965; DI49_2966 | | YJL050W |
| DI49_2968; DI49_2967 | DI49_2968; DI49_2967 | | |
| SPGP0Mtrna14Y; DI49_2382; DI49_3070; DI49_2383; SPGP0Btrna1Y; DI49_1668; DI49_4369; SPGP0Ftrna12Y; DI49_4037 | DI49_2382; DI49_3070; DI49_2383; DI49_1668; DI49_4369; DI49_4037 | SPGP0Mtrna14Y; SPGP0Btrna1Y; SPGP0Ftrna12Y | |
| SPGP0Mtrna30R; DI49_4323; SPGP0Mtrna31R; DI49_2981; SPGP0Ptrna15R; DI49_3226; SPGP0Qtrna12R; DI49_1385; SPGP0Wtrna4R; SPGP0Etrna36R; SPGP0Etrna37R; DI49_2082; SPGP0Ftrna26R | DI49_4323; DI49_2981; DI49_3226; DI49_1385; DI49_2082 | SPGP0Mtrna30R; SPGP0Mtrna31R; SPGP0Ptrna15R; SPGP0Qtrna12R; SPGP0Wtrna4R; SPGP0Etrna36R; SPGP0Etrna37R; SPGP0Ftrna26R | |
| DI49_2975; DI49_2974 | DI49_2975; DI49_2974 | | |
| SPGP0M01630; DI49_2976 | DI49_2976 | SPGP0M01630 | |
| SPGP0M01670; DI49_1172; DI49_1171 | DI49_1172; DI49_1171 | SPGP0M01670 | |
| SPGP0Mtrna25V; DI49_4248; SPGP0Ntrna4V; YJL037W; SPGP0Ptrna8V; SPGP0Ptrna9V; DI49_3302; SPGP0Qtrna5V; DI49_1558; SPGP0Qtrna6V; DI49_1540; SPGP0Ztrna18V; SPGP0Ztrna19V; YKL069W; SPGP0AKtrna24V; YGL225W; SPGP0AMtrna35V; SPGP0Ctrna3V; DI49_3783; SPGP0Dtrna17V; YOR176W; SPGP0Etrna31V; DI49_2123; DI49_4247; SPGP0Ftrna22V; DI49_4138; YMR163C; SPGP0Gtrna28V; YMR049C; DI49_5011 | DI49_4248; DI49_3302; DI49_1558; DI49_1540; DI49_3783; DI49_2123; DI49_4247; DI49_4138; DI49_5011 | SPGP0Mtrna25V; SPGP0Ntrna4V; SPGP0Ptrna8V; SPGP0Ptrna9V; SPGP0Qtrna5V; SPGP0Qtrna6V; SPGP0Ztrna18V; SPGP0Ztrna19V; SPGP0AKtrna24V; SPGP0AMtrna35V; SPGP0Ctrna3V; SPGP0Dtrna17V; SPGP0Etrna31V; SPGP0Ftrna22V; SPGP0Gtrna28V | YJL037W; YKL069W; YGL225W; YOR176W; YMR163C; YMR049C |
| DI49_2984; YJL036W; DI49_2985 | DI49_2984; DI49_2985 | | YJL036W |
| DI49_2987; YJL033W; DI49_2988 | DI49_2987; DI49_2988 | | YJL033W |
| DI49_2990; DI49_2989 | DI49_2990; DI49_2989 | | |
| DI49_2993; YJL026W; DI49_2994 | DI49_2993; DI49_2994 | | YJL026W |
| DI49_2999; YJL019W; DI49_3000; YJL020C | DI49_2999; DI49_3000 | | YJL019W; YJL020C |
| DI49_3002; DI49_3001 | DI49_3002; DI49_3001 | | |
| DI49_3004; DI49_3003 | DI49_3004; DI49_3003 | | |

Create PDF in your applications with the Pdfcrowd HTML to PDF API PDFCROWD

| | | | |
|---|---|---|---|
| SPGP0Mtrna9W; DI49_3006; DI49_2157; DI49_2156; DI49_1855 | DI49_3006; DI49_2157; DI49_2156; DI49_1855 | SPGP0Mtrna9W | |
| DI49_3007; YJL010C | DI49_3007 | | YJL010C |
| DI49_3008; YJL008C | DI49_3008 | | YJL008C |
| SPGP0Mtrna15M; DI49_4243; DI49_3010; SPGP0Ftrna13M; DI49_4242 | DI49_4243; DI49_3010; DI49_4242 | SPGP0Mtrna15M; SPGP0Ftrna13M | |
| DI49_3012; YJL005W; DI49_3013; YJL006C | DI49_3012; DI49_3013 | | YJL005W; YJL006C |
| DI49_3015; DI49_3014 | DI49_3015; DI49_3014 | | |
| DI49_3016; SPGP0M01930 | DI49_3016 | SPGP0M01930 | |
| DI49_3018; DI49_3017 | DI49_3018; DI49_3017 | | |
| DI49_3020; YJR005W; DI49_3021 | DI49_3020; DI49_3021 | | YJR005W |
| DI49_3023; YJR007W; YJR006W | DI49_3023 | | YJR007W; YJR006W |
| DI49_3024; YJR008W | DI49_3024 | | YJR008W |
| DI49_3025; DI49_2159; YJR009C; YGR192C; YGR193C | DI49_3025; DI49_2159 | | YJR009C; YGR192C; YGR193C |
| DI49_3026; YJR010W; DI49_3027 | DI49_3026; DI49_3027 | | YJR010W |
| DI49_3029; DI49_3028; YJR011C; YJR010C-A | DI49_3029; DI49_3028 | | YJR011C; YJR010C-A |
| DI49_3031; YJR014W; SPGP0M02100 | DI49_3031 | SPGP0M02100 | YJR014W |
| DI49_3033; DI49_3032; YJR016C; YJR015W | DI49_3033; DI49_3032 | | YJR016C; YJR015W |
| SPGP0M02140; YJR021C; DI49_3035; YJR019C | DI49_3035 | SPGP0M02140 | YJR021C; YJR019C |
| DI49_3037; DI49_3036 | DI49_3037; DI49_3036 | | |
| DI49_3039; DI49_3038 | DI49_3039; DI49_3038 | | |
| DI49_3041; YJR033C; DI49_3042 | DI49_3041; DI49_3042 | | YJR033C |
| DI49_3044; YJR036C; YJR035W | DI49_3044 | | YJR036C; YJR035W |
| DI49_3045; YJR039W | DI49_3045 | | YJR039W |
| DI49_3046; YJR040W; DI49_3047 | DI49_3046; DI49_3047 | | YJR040W |
| DI49_3049; DI49_3048 | DI49_3049; DI49_3048 | | |
| DI49_3053; YJR046W | DI49_3053 | | YJR046W |
| SPGP0Mtrna37S; DI49_4419; SPGP0Qtrna14S; SPGP0Wtrna6S; SPGP0Btrna1S; SPGP0Btrna3S; DI49_0593; DI49_0694; SPGP0AItrna24S; SPGP0Ftrna34S; DI49_4865; SPGP0Htrna39S; SPGP0I02800 | DI49_4419; DI49_0593; DI49_0694; DI49_4865 | SPGP0Mtrna37S; SPGP0Qtrna14S; SPGP0Wtrna6S; SPGP0Btrna1S; SPGP0Btrna3S; SPGP0AItrna24S; SPGP0Ftrna34S; SPGP0Htrna39S; SPGP0I02800 | |
| DI49_3055; DI49_1358; DI49_3056; YJR047C; YEL034W | DI49_3055; DI49_1358; DI49_3056 | | YJR047C; YEL034W |
| DI49_3058; YJR050W; DI49_3059; YJR049C | DI49_3058; DI49_3059 | | YJR050W; YJR049C |
| DI49_3061; YJR052W; DI49_3062 | DI49_3061; DI49_3062 | | YJR052W |
| SPGP0Mtrna32R; DI49_3064; DI49_3065 | DI49_3064; DI49_3065 | SPGP0Mtrna32R | |
| DI49_3072; YJR059W; DI49_3073; YJR058C | DI49_3072; DI49_3073 | | YJR059W; YJR058C |
| DI49_3075; YJR062C; DI49_3076 | DI49_3075; DI49_3076 | | YJR062C |
| DI49_3078; YJR065C | DI49_3078 | | YJR065C |

Create PDF in your applications with the Pdfcrowd HTML to PDF API PDFCROWD

| | | | |
|---|---|---|---|
| DI49_3079; YJR066W; DI49_3080 | DI49_3079; DI49_3080 | | YJR066W |
| DI49_3082; YJR069C | DI49_3082 | | YJR069C |
| DI49_3083; YJR070C; DI49_3084 | DI49_3083; DI49_3084 | | YJR070C |
| SPGP0M02600; DI49_3085 | DI49_3085 | SPGP0M02600 | |
| DI49_3087; YJR076C | DI49_3087 | | YJR076C |
| DI49_3088; YJR077C; DI49_3089 | DI49_3088; DI49_3089 | | YJR077C |
| DI49_3091; DI49_3090 | DI49_3091; DI49_3090 | | |
| DI49_3093; DI49_3092 | DI49_3093; DI49_3092 | | |
| DI49_3095; DI49_3094 | DI49_3095; DI49_3094 | | |
| DI49_3097; DI49_3096 | DI49_3097; DI49_3096 | | |
| DI49_3099; YJR091C; DI49_3100 | DI49_3099; DI49_3100 | | YJR091C |
| DI49_3102; DI49_3101 | DI49_3102; DI49_3101 | | |
| DI49_3104; YJR095W; DI49_3105 | DI49_3104; DI49_3105 | | YJR095W |
| DI49_3107; YJR098C; DI49_3108; YJR097W | DI49_3107; DI49_3108 | | YJR098C; YJR097W |
| SPGP0Mtrna33L; DI49_3110; DI49_3111; YJR100C; SPGP0Ttrna29L; YLR227C; SPGP0Ctrna9L | DI49_3110; DI49_3111 | SPGP0Mtrna33L; SPGP0Ttrna29L; SPGP0Ctrna9L | YJR100C; YLR227C |
| DI49_3113; DI49_3112 | DI49_3113; DI49_3112 | | |
| DI49_3115; YJR105W; DI49_3116 | DI49_3115; DI49_3116 | | YJR105W |
| DI49_3118; DI49_3117; YJR107W; YJR106W | DI49_3118; DI49_3117 | | YJR107W; YJR106W |
| DI49_3119; YJR109C; SPGP0N00510 | DI49_3119 | SPGP0N00510 | YJR109C |
| DI49_3120; YJR110W | DI49_3120 | | YJR110W |
| DI49_3121; YJR111C; DI49_3122 | DI49_3121; DI49_3122 | | YJR111C |
| DI49_3124; DI49_3123 | DI49_3124; DI49_3123 | | |
| DI49_3126; DI49_3125 | DI49_3126; DI49_3125 | | |
| DI49_3128; YJR118C; DI49_3129 | DI49_3128; DI49_3129 | | YJR118C |
| DI49_3131; DI49_3130 | DI49_3131; DI49_3130 | | |
| DI49_3133; YJR124C | DI49_3133 | | YJR124C |
| DI49_3134; YJR125C; DI49_3135 | DI49_3134; DI49_3135 | | YJR125C |
| DI49_3137; DI49_3136 | DI49_3137; DI49_3136 | | |
| DI49_3139; DI49_3138 | DI49_3139; DI49_3138 | | |
| DI49_3141; YJR133W | DI49_3141 | | YJR133W |
| DI49_3142; YJR134C; DI49_3143 | DI49_3142; DI49_3143 | | YJR134C |
| DI49_3145; YJR136C; YJR135W-A | DI49_3145 | | YJR136C; YJR135W-A |
| DI49_3146; YJR137C | DI49_3146 | | YJR137C |
| DI49_3147; YJR138W | DI49_3147 | | YJR138W |
| DI49_3148; YJR139C; DI49_3149 | DI49_3148; DI49_3149 | | YJR139C |
| DI49_3151; DI49_3150 | DI49_3151; DI49_3150 | | |
| DI49_3153; DI49_3152 | DI49_3153; DI49_3152 | | |
| SPGP0M03300; DI49_2608; DI49_2609 | DI49_2608; DI49_2609 | SPGP0M03300 | |

Create PDF in your applications with the Pdfcrowd HTML to PDF API PDFCROWD

| | | | |
|---|---|---|---|
| DI49_3156; DI49_3155 | DI49_3156; DI49_3155 | | |
| DI49_3158; DI49_3157; SPGP0M03360; DI49_3160 | DI49_3158; DI49_3157; DI49_3160 | SPGP0M03360 | |
| SPGP0M03390; DI49_3161 | DI49_3161 | SPGP0M03390 | |
| DI49_3163; DI49_3162 | DI49_3163; DI49_3162 | | |
| DI49_3165; DI49_3164 | DI49_3165; DI49_3164 | | |
| SPGP0M03450; SPGP0AQ00540; SPGP0Q02660; DI49_1326; DI49_2619 | DI49_1326; DI49_2619 | SPGP0M03450; SPGP0AQ00540; SPGP0Q02660 | |
| DI49_3166; SPGP0BG00110; SPGP0M03470 | DI49_3166 | SPGP0BG00110; SPGP0M03470 | |
| YGR286C; SPGP0IL00100 | | SPGP0IL00100 | YGR286C |
| SPGP0EP00100; YHL007C; DI49_4842; YHL006C; DI49_4843 | DI49_4842; DI49_4843 | SPGP0EP00100 | YHL007C; YHL006C |
| SPGP0ER00100; YOL156W; SPGP0FF00100; YPR011C | | SPGP0ER00100; SPGP0FF00100 | YOL156W; YPR011C |
| SPGP0ES00100; DI49_2501 | DI49_2501 | SPGP0ES00100 | |
| SPGP0N00100; YHR211W; YJR150C; DI49_3472; SPGP0P00110; YFL051C; YHR213W-B; DI49_0077; SPGP0AP00560; SPGP0AP00570 | DI49_3472; DI49_0077 | SPGP0N00100; SPGP0P00110; SPGP0AP00560; SPGP0AP00570 | YHR211W; YJR150C; YFL051C; YHR213W-B |
| YJR148W; YJR149W | | | YJR148W; YJR149W |
| YJR145C; SPGP0AP00480; YJR144W; YHR204W | | SPGP0AP00480 | YJR145C; YJR144W; YHR204W |
| YJR142W; YJR143C | | | YJR142W; YJR143C |
| YJR140C; YJR141W | | | YJR140C; YJR141W |
| YJR132W; DI49_3140; YJR131W | DI49_3140 | | YJR132W; YJR131W |
| YJR129C; YJR130C | | | YJR129C; YJR130C |
| YJR126C; YJR127C | | | YJR126C; YJR127C |
| YJR123W; DI49_3132; YJR122W | DI49_3132 | | YJR123W; YJR122W |
| YJR119C; YJR121W | | | YJR119C; YJR121W |
| YJR117W; DI49_3127; YJR116W | DI49_3127 | | YJR117W; YJR116W |
| YJR113C; YJR115W | | | YJR113C; YJR115W |
| YJR112W; YJR112W-A | | | YJR112W; YJR112W-A |
| YJR104C; DI49_3114; YJR103W | DI49_3114 | | YJR104C; YJR103W |
| YJR101W; YJR102C | | | YJR101W; YJR102C |
| YJR094W-A; DI49_3103; YJR094C | DI49_3103 | | YJR094W-A; YJR094C |
| YJR092W; YJR093C | | | YJR092W; YJR093C |
| YJR090C; DI49_3098; YJR089W | DI49_3098 | | YJR090C; YJR089W |
| YJR086W; YJR088C | | | YJR086W; YJR088C |
| YJR084W; YJR085C | | | YJR084W; YJR085C |
| YJR082C; YJR083C | | | YJR082C; YJR083C |
| YJR078W; YJR080C | | | YJR078W; YJR080C |
| YJR075W; DI49_3086; YJR074W | DI49_3086 | | YJR075W; YJR074W |
| YJR072C; YJR073C | | | YJR072C; YJR073C |

Create PDF in your applications with the Pdfcrowd HTML to PDF API PDFCROWD

| | | | |
|---|---|---|---|
| YJR068W; DI49_3081; YJR067C | DI49_3081 | | YJR068W; YJR067C |
| YJR064W; DI49_3077; YJR063W | DI49_3077 | | YJR064W; YJR063W |
| YJR061W; DI49_3074; YJR060W | DI49_3074 | | YJR061W; YJR060W |
| SPGP0Ntrna5Y; SPGP0Dtrna11Y; YJR056C; SPGP0Atrna8Y; YOL020W | | SPGP0Ntrna5Y; SPGP0Dtrna11Y; SPGP0Atrna8Y | YJR056C; YOL020W |
| YJR054W; DI49_3063; YJR053W | DI49_3063 | | YJR054W; YJR053W |
| YJR045C; DI49_3052; YJR044C | DI49_3052 | | YJR045C; YJR044C |
| YJR041C; YJR042W | | | YJR041C; YJR042W |
| YJR032W; DI49_3040; YJR031C | DI49_3040 | | YJR032W; YJR031C |
| YJR025C; YJR030C | | | YJR025C; YJR030C |
| YJR013W; YJR012C | | | YJR013W; YJR012C |
| YJR004C; DI49_3019; YJR003C | DI49_3019 | | YJR004C; YJR003C |
| YJR001W; YJR002W | | | YJR001W; YJR002W |
| YJL002C; YJL001W | | | YJL002C; YJL001W |
| YJL004C; YJL003W | | | YJL004C; YJL003W |
| SPGP0Ntrna1W; SPGP0Gtrna10W; YGL119W; YKL072W; SPGP0AStrna6W; YPL257W | | SPGP0Ntrna1W; SPGP0Gtrna10W; SPGP0AStrna6W | YGL119W; YKL072W; YPL257W |
| SPGP0Ntrna9K; DI49_5468; YJL011C; SPGP0Ptrna15K; DI49_3250; DI49_1538; SPGP0Qtrna11K; SPGP0FXtrna35K; YDR444W; SPGP0Atrna19K; YDR363W; SPGP0Vtrna12K; SPGP0Vtrna13K; YER134C; SPGP0Wtrna4K; DI49_2750; SPGP0Ztrna27K; YKL128C; SPGP0ABtrna31K; SPGP0Btrna2K; DI49_0613; SPGP0AEtrna5K; YFR035C; SPGP0AHtrna23K; YHR150W; SPGP0BEtrna28K; DI49_1599; DI49_4194; YMR106C; SPGP0Gtrna36K; YMR319C | DI49_5468; DI49_3250; DI49_1538; DI49_2750; DI49_0613; DI49_1599; DI49_4194 | SPGP0Ntrna9K; SPGP0Ptrna15K; SPGP0Qtrna11K; SPGP0FXtrna35K; SPGP0Atrna19K; SPGP0Vtrna12K; SPGP0Vtrna13K; SPGP0Wtrna4K; SPGP0Ztrna27K; SPGP0ABtrna31K; SPGP0Btrna2K; SPGP0AEtrna5K; SPGP0AHtrna23K; SPGP0BEtrna28K; SPGP0Gtrna36K | YJL011C; YDR444W; YDR363W; YER134C; YKL128C; YFR035C; YHR150W; YMR106C; YMR319C |
| YJL013C; YJL012C | | | YJL013C; YJL012C |
| YJL016W; YJL014W | | | YJL016W; YJL014W |
| YJL024C; DI49_2995; YJL025W | DI49_2995 | | YJL024C; YJL025W |
| SPGP0Ntrna6M; YJL007C; YJL029C; SPGP0Atrna9M; YDR349C; SPGP0AUtrna2M; YMR158W | | SPGP0Ntrna6M; SPGP0Atrna9M; SPGP0AUtrna2M | YJL007C; YJL029C; YDR349C; YMR158W |
| YJL031C; YJL030W | | | YJL031C; YJL030W |
| YJL034W; DI49_2986; YJL035C | DI49_2986 | | YJL034W; YJL035C |
| YJL039C; DI49_2978; YJL041W | DI49_2978 | | YJL039C; YJL041W |
| YJL043W; YJL042W | | | YJL043W; YJL042W |
| SPGP0Ntrna6Y; SPGP0Gtrna17Y; YJL046W; SPGP0HYtrna2Y; YIL004C; SPGP0AE00400; SPGP0AEtrna4Y; YFR011C; YMR283C; SPGP0Gtrna16Y; YML054C | | SPGP0Ntrna6Y; SPGP0Gtrna17Y; SPGP0HYtrna2Y; SPGP0AE00400; SPGP0AEtrna4Y; SPGP0Gtrna16Y | YJL046W; YIL004C; YFR011C; YMR283C; YML054C |

Create PDF in your applications with the Pdfcrowd HTML to PDF API PDFCROWD

| | | | |
|---|---|---|---|
| YJL048C; YJL047C | | | YJL048C; YJL047C |
| YJL052W; DI49_2963; YJL053W | DI49_2963 | | YJL052W; YJL053W |
| YJL055W; YJL054W | | | YJL055W; YJL054W |
| YJL060W; DI49_2955; YJL061W | DI49_2955 | | YJL060W; YJL061W |
| YJL062W-A; DI49_2952; YJL063C | DI49_2952 | | YJL062W-A; YJL063C |
| YJL066C; YJL065C | | | YJL066C; YJL065C |
| YJL069C; YJL068C | | | YJL069C; YJL068C |
| YJL076W; YJL074C | | | YJL076W; YJL074C |
| YJL080C; DI49_2938 | DI49_2938 | | YJL080C |
| YJL087C; DI49_2933; YJL088W | DI49_2933 | | YJL087C; YJL088W |
| YJL109C; DI49_2911; YJL110C | DI49_2911 | | YJL109C; YJL110C |
| YJL117W; YJL116C | | | YJL117W; YJL116C |
| YJL121C; YJL118W | | | YJL121C; YJL118W |
| YJL123C; YJL122W | | | YJL123C; YJL122W |
| YJL133C-A; DI49_2889; YJL134W | DI49_2889 | | YJL133C-A; YJL134W |
| YJL137C; YJL136C | | | YJL137C; YJL136C |
| YJL139C; YJL138C; YKR059W; YKR058W | | | YJL139C; YJL138C; YKR059W; YKR058W |
| YJL143W; DI49_2882; YJL144W | DI49_2882 | | YJL143W; YJL144W |
| YJL146W; DI49_2879; YJL147C | DI49_2879 | | YJL146W; YJL147C |
| YJL149W; DI49_2876; YJL151C | DI49_2876 | | YJL149W; YJL151C |
| YJL154C; YJL153C | | | YJL154C; YJL153C |
| YJL162C; DI49_2865; YJL163C | DI49_2865 | | YJL162C; YJL163C |
| YJL168C; DI49_2859; YJL170C | DI49_2859 | | YJL168C; YJL170C |
| YJL177W; DI49_2852; YJL178C | DI49_2852 | | YJL177W; YJL178C |
| YJL184W; YJL183W | | | YJL184W; YJL183W |
| SPGP0N03100; SPGP0AH00930 | | SPGP0N03100; SPGP0AH00930 | |
| YJL193W; DI49_2838; YJL194W | DI49_2838 | | YJL193W; YJL194W |
| YJL197W; YJL196C | | | YJL197W; YJL196C |
| YJL201W; DI49_2831; YJL203W | DI49_2831 | | YJL201W; YJL203W |
| YJL210W; YJL209W | | | YJL210W; YJL209W |
| SPGP0N03290; YJL212C | | SPGP0N03290 | YJL212C |
| YJL214W; SPGP0O02530; YJL216C; YHR092C; YHR093W; YDR342C | | SPGP0O02530 | YJL214W; YJL216C; YHR092C; YHR093W; YDR342C |
| YJL218W; YJL217W; DI49_1320; DI49_1319 | DI49_1320; DI49_1319 | | YJL218W; YJL217W |
| YJL221C; YGR287C; SPGP0GV00100; YBR299W; SPGP0E00100 | | SPGP0GV00100; SPGP0E00100 | YJL221C; YGR287C; YBR299W |
| YOL158C; DI49_1601; YOL159C | DI49_1601 | | YOL158C; YOL159C |
| YHL046C; YDR542W; YEL049W; YEL048C; YHL044W; YGL258W-A; SPGP0BM00110; YCR086W | | SPGP0BM00110 | YHL046C; YDR542W; YEL049W; YEL048C; YHL044W; YGL258W-A; YCR086W |
| SPGP0EVtrna4S; DI49_2374; SPGP0EW00100; | DI49_2374; DI49_3831 | SPGP0EVtrna4S; SPGP0EW00100; SPGP0Atrna21S; | YDR339C; YLR327C |

Create PDF in your applications with the Pdfcrowd HTML to PDF API PDFCROWD

| | | | |
|---|---|---|---|
| SPGP0Atrna21S; YDR339C; SPGP0Ttrna36S; YLR327C; SPGP0AFtrna30S; DI49_3831; SPGP0Ctrna10S | | SPGP0Ttrna36S; SPGP0AFtrna30S; SPGP0Ctrna10S | |
| YIL007C; YCR090C | | | YIL007C; YCR090C |
| YBR298C; SPGP0AD00100 | | SPGP0AD00100 | YBR298C |
| SPGP0O00100; YGR289C; SPGP0E00120; YGR288W | | SPGP0O00100; SPGP0E00120 | YGR289C; YGR288W |
| DI49_2256; DI49_2255 | DI49_2256; DI49_2255 | | |
| DI49_2258; YOL151W; YOL152W | DI49_2258 | | YOL151W; YOL152W |
| DI49_2259; YOL149W | DI49_2259 | | YOL149W |
| DI49_2260; YOL148C | DI49_2260 | | YOL148C |
| DI49_2261; YOL147C; DI49_2262 | DI49_2261; DI49_2262 | | YOL147C |
| DI49_2264; DI49_2263 | DI49_2264; DI49_2263 | | |
| DI49_2266; YOL142W; DI49_2267 | DI49_2266; DI49_2267 | | YOL142W |
| DI49_2269; YOL139C; DI49_2270 | DI49_2269; DI49_2270 | | YOL139C |
| DI49_2272; YOL136C; DI49_2273; YOL137W | DI49_2272; DI49_2273 | | YOL136C; YOL137W |
| DI49_2275; YOL132W; DI49_2276; YOL133W | DI49_2275; DI49_2276 | | YOL132W; YOL133W |
| DI49_2278; YOL129W | DI49_2278 | | YOL129W |
| DI49_2279; YOL128C | DI49_2279 | | YOL128C |
| DI49_2280; YOL127W | DI49_2280 | | YOL127W |
| DI49_2281; YOL126C; DI49_2282 | DI49_2281; DI49_2282 | | YOL126C |
| DI49_2284; YOL123W | DI49_2284 | | YOL123W |
| DI49_2285; YOL122C | DI49_2285 | | YOL122C |
| DI49_2287; YOL119C; DI49_2288 | DI49_2287; DI49_2288 | | YOL119C |
| DI49_2290; YOL115W; YOL116W | DI49_2290 | | YOL115W; YOL116W |
| DI49_2291; YOL114C; DI49_2292 | DI49_2291; DI49_2292 | | YOL114C |
| DI49_2294; DI49_2293 | DI49_2294; DI49_2293 | | |
| DI49_2296; DI49_2295 | DI49_2296; DI49_2295 | | |
| DI49_2303; DI49_2302 | DI49_2303; DI49_2302 | | |
| DI49_2305; YOL101C | DI49_2305 | | YOL101C |
| DI49_2306; YOL100W | DI49_2306 | | YOL100W |
| DI49_2307; YOL098C | DI49_2307 | | YOL098C |
| DI49_2308; YOL097C | DI49_2308 | | YOL097C |
| DI49_2309; YOL096C | DI49_2309 | | YOL096C |
| DI49_2310; YOL095C | DI49_2310 | | YOL095C |
| DI49_2311; YOL094C; DI49_2312 | DI49_2311; DI49_2312 | | YOL094C |
| DI49_2314; YOL091W; YOL092W | DI49_2314 | | YOL091W; YOL092W |
| DI49_2315; YOL090W | DI49_2315 | | YOL090W |
| DI49_2316; YOL089C; DI49_2317 | DI49_2316; DI49_2317 | | YOL089C |
| DI49_2319; DI49_2318 | DI49_2319; DI49_2318 | | |

Create PDF in your applications with the Pdfcrowd HTML to PDF API PDFCROWD

| | | | |
|---|---|---|---|
| SPGP0O00770; DI49_2320 | DI49_2320 | SPGP0O00770 | |
| DI49_2322; DI49_2321 | DI49_2322; DI49_2321 | | |
| SPGP0O00810; DI49_2323 | DI49_2323 | SPGP0O00810 | |
| DI49_2325; DI49_2324 | DI49_2325; DI49_2324 | | |
| DI49_2327; DI49_2326; YOL078W; YOL080C | DI49_2327; DI49_2326 | | YOL078W; YOL080C |
| DI49_2329; YOL076W; YOL077C | DI49_2329 | | YOL076W; YOL077C |
| DI49_2330; YOL075C; DI49_2331 | DI49_2330; DI49_2331 | | YOL075C |
| DI49_2333; DI49_2332; YOL072W; YOL073C | DI49_2333; DI49_2332 | | YOL072W; YOL073C |
| DI49_2335; YOL069W; YOL070C | DI49_2335 | | YOL069W; YOL070C |
| DI49_2336; YOL068C; DI49_2337 | DI49_2336; DI49_2337 | | YOL068C |
| DI49_2339; DI49_2338 | DI49_2339; DI49_2338 | | |
| DI49_2341; DI49_2340 | DI49_2341; DI49_2340 | | |
| DI49_2343; YOL061W | DI49_2343 | | YOL061W |
| DI49_2344; YOL060C | DI49_2344 | | YOL060C |
| DI49_2345; YOL059W | DI49_2345 | | YOL059W |
| DI49_2346; YOL058W; DI49_2347 | DI49_2346; DI49_2347 | | YOL058W |
| DI49_2348; YOL055C; YOL056W | DI49_2348 | | YOL055C; YOL056W |
| DI49_2353; DI49_2352 | DI49_2353; DI49_2352 | | |
| DI49_2355; YOL051W; DI49_2356 | DI49_2355; DI49_2356 | | YOL051W |
| SPGP0O01150; SPGP0O01140 | | SPGP0O01150; SPGP0O01140 | |
| DI49_2358; DI49_2357 | DI49_2358; DI49_2357 | | |
| DI49_2360; DI49_2359 | DI49_2360; DI49_2359 | | |
| DI49_2362; YOL040C; YOL041C | DI49_2362 | | YOL040C; YOL041C |
| DI49_2363; YOL039W; DI49_2364; DI49_0986 | DI49_2363; DI49_2364; DI49_0986 | | YOL039W |
| DI49_2366; YOL034W; DI49_2367; YOL036W | DI49_2366; DI49_2367 | | YOL034W; YOL036W |
| DI49_2369; DI49_2368 | DI49_2369; DI49_2368 | | |
| DI49_2371; DI49_2370 | DI49_2371; DI49_2370 | | |
| DI49_2373; DI49_2372; YOL027C; YOL028C | DI49_2373; DI49_2372 | | YOL027C; YOL028C |
| DI49_2375; SPGP0Otrna17S | DI49_2375 | SPGP0Otrna17S | |
| DI49_2377; YOL023W; DI49_2378; YOL024W | DI49_2377; DI49_2378 | | YOL023W; YOL024W |
| DI49_2380; YOL021C; DI49_2381 | DI49_2380; DI49_2381 | | YOL021C |
| DI49_2385; DI49_2384 | DI49_2385; DI49_2384 | | |
| DI49_2387; DI49_2386; YOL016C; YOL017W | DI49_2387; DI49_2386 | | YOL016C; YOL017W |
| DI49_2389; DI49_4051 | DI49_2389; DI49_4051 | | |
| DI49_2390; YOL013C | DI49_2390 | | YOL013C |
| DI49_2391; YOL012C | DI49_2391 | | YOL012C |
| DI49_2392; YOL011W; DI49_2393 | DI49_2392; DI49_2393 | | YOL011W |
| DI49_2395; DI49_2394; YOL009C; YOL010W | DI49_2395; DI49_2394 | | YOL009C; YOL010W |
| DI49_2397; YOL006C; DI49_2398; YOL007C | DI49_2397; DI49_2398 | | YOL006C; YOL007C |

Create PDF in your applications with the Pdfcrowd HTML to PDF API PDFCROWD

| | | | |
|---|---|---|---|
| DI49_2400; YOL003C | DI49_2400 | | YOL003C |
| DI49_2401; YOL002C; DI49_2402 | DI49_2401; DI49_2402 | | YOL002C |
| DI49_2404; YOR002W | DI49_2404 | | YOR002W |
| DI49_2405; YOR003W; DI49_2406 | DI49_2405; DI49_2406 | | YOR003W |
| DI49_2408; YOR006C | DI49_2408 | | YOR006C |
| DI49_2409; YOR007C | DI49_2409 | | YOR007C |
| SPGP0Otrna16R; DI49_2410; DI49_2411; DI49_1570; SPGP0Atrna18R; SPGP0Utrna22R; SPGP0Btrna2R; YER161C; SPGP0Ctrna5R | DI49_2410; DI49_2411; DI49_1570 | SPGP0Otrna16R; SPGP0Atrna18R; SPGP0Utrna22R; SPGP0Btrna2R; SPGP0Ctrna5R | YER161C |
| DI49_2413; DI49_1406; DI49_3279; DI49_3278; DI49_1405 | DI49_2413; DI49_1406; DI49_3279; DI49_3278; DI49_1405 | | |
| DI49_2414; SPGP0D03030 | DI49_2414 | SPGP0D03030 | |
| DI49_2417; YOR014W; YOR013W | DI49_2417 | | YOR014W; YOR013W |
| DI49_2418; YOR016C | DI49_2418 | | YOR016C |
| DI49_2419; YOR017W; DI49_2420 | DI49_2419; DI49_2420 | | YOR017W |
| DI49_2422; YHR015W; DI49_2423 | DI49_2422; DI49_2423 | | YHR015W |
| DI49_2425; YHR018C | DI49_2425 | | YHR018C |
| DI49_2426; YHR019C | DI49_2426 | | YHR019C |
| DI49_2427; YHR020W | DI49_2427 | | YHR020W |
| SPGP0O01810; YHR021C; YHR022C | | SPGP0O01810 | YHR021C; YHR022C |
| DI49_2429; YHR023W; DI49_2430; YHR024C | DI49_2429; DI49_2430 | | YHR023W; YHR024C |
| DI49_2432; YHR026W | DI49_2432 | | YHR026W |
| DI49_2433; YHR027C | DI49_2433 | | YHR027C |
| DI49_2434; YHR028C; YHR029C | DI49_2434 | | YHR028C; YHR029C |
| DI49_2435; DI49_4016 | DI49_2435; DI49_4016 | | |
| DI49_2436; YHR030C; YHR031C | DI49_2436 | | YHR030C; YHR031C |
| DI49_2437; DI49_4030; DI49_4029 | DI49_2437; DI49_4030; DI49_4029 | | |
| DI49_2438; YHR032W | DI49_2438 | | YHR032W |
| DI49_2439; YHR033W; DI49_2440; YHR034C | DI49_2439; DI49_2440 | | YHR033W; YHR034C |
| DI49_2442; YHR036W | DI49_2442 | | YHR036W |
| DI49_2443; YHR037W; DI49_2444; YHR038W | DI49_2443; DI49_2444 | | YHR037W; YHR038W |
| SPGP0O01990; YHR039C-A | | SPGP0O01990 | YHR039C-A |
| DI49_2446; YHR040W; DI49_2447; YHR041C | DI49_2446; DI49_2447 | | YHR040W; YHR041C |
| DI49_2449; DI49_2448 | DI49_2449; DI49_2448 | | |
| DI49_2451; DI49_2450 | DI49_2451; DI49_2450 | | |
| DI49_2453; DI49_2452 | DI49_2453; DI49_2452 | | |
| DI49_2455; DI49_2454 | DI49_2455; DI49_2454 | | |
| DI49_2457; YHR052W; DI49_2458; YHR055C | DI49_2457; DI49_2458 | | YHR052W; YHR055C |
| DI49_2460; DI49_2459 | DI49_2460; DI49_2459 | | |

Create PDF in your applications with the Pdfcrowd HTML to PDF API PDFCROWD

| | | | |
|---|---|---|---|
| DI49_2462; YHR059W; DI49_2463; YHR060W | DI49_2462; DI49_2463 | | YHR059W; YHR060W |
| DI49_2465; DI49_2464 | DI49_2465; DI49_2464 | | |
| DI49_2467; YHR064C | DI49_2467 | | YHR064C |
| DI49_2468; YHR065C | DI49_2468 | | YHR065C |
| DI49_2469; DI49_0490; DI49_2470; DI49_0491 | DI49_2469; DI49_0490; DI49_2470; DI49_0491 | | |
| DI49_2472; YHR069C; YHR070W | DI49_2472 | | YHR069C; YHR070W |
| DI49_2476; DI49_2475; DI49_4603; DI49_4602 | DI49_2476; DI49_2475; DI49_4603; DI49_4602 | | |
| DI49_2478; DI49_2477 | DI49_2478; DI49_2477 | | |
| DI49_2480; YHR075C; DI49_2481; YHR076W | DI49_2480; DI49_2481 | | YHR075C; YHR076W |
| DI49_2483; YHR077C; DI49_2484; YHR078W | DI49_2483; DI49_2484 | | YHR077C; YHR078W |
| DI49_2486; DI49_2485 | DI49_2486; DI49_2485 | | |
| DI49_2488; DI49_2487 | DI49_2488; DI49_2487 | | |
| DI49_2490; DI49_2489 | DI49_2490; DI49_2489 | | |
| DI49_2492; DI49_2491 | DI49_2492; DI49_2491 | | |
| DI49_2494; YHR087W; DI49_2495; YHR088W | DI49_2494; DI49_2495 | | YHR087W; YHR088W |
| DI49_2497; YHR090C; DI49_2498; YHR091C | DI49_2497; DI49_2498 | | YHR090C; YHR091C |
| DI49_2500; YHR094C; DI49_2502; YHR096C; YHR097C | DI49_2500; DI49_2502 | | YHR094C; YHR096C; YHR097C |
| DI49_2504; YHR099W; DI49_2505; YHR100C | DI49_2504; DI49_2505 | | YHR099W; YHR100C |
| DI49_2507; YHR102W; DI49_2508; YHR103W | DI49_2507; DI49_2508 | | YHR102W; YHR103W |
| DI49_2510; DI49_2509 | DI49_2510; DI49_2509 | | |
| DI49_2512; YHR107C; DI49_2513; YHR108W | DI49_2512; DI49_2513 | | YHR107C; YHR108W |
| DI49_2515; DI49_2514 | DI49_2515; DI49_2514 | | |
| DI49_2517; DI49_2516 | DI49_2517; DI49_2516 | | |
| DI49_2519; YHR114W; DI49_2520; YHR115C | DI49_2519; DI49_2520 | | YHR114W; YHR115C |
| DI49_2522; DI49_2521 | DI49_2522; DI49_2521 | | |
| DI49_2524; YHR119W; DI49_2525; YHR120W | DI49_2524; DI49_2525 | | YHR119W; YHR120W |
| DI49_2527; DI49_2526 | DI49_2527; DI49_2526 | | |
| DI49_2528; YHR124W | DI49_2528 | | YHR124W |
| SPGP0Otrna10F; DI49_2529; DI49_2530; DI49_1938 | DI49_2529; DI49_2530; DI49_1938 | SPGP0Otrna10F | |
| DI49_2532; YHR128W | DI49_2532 | | YHR128W |
| DI49_2533; YHR129C; DI49_2534; YHR131C | DI49_2533; DI49_2534 | | YHR129C; YHR131C |
| DI49_2536; YHR132W-A; DI49_2537; YHR133C | DI49_2536; DI49_2537 | | YHR132W-A; YHR133C |
| DI49_2539; DI49_2538 | DI49_2539; DI49_2538 | | |
| SPGP0O02950; DI49_2540 | DI49_2540 | SPGP0O02950 | |
| DI49_2542; DI49_2541 | DI49_2542; DI49_2541 | | |
| DI49_2544; YHR142W; DI49_2545; YHR143W | DI49_2544; DI49_2545 | | YHR142W; YHR143W |

Create PDF in your applications with the Pdfcrowd HTML to PDF API PDFCROWD

| | | | |
|---|---|---|---|
| DI49_2547; DI49_2546 | DI49_2547; DI49_2546 | | |
| DI49_2549; DI49_2548 | DI49_2549; DI49_2548 | | |
| DI49_2551; YHR148W | DI49_2551 | | YHR148W |
| DI49_2552; YHR149C; DI49_2553; SPGP0IP00100 | DI49_2552; DI49_2553 | SPGP0IP00100 | YHR149C |
| DI49_2555; DI49_2554 | DI49_2555; DI49_2554 | | |
| DI49_2557; YHR154W | DI49_2557 | | YHR154W |
| DI49_2558; YHR155W; DI49_2559; SPGP0AW00160 | DI49_2558; DI49_2559 | SPGP0AW00160 | YHR155W |
| DI49_2561; DI49_2560 | DI49_2561; DI49_2560 | | |
| DI49_2563; DI49_2562 | DI49_2563; DI49_2562 | | |
| DI49_2565; DI49_2564 | DI49_2565; DI49_2564 | | |
| DI49_2567; YHR164C | DI49_2567 | | YHR164C |
| DI49_2569; YHR165C | DI49_2569 | | YHR165C |
| SPGP0FDtrna1A; DI49_3564; SPGP0Qtrna8A; SPGP0GLtrna2A; YNR032C-A; SPGP0Utrna18A; YLR036C; SPGP0Vtrna9A; YER076C; SPGP0Ytrna17A; SPGP0AItrna15A; SPGP0Ctrna3A; DI49_3565; SPGP0Dtrna22A; YOR289W; SPGP0Etrna34A; DI49_2114 | DI49_3564; DI49_3565; DI49_2114 | SPGP0FDtrna1A; SPGP0Qtrna8A; SPGP0GLtrna2A; SPGP0Utrna18A; SPGP0Vtrna9A; SPGP0Ytrna17A; SPGP0AItrna15A; SPGP0Ctrna3A; SPGP0Dtrna22A; SPGP0Etrna34A | YNR032C-A; YLR036C; YER076C; YOR289W |
| YKL221W; SPGP0JI00100 | | SPGP0JI00100 | YKL221W |
| DI49_3470; YKR100C | DI49_3470 | | YKR100C |
| DI49_3469; YKR099W; DI49_3468; YKR098C | DI49_3469; DI49_3468 | | YKR099W; YKR098C |
| DI49_3466; DI49_3467 | DI49_3466; DI49_3467 | | |
| DI49_3464; YKR095W | DI49_3464 | | YKR095W |
| SPGP0P00200; YKR094C | | SPGP0P00200 | YKR094C |
| DI49_3463; YKR093W; DI49_3462; YKR092C; YKR091W | DI49_3463; DI49_3462 | | YKR093W; YKR092C; YKR091W |
| DI49_3460; DI49_3461 | DI49_3460; DI49_3461 | | |
| DI49_3458; DI49_3459 | DI49_3458; DI49_3459 | | |
| DI49_3456; YKR086W; DI49_3455; YKR085C | DI49_3456; DI49_3455 | | YKR086W; YKR085C |
| DI49_3453; DI49_3454 | DI49_3453; DI49_3454 | | |
| DI49_3451; YKR081C | DI49_3451 | | YKR081C |
| DI49_3450; YKR080W; DI49_3449; YKR079C | DI49_3450; DI49_3449 | | YKR080W; YKR079C |
| DI49_3447; DI49_3448 | DI49_3447; DI49_3448 | | |
| DI49_3445; DI49_3446 | DI49_3445; DI49_3446 | | |
| SPGP0Ptrna14K; DI49_5564; DI49_3442; SPGP0Ctrna6K; SPGP0Htrna38K; DI49_4901 | DI49_5564; DI49_3442; DI49_4901 | SPGP0Ptrna14K; SPGP0Ctrna6K; SPGP0Htrna38K | |
| DI49_3440; YKR070W | DI49_3440 | | YKR070W |
| DI49_3439; YKR069W; DI49_3438; YKR068C | DI49_3439; DI49_3438 | | YKR069W; YKR068C |
| DI49_3436; DI49_3437 | DI49_3436; DI49_3437 | | |
| SPGP0P00490; YKR064W | | SPGP0P00490 | YKR064W |

Create PDF in your applications with the Pdfcrowd HTML to PDF API PDFCROWD

| | | | |
|---|---|---|---|
| DI49_3434; YKR063C | DI49_3434 | | YKR063C |
| DI49_3433; YKR062W | DI49_3433 | | YKR062W |
| DI49_3432; YKR061W | DI49_3432 | | YKR061W |
| DI49_3431; YKR060W | DI49_3431 | | YKR060W |
| DI49_3427; DI49_3428 | DI49_3427; DI49_3428 | | |
| DI49_3425; YKR054C; DI49_3424; YKR053C | DI49_3425; DI49_3424 | | YKR054C; YKR053C |
| DI49_3422; DI49_3423 | DI49_3422; DI49_3423 | | |
| DI49_3420; DI49_3421 | DI49_3420; DI49_3421 | | |
| DI49_3418; YKR046C; DI49_3417; YKR045C | DI49_3418; DI49_3417 | | YKR046C; YKR045C |
| DI49_3415; YKR043C; DI49_3414; YKR042W | DI49_3415; DI49_3414 | | YKR043C; YKR042W |
| DI49_3411; YKR039W | DI49_3411 | | YKR039W |
| DI49_3407; DI49_3408 | DI49_3407; DI49_3408 | | |
| DI49_3405; YKR034W | DI49_3405 | | YKR034W |
| DI49_3404; YKR031C; DI49_3403; YKR030W | DI49_3404; DI49_3403 | | YKR031C; YKR030W |
| DI49_3401; YKR028W; DI49_3400; YKR027W | DI49_3401; DI49_3400 | | YKR028W; YKR027W |
| DI49_3398; YKR026C; DI49_3397; YKR025W | DI49_3398; DI49_3397 | | YKR026C; YKR025W |
| DI49_3395; YKR023W; DI49_3394; YKR022C | DI49_3395; DI49_3394 | | YKR023W; YKR022C |
| DI49_3392; DI49_3393 | DI49_3392; DI49_3393 | | |
| DI49_3390; YKR018C; DI49_3389; YKR017C | DI49_3390; DI49_3389 | | YKR018C; YKR017C |
| DI49_3387; SPGP0AD00940; YKR015C | DI49_3387 | SPGP0AD00940 | YKR015C |
| DI49_3386; YKR014C; DI49_3385; YKR013W | DI49_3386; DI49_3385 | | YKR014C; YKR013W |
| DI49_3383; YKR010C | DI49_3383 | | YKR010C |
| SPGP0Ptrna14L; DI49_5686; DI49_3381; SPGP0AGtrna45L; SPGP0Btrna3L; SPGP0NZ00100; SPGP0Ctrna10L; DI49_2147 | DI49_5686; DI49_3381; DI49_2147 | SPGP0Ptrna14L; SPGP0AGtrna45L; SPGP0Btrna3L; SPGP0NZ00100; SPGP0Ctrna10L | |
| DI49_3379; DI49_3380 | DI49_3379; DI49_3380 | | |
| DI49_3377; DI49_3378 | DI49_3377; DI49_3378 | | |
| DI49_3375; DI49_3376 | DI49_3375; DI49_3376 | | |
| DI49_3373; YKR001C | DI49_3373 | | YKR001C |
| DI49_3372; YKL001C | DI49_3372 | | YKL001C |
| DI49_3371; YKL002W; SPGP0P01100; YKL003C | DI49_3371 | SPGP0P01100 | YKL002W; YKL003C |
| DI49_3369; YKL004W; DI49_3368; YKL005C | DI49_3369; DI49_3368 | | YKL004W; YKL005C |
| DI49_3366; YKL006C-A; DI49_3365; YKL007W | DI49_3366; DI49_3365 | | YKL006C-A; YKL007W |
| DI49_3364; SPGP0P01170 | DI49_3364 | SPGP0P01170 | |
| DI49_3362; YKL010C; DI49_3361; YKL011C | DI49_3362; DI49_3361 | | YKL010C; YKL011C |
| DI49_3359; DI49_3360 | DI49_3359; DI49_3360 | | |
| DI49_3357; YKL015W | DI49_3357 | | YKL015W |
| DI49_3356; YKL016C; DI49_3355; YKL017C | DI49_3356; DI49_3355 | | YKL016C; YKL017C |
| DI49_3353; YKL018C-A; DI49_3352; YKL019W | DI49_3353; DI49_3352 | | YKL018C-A; YKL019W |
| DI49_3350; YKL021C | DI49_3350 | | YKL021C |

Create PDF in your applications with the Pdfcrowd HTML to PDF API PDFCROWD

| | | | |
|---|---|---|---|
| DI49_3349; YKL022C; DI49_3348; YPL055C | DI49_3349; DI49_3348 | | YKL022C; YPL055C |
| DI49_3346; YKL024C; YKL023C-A | DI49_3346 | | YKL024C; YKL023C-A |
| DI49_3345; YKL025C; SPGP0P01380; YKL026C | DI49_3345 | SPGP0P01380 | YKL025C; YKL026C |
| DI49_3343; YKL027W | DI49_3343 | | YKL027W |
| DI49_3342; YKL028W; DI49_3341; YKL029C | DI49_3342; DI49_3341 | | YKL028W; YKL029C |
| DI49_3338; YKL033W; DI49_3337; YKL033W-A | DI49_3338; DI49_3337 | | YKL033W; YKL033W-A |
| DI49_3336; YKL034W | DI49_3336 | | YKL034W |
| DI49_3335; YKL035W; DI49_3334; YKL038W | DI49_3335; DI49_3334 | | YKL035W; YKL038W |
| DI49_3332; YKL040C; DI49_3331; YKL041W | DI49_3332; DI49_3331 | | YKL040C; YKL041W |
| DI49_3329; DI49_3330 | DI49_3329; DI49_3330 | | |
| DI49_3327; YKL046C | DI49_3327 | | YKL046C |
| DI49_3326; YKL047W; DI49_3325; YKL048C | DI49_3326; DI49_3325 | | YKL047W; YKL048C |
| DI49_3324; SPGP0P01590 | DI49_3324 | SPGP0P01590 | |
| DI49_3322; DI49_3323 | DI49_3322; DI49_3323 | | |
| DI49_3320; YKL054C; DI49_3319; YKL055C | DI49_3320; DI49_3319 | | YKL054C; YKL055C |
| DI49_3317; YKL057C | DI49_3317 | | YKL057C |
| DI49_3316; YKL058W; DI49_3315; YKL059C | DI49_3316; DI49_3315 | | YKL058W; YKL059C |
| DI49_3313; DI49_3314 | DI49_3313; DI49_3314 | | |
| DI49_3311; DI49_3312 | DI49_3311; DI49_3312 | | |
| DI49_3309; DI49_3310 | DI49_3309; DI49_3310 | | |
| DI49_3307; DI49_3308 | DI49_3307; DI49_3308 | | |
| DI49_3305; DI49_4134; SPGP0Q00630; SPGP0Ftrna9H; DI49_4133; YMR044W; YMR043W; SPGP0Etrna14H; DI49_4826 | DI49_3305; DI49_4134; DI49_4133; DI49_4826 | SPGP0Q00630; SPGP0Ftrna9H; SPGP0Etrna14H | YMR044W; YMR043W |
| DI49_3300; DI49_5211; SPGP0Ptrna3W | DI49_3300; DI49_5211 | SPGP0Ptrna3W | |
| DI49_3298; DI49_3299 | DI49_3298; DI49_3299 | | |
| DI49_3296; DI49_3297 | DI49_3296; DI49_3297 | | |
| DI49_3294; DI49_3295 | DI49_3294; DI49_3295 | | |
| SPGP0P01890; DI49_3293 | DI49_3293 | SPGP0P01890 | |
| DI49_3291; DI49_3292 | DI49_3291; DI49_3292 | | |
| DI49_3289; YKL086W; DI49_3288; YKL087C | DI49_3289; DI49_3288 | | YKL086W; YKL087C |
| DI49_3286; YKL089W; DI49_3285; YKL090W | DI49_3286; DI49_3285 | | YKL089W; YKL090W |
| DI49_3283; YKL092C; DI49_3282; YKL093W | DI49_3283; DI49_3282 | | YKL092C; YKL093W |
| DI49_3280; YKL095W; YKL096W | DI49_3280 | | YKL095W; YKL096W |
| DI49_3276; YKL099C; DI49_3275; YKL100C | DI49_3276; DI49_3275 | | YKL099C; YKL100C |
| DI49_3273; YKL103C | DI49_3273 | | YKL103C |
| DI49_3272; YKL104C; DI49_3271; YKL105C | DI49_3272; DI49_3271 | | YKL104C; YKL105C |
| DI49_3269; DI49_3270 | DI49_3269; DI49_3270 | | |
| DI49_3267; DI49_3268 | DI49_3267; DI49_3268 | | |
| DI49_3265; YKL112W | DI49_3265 | | YKL112W |
| DI49_3264; YKL113C | DI49_3264 | | YKL113C |

Create PDF in your applications with the Pdfcrowd HTML to PDF API PDFCROWD

| | | | |
|---|---|---|---|
| DI49_3263; YKL114C | DI49_3263 | | YKL114C |
| DI49_3262; YKL116C; DI49_3261; YKL117W | DI49_3262; DI49_3261 | | YKL116C; YKL117W |
| DI49_3257; YKL121W; DI49_3256; YKL122C | DI49_3257; DI49_3256 | | YKL121W; YKL122C |
| DI49_3254; YKL125W; DI49_3253; YKL126W | DI49_3254; DI49_3253 | | YKL125W; YKL126W |
| DI49_3248; DI49_3249 | DI49_3248; DI49_3249 | | |
| DI49_3246; DI49_3247 | DI49_3246; DI49_3247 | | |
| DI49_3244; DI49_3245 | DI49_3244; DI49_3245 | | |
| DI49_3242; YKL138C; DI49_3241; YKL138C-A | DI49_3242; DI49_3241 | | YKL138C; YKL138C-A |
| DI49_3239; DI49_3240 | DI49_3239; DI49_3240 | | |
| DI49_3237; YKL142W | DI49_3237 | | YKL142W |
| DI49_3236; YKL143W; DI49_3235; YKL144C | DI49_3236; DI49_3235 | | YKL143W; YKL144C |
| DI49_3233; YKL146W | DI49_3233 | | YKL146W |
| DI49_3232; YKL148C; DI49_3231; YKL149C | DI49_3232; DI49_3231 | | YKL148C; YKL149C |
| DI49_3229; DI49_3230 | DI49_3229; DI49_3230 | | |
| DI49_3224; YKL156W | DI49_3224 | | YKL156W |
| SPGP0P02560; YKL157W; DI49_3223; YKL159C | DI49_3223 | SPGP0P02560 | YKL157W; YKL159C |
| DI49_3221; DI49_3222 | DI49_3221; DI49_3222 | | |
| DI49_3219; DI49_3220 | DI49_3219; DI49_3220 | | |
| DI49_3215; YKL165C | DI49_3215 | | YKL165C |
| DI49_3214; YKL166C; DI49_3213; YKL167C | DI49_3214; DI49_3213 | | YKL166C; YKL167C |
| DI49_3211; DI49_3212 | DI49_3211; DI49_3212 | | |
| DI49_3209; DI49_3210 | DI49_3209; DI49_3210 | | |
| DI49_3207; DI49_3208 | DI49_3207; DI49_3208 | | |
| DI49_3205; DI49_3206 | DI49_3205; DI49_3206 | | |
| DI49_3203; YKL179C | DI49_3203 | | YKL179C |
| DI49_3202; YKL180W | DI49_3202 | | YKL180W |
| DI49_3201; YKL181W | DI49_3201 | | YKL181W |
| DI49_3200; YKL182W | DI49_3200 | | YKL182W |
| DI49_3199; YKL183W; DI49_3198; YKL184W | DI49_3199; DI49_3198 | | YKL183W; YKL184W |
| DI49_3196; DI49_3197 | DI49_3196; DI49_3197 | | |
| DI49_3194; DI49_3195 | DI49_3194; DI49_3195 | | |
| DI49_3189; DI49_3190 | DI49_3189; DI49_3190 | | |
| DI49_3187; YKL194C; SPGP0P02910; YKL195W | DI49_3187 | SPGP0P02910 | YKL194C; YKL195W |
| DI49_3184; YKL197C; DI49_3183; YKL198C | DI49_3184; DI49_3183 | | YKL197C; YKL198C |
| DI49_3181; YKL203C | DI49_3181 | | YKL203C |
| DI49_3180; YKL204W | DI49_3180 | | YKL204W |
| SPGP0P02980; YKL205W; DI49_3179; YKL206C | DI49_3179 | SPGP0P02980 | YKL205W; YKL206C |
| DI49_3177; DI49_3178 | DI49_3177; DI49_3178 | | |
| DI49_3175; YKL209C | DI49_3175 | | YKL209C |

Create PDF in your applications with the Pdfcrowd HTML to PDF API PDFCROWD

| | | | |
|---|---|---|---|
| DI49_3174; YKL210W | DI49_3174 | | YKL210W |
| DI49_3173; YKL211C | DI49_3173 | | YKL211C |
| DI49_3172; YKL212W; DI49_3171 | DI49_3172; DI49_3171 | | YKL212W |
| SPGP0P03080; YKL215C; YKL214C | | SPGP0P03080 | YKL215C; YKL214C |
| DI49_3169; YKL216W; DI49_3168 | DI49_3169; DI49_3168 | | YKL216W |
| SPGP0P03120; DI49_3167 | DI49_3167 | SPGP0P03120 | |
| DI49_2820; YOL155C; DI49_1595; SPGP0D04550; DI49_0839 | DI49_2820; DI49_1595; DI49_0839 | SPGP0D04550 | YOL155C |
| DI49_1593; YER183C; DI49_1592; YER184C | DI49_1593; DI49_1592 | | YER183C; YER184C |
| DI49_1588; DI49_1589 | DI49_1588; DI49_1589 | | |
| DI49_1586; YER177W; DI49_1585 | DI49_1586; DI49_1585 | | YER177W |
| SPGP0Q00210; DI49_1584 | DI49_1584 | SPGP0Q00210 | |
| DI49_1582; YER172C | DI49_1582 | | YER172C |
| DI49_1581; YER171W; DI49_1580 | DI49_1581; DI49_1580 | | YER171W |
| DI49_1579; SPGP0Q00260 | DI49_1579 | SPGP0Q00260 | |
| DI49_1577; YER167W; YER168C | DI49_1577 | | YER167W; YER168C |
| DI49_1576; YER166W | DI49_1576 | | YER166W |
| DI49_1575; YER165W | DI49_1575 | | YER165W |
| DI49_1574; YER164W; SPGP0Q00330 | DI49_1574 | SPGP0Q00330 | YER164W |
| DI49_1572; DI49_1573 | DI49_1572; DI49_1573 | | |
| DI49_1566; YER157W; DI49_1565 | DI49_1566; DI49_1565 | | YER157W |
| DI49_1563; YER154W; DI49_1562 | DI49_1563; DI49_1562 | | YER154W |
| DI49_1560; YER151C | DI49_1560 | | YER151C |
| DI49_1556; YER148W; DI49_1555; YER149C | DI49_1556; DI49_1555 | | YER148W; YER149C |
| DI49_1553; YER145C; DI49_1552; YER146W | DI49_1553; DI49_1552 | | YER145C; YER146W |
| DI49_1550; YER142C; YER143W | DI49_1550 | | YER142C; YER143W |
| DI49_1549; YER141W; DI49_1548 | DI49_1549; DI49_1548 | | YER141W |
| DI49_1546; DI49_2906; SPGP0Q00580; SPGP0JD00100; YCR063W | DI49_1546; DI49_2906 | SPGP0Q00580; SPGP0JD00100 | YCR063W |
| DI49_1535; YER131W; DI49_1534 | DI49_1535; DI49_1534 | | YER131W |
| DI49_1532; DI49_1533 | DI49_1532; DI49_1533 | | |
| DI49_1530; YER126C | DI49_1530 | | YER126C |
| DI49_1529; YER125W; DI49_1528 | DI49_1529; DI49_1528 | | YER125W |
| DI49_1526; YER122C | DI49_1526 | | YER122C |
| DI49_1525; YER120W; DI49_1524 | DI49_1525; DI49_1524 | | YER120W |
| DI49_1522; DI49_0101; DI49_1521; DI49_0102 | DI49_1522; DI49_0101; DI49_1521; DI49_0102 | | |
| DI49_1519; YER114C; DI49_1518; YER115C | DI49_1519; DI49_1518 | | YER114C; YER115C |
| DI49_1516; YER111C; YER112W | DI49_1516 | | YER111C; YER112W |
| DI49_1515; YER110C | DI49_1515 | | YER110C |
| DI49_1514; YER109C | DI49_1514 | | YER109C |

Create PDF in your applications with the Pdfcrowd HTML to PDF API PDFCROWD

| | | | |
|---|---|---|---|
| DI49_1513; YER107C; DI49_1512 | DI49_1513; DI49_1512 | | YER107C |
| DI49_1510; DI49_1511 | DI49_1510; DI49_1511 | | |
| DI49_1508; DI49_0112; DI49_1507; DI49_0113 | DI49_1508; DI49_0112; DI49_1507; DI49_0113 | | |
| DI49_1505; DI49_1506 | DI49_1505; DI49_1506 | | |
| DI49_1500; YER094C; SPGP0Q01000 | DI49_1500 | SPGP0Q01000 | YER094C |
| DI49_1498; SPGP0Q01010 | DI49_1498 | SPGP0Q01010 | |
| DI49_1496; YER090W; DI49_1495 | DI49_1496; DI49_1495 | | YER090W |
| DI49_1492; DI49_1493 | DI49_1492; DI49_1493 | | |
| DI49_1490; YER086W; DI49_1489 | DI49_1490; DI49_1489 | | YER086W |
| DI49_1487; YER082C | DI49_1487 | | YER082C |
| DI49_1486; YER081W; DI49_2710; DI49_2711 | DI49_1486; DI49_2710; DI49_2711 | | YER081W |
| DI49_1485; YER080W | DI49_1485 | | YER080W |
| DI49_1484; YER079W | DI49_1484 | | YER079W |
| DI49_1483; YER078C | DI49_1483 | | YER078C |
| DI49_1482; YER077C | DI49_1482 | | YER077C |
| DI49_1480; YER075C; DI49_1479 | DI49_1480; DI49_1479 | | YER075C |
| DI49_1477; YER073W | DI49_1477 | | YER073W |
| DI49_1476; YER072W | DI49_1476 | | YER072W |
| DI49_1475; YER071C | DI49_1475 | | YER071C |
| DI49_1474; YER070W | DI49_1474 | | YER070W |
| DI49_1473; YER069W; DI49_1472 | DI49_1473; DI49_1472 | | YER069W |
| SPGP0Q01280; DI49_1645 | DI49_1645 | SPGP0Q01280 | |
| DI49_1469; YER065C | DI49_1469 | | YER065C |
| DI49_1468; YER064C; DI49_1467 | DI49_1468; DI49_1467 | | YER064C |
| DI49_1465; YER061C; YER062C | DI49_1465 | | YER061C; YER062C |
| DI49_1464; DI49_1459; DI49_1463; YER056C; DI49_1458; YER060W | DI49_1464; DI49_1459; DI49_1463; DI49_1458 | | YER056C; YER060W |
| DI49_1462; YER059W; DI49_1461 | DI49_1462; DI49_1461 | | YER059W |
| SPGP0Q01390; YER056C-A; YER057C | | SPGP0Q01390 | YER056C-A; YER057C |
| DI49_1456; YER053C-A; YER054C | DI49_1456 | | YER053C-A; YER054C |
| DI49_1455; YER053C | DI49_1455 | | YER053C |
| DI49_1454; YER052C; DI49_1453 | DI49_1454; DI49_1453 | | YER052C |
| DI49_1451; YER049W; DI49_1450; YER050C | DI49_1451; DI49_1450 | | YER049W; YER050C |
| DI49_1448; YER048C; DI49_1447 | DI49_1448; DI49_1447 | | YER048C |
| DI49_1445; DI49_1446 | DI49_1445; DI49_1446 | | |
| DI49_1443; YER044C; YER044C-A | DI49_1443 | | YER044C; YER044C-A |
| DI49_1442; YER043C | DI49_1442 | | YER043C |
| DI49_1441; YER042W; DI49_1440 | DI49_1441; DI49_1440 | | YER042W |

Create PDF in your applications with the Pdfcrowd HTML to PDF API PDFCROWD

| | | | |
|---|---|---|---|
| DI49_1438; YER039C; DI49_1437 | DI49_1438; DI49_1437 | | YER039C |
| DI49_1435; YER036C; DI49_1434; YER037W | DI49_1435; DI49_1434 | | YER036C; YER037W |
| DI49_1432; DI49_0132; DI49_1431 | DI49_1432; DI49_0132; DI49_1431 | | |
| DI49_1429; DI49_1430 | DI49_1429; DI49_1430 | | |
| DI49_1427; DI49_1428 | DI49_1427; DI49_1428 | | |
| DI49_1425; DI49_1426 | DI49_1425; DI49_1426 | | |
| DI49_1423; YER025W | DI49_1423 | | YER025W |
| DI49_1422; YER024W; DI49_1421 | DI49_1422; DI49_1421 | | YER024W |
| DI49_1419; YER021W; YER022W | DI49_1419 | | YER021W; YER022W |
| DI49_1418; YER020W | DI49_1418 | | YER020W |
| DI49_1416; YER019W; DI49_1415 | DI49_1416; DI49_1415 | | YER019W |
| DI49_1413; DI49_1414 | DI49_1413; DI49_1414 | | |
| DI49_1411; DI49_1412 | DI49_1411; DI49_1412 | | |
| DI49_1409; DI49_1410; YER014W; YER015W | DI49_1409; DI49_1410 | | YER014W; YER015W |
| DI49_1403; YER008C; DI49_1402; YER009W | DI49_1403; DI49_1402 | | YER008C; YER009W |
| DI49_1400; DI49_1401 | DI49_1400; DI49_1401 | | |
| DI49_1399; YER005W; DI49_1398 | DI49_1399; DI49_1398 | | YER005W |
| DI49_1396; DI49_1397 | DI49_1396; DI49_1397 | | |
| DI49_1395; SPGP0Q02010 | DI49_1395 | SPGP0Q02010 | |
| DI49_1393; DI49_1394 | DI49_1393; DI49_1394 | | |
| DI49_1391; DI49_1392 | DI49_1391; DI49_1392 | | |
| DI49_1389; YEL007W; DI49_1388 | DI49_1389; DI49_1388 | | YEL007W |
| DI49_1383; YEL020W-A; DI49_1382; YEL020C | DI49_1383; DI49_1382 | | YEL020W-A; YEL020C |
| DI49_1380; DI49_1381 | DI49_1380; DI49_1381 | | |
| DI49_1378; DI49_1379 | DI49_1378; DI49_1379 | | |
| DI49_1376; DI49_1377 | DI49_1376; DI49_1377 | | |
| SPGP0Qtrna6Q; DI49_1374; SPGP0Vtrna7Q | DI49_1374 | SPGP0Qtrna6Q; SPGP0Vtrna7Q | |
| DI49_1373; YEL012W | DI49_1373 | | YEL012W |
| DI49_1372; YEL011W | DI49_1372 | | YEL011W |
| DI49_1370; YEL022W | DI49_1370 | | YEL022W |
| DI49_1369; YEL023C | DI49_1369 | | YEL023C |
| DI49_1368; YEL024W; DI49_1367 | DI49_1368; DI49_1367 | | YEL024W |
| DI49_1365; YEL027W | DI49_1365 | | YEL027W |
| DI49_1361; YEL031W | DI49_1361 | | YEL031W |
| DI49_1360; YEL032W; DI49_0695 | DI49_1360; DI49_0695 | | YEL032W |
| DI49_1357; YEL036C; YEL035C | DI49_1357 | | YEL036C; YEL035C |
| DI49_1356; YEL037C; DI49_1355 | DI49_1356; DI49_1355 | | YEL037C |
| DI49_1353; DI49_1354 | DI49_1353; DI49_1354 | | |

Create PDF in your applications with the Pdfcrowd HTML to PDF API PDFCROWD

| | | | |
|---|---|---|---|
| DI49_1351; DI49_1352 | DI49_1351; DI49_1352 | | |
| DI49_1349; DI49_1350; YEL043W; YEL042W | DI49_1349; DI49_1350 | | YEL043W; YEL042W |
| DI49_1347; DI49_1348 | DI49_1347; DI49_1348 | | |
| DI49_1345; DI49_3566; DI49_3567 | DI49_1345; DI49_3566; DI49_3567 | | |
| DI49_1341; YEL052W; YEL051W | DI49_1341 | | YEL052W; YEL051W |
| DI49_1340; YEL053C | DI49_1340 | | YEL053C |
| DI49_1339; YEL054C; DI49_1338 | DI49_1339; DI49_1338 | | YEL054C |
| DI49_1336; DI49_1337 | DI49_1336; DI49_1337 | | |
| DI49_1334; DI49_1335 | DI49_1334; DI49_1335 | | |
| DI49_1332; YEL061C; DI49_1331 | DI49_1332; DI49_1331 | | YEL061C |
| DI49_1329; YEL064C; DI49_1328; YEL063C | DI49_1329; DI49_1328 | | YEL064C; YEL063C |
| SPGP0Q02650; DI49_2618; SPGP0M03440 | DI49_2618 | SPGP0Q02650; SPGP0M03440 | |
| DI49_1324; YLL062C | DI49_1324 | | YLL062C |
| SPGP0FYtrna10H; SPGP0Gtrna12H; SPGP0Rtrna6H; SPGP0Vtrna3H; YER026C; SPGP0Vtrna4H; SPGP0Xtrna7H; YHL022C; YKL068W; SPGP0AKtrna11H; YGL203C; SPGP0Ztrna8H | | SPGP0FYtrna10H; SPGP0Gtrna12H; SPGP0Rtrna6H; SPGP0Vtrna3H; SPGP0Vtrna4H; SPGP0Xtrna7H; SPGP0AKtrna11H; SPGP0Ztrna8H | YER026C; YHL022C; YKL068W; YGL203C |
| YGR081C; SPGP0R00320 | | SPGP0R00320 | YGR081C |
| SPGP0FZ00110; YGR080W; DI49_2037 | DI49_2037 | SPGP0FZ00110 | YGR080W |
| YMR006C; DI49_4094; DI49_4093 | DI49_4094; DI49_4093 | | YMR006C |
| SPGP0GCtrna21K; SPGP0Dtrna26K; SPGP0GDtrna22K; SPGP0Rtrna16K; SPGP0Ttrna32K; YLR377C; SPGP0ACtrna29K; YPR119W; SPGP0ADtrna37K; YKR072C | | SPGP0GCtrna21K; SPGP0Dtrna26K; SPGP0GDtrna22K; SPGP0Rtrna16K; SPGP0Ttrna32K; SPGP0ACtrna29K; SPGP0ADtrna37K | YLR377C; YPR119W; YKR072C |
| SPGP0Rtrna16L; DI49_2066; YGR106C; DI49_2065 | DI49_2066; DI49_2065 | SPGP0Rtrna16L | YGR106C |
| YGR104C; YGR105W | | | YGR104C; YGR105W |
| YGR102C; YGR103W | | | YGR102C; YGR103W |
| YGR100W; DI49_2059; YGR099W; DI49_2058 | DI49_2059; DI49_2058 | | YGR100W; YGR099W |
| YGR097W; DI49_2056; YGR096W; DI49_2055 | DI49_2056; DI49_2055 | | YGR097W; YGR096W |
| YGR094W; DI49_2053; YGR093W; DI49_2052 | DI49_2053; DI49_2052 | | YGR094W; YGR093W |
| YGR091W; YGR092W | | | YGR091W; YGR092W |
| SPGP0Rtrna17T; YPR108W-A; YGR089W; YPR109W | | SPGP0Rtrna17T | YPR108W-A; YGR089W; YPR109W |
| YGR084C; DI49_2041; YGR083C; DI49_2040 | DI49_2041; DI49_2040 | | YGR084C; YGR083C |
| YGR079W; DI49_2036; YGR078C; DI49_2035 | DI49_2036; DI49_2035 | | YGR079W; YGR078C |
| YGR076C; YGR077C | | | YGR076C; YGR077C |
| YGR074W; YGR075C | | | YGR074W; YGR075C |
| YGR071C; YGR072W | | | YGR071C; YGR072W |
| YGR068C; DI49_2027 | DI49_2027 | | YGR068C |

Create PDF in your applications with the Pdfcrowd HTML to PDF API PDFCROWD

| | | | |
|---|---|---|---|
| YGR067C; DI49_2026; YGR066C; DI49_2025 | DI49_2026; DI49_2025 | | YGR067C; YGR066C |
| YGR063C; YGR065C | | | YGR063C; YGR065C |
| SPGP0R00490; DI49_2021 | DI49_2021 | SPGP0R00490 | |
| YGR060W; DI49_2020; YGR059W; DI49_2019 | DI49_2020; DI49_2019 | | YGR060W; YGR059W |
| YGR057C; YGR058W | | | YGR057C; YGR058W |
| YGR055W; DI49_2015 | DI49_2015 | | YGR055W |
| YGR054W; DI49_2014; YGR053C; DI49_2013 | DI49_2014; DI49_2013 | | YGR054W; YGR053C |
| YGR049W; DI49_2011; YGR048W; DI49_2010 | DI49_2011; DI49_2010 | | YGR049W; YGR048W |
| YGR046W; DI49_2008; YGR044C; DI49_2007 | DI49_2008; DI49_2007 | | YGR046W; YGR044C |
| YGR042W; YGR043C | | | YGR042W; YGR043C |
| YGR040W; YGR041W | | | YGR040W; YGR041W |
| YGR036C; DI49_1999; YGR035C; DI49_1998 | DI49_1999; DI49_1998 | | YGR036C; YGR035C |
| YGR033C; DI49_1996 | DI49_1996 | | YGR033C |
| YGR032W; DI49_1995; YGR031W; DI49_1994 | DI49_1995; DI49_1994 | | YGR032W; YGR031W |
| YGR027C; DI49_1989; YLR333C; DI49_3836 | DI49_1989; DI49_3836 | | YGR027C; YLR333C |
| YGR026W; DI49_1988; YGR024C; DI49_1987 | DI49_1988; DI49_1987 | | YGR026W; YGR024C |
| YGR020C; DI49_1983 | DI49_1983 | | YGR020C |
| YGR019W; DI49_1982 | DI49_1982 | | YGR019W |
| YGR017W; DI49_1981; YGR016W; DI49_1980 | DI49_1981; DI49_1980 | | YGR017W; YGR016W |
| YGR014W; YGR015C | | | YGR014W; YGR015C |
| YGR012W; YGR013W | | | YGR012W; YGR013W |
| YGR009C; DI49_1974; YGR007W; DI49_1972 | DI49_1974; DI49_1972 | | YGR009C; YGR007W |
| YGR005C; DI49_1970 | DI49_1970 | | YGR005C |
| YGR004W; DI49_1969 | DI49_1969 | | YGR004W |
| YGR003W; DI49_1968 | DI49_1968 | | YGR003W |
| YGR002C; DI49_1967; YGR001C; SPGP0E02720 | DI49_1967 | SPGP0E02720 | YGR002C; YGR001C |
| YGL002W; YGL001C | | | YGL002W; YGL001C |
| YGL004C; YGL003C | | | YGL004C; YGL003C |
| YGL006W; YGL005C | | | YGL006W; YGL005C |
| YGL009C; DI49_1958; YGL010W; DI49_1957 | DI49_1958; DI49_1957 | | YGL009C; YGL010W |
| YGL012W; YGL011C | | | YGL012W; YGL011C |
| YGL014W; DI49_1953; YGL015C; DI49_1952 | DI49_1953; DI49_1952 | | YGL014W; YGL015C |
| YGL017W; YGL016W | | | YGL017W; YGL016W |
| YGL019W; YGL018C | | | YGL019W; YGL018C |
| YGL021W; DI49_1946 | DI49_1946 | | YGL021W |
| YGL022W; DI49_1945 | DI49_1945 | | YGL022W |
| YGL023C; DI49_1944; YGL025C; DI49_1943 | DI49_1944; DI49_1943 | | YGL023C; YGL025C |
| YGL027C; DI49_1941 | DI49_1941 | | YGL027C |
| YGL028C; DI49_1940 | DI49_1940 | | YGL028C |

Create PDF in your applications with the Pdfcrowd HTML to PDF API PDFCROWD

| | | | |
|---|---|---|---|
| SPGP0Rtrna11F; SPGP0MHtrna5F; YGL029W; SPGP0Atrna14F; SPGP0Xtrna12F; YHR072W; SPGP0Xtrna13F; YHR126C; SPGP0AEtrna6F; YCL064C; YMR041C | | SPGP0Rtrna11F; SPGP0MHtrna5F; SPGP0Atrna14F; SPGP0Xtrna12F; SPGP0Xtrna13F; SPGP0AEtrna6F | YGL029W; YHR072W; YHR126C; YCL064C; YMR041C |
| YGL031C; DI49_1936; YGL032C; DI49_2112; DI49_2111; DI49_1935 | DI49_1936; DI49_2112; DI49_2111; DI49_1935 | | YGL031C; YGL032C |
| YGL035C; YGL033W | | | YGL035C; YGL033W |
| YGL037C; YGL036W | | | YGL037C; YGL036W |
| YGL039W; YGL038C; DI49_1929 | DI49_1929 | | YGL039W; YGL038C |
| YGL040C; DI49_1926; YGL041W-A; DI49_1925 | DI49_1926; DI49_1925 | | YGL040C; YGL041W-A |
| YGL044C; DI49_1923; YGL045W; DI49_1922 | DI49_1923; DI49_1922 | | YGL044C; YGL045W |
| YGL047W; SPGP0Rtrna10V | | SPGP0Rtrna10V | YGL047W |
| YGL049C; DI49_1918; YGL050W; DI49_1917 | DI49_1918; DI49_1917 | | YGL049C; YGL050W |
| YGL056C; DI49_1912; YGL057C; DI49_1911 | DI49_1912; DI49_1911 | | YGL056C; YGL057C |
| YGL059W; YGL058W | | | YGL059W; YGL058W |
| YGL061C; YGL060W | | | YGL061C; YGL060W |
| YGL063W; YGL062W | | | YGL063W; YGL062W |
| YGL065C; DI49_1904 | DI49_1904 | | YGL065C |
| YGL066W; DI49_1903; YGL067W; DI49_1902 | DI49_1903; DI49_1902 | | YGL066W; YGL067W |
| YGL070C; DI49_1900 | DI49_1900 | | YGL070C |
| YGL071W; DI49_1899 | DI49_1899 | | YGL071W |
| YGL073W; DI49_1898; YGL075C; DI49_1897 | DI49_1898; DI49_1897 | | YGL073W; YGL075C |
| YGL077C; DI49_1896 | DI49_1896 | | YGL077C |
| YGL078C; DI49_1895 | DI49_1895 | | YGL078C |
| YGL079W; DI49_1894; YGL080W; DI49_1893 | DI49_1894; DI49_1893 | | YGL079W; YGL080W |
| YGL082W; YGL081W | | | YGL082W; YGL081W |
| YGL084C; DI49_1889 | DI49_1889 | | YGL084C |
| YGL085W; DI49_1888; YGL086W; DI49_1887 | DI49_1888; DI49_1887 | | YGL085W; YGL086W |
| YGL089C; YGL087C | | | YGL089C; YGL087C |
| YGL091C; YGL090W | | | YGL091C; YGL090W |
| YGL093W; DI49_1882 | DI49_1882 | | YGL093W |
| YGL094C; DI49_1881; YGL095C; DI49_1880 | DI49_1881; DI49_1880 | | YGL094C; YGL095C |
| YGL096W; SPGP0Rtrna15E | | SPGP0Rtrna15E | YGL096W |
| YGL098W; DI49_1874 | DI49_1874 | | YGL098W |
| YGL099W; DI49_1873 | DI49_1873 | | YGL099W |
| YGL100W; DI49_1872; YGL101W; DI49_1871 | DI49_1872; DI49_1871 | | YGL100W; YGL101W |
| YGL104C; YGL103W | | | YGL104C; YGL103W |
| YGL106W; YGL105W | | | YGL106W; YGL105W |
| YGL108C; YGL107C | | | YGL108C; YGL107C |
| YGL111W; YGL110C | | | YGL111W; YGL110C |
| YGL113W; YGL112C | | | YGL113W; YGL112C |

Create PDF in your applications with the Pdfcrowd HTML to PDF API PDFCROWD

| | | | |
|---|---|---|---|
| YGL115W; DI49_1859 | DI49_1859 | | YGL115W |
| YGL116W; DI49_1858; YGL117W; DI49_1857 | DI49_1858; DI49_1857 | | YGL116W; YGL117W |
| YGL121C; YGL120C | | | YGL121C; YGL120C |
| YGL123W; DI49_1851; YGL124C; DI49_1850 | DI49_1851; DI49_1850 | | YGL123W; YGL124C |
| YGL126W; YGL125W | | | YGL126W; YGL125W |
| YGL128C; DI49_1846 | DI49_1846 | | YGL128C |
| YGL129C; DI49_1845; YGL130W; DI49_1844 | DI49_1845; DI49_1844 | | YGL129C; YGL130W |
| YGL133W; YGL131C | | | YGL133W; YGL131C |
| SPGP0R02200; YPL220W | | SPGP0R02200 | YPL220W |
| YGL136C; DI49_1839 | DI49_1839 | | YGL136C |
| YGL137W; DI49_1838; YGL138C; DI49_1837 | DI49_1838; DI49_1837 | | YGL137W; YGL138C |
| YGL140C; DI49_1835 | DI49_1835 | | YGL140C |
| YGL141W; DI49_1834 | DI49_1834 | | YGL141W |
| YGL142C; DI49_1833; YGL143C; DI49_1832 | DI49_1833; DI49_1832 | | YGL142C; YGL143C |
| YGL145W; YGL144C | | | YGL145W; YGL144C |
| YGL151W; DI49_1825; YGL153W; DI49_1824 | DI49_1825; DI49_1824 | | YGL151W; YGL153W |
| YGL155W; YGL154C | | | YGL155W; YGL154C |
| YGL157W; YGL156W | | | YGL157W; YGL156W |
| YGL161C; YGL160W | | | YGL161C; YGL160W |
| YGL163C; YGL162W | | | YGL163C; YGL162W |
| YGL166W; YGL164C | | | YGL166W; YGL164C |
| YGL169W; YGL167C; DI49_1809; DI49_1808 | DI49_1809; DI49_1808 | | YGL169W; YGL167C |
| YGL170C; DI49_1806 | DI49_1806 | | YGL170C |
| YGL171W; DI49_1805 | DI49_1805 | | YGL171W |
| YGL172W; DI49_1804 | DI49_1804 | | YGL172W |
| SPGP0R02540; SPGP0E04280; SPGP0BH00100 | | SPGP0R02540; SPGP0E04280; SPGP0BH00100 | |
| DI49_4698; SPGP0Strna17P | DI49_4698 | SPGP0Strna17P | |
| DI49_4696; DI49_0162; DI49_4695; DI49_0164 | DI49_4696; DI49_0162; DI49_4695; DI49_0164 | | |
| DI49_4693; DI49_4694 | DI49_4693; DI49_4694 | | |
| DI49_4691; DI49_4692 | DI49_4691; DI49_4692 | | |
| DI49_4683; DI49_4684 | DI49_4683; DI49_4684 | | |
| DI49_4678; DI49_4679 | DI49_4678; DI49_4679 | | |
| DI49_4676; YNL054W | DI49_4676 | | YNL054W |
| DI49_4666; DI49_4667 | DI49_4666; DI49_4667 | | |
| DI49_4664; DI49_1828; DI49_4663 | DI49_4664; DI49_1828; DI49_4663 | | |
| DI49_4657; DI49_4658 | DI49_4657; DI49_4658 | | |
| DI49_4655; DI49_4656 | DI49_4655; DI49_4656 | | |

Create PDF in your applications with the Pdfcrowd HTML to PDF API PDFCROWD

| | | | |
|---|---|---|---|
| DI49_4653; DI49_4654 | DI49_4653; DI49_4654 | | |
| DI49_4632; DI49_4633 | DI49_4632; DI49_4633 | | |
| DI49_4594; DI49_4595 | DI49_4594; DI49_4595 | | |
| DI49_4592; DI49_4593 | DI49_4592; DI49_4593 | | |
| DI49_4590; DI49_4591 | DI49_4590; DI49_4591 | | |
| DI49_4588; DI49_4589 | DI49_4588; DI49_4589 | | |
| DI49_4586; DI49_4587 | DI49_4586; DI49_4587 | | |
| DI49_4584; DI49_4585 | DI49_4584; DI49_4585 | | |
| DI49_4582; DI49_4583 | DI49_4582; DI49_4583 | | |
| DI49_4574; DI49_4575 | DI49_4574; DI49_4575 | | |
| DI49_4569; DI49_4570 | DI49_4569; DI49_4570 | | |
| DI49_4563; DI49_4564 | DI49_4563; DI49_4564 | | |
| DI49_4546; DI49_4547 | DI49_4546; DI49_4547 | | |
| DI49_4544; DI49_4545 | DI49_4544; DI49_4545 | | |
| DI49_4542; DI49_4543 | DI49_4542; DI49_4543 | | |
| DI49_4534; DI49_4535 | DI49_4534; DI49_4535 | | |
| DI49_4529; DI49_4530 | DI49_4529; DI49_4530 | | |
| DI49_4527; DI49_4528 | DI49_4527; DI49_4528 | | |
| DI49_4525; DI49_4526 | DI49_4525; DI49_4526 | | |
| DI49_4523; DI49_4524 | DI49_4523; DI49_4524 | | |
| DI49_4515; DI49_4516 | DI49_4515; DI49_4516 | | |
| DI49_4510; DI49_4511 | DI49_4510; DI49_4511 | | |
| DI49_4508; DI49_4509 | DI49_4508; DI49_4509 | | |
| DI49_4502; DI49_4503 | DI49_4502; DI49_4503 | | |
| DI49_4488; DI49_4489 | DI49_4488; DI49_4489 | | |
| DI49_4486; DI49_4487 | DI49_4486; DI49_4487 | | |
| DI49_4474; DI49_4475 | DI49_4474; DI49_4475 | | |
| DI49_4472; DI49_4473 | DI49_4472; DI49_4473 | | |
| YAL068C; SPGP0CU00100; YDR541C | | SPGP0CU00100 | YAL068C; YDR541C |
| YDR539W; YDR540C | | | YDR539W; YDR540C |
| YDR536W; YDR538W | | | YDR536W; YDR538W |
| YDR533C; YDR534C | | | YDR533C; YDR534C |
| YDR531W; DI49_0690; YDR530C; SPGP0B06190 | DI49_0690 | SPGP0B06190 | YDR531W; YDR530C |
| YDR528W; DI49_0687; YDR527W; DI49_0688 | DI49_0687; DI49_0688 | | YDR528W; YDR527W |
| YDR524C; DI49_0683; YDR523C; DI49_0684 | DI49_0683; DI49_0684 | | YDR524C; YDR523C |
| YDR520C; YDR522C; DI49_0681; DI49_0682 | DI49_0681; DI49_0682 | | YDR520C; YDR522C |
| YDR518W; YDR519W | | | YDR518W; YDR519W |
| YDR516C; DI49_0678; YDR515W; DI49_0679 | DI49_0678; DI49_0679 | | YDR516C; YDR515W |
| YDR513W; YDR512C; YDR511W | | | YDR513W; YDR512C; YDR511W |

Create PDF in your applications with the Pdfcrowd HTML to PDF API PDFCROWD

| | | | |
|---|---|---|---|
| YDR508C; DI49_0671; DI49_0672 | DI49_0671; DI49_0672 | | YDR508C |
| YDR507C; DI49_0670 | DI49_0670 | | YDR507C |
| YDR506C; YHR145C | | | YDR506C; YHR145C |
| YDR505C; DI49_0667; YDR503C; DI49_0668 | DI49_0667; DI49_0668 | | YDR505C; YDR503C |
| YDR501W; YDR502C; DI49_0665; DI49_0666 | DI49_0665; DI49_0666 | | YDR501W; YDR502C |
| YDR499W; DI49_0662; YDR498C; DI49_0663 | DI49_0662; DI49_0663 | | YDR499W; YDR498C |
| YDR496C; DI49_0659 | DI49_0659 | | YDR496C |
| YDR495C; DI49_0658; YDR494W | DI49_0658 | | YDR495C; YDR494W |
| YDR492W; YDR493W | | | YDR492W; YDR493W |
| YDR489W; YDR490C; DI49_0654; DI49_0655 | DI49_0654; DI49_0655 | | YDR489W; YDR490C |
| YDR487C; DI49_0651; YDR486C; DI49_0652 | DI49_0651; DI49_0652 | | YDR487C; YDR486C |
| YDR484W; DI49_0648; YDR483W | DI49_0648 | | YDR484W; YDR483W |
| YDR481C; DI49_0645; YDR480W; DI49_0646 | DI49_0645; DI49_0646 | | YDR481C; YDR480W |
| YDR478W; YDR479C | | | YDR478W; YDR479C |
| YDR476C; YDR477W | | | YDR476C; YDR477W |
| YDR473C; YDR475C | | | YDR473C; YDR475C |
| YDR471W; DI49_0636; YDR470C | DI49_0636 | | YDR471W; YDR470C |
| YDR468C; DI49_0633; DI49_0634 | DI49_0633; DI49_0634 | | YDR468C |
| YDR466W; DI49_0632 | DI49_0632 | | YDR466W |
| YDR465C; DI49_0631; YDR464W | DI49_0631 | | YDR465C; YDR464W |
| YDR462W; YDR463W | | | YDR462W; YDR463W |
| YDR461W; YDR461C-A | | | YDR461W; YDR461C-A |
| YDR459C; YDR460W; DI49_0626; SPGP0B05530 | DI49_0626 | SPGP0B05530 | YDR459C; YDR460W |
| YDR457W; DI49_0623; DI49_0624 | DI49_0623; DI49_0624 | | YDR457W |
| YDR456W; DI49_0622; YDR454C | DI49_0622 | | YDR456W; YDR454C |
| YDR452W; YDR453C | | | YDR452W; YDR453C |
| SPGP0A00930; YML026C; DI49_0617; DI49_0618; SPGP0F03290; DI49_4062; YML027W | DI49_0617; DI49_0618; DI49_4062 | SPGP0A00930; SPGP0F03290 | YML026C; YML027W |
| YDR449C; DI49_0616 | DI49_0616 | | YDR449C |
| YDR448W; DI49_0615 | DI49_0615 | | YDR448W |
| YDR447C; DI49_0614; YDR446W; DI49_4065; SPGP0G03220 | DI49_0614; DI49_4065 | SPGP0G03220 | YDR447C; YDR446W |
| YDR441C; YDR443C | | | YDR441C; YDR443C |
| YDR439W; YDR440W | | | YDR439W; YDR440W |
| YDR437W; DI49_0605; YDR436W | DI49_0605 | | YDR437W; YDR436W |
| YDR434W; DI49_0602; DI49_0603 | DI49_0602; DI49_0603 | | YDR434W |
| YDR432W; DI49_0601; YDR430C | DI49_0601 | | YDR432W; YDR430C |
| YDR428C; YDR429C | | | YDR428C; YDR429C |
| YDR425W; YDR427W | | | YDR425W; YDR427W |
| YDR423C; YDR424C | | | YDR423C; YDR424C |

Create PDF in your applications with the Pdfcrowd HTML to PDF API PDFCROWD

| | | | |
|---|---|---|---|
| YDR421W; DI49_0594; YDR420W; DI49_0595 | DI49_0594; DI49_0595 | | YDR421W; YDR420W |
| YDR419W; DI49_0591 | DI49_0591 | | YDR419W |
| YDR418W; DI49_0590 | DI49_0590 | | YDR418W |
| YDR416W; DI49_0589; YDR415C | DI49_0589 | | YDR416W; YDR415C |
| YDR412W; DI49_0586; YDR411C; DI49_0587 | DI49_0586; DI49_0587 | | YDR412W; YDR411C |
| YDR409W; YDR410C | | | YDR409W; YDR410C |
| YDR407C; DI49_0582; SPGP0B05080 | DI49_0582 | SPGP0B05080 | YDR407C |
| YDR406W; SPGP0B05060; YDR405W | | SPGP0B05060 | YDR406W; YDR405W |
| YDR403W; YDR404C | | | YDR403W; YDR404C |
| YDR400W; YDR402C | | | YDR400W; YDR402C |
| YDR398W; YDR399W | | | YDR398W; YDR399W |
| YDR395W; DI49_0572 | DI49_0572 | | YDR395W |
| YDR394W; DI49_0571; YDR393W | DI49_0571 | | YDR394W; YDR393W |
| YDR391C; YDR392W | | | YDR391C; YDR392W |
| YDR390C; SPGP0Atrna18G | | SPGP0Atrna18G | YDR390C |
| YDR388W; DI49_0564; YDR387C | DI49_0564 | | YDR388W; YDR387C |
| YOR133W; YDR385W | | | YOR133W; YDR385W |
| YDR384C; DI49_0560; YDR383C | DI49_0560 | | YDR384C; YDR383C |
| YDR381C-A; YDR382W; DI49_0558; DI49_0559 | DI49_0558; DI49_0559 | | YDR381C-A; YDR382W |
| YDR380W; YDR381W | | | YDR380W; YDR381W |
| YDR379W; YDR379C-A | | | YDR379W; YDR379C-A |
| YDR377W; YDR378C | | | YDR377W; YDR378C |
| YDR375C; DI49_0552; YDR374W-A; DI49_0553 | DI49_0552; DI49_0553 | | YDR375C; YDR374W-A |
| YDR373W; YDR374C | | | YDR373W; YDR374C |
| YDR371W; YDR372C | | | YDR371W; YDR372C |
| YDR369C; YDR370C | | | YDR369C; YDR370C |
| YDR367W; YDR368W | | | YDR367W; YDR368W |
| YDR364C; DI49_0542 | DI49_0542 | | YDR364C |
| YDR363W-A; DI49_0541 | DI49_0541 | | YDR363W-A |
| YDR361C; DI49_0537; YDR359C | DI49_0537 | | YDR361C; YDR359C |
| YDR357C; YDR358W | | | YDR357C; YDR358W |
| YDR354W; DI49_0532 | DI49_0532 | | YDR354W |
| YDR353W; DI49_0531; YDR352W | DI49_0531 | | YDR353W; YDR352W |
| YDR350C; DI49_0528; DI49_0529 | DI49_0528; DI49_0529 | | YDR350C |
| YDR347W; YDR348C | | | YDR347W; YDR348C |
| SPGP0A01900; YDR345C | | SPGP0A01900 | YDR345C |
| YDR341C; DI49_0521 | DI49_0521 | | YDR341C |
| YDR337W; DI49_0517; YDR336W | DI49_0517 | | YDR337W; YDR336W |
| YDR334W; DI49_0514; YDR333C | DI49_0514 | | YDR334W; YDR333C |
| YDR331W; YDR332W | | | YDR331W; YDR332W |

Create PDF in your applications with the Pdfcrowd HTML to PDF API PDFCROWD

| | | | |
|---|---|---|---|
| YDR329C; DI49_0509; YDR328C; DI49_0510 | DI49_0509; DI49_0510 | | YDR329C; YDR328C |
| YDR325W; DI49_0506; DI49_0507 | DI49_0506; DI49_0507 | | YDR325W |
| YDR324C; DI49_0505; YDR323C | DI49_0505 | | YDR324C; YDR323C |
| YDR322W; DI49_0502 | DI49_0502 | | YDR322W |
| YDR321W; DI49_0501 | DI49_0501 | | YDR321W |
| YDR320C-A; DI49_0500; YDR320C | DI49_0500 | | YDR320C-A; YDR320C |
| YDR318W; YDR319C | | | YDR318W; YDR319C |
| YDR316W; DI49_0494; YDR315C | DI49_0494 | | YDR316W; YDR315C |
| YDR313C; YDR314C | | | YDR313C; YDR314C |
| YDR311W; YDR312W; YHR066W; YHR067W | | | YDR311W; YDR312W; YHR066W; YHR067W |
| YDR309C; YDR310C | | | YDR309C; YDR310C |
| YDR307W; YDR308C | | | YDR307W; YDR308C |
| YDR306C; SPGP0APtrna34V; SPGP0Atrna12V; DI49_2588 | DI49_2588 | SPGP0APtrna34V; SPGP0Atrna12V | YDR306C |
| YDR304C; YDR305C | | | YDR304C; YDR305C |
| YDR302W; YDR303C | | | YDR302W; YDR303C |
| YDR300C; DI49_0477 | DI49_0477 | | YDR300C |
| YDR299W; DI49_0476 | DI49_0476 | | YDR299W |
| YDR298C; DI49_0475 | DI49_0475 | | YDR298C |
| YDR297W; DI49_0474; YDR296W | DI49_0474 | | YDR297W; YDR296W |
| YDR294C; DI49_0471; DI49_0472 | DI49_0471; DI49_0472 | | YDR294C |
| YDR293C; DI49_0470 | DI49_0470 | | YDR293C |
| YDR292C; DI49_0469 | DI49_0469 | | YDR292C |
| YDR291W; DI49_0468; YDR289C | DI49_0468 | | YDR291W; YDR289C |
| YDR287W; DI49_0465; YDR286C; DI49_0466 | DI49_0465; DI49_0466 | | YDR287W; YDR286C |
| YDR284C; YDR285W | | | YDR284C; YDR285W |
| YDR282C; YDR283C; DI49_0461; DI49_0462 | DI49_0461; DI49_0462 | | YDR282C; YDR283C |
| YDR280W; DI49_0458; YDR279W; DI49_0459 | DI49_0458; DI49_0459 | | YDR280W; YDR279W |
| YDR277C; DI49_0455 | DI49_0455 | | YDR277C |
| YDR276C; DI49_0454; YDR275W | DI49_0454 | | YDR276C; YDR275W |
| YDR272W; YDR273W | | | YDR272W; YDR273W |
| YDR268W; DI49_0449; YDR266C | DI49_0449 | | YDR268W; YDR266C |
| YDR264C; YDR265W | | | YDR264C; YDR265W |
| YDR262W; YDR263C | | | YDR262W; YDR263C |
| YDR259C; YDR260C | | | YDR259C; YDR260C |
| YDR257C; YDR258C | | | YDR257C; YDR258C |
| YDR255C; YDR256C; DI49_0436; DI49_0437 | DI49_0436; DI49_0437 | | YDR255C; YDR256C |
| YDR253C; YDR254W | | | YDR253C; YDR254W |
| YDR251W; YDR252W | | | YDR251W; YDR252W |
| YDR248C; DI49_0429; YDR247W; DI49_0430 | DI49_0429; DI49_0430 | | YDR248C; YDR247W |

Create PDF in your applications with the Pdfcrowd HTML to PDF API PDFCROWD

| | | | |
|---|---|---|---|
| YDR245W; DI49_0426; DI49_0427 | DI49_0426; DI49_0427 | | YDR245W |
| YDR244W; DI49_0425; YDR243C | DI49_0425 | | YDR244W; YDR243C |
| YDR240C; DI49_0422; YDR239C | DI49_0422 | | YDR240C; YDR239C |
| YDR237W; YDR238C | | | YDR237W; YDR238C |
| YDR235W; DI49_0417 | DI49_0417 | | YDR235W |
| YDR234W; DI49_0416 | DI49_0416 | | YDR234W |
| YDR233C; DI49_0414; DI49_0415 | DI49_0414; DI49_0415 | | YDR233C |
| YDR232W; DI49_0413 | DI49_0413 | | YDR232W |
| YDR231C; DI49_0412 | DI49_0412 | | YDR231C |
| YDR229W; DI49_0411 | DI49_0411 | | YDR229W |
| YDR228C; DI49_0410; YDR227W | DI49_0410 | | YDR228C; YDR227W |
| YDR224C; DI49_0406; YDR223W; DI49_0180; DI49_0181 | DI49_0406; DI49_0180; DI49_0181 | | YDR224C; YDR223W |
| YDR221W; DI49_0403; YDR219C; DI49_0404 | DI49_0403; DI49_0404 | | YDR221W; YDR219C |
| YDR217C; YDR218C | | | YDR217C; YDR218C |
| YDR214W; YDR216W | | | YDR214W; YDR216W |
| YDR212W; DI49_0397; YDR211W | DI49_0397 | | YDR212W; YDR211W |
| YDR208W; DI49_0394; YDR207C | DI49_0394 | | YDR208W; YDR207C |
| YDR205W; YDR206W | | | YDR205W; YDR206W |
| YDR202C; DI49_0389 | DI49_0389 | | YDR202C |
| YDR201W; DI49_0388; YDR200C | DI49_0388 | | YDR201W; YDR200C |
| YDR197W; YDR198C; DI49_0386; DI49_0387 | DI49_0386; DI49_0387 | | YDR197W; YDR198C |
| YDR195W; DI49_0383; DI49_0384 | DI49_0383; DI49_0384 | | YDR195W |
| YDR194C; DI49_0382; YDR192C | DI49_0382 | | YDR194C; YDR192C |
| YDR190C; DI49_0380; DI49_0381 | DI49_0380; DI49_0381 | | YDR190C |
| YDR189W; DI49_0379 | DI49_0379 | | YDR189W |
| YDR188W; DI49_0378 | DI49_0378 | | YDR188W |
| YDR186C; DI49_0377 | DI49_0377 | | YDR186C |
| YDR185C; DI49_0376; YDR184C | DI49_0376 | | YDR185C; YDR184C |
| YDR182W; YDR183W | | | YDR182W; YDR183W |
| YDR180W; DI49_0371; YDR179W-A; DI49_0372 | DI49_0371; DI49_0372 | | YDR180W; YDR179W-A |
| YDR178W; YDR179C | | | YDR178W; YDR179C |
| YDR176W; DI49_0366; DI49_0367 | DI49_0366; DI49_0367 | | YDR176W |
| YDR175C; YDR174W; DI49_0364; DI49_0365 | DI49_0364; DI49_0365 | | YDR175C; YDR174W |
| YDR173C; DI49_0363 | DI49_0363 | | YDR173C |
| YDR172W; DI49_0362; YDR171W | DI49_0362 | | YDR172W; YDR171W |
| SPGP0Atrna11Q; DI49_2421; SPGP0Atrna12Q; YDR034C; SPGP0Vtrna8Q; YER048W-A; SPGP0Xtrna10Q; SPGP0Btrna3Q; DI49_0361; SPGP0AGtrna18Q; SPGP0AHtrna14Q; SPGP0C02130 | DI49_2421; DI49_0361 | SPGP0Atrna11Q; SPGP0Atrna12Q; SPGP0Vtrna8Q; SPGP0Xtrna10Q; SPGP0Btrna3Q; SPGP0AGtrna18Q; SPGP0AHtrna14Q; SPGP0C02130 | YDR034C; YER048W-A |

Create PDF in your applications with the Pdfcrowd HTML to PDF API PDFCROWD

| | | | |
|---|---|---|---|
| YDR170C; DI49_0359 | DI49_0359 | | YDR170C |
| YDR169C; DI49_0357 | DI49_0357 | | YDR169C |
| YDR168W; DI49_0356; YDR167W | DI49_0356 | | YDR168W; YDR167W |
| YDR165W; DI49_0353; YDR164C; DI49_0354 | DI49_0353; DI49_0354 | | YDR165W; YDR164C |
| YDR162C; DI49_0350; YDR161W; DI49_0351 | DI49_0350; DI49_0351 | | YDR162C; YDR161W |
| YDR159W; YDR160W | | | YDR159W; YDR160W |
| YDR156W; YDR158W | | | YDR156W; YDR158W |
| YDR153C; DI49_0343; YDR152W | DI49_0343 | | YDR153C; YDR152W |
| YDR150W; YDR151C | | | YDR150W; YDR151C |
| YDR147W; YDR148C; DI49_0340; SPGP0B02670 | DI49_0340 | SPGP0B02670 | YDR147W; YDR148C |
| YDR145W; DI49_0337; YDR144C; DI49_0338 | DI49_0337; DI49_0338 | | YDR145W; YDR144C |
| YDR142C; DI49_0334 | DI49_0334 | | YDR142C |
| YDR141C; DI49_0333; YDR140W | DI49_0333 | | YDR141C; YDR140W |
| YDR138W; YDR139C | | | YDR138W; YDR139C |
| YDR135C; DI49_0328; DI49_0329 | DI49_0328; DI49_0329 | | YDR135C |
| YDR134C; YLR110C; YDR132C; SPGP0C01720 | | SPGP0C01720 | YDR134C; YLR110C; YDR132C |
| YDR130C; YDR131C | | | YDR130C; YDR131C |
| YDR128W; YDR129C | | | YDR128W; YDR129C |
| YDR126W; DI49_0320; YDR125C | DI49_0320 | | YDR126W; YDR125C |
| YDR123C; DI49_0317; DI49_0318 | DI49_0317; DI49_0318 | | YDR123C |
| YDR122W; DI49_0316; YDR121W | DI49_0316 | | YDR122W; YDR121W |
| YDR119W-A; YDR120C | | | YDR119W-A; YDR120C |
| YDR118W; YDR119W; DI49_0312; DI49_0313 | DI49_0312; DI49_0313 | | YDR118W; YDR119W |
| YDR116C; YDR117C | | | YDR116C; YDR117C |
| YDR113C; YDR115W | | | YDR113C; YDR115W |
| YDR110W; DI49_0305; YDR109C; DI49_0306 | DI49_0305; DI49_0306 | | YDR110W; YDR109C |
| YDR107C; DI49_0303 | DI49_0303 | | YDR107C |
| YDR104C; YDR105C | | | YDR104C; YDR105C |
| YDR101C; DI49_0297; YDR100W; DI49_0298 | DI49_0297; DI49_0298 | | YDR101C; YDR100W |
| YDR098C; YDR099W; DI49_0295; DI49_0296 | DI49_0295; DI49_0296 | | YDR098C; YDR099W |
| YDR096W; DI49_0291 | DI49_0291 | | YDR096W |
| YDR093W; DI49_0290; YDR092W | DI49_0290 | | YDR093W; YDR092W |
| YDR090C; YDR091C | | | YDR090C; YDR091C |
| YDR088C; DI49_0285; YDR087C | DI49_0285 | | YDR088C; YDR087C |
| YDR085C; DI49_0282; YDR084C; DI49_0283 | DI49_0282; DI49_0283 | | YDR085C; YDR084C |
| YDR082W; YDR083W; DI49_0280; DI49_0281 | DI49_0280; DI49_0281 | | YDR082W; YDR083W |
| YDR080W; DI49_0277; YDR079C-A; DI49_0278 | DI49_0277; DI49_0278 | | YDR080W; YDR079C-A |
| YDR078C; YDR079W | | | YDR078C; YDR079W |
| YDR076W; DI49_0272 | DI49_0272 | | YDR076W |
| YDR075W; DI49_0271 | DI49_0271 | | YDR075W |

Create PDF in your applications with the Pdfcrowd HTML to PDF API PDFCROWD

| | | | |
|---|---|---|---|
| YDR074W; DI49_0270; YDR073W | DI49_0270 | | YDR074W; YDR073W |
| YDR071C; YDR072C; DI49_0268; DI49_0269 | DI49_0268; DI49_0269 | | YDR071C; YDR072C |
| YDR069C; DI49_0265; YDR068W; DI49_0266 | DI49_0265; DI49_0266 | | YDR069C; YDR068W |
| YDR066C; YDR067C; DI49_0263; DI49_0264 | DI49_0263; DI49_0264 | | YDR066C; YDR067C |
| YDR064W; DI49_0260; DI49_0261 | DI49_0260; DI49_0261 | | YDR064W |
| YDR063W; DI49_0259 | DI49_0259 | | YDR063W |
| YDR062W; DI49_0258 | DI49_0258 | | YDR062W |
| YDR061W; DI49_0257 | DI49_0257 | | YDR061W |
| YDR060W; DI49_0256; SPGP0A04500 | DI49_0256 | SPGP0A04500 | YDR060W |
| YDR056C; YDR057W | | | YDR056C; YDR057W |
| YDR054C; DI49_0248 | DI49_0248 | | YDR054C |
| YDR052C; DI49_0247 | DI49_0247 | | YDR052C |
| YDR051C; DI49_0246 | DI49_0246 | | YDR051C |
| YDR050C; DI49_0245; YDR049W | DI49_0245 | | YDR050C; YDR049W |
| YDR046C; YDR047W; DI49_0243; DI49_0244 | DI49_0243; DI49_0244 | | YDR046C; YDR047W |
| YDR044W; DI49_0240; YDR043C; DI49_0241 | DI49_0240; DI49_0241 | | YDR044W; YDR043C |
| YDR041W; DI49_0237; YDR040C; DI49_0238 | DI49_0237; DI49_0238 | | YDR041W; YDR040C |
| YDR036C; DI49_0234 | DI49_0234 | | YDR036C |
| YDR035W; DI49_0233 | DI49_0233 | | YDR035W |
| YDR032C; DI49_0229 | DI49_0229 | | YDR032C |
| YDR031W; DI49_0228; YDR030C | DI49_0228 | | YDR031W; YDR030C |
| YDR027C; DI49_0225 | DI49_0225 | | YDR027C |
| YDR026C; DI49_0224 | DI49_0224 | | YDR026C |
| YDR022C; DI49_0220 | DI49_0220 | | YDR022C |
| YDR021W; DI49_0219; YDR020C | DI49_0219 | | YDR021W; YDR020C |
| YDR018C; DI49_0216; YDR017C; DI49_0217 | DI49_0216; DI49_0217 | | YDR018C; YDR017C |
| YDR014W-A; YDR016C | | | YDR014W-A; YDR016C |
| YDR013W; YDR014W | | | YDR013W; YDR014W |
| YDR009W; YDR011W | | | YDR009W; YDR011W |
| SPGP0A04960; YDR007W | | SPGP0A04960 | YDR007W |
| YDR004W; YDR005C | | | YDR004W; YDR005C |
| YDR002W; DI49_1055 | DI49_1055 | | YDR002W |
| YDR001C; DI49_1054; YDL001W; DI49_1053 | DI49_1054; DI49_1053 | | YDR001C; YDL001W |
| YDL003W; YDL002C | | | YDL003W; YDL002C |
| YDL005C; DI49_1049 | DI49_1049 | | YDL005C |
| YDL006W; DI49_1048 | DI49_1048 | | YDL006W |
| YDL007W; DI49_1047 | DI49_1047 | | YDL007W |
| YDL010W; DI49_1042; YDL012C; SPGP0I02840 | DI49_1042 | SPGP0I02840 | YDL010W; YDL012C |
| YDL014W; DI49_1040; YDL015C; DI49_1039 | DI49_1040; DI49_1039 | | YDL014W; YDL015C |
| YDL020C; YDL019C; DI49_1035; DI49_1034 | DI49_1035; DI49_1034 | | YDL020C; YDL019C |

Create PDF in your applications with the Pdfcrowd HTML to PDF API PDFCROWD

| | | | |
|---|---|---|---|
| YDL022W; DI49_1032 | DI49_1032 | | YDL022W |
| YDL028C; DI49_1027; YDL029W; SPGP0I02990 | DI49_1027 | SPGP0I02990 | YDL028C; YDL029W |
| YDL031W; DI49_1025; YDL033C; SPGP0I03020 | DI49_1025 | SPGP0I03020 | YDL031W; YDL033C |
| YDL036C; YDL035C | | | YDL036C; YDL035C |
| YDL042C; DI49_1021; YDL043C; DI49_1020 | DI49_1021; DI49_1020 | | YDL042C; YDL043C |
| YDL045C; DI49_1018; YDL045W-A; DI49_1017 | DI49_1018; DI49_1017 | | YDL045C; YDL045W-A |
| YDL047W; DI49_1015; SPGP0I03140 | DI49_1015 | SPGP0I03140 | YDL047W |
| YDL048C; DI49_1014 | DI49_1014 | | YDL048C |
| YDL049C; DI49_1013 | DI49_1013 | | YDL049C |
| YDL051W; DI49_1012; YDL052C; DI49_1011 | DI49_1012; DI49_1011 | | YDL051W; YDL052C |
| YDL054C; YDL053C | | | YDL054C; YDL053C |
| YDL055C; DI49_1007 | DI49_1007 | | YDL055C |
| YDL056W; DI49_1006; YDL057W; DI49_1005 | DI49_1006; DI49_1005 | | YDL056W; YDL057W |
| YDL059C; YDL058W | | | YDL059C; YDL058W |
| YDL061C; DI49_1001 | DI49_1001 | | YDL061C |
| YDL063C; DI49_1000 | DI49_1000 | | YDL063C |
| YDL064W; SPGP0I03290; YDL065C; DI49_0999 | DI49_0999 | SPGP0I03290 | YDL064W; YDL065C |
| YDL067C; DI49_0997; YDL069C; DI49_0996 | DI49_0997; DI49_0996 | | YDL067C; YDL069C |
| YDL072C; YDL070W | | | YDL072C; YDL070W |
| YDL074C; DI49_0992 | DI49_0992 | | YDL074C |
| YDL075W; SPGP0I03380; YLR406C; YLR405W | | SPGP0I03380 | YDL075W; YLR406C; YLR405W |
| YDL076C; DI49_0990 | DI49_0990 | | YDL076C |
| YDL077C; DI49_0989 | DI49_0989 | | YDL077C |
| YDL078C; DI49_0988; SPGP0I03420 | DI49_0988 | SPGP0I03420 | YDL078C |
| YDL080C; DI49_0987 | DI49_0987 | | YDL080C |
| YDL082W; DI49_0985; SPGP0F01690; DI49_4228; YMR142C; YMR140W | DI49_0985; DI49_4228 | SPGP0F01690 | YDL082W; YMR142C; YMR140W |
| YDL083C; SPGP0I03470 | | SPGP0I03470 | YDL083C |
| YDL084W; DI49_0984 | DI49_0984 | | YDL084W |
| YDL085W; DI49_0983; YDL085C-A; DI49_0982 | DI49_0983; DI49_0982 | | YDL085W; YDL085C-A |
| YDL087C; DI49_0980; YDL088C; DI49_0979 | DI49_0980; DI49_0979 | | YDL087C; YDL088C |
| YDL090C; SPGP0I03550 | | SPGP0I03550 | YDL090C |
| YDL091C; DI49_0977; YDL092W; DI49_0976 | DI49_0977; DI49_0976 | | YDL091C; YDL092W |
| YDL095W; YDL093W | | | YDL095W; YDL093W |
| YDL098C; YDL097C; DI49_0973; DI49_0972 | DI49_0973; DI49_0972 | | YDL098C; YDL097C |
| YDL100C; DI49_0970; YDL101C; DI49_0969 | DI49_0970; DI49_0969 | | YDL100C; YDL101C |
| YDL103C; DI49_0967; YDL104C; DI49_0966 | DI49_0967; DI49_0966 | | YDL103C; YDL104C |
| YDL106C; DI49_0964; YDL107W; DI49_0963 | DI49_0964; DI49_0963 | | YDL106C; YDL107W |
| YDL109C; YDL108W | | | YDL109C; YDL108W |
| YDL111C; DI49_0959; YDL112W; DI49_0958 | DI49_0959; DI49_0958 | | YDL111C; YDL112W |

Create PDF in your applications with the Pdfcrowd HTML to PDF API PDFCROWD

| | | |
|---|---|---|
| YDL114W; DI49_0956 | DI49_0956 | YDL114W |
| YDL115C; DI49_0955; YDL116W; DI49_0954 | DI49_0955; DI49_0954 | YDL115C; YDL116W |
| YDL119C; DI49_0952; YDL120W; DI49_0951 | DI49_0952; DI49_0951 | YDL119C; YDL120W |
| YDL122W; DI49_0949; YDL123W; DI49_0948 | DI49_0949; DI49_0948 | YDL122W; YDL123W |
| YDL126C; DI49_0946; YDL127W; DI49_0945 | DI49_0946; DI49_0945 | YDL126C; YDL127W |
| YDL129W; YDL128W | | YDL129W; YDL128W |
| YDL130W-A; YDL130W | | YDL130W-A; YDL130W |
| YDL132W; DI49_0940 | DI49_0940 | YDL132W |
| YDL133W; DI49_0939 | DI49_0939 | YDL133W |
| YDL134C; YDL188C; YDL189W; DI49_0938 | DI49_0938 | YDL134C; YDL188C; YDL189W |
| YDL135C; DI49_0937; DI49_0890 | DI49_0937; DI49_0890 | YDL135C |
| YDL137W; YDL192W; YDL138W; YDL193W | | YDL137W; YDL192W; YDL138W; YDL193W |
| YDL140C; DI49_0934; YDL141W | DI49_0934 | YDL140C; YDL141W |
| YDL143W; YDL142C | | YDL143W; YDL142C |
| YDL145C; YDL144C | | YDL145C; YDL144C |
| YDL147W; YDL146W | | YDL147W; YDL146W |
| YDL149W; YDL148C | | YDL149W; YDL148C |
| YDL153C; YDL150W | | YDL153C; YDL150W |
| YDL155W; YDL154W | | YDL155W; YDL154W |
| YDL157C; YDL156W | | YDL157C; YDL156W |
| YDL160C; YDL159W | | YDL160C; YDL159W |
| YDL161W; YDL160C-A | | YDL161W; YDL160C-A |
| YDL165W; YDL164C | | YDL165W; YDL164C |
| YDL167C; YDL166C | | YDL167C; YDL166C |
| YDL169C; YDL168W | | YDL169C; YDL168W |
| YDL171C; YDL170W | | YDL171C; YDL170W |
| YDL174C; YDL173W | | YDL174C; YDL173W |
| YDL176W; YDL175C | | YDL176W; YDL175C |
| YDL178W; YDL177C | | YDL178W; YDL177C |
| YDL180W; YDL179W | | YDL180W; YDL179W |
| YDL183C; YDL182W | | YDL183C; YDL182W |
| YDL186W; YDL185W | | YDL186W; YDL185W |
| YDL195W; YDL194W | | YDL195W; YDL194W |
| YDL198C; YDL197C | | YDL198C; YDL197C |
| YDL200C; YDL199C | | YDL200C; YDL199C |
| YDL202W; YDL201W | | YDL202W; YDL201W |
| YDL204W; YDL203C | | YDL204W; YDL203C |
| YDL206W; YDL205C | | YDL206W; YDL205C |
| YDL208W; YDL207W | | YDL208W; YDL207W |
| YDL210W; YDL209C | | YDL210W; YDL209C |

Create PDF in your applications with the Pdfcrowd HTML to PDF API PDFCROWD

| | | | |
|---|---|---|---|
| YDL213C; YDL212W | | | YDL213C; YDL212W |
| YDL215C; YDL214C | | | YDL215C; YDL214C |
| YDL217C; YDL216C | | | YDL217C; YDL216C |
| YDL219W; YDL218W | | | YDL219W; YDL218W |
| YDL222C; YDL220C | | | YDL222C; YDL220C |
| YDL224C; YDL223C | | | YDL224C; YDL223C |
| YDL226C; YDL225W | | | YDL226C; YDL225W |
| YDL232W; YDL231C | | | YDL232W; YDL231C |
| YDL234C; YDL233W | | | YDL234C; YDL233W |
| YDL236W; YDL235C | | | YDL236W; YDL235C |
| YDL238C; YDL237W | | | YDL238C; YDL237W |
| YDL240W; YDL239C | | | YDL240W; YDL239C |
| YDL245C; YEL069C; SPGP0A07150; YJR159W; YEL066W | | SPGP0A07150 | YDL245C; YEL069C; YJR159W; YEL066W |
| SPGP0A07180; YEL073C; YEL072W | | SPGP0A07180 | YEL073C; YEL072W |
| YDL248W; SPGP0BG00140 | | SPGP0BG00140 | YDL248W |
| YLR410W; DI49_3908; DI49_3909 | DI49_3908; DI49_3909 | | YLR410W |
| YLR409C; DI49_3907; YLR408C | DI49_3907 | | YLR409C; YLR408C |
| YLR403W; YLR404W | | | YLR403W; YLR404W |
| YLR399C; YLR401C | | | YLR399C; YLR401C |
| YLR397C; YLR398C | | | YLR397C; YLR398C |
| YLR395C; YLR396C | | | YLR395C; YLR396C |
| YLR393W; YLR394W | | | YLR393W; YLR394W |
| YLR390W-A; YLR392C | | | YLR390W-A; YLR392C |
| YLR389C; YLR390W | | | YLR389C; YLR390W |
| YLR387C; DI49_3887 | DI49_3887 | | YLR387C |
| YLR386W; DI49_3886; YLR385C | DI49_3886 | | YLR386W; YLR385C |
| YLR383W; DI49_3883 | DI49_3883 | | YLR383W |
| YLR382C; DI49_3882; YLR381W | DI49_3882 | | YLR382C; YLR381W |
| YLR378C; DI49_3879; SPGP0C04060 | DI49_3879 | SPGP0C04060 | YLR378C |
| YLR375W; DI49_3874; YLR373C; DI49_3875 | DI49_3874; DI49_3875 | | YLR375W; YLR373C |
| YLR371W; DI49_3871; YLR370C; DI49_3872 | DI49_3871; DI49_3872 | | YLR371W; YLR370C |
| YLR368W; DI49_3868; SPGP0T00490 | DI49_3868 | SPGP0T00490 | YLR368W |
| YLR363W-A; YLR364W | | | YLR363W-A; YLR364W |
| YLR362W; YLR363C; DI49_3863; DI49_3864 | DI49_3863; DI49_3864 | | YLR362W; YLR363C |
| YLR361C; YLR360W | | | YLR361C; YLR360W |
| YLR359W; DI49_3858 | DI49_3858 | | YLR359W |
| YLR357W; DI49_3857; YLR356W | DI49_3857 | | YLR357W; YLR356W |
| YLR354C; DI49_3854; YLR353W | DI49_3854 | | YLR354C; YLR353W |
| YLR351C; DI49_3851 | DI49_3851 | | YLR351C |

Create PDF in your applications with the Pdfcrowd HTML to PDF API PDFCROWD

| | | | |
|---|---|---|---|
| YLR350W; DI49_2001; YLR348C; DI49_2000 | DI49_2001; DI49_2000 | | YLR350W; YLR348C |
| YLR346C; YLR347C; DI49_3848; DI49_3849 | DI49_3848; DI49_3849 | | YLR346C; YLR347C |
| YLR344W; YGR034W; SPGP0C03720; DI49_3846 | DI49_3846 | SPGP0C03720 | YLR344W; YGR034W |
| SPGP0Ttrna28R; DI49_3845 | DI49_3845 | SPGP0Ttrna28R | |
| YLR343W; DI49_3844 | DI49_3844 | | YLR343W |
| YLR342W; DI49_3843; YLR341W | DI49_3843 | | YLR342W; YLR341W |
| YLR337C; YLR340W | | | YLR337C; YLR340W |
| YLR335W; YLR336C | | | YLR335W; YLR336C |
| YLR332W; DI49_3834; YLR330W | DI49_3834 | | YLR332W; YLR330W |
| SPGP0T00820; YLR329W | | SPGP0T00820 | YLR329W |
| YLR325C; DI49_3827; YLR324W | DI49_3827 | | YLR325C; YLR324W |
| YLR321C; YLR323C | | | YLR321C; YLR323C |
| YLR319C; DI49_3822; YLR318W | DI49_3822 | | YLR319C; YLR318W |
| YLR315W; YLR316C | | | YLR315W; YLR316C |
| YLR313C; YLR314C | | | YLR313C; YLR314C |
| YLR312C; YLR312W-A | | | YLR312C; YLR312W-A |
| YLR309C; DI49_3815 | DI49_3815 | | YLR309C |
| YLR308W; DI49_3812; YLR307C-A; DI49_3813 | DI49_3812; DI49_3813 | | YLR308W; YLR307C-A |
| YLR306W; YLR307W | | | YLR306W; YLR307W |
| YLR304C; DI49_3809 | DI49_3809 | | YLR304C |
| YLR303W; DI49_3807 | DI49_3807 | | YLR303W |
| YLR301W; DI49_3805 | DI49_3805 | | YLR301W |
| YLR300W; SPGP0C03340; YLR299W | | SPGP0C03340 | YLR300W; YLR299W |
| YLR297W; YLR298C | | | YLR297W; YLR298C |
| YLR293C; DI49_3800; YLR292C | DI49_3800 | | YLR293C; YLR292C |
| YLR290C; DI49_3797; DI49_3798 | DI49_3797; DI49_3798 | | YLR290C |
| YLR289W; DI49_3796 | DI49_3796 | | YLR289W |
| YLR288C; DI49_3795; YLR287C | DI49_3795 | | YLR288C; YLR287C |
| YLR285W; YLR286C; DI49_3793; DI49_3794 | DI49_3793; DI49_3794 | | YLR285W; YLR286C |
| YLR283W; YLR284C | | | YLR283W; YLR284C |
| YLR278C; DI49_3788; DI49_3789 | DI49_3788; DI49_3789 | | YLR278C |
| YLR277C; DI49_3787 | DI49_3787 | | YLR277C |
| YLR276C; DI49_3786; YLR275W | DI49_3786 | | YLR276C; YLR275W |
| YLR273C; YLR274W | | | YLR273C; YLR274W |
| YLR272C; DI49_3781; YLR271W | DI49_3781 | | YLR272C; YLR271W |
| YLR268W; YLR270W | | | YLR268W; YLR270W |
| YLR266C; YLR267W | | | YLR266C; YLR267W |
| YLR264W; DI49_3774; YLR263W; DI49_3775 | DI49_3774; DI49_3775 | | YLR264W; YLR263W |
| YLR260W; YLR262C | | | YLR260W; YLR262C |
| YLR258W; DI49_3769 | DI49_3769 | | YLR258W |

Create PDF in your applications with the Pdfcrowd HTML to PDF API PDFCROWD

| | | | |
|---|---|---|---|
| YLR257W; DI49_3768 | DI49_3768 | | YLR257W |
| YLR256W; DI49_3766; YLR254C | DI49_3766 | | YLR256W; YLR254C |
| YLR251W; DI49_3762; YLR250W; DI49_3763 | DI49_3762; DI49_3763 | | YLR251W; YLR250W |
| YLR248W; DI49_3759 | DI49_3759 | | YLR248W |
| YLR247C; DI49_3758 | DI49_3758 | | YLR247C |
| YLR244C; DI49_3754; DI49_3755 | DI49_3754; DI49_3755 | | YLR244C |
| YLR243W; DI49_3753; YLR242C | DI49_3753 | | YLR243W; YLR242C |
| YLR240W; YLR241W | | | YLR240W; YLR241W |
| YLR238W; DI49_3748 | DI49_3748 | | YLR238W |
| YLR237W; YOR071C; YOR192C | | | YLR237W; YOR071C; YOR192C |
| YLR234W; DI49_3746 | DI49_3746 | | YLR234W |
| YLR233C; DI49_3745; YLR231C | DI49_3745 | | YLR233C; YLR231C |
| YLR229C; DI49_3742 | DI49_3742 | | YLR229C |
| YLR228C; DI49_0398; YDR213W; DI49_0399; DI49_3741 | DI49_0398; DI49_0399; DI49_3741 | | YLR228C; YDR213W |
| YLR225C; DI49_3738 | DI49_3738 | | YLR225C |
| YLR224W; DI49_3737 | DI49_3737 | | YLR224W |
| YLR223C; DI49_3736; YLR222C | DI49_3736 | | YLR223C; YLR222C |
| YLR220W; YLR221C | | | YLR220W; YLR221C |
| YLR218C; YLR219W | | | YLR218C; YLR219W |
| YLR215C; YLR216C | | | YLR215C; YLR216C |
| YLR213C; YLR214W | | | YLR213C; YLR214W |
| YLR211C; YLR212C | | | YLR211C; YLR212C |
| YLR209C; DI49_3723 | DI49_3723 | | YLR209C |
| YLR208W; DI49_3722 | DI49_3722 | | YLR208W |
| YLR207W; DI49_3721; YLR206W | DI49_3721 | | YLR207W; YLR206W |
| YLR204W; DI49_3718; DI49_3719 | DI49_3718; DI49_3719 | | YLR204W |
| YLR203C; DI49_3717; YLR201C | DI49_3717 | | YLR203C; YLR201C |
| YLR199C; YLR200W | | | YLR199C; YLR200W |
| YLR196W; DI49_3712; YLR195C | DI49_3712 | | YLR196W; YLR195C |
| YLR193C; DI49_3709; YLR192C; DI49_3710 | DI49_3709; DI49_3710 | | YLR193C; YLR192C |
| YLR190W; YLR191W | | | YLR190W; YLR191W |
| YLR188W; YLR189C; DI49_3705; DI49_3706 | DI49_3705; DI49_3706 | | YLR188W; YLR189C |
| YLR186W; DI49_3702; DI49_3703 | DI49_3702; DI49_3703 | | YLR186W |
| SPGP0T02090; DI49_3701; YLR183C | DI49_3701 | SPGP0T02090 | YLR183C |
| YLR181C; DI49_3698 | DI49_3698 | | YLR181C |
| YLR180W; DI49_3697; YLR179C | DI49_3697 | | YLR180W; YLR179C |
| YLR177W; DI49_3694; YLR176C; DI49_3695 | DI49_3694; DI49_3695 | | YLR177W; YLR176C |
| YLR174W; YLR175W; DI49_3692; DI49_3693 | DI49_3692; DI49_3693 | | YLR174W; YLR175W |
| YLR172C; DI49_3689; YLR170C; DI49_3690 | DI49_3689; DI49_3690 | | YLR172C; YLR170C |

Create PDF in your applications with the Pdfcrowd HTML to PDF API PDFCROWD

| | | | |
|---|---|---|---|
| YLR167W; DI49_3686 | DI49_3686 | | YLR167W |
| YLR166C; DI49_3685; YLR165C | DI49_3685 | | YLR166C; YLR165C |
| YLR163C; YLR164W | | | YLR163C; YLR164W |
| SPGP0HE00100; SPGP0HD00100 | | SPGP0HE00100; SPGP0HD00100 | |
| YLL061W; DI49_1325; YLL060C | DI49_1325 | | YLL061W; YLL060C |
| YLL057C; YLL058W | | | YLL057C; YLL058W |
| YLL055W; YLL056C | | | YLL055W; YLL056C |
| YLL053C; DI49_3484; YLL051C; DI49_3485 | DI49_3484; DI49_3485 | | YLL053C; YLL051C |
| YLL049W; YLL050C | | | YLL049W; YLL050C |
| YLL046C; YLL048C | | | YLL046C; YLL048C |
| YLL043W; DI49_3490; YLL042C; DI49_3491 | DI49_3490; DI49_3491 | | YLL043W; YLL042C |
| YLL040C; DI49_3493 | DI49_3493 | | YLL040C |
| YLL039C; SPGP0C00310; YLL038C; DI49_3494 | DI49_3494 | SPGP0C00310 | YLL039C; YLL038C |
| YLL035W; YLL036C | | | YLL035W; YLL036C |
| YLL033W; YLL034C | | | YLL033W; YLL034C |
| YLL031C; DI49_3500 | DI49_3500 | | YLL031C |
| YLL029W; DI49_3501 | DI49_3501 | | YLL029W |
| YLL028W; DI49_3502; YLL027W; DI49_3503 | DI49_3502; DI49_3503 | | YLL028W; YLL027W |
| YLL025W; YLR037C; YLR038C | | | YLL025W; YLR037C; YLR038C |
| YLL024C; DI49_3507; DI49_0055 | DI49_3507; DI49_0055 | | YLL024C |
| YLL023C; DI49_3508; YLL022C; DI49_3509 | DI49_3508; DI49_3509 | | YLL023C; YLL022C |
| YLL019C; DI49_3510 | DI49_3510 | | YLL019C |
| YLL018C-A; DI49_3511 | DI49_3511 | | YLL018C-A |
| YLL018C; DI49_3512; YLL016W; DI49_3513 | DI49_3512; DI49_3513 | | YLL018C; YLL016W |
| YLL014W; YLL015W | | | YLL014W; YLL015W |
| YLL012W; DI49_3517 | DI49_3517 | | YLL012W |
| YLL011W; DI49_3518 | DI49_3518 | | YLL011W |
| YLL010C; DI49_3519; YLL009C; DI49_3520 | DI49_3519; DI49_3520 | | YLL010C; YLL009C |
| YLL007C; YLL008W | | | YLL007C; YLL008W |
| YLL005C; DI49_3524 | DI49_3524 | | YLL005C |
| YLL004W; DI49_3525; YLL003W; DI49_3526 | DI49_3525; DI49_3526 | | YLL004W; YLL003W |
| YLL001W; DI49_3528; YLR001C; DI49_3529 | DI49_3528; DI49_3529 | | YLL001W; YLR001C |
| YLR003C; DI49_3530; YLR004C; DI49_3531 | DI49_3530; DI49_3531 | | YLR003C; YLR004C |
| YLR006C; DI49_3533; YLR007W; DI49_3534 | DI49_3533; DI49_3534 | | YLR006C; YLR007W |
| YLR009W; DI49_3536; YLR010C; DI49_3537 | DI49_3536; DI49_3537 | | YLR009W; YLR010C |
| YLR011W; SPGP0Utrna27S | | SPGP0Utrna27S | YLR011W |
| YLR013W; YLR012C | | | YLR013W; YLR012C |
| YLR015W; DI49_3543; YLR016C; DI49_3544 | DI49_3543; DI49_3544 | | YLR015W; YLR016C |
| YLR018C; YLR017W | | | YLR018C; YLR017W |
| YLR021W; YLR020C | | | YLR021W; YLR020C |

Create PDF in your applications with the Pdfcrowd HTML to PDF API PDFCROWD

| | | | |
|---|---|---|---|
| YLR023C; YLR022C | | | YLR023C; YLR022C |
| YLR025W; YLR024C | | | YLR025W; YLR024C |
| YLR027C; DI49_3555; DI49_3556 | DI49_3555; DI49_3556 | | YLR027C |
| YLR028C; DI49_3557 | DI49_3557 | | YLR028C |
| YLR029C; DI49_3558; YLR030W; DI49_3559; DI49_4210; YMR121C | DI49_3558; DI49_3559; DI49_4210 | | YLR029C; YLR030W; YMR121C |
| YLR033W; DI49_3561 | DI49_3561 | | YLR033W |
| YLR034C; DI49_3562 | DI49_3562 | | YLR034C |
| YLR035C; DI49_3563 | DI49_3563 | | YLR035C |
| YLR040C; YLR039C | | | YLR040C; YLR039C |
| YLR043C; DI49_3571 | DI49_3571 | | YLR043C |
| YLR044C; DI49_3572 | DI49_3572 | | YLR044C |
| YLR045C; DI49_3573; YLR046C; DI49_3574 | DI49_3573; DI49_3574 | | YLR045C; YLR046C |
| YLR048W; DI49_3576; YLR049C; DI49_3577 | DI49_3576; DI49_3577 | | YLR048W; YLR049C |
| YLR051C; YLR050C | | | YLR051C; YLR050C |
| YLR053C; YLR052W | | | YLR053C; YLR052W |
| YLR055C; DI49_3583 | DI49_3583 | | YLR055C |
| YLR056W; DI49_3584 | DI49_3584 | | YLR056W |
| YLR057W; DI49_3585 | DI49_3585 | | YLR057W |
| YLR058C; DI49_3586; YLR059C; DI49_3587 | DI49_3586; DI49_3587 | | YLR058C; YLR059C |
| YLR061W; DI49_3589; YLR063W; SPGP0AF01020; DI49_3590 | DI49_3589; DI49_3590 | SPGP0AF01020 | YLR061W; YLR063W |
| YLR065C; YLR064W | | | YLR065C; YLR064W |
| SPGP0U01280; YLR066W | | SPGP0U01280 | YLR066W |
| YLR069C; DI49_3596; YLR070C; DI49_3597 | DI49_3596; DI49_3597 | | YLR069C; YLR070C |
| YLR072W; DI49_3599; YLR073C; DI49_3600 | DI49_3599; DI49_3600 | | YLR072W; YLR073C |
| YLR075W; DI49_3602; YLR077W; DI49_3603 | DI49_3602; DI49_3603 | | YLR075W; YLR077W |
| YLR079W; YLR078C | | | YLR079W; YLR078C |
| YLR081W; DI49_3607 | DI49_3607 | | YLR081W |
| YLR082C; DI49_3609 | DI49_3609 | | YLR082C |
| YLR083C; DI49_3610 | DI49_3610 | | YLR083C |
| YLR084C; DI49_3611 | DI49_3611 | | YLR084C |
| YLR085C; DI49_3612; YLR086W; DI49_3613 | DI49_3612; DI49_3613 | | YLR085C; YLR086W |
| YLR088W; DI49_3615 | DI49_3615 | | YLR088W |
| YLR089C; DI49_3616; YLR090W; DI49_3617 | DI49_3616; DI49_3617 | | YLR089C; YLR090W |
| YLR092W; DI49_3619; YLR093C; DI49_3620 | DI49_3619; DI49_3620 | | YLR092W; YLR093C |
| YLR095C; YLR094C | | | YLR095C; YLR094C |
| YLR097C; YLR096W | | | YLR097C; YLR096W |
| YLR099C; YLR098C | | | YLR099C; YLR098C |
| YLR100W; DI49_3627; YLR102C; DI49_3628 | DI49_3627; DI49_3628 | | YLR100W; YLR102C |

Create PDF in your applications with the Pdfcrowd HTML to PDF API PDFCROWD

| | | |
|---|---|---|
| YLR104W; DI49_3630; YLR105C; DI49_3631 | DI49_3630; DI49_3631 | YLR104W; YLR105C |
| YLR107W; DI49_3633; YLR108C; DI49_3634 | DI49_3633; DI49_3634 | YLR107W; YLR108C |
| YLR113W; DI49_3638 | DI49_3638 | YLR113W |
| YLR114C; DI49_3640; YLR115W; DI49_3641 | DI49_3640; DI49_3641 | YLR114C; YLR115W |
| YLR117C; DI49_3643 | DI49_3643 | YLR117C |
| YLR118C; DI49_3644; YLR119W; DI49_3645 | DI49_3644; DI49_3645 | YLR118C; YLR119W |
| YLR121C; YLR120C | | YLR121C; YLR120C |
| YLR126C; YLR125W; DI49_3649; DI49_3650 | DI49_3649; DI49_3650 | YLR126C; YLR125W |
| YLR128W; YLR127C | | YLR128W; YLR127C |
| YLR130C; YLR129W; DI49_3652; DI49_3653 | DI49_3652; DI49_3653 | YLR130C; YLR129W |
| YLR132C; YLR131C | | YLR132C; YLR131C |
| YLR137W; YLR136C | | YLR137W; YLR136C |
| YLR139C; YLR138W | | YLR139C; YLR138W |
| YLR142W; DI49_3664 | DI49_3664 | YLR142W |
| YLR145W; YLR144C | | YLR145W; YLR144C |
| YLR146W-A; YLR146C | | YLR146W-A; YLR146C |
| YLR148W; DI49_3673 | DI49_3673 | YLR148W |
| YLR149C; DI49_3674 | DI49_3674 | YLR149C |
| YLR150W; DI49_3675; YLR151C; DI49_3676 | DI49_3675; DI49_3676 | YLR150W; YLR151C |
| YLR153C; DI49_3678; YLR154C; DI49_3679 | DI49_3678; DI49_3679 | YLR153C; YLR154C |
| R0020C; YIL001W | | R0020C; YIL001W |
| YEL070W; YEL071W; DI49_5679; DI49_4719 | DI49_5679; DI49_4719 | YEL070W; YEL071W |
| YEL060C; DI49_1333; YEL059C-A | DI49_1333 | YEL060C; YEL059C-A |
| YEL057C; YEL058W | | YEL057C; YEL058W |
| YEL055C; YEL056W | | YEL055C; YEL056W |
| YEL046C; YEL047C | | YEL046C; YEL047C |
| YEL040W; YEL041W | | YEL040W; YEL041W |
| YEL038W; YEL039C | | YEL038W; YEL039C |
| YEL030W; DI49_1362; YEL029C | DI49_1362 | YEL030W; YEL029C |
| YEL026W; DI49_1366; YEL025C | DI49_1366 | YEL026W; YEL025C |
| YEL021W; DI49_1384 | DI49_1384 | YEL021W |
| YEL018W; YEL019C | | YEL018W; YEL019C |
| YEL017W; YEL017C-A | | YEL017W; YEL017C-A |
| YEL015W; YEL016C | | YEL015W; YEL016C |
| YEL009C; DI49_1387 | DI49_1387 | YEL009C |
| YEL006W; DI49_1390; YEL005C | DI49_1390 | YEL006W; YEL005C |
| YEL003W; YEL004W | | YEL003W; YEL004W |
| YEL001C; YEL002C | | YEL001C; YEL002C |
| YER002W; YER001W | | YER002W; YER001W |
| YER004W; YER003C | | YER004W; YER003C |

Create PDF in your applications with the Pdfcrowd HTML to PDF API PDFCROWD

| | | | |
|---|---|---|---|
| YER006W; SPGP0Q01960; YER007W | | SPGP0Q01960 | YER006W; YER007W |
| YER011W; YOR010C | | | YER011W; YOR010C |
| YER016W; SPGP0V00960 | | SPGP0V00960 | YER016W |
| SPGP0V00990; YER017C | | SPGP0V00990 | YER017C |
| YER028C; YER027C | | | YER028C; YER027C |
| YER030W; YER029C | | | YER030W; YER029C |
| YER032W; YER031C | | | YER032W; YER031C |
| YER034W; YER033C | | | YER034W; YER033C |
| YER040W; DI49_1439; YER041W | DI49_1439 | | YER040W; YER041W |
| YER046W; YER045C | | | YER046W; YER045C |
| YER066W; YFL009W; YER067W | | | YER066W; YFL009W; YER067W |
| YER074W; DI49_1478; DI49_2715; YIL069C | DI49_1478; DI49_2715 | | YER074W; YIL069C |
| YER083C; DI49_1488; YER085C | DI49_1488 | | YER083C; YER085C |
| YER087W; DI49_1491; YER087C-B | DI49_1491 | | YER087W; YER087C-B |
| YER091C; DI49_1497; YER092W | DI49_1497 | | YER091C; YER092W |
| YER093C-A; YER093C | | | YER093C-A; YER093C |
| YER095W; DI49_1501; YER096W | DI49_1501 | | YER095W; YER096W |
| YER100W; YER099C | | | YER100W; YER099C |
| YER103W; DI49_1509; YER104W | DI49_1509 | | YER103W; YER104W |
| YER106W; YER105C | | | YER106W; YER105C |
| YER117W; YBL087C | | | YER117W; YBL087C |
| YER118C; DI49_1523; YER119C | DI49_1523 | | YER118C; YER119C |
| YER123W; DI49_1527; YER124C | DI49_1527 | | YER123W; YER124C |
| YER127W; DI49_1531; YER128W | DI49_1531 | | YER127W; YER128W |
| YER130C; YER129W | | | YER130C; YER129W |
| YER132C; DI49_1536 | DI49_1536 | | YER132C |
| YER136W; DI49_1542; YER137C | DI49_1542 | | YER136W; YER137C |
| SPGP0HX00100; DI49_3953 | DI49_3953 | SPGP0HX00100 | |
| DI49_2624; YIL162W; YIL164C; SPGP0OF00100 | DI49_2624 | SPGP0OF00100 | YIL162W; YIL164C |
| DI49_2625; YIL161W | DI49_2625 | | YIL161W |
| DI49_2626; YIL160C | DI49_2626 | | YIL160C |
| DI49_2627; YIL159W; DI49_2628 | DI49_2627; DI49_2628 | | YIL159W |
| DI49_2630; DI49_2629 | DI49_2630; DI49_2629 | | |
| DI49_2632; DI49_2631 | DI49_2632; DI49_2631 | | |
| DI49_2634; DI49_2633 | DI49_2634; DI49_2633 | | |
| DI49_2636; YIL151C | DI49_2636 | | YIL151C |
| DI49_2637; YIL150C | DI49_2637 | | YIL150C |
| DI49_2638; YIL149C | DI49_2638 | | YIL149C |
| DI49_2639; YIL148W | DI49_2639 | | YIL148W |
| DI49_2640; YIL147C | DI49_2640 | | YIL147C |

Create PDF in your applications with the Pdfcrowd HTML to PDF API PDFCROWD

| | | | |
|---|---|---|---|
| DI49_2641; YIL146C; DI49_2642 | DI49_2641; DI49_2642 | | YIL146C |
| DI49_2644; YIL143C; YIL144W | DI49_2644 | | YIL143C; YIL144W |
| DI49_2645; YIL142W; DI49_2646 | DI49_2645; DI49_2646 | | YIL142W |
| DI49_2648; DI49_2647 | DI49_2648; DI49_2647 | | |
| DI49_2650; DI49_2649 | DI49_2650; DI49_2649 | | |
| DI49_2652; DI49_2651 | DI49_2652; DI49_2651 | | |
| DI49_2654; DI49_2653 | DI49_2654; DI49_2653 | | |
| DI49_2656; DI49_2655 | DI49_2656; DI49_2655 | | |
| DI49_2658; YIL128W; DI49_2659 | DI49_2658; DI49_2659 | | YIL128W |
| DI49_2661; YIL125W | DI49_2661 | | YIL125W |
| DI49_2662; YIL124W | DI49_2662 | | YIL124W |
| DI49_2663; YIL123W | DI49_2663 | | YIL123W |
| DI49_2664; YIL122W; DI49_2665 | DI49_2664; DI49_2665 | | YIL122W |
| DI49_2667; DI49_2666; YIL120W; YIL121W | DI49_2667; DI49_2666 | | YIL120W; YIL121W |
| DI49_2669; DI49_2668 | DI49_2669; DI49_2668 | | |
| DI49_2671; YIL115C; DI49_2672; YIL116W | DI49_2671; DI49_2672 | | YIL115C; YIL116W |
| DI49_2674; DI49_2673 | DI49_2674; DI49_2673 | | |
| DI49_2675; YIL110W; YIL111W | DI49_2675 | | YIL110W; YIL111W |
| DI49_2676; YIL109C | DI49_2676 | | YIL109C |
| DI49_2677; YIL108W | DI49_2677 | | YIL108W |
| DI49_2678; YIL107C; DI49_2679 | DI49_2678; DI49_2679 | | YIL107C |
| DI49_2681; DI49_2680 | DI49_2681; DI49_2680 | | |
| DI49_2683; DI49_2682; YIL103W; YIL104C | DI49_2683; DI49_2682 | | YIL103W; YIL104C |
| DI49_2685; YIL101C; DI49_2686 | DI49_2685; DI49_2686 | | YIL101C |
| DI49_2688; DI49_2687 | DI49_2688; DI49_2687 | | |
| DI49_2691; YIL095W | DI49_2691 | | YIL095W |
| DI49_2692; YIL094C | DI49_2692 | | YIL094C |
| DI49_2693; YIL093C; DI49_2694 | DI49_2693; DI49_2694 | | YIL093C |
| DI49_2696; DI49_2695; YIL091C; YIL092W | DI49_2696; DI49_2695 | | YIL091C; YIL092W |
| SPGP0Wtrna2E; DI49_2698; DI49_2699; DI49_0457 | DI49_2698; DI49_2699; DI49_0457 | SPGP0Wtrna2E | |
| DI49_2701; DI49_2700 | DI49_2701; DI49_2700 | | |
| SPGP0W00880; YIL083C; YIL084C | | SPGP0W00880 | YIL083C; YIL084C |
| DI49_2706; YIL077C | DI49_2706 | | YIL077C |
| DI49_2707; DI49_4178; DI49_2708 | DI49_2707; DI49_4178; DI49_2708 | | |
| DI49_2713; DI49_2712 | DI49_2713; DI49_2712 | | |
| DI49_2716; YIL068C | DI49_2716 | | YIL068C |
| DI49_2717; YIL067C | DI49_2717 | | YIL067C |
| DI49_2718; YIL066C; DI49_2719 | DI49_2718; DI49_2719 | | YIL066C |

Create PDF in your applications with the Pdfcrowd HTML to PDF API PDFCROWD

| | | | |
|---|---|---|---|
| DI49_2720; YIL063C; YIL064W | DI49_2720 | | YIL063C; YIL064W |
| DI49_2721; YIL062C; DI49_2722 | DI49_2721; DI49_2722 | | YIL062C |
| DI49_2725; YIL056W | DI49_2725 | | YIL056W |
| DI49_2726; YIL055C | DI49_2726 | | YIL055C |
| DI49_2727; YIL053W | DI49_2727 | | YIL053W |
| SPGP0W01130; YIL052C | | SPGP0W01130 | YIL052C |
| DI49_2728; YIL051C; DI49_2729 | DI49_2728; DI49_2729 | | YIL051C |
| DI49_2731; YIL048W | DI49_2731 | | YIL048W |
| DI49_2732; YIL047C | DI49_2732 | | YIL047C |
| DI49_2733; YIL046W | DI49_2733 | | YIL046W |
| DI49_2734; YIL045W; DI49_2735 | DI49_2734; DI49_2735 | | YIL045W |
| DI49_2737; DI49_2736 | DI49_2737; DI49_2736 | | |
| DI49_2739; DI49_2738 | DI49_2739; DI49_2738 | | |
| DI49_2741; YIL038C; DI49_2742 | DI49_2741; DI49_2742 | | YIL038C |
| DI49_2744; YIL035C; YIL036W | DI49_2744 | | YIL035C; YIL036W |
| DI49_2745; YIL034C | DI49_2745 | | YIL034C |
| DI49_2746; YIL033C; DI49_2747 | DI49_2746; DI49_2747 | | YIL033C |
| DI49_2752; DI49_2751 | DI49_2752; DI49_2751 | | |
| DI49_2754; DI49_2753 | DI49_2754; DI49_2753 | | |
| DI49_2756; YIL021W; DI49_2757 | DI49_2756; DI49_2757 | | YIL021W |
| DI49_2759; SPGP0AF00380; DI49_2760; SPGP0AB00230 | DI49_2759; DI49_2760 | SPGP0AF00380; SPGP0AB00230 | |
| DI49_2762; YIL015W; YIL016W | DI49_2762 | | YIL015W; YIL016W |
| DI49_2766; DI49_2765 | DI49_2766; DI49_2765 | | |
| DI49_2768; DI49_2767 | DI49_2768; DI49_2767 | | |
| SPGP0W01530; DI49_2770; DI49_2771; DI49_1642 | DI49_2770; DI49_2771; DI49_1642 | SPGP0W01530 | |
| DI49_2773; DI49_2772 | DI49_2773; DI49_2772 | | |
| DI49_2775; DI49_2774 | DI49_2775; DI49_2774 | | |
| DI49_2777; YIL002W-A | DI49_2777 | | YIL002W-A |
| DI49_2781; SPGP0QF00100; DI49_2782; SPGP0QG00100 | DI49_2781; DI49_2782 | SPGP0QF00100; SPGP0QG00100 | |
| DI49_2784; DI49_2783 | DI49_2784; DI49_2783 | | |
| DI49_2787; YIR007W | DI49_2787 | | YIR007W |
| DI49_2788; YIR008C; DI49_2789 | DI49_2788; DI49_2789 | | YIR008C |
| DI49_2791; YIR011C; DI49_2792 | DI49_2791; DI49_2792 | | YIR011C |
| DI49_2794; DI49_2793 | DI49_2794; DI49_2793 | | |
| DI49_2796; DI49_2795 | DI49_2796; DI49_2795 | | |
| DI49_2798; YIR018W; DI49_2799 | DI49_2798; DI49_2799 | | YIR018W |
| DI49_2801; YIR022W; DI49_2802 | DI49_2801; DI49_2802 | | YIR022W |

Create PDF in your applications with the Pdfcrowd HTML to PDF API PDFCROWD

| | | | |
|---|---|---|---|
| DI49_2804; DI49_2803 | DI49_2804; DI49_2803 | | |
| DI49_2806; DI49_2805 | DI49_2806; DI49_2805 | | |
| DI49_2808; YIR029W; DI49_2809; YIR030C | DI49_2808; DI49_2809 | | YIR029W; YIR030C |
| DI49_2811; DI49_2810 | DI49_2811; DI49_2810 | | |
| DI49_2813; YIR034C; DI49_2814; YIR035C | DI49_2813; DI49_2814 | | YIR034C; YIR035C |
| DI49_2816; DI49_2815 | DI49_2816; DI49_2815 | | |
| YHL040C; YHL043W | | | YHL040C; YHL043W |
| YHL038C; YHL039W | | | YHL038C; YHL039W |
| YHL035C; YHL036W | | | YHL035C; YHL036W |
| YHL033C; YLL045C | | | YHL033C; YLL045C |
| YHL032C; SPGP0H00180 | | SPGP0H00180 | YHL032C |
| YHL031C; DI49_4816 | DI49_4816 | | YHL031C |
| YHL030W; DI49_4817 | DI49_4817 | | YHL030W |
| YHL029C; DI49_4818; YHL028W; DI49_4819 | DI49_4818; DI49_4819 | | YHL029C; YHL028W |
| YHL026C; YHL027W | | | YHL026C; YHL027W |
| YHL024W; DI49_4823 | DI49_4823 | | YHL024W |
| YHL023C; DI49_4824 | DI49_4824 | | YHL023C |
| YHL020C; DI49_4828; YHL019C; DI49_4829 | DI49_4828; DI49_4829 | | YHL020C; YHL019C |
| YHL017W; DI49_4831 | DI49_4831 | | YHL017W |
| YHL016C; DI49_4832 | DI49_4832 | | YHL016C |
| YHL015W; DI49_4833; YHL014C; SPGP0H00360 | DI49_4833 | SPGP0H00360 | YHL015W; YHL014C |
| YHL012W; YHL013C | | | YHL012W; YHL013C |
| YHL010C; DI49_4838; YHL009C; DI49_4839 | DI49_4838; DI49_4839 | | YHL010C; YHL009C |
| YHL008C; DI49_4841 | DI49_4841 | | YHL008C |
| YHL003C; DI49_4845 | DI49_4845 | | YHL003C |
| YHL002W; DI49_4846 | DI49_4846 | | YHL002W |
| SPGP0X00490; DI49_4847; YHR001W; DI49_3367; DI49_4848 | DI49_4847; DI49_3367; DI49_4848 | SPGP0X00490 | YHR001W |
| YHR002W; DI49_4850 | DI49_4850 | | YHR002W |
| YHR003C; DI49_4851; YHR004C; DI49_4852 | DI49_4851; DI49_4852 | | YHR003C; YHR004C |
| YHR005C-A; YHR005C | | | YHR005C-A; YHR005C |
| YHR008C; DI49_4858 | DI49_4858 | | YHR008C |
| YHR009C; DI49_4859 | DI49_4859 | | YHR009C |
| YHR010W; SPGP0H00600; YHR011W; DI49_4860 | DI49_4860 | SPGP0H00600 | YHR010W; YHR011W |
| YHR013C; DI49_4862; YHR014W; DI49_4863 | DI49_4862; DI49_4863 | | YHR013C; YHR014W |
| YHR017W; DI49_2424 | DI49_2424 | | YHR017W |
| YHR044C; YHR042W | | | YHR044C; YHR042W |
| YHR046C; YHR045W | | | YHR046C; YHR045W |
| YHR048W; YHR047C | | | YHR048W; YHR047C |
| YHR050W; YHR049W | | | YHR050W; YHR049W |

Create PDF in your applications with the Pdfcrowd HTML to PDF API PDFCROWD

| | | | |
|---|---|---|---|
| YHR057C; YHR056C | | | YHR057C; YHR056C |
| YHR062C; YHR061C | | | YHR062C; YHR061C |
| YHR073W; YHR072W-A | | | YHR073W; YHR072W-A |
| YHR079C-A; YHR079C | | | YHR079C-A; YHR079C |
| YHR081W; YHR080C | | | YHR081W; YHR080C |
| YHR083W; YHR082C | | | YHR083W; YHR082C |
| YHR085W; YHR084W | | | YHR085W; YHR084W |
| YHR098C; DI49_2503 | DI49_2503 | | YHR098C |
| YHR105W; YHR104W | | | YHR105W; YHR104W |
| YHR110W; YHR109W | | | YHR110W; YHR109W |
| YHR112C; YHR111W | | | YHR112C; YHR111W |
| YHR117W; YHR116W | | | YHR117W; YHR116W |
| YHR122W; YHR121W | | | YHR122W; YHR121W |
| YHR135C; YHR134W | | | YHR135C; YHR134W |
| YHR137W; YHR136C | | | YHR137W; YHR136C |
| YHR139C; YHR138C | | | YHR139C; YHR138C |
| YHR144C; YHR143W-A | | | YHR144C; YHR143W-A |
| YHR146W; SPGP0Xtrna9P | | SPGP0Xtrna9P | YHR146W |
| YIR004W; YPL245W | | | YIR004W; YPL245W |
| YGR109C; DI49_2069; DI49_2068 | DI49_2069; DI49_2068 | | YGR109C |
| YGR112W; DI49_2073; YGR113W | DI49_2073 | | YGR112W; YGR113W |
| YGR117C; YGR116W | | | YGR117C; YGR116W |
| YGR119C; DI49_2078 | DI49_2078 | | YGR119C |
| YGR120C; DI49_2079 | DI49_2079 | | YGR120C |
| YGR123C; DI49_2084 | DI49_2084 | | YGR123C |
| YGR124W; DI49_2086 | DI49_2086 | | YGR124W |
| YGR125W; DI49_2087; YGR126W | DI49_2087 | | YGR125W; YGR126W |
| YGR128C; YGR127W | | | YGR128C; YGR127W |
| YGR130C; YGR129W | | | YGR130C; YGR129W |
| YGR132C; YGR131W | | | YGR132C; YGR131W |
| YGR134W; DI49_2096 | DI49_2096 | | YGR134W |
| YGR135W; DI49_2097 | DI49_2097 | | YGR135W |
| YGR136W; DI49_2098 | DI49_2098 | | YGR136W |
| YGR138C; YPR156C | | | YGR138C; YPR156C |
| YGR140W; DI49_2101; YGR141W; DI49_2100 | DI49_2101; DI49_2100 | | YGR140W; YGR141W |
| YGR143W; DI49_2105 | DI49_2105 | | YGR143W |
| YGR144W; DI49_2107 | DI49_2107 | | YGR144W |
| YGR145W; DI49_2108 | DI49_2108 | | YGR145W |
| YGR146C; DI49_2109; YGR147C | DI49_2109 | | YGR146C; YGR147C |
| YGR149W; YGR148C | | | YGR149W; YGR148C |

Create PDF in your applications with the Pdfcrowd HTML to PDF API PDFCROWD

| | | | |
|---|---|---|---|
| YGR152C; DI49_2116; YGR153W | DI49_2116 | | YGR152C; YGR153W |
| YGR155W; DI49_2118; DI49_2117 | DI49_2118; DI49_2117 | | YGR155W |
| YGR156W; DI49_2119 | DI49_2119 | | YGR156W |
| YGR157W; DI49_2120; YGR158C | DI49_2120 | | YGR157W; YGR158C |
| YGR161C; YGR159C | | | YGR161C; YGR159C |
| YGR162W; DI49_2126 | DI49_2126 | | YGR162W |
| YGR163W; DI49_2127 | DI49_2127 | | YGR163W |
| YGR167W; YGR166W | | | YGR167W; YGR166W |
| YGR169C; YGR168C | | | YGR169C; YGR168C |
| YGR170W; DI49_2135; DI49_2134 | DI49_2135; DI49_2134 | | YGR170W |
| YGR171C; DI49_2136 | DI49_2136 | | YGR171C |
| YGR172C; DI49_2137 | DI49_2137 | | YGR172C |
| YGR173W; DI49_2138 | DI49_2138 | | YGR173W |
| YGR174C; DI49_2140 | DI49_2140 | | YGR174C |
| YGR177C; DI49_2141; DI49_2142 | DI49_2141; DI49_2142 | | YGR177C |
| YGR178C; DI49_2143; YGR179C | DI49_2143 | | YGR178C; YGR179C |
| YGR184C; DI49_2150; DI49_2149 | DI49_2150; DI49_2149 | | YGR184C |
| YGR185C; DI49_2151 | DI49_2151 | | YGR185C |
| YGR186W; DI49_2152; YGR187C | DI49_2152 | | YGR186W; YGR187C |
| YGR189C; YGR188C; DI49_2154; DI49_2153 | DI49_2154; DI49_2153 | | YGR189C; YGR188C |
| YGR191W; DI49_2158 | DI49_2158 | | YGR191W |
| YGR195W; DI49_2162; YGR196C; DI49_2161 | DI49_2162; DI49_2161 | | YGR195W; YGR196C |
| YGR198W; DI49_2165; YGR199W | DI49_2165 | | YGR198W; YGR199W |
| YGR201C; YGR200C | | | YGR201C; YGR200C |
| YGR203W; YGR202C | | | YGR203W; YGR202C |
| YGR205W; DI49_2172; YGR206W | DI49_2172 | | YGR205W; YGR206W |
| YGR208W; DI49_2175; DI49_2174 | DI49_2175; DI49_2174 | | YGR208W |
| SPGP0Y01010; DI49_2176; YGR211W | DI49_2176 | SPGP0Y01010 | YGR211W |
| YGR213C; YGR212W | | | YGR213C; YGR212W |
| YGR215W; YGR214W | | | YGR215W; YGR214W |
| YGR217W; DI49_2184 | DI49_2184 | | YGR217W |
| YGR218W; DI49_2186; YGR220C | DI49_2186 | | YGR218W; YGR220C |
| YGR222W; YGR221C | | | YGR222W; YGR221C |
| YGR224W; YGR223C | | | YGR224W; YGR223C |
| YGR227W; YGR225W | | | YGR227W; YGR225W |
| YGR230W; YGR229C | | | YGR230W; YGR229C |
| YGR232W; DI49_2196; YGR233C | DI49_2196 | | YGR232W; YGR233C |
| YGR235C; YGR234W | | | YGR235C; YGR234W |
| YGR237C; YGR236C | | | YGR237C; YGR236C |
| YGR239C; YGR238C | | | YGR239C; YGR238C |

Create PDF in your applications with the Pdfcrowd HTML to PDF API PDFCROWD

| | | | |
|---|---|---|---|
| YGR241C; YGR240C | | | YGR241C; YGR240C |
| YGR244C; YGR243W | | | YGR244C; YGR243W |
| YGR246C; DI49_2209; YGR247W | DI49_2209 | | YGR246C; YGR247W |
| YGR249W; YGR248W | | | YGR249W; YGR248W |
| YGR251W; YGR250C | | | YGR251W; YGR250C |
| YGR253C; DI49_2217 | DI49_2217 | | YGR253C |
| YGR254W; YHR174W; YGR255C; YHR175W | | | YGR254W; YHR174W; YGR255C; YHR175W |
| YGR256W; DI49_2221; YGR257C | DI49_2221 | | YGR256W; YGR257C |
| YGR260W; DI49_2224 | DI49_2224 | | YGR260W |
| YGR261C; DI49_2225; YGR262C | DI49_2225 | | YGR261C; YGR262C |
| YGR264C; DI49_2228; YGR266W | DI49_2228 | | YGR264C; YGR266W |
| YGR268C; YGR267C | | | YGR268C; YGR267C |
| YGR271W; DI49_2233 | DI49_2233 | | YGR271W |
| YGR271C-A; DI49_2234; YGR273C | DI49_2234 | | YGR271C-A; YGR273C |
| YGR275W; YGR274C | | | YGR275W; YGR274C |
| YGR277C; YGR276C; DI49_2238; DI49_2237 | DI49_2238; DI49_2237 | | YGR277C; YGR276C |
| YGR279C; DI49_2240; YGR280C; SPGP0E00220; YMR305C; DI49_4389 | DI49_2240; DI49_4389 | SPGP0E00220 | YGR279C; YGR280C; YMR305C |
| YGR282C; DI49_2243; YGR283C; SPGP0E00190 | DI49_2243 | SPGP0E00190 | YGR282C; YGR283C |
| YGR285C; DI49_2246 | DI49_2246 | | YGR285C |
| YKL043W; YKL042W | | | YKL043W; YKL042W |
| YKL050C; YKL049C | | | YKL050C; YKL049C |
| YKL052C; YKL051W | | | YKL052C; YKL051W |
| YKL061W; YKL060C | | | YKL061W; YKL060C |
| YKL063C; YKL062W | | | YKL063C; YKL062W |
| YKL065C; YKL064W | | | YKL065C; YKL064W |
| YKL067W; YKL065W-A | | | YKL067W; YKL065W-A |
| YKL071W; YKL070W | | | YKL071W; YKL070W |
| YKL074C; YKL073W | | | YKL074C; YKL073W |
| YKL077W; YKL075C | | | YKL077W; YKL075C |
| YKL079W; YKL078W | | | YKL079W; YKL078W |
| SPGP0Z00620; YKL080W | | SPGP0Z00620 | YKL080W |
| YKL084W; YKL082C | | | YKL084W; YKL082C |
| YKL107W; YKL106W | | | YKL107W; YKL106W |
| YKL109W; YKL108W | | | YKL109W; YKL108W |
| YKL130C; YKL129C | | | YKL130C; YKL129C |
| YKL133C; YKL132C | | | YKL133C; YKL132C |
| YKL135C; YKL134C | | | YKL135C; YKL134C |
| YKL140W; YKL139W | | | YKL140W; YKL139W |
| YKL151C; YKL150W | | | YKL151C; YKL150W |

Create PDF in your applications with the Pdfcrowd HTML to PDF API PDFCROWD

| | | | |
|---|---|---|---|
| YKL161C; YKL160W | | | YKL161C; YKL160W |
| YKL162C-A; YKL162C | | | YKL162C-A; YKL162C |
| YKL164C; YKL163W | | | YKL164C; YKL163W |
| YKL170W; YKL168C | | | YKL170W; YKL168C |
| YKL172W; YKL171W | | | YKL172W; YKL171W |
| YKL174C; YKL173W | | | YKL174C; YKL173W |
| YKL176C; YKL175W | | | YKL176C; YKL175W |
| YKL186C; YKL185W | | | YKL186C; YKL185W |
| YKL188C; YKL187C | | | YKL188C; YKL187C |
| YKL192C; YKL191W | | | YKL192C; YKL191W |
| YKL208W; YKL207W | | | YKL208W; YKL207W |
| SPGP0AA00120; YMR186W | | SPGP0AA00120 | YMR186W |
| YPL237W; DI49_5231; YPL236C | DI49_5231 | | YPL237W; YPL236C |
| YPL233W; DI49_5235 | DI49_5235 | | YPL233W |
| YPL232W; DI49_4266; DI49_4265 | DI49_4266; DI49_4265 | | YPL232W |
| YPL225W; DI49_5243; YPL224C | DI49_5243 | | YPL225W; YPL224C |
| YPL222W; YPL223C | | | YPL222W; YPL223C |
| YPL219W; DI49_5250; SPGP0AA00330 | DI49_5250 | SPGP0AA00330 | YPL219W |
| YPL211W; DI49_5257; YPL210C | DI49_5257 | | YPL211W; YPL210C |
| YPL208W; YPL209C | | | YPL208W; YPL209C |
| YPL206C; YPL207W | | | YPL206C; YPL207W |
| YPL203W; DI49_5264; YPL202C | DI49_5264 | | YPL203W; YPL202C |
| YPL187W; DI49_1885; YPL186C; DI49_1884 | DI49_1885; DI49_1884 | | YPL187W; YPL186C |
| YPL181W; YPL183C | | | YPL181W; YPL183C |
| YPL161C; YPL162C | | | YPL161C; YPL162C |
| YPL159C; YPL160W | | | YPL159C; YPL160W |
| YPL157W; YPL158C | | | YPL157W; YPL158C |
| YPL155C; YPL156C | | | YPL155C; YPL156C |
| YPL147W; YPL148C | | | YPL147W; YPL148C |
| YPL145C; DI49_5320; YPL144W | DI49_5320 | | YPL145C; YPL144W |
| YPL141C; DI49_5323; YPL140C | DI49_5323 | | YPL141C; YPL140C |
| YPL128C; YPL129W | | | YPL128C; YPL129W |
| YPL125W; DI49_5338; YPL124W | DI49_5338 | | YPL125W; YPL124W |
| YPL122C; DI49_5341; YPL121C | DI49_5341 | | YPL122C; YPL121C |
| YPL119C; YPL120W | | | YPL119C; YPL120W |
| YPL117C; YPL118W | | | YPL117C; YPL118W |
| YPL110C; DI49_5353; YPL109C | DI49_5353 | | YPL110C; YPL109C |
| YPL107W; YPL108W | | | YPL107W; YPL108W |
| YPL101W; YPL103C | | | YPL101W; YPL103C |
| YPL099C; YPL100W | | | YPL099C; YPL100W |

Create PDF in your applications with the Pdfcrowd HTML to PDF API PDFCROWD

| | | | |
|---|---|---|---|
| YPL097W; YPL098C | | | YPL097W; YPL098C |
| YPL096W; YPL096C-A | | | YPL096W; YPL096C-A |
| YPL093W; DI49_5369; YPL092W | DI49_5369 | | YPL093W; YPL092W |
| YPL087W; YPL088W | | | YPL087W; YPL088W |
| YPL085W; YPL086C | | | YPL085W; YPL086C |
| YPL083C; YPL084W | | | YPL083C; YPL084W |
| YPL078C; YPL079W | | | YPL078C; YPL079W |
| YPL076W; YPL077C | | | YPL076W; YPL077C |
| YPL066W; YPL067C | | | YPL066W; YPL067C |
| YPL064C; YPL065W | | | YPL064C; YPL065W |
| YGR086C; YPL063W | | | YGR086C; YPL063W |
| YIL003W; DI49_2776; YIL006W | DI49_2776 | | YIL003W; YIL006W |
| YIL011W; YIL010W | | | YIL011W; YIL010W |
| YIL014W; YIL013C | | | YIL014W; YIL013C |
| YIL019W; DI49_2758; YIL020C | DI49_2758 | | YIL019W; YIL020C |
| YIL022W; DI49_2755; YIL023C | DI49_2755 | | YIL022W; YIL023C |
| YIL026C; YIL024C | | | YIL026C; YIL024C |
| YIL030C; DI49_2748; YIL031W | DI49_2748 | | YIL030C; YIL031W |
| YIL039W; DI49_2740; YIL040W | DI49_2740 | | YIL039W; YIL040W |
| YIL042C; YIL043C; YIL044C | | | YIL042C; YIL043C; YIL044C |
| YIL049W; DI49_2730; YIL050W | DI49_2730 | | YIL049W; YIL050W |
| YIL070C; DI49_2714; YIL071C | DI49_2714 | | YIL070C; YIL071C |
| YIL073C; YIL072W | | | YIL073C; YIL072W |
| YIL075C; DI49_2709; YIL076W | DI49_2709 | | YIL075C; YIL076W |
| YIL078W; DI49_2705; YIL079C | DI49_2705 | | YIL078W; YIL079C |
| YIL087C; YIL085C | | | YIL087C; YIL085C |
| YIL089W; YIL088C | | | YIL089W; YIL088C |
| YIL098C; YIL097W | | | YIL098C; YIL097W |
| YIL106W; YIL105C | | | YIL106W; YIL105C |
| YIL113W; YIL112W | | | YIL113W; YIL112W |
| YIL118W; YIL117C | | | YIL118W; YIL117C |
| YIL126W; DI49_2660; YIL127C | DI49_2660 | | YIL126W; YIL127C |
| YIL129C; DI49_2657; YIL130W | DI49_2657 | | YIL129C; YIL130W |
| YIL132C; YIL131C | | | YIL132C; YIL131C |
| YIL134W; YIL133C | | | YIL134W; YIL133C |
| YIL136W; YIL135C | | | YIL136W; YIL135C |
| YIL138C; YIL137C | | | YIL138C; YIL137C |
| YIL140W; YIL139C | | | YIL140W; YIL139C |
| YIL152W; DI49_2635; YIL153W | DI49_2635 | | YIL152W; YIL153W |
| YIL155C; SPGP0AB01460 | | SPGP0AB01460 | YIL155C |

Create PDF in your applications with the Pdfcrowd HTML to PDF API PDFCROWD

| | | | |
|---|---|---|---|
| YIL156W-B; YIL156W | | | YIL156W-B; YIL156W |
| YIL158W; YIL157C | | | YIL158W; YIL157C |
| YIL167W; YIL166C | | | YIL167W; YIL166C |
| R0010W; SPGP0AB01590 | | SPGP0AB01590 | R0010W |
| YKL222C; SPGP0KK00100 | | SPGP0KK00100 | YKL222C |
| YPR025C; YPR024W | | | YPR025C; YPR024W |
| YPR028W; SPGP0ACtrna19F | | SPGP0ACtrna19F | YPR028W |
| YPR030W; YPR029C | | | YPR030W; YPR029C |
| YPR036W; DI49_5494; YPR036W-A | DI49_5494 | | YPR036W; YPR036W-A |
| YPR040W; YPR037C | | | YPR040W; YPR037C |
| YPR043W; DI49_5500; YPR045C | DI49_5500 | | YPR043W; YPR045C |
| YPR047W; DI49_5503; YPR048W | DI49_5503 | | YPR047W; YPR048W |
| YPR051W; YPR049C | | | YPR051W; YPR049C |
| YPR056W; DI49_5510; YPR057W | DI49_5510 | | YPR056W; YPR057W |
| YPR072W; DI49_5524; YPR073C | DI49_5524 | | YPR072W; YPR073C |
| YPR075C; YPR074C | | | YPR075C; YPR074C |
| YPR079W; YPR078C | | | YPR079W; YPR078C |
| YPR081C; DI49_5531; YPR082C | DI49_5531 | | YPR081C; YPR082C |
| YPR084W; DI49_5534; YPR085C | DI49_5534 | | YPR084W; YPR085C |
| YPR089W; DI49_5538; YPR091C | DI49_5538 | | YPR089W; YPR091C |
| YPR101W; YPR100W | | | YPR101W; YPR100W |
| YPR105C; DI49_5549; YPR106W | DI49_5549 | | YPR105C; YPR106W |
| YPR108W; YPR107C | | | YPR108W; YPR107C |
| YPR111W; YPR110C | | | YPR111W; YPR110C |
| YPR124W; YPR122W | | | YPR124W; YPR122W |
| YPR131C; YPR129W | | | YPR131C; YPR129W |
| YPR145C-A; YPR145W | | | YPR145C-A; YPR145W |
| YPR148C; YPR147C | | | YPR148C; YPR147C |
| YPR151C; YPR149W | | | YPR151C; YPR149W |
| YPR153W; YPR152C | | | YPR153W; YPR152C |
| YPR155C; YPR154W | | | YPR155C; YPR154W |
| YPR157W; DI49_5599; YPR158W | DI49_5599 | | YPR157W; YPR158W |
| DI49_0079; YJR160C; SPGP0BF00120 | DI49_0079 | SPGP0BF00120 | YJR160C |
| SPGP0B00110; DI49_5680 | DI49_5680 | SPGP0B00110 | |
| DI49_5681; SPGP0H03940; SPGP0NU00100 | DI49_5681 | SPGP0H03940; SPGP0NU00100 | |
| SPGP0B00130; DI49_5682 | DI49_5682 | SPGP0B00130 | |
| DI49_0085; YBL104C; DI49_0086 | DI49_0085; DI49_0086 | | YBL104C |
| DI49_0088; DI49_0087 | DI49_0088; DI49_0087 | | |
| DI49_0090; YBL099W; DI49_0091 | DI49_0090; DI49_0091 | | YBL099W |
| DI49_0093; DI49_0092 | DI49_0093; DI49_0092 | | |

Create PDF in your applications with the Pdfcrowd HTML to PDF API PDFCROWD

| | | | |
|---|---|---|---|
| DI49_0095; YBL092W; DI49_0096 | DI49_0095; DI49_0096 | | YBL092W |
| DI49_0104; DI49_0103 | DI49_0104; DI49_0103 | | |
| SPGP0B00370; DI49_0105 | DI49_0105 | SPGP0B00370 | |
| SPGP0B00430; YBL075C; DI49_0111 | DI49_0111 | SPGP0B00430 | YBL075C |
| DI49_0120; YBL061C; DI49_0121 | DI49_0120; DI49_0121 | | YBL061C |
| DI49_0122; SPGP0B00550 | DI49_0122 | SPGP0B00550 | |
| DI49_0124; DI49_0123 | DI49_0124; DI49_0123 | | |
| DI49_0136; DI49_0135 | DI49_0136; DI49_0135 | | |
| DI49_0142; YBL037W; DI49_0143 | DI49_0142; DI49_0143 | | YBL037W |
| DI49_0145; DI49_0144 | DI49_0145; DI49_0144 | | |
| DI49_0147; DI49_0146 | DI49_0147; DI49_0146 | | |
| DI49_0149; YBL030C; DI49_0150 | DI49_0149; DI49_0150 | | YBL030C |
| DI49_0156; DI49_0155 | DI49_0156; DI49_0155 | | |
| DI49_0159; YBL022C; DI49_0160 | DI49_0159; DI49_0160 | | YBL022C |
| DI49_0170; DI49_0169 | DI49_0170; DI49_0169 | | |
| DI49_0172; YBL009W; DI49_0173 | DI49_0172; DI49_0173 | | YBL009W |
| DI49_0183; DI49_0182 | DI49_0183; DI49_0182 | | |
| DI49_0195; DI49_0194 | DI49_0195; DI49_0194 | | |
| DI49_0197; DI49_0196 | DI49_0197; DI49_0196 | | |
| DI49_0199; DI49_0198 | DI49_0199; DI49_0198 | | |
| DI49_0201; DI49_0200 | DI49_0201; DI49_0200 | | |
| DI49_0203; DI49_0202 | DI49_0203; DI49_0202 | | |
| DI49_0209; SPGP0J02600; DI49_0210 | DI49_0209; DI49_0210 | SPGP0J02600 | |
| DI49_0212; DI49_0211 | DI49_0212; DI49_0211 | | |
| DI49_0214; DI49_0213 | DI49_0214; DI49_0213 | | |
| DI49_0222; YDR023W | DI49_0222 | | YDR023W |
| DI49_0226; YDR028C; DI49_0227 | DI49_0226; DI49_0227 | | YDR028C |
| DI49_0230; YDR033W; DI49_0231 | DI49_0230; DI49_0231 | | YDR033W |
| SPGP0B01640; DI49_1093; DI49_0236; DI49_1092 | DI49_1093; DI49_0236; DI49_1092 | SPGP0B01640 | |
| DI49_0249; YDR055W; DI49_0250 | DI49_0249; DI49_0250 | | YDR055W |
| DI49_0252; DI49_0251 | DI49_0252; DI49_0251 | | |
| DI49_0255; YDR059C | DI49_0255 | | YDR059C |
| DI49_0273; YDR077W; DI49_0274 | DI49_0273; DI49_0274 | | YDR077W |
| DI49_0276; DI49_0275 | DI49_0276; DI49_0275 | | |
| DI49_0287; YDR089W; DI49_0288 | DI49_0287; DI49_0288 | | YDR089W |
| SPGP0B02160; DI49_0289 | DI49_0289 | SPGP0B02160 | |
| DI49_0292; YDR097C; DI49_0293 | DI49_0292; DI49_0293 | | YDR097C |
| DI49_0300; DI49_0299 | DI49_0300; DI49_0299 | | |
| DI49_0304; YDR108W; SPGP0B02300 | DI49_0304 | SPGP0B02300 | YDR108W |

Create PDF in your applications with the Pdfcrowd HTML to PDF API PDFCROWD

| | | | |
|---|---|---|---|
| DI49_0308; DI49_0307 | DI49_0308; DI49_0307 | | |
| DI49_0310; DI49_0309 | DI49_0310; DI49_0309 | | |
| DI49_0315; DI49_0314 | DI49_0315; DI49_0314 | | |
| DI49_0321; YDR127W; DI49_0322 | DI49_0321; DI49_0322 | | YDR127W |
| DI49_0324; DI49_0323 | DI49_0324; DI49_0323 | | |
| DI49_0326; DI49_0325 | DI49_0326; DI49_0325 | | |
| DI49_0331; DI49_0330 | DI49_0331; DI49_0330 | | |
| DI49_0335; YDR143C; DI49_0336 | DI49_0335; DI49_0336 | | YDR143C |
| DI49_0342; DI49_0341 | DI49_0342; DI49_0341 | | |
| DI49_0344; YDR155C; DI49_0345 | DI49_0344; DI49_0345 | | YDR155C |
| DI49_0347; DI49_0346 | DI49_0347; DI49_0346 | | |
| DI49_0349; DI49_0348 | DI49_0349; DI49_0348 | | |
| DI49_0369; DI49_0368 | DI49_0369; DI49_0368 | | |
| DI49_0374; DI49_0373 | DI49_0374; DI49_0373 | | |
| DI49_0390; YDR204W; DI49_0391 | DI49_0390; DI49_0391 | | YDR204W |
| DI49_0393; DI49_0392 | DI49_0393; DI49_0392 | | |
| DI49_0395; SPGP0Ctrna9I; DI49_0396; DI49_3744 | DI49_0395; DI49_0396; DI49_3744 | SPGP0Ctrna9I | |
| SPGP0B03260; DI49_0400 | DI49_0400 | SPGP0B03260 | |
| DI49_0402; DI49_0401 | DI49_0402; DI49_0401 | | |
| DI49_0408; YDR226W; DI49_0409 | DI49_0408; DI49_0409 | | YDR226W |
| DI49_0418; YDR236C; DI49_0419 | DI49_0418; DI49_0419 | | YDR236C |
| DI49_0421; DI49_0420 | DI49_0421; DI49_0420 | | |
| SPGP0B03490; YDR242W; DI49_0424 | DI49_0424 | SPGP0B03490 | YDR242W |
| DI49_0432; DI49_0431 | DI49_0432; DI49_0431 | | |
| DI49_0434; DI49_0433 | DI49_0434; DI49_0433 | | |
| DI49_0439; DI49_0438 | DI49_0439; DI49_0438 | | |
| DI49_0441; DI49_0440 | DI49_0441; DI49_0440 | | |
| DI49_0445; DI49_0444 | DI49_0445; DI49_0444 | | |
| DI49_0447; DI49_0446 | DI49_0447; DI49_0446 | | |
| DI49_0450; YDR270W; DI49_0451 | DI49_0450; DI49_0451 | | YDR270W |
| DI49_0453; DI49_0452 | DI49_0453; DI49_0452 | | |
| DI49_0464; DI49_0463 | DI49_0464; DI49_0463 | | |
| DI49_0478; YDR301W; DI49_0479 | DI49_0478; DI49_0479 | | YDR301W |
| DI49_0481; DI49_0480 | DI49_0481; DI49_0480 | | |
| DI49_0483; DI49_0482 | DI49_0483; DI49_0482 | | |
| DI49_0485; DI49_0484 | DI49_0485; DI49_0484 | | |
| DI49_0487; DI49_0486 | DI49_0487; DI49_0486 | | |
| DI49_0489; DI49_0488 | DI49_0489; DI49_0488 | | |
| DI49_0493; DI49_0492 | DI49_0493; DI49_0492 | | |

Create PDF in your applications with the Pdfcrowd HTML to PDF API PDFCROWD

| | | | |
|---|---|---|---|
| DI49_0495; DI49_1939; DI49_0496 | DI49_0495; DI49_1939; DI49_0496 | | |
| DI49_0498; DI49_0497 | DI49_0498; DI49_0497 | | |
| DI49_0503; YDR322C-A; DI49_0504 | DI49_0503; DI49_0504 | | YDR322C-A |
| DI49_0512; DI49_0511 | DI49_0512; DI49_0511 | | |
| DI49_0515; YDR335W; DI49_0516 | DI49_0515; DI49_0516 | | YDR335W |
| DI49_0518; YDR338C; DI49_0519 | DI49_0518; DI49_0519 | | YDR338C |
| DI49_0523; YDR346C; DI49_0524 | DI49_0523; DI49_0524 | | YDR346C |
| DI49_0526; DI49_0525 | DI49_0526; DI49_0525 | | |
| DI49_0533; YDR356W; DI49_0534 | DI49_0533; DI49_0534 | | YDR356W |
| DI49_0536; DI49_0535 | DI49_0536; DI49_0535 | | |
| DI49_0538; YDR362C; DI49_0539 | DI49_0538; DI49_0539 | | YDR362C |
| DI49_0543; YDR365C; DI49_0544 | DI49_0543; DI49_0544 | | YDR365C |
| DI49_0545; SPGP0B04650 | DI49_0545 | SPGP0B04650 | |
| DI49_0547; DI49_0546 | DI49_0547; DI49_0546 | | |
| DI49_0549; DI49_0548 | DI49_0549; DI49_0548 | | |
| DI49_0551; DI49_0550 | DI49_0551; DI49_0550 | | |
| SPGP0B04760; DI49_0554 | DI49_0554 | SPGP0B04760 | |
| DI49_0556; DI49_0555 | DI49_0556; DI49_0555 | | |
| SPGP0B04800; DI49_0557 | DI49_0557 | SPGP0B04800 | |
| DI49_0561; DI49_4969; DI49_4970 | DI49_0561; DI49_4969; DI49_4970 | | |
| DI49_0562; YDR386W; SPGP0B04870 | DI49_0562 | SPGP0B04870 | YDR386W |
| DI49_0565; YDR389W; DI49_0566 | DI49_0565; DI49_0566 | | YDR389W |
| DI49_0568; DI49_0567 | DI49_0568; DI49_0567 | | |
| DI49_0570; DI49_0569 | DI49_0570; DI49_0569 | | |
| DI49_0573; YDR397C; DI49_0574 | DI49_0573; DI49_0574 | | YDR397C |
| DI49_0576; DI49_0575 | DI49_0576; DI49_0575 | | |
| DI49_0578; DI49_0577 | DI49_0578; DI49_0577 | | |
| DI49_0580; DI49_0579 | DI49_0580; DI49_0579 | | |
| DI49_0584; DI49_0583 | DI49_0584; DI49_0583 | | |
| DI49_0596; SPGP0B05210 | DI49_0596 | SPGP0B05210 | |
| DI49_0598; DI49_0597 | DI49_0598; DI49_0597 | | |
| DI49_0600; DI49_0599 | DI49_0600; DI49_0599 | | |
| DI49_0606; YDR438W; DI49_0607 | DI49_0606; DI49_0607 | | YDR438W |
| DI49_0609; DI49_0608 | DI49_0609; DI49_0608 | | |
| SPGP0B05370; DI49_0610 | DI49_0610 | SPGP0B05370 | |
| DI49_0620; DI49_0619 | DI49_0620; DI49_0619 | | |
| DI49_0628; DI49_0627 | DI49_0628; DI49_0627 | | |
| DI49_0630; DI49_0629 | DI49_0630; DI49_0629 | | |

Create PDF in your applications with the Pdfcrowd HTML to PDF API PDFCROWD

| | | | |
|---|---|---|---|
| DI49_0637; YDR472W; DI49_0638 | DI49_0637; DI49_0638 | | YDR472W |
| DI49_0640; DI49_0639 | DI49_0640; DI49_0639 | | |
| DI49_0642; DI49_0641 | DI49_0642; DI49_0641 | | |
| DI49_0644; DI49_0643 | DI49_0644; DI49_0643 | | |
| DI49_0649; YDR485C; DI49_0650 | DI49_0649; DI49_0650 | | YDR485C |
| DI49_0657; DI49_0656 | DI49_0657; DI49_0656 | | |
| DI49_0660; YDR497C; DI49_0661 | DI49_0660; DI49_0661 | | YDR497C |
| DI49_0674; DI49_0673 | DI49_0674; DI49_0673 | | |
| DI49_0676; DI49_0675 | DI49_0676; DI49_0675 | | |
| DI49_0680; SPGP0B06060 | DI49_0680 | SPGP0B06060 | |
| DI49_0686; DI49_0685 | DI49_0686; DI49_0685 | | |
| SPGP0B06210; SPGP0B06200 | | SPGP0B06210; SPGP0B06200 | |
| SPGP0AD00110; SPGP0HV00100; YKR101W; YPR196W; YPR194C | | SPGP0AD00110; SPGP0HV00100 | YKR101W; YPR196W; YPR194C |
| YKR096W; YKR097W | | | YKR096W; YKR097W |
| YKR089C; YOR081C; YKR088C; YOR080W | | | YKR089C; YOR081C; YKR088C; YOR080W |
| YKR083C; YKR084C | | | YKR083C; YKR084C |
| YKR077W; YKR078W | | | YKR077W; YKR078W |
| YKR075C; YKR076W | | | YKR075C; YKR076W |
| YKR066C; YKR067W | | | YKR066C; YKR067W |
| YKR056W; YKR057W | | | YKR056W; YKR057W |
| YKR051W; YKR052C | | | YKR051W; YKR052C |
| YKR049C; YKR050W | | | YKR049C; YKR050W |
| YKR036C; YKR037C | | | YKR036C; YKR037C |
| YKR020W; YKR021W | | | YKR020W; YKR021W |
| YKR007W; YKR008W | | | YKR007W; YKR008W |
| YKR005C; YKR006C | | | YKR005C; YKR006C |
| YKR003W; YKR004C | | | YKR003W; YKR004C |
| YKL009W; DI49_3363 | DI49_3363 | | YKL009W |
| YKL013C; YKL012W | | | YKL013C; YKL012W |
| YFR054C; YFR055W | | | YFR054C; YFR055W |
| YFR052W; YFR053C; DI49_1714; DI49_1713; YGL253W; YGL252C | DI49_1714; DI49_1713 | | YFR052W; YFR053C; YGL253W; YGL252C |
| YFR050C; DI49_1711; YFR049W; DI49_1710 | DI49_1711; DI49_1710 | | YFR050C; YFR049W |
| YFR047C; YFR048W | | | YFR047C; YFR048W |
| YFR045W; YFR046C | | | YFR045W; YFR046C |
| YFR043C; YFR044C | | | YFR043C; YFR044C |
| YFR041C; YFR042W | | | YFR041C; YFR042W |
| YFR039C; YFR040W | | | YFR039C; YFR040W |
| YFR037C; DI49_1698 | DI49_1698 | | YFR037C |
| YFR036W; DI49_1697; SPGP0AF00310 | DI49_1697 | SPGP0AF00310 | YFR036W |

Create PDF in your applications with the Pdfcrowd HTML to PDF API PDFCROWD

| | | | |
|---|---|---|---|
| YFR033C; DI49_1693 | DI49_1693 | | YFR033C |
| YFR032C-A; DI49_1692; YFR032C; DI49_1691 | DI49_1692; DI49_1691 | | YFR032C-A; YFR032C |
| YFR031C; DI49_1689 | DI49_1689 | | YFR031C |
| YFR030W; DI49_1688; YFR029W; DI49_1687 | DI49_1688; DI49_1687 | | YFR030W; YFR029W |
| YFR023W; DI49_1680; YFR022W; DI49_1679 | DI49_1680; DI49_1679 | | YFR023W; YFR022W |
| YFR020W; YFR021W | | | YFR020W; YFR021W |
| YFR019W; DI49_1676; YFR018C; DI49_1675 | DI49_1676; DI49_1675 | | YFR019W; YFR018C |
| YFR014C; YFR015C | | | YFR014C; YFR015C |
| YFR012W-A; YFR013W | | | YFR012W-A; YFR013W |
| YFR009W; DI49_1672 | DI49_1672 | | YFR009W |
| YFR006W; DI49_1659 | DI49_1659 | | YFR006W |
| YFR005C; DI49_1658 | DI49_1658 | | YFR005C |
| YFR004W; DI49_1657; YFR003C; DI49_1656 | DI49_1657; DI49_1656 | | YFR004W; YFR003C |
| YFR001W; YFR002W | | | YFR001W; YFR002W |
| YFL002C; YFL001W | | | YFL002C; YFL001W |
| YFL007W; DI49_1647; DI49_1648 | DI49_1647; DI49_1648 | | YFL007W |
| YFL008W; DI49_1646 | DI49_1646 | | YFL008W |
| YFL010C; DI49_1644; YFL011W | DI49_1644 | | YFL010C; YFL011W |
| YFL013C; DI49_1641; YFL014W; DI49_1640 | DI49_1641; DI49_1640 | | YFL013C; YFL014W |
| YFL017C; YFL016C | | | YFL017C; YFL016C |
| YFL018C; DI49_1636 | DI49_1636 | | YFL018C |
| SPGP0AEtrna5P; SPGP0AJtrna14P | | SPGP0AEtrna5P; SPGP0AJtrna14P | |
| YFL020C; SPGP0AJ00200; YAR019C | | SPGP0AJ00200 | YFL020C; YAR019C |
| YFL021W; DI49_1633 | DI49_1633 | | YFL021W |
| YFL022C; DI49_1632; SPGP0AE00850; DI49_1631 | DI49_1632; DI49_1631 | SPGP0AE00850 | YFL022C |
| YFL024C; DI49_1630 | DI49_1630 | | YFL024C |
| YFL025C; DI49_1629; YFL026W; SPGP0AF00940 | DI49_1629 | SPGP0AF00940 | YFL025C; YFL026W |
| YFL028C; YFL027C | | | YFL028C; YFL027C |
| YFL030W; YFL029C | | | YFL030W; YFL029C |
| YFL033C; DI49_1623 | DI49_1623 | | YFL033C |
| YFL034W; DI49_1622; YFL034C-A | DI49_1622 | | YFL034W; YFL034C-A |
| YFL036W; DI49_1620 | DI49_1620 | | YFL036W |
| YFL037W; DI49_1619; YFL038C; DI49_1618 | DI49_1619; DI49_1618 | | YFL037W; YFL038C |
| YFL040W; YFL039C | | | YFL040W; YFL039C |
| YFL042C; YFL041W | | | YFL042C; YFL041W |
| YFL045C; DI49_1612; YFL046W; DI49_1611 | DI49_1612; DI49_1611 | | YFL045C; YFL046W |
| YFL048C; YFL047W | | | YFL048C; YFL047W |
| YFL050C; YOL130W | | | YFL050C; YOL130W |
| YAR071W; SPGP0BW00100; DI49_4805 | DI49_4805 | SPGP0BW00100 | YAR071W |

Create PDF in your applications with the Pdfcrowd HTML to PDF API PDFCROWD

| | | | |
|---|---|---|---|
| YHR216W; YLR432W; YCR095C; DI49_3928 | DI49_3928 | | YHR216W; YLR432W; YCR095C |
| DI49_1718; DI49_1719 | DI49_1718; DI49_1719 | | |
| DI49_1708; DI49_1709 | DI49_1708; DI49_1709 | | |
| DI49_1706; DI49_1707 | DI49_1706; DI49_1707 | | |
| DI49_1704; DI49_1705 | DI49_1704; DI49_1705 | | |
| DI49_1702; DI49_1703 | DI49_1702; DI49_1703 | | |
| DI49_1700; DI49_1701 | DI49_1700; DI49_1701 | | |
| DI49_1695; SPGP0AF00330; DI49_1694 | DI49_1695; DI49_1694 | SPGP0AF00330 | |
| DI49_1685; YFR028C | DI49_1685 | | YFR028C |
| DI49_1684; YFR027W; SPGP0AF00440 | DI49_1684 | SPGP0AF00440 | YFR027W |
| DI49_1671; YFR010W; DI49_1670 | DI49_1671; DI49_1670 | | YFR010W |
| DI49_1666; DI49_1667 | DI49_1666; DI49_1667 | | |
| DI49_1663; DI49_1665 | DI49_1663; DI49_1665 | | |
| DI49_1661; DI49_1662 | DI49_1661; DI49_1662 | | |
| DI49_1654; DI49_1655 | DI49_1654; DI49_1655 | | |
| DI49_1652; DI49_1653 | DI49_1652; DI49_1653 | | |
| DI49_1638; DI49_1639 | DI49_1638; DI49_1639 | | |
| SPGP0AFtrna13P; DI49_4914; SPGP0AItrna11P; SPGP0Ctrna6P | DI49_4914 | SPGP0AFtrna13P; SPGP0AItrna11P; SPGP0Ctrna6P | |
| SPGP0AF00870; DI49_0075 | DI49_0075 | SPGP0AF00870 | |
| SPGP0AF00880; SPGP0AI00800; SPGP0AI00810; DI49_4789 | DI49_4789 | SPGP0AF00880; SPGP0AI00800; SPGP0AI00810 | |
| DI49_1627; DI49_1628 | DI49_1627; DI49_1628 | | |
| DI49_1625; DI49_1626 | DI49_1625; DI49_1626 | | |
| DI49_1621; YFL034C-B | DI49_1621 | | YFL034C-B |
| DI49_1616; DI49_1617 | DI49_1616; DI49_1617 | | |
| DI49_1614; DI49_1615 | DI49_1614; DI49_1615 | | |
| DI49_1609; DI49_1610 | DI49_1609; DI49_1610 | | |
| DI49_1607; DI49_2277; YOL131W | DI49_1607; DI49_2277 | | YOL131W |
| DI49_1606; DI49_3482; YJL078C; DI49_3483 | DI49_1606; DI49_3482; DI49_3483 | | YJL078C |
| SPGP0AG00110; SPGP0BH00120; SPGP0AG00120 | | SPGP0AG00110; SPGP0BH00120; SPGP0AG00120 | |
| SPGP0AG00140; DI49_0699 | DI49_0699 | SPGP0AG00140 | |
| DI49_0701; DI49_0700 | DI49_0701; DI49_0700 | | |
| DI49_0703; YCL057W; DI49_0704; YCL056C | DI49_0703; DI49_0704 | | YCL057W; YCL056C |
| DI49_0706; YCL054W; DI49_0707; YCL052C | DI49_0706; DI49_0707 | | YCL054W; YCL052C |
| DI49_0709; YCL050C; DI49_0710; YCL049C | DI49_0709; DI49_0710 | | YCL050C; YCL049C |
| DI49_0712; DI49_0711; YCL048W; YCL047C | DI49_0712; DI49_0711 | | YCL048W; YCL047C |
| DI49_0714; YCL044C | DI49_0714 | | YCL044C |
| DI49_0715; YCL043C | DI49_0715 | | YCL043C |

Create PDF in your applications with the Pdfcrowd HTML to PDF API PDFCROWD

| | | | |
|---|---|---|---|
| DI49_0716; YCL040W | DI49_0716 | | YCL040W |
| DI49_0717; YCL039W; DI49_0718; YCL038C | DI49_0717; DI49_0718 | | YCL039W; YCL038C |
| SPGP0AG00350; YCL036W; DI49_0720; YCL035C | DI49_0720 | SPGP0AG00350 | YCL036W; YCL035C |
| DI49_0722; DI49_0721 | DI49_0722; DI49_0721 | | |
| DI49_0724; YCL031C | DI49_0724 | | YCL031C |
| DI49_0725; YCL030C | DI49_0725 | | YCL030C |
| DI49_0726; YCL029C; DI49_0727; YCL028W | DI49_0726; DI49_0727 | | YCL029C; YCL028W |
| DI49_0729; DI49_0728 | DI49_0729; DI49_0728 | | |
| DI49_0731; DI49_0730; YCL025C; YCL024W | DI49_0731; DI49_0730 | | YCL025C; YCL024W |
| DI49_0736; YCL018W | DI49_0736 | | YCL018W |
| DI49_0737; YCL017C | DI49_0737 | | YCL017C |
| DI49_0738; YCL016C | DI49_0738 | | YCL016C |
| DI49_0739; YCL014W; DI49_0740; YCL012C | DI49_0739; DI49_0740 | | YCL014W; YCL012C |
| DI49_0742; YCL010C | DI49_0742 | | YCL010C |
| DI49_0743; YCL009C | DI49_0743 | | YCL009C |
| DI49_0744; YCL008C; DI49_0745; YCL007C | DI49_0744; DI49_0745 | | YCL008C; YCL007C |
| DI49_0747; DI49_0746 | DI49_0747; DI49_0746 | | |
| DI49_0749; YCL001W; DI49_0750; YCL001W-A | DI49_0749; DI49_0750 | | YCL001W; YCL001W-A |
| DI49_0752; DI49_0751 | DI49_0752; DI49_0751 | | |
| DI49_0754; DI49_0753; YCR004C; YCR005C | DI49_0754; DI49_0753 | | YCR004C; YCR005C |
| SPGP0AG00710; SPGP0AG00700 | | SPGP0AG00710; SPGP0AG00700 | |
| DI49_0760; DI49_0759 | DI49_0760; DI49_0759 | | |
| DI49_0762; YCR012W; DI49_0763; YCR014C | DI49_0762; DI49_0763 | | YCR012W; YCR014C |
| SPGP0AG00810; DI49_0767 | DI49_0767 | SPGP0AG00810 | |
| DI49_0770; YCR019W; DI49_0771; YCR020C | DI49_0770; DI49_0771 | | YCR019W; YCR020C |
| DI49_0773; DI49_0772 | DI49_0773; DI49_0772 | | |
| DI49_0775; YCR023C; DI49_0776; YCR024C | DI49_0775; DI49_0776 | | YCR023C; YCR024C |
| DI49_0778; DI49_0777 | DI49_0778; DI49_0777 | | |
| DI49_0781; YCR028C; SPGP0AG00930; YCR028C-A | DI49_0781 | SPGP0AG00930 | YCR028C; YCR028C-A |
| DI49_0786; YCR034W | DI49_0786 | | YCR034W |
| DI49_0787; YCR035C; DI49_0788; YCR036W | DI49_0787; DI49_0788 | | YCR035C; YCR036W |
| DI49_0789; SPGP0AG01010 | DI49_0789 | SPGP0AG01010 | |
| YCL061C; YCL063W | | | YCL061C; YCL063W |
| YCL058C; YCL059C | | | YCL058C; YCL059C |
| YCL033C; YCL034W | | | YCL033C; YCL034W |
| YCL026C-B; YCL027W | | | YCL026C-B; YCL027W |
| YCL004W; YCL005W | | | YCL004W; YCL005W |
| YCR003W; SPGP0CF00110 | | SPGP0CF00110 | YCR003W |
| YCR010C; YCR009C | | | YCR010C; YCR009C |

Create PDF in your applications with the Pdfcrowd HTML to PDF API PDFCROWD

| | | | |
|---|---|---|---|
| YCR018C; YCR017C | | | YCR018C; YCR017C |
| YCR020W-B; YCR020C-A | | | YCR020W-B; YCR020C-A |
| YCR026C; YCR024C-A | | | YCR026C; YCR024C-A |
| YCR038C; YCR037C | | | YCR038C; YCR037C |
| DI49_0006; DI49_0005 | DI49_0006; DI49_0005 | | |
| DI49_0008; DI49_0007 | DI49_0008; DI49_0007 | | |
| DI49_0010; DI49_0009 | DI49_0010; DI49_0009 | | |
| DI49_0012; DI49_0011 | DI49_0012; DI49_0011 | | |
| DI49_0014; DI49_0013 | DI49_0014; DI49_0013 | | |
| DI49_0016; DI49_0015; YAL046C; YAL047C | DI49_0016; DI49_0015 | | YAL046C; YAL047C |
| DI49_0018; YAL043C; YAL044C | DI49_0018 | | YAL043C; YAL044C |
| DI49_0019; YAL042W | DI49_0019 | | YAL042W |
| DI49_0020; YAL041W | DI49_0020 | | YAL041W |
| DI49_0021; YAL040C; DI49_0022 | DI49_0021; DI49_0022 | | YAL040C |
| DI49_0024; DI49_0023 | DI49_0024; DI49_0023 | | |
| DI49_0026; YAL035W | DI49_0026 | | YAL035W |
| DI49_0027; YAL034W-A | DI49_0027 | | YAL034W-A |
| DI49_0028; YAL034C | DI49_0028 | | YAL034C |
| DI49_0029; YAL033W; DI49_0030 | DI49_0029; DI49_0030 | | YAL033W |
| SPGP0AI00400; DI49_0031 | DI49_0031 | SPGP0AI00400 | |
| DI49_0033; DI49_0032 | DI49_0033; DI49_0032 | | |
| DI49_0035; YAL026C; DI49_0036; YAL027W | DI49_0035; DI49_0036 | | YAL026C; YAL027W |
| DI49_0038; YAL023C | DI49_0038 | | YAL023C |
| DI49_0039; YAL022C | DI49_0039 | | YAL022C |
| DI49_0040; YAL021C | DI49_0040 | | YAL021C |
| DI49_0041; YAL020C | DI49_0041 | | YAL020C |
| DI49_0042; YAL019W; DI49_0043 | DI49_0042; DI49_0043 | | YAL019W |
| DI49_0045; YAL016W; DI49_0046 | DI49_0045; DI49_0046 | | YAL016W |
| SPGP0AI00570; YAL013W | | SPGP0AI00570 | YAL013W |
| DI49_0048; YAL012W | DI49_0048 | | YAL012W |
| DI49_0049; YAL011W; DI49_0050 | DI49_0049; DI49_0050 | | YAL011W |
| DI49_0052; DI49_0051 | DI49_0052; DI49_0051 | | |
| DI49_0056; SPGP0AJ00320; DI49_0057 | DI49_0056; DI49_0057 | SPGP0AJ00320 | |
| DI49_0059; DI49_0058 | DI49_0059; DI49_0058 | | |
| DI49_0061; YAR003W | DI49_0061 | | YAR003W |
| DI49_0062; YAR007C | DI49_0062 | | YAR007C |
| DI49_0063; YAR008W; DI49_0065 | DI49_0063; DI49_0065 | | YAR008W |
| DI49_0068; YAR014C | DI49_0068 | | YAR014C |
| DI49_0069; YAR015W; DI49_0070 | DI49_0069; DI49_0070 | | YAR015W |

Create PDF in your applications with the Pdfcrowd HTML to PDF API PDFCROWD

| | | | |
|---|---|---|---|
| DI49_0072; DI49_0071 | DI49_0072; DI49_0071 | | |
| SPGP0AI00840; YAR035W | | SPGP0AI00840 | YAR035W |
| DI49_0076; YAR042W | DI49_0076 | | YAR042W |
| SPGP0MW00100; YPR204W; SPGP0C00100; SPGP0PU00100; DI49_2575 | DI49_2575 | SPGP0MW00100; SPGP0C00100; SPGP0PU00100 | YPR204W |
| SPGP0NC00100; SPGP0G04210; SPGP0AJ00100 | | SPGP0NC00100; SPGP0G04210; SPGP0AJ00100 | |
| YAR031W; YAR027W; YAR029W; YAR028W | | | YAR031W; YAR027W; YAR029W; YAR028W |
| YAR002C-A; DI49_0060; YAR002W | DI49_0060 | | YAR002C-A; YAR002W |
| YAL002W; YAL001C | | | YAL002W; YAL001C |
| YAL007C; DI49_0053; YAL008W | DI49_0053 | | YAL007C; YAL008W |
| YAL010C; YAL009W | | | YAL010C; YAL009W |
| YAL014C; DI49_0047; YAL015C | DI49_0047 | | YAL014C; YAL015C |
| YAL017W; YOL045W; YAL018C; SPGP0D03540 | | SPGP0D03540 | YAL017W; YOL045W; YAL018C |
| YAL024C; DI49_0037; YAL025C | DI49_0037 | | YAL024C; YAL025C |
| YAL030W; YAL029C | | | YAL030W; YAL029C |
| YAL032C; YAL031C | | | YAL032C; YAL031C |
| YAL036C; DI49_0025; YAL037W | DI49_0025 | | YAL036C; YAL037W |
| YAL039C; YAL038W | | | YAL039C; YAL038W |
| YAL049C; YAL048C | | | YAL049C; YAL048C |
| YAL053W; YAL051W | | | YAL053W; YAL051W |
| YAL055W; YAL054C | | | YAL055W; YAL054C |
| YAL058W; YAL056W | | | YAL058W; YAL056W |
| YAL060W; YAL059W | | | YAL060W; YAL059W |
| YGL257C; YGL258W | | | YGL257C; YGL258W |
| YGL255W; YGL256W | | | YGL255W; YGL256W |
| YGL250W; YGL251C | | | YGL250W; YGL251C |
| YGL248W; YGL249W | | | YGL248W; YGL249W |
| YGL246C; YGL247W | | | YGL246C; YGL247W |
| YGL244W; YGL245W | | | YGL244W; YGL245W |
| YGL242C; YGL243W | | | YGL242C; YGL243W |
| YGL240W; YGL241W | | | YGL240W; YGL241W |
| YGL237C; YGL238W | | | YGL237C; YGL238W |
| YGL234W; YGL236C | | | YGL234W; YGL236C |
| YGL232W; YGL233W | | | YGL232W; YGL233W |
| YGL230C; YGL231C | | | YGL230C; YGL231C |
| YGL228W; YGL229C | | | YGL228W; YGL229C |
| YGL226C-A; YGL227W | | | YGL226C-A; YGL227W |
| YGL223C; YGL224C | | | YGL223C; YGL224C |
| YGL221C; YGL222C | | | YGL221C; YGL222C |

Create PDF in your applications with the Pdfcrowd HTML to PDF API
PDFCROWD

| | | | |
|---|---|---|---|
| YGL219C; YGL220W | | | YGL219C; YGL220W |
| YGL215W; YGL216W | | | YGL215W; YGL216W |
| YGL212W; YGL213C | | | YGL212W; YGL213C |
| YGL210W; YGL211W | | | YGL210W; YGL211W |
| YGL208W; YGL209W | | | YGL208W; YGL209W |
| YGL206C; YGL207W | | | YGL206C; YGL207W |
| YGL200C; YGL201C | | | YGL200C; YGL201C |
| YGL197W; YGL198W | | | YGL197W; YGL198W |
| YGL195W; YGL196W | | | YGL195W; YGL196W |
| YGL194C; YGL194C-A | | | YGL194C; YGL194C-A |
| YGL190C; YGL191W | | | YGL190C; YGL191W |
| YGL187C; YGL189C | | | YGL187C; YGL189C |
| YGL185C; YGL186C | | | YGL185C; YGL186C |
| YGL183C; YGL184C | | | YGL183C; YGL184C |
| YGL180W; YGL181W | | | YGL180W; YGL181W |
| DI49_4743; DI49_4742 | DI49_4743; DI49_4742 | | |
| DI49_4745; DI49_4744 | DI49_4745; DI49_4744 | | |
| DI49_4752; YNR017W; DI49_4753 | DI49_4752; DI49_4753 | | YNR017W |
| DI49_4758; DI49_4757 | DI49_4758; DI49_4757 | | |
| SPGP0AL00330; DI49_4759 | DI49_4759 | SPGP0AL00330 | |
| DI49_4761; DI49_4760 | DI49_4761; DI49_4760 | | |
| DI49_4763; YNR031C; DI49_4764 | DI49_4763; DI49_4764 | | YNR031C |
| DI49_4766; DI49_4765 | DI49_4766; DI49_4765 | | |
| DI49_4768; DI49_4767 | DI49_4768; DI49_4767 | | |
| DI49_4770; DI49_4769 | DI49_4770; DI49_4769 | | |
| DI49_4772; DI49_4771 | DI49_4772; DI49_4771 | | |
| DI49_4774; DI49_4773 | DI49_4774; DI49_4773 | | |
| DI49_4778; DI49_4777 | DI49_4778; DI49_4777 | | |
| DI49_4790; YNR055C; DI49_4791 | DI49_4790; DI49_4791 | | YNR055C |
| DI49_4793; DI49_4792 | DI49_4793; DI49_4792 | | |
| SPGP0AL00690; DI49_4794 | DI49_4794 | SPGP0AL00690 | |
| SPGP0AL00710; DI49_4796 | DI49_4796 | SPGP0AL00710 | |
| DI49_4798; DI49_4797 | DI49_4798; DI49_4797 | | |
| DI49_4800; DI49_4799 | DI49_4800; DI49_4799 | | |
| YER139C; DI49_1547; YER140W | DI49_1547 | | YER139C; YER140W |
| YER152C; DI49_1561; YER153C | DI49_1561 | | YER152C; YER153C |
| YER155C; DI49_1564; YER156C | DI49_1564 | | YER155C; YER156C |
| YER158C; DI49_1569; YER159C | DI49_1569 | | YER158C; YER159C |
| YER163C; YER162C | | | YER163C; YER162C |
| YER170W; YER169W | | | YER170W; YER169W |

Create PDF in your applications with the Pdfcrowd HTML to PDF API PDFCROWD

| | | | |
|---|---|---|---|
| YER173W; DI49_1583; YER174C<br>YER176W; YER175C | DI49_1583 | | YER173W; YER174C<br>YER176W; YER175C |
| YER178W; DI49_1587; YER179W | DI49_1587 | | YER178W; YER179W |
| YER180C-A; YER180C | | | YER180C-A; YER180C |
| YER186C; YER185W | | | YER186C; YER185W |
| SPGP0AM00580; YER187W | | SPGP0AM00580 | YER187W |
| SPGP0AM00600; YPL283C | | SPGP0AM00600 | YPL283C |
| DI49_3477; DI49_3476 | DI49_3477; DI49_3476 | | |
| DI49_3479; DI49_3478 | DI49_3479; DI49_3478 | | |
| DI49_3481; DI49_4140; SPGP0C00170 | DI49_3481; DI49_4140 | SPGP0C00170 | |
| DI49_3487; DI49_3486 | DI49_3487; DI49_3486 | | |
| SPGP0C00250; DI49_3488 | DI49_3488 | SPGP0C00250 | |
| DI49_3496; DI49_3495 | DI49_3496; DI49_3495 | | |
| DI49_3498; DI49_3497 | DI49_3498; DI49_3497 | | |
| DI49_3515; DI49_3514 | DI49_3515; DI49_3514 | | |
| DI49_3522; DI49_3521 | DI49_3522; DI49_3521 | | |
| DI49_3539; SPGP0Ctrna9S | DI49_3539 | SPGP0Ctrna9S | |
| DI49_3541; DI49_3540 | DI49_3541; DI49_3540 | | |
| DI49_3546; DI49_3545 | DI49_3546; DI49_3545 | | |
| DI49_3549; DI49_3548 | DI49_3549; DI49_3548 | | |
| DI49_3551; DI49_3550 | DI49_3551; DI49_3550 | | |
| DI49_3553; DI49_3552 | DI49_3553; DI49_3552 | | |
| DI49_3569; DI49_3568 | DI49_3569; DI49_3568 | | |
| DI49_3579; DI49_3578 | DI49_3579; DI49_3578 | | |
| DI49_3581; DI49_3580 | DI49_3581; DI49_3580 | | |
| DI49_3592; DI49_3591 | DI49_3592; DI49_3591 | | |
| DI49_3594; DI49_3593 | DI49_3594; DI49_3593 | | |
| DI49_3605; DI49_3604 | DI49_3605; DI49_3604 | | |
| DI49_3622; DI49_3621 | DI49_3622; DI49_3621 | | |
| DI49_3624; DI49_3623 | DI49_3624; DI49_3623 | | |
| SPGP0C01600; DI49_3625 | DI49_3625 | SPGP0C01600 | |
| DI49_3636; DI49_3635 | DI49_3636; DI49_3635 | | |
| DI49_3647; DI49_3646 | DI49_3647; DI49_3646 | | |
| DI49_3655; DI49_3654 | DI49_3655; DI49_3654 | | |
| DI49_3660; DI49_3659 | DI49_3660; DI49_3659 | | |
| DI49_3662; DI49_3661 | DI49_3662; DI49_3661 | | |
| DI49_3668; DI49_3667 | DI49_3668; DI49_3667 | | |
| DI49_3670; DI49_3669 | DI49_3670; DI49_3669 | | |
| DI49_3672; DI49_3671 | DI49_3672; DI49_3671 | | |
| DI49_3683; DI49_3682 | DI49_3683; DI49_3682 | | |

Create PDF in your applications with the Pdfcrowd HTML to PDF API PDFCROWD

| | | | |
|---|---|---|---|
| DI49_3687; YLR168C; DI49_3688 | DI49_3687; DI49_3688 | | YLR168C |
| DI49_3699; YLR182W; DI49_3700 | DI49_3699; DI49_3700 | | YLR182W |
| DI49_3708; DI49_3707 | DI49_3708; DI49_3707 | | |
| DI49_3713; YLR197W; DI49_3714 | DI49_3713; DI49_3714 | | YLR197W |
| DI49_3716; DI49_3715 | DI49_3716; DI49_3715 | | |
| DI49_3724; YLR210W; DI49_3725 | DI49_3724; DI49_3725 | | YLR210W |
| DI49_3727; DI49_3726 | DI49_3727; DI49_3726 | | |
| DI49_3730; DI49_3728 | DI49_3730; DI49_3728 | | |
| SPGP0C02630; DI49_3731 | DI49_3731 | SPGP0C02630 | |
| DI49_3733; DI49_3732 | DI49_3733; DI49_3732 | | |
| DI49_3735; DI49_3734 | DI49_3735; DI49_3734 | | |
| DI49_3739; YLR226W; DI49_3740 | DI49_3739; DI49_3740 | | YLR226W |
| DI49_3747; DI49_3867; DI49_3866 | DI49_3747; DI49_3867; DI49_3866 | | |
| DI49_3749; YLR239C; SPGP0C02810 | DI49_3749 | SPGP0C02810 | YLR239C |
| DI49_3752; DI49_3751 | DI49_3752; DI49_3751 | | |
| DI49_3760; YLR249W; DI49_3761 | DI49_3760; DI49_3761 | | YLR249W |
| DI49_3765; DI49_3764 | DI49_3765; DI49_3764 | | |
| DI49_3770; YLR259C; DI49_3771 | DI49_3770; DI49_3771 | | YLR259C |
| DI49_3773; DI49_3772 | DI49_3773; DI49_3772 | | |
| DI49_3777; DI49_3776 | DI49_3777; DI49_3776 | | |
| DI49_3779; DI49_3778 | DI49_3779; DI49_3778 | | |
| DI49_3785; DI49_3784 | DI49_3785; DI49_3784 | | |
| DI49_3791; DI49_3790 | DI49_3791; DI49_3790 | | |
| DI49_3801; YLR295C; DI49_3802 | DI49_3801; DI49_3802 | | YLR295C |
| DI49_3804; DI49_3803 | DI49_3804; DI49_3803 | | |
| SPGP0Ctrna11L; DI49_3806 | DI49_3806 | SPGP0Ctrna11L | |
| DI49_3810; YLR305C; DI49_3811 | DI49_3810; DI49_3811 | | YLR305C |
| DI49_3816; YLR310C; DI49_3817 | DI49_3816; DI49_3817 | | YLR310C |
| DI49_3819; DI49_3818 | DI49_3819; DI49_3818 | | |
| DI49_3820; SPGP0C03480 | DI49_3820 | SPGP0C03480 | |
| DI49_3821; SPGP0C03500 | DI49_3821 | SPGP0C03500 | |
| DI49_3823; YLR320W; DI49_3824 | DI49_3823; DI49_3824 | | YLR320W |
| DI49_3826; DI49_3825 | DI49_3826; DI49_3825 | | |
| DI49_3828; YLR326W; DI49_3829 | DI49_3828; DI49_3829 | | YLR326W |
| DI49_3833; DI49_3832 | DI49_3833; DI49_3832 | | |
| DI49_3840; DI49_3839 | DI49_3840; DI49_3839 | | |
| DI49_3842; DI49_3841 | DI49_3842; DI49_3841 | | |
| DI49_3852; YLR352W; DI49_3853 | DI49_3852; DI49_3853 | | YLR352W |
| DI49_3855; YLR355C; DI49_3856 | DI49_3855; DI49_3856 | | YLR355C |

Create PDF in your applications with the Pdfcrowd HTML to PDF API PDFCROWD

| | | | |
|---|---|---|---|
| DI49_3859; DI49_3860; DI49_3861 | DI49_3859; DI49_3860; DI49_3861 | | |
| DI49_3869; YLR369W; DI49_3870 | DI49_3869; DI49_3870 | | YLR369W |
| DI49_3881; DI49_3880 | DI49_3881; DI49_3880 | | |
| DI49_3884; YLR384C; DI49_3885 | DI49_3884; DI49_3885 | | YLR384C |
| DI49_3888; YLR388W; DI49_3889 | DI49_3888; DI49_3889 | | YLR388W |
| DI49_3891; DI49_3890 | DI49_3891; DI49_3890 | | |
| DI49_3893; DI49_3892 | DI49_3893; DI49_3892 | | |
| DI49_3895; DI49_3894 | DI49_3895; DI49_3894 | | |
| DI49_3897; DI49_3896 | DI49_3897; DI49_3896 | | |
| DI49_3899; DI49_3898 | DI49_3899; DI49_3898 | | |
| SPGP0C04280; DI49_3900 | DI49_3900 | SPGP0C04280 | |
| DI49_3903; DI49_3902 | DI49_3903; DI49_3902 | | |
| DI49_3905; DI49_3904 | DI49_3905; DI49_3904 | | |
| DI49_3911; DI49_3910 | DI49_3911; DI49_3910 | | |
| DI49_3913; DI49_3912; YLR414C; SPGP0PE00100 | DI49_3913; DI49_3912 | SPGP0PE00100 | YLR414C |
| DI49_3915; YLR419W; YLR418C | DI49_3915 | | YLR419W; YLR418C |
| DI49_3919; YLR422W; YLR421C | DI49_3919 | | YLR422W; YLR421C |
| DI49_3920; YLR423C | DI49_3920 | | YLR423C |
| DI49_3921; YLR424W | DI49_3921 | | YLR424W |
| DI49_3923; YLR425W; SPGP0C04490 | DI49_3923 | SPGP0C04490 | YLR425W |
| DI49_3925; YLR429W | DI49_3925 | | YLR429W |
| DI49_3926; YLR430W | DI49_3926 | | YLR430W |
| DI49_3927; YLR431C | DI49_3927 | | YLR431C |
| DI49_3929; YLR433C; DI49_3930 | DI49_3929; DI49_3930 | | YLR433C |
| DI49_3932; DI49_3931 | DI49_3932; DI49_3931 | | |
| DI49_3934; DI49_3933 | DI49_3934; DI49_3933 | | |
| DI49_3936; DI49_3935 | DI49_3936; DI49_3935 | | |
| SPGP0C04640; DI49_3937; YLR441C; YLR440C | DI49_3937 | SPGP0C04640 | YLR441C; YLR440C |
| SPGP0C04660; DI49_3938 | DI49_3938 | SPGP0C04660 | |
| DI49_3940; YLR447C | DI49_3940 | | YLR447C |
| DI49_3941; YLR448W; DI49_4017; YML073C | DI49_3941; DI49_4017 | | YLR448W; YML073C |
| DI49_3942; YLR449W | DI49_3942 | | YLR449W |
| DI49_3943; YLR450W | DI49_3943 | | YLR450W |
| DI49_3944; YLR451W; DI49_3945 | DI49_3944; DI49_3945 | | YLR451W |
| DI49_3947; YLR454W; YLR453C | DI49_3947 | | YLR454W; YLR453C |
| DI49_3949; YLR455W; DI49_3950 | DI49_3949; DI49_3950 | | YLR455W |
| DI49_3952; DI49_3951 | DI49_3952; DI49_3951 | | |
| DI49_3954; SPGP0BG00130 | DI49_3954 | SPGP0BG00130 | |
| DI49_3955; YOL159C-A | DI49_3955 | | YOL159C-A |

Create PDF in your applications with the Pdfcrowd HTML to PDF API PDFCROWD

| | | | |
|---|---|---|---|
| YPL003W; YPL002C | | | YPL003W; YPL002C |
| YPL005W; YPL004C | | | YPL005W; YPL004C |
| YPL007C; YPL006W | | | YPL007C; YPL006W |
| YPL032C; YPL031C | | | YPL032C; YPL031C |
| YPL034W; YPL033C | | | YPL034W; YPL033C |
| YPL049C; YPL050C | | | YPL049C; YPL050C |
| YPL054W; YPL053C | | | YPL054W; YPL053C |
| YLR456W; YLR457C | | | YLR456W; YLR457C |
| YLR446W; DI49_3939; YLR445W | DI49_3939 | | YLR446W; YLR445W |
| YLR443W; SPGP0AO00240 | | SPGP0AO00240 | YLR443W |
| YLR438C-A; YLR439W | | | YLR438C-A; YLR439W |
| YLR437C; YLR438W | | | YLR437C; YLR438W |
| YLR435W; YLR436C | | | YLR435W; YLR436C |
| YLR427W; DI49_3924; YLR426W | DI49_3924 | | YLR427W; YLR426W |
| SPGP0PL00100; SPGP0PF00100 | | SPGP0PL00100; SPGP0PF00100 | |
| YHR167W; YHR166C; DI49_2570; DI49_2571 | DI49_2570; DI49_2571 | | YHR167W; YHR166C |
| YHR169W; DI49_2573 | DI49_2573 | | YHR169W |
| YHR170W; DI49_2574; YHR171W; DI49_5609 | DI49_2574; DI49_5609 | | YHR170W; YHR171W |
| YHR173C; YHR172W | | | YHR173C; YHR172W |
| YHR177W; YHR176W | | | YHR177W; YHR176W |
| YHR179W; DI49_2581; YHR180W | DI49_2581 | | YHR179W; YHR180W |
| YHR181W; DI49_2583; YHR182W; DI49_2584 | DI49_2583; DI49_2584 | | YHR181W; YHR182W |
| YHR184W; YHR183W | | | YHR184W; YHR183W |
| YHR186C; DI49_2589; YHR187W; DI49_2590 | DI49_2589; DI49_2590 | | YHR186C; YHR187W |
| YHR189W; YHR188C | | | YHR189W; YHR188C |
| YHR191C; YHR190W | | | YHR191C; YHR190W |
| YHR193C; DI49_2596 | DI49_2596 | | YHR193C |
| YHR194W; DI49_2597; YHR195W; DI49_2598 | DI49_2597; DI49_2598 | | YHR194W; YHR195W |
| YHR197W; DI49_2600; YHR198C; DI49_2601 | DI49_2600; DI49_2601 | | YHR197W; YHR198C |
| YHR199C-A; YHR199C | | | YHR199C-A; YHR199C |
| YHR201C; YHR200W | | | YHR201C; YHR200W |
| YHR206W; DI49_2611; YHR207C; DI49_2612 | DI49_2611; DI49_2612 | | YHR206W; YHR207C |
| YHR209W; YHR208W | | | YHR209W; YHR208W |
| SPGP0AQ00120; DI49_2576 | DI49_2576 | SPGP0AQ00120 | |
| DI49_2578; DI49_2577 | DI49_2578; DI49_2577 | | |
| SPGP0AQ00160; DI49_2579 | DI49_2579 | SPGP0AQ00160 | |
| DI49_2586; DI49_2585 | DI49_2586; DI49_2585 | | |
| DI49_2592; DI49_2591 | DI49_2592; DI49_2591 | | |
| DI49_2594; DI49_2593 | DI49_2594; DI49_2593 | | |
| DI49_2603; DI49_2602 | DI49_2603; DI49_2602 | | |

Create PDF in your applications with the Pdfcrowd HTML to PDF API PDFCROWD

| | | | |
|---|---|---|---|
| DI49_2605; DI49_2604 | DI49_2605; DI49_2604 | | |
| DI49_2607; DI49_2606 | DI49_2607; DI49_2606 | | |
| DI49_2614; DI49_2613 | DI49_2614; DI49_2613 | | |
| DI49_2616; DI49_2615 | DI49_2616; DI49_2615 | | |
| DI49_4422; YNL330C; DI49_4423 | DI49_4422; DI49_4423 | | YNL330C |
| DI49_4431; YNL320W; DI49_4432 | DI49_4431; DI49_4432 | | YNL320W |
| DI49_4434; DI49_4433 | DI49_4434; DI49_4433 | | |
| DI49_4436; DI49_4435 | DI49_4436; DI49_4435 | | |
| DI49_4438; DI49_4437 | DI49_4438; DI49_4437 | | |
| DI49_4440; DI49_4439 | DI49_4440; DI49_4439 | | |
| DI49_4442; DI49_4441 | DI49_4442; DI49_4441 | | |
| DI49_4453; DI49_4452 | DI49_4453; DI49_4452 | | |
| DI49_4460; YNL287W; DI49_4461 | DI49_4460; DI49_4461 | | YNL287W |
| SPGP0AR00530; SPGP0F02560; SPGP0AS00100 | | SPGP0AR00530; SPGP0F02560; SPGP0AS00100 | |
| YPL276W; YPL282C | | | YPL276W; YPL282C |
| YPL270W; YPL271W | | | YPL270W; YPL271W |
| YPL268W; YPL269W | | | YPL268W; YPL269W |
| YPL266W; YPL267W | | | YPL266W; YPL267W |
| YPL264C; YPL265W | | | YPL264C; YPL265W |
| YPL262W; YPL263C | | | YPL262W; YPL263C |
| YPL259C; YPL260W | | | YPL259C; YPL260W |
| YPL256C; SPGP0AS00300 | | SPGP0AS00300 | YPL256C |
| YPL254W; YPL255W | | | YPL254W; YPL255W |
| YPL252C; YPL253C | | | YPL252C; YPL253C |
| YPL249C-A; DI49_4277; YPL249C; YMR194W; DI49_4276; YMR193W | DI49_4277; DI49_4276 | | YPL249C-A; YPL249C; YMR194W; YMR193W |
| YPL247C; YPL248C | | | YPL247C; YPL248C |
| YPR201W; YPL246C | | | YPR201W; YPL246C |
| YPR199C; YPR200C | | | YPR199C; YPR200C |
| YPR192W; YPR193C | | | YPR192W; YPR193C |
| DI49_5634; YPR191W | DI49_5634 | | YPR191W |
| DI49_5632; DI49_5633 | DI49_5632; DI49_5633 | | |
| DI49_5630; DI49_5631 | DI49_5630; DI49_5631 | | |
| DI49_5628; DI49_5629 | DI49_5628; DI49_5629 | | |
| DI49_5626; DI49_5627 | DI49_5626; DI49_5627 | | |
| DI49_5624; DI49_5625 | DI49_5624; DI49_5625 | | |
| DI49_5622; DI49_5623 | DI49_5622; DI49_5623 | | |
| DI49_5620; DI49_5621 | DI49_5620; DI49_5621 | | |
| DI49_5618; DI49_5619 | DI49_5618; DI49_5619 | | |

Create PDF in your applications with the Pdfcrowd HTML to PDF API PDFCROWD

| | | | |
|---|---|---|---|
| DI49_5616; DI49_5617<br>DI49_5611; DI49_5612 | DI49_5616; DI49_5617<br>DI49_5611; DI49_5612 | | |
| YOR344C; DI49_5156; DI49_5157 | DI49_5156; DI49_5157 | | YOR344C |
| SPGP0AUtrna4P; SPGP0Gtrna21P; SPGP0Dtrna15P; YOL013W-A; YOL014W | | SPGP0AUtrna4P; SPGP0Gtrna21P; SPGP0Dtrna15P | YOL013W-A; YOL014W |
| YOR346W; DI49_5159 | DI49_5159 | | YOR346W |
| YOR347C; DI49_5160 | DI49_5160 | | YOR347C |
| YOR348C; DI49_5161; SPGP0AU00140; DI49_5163 | DI49_5161; DI49_5163 | SPGP0AU00140 | YOR348C |
| YOR351C; YOR350C | | | YOR351C; YOR350C |
| YOR353C; DI49_5167; YOR354C; DI49_5168 | DI49_5167; DI49_5168 | | YOR353C; YOR354C |
| YOR356W; DI49_5170 | DI49_5170 | | YOR356W |
| YOR357C; DI49_5171; YOR358W; DI49_5172 | DI49_5171; DI49_5172 | | YOR357C; YOR358W |
| YOR360C; DI49_5174 | DI49_5174 | | YOR360C |
| YOR361C; DI49_5175; YOR362C; DI49_5176 | DI49_5175; DI49_5176 | | YOR361C; YOR362C |
| YOR365C; YOR363C | | | YOR365C; YOR363C |
| SPGP0RM00100; YOR367W; DI49_5180; DI49_5181 | DI49_5180; DI49_5181 | SPGP0RM00100 | YOR367W |
| SPGP0RU00100; SPGP0RR00100 | | SPGP0RU00100; SPGP0RR00100 | |
| YKL218C; YKL219W | | | YKL218C; YKL219W |
| DI49_3176; SPGP0Ptrna13T; SPGP0RV00100 | DI49_3176 | SPGP0Ptrna13T; SPGP0RV00100 | |
| YHR152W; YHR151C | | | YHR152W; YHR151C |
| YHR157W; YHR156C | | | YHR157W; YHR156C |
| YHR159W; YHR158C | | | YHR159W; YHR158C |
| YHR161C; YHR160C | | | YHR161C; YHR160C |
| YHR163W; YHR162W | | | YHR163W; YHR162W |
| YOR342C; DI49_5154 | DI49_5154 | | YOR342C |
| YOR341W; DI49_5153 | DI49_5153 | | YOR341W |
| YOR340C; DI49_5152; YOR339C | DI49_5152 | | YOR340C; YOR339C |
| YOR337W; DI49_5149; YOR336W; DI49_5150 | DI49_5149; DI49_5150 | | YOR337W; YOR336W |
| YOR334W; YOR335C; SPGP0H03470; DI49_5148 | DI49_5148 | SPGP0H03470 | YOR334W; YOR335C |
| YOR330C; DI49_5146; YOR329C; DI49_5147 | DI49_5146; DI49_5147 | | YOR330C; YOR329C |
| YOR327C; DI49_5143 | DI49_5143 | | YOR327C |
| YOR326W; DI49_5142; YOR324C | DI49_5142 | | YOR326W; YOR324C |
| YOR322C; DI49_5139; DI49_5140 | DI49_5139; DI49_5140 | | YOR322C |
| YOR321W; DI49_5138; YOR320C | DI49_5138 | | YOR321W; YOR320C |
| YOR317W; YOR319W | | | YOR317W; YOR319W |
| YOR315W; YOR316C | | | YOR315W; YOR316C |
| YOR311C; DI49_5130 | DI49_5130 | | YOR311C |
| YOR310C; DI49_5129; YOR308C | DI49_5129 | | YOR310C; YOR308C |
| YOR306C; DI49_5126; YOR305W | DI49_5126 | | YOR306C; YOR305W |

Create PDF in your applications with the Pdfcrowd HTML to PDF API PDFCROWD

| | | | |
|---|---|---|---|
| YOR304W; DI49_5123; YOR303W; DI49_5124 | DI49_5123; DI49_5124 | | YOR304W; YOR303W |
| YOR301W; YOR302W | | | YOR301W; YOR302W |
| YOR298C-A; YOR299W | | | YOR298C-A; YOR299W |
| YOR297C; YOR298W | | | YOR297C; YOR298W |
| YOR295W; YOR296W | | | YOR295W; YOR296W |
| YOR293W; DI49_5112; YOR292C; DI49_4314; YMR230W | DI49_5112; DI49_4314 | | YOR293W; YOR292C; YMR230W |
| YOR290C; DI49_5109; DI49_5110 | DI49_5109; DI49_5110 | | YOR290C |
| YOR287C; YOR288C | | | YOR287C; YOR288C |
| YOR285W; YOR286W | | | YOR285W; YOR286W |
| YOR283W; DI49_5101; DI49_5102 | DI49_5101; DI49_5102 | | YOR283W |
| YOR281C; DI49_5100 | DI49_5100 | | YOR281C |
| YOR280C; DI49_5099; YOR279C | DI49_5099 | | YOR280C; YOR279C |
| YOR276W; DI49_5096; YOR275C; DI49_5097 | DI49_5096; DI49_5097 | | YOR276W; YOR275C |
| YOR273C; DI49_5093; DI49_5094 | DI49_5093; DI49_5094 | | YOR273C |
| YOR272W; DI49_5092 | DI49_5092 | | YOR272W |
| YOR271C; DI49_5091 | DI49_5091 | | YOR271C |
| YOR270C; DI49_5090; YOR269W | DI49_5090 | | YOR270C; YOR269W |
| YOR266W; YOR267C; DI49_5087; DI49_5088 | DI49_5087; DI49_5088 | | YOR266W; YOR267C |
| YOR264W; DI49_5084; DI49_5085 | DI49_5084; DI49_5085 | | YOR264W |
| YOR262W; DI49_5083 | DI49_5083 | | YOR262W |
| YOR261C; DI49_5082 | DI49_5082 | | YOR261C |
| YOR260W; DI49_5081 | DI49_5081 | | YOR260W |
| YOR259C; DI49_5080; YOR258W | DI49_5080 | | YOR259C; YOR258W |
| YOR256C; YOR257W | | | YOR256C; YOR257W |
| YOR254C; DI49_5075; YOR253W | DI49_5075 | | YOR254C; YOR253W |
| YOR251C; YOR252W | | | YOR251C; YOR252W |
| YOR249C; YOR250C | | | YOR249C; YOR250C |
| YOR246C; YOR247W | | | YOR246C; YOR247W |
| YOR244W; DI49_5066; YOR243C; DI49_5067 | DI49_5066; DI49_5067 | | YOR244W; YOR243C |
| YOR241W; DI49_5063; YOR239W; DI49_5064 | DI49_5063; DI49_5064 | | YOR241W; YOR239W |
| YOR237W; YOR238W | | | YOR237W; YOR238W |
| YOR234C; YPL143W; YOR233W | | | YOR234C; YPL143W; YOR233W |
| YOR231W; DI49_5056; YOR230W; DI49_5057 | DI49_5056; DI49_5057 | | YOR231W; YOR230W |
| YOR228C; DI49_5053 | DI49_5053 | | YOR228C |
| YOR227W; DI49_5052; YOR226C | DI49_5052 | | YOR227W; YOR226C |
| YOR223W; YOR224C | | | YOR223W; YOR224C |
| YOR221C; DI49_5047 | DI49_5047 | | YOR221C |
| YOR220W; DI49_5046 | DI49_5046 | | YOR220W |
| YOR219C; SPGP0H02430 | | SPGP0H02430 | YOR219C |

Create PDF in your applications with the Pdfcrowd HTML to PDF API PDFCROWD

| | | | |
|---|---|---|---|
| YOR217W; DI49_5045; YOR216C | DI49_5045 | | YOR217W; YOR216C |
| YOR214C; YOR215C; DI49_5043; DI49_5044 | DI49_5043; DI49_5044 | | YOR214C; YOR215C |
| YOR212W; DI49_5040; DI49_5041 | DI49_5040; DI49_5041 | | YOR212W |
| YOR211C; DI49_5039; YOR210W | DI49_5039 | | YOR211C; YOR210W |
| YOR208W; YOR209C | | | YOR208W; YOR209C |
| YOR206W; YOR207C; DI49_5035; DI49_5036 | DI49_5035; DI49_5036 | | YOR206W; YOR207C |
| YOR204W; DI49_5032; YOR202W; DI49_5033 | DI49_5032; DI49_5033 | | YOR204W; YOR202W |
| YOR198C; DI49_5029; YOR197W; DI49_5030 | DI49_5029; DI49_5030 | | YOR198C; YOR197W |
| YOR195W; YOR196C | | | YOR195W; YOR196C |
| YOR193W; YOR194C | | | YOR193W; YOR194C |
| YOR191W; DI49_5022 | DI49_5022 | | YOR191W |
| YOR190W; DI49_5021; YOR189W | DI49_5021 | | YOR190W; YOR189W |
| YOR187W; DI49_5019; YOR186W | DI49_5019 | | YOR187W; YOR186W |
| YOR184W; YOR185C | | | YOR184W; YOR185C |
| YOR181W; DI49_5015; DI49_5016 | DI49_5015; DI49_5016 | | YOR181W |
| YOR180C; DI49_5014; YOR179C | DI49_5014 | | YOR180C; YOR179C |
| YOR174W; YOR175C | | | YOR174W; YOR175C |
| YOR172W; YOR173W | | | YOR172W; YOR173W |
| YOR168W; DI49_5003 | DI49_5003 | | YOR168W |
| YOR167C; DI49_5002; YOR166C | DI49_5002 | | YOR167C; YOR166C |
| YOR164C; DI49_4999; YOR163W | DI49_4999 | | YOR164C; YOR163W |
| YOR161C; YOR162C; DI49_4997; DI49_4998 | DI49_4997; DI49_4998 | | YOR161C; YOR162C |
| YOR159C; YOR160W; DI49_4994; DI49_4995 | DI49_4994; DI49_4995 | | YOR159C; YOR160W |
| YOR157C; YOR158W | | | YOR157C; YOR158W |
| YOR155C; YOR156C | | | YOR155C; YOR156C |
| YOR153W; DI49_4987; YOR152C; DI49_4988 | DI49_4987; DI49_4988 | | YOR153W; YOR152C |
| YOR150W; DI49_4984 | DI49_4984 | | YOR150W |
| YOR149C; DI49_4983; YOR148C | DI49_4983 | | YOR149C; YOR148C |
| YOR145C; DI49_4980; YOR144C | DI49_4980 | | YOR145C; YOR144C |
| SPGP0Dtrna30G; DI49_4977; DI49_4978 | DI49_4977; DI49_4978 | SPGP0Dtrna30G | |
| YOR142W; DI49_4976 | DI49_4976 | | YOR142W |
| YOR141C; DI49_4975; YOR140W | DI49_4975 | | YOR141C; YOR140W |
| YOR137C; YOR138C | | | YOR137C; YOR138C |
| YOR134W; YOR136W | | | YOR134W; YOR136W |
| YOR132W; DI49_4968; YOR131C | DI49_4968 | | YOR132W; YOR131C |
| YOR130C; DI49_4965; YOR129C | DI49_4965 | | YOR130C; YOR129C |
| YOR127W; DI49_4962; YOR126C | DI49_4962 | | YOR127W; YOR126C |
| YOR124C; DI49_4959; YOR123C | DI49_4959 | | YOR124C; YOR123C |
| YOR120W; YOR122C | | | YOR120W; YOR122C |
| YOR118W; YOR119C | | | YOR118W; YOR119C |

Create PDF in your applications with the Pdfcrowd HTML to PDF API PDFCROWD

| | | | |
|---|---|---|---|
| YOR116C; DI49_4952; YOR115C | DI49_4952 | | YOR116C; YOR115C |
| YOR113W; DI49_4949 | DI49_4949 | | YOR113W |
| YOR112W; DI49_4948; YOR111W | DI49_4948 | | YOR112W; YOR111W |
| YOR109W; DI49_4945; DI49_4946 | DI49_4945; DI49_4946 | | YOR109W |
| YOR108W; DI49_4944 | DI49_4944 | | YOR108W |
| YOR107W; DI49_4943; YOR106W | DI49_4943 | | YOR107W; YOR106W |
| YOR103C; DI49_4940; DI49_4941 | DI49_4940; DI49_4941 | | YOR103C |
| YOR101W; DI49_4939; YOR100C | DI49_4939 | | YOR101W; YOR100C |
| YOR098C; YOR099W; DI49_4937; DI49_4938 | DI49_4937; DI49_4938 | | YOR098C; YOR099W |
| YOR096W; DI49_4934; DI49_4935 | DI49_4934; DI49_4935 | | YOR096W |
| YOR095C; DI49_4933; YOR094W | DI49_4933 | | YOR095C; YOR094W |
| YOR092W; DI49_4930; YOR091W | DI49_4930 | | YOR092W; YOR091W |
| YOR089C; YOR090C | | | YOR089C; YOR090C |
| YOR086C; DI49_4924; YOR085W | DI49_4924 | | YOR086C; YOR085W |
| YOR083W; YOR084W | | | YOR083W; YOR084W |
| YOR078W; YOR079C | | | YOR078W; YOR079C |
| YOR076C; YOR077W | | | YOR076C; YOR077W |
| YOR074C; SPGP0H01130 | | SPGP0H01130 | YOR074C |
| YOR073W; DI49_4915 | DI49_4915 | | YOR073W |
| YOR070C; DI49_4912 | DI49_4912 | | YOR070C |
| YOR069W; DI49_4911 | DI49_4911 | | YOR069W |
| YOR067C; SPGP0H01090 | | SPGP0H01090 | YOR067C |
| YOR066W; DI49_4909 | DI49_4909 | | YOR066W |
| YOR065W; DI49_4908; YOR064C | DI49_4908 | | YOR065W; YOR064C |
| YOR062C; YOR063W | | | YOR062C; YOR063W |
| YOR060C; DI49_4902; YOR059C; DI49_4903 | DI49_4902; DI49_4903 | | YOR060C; YOR059C |
| YOR058C; DI49_4899 | DI49_4899 | | YOR058C |
| YOR057W; DI49_4898; YOR056C | DI49_4898 | | YOR057W; YOR056C |
| YOR052C; DI49_4895; DI49_4896 | DI49_4895; DI49_4896 | | YOR052C |
| YOR051C; DI49_4894; YOR049C | DI49_4894 | | YOR051C; YOR049C |
| YOR047C; DI49_4891 | DI49_4891 | | YOR047C |
| YOR046C; DI49_4890; YOR045W | DI49_4890 | | YOR046C; YOR045W |
| YOR043W; DI49_4887; YOR042W; DI49_4888 | DI49_4887; DI49_4888 | | YOR043W; YOR042W |
| YOR039W; DI49_4884; DI49_4885 | DI49_4884; DI49_4885 | | YOR039W |
| YOR038C; DI49_4883 | DI49_4883 | | YOR038C |
| YOR037W; DI49_4882; YOR036W | DI49_4882 | | YOR037W; YOR036W |
| YOR034C; YOR035C | | | YOR034C; YOR035C |
| YOR032C; YOR033C | | | YOR032C; YOR033C |
| YOR030W; YOR031W | | | YOR030W; YOR031W |
| YOR027W; YOR028C | | | YOR027W; YOR028C |

Create PDF in your applications with the Pdfcrowd HTML to PDF API PDFCROWD

| | | | |
|---|---|---|---|
| YOR025W; YOR026W | | | YOR025W; YOR026W |
| YOR022C; YOR023C | | | YOR022C; YOR023C |
| YOR020W-A; YOR021C | | | YOR020W-A; YOR021C |
| YOR019W; YOR020C | | | YOR019W; YOR020C |
| YOR009W; DI49_2412; YOR008C | DI49_2412 | | YOR009W; YOR008C |
| YOR005C; DI49_2407; YOR004W | DI49_2407 | | YOR005C; YOR004W |
| YOR001W; DI49_2403; YOL001W | DI49_2403 | | YOR001W; YOL001W |
| YOL004W; DI49_2399; YOL005C | DI49_2399 | | YOL004W; YOL005C |
| YOL019W; YOL018C | | | YOL019W; YOL018C |
| YOL031C; YOL030W | | | YOL031C; YOL030W |
| YOL033W; YOL032W | | | YOL033W; YOL032W |
| YOL044W; YOL042W | | | YOL044W; YOL042W |
| YOL048C; YOL047C | | | YOL048C; YOL047C |
| YOL052C; DI49_2354; YOL052C-A | DI49_2354 | | YOL052C; YOL052C-A |
| YOL062C; DI49_2342; YOL063C | DI49_2342 | | YOL062C; YOL063C |
| YOL065C; YOL064C | | | YOL065C; YOL064C |
| YOL067C; YOL066C | | | YOL067C; YOL066C |
| YOL082W; YOL081W | | | YOL082W; YOL081W |
| YOL084W; YOL083W | | | YOL084W; YOL083W |
| YOL086W-A; YOL086C; YMR303C | | | YOL086W-A; YOL086C; YMR303C |
| YOL088C; YOL087C | | | YOL088C; YOL087C |
| YOL102C; DI49_2304; YOL103W | DI49_2304 | | YOL102C; YOL103W |
| YOL105C; YOL104C | | | YOL105C; YOL104C |
| YOL107W; SPGP0Dtrna23T | | SPGP0Dtrna23T | YOL107W |
| YOL111C; YOL110W | | | YOL111C; YOL110W |
| YOL113W; YOL112W | | | YOL113W; YOL112W |
| YOL124C; DI49_2283; YOL125W | DI49_2283 | | YOL124C; YOL125W |
| YOL140W; DI49_2268; YOL141W | DI49_2268 | | YOL140W; YOL141W |
| YOL143C; DI49_2265; YOL144W | DI49_2265 | | YOL143C; YOL144W |
| YOL146W; YOL145C | | | YOL146W; YOL145C |
| YOL154W; YOL153C | | | YOL154W; YOL153C |
| DI49_4721; YNL016W; DI49_4722 | DI49_4721; DI49_4722 | | YNL016W |
| DI49_4726; YNL009W; DI49_4727 | DI49_4726; DI49_4727 | | YNL009W |
| DI49_4733; YNL002C; DI49_4734 | DI49_4733; DI49_4734 | | YNL002C |
| YOR380W; DI49_5191; YOR378W; DI49_5192 | DI49_5191; DI49_5192 | | YOR380W; YOR378W |
| YOR375C; DI49_5188; DI49_5189 | DI49_5188; DI49_5189 | | YOR375C |
| YOR374W; DI49_5187; YOR373W | DI49_5187 | | YOR374W; YOR373W |
| YOR371C; YOR372C | | | YOR371C; YOR372C |
| YOR369C; DI49_5182; YOR368W; DI49_5183 | DI49_5182; DI49_5183 | | YOR369C; YOR368W |
| YIR017C; DI49_2797; YIR014W | DI49_2797 | | YIR017C; YIR014W |

Create PDF in your applications with the Pdfcrowd HTML to PDF API PDFCROWD

| | | | |
|---|---|---|---|
| YIR012W; YIR013C | | | YIR012W; YIR013C |
| YIR010W; DI49_2790; YIR009W | DI49_2790 | | YIR010W; YIR009W |
| YIR006C; DI49_2785; YIR005W | DI49_2785 | | YIR006C; YIR005W |
| YCR066W; YIR003W | | | YCR066W; YIR003W |
| YCR068W; YCR067C | | | YCR068W; YCR067C |
| YCR071C; YCR069W | | | YCR071C; YCR069W |
| YCR073C; YCR072C | | | YCR073C; YCR072C |
| YCR075C; YCR073W-A | | | YCR075C; YCR073W-A |
| YCR076C; YCR075W-A | | | YCR076C; YCR075W-A |
| YCR079W; YCR077C | | | YCR079W; YCR077C |
| DI49_4710; DI49_4709 | DI49_4710; DI49_4709 | | |
| DI49_4712; DI49_4711 | DI49_4712; DI49_4711 | | |
| YIR023W; DI49_4717 | DI49_4717 | | YIR023W |
| YIR025W; YIR024C | | | YIR025W; YIR024C |
| YIR027C; YIR026C | | | YIR027C; YIR026C |
| YIR032C; YIR031C | | | YIR032C; YIR031C |
| YMR314W; YIR033W | | | YMR314W; YIR033W |
| YMR312W; YMR313C | | | YMR312W; YMR313C |
| YMR310C; YMR311C | | | YMR310C; YMR311C |
| YMR308C; YMR309C | | | YMR308C; YMR309C |
| YPR005C; YMR307W | | | YPR005C; YMR307W |
| YPR007C; YPR006C | | | YPR007C; YPR006C |
| SPGP0BG00120; YIR039C | | SPGP0BG00120 | YIR039C |
| DI49_2245; YGR284C; DI49_2244 | DI49_2245; DI49_2244 | | YGR284C |
| DI49_2235; DI49_2236 | DI49_2235; DI49_2236 | | |
| DI49_2232; YGR270W; DI49_2231 | DI49_2232; DI49_2231 | | YGR270W |
| DI49_2229; DI49_2230 | DI49_2229; DI49_2230 | | |
| DI49_2227; YGR263C; DI49_2226 | DI49_2227; DI49_2226 | | YGR263C |
| DI49_2223; YGR258C; DI49_2222 | DI49_2223; DI49_2222 | | YGR258C |
| DI49_2216; YGR252W; DI49_2215 | DI49_2216; DI49_2215 | | YGR252W |
| DI49_2213; DI49_2214 | DI49_2213; DI49_2214 | | |
| DI49_2211; DI49_2212 | DI49_2211; DI49_2212 | | |
| DI49_2208; YGR245C; DI49_2207 | DI49_2208; DI49_2207 | | YGR245C |
| DI49_2205; DI49_2206 | DI49_2205; DI49_2206 | | |
| DI49_2203; DI49_2204 | DI49_2203; DI49_2204 | | |
| DI49_2201; DI49_2202 | DI49_2201; DI49_2202 | | |
| DI49_2199; DI49_2200 | DI49_2199; DI49_2200 | | |
| DI49_2197; DI49_2198 | DI49_2197; DI49_2198 | | |
| DI49_2195; YGR231C; DI49_2194 | DI49_2195; DI49_2194 | | YGR231C |
| DI49_2191; DI49_2192 | DI49_2191; DI49_2192 | | |

Create PDF in your applications with the Pdfcrowd HTML to PDF API PDFCROWD

| | | | |
|---|---|---|---|
| DI49_2190; SPGP0E00700 | DI49_2190 | SPGP0E00700 | |
| DI49_2188; DI49_2189 | DI49_2188; DI49_2189 | | |
| DI49_2183; YGR216C; DI49_2182 | DI49_2183; DI49_2182 | | YGR216C |
| DI49_2180; DI49_2181 | DI49_2180; DI49_2181 | | |
| DI49_2178; DI49_2179 | DI49_2178; DI49_2179 | | |
| DI49_2171; YGR204W; DI49_2170 | DI49_2171; DI49_2170 | | YGR204W |
| DI49_2168; DI49_2169 | DI49_2168; DI49_2169 | | |
| DI49_2166; DI49_2167 | DI49_2166; DI49_2167 | | |
| DI49_2164; YGR197C; DI49_2163 | DI49_2164; DI49_2163 | | YGR197C |
| DI49_2145; YGR180C; DI49_2144 | DI49_2145; DI49_2144 | | YGR180C |
| DI49_2132; DI49_2133 | DI49_2132; DI49_2133 | | |
| DI49_2130; DI49_2131 | DI49_2130; DI49_2131 | | |
| DI49_2121; DI49_2122 | DI49_2121; DI49_2122 | | |
| DI49_2103; YGR142W; DI49_2102 | DI49_2103; DI49_2102 | | YGR142W |
| DI49_2095; YGR133W; DI49_2094 | DI49_2095; DI49_2094 | | YGR133W |
| DI49_2092; DI49_2093 | DI49_2092; DI49_2093 | | |
| DI49_2090; DI49_2091 | DI49_2090; DI49_2091 | | |
| DI49_2088; DI49_2089 | DI49_2088; DI49_2089 | | |
| DI49_2074; DI49_2075 | DI49_2074; DI49_2075 | | |
| DI49_2072; YGR111W; DI49_2071 | DI49_2072; DI49_2071 | | YGR111W |
| DI49_2063; DI49_2064 | DI49_2063; DI49_2064 | | |
| DI49_2061; DI49_2062 | DI49_2061; DI49_2062 | | |
| DI49_2050; DI49_2051 | DI49_2050; DI49_2051 | | |
| DI49_2045; DI49_2046 | DI49_2045; DI49_2046 | | |
| DI49_2043; DI49_2044 | DI49_2043; DI49_2044 | | |
| DI49_2038; DI49_2039 | DI49_2038; DI49_2039 | | |
| DI49_2033; DI49_2034 | DI49_2033; DI49_2034 | | |
| DI49_2031; DI49_2032 | DI49_2031; DI49_2032 | | |
| DI49_2029; DI49_2030 | DI49_2029; DI49_2030 | | |
| DI49_2024; SPGP0E02170 | DI49_2024 | SPGP0E02170 | |
| DI49_2022; DI49_2023 | DI49_2022; DI49_2023 | | |
| DI49_2017; DI49_2018 | DI49_2017; DI49_2018 | | |
| DI49_2005; DI49_2006 | DI49_2005; DI49_2006 | | |
| DI49_2003; DI49_2004 | DI49_2003; DI49_2004 | | |
| DI49_1978; DI49_1979 | DI49_1978; DI49_1979 | | |
| DI49_1976; DI49_1977 | DI49_1976; DI49_1977 | | |
| DI49_1965; DI49_1966 | DI49_1965; DI49_1966 | | |
| DI49_1963; DI49_1964 | DI49_1963; DI49_1964 | | |
| DI49_1961; DI49_1962 | DI49_1961; DI49_1962 | | |

Create PDF in your applications with the Pdfcrowd HTML to PDF API PDFCROWD

| | | | |
|---|---|---|---|
| DI49_1955; DI49_1956 | DI49_1955; DI49_1956 | | |
| DI49_1950; DI49_1951 | DI49_1950; DI49_1951 | | |
| DI49_1948; DI49_1949 | DI49_1948; DI49_1949 | | |
| DI49_1931; DI49_1932 | DI49_1931; DI49_1932 | | |
| DI49_1928; DI49_1820; DI49_1819 | DI49_1928; DI49_1820; DI49_1819 | | |
| DI49_1920; SPGP0Etrna32V | DI49_1920 | SPGP0Etrna32V | |
| DI49_1909; DI49_1910 | DI49_1909; DI49_1910 | | |
| DI49_1908; SPGP0E03250 | DI49_1908 | SPGP0E03250 | |
| DI49_1906; DI49_1907 | DI49_1906; DI49_1907 | | |
| DI49_1891; DI49_1892 | DI49_1891; DI49_1892 | | |
| SPGP0E03530; DI49_1883 | DI49_1883 | SPGP0E03530 | |
| DI49_1878; SPGP0Etrna29E | DI49_1878 | SPGP0Etrna29E | |
| DI49_1876; DI49_1877 | DI49_1876; DI49_1877 | | |
| DI49_1869; DI49_1870 | DI49_1869; DI49_1870 | | |
| DI49_1867; DI49_1868 | DI49_1867; DI49_1868 | | |
| DI49_1865; DI49_1866 | DI49_1865; DI49_1866 | | |
| DI49_1863; DI49_1864 | DI49_1863; DI49_1864 | | |
| DI49_1861; DI49_1862 | DI49_1861; DI49_1862 | | |
| DI49_1853; DI49_1854 | DI49_1853; DI49_1854 | | |
| DI49_1848; DI49_1849 | DI49_1848; DI49_1849 | | |
| DI49_1842; DI49_1843 | DI49_1842; DI49_1843 | | |
| DI49_1830; DI49_1831 | DI49_1830; DI49_1831 | | |
| DI49_1822; DI49_1823 | DI49_1822; DI49_1823 | | |
| DI49_1817; DI49_1818 | DI49_1817; DI49_1818 | | |
| DI49_1815; DI49_1816 | DI49_1815; DI49_1816 | | |
| SPGP0E04190; DI49_1814 | DI49_1814 | SPGP0E04190 | |
| DI49_1811; DI49_1812 | DI49_1811; DI49_1812 | | |
| DI49_5607; DI49_5608 | DI49_5607; DI49_5608 | | |
| DI49_5605; DI49_5606 | DI49_5605; DI49_5606 | | |
| YGL178W; SPGP0BJ00150 | | SPGP0BJ00150 | YGL178W |
| YGL175C; YGL176C | | | YGL175C; YGL176C |
| YGL173C; YGL174W | | | YGL173C; YGL174W |
| YCR051W; YCR048W | | | YCR051W; YCR048W |
| YCR053W; YCR052W | | | YCR053W; YCR052W |
| YCR057C; YCR054C | | | YCR057C; YCR054C |
| YCR088W; YCR087W | | | YCR088W; YCR087W |
| YGL262W; SPGP0BO00120 | | SPGP0BO00120 | YGL262W |
| DI49_4387; YMR300C; DI49_4386; YMR299C | DI49_4387; DI49_4386 | | YMR300C; YMR299C |
| DI49_4384; YMR297W; DI49_4383; YMR296C | DI49_4384; DI49_4383 | | YMR297W; YMR296C |

Create PDF in your applications with the Pdfcrowd HTML to PDF API PDFCROWD

| | | | |
|---|---|---|---|
| DI49_4381; DI49_4382 | DI49_4381; DI49_4382 | | |
| DI49_4379; DI49_4380 | DI49_4379; DI49_4380 | | |
| DI49_4377; YMR290C; DI49_4376; YMR289W | DI49_4377; DI49_4376 | | YMR290C; YMR289W |
| DI49_4374; YMR287C; DI49_4373; YMR286W | DI49_4374; DI49_4373 | | YMR287C; YMR286W |
| DI49_4371; DI49_4372 | DI49_4371; DI49_4372 | | |
| DI49_4367; DI49_4368 | DI49_4367; DI49_4368 | | |
| DI49_4365; DI49_4366 | DI49_4365; DI49_4366 | | |
| DI49_4363; DI49_4364 | DI49_4363; DI49_4364 | | |
| DI49_4361; YMR275C; DI49_4360; YMR274C | DI49_4361; DI49_4360 | | YMR275C; YMR274C |
| DI49_4358; SPGP0G00400 | DI49_4358 | SPGP0G00400 | |
| DI49_4357; SPGP0Gtrna17Q; SPGP0F00410; YMR271C | DI49_4357 | SPGP0Gtrna17Q; SPGP0F00410 | YMR271C |
| DI49_4355; DI49_4356 | DI49_4355; DI49_4356 | | |
| DI49_4353; YMR269W; DI49_4352; YMR268C | DI49_4353; DI49_4352 | | YMR269W; YMR268C |
| DI49_4350; DI49_4351 | DI49_4350; DI49_4351 | | |
| DI49_4348; YMR264W; DI49_4347; YMR263W | DI49_4348; DI49_4347 | | YMR264W; YMR263W |
| DI49_4345; YMR261C | DI49_4345 | | YMR261C |
| DI49_4344; YMR260C; DI49_4343; YMR259C | DI49_4344; DI49_4343 | | YMR260C; YMR259C |
| DI49_4341; YMR257C | DI49_4341 | | YMR257C |
| DI49_4340; YMR256C; DI49_4339; YMR255W | DI49_4340; DI49_4339 | | YMR256C; YMR255W |
| DI49_4337; DI49_4338 | DI49_4337; DI49_4338 | | |
| DI49_4335; DI49_4336 | DI49_4335; DI49_4336 | | |
| DI49_4330; DI49_4331 | DI49_4330; DI49_4331 | | |
| DI49_4328; YMR243C | DI49_4328 | | YMR243C |
| DI49_4327; YMR242C; DI49_5131; DI49_5132 | DI49_4327; DI49_5131; DI49_5132 | | YMR242C |
| DI49_4326; YMR241W; DI49_4325; YMR240C | DI49_4326; DI49_4325 | | YMR241W; YMR240C |
| DI49_4322; YMR238W | DI49_4322 | | YMR238W |
| DI49_4321; YMR237W; DI49_4320; YMR236W | DI49_4321; DI49_4320 | | YMR237W; YMR236W |
| DI49_4318; YMR234W; DI49_4317; YMR233W | DI49_4318; DI49_4317 | | YMR234W; YMR233W |
| DI49_4315; YMR231W | DI49_4315 | | YMR231W |
| DI49_4313; YMR229C | DI49_4313 | | YMR229C |
| DI49_4312; YMR228W | DI49_4312 | | YMR228W |
| DI49_4311; YMR227C | DI49_4311 | | YMR227C |
| DI49_4310; YMR226C; DI49_4309; YMR225C | DI49_4310; DI49_4309 | | YMR226C; YMR225C |
| DI49_4307; DI49_4308 | DI49_4307; DI49_4308 | | |
| DI49_4305; YMR221C | DI49_4305 | | YMR221C |
| DI49_4304; YMR220W; DI49_4303; YMR219W | DI49_4304; DI49_4303 | | YMR220W; YMR219W |
| DI49_4301; YMR217W; DI49_4300; YMR216C | DI49_4301; DI49_4300 | | YMR217W; YMR216C |
| DI49_4298; YMR214W; DI49_4297; YMR213W | DI49_4298; DI49_4297 | | YMR214W; YMR213W |
| DI49_4295; DI49_4296 | DI49_4295; DI49_4296 | | |

Create PDF in your applications with the Pdfcrowd HTML to PDF API PDFCROWD

| | | | |
|---|---|---|---|
| DI49_4293; DI49_4294 | DI49_4293; DI49_4294 | | |
| DI49_4291; YMR207C; DI49_4290; YMR206W | DI49_4291; DI49_4290 | | YMR207C; YMR206W |
| DI49_4288; YMR204C | DI49_4288 | | YMR204C |
| DI49_4287; YMR203W | DI49_4287 | | YMR203W |
| DI49_4286; YMR202W; DI49_4285; SPGP0G01090 | DI49_4286; DI49_4285 | SPGP0G01090 | YMR202W |
| DI49_4283; YMR199W; DI49_4282; YMR198W | DI49_4283; DI49_4282 | | YMR199W; YMR198W |
| DI49_4280; YMR196W; DI49_4279; YMR195W | DI49_4280; DI49_4279 | | YMR196W; YMR195W |
| DI49_4274; DI49_4275 | DI49_4274; DI49_4275 | | |
| DI49_4272; YMR189W; DI49_4271; YMR188C | DI49_4272; DI49_4271 | | YMR189W; YMR188C |
| DI49_4268; YMR185W; DI49_4267; YMR184W | DI49_4268; DI49_4267 | | YMR185W; YMR184W |
| DI49_4263; DI49_4264 | DI49_4263; DI49_4264 | | |
| DI49_4261; DI49_4262 | DI49_4261; DI49_4262 | | |
| DI49_4259; YMR176W; SPGP0F01380; YMR175W | DI49_4259 | SPGP0F01380 | YMR176W; YMR175W |
| DI49_4257; DI49_4258 | DI49_4257; DI49_4258 | | |
| DI49_4255; YMR171C; SPGP0F01430; YMR168C | DI49_4255 | SPGP0F01430 | YMR171C; YMR168C |
| DI49_4251; DI49_4252 | DI49_4251; DI49_4252 | | |
| DI49_4249; YMR165C; SPGP0F01480; YMR164C | DI49_4249 | SPGP0F01480 | YMR165C; YMR164C |
| DI49_4245; YMR161W; DI49_4244; YMR160W | DI49_4245; DI49_4244 | | YMR161W; YMR160W |
| DI49_4240; DI49_4241 | DI49_4240; DI49_4241 | | |
| DI49_4238; DI49_4239 | DI49_4238; DI49_4239 | | |
| DI49_4236; DI49_4237 | DI49_4236; DI49_4237 | | |
| DI49_4234; DI49_4235 | DI49_4234; DI49_4235 | | |
| DI49_4232; DI49_4233 | DI49_4232; DI49_4233 | | |
| DI49_4230; YMR145C | DI49_4230 | | YMR145C |
| DI49_4229; YMR144W | DI49_4229 | | YMR144W |
| SPGP0F01680; YMR143W | | SPGP0F01680 | YMR143W |
| DI49_4226; DI49_4227 | DI49_4226; DI49_4227 | | |
| DI49_4224; DI49_4225 | DI49_4224; DI49_4225 | | |
| DI49_4222; DI49_4223 | DI49_4222; DI49_4223 | | |
| DI49_4221; SPGP0F01770 | DI49_4221 | SPGP0F01770 | |
| DI49_4219; DI49_4220 | DI49_4219; DI49_4220 | | |
| DI49_4217; YMR128W | DI49_4217 | | YMR128W |
| DI49_4216; YMR127C; DI49_4215; YMR126C | DI49_4216; DI49_4215 | | YMR127C; YMR126C |
| DI49_4213; YMR124W; DI49_4212; YMR123W | DI49_4213; DI49_4212 | | YMR124W; YMR123W |
| DI49_4209; YMR120C; DI49_4208; YMR119W | DI49_4209; DI49_4208 | | YMR120C; YMR119W |
| DI49_4206; SPGP0Ftrna28L | DI49_4206 | SPGP0Ftrna28L | |
| SPGP0F01940; DI49_4205 | DI49_4205 | SPGP0F01940 | |
| DI49_4203; YMR114C | DI49_4203 | | YMR114C |
| DI49_4202; YMR113W; DI49_4201; YMR112C | DI49_4202; DI49_4201 | | YMR113W; YMR112C |

Create PDF in your applications with the Pdfcrowd HTML to PDF API PDFCROWD

| | | | |
|---|---|---|---|
| DI49_4199; DI49_4200 | DI49_4199; DI49_4200 | | |
| DI49_4197; YMR108W; DI49_4196; YMR107W | DI49_4197; DI49_4196 | | YMR108W; YMR107W |
| DI49_4192; DI49_4193 | DI49_4192; DI49_4193 | | |
| DI49_4190; YMR101C; DI49_4189; YMR100W | DI49_4190; DI49_4189 | | YMR101C; YMR100W |
| DI49_4186; YMR098C; DI49_4185; YMR097C | DI49_4186; DI49_4185 | | YMR098C; YMR097C |
| DI49_4183; DI49_4184 | DI49_4183; DI49_4184 | | |
| DI49_4181; YMR093W; DI49_4180; YMR092C | DI49_4181; DI49_4180 | | YMR093W; YMR092C |
| DI49_4177; YMR089C | DI49_4177 | | YMR089C |
| DI49_4176; YMR088C; DI49_4175; YMR087W | DI49_4176; DI49_4175 | | YMR088C; YMR087W |
| DI49_4173; YMR085W | DI49_4173 | | YMR085W |
| DI49_4172; YMR083W; DI49_4171; YMR081C | DI49_4172; DI49_4171 | | YMR083W; YMR081C |
| DI49_4169; DI49_4170 | DI49_4169; DI49_4170 | | |
| DI49_4167; DI49_4168 | DI49_4167; DI49_4168 | | |
| DI49_4165; YMR076C; SPGP0F02320; YMR075W | DI49_4165 | SPGP0F02320 | YMR076C; YMR075W |
| DI49_4162; DI49_4163 | DI49_4162; DI49_4163 | | |
| DI49_4160; YMR071C; DI49_4159; YMR070W | DI49_4160; DI49_4159 | | YMR071C; YMR070W |
| DI49_4157; YMR068W; DI49_4156; YMR067C | DI49_4157; DI49_4156 | | YMR068W; YMR067C |
| DI49_4154; YMR065W; DI49_4153; YMR064W | DI49_4154; DI49_4153 | | YMR065W; YMR064W |
| DI49_4151; DI49_4152 | DI49_4151; DI49_4152 | | |
| DI49_4149; DI49_4150 | DI49_4149; DI49_4150 | | |
| DI49_4147; YMR058W | DI49_4147 | | YMR058W |
| DI49_4146; YMR056C; DI49_4145; YMR055C | DI49_4146; DI49_4145 | | YMR056C; YMR055C |
| DI49_4143; YMR053C; DI49_4142; YMR052W | DI49_4143; DI49_4142 | | YMR053C; YMR052W |
| DI49_4136; DI49_4137 | DI49_4136; DI49_4137 | | |
| DI49_4131; DI49_4132 | DI49_4131; DI49_4132 | | |
| DI49_4129; DI49_4130 | DI49_4129; DI49_4130 | | |
| DI49_4127; DI49_4128 | DI49_4127; DI49_4128 | | |
| DI49_4125; DI49_4126 | DI49_4125; DI49_4126 | | |
| DI49_4123; DI49_4124 | DI49_4123; DI49_4124 | | |
| DI49_4121; DI49_4122 | DI49_4121; DI49_4122 | | |
| DI49_4119; DI49_4120 | DI49_4119; DI49_4120 | | |
| DI49_4117; DI49_4118 | DI49_4117; DI49_4118 | | |
| DI49_4115; YMR027W; DI49_4114; YMR026C | DI49_4115; DI49_4114 | | YMR027W; YMR026C |
| DI49_4112; YMR024W | DI49_4112 | | YMR024W |
| DI49_4108; YMR021C | DI49_4108 | | YMR021C |
| DI49_4107; YMR020W; DI49_4106; YMR019W | DI49_4107; DI49_4106 | | YMR020W; YMR019W |
| DI49_4104; DI49_4105 | DI49_4104; DI49_4105 | | |
| DI49_4102; YMR015C | DI49_4102 | | YMR015C |
| DI49_4101; YMR014W | DI49_4101 | | YMR014W |
| DI49_4100; YMR013C | DI49_4100 | | YMR013C |

Create PDF in your applications with the Pdfcrowd HTML to PDF API PDFCROWD

| | | | |
|---|---|---|---|
| DI49_4099; YMR012W | DI49_4099 | | YMR012W |
| SPGP0F02950; YMR011W | | SPGP0F02950 | YMR011W |
| DI49_4097; YMR010W; DI49_4096; YMR009W | DI49_4097; DI49_4096 | | YMR010W; YMR009W |
| DI49_4091; DI49_4092; YMR004W; YMR003W | DI49_4091; DI49_4092 | | YMR004W; YMR003W |
| DI49_4089; YMR001C | DI49_4089 | | YMR001C |
| DI49_4088; YML001W | DI49_4088 | | YML001W |
| DI49_4087; YML002W | DI49_4087 | | YML002W |
| DI49_4086; YML004C | DI49_4086 | | YML004C |
| DI49_4085; YML005W | DI49_4085 | | YML005W |
| DI49_4083; YML006C | DI49_4083 | | YML006C |
| DI49_4082; YML007W | DI49_4082 | | YML007W |
| DI49_4081; YML008C | DI49_4081 | | YML008C |
| DI49_4080; YML009C | DI49_4080 | | YML009C |
| DI49_4079; YML010W; DI49_4078; YML011C | DI49_4079; DI49_4078 | | YML010W; YML011C |
| DI49_4076; DI49_4077 | DI49_4076; DI49_4077 | | |
| DI49_4074; DI49_4075 | DI49_4074; DI49_4075 | | |
| DI49_4072; YML017W; DI49_4071; YML018C | DI49_4072; DI49_4071 | | YML017W; YML018C |
| DI49_4069; YML020W | DI49_4069 | | YML020W |
| DI49_4068; YML021C; DI49_4067; YML022W | DI49_4068; DI49_4067 | | YML021C; YML022W |
| DI49_4064; YML025C | DI49_4064 | | YML025C |
| DI49_4060; DI49_4061 | DI49_4060; DI49_4061 | | |
| DI49_4058; YML031W | DI49_4058 | | YML031W |
| DI49_4057; YML032C; DI49_4056; YML034W | DI49_4057; DI49_4056 | | YML032C; YML034W |
| DI49_4054; DI49_4055 | DI49_4054; DI49_4055 | | |
| DI49_4052; YML038C | DI49_4052 | | YML038C |
| DI49_4050; YML041C | DI49_4050 | | YML041C |
| DI49_4049; YML042W; DI49_4048; YML043C | DI49_4049; DI49_4048 | | YML042W; YML043C |
| DI49_4044; YML048W; SPGP0F03470; YML049C | DI49_4044 | SPGP0F03470 | YML048W; YML049C |
| DI49_4041; YML051W | DI49_4041 | | YML051W |
| DI49_4040; YML052W | DI49_4040 | | YML052W |
| DI49_4039; YML053C | DI49_4039 | | YML053C |
| SPGP0F03540; SPGP0G03490 | | SPGP0F03540; SPGP0G03490 | |
| DI49_4035; YML057W; SPGP0F03560; YML058W | DI49_4035 | SPGP0F03560 | YML057W; YML058W |
| DI49_4032; YML059C; DI49_4031; YML060W | DI49_4032; DI49_4031 | | YML059C; YML060W |
| DI49_4027; YML064C; DI49_4026; YML065W | DI49_4027; DI49_4026 | | YML064C; YML065W |
| DI49_4024; DI49_4025 | DI49_4024; DI49_4025 | | |
| DI49_4022; DI49_4023 | DI49_4022; DI49_4023 | | |
| DI49_4015; YML074C | DI49_4015 | | YML074C |
| DI49_4014; YML075C; DI49_4013; YML076C | DI49_4014; DI49_4013 | | YML075C; YML076C |
| SPGP0F03780; YML078W; DI49_4011; YML079W | DI49_4011 | SPGP0F03780 | YML078W; YML079W |

Create PDF in your applications with the Pdfcrowd HTML to PDF API PDFCROWD

| | | | |
|---|---|---|---|
| DI49_4009; DI49_4010 | DI49_4009; DI49_4010 | | |
| DI49_4007; YML082W | DI49_4007 | | YML082W |
| DI49_4006; YML083C | DI49_4006 | | YML083C |
| DI49_4005; YML085C; DI49_4004; DI49_3969; DI49_3968; YML086C; YML124C | DI49_4005; DI49_4004; DI49_3969; DI49_3968 | | YML085C; YML086C; YML124C |
| DI49_4002; DI49_4003 | DI49_4002; DI49_4003 | | |
| DI49_4000; YML092C | DI49_4000 | | YML092C |
| DI49_3999; YML093W | DI49_3999 | | YML093W |
| DI49_3998; YML094W; DI49_3997; YML095C | DI49_3998; DI49_3997 | | YML094W; YML095C |
| DI49_3995; DI49_3996 | DI49_3995; DI49_3996 | | |
| DI49_3993; YML099C; SPGP0F03980; YML100W | DI49_3993 | SPGP0F03980 | YML099C; YML100W |
| DI49_3990; DI49_3992 | DI49_3990; DI49_3992 | | |
| DI49_3988; DI49_3989; YML103C; YML104C | DI49_3988; DI49_3989 | | YML103C; YML104C |
| DI49_3986; YML106W; DI49_3985; YML107C | DI49_3986; DI49_3985 | | YML106W; YML107C |
| DI49_3983; YML109W; DI49_3982; YML110C | DI49_3983; DI49_3982 | | YML109W; YML110C |
| DI49_3980; DI49_3981 | DI49_3980; DI49_3981 | | |
| DI49_3978; DI49_3979 | DI49_3978; DI49_3979 | | |
| DI49_3976; YML116W | DI49_3976 | | YML116W |
| DI49_3975; YML117W | DI49_3975 | | YML117W |
| DI49_3974; YML118W; DI49_3973; YML119W | DI49_3974; DI49_3973 | | YML118W; YML119W |
| DI49_3971; YML121W | DI49_3971 | | YML121W |
| DI49_3970; YML123C | DI49_3970 | | YML123C |
| DI49_3966; YML127W | DI49_3966 | | YML127W |
| DI49_3965; YML128C | DI49_3965 | | YML128C |
| DI49_3963; YML129C; SPGP0F04270; YML130C | DI49_3963 | SPGP0F04270 | YML129C; YML130C |
| DI49_3961; DI49_3962 | DI49_3961; DI49_3962 | | |
| YCR042C; DI49_3960 | DI49_3960 | | YCR042C |
| YCR044C; YCR043C | | | YCR044C; YCR043C |
| YIR021W; SPGP0BU00100 | | SPGP0BU00100 | YIR021W |
| SPGP0BT00110; SPGP0BU00120; YCR081W | | SPGP0BT00110; SPGP0BU00120 | YCR081W |
| YCR083W; YCR082W | | | YCR083W; YCR082W |
| YIR037W; YIR036C | | | YIR037W; YIR036C |
| SPGP0CA00100; YFR017C; YMR302C | | SPGP0CA00100 | YFR017C; YMR302C |
| YMR294W; YMR295C | | | YMR294W; YMR295C |
| YMR292W; YMR293C | | | YMR292W; YMR293C |
| YMR284W; YMR285C | | | YMR284W; YMR285C |
| YMR281W; YMR282C | | | YMR281W; YMR282C |
| YMR279C; YMR280C | | | YMR279C; YMR280C |
| YMR277W; YMR278W | | | YMR277W; YMR278W |
| YMR266W; YMR267W | | | YMR266W; YMR267W |

Create PDF in your applications with the Pdfcrowd HTML to PDF API PDFCROWD

| | | |
|---|---|---|
| YMR252C; YMR253C | | YMR252C; YMR253C |
| YMR251W; YMR251W-A | | YMR251W; YMR251W-A |
| YMR244C-A; YMR246W | | YMR244C-A; YMR246W |
| YMR223W; YMR224C | | YMR223W; YMR224C |
| YMR211W; YMR212C | | YMR211W; YMR212C |
| YMR209C; YMR210W | | YMR209C; YMR210W |
| YMR200W; YMR201C | | YMR200W; YMR201C |
| YMR191W; YMR192W | | YMR191W; YMR192W |
| YMR182C; YMR183C | | YMR182C; YMR183C |
| YMR180C; YMR181C | | YMR180C; YMR181C |
| YMR178W; YMR179W | | YMR178W; YMR179W |
| YMR173W; YMR174C | | YMR173W; YMR174C |
| YMR166C; DI49_4250 | DI49_4250 | YMR166C |
| YMR156C; YMR157C | | YMR156C; YMR157C |
| YMR154C; YMR155W | | YMR154C; YMR155W |
| YMR152W; YMR153W | | YMR152W; YMR153W |
| YMR149W; YMR150C | | YMR149W; YMR150C |
| YMR147W; YMR148W | | YMR147W; YMR148W |
| YMR138W; YMR139W | | YMR138W; YMR139W |
| YMR136W; YMR137C | | YMR136W; YMR137C |
| YMR134W; YMR135C | | YMR134W; YMR135C |
| YMR132C; YMR133W | | YMR132C; YMR133W |
| YMR130W; YMR131C | | YMR130W; YMR131C |
| YMR116C; YMR117C | | YMR116C; YMR117C |
| YMR110C; YMR111C | | YMR110C; YMR111C |
| YMR104C; YMR105C | | YMR104C; YMR105C |
| YMR095C; YMR096W | | YMR095C; YMR096W |
| YMR090W; YMR091C | | YMR090W; YMR091C |
| YMR079W; YMR080C | | YMR079W; YMR080C |
| YMR077C; YMR078C | | YMR077C; YMR078C |
| YMR073C; YMR074C | | YMR073C; YMR074C |
| YMR062C; YMR063W | | YMR062C; YMR063W |
| YMR060C; YMR061W | | YMR060C; YMR061W |
| YMR047C; DI49_4135 | DI49_4135 | YMR047C |
| YMR039C; YMR040W | | YMR039C; YMR040W |
| YMR037C; YMR038C | | YMR037C; YMR038C |
| YMR035W; YMR036C | | YMR035W; YMR036C |
| YMR033W; YMR034C | | YMR033W; YMR034C |
| YMR031C; YMR032W | | YMR031C; YMR032W |
| YMR029C; YMR030W | | YMR029C; YMR030W |

Create PDF in your applications with the Pdfcrowd HTML to PDF API PDFCROWD

| | | | |
|---|---|---|---|
| YMR017W; YMR018W | | | YMR017W; YMR018W |
| YMR008C; DI49_4095; YMR007W | DI49_4095 | | YMR008C; YMR007W |
| YML013W; YML012W | | | YML013W; YML012W |
| YML015C; YML014W | | | YML015C; YML014W |
| YML029W; YML028W | | | YML029W; YML028W |
| YML036W; YML035C | | | YML036W; YML035C |
| YML062C; YML061C | | | YML062C; YML061C |
| YML067C; YML066C | | | YML067C; YML066C |
| YML069W; YML068W | | | YML069W; YML068W |
| YML081W; YML080W | | | YML081W; YML080W |
| YML088W; YML087C | | | YML088W; YML087C |
| YML097C; YML096W | | | YML097C; YML096W |
| YML112W; YML111W | | | YML112W; YML111W |
| YML114C; YML113W | | | YML114C; YML113W |
| YML126C; DI49_3967 | DI49_3967 | | YML126C |
| YMR317W; YMR318C | | | YMR317W; YMR318C |
| YCR002C; SPGP0AH00640 | | SPGP0AH00640 | YCR002C |
| DI49_4809; DI49_4808 | DI49_4809; DI49_4808 | | |
| DI49_4811; DI49_4810 | DI49_4811; DI49_4810 | | |
| DI49_4813; DI49_4812 | DI49_4813; DI49_4812 | | |
| SPGP0H00170; DI49_3489 | DI49_3489 | SPGP0H00170 | |
| DI49_4821; DI49_4820 | DI49_4821; DI49_4820 | | |
| DI49_4836; DI49_4835 | DI49_4836; DI49_4835 | | |
| DI49_4854; DI49_4853 | DI49_4854; DI49_4853 | | |
| DI49_4867; DI49_4866 | DI49_4867; DI49_4866 | | |
| DI49_4869; DI49_4868 | DI49_4869; DI49_4868 | | |
| DI49_4871; DI49_4870 | DI49_4871; DI49_4870 | | |
| DI49_4873; DI49_4872 | DI49_4873; DI49_4872 | | |
| DI49_4875; DI49_4874 | DI49_4875; DI49_4874 | | |
| DI49_4877; DI49_4876 | DI49_4877; DI49_4876 | | |
| DI49_4879; DI49_4878 | DI49_4879; DI49_4878 | | |
| DI49_4881; DI49_4880 | DI49_4881; DI49_4880 | | |
| DI49_4892; YOR048C; DI49_4893 | DI49_4892; DI49_4893 | | YOR048C |
| DI49_4905; DI49_4904 | DI49_4905; DI49_4904 | | |
| DI49_4907; DI49_4906 | DI49_4907; DI49_4906 | | |
| DI49_4916; YOR075W; DI49_4917 | DI49_4916; DI49_4917 | | YOR075W |
| DI49_4919; DI49_4918 | DI49_4919; DI49_4918 | | |
| DI49_4921; DI49_4920 | DI49_4921; DI49_4920 | | |
| SPGP0H01210; SPGP0H01200 | | SPGP0H01210; SPGP0H01200 | |
| DI49_4923; DI49_4922 | DI49_4923; DI49_4922 | | |

Create PDF in your applications with the Pdfcrowd HTML to PDF API PDFCROWD

| | | | |
|---|---|---|---|
| DI49_4926; YOR087W; DI49_4927 | DI49_4926; DI49_4927 | | YOR087W |
| DI49_4929; DI49_4928 | DI49_4929; DI49_4928 | | |
| DI49_4931; YOR093C; DI49_4932 | DI49_4931; DI49_4932 | | YOR093C |
| DI49_4950; YOR114W; DI49_4951 | DI49_4950; DI49_4951 | | YOR114W |
| DI49_4953; YOR117W; DI49_4954 | DI49_4953; DI49_4954 | | YOR117W |
| DI49_4956; DI49_4955 | DI49_4956; DI49_4955 | | |
| DI49_4958; DI49_4957 | DI49_4958; DI49_4957 | | |
| DI49_4960; YOR125C; DI49_4961 | DI49_4960; DI49_4961 | | YOR125C |
| DI49_4963; YOR128C; DI49_4964 | DI49_4963; DI49_4964 | | YOR128C |
| DI49_4972; DI49_4971 | DI49_4972; DI49_4971 | | |
| DI49_4974; DI49_4973 | DI49_4974; DI49_4973 | | |
| DI49_4981; YOR147W; DI49_4982 | DI49_4981; DI49_4982 | | YOR147W |
| DI49_4985; YOR151C; DI49_4986 | DI49_4985; DI49_4986 | | YOR151C |
| DI49_4990; DI49_4989 | DI49_4990; DI49_4989 | | |
| DI49_4992; DI49_4991 | DI49_4992; DI49_4991 | | |
| DI49_5000; YOR165W; DI49_5001 | DI49_5000; DI49_5001 | | YOR165W |
| DI49_5004; YOR171C; DI49_5005 | DI49_5004; DI49_5005 | | YOR171C |
| DI49_5007; DI49_5006 | DI49_5007; DI49_5006 | | |
| DI49_5009; DI49_5008 | DI49_5009; DI49_5008 | | |
| DI49_5013; DI49_5012 | DI49_5013; DI49_5012 | | |
| DI49_5018; DI49_5017 | DI49_5018; DI49_5017 | | |
| SPGP0H02170; YOR188W; DI49_5020 | DI49_5020 | SPGP0H02170 | YOR188W |
| DI49_5026; DI49_5025 | DI49_5026; DI49_5025 | | |
| DI49_5028; DI49_5027 | DI49_5028; DI49_5027 | | |
| DI49_5038; DI49_5037 | DI49_5038; DI49_5037 | | |
| DI49_5048; YOR222W; DI49_5049 | DI49_5048; DI49_5049 | | YOR222W |
| DI49_5051; DI49_5050 | DI49_5051; DI49_5050 | | |
| DI49_5054; YOR229W; DI49_5055 | DI49_5054; DI49_5055 | | YOR229W |
| DI49_5060; YOR236W; DI49_5061 | DI49_5060; DI49_5061 | | YOR236W |
| SPGP0H02610; DI49_5062 | DI49_5062 | SPGP0H02610 | |
| DI49_5069; DI49_5068 | DI49_5069; DI49_5068 | | |
| DI49_5071; DI49_5070 | DI49_5071; DI49_5070 | | |
| DI49_5073; DI49_5072 | DI49_5073; DI49_5072 | | |
| DI49_5076; YOR255W; DI49_5077 | DI49_5076; DI49_5077 | | YOR255W |
| DI49_5079; DI49_5078 | DI49_5079; DI49_5078 | | |
| DI49_5104; DI49_5103 | DI49_5104; DI49_5103 | | |
| DI49_5106; DI49_5105 | DI49_5106; DI49_5105 | | |
| DI49_5113; YOR294W; DI49_5114 | DI49_5113; DI49_5114 | | YOR294W |
| DI49_5116; DI49_5115 | DI49_5116; DI49_5115 | | |
| DI49_5118; DI49_5117 | DI49_5118; DI49_5117 | | |

Create PDF in your applications with the Pdfcrowd HTML to PDF API PDFCROWD

| | | | |
|---|---|---|---|
| DI49_5120; DI49_5119 | DI49_5120; DI49_5119 | | |
| DI49_5122; SPGP0H03190 | DI49_5122 | SPGP0H03190 | |
| DI49_5127; YOR307C; DI49_5128 | DI49_5127; DI49_5128 | | YOR307C |
| DI49_5134; DI49_5133 | DI49_5134; DI49_5133 | | |
| DI49_5136; DI49_5135 | DI49_5136; DI49_5135 | | |
| DI49_5144; YOR328W; DI49_5145 | DI49_5144; DI49_5145 | | YOR328W |
| DI49_5165; DI49_5164 | DI49_5165; DI49_5164 | | |
| DI49_5178; DI49_5177 | DI49_5178; DI49_5177 | | |
| DI49_5185; DI49_5184 | DI49_5185; DI49_5184 | | |
| DI49_5684; DI49_5685 | DI49_5684; DI49_5685 | | |
| YMR304W; DI49_4718 | DI49_4718 | | YMR304W |
| YCR094W; YCR093W | | | YCR094W; YCR093W |
| DI49_1311; YBR288C | DI49_1311 | | YBR288C |
| DI49_1307; DI49_1308 | DI49_1307; DI49_1308 | | |
| DI49_1301; DI49_1302 | DI49_1301; DI49_1302 | | |
| DI49_1295; DI49_1296 | DI49_1295; DI49_1296 | | |
| DI49_1293; DI49_1294 | DI49_1293; DI49_1294 | | |
| DI49_1287; DI49_1288 | DI49_1287; DI49_1288 | | |
| DI49_1282; DI49_1283 | DI49_1282; DI49_1283 | | |
| DI49_1277; DI49_1278 | DI49_1277; DI49_1278 | | |
| DI49_1275; DI49_1276 | DI49_1275; DI49_1276 | | |
| DI49_1254; DI49_1255 | DI49_1254; DI49_1255 | | |
| DI49_1253; SPGP0I00770 | DI49_1253 | SPGP0I00770 | |
| DI49_1243; DI49_1244 | DI49_1243; DI49_1244 | | |
| DI49_1237; DI49_1238 | DI49_1237; DI49_1238 | | |
| DI49_1227; DI49_1228 | DI49_1227; DI49_1228 | | |
| DI49_1222; DI49_1223 | DI49_1222; DI49_1223 | | |
| DI49_1220; DI49_1221 | DI49_1220; DI49_1221 | | |
| DI49_1218; DI49_1219 | DI49_1218; DI49_1219 | | |
| DI49_1212; DI49_1213 | DI49_1212; DI49_1213 | | |
| DI49_1210; DI49_1211 | DI49_1210; DI49_1211 | | |
| DI49_1208; DI49_1209 | DI49_1208; DI49_1209 | | |
| DI49_1206; SPGP0I01220 | DI49_1206 | SPGP0I01220 | |
| DI49_1204; DI49_1205 | DI49_1204; DI49_1205 | | |
| DI49_1202; DI49_1203 | DI49_1202; DI49_1203 | | |
| DI49_1188; DI49_1189 | DI49_1188; DI49_1189 | | |
| DI49_1176; DI49_1177 | DI49_1176; DI49_1177 | | |
| DI49_1159; DI49_1160 | DI49_1159; DI49_1160 | | |
| DI49_1151; DI49_1152 | DI49_1151; DI49_1152 | | |
| DI49_1139; DI49_1140 | DI49_1139; DI49_1140 | | |

Create PDF in your applications with the Pdfcrowd HTML to PDF API PDFCROWD

| | | | |
|---|---|---|---|
| DI49_1133; DI49_1134 | DI49_1133; DI49_1134 | | |
| DI49_1131; DI49_1132 | DI49_1131; DI49_1132 | | |
| DI49_1120; DI49_1121 | DI49_1120; DI49_1121 | | |
| DI49_1118; YBR085W | DI49_1118 | | YBR085W |
| DI49_1103; DI49_1104 | DI49_1103; DI49_1104 | | |
| DI49_1096; DI49_1097 | DI49_1096; DI49_1097 | | |
| DI49_1090; DI49_1091 | DI49_1090; DI49_1091 | | |
| DI49_1088; DI49_1089 | DI49_1088; DI49_1089 | | |
| SPGP0I02630; DI49_1063 | DI49_1063 | SPGP0I02630 | |
| DI49_1061; SPGP0I02640 | DI49_1061 | SPGP0I02640 | |
| DI49_1059; DI49_1060 | DI49_1059; DI49_1060 | | |
| DI49_1057; DI49_1058 | DI49_1057; DI49_1058 | | |
| DI49_1051; DI49_1052 | DI49_1051; DI49_1052 | | |
| DI49_1037; DI49_1038 | DI49_1037; DI49_1038 | | |
| DI49_1028; DI49_1029 | DI49_1028; DI49_1029 | | |
| DI49_1023; DI49_1024 | DI49_1023; DI49_1024 | | |
| DI49_1022; YDL040C | DI49_1022 | | YDL040C |
| DI49_1009; DI49_1010 | DI49_1009; DI49_1010 | | |
| DI49_1003; DI49_1004 | DI49_1003; DI49_1004 | | |
| DI49_0994; DI49_0995 | DI49_0994; DI49_0995 | | |
| DI49_0961; DI49_0962 | DI49_0961; DI49_0962 | | |
| DI49_0943; DI49_0944 | DI49_0943; DI49_0944 | | |
| DI49_0942; SPGP0I03900 | DI49_0942 | SPGP0I03900 | |
| DI49_0935; DI49_0936 | DI49_0935; DI49_0936 | | |

Create PDF in your applications with the Pdfcrowd HTML to PDF API PDFCROWD
